# Function and phylogeny support the independent evolution of acid-sensing ion channels in the Placozoa

**DOI:** 10.1101/2022.06.28.497943

**Authors:** Wassim Elkhatib, Luis Yanez-Guerra, Tatiana D. Mayorova, Mark A. Currie, Maria Perera, Anhadvir Singh, Julia Gauberg, Adriano Senatore

## Abstract

Acid-sensing ion channels (ASICs) are proton-gated cation channels that are part of the Deg/ENaC ion channel family, which also includes neuropeptide-, bile acid-, and mechanically-gated channels. Despite sharing common tertiary and quaternary structures, strong sequence divergence within the Deg/ENaC family has made it difficult to resolve their phylogenetic relationships, and by extension, whether channels with common functional features, such as proton-activation, share common ancestry or evolved independently. Here, we report that a Deg/ENaC channel from the early diverging placozoan species *Trichoplax adhaerens*, named *Tad*NaC2, conducts proton-activated currents *in vitro* with biophysical features that resemble those of the mammalian ASIC1 to ASIC3 channels. Through a combined cluster-based and phylogenetic analysis, we successfully resolve the evolutionary relationships of most major lineages of metazoan Deg/ENaC channels, identifying two subfamilies within the larger Deg/ENaC family that are of ancient, pre-bilaterian origin. We also identify bona fide Deg/ENaC channel homologues from filasterean and heterokont single celled eukaryotes. Furthermore, we find that ASIC channels, *Tad*NaC2, and various other proton-activated channels from vertebrates and invertebrates are part of phylogenetically distinct lineages. Through structural modelling and mutation analysis, we find that *Tad*NaC2 proton-activation employs fundamentally different molecular determinants than ASIC channels, and identify two unique histidine residues in the placozoan channel that are required for its proton-activation. Together, our phylogenetic and functional analyses support the independent evolution of proton-activated channels in the phylum Placozoa. Spurred by our discovery of pH sensitive channels, we discovered that despite lacking a nervous system, *Trichoplax* can sense changes in extracellular pH to coordinate its various cell types to locomote away from acidic environments, and to contract upon rapid exposure to acidic pH in a Ca^2+^-dependent manner. Lastly, via yeast 2 hybrid screening, we find that the *Trichoplax* channels *Tad*NaC2 and *Tad*NaC10, belonging to the two separate Deg/ENaC subfamilies, interact with the cytoskeleton organizing protein filamin, similar to the interaction reported for the human ENaC channels.

## Introduction

Degenerin/Epithelial Na^+^ Channels (Deg/ENaC channels) are a large family of metazoan cation channels that exhibit a remarkable diversity in their mechanisms for activation, gating, and physiological functions. Mammals for example possess three major types of Deg/ENaC channels: Acid Sensing Ion Channels (*i.e.*, ASIC channels), which are activated by extracellular protons and serve as major extracellular pH sensors in the central and peripheral nervous system (1), Bile-Acid Sensitive Ion Channels (BASIC channels), which are expressed in the brain, liver, and intestinal epithelium and are activated by bile acids (2), and Epithelial Na^+^ Channels (ENaC channels), which conduct Na^+^ leak currents in epithelial cells of the lung and kidney important for Na^+^ reabsorption and homeostasis (3). Despite these distinctions, ASIC and ENaC channels also share some overlapping functional properties, in that both are modulated by mechanical stimuli (4, 5), and the ENaC-δ subunit, which is expressed in the brain, is proton-activated similar to ASIC channels (6). All Deg/ENaC channels are thought to form hetero- and/or homotrimeric holochannels, with each subunit comprised of two membrane spanning helices and a large extracellular domain (7). Nonetheless, during evolution these channels underwent extensive sequence divergence and genetic expansion/loss in different animal lineages, in several cases obscuring their phylogenetic relationships. Although several recent phylogenetic studies have provided some important insights (8–10), there are still many unanswered questions about the evolutionary origins ASIC, BASIC, and ENaC channels, and their relationships with the many divergent channels identified in invertebrates.

Like in mammals, invertebrate Deg/ENaC channels show striking diversity in their mechanisms for activation and physiological functions. In the nematode worm *C.elegans* for example, the subunits MEC-4 and MEC-10 form mechanically-gated heterotrimeric channels that are required for sensory mechanotransduction (11). In addition, the channel ACD-1, which is involved in acidic pH avoidance behavior (12), is a Na^+^ leak channel that is blocked by external protons (13), while the channels ACD-2, DEL-9, and ASIC1 are activated by external protons (14), making them similar to ASIC channels. Insects like the vinegar fly *Drosophila melanogaster* also possess a diverse set of Deg/ENaC channels (15), including the proton-activated channel Pickpocket 1 (PPK1) which is expressed in proprioceptive and nociceptive sensory neurons where it also thought to respond to mechanical stimuli (16, 17). In molluscs and annelids, Deg/ENaC channels act as neurotransmitter receptors activated by the secreted neuropeptides FMRFamide and Wamide (9, 18–20). Peptide-gated channels are also found in the cnidarian species *Hydra magnipapilatta* (a hydrozoan), in the form of *Hy*NaC channels that are activated by RFamide neuropeptides (21, 22). In the cnidarian *Nematostella vectensis*, an anthozoan, Deg/ENaC channels are reported unresponsive to neuropeptides, but rather, proton sensitive with the channel *Ne*NaC8 being blocked by protons, and the channels *Ne*NaC2 and *Ne*NaC14 being proton-activated (8). In this species, the ASIC-like channel *Ne*NaC2 contributes to cnidocyte discharge, a unique cellular process in cnidarians that leads to expulsion of a venom-laced barb for defense and prey capture (8, 23). To date, the most early-diverging Deg/ENaC channel to be functionally characterized *in vitro* is *Tad*NaC6 from the placozoan species *Trichoplax adhaerens*, which forms a Na^+^ leak channels that is blocked by external protons and Ca^2+^ ions (24). Placozoans are an intriguing group of animals that lack nervous systems, and yet possess a large complement of genes involved in neural and synaptic signaling.

Combined, the unclear phylogenetic relationships of Deg/ENaC channels, coupled with their diverse but sometimes similar modalities for gating (*e.g.*, mechanical force, peptides, and protons), makes it difficult to infer whether these functional features are evolutionarily conserved, or evolved independently.

Here, we report the functional properties of a second Deg/ENaC channel from *T.adhaerens*, *Tad*NaC2, which forms a proton-activated channel *in vitro* with biophysical features that are a striking hybrid of the unique features reported for the mammalian ASIC 1 to 3 channels. Specifically, *Tad*NaC2 macroscopic currents have reduced sensitivity to proton activation like those of ASIC2a, are biphasic like those of ASIC3, and exhibit rundown or tachyphylaxis like those of ASIC1a. Furthermore, the biphasic currents exhibit different ion selectivities, with the early/transient current being non-selective for monovalent cations but highly Ca^2+^ permeable, and the late/sustained current being more selective for Na^+^ over K^+^ but highly impermeable to Ca^2+^. *Tad*NaC2 is relatively insensitive to the general Deg/ENaC channel blocker amiloride, which interestingly, appears to selectively block the early transient current.

Given the uncanny functional similarities between *Tad*NaC2 and the mammalian ASIC channels, we sought to better resolve their phylogenetic relationships. Through combined cluster and phylogenetic analysis, we generated a phylogenetic tree that resolves numerous relationships within the Deg/ENaC family, including the identification of two ancient subfamilies of Deg/ENaC channels, one bearing the mammalian ASIC and BASIC channels (*i.e.*, the ASIC subfamily), and the other bearing the ENaC channels (the ENaC subfamily). Within these two subfamilies, we identified numerous previously undefined clades of Deg/ENaC channels and, within the ENaC subfamily, bona fide Deg/ENaC channel homologues from single celled eukaryotes. Importantly, we find that within the ASIC subfamily, *Tad*NaC2 and most other *T.adhaerens* Deg/ENaC channels form a strongly supported clade with bilaterian BASIC and BASIC-related channels, separate from ASIC channels, while the singleton *Tad*NaC10 channel falls within the ENaC subfamily. Thus, we propose that *Tad*NaC2 is not phylogenetically related to ASIC channels, but rather, is a BASIC-related channel. Furthermore, our phylogenetic analysis establishes that additional proton-activated channels that have been identified in mammals (ENaC-δ), cnidarians (*Ne*NaC2 and *Ne*NaC14), and *C.elegans* (ADC-2 and ASIC-1) belong to phylogenetically distant clades, suggesting that proton activation evolved numerous times independently within the ASIC and ENaC subfamilies.

To better understand the functional similarities and differences of *Tad*NaC2 relative to ASIC channels, we conducted structural and functional analyses, focusing on key extracellular regions of ASIC channels that bear molecular determinants for protons activation. We find that *Tad*NaC2 lacks all major determinants for proton activation of ASIC channels, including the critical H73 and K211 residues that are common to all ASIC channels (10, 25), and the acid pocket, a region of secondary importance for proton activation (26). Instead, we identify two histidine residues, H80 and H109, that are unique molecular determinants for *Tad*NaC2 proton activation as revealed by mutation analysis. Altogether, our analyses provide compelling evidence that proton activation of *Tad*NaC2 is fundamentally different from that of ASIC channels, which together with our phylogenetic analysis, strongly suggests that the ASIC-like features of *Tad*NaC2 evolved independently.

Our discovery of proton-sensitive Deg/ENaC channels in *T.adhaerens* prompted us to explore whether the animal is capable of sensing changes in extracellular pH and generate corresponding behavioral responses. Remarkably, we find that despite lacking a nervous system and synapses, *T.adhaerens* can sense pH gradients in their environments to undergo chemotaxis away from acidic environments (*i.e.*, acidic pH aversion behavior), and to rapidly contract in a Ca^2+^ dependent manner upon rapid perfusion of acidic saline. Lastly, we conducted a yeast 2 hybrid screen using the C-termini of *Tad*NaC2 and *Tad*NaC10 as bait, finding that both channels interact with the actin-binding protein filamin, a cytoskeleton reorganizing protein which in humans interacts with ENaC channels to regulate their function (27, 28), and is independently implicated in mechanotransduction (29).

## Results

### The placozoan DEG/ENaC channels fit within the two distinct metazoan superclades of ASIC-related and ENaC-related channels

To better understand the phylogenetic properties of the *T.adhaerens Tad*NaC channels and DEG/ENaC channels in general, we conducted a CLuster ANalysis of Sequences (CLANS) (30) with gene sequences from representative species spanning the major animal groupings, extracted from high-quality gene datasets with high BUSCO completeness scores (31). The CLANs analysis revealed two distinct clusters of metazoan DEG/ENaC homologs, hereafter referred to as the ASIC and ENaC superclusters given the respective presence of these channel homologues in these two groups (Fig. 1). Most of the *T.adhaerens Tad*NaC sequences associate with the ASIC supercluster, while the singleton *Tad*NaC10 and its *H.hongkongensis* orthologue associate with the ENaC supercluster. In addition to DEG/ENaC channels, we also included P2X receptors and proton-activated chloride (PAC) channels in our analysis, both of which are structurally similar to DEG/ENaC channels in their trimeric quaternary structures, but lack apparent homology at the protein sequence level making it altogether unclear whether these are evolutionarily related to DEG/ENaC channels (32–34). Accordingly, both the P2X receptors and PAC channels form distinct and tight clusters that are separate from all DEG/ENaC channels. Notably, several sets of *bona fide* DEG/ENaC channel homologues from the phyla Annelida, Platyhelminthes, Arthropoda, and Ctenophora also formed separate clusters that did not associate with other DEG/ENaC channels, which likely represent lineage-specific variants that underwent extensive sequence divergence. This includes the pickpocket (PPK) channels from *D.melanogaster*. Lastly, and consistent with a previous report (35), we identified DEG/ENaC homologues in the gene data for unicellular eukaryotic species from the clades Heterokonta (*i.e.*, from the SAR supergroup, for Stramenopila, Alveolata, and Rhizaria) and Filasterea, most of which associated with the ENaC supercluster.

**Figure 1.**
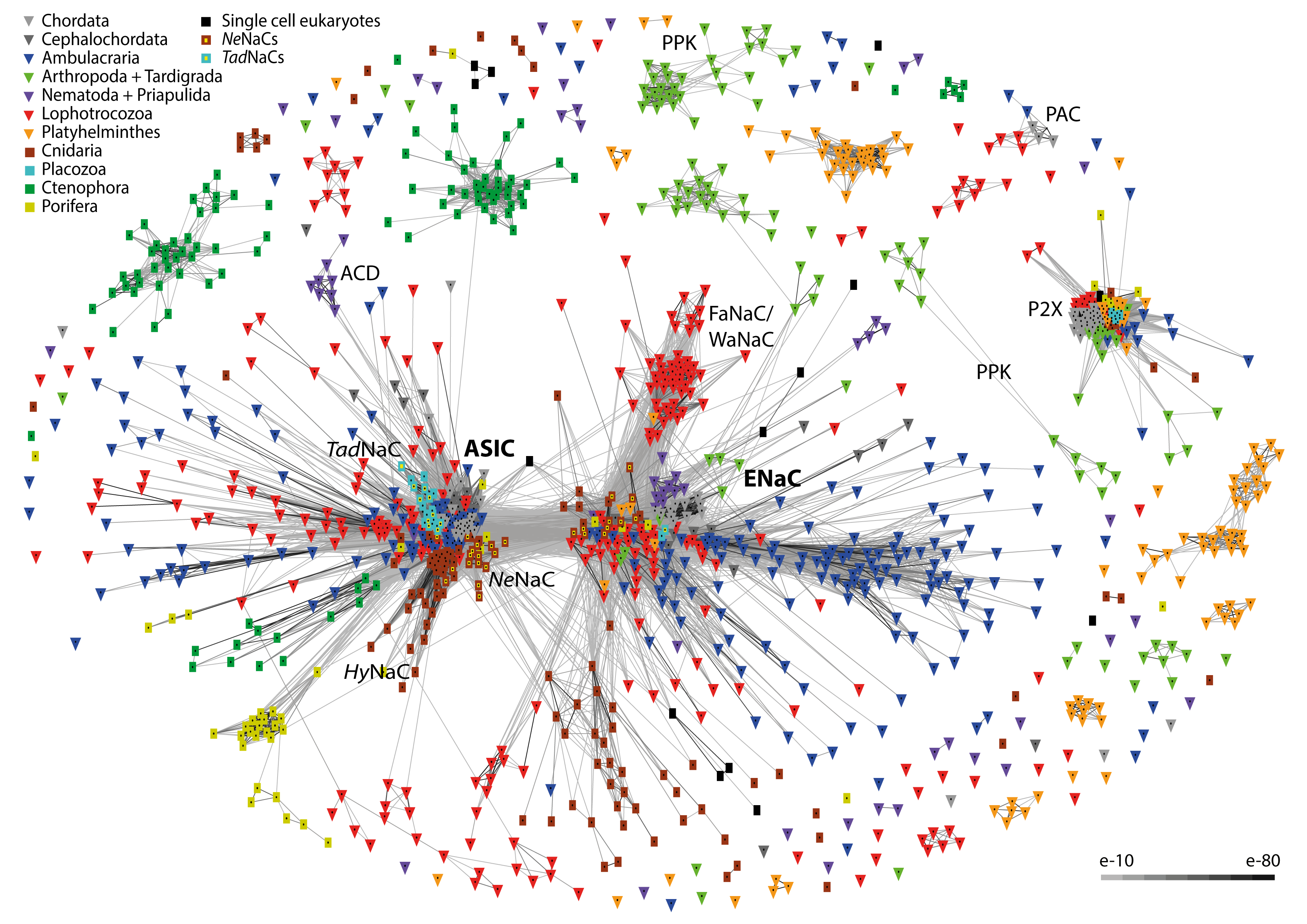
BLOSUM62 cluster map of PAC, P2X, and Deg/ENaC channels revealing two superclusters of Deg/ENaC channels. Nodes correspond to individual channel sequences and are coloured by taxon as indicated by the key. Edges correspond to BLAST connections of a P-value >1E-40.

A phylogeny inferred from aligned protein sequences selected strictly from the two DEG/ENaC superclusters from the CLANS analysis, rooted on a subset of the divergent PPK sequences from arthropods, revealed strong phylogenetic support distinguishing the ASIC and ENaC supercluster sequences, forming two corresponding superclades (Figs. 2 and S1). However, despite weakly associating with the ASIC supercluster in the CLANS analysis, the *C.elegans* “degenerin” channels ACD1 to 5 and FLR grouped separately from both the ASIC and ENaC superclades on the phylogenetic tree, most likely due to strong sequence divergence (Fig. 1).Within the ASIC superclade, *Tad*NaC channels 1 to 9 and 11 form a subclade with a sister relationship with ASIC5/BASIC channels. Together, these *Tad*NaC channels and BASIC channels form a sister clade with true ASIC channels (*e.g.*, vertebrate ASIC 1 to 4 channels). Chordate and urochordate BASIC channels fall within a larger subclade of uncharacterized BASIC- related channels found in cephalochordates, ambulacrarians (*i.e.*, echinoderms and hemichordates), and lophotrochozoans (*i.e.*, molluscs, annelids, brachiopods, and phoronids). As reported previously (10, 25), true ASIC channels form two distinct subclades, groups A and B, with chordates possessing only group A orthologues, cephalochordates possessing group A and B orthologues, and both ambulacrarians and lophotrochozoans both possessing only group B orthologues. Also within the ASIC superclade are the neuropeptide-gated *Hy*NaC channels from the cnidarian species *Hydra magnipapillata* (21), which form a separate clade from ASIC/BASIC/*Tad*NaC channels, together with the *Ne*NaC channels from the fellow cnidarian *Nematostella vectensis* (8). These form a sister clade with another group of *Ne*NaC channels which includes the proton-activated *Ne*NaC2 channel (8). Interestingly, these cnidarian channels form a sister clade with a large subclade of *Hy*NaC/*Ne*NaC-related channels found in ambulacrarians, lophotrochozoans, and a single representative from the early-diverging ecdysozoan worm *Priapulus caudatus* (phylum Priapulida). Also interesting is that this large subclade of *Hy*NaC/*Ne*NaC-related channels includes two subclades of ctenophore channels, and one comprised entirely of channels from the poriferan species *Amphimedon queenslandica*. Altogether, our analysis suggests that channels within the ASIC superclade have deep evolutionary origins, present even in the earliest diverging animal phyla of Cnidaria, Placozoa, Ctenophora, and Porifera. Furthermore, the observed sister relationship between *Tad*NaC channels and bilaterian BASIC channels, coupled with the absence of true ASIC channel sequences in placozoans, cnidarians, ctenophores, and poriferans suggests that ASIC channels evolved strictly in bilaterians as previously suggested (10). By extension, it is apparent that the *Tad*NaC channels present in the ASIC superclade are not orthologous to true ASIC channels, but rather, belong to a newly defined subclade of BASIC-related channels found broadly in Bilateria.

**Figure 2.**
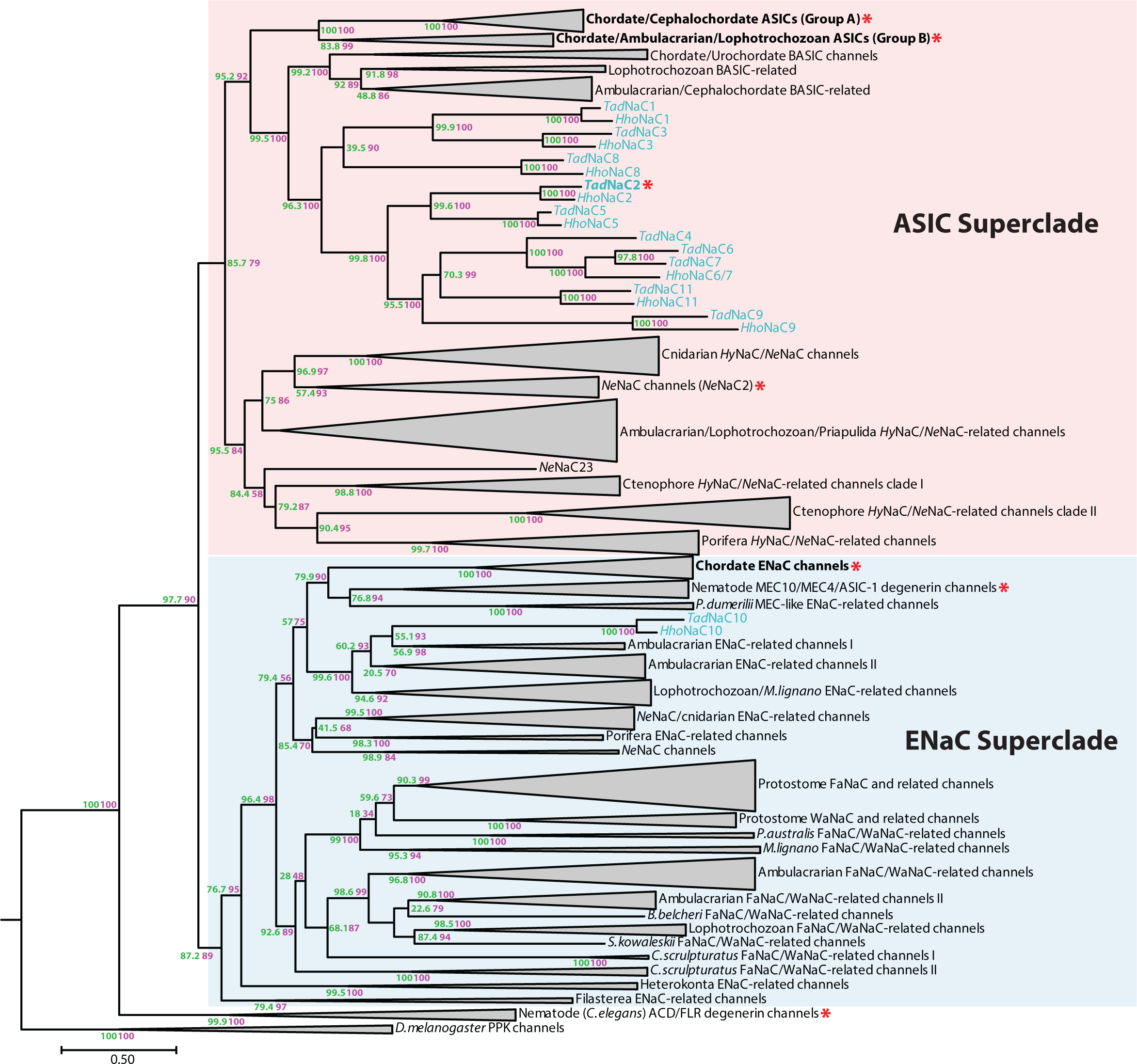
Maximum likelihood protein phylogeny delineates the ASIC and ENaC subfamilies within the Deg/ENaC family of ion channels. The tree was generated with the program IQ-TREE 2 with the best-fit model LG+F+G4, and rooted with the divergent PPK channels form *D.melanogaster*. Node support values are for 1,000 ultrafast replicates using the aLRT-SH-like method (*green*) and aBayes methods (*purple*). The asterisks indicate clades or single channels that have been identified as proton- activated.

The ENaC superclade subdivides into several distinct subclades, one of which bears the namesake ENaC channels that are unique to chordates (3). The chordate ENaC channels form a sister clade with the mechanosensory MEC4/MEC10 and the H^+^-gated ASIC-1 channels from *C.elegans* (14), as well as several undescribed homologues from the annelid species *Platynereis dumerilii*. The singleton *Tad*NaC10 forms a subclade with a large set of uncharacterized homologues present in ambulacrarians, lophotrochozoans, and the platyhelminth species *Macrostomum lignano*, which in turn form a sister clade with the ENaC/MEC4/MEC10/ASIC-1 subclade. Interestingly, all these described channels form a sister clade with a group of early diverging homologues from cnidarians and the poriferan species *Sycon ciliatum*. A distinct group of channels within the ENaC superclade contains the FMRFamide and Wamide neuropeptide-gated sodium channels (*i.e.*, FaNaC and WaNaC channels), recently shown to be conserved between the lophotrochozoan lineages of molluscs and annelids (9). These form a sister clade relationship with several channels from the phoronid *Phoronis australis* and *M.lignano*. Furthermore, our analysis identifies an additional group of FaNaC/WaNaC-related channels that are present in lophotrochozoans, ambulacrarians, the cephalochordate *B.belcheri*, and the arthropod *Centroides sculpturatus*.

Interestingly, the unicellular eukaryote DEG/ENaC channel sequences that we included in our analysis formed two separate clades on the phylogenetic tree, both forming a sister clade with ENaC channels (Fig. 2). This is mostly consistent with the CLANS analysis, except for a single sequence from the filasterean species *Tunicaraptor unikontum* which associated with both the ASIC and the ENaC superclusters (Fig. 1).

### TadNaC2 conducts desensitizing proton-activated cation currents in vitro with features that resemble mouse ASIC1

Previously, we found that the *T.adhaerens Tad*NaC6 channel conducts constitutive Na^+^ leak currents *in vitro* that are blocked by external protons and Ca^2+^ ions (24) (Fig. 3A). Here, we set out to characterize the *in vitro* properties of a second *T.adhaerens* DEG/ENaC channel, *Tad*NaC2. Whole-cell patch clamp recording of Chinese Hamster Ovary (CHO)-K1 cells transfected with the *Tad*NaC2 cDNA revealed robust inward macroscopic cation currents that could be elicited by perfusing a pH 5 solution over the recorded cells. No such currents were evident in untransfected cells, but we did observe a small endogenous inward current in these cells that became activated by solutions with a pH of 4 or lower (Fig. 3A). We similarly expressed the mouse ASIC1a splice variant channel in CHO-K1 cells, which has been subject to extensive *in vitro* experimentation, which also produced robust inward currents at pH 5 but with a noticeably faster desensitization compared to *Tad*NaC2. Remarkably, *Tad*NaC2 whole cell currents were quite large in amplitude, reaching upwards of 5,000 picoamperes (Fig. 3B), despite the cDNA not being codon optimized as was required for efficient expression of the cnidarian *Hy*NaC channels in mammalian cells (36).

**Figure 3.**
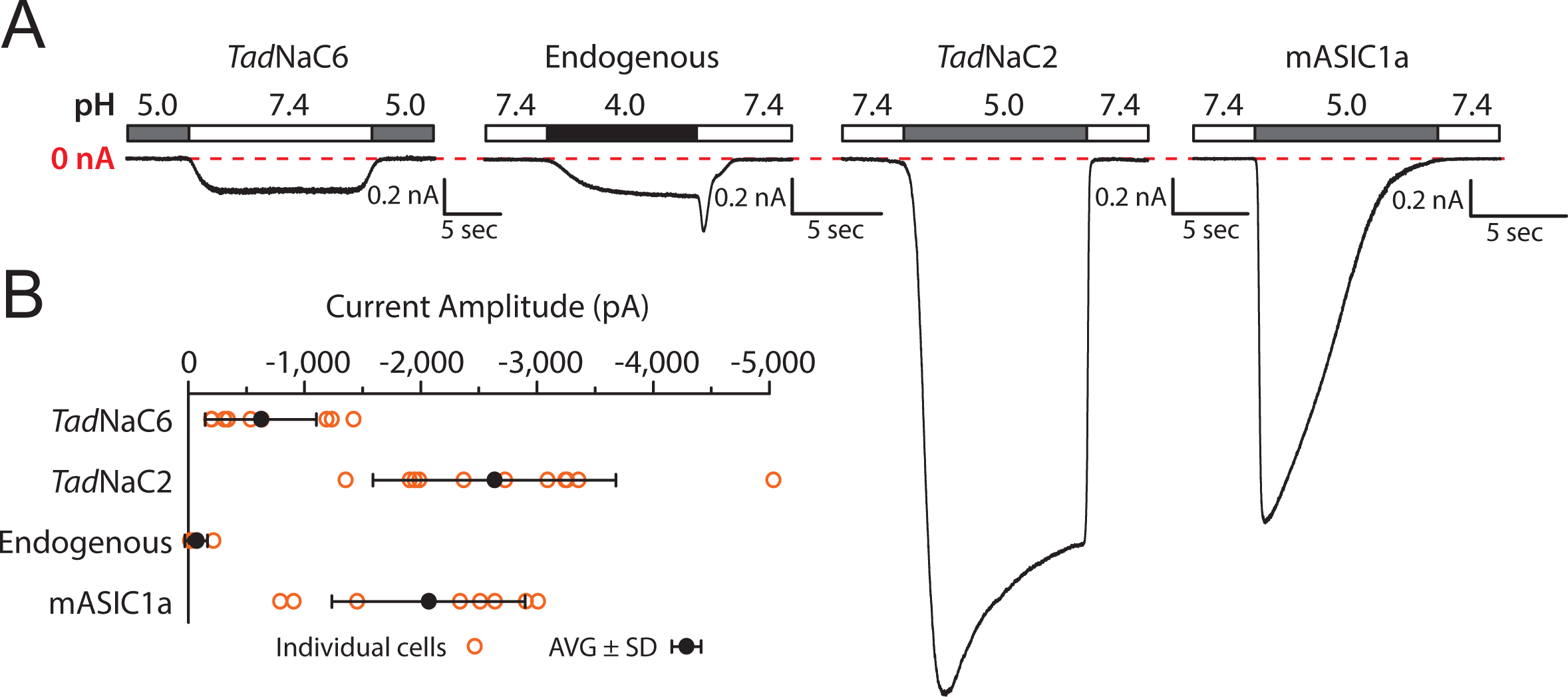
*Tad*NaC2 conducts robust proton-activated currents in Chinese Hamster Ovary (CHO- K1) cells. *(A)* Sample whole cell currents recorded for the previously characterized *Trichoplax* Deg/ENaC sodium leak channel *Tad*NaC6 that is blocked by extracellular protons (24), a newly identified endogenous current in CHO-K1 cells that becomes activated upon perfusion of strongly acidic solutions below pH 4.0, and large, prominent proton-activated currents conducted by the *in vitro* expressed *Trichoplax Tad*NaC2 and the mouse ASIC1a (mASIC1a) channels. *(B)* Plot of average peak inward current amplitude (in picoamps or pA) for currents shown in panel *A* ± standard deviation. Orange symbols denote values for individual cells/recordings.

Next, we sought to compare the general properties of *Tad*NaC2 and mASIC1a proton-activated currents. Perfusion of external solutions ranging in pH revealed that *Tad*NaC2 begins activating at pH 5.5, with current kinetics that accelerate from a slow onset current at pH 5.5, to a faster transient and desensitizing current at pH 4.0 (Fig. 4A). These currents are markedly different from those of mASIC1, which began activating at the much more basic pH of 6.5, with much faster activation and desensitization evident across all tested values of pH. Dose response curves generated from these experiments revealed that *Tad*NaC2 is considerably less sensitive to external protons than mASIC1a (Fig. 4B), with a pH_50_ of 5.1 ±0.1 vs. 6.7 ±0.1, and a Hill coefficient (*n*_H_) value of only 1.7 ±0.4 vs. 8.4 ±2.7. Notably, these values for the mASIC1a channel are closely in line with those reported for the human ASIC1a channel recorded in *Xenopus* oocytes (25, 26). Together, the lower pH_50_ and *n*_H_ values observed for *Tad*NaC2 indicate a lower binding affinity and reduced cooperativity for extracellular proton binding, more inline with the sensitivity reported for the rat ASIC2a channel (37–39).

**Figure 4.**
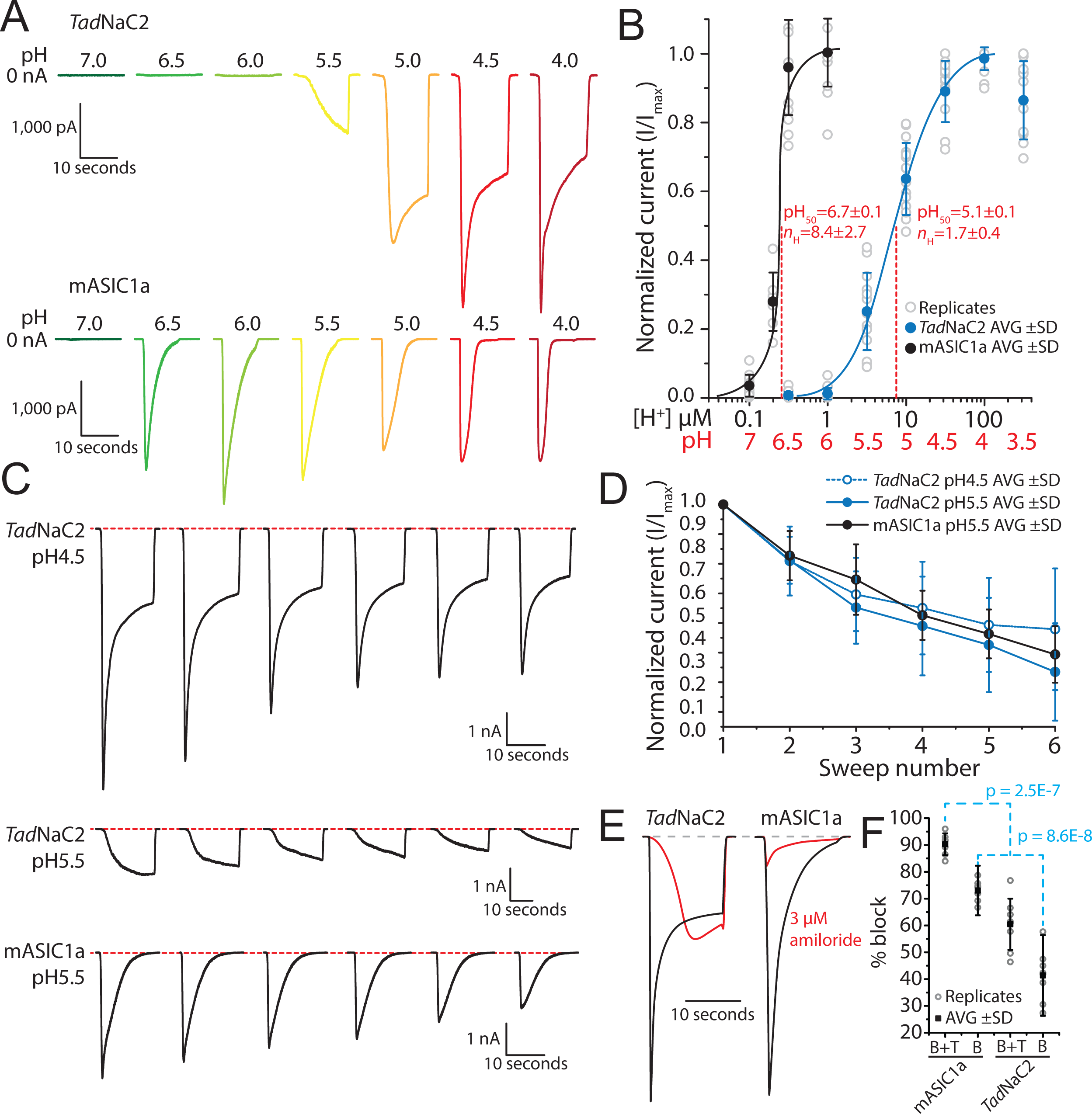
General electrophysiological properties of *Tad*NaC2 compared to mouse ASIC1a. *(A)* Sample recordings of *Tad*NaC2 currents (*top*) and mouse ASIC1a currents (mASIC1a, *bottom*) elicited by perfusion of solutions with decreasing pH. *(B)* pH dose response curves for *Tad*NaC2 and mASIC1 revealing a right shifted pH_50_ for the *Trichoplax* channel relative to mASIC1a, and a smaller Hill coefficient (*n*_H_). The values observed for mASIC1 are consistent with previous reports (26). *(C) Tad*NaC exhibits current rundown or tachyphylaxis at pH 4.5 and 5.5 similar to mASIC1a (40). *(D)* Plot of average peak current amplitude ± standard deviation through successive sweeps for the *Tad*NaC2 channel (pH 4.5 and 5.5) and mASIC1a (pH 5.5), showing statistically significant decaying peak currents (tachyphylaxis) that are statistically indistinguishable between the two channels after two-way ANOVA (p < 1.0E-14, F = 47.1 for the tachyphylactic decay of currents). *(E)* Sample current recordings for *Tad*NaC2 and mASIC1a before (black traces) and after (red traces) perfusion of 3 µM amiloride, revealing a nearly complete block for mASIC1a (at pH 5.5) and only ∼50% block for *Tad*NaC2 (pH 4.5). *(F)* Plot of average percent block of inward current ± standard deviation for *Tad*NaC2 and mASIC1a before and after perfusion of 3 µM amiloride. Individual replicates are included as *grey* circles. B+T indicates the total decay in average current for a successive sweep, which includes the effects of drug block (B) and tachyphylaxis (T), while B indicates the isolated component of drug block alone, obtained by subtracting the average decline in amplitude caused by tachyphylaxis. Denoted P values are for post hoc Tukey’s tests after one-way ANOVA (p = 1.7E-11, F = 56.1).

Interestingly, repeated activation of *Tad*NaC2 by perfusion of acidic saline resulted in a continuous decay in current amplitude, at both pH 4.5 and 5.5, resembling the unique decay of ASIC1 channels referred to as tachyphylaxis (40) (Fig. 4C). A plot of average normalized current amplitude for decaying currents of *Tad*NaC2 at pH 4.5 and 5.5, and mASIC1 at pH 5.5, revealed that the rate of tachyphylaxis was statistically indistinguishable between the two channels, irrespective of the tested values of pH (*i.e.*, P values for 2-way ANOVA were greater than 0.1; Fig. 4D). Lastly, we sought to test the sensitivity of *Tad*NaC2 to the general DEG/ENaC channel blocker amiloride, finding previously that the *Tad*NaC6 channel was potently activated by this drug (24), a rare feature also reported for ASIC3 channels (41, 42). Application of 3 µM amiloride almost completely blocked mASIC1, but only partially blocked *Tad*NaC2, slowing channel activation but maintaining a sustained current (Fig. 4E). This suggests that amiloride block of *Tad*NaC2 is more selective towards the early transient component of the macroscopic current, rather than the slower sustained component. Given that *Tad*NaC2 and mASIC1 currents respectively decay by 19.1 ±11.6% and 17.3 ±8.3% upon successive activation, we reasoned that a component of the attenuated current amplitude in these experiments was attributable to tachyphylaxis. Subtracting the effect of tachyphylaxis to isolate the amiloride block of both channels reduced the decrease in average peak inward current from 90.3 ±4.1% down to 73.0 ±4.1% for mASIC1, and from 60.5 ±9.5% to only 41.4 ±9.6% for *Tad*NaC2 (Fig. 4F).

### TadNaC2 conducts biphasic currents that exhibit differences in ion selectivity

As noted, the kinetics of *Tad*NaC2 macroscopic currents accelerate significantly with decreasing pH (Fig. 4A), such that with acidic solutions below pH 4.0, two separate components become apparent: a fast activating and desensitizing current, followed by a current with much slower activation and desensitization. These two separate components of the *Tad*NaC2 current become particularly evident at pH 3.5, where two separate peaks can be observed in the macroscopic current (Fig. 5A). Given that the whole-cell patch-clamp technique permits complete control of the intracellular and extracellular ionic compositions, we sought to determine whether these two current components exhibit differences in their ion selectivity, reflecting two fundamentally distinct conductance states for the *Tad*NaC2 channel. We thus employed the bi-ionic reversal potential technique, which compares the change in current reversal potentials (*i.e.*, voltages where currents reverse from inward to outward) under different ionic conditions (43), to see if the reversal potentials of the two currents differ. In this technique, cells expressing the channel are recorded with internal and external saline solutions each bearing 150 mM Na^+^, which typically results in a reversal potential of zero. Subsequently, perfusion is used to exchange the external Na^+^ solution with one bearing equimolar K^+^, which causes the reversal potential to shift leftwards according to the channel’s preference for Na^+^ over K^+^ ions (*i.e.*, a greater shift indicates greater Na^+^ selectivity). Application of a series of voltage steps (Fig. 5B, *top*) while perfusing 150 mM Na^+^ external saline at pH 4.5 produced slow-activating currents that lacked prominent desensitization (Fig. 5B, *top traces*). These currents were markedly different from those observed when the major internal cation was Cs^+^ (Fig. 4A). Plotting the early and late components of these currents as a function of holding voltage revealed a linear current-voltage relationship for both the early and late components (Fig. 5C), with expected average reversal potentials around zero (i.e., 0.9 ±0.9 mV for the early component, and 1.1 ±2.3 mV for the late component; Fig. 5D). Exchange of the external Na^+^ ions with equimolar K^+^ ions caused a dramatic change in the macroscopic current waveform, uncovering a distinct early transient current, and a sustained non-desensitizing late current (Fig. 5B). Both the early and late currents in these conditions exhibited linear current voltage relationships with reversal potentials that were left-shifted relative to the equimolar Na^+^ recordings (Fig. 5C), which was considerably more prominent for the late current compared to the early current (i.e., -60.4 ±2.3 mV vs. -49.0 1.6 mV; Fig. 5D). Calculation of the Na^+^/K^+^ permeability ratios of the early and late currents using the bi-ionic reversal potential equation revealed that the early current is less selective for Na^+^ over K^+^ ions compared to the late current, with pNa^+^/pK^+^ ratios of 7.2 ±0.5 and 11.6 ±1.4 respectively. Thus, the late current is more Na^+^ selective compared to the early current.

**Figure 5.**
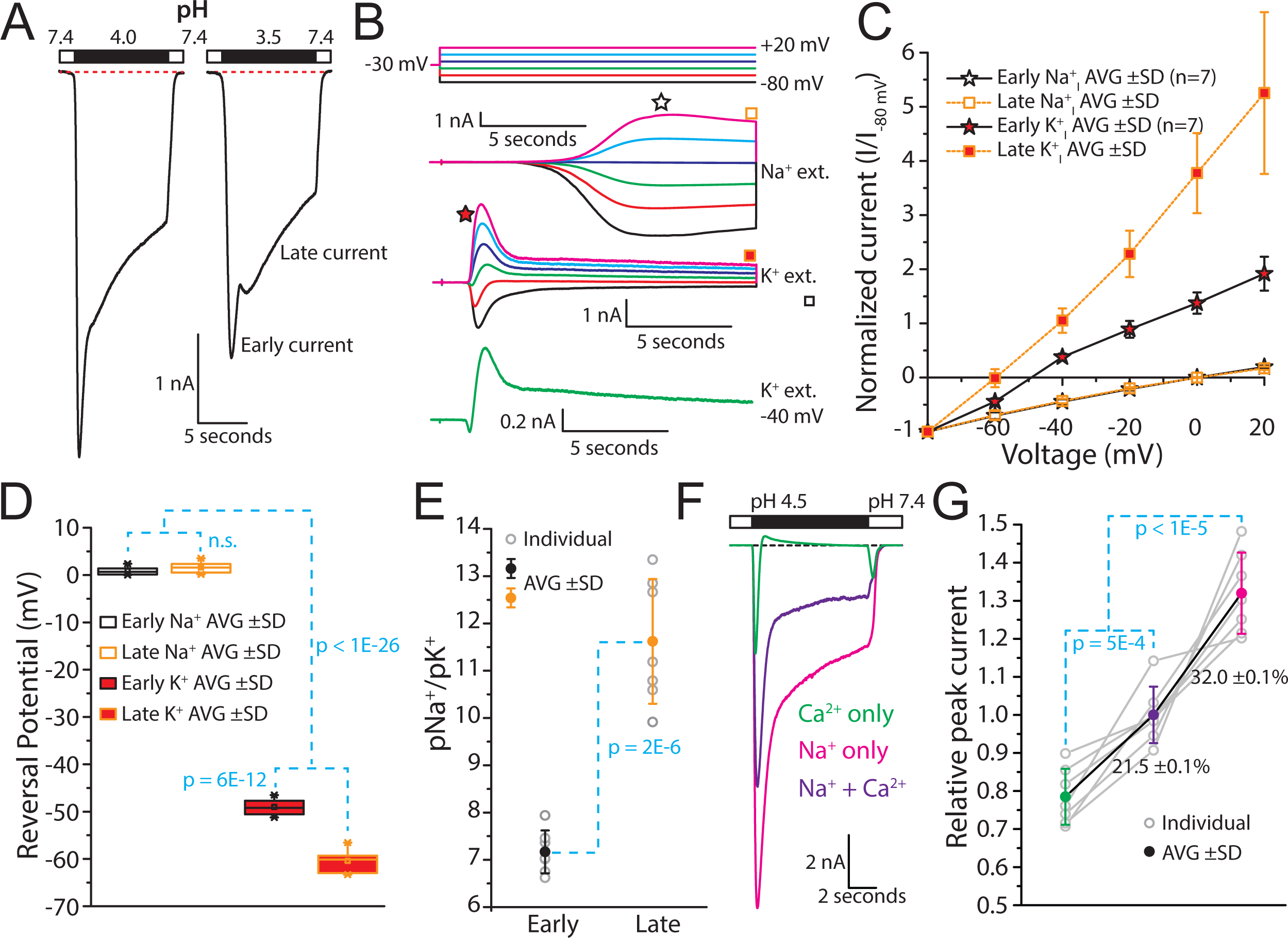
*Tad*NaC2 conducts biphasic currents *in vitro*. *(A)* Sample current recordings of the *Tad*NaC2 channel at pH 4 and 3.5, revealing a biphasic current with a fast transient component (*i.e.*, early current), and a slower, sustained (late) component. The biphasic current is particularly evident at pH 3.5. *(B)* Sample proton-activated *Tad*NaC2 currents recorded at different voltages (voltage protocol on top), under bi-ionic conditions of equimolar intracellular Na^+^ and extracellular Na^+^ (Na^+^ ext.) or extracellular K^+^ (K^+^ ext.). The star and square symbols denote portions of the currents that were considered as early/transient and late/sustained, respectively. The green trace (*bottom*) illustrates the transition of an inward early/transient current, to an outward late/sustained current, indicative of different ion selectivity for the two current components (at -40 mV). *(C)* Plot of average reversal potential data (± standard deviation) for the bio-ionic reversal potential experiments, revealing a significant leftward shift when extracellular Na^+^ was replaced with K^+^, particularly for the late current, indicating that the late current is more Na^+^ selective than the early transient current. *(D)* Box plot of average reversal potential data, showing statistically significant differences for both the early and late currents when extracellular Na^+^ was replaced with K^+^. Notably, whereas the early and late current reversal potentials were not statistically different from each other in the Na^+^ ext. condition, replacement of extracellular Na^+^ with K^+^ (*i.e.*, K^+^ ext.) caused a more significant leftward shift in the reversal potential for the late current, indicative of higher Na^+^ selectivity. The denoted P values are from Tukey post hoc tests after two-way ANOVA (p > 5.5E-4, F > 15.6 for all comparisons). *(E)* Na^+^/K^+^ permeability ratios calculated using the bi-ionic reversal potential data, revealing that the late current exhibits a significantly higher preference for Na^+^ over K^+^ compared to the early current (P value shown is for a two-sample t-test). *(F)* Sample *Tad*NaC2 currents elicited by extracellular protons (pH 4.5), with extracellular solutions containing Na^+^ only, Ca^2+^ only, or both. The largest current amplitude was observed when only Na^+^ was present (*pink* trace), while the amplitude was diminished when Ca^2+^ was included in the extracellular saline (*purple* trace). Perfusion of extracellular saline containing only Ca^2+^ (*green* trace) produced a fast inward transient current, which reversed to an outward monovalent Cs^+^ current, indicating that the late current is highly impermeable to Ca^2+^ ions. *(G)* Plot of average relative peak current amplitude for *Tad*NaC2 currents (± standard deviation) recorded with external Na^+^, Ca^2+^, or both, revealing statistically significant differences with the largest current occurring when just Na^+^ is present, and the smallest when just Ca^2+^ is present. P values are for Tukey post hoc tests after one-way ANOVA (p = 4.0E-9, F = 68.2).

Next, we sought to determine whether the early and late currents exhibit different permeabilities to Ca^2+^. Perfusion of a pH 4.5 external solution containing 123 mM Na^+^ and 10 mM Ca^2+^ ions, with an internal recording solution containing 143 mM CsCl, elicited standard *Tad*NaC2 macroscopic currents with a fast early component and a sustained late component (Fig. 5F; purple trace). Replacing the 10 mM CaCl_2_ in the external solution with the large impermeant cation NMDG^+^ (*i.e.*, 15 mM NMDG-Cl) resulted in a 32.0 ±0.1% increase in maximal inward current (Figs. 5F and G), affecting both the early and the late components. Thus, like *Tad*NaC6 and other Deg/ENaC channels, external Ca^2+^ ions appear to partially block inward Na^+^ currents through *Tad*NaC2. Instead, replacing the 123 mM Na^+^ in the external solution with equimolar NMDG^+^ resulted in a 21.5 ±0.1% decrease in peak current amplitude. This decrease was accounted for exclusively by the early current, while the late current reversed to a small outward current, attributable to an outward monovalent Cs^+^ current. Notably, switching the Na^+^ free solution to one containing 123 mM Na^+^ ions but at pH 7.4 produced a transient inward “tail” current, attributable to Na^+^ influx through *Tad*NaC2 channels caught in an open state, before undergoing deactivation. Thus, whereas the early transient current is quite permeable to Ca^2+^ ions, the late current is highly selective against Ca^2+^, preferring instead to conduct outward Cs^+^ despite the negative holding voltage of -30 mV used in these experiments which would create a significant driving force for Ca^2+^ influx.

### TadNaC2 lacks core molecular determinants for proton activation of ASIC channels

Deg/ENaC channels including ASIC channels are homo- and/or hetero-trimeric in nature, with each separate subunit forming a “ball in hand” tertiary structure comprised of wrist, palm, thumb, finger, knuckle, and β-ball regions (Fig. 6A). Cumulative efforts have uncovered the core molecular determinants for proton activation of ASIC channels, namely a critical histidine residue in the wrist region (H73 in mASIC1), and a lysine in the palm region (K211) situated at the extracellular interface between subunits (Fig. 6A) (25, 44–47). A protein alignment of several regions bearing these and other determinants for proton-activation of ASIC channels, including the group A ASIC channels from mice (*i.e.*, ASIC1 to 4), selected group A and B channels from *B.belcheri* (25), and the singleton group B channel from *L.anatina* (10) reveals near complete conservation of the H73 and K211 residues. The only exception are the proton-insensitive ASIC2b splice variant which lacks H73 (48), and ASIC4 which is also proton-insensitive and lacks K211 (49). The mouse ASIC5/BASIC channel, which as noted is phylogenetically distinct from ASIC channels (Fig. 2) and is not activated by protons, lacks both the H73 and K211 residues. Notably, these residues are also absent in other Deg/ENaC channels shown to be sensitive to external protons *in vitro*, including the proton-inhibited *T.adhaerens* channel *Tad*NaC6 (24), and the proton activated channels *Tad*NaC2, the ENaC-δ channel from human (6), the channels ACD-2, DEL-9, and ASIC-1 from *C.elegans* (14), Pickpocket1 from *D.melanogaster* (16), and *Ne*NaC2 from the sea anemone *Nematostella vectensis* (8). Notably however, *Tad*NaC2, as well as the mouse ASIC 4 and BASIC channels, possess a cationic residue just one amino acid upstream of the K211 position (*i.e.*, R201 in *Tad*NaC2). *Tad*NaC2 and its *Hoilungia hongkongensis* orthologue *Hho*NaC2 also possess a conserved lysine one position downstream (K203 in *Tad*NaC2; Fig. 6B). Furthermore, all of the ASIC channels except for the non-functional ASIC4 isotype bear a conserved aromatic residue 2 positions upstream of H73 (*i.e.*, Y71). Y71 forms an aromatic bridge with a conserved tryptophan (W287) in mouse ASIC1a, shown to be important for coupling conformation changes in the extracellular domain with gating of the pore helices (50). This aromatic residue is notably absent in all included non-ASIC channels except for *Tad*NaC2 and *Hho*NaC2 which bear phenylalanine and tyrosine residues at this position, respectively (*i.e.*, F70 in *Tad*NaC2), as well as a tryptophan corresponding to W287 (*i.e.*, W276 in *Tad*NaC2). Also notable is that *Tad*NaC2 possesses several protonatable amino acids that are in proximity of the ASIC H73 position, with aspartate and glutamate residues at positions 75 and 77, and a histidine at position 80 that aligns with residues in the palm region placing it proximal to the noted R201 and K203 residues.

**Figure 6.**
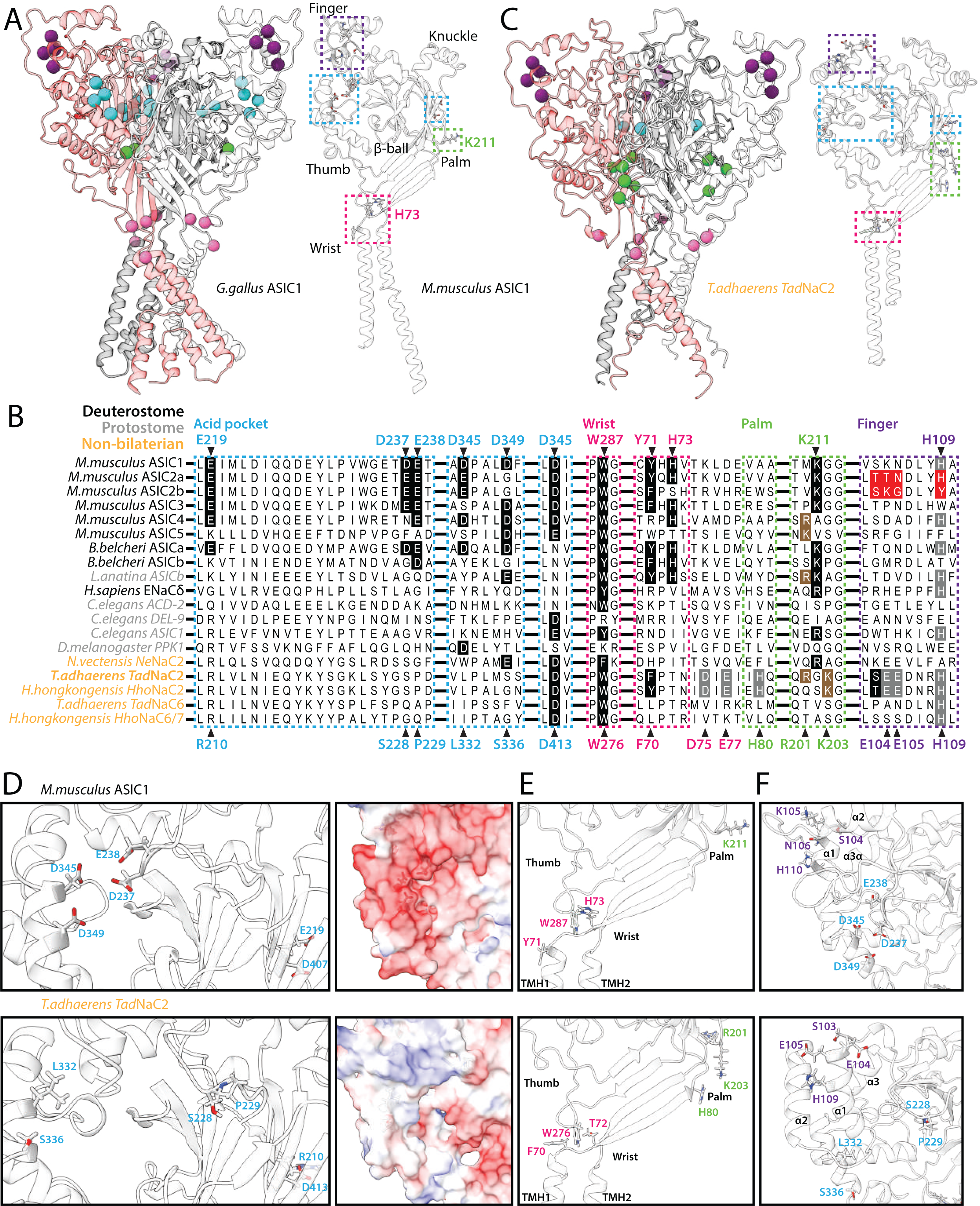
Structural features of the *Tad*NaC2 channel. *(A)* Ribbon diagrams of the chick ASIC1 homotrimeric channel crystal structure (*left*, PDB number 6VTK), and the AlphaFold-predicted tertiary structure of the mouse ASIC1a subunit (*right*). The three separate subunits of the homotrimeric channel are colored in red, white, and grey, and the colored circles denote the β carbon atoms of critical residues corresponding to the back-colored residues of the mouse ASIC1a channel in the protein alignment shown in panel *B* (*i.e.*, atoms in *blue* are within the acid pocket, *pink* are within the wrist, *green* are within the palm, and *purple* are within the finger). The dashed boxes denote structural regions of the single mASIC1 subunit structure bearing these same critical residues. *(B)* Protein sequence alignment of the acid pocket (enclosed by *blue* dashed boxes), wrist (*pink*), palm (*green*), and finger (*purple*) regions of select proton- activated Deg/ENaC channels from cnidarians and bilaterians with *Tad*NaC2 and *Tad*NaC6/7 channels from the placozoans *Trichoplax adhaerens* and *Hoilungia hongkongensis*. Residues that are back-colored in black represent conserved residues for proton activation of ASIC channels, while those back-colored red denote key residues that render the ASIC2b splice variant insensitive to external protons. Residues that are back-colored in grey denote protonatable amino acids in *Tad*NaC2 within these key structural regions, some of which are conserved in cnidarian and bilaterian homologues, while those back-colored in brown denote cationic residues in *Tad*NaC2 that flank the critical K211 residue of ASIC channels, also found in several other channels. Notable is the complete conservation of the critical residues H73 and K211 in all true ASIC channels, which are absent in most non-ASIC proton-activated channels including *Tad*NaC2. *(C)* Homology model of the homotrimeric *Tad*NaC2 channel structure (left), and AlphaFold- predicted structure of the single subunit, with a similar annotation as described for panel *A*. *(D)* Left panels: Close-up view of the acid pocket region of mASIC1 (top) and *Tad*NaC2 (bottom) within corresponding AlphaFold-predicted structures. The six rendered residues in the *Tad*NaC2 channel correspond to residues that align with the six acid pocket residues in mASIC1a as depicted in panel *A*. Right panels: Surface rendering of the acid pocket region of mASIC1a (*top*) and *Tad*NaC2 (*bottom*) reveals a stark difference in the electrostatic potential between the two channel subunits. *(E)* Close-up view of the wrist and palm regions of mASIC1a and *Tad*NaC2. Apparent in the wrist region is the absence of a critical H73 proton-sensing residue in *Tad*NaC2, but conservation of the aromatic amino acids F70 and W276, which in mASIC1a (*i.e.*, Y71 and W278) form an aromatic bridge that is critical for channel gating. Instead, *Tad*NaC2 bears a putative proton-sensing amino acid (H80) at the opposite end of a β strand that projects from the first transmembrane helix of wrist region (TMH1) to the palm domain, placing it near the R201 and K203 that flank the critical K211 residue of mASIC1a. *(F)* Close-up view of the finger and acid pocket regions, with rendered amino acids corresponding to the positions in the ASIC2b splice variant that makes the channel insensitive to protons. Also labeled are the equivalent acid pocket residues, and the predicted α1 to α3 helices in the finger region.

Another region associated with proton activation (and desensitization) of ASIC channels is the acid pocket, comprised of a cluster of four acidic amino acids located between the finger, thumb, and β- ball regions of the subunit monomer, and another pair in the β-ball region close to K211 (Fig. 6A and B). In the trimeric channel, the four acidic residues from one subunit and two from an adjacent subunit combine to form a namesake pocket-like structure where protons are thought to bind and affect channel conformation and gating (26). Of note, mutation of these glutamate/aspartate residues does not completely disrupt proton activation, and instead, these appear to be more important for channel desensitization (26). Accordingly, the group B ASIC channels from *B.belcheri* and *L.anatina* lack most glutamate/aspartate residues in the acid pocket (Fig. 6B), while they are largely conserved among the group A channels. Furthermore, the two *Tad*NaC channels, as well as the various other non-ASIC Deg/ENaC channels included in the alignment lack most if not all acidic residues at equivalent positions of the acid pocket.

A third region of interest with respect to proton activation is the finger region, where a motif of four amino acids distinguishes the proton sensitive ASIC2a mRNA splice variant from the insensitive ASIC2b variant (Fig. 6B). Specifically, ASIC2a bears a motif of TTN-XXX-H and is proton-activatable, while ASIC2b bears an SKG-XXX-Y motif and is not (51). Moreover, introducing the SKG and Y elements of the ASIC2b motif into ASIC2a, together but not separately, abrogates proton activation, and insertion of the finger region of ASIC2a into ASIC1 causes a significant reduction in proton sensitivity making the latter less sensitive to protons similar to ASIC2a channel (52). Notably, a histidine residue within the finger motif of ASIC2a (H109) is conserved among many of the included Deg/ENaC channels including *Tad*NaC2 (Fig. 6B), however, its mutation in ASIC2a does not exert noticeable effects on proton sensitivity (46). In this region, *Tad*NaC2 also bears two protonatable glutamate residues (E104 and E105).

To better infer how the structure of *Tad*NaC2 compares to the well-studied structures of ASIC channels, we generated a homology model of the homotrimeric channel using the crystal structure of chick ASIC1 as a template (Fig. 6C; left panel) (53). We also predicted the tertiary structures of the monomeric mouse ASIC1 and *Tad*NaC2 channel subunits with AlphaFold (Figs. 6A and C; right panels) (54). Labelling the β carbon atoms of the Y71, W287, H79, K211, acid pocket, and ASIC2 finger motif equivalents in the homotrimeric structure of the chick ASIC1 channel (Fig. 6A), and the F70, W276, H80, R201, K203, D413 (single acid pocket residue), and finger motif equivalents in the model of the *Tad*NaC2 homotrimer (Fig. 6C), illustrates the general absence of acid pocket residues in *Tad*NaC2. Also evident are the noted differences in the wrist region, where *Tad*NaC2 lacks a critical H73 equivalent, and in the palm, where interestingly, the residues H80, R201, and K203 in *Tad*NaC2 are arranged in a triangular cluster at the interface between subunits, in similar position as the K211 residue in ASIC channels. Furthermore, the aromatic residues F70 and W276 residues in *Tad*NaC2 are in proximity to each other, suggesting that like Y71 and W287 in ASIC1, these can form hydrophobic interactions.

The predicted structures of the mASIC1 and *Tad*NaC2 monomers also highlight key differences and similarities. First, whereas the cluster of four acid pocket residues in mASIC1 (D237, E238, D345, and D349) are arranged in a tight cluster, the equivalent residues in *Tad*NaC2 are not (S228, P229, L332, and S336) (Fig. 6D, left panels). Rendering the electrostatic potential on the surface of the two channel subunits also illustrates a stark difference in the acid pocket region, with the acidic residues of mASIC1 contributing to a highly electronegative surface, while those in *Tad*NaC2 contribute to a surface that is slightly positive and hence unlikely to attract and bind H^+^ ions (Fig. 6D, right panels). In the wrist region, the W276 residue at the base of the thumb of *Tad*NaC2 is in a similar position as W287 in mASIC1, between the Y71 and H73 equivalent residues F70 and threonine 72 from the wrist region (Fig. 6E).

Lastly, both the SKN-XXX-H and SEE-XXX-H finger motifs of mASIC1 and *Tad*NaC2 are within a short loop and adjacent descending alpha helix, consistent with the α1 helix identified in the crystal structure of the chick ASIC1a finger region (7). However, this helix is predicted to be two helical rotations longer in *Tad*NaC2, with a short loop connecting it to the downstream α2 helix that is also longer than its predicted counterpart in mASIC1 by one rotation (Fig. 6F). Lastly, it is notable that the finger regions of the two channels are positioned above the divergent acid pocket, suggesting that any structural alterations taking place in the finger region would be differentially transferred to the thumb and pore regions that mediate gating.

### Key residues in the palm and finger region of TadNaC2 are required for proton activation

Despite lacking known deterministic residues for proton activation, the similar predicted structure of *Tad*NaC2 compared to mASIC1 prompted us to examine whether corresponding structural regions in the placozoan channel bear proton-sensitive elements. Thus, we performed site-directed mutagenesis on selected aromatic or protonatable residues in the wrist region (F70, D75, and E77), protonatable residues in the finger region (E104, E105, and H109), and protonatable or cationic residues in the palm region (H80, R201, and K203) (Fig. 6A). To assess changes in H^+^ sensitivity, we tested each mutant with a series of perfused solutions to generate pH dose response curves of recorded macroscopic currents (Fig. 7A to C).

**Figure 7.**
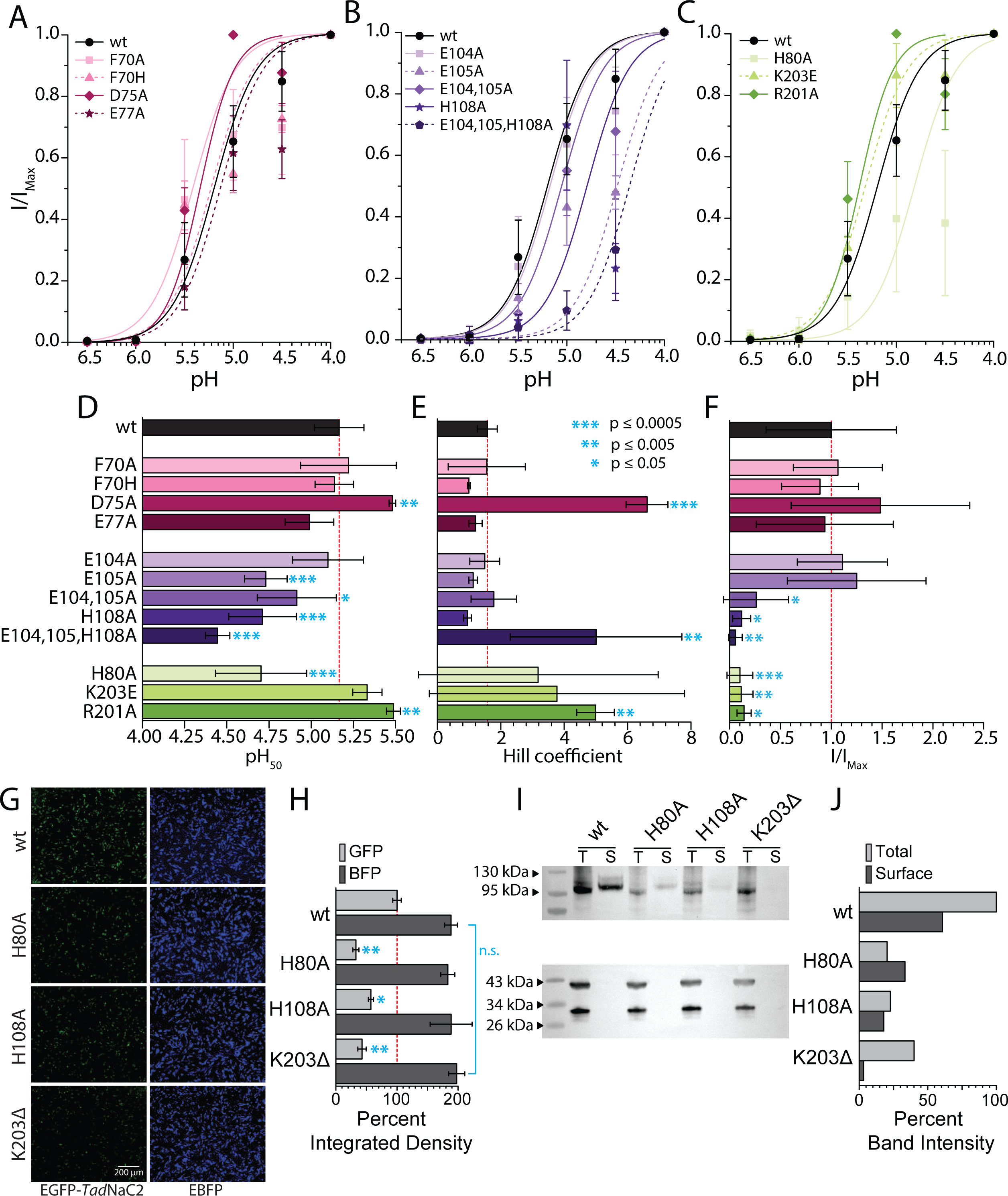
Unique residues are required for proton-activation of *Tad*NaC2. *(A* to *C)* Average pH dose response curves ± standard deviation for wildtype (wt) *Tad*NaC2 and various variants bearing amino acid substitutions within the wrist (panel *A*), finger (panel *B*), and palm (panel *C*) regions. (*D* to *F*) Plot of average pH_50_ (panel *D*), Hill coefficient (panel *E*), and normalized peak current amplitude ± standard deviation (I/I_Max_; panel *F*) deviation for wildtype (wt) and the various point mutated *Tad*NaC2 channels. Cyan-colored asterisks denote P value thresholds for Tukey post-hoc means comparisons between wildtype values and those for mutant channels, after corresponding one-way ANOVAs (pH_50_: p < 1E-14, F = 26.0; Hill coefficient: p = 2.5E-11, F = 9.2; current amplitude: p = 1.4E-15, F = 11.0). *(G)* Representative fluorescence micrographs of CHO-K1 cells co-transfected with pEGFP-*Tad*NaC2 fusion vector (*left* panels) and an empty pIRES2-EBFP vector (*right* panels). *(H)* Plot of percent average integrated density ± standard deviation, quantifying the emitted fluorescence of pEGFP-*Tad*NaC2 wildtype (wt) and mutant channels, normalized to the average integrated density of wildtype *Tad*NaC2 (n=3 for each transfection condition). EBFP fluorescence was also quantified to determine tranfection efficiency. Cyan-colored asterisks denote P value thresholds for Tukey post hoc means comparisons of fluorescence signals between wildtype and mutant channels after one-way ANOVAs (EGFP: p = 5.6E-11, F = 73.6; EBFP: not significant). *(I)* Top panel: Western blot of select EGFP-tagged *Tad*NaC2 channel variants in CHO-K1 cell lysates using anti-GFP polyclonal antibodies, comparing total channel protein content (T) with membrane/surface expressed channel protein content (S) for each variant. Bottom panel: Western blot of the lower half of the membrane used in the top panel, using anti-GAPDH (*top* bands) and anti-EBFP (*bottom* bands), polyclonal antibodies. *(J)* Quantified band intensity (mean grey area) of *Tad*NaC2 bands in panel *I*, relative to the wildtype EGFP-*Tad*NaC2 total protein band, revealing decreased total and surface protein expression of *Tad*NaC2 channels bearing mutations, and a near complete absence of membrane expressed variants harboring a K203 deletion, consistent with our inability to record current for this channel *in vitro*. Bands for each channel variant were also normalized to the intensity of EBFP present in corresponding total protein lanes.

In the wrist region of rat ASIC1a, mutation of the Y71 aromatic residue to a histidine imposes a ∼70% reduction in elicited current amplitude, while mutation to alanine completely disrupts proton activation (50). In contrast, analogous mutations of the F70 residue in *Tad*NaC2 had negligible effects on the pH_50_ and Hill coefficients (Fig. 7D and E), and no effect on average peak inward current amplitude at pH 4.0 compared to the wildtype channel (Fig. 7F). Thus, this residue in *Tad*NaC2 does not likely form an analogous aromatic interaction with the conserved W276 residue in the thumb region, akin to the Y71- W287 interaction in ASIC1 channels. As noted, *Tad*NaC2 bears two protonatable residues in the wrist region (D75 and E77), within a predicted β strand that in ASIC channels projects from the H73 residue in the wrist towards the K211 residue in the palm (Fig. 6A and B). Notably, the E77 residue in *Tad*NaC2 aligns with D78 in ASIC1a and ASIC2a, which when mutated to asparagine in the rat channels significantly disrupts proton sensitivity (45, 47). In contrast, alanine substitution of E77 in *Tad*NaC2 had no noticeable effect, while mutation of the D75 residue two positions upstream caused a significant *increase* in both the pH_50_ and Hill coefficient, indicative of enhanced proton binding and cooperativity (Fig. 7D and E). Furthermore, neither the D75A nor the E77A mutation caused a change in peak current amplitude (Fig. 7F). Overall, it is apparent that *Tad*NaC2 is fundamentally different from ASIC channels in lacking known molecular determinants in the wrist region involved in proton activation and gating.

In the finger region of *Tad*NaC2, alanine substitution of protonatable residues produced more striking effects, with the single mutations E105A and H108A both causing a decrease in pH_50_, but no effect on Hill coefficient, while E104A was indistinguishable from wildtype (Fig. 7B, D, and E). The E104A/E105A double mutation also caused a decrease in pH_50_, but not relative to the E105A single mutation, suggesting that the E104 residue plays a minimal role in proton activation. The most dramatic effect was observed for the E104/E105/H108 triple mutation, which severely diminished current elicited by pH 5.5 and 4.5 and required a much more acidic pH of 4.5 and 4.0 (Fig. 7B and F). This resulted in a pH_50_ value of 4.4 ±0.1 compared to 5.2 ±0.1 for the wildtype channel, and an increased Hill coefficient of 5.0 ±2.7 compared to 1.6 ±0.3, attributable to the sharp rise in elicited current at pH 4.0 (Fig. 7D and E). Lastly, the E104A/E105A double mutant, the H80A single mutant, and the E104A/E105A/H108A triple mutant all had diminished maximal current amplitudes at pH 4.0, which was most pronounced for the triple mutant (Fig. 7F).

In ASIC1a, deletion of the K211 palm residue produces a strong decrease in proton sensitivity, while mutation to glutamate causes a more moderate effect (25). In *Tad*NaC2, mutation of the two cationic residues that flank the K211 position, R201 and K205, produced different effects. Specifically, mutation of R201 to alanine caused an increase in proton sensitivity, with a pH_50_ of 5.5 ±0.04 and a Hill coefficient of 5.0 ±0.6 (Fig. 7C, D, and E). Instead, mutation of K203 to glutamate did not significantly impact proton sensitivity, while its deletion (K203Δ) completely disrupted channel function, as we were unable to record currents for this channel even with very acidic pH. Interestingly, alanine substitution of the unique protonatable H80 residue, which as noted is positioned proximal to the R201 and K203 residues in our predicted structures (Fig. 6), produced a strong decrease in pH_50_ (4.7 ±0.3). However, neither the H80A nor the K2034E mutations had an effect on the Hill coefficient. Like the E104/E105/H108 triple mutant, all tested mutations in the palm region caused a significant decrease in maximal current amplitude at pH 4.0 (Fig. 7F), especially the K203Δ variant which as noted did not produce currents *in vitro*.

Next, we sought to determine whether the noted decrease in current amplitude caused by select mutations was due to reduced functionality or a reduction in channel membrane expression. Hence, we N-terminally tagged the wildtype channel with enhanced green fluorescent protein (EGFP), as well as the mutants H80A, H108A, and K203Δ which respectively caused moderate, strong, and severe effects on current amplitude. This permitted inference of the total channel protein levels in transfected CHO-K1 cells via EGFP fluorescence quantification, relative to a co-transfected blue fluorescent protein from the empty vector pIRES2-EBFP. Of note, we tested whether tagging the wildtype *Tad*NaC2 channel with EGFP disrupted its function, finding it to conduct proton-activated currents that were visually indistinguishable from the untagged channel (data not shown). Fluorescence micrographs of transfected cells reveal a noticeable decrease in EGFP fluorescence of all three mutant channels relative to wildtype (Fig. 7G), with respective normalized average integrated density values of 67 ±5%, 43 ±4%, and 57 ±7% for the H80A, H108A, and K203Δ mutants (Fig. 7H). Notably, average integrated density measurements for the co-transfected EBFP were statistically indistinguishable for all transfections, indicating that the differences in EGFP fluorescence were not due to differences in transfection efficiency. Thus, all three of the tested mutations cause a decrease in total protein expression *in vitro*.

We also sought to characterize the effect of the mutations on total protein and membrane expressed protein levels in transfected cells using a cell surface biotinylation strategy. A Western blot probed with anti-EGFP antibodies reveal a significant reduction in both total protein and membrane expressed (surface) proteins levels of mutant *Tad*NaC channels relative to wildtype (Fig. 7I).

Measurements of the mean gray value of the different bands on the blot reveals similar reductions in total protein levels for all three mutants, and notably, extreme reduction of membrane expressed K203Δ (Fig. 7J). Altogether, this data is consistent with our current amplitude measurements and inability to record currents for the K203Δ variant of *Tad*NaC2.

### T.adhaerens exhibits chemotactic and contractile responses to extracellular pH changes

*T.adhaerens* is a simple invertebrate with only six identified cell types, which locomotes across surfaces through the action of beating cilia on its ventral epithelium (55). Remarkably, *T.adhaerens* lacks chemical/electrical synapses and a nervous system (55) but is nonetheless able to coordinate its various cell types for directed locomotion such as feeding (56), chemotaxis towards food and glycine (57, 58), and gravitaxis (59). Given our discovery of proton-sensitive Deg/ENaC channels in placozoans and the general prevalence of pH sensing behaviors in animals (60–62), we asked whether *T.adhaerens* is able to respond to changes in extracellular pH. First, we sought to determine whether *T.adhaerens* exhibits pH-sensitive chemotaxis by lining the base of a chamber with an array of highly buffered agar strips of pH 5.0, 7.5, and 9.0, all having the same combination of buffering agents (*i.e.*, 165 mM each of MES, HEPES, and CHES). The pH indicator phenol red was also included, revealing that the pH of each strip remained stable for over the course of three hours, despite the buffering capacity of the artificial sea water that was subsequently added to the chamber (*i.e.*, attributable to calcium carbonate and sodium bicarbonate). Placing a total of 47 animals into the central part of the chamber bearing the pH 7.5 strip, as separate groups in a total of 12 experiments, revealed a collective tendency of tracked animals to avoid moving into the area of the dish lined with the pH 5.0 strip, with only three animals ever entering this area (Fig. 8A). Instead, most animals stayed within the pH 7.5 strip or migrated towards the pH 9.0 strip, such that the average horizontal displacement approached the interface between these pH 7.5 and 9.0 strips, consistent with the pH of artificial seawater which is roughly 8.2. Therefore, *T.adhaerens* can sense external pH and respond through directed movement, which manifests as positive chemotaxis towards neutral pH, or alternatively, negative chemotaxis away from acidic pH.

**Figure 8.**
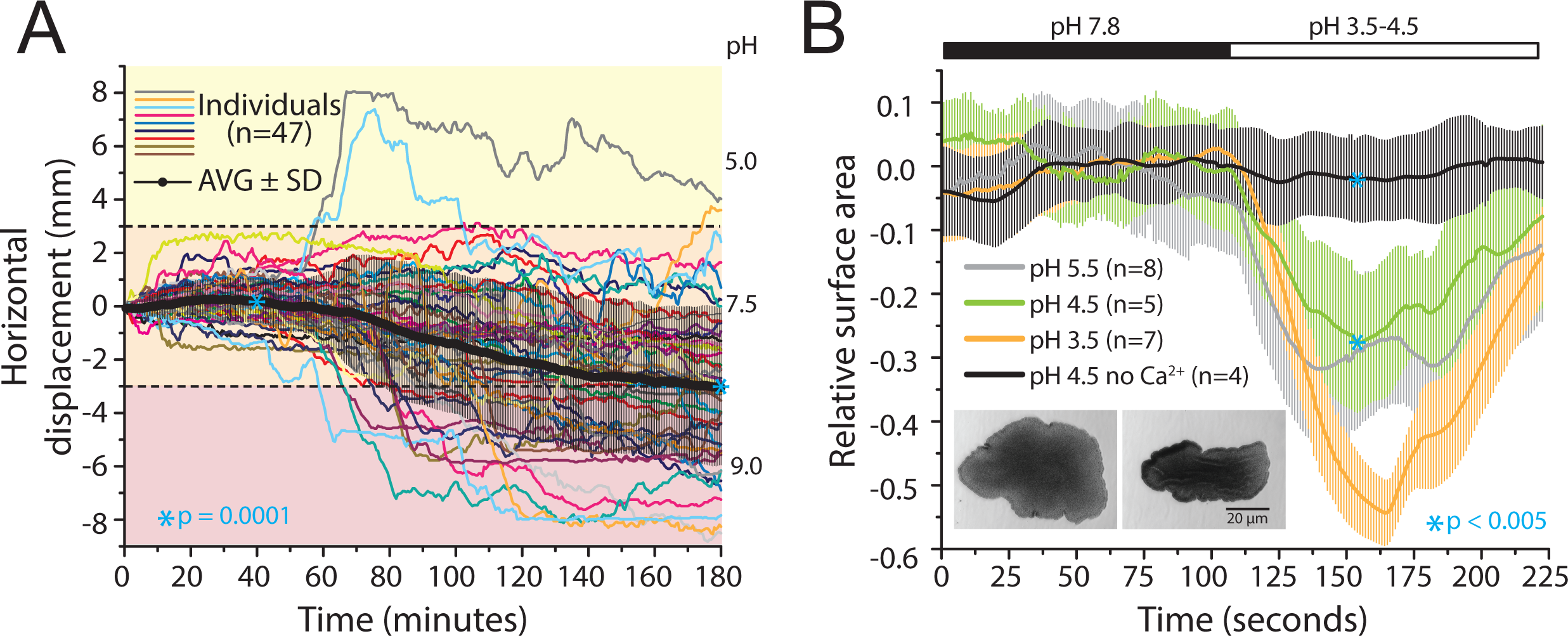
*Trichoplax* exhibits negative chemotaxis from an acidic environment, and contracts upon transient exposure to acidic pH in a Ca^2+^-dependent manner. *(A)* Plot of horizontal displacement over time of tracked *Trichoplax* specimens in a dish bearing strongly buffered agar strips (individual colored lines), revealing general avoidance of a pH 5.0 acidic environment, and a net displacement away from this environment towards the interface between pH 7.5 and 9.0 (*i.e.*, average displacement standard deviation). *(B)* Plot of average relative surface area of video-recorded *Trichoplax* specimens over time (± standard deviation), revealing strong contraction upon rapid perfusion of acidic saline solutions between pH 5.5 and 3.5. Contraction at pH 4.5 was not observed when Ca^2+^ ions were excluded from the perfused saline. Inset: Screenshots of a recorded representative animal shortly before (*left*) and after (*right*) perfusion of a pH 3.5 solution. For panel *A*, the *cyan* asterisks denote statistical differences between the mean displacement at time = 40 and time = 180 minutes, and for panel *B*, between the normalized animal surface area upon perfusion of a pH 4.5 solution bearing Ca^2+^, compared to the same solution but without Ca^2+^ (P values are for two sample t-tests).

Next, we sought to determine whether fast perfusion of seawater solutions at different pH could elicit behavioral responses. Interestingly, switching perfusion from a minimal artificial sea water solution at pH of 7.8 to ones at pH 5.5, 4.5, and 3.5 all caused *T.adhaerens* animals to rapidly contract over the course of ∼30 seconds, most pronounced for the pH 3.5 solution (Fig. 8B). Similar contractions have been reported upon the application of various endogenous neuropeptides (63, 64), as well as glycine (58), likely mediated by the dorsal epithelial cells that are highly contractile (65). However, whether any of these contractions involve membrane influx of Ca^2+^, consistent with excitation-contraction coupling, has yet to be determined. We therefore repeated these experiments at pH 4.5 but without Ca^2+^ in the solution, revealing a complete disruption of the contractile responses observed when Ca^2+^ was present (Fig. 8B).

### TadNaC2 and TadNaC10 interact with filamin at their C-terminus, like human ENaC channels

The C-termini of Deg/ENaC channels can mediate important protein-protein interactions that serve to regulate channel function. For example, both the α- and β- subunits of the human heterotrimeric ENaC channel interact with the actin-binding cytoskeletal protein filamin, which regulates their membrane expression and modulation by intracellular signalling pathways (27, 28). Also, both ASIC1a and ASIC2a channels have C-terminal ligand sequences that are bound by the PDZ domain of the scaffolding protein PICK1, which regulates their sub-cellular localization in central and peripheral neurons (66, 67). We therefore set out to determine whether *Tad*NaC2 and *Tad*NaC10, which fall within the ASIC and ENaC superclades in our phylogenetic analysis (Figs. 1 and 2), interact with any endogenous *T.adhaerens* proteins in a yeast 2-hybrid screen. Interestingly, using the 70 most distal amino acids of both channels as bait, we found that the C-termini of both channels interact with *T.adhaerens* homologues of the proteins filamin and U5 small nuclear ribonucleoprotein helicase (Table S1), with filamin being the top hit for *Tad*NaC2. *Tad*NaC2 also interacted with numerous other proteins including homologues of the scaffolding protein VAC14 and the cell adhesion protein contactin-6 (Table S1).

We sought to better understand the phylogenetic properties of *T.adhaerens* filamin homologues relative to those in humans (*i.e.*, filamins A, B, and C), *D.melanogaster* (Cheerio and Jitterbug) (68), *C.elegans* (FLN-1 and 2) (69), the cnidarian *N.vectensis*, and the fellow placozoan *H.hongkongensis*.

Interestingly, we identified two filamin genes in the gene data for *T.adhaerens* (70, 71) and *H.hongkongensis* (72). One set of orthologues is longer at 3,838 and 3,835 amino acids in length for *T.adhaerens* and *H.hongkongensis* respectively. Each bears a predicted N-terminal actin-binding domain comprised of two calponin homology (CH) domains, followed by 36 tandem immunoglobulin-like filamin domains. The other set of orthologues is shorter at 1,253 and 1,250 amino acids in length respectively, with three acting-binding CH domains and only 9 filamin domains (Fig. 9A). The longer placozoan filamin orthologues form a well-supported clade with the three human filamin proteins on a phylogenetic tree, along with Cheerio from *D.melanogaster*, FLN-1 from *C.elegans*, and one of the two filamin homologues identified for *N.vectensis* (Fig. 9B). Given their similar secondary structure and proximal phylogenetic relationship, were hereafter refer to this group as type 1 filamins or filamin-1. The two shorter placozoan orthologues on the other hand, which correspond to the filamin identified in our yeast 2-hybrid screen, form a clade with Jitterbug and FLN-2 from *D.melanogaster* and *C.elegans*, and a particularly short filamin homologue from *N.vectensis* which differs from all other examined sequences by possessing a predicted thioredoxin-like domain towards its C-terminus. Outside of this sequence, all other filamin homologues in this clade are distinct from type 1 filamins in possessing three tandem CH domains at their N-termini, a somewhat atypical feature relative to other actin-binding proteins (73). Hence, we refer to this clade as type 2 filamins or filamin-2. Interestingly, mapping the partial filamin prey sequences identified in the yeast 2-hybrid screen revealed that all interactions, for both *Tad*NaC2 and *Tad*NaC10, occurred within the C-terminus of *T.adhaerens* filamin-2, within the last two filamin repeat domains (Fig. 9A). This is notable because all three of the human filamin paralogues were found to interact with α- and β-ENaC channels via their C-termini (27), within the five distal filamin repeat domains (Fig. 9A).

**Figure 9.**
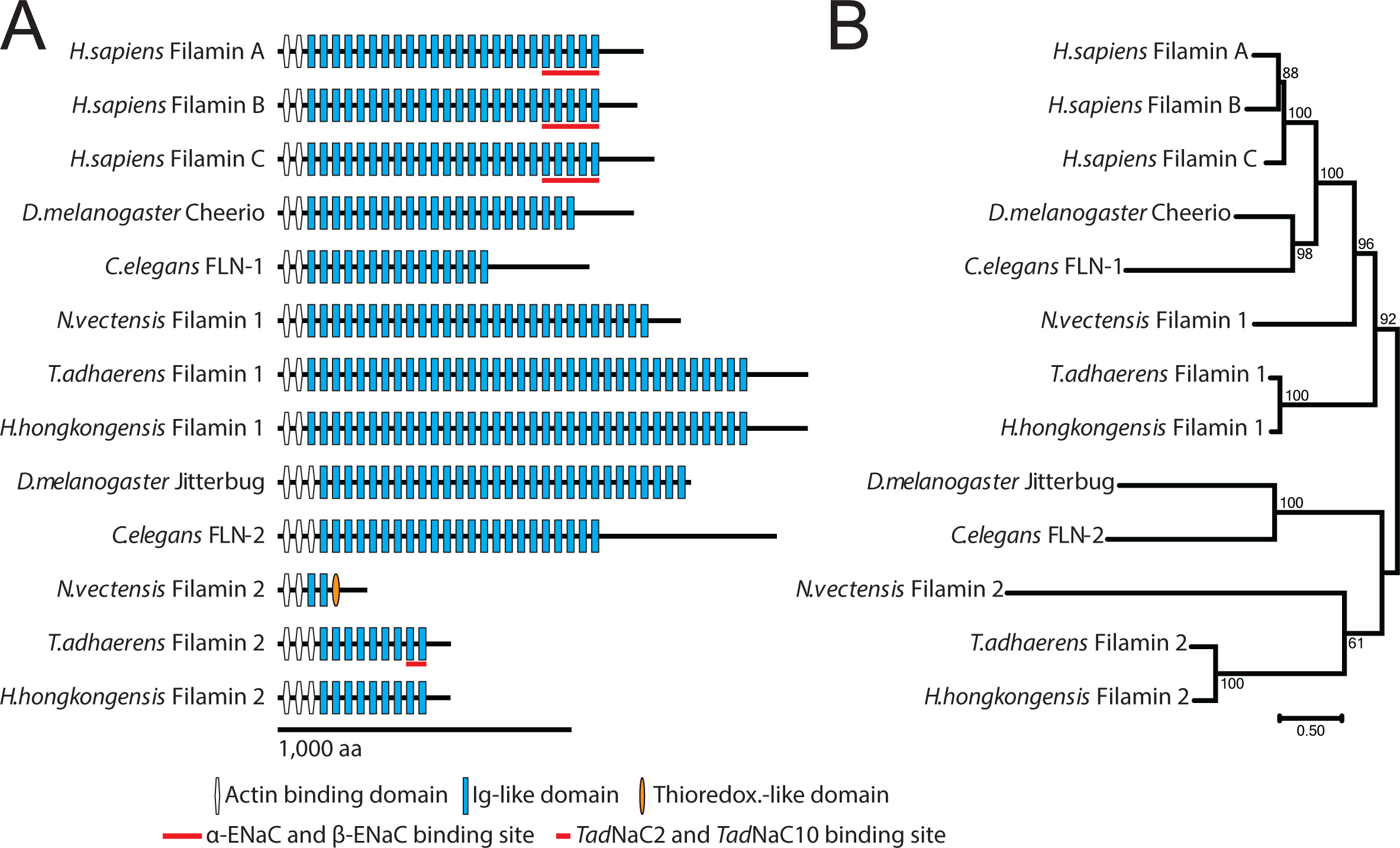
The C-termini of *Tad*NaC2 and *Tad*NaC10 interact with a jitterbug-like homologue of the actin-binding protein filamin. *(A)* Illustration of the relative amino acid sequence length and InterProScan-predicted domains for each protein, revealing conserved N-terminal actin binding calponin homology domains, followed by numerous immunoglobulin-like filamin repeat domains (Ig-like). The red bars underneath the three human filamin 1 paralogues (filamin A to C) denote regions found to interact with the C-terminus of α- and β-ENaC channels (27), while the red bar underneath the *Trichoplax* filamin 2 homologue denotes the region found to interact with *Tad*NaC2 and *Tad*NaC10 C-termini in a yeast 2-hybrid library screen. *(B)* Maximum likelihood protein phylogeny of aligned filamin homologues from humans, the vinegar fly *D.melanogaster*, the nematode worm *C.elegans*, the cnidarian sea anemone *N.vectensis*, and the placozoans *T.adhaerens* and *H.hongkongensis*. Notable are two separate clades of filamin proteins (*i.e.*, filamin 1 and 2 clades), distinguished by the presence of two vs. three actin binding domains at the N-terminus, with the exception of the filamin 2 homologue from *N.vectensis*.

## Discussion

### New insights into the phylogenetic groupings of Deg/ENaC channels

Combined, our CLANS and phylogenetic analysis provides strong evidence for the existence of two major clades of Deg/ENaC channels in metazoans (Figs. 1 and 2), which we propose as the ASIC and ENaC subfamilies. Both ASIC and ENaC subfamily channels are represented in most animal phyla examined including the placozoans *T.adhaerens* and *H.hongkongensis*, the poriferans *A.queenslandica* and *S.ciliatum*, and cnidarians including *N.vectensis*. The only exception was the ctenophores, for which we could not identify ENaC family channels. However, it is worth noting that ctenophores possess two separate sets of Deg/ENaC channels that formed distinct clusters in the CLANS analysis, representing highly divergent channels with obscured phylogenetic histories. Since most of the recent phylogenomic studies point to either ctenophores or poriferans as the earliest diverging animals (74–76), our phylogenetic analyses suggest that the ASIC and ENaC subfamilies were established at the very early stages of animal evolution. Indeed, these two subfamilies could have been present in the last common ancestor of all animals, and ENaC channels were subsequently lost in ctenophores or diverged beyond recognition by the cluster-based and phylogenetic analysis methods. Alternatively, if ctenophores are the earliest diverging, ASIC channels emerged first, followed by ENaC channels after ctenophores separated from other animals.

Our CLANS analysis also revealed numerous additional lineages of Deg/ENaC channels that formed separate clusters, indicative of strong sequence divergence. This permitted us to exclude these problematic sequences from our phylogenetic analysis, as they would provide little phylogenetic information and interfere with the phylogenetic inference. As a result, we were able to produce a tree with strong support for most nodes, including the key node resolving the ASIC and ENaC subfamilies. Of note, a recent comprehensive phylogenetic study similarly delineated two major superclades of Deg/ENaC channels, referred to as clades A and B which somewhat correspond with our ASIC and ENaC superclades, respectively (8). Nevertheless, there are some notable differences. For example, because we included a large set of sequences from bilaterians, ctenophores, and sponges, all cnidarian channels (*Hy*NaC and *Ne*NaC) within the ASIC subfamily formed a clade that was separate from ASIC, BASIC, and *Tad*NaC channels. In contrast, the other study found a subset of *Ne*NaC channels to form a clade with ASIC, BASIC, and *Tad*NaC channels, including the proton-activated channel *Ne*NaC2 (8), although the node support for these relationships was weak. A separate study found that ASIC, BASIC, and *Hy*NaC channels form a clade with each other, however also with poor node support. Instead, our tree resolves the elusive phylogenetic relationship between ASIC, BASIC, and *Hy*NaC channels (10), revealing that both sets of cnidarian channels within the ASIC subfamily, *Ne*NaC and *Hy*NaC, form a distinct clade with uncharacterized channels from ctenophores, poriferans, and bilaterians. Also, whereas in our analysis we excluded the *C.elegans* channels DEL9 and DEL10 because of their strong divergence evident in the CLANS analysis, these formed a sister relationship with BASIC channels in the other study (8), perhaps because of long-branch attraction. Furthermore, whereas we found both the *D.melanogaster* PPK channels and the *C.elegans* ACD/FLR channels to be highly divergent and fall outside of the ASIC and ENaC superclades, these formed a clade with each other within the ASIC superclade in the other study (8), perhaps also because of long-branch attraction.

Within the ENaC subfamily, we identified two major subclades of Deg/ENaC channels. One is comprised of the namesake chordate ENaC channels, nematode channels including *C.elegans* MEC10, MEC4, and ASIC-1, a set of channels from the annelid *P.dumerilii*, the singleton placozoan *Tad*NaC10 and *Hho*NaC10 channels, and channels from cnidarians (including *Ne*NaCs) and the poriferan species *S.ciliatum*. The other major ENaC subfamily clade was restricted to bilaterians, comprised of the protostome neuropeptide gated FaNaC and WaNaC channels, and several clades of newly identified FaNaC/WaNaC-related channels that are found broadly in bilaterians. Thus, while the ENaC subfamily itself is likely of pre-bilaterian origin, the FaNaC/WaNaC subclade appears to be a bilaterian innovation. Notably, the phylogenetic relationships within the ENaC family also differ between our tree and the one in the described recent study (8). First, whereas we find the *Ne*NaC channels to form a clade with ENaC, MEC, and *Tad*NaC channels, along with sequences from poriferans and placozoans, in the other tree these form a separate clade from ENaC, nematode, and FaNaC/WaNaC channels, along with *Tad*NaC10.

Furthermore, although the other study also found FaNaC/WaNaC channels to falls within the ENaC superclade, their relationship to other channels within the ENaC subfamily was not resolved.

In this study we provide the first robust phylogenetic evidence that Deg/ENaC channels are not unique to animals, present in unicellular eukaryotic lineages of Heterokonta and Filasterea. Although our search was not exhaustive, we were unable to find Deg/ENaC sequences in the intervening opisthokont lineages between Metazoa and Filasterea, including choanoflagellates, or between these lineages and Heterokonta. This indicates that there was either extensive gene loss of Deg/ENaC channels in eukaryotes, or alternatively, lateral gene transfer between these three groups. Interestingly, similar phylogenetic gaps are apparent for other major cation channels shared between animals and unicellular eukaryotes. For example, the Na^+^ leak channel NALCN and its ancillary subunit FAM155 are conserved between animals and fungi (77), but are absent in all intervening eukaryotic lineages (78). Similarly, Ca_V_3 voltage-gated calcium channels, conserved between animals and choanoflagellates, are also found in algae but not in the vast lineages of eukaryotes that fall between these organisms (78). Hence, it is conceivable that Deg/ENaC channels arose in the metazoan ancestor through lateral gene transfer, perhaps giving rise to the ENaC family, and that the ASIC family evolved thereafter. Alternatively, Deg/ENaC channels first evolved in animals, and these were subsequently transferred to select eukaryotes. This is also conceivable if one considers the P2X receptors that are clearly of premetazoan origin (79) (Fig. 1). These channels have remarkably similar secondary and tertiary structures compared to Deg/ENaC channels (32), and thus, could have given rise to Deg/ENaC channels through extreme sequence divergence but retention of a similar protein architecture. Although this is speculative, it is interesting that numerous clusters of bona fide Deg/ENaC channels formed distinct clusters in our CLANS analysis, perhaps reflecting emerging lineages of divergent channels on the path to having obscured ancestries. Perhaps, the proton-activated PAC chloride channels, which also have a similar secondary and tertiary structures but are restricted to deuterostomes, represent a current example of such specialization, in that these channels do not resemble Deg/ENaC channels (or P2X receptors) in protein alignments, but have very similar trimeric architectures (33, 34).

### On the independent evolution of proton-activated Deg/ENaC channels and the unique functional features of TadNaC2

As noted in the introduction, whereas ASIC channels were once thought to be unique to chordates (44, 80, 81), more recent studies indicate a much broader presence, with the identification of group A and B ASIC channels thought to have emerged in an early bilaterian ancestor (10). In our sequence alignments, all included group A and B ASIC homologues that are proton-sensitive bear the quintessential H73 and K211 residues, considered core determinants for ASIC channel proton-activation (Fig. 6) (25). Also fully conserved are the pair of aromatic amino acids, W287 and Y71 in mouse ASIC1a, which form a hydrophobic bridge between the thumb and wrist regions essential for channel gating (50). Instead, the protonatable glutamate and aspartate residues that make up the acid pocket are absent in group B ASICs, indicating that the acid pocket is a unique feature of group A channels, altogether consistent with the dispensability of the acid pocket for proton activation (26).

Importantly, numerous non-ASIC Deg/ENaC channels have been identified that are also activated by extracellular protons but lack most key residues involved in proton-activation of ASIC channels. This includes *Tad*NaC2, the human ENaC-δ channel (6) and the *C.elegans* channels ACD-2, DEL9, and ASIC- 1 (14), which form homotrimeric channels *in vitro* that conduct slow onset proton-activated currents with minimal desensitization, the *Ne*NaC2 channel from *N.vectensis*, which conducts moderately desensitizing currents *in vitro* (8), and the *D.melanogaster* channel PPK1, which conducts transient, fast desensitizing cation currents in sensory neurons (16). Indeed, all of these non-ASIC channels lack an H73 equivalent residue, and most lack K211 and other important residues within the acid pocket, wrist, and palm regions (Fig. 6). Within the ASIC subfamily, *Tad*NaC2 forms a strongly supported clade with BASIC channels (Fig. 2), which are activated by bile acids but are mildly blocked instead of activated by extracellular protons (82–84). Similarly, *Ne*NaC2 falls within a strongly supported clade of cnidarian channels that includes the *Hy*NaC channels, which are activated by RFamide neuropeptides but are insensitive to protons (21, 22). Within the ENaC subfamily, proton activation of ENaC-δ is a unique feature not observed for other ENaC subunits (*i.e.*, α, β, and γ) (85), while the *C.elegans* channel ASIC-1 forms a subclade with other nematode channels including the mechanically-gated channels MEC4 and MEC10 (86). It is more difficult to make inferences about the evolution of the *C.elegans* ACD-2 channel, which formed a clade that fell outside of the ASIC/ENaC subfamilies, and the *C.elegans* and *D.melanogaster* channels DEL-9 and PPK1, which were excluded from our phylogenetic analysis due to strong sequence divergence. Taken together, these molecular and phylogenetic observations suggest that proton-activation evolved numerous times independently within the Deg/ENaC superfamily, mirroring to the apparent independent evolution of neuropeptide gating of FaNaC/WaNaC and *Hy*NaC channels.

Here, we provide a detailed functional characterization of *Tad*NaC2, revealing a striking functional resemblance to ASIC channels and a conglomeration of isotype-specific biophysical features. With respect to gating, *Tad*NaC2 more resembles ASIC2a in having a lower pH_50_ value of ∼5 compared to ASIC1a and ASIC3 channels, indicative of reduced proton-sensitivity (Fig. 4A and B) (39). Instead, the biphasic macroscopic currents of *Tad*NaC2, consisting of early transient and a late sustained components (Fig. 5A), resemble those of ASIC3, whose biphasic currents are thought necessary for this channel’s dynamic contribution to nociceptive signaling in peripheral neurons (87). Interestingly, these two current components of *Tad*NaC2 appear to be differentially sensitive to amiloride, with 3 µM amiloride causing a sharp reduction in the early current, but no apparent block of the sustained current (Fig. 4E). Lastly, the observed rundown of *Tad*NaC2 currents in response to repeated activation, or tachyphylaxis, is a unique negative feedback feature of ASIC1a channels with an undetermined physiological function (40, 88). Taken together, if we consider *Tad*NaC2 and ASIC channels to have evolved proton-activation independently, then it is remarkable that this placozoan channel converged upon these several distinct and defining properties of the mammalian ASIC channels. In this respect, *Tad*NaC2 provides some unique opportunities for exploring molecular mechanisms for functional convergence of these various biophysical attributes.

Another interesting feature of *Tad*NaC2 is its differential ion selectivity, having an early current that is highly Ca^2+^ permeable and poorly selective for Na^+^ over K^+^ ions, and a late/sustained current that is highly *impermeable* to Ca^2+^ but more Na^+^ selective than the early current (Fig. 5). Dual cation selectivity has also been reported for the early and late current components of the mammalian ASIC3/ASIC2b heteromeric channel (48), as well as the homomeric shark ASIC1b channel (44). However, the monovalent selectivity profiles reported in these studies are reversed relative to *Tad*NaC2, with the transient current being more Na^+^ selective than the sustained current. To the best of our knowledge, here we provide the first report of dual Ca^2+^ selectivity for a Deg/ENaC channel carried by separate early and late current components. This particular feature instantly evokes questions about physiological relevance, where perhaps, an early Ca^2+^ current at the onset of channel activation permits transient cytoplasmic Ca^2+^ signaling in response to extracellular acidification, while the sustained current permits prolonged signaling in the form of membrane depolarization, without excess Ca^2+^ influx which can be cytotoxic when concentrations remain elevated for extended periods (89).

### Unique molecular determinants underlie proton activation of TadNaC2

Our work provides a first account of molecular determinants for proton activation of a non-ASIC Deg/ENaC channel. Focusing our attention on channel regions that are critical for ASIC channel gating enabled us to uncover fundamental insights into how *Tad*NaC2 operates and differs from ASIC channels. In the wrist region, *Tad*NaC2 and *Hho*NaC2 channels bear conserved aromatic residues W276 and F70 which align with the W287 and Y71 residues of ASIC channels (Fig. 6B). Nevertheless, alanine mutation of the F70 residue in *Tad*NaC2, which renders ASIC1a channels non-functional (*i.e.*, Y71A) (50), had no effect on proton-activation (Fig. 7A, D, and E). In ASIC channels, the aromatic interaction between W287 and Y71 is thought to couple conformational changes that occur in extracellular regions such as the thumb and acid pocket, with the first transmembrane helix of each subunit that contributes to the pore in the holomeric channel. This, combined with the absence of an H73 equivalent in *Tad*NaC2, indicates a fundamental difference in gating between *Tad*NaC2 and ASIC channels. This notion was also supported by the E77A mutation in *Tad*NaC2, which unlike mutation of the aligned residue D78 in ASIC1a and ASIC2a channels (45, 47), did not affect proton activation, and the upstream D75A mutation, that enhanced activation by increasing both the pH_50_ and Hill coefficient (Fig. 7A, D, and E). Notably, the cnidarian channel *Ne*NaC2 also lacks an H73 equivalent, but bears a histidine two positions upstream, aligning with residue Y71 in ASIC channels (Fig. 6B). However, mutation of this residue does not disrupt proton activation (8), indicating that like *Tad*NaC2, this channel differs from ASIC channels in lacking core determinants for proton-gating within the wrist region.

In the palm region, we found that the H80 residue in *Tad*NaC2 plays an important role in proton activation, wherein its mutation caused a significant reduction in pH_50_ (Fig. 7C and D). An interesting feature of the H80A mutant channel was its plateaued activation between pH 5.0 and 4.5, which somewhat masked the measured severity of the mutation by imposing a leftward shift in the pH dose-response curve and increasing the variability of the Hill coefficient (Fig. 7C and E). This residue, which is conserved in *Hho*NaC2, is notably absent in all other channels that we included in our analyses (Fig. 6B), representing a unique molecular determinant for proton activation in the placozoan channel. Based on our structural modelling, which of course, should be interpreted with caution, this H80 residue appears proximal to the residues R201 and K203 in *Tad*NaC2, all clustered within the same structural region that harbors K211 in ASIC channels (Fig. 6). However, the contributions of these various cationic residues appear to be different between *Tad*NaC2 and ASIC channels. Alanine mutation of R201 in *Tad*NaC2, which is notably absent in *Hho*NaC2, enhanced proton sensitivity evident in increased pH_50_ and Hill coefficient values (Fig. 7D and E). Furthermore, a K203E mutation did not affect on proton sensitivity (Fig. 7C), while deletion of K203 resulted in channels that did not traffic to the cell membrane and were thus likely non-functional (Fig. 7G to J). This contrasts with ASIC channels, for which a K211E mutation, expected to moderately disrupt molecular interactions with several residues in the adjacent subunit, causes a moderate reduction in proton-sensitivity, while its deletion imposes a more marked effect but does not completely disrupt channel function (25). In ASIC channels, a hub of hydrophobic residues that lies adjacent to K211 (*i.e.*, residues F87, F174, F197, and L207), is thought to functionally couple conformational alterations between the palm and wrist regions during gating. Notably, these hydrophobic residues are strongly conserved, with *Tad*NaC2 having four equivalent hydrophobic residues in corresponding positions in a protein alignment (*i.e.*, residues F86, W165, F188, and L198; Fig. S2).

Altogether, *Tad*NaC2 and ASIC channels exhibit significant structural and functional differences in the palm region, in that the non-homologous H80 residue in *Tad*NaC2 fulfills important functions in proton gating similar to K211 in ASIC channels, with perhaps, a similar role for hydrophobic hub residues coupling this region to the pore.

In the finger region we identify the residues H108 and E105 as important for proton activation of *Tad*NaC2, with single mutations causing moderate reductions in pH_50_, and the triple mutation of E104A/E105A/H108A having the most severe effect of all analyzed channel variants (Fig. 7B and D).

These residues are fully conserved between *Tad*NaC2 and *Hho*NaC2, while H108 is also found in several other Deg/ENaC channels including several ASIC channels, and the E104/E105 doublet is found in *Ne*NaC2 (Fig. 6B). As noted, the finger region is of particular importance for ASIC2 channels where four amino acids differentiate proton-sensitive ASIC2a channel variants from non-functional ASIC2b variants (51). Alanine substitution of the H108 equivalent in ASIC2a (*i.e.*, H109A) produced different effects in separate studies, two describing no effect (46, 51), and the other a complete loss of proton-activation which may have been due to complete loss of membrane expression (47). To the best of our knowledge, whether this equivalent residue in ASIC1a, or a histidine residue one position upstream in ASIC3 (Fig. 6B), contribute to proton activation has not been explored. Altogether, it is difficult to infer whether the finger region might play a broad role in the activation of ASIC and non-ASIC proton-gated channels. However given the importance of the E105 and H109 residues in *Tad*NaC2, and the presence of several protonatable residues in this region of other channels (Fig. 6B), it may be an area for further exploration.

Lastly, during our experiments we noticed that several *Tad*NaC2 mutations caused significant decreases in current amplitude, most marked for the K203Δ variant. We therefore selected the H80A, H108A, and K203Δ variant channels, which produce moderate to absolute reduced current amplitude (Fig. 7F), to explore whether these specific mutations are affecting total protein expression and/or targeting of the channels to the cell membrane. Interestingly, and corroborating the electrophysiological recordings, all three mutations caused a reduction in total and surface/membrane protein expression, with the K203Δ variant being almost undetectable at the cell membrane (Fig. 7 G to J). Thus, in addition to affecting proton sensitivity, all three these mutations also affect total protein expression, perhaps reflecting decreased translation or stability of the single subunit protein. Instead, only the K203Δ variant had a noticeable effect on the apparent ratio of membrane vs. total expression (Fig. 7J). This suggests that this particular mutation imposes instability in homotrimeric assembly, assuming that membrane trafficking requires holochannel formation. Importantly, despite the observed changes in current amplitude, our ability to record current for these variant channels and detect their diminished pH sensitivity indicates that these mutations also affected ligand activation. Indeed, future studies of *Tad*NaC2, including protein misfolding analysis and direct structural analysis, will shed more light on the unique functional features of this intriguing ASIC-like channel.

### T.adhaerens conducts pH sensing behaviours

Our discovery of the pH sensitive *Tad*NaC6 (24) and *Tad*NaC2 channels prompted us to explore whether *T.adhaerens* exhibits behavioral responses to changes in extracellular pH. To our surprise, we found that animals will chemotax away from acidic environments (Fig. 8A), consistent with a pH aversion behavior commonly seen in other animals (60–62). As noted, *T.adhaerens* locomotes through the action of beating cilia on its ventral epithelium (55), which allow it to glide over substrates in the aqueous environment.

Placozoans similarly exhibit negative locomotory responses to light and gravity (59, 90), and positive chemotactic responses to food (algae) and glycine (57, 58), as well as start and stop behaviors (56, 63, 64). A central question is how an animal that lacks body symmetry and a nervous system can generate directed locomotion, especially given that ciliary beating is asynchronous, and that ciliated cells lack overt functional coupling or coordination between them (56, 57, 91). A model that has been put forward is that individual ciliated cells along the ventral epithelium autonomously detect different concentrations of signaling molecules present along a gradient, generating proportional (graded) responses in altered ciliary beating, such that cells located closer to the ligand source would elicit different responses than those that are farther away. This asymmetry would result in a collective semi-directional propulsive force generated by the cilia, that would eventually bring the animal to the source (57). If so, this would require receptors on the ciliated cells that can detect relevant external ligands (*e.g.*, protons) to elicit graded intracellular responses that alter ciliary beating. In other eukaryotic cells including those from animals, ciliary beating can be regulated by various ligand-dependent signaling pathways including those linked to G-protein coupled receptors (GPCRs), as well as calcium signaling through the influx of extracellular Ca^2+^ or release from internal stores (92). Indeed, both *Tad*NaC6 and *Tad*NaC2 can generate sustained currents with graded responses to extracellular pH changes (24) (Fig. 4), and as such, these channels could conceivably contribute to cellular excitation that regulates ventral ciliary beating and chemotactic behavior. A role for Deg/ENaC channels in chemotaxis has been demonstrated in *C.elegans*, in which the proton inhibited channels ACD-1 and DEG are important mediators of acid avoidance behavior (12).

We also discovered that *T.adhaerens* will contract upon fast perfusion of acidic solutions, but not when Ca^2+^ ions are excluded from the external solution (Fig. 8B). Evoked contractions have also been reported for applied glycine and endogenous regulatory peptides (58, 63, 64), however, to the best of our knowledge ours is the first report that extracellular Ca^2+^ is involved in elicited contractions, in this case consistent proton-dependent Ca^2+^ influx, and Ca^2+^-dependent excitation-contraction coupling as occurs in myocytes. Of note, the homology of placozoan cell types to those in other animals, including of myocytes, is still unclear (55, 93). *T.adhaerens* possesses dynamically contractile cells along its dorsal epithelium which mediate contractile behaviors, for which drug-induced rises in cytoplasmic Ca^2+^ have been shown to elicit contractions (65). However, these lack obvious contractile actinomyosin arrangements in electron micrographs (55), and the molecular mechanisms that mediate contraction in these cells is unknown. Nevertheless, our observation that Ca^2+^ influx is required for proton-elicited contractions in *T.adhaerens* suggests ionotropic signaling is important for elicited contractions. By extension, *Tad*NaC2 would be a good candidate to mediate such an ionotropic process, with its early current providing a direct Ca^2+^ signal at the onset of a pH pulse that could in turn trigger the release of internal Ca^2+^ stores via calcium-induced calcium release.

Of course, an important question is whether *Tad*NaC channels are expressed in ventral ciliated cells and/or contractile dorsal epithelial cells, and directly contribute to the described proton-sensing behaviors. Previously, using fluorescent RNA probes, we found *Tad*NaC6 mRNA expression to be enriched along the outer edge of the animal (24), in a region bearing neuroendocrine-like gland cells thought to secrete ligands that regulate cell activity and behavior (55, 94). Instead, *Tad*NaC2 exhibited a more centralized expression, consistent with the location of ciliated cells and digestive lipophil cells located in the ventral epithelium, or contractile cells in the dorsal epithelium (24). Furthermore, a single cell transcriptome study of *T.adhaerens* identified expression of *Tad*NaC2, together with *Tad*NaC3 and *Tad*NaC8, in lipophil cells (93). Furthermore, *Tad*NaC10, for which we have so far been unable to record currents *in vitro*, was detected in ciliated epithelial cells. Notably, although an important step forward, the sequencing depth in this study was limited to roughly 100 genes per cell type, precluding a complete assessment of *Tad*NaC channel cellular expression. As the depth of single-cell sequencing improves, we will hopefully develop a clearer understanding of the unique cellular expression patterns of all *Tad*NaC channels, including their cellular co-expression. This, in turn, would facilitate the study of heteromeric *Tad*NaC channels that might be active *in vivo*. Another crucial step forward will be the development of tools that permit manipulation of *T.adhaerens* genes *in vivo*, such as gene knockout, gene editing, and ectopic overexpression, required for assessing the direct contributions of the different *Tad*NaC channels to physiology and behavior.

### C-terminal interactions with filamin point to ancient and conserved associations with the cytoskeleton

Our yeast 2 hybrid screen revealed that the C-termini of both *Tad*NaC2 and *Tad*NaC10 interact with the shorter of two filamin paralogues possessed by *T.adhaerens in vitro*. Both also interacted with a U5 small nuclear ribonucleoprotein helicase, while *Tad*NaC2 interacted with additional proteins including the scaffolding protein VAC14 and the cell adhesion protein contactin-6 (Table S1). Interestingly, both *Tad*NaC2 and *Tad*NaC10 interacted with the C-terminus of filamin, like the human ENaC channel α and β subunits that interact with the C-terminus of human filamins A, B, and C (27). Of course, the interactions reported here should be interpreted with some restraint since we did not verify cellular co-expression of these channels with the short filamin paralogue, nor the *in vitro* interaction itself by other means such as co-immunoprecipitation. Nonetheless, it is worth noting that we have previously used our *T.adhaerens* yeast 2 hybrid library to screen numerous other proteins, including other ion channel C-termini, and have never encountered any of the hit sequences identified here, including filamin.

Furthermore, both *Tad*NaC2 and *Tad*NaC10 interacted with the last two immunoglobulin-like repeats of the short filamin paralogue, suggesting that this region of the filamin protein bears specific binding motifs for interacting with these two channels. For mammalian filamins, the C-terminal immunoglobulin-like repeats are a hotbed for interactions with various ion channels and receptors (95), which in addition to ENaC channels, includes the voltage-gated potassium channel K_V_4.2 (96) and the D2 dopamine receptor (97). Lastly, the interaction between *Tad*NaC2 and filamin was the single most frequent hit in our screen, suggesting that this interaction is robust.

Based on our phylogenetic tree, the *T.adhaerens* short filamin paralogue is orthologous to the *D.melanogaster* filamin Jitterbug, and the *C.elegans* filamin FLN-2 (Fig. 9B). This clade, which we refer to as type 2 filamins, includes a filamin homologue from *N.vectensis* that has a particularly long branch and hence questionable relationship to the other type 2 sequences. This notion is also supported by the absence of three N-terminal calponin homology domains that are present in all other type 2 orthologues (Fig. 9A), an atypical structural feature not seen in other actin-binding proteins including type 1 filamins (73). All four of these invertebrate species also possess a type 1 filamin paralogue, which on our phylogenetic tree form a clade with the human filamin A, B, and C paralogues. Although our tree is by no means exhaustive, the presence of both type 1 and type 2 filamins in placozoans and ecdysozoans suggests these paralogues predate bilaterians, and that somewhere along the human lineage, filamin 2 was lost and filamin 1 underwent expansion via gene duplication (98). By extension, although Deg/ENaC channels from both humans and placozoans interact with filamins, the interaction for human ENaC channels is with the type 1 filamins, while for the *Tad*NaC channels it is with type 2 filamins. Nonetheless, since *Tad*NaC10 and *Tad*NaC2 belong to the two separate subfamilies of Deg/ENaC channels, it is possible that associations with filamins have ancient origins, perhaps predating the emergence of the two filamin paralogues.

In cells, filamins form dimers with each other at their C-termini, and bind actin filaments at their N-termini, where their general function is to dynamically remodel the actin cytoskeleton through the formation of orthogonal actin networks, and tether the cytoskeleton to various receptors and ion channel expressed at the cell membrane (95). In various cellular contexts, filamins also contribute to cytoskeletal complexes capable of sensing forces acting on the cell membrane and cytoskeleton, effectively acting as mechanotransducers (29, 99). In *D.melanogaster*, Cheerio (type 1 filamin) is involved in mechanotransduction during egg development (100), and tissue remodelling of the epithelium during development (101), while Jitterbug (type 2 filamin) regulates axon migration and targeting during neural development (102), and the formation of actin-rich protrusions (bristles) on the epithelium (68). In *C.elegans*, FLN-1 is involved in actin remodelling in the spermatheca during reproduction (103, 104), and FLN-2 is involved in biogenesis of multivesicular bodies that associate with lysosomes (105), and interestingly, its loss of function increases life span in animals that experience pharyngeal infections (106).

Indeed, the interaction of *Tad*NaC and ENaC channels with filamin is tantalizing because interactions of Deg/ENaC channels with elements of the cytoskeleton can render them mechanically gated. Indeed, Deg/ENaC channels were first discovered in *C.elegans* (i.e., MEC-4 and MEC-10) as mechanically-gated channels that interact with cytoskeletal and extracellular matrix proteins in mechanosensitive sensory neurons (1). Interestingly, both ENaC channels and ASIC channels are also mechanically sensitive. *In vitro*, mammalian ENaC subunits form homo- and heterotrimeric channels that are activated by shear stress (*i.e.*, flow of fluid over the cell membrane) (4). *In vivo*, the mechanosensitivity of these channels is implicated in dynamic contractility responses of vascular smooth muscle and endothelial cells that control blood pressure (107, 108). Similarly, ASIC channels currents that are elicited by proton and non-proton ligands are potentiated by shear force *in vitro* (5), and various lines of evidence indicate that ASIC channels act as mechanotransducers *in vivo*, both in tactile and nociceptive sensory neurons and in vascular tissues similar to ENaC channels (1). Currently, it is unknown whether the observed mechanosensitivity of ASIC and ENaC channels requires association with the cytoskeleton, similar to the *C.elegans* MEC-4/MEC-10 channels (1, 109). However, given that both filamin and ENaC channels are directly involved in mechanotransduction, it is certainly tempting to speculate that their physical association is of relevance in this regard. Whether this might also be true for the placozoan *Tad*NaC channels, and other metazoan Deg/ENaC channels for that matter, is certainly an interesting and worthwhile question to consider.

## Materials and Methods

### Transcriptomic resources

To identify in which phyla to search for ASIC and Deg/ENaC channels, we performed an initial BLASTp analysis in the NCBI databases in the following taxa: Picozoa (taxid:419944), Ancoracysta (taxid:2056028), Rhodophyta (taxid:2763), Chloroplastida (taxid:33090), Glaucophyta (taxid:38254), Palpitomonas (taxid:759891), Katablepharida (taxid:339960), Cryptophyta (taxid:3027), Centrohelida (taxid:193537), Haptophyta (taxid:2830), Telonemida (taxid:589438), Discoba (taxid:2611352), Metamonada (taxid:2611341), Malawimonadidae (taxid:136087), Collodictyonidae (taxid:190322), Mantamonadidae (taxid:1238961), Breviatea (taxid:1401294), Amoebozoa (taxid:554915), Apusomonadida (taxid:172820), Rhizaria (taxid:543769), Alveolata (taxid:33630), Stramenopiles (taxid:33634), Fungi (taxid:4751), and Holozoa (taxid:33208). The ASIC and Deg/ENaC receptors from human, *Platynereis dumerilii*, *Caenorhabditis elegans*, and *Aplysia californica* were used as queries (see file S1 for the list of these queries).

Candidate sequences were identified in holozoans (multiple species), Alveolata (one species) and Stramenopiles (one species). Thus, whole transcriptomes of predicted proteomes from different classes of holozoans, including: metazoans, choanoflagellates, one filasterean, and *Tunicaraptor unikontum* were obtained. The more distantly related eukaryotic species, *Symbiodinium sp KB8 (*Alveolata) and *Cafeteria roenbergensis* (Stramenopiles) were also added to the list. The transcriptomic databases were translated to protein using the tool TransDecoder (http://transdecoder.github.io/) with a minimum length of 75 amino acids. The databases that were available as predicted proteins were used directly. For completeness assessment of the transcriptomes, we ran BUSCO v5.2.1 (110) in protein mode and with the lineage set to ‘eukaryote’ with the database ‘eukaryota_odb10’ (database creation: April 2022; number of BUSCOs: 255). The source of the database used for this analysis, and the results of the completeness analysis is available in Table S2.

### Clustering and phylogenetic analyses of Deg/ENaC channel protein sequences

Multiple-species sequences of the families Deg/ENaC (PF00858) and P2X (PF00864) were obtained from the PFAM database (https://pfam.xfam.org). For the proton-activated chloride (PAC) channels, we identified chordate sequences in the NCBI non-redundant database using a BLASTp search with the human PAC protein as query (UniProt accession number Q9H813). These sequences were aligned using MUSCLE, and automatically trimmed with trimAl (111) using the gappy-out mode. The trimmed sequences were used to produce Hidden Markov Models using HMMR3 (112). The subsequent search for receptors in the database obtained as described above was performed using HMMER3 with an expect value cut-off of 1E-10 (113). All identified sequences are provided in file S2.

The relationship between these protein families was analyzed first using a cluster-based strategy with the CLuster ANalysis of Sequences CLANS algorithm (30). The sequences were clustered with an expect value of 1E-40. The ASIC and Deg/ENaC receptors are part of an easily identifiable supercluster that contains the ASIC and ENaC channels, different from the P2X receptors. Thus, the ASIC and ENaC superclusters were pooled and obtained for the phylogenetic analysis of sequences. These selected sequences were first analyzed using Phobius (114) to predict the transmembrane domains (TMDs), and sequences with a minimum of one TMD were kept in order to remove sequences that are too fragmented to contribute to the phylogenetic analysis. Selected and filtered protein sequences were aligned with MAFFT version 7, with the iterative refinement method L-INS-I (115), and alignments were trimmed with ClipKIT automated mode (116). The maximum-likelihood trees were generated using IQ-TREE 2 (117) with the best-fit model LG+F+G4. To calculate branch support, we ran 1,000 replicates with the aLRT-SH-like and aBayes methods (117) (https://paperpile.com/c/LYvsPL/D2o0G). The tree was coloured and analyzed in Figtree v1.4.4 (118). The sequences used for the phylogeny, and aligned- trimmed sequences are available in files S3 and S4.

### Cloning of Deg/ENaC channel cDNAs for in vitro expression

The *Tad*NaC2 and *Tad*NaC10 cDNAs were cloned from a cDNA library prepared from whole-animal total RNA, as reported previously (24, 71). Briefly, corresponding cDNAs were amplified by nested PCR using two sets of primers for each channel, with the second forward primer incorporating an *Xho*I restriction site followed by a Kozak translation initiation sequence flanking the channel start codon, and the second reverse primer incorporating a *BamH*I restriction site followed by a stop codon (Table S3).

PCR reactions were done with the Phusion high fidelity DNA polymerase (New England Biolabs Canada, Whitby Ontario) using standard protocols. The included restriction sites permitted cloning into both the pIRES2-EGFP and pEGFP-C1 mammalian expression vectors (Clontech), the former a bicistronic vector expressing enhanced green fluorescence protein (EGFP) from the same transcript bearing the *Tad*NaC channel coding sequence (through an internal ribosome entry site), and the latter expressing the channel with EGFP fused to its N-terminus. These plasmid vectors were respectively named p*Tad*NaC2-IR-EGFP and pEGFP-*Tad*NaC2. The various mutations of *Tad*NaC2 were generated via site directed mutagenesis using the primers listed in Table S3, using a modified PCR protocol of only 15 cycles, after which the PCR reactions were each treated with 1 µL of the restriction enzyme *Dpn*I, then transformed into the *E.coli* NEB Stable strain from New England Biolabs (NEB, MA, USA) for plasmid isolation. All generated plasmid vectors were verified through diagnostic restriction digestion followed by Sanger DNA sequencing. The mouse ASIC1a subunit was synthesized by GenScript (new Jersey, USA) with *Sac*I and *BamH*I restriction sites flanking the start and stop codons, and a 5’ Kozak sequence flanking the start codon, which was cloned into the bicistronic vector pIRES2-EGFP to provide the construct pmASIC1-IR- EGFP.

### In vitro expression and electrophysiology

Chinese Hamster Ovary cells (CHO-K1; Sigma) were cultured and transfected in vented T-25 flasks at 37°C in a 5% CO_2_ incubator. Cells were grown in a DMEM/F12 and DMEM/Ham’s F-12 50/50 MIX media supplemented with 10% fetal bovine serum (both from Wisent Inc., Saint-Jean-Baptiste, Quebec). For *in vitro* expression, 3 μg of the p*Tad*NaC2-IR-EGFP vector, or 2μg of the pmASIC-IR-EGFP vector, were transfected into CHO-K1 cells grown to 80-90% cell confluency using the transfection reagent PolyJet^TM^ (FroggaBio, Concord Ontario), according to the manufacturer protocol. Cells were incubated with the transfection mix for 6 hours, and then washed three times with serum-free media, followed by an overnight incubation at 37°C. For electrophysiological recordings, cells were briefly treated with a trypsin solution (Sigma-Aldrich), plated onto glass coverslips in 35-mm tissue culture-treated dishes (Eppendorf, Mississauga Ontario), and then left to adhere for a minimum of 6 hours before recording.

Before performing the patch-clamp electrophysiology experiments, cells on glass coverslips were transferred into 35-mm cell culture dishes (Eppendorf) containing ∼3 mL of extracellular recording solution. For all pH-dose response curve experiments, the external solution contained 140 mM NaCl, 4 mM KCl, 2 mM CaCl_2_, 1 mM MgCl_2_, 5 mM HEPES, and 5 mM MES (pH 3.5 to 7.5 with HCl/NaOH; osmolarity set to 320 mOsM with glucose), while the internal solution contained 120 mM KCl, 2 mM MgCl_2_, 10 mM EGTA, 10 mM HEPES (pH 7.2 with KOH; osmolarity set to 300 mOsM with glucose).

For the Na^+^/K^+^ ion selectivity experiments, the external solution contained 150 mM XCl (where X=Na^+^ or K^+^), 10 mM TEA-Cl, and 10 mM of MES buffer (pH4.5 with TEA-OH). The internal solution contained 150 mM NaCl, 10 mM EGTA, 10 mM TEA-Cl, and 10m M HEPES (pH 7.2 with TEA-OH). The Na^+^/Ca^2+^ permeation experiments were done using three external solutions: 1) 10 mM CaCl_2_, 123 mM NMDG-Cl, 10 mM TEA-Cl, 10 mM MES; 2) 10 mM CaCl_2_, 123 mM NaCl, 10 mM TEA-Cl, 10 mM MES; 3) 123m M NaCl, 15 mM NMDG-Cl, 10 mM TEA-Cl, 10 mM MES (all solutions were set to pH 4.5 using TEA-OH). The intracellular solution contained 143 mM CsCl, 10 mM EGTA, 10 mM TEA-Cl, and 10 mM HEPES (pH7.2 with Cs-OH). Pharmacology experiments with amiloride were done by first perfusing standard pH 4.5 or pH 5.5 solutions for *Tad*NaC2 and mASIC1a respectively, followed by the same solution but containing 3 mM amiloride. Whole-cell patch clamp recordings were done on an inverted fluorescent microscope, with recording electrodes coupled to an Axopatch 200B amplifier interfacing with a Digidata 1550A data acquisition digitizer controlled on a personal computer using the pClamp 10 software (Molecular Devices, California USA). Patch pipettes were made from thick-walled borosilicate glass (1.5 x 0.86mm outer and inner diameter, respectively) pulled using a P-1000 micropipette puller (Sutter, Novatto California) with a resistance in the bath solution of 2 to 6 megaohms. All recordings were sampled at 2000 Hz and filtered at 500 Hz using the pClamp10 software (Molecular Devices). Perfusion of the various solutions was achieved using a Valvelink8.2 gravity flow perfusion system (AutoMate Scientific, Berkeley CA). Dose-response curves were fitted using monophasic exponential function using the program Origin 2017 (OriginLab), which produced pH_50_ (EC_50_) and Hill coefficients (*n*_H_) according to equation 1 below. The relative permeability of *Tad*NaC2 to Na^+^ and K^+^ ions (i.e., pNa^+^/pK^+^) was calculated using equation 2 below, where *E_Rev,K_*and *E_Rev,Na_* denote the reversal potentials observed with K^+^ and Na^+^ present in the external solution, respectively.

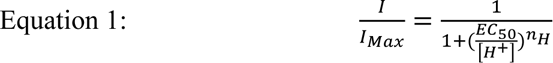

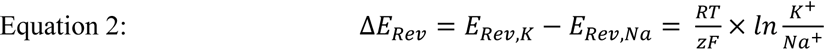

### Structural analyses

For prediction of the *Tad*NaC2 homotrimeric structure, a monomeric structure was first generated using Phyre2 (119) run in intensive mode. The ternary complex was assembled using the three-fold rotational symmetry of the *Gallus gallus* ASIC1a channel structure in its resting state (PDB number 6AVE) (53). Energy minimization, to minimize steric clashes, and geometry optimization were performed using Chiron (120) and Phenix (121), respectively. Prediction of the *Tad*NaC2 monomeric structure was achieved with the AlphaFold algorithm (54) available on the Google Colab webserver. All structural annotations were prepared using the visualization software ChimeraX version 1.3 (122).

### Imaging and quantification of EGFP fluorescence

CHO-K1 cells were transfected and imaged in 6 mL vented T-25 flasks on an Axio Observer 3 inverted microscope (Zeiss Canada, Toronto Ontario) bearing an LED light source and appropriate fluorescence filters. Micrographs were acquired through a 10x objective using a mounted Axiocam 506 camera with the ZEN lite software (Zeiss), selecting fully confluent areas of each dish. Fluorescence quantification was done via integrated density analysis of the acquired micrographs using ImageJ (123).

### Quantification of total and surface expressed TadNaC2 channel protein in vitro

The Pierce^TM^ Cell Surface Protein Biotinylation and Isolation Kit (Thermo Fisher Scientific, Massachusetts USA) was used to quantify total and cell surface expression levels of N-terminal GFP-tagged *Tad*NaC2 protein expressed in CHO-K1 cells from the pEGFP-*Tad*NaC2 vector, as well as GPG- tagged *Tad*NaC2 variants bearing H80A, H109A, and K203Δ mutations. Briefly, for each *Tad*NaC2 variant, two T-75 flasks bearing CHO-K1 cells at ∼80% confluency were each transfected with 9 μg of DNA using the PolyJet^TM^ reagent. The flasks were the washed with 10 mL of phosphate buffered saline (PBS; MilliporeSigma, Ontario Canada), then 5 mL of a PBS solution bearing 0.25μg/ml EZ-Link^TM^ Sulfo-NHS-SS-Biotin was applied over the cells and these were incubated on ice for 30 minutes. This solution was removed by aspiration, then cells were washed twice with 10 mL of ice-cold Tris Buffered Saline (TBS; 150 mM NaCl, 50 mM Tris-Cl pH 7.5) and suspended in 5 mL TBS using a cell scraper. For each *Tad*NaC2 variant, corresponding sets of suspended cells were combined and lysed with 500 µL of the provided lysis buffer supplemented with Protease Inhibitor Cocktail I (MilliporeSigma). The protein lysates were clarified by centrifugation at 10,000 x g for 2 minutes, transferred to a new tube, then quantified using the Pierce™ BCA Protein Assay Kit (Thermo Fisher). The volumes of all samples were adjusted with lysis buffer to dilute them to the same concentration (5 µg/µL), then 50 µL of each lysate was set aside, representing the total *Tad*NaC2 protein component. Membrane expressed *Tad*NaC2 proteins were isolated by applying 600 µL of the clarified protein lysates to columns loaded with the 250 µL of the provided NeutrAvidin^TM^ Agarose slurry, followed by a wash using the provided buffer, and elution with 200 µL of the provided SDS-PAGE sample elution buffer. For each experiment, 40 μg (8 µL) of the total lysate and 40 μL of the isolated surface proteins, which corresponds to a 5-fold enrichment of surface relative to total protein fractions, were loaded onto a 4-12% Bis-Tris gel for Western blotting.

For Western blotting, protein samples were first electrophoretically separated using a 4-12% Bis-Tris acrylamide gel (Invitrogen, Massachussets USA), and transferred onto nitrocellulose membrane using a wet transfer system at 25 mV overnight. Membranes were then washed with TBS containing 0.05% v/v Tween-20 (TBS-T), then incubated for 1 hour in TBS-T with 5% w/v skim milk powder at room temperature. The membrane was then washed with TBS-T and incubated at 4°C overnight in TBS-T plus 5% milk and rabbit monoclonal anti-EGFP antibody (1:1,000 dilution) and rabbit monoclonal anti-GAPDH antibody (1:50,000 dilution; both from Cell Signalling Technology, Massachusetts USA). The membranes were then incubated for 2 hours at room temperature in TBS-T containing 5% milk and a 1:3000 dilution of goat anti-rabbit secondary antibodies conjugated to horseradish peroxidase (Cell Signalling). Imaging of the membrane done by applying 5 mL of Clarity Western ECL Substrate (Bio- Rad, Califormia USA) using an IBright^TM^ FL1500 Imager System (Thermo Fisher). Protein band intensity was quantified via mean gray area analysis with the software ImageJ.

### Behavioral analyses

To analyze the locomotor activity and pH preference of *T.adhaerens* animals, we first prepared buffered solutions bearing equimolar (165 mM) MES, HEPES, and CHES buffers (Sigma-Aldrich, St. Louis, MO, USA) supplemented with a 1:1,000 dilution of the pH indicator phenol red (GIBCO, Waltham, MA, USA). The buffer mixture was separated into three portions, each of which was pH-adjusted to one of the following values: 5.0, 7.5 and 9.0. These pH values were visualized by phenol red coloration as yellow, peach, and purple, respectively. Each buffer mixture was then mixed with SeaKem®GTG agarose (2% m/v; Life Technologies, Gaithersburg, MD, USA) and gelled in a Petri dish so that the thickness of the gel layer was about 1 mm. Behavioral tests were carried out in one well Lab-Tek II coverslip chambers (Thermo Fisher Scientific, Waltham, MA, USA). We cut out strips of each of the three buffered gels with lengths equal to the chamber length, and widths equal to one third of the chamber width. When placed inside the chamber, three strips covered completely the bottom of the chamber but did not overlap (with the pH 5.0 strip at the top along a horizontal axis, pH 7.0 in the middle, and pH 9.0 at the bottom). Once the buffered gel strips were in place, we poured ASW (*i.e.*, 32 ppt Instant Ocean salt, Instant Ocean, Virginia USA) with low melting point agarose (2% m/v; FMC BioProducts, Rockland, ME, USA) to create a ∼1 mm layer of gelled ASW on top. 10 ml of ASW was then added to the chamber, and 4-5 *T.adhaerens* animals were placed along the midline of the chamber (12 replicates), such that all of them were situated over the pH 7.5 strip at the beginning of the experiment. Once the animals attached to the gelled ASW substrate, time lapse recording was initiated at 1 shot every 30 sec, using a Canon EOS 5D Mark III 22.3 MP Digital Camera equipped with a Canon MP-E 65mm f/2.8 1-5x Macro Photo lens. EOS Utility 2.13.40.2 software was used to control the camera and collect images. Lighting of the chambers for imaging was achieved using a precision light box of adjustable intensity and heat control (Northern Light Technologies, Inc., Canada). Animal movement was recorded for a total of three hours, and the obtained images were transferred to ImageJ software and processed with the Manual Tracking plugin that registers the location, velocity, and travelled distance of each individual at each time point (123).

For the *T.adhaerens* contraction experiments, we used solutions consisting of 433.2 mM NaCl, 9.6 mM KCl, 8.7 mM CaCl_2_, 28.9 mM MgCl_2_, 15.4 mM MgSO_4_, 20 mM HEPES, and 20 mM MES, with different pH set to 7.8, 5.5, 4.5, 3.5, 4.5 with HCl. We also prepared a pH 4.5 Ca^2+^-free solution in which the 8.7 mM CaCl_2_ was replaced with 13 mM NMDG-Cl. The osmolality for all solutions was adjusted with a 1 molar glucose solution to a range of 953 to 990 mOsm, to roughly match the osmolality of the 32 ppt Instant Ocean ASW solution. *T.adhaerens* animals were placed on 35 mm culture dishes (Eppendorf) containing the pH 7.8 ASW solution, and imaged using the same microscope system used for imaging CHO-K1 cells as indicated above, but with a 2x objective. Animals were left to settle for 30 minutes, after which the pH 7.8 solution was perfused over the animals for 110 seconds, followed by the various acidic solutions for another 110 seconds. Captured videos for each pH application were separated into individual frames using FFmpeg tool (124), and the surface area of each animal was quantified using ImageJ (123).

### Yeast 2-hybrid screening

A yeast two-hybrid prey cDNA library of *Trichoplax* whole animal RNA was prepared in the pGADT7- AD plasmid vector using the Make Your Own “Mate & Plate™” Library System (Takara Bio, California USA) according to the manufacturer’s instructions. To extract *Trichoplax* RNA, 35 mg of animals were transferred into a 1.5 mL tube and total RNA was extracted using a Nucleospin® RNA Plus Kit (Macherey-Nagel, Düren, Germany). The prey cDNA library was screened for interactions with the 70 distal C-terminal amino acids of the *Tad*NaC2 and *Tad*NaC10 channels expressed in yeast from the bait vector pGBK-T7. To generate the bait plasmid constructs, the selected coding sequences of each channel were PCR amplified using the corresponding pIRES2-EGFP plasmids as template and corresponding forward and reverse pGBK primer sets (Table S2). These primers incorporated an *Eco*RI and restriction site at the 5’ end of the PCR product, and a *Sal*I flanking the 3’ stop codon, permitting cloning into the empty pGBK-T7 plasmid using the same enzyme sites. These constructs were then transformed into the yeast strain Y2HGold, and cells were plated on SD/-Trp selection plates. Toxicity and autoactivation of the bait proteins was tested on SD/-Trp/X-α-GAL and SD/-Trp/X-α-GAL/Aureobasidin A (AbA) plates, respectively. For all media, X-α-Gal and AbA were at respective concentrations of 40 µg/mL and 125 ng/mL. Yeast 2-hybrid library screening was conducted using the Matchmaker Gold Yeast Two-Hybrid System (Takara Bio USA) according to the manufacturer’s instructions. Briefly, single colonies of each transformed Y2HGold bait strains were innoculated into 50 mL of SD/Trp- liquid medium and grown overnight at 37°C, and these were concentrated by centrifugation and resuspension in 5 mL of SD/Trp- liquid media. Concentrated yeast cell were then each combined with a 1 mL aliquot of the *Trichoplax* cDNA prey library and 45 mL of 2X yeast extract peptone dextrose liquid media with adenine (*i.e.*, 2X YPDA broth), and allowed to mate overnight at 30°C. Diploids were selected on plates bearing SD/-Leu/- Trp/X-α-GAL/AbA (DDO/X/A) medium, permitting growth of blue colonies representing transformants with putative bait-prey inetractions. Positive clones from the DDO/X/A plates were then streaked on maximally selective SD/-Leu/-Trp/-Ade/-His/X-α-GAL/AbA (QDO/X/A) plates to confirm true positives. Colony PCRs were conducted using primers flanking MCS of pGADT7 to amplify the prey sequences from the positive colonies on QDO/X/A plates. PCR products were then subjected to DNA electrophoresis, from which amplified DNA products were purified using the Monarch DNA Gel Extraction Kit (NEB) and analyzed via Sanger DNA sequencing.

### Bioinformatic analysis of filamin protein sequences

The Human, *C.elegans*, and *D.melanogaster* filamin protein sequences were manually extracted from the UniProt database, and these were used to find corresponding entries in the NCBI Reference Sequence database. The *D.melanogaster* Cheerio and Jitterbug filamin sequences (68) and the human filamin-A protein sequence (69) were then used to research through transcriptome databases available for *T.adhaerens* (71) and *H.honkgongensis* (72), and a protein database available for *N.vectensis* (125), using appropriate BLAST algorithms available in the BLAST+ suite (126). Candidate filamin sequences where then verified as true filamin homologues via reciprocal BLAST against the UniProt database, Smart BLAST analysis (127), and domain prediction with InterProScan (128). The maximum likelihood phylogenetic tree was inferred with the program IQ-TREE2 (117) from a MUSCLE protein alignment (129) that had been trimmed with trimAl using a gap threshold of 0.6, and a best-fit model of LG+F+R4. Node support values were generated though 1,000 ultrafast bootstrap replicates. All sequences used in the analysis are available in file S5.

## Acknowledgements

We would like to thank Dr. Carolyn Smith for early discussions about behavioral pH experiments on *T.adhaerens*.

## Author Contributions

Conceived the study: A.S and W.E; molecular biology: W.E., A.S, M.P., A.S., and J.G.; electrophysiology: W.E.; Bioinformatics analyses: L.Y.G, A.S.; Structural analyses: M.C., W.E., A.S; Original draft of manuscript: W.E. and A.S.; revision of manuscript: all authors.

---

Figure S1. Expanded phylogenetic tree.

Figure S2. Protein sequence alignment reveals conserved aromatic residues that contribute to the hydrophobic hub in ASIC channels.

