## Supplementary figures and images for "Function and phylogeny support the independent evolution of acid-sensing ion channels in the Placozoa"

### Figure S1

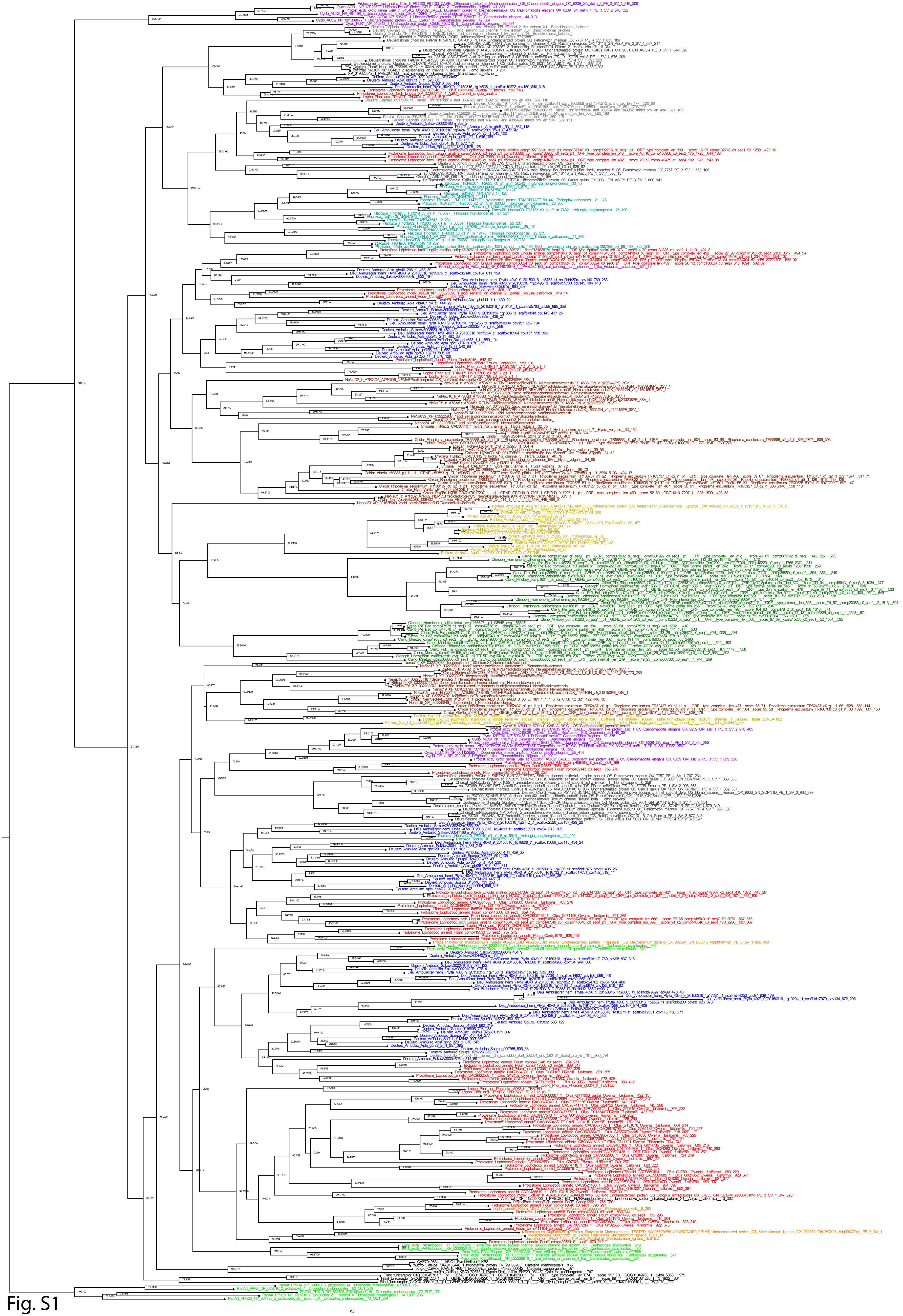

Fig. S1

### Figure S2

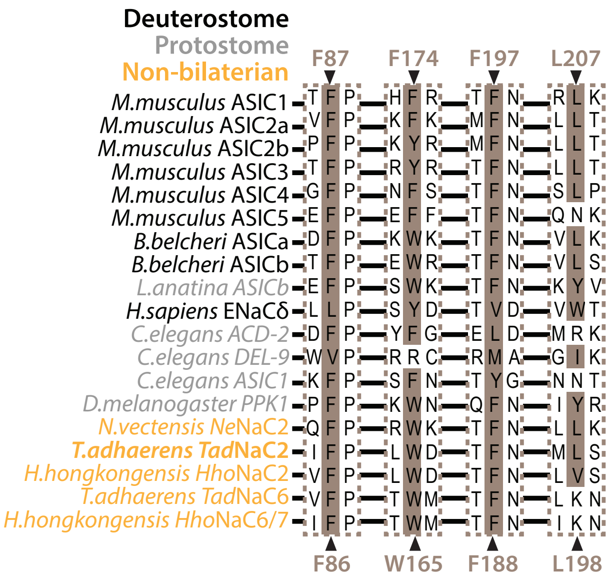
