## Supplementary material for "Function and phylogeny support the independent evolution of acid-sensing ion channels in the Placozoa": FIle S1

>Chordat_hENaCalpha_NP_001029_1_amiloride_sensitive_sodium_channel_subunit_alpha_isoform_1__Homo_sapiens_

MEGNKLEEQDSSPPQSTPGLMKGNKREEQGLGPEPAAPQQPTAEEEALIEFHRSYRELFEFFCNNTTIHGAIRLVCSQH
NRMKTAFWAVLWLCTFGMMYWQFGLLFGEYFSYPVSLNINLNSDKLVFPAVTICTLNPYRYPEIKEELEELDRITEQTLF
DLYKYSSFTTLVAGSRSRRDLRGTLPHPLQRLRVPPPPHGARRARSVASSLRDNNPQVDWKDWKIGFQLCNQNKSDCF
YQTYSSGVDAVREWYRFHYINILSRLPETLPSLEEDTLGNFIFACRFNQVSCNQANYSHFHHPMYGNCYTFNDKNNSNL
WMSSMPGINNGLSLMLRAEQNDFIPLLSTVTGARVMVHGQDEPAFMDDGGFNLRPGVETSISMRKETLDRLGGDYG
DCTKNGSDVPVENLYPSKYTQQVCIHSCFQESMIKECGCAYIFYPRPQNVEYCDYRKHSSWGYCYYKLQVDFSSDHLGC
FTKCRKPCSVTSYQLSAGYSRWPSVTSQEWVFQMLSRQNNYTVNNKRNGVAKVNIFFKELNYKTNSESPSVTMVTLLS
NLGSQWSLWFGSSVLSVVEMAELVFDLLVIMFLMLLRRFRSRYWSPGRGGRGAQEVASTLASSPPSHFCPHPMSLSLS
QPGPAPSPALTAPPPAYATLGPRPSPGGSAGASSSTCPLGGP

>Chordat_hENaCbeta_NP_000327_2_amiloride_sensitive_sodium_channel_subunit_beta__Homo_sapiens_

MHVKKYLLKGLHRLQKGPGYTYKELLVWYCDNTNTHGPKRIICEGPKKKAMWFLLTLLFAALVCWQWGIFIRTYLSWE
VSVSLSVGFKTMDFPAVTICNASPFKYSKIKHLLKDLDELMEAVLERILAPELSHANATRNLNFSIWNHTPLVLIDERNPH
HPMVLDLFGDNHNGLTSSSASEKICNAHGCKMAMRLCSLNRTQCTFRNFTSATQALTEWYILQATNIFAQVPQQELVE
MSYPGEQMILACLFGAEPCNYRNFTSIFYPHYGNCYIFNWGMTEKALPSANPGTEFGLKLILDIGQEDYVPFLASTAGVR
LMLHEQRSYPFIRDEGIYAMSGTETSIGVLVDKLQRMGEPYSPCTVNGSEVPVQNFYSDYNTTYSIQACLRSCFQDHMI
RNCNCGHYLYPLPRGEKYCNNRDFPDWAHCYSDLQMSVAQRETCIGMCKESCNDTQYKMTISMADWPSEASEDWI
FHVLSQERDQSTNITLSRKGIVKLNIYFQEFNYRTIEESAANNIVWLLSNLGGQFGFWMGGSVLCLIEFGEIIIDFVWITIIK
LVALAKSLRQRRAQASYAGPPPTVAELVEAHTNFGFQPDTAPRSPNTGPYPSEQALPIPGTPPPNYDSLRLQPLDVIESD
SEGDAI

>Chordat_hENaCgamma_NP_001030_2_amiloride_sensitive_sodium_channel_subunit_gamma__Homo_sapiens_

MAPGEKIKAKIKKNLPVTGPQAPTIKELMRWYCLNTNTHGCRRIVVSRGRLRRLLWIGFTLTAVALILWQCALLVFSFYT
VSVSIKVHFRKLDFPAVTICNINPYKYSTVRHLLADLEQETREALKSLYGFPESRKRREAESWNSVSEGKQPRFSHRIPLLIF
DQDEKGKARDFFTGRKRKVGGSIIHKASNVMHIESKQVVGFQLCSNDTSDCATYTFSSGINAIQEWYKLHYMNIMAQV
PLEKKINMSYSAEELLVTCFFDGVSCDARNFTLFHHPMHGNCYTFNNRENETILSTSMGGSEYGLQVILYINEEEYNPFLV
SSTGAKVIIHRQDEYPFVEDVGTEIETAMVTSIGMHLTESFKLSEPYSQCTEDGSDVPIRNIYNAAYSLQICLHSCFQTKM
VEKCGCAQYSQPLPPAANYCNYQQHPNWMYCYYQLHRAFVQEELGCQSVCKEACSFKEWTLTTSLAQWPSVVSEK
WLLPVLTWDQGRQVNKKLNKTDLAKLLIFYKDLNQRSIMESPANSIEMLLSNFGGQLGLWMSCSVVCVIEIIEVFFIDFF
SIIARRQWQKAKEWWAWKQAPPCPEAPRSPQGQDNPALDIDDDLPTFNSALHLPPALGTQVPGTPPPKYNTLRLERA
FSNQLTDTQMLDEL

>Chordat_hASIC1_NP_064423_2_acid_sensing_ion_channel_1_isoform_a__Homo_sapiens_

MELKAEEEEVGGVQPVSIQAFASSSTLHGLAHIFSYERLSLKRALWALCFLGSLAVLLCVCTERVQYYFHYHHVTKLDEVA
ASQLTFPAVTLCNLNEFRFSQVSKNDLYHAGELLALLNNRYEIPDTQMADEKQLEILQDKANFRSFKPKPFNMREFYDR
AGHDIRDMLLSCHFRGEVCSAEDFKVVFTRYGKCYTFNSGRDGRPRLKTMKGGTGNGLEIMLDIQQDEYLPVWGETD
ETSFEAGIKVQIHSQDEPPFIDQLGFGVAPGFQTFVACQEQRLIYLPPPWGTCKAVTMDSDLDFFDSYSITACRIDCETRY
LVENCNCRMVHMPGDAPYCTPEQYKECADPALDFLVEKDQEYCVCEMPCNLTRYGKELSMVKIPSKASAKYLAKKFNK
SEQYIGENILVLDIFFEVLNYETIEQKKAYEIAGLLGELLMTPVPFSCHGHGVAPYHPKAGCSLLSHEGPPPQRPFPKPCCL

GDIGGQMGLFIGASILTVLELFDYAYEVIKHKLCRRGKCQKEAKRSSADKGVALSLDDVKRHNPCESLRGHPAGMTYAA
NILPHHPARGTFEDFTC

>Chordat_hASIC2_NP_001085_2_acid_sensing_ion_channel_2_isoform_MDEG1__Homo_sapiens_

MDLKESPSEGSLQPSSIQIFANTSTLHGIRHIFVYGPLTIRRVLWAVAFVGSLGLLLVESSERVSYYFSYQHVTKVDEVVAQ
SLVFPAVTLCNLNGFRFSRLTTNDLYHAGELLALLDVNLQIPDPHLADPSVLEALRQKANFKHYKPKQFSMLEFLHRVGH
DLKDMMLYCKFKGQECGHQDFTTVFTKYGKCYMFNSGEDGKPLLTTVKGGTGNGLEIMLDIQQDEYLPIWGETEETTF
EAGVKVQIHSQSEPPFIQELGFGVAPGFQTFVATQEQRLTYLPPPWGECRSSEMGLDFFPVYSITACRIDCETRYIVENC
NCRMVHMPGDAPFCTPEQHKECAEPALGLLAEKDSNYCLCRTPCNLTRYNKELSMVKIPSKTSAKYLEKKFNKSEKYISE
NILVLDIFFEALNYETIEQKKAYEVAALLGDIGGQMGLFIGASILTILELFDYIYELIKEKLLDLLGKEEDEGSHDENVSTCDT
MPNHSETISHTVNVPLQTTLGTLEEIAC

>Chordat_hASIC3_NP_004760_1_acid_sensing_ion_channel_3_isoform_a__Homo_sapiens_

MKPTSGPEEARRPASDIRVFASNCSMHGLGHVFGPGSLSLRRGMWAAAVVLSVATFLYQVAERVRYYREFHHQTALD
ERESHRLIFPAVTLCNINPLRRSRLTPNDLHWAGSALLGLDPAEHAAFLRALGRPPAPPGFMPSPTFDMAQLYARAGHS
LDDMLLDCRFRGQPCGPENFTTIFTRMGKCYTFNSGADGAELLTTTRGGMGNGLDIMLDVQQEEYLPVWRDNEETPF
EVGIRVQIHSQEEPPIIDQLGLGVSPGYQTFVSCQQQQLSFLPPPWGDCSSASLNPNYEPEPSDPLGSPSPSPSPPYTLM
GCRLACETRYVARKCGCRMVYMPGDVPVCSPQQYKNCAHPAIDAMLRKDSCACPNPCASTRYAKELSMVRIPSRAAA
RFLARKLNRSEAYIAENVLALDIFFEALNYETVEQKKAYEMSELLGDIGGQMGLFIGASLLTILEILDYLCEVFRDKVLGYF
WNRQHSQRHSSTNLLQEGLGSHRTQVPHLSLGPRPPTPPCAVTKTLSASHRTCYLVTQL

>Chordat_hASIC4_NP_878267_2_acid_sensing_ion_channel_4_isoform_2__Homo_sapiens_

MLSGAAGAARRGGAALAPSLTRSLAGTHAGADSCAGADKGSHKETIEERDKRQQRQQRQRQHQGCGAAGSGSDSP
TSGPHPVPVLFPLALSLEEQPLPPLPLGRAPGLLAREGQGREALASPSSRGQMPIEIVCKIKFAEEDAKPKEKEAGDEQSLL
GAVAPGAAPRDLATFASTSTLHGLGRACGPGPHGLRRTLWALALLTSLAAFLYQAAGLARGYLTRPHLVAMDPAAPAP
VAGFPAVTLCNINRFRHSALSDADIFHLANLTGLPPKDRDGHRAAGLRYPEPDMVDILNRTGHQLADMLKSCNFSGHH
CSASNFSVVYTRYGKCYTFNADPRSSLPSRAGGMGSGLEIMLDIQQEEYLPIWRETNETSFEAGIRVQIHSQEEPPYIHQL
GFGVSPGFQTFVSCQEQRLTYLPQPWGNCRAESELREPELQGYSAYSVSACRLRCEKEAVLQRCHCRMVHMPDSLGG
GPEGPCFCPTPCNLTRYGKEISMVRIPNRGSARYLARKYNRNETYIRENFLVLDVFFEALTSEAMEQRAAYGLSALLGDL
GGQMGLFIGASILTLLEILDYIYEVSWDRLKRVWRRPKTPLRTSTGGISTLGLQELKEQSPCPSRGRVEGGGVSSLLPNHH
HPHGPPGGLFEDFAC

>Chordat_hASIC5_NP_059115_1_acid_sensing_ion_channel_5__Homo_sapiens_

MEQTEKSKVYAENGLLEKIKLCLSKKPLPSPTERKKFDHDFAISTSFHGIHNIVQNRSKIRRVLWLVVVLGSVSLVTWQIYI
RLLNYFTWPTTTSIEVQYVEKMEFPAVTFCNLNRFQTDAVAKFGVIFFLWHIVSKVLHLQEITANSTGSREATDFAASHQ
NFSIVEFIRNKGFYLNNSTLLDCEFFGKPCSPKDFAHVFTEYGNCFTFNHGETLQAKRKVSVSGRGLSLLFNVNQEAFTD
NPALGFVDAGIIFVIHSPKKVPQFDGLGLLSPVGMHARVTIRQVKTVHQEYPWGECNPNIKLQNFSSYSTSGCLKECKA
QHIKKQCGCVPFLLPGYGIECDLQKYFSCVSPVLDHIEFKDLCTVGTHNSSCPVSCEEIEYPATISYSSFPSQKALKYLSKKL
NQSRKYIRENLVKIEINYSDLNYKITQQQKAVSVSELLADLGGQLGLFCGASLITIIEIIEYLFTNFYWICIFFLLKISEMTQWT
PPPQNHLGNKNRIEEC

>AcFaNaC_XP_012938733_1_PREDICTED__FMRFamide_activated_amiloride_sensitive_sodium_channel
_isoform_X1__Aplysia_californica__tested

MWGRGKRQRNKNYPSGSGGGGGAFRSPAMRNDNELEGFVSILHTSGDNYVPIRDSSADHMKYTSVSAKSGMVPEH
RYTMVRSRHHGRHHHHHSYQEYNTQRSAISLIAELGSESNAHGLAKIVTSRDTKRKVIWALMVIIGFTAATLQLSLLVRK
YLQFQVVELSEIKDSMPVEYPSVTICNIEPISLRKIRKAYNKNESQNLKDWLNFTQTFHFKDMSFMNSIRAFYENLGSDAK
KISHDLRDLLIHCRFNREECTTENFTSSFDGNYFNCFTFNGGQLRDQLQMHATGPENGLSLIISIEKDEPLPGTYGVYNFE
NNILHSAGVRVVVHAPGSMPSPVDHGFDIPPGYSSSVGLKALLHTRLSEPYGNCTEDSLEGIQTYRNTFFACLQLCKQRR
LIRECKCKSSALPDLSVENITFCGVIPDWKDIRRNVTGEYKMNQTIPTISLACEARVQKQLNNDRSYETECGCYQPCSETS
YLKSVSLSYWPLEFYQLSALERFFSQKNPTDQQHFMKIAQDFLSRLAHPQQQALARNNSHDKDILTTSYSLSEKEMAKE
ASDLIRQNLLRLNIYLEDLSVVEYRQLPAYGLADLFADIGGTLGLWMGISVLTIMELMELIIRLFALIFNAEREVPKAPVHSS
NNGGGGGGDGQHNFANGDVEHERDTHFPDLGSSDFDFRRGGGIGAESPV

>Lopho_annelid_Pdum_MGIC_AWC68057_1_MIP_gated_ion_channel__Platynereis_dumerilii_

MALRALMQEFAGGTTMHGIPKAIRSRSISARIFWSIVCICAATMFCVQFAQLISKFYAFPKKVTIEIVPAMVPFPAISLCN
MRNLDIMVLNTLNSIFKNATDPLTWTNITEDPFINAYMMTVAKYHPMFVRNDTDMKIFQTILTRTLIATNVDRHLVQK
AGVPFKEFIVTCRYGGLACNRSEEFTQFFDPYYYNCFTYTAPELMYADTTLAEGLENGWSTVVLTGSGMLDQNDELRIIP
GTHEKFSPMASNEGVRLVIHPPHTEPFPHTEGFDVPPGFSVSLGVKARLNLRIGPPHGNCSHIDPFGQGKSREYRLISCQ
KKCLQREIVKECGCKEISLPNHEKYDNLKYCTQDDDLPDSCSVGATPECFERLYQVYDRFLCVQNTTARLTRNMTFAGQ
CKCFPPCREVSYDVTYSLSKWPAESFDGEEAYVDIFETEAYPVRFMGPDDYKKFELYANYFDMSNRKRAMKDFARLNV
YIADSNVLKTEESQDYTQSQLLSDIGGQLGLWVGISVITLAEVLELIIDLCKFIASNHGPYSKGRTFNKRNNKYSAPNDEPV
PNCRSCRLYGQMNGTIPLTAVPEPMDPSHMV

>Cyclo_ACD_1_NP_491295_2_Uncharacterized_protein_CELE_C24G7_2__Caenorhabditis_elegans_

MEPTLSPNYRNEAFEHDDSYLVNFVAGSSSGESSTPPPSFVPKCNFRYNQSRSQMIIEVPVAQLKKLRKLEGTVSIKRET
QHFCETTTMHGPKRIFQGKRWATLFWLIMVSCSLGLLITQVFILASEYLSKPTVSDVSFLINEDGMDFPLITICNLNPIRKT
YVNEINKTGEVSPPMINYMMKWFTEIPTLIGGADRPTLHEGNEELKLYMKNHLNFTVDSFFMNSGFSCPDIFKLCSFQG
EIFDCCTLSTEVLTPLGKCFTLDLSSSTKASMHKQTEPGIQAGLAITLDAHLEEQFDGSNGMDALFTNSFVNGFRYFVHP
PNTIPHLSSDEFTVSPNTVAFSAISSDRYVLLPTHQWGNCTENFPDGIQSNLSYSSGNCLSLCKAKFYMENCGCTPALYNI
ENNLKECTPYETTTCLDNILAKPNKETGKIEFQTPNCKACAQQCNSLVYRAYNSYGSQFSAGAFHYLKSINPEWTDGHM
RANFQMINIFYRDMSYTEYNQVQDASVTQLLSDIGGNMGMFLGMSVITITEICLFFSKMFWLGFSKKRRDYMYSKRV
NEKTHEREVCETVEKMKAIASQGNLSSIAGTTSAKNSIPNDNVEFRINLKDLADQLDSDSGYSQNQPDSRNASFQKY

>Cyclo_ACD_2_NP_001309477_1_Uncharacterized_protein_CELE_C24G7_4__Caenorhabditis_elegans_

MHLEDGPSTKPPDFENEKTQETSLSGEEFENNSTLGTMRDAKSWAAANKQFANQLVIQVPVNSFKNGKKIKGVGSAF
RETKHFSSTTTMHGPKRIFYGKGVARAFWMLIVGLALAMLCFQIFILLQMYFSKPTLSQVSFIVNEGGMDFPAVTVCNF
NPIKKSYVRELNVSGDLTGETLEYLLQTNMDAMFLFSNLDRHNLKETHDEAETYFQNHTDFQIIKFLRTAGYDCGEMFM
TCYFGGRRFDCCKYMKQKVTSLGKCWELDLRNLAPEWMRKQISPGSEAGLQIVVDAQLEEELKGENDDAKAIFSDIYE
NGFRYFIHPPGTNAQLTSEGISVSPSRTVYSAIKTVTHNLLNRGNWGNCSENWPEGYNTFLSYSASACRALCIAQFFNDT
CGCAPFTYNVDGRKKICAPYESITCMDNHMLKKVNGTDYLELPDCEECHMECQSTSYTSYNSYGDGFNRGSLEWLKKIS
NKSETHIKNNVAVINIFFLEMFYTSYSQVQATSLTEILSDIGGNMGMFLGMSVITITELSLFFSKIFWIMVSKRRRQYMYS

KKTHEKEKEHQLDEAVKEFQERRSRRNSRENISALGHYSNRITPVDDFQTKFGYKNAFSEGNMSNSSLDSVMELKFDIN
ELRRQLNQPSTDGIARIRLPHQTSRQNSTENYYSSSPPIFTIEPMSRKQSKTSLPSSLSPR

>Cyclo_ACD_3_NP_001257250_1_Uncharacterized_protein_CELE_C27C12_5__Caenorhabditis_elegans_

MTETSNCSSSSSEYEEEEERIVLHVYDDESKEFTSLTTYHGMIRIYTSETWPSRIFWGVVVVTCVTLFMIQGGVLLEFYNS
HPTATKIDEYRLPTSFLPSISICPYGFKTDDNLFYLITQGDKDVYIPPDYWKNSSKQLLKRLSYKCEDVVESIMINPNQIIDFC
ANSRTQITEIGKCFTFENWREFETNTLKIKLKSDFTKMYTAHIHSEYYEVSRSSTQAWLKPGSHAKLSFRIEEQHNLPQNN
WGTCKVQTGEIYNHLGCLEQCLVAGYDQSCHCSPFFNRFTRFHCSIDELLNCPKLKEVPCDCPMQCYSQNYVLQPVSLL
KSRSNISTVTFHLNSNLLRSHQQYKRFKQIDLMSYIGGVMGLFLGMSCVTLLEVFIYLFKTIFGTLNSTRHKAFIERLLSNED
GSIHGSHEEIIITQKIEKTVQKETLEQPLEQVADTRPLAERRFSLMPNNQLGVKVQFHRPNHLLKRNSVYLGNCDF

>Cyclo_ACD_4_NP_505230_1_Uncharacterized_protein_CELE_F28A12_1__Caenorhabditis_elegans_

MNRKRKLSCFVSVKFPVDSVKKLRKTEGVGSVYQETQHFSTITTVNGPRRIFYGKRSAQIFWILVVISILAFLVYQIVILIQY
FYSKPTLSQINFITNEGAVYFPSVTVCNLNPVKTSFIKKLNSSGDLSEELLNYLLATKTNSMYMFNNANIFELKRAHLNALV
YLANHPDFEIVNFLNSAQFDCDELFETCFYGGKQFNCCKYMTQSITSLGRCWELNLRNETDAWLTKKGRSGTSPKTGLQ
IIANARQSEQFINFHYSSFQENGFRYFIHPPHVSPDLAAEGITVSPSRVVNSAIKTVLHDLLNHENWGNCTSSWPEHYNT
NLPYSSSACQALCVSNYFKKLCGCSPYSYNIDNNTQVCLPYEEVICMMEKMTKSDSNGTVSLDFPFCAECHLACQKTSYS
SYTSYGDGFNYNSMKWLTRETNRSASYIRQNIAIINIHFMELFYTSYSQVKATTILNTFNKIFGLNGLWFGMSVVSLTELIL
YFTKISWIAVSSKRRQYLFEKKMSEKRKERNIEEAVQEAEISRSRRSSAANLKFLDIESLDDEDYWYRSSSQLSDSHLDNVI
QLAIDFEKPLQRPSAISLPRISECCEDFEEEDEDENNDLGIIIKL

>Cyclo_ACD_5_NP_491196_3_Uncharacterized_protein_CELE_T28F2_7__Caenorhabditis_elegans_

MRRVRNLSLLYNDGPMGRFADQENPVENRNKKETVHFQSGSYDDDMSNSPSSSCSTVGDMPNIKPSASKGSFLSELK
PFSKRASQLIVDVPVAHLRKIKNTEGVSSITRESEHFSNTTTLHGPKRIYNGKGWSCVFWVFIWISSMIMLLTQVTSLISM
YISKPTVSQVSFLLSEGGMQFPRVTVCSFNPIKRTTVEALNSTKDLSDDLLDYLMMFNSDAMTLYGRADAASLHSGDNV
FKHYVSSHPNFTADNFFMDAGFSCGDMFKMCSFGGRRFDCCKYATPIFSDLGKCFTLNLQGSDKSWMKMQTEPGIA
AGLQIILDSHLEEQFDSETDGVTPVFSSAFENGFRFYIHSSEEIPFLASEGIAVSPDSVVYSALSSSKYILLSSNAWGNCSDS
WPRGYDYSFPYTSAMCSTMCKAQYFQNLCGCSPSIYNHLNRFNDCTPYETFICMDTKMKKVVNQSFNIEMPTCEECK
VECKSQVYHSFNSYGKGLSRGALMWLTKQGKQETWTIPHMKLNFQVVNVFFRDMSYTEYIQKRGMSLTELLSDIGGN
MGMFMGMSVFTIIELFLFLSKIGWIGFSRKRRDYMYSKKKNEEMHEKELEDVVTGFKLFRHRKSGKDMSHLREKIKGLS
MHRVTSEQLNVCKLAWENEPDIERRLASVTRQNSALKEHKDYKQPTILPFDLKDIKDQITRGRAASMFRSRRSRSETAP
AVIHEA

>Cyclo_DEL_1_sp_Q19038_1_DEL1_CAEEL_RecName__Full_Degenerin_del_1

MARKYIDILKKSKMMLFQDVGKSFEDDSPCKEEAPKTQIQHSVRDFCEQTTFHGVNMIFTTSLYWVRFLWVVVSLVCIC
LCMYSFSHVKDKYDRKEKIVNVELVFESAPFPAITVCNLNPFKNHLARSVPEISETLDAFHQAVVYSNDATMDELSGRGR
RSLNDGPSFKYLQYEPVYSDCSCVPGRQECIAQTSAPRTLENACICNYDRHDGSAWPCYSAQTWEKSICPECNDIGFCN
VPNTTGSGNIPCYCQLEMGYCVFQPESRVRRIWEFQGNKIPEKGSPLRKEYMEQLTQLGYGNMTDQVAITTQAKEKMI
LKMSGLHPQRRAALGYGKSELIKMCSFNGQQCNIDTEFKLHIDPSFGNCYTFNANPEKKLASSRAGPSYGLRLMMFVN
SSDYLPTTEATGVRIAIHGKEECPFPDTFGYSAPTGVISSFGISLRNINRLPQPYGNCLQKDNPQSRSIYKGYKYEPEGCFRS
CYQYRIIAKCGCADPRYPKPWKRSAWCDSTNTTTLNCLTTEGAKLSTKENQKHCKCIQPCQQDQYTTTYSAAKWPSGSI
QTSCDNHSKDCNSYLREHAAMIEIYYEQMSYEILRESESYSWFNLMADMGGQAGLFLGASIMSVIEFLFFAVRTLGIACK
PRRWRQKTELLRAEELNDAEKGVSTNNN

>Cyclo_DEL_4_NP_492230_2_DEgenerin_Like__Caenorhabditis_elegans_

MGVFWTGLKYVFTDFSCWTSTHGVPHIGMANARWLRAFWILVVVVSIALFIWQFITLLTNYLSFSVNTETTLQFAERTF
PTVTICHLNPWKLSETKSVDPDMSALIDAYNSDSSSAQFGLPASLTADRQQQASKWTLMYSERLNEKQYNDAADIAYS
YDDMVVSCTYNAKTCNITDFNDFYNPSYGNCLQFNTDGMYSSSRAGPLYGLRMVMRTDQDTYLPWTEASGVIIDIHM
QDEIPYPDVFGYFAPPGTASSLGVSYVQTTRLSKPYGSCTTKTKLKTTHYTGTYTVEACFRSCMQEKIIASCGCYYPAYSH
ASNTTQYVSCDNGVQTLSNLNCVDLINSADSTEFDVLTDCDCPQPCEIDSYGVTVSTAQWPSDSYVPTECNPGGPSGP
WDASGESCLDWYKANTILIEIYYERMNFQVLTESPAYTFVNFISDVGGQVGLFLGMSIISAIEYLVLIFLVFFYCCTHKSRR
AEIEQLEMDIKKAKDDVDQVAEKRKKHQKANAELYEMDTAHDIVPPKPHSND

>Cyclo_DEL_7_NP_501276_4_DEgenerin_Like__Caenorhabditis_elegans_

MNCSCGHQTDRERVTHAARNMYIMKPPPENEVFISPITLASVNTTQYEEALKIAQFHCNCKYLWLRELHGLSAFMMS
NSLMSKVFWAIVIMACAGWSIGNTISILKQYGDEATTTLLTILPTKQLKFPTMIFCPRNPDFLNYYNVLEDMYNHLGYM
ENTTNFHILQYAMTGFGFDNANGDTFNETYREQIHIYYLKWRGERTQYEMFDFMFNKNGYTCTDLFQTCYGGSQTYN
CCDIFQPTYAMLRGRCFRLIDSYYQNDTDEVAKLSIFFNNMTSPILNTGVLPQLVLYNGDSNVEIGIYPRYYLNSNDWNR
VRFYQKSMILLPKSDGCSTDPIYQGKFTCFVYKWLMQLIEQYNCTVPYYKYTLSYLKDVPICEPDVIVNNFENISLTPSTIG
YKCTSACSRIENTVTLATSIDTDPDPSYMFRIEASFTYLEYEQYKEIRTTSTAGFISELGGQAGLFVGSSVMSFVQLFNSVFI
QIYKILRNYCNKKGIRVRIGLYQTHPSEPADKYYERDAALSDEPPYPLDTILEVEEPNTMDADGMLLEAGDPISEVDAIELE
TWSSVSEIESFDSSNYLPTPSFSSSEQHSTICEEPEDIRTAKLIWIRENREMDL

>Cyclo_DEL_9_NP_508622_2_DEgenerin_Like__Caenorhabditis_elegans_

MYMNGNFPETTVVRSHSLTQSQSLSDSAESAIRRLKKENHCLKKTLSRRWSGRSRASTRSHDPTDASSLSDSSDDNDDS
LYLLRETSTLHGLRDVMLSSSGRLRMVWVLIVILALFMTFQGCYQIMDEYSMRRIVVSYFIQEAQSIWVPDVVVCPYNR
LNRSFIEANNVSFELAQFLELSFPSMDLPFENIQQQQEEIISKIDTLDFQIEILLEQNNMTYAQFLRKASLNCEAFFEDSRRC
ANTTEIMTSAGKCFRMAGIKQEIAGFGNGDRYVIDLPEEYYNPGINQMINSGVIIKLAERGQGIDNDLTFLPAGVHAIM
PLLGTQFEFMNDPPRYECEEDPHGNYSRVHCFEDCLTLDAQQTCQCSPAAAQNPAYPDKLCTATQLYHCFFTKLFPED
SNLSKAIVDACKKECKAPCHAWNYNKQVSYSSIPSEASKKLLPREEWEKMKRKIILDIYYSELDYTIIKHVIAMPLSSLIAQI
GGQFSLFAGGSLISLCQIVIYSVRYVMHKFCGLKSQRRRCRETREAHEARQRRDMRRRRRNSTSHSRSTPKPHNGNGN
GKVVNVEMVTTHTSETTPPI

>Cyclo_DEL_10_NP_495302_3_Degenerin_like_protein_del_10__Caenorhabditis_elegans_

MVRMAERLAENFIPEANQRNENEPAYSRYKRVGQNRSRLNSRASLGSSMGRRITLIETDSGVIEVESDKQFLDAFKDAN
MDAVHHLNAASPVTRGLWCMIIIAFVILVLVQCYSQIKLYISEPVATNIEAEYPSKISFPTVAICNNNQFRLTYLTGGRIMN
RRSKSISGSLLSTGHDVESVFDTVLRKSWDMDAVKFLRSAAHWKSRMILGCTWPNGTSCKLSDFKAVWTTTGLCWAI
NTDPHNPYEVTGSGEGHGLRLLLNVESYERVDACTKHFRTKTLPGLKILIYNQTDIPDSSMNGVNVPSGYSMDIPFKMQ
HRSKLTGVHCIEENDEQIEASTDFNNPENIRTCTLRRYMTEVENSCHCTLRRAYTSNSTDVKMKACNVDQYFGCAQKA
MQRIREEGTASTCLPPCKSIDYTAWQDMNRLPQNLMPALIEEQEEDDEDDVEQEELDENVSFSTVSGGETFSCEDSAY
LDDKQVMRIKRDAHRAYEMQARHQEDIFLRSRRLIARLRNAINSIERYKWGWHYDTFSGVADRLSNLTCFSNFSERHR
DIISILESRPITSEEKKANQMFFLLDETAFNRNATRYMSVGDLKSRYGDKVDDVAEEIAVILRIMEKLWHVFMPDSYIRT
MTGDFSRMDRIIELMNQYELNKLQRRAWAEKMQSRQMKHFFEDDFYESYYQPLIKDLDTTLVKQIDEVEADWPKVEY
YLQRGSAGKTGAIMFFGDGNKDNRQKFEKLIVEMHECASGKMRKEAGKMLSSFKKSYRELQAAYGKLFKEELPDYLEN
FQFGNKFVGDNFAMVNIFLHRMNLEVWSQDRTYGFWSLACDIGGALGLFLGASLLTIIEIVYLCIQYGLCGKRARNMKC
IPMDALTRQMKKVATCSCCKKPIEKSREPIYKKKSQSYQRRFTADDEDQGDKFRSRASSEESKRRKNIWAQMNDPSGN

STLTPSEIKNFLDQVQRNSQPPSYHDDHHPEDHYYYNDPNYLTISPPSDRPEDNREGSTSGYYSSPRAPPQANPRDESPI
PDTEPISVSEFSDALAPPLFKTPPRERRTRKEQKIDEEDDDKHSYV

>Cyclo_MEC_10_NP_509438_1_Degenerin_mec_10__Caenorhabditis_elegans_

MNRNPRMSKFQPNPRSRSRFQDETDLRSLRSFKTDFSNYLASDTNFLNVAEIMTSYAYGESNNAHEKEIQCDLLTENG
GIEIDPTRLSYRERIRWHLQQFCYKTSSHGIPMLGQAPNSLYRAAWVFLLLICAIQFINQAVAVIQKYQKMDKITDIQLKF
DTAPFPAITLCNLNPYKDSVIRSHDSISKILGVFKSVMKKAGDSSSEALEEEEETEYDMNGITIQAKRKKRGAGEKGTFEPA
NSACECDEEDGSNECEERSTEKPSGDNDMCICAFDRQTNDAWPCHRKEQWTNTTCQTCDEHYLCSKKAKKGTKRSEL
KKEPCICESKGLFCIKHEHAAMVLNLWEYFGDSEDFSEISTEEREALGFGNMTDEVAIVTKAKENIIFAMSALSEEQRILM
SQAKHNLIHKCSFNGKPCDIDQDFELVADPTFGNCFVFNHDREIFKSSVRAGPQYGLRVMLFVNASDYLPTSEAVGIRLT
IHDKDDFPFPDTFGYSAPTGYISSFGMRMKKMSRLPAPYGDCVEDGATSNYIYKGYAYSTEGCYRTCFQELIIDRCGCSD
PRFPSIGGVQPCQVFNKNHRECLEKHTHQIGEIHGSFKCRCQQPCNQTIYTTSYSEAIWPSQALNISLGQCEKEAEECNE
EYKENAAMLEVFYEALNFEVLSESEAYGIVKMMADFGGHLGLWSGVSVMTCCEFVCLAFELIYMAIAHHINQQRIRRR
ENAANEY

>Cyclo_MEC_4_NP_510712_2_Degenerin_mec_4__Caenorhabditis_elegans_

MSWMQNLKNYQHLRDPSEYMSQVYGDPLAYLQETTKFVTEREYYEDFGYGECFNSTESEVQCELITGEFDPKLLPYDK
RLAWHFKEFCYKTSAHGIPMIGEAPNVYYRAVWVVLFLGCMIMLYLNAQSVLDKYNRNEKIVDIQLKFDTAPFPAITLC
NLNPYKASLATSVDLVKRTLSAFDGAMGKAGGNKDHEEEREVVTEPPTTPAPTTKPARRRGKRDLSGAFFEPGFARCLC
GSQGSSEQEDKDEEKEEELLETTTKKVFNINDADEEWDGMEEYDNEHYENYDVEATTGMNMMEECQSERTKFDEPT
GFDDRCICAFDRSTHDAWPCFLNGTWETTECDTCNEHAFCTKDNKTAKGHRSPCICAPSRFCVAYNGKTPPIEIWTYLQ
GGTPTEDPNFLEAMGFQGMTDEVAIVTKAKENIMFAMATLSMQDRERLSTTKRELVHKCSFNGKACDIEADFLTHIDP
AFGSCFTFNHNRTVNLTSIRAGPMYGLRMLVYVNASDYMPTTEATGVRLTIHDKEDFPFPDTFGYSAPTGYVSSFGLRL
RKMSRLPAPYGDCVPDGKTSDYIYSNYEYSVEGCYRSCFQQLVLKECRCGDPRFPVPENARHCDAADPIARKCLDARM
NDLGGLHGSFRCRCQQPCRQSIYSVTYSPAKWPSLSLQIQLGSCNGTAVECNKHYKENGAMVEVFYEQLNFEMLTESE
AYGFVNLLADFGGQLGLWCGISFLTCCEFVFLFLETAYMSAEHNYSLYKKKKAEKAKKIASGSF

>Cyclo_FLR_1_NP_510243_1_Uncharacterized_protein_CELE_F02D10_5__Caenorhabditis_elegans_

METETESERIYLQLYDYETKEFSGLTTYHGLVRIYNSNTWPSRIFWVVVVVLSCLSLFMIHSGYLLLGYHAKPTLFQTNTIVP
MNGLLFPEVTICNLNPLNTTKLEELNISKSTWTYIFGYFDEITTSEHKSTKLGEEQFLEIMNNYQELTKQEFNVKNFLKSVS
KSCEETFISCSFGREKLHNCCEHVTTEMTEVGVCFRLSNVNKKYRQWYSGNGFGWEFVLNGNNEIDDHADSLDFEPDR
GFLIMVHESEKYPKINSYGVAVSPDSQLHAAISMKNISLLDKANWGSCSKGWNRNDTDVPYTATHCEIDCKLRKVRNLC
GCSPLAYSARESGSNDTICTPYQIQQCFRKVRGLDNRWEDECDCPSECNMLEFDVTNSYSDLDGRSRGLSSSKVESDIS
HVSLYFSHVAYERIEQQKQLQTADLLSNIAGSMGLFLGMSTVTLLEIFIYLFKSVWGTVNSTRQQQFVDAVAEEEKERSE
SIVIIQNGRNDDMDDQKPSRFPGADRKLSGNSIHIHLDRRNSRMIRGGDLSAASRGSVSIPSQLLSPLSRHNRQSISYGQ
LGRKVSAIGIPLQPNHDTVESGTSLMPPKSPIRRCSTSTTPSMLTRKLSFASQQSDPAQPAHQSRKVSTSSIFKSQLI

>Cyclo_DEL_3_NP_492135_1_DEgenerin_Like__Caenorhabditis_elegans_

MWLRGLFGGLFFLFLLCSCVALFVLHSLYFIFRTASQTEKTESKTVHDDQLFIPALVICNRMPFSQDGLNNVNVNLRQDS
ALRYLLEWTNPSLREAADYVAPSADLMNQGQNTVLQYITQSTRNQTIQNMQYQCQSVINSCTYQGIQLSSFDCCRNLL
SMIPSTNGLCWVWKDSTMWQNSTGINRQFSITFQMTRNSWFSTYVPTHPGVDIYLREDGNDVMRMATELENPIRLL
DKRGVRLQMRKTKKADIRRTSCGYALGDARRSDEHAFKNNHTNYLMCNMLVAMRYCSCHPLMAELIHYDPSSYRDFL
LRVTQTSVCSVDAYDNCARRYIDLTRIENWEEDIPKDLPGYEDAKKCRNDNQRSCLITSYPGTIEGYDLPEEYRTTQDYVS

RLVLEYSTMRTTEILVSKDPNIYELLSFIGYNMALWFTVGHILWSMFWYATGLCCPKRQSSNRISPEIRRKRSVSSPEPIVV
EHRPSQDVASGDT
