## Supplementary material for "Function and phylogeny support the independent evolution of acid-sensing ion channels in the Placozoa": File S2

>Cyclo_NP_491214_3_Degenerin_like_protein_asic_1__Caenorhabditis_elegans_

MNPYKNSAARELGAIRNAIEAFELAIDKSDGNAHSKRKKRSANSKMVPIDLLCKEEHGMFTAHDYGHVECTCVTFEDM
SKVGDTDDDEIFWNCHQRKDWTHKICHLAEGSNQLKTCKCFEDTCVSDEVTKQLVWPLQLSKNGTKLCISPESSGPRY

CASAQKFQVSTCSNCDWLGKCEESDDMDLEEEIDSKTCICHHGNCFQIKGNVKKRKRRTPERKVHERLLSRYEGLLAVY
SHCNCTKQHGCVSTSVPDMDLENSNKTCLCFYNKKNEQIWPCYKEPEWEERKCSRCNTMGDCVYSDKPKKQTISCLC
ATPIKMCVRIDPPQTNDTSLDDRVVKFWDIQPSTTMSPIVKKKEERDKAYGYTGVKDRIALRAKAMENMIFAVDALTEE
EKWKISYNKSDFIMKCSFNGRECNVKHDFVEYLDPTYGACFTYGQKLGNNTNERSGPAYGLRLEVFVNVTEYLPTTEAA
GVRLTVHATDEQPFPDTLGFSAPTGFVSSFGIKLKSMVRLPAPYGDCVREGKTEDFIYTQKAYNTEGCQRSCIQKHLSKT
CGCGDPRFPPYRESKNCPVDDPYKRECIKNEMHVATRDSKKLGCSCKQPCNQDVYSVSYSASRWPAIAGDLSGCPLG
MAAHHCLNYKREQGSMIEVYFEQLNYESLLESEAYGWSNLLSDFGGQLGLWMGVSVITIGEVACFFFEVFISLISSNRTK
RRPARKSFSSSLRCSTDYNLNKDGFNLDN

>Cyclo_tr_K7H9J0_K7H9J0_CAEJA_Uncharacterized_protein_OS_Caenorhabditis_japonica_tested

MLCQSNVNVIANIIMKEEVEVEVEAKKSCCNTLGDCVYTEKPKKKTISCLCATPIKMCVRIDPPQTNATTLDDRLVKFWH
IQPSTTVSPTIKRKEERDKAYGYTGVKDRIALRAKAMENIIFAVDALTEEQKWKISYNKSDFIVKCSFNGRECNVKHDFVE
YLDPTYGACFTYGQKLGNITNERSGPAYGLRLEVFVNVTEYLPTTEAAGVRLTVHATDEQPFPDTLGFSAPTGFVSSFGIK
LKSMVRLPAPYGDCVKEGKTEDFIYTKKAYNTEGCQRSCIQKHLSATCGCGDPRFPPYRESKNCPVDDPYKRECIKSEM
HVATRDSKKLGCSCKQPCNQDVYSVSYSASRWPAIAGDLSGCPLGMSAHHCLGYKREQGSMIEVYFEQLNYESLLESE
AYGWSNLLSDFGGQLGLWMGVSVITIGEVACFIFEMFISIFSAKRVKRRPARKSFSSSLRCSTDYNLNKDGFNLDN

>LsFaNaC_AAK20896_1_FMRFamide_gated_and_pH_modulated_sodium_channel__Lymnaea_stagnalis
_

MYRSRVHSQNEAYARNGYISTRTRSETSSGHMKYTNIGSKPGMRPDQYSMGKPRGQPRHTALGIIAELGSESNAHGLA
KIVASHDTKRKVIWALLVIIGFTAATLQLSLLVRKYLQFQVVELSEIKDSMPVEYPSVTICNTDPISLRKIRRSVFSNESLLLRS
WLTFIETFKFEQSAIMQSIRAFYENLGTEAKKISHDLQDLLIHCRFNREECSVSNFTYSFDGNYFNCFTFNGGQLKEKLQM
HATGPDNGLSLIISVEKDDPLPGTYGIYNFENNILHSAGIRVVVHAPGSLPSPVDQGIDIPPGYSSSVGLKALLHSRLSEPY
GNCTEGTLQGMHTYRNTFFACLQLCKQNLIIRRCGCKSSALPDLPKENVTFCGVIPNWEKILRNESGDAFHGMTIPTPN
LECEEKVHRELNNDRAYESECQCYQPCSETSYLKSVSLSYWPVEFYQLCVLQKFFNATDPDHFMKVAFYNLDKVAREGQ

AEMASDDNRKFNKSYSDEQKKMTKEASDLIRQTMLRLNIYLEDLSVVEYRQLPAYGLADLFADIGGTLGLWMGISVLTI
MELMELIIRLVGLVFSERDEPRRQTNETFTADNRRTHPKTSNAPQESNYPDPFGAPEFNFRRGVEAPPI

>AcFaNaC_XP_012938733_1_PREDICTED__FMRFamide_activated_amiloride_sensitive_sodium_channel_isoform_X1__Aplysia_californica__tested

>Cnidaria_HyNaC7_CDG50528_1_Hydra_sodium_channel_7__Hydra_vulgaris_

MRIRLKERMRKIVQERLTVKQIFKRYIESSTLHGFCYVCMDTFLGRRLIWAVLMILGAIYFIFKLRYGIKEYFDYPFSTLSTVE
YVDDLLFPAVSVCATNSYIASQVYTNQLNTMYKEGRLPLDNNQSIPEYNIPGDELVKTLKNSSLTIESLLKYCDWIMQDTS
HPLVTPNNCGALNFTSYFNYKGEQCHTLNSGAKGHELLKVSDVGISHGYELVFDLQTNEVIKNYQLSGMRIVIHDQVFP
PQLVDGFFISPGFKTYIKLGITQSQSLPPPYSTECGQKKLKYYAIYSQRLCLLETLTDFTGDLCGCRDVFMPENGLPFCSLQ
ELYSCMYPAKESFSEFTMRKECPSDCEERTFSYELSEARYLHNPPIGLSLSRLDKLQLDKSLPSEAHLSKLAKSLTPKELDAYI

ESNIISVILFFGDTRIDYHEQEATNDFFQFLGNMGGEFGLMLGASLLTFVEFIDLFIVLLYHQMLRLYNLRKIPDIFASKRNR
NKEKTKKCENV

>Cnidaria_HyNaC11_NP_001296594_1_acid_sensing_ion_channel_1_like__Hydra_vulgaris_

MLNFKDIAQITVEAIQETNEVTNKDEKNIQTIQDLRNKKIREHISYMIDNSSFHGLSYIFDKRHSVRRTIWFFITIAAFAYA
MQKVYESTMNYFSYPFYTVRMRMYVNQIDFPAISFCNLNDIKFSAMNGTIVDDAVVTQNHEANITGEEYRSYNQAAR
HTLNEMLVDCDFEGKKCSHKNFTEFSWMQGESCFTFNSGKPPHTLLKVKGAGINRSLKLTINVQHYDYYRDKMDSGIR
LILHGQDETPVKMSGLTVPPGFTTYIQIEKKTIINLEAPYKTKCGSVKLKYFDSYSMHTCWLEQLTDYVYKTCNCKDYFMP
GDIPICSFDLLYNCAWPEWETFDKQKLYQCPLPCKIDSYEVSLSRALFPTGLYASSLANDLRKYQQVPIALKSKTDELIFMR
ENLLRLVIYYDDLAYELVEQKPSYNTLLWLGDVGGQIGLFIGAGVMSYFEFIDCLAMVIYTRFFEKISLKNPTTV

>Cnidaria_HyNaC12_NP_001296710_1_amiloride_sensitive_sodium_channel_subunit_alpha_like__Hyd
ra_vulgaris_

MDYLNLLERIRSSKVRRINKTSTDKEFSVYDNLSIRENFQDKMKTIRTKLHSSVTSAHGCLMIFKPKTKLGKLMWFFVILC
CFFFCIVNLVKITRDYSSENPEVKILELYEKSPTFPVVVICNENALNSSIKEHIISLTGKKLDQFDSNLKKYITHKNFNSRLFNE
FGNRLNSTLVSCTIGVDDCSTLLSWYQIWHLEYGLCFAFNSGFNIEGNEVTTRRVFLPGYSHGLNLNLKLEKNEYNDKSK
TTALRVFITHQGEYLFPPNEDLLLTSGFYYSISFQKRLMNREYGSNKMCRKDKIIKLSNFSLSQPLITKYTQNFCNLNCLSEE
IARECMCVEYDRPVLEQHAKIPKCNVDASTQSCIKQQYSYWKSGSSECLQKCKLECFEVKYEPRIHLQKYCSNDLCEGDL
KLSVNFRSFNYFFYQVHINHVLAEFLGHAGGIITFLTGFSIISFIEASFITMQLVITILYMRCKSFKKSCDKTITSRKSFVTCQA
SLEV

>Cnidaria_HyNaC8_NP_001296668_1_acid_sensing_ion_channel_5_like__Hydra_vulgaris_

MLELKCEISKRRRKRYQNAISYLPGANHDTVDELRNQSINDYLSDMAVNSSFHGINYICDSTYKVRRIIWIVVTLTAMLYA
MREVYESTRKYLNYPVSTVRMKIYVDDLEFPAVSLCNLNDVRMSIVNGTSFDNALIDQNAQNISADDALMIAREARHN
LEDMLLECKFNGRSCSAKNFSEFNWMQGDRCFTINSGKPGHSRLSVKGTGIKRNLELILNLQHYEYYRDEMESGIHFILH
SQEETPVRMRGPIVSPGFTTYFRINKIKTKNLKYPYKTRCGSLNLKFFKGYSKQLCWLDQLTDYVNSKCGCKDFFMPGNI
SICTFSTAFGCMWPAWEEFEKKKISNCPLPCDADTYFGQTVSRALFPSNEYKKSFIKNLMHIQQFRDVYGNKKKELQFM
RDNLLRIVLYYDDLSYELLEQKPSYDLLSWLGDIGGQIGLFVGSSAMSYFEFLDCLIMIIYAKYFKQYK

>Cnidaria_HyNaC4_CAL36113_1_hydra_Na_channel_4__Hydra_vulgaris_

MNFEELAKIVVEAVQENNNQPINTNKIKTPRELRNNKTKEYVSEMIDNSSFHGISYIAGKENHFIRRTIWLLITMTAFGYA
AQKVYESTVNYFSFPISTTQMRIYVNEIDFPAVSFCNFNEFRLSKMDGTKVDQAILNPKLQGLVTAEEYRNVTFGAMFDL
KEMLVDCEFNGIPCSDENFTMFSWMQGERCFTFNSGKFPHKLLKIGGAGMKRSLKITINIIHYEYYKDEMDAAIHMIVH
GQQDTPLKMRGPTLSPGFTTYIQLEKKMIINLEAPYKTKCGSLKLKYFDSYSLDTCWLEQLTDHVHSVCKCKDFFMPGDI
PICSLNDAMSCMWPEWARFDQLKMQNCPLPCVIDSYKLSLSRALFPSPQYADSLTNSLQQPPHIINSIKNISDKVQFAR
DNLLRFVIYYDDLSYELIEQKPSYDTLVWLGDIGGQIGLLIGAGAMSYFEFIDCLFLVIYNQFFKTYR

>Cnidaria_HyNaC5_CAX62741_1_hydra_Na_channel_subunit_5__Hydra_vulgaris_

MPQKKKLKEIAEERLNAKEIFSRYIEASTLHGYAYVCSDTFYIRRVLWAVLMLLGGIYFVFKLKVGIIEYFQYPFSTLSTEDFP
EVLTFPAISICPINNFNISRVQNSILKQLYDADRLPLDKNWIDPGFDIPGNELADILKRSSLQIDKMHFYCDWISRNTTHPYI
GPNPCGIENFTEYFNFKDQLCFTLNSGKKGFNLLSVDHEGLTYGYELGLDVHTNEIISSYPYNGIKLIIHNQQESPVMSDG
FVISPGDRTFIKMATVEKKSLQPPYSTECGSVELKYHSVYNRRACLLELLTEYLGLKCGCRSLYMPDNGMQYCSLKETFTC
MFPAQESFNEYEMGKRCPVECNSITFKHELTYSRYQPQSVSVPLMETLSTKNITRPVLKLMKEKNMGYNEIINYIRENIIF
VIFFFGDMKKEVHEQEATFDFYQFLGDMGGEIGLMLGASLLTFVEFMDLLIFFVYHQLLRIQS

>PanArt_PPK4_NP_001334723_1_nach__isoform_D__Drosophila_melanogaster_

MGHQEELKPEQVDLKVTPFVGYLRRTWSDFCATSSIHGLKYTRDEDTNKIVHLVWLLISVVMFICAVVMARTFYMDYR
SSPTRMNVESDNTPVNRLYFPPVTICPDVLFNMQKSEAFLNTLRLPKGAELRGILRKLHIFYGFMLDDERYSAEDIEQME
ALLFLNNLTIPEFVEHLRWNCDEILYRCRFNGEIMDCSKIFQLSKTFFGHCCSFNLRQKGWVNNKLNNLESFKVFHLNSL
NFTAQRAIGGLRYGLSVVVRYKDDNYDPLQSYSYGVKLLIQEADAFPSAHSAAKFIAFNSETFAAVRPQETFCSSAVKALII
EERNCVFQNEFPMRYFSDYVYPNCELNCRVTNMVKFCGCHTYFFDFNRTSDRICTFRDIPCLVDNFANIITRKKSTQCYC
PLTCEHIDYDVQLTNFPLELNMPVADKFYSGLAKNDGVLHVFINSFSYRRLRRDLLSNMVTLVSNLGSAFSLFVGMSML
SVVEIIYYFSVILRKNYKLECETRSQMLHKKPKFAWPKANDTHSKEQKSVFIIHKS

>PanArt_PPK5_NP_996138_2_pickpocket_5__isoform_B__Drosophila_melanogaster_

MEMLKYVISDNNYSWWIKLYFAIIFALVLFVAVNLAVGIYNKWDSTPVIIGISSKMTPIDQIPFPTITVCNMNQAKKSKVE
HLMPGSIRYAMLQKTCYKESNFSQYMDTQHRNETFSNFILDVSEKCADLIVSCIFHQQRIPCTDIFRETFVDEGLCCIFNV
LHPYYLYKFKSPYIRDFTSSDRFADIAVDWDPISGYPQRLPSSYYPRPGVGVGTSMGLQIVLNGHVDDYFCSSTNGQGFK
ILLYNPIDQPRMKESGLPVMIGHQTSFRIIARNVEATPSIRNIHRTKRQCIFSDEQELLFYRYYTRRNCEAECDSMFFLRLC
SCIPYYLPLIYPNASVCDVFHFECLNRAESQIFDLQSSQCKEFCLTSCHDLIFFPDAFSTPFSQKDVKAQTNYLTNFSSEYM
RKNLAVVNFFHTDNYFRSSVRTSYTGPTEYMASTGGIMSLMIGFSVIFLAEIIYIILFVSTLLLILIF

>PanArt_PPK6_NP_611461_3_pickpocket_6__Drosophila_melanogaster_

MIESGKWPTPGKKNSKGDGSRKRGKDPSLIRTAILATFWEYTERTKVSGMWLMRRNRTYGLSRFIWSSVLLQLLLLSIYL
TLLLWLKFYSYPILNTISNDLSITDVAFPGVTICSPKVVNSERVDRYVKTLKIPKEYDMAEVIAGFDFLNAFTDQSFEPPGH
DSYRATDAVLRLNNVSIWEAAMAVSPGCFDYVKRCFWGHTEFQCNQSHEYLSFIPTTAYLGPCCSFNYNPRNASFVPF
SANIFGMDGGLTFVGAEGSERNLNTGLIVLVHHPMDYVTEAAASVTITAQSESFVEVSPTVQSSSVEVLELSERKRDCLIS
GDLQLSNYRQAACLLACQTEAIVKKCGCHPYLLPIVGNKFKECNLNDTFCYSANYDNFKSVRCDQCLPNCYDVTYSTLSY
KTDLNQHKYSVSRFYSPELLNNDSFVLRVYLAKQVVPVIRKVTVMSWIGLLSDLGGIFNLCLGLSMISVVEFFYYCTYRLYI
NYQLQKVQQPRKAWQ

>PanArt_PPK7_NP_609016_2_pickpocket_7__Drosophila_melanogaster_

MTLVYFPPSKLQQQQQPSRSSRLAQQLAQSSWQLALRFGKRTTIHGLDRLLSAKASRWERFVWLCTFVSAFLGAVYVC
LILSARYNAAHFQTVVDSTRFPVYRIPFPVITICNRNRLNWQRLAEAKSRFLANGSNSAQQELFELIVGTYDDAYFGHFQS
FERLRNQPTELLNYVNFSQVVDFMTWRCNELLAECLWRHHAYDCCEIFSKRRSKNGLCWAFNSLETEEGRRMQLLDP
MWPWRTGSAGPMSALSVRVLIQPAKHWPGHRETNAMKGIDVMVTEPFVWHNNPFFVAANTETTMEIEPVIYFYDN
DTRGVRSDQRQCVFDDEHNSKDFKSLQGYVYMIENCQSECHQEYLVRYCNCTMDLLFPPGQYRSCRAQDLLCLAEHN
DLLIYSHNPGEKEFVRNQFQGMSCKCFRNCYSLNYISDVRPAFLPPDVYANNSYVDLDVHFRFETIMVYRTSLVFGWVD
LMVSFGGIAGLFLGCSLISGMELAYFLCIEVPAFGLDGLRRRWKARRQMDLGVTVPTPTLNFQQTTPSQLMENYIMQL
KAEKAQQQKANFQNWHRITFAQKHVIGK

>PanArt_PPK8_NP_001036260_3_pickpocket_8__isoform_H__Drosophila_melanogaster_

MRVPRNRQRAFFASQQKRSRPQHLQITLSTWRRILSRNTDEFCRNTTIHGLKYINNSKLRSSDRLFFGIALLVVLSLAIYLI
QDAFDKWNTNPVIVGIDPELTSIANEPFPAVTICNLNQALAERVEHFSNDSVEFAMLQLLCRRKVDVELVKTNDSRWEE
FILNISQPCNSMVIHCRFGADDYECARLFHPIVTDEGLCCVFNMLHPRFMYRKRVPYSHRNISLPEGFHAVNWHAELGY
RKRGFQPDGDNPLYPRRAQGTGESLGLSLTLDVQADAYYCSSSSSIGFKIALHSPNESPNVRETGVLLAPGMETKLRIDPS
KILTEKHLRNVDRRSRRCLFHNELKLRWFAHYTQRNCVAECLSGWLIRHCGCVTFYMPRLNANDTICPLHKRECVELIRF
RTIIAMESCLDECLPSCFDLSFSAIAYSTRISLDGFRETPSNGGWNFTDAYVERSVAVVNMYFKDPTFRANKQTEFIGFSD
FLSGVGGLMGLFLGFSFLSIAECVYFALIRPCRTCSEIRQLNQPRVHQMNLKRSPTPGNIRYISAADWFQAELAHQQPN
NRQHFNPA

>PanArt_PPK9_NP_611622_2_pickpocket_9__isoform_A__Drosophila_melanogaster_

MQPRIDDSSPSAAARTLTSSSSRSSWAENEGSASGSDSLLTSSTASLLPPQDNNRRLGPGPGLLIFPGTGQRHQHNPNG
STGSKTIAKANDGNAAINQPGPARSIRSFRRPPLWQPPPQDHAQDGQEMEVLDRQKRKDCRPACLVKGQAFIIEFLQG
SSIHGFIYLAKLGLSFVERVLWLAFICVALFSIISLSKRTWHRFQTSPMVISMDRNKLVWNTSFPSLTVCPHKRIDELKVEEY
ILAHPDQFLSEEDQEDFRDFIVKLASLTYDNLETLPLNKSYGIPSTKYLELLYELKWAFEPEISSGAAVKMFIYETQTEFGICH
SVNSMVARYNSFDYWRSGDWSLMDHGDRVTVHPLDGEIYAQIINLSTAYDVYFHGAGDVPSISKQRYTFPESDYTTVE
LIALEIFTNEEARATKQRSCRFNYEAEEMMTVPIYSFGLCLSECRMFFALRVCGCVPHFYRNRCGYNHIFGTQMRNGRR
LPVCGLEGIGCLVKIKREIISLKSDKYKINCNCLANCDDSNFFVQSYRSRVWFLGANLQWGILEHPKMQLRREVLFSFADV
LVYIGGLVGFFLGCSALSFTEIIYYFTARFVRRMFFD

>PanArt_PPK10_NP_001033894_1_pickpocket_10__Drosophila_melanogaster_

MVAKNSKYRNGLKIVINYFVLYFNNCCIHGFRYLTDSMLILFEKFLWLILLIASIYFCIIVCLSSIDRYYTKSTHIGIERNYIFWN
TTLPSVTVCPVDRLNITYFADYCRNNGIKGPQRDILWDFLENLANSTYINFQNIPQNEQIDQIIEDIGLKPEHYTELIYNLTY
DRTYEPNFNERIRCMDGAMFIHVRQVLTEWGLCYLGNSKLTEEYSSRYFIFGKYPEYNKYEYENIRLPYQVGSFFQKDTQ

YALLGFKGPAIIAFAHSAFEVMKVDSNSDYAYDGILYDLSTEEITAEDNLEQDTTVAMRRCRFPHESNLTHFPFYTRNICQ
QECRINLAYKICKCIPHFYPNRIANPKPVCDYKTLRSCFPHHASFFLKLYEENGKHENPAICYCEQNCHDSVVTMKSMNP
MSGAKQLLGGIGSAVSVKTWPQSRLRRQVIFSLTDLLVSIGGTAGLFLGFSVLGFVEVIYFFTIRIVFQILGYTL

>PanArt_PPK11_NP_001334672_1_pickpocket_11__Drosophila_melanogaster_

MSDVPGEDSPTHFYPVNFENYLRPKQSIKCQPLQRFKKPNERATNLYRNLKRLKILRWYNRVSKRFEEFPLPKFLGFLQA
RNDDGLCKRKTGFEIYCEMASIHGFHIFVGAKTWQRILWWLLICNAVLLSFTLVIMSLSMSKETPTIRFIDTMMKPTAEV
PFPAVTICGFNTKEWMNSSQIVNQRNASWLELLEDLALPICPQIKICQWDNRMVNCLDQLQPIWTLDQRLCCSFNYN
KQLFSSYLGVSFVLRSNDEILQSSKSAGFEVLIHESHEIPNGATPRVFVPGESDAHIMLRPYINRFTKNLKGLSLQKRGCYF
STERRLILSDVYNQINCLAECRTESILKSCGCIPPKSPIEKSWLICDLKQMQCVIDFDHDEIISGEQKNCDCLPPCEFNRYEF
QSDIRFIKGMINNSIVNTSNQETTNEVRVRVYYDSAIAEELLLDVYENWLTFIGTFGGITGLFMGCSFVSVFELIFFSCVRP
TCNWLTRQQILWRRRRNQRVGITESRSLGPAN

>PanArt_PPK12_NP_611672_1_pickpocket_12__Drosophila_melanogaster_

MEPSPSAGTERQGSSLFLLKSFVRSLRQFLNQTSLHGLKFVGDSGLSSWERSFFLGSFVTALIITVHLISNIYVKWDSTPVII
GISPQATSILKVPFPAITICNMNQVQRSLVANYREGSNESALLKLLCESDSWESSEADEEFSASNFVTNNLKISEFVSNHS
QSCERMLLFCRFSAVERNCSQLFQQILTDEGLCCVFNFQPPEYLYKPFANNNRNLTNSDGFESVMWDPESGYPEQLPP
KFYPATASGTGITLGFTAVLDAEMGEYYCSSTNGPGFKVYFHNPIEVPKVKEAGLISAIGYETNYRIEMVRAEAVPAIRSIS
RDGRQCLFKNEKELIFYRIYTRLNCENECLAAFLYDTCSCIPFDHPLIYSNASICSMGDTSCVRRAQRASNRPGWAKCRQ
QCLPSCFDLNYLASGFSFPLASNNFQLANALVESFNKSYLSKNIAVINVYFRESVYYGNTKNAYVGLTEFLSNVGGVMGL
FMGFSVISLAEILYFLILKPLAELFVWKRSSHVDSEKSLKHNAFGIEKGQSSDNPSFWHSKELYPKGVSISAYENAKSTKGL
NYQ

>PanArt_PPK13_NP_001014495_1_pickpocket_13__Drosophila_melanogaster_

MRFSRVIEEYLKNSTLHGARFIVDKDASWIERIFWVICLVASWYASWLLIKASLSAFENNAISFVVESSFRDWNTNFPAII
VCESKNMDRIQEVAEQLWGADHDFTLEEVLSEIAFFRGESYHTVHECSGEEVTASCFYSNFSYYAELVRSSCVDSISNCE
WNNKPFECCKYFHRMETELGICYAINSMQAGKPKMPKLNMYSNRGSGPGALKMELHTEATVFILGTEEVPTLVTPKTD
FLVVGPYISFVRYISKRDIENDEEIRQTSVHQRNCRFADENILDVHKFYSYSACTVQCRKDRQMELCNCSSHLSPNSPDW
QLCNMEGLECLNRNYEDLSVIIAKWSKLRGRKGLVCDCLPSCTEVDITTVYDSKENMAGSQNAISRIEVGLIELPTERYKR
NLVRGKLDLVVSIGGTTGLFVGASLLSFVEIFYYITIRPYTTYINEKRRLKRLIIPKRFQ

>PanArt_PPK14_NP_609017_2_pickpocket_14__Drosophila_melanogaster_

MFVRSTEKETRVVADRIRRQDQNPLAPVNTKSEIQRAWTLLIDSYISRSHIHGLYLLFLPSMRRRMRVLWALALICACTV
LFHVSYLLGDRYHNKQFQTIVAHAHASIHHIAFPVVIICNKNRLNWSRLPEIKSLYNITPSQDELFDRILTAYDGFSFHKFN
AFDSLLGESLDELNHLNFTEIVIQMSWRCDEILRDCHWQTASRDCCKLFRPRRLPLGYCLAFNELEKRRGTETGINTGLLL
RLLLREGQHAPGNSGLKGFWLTVVESSVWFGFPIEVVPHSRTNVAVTAVYHYFDESTLSLPSSWRHCVMDYEEESEHF
RTLEGQKYMLENCQAECQQRYLLRYCNCTVDLFYPPSNYPACRLKDLPCLAAHNHLLQNFEQPGEHPYVHREESGLVC
ECLHNCKSLTLLTDMRKSVQQPWLQPNSSAIESMWLNVYFKKPSMLVYKTNLIYTWVDLIVSFGGICQLCLGCSIISLIEF
VFFALYKVPQLYWERFSNEHRSNK

>PanArt_PPK15_NP_001097937_1_pickpocket_15__Drosophila_melanogaster_

MGGTENAKGQVLRKKRGFGFVTNIKDYCTNCTLAGFAYIANSRLHFMERIFWLICVFMSSLGCYQLIMGYQRSFPTRAV
SIVYESLPPFSKWKFPAVSVCELAYRGNLFPKFEEYITSLGVDVTGDYPYDVETGVSILLFPALYNENGLKGKCGTVHKNCT
DACAKCPSDNYRQILTWYGANCSDLFVECKLSHEPFDCCRHFLPLLTPFGRCYMLNSLLNNEPGSKHWLPNELDPAHQ

KAVINVITRLDVQISVINAEDIPHTAFFPPGIPLITEGLSKYMQFNQVAMKNDPDVKDIDPKIRSCFFPEEIPADSLYKSYSF
SVCITECIRRLQMKACNCTSFLYNPNADPRYPDCDLEGFLCLEKTRMIKPDSRVLVNNNKGNNASCGCLPSCNDGDITTI
YEPLLFVRNPNKYYNGTLDMPFLPTDQYRRQSLRTPLDVVVSMGGMLGLFLGASILSAIEFVYYFTVRPLSNMLGARAA
RP

>PanArt_PPK16_NP_001334673_1_pickpocket_16__Drosophila_melanogaster_

MAFKKRRIFDLRHVQAQQMATTAIHRRSHPMHNHSHNPIQPQHRFRQIGEWFTENMRNYCQTTSLHGFSYITRQDIS
RHERWFWLVVVILAIITSIVLVVVSWYWSQETPTVTVIESSHFPTWNIPFPAVTICNFNKISKSKALSLLDQMQVPVGINR
SELHNLFNLTLLPVDTMISNDSLQKYDRILSLNNLTLNRLTQQLSPDCIEMISSCIWKGINTRCESLFQRIDTMEGQCCTF
NYFGGISNNFPEKIAYQVPKRPYRVTGCGYPTGLSVLLNPMISDYYGTFFSGFGFRLLLHDAYNFPDENSETKVVTTTRES
FVRINPESTYATNDIRRMDLSLRNCLFGSEMTMHGLRRYSFINCMFECRVRMTVDLCGCLPPYVYNNGSYKVCGVLQT
NCIIHSKRLFSHALANLNFSLSIVRETDSFPCGCLPDCQSNHYVSESTTGRLDISYFANRPTFNNATDRILLHVFFSDLMSTR
YRTDIFQNWLSALASFGGLLGLIMGFSIVTAFEFIYFLTFRPVFNYINRE

>PanArt_PPK17_NP_001285988_1_pickpocket_17__isoform_B__Drosophila_melanogaster_

MGYWSETRAWLFRDPAKLIRYLVLFACCIVVIVQLYECFAKLYNPPISTHSYYSLNETIEMPSVTICREPPYKEEVLTRLSGG
ACPHPKYATCWMKYPFGEISLDEFFENSTHDSGDTFVFYGLNEDKNNVVMNSSLHFYMGRCYTLRPKESAKRVSKAVG
YSIMLEHSMLTTSVSDVDTGSVGWHVFIHDKKENFTEINMKGSGRVEYVFVGVNEEIEIKLQTQYFSNVQTREEACSDD
ENYSDLKCGEQCIWQDLADNMQCSGPWMHEIASEPCNDSLSMRKLISDYKDVYENEDDFDCDCVQPCQSRIYTTFIQ
NRKAFNQPEPRTQIYIYYTTKLISMIEERPSYDTTQFIADVGGSLGFLLGLSVLGLIGILEHMMLFFCGGFIKRMQQKEQA
KLEANSEDGQSQTSDETIDVEIAYKKKEKQPKY

>PanArt_PPK18_NP_609308_3_pickpocket_18__Drosophila_melanogaster_

MARVEESYYRRLFREFLRQSYINGLHPFLYHTPVRYAKAIWLAFLTAIMIYTHIVIADLILEYLVQPTEIHMAPDLVHVANS
PFPAVGVCTSNKINGRLLRSYAEKLFHHQGNSSLQAVALDLDDMANRLLVLAQFYLNPLESDHERWNATELDRLHRLLV
AYHNGAGYSVRDILRDLSPGCGDLVIRGIVFGLPVNASQLFLKRPTSSNVCCIFNYRRPSYSQFLTKRAEEDSEMRAVPM
VVFESNSILNSVQFVLQDTPKEDITNTFHSNSAFQLIIFPQEDYATIQSTTLGEVLVDHYSIVEIPIQPMFFSSGDGVRGVP
PHIRRCYYPEEGMRLLNQSYYSLDECLLVCRMKSMIDHCGCVSPPLTGSSMDLDYCMLPDLPCLMRWKSIWYGYSDFA
YLQNKEENLVQRQCEHCLPTCNGVSMSITTNVAPLRRTYNSSGYFTGLLKGLTNERPLAIIKLFFKLRFAQATKSQMVSS
WVVLLNRFGGILSLMYGFSIVSYIEILYYLTGKGFVYIYRSFLLDKEPINHNYSLYWRELRSNLPRQEQLRK

>PanArt_PPK19_NP_651708_2_pickpocket_19__Drosophila_melanogaster_

MLLYTKELVIPRPRPGLLRFRNNPRGIKFREKLCNSFAHSNIHGMQHVFGEQHLWQRCLWLAIVLGAVITGFSLYTVLM
HRHSEQLLVSLIETTQLPVYHIDFPAVAVCPWNHFNWQRAPSAFIRFLPRHPNAELRETFRQLLASMDIMNFSNFNRIRI
LTKRNLTGISYLKMTDLMNFMTYRCDELFVADSCVFDETPYDCCKLFVREQTVKGQCLVFNSMISENSRKKHLINQFYP
HKLSTAGEDSGLKFTINASYSFMNNIDALTPFGMNLMIKEPRQWSNEMMYHLYPDTENFVAVHPLVTETSPNTYEMS
PKKRRCYFDDEKNPTFQNTSLTYNRENCLVVCLHLVVWKTCQCSLPAFLPPIDGVPECGINDAQCLGNNSDIFTYVKMG
DQEKYINDSRQGHFCDCPDNCNSRLYEMSLNVRKLDYPKNSTDQLIKAQVYYGQRVMTKIITKLKYTNIDLLANFGGIISL
YIGASVMSFIELLFVLGKLMWGFIRDARIKLKEYTK

>PanArt_PPK20_NP_651705_2_pickpocket_20__isoform_A__Drosophila_melanogaster_

MAKGDNSVKPTEAAGFGDNERCLADLLKVHFRSYCEKSTIHCVRYLYDSHLHNLERFVVYPLFISTFNFKLYIIIVFRIIWS
MLLIISIGLSFFFYLLLSERFVSQKLQTVVHDPQFPVFLVPFPAVGICTDNRINWNKLEAAKEQFLPTNASVELVESFTVLVS
RMETLRFGSYLSSLAELEDDNLEAVGFVNLTELAIFMTLQCSDIMVPKSCLWRSSSFNCCEYFVLEKTEFGFCLVFNSEVS

PRSKAIKQKEGNNFYPRHNAKAGQSTGLNFDLILNESFRRPDSQANNNVYVSICQAPDQLNNVVYSITQNTETYVTVRP
GLTWTDNTTRSIPPERRNCLFADEQGELDANDSAKNFGKPFQLSNCLNRCHESYLIQLCNCSLPIFFLYNHRVPDCNAVS
LRCLARHNDIFSYDKRRDEDALFSATKLGMTCSCLVDCYLLDYYTSTTTLPLSAHKLPKDPHQKLFRVDVHYQVETTPLYR
TSLEFTIIDLIANLGGIFGLCLGASMVSAFELIYYLTVGLAMHLYDHQYYGVLFKHLKAKWVNLKGYLRNEVGHLAENPAA
HKDTNDRKLRHPFNYRKNVW

>PanArt_PPK21_NP_651704_2_pickpocket_21__isoform_A__Drosophila_melanogaster_

MLYPLELPRARRPLYRDYGGKSGLIQTQRDARKNSRLGKLWHFMLPYLKDYAAESSVHGIRYLADPKMRNYLNIRVIWL
LILLTTSIGAIVVYVDLNELYQTVRIQTTIKNTMLPIFRIPFPSIGLCPRNRLNWKILETEAVDHFLGANVSAAQKDLFVKFFT
AAGDPHLSRLNEMSNFFGNKTLTDELHMLDHLDLREVYKFIQFRCQDLFHTCRWRGNPVNCCEVIEYQFTEAGLCFVF
NTEISPASRQKAREDKYYPLRTPHYGEGSGLDLFLRLNRSFIRPGKRGINVMIKQPQQWSDVVRHVPHEAHTRISITPRF
TVTDERTRTVTPEIRRCIFGDEVDNPHYKNFPDFEYWVGNCRSRCHQEHVLNLCKCSPSIFFPISDKDNFTACKASDFKCL
YDNRFTFSIERHPEEDDFVKNPFKESMICDCFTSCSQLVFDRVFTTTTLDNNETDTEAGTMRLDIFYQSGWFIQYQTNM
RFTFVELLASFGGIIGLFLGASLLSAFELAYYFSIGLYLYIHGKRKLRKPEPGVLTIQFGQRKITPIKF

>PanArt_PPK22_NP_733051_2_pickpocket_22__isoform_B__Drosophila_melanogaster_

MVKLASTSNAIWWIKNPKAAGEHNSKGKRKESLGSAFCLDMADLVRNISLQGYNKLLSPDLTLGQRLIWLLVHMATTV
SLIVVLSLTWEQFVAQSFVTNLKDPLFPVENVPFPAVSICPNNRISRQAVIQYAEELRLNSPVIRPVEYFLERLRFFREFYTH
VGVVVDTDDFITFQTFLDVFGTWNNETFFDTRRIMKMLTPRCQGFVLKCTVANVEVPCFSKDAFQDSLTMYGPCCTF
NMENKLKKRHFKNRLASSELGLKVVLNDSHVDYFAPILNTNGYIVMIHNAENYASVYSSNVLEMFPGQGEDSYIAVCAR
VVDTDDSLKSFSPFSRRCYFEYEAQNPIHEQLMNTYSLSYTFPNCITRCRIRSIIALCRCLPFQMPLQLVENLDGVVYCTLG
HVSCLNQYIFKWRNILTERHIVNGLEREIEEALYCPQCLPSCRDVQYEVSMSALPIDNYLATLKLDENNETEFGTDISVLRV
YFGDPHAQYYIRLLNNTWFEVFSTIGNIMSIFVGFSMVAIFEILFFVTKYIYKGCNRMVEQNIMDRKAKELETKKLYICP

>PanArt_PPK24_NP_651860_2_pickpocket_24__Drosophila_melanogaster_

MPVEMESLDQKPFRLAVGRAAWWIPNPGRVARYPPIRTLAPNHLPGTSAEDTRKLSFAAALKDLLQNLSFHCYSKLVES
GRRIQERFFWFVFHITALTVLIAFLWGTYTKEQEALVTTMYHPMYPIWKVEFPAISVCSLNRISQRAAWQYAHNLSGKD
PKQRNASHFYDQLKAFMYMYYDPSDLMDIDNALRFQSFLDRFDTAKEELFFNTRNRMSALTPNCSDMFVSCRIAGRLF
DCMDKFETTLTSHGFCCTFNYDGRYVNKRDFRQRYFGPDMGLVLTLKTDPSDNFYKINGHNGYPDPFSGGVAERVAE
TGFNTLLPVRAKIFETLPEARSMSPSVRKCLFENEMPWIFARHYTFSKCISACRAQSVVSLCECVPFSLPHRYIDGSEKRIY
CTLQHLACLKRYEFKWLNVITSRENVTGLEHELQDALYCPLCLASCTETRYSVRGAMTLGLPTPSQIKGPARSNPGPRAN
NSYGPASSSGSAKGPVHSSSAPAELAVVRIYFAETHIQYFRQIIKSAWYETFSTIGNICGIIAGFSLIGICELLFFLAKQLWQA
CRAELRAELAHIHVQIRNAQVAEPEAERPMKLFILP

>PanArt_PPK25_NP_995766_1_pickpocket_25__Drosophila_melanogaster_

MESKMSRPKQPDWLAELCQESSIHGMPYIARRDLHWAERLFWTFIILGSAYYAISSCLNQWYRFRDNPIVYEYEYLFGL
RIFPFVGITLCPRYHDETEIPRLINQTWGVDPSEDKEKAVYYRKFLLAINGLRYSTLETLEPFENDTTLDNVNYLNILLTLQK
KVIAVKIPPELAPIITEVGLCQTSSQLNRYGNPYGKLETQDMEPMKQCGYLSNCITSLKPINSIVAPIFMYLHDVEEMMLP
DDMRTPSFDAKDIESKDLDIMLHTTSAESEVRNLPVAYRKCRFSDENNLQYYSPYHPSLCRLECRIKWALSLCNCKPYFYV
AAPEVPICTVSGMLCLARSKWLERPCDCYPSCREETFTIFKVSDQTGGDDNYSGERFERTLIINLQISRMGINRRVVFSTD
QLIMSFGGAIGLFLGASFMTIYGVVYFFLTFIAYTCKNRFCKRFFF

>PanArt_PPK26_NP_648125_3_pickpocket_26__Drosophila_melanogaster_

MSTDSTQPHTPSHEVAVRPPPCAANQWKRAALMARMAPAFRETVSDDSEDEVDTKKQIKYGSGFQAAKAIIHEYCDY
TTIHGIRYLGEKKRPLLERLFWISVLVLSVFTCVKLTLNIWDKWNNNPVIVSFAEKSTPVWQIPFPAVTVCPETKTRREIFN
FTDSYHQVRDFQSNVSGIVDLSDKQKGLYGAVSQVCEPHLHDVTLGNKTRRGMEIIDALTEVSPHFDDTYLNCKWRNS
PVKCSDIFHKFVTEDGVCFSFNSLSPAEIFRAEGIIPDFIFREENRLSMDWNVEDGYSASADTSPYPNRVLGPGARAGLYL
FMGGAEIDFDDMCRGPVQGFKILLHTPGDVAQVSKQYFRIPFDQEVLISIRPKIITTSDGLKHYEPNRRQCYFQKERELRY
FNIYSQSNCELECLANFTLTKCGCVKFSMPRNVNMPVCGDASLKCYNQAEDELLLREFTQGLVNAGENTRGETECNCLP
SCTSIAYEAEISQADFDYKTVINTDSPEGKEEQSKRQGMKMSRVSIFFKEAQFLTSRRSELYGTTDFLANCGGLLGLFMG
VSMLSIVELIYFCTVRLISNLRMRRKTRKELLKAVNKDA

>PanArt_PPK27_NP_647826_2_pickpocket_27__Drosophila_melanogaster_

MSRIVTAFNKTVVEYFRKTSLNGFGLLYFIRKRRIQRIFWFLFISFGILFASYAVFSMVLEFLSYSTIADLSELKVLEDEIHFPEL
KICSGYKFSYRNMLASAHDLVSSQNKSLDYWLNKLSLLSGYFDALSVKAENVDDLNSLLDIKNISSFLLALTPACESLILKCK
LNNIPANCLKLFTLKAYNDGNCCVLRNSNLTGELTLFMDSSQIDEYPLNGNLPGFSLHVPSWQGRVSINPGEMAAVEIE
VMELQGNSQLNEYAVEKRACYFSQEGESREKCLHECRIKATLINCQCVPYPFEFRTQKFGYCTLENIRCLQLVERNWSPA
QCPQCLPLCNQLFYRLNKQILGHLHPWRSELNFKFKTPHRQRYKTNILYHWYQMLSNVGGVLGICIGCSFISGFELIYFLV
FRLWTNYLRQPET

>PanArt_PPK28_NP_573169_2_pickpocket_28__Drosophila_melanogaster_

MRTLTESRRRQSGSSGCKKDSESDDDENTICSRAAIKRSVVYYLKNSTLHGLKYIAEESITIPERIFFGLAFVLVVILSVFFISN
VYVKWSASPIIISTSAKQKLTSNMPFPAITICNLNQALLSKVDRIGRTSTNFSLLMGLCDQGGDTTISYIGTWKYFKAILVD
VAQPCEKMLLYCSFGSREEDCSWLFTSILTDDGLCCNFNALHPSYLIRNYSDDVRLETAHPNTRYELIDWTPEKGYARNL
PEFYFPRTSGGTGIRMGLTVVLNASIAEYYCTKSMSVGFKVLVHNPAELPKVSNYGFVVTAGREARIPIEPVYEDALPTIR
SIKKSVRRCLFSDENDLAYYRTYSRKNCELECEAKLLLRECSCVLYYLPRIDPLARVCGPNDNQCTDRVQTEIESSLTNLSCE
NCWPGCFELTYRATLSTASIVSDPRFQAGENLPEYIFHGPYSNASEISILHFYYMTNIFRSTTKSEMFGFTEFLSNTGGLLG
LFMGFSIFSVIEIFFYITVRPYCASRTLRQRHKRRLEQLSWLTPIRMPVRRALRRNRGGLLRNPPPPAYSDLQKFRGKLDKP
ETCKRKLWRTLQVRPVVDRADEELPTYPYLD

>Cnidar_AlaAla_agalma_gene_36048_1_isoform_1_p1__GENE_agalma_gene_36048_1_isoform_1__ag
alma_gene_36048_1_isoform_1_p1__ORF__type_5prime_partial__len_128__score_12_61__agalma_ge
ne_36048_1_isoform_1_2_385_

ISQTKWPSQADLPLYKQLFAYALGVKTSDLTDDFVKNNFLQVNIYFDEMSYQDVLEKREWTAEKLISDIGGQIGVWIGA
SVLSLFEIAFFIVKLLSTIYSKRKKKNKKVDPKCKPSENSLPKAEIYK

>Cteno_Puk_Fal_comp1688_c0_seq1__p1__GENE_comp1688_c0_seq1_comp1688_c0_seq1__p1__ORF
__type_5prime_partial__len_128__score_29_17__comp1688_c0_seq1__3_386_

KFYAENKKQAQLPALVNAFRLNEGIDEPYDLEDTATVYENIMVLQIIFTSLQESNYKEYVKYDYTQLIGDVGGQMGMLL
GASLFTIIEFIHFFVKAAWRYCIRRGRRAVAGQESGQPTYEMHQDQEE

>Cnidar_AlaAla_agalma_gene_18006_1_isoform_1_p1__GENE_agalma_gene_18006_1_isoform_1__ag
alma_gene_18006_1_isoform_1_p1__ORF__type_3prime_partial__len_129__score_14_41__agalma_ge
ne_18006_1_isoform_1_2_385_

MFPDNSTVVKSSEKKTLKSVRKEYFLSATTHGLGRIYEATSQFTRAFWIIAFLTMFTLLSLTVTGIITKIYDKEIITVTRAILKT
PMNFPAVTVCNANPFHPAVEKLLSGNASTVSKISTGHRWIILQ

>Porifera_Syc_Cil_scpid86841_scgid1215_

MSGLFPLQCWDMDGLGMPCVRVLPCGFFCRRNYVHLNMYYEELNYQKVEQVIAYDWLSLVSEIGGLAGILLGASVLT
FLEFLDFVTGSILCAICDVSHRRRYNAQRKSSFQGNVELTTNWSLEPPPNT

>Protost_ecdy_cyclo_PriCa_ecdy_XP_014677754_1__PREDICTED_acid_sensing__ion__channel__3_like
_Priapulus__caudatus_

MSSHVAPSVRWSKVLNRSLAESRQNLVALEVHYSSVANRTVWNEPSYSAEELICDIGGLMGAFAGASLLTILELLHFICN
RRLSVPPSGGGGGFDGYYGGDGGGGGSGRYNVRKIFVVSDADKLQPSRM

>Protostome_Lophotroco_brch_Lingula_anatina_comp134404_c2_seq3_p1_comp134404_c2__comp134404_c2_seq3_p1__ORF_type_internal_len_131___score_14_29_comp134404_c2_seq3_2_391_

ICNNNMFKLPYLIANGHLEQANYFDIVFNHPNFTNIEQYNWTDFHTRNQYKMADLMPSFCLFNQVICTDADFSMIETEMGSCIQFNAEGELKSVETEGSVFGLKLYLFAQQSDYASFTTISGFTVLMHER

>Deu_Ambulacrar_hemi_Ptyfla_40v0_9_20150316_1g20700_t1_scaffold21422_cov88

MKLKILTDLTQKPISIKPLNMMSRDAWKFVTKFATGHNVASPFGKTSTNLFFWSIFEPKIYHSMNLASLVIYFDALNYELISENPDIPVESLFSDIGGTLGLYIGLSIITVFEFFEFMYEIGIFLATRVYSP

>Cnidar_Rhopilema_esculentum_TR52801_c0_g1_i1_p1__Rhopilema_esculentum_TR52801_c0_g1__Rhopilema_esculentum_TR52801_c0_g1_i1_p1__ORF__type_3prime_partial__len_134__score_25_44__Rhopilema_esculentum_TR52801_c0_g1_i1_3_401_

MTDKMDLQDSAASASMHGFGRIFITKSKVLRLVWLCFMILFVALLAYFLTPVLKRYFSNPSVTKEETVIAPRIRFPAITICGAGMSKEKLDKFGKLHNKSFEMNDDVEGIGFSRDMHEEDIFQKLGTHENIKG

>Cnidar_Rhopilema_esculentum_TR14902_c0_g1_i1_p1__Rhopilema_esculentum_TR14902_c0_g1__Rhopilema_esculentum_TR14902_c0_g1_i1_p1__ORF__type_3prime_partial__len_135__score_16_69__Rhopilema_esculentum_TR14902_c0_g1_i1_247_648_

MELKESAADLTMHGLGKIASSTSKTMKFLWINLMTFSLGTLIFFFQKMTIEYLAHGTLTNHQTIIKSKIPFPAITVCGPTVSKAKMQKFFKESNKTFPFDMDSLHLGLGRYFNSEKILHQLETRENFQKVIPDE

>Ctenoph_Hormiphora_californiensis_evg14407__p1_GENE_evg14407_evg14407__p1__ORF_type_internal_len_135___score_16_03_evg14407__3_404_

FSQIVNGKAGGFSVDLDTKTDDYLPTTASRGFVVFIRDQDETMMLNQGGYVVGTGVETFMKISVNDVKRLGPPHGTCKDVPSKFSKYGTRAESVRECMQRQFIEFFLDGCGCIPWYVVARMIDLEKESVLDEMV

>Deutero_Ambulac_Spurpu_008248

MSYRRAKSASIWFGTQDLPVTIPQLWFTLPLQRCRQLIQSLNDEGKLDCYCPNACTKNLARIRIYFEELNFQSVEEIPAYTFGSFLGGVGGLLGLYIGFSVVTISEIILLFLFLLKHTFCAAFCNTKVQPVHT

>Porifera_AmpQue_tr_A0A1X7TEY9_A0A1X7TEY9_AMPQE_Uncharacterized_protein_OS_Amphimedon_queenslandica__Sponge__OX_400682_GN_Aqu2_1_13198_PE_3_SV_1

MSAEYFKDNYLSIDVFFDDFQVTTTTTKYTYGVEALLGEIGGLLGLFIGVNIINFFELLMLCVDGLKNCFQLKRKKTNLGGSAKSNSNSFCLTIAHIPSIESGTPPAESEMPPTESETPPAESEMPPAESETPPD

>Cteno_Ple_Bac_comp46220_c0_seq1_p1__comp46220_c0__comp46220_c0_seq1_p1__ORF__type_complete__len_137__score_9_52__comp46220_c0_seq1_2227_2637_

MHPHILRSYTNTSSLGFFMSPGSIYKVDLGLVREFREPPPAGLCDSTLTTNTYGRYEENSCIAECRDNAIIRACGCVHVVPPHPDNNPLVEYLPCTLEQWAKCGMREYKKWGESNSNRHFKARKILSKLLETQKTI

>Deutero_Urochord_tr_F7B3I7_F7B3I7_CIOIN_Uncharacterized_protein_OS_CioInt

AKAEAVVTPQNCNCAYPCKDIKYQTTLSYSTFPHVNGARQAEALFNFSKGHYHTNVVGLSVYYESLTITKIEESQSITGSALLSDIGGQLGLWIGISVVTLFEFFQFIVRFVHNEIISLRKPKVTKTQVLDIQLHG

>Cteno_Puk_Fal_comp27722_c0_seq11__p1__GENE_comp27722_c0_seq11_comp27722_c0_seq11__p
1__ORF__type_internal__len_138__score_19_61__comp27722_c0_seq11__2_412_

FQYFNVKNYYRDSGQRLENFIERPVFLEEPIDVSEAFVEKFTTHGLCWTFNDKGDKFVNRTGMDFGLSFNMNINQDGY
PDFITTAGVKVMFHNFYEPALIDEYGMALAPGTENYISLSYERVTACTSLCQSSHNVVS

>Cnidar_AlaAla_agalma_gene_74195_1_isoform_1_p1__GENE_agalma_gene_74195_1_isoform_1__ag
alma_gene_74195_1_isoform_1_p1__ORF__type_internal__len_139__score_8_11__agalma_gene_741
95_1_isoform_1_1_414_

QSCKASCLFTRTLARCGLAEATSMLFLNQTGTNTVQNFNESTQKCVLDNMRKAIKENSEDCVCPVPCKEVQYIPTISQT
KWPSKADLKYYKHAFAYAIGKDASTMSDDFIYDNFIYLNIYFPELNYREINETRAWTIA

>Cnidar_Rhopilema_esculentum_TR51305_c0_g1_i1_p1__Rhopilema_esculentum_TR51305_c0_g1__R
hopilema_esculentum_TR51305_c0_g1_i1_p1__ORF__type_5prime_partial__len_139__score_19_66__
Rhopilema_esculentum_TR51305_c0_g1_i1_373_789_

DNFRKHGNKCVCPDPCESVEYTPYLSYAYFPSDVYAVEVANHLVNIGISKSNESALIYFRKNVIELDVFFQQMSHIVIRQS
PAKTLRDIYSELGGLLGASLGASFLTFIELFDFIWTCILRKFEKRREKKTHIESLSQ

>Protos_Platyhelmin_Macrostomum_lignano_A0A267F1U0_A0A267F1U0_9PLAT_Uncharacterized_prot
ein_OS_Macrostomum_lignano_OX_282301_GN_BOX15_Mlig013326g1_PE_3_SV_1

MSELEQRVTKIDVLAESYLADLEVEELAYTMEKLLSDLGGNLGLWIGVSVLTVLEILDLLLSYIRLAVGQQFARRCGCRSR
GCGESSNRRSTVLMELSAGSYRQRRASVYPSCYLHQDSAGSNNNGGSARWQPETHYF

>Cnidar_AlaAla_agalma_gene_10114_1_isoform_1_p1__GENE_agalma_gene_10114_1_isoform_1__ag
alma_gene_10114_1_isoform_1_p1__ORF__type_internal__len_140__score_13_22__agalma_gene_10
114_1_isoform_1_3_419_

DLFGLQIILFVADGYSNSLLSGGCGIQVIIHDPDEYPLIPLGRGLYVMAGYETHIITKANMKVRKPSPYPSKCADGAPGKQ
SLFPGKYTTNKCAVDCYYRKLYDDCGDLPDIVRMLIGSSMLPVKGNATMQCYTESTEA

>Ctenoph_Hormiphora_californiensis_evg191620__p2_GENE_evg191620_evg191620__p2__ORF_type_
5prime_partial_len_140___score_34_13_evg191620__89_508_

EYARENIVRIHIYHDDIRRETITQEAAYDVASFIAEMGGIGDLFIGFSFFTMFQLVEIIIALIARSSYTQWLKWKERKRREQD
PVNRGNKDIDNHSGNHGKNIGNHDNNGNNSNNGNGGIGGDIGRSGNHGSFGNTTLV

>Deutero_Ambulac_Sakowv30016344m

MLFMFNWNVLACKCPQSCYEVDYGMKISLSKWPSDVYLNNLLKTLHVKNSKTRNINDNIAASNNLARLEIYFDKLSYS
MTSQYKAYTWEDLLSDIGGTLGLYIGTSLITVIEITTFIIAICQHLCKKRSNEANNKVASG

>Cteno_Ple_Bac_comp46220_c0_seq29_p1__comp46220_c0__comp46220_c0_seq29_p1__ORF__type
_3prime_partial__len_142__score_19_04__comp46220_c0_seq29_2008_2430_

MFFADLNDYTAPAKNQAQYGYTVAFHDHETYSSTIPSGFFMSPGSIYKVDLGLVREFREPPPAGLCDSTLTTNTYGRYEE
NSCIAECRDNAIIRACGCVHVVPPHPDNNPLVEYLPCTLEQWAKCGMREYKKWVKRYTDAT

>Cteno_Puk_Fal_comp49951_c0_seq1__p1__GENE_comp49951_c0_seq1_comp49951_c0_seq1__p1__ORF__type_internal__len_142__score_16_39__comp49951_c0_seq1__1_423_

NAEYGVSSKPWFFRTLNRGERTPKMIPEYRPVGDRPTVGFTSPTPSSNKTTTSWVRDFKSFAESTTAHGVSKIFAGGNKAVRLFYLLAWIACVIYTMYMIITSIITFINKPTGTKYEVIVDNVDGLKPQSIKFPTISICSH

>Cnidar_AlaAla_agalma_gene_91159_1_isoform_1_p2__GENE_agalma_gene_91159_1_isoform_1__agalma_gene_91159_1_isoform_1_p2__ORF__type_3prime_partial__len_143__score_22_99__agalma_gene_91159_1_isoform_1_483_908_

MFSTLKSTFGEATVKEKYNLVDSYSNNDCFEKISDKFGKGEIQCPCPVVCEEDIYRSTVSSAKWPSKPLASAILEELQGSSKRYENWTLETISENLLSVKIFFDQFFTYETKQKPAYDKTSFISDLGGQLGLWIGASVFSVL

>Deu_Ambulacrar_hemi_Ptyfla_40v0_9_20150316_1g33947_t1_scaffold1162086_cov166

MASRENNGLVGKDEMKNKYKVLLCEVMSNSTVHGLPNIRRAKSMPKRIFWSISFLTAVGMFVGFSSGLIMKYYEYEVNVGVQVNFQRNMTFPAITICNLNPLRKSFLRENHSARLRKIVQRAANTTIYDIKATPDKVTNGTEN

>Deutero_Ambulac_Spurpu_025729

MDSTARSITRDSHHSQNGQRRALTKNLVRVEIFYEKLNYEAYTQMPKYTFGSLLGGIGGIMGFFAGMSLITVFELCGFIIQLLGLLCGRLTTPMPEEATMEGEHPPPHPESRIGRVGRRFTKVFSHHPHGHGGSARARQTDL

>Protos_Platyhelmin_Macrostomum_lignano_A0A267GI52_A0A267GI52_9PLAT_Uncharacterized_protein_OS_Macrostomum_lignano_OX_282301_GN_BOX15_Mlig029354g1_PE_3_SV_1

MRARLRTVTGAAQEVAGITRTHGLLHIFLSRSTARRIFWLLLFVAALIGCTVHLAKLVQKFTDRSVESQLKLRSERAQFPDVTICNFKPASTGLMEGFFLNDTWKSSGATSPTGSRRTPRSARKNQRKATKIDLRRCVRSSWT

>Protost_ecdy_cyclo_PriCa_ecdy_XP_014678934_1__PREDICTED_acid_sensing__ion__channel__1_like_Priapulus__caudatus_

MYFLLTSISAFTSSTTPHGVNHACGSRLLPARAVWAAITLAFLAYQHSLVLRRYFSYPVTVKVEMGYSRLPFPAVTICNKNPVHMSRLPSHVVHLVTSGYGNVLHTHESAFDEFRRIVSGNESQTTEGSQATATPPMSLKGIV

>Deu_Ambulacrar_hemi_Ptyfla_40v0_9_20150316_1g29543_t1_scaffold132617_cov79

LKLTVFAEQDEYVGPASPSAGFRVVVHSPQSMASPEYEGVSVHPGTSTSLGVNLVSFARMNEPYNYCPDDENLQLFTEYNYTLKTAHLWANIGGSLGLWIGISVVTIVEVLEYFIDLAVHIKRRGIQCQLLPDDETTLRSPIRR

>Cnidar_Rhopilema_esculentum_TR3920_c0_g1_i1_p1__Rhopilema_esculentum_TR3920_c0_g1__Rhopilema_esculentum_TR3920_c0_g1_i1_p1__ORF__type_internal__len_146__score_20_30__Rhopilema_esculentum_TR3920_c0_g1_i1_1_435_

EIKKTEIQRLKPPYPSKCTNGQNMKLLFKGEYTISNCRLSCFLSKVVHKCKTIYHFYNLLLPKKMRKPPSSREDDFLCQDNIEETLRSEKYASCGCRLPCNEIRFRKSASFSKWPSTQDIAWERVSLGFEIGLNKSNLTEKFVRD

>Ctenoph_Hormiphora_californiensis_evg193137__p1_GENE_evg193137_evg193137__p1__ORF_type_internal_len_146___score_26_51_evg193137__1_435_

QCLDIGTCGISQQIQCENLIMNSLTSGAVTVPACYEPCSYEKYSFTMTTENFPTEMYWDDALLSTNANYSYEEAKENLLQLVFYTDLIMETRDIQTASYEIQNFIAELGGLSDLFIGVSFYTAFQVMEWFIVRVVHRETKRLVHQ

>Protost_ecdy_cyclo_PriCa_ecdy_XP_014669085_1__PREDICTED_amiloride_sensitive__sodium__channel__subunit__gamma_like_Priapulus__caudatus_

MTVGEIVTAFTSSTTAHGVKRACGSRSLPARAAWAAITLCALAFLAYQHSLVVRRYFSYTVTVKVEMEYSRLPFPAVTICNKNPVRMSRLPSHVVNLVTSGYGNVLHTHESAFDEFRRIVSGNESQTTEGSQATATPPMSLKGIV

>Protost_ecdy_cyclo_PriCa_ecdy_XP_014679204_1__PREDICTED_acid_sensing__ion__channel__5_like_Priapulus__caudatus_

MQVLRDFVDSTSAHGVYRAATPMSLFRRTVWSLLFVLLFGFAIVQAYNILEKYLSYPKLVDVDVIYAKELMFPAVTICNQNAYAGYKLDAYNISVDTVQEAKICDPFDYEYSYENDDDSCDDAEVGEQLRRQLLLKVLSSLTWQE

>Porifera_AmpQue_tr_A0A1X7TP70_A0A1X7TP70_AMPQE_Uncharacterized_protein_OS_Amphimedon_queenslandica__Sponge__OX_400682_GN_Aqu2_1_16668_PE_3_SV_1

MELSDAIVANASPQDQGEAVPAVQPRKGPLTILKSRLKRGQHHFNKWLDNTSIHGVVHIFKGKSRLRRILWLLVFLFASITCAGIIIFYIILWAKDPTSTSISFISDDEGQSFPAITVCNLNAVKKDYADQNVVDLLNNYANPTSGF

>Protost_ecdy_cyclo_PriCa_ecdy_XP_014663983_1__PREDICTED_FMRFamide_activated__amiloride_sensitive__sodium__channel_like_Priapulus__caudatus_

MIAKWPSNAFLSTVAGRYNRQHSAMKLLRQQGTDGNDGPVVSNYGVVILDNIAKIQIYFSDLSVEHIQQSAYIKFTPLVSNIGGTLGLWVGISILALFEFLELVYDLAVYGCNSLCGKKKSQNGEQQETHKTTMTKVYPLDLKSTKF

>Cnidar_Rhopilema_esculentum_TR38848_c0_g1_i4_p2__Rhopilema_esculentum_TR38848_c0_g1__Rhopilema_esculentum_TR38848_c0_g1_i4_p2__ORF__type_complete__len_149__score_24_27__Rhopilema_esculentum_TR38848_c0_g1_i4_1899_2345_

MKEEGKNGHFVTEESKMEEVNSLTIHGVPKIFTTKSYFSRVLWALLCTSTVVFLIYFMLDAIRKHLNFEIYIKEERIPKHEMSLPAITICNTNFYSGDDDDYAVNQVLPKNCSYTDAKYFKTNTSRENFLDICRWFYGGIWASSTSME

>Cnidar_Rhopilema_esculentum_TR85881_c1_g1_i1_p1__Rhopilema_esculentum_TR85881_c1_g1__Rhopilema_esculentum_TR85881_c1_g1_i1_p1__ORF__type_3prime_partial__len_149__score_20_75__Rhopilema_esculentum_TR85881_c1_g1_i1_66_509_

MSKQDRIKTLLRFYGENATVHGIAAIHNSLSILRKVIFAFLFTSLFGAVIYAVYSSFYSFSLRQVYTSVEKESRRNIEFPSVTICRNGKATKSSLPVNIVDKTMAKLEGIKNGNASLSALHKFVQNASLEVRAAFENSNMSVPEHPPM

>Deutero_Ambulac_Sakowv30034301m

MEEVHGELSTITQSLWPSNMYVNGLIQVLRHVKPKVRDIVVDSTTARENLVNLEVYFEELNYHNTIETISYTIEELLGDVGGLMGLYIAAIEYERLKLAKEDNGLDNMIENWDWKRGILSQSLPVVCTTVPRSYNYLEIKCGSPVIKQ

>Protostome_Lophotroco_annelid_Pdum_comp1073202_c0_seq1_

GVSRDLRKSIGHLKNVSVLNCVFNGRQCDTGLFSDFTNIEFGNCWTFNSHFIQPSAFKTRFTGVDHGLTLHLFLESSGYLPFTEGNGFRVVIHPFGSKPFPLQEGLSITPGKETNFALSYTNVTRLGGKFGKCNGDDYFKETFNVSYS

>Cteno_Ple_Bac_comp48643_c0_seq2_p1__comp48643_c0__comp48643_c0_seq2_p1__ORF__type_complete__len_150__score_12_32__comp48643_c0_seq2_628_1077_

MGFPVNKGSVQQNNIFSKFYANDKAISIDGQPAQLARIVDGYGLNHGIEETYNISKETVYENLMVLQVLFTSIQESNYKE YVKYDYTQLIGDVGGQMGMLLGASLFTIIEFVHFFIKVAWRYCVKKATRRAGEAAERPAYEMHRDQVEE

>Protost_ecdy_cyclo_PriCa_ecdy_XP_014679206_1__PREDICTED_degenerin__unc_8_like_Priapulus__caudatus_

MVREGSPAVHKTLAMWLNTSESALKSGAECLIEKSNCTDDSYDFIDTSDDSTYFLAVAAGRSNFLKISVYYKDMKFKSTK LQKAYTWVSFLAELGGMFGLCIGFSLLTFVEVLELPVLLLHIWVCGGANKVGEKPTTIQINVKNNVGEV

>Protostome_Lophotroco_annelid_Pdum_comp400297_c2_seq1_

MTYKESTEKINEPLTYKDATNKFADNTNCNGVSSIKRSDHVVRKAFWIILVLTGAGLTIYQVWEVFNKYSSYPVAINVNV TYANTVIFPAITLCNNNMVRKSKLELMNGTDTLDLLTYTVGGNADISKWANSKGRAELRQKFMNSWSQL

>Proto_ecdy_PANarthopod__XP_023234682_1_uncharacterized_protein_LOC111634194__Centruroides_sculpturatus_

MDEDNKTKHIKRPNAHQRHAYLENLLKNTVMTGLPQIATEHNLKRKVFKILVFFACLFGFFYQLTEFLMLYFTYPTSLEVL VKITTEDYLPKPVITMCDLNGIRRTVFCKDFPNLCEKPKNVSLFCIKNSQFCDKENFTDDDLMVSLVVN

>Deutero_Ambulac_Sakowv30034166m

MDSFDSNSKMNLSRQSLTAINLKERRRPLTPRMSDHEDKSVKDVVEEFGLNTTAHGVSHIVSADNGYVRLAWISIVLTA TASLLIQVALLVHQYFEYNVNVQIAVVSSNELMFPSVTVCNTNKLRRSAVRDTNYSDLLVVDTSLVLPYYGK

>Cteno_Puk_Fal_comp42050_c0_seq1__p1__GENE_comp42050_c0_seq1_comp42050_c0_seq1__p1__ORF__type_5prime_partial__len_153__score_18_78__comp42050_c0_seq1__2_460_

CTIREMGECGLMAYYKFVLNESATGPNQRTHECGCWVPCEHFDYELSVATTAIPREYAETKVSGGEGEKFNFTVNDFL QNNVILEVYFRDMYAEQVFQVPEYNFWNLLADIGGMLGLFLGASIFTIIDFIRYSLDFGWRKIRSFRCYIRDRS

>Deu_Ambulacrar_hemi_Ptyfla_40v0_9_20150316_1g34047_t1_scaffold1165291_cov166

MESRHNRRLDEGGATKDKYRVLLSDVIANSSVHGLPNIRRAKSVPKRVFWLVSFLTAVGTFIGLSSNLVEKYFDYGVAVG LQVDVERDLTFPAVTICNMNPLRKSLLNEDQLSHLRSIVAGAEGENEANKPVKTTDETPSNDTRRNSSDQVR

>Protos_Platyhelmin_Macrostomum_lignano_A0A267H5P1_A0A267H5P1_9PLAT_Uncharacterized_protein_OS_Macrostomum_lignano_OX_282301_GN_BOX15_Mlig011411g1_PE_3_SV_1

MRARLRTVTGAAQEVAGITRTHGLLHIFLSRGSARRLFWLLLFVAALIGCSVHLAKLVQKFTERSVESQLKLRSERAQFPD VTICNFKPASASLLISLNSTLRMSLSLRCLTTLKSSGATSTTGSRRTPRSARKNRMKAMITDLRRCARRSWT

>Protostome_Lophotroco_annelid_Pdum_comp333774_c0_seq1_

MIIHEVTMSIKKYYSYPVQTNIIPQTALSQPFPAVTICNQNSMMTRAINRADNDIVQLLKVNPANASSRAVLQEEILRSY AQLPLDARMRLGHTMSTMLLDCSFGNTACTHNDFLLFNNFYYGNCYTFNSGSDNRSTLIAENKGPLHSFQLALF

>Cnidar_AlaAla_agalma_gene_91159_1_isoform_1_p1__GENE_agalma_gene_91159_1_isoform_1__agalma_gene_91159_1_isoform_1_p1__ORF__type_5prime_partial__len_156__score_8_97__agalma_gene_91159_1_isoform_1_2_469_

FDQFVLNAVPTTFNSVPFTNPKDFKVITSFLGNCYTFNYNGTLKQQRPGHIFGLSININLNVNQDEVKSDTMMQDIVGAL
VIIHRPEIIPTTLSSTIVVAPGQQSRIMLKHERLIRLPDPYPDHCKPSNTRIMGDVRYSANLCNIKCVADQQLKKLR

>Porifera_AmpQue_tr_A0A1X7T6E0_A0A1X7T6E0_AMPQE_Uncharacterized_protein_OS_Amphimedon
_queenslandica__Sponge__OX_400682_GN_Aqu2_1_09807_PE_3_SV_1

MIACITLISISFRRFINKPTSSTITVVTENTKSGVEFPAVTICNHNLEYNVSDLVIRNTYHLMNYLFNADKNFHLTGSNSSSVI
KQCQALVGDALSEILNATIWNIQNPSKTIDELIHYCGFIEGVNGEVEPCKDAFKPVLTSAGIYFTLNGSDH

>Protost_ecdy_cyclo_nema_Cele_tr_Q7YTI3_Q7YTI3_CAEEL_Uncharacterized_protein_OS_Caenorhabdi
tis_elegans_OX_6239_GN_WormBase_WBGene00013333_PE_3_SV_1

MSEVRQRKSSIIDSDLDFSDSDGEFKEIIKDIENDQWKDKPVDETRWSKTKSAVHEWGLSCSWHGIPHMAQSLSWPTI
LLWTTLLIISAVLFVYLITVTVRQYFSFQKLVNLNIGMVESNFPSITFCNTNPYKLSAVRAIPELEALLTVYSQAYT

>Cteno_Ple_Bac_comp31243_c0_seq1_p1__comp31243_c0__comp31243_c0_seq1_p1__ORF__type_in
ternal__len_157__score_25_91__comp31243_c0_seq1_1_468_

QVAAAIYSFHNITIFIGRYFDRPTTTQITSVAPVNDVIDFPTISICNMNNIEKSFLRLNPLLHEVWNRIEDVKLINWKNDSIR
EIGSLTYPDVYSKSFNWKHVVDSCKYGGFRDCENLPYHDDELFGEYVGSAGKCYLFNPKGSIYSTGIGENGCF

>Protost_ecdy_cyclo_PriCa_ecdy_XP_014678701_1__PREDICTED_acid_sensing__ion__channel__5_like
_Priapulus__caudatus_

MPHGFKECDIIKYFTCYAPNYYKYNLDKFGPPDKSCPDNCIVDYYNFQTSFNVAPSVVMSQKFNKSLDAQRKNMAILEV
YYADIANTTVWTEPSYTPQDLICDIGGLMGAFAGASLLTILEVLIFIRPKRHLVGDSGTTQTIDVERGKQDTTEQAI

>Cnidar_AlaAla_agalma_gene_14319_1_isoform_1_p1__GENE_agalma_gene_14319_1_isoform_1__ag
alma_gene_14319_1_isoform_1_p1__ORF__type_internal__len_158__score_18_70__agalma_gene_14
319_1_isoform_1_1_471_

VARASEMNFPAVSICNLNSVRASVMNGSELDKVLRNKVSRWRARWTMNASAFDIVEEARHEMKDMLIRCRHRRTRC
TEKDFETFTYRFYDICYTFNSGKHGKRLFTQVAPGRYYGLKLTLNIEHEKYYIGTPGAGFRVIVHQQGDVPVRKAGILVSP

>Deu_Ambulacrar_hemi_Ptyfla_40v0_9_20150316_1g31517_t1_scaffold334992_cov137

VDNNLAHAQDEKYWGYQDPERAPGKTGDDSKDLDKSKNAKSFGSIVRKFGLETTAHGIPRIAGAKSTLGRLIWSTLFLIA
AGAFARQFIILVDLYYRERPVTVHMKVVSKDSLDFPSVTVCNTNMLRNSAVRNSVHNELTKAVEGTEILPYYGKLIEH

>Protost_ecdy_cyclo_PriCa_ecdy_XP_014671065_1__PREDICTED_acid_sensing__ion__channel__5_like
_Priapulus__caudatus_

MTGDTLRSSPAAADAPSLSSRLKRTTREWARNTTLHGVKHAADSDNIILRIIWTLSIVGCWSFLIWLMHVRVSTWLSSP
YISIINYEYEKEGIEFPAVTICNNNKLNKTKMEKFYPDVAKIGMVRLHRFSIFRYISAINFVCGFYSRNLAIYYCVIGW

>Deu_Ambulacrar_hemi_Ptyfla_40v0_9_20150316_1g25472_t1_scaffold44668_cov97

MFKPILIRVSRYRYRSTIYSMNTAGNSTYCSPYEMQNCYFSAMDDFALKVDPCKHCEEPCVQTSTHSSDDLMDIATNM
ANFVLEHYYSLGANGQNISAFIPSPEFHAVILKALTEGPFTDVISEVPWVGLFFGASIVTFVEIIDFFAVQFWLMKVSGNK

>Protostome_Lophotroco_annelid_CAC9668827_1__Ofus_G161665_partial_Owenia__fusiformis_

INLSQWPHEKYYQGLDKRRFFELINGNPVSKKYFEEKLDTDKLDQYLADTTKFTEETNKDIYGHTMFTKYQLANKLIEKNF
IKLNVYFQSLEVEIVSSSKAYQLDQVISDIGGVFGFYIGISIVSIFELGSLLMKIGKITMCARKPEEMKKRQKVVHLI

>Cteno_Ple_Bac_comp46220_c0_seq6_p1__comp46220_c0__comp46220_c0_seq6_p1__ORF__type_c
omplete__len_161__score_17_01__comp46220_c0_seq6_2227_2709_

MHPHILRSYTNTSSLGFFMSPGSIYKVDLGLVREFREPPPAGLCDSTLTTNTYGRYEENSCIAECRDNAIIRACGCVHVVP
PHPDNNPLVEYLPCTLEQWAKCGMREYKKWVKRYTDATSLYPECHCPPPCQETHYNAQLSSSALNKFYAESKTSQLPY
G

>Porifera_AmpQue_tr_A0A1X7TYE2_A0A1X7TYE2_AMPQE_Uncharacterized_protein_OS_Amphimedon
_queenslandica__Sponge__OX_400682_GN_Aqu2_1_20432_PE_3_SV_1

MEVNELIHYCGFLHGADSDVVTCKDLFEPVLTSAGICFTFNGTNKLANSTGIRYGLKLVLNIQQKERPSFNGKLGVKLVIH
NGRDIARPYLYGISIPPGFAVDVGVRKMATQDETSEAKCIDGMNLPFFPSDKFEYSQFACRENAVAENIAKRSKCNCVV

>Cteno_MneLey_comp19091_c0_seq1__p1__GENE_comp19091_c0_seq1_comp19091_c0_seq1__p1__
ORF__type_3prime_partial__len_162__score_22_03__comp19091_c0_seq1__3_485_

MTGFCLYIVVQLLSGYYQYDSYNLEYSRVAQKYTLPAITICNVNPLNYTKLKERGIYPAYKDWINVVKKRNSTEDEEKTEP
NGDLDDIDIHTQFKMDIKASVIDSLTSFAREPTTYKNFTSFAYTSNGDCMEMNDNEVLVQRVNGAIGGLTLVLDTLTGD
Y

>Porifera_AmpQue_tr_A0A1X7TNW0_A0A1X7TNW0_AMPQE_Uncharacterized_protein_OS_Amphimed
on_queenslandica__Sponge__OX_400682_GN_Aqu2_1_16667_PE_3_SV_1

MVSRYNKSCNDSLNDISSESKEETLLEAFANGGHKIDDFVVYCGFAGSDGNIVRCEESLEPVFTSLGVCFIFNSAYNGQP
DRKVTLAGPQFGLNLVLNVSVSEYSRLSSSNVGVKVSVHDRNSIPVPHQVGVTLSPGKNSYLAIETVKHVDKTRESKNLR
Q

>Ctenoph_Hormiphora_californiensis_evg15555__p1_GENE_evg15555_evg15555__p1__ORF_type_inte
rnal_len_163___score_30_02_evg15555__3_488_

VYLICAPGRSKIVRLFYLLIWLTCVTYTCYNISLAIRTYINKPTGTKWEVIVESDDAGKQPYFIKFPTVSVCSNNIMKRSYLER
DGHDPIRRLWHLFDTVQHNMEDPKDIAALNFTQLISSFSGEPVIRNKTLEDVFSQGGPDDNRLLKCVQRNKKCSKIAS

>Proto_ecdy_PANarthopod__XP_023226920_1_degenerin_like_protein_asic_1__Centruroides_sculptur
atus_

MNVSSVRVSRNIGTLSGLTLVLNLELEEYIYKTQSVGARIVIHSSDEFPDTEGEGINILPGVETSIAINKKTTKRLPFPYKDRC
RDYKKETTQSAAILNGKDCVIQCLQDISMQRCNCTDPTLIFTKSRKKCNLENEIEVCCLDEVFEEAFSNISKMCDCPISCS

>Cteno_Puk_Fal_comp16663_c0_seq1__p1__GENE_comp16663_c0_seq1_comp16663_c0_seq1__p1__
ORF__type_internal__len_167__score_53_44__comp16663_c0_seq1__3_500_

NAEYGGREVQELLSTCNEPCYYREWSVDMTSTSFPPTAPYLEQVMEFRAGYTVPTFEYARENLVRIHVYYDDLRREEVK
QEAAYDVASFIAEMGGIGDLFVGFSFFTMFQLVEILIALVARGSYAQWAKWKERRRRRREDEGEVVFANQDNGSTG
GHHDGYHGDD

>Ctenoph_Hormiphora_californiensis_evg197868__p1_GENE_evg197868_evg197868__p1__ORF_type_
3prime_partial_len_168___score_33_29_evg197868__69_569_

MSDHEASVRRPPISYLHPSSERLIGSVLQRSPAAASDPQQSQPRLDREKEEEELQEKEKQTDKKRAWIRTVRATKWLFYIAWALALSYTLLNISKSVQKYLDKPVSTKITVFHDDNGTGSMQFPTISLCNLNRVSKSYLEKEELLKETWESLSQGKEIDWSKKGMDR

>Deu_Ambulacrar_hemi_Ptyfla_40v0_9_20150316_1g20851_t1_scaffold21886_cov93

MILLFRKTNDSVTVFTLMGIQISPDCRRVLDLTKSKLNRQTYPFGNCTNQVNSKGMDERVAAMYMSGQYTLANNLLKLVQISNKKTKNLHNLESARANLVRLKVYFETLNYDLTSELPAYTWEDLLSDIGGTLGLYVGFSVITVCEFVTLLLNLCGKVRRRFKKKTT

>Proto_ecdy_PANarthopod__XP_023229491_1_acid_sensing_ion_channel_1_like__Centruroides_sculpturatus_

MSFDHRVKVRTAYCYNCPKHGYARQAARFSNSSSSYMKNEEEEEDFWNHCSVILGSSSINGVSQIASTKQIYRKILWVIVFSGCMAGFLFQTISFLIHFSNYPTTVEVLVLNEGVLEFPSVTICNSNRIRKSQYCKYLESKPNVNKTCPDVMYFGYYDVILYQTKKL

>Protostome_Lophotroco_annelid_Pdum_comp286272_c0_seq1_

MINDIFGGNQLFTVAVHTKAYLSALWNELDTSKQEINRAKKNSLLMLNELSVNLSEFRKINEINEDFVKKNFLTVDFYYSVLKYTLVRESLKYTYTNYLGEIGGYMGLLLGCSVLTILEFLDLVMYNLFTKFVDHRHQRRSTATVKNTCAGDVENEPSKGAVLSTNE

>Cnidar_Rhopilema_esculentum_TR25170_c0_g1_i1_p1__Rhopilema_esculentum_TR25170_c0_g1__Rhopilema_esculentum_TR25170_c0_g1_i1_p1__ORF__type_5prime_partial__len_169__score_20_27__Rhopilema_esculentum_TR25170_c0_g1_i1_75_581_

SDVEDVLECRNRILMELYEHGFHEKCNCPPPCVDEVYHHIVSMTKWPALVDIPLYRIAFAKALSLEPNKLTREFIRSNFLQIRVYFDELGYKHVYEQCKISLAELFSNLGGQLGLWIGYSLFSIIELMAMLVTVAYLFVKQKCTNRTNVKSNGISQFEEKKSDDENAQ

>Protostome_Lophotroco_annelid_Pdum_comp347366_c0_seq2_

ANNALSTASPNNIVHDFAAGKNVTTAASMLTSQFIPTTSELNFHTSNPSSRKKRQLQNDNRLSELQVPEKAHIENVSDSGDLPKPAYGEEAIENMDPFYADAGHQMSDIIYRCKWDKKSCQSVGFKSHVTDLGYCHTFNVDPETVLNVSKTGSGFGLSLLVNIEQYEY

>Ctenoph_Hormiphora_californiensis_evg1106631__p1_GENE_evg1106631_evg1106631__p1__ORF_type_5prime_partial_len_170___score_47_55_evg1106631__2_511_

AHPLASAPAPPTSDPASPVTTTTWPPTSIPFISADEKLNCSSSCPVPCHKTYFDIMVSTSKVSARHANMLAKQIRAKKRNPNITAEYLLQNHVLMEFYIQDMFTEQITNVPAYGLWHLLSDIGGVTGLFLGASLFSFIVFVKSSVLRVYLYVKNSDGPWNAIKRNLKKP

>Cteno_MneLey_comp9400_c0_seq1__p1__GENE_comp9400_c0_seq1_comp9400_c0_seq1__p1__ORF__type_internal__len_171__score_23_27__comp9400_c0_seq1__1_510_

HVTSQRDHVTPQRDHATTGGYSLTTFEQKPLEGGLKRRGKPVDYDLVLASRAAAAAKETTNYCEIGTGSEVDRESLVPDEEEMSGVEVESCSEFMQSITASFIRFIFESDSPPFRRILWFLFWVTCCGYALSNIKAAFTQYFSYETASQLSFIQSLGEELTMPAITICNN

>Deu_Ambulacrar_hemi_Ptyfla_40v0_9_20150316_1g12295_t1_scaffold7626_cov144

MDEYRYGNGVTGMNTFTEDIRDSGPAKADASGYQEVKLKELGSELLENTTCHGVPRIQSARNAYRRIFWSMVFLAALS LFTWQGDDFDYMNWDTMDIDYEARDSGFHTILLIYDYIASLNYSEKLSLGHQLSDLLIECTWQGYPCGPKYASSITHLPF DCTRFVGQAAAD

>Deutero_Ambulac_Sakowv30041594m

MLEFCGCVDTMLRDEPRCMLLNRTQDSCKQLIHYLARKEILDCSCSQSCTESWYTTTTSQSVWPSNTYLKHLLKQIHYN NAKTNVINDLESVRQNLVRLEVYFEELNYESIEEIPAITEENLLSDIGGTLGLYCGFSFITIVEFLQFGFELLRVIFLRSTKVAPS NGSHA

>Ctenoph_Hormiphora_californiensis_evg1102714__p1_GENE_evg1102714_evg1102714__p1__ORF_type_internal_len_172___score_14_83_evg1102714__3_515_

KRNEPGEDLRMKIAHQARSRRSSDAFVGNDFTQYMCPFVVAKTCHFLINYYKRNGDIVYQPCSEPCSYRSWNVELTQS QFPSTESYFDTFVKDSVYVFPTPTYAYAQNNMARLHIYYDELKENHVDQVKAYEIQNFVAEFGGTVDLFIGFSFFTIFQLI EIAIAWCVLKCC

>Cteno_Ple_Bac_comp29868_c0_seq1_p1__comp29868_c0__comp29868_c0_seq1_p1__ORF__type_complete__len_184__score_26_16__comp29868_c0_seq1_53_568_

MHNGESMERELSMRGYCYRINPNGKLMGKGGDYGKLSIKLFSDLNQYSSTGKKLAQVGYSVQFHDHEQYSGSIPNSY WMSPGLIYKADLSLTRETRLQPPAGRCNDTVQYNTYGRNEENSCLAQCRDDAMMRSCGCVLVPPPFNPDNYYTPCTL KQWTQCGLAAYDNWCEY

>Deutero_Ambulac_Spurpu_015718

MHDVIEIAYVDFRKSLRSCFVQVFDALLLSYNPGLHLTLMVEQPEYIGILSPDSGVKVAINDPRIHAFPEDDGFMAAPGIA TSIGLKKTTITRLPKPFGDCKSDESSEYYEPEKYDFNRRSCMKLCLQKIMKQNCSCITDILIDGDMCGAIDKEQVKLCLTRQ YRHHDGQV

>Deutero_Cephalo_088090F_t1__name__chr_scaffold187_start_382926_end_384070_strand_pro_len_173_

MMGLFLYWSITHILVYFSYPAVTHIDEGDESQFFPAVTFCNVNKYRLSALTNQDFYHLGRSLGYLHPNRSLKAPSNRDLD YNRLEAIQRLADFSQLGNVTNGSLTELQNRTGHQLTDIRKEVQVSGQAVLRGHVHYQSQMEKTEKKLLQESKKNVLSV FLTLCEAPSGPKDP

>Protostome_Lophotroco_annelid_Pdum_comp382617_c0_seq1_

FYTFQSTRVLKFHVGDFFGELSDRLDAFEKEYQVLFTRMSTSFQSLNKSMAKVQQSSVIDEQFILNNFLKLDVYYGELTH ELIEQTPAYTLPNFFSDMGGYLGLLLGASVLTIFELLDVIFYNSALKCSEGRKMKSKDEINEDLSDITVPKCWDEKTVKSLPL DVNKVSPYN

>Protostome_Lophotroco_brch_Lingula_anatina_comp135130_c1_seq3_p1_comp135130_c1__comp135130_c1_seq3_p1__ORF_type_5prime_partial_len_173___score_9_94_comp135130_c1_seq3_2_520_

GLGTGSALGLSSTGTSSSGHDYLSKSSRTQSLDQFQYQFAKLNDTERMAIGHNMTTFLVDCSWQGYPCDKSNFTEFSD YKYGNCFIFNSGMNKTVRSAARGGHTYGLSLELYIEQDEYIMTTESAGVRVTIHEQNSMPFAEEHSIVVAPGKHTSIGISK VGLDYPVPGWQVL

>Protostome_Lophotroco_brch_Lingula_anatina_comp149479_c1_seq6_p1_comp149479_c1__comp149479_c1_seq6_p1__ORF_type_3prime_partial_len_173___score_3_67_comp149479_c1_seq6_91_606_

MSFVRGGFDFGNFTRNIGWELNNETMPLCLFRMKQCFPENFTHAFQSEYGNCYTFNSHHDEDSLSLQQTRPGAQQGLYILFDIQQAEYTELLNLNGALGVGIAFQIHSWYERPNVAQHGLGAGIGTAALVSMYQNELQNLGHPWGNCSNDRTLKHFPRYSRSGCEVECYLEK

>Proto_ecdy_PANarthopod__XP_023234672_1_amiloride_sensitive_sodium_channel_subunit_alpha_like__Centruroides_sculpturatus_

MYSNHKIVGDVVITFNSKELFRPTVPIGAKVSVHSKDEIINPFIDGFLVRPGNRYIVQLKVIEEDLLTNPYSTNCSYYEQSKEKSFDELYTRQRCILNCQKEKWLESCGCISGDYPFETDGIFCSEHKLRCIHETLGPAFWTVQTSRNPIESTSRFTSFGVNNGTQYYLLLRDK

>Protostome_Lophotroco_annelid_Pdum_comp419518_c0_seq4_

MADHKFAGTNNRGYYDNVKLTDFNEPPPKKKRKKRDNVKKVCTTFFREEASLHGIRYVFLKSPWFCRRFIWFCILITAIIIFIYLNSLNLQKYLSHPSQVSIDVRVKPSLDFPAVTICNYNRHRFSRVADLGLFDSLFEKAESEIASRSVNGAWYKYRSDVCIDDRDVHTSIIR

>Cteno_Puk_Fal_comp12245_c0_seq1__p1__GENE_comp12245_c0_seq1_comp12245_c0_seq1__p1__ORF__type_internal__len_176__score_12_40__comp12245_c0_seq1__2_526_

QRRVRGIGVLPPLNISGYTPCTLQEWANCGLPTYKKWYATLTDPTRKTPVCDCKIGCKRKFYISKVSGSEIHDPFASALLTSDNILEEMSNKEFGIEYNTTRDLIDNILILDVQMASAQVPVTQQIVTYTFGTLLGDVGGVLGLFLGASVFTVMEFIRLVVCWPCNRVEREKEEE

>Cteno_Puk_Fal_comp28403_c0_seq19__p2__GENE_comp28403_c0_seq19_comp28403_c0_seq19__p2__ORF__type_5prime_partial__len_176__score_35_15__comp28403_c0_seq19__2_529_

QRRVRDSQHTYNNYLSPTLHHPFAGCNFSSNCFSKLVLDSLRQIMSEEGKGDPPEKQFYSYKDFGSDIFQVLAVERNRPKWCKALWFLVGLAFFGAAAYLVYQLMTEYFEYKSYNKVSTAFTTNETLPAITICSKNPYNYTQLKSLLTEGEVGMLPWYLGSASIFHQHHSAPPLS

>Protostome_Lophotroco_annelid_Pdum_comp422306_c0_seq1_

MRSIKEEAKRFCQTTSIKGVPRIMRSQSTLLCLIWTLAVISFLSVALYQTVYLVINYFQYSTITIVKEKSIDLSSAPDIAICNSHPLSSLSEEAKNNMSSLREFHWHLNALTNCSNCSTAEMKLRKNMQQRLSSPYGYYQFIGKEAAKAASHLESDILVDCSLVVLRGSSFEHVPC

>Deutero_Ambulac_Spurpu_019480

MKYAFNPAEVKWRRFSWLLLVMAAFGALTYQVVDRVQHFASHPKSVDVSIEYTDSIIFPTVAICNYNTFRSSQLTGTELGDFLTIASTDPTSLNMTKYYDVLTSTNMSYLMTEYGHELDLSVVFATYSRVDLEKKANFSVKLTEWGPCFVFNDADNGYPPVTVQRAGRYQRGRSYL

>Ctenoph_Hormiphora_californiensis_evg17735__p1_GENE_evg17735_evg17735__p1__ORF_type_internal_len_179___score_18_12_evg17735__2_535_

LQTKKEFRQPPPGGTCNASLTQTNYGNYTDDACIADCRDRFIYEECGCVYVVPPINNDNYTSCTLVQWSECGVHAYFE
WYDLYTDIESDDPICTCSIPCEEITYEARVTSGELSQQYVERVFKDPIIHGWLANISDPSLNISYNTSEDIMKNVLAIDIQFT
TSQVSEIHEIVTYTFSN

>Protostome_Lophotroco_brch_Lingula_anatina_comp130423_c2_seq1_p1_comp130423_c2__comp13
0423_c2_seq1_p1__ORF_type_internal_len_179___score_19_73_comp130423_c2_seq1_3_536_

TVLDTSQEKQHHGNGKKTETNANLPLSIYLTNFANEVSILGLKQAINPKFSLFRRFIWGLCVMGGMICMFYHIYNRFATF
AGNPVSVNIAVHHKDSLQFPTVTICNNNVFNASSLSANNESVVAYMYPPHRLTPTPFASSVNWTDFHIRNGYEFENIYG
GFCAWGASSGSVSSCSAAN

>Protostome_Lophotroco_brch_Lingula_anatina_comp137912_c0_seq2_p1_comp137912_c0__comp13
7912_c0_seq2_p1__ORF_type_complete_len_204___score_29_30_comp137912_c0_seq2_78_614_

MSGCVAECKLGHMMNRCGCKLMHLPGPGATCSPEVMLTCIRPLMGEIKRSFSEMCECEVPCSSVLFETSLSYAAIPSES
TRAELALELGTTVENARKNYVIIDVYFESLNYVESVQMQAVEVTGIISDIGGQFGLFMGFSLLTIVEFLEFGVLRIISLLSKKP
NKKVNVLKINTNKNP

>Protostome_Lophotroco_molsk_AplCal_XP_035826425_1_uncharacterized_protein_LOC101861515__
Aplysia_californica_

MSVHCQVSETPIGRCFTFNGPEYVKRHGVRQATYDGRYSGLQVYVNLHQEQYFIYDDITAGLRIFISQHPETARLFKDGI
DVHPGTSTRLALSPVKFTFLPAPYNSYGSDTCVDTSVNDLQPKMVNASFYSHPECIEQCLNLRVSHICQCYTHLLYVPGL
RECTFGEYLLCSRPLKDT

>Deu_Ambulacrar_hemi_Ptyfla_40v0_9_20150316_1g25771_t1_scaffold47285_cov141

MRRGVSTQIRRLADSSTGVTFGYVGSERKESNPFAGNLGDGPKKSSNGDEEGVSTAREKRAMTLKDISRDFALRTTSHG
VGRIAEAGNGWAKTLWAAVVVFALCAFTAQLVTLIIRYLDYPVSISIGVVTNSSLQFPAVTICNNNKVRKSAILESVYDELL
SIDDAGSRRGYLTYIGKL

>Deutero_Ambulac_Spurpu_010266

MKINAPQLLCRYLLLRHLLLVHHRGDVPLVSDIGVAIMPGVRTLIAIKKHTERNMSHPYKTDCRDVPLEYYPFYSEEACLT
ESATKQLTEACGCRFPYMPGPSPVCTVRQIIDCGLHDFEVGDLGGLIGVHLGASLLTVFEFVDYTFVEQAHKILYIRFYWR
KLKSFCSRDKKEKETST

>Deuterostome_chordata_Ggallus_tr_A0A1D5PPP3_A0A1D5PPP3_CHICK_Uncharacterized_protein_OS
_Gallus_gallus_OX_9031_PE_3_SV_3

MLNIQQEEYLPVWGDTDETSYEAGVKVQIHSQQEPPFIDQLGFGVAPGFQTFVSCQQQRLVYLPPPWGDCKATPIESD
FFTNYSLTACRLDCETRYLAENCNCRMVHMPGNANVCTPEQYKECADPALDFLVKKDSEYCACRTPCDTVRYGKELSM
VKIPSKASARYLARKFNKTEQYIA

>Cnidar_HydraVulSc4wPfr_248_2_g2926_t1

MEFVKSKYRANLTSEHLLDLLRIPAKSLKPGNIPYCSLPELQNCTWIEWAKFNKDKMYKCPLPCKIDLYGVSLSRALFPTT
QYSSILAEQFRKQPHVLSIVHNITDELLFMRDNLLRFIIYYDDLSYEVLEQKPSYETLVWLGDIGGQIGLFIGAGVMSYFEFL
DCLAIVIYTRFFQKFTSS

>Deutero_Ambulac_Spurpu_010858

MLIDDFKNVKKQISSICPKSVSQNSDVDKFDWFLSQLIFRLIWMCIMLASYIACLITCKELRTKYDARLTIEEFSTTRKSSIAF
PAVTICNSNLVKASLIDNPTSKAIVQVLVDGSTELGLNATDEAFETQLDEILLAHAHQASDMILECAWKRGSNGCGAAN
FTTKITSAGVCYTFNDGE

>Protostome_Lophotroco_annelid_CAC9672764_1__Ofus_G22305_Owenia__fusiformis_

MKKDVSTQELGKANEDGDSADQTDSNITNISKSFAESTTLHGVSKVFGSRSNVRKCIWGLIFIGSVIGFIAQSVTLFVRYL
DYPNDVSVGIINPNYLEFPAVTICNHNAMKLSKLKDTRFQEFVDLQSQFNSKKVKPSTQTPVSTPSPDWTDSLTTSAAD
GFTSATSKFDKFIRILSDIRPR

>Cteno_MneLey_comp614729_c0_seq1__p1__GENE_comp614729_c0_seq1_comp614729_c0_seq1__p
1__ORF__type_internal__len_184__score_30_14__comp614729_c0_seq1__1_549_

NFISLSYERVRNLKQPYGDCRDDAMPLYRNYSISGCERQCFTEIMIKECNCSSYYLKAMPGGKDVECNLYQEKNCVKELI
RRYHAGEFINSSECRCPDPCVSESYLATLSTAEYPAKNHGEALQRYNEANRGPNDVVQDVTFYRENLMKVHIYFEILSIKE
VEKVGSYKWSNLMGDVGGQLGL

>Porifera_Syc_Cil_scpid91493_scgid1215_

RCTHVHGWVHCTSMCIIMCMLSSPSAAIIQGTNQTFDCAAQQNCVQPCEYSAYSYVLSSATYPSTRSVISPALTQAEEA
ALLSTKRRNYVHLNMYYEELNYQKVEQVIAYDWLSLVSEIGGLAGILLGASVLTFLEFLDFVTGSILCAICDVSHRRRYNAQ
RKSSFQGNVELTTNWSLEPPPNT

>Protostome_Lophotroco_annelid_Pdum_comp368911_c0_seq1_

MMKTLELCQCFSAMDIDDLPLNAMDTSSICKSDAQLDCMENVTFNVAMGNIGCECHQACRDQIFHTEASSHSWPEV
NQFDDFRKAICSKLPFTQESCQRMREIKNEEYMKDWFLKTDFLSVRVYFKTLTFTKLKEIASYPLANFASDIGGSVGLWL
GLSVYSGVELLNLVLQLFLYCCRKYSKC

>Deutero_Ambulac_Sakowv30018851m

MNHMLLYHMLKWQRMGESEELYSQFYNTTAENLSERKLEKAHLKAIYSAVYTYSNTVGISPETVPVTTLYDEFYVPLFYE
MILNIYVTLKRVKDTTGVELLAREDSGYVCKRYMRDNFSKTTVYFEDLKIQSINQQRGYEPFKLVCDIGGTLGLFFGASM
VTFLEIIDFFILQFWRTKLRSGRIQT

>Proto_ecdy_PANarthopod__XP_023234400_1_amiloride_sensitive_sodium_channel_subunit_beta_lik
e__Centruroides_sculpturatus_

MDKNSFNETINQDRCIIECIKENWKKTCECISNSYPFRFEGEFCSSVDCIQKVNNDECYQKCRRSCRFTVFEYDINEFSND
FYYFLYPKNLGKEYYDIPYSKWLTEMTFLYKRLELHTYRYKAQYEMIELFGYVGGYIGVWLGISLIAVFDLLEIIIVYIYNKLY
VDITVKTEEKSNSSSKQVTKF

>Cnidar_AlaAla_agalma_gene_62598_1_isoform_1_p1__GENE_agalma_gene_62598_1_isoform_1__ag
alma_gene_62598_1_isoform_1_p1__ORF__type_internal__len_188__score_21_27__agalma_gene_62
598_1_isoform_1_3_563_

FMDDSEKYKRYVQYYWNVSESTHGDFVTPPYLEYAVISCSFNSKNCSGNIALTNFSSWSLGKCYTFNHNPNVRVTQIRQ
GYLHGLSIVIYLNQNDYDIESPWVIPTAGILVDVFTKGLEIPSAKGLMVAPGEYTRLAIKKSTMKRLEHPYKDNCTHGNHE
ALLFKGGYSTEGCILSCFMEQQIKRCG

>Cteno_MneLey_comp671018_c0_seq1__p1__GENE_comp671018_c0_seq1_comp671018_c0_seq1__p1__ORF__type_internal__len_188__score_39_00__comp671018_c0_seq1__3_563_

PNETLLIDDGGILVSPGTETFISLSKSKVTRLPPKYGSCQNVTSPFAPTVSRSVRECIQDHKLHLAVKNCSCVPWYLRDKLMKLNNTNYTNYWYNMLMTSAEVAEEDVDNDQICGFATQSMCDVQVLEDLVGTIEGNGTVCPEPCSFNQYEWTINKGDFPPTEDYWSMMLKDSVVYEDEVKDKTFSF

>Protostome_Lophotroco_brch_Lingula_anatina_comp141642_c3_seq5_p1_comp141642_c3__comp141642_c3_seq5_p1__ORF_type_3prime_partial_len_188___score_13_41_comp141642_c3_seq5_166_726_

MELDNLAEVKKKLNETYNYGVNVIDTDIKGNEEESEREPTWKSAFATFSTATTFHGVGKIVEDTPFMLRRVIWIFLVLLGAGLFLMQLIGCVVYYRSWPVTVNVKINYADKINFPAVTICNLNAFKMSKAAENGHYEIINELYSSGLSGNHTDRTGNGTIEDLFQLNAHTKEDMILSCRWNNELCSD

>Cnidar_Rhopilema_esculentum_TR103753_c0_g1_i3_p1__Rhopilema_esculentum_TR103753_c0_g1__Rhopilema_esculentum_TR103753_c0_g1_i3_p1__ORF__type_internal__len_189__score_20_84__Rhopilema_esculentum_TR103753_c0_g1_i3_1_564_

IWLLLMLFFVSLVILTVTPTVRRYFKSPSLTKVETVRVSEMKMPAITVCGPSFSKAKVTNFAKEKNLTVKYNRIGLIDNIDDILGRASSLEDLAFDNNTMFIINPDENDIAIERVLVANQGYCYKVNANGTAIQKSVDLGFTFEIVLFVNLSNGVHLFDSPTDAVHVIIQDHRQFPSLDSGRIFAPIG

>Cteno_MneLey_comp1086195_c0_seq1__p1__GENE_comp1086195_c0_seq1_comp1086195_c0_seq1__p1__ORF__type_internal__len_190__score_29_76__comp1086195_c0_seq1__2_568_

KLLWAVLFLGFFVYSFQCCLKSFNTYFSMPTSTKYNMYFEPNKLLRFPSITICNMNPHNTAYLDKEENNPMKAYLLSESRVPWDTSNLTPDVSETDYRNFNVKKYYRQAGQKLNDFISEVVFFEEEIETSDAFREQFTTHGMCWTFNADGSRQVNRTGMDFGLSFKLNINQSSYPNFVTTAGVKVMFHN

>Deutero_Ambulac_Spurpu_007183

MAEGLNKSFPPKIGASIAQSVDFGVAGASLAENAHQVEPPEIKKSRWYLLSSSFPHAIGVAGIKFIFNGDEMKIRRYVWGILLLICLSAMVYQTGTQCVRYAMKPVNTLLDVEYVDRLDFPSVVLCNYNAYRKSYIEDNEAEFAALLEHLANRTSRRHQDIDATFYDGNLKYTNLTELRLNAAHQLDTIV

>Deu_Ambulacrar_hemi_Ptyfla_40v0_9_20150316_1g32037_t1_scaffold442052_cov104

MWASKTKIHAINVGEDSKSNANEGLQEPIQNLDLKRDVEDEDAKAYKKETGCRKILFSNFADDFGISGVQVIVGRAVVPLRRLLWIVVLLAGMGVMTYQIAERVAYFASFPSHVSVKLHYVDHLTFPAVAICNFNMFGYTATENSNLTHLLYDIYGNSEAMHVIDFDAYDINATASEEFYISHAHALSEMIL

>Cteno_MneLey_comp601992_c0_seq1__p1__GENE_comp601992_c0_seq1_comp601992_c0_seq1__p1__ORF__type_complete__len_212__score_42_61__comp601992_c0_seq1__142_726_

MLNKTCGCLNSTPPLNDVFNYKECTIKQVATCSLKAYYQYVLDFADVKSDYEECGCDVVACEYTLFRTTVTATPLPDTEIEHKLKAWNSPRHFDRFQRFSNYTAEDLAKNMARIEFFFGSSSITKVEEIISYDFDNLLGDIGGVMGLFLGASIFTTMEFCTLVVNLLSRLWQWKIQPWLRSETRDRKIISETNL

>Deutero_Ambulac_Spurpu_001740

MAQQAKEAWYDAPSKPVTAFKSLDKESATPIEQPSFNMSKEEENSVLFLVGNRMASTGAHGIPNIQRASNIWRRLIWL
LLFIGGVTVVVWQLTTIINNFFAYEVDVNIDFVRQRAIKFPAITVCNLNPVPMSKADVLWNIASNEEDATSLRDFLPAQK
PSGVMNATQNQNDAGASMRVIFFSYFHIKFHLSKC

>Deutero_Ambulac_Spurpu_001741

MDGPTDPQAEDPFNATSPGMVSAIFGDKHLGRGVPKQKQQHPVTQGVAISALINSRLATTGAHGIPNIMRASNLVRR
ITWAALFLCSVTIAMSQWQVLIIKFFEYDVATNVDLVRKRETVFPAITLCNMNPIANPSAVPSLNDYLDSVGASATSPTG
GGGGGDGGGTGSGQSGNSAGSNGGRKVRPLENSHDLNT

>Deutero_Ambulac_Spurpu_017119

VFLRSTRRETMPTQQVKVKPAGNGASGKPPLATRNSKIFTVSQPPRKESFIGSDIEKGKDKVVELVNGKLNRNAKVNGA
VNAKDNGDASSSPRPFNRFMYRDRMSWIAVLYTSIPEAIGLSGIKYAINVGENKLRRLFWILLILAGFGLTCFQIVDRIAFY
LKYPTNINIAINYTDELPLPAIAICNVNPYRYVT

>Protos_Platyhelmin_Macrostomum_lignano_A0A1I8GKY4_A0A1I8GKY4_9PLAT_Uncharacterized_prote
in_OS_Macrostomum_lignano_OX_282301_GN_BOX15_Mlig021706g1_PE_3_SV_1

MDILRSSAKAILARINGTGLGLRRLETLRDHPYASLNQKANLTANNLASTGDFISVLDVYPVSTKASVFTESFAYPLRNLFS
DLGGILGLWVGVSIITLLELLELVIQLVQLAALRLKTVSEMPNQKVHSWQEDKHEGPASDIRHSTIVPKLPVDQTADLQL
HVALPPKPPPGSTEQPCPDFVVRQPSWFERIFSF

>Protost_ecdy_cyclo_nema_Cele_tr_Q7YX13_Q7YX13_CAEEL_Uncharacterized_protein_OS_Caenorhab
ditis_elegans_OX_6239_GN_WormBase_WBGene00010278_PE_3_SV_1

MVSVGHQIFSLLSFTLFSHFIFGNPFLILHFSVPVKMALPSSEGLRKRKDSLSDEFDSDLDDNEMKAIVSDIEQDKLHDETP
DTRWIQFKNHLKNWGETATIHGVPHMAQAHTVIAIIVWSIILIVSAVAFVYMFYSIAASYLAFNVVVNLNTGLDSEPFPS
ITFCNTNPYKLSEMVNVPELNALLTVYQASADGSLS

>Cteno_Ple_Bac_comp46220_c0_seq11_p1__comp46220_c0__comp46220_c0_seq11_p1__ORF__type
_complete__len_202__score_27_04__comp46220_c0_seq11_1766_2371_

MLCSDVLKDNSFTREASVNGNCFRVNPEGKLKGKGGEYGRLRLMFFADLNDYTAPAKNQAQYGYTVAFHDHETYSSTI
PSGFFMSPGSIYKVDLGLVREFREPPPAGLCDSTLTTNTYGRYEENSCIAECRDNAIIRACGCVHVVPPHPDNNPLVEYLP
CTLEQWAKCGMREYKKWGESNSNRHFKARKILSKLLETQKTI

>Deutero_Ambulac_Spurpu_018816

MGLGPFVSCATPSRAASACKSYGLELMLDVQQWEYFFSPFTRQSAGFQILLYDYGSVPLIEDQGFGISPGTHTHVGVEV
TDVNNLKPPFGSCEDKELKYSDHYGYAECDQECKADFYIDTCGCRPYAAPGNNDDDTGDDDDDDDDDDNDDDDATT
TTSTTTTTTTTLTTNYNNDDNNNNNNHNHNHDNNDTNDSNYNNDN

>Protost_ecdy_cyclo_PriCa_ecdy_XP_014676234_1__PREDICTED_acid_sensing__ion__channel__1_like
_Priapulus__caudatus_

MATWTKRARRSISACSRQTTLHCVHWAFDSRIPLLLRLGWIIAILSAWSYMFYLTYKRVTYLSSQPMKTIIDEQFSARGLE
FPGVTICNNNLVKRSAARQLKEYAELERFLSDYADFTWRYGARAAVNFSTSGYKLAHVTPNDLYRLGRRDSAAAILRCSY
GGENYRNHLQLMTRSHGNCFTFNSQEFKLSHDTIDPGCPG

>Protostome_Lophotroco_molsk_OctBim_tr_A0A0L8GE48_A0A0L8GE48_OCTBM_Uncharacterized_protein__Fragment__OS_Octopus_bimaculoides_OX_37653_GN_OCBIM_22035600mg_PE_3_SV_1

LTLLGYPYHPIEEDCVSVEKQSLLDNTSPYSQVICEEKCRVDFIIRSCNCSPSFAGGNNRRCTFREYITCERSATDEFFSNKS
HLNMCNCIRSCSFTSYSISTSVAYYPRPAHKLPTKFKIHLYYESLILKEMSEQEMFTSEAFLAQFGGQMGLFLGASLITALEL
VELIIMVLWTQFRCLPSQAHGAGFPVSMAYVFPPS

>Deutero_Ambulac_Sakowv30017787m

MKYFDHYSMSACRIECETYAVEEMCGCRLVEQPGDGTWPVCSANQTHLCAHPSLVDYNNGHLDINCSCREPCDVEDY
PHVVTMSKLRPAFIEKLFHNTKHNMSETYIEQNVAVLSIFYEALNMETITQLPDTDGIILLSQIGGNMGLFLGASILTVVQ
LGEFLFDFMRAKMRMSGQEEYSGRAEKQIKNAWDINSTDTSLDTITT

>Cnidar_Rhopilema_esculentum_TR38848_c0_g1_i1_p1__Rhopilema_esculentum_TR38848_c0_g1__Rhopilema_esculentum_TR38848_c0_g1_i1_p1__ORF__type_complete__len_207__score_33_00__Rhopilema_esculentum_TR38848_c0_g1_i1_410_1030_

MKEEGKNGHFVTEESKMEEVNSLTIHGVPKIFTTKSYFSRVLWALLCTSTVVFLIYFMLDAIRKHLNFEIYIKEERIPKHEM
SLPAITICNTNFYSGDDDDYAVNQVLPKNCSYTDAEYFETETNRKYFLDICHWFHGGIMGEFSPKNISFPQNASFLPNFW
PCVTLNKNKTMVQKRESLNSGLKLLLIYGKGQIRQFLGMGMKFL

>Cteno_Puk_Fal_comp37466_c0_seq1__p1__GENE_comp37466_c0_seq1_comp37466_c0_seq1__p1__ORF__type_internal__len_207__score_26_70__comp37466_c0_seq1__3_620_

NDDPFMFGFQDLASDYYQVLRERARPKICKILWAILVCGLLAMTIFMVYKLLSDYFKYDSYDKISTEFKSSLVFPAVSICNL
NYVNNSRLEGELTDDILDVYGEYLKAMEDLKYSPDPITQFEEYEELFVTLESYETDNFGILDYYFKLNLYDTFVGSYDYVLG
GQGFVIPYDKWPFYDGYTKFGSCLEINDNSTFKQSIEGKQGG

>Cnidar_Rhopilema_esculentum_TR39512_c0_g1_i1_p1__Rhopilema_esculentum_TR39512_c0_g1__Rhopilema_esculentum_TR39512_c0_g1_i1_p1__ORF__type_3prime_partial__len_208__score_17_84__Rhopilema_esculentum_TR39512_c0_g1_i1_1_621_

MANTVHPKQDQKVHTKLAARFKQFSEVATVHGVHKVANSTSRSRMIAWSVLCLTLVGHAVYSISSSVKEYMSYPTLTYI
KKETSNKMDFPSITICNANNYRLSALQSMSSNDTDIEQLLASYLQTSLQTKNVLPLSRSDVAMKYSSALLNSTTIKTLSSSI
EWMKWNFKDWCKFSFSIKCESPEDFTDYFYHSNTGVCQTFNYDGKY

>Deutero_Ambulac_Spurpu_025391

MALNTSLGATKISPSVQNTGKGTQEEEMAGVKGVRHEKYREDGRELQRPRRRTKSSGEDQELGGMNAYILIGSAIQNT
GAHGIPNIARANSMSKRIFWSVLFVGAMAVFFWQMVMIIDAYTDYEVITSLDVLYASKIGFPAITVCNLNSYASSKASVF
SKEFAEAVGVTYNANGNDGVNATSVVPGPTGTATQASSTERVRGLHFD

>Deu_Ambulacrar_hemi_Ptyfla_40v0_9_20150316_1g31105_t1_scaffold269182_cov104

MDEAREELYTQPSQFWNMTTIFTQIYSSLNASEKQDMGHQLKDFLINYTQASTVPAYAHCIQGHVDYKFDYLIRSRKXX
XXXXXXXXXXXXXXXXXXXXGLTLQLNIEQDDYLEEYSEIAGVRVLIHAQDVAPMPEDDGFDVNPGVLTSIGIRRTEITRL
GDPYSNCVSKTGDIGQYSNVFTDMFNANYSQEVPQAEFNRGSGFLQHP

>Deu_Ambulacrar_hemi_Ptyfla_40v0_9_20150316_1g3256_t1_scaffold1378_cov105

MKSGQYEYVRQPAPYGNCTSGDEVYMVNQQFNYSVVLCMKECLQKNLIERCGCVTDVLIDYDKCSVLNTTQQRCRQVVEALYEEDDLNCTCKNPCHEVTYAAQVSSSLWPSERYETHLYGVLADKSIKAAITLQDVTESRRNLVRLKVFFEELNFQTIIQTKAYKIENLLGDIGGTLGLYIGLSLITVFEIIVFFLHTCRFCCRRIWS

>Deutero_Ambulac_Sakowv30012023m

MSTGVAFDTVAVERLPLPYGDCYDTAYPENDGNYSAKICEEKCFQNHLLDKCGCVTSVVLDGETMCSALDEHQDMCVQMMTHFRNNGNLFCRCPPPCRERDYKITTSDSNWPSKVYMEHFARYVKTVNDDNEDMDEDRIRENIVSVNVYMSDMSTMVFVEVPSFTLVQMLADLLAAFGGWIGMAVFINVIANMCWSRKVRTHVGGSPSSQPI

>Protost_ecdy_cyclo_PriCa_ecdy_XP_014679704_1__PREDICTED_degenerin__del_1_like_partial_Priapulus__caudatus_

DDAEVGEPLRRQLLLKVLSSLTWQEMINASTEMDTMIISCYFNGIKCNMEKDFKLHLSTDYGNCFTFNYASDNLRYLVKQGGATRGLTLTIFVDQLNYVSGLTEAAGARISVHEPGHSPFIVENGIDVPVGTRTSIGVTQGKIQRLKYPYDSNCTDGDEPEDEKSYFGEEFAYSLKGCVFACAKSRIMDECDCIPYQYVDLDDTSSRLCNTA

>Protostome_Lophotroco_annelid_Pdum_comp342860_c0_seq1_

TAAALSSHTYLAVKLVESISLPDPYGTCVSEKLEMFPDQNYTQFKCNRQCLTKAKVNLCQCRDQYMPGNASYCTLGKHLSCHANLAELNCNCSNACHVTAFASQASFSSLSKNGVSTLLQDNYKYRENYHKVLNYRYRIQPKHTERIISALGRFSKAVENLFVFLKRETSFGETSTIYLMNAAIKAMDEIAKQDVKEAFSFLPDLVEAFSCA

>Protostome_Lophotroco_brch_Lingula_anatina_comp141642_c3_seq1_p1_comp141642_c3__comp141642_c3_seq1_p1__ORF_type_3prime_partial_len_213___score_11_60_comp141642_c3_seq1_35_670_

MNGPIVSQFGTGTDANSNGIIRSGKMELNNLPEVKKNLNGNFSYGVNVIDTDFKAKEEESEEEPTWKSAFATFSTATTFHGVGKIVEDTPFMLRRVIWIFLVLLGAGLFLMQLIGCVVYYRSWPVTVNVKINYADKINFPAVTICNLNAFKMSKAAENGHYEIINELYSSGLSGNHTDRTGNGTIEDLFQLNAHTKEDMILSCRWNNELCSD

>Deutero_Cephalo_088070F_t1__name__chr_scaffold187_start_346867_end_347508_strand_pro_len_214_

MEGGSQNHDVPRQSDINVFAASASMHGLAHIFTEGKFTVRRILWAGVFCGCVAVLLVQSVDRVQYYLSNPHATKLDEITAVNGLNFPAVTICNMNSFRFSQVNQQDLFYAGPDILDLIDHTLPRPKLKISEEFLNHMNENPSDHQEMRQKMERIRELTNFDGFVPGKFSMSDFYDRTGHQIEDMLLDCKYKGEPCSAKNFTTVSLVLLLFLKS

>Porifera_Syc_Cil_scpid76710_scgid1215_Acid_sensing__ion__channel__4_Acid_sensing__ion__channel__4_2_Amiloride_sensitive__cation__channel__4_B_ZASIC4_2

MLVIDQTYMNRACADLFYSLVIVCCLLVGPLCSANDNSTVCNPIQMFRCIDEVKTAIIQGTNQTFDCAAQQNCVQPCEYSAYSYVLSSATYPSTRSVISPALTQAEEAALLSTKRRNYVHLNMYYEELNYQKVEQVIAYDWLSLVSEIGGLAGILLGASVLTFLEFLDFVTGSILCAICDVSHRRRYNAQRKSSFQGNVELTTNWSLEPPPNT

>Deutero_Ambulac_Sakowv30015883m

MHAPWGNCTYGKTEQVFNIDGFQNEYSLMACERACIMNVLEQYCGCTNSIHRNGSMCTILDEYEDRCNNLIQFFYRNDLLQCDCRQPCSETQYDLTISQTLWPSDIYLNHFLQNVHTKNSKTRTITDASSARSNFVHLEVYFQRLSYETTSEYKAYSWEDLLSDIGGTVGLYVGLSVITVCEFLKLFFELFKLCTAKKRKKKQTYDTISVKNTAV

>Cnidar_AlaAla_agalma_gene_74369_1_isoform_1_p1__GENE_agalma_gene_74369_1_isoform_1__ag alma_gene_74369_1_isoform_1_p1__ORF__type_internal__len_217__score_15_73__agalma_gene_74 369_1_isoform_1_3_650_

ASATNGQHTQRRPSVLEKFRRAGMKAKLVQRMEKIIEDRLTAKQVFSRYVQSSTLHGFRFIFTRTFYIRRFAWLVVTLT MAGLFLKELRDSINLYYAHPFSTTSTVEYVNKIKFPAISVCNLNSFKYSGILGSEFGKRLINSTKPLERDWGMPKIDISGPKL AKTMKESSHDIKTMLLGCEWIARDTAQNRPNFCGWQNFTRFYQYYGQSCFTFNS

>Protostome_Lophotroco_annelid_Pdum_comp414035_c0_seq2_

MVALTQKDSLGLAVEKSGWVETKRPMESWDLELSIPGTKYLLEKKTSKFHRFIWFIILLGAFIGMCFVISNRIIRFLQSPKS VNVEINYNPSVVFPSITICNKNLFSASSVYNYEKRFEAENRTAGKAFGTFKQMFVFQFIADYFTHLQGKHSYDAATLEKFE IEYKDIINGQADFVHIYNVSTPDVKNMILECSWKGKPCGPQNFTKLMTDFGFC

>Porifera_AmpQue_tr_A0A1X7TPC1_A0A1X7TPC1_AMPQE_Uncharacterized_protein_OS_Amphimedon _queenslandica__Sponge__OX_400682_GN_105314570_PE_3_SV_1

MTRESNCVRSNSNTRSLDFFPGLPYTASTCHIDAEYRNYADTNKCGCSPEVNPSSNVNTRNCTIADICCLSQERRTFNPS TNCPLSCDHLTYNVEPSYSKLFSGDSSESAAESFGTAIEEVEASLLSVYIYFDEMLATQLFTDYSYTFANLLADMGGLLGLF TGASVISLLEVCILLFDVLKNLLLTREMKKAVEKLESKINLPNVDAITKEEIEDD

>Protost_ecdy_cyclo_PriCa_ecdy_XP_014669620_1__PREDICTED_amiloride_sensitive__sodium__chann el__subunit__gamma_like_Priapulus__caudatus_

MVTVLRARYTSRGDADEAAVADDGGEGQTTAATPSGSETDQTIRNRLYHFIASLYGEQATQGAENIAHYGHQLTDLIQ ECNWVGMPCNMTTDWDHSQHPVYGNCYTFNSGEHQPVKMIGRTGSLHGLSLTLYIEQDEYIGRYAQSAGVVMSLH PANTHGFPEQYGIDLAPGLETSVGLHVKKVFRLSHPYGNCSHDGHNVEYFNSFHNMQYTLEVNP

>Protostome_Lophotroco_brch_Lingula_anatina_comp136149_c7_seq1_p1_comp136149_c7__comp13 6149_c7_seq1_p1__ORF_type_5prime_partial_len_219___score_18_23_comp136149_c7_seq1_2_658 _

ETRGLGVSPGFHAFAGLKRLQTVNIEPPWGKCNKSLRLDHYPQYTFSGCIIECKLQKAVEKCGCKPLEYPGPYPVCSPREI QECVQEEMRIINQDFSKHCSGCTVPCNSTNYNIQLSYVALRNATMHHKYGQHIIPSGYSGDYVQDNLLALDVYYEELNL ESMEQKKAVEESGLLSDIGGQLGLFMGFSALTFIEFFEYIILKFRRLTRRRKLIEPIV

>Cteno_Puk_Fal_comp26472_c0_seq1__p1__GENE_comp26472_c0_seq1_comp26472_c0_seq1__p1__ ORF__type_5prime_partial__len_221__score_43_26__comp26472_c0_seq1__674_1336_

NAEYGDPVLEQDYDPDGYTIAKCVMNNRARYTIQKCNCIPFYMEVPRDLSSTSPAVCNLYQEWDCVEPLLEDLALGTLT MGGQLVQDPDCPSPCKYLTYSYTISQSEYPSQSVEDQVLSRVGASRGAGWNIERIRDNYLKVHIYFEELSTLEMERHPSY MIANLIGDVGGQLGLLLGMNICSLVQFTDYIVRFSFFKGIMKLVRMSKRNNKRSVGGRSRL

>Protostome_Lophotroco_molsk_AplCal_XP_035824945_1_amiloride_sensitive_sodium_channel_subu nit_beta_2_like__Aplysia_californica_

MFGGKYSEPACRKTCYQQEMYSSCHCCDIQYPCLEEKLVKITGINPTGGYIRFCNVSDTYQTDCAENVHEDFMAGKLG CSKRCPPPCDEEQFQYMVSSGTWPSVTFSATFIDKVNAAAQGGMGKYLDFAKDPLFVRNNFLRLVVYDEAMTFQVLE SQPLYDWSRLLSDIGGQLGLWLGFSLMTAVEWLELLYDVVLNIGCKLCNRHPNKPHPEGDTLSVS

>Cnidar_Rhopilema_esculentum_TR51234_c0_g1_i1_p1__Rhopilema_esculentum_TR51234_c0_g1__Rhopilema_esculentum_TR51234_c0_g1_i1_p1__ORF__type_5prime_partial__len_222__score_25_64__Rhopilema_esculentum_TR51234_c0_g1_i1_840_1505_

PIHKVITKRLKSPFKTNCTNGDGKFLIFPGNYTVGNCQYSCFFKKALEECGLIEPAYKFHKPREFQESLKRFQENNSSKADLSSCFHHLKMNEDDCKCPPSCYEEKYRTKISYLQWPTPANIPYYRSLIANFTGREKITDADVHDNIVFLKIYFDELSYDIIAEQAVHLWPRFFGTVGGQLGLWMGASAFSIIEFFLFIADAVLEMCRSKVRNDDSSNTSA

>Cnidar_AlaAla_agalma_gene_93708_1_isoform_1_p1__GENE_agalma_gene_93708_1_isoform_1__agalma_gene_93708_1_isoform_1_p1__ORF__type_5prime_partial__len_223__score_15_94__agalma_gene_93708_1_isoform_1_482_1150_

ISGILVSPGTLNHFVLNLKKVNRLPVPYQSGCGSRKLKYFKTYTTDNCWTETLSAKVIKKCNCKLSFMKGTAKMCEGRTLMRCTFSVWSKFNKDQEFDKMCPLPCNFEKYDVTASSALFPSNSDTGRIASMFNVSGTHEEQRTYLRENFIKLNIFFEDFHHEIYYEEPSFRMYTLISLICAMIVFYFGASCLTIYEFFDLLITIIIRKIKKKRRTSNYPSRM

>Cteno_Ple_Bac_comp44341_c1_seq10_p1__comp44341_c1__comp44341_c1_seq10_p1__ORF__type_5prime_partial__len_223__score_28_26__comp44341_c1_seq10_1_669_

ETTRECLQSVWLNISAEYCGCVPWFLIDKLEELERKEQIEKWISISLDEDEVFNWTAYADGEVAEDDYMCGYSTYSLCEAYVMDKFNNATWLASSPFKTVCNEPCKYYKYESSLDYSNFPVSEEHFNTMMGDFSKNYTYVSENFLRVHIFYEDLKLTTITQSASYEIFNFIGKWNDILAPSVWRAQSGSVAPDIAPLIIVIPAAVSIRQECRARHSDRARAG

>Proto_ecdy_PANarthopod__XP_023243711_1_uncharacterized_protein_LOC111641737__Centruroides_sculpturatus_

MFPSDVDYEYNRSIIDKYGIREEDILHDFAYNNIAFRTIQTDELLRQRNCYSIDPDDYVKKNMNRQEYASIEMYNIDIRLIGSIRIKFNPEETIWPSNPIGIVAAVHSKDDIINPFISGIPMTPGNRYIVQLTPILKTLLPSPFSTNCRNYINSSNNSFLFTRNYCFAECRKKLWLEKCGCLSPEYPFVTDEEICDISTYVYECLSSINLTLCLDKCRKRCK

>Deu_Ambulacrar_hemi_Ptyfla_40v0_9_20150316_1g12296_t1_scaffold7626_cov144

MVDLGSKDTFSRSSPSGLYISDVTLSTNLFPLVLFSNFTTFFNPVYGNCYTFNSGRDGVRLQTSRPGPFYVKCLMSVENAYTTNTLTPSCATECTERCSETTFRTTVTTALWPNQMYESTIIDKVMTISKDLRSRVASSPEFAENNLCKLNIYYQEFNYESISESPKYEIEDFLSDMGGQMGLWIGMSFCTIFEFVEFCFDVIRWIRGKLSARARIHNNPSHA

>Deutero_Urochord_tr_H2XR32_H2XR32_CIOIN_Uncharacterized_protein_OS_CioInt

MSKNCTVSSFDQAKQNRTHYICQDDQSTGTAIADRCNAYFAYKTSVQIKDVFVNEIPLPAITVCNFNRYFGIDNPIEQLAMDQLSTVLDNRYNIESVLYHRHNESVNYFNWTKPGFSLRNYMMDKGWNFVNGYNQGIYFPTVFYCTLGKTVACSHANFTAVLTNHLGVCYTLNVNTSQTIAGKMHGFTVVLDTYQNQYTENIIHGNWEGGILFQVHSPYEPPE

>Ctenoph_Hormiphora_californiensis_evg14410__p1_GENE_evg14410_evg14410__p1__ORF_type_internal_len_225___score_36_04_evg14410__1_672_

EELSATGKCFRFNPNGILYSKAGDYGSVYLRLNANLRDYTDTSGPIGFVMAVHHHTQYSSTRNTGIIMSPGLKYRLTVKRIERTELPAEEGGKCNSALRSTSYGAYDLESCHTECRDKHLNASCGCIPVLPPHNVANYTACTLRQMDACGNNAYLQFVLRESDTDPSSARHTCNCWLPCDHFDYELNVATTAIPREYAQAKVQGEEGKRLNFTLDDFLANNVML

>Cteno_Puk_Fal_comp20056_c0_seq1__p1__GENE_comp20056_c0_seq1_comp20056_c0_seq1__p1__ORF__type_5prime_partial__len_225__score_62_77__comp20056_c0_seq1__184_858_

AEYGASLTTTTYGDYSEGSCIHECKDRWLMEKCGCVNMVPPRNYLNYTSCTLRKWADCGLEAYLEFSRKYMDPSLEEG
NICPCNSPCEETHYEAQLSSSSLSPAYTDAIFNKEGIKKYLNASSIPDLGIYYNTSTDLNKNLMVLEILFSSMVTSEIKQVVTY
TSANLLGDIGGVLGLFCGASVFTIVEFLQHFLSLLEKYCLGGLCGGRKRKKENGTTGHESEL

>Ctenoph_Hormiphora_californiensis_evg14406__p1_GENE_evg14406_evg14406__p1__ORF_type_3prime_partial_len_227___score_48_91_evg14406__1_678_

MVVLNVKPWEEEGGDHKKDEGSTANVDNEFLFGFQDLASDYYQVLREKSRPRIFKILWAIITTVLIAVTIYSVYRLLREYL
HFDSFDKTSTVFLTSLTFPAISICNLNYINYTAAHIEMMKLPGSEELFSKFLVNLQAMEYVDESPELNFTLFDTLVHIEADY
VNGSIDYYFRTNLYDMLVGEYDYIFGGVGQNIPLSNWSYYSGSTDLGQCFEMNDDGGFSQIVN

>Protostome_Lophotroco_annelid_Pdum_comp377163_c0_seq1_

LSEKINKTVIQTMGETIGKIVKNLSIYSIDPLKQEGSDIIGFLPAAYHESLDGVSKIEEYYTQLSSTKDRTRDLLVWKGPMAK
LDTEHILVNKYETESWKAWPTSYTLKDFVQAQDALKFISSITKDYVKVVNDLLHELEGGLESARTDAFTAMGSLVDLLEEF
RKKAQISPAFVLENFLKIDVYFPSLRYELWEQTEAYGFQNFLSEAGGYMGLLIGGSALTLCEILD

>Cnidar_AlaAla_agalma_gene_723_1_isoform_1_p1__GENE_agalma_gene_723_1_isoform_1__agalma_gene_723_1_isoform_1_p1__ORF__type_5prime_partial__len_231__score_31_89__agalma_gene_723_1_isoform_1_79_771_

GFHTFIEMGVQKVQNLPPPHDTKCGSIELSYFPGYKRSKCHLEQLTNIVAKECQYRGLFMPDESLPLCSLKQTVCCLYPVI
ARQDKKAVDQCPVDCHTMQYSYRQSYARFISKPPIGAGNIRYLFEDKRFKNLTLEKLQNEIRDNVVVMTFFFQDLKVED
IRQTPSYDFYKLVGDVGGQLGLMLGASVLTLVEFIDLVAFIIYHQVLRLSRRKKRAKEDEAMTAAIHYKR

>Cnidar_Rhopilema_esculentum_TR9696_c0_g1_i1_p1__Rhopilema_esculentum_TR9696_c0_g1__Rhopilema_esculentum_TR9696_c0_g1_i1_p1__ORF__type_internal__len_232__score_16_73__Rhopilema_esculentum_TR9696_c0_g1_i1_2_694_

ESEANLPKVTPDEDQLFMKNLSGPGLANRYLYSMGTQRIPVMKRGYCYKINPNGTFFQKEAGSDEALSAFFFLNVTDAI
SGAKSDTGDSIEIIVQDHQEHPFLDSGSVFAPVGYLTRIEIRKTVIHRLRAPYHSKCTNGEDMQLLFRGKYTASNCQQSCF
LARASRNCKALSFLYNLHLPENLKKPFPNTELESACSQGTLQELIDTDFSSCNCALPCFETKFYKSASYSK

>Cnidar_Rhopilema_esculentum_TR51440_c0_g2_i1_p2__Rhopilema_esculentum_TR51440_c0_g2__Rhopilema_esculentum_TR51440_c0_g2_i1_p2__ORF__type_5prime_partial__len_234__score_26_81__Rhopilema_esculentum_TR51440_c0_g2_i1_1_702_

IGFNSVDIIHLSPGYYTEVAVKKVVRKRLKSPYSSTCSSKSDVRTIFEGNYTVIGCLASCFTMNMYEKCGDVLPPGRVFLPK
EKFPKTPKTANLDKCLFEIYQTSGQLDCECPLPCYEEEFVTEVQQTPLRITSTTPRMNKNLARALNLSEEEISLEFYKKNIVR
LNLFYKNFMVETIREQPLYDVESLLGDFGGLMGLLIGASTISLIEILWVLAASISNLIRKKTRVMDIS

>Cteno_MneLey_comp353714_c0_seq1__p1__GENE_comp353714_c0_seq1_comp353714_c0_seq1__p1__ORF__type_5prime_partial__len_234__score_32_15__comp353714_c0_seq1__189_890_

NTDTENTDTENTDTENTDTDNTDTENTETENTDTENETVEESPTTDSPQPPTTTPWYFVTNEPLHETDYVNLICGFVEM
YICDKKLETAEMEGEIEFEPCPEPCEYHEWNVEIASTLFPPSEEYFDKFLRDQLYLPNTPTYDSIRENMARVHIYYDELKVN
QMVQTKAYEPQNFVAEFGGTVDLFIGFSFFTIFQLIEISVAMCLTKLWQRRRKTRKKEVNTSNSVLANPSVM

>Proto_ecdy_PANarthopod__XP_023232914_1_uncharacterized_protein_LOC111632651__Centruroides_sculpturatus_

MMNHKKAFRKKITVRNYVIALLKQSILTGVPQIFFATNFRRYWKILVLFSCIIGFCYQMGTFLRIYWKYPIMEEADNYFFK
DIALPAITICNYNGLSRSKYCKDYPDNCLTLEKNSDFCKRFPNSCLLKENITSIQIPKPEYSVMEKNAPREIVLKYGQQKEEL
LRRCTLTVSDKSMACPNITVSFVNFYEDIPRICHTINGRYRNAKKLPILKVNIDISGMNVSEESSKILIFG

>Protost_ecdy_cyclo_PriCa_ecdy_XP_014676231_1__PREDICTED_uncharacterized__protein__LOC1068
16184_Priapulus__caudatus_

MASLKLTIKNDKFPGRHPDLLGMSYKFYENEFLAPKWIAIMSIELSDHIVAAKTLSYREASSLPIPWLLNLHPCTNSTVENS
LVVNVAPGFRTTIGLTYRRVYMKGTPTAPKCESPREQSLKRYLNATSSYNKAGCMHTAHLTRAEKKCKCVMKASETLFD
DREECSILTYFTCFLPYLQEFKLSHDTIDPGCPDDCIVDTYEYQMSSHVAPSVRWSKVLNRSLAESRLEVIFSR

>Protost_ecdy_cyclo_PriCa_ecdy_XP_014678557_1__PREDICTED_acid_sensing__ion__channel__1_like
_Priapulus__caudatus_

MSSKAEQTTLHGVHWVFDSRLPLLLRLGWIVAILSSWSYLLYLTYSRVTYLSTHPKKTIVDVKYDPGGLEFPAVTICNNNL
VKRSTSHKLPQYEELERFLNDYTTYRWRHGLAAGENFPAADYNLSHLTMNELTALSEKDEGAGSVIGCHFDGRSCMGS
LRHITRLYGHCYVFNSQVSVPKTQDHDGWDPRPPLYSSQTTSDHGLTMVLWADADEYFYGYKGDMVGFTVSISFVVK

>Ctenoph_Hormiphora_californiensis_evg166283__p1_GENE_evg166283_evg166283__p1__ORF_type_
complete_len_264___score_29_51_evg166283__1291_2004_

MDIRSEIVGNIRFAKTTLAFESIGNVESTERGSCLELNDDNTLIQRVNGPTGALVITLDAETAKYLSGTTSAGFSVILRQAN
ETVISKEFGFSVAPGKRYMIDLKTTFTSRIEPPWGNCTNEVDILDMEGYFTLKECFLNQILYMYPKLEEQCACFPWFFVTR
FISDEKSKKYDPYSRIRMMRYWTEGISDEEQRLSFKGVGGSFRRGSDIPATGSCLRKNRDITGPVLQRAQLNAV

>Protostome_Lophotroco_annelid_Pdum_comp405647_c0_seq2_

KWYSDVLLSVNQLLKHFDDPTELDRHIQQMQVWRGFEAILEVSPVLQFKHSKEDIWHYWPTTRSALQFIESGKFQKPIS
DDATKFTKTLSKTLYESILQLETIMGDIYGGLDALNTDLKAIDEQRADYIIQNILKLDIYYSSLESQSITQSKAYTFVNFISEVG
GYMGLFLGCSVMTLCEFLDLVVYNFFLKAANSRQKKKATAPLNEKKFPQNYQDNWNTDLQNEKAKNVVYSSHT

>Cnidar_Rhopilema_esculentum_TR103753_c0_g1_i1_p1__Rhopilema_esculentum_TR103753_c0_g1__
Rhopilema_esculentum_TR103753_c0_g1_i1_p1__ORF__type_internal__len_240__score_28_34__Rho
pilema_esculentum_TR103753_c0_g1_i1_1_717_

YKVNANGSFVQRSTDIEKFLFIGLFLNISDGVYLRRSTTDSVEVNIQDHRQLPSRDSGTVLAPVGHLTSIEIKKTEIKRLKPP
YPSKCTNGENMKLLFKGDYTVSNCRLSCLLSQVSQKCSYINAWHKHLLPKAMKKQLSRNQSEASCSLKISKKLIESNYAS
CNCELPCTETKFRKYSSTAKMTSENPRILKHTVVITVFFGDMEYEKITELKMYTTERFVGEIGGQMGIWIGASIFS

>Protostome_Lophotroco_annelid_Pdum_Contig8512_

MRMNDVKRLNAPYSNCTEKNDTSFRNAYEHYYTKEACLVTCYYEALVHRCGCIKTPYEVYHRPSHQLSYPKCQDADDT
CVTALDEQMKIGNVTECIDCLPACREESFDLQISSALWPEPSLSEYYSNKLINANSRYWENFAQVQGSLMGVSEETVES
NLLQVNIFYRDLNYQQIEEKPDYEVAAFASDIGGTLGLWLGISAVTVVELLQLASQLVWNCRRNNKMDDRMKNRSWR
RPQAK

>Cteno_Puk_Fal_comp4629_c0_seq1__p1__GENE_comp4629_c0_seq1_comp4629_c0_seq1__p1__ORF
__type_5prime_partial__len_241__score_34_98__comp4629_c0_seq1__89_811_

PQPPLGFFMSPGSIYKVDLGLVREFREPPPAGLCDSTLTTNTYGRYEENSCIAECRDNALLRACGCVHVVPPHPANNPLV
DYRPCTLEQWAKCGMREYKKWVKRYTDATSMYPECHCPPPCQETHYTAQLSSSALNKFYASSKISHLPKGYNTTQDV

MENLLIIDILFTSMQVSEIREIVTYGWGNFLGDVGGVLGLFLGASAFTIIEFCQFFTFAIAKWCFNTNLATGKPPSNRTLFRV

>Deuterostome_chordata_Ggallus_tr_F1NBJ4_F1NBJ4_CHICK_Uncharacterized_protein_OS_Gallus_gallus_OX_9031_PE_3_SV_4

MDLKDSTSEGSMGSLQPSSIQIFANTSTLHGIRHVFVYGPVTIRRLLWTLAFVGSLGLLLVESSDRVAFYFSYQHVTKVDEVVANSLVFPAVTICNLNEFRFSRLTTNDLYHAGELLALLDVNLQIPNPHLADPAVLAILQEKANFKQYKPKVFNMQEFLARVGHDLKDMMLYCKFKGQECNHEDFKTVSMSRCSAFLHSGYMDGGGAGPGQPWMSLAVLRGHCVTSTGCGTAEEEQAAC

>Cteno_Ple_Bac_comp46220_c0_seq48_p1__comp46220_c0__comp46220_c0_seq48_p1__ORF__type_3prime_partial__len_242__score_38_17__comp46220_c0_seq48_1236_1958_

MIQTLFLFMWLASIGYAVYVIASSTMSFIGKPTGTKFQVIASQLDHQENVNLQSAVQFPTISVCSHNQVSLKWSSERPGLEKLLEKFDKWNPVLAANIDWETPQFNKWRNATYEEVIKKGGPDNRTFIQCEHGIMLCSDVLKDNSFTREASVNGNCFRVNPEGKLKGKGGEYGRLRLMFFADLNDYTAPAKNQAQYGYTVAFHDHETYSSTIPSGFFMSPGSIYKVDLGLVREFREPPPAG

>Protostome_Lophotroco_annelid_Pdum_comp41273_c0_seq1_

IIPRTTLHGVRYITLNTSRIRRSLWLCVVLLAMSIFIYLFAARLAHYFKYPRAVNMELTYKTDLPFPAVTICNQNRFRLTKVQEYGLYDILTENLQKPVDKKSSIRDKRQAVTKECKDQYVVPKDPHTCFNSNYQTNHTFAYNEVTLNGCKYLCSHLHDLTCSTVLFMLTNRSCLLIPRINLDSIARNSQCVFTEIYQRQRCLNDSYGDISCNYEQSPCPLIQDTTDDEDWKRVKAHLTPS

>Cteno_Puk_Fal_comp16196_c0_seq1__p1__GENE_comp16196_c0_seq1_comp16196_c0_seq1__p1__ORF__type_5prime_partial__len_243__score_44_61__comp16196_c0_seq1__1_729_

VNLDMMNVKNHKKHCREGDYISGYGRYDQATCLLDCRDRILYEKCGCLMTVPPNAAQAGLNYTSCTLQQVSQCALRTYVDYVKRYADVTADQTVANDCDCYITACSYTLFTSSVTATPIPDLELKYKVDRWRDQKRFARFRNYTYEDLGKNIAIIEVYFGTSSITSIEEVVAYDFDNLLGDIGGVMGLFLGASMFTAMEFITLLISLVMRLWKSFFQARNQRNHQPLPLKVPQVASVSEVNL

>Protostome_Lophotroco_annelid_Pdum_Contig3183_

MGSKVHIAIWIFFTILALFVLLYTLANNIMRYLEYPMKSEYKILFKDKRHWPEITFCSLKPLDVSSVTIMNDVIKGTNAMSGQQPSWKPKFMDFEKNAKVFVNAYSKMGQNNTGATEMMKKQVNQVFSVQGLIKNIDEEFAQAVVAKNFVVGCDFDGEKCSVTPKFSKEDGLCTSIHLKNKNSMQAAGDQVLTITAFAGETAEMDMMDMNQGNQHGFKVYLKQNKMEEHGDEEGEENGEENG

>Protostome_Lophotroco_brch_Lingula_anatina_comp131367_c0_seq1_p1_comp131367_c0__comp131367_c0_seq1_p1__ORF_type_3prime_partial_len_243___score_7_97_comp131367_c0_seq1_219_944_

MGTEATTEEEHTPEYEFATNTSAHGFARVADNKYTLSRVIWVLLILTGFGVATWQITMRFIAYYKYDTTTKITMTFSKTLQFPAVTICNVNKYLKSQLDQTDIDFLDQATSMLNPYDYAYNPYSYEDYSYYDYGGSGDAAYGVPPAAPSGNTGSVNIDDLMANISLWYPGGFDFANFTKEKGWQLTDTNIHCEFKGVKCNISKSFTPVLTEYGLCYTFNAVPNSKPRVFQDQPGIGNGLKLI

>Protostome_Lophotroco_brch_Lingula_anatina_comp140408_c2_seq1_p1_comp140408_c2__comp140408_c2_seq1_p1__ORF_type_complete_len_257___score_35_99_comp140408_c2_seq1_43_771_

MDAMQHEYFRGFAGIGAGYKVMVHDPREPPLMANLGFSVGVGMHTLAALKMYNISNLDTENWGKCRRNMTLKYYDYYSLSACRLECLTDFIINKCKCQFINMRAPDVPICDPKQIFDCYEPNRKLFIKEKGQCQQECPIPCDSVTYDVSLSQAPLSDYHTGTFTVNNVKPDYIKNNFVQLDIFFQGISYERLEQYPAYNPFALGCDVGGAMGLVLGASALTLCEIVDFVIVWALRGKGKRLC

>Protostome_Lophotroco_molsk_AplCal_XP_005111319_1_acid_sensing_ion_channel_1__Aplysia_californica_

MFEEIVESLFTTYFSFNKHSQVTLGFNDLPFPAITLCNINPMKHSKRHLASDELQSIIEAADPDAIKRLAGEWKPEFHDDADDSDDRLRTGNFLDDMNVTFNFSNPRLDADNVLENLDLDGYSDSWEGEAEKSTFSSVQEDFIEYFNAENANFTLHQSATYGNCYTLSYSKFIARKSGPAKGLELILFLETDDYLHGLTNGKGLQLVVHEQGTLAYPDEQGIAIDAGTHTLIGLRQNEKECSE

>Cnidar_Rhopilema_esculentum_TR53675_c0_g1_i1_p1__Rhopilema_esculentum_TR53675_c0_g1__Rhopilema_esculentum_TR53675_c0_g1_i1_p1__ORF__type_5prime_partial__len_245__score_35_71__Rhopilema_esculentum_TR53675_c0_g1_i1_1_735_

QISIYKSETISLPYPFRTNCSDGEGLTNYFPGRYSDQSCLVTCFYKHVFEKCGYVNPTIGGKIKPKKKSKVTIDMLKCELEMRDQFLEGDSLAHCDCPKACKEVIYDSKTSSSKWPVDGDLPLYRHLAKSILGFDPINASDDFSSKNFLKVRIYFDELSVMRYVTKELIGELDLISNIGGHLGFWCGASVFSVFEAIALIASILFMAIFGKSGIKVNNDKDLDMEAFEKSVVKVKSDKDLDKDA

>Cteno_Ple_Bac_comp32779_c0_seq1_p1__comp32779_c0__comp32779_c0_seq1_p1__ORF__type_internal__len_245__score_29_27__comp32779_c0_seq1_2_733_

TFFFFGLTANGHDTRDPFVIINETAFMLLEEWEGNLNESVSLAFRNRLYDMMIGKYDYVFGGAGMYIPSSNWSLYDGYTKLGNCLEMNDDGSMVQIIHGKDGGLTVDLDTKSDNYIFTAGSRGFVAFIRDQDETVLINQGGTYIQPGTETFLKISKKKISRLGDPYGTCKDTYSKFSKFATPKAETVRECIQRQFIAFYTTGCGCIPWYVVQRMGNLGKFNVLDEMVLDVIERSDEVEEPGRNG

>Cnidar_HydraVulSc4wPfr_248_2_g2932_t1

MGKACRVKKCTQTHKKWKRFMENNQNSVNTVIVSDSTNIELPTINPSAPNITCVTEPIFENPPNLSASAHKIENLKDKIISKPAISGYLIIYVAVLMSILSSVLCPECGCSTISLTRFDQLKMQNCPLPCVIDSYKLSLSRALFPSPQYADSLTNSLQQPPHIINSIKNISDKVQFARDNLLRFVIYYDDLSYELIEQKPSYDTLVWLGDIGGQIGLLIGAGAMSYFEFIDCLFLVIYNQFFKTYR

>Cteno_MneLey_comp860380_c0_seq1__p1__GENE_comp860380_c0_seq1_comp860380_c0_seq1__p1__ORF__type_internal__len_249__score_49_23__comp860380_c0_seq1__1_744_

FPEINTELGICYTFNADGKKRVKRTGNDFGATFTININQSEYAETQESAGIKVMLHHFKEPPLIKEYGLAIPPGSEAYISTRTLQLNNLREPYFNCTDRKLRTTDYYSVVACTLECHAKILDEECQCKQSHMPLENITICSLYTHRMCTENAIKKIHNGSYADRISDCWCIDACKDTYYEYSLSRAQYASINAAKKILEVNRNVSEYSTISDLRENFIRLHVYFETLAIENVEKVPAYTTMNLLGDVG

>Deu_Ambulacrar_hemi_Ptyfla_40v0_9_20150316_1g22491_t1_scaffold26765_cov105

MSSVFNDIVSITLDIQPSQKPISVDFLTLCTHTSFRISQVLTSHQRPQNNSIYSLHTNSTQCFMRESFGFSVMVDAVQAEYNIGPRFSVGFQVMLYSQGDVPLVGDLGFAVSVGQEVRIGIDATNETLMYVTSNSLFIVAKQRQILDRIAYFAGFPVNVN

VRMKYVEELTFPAVAICNFNLHRRSAIAKANLTGLFDTMYTSSNPNFTQYDISTTATEQFYRESAHHRYTMFFERVHYIRRTRYRGQYD

>Ctenoph_Hormiphora_californiensis_evg190294__p1_GENE_evg190294_evg190294__p1__ORF_type_internal_len_253___score_30_09_evg190294__2_757_

LMAEYSPANRRVSDGGRSSATSRSDTPSSWVKGFQLFAETTTAHGVNQIFTGSNRAVRLFYLLAWIACVIYTMYMIITSIVTFINKPTGTKYEVIVNDLGGLKPQSIKFPTVSICSHNKVKKSYLRKEGNEPIRDLWSKLDMYDQAEADKINWTNIGTASIQHMTLEDIISEGGPDDDRLLSCTQRNKNCQDVLGKNESYSVRENTPSGKCWRINPEGKIFGKSGDWGKMQLIVFADLQDYSKRSADQPQYG

>Deutero_Ambulac_Spurpu_027130

MDNTKYYWFGPEKSISENLFVSSIFRSILGLDKTPPRKVRGPWSPAPPPGSATDLGYLNHVDRQWCSPGAYDVQVNIMEYGQNDYVYNSEKQRMSVGLGVPPGDDMVGVETKALPPSPRSPMATPFQEPPKRATYGSVLNDFADSTTAHGLPRVISEGKWYWKIFWFLVFSGATYVFCRQAWDILTDYRSNPVNTNIKIQTRTYVDFPAVTVCNMNRMRRSKMVGTRFEGRSNIPLLQKATDKLLSCQSYYVD

>Protostome_Lophotroco_annelid_Pdum_comp409606_c1_seq5_

MESKLEHEEKEKELEVIGDGFSYAERTTCHGPNIIASSQSWVRRISWSVIFTTILGLLLYQVIAQLLAYNEKAVYINTKYRYPQNITMPSFTICNFNRFKKSVLASLAEKYQVTENALWHYMATALSSPAFSDISPEDAEPLFDLLNKTEQAIIAVDGTVNIMDLLTELTFTLEDFQRNPLPLRLRCYMIDSRRRKCSPEMFGTILTDHGNCVTFQPKHSGPYVQTQSSNANGGVHFHFYLNTNESSEMFGSYI

>Deutero_Ambulac_Spurpu_015924

MAEGLDKSFPPKIGTSVAQSVDFGLAGASLTENADQVEPPEIKKSRWYLLSSSFPHAIGVAGIKFIFNGDEMKIRRCVWGILLLICLSAMVYQTGNQCVRYAMKPVNTLLDVEYVDLLDFPSVVLCNYNAYSATFGNEKLTVDNFSLALFEFGVCFTFNLDGTHRLQVRSPEDFNSKATAACPCPRPCDVVTYATTVSQAKFPSDFYSKFLAETLTERRNISLTAAYFRHMTRCENSVAREAASYGLIILLQLN

>Proto_ecdy_PANarthopod__XP_023219715_1_acid_sensing_ion_channel_5_like__Centruroides_sculpturatus_

MAKSEIDSNYCANCQVHGWKNPSRSINSISSSPSKQQLSVSKYCLNTFKGSSIAGVSNIAAAKGKFERFIWIFIVICCLTGFIYQVTLFLQQYYDYPTRVEVVVTTEGLMEFPSVTLCNNNRIKKSEYCKYNNGFCPNNSYELEHIKLKENDIPKIQLQLLVEDKPNIRRILGHNISDMIISCTFTGTPPMNSNECVKRFEYFHDPDFGNCYSLESVGDNDTLKASESDVWEELSGHATEGIFTVKYRLYMDIVT

>Protostome_Lophotroco_brch_Lingula_anatina_comp131367_c2_seq1_p1_comp131367_c2__comp131367_c2_seq1_p1__ORF_type_complete_len_272___score_27_15_comp131367_c2_seq1_50_817_

MERGDPIEVGLLFQVHNQSEPPMMETKSLAVPPGFHAYAGVKRTDSAEMEPPYGICDRASPLKYYSVYTMSACVQECKLDHMMGNCTCKLLNYPGPGEVCSPKQIVECVKPILLEIKKNFTHMCPCKIPCQSVNYETYLSYAKLPSDETAPEVAMEYGVAQSDIEKNILVLDIFFQDLNYVISEQLPAIESSGLISDLGGQFGLFMGFSLLTLVEFIEFGFTRLILFICRPKKKKRTITPVLPKEQSNSKSAFHY

>Cteno_Ple_Bac_comp44341_c1_seq21_p1__comp44341_c1__comp44341_c1_seq21_p1__ORF__type_5prime_partial__len_257__score_40_50__comp44341_c1_seq21_1_771_

ETTRECLQSVWLNISAEYCGCVPWFLIDKLEELERKEQIEKWISISLDEDEVFNWTAYADGEVAEDDYMCGYSTYSLCEA
YVMDKFNNATWLASSPFKTVCNEPCKYYKYESSLDYSNFPVSEEHFNTMMGDFSKNYTYVSENFLRVHIFYEDLKLTTIT
QSASYEIFNFIAELGGTVDLMIGFSFFTCFQLLEIFIAYLLFRCCRRGPPPPGPRDSERGPPPPPGGRRPKRGDIELSRSDNG
IMRSPTGFLGFKF

>Filast_tunicaraptor_GIQG01084969_1__p1__GENE_GIQG01084969_1_GIQG01084969_1__p1__ORF__
type_internal__len_257__score_7_54__GIQG01084969_1__3_770_

PCNVTTEAGSHWSSFLSLEYGSRCFTFNANNNPSLKVSRSGSTFTVDITVFLDQLSFLSNTESAGLIVAIHDQLDRPDFDR
GAVLAPGKQHVVAFEKIVREDLPDPYGGCREDPFLDVDLPYHRGLCQINCIADQVVSDCGCRYNLDSRPSATSYDVCTY
QEMEDCGDAAVEVVLVGNDCECPDLCSRTRYNYQLSGGNWPTHPVVKDILTTLRAAFEERNPGRNGTETIDASVITEN
FVNVVLYYSTLEVTTMQE

>Protostome_Lophotroco_molsk_AplCal_XP_035824940_1_amiloride_sensitive_sodium_channel_subu
nit_gamma__Aplysia_californica_

MATSSKQLFTEFSESTSMHGLQRAVNSVSKWRRPIWAVLVICGAAVATYNLAKTLQDFVNYPTSTKISETYKTSLDFPAV
TICNLNPARMSLFNRTGKWTDLEASWNEDRLNISKRQRELATVLRETDEGVRTEQSHKIKTMIVDCQFDNRKCTHADF
QEMPSSKYGSCYTFNGDQFGESRNTTREGQDYGLTIEVDVEAMEYLPLTSTAGLKVLVHPMGITPFPEDEGIVVGTGAS
TSIGLRRFPVAVLRQPGIP

>Cnidar_Rhopilema_esculentum_TR116928_c0_g1_i1_p1__Rhopilema_esculentum_TR116928_c0_g1__
Rhopilema_esculentum_TR116928_c0_g1_i1_p1__ORF__type_5prime_partial__len_258__score_31_27
__Rhopilema_esculentum_TR116928_c0_g1_i1_2_775_

WALSSEGDSVYVDIKEFDEPVFHGSGSILASVGQLTRIEISKTVVTRLKSPFHSNCTNGENKPLLFEGTYTIENCHITCALN
MAHKQCKSIDFHNNMFLPEHMRKNSTNISEAVACSTSIYNYLIDTNFESCHCQLPCMETKFHKTVSTAKWPPKNDLPA
YKALASAAFGLNKSEVTDEFIRNNFLKIDVYFGELSFQKITEVEEFTTVKLISDIGGQMGIWVGASVFSMVELLFLLWQSIK
YFVNRNNITEEHPDN

>Protost_ecdy_cyclo_PriCa_ecdy_XP_014670797_1__PREDICTED_acid_sensing__ion__channel__1_like
_partial_Priapulus__caudatus_

MSSKAEQTTLHGVHWAFDSRLPLLLRLGWIVAILSSWSYMLYLTYSRVTYLSTHPKKTIVDVNYDPRGLEFPAVTICDNN
LIKKSTARHLAEYEDLKRFLADYTAYRWSHGGENFPAADYNLSHLTMSDLARMSHRGHDFVGGCQFASRRCRDDLLYT
MRTFGHCYTFNALFPTERQTYKSDWSPGPPLYTSQTTSNHGLSMLLWADEEEYFYGYRSDTVGFTVAVHSPHTMTTID
NAVVIRIAPGFLTSIGLSYKR

>Deu_Ambulacrar_hemi_Ptyfla_40v0_9_20150316_1g15999_t1_scaffold12160_cov117

MDTIRSGQKHSNKPYPILSMDAVKIEERDMYTSSKTRLFAGQGQTNQSIPDYKSASETSWPDGEKDGEKDDSIKDMLV
DFSKETTAHGLVHITSAKSSLTRSTWLAIVLGASSFMLLQMTLLLFQYFEYNVSVTVALVSEKALEFPSVTVCNTNKLRRS
AVRDSNYSEMLMLESNFVPPYYGDFLCRSGKHCIKPYLVCDGVNHCGDMSDERQCNYHDCGDNQFRCVSGSELGICIS
NDLLCDKISHCYGGEDEDNCG

>Deutero_Ambulac_Spurpu_017120

MVAQFAWDPFSFDITSYEAALNGTNMTQLLLDLKHPLNETVLQVKLGLEYVSVSNFATRFTDYGICYTFNSGEDGQPPI
YIKASGPSFGISIILHTKQEWYFFEERNRYSAGFHVILYDRGTEPVEDLGFSIKPGQFLEQRSSACDCPTPCFQRKYDREMS

YSSFPSNYWADLLANAYFKDQNVTAGYFTQNLAYINVFFKDLTIEKVDQIKAYEFFSLMCDIGGSLGLWLGGSILTIFEFV
DLFGHSIYVYSRKAR

>Protostome_Lophotroco_brch_Lingula_anatina_comp141642_c0_seq2_p1_comp141642_c0__comp14
1642_c0_seq2_p1__ORF_type_5prime_partial_len_259___score_16_43_comp141642_c0_seq2_3_779
_

KAQDYIKHDNISKTEVAVSFTSPAVEEELLNMQRFFGAARSRGRELLDSLHQYQFAHTAMWQNMMDEVILDNFYQKI
HQDVRDIFSNTSDSAYLLDIVRRMLHYPRGSLDNISMPELQVLTNADFAEIDMQNKSAEIDIKLGRLIQQADLDMSLGN
KDELFLSAFFKLQEQLGSFQRKNTIDATFFMENFLQLDVFLKELSYEEIKQQIAYDTNSLWSDIGGSLGLFVGASLITLLEFV
DVIIHTLLLKRRIRNKKRH

>Cteno_MneLey_comp363063_c0_seq1__p1__GENE_comp363063_c0_seq1_comp363063_c0_seq1__p
1__ORF__type_internal__len_260__score_43_05__comp363063_c0_seq1__1_777_

DQFNQLMEDFVKSSTIENGHLDISLDKFEDFNAFEHQNGSLVLRFGPHLYNMVFGQYDYVFRGVGYFLDYFEWNKYS
QMTELGVCLELNDDGKLKQTVGGVNGGLVIDLDANVDHYLFTTMTKGFVVFIRDQDETIMLNRGGYVVSPGVENFIKL
TTTTHTRLGKPHGTCKNVPSKFGKYDHHYESVRECVQRQRMEVMVDMCHCIPWYLADRLFTLNKTDILDTYVERLQL
HQPARERGRRKRQSEDGTTNTDTENTD

>Porifera_AmpQue_tr_A0A1X7V3J9_A0A1X7V3J9_AMPQE_Uncharacterized_protein_OS_Amphimedon
_queenslandica__Sponge__OX_400682_GN_Aqu2_1_34379_PE_3_SV_1

MVNLIVSVSEYSRLSSSNVGVKVSVHDRNSIPVPHQVGVTLSPGKNSYLAIETVKHIDKTRESNCVRHDRSLDFFPGYSYA
VSTCRIDAEYRNYADVNKCGCSPEVNPSSNVNTRNCTIADICCLFQERRTFDPSVNCPPSCEYLSYDIQPSYSDLYTDGES
GRINLISLYVYFDTMLVTQLFTDYSYTFANLLADMGGLLGLFTGASVISLLEVCILLFDVLKSLLLTPRVNKVVAELENKIHLP
EVEIEEPESDDSL

>Cnidar_Rhopilema_esculentum_TR112468_c0_g1_i1_p1__Rhopilema_esculentum_TR112468_c0_g1__
Rhopilema_esculentum_TR112468_c0_g1_i1_p1__ORF__type_complete__len_261__score_40_07__Rh
opilema_esculentum_TR112468_c0_g1_i1_422_1204_

MLLCQTDLRITIDMKKSEYIPSLTLQEGIRILIEDQDMPLYPFEKALSIAPGMSASISIKKSVVEREDVHGNGSCRAAGSETR
KYSVQDCVAKCKGNTQIRLCNCTDVQYSQLVESSTCKTMEEKQCIMFVHGNFSRGLIECNSKCPQPCREIFFAKSISQAN
VYPTGFLKIHSIKNMTTAYVKENIILIEIKYRDQTVQTVATKAYYKFDNLMSDIGGQLGLWIGVSAVTCIEFVALLWNIVAY
IMHRRKLNKIKQSNT

>Deutero_Ambulac_Spurpu_025390

MALNTSLGATKISPSVQNTGKGTQEEETWLSSDDRDDPSLADLTSTLLSHKWYHNPCNGRQGGGVEMAGVKGVRHE
KYREDGRELQRPRRRTKSSGEDQELGGMNAYILIGSAIQNTGAHGIPNIARANSMSKRIFWSVLFVGAMAVFFWQMV
MIIDAYTDYEVITSLDVLYASKIGFPAITVCNLNSYASSKASVFSKEFADAVGVTYNANGNDGVNATSVVPGSTGTATQA
SSTERVRGLHFDRALWSTNNLVASTP

>Protostome_Lophotroco_annelid_Pdum_comp401803_c0_seq2_

EDDGISVGPGQSATIGLRMIKNKKYGGKYTECFDESQQTASSVFAQKYPWLGYSIGACYITCYQRELQKLCGCSDPSYPK
FSEDGNVTGIETCTINTTMRMPSFPQTTNVYIDYGNACYETSSLKLNLQNCENECKLPCDQTSYTPSVSYHGFPSNTYLE
YMMQDLKQQGRNVTKEYIKDNFLEVSIYFEEMGYQEISESPAVQITEMLSDLGGNLGLYIGASLFTFFEFIQFFIDLFMLL
CWRVNSCSPKVTNVEPVLN

>Protostome_Lophotroco_molsk_AplCal_XP_005106187_1_amiloride_sensitive_sodium_channel_subunit_alpha_like__Aplysia_californica_

MAVDVTKVETERESPPFGTCEKVDKKRHESISFYADRYSYSDTACQKSCLQVTTIKRCGCCFVDFPCNPLNKTHVFDNLDLGPKVHFCNTTEYQACLYELDTSRASKFLAECYERCPPACSDINYDITLSMALWPSENRKDQYLQSFDNRHNATGSQEQFQALRQNLLLLSVYPETLKVDKYVTNAKYDWSILLSNFGGQLGLCIGFSILTALEIFELIFDIFTHLVFFFFPRIQSKVRVKVSDGNSRENPGRSNTTSSGQP

>Ctenoph_Hormiphora_californiensis_evg1639__p1_GENE_evg1639_evg1639__p1__ORF_type_internal_len_264___score_48_35_evg1639__2_790_

HGVKYVFHSNHRVIQALFLIMWFVSIGYAVYVIASSTISFIGKPTGTKFQVIASQLDRNNVNLQSAVQFPTISVCSHNQVSAKWSRQRPGLEKILERFDTWSPGITEAINWDRPEFNEWRNSTFETIIKEGGPQDSTFIQCEQGIQLCEEVLGAKYVTREPSVNGNCFRVNPEGQLRGKGGDYGRLRLMFFADLNDYTVPAKNQAQYGYTVAFHDHETYSSTIPSGFYMSPGSIYKVDLGLVREFREPPPAGLCDSTLTYNTY

>Protost_ecdy_cyclo_PriCa_ecdy_XP_014675642_1__PREDICTED_amiloride_sensitive__sodium__channel__subunit__beta_like_Priapulus__caudatus_

MVHMETATFVAPETSTHVSFQRTLIERKKRPEQDCIMAEEVNKAGVYHDVFAEVYGTDYTVRSCLKTCSQKKAVKICDCYMLSTEIPLTDAMLGNARSAASVSRCNGTELNACVLRIINGTYDSDPCLCPPPCHEVQTEAVVTSAPMTTLVSASENEQLLELLKTAGVSVDVMERYEARGIVVSSPAFAVDTYSRLYIRFRDTFDREIREEFAFTSSDLGANIGGLLGLYIGLSILTVYEVIEFLYDICVLLVRKAKPKSVKI

>Protostome_Lophotroco_annelid_Pdum_comp417613_c0_seq11_

DMAHRYNVSQEALWEYLALLLLNSREVQFDQTLFSISMEELRKVYDAIVLKDGIVIPEKLIDQVTFGYSDFFVRGLRSKFVTCGWSGHNHRVCDASMFSDIFTRFGRCLVFNSLKNAENEKLIQVMEGPDNGFSGIFDLNMNESSEIMTLTVSGHLSSGYLVAVHPFGTLPDMNLRSGTANAGTHMLASLTYAKMINYNSSRNPCVNTGRVTLMWQYSFNTCFQECIASVTLTECGCVPFNTYNVAVEGMPFCKNISDFSCMW

>Ctenoph_Hormiphora_californiensis_evg16615__p1_GENE_evg16615_evg16615__p1__ORF_type_5prime_partial_len_265___score_62_43_evg16615__102_896_

AFHDSVYHGVTLNSGYYMSPGSIYKVDLSLKKETREPPPGGNCNASLTMTTYGAYSEGSCIQECKDKDLMKRCGCINMVPPQNYLNYSSCTVRQWAECGLSAYLNFSERYRDPNLTDDICPCNSPCSETHYEAQLSSSSLSPAYTENLFASKYIKGYLSKVSQPQYGIIYNSSDDILKNVMVLEILFSSMVTSEIQQVVTYTSANLLGDIGGVLGLFCGASVFTIVEFLQLFLSSIQKHCLTDLCSSDEDTVEAADDKRGDKEL

>Protos_Platyhelmin_Macrostomum_lignano_A0A267E0A8_A0A267E0A8_9PLAT_Uncharacterized_protein_OS_Macrostomum_lignano_OX_282301_GN_BOX15_Mlig014185g1_PE_3_SV_1

MATASQAPRIKQEQQQQQQRREQERRNTLRFIINEFNDYGHRTTVHGLGHMYQSTDTCRKLFWLCITLVSCGACAVHIYFIVANYMETPVNSVILQGRIRQEFPDVTFCNMYPISESVQYHAAKEIHGHISNRWKYFSGFIRAGNFSANDKVGRLKVARTFMQIFWASEDTRDLAHDDDLFIVQCSYKNRQCSNKQFKMVQNLRYWNCYTFAPKFDGGHEDRQVYSSNEDEGLSLILYTNSHLRNVHPRTASPRFETFTTTTRTKS

>Deutero_Ambulac_Sakowv30018219m

METVLHSTHQLKTENCIQLRKLANKTSEYLTLLSEGVGFVLSVHNPNNHPFPEDESISVVPGYETSIGISISETKRLGQPWY
NCTKEDKTPYGNYPYSTSACRKECLQTFLRGKCGCENAISKNGYKKCNMLNKTQDLCHQMIRYLHNHGKLTCNCPEPC
SRYSYSMTTSLAVWPSDGYSERLLHLLHVMNNKTKAIKTSTLRGNIALVQVYIDSLHIETVSQEKAYDMFDLFADIGGAL
GLWIGMSAVTLLEVTELIFGILTILCT

>Deu_Ambulacrar_hemi_Ptyfla_40v0_9_20150316_1g11967_t1_scaffold7374_cov130

MAKPETHYMSCLHATDLKPISDLLLTYLRDYSCCDNNISKDIQYEYMSGLLNATLPVFYEKIYYELNLIVTLKMYQDAYDM
VKENDTSILIVGGEIYNRTYDAAYEHLIDRKWYEVLLGRTIVTTVENPQSSVGNMTIDGAVIHDFFLVALMKSYDEVFNLI
YDSYRDCFEQRFADLMEAMSSWSGEEICFQYARQNVVKVNVYFEDMVVNVITQRADYRVFSVICDIGGVLGLFFGAS
MLSVFEIFDLLIFKRFCKKTVRPSTNMAV

>Porifera_AmpQue_tr_A0A1X7V3I9_A0A1X7V3I9_AMPQE_Uncharacterized_protein_OS_Amphimedon_
queenslandica__Sponge__OX_400682_GN_Aqu2_1_34369_PE_3_SV_1

MDVQESRAGSSSHNFDQADQPDADETIDEDKVNILDPSPQKKSFKLRLKRGQHHFNKWLDNTSIHGIVHVFKGRSRLR
RILWLLVFLFASTTCAGIIIFYIVLWAQDPTSTSISFITDDEGQSFPAVTVCNLNAVKKDYADQYVVDLLNNYANPTSGFIQ
NWPFSNISRSCTDSLNSITIEDQQEKLFDAYTNGGNKIDDFVVYCGFAGSDGNIVRCEESLEPVFTSLGVCFTFNSAYNG
QPDRKVTLAGPQFGLILFSMSVYQSIQG

>Deutero_Urochord_tr_F6TDC8_F6TDC8_CIOIN_Uncharacterized_protein_OS_CioInt

MAEKEDIKSVSDSFCGPEKYFASKTTCGGVSYWCGGIHKTQKFAWVIIVLGLTAGLAFQVFLRISAYLSYAATRQTTVKFT
SEIEFPAVTICNFNRYFGLISAREVSAVHTLNRLTSPQVYSDLANSSTGFDSSPLSYFNDSFSVTSYLTQKGYPLDSSPILIHC
MFQSKPCTAADFTPVITRMGKCYTFNKSRSIQNVAGSGQGLLFILIVLHSLYTEEPLHGEIEAGIRFQLHSKFEPPEVHKYG
VGIPTGMKAFSTIQRTETSLVE

>Cteno_Ple_Bac_comp46643_c0_seq4_p1__comp46643_c0__comp46643_c0_seq4_p1__ORF__type_5
prime_partial__len_272__score_55_68__comp46643_c0_seq4_3_818_

DDYFRTQDVAGFKVLLHNSYEPPMIEEYGFALRPGSESYVRIRLQKFRDTERPLGYCDPVLEQDYQPAGYTIAKCVMNN
RARYTIQKCNCIPFYMEVPRDLSSISPKVCNLYQEWDCVEPLLEEIALGTLTMGGELIQDPDCPSPCNYVTYSYTISQSEYP
SQSVEAEVKSKVVGARGAGWNIDRIRDNYLKVHIYFEELSTLEMERHPSYMISNLIGDVGGQLGLLLGMNICSLVQFTD
YIIRFSFFNGIMKLVRMSHRNNKRAAGRSRL

>Cnidar_Rhopilema_esculentum_TR38848_c0_g1_i2_p1__Rhopilema_esculentum_TR38848_c0_g1__R
hopilema_esculentum_TR38848_c0_g1_i2_p1__ORF__type_5prime_partial__len_274__score_35_70__
Rhopilema_esculentum_TR38848_c0_g1_i2_230_1051_

QKRESLNSGLKLLLVYGERPNITVPSYANEIFEDERQGLFVTAHDQREVIPAFSGIFLPPGYHTIVKLRKKVIRRKESPYPSN
CIKSGQKTGFNVFPGKAGRDNCLFSCLALRGYQICHGVDERWRPFMNKKEYPNTLNYTDKKMFDCMFEVVQSVDAV
NCKCNVYCDEEVYETQTSRVPWPQQWKAEELTRIFPNLTSSTIRQTFIKLTIYYEEISEFRYTEEEKYSIMNVWTDFGGQ
MGLFIGASLISFVEILLLLLGCLKKCWKKKINVV

>Deu_Ambulacrar_hemi_Ptyfla_40v0_9_20150316_1g3606_t1_scaffold1495_cov133

MEIVSNSTMDEGMRLLNGHETGRYLRYVNTDEYIGIYGHDAGVRVAIEPSEYVAFPLDAGITVQPGTVTSIGLREIRVSK
KPDPFGNCTLDDSNDSIFGNHYQYSALDFRSSSVEWKTDLQIDLSRLHLLTKFIDISRQSKRSEGYGKHLLKTIHVINPKTN
NVNDLRSVKKNLARLEVYFEELTYQSSVDVEAYGREDLLSDIGGTLGLYIGLSVITVAEFIELLIVVIKFVGRRRGALRTAKLS
PSSDTEISHRNDKITDHPEAAKQAWHP

>Protostome_Lophotroco_annelid_Pdum_comp409606_c1_seq13_

QSSNANGGVHFHFYLNTNESSEMFGSYILGHFNDGFRVAVHDRNTPPIMRRNSAVINAGVHAYLSITLEKWLYLKEVYP
CNSNEDYSYGSCLWKCGVEAHLKVCDCVTVSLLPFMENNRTICATAEELSCRLRNAEKNRMWEDIRGSCAEKCVHECV
QLMYTPTVNSVDIPTPSVGRVLTKEEVNTVKENLIILNVYYESLNYRVNQQVRKETPLDLLSSLGGQMGLMLGASVITCL
ELLEFLAAAIINCFGKRHGVTNIFVGQKIKSNPPSA

>Deu_Ambulacrar_hemi_Ptyfla_40v0_9_20150316_1g22275_t1_scaffold26004_cov112

MRSKRRRRIVIRRMWASKTKIHAINVGDDGISNTEEGLQGPIPNLDVKKGVEDKDAKANRKETGCRKILFSNFADDFGIS
GVQVIVGRAVVPLRRYTATENSNLTHLLYDIYANSEAMHVIDFDAYDINATASQEFYISHAHALSEMILEPTWKKTSIDR
DAFQTILTDFGVCYAFNTGDDNYGIREVDDTAGKTFGLGMLLDTYHRNYNVGPESGVGFQLMLYPQGDVPLVTDLGF
AIAVGEEVRIGVEVTRVCMAKCELITLNKILLTHLMK

>Deutero_Ambulac_Sakowv30031449m

MLDESHHEDTEDSLQACSTSPASNTEDKITWKDVVKEFGMNTTAHGIPRITTSTNRCGRLAWITIVSIAFGVLTLQAGLLI
NQYFDYNVNVRVDAVSATQLPFPTVTVCNTNKLRRSAIRDSNYSDLLVVDTDLVLPYYDECGRDEFTCEHGSERGVCIP
ANLVCDKNIDCYDGQDEDKCGLLDDGIPSIVRHTFNHSASHDELLRAARKAEAEINNRKPTLTKHRTARTHLATRAGMQ
SQSDTDGNDGVITSSSDTDKQLSMMIKAKLMKEQS

>Protostome_Lophotroco_annelid_CAC9605582_1__Ofus_G07899_partial_Owenia__fusiformis_

MMGPDIEPNVNDSYNSDSVDMEKQEQLRPPDVISNQPFGSLAFVEETQPKDMDNQDWENVASSKSSLKESINEKSV
KYKVDEFCNDTSINAITRAYKSKSALRRYTWIVMFITSVIIVVVQSSALVQKYFTWPSEVKVDILYDAVIEFPSITVCNLNPL
RRSQLRDDDDQRKGFKNFATNSDMDEEDELYSLYSKAKLELQQTLENGLNIGEPPPEPTSIPGDLGPPSAPAGPTSGGP
GPISAPVGPTSGVPGATSAPAGPTSGVPGATSAPAGP

>Protostome_Lophotroco_annelid_CAC9646860_1__Ofus_G10414_partial_Owenia__fusiformis_

YVDFPAVTVCRTNGAHGFARLQDISKKNVADTVKDKTDLWAMLNPNIMTRMMDETDNIKKLTRAQAISQHSPKMA
QYLAPKYETFIVSCRFKGKKCNKTNFIKRMNGEYFKCFTFNNSATIKNVGPRWGLSLTFYLDRIYGIDLNYDMDAMDITE
YGVKVSIHEPNTIPDMVNNAITISPGYNTKMALRQTVFGKMEHPWGQCDKFRVKLVDIAYEATNIAYNQDTCLKICQQ
RYIQGKCGCLNIMRPVPNDLLNETYCSDISDNNESTIFLNDL

>Protostome_Lophotroco_molsk_OctBim_tr_A0A0L8GCR6_A0A0L8GCR6_OCTBM_Uncharacterized_pro
tein_OS_Octopus_bimaculoides_OX_37653_GN_OCBIM_22035601mg_PE_3_SV_1

MRGNMNFRWPSFIRNDKTGDTTQGKSWIDYISEKIHFDKKQHVSSLGNELYKKEKSYTNSQNNLIQEEKKEKDATKSLL
MHFSNNTSLHGVRNIADSNNTLLRLFYIGIVASFMILVSYYLIKLFMIYNSHPVNRQYTASLQEKLMFPGITMCDFHPYNL
RNQSIFVTLIEGLDAKNYTNPLFSLEEETIYTLLYSDQNDDLESKFNSNELCADDWSFRWMVIDNDILKCYTFNTVNKTEP
RITNKPGIKYGLEIVRNFSDLYRDYTTGVRVSNF

>Protost_ecdy_cyclo_PriCa_ecdy_XP_014672232_1__PREDICTED_degenerin__mec_10_like__isoform__
X1_Priapulus__caudatus_

MSYDRREFPAVTVCNSNPFTATRMAYDVNFTSFWFLPPNLTEGSFYDDFQELDNQLEAAYSTGNVAEYIATLKELETLV
VQQNISSREQQAYFMMELYGRDLNDTEMVGLSQPLNELVLSCEWQGKPCQMSKHFYAFFNPTYGNCFTFNSGLNPN
ASELETVEQVGEDFGLRLMLFVNQDEYIGPIAKSAGIRVAVHEARQYSWPLNEGFFISPQAETSVAISRKLVTRLESPYSD
CVYEDVGKYSYLTYFPEYSYSMTGDALPALGDACSLASCCL

>Protostome_Lophotroco_annelid_Pdum_comp406638_c0_seq8_

MDSKGSKAGIDSMDFPSDTTCHGIHHVAHAPNRAFRAVWFLCILGSVSFLMWQLALLGKGYFAYVALTEVTVHYENM
KLPWGNCTNLPLKHFPVYSKPACKAECMVDFVQEKCGCHIISLPLSPSDNTRPCFPYDMLTCVPNKLGELQNMKGGSC
VVKCSTACDVTEFKTMVDEVDLSTIGATQSFANTLDVDGTSFLESGTSVRVYYRSLQETQYIQKQGTTVTELISYIGGQLG
LALGISLITITEFVIYGFVHIYKKLFSKRNIIQVQTVKLPV

>Cnidar_HydraVulSc4wPfr_824_g11301_t1

MNFEELAKIVVEAVQENNNQPINTNKIKTPRELRNNKTKEYVSEMIDNSSFHGISYIAGKENHFIRRTIWLLITMTAFGYA
AQKVYESTVNYFSFPISTTQMRIYVNEIDFPAVSFCNFNEFRLSKMDGTKVDQAILNPKLQGLVTAEEYRNVTFGAMFDL
KEMLVDCEFNGIPCSDENFTMFSWMQGERCFTFNSGKFPHKLLKIGGAGMKRSLKITINIIHYEYYKDEMDAAIHMIVH
GQQDTPLKMRGPTLSPGFTTYIQLEKKMVILFLKIFINKI

>Cnidar_Rhopilema_esculentum_TR91560_c0_g1_i1_p1__Rhopilema_esculentum_TR91560_c0_g1__R
hopilema_esculentum_TR91560_c0_g1_i1_p1__ORF__type_5prime_partial__len_281__score_42_85__
Rhopilema_esculentum_TR91560_c0_g1_i1_154_996_

GPDFSLQVFLDLDVSNSISLSTLHAGDSVEVIIQDHRQLPSLDTGTVLAPVGHLTQIEIHKTEIQRLKPPYPSKCTNGENMK
LLFKGEYTISNCELSCFLRKVAQKCTHTTFLYNHLLPKEMKMPIAKTERENECTDRILEDLQKTLFASCNCGLPCRETKFRK
YSSQSKWPSTEYIDEYKEMVGRLRSINGSKETEDYIRNNLIAVNVFFGEKQYKKVTEMPMYTAERFVGEIGGQMGIWL
GASIFSLLELVYLLAQFVGVSLRKKEKKVTDLEESGVN

>Proto_ecdy_PANarthopod__XP_023239326_1_amiloride_sensitive_sodium_channel_subunit_alpha_li
ke__Centruroides_sculpturatus_

MLSIRQTNVLRLWIDFQYNEYTPLQFRTSGKVVIHEATNLPNPEWSGLNIEPGFFYKIKIQQLTKELLPAPYQTNCIDYSKF
PRDMRSNAKLDQESCVDECLLNHTINICGCTFMRLALSYNRKICESEESLKGSSCEKNDKFCVCYAKKEAAKHCPHLCKLS
CKQVYYEEKGLEILNWPGSIKKISNYYNYTNLTEARKTVASVQIFFTTMETTTRKYQPKYHFVEIFSYLGGFVGLWLGVSLL
NIFEYIKDEALCLARCVRKIKKKSVKHGDISAHKE

>Deutero_Ambulac_Sakowv30045927m

VIWSIIVLSMLVVFTYSTINSFVRYGKYNVSTVIQINYVSDLTFPAVTLCNYNQYRKSVVNSSGEEFLKLLFPTTGNSSLTEN
IDWSFFNFTGFNMTQTAIASAHQIEEMLVWCKWRNSIECGPGDFKQILTDFGICYTFNGGSNPLKIQQTGSTNGLYLRL
NVQQDEYTNNENTGAGFKVLVHPQGQDPMVKLLGFAVSPGFETLVSMRYTKLHSLGDPYPSNCSEKKLKFFDTYTRQ
ACIAECFSDRVVAECGCMKHTQPGAVESCNPKEWVQCADPVI

>Protos_Platyhelmin_Macrostomum_lignano_A0A267H346_A0A267H346_9PLAT_Uncharacterized_prot
ein_OS_Macrostomum_lignano_OX_282301_GN_BOX15_Mlig024348g1_PE_3_SV_1

MQSTLHFPMASRSDCINDFIVKKFIEHCGCLPSGLPVQLKHFDNHTYCFNLTHSQQLLAIESCYQAQVASMPQYETEAH
RSCLPQCETQRYSIRQSSHPWPNLRRETAVKFLKVLAELHGYPDEMRSTWGESALYFMYERAAQLVDQKSSEKSAEDW
INFSRQLEALQDQLAMVSVQGRQPFGEETFEEEAYPTKNLLADVGGALGLWSGVSVLTVCELVELCIYMVAAVRYRIAH
PQTSTTADPYNEVDSPLRSSGHAADKEAVKMFAAEGGILNADTDV

>Deutero_Ambulac_Apla_gbr54_22_t1

MAKTVADQFPRPSLEREFAESTSLHGIAKVFQASRLLVRVFWIAILLTCLCVCAWQISDRFRRFLQYETNTEISVEYVGHL
SFPAVTVCNFNRYRAGALTQDDEDVLAYIWQLDNYDFYHDYYDDDDAIDEIDSPHPDFNYTEFTLRAGFRMDDVTLLS

CQWYGERNSCSEASEYTETEHVDAGLKIQVHDQATPPTVESSGFALSPGEHAFLAIRRQQAASTFLRSAQSSLYVLHCM
AGFFLKEPARRVGYLGLSPSNHKKSRDCHIERICAYSSKQSNTRKE

>Deutero_Ambulac_Spurpu_027071

MNRLDRSNKVESISGKVTCGNHVQPFVDTDIDVHSVDEEEDVQAPEKKDSGDDLDSTLHGIKYVFNKKYNVIRRLSW
VCVMLASYSALFWTGNTLRMKYTAYMTKEVSLTKRTSSIAFPAVTICNSNLIKASRINNPISKAIVQVLVDGSPEREWDTF
EFNAIDHALETQLDEVFWVHAQQASDMILECAWKKGSKACGAANFTTKITSAGVCYTFNDGEFEGDDDVEGDTGLEV
TRSGTRQSLLVRLNAEVDEYLSALFSEGIGFKASSLKKKKIYIITMWNF

>Protostome_Lophotroco_annelid_Pdum_comp400297_c0_seq5_

EYIPELTESAGVVVTVHDQDVMPFPEDDGILIAPDVQSNLAMVVEQIQRKEQPYPSDCLDVSKKGAWDYSAYSEMYPV
RYTHRACYRTCYQAHLFKECNCGDPAIPLEGSALYNFVDDKENIRNICTDDQEECVVRVYQQYLDNELKCNCTVACEDTI
YDMTVSTLHWPSDNYKELYYNHLATRNPHLESVMKSHSENATNPGNHLVKLFVYYSEFNFKSIEEYASYPLTSLLSNLGG
SISFYLGISVIAVFEFFEYFFDLFLIACTRPREKREEIQVNQVKNIEE

>Protostome_Lophotroco_brch_Lingula_anatina_comp138624_c0_seq2_p1_comp138624_c0__comp13
8624_c0_seq2_p1__ORF_type_complete_len_288___score_20_95_comp138624_c0_seq2_1394_2257_

MKNFTPVYTDMGICYTFNSGYSGHRLTSDSSGRLFGLQLTLDVDQRNYFRGFGGYGAGFKVRVHARDPPMMSNMGF
SVGAGMHTLVGLKMINVSNLDVTQWGSCRKDITLKFHPEYSQSACRRECLYDFMIEKCQCVKFLKFGKIRQSTGRFCNP
KELFYCYGPNYDVFIQKKGQCKHVCPVACESIKYEVSLSQVPFADYYAEEMINDSGYRDVIDFKKNTARLDIFFMEISYER
REQYQAYNAFTLGCDIGGALGLILGASVLTLFEVGDFIILMVLKGKGKKLC

>Protostome_Lophotroco_brch_Lingula_anatina_comp141843_c0_seq1_p2_comp141843_c0__comp14
1843_c0_seq1_p2__ORF_type_complete_len_288___score_27_42_comp141843_c0_seq1_2_865_

MVVFCLWKTKECKSNAFRPVLTEFGLCHQFNGGEEIHQVDMEGSRNGLELYLDTQQQQEYTQNVLSNGALILLHDRH
DHPDMLSFAIHTVPGESAFIALKLKKSVNLPAPYGNCGEKKLAYYKKYSKVNCRAEYLHNMTLKTCGCRMVYYPEVKGY
RDCSPRDIITCYFPLSENTTQIKRQCDCIVPCEILSFDYSISSSKISAKPVSLEFNRTAESVEKDFVSVALYYTDMTYEEVVQQ
EAYSTLTLFADVGGALGLVLGSTLITFAEIIDFIISLCIWKFLRVKV

>Protostome_Lophotroco_molsk_OctBim_tr_A0A0L8GPS6_A0A0L8GPS6_OCTBM_Uncharacterized_prot
ein__Fragment__OS_Octopus_bimaculoides_OX_37653_GN_OCBIM_22030190mg_PE_3_SV_1

MPDVKIDFISFPTSSSYSNESRNIERGNCNTKQSGSKTDYLPNWIRSFSNQTEFMKNCTLHGVRYLSLSNTFIIRKLLWLLL
VLLCLGSMSYQIIDRIIYYYNYPVSVNVHVNYNRTLIFPSVTICNENAFRATKAVELGRYELLEYIYHDMRYVNSSELERFGY
NNMTMEELFKSIEHQKEDMIISCMWGSEPCSFKNFKQTYTDHGVCYTYSQLQAGNKYQKALSTGADHGLRLILNVEQ
YEYMPGPNNAAGIKILMHNEDEFPKVHELGLAIPTGSHAFVGLKLVS

>Protostome_Lophotroco_annelid_Pdum_comp420047_c0_seq12_

LGGLTNNSVINTLGGLTNNSLTNTLGGLTNNSVTNTVGGLTNNSVTNTLGGLTNNSLTNGLTNPLGSLGNTFGSLSGVA
ASTGQGTMGLPTGPVAASLSSINPSSYAAWNDPNNYTQGTGSTNVVTYDYSQTTNYSVPKSGEKDHNMSTFYSTIGH
QLEDFIYDCQWGYTKCVKVKSVIADLGNCITFNSEKEGAHTTEKIGSNFGLRMKLNVEQYEYIRSPNPGAGVKVYLHQQ
GDVPRTKDLGFAVYPGVHALVSAKLNRATGLKDPYGKCGTKPLNLYGDVEYNRAK

>Proto_ecdy_PANarthopod_Tcas_tr_A0A139WIQ3_A0A139WIQ3_TRICA_Pickpocket_protein_28_like_p
rotein_OS_Tribolium_castaneum_OX_7070_GN_TcasGA2_TC034647_PE_3_SV_1

MAQLAQPCEQMISDCFWNSKQSNCSDIFTLVKTTSGYCCSFNYKGIKTDHPNFREIYANGVGPSFGLSVRLHAEENQYL
SSVTKTYGIDVSVHENIKYPDLSIYSTTIRAGSEVSLPVRPEIIQSEDSVKDTDITTRGCAFSDEIKLSWSHNYSYATCINQCK
ARAIKKLCGCVPYYYYPVFGKSKYCTLLDITCVKTNHVRYKNSEIELQTQKGHHAFVKCQCYPQCNEISYSYQLDQEQLKIS
ENEPKSNESVLQVYFEKASCTKLQRSVLITWELILGTKILVNCGRKY

>Protostome_Lophotroco_annelid_Pdum_comp401897_c0_seq2_

MTVMHNKVKKLDNPAPLATSGSVLLKNLTDNTTAHGIPHIAKSKEFCRKLCWFVITLVLISFWLYHSQYTISQYLKREVNI
RTKPVVVNELSFPAVTVCNVNPFLNSSLIDFQAKLFGRLYYEVFQGTYKASNHPAWVSYYYDNYLSKLGAITSSNSSNSSR
GTSSQNGQINFLEFGEDYAQLMTDTIISYGSAHNYSLHNFSLGFSDFVDTCYYEKVFCRKRGIVIKHIFNWLYGKCYVINA
LPDILKSTKTGPRFGLRLSIDIRQDEYSSVNPMAGVRITIHDPKVMPFPE

>Deutero_Ambulac_Spurpu_010265

MNRLDRSNKVESISEKVIYGNHVQPFVDTDIDVRSVDDDEEEEEGVQAPEKKDSGDDLDTLSTIEEGSKLLLNDFLGITTL
HGIKYVYNKKYSVIRRLSWVCAMLVSYVFLIMTCRELVVKYKASLTKEEFSTIRKSSIAFPAVTICNSNLAKASLIDNPTSRAI
VQVLVDGSPERGWDTFGINATDDAFETQVDELLLAYAHQASDMILECAWKRGLKACGAANFTRKITDAGVCYTFNDG
EFESDDDVEGDIRLEINRSGTRQSLWVRLNAEVDEYVSALSSEGIGFKASS

>Proto_ecdy_PANarthopod__XP_023224595_1_uncharacterized_protein_LOC111625622__Centruroide
s_sculpturatus_

MKHEANMILPSVTVCNNNRYKRSKYCPENIRNCQMNINESFSSRCSYICKTLNSHRLSTHNALKFCSVCNRSQLILYGHS
FHSLVRKLEIIQADNHSYELMDDFLHVTLSQGSTCQRNCYVLNSVVDKMNFTLKNASLRTETRQIMKLFLDLEPEEYIDP
TERVEAEIAVHSPFEIVNPFIKGFRAKPGYVNYVNLNVFQKTLLPSPYSTNCINYLQKWKEEGRVGPYTWEVGTISFTGSK
QHDSRMAEQSGLQCTVQYHSEGSAIEKLQEFFQRWFLTGCLILSPIEPIMGLLK

>Ctenoph_Hormiphora_californiensis_evg16414__p1_GENE_evg16414_evg16414__p1__ORF_type_inte
rnal_len_297___score_76_10_evg16414__3_890_

APVKKKNEFRPYPAPYHVPPQQESASAYQSVNEYQAPSERERNPVKRSQSDSSTAGSFRAAPWLNGQTKILAPKYPVM
SSVAATPTNSLKMRSMGGGGGRTVLPPPPTWHDSLLGEDRPDQVIQDREEETTVKSSGDKEPTKSESFADFTGDMTL
NYLRYVWNSPEYIRKLIWAVLFLGFFIYSFSCCYRSIAHYLARPTSTKYNMFYEPNKQLRFPAITICNMNPHNKTYLDQDE
QIAMRSYLRKTWKTPWAKDLPEAKEELMQNFNVKGYYRNSGQRVNDFIRKAVFIEEEQVV

>Ctenoph_Hormiphora_californiensis_evg15900__p1_GENE_evg15900_evg15900__p1__ORF_type_5pri
me_partial_len_299___score_42_58_evg15900__2_898_

EEQVVEENFVERFTTHGLCWTFNAKGDKFVNRTGMDFGLSFSMDINQGGYPDFVRTAGVKVMFHNYYEPALIDEYG
MALSPGTENFISLSYERIRNLDSPYGNCTKDPSPPFANYSVSGCERICFTRALEGDCNCTSYYLPVGEKKECTLFDEDVCVE
KVIKRYHAGLYSDRPYCQCQDACVSENYIASLSTAEYPSKNYGEALEAHNKETLNDTVRDVTFYRENYIKVHIYFEILSIKEV
EKVASYVWSNLMGDVGGQLGLFLGANIVTIFEFVDFFLRVFWYRLLTPYIFKRFRGN

>Deu_Ambulacrar_hemi_Ptyfla_40v0_9_20150316_1g18820_t1_scaffold17133_cov166

MSTAWCYTIDRDVEYELCDVKFDREQCITGENITYRTVTEQFLDINGVTPEMLSEFENGYYKDNGFSGVVKGEVPPDCD
FSTFQDDHYGNCFRFNHGRSGTVIRDANRQGALYGLRLTLFLEQNEYISIYGRNAGARVNINPLNASVLPQDEGITIMPG
TVTSIGIRQKHTFRRTEPYGKCTDGTNTDNLFGHNRVYSVLVGIEVFANLKRSLPIFKTEDFLSDIGGTLGLYIGLSVITVAE
FIELVVSVIGYIYCGKRDMNSQGESENKETKEHFSGAVAFPGFRSFDAINDRNTAT

>Deu_Ambulacrar_hemi_Ptyfla_40v0_9_20150316_1g22276_t1_scaffold26004_cov112

MWAEKTKIHAIDLTGENDKKSKDKKGIAEPIINDEIKKSIDTEDFRVSKKKSGCTAALFSGFADDFGIVGVKYVVGKSVAV
LRRKDATDATNYTDLIYGMYGTYSPDFSGYDLEFTASEEFYKLAAHQPETMLLMPTWKSVYISQDSFKTIITDFGVCYAF
NTGEDEFGVRTSDNAGKSYGFSIILDASQWNYNIGPQIGVGFQVMLYKQGDVPLVGEMGFAVPVGEEVRVGVEVTNS
HVQTKPITCKVSIRYLFIFLVVVIRNAIVNVFVAFVPDTIEFTKFPARTIRPTLSLKWSNYR

>Deutero_Ambulac_Spurpu_028125

MNTLPISNGPNTISQKMSDYDNDDDDVPKSGTKTIVEDFVDNASTHGIPRVLNSSRSSISRLFWCLVTVGLMAALLYQG
SKLVISFIGRPTTTKISLVTKSKLEFPSVTICNLNMMRRSQLNGTRFEALADLDKASDIDGIGHDDSDYDWFFSSDDASPSS
PGGDPSTTPQPGRRRRRRAVSNSDNSYNYDYNYDTWDSIEDPNDWERFYESSTASDFSDFQAVVRPTEEELKKYGHQ
AKDFILQCTFDTQPCSYKNFTVIQNVNYGNCFVFNNAHKLNRSTTINTTRTGTQYGMYLPRP

>Cnidar_Rhopilema_esculentum_TR99659_c0_g1_i1_p1__Rhopilema_esculentum_TR99659_c0_g1__R
hopilema_esculentum_TR99659_c0_g1_i1_p1__ORF__type_5prime_partial__len_303__score_41_16__
Rhopilema_esculentum_TR99659_c0_g1_i1_325_1233_

YKGYCYKFNPKGSFIQARHGRDEAISIILFLNVSDFLPWPQYDSGESVDVFVNDYKEFPFVYTGPVLAPAGHITQIEIRKSE
LKRKKAPYSSDCTNGENIKLYFDGVYSVINCQYSCFLSKISQQCKTHDFYTNMILPKDKRKPFFSFPPTDFCVQKVYEKLLE
ENYASCNCKLPCNEVKYDKTVSYSNWPSDADIPIYKQLFSKALDLNASEVTDDFVRRNFLRIDVYYGDMMYRMITEEES
YSFGKLIGDIGGYMGLLVGASVFSLLELIVFLIQAVHFATKRNKTNDVKTIAITVDELS

>Ctenoph_Hormiphora_californiensis_evg1113587__p1_GENE_evg1113587_evg1113587__p1__ORF_ty
pe_internal_len_303___score_50_98_evg1113587__2_907_

NYTMLKELITEEELESLEYILYEMNVYNGTGDLGSSGNFFLDVSNVTDVIDYLDSMGDYTGDNYLDYILDRISFFPEDFVV
GYYYDFKMGNTWKYLNMTSTNASNVIPHMLGYCITVNDDESLVQIVPGIHGGFTIMLDAYIDHYMPNVAEKGFEVFF
RTPYEEVMVTRGGITVEAGKETNIMISKKKMTRLGSHWGTCESNPSLYHPEKKGIYQSQRECYQEAIVSISQSECNCTP
WYYEERVNYYLGLNMTWTDSETGEAIESCTFANEIVCDTVIYQLMEQWIKSNEIKCMNERCSYNA

>Protos_Platyhelmin_Macrostomum_lignano_A0A267FKN2_A0A267FKN2_9PLAT_Uncharacterized_prot
ein_OS_Macrostomum_lignano_OX_282301_GN_BOX15_Mlig022345g2_PE_3_SV_1

MQARLRTVTGAAQEVAGITRTHGLLHIFLSSSTARRLFWLLLFVAALIGCSVHLAKLVQKFTERSVESQLKLRSERAQFPDI
TICNFKPASASLLTLGDFFEFHLKHKKFKFKVLQYFENFWGNFVNWFKEDTQRRPQESDESYDSRLEKVCKKVLDVMW
VSGDTRDIFQLDETMLLSCRYNSEPCSHRNFSLVQTSRFWNCYTFHPDPRQSSSGPGGRGAELDMVLFTDGNEKYPD
WHDTQPDGCISRKFSRSKAVNRLLQSNEPQSNGFGIFIHEEKSYPMSRALFCFATREFDFFCSKM

>Deuterostome_chordata_Ggallus_tr_A0A3Q3A9Z6_A0A3Q3A9Z6_CHICK_Uncharacterized_protein_OS
_Gallus_gallus_OX_9031_PE_3_SV_1

MAWVAKAHSAHPAPTPCYVQGRQPAAQAAQSHIQPGLECLQGWGIHTSLGNLFSASPRCEQHGPAWHSPPHAQM
HQHRARFATRHCMAQHCMARPYWCTEQTSQHGTARRSLAWHDPQPTQRTQSHRTAHHSAAQPPTLAPPAPPPPP
CRDNVLVLDIFFEALNYEMIEQKKAYEVAGLLGDIGGQMGLFIGASLLTILEIFDYLYEVSPPTPPAPAPRRVGALTRGPPG
VPGQAHRLLQGQEADAARQQHHPGASRRAGQPRRHAPTQNAHRALRGHADGVTFAPHLLPRHSAVGTLEEFSC

>Protostome_Lophotroco_annelid_Pdum_comp417613_c0_seq6_

NGKEKRLIQVMEGPENGFSGIFDLNDNESSEIMTPTIRGHLSSGYRVVIHPFGTLPDINQRSSTANVGTLMLAALKYAKV
TSYNSLPNPCVETARRSKIWLYSINTCIEECIANVTLAECGCVPFNTYNVAVDGIPFCKTIDDFKCMWLNNKAIYRIVNKT
TECCLDCKNLCEQEIYDMRRTSMSYPNPSIVPTITGRDGPDVHYMRQHTAAINIYLDSFVTKVTTHKQKTTLSDVFTAFG
SQMGLMLGASVLTILEFIDFVISSVITSMAKRKQKSSTKDLEQTNTSYSGSDSKPSNFKNNF

>Protostome_Lophotroco_annelid_Pdum_KN_1127_B_01_1_N11_SP6_abi_

SGFPGSVIHMIYRGLAHRILPILRETWKDILADYSQDTTMHGIRYTTLPGKKQRFRRFLWVVLVLTVTGIFLYLFTSRLQHF
FTWPKSVSVDVTYSDTLPFPAVTVCNQNQFRLSKIQEHGLYDIFKEKYVSPGKNKTINGPESKYKFIRTGCIRRNYWTKN
VHRAYKMTLSSCFALCESVLHTMDCSYVIYLPVKQTCYKLPLIKVSEPKGHCDLMEVYHRIRNPRENFSISCDFDGAPCPL
VQDEENVINFAEVKAAELNLQQTDHTTESGRGGLMYIPAMVELLTTNSEMYTDYFQSDGKC

>Cteno_Ple_Bac_comp39293_c0_seq1_p1__comp39293_c0__comp39293_c0_seq1_p1__ORF__type_complete__len_307__score_10_40__comp39293_c0_seq1_54_974_

MEFEIENRFKTELQNHRSRRWKAYWVIVLLLMSGVCGQMVYQIITNYLAYDSFNLYYTNLGSEYTIPAITICNLNILDRTK
LEKASVSVQGKNITYAEIFTEILETYSSRDSTEKGANIIDGAGDLEDAISVTELYQIYGMDIENEIVGNIQFAKKDFGFRSST
MGSVASTENGNCLELNDNEMLVQRVSSFIIEVSQTLIFNIRNILKISPSGVKQNGPNTSYHELVSYILLLKKEKTRYICVLRN
TYYSTIQLSFPQVSGPSGGLVITLDAEVAKYLDIQQSSGFRVILRQTNETVRRLKNNGS

>Deutero_Ambulac_Sakowv30042265m

MDMFGRHTEDISTSVETPKKSYLAESENSPPGDGNSKIGELGKNLMETTTCHGIGRIYGGTNCYRKSFWTLVFMVATS
MFLWQFSELLINYYDRDVTVSLDIKFDRNLNFPAVTICNLNPIKLSIMAMDENLREMFGFEAEYEPTLAPAPGPPGTGPP
LGTMAPTPSSRKRRHLQREKRRMRFGHGLKTFLVDCTWKGYPCGPGNFTKFYNYMYGNCFTFNSGEDGTTQATNNP
GPLHECDAFPGRQWFDFTAWHGDKCRSTEGRYLPSTISVRRLRNIDESDVRQQHIHQFLWNGVQPADVLRRY

>Cnidar_Rhopilema_esculentum_TR43793_c0_g3_i1_p1__Rhopilema_esculentum_TR43793_c0_g3__Rhopilema_esculentum_TR43793_c0_g3_i1_p1__ORF__type_5prime_partial__len_308__score_43_44__Rhopilema_esculentum_TR43793_c0_g3_i1_708_1631_

TILVVERGYCFRINPNGTYFQSRAGPGHGLAITFFLNTSDKVPWTKSNNGDSVEAIIGYHDEYPFPSIGSVLIPVGHSTQV
ELGKTEIERLPPPFPSKCTNGDNMPLLLYPGEYSIQNCQESCFAFKVSHKCDFLDFYTNTSLPKELKKPFPKTEEDFLCGFRI
YENLEKTNFASCKCPLSCFETKFTKTISYSRWPSAADMPIYKKLFSKALGVNISQMTDKFVERNFMRLNIFFGDMTYHKV
KETKMYTADKLISEIGGLMGIWIGASMFSLLEILFLLVQVIQYGFGSKKLRKENDLTLENTKT

>Deutero_Cephalo_132380R_t1__name__chr_scaffold26_start_1779766_end_1783909_strand_pro_len_308_

MPPCSDPXXXXXXXNCFTFNSDKSNMLWQTMEGHGNGMMVFIDIRAEEYTENFFSYGNSEVGLKLLVHDQDEPPQ
MDAKGIALAPGNHAFITVKRTVYENHGPPWGNCQDLQLEYYDTYTLPACYLECRSKHVVANCSCRPFFLPGTAPFCDP
TQIVQCVIRVMAKVTSGELKCNCPVPCSMTKYSTSVSYSGWPNKLTKDYIVRTYSQTKDYMRENGVVFSVFYEQLNYER
VRELKAMDGGQLASNIGGMMGLFIGASLLTILEVWEYLWMRLLGCLSKARRPRTRIIQVQVSEKNGGLNNGAFRS

>Deu_Ambulacrar_hemi_Ptyfla_40v0_9_20150316_1g22274_t1_scaffold26004_cov112

MVVDSYNVLYNVYCKSGIFHRRHIHMALPEDLVQTAFSPVGEKGVTWDAIKLTLNMYTIPAFVSALLALINLIIVLVLFKE
YRISDTADADDPESKKEDDEAGSIDKVAVVFSNVIFFVVLFVFSLNETRSTTDPENLTDLMTQMFGGFSQSVDFHLHDLD
ATASEDFYIRHAHNKSMLYWAFWKGVTMTKDVFKTILTDFGVCYAFNTGEDGHQLRKVYGTGKSFGLQMILLTQISEY
NIGQESGAGFQLMLYSQGDVPLVADLGFALSPGEHVKIGIEITNCLCNDDVVENMLIICGSRVHIRILK

>Cnidar_Rhopilema_esculentum_TR38848_c0_g1_i4_p1__Rhopilema_esculentum_TR38848_c0_g1__Rhopilema_esculentum_TR38848_c0_g1_i4_p1__ORF__type_complete__len_310__score_43_88__Rhopilema_esculentum_TR38848_c0_g1_i4_997_1926_

MGEFNFNGIEFPQNASLLPHVWPCVTLNQNSSLAQKGEKQRSGLKLIMVFDERRRSKNPFYLDPMYEDERDGLFVAVHDPREVIPSFSGISLPPGYHTIVKLRKKVIQRKEAPYPSKCLHSGPNKTFNLFPGKDRRDKCLYSCFGLRLYQKCNGTEVTMRQFMNKKEFPNILNMTDGKTIKCVMEVAESVDAVNCKCGIFCDEEVYETQTSRVPWPQQWKAEELTRIFPSLTSSTIRQTFIKLTIYYEEISEFRYTEEEKYSIMNVWTDFGGQMGLFIGASLISFVEILLLLLGCLQEFWKKKKIEVSC

>Cteno_Puk_Fal_comp27371_c0_seq1__p1__GENE_comp27371_c0_seq1_comp27371_c0_seq1__p1__ORF__type_5prime_partial__len_312__score_53_58__comp27371_c0_seq1__3_938_

AGVRYCTQIRHDNKPIYTEDFGPTGRCFRFNADGWIKSKAGDYGSVYLRLNVNRREYLNLDEIYFTVAIHHHTQFSVTRNTGIQISPGFKYKVGVRTTERFELDKTEGGDCNRSQGQNSYGAYDLESCHMECRDRYLYHKCNCVLWLPPIDMFNYTVCTLKQVHNCGYPAYVDYIHAITDTDHENEYHCDCPVPCNNTIYNLQTTTTALSKEWARKKWQYEGERFNYTVQDILDNYVVLEVYFSDMYIDKVEQIPAYSFWNLLADVGGMMGLLLGASVFTLLEAVTQASTKIYNCSKKNSASVSTRSNGTA

>Proto_ecdy_PANarthopod__XP_023236075_1_amiloride_sensitive_sodium_channel_subunit_beta_2_like__Centruroides_sculpturatus_

MSRQDNSYSFCRLYRLLGFLCLTAGCLYQSFSFMAMYLEYPTTVEVNVMHLPALKFPAVTICNSNRGLRMKLGHQKQDFIIYCDYDGLNCFQQFYDYEYTNCYTFNAEWGKSDTELPMATLFNPISGKSSELLICMNLEKDEYTKMDKTVRARMTIHDSQTIPNPRYDGVSLLPDTFYSYGVRKNTIKLLEAPYQPGCRNYKSEKEKRVGALMSQEDCLAECLFMATLKKCNCTYHRFALIYEDIPCKKEKENCLKNNFTVRREACYKSCGMPCEKTTYDYIAVDSHKLQTASFQYDDKVLPFIKNKKEEGL

>Protos_Platyhelmin_Macrostomum_lignano_A0A267GXA7_A0A267GXA7_9PLAT_Uncharacterized_protein_OS_Macrostomum_lignano_OX_282301_GN_BOX15_Mlig022345g1_PE_3_SV_1

MRARLRTVTGAAQEVAGITRTHGLLHILLSRSTARRLFWLLLFVAALIGCSVHLAKLVQKFTDHSVESQLKLRSERAQFPDITICNLKPASASLLTLRDFFEFHLKHKKLKFGAFYYFDNFWGNFVNWFKEDTQRRPQESDESYDSRLEKVCKKVLDVMWVSGDTRDISQLDETMLLSCRYNSEPCSHRNFSLVQTSRFWSCYTFHPDPRQSSSGPGGRGAELDMVLFIDGNEKYPDWHGIWLNEALHDGCMSRKFSRSKAINRLLQSDEPQSAGLRIFIHEEKSFPMSRALFCFATREFDFFAAKCSPKLNKLVA

>Deu_Ambulacrar_hemi_Ptyfla_40v0_9_20150316_1g9916_t1_scaffold5519_cov135

VCNRNSCLHNSSCITVVVQNYYGFNDEWWDCQCLDGFHGSNCQYAERILRSVEARSYERGMVELTFYLDWPWSSYEIDYEVMYWNDSNGNDTEIITANSQPINLTSLRESTTYSISVTVTLTYFGLRERLKLNYSDPCVVDLCQNNGSCISYVKQYYYEDYYEWWNCDCLDGFVGEYCETGLRLTLFLEQDEYISIYGRNAGARVAINPINDTVLPQDESITIMPGTVTSIGIREVEDFLSDIGGTLGLYIGLSVITVAEFIELVASIVRYIYYNQKDSKSRSIESRNKVTKERLSRGRAYPSSERVNYNPDISVAKM

>Protostome_Lophotroco_RotSoc__A0A817WV45_A0A817WV45_9BILA_Hypothetical_protein_OS_Rotaria_socialis_OX_392032_GN_FME351_LOCUS5404_PE_4_SV_1

MRHMLFPGVTVCSANPYREDRVKEAIDVYARSHSSDANETDRETLFVFMLIDLFNRNESDELVHLGFQKSDMLLECSYNGISCSSNFIHSLSLVFGNCFTFNWKDSSHKLYSLAELGSTLMPYKGLSMAFYVPSHLNYPLNDSEDGLILFLHDNNAIPFFAKNTERLRPGLAHTIAYRKSQTIFLPKPYTNCTTIATGVYSRIHALKARQDLAVNETLMAVWCSRCVPQCVHTHFSIDVSALSAPNVKQKEYWSKRLLRNDLNISLPNDFAVNHNKYVDSNYLRVTVTCGSAYVTVHKQAAKITLIHAFSAVGGQTGL

>Cteno_Puk_Fal_comp26850_c0_seq10__p1__GENE_comp26850_c0_seq10_comp26850_c0_seq10__p1__ORF__type_5prime_partial__len_320__score_71_31__comp26850_c0_seq10__394_1353_

RRVRGHEVIPEPEVNFFTREVSLTGNCFRFNPNGVLRGKGGDYGKMQLMFFTDLNEYSDLTRREPQYGYTVVFHDHES
YSSTIPSGFWLSPGSIYKADLSLSQEFREGPPAGTCDPTRVNNTYGRYEENSCIAQCRDDALMAECGCVHVAPPLPENEP
GKDYRGCTLQEWATCGLRAYKEWVKKYSDVNRVSMGCNCPPTCFETEYKAQISSSSLSKFYAESKVGRLPPGYNTTQD
VMENLLIIDILFTSMQVSEIREIVTYGWANFLGDVGGVLGLFLGASIFTIMEFLQFFVLAIWDALCGLPGRKKAPNHDTVS
LT

>Protos_Platyhelmin_Macrostomum_lignano_A0A267H5N2_A0A267H5N2_9PLAT_Uncharacterized_protein__Fragment__OS_Macrostomum_lignano_OX_282301_GN_BOX15_Mlig013723g1_PE_3_SV_1

SDKTRFTVKNSESAILHVFSMERNRINQADSPCSSRPLGTIDYVTNFGIGPDNQKRRDFTESVEDVLNDRIQDLIYQRCG
CYSHRMPFRVGKSLLCYYAPLQYRKQPSSELLTQIYCHDYWHQKFIGISELDDVSTVVRPCHETYREVRYKSTRWPTDED
LVSVTILYFLHRLANGTELSGPNDPHDHRIFNFSDAEVVAAATKNGTIKLSSRENVAIRSLLLPRCAGIHEFHEGNIAETAA
CLNKDIVRRNLLRINVRLASPAATVYNESYSYTGIEFLSEIGGVLGLWFGVSLLTIFEFFELIYVIMRSKQRIDKKF

>Protostome_Lophotroco_annelid_Pdum_comp421915_c0_seq9_

MADANHYDATKVAEKESSTEPWKTLLFDFTQNTTLHGIRYIFEPTKWLLRRIVWFLSVSAALTGFLLINYFNVKQYRAHP
KKVNVEVSVKDNLTFPSVTICNYNFHRFSKVVEMGLYDTFYDVYSPDAKQSIGDDTYSYSFCLPTEIEVGLSICPSQNNSF
ECLVQAPVYHDISSVACERLCSLVHSKTCNGFLYQRSKRKCYIWKHIVTVEVTYEDFGPPPLPPPLPPPDFIPPIPGSSNTP
ELGSIMRGSSSPNDSLAKPEGAGISQTSSDLTSAATGNEEPTAEPSNLEPEAVAMISGSREIAPGPQPPAPPPPHD

>Deu_Ambulacrar_hemi_Ptyfla_40v0_9_20150316_1g24775_t1_scaffold38646_cov86

MSLFFVYFPYLYAIFSSDGEAISEFGDVRVPEDFKYGEGHTQLSFMEESAHDVEDIVRSNCVENVEGNLKYFDGTYSLSKC
WMECETDYAIEQCGCRYFYMPGNSTYCSPYALQNCYFGAMVKALREGMLTDFFRDLTFEVLKYTLTLIYKNGTDDVGS
DFYVQTKKTLQQKYMNLSVNPWASHNTTMWNMFASANHTSDTIQYLIDTLFQDFYLPLFDETFLYLYSELKSWRDDKT
EEYPRKKLRDICKAHMQENFSKVTVYFEDLNVNSIIQQPGYHSFNLICDIGGSLGLFFGASIVTFVEIIDFFIVQFWLMKISG
NK

>Cteno_Puk_Fal_comp25938_c0_seq2__p1__GENE_comp25938_c0_seq2_comp25938_c0_seq2__p1__ORF__type_complete__len_359__score_71_32__comp25938_c0_seq2__106_1077_

MSSRWMGIDYVMDVIPHMLGYCLILNDDQRLVQVVPGIQGGFSILLDAVVADYMPNVGNKGFEVYLRTPHEEVLINR
GGITVEPGRETNIRIEKRQVTRLSGNWGTCHSKQSMYHPDKAGVYESQRECYQESIVALTREACKCQPWYYLSRMESL
GLNREDFLEDDIEECTFARQFICDQVIYQMLETDKNLSESLSKNWTCSRERCAYDTWSYTITSSEWPSQDAYYDLALLPH
MLNDYDAAYASKNIAKVNIFYEDLKEIRVEQKKAYEIFSFIAEFGGGVDFFIGVSFFTVFQALEILVAYTCCRACRKRRLAA
GGEESA

>Deuterostome_chordata_PetMar_tr_S4RYH7_S4RYH7_PETMA_Uncharacterized_protein_OS_Petromyzon_marinus_OX_7757_PE_3_SV_1

FGSRTGLHGVRHVLAARRTRTYRLLWLLACVAALALLLASSADRVHHLLSRPSNSKVSTVWPRGLTFPAITLCNNNLVRF
SRLTRTDLHALGIWLGLLTRLANGPSTIVNLTADPDVLASLAESQRQWFLRLSDFSTFLPPRDPAGLSRTLLERLGHPLGE
MMLTCRFGGQQCGPDDFTVVYTRYGKCYTFNSGKDGRPLLTSMKGGMGNGLEMMLDIQQDEYLPVWGETDETSFE
AGIKVQIHSQDEPPFIDQLGFGVAPGFQTFVSCQEQRLTYLPYPWGDCKDTPPESEFFDTYSIAACQIDCETRYLVENCN
CRMVHHA

>Proto_ecdy_PANarthopod__XP_023219714_1_amiloride_sensitive_sodium_channel_subunit_alpha_like__Centruroides_sculpturatus_

MRNRPSFRCYAFEERQKNRKWISQLDLILNVEEEEYLEPTRDVLIVLVVHATDVYPDPYSEGHEIYPGFSYTFGVQKKAIE
LLPRPYKTNCTDYHSLEWEHNNPRMSNTRVCTAKCSQSIQLKQCNYITDELSLFFDDVPWKDGEKDENKIRCAKKVAN
DTKEYCRALCRVPCRQTSYDVDSDTTLWPRPKQVTEIDQLGSWRNRTYEDITNNVARIRIFYSSMDYTIYKHTPIIESLELF
SHIGGLIGIWLGTSLIAVCEFFQKLISVLCYVAQKKQKTKGKFRKVSHFNRVVPLPIRKERSIQLNNNNHKRERPRWYLATI
D

>Deutero_Ambulac_Spurpu_016036

MTTNDDDDYDNGDGEDDEDECNGHNRIIWCLILVAATSIMLYQIIYQCIFYSTLPINTRVDIEFAENLDFPSVAICNYNVY
RATFNNEVITRENFSISLLEFGACFTFNLGAANRLKVHAPGISHGLQVVLDTRQWDYFYATSSVGSAISAGFQIQAYDSN
TSSAQVNEYGIAVGPGTRTLIGLTLFEVAYLMKVLMEVVMMVVMAGGGGGGGGGGGGGGGGGGGGGGGGGVDGD
DDGCGGGGVDGDDSDGGDDGGGLWWLLCCWFFWGVVMMVLVEEVLMVIIVEVVVVGGGGEGGWVLWWWG
GGSGGVVVVMMMRMKLMTLMVK

>Protos_Platyhelmin_Macrostomum_lignano_A0A267ECD9_A0A267ECD9_9PLAT_Uncharacterized_protein_OS_Macrostomum_lignano_OX_282301_GN_BOX15_Mlig024322g2_PE_3_SV_1

MRTSTLIKYKAVKVQKLDYPAGECADSARLPDILDDDSRYKYTFAACLSSQIGQLIRRQCNCTSAFHVRPPAAGPAAPYC
GHYSAENPLDFIRRLDCVRNLNLSNLMASVMHSTCQKPCTTFHYESHLSSSNWTPQVWQIHWMMHISSALKQVMD
NPFDQHFSEQLQRLINFTDLESVAGAPEVEALKVFESELAEKMALVTVTRMNYNTLVKQEALQITPSALMSRLGGLGSLT
IGVTLTCAFELVELIYLTARRLRRLDQTAEDGGGGGGVDSNGSAAGANQRAVAAATVELLLLQQQAGALASSERQANH
VQQGSYYDNCPSG

>Cteno_Puk_Fal_comp27722_c0_seq1__p1__GENE_comp27722_c0_seq1_comp27722_c0_seq1__p1__ORF__type_5prime_partial__len_329__score_62_50__comp27722_c0_seq1__161_1147_

FQYFNVKNYYRDSGQRLENFIERPVFLEEPIDVSEAFVEKFTTHGLCWTFNDKGDKFVNRTGMDFGLSFNMNINQDGY
PDFITTAGVKVMFHNFYEPALIDEYGMALAPGTENYISLSYERIRNLQSPYGNCTKDSSNLPFANYSVSGCERMCFTLALK
EECECTSYYLPPSNVKECTLFDEQECVEKVIKKYHAGEYADKPFCKCQDACVSENYIATLSTAEYPSKNYGEAIERENKKN
ANGSDVRDILFYRENMLKVHIYFEILSIKEVEKVKSYIWSNLMGDVGGQLGLFLGANIVTIFEFVDFFLRVIWYRFLTPYVF
KRFQRH

>Proto_ecdy_PANarthopod_Tcas_tr_D6WCA7_D6WCA7_TRICA_Sodium_channel_protein_Nach_like_Protein_OS_Tribolium_castaneum_OX_7070_GN_TcasGA2_TC001349_PE_3_SV_2

MLEANKKIFGSISQQFPKLETFVFFDGSIRHTQVCQPNGALRDKCLLTNYSQYMHKFRSTCSEAISSCFYNNEEFPCCDEF
LPIDTYHGPCFGFNSLVSGKYTSREDLPFKITRDSNFGYGNLSIELYHDGFIVYGLSQDSVPTKESSSEFVLLDSRTSSGIVAK
TAYKAFLSVKETFTDEAARLMKIANRGCRFYDENNLQHAKLYSTSACVNECLVEAQREICNCTYHILPHNDQVKDCDFE
GLLCTIDNKKVKEKTTSCSCTSSCSDVEVVNIGFTKTPIHSSQFRNDKSQFQLVVLLVCLWVQVCSQWSRLFIIFSCVRLAT
NT

>Protostome_Lophotroco_annelid_CAC9662881_1__Ofus_G12679_Owenia__fusiformis_

MDVKIESHNQQKDKLPTFKERIYDFADNTTLHGLPRSFKTSVHLFRRLVWLGIFFGCSGYFTFQVIELINDYSTWPILIKSE
MKLQTFMKFPAVTICNMNMLRKSKIKGTAFQGLIDLDEVTVPNLAQTVSPDQTITENKPTATDQTMSMHPTTNPPNN
TETADGTITPSDIATKPVTTPYAAITTTDHITIHDEMTASAYQTTITDEATTTTDINWIADTTTTTADKNTSIDVATATTVE
NRNPETTSARVTPTLDNSSCTCATCQDPCQSNTLSDSTLAPTATNADPVGETYLSEVQNYGYTPKMQSDFRRRFDAKF
RCRFGTKWTD

>Deu_Ambulacrar_hemi_Ptyfla_40v0_9_20150316_1g24895_t1_scaffold39397_cov108

MGKMGYQCSMQPGLAAIMRYFNEPPRRREYDLVIGEEFASSRNMFAAICKNLKNNGKGNVVHKPAILPGDLDKLSTY
FQGAYNNPKVLLHRVWLAVMLYFCRHGREGQRELQVDNFQVKTDDRGVKYIFQVGSEVTKNHQVDDEGVTGGIMY
ANDSNPLECPVRAFELYASKTNKGXLKLTLFLEQSEYISIYGRDAGARVIIVPYDVPSFPFMEGITIKPGTVTSLAINEVGTYF
CLSITAENQSRCSYKHVVTRLPDPYGDCEGDRSTMQGDADTGSGYRYKVEACQDECLRRKIALECKCYQQFRFEELPPC
SVLNKTQGELTPGVE

>Proto_ecdy_PANarthopod__XP_023229573_1_amiloride_sensitive_sodium_channel_subunit_beta_2_like__Centruroides_sculpturatus_

MILYCDFKGDPKLSKAECVDRFAHFYDPDFGNCYSFDSSQDDNGDYLKATESDVWQELSELILILNTEEDEYLESSRDPG
FLITFHTARIYPDPFSDGFQLRPGFSYNFGLQKFTNELLTLPYKTNCTDYDKLPWKRRIEDRKLSTRMCTAECSQHEQLVQ
CDYVTNNIRLFFDDWSDKINDPEKEECAEKVSNLTMDYCRSLCRVPCKQTSFKVTSDSSAWPRKDKNLPLELNTWRKEK
IEIITKNLVRVRIYFSTMEHTIYKNYPKYRPLELFSYLGGYVGIWLGISLVAFYEFLERLILILMFPLNKKRRKRKSSKVRHIMA
SQIHR

>Protos_Platyhelmin_Macrostomum_lignano_A0A267FLQ9_A0A267FLQ9_9PLAT_Uncharacterized_protein_OS_Macrostomum_lignano_OX_282301_GN_BOX15_Mlig009645g3_PE_3_SV_1

MTVGSNAGGSVRRRWEVRKHWNDFLVTSSVRGLKRTTTSETRFLRLFWVVFLLGSLALMLYQISSVVDTYLHQKDSTA
MQSKQLDIDVPFPAVSVCREATYSSHKQMRQAMERHNASVTTPQSYSRRVADLLYRHFRNATNMTEDELEEAKMAL
LRFDTARSYQAQIPRAAIKEVALPLIRQIATARANFANGSSTSLASNMDIIGMDVETDDYPSSSYDAELWGMRVMDFYH
WRYFLCQTLEITLSREEMNEMRSLDLLLWYAAMTSTPASRVSRIEPLPCAASGRLTASDWHCIRQGSMARCTSTTAPR
WCGAAGIRRSGSDPASPGT

>Protostome_Lophotroco_annelid_Pdum_comp409579_c0_seq9_

TFVTRDNSDIAELLRTNPGNASSRAVLQEEILRSYAKLPLHQRIKLGHSMSTMLLDCSFGNTPCSYNNFSLFNNFYYGNC
YTFNSGYDNHSTIIAENKGPLYSFQLALFVEQIQYIGDLADGAGFKVVIHNKSRMPFPEDEGFTIAAGQITYVGLIRKEVSR
APPPHGECQDDSKAYNAYERLLPSVGYSKLACMKSCLQLHVLTECGCGDSILEMEYFRSLFANITVCSNKSEISCNKQIRD
KFEGRELNCTTECLSKCSEVTHLSSISTALWPGDNAMPDVITQLSTGKLARELERMNDAETETFVRKNTLKLQIYFKSFEY
DLVSTSA

>Deutero_Ambulac_Sakowv30032757m

MSTQLQRPLPVNGPSTDLLQDYRSAHEEGITPMEEPPKEVIMKGDNPCDAIIGNFASSTTVHGLPHILSSTSLRAKLVWA
AVCLVAICVFAYQSSELIHLYLRFEVRMKIETGTAESLSFPAVTICNTNTIRASEAVRSGFAEVVKTDPQHPESIHRKLIYKRL
LSRSTSPDYSDLQDMLKLSAEEVAKFGHQGEDFILRCTFDEKYCNSSQFSVFQNDNYGNCYTFNDEKLINGTIRRSSRPG
ARFGLKITLYLEQNEYIPIFGQTAGVRVLIHPPDVAPFPENNGITVPPGFKSSFAIRRDITNGVGYPYSNCTYRQDRQTAY
GNDYAYTQL

>Protostome_Lophotroco_brch_Lingula_anatina_comp113654_c0_seq1_p1_comp113654_c0__comp113654_c0_seq1_p1__ORF_type_3prime_partial_len_335___score_38_56_comp113654_c0_seq1_158_1159_

MSCCCAPKAPENDDREDDDATFGHFASSTSAHGFARVVGSRYLIFKIFWALLILAGFSIASINIRDRFLSFLRHDTTTKMS
MIFPKKLQFPAVTICNANKYNQKLLTENDEKFLNSTASILFSLEYGDDYGDDYDYAEYGDEDYSDDSYPYSDQIYADGAR
TLEQELNALMADINQWYPDGFDMEKFTKEKGWTLNDENLVCSFRGFPCNASEDFTHVFTQFGNCYTFNGDGALYRY

QDQAGAGRGLTLGIDIEQDSYSNKLSRGDRMDAGIFFQVHSQFEPPSVEAYGRAVPPGFHAYAGLIRKDSSTMEPPYG
DCNASSTLDYHAVYTMSGC

>Protostome_Lophotroco_annelid_Pdum_comp409123_c1_seq3_

MGLGGRHGDFYDNRGFEGPPSQEQRRLARMEMIDSRSMDRHTGGPFGLTRDWDYNDTPSRMSFPARSSSSSGLEP
YRMHPKRPAQWMEEWKEDRKEKHRRKTSKDVVANWADQTTLQGVIFIKNSNRWWSTGFWALCFLAATVMSLIQL
VAIFQDYYARPYSTLVSLKYTFLAFPSVTVCNMNPIRISQLQNTSEELVAFFDEYGPDNRIGIGSGPPNNGNNNGGPGNA
GQGSFNPGQGGFNAGLGSFNPGQGGFNQGPDFNPGGGEGEPDGGVPPGGVPPGGGEPPGGGEPPGGEQSHHRG
KRAPRPPPRPPPQGGFLGAHPGVCSVYILFSPL

>Cteno_Ple_Bac_comp45500_c0_seq7_p1__comp45500_c0__comp45500_c0_seq7_p1__ORF__type_3
prime_partial__len_337__score_42_88__comp45500_c0_seq7_171_1178_

MVILKVFPWTKERDDEKNEKDIQSTAPGTAPGGDPFLFGFQDLASDYVQVLRENARPRWCKILWWVLTAALGILTIYM
VHELLSDYFKYESFNTASTIWTAKLVLPAITICSTNFINYTALEPDLLKYGDTGQFKYIVTQIEKRVAQGRDLSKLKTLAEYK
NVSAFEEYVGSIDYKYANELYGTFVGEYDYTFAGNGKIISHSEWNKLQGMTQLGRCLEINDDKKFYQTYYSDSAGFTIDA
SANLADYLFSTRTQGFKVFLRSHDETVMLDQGGFLVQPGTETFFKISAKEATRLGSHWGTCRNVPSAYSKYGNDSNPH
YETVRECLQRQRIELYA

>Porifera_Syc_Cil_scpid76368_scgid11417_Acid_sensing__ion__channel__5_Amiloride_sensitive__catio
n__channel__5_Brain_liver_intestine__amiloride_sensitive__Na_channel

MSNAGQTNSNRPLIGNGMNGNLLHPAAHSTSSLSAFIDDTVASGRRDLDDVDGARSRYRHRAHGAGGDRSRFERFA
CDESGMHAMPMIGNPKSPIFRRIIWLLLVLAAAGILTYMIQDRVGKYRRFETTTNVVLKYEKNLSFPAVTVCNFNPFYQS
LVNDTIDFAVLSLIRSNLVDSSVRELGRATARTIPRDEIRQRPSDNTRNTGGPLAQNSPALGGDAQQRNRRQAATGRAP
TAATTKKASVTTKQATAAATTVKQATSKATTKSPGTAGKTTKPRNAGAASATTAAAAVTSRAAAAATTRAKATGRQTD
ASTQGQGRSTPTPIPTQQQQQQQL

>Porifera_AmpQue_tr_A0A1X7TFM5_A0A1X7TFM5_AMPQE_Uncharacterized_protein_OS_Amphimedo
n_queenslandica__Sponge__OX_400682_GN_Aqu2_1_13197_PE_3_SV_1

MCFSQKKKEGCLVLIGLSIKRFGDKPTASTITVVSSHESGLPFPAVTICNLNLKKNDSDFLLNTTYQAMNFLYNPDKSYQF
NSTKKKNHLLNTCTAPHSNSIQNATIWDIVNPDRAVNELIQYCGFLHGADSDVVLCKDLFEPVLTSAGICFTFNGTNKLA
NSTGIRYGLKLVLDVQKKERPSFNGKLGVKLVVHDGRDIARPNLYGLTVPPGIAVDVGVRKVITRDETNEAKCIDGMNL
PFFPSDKFEYSQFACRANAVAENIAKRSKCNCVVQPDRPPGFYTSTPNCTFSKACCLLQEQYTFHPLKKSTVLCHATFNI
MSILLVFPVFLMDNTCNY

>Cnidar_Rhopilema_esculentum_TR85881_c0_g2_i1_p1__Rhopilema_esculentum_TR85881_c0_g2__R
hopilema_esculentum_TR85881_c0_g2_i1_p1__ORF__type_3prime_partial__len_340__score_32_55__
Rhopilema_esculentum_TR85881_c0_g2_i1_1_1017_

MYRKDRIRSALKFYGENATIHGIGAIYNSLSALKKFVFVLLFICLLVTVTFAVYDSIYVFTQREVYTKIEKLTHSRIEFPSVTICR
NGKATKSSLPIDIVNKITRKLDEMKTGNLNISDLLDFIENTSLEVQAVSENSNISVPEHPPMLLRHIKDACLFASNKYCNVT
RDFRSTYGTDACYTFHGPGPNTYFQAENGPFFGLQLILYVNESDYLPLIGTQNGAGFTIYIHERDTIPLFEIGGILVAPGTST
HISFFKSETISLKYPFHTNCTDGEGFTNYLPGRYTYLSCLATCVYKNVFEKCSYVNPLIRGSFKPKNKSNVTLDMLKCELKM
RNKYLE

>Cteno_Ple_Bac_comp45500_c0_seq4_p1__comp45500_c0__comp45500_c0_seq4_p1__ORF__type_3prime_partial__len_340__score_29_57__comp45500_c0_seq4_346_1362_

MVVLKVSPWSTGEIDSPISPPSTHKELFDLSQTGTDSTFLFGFQDLTSDYIQVLCENARPRWCKILWWILTAMLGLFTVF
MVYELLREYFEYESFNKATTSWTNKLILPAITFCSTNFINYTALKSDLATYGDTGEFKYIVAQIQKEVAWGRDPSKLKSLDE
YRNVSAFEEFVGSIDYKYANELYGTFVGEYDYTFAGNGKIISHSEWNKLQGMTQLGRCLEINDDKKFYQTYYSDSAGFTI
DASANLADYLFSTRTQGFKVFLRSHDETVMLDQGGFLVQPGTETFFKISAKEATRLGSHWGTCRNVPSAYSKYGNDSN
PHYETVRECLQRQRIELYA

>Proto_ecdy_PANarthopod__XP_023233835_1_degenerin_mec_10_like__Centruroides_sculpturatus_

MLIKEVCENFDYYCKKENFTEEDLKFPYDFWHEYNRSIIDEYGIREEDILPFLSDNHSTLRTIQLDDLLRQRNCYSIQPEVY
MERYKMHDIIDNDTKAIGSLFIRFNPEETTYPGKPISVVASFHSKDDIVNPFISGITMTPGNAYIVQLTPIVKTLLPSPFSTN
CENYFDSQNNSFSFPRYRCLAECKKELSLKKCGCILSDYPFVTDEVCDSLEFLNFPCSINMKFCTDKCPKSCEFTVFEYSVR
EKTFDSKMFNEKVLKSRDEYARIHLIFKTRELYTYSLKPKFQPIELFGYVGGYIGIWLGISLIAVCDFLEAIFIYFYHIITRRIPNI
KLFSL

>Proto_ecdy_PANarthopod__XP_023233850_1_degenerin_mec_10_like_isoform_X2__Centruroides_sculpturatus_

MLIKEVCENFDYFCKKENITEEDLKFPLEIWDVYNRSIIDKYGIRKEDILRDLSDNNSTFRTFQLDEVSRQRNCYSIQPEVY
MEQKFKMYDISNVDIKAFGSIQIKFNPDEMTYPGNPISVVAAIHSKDGIVNPFTSGIAMTPGNRYIVQLTPIVKTLLPSPF
NTNCENYFDSQNNSFSFPRYRCLAECKKELSLKKCGCILPDYPFVTDEKDCEYRDSLNCLSSINLKFCSDKCPNSCEFTVFE
YSVRKRPYSYKMFHEKVLKLRQEYATIHLIFKTRELYTYSLKPKFQPIELFGYIGGYIGIWLGISLIAVCDFLEAIFIYLYCITSR
MPEFSTMFRF

>Deutero_Ambulac_Spurpu_019949

MYESYGFGMNFSKYYESGLPNVNTTEFYFNVSHKIEDSFVYGFWGTEELTAANFSFDIFDFGVCFTFNTGTDGTDLLMV
ETSGKLHGLGMILDTQQWDYFYSSDFHRVSSGFQVMVYDRNEVPLVSELGFAVGPGTENLVGLEITEVNNLPPPHGVC
GHKNLKYYDHYSTSGCRDECHTDYVIQRCGCKEPYMPGNTSVCNPVELYDCAATATIDFIKTAAHDCECPVPCHSVTYE
PVVTTAGFPSHFWSASLSDIYQRVYNITLSPEYFSENMVFINIFFKELSIEKITQQTAYTFFSLLCDIGGAMGLWLGGSILTI
FEFFDLLSNSTHFYYTRLARRK

>Protostome_Lophotroco_annelid_Pdum_comp417550_c0_seq5_

MTVTVAPASKMSSMDTTEDLRREENESTTFQDEFCEGVGLSGVKFVCGVSVNALRRFLWTLLLLAGIGATLYQIGVRV
QSFMKDPKVVDLDFTQKTRVPFPSVTICNQNTYRMSMTYNFSRSADLVITPVMVAEKTIYQMVSDSLKIKNEHGPSFKE
KVAAYEKEFRSLFSDISLYPQTINQFYERSYHIPNSRGMLINCLWNGKQCDDTLLSSSMTNMGSCLLFNPGPAELITTIPG
PSNGLELLLDVEYLDLLYGVEAIKGFRVHVHEQNDTIKLMTDLGSSIQTGAITSISLKRNKLKSLSSDRCSDTSDMDSCMLK
CISEHVSKTCGCRENYMKGDAI

>Cteno_Ple_Bac_comp48643_c0_seq3_p1__comp48643_c0__comp48643_c0_seq3_p1__ORF__type_5prime_partial__len_344__score_41_94__comp48643_c0_seq3_3_1034_

LSSLYFRVPKFTLFQEVSLSGSCYRINPKGSLLTQVGDYGVLKLTFFTDLNDYSKFSKKDPMYGYTVVFHDNGTYSSTMTS
GFYMSPGNVYKADMRKYKEFNIGPPYGRCNPHLKNNVYGTYSSSSCIARCRDQYVKDMCGCIQILPPHPEYEPELDFRG
CTLKEWVECGARATLNFTDRFVDLKEEAMCECTPACMETWYDATISSSRLSKFYANDKAISIDGQPAQLARIVDGYGLN

HGIEETYNISKETVYENLMVLQVLFTSIQESNYKEYVKYDYTQLIGDVGGQMGMLLGASLFTIIEFVHFFIKVAWRYCVKK
ATRRAGEAAERPAYEMHRDQVEE

>Cteno_Puk_Fal_comp28008_c0_seq1__p2__GENE_comp28008_c0_seq1_comp28008_c0_seq1__p2__ORF__type_5prime_partial__len_344__score_55_04__comp28008_c0_seq1__1998_3029_

TQSTGGKQGGLTIDIDIKLDDYLPTVATQGYAVFIRDQDETVFLEHGAFYLSAGSETFMKLTKTEVKRQPPPYGTCKDVQ
SKYSKYGNKSETVRECIQRQTIEQYLNVCGCVPWYFGKRMIELGKYDVLDDLIEKMESKKRSTNTVRDRLIKKQKDSSSE
GGRTKRDLDDPFLNNNYTTYMCSMTIEKTCEFFYNAAVRSGELVLQNCSEPCSYNSWDVEIDQVKFPSTESYYEMFMR
DGLWIYPSPSFEYAQQNLARLHIYYPELKKVVKTQEKAYEVQNFIAEFGGTVDLFIGFSFFTVFQLMEIATAWVVYRCCK
RGRNNRNTEEADKPGKSTVVENTAI

>Deutero_Ambulac_Spurpu_018815

MENGNRTYPHKHGPNTVSGDWVQQEADSVPAMNTASYRDRVSWWAVFYKSVPNAIGIGGMKYAFNSSEVKWRRL
FWLLLILVAFCLLLYHVTNRAQYLATYPTSVDIYTNYTDVLTFPTVAFCNYNAYRLPRDSVWASFTPPSKEQGLELGLNVE
QYESFIDSTARYSTGFQVLLYDYGTEPLITDQGFAIAPGTHTHVGVEVTETSNLEPPYGLCQDKALKYFNNYGYPECFIECR
SDYFYNACGCKFFALPKGFVKNHTCDCPVPCHVRSYKPSISSTAYPSAYIDSRYSSLFQFPDGFMHLLTRPRDSGYFPRQS
AAEMSLAGDEERSRDSGERVIEWR

>Proto_ecdy_PANarthopod__XP_023234691_1_uncharacterized_protein_LOC111634203__Centruroides_sculpturatus_

MDEDNKTKHIKRPNAHQRHAYLENLLKNTVMTGLPQIATEHNLKRKVFKILVFFACLFGFFYQLTEFLMLYFTYPTSLEVL
VKITGKDYLPKPVITTCDVNGIRRTVFCKDFPNLCEKPKNVSLFCIKNSQFCDKENFTDDDLMFPLPLVYLDIDFNRSIIEDY
GTMAEDILYIYKNFTKRIFYANELWKIRSCYSIDLDLIDPNITENNFPLLKMYSNTKTAGDVVITFNSKELFRPTVPIGAKVS
VHSKDEIINPFIDGFLVKPGNRYIVQLKVIEENLLTNPYSTNCSYYERSKEKSFDELYTRQRCILNCQKEKWLESCGCISGDY
PFETDGTFCSERN

>Deutero_Cephalo_088100F_t1__name__chr_scaffold187_start_384564_end_386936_strand_pro_len_349_

MGLKHRGLGWLRFNSGDLDTTVIAPPAAFRDSALPADDMKNCLIMAAEADKEKAETSQKKDMKNCLIMTKAEADTS
HETNPKTSWKQQTEPRQQLTKRLDQTVLTKRSSLTHFANNCSLHGLPRVFPEESKGWDERRSLWGMTVLVMMLVFL
YWAAVHLWAYFSFPISTKLHLQSAIELFPAVTICNLNTYRVSQLTSDDLYTWGHKLGFLHKNKTLKPHGLPYLDHREEAV
LELAKFSPEGHVNVSMSDMLNRTGHDLKDMVKKCTYQGRACNDSMFTPDMVKKCTYQGRACNDSMFTPVSNIKHV
HTHNLAACRVISCWVRVVFLSGHISDTRVVGGLAASIIVP

>Deutero_Ambulac_Sakowv30000480m

MLQHYGITLELLQRDLERKFPDGLDVENFTRAIGWQLNDFTLPLCQWRGHKCFPQNFTHTFTRFGNCYTFNGGDEYID
QRIPGAGHGLSVVLNIQQSEYTESLYGNIEAGIKFLVHPKEEPPQIGSQGYALQPGTRAFADIRQVRYISLEPPWGQCDN
NAALDYFTSYSLSGCSLEYLKNLIYKDCGCRPLFIPGNEPVCSLNQSSTCVQQISAEWASNAPELPCTVPCNYTLYPTSLSY
TTFPSTNVAEQFEILLDITLEDMRENYVYLDVYYSQIQYEQYKQTKAMTSSALLSDVGGQFGLFIGISVITFVEIIEYIAKKIT
GKCCRAEKTMKVNSYNGEPHSTRN

>Protostome_Lophotroco_molsk_AplCal_XP_005109081_2_degenerin_unc_8__Aplysia_californica_

MLLGCRYNDNQCGVEDFYWSYSNEFGNCYTFNFHLHENGTKRDKLHWTSYTGPGWGLRLILDIHLEDYEESDVAGVR
VNVHHNDQPPFPEERSVLASPGFFTSIGFRKIKTGRESPPFGTCQDVGKTRHEGISFYADRYSYSDTACQKSCLQVTAIK

QCGCCLTNYPCNPLNKTRVFDNLHLKPKVNFCNTTEYSSCVSEFDTLRALKFLAECYDRCPPACTEISYDITLSMAMWPT
ENKKDDYFRLFNDDRHKPAGSQEQFQVLRQNFLQLAVYPETLKVDKYVTNAKYDWSILLSNFGGQLGLCIGFSILSALEIF
ELIFDIITHLVFFFIYHTRSKVRVRVSDSSSRE

>Cnidar_HydraVulSc4wPfr_248_2_g2925_t1

MGRYFPSQPTPKVEDKIKSPQEQKRRTIKQYLAYAGENISIHGLSHVFDKRENFICRTVWLLITIAAFGYAVQKVYESTLNY
FSYPFSTARMKIHVNQMDFPAVSFCNLNDFRYSATKGTKLDKAILSNDKKISGEEYLNITLKARHKIEEMLVDCEFDGNK
CSVHNFTEFNWNQGELCFTFNSGKFPHSLLKVNGVGMKRSLVLTINVQHYEYYGDELDAGIHLILHDQEETPIKKRGPVI
PPGFTTYIQVEKKTILNLESPYKTKCGSIKLKYFDSYSMHTCWLEQLTDHVYKICKCKDFFMPGDVGGQIGLFVGAGVMS
YFEIIDCLVLIIYTRFFQKFKTINSIRNHSL

>Deu_Ambulacrar_hemi_Ptyfla_40v0_9_20150316_1g20665_t1_scaffold21274_cov123

MAEGHESRRFHRAFRILWLVLIVIVAVLFGYSAFTYVTRYLQYPVTDEVTMVRSRTQQFPAVTLCNTNTFRRSALGESSY
KDLLLLDRPGALSNLMPYKGPCFEGDFSCGESSTECVKRYMKCNGVEECSNAADEDNCSGGPAPVSDDDEPCALDNTF
QCNNSRCISSELTCDFIDNCGDFSDEEKCKAEMRLVAFTPRFLNCASLWYCWKDQNNGFRKDPTLRLTGVPTLLRVGK
VNVLALSICTREAVKQMMERNIDDGHIIHINSTLSWSPTTDLMANFYAATKCMVTFLTEGLRRELRELGSHIRVTSISPAA
VRDTEITLRASGGDEEFVRTKFASKPINGIIVRSIAVKT

>Protostome_Lophotroco_brch_Lingula_anatina_comp135130_c1_seq2_p1_comp135130_c1__comp13
5130_c1_seq2_p1__ORF_type_complete_len_410___score_21_83_comp135130_c1_seq2_161_1231_

MTTFLVDCSWQGYPCDKSNFTEFSDYKYGNCFIFNSGMNKTVRSAARGGHTYGLSLELYIEQDEYIMTTESAGVRVTIH
EQNSMPFAEEHSIVVAPGKHTSIGISKKRFVRIDSPYGDCKNYTEEENYELNAYSEYYNLIVNYSREACFKTCLQRQIIRDC
GCAHPSYPANATALQASWNIAYNKNLTNSRDDPVLCNIANSTTHIECVDNIVDMYKKGQITCDCPPKCVEDTFEASVSS
MRWPSDVYQMEMLQTLSAYVNTDLDLLKMRRNVLKVEIFYRDMNLETVTTNVAYSLTDLGSDLGGNLGLWLGVSVL
TACELFNLILDVIQLALRNHKKNSKNNHVSPVHKITTKGF

>Protostome_Lophotroco_molsk_OctBim_tr_A0A0L8HEL1_A0A0L8HEL1_OCTBM_Uncharacterized_prot
ein__Fragment__OS_Octopus_bimaculoides_OX_37653_GN_OCBIM_22016435mg_PE_3_SV_1

MSYQIIDRIIYYYSWPVTVNVDINYNKTLIFPSVTICNQNAFRDSQNIATRDSQNIASRGSRNIATRDSQNIATRGSRNIAT
KDSQNIASRGSQNIATRDSQNITTRGSRNIATRGSRNIATRGSQNIATRGSQNIATRGSRNIANRDSQNIATRGSQNIAT
RGSRNITTRGSRNIATRGSQNIATRDSQNIVIIKVVVIQTPMMEYIYIFNRATKAAELQRYRLLEYIYNNVRYVDSSELERF
GYNNITMEELFKSVAHQKEDMIISCMWGSEPCSFKNFKQTYTDHGVCYTYSQLQAGNKYRKALSTGRDCRYHCFCGSF
QMLSRSCRFNVYGVDSFLLQFNHIFQVLESVLVW

>Cnidar_AlaAla_agalma_gene_38098_1_isoform_1_p1__GENE_agalma_gene_38098_1_isoform_1__ag
alma_gene_38098_1_isoform_1_p1__ORF__type_3prime_partial__len_358__score_25_88__agalma_ge
ne_38098_1_isoform_1_1_1071_

MAQKKSVRKVVKDFAEETSTHGIRYIILRKSVTIRMVWFILFGLSSGFFIYQTHKLLSAYYNRPTLISIEKKAVKAMKFPSITI
CKGSTLSRSAGVQEHISLRQPAFIQHFLSFNHLLGFRAFLASEYTRQFHTLEDAKKAAGNPHNLFLTKLDSSCIFGIHEYCS
YPINISDVITSSYGGICYEFNQNGQLRQRHAGSYFGLTMIMFVNQSDFVPFTTIDFGSGIILSLHDSGEFSFPDLNGIPISPG
FLTRISLKKTIYKRLKDPYPSNCSDGKNVDLLYPGSYTISNCKRSCLVRDSLKSCGSSDPLTMMLLNQTGKYKYKPINDTTQ
QCIINAAIATTSQNLEKCMCPVPCTY

>Deu_Ambulacrar_hemi_Ptyfla_40v0_9_20150316_1g26587_t1_scaffold56434_cov87

MILTIIKRKDRQRPQKEIFILKGPAPADIDSSQVELSCRTEQTSYTPISDLGCEENRGSHDTENNEEETLLDDWEFPRDSLQ
LCGELLGEGQFGEVRKAKAVNIRGRDGVTEVAVKSLRGDINDNDVADFHREISLLKSFAKGHPNIVAFLGCCTEQVDFLL
FLIGLSLHVSCVLTENSTRYAVPAMDQVLGNDKNVWLLWLIIVLAALSGLLFQVSTMITSFTGNPVNVNIEYVEPSQKPD
FPAITICNNNMFNYRSIGRQSRIEIKALKETHRADAIELLTDVFWRHSPMLKHLGASYECVKHLKEIAVNCSIRDGVGTVA
IDREINKIAGVSTVFLKTHERDGSEDYGDSRLLRP

>Cnidar_HydraVulSc4wPfr_412_g24358_t1

MLFSLCIIIYGISTLVIKYKRNDVYIQFYATEYSEADTPIFTVCPYEVDKRGTCNLTNNCRKNVTFSQLFIPKNSSYSWHNDII
SIVVDDYSKSSVKAGVQISDYIKKTYHNCVRLSLNFFNTLSSTSNGIRVYTIYPENYLEIFVHDSDEEYPFLQSKPVFIGSYES
DDINIELKKYKRLKQPFASNCSQSSDSMIFPGKYSKLKCIESLKCINSFQACGNSYDFCKNFLPKNIFDNNSSSVYDINNLTE
CLKLEYEKKYDHECALPCDENVYTVISSFTSNCDWLKNNAREIWIDYPITPYYHVFIEAPLYPFIQFISECGGLLGFLTGSSLI
SFIEVLVFFSLIVLKKIINIKSKKEE

>Deu_Ambulacrar_hemi_Ptyfla_40v0_9_20150316_1g29954_t1_scaffold158390_cov89

MVTANRVLVLETEILGRYTGYDVAQCEFDVDERSAFLVNVGLVSVCCIKASERRVDPQYIETLENGDQRKQRFCGQYSD
LRIISASSTMELIFHSDKVVSAKGFNATYEFVPCGTWFASPWGECTSACGMGTQVRRVRCIGRTQNAQINDEFCQGVK
PATQKECITACPDYKRPEVPTNDLESLMNKHGHINNDVQLYEDFLHNYYMPNSFHRVENDSEWTEYLTYSSSPDYSDL
QNVLQLTREDVAKFGHQAEDFILQCSFDESYCDHRNFHAFENDHYGNCFTFNSWQLNGQRLLNSTRRGSRYGLKLTLF
IEQDEYIPLYGQEAGVRVLVHPANVTPFPEDEGITVAPGLKTSIGLRMV

>Cteno_Ple_Bac_comp39160_c0_seq1_p1__comp39160_c0__comp39160_c0_seq1_p1__ORF__type_5
prime_partial__len_365__score_60_65__comp39160_c0_seq1_1_1095_

EYVGSAGKCYLFNPKGSIYSTGIGENGCFRFLVNAHANEYLPQVDEIGYTVDIHHFTFFQGGNKIEMSPGFKYRVGIKKRI
REELPIEEGGECDPTKVSNSYGSYDTATCEYECRDKYINATCGCVMSAPPNNIHDYKTCTIRQMQNCGWLAYYNFVTR
DEITPQHQETDDDEYAHDLLNEDEGGNPDHLVVNENEVHTRASKLPDPTTRPEQLATTTWPPTSMPFVSADDKLNCS
SSCPVPCHKVYFDIMVSTSKISKRYATSLAKQLRSQKTDSNATAEYILKNHVLMEFYIQDMFTEKITNVPAYGLWHLLSDI
GGVTGLFLGASLFSFIVFFKSTVLRVYLYIRNSDGPWNAIKKNLKKP

>Deutero_Ambulac_Sakowv30045925m

MYCADDYFQCDDQRKCIPPSWQCDDYSDCDDNTDELDCATQSAGETGIVVCSNTEFQCDAANCIPLSWSCDWYLDC
LSGNDEENCDDSATNDCDSDLFMCDDNTCIVSYWVCDGVADCLDQTDEAEYNCGVVSEDPPDWYGFIAYSNTPDYS
DLENVLKLDVNETAGYGHQLSDLVLECSFDATSCNVTRDFQEFYDDRYGNCYQFNPFNDNQDDERYSTKTGAKYGLK
MTLFTEQDEYISVYGRDSGARVVIHPPHVPAIPWSEGITIPPGMITSIGMKETRIKRQPHPYGNCTEQTKFESMYGTHYT
QTICEEECLQAKMLEFCGCVDTMLRDEPRCMLLNRTQGRAGTQQQPSSQYGHLIHI

>Proto_ecdy_PANarthopod__XP_023233842_1_uncharacterized_protein_LOC111633492_isoform_X1_
_Centruroides_sculpturatus_

MLIKEVCENFDYFCKKENITEEDLKFPLEIWDVYNRSIIDKYGIRKEDILRDLSDNNSTFRTFQLDEVSRQRNCYSIQPEVY
MEQKFKMYDISNVDIKAFGSIQIKFNPDEMTYPGNPISVVAAIHSKDGIVNPFTSGIAMTPGNRYIVQLTPASDCSIRNK
YFYFRLNLYLILLLQIVKTLLPSPFNTNCENYFDSQNNSFSFPRYRCLAECKKELSLKKCGCILPDYPFVTDEKDCEYRDSLNC
LSSINLKFCSDKCPNSCEFTVFEYSVRKRPYSYKMFHEKVLKLRQEYATIHLIFKTRELYTYSLKPKFQPIELFGYIGGYIGIWL
GISLIAVCDFLEAIFIYLYCITSRMPEFSTMFRF

>Deutero_Cephalo_088110F_t1__name__chr_scaffold187_start_387182_end_389737_strand_pro_len_368_

MKKVLRRLLGRRPQVSTTTVAVQPGENFLPFGAPLTSDLTSNTLAVFAHDPDIDDPAGLHGDGYRGDPAQLDSLLATM
MAKLDAPEEFSDSESVRGKNDKRQEPDGCHANGCHADSPSSYDGDESCNVSLSSRHYSNTMATLNNNTIQPAERDGR
ERKSHISQFAGSTTAHGFWHVFPPGYGPWTVQRAVWAVLFTWGLIMFLYMKIIHIQMYFDYPHSTMIDEGAFHTDRF
PAITICNLNKYRLSRLRTSDFWHLGQSLNYLDADRSLRQEPVINRTLPNPTAAFTDPQNLAVLSNLSSFPTAWQENPPPF
CLLDAQDRAGHQLEDMVLDCHYRGRKCCKDWFKPVEMELHMYGILLRETGYGIGG

>Protostome_Lophotroco_brch_Lingula_anatina_comp120979_c0_seq1_p1_comp120979_c0__comp120979_c0_seq1_p1__ORF_type_3prime_partial_len_369___score_33_21_comp120979_c0_seq1_206_1309_

MKMKTAKKVREVSCEFLENTSAHGLPNAHRATSVPRKVIWSLIFLGSLAGFTTQSYLLLDAFFKYPHNVDIRREFGKKVT
FPAVTICNLNPIRLNKIREKLESGNVSSIVTSLTDSRWPEQLVPEEFVQEAQGILSQMDEIQEDLLSPPPPLRRKRSLLSHM
GSRINGSISSPSSSSTLSSPSSSISTSPSQTTDTLYTTESSNSNSLSQYWDAREGTGNYFKDKPNDMKTSELFEQLYTKMDD
QDRAYFGHDLKNMLVSCTYGGSSCSPSNFTKSFHYKYGSCYTFNSVKVDGSYLTSYRPGPLFGLSLQLDVEQEQYLGGLS
NEAGVRVLVHSQGNIAFPEDEGFTVSPGFATSVGLKLVSTSRLT

>Deu_Ambulacrar_hemi_Ptyfla_40v0_9_20150316_1g3003_t1_scaffold1254_cov111

MLSSQKASKVRVKMSWIAKLQVIHKQLERKLLVLIGRTFRPIPMQKDPDSEERQDDSAASAPLDSNAVMNVDSFDSLK
EGMLVAVYIPQLKHKPQIGESFGFSVMVDAVQAEYFIGPRFSVGFQVMLYRQGDVPLVGDLGFAVSVGQEVRIGIDVT
NGEVPLVGDLGFAVSVGQEVRIGIDVTNIWDRISYFADFPVNVNVRMKYVDELTFPAIAICNFNLHRRSAIAEANLTGLY
DMLYTTDNVDFKQYNVSTTASEQFYIESAHDVYTMLYQVLWLLVSSAGLGLVTFQILDRIAYFASFPANVNVRMKYVDE
LTFPAVAICNFNLHRPTWQGNFLSKDDFTPMLTDFGVCHTFNAEDANDTFKGQKH

>Protos_Platyhelmin_Macrostomum_lignano_A0A267ECQ9_A0A267ECQ9_9PLAT_Uncharacterized_protein_OS_Macrostomum_lignano_OX_282301_GN_BOX15_Mlig026396g1_PE_3_SV_1

MDIYLNVPGLFDNYNISHLPYNEDDTTGARVCVHTPLSWPYTEEIGIDIEPGTKTMIDVSPTKLKLLPNYGGQVCTEHNL
LEEVMEAYTDMGSTKFAYTSKFCKDLHLQNLTVDRCNCIDPRKPYPPSLHKYQFCGKMNGSDDKSLNETLHQLRCADN
IELDNSYSASQHDCALSCETCQYLYQTSQSLWPTKNKEVEYYQQMLKPLPKLREKYKAYDHIQNLSRVNAIKSRKLLMKE
DDIRNNFVRLHIFRSDFSASEKSMALKTTKESMVSQVGGIFHLLTGITFACVVEVLECLYYLLKKTLLPRSNGPNHGVLLQ
RLRRRPHRNDSANNCQGVDGSAYEMQKLQTPEPKAEEFNHHAIAEDGAASA

>Proto_ecdy_PANarthopod_Tcas_tr_A0A139WHT8_A0A139WHT8_TRICA_Pickpocket_protein_28_like_Protein_OS_Tribolium_castaneum_OX_7070_GN_TcasGA2_TC033005_PE_3_SV_1

MGATDREPDCHIFRKCDFSSLRGAISCDWCLSKNTRYDIETIKEYMQALGSFYDFDHTKTPKLLTLQKILEQFPQIEHFRP
GNLIKWLAPSCDDLIINCSWKNEQRNCSELFQQRLSFEGSCCMFNYVKNTVENISDNFRNRKIMSGDAGINYGLTIELN
NQIDDYFYTQHASLGANIHIFNPDDYPDKNSGGLTEQLVGLSREFYMQLETIGFRSVPEVIDYSVDKRMCYFEFEKTTAF
GLYSESDCLVDCRIKSMIGLCGCVPFMIPQNRSRVCGFSDMHCLNKFRNKWESLAPPDGFQEASLLREAHHSLRCERCY
PVCESFLYNMATNNEFFPSFGNNSLLHIYYVDRFSTIYKQDIQNYWFEIISK

>Protostome_Lophotroco_annelid_Pdum_comp420350_c0_seq13_

MSDTGKSEAPPSWPTLIDDFTQDTSFHGWRYLHTGTQYNIRRFLWLVVVGCCTGLFLWVVYGRVEKYRQHEKNVNVE
VESSDKIKFPSVTLCNQNQFRLTKADETGVLDLLEFHFQKDIQGRTLTDPNDYANVSVTNGIQITHDLESLKGQVQTFVK

VSLPVCSTTCCSEFFRKECKSIQFDFATKTCTIQPYTTSTPGVNITANANMTYHEFHKCGKPQYTDKFRPVSNADYEISLY
DFSLAAAHTKEDMIQGCSFEGKPCSADNFTRTITDYGVCYTFNGAPDRTLTTRQTGSSSGLRIIFNIERDEYLSGPNHDM
GLKLYLHEANELAMVRDLGMALDVGKHSYIPVKATILKNEKPPVGKCGSQQLQF

>Protostome_Lophotroco_brch_Lingula_anatina_comp131826_c1_seq2_p1_comp131826_c1__comp131826_c1_seq2_p1__ORF_type_5prime_partial_len_373___score_4_70_comp131826_c1_seq2_1_1119_

IDYYWSNPVSVNIELHYVDFIPFPAVTICNNNKLRVLNLYHTGEIDVGYLFTSPTATTTKNLSSYNWTEFITEYGHDAPTM
LNAPGSCEWNAQPCSVDNFTRILTEMGVCYTFNHGDTNLKTELLGSTFGLKLILNAEQYNYLATTHSAGFTVLLHHPDD
VPYIENLGFQVAPGESISVGISQTKVTNLPSPYGNCAEKNLAYYDSYSTLNCIAECTLNFTMDNCGCRAYYMINVPGVPV
CSPTQWWECLDELRANTSAALQACACPVPCEEVHYGYTISHSKMSNAITAALMAGYPGTTEDYFRDNYLILQIFYKELN
FQGIEQQPAYSGLALFSDIGGALGLVLGSTLMTAAEIIDFAVSLCLWVAFRRK

>Cnidar_Rhopilema_esculentum_TR36173_c0_g1_i3_p1__Rhopilema_esculentum_TR36173_c0_g1__Rhopilema_esculentum_TR36173_c0_g1_i3_p1__ORF__type_internal__len_374__score_41_07__Rhopilema_esculentum_TR36173_c0_g1_i3_3_1121_

SSVIIALFAASVANIYNLAAEYFRRRTYLLVEEHANHTAKFPSVTICNANNFQRSKVGNLAVDPEARFQHFLNISLDKGAV
VNLLNFPNAKSFMDQGSFSTAKATQSASDMLFLESIEGWCRFGHFKTCNQSDFKDYFYGLESGFCKTFNPDGKFTQVV
VGPHGGLLLTLYLNQDDYMRLLPFDNGAGVFVKVHPQDVFPYPSNDWLAVEPGTQTHISIEQKIMLREQKPYKSECVN
RDEDLIYPGRYHVQNCLKSCFHLKLYESCGIVEASVLYHLEQKGKKFPSLPRNISKDPAACGREFYANLTANKVFCDCPLP
CYEETFKLTATSSQWPSEADMQYYKQVLSKVLNRTNLSKDFISSNFLQLKVYFKS

>Proto_ecdy_PANarthopod_Tcas_tr_A0A139WEM9_A0A139WEM9_TRICA_Sodium_channel_protein_Nach_like_Protein_OS_Tribolium_castaneum_OX_7070_GN_TcasGA2_TC033776_PE_3_SV_1

MILSRLKYYGKNTTLHGLKYVADEKRRPVIRLTWIAIFILSFYGMILVFKASYKSYTENAISFVTETAYLNWNTSFPAVTLCEI
TSSEVAWNGDKEEAEAVQQFIADLIFFTGSCYSCHTPCSICEELDFGDIVEKFRKKCSQLLSKCKWNETPFDCCEKFLPLET
EYGICYSFNSLHTKKPSQHASDFYFFSNRKTGPGALFIQTIEDVRLYFHAREDVPFINSDPDQRKDVMLGETYNISINIIEIE
NDENVKLLPVEKRKCRFPEEKPGDLIVHDYYSYSTCVVQCHADAHIDLCNCTHHLMPVLNKEKYCDIEGLKCLTDNFETV
NRLHAKGFDKPGLVCDCLPSCIEPEYKVVADTKGVKSSHGLRWT

>Deutero_Ambulac_Sakowv30023603m

MTKDNHPSTENGDTTVRRITRRFALETSAHALPKSAKSSHTYTQCMWVVIFCAACSVYLWQIGDLLSLYLSVPKRTENIL
VSANTLPFPAVTLCNTNKVRRSMIQQSGYSSLLDSLEINTLIYYLKSEDPPDWRAFMSFSRDPSWTPLRKVLKFSKDELKV
LGHQLEDLILECSFDGNECDMEKDFHTFQHDEYGNCFTFNHGMHGVPLRNSTKAGGDYGLRLTMNVEQPEYITFYGK
DAGVLMTIHSPEDTPFPEDHGMVLRTGSKTSIGLYKEDDILIPEPNRECENVDTFYGIYGDYKYTTLACDKSCLNRYIQYY
CGCVDTIMIEGVPRCMILNDTQEVCQQLMNYLYYRGLLVCRCPRRCRQVHKNTMR

>Filast_tunicaraptor_GIQG01084212_1__p1__GENE_GIQG01084212_1_GIQG01084212_1__p1__ORF__type_3prime_partial__len_375__score_43_44__GIQG01084212_1__328_1449_

MSDHKKDSTDLGVREPYGEGTSQSLRQFAAGCMGETTVHGIPHAFPPDGRSRLRRTLWAIWFVASIVGLVAFIAGRVE
FFYTRPINSVVSIEYSPILAFPAITVCNLNGYKSSAFVAWSEEVAAKVMMALLPELAEYPIAQIIEYLAAKGLLDSGTPEGKA
ADRRRRDADQGVRTEGRWRRIHERHERRHGRRDAADDLYSELRALQAERKPEVSAPEVSVGAASTAWEICSVVVAAA
EAGEPAPTTVEYAGADVPDRLRDTNWAPLCTKYLPLLADEVFVRLADAVKAAIDGGDAEGEAMVAVLRSDAVSSSKNS
LVVAVVAELNELFDLAPATDDRGDGEDAGTGEAEGGGGGGGFAGGDDGEGGDGDGDGD

>Cteno_Ple_Bac_comp45633_c0_seq3_p1__comp45633_c0__comp45633_c0_seq3_p1__ORF__type_complete__len_379__score_46_26__comp45633_c0_seq3_144_1280_

MPTTQHYPGVKEIVTWLEDIGDYNGTDYTGWILDQVAFWAQDYVASINYDYKLGSKWKFLNTDDYGSTNVIPHMLG
YCIIVNDDQSLVQIVPGKPGGFSILLDARISLYMPNVDTYGFEIYLRTPYEQVMINKGGISAEAGRETSIQINKNKMTRLAS
HWGTCQSKASPYYPAKMKDGVFESQRECYQEAIVEIAQYECNCTPWYYVYRMNELDLTIVGAGGGNYTQCSFARQVI
CDGVIYQLMENWLEEDENNINCGHERCKYDEWTYQVQQSDWPAYEAAFDLSIKDILTSDKNYDYAKKNIAKVNIFYGD
MKENTVSQEKAYEVFSFISEFGGGVDFFIGLSFFTAFQAIEIFIAYTCCRRLRGRGGTIANSGAK

>Cteno_Ple_Bac_comp45500_c0_seq11_p1__comp45500_c0__comp45500_c0_seq11_p1__ORF__type_internal__len_382__score_55_56__comp45500_c0_seq11_3_1145_

SKLKLMDEYTFVNAFEDHVGSIDYVYANHLYGTFVGEYDYTFSGSGNVISTGEWDKLQGMTQLGRCLEVNDDKSFYQT
YYSDSAGFTIDASANIADYLFSTRTQGFKVFLRTHDETVMIDQGGFLVQPGTETFFKISAKEATRLGSHWGTCQNVPSAY
SKYGNDSNPHYETVRECLQRQRIELYATKCSCMPWYFVSRMKTQEKYEVLDELVMQILMTEYSNTTDDHDHDHDRKR
RQTKFRNTVRVPKHHGKKSAVKADTADRVKRDTVTTEHTTPWWDFFTTDQPTSQTPYYADTSYTNYTCSYTELLICEAF
IEEEISNGHITLEDCPEPCEYLEWGAEVASSSFPPSEKYYNEYIKDDWYIETNTSYSYAQENFARIH

>Protostome_Lophotroco_annelid_Pdum_comp418513_c0_seq7_

MDGEIGSKKPDNWTSALYRFSQTTTVDGVKQITEPTPFCARRLVWIALLLGGIALVARQIADSVLYYHSRPVTVNLNIKY
NKTLKFPAVTICNQNNFRATVASTRGQFDLISDIYVRNSSLEDKTYEDDFYSESVAGSMFALKHDKHDLIHSCKWDNDY
NCSADDFITSPTDHGICYSFNSGRDDKPIRYISQTGADSGLKMKLNIEQYEYMQGPNHGAGLKILVHDPSDVPMVKDLG
LAVKPGTHTYMDINIIEVHNLPPPHGHCEEKNLTLFSKYAMSTCLLECKTLHITSLCGCIDYYMPWKTSGGPPFCTIRQFL
ECVKPSLEAFNVADECDCPVPCNMTILNPSITLLANSDFDVEKMLHQNHSELQEKYTYAQEV

>Cnidar_Rhopilema_esculentum_TR106686_c0_g1_i1_p1__Rhopilema_esculentum_TR106686_c0_g1__Rhopilema_esculentum_TR106686_c0_g1_i1_p1__ORF__type_3prime_partial__len_383__score_42_81__Rhopilema_esculentum_TR106686_c0_g1_i1_3_1148_

MTIVPNEPTVNAEPLTIAKSWDGFVSSSSVHGIRFTSPGFSRIRRFTWALMLVTGFSVLLGSITSTVVEYFRYEVNTVITM
VSEREVSFPAVTICNLNTLRRSKFNEASKDANYSQMFNLAGILSMAGLKGSQRNLTIPYISGKKVQDMYKYFGHNLNSY
EEGGMLLNCTFNEISCNKSYFKPVLTNSGQCYTFNPGGQFGRERFGKEVLTVKQPGSGFGLKIRLVVQANEYVLMPTHE
FSAGWKMLIHDQKELPLVRDYGFALSPGTHTLVGLVKRKVLNLEPPYTTACRKEPLETGSAYTLNGCLRICQEKYIYAQC
GCYAPINFVDGKDFELCDLEKSIKCVLPALDNFRKHGNKCVCPDPCESVEYTPYLSYAYFPSD

>Cteno_Ple_Bac_comp47578_c1_seq3_p1__comp47578_c1__comp47578_c1_seq3_p1__ORF__type_complete__len_384__score_68_54__comp47578_c1_seq3_141_1292_

MELNTSPGGMKKLVTDVKDLAILMNYRNFKEGVYPGVEAFTPYSKYKQDESAFDMTGRGERTSYYPYLEHYQPSYPRP
PRGGKPPPPTGAPVPNNHRTSDQNGKIKYPPNHRDTGSPTELEDRDGEDCEEFMPPPKRKKKKKRRANRAALEQRRE
EMEPDEMESFKEFASTVTADYLAEMFAKASNGRRAIWVSLWLASFVYAWYNIALSVGVYLSKPTATKLNFYEAPPGGV
AFPTVTICNFNKYNKSFFEREGDSVNSQLRDFLVATTPIWDHKERLDQDQWTSKYPNIMSLNLKELYANGSHLITNTVEF
CRFGGKPCKDTANRETSIPSEAFTKVYTEHGLCYAFNLDGKLNMTRTGAQYGLSLRLHVAQDDYFRTQVT

>Protos_Platyhelmin_Macrostomum_lignano_A0A267EYP1_A0A267EYP1_9PLAT_Uncharacterized_protein_OS_Macrostomum_lignano_OX_282301_GN_BOX15_Mlig031432g2_PE_3_SV_1

MHPPRVLKDYSSPFPSVTLCNMRPLSSEGRQQLRQQGWKYPRDFSESLELFARNLYFYNESTKYVYYNFSGSMGFPAYL
ESLPRSASRDIGHTLQDFILDCRAVIRNGSARNLKFCSPEYVWGGKGSDMFAGNFHAFLHGNYMNCYTLDLHEELRDN
TTMVEMAVYLDNEWNLDNFNCVDCYTMDISTQLSGAVLTIHSGDTYPDVNHEAIHLKAGSLTEVKLKHVLNVQKQAP
YGRCSQDTQRNLNLFGREYKYSEDACRQATIQRDILNTCGCQAIEYPLVRESNFTYCTGLEEFLRSDGCNWATAENDQK
RLADCSATLQKVYGYIRCKNDVISKYSKDVVESCTLPCSFTATRPSGRHPTGRPGRSSCACSTAGSTAAC

>Proto_ecdy_PANarthopod_Tcas_tr_D2A5I2_D2A5I2_TRICA_Sodium_channel_protein_Nach_like_Prote
in_OS_Tribolium_castaneum_OX_7070_GN_TcasGA2_TC015541_PE_3_SV_2

MANLDDYHFVKENKLWANRTSIKFSEAFKLAKFFGNFYDFDFNIADENHYKHLQGIVDVLELDIATLMKQFTTRCPDLLL
NCRWKGVEKKCNELFTMKLTYDGFCCTFNYFKESLMNGSVSILNGTDGLTRGLSVTVMNDLDDYFYTPLESQGVTVQI
FVPNDYPDKPSGSLSEVVVNVGTENYVKITPTTIFSTKEVMKYAVAKRQCLFDEERPTTFEGLYSQSDCNVDCRIASALAL
CHCLPFTIPLSEGESVCNFQDLKCLNQYKNKWNTLYPDNDDFGILQHEQHDSLRCEQRCYPSCDSTLYEVFPDSSPFHKK
GNFAIIHIYYTNRFSRLFKHDVVYNWFEILSNFGGTFGIIVGFSVISVIECLVFVLQFTKKKCFT

>Proto_ecdy_PANarthopod_Tcas_tr_A0A139WGL3_A0A139WGL3_TRICA_Pickpocket_protein_19_like_
protein_OS_Tribolium_castaneum_OX_7070_GN_TcasGA2_TC034728_PE_3_SV_1

MITTQKIIFRANQTKKSPEEVMAFLHFTGQLYDFDFQPDTVSKLLLFQNILDSLNIKIDDLAREMVTPCEDLLSSCLWRN
MEVNCSTIFYQRLTYDGFCCVFNYVKFTQELTYEIDQKPVTTDAGVENGLFVAVNNNLQDYFYTTISAVGVIIHIFVPTDY
PDKSSGGLTEILTHPNSENFVEIVPTTLTSVRDVADYSLEKRQCLFDHERYTSFGTTYSQSDCNIECRTMSIYAMCQCIPFLI
PLSNSSNVCTLADVACLNRYKEKWTSLFPDDLEPGPYLQKERQDSLRCDNKCYSSCAETVYEAHANSFPILNKTDNNVT
RIFVYYSGRFSKLYLQDVVYYWFEVLSNFGGIFGIFVGWSLIFIIQLSAYYTRKILEKVLKT

>Proto_ecdy_PANarthopod__XP_023243708_1_uncharacterized_protein_LOC111641733__Centruroide
s_sculpturatus_

MIEYIKNPSSKDMRRYLRRMLENVIMTGIPQIVTEPKIKRKIFKLFIFFGALFGFFFQLTEFLLLYFTYPTNLEASIKFMDVKD
LPKPIITVCDNNGVRRTTFCRDVPRSCEKTEFIKEVCENFDYFCKKENFTEEDLKFPYNFRQECNRSIIDEYGIREEDILPDLS
VNNITFRTIQLDELLRHRNCYSIYPEVYMERKYDIYDMHNKDTKSIGSVQIKFNPEETTLPGNPISVVAAIHSKDVIVNPFIS
GITMTPGNRYIVQLTPATQTAKTTSIRKTIHFHSLAIVVWRNVKKKLSLKTCGCISAEYPFVSDEKVCYSWDDLQITLRGRE
AFQETIEEEVEKYKRELISLQKDLEYKEMFEYSVRERPFIKQLTLYDSVEHG

>Protos_Platyhelmin_Macrostomum_lignano_A0A267H3M8_A0A267H3M8_9PLAT_Uncharacterized_pr
otein_OS_Macrostomum_lignano_OX_282301_GN_BOX15_Mlig029354g2_PE_3_SV_1

MEIFWGNFTNWFQEDTKKRPQESEEGYENRLEKVCKKLLDVMWVSGDTRDVSHLDKTMLLSCRYNNEPCSHKNFSL
VQTSRFWSCYTFHPDPGQSSSGPGGRGAELDMVLFTDGYHKVSMEDNLNENDLEYLSQSCSTRLFKCSNDVNRLLQN
NELQSTGLRVFVHEAKSYPMSQTEFVDVASATSTSIKLRPVHNRLMSKPSRRCSEPPIETINYVRHFSNASLNVVTKAYSK
SVSDFVVEAQQSILHSNCGCYSHLLPFSVNTSDLCYFAPPQEWITPSDAVLKRIDCHDHWFEHAQRQSQELAKEFQHR
MWCQFTQQRPWQQSRRWPPFSAIQEIWENLMVPQVRHGVEFLPEDGKVHHIYRNNTGRRNSGTLARHQRREHI

>Deu_Ambulacrar_hemi_Ptyfla_40v0_9_20150316_1g27486_t1_scaffold69225_cov128

MARKDTDERSFRQLFVELVQNSSAHGIPNAGRAKTKFRTVVWSIIFVIGVAGFLFQFSELFIRFIDFPVTTNVDVTSNRSL
VFPAVTICNQNPVRVSALEYASDPKLRENFDDTYEPSTSSQAPQDVTGNTAFTSPDMASALPIGNPPPPQGRKRRATDY
LERQGNKQVEITRKPYPYSNCVKKYDSAYSNGYSELYGVGYSVQACMKDCYQRHVIERCQCADPYYPFDADQYQPCSL
EDSEQMYCKAVVEYEYSTQSSDVFCDCPQPCSENLLRIDVYYQELNFETIYAEAAYVHKDRSHVLDQASKLKGTAISIGED
FSKRIRIIRSILLDYRKCLIAENSKLKARVRYDKLVYTDEKGRKKIYTVNEATRQVEEREVRHYDSHS

>Cnidar_HydraVulSc4wPfr_1152_g30239_t1

MSDTNIVNKHFITNDSKGWQSPCPKIKLNNQFRKESNEFIHRNGKPYFPVLKSEKEARREKLDETIQKFVDASTCHGFKH
CFNSGSKVRQLIWMMILAASVALLVQKLYESGVKFLERPFSTKTTLTYVDQMEFPAISICNMNDMRNSKYDYYEDIKEA
GIKLIIHDQHETPVRMAGVKLSPGFSATVQIKKKKTLNLKAPYATNCGSKPLKYFDHYSTNTCWLERLTDHVVTSCNCKD
SFMPGHARVCSIPELMNCTFLKWEEFNKLKDIQCPIPCESQEFESSISFARYPSNILADKIAKDMQLPGSVQENREFIRDN
YLRVEIFYEEMSYIQVEQTPSYDLMILLGDIGGQFGLFLGSSIITYVEFFDFFAALIYTKYFRIFKPPKI

>Deutero_Ambulac_Spurpu_017929

MALISKVMNLRYIISVITSLLIFCSPSHGSGIESITVQNSPSYINISWATAAGSSWDYYTVDVIKSSQRSVGGYPTNTIDTTH
LITGLKPEEQCTIVVTPWTTEGAGRSLNITVTTEAIAEGEILVSDVTSSSAFIQWKPVAGLFFYGMEAEPSDAIEDTGGIFE
AEVGLTILLPGTVFTVTLRGGSPVMDLASTQIRSAPESPVNVSLKEIGTTEATLSWTPPTSPHDAYEFYVINNSTNERSPTT
TKQRGDVSDNEVTLTGLSPATIYSFGVGSVLNAMDSFGLQRSGQDDNPTVYGRTEDFNSKATAACPCPRPCDVVTYAT
TVSQAKFPSDFYSKFLAETLTERRNRSLNAAYFSNSMCLINIFFNELSRQTNTQQEAYGFYSLLC

>Protostome_Lophotroco_molsk_AplCal_XP_035825598_1_acid_sensing_ion_channel_2__partial__Apl
ysia_californica_

MNNTAHHNSFEPRPFNTQYDNNIPESISMKPYAKDGVTSSSLDPDVISPQKKKGSLTWKEALFDFTQNTTLHGIRFIFM
NDVFILRRLLWLALFLTCSVLMSVQIVERIVFFYSYPVTVNVHVNFNKTLAFPAFTVCNQNAFRASAATDRHLYRLIERLH
SGDTSALLSQSPDPLPSNLSLDELYLTTAHRKEDLIVRCEWQNKPCGPENFTLVLTDHGVCYNFNDNPSEPLWVTSTGA
EYGLKLTLNVEQYEYMPGPHDAAGIKILLHDGKEFPKVAELGLSIPTGTHTYVGIQLLKIQNLPAPHGTCRSERSPYYERYS
PDACQLACLTKYVSEQCHCRHFYMPHIDGSPPVCTLGEYLSCYERIIDQVKDRVRAECDCPVPCDFLIYD

>Protostome_Lophotroco_annelid_Pdum_comp406703_c0_seq2_

MRSLQKYIFTIKVTVRSNFIRELQKKLIIKKKSNIPLNVAGAIFTSSSQPTTGVSKAPSFGEAELFSHLSSVFSPTSQEPVSFG
HFLRTDPRWELVEEGYGDLPPMGAAPGFGPQNTRRDLSDQILVNEVFKKTAPEQDTFIHSCKFNGKYCNGDDIVTTAT
DFGTCYTFNPRGERYTVTPGTRNGLQLIMNAETYENMRSKNTDAGVKVHVHDKDDIPYVRYVGTAAALSSHTYLAITL
VESISLGDPYGNCSSNKLDLYPDQNYSQFKCNQQCVTKAKADRCHCRDHYMPGNASFCTFQEHLHCKVDLGELNCNC
PSSCHVTAFSSQASFSSLSKDGVSALLHNKVLFESNYRHALNYHYRMDPKLAEKVIFHLGGFSESVEKLFVFLKRE

>Proto_ecdy_PANarthopod__XP_023224506_1_uncharacterized_protein_LOC111625567__Centruroide
s_sculpturatus_

MTSYDFVHRTRYCKENPQKCKPVENITNFCKTFQDFCENLENFSSLENLMIPEKPLFAAYLSEYKKAAVQENELIPNCTLV
LGDLSTKSCTADYFLMNDKLPSSEGTCYVFNIRFTRASWDDPLISTRVAPMYSAEFFIDFLPHEYFFPHLHVEGNFAIHSP
FSIEDPSIYGKNLDPGYIYTVLIKPVKKHLLESPYSTNCFDYVSQWKKNNYQGPLSMASCLAKCDVKEQKEQIKCILRDNFY
PHEYNLCVNDEEFIDSLDCYSFCSRPCYSLEFEYEVEEREVSTASIEEVEKLIKDLKYPNYNLSQIRKRFSLIQLYIRKPEVLVY
KYSPKFEAIEISSYLGGYLGMWLGLSFFNVYDAFVAFIISLKNRKNKKQSISNTRLSNAWIVQKV

>Protostome_Lophotroco_annelid_CAC9660887_1__Ofus_G111053_partial_Owenia__fusiformis_

IGCTFKGEQCNRSMFVLQKDSHYHNCYTFNGIGSNNFNKNVSKTDPGSGLSLVVFTDTYTDTLTSDPGVIYNPSDPTSG
NNGIRVIVHSPKTRPAPVDKGFDVPNGFSTTVALRATRRTLLTEPHGDCTTDASNKGTEYRYTTDICYEQCHQDIITQKC
GCKSSHFIPPRNETGFQYCGKLNITNLLSGDDTSKESILEDFTHLECEKSVIDTFWSNVTMVSKCNCRPPCNSTYYIKTTSQ
AVWPNEFTQQSFYKTYVNITDSTARPNILFQGLTPKEIKDRGLIQKNFLRLNIYFEDLQVEETSQEADYPITQVISDIGGN
MGFYVGVSVITLVEFLSLIGALLLIFFKSCIKAGGKLREAKVSDVNIQPAHSLENKLGAQYTYEYKSYNDMISR

>Protostome_Lophotroco_brch_Lingula_anatina_comp132718_c0_seq1_p1_comp132718_c0__comp132718_c0_seq1_p1__ORF_type_complete_len_399___score_16_97_comp132718_c0_seq1_93_1289_

MDRLYLNGATLYLAALSSKNSRAYAHINLDNIKNLNMTAWYGSGYTFANFTQDKGWLLNERNARCTFRGEDCNITAD
FKHVFTEFGNCYTFNSGDDPSRNLFQDQAGIGNGLRIEINIQQEQYTNLILRGDAPDAGILFHVHNQSEPASVETDGRA
VGPGLHAYAGLTRSDFATLHPPYGQCNETASLEFYPVYTMSGCVVECKRKHILRHCNCRLMEHPGYEPVCGLIRILTCVK
PLLVNLTQDFTGMCSCSVPCSSVVYQTALSYSLVPSESTKVEASNTLGVAVDDARKNHIILDVYFQSLNYQQSEQLPAVE
WTALISDIGGQFGLFMGFSLLTVVEFIEFAIMSVVTLTAAKRSQNKVKGHDTIHGKGDKVTKSSKTGRISMQGDFVQVH
EIF

>Cnidar_AlaAla_c59883_g1_i1_p1__GENE_c59883_g1_i1__c59883_g1_i1_p1__ORF__type_5prime_partial__len_400__score_24_40__c59883_g1_i1_3994_5193_

IWSLLILTALGLLVEKLYESTMHYFSYPFSTTTTIKYAGKMKFPAVSFCNLNDLRMSKLNGTKLHTAILQSKNLLTTLTGEEY
KNTTKNANHRIHDMLDTCQFVHDNCSTFNFTQFFHNQGDRCYTFNSGKNNHPVLEVDNTGVGHSLQLVINIQHYDYY
MDTEKSGLHLILHGQNETPVKMQGVIVSPGFITYVDIRRRKLKNLPYPYKTNCGSKPLHYFSGYSMHLCWLESLTDHVV
EKCNCRDWFMPGSHKVCSLNESMNCMWPEWAAFDKFKMYNCPLPCEIETYSSRLSFAQFPSNSHADVMAKRLNLH
GSPIENRLFIRDNYLKIVIYYSELSYDHMEQVPSYDMMVLLGDIGGQLGLFLGSSILTYLEFFDCLVMLIYSRFFEKIKPTSVP
V

>Cteno_MneLey_comp19091_c0_seq2__p1__GENE_comp19091_c0_seq2_comp19091_c0_seq2__p1__ORF__type_5prime_partial__len_400__score_77_42__comp19091_c0_seq2__234_1433_

GLTLVLDTLTGDYLPETETAGFLVTLRMSNETVLSQEFGFLVAPGKQHNINLRTRKVTRLPEPWGNCEETVDHFDGKGF
LTFKECYLNQMLWMYPKLRDLGCGCYPWYFYERYIQDPEDSKYEPNIGWKLIKYWTEVLPEQFRFNFTDTCKYEESSYK
PNSGQIREESGVVSMEECVEICRDDTNCRYFLFTDTTNDILEANKCVLYKRGAEMEIIPSKNIVAGLVWGDTQCLNMEK
CSVSAGAACVNKIMNELMGAEEEDLDDNGKSKTPPTCFEPCSYMEHSYDVSVGDFPTRKYWDDYLSKEVSNYDNYDT
ARLNLLKLVFYTDQMSSIELEQTPSYEITSFIAELGGLTDLFIGFSFFTAYQFFEWLVTRVVRKLHTERRTIDSSDNVNVEDA
NL

>Deu_Ambulacrar_hemi_Ptyfla_40v0_9_20150316_1g4290_t1_scaffold1846_cov106

MFIVSVAYTVPRYLEYPVSYDVAMVTSQKLPFPAVTLCNTNRFRKSEIEDSRFRDLLLLDRPAGLSNLMPYIDCGSDSFQC
SSNEGEVHCLPGYLRCDRIKHCRNGEDEDGCEYPTNTGFFDVFAENWTHPYHDITTASRYYDKFREDVYLPNLIDFVSN
DTSPSLIPLLLASTSPTFDDVRAALMMTAGEMHQFGHQADDAILECSYNGMECETEDFTVSQHENFGNCFTFNNKEK
GRLKVEKPGPAYGLELLLSAELYEYVGVYGGRSGFKVVIHDRDVMPFPDNEGFYASPGTFISASLESSVISRLSAPYGDCSS
DFSSVKMSALMMDVGLFLAIWMTLALLVVVLGYVCCGPKGETGQEADEVKVNHVDNVKNANSVNVSLDEITEQDTM
DTKL

>Protostome_Lophotroco_annelid_CAC9477573_1__Ofus_G013340_Owenia__fusiformis_

MIMPQMRMKLYLLGMTWKTFCFHVQLLAMTNMSQIESFINPYFFKCYTMNPKDFNDSSIEPLVHSTGAINGLSALFFL
DIGTAKRIYNPTSPLGGSSGLLVVIHPPGTQPDPLQNGFNIAPGFSTSVELQNKNRELLGHPWGDCENREKLSDFPYKYD
KSSCERQCQQKFIMDQCGCVTSIQPIPDNLKNMSMCGKMDLKNLNSDSPNISQIAEEIDKLECEHAKKRVLKKDYPTEN
KCDCPRACHNTIYDYTISQSEWPSEGVMLDFYKHAITLRDMQGSGYVHSTMYNLLDSKVNSNNTKENRANQEMIRKN
FLRLNVFFSSLNTEITKQVVEYTFTDLISGVGGGFGVYVGFSIVTICEFCVLFAHLVKAMMNNRKNAVNDEPWAFKRKS
MEIPSSK

>Protos_Platyhelmin_Macrostomum_lignano_A0A267FME3_A0A267FME3_9PLAT_Uncharacterized_protein_OS_Macrostomum_lignano_OX_282301_GN_BOX15_Mlig017366g2_PE_3_SV_1

MATARKDLMQRAVKPVSLQDEFVRFCMLSSVRGVPRILRSRSLCWQAVWLVSVFGFLGISLLSITLLVNTYLSYQTVLEVTTRHYTQTVPFPSVTICNLQPFSTDMVKRFIRNPNVNVTDMRFLNELKKMKNHPNYTSSFKALNEKSFLFQNWGIETVRKIGLTFHEFVLDGTCNFSFVQAAKFQQRECNLRTDVLLIPNPQFFTCFMISPTPDIANTTYSLEMDLYLDTEQLFDCQQTTECHDALFDLDDSIGAHVLVHPAMTYPRISQLGIHVLPGTKSVIEVVETRSLEKYQQSPKETCTNKQLIDLFIGAKERTTAFPFAYESFTCRQLMVQEKILSNCGCTDPDYPLTADMKDAPFCAELKKPGQSTDAFLKNSAAWLLKFAATIAPRAWSWTPET

>Deu_Ambulacrar_hemi_Ptyfla_40v0_9_20150316_1g5442_t1_scaffold2520_cov137

MPSEDSLTYKALIQNVLRNSSAHGLPNVQRATSLPMRLFWLLAFFTAVGIFVWMSSGLIAEYLRYDVDVNLQIQFSRDLTFPAVTICNLNPLRKSKLSEFGHDILQRRPLGRMEGRRRQERQTTERSETSNDTLNDDQERTFEEYNYWDQIPTNYHASPSSSWSIIERIEHYISDIPAVQRAEIGHQLKDLVVDCQWNNIKCSPRNFTTFTNVMYGNCFTFNGEHNNVMPLSTHYSGSVFGLTLILFVEQAEYMDDVIDSPGVRVTIHSQDDTPFPEDSGFDIQPGRATSVGILMGRTQRLPKPYTNCISDNIPTYDTAFGDIYSVKDYIREFLKSRSSKLRTLMEQEERTGLDLLSNANRDTSKCYSEHHRKRSVATFDPLRFICAALLDSNDRYNLRTSPDDC

>Deutero_Ambulac_Apla_gbr414_1_t1

MSDAVGIPGIRYVVSSNFHILRRLLWTVALLGGMAAASYQVIDRATFFFSNPKSVNVVIENAEPMLFPAVTFCNTNRFRDVNGQLSTHPFGQFLAATYFHGGNGTGFTSTLGGANTTALYLEFAHQMEYGQMFIRGALGGKNITSADFRRVLTDAGVCFTFNSGLNTELRGQSFPGRAHGLYLFLNVEQYYHYYAYEFSGAGILVAVHDQTEIPDMDSMGFGIPPGRHAIVSVGKKLTSRKDGGCVEKQLKYFSKYTKSMCLRECVIDLLLQNCGCVEPYMTDKRRYVDMCDCPLPCSTVKFTTSVSYTLFPGNFYSWLLAEQFSTDEFTLPHDFYQENFVIIQMFFEELTVEHREQHSDYTFFALLCDIGGALGLWLGGSILTIVEILDHFTSTTVIPGSTSHQRG

>Protostome_Lophotroco_annelid_CAC9608250_1__Ofus_G071455_Owenia__fusiformis_

MFIHKQHLNCEEWLLSDIGNYVRPETFGSSLRKAPDSGFTPSVDYNNAPWLAENSESLTYSTRQALAEAMSSLSQQQRIDMGHDANNLIQECSWNGKTCSPANFSGFNNAYFGNCFTFNHDDLPTVLSTSKTGPLYGLSLTLYVETNEYVPALTSAVGVRVLVHVPDKQPFPEDEGIDVHPGTKTSVGLKYIKVTRETYPYPSDCTDGSDIYLLYDKTYSIQACIRTCKQEAVIDECGCANALEEAPENETICKSDKLDCVNSVLEQFERKELTCTCPQSCLDQSFSMTVSSSVWPANSYGSTLLDAFGGRDGIPDFESVEKLRDNMIKLDIYYQELNEQTFKEEPAYLSENLLGDLGGQMGLWMGISVLTIAELIELVSTIIYTVCKKLRHPEKVHVKNDLPLVEKY

>Deutero_Ambulac_Sakowv30036990m

MKSPSNADIIRNVNRRRPTPTSTSFDMSNTEKFTSTSSLSLDGGNVRRRGAVYIFSRNFRQFLSETTLHGARYTANNEYHVVRRLDVKYNINFDWSLLENTTAILNRTQFELEAAHQKESFIVACNYQGDGVERLCGPNNFTTTFTEFGVCYTFNNDPENQLFVTKFGSNSGLRVRLFTDENEYTFGKQTGSGFKVLLYSPGDVPLVQQLGFAVPTGVDALVAIRLEKSINLPPPFPTKCSNAPLKYYDGNYTYVWCALEKITDFVVSACGCREPYMPGSKRVCNLKESIVCVIPVMDDSTERSVTYCGVGCDVTRFDSRVTYADFPAKPILEHMSSTTNESSEYFRRNYADISFFAEDMTFKLTKTVPVMTGATLVSNFGGLMGLCLGASLLTAVEFIDFILFCFCKS

>Proto_ecdy_PANarthopod__XP_023234316_1_uncharacterized_protein_LOC111633896__Centruroides_sculpturatus_

MKDKMDEVNNRRNKKPTWKEKWLYFLNLLENVLLTGVPQIVTERNTRRKTFRTCVFFLCLTGFFYHLISFIMLYLTYPTT
LEVLVKYQNESTLPRPLVTVCDINGIRRSSYCEAFPDNCEKVEDPDDVKALCKHFEYFCKNESSVGDDLMFPTYETFIFSD
FDRNVVEEYGVRKEDILNIYSNETNHIIFGNEFGKLRNCYSVDLHKFDTTNSLEDSRHRFYEMHMIQGTTIGNLIITFNPEE
SFLPNVPIGAKISIHSHHQLINPFVDGFLVKPGMRYVVQLKAIEENLLASPYSTNCTIYEVPEGKEQTENNHERCLIECKKE
KWIKECGCVSGEYAYHFDGKFCLNQSEVNCITNAAVYDCYDKCKVSCKSMFFEYEIDEVPLENKELLLLMLEELNVSDFN
KYKPEK

>Deutero_Ambulac_Sakowv30000474m

MHVAYTTSGFLFKANLIDLDKCWPIVLLISLRMYRKSALTDEEVRYVGYLTQSQSTRDYDTTYSMYDDNMYDDFVNSTG
LSYPDEFDFAAFTRRVGYQFTMHDDLLHCVWKGGQLCTSDDFVHVFTRYGNCYTFNAPAKHQYLTQDHVGAGNGLV
MFLNVKQEEYTETAFGNYAAGLKIYVHNKDEPPLIESQGHAITPGTLSYVDITQHRSLNLPPKWGQCNPSQQLEFYDSY

TIPGCLLEWYSKIMTSVCGCRPFIYPGRARHCTLWETFECVQPTEAAIQSGEYGDMPCPVPCNHTTYPTTMSYANFPSK
SVASSMAQLFNISAEYASENYVAVDIFYGRMSYQLSKQTYAMTASALVSDIGGQMGLFLGASVITLMEIAEYIGNKMKH
LFCFKKKKTRANNSSNAIHTEQ

>Deutero_Ambulac_Sakowv30036989m

MSSSELNELTYRRDSIRTDIVVTEGNDGNKSTAGLNTTTAMNTQTAFSAKLSQFMGRTTLHGVAFTVGPNNSWKRRS
VWIFLVIVSVVFLCLCLFSLVGLYLTYPVDTMTSLQFTQSLTFPTVTICNKNRYRKSVINGTAFEEFLWSIYPLAGFGPQLHV
NYNWSLLENDPALQNRTEFELTAAHQLEDTILKCQFVNSDDKHECGRENFTTVFTEHGVCYSFNNGLDNILKATSTGSS
TGLHMLINVQRIEYTIGPRTGIGITVILSEPGADITGENTALSIAPGVEALVAMPMDKYLFLKAPYQTNCCTEGTSYYPVYS
YHGCMNEKMSNEMAEKCNCKEIELPGPVRTCTLQESVECALPLKVSNALTNIDCNVPCNFTLYNPRVTYGHFPSPSIIQR
LSDMYGISEESVK

>Protostome_Lophotroco_molsk_AplCal_XP_012942491_1_degenerin_mec_10_like__Aplysia_californic
a_

MSKVVKLGNVSSPAVTVGDTLLEEMDSFIYLGSVIDDKGGTEADVPCGIGDFLTIFNYKYGRCFIFNDMLQKHNDISSQR
YTTQPGPDNGLILELDVEQSEYLDTTESVGFKIVVNPQDTIAFPEDQGLIVMPGTSTNIGLKKVGFRFQSSSKLLQTLKTLV
QHDLKVIKDIIVDLRESSLLYFQVEMSRAPPPYSACVELTDELNVELNVYSFTEGTRYTAEACRKTCYMRELGYQCGCCDL
HFICPPEFYNISFCNATNTDEFVCKKQVLKMFEKGELDCLSRCLPPCEEVAYDVAHSTALWPSNNAILNEIVNNGWDKA
DYNSIKSSTRENVLRVVIYFEDLAVRRVESRPSYDWNRLLSDIGGQLGLWLGFSILTAMEWLELAYDAGIRVATRKSQGA
HKTRVHVQESGNNY

>Cnidar_HydraVulSc4wPfr_116_2_g12647_t1

MKGKKNEKQFCFVDIYDEKNETVPKKRKVSALRYKSKEYCSSFTVHGLTKVFTGNWFEKLFWGFTILLSLCTVVYIVLTLAI
KYNKRDVYTKVHTKEYLASDMPVLTICPADKKEKCNLTQNPNTNCFINNIPKVSDNYSEWSNENMYLYVYYLMKKGYF
VGDFVKPSYFNCIRMDLSFLDTSSKAFISFTTAYPNDAIQIYVHDADELYPFFRQKPVIILRNSTSEISIEMKQYKRLPHPFPS
KCVQNFDSLIYPGKYSKLKCHESLRCLNALKTCGASFNECLGFLPEKAKVTHAETIRNNLDNVTTCLNSKYVQNYESKCML
PCDERVFTTIVSNSMFNSLFSEEYNMDFYNLFIDYPLNPYYLVYEESELYAFEQLLSECGGLFGLFTGSSAISCVEVVFFISLII
LRKWWKNYN

>Cnidar_HydraVulSc4wPfr_2261_1_g31471_t1

MKQKQKNKQSYFIDIYDGKNRAVPTKRRGNTIRYKSREYSSSFTVQGLTKVFNGILFEKLFWGFTTLFSLCTVVYIVLTLSV
KYNKRDVYTKVHTKEYLASDMPVLTICPSDKKEKCNLTQNPNTQCFINNIPKVSDNYSEWSNENMYLYVYYLMKKGYF
VGDFIKPSYFNCIRMDLSFLDTSSKAFISFKTAYPNDAIQVYVHNAEELYPFFQRKPITVLRNRTSEISIEMKQYKRLPHPFP
SKCVQSFDSLIYPGEYSKLKCYESLRCLNALKTCGATFNECLGFLPEKAKVTHAKTIENNRDNVTTCLNSSFVQNYESKCMI
PCDERVFISTVSNSNFNLLLSSNKTVDSYNIVIDYPLNPYYHVYEESELYAFEQLLSECGGLFALLMGSSVILLVKVFVFICLD
ALRKWWKNNN

>Protostome_Lophotroco_brch_Lingula_anatina_comp147337_c2_seq1_p1_comp147337_c2__comp14
7337_c2_seq1_p1__ORF_type_complete_len_421___score__4_96_comp147337_c2_seq1_410_1672_

MFYKFFEYDVGVKLEITSNSTLKFPAVTVCNENAFRKSALLSSPSKLPTLDAFVTNGTVPTTGFNSSLIDSELGGRSNRATLI
DKTLEDISELTTGEKQALGHQRSDFILDCEYGGYSCNGKTFESFYSTYNGNCFIFNSGWNNSSILLATESGPNFGLSLTFNI
EQSEYVGDLTQTAGVRVLVHDQDVMPFPEDQGFLAAPGEMTFVGIKMVEIDRYGGRYTTCKKTDTFNITENMYQALY
PSVGYSQKACKKTCYQRTVIETCKCALSIYPRVETLFDSTTSVRTCDSLNATDVACTVKVQTQFINGELNCSCIQPCSATSF

SIDTSSAYWPSDEYATEFLSNYNSKSSIVRKIYNTSGSAGVMKNLAKVHIFYKDLNYEYISEYVYYDVSIIGIASCILVVFLLN YEKRTLCTQLTA

>Deutero_Ambulac_Apla_gbr467_14_t1

MGIRVKPIEDDISIISFNTDEVKIDKRERDKETTGDSLKTGRAGPQKGSKYHSSRCTKFLATWSSTLPDAVGITGIQYIVSN QDHILRRFVWLLALFGGFAAASYQTIDRTIYFLSNPKSVDYEIKYISPLEFPAVTVCNYNSYRAYWDGWGSVTSANFTRVI TDFGVCYTFNAGQAGQELLKQKVAGKGHGLSLMLDAQQYFYFYSSKALQVSAGFVVAIHNQSEVPQVDSLGVGVAPG TEVRIGLKRKEAINLEPPHGECGSKELKYFSSYSINSCRQECLTDFVLEGCGCVEPYMAERFNSEGVCECPIPCQQTTYTTS LSFATFPSDFYIQTLVNLYSDFNVTPEYFSSNTLKIEIYYEELSVESMEQQEAYTFFALLCDLGGALGLWLGGSILTFVEILDH FGHTAFLRGTAFQHS

>Proto_ecdy_PANarthopod_Dmel_tr_A8JPJ8_A8JPJ8_DROME_4Fe_4S_ferredoxin_type_domain_containing_protein_OS_Drosophila_melanogaster__Fruit_fly__OX_7227_GN_ppk31_PE_3_SV_1

MSMILYLVTFYHTLPFFANVSLSLLNSYLSSSVSFNLDTIYLNWNSTFPAITVCEIYNAERIWDLSDSNFGVEHDMHVDDF ISEIVFYRGVCSSCEDCSRMRCPSNFTHLVDVFRTKCHQLLVNCTYLNKLFDCCDQFLPLPTEYGLCFSFNSHQARKVSHL QFTNNRMTGPGHLSFHAAADIQLYVHAPIDVPYQFSEGMIRETVLLGHYKELILNVIEVHNDESVQDLSMEQRRCRYG HEHVPERQGIYDFYSYSGCVVECTVLLQLDNCNCTSHFMAIPGQNYLPVCDVRGLICLTRIRDKIMTERKSCECMSSCEE PEYNIIYNSADEDDEASDEVSEIRVALVELPTQRYVRRVTKTILDFLISLGGLVGLFFNTSALRIVETIVICLRYRKTIVKWVKR FFLGTVQYLQILDQST

>Cnidar_NemVecNVEC200_012401_1_1_protein_AED_0_06_eAED_0_06_QI_181_1_1_1_0_72_0_58_12_321_422

MSRSGPSVRALLRDFSDRTSCHGIGQINGSHSPTWRIFWLLTFLAGLGMVLFQCITLLGIYLDKPTATSVDVTYDEVTNF PAVTICNLNMIKKKNLANFTQSKKIFDDFEAFVSSNSSMDSSAFLGSKMETVLKDRLSMDSNDGSNSVSLDDTSMDTEL YVEDMLIRHMAMVDDKDLIKAGHEFDELVFRCVWNGFTCNKGGFMKFWRRFWHWRYGNCYIFNQGVDENGTLLA HLTSSKPGPMYGLTLDLFIDQEQYLIPLSQEAGVKVLLSDQRNVPFPFTDGFSVQPGVSASVGIRKLVINRIDPFNNGSCY SGDGLEKDNIYSKHKSLKYSVQGCMSSCLANSEFSICNCTEGKFRVKGRPCMSESEVKCLNTVNKMYEKGTLGCTRKCP QPCSHFSFRRTISQSQWSESYEQTFQKWS

>Ctenoph_Hormiphora_californiensis_evg134570__p1_GENE_evg134570_evg134570__p1__ORF_type_5prime_partial_len_424___score_58_64_evg134570__2_1273_

NYTMLKELITEEELESLEYILYEMNVYNGTGDLGSSGNFFLDVSNVTDVIDYLDSMGDYTGDNYLDYILDRISFFPEDFVV GYYYDFKMGNTWKYLNMTSTNASNVIPHMLGYCITVNDDESLVQIVPGIHGGFTIMLDAYIDHYMPNVAEKGFEVFLR TPYEEVMIDRGGIIAEAGKETNIMISKKKMTRLGSHWGTCESIPSPYYPNKKNVHRSQRECYQEGMIYESQLSCNCTPW YYVERVNMELNLNWTSSLNVSTCSFASEIICDTVIYQLMETYNKDEWFQCQNERCSYNTWDYSLQAFEFPPTEDYYDLL ISPILKNEMYTMEYAQKNIAKINIYYGDLKETTVIQEKAYEVFSFIAEFGGGVDFFIGISFFTAFQALEIAIAYTCCRGLRSRKL TEEERKNNYKENNVNSVDTPK

>Deutero_Ambulac_Spurpu_014120

MKAPKVDGQEGSWGTAPEREEKSLRTILNSRMENSSAHGVPNIQRSSGPVTKLAWSLVFLTGISVMTWQAVILFQTYF EWNYSVNLEVRFNRTQSFPAITICNANPIKRSELETRDALFQALFDVHYVPSMPDLPDQQPLPDEQPLPDVPGLSLELFV QQDEYVEGMTEVAGFRVSVHHPSIMPFPEYNGLLVSPGFATDIGLRVLEVDRLPKPYGDCKADLTQGIEDDIFHQHYNI TYNRKTCEVSCFQNEVISRCDCFDATYPNSLKVNRTVYPCEYINDVETQCMADIEMEHARDELECNCPLACRETTYLTSV

SSSIWPSDAYESTLFQKMVKYNAEIRRNVVGENASDWTRRNMAKVEIFYDEFNYEHIRQDAAYTISTIMETRYLHPFFIH
TNSIAYISETVHNLKQTFNVLSKSI

>Protostome_Lophotroco_RotSoc__A0A818Q233_A0A818Q233_9BILA_Hypothetical_protein_OS_Rotaria_socialis_OX_392032_GN_FME351_LOCUS4621_PE_4_SV_1

MNAALLAFSEKNRVNLTETNAESLAYSLLADLFNQNETAELVDLGFQLDNMLLSCSYNGIECSHDFVLSISSVFGNCFTF
NWNAFMKKLFTLGDMNSTLVIFQGLTLTFYVPSQLNFPSTLFDEGLVVLLHDNDELPRTTEKGLRLKPGLAHTINYEKSQ
TIFLPKPYTNCTSSVELDLIALYKTMFNSIAIGQVAYSEVVCYQLCEQAFIFSVCSCILPIPLYTRYVFNREHDNLIFTNVCHP
YTEEQTCALTTAQRFVSSPTTIAAWCSHCVLQCTYTYFPTELSANLAPTTAQKAYWAPILLSNTSNWTNLSLPNDFAANY
DTYMSSNYLRVTITCSKRYVTLNVQEAKLTIVDTFAAIGGQTGLWIGLSVLSFIEVCELIYWQTVKWILAFRSWHQVKPST
EIDLEKVESCSDQVDTKK

>Cnidar_HydraVulSc4wPfr_417_3_g31877_t2

MNKKGFVKSKYIFEKNGTVTHSCEKKDSPSKKINDHSRHFSALIINRILLGCYQEKLFWSFVLILCCALVALPIYFRVQEYLS
YESSQIISTIVTPKNLFPAVTICKERKSMFLYCDLDIYDVNSNYNVPCPRDKQTVEERSNVVGTGSYRTNIQPFEFYCDELR
RKCSPSDLQSQFFKTVRNERGECVTWNGNGTYTNANNIIELIVKVVKVERYFEVIEVIVHEPQVDGTLQDLTFRVTTAKK
YEISFTKVVSELLPAPYSNCINKKQNDIFPGKYTLENCVNSHNCISTFKQCGDTFDYCRKYIPESIAIHYGRNNTISRNINCLK
KQNETLPPICLLPCKQTDFEVSIFSEVDTRLNNNQFVFVMKNKNRNTYIEKKDKLDKTNNWTRLASEIAGLFSLVTGSLLI
CIIELFVYAVLSCF

>Cnidar_Rhopilema_esculentum_TR103753_c0_g1_i2_p1__Rhopilema_esculentum_TR103753_c0_g1__Rhopilema_esculentum_TR103753_c0_g1_i2_p1__ORF__type_5prime_partial__len_425__score_38_48__Rhopilema_esculentum_TR103753_c0_g1_i2_1072_2346_

SAASVTMHGFGRLFSTKSKAMRCIWLLLMLFFVSLVILTVTPTVRRYFKSPSLTKVETVRVSEMKMPAITVCGPSFSKAK
VTNFAKEKNLTVKYNRIGLIDNIDDILGRASSLEDLAFDNNTMFIINPDENDIAIERVLVANQGYCYKVNANGSFVQRSTD
IEKFLFIGLFLNISDGVYLRRSTTDSVEVNIQDHRQLPSRDSGTVLAPVGHLTSIEIKKTEIKRLKPPYPSKCTNGENLRLLFK
GDYSISNCRLSCFLIQVSQKCSRIHFFDSILLPKAMRKPLARNESELSCTYSIHESLYNSNYASCNCELPCTETRYNKYSSSAK
LTSSDPSIHKHIVAINVFFGGMEYEKITELKMYTTERFVGEIGGQMGIWIGASIFSLFELIYLFVQFAGVLCRKKRRKSSNLD
KGMSKEANDSVL

>Deutero_Ambulac_Spurpu_007150

MENKHHTTTDPRGFVTVELGMEDRGNMQHAGAKRLDRSLSGVQLVSSWWLALSHDAPEGIGIAGVKYVLMSRESRL
MRLGWLVILMVAMTIVSYEIVDRGIYFSTGPLSVNIAVNQSEKELTFPKVVVCNYNIYVKSKMQALPAVDKVLRNYEYH
HYDPNRTIALTPEEVSEVKATDTARLLEQVAHEKEHMFLRANFAGKPINLDDIVPVFTKEGRCFEINDGLNGTELLKSKGI
GRAFGMTLVLNTEQDEYYYNKYLNDAAGFIIQLLDQDEVPQVLEQGFDVGPGTLTSVAIMATQHIYEKPPYGVCGEKKE
KFYEKYSQQKCDLECTTDYVVGECGCRTAYMPGLLFAKDDARLSHWVTQYSMRHCRVTQQFVTRLPPLPVRHLLLVSR
LGICEIFVSAHTKHTHTRTHTCVYIYIYIYI

>Proto_ecdy_PANarthopod_Dmel_tr_A8DYP1_A8DYP1_DROME_Pickpocket_29__isoform_D_OS_Drosophila_melanogaster__Fruit_fly__OX_7227_GN_ppk29_PE_3_SV_2

MWRKSVMNFRGFIRSFVENKGLLWIFVIGISGWSTVSILVLLKTRYETDSTTIGVSTAYSRWINTFPSIGICLTKSRAFNEF
KAMMREYFQEDFAFSFTRMIYEYAFLNPNNIFTKEPTKNTSYPYNFNILDIRRKMFPTNCTECFKEIYFRGELVTDCEEIFK
FHVTEMGYCFLANNLLDYDSIEEMPLRYSSLDNNRSLRLYMRSSVMYKYEMYVNSPEDLPFFNSLTYTISTDPTTYAFNV
EEIHNHEGVIDEPISQRKCKFPSESSIEGFPYSFSACMSIIRSEFEMKTCDCSLFNPKDRNESLYCGLQHADCLIKEGFATRV

KEYVGSSTVCLPSCVEQQISLVGVITENGTLYNNNTQITEIQIASPPTVRYERKVTQTKLDLIVGIGSVAGLFFGASLLNLLEI
ISYFIKKLKTMIFG

>Protostome_Lophotroco_annelid_CAC9668426_1__Ofus_G16866_partial_Owenia__fusiformis_

VGAPLKRNLSSDALGYGYFNKPSRPLPPGYDETKVEPEADSETWGSLLYDFTQNTTFHGVRYIPQPTVFLVRRILWIAVV
LTSTCFFCNIVYNRLFTYFEYNTIVNVQVESASSLKFPAVTICNHNFLRGSEVYKTGMFDEISSRFKAIQAGYSEVRPCTVQ
NPFTLLPNKTVSISNFVELEKHEVATFIGLTLEVCKYLCVKDTKLKCKSIFYSKAQASCSITSIETDDFELAESSTEDIFFRCRVC
GKDEFECDNGDCIPESLVCNGMSNCDDGSDEINQCWEDCEFNYDNLCGFSIEPKQSELYTWMLMDEDKENEIFKDDI
MCNVIGSESNKYGLECVSYDYDDSSYYSSPYASTTSPYGASGESYGECCSSYPTDPSYENAGSSGGYPPPEDQTSYSVIVD
VFFTPPFPETTNTGATGST

>Deutero_Ambulac_Spurpu_025196

MNGNNAQTSPADDVSVELGMVSNEGVMFDKKHSRIHLVSSWWLALSRSAPEGIGVPGIKYILVSRESRCRRKSKTQAR
PAMDKLLRIYENTHYNSNETILLTQEELTHLRNVNTTQMFEELKHQKEDMFIRVVFAGVPVSLDDIIPVATTFGRCYQIN
DGQNGRQLLNSSVAGVSFGLSLILNTEQDEYYYNTYLKDSAGFSIQLLDQDETSDLHVRGFQVGPGTSADVSITAKQQN
YAKPPYGGCGERKLQYFDKYSRIKCLLECMTHTVIEACGCRSPALPEEITSTHHAELCQCPMNCEERDFPVSVSTAKYPSA
FRAGLLEKYISEGLLDNISFKSANLSVIEQNFVAVNIYFSDISTTVVTHRPAYTVSSFIVLLIENDERIEHQCEGRKEGKPGQG
GVSQRSQVELNSKLDLNIMESHL

>Protostome_Lophotroco_annelid_CAC9627304_1__Ofus_G09267_partial_Owenia__fusiformis_

DDDDDDDDDYQLNDALKDFHIHIEEPATFFSFLTSKERFDISHKKKDMITDCRKLHRFAKKNVYLDIECGENLFTKYNDM
DNMNCHTIQLQNIPNVTLETKGISLVVNLNHYYEDVYVKKKIPITTSSERGLKVYIHEHNTMAYAQNHIKVTPGHLIDIEI
DVSHHSKLGEPYGPCTKTSTVEFRTPSGTKYAYSEGSCIAACIQQKILKKCDCILQEFPLLENKKVCMSIRNNKSTYLKQLS
CLDEFWKSSTSHNRICNQQCGPKCKGYSYTTTVSQTRWPNEDRHLWFYNKYISGSTFQDFYEDYAKLAEEHMDAESTE
EENENFKITDTIDKLRNMTLLRDNFVRLDVSVTTQKTIYREVELVSTFTFYNTIASMFIFWIGCTFFSCLEIIEAVALWIRGR
VTSANKHPITKVIPLDKREVKM

>Cnidar_HydraVulSc4wPfr_283_g24324_t1

MSISCNGLPKSFRKNPIIKVKITRDYSSENPEVKILELYEKSPTFPVVVICNENALNSSIKEHIISLTGKKLDQFDSNLKKYITH
KNFNSRLFNEFGNRLNSTLVSCTIGVDDCSTLLSWYQIWHLEYGLCFAFNSGFNIEGNEVTTRRVFLPGYSHGLNLNLKL
EKNEYNDKSKTTALRVFITHQGEYLFPPNEDLLLTSGFYYSISFQKRLMNREYGSNKMCRKDKIIKLSNFSLSQPLITKYTQ
NFCNLNCLSEEIARECMCVEYDRPVLEQHAKIPKCNVDASTQSCIKQQYSYWKSGSSECLQKCKLECFEVKYEPRIHLQK
YCSNDLCEGDLKLSVNFRSFNYFFYQVHINHVLAEFLGHAGGIITFLTGFSIISFIEASFITMQLVITILYMRCKSFKKSCDKTI
TSRKSFVTCQASLEV

>Deutero_Ambulac_Sakowv30026092m

TTAHGVGRFTAATSLVVRLIWLVICFLMWGIFLWQGSLEVKKYFNYDVTYDIAVVFRELPFPAVTICNLNPLRQSKLGDL
GTEYLNELGFLDFDTTDDNQTSEFDWNAEIPVDSDWDDDSNFFDGFDDEYAAEINLTRIMSILDPSLRKSVGHQSKDFI
QECTFNGGKCSTANFTLVLNSKYGNCFSFNTGINDSEILSTVRPGPTRGLTLTLFIEQDEYLTDYTENAGAVVLVHPHSIV
AFPEEQGIEVPPGFSTSIGVRLSEIIRKGGDYSDCTDGTDIDMLYDDDYSYSREACLSSCLQQALIDTCLCYEPTLPHSADV
EMCSKVSGGTTCQAGVTNQHNKNQLGCTDTCPPSCNDRTYETVVSMSQWPSEAYDDTLFRRLSKRSSKIQDQLGSIE
DSRRNLIQLKVYYQELNYQSITEKPALD

>Deutero_Ambulac_Apla_gbr500_4_t1

MRASEKPADTPRFVCSVLERIGAHGIPNIGRAGTQQRRLAWTGLVLAGLGFLVWQGTLLVTRYLQYDVKVSVGLGYEA
LTTFPAVTVCNLNRIPRTAMHTNSKLAELEGYIEYILETQVYQAWSDEMRYTNQVADFGITTESFDIMADIPYANRSSAG
HQLSRMLLYCTFNGYPCSPLNFTHFYNYFYGNCYTFNFDSRLARRVAQAGPFYGLTMELFVEQTEYVRGLQNSAGLKVF
ISGQNQVPFPEDRGIIVSPGRETSIALHKACEKSCVAQEILEMCHCADPREVEFEETVSAASWPNENYKFALENIITRLSA
DFSSAIEEDDRFIEKNVVKLNVFLSSLQYQYIYQTPDYSQENLISDLGGQVGLWMGFSILTFFEIAELVLDLFVYTRGKSAN
RVSREAKVGEGLVGMSLSPSVNTLTPIG

>Porifera_Syc_Cil_scpid69199_scgid1215_Acid_sensing__ion__channel__4_Acid_sensing__ion__chann
el__4_2_Amiloride_sensitive__cation__channel__4_B_ZASIC4_2

QQQQQQQSTPRATRNGNQAATASPGQGQGQQQGGQGQGQGGQQQQGSDGQQSRLSSELRQILQNESALVSLLE
PVDVGQYTLEKGMQLDRLLESCTFRGTECNASDFQHVFTTFGSCYTFNPDETRQVSQAGQGHGLDLILNIEEDEYISGV
DQSVGVRVLLHSSGESPHTTVDSLGFAAAPGETTLVGISRRQVTNLDRPYNANGCRSNTGEDKLKYFDQYTLSNCALEC
FADLSELVCGCRAPYMPSNDNSTVCNPIQMFRCIDEVKTAIIQGTNQTFDCAAQQNCVQPCEYSAYSYVLSSATYPSTR
SVISPALTQAEEAALLSTKRRNYVHLNMYYEELNYQKVEQVIAYDWLSLVSEIGGLAGILLGASVLTFLEFLDFVTGSILCAI
CDVSHRRRYNAQRKSSFQGNVELTTNWSLEPPPNT

>Deu_Ambulacrar_hemi_Ptyfla_40v0_9_20150316_1g25474_t1_scaffold44668_cov97

MNSGSSSSDSGSLSGSSQSSDSNNDVTDIHTARQRRFSEREPAKTKTYAMVDPEADKTAWSLPPNLQSRMGSSPSDQ
PETIKPVHTLPYEHFKIEGLHILKDLLREGDWICKVDLKDAYLSIPVSNLGAPYGNCVENVQNKLKYFDGAYSLSKCWMEC
ETDYAVEQCGCRYLYMPGNSTYCSPYVLQNCYFSAMDDFALNEDPCKQCGERCVQTKYETKLSYDSYPSSIYSAMLSVT
GRWPCNEALAFYAMDHSSNYLIEIAANMKNFVMEHYYTVEANGQNISAFIPSPEFHDIIVTALTERLFPKVLRDLNFDVL
KYFPTFIFKNGTDDVGSDFYVETKETLKQKYMNLSVYPWTGHNTTMWNLFASANHTSDSIQYLIDTLFQDYYLPLFDET
FLYLYSEVKSWRDNKTEEFLSEMTPDICRAYMQ

>Deutero_Ambulac_Apla_gbr243_2_t1

MEGNRSKLVADFKKFGDQTTFHGLRYVTNNTYHSVRRLIWLAVVLGMTTWLVINIIRAVVYMFQYPETSSISVNYVPNI
TFPAVTLCNFNQYRKDALDARAIKILAKVFGNPALRESIDLDVSDVKLFPTTNQANVTATTVQATHQIEDMLVDCHWRT
EPCSHLNFTQRLTDYGVCYTFNDDLAGLGDVLTIQNPGASNGLHLRLNIQQDLYTFGESTAAGMKVLLHPQGEFPIMKE
FAFSLSPGFETSVAVRKETVTSLKAPYKSDCIDGSLKEFPYKYSVAACQLECRALYVIDKCGCRDIRWPEEFLSRGERCHCP
MACETTTYQSKLSLAYWPAGYLTAELQAKRNLTEDFIRKNYIDVYIYFEEIMYMAIEQKMAYTIDNVQSDIGGYLGLLCG
MSLITLVEWADFILITLYKRCRRLASRRIST

>Porifera_AmpQue_tr_A0A1X7V291_A0A1X7V291_AMPQE_Uncharacterized_protein_OS_Amphimedon
_queenslandica__Sponge__OX_400682_GN_Aqu2_1_34370_PE_3_SV_1

MAGQAEGSSQSFDLPVAVDGDKAIILDPSLRKKSFRSKIKKGQHDFNKRSLLAPRSRLKRILWVVVILFVFLFFPQMMIFS
MVLGIAKVQYLTSTSISFISDDEGQSFPAVTVCNLNAVKKDYADQNVVDLLNNYANPTSGFVQDWPFSNTSRSCTDSLN
SISIEDRQEXXXXLAGPQFGLNLVLNVSVSEYLRLGSSNVGVKVSVHDRNSMPVPQVGVTLSPGTNSYLAIETVKHVDKT
RESNCVRHNRSLAFFPGYSYAASTCRINAEYRNYADANKCGCSPEVNPSSNVNTRNCTIADICCLSQERRTFNPSTNCPL
SCDHLTYNVEQSYSKLFSSASSESAAESLGKTVEASLLSVYIHFDDILITQLLTRYSYTIANLIADMGSLFGLLLMNLLIVSIM
RRAVKKLESKIMALPEVNLDGPIDDA

>Protost_ecdy_cyclo_PriCa_ecdy_XP_014665185_1__PREDICTED_FMRFamide_activated__amiloride_se
nsitive__sodium__channel_like_Priapulus__caudatus_

MVSKRARLAVINAHDARKENKRDLTPPPRQRHGGVRYVKHHGPGDAIHAFMRRTPAHGFGALLQGKSTAGRIFWML
CTLLFYVFGIVALGMVISRYREYLYRQNISVRNTAVDFPTVTICNINPARVSQRHLGGAQFEAEVAELESNVQSKFQSQEY
VDAVSSLKAGVTSPLHVGRESFAIALSKLNASEKVAMGHQLEDMVVSCTFADEDCDLDKLLLNVEEGEYSQWLHDGTA
LRVAVHERGSMPQPDVAGVVVSPSSQLTVRLTTTSFERLGSPYRAADCATENHFTATTFTENFGFDYTKSSEIEYGCGCY
DARLQPTLASHPCEEQNREDLSCMMARTSEYEKGQSSCNCVDECRLASVEVHFMDVAQTQVKEVPVMEVPEALAQV
GGMLRFFTGSTLLTGVEMATLFVNVFGLVCSSGSYVSVD

>Protostome_Lophotroco_RotSoc__A0A818JTG6_A0A818JTG6_9BILA_Hypothetical_protein_OS_Rotaria_socialis_OX_392032_GN_FME351_LOCUS19348_PE_4_SV_1

MVVIFQAYFECPLSMNSDAIQQQPQYFPAFSFCNVGELRYDQFIDPFLNYTNANNVTSSNDTTTITRNQASYIPNFLLEQ
LNQNKSLEQYFFSLSPMLYQCSFDSKPCSAADFISFTEAGFGSCYTFNAQLKNTTAQTPRYAILGGTGLQLGLYVYSQQY
VPYVSDGIGMISLLHNNTQPPFLDLTGMYLSPGRKYKISYTTTITNYLATPNSQCTDDVPQSMKGILSNYPSADYGYSQA
LCIENNVQIQTYQQCGCVSPYAWDKRAITLPGTSNIFIAPLCNITDTCYLQAVSAYLTAPISSNSNNFTCPKRCSNVNFPV
EKSELSTPLEWQMSDIKSFVENSSVPLPANWSVMWRDYIRMNYLSINVVCETNVIDAKQQTLVYTTNTLISNIGAQAGL
WLGLTVLVFAELIELLYHLLRFTCQKIRSKM

>Deu_Ambulacrar_hemi_Ptyfla_40v0_9_20150316_1g8126_t1_scaffold4191_cov132

MSSSEFAWDGKKICEHETAKNTLSVIFDELLKNSSSHGLPNIQRSVSAVGKLSWALLFFVAVIMLLIQTVGLLQTYFSYPV
AVSLTMEFDKKLFFPAVTVCNANPVRMSELRAASDYRLRSQFDPSFSPSVKVEVEEVTDSGKPLITEGGSNTTPVGKLKF
SRTTDVETTLAQPTTTENSSEPIMSWDDRVGGDFYRKPSNGFSKANSLSVLMANQSTQTKMTMGHQLSTMLLDCFY
NYKYGNCFTFNSGRNGEALTINKPGPMYGLSLELYIEQIEYMIGTTEAAGVRVLLHHQDEMPFPEDEGFSVAPGQAASV
GVRQMELLRKSAPYGDCKDMENLSQEEIDNNIFMSRFDVGYSVQVCGRNCYQYEVIEECGCFDSNYPIPVSAEGRYDA
CDIVSDEKAITVMNLLPQTLIETYKLLPNMEWNMPLLC

>Proto_ecdy_PANarthopod_Tcas_tr_D7EHU3_D7EHU3_TRICA_Pickpocket_protein_11_like_Protein_OS_Tribolium_castaneum_OX_7070_GN_TcasGA2_TC002260_PE_3_SV_2

MAGRKVRGKMLIEFIKMFFAQSSIHGFPHMSARKRHPIELIIWIVIVGAAIYGAVVLSSITLRRYTENPTVISMERDRFSW
NTTFPAATICPTYKINEPLLELYVQSSTEKNKTLLREFLVTLAGATYTNFDTVVEYEGIAEENYLDILLDLQFEFKPAVSNSG
GNDKMYTLQKIVSEMGICYSFNSKLAIYNSPDYWDEGNWDLVEGNETFYVNPLDGEVFANVMNMSTGFDIYVHGPY
EVADIASKKLLHLMAISYKCLYLQRKCRFYDESNLKHSPAYSYVLCRMECRAALAKRLCGCVPHFYRKLEGEKICNVPGLH
CLAKHKEKLILLKGECSCLPNCDEVNYIIEDMDTREWFLGSNLQWGLKEYPKMRLRRDIIFGFSDLLVYIGGMAGLFLGCS
VLSFIEIVYFFTLRLYWFVIKYGKETRERF

>Deutero_Ambulac_Apla_gbr40_206_t1

MRVDKLVKILGWEVHGDPNSAAVSIPGEVLDQTLCAEAPREVFHKPTSGEVPKTRQPAWHVCSRMDRVWTKEEKRR
TSHGASLRLKTALKDDTSVVGIKNVVDEKVQTGRKKCAVEKYGPEFVDYIDLAYPVGMMNNKQTPPDFMVFKDLDLK
QFYVDTGHLKDDMILECSWQGQPCGHDDFISTLTDFGMCHTFNSHTKGMAVRRIKHRGSRFGLRLRLNVETDQYMP
GPRDSVGIKVALHHQNDIPKGMSKDLGFALSPGTYNLAEVTMSQITNQPMPFGKCATKQLKHVEMYSVMACILDCQT
DYVVEKCHCKDAFMPESFISSFETNCSCPVPCERTEYSTMLSSGIFPARHVAMALDKELNMSMDYSRNNLLELIVYFEEL
KYTRITQQPAYSLESLLSDIGGSMGLFMGVSILTLFELMDVLLRTVY

>Cteno_Puk_Fal_comp22820_c0_seq4__p1__GENE_comp22820_c0_seq4_comp22820_c0_seq4__p1__ORF__type_5prime_partial__len_436__score_74_28__comp22820_c0_seq4__1091_2398_

EDVKYDDIGHTEYTERGKCLELNDDANLVQRISGPSGALTVTLDADVAHYTEFQETAGFAVVLRSQDETVISQEFGFSVA
PGRKYLVNLRTTDVTRIGAPWGNCSNQADIMDDEGYFTLKECFLNQILYSYPNLKDDCSCFPWFFVTRFLQDGSSRKYD
AYTKMRMIYYWTEQISDVSQRLSFNDTCLYDTSPVEFTSNKMETIENVANVTSCKTECIDSPDCRYFMYSSHDNDYQQ
KGECALFKAGASYTELTESKVGRVNTTGLTPGDTKCINIDSCGISQQIECENKIMNQLMAGTMSVPTCYEPCTYQKFEHT
ISSEKFPTESYYKYSGFEDETGYTFDQASENLVQLVFYSDMIVEQKDVQTASYEIQNFIAELGGLSDLFIGVSFFTVFQFLE
WFIVKIVHRETRAPAGGGPTTQVNPNVAMTTIGTDN

>Deu_Ambulacrar_hemi_Ptyfla_40v0_9_20150316_1g28005_t1_scaffold79692_cov99

MFSESPDLSQLTGVLQLSVDEISELGHQAEDFILQCTYAGEKCDANETTYDMTISQSQWPSHAYLNNLLKLVQVSNKKT
QNMHNLESARANLVRLKIYFETLNHDLTSELPAYTWEDLLSDVGGTLGLYVGFSVITVCEFVTLLFNLCGKSPHLDHTKKE
EQPLHERVSDTNEGTNEFSDKTNVQHDFFKHSDDTGKQHDESVKHVLEDFGRETTAHGIVHITTAKSNVTRSTWIMIV
LVAACVMFVQMTLLLVQYFEYNVNVKVTLVSEKALHFPSVTICNTNKLRHSAIRSSKYSEMLMLERNFVSPYYAPCIEGD
FQCKNGIHCIKPYLICDGVNHCGDMSDEHNCAYPECGPDQFRCDSGSELGVCIPNDKFCDRISHCYGAEDETSCGECSE
DEFQCITSRECVASLKTCDQLFDCRDGSDESASICGKQ

>Proto_ecdy_PANarthopod_Tcas_tr_D6WY25_D6WY25_TRICA_Uncharacterized_protein_OS_Tribolium
_castaneum_OX_7070_GN_TcasGA2_TC005481_PE_3_SV_1

MAFINVKDCLTDPRQLIKLILLVVCVSITIYQLSECVNKIISPPITSHYRFRYNQSIKYPAITVCRRPSFKTFLFPQYGIRASTLD
LAANNLFEHFPFNTTTLEHFLNATTYDFKEVVPLYGYNETGMGENLWFRKIFFTKRGLCHTMIPKEMSQNFAIGGGFW
LYLKHTAKDKTKDTNGLSTTGFEIHIHDEFEVIKEEDGSNTGFVYVETGEDLHVTLTVQSYSQVSTKQQPCVKDPTYGKSR
CEQKCVHNFAARKVGCRVPWLRDLDQDYPLCSDYVSINEITNAMTEEFSSRALSECNCQGRCNNTLYKYQIVHRRDVD
VIQGSSSTIAIYFDSNLVTELIDVISYDWNIFLADIGGSLGFLLGLSVIGFVNVIEEIVKLIFRMREKTEDFPELELEGKGEGKE
GESLKSVMEFVANCDMYEEKHGEKITKY

>Deu_Ambulacrar_hemi_Ptyfla_40v0_9_20150316_1g4291_t1_scaffold1846_cov106

MMKIYVQQRSCGNPASDIKVSIKSLLLHSLSKVSRFSPNPQGGTRPLDLVKQNLSRSSGKDVAKAFIEYYYGDNDVTESII
AACYDRDIRTELVNHQQKHRLKLEYNSLMKEPGVRRNIMKICLCGYGGKGKTTLKYALQRAYQNITKVERHYDDFREKV
YKPKPMPSLIPLLLASTSPAFDDVRDALAVTKEEMRVLGHQQDDAILKCSYNGMECGSGAFTMSQHRRYGNCFTFNG
DASAALKAAKPGPAHGLKLLLSAQQNEHVGIYGGETGFKVVIHAPDVMPFPELNGFYASPGTFTSVAVEKISAIRLPEPY
GSACSTEYRPVEVMTFMMDLGAVLATWMSLALLLVVIVYICCCRALSGKSGKASRYQVHTPRIFCCGSTRDWRRYLLH
RPLSGFFGELNVHFVPCPCRTDDTGVNRPNSREDVIEYCERW

>Deu_Ambulacrar_hemi_Ptyfla_40v0_9_20150316_1g5424_t1_scaffold2509_cov138

MACDDVDKEFFTGTTIHGVGRIVERSRLVFKILWFFIVLVAFGFCAWQITDRFVRFFEFNVNTEITLKFESSIEFPAITICNF
NRYYSSAITASDQAIVNQLLYAMDYDYDYSDSEYSSFDWDNWYSNLSLAADFDYANFTRRVGMQLDDTTLLSCEWRG
RSSSCSAREEYTESASGNVQAGVKFLVHDKNTPPLMDSQGSAVAPGTYAFVAVQQVRTVSLSKPYGSCVSSSTLQYYDE
YSLSACLIEWRLELIVQECGCKPFRYPGPARDCTVTESATCAKTTLGRIKNGDFGQSSCYVPCNMTSYPTSVSYAQYPSQS
VAPEMASFYNVSESYGRNNLIYLDIYYDEINYEELKQTEAMTPSALVSDIGGQLGFFLGASIITLCEFIEYLIMKCHGACRTK
SKKVWDSNDEKEKKVDVAANGDTELNFGPRFAFS

>Deu_Ambulacrar_hemi_Ptyfla_40v0_9_20150316_1g18731_t1_scaffold17031_cov102

MDDEQFYETMDNSHVLKLLLRQHLAKKNESERVGMGYDIRRMLVDCIWKGYPCSPRNFTRIPNAIYGNCYMFNTVKN
DVALKTNRPGMAYGLTLELFVDQDDYIPDITETAGIRVAIDPQDMVPFPEDNGISVSPGFSTEIGLRLIQIERQPDPYGDC
IDINNIPNKYYKNNIYMTKFKVGYSVQSCEKSCYQDTLIEHCSCYDTDYPPTDEAMEQGIKPCNILGNYTVKCANNHDDR

YRVSHEQTHKIRILLRYQTVKYVQRSTKDVQDFPDLGFSINHGKNFTRISNAAYGNCYMFNAVNNDLALKTNRPGMTH
GLTLELYINQNNYVPDITETAGIRVAVDPQQIVPFPEDNGISVSPGYSTEIGLRLVKIERQPDPYGDCIDVNNIPNKYYKNN
IYMNRFNVGYSVAKDIATSMSMLYSMRINLGATALYRASE

>Deu_Ambulacrar_hemi_Ptyfla_40v0_9_20150316_1g15842_t1_scaffold11883_cov108

MQENSCKTLDAADIHGVKHITATDIGLCISSIIYYVERHHVTKTDVNYTSELEFPAVTLCNFNKYRESAFTERDIGEVGIHL
GIYETGNEHTNTNGFIDNNSSLIESNYLRLKCEYSGDECFYSNFSHVNTHHGNCYTFNKWEEGRGPLISKRAGAGNGLKL
ILDIELEEYTPSNDLDGDSMDAGLKILIHPTNEPPYVKELGFALTPGYHSFISMKKELITGLPVPYGECLNDGNIQYHDHYS
LSACRIECETLAVQTECGCRLVEQPGDLPVCTASQTHHCAHQTLVDFVDGHLDVDCSCADPCTVEDYPHVITMAKLRP
AFIEKMFHDTGHNMTEEYIQENVIVLSIFYEGLNLQTITQLPDTDVIVLVSQIGGNMGLCLGASILTILELLEFALDYIASKC
KLVVTRYQKRKIIHVKPAVEVNEEKKEIRPTVVN

>Deutero_Ambulac_Sakowv30044075m

MPTTLRASPIKDIVNIHSLQERYQELTQDTRLFDEFRVNYYQGNHFYQSDLANNDTPDWKRYLTYSSSTDYSDLRDVLTL
NMSDVFKYGHQFEDFVLQCAFNENKCDQSDFSTFQNDRYGNCFTFNNVRNDALTRRADKMGSQYGLKLTLYIEQDEY
IPLYGQEAGVRILVHPSDITPFPEDDGITVAPGRKASIALREELHSSVAHPYSNCENNPQIDSIFGSQYNYSVQACRKTYLH
QKIQQKCNCRDTLWLSKDRPACKITNTAQEVCRRLINFLDRRGVFENNCKEPCRKTTFKKTVSQSMWPSNKHLGGLLK
VLRPINTKIRDKVVDEKSARDNLVLIEVFFEVPNYRSILNTPAYTVDQLFADIGGLMGLYVGISLITVVEFIELVYNLYNYFSL
NREEKLAKRAKKMESKNMNHKYDPIPCNLPQSMIGTSV

>Deutero_Ambulac_Spurpu_024200

MAQQAKEAWYDAPSKPVTAFKSLDKESATPIEQPSFNMSKEEENSVLFLVGNRMASTGAHGIPNIQRASNIWRRLIWL
LLFTGGVTVVVWQLTTIINNFFAYEVDVNIDFVRQRAIKFPAITVCNLNPVPLSKADVLWNIASNEEDATSLRDFLPAQK
PSGVMNATQNQNDAGASMRGYQDWVNVDFNQLNTRVSNLEVAQSIIGSTAYENRSETGHDLDDMLLDCNFENYPC
APDNFTHFYNYMYGNCYTFNSGQTGKPLEVSTVGPLYGLSLELYIQQSEYISDLQPSAGLRLLVHDQNEMPFPEDRGINL
APGAHTSVGLSLVYLERKESPYTNCSLEFKPGNVFKEQFDHLEYSRSIYSLISDLGGQIGLWIGVSILTIFEFIELIYDLIKLICLK
LMDPRMKKKRSSQSQTKQSNVMALDGQQGRTNSLFADR

>Cteno_Ple_Bac_comp48680_c4_seq16_p1__comp48680_c4__comp48680_c4_seq16_p1__ORF__type
_3prime_partial__len_441__score_80_94__comp48680_c4_seq16_2_1321_

MFTVFLVYKLISDFFRFASFDQTSTVWSTAVPFPALTICNTNKINYTALRERFRASEPGLDAKFEHLLSDMGESAARGEYV
PESIDTSLVIDVLGWAGLEGVSLTEIAEARVVTVGRYDYVFGGLGYNVTNTSHAPTEMGTCIEVNDDGGMVQVIPGES
GGFSITLDARTHDYMFTTDTRGFLVFIRDQGEVVLTNEGGYLAPTGMETRLRLTEERYKRLGKPHGTCQNVPGWFQGR
AFETVRECLQRQRLELHFQHCQCIPEFLAKRWYEAGDFDNLEKAKKMLKTYDFDEMAMGRKRANGGGGYETYSDATC
NLIYSQLCHYNIESLLRNGTVGYTGCHEPCDYRCWHADQNLSPFPPTKQYFAAHLAEHGEHRTLNYTYAKNNFARIRVF
YPELKRKTTTQVKVYSIQNMVAELGGTVDLFVGFSFFTVFHPAVRPL

>Protos_Platyhelmin_Macrostomum_lignano_A0A267GWZ5_A0A267GWZ5_9PLAT_Uncharacterized_pr
otein__Fragment__OS_Macrostomum_lignano_OX_282301_GN_BOX15_Mlig024322g1_PE_3_SV_1

HLKKDPQPIGRAREPLAAAFIWPAALLRCPAQPRGFPAAKPPAAVAMTSEAARRERSFCSFLLATFIDNTSMRGFRKCC
GSRYAALRLHWLFYIASMTVILAYGSFNLFAEFFTYNKSYQIEEVLNDPTPFPSLTVCSLSPFSPEAASLWRSGEVPSPARF
SRDLRHGMGRMVRLGRYADAASLFRNDKMATYFQFISESQATQLGHQLAELVPFCVMRHSNNANPSVAHEEKCNFK
VTGARLHHPEYLVCYDLELDDAESIYVSSLSLILSNAHLNEYLLHNISSTAGGGVRVSRDPADKEHVFDIFNQGAGFRIVLH

EPGTHPLADTDGFSVQMRTSTLIKYKAVKVQKLDYPAGECADSARLPDILDDDSRYKYTFAACLSSQIGQLIRRQCNCTSAFHVRRQRRGRPRPTAATTAPRTRLTSSAGWTASGTSTSAI

>Deutero_Cephalo_070750F_t1__name__chr_scaffold16_start_2515230_end_2516754_strand_pro_len_444_

MAGRKTPAAFWVCFFILLIPVLGICIYQIADRISFYVSYPVTRESREIAVQSQPFPSVTVCNNNPIRKAALSGTTHASLLDTDEQVRQGVENLLASTSSGAVNPSAAREILNTAYQDRRTARATGDLVGMSRDDIANLGHQLSDFVMSCSYNGRQCVSSDLSSVFTTFQSAKFGNCFTFNANFKVSSPNKGQGLTMTLYTDVAEYVGLFGRQPGVTVTVHPANTTAFPEDDGIVVRAGESAAIGLRKTVNNRMLAPFGDCQQDTDAESLYPGGYSLAACRKECLQKTLLQNCGCLDDPRTTSGNLCTASEESCRSRYYEQGATWRDNCDCKRPCREESYVTAMTSSPWAPENVLIPELDYVTPLPDVGGIIKQNIVQVQVYYLDNTSQQVDERAAYPVNKLMRDIGLDIAVWLALLLILIGIGYLVVLIIRACCGEGGGNKAI

>Deutero_Urochord_tr_F6ULW5_F6ULW5_CIOIN_Uncharacterized_protein_OS_CioInt

VMSSLSDLIYEFAQRTTCHGLSQILQTKRKTIYRCLWFMMVSGLCGGATWQLTQAVQEYLSYKTSTDIQEMYTSELPFPAVTICNINSFSHLVKKFLNVLSDKLFSVWPSPEFEMIAFNNFDNWWRREVELYYEKLNERGIDKFYLSDEILLKGWTLNETTLSHCTFRQKPCSIQNFTQVVTPMGLCYTFTSNESMYQNVPGSGHGLVVHLNIDQKRYSEHPKYGNPNAGVKVHVHNHNDPSEVQNYGIGVPSGFGAHIVIRKTERILMYKPWGVCSEKTKTLLHHDHYSTAGCLRECRAEYVYSKCGCKPYWALNMANVPECDALNILNCSGSADAVFSTKFSLRTCECVEPCNFVVYDSSTSYSTFPSIEVGEFFALHAGKNLRKYRENHLMLDIFYDSLSYVRMKQSKAVTEISLMSDMGGLLGLWLGISLVTVSEFVQLFF

>Deutero_Ambulac_Sakowv30035231m

MDASNSSRNIMETVFQTTNDVHPSSDEDMSHQRNCAPRCRDRLREFTQETTLHGIRFTTDSKSNIYRRLMWVLFVSLGTAGVVITLKDSAIRFIAQPVTTVVEIHHPNAVPFPAVTVCNYNKYRQSVLAGTWAEDLLYKMYGSPVQGEIGEVDWSNYTDVITNLNRTAFEMEAAHQIEDMLIACTWSSETACGPENFTNVMTNYGVCYTFNGDVKNGLTVKNRGSMFGLYMVLDIEQHEYMPGVSKGAGVKIMTHTQTDVPLVNEVSMSFGPGMDSTIALRLHQTKLLRCTNQPLQYFDVYSRANCEMESFTLAATKSCTCREPYMPEESELSCESACQEVIYDSTISYGSYPSLQILKKIQLMRNITEAEIRENYLAISVFIEDLTVLSTEEEYSYTYDAFLGDIGGQFGLCLGASLLTLLEFFDFLMMSLCGACRRWYNRSRKQ

>Protostome_Lophotroco_molsk_AplCal_XP_035827487_1_amiloride_sensitive_sodium_channel_subunit_gamma_2_like__Aplysia_californica_

MSPMTPQEDSEEDKKTKRFGDIVAKFISSTTLHGIPHISDTSKFYNVFFWKNVIWSLCVAAGCAMAAGFFPQLAPEYYRLRYRRMSPDERWARSHNLDEMMVGCKFGNKDIAHFKRLVHHRKGRRKRNPTTKTGEEYGLQLLLDVQQDLYMPYTEATGVRIILHSPEVAVFREDTGFKVPTGFECTAPIKRLDMLPSQAEYPISMDENQKVTRPTRPHGGCEVISDSDNKQMNALAQDYGYSALACSKTCLQRNILTKCHCCTPEVPCPSSLTTLVFGTNQSEETMPPICGRESNVTMNFCSIQVKGNFFSDQLGCTEMCPELCEEMFYGTNPSFAVWPGKAYHSTIVEKLKITHGAALTKQELNSIVEDSLLKVDIFYPSFEYQQFTISPKYDWSVLLSNIGGSVSLWLGFSLLTALELAELILDSVLFVFSKVFMPRKSQINPR

>Deu_Ambulacrar_hemi_Ptyfla_40v0_9_20150316_1g21584_t1_scaffold23944_cov133

MADIDRSRNESFGVLLNNLLQNSSAHGLPNISRASSIPQRLMWFILFWGAVSAFVIISVDLVERLLANGVKVDYDMKFSRNLPFPAVTICNLNPLRKSAIEKSGNDDLREAFDKNWTPPPRPTRQYYTLHARTNPPREPSTTVRIRPTASDIGPIRDVSGDQGGERTNISGSRPLDSAQSVANEGTQTFDILNTDEENSEGESSWESGEYDDWDMDEAREELYTQPSQFWNMTTIFTQIYSSLNASEKQDMGHQLKDFLINCQWNNVRCSTQQVTYTTEFSTAYWPARNSQRYVNDWLTQRNRELHRMVMKQEDLNEDLVRLNILKLRVYYRDLNYETITQSPSYTRVLKYQKDGLATLDSASYVGQNTMQSIRNGIHEKNQDLKRLCIERIPRLTLDNIDTSMDLFLKLKDQGMISKDNKAFLTDLLNAVDLHELVQKYMPAYNPAS

>Proto_ecdy_PANarthopod__XP_023225242_1_amiloride_sensitive_sodium_channel_subunit_gamma_like_isoform_X1__Centruroides_sculpturatus_

MGFCLFHSIAYLREYFQYPTVLNVQVNTESKLDFPAVTVCNYNRIRSNALDSFCCTSGLLLKASSLCHVLNVTCSQTNNEEASSKRLKTLIGSPPLSASTAFATCPRNRTNNTVSYEGSIMKSFSNAYVSLPLSKKIQMGHQVQKFVHQCQFMGQPCDHLNFTTFHTFLYGNCFTFNSGQNGKDILRATTDGSMSGLHLELNLETDEYIFDLTNNIGARLIIHPSNVKPLPENEGIAITSELETSISIEQVNIHRSKPPYPDNCIDYPNDGEGFLYTRNTCLRDCFQQLSQNKCSCADPTWPLPRNSTSCDLRDVTQVCCLDDVRQLLRNDENICICPLPCCEIQYKLSVSSTKWNPLKSKINKEMYSSILPSDIAKVRIYFQTLDHVVLKSHPKYQVQDIFSNLGGQMGMWLGISLMTLLHYFETILGSVFCKQKQTIHQSPIGINP

>Cnidar_NemVecNVEC200_006432_1_1_protein_AED_0_27_eAED_0_27_QI_414_1_1_1_1_1_6_1486_449

MIMAAAKKLLDGRKKRAKVKELWENFLGGCTLHGFHYCFAGNPPLRRLIWSLLLLGAFAMFFEKCTESFINFFDYPFTTTTLLVYDKRLPFPAISMCNYNDARMSKMNGTLMNEIFVASKLEGRNTSHLQSQLTGELMQRTLKEAAHRLPDMIKECSWQKHGKCSWKNFTSFKSADGDTCYTFNSGRKDPILSMSNVGEENGLRLVIDTQHSEYYYDVKNAGFKVILHDQGETPVKMQGLSVSPGFTSYMELKRTKVTNLPFPYKTMCGMPELKYFNSYSKSKCFLDKLTQYVVTLCGCRDWFMPGQGKIPVCDYETAASCMWKAWAYFEENKLDQCPVACNSVEYSAQLSYARFPANNYAKMLAKEYGLKGSDEENRQYLRDNLIEIKIYYEDLTYFDVQQVPSYDLYSLLGDVGGQIGLFLGASLLTVVEYLDLLGMVAYTSFKYRN

>Proto_ecdy_PANarthopod__XP_023226822_1_amiloride_sensitive_sodium_channel_subunit_gamma_like__Centruroides_sculpturatus_

MLRQVNKLKLFWILCCLGGFLYQAWSFLEIYLAFETVVDVEIFHPESIEMPAFTFCSWNGIRKSKYCEKFPDRCNGNDTIISVCSTYPSMCKYGASQNLHLSKRIAEYEKMEHEASREIMKEFGHQKSIINDCDYSWKSKTSCKKLRYQYGVSFEGPRNCYTSNSRIWKNLTEPEMIPVLQGSNVKNKYAKMNYFISFDPEEYMVPERQIATQFDIHPADELVDPFRTGISLFPGYNYQILMSVSEIHLLPYPYNTDCFNYTKDYLEKKGTTPWRRSMCMEKCYVDVYIKKCNCVPTDMKYVHDTQYCNFSSYGCTEEFDVTECLKKCKKDCRIIKYDYEVKKVPLLPIVKYFLTDPALKGLHKFTYVTVTLKQEDVIVFNHKPKYEKIELFSYVGGFLGMWLGFSLVTISEVVKHFFVARKYFKRNNSMKLMKHRTRISVIYPKDYKF

>Proto_ecdy_PANarthopod__XP_023234703_1_uncharacterized_protein_LOC111634211__Centruroides_sculpturatus_

MADDSGSKRRDKSSCKEKRIFFMKMLEDVLLTGVPQIVTESDGRRKIFKIIVFFACLSGFLYHLISYIILYLAYPTTLDVLVEFKETEQLPKPVLTICDLNGLRRIPLCKDFPEECEKPKNMKALCERFEYFCKDSDVDNLMFPTHNLSVVSDYLRSIVDIYGPTVDDILKNYSNLTTPVFYTNTYRKLRKCYSLDLAKFDNSYYTVVYDKFNAFISEGELVGTILVTFNPKEISNPSTPVGAKVSVHNRDQLVNPFLDGFQIKPGRTYNVKLKAVQENLLDSPYNTNCTDYEKQKDKGFNGIYSRDFCIIECRKELWLENCNCVLGEYNFPFTGNFCDEEEEFQCISDVDVDKCYKQCRRGCKIWYSTISVFYKRPEILVYKYKPQYELIEFFSYIGGYIGIWLGISLIAVFDFLEMLFVYFCCIIRGKIPTNKETNSNILLESKPSA

>Protostome_Lophotroco_molsk_AplCal_XP_012942490_1_degenerin_deg_1_like__Aplysia_californica_

MGLFQEFSESTSMHGLQRAVSSESKWKRPLWAALVLAGACLATFNLSQTLKDFLHNPTTTQTSVHYNPDMEFPVITICNLNPFRSSLINAVTDSDFIENLWRNPGNFTYEDVVDLTWQYYKMKNVSVTHQLEATVMTCKFGTGPRGYGKFLPSFNLKYTTKPGPSNGLILELNVQQSEYLSATESAGFKIAVRPRDTLAFPEDQGLIVMPGTNTNIGLKKVEMSRTPPPYSACVELTDEPNVELNMFSFTEGTRYTAEACRKTCYMRELVRQCGCCDLHFICPPEIYNNISFCNATNTDEFVCKKQVLKMFEKGEL

DCLSRCLPPCEEVVYDVALSTAIWPSNNAIPYKMTALGFDQANLNTAMSEMKRNFLRVAIFYEDLAVRRVESRPSYDW
NRLLSDIGGQLGLWLGFSILTAMEWLELAYDAGIRATTRMSLGAQTTRAHGQEQGP

>Cteno_Puk_Fal_comp18295_c0_seq1__p1__GENE_comp18295_c0_seq1_comp18295_c0_seq1__p1__ORF__type_internal__len_452__score_82_45__comp18295_c0_seq1__1_1353_

DPFLFGFQDLASDYVQVLQEKHRPRWCKILWWIITAGLAAMTIFMVYELLSDYFAYDSFNTASTAWDSKLVLPAITICST
NFINFTELQPELSDFGDTGEFQYVVTRIEKMVAQGRDPMKLKDTSAYINVTQFEDWAGSIDYNYANHLYGTFIGEYDYS
FAGQGKMISSGEWSKLEGMTQLGRCLEINDDMKFYQTFYSDSAGFTIDATANLDDYMFSTRTQGFKVFLRSHDETVM
IDQGGYLVAPGTETFIKISAKKAQRLGSHWGTCKNVPSAYSKYGNETHPHYETVRECLQRQRIELYARKCSCLPWYFVSR
MQTLEKLSVLDELIEDIKANAGNSTDGSTRNARNSSHLKVQRKQHYRKIRRNQDTVRRQRREAEDRMRRDAEDATSP
WWEYSSTGPTTPTSAPPFWSDKSKYNSYTCSYTDLIICEAHIEEEIAKGSISLEGCPE

>Proto_ecdy_PANarthopod_Tcas_tr_A0A139WCP8_A0A139WCP8_TRICA_Pickpocket_protein_28_like_Protein_OS_Tribolium_castaneum_OX_7070_GN_TcasGA2_TC034126_PE_3_SV_1

MKMEENTKFNQKLRNHLVQYSSSTSIHGFKYLTEKRSKIEKTIWSLMLLILLSGCMFMIYKVYHKYESCPVVVSFSTKDTP
LYQIPFPAVTICPESKYSRKEFNYSKVYFEIMQDKEINETELKNFYYLTLLCDPDLDISKRANKTVDDQFFQVLDQVKLNVE
SILHFCNFMGKFYSCDKLFTPILIDQGVCYSFNNLGRDELFKKHVFHYSDYYHVPNNSHNAYDTEKGYETNSGVDTHPKR
ALMSGADNSLTVFFNYDTADTDFICNSFLQGFRVLIHNPWDVPRLTKHYFRIPVSKVVVAAIEPEIVKTSIEVRKFPPEKRK
CYMNNERPLKHFKNYTQPNCYLECLTNYTLHECGCVQFFMPRDNLTAICGSGSGSCLIRAEEKLKLKDLEFKILGQTFCD
CKPSCTNLRFNMETSHSDFYYKEYFEAHLNMKIDDDGFGDISSPTFG

>Protos_Platyhelmin_Macrostomum_lignano_A0A267FSS2_A0A267FSS2_9PLAT_Uncharacterized_protein_OS_Macrostomum_lignano_OX_282301_GN_BOX15_Mlig024294g1_PE_3_SV_1

MRTRLRTVTGAAQEVAGITRTHGLLHIFLSRSTARRLFWLLLFVAALIGCSVHLAKLVQKFTERSVESQLKLRSERAQFPDI
TICNFKPASASLLTLGDFFEFHLKHKKFKFKVLQYFENFWGNFVNWFKEDTQRRPQESDESYDSRLEKVCKKVLDVMW
VSGDTRDISQLDETMLLSCRYNSEPCSHRNFSLVQTSRFWNCYTFHPDPRQSSSGPGGRGVELDMVLFTDGNEKYPD
WHDTQPDGCMSRKFSRSKAVNRLLQSNEPQSAGLRIFIHEEKSFPMSETEFVDVASATSTSIKLRPIHNRLMSKPSRRCS
EPPIETINYVRHFSNASLNVVTKAYSKSISDFVVEAQQSILHSNCGCYSHLLPFSVNTSELCYFAPPQEWITPSDAVLKRIGC
HDIWFQHAQSRGQELAKQFQHRMWCHFTQQRPWQQSRRWPPGSAPFKRSGRT

>Deutero_Ambulac_Spurpu_015893

MGSDDLQNETVIGSVNTFGNETTVHGVRYMIARGHFLFRLCWLAIIVLAFVLFVIQAFNVYGDYASSPYSTKIDIVQQTF
KAFPAVTVCNANRIRRSMLYDTNYEGLIDIDDNNRSESEVEDGEAERSEEIWRKVNGLYDWWGFYSASVADDFSDLIN
VVNPSQEQLMGYGHQLEKFLIQCTYDQQPCNMSSDFRVWQSRYYGNCFTFNFGMMETAEAITTSKTGALNGLHLTL
WVEEDEYMGLLSPSRGAKVTIHPQNTLPFPEDEGINVGTGMATSIGIREEVIERISGMTNCVEEDGAGTNFTVTRTNTIY
TVAYCLKLCFQQSLINICGCVDGILLDYTHCDVLNSTQRACVSMVHQLYINNGLYCNCPLPCERNLIRLQIYYETLNNEVV
EQVPRYTIPNILGNWGGLMGLFTGMSFISVFEVIFLLLRVSKISCTKLFFPNRVQPVKT

>Protostome_Lophotroco_brch_Lingula_anatina_comp141843_c0_seq1_p1_comp141843_c0__comp141843_c0_seq1_p1__ORF_type_complete_len_456___score_27_90_comp141843_c0_seq1_2150_3517_

MNRNRVTPTDIPKGKIGDEATKDSEIDQTSSTQEKLATFAGEVSILGLKQSTNPKYSPIRRLIWLCFVLIGFGFLIFHLKNRF
DYYLSNPATLNMEVVPNDTLVFPAITICNSSPLKASNLAANNESELGNIFSPFVNVDWSQYNLSHINWTDFHMRNGFQ
LEELLFWCTWKKTNCTKHDFTPIITEIGLCYQFNAGGEEQLHLETQGSGYGFGMTLLAKQRDYVSYAYPAGFLIIVHSPG
DHPDINLGTLISSGGFTGVSLKLKRLINLPPPHGKCGEKPLVYYNKYSKANCHAECFHNFTLTKCDCRLTHFPRVPGYREC

SPEDSRVCVDNLKLIITESVYLRECGCKTPCDSFKYDVSLSSARLSETAMRTTAAILRTTQEEISQDFLSVFIYYSEMSYEEVV
QQKAYSALALLADIGGALGLVLGSTLMTLAEVVDFFFGLCFWKLFGVEM

>Protostome_Lophotroco_molsk_AplCal_XP_012942493_1_degenerin_del_1_like__Aplysia_californica_

MGLFQEFSESTSMHGLQRAVSSESKWKRPLWAALVLTGACLATFNLSQTLKDFLHNPTTTQTKVYYNPKMEFPVVTLC
NLNPFPVTNLVAEMDFMTMSNFWEANNFKLEDVMESIVQFQKSQNVPITFKLGDLVTDCQFDMVPCRADDFLTVFN
YKYGDCFVFNKQIKEDSNQEYMHSSEAAGFKVVVHSQDTIAFPENQGFMVMPGASTSVGLKRTEMSRAPPPYSTCIEL
EDEYNIDLNVFSFTDGTRYTGEACRKTCYIREVVSQCGCCDLYFVCPPEVYNVYVVNETIPYCNATNIDDFFCKKRVLEAF
EKGELDCLSRCLPPCDEVVYDVALSTALWPSDSSIYFDMESNDGSDIAISEFEEFKSHARDNFLRVAIFYEDLTVQRVESRP
SYDWNRLLSDIGGQLGLWLGFSILTAAEWLELAYDAGIRAATRKAQGAHTTKVHVQEEGK

>Deutero_Cephalo_091770R_t1__name__chr_scaffold19_start_3668070_end_3674027_strand_pro_le
n_457_

MTSRCCRKARAALCGGPLDYEFATNTTCHGAAKIANAHNRVHRLLWVLVFLGSFSVCVWWIADRVGNYYGFRTNTEI
SRRFEDKLIFPAVTVCNFNRYRASKVSNDTLEHLWELLNLHTAVVNKDNATYNASLWNTPPINATEFADLTKQAGFRLD
EDTLIMCKWRDEDCHQEDFEHHFTPYGNCWTFNWNTSSARAQPTAGEGNGLRMLINIHADEYTEPANLDSMDVGL
KFLVHDQHDPPKMDSEGLAVAAGTHVYTSIVQQKYINERKPWGQCVPEKTLQYFDRYSLTGCLLECRARRLMNSCNCL
PASLPGPADMPHCSPIQLGTCAYEELAKFDQGEEEVGLCNCSVPCEETKYQVSLSYSTWPNKDAPRLFNRSLQDLRENY
VSMDIFYQDLNYQTFQQFQTVTVPGMISDIGGQLGFFLGASIITLSEFLEYLILKLIVHRFYESEVDF

>Proto_ecdy_PANarthopod_Dmel_tr_Q9VAJ5_Q9VAJ5_DROME_Pickpocket_30__isoform_A_OS_Droso
phila_melanogaster__Fruit_fly__OX_7227_GN_ppk30_PE_3_SV_2

MSATAWKPEPSSEDNDDDHKESKKKLKKEGAGGWASTRRYARIPLKIMAALATVYVSILSSERYFYYWVQTSIERTDM
HVSEIDFPAVTIIPINLSILKAEKLSKAYNLVQSVVWQTPMSARLTDENFTEFSELENWNAQSWGIYQALQMNCQHFFT
ECQWRRKAMNCCDLFRPTKTFNGFAFEFNSLVSSGRDETWPWSVASCGSYSGLNVKIKRQQGLYTLNTMGVIVHEPT
QLLGMSIDYSSEDRIVVPVEPLHFTAELDVRARPVQMRRCYFENEIPTGKSRSECIYKCHVNYIISKCNCSLELPVKATQDE
DNIAAAKESNGRRICGVKDLACFNQHRLSLFSMSNIIEESRDNVFSTVDCGCFPQCGHTQYHTSTYTEKMSAHTNLAAA
IEIDVYFQEETLFSYRSMLRFTLIDLMVSYGGIAGLIMGISVLGCINELLDRFACCRIPHGS

>Cnidar_Polpod_Hydrif_GBGH01017297_1__p1__GENE_GBGH01017297_1_GBGH01017297_1__p1__O
RF__type_complete__len_459__score_52_80__GBGH01017297_1__223_1599_

MQVSRGAKPNTLVEVSSKKQTAPTTLVRKHLSNFFGQSTIHGLHYCFDGNLPMRRVVWCCLVLGALALALQKISESATE
YLSYPISTVNTIRYEETMVFPAVTLCNMNDWRLSVTVNTSLFYTVTSFYNTGEKKEVSAEELLATQRRANHRLENMLLSC
KWKGQSCGLQNFSEYFTVQGERCYTFNSGRDNHSTLYAETASIIDSLELILNVEYYDYFIDNFVSGIRIILHSQEELPVTSQ
GYVLTAGDKTYIKVKKTKARNLPYPYKTNCSSIPLKYFNKYTADMCWLEQLTDHVTSKCGCRDVCMPGNASVCPLSVM
DACGWAEWDMFNIGKLAKCPLPCEVDKYSALMSHAHMAPSIVSTTLTESLRIERPTMAHETFVGENVIRAVIYYEDLSY
VLIEQEPTYTLMKLLGDIGGNLGLLVGASVLSYMEIIDCIILIVITLIKNSRSSSKKQKS

>Cnidar_Rhopilema_esculentum_TR36173_c0_g1_i2_p1__Rhopilema_esculentum_TR36173_c0_g1__R
hopilema_esculentum_TR36173_c0_g1_i2_p1__ORF__type_3prime_partial__len_459__score_45_96__
Rhopilema_esculentum_TR36173_c0_g1_i2_2_1375_

MEHNTKKYEVNSNVKSQLTEMALRSYLANSTIHGLSKIEASKHWSLKLLWSSVIIALFAASVANIYSLAGEYFKYRTYLLIEE
CANHTAKFPSVTICNANNFQRSKVGNLAVDPEARFQHFLNISLDKGAVVNLLNFPNAKSFMDQGSFSTAKATQSASD
MLFLESIEGWCRFGHFKTCNQSDFKDYFYGLESGFCKTFNPDGKFTQVVVGPHGGLLLTLYLNQDDYMRLLPFDNGAG

VFVKVHPQDVFPYPSNDWLAVEPGTQTHISIEQKIMLREQKPYKSECVNRDEDLIYPGRYHVNNCLQSCYNLKLYESCGI
VEASVLYHLEQKGKKFPSLPRNISKNPAACGIEVSTNLTANKVFCDCPLPCYEETFKLTATSSRWPTEADMQYYNPVLSK
VLNRTNLSKDFINSNFLQLKVYFKSFSYQKVMEVPAMNFFSLVAGIGGSLGLFIGASVFS

>Cteno_Ple_Bac_comp47578_c1_seq5_p1__comp47578_c1__comp47578_c1_seq5_p1__ORF__type_5prime_partial__len_459__score_71_82__comp47578_c1_seq5_141_1517_

TDVKDLAILMNYRNFKEVQLVVPATLSVPAPCMHQNGRPKDYIFTEIEQPLSANGFFKRVPVPNRTTGPGFHGRGGPP
PREPAFPIYDNLLDRDDVLSSNCPSTVPGVYPGVEAFTPYSKYKQDESAFDMTGRGERTSYYPYLEHYQPSYPRPPRGGK
PPPPTGAPVPNNHRTSDQNGKIKYPPNHRDTGSPTELEDRDGEDCEEFMPPPKRKKKKKRRANRAALEQRREEMEPD
EMESFKEFASTVTADYLAEMFAKASNGRRAIWVSLWLASFVYAWYNIALSVGVYLSKPTATKLNFYEAPPGGVAFPTVT
ICNFNKYNKSFFEREGDSVNSQLRDFLVATTPIWDHKERLDQDQWTSKYPNIMSLNLKELYANGSHLITNTVEFCRFGG
KPCKDTANRETSIPSEAFTKVYTEHGLCYAFNLDGKLNMTRTGAQYGLSLRLHVAQDDYFRTQVT

>Protostome_Lophotroco_brch_Lingula_anatina_comp130467_c0_seq1_p1_comp130467_c0__comp130467_c0_seq1_p1__ORF_type_complete_len_459___score_26_84_comp130467_c0_seq1_263_1639_

MQKEVELHGTPKETTETRNASSVWMTFFEETTLHGARFLAKRSLIRRIVWAIFLLMATVTCLVMITYSVIDYYHYPVQTK
VGIEYRRSLRFPAVTICNVNRYKKSKIIKNQTLMSLLHYISPLPASLKNDINRSDPLLPIVLNNTKTDSLISMAGYSVDDMF
WSCFWKNHAINCSEAFTTTFTSMGRCFTFNSNGSLTAERTGSSSGLWLRMKLQHEEYTPGFALSAGVKALLHEPYEVPL
VHEQGFAVSAGYEALVAIRMTQIIGQALPYRGIDCIDTKASTFRNPLKFYPTYSLSGCIYECQARYVNDVCGCTLFYHPGG
YRCCNVWEALVCASKAMVSFMQNTMIETSCGCKPPCYQTLYSSQLSHLSYPNEPTLQQLKKFNYTEEYLKKEFLEMRLF
YQQMNYDIIEQSPKYYFAQLIGDIGGLMGIFVGSSFLTICEFGDFLFLLLKRCKFRK

>Protostome_Lophotroco_brch_Lingula_anatina_comp138624_c0_seq6_p1_comp138624_c0__comp138624_c0_seq6_p1__ORF_type_complete_len_459___score_26_12_comp138624_c0_seq6_318_1694_

MISPSRLPRKTDNSSDQETPVIEKFAERVSCQALQEIVSRESGYIRRLIWLVCFLGGLILMAFQVYERVTFYAGHPVRFNT
YTVTPLSVMFPAVTICNYNAISKMRARELDLLEEFERYFPLKVGSENFTGLGDDWFDWDSVLNISKPFVKECTFRGISCS
MKNFTPVYTDMGICYTFNSGYSGHRLTSDSSGRLFGLQLTLDVDQRNYFRGFGGYGAGFKVRVHARDPPMMSNMGF
SVGAGMHTLVGLKMINISNLDTSQWGKCNRDIKLKHHVVYSQSACRLECLYEYMEKKCQCVRFLKFKHHQGPRARLC
DPEQLFNCYTPNYNLFIQEKGQCSHTCPVACESIKYEVSLSQAPVADYHAYNSATTTNETKVNKYRKNFVQLDIFFQEISY
ERREQYQAYNAFTLGCDIGGALGLILGASVLTLFEVGDFIILMILKGKWRRQTQVTSVSVHKK

>Cnidar_Rhopilema_esculentum_TR119850_c1_g1_i1_p1__Rhopilema_esculentum_TR119850_c1_g1__Rhopilema_esculentum_TR119850_c1_g1_i1_p1__ORF__type_complete__len_461__score_57_27__Rhopilema_esculentum_TR119850_c1_g1_i1_1311_2693_

MEDKEKEQVTNDIGSLTIHGLPKIVSSKTNISRIIWLVICVSALVFLVITATRSLIKYYRHEVFIAVGTKPTNHMHLPAVTFC
HTNFYNPMLYGEEAPMFQHFPKNCSFMDDKYFVNDMNKHMFYFACRLFIGIFDAKTSGVGQELPEYFRFPEKFSFLPN
IYPCVTLNRNSTLEQKASGEKYGLQMILYNEEMADETFTYAKKAEPLTERRQGVIVHIHDPKQHVPISDGIPIPEGFHTHV
AIRRRIIKLKPSPYPSKCVRGDEQDKESIYPGKNTQMQCYSSCFFKQTYKQCPGVLLEMRGFMGPPKYPKVANYSDPQF
LECIMKSIVKYDYQKCNCSPICEEEAYEVKVNRNPWPQSWQASSVSHIVSAMKGVESVSPDEVRERLIRLSVYYDEMVE
STFEEKEMYDLSAFTSDLGGQMGLFLGASLLSLIEIFALLFSFLSNKFFGSNKVKTISVK

>Cnidar_Rhopilema_esculentum_TR92117_c0_g1_i1_p1__Rhopilema_esculentum_TR92117_c0_g1__Rhopilema_esculentum_TR92117_c0_g1_i1_p1__ORF__type_5prime_partial__len_461__score_66_22__Rhopilema_esculentum_TR92117_c0_g1_i1_1_1383_

VNFGQSLKRRLFWGALVLVGTALFASQFIETVGNYFKFKVNTVVYLSNDGNTKFPAITICNQNALRKSKLEANKHDAIIQ
SLLHMTRKFLDNSTSNKSTPDGHITGQQLRDLYFKYGHSMEHIDDGGMLEFCRTPDGSLCSAKDFVRTLTFAGLCYTLN
SMRTAQMKTAYSSLSGNLASIRLILSTQVDEYSEAIGKSLSTGIRLLVHDQLDPPLADTHGMTIGPGVAASIGIRKQKIKSK
KYPFESECDDKKLGPGVPYSRLGCLRRCQEKFVVKKCGCNNIVDTILNQTGIKRCDMDQTFMCALPAATETFNNQKIKC
SCPFACESTEYIPTLSYSPYPSDIEGRKLAIKHLTKMKKNTTEEAIQEFIKEERRNKIIDIFYHQLDYLTMMHVPATTGGGLI
AEIGGLLGISLGASFLTIIELLDFLFFYLFQKISKEKKSQTSVSSKETTSSPMAAI

>Deutero_Ambulac_Spurpu_009765

MGDLPRDTKTSPEKMADYSEKAYGRPDYDESPKEGTRKIVNEFVDNATTHGIPRVLNASRPWQSRLFWCVVTLIFAGV
FLFQGSKLVQSYIARPTTTKISLITKSRLEFPAVTICNLNMLRRSMLKGDEYDYSGIEDIEDPNDWEALYNLSKGSDYNNFK
NFVKPTKEELRTLGHKAKDFILQCSFDTEACSYENFTVIQNAEYGNCFVFNNAHKLKRGKRTTTSRTGSQYGLQLTLMVE
QPEYIGILSPNSGVKVAIKDPRIYAFPEDDGIEAAPGFATSIGLTKTSISRLPEPYGNCIRKHDSFYAPEKYDFSQRSCLKLCL
QETLNETCSCITDVLIDGTMCEVTNREQGNCRNSVFKDFLKNKLNCDCSNACRSRSSSNSSSSSSSSSSSSSRSSSSSSSSSIS
SIFSSISSGSRSSRSSSIGSSSRGSSSSSSSSSSSSNIRSSIIVVAVVEVVQ

>Cnidar_NemVecNVEC200_002361_1_1_protein_AED_0_18_eAED_0_18_QI_193_1_1_1_0_83_0_71_7
_1153_461

MESFNQNTKTDIKDCKEDEKREPQPTMLQDFAGYTTLHGFHFLIHPGSPFRRFIWLLMLLGCWVALFYQLYNSVDRLLA
HRIVMSRGVEQPDEIDFPAITICNQNIMRMSKINGTEAQKYLDELDDVFRKDIKKENVSYDAETFVNKYGHDWENMFE
NIPYSCMFQRFFICSAKNFTSFLSFTRGLCYTYNSGVNRSYVQRVSEAGRNNRLEFHLEAHPEEYYGPYSYEGIGFKIAVH
DQSYVPNMDQEGYDITAGFYTNVRVKRYKEKSLPHPYKTNCGERKLEYYERYSRSACILECQAREFVRKTKCRIIGFPPIK
VIRDVPFCSVLKIETGLTYMYYNWNTNQCDCPKPCVEISYSAQMSLLQYPTPSLVREIRKSFNDTEDYINNMRANSVIVSI
FYETLLTDVFEEKNDYDISRFGSDLGGNLGLFLGCSLLTLVEFFDLGIRWCLGRKDKVQRS

>Deu_Ambulacrar_hemi_Ptyfla_40v0_9_20150316_1g1255_t1_scaffold573_cov135

MELTTVEGNGSVTFDRNTTASNRQDQGHADKQDGNRNKQSIKDAAYEFSQNTTAHGIPRAVAAESIPVRIIWLLICGL
CWGVFLWQAATSVKNYFEYKVTYDISVEFRVLKFPAVTVCNINPLRQSKLGELGEEYRDQLGFLDFDYETDGNETSFDW
NEDVADEHIDETWWDSEEEGDYDEYDTEFASEVNLTTIMSLLPDDVRESIGHNKTDLIQDCTFNGVKCSPHNFSLVLNP
KYGNCYSFNVGANDTEIASSSRPGPSKGLQLTLFIEQDEYLYGFTESAGAVVLIHPHKIVPFPEEQGNEIPPGFSTSIGVRL
ACRRTCFQREVMESCNCYDPTMPHDGDNLLACSEETDTTYDFKLFSELSKRSRKINEQLDSLDESRRNLVRLKVYYQELN
YESIREKPALDTVALLSNLGGLIGLYIGASVISVFEFLEFLVTIFFLATKKARKAGQISNMPT

>Deutero_Ambulac_Apla_gbr40_182_t1

MILDSEARPGSDRGAGNYRHGISLTMMTEATAGGTSPSGTASTTEQPSEECENSSLKERIATFGDETTFHGVRFVTNHQ
LHFLRRLLWFCIVGGMTGYLIYGVSVSFITFYQRPVTSVVKINYVRQIPFPTVTFCNYNQWKKSRVPPEFYDAVRSLNPA
NKKPVDMTYLKAHDVTNLTDKLINDTHQIEDMLLECTFKEANCSAENFTQIITDFGVCYSFNDDPSNVHYVHQSGSQN
ALFMKITVEQEEYIFGENNAAGIKVVNLKYPYESNCTDRGNVKYSAEYTVPLCQYEHKIETVKAKCGCKDMRYPANFTSE
DKSFNCPVPCDLTIYDTRMSFAYYPGQHHLEEIMQQNNLSENSIRKNVLDLRIFFEELSFQKIEEIPSYTFYSLQSSIGGYM
GLLIGASLITLFEFLDFILVTTTSCLLKHSVLARPRGNRVRGAARQRDRSSHGEAKPCFIKK

>Proto_ecdy_PANarthopod_Tcas_tr_A0A139WNY4_A0A139WNY4_TRICA_Uncharacterized_protein_OS
_Tribolium_castaneum_OX_7070_GN_TcasGA2_TC004432_PE_3_SV_1

MKTHLFNLIQNFAKQTTIHGLHIIGIPKLHVLERIFWVLVVTMAAGGAIAITYSNWMRYLANPTVVSIQKDFRNWLNPF
PAVTGCFVDKTDKKKADKYIQEKWGINSSDPKYAYYMDFIKIVSNITYKNLDQLDKFATDSKLQNIDLIDLTAKVHPDLSG

TLVTFDTKHKTKWDLIMSELGMCFSVNSLFVDLLTLNPNKTSVVTQRILRCHYLNGLCYARYDSDPDLPIKYYTHSYLEVA
HATIDPPLIVHESEELEINYRMQETESSLALRDLSPLQRKCRFYDEPISDELPVYSSSLCYMECRYKLALKLCGCRPIFYHNF
GGKVCDIKGHLCLAQHSDYITKRPSQIGCDCPQPCDIIIYFPQVPKFTKWEYGYFEQRMTFRWALLAPTTKLHRDVLFGF
EDLVVSFGGTLALFLGISFISAIEFVFLLVENIYALYKNQRKTNVIVVGPRRKIHHF

>Protostome_Lophotroco_annelid_Pdum_comp416029_c0_seq12_

AKASEVTQLVNDVTFGMMAFPSVAYNVSLVANYFKWNVTQYSFFPDAIIKGNSCIITFDSYANLSNKMKIDIQGIANLV
NGSSGNALDEAETQNAFGNILRSMSSFQDFEIAFRANCLNTFDNTIAGYVERRDNIKYEVNRRISGILVSYYYDMEIVNV
HQDKNDLYAKAGGFLGDSSITAFEVASSLGQDTTTSMTSKVMDFISKTTLQTSSSLRPAYFDYRNSIVNAYIEMLLELCVY
RDYVSTAFVESRISQLALLKKPLGNVDNLSITFTENGINSTLFGLMKQICTNISVVPLVRQEISDAIDHSFLKFGTTLMTYDV
NLTKEQGTLTDLIKEMGTSMAFITNGYFAIEDTFMNENFGSFDIFFSELRMEYINQSPAYTASDALSDIGGFMGLLLGAS
VITVCELVDLFVYNCVRKLLYKKKSNGSTTPVTSVTALTEKQNAEKDFLNEALFAPRR

>Proto_ecdy_PANarthopod_Tcas_tr_A0A139WEV4_A0A139WEV4_TRICA_Pickpocket_protein_28_like_
protein_OS_Tribolium_castaneum_OX_7070_GN_TcasGA2_TC034797_PE_3_SV_1

MLKLKNNNLKAANFRNKLLWPFLVYMVFTGFTIFLVVSLWNRFLANPTLTSLESSVYSVGNIPFPGVSLCNLNKISKKRA
QEMAEYIVKQTGQNFTRIMSSFKFLGFLHDSNYDIRFKTEIEEFQSILDSLQLKITALIRNLTTPCDDLLQMCRWKGHKIN
CSNIFRLRLTFEGYCCVFNYVKNTSEWWMRRARDETLVLEYGSPGISNGLSVELNIDIDDYFYTMMSSTGVNVHIFIPSD
YPDKSSGDLIENIADIGTENFVEIIPTTVRAVRDVMNYAVDKRECLFEVEQRTQFGVYSQSDCFVDCRVKSMTTLCECIPF
TVPLSDDYTVCTLRDLPCLARYKSKWSSLVPDKWHEYNIRELEDALLCSKTCYPSCDSTLYRVGASAVQINDSRFSNITAIH
IFYRDRHHSLYKQDVVYYWFEILSNYGGVFGISLGEPGGVSKLTTDSISRLQSLSMKL

>Proto_ecdy_PANarthopod__XP_023228162_1_amiloride_sensitive_sodium_channel_subunit_gamma_
like__Centruroides_sculpturatus_

MTENKNETQFCRSFVKIFVFFLCLGGFLYQSVTFLNIYWTYPTVVDIRIEYKLNMTMPSITVCNNNRLRRSVYCKARPEN
CEEKSSMEFCSDYPHLCGNGSSIYTAKREVFLESNKCNHSELQLYGHSITPLVTRFEILDSENYDFVEKKFNYIRISLSEGITC
PRNCYALNSVVENSNFDFVHISLQTEERKIMKLFLDLEPEEYLDLSSSAEGYLAVHSPFKVLNPFIRGFKLQPGNIYYVDLF
MEEKLSLQSPYRTNCRDYLKIWKEKNITGPFTWEMCVHHCKLTKTSTNCGCVQKRIIYPHNFPFCHSDDTCVSKLDLEA
CYSSCRQECISHSFKVSVTHKRIPDDLLRIYKEFYPQNVYNTLRNRCIILIFSVERPEFLLYQYLPKYEIIEVFSFIGGYLGMWL
GISLFAIFNFLEDAAIKIFTKYENQKKHKKQNVWTKVLCARPKPFIRRKILA

>Protos_Platyhelmin_Macrostomum_lignano_A0A267GN24_A0A267GN24_9PLAT_Uncharacterized_pro
tein__Fragment__OS_Macrostomum_lignano_OX_282301_GN_BOX15_Mlig008995g1_PE_3_SV_1

VQKGPGRDVGVGRHQRHLPARRDHAAELPLQQRAVQSQKLLACPDQPVLELLHVPPGPTAVVQRPGAGEPSWTWC
SSPMATISFPTGMEIGEMKHCATAACFASSHAAKPSTDCCRATSCSPLDCAFSFTREKSFPMSQTEFVDVASASSTSIKLR
PIHNRLMSKPSRRCSEPPIETINYVRHFSNASLNVVTKAYSKSISDFVVEAQQSILHSNCGCYSHLLPFSVNTSDLCYFAPP
QEWITPSDAVLKRIDCHDHWFQHATEEKPRASEAVPAPDVVPLHPATALAAEPQLAPVQRHSRDLGEPDGAASQTWS
RVPAEDGKVHHIFRNSTETRQEIQLYMLARGYEGCNFNVLGKFNVTDLSSLEDYRRKIVESAASCIPRSELAKELAAVRISL
QSPAADAYEEMYKYHWTEALSEIGGTLGLWLGISVVSTFELLEFIYILVQKCRRNDDDSDIEEES

>Deutero_Ambulac_Sakowv30004505m

MDVVVPLSQMDNTDHQTKKAEDDTASQSQEPEKPSRLTLAVIYAQNTTAHGLANTVNAGTLHKRLFWILTFIICMVTL
VVQAQSVIRLYLSYPKAVSVEIKSRSKLDFPAVTVCNLNMARVSRWNGTRFQELLDVDRMQFDIDEWFSENLGSSDAP
EPDTSTQADVPINATSSGDAPVNTTSSGDAPVNTTSSGDAPVNTTSSGARKRRAVVAKHPVTDLLGQGTLSEFLKTLGP

GLQSSWIRQYAKENVITRPKRNSGNDSETSETDGSDDTDNGNSEGSEMGSERGSMGFYASDDYDDFSSNAFMDYGL
GGTNPNDWSTFYTKSSTDDFSDLRAALRMTQAEIEEYGERAEDFIVQCSFDRSSCSSKDFTVSQNDVYGNCFTFNGITG
NKTRRTTKAGEQYGLKLTLYVDADEYVGLFTQQEGARVVVHPGSLYPVPESVGLTAGVGQSLSIGLKKVKPKY

>Proto_ecdy_PANarthopod_Tcas_tr_D6WYQ4_D6WYQ4_TRICA_Sodium_channel_protein_Nach_like_Protein_OS_Tribolium_castaneum_OX_7070_GN_TcasGA2_TC006095_PE_3_SV_1

MSEKTSPLRYYLLNTSFHGFRYISEAGRHWTERVFWALCCLLSWTATLLLIKSSWEDFQTNAISFVVDTSYLPWDTHFPSI
SVCESDNQKRIAEITDRIYGDPHDYNLDEIVKELVYFRGLSFYTLQICGPEAQVPSEDCAKGDLGHFSQLVRSNCSQIFRKC
RWNEEEFDCCAHFGALDTEMGVCYAINSIQEDKTKRLKMVSNIKTGPGKLYLEINGFANVYILGEQEVPSMTTLTTDVI
QVTPHIHYRRFLAVKEIDNQPEVKDVSIRQRKCRFSDESDLNVYRYYSYSACCVQCRKDAQLKICNCAHHLMPNTPPKN
KCNITGLYCLSKHYNELSVLKPYWANRTGLVCDCLPSCTEMELAVVRDDKIGIPEDYAIVELALERLPTERYKRNVVRGKL
DLVVSMGGSTALFLGASILSFVEVLYYFFVRPVSDSLITKQKMKENQKFRRFKDQSRLPTIS

>Cnidar_HydraVulSc4wPfr_875_1_g17144_t1

MFGSRLIRLSDSSDNYQYDYDYLQMEFAGMDAETGYQNAFLFPTTSPMALEDNRFNDKGIIKGIAPEKRIEQYMKSFSL
HGLTKIFVARSRFESLFWLIALLGGISLSMYVSVNLVIKYLNYEVYTDAKLEITDKNYFPAVTICERDLLQSFYFSYCGLTSDN
QSSFIPCQNQKKIHITTDNKTWWSNGLFNVTLCTTWGAKQCVKNDFFLSLSQYNHACIRWNYNGDLYDMYSHVDIEF
TCISDIDQPPPNIILISHDPLVSEIDFTNAISLEAKQYSINMEKTIIELLPSPFPSKCKDDKDNDMFPGKYTRRSCLETQNYIG
MFKECGDIFDYMKQYIPKGILLKYKKNDTKSNFFNCMYSYSMREAKSSSVLCPFPCSETQISMVVSSRNYFKNDQRRDS
KIYSIGIQYSLVDSYRVVKEKQLYYIHQLACEIGGLIGLMVGFSLISAIEIVTCSVLWLICKIRKLNKF

>Cnidar_Rhopilema_esculentum_TR102937_c0_g1_i1_p1__Rhopilema_esculentum_TR102937_c0_g1__Rhopilema_esculentum_TR102937_c0_g1_i1_p1__ORF__type_complete__len_475__score_61_96__Rhopilema_esculentum_TR102937_c0_g1_i1_837_2261_

MPVDKNGISVAKDHKKNTTKEDDSVLNAAESLTVHAFPRIFNSKNWVFKVFWFLILSGALGVLIWLIYQRVTDYKKYEV
YIKRESNVVQPLPFPAVTVCNSDRFSGIPVKQFPMDNKICSDATTSKSTEHQDIEFNKACRMFLSGFKDTFRFGGKDLPG
FPDQFMSSNGAYPCFTFNKNGKAQQNVSGGGKGLDMIFFSDPNDIYHSEMVNLSRFDDQRRGILVQIHDPNTEVSLD
QSSSITISPGYLTEIVIKRVTTSRKKHPFPSKCHTSETTKYKLIGGRYTRSNCISSCFQLKLQEKCGDIMKPNKSATQILCESD
MYRRQPIEDCQCPQPCYEVAFPRMVSVSVWPRMFQFESLKDDFSKMANTSDNMTLQYMQKRFSQVKIYYQDLLETK
LSEEELYGITNLLSEVGGLMGLLIGSSVISVVEIIWLSGIIIKKLFGSAVSHTRKEPKKTNTIKNSTMQLYPVKFYSK

>Deutero_Ambulac_Sakowv30036986m

MNTLGATVSPEEASSTAKIYAMTGIKEGKDDIGQGDRSRKHGCTCEYTKERIRKFTQETSMHGVRYTGMTDIHSVRRII
WTVLVLWSLGWLTYFLYTSVVKYRAFNVNTLLSMKYTDVLLFPAVTVCNYNRYRKSYIQGTTFAEWQQTLYHPINLIHG
STTINWTTDAAIEELAKNRTYLELSAAHQKESMILECKLGRASSGCNAVDNFTTTLTDYGVCYTYNNALSNHMLVTESGS

RHGLSLTLNLEQEEYTYGQNYGAGFKILVHFPGDVPLVTDFGTAVSPGTEVFMPVRLVVEKNKESPYRTNCTNVSLEYYA
HYTKSNCIMEKVTQMVVSKCGCREPYMPGNVRVCNYQEKEECVQIAVNAANCPTACDIETFQTSSSFATYPSYHMND
YLEQTYNMTIANIRENVAVLHVFYEELSFMSIEQSAAYNGMDFLADFGGQMGLCVGASLLTVCEAFEFVFIGIWKRLCR

>Deutero_Urochord_tr_F6VJJ2_F6VJJ2_CIOIN_Uncharacterized_protein_OS_CioInt

NNHTEQNGLSVLKRLNNRIRNNLRMKPKLGLRTKQKKLAVTFSEETSCHGILQIKNNAKIWLKVFWALLVLAAFSVLIW
QIIDRINAYNKHETNTQISEEYVQSVHFPAVTICNSNRILYEAIQRIKKYFYFSSDEERLAMLTLSKAFDTGVPQNASVVQN
LGLASNFSLREFMMAKGWRLDVGGALAFCGVSKTEVCSHVSNSLGLCWTLNAQRRQSIPGYPHGIHLIFITHQSDSTED
PISGNSETGIKFQVHSQDEPPQVESLGHSVGVGLWASVQMHKQEKSNMYSPWGACRPSNENLRFYSKYTLNACLKQC
YDTLIQQTCNCRGPLMEYNYEDVVECTPIQVLNCTAPVRYAIESSYTPQKCGCYQPCNEVKYQTTLSYSSFNPKHSEFFEK
GMNLSSGYLSENSVGISVYFDALSVTKHVESKAITESALLSDIGGQLGLWVGVSVVTLFELLQFLTSNIYCILIYL

>Protostome_Lophotroco_molsk_AplCal_XP_012942494_1_amiloride_sensitive_sodium_channel_subunit_gamma_like__Aplysia_californica_

MGLFQEFSESTSMHGLQRAVSSESKWKRPLWAALVLAGACLATFNLSKTIKDFLHNPTTTETSVHYSPDMEFPVVTICN
LNPFSYGKLTGEMDFMAVIKNLVENGFTEEDDIFEYMAQFQRSRDVAITYSLKDLLIECQFDMMPCSADDFLTVPNYKY
GDCFVFNKQIPENFTHSNPRYTTKPGPDNGLIMKLDVQQMQYLYTTEDAGFKIVVHSQDTTAFPEDEGFLVMPGTRT
NVGLKRTEMSRAPPPYSTCIGLADEYNVDMNVYSLTDGTRYSGEACRKTCYIREVVRQCGCCDILFVCPSEIYNTYEVNE
TNPYCNATNTDQFFCKKRVLNAFEKGELDCLGRCLPPCEEVVYDVALSTALWPSENWVFVPPGTSMQDAKNIINYYRE
NMLRVAIYFEDLTEKHVESRPSYDWNRVLSDIGGQLGLWLGFSILTAMEWLELAYDAGIRAGTRKSQGAQAAKVHVQ
QQGQ

>Cnidar_AlaAla_agalma_gene_23162_1_isoform_1_p1__GENE_agalma_gene_23162_1_isoform_1__agalma_gene_23162_1_isoform_1_p1__ORF__type_complete__len_476__score_44_57__agalma_gene_23162_1_isoform_1_424_1851_

MEVSWSGFSSSASVHVIRYTSSEFSWRRRVFWIIILCMSFGFFIFFAYTTLTQYFQYDVNSVVRITFENRSTFPAVTICNFN
AIRKSKVSEANENIRAFVEVERQIAIDTALNKDETINYTGPNLSGQEVLRVYHDSGHTMDYFIDGGMLLTCSFHGRKCDK
NNFTSVITRRGLCHTFNSGKDGHLKSLSQPGSRFGLRLLLSVQKHEYMLFGDNGISVGFQVYVHDQLQIPLMHLYGESV
APQTHTEISVTKEKVENLEPPYKTQCIRHIRGNVPYRRTNCMLHCEEDFIMKRCGCLPPTYLGVGMDFKYCDLKTMINC
ATPASNLFTKEDRSCKCSEPCEDTQYIRSLSSSQFPSDEFVKTVSTIISEANKTASPEQVNRYIRENLVQIDIYFKQMNYIIIK
QTPAKTISDVYGDLGGILGVFLGASFLTALELIDFLIAYATNIFKLRNGFTAVKDEKETQSNNDHVHVASL

>Deutero_Ambulac_Apla_gbr173_7_t1

MWPGAAGVVRRLIWAVVTLAALGFFLYQTALSFISFFEWNHVTKVDINYADELEFPAVTLCNFNKYRESAWTERDLKN
VGTHLGIIDEDHNLINPHLYTDEFKQNLAAVNWTALEVEEDYDMTEFTNRTGHQLDEMIVECTWKGSPCNSQDFHHV
FSHLGNCYTFNHVELDQDRHSTISAGAANGLKLTLNIQEEEYTPSNDLDGAAEDAGIKWMLHHPSEPPYVKELGFGAG
PGHHTFVATRHEQISSLPFPYTMCEEDGSLKYFDHYSIQACRIECETDLVVDQCGCRLVEQPGTAPVCNPAESHECAHV
VLVQAVTGETESSCDCKSPCELEVYPFSTTNVKLRANFIETIYSNSTTHNFTAEYIENNLVLLSIYFEALNHEVIEQLPEMTV
FTILATLGGNFGLFLGGSVLTVMQLLEYLFDEVTASCASKGSSNRVGDSRANLTGKEKDVENAEDGISIIDMGKDKSKDR

>Porifera_OscCar_comp36352_c1_seq9_p1__comp36352_c1__comp36352_c1_seq9_p1__ORF__type_5prime_partial__len_476__score_66_85__comp36352_c1_seq9_2_1429_

SWILQKVTAVFLYSISLWPTLNEVLCFADESESGRDPFANIREHCESTTMHGVRFFFIGAYWRRLIWGLLVAASLAGFLRL
AIGNIIQFTDLPTTTKVSVVYPRAIEFPVVTICNLNTFIKSKVLSIGGIDLVKALESVNLYAQTIAAENETDRANKIKFLEEAG

GLEKLVMDAGIDESFFGDCLWNGLPCHLYDDAFTPSLTPFGKCYSFNKRAVPYNSSGGGSLSGLTFIVDIQQNEYLNPGF
FWSAGVQILVHSQAETSASILTSGFNANPGTHTSIQIRETRTKNLDQPYGDCNASNPADYSVNNCANLCYYNVLKEKCG
CKELYMKEFYNDNVRICSFRDFSCINLAQDDFIVDSTCSCPPPCEERLFGIQLSQSQFPSNEGAADWTKRFNASESYTKSS
LIWLDFFYPQIQYTQNEQQKSYTIANLFGDIGGNLGLFIGASILTMGEFAQYFIKEIYDACKRRRRRQKV

>Deutero_Cephalo_194160R_t1__name__chr_scaffold41_start_1865858_end_1873272_strand_pro_le
n_477_

MSPGEEKESPEKEFAGNTTLHGLNRIVIAPSGYFRVLWVVAILGSYSGFAYMFSSMIIDYFSYRTITDTKLEFAEDLTFPAV
TICNMNKFDESKLKIIDFSYLSLFLYGVQLDVDAILASPGAYPDETVNSTLEGVTVGDLLLEKGFNLQWPRMPLCKWKGN
PCTQQDFNHSFGYYGNCYTFNADANSPLKQTIQGSGNGFVAFVDLKVNDYTESYYASGNSEVGLKLLLHDQNEPPMM
DTQGIALAPGSHAFIAAKRTLYENHVPPWGVCQDLQLEYYDTYTLNGCYLECRSKHVVRNCSCRPYDLPGTAPSCDPNT
MFSCVRAVLDQVTRGELECNCPVPCSMTAYSMSVSYAGWPNKLTGEYISPLWGSTGEYLELNGVMFSVYYEKLNYQKI
TELKAMDGGQLASNIGGMMGLFIGASLLTVLEVWEYLWQRVLGLLGKQRRPRVIEVKSPPDDMERGIRNVAFPMENT
KIN

>Cnidar_AlaAla_c57327_g1_i1_p1__GENE_c57327_g1_i1__c57327_g1_i1_p1__ORF__type_complete__l
en_478__score_37_33__c57327_g1_i1_249_1682_

MQKFKDIAVKQYKKIKLKKIIEEFFSSTTIHGLGRIYGASNHFTRCFWILAFLGILAYLIYNVLCIVQKLNGNNIVTVTQATSN
TLMNFPAITVCNANPFRKSEVKKVLSQHMVNESNTDDLERRKTLGSIVATMSFSQLFQMGHLPDEFLVGNQSCSFGDT
PCSYPTDFQVSSTSINGNCFTFNSNNTIKQERPGQFFGLTFTLNVNQDEYDLTSKLTNVVGAFVTVHHPSSPATLMSTSV
LAGPGQLTRIALRHERVTRLPHPYPDNCTKSSTKIMEGINYNSKVCFLKCFIDQQHKFCGVVDSGVISYFKGTFGRIPRDV
NYTIATKASDFSCIWRYEQQFKRNIFKCSCPATCKEDIFRTTVSSARWPSRPHANKLLQQLKQMSTRYSNWTSETMYEN
LLSIEIFFEEFSLFETKQKPEYDSNNFLSDLGGQLGLWIGASVFSVLELMAFISHLTLHGFDCHSEQNKVHKIVT

>Proto_ecdy_PANarthopod_Tcas_tr_D6WHS4_D6WHS4_TRICA_Pickpocket_protein_28_like_Protein_O
S_Tribolium_castaneum_OX_7070_GN_TcasGA2_TC003563_PE_3_SV_2

MTIMSMCGSERGDLELTDKENCGKNENGKSSYNCGQNLLNYVVEFCNNGTIHGIRYFGEYRTWIEKLWWGVSISLSLY
FCVSLIIQTYIKWETSPVIVSFATKETAVWQIPFPAVTICPESKTRASVFNYTYYFRKKRNNEPLTEEEERRFGYLSFICDTKP
NIEYKAPKFAGEEMLEFLDEAQPKFLENSFDCRFMGIEYNCSELFVPIFTDDGICYTFNIVDRTQLLHDIVIHYKNFHKTKV
AHDWTLEDGYVSSAGIDAYPRRALRAGVPFSLEIKMSVDVDDLDYTCGKTSVKGFKVQISHPQRMPRVRHQHFVVPLE
HLAKARIAPDMMTTSAEVKQYKPKKRNCHFPSERGLKYFKKYSKSNCALECFANFTMIRCGCVPYYAPREKGMPLCGG
GSLRCLEEAQKYQLRGNLEKKFDALNNRKTHSDIPNCRCYPICTSMYYDIENTQTSWDWRTTYKYNDDERVNISDSK

>Cteno_MneLey_comp13853_c0_seq1__p1__GENE_comp13853_c0_seq1_comp13853_c0_seq1__p1__
ORF__type_3prime_partial__len_479__score_87_11__comp13853_c0_seq1__3_1436_

MLLNNDTKPPDVPKKQSFFEDFKEFTTETTSHGIKYVFHAAYKWVRVIYLILWLAAVGYTVSIVFISCKKYFSMETGTKIEE
KHTPKHSSNEVAITHPTITVCPNNLISKSYLAKHPNLDELMTKLVTFNENETPSLFDDPKYAGYGDMTYHDLMWQGRP
RVGLLQCTYFAHKCGAVDPFNSEDYETKTYHWDKTYNGSNHKFYEWEVSLSGSCYRINPRGNLWAKVGDYGEIKLTFF
ADLNDYSSASKEEPEYGYLVTFTDSGTYSSTATSGFFMSPGNVYKADLRKYKEENLGPPAGKCDIESKTNVYGAYSTNSC
NKLCRDQHVMEKCGCVHILPPHPENDPDTKFVGCTLKQWAECGLRESTKFTEEFVNMNDDKVHCEHCNPACMEIWY
EATISSSRLSTFYAEAKVSELDKLITGYQLNEGNKTTNYTIDKDTVYENLMVLKVMFTSIQESTIKEYYKYDYTQLIGDVGG
QM

>Deutero_Cephalo_131190F_t1__name__chr_scaffold26_start_146491_end_151392_strand_pro_len_
479_

MAEEEDEQSPEREWASGTTLHGPAKVVDGPNLVVRLLWLALFLVSLGIATYLITMSFIDFFSWGTVTDVKLEFTDQLEFP
AVTICNFNKFEKDLLCMDQRAVEYLRPLIGNDAICETDNSTSPTPGPTYYPMSLAGITAYTGFQMDRYALKECTWRGET
CTWQNFTHVFGEYGNCYVFNQKTSGTPLQQIIPGQGNGLHMVLDVHEAFYTENPATGGSSDVGVKVHIHPQGEPAK
MDIQGIPIGPGSHAYIGISQVTYQNEVPPWGLCEDLDLEYYDTYTLTGCLMECMAHWVESECGCTALELPGNLSYCEPQ
LIVDCVLGVKDQITTGVLACNCPTPCRIQTFDYFVSYAGWPNHIVEVYLGEENNVDMEYMQSNWASIDVFYEELNFEQI
TQKKAMTPGDLLSGLGGQLGLFVGASVITIMEFVEYLLKRLTYCCRKKKKNKVSASTAQVWTADKVDPLALQKVNMLQ
ETKMYKE

>Protos_Platyhelmin_Macrostomum_lignano_A0A267DAD5_A0A267DAD5_9PLAT_Uncharacterized_pro
tein_OS_Macrostomum_lignano_OX_282301_GN_BOX15_Mlig009645g2_PE_3_SV_1

MTVGSNAGGSVRRRWEVRKHWNDFLVTSSVRGLKRTTTSETRFLRLFWVVFLLGSLALMLYQISSVVDTYLHQKDSTA
MQSKQLDIDVPFPAVSVCREATYSSHKQMRQAMERHNASVTTPQSYSRRVADLLYRHFRNATNMTEDELEEAKMAL
LRFDTARSYQAQIPRAAIKEVALPLIRQIATARANFANGSSTSLASNMDIIGMDVETDDYPSSSYDAELWGMRVMDFYH
WRYFLCQTLEITLSREEMNEMRSLDLLLWYAADDFNTSISGVADRAFAMRGFWSADGIRLALHPAGEHGKVHIDNGA
TLVRSGGYSEIRFRPSITRYLNCSDRELQHGDQGGYSYNMNDCKYLNYDEELRRQCNCTMDPLAWDYHQCIRINWTDL
DMSDWAAAWAESSGTASPAAARPALTPNIGTRPRSRPGTSTTSNGGGLSTCCSIETKPCRSIWPCRRWARRRAVSTIA
SRRRFFSELKTT

>Protostome_Lophotroco_brch_Lingula_anatina_comp134404_c2_seq5_p1_comp134404_c2__comp13
4404_c2_seq5_p1__ORF_type_complete_len_479___score_28_84_comp134404_c2_seq5_272_1708_

MSSVHPFSFEEKGTEFIKQYNNMHGDFSNKSPHRGTNITKRKPDQPKDETIPSLKDEISDFANNVSILGMKQVANPKFS
PVRRLLWLAFVLCGSGFLIFHLWNRISYFLENPASVDIKFHHNDSIHFPTVTICNNNRFKLPYLRANGHLEQANYFDMFN
FNLTNIEQYNWTDFHTRNQYKMADLMPSFCLFNQVICTDADFSMIETEMGSCIQFNAEGELKSVSRAGSLSGLQMWL
FSHQDDYLNFTTKAGYTVLMHERGEFPHMIGLGLKVSPGESVDIAMKQRRRTNLPPPHGQCKERTLKYFSKYTKLNCD
AECSFIAMQACGCRLFHQPKTGNQTIDGQRTCNLKDTMFGCFNISKEPMAGYSSCACMEACQSTLYTHSISHTRMSEV
FIKRLIAMYNNTDTSFFRENIIMLNIFYSDISVEEVVQQEAYSALTLFADIGGALGLVLGSTLMTAAEIVDFFLGVSLWKLFR
IRV

>Proto_ecdy_PANarthopod__XP_023240338_1_acid_sensing_ion_channel_4_like__Centruroides_sculp
turatus_

MKSYEKRSNTKPPYIIAWENSSKNKKKLSRKKCSLSGFTYSSYKFLTNFFEYPVVVNLEVENEGELFFPAVTICNSNRMRV
SELHKDVCKSKRKSCEYLRIIHESSSDIENKPEFVFKFATCSNSSKREMTLYHQKLLAFTNTFSRFEDWKIKKMSHQIEDMI
EECTFDGEKCDASSFTPLESFTYGSCYTFNGRWMNSRGQEDMGRRKRKKFIDDGNEKLISRSTNPFSGLSLTLNVQIEEY
LSITSKMGVHVIIHSPDERPYPEETGIDISTGFDTAFAIQENLYRRMSKPFKDKCVLYGKDDFKKIKSQKECLSLCTRDLIKK
ECGCLDPTLKFYRNVTFCDLTKEEDACCMMKMRKVNISYCQCPLPCHEITYDVSLYSTIWPSESEFKLRSLKNLSKQITYE
EYRKGHLQMNVFFDRMERFIYNQHPVYEKDEFFSHLGGQLSLWLGLSLLTLFEYLEKFSLFCIQVYKNFSRK

>Protostome_Lophotroco_annelid_Pdum_comp417182_c0_seq3_

MTQELHFADETSCHGVRHVFYAANIIARILWILCILGSLGFFIWQFHSLMSSYTKYEAVTEVTVKTHPAVPFPALTICNYN
RLKKSVLEQYAERYNLPAVQIWQYLILAMNIHVSMSLSGPGSPKPQNLSMHSKATSDRIRQVDGAIDLDGMIRNASFG
VYELFAATSFAEKYTKKLIDDMMSGVFTSSGVCVTFNPREESSLVQGSVGLTSGLFTGLTTNSHENSENMTDYLGGNFA

VGFQIMVHQHGTWPSLVPAVSISPGKQAFVSIEKMVYENAEKPWGECVTRQLKYFQVYTEESCRTECLIEYLHQQCKC
HPIVYPRNVSDNVSLCLAEDMVSCVISKSESYKNQESSCTSNCKVACKYEEYRVSVSDAGTGEDGPSKTLAIMLNMTVE
HFIKEYALVNVYLKELKSLHFVQKEGKNFSDLVADIGGQLGLAMGISLITLAEFIIYSVNFIGRVVTSKSKQNQVREMEKPP
LD

>Cnidar_Rhopilema_esculentum_TR74840_c0_g1_i1_p1__Rhopilema_esculentum_TR74840_c0_g1__Rhopilema_esculentum_TR74840_c0_g1_i1_p1__ORF__type_complete__len_481__score_64_39__Rhopilema_esculentum_TR74840_c0_g1_i1_867_2309_

MEASKANAFGEFAERTSLHGVGRIVETKTKWGKISWAALLVGFILFLCIMLFNVFTSYMSNNIYIVREKIPVTKMQFPGIS
FCPTTLLRRSGFSSPNTALIDAVFHRFKGREKDIEINNLRIANLHRHEDLMKGFESIRRLTSNADSLFIAGMNSACRFGLNT
PCHYPMRFKETPVSLFEGICFKFNPNGTFFQVGEGSYYGMSIILFVNQSDTDPFSGFDEGAGIKLVIQSHDTFPFPLENGI
LVKPGAYTRIIVEKQVYRRLPHPYPSKCKSQGDGIFPGKYTPRNCKRTCFVRYAKKACGGTDAFVQYFTGEKRDPPLNITE
LQCFYNKQIEAAWPHNAKKCNCALQCHEETFHPMVSQSLWPSPSDLPYYKQIFAAALGLNHSTMTDAYVYNNFLKINI
FFSELAYEQVSEMPEWTSQKLVSDIGGQMGIWIGASLFSVLELIVFLFYFAKSKAHVWMLKRNVIVVEPAPTKFAVK

>Deutero_Ambulac_Apla_gbr256_12_t1

MDPRVIIETTPRNGFNERRVMGKDNLDELNIIKNDRNRHTQDDDANRTFAERFQDFGNSTTLHGISYVINLNYKKQRRL
LWLLLVLVMSIWLVYSIVEAFITYFNYPMTSAISIHYMDTLTFPAVTLCNFNQFRRSALTPGQVMVMNEVYGPNPPESID
FGPFNEITNYDFSEDHLANISHHLDRMLVKCKWRGTQPCTGQNFTRRFTDHGVCFTFNDPKDASHRLEVSNAGRLDGL
FLRLFAETDEYTFGENSAAGFRILLHPQGLMPLVKELGISVSPGFESSISIRQNVFKSLPWPFESNCTSSKSKNSTTYSVPTC
KYECKVQYVVERCGCRDYRWTGSAPLCPPEQHFNCIDKYKAEYQKQEKYRDCHCPVSCELKTYDSRVSFGYWPSSYVSE
NLPLNSTSEEYIRKNYLDVYIFFEELVYLQIEQKSAYDFAALQSGVGGNMGLLCGMSLITVVEWADFIVISLLRKFKARRI

>Protos_Platyhelmin_Macrostomum_lignano_A0A267G831_A0A267G831_9PLAT_Uncharacterized_protein_OS_Macrostomum_lignano_OX_282301_GN_BOX15_Mlig007609g1_PE_3_SV_1

MGSLVRQGRYLDASKLFHNDRVTTYFQLINESELNEVSHHLTDLMPFCIARHIGNASPSKIQETTCQFKFTRIHHPEFFTC
YNLKMLGTTEQLSEVSLILSNVRNSDLYGGSAVNKSFREMEEQAFNAFNQATGFRVVLHEPGTFPNVDSDSFTVQTGT
ATLIRYQTVRVEKENYPEGECSNPKEQTPIRDHDHVYNYTQTACVSDLLGKRVTEICNCVSAFQIISQVPSKEVPYCGHYH
ETQLEEFLKRYTCVQHLNLSSIIRHFTLNACRKPCTQYFYNSHLSVANWAPRLSQLRWMGLVSGALRHASLNASDRLAR
WALPKLLEFCNLESIGASGDWKNNPEEQALVERMALIRVSRLNYNTVLKQEKLQITLSALVSRIGGLGSLTIGVTLTFAFEF
LELLFLLLHRAVYRCQQEMDASHSRQSCPPDVRVENVDTIEELMPLPARSPMMEARLRGWSDVGFDEAALRRLSEIGN
GC

>Protostome_Lophotroco_annelid_Pdum_Contig8016_

MSKVTHYPMDDLGKSPRPLVNDMGQGHRVRKSSSMQRVLSDYSEHSSMHGVWIVANPDYPKLRRLIWMLALITTTT
VFVYAFIQQMTMFFQYKVTTVLSVEEQATLNFPAVTLCNYNQFRKSKAEQKPGVMGVLRDLFSVDAIIERQKSKLRMS
AQSLVSQDINIINELVALAHTLKDMVRHCQWCGKTIPCSDYFSQTLTEVGVCYTFNSHDVRKKIPPLEVHDAGTDCGLKL
RLNIEEYEYYYGEGSSAGIQMLIHDPEDVPLVRQHGFSAAPGTSVMAAIRLTKETYAQSPYGDCVDAKHPSFENPLRYHE
IYRYSACKLEHRVDYIYNACGCVMYYNPVGSRSVCKTVEEVLCFKDAKENHTRYEMQAMRHACPVTCQSDNYEARLS
WAAFPAPQILYDLKKNFNMSGDQVRKNLLQVTVYFEDLGIHKKEQQPAYTGNSLFADIGGMMGLCIGASLLTIVELLEL
GAQLMLSAGKR

>Protostome_Lophotroco_brch_Lingula_anatina_comp149479_c1_seq4_p1_comp149479_c1__comp149479_c1_seq4_p1__ORF_type_complete_len_482___score_29_72_comp149479_c1_seq4_182_1627_

MGKGEGESLETEFASGTTCHGFNRLAVAKKWPHRAFWTVAILWGFGMTTWMISSRIEAYFKYETSTEIRVEEKYALEFP
AVTFCNFNKYRRSAMTALDWYIYANSKLAEKYAFEADYDPEATAALRAAFEGFEANLTATYPGGFDFGNFTRRIGWEL
NNETMPLCLFRMKQCFPENFTHAFQSEYGNCYTFNSHHDEDSLSLQQTRPGAQQGLYILFDIQQAEYTELLNLNGALG
VGIAFQIHSWYERPNVAQHGLGAGVGTAALVSMHQNELQNLAHPWGNCSNDRTLKHFPRYSRSGCEVECYLEKVLEL
CGCKPFSYPGNAVNCNPVDQVLCADPTIFKVASNFQKECGARCPEACQIIEYPPTVSYAPFPNYYMAQELATMYGTNA
NYASENLVGLSIYYKDLNNRLTKKNPAMTPSGLLSDIGGQLGLFIGMSTLTLMEFGEYIIFKIIQCCRKRKPKTSQSKGVAN
NVKVNESGM

>Cnidar_AlaAla_agalma_gene_58222_2_isoform_1_p1__GENE_agalma_gene_58222_2_isoform_1__agalma_gene_58222_2_isoform_1_p1__ORF__type_complete__len_483__score_50_45__agalma_gene_58222_2_isoform_1_499_1947_

MAPEKSAKKVVKKFADETSTHAIRYIVLAKSKVVRLIWLVLFLTASGFFIYELQKLLKAYCKRPTLVSVEKKEVKELNFPSVTI
CKASTLSRSGGVQEHPFIRDPAKRDYYLRQNDITTLRAQLALEYNRYFTFKDLKEAAGDPDKLFLTDLPGGCIFGLHQHCY
YPANISRVITSSYGGLCYQFNKDGMIKQRDAGSYYGLKMLLFVNQSDFVPFTTFDFGSGILLNIHDSGEYPFPEINGLPISP
GFLTRISIKKSVYKRLKDPYPTNCTDGKGIDMLYPGSYTIANCKRTCLVRDSLKACGSSDPLTMMLLDQAGKYKYKPINDT
TQQCIFESAARSTSADREACLCPVPCREIEYTPTISQTKWPSEADLPLYKLIFSRTLGVPISTLTDDFVHNNFLQLNIFFDEM
SFQDISEKKEWTLEKLISDIGGQIGVWIGASVLSLCEVAFFVVKLVRALHAKRTKPAKETVSMDGIAENMK

>Deutero_Ambulac_Apla_gbr54_23_t1

MVREDADQSPRPSLEREFAESTSLHGIAKVFHASRLLVRVFWVIIMLTCLCVCVWQISDRFHRFLQYKANTEISVEYVGH
LPFPAVTICNFNRYRSGALAPGDEDALAYIWELSNYQNDYDYEGESNSTDSSYLDSLVDFNYTEFTLRAGFRMDNVTLLA
CSWRGKRNSCSAEDFSHVFTSYGNCWTFNSGKTGPVLTESYSGSGNGVSLLVDIQQSEYTETETIEAGLKIQVHDQVTP
PTVESSGLALSPGVHAFMAVRRQQYLNLEHPYGKCDASRTLKRYDNYTLEGCNIECRANAVWEKCRCRLIQHPGNEVE
CTPEEVRFCAREAIDGLIGGDTESCDCPVPCNFTAFTTSLSTAFLPNEKILPILLLLLNNISTSDERNVSRSISEQQEYIRKNW
VAIDTFYEDINYQKFEQTEAITPSALISDIGGQLGLFLGASFITLAEIFSYLSRKIQRLIHGPTRRPEKQRQMSHSVETLA

>Proto_ecdy_PANarthopod__XP_023226921_1_amiloride_sensitive_sodium_channel_subunit_alpha_like__Centruroides_sculpturatus_

MQEPSEKDKNTSIFEKFLGSSSVIGLSQITKSRSIVRKLLWLAVLVTGLTFCAIESHKFMREFYKYPVVVNLEIENKGALEFP
AVTVCNLNSVRKSEYQKRFVKTITIESAFDSPFLPSAARGFFIPFRKFTSQCDAVEQKSRNENNQNDFYNTYFSLNKTVRG
SLGHQKENFIMSSSFNVKQISLFDNFQMVVKPRHGPCYTFNGRNTTSQNMNVSSVRVSRNIGTLSGLTLVLNLELEEYF
DKTKSVGARIVIHSPDEFPDTEGEGINILPGVETSIAINKQTTKRLPFPYKDRCKDYKKQTIRNVSILNEKDCIIHCLQDLNIQ
RCNCTDPTLIFKKKRQKCDLENNEEVCCLDKVFEEMSSLNFETCDCRVSCSITNYEMTTSMSLWPSPEEFQYNYQYYGF
NYSSFRNTYAKLNVYYATLEETIYAQRPVYQNNEWYSHLGGQLGLWLGISLPAIFDCIETIVLLIHHAIIKYITKN

>Proto_ecdy_PANarthopod__XP_023234266_1_degenerin_mec_10_like__Centruroides_sculpturatus_

MDNNTNETRNKSTCKEKGIYFLKLLEDVQLTGIPQIVTATDKRRKVFKIIVFCTCLAGFLYHLISFVLLFLTYPSTLEVLVTFK
DTNDLPKPVLTICDLNGIRRTSFCNELPEDCEKPKNMKVLCERFEYFCEDNDTDNLVFPIHNSLRAAEFFRQTVDRYGPT
AEDMLETYSDIATPIFYTNTYGMLRKCYSIDLSELHELHYLFSYDKLSSFLTSGEVIATIFLTFNPKESSNPAAAIGAKVSVHN
KDQLVNPFVDGFRVKPGRIYNLQLKAIEENLLDSPYGTNCTRYEQLIDRGISGSYSRDFCIIECKKELWLKECNCVFGEYNL
DFTGNFCDKVEEYKCIRDVDEDECYNKCKKSCELRYFEYEIDQLEIEQYHVENFTWDKNDDWWNNERIWYSTISVYYKR
PEILIYKYKPQYELIEFFSYIGGYIGIWLGISLIAVFDFLEMLFVFCCCVIKGKIPIKREKKSNVLLQNDTNV

>Ctenoph_Hormiphora_californiensis_evg120240__p1_GENE_evg120240_evg120240__p1__ORF_type_complete_len_484___score_100_73_evg120240__313_1764_

MGWLADFKEFSDGTTAHGVKWIFHPSVHGNFKIVKLIFFIVWLAVIGYSLFTIYNAVHNYVNKPTGTKFQVIAEGPEQE
KPASIQFPTISVCSHNLIKQSYFDSNTGLEELWSDLDKWDPDVTEKLNFTDSKFARYRNMTYQEIIRAGGPDNTTFVHCR
HFINKCEDAIPNPEENFYVREASLSGNCFRINPNGVLHGKGGDYGKMQLLFFADLKEYSTMTRSEPQYGYTVVFHDHES
YSSTMSSGFWLSPGSIYKVDLSLSQEFREGPPAGTCDPTRVNNTYGRYEENSCIAQCRDDALMEKCGCVHISPPLPHNE
PGKDYRGCTLEEWATCGLRAYKEWVVKYSDVNRISMGCDCPPTCFETEYKAQVSSSKLNRFYAEEKANMGLLPPGYN
GTQDVMDNLLIIDILFTSMQVSEIREIVTYGWGNFLGDVGGVLGLFLGASIFTIMEFIQFFILAICNACCPGLTSRSKANSP
VDTYALS

>Deu_Ambulacrar_hemi_Ptyfla_40v0_9_20150316_1g7627_t1_scaffold3848_cov141

MNRIVNSKYAWAKVLWFLIVGSSTGVFVWQASKLVTEYFSWPVDVSIDVISKKDVAFPAVTVCNMNRIRRSEMEGTR
FEGILDIDGGAGSNIWDYGGFEWYFYSDMYSDGSSATSEPATGAPSSSAGGVGRRRRRAADNVDNQEPSLDEFHPES
DQARPLQRENLGNDSHHPKDSPKRKSKHKHGKKMHHSEVDFEPEYPDHEDYPGPREKPASRQRHPIDRKKYADAVHK
LLAPEIQAKNEKRRRRLEEMKQKIEHLKAEGKLPQKMKRKKRAASSSSSSQSYSWSLDYSWFDDFDWDFDMSDFEYYD
YGWDGVTGEDDWAGFYANSKMEDFSDLADVANPTDDELKDMGHQAVDLILQCTFDKRPCNYTEVTYAAQVSSSLW
PSERYETHLYGVLADKSIKAAITLQDVTESRRNLVRLKVFFEELNFQTIIQTKAYRIENLLGDIGGTLGLYIGLSVITVVEIIVFF
LHTCRYCCRRIWS

>Deutero_Ambulac_Apla_gbr658_1_t1

MNRPCDGTDHNFDMETDDLRDGRERRRPSVTRRKRGFKRKKSKDTCKNILAARIQQFGDDTTFHGLRYVTNSTYHTFR
RFVWLLVVTGMTALLIFDIIRAIMLMFEYPEMSSISVQYPPNITFPALTLCNINMLRKDALDASEVQLLSDIFANPATPPD
TNALVDSKLFNSSSNEELMGRMIDASYRIEDMLVKCRWVSETCSYRNFTRSITDHGVCYTFNDPLSDHEALIVRHPGSR
HGLFMRLNVQQYLYTYGESTAAGLKVLLHAQGEYPMIKDFAFSLSPGFETSIVVRQKVMINQKAPYKSNCTDGNWGR
HRFKYSAASCKYACKILYVTRKCGCRDYRWSIRDIPICSLEKQVHCIYRYEDEFVLSQENQCHCPIACKAITYGRETSQAYW
PASFLSKELQAEMNVTEDFIRENYLDVYIYYDEIMYTRIEQIGAYTVDNLQSDIGGYLGLLCGMSLMTVVELLDFILVALCS
RLKR

>Deutero_Cephalo_298690R_t1__name__chr_scaffold94_start_933624_end_940952_strand_pro_len_484_

MITSDEKTPEREFMDNTTLHGFNKIHHPNRAIRTVWIVAMLAAYAGFIYMAVNMAQTYFSYDTVCDVKLSFESKMTFP
AVTICNYNRIDSHKLTWEEWSILSTLLYGRAMDVPTLQAFLGPTFTDRNRTTNGTVTNATTMDMAAAVRMKGFDISP
ARMYLCSFAGDSCTELNYTHSFSNYGNCYTFNADAANKLNQTIPGSTMGFQIVVNVMADSYTETMAVGGHEEVGLKF
LIHDQNEPPMVETHGLAIKPGIHSFVSVKRTEYYNHVPPWGDCHDKALTYFDAYTLPGCNAECRGKHVQDRCGCKAYY
LPGSAPPCAADVLFTCVQEVLAEVNEGKLTCDCPLPCTMVKYSPTLSFSAWPNQLTEDYYSRTYNLSRGYLTLFMLFPRD
NVVALTVYYEDFSYMKISQLKAMDSGQLGCNMGGMMGLFIGASVLTLVELTEYLVARLSISWGKKKRPLHVQPAPDA
EMSDKKQPPDSLVFHK

>Deuterostome_chordata_PetMar_tr_S4RS34_S4RS34_PETMA_Acid_sensing_ion_channel_subunit_family_member_5_OS_Petromyzon_marinus_OX_7757_PE_3_SV_1

STGSPAGPTDSAKETIAFSSSTSLHGVARATSAQSRAQRAFWMMVVVAGVAMAGWQVALRLAAYYAWPTSTSLTLH
YVRDLSFPAVTFCNYNRYRRSSLHGTTDLLAALSILSAGIKTPHPEYLSGFLWRITLRGYSSPPITQTYSARNQDLSTTILEID
DTYCENSCTPQDFQHVFTEYGNCYTFNGGFSEEQRSLKQKTAGAGKGLQVLFDVQQKSHLWDYWENPELGFTGVGIR

LVVHDPDEPPQVEALGLSIGVGTQAYVSVQHTKSINQQYPWGDCDPSKKLQSPAPYSMARCLQECEADLVERICGCKP
FNFPGDIKECSLYQHYHCARDALVIFLAQGMCTGGSHLSQCPVPCLHKNFPSSVSYATFPSLQAAQRYERSLNYSLVYM
RENLLSIDVAYTDLNYALIQQQKTLPESVLLSDIGGQLGLFVGASVITIIEVLEFVARSAARALNAAASFLSNRAATCTKAAL
SPPVQG

>Protostome_Lophotroco_annelid_Pdum_comp406059_c1_seq1_

MNLILTGVFIFHTFTTVEAYLDFKVNIHMKVNNSNQIEFPAITICNQNSYKSNESQAFMERLITAYLTDPSKVGMPLWLR
SFVQPGLDPSLLGRELSKVDDPTTLSALTVETVTVYGPQIGFTMENITHRTEDLLLDCQFNKEQCNYSSIRHVYNWLYGR
CITLNSDEKWISYEPGPRQGLDLLLNIETMTHIGIAESSGVRVLVHSPSEMPIVDEDGFDVAPGYAASVALTAQKQAHLG
GRYGDCFSAEQGEKVSNFYRDHYNWTSYSVKSCMSTCMQETVIRKCNCFYGKYATPKAEGLNNLKSCSDNSTISAYDD
GSPCWMRLKESYEEFDYCDQKCWQPCSSIKYTTSLSFSKRTTLYQQAQGSAKAIYNDSLSEFANSFPESETAGQIIRSNLE
DQNFVRVRVYFETFSHQMMSEVPAIQASVLLCIKYCHNMVISKWNYMICRPAKDVLVRKTVAASFYSEIQQPGLAPFV
KSSQTN

>Protostome_Lophotroco_brch_Lingula_anatina_comp131676_c0_seq1_p1_comp131676_c0__comp13
1676_c0_seq1_p1__ORF_type_complete_len_484___score_23_39_comp131676_c0_seq1_234_1685_

MSQVDPFSVDYRTSERKQAFTDDGLISFNVYNRGKQKQNSRDIVKQFKAEKDGNDEFSLSSELSSFANDVSILGMKQIV
NPKFSLFRRATWLCFVLAGTGFLIYHLWNRLDYYFSNPASVNIRVNYDGSLLFPTVTICNNNVFLKSKLAANNETYIGDLF
YPSFDSYPNWSAYNFTGFNWTDFYYRNGYSVMDVFGGFCFWKSSPCQANVLKPVITEHGVCYQFNSGDEGKRSDVE
GSLYGLMISFMTKQEEYLVGSYIAGYTILLHNSKDHPNVVTQGFQVAPGGSANVAIKQKYVKNLPQPHGQCADKQLAY
FPHYSKINCQAECLHNKTIEKCGCRLVYYPEVPGLRECSPIDVNTCWGPLLTNVSQIMKNCACTEACETFYFEYSVTTGLV
SDVSAQAAAVFLNTTQTYVEKNYLAVYLYYKEMSYEEVIQQEAYSTLTLLADIGGALGLILGSTLMTVAEILDFFLGLSIWK
FFGVKL

>Cnidar_Rhopilema_esculentum_TR36932_c0_g1_i1_p1__Rhopilema_esculentum_TR36932_c0_g1__R
hopilema_esculentum_TR36932_c0_g1_i1_p1__ORF__type_complete__len_485__score_76_67__Rhopi
lema_esculentum_TR36932_c0_g1_i1_1028_2482_

MSSDNTRVVANTWNGFANSSTVHGIKFTSHDFSRTRRIAWGILLFTGLSLLLGFTVSAVVEYFQYKVNTIVTVGSEREAA
FPAVTLCNLNALRKSKFEESKNNSKYSQLFMLAKILSMMDLTGSNENVTIPSFSGENVREMYRYFGHNMSPFSKGGML
LNCTYRNTPCNESFFKPVLTNLGQCYSFNPGGDFGKERYGKEILTATQPGSSFGLGMRLTVQSDDYVLMPTQEFSAGW
KIFVHDQRELPLVADYGFALSPGTHTFVGLVKRKVVNLEPPYTTACRKQPLATGHPYSLKGCIRVCEEMYVRKQCGCNS
PIGFMDDTASELCDLKKSVNCVLPALAMFRKYGNDCVCPDPCESVEYTPYLSYAYFPSDVYGVEVADHLVSLGVASSNE
KALSYLRKNIIELDIFFQQMDHIIVKQVPAKTLRDIYSELGGLLGASLGASFLTFIELVDFLATRFLRRLEKKRGQIIRVASVG
NEGPSS

>Cnidar_Rhopilema_esculentum_TR91732_c0_g2_i1_p1__Rhopilema_esculentum_TR91732_c0_g2__R
hopilema_esculentum_TR91732_c0_g2_i1_p1__ORF__type_complete__len_486__score_46_22__Rhopi
lema_esculentum_TR91732_c0_g2_i1_876_2333_

MSEKLKKTLSFFLENATTHGISGIYQMKNIFVKILIMLLFLTLVSFMGYSLLATFRTFFEYEVYTSVEIETNHRLQFPAVTICS
NNAIPKSYLNSSLLQDVLNKTNDVVTGKIPFDEINEFFLNLTYELTAKLESSKTTAPKQPPMILETLNDACLFAQKQRCNLT
KDFKPTFPTEILNTCYTFNAGLENRYVQTGKGSGFGLSLVMFMNQSDFLPLIGKKRDTGFTVYIHDSDTLPMLGIRGML
VGPGFHTKIAIRKTEIVAKPSPFPSNCSNGEGIVNYIPGAYTSLACQLSCVFNSVLSECGYAQNLIRSHINTTKIIQSNASML
QCRDRVMNAVLGAQTFPCQCPRACKEIRYESTMSQSRWPNDIDLPIFKALSQKVVGVNPSSITDDFFTKNFLKLDIYFD
DMSVRKYVEKEMMGGISLLSIIGGHLGFWTGSSLFSIFEWLVALISLLMLFIQIMANKFLGERDQSVTKVESIGINPM

>Cteno_MneLey_comp6894_c0_seq1__p1__GENE_comp6894_c0_seq1_comp6894_c0_seq1__p1__OR
F__type_5prime_partial__len_488__score_56_57__comp6894_c0_seq1__85_1548_

LMITDFISYGSYNKISTVSTSSIILPSVTICSMNPMNYSSLVEDLGEDSDVVKELKELMWDAQLANDYNFRLSSDNYSNLIEYEEENGPLSFIYRNDAASLIALTSRHDYYLGKIGENIRNWLTETGSNELGSCIELNDDSSLSQNISGSRYGLTLDLDASLLTYAPFTSSVGLKVFIKDHDETVFFNEGGIVVQPGTETFMKVSLSKIVRLGPNWGTCQNVTSSVSKYDSTIESRRECLEVAKVREMLDKVEYCGCIPWFVVIRLREIGRVEILDQLLAPVTNSSSNSRKKRDSWDQANENYTEIWGDDIAEEMEKYDDFDDFFIKGLNYSEYFCDSYSQAKCDNRFPSHWASVQEKYSTICHDPCASNHYDVVVSSTKFPPTENYFNDHLKDYEGLEYTWEYAKENLIRLHVYYDKMEYLSVVQTKDYEIQNFIGELGGVVDLFIGFSFFTVFQLIEIVIAAVILRKCRRKTASDSENDERNGDV

>Deutero_Cephalo_091790R_t1__name__chr_scaffold19_start_3687569_end_3692166_strand_pro_len_488_

MGDEQNSLDHEFANYTTCHGAARIANAKNKPHMALWTLIFLAAFGLCTWQIVDRFNNYFAYRTGTEINVQLESELTFPAVTICDFNRIRGSGVTENDLAYLGALYQIAALVSDPVALRNLSESIDWDAFNASAGLVEDFPGFVRRNGFVLDNVTLAECSWRGSRCYAENFTHVFTEYGNCWTFNSEKNSPLRQTSPGAGNGLKLILDIQTHEYTEDPFIGNLEYGLVFQVHDQYEPPRPELVGTAVAPGSHTYASTFQTHVKNQQAPWGQCDPRKDQNFLKYFDKYTLAGCLLECRAEIVVRACNCRPLYYPATFLSETNEFDFCGCTVPCEYTSYSSQVSYAGYPSLKAAEFYETTFNYSRGYMSTNVVLLDVYYHELRSTSWEQFKAVEESGLISDVGGQLGFFIGCSVITLWEFLEYLALKFSAFFLFSSKPSSKTATKDGVKVDNAMVFENTAAGDAEKTEITELSTIRTTSNGEKSAVLEMT

>Protostome_Lophotroco_annelid_Pdum_comp406638_c0_seq15_

MDSKGSKAGIDSMDFPSDTTCHGIHHVAHAPNRAFRAVWFLCILGSVSFLIWQLALLGKGYLAHVALTEVTVQPKAKMEFPALTFCNYNRFSKSLLEKYAPMYNMTVAELWEYLALAITTDNLSLAMNPITTDSLERINATTKKIRQTDGKIDLVALIEKVSSFPPKFLLQAPCTFGSFYSDNRCKPEMFSSVLTPNGLCITFNGNTSTNIYQSNSKAINGLQISLSAEESDTSEIMTWLIGGNVQHGFELTVHEKHEWPVLRPQTSFSPGKRTFVAIEKNVYENIQKPWGNCVNLTLKHFPAYSKSACQLECKMDHVQEECGCHIISFPWNATDRTRSCLPYDMLTCVPETLGKMQSQTRPDPCSSRCLTACKVTEYKTAVTEVKFSTTGAGKSLSDMMNMSSSEFLARSAGVLVHYHSLQEVHYIQKKETTEAELISNVGGQLGLALGISLITITEFVIYGFVHIYKKLFSKRNIIQVQTVKLPV

>Deutero_Ambulac_Apla_gbr81_93_t1

MENTAQKELNTAKNKQKPNLEYEFATTTTLHGISRVAESNGARAKTLWVLVLLIAGCFYVVVTVDRVRAYLKFDANTNVIVEFNDTLEFPAVTICNYNRFMNNKISEDDTKYVRHLLELNEDLVDYEEDFNYTHIDDLFGEGFNYTQFALTTGFTLNESVIRCDWKGVAYSCNALNFSSYYSPSYGQCYTFNTRGDQHSHEQSQQGIGNGLQLIVDIGQSEYTETFKSGHLEAGLKFAIHPRDELPLIGTMGLSAAPGFHTYASMRRVRHINLPEPWGVCGTGETIAVTGDSRKYSRNTCLRICRRDAIIEVCGCQPFGYERAVEPDSSELVPLCDIDRFYCVAHALERYRASFSALDCTCPVACEYITYETTLSMAKYPSQSVINDILSGFPDDEASKAGAVDAEYLDENLVYLDVYFEELSTITYKQVEALPFTALVGDMGGQLGLFLGASIITGAELLDYAIRRLKLLILKVKSAKKIRSGLMRS

>Deuterostome_chordata_Ggallus_tr_F1NU43_F1NU43_CHICK_Uncharacterized_protein_OS_Gallus_gallus_OX_9031_GN_ASIC2_PE_4_SV_5

LLHPVMPSRAQLLPSSSPMARGRRVPALRTRLHGLRHMCSRRAALPRRALWVLAFCTALGLLLSWSSNRLLHWLSFPSHTQVRTEWSRQLAFPAVTLCNNNPLRFPRLSKGDLYYAGHWLGLLLPNRTLADFMDRLGHQLEDMLLSCKYRGELCGPHNFSAVFTKYGKCYMFNSGEEGRPLLTTVKGGTGNGLEIMLDIQQDEYLPIWGETEETTFEAGVKVQIHSQSEPPFVQELGFGVAPGFQTFVATQEQRLTYLPPPWGECRSSDMGLDFFPVYSITACRIDCETRYIVENCNCKMVHMPGDAPFCTPEQYKECAEPALGLLAEKDSNYCICRTPCNLTRYNKELSMVKIPSKTSAKYLEKKFNKSEKYISENILVLDIFFEALNYETIEQK

KAYEVAALLGDIGGQMGLFIGASILTILELFDYIYELIKEKLLDLLGKEEEEGSHDENVSTCDPMPNHSETISHTVNVPLQATLGTLEEIAC

>Protos_Platyhelmin_Macrostomum_lignano_A0A1I8J2R4_A0A1I8J2R4_9PLAT_Uncharacterized_protein_OS_Macrostomum_lignano_OX_282301_GN_BOX15_Mlig033047g1_PE_3_SV_1

MSDSIIQQLNDSLTEISAAIERSVRNSLKNTPGSSEADIAQQVEKLKLTASYVYPLSIPYTLLVSALAHRAEQAVLLSSALKANKNGFAEEFPVPFTQFYHQTQLRCFRWSADNNSELQSIQSRNSLEIAMFLDNLDETSAPNLTGKSVESPVYAEYRKRNTGTPVYIRMPVTDWPLALRDSLRPDRRSGVRLFLMPPGEYPSNFGTSILLEPGKYYQIELSMRQEERNADQFDCVSTNDLIQFLNPYDFEPYSHSYSYSYCYFDKFNENSLAIYNFTTPGIPTTWRLRHSQTTLNMSGPSNILDHLPAEVSKIESTPKNFDEATASACKSRAPCSSLIYDAQISSASWPTSTSLRAFIGTTVADARDTAVLKGISVPGLEFLVNASSESSSMSPAAIAFANANLLKLEIRPQRMDCDKLKFSIDYGFENFVADIGGIMGIYLGMSLLTICELLDLIAELAYTRHRSASVEEPSAVEGEPKPSCLTTQM

>Protostome_Lophotroco_annelid_Pdum_comp404514_c0_seq1_

MDTKKEEASAFALIAAMANNTGMHGLPNVGRSQHIVRKVFWMLIFLAGAGMLIWQVYEALAQYYSYPTKTSIIPKVYNKLNLPAVTICNVNPVRKSLLYQLDALTDPSDRNYENFQTAMNALNVTEKQKIGHQLKDMMLYCNYNGYTCEVGNFTTFYNSYLGNCFTFNGGNRPTVDGLGKTGPYYGLDIEFYIEQDEYEPEYSEYAGMKVVIHNPKEMSFPEDQGLFVSPGAKSQVSLKKIENTRADPPHGECRTFNEEETKARNAYKDSYDYLEYTSKACGKTCYQQYVMSKCGCYDTSYPSTGSAFENITSGKTFSSCKSLNSTYTDCNTAVYYNYTDNIISCANMCPPACTEVLYDMTTSNLAWPSVNALSEIVKKLEDKSTALNTTIQNMTEIEKESFVRKNVVSISVFYGSLEVTKIKTEPSFGLVDLLFAIGGSVGLWLGLSVISVIEIVELLYDISSILVCCCFKTKPNGNNPKTIQVDPQS

>Protostome_Lophotroco_annelid_Pdum_comp407836_c0_seq2_

MTANLESDFASETSCHGVRQVVYAPKNSIRLIWITCITASLCFFVWQFYELTTSYGQYNAVTEVKVKTHSFLDFPSVTFCGYNRFKKSILQKYAIRYNVTLDAIWEYLILSFKTENVVRGMSDKNSSNAAVNLRSLTDKIRAVEQSQMDLKKLVIDASSRFEDVVSSTPCYFADYFEPDFCQNNFKEQLTPDGLCFTFNDNINSTLKQSNIGTSDGFSTVLHANFSENSERLTELVAGKYPLGFLVKVHPKGMWPSLVSGVSFSPGKMAYLTIDKIVYENEKKPWGECVHHNLKYFPVYTEKACKEECIINHIHKECGCHPISFPAQVSENYRVCYPDDMLSCVTVKISQYKKLEQDCITNCKVACHFEEYRPTVNEIRSGSKDFINTISRMARSDYETFIKEWSMIEVYFRELQTLHYIQKEGKKMSDLIADIGGQLGLAMGISLITIAEFLVFSSGFLFGKFCQRCKKNKVNEVKRTLSTTTLTSVS

>Protostome_Lophotroco_brch_Lingula_anatina_comp141642_c0_seq1_p1_comp141642_c0__comp141642_c0_seq1_p1__ORF_type_internal_len_491___score_12_80_comp141642_c0_seq1_1_1470_

WNNGICSNSDFEKIQTDHGICFTFNGGNKPVKTTAQTGAEHGLRLTINIEQYEYMSGPFSSAGLKVLLHEQHEVPLVRDLGQAIPPGSHAFVGMSVIKVENLIPPHGKCGSRELRYYDYYSVPACDLDCEARHVVEQCGCRDFYMPEIYTGHPSVCNLSTYATCLRPSLETYERSAEACPCPVPCNRTLFDTTTTYATISNFNTETWLRSEGLQLLQKKYVKAREIRQKVDQAIRTDDEAKFKDLLEKYEDILRFFDNITIILKTVKSSISTMMEELKPRIAFHSNWGLESSVFTYHNDFIRGWNVMDERTFSHVTTGFFQTHDLFKHMMDCARSSDHNDTCFRPAVFHIVDMNLLTKLSLANRARANLTKVYEAYINGTPLLTFLRTPERHYDMTYVSLYLLHHDLKNTHEHFEYIMRYLGLYIQSLHNLRLVAKQIFENGTHNETLYQTSIDTFISMSKKFLYQKYMIYDRIVAKPKEMLENIVQTFN

>Protostome_Lophotroco_annelid_Pdum_comp414202_c0_seq1_

MSIGYRINLTLSFWTVLSLISTTFLIAKSVHSVNKFLTAKHGIMSSEEAKATDYPAITICNQNHLDREAIHAMEKILLNENCT
YAEGSDLFEVAKHFGLFVEKASSFFAPYLNFPDTSAKARQDFKAVFSNHGFLAHVGEDIALKSGVKMKDFIIACHTGQKE
CLANFTQHFDSSYYNCFTLKLNVSHLNQDNLLLLLLHTGNYEDNLLPRKNRLEQFIPESQRNIHTQKSGVKIFIHQRDRSY
QTMLPSSVNIPTGFVANVKVKITHTYHKRAALKPCNPDNDYVQKICEDRNFSDLTFKECDCVDKSVLDSNTSELYVHNE
HCLHLPYMPKECAFDMRSNKQCSVILQKWWKRMRCLEKIYDRETKPNSGCLPSCEKVTYDFDTNMVSLEHTLTEDME
EKMDYFLRGKNSSLDANNKTRPYSRFSMLMLHVDTSMLNYKEVSNYSFTDIINDICIVLFITILTVAQLITIAAKRLAICRTQ
GINFNSIKK

>Deutero_Ambulac_Apla_gbr256_11_t1

MDVWTVEKSLNRIDESTSADSIKMKGWRKDEGGVNGDGRKRSVHTQTESDRRNADNCRQIFNRRKWSEFGTVTTLH
GMRYITQPNHSFTRKVIWAIILLSFMGVLIFVVIDNTVRYFNYNVTSIVTVKYVNDLTFPAVTICNYNLLRKSYSEQAFDED
LLEAFKIISPGGGQNSVNFTALKDWNITEDLIKGAHQKEDMIIECTWRSTRNCTADNFTQVLTDFGVCYTFNNPKDPND
ALTVQQTGYDSGLYLRLNLEQYDYFYGSSKGAGFKIMLHKQGEFPFVKQLGFAVSPGFETLVTLKFTQLVNLETPYVSEC
VVAEDIQGFKYTVETCQALCKTKYILEGCGCRTTDLPAGPGAFMADLDARCNCSVPCFREVFTPRISEVFWPAEHITKAL
QDAFNSSESFLRKNIIDMVIFFEELNYEEIRQIAAYTFTDLLSDIGGYMGLMVGASLITFLEFIDFLVLGAIEYCQKHNMFKR
RGKINISQSHLHR

>Deutero_Ambulac_Apla_gbr40_81_t1

MEYSEDGEATVKPSRGLPEQTNLSDEGAQSAAAEDGDVPNKWRRKNTIWRMERKSEHLFHQFVENTTLHGVKYVTN
PVHSKKRRAIWMVLFMTMVVIVSCAIVSHFNEYFAYATKTVISYTTPNQTAFPSVVLCNCNRFRVSALNGSQWEELLP
MVVNKGDDRSEQIEKLRSGEFPLNVSDEEERRIKGANYRKLGLALSHEFEDMLLDCRWAGRQRCSAANFSQVWTDFG
VCYVFNAEDGPGAGEEGLMADNTGLDGGLWLRLNVEQSEHTSTNTHGSCFMVAIQPRGEISFVREVGRTVRSGYHSV
IGLKKHGRTSLESPYDTNCRKDGLKYFNNYSIGACRMECLTDMAMEKCGCIAPFMPGLPYCPCSTPCSEVLYDERVSYS
QAPSISVKDEVQQMFNWSREQVLENILEVSIFFEDLREKWTKQEVAYGLTALMSSIGGELGLCLGGSIITVFEFADFLLTR
EALHWRLFTKKLSRFSQRKQRNHPV

>Deutero_Ambulac_Apla_gbr54_18_t1

MTGSASGKTTDLDREFAANTSLHGVARFFDSTRLPVRLLWLSILLGCLGVCVWQITDRFQRYLLHEATTAVSVEYVGDL
DFPAVTICNFNRYRKSALTEADVAKLRAYQRYIDYDYDFYDYDSSVGGTPPPDDISDFRFTNFTLRTGFQMDEDTLLGCL
WQSNRNSCTAKNFTHVFTPFGNCWTFNSGEDGEKDVPILKQTQPGSDNGLVMTIDILQREYTQHLQSGYVEAGLKILV
HDQKTPPPIDSEGSAIAPGVHAFVGVRKIEYSNLEPPWGKCDKSRRLTYYDKYTLSGCVIECRARKIDEECKCRLFSHPGN
AVECTPTQVKDCAIPVIVKLRSGERAGCGCPLPCNYSEFRTSLSMATLPSNNLRDEIWEELRQEGINDTLPETYIRDNWIL
LDVYYETLNYEKYVQSEAITPSALISDIGGQLGLFLGASFITVTEVLHYLGCKAGRWMTTTCIKVDGKPTHESAKEVVMEE
GQQTSGSKDSTLT

>Deutero_Cephalo_045710F_t1__name__chr_scaffold13_start_1042295_end_1051096_strand_pro_le
n_494_

MADDEDISPEKEFADNTTLHGIGKVSGSEKCGLKLIWLAVFLACFGVAAWQITETVIRYFQFNTVTNIKLEFEDELIFPAV
TICNFNRFEENRLEADTQAYLSALLYGFELPEPVGVYYDPVNDVLVNPVRDNLDPVNTTLEAGWQLVKNWNFYVCQFK
GEDCDVVDFEHVFTTYGNCYTFNAVVDPDNPRRQKLPGAGNGLKLYFNIQSEFYTEDPAQGGSDDVGLKVLIHDQKEP
PKMDTQGIAVGPGSHAYIAMSKVEYINEIPPWGECQDLKLEYYENYTLTGCLLECRTKHVYAQCGCRLFYLPGGNDYCD
PRAIADCALDTLKAVTAGNFTCDCPTPCDIAHFQTSVSYAGWPNKLTAAHWLTELNLWDVWDEQSATDYMRQNWIV

LDIYYSELNYQVIEQQREMTEGDLLSNIGGQLGLFIGASVITLFEFVEYLIKRFTSCCRKKYRKDKVQAVDIKPARVDVKPA
TSRFDTAHPRHPATDGSMW

>Protostome_Lophotroco_annelid_Pdum_Contig6895_

MTRVGEPSRDPPVVIASQGGANKPTATKNPEEETLRKILDDYVEHSSMHGIWKAAKTSPFSRAVWTVVLLCMICLCLFT
FVSQLLFFLQFKTNTLVTVIEQPFLPFPAVTVCNYNSFRKSKALEKPSITKVLEQLFSLDRIFHINVYTTSTAEQSDDVDILKE
LIDLAHTLEGMVVYCFWCGMRVDCKQHFVQTMTDVGVCYTFNSKNTTGDTPMLVRDAGSDCGLKLILNIQRHEYFYG
QGDSTGVKVLVHDPKEVPLVGQQGFSAAPGTTVIASIKLQQELYASSPYGDCVDVSKRDFKNPLQYHDEYTYTGCKME
HRVNYIYSSCHCVQYFHPVGGREICTTFDQVECSRKAQENHTMYQLEQLSQDCPVTCEYNTFDEKLSWASFPAQHILD
DLQTTFNMSAQAISDNLVQLHIYYENLAVHRQEQTPAYTVGALFADTGGEMGLCIGASLLTVVELFELCGHLAAFSWKK
ITRHMFYKKPRSSLSKPI

>Protostome_Lophotroco_brch_Lingula_anatina_comp140967_c6_seq14_p1_comp140967_c6__comp1
40967_c6_seq14_p1__ORF_type_complete_len_494___score_28_99_comp140967_c6_seq14_162_164
3_

MVALKGVNKIRTPYKVWINFLKWTTAHGLPHVVRSHSSYRKVFWLLSAIMCLVGLIFQLEKIAVQFLTNPYAVSTYMEY
AVELQYPAVTLCNLNPVRTSVLRQEAKTGGRLDNLLAQLYGKGENATEPKEILANELMWSWLQFDDSTKARLGHRIED
MLLGCTLHGQTCAPENFTLLFNSKYGNCYTIKPLASQIHKPGHSHGLTVELNIQQEEYLPVIAEAGVRVVITDHKSVPFPE
DNGLSVSPGFYSAVGMSMVEISRLGPPYKSNCTNGFPTLYTGDTTAGSGYNYTVHACMKSCVQTVTIEECGCSLMNCP
NPNKTRLCAINTNSTDYECTQRMHRQLASRSYDACSCPQRCSENLYHLTVSSATWPSQSYRESAFLRKEVLSAYNSTLSP
AENLLKLEIYISDINVMKIEEHPSYLAANLFADVGGLAGFYIGASVFTAGEFFEFAMDLVIAFIYHRGSCRVPRSGQRKKGS
EEHDLEQRTTVEQYV

>Deutero_Ambulac_Apla_gbr256_13_t1

MYAWTEGTKTPNLLGDQESIFSIGSPDFTVRSTIINMEDKETKGNNCRSQTCSADVDRRSFADRFREFANSTTFHGISNV
TNSKHNGIRRLFWSLIVIGVSSWLVTGITQTVIEFFKYPVTSAISINYVDSMTFPAVTICNFNQFRRSVIPDDQVDFINKLY
GLDPEAPTDLNISDFEITGLTRDQYEDLFAAMSHQLETMLSECVWRSSETCTVQNFTRRFTDHGICFTFNDPKNGSEILQ
VNSAGTRNGLFLRLYADTEEYMFSENTAAGFRMLLHPQGVAPLVKELGFSVSPGFESSISVRQTLVESLPQPYESNCTNS
TLRYSDTYTVPSCRFECKVEYVVDKCGCRDYRWPGPAPICNPQEQFKCVYPQEAEFLRQNVECSCPVSCETKTYESRISL
GYWPAGHVTEFVEAHMNLTEEYVRKNYLDVYIFFEELMYLKIEQKEAYSTSSLQSNIGGYMGLLCGMSLITVLEWADFT
LVSIIRRFRSRKE

>Deutero_Ambulac_Spurpu_015892

MGNIEKEKLNELDDDTVVGAVRVFSNETTLHGIRYLIGRGHLLIRLCWLAIVVLAFALLVMQAQIVYQDFRSSPYSTKIDI
VSEVVKTFPAVTVCNDNKIRRSQLYNTKYEGLIDIDDGGRSTEGRSSTRDKSSDNVWASVKDDYDWQGFYSASSADDF
SDFINVVNPTKAELNDYGHTLEDFVIQCTYNQKSCNLSADFTVSQNRHYGNCFTFNFGDGDDEPPRVTSKTGALSGLHL
SLMVDEEEYMGLLAPNRGAMVVIHAPNTVPFPEDDGINLETGSAISIGIREEVIQRLPKFSQCVVDRKIGTNTSISGTNYT
VRCCLKICYQQSLLERCSCVDDILLDYRQCSVTNKSEGDHGGHGSPVYHLSIDLLYLECNVPPAYRGLKFKKGKRDVRVG
KRVLNHSMVYFRGKTYEFGANSLVLHEGGVGRTNPTSCPTVRKTTAGRSLCTEYQATELAVRYHQDHNFNLFSKNCH
MFAEWLINRLLNNKCTI

>Protostome_Lophotroco_RotSoc__A0A820XFQ4_A0A820XFQ4_9BILA_Hypothetical_protein_OS_Rotari
a_socialis_OX_392032_GN_FME351_LOCUS29310_PE_4_SV_1

MSSSKTTTEIVKTWLFNAPSHGIRPISRAKTLKTRLFWAYTFWIFTVLMCTFIYTISMNHITNPMKIHLSIKHYRDPAHFPA
ITFCNINPLQHDGFNDDIRKSILPGGDPSSRVNETEYRKAVSDYVNRILMAHVNGEVDPFVKWGFKLNDLLIRCIFNKRR
CHRNLTQLYHPDYGNCYTFDNDHDVEDQELSDIRHDWSVDDDNGQDNYKLFLELFLHQMQYNEYFENRASFRIFIHR
KNELPVLSQNSLFLAPNKYTKLSFSQSIKSFSRQCRTELTDEMKNIFRTHNIRYTQALCYKLCEKRYLEQRCHCVERTLAVF
YQFFNDKNTNNAHTTRKSSICSMGEPCLNERIHFNSSESCQECLPECEFIQYSVQSSYAHYPNMKANRHVSKRVQQHFI
RDKTNNESLSGTQLANVYMGIRDNIVAVEISASSDPTEILTESPMYTWVDLISSIGGQTSLWIGVSLISLVEIAELLYLLLNR
FYSYIFPMKKN

>Deutero_Cephalo_070040R_t1__name__chr_scaffold16_start_1701523_end_1711747_strand_pro_le
n_496_

MSEKRPSVRSTLRKYGENTSAHGIPRAVTTKSLPRRLFWTCLFLASFSYFLYQAQTLVNKYLVYPVNTDVKIEWSELEFPA
VTICNANPLRYRELIKRGSAAFQNAGFIPKPQPQGQNNPNSNGSTPATGSTTVATAPTTTQNSTTRKSDRLVHPFLFQS
DEDDDDYDYDGYHGDFEMVNSFMGLLVQLTASMRQSLGHQGRDFIQECQFDGRTCSHRNFTTFEDSTYGNCFTFNK
DKDGEVLHTATSAGPLHGLSLILYIEQDEYIPAIAEKAGARVVIHNPYVFPFPESEGFDAAPGFLTSAGLRLNCLATCHQEH
MVEICGCADVTFLQPNGTEFCNSSKSVGQDRIYRKAIGLSEWPADSYVSTVLNKLKTKRRGPGTGILDDRDEFKKNLLKL
NIYYEALNYETITESPAYEVENLLGDLGGQLGLWVGMSCLSAMELLEFLVDIAIILWKKMSNREKTTSRNNVVNVAETGK
IGTSNGHTIPIHMTEVM

>sp_Q9R0W5_ASIC5_RAT_Acid_sensing_ion_channel_5_OS_Rattus_norvegicus_OX_10116_GN_Asic5_P
E_1_SV_1

MEHTEKSKGPAEKGLLGKIRRYLSKRPLPSPTDRKKFDHDFAISTSFHGIHNIAQNQNKVRKVIWLSVVLGSVSLLVWQIY
SRLVNYFMWPTTTSIEVQYVEKIEFPAVTFCNLNRFQTEAVSRFGIIFFLWDIVSKVLRLQEISGNNTGSPEALDFVASHR
NFSITEFVKNNGFYLNHDTLVHCEFFGKTCDPKDFKHVFTEYGNCFTFNYGENVQSKNKVSVSGRGLKLLLDVHQEEFT
DNPVPGFADAGVIFVIHSPKKEPQFDGLGLSSPVGMHARVTIRQLKTIHQEYPWGECNPDIKLRNFTTYSTYGCLKECKA
KHIQRLCGCLPFLLPGNGVECDLLKYYNCVSPILDHIERKGLCTMGTHNSSCPVPCEETEYPATIAYSTFPSQRATKFLAKK
LNQSQEYIRENLVNIEINYSDLNYKITQQQKAVSVPELLADVGGQLGLFCGASLITIIEIIEYLFTSFYWVFIFFLLKILEMIQR
TSPPQTV

>Cnidar_Rhopilema_esculentum_TR59902_c0_g1_i1_p1__Rhopilema_esculentum_TR59902_c0_g1__R
hopilema_esculentum_TR59902_c0_g1_i1_p1__ORF__type_complete__len_497__score_41_10__Rhopi
lema_esculentum_TR59902_c0_g1_i1_1078_2568_

MKQMDETKDRRSFFERFKHFSEIATIHGLYRVGHSTGWIRRVGWGLLFVSLLTLVTVLTASSLMQHFSYPILTEVNKETT
SSMKFPSVTICHANNYRLGAFRVLSEGFDSDLSQLITSTIEKSLKNQSIKGLEESDVTFALDSVYLNSTVIKMVSSTVDSMK
WKFEKWCTFSMMDECKFPQDFTDYYYHSTVGFCKTFNFHGNYTQIAPGLPFGLSIKLFIDQWDKVPLLRDSVAGILLMI
HPQDVYPNPFAEAVQVPTGYETHISMRKLLFHRQKKPYKANCTNGDGIFLIYPGRYTVINCQYSCFVKNLMEKCGLVES
AYQFHSPKEFAESLKRFSARNMSATEVKKCILETVKSLKGPKDECICPAACFEESFNTKVSYSQWPNPSNIHYYRTMLANL
TNRENVTTDDVYKSVISLKVYFDELGQQVIRETPVSKWTDLFSKVGGHFGLWLGASIFSLMEMIGFTIKSIICLFKKKVYP
DRENPIKIKVKMRE

>Cnidar_Rhopilema_esculentum_TR90022_c1_g5_i1_p1__Rhopilema_esculentum_TR90022_c1_g5__R
hopilema_esculentum_TR90022_c1_g5_i1_p1__ORF__type_complete__len_497__score_69_36__Rhopi
lema_esculentum_TR90022_c1_g5_i1_126_1616_

MDNEGDEHLSSSTDESAFDEEYCDFQQEEVFKGIEWILDCDDLTESEYLRYITRFLAEMEDHARRLDKQTRQRKIHIYKR
SLISKSTVHGLRNCFDRKSRVRRFIWTILLLTAVGLLMQKLYECTMHFISHPFSTTTTVRYEDSMRFPAVSICNLNDMRTS

VMKGTKLDMILKGDRNVSLSGDEYRNTIRKANHELPDMLYKCKILDKECSVQDFIQFNKDQGDRCFTFNHGHEGQQIL
FFNKTGPTHALELILNIEEYEYYPGTDYSGIQLILHGQDETPVKMLGVMLSPGFITYVQVKRRKIKNLPPPYKSNCGSKKLK
YFSERYSKHLCWLETLTDHVIEECGCKEWFMPGGYRVCSLNESKECMWPRWAEFDKYMKYDCPLPCVMDTYTPSLSL
SQFLLDRDVDEIAQDPRFRGAKEMNRKTIRGTFVKAVIFYGELSYEYSEQVPSYDKLDFLADIGGQLGLFLGSSALTYMEF
IDCLVMIIYTKYISVKP

>Ctenoph_Hormiphora_californiensis_evg134569__p1_GENE_evg134569_evg134569__p1__ORF_type_5prime_partial_len_497___score_60_08_evg134569__2_1492_

PDIYQVTIVEKNRPWWCKMIWVVIGLAFFGAAAYMVYNLMTSYFQYSSYDKINTSFTKNVTLPAITICSKNPYNYTRLKE
YVGDDELLLLEELMYEMNVYNGTGSNGSSGNFFYDVPNVQDIVDYLDSLGDYSGDDYLKYILEMVSFFPDDFVVGYYY
DFKMGKNWKYLNMSSIADSNVTPHMLGYCIILNDDESLVQVVPGIHGGFTIMLDAYIDHYMPNVAEKGFEVFLRTPYE
EVMIDRGGIIAEAGKETNIMISKKKMTRLGSHWGTCESIPSPYYPNKKNVHRSQRECYQEGMIYESQLSCNCTPWYYVE
RVNMELNLNWTSSLNVSTCSFASEIICDTVIYQLMETYNKDEWFQCQNERCSYNTWDYSLQAFEFPPTEDYYDLLISPIL
KNEMYTMEYAQKNIAKINIYYGDLKETTVIQEKAYEVFSFIAEFGGGVDFFIGISFFTAFQALEIAIAYTCCRGLRSRKLTEE
ERKNNYKENNVNSVDTPK

>Deutero_Ambulac_Apla_gbr173_6_t1

MGCYKRCYLRQWALTDTDLHGVKHIAGEGGILRRLIWAACFLAALVAFLHQATLTLIHFVEKHHVTKVDISYRKKLDFPA
VTVCNFNKYRESALTDRDIRNVGYHLGIVDEDHNLINPYLYTEEFRRKMAAVDWSVHDIDDEYNMTEFTNRTGHQLDE
MIVECSWRDEPCSPDDFHHIFSHLGNCYTFNHVALATERHSSISAGAANGLKLTLNIQEEEYTPSNDLHGAAEDAGIKW
MLHHPSEPPYVKELGFGAGPGDHTFVAVRHEEVESLPSPYTPCMETSAGFLDHFDHYSLQACRIECETEVVVQRCGCRL
VEQPGNAPVCNPAETHECAQAALVHAVAGHGDQACDCSSPCSVESYPFTTTNVRLRAKYIERIYSNTTHNFSADYIQN
NLVLLSIYYEALNAEVIEQLPEMTVPSLLAALGGNFGLFLGASVLTIVELLEYVFDELTASCTRRKTSGRISNAREILTVRAAP
ISVIDTEKELEHGRQWNR

>Deutero_Ambulac_Spurpu_006217

MAYDERMNATSVRHTTTNSTMDGRPPKSEVYAMVKPMLENTAAHGIPNIVRAESTPRRVAWSFLFVVALCCFIGLSG
NLIRKYYSFDFNVNVEVLFEPSINFPAVTLCNMNPFRQSSVVNASLELTEILGLENDPMSKWNSSTNALYGDWENLEFSS
LDAQTEILISAIQVVGNMSYEQRFDIGHDLGDMLLSCSFHGLPCAPAMETGRTSNFHRSMLKLRSSYRQSRSLATCLTN
NGLTSGMISVICYSVVRSMVYRVLQRLAIELFIDQEEYISTLQSSAGIRIIIHDPNEMPFPEDSGSTLAPGRQTSIGLTKVAV
HQLDYPYTTCTNDFPNDNIFQERFPHTIYSVLACEKNCLFKYIKQECNCADARYRYNETIETCDFDLNEAKIRTKNQFLKQ
FGQTMLIGTNVVKVIVYFSSLEYENFYQTADYTVYDLISALGGQVGLWIGVSVLTVFEFVELLYDILKFGCSKLTKLGKRTIF
AKDSLEMGPHRT

>Protostome_Lophotroco_annelid_CAC9633462_1__Ofus_G091394_Owenia__fusiformis_

MEVMTETKEKQTDPVIEHSFSTRLKEFGEDVEFHGVRHLFRDNSIVTKVTWILLFLCGCSWLTYQIHDRIIYYYQFPHITKI
DKLYVPSLDFPTITICNINTFRRHKLNEYDLLHYGTSLNILDENWTLLHPEHYDKAFDDWVYSINWTDVEIHDRSINHSME
EMYERAGHQIEDMLIYCKWKQQECSVANFTLINTHYGRCYQFNSGKDGVKHQSFKGGKANGLKLYLNVEELEYLNYM
EASDLGFKILAHDQDEPPLIQELGFGVTTGNHYFIALETERVTSLPDPYGNCEEDHKLDHYDHYSIPACRIECETLIVEEKCS
CRLVEMHGDHGIRVCTAEEYHDCALPTLESITESDTCVCQNPCELTQFQHSISSVKLRESAVEMIHGHTSSNINLTEFKSV
EFIEKNLLVVNLFFDSLNYQYIEQTVAYPGVSLLSDVGGQMGLCIGASILTVLHLVQFAIGEIIKKFQKRENKEVNTNVIHV
ASANPVDDKL

>Deutero_Ambulac_Spurpu_015310

MGCRSGRCHLRKWAADTSDLHGLKHIAGSGNLFRRLVWLCLFLTALGFCIYECYLVFVGFLSFNHVTQVDVLYSTEVEFP
AVTVCNINKYRESAFTEDDIKNVGVHLGIIDEDHNLLLPELYTDEFRDFIDSVNWTVVEEDPDYNMTDFTIRTGHQKED
MILSCLWKEEPCEESDFQHKLTHLGNCYSFNLQGANSDEADWKHSYAAGAANGLQLILNMETPEYTPTNDLDGGAM
DAGLRIMFHYPTEPPYPKELGFAVAAGDHSFISMRHEKITSLPDPYSECESDGADIITHFDHYSLQACRIECETLVVVAEC
GCRLPEMPGDDDVCGPADLHECAHPTLVEFITGNLDSAESCECFSPCEVENYPFTLSTSRLRLTYLEALFANTSTNFTAAY
IGENIAVVSMYYEALNFETIDMLPEYTVATLLAVLGGNLGLFLGASFLTLAQLGEYCFDEVIGWCICTPKKEDDDKNDGG
NKVEPVSTVTAMDAWGAQKL

>Protost_ecdy_cyclo_nema__A0A0V1BEP0_A0A0V1BEP0_TRISP_Acid_sensing_ion_channel_5_OS_Trichinella_spiralis_OX_6334_GN_Asic5_PE_3_SV_1

MPQVEVYPCKCSGNDGMTKFCEITTFHGLRDWHNADRFYLRVIWTVVLIGALSAVAFGCYSILNEYISSPLAISYLVKEA
DEDVHVPDVIVCPLNRFNGKYLKEKNVSDSLAWYMQLSFGLATKHPYMRKFMIENILSKTETLKVEAQQMLDRLGMSF
EMFVQEASFKCSDVLIQCQGPQGIFSCCYDSSNATLMTFAGNCFRIPGVLQQLPGYGMGISVVVHLPKDDFHIAPNNF
NSDGILVKLAERGTEIGHDMTFIPTGSYALLPLRATKYDFINNPPYFECKPDDPSYGRVSCFDDCIYTVAEKKCNCSHVAL
GSQRRYPPCDTDQFLNCLQVELIKQLKEQWEEIFNICRSRCLLQCVRTQYTAQVSYLKFPIDETRRFVESDEQWEEMKN
SIVLDIYFERLGFTLIKHYPSMTEQGFLANIGGQYSLWLGGSFLTLVQLILFITRCIWDQLISKCPITNSKRLANHGPTSDVH
KFYHFAPFKSGPKLQILII

>Cteno_MneLey_comp15203_c0_seq1__p1__GENE_comp15203_c0_seq1_comp15203_c0_seq1__p1__ORF__type_complete__len_500__score_87_69__comp15203_c0_seq1__52_1551_

MKKVTEMSWVKDFREFANDTSAHGVKYIFEGRYKLVKLLFLVTWLGFSIYACHVIITSIVRFVEKPTSTKYEVIQDDSEGR
PERIEFPTISVCSMNKVRKSYLEAPENEAIREYYEVVDKYNVSLVKDLAKRFKNSDDPLHSIKDMTYEELIRNGGPNPDRLL
KCTQRAKYCHELPAFNGRDVSVMENSMTGNCWRVNPEGRLMGKMGDYGAMKLMFWADVQDYSARTADVENQG
FVVAFHDNSTYGSTMTAGFLMSPGTYYKADLRRKQEIRNRDKIESCNASLTENTYGAYNEGSCALECKDEALHKACGCT
NVVPPLNNGKYKSCTLEQWVDCGLAVYKEWFHNFTDTDRADQLCPCQIQCEEVRYEAQISSSSISPAFAEKLFPSVQPII
SQPQYGYSNPDFNILYNTTQDILDNVMVLEVLFTSMRTNEIKEIISYGLSNLLGDIGGVLGLFLGASLFTILEVFQFVFFSISK
YCFDKGQPKHDSLTNGDKLSI

>Cnidar_NemVecNVEC200_006433_1_1_protein_AED_0_26_eAED_0_26_QI_422_1_1_1_1_1_6_1527_500

MALCGCCASKPSPTNSASEDESTGPESKEEILLANQNGNASTYENIRKRSAAKRDKKHAEYGKRKMREVFAYYINHCTL
HGFHYIFETKSLFRKIAWFSVLAIAGGFFFKEIKTSTTQYFKYPFSIMSTVEYPRTLVFPAVTICDFHDIRQTLVSNAKGEQAI
NISSSEKLETLARKTYKSFNETLISCSLRRGVRGARPFNIHDFKVFFTAKGQTCYTFNAAMDGKKLLEVDNVGPKFGLEIYL
NAQHWWFKDDVRESGFRFILHQQDDLPLTREGFRVSSGYVTYVDMRLEKVKNLPYPFSSDCDGRGLDLYPKYSRNNC
YMESLTKYILQQCKCRAWFMADVINTSTCSIKEALDCMWPAWEDFDNAYNYTCPVDCEERVYKTRLSSALFLPQKLLPL
TKKYKFLMRPKGIPNDTEGAVDFILENYSVINLFFDELRLDTIQQTPAYGFFRLVGDVGGQLGLVLGASVITIVEIIDLVIMY
SIYWIKSKTVPRAATS

>Cnidar_Rhopilema_esculentum_TR40535_c0_g1_i1_p1__Rhopilema_esculentum_TR40535_c0_g1__Rhopilema_esculentum_TR40535_c0_g1_i1_p1__ORF__type_complete__len_501__score_55_33__Rhopilema_esculentum_TR40535_c0_g1_i1_597_2099_

MDSKTEMATLEREWTEQYLEYYVKKYLEEYEESTSDTSEGSDYSMMDWDFQKKGLLERERLWLKAKRRKKITKHFNT
MVKYSTFHGLQFCFKKESPLRRTVWCLLLLICNGLLVQKMFESTQHYLEHPFSTTKSVKYKESLTFPAVSLCNLNDMRHS
RMVGTKLHKLIIERQQNISKQLSGNEYMNTIRQANHRLENMLYSCTMDGVKCTSEDFSLFYHKQGDKCFTFNSGRPRY

KLVKTNRIGPQHALELTINIEFWDYYDDAVQSGIHLILHGQEETPVTMEGFRISPGFITYAEVKKTQRRNLPSPYKTQCGS
LKLKYFKGYSRNLCWLEQLTDDVVGHCGCKDWFMPGDYKVCSLTELEMCLWSRWVDFDKYKKYSCPLPCVIDSFTTKT
SFSRFPTHDGSESMAKELNLNGTSRDNHKFLSDNFLKVVIYYGDLSYEYLEQKPSYDLLVFLGDIGGQIGLFVGASVLTFFE
FIDCFLMCIHARFFEIFRPRETQL

>Cteno_Ple_Bac_comp45500_c0_seq2_p1__comp45500_c0__comp45500_c0_seq2_p1__ORF__type_5
prime_partial__len_501__score_51_82__comp45500_c0_seq2_3_1505_

SKLKLMDEYTFVNAFEDHVGSIDYVYANHLYGTFVGEYDYTFSGSGNVISTGEWDKLQGMTQLGRCLEVNDDKSFYQT
YYSDSAGFTIDASANIADYLFSTRTQGFKVFLRTHDETVMIDQGGFLVQPGTETFFKISAKEATRLGSHWGTCQNVPSAY
SKYGNDSNPHYETVRECLQRQRIELYATKCSCMPWYFVIRMKTQEKQEVLNEIIERIKRGADNTTENHDRKRRKTRNTL
RLPKNQSTGKKSAVREGMNDRRRREIDTYTTSWYDTYTTATPGPTEYAPFWSDKSESSDYTCSYAQLLICEAFIEEEISNG
NITLEGCPEPCEYLEWDAEVASSSFPPSQRYYNEYIKDDWYIETNTSYSYAQENFARIHVYYDDLKTTTNEQVKAYEIQNF
IAEFGGTVDLFIGFSFFTVFQLIEIGFALCLYRWRQRTISITSVETGKGVTPEKFESENPKLVSLVPHSSKSQKIETQKVEYRY
TYLDLFVHCLKSWFIKF

>Deu_Ambulacrar_hemi_Ptyfla_40v0_9_20150316_1g17738_t1_scaffold14937_cov156

MPSAWCYTTNRDVGYELCDLKIDRKECVTDHVLNDFNVSFAPLEDLEYDQQYEQMTEAQCAAMCLEYSAFVCMFFDY
SVQNHTCYLSSNSSDQSQPSTVQQDIVRYIRILDDTEVEILHLKFSRYPNKKMIEGGLQRKVIAGSISECITACIEMDIFKCK
SLDFSPSQSTCNLYEKPSANVGGLMKEDISYIHLTRQGEYYDHRTVTEQFLDISGITPEILNDFENGYYRDNGFSGVVKGE
VPPDWYGFKTFSSTPDYSDLRSVLKLTGEEIAKYGHQAEDFILQCTYDEKTCSHSDFSTFQDDHYGNCFKFNHGLNGTV
VRDANRQGALSGLRLTLFLEQNEYISIYGRNAGARVNINPPNDTVLPQDEGITIMPGTVTSIGIREKHTFRKTKPYGKCKD
GTNEDNLYGPGNTYKVLIEDFLSDIGGTLGLYIGLSVITVAEFIELVASVIRYMYCSQKDSNPRTAESENKDRERRFSGAYA
YPSFGRVDINDTKVAKL

>Deuterostome_chordata_Ggallus_tr_F1P0L7_F1P0L7_CHICK_Uncharacterized_protein_OS_Gallus_gall
us_OX_9031_GN_ASIC5_PE_3_SV_3

KTNKSHLISEFSIIFHCLSLDKMLPRLEDRRKYRQEFASSTSFHGVYNIVQTQTRTRRVLWLLVVTGCLGIVIWQICSRFNY
YFSWPTTTTVVVQHVENVKFPAVTFCNLNRFQAHAVSNLQIIFFLWNIVSGIVQKFAMEDKYFHELNGFLLGNQNFSIK
EFTRENGFYLNGSTLLECEFFGKTCHPEDFEHIFTEYGNCFTFNYNDLPARRVSLSGRGLHLLFDVQQEQFTDDPALGYT
DAGITFVIHSPKEIPRFDGLGLLTPVGMHAQVSIQQLKSIIQEYPWGECKPDIKLQYQDTYSTNGCLKECKAWYIQDWC
GCLPFILPGNGIECDLMKYYNCVYPAIHDIEVKGLCTVGTHNSTCPAPCEETHYPTTVTYSSFGGENAIKYISAKLKKSPEYI
RQNLVIIDIKYDDLNYRITQQQKALTISELLADVGGQLGLFCGASMITIIEVLEYIFTNFFWMCLFLLLKAPEIPRWNNPSH
DQPTHVEKNKGIQEC

>Cnidar_Rhopilema_esculentum_TR106708_c0_g1_i1_p1__Rhopilema_esculentum_TR106708_c0_g1__
Rhopilema_esculentum_TR106708_c0_g1_i1_p1__ORF__type_complete__len_504__score_95_58__Rh
opilema_esculentum_TR106708_c0_g1_i1_88_1599_

MMKSSEKKENNRNEQGSVKKIFDDFANSTTAHGIQHLVSSKKTLGKAVWLIVCLAALTTNLFLCAQLVKKYLSKPVSTTV
GVRHEQTMIFPSVMVCNHNMLRKSEFQKAKRRKLINFKQDASMMSLYKEMMLKAIGMANQASNTSSVNISVNTDIIE
RRCLLEYLSNIEKPQLKGLGHQFHNFVVRCDWKGIDCKSGMLENRWYKVWNHIYGNCFIFNYGYNMERKKQEPLRLE
DVGMQNGLTLDLNIELYEYDEELTKETGVRVLLADQGVFPLPSIQGFSVPPGVHASIGIKKVEAVRVDPFNNGSCRRSDS
LDEDGDKYTMQRCTQRCHWQRQLKKCNCTGLRLSSPNRTCLTVEEHMCKAEVSTEMDALGINKICSTKCPPPCFEAKF
SHSVTYSTLYTAALLKLKTSIFRNDDDYRKNFVRLQFFFESDIVEFIETYTSYEMEQLLSDIGGQLGLWVGVSFVTCVEIFLF
LYSFSYHVLKKVWCTSFKVHNNKQTQEN

>Deutero_Cephalo_292990F_t1__name__chr_scaffold9_start_4371418_end_4385499_strand_pro_len_504_

MRNVVGHKIGGTVACQDSNSGPLGSESTTLPLRHTTPQNLFLNPSFMATCCGDDSVEREYAAETSLHGVGKIAGARHRGVRLLWAVLFLGMFGVATWQITERFVAYFQFDTVTNMKVEFRDVLDFPTVTICNFNKYRDSQITEEEEQYVSTLLEGSTGFSDYDYDADYYDDDGDGEYQYSYDWNNTGIPDDFNLAEFTLRAGFDIHTSLKHCLWRGQTCNSDNFTHIFTSFGNCWMFNAEGRMNQTISGVGNGLQVAIDIQQDEYTENAPTGNLDAGIRFIVHSPSEPPKVDTQGISVGPGIHAYASISKIEFRNEIPPWGQCDPGRALQYYSGYTKTGCLLECRADHVADECGCRTVSMPGTLDYCQPAVVTGCVKTTIAELKTGKRSCNCPTPCSATAYPATLSYGGWPSSSTMDYFTDLLNKTEQEIKDNIVLLDVYYQQLNLQTVQQRRAISTNALLGDLGGQLGLFLGASVITIIEFLEFLVKKGTSCCFRHNNRVKT

>Cnidar_Rhopilema_esculentum_TR42024_c0_g3_i2_p1__Rhopilema_esculentum_TR42024_c0_g3__Rhopilema_esculentum_TR42024_c0_g3_i2_p1__ORF__type_complete__len_505__score_64_02__Rhopilema_esculentum_TR42024_c0_g3_i2_373_1887_

MKFKVIPCTKDVEMMKRETSKERDDTEQAKSRMPVDVKFSAKPHILFKEQFYEYLNSATTHGLNKIANAPNTLARLLWSIIVACMFFYLIYAVYKLHLKLKRDEVKVSVRTNVVNSLQLPGITVCDAGPIRMLFPAEADKNTTVSNKTAKTPLEASNEHHRRMLATKDQSNYSNHSHADANRFVLKRESGEMPLCHVSGDLCSYHKHLKHFVTIHKGHCFTFNHDGKARQVLKGSDMGIFTVFHLEKKRNYTSSASTTSLHGVEIFIHEPKEKNPTHYGGLLAVPGRLTRIKLKKTIHERLPAPYPDKCRDEGDLSLHECQINCIAKNQYTKCNAVSLLLQIYLHDITKQEWNNGSTIRQPMTTAEHECVEGVTDSYADGDITCDCFLPCHEIVYDTSVSSSKWPLSEITDEILLLIKSKLGIVVEKENVDQQLVAIQVYYGDLRYEEIKQHPAYSLEEFSYDLGGLLGLVIGASVFSGIEMVMFVLYAIQSLLKNIVNRGSVS

>Cteno_Puk_Fal_comp20088_c0_seq2__p1__GENE_comp20088_c0_seq2_comp20088_c0_seq2__p1__ORF__type_internal__len_505__score_74_57__comp20088_c0_seq2__2_1513_

AEFTTAHGVNLIFVGGHKALRFLYFVIWLLCVLYAIYIILQAAVAFISRPTGTTEQLVVETDHGQDIQFVKFPTITVCSTNIMKRSFTETDKSISALWYLISRLKVMDPQYVSHVLEHPKRFFQSYCDYHGVTFEHLKNYTVKEMFERGGPDDNRLVYCKQRNIDCQDLTESYFVRETSSDGMCWRVNPNGSMYGRSGEWGRMELFFFADIQDYTSTSPVYGYKVKFHDSEYCGEGITGAHFMSPGFVYKVDLGLTKRSWIPKPIGNCDANAKKSTYGKNSEESCLKECEDRYMNKVCGCVDLLPPHNHHNYTACTIGQYIGCGFREMVHFRMGLRDASRRHGTGEDGCSCSHSCDETHYDVKLLSTSALSPVWADDLSRDKWIAEAVANFSQPSFNISYNVSSDLLKNIVILQVEFSTVFTKEFKQIVTYSYSNLLGDIGGVLGLFLGASLFSVLEFFQFTLTSLHKHFCGREDKPKTLPETEKLSSNLAFSPAIRSTPVLCV

>Protos_Platyhelmin_Macrostomum_lignano_A0A267FRW6_A0A267FRW6_9PLAT_Uncharacterized_protein_OS_Macrostomum_lignano_OX_282301_GN_BOX15_Mlig006976g1_PE_3_SV_1

MSISSKKSSAMERDSILSLLVGNTKPDMKNLVKLRRRLKLRELALAHTRLWSERTTIEGVPHLMAAHSVQRKFCWAIIMLTVTILGLVHLVRSVQDFLASGVASQVTVDAAPAEFPSVTLCSLKPASESLLSYYRPVMDRHLERHWGYFDAWFTHQRRQGKTESYRDYRLRIVKVLAKMLEVLWVSDDTRDLSLLDYSTILECQFNGQPCNKKNFSLVQHYRHWNCYMFRPHSSLRYDNPTGEHNAGLMLNIFADSNFFSSTSSSHSEYNRSRRDSNRDYRIEGLSLRQFLADDSEFRLNRLLDVNHLSSDGMQVILHPTGSYPMFYKTRFTVKNSESAILHVFSMERNRINQADSPCSSRPLGTIDYVTNFGIGPDNQKRREFTESVEDVLNDRIQDLIYQRCGCYSHRMPFRVGKSLLCYYAPLQYRKQPSSELLTQIYCHDYWYQKFIGISELDDVSTVVRPCHETYREMAHGRGPGIGDHFVLSTPVGKRHGIVRTERPP

>Deutero_Ambulac_Apla_gbr54_16_t1

MTASAPDTETNLEEEFVSNTTLHGIARVAHNSRTIFRLAWLAIMLVFLGVCIAQITDRLQRFFMYKANTAISVEYVGDLD
FPAVTICNFNRYRWSELTSSDLRSLNYILEATDYDYDSYGEASSSSLSDTYPLDDDNASSAFNYTEFTLRAGFLMDDLTLLS
CEWRGKTNSCSAANFSHVFTSFGNCWTFNSGSDGGPLLKEIQAGAGNGLRVQIDIQQNEYTEPLSGNREAGLKVLVH
DQQTPPMMDSQGFAIQPGVHAFVATRRQQFLNLEPPYGKCDASRTLNRFPNYTLEGCNIECRTEKIIERCNCRLVRHP
GTEVECTPMQTSECARDVLSKLRSGEIPACDCPVPCNYTTFSTSLSTATLPNQRATQLLVDQYDDLAQYSSNISDTTNTG
SASSAGQSNPILLDSTYMQNNWILLDVFYENINFQRYEQSEAITPSALISDIGGQLGLFLGASFITLTEILSYLGRKLGSILAS
RRPRRAATSPDTSPGWDMNGVKIT

>Protost_ecdy_cyclo_PriCa_ecdy_XP_014662731_1__PREDICTED_degenerin_like__protein__asic_1_Priapulus__caudatus_

MVLVSHIHNFASNTSAHGVARAFNSVSIRQRVFWGFVFVVGLSYSIFQTYRILEQYLRYPRNTLIDVEYVKYIEFPAVTICN
KNPYSQAVLDEVIHYLPDENDYDWSNYDYTYYDDDGEPLFEENSDFLDEVKDSSEYISLVNEQLPTLREKTIAGTAIGDFT
IECAFNGRKCDLKKDFILTLSGRYGNCYTYNYGFDKPAIRFSKTGARAGLTLTLFVNQDDYGTAYTQSAGARIVLHQPGQ
SPFPIEEGFDVAPGFETSLAVQLVELTPTSTIVSTDRSHVSQDSEGISRRDSLRVKGFADIITEFDLQSTGGTQIHETYNATT
PDYESNTDYNQSTPSTDSTSPGYESNTTASADGYDFITDSSLGNGTSQNETGTNYNTGTNMTEEIIQILDTSLNIARNNFL
KVNVYFQELIIYNITDVPAYAWENLMAELGGTFGLCIGFSLMTIMEFVELVVIIIQDVACKRRKSDETQVVISSSADIQEM
PQIAWPTDKKHQSKHMR

>Protostome_Lophotroco_annelid_Pdum_Contig1978_

FHSINRMHMMAEIGGDGNPAMSFLTRLKTLTENTGMDVLPHVGRARSKVKRLFWTLLLFVGTTMMISGVAKSFQRF
YSRPVETHLIQRVSESLPFPAITICNENPVRFSAVERAYPDFLKKLQNSSDKTFWVREELTSFLNSLPLEERINLGHNLTTML
LDCQQKGITCNESNFESMNSLYHGNCYTYKGMGDNLASNIGPHTGIGITLYIEESEYIAGVADGTGFKVVIHNGSDLPFP
EEEGFSVTPGFITFVALTKREVERAEHPYGVCHQEEDVPIDDIQYRTIGCRRRCLLRRQIAKCGCGLATFPYTEQTSNITDD
FNPPQCNTAEEKECSENVQQKYMDGKLNCNVGCPPRCKEVYHTYELSYTGWPPVCSRQLIMKIIERNTHTPLQLANKC
GKKYENIRYQTLKLVFFFNTMETTITKTASAYTCQDVLSNIGGQMGLWLGLSLCTLCEFLEFFFECFHILLRYLCKKNITTV
QSPQFGMRDHSHQNANNSSPKDKQ

>Cnidar_AlaAla_c58222_g1_i1_p1__GENE_c58222_g1_i1__c58222_g1_i1_p1__ORF__type_complete__len_507__score_37_28__c58222_g1_i1_27_1547_

MLLVMGSNKSVKKVVQTFAEETSTHGIGFILGKKSKGLRLLWLFLLLTSTGFFIYQIQKLLKAYNSNHTLISVEKKEVKAINF
PSLTICKGSTLSRSGGVQDHIFLRHPSLKHYYLTQYHLIGLRALLAQDYTRHFYTLEDLKRAAGNPRKLFLTALESSCIFGIHE
FCNYPTNISDVITSSYGGLCYEFNQNGLLTQDEPGSYFGLTMIMFVNQSDYVPFTTFDFGSGIVLKINNAGEYAFPEIDGI
PISPGHLTRISLKKVVYKRLKAPYPSNCTDGENINLLYPGKYTVLNCKRSCLVRDSLESCGSTDPLTMMLLNRTGKYKYKR
NNITTINCIYRSAAKSTSKDRKECICPVPCTEVEFIPTISQTKWPSQSDLPFYKYIFAHTLGMKVSNITDDFIYNNFLQLNIYF
NEMNYQEVREKEEWTLEKLISDIGGQMGVWIGASILSLCEIVFFFVELVKAMYLKHKKLKENMNPKNVVPLDKQKVEN
ESYANHACNNNSWFS

>Cteno_MneLey_comp16014_c0_seq2__p1__GENE_comp16014_c0_seq2_comp16014_c0_seq2__p1__ORF__type_5prime_partial__len_507__score_115_90__comp16014_c0_seq2__352_1872_

PTRKNRRIKCPRCRQHFVFNKVRKDGIMGWLQDFKEFSDGTTAHGVKYIFHPSVHGNFKIVRILFLVVWLVAIAYSLFVI
YNAVGNYVGKPTGTKFQVIASNPYKEKPASIQFPTISVCSHNLVTKSYFKSNDGLEELWSELDQWNPDTAENIDFNEGS
PAAKYKDWTYEKIIAEGGPANWTFLQCEHFINLCQEVIPQPDFFERETTLTGNCFRINPNGKLRGKGGDYGKMQLMFF
ADLNEYSDLTRREPQYGYTVVFHDHESYSSTIPSGFWLSPGSIYKVDLSLGKEFREPPPAGSCDPTRVNNTYGRYEENSCI
AQCRDDVLMEKCGCVHVSPPHPRNVPDKYYRGCTLEEWATCGLRAYKEWVVEYSNVNKVKTECNCPATCTEIAYKAQ

LSSSQLSKFYAEEKAAKLPPGYENAQDVLENLLIIDILFTSMQISEIREIVTYGWGNFLGDVGGVLGLFLGASMFTIIEFIQFI
VCAVLKGCCGLNDKESRSHSPLYNENL

>Deu_Ambulacrar_hemi_Ptyfla_40v0_9_20150316_1g15975_t1_scaffold12140_cov134

MKPRENSEDIHQEKSPSFSVQFKNTTGSSVIGLFFGDADSKKGTVQNLDNKYENLDENAAIQDFTQKATISGMKYVGD
KKSRIPRRCFWLLVVLTGAGFLTYQIIRSVNLYRSFPVNVDVKIKYAQDLPFPAITICNMNLLRKKEMEKHNEKNVAGVLK
LLYPLVTEIIEDSDALHNFTLSSVNNLTELLLRYGHRKEDMIIWCEWKGRRCSYENFTSTLTDYGLCHTFNSGSNGLPSLRV
DNTGARYALTLILNAEQWEYGRGPSQSAGFKIAIHSQYDIPMVSDLGFAVAPGTSAFIGLKMTKVSSFPEPYGTCKSRKL
KYHGNYTMSKCQLECLIDFIVEKCGCKRPYMPVAVDAPYCSPEEEYLCCLPKLDEYASHRQMCDCPVPCERTLYSASLSY
AQFPSDFMAVEYSKLLNKSDPQYPRRNGAKLNIFFEELSYEEISHRAAYELFTLFSDIGGSMGLLLGASFVTVVEIVDFACV
NIYRRWRRKTSSKIYPKTSSNDR

>Porifera_OscCar_comp39857_c1_seq3_p1__comp39857_c1__comp39857_c1_seq3_p1__ORF__type_
5prime_partial__len_507__score_68_95__comp39857_c1_seq3_3_1523_

PPPQFLIDIETFSRLKCVMEVVAKRRTRSGGPAQYSDGLSPAKKPARSPAKKTAEKKKVKKPAKAKPPPLTCVRFLFIGAP
WRRMIWVLLLVGAFAGFLSFVTGSIIQFANLPTVTKVSVEQPDKIEFPVITICNLNTFVKSKVSSMGGDPLLGAIESIDVFA
PRISANDETDRENKFKILEQAGSMEELVTKAGINRFDFLPINDNNSATCVWKGDPCYHDDFTLTLTPFGNCYSFNKSPQ
PDFVSDGGILNGLTIALNIQQEEYLDSGFFDAAGIQVLIHSQRETPASVLSYGFSVFPGTVTTVRIRETRVKNLEQPFGVCD
SGNPANYSINNCKDKCYYDALKQKCKCKEMYMRDFYQDDLPICSFRDFQCIQKLENILCRCLPPCEERLFDVELSQAQFP
SNRGVEKWTKRLNKSESFVKTNFIWLNVFYPKILFILNEQQRSYTVNNLIGDIGGHMGLFIGASLLTLAEFIQCIVTKICDA
CKKKNKIDIDTKEESSTLKP

>Protos_Platyhelmin_Macrostomum_lignano_A0A267DD09_A0A267DD09_9PLAT_Uncharacterized_pro
tein__Fragment__OS_Macrostomum_lignano_OX_282301_GN_BOX15_Mlig015081g3_PE_3_SV_1

VREREGGRGGTTKRLLNLAIPQNRDKFSAKKVGHNSLELRWRLHAKQRGPNGVQQRQQLQQKQQDDTGSIASYSQV
PRIKAIRTRDIFRALRTFAANTTAHGFYHLNHPVRFFKIGWIVLIIAAFVGIIVHLSILVVKYRDEPTSTKINPRGSVYTPPDIT
VCNRMMFAVSKLFEKSGDSYTVNPEFQPVLDSVLKVREFLEVSVKKYLRLSYGNYESDSDLLRIEDEGVYNLPNLAIRDTY
GAPFLSRFSAEPRQMLLFCTYSSIACDFLDFKEFGLSTFPSCMTFSPSNFLSRRPNISVNSLGELRLVLFTDSYHPSQLVTDT
LQLLPGDWFNMQGQGGAVVFVHQPSTYPTLGSRIEAPASRETIVDIRYQVYTSKNTRNYRCSESAESVQYLAIVNRTTQ
PERYQTQLADCYVDVTQKILLQKCGFIDPSLPLSLALSDRPRRGNLSNLLINSPSGRPLQPTDLITFIYQVFKRPTCQPLRSC
QRFLTPCLIRRSACDLPGS

>Protostome_Lophotroco_RotSoc__A0A818ZHA4_A0A818ZHA4_9BILA_Hypothetical_protein_OS_Rotar
ia_socialis_OX_392032_GN_FME351_LOCUS31822_PE_4_SV_1

MRVATVFRVDRSTIKSDRVRQGWRSIMREFALNTSTHGIPGIARSQSKKNRMFWSLSFLVFSGITLYFVIQTIREYFQYST
QTSVNLVSEWPQNFPAFTICNVAQIRYDQFIEPFLNYTNTLNITNTTDTSSFSLLQSQYVKEFFRIKINQNESIIKFFFPLSA
MLIKCVFNDRICSTSDFTYFLSSSHGLCYTFNAKLKNMTNVRKSDEYGGPGKLELSLYAHSHQYIPYAREDIGMIAMVHD
NGQFPMIEFAGRALVTGFKHRMTFSKKSIFYLSAPYSTCNDEIPLIMQTMFDNYPNVEYSYSEDLCYDLCTEVYTYGICGC
IDPKQWNARSIVLPRTKTIIVAPLCNLSDTCYSNASITLSSLSSLLTEYCPYCSQQCSITDFIVKASMWKAPAYWLTNDIKHF
VENSEILLPNDWSTDWQRHIQTNYLSIELVHESILVENYTQIPAMGPINVLSDVGGQTGLWIGISFLSLVEVAEMFYRLIR
YQYYVIRRSIRKYFVQK

>Ctenoph_Hormiphora_californiensis_evg1138204__p1_GENE_evg1138204_evg1138204__p1__ORF_ty
pe_complete_len_508___score_90_51_evg1138204__158_1681_

MGITQMTWIKDFREFAGDTSAHGVKNIFEGPTKILKLLFLITWIVCSVYATYVIFNAIVTFINKPTGTKFQVTVESTEREEG
KPAVIKFPTISVCSYNKVKKSFLRAPGNELLRRVYLTLNKYDLDEAAELAAEFEDPDSEVAQASVGDMKYEDLLADGSPDS
NRFLMCSQRAKNCDKLAVFKGNASEGIPPGHFVRENSISGMCWRVNPNGKLKGLMGDYGALELRFWADIQDYSEST
KNMATHGFIVAFHDNGTFGSTLSSGYLMSPGTYYKVDLRLRKEIRKPPPGGNCNASLTDTSYGVYTEGSCIMECKDRWL
YERCGCVNVVPPINKDNYESCNLRQWASCGLKAYLEWYDRYTDTDTANPICTCSMPCEETNYEAKVSSSDISQAYADH
MFTTKGVAGFLNSISDPSLNISYDSANDIRENMMVVEVLFSSAQVSEIREVVTYTANNLLGDIGGVLGLFLGASLFTILEFF
QFIIISVAKHCCNWIPGGGKKKYESAEMS

>Deu_Ambulacrar_hemi_Ptyfla_40v0_9_20150316_1g14166_t1_scaffold9674_cov125

MNKARARKIKMTGKKKRGLRSIISKFAKNTTAHGVANISNSTSTFGRTVWSVVCVTAFSLFLWQSSELLKDYLEYGIKIKF
DVVSVPKLSFPTVTVCNTNKIRKSEILKSEHRRILVVDEDVVHPYYALQAFKSTYNKRLSPKDAVLKTNTSLGLCALSCMHF
SEFTCLSFDYDSQRRECYINPGSKDTLVRFTLTNDDDYDYYERIYRGMECYTLDDGTDYRGHVSLSSWGERCMNWSSL
DPSKFKYTPSTAPGKGLGNHNYCRNPNDEPGTWCYVYRFNSNVPVAARCTVNPPKYGCLSQKQWEKEIVESDTFRDS
KPIPAIAVNCSADPSGQGYRGNKSTTVSGKQCQNCDFYMFQDDQYGNCFKFNDGEKEPVIRQADRQGALSGLTLTLFL
EQNEYISIYGRDAGARVNINPLNETVFPQDEGITIMPGTVTSIGIRELDDILSDIGGTLGLYIGLSIITVAEFIEFVAAILRYLY
QSRRIKKLRTVDSDIKEKKRRRKCKF

>Deutero_Ambulac_Apla_gbr129_26_t1

MADEKSGENDARSLRALFKNLMGNTSAHGLPNIDRAGNPFRRCFWSVLFVWALGMFLWQCYELVYAYFQWEVDVN
INIQYKTQIDFPAVTICNLNPVKASSLSNSRELEILLGSLDSETSGSTDGSWTAVTAVTPKVTSDQLSVNSTDGELFGGTAS
SVTKLLPATEAGLTATGDTHTLKASSERDQLTPTSTDASSPSSGVSSASPAGSASIHVVGGTQMATNAVTAGVGIPTKEV
ERRKRDTSTNSERQYQNWDNIDYTAKESDVDLFDQVADAIAKTEYGKRLNFGHSIDDMLLACSYKGYPCGPTNFSYFH
NYLYGNCYTFNAGMNSDIQTSSKPGPLYGLILELNLEETEYIPSIQQASGARVVIDQQYRMPFPEDAGINVAPGVLTSIGI
RKVEIDRKSDPYTNCTERDSDDSQTVFSDFFGANYSLEGFGLISNLGGQVGLWIGISMCTVFEFVELLYDVCKILLLKVIFC
SHAKRATAPSASDLKLDEKPSKPRLALNN

>Protostome_Lophotroco_annelid_Pdum_comp414035_c0_seq4_

NMILECSWKGKPCGPQNFTKLMTDFGFCYTFNTESLDPYTTEAAGPRGGLRLVLNAETYDSLASWDMSSGFLVGITHP
NETKQPIKDLALQVSTGAQTYMGIRVSEFHELHDRKNPKCSMEESLLSCSFKCRSKVIIDKCGCREVSLAGDDVKLCNPS
DMISCSGRAVGDLYTLKNFSSICNCLPACHRVEYGVTVTQSAFSEFKTSLYALQDGNVDQRLESLFKFQDLSYLNAIDKN
AVRSLVSQINYPVVDIIKTFIQKYSFCNDEISTILKNMVGLDQNISTCASGKVDDINTTVSSDFMKFVDGNYFTSHTLEKGY
LDDFLVLISHDLLRKKYEFMKWLTDFKDYITHVNDTASAVPSCFNWNISQRISDAVSNCTDNSVTLQNNIYALQSAYLEI
NKTVTELTAIANLPERPKTKDRMWYKENIAFLDIFLEHLTVMRVVQNEDDSFNSFICDIGGSIGLYLGGSLLTLMECVDLLI
WFCHKKDKKTKVISVAEKRPDKEMY

>Cnidar_HydraVulSc4wPfr_2449_g1062_t1

MAYFQEKAETPKKGFEVNGGEEKADTKNSFVVKDGKKKAEKPIKTIVVEDSEVQSTQPTTAKFVTYHRILLIATKNYQET
GMEAKKGEKILDFVANKDKKDAVSIANEVNDHYDKSNDYRGSFTIHGLSRALTGNFYENIFWGFIVLSSFTAIAVVAWKL
AIKYQKNEVYIKFFSKEYSKLNMPILTICPNRINKRRICNLTNDCMNNSTDISTLYIENSSYAWWNDEMSVIVIKESRKSVK
TGIELRNYIKSTYYQCIRMNLSFFDISSSSSNGIHVYTIYPKSYIEVYVHDENEQYPFFQTKPNYIEALESDEINIEIVQYNRLK
YPFSSNCSQALKDMIFPDSYTKLKCVESLKCMNSFKACGKTYDFCEQFLNDNFRVKYAKNLSALDLKILTKCLKSEYGKSY
DNECALACDESIFKIIMSYTTQCSWLQNNAREIYVNYPLGPYFHVFEEAQLYNFEQFLSECGGLIGLFTGSSAISIIELITFLC
LLLIKYIFNKSNKKTP

>Cnidar_HydraVulSc4wPfr_875_1_g17145_t1

MKNSDESISSSSYLQSSSVWHVGQNSFDKKTKKFGSRLIESQDDDYQYDHDYLQMEVIGKDVERGCVKKYNGNQRSSL
EIEDKNLYKSCQKSRNVIENIIEEYINGFSLHGLTKIFVAHTRLESLFWLIALLSGIMLSMYVSTNLVNKYLRFEVYTETKLEIT
DKNYFPAVTICERQLLLDFYYAYCKFEQPNDSYPCQHKKQKIEDNQSTTDNKTWWSNGLFNITKCSSWGAKQCANND
FLYSLSQYNHTCIRWNYNGDLYDMYGHVDIEFTINNSLLLPYQEPNIILMSHDPLIFEIDITNAVNLEAKQYNINLEKTFIER
LPFPFPSKCTDSKNNDMFTGKYTRRSCLETQNYIDMYKECGDTIDYIRQYIPKDILQKYKKIDTISNIGNCINLYSHKEVNPL
SKNCPFSCTEYQINMLVSSREYFKNDDKITTETYSVGIQYITVDSYRIIKEKQLYSIHQMACEIGGLIGLMVGCSLLSAIEIIV
CSVLWLVVKFKKIVLRIN

>Cteno_Ple_Bac_comp44971_c3_seq2_p1__comp44971_c3__comp44971_c3_seq2_p1__ORF__type_c
omplete__len_509__score_81_35__comp44971_c3_seq2_350_1876_

MGDEGLQRMRFIHPNSKKLISGVIQKKFVPKRNGNHQNIIPMAEPLDEPSTPEEEDDEAENKEGKEEGGTKKKRRWM
TVKWLFYLAWALALTYTLHNISKSVHKYLSKPVSTKINVYHNEHGSMQFPTISLCNLNRVSKSYLEEEELLEDTWNTVSD
WGSVDWTQEGMDDIGNLPYQDFYRNSPSWEDTKIFCAYAHARRCEDMDIFADPSNMVEEELSPTGKCFRFNPNGVL
HSKAGDYGSVYLRLNANQRDYIDNTGQIGFVMAIHHYTQYSSTRNTGIIMSPGLHYRVTLKRIERRELPEEDGGKCNSAL
QRNSYGAYDLESCHTECRDKQLNASCGCIPVLPPYNIYNYTACSLRQMNECGQWAYHQFVLDESQTSPEKGEERECGC
WVPCEHFDYELSVATTAIPREYAAGKIRAGEGEKYNYTVDDFLKNNVILETYFRDMYSEQVFQVREYDFWNLLADIGG
MLGLFLGASIFTIIDFIRYSVKFGYRKLRVIGIRMKERS

>Deutero_Ambulac_Sakowv30038064m

MDNTEKRECDKANNGFPKFKKSTSLLALFFRDLSEQRSQKDRDSQDSSDEEENGDEKKVEDESVIGDFAQKSTISGMKY
VSDKKSRIPRRCFWLLVVLTGAGFLVYQIVRSVNLYLDYPVNVDVKITYANDLPFPAITICNMNLLRRLELEHYEGHDAIEF
LELLYPLVTEKTSPVPENIIQNFTLTTVANLTDLLMRYGHQKENMVILCHWKSSACSSVNFTTTITDYGLCHTFNSGLEGT
APLRVDNTGARYGLTLYVDIEQGEYGRGPRESAGLKIAIHNQRDIPLVSDLGFAVSPGTHTLIGLRMTKISSLPNPYGLCK
SKKLQFYGDYTMSKCQLECLTHFIVGRCGCKQAYMPGQVRHCSPQDVYNCYLPNLGEYAGHKDMCECPVPCERTIYSS
RLSYAQFPSDFAANEYSKLLNNSDSDYIRRNGLKLEIFYEELSFKEVTHQVEYELFKLFSDIGGSMGLLLGASFVTIVEIFDFA
CLNLYRRVRSKWRPRITPIETKS

>Deutero_Cephalo_032830F_t2__name__chr_scaffold117_start_653459_end_666145_strand_pro_len
_509_

MPKIKTPMSSDEGHDWPESPEYVFATTTSFHGIGRIATAPNIPLRVLWTLATLASYIGFFILMVEMLQSYFTHGTVTDVT
LEFVPEVRFPAVTICNLNKFEDKALTVEEKHYLSYILYGVQEDIDVIAGAPVNNKYTPNSTDANVTTETPRTEAPHTHGSF
SVDSITREKGFRLGNDMHVCTWMGQTCTELDFTHSFSTYGNCYTFNASPENPINQTIPGSGYGLSVMIDIKSHLYTENP
LLPGGTADVGLKLLVHDQNEPPKMDTQGIAVAPGSHAYIAIRQIQYQNHIPPWGVCQDLQLEYYDTYTLTGCQLECRS
KYVVQNCTCRPIYLPGDAPYCEPADVAHCVRPVEAAVTSGALPCDCPVPCSLMTYRTSSSYAKWPNNKADVVYTTAFG
LSPGYMADNSVVFDVYYEELNYEIISQLKAMDSGQLASDLGGQMGLFIGASILTLLEIFEYLGLRLIAFIQRRKVRTRRVHT
SAYPMPDTLSVNQAAKNKESKEKSLMNLKE

>Proto_ecdy_PANarthopod__XP_023235401_1_degenerin_unc_8_like__Centruroides_sculpturatus_

MALVSLIYNCYDVVDEYYNYPTALIVRMEDPKEMDFPAITICVLLKPKHFHYQNQKMCVEKTQNTLQPLSDINNDNKQL
PPRTTGETFYESSKKNVNHLYRNLSIVKGSDGISNFNLKRGNENNADGGNGRRFQAEKRPLTGPRQNGMCDFEFEKHI
KYNGSAQETILICSFDLNECYYSDFKEWESEDYGKCYTFSGVRTTTKFGSTSGLHLVIKVTHTDPNIRTSGIRTIVHSPNEFP
FPEDDGINVPRCMLADIAIMKEEIERLQPPHGNCTTNTAIPHWEEMFGELPYSRQGCIKACHSQKVLDQCNCTLNIRIG

KSLRGDVIHCRQHKDKEDCSENVISGADCECNPACSEETFDTSVSYSQFVFEENCPEKISCGNSTCYTKKLPCRDIMGIHV
YYKDLTFEKKTQTPTYTTPALIANIGGTLGLCLGLSLVTIVEVIELILDMLMTCFRSSSDVKPVNLPRPKTETKQGDHPNLH
QIRVTPYSVQVRQIAMEDAPERNQQK

>Protostome_Lophotroco_annelid_Pdum_comp414620_c0_seq3_

MGCKEWFTMGKSNPPLTYTEVTNQFADNTNCNGISSIKRSNHVVRKAFWTLLVLAGAGLTIKLLVDVFTKFFAYPVSIS
VNVTYESTVQFPAVTLCNVNMIRKSQIGLMTGSSALDKLTFTVGGNPDISAWATTEGRSILRHSFVDGWSQLNDTVKK
QQGHQLEEMLLDCSYNGDVCGTNNFTHYINPNFGNCYTFNSNWENQTIQYSHRAGPFYGLMLELFIEQDEYIPELTEV
AGVIVSIHHQDVMPFPEDDGVLIPPDMSSNIAITAETITRKAKPYPSDCLNATFKEAAHYSAYSELHPVRYTHTACYRTCY
QAHLFSECGCGDPVMPLEGQALYNLVTDKQNKRTVCTSQDEITCELKVYKQYSNNELECNCSVACEDHIYQTAVTSLR
WPSANYKNLYYGHLATRNPHLATIMQSYVTNGTNPGEHLVKLLIYFNEFNFNYIEEYASYPSSSLLSDLGGAIGFYLGISVI
AAFEFIEYIFDLIFIGCKKMKNRKANEVKSID

>Protostome_Lophotroco_RotSoc__A0A820X5A2_A0A820X5A2_9BILA_Hypothetical_protein_OS_Rotari
a_socialis_OX_392032_GN_FME351_LOCUS2233_PE_4_SV_1

MYSDTIFVISNKTNKRRERVVRRRSIIRQFALNTSTHGIPGIARSESMHNRIFWSISFVVFTGITIFFVTQEIRKYFEYSTQTS
VDIVNEWPQHFPAFTICNLAPVRYDLFVQSFSNYTTTLNITNANDTNSMSLQQIKRIGDFFQMKINRNESLDEFFFPLSS
MLMTCVFNGRACSVVNFTSFSSSSYGFCYTFNAKLKNINSVHYSDEYGGPGLLNLGFYVHSHQYVPYIREGVGMIAVLH
DNAQLPMIDSAGMALATGFKHRITYTKKMISFLRSPYSTCDDKIPPMMQAMFDNYQGTEYGYTEDICYELCTQVFTYK
HCGCIDPLQWNARWVFLPETEQMILAPLCNISNKCYSEAAYVFLTSSSLLEQHCSYCPQQCFITDFSIKNSLWRTPPAWL
VDDIKRFVENSKIPLPHDWLTNWRSHIDASYLSIELVHESTRIENYTQAATLTAVGVLSNVGGQTGLWIGVSFLSLMELA
EMLYRLARQQYHTMRRNCIRTSDESKV

>Deutero_Ambulac_Spurpu_016076

MNQFDRSNKVESISGKVTCGNHVQPVVDTDIDIRSGDNDEEDDVQAPEKKDSGDDLDTLSTIDEGSREGSKLLFNDFL
GITTLHGIKYVYNGKYSVFRRLSWVCVMIVSYVALILAGKEHVTKYIARSTKEEFSTIRKSSIAFPAVTICNSNPVKASLIDNP
TSKALVQVLVDGSTERGWDTFGLNATDDALETRVDEIYVAHAHQASDMILECAWKRGSKPCGAANFTTKITSDGVCYT
FNDGEFEGESGDDDVEGDMRLEVTRSGTRQSLWLLVHRRGDVPLVSDIGVAIMPGVRTLIAIKKHTERNMPNPYDTN
CRNASLDYYPFYSEEACLTESATKQLTEACGCRFPHMPGPTPVCTVRQYIECWLPDFELGCDLEFHCQRACYTEHFQMT
MSYLKAPGKDDSQRLQNQFNWTGDDVTRNIAEVEIYFADMNEHHTESVAAYSFADLWGDLGGLIGVHMGASLLTFF
EVVDYTFVHKILQIRFLWRKIKSFCSRDKKRPALDQ

>Protostome_Lophotroco_annelid_CAC9651275_1__Ofus_G101167_Owenia__fusiformis_

MGRANMIEQDEPKPGGVRAITDEFIETTKAPGMTQIHRNTKPAGKALWALVYIVLICLFGYTLAIALMKYVQYPKRTEM
SKVHGEALEFPAVTFCMTTPFAATKKVINGSEISFPGKAAGLAKMDELLTKDAYQSIQDDILEFKERLLSQEFMNENLQS
DPGNDALYHQLDDEIIHCSLDGENCDMNNFDIFKHPRYGNCFMFNGTKQSNDSGHAIHEGLSLVIFTNSYIPGDDALDF
SMESTDATNGVKMVIHDKDIKTLLDKGLDLSPGNQFNIGLSVSHNLRLPAPYGMCQDSMNQNHDACKNECLQKHFK
EQCTCLSPFAPIPSNSDATEYCGSFDTDNIDSFKKGLKRLQCEKDILNEGHNDTGCDCMAPCSETTYSSTVRQTPWPHK
NHFKKFLKTTLEGNVAVKSLMELTRNYANVTTGDYNDTSLEKDLILENFLKVNIKFNSFEMTRIVESAAYTANALVMDVA
LNLVIILGLSLIPLVHMLVYIVHVFWALMCGRRSNE

>Protostome_Lophotroco_annelid_Pdum_comp405972_c0_seq2_

MMSGESSIKPILGSMAENTAMHGLPNVKRSISTARKIFWSIIVLAGVAMILYEVIIATIKYYQYPVQTTISINSAIVHQFPV
VTICNQNSIRRSALEDAGSDIYMLLNSTGSRQTGSSIVSLRGNILKYYSQLTPQVRMDIGHQMETMMLDCAYGAMECN

TTYFEQFSNYYYGNCYSFNNGLNYGIKTASTVGPDHAFQIVLFIQQDEYISQLSEGAGVRVIVHNRTHMPFPENEGFSAP
PGQITYVGLSRREIKRASPPYGDCSENHLTMNSYTDFLTGVDYSKMACKKSCLQYTIIETCSCADTSLVTNSTALNTFFKNI
EPCRNLTQDACISTVTQKYYNNELNCSTGCSPECFEVQHERTLSYSLWPGDNALNDTLTLLSENTNAAQLMKNMNDDE
KESFLRKNVVKLAVYFESLDYDYIATSSSYSSSDLAASYGGQMGLWLGVSVCTLFEVVEMLWEMGLVISQKMNSRKNK
TGNMETTNSDGKKMPSDKAVIVKKDSWTASY

>Cnidar_NemVecNVEC200_011922_1_1_protein_AED_0_16_eAED_0_16_QI_93_1_1_1_0_90_0_83_12
_814_510

MADQPRPSVRALVRDFVDRTTCHGIGQINGSQSPLWRVFWVVTFVAGLGMVVHQGVTLFGTFLDRPTSTTIDMTYA
PAMDFPAVTICNLNAIRKDHLSQFPDADVLLKGFSEASTPSVTVFLGKDVESFIKQHSEAGPNVTLDPELAFKDAMVEIF
AQSELKKLQMAGHGFEELVLGCTWNNIKCNKGDFLKYWRPFWNFRYGNCYTFNQGMSEKGVAIKPLTLSNTGPNYG
LTLDLFINQEQYIAPYTQEAGVRILLSDQNQIPFPDSDGFTVSPSSSSAVGIKKIFITRIDPFNNGSCYKVTKGLEEGSIYKSV
FSHDMGYSVQGCMNSCLANKQREMCNCTEGRFDMMTGLICQQIEAWRCLNRVNMMYQEGGLKCLEKCPQPCTQ
NVFSRSMSHAHWAEEYKKALSKFIPNNGNGSSVDPNELISHNILRVKIYFEELNMETITYKRNYPVESFLGDVGGQLGL
WIGVSVITCAEFAKLLIDLVLCVAKKMNRSDKVQSVCIGRGN

>Proto_ecdy_PANarthopod__XP_023234948_1_degenerin_unc_8_like__Centruroides_sculpturatus_

MTSRRTSSAFTNFSEIPKLSDLFQDILENSILTGIPQIALAKKKSRKILKVVVLFGCIAGFLYQLSAFLIIYWAYPTLVDVQITK
PLMVDMPSITICNRNGIRRTSFCKKFPSECELPNNMTTFCEQYPYVCNAAKTEEDMMKPKLEAIDREETVTRDLLLDIGH
ASYPLISQCTFGTGKDEFQICDNETFRYYAFLDIYQRSKLCHIINPRWSKKELKLPRVPLHEVLTIVVDFEPTEAFQLDRRIG
GQAALHSPYSIINPFIEGFTITPSKLYLVHLKEIIKHLLPSPYLTNCTNYKKLWKRRLGRGPLSREMCIEECLMNATLDHCGC
VKTSSIYPHYEKFCHEVDIPCTDKINYSHCFIACGPACHSISFEVNIQEENIEVGIESTNATTLWNQTAIILLFFKKSDLVIYTY
RPKYESIEFFSYVGGYIGIWLGMSLIAVFDFLEALSSLVIPIKMKIKRACANPLQTLKSASSRDCILSVTPYPYRDNMNSQLS
VRTRSMPRAWE

>Protos_Platyhelmin_Macrostomum_lignano_A0A267EWW3_A0A267EWW3_9PLAT_Uncharacterized_
protein_OS_Macrostomum_lignano_OX_282301_GN_BOX15_Mlig009331g1_PE_3_SV_1

MTSSTSPGPCATTRCSSPPSRCATRGPSPRRASSTSWTAASCCPRTTRSSSCRQCATPRRTSGASGSTTPPRRRSAPSSR
PRATRERAQGLRHRALGHTIDQTILICETTILDKDFKRSKFCETIGTWTRTYDNKFLNCYTFTADANNTQKILTVRIMAYT
NERNLTNCPDCSSFPATQVSGLRVQLHNAGNYPEVQEEGLNIRPGSLTEIRFETKEWVMKEPPPAAAPLGCAPSVLFNG
DAFTYSYQACKKKLLHEEIMKKCGCLDSTIPLPDSMRNTGLDFCGRIPPSQFSLQSEYIQRHNHSTPHDDNDDQRKRPSF
HFDADFNDFLRRLTCNTAVFKVSGHATAHQCFTPCTYYTYQTTISTTNWPTRAYHYEIVNDFKNLRDRIDNLHRRGKISR
NSSYLDPVFQRLQRIYGPLSEADSPAEAERMLRGIDYIERNFIEILINRGKSNFDIERIEEKAVISLTSLLSQIGGLLSIWVGLT
FVCIVEIVDLLYNICVSYNGSRQAA

>Protostome_Lophotroco_annelid_Pdum_comp392061_c0_seq1_

KERVPPWKSLLFDYSQDSSCHGVRYITGQSRFFIRRVVWTLIVLTCIGLFMYFFATRLKYKASSPSNVKIEVKNRNPMPFP
TVTICNQNQVRLTAVQELGLYDEVSSGLSTVDKKRATTSQVNEGCEDAYIEEKHPRCVSREFLQSFRFSHTYVEVTLSVCK
NLCQNLHSSTCSRIFYFPLARTCILTPRTEIYDIEAATEGCLMAVVYQRQRCIDATPFTCDFEQETGIASSTCHLWKNGNE
YNDEKWIRIDASLYTPFPRDHSTDSGKGGIIHLINAERPKDAKAQLQTPYKNTTGYCLEFFYIKPMEGSQLNVSVIGENFQ
SKTLLLIEESAFYWRRQFVQLPKGLHQISIQGHMKKGSAQTIAIDDVTVMSCEDFEKSCLRYKYGDEYVGTETYEGRDGR
KVEACIEWNVLKLLESSMDEFSSYENTNLCRLPYFKGFTSYYGTQPFCLQEYQQVYCDIPYCECTEGWTKCDNGQCLPKS
FFCDGEVDCADKSDEADQCASQQSLS

>Cnidar_Rhopilema_esculentum_TR89610_c0_g1_i1_p1__Rhopilema_esculentum_TR89610_c0_g1__Rhopilema_esculentum_TR89610_c0_g1_i1_p1__ORF__type_complete__len_512__score_64_81__Rhopilema_esculentum_TR89610_c0_g1_i1_43_1578_

MRFAVCDFKMSNHSSSETDNSPMKLDLKHIERPTYISQDEIQLKVRKISLKKEANNWTHKEIVRKSLSDAECEEDMVELV
REELYLRKGSQISLHGVSHLAYAKDTSKRISWLLVISIFWAFLAYMLVSVFVSYAAHDVYTVREKVQVNEMVFPAISFCST
PKLDEIFLRYNVSYQNHNDVTALPVLHRFVGDYISNRSTKNRLSVLKTMFVEGIGACQFGNKPCSFERDFKAISEFHFDG
VCFTFNPSGTFSQQGEGSKFGLSVIVFVNNTEETVDMAVRPGLSLVINSHDVFPFPLINGILVSPGELTLIRLQKQRMTRL
PAPFPRKCADNGPILFPGKYTPNNCKQSCYIQKALDACGASVFPVNKEKQNCFYSKLVEENLAHHPHACACPLPCKEDR
FIPLATQSDWPNEADLPYLKKLFATAIGKNSSTLSNEYVKKNFLRVNIFFHELAYEKVDDVAVWTQQKLVSDIGGQMGI
WVGASLFTVMEMMAYMASKIKSFLYLLCHRS

>Proto_ecdy_PANarthopod_Tcas_tr_D2A5Z0_D2A5Z0_TRICA_Pickpocket_protein_28_like_Protein_OS_Tribolium_castaneum_OX_7070_GN_TcasGA2_TC015087_PE_3_SV_2

MSRQKFWQNVCKYFTEYTNVTAIHGLLYLTEKRSKTEKIFWLFVILSSLAGCLFMIGQILEKYHNNPVLVSFGTKDTAISEI
PFPAVTICPESKCLEERFNYSDAYLRVLRSNNLNKSEQLKFKYVSTLCNDLEQNVFLKKSGKTLKDDFFDVVNELQPYELST
CYYMGESFGCDELFTPIITDEGICYSFNILDRKDMFRKNVLWYKNYQDVTSEPLKYDIDLGYLTPAVTTYPSRALLSGADN
SLQVSFQHVQQNTDFLCNKYYQGFRVLLHNPVEIPRLSKFCFCVPLDEIVTASIEPNLIQTSDSVRKYNPEKRNCYMGYER
PLKYFKIYTQQNCQFECLTNFTLNYCECVRYFMPRTEDTKICGFDSLDCVTQVDKKLKTQEMSGTASQCDCKPSCTSLSY
NVETSNSRYYFKEYLDLRNISNNFEEPDKHHFSMLLVYFKVDQFLSIQRNELYGVSDFIANVGGLFGLFLGFSLISTVEIVYF
LTMRIYYNLKLYASWSGVSNFT

>Protostome_Lophotroco_annelid_Pdum_Contig17650_

MSTTPDESKESKMGSVKGEAKRFCETTSIKGVSRVMRSQSKLLCVIWTVAVLGFLFMASYQTVILVLDYLKYGTITLVSEE
MVEYDSVALNPDLIICNSRALTSEFSNSSKQHLDKFQSHLGKLDVCDNCSLEEFYGQMAVKHELSSLKGLYQFMGKEEA
AMMSHRLDEMIVKCETIYYLGGLVYNPCPDSVVITPILSMNHLNCFKVQTGDKYALGYRFVSYIDNFPAHNYFESSRDD
PVGLFVYLLEQKTMWSSLSVKTWLHPGTITEIQYATQKRDLLPPPYGSCEEFESNSSLQLAPYKKTLGTYISLCVETEIAHA
CSCSAPTEVSFLDELERRHNLSYCGNVSLPVQKLLENMKCATETKHKVIEDCISSCRVPCSELSFSPRISSSQWPSLSNLKSF
YESYVQGRSYEHHFNEAWQEVLLGSTDKFEKLIRNNFVKFDIIPSDYRYLHMKDEPKVKIVNLVSQLGGVLNLWAGITVV
IIIELVEAVIRIFIEKNDSNNVKPFNK

>Protostome_Lophotroco_annelid_CAC9672484_1__Ofus_G21979_partial_Owenia__fusiformis_

LYFVYTTLEIYFEYNTHTKASILRINEVDYFPDITLCNIRNQANSRFFTNDGNLTIYAQRIRKYMTSALYKLHDLFDADITSDL
TTIGMYDLEKLFITTINKVGANQKNLTRSEQKVLESLTDFRSYFTMFNSSEREDMGHQKHDIIKQCRVLKWLETDEEDCN
VESIILYQDMKYINCYTVQLQKQVTNKTRGISTVLYMTPHNFAVLESIPEKIPQDSGIVVFIHQHNTMPFGAPSFTAAPGF
KTMVNIDVTNTITLPAPFTNCVQERKLNFTKYNGDEFVYDKEPCVAQCVQEQALKQCKCLNADFPPLPNASYCTDLTLD
SKLYEANIACMHTALPTAETKPCLKQCPSKCNQYIYNYKLSQSTWPQEMSHLGIYKTYIRGTQYESEFTDYEALLKENAP
NLTNKERYDKLRTLSTIKDTFVKLEIGLTDKDVIQFKDTPQLTKFDIWNTVASTFIFWIGFTFFSIMEIIECAFHVAIGYMFH
GGRTREHPDKRKQVEMGDKGDNNTKY

>Cnidar_AlaAla_c46701_g1_i1_p1__GENE_c46701_g1_i1__c46701_g1_i1_p1__ORF__type_complete__len_515__score_83_16__c46701_g1_i1_223_1767_

MDIKEKDLPENVKAVLKQFASSTSAHGLSHIASAKYSYQKLLWAFVLLLNLFMFFFMITRLITLYTSKPVSTKVEFTFEKEL
AFPAVTICNQNMFRRSKLTPSDHARLRPSETQITSTMVEKMQKKFQDELKRQGHEGTHGGSGGTTGATSGGSNSTST

DNMVTVDSSHLMRMSILESVNLMEEQEKMGKGHSFEDLIKFCRWDGVACHKGFMKQYWYRTWDWYYGNCYTFNY
GKTGNGTSLKRLKATGSGSENGLFLEIDIQNKEYFPELTLESGVRLLLTEQEVNLFPLQEGFSIPPAVSAMVGVKMSTAIR
ADPFNNGSCRANEYADGGFRYTVQNCVQNCTALAQESKCNCTLLKYAAVNSTRVTCHTEAEKYCADEVVKSLKNGTN
DCKANCPQPCSESYYSKSMSYSRLYGEGYQKDIAIRDKSNPKEKDIMENYIRLLVFFEDQNVHKIRTAVYYEVENLLSDIG
GQLGLWIGVSMVTMVELILFIYKFVEYLMEKKKAGKVIHVAH

>Deu_Ambulacrar_hemi_Ptyfla_40v0_9_20150316_1g17281_t1_scaffold14250_cov87

MLIYSYFSWQDDRPGDPYPLVSLNPTTLNGNGEGQDQNRNLRKRQGNNTDERTDEISSLDKSAGETNGEYEDSVKHM
LGDFGRETTAHGIVKITTAKSNVTRSVWSAIVLLAASVMLVQMTLLLVQYFEYNVNVKVTQVSEKALEFPPVTICNTNKL
RKSAVRASKYSEILLLEQSFVPPYYTPCIEGDFLCQDGIHCIKPYLVCDGVNNCGDMSDEDNCTYRECGPNQFRCDSGSN
LGICLTKDRVCDRIKDCYEGEDETDCGSERIYGAVVRIGNSDVIVNNTQCGEIIISDNIDSNAAVEVGYALRGRFVSIQLEN
RTDYLSLCEVEVIGEASRILGAVIRIGYYKEEITQNVPCGGQITETDISNKTIIRDCIIPILGRYVIVQNDPYREDYLHFCEVKV
SAKEFVPSEFTDPLKTRKFVMFEPQEEKTLPVNSTDVFSNLLLNKCLSECLNWKRYECHSFDFHFDSKTCRIHRQKAGMD
GLHVVHSPGTVYYQKLRMSGMKNRYSVLIVT

>Protostome_Lophotroco_annelid_Pdum_comp401362_c0_seq2_

KRLWFHKSWGLEKVKYAMEYNFVRGWQVMKERTLDHMTVGYQEITAYLEHVADLVNNNKVDDQMKAALVVQVD
RMLYEKKVFALRAQSNITVADNAYNNAEPLNSFEVSPRLWYDLAYVTPLVVKSSPFTNMNYFQRLQTRVGRFMEALD
NLFALMSNVNNGRQIDTDLLKTRNDRLQSVSRSYNFFRYMLDERVINIPIQFVDGLINEFDNLNKTLNIRYQDFELKFNSL
LMFLKKATDSWSRILEARSMADQYLTDLMAKKADLAERLTSKQFKADLETTIDFLSHVTKRSTDVILALNAYERSYNRLW
NNMLSEASLKDFYAQVSYDWKNAKLNRTYLKIWDGVAAEMLGVKLEWIANNLNSIEEIMNADFVNRSIEDQERIDHEF
METVSQFVDASECLETGNQFVKAANNLTGNMEAFIEETTLNTAYYRNNFLNLDVFMMKLSYEEVTQQAAYDMTSLW
SDVGGSIGLFIGASIITVFEIIDVIVNGFFKLHLCQAKKKKTQREESTFL

>Protostome_Lophotroco_annelid_Pdum_comp413322_c0_seq6_

MAFMEELRNFGLTTSVKGIPRALRNQSKCMTFTWASAVVIFFGIACTQTCFLILDFASFPKISTLSETVVNDSFPLELIICNS
KPISSLSQLKDNVSDIPTPKDYFMHLDKVLQCENCTKEEIRRINNVKEGLSRPPGYYQYLGKEKIRSIGHERDDLIVDCRVIL
LDGAGKSMEPCEKYFNTTLQLCTWLYNCYRLKVKATAKARILGVTLISYVDDFNTDIMERFSINSPVINSPGLVVYLVEED
VMWKDLSGQIWLEPGTFTHIQYRISQYEVLPEPYGNCTDKWPYQSEYDIPFKYSYFCCITMCFELTAAEQCLCKDLQSW
GILDKIGLREELPYCANISQGHEAILEKGMCSERIRSKNRQNCLSICKPPCNYLTVSHRLSTISWPPSYALKQFYETYVEGRR
YEERFRAAWQEASRPNKQELSVDDFMKLLRENFVKYTISLESYKVLKMIDRPKYTVTSLVSQVGSILNLWSGITVMVLIEL
MEVCVRFFGNRMTTVGAVGSKSGNQ

>Proto_ecdy_PANarthopod_Tcas_tr_D6X3F2_D6X3F2_TRICA_Pickpocket_protein_28_like_Protein_OS_
Tribolium_castaneum_OX_7070_GN_TcasGA2_TC012955_PE_3_SV_1

MEKPTNSFCKNVRTYFREYCNCTSIHGFRYFGEKRTIFERIWWIIVFLICLSACIFSIYGVYKKWDGSPVIVNFANQGTSIY
QIPFPAVTICPESKSVQNLFSFTKIMLKKLRRQNTTQAEDTKFEYMALVCKYHEKINYTNPNTFPDEFLDVLNTVKPNLIK
WNCMYLGEKYDCNQMFVKILTDEGICYTFNMLDRTDIFRDKVLQYKDYQRSLYKMSEWSVEHGYTENAQFSPYPRRA
LLAGASNGLTLNLMTPKNYLDYACRQTSLQGYKVILHTPMRIPRPSQQYFRVPLDQNVVGAVQPVMISTSDRVQMYD
PKRRGCYFPSERHLKYFKIYTSLNCKLECLTNYTLSRCKCVNFFMPRESGTNICGTGKIACMREAERFLQAKDLYLKLSKNK
SLKKTSQCDCLPICTDTTYEVETSQSDWDWKMTYSAHGTPWPNDTHLSSLTIFFKSDTFITSERNELYGPIDFLANFGGLL
GLFTGFSVLSLMEILYFLSVRIICNNRLYGFWAGPSI

>Proto_ecdy_PANarthopod__XP_023219716_1_acid_sensing_ion_channel_1_like__Centruroides_sculp turatus_

MAKFQSDLNSGIHCPVHGWKRSSFHICNSSPPSSKKCISTTKYCLNIFKESSIPGVNNIAAAKGKFERFIWIFIVICCLTGFIY
QVTLFLRHYYNYPTRVQVLVKTEGLMEFPSVTLCNNNRIRKSEYCKYNKSSCSSDPIELDEMKLKEKEIPKLQLQLLVEDRP
DIRKILGHNISDMIVSCTFTGIPPMKGRECVNRFEYFYDPDFGNCYSLESVEDDGDALRASESDVWQELSELVLVLNVEE
DEYLEPSRDVAITVVMHTTDVYPDPFSEGHEIRPGFSYKFGVQKTSIQLLPPPYKTNCTEYSKLEWEKNNDRMSNTRMC
TAECSQSIQLKKCNYITDELSLFFDDVPWKDGEKDEVKIKCAEKIANDTKEYCRKLCRVPCWQSSFEVESDATIWPRKKKY
SQLEELGDWRNRSYDDISHNLARIRVFYSSMDHTMYRHIPIIESLELFSHLGGLIGIWLGTSLIAVCELFEKTISIFTYCFRRR
RNQRKVKKVVCFNHMSSLPIRKTKLRN

>Protostome_Lophotroco_brch_Lingula_anatina_comp136986_c0_seq2_p1_comp136986_c0__comp13 6986_c0_seq2_p1__ORF_type_complete_len_518___score_46_16_comp136986_c0_seq2_175_1728_

MTEACEKMVLQTIIVEPMDKCKGLEKDKVSQDITMEEEESLEREFATGTTAHGFSRAAETKSIFRRSAWILLILGGLGMA
TWMISLRVINYFQYDTSTEITLTFEPRMAFPAVTICNFNRYMRRNLNAEDYAIISLTEDVYSGYDSGYSYGQYYYNYYYDY
SGDGESSPSTTTQPTNSNFNESWANISASYPNGFDFGNFTLEKGWKLDNISVLECTFKGIKCNLKKDFIHTFSTYGNCYTY
NPSLNSNRSRYQDQPGLGNGLHLTIDVEQSEYTESLPDGSAQTGVIFQVHPQWEPPLVESKGLGASPGFHAFGALKRQ
ETVNIEPPWGKCNKSLTLLHFDQYTFSGCIIECKLQKIETECGCKPVQYPGTYRICDPAEMMYCVKPLLKTINQAFSDHCH
MCTVPCNSTQYNVQLSYTTIPNNNNIEAAFATKYNLPTGSNYIKDNIVALDIYYEELNLETMEQKKAVEESGLLSDIGGQLG
LFMGFSALTFLEFFEYIILKFRRITQRKKRIKPLA

>Deutero_Ambulac_Apla_gbr54_20_t1

MTHEASRQRHGKSSLEREFAESTSLHGIAKVFHASRLLIRFVWVAIMLTCLCVCVWQISDRFHRFLQYKANTEISVEYTR
DLDFPAVTICNFNRYRSSALTESDKTYLPYIIWANDYDYDSFGDSQGGVEYWDMVADDGRDNDTHFNYTDFTLRAGFL
MDDITLRQCEWRRKTKSCSPANFSHVFTSFGNCWTFNSGKTGPILKETQAGSGNGVRMLIDIQQSEYTETVDGNIPAG
LKVLIHDQATPPFVESSGLAISPGVYAFIGVRKQQYLNLEHPYGKCNASKTLTRFSHYTREGCKIECRARAILEKCNCRLVR
HPGNETECSPQQTSGCAMSTLGALVGGEDNPCDCPVPCNYTTFTTSLSTASIPSNSVAAHLQGLTETSYSSDYSYSSFSV
PYPAQLDLFEEGYIPKNVILLDVFYEDINFERYEQSEAITPSALISDIGGQLGLFLGASFITLAEILSYFGKKIDFWIQRVLGKH
RRLLERQQDNRHGRDVPLKSVRIDNGNHRTLWADV

>Protostome_Lophotroco_RotSoc__A0A817V161_A0A817V161_9BILA_Hypothetical_protein_OS_Rotari a_socialis_OX_392032_GN_FME351_LOCUS3194_PE_4_SV_1

MSSTVEMKPEQEKTALNIIKKWFLNSPTHGIRRISIATSIFERVFWSLTFVVFTTLMCIFIYTVIVKYIAHPTKINLSVRQHRD
TKNFPAITFCNLNPLRSEKFIDDDLYDQRTPPNSNQFYTDDDYKNRFIQYMGRLVAKNLENKQPTIIPSGYKLDDLLVQC
TFNGRTCHRNLTSFFHPNYGNCYTFNNDYHVEAIQRNDTYHFWSIDDENTVESYKLFLELYLHQNEYIQYLDDRAAFRIF
IHRKDDIPILSQNSFFLAPTTFTKLIFSQRIISFSQQCRNDLTPGMKQIFNSHDIRYSQALCFKLCELRYIRKICQCTDQSLMV
YFQFFSQNNTKTRISRIPCSTNEECLENRATFNSRQHCPDCLPECEIIQYKVQSSFAAYPNTRSIEKVLQRVDKHFKAMGQ
SLVQSNTSACSNNRKVSLLDNIVAVEVSASPVATEILAESPMYTWVDVISSIGGQTGLWIGVSLISFVEIAELLFLLFCHCM
EKIYQSISIRWKHFCQRYISKKISSRF

>Protostome_Lophotroco_RotSoc__A0A818AWC3_A0A818AWC3_9BILA_Hypothetical_protein_OS_Rot aria_socialis_OX_392032_GN_FME351_LOCUS9994_PE_4_SV_1

MSDMTTKHVTVATRTDTEVSQSVPKRRRAGSVINEFCLTTSIQGFPGIARSESIHNWLFWTVALICFTGIMVYFITQTIRA
YFNYPTQTSVTIVVEWPQLFPAVSICNYGMIRYDRFIGPFVNYTNTLNITNTTDTANFTETQSYYINDFLIDQLNQGAPFD

NYFFPLDGMMISCAYNGQTCTVANFTSFISSAYGMCYTFNAKLKNVVDGGIRYGSENGENGLLKLVLYAHQQQYVPYT
KIGTGIVALVHDNSKVPNVDLEAVFLSPGRHHKLGFQKKKNVFLAPPYTTCTEQISPGLQVVYDGYNGTDYAYSLFYCTN
ACVQEYIYTQCGCGDAELWNIRAVVIQGTEQRINISLCNTSNPCPRKYKTFIISSDSILNGYCPQCTTECTDTEFIMKLSSLS
APPAVLLPRIKQFVESSNITLPNDWSTSWVSEIQSNYVSLEVAYETVRTELYSQQASLGPVDVLSNVGGQTGLWIGISFLS
VMEIAEMIYRLIRCLCFNRRPVVNNEIKAQEAFQ

>Protostome_Lophotroco_RotSoc__A0A818CK20_A0A818CK20_9BILA_Hypothetical_protein_OS_Rotari
a_socialis_OX_392032_GN_FME351_LOCUS12048_PE_4_SV_1

MQNTGNSRVVPTVLPTKQEARLDLTKPRRLAIIRDFALNTSTHGLPGIARSQSKHNCIFWTISFLIFTGVMIYFVTRSIQNY
FQYPTQTSVSIFVERSQVFPAVTFCNYSPARFDRLAGPFLNYLNSINATNTKDTETFTREQYFLLSSYFQHLVNTGQSISEY
FFSLDIMLIECVYNTKTCTHDDFISFISSAYGKCYTFNAKSKSTNGTSLRYTNDNGGSGKLILRLYAQSHLYSPHINEVDVSA
GMVTMIHDNTQLPLINVAGILLTPGRKHKLGYKKKQYTFLPSPYTYCTTNIPRPMRAMFDQYEGADYAYSQGVCYTLCT
QAYIYEQCGCVSPYEWSARSIVLPGTNKIAEAPLCNLTDVCYTVAGLKITKISSIWDQYCSHCHQECSTVTFALMKSSIAA
PSMPFIFATKAHIESLPIPLPANWSTNWLSEIQNNYVSLEVVCESNQVENYTEEASVTAVSVLSSVGGHTGLWIGISFLSI
MEFIEMLYRLCRYEFHSMQQIPRNRFNNNN

>Cteno_Puk_Fal_comp27454_c0_seq1__p1__GENE_comp27454_c0_seq1_comp27454_c0_seq1__p1__
ORF__type_complete__len_520__score_91_49__comp27454_c0_seq1__400_1959_

MSRYGERYPPSSGLQLFPDPDKEEEGISPVQQPCRCTCPCNTQNTRDTGSQMDVKTISTSSGKRATFYDRWFDFSDGT
SLHGFKFIFSGGRVTRAAFAVFMVLFVFKLGWQLRESVISYMEHPTSSKISVSNVEGPALVTFPTISICSHNLVSQSYLDD
NPGLDFLFEMIDTWQPEVVSQIDFTDPQFAPYADFKYSDILRDGGPPSINVLQCSHFTNDCTPDLQSEKSMERELSMRG
NCYRINPKGQLMGKGGDYGKLSIKLFSDLNEYSATGKKLAQVGYSVQFHDHEQYSGAIPNSYWMSPGLIYKADLSLTRE
TRLTPPAGRCNDTVLYNSYGRNEENSCLAQCRDDAMMRACGCVLVPPPYDPTSYTPCTLQQWSECGLTAYNGWYQD
YVDVSRTEEMCYCYPSCREVFYTVSMSSSLLSRFYARDHVQFLPKGYNGTNDVLNNLVVLDILFVNTQELEIEQVVAYG
WTNFLGDVGGVLGLFLGASAFSLIEFFFFVGHLFWTYVLCIWREEHQ

>Deu_Ambulacrar_hemi_Ptyfla_40v0_9_20150316_1g16876_t1_scaffold13442_cov91

MKIKLQMYEHLPWTLQTTQLLTVYRGLSVTLICGKEIEFPTVTVCNTNKLRRSAVRESVYKEMLIMENDKVAPYYNQCD
ETQFRCHTGSDLGICINEDLRCDQYKDCYGGEDEDDCECNPETTFQCLVSRECVDKIVVCDDVYDCRDGSDESNTCKRL
EGFNALVGVQRERFYENYPCIESIYFDYSITRASTFQFTCQPSPLVGRYAGIVLAEFEDMVLCEVEVMATDDRLLNAQIYV
LEETGLTAQTSCAEPLNYEDLSTNPVTVNCGGRKLGRYVLIKLLNTTNYLTLCEVEVKGQVLIDRLPNVALGKPTIQISTFY
DAVSDRAVDGNTTQDNSGCSSTQWEFEPWLRVDLQTVYSVYRVEIVKPVGTGFYLHGNQTSCAGGFVKCPSGECIHP
YRMCDIMNDCDNGFDESGCVNSTTTTDSPDMDDVFRYGWEENFQTITSEPSHFTYFKDNIYQDRGLGRWEDLLSDIG
GTMGLYIGISLMTICEFVSFLAVLCARCFGKYGKNAIHDAHGTA

>Protost_ecdy_cyclo_nema__A0A0V1API1_A0A0V1API1_TRISP_Acid_sensing_ion_channel_4_OS_Trichi
nella_spiralis_OX_6334_GN_asic4_PE_3_SV_1

MEMVCVNIMPANADDDTVLSVVFNIPTYKRVSLVCTKMRRNSAKKKVLKFSWIQFLETTSFEGLRNIHYSKEPCERTLW
TLLFISAAFTSVYFSAKTIASYMENPIATSFTTEWNNSMLLPNLAICHLGGPNVTKLKMMNISEELSDFIICNFLPAKWRLR
CLKNMHNSTEMELVNFLKFRNFTFDQILRQVSFTCEDLGTISNQPHICQQSRIIAHPYTGTCISLRRFAQNVPGMDGGLI
FKVTIPPKEHLYNGMRYHSGFVEAFAFEIGHPSWHSNLHTLASSMNGVLQLGLSVVVYERLESRKGGCLTSDSVSSSASC
YFLCNNKQTLAECGCLDAHFSETNFKNMTNICTPLQLFKCNERLLSTRNVTIEQLINDCQKDCQPLCYETRFDRTVTFMG
LSPKRKFNQPSAQCLKCKSKEQEHQLTIIMAYPNFIIEKIEEKTVLTLEMVIGGIGGQVGLWLGCSLMSFVHLLVYLLQFCF
HVKKDVGHNLNTHFSQARSSFKSLRHFRSKYDGDVV

>Deutero_Ambulac_Apla_gbr14_3_t1

MLKPSARIQALSSGGAVGPMPSSVEESDFVIADNASRSVDHEETTNRRDLRQTADDPVGLEVRNKRVLLFRDFMSETSL
HGIKYVTNSKYHICRRFVWLILFLAAWIGFIILTADHFRTFSEFDTTTSLSYMPVSNLTLPAITICNYNAFKKSAVIESGWESL
VPHLLPFLTTTSFPQNEVDVTSPDYENLTFPDWEAEELSTRWGHRIEDMLLECRWKQSFTCGPQDFEMTWTDFGPCF
TFNGGDDRQPGGGESPYVANDSTSEEGLWLRLNLEQHEYIATPFDTAGFRILIHSRYDWATFVNPGIIQAAPGFTTIVPL
TKRIINAKKYPYVTNCTDGGLKYYATYTPNTCLLECMADAVLEKCGCYMPWMSAPSGGDQEVCDLATVAACGFDALV
GVNCTCNMPCTEEQYMTRVSHTATPAQYAIPKLSAKYNMTAEELRNNIVDLYVYFEDMRVTVIEHKEAYDFAKLTSDIG
GALGLLLGASVLTVFEFLDYLAVYHLFGLTLLKNFCQRIMGKGK

>Deuterostome_chordata_PetMar_tr_S4RU13_S4RU13_PETMA_Uncharacterized_protein_OS_Petromy
zon_marinus_OX_7757_PE_4_SV_1

KAAHAAFVGRAKIHGVRHFLWPRHAVTRRILWLLAFMAALGLLLAWSADRIQHLLSRPAHTRVSTSWVRGLPFPAVTF
CNNNPVRFPRLTKSDLYVAGMWLGLLTHAESQQPQATFAATAANLTANPHMMAGLSEPRRQWFQRMSDFSRFLPP
REAPGISQGLLDRLGHPLDEMVLSCRFCGQDCKPQDFITVYTRYGKCYTFNSGKGGRPLLTTVKGGMGNGLEIMLDIQ
QEEYLPVWGENDETSFEAGIKVQIHSQDEPPLIDQLGFGVSPGFQTFVSCQEQRLIYLPPPWGDCKSTPISSDFFETYSIT
ACRIDCETRYLVENCNCRMVHMPGDAPYCTPEQYTECARPVLDVLVKRDNDFCLCETPCNMTRYGKELSMVKIPSKAS
AKYLAKKYNKTEAYIAENVLVLDIFFEALNYETIEQKKAYEVAGLLETGDIGGQMGLFIGASILTILEIFDYVYELIRDKILFYF
RAKKKQKTESKDTGPCEALRGHSESAGYAASMLPPPHHHHQHHAL

>Proto_ecdy_PANarthopod_Tcas_tr_D6WXM8_D6WXM8_TRICA_Pickpocket_protein_28_like_Protein_
OS_Tribolium_castaneum_OX_7070_GN_TcasGA2_TC005542_PE_3_SV_1

MGEIAPRVVTAKDKDFCHNLRKYFREFTEVTGIHGLRYVSEKRSRTEKVIWVIILLISLGGCIYMIYEVLDKYNKSPVVVSFA
TEDTPLYQIAFPAVTICPESKYSREKFNYSGVYSKLVEEESVTIDERRKFDYLSMLCDDSPDNVATTQNFGNEFYEIIHKMK
LNQSELFPLCIFDNKFTECDRIFTPIFIDEGICYSFNIQGAHEIFRDHVYNFKDYYKVSNNSEKYFDPEFGYRPSAGVHTYPR
RALRSGADSAFGIFFNYDTADTDFNCKNFLQGFRVLIHNSSNVPRVSEHYFRIPFDKVVVAAMEPVLVVTSQNVRGFRP
EKRKCFMRNERFLKHFLYYSQPNCMLECLTNYTLAKCGCVQFYMPRDNSTLICGPNSEQCTENAESEMQIKDLEYRISG
RSLCDCKPTCTNMKYNVETSHSDFYFKEYFKVHNEFNLGDGGHWSVLQLYFKDEQFMTMERNELYGISDLISNLGGLL
GLFTGFSLVTMAEIIYFCSLRVVYNRRLYGLWSGPN

>Porifera_Syc_Cil_scpid45734_scgid16157_Acid_sensing__ion__channel__1_Amiloride_sensitive__catio
n__channel__2_neuronal_Brain__sodium__channel__2

MTDSGDPSDGSTAASSDEVDYGGISGLREFFGETTMHGVQYTFRGGLLLRFVWLLGVLGCLCGFFYFFITAAINYAGFPS
NTKVDIITADSLPTPAITICNSNVALASGVAKSLRNLAFLCRTYNLTVIDDARGCPFTAQQRDTLTESDTRLPAEIIQEIGQS
KENMVLDELVRSLSTNGTVYTYENSSSFITFITALGLCHTYNADGRFPYQTTIGVNGGRRFIINVNQNEYAGITLLSAGVE
VYIHNPGELVQRGQHAIYVGPGQRANIAVKPARRHILGSPYRTPPCTNENLNRDKSLPYSPARCELDCQNQYIAEECECL
PLTVSAGVTLTLPSGRRVSRYCSVQEYFSCSSLLDYAFHNNVSVYSECRSRCLLPCEAPPNTYIYGSALSWDLFPNFIASELII
EVLTDFGLYDNPRNITTNLQLVRENFAQINVFYESLSYERIDEQVSVTLSTLFGSVGGNLGLFIGASMLTLLEFGQYITRFFI
ARFRRLRRQEKDIWQKDDASNDTALDDVVASS

>Deu_Ambulacrar_hemi_Ptyfla_40v0_9_20150316_1g15265_t1_scaffold10904_cov157

MSSTVYCDRYSDGALQFDLQPVATATLAKPKTRKSFAVTSFRDRGYHSKVEVIKEEEKKGDRESDSRSVGQKDTCDDCC
CCRTFYSERFRQFSEDTTLHGLRYAVGPHVGTSRRIFWTLLMIGMIVSSCIYLYLCWVKFLAFPVKTVVSMNYKTEISFPAI
TICNYNHYRKSYVNGTRFEDWVRQRYPTFNFSDDEPPVELDEDFLRINRTWFEISAAHDKWNMIYQCTFGRSKIPCDAS

NFTLTLTDFGVCYTFNGGKESEDNVLHISNGGSEHGLRLRLFVEQFEYTYSENSAAGFKILAHRQNDVPMIRNFGFAIPP
GTESLIGLKMIQEHNLPAPYVTRCSNEKLEYFEVYTHSNCVRELNTNAVVENCGCREPYMPGSARVCNYNESVKCAVPV
IETLHTDKDCPVACSNIGYETRVSYASFPGRHIVKDLEDRLNLTQYDIRENIADLKIYFEEISYEEIRQEKGYAFPDLIGDSGG
ELGLCLGASLLTMCEIVEFFLMWIWRKMTRFQRGVGPNLKR

>Protostome_Lophotroco_annelid_Pdum_comp395003_c1_seq1_

MQIISILHSTSFNIMSFKDELKNFGLTTSVKGIPRALKNQTKCMTFIWAMAVVVFFGVACTQTSTLILDYLKYPKITTLSEN
VLNDPFPLELLVCNSKPLSSLSRMKYNTSDIPTPRDYFLRLNDVMSCQGCTEEEKRKISIYQEELSHPTGYYQYLGREKIKSI
SHEMKDLIVECYVILLDGAGKALLPCAGYFNITQQPCNRFYNCYRLTPMPVLGIDILGVTLVSYVDNFDTEVMEHFSIHS
DIINSPGLMVSLIEKGVLWNDLSGEMWLEPGSFNHIQYNIEQRRELPAPYSDCQDKSDFNSDKGIPFTYNIFSCYSLCLEL
EVTKDCSCRDMTSRAVLDALDLNQNMSYCGDVTLSPETIIANRICSENTRHKNFESCTEKCKPPCSYQTINHRSTSVSWP
PSYALKQFYSSYIKGQRYEERFKTAWQEASAGSVDGFLKLLRKNFIKCTISLENYRALHMDDQPKYTIVNLISQVGSILNL
WSGITIMVVIELVELGVSFCSNSITKVHVMEKNKNLK

>Protostome_Lophotroco_annelid_Pdum_comp403194_c0_seq1_

MESISNVKGLSISADGKSQKTSNNRHATCVCQANEPEESSENLLEFADRTSCHGPKMIVMSRNIFRKIVWIVAFLTVVVA
LICSVVIQSFNYKEAATFQDIRTTYSANLPFPSFTICKFNRFKKSSLMKMSARYNVSHEALWEYLGLLLHNTKEYQFDDFSF
NASMPALKAVYEAIKSVDGFVNPQNLIEEATFQYDDFFIEGMNPWYRCAWFGSKYAYCNGSTFRSIFTRHGRCLVFNS
HWNGTGHRLQLKEGPDNGFSGILDLHPEESSEILTPTMRGYINLGFRVMIHSHGTLPDLNAMSLTASPGFNVMAALSY
TKVISYNSSPNPCHENPPDTSDWRYSLVNCFLTCIAGVTYNSCGCVPLNAHPVAGGKKYCKTIDEFICMSTGKSSEIIYEKI
TDCCLECQDICVRDRYTVQKGWIAFPNPSIVPMLIDGETPDAEYIRENCSALDIYLDTFFHQIVEHKRKVTVSDVFTAFGS
QLGLMLGASLLTILEFVDYIITAIAALFVKRKSKQIKNGMQEQ

>Protostome_Lophotroco_annelid_Pdum_comp415688_c0_seq3_

MKYTKCLKTGKSNEPLTYTEVTNNFADNTNCHAIGSIKRSESALRKSFWIILFLTGAALTIHQICEIFETYFSYPVSINVNVTY
EKSLVFPTIAICNYNMVRKSQLGLMKGSEALDLLTYTVGGDPDMSQWENSEGRAELRHKFLNSWSQLNESSKAAQGH
QIEEMLLDCSYNEKRCYPDDFTHFFNPKFGNCYMFNSNWKNQNVQVSHQPGPFYGLKLELFIDQDEYIPELTESAGVVI
AIHDQDVMPFPEEDGVLITPDAQSNIAIVVEKIQRMQSPYPSDCLNVSRKEAYDYSAYSELYPVRYTHHACYKTCLQAHL
FKECNCGDPEIPMEGVALYNLVDDKENIRTVCTKDQQACMEKVHKKYLNGELKCNCSVACIDTVFEKTVSTLQWPNAN
YEDLYYSRLATRNPQMKSVLKSHSQSGTNPGTNLVKLLIYYSEFNFKAIEEYASYPVSSLLSDLGGSIGFYLGISIIAAFELIEY
LFDLFLVTCQKSRKKDDNRNAKSSPESRILNDLENEREERQ

>Deutero_Cephalo_026690R_t1__name__chr_scaffold11_start_5135990_end_5141804_strand_pro_le
n_526_

MVLVDSRLLEVIEETAEERAQKIVEKHLKERDEKIKEQRKKDIKTMTKEFCTTTTAHGISRVAEAQTTKSRIIWTLILITCITL
FVYQAAILIKSYFEYPVNVDIKVVDNKKLTFPSVTVCNNNRLRKSKLSGSRHSSLLELDERTEREVRDYLRLPGGFGPSPFP
PRPGSTWDFEVADLSDENSDASSVCRVKSKQLCTLAQILRQSKDNDIMSGWSYYNRDGWAAMLSSQCTAADQQDTC
DVCYNDRILLRRIEDIPDKINSTFCCRPDSYIKLTNQMITIDPSSIICSSNDCVSNVTTYRGTVRSAGNAVCQDWTCPNRHF
DFCLEENFCRFAPDGASTDLVCPSDSRQGNLTSEPCQSIARCPISENEDEPVFPTRHKFYQQTYAAQYCEERGKQLCSLE
QLVLLNTFKDVQAIVWGWIRNHSMEAMFEEDCTNSTSEFCFKNMYRYRVSVARTPNHARLNGRSSWKMEENDNEP
WIQVDLLDSHILAGVITQGGGDVRRTRVHRGTRSELQRQAERYDER

>Protostome_Lophotroco_annelid_Pdum_comp403089_c0_seq3_

LEISHRVRVSNLQNTVEMLSNISSSLERFVRVMESMGRYALNRVNKALYALEFEINKDFNTSLHKLNEFKFAYHMFHGT
NVSLTMTLFDDLSKHVNLRILRKLGMYAPSSLYYLMDALGWAVNHEKLPPPAEHRDRVPYNLWASNATGTSKRDCKKI
VEDYRTQINTLRSMTVVRFRKATADWPKTRECLLDYEHGLDRTLVGVKSTLTVLHKFLYKEDSFDFEAIIATVKTELDNIK
GNITHFKELSILYGENKISKLELTKQVNQEVLKEMDSSIEKLVYNLELYAIRPLKTEASELNGYIPFWYQRLLQNLFELEQHF
ERKSLSNVARDLDIWRVPTARMEIPGVLEFKFRANQKWRTWSSSYSLQYIVEEGHATATISEAVQNYVKVSNELLYQLE
NDLATAKAQLLESMNKGANLLEEFRKEAEVSTGFVLNNFLVMDVYFPALRYEAWEQVEAYGVQNFLSETGGYMGLLIG
ASALTLCEILDLIIYNLVIRCSKKRYSQIAVQPAAVVMDTTNYWNQP

>Deutero_Ambulac_Spurpu_021761

MPIKKMTVEPMKTTSAKEQTISSMVDQGMMDMKRTPYPPSYNDRMSWFAVLYRFVPDGIGVAGIRYAFNPREIKLR
RLTWLLLVFIAFCMTGGQVIDRILFFASHPKDVDISIDYKEEVPFPSVAICNINQFRRRAVTGTVFEDFLPDMHFNYPSVN
LTKYRSELEAYEMTRVLREMGHDIELTLPLGMSWNGRELTHANFTRRVTDAGVCIVFNDIDNGFPQLMVKSSGRCLCS
LRLPLDLVCAFYTNPDVREWRNGNKASCATGSYVMPRLVSSGRRRTGQLSRRFQLVKYKPSTNPDTICPFITKKDRLPPP
PPPTQYSLLDNGLLEFLTQGLKYGLHLNLNAAQHEYYFQPFGKIGAGFEVLLYDYGTEPLVEKGAFGVAIGMETRVGVEI
TEYHNLKHPHGRCENNTLKYFETYGYSECVLECLSDFFLERCNCNCLSMRAPARECTAYEFFDCMTPSYDSYTQGVIEQS
CHCPVPCFNRQYKPSVSTLTSPSLHWSNYAADQLQLTPEHFSRRSRMRQP

>Protost_ecdy_cyclo_nema_Cele_tr_G5EFH3_G5EFH3_CAEEL_DEgenerin_Like_OS_Caenorhabditis_eleg
ans_OX_6239_GN_del_5_PE_3_SV_1

MTSVSFGGSDSQFQSLDPAPGYSDDASGYPDIEVIDNEDEMYIDEETYEYLTNHYFRAGHDGCDEYTNLTTFHGMIRVF
NSRNCPSLIFWCLVVTTCLVFYIMVCGTMIKTYSTQPSFMRINETKSHRAENNIELCSEKQLTCNEILKENKQMTCKEVD
AYCVSIRFSTKTKIRLKKKGLYFKHGTEEDVHYLSSKPHTHHMIRLKLIQIDRLNLGRAPCTTNWREITWIEKDSIPDYRYSL
NMCENIRFELTRKVEYLQYDFPCYPSCSEIKYQVSKSKLRHSSDSVVITFSVLPTITLMQETRKTTLIDILCYLGGASSLFMG
CSCVTLMEMFVFLFKLVAKSACSQEVPEPDDSFYNDKIEFEYSDHRNKRRYAIFDRETMEKYLISNNSIDVKSINLDKRLSV
FSVKKAPKLRALKDNRNFDKIDIFDEGISENDAKNEVALSQKDECSVEYQDEKMDTLGEIPLENRRPLRPQLRRCSTATSF
ASHTSKGSSSLTVRSYAPNQQKRRMSKAFTKNMPMNDF

>Protostome_Lophotroco_annelid_Pdum_comp408485_c0_seq2_

MSTSKEQQDGVAVCMDSCAEDSSTQCIPQIIRAKNRLRQLTWVLITVTLVTAWIWHSCFTIGKYLKFDVNVGTKIIPSRQ
LANPAVTVCNKNAYHGVKADEFLTEMGQRTKSALKNNLENASLYGPSWFLAWVEALKSNKNVYKIPSSAQFLQMFSE
FTDIYGPTYGVDYSNFTMKYEQFVQRCLYAGAPCALKGVHYQVVFNWLYGNCFTFTVDKNATGTGPLYGLQLSLFIDQ
EGYSELTTGAGVQLVVHDESQMPFPDEDGISVPPGQSAFVGIRLHEVQRLGGAFGDCFDEAKQRNSLDYFRKYSYSTSY
SLRTCERSCFQRNMMSQCQCLDNRYPPYTVKHIESSKYNNCITQYLSNFNFSNGLPEDLGCKDRLEDMFNNVELGCDI
DCNIPCNTTKYEVMVSYAKWPSEKAMNETKAFMSLKEAPEMSDEEFRRNFLEVTVFFKTLDYVITSESKAVTILQTISDL
GGNMGLYIGASVFTFFELTQLVLDFMNWLCGKKEKKKNATGEAQEGHERYEVKL

>Protostome_Lophotroco_brch_Lingula_anatina_comp147337_c2_seq2_p1_comp147337_c2__comp14
7337_c2_seq2_p1__ORF_type_complete_len_527___score_4_70_comp147337_c2_seq2_294_1874_

MTTKTKFFRYDVPSDFDEENDTLTYYVLWKGLKERTTSHIIPQVTNAKGIFKHFFWILITLATFVGLGYHLYTMFYKFFEYD
VGVKLEITSNSTLKFPAVTVCNENAFRKSALLSSPSKLPTLDAFVTNGTVPTTGFNSSLIDSELGGRSNRATLIDKTLEDISEL
TTGEKQALGHQRSDFILDCEYGGYSCNGKTFESFYSTYNGNCFIFNSGWNNSSILRATESGPNFGLSLTFNIEQSEYVGDL
TQTAGVRVLVHDQDVMPFPEDQGFLAAPGEMTFVGIKMVEIDRYGGRYTTCKKTDTFNITENMYQALYPSVGYSQKA
CKKTCYQRTVIETCKCALSIYPRVETLFDSTTSVRTCDSLNATDVACTVKVQTQFINGELNCSCIQPCSATSFSIDTSSAYW

PSDEYATEFLSNYNSKSSIVRKIYNTSGSAGVMKNLAKVHIFYKDLNYEYISEYVYYDSIQFLSDFGGALGLWLGWSVVTF
FEFFELFMDVLAFSCRCGRKRKVSRSSVNDITLKEKASF

>Deu_Ambulacrar_hemi_Ptyfla_40v0_9_20150316_1g29162_t1_scaffold115794_cov80

MADMSRLNGNNSSVLQMDNLGYEAEFSGGKRIRDEPTNSNTEQDHNQSRKKDMERPGVFRTFTQEAGVSGVKYIG
DVNVSPARRYDNLEGILGGVFEFNSIFPYLYTLFSVDGENIANFASFLTSEDDSYGEGQTQFTFIDGSAPSLENMLRSCSF
GGLPCGPENFTKVITNYGICYTFNSDKEDYVHKVRNADVENNLKYFDGAYSLSKCWMECETDYAVEQCGCRYFYMPG
NATYCSPYMLQDCYFSKMDEFALKTEPCKHCEEPCVQTHYGTKLSYSSYPSTIYATLLTDTVSWPCDEALAYYLMDRSLE
EKSDIFVDIKGFIFERYYTIEVYGQNISVLIPSAEFHSYMIKLLTVRLFVDVFRDLFFEVLLHTLASLNGNKTDGVNSAVYNVT
KDMLTAKYINFSANPWTDLNATLRGIFAAKNLTSDVIEYLIDSLFQEYYVPLFEEVFVYSYSKLTEMRDNKTAELCVIKLGA
ICKNYMLENLSKVTVYFEDLKVNTIVQQTGYHSFNLICTYFTFIKDRV

>Deuterostome_chordata_Ggallus_sp_Q1XA76_ASIC1_CHICK_Acid_sensing_ion_channel_1_OS_Gallus_
gallus_OX_9031_GN_ASIC1_PE_1_SV_1

MMDLKVDEEEVDSGQPVSIQAFASSSTLHGISHIFSYERLSLKRVVWALCFMGSLALLALVCTNRIQYYFLYPHVTKLDEV
AATRLTFPAVTFCNLNEFRFSRVTKNDLYHAGELLALLNNRYEIPDTQTADEKQLEILQDKANFRNFKPKPFNMLEFYDR
AGHDIREMLLSCFFRGEQCSPEDFKVVFTRYGKCYTFNAGQDGKPRLITMKGGTGNGLEIMLDIQQDEYLPVWGETDE
TSFEAGIKVQIHSQDEPPLIDQLGFGVAPGFQTFVSCQEQRLIYLPPPWGDCKATTGDSEFYDTYSITACRIDCETRYLVE
NCNCRMVHMPGDAPYCTPEQYKECADPALDFLVEKDNEYCVCEMPCNVTRYGKELSMVKIPSKASAKYLAKKYNKSE
QYIGENILVLDIFFEALNYETIEQKKAYEVAGLLGDIGGQMGLFIGASILTVLELFDYAYEVIKHRLCRRGKCRKNHKRNNT
DKGVALSMDDVKRHNPCESLRGHPAGMTYAANILPHHPARGTFEDFTC

>Deu_Ambulacrar_hemi_Ptyfla_40v0_9_20150316_1g2678_t1_scaffold1096_cov89

MSCTVLDENNNCPDEYRECGSGECVYQYAICDTIHNCADGSDEVNCTTTTKGENISLRTVTEQFLDISGVTPEILSEFENG
YYKDNGFSGVVKGEVPPDWYGFKTFSSTPDYSDLRRVLKLTREEIAKYGHQAEDFILQCTYDEKSCSPSEFLTFQDDHYG
NCFKFNHGRGGTVLRDAKRQGALYGLRLTLFLEQNEYISIYGRNAGTRVNISPLDASVLPQDEGITIMPGTVTSIGIREIPV
DNSLDICFKHRNRTVIAGFRARLYIGLSVITVAEFIELVASIIQYMYCGQRDMKSRVKSANEETKEHFSGAVAFPGFRSFD
VDVAHSTKKFRIVSGKIFHRSVKYGTIVNLTLYNHMAEDYTTMHWHGIYQIRTPWSDGVPYITECPIPPGGVYVYAFAAI
PAENPVDQPTFDSGCGIPHDFYGESGTITSMGYDGTTLYENNAKCEFRITASEGKKIVLNFVAFDLEEGSDFLSVNGIGG
ESIAEYTGSNIPGPLATSSSQVIVHFNTNESGQAKGFHADWEAV

>Deutero_Chord_Hsap_sp_P78348_ASIC1_HUMAN_Acid_sensing_ion_channel_1_OS_Homo_sapiens__
Human__OX_9606_GN_ASIC1_PE_1_SV_3

MELKAEEEEVGGVQPVSIQAFASSSTLHGLAHIFSYERLSLKRALWALCFLGSLAVLLCVCTERVQYYFHYHHVTKLDEVA
ASQLTFPAVTLCNLNEFRFSQVSKNDLYHAGELLALLNNRYEIPDTQMADEKQLEILQDKANFRSFKPKPFNMREFYDR
AGHDIRDMLLSCHFRGEVCSAEDFKVVFTRYGKCYTFNSGRDGRPRLKTMKGGTGNGLEIMLDIQQDEYLPVWGETD
ETSFEAGIKVQIHSQDEPPFIDQLGFGVAPGFQTFVACQEQRLIYLPPPWGTCKAVTMDSDLDFFDSYSITACRIDCETRY
LVENCNCRMVHMPGDAPYCTPEQYKECADPALDFLVEKDQEYCVCEMPCNLTRYGKELSMVKIPSKASAKYLAKKFNK
SEQYIGENILVLDIFFEVLNYETIEQKKAYEIAGLLGDIGGQMGLFIGASILTVLELFDYAYEVIKHKLCRRGKCQKEAKRSSA
DKGVALSLDDVKRHNPCESLRGHPAGMTYAANILPHHPARGTFEDFTC

>Protostome_Lophotroco_annelid_CAC9453486_1__Ofus_G01826_Owenia__fusiformis_

MILRKSSQIRHENDDVTDANEKANTCTQRVIDVLKQFCETTSIRGLPRIVKSKYMFGKIYWTLAFLLATGAMCHTLYDHI
DVYFSYSTVTKTTIVSELEIIKTFPDVTVCNLFGFAHMNETPKTALEYRSIVFDLVTKYRLPVDFFNYFEAYLSFMTEAERYE

AGHKIHDFIVDCKKINMISGFNDTCPENMFVLFTDLTYLNCYTIQLQNDAALRYITSGISLTINLNHYNLNVSLLDLIDGMP
TERGIAVYIHEHNTMPPYHTPLKISPGHAVEIQIETEDVIKLPPPYGTCMDGSDTKYTVVGSDSKNYSYTMISCVYDCVQT
LILERCKCVSREYPLFRGQPICLNGRNNVPRIINETDCLFELLEENIEFQCLSECKEPCRGTSYKYTASQSIWPNEGSQLDIY
KKYIQKSKYQHFYEEYEELLTLGKYTLFEKGKKLEQLNTLRNNMVKLDISLTSNSKIVRKEIPLINDISLFTSISGTFIFYIGFTFF
SLLEMVEYTWKILKVIFNSKKQRSDREISLKG

>Cteno_Puk_Fal_comp26884_c0_seq1__p1__GENE_comp26884_c0_seq1_comp26884_c0_seq1__p1__ORF__type_5prime_partial__len_531__score_77_18__comp26884_c0_seq1__1_1593_

QRRVRGFFICIYPSCVCFEARNVKRTAHSTGTSTIGRRMTWIKDFRDFAGDTSAHGVKNIFEGPTKLLKLIFLICWIVCSVY
ATYVIMVSIITFMQRPTGTKYTFLVENENRKLGKSAIVTMPTISVCSHNKVSKSFLKTEGNEILSEVYTTLDKYDKEDAEEL
AKRFDNSSDPLSEVADYLYEDLLADGSIPDDRLLLCQQRAKQCFKLPAFAKTPFFTRENSITGKCFRINPMGTLSALMGD
YGALKLYFWADVQDYSEASSNSATHGFTVAFHDSDTYGSTLSSGYLMSPGSYYKVDLRLRKEVREPPPSGHCDKTREFT
SYGNYSEGACVMECKDKTLNETCGCVNVVPPINDLGYRSCTLKEWATCGLKTYLDWYDAYTDTDTAQTTCKCHLPCQE
TNYEAKVSSSSISTAYSEKMYSNPDVVNWMKKISKPELGINYSTAEDIRENIMALEVMFSSSQVSQVEEFVKYTSTNLLG
DIGGVLGLFLGASLFTILEFFQFIIMSIAKYCCQWTPDGRKKLDGHEMS

>Deu_Ambulacrar_hemi_Ptyfla_40v0_9_20150316_1g19294_t1_scaffold17970_cov134

MFHTEASYAFQRKKRIIPIIVEKDYRPDGWLGLLTLPELYFDFTSDEQLEKNFDELVRQIGDHGNLNDDGKDELHISLQRS
KEGLKHHQLPLKSGAYNVERGKAYQKCQTNDNSEEDESDVDDKQQRVDSRTCGDGQERSQATLKHVLADFGKGTTA
HGIVHITEAETSVSRTVWLAIVVAAAGIMVIQMTLLLVQYFEFGVTVKVSLVSEKMLEFPSVTVCNTNKLRKSAVRNSIY
NQMLILESNFVPAYYECEEPNFRCLASSECINNLKVCDGTYDCRDGSDESGSICEPKLKNVARGKATSLTGPILENRDSNL
AVDGYTYTCVQTDWIWEPLWSVDLGKNYLVYKLKITHMATYDPKLVNAEIRVGLAPKAGPDNPICVKSIREHETQTETF
TVRCHPVPVAGRHVFIQVKGKTEDMRLCEVEVLATDIAEELENVAYGKPAEQSSTYYEHVASRAVDGNENFALETGHSC
IHSAKEAGAWWRVDLLNVYSVYKVVMINRYDCCRERASFVVGYSVLQVLSSLSYP

>Deutero_Cephalo_027180R_t1__name__chr_scaffold110_start_763085_end_774815_strand_pro_len_532_

MESKAASETPAGLWSSYREETTAHGLPKTVTAKSRFRRAVWGIIFLASFVYFLYNFSLMLQRYFSYPVTTSIDIKFASLPFP
AVTVCNVNPVRASKLPKVLREYLGLEPENVSPSIGPDHAHVREKREDGLPTPAPTDELGTTSQSNSSSEDDYYYVYEYEY
DYYDYDEGFEVEQNFTIMLAHKNHTQRKELGHQKKDFIQKCTYNGRPCSRKNFVQFLDEKYGNCFTFNGGMTKNGSR
IPIRNVTQPGPLHGLQVLLYVEQDEYVPVLTETSGVRVLINTQAERPFPAQAGFSVAPGFSSSVGLKLTEIERLGGNYGTC
TDGTGKSYDLYGSRHAYSPVLCQKTCYQRAVVRGCLCADAGLPIPEGTAVCDPAVRRTAECLKETRDALRKNLLDCDCP
EACKESVFGQTFGFSELPSATYQEQFIEDLKKNDNPKIQKMTGTDARLNFVELTIYFEGLHYDKISQSPAYEVINLLGDIGG
QLGLWMGVSVMTVLEFFEFIVDLLSWPLQKQKGDKKDDIELGPADSTVKT

>outgro_CafRoe_KAA0157466_1_hypothetical_protein_FNF29_00042__Cafeteria_roenbergensis_

MASSKNAPGSMAITGADEGFQPLQEFSDNVTAHGVRKVMYLSKERSCRKWVWVALLVGALIALVVIVSERIGYFMEN
PTSTVVVWENIEPVPFPAVTVCNENGFQQSRVAEYGLDTLNNSALLMMVQPVGNLIVRCEFAGQECFADNFSVLLDA
QFGACYTFHGLQPVTVGSEVYDANNGNRLTSARTGPDYGLRLMLDIEQSEYVGYTPKTSVQPPQQDVKAAPDNRNST
VEALFEGPESAGIRVIIHDSSVLANPTSGIGVAPGTASDLAFVRADRSRLFAPWGTCNQAWHSESQGTDPTGYAKSACE
RDCYYASVADVCECLLLPWPESLSDRPSLFVCSVEEKTCIAEQLSKFQSGKLGCSENCPDRCLETLYTATVGSQLWPSSSS
ADTILSLVRDTRQDNTISAEYLTENMMSVRVFAPSSEFQVITTVRAYTFESMLGEVGGNTGLWAGMSALTLVEILEFVFL
ALSAWLCAGCRCCSRDRAVRTVGGSELHAEMSPGSARPGPDAAASRAAFDDRELAAKASP

>sp_O35240_ASIC3_RAT_Acid_sensing_ion_channel_3_OS_Rattus_norvegicus_OX_10116_GN_Asic3_PE_1_SV_1

MKPRSGLEEAQRRQASDIRVFASSCTMHGLGHIFGPGGLTLRRGLWATAVLLSLAAFLYQVAERVRYYGEFHHKTTLDE
RESHQLTFPAVTLCNINPLRRSRLTPNDLHWAGTALLGLDPAEHAAYLRALGQPPAPPGFMPSPTFDMAQLYARAGHS
LEDMLLDCRYRGQPCGPENFTVIFTRMGQCYTFNSGAHGAELLTTPKGGAGNGLEIMLDVQQEEYLPIWKDMEETPF
EVGIRVQIHSQDEPPAIDQLGFGAAPGHQTFVSCQQQQLSFLPPPWGDCNTASLDPDDFDPEPSDPLGSPRPRPSPPY
SLIGCRLACESRYVARKCGCRMMHMPGNSPVCSPQQYKDCASPALDAMLRKDTCVCPNPCATTRYAKELSMVRIPSR
ASARYLARKYNRSESYITENVLVLDIFFEALNYEAVEQKAAYEVSELLGDIGGQMGLFIGASLLTILEILDYLCEVFQDRVLG
YFWNRRSAQKRSGNTLLQEELNGHRTHVPHLSLGPRPPTTPCAVTKTLSASHRTCYLVTRL

>Deu_Ambulacrar_hemi_Ptyfla_40v0_9_20150316_1g2755_t1_scaffold1119_cov138

MDHQDKELSHQHLYFTTAPYRSAWLYDHMQSNDTQRMATVFGAGSTLNILTPNGTRQGNAGSQEMEIGGGSVANS
GSSASKCKKRAKRWADETSSHGIARVANARSKCGRFFWTVIIICASVGLLTIITLLVREFLGYPVNTVLTIVPQNSAIFPAVT
VCNTNRVRASELKNSRFAALIEADINPNHRYYVVPIGTMMVLDEKCLMVHNENQILHTIKNVTESWCEVSFNVTPCFRC
LSSQYNRGTSECTILMESHLIGGNQLVDDENCTYYYKNLTTYCNGFSCDNKTKCLPKSTQCDFLMDCDDGTDEDGCSR
QCFDTEFACGNGQCLDPYKVCDKTEDCVNGSDEINCPENCSDIHFLCRDSLNGDQCIPKYKYCDYRQDCKNPNDEDPD
ECRHRQVLGGEDLFSEGWYDSFVNLTDNITRFETFRNNVYHDLGFERVQYEDPPDWNGFFLFSSTVDFSDLKRVLKFDE
EEVRILGHQREDMILLCKYNDQKCDVKDFLQESNDVYGNCFTFNHGIDLPTRNASRTGPAYG

>Proto_ecdy_PANarthopod_Dmel_tr_Q8MLR6_Q8MLR6_DROME_Pickpocket_3_OS_Drosophila_melanogaster__Fruit_fly__OX_7227_GN_ppk3_PE_3_SV_1

MSNERVLLWIRGAAARILRGLLALLWLILAGCGRLFGRLRDVVAQAGAETGLLALSLCVNRSIHLLERLFWLAVVVSSVIA
ALVLSRLQLERYFSSPTVISVDRDYRGWNGSLPAVTLCYYDHIDSFKANEYIQEMWNVSIIDEDYFYFMDFLYAVVNATA
GNYAELAKFAEDERFDQIDLYEMIQRVDRPFEQVISSFDAGFQVHVQRVMTERGACYAINSPMSTVLSGQPVAYELMP
QPLSCQYGKQQCYIRMDLYESTGVLDVHSPFEVSATEANIVALHKSDEITASFKVLETVASKNLRQLSVAQRKCVFNNEE
TSNLKIYSKSLCLARCRAVMALEMCNCVPFFYPYVDGPSCNPAGFECLLDFKWPIWALHICKCPSTCTEIEYTMQTVKKS
SWGVKNNEEVASSETATSSFRWDLIPPKVRMRRDVVYSFEDLVVSFGGVLALFVGVSVMGLVEMVHVLVNNLLLDVF
TLLRLMLGRLRRCWRRDDHMREANHHHHANAHAHARLHDFNERLSLAATAELPPFDYVN

>Deutero_Ambulac_Apla_gbr160_4_t1

MMYRNRSADEAHLNNSTLSGWPTIGSVYSLPTTINEGKVTATKTPAGGKNGAVGERLGGFGHASTFHGLNYVMNNSL
SRNRRLVWFIVVLGMTTLLVVNTIQAILVLLRHPETSAISLNYVPRITFPAVTVCNINLLRNSALDESVIPTLAAIYLEDRSDS
TGFDFGPVDALYHYPNGSDSLETIFNASHQIEDMLFRCRWRHEKCSAANFTRRLTDHGVCYTFNDPADERDALQVQN
PGSSNGLFMRLNIEQDLYTYGESTSAGIKVLLHPQGEFPIVKEFAFSLTPGFETSIAVRKHMVRSLRAPYKSNCTESTLREF
PFKYSVAACHFECLVHFVVEHCGCRDYRWPAVKAPICSVEKQVTCVYHYEDEFVWDSGNCSCPMSCETTTYESKISQA
QWPAKYFDRDLQEKYNLSHQYLRDNYVDVYIFMEEIMFMEIEQQEAYTLEHFEGDIGGYMGLLCGMSLITVVEWVDFI
IITIYKRFIGSKWMCSRKTRVMSSDDAGELETFGEEPSYAWRARRGASLLGLAQLSLAGT

>Protostome_Lophotroco_annelid_Pdum_comp409581_c1_seq8_

MFVIRQKPTDHQNETSEKGIIAEFCGSTTGHGYYKIYSARHQPLARFFWIFLTFFAMVCFIVHATSLGLRYVSYPLRSDQIT
QSEEVHFPDITICNIYPYDVATWVLNDLEPIKKYRSELDCLRANHSSIGNEKQWNTLKSITGIYANMKPEEFSQLGHKKED
LIKDCVYGGKPCNMDHWKLMPHPKLVNCYTFHPKHEGLVYRQKSLSATVYIQSDDISQQCLHSSNYQIGSAHGLRLSIH
PPGTAPRKKHAMNLPPGVLVLLPLTNTKSTRPGHPYSDCVTDRNLVALKEDLNYSLTACSSLCQQQLVQRRCNCSSTGE

LIPDRNDPDLCKKVIPSDLDLTFKKMTCEDEADEEAAEMDTVTECGCTWPCELWSYKVVPSTAVWPTESATLDILKTYV
SATNLEKCLEKEAGPMPPELKTIDHSSPIPAEYHSYLYKANRWIKGHFLRINIYMRDPRVVEYRYQAAYGLQEFLSDLGGS
LGLWMGFSILTIVELFELIPRFLIRFSNLTTTTQPTGGAVHNYSKENASISKISH

>Protostome_Lophotroco_annelid_Pdum_comp414467_c0_seq1_

MMTVKTVLRSFLETTSIKGVSKATKSRVRFLRISWTICILFGLSMTVYFLYNTLVEYYRYEYTTNFMEKRSAPPFPDITICKI
NPILFRGKDATDYENYKEILESHNDMNQNNSEDSQTDRSKIQGFLKDLTGRVGYFQYSNASKVLENFNKSFVIHCEIGL
WTGQNKDCHLHVEQQPLFLQPDTPLCVKISPLEADKTSIRSIETILYLDDNRCDEDTDLSVQNLLTVGADGIFVLPHGTA
VEPAKYDGIRIPPGVEMDLLVNAHEFKRLGNPYSNCSDHTLVSEEVCVLSCVQKMVIRKCKCLDPYLPTTYPQRLEYSFC
GNVSVDHENFKKNIKCYINRDWTEDEAFCTEQCFPPCLETKYDFQEAASKWPHEIFRTVLYSYIKNNPAISSDVRKEISCSI
KHDKDIKIFGDGFDDASDSDVQSADNESVNCNDLRNSCTKSSHADPIERNFLKVNIILTRRALVSSIDTIKVSISQVYGTVG
GTLNLWIGITFVTLIELFELIYNLMKDRYDKDKVVAMTEKSEKMPPSGKF

>Deutero_Ambulac_Sakowv30041176m

MTDNKLETPIRGTFCGLVQNQMSNTTAHGVPRIESSTNLLRRLYWVVVFLGGVSLFLWQCSTIISKFYDRETTVNIDMK
FKPMLSFPAVTICNLNPVKSPDATPPTLQDNSTANPIEAITNKEFTMPYKATTDTGTTADYSRNWESKHVAFNEKREGF
KLREKVQNYMASKDYEDRSAYGHSLNDLLFDCTWKGYSCSPSNFTSFFNHMYGNCYTFNSGSLEETLMTSKAGPLYGL
NLELYIEEKEYISDLQESSGVRIVIHDQNEMPFPEDDGFMAAPGFLTSVGLRMIKVTRQPDPYSTCKTDADADATTNIFS
ELFSTGYTRKACEKSCFSWAVTEHCMCTDIIYKYHSYETVCNSTDPDIVECQEMVSLLNDNGELSCNVTCAQPCEQRLYE
ATVSNVVWPNEQYKSALTEELFKFSDDVKQKLNDNNNFISENMVKIDIYYNELNYEYIQEQIAYDNGDVVSDLGGQVG
LWLGVSVITCCEFIEFMIDVCILLWKKLTKASNLKVTPSKLKETKGFSEEKSGMPKEFQMVNT

>Protostome_Lophotroco_annelid_Pdum_comp418361_c0_seq1_

MLDHSGEAHRRRMASVKEEMKRFLQTTSVKGLPRIMRNQTKLLRTIWIVAVLAFFMVAMYQTVMLFIQYFRYETSTIV
SEKILEFPSAPDVIICNMKPVSSLRNQTNNQTATLHDFNAYLNTIATCNNCSAEELKVRTHLVNKLSSPAGYYQYIGKEGA
ISVSHSLEDIFVDCRIEFIDGMLTSRYPCNSSVNITRHVSMTHFNCYKIQPSTPAALGWSIISFVDNFQTDILGQFSAHEAG
ETLAGLDVSLIEAGTLWSGIKGQIWLAPGTSNNIQFLLQKRENLPPPYGYCRNPGYEGEKYKMPYKKTMTSCISLCLETEV
VNKCFCKDPISLAMLDELGLKEELPYCASSSQPIEQMIKNLQCANSVRMPSFSLCAEECHLPCTRKLYTHTISNIPWPSEQ
NLNYFYKNFIQGRRYESLFRSAWEEANRPNGSNANLLNLVKRNFVHFSIFPLDHTFLHLRDQPKLELVNLSSQLGGALNL
WSGITLIVAIEVIELFVRLCSKDSGSSRSTVVDVKKPHFQNNQKQKFGFTETPH

>Proto_ecdy_PANarthopod_Tcas_tr_D7EL39_D7EL39_TRICA_Pickpocket_protein_28_like_Protein_OS_T
ribolium_castaneum_OX_7070_GN_TcasGA2_TC002095_PE_3_SV_2

MDEPIKSTQPQLKASYITVILSENVTPRYHFMPTMPIIEDVTYGKVQRNFIVQVYRKKKSGRRMYSEGNIIRPRKIQGWT
QPNFSSPVFRIEDEFALALPLRSGSNPTRRNHTEVVKKFLKVVLRNVENYCDKSTLHGLKYIGDTSLSLVERIFWLLAFVSA
FVFACYYISNIFDKWNSNPVLVAFNPTDATFNQIPFPSITICSMNQIKRVEAERILSEQNPVERQLLNDYCNFNNNSLETA
VQSPNWETVQNFLIRVGQTCSDSIKSCIWQNDNISCEEYFNNDLTDEGLCCSFNRLPAENIFKNPDDYTYLNHTFPKRIY
DWSPEKGFKDDNDGDMGTIPIRPSGSGTHLGLSIVVDAQTDNYFCSSTNSIGFKIVLSNPIETPKIQDYGFLVSPGVEARY
VIQPEIREATRSLRSVDVGKRQCFFQDERPLRYFRSYTKRNCRLECQSNHTLRKCGCIPYYLPKNKKVLDCGKRDEKCASQ
AKEDMELIYGNGSSCNCLASCYEMNFETSATYSKLYQMSKNQRLENVTNEYFV

>Protostome_Lophotroco_annelid_CAC9667732_1__Ofus_G151678_Owenia__fusiformis_

MKKLLEDFAGNTTAHGWGNVIKSTTKIGKAVWIVITSGCCIVALYLIINVIIRYGQFDTKDRVRVTKDDAVFPSVTICPLSP
VSRKGVVNYGKDVKDNFELSAFNNLMNTLEQYDLNISNLLDASQEVQMLYNQSFKRLNTNSGQYENFQLSQLLRYTM

THEDFIPSCKYQGNKCKEEPYRSGHHAHFNCFTYNGPSSNIPGLTSEIAGPTAGLSLLLFLDTDEDFSTSFDPDYPTSGSQ
GVRVVIHDRGTLPDPENDGFDIKAGNSVNVALSAKRRELMIPPWGECVDHNDIRYDVWKYSTKACTSICLQRMIYKTC
GCILGSLPVDDQTKNAEFCANIQAVEWLKEDPNVTILRQDLEQLNCSMQEYTWTDVLAQEGCECKENCSNNKYDITISE
SQWPMKGIEFNFYCKLIVKLDNYENSSIYDVFHQRISQQCSNNSMENIRVVNKQVGDDLREQFIRLNVYFKDLDTKITT
QTEDFTLGSMVSEIGGSLGLLVGMSVISMTEAFVLWYGIIVGLLTKFKRKSKVETLKDKMKN

>Proto_ecdy_PANarthopod_Tcas_tr_D6X1W4_D6X1W4_TRICA_Sodium_channel_protein_Nach_like_Protein_OS_Tribolium_castaneum_OX_7070_GN_TcasGA2_TC012362_PE_3_SV_2

MRFRLEKGPKKRLQNLYKLHKNLGTSWQVQTKLFFEHSTLHGVRYIAEYGRPFLERFMWFSCVGIGAVATAVIIVSLWD
KFQTNPTITGLDTDFHNWKVPFPAITVCQEKPYNLSRIEEYFEDKYSDPSLGYSVGLLTELTKLSFDSIRELTFKIKPSASNTF
DLKEVAFELMNRCEEIFESCEFKSYKSNCCEDFFPVFTEKGFCYTFNSRHYEKKLPWNDTVVPDFQMRYIKETDLAWSLK
FTVKDTSVNMPIYILNSDEQAGLDIQPQHIWDYKINKISFSVKQTYTTEDTKQLSIKQRHCVFSDELKLQTDNIYTYTACTR
QCRMDNSMRLCKCVPFFYPQIGGYKHCSLLQLSCIWENLDSIKSVDRCFCNLGCSNTVYEVEKLTEATGDSGRNELEAR
FVSWPMVRYKREVLFGWVDLLVSFGGIAGLFLGFSLLSGVEILYYFTIRACCMVVREKRELEKIQFEQKTKLPEDYDLSLM
PYCCFDPFPGNGINQVAKSYYHENLYQNKIAPLGKGKWNPSEKILPPFGIEFIN

>Protostome_Lophotroco_annelid_CAC9485189_1__Ofus_G014735_Owenia__fusiformis_

MKRIVNKFASNTSGHGWGMVGCTNSKLGKLLWVAITLGSTIAAIIWVLTIVQTYAKFETKDRVELRADVDIVFPSVTVCL
LYGFSPKKIRATLTDPNSGYVLHNSFLKTIYTNHVVLTQMTNESILENIVGQARTNRGYYENLGPSEISSLGHNIGSLIPYC
MYQESKCTEADFRKLVHHEYKNCYTFNGEDVNLTKPIITEVGAQSGLSFTLYLDHKEGARVTYEKSRAMTVASGARVIIH
EKGTLPDPDSKGFDVEPGHLINVALSVDKRELMKPPWGECADHTTLHSTDFKYTRNTCKRKCMEKMIEKQCGCWKDL
LPVDLNDTRDDDLQACGKMNREEWLKTTRANVTVLKEDIARIKCQNEKYSWSEVYEAVVGCECKLNCTYLSYNIETSQS
VWPMKGSELDFYCIGFSYDTGYNGSSIYDEFHDKIGVHCSDHFEFENVSKNWGDELRANFLRINVYFKDTETKYIIEEEN
FPLVSMISEIGGVLGLFIGVSLITILEVFVLCSSIIYTLLGRNLEETRYMKKKICQSLKPI

>sp_Q9JHS6_ASIC4_RAT_Acid_sensing_ion_channel_4_OS_Rattus_norvegicus_OX_10116_GN_Asic4_PE_2_SV_1

MPIEIVCKIKFAEEDAKPKEKEAGDEQSLLGAAQGPAAPRDLATFASTSTLHGLGRACGPGPHGLRRTLWVLALLTSLAA
FLYQAASLARGYLTRPHLVAMDPAAPAPVAGFPAVTLCNINRFRHSALSDADIFHLANLTGLPPKDRDGHRAAGLRYPE
PDMVDILNRTGHQLADMLKSCNFSGHHCSASNFSVVYTRYGKCYTFNADPQSSLPSRAGGMGSGLEIMLDIQQEEYLP
IWRETNETSFEAGIRVQIHSQEEPPYIHQLGFGVSPGFQTFVSCQKQRLTYLPQPWGNCRAESKLREPELQGYSAYSVSA
CRLRCEKEAVLQRCHCRMVHMPGNETICPPNIYIECADHTLDSLGGGSEGPCFCPTPCNLTRYGKEISMVKIPNRGSAR
YLARKYNRNETYIRENFLVLDVFFEALTSEAMEQRAAYGLSALLGDLGGQMGLFIGASILTLLEILDYIYEVSWDRLKRVW
RRPKTPLRTSTGGISTLGLQELKEQSPCPNRGRAEGGGASNLLPNHHHPHGPPGSLFEDFAC

>Cnidar_NemVecNVEC200_011693_1_1_protein_AED_0_21_eAED_0_21_QI_234_1_1_1_0_91_0_84_13_97_540

MLKLPCSEEMRRNTDPKETALEQINRFLQETTAHGFGRLGATAGSKWRLYWVMFCLAAYCVFVWQLVGLVNQYNSK
PIKTRTQLKHAQKLDFPVVTICNMNVLRASRLPPKLRTKFDQIINNTNKTSPRNSSNSRNAFVDPQDLSFEETKKFEILHA
VTAHDDYRELVSAAHQLEDILLSCNFNGVNCRNSKDPTIPTYWTQTWNDNFGNCYMFNSVKTHNGEKVDLYSSSVPG
ESNGLTLQLNLEQNEYLEGITEVAGMKVTISDQGVLPFPGQQGIRIMPGQSTGIQMTKLQTRRIDPFKNRSCESSNDMS
DKNLFFGYNMTYSIMACKYSCLNARKIERCGCTNYNTPEMQKRNIPLCNRLNSTIIDCLNKAYDTFEDGSCDRECPPSCS
EVSFDLTVSSARWPARSFEKTKLKTLQTEYGINMTKEEMFENIAQVHVYYGELDYLLVQETLAYTFMSLLSDIGGQMGM
WIGISALTCAELIELVCVILANMSNRSKKIVHINSFRPEKLGGVETIEKGGVKRIEQEVQGGVGTIEQ

>Protostome_Lophotroco_annelid_CAC9669404_1__Ofus_G17801_Owenia__fusiformis_

MTDIVETGAETNKQRYSEEHLYPSSVVETGHQDVNDFDKNRQEENKSSIEIDQQEEHTNEFSCDVQKQESTTRNDMHL
REGVLIDTLLGQESHSNKDHVENKLELSVENKIPDDPNISASTSGAESSHNKPAIMATQGEQMKSFRRKISSSLAESAEM
TSIHGSKRILKAKGKLSTIKWTIIFLGTVSMAVSLIASVVVKFYSYPVQPVEDGISVTTQPLAFPAVTLCPFIPYYMRDINPK
VRGKLKNFTEKLWARVHMLDCSYKVIFQPDENYDANTIRKIDKYERGCLPLQDIFQLRHKLISSGWLVENLPIDYETFATP
IKVEDFILECTYNNKPCESQINITRTNTVYGQCFTLTVNEPNDHVSEIGPDKGIALILYTGRANPLSSVNSNKEVRDGFTTIL
STPSDGVQVVLHKPGTMPRPYNEGFYICPGRFSSVEITQETRTRLLPPYGECTDAGYITKTTYGYSYDVCVDQCIQERIAK
SCGCISPMFITPTKHNFMKTPFCGNIQGRIWERTRGGGPNFRSEKKTSAKFWAP

>Deutero_Ambulac_Spurpu_015894

MRGKEKDTLGKSLTDFGNETTIHGVQYIVNKDNIIYRLCWVAICIGFLVVFIIQGSDIITDFLQWPYSTKIDIVGRPSLTFPA
VTVCNANMMRRSQLVDTRFEGLIALDGGVSGADYDYSWWFSSDFVNGFEQSSSSVQRKKRSPPASIDYDMFSWYDP
SDVSDFQYYADNWDTVTSGDDWTGFYMSSRADDYSDFVDVVNPTRAELTDYGHQLEDFVLQCTFDRRQCNISEDFY
VWQNRYYGNCFTFNSALSPANQTRTTGKTGGLSGLQLTLFVEQPEYLGILSPHTGAKVSIHHPDVFPFPEDDALELSTG
QETSIGIRQEYIKRLGGFYTNCTIDGDDTNFTSSEYIYSKAACKKICYQLHLSQFCGCVDNQFFDDYPTKCDVLNATQQW
CRQYVEDRYLDDELACVCPSPCAETKYVKTASSLLWPSERYEEHLLRRLNASSHSNAVRILASSELSSKNLIRVKIFYEDLNY
EFVEMVPVYTIPSVLGSVGGLMGLYIGMSFISVFEVLSLLLRICRIGFAKLFFNGERVLPIKT

>Deutero_Ambulac_Apla_gbr158_31_t1

MQAFSRRDSEQWTIDFSQRTDLGHVWAKGPDMMNSLLGVLLRFRENGIAMVGDIPKMYHSRIFTTLKDQHCHRFL
WRDMQVVQEPKTYCMRVVNIGDRPSATFSTVALKNTAELGKDQFPKATETILKYVYVDEIVESVSTPDETRICAAQTDQ
LIKPGNFAVKGWIFSGAESVGNDKKLPDDDMSEKVQGVTWSPRKDVLHIVTGANFFLKKKTRKEPGEEFYAMHVHNE
GHPGNQTTVSKMCSKFRIVNLYRMVKSVHHPCVPCKRQERDMAGQQMGQLPVERLKPSPPWTYIGVDLFGTYTIQS
EGMAGDMNYCVEFVRTQESKTGNHPWLEILIPLLYSGNLGSRDNVEPSDMDSMHLKPLRAGEAEFKYNSEDTQIESPR
KTVCATVKKFALEQTTAHGFPNIFRATTLFGRLLWTVLFLAGVVATFYQGTVLVLKFRRYPVNVEVKIVTKALLELPSVTV
CNTNKIRTSAIEKSKHRRMLVVDQTTSLPYYEPCLDDDFACKKRPVCIKKYLLCDAIRQCGVNDTSDERDCNYGKF

>Protostome_Lophotroco_annelid_Pdum_Contig14378_

MAGRKKSTVIKFMETSSMKGLPRIVRAQNPVLRIVWLVGFLFGMAMAIYYLYALLSVYFSHTSVTSISEGHSNVSFPDVT
VCNLNPISNIEGGEIRYAEHVETIGSLQDRLQGRGKTELTDMLAYLMTPQGFLQNIADNKLHSEVSSNFSVTCDWWFTE
GAAKEDRDQNCSATITHRFLSASAGHCLTFHAPKKVNGLTAIFFLDDFAYISLPDFDVSSQQVFGGGGSVVLHEPDTMP
DLEGSMAVPSGYHSEFDLYTSTRVRLGKPYSQCASPRDVHWPKQLIMDPGVKYSYYTCVQLCKQFTTIERCGCIAPELLS
TKYMRAEFEFCGKIPMSTNPDGQPNYEKLADMLEKYQCYISTDTTRPMCQCPTPCKENYYHFKYVQAKWPHPSFALPF
YLKYIKKKPYAHKFNYYDEIEKTMVKNKSAAFDMLWKEEKMQQNFFHLKVYFGTQRLDVTYEDKPAIGSETLWGNVG
GTLNLWMGVSFVTLIELVEMIYHLLLSYRTDNDNNAVDRNDSFHSSRSRNASDSGDRSKRESRNGTSSR

>Protos_Platyhelmin_Macrostomum_lignano_A0A267G1A0_A0A267G1A0_9PLAT_Uncharacterized_protein__Fragment__OS_Macrostomum_lignano_OX_282301_GN_BOX15_Mlig019094g1_PE_3_SV_1

QTSAMDSKQSDQPHTAAPSATPQSRLQQAAAEAVARSRLRRQLIEFTKETSLPGVSQAGLSSGRGLVWTGLLSLSLVAL
IVHLCIIVTVYLRYSTTTSSSYDQPFQWPDVTLCNLVPLSVSAVNEHLAEAPTSPSNSLGRFFDALPALAESISAANGPDVT
NSQSLRELMTRSRNLTPVEHKLANSLLSCNYRGVQCPTDSIKQTYVPRFHTCFTVSIGSAGGSSGTLADDDFLDMMLFV
ESGVLSYWDRRTLADFSPAGKVFVHARGSYPDVEDILDLPTGAWVMLGLRYTERHNFEGRCAKYQGEVKFSTPSGEKL
FTSPSDRRSCLESSIQAELIRQCGCISEKFRRFDAHPGQPGCHSVGNASSGSASTEDRARAACHGRVLPQALEAAGTECR
IDCLERDWAIRVSPHVWPKLDASVTALFDAYAEPAVRLRGFDPRLLLVNGTTNVTTLDAKSQLEFVQRSFLRITVNPTSTI
GLRIDESLVYTAGNLLSDLGGILGLYLGFSIVFIYEIAFLLVKLVRDSRAGGGGAVGTADSKSSC

>Protostome_Lophotroco_annelid_CAC9640639_1__Ofus_G092493_partial_Owenia__fusiformis_

VSYGYLKKNISLVRLFFPEFSMTYNMKKVVEEFADHTTAHGWGNISQRSSTLGKTVWIIFTLGCTCLAIASIIGTISKYFEYE
TKDIVELRDEPLEFPSVTVCPLNPIAHTHQLMYRQNIYYEIVDYHNLTNNILPTERKTPQHNTLYKDYINQLVSYQGYYEN
AMQILVYSHNKDDFITSCTYMENMCNRSDITTSLHHEHYNCFTYSTSRSLPDFAKTTVGPNSGLSMILYLDVSKLSMTLY
DPNYPTSGSNGARVVIHERGSLPDPEKDGSDIEPGHSVNAALSVNRRELMKKPWGNCVDYSALDAGGFVHTMNSCIK
RCQQKRVYEECGCVKASLPLEPDLENAQFCGKLDVKEWLKNEPNITILNEDLNRLECQKSAWRPCKTCAKNCTYHTYDV
SLSQSEWPTDGAVKSFYERWIMQQPNYENSTLYHRFHREMKDLKDIYSVTETTKSAIKNNFVRLNVYFKESETKVTTKA
QAFDLANLIAETGGFLGFYVGISVISIAEVMILLYNLLNFLRNKLIQQKYNNTNNAGVNIKQNGK

>Protostome_Lophotroco_annelid_Pdum_comp417802_c0_seq3_

MTGILQHIKSFCESTSVKGVPKITKSPTRGLGALWLFAVLFCAACCCIQVGDIVSRYMSHPRVTLVKEEQISILNGKNMAD
LEFPHIIVCNLNLMSADVSERQDMIDFPKRYYNEVIKWTTCPSCTPEQNELLMKLRRFYYTTHGFYSLSNLSAMQSFAIP
AGQLISSCKALVLTNLELQKLPCEQFADLSILALPDAFNCVNIKAKPPYRSFMPLVGFEFVMYLDDLGPEMPLYFNEGSTS
AYLTGGKVTIYKPGTIPLPERDGIYLSPGYYNDVKLEVEAWKRLPPPYSQCSLNKTVLDAYKFDGNTVRYNSDTCLGIHVA
KKIEEICGCRDSFSFVTVSTKYSDLGSLPQCVEPTDPKIMAERYNCTLHIRKIATSALSSRCHEGCHETYYRSFNSQTQWTS
KKYLSRIYKEHVENTSYAWRFDKNDETKLLQQIKDNFVSLKVMLSTRKYIKFQEEPAYTASYMVSSLGGALNLWLGITVIL
VVEVIDYILKVFLVSNDKKQTAPSGGDQKQTEITKNGNNPGFSNEYVSEYGMDVVKKN

>Protost_ecdy_cyclo_nema_Cele_sp_Q22851_ASIC2_CAEEL_Degenerin_like_protein_asic_2_OS_Caeno
rhabditis_elegans_OX_6239_GN_asic_2_PE_3_SV_1

MRGGGFVQIFKDFSNWSTVAVVPHVANANNKISRIFWIAIFLFVLGMFAYELYILIAKFFSYPATVNTEILFEKQIFPVVTV
CNMNPYKYSVVKSNSAFSSVNTLMTTYSDATVGTFSTDKWGLYADNDETYDLDSRAADALVLEANLISDTAKVPALYTY
ADLIQDCSFAGIPCSESDFTKFIDPVYGACYSFNEDASLNYSVSREGIQFGLKLMLTVTQTKTNGNTDSLPTTKLAGARIG
VNSRGSSPGLDSNGIDAGVGYESAVSVSLTQNVRAKKPYGTCVDREPDSSDYYKDFIYTLETCFNGCKQRDTIAKCQCA
NPRLALGSTDTACQPIKADLDCLQTLKGNQTSSTPNIDLLVECNCNPPCDESTYTPTVSLAQFPSTSYYVATSSTAGVGSC
SSTNSKFSSKSDCQKWYNNNGMIIQVFLETLSYELYTETAGYTVSNVINDLGGQAGLWLGLSVISVVEMTGLMLVMGA
FCVTGGAIKMAPDDDEIENDHRIKDVEDVKKEIDHLEKKHGEMESGSDGEVDDIENKGDEEKKKK

>Protostome_Lophotroco_annelid_CAC9618795_1__Ofus_G081188_Owenia__fusiformis_

MRTLIGEFAGNTSAHGWGKASQTTSKLAKAAWIFTSSACSIVALFLIIKVVIKYCEFNTKDSIRVSKDEAVFPSVTVCPLTPI
SRKGMATFRKELAENNSAIDSMLYYNNFLYVLGQYNLNISSLQNASEELQLLYNQSYKRLHTNAGYYENYQQSERLRYTI
KQKDFIPSCKYQSEVCSEELYHSGHHEHFNCFTFNGPLSNISSLDVDSGPSAGLSLILYLDIDLEHSSFYDPDYPTSGSQGV
RVVIHEQGTWPDPENDGFDIKAGNSVNVALSSERRELMGQPWGDCADHNDIKYDRWKYSTKACLLTCIQRLGFKRCG
CIRGSLPVDDKTKDVEFCANIQSREWLKSDPNVTLLRQDLNQLNCSLQEYSLSDILAQEGCECKENCKIDKYDIMLSDSE
WPMKGIEFHFYCDMIMKLGNYNENSSIYQAFHDRVASNCSSLAYDVERKKIINKDIGKDLREQFLRLNVYFKDSDTKVT
TQTEDFSIGSMVSEIGGSLGLLIGMSVISISEVFILCLGILAILIRKSKAQTKVETFKDKMEYP

>Protostome_Lophotroco_annelid_CAC9629795_1__Ofus_G09673_Owenia__fusiformis_

MESMNTEMTDKQHMNNTEKDKNKLKSKNKTKKELAKDFASNTSAHGFAYLVTSKYGAIHVLWVICIIASGTGMLYGIF
SSFVIYLNFPTNEIVSKYTTKQDFPAVTLCNVSPVSAYKAKLKNSVNSQSFIEYVQFLMKYDTLKSIPGFEEFKERFQSSESI
MTNVNFENWSLQSHNWTDIILSCKINSKACPHNKDFFKPIMNPSFGICVTINGNQSAEKIKISTSGPTHGLSMLVFLDTY
QEGLDNVIYDTESVIQGGSGLYVQVHPPGIQPDPVTKGVQIMPGQSTSIRIKTHVNKYLTYPHGNCTKPFFIDDIGYAHS
ETLCMGICKQALTREKCGCTSLGEPHDSRPYKSGLKMTGNLSYCGRFDIENALSPHPNITELDQLFATLRCQQNINLNFY
SENDKTSCACFPPCMDTTYEITTSASAWPEEMYMKGIYDHYINDRPDRDDLTVYNSMKHLNISSVVWKRHMKKNML
QLNIFFKDLEITLTEQVKSFTFTEMVSSIGGFFGFYMGISVLTLLEIGLFLAAIVKKTCCNQSNKDTI

>Protostome_Lophotroco_annelid_Pdum_comp420144_c1_seq15_

NVEHILPSYRKALETGHRVRVHELQRTLEMLSNISSSLERFIWVLESTGRFTLQRANDAFSHLEREIYYDFYILFQQMRDLK
NNYNQMRGTNVSLAIEMYEQTHWSVKKTLLVQGTQLVEQDSLQYLKQALNWIAIHTKVKTPKNTGTGGHFPLPQYL
WKPGFWMDRCMGQIKQQITLVDELISEYSKQSALEFLEEYQGTMDCLKTFDKEFDEFLDEIEKRAEAMKRFREKESALE
FDNVLAIVEAQLSYIKSNLTYFKDLGRMYGENQITKFQLTERINKNILEKMDDTIKKLVDYLQMYAIQPLRHEASTLIGLIPY
FYHDTMVDMSQLEEYFEKKTLSTFARRFGIWRGPTARLEKSEILEYEHAYNEIWKTWPMSFSLSDIIETSHDIDIISGSINK
YISVVNDLLYELESELDSAKTQVLHAMNSAADLLEEFRRKAQISAEFVLNNFLKMDVYFPSLKYEAWEQIEAYGVQNFLS
EAGGYMGLLIGGSALTLCEILDLVIYNMVIRLMAKRQGKVHPLRSVKSNRASVRTAFQDSIRK

>Protostome_Lophotroco_annelid_CAC9667655_1__Ofus_G151630_Owenia__fusiformis_

MKRILKDFSSNTTAHGWGNIHQRTTTLSKVIWIIICLGCTAVAIWQVITIVMRYGLFETKDRIKVEVGEIVFPSVTVCPLIP
VPHDGNLKFERDMKKGNIDVEPFLNFRLVFHSETFYINSNVSHNLHSEKLEFYNVFAEQIYSNQGFFENFKFMSDYTHQ
QGNFIPVCSYQGKPCLRSDFHTLDHPEYNSCFTFNGANKTINNPITKITGPKGGLSLVLYLDVGQLSMIYTPSYPTSGSQG
VRVVIHEKGTLPDPENDGFDIEPGHSINAALSVNERRLMKPPWGECNDQNPELKGGFTYTKKHCQLSCLQKFFYRTCG
CIKSSLPIDEETVYKEYCLKLHAKEWAKLTTDQYNKTTIVSDMENFRCQNREFLSSDVEVQDGCQCLEKCSYQTYDMTLS
QSEWPMKGVEIEFYCRTIMSMDSYINSSIYDRFHDIFPEYCDHSKDEDLLTKIQKETLRENFIRLNVYFKDLETKITNQVED
FTISSMISEIGGSLGFFVGMSIITIAEIILLCWNITCLLGKKFVISPFEQKIGSDMKHASTNTS

>Deutero_Ambulac_Apla_gbr361_5_t1

MYHQESGSQEGLHLAKRKVSPMTVEDISAEDGAAPAAPPDPAASAKQTSLLYVVSHRLAESGAHGIPNIQRANSNFRR
VAWTLLFLAGFGMFIYQGCDLIIKFYKWPYNVNIEVRTPKRVEFPAVTWCNLNPIRKDALADFTDLRSLLEGGPSSCGTA
VNATFGGYPDDWESADIDYKLEDCTSQTVDAAIDIVGAMPYGERAVTGHRLNDMLLMCTFQQKPCSPRNFTSFYNSR
YGNCFTFNSASRGVAPLKVTRTGPSYGLSMELYIQQDRYITGVETGAGIRIMVHNQTEMAFPEDIGANIAPGTESFLGLY
RAMVNRLSDPYGDCATDFTQDNIFKEQFKRLEYTKEACEKDCFFRAVLLDCNCAYINYRYNDSQAPCNSSVEAVKTCIEK
VENNFTSGELRCGHCHKSCNETRYQISFSNSKWPNREYMETVRKNIQESNKETISLIKNNSYFIEDNVLRVNVYFDSLNY
DDIFQTAAYTLGSLVSDLGGQVGLWIGVSVLTLFEFLELMIDIFVLISTRCRAGGENRRNGKTGQENSRPQA

>Proto_ecdy_PANarthopod_Tcas_tr_A0A139WHB4_A0A139WHB4_TRICA_Pickpocket_protein_28_like_
protein_OS_Tribolium_castaneum_OX_7070_GN_TcasGA2_TC034701_PE_3_SV_1

MPQKHFSFWAMLKYEGKQFCLNTALHGYRFIVLPKRILLERVFWTTIVTISLISAVVLLLLAWINFQENPTLLVTDSTHYPI
WNYPFPAVTICSFNKILKRRAVALAEELDKAYCITTEELANDLRIVFQLIYNNNKNIPQRNYSRLQDILDYNEYPIEDVMKY
LAPKCSELLQRCKWKGEEKRCESLFEPIKTTEGYCCAFNYYALKNHTFGGALANKIPAYPRRVSACGYLTALEVLVNSYPA
DYFGTQIAAIGHRVLIHNSYVYPDWGIQNLLTEKGIINLIGVTPSKTYSTESIRGIDVSKRKCLFNNESQLHNFVHYNFQNC
IIECRMNLSRQYCGCIPFYYQHETGTYPKTRTCNLRDVQCLTDHRKLLMSSVPGYDFSRIGPDHKWGTEQQICRKCLPD
CDYVEYAGEASSGVFTREYSSNQLNLYGGIVIKDQSVVRVYFNDLVATKNRRDVIFSWHTLLAFYGGLLGLFTGFSFVSAI
ELVYFFTIRLLIDVRQKNEIARKKKERFVQVKPHSDEILNPKHLPTNYNRFNLRRRSNSNS

>Protostome_Lophotroco_annelid_CAC9628722_1__Ofus_G09545_Owenia__fusiformis_

MSLTLLLDHPVDDRGTISHYMSLITTHGCQQIYQTKSKLIKICWTIFLLASVCVATYFVSVAISKYYKYRTHEETSLKDTYVD
FPAVTVCRLDDDGVSRLFSEHYTPDKGDEEIIKQEMNILMKIIDIDLDFENEDVSYLHLLLLYAEYAPRLLDRQSIPQAFPAI
AEYLASDYDRFIVSCQFKEHDCNKTHFVEKKNGQYFNCFTFNHTDTIKKEGPTSGLSLVFFLDTLKEIELQEISLGDSSLRFT
DSGIKVAIHEPNTIPDIINDGFIVAPGYSTNIALKQSVFEKMKSPWGECDKSRIKINDITNENTDILYNRETCMKICQQRYI
QGKCGCLNVMLPVPNDLNKEKYCFYIDKDNMLQNPNWTQVFHEELKHVDCESKHLLDLPDSYVGQYKCDCPLSCHYK
TYNSLISQANWPPQNGVHRFIKKYRNEQPTLLNGTDIYKAYEENNATGDLYDMVQNNFLRLNVYHRSLTVQTTTQVAA
YTVQDLVSEVGGIIGICVGMSVLSILELFQLVAIVIRSSGKKADRRGMSLKDGDMELNEEGV

>Deuterostome_chordata_PetMar_tr_S4RTA3_S4RTA3_PETMA_Sodium_channel_epithelial_1_alpha_su
bunit_OS_Petromyzon_marinus_OX_7757_PE_4_SV_1

PPTVLEFWRGSVTDFYDSYDEMFEFFCDNTTIHGTIRLVCSKRNKLKTAFWSLLFTVTVILFYYTSALVFLQYYSYTVAVT
MGLMFQQSTFPAITVCSLNPYRYEVVQSSLSQLDSMTGQALQQLYGYQPPATKAAGTAAAPGIRLDTGVVLERTGPDT
VGFKLCNATGGDCFYQSYGSGVQAVTEWYTFQYVNIMSQVPSYIKQSDDANIEDFIFSCMFSGMPCSDSEYSRFHHPT
YGNCYTFNSANSSKLWQASKPGRDYGLSLILRTEQNDYIPFLSTVAGARIMVHDQESPPFMEEGGFDMRPGFETSLGIR

MLEATRMPDPYGNCTEDGSNVPVLNLYSSAYTVQVPECSCFQLALVEACGCGYYFYPLPPNASYCSYNNTAWAGHCY
YKLYRQFISDELGCVDKCAQPCTTKRFAVTPGYAAWPDSSSEKWIFNLLSLQNNYSVTTVRNDVAKLNVYFRELNMKTI
SESAATNVIWLLSNIGSQWSLWFGSSVLSWLEVGELGIDCCIMVFVLAYRRRRSRAERRRARGTGDSEAAPPPPPSF

>Deutero_Ambulac_Spurpu_015895

MDTEEKDTVKKSVTDFGNETTIHGLQFVVNKKNIIYRLCWLGICTTFLVVFLIQGNVILKDFLRWPYSTKIDIVGRPNLAFP
AVTVCNANMMRRSQIEGSRFEDLVNLDGGVEGADYDYSWWFSSAYRNWYASSSASSYGQSSDQNSNGRSSSSSQSD
SASSSDGQSSSSNYDPSSSEESSSASSGESSGVTSDGQSSSSSYSPSSSEESSSASSGESSGGTSDGQSSSSNDGPSSSEEP
ASSSPGGSADGQSSSTNYGPPPSGSVSSFGLPNSEFPAPNWVNEWSWYDPSFFADFEFSENGWDGVSGGEDWQGF
YEASKADDYSDFLNVINPTKEELEQYGHQLEDFIVQCTFDRRPCNISRDFHVWQNRHYGNCFTFNTELSPDNKNILTGK
TGGLNGLHLTLFVEQPEYLGVLSHQTGAKVTIHHPNEYPFPEDNALSLGTGQETSIGIRQEYIKRLGGYYTNCTSDGKDT
NFTSTTELSYSSVACKKICYQLHLSQLCKCVDDQFYDGFPTKCDVLNMTHQICRKFVEDLFLDDKLPCSCPPPCT

>Cteno_Puk_Fal_comp28403_c0_seq1__p1__GENE_comp28403_c0_seq1_comp28403_c0_seq1__p1__
ORF__type_5prime_partial__len_552__score_111_86__comp28403_c0_seq1__2_1657_

QRRVRDSQHTYNNYLSPTLHHPFAGCNFSSNCFSKLVLDSLRQIMSEEGKGDPPEKQFYSYKDFGSDIFQVLAVERNRP
KWCKALWFLVGLAFFGAAAYLVYQLMTEYFEYKSYNKVSTAFTTNETLPAITICSKNPYNYTQLKSLLTEGEVTDLEDVL
WEMMYYNGTGTNASTGNFFQDELSINSLIKWLDEIKMYNNQTILSNITFWADDYIANRVYDYRLGSQWKYLDTYNYV
SANVIPHMLGYCIMVNDDQKLVQILPGVNGGFTILLDAGIQHYMPNVDSYGFEVYLRTPYEQVMINRGGITAPAGKQT
NIKINKNKMTRLDSHWGTCVSKQSMYYPNKTNVYESQRECYQEALIEIAQDQCNCTPWYWVDRLEQLNLTVSGLETY
PECSFARQIVCDGVIWQLIENWVKKDGNSIGCGHERCSYDAWDYQVSQTEFPVSEKYYEVLYLGSLLDYTYDYAKDNV
AKINIYYGDMKETTVTQEKAYEVFSFISEFGGGVDFFIGLSFFTGFQAIEIFIAYTCCRRLSRRGSAAPSDANATPVKPFNG

>Protostome_Lophotroco_annelid_CAC9666691_1__Ofus_G142267_Owenia__fusiformis_

MEANRDEQTLKQILKDFASNTTSHGYSQVKLSTTRLQRWIWIAICAVVNVVGLALFVTLMFKYFQFETREVVSVENKHL
KFPSVTICALYPLAGSKLGKYAKPLENKTSFESANLMSSTKAMATLTTFLENLNEKYVVQQNKSGHADRFSKHYNRLLSFM
FRSENSQIEMAQRSHSYEDLIVSCDFKGEPCKREMFRYEFNNEYFNCYTFNGVGTNVSSKMVWTGSLYGLNIVFYIDVS
TLNPNYTIYNPKSPSGGNSGIRVIIHPPHTQPDIINNGINIAPGFSTDIGLNLYRKERLEPPWGECKNGKRIEGFEYEYSVTS
CRGQCRQWYLYAKCHCVSAFFLVPKDLVNVQYCGKLNLKNLNSDKANISVITEDLDRLDCERNHIRHHVGSYPKGLRCD
CPMPCSHNQYTKQLSYSKWPFKAAINSYYHTQLKYIANYEHLSMYPAIENARNKSDEEFYDVFHQNFLRLNVFYDSLNE
YVTKQVKDYELVDFAAEVGGVYGICAGISIVTVCEIIQLILAVVKWHQNQYESRKVNVIEADRNMKMSNTKT

>Deu_Ambulacrar_hemi_Ptyfla_40v0_9_20150316_1g20853_t1_scaffold21886_cov93

MSKEGLKHRQLPLKSGAYNVERGKANQQCQTDDNSEEDDSDVDDKQQRVDSRTCGDGQERSQATFKNVLADFGKG
TTAHGIVHITEAETSVSRTVWLAMVVAAAGIMVIQMTLLLVQYFEYGVTVKTFFRGINICTTIFVIPIEFLRDDFKDIMKRR
MFSMFDRQVDRSLSVNSTDILPSIPLYKCFESCLRWKRYSCRSFDYNRTTLTCRLFKESAGNSGGEVVKAIGTDFYQNLRT
RIPDTVFKPGWQMPYQSIIDDPDIYKYFKENVFINPGYHRVKGEDPPDWLRFKSFSKSPDYTELADVLKVNTNEIAAMG
HQAEDFILQCTYGGEVCHPRSDFVITQDEVYGNCFHFNADPNKDTRFTRGTGTAHGLSLVLFTEQYEYLGIYGQDSATV
VSIVPRNLRPFPVDHGFFAMPGTATSVSLTESKIHRQNQPYGNCTDENSNLHFTDNMKKDSSDKMYSIVDNLLKLVKV
KNQKTRNLNDLESARSNLVRLKVYFESLNYESTSEHPAYTVRILVCFDDVTVSSLGEFRSVCLRLKVNFDFVSLLLTF

>Deu_Ambulacrar_hemi_Ptyfla_40v0_9_20150316_1g29125_t1_scaffold114812_cov86

MSSFNENNSSVLQMDNLGYEAEFPDVKRIPDEPTTSNAEQSDHQIGNKDMKRPGVFRTFTQEAGVSGVKYIGDVNVS
PARRLLWLLIVLLALSGLVYQVIRLIVSFAERPVNVNINFEEPNTRPEFPAVTICNNNMFEFDNVVGILGGLFEYTVFFPYL

YALFSVDGDKIANFASILIPEDVNYGERQNQFTFIDGSAPSLENMLRSCSFGGLPCGPENFTKVITNYGICYTFNSNKEGY
VNKVRNADEFALKSDPCRHCEEPCVQTHYGTKLSYSSYPSTTYANLLALTGSWPCDGALAYYLMDHLSDEASDIFENIQE
FILERYYTLEANGRNISVSMPSAEFHSFMVEHLMDRLFLDVFHDLHFEVLLRALASLNGNKTDGVNSAVYNETKHMLM
ATHTNFSSNPGSELNATMWGIFAQANLTADVIEYLIDKFFQEYYVPLFEEVFVYSYSKLTAMRDNKTAEITEEESGAICKN
YMLENLSKVTVYFEDLEVNTIVQQPGYHSFNFICDIGGSLGLFFGASMVTFLEIIDFFIVQFWMKKHSKPDNK

>Deutero_Ambulac_Apla_gbr412_20_t1

MGSGDQKETFLGALNTLLETSSAHGLPNIHRSKSFLSKAAWTLLFFAGVTALTIQVISLVTTYLKFEYTVSLDVRFDRSLNF
PAVTVCNINPVRKSKLERASDGFRELFDVNFAPPRPEDQTGAVNPGDNTDMSPGSTAGGTMKQAGQPLTTQGLAGA
GMGQGKGQGGSGMPTAGATMPIAIPTGITTEGPETTAGIEYSNSTSSVNNGTTEATILAWEERQTGDFFKKRDDDYLK
EQRLISHLANLTETQRMDMGHLLEDMLLDCQWQGYPCSPANFTSFYHYKFGNCYTFNSFRFGLTLELFLEQEEFMPEIT
EAAGFRLVVHDRDTMPFPEDDGISISPGSKTAITARVVTIERLGNPYGNCTKKHDEPDDGNIFRDRYGVSYSLKACERSC
YQKEVISQCGCFDPHYPNTLNDTVYPCDIDNDEAQSCMSDLEEKYKQGTLTCYCFQQCNKNVAKIEVYFQEFNFEYIKQ
SPAHTIPSLMSDIGGQLGLWLGLSILTVFELFEHCGTFLAVIVSKLCREGGNLKSSSSKVQKIKIFPTEDPGITLQNY

>Protostome_Lophotroco_annelid_CAC9661516_1__Ofus_G111721_Owenia__fusiformis_

MKRTISDFASNTTAHGWGNVTQRTTKLGRTIWIGLCLGCTGVAIWQVITIILRYGTFDTRDRIVVKNGDIVFPSVTVCPL
VGMPASGYDQLFQDMREGKQETIPLMRLTELLQNLTAYVDNNISQSAQSEKLEFYEEFHTQLYSFQGFFDNSKKSIQDY
SHKQDNFIPVCVYQGKSCLKSDFTTLDHHEHYTCFTFNGGNTTIAKPSTKISGPKGGLSLTLYLDAVEGSLIYNPNYATSGS
QGVRVVIHEKGTLPDPENDGFDIEPGHSVNAALSVNERQLMKAPWGECADHDSEAYGSFKYTKKHCLLKCLQKTIYET
CGCVQSSLPVDDDMAHTRYCLKLHLNEWVKLQSNEYNLTIIQNDLEQLRCQNNFSLSDFVGQDGCQCREKCLYQSYE
MTLSQSVWPLEGVEIDLYCRSLMNMDNYINSSIYNSFHDVFPEYCGHAKSEEENKILRDTYGIKLRKNVVRLNVYFKDLD
TKITQQAEDFTINSMISEIGGSFGFFIGMSVITIAEVIILCLNLAQVIRKQMVISPTHQEKEMARKNAWVETWPKQ

>Deutero_Ambulac_Sakowv30044073m

MNSTRELSHGTLTVESMDASPKFKKRMKRFTRRDIVNEFSRNTRCHGLPRIISAKTLPSRFVWSLVFFAALGAFVFQATK
LIKLYLNYDVTVTIEDETIASLMFPAITICNTNKLRFSEIEKSEHAVLLRTDPNHPNSLHRSLSYQGPCLKGDFECSDGIHCIK
PHLHCDGYIHCWDGLSDEVNCTYPPCGRDQFKCNGPGYYGICIDSDKRCDGLPHCLSGEDEAYCNECKSGFKCDDNDD
GQGGKCVQEEQRCDRFEDCKDGQDESKCDSVYADPCDKNLSATSMQENLYSPLYPKAYPVEKICTVTIIAPPNKSILITFL
VFDIEHNSFCDYDYLKIQDNMNSDLNVKLCGGLRPPPWISSSNVVTVTFRSDTEYTARGFHLVYEAVNSTQYTMRWEV
GPWSNCSKRCGGGVQTRAVMCNGASQTGEVTNLDECKAIGSRPVSTKVCKQEVCESTCQSRLTHCCQVIQSKGYPDQ
YTNGQNCYTNIINEGGCINITFSDFPLEQGGSCGDFVELSDHNKPTLYRRICDMETGDSNPTWGSFSGNVSVTYR

>Proto_ecdy_PANarthopod_Tcas_tr_D6WXM7_D6WXM7_TRICA_Pickpocket_protein_28_like_Protein_
OS_Tribolium_castaneum_OX_7070_GN_TcasGA2_TC006575_PE_3_SV_1

MTIIGICGGSERGDLELTDKANYVKNEEGKRSYTCGQNLLNYVVEFCNNGTIHGIRYFGEYRTWIEKLWWAISICLSLYFC
ITLIIQTYIKWETSPVIVSFATKETAVWQIPFPAVTICPESKTRASVFNYTHYLHKKQDNEVLSEEEERKFGYLSFICDTKFDI
EYKSPKYADEEMLEFLEQVQPKFLEDSFDCKFLGIPYNCSELFVPILTDDGICYVFNIVDRTQLLSDIVVHFKNYHKTQVSH
DWNLEQGYAPTAGIDAYPRRALRAGIPYALEIKMAVHVDDLDYTCGTDVKGFKVQIAHPQRMPRVKQQHFIVPLERV
AKAGIAPDMMTTSVEVKEYKPRKRNCHFPSERGLKYFKKYSQNNCAIECWANFTMSRCGCVPFYVPREESVPLCGAGS
LHCLEEAQKYQFRGNLEKKIDALSNKKTHSDIPDCRCFPICTSMYYDVENTQATWDWRTTYKYNKHKRIDVNDKKKRM
SMLQVFFKGNQFIYSERNELYGSTDLLANVGGLLGLFIGFSALSLVEIIYYLTLRIICNVAMFGRRNWSGTVN

>Protost_ecdy_cyclo_nema_Cele_tr_Q93205_Q93205_CAEEL_DEL8_DEgenerin_Like_OS_Caenorhabditis_elegans_OX_6239_GN_del_8_PE_3_SV_2

MPDKITIAGWKHVRMFLPKPQKQCDCHVNDPPPWTGEVHGLSQALMSKTLRMRIFWWIVIIICISLGAFTTVLIIIEYIQ
GPTATSTTIRLVDSLEFPAITICPKVPDSFNITGLRGDIKESLPDISDKTADDLLEFFVAGSGLENMNDVTYFNRTYLSHLNN
YYKIWSEGYSVDGFFEIIQQKYGYKCEDLFYECQLAGKILDCCNDLMVRRVVMRRGICYQTRKNVNQSEADDIGRIVFSL
KAPTSYTSVKTNYTQTQLIVYITDNHDVVTEFPRFYLYPNEWNRMRFTARLIELIQNKGVCTNHIFGRDAECLVRKWYLS
NIIIPFNCTLPYLKSITKLPPTTGVCKPDVIADNYLDKIQYVWNSAEVNEECTPGCNRWDYQTSVQQSQTLSPFEDYTFNL
EASFNDLQYEYVKEIYTTSIPGFMSQIGGQFGFFLGLSIITLIQMVLYGFHSAFMFAKKHIQRKFPFCKIHPSDDYPRSTMT
FDNSTNIYPPEKNISKEHIATLSRRVIATNPPSPVSTVIDSVDLPPHWGLRNIDQSRENPLFGNSNM

>Protostome_Lophotroco_annelid_CAC9674854_1__Ofus_G26524_partial_Owenia__fusiformis_

IQEMTVKSLLSDFSKTTTGHGWGNIHERKSMCTKVAWLIFCIASTLASVGHVTSIIYRYSLFETKDRVLVQRQDLLFPSVTI
CPLTQMSKQGILELTQDSINGEDTYLSIFALQTLLADMVKIEQNSTLLMEASSERQQLYEKFFNRLYTYNGFYENVPTVTK
YGHKKHDFISSCMYQGKPCSDEDVHSVIHSHYYKCFTFNDINTSLSNPYSNTTGPTGGLSLQLFLDTDENLYAIYNPEYPT
SGSQGVRVVIHEKGTLPDPENEGFDIEQGHSANVALSVTRRELMKPPWGDCTEHPQLDEDGFTFNTRVCQMRCLQK
MVYKSCGCKRGSLPMYPDVENVQFCNKIQYNEWIKKNPNITALLYDLEKLQCGMTEYNWDVVLDQDDCQCQDNCSF
HTYDMTLSQSQWPKKGVELDFYCRKIIHLPNYSNSSVYKAFHDQIEQLCTDIYSNASTTKWNEVKQEVGDRLRENFIRL
NVYFKDLETKIIKQEPDFALGSMISEIGGSLGIFIGVSIITIVEVFILCLSIIRVLSQKSKVMSIQDPNGVNHEKQWP

>Deutero_Cephalo_105580R_t1__name__chr_scaffold204_start_135507_end_150745_strand_pro_len_555_

MSFECMESLIYYFEYHHVTKVELQYAKNMTFPAVTICNMNKYRRSALKMRDLATVGPYIGIVDQEKFQLKNEHLYPKE
WVDFVRMHDLKTISKISRFPELSEFVKRVGHQIKQMIISCEWRGETCSEKNFTHIITHLGNCFTFNGAPGAGGTVLSSSK
PGEGNGLKVTINIENDEYMATNDVAGDAMDAGIKVMVHPQHEPPFVKELGFAVSPGFHTFVAIRKEVITTLPAPYGNC
QDEGSQKYFPHYSLSACRIECETEYVINECGCRLVEMPGNVPVCTPKMYECAYGKLLKVTEGDLCACRSPCHNVQYNVK
LSTLKLRPKSVAEVQKMDSEDDFEMSHKEIHDNIAVLNVYYEALNYESIVQIPAIETEGLLGDLGGQMGLFLGASFLTLVE
VFEYLMDELYGRVLKKCCRCKATASAHSASPSADVLSVNINGNKSKPTPLRGDDNILVLNVYYESLNYELIQQMPAIETE
GLLGDLGGQMGLFLGASFLSLVEIFEYLMDELYDYGKVLRRCCKSKAASGATNASHCADVVNLNINGSATQLPQTSRA

>Protos_Platyhelmin_Macrostomum_lignano_A0A267G3Q5_A0A267G3Q5_9PLAT_Uncharacterized_protein_OS_Macrostomum_lignano_OX_282301_GN_BOX15_Mlig003120g3_PE_3_SV_1

MATNSSEKRQSAGNSRPNKLSKMVEKMRRDGFGESPSKKLLKQFCESTSAHGFHKIFQGSSILHRIVWLLIIITAFSGFSA
HLISLFSRFNEGPVNTEISVYSTNFVFPDISICATSHFSLRRYERNPKIRSLNKPIKELVSDIRSFWDPNNESDSEINGDLLNSL
KSYLHEVNITKQSQEMILHCDFAGTICSYINFTEYIDPNYFSCFTFSPRERHLLVGGERDGGLSVIMYVGDMMNFISEEDS
LRYSKGFRAVIHEKGTRPEMLEGFYVQEHACTYVQLGYTEEVVLRKCAEEESLQPLEFVDRYGKVTQYRYSVRDCQVNLY
QNFLNESCRCVSEILPLPAKFRNITRHNITFCHHINEERLKQSRGAKSNDSSSVREKQECHDNAINEIHKQVSAKKTKKDS
YCPLKCNNTRYEMRVMQNKWPNDANETRQALWEATQRTMHEARSRGDSTEVLEEIMEHLKNDPYALQRFAKLTVY
SHSTMGNIVRENWAYLARNLASDIGGILGLWLGVSIISTFEFFELFIQLFSVRRKENQQLQAASAADAELA

>Deutero_Cephalo_036230R_t1__name__chr_scaffold12_start_2806583_end_2814836_strand_pro_len_556_

MDEKTGSILGTLRDYGENTSAHGIPRAVTTTSVPRRLFWTCLFLASFSYFCSQAYDLVNKYIDYPVTTDIKIQWSELEFPA
VTVCNANPLRFSMLQRLGDEAGLGFQNAGFIPRQAQNAQSNINRGEMPIFNGRPNQQAQATTTPTTAVPTTTTEETT

TPGYSSTADYEDYDYDYDDPHEDFRTVNTFVGLIANLNTSTRQAIGHQARDFIQECQFNGKACSPKNFSVFTDPTYGNC
FTFNKASSNATPHSATSAGPLNGLSLILYIEQEEYIPSLTEKAGARVVVHSPSRWPFPEFEGFDAAPGFLTSAGLRLTSITRL
GERYSECTDGEGYELLYPGTYTQTNCLVTCYQGHMVSECGCADPSQKVPDNTTICAVTSTNVACKERVTRQIATGVLSC
ACPASCSDKVYGKAIGLSEWPADSYLSKLLGKLGTKRAGRGPGAAVLADTDRFRRNLLKLNIYYEALNYEQISESPGYEVE
NLLGDLGGQLGLWVGMSLLSAMEVVEIFADLFAILCKKISSKEMKSSNKKVEVAPIHKKSVFELENPALTANGKT

>Protos_Platyhelmin_Macrostomum_lignano_A0A267EKX1_A0A267EKX1_9PLAT_Uncharacterized_prot
ein_OS_Macrostomum_lignano_OX_282301_GN_BOX15_Mlig014550g1_PE_3_SV_1

MGLRQDFELFCTATSLRGVAKTVKAPTRPLRCIWTAYVAAMAAFLVTCVCFIVTDYLSYRTVIQTTVVLNDPGLSWPAV
TVCGHYPFTESGRQLWQSGQILSPTNFSRMLRAHALKMFDAGNHIGAEEILASDSKRVYYQNLEPDDAEKLSHTAQQL
FIFCINATGSFMEMSSGDGCAMQLKLISLPDYFNCFQVQATPTIDSLTVVLRVTDSHHQHGADPHEQAFVQDAFQQAS
GFRVQLQEHDRFPEPTDRGIHVAPGKMVEMAYQPVRLIEVGSPTDACLPETIASSRRVLDLGRDFAYSQEGCLRQELGR
IVLKACGCHFADLLRETPPNATHPYCGDYRRGVGPVLRSRACLQSLNLPSLRANLGDTHCLKPCHYHDFEMVSSKTIWR
AADWQLHWLALQAKSFRDLAEASERLADLTASSEQRRQAEASIQSAKNSLRDFLTAKNLTAISGPQVLEPSGRDFAYITV
RRQTRNTIQKTETLSLSFSALMSQVGGLSSLTLGLTCAFAVELLELCLLKRHRKKVSKASEGCAEVADNSLVPQQLQQQR
GMI

>Protostome_Lophotroco_annelid_CAC9623000_1__Ofus_G082429_Owenia__fusiformis_

MPKLMKPKGGKDSNTKEILIKFMKTTSIRGLPRIINSESKTWRMVHIVFFLSALTLTTYMVQQTLTDYFQYHVSTKTYIIRD
GENAYFPDITLCNINPHSDKFEDYLRDVLLKALNKSDIDTSGLNDNDIFDIWKTVEKQGMNGWLNNITNSLTFVHTSLTS
RQQEMSGHKKEDFIRDCNMMAWYGEEEQACKMDHIILHRDLQKLNCYTFQLQTQQDKLQMRKTCGVSFVLFLAPTK
HFVFPEIPKYFPSSNGVIVYIHEPNTMPSKSKGFTVSPGFETRVSMSVVHSIQQPAPYGNCKEDFENIKSGDREFVYEYDT
CIAKCAQEAITKTCRCLWEEAPPFGDYSYCKNVSLPIKTLINNMKCQYDASTKLVSSYNSGQKLCNHCKDKCKKYTYLNSI
SQSVWPDVSTHLGIYKQYIQNVTKYKDFFTEYEELLIGKGKNLNMTEKYTKLRAYNRVKDSLIKLDVMLNSKDVMTYEQ
KKIWTPVSLCASLAATLIFGIGFTYFAVAEILECIFNIIKSCFKSRKNTRDFQSEPDRDDIKMENLDHTEYNTSTLS

>Protostome_Lophotroco_annelid_CAC9651783_1__Ofus_G101132_Owenia__fusiformis_

MTQTRQRKKDKNASPNEVLVKFMKTTSIRGLPRIVNSESKIWCVVHILVFLAALGLTLYMVQNILKEYFEYKVTTSVDLSR
NTEDTHFPDVTLCNINPHSKTFKDITRETLSKAWSKVDGNTAQKTKDPHKSYKKLLEEFRRMGMKDAVESKTFFHASLS
PHELEMSGHKKGDFIRDCKLKDWYGRETPCEMKQLILFRDLDKLNCYTFQLHTQELNATRLQRLRGVSFVLFLAPLAPD
VFYEIPENVPSSNGAIVYIHEPNTMPSKSKGFTVSPGFETLVSVSVIHSIQLPAPYGNCKEEFEDIKSGRREFVYDHATCFA
KCAQDVFKESCNCSWEEIPSFGDHHPLYCMNMNLPREALKKNDRCLQKARIKLTSHKQKHQCSHCKEKCSTYSYQNSIS
QSVWPDVSTHLGMYKRYIRKGTACNQLFTEYEDLLTGNGKNLTMLEIHTKLRAYNGVKDSLIKLDVMLNSKDVLTYEQK
KIVTLANLLSSLGSTLIFGIGFTYFAFAEIFECLFYFIKSCIKRENRDVQTDSICIDIKGENIKGAMSDKSTIRVTP

>Protostome_Lophotroco_annelid_Pdum_comp402494_c0_seq1_

MDNNADFNKNRNCRNVRPQEKLMYLSHRLTEETTAHGIPHIARANGFWRSLIWIIISLVMLVTWIYHSQYTISQYLKME
VNVKLEFKTAQNLPFPAMTVCNKNAFKNTQGMWLMERMIMRFNTPEGKNASYPPWLISYLKYLATFNGTDEPEAGT
VQTSNPTQLGEEMMQLLYDSIIKYGPEYNLTVDDLTYSFKDFVLTCDYERTPCEKRGVKINRIFNWLHGHCYVLNAEKAT
LLSTQTGPRYGLKVALNIDQDEYWEVAEQAGVRVLIHDPDQMPFPEDDGYNIPPGLAASLGVRLVKQSRLGGSYTPCF
NEKEQNEKHDIFRAHYGWTKYSLKTCMISCYQREVEARCQCSDIRYPAYKKGGGKCVSCTENMTMSFGAEMKTHNPN
ECLRFVQEMYRNQSLNCDKECLVPCSQNSYVSSVSYSGWPSTKSLETVTQNLKKIPSAAKLLDESIANIYKNFVEVTLYYE
DFSYNAVSERPAIRLTECLSDLGGNMGMYVGASLFTFFEFFQYLIDVFLWSCVWCCKSLTRNGDGHLKELFAATKVRKI
QVKPEH

>Deu_Ambulacrar_hemi_Ptyfla_40v0_9_20150316_1g8769_t1_scaffold4611_cov111

MEQINTEEGVQAAKPQKGNHRIRNTLADFGSNTTAHGFARLVAADSKIVKFLWMLICCLCWGVFIWQAIVFIEKYYRFE
VSYSISVDFRVLPFPAVTVCNMNPLRDSALEIIDDEYKERLAFLKYEEQSEDDADADEESIFTTTSESESVIERIGGKYSDCIK
AGEMPLLYKGYNYSVEKFRIRLSEEIENFLLRQTIEPIINDAKSDQVESLLFGTELRKHKGETSFRXMEQINTEEGVQAAKP
QIGDHRIRNTLADFGSNTTAHGFARLVAADSKIVKFLWMLICCLCCGLFIWQAIVFIEKYYRFEVSYSISVDFRVLPFPAVT
VCNMNPLRESALENIADKYKERLAFLKYEEQSEDDADADEESIFTTTSESESLSEYSLEENYDFDFRQGLNLSMMMSNLP

EDVRNATGHRKEDLIQECVFNGITCSSNGRSYKETVSMSHWPSYAFDDKLYERLSKRSSKINETLLDHNESRRNLIKLKVF
FSELNFESITEKPAFDSVDLLSNLGGLMGLYIGASVISLFEFFEFVGKVCCLIKTKWNVSDGADERVQMG

>Deutero_Ambulac_Spurpu_018446

MDAPKVNCEESSWGTAPEREEKSLRTLLNSRMENSSAHGIPNIQRSSGLVTKLAWSLIFLAGIGVMTWQAVILFQTYFE
WKYSVDIEMRFNRTQSFPAITICNTNPVKRSELETRDASFRRAFDVHYVSSTPDQQPLPDVPDGVTPSMLNNSDSGNE
TQGGSTNEQVDMSEAVNDWRSRIAIPKFYRMKSEDYGKKRIRVVTLANETLEERVSLGHKLDDMLLDCSWKGIPCSPE
NFTKFYDSILGNCYTFNSGKNGEQLTTNRPGSTHGLTLELFVQQDEYVEGMTEEAGFRVSIHHPSKMPFPQFNGLLVSP
GFATNIGLRKLEVDRLPKPYGDCEADLSKNIEDDIYHQHYSITYNRKTCEVSCFQNEVISRCDCFDATYPNSLKVNHTVYP
CEYINDVETQCIADIEMEHARDELECNCPLACRETTYLTGASSSIWPSDAYESTLIEKMLKYNAEIRGHVVGENASDWTR
RNMAKVEIFYDEFNYEYIRQDPAYTIPDLLSDIGGQLGLWLGLSIITIFEFFEGAWLVLAFFCSRSGNKTNRSEIDPGTKTT

>Proto_ecdy_PANarthopod__XP_023244170_1_amiloride_sensitive_sodium_channel_subunit_gamma_
like__Centruroides_sculpturatus_

MRKLNLEAQSQNSSLCGLCRSFAYRSSAHGVQRIASSQDNARRLMWSVVFLFAIAGCGFHSVYLILTYLSYPRMTITEEI
HADHIDFPALTVCNLNPLKKSSIREHLLETHQSVSEFYNPDKYDDDDEIEERGICFRSENEFLKKSKTMDLSDVWMSVIAT
KDSLMKCGHQAQDLITQCTYNARNCFNDSSTIVLLDQYPSPRYGLCHTIVVDKEPLRKVKKTGLSLGLRLTLNIEREDYLD
LVSPEFGARLLVHPKGTYPTLQGGGVVLQPGTKTYVGVRMRKIERLPAPYRGCYDNFQSSQLIHFLRKQDQSQAPFAVI
HNVYTYEYCQTLCRDIHLLKNCGCVEEVTPVNNKFCDPCNTTQARCRTRFYRSFTRANVDHDCQKFCLPACTDVRYDLT
VSRSEWPNVRHQNYVLKRWPNLRQSLTVFLTNKTLKNGSDELDVDNLRKNFLRVHVYIQEMNYLSVRDIPAYTLPQLF
ADLGGCLGLYIGVSAITIVEVLEHMASVMAFLYKKRKEYFPKSNSVKKSEIKKRRTNFRLSERSHVQKYSKSKDCKRGNH

>Protos_Platyhelmin_Macrostomum_lignano_A0A267DCW3_A0A267DCW3_9PLAT_Uncharacterized_pr
otein_OS_Macrostomum_lignano_OX_282301_GN_BOX15_Mlig012953g2_PE_3_SV_1

MTFHEVFQQFCSSTSLRGAGRLVKAPTRHLRAIWAGYLCLMAAALIACVCFIAKDYLQYKTIIQTTIVLNDPTIAIPAVTIC
SHYPFSKTAATLWSSGRVLSPTNFSHELRRTAFQLFNSSEFLMAENVLASDTKKVYYQSLEHNDAVNLSFSVPQLFIFCVT
GVGPFVEISNWKDCSLLLKRVSLPSYFNCYQVQAPRDIDWLTLVLRVSSAVPNEYEEAFVQDIFEQADGIRLEVHENDTY
PEPTLKGIHVEPGKLVEISYKTERWAEVDSPADACMSAEAASQRKIRDLDEEFAYSYEACLRQELGKLALERCGCHYVKYL
RSSVPNGSHPYCGDTRIGMRKMLANIDCLNRLNLDSERERLSRSVCLMPCHFHYFDTVASKTVWRATQWQLHWLMQ
HSRCFRDLDEASKRLEMGGLPPDEERRLREKLEQASVTMKTFFETANITKAASADGGTNGFDESAEEKQQTLDQVMPK
DDTNFAYVTIRRQSRNTVQKTEILSLGLSALMSQIGGLSSLFLGLTCAVAVELLELCLLRRYQSNQKRKENQQQLPQQVL

>Protostome_Lophotroco_RotSoc__A0A818HJ21_A0A818HJ21_9BILA_Hypothetical_protein_OS_Rotari
a_socialis_OX_392032_GN_FME351_LOCUS17285_PE_4_SV_1

MMVATGDIIAHVVSNNPMNPSVQDASIHHLRISILREFALNATTHDIPGITRSRSMPNRIFWAIPFMAFTSAMVYFITEA
LCAYFEYPSQTKVSITNARIQHFPAFTICNAGGIRADALLKTYNEYTAHLNLTATSSNSVILKLADFMRFYFSSLVNEDGSL
GQYVFPLESMWMSCYYVGIRCSVNDFIGFDSAEFGRCYKFNAKRKNGSLLYNHETDSTGKLLLRLYIHDHQYVPTVTEG
VAMLAMAHDNTQLPMIEEAGMTLAPGFQHRLSYVKRDIQFLSEPYSSCTNEIPCALQIMFEKYDGAEYVYSEELRNML
CSQAYMYVLSILDIHNENKYFVAFVYLLICSNDQCGCVSPLQWNARTIVLPVTDKTIATRLCNVTNPCFNKAMNDFDLT
QSIQDLYCSHCLQQCSTTEFVVKSSLSGTLPEWLIPFIREFVQNTSIPLPDDWSTRWREHIFSSYLSLELFSESSLIETYTQTP
TLQPFDIFSNVGGLSSPWIGVSFLTIMELIEMLFLLIQHQIRLSRSANPENKKSDLHAYMKLLQSTLKEDYAAIVL

>Cnidar_NemVecNVEC200_001060_1_1_protein_AED_0_01_eAED_0_01_QI_333_1_1_1_1_1_8_1063_
559

MSLDICDAYIQQETDIGKLYFVNTHILDAAITGCLSLYDLKLIAAVMSCPVAQRHFETEEDKKDEDEDDRAEDPVDENPD
DTITVSQMWQDFLHTLTLHGFRFVFERGPTIRKVLWLAILLFAVGMLMMQSKKSIQKYFDHPITTSVQVEFLEEIQFPAV
TICNFNLFPYYLINGTIGEKVMSILAPQKYIDNKEEVLFARSPIPNFLNYARKRRRSTGGTIVTDDMLQSEKDFGELDEKFD
FAEFVRTHGHRIDHMIKKCRWKSQPCGPENFTAVITEFGLCYTFNSGMKGHPLLKVQRAGVDYALRLQLSVQQDQYY
GSLRDSSGFKVMVHDQEEPPLINELGIAIQPGTHTFCGLRKEEMHNLPAPFKTACRDMQLEGFKKYTKSACLLKCRADY
VMKMCKCRSYDLKGPAPPCQPREVKNCVWPAMEIFRNESINCECPVPCEITKYQTQLSYAQTPAKHFSEVLARRKHIN
KDVMRHYLRDNFLELDVYFEEMQVTLIQQRQAYDQESLFGDIGGQVGLFLGASILTVLEFLDLLWRILIHKFKKRKNRKV
RNV

>Protostome_Lophotroco_annelid_CAC9522438_1__Ofus_G031129_Owenia__fusiformis_

MKRIINEFTSNTSGHGWGMINQTSSKFGKALWVAITLLSTIAAIIWVSTIIVRYVKYETIDRVEAKVDEDIIFPSVTVCPLYG
ISSMKIAEVYSDPNSDYISLNNFLSSSYALVDELAQITNETYLSIMTNTILVARSNRGYYANLGVSGVSTIGHEFGDFIPYCL
YQEKECSAADFQKLVHHEYKNCYTFNGGDINISKPIISSTGAQKGLSLTLYLENSENAYVAYNVNRVMTAASGARVIIHEK
GTLPDPDNEGFDVEPGHLISVALSANKRQLLKQPWGECAEHNTLHSTDYKYTRNTCRIKCIEKVIQRQCGCRYDLLPVDI
NATTELDIQPCGKFNIKEWLKATEANTTVLKEDLAKVICSKEEHQWTALAKDIPSCECEYNCTYVDYEIETSQSVWPMRG
SELDFFCQLSYLNISGSFFERFQNSIGIHCNSFMEYINVAQNHGDELRANVLRVNVYFKTLEVKYTIQDEGFSLVSMISEIG
GVLGIFVGVSIITILEIFVLCSGIVHVIINSKKVSSEINVKEIKSTSKSKLEESSNHKPDDSYYDQFKKY

>Protostome_Lophotroco_annelid_Pdum_comp416533_c1_seq1_

MKLFNTNMELESKGHHLCDIVRNALETTGVKGIPRIIKSEEKVLRFLWGFGVTFLLSMTIYQCYKLIKIYASFPSTTLISQGY
VFGPNNFSVKFPDVTVCNLNPIYVSNTKETVQEVEDFYNLVMQVTSCNNCSQRDTDKLATIRAQLLTPDGFYQYLGAKT
GNVTVKDPPQDNFVLDCSWNELQGVANVRQSCAEISIIRHWMSSSHFRCFTVSQRHINRMLVGLTLVLHLDTSDQRP
VDAFFDVTSYMGLKNGVRLAVHPAGTTPMFDLTSVEVAAGMATLANFHFQHWEKLPPPHGKCSTALPQLNDTLGRP
VAYDYSVCYANCIESRVIEHCGCRDSSSVLKPTSHAKGNDVPFCQDVRHGRSIILNLTKCMVTTRLSAMQDCVRFCPVA
CHKTSVFMTVSHSQWPSHPIPNEFYNYFLANQELRNLFPGGFSRKVNSSSYEDLVRSYTEITIVRNKFLKLQVQLDSVNYI
HLKDVVVMTIENLTSALGGVLNLWSGITAVFFIEVLDVIIRLILERCNEASLDKHECNKVQVPSIKPIKRTWLPWYDPYKG
PVI

>Protostome_Lophotroco_annelid_Pdum_comp404879_c1_seq1_

NCTCPIACNERYYAPAMSYAALSIVSVDTLLRSNTSQLKKRYRKANEIRERVMSENLIQTLSDLKDIPLKLKDYKFALETDIG
SATIRVKEAIIGLLQMAENDFEKLKRGLEQLQQDFEINIKSHLDLFRYSIEEPLLEILKKLHIPHIEEQIMESIEIHTIVKNLKNY
ENSFSPKIYPFLKRIWKTDMYLDCEDLCVKIHNDLQCFGSEKNFTLLIATASDIPRFRECMYAHKSSLDEATSSLDEALSFIN
NTISNAHKLAVGNSDALEDKYGWILSNQSYFKKLEERYSQFEIDKLGLQEHLSYNLLLSLRVLFSQIMEGCNEWKITSNYQ
IKEIGHRLIGHYYSVLGSYRHLLTNFDDRHQLSKLVSDLQIWRQPMAELGASPILSFHWSSKENSYYWPQSRKAADFIAN
GAANEIVYRNVWSYLDSLSKDLYHTSLQMNSMGNELFIFTQGVVNNYQKFAARNELDEHFIMNNFITVDIYYSALRYQE
VIQVRGYTLLNFFSEIGGYMGLLLGCSVITLFEILDLFFHNLSLKANEKRQRLGAKRRDSKKSLPQFH

>Protostome_Lophotroco_annelid_CAC9660748_1__Ofus_G11941_Owenia__fusiformis_

MAESTEGSNSEKGPMTYKLIFIDFLKTTSIRGPPRIIRTKHKLWRLLYSFIFMFVLGTNCFCIWTTIGDYLAYPVISQSYIIHS
DMVSDRVEHFPDITVCDINAPQPQYAHPNGSLTELGIEAEVFLRMILTNIDQILGGEPEKDLNKLNMYELDLLRIQKKEQ
FLIENNTQGLELMEMFYSTRGYLQQYTRSQRYQIGHRKEDFIVDCKSLDWYVSDQETCNMTRILLHQDLGFPNLNCYTI
QLQTQPFDMVKTNGVSLVLFLGPPGDFLPEDPPSPVPLDSGAVVYLHEHNTMPFQQTSFNIPAGFKTTVHLDVSKVMK
LKNPYSTCSDSIELLFKNLDGDPYKYTRDACLAENMQKCMIKNCKCIIDEVPAIENVSYCLKTQSDSNIVKENIKCMKRMK
SPKHIANCALGEICLDPCEKYIYKPRLSHTEWPQENKQLPVYNEFIRGGPHQVVYHEYEALLYENSPNMTSSERMNALK
QLNLIKESFIKLDVVLNSKDIVAFSDKPFYTGLSVLSVIASSLIFWIGFTFFSLIEILECLFHMVLSCVSQGNKKPSSVH

>Protostome_Lophotroco_annelid_CAC9574840_1__Ofus_G053378_Owenia__fusiformis_

MMKEQNKMDGKSIQERVKEFIETTSAHGCGQIPTAKFKLTKFLWGLTFLSFFTFATVSSGILIAKYFTYPTKDVVSIEFGNI
EFPAVTFCSLQPITITSLIDFRQNPPPPGHSVSNMIYVMQNIRNIIIDRDIGGLGQWYLKHFDRFLSNQFIFENGEENDME
QFIHDLRSFKLGCKFQGQVCNGTSFVLHKDGGFHNCFTFNGAGSDLSNVNIVRTDPASGLSLVLFMDAYAKGKSVSAYT
IYNPDDPTSGQTGVRAVIHSPRTRPMPFDKGFDIPTGLSTSVALKETKRELMTQPHNNCTNDEFNQGTNYSYSVDTCYE
QCQQDLLIEECGCKSSLLIPPSFDKSDLEYCGKVNITNMMLMMNQNDDITQSIAEKELENLDCEEGVMYSFGNLAILEQ
CACKDHCQTINYVKTISQAVWPNDNAQEAFYDTYVNRTDQTLRPNKLFKGYNTSEIINNDMIHKNFLRLNIYFESLYVET
TSEVEDYPAGTLISEIGGNMGFYVGISIITLMEIFCITGAIILYCFKDWLIKCGHMNKTKVSHVNVKPVELDGDSTEKIKDKY

>Protostome_Lophotroco_molsk_AplCal_XP_012941892_1_amiloride_sensitive_sodium_channel_subu
nit_gamma_like__Aplysia_californica_

METKKTDFVPNAFLADRWGSDLTSCTSSKAEEDSESKKANVGRIANEFAENTSMHGMANAVNIHKRFNGLLWKNVF
WFLLILCGIVMSIMNLVKIAEDFRSSPVTTEVKMEYQPKIEFPSVTICNMNKWRNSQFATFNLSIFVPPPRLSLVLSEMET

SFGDFHLLLYVTEKCRKGVKSILLPSLLLIQANATKDEKELYYLLSAMRVPDSQKIQWGHQLSDMLLECTFDAVSIHFIVVL
MVVLFYILVRESRHLLKTTSRPGPANGLTVVMDIQQDEYMDAVDTGGLRVVIGERDEMPFPEDEAVDVAPGFATSIAL
TKVEIRRQSPPYGPCLEMSKEKNRELNLYAATYRYTDNTCKKTCLQATTMTECKCCAAGYPCNPYGVEKVMNYTLPVG
GNLTYCDLYNDNQTNCFLHVDARQTSGNLGCEHFCPPACFETKFKKTISFGSWPSLAVQKEYSGRLNYSRFQSAYSSVE
EYQILRQNLLKLHVFYEQLNFEKIVTTPKYDWSILLSNIGGQLGLWLGFSLLTAFEVIELLLDMFLNGAFRMACCKSASAR
GKVASGGQ

>Protostome_Lophotroco_annelid_CAC9674171_1__Ofus_G24722_Owenia__fusiformis_

MDGKNMIATVKEFTENTSAHGCGQIPTSKFKLTKFLWGLIFVGFFMFATISSVNLITIYFTFPTKDVVSVNFEDIEFPAVTF
CTQQPVTATALIDFIDNQSHRDDAIGKLVLTMGSFPGMLVKPNIGGLGDWYFKYSYRFLSNQFVFENLDKKDLKTIQHD
LKSLKISCKFKGHPCKDSAFVVHQDGGFQHCFTFNGAGAQLNDTKVIKADPTSGLSLILFVDAYTTKRDLSELTLYNPFDP
TSGQSGVRVIIHSPKTRPMPFEKGFDIPTGFSTSVALKETKRELMTEPHGNCTMAKFNGGTNYAYSEDTCLEQCKQKILI
ENCRCMSSLLPTPSGELKPQYCGKVNVTNMFTMMYGKLDATMRGIAVKELEMLDCETKLMESLGNSEVLNKCECKKP
CVHTNYVKMLSQAVWPSDYNQKNFLAESINMSDPTQRATILFEGLDIGQITKNESEMIQKNFLRFNVYFESLQVKTTSQ
VEDFPASTLISEIGGNMGFYVGISIITLMEMLTLTGAIILYCFKDLIIKCGQSNTTKVPDVNIKQIKSYDNHMDEDKENINAE
IHG

>Ctenoph_Hormiphora_californiensis_evg140127__p1_GENE_evg140127_evg140127__p1__ORF_type_
complete_len_566___score_73_21_evg140127__173_1870_

MPESEPDRMVDDGKLQRHDFYKTRDLTSDYFQVMREKSRPVWCKIQWFIITMAFLTVTVYMVYRLLKSYLKYESYDSV
SLEWVSQLALPAITICNVNPMDYTSMKKKYNASILAELIEYAAEIKKYSGIDMAHFNTDETLETMKQLDAEFVAGDADIF
RENVYDIQDFIVGDGNGHPVYTFAGQWMNESGIKQVSHYYGKCIELNSEETLVQTVAGPKGGLTIDLDVKTNTYMPIT
PQSGLMVYLRSLNETVVLDNEGILITPGSETSISLQKYSVKRLKPRVGQCDNVTSDFIPSAVQTVRECSQGVWLKMVVRL
CECVPWYLIDKLEELGRGDALSGWVAMAFTDKEEEENDINVDQLEGVDEDEYMCGYSTAALCDARVLDKFRNVTWLT
TVGSENYSIECKEPCQYEDYEYDTIATTFPVSETYFNSTLFYLDQARGLNFSYTQNNLVRINIFYDQLKTTKVEQVAAYEFF
NFVAELGGTVDLLIGFSFFTCFQLLEIGLAVIIFRCCRRGGPELPPPHHHDRRVVPGRRQEKKAGDFVLTRSDQGMFRMP
DSFLGIRIGD

>outgro_CafRoe_KAA0150573_1_hypothetical_protein_FNF29_05148__Cafeteria_roenbergensis_

MAEGTVKGQRRWWGPRLAARWASSTSAHGFGRIAHYCRQPGSCLRLAGCVWFSSVVAFLVLLVVFILDRLAFYRTWP
VTTSFALKQERPIQFPAVTVCSNVFAFDSRIALVNFTDVIEHPTDAALSRLGAQPDELVIMCVFGGRKCSGLHAEEWIHP
AFGLCLTLNGGQRPFRVGNTIYDAASLPSSFSTVGEYLSAREERVDYPPLEASEIAGLSNSPKQLSVKAPRADDGLTLYLG
VWQNEYTAVASSADEIGIGWSNEPVTGVFNAESAGVRIAVHDRLKTPDMDTATMLHPGKETHVTYRKRVVTSQPPS
WGDCSPRFWNETAGQPLPSGNYSVAGCELVCAAKLVEAECGCLTIPTPRGFVPANPSMRACPLTMECPVVQQEKVLL
GQAECPDCVKDCATTEYPLTTSTLSWPAAPSQQVFLDLFDQFSQIGIGASALENISTAEDGLQRLQSNVLLARIYPSSATY
EEVTSNRAYSITSLLSDLGGQAGLWGGISILTAIEFVEFIGLGFMACFGVACARNAFTEPGDEDELTPDDQATTHTSPIGS
AHALQSKAATDAA

>Protos_Platyhelmin_Macrostomum_lignano_A0A267G6S3_A0A267G6S3_9PLAT_Uncharacterized_prot
ein__Fragment__OS_Macrostomum_lignano_OX_282301_GN_BOX15_Mlig031004g1_PE_3_SV_1

SLGVSRAATPAKMPSAVMDTLAESSAHGISRVATSRSRVMKVMWFLLFLLFLALFVRSAVTLVIRLRGGGLTKSFELDKV
PFVFPDLYFCPNNPYSGSYPHDLSQADAELINTAITNLSAEFDSRLGPTRSIELRARLVQSSLPVHLSYRLLAQRRAQSVLQ
TTATPTMGPQSSFPIFPRVLPFSHPVSLVCFQLRLTDNDRRKIVAENWQVNLRLSTDPFFAEFNYSGLSVDPIRSPATGEE
VPLQDWSISVGRERLQEAGFTMLIVPVGSYPSFAHEAKFQLAPSNSYSVALSMEETEYSEELRGQQCQGDSEAVDFSDH

LTGRKLTFAYSPAACYYENLVRLVTQAYGCMVFSFPLTWSLRNTTRCFNMSETGRDQFNVVQTTKTNWSDLDRQVRE
ACRSRQPCRYQRYSAEFSFSRWPSIGSSVVAHFAEMGSSPYSTPATRRLIDMARSLRTRNHSAHTEAQFRAYLSESLINV
RIVARSTTSPQVVISPAYSVTAFLADTGGLLGLYVGCSLLTLCEFFELFYRLLSGALLKHRGGARPAEPEAPQGDSQPDAK
NDTSEA

>Protostome_Lophotroco_annelid_CAC9602553_1__Ofus_G07376_Owenia__fusiformis_

MEKLKEQLKHFAESTTAHGLARIPTSRTKIVKFIWIIIILGCGITSFVFLYQSFAKYLAFKPKDVVSISREITVKFPSVTVCPLYP
IATGKNFNIYDYLDNNSTYVYKLNQIISGFYEKFDEPEDPLYAKYIRHKNRVVAWQWIYENNPNITNASHSWPDLVPICS
FKGKPCKAEHIEKFVDPNYYNCYTFNGMNSSTLNGENHTSTEDEMIVESIGPTSGLSLIMFLDLHEEQRSMFNPMIPTS
GSAGIRVVIHETGTVPDPTANGFDIPPGFSTNVPLRLIQRNHMEEPWGTCDKKADPYLKGSRYTYSRSSCKRQCVQGLL
KAKCGCISSLWPINNDADDSKYCGTFNIDEWIKQESNRSILEEDLLRLDCEEDLLSHLTEISEFIQSPDCQSNPDCHDCDCR
PACEHYKYYQDISMSYWPMESAQMTFYESLMTMPGYNDSTIYKLLHKPHDDLGTADNSSQVNKDLIRKNFLRLNIYYK
TIETEVISMYAEFTSGELVAEVGGTLGIFLGVSFVTLCEVVGLLSNLISSLLCNKNRKIKDTHDLSEKISKYDANDVKIVNQ

>Protos_Platyhelmin_Macrostomum_lignano_A0A267GWM4_A0A267GWM4_9PLAT_Uncharacterized_
protein_OS_Macrostomum_lignano_OX_282301_GN_BOX15_Mlig026138g1_PE_3_SV_1

MAKKLAETIASSSAHGISRAAASENKLLRVMWILLFLICLAVFLKSGINLVLKLQKSEMTKKYQLDQATFVLPDLYFCPINP
YSGSYPFTLTKEEADLVNSRIKQLIDRFGELYDVPKSILFHAFLWTPQVQLSLPFDLLAKYLLVRKEESVQFAAITDDFVAH
NVDVEAFYNYENLVCFRLILSDKLKKMIMNPGSTLKLRLSTDPYLAELNFTGHPIQPIADSNGNSVPFLEWHPYRYYEQL
RTSGFTMLIVPPGAYPSLRFEESFHMSPGFEYSVQLKMTESTCSEELHDKFCLLETPPVEFMNSLSGKRERFNYSNAACY
YRNVNVQVVDEIGCQLSQRPSLWEFRDYPHCSNMSEMTAATVKRMRKAGSGRRIELETLTQKYCKTRIPCSYPIYSTSY
SFGKWPSAGSSILRYFAEYFNRTSVSKAEAPALARVSDIAKQLQMGDHSAHLEGIFHSYVSESMLSVQITIDSLLSPQVML
AWSYTFADLLADTGGLLGLYIGCSVLTVCEMFELIYICVSLLLKRRSAKRSSSRTGDQETAGCDVGEKQALGGENAEEIEL
RS

>Proto_ecdy_PANarthopod_Dmel_tr_A0A0B4K691_A0A0B4K691_DROME_Ripped_pocket__isoform_B_
OS_Drosophila_melanogaster__Fruit_fly__OX_7227_GN_rpk_PE_3_SV_1

MTISDSELDSSKGIDLTFRRRRKAGSVACEGFITTYHEYCRNTSIHGVQYLGEQERPFRERIFWLFVFLISIYGCSTLIQSAYT
KWTETPVIVSFAEKSTPVWNIPFPAVTVCSETKRVLKQKGKETTYADLYSQFSEDMRASRVFRPENVSALEMEEFRTLLH
VCNTQIIEEDIPLIAGDDLDYFDVLQRMLPQFDRYFFYCRWLSRFGECETFFRKTLTEEGICYTFNGLRATEIYRDDTYQYQ
HSGEPLEMENISSQHTAWTLETGYALDSDVETFPARVLSAGARSGIFLALQSFKQEVDYACRGPVQGFKVGKFENVLLH
APDDVPQVSKQFVRIPMGKEVLIAVKPNMITMSSGIAEYHPVRRQCFLSHERSLRFFKVYTESNCQLECLANFTLTKCGC
VKFSMPRNVDMPVCGEDKIHCYDRAERELLVREFKRVKALNAGRENSRSVESACNCMPACTSLVYNTEISQANFDLEE
MLVAEGDTEFLKEYPGSQMSRLSIYFKQSQFITSKRSELYGMTEFLANCGGIFGLFMGFSILSLVEMIYHFTLRLFTNLKRL
VKG

>Proto_ecdy_PANarthopod_Tcas_tr_D6WXM6_D6WXM6_TRICA_Pickpocket_protein_28_like_Protein_
OS_Tribolium_castaneum_OX_7070_GN_TcasGA2_TC006574_PE_3_SV_2

MSASPKPSPNRGPLDNVVFVKSPKKKKFPGFRKNIYDYFTEFSSNTGIHGFKYMGEQERSILEKLYWVVLFAISLYICVSLII
QTWVKWDQSPVLVSFAQSPTPVWQVPFPAITICSETKARQRAYNFTYYYHKVIEQLTTGDGNLTQEELQRFSDSSLICD
NHLYQQGNKTTSFETIEYLMSIAPPFSEVFFACKWTSLNETCSNFFSPMLTEDGMCFTFNMLDRSELFTNAVYLHGDYM
GHEKRSEGWTLENGYPKTTGKDTFPRRAMSAGSKAGLFLLLRAYEQDLDYVCRGPVQGFKVLLHHPAEVPRVGTQYFR
APLNQEIVVAVKPDMMTTSQGLRNYDPHRRQCFFAEERQLQFFQNYTQQNCQVECVANFTLARCGCVAFHMPHAE
STKICGSGSNVCIFEAEQELLAREVESGLSRYDDNADDDLPTNCDCLPACTSITYNAETSQADFNWPKVFEAFKANFSEF
PGVQMTRLTIFFKDMQFITSERNELYGQTDFLANCGGLLGLFTGFSFLSIVEIVYFLSLRLICNVKKYGIHFWSGSKDLLND
DAYLHKEQKDLS

>Deu_Ambulacrar_hemi_Ptyfla_40v0_9_20150316_1g20263_t1_scaffold20289_cov133

MTSQEERTSKPENNTKSLNQLEQGELLAGKKMAFRQKKAKRKRQGRERLRNIVKEFGNRTSCHGLPRILGAKSLIGRIL
WLIIFVGLCSLFFYQLHALVVLYLNFDVAVRIEMGTAKSLNFPAVTLCNTNKLRLSEIEQSEEHKSLALTNPSHPDTIYRSLS
YETPCLEGDFPCGDGERCIKSHLVCDGYIHCWATLRDEVNCTYPPCGEKQFDCGQAGLKGVCILDSLTCNGEWDCSDG

RDELNCVCNDESSSTPLDTIRFEDNGMLYSPLYPMHDSAQSYCEVVISPAHVVFWDCVQLTFFDIDLKYTEGCEDEYVM
CSATCGSGVRKRTIGCYSARGIRKSSRSCRAYGHKPRDYEICNMEPCYGAVYCDQTFNTSRRIIMSPKYPLNYPANAHCEI
DIKNPESDNCILITFDDFYLDSSQDCNDDNVNGDGKNHIMISIERKTYPYGNCSNVKDFVTIYDDFYNYSSLSGLIQELRHI
NPKVTRTVVDSRSARENLVNLEVYFEELNYHSTLETVAYTLEELLGDVGGLMGLFIGLSAVTVFEIIEFLLEIVYYVTLRFRLI
GQVY

>Protos_Platyhelmin_Macrostomum_lignano_A0A267G058_A0A267G058_9PLAT_Uncharacterized_protein_OS_Macrostomum_lignano_OX_282301_GN_BOX15_Mlig029271g2_PE_3_SV_1

MLYQYLIALHVFFLSLDARFKKLRKSQVPHTPSRHQIVKSDTPVCFVMINQRSAILGACRVDACSRVTQSIPGLVRDQVG
NALLASRFGSSIGIGFTGCSFGNQACTMEDFTTLLHPELGLCHRFNQEKFVGLTKDLKATESLLSFNYFTDSTDSKGRRLP
GLSRMAKVIFPGLVDANDDTLSAMKVILVPKGQFPWRENAITASPGQHLQLNIRFDKLNRYSKPPTQECRPPAGPTVYR
IPSWRTSWNYSSPELDSFNMTTTISSLNWSAKLQTLRGRLRTVNSSISPMSENLTFEMQDQRDCVNLYAVKKFIEHCGC
LPSFLPVSVQLFQNYTYCFNQTDMESIKRMEKCYTERMSSISRYEMEARQVCLTMCNTRFYTFRPITYPWPGLTIETAVD
FMQNSQGDWSYPYNFLFSKGIANLFYLDIRSSEMYNRSTMLSASELLDFRNKLEVLQESLVKVSIKPVSDFGQEMTEEPS
YTIKNLLADLGGALGLWSGISVLTACELIELLVYLVAATRRTEQPAQPSGPATGNAEQQDSGEMKADDSATEVEMTDK
MTDIAAPEVSISVH

>Protostome_Lophotroco_annelid_CAC9661806_1__Ofus_G112085_Owenia__fusiformis_

MHRLVCSQKIGSFTFDDANGYAITVNIKLYFVIPIEFLGTHARADTEEDIPKGKKTPMSYYYLLGNDYQNCSDHEVRPMG
LMRGLAENTSAHGITEIYYSKGFFKITCWVLLTLAAIAVMILHMTTLFIDFYSYETTMSISVQNKRSLPFPAVTICNVNPIRA
SRLSQSTILQSTIDGRDNITGATRDDKQTGLGAMSNEFLIEELIEETIADIDTDTKIAMGHQYSDFILDCQYDGYYCDEGEF
LAFYNYKYGNCFTFNSGSNSDVFLSGRAGPLHGLKLVLDIEAAEYIGDMSPAYGVRVLVHSQGDMPFPEDQGITIGPGQ
ATVIGTNMLNIQLAGDKYSDCTNSSFSNVNINVYEEHYPGTSYTQTACMKTCYQSHLLSGCQCGDPSVPLDGKAFPSAF
HTEPATSCNSDNQAQVTCEENTYASYVNGSLSCTCYPECNQTTFEAHVSSTLWPTDQYISRLIQDLSPKLNQYTNDSAG
LRKNVAHLQVYYEELNLQMIQEQPSYSTVQFASDVGGTVGLYVGASLLTAFEFGEFFLDLLVYFIRKPFHKNKVSEVKME
QTKSNVQFS

>Deutero_Ambulac_Spurpu_014519

MTTGYTFEWYDCLLTQATLLLGRVKRDSLFLMRTDKDLPHLNSVNAVLIDYSGSTSAHGIPRIITSRSVKSRLFWSFVTLV
CLGAFLWQGSLLLLDFSKYPYTTQIDVVARTELQFPAVTVCNMNKMKRSAMVHTRFQSLIQADLGVNGGDADYSWW
FDWSSQWSQEEEEQEEQEEQELASASAPDTGSSSSDWSSDAREHYSEQAPALVTGSSTSSSDSSSDAQEHYSEQYEW
WDPGWDTDGFDFHEYDWTNVSDDDDWEGFYKQSTSDDFSDLLEVVNPTREELKVMGHQAEDFILQCTFDRHQCNY
TDFHQFQNKYYGNCFTFNREVGNSTRARSTGKTGAQYGLHLTLFTEQPEYVGLFAQEAGVRVAIHPPNVFPFPEDDGV
VASTGQATDIGMRQSYFNRLPHPHGNCTEGTRTIFMSEEYAYTTRACVKSCVQQQLFDNCGCVTDIIMNETMCGARN
KSQQVCRQAIEHFHEDQSSCNCPIACDRNLARVRIYFEELNFEQMIQKPKYTIESLLGGIGGLLGLYIGFSVITICEVGVLVV
DLVKFLLRKAYNRREKIVPIELKC

>Protostome_Lophotroco_annelid_Pdum_Contig13621_

MAGKRVLKSKSNIAHLFWLLVCLAAIAMFCMQMAEVLQRYFSYPKKVTVEVISAQVPFPAVSLCNMRNMDVHVLNTL
NRKFIEDHSPINHINNSDNDFVREYMKLSAKYGPLWYKYQGKYPLAFQEVFSRTTYSANIPQHIVSSAAVQLDEFVVSCY
FGEYHCNTTNDFTTFFDPYYFNCFTYNMADPDQASSSEGIENGWSSVLYSGSGILDRNKDIRVLPGLHESRSAVSANEG
VRVVIHPPGTIPFPLTEGFDVPPGFSASFGIRPRQIKRIGPPHGNCITSNPFGDSSSQYRVLSCQKMCLQRYIINSCGCQDK
TLPDVPDLKATPCRNATDFPDSCMTDASDECLKQFFHLYDRIQCVRSTKDRFMKNSSLLTDCGCFPQCDEISYDVSYSLS
KWPASGYEGDAAFFDIFYIEGFRERFLNTPKYNMVMSYFHEDTREKTMQDFARLNVYIADSNVIITQEMEDYTTTQLVS

DIGGQLGLWIGISIITLAEVLELFIDTCRMLFSKRRIMVRNARRKETDRYRSHSGIAGLVPIKKTNCNNIFMSPLDRSETHAMILRRQNGDSSSIR

>Protos_Platyhelmin_Macrostomum_lignano_A0A267FKQ4_A0A267FKQ4_9PLAT_Uncharacterized_protein__Fragment__OS_Macrostomum_lignano_OX_282301_GN_BOX15_Mlig033565g2_PE_3_SV_1

ILMSSIRLLKKFYSTVERTAKMRRALVESLASSSAHGISRAVSVRSRQLKVIWLLLFLAFTALFCKDALELVGKLINASIVTTSEVDSVPFEFPDLFACPRHPISGSIPYALSESEAATANEEIFRLELLYERLKDEKGEPKYLSLETRSRIILNSLPTELLNKIILSRSQSIIMKSNNSGVDNETALFSEMKVLCFKIPAKNMGRSGLPPELIKLDTLELELFTDPYDSHFNYTGSPIREIMNLRNTSRRAPVSHWDLEKGLRDPVSDSGFSLLLVPRGTYPSVLIEPSVSLAPGHSYDISLSMEQVTYSEELQRKSCTIKEPKLQFVDLLNGGMLEYEYSYAGCLIDRLSRKKEKDSGGYVWPSYQVPWDLRDSLRWETLPDILVEMGRSPAEFFGNPNELPNCQQPCQRLRYKASISATNWPTFNRNYSHGFWNVSAGKQPALAQLANMAKQLALDNHSSHLEATFRSRVANSMLRIKISADSPMSTRLVIAKSYSTTAFLADVGGVLGLYIGCSLLTVCEVIELIYHLVASIVGSRPGPKQQEEIKEADAADVYTLEEKNDIPNTVTENS

>Cnidar_AlaAla_c48440_g1_i1_p1__GENE_c48440_g1_i1__c48440_g1_i1_p1__ORF__type_complete__len_576__score_42_88__c48440_g1_i1_504_2231_

MEVPLEEPGLVRSTKFLDLVTRLAVLSYILKKRKSKTRKIEQSVSDKDKRHSVDLPQYNELWYNFLHNFTLHGFRYVFERGAIVRRFLWLVVLVTAIGAVSFQTERSIRKYFSRPISTTVSLEYTNEIVFPAVTLCNFNFFPHYLINGTIGEKVVSILEPPEFVRSHQENILISPYTENIMDYRSHIPHPFPPDDGISDTFYYEDDNEIYRSYRQTEGSHAKESSKKKSTNPKFLKIFKNFNSNSQKRQNGPPNLMDNTFDFAEFVKKHGHRIGHMIKRCHWKAGKCGPHNFTSVLTNFGLCHTFNSGRAGYPLLKVHRAGVDFGLRLQLNVQQDQYYGIFRDSSGFKVMIHDQDEPPLINELGFAIHPGMHTFCGVRKTKIINLPKPWNTACQEQRTDSARRYTKAACLLKCRAEYIVRACKCRSFEYEGDTPLCLPQQIRDCVRPSVAAFMNESDHCQCPVPCEIMKFQPQLSYAQLPAKHFSEALAKIKHIDKHQMRHYLRDNLLELDVYFEEMSTQVIEQVPAYDEESLFGDIGGQVGLFLGASILTILEVIDLLGRVLVIRLRKSPQLTV

>Protos_Platyhelmin_Macrostomum_lignano_A0A267G7E7_A0A267G7E7_9PLAT_Uncharacterized_protein_OS_Macrostomum_lignano_OX_282301_GN_BOX15_Mlig023951g1_PE_3_SV_1

MATARKDLMQRAVKPVSPQDEFVRFCMLSSVRGVPRILRSRSLCWQAVWLVSVFGFLGISLLSITLLVNTYLSYQTVLEVTTRHYTQTVPFPSVTICNLQPFSTDMVKRFIRNPNVNVTDMRFLNELKKMKNHPNYTSSFKALNEKSFLFQNWGIETVRKIGLTFHEFVLDGTCNFSFVQAAKFQQRECNLRTDVLLIPNPQFFTCFMISPTPDIANTTYSLEMDLYLDTEQLFDCQQTTECHDALFDLDDSIGAHVLVHPAMTYPRISQLGIHVLPGTKSVIEVVETRSLEKYEQSPKETCTNKQLIDLFIGAKERTTAFPFAYESFTCRQLMVQEKILSNCGCTDPDYPLTADMKDAPFCAELKKPGQSTDAFLKNYDCASRLELDPRNLIKENLNDQCATPCQSISFKTTAYQSLWPPKTNELDFYRLRVQGKPLFGRKFDAYEQAENFTRLGQHTEAARALKSPDANAIRDNFLRLRVFRPDLSLTVTKEKLQITPETLISQIGGSFSLWGGLTVITTVELLEFVYYMAKRFVCSGYGGDSQSEQSGEAGEAGDVAMPLAVYASVRGPAAS

>Deu_Ambulacrar_hemi_Ptyfla_40v0_9_20150316_1g8763_t1_scaffold4594_cov140

YGVMAGKSLTEFGQETSIGGVKYVTDVSSRNLRRFFWFLVVCTALGALTFQIVNICTAYIARPVSVNREYIPMGQMKFPAVTICNYNQIRKTGLLEAFGYSHGDGVIKTAHAIMSPSPETMDQDVNLVGLNANYDFEYLMTTVGHKKDEMIMRCKWPGRQCSADNFTTTYARERYGLTVYLDVESQEYVDNWQNFVGFRVIVHDQDDKPNMKDKGFNVAPGTYTAAALKKLSVMHLFAMSYSFTIAMCQRWYRFLPPTAKFTDVQGLCSHCKTPCYEESYGEHLSYALFPSGPSVEELAEKAEKPCPKAMALYYKEKITAYVFASLSMANLSKILITTKDKLLDAIFLGETHISIDRIINDITASAPEGGLLPSVDVTDSPIENDQLTVGKLYEILTEALQSYYDQIYSEISIPLSSNLYRYIHSERQSSVEVGGDIYRKVYETMINSSASARRLKKIAPDLAVQGTFRAALNVTDSYGA

LVNDTKYALSDAFTDIFDLIYTTVLDTFEENRITALENHTMYQICMEYMSMCPLDRLPGPSHVFQAFGGGNGNDTERSA
IAPKKLILEVGEPMMLL

>Deu_Ambulacrar_hemi_Ptyfla_40v0_9_20150316_1g8782_t1_scaffold4615_cov164

MEDNINSKNESTKGNIRDLAKGFALRTSSHGISRIAEANTRFRRAAWIIVVTASFIGYSVHVGILVSRFLRYDVSIKMDVIT
GDSVLFPAVTVCNSNKLRLSEVKKSPYSDLATVNEPHFSFTYRGPCVPGDRACDEDNRICIKRYMWCNGIVDCESGIDET
DCNRVPCQYGEFQCEETRQCLPLYVLCNDRLDCADGSDEPDGQFCPFYRVGLCNYDVFTGERSCSGPPLQVSYCPGGF
RFMHCGGSETKICLPERNMCDGNVDCQIGEDEVECQNYQCPVGTIKCTGSHLCVDNSKICDGSYDCAYSSVFDLTDEV
NCQENKCPEGYFRCRSGQCIRHHLYCDEPTRPDCADGSDEIFCNVCGSSQYTCRSGFEEGNRIVVKCIPADRKCDRFVD
CHFSDDEYNCEYGLKLTLFTEQAEYIGIFGQETGVRVVIHDPHSTPFPEDEGIDAKTGTATSVIIKEVSLNFSFAFGICYLKR
NNLARLKIYYESLNLELMSEVEDYPEYVLHELENQFDKNVSNDRQEQRYAKSWMTSATNDSARIIEIRIPTGGKDSRRST
ASFVYFHGNGENSGTF

>Deutero_Urochord_tr_F6SR69_F6SR69_CIOIN_Uncharacterized_protein_OS_CioInt

MHSIHNRDVRVSSLSAHFEPVENCVVHSDDVTVADDVSYTSSSMSSDLESMKPSNIHVFSSSSTLHGFNHIFALHHSSLR
RFAWTVAFLTSLSILLYQSSNRAIYYLSYPHVTRLDEVAAGNLTFPAVTICNLNEFQFSKITKNDMYHVGELLGFLDKNYAL
CCNTIVPERQTLNFLNIQRSSFRHFNPRGTFSMKEFYERVGHNLSDFLLACNYRGRFCNADNFTTVYTRYGKCYTFNSGV
KQPPLKTLKGGVDNGLELLLDTQQNEYMPVWKETDEITIEAGFKVQIHVQSEPPFIHELGFGISPGFQTLVSTQEQRITFL
PNPWGKCRNGFDKSEHYHFPNYSISACRISCETLYVYLHCNCRMIHMPGKERYCTPEEYKTCADKALDFLVMTDDQLC
VCETPCEVIRYNLEMSTLQLPSTQASSYLAYKYQVSEDYVRKNFAKLNIFFEALNYETIEQKVAYEIPGLFGDIGGQMGLFI
GASILTILELVDYFYEVVKDRTWGRKLRKNTDRSPTREEANNKTVSVMVREQAIFPVITNLFLGVVCCLYIYIILVVYKCCN
VPHILTTESGS

>Protost_ecdy_cyclo_nema_Cele_tr_Q8MPW0_Q8MPW0_CAEEL_DEgenerin_Like_OS_Caenorhabditis_
elegans_OX_6239_GN_del_6_PE_1_SV_2

MGAKVKDQTSVEMHDIPKNQAHPEDDKPHSWKDALFGRRARKYAFVFIVAFLAVLTVKDVFDLFEEYFDYPKESDINIV
FNESMTMPNVTFCMSRQQAWSHFKLNLTAPADEWDAVVDENLANMTDHDAFMKTPWDYRLVMEAYDMIATYSS
LERETTAHGSARSIHVFKNAPRLAAKRKTFKKWRDILDSRGITFDEFTQKTGIEVLRRSMQRFRRRTFDDDETVIKTKLRIS
WISQMQICFQPEFDQDNFKTIDDQGVFFDMLLSHNAENTEGQKIDCMSVDFHGRPSSLNRFMEGKGRSRDGYVDEV
CLGQRHEVTAHVTALYQMLENDEQGTRCRDVEDGEDSEFNCRSRCRMEMIRDACHCTPLSLSYLAKKEDMEIFPLCDY
TQCTVDVQKGNYSDTECANKCFPDCRQIRFEVDHSVKGRMLRPDLTLVELSWGPFEYLTMEQQWKYSATSFIAALGGS
IGMWLGLSILSLIQLVTYSYTYFTKKIVNERILKKGNTFDKKDEGDEDGDFTIDNGTERKKMSLAENPFAEFFPAGSPEQP
KKMGSKPNSRQASLDKDGGNAQGYSSNPRA

>Cnidar_NemVecNVEC200_011692_1_1_protein_AED_0_06_eAED_0_06_QI_233_1_1_1_0_91_0_84_1
3_1484_578

MLKLPCNKEMLRNTDPKETALEQINQFLQETTAHGFGRLGATAGSKWRIYWVMFCLAAYCVFAWQLVGLVNQYNSK
PIKTRTQLKHAQKLDFPVVTICNMNVLRASRLPPKLRTKFDEIINNTKKTSSRKSSNSRNAFVDPQDLSFEETKKIEILHAVT
THDNYRELVSAAHQLEDILLSCNFNGVNCRNSNDPTIPTSWTQTWNDNFGNCYMFNPAQTHNGEKVDPYSSSIPGES
NGLTLQLNIEQNEYLEGITEVAGIKVSISDQGVLPFPGQQGIRIMPGQSTGIQMTKLQTRRIDPFKNRSCENSNEMSDK
NLFFGYNMRYSKMACKYSCLNAKTIERCGCTNYNTPELQKRNISLCNRLNNAIIDCLNKAYDTFEDGSCDRECPPSCSEV
SFDLTISSAKWPAKSYEKTVLKTLQTEYGINMTKEEMFENIAQVHVYYGELDYLLVQETLAYTFMSLLSDIGGQMGMWI
GISALTCAELVELVCVILANMSNRSKKIVHISSNIGCFRPAKLGGVEAIEKVGGVKTIEQEVQGGVGIIEQEVQGAVGTIEQ
EVQGGVGTIEQEMQGRVGTIEQ

>outgro_CafRoe_KAA0154385_1_hypothetical_protein_FNF29_02605__Cafeteria_roenbergensis_

MPGGGRGEGRVRERTKLVWLPADADLWLDRLIAAVALTGFIILTSWMTSVLIRRTGEMQHEQPTTAVHMRAAENLPF
PAITVCNMAPHVPLERVWCGSYVSNATTRCAAVVPPGMPHCLAFNNHLNGSLYTTEKTGLADTLMIMLRIHRDKYPP
TSRFVGVHVSLHPQCEHGKTCPQELSNVLLASPGQPKFFRMRRLTQTFLNGTSDSQFEARDTVSGVSNFSEAFGDPDQ
TVVMGFHYGSMTETVISELPRYSLLDLGAELGGIMGLVFGTGAFDVMMHLLRWCAGRGSLYHDLTGRHPSQYRPWG
KGGKGGGDGEDSRGARGAAFEEDEEAAAAGAAAAEDEDEEGSTVAAGRAAAAGAPPSAGGIPWPAGGASGRWWT
RRYAAWLRRGASVPRACAAMDECGDACCPRQCYASRATALAEAEADRRAGLTGGPLPPSAAHAEGTATVRGRDADIL
TLGPGAERAGWWPLLCCCAVQRVAELEPRAPGLRGAGGVASGGASPAQPLLRRGHGDTDGYSHASDGVFSAEPREP
GLGDAGRIGHEAGPEGMANAAGDEGGVARRGAWGRDAAGEATG

>Cteno_Ple_Bac_comp42060_c0_seq1_p1__comp42060_c0__comp42060_c0_seq1_p1__ORF__type_internal__len_580__score_119_70__comp42060_c0_seq1_2_1738_

FSIHDLANDYVQTFSEERRPRICKIIWSVIIVTVMSISVYMVYKLVQDYIQFKSFNKSTTVWLENIALPAITICSTNYVNNTA
FRESLPTNETHLEEKFNRFLRDVAHFNKRGDILRNINISEALDLIAWEKEQGLVNLRFGNQFSSLLVGEYDFEFRGAGRTI
WNMDKKEMPTELGMCLEMNDKGSLVQTSGGVDGGLTFNLDATVTDYLISTKTKGFVVFIRDQDEIVMYNQGGYVIA
PGTEAFIKLNAKQIKRLGGKYGTCENNKSQYSRFNMHFETVRECIQRHKISMALKHCGCVPWYVAERLYKSNKTDTLEG
FIEEIDRWAVEGEQRRRRRSDTSGSHGDETFQDFLERSKREVEQDSVSATDSPPTTTVYTTTGETVTDWDKTPREPLYD
TPYVDHVCGYVMQTACDKYLSGSVGSEVQELLSTCNEPCTYKEWGVEMTSTAFPPTEGYLSQVMEFREGYTVPSWQY
ARENLVRIHIYHDDIRIENINQEAAYDVASFIAEMGGIGDLFIGFSFFTMFQLVEILIALVARSGYGQWLRWKERKQKNN
DQLVVISNSVTNRGNRGDGHSALI

>Deu_Ambulacrar_hemi_Ptyfla_40v0_9_20150316_1g12811_t1_scaffold8178_cov112

MTEGRESRRFHRTFRILWMVLIVIVAVLFGYSAFTYVTMYLQYPVTDEVTMVRSRTQQFPAVTVCNTNTFRSDILTLKSK
ENPNNVAYTGFKLGLHTDFSHHYHTPSITIFHCIKQSTIGGESLFADGFKTALQLKEENPEAFQILCDSHCDFIQKNKEKIEY
YTRYAWPTFRSILKVSGGVRCIRNVRLSRKSTQMLSRTSLASRSNDGSNLCTQNQRAFSGDGLGLQSQFNQHAARGAK
PLIKQAKICGSSKMVLVGWDDGTVSRYPFVWLRANCRCNTCWHHDAYGRISLMEDLDIDIEPTGVKVSTQGSEIEIHW
PDNHLSPFEASLLKDVTFKEDVRKKNIRLWGSELLDELPTHQFDEIMSDDTKLLNFLEDLRDVGLTLMKGVPTETGNVEK
IAERISYIKRVNNGHSVGPTLLLLPFQQITALHCIQQSTIGGESMFADGFKTALQLKEENPAAFQILCDFHCDFTQKNKHK
TEFYTKCAWPTFRMPGQYECDGVTFYDFVNPHVVRDPEIVKCHEDDVFVVSYPKSGVIGDWQNFFTVAKNREFEKNC
TQNGWKELTLTLSGESHAVTEK

>Filast_tunicaraptor_GIQG01095491_1__p1__GENE_GIQG01095491_1_GIQG01095491_1__p1__ORF__type_complete__len_581__score_92_90__GIQG01095491_1__198_1940_

MTARGNNWSLELDEANPGNAPEAVGAHREGMGRARNHDKPFDEDEETKMFENAYGCMPLTRFFMHTTTIHGLPR
VFEKGHHWFRTALWAALFCASFGMFVYVAQFRFDEFYAYNTSTSVDISFQPRVQFPTVTICNSNMYRRSALSYDVLEAL
WDDEGLTNQEIVEFVAVTINELNSDSSLPTLTREQLFAIAVDELGKEGEDDQELITDDLIELLADRYHFDDDDFYDLLLEKY
LRSNRNGGTDAQVLASLFSDEQIGEIGHQLDDLVAFGYAKFKGESIDIATEFERTISHPYGNCYSWNSNGNYVTTRPGPD
LAFDLIINLEQQEYLSSSLSGTGKGIDDVSIIPAGLRVHIHDKSEPPFPDAGLTVAPGTQAFISLQQQTITKMTPPHGECVE
RTDTAAYSVESCFKGCFEKAVFTQCGCRYPAAHAVSRTVGTCSTSDFTDCVDDIEDAFSTDVASFSCGCDDPCNRITYRK
NLSHGRWPSVAVSGIISDAYNISTREVGDSFLRVTIYYEELNVIAMTQEEKYSMSALGAELGGLLGLFLGVSVMTVFEFCE
YMTFFFRPRMYRKKSVKVTPQS

>Proto_ecdy_PANarthopod__XP_023226919_1_acid_sensing_ion_channel_4_like__Centruroides_sculpturatus_

MRELNNKSKEDGGENEEEFSIQTFFGSSAVIGLPQIASSQHILRKLLWCAVLITGITFCALESYKFMTEFYKYPVVINLEIEN
KGILEFPAVTICNINRVRKSEYHVINPENVNNRTTFINSTPRDSYCSRERLVDERNQAEFYSKYFFLNKTVRNKIGHQYHKF
IIMKSLRNMNFASHFFTSITKPNYGACYTFNSKINTSYKDVPMSTHIGPKSGLAMTLNLELEEYLDDVKTVGARVVIHSRE
EFPDTEGEGINIPPGMETSLAITKKSIRRLKSPYKDKCRDYKGEYYANKGPAILNQKDCRIYCLQQKNLKTCKCTDPLQVFT
ANKKCNLKNLNKMCCLDNVFDKMIHSQYAECDCPLECLTTQYEPIISSTAWPAPENFKRLELQNYSEFRKTYSKVNIFYS
TLEETIYVQRPAYENSEWYSHLGGQLGLWLGISLPAIFDCIESIILLLCYICLPKYRNKKKFKNDFETIVKPVNGPCYTFNGR
NTTSENMNVSSVRLSRNIGTLSGLTLVLNLELEEYIYKTQSVGARIVIHSSDEXVTNFYICSSVTSKWLSNQDMGRVIHLM
VETRLVKI

>Protos_Platyhelmin_Macrostomum_lignano_A0A267FWM6_A0A267FWM6_9PLAT_Uncharacterized_protein__Fragment__OS_Macrostomum_lignano_OX_282301_GN_BOX15_Mlig032051g1_PE_3_SV_1

KHDCCGCCPAKLCNSRTTAALSGTVLTILAGSTAHGISRAAASHSTPMRAMWLLLFLVFLGLFFNNASTLVRRLSENKISK
SYELDKVPFVFPDLYFCPINPYSGSFPFVLSTEEARQVNQEIRRLGKLYDDVFGNSSGEMHMSLKSRASFIQNSLPVDLITK
YLTHRKSQSILRCNISSLHETFSVAVEEFSHPENLICFKLVLTEEQRHQMETKMYKLEVQLSADPYFKDFSYSGHHIKPMV
DPDSGEEILYQDWHPTVGIEKLKSSGFTMLIVQNGTYPSFDHETSFLLAPGKSYTVSLEMEENFFSHDLHGTNCSVNNP
NVTFKDYLTDEKVNFTYSYAACYYHEMTKQLLKELDCVFSHYPIIWSMRHAKRCFNMSESSKRSIQALKNKPIDWKQIEA
DVKRICRLSQPCQHPEYSASFSFATWPSVGSSAVSYFARMPLLNDSEHLAPAIAELIKIAKQLDSGEHPAHLEGVFRSYIA
ESMVSVTIAPSSTVSPQVVISRSYSFTAFLADTGGLLGLYIGCSLLTVCEIFELFYSLVEKLRSSGGKKTPANCQTELEMQAD
GAAESLVRPPPSA

>Deutero_Ambulac_Sakowv30035415m

MIADYDNMVEDYENMIADCDSIVADYDNMVADYDSMVADYDNMIADYNNNLVVFKELGECGSKANRYRIMASSYS
FYGDRYSDGALQFSQPIPHPCKTKTRNTIALSSFSDGRFRSKVNSIKEDEEKKVQLQKDPPTATSKGDVNACGDCCCCRV
FYSDRFREFSSDTTLHGLRYAVAEGVEPWRRFLWTVLLVIAVSATAVYMYLCWYKFFSFPVNTVVTLTYKSKLRFPAITIC
NYNQYRKSVVKGTRFEEWVRQQYPLFNFSDDDVQVELDESILEANRTWFDQMAAHSKWSMIYQCTFGSVKIPCSASN
FTETFTDFGVCYTFNGADREGGALLVGHSGSEHGLRMRLFVNQSEYTFGEHCGAGFKILPHPQDEVPLVRNFGFAVSP
GTEALVGLKMIQEHNLPAPYASQCSNETLQYFAKYARSNCVRERQTDAVVAKCGCREPYMPSDARVCNYNETRVCVR
PELERIQVQCPMACESSTYAPRVSYAMFPGRHIVEDLQMRFNLSYVDIRENVVDVKIYFEEISLEEIYQEDGYPISELIGELF
YNIDDARAQDNDVINITVRIIRYDDCSIPH

>Protos_Platyhelmin_Macrostomum_lignano_A0A267DFK0_A0A267DFK0_9PLAT_Uncharacterized_protein__Fragment__OS_Macrostomum_lignano_OX_282301_GN_BOX15_Mlig021913g2_PE_3_SV_1

FNFPFVLKFLKPQQPEEMKPLPGEADCPVFTLSEGQQAKAREAWAAAADTASSTDSPGRSSVYSASSAKMRLAVTQW
ARTTSMHGIGKLTEDEADAPSAMRRIVWTCLLVCSVGFFAYQISQLAIKFSQFPVATRRYEEDVTSVQFPAVTICPINGLS
NFSTQANMKALTEVMGYIFSTAPPGSASTNSSSLPGNMTCEEKWRLTTSNVPSKENFLMGSYAMGINQVNDSLLAGV
SVRLQRILLDCSYLGMQCTPKDFVTFRNAAFGFCYTINLNGSWRMASYGPFNSLQMTLALNQSDYVDNGYGDAGFRIL
IHEPHTYPFAEAAGFHLSPGSSAQIGLSFHRIDTLPPPYGTCKIYTDEENRQLNAYVNASVAVGAESPPAAAEVGYSHQA
CLRTCMQQAALSTCGCYLSCSYASNSLRNSKRPCELRSAMLIRGNVTEKSCYEKLLTGGVDCLERCKFPCSRRVYRATSSY
GQFPALSYLPRLIELATASGYANDVAKAKADNMLLDFARGNFLRVSIFAEHLKVYRVEDSPSYELKDLLSDCGGQGGFW
LGLSILTMFELADLLSQLMRSCCRGLGE

>Protostome_Lophotroco_annelid_CAC9665778_1__Ofus_G14632_Owenia__fusiformis_

MWQLNTNHTNNEKLKEENSKDGKFNDDLTNDSIIIKQKFPVESKNGTISLKEYLYDWVDNCVLHGIKNIRASKNTFHGL
FWLIAVLNFTVCFVFFGKHQVTLFISKPVKQIQTIEKSQIEFPTITICPINTYSQRNMDRIRGNSSTTTSTFFENIQKMPFSQ

DDAAFSSSSVGTTRQFAYTNIYYSELQQSGHQAKDFIVSCRYKNRGCTYKDFKLLESVMYQNCYSFKANDSVDIAVETM
GLSLVLFTEDEVHSDVMKGGGKYKRNAYSEAGKPSDYGAGVKVFIHPPGTPIDVAAGGITAAVGMTTDIGVNVENIKW
LDDPYPNQACVPSTEIEHHHNDTYSQETCFSNCVQDFLKETCGCGSISYAGTPAESPPEEMCGNFANISALTFYAYATGI
LEWVCLLKNQEQFNASSKAECNCPPPCDQITYKPTISQGVFLPSSYQYSFLERLTGTACPEDSSDLPKYYSLLMGAVMDE
NGKKVDCNKALETILAEELVSKNFLRVNVKFESTLVSTTTMTPDYTGSDLFSDLGGTLGLCLGGSMVGVVEICELLVGLFL
KTLRKIKVSTVKPINKAIKTAW

>Protos_Platyhelmin_Macrostomum_lignano_A0A267E6Y3_A0A267E6Y3_9PLAT_Uncharacterized_protein_OS_Macrostomum_lignano_OX_282301_GN_BOX15_Mlig009645g1_PE_3_SV_1

MTVGSNAGGSVRRRWEVRKHWNDFLVTSSVRGLKRTTTSETRFLRLFWVVFLLGSLALMLYQISSVVDTYLHQKDSTA
MQSKQLDIDVPFPAVSVCREATYSSHKQMRQAMERHNASVTTPQSYSRRVADLLYRHFRNATNMTEDELEEAKMAL
LRFDTARSYQAQIPRAAIKEVALPLIRQIATARANFANGSSTSLASNMDIIGMDVETDDYPSSSYDAELWGMRVMDFYH
WRYFLCQTLEITLSREEMNEMRSLDLLLWYAADDFNTSISGVADRAFAMRGFWSADGIRLALHPAGEHGKVHIDNGA
TLVRSGGYSEIRFRPSITRYLNCSDRELQHGDQGGYSYNMNDCKYLNYDEELRRQCNCTMDPLAWDYHQCIRINWTDL
DMSDWGGCMGRVEWNRESCSRPACTHTEYRYEASFAPWNINDFQWRWIVDLLLNRDEALPLYLAMSQVGEAESGI
DYRESSALFQRIKNNLTSLTISRQSGLVAYQQEVRLTSIANMLSSCGGALSLFMGITLTLMAEFIEFLYKLFRPAFQRSKTP
AKQQDAASDNEGQADRKEGDSKNESVTLDSLQVVEQEV

>Protostome_Lophotroco_annelid_CAC9588452_1__Ofus_G061690_Owenia__fusiformis_

MKEEETKCKATVGEILNTFADNTTAHGCGSISRSKHKLGKLVWSVIFLGCICTACQGLVAIFVTYFGYPTTDTVTLERRELE
IPSVTICSLFPFTISSQMTLPVPGADTNLAKLLTLFHVYHYGTFYGEHKDEYRKHKNRINSYAALFENVEDDQKLLYTHDLS
NMLEQCFFNGKPCDDSFFSEIENANYQKCFTFNGAGLNLTDMTALATDPNNGLSMVLFLDAYKANYTLPGGMIYNPE
NPNSGTTGIHVTIHSPQTMPFPTIEGFDVPPGYSTTIGLTPVARERLPEPYGVCTNQEYIEGTQFKYSELGCLAQCVQKE
MMDQCGCITAYLPYPNDTQGMLYCGDLNLDNFLNPLGPNITLMDDDFNSLECEDRLMMLGFTDNMTERCPLCLPAC
NAMKYEQTIAQALWPFNYVQGNFLTNTLVYGPDNNQTRRSFKLFEDLLADWPAQMDYIINTGMVRQNFLRVNIYFK
DFTVETTKQTVAYGLNNMVSDIGGTLGFYVGISVVTLCEFLNLFWMIICACLRRCKSAGARRSDNVEVIQVKEANQSTD
GKTQYDIEFTKNLAHLHNDTMPRNDFSSAFKY

>Protos_Platyhelmin_Macrostomum_lignano_A0A267G0V5_A0A267G0V5_9PLAT_Uncharacterized_protein__Fragment__OS_Macrostomum_lignano_OX_282301_GN_BOX15_Mlig024322g3_PE_3_SV_1

HLKKAPQPIGRAREPLAAAFIWPAALLRCPAQPRGFPAAKPPAAVAMTPEAARRERSFCSFLLATFIDNTSMRGFRKCC
GSRYAALRLHWLFYIASMTVILAYGSFNLFAEFFTYNKSYQIEEVLNDPTPFPSLTVCSLSPFSPEAASLWRSGEVPSPARF
SRDLRHGMGRMVRLGRYADAASLFRNDKMATYFQFISESQATQLGHQLAELVPFCVMRHSNNANPSVAHEEKCNFK
VTGARLHHPEYLVCYDLELDDAESIYVSSLSLILSNAHLNEYLLHNISSTAGGGVRVSRDPADKEHVFDIFNQGAGFRIVLH
EPGTHPLADTDGFSVQMRTSTLIKYKAVKVQKLDYPAGECADSARLPDILDDDSRYKYTFAACLSSQIGQLIRRQCNCTS
AFHVRPPAAGPAAPYCGHYSAENPLDFIRRLDCVRNLNLSNLMASVMHSTCQKPCTTFHYESHLSSSNWTPQVWQIH
WMMHISSALKQVMDNPFDQHFSEQLQRLINFTDLESVAGAPEVEALKVFESELAEKMALVTVTRMNYNTLVKQEALQ
ITPSALMSRLGGLGSLTIGSPSPALSSWLNSST

>Protostome_Lophotroco_annelid_Pdum_comp417550_c0_seq1_

VDLDFTQRTRIPFPSVTICNQNTHRMSMTYNFSRKGEPVITPASVPPMTIYQMLSDALKIKNDRGPIPAKKVVAHEKEFR
VLFQGIDLAPGMAKEFYKDSYHLPDRRGMIKDCLWNGKPCNDTLLSTSITNMGSCLLFNPGPAVLNTSVPGPSNGLDLL
LDVEYLDLLYGVEAIKGFRVHVHEQNDTIKLMTDLGASIRTGAVTTISLKRNKHKSLSSDQCSDTWDMESCMLKCISDHV
SKTCGCRENYMKGDAILCDPLNFINCSVPQTKYIMTQGIDDQCKCTAACERVTYDTNVVKVPFSQEYIWQIWLASSHQ

NQSDSALKKLSLLRDWAILDAWNRRLTADIRTIAMTFNEIYMVHFLDNNAESISRVLDESISAANESISCTNSTKKNVSDS
YQAFRTNFLKSPFISFIFDDYIVEDFQAVLASDILRNKTVFSPFLQQTLTHINSLWDAITRFQTCLNISAIPEFSKFNSTKYRE
SLADALSIHSRVSAATLGSTLMKFIPKPFETKMPDHYEKSYAMLRVSYEDMTVTTITQRKADSLTSLICDIGGSFGLWIGA
SVLTICEVIDIIGFYCVCRKSKK

>Protos_Platyhelmin_Macrostomum_lignano_A0A267DVT2_A0A267DVT2_9PLAT_Uncharacterized_protein_OS_Macrostomum_lignano_OX_282301_GN_BOX15_Mlig012217g1_PE_3_SV_1

MRDRAARANKPQTVGDEFLRFLTFSSVRGVPRLLRARSLGWSVIWAVSVLGFLTVSFYFTIRLVMQFSTYQTVYEIKNEF
NSQKAPFPSVTICNLQPFSATLLDRIFASTSPLDNVTADMYNDAFLRLQASSEDFRSLTWKSFFLRNWGRKASEKTGLKF
YEFVLNNTCNFTYSSNSKLRSMRCDPTRDVQELFSSRYLTCFMLSPPQNISNSTFSVEMDMYLDTESSLFNRSHLGDESK
AALFTLEDSIGAFVMVHTARTYPEGYHGVHVLPGTKSVIEITSVNKEVKLEHFAKDPCTRKLDQDILALYQEIRPFAYKSTI
CRRLMIQRMIARSCNCTDPNHPVPEELASQGFCGEVSQDAFSSVESVRNAALNATIKKLVCGLKIELSSDPTSSHGETQC
PQPCEEISYRTTAYQSLWPPKSNEIDFYRYHIKGNSHLARKFKVYAEASALIDSGRLSEATALLDNLGTDLIRENFLRLHVFR
PDIMLTISEEKLQITLEALISQIGGAFSLWAGLTIITTVELMELLYYVAKRVVVSLPGARPATQSDDGAEAAPRKQPEPGGS
SDLISGGGGGGFATQTFSKL

>Protostome_Lophotroco_annelid_Pdum_comp418513_c0_seq4_

LANSDFDVEKMLHQNHSELQEKYTYAQEVRQRVDASVSEVDQKMAADLKGTVKDFNEMFDGILASLDSIDEEVKRISD
TMDLRTKFHYDTALDSVHYVVRHNFIRGWEIIDERTFPYISGSFFQLRSSILQLFTNIAQKDELTKRVEWQVLFELIQNKLH
LCQRAKENLTKVYNAYKTGDPLLNFKKTKDGRYDLDFISLELLHGVIEDGGRETYYSDALQAVSAIEDSLKKVDLAMRLPE
DGMVLALEAIDEFVDACKTYNLNIFLFKERIVEAPLTEAERRMSEFQLKNKAFKDNARLVLDMTFELCTSINAFLEITWRPI
LESVDMFVDYVDGKRISKSEMAETFRSRGILMSLLTVEQYFDTLHLRLVDIKDLITKFGQSYLKVYGNMLSEPSTRKFYSRL
GEELKELEINGSRRGKMDRYYAHLLGVKRREISRIPTQDLYTLMNADFSMTNWEILENNTKVIINRLLLKSDHDAALGKR
VTAFINTVGDIVQSLKDFEGKSQIDHNFYRENFIHLDIFFHELSLEVVDHQIAYDLTNLLSDIGGLIELFLGASILSFFEIFDFIF
CTFVCRGKKSKTCCNK

>Proto_ecdy_PANarthopod__XP_023234660_1_uncharacterized_protein_LOC111634182__Centruroides_sculpturatus_

MGDESIECIKIPMRKKKRKPIRLRELLENVTMTGIAQIARERNCRRMILKLFIFIACLIGFIIQLTEFFLFFFTYPTTLNVLIEHK
DTKDLPKPMVTICDMNSIRRSSYCRQFSKQCAKLKNMKMFCGRFHRFCEIHNITEEDLVFPGNYFYEMAQYQRSIINEF
GIRSEDILPLILERNDIIRTIALDVNLKYRTCYSIDLGNPLSENYTIESTVQHVDGHLYLKDSRTVGEIFVQFNPADIFCPTYPL
GAMVAIHSKDYIINPFKNGFPIYPKNRYIIQLKPLRKSLLPAPYGTNCTQYRKTDNVSFDKILSRTRCLFECTKRLWIKNCGC
ISPEYPFIIDEKFCDILESYRCLNETNVESCYSECQRDCILTIFEYDIQNLPIHNFHIEKNLLTNPYSTNCSYYERSKEKSFDELY
TRQRCILDCQKEKWLESCGCITGDYPFETDETFCSGRAYMCLHSINTDHCHDKCQVACQATVFEYEVDDTPIEFYDFDV
PLETTDEDKNKEYSRKQEQVAVMNFFFNRPELLIYKYKPKFQSIELFSYIGGYIGMWLGISLIAICDLLEVITVYIYHIITRDIP
NIENY

>Protost_ecdy_cyclo_nema__A0A0V1B2K5_A0A0V1B2K5_TRISP_Acid_sensing_ion_channel_1__Fragment__OS_Trichinella_spiralis_OX_6334_GN_asic1_PE_3_SV_1

LKCHRNMQQANGTRKLDDDNMNNKNKHNISKSNTAPELSEYKGRKRRSTAASLENRTGAMSKIDESIMDVNLSGLHE
FSNATTTYGKMLWAILIALAIVGSCYEVYKILYNYYNCPIVTEWSVKATRAIDFPKVEICLTSSIGQSAIDEYPNAAAVAAT
MKKRLLFTDSNRPNKPTKERSKRQVHQHVPPQFWDIAPEWHPSKQKRDTTTATTLDSNNKNNNNNNTDLSNMTKLF
DDFMRYSFSVDDVFLDCVFRDTMMRQLNCNDVITEVLNPNWGKCFVVSVGDRKQHVPGQGLVLVMNAQNHNYPT
EKSLMPKTAGMIVSIHHDYNPYSFNYINVPTKSYTRMGLYAEYHEYLDVETGPKAQPCISEEEKHFSVLDLEYNVAACQ

MDCFQNLTLTKCDCIILQDVSFVKADIRDSVHFCSPEEITNCVLPEVTNSNETQTLYESCCDKCHTSCSIWHYNVQSSSM
GLHEPAFNFLQKYNISLDNLVYLHIAHTSLEYTEVGQEWATEIDNLIADFGGQVGLWIGGNLITFIQIPIMLLLIFFVSMYE
KFVGVRKIRRVVDQKRNMWSTWSTIIQAGRDGQGSPLP

>Protostome_Lophotroco_annelid_Pdum_comp411104_c0_seq3_

MEVIKEFGDTTTMHGVQKIAKAQYKRLRVFWIVAVICAIGMFIFQLYLLISQYLKWPTRTTMELSRDSIRFPDVSVCNMR
NIDVDVLYSLVKQFSLNKHPFELLTWTNITRASPEFEVKFLQLTGEYYLFYVYHYFAYPEVFSTLFSRSNLVGNFGKDTMIK
GAVPPWELTLRCSWQGAPCYSSNFTTFYDTYYSNCVTFKAPVEKIGSEGAESGLSLVALVGSGMIDWANISAKKVLIPGL
QENNYPLAGDGGLRVVIHPPGSHPLPSIEGYDVPPGFSGSFALKVTNNSLLGKPYGNCSTDYVEQDGSSYRVLNCLRKC
MQKQIIEKCKCVDARLPFSDDMIVNTDFCGKVAPLTAACKMSSSVSCMGPLFEAANNITCAKQVAGEMISKDTLVSDC
NCFPPCKETLYDLTYGLAKWPADHETDFVYQELLLQDRFFEKLNASNTSALKLDLFHQYFQFQNREKNLKEFAKINVYFA
DLTITKTKQVPDYNLIDLMSDIGGNMGLWLGMSLLTWTEFLQLALDLCIHAKSANSNKTKVQKMKNVTKVSVLPADD
AKYDFDYKKRTTSRENLIKQAMETDGLYYINR

>Porifera_Syc_Cil_scpid47944_scgid21366_Acid_sensing__ion__channel__3_Amiloride_sensitive__catio
n__channel__3_Dorsal__root__ASIC

MDTSATDQANDSSSSNNSAELNADSIFVDVSAHTDKVTPKKRKPIWPLSRFLHGKPDSSTAYEYHVDDDSDFPLKEYTR
NGGGHAGNGGTSNGHIGQQKQTFEQTDEADTPAHHSGIDWRERKAMALKFTYTYVYLAACHGIARAMQNKASVAR
RLFWLLAFSGAVSMFFYQISLLVKQFTDYEFQVDVEFSFNRQLAFPAVTFCNLNPMNVKLLNCSRFHDECLQLAGANQS
ADGQLIKTDDAIGYAHEYNFSGEPITRAEYLQYGFLPEDMIITCTWNGEPCSARNFTISQNPFYGNCFTLNGDPKNPLQT
NYPGREAGLALELYMNPDEGIGGETSTSSGVRMSLHQPGVKPFPDEDGFSLATGEATFIGVRQLEVSRLGSPYSNCTQE
NDNWKSLYPGYSYSRTACVQSCLAQEMFDRCGCVEIFLVDKPICNNANVNKEQADCAAGVRRDYRLDRLNCDLCYNP
CVDRIYLHTLSKATWPAKAYQYSFRETLKELNFSDGNSEDSNAEEIYREKFLRAVVYFEELNYQRIVQKPAYPAENLFADF
GGQMGLWAGLSFLAFVELFDYVGQVLLIALGLKKVVVLP

>Deutero_Cephalo_107760F_t1__name__chr_scaffold21_start_1172790_end_1185891_strand_pro_le
n_591_

MSPEDDKESPEKAFAGNTSLHGFNRIASSPGGSPRVLWVIAILGSYTGFGYMFGSMMASYFKYDTITDTTLEFADELLFP
AVTICNMNKFDSDKLQVADWSYLSLFLYGIQADIPTILALGVPLDETVNSTMTGVKIEDLVLAKGFNVSPNRMSLCWW
RGTNCSERNFTHSFGHYGNCYTFNADADNPLKQTMPGAVNGFMAFVDIKEDKYTESFLVSGNAEVGLKLLVHDPREP
PMMDTQGIALAPGNHAFIAIKQILYENHVPPWGVCKDLQLEYYDTYTLNGCYLECRSKHLVRNCSCRPYDLPGTAPSCD
PRTMFTCVRAVLAQVITGDLKCDCPVPCRMTSYSTSLSFAGFPNKHTREYLSPLLGMEPSYMGDNGVVFSVFYEKLNYQ
KIRQLKAMEEGQLASNIGGMMGLFLGASVLSLLEVCEYLLKRPLGFLGRTRHAKVVHVQPQETKNDVIQHVPVLRGISQ
QTHKGIPLPAAGDGAKLHGMFSFVTYSIRFRDNGVVFSVFYEKLNYQKIRQLKAMEEGQLASNIGGMMGLFLGASVLS
LLEVCEYLLKRPLGFLGRTRHAKVVHVQPQEPKSATALHK

>Proto_ecdy_PANarthopod_Dmel_tr_A8JV41_A8JV41_DROME_Pickpocket_23__isoform_C_OS_Drosop
hila_melanogaster__Fruit_fly__OX_7227_GN_ppk23_PE_3_SV_1

MPQEKRHPPESRTQRFLETLVIFRRSLIYQTKEFFQNSTLHGVRYIAESGRPIGEKFMWFCFTSIGAVTALVIIMSLWEKF
QTNPTITGLDTDFHNQNVVFPTTVVCPEAAFDHDKTYEKVYNTLANYDEAQAQMYTPFLRILTSLNFENVRDAKVLSQS
IPQNLLDAHTIREWAFEGHIDCKNVFVSCKYRDEDIPCCDHFEPIYTEHGFCYAFNSRFKSTPTEDVKTGAPHDLYETDKK
WALFFIPNSTSRIFIFSNEEYFGSDFNAQIDWSEPQLVEVRISKKNTYTTDDARQLSIGQRKCIFSDEVKLNYFPDAYTFSSC
MKQCRMNKAIKLCKCNPPFYKPIRELSCVINIFTNLIAYILLLYLTPKANVPMCSIKDFDCLDEFKSNITNIKDCLQCELSCSK
TVFNIDKLIKMSDRPESLGVLVEFLTWPIIRYKREVLFGWVDLLVSFGGIASLFLGFSLLSGVEIIYYFTLRACCMVYKNRQE

LYEIEEKIRQEPPPKIDLKLSLKSHNPRIGDTPTSAVLKVKPAEESAGPNPLGGYQRKHEKDYSLNTKNYYKTPKVVPDYGY
TSKTKNSWTTYDHSQYMP

>Protostome_Lophotroco_annelid_Pdum_comp414793_c0_seq1_

MGKLQGLMDDYAQSMTAHGVGRISRASSFKAKIFWAVIWTGMIAMFFLQATTLMRRFFAYPKSVRLEMANQPVPFP
AVSLCNLRPLDVFMFKDAIFKDNVTMRWIPGNYSYTGRSGASDFERFLIKYGNVYMSYRSYIEVLLEKEQVDQDWFWE
ARRALFSRLTLAANFNEKAAQQGGIQADQFIAQCEYAGKKCSFENFTYFLDPAYFNCYTFDMNQMGWESQTIYEGPD
HGLSVVLFIPSLPSTDLSPEAYRLIAFTQELSFGGEGIRLVIHEQNTVPYPLTDGLDIPRGVSASIGVQLNHNMRLSPPYGN
CTDKNTLEGTLVNYTYTMASCKKTCLQGLIADTCGCTDIGLPLSENFEEGSCSRFDALPTECQQKGDIRMNLDKCKTFFE
PWFERNLCKRTVRTNISKNLSAWNLCKCFPRCYDVAYVTSTSQSDWPTIESSVYLLQDILTAGGFASKFPEEKAKQYFGP
LLGAETPTFQEARDHILAKNWLRINLYISDTSVVKIEETEDYGISQLISDIGGQLGLWIGVSIISIIEIFDLIYQICKYLLDPERQ
QKRHRGSTIRETKQNGSNDLSYAQSNGNPLRF

>Protostome_Lophotroco_annelid_Pdum_comp418040_c0_seq1_

MMVLTLLLNLRFKSKMALRDVLGNFAVSTTMHGVPKVINAKSAMARLFWSIVCLAAGSMFCLQMSEVLTRYYSYPKK
VTVEVVPTPVPFPSISICNMRNLDVHILNTLNAAFLRDDNPINHINKSDNAFINSYMYNVGRLAHLFWKYQDAHPEVFQ
EVFSRTTFSANIPEDIIALAAVQLDGFVVNCHYAGHRCNKSRDFLRFFDPYYFSCFTYKAPEPSELDDSLSEGIENGWSSIL
LSGSGMLDKNDEIRMLPGLHEWRSAVSASEGVRVVIHPPNTEPYPFTEGYDVPPGFSASFGIKPRRNIRIGPPHGNCTN
KNPFDQATERYRLISCQKMCLQSAIIKSCNCSDVGLPRLNHMDVPLCRSSESFPESCFDDATDECLEALMNMSSLIKCVR
VTKAKLTKNTTQMEECQCFPPCDEVAYDVSYSLSKWPASGYEGDAAYFDVFGIEGFNERFNNSETKGKYELFAKYFNVS
NREETMKNFARLNVYIADSNVVKTQESADYTRNQLVSDIGGQLGLWVGISVITLTEVLELLSDLVRFFTSSYSRISRHDKT
YRNNNMNHHHNIIPLKRPMLHDHDLQFRMENTTL

>Cnidar_Rhopilema_esculentum_TR96298_c0_g1_i1_p1__Rhopilema_esculentum_TR96298_c0_g1__R
hopilema_esculentum_TR96298_c0_g1_i1_p1__ORF__type_complete__len_593__score_56_12__Rhopi
lema_esculentum_TR96298_c0_g1_i1_111_1889_

MLFSVNLKMLSLISFKLRNLLQEAADRSFYLGLNRSEFMRKSAKDTFNRINNCSSTITNFSDQQQLIKIETELGMEKHTKK
SERESSTREKVIDSEANLRSQQKESYLTKSTIHGLSKIEESKHWSLKLLWSLAILALFVGSVTNIVFLVAEFLEFRTYVLVEEH
TNDTAKFPSVTICNLNCYQKSKFNSLVTDMERRFQSFLNKSMERKKVLSMLKHPGFFNFMIHSNITLTKATQFASDKLFL
DGIEDWCRFGSFQDCNRSDFTDSFLNHHSGFCKTFNVNGKFTQTLSGVYGGLNLVLYVNQDDYMKLVPYDNAAGVLV
MVHPQDVFPRPDNDWFAVQPGTQTLISIEQKFIFRKEKPYTSKCVDREGDPVYPGRYGVNNCLQSCHHLKLYERCGIIA
NSALYHLSRRGKKLPNPPKNVSGDAGTCLLEFYGDLSSKKGTCDCPLPCYEETFKLTGTSSRWPTNVDMQYYRPVLSKVL
NRTNVSEDFIRSNFLQLNVYFKSFSYQKAMEAPAMNFFSLVAGIGGSLGLFIGASFFSILEFLTFIITALFGRYHRRIGHQG
AWLRDKGLTAEQFTGTRRSRAIGMVENFK

>Cteno_Puk_Fal_comp23896_c0_seq2__p1__GENE_comp23896_c0_seq2_comp23896_c0_seq2__p1__
ORF__type_complete__len_594__score_25_80__comp23896_c0_seq2__92_1873_

MKFVEVIDNRFITTITNVSSHIIRRLCWTVIAIVCLAVTTWMTFLLLSDYFGHPSYTLIQSSYTDRYVLPAITICNVNFLNRT
MMEEKTLEDGTSIYEVYKNLVGELGLTNDTMRGERERLEKEYARIDQQNIYVLYKWNIRQTLADGLVEFSKARLAHYFD
KDDVAGTLRTSFTEMGNCLDINDNQILVQRVNGPIGGLSLVLDAQTDHYLESTQSEGFYISLRMPNETVLKKEYSFMVA
PGTETYVQLSTSEVERLEHPWGNCKNMENLYNIEERNFLSWKECFATKVVWYYLNAPDCQCFPWYFHTRYLSNPATR
DPSQEILSRLRTAWKGDALFREPCVNYTTTELVYEGLDLVETTPNTPSAKDCLTLCNSTEGCQFIQYVLNSTLAGDNAAD
CEMYRGDAYDLVENQRGYQIRIGALQCEVERTDCLLSKQMYCDNVVAENIYHEFNKSKKIDAITETDFECSEPCTYSRTT

YQLSSSVFPSQKYWNSGLKKHFRHYSTLLDARQNLVKIHFYQETLHKTKEVQTKAYELENLIAELGGLSDLLVGISFFTFFAI
VDWLMVRMCSGCPRGRRDPPSNNSDKKANDAV

>Deutero_Cephalo_088030F_t1__name__chr_scaffold187_start_204160_end_230405_strand_pro_len_594_

MDKGVKSAISDCFVCVDEGLKSAISDCPVCVGKGVKSAISDCLVCVGKGVKSAISDCPVCMDYNKRYSRIVVRSGPRLSA
VCFSFVLKVVAFTPEVIILPLTEKKQSQKPPLEYFISDSGDVLYLVTDGNHLVTGGNHPVAMMNRDPTMSSLSDIDEVAP
KPSDIEVFAGNSTFHGINHVFTNGRCSIRRILWGLAFLTSFGLLVNQGVDRVLYFFQYPHVTKLDELEAQMMSFPAITIC
NLNPFRFSQITKNDLYHMGHAFEMLKPDKTLAHSNYIDPEHLRTLKMLSDFDNNFTPTKSFSMTEFYDRAGIQIDEMLL
KCSYRGEDNCGPRNFSTIRSGKTAKLEPQYGINAAEFRDPAEFRPGYRYREAGSMSSYEENEPKPSDINVFAGNATMHG
ISHIFTEGSFSFRRFLWAAAFLICFGMVSYNGYDRVIYYLSYPRVTKLDELEAQMMNFPAITMCNLNPFRFSKLTKDDLW
QVGHALEILDENKKLAHSEFVDPEHLSTLRRLAEFGPGYVPKNKDSFNMKEFYNRTAHQIQDMVLECSYRGTKCSWAN
FTVTIFPSTPVVDMQGFSPVWIQQLWTDPAQSHKSDE

>Deu_Ambulacrar_hemi_Ptyfla_40v0_9_20150316_1g21914_t1_scaffold24959_cov85

MTINMDKAHVRRIKMTGKKKRGLRSIISKFAKNTTAHGIANISNSTSTFSRTVWSVVCVTAFSLFLWQSSELLKDYFEYGI
KIKFDVVTLPKLVFPTVTVCNTNKIRRSEILKSEHQRILIVDEDFVHPYYALKAFERTYDKRLSPKDAVRKTNTSLRDCALSC
LHSNEFTCLSFDYDSQRRECYINPGSKDTLIRFSLTDDDEYDYYERIYRVFSVLDSNTALKFERFRNVRIPSDYPPQKTTTEE
CAQLCLDEVFFDCLAFTYEPFTSRCWLHSESHSLDSIPLIKDSTCDYYDRRGIECYALDDGTDYRGRMSVTRYGSHYKECY
TSQNGSDYHGFAYVTVSGNVCKSWLDVQSQNDLVFNSSDGIGDHNYCRNPNNADAPWCYTSVSGDSYESCNVGKS
QKKCPPPTKHTTNLKECYSDPKGLDYRGTISTSVSGYTCKNWTYFNTYPVFLKSYYGLGDHNYCRNPDHDRGPWCYTG
RGEKPWEFCDVGERNISCNVLGKPASLYIEKAFCPCINDRVRFFPRMVWDVQYCYSVKNMSHARMLKGVLLVNSLSLH
VRNFLSIFDMLTSYDGVMYSFHYSFVHFIRSKSILR

>Protos_Platyhelmin_Macrostomum_lignano_A0A267F1U6_A0A267F1U6_9PLAT_Uncharacterized_protein_OS_Macrostomum_lignano_OX_282301_GN_BOX15_Mlig031795g1_PE_3_SV_1

MFKESSDSSNKRVTSLEKSSDDEKADSSRTSGLPDYGATKSTDEDSNSGKFNGRLIREKLRPKRRTREEQTLYENNIKDFL
DALLSNTSLLAISNIVKRSHPVFKTLWAIAFTVCLILTIRQLYSLSRTYFSDPKSTIIQLSYKPFEFPSVTVCNHNPLRRSMVSL
FGQPDLQRFIAGLETKNYSVVDNGTQLMTQQVDWMRFHRLISNHTPEQRLRMGYQMSDLILQCSFAGQRCNTKHFS
VSLTKTYGSCFTFNGWQSAMAGRPVIGKNGREHGLILTLYLELEEYLQNFIPSSGIKVHIHEFGSLAFPETTGFSVPPGFQT
SIGMSLRMINRINRPKGSCHDGEGFLQNHGFKYTYNACTMLCLYDRVHKLCGCYDRANAVYYSNLKARLKPCQTEEQL
SCLMNTVYASDQGDLKCDCQLPCEESHFTFKQSLSEWPTDSYEMRLFGDLVDVKPKVADLKNQTKLAARRNLLNLVVY
VDGVMAEHIIEKQAYAIDNYLSDIGGTLGLFLGFSVLTGFELLEASALIMAYLCGKIGYTSDEDDDQPAASEDSDESGDAD
RRQSISASNEALTGRRASQILTLYGSNPTIHLHSN

>Protostome_Lophotroco_annelid_CAC9486619_1__Ofus_G015124_Owenia__fusiformis_

MHSESNNKLEFPPISGYDFEVTTTPKEKRYSEFDFTATTPRSKGLDGIDEKNESPKRKEITPEITPLFREFCDNASVHGVNH
LRTERVKVKRWVWTIFVILALGFNLFHCSLLIDKYLGFPSEETQFVDQSWIEFPSVTICNINAMSRITRKKMLEDNSTLLYK
WHDYVNNRFEALVSDLADGYSLEGSNLLDTLELLYNRVHLPTGYLENIGEEALEVGHKLNDLVVDCTFGITKCHAVNFTS
FFESTYYNCYTFNGGNVSTSHNNLITRTTGPQEGLSIIMYLESDNGNINENGSYLTMSKLNNAAGARVMIHAPNTMPSP
TDQGFDIPPGFSSSVGVSVSTRTRLGEPYAKCSKREINIGTDYLYSDNICLKLCQQRYVMGNCNCISSLLPFDRSTDLYFCG
HMDYNNNASFFENMACESRVLEEFVHNEDVRRECGCHPPCHNYIYNYQTSQSYWPLEYYQADFYDLYILSDPNKENLK
AYQNLKHHNVPQLIEKGLIRKNFLRLNVYLKDLTIVQHIEKRSYAIENLFSDVGGTFGLWAGMSILTICEFSELFFRVMAIL
VSKFLSIFGPMSTRAVVEDSSSNSHTPVRRT

>Protos_Platyhelmin_Macrostomum_lignano_A0A267E3E4_A0A267E3E4_9PLAT_Uncharacterized_protein__Fragment__OS_Macrostomum_lignano_OX_282301_GN_BOX15_Mlig000966g1_PE_3_SV_1

PVSQSLTKNSLQLFSTMSTRSIVAAAIAESSAHGIGRAVGARSIALKIMWTVLFLVFLVLCCRSSWSLIEKLINITIDQTYEA
EKVPFEFPDLYLCPSLIYSGSIPYTLQENEAKIVNDEVERLNRLYDRLYCENDSKLCTGPHETLPRVSKIQMSMPVHILQKYL
LQSRLQAFLIPPAVMHSSLSQSKFQTGITTFLHPNNEVCFKVNLPEEEYDNLNYRRSNVLAWPFVDPHNSDFNYGGAAI
NPITNPFNNESVPFEDWQTTDGYYGLSSLGYNLQVVPRGAYPSPLHETKVSLAAGRQYEISLEMEDKEFSSELHGTYCN
PSQAKPEFQDQLTGAQKEYKNSRGACFFIHMSTLLENRFKVLVSQYPKPWKWKDVPMIENMTEIAMSNISSAKTSIEEK
MKDFIDKHTQQVIDHCTKHQPCEYKKYTVQTTSMKWPRSSKGLARFILKFFGSYIASFKKCCPGTARLIELAKNFSSGNH
SAHTESFFRHYVSQSMIQVMLSVKSPMSPKITMAYSYSWVAFLADMGGTLGLYIGCSLLTLCELLELAYHLVMGAVAS
WAKSGDPSDKDIIDREAGIAMTAEVKSEQAEDELVA

>Protos_Platyhelmin_Macrostomum_lignano_A0A267EYX5_A0A267EYX5_9PLAT_Uncharacterized_protein_OS_Macrostomum_lignano_OX_282301_GN_BOX15_Mlig003382g1_PE_3_SV_1

MTRKSSGASKVRRGARTAYFVLEIYGKNSSLHGLYHLFQSKDMTRRAFWLIVLLAATTAGVYHTVMSIKRYSAMDTFVQ
IQDEAVPAEFPDITLCRVKPFSESLLIHYDKVLDRYFDTKWSYFNNWFKEERMQKPNESNIAYKHRLSVIFEKFMKIVWA
SEDFRDLGMFIDSALLSCSYEGQKCDHRNFSLIRDPDGWYCYSFSPSGDVAVGTEGLKLDLFTDTSFYHPVFEGTIVQFY
GRDVKIHGSENMSDWNSLDFFLNTHWRSLNQIAEPIITQSNGFRVILHDSGTHPMMSLDSLEFEAGVSAAVNIGQMF
VIRQQRLGVSDCIKKPFEQVRYVTNFDNRGKLEYRSYDKQQDEQVIQKKQEELMSQCGCYSHKYPFESGMETLCYHVP
DSVKRTPTKEVMQRINCHDRVVAEVDKDTYQLLRKYSKFEPCRSAKLDFFNNVYEWPKTEDLPSVVNKYVLETLYERTD
LLPGDGEHHSILNFSDPDIVRLVGKDKIVSRQTATLQQLFHGAGKFSRSSWRLVNNTRLTSNLLRLNIRLMKTKSKLIQEH
PSLLEIAYLFFSANSDLQQDNDSSSQGVTADAKLGSEVESTQI

>Cnidar_HydraVulSc4wPfr_230_g26226_t1

MSSKVSSKENMTEVGIETPTECRFDNNDMNQNEKDNTSINKINHFHIKSIEYRGSFTVHGLSRALTGNVFEKIFWGFIILL
SIGIIVRIAWSLIIKYNKNDVYIRFYSNEYTEEETPVLTICPIQVDKRGICKLDNNCTTTSNISQLPYSENSSSIWLNDKISVEA
PDSMVEGSQIEVGIETPTECRFDNNDMNQNEKDNTSINKINHFHIKSIEYRGSFTVHGLSRALTGNVFEKIFWGFIILLSIG
IIVRIAWSLITKYNKNDVYIRFYSNEYTEEETPVLTICPIQVDKRGICKLDNNCTTTSNISQLPYSENSSSIWLNDKISVEAPG
SMVEGSQIGDYIESTYYDCVRVNPRFFDTSTYTENSLRIYTIYPQEKIELFVHDAKDLYPFFQTQPDYIEVNESDVISIELKHY
NRLERPYASNCSQDSEFLIFPGNYSKIKCTESLKCISSYKVCGGSYDFCEQFIPKAFKAVQANFSQNEAENLTECLLQEYKK
KYDRECPLPCYENVYTFLPSFSFNCNWLDNSREIVLFFPISPYYHVFEEVPLYSFEQLLSECGGLIGLFTGSSLISFIEVMAFLS
LILLKMFNKNKNKNET

>Protos_Platyhelmin_Macrostomum_lignano_A0A267EZH7_A0A267EZH7_9PLAT_Uncharacterized_protein_OS_Macrostomum_lignano_OX_282301_GN_BOX15_Mlig010245g2_PE_3_SV_1

MRARLSTVTGAAQEVAGITRTHGLLHIFLSRGTARRLFWLLLFVAALIGCTVHLAKLVQKFTEHSVESQLKLRSERAQFPD
VTICNFKPASASLKRSKLYHDYLNESWGNFDNWFQEDTKKRPHESAESYENRLEKVCKKLLDVMWVSGDTRDISHLDE
TMLLSCRYNSEPCSHKNFSLVQTSRFWSCYTFHPDPGQSSSGSGGRGAELNMVLFTDGYTGASVHYSEFLHSDSVGVC
NKRLILGQAATNTLLGGSDPQSTGLRVFIHAASSYPMSQTEFVDVASASSTSIKLRPIHNRLKSTPSRRCSEPPIETINYVR
HFSNASLNVVTKAYSKSVSDFVVEAQQSILHSNCGCYSHLLPFSVNTSDLCYFAPPQEWITPSDAVLKRIDCHDHWFQH
AKRKSEELTKEFQHRMWCQFTQQRPWQQSRGWPPFSAVQEIWENLMVPQVRHGVEFLPEDGKEHHIFRNMNQSR
QFYQFNVFLTQDDNCLSFFCRNSSIKYYECNSSLMLQHLENASYCIYQKSLARSLASVSISLQSPAADVYEEKAGYHWTET
LSEVGGTLGLWLGISVVSTFELLEFFYILAKRCRGGTTGSESEGSRD

>Protos_Platyhelmin_Macrostomum_lignano_A0A267GW51_A0A267GW51_9PLAT_Uncharacterized_protein_OS_Macrostomum_lignano_OX_282301_GN_BOX15_Mlig013503g1_PE_3_SV_1

MMHQQQQQQQQQQQQQQQQQQQQQQQQQRVVASKPPQEKSEVRSLCLDCLKRTSVRGLPRILSAKSAGLRTLWFVAVLAFFLLLLYNVVKLTFVYIRGDVILAFEPWTPRPAFPNITICDASPMTYSGMRRFNLTPRAYSRHVKELAAATQSRVQAAALSKLLSPVAYHDHLRLTEFQAALHEDPLVLSESCSVHLHQNGVLINQACQRVEKLQVPGYFSCSTITVAESAERVTFHLVLDSETDSLECSKCIPDDPETLNTGAILFLHERDSYPVSIDNGLRVMPGTQNFVLVKATRSMQRRSCSESTLEDVEFQYSGLSLSTKYSQSLCEMLHRHDKYQELCNCSVDSMPHAIEDDETHGFESALPVCHRQKPLADRQRYYGVADVGLQHLDRVDCVLQVDGRLPVPPEHRERFKDLESSRNCPKPCQQVQFSVARSQSKGIDKNSVLDVFERYIWNTSLYGHMEDRLSVYSRAIQLREAGRKDEATELLRSTTYLEENFMTVILQFGPDQAKVQSENLILGLSAFISQIGGLLALFSGITVVCVAEVAELLWLLAVRLKLRRTAPCCLREADDSGGEGKTEVAVDSRGRDGQEAVAML

>Protos_Platyhelmin_Macrostomum_lignano_A0A267EQY0_A0A267EQY0_9PLAT_Uncharacterized_protein_OS_Macrostomum_lignano_OX_282301_GN_BOX15_Mlig025217g1_PE_3_SV_1

MSGSTTEAEPKPKPEDAAAEPTAEQTPQRMTRYGAISTKEEPLPQPLPPPQPPKPPPLPSKSSRSEGIAGALRSSALPKRRRTKEEKQIYEDSIKGFMDALLSNSSLLAISNIVKRNHPVLKTLWAIALCLCLILSIRQLYSLTRTYLGHPKSTTIELSYKPFDFPSVTFCNHNPLRRSRVELNGPSDLQSFFSDLLSRNYSVADNGTQLMTQQSDWLKFEKLISNISVLDRLSMGYQHGDFILQCNFAGKKCDPDMFSVSLTKTYGNCFTFNGWQSVRGFERPVAKNGREHGLIMTLFLELDEYLPNFIPSAGVKVQLHEPGSMGFPEYSGFSVPPGFQSSIGVSLKKITRLDHQNDHCEDGEKFYATYGIKYTYSACMARCKHDRVRQKCGCYDLANKLFYANHVRHMQPCQSEKELSCMIDVEYSGEEGNLNCGCLLPCQESSYKYGQSLSEWPTDSYEARLFGDLMDVKPKISSLKNETKILARRNLLNLLVYMDGLMTEHVIEKNSYAVDNYLSDIGGTLGLFLGFSVLTGFELLEASALVLAYVCGKIIYTEEEDDETSAFADDDSLTPPGSPRGAHRSSRIASLAGTTPHLHPG

>Cnidar_Rhopilema_esculentum_TR55888_c0_g2_i1_p1__Rhopilema_esculentum_TR55888_c0_g2__Rhopilema_esculentum_TR55888_c0_g2_i1_p1__ORF__type_complete__len_604__score_83_96__Rhopilema_esculentum_TR55888_c0_g2_i1_896_2707_

MAATNTRGLFLRAQRHIMDRELSKDENEEEPSQPYSSADESKESLLLQNFGMKYGHAFKVKPKKTEPEQKNGNTELPQRKVSPEIETNKPAQVTVNPAFKNVALDRFRRAGMKARLVRRVNKIVEDRLTAKQIFARFTKESTLHGFRFIFTKTFYIRRFIWFVITITMAAMFLKELTDSINLYFQHPFSTTSTIEYVNRLTFPAISFCNLNDFRFSKINGSDLQDVFMHEKGKFYLHRNSSFDLDGKKLGERLEDASHRITDMFMKCVWLFSQTAAGQPVPCNYTNITTYYGLNGQTCYTFNPGDRGHRLLTLNETGLFHAFELQLDLETHEYLKDIQEGGVRVHIHDQNETPFSSAGFAVPPGFKTFVSLNVQKIENLPPPYTTQCGSKKLKYYSSYKKSKCYLEELSKFVTKKCKCRGRFMPDASYPLCSLKQTVECLRPTIRTFDHRAGGACPEDCLSTQYNAQLSYARYVSRPPIGTSGITIPERFQKANLTNSQLEAVIRDNIVYLIFFFPELRVENVKQTVTYDFYKLVGDVGGQLGLMLGASVLTLVEFIDLIAFIIYHQILRLTRLKKKKKEETEQLKLREMNGNGAREGRFINF

>Protostome_Lophotroco_annelid_CAC9670605_1__Ofus_G181373_Owenia__fusiformis_

MVYCSSSIIKLNMAKRIHDVVQRFASGTTIHGVPKLIRAQTIPGKIFWSLICLAALGVFIFELALLLQKFFGYPKSVDVKVVQEPVAFPSVTVCSVHAMDPFVIHKIYKLRQEQMEHRMRRKPWVISKNVSDFEKWYFDTFLKKSMSYVLEPSYLEGLIDRHDLKTEFFPSLSSRLTLAANLNKSITFPGVEPAELIAFCRYARQECSFEHFKTVFDPYYFNCFTFNASVIAGSLKTLAEGIENALSIVMYIPKLHNEIKIGTKVKLPGIVEHDIRDPLAGSGGVRVVIHPPDTQPHPATEGFDIMPGYSVSIGVKTTENTRLGRPYGNCTETTKGVFSDKASYRYTMTSCRKKCLQNLLSREVKDGCGCLDVVLPTFPEVNGVDFCAKFDDIPRKCIFGPFARRNESCKELKARWLKRMICMRDIETSGSHEHIAITENCNCHPPCKDLSYEFFYSNSLWPGQQHKQDVYRDLFISRRFERRFQEGQRTYYFGAENAQALNVTEQEKTVDRFARLNVYLYDTNVVKITETKDYTGIQLISDVGGQLGLWLGISIITLTEVFELIADIVHLCLKRKRKQSIREGNCTPAEITALNRRNGSTTV

>Cteno_MneLey_comp6738_c0_seq1__p1__GENE_comp6738_c0_seq1_comp6738_c0_seq1__p1__OR
F__type_complete__len_605__score_106_06__comp6738_c0_seq1__64_1878_

MGRGGELIEGIDNDYLNSIASYKHILPKLIWTVLFSGMLTLGIYCSIELVRDYMDYPSYTEISSQFTTEFHFPAITICNINRLN
RTLMEESTIGNTNTNVYDLYSELMEETSFRSKSKGGVKNPKSKSKGRNKRNEPKISYEVINNMNVYRNFKWNIYSTLDK
KTLTFAKDNLANLGDTFKVDLTEMGNCLEINDNQALVQKVNGPVGGLSMILDAQTDHYVRSTESEGFYIVLRMANESV
ISKEYAFAVSPGKETFIQLETTEVTRLRPPYGSCQDTSDLFHVKGEEETPGKISNPMTIKECFTGQVLWQFMKEPQCKCY
PWYIYSRHIDRTVGKPTRNIDLEAQLYRYFTSDLPQEKRIEFDNTTCYFQDELSYSGTTLATFESVQRVEGCADRCKTTEN
CVYFNWADALEQCDLFGEGAELTADLDYRVTRGNVSCSSSAMEFHECSLTVEAKCNNLMIDLMMSGEDTQKPMECH
EPCTYNKTTYTLSSTNFPSKGIWDSSLAAAFPHYKTFEDAKRNLVKLVFYQELMTRTEEVQTAAYNWRSFVGELGGVLD
LFVGISVLTGFRIIEWLLCLVWKPRKRVGSESKLDVMNMNSSPTAIS

>Cteno_Puk_Fal_comp28008_c0_seq2__p1__GENE_comp28008_c0_seq2_comp28008_c0_seq2__p1__
ORF__type_5prime_partial__len_605__score_101_68__comp28008_c0_seq2__3_1817_

TQSTGVGLVVAMVVLNVKPWEEEGTHREAEENFSDDPILFGFQDLASEYFQVLREKGRPKFCKLVWAVLVCGLMSLTL
FMVYQLLDDYFQYDSFNKISTKFETSMVFPAISICNLNYINYTRLTASLQDDPELLDTLNDFLYHVTVDSEDNVDPLESTN
RTFLSIVDTWERENITSISVHFKNHLYDMLVGRYDFVFGGEGKYIPPRNWSLYDGYTKLGNCLEMNDDGQMKQIIHGK
DGGLSIYLDTKSEDYIFTAESRGFVVFIRDQDETILINQGGTYISAGTESFLKLTKRRVSRQGEPYGTCKDIPSKFSKYSTTKA
ESVRECIQRQYIEFYTQGCGCIPWYVVERMIGLRKFHVLDEMVVEVNRTRDDIPVDGEERRRGRRSPIRRMRSLASKLR
RNRSLPDDLSSSDNVAPSEEPTSYTTLPPPGDLSTTNYTQYTCPFTLDQMCNSIINYLTRSGHLEYVSCSEPCNYEAWDVE
LDQIEFPSTESYFDRFLKHELYVFPTPSFEYARSNLARLHIYYSELKEEVIAQEKAYEVQNFVAEFGGTVDLFVGFSFFTVFQ
LIEISVACCLWKCCRKSRDEDNLPSYEDKNYPTVYENSAI

>Deu_Ambulacrar_hemi_Ptyfla_40v0_9_20150316_1g24013_t1_scaffold33951_cov84

MARKDTDERSFRQLFVELVQNSSAHGIPNAGRAKTKFRTVVWSIIFVIGVAGFLFQFSELFIRFIDFPVTTNVDVTSNRSL
VFPAVTICNQNPVRVSALEYASDPKLRENFDDTYEPSTSSQAPQDVTGNTAFTSPDIASAPPIGNPPPPQGRKRRATDYL
ERQGNKQPPSPDSYNGPSAGLAQVSLLTELMSKVSIEERIELGHQAQDFIVNCTWQGERCSSDNFTVFNNPVYGNCFT
FNGKDVADEDVLLTNFPGPLFGLSLQLFIEQGEYLVDYTEIAGVRIVIHDRDKMPFPEDTGYTVPPGDSTSLGVEITRKPY
PYSNCVKKYDSAYSNGYSELYGVGYSVQRHDLRERSISGSVAGKDFNGLTLQLNIEQDDYLEEYSETAGVRVLIHAQDVA
LMPEDDGFDVNPGVLTSIGIRRTEITRLGDPYSNCVSKTGDIGQYSNVFTDMFNANYSQETCMKACYLTRVAERCDCY
DGRYPKPNSTEDLDFCNTHNNANYRCVKNIEEEVRTNSTSFRCDSPQPCEQVTYTTEFSTAYWPARISQRYVNDWLTQ
RNRELHRMVMKQEDLNEDLVRLNILKLRVYYRDLNYETITQSPSYTDAM

>Deutero_Cephalo_132390R_t1__name__chr_scaffold26_start_1783973_end_1797692_strand_pro_le
n_606_

MNAPSMGQLCGGTLTSSGGEVTSHGFPNFYLHNQDCTWVIEARGGRASVTFLDFELEQPAGGAFGTCRYDYVEILSSG
ESIGTYCGNMNPGSASSDDRIEVRFVSDATFDARGFRFTYKVQGAIEEPSCGGILNDTSGVVTSPGYPGEYDNFLDCTW
TLPATGVPVTVTFTDFQTETSFTGLTTCLYDWVDMYEDGTQLGRWCGTEKPDDVTSTKSLRIRFRSDGSVGDRGFRYP
EEYDNFLDCTWTLPATGVPVTVTFTDFQTETSFTGLTTCLYDWVDMYEDGTQLGRWCGTEKPDDVTSTKSLRIRFRSD
GSVGDRGFRFVYEYRQPDETTTSSPSKLPELEPTQIPSEETTSKPSAMSPEEDKQSPEKEFVGNTTLHGLNRIVIAPSGYFQ
ALWVLVILGSYAGFAFMFSSMIMDYFRYDTITDTKLKFADAIPFPAVTICNMNKFDSTKLKLVEWSYLSPLLLGKEYSTSEL
LTLGLQPDETINSSIANISIKDFVLENGFDINQDRMALCYWRNEACSYLNFTHSYTFFGNCFTFNSDKSNMLWQTMEGH
GNGMMVFIDIRARAAGDLVTLKFRLDYASPPGAALVRQLKTTAVGDLTL

>Proto_ecdy_PANarthopod_Dmel_tr_Q7KT94_Q7KT94_DROME_Amiloride_sensitive_Na__channel_OS_Drosophila_melanogaster__Fruit_fly__OX_7227_GN_ppk_PE_2_SV_1

MAEIREDEEEKKSGISILPGPELLALPGFDTRASIASAALSDVPSDVIIKSRIRYGSPLSACKGLLLEYAKSTTIHGIRYIFEVHR
PIYEKLYWLFFTCISVYFAVSLIWDTYLKWQESPVILGFDETLVPVHKIPFPTITICPEIKMERNVFDYTNVSRQLWEEYKQ
NGNISDLDDEDLARMAVAMHICDSEVVQRFTPLLSQLNPPNVDVTQTLIDLSISKNETGPFCKWNGRFYFCDKIFDFVA
TDEGICYQFNGLRPKDIYRDEKFISYVDPDVVDFNKYFDVDLPPWNNITGNWSLDTGFVDQGQNAYPQRTVFSSVKNG
FFAFLQGLQHNFDYDCRSFKQGYKVFLNSPESVPLTTGNYILVPHGDEVLVSVLPAYVVSTDNLHEITPEKRQCLFDDERS
LRFFRSYSQSNCQTECLANYTVSKCGCAKFWMPKPLGTPVCGLKDINCYTSAQDELYTLMQNQTMAKSIDESVDITCN
CMPACTSLEYNFEISRAKYDVAKTIRAFREEYEHTDAIGSRLSVYFKEHQFTAIKRTILFGVSTLISNCGGICGLFMGISCLSF
LELIYFFCMRICGSCRDRRKHKIQQQNSVDLPEEKSEN

>Protos_Platyhelmin_Macrostomum_lignano_A0A1I8J0N4_A0A1I8J0N4_9PLAT_Uncharacterized_protein_OS_Macrostomum_lignano_OX_282301_GN_BOX15_Mlig007166g1_PE_3_SV_1

MRARLRTVTGAAQEVAGITRTHGLLHIFLSRSTARRLFWLLLFVAALIGCTVHLAKLVQKFTEHSVESQLKLRSEQAQFPE
VTICNFKPLSASVGPNILVSKYFPTFWGNFVNWFKEDTPKRPQESDESYESRLETVCKKVLDVMWVSDDTRDISHLDET
MLLRCRYNNEPCSYKNFSLVQTSRFWNCYTFHPDPKQSSSGSGGRGAKLDMVLFTDGYIGSSSFTYRPEFSNIISKLVCL
GRLAFNMIAISRLLNNNQPQSNGLRIFFHEAKSYPLTDAEFVDVASARSTSIKLRPIHNRLVSTPSRRCSEPPLETINYVRYF
SNASTNVVSKEYFKSISDFVFEAQQSILQSNCGCYSHLLPFSVNTSDLCYFAPPQNWITPSDAVLKRISCHDLWFEHAQR
RGQELVKEFQHRMRCDFTQQRPWQQSRSWPPLSAIQEISEELIAPLVRHGMKFRRSRDGKKQHMIKKFLLFSRKEHVL
MHLLEQDYWNCTAWFSYNRTIDNTNTSTLNELLEEYLESAPYCLQRRSLSKNLAAVSINLQSPAADAYEEKYKYHWTEA
LSEIGGALGLWLGISVVSTFELLEFIYILVQRCRRNDSDSDSEEES

>Protostome_Lophotroco_annelid_Pdum_comp420310_c0_seq1_

AAALLEGLHGLKMSSDYGNKFDKSPPWQDLLYDYTQNSTLHGVRYVTSNTRYRLRRFIWVCVLLTYVSIFLYFFIHRLIYF
FSHPTTVNIEVAYNETMLFPAVTICNQNEFRLTAIQERGLYDVFAKKFSRASDNRTDLEQSSGCVDKIEKYPGPMKCIPIA
TLRKVENIHSYSEITLSGCEKLCYNLHNWKCSRIYFFPKNRTCVLTPRINFNFTHESEKCEGVVIYNRTRCLEDKKITCDFET
PTCEFTDDSADNEDWNKVFAALYPDRPLDHTTDTPKGSIIHMLKQERVRSAVAIIQTPYLRTDGMCMEFFYIMPDANS
SSLIVSVRGENFVKKPLLEVRTVAFHWRRKFIKLPSGVFRIMIEGNVSQGVSASIAIDDVSVWPCQDFVSECRRFSKGDD
YIGKKTVPVEGTKCMAWSIALDRNGQKTAEFTDSSVATAKNYCRRPYFQDKNLGELNLMCLVAPGTHRNCDIAYCKCP
NHLKACSNGQCLPESFFCDGQEDCEDKSDEKDFCGRFVSVNCTFDHAFVCGYSYSGDPSTSPVWTRGGYNDKTFMKV
QGNLIKGTTTSMVSPEIIGQTDYCLKLNYGLNDEEASAQLNIYQRYGK

>Porifera_Syc_Cil_scpid46263_scgid5448_Acid_sensing__ion__channel__1_Acid_sensing__ion__channel__1_3_Amiloride_sensitive__cation__channel__2_C_neuronal_ZASIC1_3

MTIATTMEPFEDASAKLGNRGGRRIWRNGSVGKRRAVEIPTNGVTTQSNGHSDDHVRSLSNASSFLKTSTLHGLRFWP
TYPNQPRFWFIVQLLWILLTLGALGAVTYFSYDAVNKYLQYKTTTHYDVQRASTLPFPSVTVCNFNPLRFTDILGGLKEDT
SLSTLPNLYLLCTLYHLSDKFCEESLVSDKIAADNRTLGEIYAEFGHKKWQMIPDIGPPWWYSTGCKFAGRPCSGANFST
TISEYGNCYTFNGDQNSQLVSRAQGQKAGLEVVLNVEQDEYFSRGTTAVGMAVFIHPPHRPLLAYHEALALTPGTENFL
AIRARKTVREPKPYNGPDFCITPDEGKAIVQYYQDYSLAACKKECAARFVYDRCGCRMPEMPPLLVDGQVMKVCKLFD
TIACGEKERDYFESNYEAQLEYCVNCKEPCNETGFQYTYSLTSSNFLEHSNSLQMIQALDAIGIDLGLLKEDSTNDTSDTY
RTSLTGARENLLKLVVYFDDLNFEERKEIPVYPLVSMLGDLGGNLGLFLGASILTIIEVIQFALTYTHRKAIVALKKREVVQRI
SRAASRRQLVTWTDGLAAEKQNAGDHDATVVEYNRSSVALAAENRV

>Protost_ecdy_cyclo_nema_Cele_tr_P91103_P91103_CAEEL_DEgenerin_Linked_to_Mechanosensation_OS_Caenorhabditis_elegans_OX_6239_GN_delm_2_PE_3_SV_1

MIPTISKPKTNQSTKSARKSSNESPYPSVTFRPNSSLSHLIIEVPVAQLKKFKHAGETVSVERETQHFCETTTMHGPKRIFQ
GKRWATLFWLIMVCTSIGLLITQVCILASNYLSKPVVSDVSFLINEEGIQFPQITICNFTPIRKTFVNEMNKTGQISPNMIN
YIMHWFTEVPILIGSSNWQLLHEGNKDLQEYQKNNPNFTVQGFFIDAGFSCSDIFKLCSFQGETFDCCSISTPVLTPLGKC
YTLDLLSSTKPSMHKQTEPGIQAGLAITLDAHLEEQFDGSNGMDALFTNSFVNGFQYFVHPPNTIPHLSSDEFTVTPNSV
AYTAISSERFELLPTNKWGNCTEHYPSGIKSDLPYLTGNCVSLCKAKFFMENCGCTPAVYNNERNLKECTPFETLACVNN
HNGSNNATGKLEFKLPRCAQCAQQCNNLIYRAFNSQGNQFSARAFEWFRTNNSNWTMAHIKTNFQIIHIFYQDMSN
TEYNQVQDASISDLLSNIGGNMGMFLGMSLITITEICLYFSKIIWLGISKKRREYMYSKRVHEKTHKKEIVETVEKMKVIES
KRKLSSPIGFENQISWNSNDNVEFRINLNNLEREFEAVGSRSVLI

>Protostome_Lophotroco_annelid_Pdum_comp405647_c0_seq1_

MKPENSDPDRPFYEMLPVMNDSYGLAMVDDVGKFMKQAGHRRSDFVQQCFWKGRTCISDFIQDTETDMGKCIVIN
ADDEEVVSQTGSGFGLTLTLNVEQYEYMPGANTDAGVKIFMHSNDDVPQMMDFAYSIAPGQHAFIDIFNTETKNLKN
PFGICDEKPYKYYNKSSVYTWSKCQTECASDLVMEKCECILPYMPGDAPVCNLTSYLYCILPLLYIDGLKSDECKCPTACLS
STFAGRISYAEFAVTSVKDMLSSESQSFDLDDASAFEIRNRVNNAKFLEIIELLDIMHNDIERFLLSVSGTLGNTIGRVQQG
EDQLYSMVLDDMLYLSNVLHQLKKEVTPEYESLLNMYRYVFRDQLQRFEYFRTYVRLGMNITEEFRADVFLTVKLGRMF
RDTMNTFFDSYNGYVLSSGVELSSEYCHYVVVPNLAEAVKNIGEAIYSTEKEPEQLEDVFTKIIDMTPQFDECMFGFEKSL
NAAVAMVDATVELINLQIKLAKERKLNLNEELSIDVAYLASNASFYAELGEQYRENKTRKIDFISFITQEHMTSLRTGLQH
MTNKAYAKGVVEIDYRLGILPENVIKWYSDVLLSVNQLLKHFDDPTELDRH

>Protos_Platyhelmin_Macrostomum_lignano_A0A267GGG8_A0A267GGG8_9PLAT_Uncharacterized_protein_OS_Macrostomum_lignano_OX_282301_GN_BOX15_Mlig025378g2_PE_3_SV_1

MLEKPKRKMRARLRTVTGAAQEVAGITRTHGLLHIFLSRSTARRLFWLLLFVAALIGCTVHLAKLVQKFTEHSIESQLKLR
SKRAQFPDVTICNFKPASSSLTRFDVYHDYLKKYWGKFDNWFQEDTKKRPQESDESYEDRLETVCKKLLDVMWVSGDT
RDISHLDKTMLLSCRYNNEPCSHKNFSLVQTSRFWSCYTFHPDPGQSYSGPGGRGAELDMVLFTDGYTVSSEPHNDFL
NSDSVGVCFKRLIARQGATNRLLKSSDPQSKGLRLFIHEAHSYPMSQTEFVDVASATSTSIKLRPVHNRLISTPSRRCSEPP
IETINYVRHFSNASLNVVTKAYSKSVSDFVVEAQQSILHSNCGCYSHLLPFSVNTSDLCYFAPPQEWITPSDAVLKRIDCH
DIWLEHAQRKSEELAKEFQHRMWCQFTQQRPWQQSRRWPPFSAVQEIWENLMMPPQYRHGVEFLPEDGKEHHIF
RNMNESRQLGQILVFLTQDYNCFSFFCRNSTRSYYYECNISLMLQHLKNMSYCIYQKTLADTLASVSISLQSPGADVYKE
KAGYHWTEALSEVGGTLGLWLGISVVSTFELLEFFYILAQRCRGGTTGSESEGSPD

>Protos_Platyhelmin_Macrostomum_lignano_A0A267GWB9_A0A267GWB9_9PLAT_Uncharacterized_protein_OS_Macrostomum_lignano_OX_282301_GN_BOX15_Mlig003720g1_PE_3_SV_1

MQDRLRTVTGAAQEVAGITRTHGLLHIFLSRGSARRLFWLLLFVAALIGCSVHLAKLVQKFTERSVESQLKLRSERAQFP
DVTICNFKPASASLLSIINFLEFHSKNEFKFEVFDYFEIFWGNFDNWFKEDTQKRPQESDESYDNRLEKVCKKVLDVMWV
SGDTRDISQLDETMLLSCRYNSEPCSHKNFSLVQTSRFWNCYTFHPDPRQSSSGPGGRGAELDMVLFTDSNEKYPDLYE
IYPNGALPDGCITRKFLRSKAVNRLLQNNEPQSAGLRIFIHEEKSFPMSETEFVDVASATSTSIKLRPVHNRLMSKPSRRCS
EPPIETINYVRHFSNASLNVVTKAYSKSVSDFVVEAQQSILHSNCGCYSHLLPFSVNTSDLCYFAPPQEWITPSDAVLKRID
CHDHWFEHAQRQSEELAKEFQHRMWCQFTQQRPWQQSRRWPPFSAIQEIWENLMVPQVRHGVEFLPEDGKEHHI
FRNSTKTRQEIQLYLLARGSDYCRFNFLRKFNFIDRLLGKDYKKKIIESATSCVSRSELAKELAAVSISLQSPAADAYEEKYKY
HWTEALSEIGGTLGLWLGISVVSTFELLEFIYILVQKCRRNDDDSDTEE

>Protostome_Lophotroco_annelid_CAC9484202_1__Ofus_G014375_Owenia__fusiformis_

MPQKYYDKNMKLPGVLQVQPTAEGYEEYRDVGKDNTKDKTPQNYYNYLLQEKDKNLLNDEVKLTHLYRGLVEHTSSL
GITQISYSKGPVKIGFWVALTLAMSVLTIWNVSTVFMEYLKFDVDMTIVVEQRATLDFPAITVCNMNSIRLSKYLSNPML

NATLNGDPGSDGANATATSTTTTTTTTTTTVAPAITTAASKGNGKNSKTKGGTNTDKGNGKTDKKPINGNGNKQTND
NLGEVDPMFAFQERVSEMLANMGESEKVPMGHQREDFIIDCKYNGYECFIDQFATFYNPKHGNCYIFNAGWNNSQS
RFTSSRPGPFYGLQLTFNIEQSEYIGKLTSTAGVRVQVHHQNVMPFPEDEGINIVPGQSTSVGIQMLNLQKQGGKYSDC
FDDDVRNERMNVYEEYYNVKYSNPACMKTCFQRHLLESCGCADKQYPMTGVAFNNANATVETCNSNDISQSRCIQNI
TDQHIGGKLRCSCYSACNETLYKTEVSSVLWPADAYKDDLLANLALRSPSASAVVSDADTDAYKRNFAHLQIYYSEFNFQ
SIVENIAYDEGALASDLGGAFGLWLGASILTICEYIDFIMDVCVWALRKLKKKMTVNKVDKHPS

>Ctenoph_Hormiphora_californiensis_evg120547__p1_GENE_evg120547_evg120547__p1__ORF_type_5prime_partial_len_612___score_109_77_evg120547__2_1837_

DLVDHHGMSPNRGGEEEEEGNYRPTLTDIDNEYLSELLRKRSKLWKLFWLLIFLLVTGFATWLLTLLIQDYLTYSSYNLYY
TNLGSSYTLPAITICNLNSLDRTKLENEFVTTNSSKNVSYYSIYKALQSEYIDRNGGGDNDNVEGLIEFEQSLEAYELYVNYS
MDISSEVVGDIRFAKQSYSLDDIGEIESTERGSCLELNDDNELIQRVNGPTGALIVTLDAETAKYLSLTSSAGFSVILRQANE
TVISKEFGFSVAPGKRYMINLATTSVTRIEEPWGSCTNEPDIMDNEGYFTQKECFLNQILYMYPKLVQENCTCFPWFFV
QRFLNDETNKKYDPYTKIRMIRYWTEEISEESKRLRFNETCYYDNSPLSYTSKHIETLTSVSSSVECAGLCSNNTDCRYFMY
SSKSNGYQNANECALYKSGAEYEELDAKANTTGLVPGDTQCKDLAACSISQQITCENHIMNNLMNGAISPPSCYEPCSY
NKYSFSMTTEDFPTELYYNEGEIKTKSNYSYEDAKLNLLQLVFYTDLIMETQDIQTASYEIQNLIAEMGGLSDLFIGVSFYTI
FQILEWMIVKIVHREYRKNKATSPADGVTTGNTGVALSSLE

>Protos_Platyhelmin_Macrostomum_lignano_A0A267EZS4_A0A267EZS4_9PLAT_Uncharacterized_protein_OS_Macrostomum_lignano_OX_282301_GN_BOX15_Mlig028584g2_PE_3_SV_1

MSIRRRQNRLESCPDRRDIQSSAAGVELRREFIRFCQNTTIRGLPRIVKSDSRRIRTLWLVSVLALFVSCLLCLTFLVNQFLE
YDVMHPPRVLKDASSPFPSLTICNMRPLSTPGKDLLSQLGWKNPRQFSRSVEGLTRHLYYSNESTKHIYYSFSGSFGLAA
FLESLPRNDSLNLGHRLQDFVLDCRAVIRNGSARNQFFCSPDYGPGRADVSAGTFHAFRHASFLNCFTLDIAESLRNDTT
LVEMSVYLDNEWGPDNFECVDCYLQDINTQLSGAILTIHDGDTFPDVNHDAVHLKSGSLTEVKIKNVLNVQKRPPYGR
CSADTLKSLDLFARRYKYIEGACRQATIQRDIRDTCGCQAAEYPQIGDPDEVRFCTGLPEFLPANACSDGDSDLGVCRAA
LERAHRFIKCKQAVISKYFKDVVDSCTLPCSFYSYQTERSTSDWPTKSWQLQLLQSSGFGRLRRRKEMAVYDRAIRLAKA
GDEMGAYTLLRNVSLLERNLLSIQLIRPNFDLHKVAEKEVLSLTSFLSQVGGLFSIFIGLTMISVVELIDFCIHCYGVCAMRR
GRHRDSGNNTAQSEASGTAAGTDLAGTADENLLEELPKPVWEKNDGV

>Protos_Platyhelmin_schman_A0A5K4FE55_A0A5K4FE55_SCHMA_Uncharacterized_protein_OS_Schistosoma_mansoni_OX_6183_PE_3_SV_1

MLPNSLDDILERTQMNGINTTNNNNHNDNNNINQFNNEHPFHIGITRLLNHSTIHGLSHISTATNHITRLIWFTLWFLSI
IGFLINLSYIIEHYLSHPILTTYINDYEPFQWPDIILCNPMAPYNVYTKDKYQLWNDYVQLARNLSIRLPKDLFIHIDELDTNE
IMIQSLIHSTISYWQFNIGNVEDMFQFIAMDNGYEGIRLSIGLDNNLVMVPMPLSRYFYTQILQKRYNIPCFNFQISRYLN
QTIYQRIEKLVFGIKFNYQNYLIINSSYTSHQIDLYITLPNHFPNINSIELIPGYIHHIKLEMLRIKRIKHHPHYHPYHPHHHHS
YQSLCNDKKFSTILYDGDFIHKRLIYGSNELCHRILSQYLYLNECHCYNPFLPLGYYGNNNNDNQTNEKPILCLNMSIFNEK
QLINNIDCMYRVYQKYNNDNLYMSLMEAKQCDVYGDLPYCDDIYYQHLGINRIIMNELWSNTYNEARISFLMNTFYHV
FTNTTTTTTTTTTDNTTYNTTNDNTDKTTDNIISSQNLSTKTILSELRNNFGLLTIERNSPHGRLIREEQEYSLAQLLSDIGG
NMGLWIGISVIGLFEFIELISFILYTLFNYIMKLCRKN

>Proto_ecdy_PANarthopod__XP_023234651_1_uncharacterized_protein_LOC111634174__Centruroides_sculpturatus_

MCLAVFKFLVVKSEGNKYTLKPTNEEKRVYLVHLLGKVLMTGIPQIVTESNNKRRIFKLFVFVACLIGFLYQLGSFLSFYFTY
PTSLEILVKYKSIKNLPKPMVTICNLNEVRRTMYCKDFPDNCERPKNIRNFCQHFNHFCQKENYNNEDLLFPLPHYYYYTS

HNRTLIEKYGDEVEDMLNPYRNFSNRVILANDGIEIRKCYTVDLSKVYSNEFAKENYIHYSKRLVTESNIFVEMNVTFNPE
ETFYPTIPIGARLSVHSRTQLINPFINGVLIRPGRKYGIQLRIIEHNLLPAPYRTNCKMYDRSHGKTRHILYSRDRCLIECSKIN
WIKKCGCVGEEYPFRFEGKFCDDFDSLQCVRNLNNEDCYQKCRINCRYYIFRYSVDESPIECNNCTDVLKNPESVRKHGN
RIKHSLLSFFLKSPEILIYKYKPQFQVIEVFGYIGGYVGIWLGISLIAVFDLLETLFVYFSYRLKGKIPKFSIKSFLGHVQYNRSFL
DQFGIRLEDILPSHMQNIACMNAKEKFGCVSTEYPFLTNEHFCDATADFYSTNEINVTSCYNECQKEFSELFGYVGGYIG
MWLGISLIAVCDFSEVICIYLYRITRERINYLNAK

>Protos_Platyhelmin_Macrostomum_lignano_A0A267GJX2_A0A267GJX2_9PLAT_Uncharacterized_protein_OS_Macrostomum_lignano_OX_282301_GN_BOX15_Mlig008995g3_PE_3_SV_1

MRTRLRTVTGAAQELAGITRTHGLLHIFLSRSTARRLFWLLLFVAALIGCTVHLAKLVQKFTERSVESQLKLRSERAQFPDI
TICNLKPASASLLSLKDFYESHFKFRFKFEVFDYFENFWGNFVNWFKEDTQRRPQESGESYDSRLEKVCKKVLDVMWVS
GDTRDISQLDETMLLSCRYNSEPCSHRNFSLVQTSRFWNCYTFHPDPRQSSSGPGGRGAELDMVLFTDGNDKFPDWH
GNWRNEALRDGCMFRKFSRSKAINRLLQSDELQSAGLRIFIHEEKSFPMSQTEFVDVASASSTSIKLRPIHNRLMSKPSR
RCSEPPIETINYVRHFSNASLNVVTKAYSKSISDFVVEAQQSILHSNCGCYSHLLPFSVNTSDLCYFAPPQEWITPSDAVLK
RIDCHDHWFQHAQRKSQELAKQFQHRMWCHFTQQRPWQQSRSWPPFSAIQEIWENLMVPQVRHGVEFLPEDGK
VHHIFRNSTETRQEIQLYMLARGYEGCNFNVLGKFNVTDLSSLEDYRRKIVESAASCIPRSELAKELAAVRISLQSPAADAY
EEMYKYHWTEALSEIGGTLGLWLGISVVSTFELLEFIYILVQKCRRNDDDSDIEEES

>Protostome_Lophotroco_annelid_Pdum_comp408841_c0_seq5_

MAHPLHRTMYLGASHTYGADMRNHYHNHSPRPLKPVRPNRKTMDVHRVFREFADSTSMHGVPRIINARSLAARVF
WSITCICAFGIFLWQCIILLQRFYSYPKKVNVEVVQRPVRFPSVSFCNTDHLDLVVVKRLEEMLLESDNITYGNDTHFMKF
KEAYQAFWDSSSFFFQPYLQHVKPYEKAMTHMLAAYSRLGLIANLGVDLASQGGIEMRDFIVNCRFMGEPCDIEKSFV
KFFDPYFFNCFTFDPSTILSSKTTRLQGAEYGLTIILFTGSAGQLTKKSEFEYVIPGMEEADGVLASGRGARVLVHSPGTAP
RPASSGFDAPPEFSVSLGVRARENVRIDKPWGNCSFGNMDSKFKYTLEDCQNTCLQRNIMQKCGCIDNKISIPSYTKGL
PFCLTLPTIPYTCYDMPLPEKCAKIMDEWTFRMDCRKEVYENLTMKDPDAMDNCGCFPPCNDIIYEASYSLSTIPEQTEE
NSAFYSIISNFLSSLAEPKKKLLNDLYKLNKKDGRIRGYIGRINVFIADSNVVKTTEAPDYEAIRLISDIGGQLGLWIGISVMT
LFEVMQLTCDICRFLSASGRQKSRDRRRARPTRVDRVADRDVEIHFEVGDKLTAV

>Protos_Platyhelmin_Macrostomum_lignano_A0A267DUE3_A0A267DUE3_9PLAT_Uncharacterized_protein_OS_Macrostomum_lignano_OX_282301_GN_BOX15_Mlig022931g1_PE_3_SV_1

MERRKNGVMLFCECSSIRGMPKIARSSSKFLRKMWLAFVVGMSLALIVSVSFSVHRYLNYDVIVHISDKQKFNSKFPAIT
VCLHQPFSRSAFPLWSNKSVMSPRSYNRLLRDLALKAINRTGDFSAISNDNMDPIEYALYYDSVEVYYQNIPYESIIRLSH
AQSVIRKCIVRQMGVIFFKDPCDLSEHGFDIKLFSHPKHLNCYTVEPREQKTSLETYSLGLAVDLGPKPDYANHQQGFLN
DVFDQAAGLRVVIHEPNRYPDVEHQGMNIEAGKMTELAYTPTIWNRSNTPVNPCIEGEGESYMDLGRNFDYSQKECI
KLNRQQLVMENCNCILGVPGYPRPRVPTKENAYCAAFSEAFNKTFKDFIDRVSCVGEIMSRPEPPCLSRCVLFEYETKISV
TEWGATDWQVDWSGRYVNDIETLVKLIEARETTPEDSRSPGWRKNFDNVKQRLLRYLEEDNLTSTAQTNDRMLESKS
FAYVLLKRKGDNLILKEEKQVLTLNALMSQIGGLCSLTIGLTCAVAVEIVEFVYIYLTSRRNDSTENSANATAAAAVTTAAA
TAESQAAAAATSMSADAMRLRAANSGSDGDDMDVDADGDGSDQLPMNCHRRADVSRL

>Protos_Platyhelmin_Macrostomum_lignano_A0A267DWC3_A0A267DWC3_9PLAT_Uncharacterized_protein_OS_Macrostomum_lignano_OX_282301_GN_BOX15_Mlig016990g1_PE_3_SV_1

MSKTLAFPKFLVSTFIENTTIRGIKKCYTTKYSALRLLWLLYLISMTVLLGYGTCDLFSEFFEYHKSYQIEKRLNDPTPFPSLTI
CSMSPFSARALKLWREGRIASPAEFSRQLRRGMGDFLRDGRYKEAASLFHSDKVSTYFQFINESSAKQLAYSFNDLMPF
CIMRHSANGNPSVAHEEDCEFQESSSFHHPDFFFCHDLELLQPDSSYVSSLSILLSNAHQNEMLMNDFNLTESDLEDSSS

EDYAFNVFNEASGFRIVLHEPGTYPLIDTQSFSVQMQTSTLIKYRPVKVEKQNCPEGECSEASQLPIIQDDDSKYLYTFDA
CLSSHIGQLITEKCNCTSTYHVRPLASDAQEKAPYCGFYSAEHPMDFIRRLNCVRNLNLSELTNSVSQTECPKRCSFFHYE
SQLSAADWFLRLWQIHWMSDISSALNALSERPVSRKAVEKLEQLLSFSQLELIATAPQVTRLRRLEKRLSKKMALVTVSR
FDYTTQVNRETLQISLSALLSRLGGMGGLTLGVTVTTAFELLELLYLSSKRLLQSCGFTAKQPAAAGAGATEEADENFARA
LGSLPRSRQNGDFRTAANDIEDAETQLPLLILSKPLPSFYDNCPTECVR

>Protos_Platyhelmin_schman_A0A3Q0KGD7_A0A3Q0KGD7_SCHMA_Putative_amiloride_sensitive_sodium_channel_OS_Schistosoma_mansoni_OX_6183_PE_3_SV_1

MEQPKFLTGWTPVVRTFVNSNRAVKALWALFATCMVIALVSSVSVVIHKYLTYVTVVRLDQRGPKDGVPPPAVTICLTE
HQTKYLHSEYYRTNSKLNIVQGASKSCQLKPGIIWDSRVDAISKFYNYTKHFQVTFRTEPPGCYSLNVMEQVPWPPDSIC
TTFQLVQVGLLNTTFWKIFHTDVRLTEVNQTLLGRPLIITHETNSFPFDNFGRYLYEYMNPGSLLNIFYSKSITKRLNTRRN
PCTSKPINLLDNSFDYDQITCQWHTVCARYNKECNCTCPLELLRLRLDREIKNKASNKSFFMNVDRNRQHYRSQHYCPP
NCQLDYRMAFVNNSICPLACRTVTYKKLNEVLDNITDRSAVQLELIQAEGLTEMTEEALYSLPKLFSEIGGLSSFCFGFSCIL
FFELLELAFWLRCTYRSQFLERVTKMLGQHMEIIDEPNDKTFKNLQNATNKLSFTENVWQIQNKRRNKLSNRNISPQRQ
KQQKNESFRPVTSISKWSSHDQPKGKSYKYSLICTPSRLKSRTSNRIELNPLKMYKVDGTRLSEYPKDQDSPLIGIKSHKTF
LSEKPTLGIHRNSSNSIQYSINLSAFNDKVQILTVPVILNNHLFQVAIDYEIAL

>Deu_Ambulacrar_hemi_Ptyfla_40v0_9_20150316_1g34414_t1_scaffold1171760_cov98

MTLYPPVSEVPKIHNFPQFTWENTLKHLNVNQNVRNNVNQAEIPQRLDKHGNCGHAEPMERSQTVNDEYHVGNTR
SVAAQADEDISRSPTRQKIREKSNTTNAARLPNWTIDKQGHELKYSEELWDRASLRTITTSRARTLCDSGRLRGHISLLNR
VDNSNRKLMWLAVFSIAVAVFAFQAYELVALFLNYDVSVNIEIGTATSLAFPAITICNTNKLRLSEIEKSEQHQDLAKTDPE
HPDSVHRLLSYEATCRDGVTCDFENDGRTFTCVPRKNQCDRFPDCEDGTDEKNCDLNDQCPTHTVYIWNSPEVLFSVN
YPSNYDDDYRCSWILSAYSSYCILLSFLDLDIEPSEDCSKDYLEISDVGNLSIKMRYCGSRIPPSWRSSSNTVKIEFVSNDKIT
GKGFRMEYFSVVCSSSQISDVTSWSECSVSCGWGVRKRNIKCSVVHEESLSNSDTGSGYQPHYGSGESPDSNDHDDED
YLSDAESMPSSDMVNHYNRTNVTYLNPNDDMLXLKLTLFIEQNEYIPLYGQEAGVRVLINPQDITPFPEDEAITVAPGLK
TSIGIRKDCHKSMLQQYIRHYCDCVDTLHLKGRYCNIANQEEGKRMNMFFVVINHLLFDNTCSN

>Protos_Platyhelmin_Macrostomum_lignano_A0A267EL21_A0A267EL21_9PLAT_Uncharacterized_protein_OS_Macrostomum_lignano_OX_282301_GN_BOX15_Mlig010186g1_PE_3_SV_1

MLSDVSQLQAGDGESGERFRRVLAACLKASLAESTAHALPRLVSGKRPARVRVVWLVLFGLLACVFGYQAYSFVLKLID
RPETTGIQQDSVPMVFPDLFLCPQPAMSGSYGNYVSNETADQVNGFIHAIARQLNFTPQTDKETETFIRILLSTMPYRLL
SDEFAHQFQQAILYDLYSDNRAMELAGNSTGSLLVFHSPSRLNCFHIRLTAERRAFLEDPRNFLTVYLFSDAPMAGLYPT
HPRLGRVPYALIKDGVGFSMWDPEYGMSIRSGAAMTLQMVPPGHYPTDAAAAVLIEPGSECSINLRMSQLAMQDG
MECVAESPKARIVNPYTEQVEEFEYSDHLCWIEFKELMRYKKLGCIEPTAPRLKAFSYLPRCMDARGYLNASQQDRRSFL
QASLAMKRQHFQYDLEAVRSACNPRRPCRQNHYTATTHSIRWPSFNGYQDAVLRRMFSDIASTKRRTMGNFTYPGY
DVLTKLLDEMEANYTAASRARIASFASIMVSKVTIKTAAAIGSVVTISRAYTVFDFVSNVGGILGLYLGCSLLTLFELFELLHS
VCLSLRLCDCRSRRRPEGSGDAEAAAATQAGAAKSAVPMESTESRRPPPPQEWGGFVQTLRLRVLDLE

>Protostome_Lophotroco_annelid_Pdum_comp417306_c0_seq28_

MMNPEKVPPPMTPVSAFSDPPQYNDSFKSVTLEYLDTTTAHGLPSVITKRRKIQKVLWFLIFWALMGYAVYQLFGIVNE
FNEYQVTVKTLKKHRNLAEFPAVTICNENKLKKSKLGGTNYASIIELQQKYSEEVFGPSAPTTSNTAGSSLPTNVANSTAL
QNVGTASLLNTTTNGTSSTNLTNTISNAVSNLTNNGTNVPNLSNTISNALTNLTTNSTSLTNLTNTISNAMSNLTTNGTS
LTNLTNTISNALSNLTTNATGLTNLTNTISNALSNLTANGTSLTNLTNTISNAVSNLTSPLTNSTGNTNQTGSTGLNLPSIP
LGRKKRQILSGNTLSPRVPPGESGTNNLLSMELKGDHDYETLFASSKTSDFSDVVSMLRPSMTDLDKYGHQFDDFVLM

CTFDGENCTSDDFVRVYNEVYGNCFTFNRQTNGSTVRTTTKFGSQYGLKLSLNIEKDEYLGLFSHQYGARVTVHSNNIT
AFPQDNSVSIPVGYYASVAVDTEVMDSKERPYETNCTHGKSADLFYEGLYSTDNCLNSCLRKMVKEVCSCVETVLVTGN
ESRCSISNTTEVACRLEVYANFRDNAYSCQHECNEPCLEVTYRTSVSYANWPTDGFNALVGRQL

>Deu_Ambulacrar_hemi_Ptyfla_40v0_9_20150316_1g14838_t1_scaffold10372_cov150

MEMKDNNYDDANSRKNNNVSPDSISAAIARQSYIDSPETPGVDIDSTPSGRIRIWAAQISDIHGVKHIVGERSRLRKLL
WALVVLASFGILLQQCIKAAINYGEFHHVTKVDVEYVQHMPFPAITICNFNKYRKSAITPADMVHVGKPLGLMDEDGN
LNSDLFSEEFVRQMKDLDWDSEEKKRFNYTEFTYRVGSQVKETIVECTWNGHKCTEHDFVKVFTHYGICFAFNKYHRD
TEARHAGKPGADNGLRVVLNAQTSEHLPTADLEDSFINVGFKLMIHPPTEPPYPKELGFAVGPGSHIFLAITRQEIKRLSK
PYGECDMKSVGSKYFDHYSMSACRIECETALLLEMCGCRLVEQPGNGPVCTPKIVKECAHVKLLEYIEGHIEFDCPCHIP
CDSEVYSVTPSSSRLKPDRSGKSPAMSNYTQEYIDSNVLVLTIFYEELNFETITQLPETSLVGLLGQLGGNMGLFLGASILTL
IQIIEYFVDECIHCFRPMAPKKPKRTYRGEDKDVNTPLSVQHWQGSHPVRNTTLMIVQHDCEIVSVEHFIPLYEILTGSAN
TGLFVFPRCLRTRAANMLRFTGPVWRQRLSFELLRNAENQFITFLRHDAWYSCLTYKMSSKLVR

>Protos_Platyhelmin_Macrostomum_lignano_A0A267H0F5_A0A267H0F5_9PLAT_Uncharacterized_prot
ein_OS_Macrostomum_lignano_OX_282301_GN_BOX15_Mlig018909g1_PE_3_SV_1

MTRKSSGASKVRRGARTAYFVLEIYGKNSSLHGLYHLFQSKDMTRRAFWLIVLLAATTAGVYHTVMSIKRYSAMDTFVQ
IQDEAVPAEFPDITLCRVKPFSESLLIHYDKVLDRYFDTKWSYFNNWFKEERMQKPNESNIAYKHRLSVIFEKFMKIVWA
SEDFRDLGMFIDSALLSCSYEGQKCDHRNFSLIRDPDGWYCYSFSPSGDVAVGTEGLKLDLFTDTSFYHPVFEGTIVQFY
GRDVKIHGSENMSDWNSLDFFLNTHWRSLNQIAEPIITQSNGFRVILHDSGTHPMMSLDSLEFEAGVSAAVNIGQMF
VIRQQRLGVSDCIKKPFEQVRYVTNFDNRGKLEYRSYDKQQDEQVIQKKQEELMSQCGCYSHKYPFESGMETLCYHVP
DSVKRTPTKEVMQRINCHDRVVAEVDKDTYQLLRKYSKFEPCRSAKLDFFNNVYEWPKTEDLPSVVNKYVLETLYERTD
LLPGDGEHHSILNFSDPDIVRLVGKDKIVSRQTATLQQLFHGAGKFSRSSWRLVNNTRLTSNLLRLNIRLMKTKSKLIQEH
PSFLYPEALSEIGGVCGLWLGMCVLTLFELLEIAYLFFSANSDLQQDNDSSSQGVTADAKLESEVESTQI

>Protos_Platyhelmin_schman_A0A3Q0KFY2_A0A3Q0KFY2_SCHMA_Amiloride_sensitive_sodium_chann
el_putative_OS_Schistosoma_mansoni_OX_6183_PE_3_SV_1

MLPLENNNNSHNTLINDQRSATDKLAYNCPHIIQMKEPSKIVKKVSILEIICDTFSIRGVARMKRGPPFLRGLWFCFVLIM
TVGLLLTTYLLVQDYLVYDVSVNIHVELDAKSPFPALTICHHHPFSQNAYHLWRNGDVMSPSIFNKHMRNLTHKFLMD
NDVDAAETLYQYDSLSIYYQNLKAIDAYRLGHDTTAFLNCMRRVNHTMQIEDNCMHMDGFQIRKFSHHTYFNCHTFE
PKTKVEAEDTDTIALIVSLGTDPNYSNQEQAFLVDLFEMARGLRVVVHEPGTYPDLEREGLHVEPGKLNEINYQPVLWK
RLNTPQNPCRAEFIDRNPELASITDHKLPRLTGLDFAYAYNDLNVSYRYTQSQCILFHQQKDIIKECGCQYIYNPRPQYPS
SNLPYCGSILEGDFDILALAKRIQCLTSGPLNVTVKSQYESTLCHPRCRYYTYESTISVTTWRAVDWQLHWLRQLNRAFKS
MQYDVKINGEDWNITQSNGFHRWKDIENELSLSNDFQISVNAINDLNLQGDNFAYVVLKRRASDVRVSQEKLVLSISVL
LSRIGGLCSLSIGLTAAFIVELIEFTYRLATTERSNIRRNKSIENKTHSNSVSDGEYKNKLKRSATCI

>Deutero_Ambulac_Sakowv30037801m

MAIGERKFHHFHHFYRALWTVFLVIIALISLIEIVFIVTTYLSYPSRTEVVMVTREQAPFPSVTVCNVNRLRRSAVANSQYE
QLLSLDSLGSISHLMPYMGPCLSGDFECGGECVKWYMQCNGVEECRNGEDEINCASDTVIPYGGSRLNCDGGDAVFA
CDNSECVSSVQQCDYFDNCGDGSDERNCTYYDQDCGSDGVTCFESDTSYCVPADLVCDRIKHCRTNGEDENNCDYPD
IGSNDLFSANWEDSYLHLTDSDDFYEDFRENIYREPVYGDDGKLSLINTLLLSRSSRVNDVIEQTVKFNTEELKTLGHQFE
DMLIKCQYNGGKCGHSDFTVFQDNKLGNCFTFNSPSPVNVAGTQNGLKLTLSLEQDEYLGLFTSGEVGVKVLIHDPDV
KPSPETDWFYARSGVSTSKEIKRVEIKRLSEPFGRCSAEYSSVNECREKCLRYEMLEYCGCTDSLIANETAPCDVTDIRQDL

CLQLLRHIHRTGVVTCDCPLPCSETKYKVEMTSEATWPSSPYVPHLLEELRYSNVTSQVLNDETAIRRNLLLLETYYGSMD
SENVASFPEYTMQKLMKDIGVSIAIYLSLFMIIALLVYVCCGPKCDTQKKKDVEITSKKNSIDATTNF

>Deuterostome_chordata_Ggallus_tr_A0A3Q2UMV1_A0A3Q2UMV1_CHICK_Uncharacterized_protein_
OS_Gallus_gallus_OX_9031_GN_ASIC4_PE_3_SV_1

>Cteno_Ple_Bac_comp45500_c0_seq1_p1__comp45500_c0__comp45500_c0_seq1_p1__ORF__type_c
omplete__len_630__score_54_98__comp45500_c0_seq1_346_2235_

MVVLKVSPWSTGEIDSPISPPSTHKELFDLSQTGTDSTFLFGFQDLTSDYIQVLCENARPRWCKILWWILTAMLGLFTVF
MVYELLREYFEYESFNKATTSWTNKLILPAITFCSTNFINYTALKSDLATYGDTGEFKYIVAQIQKEVAWGRDPSKLKSLDE
YRNVSAFEEFVGSIDYKYANHLYGTFVGEFDYNFAGNGHTIGAYEWDKLQGMTHSGRCLEINDLKQLYQTHTTDSAGF
TIDVSGNVADYLFSTRTKGFKVFLRSHDETVMMDQGGFLVQPGTETFFKITAKRALRLGYPWGTCQNVPSAYSKYGTH
FETVRECRQRQRIELSAARCGCLPWYFVSRMETQLKRDVLDKLIERMGHANTTENYGRKRRHIPLHSTDRIPERQRGIKE
MKWAEEIVPPMPPLYPPFWSQDSKYNNYTCSYLEMLVCEAFIEKEMSNGHITLENCPEPWEYLEWDAEVAPSSFPPSE
EFYNEYIKDDWFIAANTSYNYAQENFARIQVYYDDLKTTTNEQIKAYEVHNFIAEFGGTVDLFIGFSFFTVFQLIEIGFALCL
YRWRQRTISITSVETGKGVTPEKFESENPKLVSLVPHSSKSQKIETQKVEYRYTYLDLFVHCLKSWFIKF

>Protos_Platyhelmin_Macrostomum_lignano_A0A267GRS7_A0A267GRS7_9PLAT_Uncharacterized_prot
ein_OS_Macrostomum_lignano_OX_282301_GN_BOX15_Mlig020704g1_PE_3_SV_1

MEMNQQQNQQQQQPQKEDEKEPLNSDPEAGTATAGSTKKACRQPRKRIAQVKAAFQENSIPALRKMARSADVCHR
LFWAVVFLATFTGFVIHLVSITIRYQSTPVSTETRVERVDRMPSILLCPQNYLSGTRIRSMLNVPNPRLRNVTFEMIYNEIL
PMFWNFIYKIRDNNVDRKMVGMKIKNNFLWYLIHMWRERGQIQHEKEDVFVHIGLQDDDEPAMKDLGTTFDSQRN
RCFSWEPQEHVTSPLTAVINVPDGWYFYAEQDAFELVSDSGLWLYICFNESFGCHDSIRYFLRTGHYTKIDLRKMVVSQ
LPVVADNKTSCIAENKFYQIKEDLFVSSSRTLNVAYTSALCTDVTLQRQIYEQCGCLSALLPTEIDLYNSALSSLGDTRNLFC
NLPPDDAFTHQQFPRECREDPRNDTLCMVRGLDGVSTMPFSQLLKHDDIAQFLKRDECHDHEFRSRVSVDKCPKVCR
RQQFSSVHSIVPWPESQLSRKHPSTFARIFVNTIKPFLDKAIVQLNSTSVGARVLETMRAAFLKGMNYSNPQPKPLYEAG
LEYMQERLLLLEIGYASADMQVMSESEAYPLKSFLSDIGGIMGLWIGMAIVSICEFLLIFGRLVYFVGHQCARRGPKT

>Protos_Platyhelmin_Macrostomum_lignano_A0A267GVL0_A0A267GVL0_9PLAT_Uncharacterized_prot
ein__Fragment__OS_Macrostomum_lignano_OX_282301_GN_BOX15_Mlig005363g1_PE_3_SV_1

SSGLTHHRLSAQQSLNCVTMLRSAINSRPDVRTAQAAFGNEDSIEDQQPTKQQQQLASSHGIFRLSQGSGVQRVLWA
LVVVGVCVGFILHLSMLVQQYTSYPVSTELEARTDFFEFPDVTICSTNYLSVTYGYDPVPSPVAVPGRARGLVDMIYDVS
RFFHLLNQTDMNASVPVNPYSSYRHGKLALNKIAYRSKQLPDPYEIVLLCRFNSIPCSYLNFTIYKDDELFKCMTFAPLNN
TVVSAGPGNGLYLLMYTYSSSFFTDEEEIDDIPGMKVVLHPRGVKPNMNGPHVMSPLKYKTEAVIDTSIQEKVDRSSYR
CLESLPNATYITDYSQLTGPENETFRGSTEDCRVRIMQQEYLDTCGCLATHLPKPADLYYMAQPGVCHDINEHVLFRDIF
YRRPTSSYTYYTIRNDSWYKYPLMDYVATNYDMYNGTIARLDCYQRVLDRQNREGVDTHGCPVLCRVKSHEVMPRYY

QWPNSVKYSGKAVSNLATEALNEAKARGLDVRGLKDLVSNYESRVKYYAGVLDLYPRDLIVQMSIESWAYPLRNFFSDL
GGILGLWIGASIVTLVEILELAMYGALLVHKKLADRKRNQVIVIKPQEQQQEEKPPGYNHSELETDQQMPEPKVEL

>Protostome_Lophotroco_annelid_Pdum_comp417306_c0_seq24_

MMNPEKVPPPMTPVSAFSDPPQYNDSFKSVTLEYLDTTTAHGLPSVITKRRKIQKVLWFLIFWALMGYAVYQLFGIVNE
FNEYQVTVKTLKKHRNLAEFPAVTICNENKLKKSKLGGTNYASIIELQQKYSEEVFGPSAPTTSNTAGSSLPTNVFNSTAL
QNASPPVVGTGSLLNTTTNGTSSTNLTNTISNAVSNLTNNVSNLTSPLTNSTGNTNQTGSTGLNLPSIPLGRKKRQILSG
NTLSPRVPPGESGTNNLLSMELKGDHDYETLFASSKTSDFSDVVSMLRPSMTDLDKYGHQFDDFVLMCTFDGENCTSD
DFVRVYNEVYGNCFTFNRQTNGSTVRTTTKFGSQYGLKLSLNIEKDEYLGLFSHQYGARVTVHSNNITAFPQDNSVSIPV
GYYASVAVDTEVMDSKERPYETNCTHGKSADLFYEGLYSTDNCLNSCLRKMVKEVCSCVETVLVTGNESRCSISNTTEV
ACRLEVYANFRDNAYSCQHECNEPCLEVTYRTSVSYANWPTDGFNALVGRQLVGKMARNEIKEYIENNILRVNIFFRTL
TYSHQQVFPTYTWETLLSNIGGTWGLFVGFSVCTFLEAAEYFLELICVCCGLSSKKKKEKVAPEGFNYAKGLS

>Deutero_Ambulac_Spurpu_025728

MEVPLDEEEKPLSPEKPSSPEKPKAVLEKPKALVREMTNDILVKSWRTVVQSFCKSTTMHGMSRVIDSTRAIFRLAWLLT
VLTFLCVLMWQAYRLVDEYRGNPTTTTIQMVTNNKLSFPAVTVCNMNRLRRSKLVGTRFEPLLDIDRHVTILNLGLDDG
EIEVDDHAEPSPSLTPATTTSTITTTTASQVTTKSTSVPPNDEDLVSSTTVEGLEMSSTKDTPLVVDVDNPDARRRKRSVN
WQTTNKRLSSLRQMNNVPAPAIPLIGAPVENDDSNWGYLLQLSESDDFQDFIGSVNPSKGELRKLGHQAEEFILQCSF
NQHYCDYRNFTTTHNSQYGNCFTFNRPAENNSVLQTGKIGSRFGLHLTLFTDQSEYIGLLSHQSGVRVAIHEPNARPFP
EDEGITASTGTLTSIGLRLRNITRLSGRYSDCRKKNRGDPSEFTHDFDYSLRACLNACYQGKLRYECGCVNDVMGNTTICS
TLDRRQEACQRRVDQMAIDDKLGCPCVSPCRKNLVRVEIFYEKLNYEAYTQMPKYTFGSLLGGIGGIMGFFAGMSLITV
FELCGFIIQLLGLLCGRLTTPMPEEAATEGGEPAPPHPESRIGRVGRRFTKVFSHHPHAHGHGGSARARQTDL

>Cnidar_NemVecNVEC200_012234_1_1_protein_AED_0_06_eAED_0_06_QI_105_1_1_1_1_1_14_954_
633

MGKEKKKIVTKIKDFLGYTTAHGFARLVESKNPVQRAFWVLACLGAFVAFGLQVHKLFSKFNRKPVDTYITVKHEMNLE
FPAVTICNLNPIRKKHLPESLREYFKYSLGDSSTKEPTSKPTTSSLLGKAFSSVDSLLTKPALLSNKTLPSLVNSSLDTLGGAL
NDSLSSVNKLKDSGESLVNSTLGLVRGLLSRRRRGLLGDVLDVDGVLPEKEKDGGKPGLLGNLLDGVLPEKEKDGGKRK
DGERPAPEELCESARRRKYLYWRIAETPTETLIQSGHQFLDMVQECSFAGVSCLGYNWTQFWNHLYGNCFVFNTAMI
GSNRERNNISYTSSIPGPSDGLLLRLRIEGEEYLGNFSTVSGVRVHISDRHEMPFPGDKGFSVSPGFETSVGMRKVFIHRE
DPFGNNSCISDVGKPVVRPDYYMENFGVNYSTIACRSSCVANLQKTMYGCVESAYAITDELRKIVCNALNTTVFRCIVKA
NETAEDSDGKSCLTKCAIRCDENIYKLTISSAVFPSPSYQNISSQDLKSDSGNEEQIRANLLSLRIFFEEHNYETIEQRKSYELI
DFLSDIGGQMGMFVGLSVLTCAELIELLALIIMHLFNRDHKRGAVGADTVTPVIELKKKPLVRLPRVE

>Deutero_Ambulac_Sakowv30039242m

MSHIGVDNLVSPGSSNCKEKQQTFSSILGSCFVQSSAHGLPNISRAGNAPRRSLWIISLGGTVAIFCLISATLIVRYFDYDV
NVSVAMQFSRELTFPAVTICNLNPLRKSKTQMDGPSDPGQQRAEEAERNSQTSQGTNIKTETTQHPNHIAVSEDIQRN
PFTQVPTTTKIYENRQYSRFTHQTYHVEQDQQSIYTHENNTDGQSLDVTNSDWNGRTLDDTVTNSEENGRPPPETSTG
GAPELVNVLLNETNDTEAEVYDDATGDTEAEVDDDATGDTEAEVDDDETSDTEAKVDDDETGDTEAEVDDDSEASE
WDYFDWDDLDTKTEFYNEPTQYWNKSASLNQWLASMPTQQKEEWGHQLKDFLLDCQWNDVMCSPKNFTKFVSSR
YGNCYTFNSGANNSTVVKTNYAGPYYGLTLELFIEQNEYLDDYPDFAGVRVAIHSQKTMPFPEDDGFNVEPGRVTSVGI
RRTCMKSCYQKRVAERCRCLDGRYPPPIVVIHRTLNACFIGANMAYQYDVMSYLFQDYVNVQLKKKSQQLKLMIEEEE
KGGNEFVRENIVKLQVFYQDLNYEGITQSIAYTEESLASDLGGQVGLWIGISFLTLIEFIELVYDLFKLWIRRLCCMATKNT
GVTNVI

>Deutero_Ambulac_Spurpu_020954

MKAPKVDGEKSSRGTTPEREEKSLRTILNSRMENSSAHGVPNIQRSSGLVTKLAWSLIFLAGIGVMIWQVVTIFQAYYE
WNYSVIIEVKFNRTQSFPAITLCNANPMRKSKLKTKNASFQETFDVNYVPSMPDLPEQQPQPDVPALAPTCPVGLGHY
RPADQQPLPDMPDGVTPSMLNNSDSGNETQGGSTNEQVDMSEAVKDWKSRIASPSFYVMESVDYEKRRIRVDALA
NETLEERVSLGHKLDDMLLDCSWKGKPCSPENFTKFYDSQLGNCYTFNSGQNGEQLTTTRPGSKYVNGPIDCLVLKPGL
SLELFVQQDEYVEGMTEEASFRVSVHHPSIMPFPADDGVLVSPGFATAIAFIKLELDRLPKPYGDCKADLSTDIEDDIYHQ
HYNITYNMKTCEVSCFQNEVISRCDCFDATYPNSLKVNRTVYPCEYNNRVACPYCQQVLHLIIIKKLKSWESFFSHFCLRF
EITDQFFAHSTLFEKMVEYNAEIRRYVNRENTAEWTRRNMAKVEIFYDEFNYEHIRQEPAYMVCKKTRQIASYDVHKPP
PPPPHSENGRTPCVLPKCPFRNKRKRSTRIDFTHRTFSEKGGAEKGSYTKEKGRAVLTLHTGLLLKKGGGRLEHAYAYPK

>Protos_Platyhelmin_Macrostomum_lignano_A0A267E617_A0A267E617_9PLAT_Uncharacterized_protein_OS_Macrostomum_lignano_OX_282301_GN_BOX15_Mlig030622g1_PE_3_SV_1

MSGPSSPAHPAENGSLRLSTCRVPGAPRPLRFARSRRSVWQPRPQQRQASSRCRHGRSCHQWPQAGLSAIFKDFAG
NTTAHGIRQLAQANGWCRRIVWAAILTVVTAGVATHLTVEVMHYFSYPISVQTSFILGGYAFPDVIICDPIGPINPRLTLR
RNASSGPAGDILIGTEDAIVLPDSGELQQKDKFRFFLSHFIDLPASGLQPADYIIFCRYQKLPCTFLNFSVYYHHDFMNCFV
FRPPTRLVAAAGPDSGLTLVLNVPQRSSVESFSEFGERYEEALLPLGLRVSLQQADTVPRPSEGGWMLPKGMLTEISTW
HFRSALANTPVAPCSGESSNLRQRNHATGRIEEFYYRKEDCEYNRLVERAGQVCNCSVHADYPLGGSGNSSSDDGGRD
RFVGCHEPFRITRRLFDMILRLRKAGHSHIGDQAAQPAMEILEDAASPLNHIIDCFQEARRQPVSHSDCPNPCEFDNYVT
AVSQALWPQQELLSRIKELLLDSIHQEYFNTVRRRGLQDSLIHPLKPDFWDQFAALEGCTAPDCFSRINRQIFSFMEENF
LELRVFPRSLDIQYTEERCNFPLSSLLSQLGGLSGLWLGISIVTVCEFIDFLGALVRRLFSSTFRERLTDTELVTELSV

>Protostome_Lophotroco_annelid_Pdum_Contig8214_

GTALQICQHTKMAKKDKGLWKEFSQNTTIHGLNKVAEENGLTFRRLFWITLILGGTALFAYQVVERSLYYIKWPVSVNY
KFVYQSEMPFPTVTICNQNAFKLSTSVQSGLHDVITSIFSKRNSIPKYSRNNISSELEDIFGNSSSLKTDDVWQLLAHAKED
MIYSCSWQEDPCSPDNFTTLLTDHGVCFQFNSASNSHPPLKVTQSGSQQGLSLVMNVEQYEYMTGPHDSVGLKVLLH
DSSHVAMVRDLGEALPTGSHAFVAVTLSKIQNLKHPYGECVNNVSLDYYDTFSLSTCRRDCETRKTVKDCGCRDAYMP
SYTDGTPQVCTLQEYSQCIQANKDQQSGIAGDVCSSNCPIPCLTTSYKSSTTFSTAFENYADDIDKESYWNRTRQHYILA
REISQKVNQGLRRKELEILSRLETEWQVITQIINEVKETVKHSKRFKADILDEMEPRTQFHTFWGLEKIKFVVDYDYVRIW
DVMEERTMAYATIGFYQTAHLYRNMVDKFLLEPELTVEKVAIFTSTMRDLDVRLQLTMSTQINLTQVCDAFANGTLRW
KYLTSPKRDNFVKVNIFMKDLYLEDMESNEAYSLIALLSDIGGSMGLFIGGSVMTILEVLDILVIACMRKSKKKNETNET

>Cnidar_HydraVulSc4wPfr_391_g24657_t1

MEVSTRDPELEINYLSCMLELKCEISKRRRKRYQNAISYLPGANHDTVDELRNQSINDYLSDMAVNSSFHGINYICDSTYK
VRRIIWIVVTLTAMLYAMREVYESTRKYLNYPVSTVRMKIYVDDLEFPAVSLCNLNDVRMSIVNGTSFDNALIDQNTQNI
SADDALMIAREARHNLEDMLLECKFNGRSCSAKNFSEFNWMQGDRCFTINSGKPGHSRLSVKGTGIKRNLELILNLQH
YEYYRDEMESGIHFILHSQEETPVRMRGPIVSPGFTTYFRINKIKTKNLKYPYKTRCGSLNLKFFKGYSKQLCWLDQLTDY
VNSKCGCKDFFMPVKMQRKNKKISPKHRGMVKTEEKNVYKYNTKADLETKAQSNIIDLAVSNNIMVVALEGCQVKRIE
LSNPFEVDTVVISKRSDDQIDKIFLDILGCHLLISMKSSESFYLGRNNKKPRLLQKLKNHKIEAVAWNKFSMNEITTGLILIG
TSKXXXXXXXXXXXTPNXXXXXXXXXXXXXXXXXXXXXXXXXXXXXXXXXXXXXXXXXXXXIQLIQRLVCETEIDSEEKFLSM
GSEKNVKVIFNVNENRDIEEPLTGMHMEKMNDIENRYVVILTTPSRMYQFVGNASIEVASFQNVLSVFNPDPSK

>Protos_Platyhelmin_schman_A0A5K4FAT6_A0A5K4FAT6_SCHMA_Uncharacterized_protein_OS_Schistosoma_mansoni_OX_6183_PE_3_SV_1

MHRNKNLEKSIKYKQSNKCHNLNIRPPAPPVPPPAPPPPPIHHHHPCGYCSLKYAFNLFCNCTSVRGMSKLCRGPKVLRI
MWLIFVLSMTFLLLSSTALLIKDFLNYDVTVNTQLILDNISPFPSLTVCHQPPFSYKAYKLWNQSKILSPSQYNLYMRKLIY
NSLYKNDIYLASSIWSYDSLSVYYQNINYKDTILLGHNITIFFNCMRIYSHISIFEDNCTLLNGYRIKQLSHHQYLNCYTFEPI
NMIAANETTFLSLIVGLGPKDKDIHIQQAFLPDIFEQARGLRVVIHEIGTMPDLEKNGIHVEPGKLNEINYEPIRWKRLNT
PKRPCITPNESHQYLDLNIKYNYTHTQCLLVHQQIEIMNNCHCIYVMHPRLIIPSNTLPYCGQLWPKLNQTEFIEKILCLSK
YLNNTIKQKYDGHMCFPRCEFYDYESTTSVTKWRAYPWQLYWLRVQNQATQSLLNYYEKYPNHNLTQTSAFKLWEYY

LNYPDLTRLPNKTELLRNHSNNTLSDISKLLPSWPPENLDLDSEDFAYVILKRKSQSTIEKTEKLVLSLYVLTSRIGGLCSVTI
GLTAAFFVELLEFSYLLYKNGSNDNNSNIKSTIIVQTEKCIQLNEIQQTEQKNSSLQYNITTNTTSNTSVEH

>Protostome_Lophotroco_annelid_CAC9669406_1__Ofus_G17804_Owenia__fusiformis_

MDKIQDDTNISASTSGAESSQNKTIGTQGGQMESLQSNIRPTLVDFAETTSINGPKRILKAKGKFSTITWTIIFFGMVSM
AVNLIASVVVKFYSYPVEDGLSVTTQPLAFPAVTLCPFIPYHMADINPKVRRKIKGFTGELWTRVHMLDCSYKVIFQPEE
NYDAHTIRKIDKYERGCLPLQDIFQLRHKLISSGWLVENVPIDYETFATPIKVEDFILECIYNNKPCESQMNITRTNTVYGQ
CFTLTVNEPNDHVSEIGPDKGIALILYTGRANPLSSTVSNWEVRDGFTTIFSTPSDGVQVVLHKPGTMPRPYNEGFYIFP
GRFSSVGITQETRTRLLPPYGECTDAGYITNSTYGYSYDVCVDQCIQERIAKTCGCISPMFITPTKHNFTKAPFCGNMSEL
LLKVFVKPTWPNCKISGDIKELFKISNSIIERLRCEQRLMSERTSCKCKRPCKENTYTHLINTIPWPQKKVVERGHHRVLFK
FLQRHPKTKNLLYKLLDTSSRFYRTSHSCENAPLELIEFCFHVATRAQRYRIYIPPPTHESMRYTFGNSRFTLTNALDKVMD
QNFMKLNLHFESLDVRTVFEERTYTYDNVMTDIGNVFGFYLGMSVVSVVEALIIIGVMIKLIIWRLTCARCNYTR

>Protostome_Lophotroco_annelid_Pdum_Contig18637_

MAGQYIKDPIKRDINKGHTLIICHEFATETSAHGMSHIIRARGTCRKLFWLVVTLGLIAIWLLQSQDTIRKYLKNEVNVQV
EIKASRELPFPAVTICNKNPYKGDDTIRFIELMNHTASLAISRNQSLEDIAPSWFVPWVRSLTGSDDGTFETGISSAKFGE
QMLQLLAELVEVYGEYHNFSLAELSVDPDNFILNCMYERVHCDMRNDIEMTSIFNWLYGGCIVYIIKQNTTRTGPRFGL
ELTLNIEQSDYMDLTTQAGALVLVHPPEEMPFPEDDGISIPPGQAALIGVKVKRTERLGGKFGDCAHANKLPNFTNYFQ
AENPWTNYSLKACQRSCFQWNVRQMCQCLDVRFPPFDEYKNFTSCVNKYFSTFNFSQGFPESVGCLDEVTKKFNAHQ
LGCDKLCHVPCKQTYYEPFVSYAAWPSDAALNVTLENLKGSSASADEILEVGHDNFRKNFLQLTVYFQDLNFEFITEKEA
VELTQMLSDLGGNTGMYVGASLFTMIEFVELFGDLWLWVCCHRMRRKRPKYAPPTEEDLDLYFITPNPSKQNNNHTN
NYQNRQGHAPNQQNRHGHANNKQNRQPHPNKHDRRHSKNNGRAPGRGQDNNGYHFNYNGHLERPRKSLPRTPL
PMDMAEVQDTMVGLY

>Deutero_Ambulac_Apla_gbr2_220_t1

MMSKNTMTTGIPAGYDDAFVVMPSESIPLPSYHNKAPTSVEAKAAEPEKENHPVTCGSIVTNFADSTTAHGVARIANA
SSCFASFLWLVILCVAFGGFFFQGTNLVLDFFSWPYGTTIDIITNTSVDFPAVTVCNMNRLRRSKLPGTRFEGVIAIDGGIS
GGDNDYSWFFEWSSAGDFYNQFVSASAGGGSSGGGGGSSAGGGNSSSVGGGSSAGGGNSSSVGGGSSAGGGGSS
AGGGGSSAGGGGSSAAGSSYSYPPASSNYYYWETEFGDDFNFEYDNYYDFGTVTGESDWDGFLANSKSEDFSDIINVA
NPTQDEMDELGHQAEDFILQCTFDRRKCNYTDFYKFQNSHYGNCFTFNHGRNETVRTTSKSGFQYGLHLTLFIEQPEYV
GLFSPESGVRVSINHWQTTPHPEDSGITATTGQATSIALRKNFIKRLGGWYSNCTYGTETNFTSDFFTYSPLTCKKQCMQ
RNLKDRCDCVTDLLLDGEKCSYLNSTQQRCRQLVEALYEEDKLDCLCPVACEENVFKTSASVSIWPSERYEEHLYSRLSSK
NAVAARMLQDVETTRKNLARVRIYFEELNYQKVEQIPQWTIESILGAVGGLMGLYVGISSITLMEIIVFVFSLLKQCCKSVI
CLNKVQPSN

>Protos_Platyhelmin_Macrostomum_lignano_A0A1I8H6N9_A0A1I8H6N9_9PLAT_Uncharacterized_prot
ein_OS_Macrostomum_lignano_OX_282301_GN_BOX15_Mlig002809g2_PE_3_SV_1

MLPETARIFLETSSIRGVGKVFKTQSPLLSRMWGIFVLGSTSFLFYSVFKITLDYLQYDVNIQTYQEIDDETPFPAVSICNN
QPFSVNAYQRWSNKQSMSPTEFNKYLRQQALSLVSPDLMRDKYRDLQAARDRKDSKAFRQIHQSMFDSSVAAMTYS
LIYDNLRMYYQNMLADEAMELGHTRNESIVDCIFKYGSIQVLGTRGCHDDMVKVKTFSDPNFFNCLSVEFTPEYAMTII
SITLIVWLGPDDSYDIRHRQSFLNEVYEQAYGLRVDVHEPKTQPRVLEQGIQTEPGKMNEIKYQPIRFERKDTPKNPCISL
DAAEKQKFRDLRMEYNYDFELCLNSKVQEVVLNKCGCMYIELPRIAMPNKTHPYCGFMDEKFKERSQCVKSLDMRTIR
KSFEGKDCFPKCVDLSFEKELSTTKWRETKWMLHWIKELCERWDKQFKQLSGSNASEFEESLTGVTDYLTRDNITDLRSL
GNSTSKASFDFDDRFSFIVVKRQSNNTFVKTEKIVLTLNALFSRVGGLCSMYIGLTLTFIMEFIEFIFRLASCDKSNDKTGG

NGGDDGGRSEAKSNQSTRNSCRVVEVTSQDNGAPENQSLIGQLAGDTELSDSGSVMLPGSEHRVLPDNRPDFRHDQ
VWVKQADSRV

>Protos_Platyhelmin_Macrostomum_lignano_A0A267EAF9_A0A267EAF9_9PLAT_Uncharacterized_prot
ein_OS_Macrostomum_lignano_OX_282301_GN_BOX15_Mlig011397g1_PE_3_SV_1

MPTVRSVVRNFHEYGHRTSMHGLGHMYQSGDLCRKAFWFVLTIAGTCACFFHVIMTVQNYLETPVTSILIDDAAMAE
FPDVTLCNLKPMSESLGEYYKDVVHQHFEDNWGFFEQWWPNERKRQVNESRVDYQRRLRKIFERYLQIFWVSDDSR
DLGMMDKVMLLHCSFNGIPCGHKNFSLVQNYRYWNCYTFSPSPQFRFASSGHTEDTGLNLILYTDSNFHESIYDQFAA
YSRQKFDEEEMTHPYSLKEYITTGGRTFNRILSGGRYDIEGIRIWLHMRVDHPMMSSIYNDVQNGKSALVHVDQMQY
VTLNQPGERQCTNDPQEIIEYTTSFGHNPSTERKRRYTKTDREFAVEAKQQFVFDNCGCYSHHFPFHVNLTQLCYFVPL
ALRRHPSKKLLDRVACHDEHLRIAEENSYNFIQDYLKYKPCEYRKFTTFKTISTWPRSDEHIDFVNEFLIPKMQSGKEFIGK
GDAHVHHIFNFTNENSAALLNKPILSHEELALSRLLNQDCHPNKSAEDSIRDMWCLDKEKMAYNVLQVIIRLTSPAAAK
YEEKFSYEWPEALSEIGGIFGLWVGMSLLTFFEFIELFYVMVVSRDSRPQQQPSGDRESLPNSSSMQHQMELLLSRLME
EETIRRAPGPVQMTAGRR

>Protos_Platyhelmin_Macrostomum_lignano_A0A267FGP1_A0A267FGP1_9PLAT_Uncharacterized_prot
ein_OS_Macrostomum_lignano_OX_282301_GN_BOX15_Mlig027426g1_PE_3_SV_1

MTSDLQRSRRAMLRGYVHNFTSGTTAHGFSRINSGQNWVRMVWIGIVSCALLGAIIHATKQVTTYLQYPVSSTNRQE
PNSFEFPDLTFCDPLNKRFFHNSSAELAKQADRVDERVITYSLVYSHFGTENDWRKERKAIEMIYTTWEHLLNITNVNIR
PMDMVLYCTFDGEPCSHHNFRVFYHRLYTNCFTFKPPARHLRSSGMDRGLFLLLYVPSADPSFKLDSLARLASGLENNG
IRFQIHQRDTIPHPLEYGIMAPTGTLTAVGLEQVKTSLANTPTHPCVPDVQLELFDNQFNFTRSYERQLFDCIKAEYHQE
VYKLCTCLLESHVILKPGFNSTTESFCHDLIGVTTPGRVAAVKRLYRKALDSLEGPPQGLGDYIKIGLLMRQMDPSLDVPL
TRMMCADNVSENAVKPESCYDSCNYNKYSYSLSQCPWPEDSFKVRNAQIQMIKIAQMLDEYERQHGDEMPVSALAR
SMQHHKLNTNSCQTYDTQDMTVPEPKECHRVQLFVRRSIVQLRVYPETLTVRHIIEERSYELVNLCSDMGGVLGLWIGF
SIVTLFEFAELLMVCASYYWYELIGSFTRSKRNPMRIPKLLAGSSGRPDLRDVELFMNASMCTSPSVLSAQSRSTAMLGR
SGSRQDGQLLQTAS

>Porifera_Syc_Cil_scpid49206_scgid6888_Amiloride_sensitive__sodium__channel__subunit__alpha_Al
pha_NaCH_Epithelial__Na_channel__subunit__alpha_Nonvoltage_gated__sodium__channel__1__subu
nit__alpha_SCNEA

MDVDQDALIRAEADNLFEELASTPEHRRPLPHMPSSSQFKRQTSLSFSTSQSSSLQRWYGEKNADEVASEAKEELDDDL
VGDMPPNWESKKTKRRTSAQSAWSTSSSSSKGSRESNVWSRDSLFSFGPKETPQITDRDVADIADEGAVKGLQRKRV
GFIDFTMRFVLDTAGHGMPRIVEKDIWLIRRIIWGCIVLGAAVGFIIQTHALVSKYLDKDFKVDVKYKFNKEIGFPAVTLC
NLNPLNRSKLECSRFDTSVHKGALVCQQTASGNHTTVQVTLAQILQARINALLKSDLQQAKERAFQYNFTHQEAMTAA
EIAKYGYDLSKLMVSCTFRGQPCGIENFTRFESARYGNCYTFNNNISTQIQISQPGPAFGLAIDLAYDPLEEIGLLSHGAGF
QVAVHEPRSKVFPEEDGMSISPATATSIGIRQAVVERLGEPYGNCTNITGYDWGNLYSGYKYTRTICLKECFAQAMLDA
CGCVSENILVGKRLCLPATLNKTEADCRAEIEELYRIYKLPCLQCFNPCKERIYRTTVSTASWPSFEYQREFKQLLKQKGFQ
VDHFDVVRFRDQYARVEVFYEELNFENIVQKPAYEPENLLADMGGQLGLWLGFSVLAILEIIEYLIYMLLVALGKVKLPDE
EDYLKARDD

>Deuterostome_chordata_Ggallus_tr_F1NW62_F1NW62_CHICK_Uncharacterized_protein_OS_Gallus_g
allus_OX_9031_GN_SCNN1G_PE_4_SV_2

MAPGKITARIKKTLPVRGPQAPTLRELMRWYCLNTNTHGCRRIVVSRGRLRRFIWILLTLSAVGLILWQCAELLLNYYSAS
VSVTVQFQKLPFPAVTICNINPYKYSSMKDYLSELDKETKKALETFYGFSEGKTKVRRAAGDWNGTESLFFRHVPLLRFEN

SFRAATDLRSGRKRKVEGSVFHKDSSIVNSGDSNDIIGFQLCDANNSSECALYTFSSGVNAIQEWYKLHYMNIMAQIPLE
TKEELSYSADDLLLTCFFDGLSCDKRHFTRFHHPLHGNCYTFNSGENGTVLSTSTGGSEYGLQVVLYIDEADYNPFLVTST
GAKIIVHDQDEYPFIEDIGTEIETAAATSIGMHFTRSRKLSKPYSDCTETGADIPVENLYNKSYSLQICLHSCFQKAMVESC
GCAQYAQPLPNGAEYCNYKKNPNWMYCYYRLHEKFVKEQLGCQQICKDACSFKEWALTTSIAQWPSTVSEDWMLRV
LSWDKGQKINKKLNKTDLANLMVFYKDLNERFISENPANTLVILLSNFGGQLGLWMSCSVVCVIEIIEVFFIDSFSIVMRR
QWQKAKKWWNHRKRDETGKPPEVGDAEQQGHDNPACSDEDLPTFNTALRLPLPQEGHPPRTPPPNYSTLRLETAFT
EQLPDTLEAGQH

>Protos_Platyhelmin_Macrostomum_lignano_A0A267EUK8_A0A267EUK8_9PLAT_Uncharacterized_protein_OS_Macrostomum_lignano_OX_282301_GN_BOX15_Mlig024916g1_PE_3_SV_1

MGVLRMLAETFVFNTSMRGFMKLYKTPHRFLRCLWALYAISMLSLWITACYFLIEHYSGFYVTVQQSERIQDITPLPSITL
CSQAPMGDHAASLWSNGTIMSPMHLHAAMKRSMRNAFGNGTSRRLSQFYAAKDVFYADKISTYFQNIDITEARRIG
MRPEDLLVFCLMKHTPNNNPSVSHDEICRLSVKWVRHPLYLNCYTVTVDQEDTLDVSSVSLILRSGVAQPLPLQGHVFD
LFSQSQGLKISLHEQGSHPNIDESAISVGMGRNTEIRYTLVQHRRLNLGNQRSNCLDSEKTAKQRVLRDFDETYNYTFDA
CMSQAIGQLILGECNCVSAFHLVTQVPTRNTPFCHYHEGENTSSLLSRLDCVRNIKLPHRIAQLRTTTCLRRCNFNQFDM
QVSSAQWMPTRWQLKLVLQLAPAWDIMIDHHSSEAEVAEARRRLMSFIDSPNEQRQLLSNTSEPDISRLTLVTVSRAT
YNTILKTETLQVTLSTLLAQIGGLGSVTIGLTITVLFEACEYFYLTTFNIYKHLRDKRRKRKNPAALEEDEKSDDDGGSGAGE
GVCDGDCEEDAIGDVFGDDNDDDDDNKSGVKQDDSAGDHAHNSDQGETLPLNSLNRASGRNSGRHSLGDNEDEDA
SSESWKAEARIHRPLALA

>sp_P37091_SCNNG_RAT_Amiloride_sensitive_sodium_channel_subunit_gamma_OS_Rattus_norvegicus_OX_10116_GN_Scnn1g_PE_1_SV_2

MAPGEKIKAKIKKNLPVRGPQAPTIKDLMHWYCMNTNTHGCRRIVVSRGRLRRLLWIAFTLTAVALIIWQCALLVFSFY
TVSVSIKVHFQKLDFPAVTICNINPYKYSAVSDLLTDLDSETKQALLSLYGVKESRKRREAGSMPSTLEGTPPRFFKLIPLLV
FNENEKGKARDFFTGRKRKISGKIIHKASNVMHVHESKKLVGFQLCSNDTSDCATYTFSSGINAIQEWYKLHYMNIMAQ
VPLEKKINMSYSAEELLVTCFFDGMSCDARNFTLFHHPMYGNCYTFNNKENATILSTSMGGSEYGLQVILYINEDEYNPF
LVSSTGAKVLIHQQNEYPFIEDVGMEIETAMSTSIGMHLTESFKLSEPYSQCTEDGSDVPVTNIYNAAYSLQICLYSCFQT
KMVEKCGCAQYSQPLPPAANYCNYQQHPNWMYCYYQLYQAFVREELGCQSVCKQSCSFKEWTLTTSLAQWPSEASE
KWLLNVLTWDQSQQINKKLNKTDLAKLLIFYKDLNQRSIMESPANSIEMLLSNFGGQLGLWMSCSVVCVIEIIEVFFIDFF
SIIARRQWHKAKDWWARRQTPPSTETPSSRQGQDNPALDTDDDLPTFTSAMRLPPAPGSTVPGTPPPRYNTLRLDRA
FSSQLTDTQLTNEL

>Protos_Platyhelmin_Macrostomum_lignano_A0A267H0I3_A0A267H0I3_9PLAT_Uncharacterized_protein_OS_Macrostomum_lignano_OX_282301_GN_BOX15_Mlig017198g3_PE_3_SV_1

MLEPQDIDTLASTPRNSVSDDCARCEQLAGPMGHCAQHTTVEGLPNLLTAGSLWRKIVWSVILLMALACCTTHLILLIID
FHNSGVASTMMLNENVVEFPDLTICGLKPISESFLAQSFERNWTDLLLHYTRRLDGWFNDESSLQRRNESITDYMERM
KALYYKMFEIVRLDKTFLDSWLSMHYQSSIFSCRFKRLPCEPSNFSLIQDHHHFDCLTFSPSESNKMADHGSKFNSILLINL
FTDSSIFESHYQKHAAYKASSKTYYDKGLSLTEYLTTNPEEFPLNQILANNEQPNGISAILHPRNTYPFNGYQSYIISSGTSA
TFRISTLEKHKLNYKGRERCSDKPPDEVVYRTVRSVVNGTSESRAFVRSPEDDLYTWAQAELFKKCGCRSHRSPTGASSS
SLCYHVQPDEDGAPRPTLPKEALDRIRCHDRWWSKLLNSAPNHVTMWKPCHSVSREVQLESLDWPSDEDLMPIAVSF
IKMRQRNAQHRDEYAVPSWNSSLTARTTYLRDYSAVHSLSRDSESLTKLLKESMPKAAKALNTQKVKENVVRLVVRLES
PYSIAYTESFTYQWFEILSEIGGILGLWFGVSLITGFELIDLIWLIAIWDSQVGHVKRSDREDVKQLEKLGSALQNERVPEFF
TSGCSMYIE

>Deuterostome_chordata_PetMar_tr_S4RY81_S4RY81_PETMA_Sodium_channel_epithelial_1_beta_subunit_OS_Petromyzon_marinus_OX_7757_GN_SCNN1B_PE_4_SV_1

MKIRKYLTRSLHRLQKGPVASVSELLYWYCMNTNTHGCKRIVVYGKKKRVLWFLITIIMLGVVLWQWVLLFQAYLSYGV
SVSVNMGFQRMNFPAVTVCNLNAYRYSSMKDKIKDLEAYTRVALQTLYNYTDSSTPSAYDTSYAVGPWQEIPLVLIDRR
DPNRTVVTEVMCSRAAVGIETHTVDNRVFHIGFCKSALGVCCDSAGDKCFYSEYLSGMTAVKQWFHFNLLSLLGNLST
EEKSNLSSSGDELIRSCLFSSDTCSATNFTTFFHPMYGNCFIFNWGENETVMQVSNPGVEYGLKLVLSIDQDEYIPFLTTIA
GAVIMVHDQNTYPFLSDLGVFVKTGVETSVGIEVGQLQRQGAPYSDCTMDGTDLPITNLYNGTAYSVQACLRSCFQTK
MIEMCGCGYYLYPLPPGEKYCQNQNFTGWRYCYYKLYEQFVEEDMDCYTICKQPCIESEYKMSISMSDWPSQSSEDWI
FHILSKERKHNVSRIFNRKQDIIKLNLFFQEFNSMTISESPAQTIVTLLSNLGGQFGFWMGGSVLCIIEFIEIIIDCVWIGMIK
ASNDVRERRKTSRKPRYSDEPPTLSSIVQGQGNSGFEMEERGPPGEAPEANGSAAQPPAAAAEQQPDVPGTPPPHYD
TLRISKTELHDEINSDDDGEFV

>Protos_Platyhelmin_schman_A0A3Q0KJ87_A0A3Q0KJ87_SCHMA_Putative_amiloride_sensitive_sodium_channel_OS_Schistosoma_mansoni_OX_6183_PE_3_SV_1

MEINFDKYDETKEVESQCQEERQPNVLNELTDLTNMKCTHSSSLIQPQSVKTNHKNIKAPKSNYNNRVKNKHLDQCSN
NELNKTSNEFTSTSLSPLSNEYNPSLLTQIKRQFRQFCETTSLRGVPRIVSTKDSKRRFLWLIFVISFFIGCITCLTFIINQYLEF
DVIHQPKKLYDQPRPFPSVTLCNLRPFSTRDIRKLKKAEVLPVDMYFEMLKRFMISVPVEYRRYLTMLWDFSTYLANLPD
LVDANNLGHTLEDIVKRCFILYKTGSVTRGTACEKSGYWTKTQDRVFHNCYTYTINPNKTNTALNMELVVYLDNLIEQTD
CFDCQERVMASQLTGARLLLHPGASYPRVAEEGVNLMPGTMTDIRFTVYEWSMMEPPHGRCSKNTPQFISFNNVNY
TFSEEACHAHMVQEKVASNCQCLTQHLPIPTHLLGKDLRKCQAFKLPATYQMLDNFNESKQLYFNNEAVSTYARFAEC
AEKTGTAMDSYQIGCRPACVRYTYKPSITAAQWPTKTFLTWFVTKFIKEMELSGYSRLKSTNNETLSIFNERMEEIRKPLQ
PYEIIANLTKEGRLQEAIKMLMQTQIFEQNFLSIIISRPNFDLERVEEKAVVSLTSLFSQIGGLLSIWIGLTFVCIVELVELGLN
VADAVIHYKKLKS

>Protos_Platyhelmin_Macrostomum_lignano_A0A267GSY3_A0A267GSY3_9PLAT_Uncharacterized_protein_OS_Macrostomum_lignano_OX_282301_GN_BOX15_Mlig015385g1_PE_3_SV_1

MNNNDSKQIKANSLGKGNDLYCQVSGFCQRTTIHGLSHMYQSKTSKRKIFWLTVFVLGLVATIVHLTFHAMQFLESPTII
TLDHEGSEFEFPDVTICSQTPTTSRFYERGTEKFPNIQNGFTAMDLLLYPVYNQTYDSVHLLPYKEEILLHKRLIQHQYAS
MPTKHKGFALDELVIKCSFHKSDCRLQFTLLQNPYYLNCYTFQPPERNLKTTRPEDGLLLLLYIGNAQKNNIFPGKNATITS
RSRPTSQKRETGNDWFARILLPEYEPAGVKIVFHEKDAFPALRAAGSIVGAGQFSEIELGMIQERKLNKGSSPCVDTEPK
VWFLTSLDESGWPVFREFNAEPGDCIYEFQQRDLYKECGCTSHRVPLQRDHKMFNSIDRGDQVGWCYNSTSLLLDLGL
DVNQIVKSVKGDPAKLVNISKTAIQSSHANSRLVERYLRRVNCHDSLLDRFSDPHYNSIQQCTDGGEKALYACQKNEYKS
VHSTASWPRYSDIQEIIEEDILPALLIPKNNLSDGNKTKVEYRPLAHDLFDFDSNGQWTAQVELVKENFVKLQLHPVEVL
VNVHREEPAYSIWEALGELGGIIGLWLGMSFLGVIEVCEFLINLTSAARRGKTLQLVKRQEPDAVSGPYGEAESAAPELLL
SEISNGVDVDQPTQNC

>Protos_Platyhelmin_Macrostomum_lignano_A0A267E4G4_A0A267E4G4_9PLAT_Uncharacterized_protein_OS_Macrostomum_lignano_OX_282301_GN_BOX15_Mlig025621g1_PE_3_SV_1

MSNVKAAATVLTEYGHRSTLHGLSHMFQSGDLWRKAFWLVLTFAASGACVYHIYATVSNYLESPITSIYRDEGPQALFP
GVTICNLKPVSEALLHAEEEKAALFHHLDQVWNFFDNWLFGLRSRKHSETRVEYKERQRELLKTYTDIFWVSDDTREFG
LSDRHTLLMCLYKNKHCNASNFKLVQHHRYWNCYTFQPLDGDRTIDNSDQDSGLRLILFTDSNFDSNRLDDSVNYLRS
MTREDEITNPYSLRTYLGRGNRTLNQVLTGNKWEAEGLRIFIHNESQFPTTSDIAFDVSSGASTTIEVNTKQVNTLNRPP
SHVCNAEEPETISYVRYFGPNVTGLRSYRLTQPEAVMEAQQKMIFDKCGCYTHLLPFKFNTSLLCLYADPQHRIHPSMEF

MQRLACHTNSFKLAEQKTSELLQQKDEYKPCTSWKLQATKVQVPWPRSDELIDFVNYVFMPSYIAELQNEDQVMHLL
KSRNSRAEAIIRHLFNPDCRKENGTDAPLVDSWCISRAKLKANVAQIWIKPASTHAGVYMEEYSYDWPEALSEVGGIFG
LWMGISLLSVMEFIELLCILMSVCTSRDAGKRSSNVGSAAASRNRMPQDFEAILRYLASLEDEEMREPVRQLQYQRQQ
PQQQQQPQQQQQLNSARAAREPSGRLGLQLVP

>Protos_Platyhelmin_Macrostomum_lignano_A0A267E4J1_A0A267E4J1_9PLAT_Uncharacterized_prote
in_OS_Macrostomum_lignano_OX_282301_GN_BOX15_Mlig033810g1_PE_3_SV_1

MHIGESSFLEAFSSGPYCFFLDSERLNAGLCDVPYCASSEKFDCVKSNSRASLAHYTGQMNRSRSGKDCMPWSRLWNV
QEDMMYFHGIELFGKMVRLLIQTPDRIEDTSDNFCRPASIVIDVLETDSGAFKRQIVSGSVPVCFPAAAQLHMSSGRISL
QNFFLIGVEKCNIPVCSSKTVVRGAFTDSVGSHLLARQFSNDLSIGLADCTFAGQKCTKEHFVNIDHPQLGMCHQFLKQ
SFIANMSSTQLAGSQLSITYFTDSTDCKGKTVEEIPEMNCAFRNGMSTDTGSLAFKAVIVPEDSFPWSSKGIDVAASQR
MDVQLRFERYSRLSTASRPCLETTEPIVYRLPTWEKSMSNEDPKVWDAADWMAVAKDWKEMIARFRAMLLLPNSSS
VLTATERQFNVSDREDCINSYIVQQFIENCNCLPSILPVKVDLFDQYSYCFEVKDIRNLSIDLETRYNEQVSRINQYEFEAM
KRCQSPCRTDGYSVRYVTYPWPNLKTARSIPLLEKLASNYGYPTSISELRGVANFYQVYLKSESIKRVPFANQSAKDILSFE
KELESLQNRLARLTVQAWTSSSQEITEEEAYPTKNLVADIGGALGLWSGISILTLCELLELLIYISRGLARRTQQNPSRERQ
QTAAGGVSQLDPEAVGLESGKIAL

>Deutero_Cephalo_088140F_t4__name__chr_scaffold187_start_416588_end_437049_strand_pro_len
_664_

MHVKLTCSVELDCCPCCDCGQCCCGSSTAGSEPRDAGSELGDGTDAGEQTYYRKQAQTISDFAAGSTLHGLPHIFPDA
PLSIRQVAWALAFIGSLSVLLYQCSDRVKFYFQYPHITKLDMVLAERLDFPAITICNMNMFRWKQFTQNDLWHMGKGL
NILDENNNLRCSEYATPGDMEALKNKANFDSFKPSPFSIMEFANRTGHQIESFLLDCKWKNFTCGPEYFTPYRGLLWSIT
TVGSQNGCLSGYLCVYLSVYLSGYLSGYQSGYLCVYLSVYLSVYLSGTRVVGYLSVYLSVYQSRYLSKYQSRYQSRYQSKYQ
SRYQSSVHWVKLTVFSSLPPTLKLLQDCQRKDLGCWEVFTRYGKCYTFNSGSPNQPVLKTLKGGIGNGVEFFLDVQQE
DYMPAWGESDEVTFEVGFKIQLHTQEEPPFIHELGFGVGPGMQYYVSTQEQRITYLPAPWGQCKAENDLTFDFAKYTT
SACRIDCETKFVVSQCGCKMVHMPGNFPICTPDIYVECADQALDFLVKSDNKKCVCDTPCNTTRYNLFMSHVKFPSEQ
AVKYLARKYGKPEDYFRKNTLVLNVFFEALNYETIEQQKAYEVASLLGDIGGQMGLFIGASILTILELFDYLYEVLKDKCTER
RHQPRSESNAVSVNLEDCKRDNSRAPLS

>Protos_Platyhelmin_schman_A0A3Q0KSP8_A0A3Q0KSP8_SCHMA_Putative_amiloride_sensitive_sodiu
m_channel_OS_Schistosoma_mansoni_OX_6183_PE_3_SV_1

MTKKSKKLITKEELQLRKGLKKEFLYFCQTTTIRGVTRVVNARNKTLQILWIGSVILFFGGLFICMFILTRQYLEYNVIHPPQ
VLRDTPSPFPSLTLCNLRPLSSEANIILNKYKLKSPRQFAIDLNNFAAKAYYYSNDIHSYQQVTSAVSMGGYLESLPQIVVP
KLGHNLSHTVIHCMVIHAEGSKRVMNECIQVGRWRRFVHAQYLNCYTYDIHEIYRNHVRTIELFVYLDESMNITSCSDC
FSSEIKSQLSGAVVTVHNAETYPDINQEGINIQPGSLTEIKVKTIKHTQKTPPYGRCSPDTPTKINLYGSEVYAYSEHACRM
STIQAKINSLCGCNAIEFPYINESLPFCLEMSSFVKIGNCDIQHPIIQNYTIHNSPSNQNYTLLSSVNQNQKTKPHEMPFNF
ECLSELEAVKNRVLCKSEVTKHYQGDVVLSCTLPCAFFAYETDRSTSTWPTKSWQLSWLSTRAGKKVANRPDLVGYRLA
KEKLDSSEGEKEAEEFLSKSNVLERNLLAIMLIRPNFNVHKVDEKEVLSLTSLLSQSGGLFSIWIGLNMMSVIEVVELIFHLI
TRCRQFSKRKRLRRTQKKQTTCDTPKSLTNTNSETGVVQDQITNQGVVKQTEKYRKDDMTINNGPIQIHPRKRSPSAP
NQMVSGNVSLYLPQI

>Deutero_Cephalo_088030F_t2__name__chr_scaffold187_start_203431_end_230405_strand_pro_len
_665_

MKKLNPFKKWQEEKEEEEKKKKKHQKKQYGFTTPLREKCALVLLFLIMDKGVKSAISDCFVCVDEGLKSAISDCPVCVGK
GVKSAISDCLVCVDYNKRYSRIVVRSGPRLSAVCFSFVLKVVAFTPEVIILPLTEKKQSQKPPLEYFISDSGDVLYLVTDGNH
LVTGGNHPVAMMNRDPTMSSLSDIDEVAPKPSDIEVFAGNSTFHGINHVFTNGRCSIRRILWGLAFLTSFGLLVNQGV
DRVLYFFQYPHVTKLDELEAQMMSFPAITICNLNPFRFSQITKNDLYHMGHAFEMLKPDKTLAHSNYIDPEHLRTLKML
SDFDNNFTPTKSFSMKEFYVRAVIQIDEMLSFSMKEFYDRAGIQIDEMLLSFSMTEFYDRAGIQIDEMLLKCSYRGEDNC
GPRNFSTIRSGKTAKLEPQYGINAAEFRDPAEFRPGYRYREAGSMSSYEENEPKPSDINVFAGNATMHGISHIFTEGSFSF
RRFLWAAAFLICFGMVSYNGYDRVIYYLSYPRVTKLDELEAQMMNFPAITMCNLNPFRFSKLTKDDLWQVGHALEILD
ENKKLAHSEFVDPEHLSTLRRLAEFGPGYVPKNKDSFNMKEFYNRTAHQIQDMVLECSYRGTKCSWANFTVTIFPSTPV
VDMQGFSPVWIQQLWTDPAQSHKSDE

>Protostome_Lophotroco_brch_Lingula_anatina_comp148540_c0_seq7_p1_comp148540_c0__comp14
8540_c0_seq7_p1__ORF_type_complete_len_666___score_77_65_comp148540_c0_seq7_79_2076_

MSTGAQTMYPSEKNQFDFFKNSLVVDHNSQDPEKSAREESLIALAYDWASNSPVNGIPNVVRSQNLIKRFFWLVALLG
CFGVMGYQTWELVAKFYRFPVDTTLTYTHSKEVYFPAVTVCNVNPLRRSMLSSAGDDVYALLGPPKSQMGSSPGGAQ
APAGPAGSFASQSQTTQAAAGPPGPASSSITTQAPAAPGPKGSGSGGTTQAPSASQTTAGAPAGGGGGTTQAPAGP
PAPPSGRKRRSTGTNSTSTSRRKPGQMNERFEMRKRFGNIWAKLNYTTRETVGHELDTMIISCTFNGVTCSSANFTRF
NNHQFGNCYTFNSGWDSSIPVETSSNAGPLYGLSLEMYIEQSEYIGDLSESAGVRVQIHSQRSMAFPEDEGFNIAPGYL
TSMAMTRVEITRRPHPYPSKCRNFTTEESKQKSVFTSVHNVDYSVSGCMKTCYQRYVISECACGDPAYPFSLDLEAFSDL
KNNINGTTAESVDPCVSDADDECVAGIKRRFADDDLSCDCPLTCVDVEYSGVPSLAKWPSKQYMSTLHATLASAGAHL
DNIINGKNGMAAEPGDNLLKLEVYFQELNFQKISENVAYDVFALLADIGGQIGFWVGLSIMALFEVVELILDVFRLLLFRV
GSFCFKKQQKRPPSRQVKVQPASEENVLTPRLEKVRFS

>Protos_Platyhelmin_Macrostomum_lignano_A0A267ESF4_A0A267ESF4_9PLAT_Uncharacterized_prot
ein_OS_Macrostomum_lignano_OX_282301_GN_BOX15_Mlig013241g1_PE_3_SV_1

MVNHQADASVDRIREVRRRLVQDLLNGIEPDPRALDPTADIEDNRQFSSPILQKLYTTVEQFGARSTIHGLQHIFHSREL
TRRIFWLVLMLVALCLGLAHMHATAMEFLSNPLTSQLKLMSEQVKFPDITLCNLKPASESMSLAPEFRQYIRDKWGYFD
NWYVEERKRRGNESKVNFYERARLVFGRLFQILWVSGDSRDIGVLDVTLLLDCSYNSMKCHQKNFSLRQNYRFWNCYT
FFVHKDNLVTGKGGKGAELDLVLFTDNHNTVRLEDLDGSARQKNLSTGIYKRSRQTEEQEPISLKELLNEATGLNHLLTE
GSYEPGGFRVFIHERKLYPMLSREFVDVQSGTKSIIKLRPLQHSLLSSAAHQCTPRSAAENVSYVRYFAHGSKHIGHKKYQ
KTDSDFVAEEKQNLVHENCGCYSHRYPFTVDNSELCYYVPPSSWKTPPEQVLRKIRCHDRWILDAEKRTYELMQKFHSK
RFCEFTLYRPLEQSSSWPRFEHVQDIWTDFIAPKIASGKEFTMECGHNHHIFNRTETAAFNRSTSRQELVYYPLLLTDQLC
LEQFCNGTDTSCDLNLLRRLYPTATNCLQKAVMTKNMALIGIHLLSSAADSYVESVSYEWTEALSETGGILGLWLGISLLS
VLEIFEFFYVLARNCFDQRLALRNRDA

>Deuterostome_chordata_Ggallus_tr_F1NE95_F1NE95_CHICK_Uncharacterized_protein_OS_Gallus_gall
us_OX_9031_GN_SCNN1B_PE_4_SV_4

MNLKRYFVRALHRLQKGPGYTYKELLVWYCDNTNTHGPKRIIKEGPKKKVMWFFLTLLFASLVFWQWGILINTYLSYNV
TSSLSIGFKTMKFPAVTVCNANPFKYSEVRPLLKELDKLIEAALERILQPTHGDPISPLLLNNSNATEGLDLDLWNQIPLVLI
DEQDKDNPVIVEIFETNQSAAGNQTAAPPAPANVTSEEKKYKLAVKLCSHQGSNNCTYRNFTSAAQAVTEWYILQSTSI
LSKVPLQERIRMGYQAEDMILACLYGAEPCNYKNFTQIYHPDHGNCYIFNWGMDKEALNSSNPGAEFGLKLILDISQQ
DYIPYLSSAAGARLMLHQQKSFPFLKDQGIYAMAGTETSIGVLVDELERMGYPYSDCTANGSDVPVKNLYSEYNTSYSIQ
ACLRSCFQNHMTEICGCGHYMFPLPEGVTYCNNEDNPGWAYCYSSLRSSIRHRQICIDSCKETCNDTQYKMTISMAD
WPSEASEDWIFHILSYERDMSTNVTLDRNGIIKLNIYFQEYNYRTISESAATTIVWLLSSLGGQFGFWMGGSVLCLIEFGE

IIIDSLWITVINIISWCKGLKQKRVRARYPDTPPTVSELVEAHTNLGFQHEEAGTETQGEALPPEPGTPPPNYDSLRVQPS
HNPGTDSDIECEEQRPAANHHGDASVWAE

>Protos_Platyhelmin_Macrostomum_lignano_A0A267G692_A0A267G692_9PLAT_Uncharacterized_protein_OS_Macrostomum_lignano_OX_282301_GN_BOX15_Mlig031432g4_PE_3_SV_1

MKRRSRQDAELAVAIAAPDSAAMNSAVTLPLHKYTVQTNNRAVEADADPTDADADAAASAVASFGAATAGQSDLAP
VRSALHRQFICFCENTTIRGIPRIIKSESRRIRLLWIFFVVALFVACILCLSFLTLQFIEYDVMHPPRVLKDYSSPFPSVTLCNM
RPLSSEGRQQLRQQGWKYPRDFSESLELFARNLYFYNESTKYVYYNFSGSMGFPAYLESLPRSASRDIGHTLQDFILDCR
AVIRNGSARNLKFCSPEYVWGGKGSDMFAGNFHAFLHGNYMNCYTLDLHEELRDNTTMVEMAVYLDNEWNLDNFN
CVDCYTMDISTQLSGAVLTIHSGDTYPDVNHEAIHLKAGSLTEVKLKHVLNVQKQAPYGRCSQDTQRNLNLFGREYKYS
EDACRQATIQRDILNTCGCQAIEYPLVRESNFTYCTGLEEFLRSDGCNWATAENDQKRLADCSATLQKVYGYIRCKNDVI
SKYSKDVVESCTLPCSFYSYQTERSTSDWPTRSFQLRLLNSRVYSRLLNRKEMKAYRKAIEKLNNNGSIQEAYEMLKHENL
LERNLLSIQLIRPNFDLHKVEEKEVLSLTSFLSQIGGLFSIFVGLTMISVVELIDFFIHCYEICTRQSRGVGGASAVRRCDGAA
SRSEADVLLPAVAAGPRPHQKLPNADS

>Protostome_Lophotroco_annelid_CAC9584386_1__Ofus_G061163_Owenia__fusiformis_

MNEKDDLDNMTVIDTKEEEIKGASDAKTVTDEFMGSFGAHGLPRVWSSASPVKKIVWLVLFLAASGYCLNIIIRVGIKFG
TFPSGINSKVYPRSIVDFPAVTICNLNMLKATSISNNNVEKYINLLSVQSEAEASMNSWITWALNYTDSVTTSFGPTTDST
SRTNSSNAGTTDHTTVADRRKRAVEEFPFNEDLLEDELLLNLRADHVNDLTRLIHMPEVVKMATLRLLKKRHGFDMRE
QRSAPPIPHRQKRDNQAAETNSSDYYDDNFYYYDNYDYSYYDDYGFGDVSENDFRTLLTKSKTDDYQDLMGVLKPTKT
ELELYGHSAQDMIVSCTFDSKKCNYTLFKTFQNSYYGNCFTFNYDDGNSTTQEVVFNTTKKGSRFGLKLTLNIERQEYIGL
FAHGSGVRVAVHPRNATPFPEDYGISAPTGWETAIGVRENRVTRLTKPFESNCSHGNHDPSETYTGLYTLLECNQRCLQ
KQIRSKCKCLDEIMDDQTWESVYASDSATSMLNPCSILNSTEEKCRQRMYYLSQTNRLGCDCPLPCLELGFEESLHSSE
WPSAQYYPLMMQKLTDSNIISEGQLKIMNDLKTTRENFVRLHVYYKDMILESLEEFEAYTVDSLIGDIGGILGLFIGMSVL
TIAEAVEYLIDLCLLRCKTNNKRNSVVALK

>Protos_Platyhelmin_Macrostomum_lignano_A0A267GU61_A0A267GU61_9PLAT_Uncharacterized_protein_OS_Macrostomum_lignano_OX_282301_GN_BOX15_Mlig030616g2_PE_3_SV_1

MSNVKAAATVLTEYGHRSTLHGLSHMFQSGDLWRKAFWLVLTFAASGACVYHIYATVSNYLESPITSIYRDEGPQALFP
GVTICNLKPVSEALLHAEEEKAALFHHLDQVWNFFDNWLFGLRSRKHSETRVEYKERQRELLKTYTDIFWVSDDTREFG
LSDRHTLLMCLYKNKHCNASNFKLVQHHRYWNCYTFQPLDGDRTIDNSDQDSGLRLILFTDSNFDSNRLDDSVNYLRS
MTREDEITNPYSLRTYLGRGNRTLNQVLTGNKWEAEGLRIFIHNESQFPTTSDIAFDVSSGASTTIEVNTKQVNTLNRPP
SHVCNAEEPETISYVRYFGPNVTGLRSYRLTQPEAVMEAQQKMIFDKCGCYTHLLPFKFNTSLLCLYADPQHRIHPSMEF
MQRLACHTNSFKLAEQKTSELLQQKDEYKPCTSWKLQATKVQVPWPRSDELIDFVNYVFMPSYIAELQNEDQVMHLL
KSRNSRAEAIIRHLFNPDCRKENGTDAPLVDSWCISRAKLKANVAQIWIKPASTHAGVYMEEYSYDWPEAQRGWRHL
RPVDGNFAAQRHGVHRAAVHPDERLHQPRRGQKVFKRWQCSSLSKQDAARFRSHTKVPGESGGRRDEGAGASAAV
PAATTSTAAAAATAATVELSQSGKGAIRQAWIAADTVESLNRLLR

>Deutero_Ambulac_Sakowv30002567m

MADISVSPEVNKQALRGWSAQRKKDIRNDGYKTWKKFSQETSLAGVKFVGDDSTIYSRRIIWLLIVLAGAAGFVYQVYR
MVDTFAEMPVTVNYRYTYPEYQRSFPAITLCNNNLFSYVSMEKYYGILGITEYLFLLFSFIPSKGLEEMAGFIPLEEVPENF
GDNLTAFEYMDMHNHDIKDMLISCDFGGIYCDAHNFTSTVTNGGICYTFNGGQGNEDLLKLTGTGLTHGLNIVLDANK
NDYLAPSDTVGFQFAVHDQKDIPNIRDKGIGIPTGMHSRIALSTTAISNLGSPHSDCVTDHTRTLNYFPGEYTESKCLME
CEEDFAVRQCECRYYYMPAYAFRTNANQNFILENFTVHTSTICDCPERCNTISYNWKLSYNTYPTNVISIFKADMGYPPC

DRAVGNYVFFATTANESYTSGALPLIDAIINNTISAQDLTRLSPSRGIPDSVFRDVTYWAALEWMIDLNYEICFNSLFYILY
VQKGNYSFYDGKETSYHNQTIALINEKYSSQEKLDQIFTLVLKNNNISLLKEEFVNVSRVFYENYYLQILRQALEVTYGYFW
VERNIVIDYLFRYRRHSDICEKFMRENFAKVSIFFEDLKFENITQKADYEPFQLVCDFGGSLGLFFGASLISFLEIFDFLLSWT
FARFKKKNQMNPDEELSETPGTIVK

>Protos_Platyhelmin_Macrostomum_lignano_A0A267EDH7_A0A267EDH7_9PLAT_Uncharacterized_protein_OS_Macrostomum_lignano_OX_282301_GN_BOX15_Mlig004551g1_PE_3_SV_1

MMARFRAQASREDGALEDFEMQGIDEDYRKAFLEDEAEQQQPLTQQKSSAVSAKSKLPSNNNMEGRKTSAMLFCEC
SSIRGMPKIARSSSSFLRRMWLSFIIGMTLVLIVSSSLSVRSYLQYDVIVHTSDIPEFPSKFPAITICLHRPFSQRALSLWANG
SVMSPRGYNQRLRRWAQEAINKTESNFSRIKQKDAVPIFYALYYDMLEMYYQNVRYEDTTLLSHDPSVIRKCIIMTKWG
ESHFKDPCNLTDHNFNIKLFSHPKHFNCYTVEPNSEASSLDTIFLGLAVDLGPKPDYLNLQQGFLNDVFDQATGLRLVIHE
PNRYPDVEQFGKNIEAGKMTELAYTPTLWNRSNTPVKPCIEGDAEPFEDLGRRFDYSQRECTRMKTQRAAMERCGCII
CLPGHPRPRIPGPDVAYCGTFTDAYNRTTDDFIRFSNCYVEDVQPSEDHLKPCLSRCVFSEYETKLSVTEWGATDWQVY
WSGKYLNDIEELQRLMNQRDSRPEEEQQGLAPRFDDVKKRIMHYLTKENLTSVQASADQMVESKNFAYVVLKRKGEN
LILKQEKQVLTLNALMSQIGGLCSLTIGLTCACIVEVVEFFYIYLTAFRTNGQAGTAAINNDGGGGGVGASESSAGVGTH
GQRTVLLPGTAPTSLDLDNAGNHGDQRIEHRQLHQRSSQI

>Deu_Ambulacrar_hemi_Ptyfla_40v0_9_20150316_1g10161_t1_scaffold5763_cov99

MWSSNNTVQPNGLFKDTTGEKNGTELTDFGSNLRLTREPGRMDHGAAGKRPGCMGILLSGFADDCGISGMKFLAGK
SVVPLRRIVWLLVLLTGIGLLSYQILDRVSYFAIYPLNVNVQLNYVEEVTFPAIAICNFNMYSLQYTNELNLTGLFYEMYSTY
NPNFTNYDIELTASEQFYRDAAQQPSAMFYRATWQWRPMEIDAVQSILTDFGACYAFNTGNDKYGIRKVRNKGTSHG
LSVYLDVNQWDYNIGPRVGAGFQLMLYEQGDVPMVSDLGFSVAPGEDVRVGIEVTKVTNLPPPHGVCAEKSLQYYDR
YSVNACQLECKTDYVVERCGCRMYYMPGNATVCDLQQFSFCVQTTLWYMNQTCDCPVPCTQVMYKPSLSHAKYPS
DAYATYLENTYGFPKDVFSLTYTNEVNLTGILYEMYGTFNPNFDNYDIEITTSEQFYRDAAQQPSSMFYTATWQWRPM
ELDAVQSILTDFGACYAFNTGNDKYGIRKVRSKGTSHGLSVNLDVNQWDYNVGPRVGAGFQLMLYEQGDVPMISDL
GFAVAPGEDVRVGIEVIKVVNLPPPHGICGEKSLKYYNRYSVNACQMECKTDHVVEHCGCKMFFMPGNANICDVKHY
YNCLQYAEFSFTNVTCDCPVPCTQVIYKPSLSHAKYPSDAYAMHLENAYGFPKEVFS

>Protostome_Lophotroco_annelid_CAC9662587_1__Ofus_G12132_Owenia__fusiformis_

MSEKNVEPHDASVKDLTQGFADATSLHGLPRVYSSKSLTRKIIWGLVFFGCFVGFCWQVTALTLNYFEYPISVTTEVKTR
LKVDFPAVTVCNMNMLKKSLLMGTRFENLTKVDKRLSDIYGTSATSNSQTTPPAEMPQDPENEDPLSGADFNEGSTTE
MDTEDDGTSTIEAEDTTDGITYVTTEDPYDETIDTTQEDIATEFITTEASAGDATTDYDYLTTETSRKKRSVGKRRPKENE
HKLPEQLHKQMKPLDGSAHKHAHPYSNNAKGESRRRKRQADSYYDDNYEDDGDEYDSDPYDDYYDNYYDNGDDYYP
EERFEFEDTGFSEIEPNDYFGFLQNSQTQDLSDLVGIVVPTTDEMDSYGHTFEDFVLQCSFDKKNCSNSDWKKLYNSQY
GNCYTWNFGYNNTIKSTSRFGSRYGLRLTLNAQADEYIGLLSHTVGARVTVHSHNVMPFPEDQGVSASVGRKTGIGVQ
MQNIKRKPHPFPTNCTYGTHLQSNYEGDYSVLSCMMSCLQNKIKTNCKCVDKIVHNQTACDITNTTQEKCRQRMYHL
FDEAKLGCDCPQACEELVYGTTISGSEWPSNQYSPYLLQKLEGGSPVSDIKQNLVRVHVYFETLAVHTIEEVPSYTWDNL
LADIGGTMGMFIGISICTAFEIVELLMEIGKLVVGKMTNKNKVTSLK

>Protostome_Lophotroco_annelid_Pdum_comp412166_c0_seq4_

MAATAHPAPKKPAANSRIKENVKPKFREEFQDAVGLSGISFVCGTSVHSIRRCFWGLLLFLGLGATGYQIWMRVVHFLD
DPRVVDLSIKQRSQLQFPAITICNLNRYRMSGVYNFSRYLSSNSPFQETRRTPMQMVTDMLEILRMYRMGENNSRLKA
KIEKFYHDHKTLIDQFSTDQNLLREFEELSAYKIYDTGMFLLCFWKKINCSREWLGESQTNMGKCLTFNMQGNLKSDIA
GSMGGARMILDVQDYERMFRGSMTTGLKVFIHDQNVSNTLMSNAGISVRPGVMTAIGLMRTELTSLPKKGDSESCTA

DSSGECMLECLSEYVVKSCKCRESYMPGDADLCHPHQMINCSFPATAYILTHSKECNCRPACRRVTFGTKISQIPLSEQF
VWQYWVDMMKRGDSSQAFEKLSKVRAWASLEIWNRSLTDQIFEIGQQVKIIYDALIIEDLRDNLVSYAEDLFVGADSV
STAKPFQCRNESESNLHEAYKALRSTFLDSPFFSYLTDCNRVEDFNAILANDVLQRDDEPSVFMDWLKQVEVGISNLTET
IDLLKACGNKQNTSSPLDAYKSNRFKATFREITKLFSSLISLISDASFFLHNLPKPTIRITPDFFRNNYAIIEVYYEDLTLKNITQ
LPGDSLSSLVCDIGGSLGLWIGASVLTIFEIFDTIMWFCVGRKNK

>Protostome_Lophotroco_annelid_Pdum_comp420047_c0_seq10_

GDVPRTKDLGFAVYPGVHALVSAKLNKATGLKDPYGKCGNKPLNLYGDVEYNRAKCELECLTKQLEKQCKCKMAYMP
GDIPICNLNQYLDCVVGVLYNQGLQNPNCSDQCIDACEIYSYSADVSYGGISEESIGSLVSGDKTVLAQKYVTATEISERV
KTQKLEDTVTYMKTVSKSIQVFTTYNLQSADASCNNIAKATRVMLDFFKQDLETVQNKVKLYKDDYNAKIGYLIPDFLYH
VDVALPQLKYIIASNMSRARDDIKKTDLTFTITKISEYTKLIAAAMDRQKILQNKGNMPASFHNIEGINCKNTVMLVSNM
EKTALQVNQTIINGTANLKQHAIELQTAIVSVQKCFTAYYSTLTSAEEELLKMSTATSALKEQLQFNFQTVVESYKTELGHL
NTNVTWLEKLWLAYSKNESLKLDLARQIDEGRMKMINTDATDITGKIEKTAVGFLSDRIGKLSEFTANNYADIIKIMAGL
STYYNDTRWEYFARQLQIWQVPIADIDTPYILRTTVKKQEIWRTWPRTTNLAAMNASAKSNVDRVLKVYILGLMSELYE
AEVKLYSAKTKIEVALQNLVKHLQEYKNFLQINKDFILLNFIQMDVYFAVLKYEHVSRQVAYEIVNFLSEIGGYMGLLIGGS
CLTLCEILDLITFNGCKKAKNGCKKSRPKVEAWEEKNGKIIA

>Deu_Ambulacrar_hemi_Ptyfla_40v0_9_20150316_1g19295_t1_scaffold17970_cov134

MQDPTIERDCMIPILGRYVALRNSPNRDDFLQICEVKVYAKEFLRDNFKDIMKRRMFSMFDIQVDRSLSVNSTDILSSIPL
YKCFESCLRWKRYSCRSFDYNRATRTCRLYKESAGDNGGGVVKAIGTDFYQNLRTRNPDTVFKPGWQEPYQSIIDDPDI
YKYFKDNVFINPGYQRVKGEDPPDWLRFKSFSKSPDYTELADVLKVNTDEIAAMGHQAEDFILQCTYGGEVCHPRSDFV
ITQDEVYGNCFHFNADPNKDTRFTRGTGTAHGLSLVLFTEQYEYLGIYGQDSATVVSIVPRNLRPFPVDHGFFAMPGTA
TSVSLTESKIHRQNQPYGNCTDETSNLRFTDNVKKDSSDKMYSIVSCEKACFRKVLLDYCGCTDSVNLNAPSCRILNDTQ
GQKITDENQAIQFLREVSDQAEIVRYHSVMANWIYNTNITDYNEQKMLEASSVLAEFDKEARHNASLFDISGFDEDTIR
QFEFLLYIGSAALSSEDKLEYDTIISKMKRTYSTAKVCGRPGHNPSECFSLEPDLTRILADSRDYDELLWCWEGWRDEAGK
PTRDDYTQFVELANKAAKANDEKFDDFGDYLRYSYDVDDLQADVLRLYAQLEPLYKKLHAYVRRRLYQHYGDDKIDLKG
PIPAHLLGNMWAQQWNNILDLVEPFPGKPTVDVTDRLQELVNISSLF

>Protos_Platyhelmin_Macrostomum_lignano_A0A267DAJ5_A0A267DAJ5_9PLAT_Uncharacterized_protein_OS_Macrostomum_lignano_OX_282301_GN_BOX15_Mlig015178g2_PE_3_SV_1

MSNQTQQLKQGRSSGGGDHSSPCVPGSTSFSDCLQPRSPTSAQQIGQMVLAFSKRSTIHGLNHIAQARSARRRIFWM
LIFSAASCGFLTNLYHIAVKYIQVPVLTNVKHDHLSFKFPDITVCNINPIFFPPKNSQHYEDLLRVIKQYDQDKKIYKQRAAK
LKNWKRGVFTSVINGVYISDYLYMQRNITLYSIPHEIFVQDCRYMGTHKDSCSSSWITHRRWPYGACYQLNITKLGRDAI
SRSNKKAFSSGFRPDLFLLLYKGVEYPDEFKYLDAHGLIGMPSGVLLMLHEPGTYPQIGESFLLDKYAQVDLHMTRKQHL
RKLKLCESHRDIYDYVDVETGERRFYKETRTDCVENITQTAIKQQCKCVGHEYPLKYDDEKYPFCRDFKWFDQHFNDSD
CANRMHRQVKEANPLKMCYKDLCEHTTYSTVISQANFPAVIERKTHRGWLRLLSMMERDELKKYGTTDLASKALKLDS
TTEPMNISEVLTNGQFELLNVDFVERNFMYLRIVPSSLFADEMLEDEEYPLSRFLTDVGGCVGLWVGASLITVFEFCDLLL
DFLGLDAIFRRKRSVRQNGHYYGQLRRALSPHEEVTRLVEYNRRTPLPPPPSPQMTRNSPADINSAKGGQDDEAGSNS
VATAAAAAATENRLNSGITMSASTLSCSSAGGDPVKRPRAVQIGMVSSV

>Protos_Platyhelmin_Macrostomum_lignano_A0A267EVJ3_A0A267EVJ3_9PLAT_Uncharacterized_protein_OS_Macrostomum_lignano_OX_282301_GN_BOX15_Mlig018669g1_PE_3_SV_1

MPPNLKAAVSVLHDYGQRSTLHGLSHMHQSGDPWRRAFWVILTFGGCAACFYHTYATISNYLESPITSIFRDEGVSATF
PDVTICNLKPVSETLLHTPSERSLLFQHLENNWNFFRNWLPEKRKRRFNETRVDYKLRQRDLFKTYMDVFWATDDTRD

FGQSDLHMLLSCVYKGRQCNSDNLKLIQHNRYWNCYTFLPTNGDRSIDSSDQNSGLKLILFTDSNFDDDRLNQQTLYER
KKSADDEMHRPYDLEEYLSVGDRTMNQVLTGNSWEAEGLRIFVHREGQHPMTTGLAYDISSGASTTIEFSEVQVNALN
IPPEHVCQEKTPDEILYVRFFGDNSTGVGFKNYSLTQSLAVIEAKQRMVYETCSCYSHLLPFRHNTSTLCMFAPHGERHE
PSRALLDRMNCHDAAYQEAESRTADLMQTMEGYTPCNNWKLHATKTEVSWPRNDELIDFVNYIVLPRFTEPKEYKSS
RLNHLLGSSNSSRGQSVLKNLFNATCKPGGLNSSLEDAWCLSKHKLITNVARVWIKPDSPRAAVYMEDYSYDWPEALS
EIGGIFGLWMGISLLSVCEFVELVFILFKACSTKDSDDRRLMRLPGVCSGGAGAASGLTAGELENLLRQVLVNQSLEEEQ
ELANFWAARGFKQQQSLDESAELQTTAAEYRNSPTYRRPMQFSPPQQQQQQQGKQQQN

>Protostome_Lophotroco_annelid_Pdum_Contig1928_

MPLSLSVGVPSCDITKYLTCVHPILETHKFSLTSCECPVSCDEAIYDPVLTHARISNYDLDDFLRKSDAKAIEERFVEAREAS
ERIMAPLVNDNTQQIVDMAYALTEVTIAMEKVENQLTLARTFCANLTSEMTPRTDFHLRTSLDGVKYVVHNNFVRGW
EVIDERTMQHVTTDFYEIINAYDRTLSQLRTLSDSDPDTRKILFLMMDKDLSAKLELAIRSIDNITSAYYNFLNASPMFSYQ
YTPGRRYDRALVPVDLLQNLTTEHKSYYENISTHLTGYMDCILTLKATLKQVYKTGNLRNQTLVTIAKEQFIHHSKLINYFR
FLFSSRIVLRSEEKVDETIRNFQKSDYLFMKAVSDVQGLVTGLHEQLSVAYNNMTGKLASLSSTATRYLHNSTLTKSGLSK
WIQSREIQDIVHSVKVLFRQIRSKANDLRDAWLRMRDAYIAIWRSILLEESTREFYRKLHQDVFDMGREPMRRDFYRRV
FSDPVMLNLNDSTSATMTIDDFVLRMNADMETLNPSVKLADINTLFLTVMQKTNLEATMGNTDYKFLAAFNDLEKSLK
MFSKGNKIDAKFLRENFLRLDLFMRELSYEEFEQHCAYDLVKLLGDIGGCMALFIGASLLSLIEVFDICCHTACLRRHHDR
KRKEAAPSADTSHRKKVTFDEKNLKNHKNPNGRTYIRYTQDTDV

>Deutero_Cephalo_088140F_t1__name__chr_scaffold187_start_416588_end_437049_strand_pro_len_689_

MHVKLTCSVELDCCPCCDCGQCCCGSSTAGSEPRDAGSELGDGTDAGEQTYYRKQAQTISDFAAGSTLHGLPHIFPDA
PLSIRQVAWALAFIGSLSVLLYQCSDRVKFYFQYPHITKLDMVLAERLDFPAITICNMNMFRWKQFTQNDLWHMGKGL
NILDENNNLRCSEYATPGDMEALKNKANFDSFKPSPFSIMEFANRTGHQIESFLLDCKWKNFTCGPEYFTPYRGLLWSIT
TVGSQNGCLSGYLCVYLSVYLSGYLSGYQSGYLCVYLSVYLSVYLSGTRHSGSIPASSAFPGASLALGKKRSAADVTSSRQI
PDEHVDTVDVFTRYGKCYTFNSGSPNQPVLKTLKGGIGNGVEFFLDVQQEDYMPAWGESDEVTFEVGFKIQLHTQEEP
PFIHELGFGVGPGMQYYVSTQEQRLNSDLLLSAEPTLLLPPPMCKAGRNQVELLFGWTKYHETITYLPAPWGQCKAEN
DLTFDFAKYTTSACRIDCETKFVVSQCGCKMVHMPGNFPICTPDIYVECADQALDFLVKSDNKKCVCDTPCNTTRYNLF
MSHVKFPSEQAVKYLARKYGKPEDYFRGVGLKYGKPEDYFRKNTLVMNIFFEALNYETIEQQKAYEVASLLGDIGGQMG
LFIGASILTILELFDYLYEVLKDKCTERRHQPRSESNAVSVNLEDCKRDNSRAPLS

>Protos_Platyhelmin_Macrostomum_lignano_A0A267F391_A0A267F391_9PLAT_Uncharacterized_protein_OS_Macrostomum_lignano_OX_282301_GN_BOX15_Mlig006075g1_PE_3_SV_1

MATASQAPRTKQEQQQQQQRREQERRNTLRFIINEFNDYGHRTTVHGLGHMYQSTDTRRKLFWLCITLVSCGACAV
HIYFIVANYMETPVNSVILQGRIRQEFPDVTFCNMYPISESVQYHAAKEIHGHISNRWKYFSGFIRAGNFSANDKVGRLK
VARTFMQIFWASEDTRDLAHDDDLFIVQCSYKNRQCSNKQFKMVQNLRYWNCYTFAPKFDGGHEDRQVYSSNEDEG
LSLILYTNSHLRNVHPKNGLTTIRDVYYDYEDEILNHLGDQSNFLERYLENYNLSLNQLLAGDKLEPDGMRVILHEKDTYP
HMDMNSVIVGNGDFSNINFYMERKLSKNRPEHPCRVETSGTEDRMEYMAYYPGSNIRYPGPRPPPNFKKFKRARLDYI
VDKSQALFWSACHCYTHIMPFQCFNTELCYHCRDGTDPAEIERTLRHIWCHDRVYEHRLKHKIFDYFRDSQNFQPCSV
NNYNVKMGSSAWPSQRDIQFLINQYVLPAFQKETVGARSYNHILKNTSVAPIHDIISQESSGHGVNWTAAEFLVRKNFL
KVAIHADSPTGTLHEEVESYSFAEALSEIGGIFGLYAGVSILSFFEFVELVYLMSTKPKEARPLRCQQHQLPGSLDRRRSPLR
PSSSSMMHAAASIFLKEAAESGELAPDELDGCRHLLGLPAADAPAAAADAQEAATLA

>Deu_Ambulacrar_hemi_Ptyfla_40v0_9_20150316_1g2128_t1_scaffold965_cov108

MTNSFEEIDKFSHISPDIVPNQRLDVIVTHVSPSIIMVTLDIHSDYDHGHTALPSMYTLDIEPDIGDAFPINITVDKESETEY
TFSENELGCFIGRSCQDLLNPCDGNPCGEHGLCFRDSQSCFQYACVCNECHAGKNCQIHVNDPCQFFSPCRNGGLCIPS
PDSCMEYTCECSQCFTGKFCEQVPCEPNDFQCNSGIHCVKKYLVCDGIYHCPDTSDEFGCDYHSYECSRNQIKCETGGP
SGICIDDSKQCDGVEDCYQGGDEANCDTYGEKNTHICTGFQCDDGLCHPLEDRCNYFTDCNDATDEEDCKFRDCQPW
EFSCDNHRCIDADSACDWKDDCGDNSDEVACDYRECTESEFKCVSNSQCIAGWKRCNSYSECEDQSDELECTTPGNS
HPGREMLFRGNWTNIYANVTTNQTYFDDFQYHYFTDPGFTRVKREKPPDLNGFVTFSSTPDFSDVRRVLKLTADEIEEY
GHQAKDFILQXLKLTLFVEQDEYLGVFGQSAGVRVTVHPNNQLPWPEDVGMTAKTGAATSFAIKQTCKKRCVQDYMI
RYCRCTDTFDSDHAQCPILNILQEACRQIIHYFYQKALLACDCPPLCKKNLALVSVYYETLSSDLVKESPGYGGEDLTSDLG
GLLGLYIGVSVITTIECVIFVKGVIVIAFRSMRDNNKDSRDITKDEIDESASDDSR

>Ctenoph_Hormiphora_californiensis_evg1166627__p1_GENE_evg1166627_evg1166627__p1__ORF_type_complete_len_698___score_158_10_evg1166627__48_2141_

MQPNGRPKDYVFTDIDQPLSANGFYKRIPAPPRPYRPTPAFPIYDNLLDREDVMSSNCPSTIPGVYPGVEAFTPYSKYKQ
DESAFDMTGRGERTSYYPYLEHYQPSYPRPARPKTTANPTTTTANHRAEPNGKIKYPPNHHNRDTDSSGSPTEMDDRD
GEECEEFMPPPKRKKKKKRRANRSLENRKNLRLRRRKKAALEQRREDMEPDEMESFKEFASTVTADYLAEMFAKASNG
RRAIWVTLWLASFVYAWYNIALSIGVYLSKPTATKLNFYEAPPNGVAFPTVTICNFNKYNKSFFEREGDSVNAQLRDFLV
ETTPIWDHKERLAKDQWLSKYPSIMNLELKQVLTAATHRLEDTVEYCRFGGKPCLDSQYRETSIPSEAFTKIYTEHGLCYS
FNQDGKLNMTRTGAQYGLSLRLHVAQDDYFKTQDVAGFKVLLHNWYEPPMIEEYGFALRPGSESYVRIRLQKFRDTER
PLGYCDPLLEQDYGPGSYTIAKCVMANRASLTIKKCGCIPFYMDVARSAFAVPPKYPSSTPVCNLYQEWDCVEPLLEDIA
LGTLVIDGEPIGDPDCPSPCNYLTYSYTISQSEYPSQSVEQEILDNVLGARGKSWNIEKIRDNYLKVHIYFEELSTLEMERH
PSYLISNLIGDVGGQLGLLLGMNICSLVQFTDYIIRFSFFNGIMKLVRISQRNSKRRAKL

>Protostome_Lophotroco_brch_Lingula_anatina_comp148540_c0_seq4_p1_comp148540_c0__comp148540_c0_seq4_p1__ORF_type_complete_len_699___score_91_82_comp148540_c0_seq4_170_2266_

MQQDKYEQEQRTTSQRSLLNSQKNGGTGFDSEQSENGKYSSCSKTASFEGGGLFVRQAQDQLPSAAEQQDETSMLLL
CKNWGDNSPINGVPNIARSTSGVGKVTWTLLLLMCFGVMGYQTWELVAKFYRFPVDTTLTYTHSKEVYFPAVTVCNV
NPLRRSMLSSAGDDVYALLGPPKSQMGSSPGGAQAPAGPAGSFASQSQTTQAAAGPPGPASSSITTQAPAAPGPKGS
GSGGTTQAPSASQTTAGAPAGGGGGTTQAPAGPPAPPSGRKRRSTGTNSTSTSRRKPGQMNERFEMRKRFGNIWAK
LNYTTRETVGHELDTMIISCTFNGVTCSSANFTRFNNHQFGNCYTFNSGWDSSIPVETSSNAGPLYGLSLEMYIEQSEYI
GDLSESAGVRVQIHSQRSMAFPEDEGFNIAPGYLTSMAMTRVEITRRPHPYPSKCRNFTTEESKQKSVFTSVHNVDYSV
SGCMKTCYQRYVISECACGDPAYPFSLDLEAFSDLKNNINGTTAESVDPCVSDADDECVAGIKRRFADDDLSCDCPLTCV
DVEYSGVPSLAKWPSKQYMSTLHATLASAGAHLDNIINGKNGMAAEPGDNLLKLEVYFQELNFQKISENVAYDVFALL
ADIGGQIGFWVGLSIMALFEVVELILDVFRLLLFRVGSFCFKKQQKRPPSRQVKVQPASEENVLTPRLEKVRFS

>sp_P37089_SCNNA_RAT_Amiloride_sensitive_sodium_channel_subunit_alpha_OS_Rattus_norvegicus_OX_10116_GN_Scnn1a_PE_1_SV_2

MLDHTRAPELNIDLDLHASNSPKGSMKGNQFKEQDPCPPQPMQGLGKGDKREEQGLGPEPSAPRQPTEEEEALIEFH
RSYRELFQFFCNNTTIHGAIRLVCSKHNRMKTAFWAVLWLCTFGMMYWQFALLFEEYLSYPVSLNINLNSDKLVFPAVT
VCTLNPYRYTEIKEELEELDRITEQTLFDLYKYNSSYTRQAGARRRSSRDLLGAFPHPLQRLRTPPPPYSGRTARSGSSSVR
DNNPQVDRKDWKIGFQLCNQNKSDCFYQTYSSGVDAVREWYRFHYINILSRLSDTSPALEEEALGNFIFTCRFNQAPCN
QANYSKFHHPMYGNCYTFNDKNNSNLWMSSMPGVNNGLSLTLRTEQNDFIPLLSTVTGARVMVHGQDEPAFMDD
GGFNLRPGVETSISMRKEALDSLGGNYGDCTENGSDVPVKNLYPSKYTQQVCIHSCFQENMIKKCGCAYIFYPKPKGVE
FCDYRKQSSWGYCYYKLQGAFSLDSLGCFSKCRKPCSVINYKLSAGYSRWPSVKSQDWIFEMLSLQNNYTINNKRNGV

AKLNIFFKELNYKTNSESPSVTMVSLLSNLGSQWSLWFGSSVLSVVEMAELIFDLLVITLLMLLRRFRSRYWSPGRGARG
AREVASTPASSFPSRFCPHPTSPPPSLPQQGMTPPLALTAPPPAYATLGPSAPPLDSAAPDCSACALAAL

>Deutero_Ambulac_Spurpu_018442

MDKSRRPGASYQSLCYRASPIIAKENSFFPPEYDDIHPMKTDSVKAVLNEYSGVTTAHGVPRIITSKSILSKLFWACVTLAA
LGAFLWQGSLLLFDYKGHPYTTQIDVVTQTEVQFPAVTVCNMNKMRRSAMVGTRFESLIEADGGAMGGDVDYSWW
FDWSSEWWLKYEDSSSSREESSKEDIGPSAEGSVISGTSQSVVDMSGDSATDGSSFNTMANFTDIPDGNSSTYPSSTPQ
NQEYSTQSDVDTASATSDGTESFVDEEWSSTDEVPPLRKRRSGLRLIPDPTTGKLDQRRFVRRKRQTNTGPSSSDPSPS
DSSEDPERFDWWEPEWETNMFDYQAYDWGDVTSDNDWEGFYRQSTADDFSDLLDVINPTREELEIMGHQAEDFIL
QCTFDRRPCNYTNFRQFQNKYYGNCFTFNQEVGNSSNARRTGQTGAQYGLHLTLFTEQPEYVGLFAQEAGVRVAIHP
PNVFPFPEDDGVVASTGQATNIGIRQSYFERLTKPHGNCSDGTQTNFTSEEYTYTTRACVKSCVQQHIFNKCGCVTDIM
MNDTICSPRNRTEQMCRQAIEQFFHEGQLECFCPIACNDLNSLDHVITFQTHLKSRLTNEKATRILQNIEQTRKNLARVR
IYFEELNYEQMIQKPQYTFESLLGGIGGLLGLYIGFSVITICEVGVLVLDIVKYLFRKAYSHDRVVPVDFKT

>Protos_Platyhelmin_Macrostomum_lignano_A0A267E2M2_A0A267E2M2_9PLAT_Uncharacterized_protein_OS_Macrostomum_lignano_OX_282301_GN_BOX15_Mlig015081g1_PE_3_SV_1

METPCQAAGPNGDQQRQQLQQQKQQDDTGSIASYSQVPRIKAIRTRDIFRALRTFAANTTAHGFYHLNHPVRYFKIG
WIVLIIAAFVGIIVHLSILVVKYRDEPTSTKINPRGSVYAPPDITVCNRMMFAVSKLFEKSGDNYTVNPEFQPVLDSVLKVR
EFLEVSVKKYLRLSYGNYESDSDLLRIEDEGVYNLPNLAIRDTYGAPFLSRFSAEPRQMLLFCTYSSIACDFLDFKEFGLSTFP
SCMTFSPSNFLSRRPNISVNSLGELRLVLFTDSYHPSQLVTDTLQLLPGDWFNMQGQGGAVVFVHQPSTYPTLGSRIEA
PASRETIVDIRYQVYTSKNTRNYRCSESDESVQYLAIVNRTTQPERYQTQLADCYVDVTQKILLQKCGFIDPSLPLSLALSD
RPRRGNLSNLLINSPSGRPLQPTDLITFIYQVFKRANVSTAAELPAILDTLLDQEICLRFARVIASNDSQNQLCLDSCQAKV
YSTSVMPNLWPVGSVEFLQPQVQRWLQSNVFYLNGANKANLSVNEEFIRRTNWNLYDAYKRLNGSGANSNNFLALR
FRIGDSQGLGVSTDESAAYELSDLFSDIGGILGLYMGISVLTLCEMAELCLHIVTMLGMLLHRNRTRAAALPRSVAQMQ
ARKKSVARVAARTNSGFDLEPDEHQLQNHHQHQHQHQHQHQHWQRPQYPVQMPARISRV

>Protostome_Lophotroco_annelid_Pdum_comp410400_c0_seq10_

PLQVTQTGSHNGLSVRLNVEQYEYMRGPNSGAGVKILVHGRSDIPQMRDLGQVIYPGAHAFLGIQLKQMTNLEPPFG
ECTTKDLQFFEIYSQGACRMDCLESFLSKTCNCRLYHMPEGRNTTDDTDNPEVCSMDQFINCVPYALDKFHTENMKFC
VCPVPCKKVIYEASVTYAATSQFGTDQVLQNVDTAYLKIKYIAAQEVAQRVDKEIARRDKALVDNLVAASKRVDDSLEE
MGRRLAETNHYKEAIDSEMYHRSIFHKHAMHRIIQAYQYWFAQGWELTSEKIDELIGGNAFFIRIYNNTMRALAEKNR
RESFRGILKFKIVHTTSDRFNLAQEVMAMIQNVQNALITLKPIDDYTINFPQKYLTYLLPAIAISQGLNQSAEIYNSLLGFTE
YIVGNMSLLTEETKQVDVSRSNDFDIDSFNNVSSLLSEDLESINHWQEQYTKEVIDFIGNILRKSENGFLSIMDELGEQSN
DLLNTLKKTDSELNDIHDDWKAVKGFIAESEEYVSSRGKGKLEMSLASNKIYREVENMRTFFTNMRSRAQNIEHDWSK
VIDMYRIAMTHIATDNATKKLYHRLHHDVSQMVAISDQEQKTTIWDNLPQPPPHGTKVWLGKDVWQYMKYSNADT
ALMTKQHVELMVQLLDEQVNNSAIYQQGSNFFAAFDDWRAVLDKFNASTNVDSNFFKENFIDMDIYFTELY

>Protos_Platyhelmin_Macrostomum_lignano_A0A267EHR1_A0A267EHR1_9PLAT_Uncharacterized_protein__Fragment__OS_Macrostomum_lignano_OX_282301_GN_BOX15_Mlig022967g1_PE_3_SV_1

IPIPEKIGQSHQLSNNIPNVSAMDSETESVGIISNRLQKYEMIESTHCLLPDASQPPSQQSNGDGPARDGQAHLQQQFQ
RDAEGLATAVASPAAFQYPNPWVRRSHPTRPPSASQSEGQQLTVAAGGTPSLFEIVRQNFISYCQTTSLRGVPKIVKTR
DRLLRHIWILFTCTFFLGCFTCLALIIKQYLAYDVIHQPRTVRNHPVQFPSFTVCNTRPFSQAGIQYILDRGLVLPKNYSQFV
MQAMRNTAENQRGVGFNDTTKEALSTIFETKGYFENVPRGFDIARLGHTIDQTILICETTILDKDFKRSKFCETIGTWTRT
YDNKFLNCYTFTADANNTQKILTVRIMAYTNERNLTNCPDCSSFPATQVSGLRVQLHNAGNYPEVQEEGLNIRPGSLTEI

RFETKEWVMKEPPHGRCSTRVPPSVLFNGDAFTYSYQACKKKLLHEEIMKKCGCLDSTIPLPDSMRNTGLDFCGRIPPS
QFSLQSEYIQRHNHSTPHDDNDDQRKRPSFHFDADFNDFLRRLTCNTAVFKVSGHATAHQCFTPCTYYTYQTTISTTN
WPTRAYHYEIVNDFKNLRDRIDNLHRRGKISRNSSYLDPVFQRLQRIYGPLSEADSPAEAERMLRGIDYIERNFIEILINRG
KSNFDIERIEEKAVISLTSLLSQIGGLLSIWVGLTFVCIVEIVDLLYNICVSYNGSRQAA

>Protos_Platyhelmin_Macrostomum_lignano_A0A267DZS2_A0A267DZS2_9PLAT_Uncharacterized_protein_OS_Macrostomum_lignano_OX_282301_GN_BOX15_Mlig031609g1_PE_3_SV_1

MKKSAVLASAAKVGKPPSQRLDSYNRVPGIQTIRASDLISTLRVFAACTSAHGMARLNHPVKSCQIGWALVITAAFVAL
TVNMSSLVTKYRDEPITTKINARGNKYIPPDITICNRMMFAISRIFNSTLAANASAPDQTEFRHDIAPEFRPVFDSVQKIR
QVLGLSARRYLMLNYGAYETDSNLLQIEDESAYNLPNLVIRDTWGDRFLSQFRADPSQGLLSCTYGTRACDINDFQLRSL
PAFPSCATFSVQRFSQRRPNVTKNSLNGLRIGMFVDSHHSSPAVTDALLKLPGDWFNMQKPGGAVVFVHQPGTYPT
QENQIEAPVNQETVVSIRYLEYTSKSTRKYRCMSATDESNAQVQYMVQKDQSPQPETFQAQLADCLVTLTQEVLLERC
GFIDPNLPLTVSMFSQVRRGNLSHLLSGIGLARRDPASLIDYVYQTFLGSNASELPAIIDTLLDNEICLRRALVAARNDSRM
HQRCINRCQAKIYETSVMPIAWPEGPEIGDFLGPMLQRWLHSNSYYPNNGSIVDEAFLQHSNWNLYQSYVLISSSRELR
NANSSLFGRIANKNFLAIRFDVADAQGRDIKTDESFAYELYDLLSDMGGILGLYIGISVLTVCEVAELLLQLVVMLNMIAH
RRRMGNRTEQSDNNAERIRRVSIMSATAAAAMAKGSRRHSEIEGRLRPSVSVGASQIWQQLPVRVTRV

>Protostome_Lophotroco_annelid_Pdum_comp418207_c0_seq2_

AKEGAPCPWKPELSITGIKYILEKKTSKFHRLIWLLVLLASLACMLIVIGIRIQRITSNPKSVNVIISSNNSVPFPRITICNKNL
MRSSYIYQYMKYLAKTSTSEDMADKKPYSESPFAIPSQFLVLQMVSVFYTEFYRNPGEPRMTIKEFERHFEAILDGPEDFL
EFYRDAAHDIKNILVECSWSGKPCSSKDFRGTTTDYGYCYTFNHNISNPLMTHTQGPNGGLRLVLNAETYDYTPSWDM
SAGFLVSIKHQDDSIEFIKDVGLQTSTGMQTYMAIKSTEFKEIYEKNDRKCSINETLNECLLRCQSAYIIEKCGCKDMYMY
QNDTKEVKTPICSPLKMVACSLKNLDGMVLLPNYTSICPCLPACQRVSYDVTVSQSAFSHKGAMVYALRGEKLTERQKE
LVRLQDLGYLDIFNRSIATQVDHEINTQAFYMHYEFLGTYQQCLQGLSDTVDFMTTAGLDTTCLAQKVTDLAKSLQNYQ
EWLANDHFTSHTYGKNVDDFLVLVVDDILREKSKFPGWLDNFSRQISTMNSSLTNFTICTNNTLPGVITAVTGCQQAGY
PLVETVETIRAAYFSIIQELRRQELFSRLPKPQLVKDANWYRENYAFLDIFLEDRSVRGVKQNKDDSFNSFICDIGGSIGVFL
GGSFLTLIELAEVIAWTCCRGKKDRKIKGSAEANFQNADYITPITQVHLPKVHLPKVQSS

>Protostome_Lophotroco_annelid_Pdum_comp420350_c0_seq1_

YIPVKATILKNEKPPVGKCGSQQLQFSDGYSWFRCVRECETVQTIKTAGCRDIYMAPHATGIPGFCNIDNYTRYLVDEQA
ETLYDGCKCIEPCERTTFEPSISSIGMKASKLESLLTTNMDKFIEKTRKVKNLRFRVDSEYFQLLAGSVMDVVIGYIDFMFQ
TKNLTDSFMKEAEASRNLATAIYTQVVEHDLNEVLFGGLTEAKEVVNALMGPLKTSLLSLLERINVQANILVMLVWQST
NLNMTSLHPKIIEKSTEVSQLLNDITYAMEIYPAAAYNITFLTEYFQWNVSDLKFFPNSTVQYSACQDTFMMYQNTSTA
MQEDIKKMTELINGTVATEAAVQKASAQSMRAIYSFQKFQKAFTANCLDTFDVSVNAYIEKKENIQEELVRRLRLISVSF
YYEIEIKPIIVAQERILKYGAWVLTWANVTVYNMTHAFGTETTTTMLSQLDDFISKVKTHTSGALKTAFFDYQNSVANAY
VEMMTELCNFKGYTSDAFIESKLSQLTILKGLKGSIEDKNMTFIRETPDSVLQSMFSQICKNKSSIPDINAQIKKLTEGAAT
GLKTDIFSFELNIVQQGSSLQDKIKRMRDMINLLSFSYFSTDENFVGDNFGSLDIFFSELRVQYINQSVAYTISDALSDVGG
FMGLLLGASVITVVELVDLFVYNSVRKISYSRKKRSPVTSPVKSMSNFDDREKKHSHLSSSFHS

>Deutero_Cephalo_109370F_t1__name__chr_scaffold21_start_3101811_end_3109315_strand_pro_len_710_

MPRTTDNKVASALLEFADTTTTHGVPRAVGSSSLLRKICWTVAFVASLGYFLYQASMLFNKYFDYPVATDISIKFATIEFP
AVTICNLNPVRLSKLNTAGGEFSSYIISDVGTTAPPSPGPRRKRGVDTSQDNRPEYDVDWEDPYPPREESAEEHELLRER
RETGSSAGSADSQGSSDSSDSSDSSDSSDSSDSSDSSDSSDSSDSSDSSDYASYMYGGTWSSHYGQDGSSSSDYMSEDY

HHEFELVQNFSSSILGLNRTSRRTMGHQYQDLVLECAYDGRSCSRTDFGRILDEKYGNCYTFNSDKVLQRKVRDPGPAH
GLQLTLYIEQDEYVPAVTPAAGVRVVIHQPGEWPFPAEEGFDVGPGYSTSIGLQVTTIRRLGGKYGNCTDGRDKDNLYR
SKYSTKTCLHTCFQRLLVEKCGCGSRFIPLPKKVPACPVPINSYEKSLRQKLSKLDEESQEKLQTSHDARRNLLKLSVYFEQL
NQQTISESPAYRVENLLGDLGGQLGLWVGVSVMTILEIVAANTVCSAENQGLTCLNGVPTGFDQSVTGISLNRLFNLTT
LTRHHIPPLPNLATLVIANSTIQAIGKGAFSSVPAITVIAIRCSRLRHIGDFSFHSLPALSAIYLDHNIIGTLSPKAFTGLISLKRI
DLNSNQLKAVPYEVVFMISRSQTVANILVNLIDNQISTVLEERWNKIPASDVSFLLLPSLCL

>Proto_ecdy_PANarthopod_Tcas_tr_D6W9K7_D6W9K7_TRICA_Uncharacterized_protein_OS_Tribolium_castaneum_OX_7070_GN_TcasGA2_TC000562_PE_3_SV_2

MSAVCSEVPFNVRKLLQVWTVLLTCSPVMAGYTWTNGVKSAPDHNAGSSPVFDDFTIRELEATEGQTLLLPCTVRYLG
DKVVSWIRSRDLHILTSGAATFSSDSRFEAVPSPGGDFWGLRIRGVHLSDSGQYECQVNTDPKMSLAFNLSGARKLEAS
KWLGQALIKGPREVYVRDGSLVTFTCEISPLSTPQSGGFMSAKPKVRWLHDNKELLFEPFRSHISTKTFFEESSQKVIGILK
ISKASWRDSGQYTCLQMSVKSDSIKMFVVESEHSEAMQRDYPVQSCASKCLLHLGLVLLFTLFLWIGLVIGAFSVATYQA
LQAWNRYDTSPVVITIEKDFFNWETEFPAVFFCARAKLTTAYKFLKIFRNKYNISASVEQLEGYVDDLLFTTYENYSSLEKY
ANGSQDDFVPTDLYLELVNFTSFRIPKNSNKTKIKYSEWPKNVNGEYSFVLTEFGFCQCFHSKVAGFFNIDNFTIDGATN
TTFLPITKKYKEPALTLSQSDFSQTNFLFVIGTNEIIDITSQYFVVTPQTLGDVFLDIVDIVGDEDIKMLSPKQRNCRYWNE
ADDLGHSRVYSYNLCRRECRIQLCLKFCYCTPYYYKRIDNEPICNMSGMFCLSKYPEILTLLDPKTGEDLCPCFQSCDFQRI
TAELTIKATWFFGSKVTVNLERFPRERELEKNPLHNFARPSINELTKPLKVASPEKIDQTVNKPK

>Protos_Platyhelmin_Macrostomum_lignano_A0A267F762_A0A267F762_9PLAT_Uncharacterized_protein_OS_Macrostomum_lignano_OX_282301_GN_BOX15_Mlig020801g1_PE_3_SV_1

MTSEEEDEEMSEPYHGLFQSGQQQQQQQQQQQPECIQLEQPSANGAAMLASRRKPTTKSQWSYLAHRLTVYGRTT
SMPGVTRLTEASSTASRFYWCLLGLMASSFCIYHVVWLIYAFETSPVSSVLRQGATPFIFPDVTVCSMTPTDATLVATHL
NANSKLHYFYSNLIIPPRWMQLPKIREVLNSSETRPDVRQKIMRYLQLQIPSVPKFMMNVSTSERNSTAEYKTESLRQHF
KMLRGTFNMYAAGYTLDSLIIRCKFEGRNCKKSEMRLVVDNHLWNCYTMRPTERNVTSMEGLDLFILTHQQHYTESES
QSSADTNCSFAHKEYDWATLKISSNWSTSNFWKQENFKGLITNALLPRYDSNGIRVAIHEENTFPHEGQSSFTIDSGKQ
ANMEIRMLELRKLDNPKSSCVKNRKNIKYRQASMNDLENFNDTPFLSTPEDDIVHEYQRLLLGKCGCYSHEVPVRSDLP
ASRLCFNVSTTLQVMLDVLNRKFNWTIRYDRLETYGDLEKFNSIEQLLNSTFPVLLRSHQFLDAAIATSEGMLCYETHSR
QMWRLARQMTEYPPNCYYREYVTSYSVTDWPRLKDIPYLVEDEILTNREDNCMVKKNLFVHYENRTSWVTTAKRVRE
NVIRVSVRPVSLSSDVYFEEQSYSVWQVVGELGGVLGLWSGLTFTGICELLELLAEVSSSLLLIFMQKLRQGLTGGVSDGL
W

>Protos_Platyhelmin_schman_A0A3Q0KNT0_A0A3Q0KNT0_SCHMA_Putative_fmrfamide_gated_Na__channel_OS_Schistosoma_mansoni_OX_6183_PE_3_SV_1

MIYPVKENCLSGYNESVKCIHCNQSEHESSCITCSTRTHQNLSIPNDNFISTRRNEPKGKLELFCVLSSIRGVQRIYSSQIPFI
RILWILFVIIMTCVLMVSSGFLISDYLCYSTAWNTRSLLDDKSEFPAITLCAHNPFSLNVNGLWKENIVPSPRQIKSRLRQL
LVTMLDNGITDYTGDLAFSDSIEFYYTNLNLDDTIKVGHDKDMLYHCMLYEEGSTVVAEKCDLELETSVRIRRFTHPKYF
NCWTIDGKPNASTSDVTRLIILAHLIPLKKIEGANTSFIMDSFTRGEGIKVVIHEKGTYPEIEKHGVNIQPGRMNEINYETLF
WKRLHTPVAPCTNENISIEDLGVHYTYKQSQCLNRMLQNELYKRCGCLNSEWPRSIKLLDQQKFLPYCQKLPINVNDIN
GAEEMKTRLDCFGTLLNDLKQLKEKAVQQGVCIPRCGYYTYDTKLSVTSWDPDPYKLALYTKGYRHVDKFLNEPNQRK
HIDELLENFDASIDPIKSKSQLTTIRTLFSNDYQRFDDGTHTYISLIRQDFDTIMKEERLVFDIYILISRIGGLCSLCIGLTMAFI
VELVEFIYLLCCKNNMKLTKSNKNLKLMKTNRTQQGSYEKEEMEHNRYNVNEFKLSAIPNNHHCHYHHHQYQQQQC
QKFWQIHCDNNNNSNNNNSNDNNNNNEEISTNEIIPNCFTLVNNNENDNQSIMTNITQTPNEN

>Protostome_Lophotroco_annelid_CAC9509085_1__Ofus_G024052_Owenia__fusiformis_

MAKQVECKKNVIALFKATLHTSPKITQQSVLTDGYVIIVQNKDDENQIAIEMKEMIKCDKDTFAPSVKQDTKCIGNGYV
KADSSIDSYTLYDLENVVSVITDVQGDKEGSINPKVLNEQTDDSSQNGTQTVKKSLKVQIRSALEEFSNTTSIHGPKRILKA
NGKYSTVGWTFVFIGVIILTVTLVTGVVVKYYSYPVEDGISITTPPLGFPAVTLCPLIPFDIDQILVDLENEAIVTPDHVYDLI
RYFFLRSEHFTPDIECISDVNCEVEADDNILQSWLELSSWLHRKLNPTYRWQVENIPLNETIVSSMLPTKTEEFILECTYNE
RQCDEQIEVTRILTAYGKCFTLNIKNQAEHINEIGPNKGITLILYTGKHNPIQGIIGAEFSTMGFTSTYSQPDDGVQVVIHN
PGTMPQPFREGFHVTPGRFSSIKLTKTERTRLLPPYGECTDENYLFNSEFKYSYEMCVEQCLQERIMQECGCVSPMYIIP
RNHNFNQTQYCGNISSVLSDKMLINAMFFRNSNNKLIKALKGVIDNLKCEKKLSKIGKELCQCKRPCKENTNIHLINTIPW
PEKSVVNTGGQRGFVKFKKKHKNVSKLIKQLIKYTGNLHSLNDSIDDMIRDFLTHIDHVNMSKPEYNRDSLAIDVVETNF
MKLNIHFESLDVRTVYEERTYTYADVMKDIGNVFGFYLGMSAVSVAEGVYIIGVMIKLIISRILC

>Protostome_Lophotroco_annelid_CAC9647536_1__Ofus_G10512_Owenia__fusiformis_

MVSTMTNIGMKDTINEFVNTTSIHGIPHTNDRRPIWLKVTWALLTLGGFSVALWQIIMVGLAYKKYPVNVLLDIKHSKQ
ISFPAVTICNSNQLRKSHVYASSSPAAQNLKAAILEHQQHYYNRSVEHNIMNSTCFAGSPTYDEITSCLWARLESDKTGIS
PATSVEVNDRLNDALKAMHISEKEPMGHTLEGMLLDCSFEKAFCSPRDFKRFYDVFEYGNCYTFNGIDDDATDSIRNTT
KPGKEHGLHLELFLEDDEYIPELTPSLGIRVVIHDQHDVPFPTENGFFVSPGFDTSVALKKIEYERLGEPWYGCIAPCSEEF
NVYQDIANCSYSISVCIHTWLQKHVMEICQCTSPYIPTAGLPGENTLASCDIHNETQEDCLEETEYKFNIGVLKSNCSPSC
KETKFDSRISTSVWPTSGYEDILEEKYASHVTPKVINILENKTSSLRNHILKVDIFYQELNNERIYQEAAYTWLSLIGDLGGQ
LGLWIGISILSKSSKWKSKFRDGDFSSDYQPSPWPQTFMNHHVNAGRKQSEMLAMAKYALEFFNIQFGIDVDNVSDA
KVIAGEPVTDEAQGLIFAPGILPKRGKLRLDVAISADGSKRYKEAYISDAGWSVTAIKPFTCAGRFKGILKPGWSVVYGDY
VINCDDHSDPLVISYKSKCPIGNVELSTSGDTFSYRVELFSELFGRGVGQVTCTDVPPKTFTTAVMTFPPILHID

>Protostome_Lophotroco_annelid_CAC9657286_1__Ofus_G102358_Owenia__fusiformis_

MVDKKTFSSRCDRTSPQQMITILTYDGDQGKCAKAYVMTSDDEDFIALRPLAYQRGAKPKFVVKNAEDIDIDKLSDDSD
FDDKSLKDIAVEFANETTLHGVPKIIKDRSCGTRLFWSCVCLSALCMFLIQASTLMGKYLDYKKKVDIEVLDDPAPFPSISL
CNIRHIDAILFEEIQGYSQQHYMKVLEVMEASKNPNYVRNKTVEEDKPFYWTSLETSINVQGYLSTKPELVKELGFDNIN
DLTEIMFTRETPAANIDKDKVTQFGIEAIEFIITCVYSGDRCNYSDFKRFFHPFFYNCFTFNTSSFLTDNSTTGRSFSLIAFLG
KMFSKESTKLDRGDTFGDPVYNSDGLRIVIHSSNSEPDPIQDGFNIPAGFSASIGVKATQYERIDYPYGNCSEQGSLSMD
NGIYDYTLISCQNLCLQNEIIEHCQCIDIALPIPENVNVSFCQDIENPPIFDCLEDKANEWKCNDAIRRSWNKFKCMRSVK
GRVSITQLAKQKCRCYPPCHEITYGVFHSLTSWPSMDQTIPTMEKVITGDFMQRFTTDHERNLVWENYFPNYSNDTDL
VKLYLETDNLTVQHDLTKVCARFLKDFSHVYVYIADDNVVKITESEFYSGVQLVSDIGGQLGLWVGISVVTLAELLQLCAT
LCGFLCKNKHRRKLKKEFERRSMRKKQRGASVRTKRYNHHHHRPRSRSLSNTRQNGTPTRNHLLKNQNHHVNV

>Protos_Platyhelmin_Macrostomum_lignano_A0A267GWK9_A0A267GWK9_9PLAT_Uncharacterized_protein_OS_Macrostomum_lignano_OX_282301_GN_BOX15_Mlig007558g1_PE_3_SV_1

MHNFSAEQLAMTDQATDSAAVDAAMTSSASIAGRQVRYCMLRPPGHADLSAAHKHRSGSGGGGGAHSRHRSDNG
ELQSVNNSIAGTAVPAGSPWVRMELSAGPASQRTSMTRAEQQDPLLIGEVASAPSVGAKHLSVMQIISMNFVRYCQT
TSLRGVPKIVKTKEKFLKVVWASFVLAFFLGCLTCLALIINQYLAHDVIHQPKTMRNYSVEFPSFTVCNTRPLSQAGIAYIR
SHGLMLPKEYSQHYIRAIRNTARNMNGLKTELFNLTVKDALSTIFDTKGYYENMPAGIDMNQFGHALEQTILLCESTILM
NSFKRSKICDSIGHWTQTYDNKFLNCYTFTADANHTKSILTVRIMAFTNERNLTYCPDCSNFPATQVSGLRVQLHNAGN
YPEVQEEGLNIKPGSLTEIRFDTKQWMMKEPPHGRCSETVPPTLQFNGTNFTYSYQACKNKLLHEQIHRQCGCLDSTIP
LPDSLQHSKADYCGRIPDPQFSLVKNATVLSNSLTSETRHRLAKFYGVQTDSIQLPSFISDAEFKAFLSRVTCNMEVFIKAD

EVTIDKCFTPCTFYSYDTTISTTNWPTKAYLYEFVNDFAAIKSRIESRQVRRSSASYLEPIYQDLKRLYEPVYRMAQTNQTE
ANRMLREIDYVQNNFIEILINRGKSNFDIERIEEKAVISLTSLLSQIGGLLSIWVGLTFVCIVEVIELLYNIVVMCTSRRLPET

>Protostome_Lophotroco_annelid_Pdum_comp411065_c0_seq3_

MKSAVSLSWLELKEDSTVMVAPATPGPWKAALYQYTQETTVHGVRFITLQGQSKLRRVVWICSFFTAVAFLLYAVIERL
EYMLSDPRTVNVEIRQVETLPFPGITICSRNQWRLTALQREGLTDTVNEYTKQISYDNENSTFVKKGKICTTEKELLNVFH
NSASVIPEAVLIYAEITIEVCTKLCSSGKRQGLCSMVVYFHKTRSCYSGPLIDFRRINASSDCHYAHVYRRIRKNLTNEIYCN
FEVGTTCHTIGIDGSDVPWKRIVPASVNRPGGLMKITDHTTGGEFGSVYSVGGLRTTEETVTLSNFSLPFIETTNKCLEVY
YAAVGSQIQVSYFREDLTEDVILPFKTAENYKWNRLFYELPAGLNMIMFSSKRSSEKYRLAVDDLRVMDCDTFFNSECKR
TIDGIDYMGNLSSNKLKSECMKWSDVNPQFPSHNFCRYLPMNMPWYNAKNLGPWCIGIGSKDATFCDVPFCDCDAG
MFQCMDKFGNKGPCLPKSMLCNGKKECPDDWESEENSDHCGNRLSITCSFIDDYICGYESARSYSNNTWIRKKDCLNK
KACMYLNRVSSNPGVVERLKSPFQLLEVEAEYSLTVEYNISQSNRSWMTVYVSDSTYSPSGKLLTRLNPAQTRKKVTFRP
IDHGFYNSSGYLDILFLIEAELAMFQDDSFEVYSVMLTRTSKPLLCPAGNFSCNSTGKCVVQSDVCSRKENCPGLYSVMH
C

>Protostome_Lophotroco_molsk_AplCal_XP_035828247_1_amiloride_sensitive_sodium_channel_subunit_gamma_2__Aplysia_californica_

MGKRKRRSSGAKWEEPGASGVKWEESGASGIKWEEPGASGIKWEEPGASGVKWEESGTSGVKWEESGAGLKRSKRF
IDGFEVSESNPRYNDNNGPPNDQGLFTGNKSALRAIEDEFKFIFSTMPKNTREFIGHQIDDMLMECTFRGIECFSEQFRL
LQTAEYGNCFTLENEQFVSTVGGPNDGLTLMFFIEDNEYLSGITSSLGFQVHVHAPGSVPNPYEEGLSVSAGMETFVGL
KQVNIVREKDPYGTCTDGDFITKLYNITYTQKTCQQFCVQSSIVTRCGCFDPEDEEIFKRLNETDTDISFCRSANETFCLVEI
LSEYNSGTLQCDCPSPCEETLYSKALSTRTWPTSEYATILVEYLCAHLNDTVCQEYASKSRSDLAASFLKIVVYFEDLNYENI
TEQPDYEPFQFVSDIGGTIGLWIGLSLISVFEVFQFFYELLHFFIYTKPKQDRKMKQKKKLRARKHGSRVAMKFIPGLSTD
PSLQTASSKLAALRSLDQKYLPGDQSRSVSLPRTRSSRSSDGTEENRLIGISDIPGSDPRKVGFSDAVNHADVNWEINEK
QSPIPERSIGNGLSLGAAVFYTGGKNRDRRNPHRDNTKDGTGKDYNDARNDIMYPGPKPEANNLSSDKEQYYEIPADV
RQTGDRGGDYSRSAKNGRILFRGHHSGWPVNDRSFHGNHHHANHGNQQHANHSNHHQANHGYENPAFQGRMK
DSSVLHLNNHMSMHDTSGTA

>Protostome_Lophotroco_annelid_Pdum_comp419518_c0_seq2_

MKGLKSDDGVKIHIHDPNDIPYIQYVGKAAAVSHHTYFEISKQDMYSMREPYGTCDSKELRLYHDQNYTRFRCNQQCE
ALAWQKNCGCIDHYMPGDGRYCKYHEWEKCRTIVDTKSYGRSDNCTGNCPTACHTVSYPSQSSSSSLAYFGVAKYLM
EEKKIKRIQERVEYSENMNYRMLPDAFKSILLKQDEIYLTMQKIYDFVNAQAVDKETSKFSLLDRALSKFHEHVQDDIGRL
KPILQKLRKESDCLKWLHGSFEDGISEAKKNIMYLKQKTMPKSSDVHPTDDLSDDMQKSILSTKSQQIIAFIASYLEMDG
HVCYGNKEAEDWLPVARQMFLDPIYQAKWSYYSKTTVDRKVTGPILEAMKLIVPGGSITYEEYIANSTVRSAVDQLYQYI
NVNATTGLKAFNHYLETLDMASKDVDDAMTFKHVEIKSPDYESEGLIQEKINELQSQFYDKMYSKFRNNKITIVEMARD
LGQGKSDEVIDVMKKSKAMMQKRILQGLSAEIYKHGPVASYHAYMKAMSGVLKIFSYMSETVQKKFTDTLRNMSIWH
KPTANIDSDDFVEYDSNKKILSRIWPNTIAFPDFFTTAASEIIEKNVNGFYEKLEEAIYVLEVEMTEQIEAFENSYKSFNDTL
RSILRKTDVNEELLMENFLKVDVFYGELVHEVIEASAAYPLTEFFNGIGGYFGLLLGASCITLIEFLDLIVYNFYWQLVKFCN
KKKRPEENGNQKAFV

>Deu_Ambulacrar_hemi_Ptyfla_40v0_9_20150316_1g1519_t1_scaffold723_cov156

MQNIDRITHFTFVDISENSSKCDADDFKCGDESCISHYKVCNTIIDCPDGSDEHGNCSRQGHGNESIFNDGWYHQFIDIT
QDTELFKDFRESQYRDPRFDRVKGLILTLFMEQPEYLSLYGRDVGARVTITPHDIISHPATEGFTIMPGTRTSIAVSEITLVT
KTSLTFPTVTVCNTNKLRRSAIADSPYEDMLLIDDSATLPYYGPCMEGDFMCDDGILCIKPFLKCDGVRNCLGDGSDELG
CEYGTCGSEHFRCANGSQYGFCIEKHRYCDRKKDCYEGEDEIGCECKRDEFQCADKSACITKKLRCDIHFDCDDESDEKD
CPWICNDEMFRCADGKCVPQYWKCDGVQDCSDGSDEDLHICDPLPEQVDCDKFLSEPNGTITSLGYPNEYPGHLMCT
MTIDNRDLGIGSIYLYFSDFDLEEAYNYTTCFYDYVQPAMDIPVIVDRNVFPMHGYATGNRTATTEEMKQTVLLIVFYN
DVNEPIKETQRSCGKRPPESLTIEFFILMVTFVSNSNITAKGFRLKYEQVEPENISKCDADDFKCEDESCISRYKVCNTVIDC
LDRSDEHGNCSRQDFSDLEDVLKLDKEAIQKYGHQLEDFVLQCVFNGHPCNTTESCNSSLYLSYEKYSSNPNRTGFRMT

SPLTRLYLMIGSMADVLERIVRLGGLYGNCHKEVDGYNSLYGPFYRYDVQIENLFSDIGGSLGLYIGLSIITVCEFFEFVLET
LRVCLRKDNGRQP

>Protos_Platyhelmin_Macrostomum_lignano_A0A267GQB4_A0A267GQB4_9PLAT_Uncharacterized_pr
otein_OS_Macrostomum_lignano_OX_282301_GN_BOX15_Mlig007222g2_PE_3_SV_1

MDRDINKSSDRYRRRVLFANIHSFASRTSAHGLSRFFSQQIPWLRLLWIVVVMGALTGLTMHTYKQISLFRDYPVSSTTR
LLTNKFTFPDVTICDPLMKKFLRKQSSDLRRSNESILNYFVLSYMMAMQFANNKNSNQSDEAISVQATKMAYASFEHA
VHTWPIFNMTEVSARMLDMLVFCQYNQETCSEKDFKVFYHRDYGNCFSFKPHNRLMKSSGVEKGLVLVLYSPSVDHI
QNNDWYSNIVTTGLETGGLRVVVHQKDTIPYPSEYGINSPSGMQTVLALNQVQTSLANTPTKQCEPRTRFHAYDHILG
NTQSYEKQLTDCTTFEHMKRVRDKCNCTIEGPVPTETYGSGERRINFCHDIMGAQNPERTIKLIKNMVEFLRKNRNDLI
NPRNNQLNMLAYNNLPMHEKQAISAALEHEFHNEVKQRLKSLRCALPELRSGRSQSKCLNMCKYTTYDFSVFQSRWP
DLSLEIRGANIRLARIAQMLKNRVDSLNRNITNNHALNHSIASHNLNLSHCMVNELNLKSNKFRNDTYCNQVHNIIRQC
FLQLRVYPKTLTVRQLFEERSYELVNLFSDLGGIMGLWIGVSMVTLCEFAEIILIILSYNIRALLAKLPSLTRSRRACGATRRE
TWSCVAEAEGQSLTCPIGCKRHGELAASIDSRQSPSRSKSYDHSVNPNLYRQPRLLTGNNSQEDAAIATVRQLLQSGLR
GNAAGDPGNGLLNSSTGASIGKETRL

>Protost_ecdy_cyclo_nema__A0A0V1BKZ2_A0A0V1BKZ2_TRISP_Degenerin_mec_10_OS_Trichinella_spi
ralis_OX_6334_GN_mec_10_PE_3_SV_1

MSNRVTFQRDTTTKRRFKNLLLNFSDKTTAHGVPRIGMATTKREGAFWTFVCVTSLTVFLFQLSLLIARFLRFETTTQVEL
KFETSLFPVVTLCNLNPYRNSLLKQVDQMKLMFDAFDNKPLSTPKTSTLLARKRRASSADEKQYRYTPLYARCSCPNEES
DCVPFTSAPLEDGEEICICYFDHVTKSPWPCYSEKQWRSEWCSRCSPNGNCLQILKKRQLSLYEDVKQCVCHSNGKTCL
VADKNKPERIWDPAYRDVRLDRYTTSTTTTTTTTTTTTTTTTTTTTTTSTATTTTTPIPTTPSEVQEYVDALGFEGITDETAK
VSRAQENTLFIMAELPPETKKKVSHQKKDFLLQCSFNGRQCDIENDFELLMDPVYGNCYMFNSNSTINRTTSRAGPIYG
LSLVIYVDADDYLPTTKASGVRLLVHSQEEYPFPDTNGYNAPTGLLSSFGIRMKKIERLPQPYGDCIKEGKTKDYIYKDQIY
SLEGCYRSCFQMEMIKSCGCGDPRFPVPEGRRHCRVKERKARECLENVIQKSGGLHGSFSGKCDCRQPCVQYVYDMSF
SAAKWPTPSIELDDCNDTAEECIRKLKKNALSLEIYFEQLNYEVLKELQAYQWVNLMADFGGQLGLWMGVSVITIIEVLV
LIYEVIKICCTRRKKPTMNKHHRRSIYDDSTVEMTIKIHPPNQNNPKDSIATAHSENDDECSQNSYAMENHLHRHDAEIP
NSVITFCAHCSLNIKQ

>Protos_Platyhelmin_Macrostomum_lignano_A0A267FBU7_A0A267FBU7_9PLAT_Uncharacterized_prot
ein_OS_Macrostomum_lignano_OX_282301_GN_BOX15_Mlig030741g1_PE_3_SV_1

MILQPIMPDGNSSNAEEASDHRSQQASQQASQISPQSTAQQPQEGIIQQQQPAAQPAAYRPQQARMDQRKKKKSS
ETVKLLISEFNDYGHRTTVHGLGHMYQSTDVRRKLFWLLITIVGCFACGIHIYFIVANYNETPVNSVILNGGVHVNFPDVT
LCNMYPISESVQNHGAGEIRGHIRKYWRFFRGFIKASAGMDNSTKDRFKMARKFFQIFWASEDTRDLAHDDDLFLVYC
RYKSRPCSNKQFKMVQNIDYWNCYTFTPKFDGEEEEDRRVYSGNEDEGLNLIMYTNSHLRNIHASNALTTTKGFYDEE
NNGKDSSNGDKSNYLERYLVNYNLSLNQLLAGDKMEPDGIRVILHEKGTYPHMDMNSVIVGNGDYSKVVFSMERKISK
NRPDFPCRETSEDAADEEVEYVSYYPGVDVRYTTKRPPPNFKKFKRDRLDYVVTRVQDLFWSSCSCYTHVVPFSHFNSK
LCYFCRNGRDPVEIRRTLERVHCHDRVYQNQLKKNILTYFSDSQRFQPCTGDEYDVKMGSSAWPSQRDIMFLVDQYVI
PAFLHESVGNRSYNSILKNTRVYPIRDIIGTTPEQAIGSKTIKSFNWTVDEFLIRKSFLKVAIHADSPTGTLNEEVESYSFAEA
LSEIGGIFGLYAGVSLLTFFEFVELAYLMATKPKEVTRHEKDGKSKFAMLQQTMEDGELPASEVLKYKQLFDRGNCCALP
TASASQQSRSFIMQAEREGSVRRPVIVS

>Deutero_Cephalo_105620R_t1__name__chr_scaffold204_start_174789_end_189091_strand_pro_len
_745_

MGWVVSPGAFRVELRVRTDVTISFPALTRARSTQGKSCCRASPPPSSRSQPIPTRSRVEVSTVAASIAMELANRKDMGK
EDSFTPLALAATPDVREVKPVQCVPTSTVGTGYTSYRKPSRFRRFALDTDIHGMKHLIKRGPIRRTAWVLMLLLSLGLMS
FECMESLIYYFEFHHVTKVELQYTNNMTFPAVTICNMNKYRRSALGMRDLATVGPYIGIVDRQQFTLQQEHLYPKAWV
DKVKNENLKALSKMTTGQQNFAVFVERVGHQKEDMIVNCEWRGSPCSAKNFTHTFTHLGNCYTFNGMTLSRKVLSSS
KPGEGNGLKVTINIENDEYMATNDVAGDAMDAGIKVMVHPQHEPPFVKELGFAVSPGFHTFVAIRKEVITTLPAPYGN
CQQRGGQKYYPDYSLSACRIECETEHVVHECGCRLVEMPGEHVPVCTPERYQCAYEKLGIVDRQQFTLQQEHLYPKAW
VDKVKNENLKALSKMTTGHQNFAVFVERVGHQKEDMIVNCEWRGSPCSAKNFTHTFTHLGNCYTFNGMTLSRKVLSS
SKPGEGNGLKVTINIENDEYMATNDVAGDAMDAGIKVMVHPQHEPPFVKELGFAVSPGFHTFVAIRKEVITTLPAPYG
NCQERGGQKYYPDYSLDNIAVLNVYYEALNYESIVQIPAIETEGLLGDLGGQMGLFLGASFLTLVEVFEYLMDELYGRVLK
KCCRCKATASAHTTSPSADVLSVNINGSATQLPQTSRA

>Deutero_Ambulac_Sakowv30041168m

MSVSRRKILLTSDVEEESESECHGRSFEDGEIRLVDMSLLNGALEKEICQFCKVGNISIVERRRHGLGSCIRSKLSQPSFPSS
RKKGAVFLINKKAAIGMRAIGKGTVRSGETVKLLGVTSTSNGSALSPEYPTNDFYLAGMENVSLRRESKQVALRKLFGR
MVNNSTAHGIPNAARAESLPRRLFWSVLFMVAVGMFLWQFSDQLVRFIDRPVNVKLNISFSRELTFPAVTVCNQNPIR
QSAIDFNQEPQSSPPTQNLNQTLNDTALDEDLGNHGGTNNNTNSPPYNTRRRRQTAQTSGDQSGQESKNDPPHPDG
DNGPSAGNRDMIRITEILTALPTSERITLGHQADMFIVNCTWQGEQCYASNFTTFSNSMYGNCFTINGPSGYIDPYWTT
SFSGPIYGLSLQLFIEQDEYIKGYTAIAGARIVIHDQDKMPFPEDNGFTIAPGAATSIGIRKVLISREPDPYSDCVEEGDSDY
TNIYMDSYDVGYSVQACMKACYQSEVINNCGCADPLYPFPYGVYEACKTSNSTQMYCKALVEYDYSTDGLECDCPQAC
SDVGFTTETSSAMWPANSYKETLLNYLKGSNKKLESLVGDDLKNGTNSVSENLLRVDIYYQELNYEVIEQEPAYLFSDLGS
DFGGLIGLWIGVSILTCFEFLELIFDFCHVFCSKGALAVNRTTDISAVSGGDIFFRDKYGNDPALTLNDRSSNKPRMNINN
RTTYVPETNAQSLYIPQYSMKIYEPGLLNTS

>Deu_Ambulacrar_hemi_Ptyfla_40v0_9_20150316_1g7150_t1_scaffold3457_cov142

MELDELRYDPPKRPSSPPAGAVDRDVGDTGFLALLYQFGDTVSAAGLPRVFTERSSLWSRLIYAIVFVASFGTGVYFSVKI
VQEFLEYPVIVTTAFTAESRMDFPSVTICNTNRVRYSAIIESKHSDLEYLLTDRSLGGLYYPPCLEDDMLCDTNECLRRFKV
CDGIRDCDGGEDEVNCYYGDCGYRQHRCETGGDHGYCINHNDYVCNGEQDCYDNSDESDCDNVFFGLNSYVDDGD
CLGFRCTNSSDCISQTEVCDGTYDCSDGSDELNCTSAFTCMTGMVKCATIDYCIYPSWVCDNYADCSDRSDEAPLATCP
SSSAARKRRSSSSGVSPTSSSGCYWCSAKYGSDNTTLQCFPLWYECDGYEDCVGGTDESNCDNINYHCPDNFFACTDG
ECIHEQFECDGWDDCASGSDEDNCEEPTRSSNSSTISDLFQDGWYSKYLHITDEEWFYEDFEDNYLQEKYNVGFERVSS
EDPPDWHGFATFSATGDYTDLEDVVKLTKAEMADLGHQNEDFILQCTFDQTKCHYEDFEVIQDDKYGNCFTWNHHS
RNMSLRSTGPGSTHGLKLTLFTEQSEYISIYGQDSGVRVSIHEMGTTPFPEDDGFTVAPGKATGVGLKQLIINREDPPHG
ECSNSTSFESMYGDGYSVAYHLLKTIHSINDKTKDINDFEQAQSNLVRLEVYFEEMTYETMSETPKYEGVYNLLAEIGGVI
GIYVGLSFITVVEFVEFFVAVVRYLFRRIKRSMKVEALK

>Protos_Platyhelmin_schman_A0A3Q0KIM0_A0A3Q0KIM0_SCHMA_Amiloride_sensitive_sodium_chan
nel_putative_OS_Schistosoma_mansoni_OX_6183_PE_3_SV_1

MSDTTDNNINNENGNIENPERHNLHSICPLCCIFIQEIRKCYEQTSKSLSMDNTNHCHINLNQNNTVHNLTGSIKVNKE
NSGRSSSSSSSSSQPPPPSSSSPPPPPPSSSSPLKHDNISVIILDELKRFCENTTIRGLPRIVRAHNRYLRSLWLILVCILILGCFI
CMFFLARQYLEYNVIHPPRVLRHTISPFPSVTICNFRPISPDGLKYLQMKGWKTPRHFVLDIATYAQYIFYDQQKIEEYNA
ASAAFSLAGFLESLPNNEERRKLGYQHNKLIIKCQITYLNGSKPVAAPCERKGIWRKFIHAQYLNCATFEPFSYLRSDITNIE
MYIYLDEMLNRADCIDCFHGEVKSQLSGALISIHGADTLPNINDKSINLKPGTLTELRLSIIENIQQMPPYGRCSINHPLNIT
LYDMIYNYSEYACRVATIQNNINQQCGCYTMEYPYNENKGYKPCTDLSQFINQSNCNRNNFNLTNLCINNFKNFISRIM
CKRSVIENYLHGELPNCKMPCSFYSYESEQSISTWPTKSWQLTWLNSSYNKKLGIFNNTEFIPYLKAQQLLELGNHIDAV

NILDKIDVLERKLLAILINRPNFDVRKVEEKIVLSLTSLLSQTGGLFSIWIGLSMISLGELFELFIRCYIRLKHSNHNNNNNTN
NSNSNDNNTNNNNKVIQHNNNSLKQNNYYPLSNYNDHILHNLLLLNQLHNNNNINNNNINDITNKINKILINEENNLH
NLLLLKKNENDQSINQ

>Protostome_Lophotroco_brch_Lingula_anatina_comp140230_c0_seq1_p1_comp140230_c0__comp14
0230_c0_seq1_p1__ORF_type_5prime_partial_len_751___score_30_91_comp140230_c0_seq1_1_225
3_

AAGVKILLHDSSDVPLVKDLGLAIPPGSHAFLGLFMLKVKNLPPPYGQCDDSIPLNYFDHYTSHTCSVDRKLRAMMDVC
GCIDISMEFSPPVKLPICNLTQYISCVVPLLETYRRDTMESPCPVPCDEHIYDPSVSYASISNFQVETLLNDKEVTKLQRRYI
AARDIFQKKNQNTFRADSETILKVSGAADAIQTMLEQMSKILQHLRHEIGQLLATIAPRVNFQDVALHKLWYSIHNGFV
RGWEVMEERTFSHVTSGYQETLGAFEDSAIRSTTATPEQEKCCLRTVLFEIIDMDLLKKQYLALRALDNITAVHASYTNG
TPIVNYIVTLPDERYDDTFNNPEMLQYHPYKQEYLMYLSSATQSYIDSIIALRNLSKNVLDNRYFNRTRFEECKVQIARNA
RAFNYRKYLLHDRLVKRSEDIVNEKRERFNNSVTNLIKVRDSLYFLLKIVNNSLESKALVSWADVYSLVRYAKDYISNKSLT
KTNLASTFLNESHVTSFQELRKALNGLHSRSRDITDSLEVFRKNYIAVWKTMLEEGSLKKFYQKLNSDVMLVSSHNVTSS
EKRKILEHIAHLLNTSPERVNALSRNHSVLLSRLNGDFSDISLNETCSRLNRIFKNLQSQSDLSGVMGTTDKQFLDAFVVLK
NQLEEYLKGNGIDAAFFRENFLQFDVFLKELQIEEIEQQKAYDINKLWSDIGGSLGLFVGASFLTLFEIADMLLHTALRRKL
VIKPKTPAVPNGIVTGRLQTMI

>Proto_ecdy_PANarthopod_Tcas_tr_A0A139WKN8_A0A139WKN8_TRICA_Eukaryotic_translation_initia
tion_factor_6_OS_Tribolium_castaneum_OX_7070_GN_EIF6_PE_3_SV_1

MVTRVGPKKDPTLCLNACRYLREYCETCNIHGVHYLTERRPKAEKIFWFLTLLLSISGCSFMIHEIWIKYQKNPIVVSFANE
VKSLYEIPFPAVTVCPESKCSQSKFNYSRVYGKLLNNESLSHFESHAFFYLRMVCKNLGNEFPNLLVNKTVGDDFFLLIEEL
KPNIELSCSFLEHFTCDESSYTPIITDVGICYSMNMLEKSEIFRPNVIYDEKLNFTKSNYSSGFDGETGFTESAGVHSYPYRA
FKSGVDNSLDLVLVDQVDYDEKCSAVQGYRLLIHRPQDIPRMSKNYIRVSHDKETTVAIEPALILTSPEVKNINPKKRRCYL
KGEKVLKYFLVYSQSNCQLECYTNFTMKTCGCVFYFMPREGETPVCGAGSYNCLELVKVQFKLEEVKIKRSDPSEYACGC
RPSCNNLHYNVEVSQTDVSYQTFSENVDRNASEKRRYAQVKIYFKHDEFMSLERKELYVFLQLELAVNCSVVESSSNSA
MSSDNTNSDIIPTRNYTSSGEYGNWWLLKRISKDSDRLQGDPWLLSKQEFRMRKSWLSRSPKAAFRIQYGNSNEIGAFI
RLTNSYCLMGGEPQQYEKIQRIIQSDLGEIPAINTSLAGCKVVGRLCVGNKNGLLVPELTYDTELQRLKEELPESLKISKIED
KLSALGNVIVCNDSVALIHPDLDRETEEVIADTLQVEVFRHMISNNPLVGSHCTLNNLGGLVNSDIGQNELENLSSLLQLS
LVVTWNCKSREQTYWLWCSM

>Protos_Platyhelmin_schman_A0A3Q0KRL7_A0A3Q0KRL7_SCHMA_Putative_amiloride_sensitive_sodiu
m_channel_OS_Schistosoma_mansoni_OX_6183_PE_3_SV_1

MHSNSTFPGSSCKEVLKSFCRWTSVRGLQHLLRSEHPLLRTIWAIFVALMCLANIGLVTLLLIHYYQFNTIENMRTLRGIH
VAFPHITLCNVDPVNSYRVYCLQNPNSKYCNINPKYSNILTKYQEANRYFKTYKGSTIWHKLEDPSMVAIFYQLIGMEAAI
QIGHQIDDFIIPIFCEVTTREKGGILMKRKCEEAGIQGLHLITYKYSNCYTLSMENSHISHNAVRLSMILYLDEEEDLNCRPY
CTHEFTEWAGSKIVVHSPSSYPDIEASALNLLPGASNQILIDGMHRIEKKNMESKPCENNERNFSIVYFDHAQQKFNQR
TMGYTDTLCIQLLIQSAIMEECQCIDFLMPIPGTQMELVNQLPFCGNLSRSTVNESLDCSQQVRQKNSKHFRLLCPIPCV
QLQYNYELTQLRWPQKPRILNYYAQLKDRFDYNRKFDIYEKIEEISMINATQALTMLQDTDIFEKNFLHVDVNRPNFDTL
MSYRENEEYTLPTLLSQIGGICSIFLSFTCVTVLELIELLCRLIIIAFPRFLTVSKNTIHYLNRKTLKLNHPTFTNSTKNKSNRTIV
ETTKNLSTLSYKKDQNHKPCDNCRNYTKSTIKNIMVDNNDNEINYLKEVLLTTNTNHSSSSNNINNNNIKLRGQSYVGLT
NYQIKEYITDNFIQIHEPDDNYILHKPRIKFFNPNQTIYTLDNSFNHPENILNQPNNEFINYNNLLNNLNYYKSMTNQNLV
TMTTTNIIKSKSFPDLFSLH

>Deutero_Ambulac_Sakowv30040445m

MGTCGNDQKNQSVKDVFSDLLANSSSHGLPNIHRSENLVGKLLWSMVFFTGLGVFCWQVSELARAFRNRDVSVQLK
VHFNTSINFPAVTVCNINPVKLSALLADPELLNALIPELNELNDGEGQGNTHASGVGDGGASVRKRRDVPSNITTIDPSS
SVTHVTPGGTTGFSTTSGTTTIDPSSSVTYVTPEGTTGFSATGGITTNDPSSSATHESPEGATEFATTSGNTTSDLSSSVTN
VTPEVTTGFSATGGITTNHPSSSATHESPEGATKFAITSGITTSDPSSSVTHLTPEGTTGFSTTSGTTTIDSSSSMAHVTPEG
TRGFLTTSGTTTTEFSTTSGATTINSFSSTAKEESTVGSTEGTKYTTTQFDWDSKMQDEHYYEQMDDSYKYKKLLRHYLS
RKTAEEKVELGHRLEDMLVNCVWRGYPCTPARALFVAENLAIRTLLRLSMISSYRFIYHLYFPDHRLLTLVRIHSESYNVPL
KFQDFGDTQGLCALRSAHLTVRSLVLRLAGSRLKCHGVPSHKRPVTSNESGILIGYLRPTYHSQRTPGTSEVLRNFTPIAS
GAYGNCYVFNSDSISPLKTNRPGAMYGLTLELDIDQDNYLPHITETAGIRLAIHPQNIYPFTEDNGISVPPGFASEIGLRLV
TIEREPQPYGNCVEMESGGQGQYYANNIYRRKFGVDYTVQACEKSCYQNYLIATCRCSDPDYPVLVINETMREILLPCD
NYDNGSGMLYFFVFTCTLSSVSLDRRHI

>Deutero_Cephalo_298710F_t1__name__chr_scaffold94_start_964637_end_983448_strand_pro_len_
755_

MVVYSWACVCCIAIAALSCSTTCRQHDWDALNGMRAMHYKTFDCTPASIGLHAVNIFLAVLELILGFVVSILSCCGLCT
MRRPAAQTVIYAAGPVPGGAEGNIVIMQGITGAQGSQPQMFLAQNPGAAGAPVQVFYPAQQAPPAYQVDPVVMA
CAYEPRVVRGLGWTLVVLGCLSVTLGVAADIAFADHRTPFHFFSAPVWSGLLVLTAGILGILSAKKPTNKCLVVAFLVVG
VIVIVMCVLCIAFAALGIVLNHHVPHLCGHFDWDAYNATRALNVSYPNPFLDPKFGDYQTFDCTPHSIGLHAVNIFLALL
EIILGFVVSILSCCEMCTIRRLAAQVAAVICAPCAARWHRLQDYFRYHTITDMTLEFADKITFPAVTICNYNRVDLDLATM
EELQYLSPMLYGSSMDEAALRAWLDLPAADNTSSSGFNVADIIRRKGFDLGIMSRAWPRMVGCTFKGQLCNSENFTH
SFSVYGNCYTFNGDRTSPLQQTMEGTGFPSSNGLQIILDVMDVWIQFSTETVSTGGHGEVGIRVLLHDQTEPPVMDSQ
AVALAPGNHAFMAVRQTQYRNHEPPWGKCGELQLTHYDPYTLPACLLECRSRHVEQACQCTPYFLPGTATPCEPPTIF
QCVNQVLTQLVSGELKCDCPIPCTMTKYGASVTYAGYPNRLTKKSYNAAYNLTPDYIRENGVVLDIYYDTLNYQKISQLK
AVDSGQLASNMGGMMGLFIGASVMTILEVAEYLGLKLLHQEAAPPRD

>Protostome_Lophotroco_annelid_Pdum_comp410722_c0_seq11_

RCKFNATHCNASDMVQDLTDFGTCYTFNANGSLSSRTPGPSHGLQLVINAETYEYMWNKNTNAGVKVHVHDYEEIPF
VRRLGKAAPLSSHTYLSVRHYTTRNVGYPYGKCREGTLKYFDHDPRYTEHKCIREFLTDRQVDQCNCRNHYMPGYAQY
CTLKEYLNCKVPDGLNHSCPTACHVWGYTTQASYASLSKDGVTDLLRNKAAFEPNYHNALKHRRRMDPKLAEKDILDL
GKFSDAVEKLFVFLKRETSFGETSVISLTNAAIKAMDRIAKEDVKEAFTFLPDLVDAFSCADKFLKSRETELEVAQMKAHR
FEDLMNKLLSEKPNEMATDETLIISATADLKETYKMLSVPFDYDSKCYGVDSAMKRYHIPKSLYKNKVMEKYCNESITNV
WTKLQSADVDMLTPNRGEPITDFWSRVKDTVLTLQGVDDLLTLNRNCTKSYSELLQEIQNWLALNLVEDDLRSVGSDFI
YEKEILQASGIANWYNHELIEDYKLGKKDVINLANSITSEKFHEFFGEITRFKKKLDDTYVTYLSQDVNINGPRRIKTIYKGLL
KYMSDLKPYVAKALGLKRHARAMKIWRKPKVNLDGTELVEYSYDNDDVWRAWPPTVNVKNFYTFQSQRVLEFHVGD
FFHDLAARLDTFEMDYLGLYSKTMQSFQSLNKSMAEVQRSSVIDEQFILDNFLKLDVYYGELTHELIEQTPAYTLSNFFSD
MGGYLGLLLGASVLTICELLDVILYNTALKCHEGRKRKQDKTYV

>Deu_Ambulacrar_hemi_Ptyfla_40v0_9_20150316_1g8260_t1_scaffold4268_cov134

MEMQAQNGTTERNLRMASMEASEDPDQKNCSILVTNFGQSTTCHGLQQILIGDSPFRKLIWLAVFLTAMGVFTFQAY
ELVSLFLTYDVRINMEVGTASSLPFPAVTICNTNKLRLSEIERSVHHELTKTDPEHPDSIHRQLSYETPCGSEEFNCGQAGF
HGICISNEKRCDGVRDCFDGRDEMNCTVCYGGRQCDVGLDSHASKCIPEDNICDRFPDCIDGADEESCVLTDQCEMG
VLFARSDPSVLYSRNYPKKYDTDYTCELRLTAETGFDGLQSSDCISLAFLELDIERGKNCQRDYIECSKTCGGGVRKRNVTC
SVIESDDPSMDFADMSLDSGENFDNPLVEGSNGKMVVARGTRGHYDSPDTGDYFADEGDVSRRCTSAGSSPERVESC

NNTPCSGGPGGIQILDKLFEKYGHISDDKKIYENFELNYYKSNNFDRVQTFPDPNWKGEFSFVKTTSPDYSDLQKVLHLD
RDEVARFGHQAEDFILQCSFDEVYCDHSNFYRFENDVYGNCFTFNSLQQNNQTTVTSSRPGSRYGLKLTLFIEQDEYIPL
YGQEAGVRVLIHPQDITPFPEDEAITVAPGLKTSIGIRMDTIKSLSEPYTNCSNDEDFESVYGEGYKYSDMGGVIKVLRPIN
EKVRNIVVDEQSARIYILQGGVIKVLRPINEKVRNIVVDEQSARENLVRLAVYYEELNYQKITKLPAYTIEELLADLGGILGLY
IGMSLITAFEVLELFLNALRHLAKKFCKPKPEDDGPTYL

>Protos_Platyhelmin_Macrostomum_lignano_A0A267FQU6_A0A267FQU6_9PLAT_Uncharacterized_protein_OS_Macrostomum_lignano_OX_282301_GN_BOX15_Mlig021130g3_PE_3_SV_1

MKAEQDEKKQKQQEFPCDPPAVTIASRQQRYHMDELNRLLPLQNVVRRNGDLGVSGGSGGAGGGASSGSVGRQFQ
SRPDFYPNPWVRRSERDFASGYRSPGPTKRNISRCFTEPPQPGDSSEFMEMSTELNQLQTTSVPSRDSQCLLGKEEQQ
QLELQQQPQQKQTVANGELLLQIIKRNFINYCQTTSLRGVPKIVKTNERLLKYIWIIFTFAFFFGCFTCLALIINQYLAYDVI
HQPKTIRNFPVQFPSFTVCNTRPLSQDGIHFIHSHSMMLPKQYTSNVFRAIRNTQANMKMNDASSEFNETIKDALSTIF
DTKGYYENVPAGINTTQFGHTLDQTILICESTILIKDFKRSKFCETIGTWTRTYDNKFLNCYTFTAGANYTNRILTVRIMAY
TNERNLTYCPDCSEFPATQVSGLRVQLHNAGNYPEVQEEGLNIKPGSLTEIRFDTKEWMMKEPPHGRCSKAVPPSMLF
SHSNFTYSYEACKKKVLHEQIMRKCDCLDSTIPLPDSMLDTGLHYCGMIPPRQFSLYTNSSVIAAQDTPSARQTLARYYNI
SADRVQLPSFTNNAEFKDFLRRVTCNMEIFKVAGNVTDKECFTPCTYYTYGTTISTTNWPTKAYLYEILRDFEQMMLRID
SLYRKGIMARNSSYLDPVHRRLKQIYDPVYTAVKDNPQKAEQMLGDIDYIQKNFIEILINRGKSNFEIERIEEKAVISMTSL
LSQIGGLLSIWVGLTFVCIVEIVDMLYNIVTMYNQQRRLGIVS

>Protos_Platyhelmin_Macrostomum_lignano_A0A267FRQ2_A0A267FRQ2_9PLAT_Uncharacterized_protein_OS_Macrostomum_lignano_OX_282301_GN_BOX15_Mlig000989g3_PE_3_SV_1

MYSNDRIISGGQWSQSQQQQRRRRQRQQAEQAAASIAQRLHRYRMLDDVGDVPGLQPADAPFRSSVSPALSPLLGD
SPRRADRANGLRTQCPAAAAAAKAATAALSSPDSAEAACVYPNPWVRRSDAGDMLELRPVTPAEQPPQPSRKRQQP
ASPAELQDLLAPQQPQALKPQPSFLEIVRVCLIKYCQTTSLRGVPKVVKTKEKLLKGIWATFVLAFFLGCLTCLGLIIKQYM
AFDVIHQPKTVRNFSVQFPSFTVCNTRPLSQQGIEYINQHRLLLPKKFSERVTRSLVNTRSNMAAINQTFNETVRDALSTI
FDTKGYYENMPPDIDVTQFGHTLDETIILCETTILIKDFKRSMFCDKIGSWTQTYDNKFLNCYSFTAHANFTKQILTMRIM
AHTNERNLTFCPDCSSFPATQVSGLRVQLHNAGNYPEVQEEGLNLKPGSLTEIRFETKEWSMKTPPHGRCSEEVPPSM
KFRESRFTYSYQACKKKILHEQIHSQCGCLDSTVPLPDTLSHPGSIYCGRIPERQFSLVHNASTLTSDSQLAAAARAVSDFY
GLNASSVALPSFLFNEEFQDFLRRVTCNMEIFKTAGQVTDDQCFTPCTYYTYETTISTTNWPTKAYLYEIVNDFQQLKSRI
DTMYSRGTTGRNSSYLVPILAKLQHIYGPVYREARRDPAKAEQMLKEIDYVQSNFIEILINRGKSNFDIERIEEKAVISLTSL
LSQIGGLLSIWVGLTFVCIVEIIELLYNIVTMCAGRAPLPGG

>Protos_Platyhelmin_Macrostomum_lignano_A0A267EP12_A0A267EP12_9PLAT_Uncharacterized_protein_OS_Macrostomum_lignano_OX_282301_GN_BOX15_Mlig009517g1_PE_3_SV_1

MDPKSDMLERALEEADDSDWRTDLGGDPVEQRRQLGSKSKSRHNYRRRMLCANLQSFAHRTSAHGLGRFFSQQVP
WLRGLWALVVFGAMTGLTLHTYKQISLFRAYPVSSTTRLLTNRFTFPDVTFCDPLMKKFFTIGSRQLRSDVNRNNYFVA
NYMLGYLVSDEERDDEASSRVKAIKLAYATFEHAVHTWPKFNMTAASARIFDMLLYCQYNQERCTGKNFTVFYHRDYG
NCFSFKATNRFIRSSGVEKGLVLVLYSPSVDVKQSIDYYSDTVTTGLETGGLRVAVHQQDTIPYPSEYGINSPSGMQTVLA
LNQVQTSLANTPTKRCEPKGRFHAFDHLLGSVTSYENQLADCTTFEYLKLIRDNCKCTVEGLVKSEPDGQPQLEYCHDLL
GAQNKPDLLSLLQRIRKFFETDPSIANYSLLDRSTGRMRTDIFDRLHLPDRFDAAQLLADRFPEIQRRLNRLKCAMPYLKS
GRKQSSCLNICNYTTYDFSVFQSRWPDHNFQIRGARIRLARIAGMLKSRVDEINKMMQRKLDNYASTEHALNFSITHHR
VNLSRCLDTELDRKSDLFEDDEHCVQVHNIIKQSFLQLRVYPKTLTVRQLFEERSYELVNLFSDLGGIMGLWIGVSMVTLC
EFAEISLILCSYYALSMVRRVSNALSPRRTNEEDEDDDYPTDCHGNHCDAADKGQQRSSAAANSDPWTELQSPLLIARP
TAEGRAASQSPSDRSDLHRHDGASEPPLQQSAASAPSSRAASPNSLSDGKETRL

>Protostome_Lophotroco_annelid_Pdum_comp417651_c0_seq7_

LMGNPCPDNLIEETVTDMGHCLVFNNKTHFNTTKTGSAFSLSLHINTEQYEYIRGPNIDSGIKVYLHGASVVPEMTQMA
FAVAPGLHAGVDIGMTKTEYLPPKYSDCAEAPLDSSSSAEVYSTSLCRQQCNTVNHQEQCGCILPYMSGNYSQCNLSRF
FECSMSILYSSEVTLQPCDCLSPCHFTAYQPQISYAGLSTLSVTNMMQAHTLQTVKEKLMIANEIRSRLDINQLMELLSSL
KLIKSNIQAFKTMVLVDLRAVMQRVKGALMELFGMLEADLGLLKTGLDTYKRKYDTYCRPCIEHYRKSTHEKRQIFAYYS
EQLKNNNTALNVDKSEMLYDLKYARYFFSCSANDPLIYGKNDIPSFPDTWSSPENSHCLHVVTSTSTLLDQFDVYRMLG
NVYVSNFFMMIARDVPLLSRCLFTYETRLEEMSNTFINILNEVSRTKSEISALDLDLDQILEEEMAEVERYLRSFENDITKFR
NLEITQYELEDNLNGTIISQYELLLNHITGKLDREVLHPLMSQLTKGETMIWSFYKDLFQVSNKMLDSFKHPSSFNSLIPSL
DVWRRPKAILNVSPMLNFEWSKDKKTFYWSKSLSVSSLIKDGQAQKKVAMHLKPYLATLRDALDISAQKVNQARKDSL
VILKELSLDLEEFREINEVNEDCVQKNFLTIDFYYSVLKYTNVRESLAYTYINFLGEIGGYMGLLLGCSVLTILEFLDLLIYNLFT
KIEDKHHLYRSTTPVSKMMPGDFDKCEKGMPNQVMYSSHQ

>Cnidar_NemVecNVEC200_012211_1_1_protein_AED_0_36_eAED_0_36_QI_180_1_1_1_1_1_14_153_
768

MSQNEEKCRQTVSGLIWDFFGQTTAHGFGKIANAKSRLRRMFWVCLVMAAFGMFSLQIYDLLMIYLSRPTGTQIWM
KHAPSIAFPAVTLCNFNIVRRSQITSELAGSFENFITPNISDVDDARSLRKRRQSVDGCSCSCYLKCRNFHDPHRNTPEPA
TTCAPVPTTIPTTATTTEPTTVPTTIPTTIPTTLPTTEPTTAPTTPEPTTLEPTDPPTTEAPTTLPPGVTPAPTEKPLTIEEKKE
QAFTGEVSQPDPATLDKQTLKTEELVLSLATKPEDMLMEVGHQFNDMVLSCTYRGIPCTNYTENLWSRFWHYRYGNC
YVFNSGKDWQGEPKPILRSNKPGPASGLTLELNAEQHEYVGQLSHEAGMRVLISTQGEMPSPLEKGISVSPGFSTATGI
RLTNILRADPFNNKSCLSSDDIDPENLYRKRYNVSYSRTTCMDSCLAHKQVDMCGCLEYRFPNIDNRTVCDILDIDVIRCL
NKVQSLYQDNKLGCSEKCRPPCNEEVFKLSISSARWPTDDYEAHFLDELKEEKGFVLPANADARAYFVKLQIFYEELNHE
VIEEYRSYELVNFVSDIGGQLGLWIGISALTASEFIELLIVIALHLLRKVTRIGLIDNSAEQPADTNPCYLDDITLTGDLKERRG
ALTESNTSLTVRYLAEVARKRNKKGHKEKREASWHIDEEAFPCYRDQATIHPNVESNTSLTVRYLDDIVQKKSKHRKRSK
GDTSMVQKGDGHRGDLSGRRGELTESSGSLTVRYLAEVASKRKR

>Protostome_Lophotroco_annelid_CAC9605609_1__Ofus_G07900_Owenia__fusiformis_

MSLDTPVTSRRSKLGVSLINIGYQNERTPELDARSFNSSIAFGSKNTLMSDDDEEKDEKSMKYKLEEFCTESSVNAITRAY
KTKSKLRRYTWLTMFLFSVTLVFAQSRTLVTKYFTYPSEVKLDMIYEPELELPSVTICNLNPIRRSKLFHNEKFEMFATNSD
TDEEDELYKSYKNKLNVNNYEEPDEENDDVENGSFLNDETQDGTEQTEAQDTTTTTQDLTTETVLSNDANEETVNNPG
ESFDEPGETNNNPVEFNDGPGEPFGYPVGPNAYPGESFDPAEPSEYPEETIDDPEETIDDPEESIDDPEESIDYPSEGTAT
TDAPTLSWEEEEEEEEEEDDFNKWTYLANQGSSFYSQENSRHKEATKMMQMISEADEETMGEIGHQLEDMLIECSW
AGYQCSPQNFTRLQNDYYGNCFTFNAGDMGNAYNLSWTNETTEPLTVSKGGPLYGLTLELFIEQHEYVGDLSPGAGM
RVVVHSVSQMPFPEDEGFDIAPGYATSVGLKKTKITRLPSPHGTCDPDLDATTLYSDKYGVEYSKEACMKSCYQKNVID
VCGCAVHEVPKPDNVPMCTSVSDSITTNTTNTTTAAPTTISSNTTTDDDNSGEDHDEMGPSPNEQCMMSVEAAYKK
NELGCGEECPLRCLDYTYDMLISNAEWPSMDYSRKLTRKMMQSSSSMKGVTGNMKANLVKLEVYFQEMSVEHIEQQ
KSYEIENLLSDLGGQLGLWLGLSAITACEFIEFVIDVIKLIITKCSSKRNERSTTPIEPFKS

>Protostome_Lophotroco_annelid_CAC9521433_1__Ofus_G03942_Owenia__fusiformis_

MIMDHPMINTGTMADGENVTKTVKADVEINADNTNIEGTENVAQTGKVDDEKNTDDVNRADIDRVTQAVNVDESN
DIEMGTFKRIGNTGVVIELNQNVGIVPVDEPNIERATEDMITEHTQNNEHIVNERMAIDELISDTPIVTSQTQHHDIIAD
ALEFKYNSLKNQISKFALITSIQSLLQVRNAKSVLGRLIWFAITLFVCSLAIIGVHEITRKYLDHPSEDILYSREELVTFPSVTIC
GVKPVVFTDEHLKKKLSSRANTGDTINTWIYHDYRLKMLESRAFSDEREKDHERINAVMRRIVSEERYIENLNEWRDYRL
RMYDDLILECRFKGKPCGKTDFKQVMLGRYGNCFTFQPNSKDDTNILSEPGPDFGLNLMLFTNSFSPLDEFTDSDTFNYI

KLIGSSDSAILNEFNSDFSTASDGIRLSIHAPGTMPDPEKDGIDLSTGTFSSIGLSQDVRTLLGPPHGNCTEERLNTHTTYTY
TQNMCLRQCQQIETIKRCKCMNPKLPLPPLTEKPHKIHYCGYEMRQEYLKEKEREMTYGNVFPLENVQFDWKIFIENFK
LYECEKEVTELFSREGEKCGCKRECKSTAYYYLKHSIPWPNERYYKGLKHRNFISFLRNHPKADQFLNKTYWSGNRLNRL
GICMSGDNIFSSYSSAVVKGAYPYLTCPCNIANSTDQPDRCAPIINWYRFINKFVETNFLKVNVFYSTLNVNRIAAEPSYP
FNKWLSDFGGIIGMNMGMSVVSLVELGYLMVSLAFILYCKHKRDN

>Protostome_Lophotroco_annelid_CAC9668492_1__Ofus_G161027_Owenia__fusiformis_

MNTCAKIEEQIKVSVIGYAEEHVLNESHEAETSGDTKRDTNLEVDCVSHDELELSIEKTNDGDDNYGHINKETSEQCLNI
DCDARDEPQVRQENKSCNISKRKAMCSIEDDHDDPIVDEAQTAKENENYDVCKGTTSYKTDTEDDNEDAIINEASSDK
NNDHNTNDDIEDPFDSWIWKYNQLKERLSYMASITSIQGCINIRNSQTMFGRIIWFSIALALGFTTVIKVSNIVVQYLSYP
SADALYSREEDVTFPSITICGARPIVFSENFHRSCMNSNCMPKTMDVQDGRLNQGILLLMNESGYESQTETLDFLRKRLL
SPERYVENVDDFPYSRLRYLKKNDLILACTFQGKECPDALKFKEVKFGRYGNCVTIDSNNTMKESGPDSGLSLILFSNSYS
PLDEFQKAMNLFIYLVSGVSEFDSTHATSSDGIRLAIHAPGTIAEPEHDGIDLSPGTFNTIGLTQDVRTLLEPPYGDCTKKIF
NEKTTYKYTKDLCLDKCMQEEMVAKCGCMTPKAQASLGKNGYDVSYCGRYYNTIKPKGYPQYLKKPNWRRRLRSQLL
QFLRRLKCESDVLRLYSRDSSNMCTSKCKTECTKIVYDPLTHATPWPNEKYYKQLNQRPFITFMKSKDNVAKLLEYWFD
VRRSVLDSYCELLHGDLRKNLLKRRVTGHVFKPLFADVNFPFVTCSENITAHINRYNIINKVIEKNFLKVNIFFRSLNVNRIA
AEPNYPLNKFLSEFGGIIGMYLGMSAVSLVELGYIMVAMVFILLFKYKR

>Deu_Ambulacrar_hemi_Ptyfla_40v0_9_20150316_1g14067_t1_scaffold9635_cov109

QLSSTLQSVVKSTFEVVRRPLVFKRSSREGKIYEKTSDIQNICSDIDKIDKSYQNSITLVTKTSLTFPTVTVCNTNKLRRSAIA
DSPYNDMLAIDDSATLPYYGPCMEGDFMCDDGLLCIKQFLKCDGVRNCLGDGSDEFGCKYGTCGNEHFRCANGSQY
GFCIEKHRYCDRKKDCYEGEDEIGCECKGNEFQCADKSACIAKKLRCDVHFDCNDGSDEKDCPIDCDKFLSELNGTITSLG
YPNEYPDHLMCTITIDNRNMGIGSIYLYFGDFDLEEAYNYTTCFYDYVQPATDIHVIVDRNVFPLHKYATGNTTAEMEE
MKQTALLVVFYNDEFEPIKETQRSCGKRPPETLTIEFFLLMVTFMSNSNITAKGFRLRYEQVEPENISKCAADEFKCEDES
CISRYKVCNTVIDCPDRSDEHGNCSRQGHDRQSIFNDGWNSQFLDVTQDTELFEDFRENQYRDPRFERVKGEDPPDW
QGFMTFSSTPDFSDLEDVLKLDKKEIQKYGHQLEDFVLQCVFNGHPCNTTQQFYTFQHDVYGNCFKFNHGKNGQPLL
NATNAGFSSGLILTLFMEQSEYLSLYGRDVGARVTITPHDMLSHPATEGFTIMPGTRTSIAIENLFSDIGGSLGLYIGLSIIT
VCEFFEFVLETLRVCLLFSVECPDRLDFDTPLDVCFQADSGHPSQRCMSGYVSSEWHYLRLQSKSLRSFATQYLHSFNDA
QLKLYDHLINKPSNDWDISHLTVGNKPTPEEYNTEVMDLLKKFAKNEKMDKAARS

>Protos_Platyhelmin_Macrostomum_lignano_A0A267FFW2_A0A267FFW2_9PLAT_Uncharacterized_protein_OS_Macrostomum_lignano_OX_282301_GN_BOX15_Mlig020199g1_PE_3_SV_1

MKLGSDPPMSIRPHRKSTMNPLTESFGLRCDGGDCTSSFVDSTTNDCERKLRHHHRQRMLYTNLHGFAGRTSAHGLG
RFFAQQVPWLRLLWGCVVLGAVTGLTLHTYKQISLFRAYPVSSTTRLLTNQFTFPDVTFCDPLMKKFLTLNSDSLRTESR
EISDYFTASYMLGYGFASTPDSSGPHPKQEESAGRVKAIKIAYASFEHAVHSWPKFNMSAASARIYDMLIYCQYNQERC
SEANFTVFYHRGYGNCFSFKADSRSIKSSGVEKGLVLILYSPSVDVKQSIDYYSEIVTTGLETGGLRVAVHQKDTIPYPGEY
GINSPSGMQTVLALSQVQTNLADTPTKRCEARTNFQAFDHLLDTVQVYDRQLADCTTFEHLKSVQEKCGCTIEGLVKTH
SGRRPDEDYCHDLLGAQDTKQLPKLLSEINQFFQRDLLVSNMSLIEKSGRLSLRTFSRLAYTDRLRVAELLSSKFPVIGRRL
NRLKCALPHLRTGRKPSSCLNICNYTTYDFSVFQSRWPDPNFQIRGARIRLARIAAMLKSRIKEIDTNLLRLRAVNPAFDT
NHALNHSIAYHGVNLSECLDDRLNKSLDRFDSDEHCQQVHSIIRQSFLQLRVYPKTLTVRQLYEERSYELVNLFSDLGGIM
GLWIGVSVVTLCEFGEISLIVASYYVRLLIRRLALALGTRALLWAEEKPELNQKSQQNRLSKRQRERQKPQKADNQPFRQ
QRLGSSGRQKLFSNRDPNRYWQPRLRHGLDRSLNSVKAEVDNEEVSKRFSIIRLAL

>Proto_ecdy_PANarthopod_Tcas_tr_A0A139WFX5_A0A139WFX5_TRICA_Uncharacterized_protein_OS_
Tribolium_castaneum_OX_7070_GN_TcasGA2_TC033556_PE_3_SV_1

MIDGRKNRKDFTSIVYETFKDFAENTSIHGLKYAVRRDIVIYEKVFWVFVVLTGLGGAGYMTVLFWSRYTSNPTRTSILTA
YAPNTAVPFPAVTICNINRIMADQVEDFTEKLTIRAEDVETVRRAFPQLLAFVAIDTKVYNMSELLVLHNVLVENNYNDI
GVVMKKITQPCKDMILKCEWEFETVPCEQLFQETLTTDGNCCSFNYIREYIDMSRISEQGPSVGSSRRRPIYTPFNGFNG
GLKLHLDPEIQAVQYSSLHSSGLKVMVHSPWDYPGSGSIYKIVSAGRQSYLQVSASKIVCSSDFRQLSIQQRQCAYSDEV
KLRYFKLYSDTNCLTECEEKYLYAKCGCVPFYYPFSSQSDFGKWSRTKINLILTLGTTLVNITKACDCPSQCEDIFYNVLSST
AALGREEAAPTSDFFEDQNVTGNSIALYVYFQGLKQTILYKDIITSTIYLLSSFGGVYSLFLGCSFISVVEIVYYSTFRLLVNM
KLIGRIDKADARPRKDEAKQPVDELQKGGNMVQEGFLIKIAIVSMFISTGWSRPKTKDDSDSPETEAEIEPLIIDRGPYFD
KTASKNVTALVGKTTYLNCRIRNLGNRTVSWIRHRDLHLLTVGRSTYTSDQRFTSIHNPMTEEWTLQVRYPQRRDSGIY
ECQVGTTPPIGFSMSLSVVEPITTLLGGPEMYINKGSTMNLTCIVKHSPEPPPTIYWTHNSEEINYDSPRGGVSVITEKGD
ITFSYLLVQRAKDSDSGKYTCNPSNANPKTVIVHVLNGEHPAAMQHGGQLRLEYPFSACLLSVLVAVAGP

>Cteno_MneLey_comp19805_c0_seq2__p1__GENE_comp19805_c0_seq2_comp19805_c0_seq2__p1__
ORF__type_complete__len_802__score_191_14__comp19805_c0_seq2__85_2490_

MSNRVITTDLRLACQTALTPNTMPETKIKIGLTRGLVDGGEIFAQFSLSSTAHYQHLLTYISAICSREKVLAIYWFNRHDRE
FVLIHDNASMDYCLVQSGLHEILYIITETGGKPAGYNSKHKPEPIQLKTSSPSSGPSPDTANLERSSDEDRPGGTVQREDH
IPLRKMKKSASLEWSYDNEQPPSVQEISSRDESAFDMTGRGERPTYYPYLEHYQPSYPARPKPHRTPTRNKRPEPKVNN
KYHTHSDQSGSLSDSDDDRNAKEVDQFMEPEPPKRKKKAKRRANRAALEQKRQEIEMETFKEFASTVTADYLAEMCSK
ASNGRRLIWVSLWLASFVYAWYNIALSIGVYMSKPTATKLNFYEASSDGVTFPTVTICNFNKFNKSYFDTDDITTKKGQL
KDFLKITTPIWGHKQELDSASWDTKYDKIVQQSMKDALQTASHSLEGTVEFCKFSGKPCLPADASPGDVADVEDADPT
AAFNPVFTEHGLCYSFNQNGNLTMSRTGALYGLSLRLHVAQDDYFETQDVAGFKVLLHNSYEPPMIEEYGFALRPGSET
YVRIRLQKDVYPQYKDTPRPLGYCDPYLEDQFSTNYTISVCVMQYRAINMLKVCGCITFYMDRPDIVLSEGKPYMKPKIC
NLKEERTCADNVLEGLALGTLKIDGEKVPEPDCPSPCVYTTYTYTVSQSEYPSSAVKDKVVEEVNRQKGTSPGEDDASS

WTIERIRDNYLKVHIYFEELSTLEMERHPSYLISNLIGDVGGQLGLLLGMNICSLVQFTDYIVRFSFFKGFMSLVRLYKARRRNK

>Deutero_Chord_Hsap_sp_P51172_SCNND_HUMAN_Amiloride_sensitive_sodium_channel_subunit_delta_OS_Homo_sapiens__Human__OX_9606_GN_SCNN1D_PE_1_SV_3

MRAVLSQKTTPLPRYLWPGHLSGPRRLTWSWCSDHRTPTCRELGSPHPTPCTGPARGWPRRGGGPCGFTSAGHVLCGYPLCLLSGPIQGCGTGLGDSSMAFLSRTSPVAAASFQSRQEARGSILLQSCQLPPQWLSTEAWTGEWKQPHGGALTSRSPGPVAPQRPCHLKGWQHRPTQHNAACKQGQAAAQTPPRPGPPSAPPPPPKEGHQEGLVELPASFRELLTFFCTNATIHGAIRLVCSRGNRLKTTSWGLLSLGALVALCWQLGLLFERHWHRPVLMAVSVHSERKLLPLVTLCDGNPRRPSPVLRHLELLDEFARENIDSLYNVNLSKGRAALSATVPRHEPPFHLDREIRLQRLSHSGSRVRVGFRLCNSTGGDCFYRGYTSGVAAVQDWYHFHYVDILALLPAAWEDSHGSQDGHFVLSCSYDGLDCQARQFRTFHHPTYGSCYTVDGVWTAQRPGITHGVGLVLRVEQQPHLPLLSTLAGIRVMVHGRNHTPFLGHHSFSVRPGTEATISIREDEVHRLGSPYGHCTAGGEGVEVELLHNTSYTRQACLVSCFQQLMVETCSCGYYLHPLPAGAEYCSSARHPAWGHCFYRLYQDLETHRLPCTSRCPRPCRESAFKLSTGTSRWPSAKSAGWTLATLGEQGLPHQSHRQRSSLAKINIVYQELNYRSVEEAPVYSVPQLLSAMGSLCSLWFGASVLSLLELLELLLDASALTLVLGGRRLRRAWFSWPRASPASGASSIKPEASQMPPPAGGTSDDPEPSGPHLPRVMLPGVLAGVSAEESWAGPQPLETLDT

>Protos_Platyhelmin_Macrostomum_lignano_A0A267EDL7_A0A267EDL7_9PLAT_Uncharacterized_protein_OS_Macrostomum_lignano_OX_282301_GN_BOX15_Mlig014011g1_PE_3_SV_1

MASSVGCGLFVSSSSSRVVCSISEHLPVQQQQQPTTAARSFIEPFRQGLLQQRRQKRNRTSVMEPFRQTPQLPPPPPPPPPPPPPPSVSYQVHPPPSVSEIARSSASANTGGLFEAVAAFLQQADDFDGNDEVSPGSWNLQESSEQQQKQKSTMTRQKTEPSEFSSETRKSFVFSSHRLRTAKAFLDNSTFHGLWRVQPSANKQNLLSRKMHLLWTLIVIATISCLWLHVYKQVSRYFKYPLQSSTREILNDFHFPSVTICEAWQMKFFKDNYKSVSLRLANEPEDYLNMTYWLFDVPKSENTLQYLSNSTAYGLSVLDLLIDCKLNGLPCDKDSFSAFYHRQYGNCFTFDAEVHHDEKLGHSGSRVHEGLSLILYSPSSDPYFRNFSDPVPLDSSGVSGLEIDGVLVTAHEANTVPRVSEDSIIAATSHETKIGLKEVRTTLVNTPVRPCHESSNFRVHDHLVPIERTYKTSLPDRLEWEDIVQVLKSCNCLASSPVGVPTHLANYSGVRQCADLFDALRGDFPKYRNFRNHECQEKFNRGKFSCREKLLKEFPALARNWEEFQCYQRLTNQSFHESDAQVLEADDQCSSISYQFQIFTSELPDSQSASWQFGRYDARLIFQTFSTILDRYLKRFENSREDELPAVVRSVRSKGLRPGDCTTMIDRKFKDEENCSDVQEFISRHFLRVQVYPFSLSVVHIAEERSYSIVDLFSHLGGILGLWVGSSIVSLFELLELVTFLSASSLSFSLNKRLQKSNSGAAVLRYPVPSKKPPAAAAAGLSPMAPSVQVSLATEFDEASAN

>Deu_Ambulacrar_hemi_Ptyfla_40v0_9_20150316_1g21852_t1_scaffold24782_cov84

MRFDDVDRPTWRRSASMHVSFVKEASFSTQHIVISVRQSVPFARVVHTGTRCTVAICGAISHFSKIPNTNMQAYDADVIENVTLEVCASVCFRYEPWECMSFDYHETFRVCRINPGKADVFVRLKNGKSSYDHFAKTTYGIECYAELDSSDYRGHVSVTRNGNICQPWNVQVPHVHYLYGRXXXXXXXXXXXXXXXXXXXXXXXXXXDPNIRFELCDVGQPRVGCLSKVEPSARECFTSANGDDYRGTVSTTESGYICQPWTSQLPYSHRYTPEIYPNAGLGDHNYCRNPDGWSATWCYTLDPLVRWESCDVGSPQTTCQNTDMPIHEFFRTTNQTQRGFDSMTGNVTVEECASSCLNSRKFTCLSFDFSNEDGICYHSDANESVSKPSTVSGILVDHYDRRFEDEERLLLDFWKFPDYRIPSDHVALLTNVTAHGCAAKCLDEITYKCQSFDYKPSLAKCYLYDKPFLSVIADTPEKDISYIHYELKVASSNGCPSDYRQCGSGECIHQYSFCDTIINCADESDEDNCYGNHRAIPDMDDIPGVDWHDHPDKATLMADFLSDYYHDNGFASVKAEVPPDWYGFKTFSSTPDYSDLQSVLKLSRDELRRYGHQREDFVLQCTYNEEDCVQNSDFYTFQDEKYGNCFKFNHGRGNETVKMATKQGATFGLKLTLCLEQDEYIGIYGHDAGVRVAIEPSKYVAFPLDAGITVQPGTVTSIGLREIRVSKKPHPFGNCTLDDTQDSIFGSHYQYSVLREDLLSDIGGTLGLYIGLSVITVAEFIELLIVVIKFVGRRRGALRTAKLSPSS

>Protos_Platyhelmin_Macrostomum_lignano_A0A267FQ21_A0A267FQ21_9PLAT_Uncharacterized_protein__Fragment__OS_Macrostomum_lignano_OX_282301_GN_BOX15_Mlig000989g5_PE_3_SV_1

GDLSTSEISAPRRPQHLGDLSNLGDLSTSETSATSEISAPRRPQQPRRPQQPRRSQHLGELRISETSATSEISAPRRSQHLG
DLSTSENSAPRRPQNRGELADFVPVKRSPAQFAALPPLLGDSPRRADRANGLRTQCPAAAAAAKAATAVLSSPDSAEA
ACVYPNPWVRRSDAGDMLELRPVTPAEQPLQPSRKRQQPASPAELQDLLAPQQPQALKPQPSFLEIVRVCLIKYCQTTS
LRGVPKVVKTKEKLLKGIWATFVLAFFLGCLTCLGLIIKQYMAFDVIHQPKTVRNFSVQFPSFTVCNTRPLSQQGIEYINQ
HRLLLPKKFSERVTRSLVNTRSNMAAINQTFNETVRDALSTIFDTKGYYENMPPDIDVTQFGHTLDETIILCETTILIKDFKR
SMFCDKIGSWTQTYDNKFLNCYSFTAHANFTKQILTMRIMAHTNERNLTFCPDCSSFPATQVSGLRVQLHNAGNYPEV
QEEGLNLKPGSLTEIRFETKEWSMKTPPHGRCSEEVPPSMKFRESRFTYSYQACKKKILHEQIHSQCGCLDSTVPLPDTLS
HPGSIYCGRIPERQFSLVHNASTLTSDSQLAAAARAVGDFYGLNASSVALPSFLFNEEFQDFLRRVTCNMEIFKTAGQVT
DDQCFTPCTYYTYETTISTTNWPTKAYLYEIVNDFQQLKSRIDTMYSRGTTGRNSSYLEPILAKLQHIYGPVYREARRNPA
KAEQMLKEIDYVQSNFIEILINRGKSNFDIERIEEKAVISLTSLLSQIGGLLSIWVGLTFVCIVEIIELLYNIVTMCAGRAPLPG
G

>Deu_Ambulacrar_hemi_Ptyfla_40v0_9_20150316_1g10746_t1_scaffold6363_cov125

SVIERIGGKYSDCIKAGEMPLLYKGYNYSVEACVKTCYQNTLVNDCKCYDTTLPNNKPIPACVDGTKRTMEDHTKRRCR
CLIGHPMHLMINFMDVSAREVRKSKKHYWIITNRGMEQINTEEGVQVVKPQKGNHRIRNTLADFGSNTTAHGFARLV
AADSKIVKFLWMLICCLCWGVFIWQAIVFYSISVDFRVLPFPTVTVCNMNPLRDSALEIIDDEYKERLAFLKYEEQSEDNA
DADEESIFTTTSESEDLADYSSVENYDFDFRQGLNLLMMMSNLPVDVRNATGHQKEDLIQECVFNGITCSSKNFTLVLN
PKYGNCYKFNTGANTSKPGPPTGLRLTLFIEQDEYLDDITENAGAVVLVHPPDIVPFPEEQGGRSYKETVSMSHWPSDA
FGDKVYERLGKRSKEINETLQDRNELRISGIFFLDIIRINNLRYLLIKQRSHSFFSASLPLDQSDCTYGDGLHKSYSKSKLRGL
MEGMDLMHKEEGVQVEKPQKDDHRIRDTLADFSSNTTAHGFARLVAADSKIVKFLWILICCLCWGVFIWQAIVFIEKY
YRFEVSYSISVDFRVLPFPAVTVCNMNPLRQSALENIDDKYKDLLEFLKSGAQSEDNAHADEESNSATSESEGFSDYSWE
EPVDFDFDFTRELNLSMIMSHLPSEVRNKTGHQKEDLIQECVFNGIKCSSNTETATLSTRERMVPISFIRRNQAPRRSVID
RIGGKYSDCIKAGEMPLLYEGYNYSVEACAKSCYQNKLIDECKCYDVTLPNNKSFPACIDGTKEAPSEVSAAEWGTQLRT
VYRANAQHLYRNF

>Deu_Ambulacrar_hemi_Ptyfla_40v0_9_20150316_1g18537_t1_scaffold16732_cov160

MVTPTAIGISWRSPLQLLHPEDRFVLTLDSPDGNSDFPLHIPANTTEYLCQDLVPNTEYNISLVIPTQVPIIANYTQKTGEN
PCAESPCENSGTCRAVVGQLSAYTCQCQPCYEGPHCEIAIDACASNPCQNHGTCTSDAATCTEYTCECLNCYVGRNCET
YIDPCESSPCKNDTAVCQPMQHLDSCYDYKCFCNDSVRGCFVGDYCQQRLDSANEERAFVLPFAIVLTVLLAHIVKRCW
IRVVQTRAMGEGVSNRVDLVISFTANVQSVILENHVKHESTHVTQQIAKTTVSVFLPMIRVWNTHAAVDTAIPGTTVN
NDNVVHRGDRHRAASDDVFTREATEPGRSLSVVDPREDSADVTAVSSSTTGNENEGKTPTCQVIAKAFFESTTWHGLP
RISQTPYRSRKLFWTVLFLTSCTLLFIQLQAVIARFWSFPINIKVEEVARSSVSFPAVTVCNTNKLRSSALAKSRYSPLVEVM
KDAKVYPYYVPCLGGDFLCNDGIDCIKSYGVCDGIPNCPDGSDELHCDHNYSSRGGESDDDTCLSGFKCDHGICHPNIH
LCNHVQDCVDGSDETDCTQRGCTEDEFRCDMGQCIAKWKVCNSHVDCRDKSDEKQNCTNILIAQNGACPAGYFRCT
SGHCIENIQFCNNERECNNGEDEDWPRCIYTESSNLTSAFNDGWYERYTNITDRFVNFTEFRQYYYSDPGFDRVPREEP
AAWKTFMSFSSTPDFSDLTNVLKAETIRIGDPYGECKPEMSDEYFYIYSKAREDLTSDIGGLLGLYIGISAITLFESLEFFIDV
FRLFVRKIIIRIQRMKH

>Cnidar_HydraVulSc4wPfr_2074_g13182_t1

MYLQFVNHFTTSSGKVWYIKTSISCNSKNIIYFLSCNRCDGKTTYIGKTNSLSFLKKMSSIARFANIAKVSSRHVDDVSKKE
SEPLIEKKEEEIDTLLKNKDAIFNNETKHENKKSFKRWSLLKEHPNYSQIINEVVEVPDFQETDTNKELISQGTDTKEVPNF

QETSNKEITNFQGTESKNDEASDKIVLPADNQSVDSEVTLNLSGNTKWAALTKKNAKKIILESDKLKQPVLSLSSDSTSQT
DNKILNNKNSLGKSKWNAVRERTNSIKNTKDDVNSNEKPSLPKLLQVVNTATATKTAASKFLEASQMRVNLKARMRKI
VQERLTVKQIFKRYVESSTLHGFCYVCGDTFLVRRVLWALLMILGAIYFIIKLRYGIEEYLNYPFSTLSTVDYVHELFFPAMS
LCATNSYLASMVSKNQLKLLYDEGRLPLDNNQTNPSYNMSGNELVKAIQESSLSIESMLAFCDWIQQDTNNPDIPPNPC
GPQNFTTYLNYKGEQCYTLNSGLEGHKLLKVDTVGLTYGYELIFDLQTNKVIKNNQFSGMRVVIHNQDVPPQLADGFIIP
PGFKTFVKMGTVQSKSLPPPYSTECGTKKLKYYNTYSQRFCLLETLTDFTGKLCGCRDVFMPENGLPFCSLKELYSCMYP
AKESFSEFTMRKECPADCEERTYPYELSEARFIHNPPIGLSIQGLANLQHKKHLEEELYLSKLAQSMTSEELDAYVEDNIVS
VIFFFGDTRIDYNEQEATNDFFQFLGNMGGEFGLMLGASLLTFVEFVDLFIFLIYHQMLRLHTLKKVPDIFGRKRSRINEYI
KKREKTRV

>Protostome_Lophotroco_annelid_CAC9484044_1__Ofus_G014320_Owenia__fusiformis_

MTELTSISIPSGKQSVKKGDCLIVESSVDISENTAVKEFLEDKSLCKINENDEISENNPENTLLDEEKTIKAEISGILKQIVESP
VDKSENAAALELNRNDNLFRIKKVNGNNEEKISIALCQDIIHTTKTELEKCEKLQKEVKDHDVPVETNEFIRDVNEQKTLS
DKTSDSKTRSTGNEDTNCISPKKDVESEIDIEASHIPETYESISDACLRKYNSLKKKLSSFANVTSIQPFINIRNADTMSARLV
WFLIAFCVWSIVLNGVTEITSKYLGFPTENSLYNREELVTFPSITLCPVLPVIFNNTWRLSFEDMLFQLHSQNIETYFSDNR
SVHENLIAPIMDRVVRKEGYLDLVGDKVRLRTYSDLIIECRFNGELCNETDFINVTVGSYGNCYTFDTNSTGIFDVSETGP
DFGLRLVLFLNSYSPLDELISRDDFKSELREFDSNYATLSDGIRMAIHAPGTMPDMDIEGIDLSPGKFITIGLAQDVRTLLE
PPYGECTNEQNNKDTPYAYTVDLCLRQCQQELMIAKCGCMTYNGPVPPNTNVSQTPYCGQKVKSMYFQVVSNVPYL
VTNNKFEWDEVIKWSDLEEILQVINCEKNVTKMVSVGNQTCGCKKQCVDIVYDPLVHSVPWPNEKYYKGINRRNFIRF
LRQQPKIARILNENLFHNDIDSIDSQREFCDIVNPFNHVKNAEYREPPLFYTETWPYLDCIFNLGNPSNLYDYGYYYWYH
NLYDGDPTINKYNIVNKIIEKNFLKVNIYYRSLTVNTIAEEPSYPLNKYLSELGGIIGMYAGMSAVSIVELASLLVPLLLVLCR
YRSRF

>Protost_ecdy_cyclo_nema_Cele_sp_O01635_ASIC1_CAEEL_Degenerin_like_protein_asic_1_OS_Caeno
rhabditis_elegans_OX_6239_GN_asic_1_PE_3_SV_2

MGKNSLKRALELDVVDFAEHTSAHGIPRAYVSTGWRRYMWLLCFLFCLSCFGHQAYLIVERFNRNDIIVGVEIKFEEIKFP
AVTICNMNPYKNSAARELGAIRNAIEAFELAIDKSDGNAHSKRKKRSANSKMVPIDLLCKEEHGMFTAHDYGHVECTCV
TFEDMSKVGDTDDDEIFWNCHQRKDWTHKICHLAEGSNQLKTCKCFEDTCVSDEVTKQLVWPLQLSKNGTKLCISPES
SGPRYCASAQKFQVSTCSNCDWLGKCEESDDMDLEEEIDSKTCICHHGNCFQIKGNVKKRKRRTPERKVHERLLSRYEG
LLAVYSHCNCTKQHGCVSTSVPDMDLENSNKTCLCFYNKKNEQIWPCYKEPEWEERKCSRCNTMGDCVYSDKPKKQTI
SCLCATPIKMCVRIDPPQTNDTSLDDRVVKFWDIQPSTTMSPIVKKKEERDKAYGYTGVKDRIALRAKAMENMIFAVDA
LTEEEKWKISYNKSDFIMKCSFNGRECNVKHDFVEYLDPTYGACFTYGQKLGNNTNERSGPAYGLRLEVFVNVTEYLPTT
EAAGVRLTVHATDEQPFPDTLGFSAPTGFVSSFGIKLKSMVRLPAPYGDCVREGKTEDFIYTQKAYNTEGCQRSCIQKHL
SKTCGCGDPRFPPYRESKNCPVDDPYKRECIKNEMHVATRDSKKLGCSCKQPCNQDVYSVSYSASRWPAIAGDLSGCP
LGMAAHHCLNYKREQGSMIEVYFEQLNYESLLESEAYGWSNLLSDFGGQLGLWMGVSVITIGEVACFFFEVFISLISSNR
TKRRPARKSFSSSLRCSTDYNLNKDGFNLDN

>Protostome_Lophotroco_annelid_CAC9572753_1__Ofus_G053069_Owenia__fusiformis_

MEPRSSTKRLNRYKAARDDNVYHNGSSSQSSQHPGNGHEKWENNNEQNWGHFGGYDNGFIAGMNYRKDHDNLFS
EDRVSMMSYHESETFQLPKTFRNVISSCANRTSVQGVPNINNSKSMVQRVGWSLLFIAACVVTVEQLYEVFDKYYSYD
KNTVISLEYQHIRFPSITICNQNAIRKDKLHLARPELKAFLENYMKEKPFGNRKKRDVGPQPLGPSSTPSDSASIQSATSAT
PNSTFGPNGLNIIPTNGPNGQEVTNDSNTQMPTSDTSTQRLTNNPNIQGDINNPNVQGDINNPNVQGDINNPNVQG
DINNPNIQGDINNPNVQGDINNPNIQGDIINPNVQGDTDLGQSSSTFPGQRQRVSNTTFLSDKFNSFRF
EEDGFEGLTEDAARDLFEEYQYLVHEESDETRQQMGHQGNSFIQSCSFNGYECEASNFEEFLNTEYGNCFTFNNKKSRK

GESLRSTKAGYQHGLNLVLYLDGRNYMPGITQGSGAILVVHEFDTTPYPSQAGIAINVGTETRLALKMVDIKRLGGIYGD
CTFDSTWEKDYGALYTQQSCIETCQHRGIMELCGCHVHGYKEYYPIHFRSNGTLCETDAELLCAFKLKRDYLKGNYTCDC
NQPCDERIYEYSTSSNDWPHTAYEDTVKELGCSDDVDDEKICEDVSSPQFNVRDDYLSLRLFYQTLNYESIEEELDYSIAN
FLSDIGGTVGLWIGASVLTAVEGLEMVALLFVFFYRKCKRWYRADQEYERKVQEAKRKKREEEEQAKREEKIKQEKRQS
KRSSQPSFANSNGLSPMGSTFRPPLRHTPSYSSARY

>Protostome_Lophotroco_annelid_CAC9668428_1__Ofus_G16870_Owenia__fusiformis_

MNGFFIDETILYKADCRNPLSEDELIRNDALNSSVENQGLEELLNYESKTMFFYSHHPDHFALFTPDAHWHQGEGLNTD
GEHITDWGAIVQQTGHKASDMISSCVYDGESCKDNITMKLTDFGMCYTFRTNRNCTRPGATYGLQLLLNAEIYEYMA
GPRTDAGIKVLMHSPDDEPMVKGLGFSVAPGLHSLIAIKASETYSIPKPWGSCASKDLTYFDKYTQVNCMRECLTKNVV
DLCGCRTEFMPYASENDPPVCDMHSYFDCIIPNIDTANITTTNCQEDCNEQCEKFVYEPMLSFGAMSELSVDEVLNKDI
SGELKDKYYNAINVQTKTNKYKFLNVQRVFRDLSTSMKVFQQFLDKWVFAETNIFKKVEDGASAMNEYFKNDAMLTN
TDRTHFNIDNILLDFRNNILVFTESLFSSLNEVEQYMHGLSAGYKSDWVQFRNLTKTVNTFVELKAHEFGNFVVRQTLSL
PSCNQFSNLKDLLDDLTGAYYVWNLNTEGFRKNKITLKTCLDNLHTNLTNVVLLKEQLESMYNASFIIEAKDQYPDLSSS
HSVISEKAGDIFDIVELFLENNPSYDFMITITEIEELIAPVQGELNILTDTILLKYMTISSEILSSYDKIGRENYARLLQVMLGV
QQYFPSSDVFRYDEMNIWDEPYVVFDRDQHEYETRFRPIKSEIKQAFIKTKGKEKSTAIIKSFFESLKPQLDVFEKEYRAAK
INIEEALDDVENLIEDSKKHGKLDDQFVKRNFLELDIYYSKLAYQNIQQMKALDEESVLSDIGGFMGLLLGASVLTVFEILD
LIFYNIGKKLRSKRNFTPIKVSPLSTERI

>Protostome_Lophotroco_annelid_CAC9662918_1__Ofus_G12800_Owenia__fusiformis_

METKISTNDIKIQQENEPTTFKGLLTDFANNTTLHGLPRVITSAQLIRKFVWFCIFFGSFGYFSFQVAELINDYYKWPVLIR
SEMKLRTFMEVPAVTICNMNMLRRSKIEGTMFNSLVELDNDTANEIIEKQDTTENEINNQNQIQNSDKTSRKKRNINM
IQKSIQRPMSKENNALKGHENKFEISRVKTKSMESVNNILAFEHEGSGSTKIEHTFFDGSLPENEADDPHSLNDEYIALED
GHSVEPIDDTNDNGRRVRREVAGGSGGEATDSYVSDPTDTTTDSYDTYTDPTDTTTDSYDTNTDPTDTTTDSYDTNTD
PTDTTTDSFDTNTNPTDTTIDSYDTNTDPTDTTTDSYDTNTDPTDTTTDSYDTNTDPTDTIEDSFDTNTDPVSTNTYPIYT
DSYPTDPNADPMDFDTYPTESYTDPNFQSSDYYESYEYSEYESSEQEMDQESGEDPGYDVPFQFEQSKLMGIKDPNNY
AEILRLAEAEDLSDIFGLASPTVEQLDEYGHNLSDLVALCTFDGENCNRSSDHWLETYNKYYGKCYTWNSVINKPRGERP
LVTTNFGSRYGLRLTLNVERDEYTGLFSPTYGTRVAIHDSSIISYPENDGITASVGNEMVIRLRLKKYERETEPYPSDCVKD
DDLVKQFGGYTVIGCMRECLEERLVKRCKCLDRIVEKDEHRCSYFNRTEELCRQRVYYAYNSGKLGCKCRARCRENKYDI
STTVSTWPSKHHDYYLLNRLKERGIQTLEEIEDNILRVHIYFEDFNVETIKEVPAYSVISLLSNIGGTMGMFVGLSICTLGEF
LELIWEMIMLLRRKISRRHNMVNASSA

>Deu_Ambulacrar_hemi_Ptyfla_40v0_9_20150316_1g11917_t1_scaffold7296_cov167

MYGRHARYFERGERTKYGGYAEGLAKPTDTDIDRQIRNQTFSVSRSVNMYSEPTLDSPQRTDEKASKRSIIAWKVVKEY
GLNSTSHGIPRIFGAKSRVLGFFWTVLTLTAFGAFLWQGSELIMQFKRFDVITKVEVVTEKRLVFPSVTVCNVNKLRKSAI
ASSSYKKMLIVDEQIVLPYYAPCIPGDFACANRIYCVKEYLRCDGINHCKDFSDEDGCVYELHKSYVMKTGGHKLKTCQR
LATKSTYFNDNKLRNQRCSLRMPFMTVCEEQAVMRSKSPVTDNRQIPVKPGNCGENQFKCINGSDFGFCISKKYECDR
KSHCYDGADEDNCRACGSYDQFTCDDGVCFPAWYQCDNYEDCADGSDERDCDCTFSCDNGTLCIPDFWICDGVQDC
MDKSDEPDYCNYDAIPDSGDETSQMLFPSDDSDVLTMPSDEEGDLFSFSLLHDDAMSVASPSNEDLTTEAENQTLKYN
CEGFRCFDGQCVELYSDCDTIVHCEDSSDEDDCIPSDDRYGNCYTFPGSRNNEKPLYVSRSGQSHGLKLTLFTEQDEYISI
YGRQAGIRVTITAPENSESIHTQDGITIQPGTETHIGLREKNLLRTIHSLNNKTLNINDKYSTRGDRTQYGGFTEGLAKPTD
TDIELPIRDHTLSVSCSTNMYSEPTLDSPQRTGEKVSKRSTIAWKVVKEYGLNTTSHGIPKIVGAKNKVLGFFWTVLTLTA
FGAFLWHGSELIREFQRFDVITKIEVVTEKRLVFPSVTICNVNKLRKSAIASSPYKEMLIIDEQIALPYPAPCIPGDFVCANGI
YCVKQYLRCDGIDHCKDFSDEDGCVYGELKE

>Cnidar_NemVecNVEC200_012212_1_1_protein_AED_0_36_eAED_0_36_QI_113_1_1_1_0_92_0_86_1
5_153_835

MSQNEGKTAKTVLGFIWSFFDQTTAHGFGRIANTISKYNRLFWASIIMVAFGMFAVEISDLLIIYLSRPIGTQIWMKHAA
SIAFPAVTLCNFNIVRRSQITSELAGSFENFITPNITKDGDGGGLRKRRVAVDWCPCTYYLCCRNKHDLVQPTEPPVTTPV
PTTESTAVPTTAPTTEPTTVPTTIPTTIPTTVPTTEPTTAPTTPEPTTLEPTDPPTTEAPTTLPPGVTPTPTERPLTMEEKKEQ
AFTGEVSQPDPATLDKQTLKTEELVLSLATKPEDMLMEAGHQFNDMVLSCTYRGIPCTNYTENLWSRFWHYRYGNCY
VFNSGKDWQGEPKPILRSNKPGPASGLTLELNAEQHEYVGQLSHEAGMRVLISTQGEMPSPLEKGISVSPGFSTTTGIR
MTNILRADPFNNKSCLSSDDIDPENLYRKRYNVSYSRTTCMDSCLAHKQVDMCGCLEYRFPNIDNRTVCDILDIDVIRCL
NKVQSLYQDNKLGCSEKCRPPCNEEVFKLSISSARWPTDDYEAHFLDELKEDKGFVLPANADARAYFVKLQIFYEELNHE
VIEEYRSYELVNFVSDIGGQLGLWIGISALTASELVELLIVIALHLLRKVTRTGLIDNSAEQPADTNPRYLDDITLTGDLGERR
GALTDSNTSLTVRYLGEIARKRKMKGHKEKRETSWHIDNSADQPADTNPHYLDNITFTGDLGERRGALTDSNTSLTVRY
LAEVARKRNKKGHKEKREASWHIDEEAFPCYRDQATIHPNVESNTSLTVRYLDDIVQKKSKHRKRSKGDTSMVQKGDG
HRGDLSGRRGELTESSGSLTVRYLAEVASKRKR

>Deutero_Ambulac_Sakowv30020102m

MEMQNSQVTEKDDTQRKKDDKESYKALVNHFISTTNAHGLPKAFEGRGIFVSSFWAIVFLAALTGAALQISELLVQYLE
WEVKIKMKVVSESYLQFPGISICNTNKLRKSAIEKSEHKGLNGVDDDIVFPYYDEGGCMANDFACNNTGCIKNYLHCDG
YDNCRDRSDEMGCKYGRCGNTEFRCPSGSDMGVCIKLKFKCDRHLDCYDGEDENECVCKEKKEFSCLDNGRCVPVNK
LCDGNNDCADGSDEKECSEDGNYCPSDYFKCDNGRTCIPAIFKCDGGADCSDETDEMSCPTQNPGCADDQFHCDESY
CIDNSWLCDGEYDCMDGTDELEMNCITSSGTNSTNTNPCQDWEFYCGDDKCINGDWECDTESDCPGGQDEVNCLT
TRAPTSTEVGSFMCELSGKLVERQYVCNNVVDCDDASDEYNCHNDASPTYIPDAEELLRNYSTLSTDLSLFEDFVSRYYID
NMFGRVVRENPPDWPGFIAYSSSPDYSDLAYVLKLRTDEMHKLGHQLNDFVLHCTFDSVKCDMEKDFLTFYDDRYGN
CFQYNYKHDHDSEQLLSTKTGPRYGLKLTLFTEQDEYISVYGHDSGARVVIHPSYMRAMPWSEGFTIAPGKIAFIGIKETR
VDRQPAPYGECATNIYQETVYGKYYKESTCEESCIQDRMMEYCGCVDTMLRNATRCMLLNRTQDTCRQLIYYSIQQKL
LGCDCPQSCKERYYETSLSQSLWPSNTYLKHLLKQIHAENPKTLNINNLETSRQNLIRLELYYDDLNFQQITEKPEVSEEAL
LSSIGGSLGLYCGFSFLTIVEFIQFAFDLVKLTYVRVFLRRLKPVAVSA

>Protostome_Lophotroco_annelid_Pdum_comp414816_c1_seq14_

STREISFLAWSVNSHTMRNADNIADDTADNISAKCEEQGNGEDQYDYPNSDEDNQALDKQIKEAEERLDQIKSKEQLE
EKRRRLRELQNEIDREEERQRSGQKKSHREETVPRSRGERDETRGTAQQKDNWLLDEDSRPAEEHTSRDKWSNLFHDF
TDSTTLHGLRYVFMRSAWMLRRILWFLVLAAAITGFVYLNAINVERYLAKPKKINMDVSVRKNLTFPAVTICNNNFHKF
TSVVDMGLSDTFYSYYDQRASESVKGDFYQYQDDICLYDELLVKKIISCGTAFCQLLWKQFSPLATAVSSTVCEHLCSDRS
ACTGFFYDRITKKCSLHSGLIAANLSALVENTDWSDYEQPPIQVIRPDKNEICRKESIEYYLRIRMLEKPSFLCTFEMGDCP
PLKNSTDNYRAEKLFISNAIDPASDFTFGSDRGHYLFLPERKRLEYSLAELDTNYFTMKDSCLEFYYYIVGDVQTKVEVSLF
GKTYGNGVMKLAKFQGNQSEHWQRLYRRIPDGEGFVKIRMKRGKEGGTAGFAIDNFAIQPCLKYEVNCLQDPRGETY
AGDISLTGTGKNCSGRISGSPTRCRNENGDIRPWCYTEETWEYCDIPHCGCSAEEFWCRKSERCIPRGFLCDGWNDCG
DRQDETEPCAALMTLYCPFNKKFICGYQSVMVRDGLQFKHFTEIINSNWPTSLPFNDYTFVSVVTDFAAEGDRTILRTPP
FNLLENKDTCLLLNIFSLTHVYGTQEQLEVLLVPMDQTEYRNVTLKVFNLPKESRKTYGEEPRLRSFLSKLPQDDGRNMSF
FIDLVYTLGIPTKSKFSFFFIITMGCNQEEKKTLQYLSKGFNQNPI

>Protost_ecdy_cyclo_nema_Cele_tr_Q19777_Q19777_CAEEL_DEGenerin_related_Touch_involved_cha
nnel_OS_Caenorhabditis_elegans_OX_6239_GN_degt_1_PE_3_SV_3

MPRKRRSEDVDAMVIATSPIKIKVEQFIEETSSMDVYIPALDRFAEDTSMLGFRYLHTRYKTWFRVLWGFVVVFFIGLTF
YQVFERVTYYFIKNPLTTRRSYETLPNMYFPTIGVCNKMQIKASSVASKNPDLLRGMCSVLDESSSNSSRFDELDKFDDV
DILSLYRNSFQSADDLFVSCEFGKSGSCQDEIRPIYTPFGLCYSVSPNKTILRPGPETTLSLVLNLEVHEIIPGTVVEPGVVLSI
YDGASSLSHYSEGIHLEAGKVVTIPVNEVRKLRLHESSCGSTKMESFSEKEYSKAACEWSVSVKQIEKECGCIPIRNPIYRG
VFDNKNDPLNNSTEIPKKKYKKWKKRNIPRCTLRQEIECVQEKLNIRPHIDDSICPDDCDDISFSSIVFGGKLSSSEIVSMLP
SDWEDTKEKRVADYQKVLEVIPNKMIPVVRNVQQLADELQLFVKEASEIFGVSDNFHDVKCLSLDGRTYESFINQFYSYE
PTWERITTYLQHSLARELNSTALCLGLFLTEKGEIDDTVTPVVNMTLASISLLQLGQIENSLGKANFNYGLTMMHESTRR
VVLGLAHPLITEVRDCVTKMYENLERVEEIAEDCRIIFKNRYTPLIEASNVHTKTNPNSESFKLYTEAMKKVLAKLQILKTRV
RMSEWKDFKIDLKEFESSYREIAKDHVEIEEMLKLRKTMVTDIPALANELSETFLAATESRRKFAVLTGIAADESFSDKFSN
FSKCLDELHEKAPNLKKTRFIRGEWLSRLRNQVLMAQSYSPTPQYDVVNLLHIKFYFAHFKQETILQERSYNLFLLLAEIGG
TIGLYVGATLLTAAETLVFLCERKKSNIFLKPQFI

>Filast_tunicaraptor_GIQG01049703_1__p1__GENE_GIQG01049703_1_GIQG01049703_1__p1__ORF__
type_complete__len_874__score_117_73__GIQG01049703_1__2484_5063_

MTGAASDERLEKLQAQLDVLAAQLRAARKEADNGGEYDDAALGSPAEDPYGAGSGESCLSMLTGCMKESTVHGIPHA
FAFNRTIWRRAMWVTYFTAAMCGLLAFCVGRVRYYYERPFSSSVSLEYAPNLAFPAVSVCLTNGYKNSGYVAWFRHA
AAEAVREHVPGAAAFDNAEVQQHLESLLFYGDTLASPNGASGGTSVARRVRQSPAMRKVRAEKVAAAAAVAAAAAR
VAREAEDGGACPRPLSDTPGDVAFAELIALLHDSGLASSALSFNVDDTVETACAAWADAYYEGEETPTVVSYGESAVNF
EPMCAKYVLLGDDPVYHEFKTALTAAGAPAHNSTAAWSDALAAEGVASSQNALVLPLRCEVIASLSELGVAPPTPVPSS
TGAAPTPTARAAAPSATPSAGSAHISTLIHPAVLSAKESALIAFISDELDEEELEKMAYTAEEVILGCLWKGGALPCSHRNF
TTFFANEWGNCFTFNNPQTLRNDVLHSAAPGPKGGLVVTLFIHQDEYEALPAGSITEDNFAALAIEGASVKLLMHDQR
DHPEMDSALDVAPGRYHLAAMRKLVVNNLGQPYSDCQEDVAAALTSTDGVVLSTIPGRTRYNPKTCFKECTVREIFKE
CACLPPTSRVARGKRVDRCLSDEQVACQDAVEGRLSNDPNLQTTCACVKPCESVTFPVSLSAAQWPSEVNRDVFLADI
EGVLGMQVPPTFFDVNIVNLNVFYSDLNFERIEQQGVMTVGSLLAEIGGMLGLFVGMSVLTLAEFIEFLFFGSRRQVRK
CKQGGDPIGPKNGVPSDACGGADRVDVVLEEPNAMNMQGLPLSKEATAAGAGRGADPVRILKLQRADSVNSVHSAV

>Protos_Platyhelmin_Macrostomum_lignano_A0A267DEH5_A0A267DEH5_9PLAT_Uncharacterized_pro
tein_OS_Macrostomum_lignano_OX_282301_GN_BOX15_Mlig002193g1_PE_3_SV_1

MVASQLCPSTNVLEKKNSYRTSASRAPCRTWATRIAPSGTELSHQAGLILALNLSESDNKCRAPPGVPLTQWWNGSYR
PLQIELGCYFKAAASAPYKCAINDFAGMTSQMCRECCYGKGYVYMGLTSPDCYCITKSDGDNMKMFGSASNCDATCN
GEKRVCGSNDLLSLSLLQIDFDDSRPMRIPLGNSLTNSMQGPYCFYFDSTQLSAYYCGVPFCGANENHDCVPSQSEKVA
PDQTELAHYTGFRNVTVTGKPCVNWRNLQNISRTIEGYVWNASRTFLQMPDSVGFSDHNYCRSVSHVQENIIHIYGSL
KKIRWTLESSLTPGCYVYTPDEENIYTFRTCSIPMCYTESPSITGAIRHSIGTDLLERQFGSNLDVSLTDCSFGGQKCTQDD
FLNMLHPQLGLCHRFKADRFVAGINNVQLANSELSITFFTDATDSFGRAFNNISKLEAGVRPNEMLADETNNALKVIILP
QGEFPDSSYAFEVDPAQLVVIRLRYDRVTKISRSELPCKKLEAPVTYRMSSWASARDKNLTKLKIDARNLVQSSANWSQ
KIAMLKAALLENNQDDGGFQQKMEFKVSGFQDCIYAYQVQKFVDACNCLPESLPIPEAYFDRYSYCFNSSNVAYLDATE
QCYVRQMARLFLYQKETFSVCESQCRIRKYVLRQATRTWPVISRVNAVKFLANLASLTGYRFNLSNSRGIASIYHTYLSAK
SVENVPRDNMSSSELLKFDYQLQAIKNRLAKLTVKASSAFGEDITEEEAYPAKNLVADIGGALGLWSGVSVLTVCELVELL
IYIGMVVGQRMRKQDVEPETGKINDNGTQEAQNEIETVDTGESEQVGNGETQHVVTSSKDSAETCL

>Protos_Platyhelmin_Macrostomum_lignano_A0A267FD09_A0A267FD09_9PLAT_Uncharacterized_prot
ein_OS_Macrostomum_lignano_OX_282301_GN_BOX15_Mlig000209g3_PE_3_SV_1

MTCRHQHGSAVRRRRRNSCRLAAVVLVWSAAAAYTVFRLVEMVLTEPSDASTSGSAAASASISAASLLASSARLGTSSA
SSAAAAASAAAAEASSLEEPEFPAVTLCNINPAKGSELVRLNTAQYNVSNAYESLVDFITHKSSLQSSFIPIKKIFQETGHTI

ADMLKGCQLSNKRQCNEYNFEHSTTILGQCYTLHPSSLSNLNSIQPHGLRERENSEVTSLKVLLDPQGYDYLLPNEGFVG
VTVIIHSPFEPVVVNLDQSLQVGPKFHTTVHVTASSKVNPAPCSNRATWKSCVEKCFASALLSKCKCAIIDLPYCTLNDLA
RCGIDISENPGINWLSKCSCGKPCVESDYRIQSVVQTPRSSLLSRLQTIKERDALARNTDMVEDANLVQQWAFLRETNR
TAFATLSRLSAILSRFSSKFAETVHGLRRMAVTWHLRYVQNQHYGSQCANIVIEISQALSNMRQKAQEVREFSGSLFFET
SFNYPNDDFLFRVVNLFFKYSPENIRAEFDLFDLTRFYQDLTAFEISFDKFKVAKCNSLENGGGAAGDQHRSLMQEDSAI
KILLSDLQSLSLEFLRIFTDIYSNNEAIRQAYPSSRSKIKQTASPKTQSSGGSQPQSKQKKSGPASRRHGSRRSDRRGSGGG
TRRLLKTNRQKQLRQQKRQRKQQEKQQKLQSKEDDEISKNLLKSFVALTVKLQFEQKMRSQPTAVRIVQTQGQDRVLF
LITSTLSISLFFTLVVHLLSRLRCRPKSKLQPPKGESSAGTCELHQRSLRGVKAVRPSHSHRTFYFGEEEDRLSCSPAGLGEM
LPASASASSSAAAAAAAAAAAACGHGTIERFNKPLGNQGESDCALTNNDYEDGWM

>Deu_Ambulacrar_hemi_Ptyfla_40v0_9_20150316_1g10747_t1_scaffold6363_cov125

MENLEIEGDVQAVNQRKGDQRIRNTLADFGSNTTAHGFARIVAANSKIVKVLWTIVCGFCWGIFIWQAIHFIVKYYRFQ
VTYSISVDFRELPFPAVTVCNMNPLRQSALDEIDDKYKEKLGFLNFEKKTQNDTDEAFDWSATVPTQSFQPSDEDFDET
LVDLDHAKGLNLSAIMSQLPGDVRAKIGHHKEDMIQECVFNGVKCSSKNFNLVLNPKYGNCYSFNTGANNTATFRTTK
PGPSTGTQTLEMERDVQATKQHRGNDHRIRDILAEFGSNTTAHGFARLVDANSKILKTLLILACCFCWGLFIWQATHFI
VKYYRFEVSYSMSVDFRELPFPAVTVCNMNPLRMSALEEIDNKYKKRLGILVDEKENHIYSDEEYNSTNTDEEYYLTDTEA
KACVKTCHQFNLIAECQCYDSTLPHNNRTIPACVETDKGNKKEVSLKVQHILEDLENAIASTAYPKATDMFEGYSDCDSR
CMIQHCLQIETLRRKSITSIRVNAALTQKPAQYRCRIVAMEKMEMEGDVKAAKPKKEDHRIRNTLADFGSNTTAHGFAR
VVAADSKILKFLWVLVCCLCWGVFIWQAIDFIEKYFRFEVTYSISVDFRELPFPAVTVCNMNPLRQSELDEIDDKYKEKLG
FLNFEQRTENETDEEFDWAAPAVTQQILNVSGEGQEEESFEDFYDLDHVQGLNLSAIMSQLPGELRARIGHQKEDLIQE
CSFNGVKCSSKNFTLVLNPKYGNCYSFNIGANDTGIIRTSKPGPSTGLRLTLFIEQDEYLDDITENAGAVVLVHPPDNVPF
PEEQGLELPPGFSTSLGVRLRIDLLSSLGGLMGLYIGASVISLFEFLEFVGKVCCLIPKKRNVLDDANTKA

>Protostome_Lophotroco_annelid_CAC9671100_1__Ofus_G19963_Owenia__fusiformis_

MANKTSPEEVDTQHDKDHPTFKQLIIDFADNSTLHGLPRSLRNSVRVFRRLIWLGIFAGCFGYFIFQVIELIKDYNKWPILI
KSEIRLRTFMDVPAVTICNVNMLRKSKIEGTTFDSLVDLDRKNFANGNTRSTTVHKKATPKPRNKRDGKVEETWERSRP
PLNSNTDVKAGRVNIRQILRGYRPQVGIGSKENGVSHSVNTLRDVKPDDPGPYNENGLQGEGIGENNYETFDNRPIEAY
NAVNDESPSRDDGLLRQHDTADDVANDKIRIKRTLDTKPSAPGATATDPTTMFDQTTTTISGQIPTKSDPVTTISDQTTT
ISGQIPTKSDPVTTISDQTTTISGQTPTKSDPVTTISDQTTTISGQTPTKSDPVTTISDQTTTISGQIPTKLDQTTTIFDQTAT
PGQIATNSDQTTTVTSVPICTCTIPVAATSAHTQTSPPGSTTVSNDTFVESDTTSSYTGNNGWSNNSSLGQGHDPNTYD
RYGPRMDTGYTAPFTFKESPFLKIKNRNNFGQILNSSTAEDLSDIFGLATPTVQQLDMYGHRISDLAALCTFDMENCKRT
SGHWLETYNKYYGKCYTWNSAINKPEGERPLVTTNFGSRYGLRLTLNVEPDEYTGLLSPTYGVRVSVHDNSVVSYPESD
GLTVSVGNEMVFRLRKKKYERQPNPYPSNCMDGWMRPSKNRNRGEYTVLGCMMECLQQRLFEICGCFDRIVDEGEH
RCNYFNRTEEFCRQKVYHAYNTGRLGCRCPAKCRESTYEVSTSVSAWPSKHHDHFIRERLNKINNISLHDIKNTTVRVHV
YFEDFNEETVIEVPVYTVSGILSNIGGTMGMFIGLSLCTIGELLELMLDVAILLYKKLSTRFNIRRVGTAVT

>Deu_Ambulacrar_hemi_Ptyfla_40v0_9_20150316_1g26495_t1_scaffold55703_cov149

MVSVVMVVMRQPLSRNLSQKITPYVLESLSVVYLGEIDITTRQRLLNVLFPGETGISMDRKIIDNILVSTPGGGLLSPNLQ
VIDLPVENNQLTVGQLYEILHNALQSYYNLIYAEISIPLSSNLYRYIHSETQSSVDVGSDIYRKVYEATMNRNTSAWHLRTT
ASHYAVEIAFRQALNGDHPYGALMNATMHAVSDAFPDIFDLIYTTVLDTFEENRLTALENITAYQICMEYMRGAGYRG
WSRNRYNIFFAISNVEEAMYVHTCKTTVLAITCRKTGLLEAFGYSHGDDVIKTAHAILSPSTETMDQDVNLVGVNANYD
FEYLMTTVGHKKDEMIMRCKWPGHQCSVDNFTTTYASDSICVTYNLGERGDIVEQVNGGERYGLTVYLDVESHEYVN
SRQHYVGFRVIVHDQDDIPNTKDRGFNVAPGAYTAAALKKSSSSYLKEPVGHTDCIDSPRGTLHYFHGNYSLAKCKVEC
ETDYAIKQCQCRYYTMPGDAPICNVMQLHNCYVPKMAKFTDVQGVCSYCKQPCYEESFTEHLSYALFPSGPLVEELAKK

AEKPCPKAMALHYKKKITPYVFASLSIVNLDEIHITTQDKVLDIIFPGETNISVHRIVENITASMPEAGLLPLIDVHDSPVKIH
QLTVGQLSEILRKALQSYYEQIYAEITIPLSLNLYRYIHSEGQSSVDIGSDRYRKVFEATMNSSASARLLRTIAPDLVVNGEF
RAALNMSYSYGALVNTTKHAVSNEFMDIFDMIYTTILDTFEENRITALENRTMYQICMEYMRKNTLELRVYFNEMREEK
MTQQQHYEPFNLVCDIGGTLGLFFGASLLSVIEIIDFLLFRRCNKVESQKNESSIALENRGSEKSATSPNEEHKT

>Protost_ecdy_cyclo_nema_Cele_tr_Q20852_Q20852_CAEEL_EGF_plus_ASC_domain_ion_channel_OS_Caenorhabditis_elegans_OX_6239_GN_egas_4_PE_3_SV_3

MLLLWFFLPLTLAQIDTPYDCGAHGYCYPANRTSATTCSECTCSDESTQTSDTTCTAPTSCSPNPCTLSNQQCNMVNDI
PTCTCAVGYTGTDCTMLTSDPCSPQPCLQNGVCSSSGGTYSCACATGFFGEQCQYSGDPCSGYCSNGGTCEMLFSDTT
PYCQCPFDYYGDKCQTTRSVQTSYVGCFDDSSASFSDYYVYSTTITTGTDCRDALISYREANPTSNYAYSTMSGTTGECLF
STANALTDPPTGGGLLVALLNALLSTCAYSNLNSGFAAVYSLNDLCTPEPCGEAAGNGKCIQTSATAYTCLCNPIKTGSAC
QSDATLTPCASIDCGVGTCGVTDDQIGYYCLCASTNQTVACVGNPCTSEPCLYGGTCSDLGNGSYSCACLNLYMDTNC
ETFDSCYINKCENGGTCVPTYNLLDSTFTCQCTPDWKGTYCEEERYYCDETPCENGGECEDIIGPPNSYNCTCTPQWTG
FNCTIDVDECVEDTTLCKTKDPDATCVNTNGSYYCVCSPNMFGKSCLFNKIIYQILNATYGDLGPDKLDEMAQELTNDP
TLVRDIVPFLIGGYSLEVRTSLSWTAADMFLWVAYEQQLIDLNSNFVQWNDKVLGNCFTFNHMNSSFKYEARSSGYPG
GLEMQMNVKQDEYLPWTETAGVMVFTSTKEEAVTSESVRINTAPHFESRIAINRVDYYRLGGRYGVCINSVSEVKSYYY
DGDYTTDGCLRSCYQDVVNGDCGCMDPRYPMPNDGISCSISQKTCIDELVDSRGDPSTWPECTCPLPCSQTVYTSKLS
RLPYVNKIVDCEEAYTNKTACYETFLDSVILRISLPKLDYMIYSETPAMDLTKFMSYLGGILSILIGVSIVSFVELFFLFVQLIVI
LLFNKRL

>Deutero_Ambulac_Sakowv30029067m

MMSETDDDANRSEEGSTSRDDENSKKQLTYSNIFLSFSSETTFHGIPRIGSAKSLQSRVLWCIIFLCALILFVWQLSTLISQY
FEYNVVVNLKVEMGYDIVFPAVTICNTNRIRKSALNGTKYENLLYLDQQDDPKYRYYAPCFLGDFACQNGICIKKFRICD
GVNDCGDMSDEECQPYADCPDGQVKCESGGPYGVCIDVNNECDRKCDCYGCSDEYLCKVEQITCSKYEYKCKSGASW
KDESLMENVQRVLGEIETSGDISPYYNGFYHGANTFCIPMLNVCDGHQDCNSGEDEEEDLCLDPCDDYEFYCLFGGIHQ
LCIPEIWKCDGMNDCYDNRDEKLCYSVNGYKCTSGDAKNGTQCDFIPDCEFGEDEYNCHFSPCKHGEFRCSNQQCIPS
SLVCNNIYDCYDNSDEDGCCDADYFICDNGGCTAQENVCDTVQQCTDGSDEQNCSRPWYVGYFGSNYTFKYMPNAL
RVDNITINITFINIQSVDCLYLTLSKEKEIMPSMYVITIPLSLKKNITIQRCLDCLEEATALPTIQWLDGLTFFLSFKAGVFEIGK
PGGMTYISWTDPYPLTIHYVGIAQLYTPAFPQLIKVTIYQDTLNAFVNQSYVDTWDYDWYAPYLTVNDNTSLYDDFKN
NYVDRKFDRVTTEDPPDIESFQRYSSRRDHSDLEDILKLKSEEVEYLGHQLTDLLLQCTFDQSECDESYFYEFQHEQYGNC
FTYNHGKNGSDIMRISRAGAQYGLILTLFTEQTEYISLFGQESGIRVVIHPFDRIPFPEDEGVTVPPGRATSIGIRMTEIIRK
PEPHGPCTDEGTSSFIFDKDVRYSTVACEKSCILSHLVDMCGCVDTMMVTGYPRCMVLNKEQERCRQLVYYMNKKRA
LNCNCREPCT

>Deu_Ambulacrar_hemi_Ptyfla_40v0_9_20150316_1g21913_t1_scaffold24959_cov85

MSPLDIHTDSATLAPRWKRWLARFENSCTAWKITDPDQKKALLLDRGGEDLFDISETLVMPTDAQYDDNYKKIVKALD
DYFLPKKKTEYEIFQFRRTVQHDTETIDQFVTRLKKLAVNCGFTNQAMADAEIKHQIIQGGRNKKVRRKALQQDMTLD
QIVSLARSLKISEREAHGIEHLQLPDTVNKTAHTPSIRQPQTQHGGYKKNRKPLQVVTFVAVNIHIKVSAQQRNHRRVTY
HLRKKVEAEIKSLEDADILEKIDGPTPWVSPIVIVPKSGTDKVRICGDYRALNMAIQRERHVTPTVQEIRTKLHGSTVFSKL
DANKHYLQFVLDQESRNITTISTHVGLRRYKRLPFGINLAAEICQNEMSLILQDIPGVESIADDIIIHGVDQEAHDKALHAC
LSRLAEYGLTLGLSKCMFSVPEIEFFGFTFDRNGMHCNEKQIKTIRELKRPENFSKVRSFLVIDEYSCFPEVEVTNSTSSASV
IPKLDRIFTSYGYPSKLKSDNGPPFNSNEFASYSKNSGFKHQKITPEYAMANGLAENFMKNLGKWTEPCIIWTAVAARK
GEKKSLAGKYVISGVKAMQSKEQQLWKNSDGGQGSPGGAVRDHRKQQRYTASPCRFFLNIVADFGLGTIEVTRKCLLF
RKRKFEELENENNTCQDEYRECGSGECVYQYAICDTLQNCADNSDEVNCNATVKDDIYDKKPVTERFLDIPGVTPETLR

DFKDGYYKGNGFSGAVKGEVPPDWVGFKTFSSTPDYSDLKNVLKLTREEIAKNGHQAEDFILQCTYDEETCSSSDFYTFQ
DDQYGNCFKFNAGLKGRVIRHADRQGALSGLTLTLFLEQNEYISIYGRDAGARVNINPLNETVFPQDEGITIMPGTVTSI
GIRKVTFTLP

>Deu_Ambulacrar_hemi_Ptyfla_40v0_9_20150316_1g16875_t1_scaffold13442_cov91

MMKKKTLMQTVTSRNPERKKCSSDEEHQSQGSGERRKQKAKKVITQQHQVTAKQPFKSVQCRGQLHPINTLKAPIQS
PGALPHDNPIDEMDVTQVIEFESDCVPGNETASVSSVANKMKRLEERLDHLKNENKELKQQNAALYMLDLQTKTEEGT
ECVVQSAQNCRHQYDGLSHTMDTSFDGVSSGSEHVASAGGCGDADPDRNIDGELNGPLVTEEEHLAGRKGTSTNHQ
EIDNPQREAVSVCDEQNHESGHPDAAPVDQHGRVGKNSKRASKTKAKGVSVELISGKQIEFPTVTVCNTNKLRRSAVR
ESVYKEMLFMENDIVAPYYNQCDETQFRCHSGSDLGICINEDLRCDQYKDCYGGEDEDDCECDPETTFQCLVSRECVD
KIVVCDKVYDCRDASDESNICNDRLFNAQIYVIEDTGITGNTSCADPLNYEDLSTNPITVNCGGPKLGRYVLIKLLNTTNYL
TLCEVEVKGKVLIDRLPNVALGKPTIQISTLYDAVSDRAVDGNTTQDNSGCSSTQWDYEPWLRVDLETVYSVYRVEIVKP
VGTEFQRSVRPSLKSMFGKQANTRLCAEPNDVFVDVSLEICLQKCYGFRRYLCRCFDYEAATHSCRLYDKCSPLGNASLV
YAYGIDNYQRLRAEDMEDVFRYGWEENFHTITSDPSHFKYFKDNIYQDRGLGRVRGETPPDWERFKSFSLSLTLYTEQN
EYLSIYGKDSGAIVTIQPRGELTFPQEQGFAAKPGTITRAALRKTEFRREGDPYGDCKIGKDDTYIDTVYGSTNDKYSVMK
KLLKLLKRRNNKTETLVTDEDARSNLIRLEVYFETLNYEETSEHPSYTWEDLLSDIGGTMGLYIGISLMTVCELVSFLAVLVA
RSFGKNDKNAMHDVNGTV

>Deu_Ambulacrar_hemi_Ptyfla_40v0_9_20150316_1g11968_t1_scaffold7374_cov130

MYMTENEKKIAISFARMAIPDDHYIPCLLATDLLSILTLLSTQLRYHGHNASKGIQYEYMSGLLNATLPVFYEKIYHELNLIV
TLIMYQDAYDMVNENDTSDLTVGGDIYNRTYDAAYEHLIHRKWYEILLSRSIPATIEDSQVFQGNATIDSAATGGVLLVT
FMKAYDEIFDSIYDSYRDCFEERFASLLETMSGEDICFLHVRRYRWCSWFVLRGQFAVRFRNFRFSHLEEMLGAWKMC
FYQLKSQSEIFTNKEQMNAQGFVLLEKCNTVLTSIYIVNNRSSCTHCQDACDLTSYDVSLSYQSFPSFPVVRDLDRNGTYL
CRDAYADNFRFLWLLIVLGAATALGIQCHKIVTTFMMKPVSINIERLEVPTMPFPALTFCNNNLVRKSLLAEDAGYLNLEI
SQQVSQTIFSREYMGIERYTDVDYRGADYGVKFGSDLLGYMEAVAHKTEDAILXMAIADDHYIPCLLATDLQSILTLLSTH
LRYYGHNISKNIQYEYMIGLLNATLPVFYENIYYELNLIVTLNMYREAYDMVNDNDTSDLTVGGDIYNRTYNAAHEHLIH
RQWYEILLSRSIPTTFEDSQVFQGNKTIDSAAASRVLLVTFMKAYDEIFDSIYDSYRDCFEERFASLREKMSGEEICFLHVR
RYRWCPWFVLRSKFAVRFRNFRFSHLEEMLLRDCAVTRASNRYAISVIHKLHITCTCRQLLTITLVGLNPGIGALCPHSTC
ETKITTDIMTMGCSASNIHLTVKSENTVTSQTTTVTRGDISIRETEEVKETHFQGLRFGEVDEIGTALNIRKQLTKFSMKQ
RSVGSIQSQCSDLSLEGSGLLQQDRYWLKKKRRFSYLNDVEMTGSMNNYYPSREGSVCSAPEYCKQGTDTCSEEGHKT
PCSFVVAFT

>Deu_Ambulacrar_hemi_Ptyfla_40v0_9_20150316_1g24341_t1_scaffold35678_cov111

MWEDLLSDVGGTLGLYIGFSVITISPCIEGDFQCENGIHCIKPFLVCDGVNHCGDNSDEHDCTYPECGPAQFRCDSGSEL
GVCIPNDNVCDRVIHCYNAEDETSCGARLFDSEVLVGSDELSPNNSVCYTRVQKHQTLAETFTLKCTPNAVAGRYVSIQ
VKGKKEDMRLCEVQVMAIDSKYWNLALSQKATQSSTFAYGSAVKAVDGEKSNNYDERSCTHTQKEYQPWWKVDLG
GKYEIHKVIITNRADCCGDRLYGAVVRVGKNDVIENNERCGQTVTTDEIYQDGEVTVECVLFGRFVSVQLENKTDYLSLC
EVQVFGEDIVDNLPNVALYRPAEQSSTYYDYEASNAVDEYVKNEVANIFNMRHFSMFDARDGFTLDVNSTDVFSELLL
HQCMYKCIKWKRYECHSFDFHYGSKACRIHNLKDGMDGVQVAKSTGSIYYQRLRLTAFIPSDNETGCPTDYIRCTSGEC
VHHYKICDVIVDCADGFDEMECVNTESNSARVFNFSPCWYASYLSIADDPAVYKYFQENIYVHHNFDRVKGEDPPDWI
RFKTFSGSPDDSELAEMILLFRKTNNSVTVFTLMGIQISRDLRQVPDLNKSKLNRQSYPFGNCTNEIINNGMDERVAAIM
SGQYTLAKNLLKVVQVSNKKTRNMHNLESASRQDGRSDNPYPLLSVCSPYFDDTEKTTQSVNGRASYTDKGNNEVSN
VQNESFKPSGEKEEQNDDSVKHVLEDFGRETTAHGIIRITNATSSVTRSTWILIVLVAACVMFLQMTLLLVQYFEYNVHV

KVTLVSEKALDFPSVTICNTNKLRYSAIRSSKYSEMLMLERNFVSPYYKPCLEGDFQCKNGIHCIKPFLVCDGVNNCGDSS
DEHDCTYLFTGTKYRLRLDYQIRFSEKSEKP

>Deu_Ambulacrar_hemi_Ptyfla_40v0_9_20150316_1g16645_t1_scaffold13124_cov152

MVDMSRFNGKGSSVLPMDNFGYETEFSGGRGTGNESANSITEQSDHQSGNNDTTWSGVFRTFTQEAGVSGVKYIGD
VKASQGRRYDNLVRLVGGASVFNFLFPYLYTLFSVDGEKIANFASFLIPEDADYGEGQNQFTFIDESAHSLETMLQSCSF
GGLPCGPENFTKVITNYGICYTFNSDREDFVHTVRNAGQQFGLQLLLFANERQYVAPRSNVGFRLMVHGQDEIPNVAS
YGLLLSPGTASTIGLKTTQISNLGSPYGNCVEDVENNLKYFDGAYSLSKCWMECETDYAVEQCGCRYFYMPDEFALKTD
PCKHCEEPCVQTNYGTKLSHSSYPSTIYANLLAYTVSWPCDEALAYYFLDHSSEEASAISENVKEFVIERYYAIEANGQNIS
VFIPSAEFHLIMTEKLIDQLFVDVFHELRFEVLLHTLVSLYGNMTDGVNSVVHNETKDILTAKYMNFLTNPWSDLNATM
WGMFALNRNLTAGVIEYLINSLFTEYYVPLFEEVFVYGYPKMIAMRDNITAILRDGEPGAICKNYMQENLSKNSNLGSPY
GNCVEDIENNLKYFDGAYSLSKCWMECETDYAVEHCGCRYFYMPDEFALKADLCKHCEEPCVQTNYGTKMSYSSYPST
IYSDLLAYSVSWPCDEALAYYLLDQSSEELSAISENMKELIMKRYYAIEAKGQNISAFIPAAVFHSIMVNALNWVCVDAFY
DVLFEVLLHALASLNGKTADGMDSAVYNETKDIFIAKYTNVSADPLSDLNATMWGMFIAGNLTSDVIEYLVDSLFRGYYI
PLIEETFVYSYSKTTEMRNNRTAKLREGETGAVCKNYMQGTMLYGTFLNKDNAIDFMNELFHLVLRTWKHGTCLVKVA
RSKHQCESGYAADDRKPEPTRYHGPITYW

>Deu_Ambulacrar_hemi_Ptyfla_40v0_9_20150316_1g6257_t1_scaffold2963_cov116

MLFNEYRILKSESERKFYVTCTENCDLPTDINVSDGQKTPLNYSRLSHTISSQDDRPGDPYPLVSLNSATFNRSDKGKHLK
RYLHKNQVNYTSEETDELSSLDKSSEEKGGNHEDSVKHMLEDFGRETTAHGIVQITTAKSNVTRSMWLAIVLLAASVML
VQMTLVIIQYFEYNVNVKVTLVSEKALEFPSVTICNTNKLRKSAVRDSKYSEMLLLEQNFAPPYYTPCMEGDFLCKDGIHC
IKPYLVCDGVNHCGDMSDEDKCNYPECGPNQFRCDSGSELGICLTNGKVCDRIKNCYGGEDETDCGEPEVRNVASHK
HTWQSGRSYWESGSEKAVDGNDSTCAHSEEQYEPLWLVDLGKTYLVYEIRITHMPTWNQLKYWNLALDQKATQSST
FAYGSAVKAVDGDNSNNYNDRSCTQTQKEYQPWWKVDLGGEFEIYSVIVTNRADYCGDRLHGAVVRIGNSDVIVDNT
QCGETITADTIDSNAAAEVGCVLRGRFVSIQLENKTDYLSLCEVEVIGEDPHYAAVKLFIPLRCLTYILLQIKLFHSFVEFIPSE
SIDILKPRMFAMFEPQEEKALPVNSSDVFSNLPLDKCLSECLNWKRYECHSFDFHYDSKTCRIHRKKVGMDSLQVVHSP
GTVHYQKLRMSAFIPSDNETECPIDYVRYNRLFHFYPGWNAPFLSITDDIDVYKYFEENVYVHHDFDRVKGEDPPDWTR
FRMFSKSPDLSELERVLKLSVERILQFRKTNDMAIVFTLTWIQKVLDNRQALDLTKSCEKACLQNALVVYCGCTDTLGIDA
QQCLILNETQVVCKTLIYYLYQKDLLECNCPPPCRANLVRLRVYFETLNYDLTSELPAHTWEDLLSDIGGTLGLYVGFSVIT
VCEFMRLILGLCRRGCSKRQETRG

>Protostome_Lophotroco_annelid_Pdum_comp416177_c0_seq3_

MTILVKEKGGSQKRPIAFESKNGEENKSFSEFAQDTSFHGVKYIFSHDISHPVRRSVWLLCWSICVGLFLYQVSVRIAVYS
SHPTNINVQVIYNTTMEFPSVTICNMNQFRLRKAANLGLYDLLDGVYKDGVINETELALHKRSNMTLKQTLTELSHQKE
DLIVGCMWDYEPCNMTEFFQEVMTDMGLCHTFNGIGQRQKPLALREPGMKGALRLTFNVEQYEHTKGANVDTGVK
VLLHDWHEFPLMSDAGFTVMPGSHTSAKIRLQQMNSLGRPYGTCNVANTAVSTCRRNCFTEQAYHSCGCRDLFMLD
TNGSTPICSLEDKLICLSDVTTKSNNLQSCRCSQPCNRRVFDTSITSGLLSRLNIGRTVIHNESIRTRVISNYHKAMEKSQRY
VLQKIKSDERLIGNAIKLTEQMMESVSRANSVMTNTSDTAEELGTPDITLFMQPWDDFLHVLHFKTREFAHFATLAEVH
GQKVFERRQALDDAFEAMVRSVCRDKIQQASCSKHFFSKIGEANKTIRIYRDSLEVWMTDTFSRAEELFRDQDVEWSN
KLDKCNDVIGKSEIIPSIIDIVSNATVMMQANTSAFEVSQFIIEKESELNFHGKINPDGKSACGSLSRDLTDESGELMKMPI
NNNLNSMTNLLTAYQASLSQLLMVETLIHDNMATPLMLKEYFSFDKTKLELTLPFSNSTFRLAIDKIMVYGKTLDQANTA
MSQAASEMHREVDVICNDLERFAIFLFTLSEIKNISVVQELSQHMTPDIRNITKVKDMVFSFKTTIKPAIKKFYGHVFANV
SYLTKDIQNFFDAMHLVMTDLAEYAERATLSDGFYMDNFGMLDVFYSEMSYEQVTQQMSFSFISLLSEIGGFLGLLLGA
SVLTVLEVVDYVLMKCRVGVMGRKNEVTQIK

>Protostome_Lophotroco_molsk_AplCal_XP_005104865_2_uncharacterized_protein_LOC101849634__Aplysia_californica_

MDQEDNWRLLTVCRRLPVCDESPSDETDLEPKVNKDAILSPTRQQLNHGKGNDVIKREDALRLLPGFKQFPLLESLVHP
FHQYHQQFKDKCTQTGSDDSVYNAARNIPNGGSSEFGTNQLCAAMSSVVTWDGRTERHDRVWLETQETIWRQSQG
TPPGGRTGKAEAPLGITSEPIETGASSNRLKTGQKETCATSFYKLYSDENRANSDMFYSDDTPSKSQKFCSDDTHTKLEKL
HLDDTPSKSERLYSDETCAKSDEFYSDENLATPYCKFHSDENRATSDIFYSDETRASNATSLSFHHTEGTEKICETSDRTLA
EDLNIMFNTENIDTNVDLSDYNNVSEMCASIKTVGKLRPSVVWKRGPLPSLGSGPKSEGGKVGDGSWPDYWTQVKLL
VNGFASETTAHGYKRTICAERSRTLRLVWLAAILGVASLVIFELVQVMRQRQTYSFSSSIRSQPEERLPFPAVTVCNLNPY
NPAQIADSLAANHSAVLRGFLREVVSLGRIQWSNQSEAALRPFTTRDLSQLTGWTVASFLKVCDWQGQALDCSNVFQ
RSVTGDQTCFTFNSDPAHPLWTDRQGASSGLGLLVNLGHQTLAQGALGISGVKVLLHEPGQLPNVREMGMLAAPGTL
TYLNVIRRQYHFLPSPFKAYGTDYCSDAPTGSGDVTAEARQSLYTNYTGYSACVADCRLRRIIELCGCVAYGGDPEVAGV
PYCSMDVYARCYLPTLKEFHVTSRAKSECGCTLPCSKVEHSAGVSTARYPLSDSDDLIVDFGDEIYQGFPSAPSNNKTMY
LELKIFFDDLVTHVEEHLPMYTWPTVLGNIGGQLGLLTGASVLTVLEFLELTALLAWLTMKRAAGGGTFRRARQRLLTLF
RRRISPGRRWWGGQRSTGVSGHRCWERGDEDLFYNIV

>Deu_Ambulacrar_hemi_Ptyfla_40v0_9_20150316_1g22725_t1_scaffold27658_cov84

MENQMSLVEDRLRVSEGQGTDVQSSGIPATNYQMADSDVSVIATIEQQPNENLTEKMHEFDVVTLPKLAFPTVTVCN
TNKIRKSEILKSEHQRMLIVDEDFIHPYYALKAFKRTYNKRLSPKDAVRKTNTSLRHCALSCLHSNEFTCLSFDYDSQRREC
YINPGSKDTLNRFSLTXXXXXXXXXXXYRVFSVLDSNTALKFERFPNAKISLDYPPQKTTTEECAQLCLDEVSFDCLAFTYEP
STSRCWLHSESHSLDSIPLIKDSTCDYYDRRGIECYALDDGTDYRGRIFVTRDAFFCKNWSSLDPAIFRYTPSTAPGKGLG
DHNYCRNPNDKECYTAQNGSDYHGFAYVTVSGNVCKSWLDVQSQNDLAFDSSDGIGDHNYCRNPNNADAPWCYTT
VSGDSYEFCNVGKSQKKCPPHTKHTTNLKECYSDPKGLDYRGTISTTVSGYTCKNWTYKHTRLHIDPVSLKSHYGLGDH
NYCRNPDHDKGPWCYTVSDEKLWEFCDVGERNISCNVLVAMTKFSLASTGIECFYNRRATDYRGYMNTTSSGRTCQK
WSEQSPHSHRYYAEFYHQRGIGDHNYCRNPTGESQTAWCYTTDRQMSTDLCDVELDREECVKDENNTCPDEYRECG
SGECAYQYAICDTLQNCADNSDEVNCNATVKDDIYDKKPVTERFLDIRGVTPETLRDFEGGYYKGNGFSGAVKGEVPPD
WVGFKTFSSTPDYSDLKNVLKLTREEIAKYGHQAEDFILQCTYDEETCSSSDFYTFQDDQYGNCFKFNAGLNGRVIRHAD
RQGALSGLMLTLFLEQNEYISIYGRDAGARVNINPLNDTVFPQDEGITIMPGTVTSIGIRKKHTFRQTKPYGKCRDGMNK
DNLFGRTKIYSTLVCQNTCVQRQMLKYCKCVDSYILGEAQCLITNKTQ

>Deu_Ambulacrar_hemi_Ptyfla_40v0_9_20150316_1g7465_t1_scaffold3698_cov112

MVSSSGSSTTARKLLVVVLLLHAHHGLNDAARKPNIIIMLMDDMGWGDLGVFGNPAKETPNLDRMTREGAQMMDF
YAPNPLCSPSRAALLSGRLPIRNGFYTTNYHARCSYTPQEIVGGIPDSEILIPEELSKAGYRSKIVGKWHLGHRPQYLPLKH
GFDEYLGAPNCHVGPEDDVKAPNIPVYRDSEMIGRYYEEFPINKKTGESNLTQLYIQEAISFIEKENNAGVPFFLYWAPDS
THSPHYASKTFLGTSQRELYGDAVRELDFGVGQILDKLRSLNIDNNTFVFFSSDNGAALSGGATGRDPGEKFKLKHLGQ
QPEHLPFNRGFDEVFSSPNCHFGPYDNKQTPNIPVYKNNAMAGRYYEDFPIDHQSGESNMTQHFIQEGLSFIESKAKA
KEPFFLYWTPDATHGPVYASKMFLGTSVRGTYGDAVRELDYGVGQILNKLKELKIEHNTFVFFTSDNGAATYAKTKVQL
LLYVDNIGSTEDFLLPYIVFYYRFPRPTHCFLVDGIDNTYTRKILTLLKGRPVLAEYSTENGKHRYGRSTILDVDYKEELTFPAI
TICNFNKYRESLITANDIVNVGVHLGIVDSNFEVIDEQLYTEEFVSRINAVNWTDYEEDPNFDMVEFTNRTGHQIDEMIV
QCEFGGQECTADDFEHVFTHQGNCFTFNGYKEGYTLHESKRAGAANGLKLVLDADTPEYTPSNDLDGDALDAGFRIAV
HHNNEPPYVTALPAPYTTCEESGNVQYYDHYSLQACRIELNFMNGQLTGVEECECQTPCDLHYYPIAVSSAKLRPTFIEK
TFENTTSSNMSIAYISDNIAVTTIYYEALNMRKIDIVAATGWFDVLSSLGGNFGLFLGASLLTIVQFLEYLVDECMYTFFKK
KKVEPDDEEAPAENGVAMKVQDVDEPEKKDDAWTTEYS

>Protost_ecdy_cyclo_nema_Cele_tr_Q9U1T8_Q9U1T8_CAEEL_EGF_plus_ASC_domain_ion_channel_OS_Caenorhabditis_elegans_OX_6239_GN_egas_2_PE_3_SV_2

MIFLLFLIFPYYSTPQTTVEYVAASPNPPHGCPNFGLCTDVTTTLVSSTNVSTTYEYGCAPENCYCVDGTTNGTEPCDTIV
DQCGDDPCGEPEYFLCTSKIESHTCACQAGYTGADCTSELGTACATSPCRSGATCVSSNSSATTEYSCICTDQQFGTHCE
YDNLCASNPCQNSGNCTMVLENSNYLCTCSPDWQGRRCEVADAVSTPPVQDGCYLYDTGSIGACTTKFTENSLTSSKC
NNYALEQSTDTVTMNYLTLCGAWCMVSEAPLCNTSWYMDGDCKKSCGGDSAEYCGRLNSRCIVYEAGTSETDANAC
TDNSTLCNANLGQGICINLSSDVTNGYQCICGPLWTDTDCETPVASACTPSPCWNGICVLNEQYNTYTCACDDGYFGD
LCQYPDVCTSSTCLYGGTCTETSSGGYTCSCLSQYFGTNCEEINRCNYADPCVNGDCQTTVDGITTNYTCTCDSGWTGE
NCDTMIDYCIPNPCSYNSTCSPYFKGFNCTCITGLTGANCSTIIDLCVPYKDSTGKWIKSPCNSKDDLANCTIEINGFNCSC
SDKWTDTLCDLNVLIKDVLLTIYGHVDLTMITLLNDLMQNPSQIKDMVPFITGLLSDSERSDLSWDVGDLFNWIAFEDQ
RLDLENDIYKWNDVVLGNCFTFNHQDQNFTYLMRRPGRHGGIQAFMKTRQDEYAPWYDTAGMLVFIHNREDYVFSE
SVRYNAQPNAQSTINIFMTRYTRLGGRYGKCVKKPSEVKNYYYPGAYTTDGCLRTCYQDRMQQECQCMDPRYPKAPN
ATSCQLSERSCVTEASDAAGDPSTWSSCVCPLPCSNQEYSVTWSKANFVNMPITCEKSSDVATCQANYVDQLMVSIVL
PKLDFQIYAETPAMDFNKFLSQLGGQLGVLMGINLVTFIEVVFLMFGVLTICCRK

>Protost_ecdy_cyclo_nema_Cele_tr_Q9XTS9_Q9XTS9_CAEEL_EGF_plus_ASC_domain_ion_channel_OS_Caenorhabditis_elegans_OX_6239_GN_egas_3_PE_3_SV_2

MIFLLFLIFPHYSTPQTIEYVAASPNPPHGCPNYGLCTDVTTTLVSSTNVSTTYKYGCAPENCYCVDGTTNGTEPCDTIVD
QCGNDPCGTPDKFNCTSKIESYDCTCQAGFTGEKCDSEIGSACATSQCREGSTCEASDNSTTGYTCRCTDKQYGTYCEN
DNLCADWPCQNGGTCSMVLSNSNWICNCTEGWQGRRCETKNSVLSSPVMGKCYKYDYVKIGTHQTKYKDDKMTSD
KCNEYAQEKSTDDKTLKYLTLCGEWCMVSEDMLVKDDVSMKDDCKKSCGGDSSEYCGKVNSRCIVYEAGEMETDPN
ACEDNSTLCGAELGHGMCINWQSDVTDGFACICEPLWTGPDCENPVASACTPNPCVNGTCVLNEQYNKYKCDCDDG
FSGDNCEYSKTCTAATCLYGGTCTETNNGGFKCDCLNLYSGNRCEEINRCNYGDPCVNGKCETTVDGITTNYTCTCDDG
WTGANCETMIDFCTPDPCKYNSTCTPKFKGYDCTCLTGLTGANCSEIIDLCVPYWDNEAWKWIKSPCNSKDDMANCT
KKINGFTCSCGEKWTDTLCDLNVLIKDVLMAIYGHVDLTMITLLNDLMQNPSQIKDMVPFITGLLTDSERSDLSWDAGD
LFNWIAFEDQRLDLNRDIHKWNDVVLGNCFTFNHRDRNFTYLMRRPGRHGGIQAFMKTRQDEYAPWYDTAAINVFI
HNRDDYVFSESVRYNAQPNAQSTINIFMTRYTRLGGNYGKCIKKPSEVKNYYYPGAYTTDGCLRTCYQDRMKEECNCM
DPRYPQAPNSTSCQLSERSCVTEASEAAGDPSTWSSCVCPLPCSNQEYSVTWSKANFVNLPITCEKSSDVATCQKQYKD
QLMVSIILPQLDFKIYAETPAMDFNKFLSQLGGQLGVLMGINVVTFIEVVFLFFGMFMVLCQKYEK

>Protost_ecdy_cyclo_nema_Cele_tr_Q9U1T9_Q9U1T9_CAEEL_EGF_plus_ASC_domain_ion_channel_OS_Caenorhabditis_elegans_OX_6239_GN_egas_1_PE_3_SV_4

MLLFLFFFPELPTPSTPQTVEYVPASINGPHGCANYGLCTDVTTTLVSSTNISVIYEYGCDPDNCYCLEGTTNGTEPCDTIE
DQCGNNPCGDPKYFLCTSKIKSYSCSCKPGFTGNDCKSELGSACATSPCRSGATCESVDNSTAGYKCICKWNQMGENC
EYDNFCANPQCQNNGNCSMVVDNANWICSCPPGWQGRRCETPDTVTTPPVWDGCYKYNFQSIGATTTKFSDNQLT
VDKCNKYALQQSSDTVTMNYLTLCGGFCMVSEGPLCNDTSNWDDDCKKKCGGNNNEFCGVLNKRCIVYEAGTAEVD
ENACDNSTFCSADLGQGLCINWQSDFTDGYACICKPLWTGRNCENPVPEACTPSPCLNGTCVLNDQYNKFTCVCDDG
FFGDKCQYSDICTSATCLYGGTCTELNNGDYKCDCLLQYFGKNCEVINRCDYGKPCNNGKCASTIDGITTNYTCTCDDG
WTGTNCDTMIDFCIPNPCSYNSTCKPKFKGYDCTCITGLTGVNCSTIIDLCVPYKDSTGKWIKTPCNSKDDMANCTKGIN
TLTCSCGDKWTDTLCDLNILIKDVLMAIYGYVDLTMISLLNDLMQNPSQIKDMVPFITGLLTDSERSDLSWDAGDLFNW
IAFEDQRLDLNRDIHKWNDVVLGNCFTFNHRDRNFTYRLRSSGRHGGIQAFMKTRQDEYAPWYDTAAINVFIHNRDD
YVFSESVRYNAQPNAQSTMNIFMTRYTRLGGRYGKCVKKPSEVKNYYYPGAYTTDGCLRTCYQDRMKQECNCMDPR
YPQAPGNVTSCQLSERSCVTVASEAAGDPSKWWDCVCPLPCSNQEYSVTWSKANFVNLPIICGKSSDVSTCKAHYIDQ
LMVSIVLPQLDFKIYAENPAMDFNKFLSQLGGQLGVLMGINLVTFIEVVFLLFGLFTLCCKKYGK

>Deutero_Ambulac_Sakowv30011650m

MDPCDPDPCSNTGWSEHQCVRLDNSCTEYTCLCRDPIRGCYTGPQCNRRHDPCLPNPCGRGVCVPDALSCNYTCTCL
GCYTGDDCDILQNPCSPNPCEHGVCYRDERSCFHYTCVCEDCYAGRVCSTYFNDPCSQFSPCRNGGTCIASSDTCMDY
SCHCPDCFEGEFCQFELDPCQLKPCQNRGTCIPVTNSCTERTCSCKDCYIGESCTIRVGGCTDNPCLHDGICTESGDNCS
DYTCQCPWTWLGTNCSWHIALVIVLSQVCVILIMWVIIMTILLFRHRRYDKQIKKSKQRRLPSPDVVRSDMGISIVEVVN
GSTTIRNGERDVRSEEDVGDERSAVVDDGDDGCVSDSTEFTHVSVASHDKKHGSLSAVTHEFCRATTWHGLARVSRST
NPLNRLLWTVVVLVACGFFISQGTTMVLRYMSSPVAVRMGEHSRSELTFPAVTICNTNTVRASALAASKYFPFVAIMTN
AKYNPYYVPCLTGDFVCNNGIHCIKPFLVCDGSWHCPDTSDEFGCDYGHYQCSSEQFRCASGSGSGICIDGKRQCDVVE
DCYAGEDEQNCDYEYEHSAVCTGFYCDDGICHPLSNKCNYITECMDGSDEEDCKYRHCQDWEFHCGNQRCITSDKLC
DWKDDCGDQSDEQDCEYRDCNEDEFKCFNGQCIYAWQRCNSYIDCIDLSDEDQCPGLELYCLDRNEECRYWASNGE
CQQNEVFMKISCPLSCGWCVGEQATVDETGQCPKGYFTCNSGFCIDEIHACNAVPDCHDAEDEFWPNCVYQSSRPEA
LDVFHGNWSDIFREISTDADVFDEFEYTVYKDPMFDRVRREEPPDWNGFVTFSSTPDFSDLRSILKLKRNELKEYGHKPE
DFILQCSYAGVKCSYSDFQWLPNREFGNCYTFNVASNGSEPRSAFNAGAEFGKCEYMPTSLP

>Protostome_Lophotroco_molsk_OctBim_tr_A0A0L8I926_A0A0L8I926_OCTBM_Uncharacterized_protein_OS_Octopus_bimaculoides_OX_37653_GN_OCBIM_22028050mg_PE_3_SV_1

MTSVRNRFEKGNVEEGPTVEVPTDDEKPSSMFLDFAMTCSLHGLKNAFSKSSKKPQKVLWLLLLMTCIAAALFQILDRIL
YFYQYPVSVLLDVNYNDSLLFPTITICNQNKFRATEAYKLGIYRMIENVNKAENRSFAFSSEFIQQAKAMNISERDLRQQI
AHTKEDMIIDCHWSSERCAPENFTTIFTDEGVCYSFNTDASNPVKVASSGTENGLRLTLNVEQYEYMSGGQKSVGIKVL
FHNSHDVPTIKDLGLASATGTNSFFGLQVVEVIGLPKPRGVCEDRKLNLFYKYSRSSCEAECITYALVETCGCRLSYMPKV
NDSVPLCSLVSFITCYIPQRDKFHSFQLKCDCPLPCNMLLFDPSISYTAHSENKVSKLIMDPRMADVKQKLINAKEVKHR
MDAGSISEFRNMLLNFNASNVAFRTVMLYKLETTIKINLAILQNISKKVEKVYASKLFLINYQKYLIEKNFERPWEAIAERTF
HHVSFDFFNYIYTLKNMFLKLDEFINNSSNQRASEMLIHNIKMNIETKLNMVEKAEGNFTEYYQSLTSGVGIFRYRYLKVP
RSHNFYAVPKQLLTIKLNQSKSDYSLRLSNTLISLKECLFNFSDMLDTRDTGFNLTKFTKVSDKFIHLSKLFNSIKSVFNGYT
TKYALGIIKSKAAKLQTSLTNIRQIINDMNNSFTSLQIEQKHLNLTSSQNVFAVSSDIIRYLTNTSVTKISLAAILHSPNHVLN
MMNLQVFMEELRERNSLLHYSWTKLNETVALLWQCIIQDRDSYAYYEYANYTKFSLPLENVTSELHDEYADYQEGSNV
AKMFGTIDRDFFYRHKTVKEYVTKFKERNTINDLFVSENILEIAFFYKQLSYEIITDQVAYDFFSLMCDTGGALGLLLGSSIL
TIFELADFAIGFSFQKLLAKLLMKKRVENL

>Deu_Ambulacrar_hemi_Ptyfla_40v0_9_20150316_1g20262_t1_scaffold20289_cov133

MHQFSFTIMTKSTPTESNSSSSPTMDTIRDANGKTIPEDLLTDSLDSPVDEFKGKEEKSHKDIIGDFAKKTSCHCLPRIIGA
KSTRSRLFWSILFCTAIILFVFQSAKLISLYASFELAVTIDDGNTANISFPAVTVCNTNKLRLSKIEASEFSELAKTDPDHPETI
HKAMSYQDGTFKLPQSIALSPNNRLLPFRKMSAADSCGQNVTASPLPASLYSRRYPEPYDRNESCEVLIISEEDTCIAISFL
DMNVETHKACENDYVRITDPKTLESVKLCGANLPDRWISTSNAASVLFRSDSAHHSIGFHLLYQTTPCPSSLMKWKAKK
WSVCSEDSCGGGMRSRDVTCEPREDVSVPPKEREKGCPGIMPDRQETCNNELCSDDICRQEIRHCCTTINKLDGYRNG
VNCAYSIVNRNGCVNITIMTMALPENEDCSLDYIKFADARKESLTKKFCGSTTATQEPALTWNSYSGIVNITFHSDGKKT
GKEDGYVLIYDFVACGVWHVSPWSQCPVTCGEGTKFRQVQCRSRATLTVEPDSYCRGPKPPTTEQCVGSQPCEDPALL
TRFSKNYDKSWCSQIAGRTIDAQICQEFFNLYRQTNKFAGVPYSSQIKWQDFLTFSSSQDYTDLRKLLKFNKNELAEFGH
QLQDFVLQCSFNDEMCDIPSDFIQTQNDEYGNCYTYNHGQDGARLKYTNGEIVGTGLKLTLYIEQDEYVPFYGQDAGV
RVLIHQPGITPFPEDYGVTVPPGKKAGIAIRRDSYIMKGSPFLDCIDQRDYRSIYGKESYYTTANAVDSSSDSFTFVEFLRTI
ASSHVDESTTRRLCHYRNGRPTSIWDNLLKLEVSYEVPNSRSTISSPRYTFDQLIADIGGLFGLYLGISLITVVEFIEFLYDMY
CYFISWKRVDERRLKKNRGNKMYDLAPSHSPDEFGTRV

>Deutero_Ambulac_Sakowv30000561m

MDEPVKNTRIGYLVQEFARRTTSHGVARIAASNTRFRKTAWILVVLGAFTGYSIHVGTLVARYLRYDVAVKIDVMSEDS
VTFPAVTVCNSNKLRLSAILNSTYKDLAKVNEPRFSFTYRGPCMPGDMSCDDDSTICIKQYMWCNGIVDCPNAIDELDC
QHGNCLSNQYKCDNGKCLPLSLQCNYRDDCGDNSDELWGGSCFIRGCVQGEYECESRECIPEYKLCDGHVGDCADGT
DEPDDCTLYREPLQGLPCYGGQIRSCHTSWQLDNMCIQSYDVCNGVVDCIDGSDELIGCEDHYWKCPYNHFKCATGH
CIMNALTCDGVMDCKDGSDELGCGDYIFSEDGRWTCEDNLTTIPISLRCNGFPNCRNGNECPSTMMKCESSNICIAESK
MCDGSNDCVVLGVPGIFDLSDEENCHQGICSSDRYRCGNGQCIDMRHRCDYHYHCLDHSDEFNCKQDDCPVNTILCE
RNSGDGQYCLDNTHVCDSIYDCYRDEDEQCSNHTCASSTFLCGLSCALLERRCNMVSDCPGGTDEVNYMCPDDSYFR
CRSGQCIPWYLYCDEPIRPDCADGSDELFCNHTCMHDEFQCSSGWCINATWKCDGVNDCGDHSDEDNCPQYVHDV
TAEPNWYAGYLDISDDPNLYEDFRKDVFVNRPIDFIPRRQPPLWEHFLVASMTPDFSDLKHVLHLRPDEIRDLGHQRED
LILQCSYDQRDCATSDFYNFQDNVYGNCYTFNHGNDGSDLKRATNTGSEYGLKLTLFTEQSEYISIFGQESGVRFMVHD
PFTTPFLGDMGIDAKTGTATSVIIKQREEKRASPPWGNCTDKAYVDSGEYHYSLLPRLLSSLLAVNPKVRTISDQDDNGIS
MNLARVKIYFESLNFERVSEVEDYPFNQLMADVGGALGLWVGLSIITVIEFGEFVYEMLRTLSQNPTQSTNYPVK

>Deutero_Ambulac_Spurpu_027370

MEAWKANAEKEEESFLKILHSRMETSSAHGLPNIHRSSKSITKLFWLLLFLTGIGVITWQVIALFKTYFAYEYTVSLEIKFNR
SQEFPAITICNMNPVMKSKLEKADDSFRELFDTNFQPPGPMDMVDQVVQGVGQVGGVFSSGVGSFMGSGGNSGG
GVEANNAKTTQRPGSSTAGGRRATDGNVEVAGSVEPPNGPSGPLSSQALCSRHQASGQPPNLREIQQRYGVRNLRG
GSMDLYKKGEMSANRGPTQKSMTSKKGVRTLCSPSLYLPLTLSHYVGFRSDLFRGRPSFGCSRANTSITPTRPTRPITDR
AKTYITRPSRESVGFEIGLKAVETWQDRQKGDFFRKQSNDYLKQQRLMVKLANKTIEERKSLGHSLDDMLLDCSWRGF
PCSPANFTNIFDPQFGNCYIFNSGLDGDTLTTNRPGPNYGLSMELFVQQDEYMEDQTEVAGFRVIIHHPSNMPFPSDD
GIFVSPGMVTAIGVRVVTAIGVRVVEIERLPEPYGLCKEDTKENIEDNIYYQNYNMSYSIQKKLFEKMIRSNEEMKRQIRG
QSPSVWTRSNMAKLEIYFDEFNYEYIRQDPAYTFATVYKALDLQHDGKIVAVKKIKLGHRSEAQDGINRTALREIKLLQEI
KHENVIGVIIKDSSVVLTPSHIKAYTMMALRGLEYLHANWILHRDMKPNNLLLDENGVLKIGDFGLAKFYGSPNRIYTHQ
VVTRWYRCPELLFGARIYGTGVDMWAMGCILAELLLRVPFLPGESDLDQLSRIFQTMGTPSEEEWPNMSALADYIEFK
KFPGTPLRDIFTAGTDDLLTLLKGLLLMDPCRRCKATEALKMPYFYTKPAPTPGHLLPRPGMVAAAANGKQQPKLGSKR
KMADDIGTAAQCYLLGQQLKRHVLRIPEGVQNTYPLADGASCLLTPTEYGSYADGMFRMPGQYRLSHTCAEVMPIAV
RL

>Protostome_Lophotroco_molsk_AplCal_XP_005111008_2_uncharacterized_protein_LOC101857070__
Aplysia_californica_

MRLFPKDNPNNSAMGELGSVYKVPDPSVSSKSKEKKSPTRRQSWKDTAQFFSQNTTLHGVRFITMDRPYTVQRLLWV
SLFLTCTVCMGVQIVERIVHYYSYPTSVNVHVNFNKTLAFPAITICNQNAFRATAASEQGVYRLLEELFSNPLSVNESTLA
QHGVLDLPMDTLYRMTSHKKEDLIVRCEWHDSECGPENFTEVLTDHGLCFSFNSDPHSQLWVNSAGAEYGLRLTLNV
EQYEFMPGPHDAAGVKLLLHDKHESPKVAELGLAVPTGTHAYVGMQLFWIQNLGHPHGPCRSEESEFYDVYSPEMCR
LAHMTRAMADMCGCRHFYMPSVNGLPPVCTLEQYLTCYLENIGQVKQEVREGVVCPVPCDFLIYDPTMSFGSTSVYA
RDRLLADTSMASLKGSLLKAREVTSRMDNLKFQHFLALEIAVRTKFRKLQEVLTGDVTSRIQAQRDNLYEIRERMSQVW
ERKKYLLEWQKYHVEKNFMRARNAMEERTMFYLMLGFQEFSFQIETRIRQMAEPPDNFTMSQDVRSSLYLDTMNTL
MGRIDLAERAHANYTQLYQAYHNGTPIFRYKFQEEKRADNLLMTPKPLLRQSLFRTSYTRKYSSRVGKDILEIKKYLEFYS
GLANTSFANWTVDEDDMYLANILFLNRGRRFYQSRSVFYVESVEYPQRVMVQKIEDLDRAWRGLGDDLRQLTSGVDT
LLTSLAVLEQSIMARLQSLIEVAQVYIDQGSVTKMQVAKDLTSDDVYEGIHDMKNFFFALRSRGQEVYDGWTRLVLSTN
QVWKAVLNDENLEEYCEAKNVVDFLRNYTVVAEETENNFTASRDAFDFRYSLGNLDSEFLQSLNLLMDDMRSYIQEAK
LDSSFIRENFLQLDVFYREKSYEQITQQEAYDKFALFCDIGGSMGLFVGASVLTIYEVIDLLVQHIITSLACGRRLSLWLKSL
TCQRAVTRTK

>Protostome_Lophotroco_annelid_Pdum_Contig12523_

MVMQMEKAREAMAQEKKEKKKNSFSWTHVLYEFSQNTTMHGISKTTEDTPFLLRRMIWILVVLGGISLFLYQLSTSIM
TYLSYPVTVNVQINHNKSLDFPTVTICNLNVFRTSKADELGYYELIHEIYDPARTHNISADKIPAGMTLERIYKELVHAKED
MIKSCWWDSHKCSAKDFKVTHTDHGVCYSFNAGGENPLQVTQTGSDHGLRLRLNIEQYEYMRGPNKGAGIKVLVYG
RSDVALVRDLGQAIPPGAHAFIGVQLKKMTNLQPPFGECNDDKHLRFFDQYSQSKCSMDCITEYLSEKCNCRLKYMPR
ERLHEHPVVCDMDQFVNCVPSALETFNVDRVHFCQCPVPCRKLIYEASVTYASTSHFDTDQVLQNSDIGALQNRYIAAK
EVHQRVDKQVAEKDKLLVKNLIETSSTIDTSIRAINQSLADAQSSMESIYKEMQNRAFFHKYVTRKIIQGYQYWFAQGW
ELTSKMIDELIGGHAGFVKTYNNTMTALTQNNHQQSFQEVLIWQIMQEASERFHLAEKVVKEFKNVQDSYTNCTPIFE
YPIEFADAYVKLLVPETVMEKRRKYSHQLFAFILEYLTNVVNRYNLLLDKVTEVSKDSSYDFDMTVFKNITDDLEEDIKSIN
LGQKTYDREVVNYPGELLGRQADNFEGAMADMAKECDDLMNTLKKTRSLLNNFERNWDGVRNFIKESNAYIDDKKL
GKATLSVRVNSIDHQVEAMRTFFTDVRSRARNLDHDWTKVFIMYENSMKEIAMEKSTKALYAKLNEDVNRMLKSDDR
TLQQEIWSNLPFPRTPGNGNADEMWPGHDMFQYQKYANLDTAFMNETDVDNLVVSEREKVNNVSDISKAIDDKDI
NFFASFDEWEKDLDFFNTSTKVDSGFFRENFIDLDIYLTELYYEEIDQQQAYDLTSLFSDIGGSMGLFIGASVITIIELIDIIFL
SLFKTKCSCRA

>Deutero_Ambulac_Apla_gbr34_30_t1

MPKFKVKPAAQGDTLPFSANLGSSADNLSQFQVADPDGEKQSEKKRRLRTSPCEIISDFALNVSSAHGLPNIFRAKTKLG
RLVWSLIFTGGIVLSIWQGALIFETYLRFQVTTEIKIITHNSLQFPSVTVCNTNKLRRSAVKSSKHRQMLVVDLVASLPYYA
PCVGDDFLCRDRPVCVKSFLICDGIRQCGEQDTSDEEDCDYDFGSCAEDEFRCVYSGSRWGLCIPEEKRCNRRKDCYEG
EDEDNCECKSSEFKCEKRGGCIPKTAVCNDVSDCLDESDEKNCSKFVDDGISDNAPDGTYQCSETSFKCGRGDVCIPKAY
LCDNHEDCPDGSDETQDICNQLSTTTEATPACYYGQFNCGDGQCISSGWMCDSYSDCANGMDESELYCGKTICDSGR
VPCDLGGGCGVPCNLESDCFNSTDERNCVYTECYTNPNGQDYRGSIMDARCLPWTDDGVAALGYTGAAYQELEGVQ
PGSTVNHAFCRNPGGEGVGPWCFVVEGEPGFTRLTIHNCTVGPPSVTCSRDHIRIYGSSSLQFQCLSGERIRAWQTCN
GFPDCQDGSDEWSKDCGDVVSECAIDEFECREKTPEGTTKCIPKKYVCDDINHCSSDIGGPGLISSDEEACATPGDVPPA
NYFRCANGHKMVHIYDVCNTVPDCYPEEDDEYACTWKNGTDGRVEDPFRDPLWYEDYATLVKANHHYLFDEFSSRYG
RSPFYRVKGEYPPDWNGFVTYSATPDYRDLVDVPKLSRDEVHRFGHQELDFIIQCSYDENKCDNRDFKEFQDDTYGNC
FTFNPNGTFIASKTGSRFGLRMTLFLEQSEYISIFGQEAGVRVAITPPNIRPNPTDNGFTVKPGAVTSIGLRYNFIKRQSDP
YGKCQPSIESSFILENGHVRHIKPGEEYDRGNIVRLEVFYEELNYESTKEIPAYESSSLLGDIGGIIGLYIGFSFITVAEFLCVIYK
LVKNAFN

>Protostome_Lophotroco_annelid_Pdum_comp421430_c0_seq2_

MGKVTVAPQQKSLAAALPRSNSVGSWSVVLKSLSQKHHESWEDFTKQTTFHGVKYIFSEGQKYPFRRALWIVCVLGAI
LLFMYQVGTRIRDYRNHSSNVNMNIQGARGGLQFPAVTLCNLNAFRLTEAVETNMFDLLDSYYDTVTNTDAAFSHYP
NTTDDTGTIFSRLAHQKTDFIEWCWWNGSPCDVEKNFDVIITRLGYCYTFNSDPRRVKKVYDTGGRRSGLKLVINVEQY
EYMKGPQDDTGVKILLHEPGEFPAILDQGFAAAPGTHTIVAVSQIRDNSLPDPYGTCGDVAIPQSDCLLSCDTAQAQTIC
GCRDLYMHAPDINGSLPYCDLRQTMECLPKAACGNDSNSECRQKCLIPCKRTMYVPSLSNSQLSQTNIERTVLKSSERKL
EITKKYHRAAETFQNVDPDASATNKRLITDLIESAEDFKSKVQYLDSLFSPGDKLADLTKSVNMRTNGSDWYDEDFTEFR
RSIMNLVSVGTRPIFRPDGWEIMDDTQPARKAFFQGYFTGISKTKLYSDLCLAYKNKLNDMSGEEGDDEFEFARDKVTF
ADSMRDDVRDELLKLYGFYEEKLTDMFGNDSSVVMANHVQGAQCTQLTTSLNLTLGLYPGAQVNAKCMKNTDFTQ
QQDVNEMNAILDPFFTQVRSSVSQDDWNLMTSTTDMLNTCDWFKQVFHENKMSFESWGNLASCQKHDVVLMNE
KFNDIATSINFIKTSMEVDLADFINNTKLYADKNVTKLQLAESFDHASVLDYVKDVKSQVSDLSRNSKSLTVKMTHFLTE
WQRITKEIIALSFPPVMNPETMRKMHFFNELSEMMGKEIDTNDYIGSLNDMLEVHDYHVSVFFGNATAFNEELVDSG
RNFVRAAGDLVENLSGHKSETAVNNNFFVHNFLTLSIHYRELTYQVVTQQKAFEFLSLLSEIGGFLGLLLGASLLTMCELID
YLVSLTSKTCQDRKDKDSDTVDILSP

>Cteno_Ple_Bac_comp47367_c0_seq2_p1__comp47367_c0__comp47367_c0_seq2_p1__ORF__type_complete__len_969__score_158_83__comp47367_c0_seq2_123_3029_

MQHSGAHYLNDYEMAPPPDLTPPTRRPPSNTRSQSFQNQETQRQISGVPDQSGGGGGDSLVIPVHPRAVQQRSSSV
GDSITTPKSSPEKEVYRSSRSRTPDRHHGGRKRTPEPRDRSQDRVPRGRTPDHSTRRSKTPDPDQSRRERRRGTCDDTG
PPPPPPPPLPAAAQSDSEPQNIHCGPHDNYPHHGGSHGSPELRPLVVQDQPSSNQGGAGDLAQISMMSEQIMVSEA
ASENHHHHHHRGRSRAYSDSDAEYYTEPYFEDSPVKQGRREFRVPPPPPPPLPVAHPYLPHPEYIYNNTQETRRFTKMP
ARVPYPNMPYRGLVPDSQDESDYDPRQQRRHHSTHRHQVFRSPPPVPPPPPTTVPALTGGPTKKKNEFRPYPAPYSVP
PTTESAYQSGNEYQTPSERDNTRSKVKRSQSDSSPSSFRTGPWLNGQSKILAPKYPVMTSVTATPTNSLKMRPVGRPNT
VPSHQPHHTWHDSLMGEDRQDPVIRDREETTTVKSGPPDNNSKSETFAEFTGDMTLNYLRYVWNSPELIRKLIWAALF
LGFFIYSFMCCFRSFGTYLSRPTSTKYNMFYEPRKEIRFPAITICNMNPHNRTYLEEFDQRPMRFYLSKTWLTPDLPKVPM
EEPEKESSMNFNVKNYYRNSGQRLESFIEKAVFLEEKINVSDAFVEKFTTHGLCWTFNDKGDKKVNRTGMDFGLSFSLN
INQDGYPDFVTTAGLKVMFHNFYEPALIDEYGMALAPGTENFISLSYERIRNLQSPYGNCTKDTTLPFGNYSVSGCERM
CFTMALKEECDCTSYYLPPSGVKECTLYDEEECVEGVIKKYHAGAYMDKPYCQCQDACVSENYIATLSNAEYPSKRYGEA
ITRENKRKSDLAKNGNSTDDVLRDINYYRENMLKVHIYFEILSIKEVEKVGSYVWSNLMGDVGGQLGLFLGANIVTIFEFV
DFFLRVLWYRLLTPYVFKRFTGH

>Deu_Ambulacrar_hemi_Ptyfla_40v0_9_20150316_1g9451_t1_scaffold5118_cov147

MQYLDFGIKIKYDLVTVPKLTFPTVTVCNTNKIRKTEILNSEHRRIRVVDEDSVLPYYGLMAFEATYSKRLSPKDAVLIRDSS
IGACAQSCLRFSEFVCLSFDFNYDKGECHINPGNKDTIVFFLTEDDDYVYYERIYHDFSALDSNTALKFERFPNAKIPSDNP
PQKTTTEECAQLCLDEISFDCLAFTYEQSASRCWLHNASHSSDSIPLIKVSTHDYYDRREKECYTAENGADYRGFTYVTIF
GNVCKSWMDIQLPDGVVFKFNSSDGIADHNYCRNPNDTDAPWCYTTISGDSYQFCDVGPPQSECPPPAKSDTKVKEC
FSDPKGSDYRGTVSTTVSGLTCQKWTVQYPHAHSRTPDLFLNAGLGNHNYCRNPDQEPSPWCYTTSPHIRWEYCDVG
ERDMSCTLLGIECFYNRRATDYRGYMSTTSSGRICQKWTEQSPHSHKYDTEFYHQRGIGDHNYCRNPTGEMSTAWCY
TTDRDFEYELCDVEFDREECVTDSETGILNLKFNMYPNTNVLEGGLQMIVTAANMSQCVDACVDVETFKCKSLDFSPSR
SACSLYEKPAANVGDLLQEDISYIHLERRDENNNCPDEYRECGSGECVYQYAICDTILNCADHSDEVDCNATVKGSIDHR
AFTEQYIDISGVTPEILSDFENGYYRDNGFSGVVKGEVPPDWYGFKTFSSTPDYSDLRSVLKLTGEEIAKYGHQAEDFILQ
CTYDEKTCSPSDFSTFQDDHYGNCFKFNHGQNGTIVRDANRQGALSGLRLTLFLEQNEYISVYGRNAGARVNINPLNDT
VLPQDEGITIMPGTVTSIGIREKHTLRKTKPYGKCKDEANEDGLYGPNRTYGVLSYLLKTIHSISSKTKDINDAHSVSLNLA
QLEVYFEELTVESTIELPAYEIEDFLSDIGGTLGLYIGLSVITVAEFIELVASIVRYLYCSQKDSKSRTVESGDRETKDRFSGAY
AYPAFGSINSTDKSSATT

>Protostome_Lophotroco_annelid_Pdum_comp421979_c0_seq3_

MANGKTMDGFPETAALQSNEIMPREKENSKSSQWKEAWFNFTQNTTLHGVNYITGETPFPIRRDYGFVEHDDCTTFR
RVIWFLIVSLGIALFARQVVESVLYYWSWPVTVNVKINYNKTITFPAVTICNQNDFRATATANLSKYHLVTDYYSNLEKHS
ITNLTEKEGNLSLSTLYYEAAHRMEDMIMDCRWDETKCNLTEFERQLTDYGVCYTFNMEQGNRTLRKVSRTGADRGLR
LHLNIEQYEYMHGPNPGAGIKLLVHDNQEVPLVRDLGVAIPPGTHAYVGVQITEIKNLKEPHGQCNDSVKLKYYKSYSV
PACVVECKEDFMVENCHCRDVYMQSEDDDNPPICRLEEYQSCALPKMEEFYDDKDRCDCPIPCRRLLFDTSISYAYASS
HDINFFLSEDLTELRESYLLARETVERKKFILRDENIRLINAFGATVDNFTQALAKVTTDLTNAGIMLSLKLTQFRKRLWFH
KRWGLEKVKYVMEYDFVRGWEVFNERTLDGVTTGYQEITAYLEHVAEIVNRGTADEYTLSALAVQVDRLLYEKKILCTR
AHDNVTEAQNAYENAEPILDYEGSPGRRYDDTWVTPLVVQASSYFKSKYFDKMLLRTRQFFNETEALEAIMANVKTGG
PIDVASLKATNVQFEKRARSYNYYKYVLEQRVISAPVEYAEKATERFEDLNTTLTRRYDDFEFKYTSLMNFLDQVTESWSR
IQQAREQAELYLADLTQRKTALAKHLTSKGLNNDLETVNEFLAQVTTRSHDLLEAVQAFERSYNKVWSNMLSEESLKDF
YALVADDWEMAGQNDTFMEYWDGKAAYMIEEKAAWVKSNMRDVIWRMNADFSNTTIEQQSQANRAFIATLQPLF

DVSECLETGQKFFKSMSNLTENMNKFLVENELDSEYYRNNFLTLDVFMKELSYEEVQQQAAYDLTSLWSDVGGSIGLCI
GASIITVFELVDVIIHNFFRLHVFSSRAKKKGEPEKEYL

>Protostome_Lophotroco_annelid_Pdum_comp405417_c2_seq1_

MGEDSIIGTTTNFGVTDYGRRNSRAVANRWEDEDDIPDEDRPEEWSSVCFHFTQSTTFHGINKVTEATPFVLRRIIWVLI
VLGGMTLFVNQMIDSVLHYMDRPVTVNIKINYNKTVKFPAVTICNQNVFRTTEAASKGHYFLIDTIFSKEKHGEVNLSEL
GAQDVTMLDLYHSTGHDKHDLIVSCLWDQSFCGPEDFVTTITDHGLCYTFNSGRNGSVRSVTQTGSDKGLSLRLNVEQ
YEYMHGPNSAAGVKILVHEQDEIPMVRQLGQAISPGSHAFVGIQMLQINNLHPPYGECRDDKELKYVQHYTIPGCLVD
CREIFLVDKCKCREAYMPEKGTDGWPPVCTLVDYQDCMPKAYNLFNAKDEKLCECPVPCQRKMYETTISYAQTSKFDT
RRHQRDANTVELQQKFVKAMETVQRVDNVIAARDFELVKRLRHTAKGIVATVNRIEQIMMNVSSSLDIFTAEMGHRI
NFHDFYGLRVVERAVMYGFVQAWDIQQQSIFQPLAAGTVEFLAYYRKAILRIINSTEEEVSFRRMLEWNLEDQISTRKEL
IRRARSEMKNIHNAFLEGRTLIDYQWGIHERYDNLFVSKQLLNHRMEYQAYLYQDLDVTLRLLDQEIVMLKNHMLDSIR
TGSFNHTLYRVARRQYSDLIRNFNLVFNVFKESIVHRTKTLIDKRRNDFKKLNETLYKESGDCKITIHQSMDLVNTMRKDT
WPIVEEVINDSRDYTGQNHLGNKTSLANMVTSETIHVKLQQMHDFFNDVRARARNFDHDWKKYMGAYIRAWKTIKT
EDTTQEFYQQLYKDFLEIRNPESNVSDELLNIFVNRTYLSTTDPDVLFTRTNADLAPINATVALKDLNDKFYKLLRSSNIAS
VVKERDDKFFEAMDDWIQSLSKFLDECKTDESFFKENILEIDVFMKELYYEQVEQRIAYDLTNLWSDIGGSMGLFVGAS
VITVFELCDAIIHNFLKMQIDKNRRRKRAAGIAKRQNGKV

>Protostome_Lophotroco_annelid_Pdum_Contig12073_

MCWEKKDSYSEEPADRCVTCGRSKADDTEPETWGAVLYEYTQTTGFHGLKQITEPTPFCFRRLIWAILLTGGIILFCIQMI
DSIIYYRSYPVTVNLKINYNQTLNFPAITICNQNSFRVTVAADKNQYSNISSLYLHQDVHHEDGHSDHDHFGDQFLKQS
MRDLFLEAGHTKSDLIVSCEWDGSDCGPYDFRTSVTDHGVCYTFNGDGMYNVSQTGADRGLRLRLNIEQYEYMQGP
NDGAGLKILIHDQNDVPMVKDLGQAIPPGSHAFVYINIVQVENLGEPHGVCNSKELEYYDRYTMSACLMDCRTKFIVSK
CRCRDYYMPKSRVFDYPPVCTVDKYIKCLRPALELFNSLRDHCDCPVPCSSVMFNPTITYAANSNFDMEKMLQADTSHL
LQQYQKAREVRQKVDTLVLQNDQLMAGKLQLTVYQLSQSFEQIVHNTQSTFDELQVMVRRLNDRIHFHYERALDGVY
YNVKHDFIRGWEIIEERAFPYLSGSFFETYSQILRFFDMDDSHDSLSKQITWLIINNSIDNRLELCNRVRANLTEINQAFTIG
KTLLNFKYTVDRRYDKDFVSLEALQSVPDYEHKDTYFRETMSAIDKIEGSLREIKTFAAGFLSSSTVDKIALRKTASNFVSGC
KSYNYRLFQFRDRIVLEPLNVVKARQQEFQHLNNTFQSKHKELTALLFGIRASITLFNHSNWMALANATNMYGKYVSG
DRITKAELADIFNSDIIEEQLITTQQYFTTLHSRTTDLKDKLKEFGKAYADVYANMLEEPSTKTFYSRLNSELKQMESNNS
MRPILDKHYTFLAMRNVSDEPISDLYRLLNADLVDTNLTGVFEFMDNVVSRMQTDTDLSSMIGQVDEQFIKNIEAITVG
FANFVESSQINFEFYRGNFMHLDVFFRELSYEQVTQQEAYDMTSLWSDIGGSIGLFVGASVLTVFEVFDVFFFTALKMQ
TKPKSKKQHRNGNSLNNHAKENNSFTNINNTTALSAL

>Protostome_Lophotroco_molsk_AplCal_XP_035829875_1_uncharacterized_protein_LOC101853999__
Aplysia_californica_

MPGVDTRWQERCARYFSSMHEAAPHYQETKKRKKDRVRKKDLFSLPKEPHPYRVTFSRKTQPCKGGVKWLSGLDLHG
TSMLLRTRSYFYFIFWVLISLAACVGFGVMVFFLVREYQDSDKIFKVNVSDTYENASPLSLTVCNTNMLKKSTLDNRTRFY
DLLLLDIDKVVNNSKTRLSTPGKSLAVLLDNTKLKNFSDSLNYNEASLIFDDLQREFEVYSGMQSDTDWAYVASRGLLET
AELFPPAFRLTRSELNELGHSMETGIIECTKNGRSCQAGELTYSQSRLLGNCATIHLSLGGSWFESLTLTVDVEPEEYLDLV
SPPPSVRALLTTSTSLRHASSVTESYVAVPGSELDLQVIKVKEINKHKDNCRADTDYKYEECVALCHASLVSDICGCSVLLD
NPLPQRCRVTDQYQFVCHRFLDILWRRGQLPCNCPARCREEEIRVGASHTIGSFSHSEINEDCNDRTGSLLHQVTFHIITI
PLTSHREKCSIFQEYITKLLAASPAAKNASTLSKSLTRIRVTSVGAQYEIWTQEEEITILGLLLQVGALSALTVGISVVTVLELL
LLAGKTLVEACRALGRSGHDNEVERLYPYEDLDAPPEFQNLDANWYTCSADLSRDTWHGTLATDNDQHLLQGGQRKY
VVSRSSGTDGVPYSVARNSGPGGLPYSVSRNSGSYGAPFSASRHSGSVGAPYAISRSSSGTGPHATTIEHSRTAHAYSNR

SFIRTNITSVPERGSLNPLLEESGDADDNTHAKDDNLATSPEALYVRPWDLSGVSRPYPKILVSSQKNSASDVPVPNAPG
RSLRDHADPENVVLEDADFVPFSQSALRLNLPDDLKNGSDDEDVADINNENLRGGHANQSANDSRYSSTIDDPVTHG
DHYNVKRWDKSRQLSDTKPIYAVSDGQLQSLDTAGLGASSKTYQSSQLQIKDEINYGISGEEGRKTGNVLDGGRSSQHR
QEWGGRDTLEPSYYIYQGRAAAETQESIFVI

>Deutero_Ambulac_Spurpu_009902

MPLQNIKVKPFASGTRSCSDGTGLKTVSGEDFRVDIPLSMKYRDKVSWWALFYKSFPGAIGIGGMKYAFNPAEMKFRR
YSLLAGTQLESFMYEFHFDHSLVESLNSSSPYFHYLSNITMSDLFSQYGHRPDETVEFAVWEKVKVLDRDTYFTTKITDFG
LCTVFNDADSGNPPLTIRRPGTNNGLQIRLNLEQYEYYFNPFYRVSAGFQMLIYEYGSQALMEDQGFGISPGTHTFVGV
DVIETSNLEPPYGLCHDKPLKYFDNYGYSECIHECHVDYYINTCGCGPPWDRNISSIRECTAWEIFTCVVTTYVIRMDGEN
VFGQLRRIARNVQNESALAQERMSRPIALRTSGDSARPVLRRMQADAHNVKLAAGDAHSSLPSSVEFSKVLESCSQLC
DAYRSKVNLLEGYLQQYGYQPQPVKVKGVPVNSIIKKEARANETQHPKGEQLLTSTYDAEEMKENVPTTPTNKPDLGSS
WKGGTPRLEDFGLSRGFLEQMTRSKHQSHQQPQEATEKPQPKAVAKLNPTTPAKKPDVVLPVAGGTPKLEDFGLSSN
FLDSVSKNKLRNQHRVESTQEGTHLNSTYNKRQMEMAQTTMFAPSQASTCPFTPVGVTTTPSARVTVTPGMFGGSEL
PDETPTKFIDLNSAMPRSHAAPEDDFGGSQGAKKSLHYEPSSKKGTASFLDSISSDLAENEDQFGTPQPPELTARWEFTS
IQAPLPSQPEFTMSSDKPPNKLDMFPSLGYQTDTPPTPEMTAKWNIRNALTSLFFPPLPLTIRTTSKAPGSLPPAEPPFV
MEEPIRLTSHEPKQTPEEPQRLYVEKPAPSLQDMPETPPGLPGKLGDSKSFGFSRFMADEMPKTPELTVSYPRYVDPST
KPANVPKPVVDSIPQKPSASLHSGIPRMVKVEESNPSHSTKPVDISYVAMVTDEEFSKQSAHMRRILPLNLVNETVSCIN
TILNIKHKQGEGVYLTDEDVQVISLGPKSKAMLLFFITVKRLSSGKRPGTSEVAYFPVLY

>Deu_Ambulacrar_hemi_Ptyfla_40v0_9_20150316_1g24774_t1_scaffold38646_cov86

MFQVSTLIMSFADRPINVNIKFEESKERPDFPAITICNNNMFSNCVENVEGNLKYFDGTYSLSKCWMECETDYAVEQCG
CRYFYMPGNSTYCSPYTLQNCYFGAMDEFALKVNPCKHCEEPCVQTKYETKLSYDSYPSSSYSQLLTLTGRWPCNEALA
YYAITHSSDDLMEIDANMKNFVLERYYTLEANGQNISALIPLPEFHAIIVKALREGLLTDFFRDLTFEVLKYTLTLIYKNGTD
EVGSDFYVQTKKTLQQKYMNLSVNPWASHNTTMWNMFASANHTSDTIQYLIDTLFQDYYLPLFDETFFYLYSELKSWR
DDKTEEYPRKKLRDICKAHMQNSIVPPDNVADETEFSDDRRTQENPIPTDEDNNNSEKDGKTWTGVFKSFTQEAGLSG
VKFVGSANVSFGRRVFWLSIVLCAFGGLLFQVSTLIMSFTDRPINVNIKIEEPKERPDFPAITICNNNMFRFNNVKNLLGG
ESMFLLYFPYLYALFSSNAEAISEFDDFRVSEEQDYGEGHTQLTFMEESAHHVRDIVRRCSFGGLPCGTENFTKVVTNYGI
CYTFNSHEQGYVNKVRNAGQQFGLQLLLFVNQREYVAPRTNIGFRLMVHGQDEIPNVASYGLSISPGTASTIGLKSKLV
SNLGSPYGNCIENVENNLKYFDGAYSLSKCWMECETDYAVEQCGCRYFYMPGNSTYCSPYELQNCYFSAMDDFALKV
NPCNHCEEPCVQTKYETKLSYDSYPSSIYSYLLTTTGRWPCNEALAFYGIIHSPNDLMEITANMKNFVLQHYYPIEANGQ
NISAFIPSPEYHAVTVKFMTESLFTDILNDLRFEVLKYTLTLIYKNGTDDVGSDFYAETKETLQQKYMNLSVNPWTGHNTT
MWNMLAFANHTSDTIQYLIDTLFQDYYLPLFDETFLYLYSELKSFRDNETDEFVNEGPGDICKAYMQENFSKVTVYFEDL
NVNSIIQQPGYHSFNLICDIGGSLGLFFGASIVTFVEIIDFFIVQFWLMKISGNK

>Protostome_Lophotroco_annelid_Pdum_comp417475_c0_seq2_

MNDKTADEKAMMKKVAWQSTIHKALEQEHEKSLAEFAKDTSFHGVKYIFDGETSSPFRRILWLLCWMTAIGLFAYQV
ATRVLEYRKHPTNVDVKIMYTDSVTFPTVTICNQNQFRFRRAVDLKMYDLLDNIYKTTNPKGINMSAYEYPKDSTVEEIF
KELAHEKLDFIAKCTWDYRECDRDADFTMTTTDMGYCFTFNAANSTANTKNVTDPGIGAGLRLIFNVEQYEYMRGPH
SDAGIKVYLHDWKEFPLVTDKGFSAGPGSHTVATVESIQVSNLNYPYGNCIKGAEPMNNCRLKCVSQKAMKVCGCRD
MFMLETLGNETIPICDMTSKLKCLTTLTSNIQKNDYCSCSQPCERQKYEASLSNAQLSRFNVKTTVIRDDQHLQKVNND
FLESMERSQREISDNVKANENVMGRARYVYKDIAINAYGLVKQTTETHIMAQMEKTLDLKNSDQMQLLEKSIEAFGDS
VGVIEEAVKKLDPTKDELEDSEDVIEIFEKMEWKNGRPVKGFRDAVQRCVNIEAALPRSSSSGQSGSSGPSSAPSPISAPV
PAPVPAPAPAAAPAPPPAGISAPAGISGPGQTSFTFTSQDPDCKELNEKIKEIFEKCTEVQEDFGPEKQVMFLQWKIKM

DNFTTELFQGTSKTPSALNQQCLTGLDAVHEKFIKVISVQANASTAQQSGADQAQIILDLADSLLQMNITDGSWLKKYG
NYDSCEWLENEFKASPEFTKIRSSADFKTKMTSLMASFDSVVDVFQNMWDKFNETMIPVFVHLDHYFDEKATKLSLVD
PFDSMEFKEATTSLFSTADTILKYAIDLESDKWYVDQFIDKFVESLVEAPVPVLKPAEYLNLSVIQAVQDLQSKRLETHLLA
LQNSENSEKNFGEVLKVLIEAFLQPVDGGGRLVGVKSKEASSTFEALVSFLQEYKKAVLMDQEFYMQNFLMLDIYFSELS
FEKINQQEAFVFLSLISEIGGFLGLLLGASILTVCEVLDYTLMYCIRCNKKRKAVDQDPLQPFNK

>Deu_Ambulacrar_hemi_Ptyfla_40v0_9_20150316_1g971_t1_scaffold462_cov140

MYLTVVDDQTGFPALITCTSEELKAPEQKHSKTSLKVQNGYIVLKLNELTATDNNTVYTWWSHSTVVGETAVTVRARDF
NSTIIDFCFTELTLTVYTKGHQAVNTTRSMSFLTIYNMQEEDSGQYSCRVTTKCGNHFESRQIRITKIQEISTFQVSTGIGM
NYSHTSMEPSSDESPIDNTYFVLLIILCIVIFLLLVVIAVITLRPRFFSTIEMSTLHDQPFEPRTYDDANSTDEAVVAVENWE
QTEASLIVGNVEERSDDSRCEVKILEPNIPLLESCDGLYEPVEYDYDHLLFCENGFLGRGEFGVVQKAVVSDKRYKCVDQE
VAVKCLKENATMDDREDSLREIEIMKSLNASHPNVISLIGFCTDPLCIIIDLAVNGNLKNYLLKNRSSSDYQNILPNSRSLRS
YDLLQFAWQIAKGMSFIASKKIIHRDLATRNILLDENRCCKISDFGFARDVSDTNAYQRKSQLRGLHHTRHYHHRKLFKN
FVAVEDSLNLYTVVKNLQISDDDGCFDKVDIANLKKQGNCGENQFKCVKGSEDGFCISKKHECDRKSHCYDESDEENCR
CTSYEFQCESKGGCIDKGKVCNGQEDCTDGSDEWSCDFYQGSGENCTGYPCDQGNRCIPESYKCDGYYDCMDQSDE
DGCQGTCGSDDHFTCDDGICIPAPFQCDYYRDCIDGSDERNCGTCESDDQFTCDDGTCFPAWFQCDNYQHCIDGSDE
RDCDCTLICDNGTTCIVDSWICDGEEDCLDKSDEPYSCYLNSNDTASSAPSPNEDTSTEPGASSEDPFSGNWREPFVTIT
KDWTKYNAFKENYYKDRGFERVKGENPPDWNGFLSFSSNPDYSDLEEVLKLQAEDIETLGHQAEDLILLCTFGQEKCTP
SDFSRFQDDRYGNCYTFPGSRNDTEPLYVSRSGQNHGLKLTLFIEQDEYTSIYGRQAGIRVIITAPEDSESIQTQDGIIIIPG
TETHIGLREKNLLRTIHSLNNKTLNINDKYSTSCTNLSAILVALLDFSSGCRS

>Protostome_Lophotroco_annelid_Pdum_comp413124_c0_seq3_

MTRSSNVISTISNFSNIEEMSQNGTADPQCKGNDNLAFETQTPDENGQSNGNTTNSNFEFPARKAPSAALRTFRRAAR
TVGVSVKVVGVWNNFAQQTTIHGVRMITEKNIYKLRREIWLFLVLTGVGLFLYQLLERSLYYNSNPVALNVKVTYNETIQ
FPAVTICNQNSFRLSKTMEKGMYKALQEMYSRPLHEFDFNKHEIGNVSLQDIFSDFAHTRDDMILSCSMNGVQCKQAF
SPVLTDHGLCYTFNGDAENIIAESSAGSDQGLRLLLNAEQYEYMAGPHEGTGVKVLVHSAQEIPSVNDAGISLQTGYSN
LIATKVIKVENLPEPYGKCQPNLHLDNFDHYTLGNCNLECTSAKSQEFVGCVLPHMVQTYDELSRSEDREDFPVCSVEEY
VTKYIARDSKMARTTCNCPLPCDSVQHEPSVTYAKILTKQVVESLDANVIENIRAKFLNSLEKQQIVKESIVQEDVMAIER
FLRMVSVCKAESASVKDVMISTADYLSEMKMTASARSQFHNKWGLEKQNILIYEGQIRGWETQDETTFQFVTRIYKEF
GSVFLNMVEILTSNQDEAVVESWKSETMRDIQDRQYMTQMAIEMFKNYHTAFEQGSLVNELEIEITPEGYFDSSYSIKN
LLVDPKIERQIYKAVWNVLNNYTSSLEKLSNFSLSYFETINFTQSQFDDIFIKFVEESETYNELRFGYKTNVVYKSFDKLDSL
QEQFQQRNSHFLVRMTSCRKAVDKVLEKLSEWRKGILPKLIEVREVSEEYIKFNGTSKRHVAEMITEPKKMKLVKDLKTS
SRDLRVLGIVITDNIKQLKGDYLALVEFIRSSPGLQGFYRDVGNVIINETDSARSNMVGDSQGFQSLNTMLNGTMTNFD
ANDTQSVLPLVNADLFIELEAFEAKLNREIQKMTEDVQTVIRFSDNFDLLTTTLSGLQERMEGYLKQTKMTNKFYGDNFI
RLNVFLPEMSFENRIQDRIYDEFALICDIGGSLGLFVGASFMTFFEIIDAGLRWLARHIKVK

>Deutero_Ambulac_Apla_gbr158_30_t1

MFPPQPKILLSRALESRTCMDSQFRCVNSGSSMGLCIPSEKKCDRHWDCYEGEDEDDCPCKTSEYKCKGRGGCIPRKH
VCDGKRNCFDGSDENEEICGTSIFSPGHCLEPWFFKCSSHKCIPKIFVCDGIADCLDKSDESEIACGSRDVVRGVVPPESV
FPTCSPGQFMCGDSVYCIQAEWTCDGSLDCSDGSDEAWDLCGRPVCPPPKQPCAEGGGCGTLCDLWPECVNGVDET
NCTCDILRNDNADEVVQKVICNGVSSRDFFYAFSECYTAPNGWDYRGTRTTPGCLPWSDPRVAGHNYTGLIYRELEGV
EQGSTVGHNFCRNPGGHRLVPWCFAENENGSLGYQDCSLTTRSEIPCHVESFRVELSQNGSDGRPTQGRVEVSFRTG
AVGTISDPDWTIDDANVVCRQLGFPAGALANVTADFWPATGRVLLKDMQCMGWEENLIDCLYSLATDQDTNHTYD
VGVACIDTNESLPAAPKRERRDLSLVAVTEAEACPFCSLERETHRTPSGCGEGSFRCGSGECIRYWQTCNKFPDCLDASD

EWLPECTEEIGVDARLLELESDAWSDGVMPVDIDGTWHSVGKPPLERGNGPCCIQTDEMVRVCGQRKFQCKEPRRD
GSVRCISQNFVCDGYNDCGQASLRRSRAVKTSDEMRCPLQNEEPPKNYYRCRGGVEFIHEYDVCNTVEDCRLGDDEA
NCPLSGRREPFKDVFRQPLWYFRYRKLVPEDHHYLFENFPAQFRVPPFGRVKGEDPADWNGFVTFSATPDYSDLVDVP
KLTKDEVSQYGHQKEDFILQCTYEQTKCNISNFETIQDDTYGNCFTFNADIEKPITASKTGARYGLQMTLFLEQDEYIGIF
GPEAGARVSINPPHIRPSPTNDGITVKPGTVTSVGLRYLCKKTCIHEAILKNCDCVDTLELDGPRCRLLNETQEVCRQLIYY
LNQHNNLSCDCPEQCSDNIARLEVFYEELNYESTKEVPAYPESSLFGDIGGIVGLYIGFSLITVCEFIALFLKLIKKSCCNRAL
RI

>Protos_Platyhelmin_schman_A0A5K4FAF1_A0A5K4FAF1_SCHMA_Uncharacterized_protein_OS_Schistosoma_mansoni_OX_6183_PE_3_SV_1

MNIHCCPKWTSVGIKKLICFLLCISVACLITYRVHVWLKHYDSLASKMIISAYDDHQSNLPFPLISICNINPARGTKLYDIQS
AESQDRGVDYEIFSDAFQGRPSENFPESKLGMPIFKLMDKASHQIGQMLRSCKVGQRHCSVLNFTKSILPNGACYTLAG
DLTGIDEIQLVLDPQSYDYLVPNQGFIGFRILLHGYGDSLWGLIPTAVYAGPTFHTMLRAVGLKKIYKQHCTSQSVWASC
MHDCMQEMLLAKCQCQLPDCTVFEMMRCGLTMNSMIESLSPVQCRCHNPCEAISYSVESITQALKSPLLFTHSLNLFT
KINQTNFELNNNFQLSTDQQQFNNKITSKNQKSATVSDDISDYDPNSNNHKNPKDLKLNQPEQFLHIYRKEIWNGLLL
QTWAFLREANRTAFEQLRQLSRLLQSLNLDLQTVERAFYKVHRKHRLNLEAHDLMIWDSERLSTNKHSSSDQSNGSCM
DSEEHSRMLERVSHLGKEFVRLFRQSILFRDITLVPGLDFHIHLVRLFFLNVVDELDHLANQSLHNDLFSDVTMLDLQRKT
HQCKQTNNNLVYNDTSWKNQLEQSQLIVAGEDITRLQAVLLATDSQFKQLRTVFSKLVLDNRELIETVPIKRASLNTKPD
SLISTEEGLVSVTIQLTRDKNRLIRLESVQAIYSPIAELLDLIVSGLLLAFLSIFLLNICVSHPNNSDSCYSECCCCCSTLVNKST
NSKSCESPKLATHTTLPTSSYINCNTDSGSSSKRLPKLNTISYRNYSDLVPTDILQKKPTMSSVYLCDSHQQDGVPISSKCII
TGLPITSNKPCAKCFGENCRHNFNICPNESSQYMTYNLPTLLGSSVTLPKWDNKVVQLDMDGFNCMDASEHSATECD
GEQSHANTNHHYLCHQQQSLPRQQQQLNHQQNSVNESCISFCDCPYSMEDNTRTDSASCLNSNKLLFCNNHLNTGR
NCSSIRKPTDINYSKNEMLNCLIITPQQQDTTTFNEVTPNIITTHQLPFSVDLLQSSKLLDNKTTLPVNTSH

>Protos_Platyhelmin_Macrostomum_lignano_A0A267GN93_A0A267GN93_9PLAT_Uncharacterized_protein_OS_Macrostomum_lignano_OX_282301_GN_BOX15_Mlig006398g1_PE_3_SV_1

MSTAILATKHGIQEPSMEQLQQQQQHDRTPKANQLRRSAIEFLGYTTAHGLGRFASQSSAIGKLIWIGFLLGTYSGFAY
HLYTLFDRFAQAPVLTTMTMSSLGFQFPDVYICPAQPVTMSRASKEIESVYISFNARDHVKPEFKANFFRFNGTNDDLY
ALSKPARAILYVLKATEAAFIAKFQLPQDEDALLDCRTSLSGEEYAGNRSFAVTYDRARIDLQRTDRSLKPYTTTKPCIRWS
EAARMLKARGAASLGDRLLMLNYSDSESRCRAPVRLPLSEFFPIGSRSLLQRVGCFAKNSSTVEENFNKTCLVRGRITTEL
CADCCDELDFTHIALRDGECHCGTWNNTQQGFCDKLSINSVARKCDCDVNVFCIRDTDSKPLRISNSSVLTALSNGPYC
FFKDDFGRLEAAFCNVPLCRMNSGYDCIFGTTDDAARQRQQAVYTGYRNVAASGEPCLAWNQIVERNKQRKQKIDPV
ANYDYKMRIHLETPDYDRLEGADNFCRPAQIIREFGFSDRVYYRNPDLISDDEAPSCVSDTFGKSNLVNLALTKCDIPACY
TKVPTVTAAIRDGIGSHLLALQFSSDLSFTLVECSFGGQSCTKDDFLSVSHPQLGMCHHFMKQRFTQNMSSSQLADSQL
SIKYFTDSTDSNGGTIPELDKLNRIVHRSVSNDERRIAFKAVIVPEGSYPWPSKGVDVATAQSLDIQLSLKKIKKLSTASRPC
VENTEPVTYRMLSWKQAKTTSSNEENIRLHFVKAASEWWQKIAVLRAQLEKQNALRPLQAMDWRYNIPSKEDYTNKF
TVQHFAKACNCLPSFLPIEVELFSNYTFCFDTNDTETVSRLRESCHREQLSKLLQYQFEAAKKALSPCEETDYDMRLISYA
WPNLSGAKSMEILKKLAHNFGLGTEYSRLRGISSIYSTYWRSLVMLESNLTESNLHDRINFENELKFLRDRLAQLTVKGLT
SYGQEISEEEAYPAKNLLADIGGALGLWSGIFVLTICELLEFFYYISIGLALSLRHRRDSLRRDSVAREIETHA

>Deu_Ambulacrar_hemi_Ptyfla_40v0_9_20150316_1g5443_t1_scaffold2520_cov137

MTRKYKVLLGDVMTNSTVHGLPNIRRAKSVPKRIFWSVSFLTAVGLFIGFSSVLVSKYLEYDVDVGVQVNVERDLTFPAV
TICNLNPLRKLLLDEGETKHLQRLIEQALSFSNDNGQGHYNHWDHIDENFYRSPSSLWLLNNQIQEYLADFEGLRRREIG
HQIDDMLITCQWSNEQCSPRNFTHFLNSMYGNCYTFNGGNNTSFLSTNFAGSTYGLTLELFVEQDEYLDGLIHSAGIRV

TIHSQDDTPFPEDSGFNVEPGKLTSVGITMGKTERLPDPFTDCVPANTSKYQTAFRSSYSVQDYVYDVMKGRSSKLRML
LEEQERTGLDLFGSTLIKHYFEYEVDVSVSVTFERDLAFPAVTICNLNPLRKSLLTEMQWTQLQGIVDYSRRNNRGEVAS
ANGTFKNETGGNSSHQQSTDYNQWDNMDETFHTDPSPEWWMNKQIEEYLAQQDVSQRVHIGHQIDDMLITCRW
SNRLCSPKNFTHHLDIMYGNCYTFNGGGKNAHISTNFAGPTHAETCNCTDSRYPKPDGFDDIEHCQFENLENRKCKSN
ADMMYQTSKLPCDCPVKCMKNVLKLQVYFRDLTHEDIIQSQAYSVSIAKVNIIITSERQQEIDLVSDIGGNLGLWIGLSVL
TIMEFVELIVDMFSILSFTRVSRGKRPQQDVEWGTDEDHNDDGVMYRSGEQELTSGDGDVYYSQRAHYNSDYGRQR
GAIEPLQSLGRYRRNLLQKPFAAMVDGQASHGVSNRNRSTTQERDWPPIRGFADKLSVEQLSVFSKGKTVRLPHPFTD
CVSADTTEHEIAFKNSYSVEACMKSCLLTHIAEACNCTDGRYPKPDDYEDVGHCQYQNEEQRLTLELFVEQDEYLDSLIH
SAGIRVTINPQDETPFPEDTGFNVEPGKMTSVGIVMGAMERLSEPFTDCVLANTTEHKTVFGGSYSVERMDLVSDIGG
NLGLWIGLSVLTIMEFVELIFDLFAILRFVQVPRAKRPEMTSGLEARVRRGSFPYATRRQGLTFRSGDVYYTNGKGYDRD
TTRHNDEISYVRRNIYLPNS

>Protos_Platyhelmin_Macrostomum_lignano_A0A267G1X8_A0A267G1X8_9PLAT_WSC_domain_contai
ning_protein_OS_Macrostomum_lignano_OX_282301_GN_BOX15_Mlig006056g2_PE_3_SV_1

MGSSSDAPEGLSSPVKRFCGYTTAHGLGRFSEQSNLIGRVIWVSFLLATYSGFAYHLSTLFERFLQAPVLTKMTVSTLQFE
FPDVYICPTNAVTMSRIFKKLPQNDVTFGEFFLKKSVEEKIGIYSFDLSSGRYDLELNDMTKALLSVLHSMEEIIMSYFQFF
DIADAAKCRRTPSGEEYSGNVSSAVVYGFTTMTVTSINTSAFTPSPYRGVKQCMRWNDAAQKLLSLGESGLANRLLAL
NFSDSESRCRSPPGLPLSEFFPIASRSVFRHAGCLPLALTNSSSFNFTCYRGGLSSLETCAECCGRRSFQYMAISRQDCFCG
SSIGGYENSSSVCNASCLHDNSAICGSETAVTVLQIRDNGSRPLHVANFSDLDAISSGPYCFFADELSDKLNAAFCEIPYC
QISRNYDCIVSSEDKLTQQQRLSRYTGRRNVTSTGRRCMPWSQLSTIEQVLKARRGSEVGFMDEVREMLIQTPDRLEG
ESDNYCRPVSVVVDSVNIHGPKDFQRFSAIDSEVPVCIPEDFWVNIPSITIKFGYLFNMIFQTCNVPLCYTEVPSVTATIRD
SVGDHLLARQFGSDLNAGLADCSFAGQKCTKDDFVNVAHPQLGMCHKFLKQRFTANMTSTQLAESQLSIKYFTDSTDS
KGQTVSNIDKLTRVVRDSHGSDATQVAFKAVIVPDSSFPWASKGVNVATAQRMDVHLRFDKYSRLSTSSRPCLETTEPI
TYRLPSWKLFTSNEDSGVNSDAQEIMRCSNWTLKLSMFRNALLKLNGSSALTVSGSQFKVSDRENCVNTYVVEKFTNT
CRCLPGMLPIQEERFKKLSYCFDMYSRGNISANTEKCYIEQVSKITQYHFEAMNECKSPCSKTDFSTRYITYPWPNLLSVQ
SLDFLKKLAADFGYRHDISLMRGASSIYYLYLKAQNLSDTPVINRSTEAIQSFESELAALQNRLAQLTVQAWQSRGHEVSE
EEAYPAKNLVADIGGALGLWSGISILTICELLELFIYLGQALAARVQAQSRMQSKNADGQPEESARPESEMVALDTDAIQ
VQQKVASS

>Protos_Platyhelmin_Macrostomum_lignano_A0A267EE85_A0A267EE85_9PLAT_WSC_domain_containi
ng_protein_OS_Macrostomum_lignano_OX_282301_GN_BOX15_Mlig028168g1_PE_3_SV_1

MATKEEDDESLMRQFCGYTTAHGLGRFGAQTGAIGRGLWICFLLGTYGGFTFHMYSLGQQFAAVPLRTSLSTSLPFEFP
DVYFCPTDLVTMSRASPQVNIKELTRALTCDTDCKKQLKFDIHWSPESGTTSAAQYGPAAQGLLHVISAVESNISQSLHY
GKNTPVSTCRTTLGGEEYIGNLSIVHLFGTAQVREPVSDMNNLPKIAPYLSTANCIGWNSAARELQAAGVPLAAQILAIG
YSEDDSRCRQPRGLPLKSFQDNSTHQMHLYVGCLDKNASAADFPHLCLQDPYGWMNHERCAECCYVANYTYMALH
GDQCLCGEEVSNVTALSLSDFACNRKCRGNIAEACGGDAAAIVYQVRSSGSNPLFIPFRASIDSFNEGPYCFYRGNSGQL
SATYCNVPFCPISPGFDCLALNSEDQVSSGEYLRYTGYKSLTNTKDSCIDWTAMRQALLNGSRTFRFDAEPSILTVNILDV
YIETPDFELYTNGRKANNFCRAARVLLDAAVTEQKTGRHYFIRTTIVDTAKPVCPISITNDEILLSSCSIDACSRKTKSIPGLV
RDRIGSALLASRFSSSMGIGFTGCTFGEHLCTIDDFTTVFHPEFGVCHRFNQEKWVQQSKDVKATDSTLTVSYFTDATDS
LGQQLPGIDRMSRVVFPGVENPNDDSFSAMQVVVVPVGQFPWRDKALTASPGQHLELRMRYDKVNRFSKPPDQAC
NQSKNDVTYRLPSWQSSWRYTPQALQSVNITELKNTADWAKKTQILESILMSLNASISEMNQTIHFSRATRRDCVNTY
AVHKFIQHCGCLPSFLPIPVEFFPTYNYCFDDKTDLLSIESTEACYMRLMASISRYEMEARQLCLDMCERRSYAIRPVTYP
WPGLTIDSATDFLHNSLGDWSYPYNFLFSKGVANLFYMDIRTKEVQARRSNLSANEMLEFTNELQNVERSLAKISIQALS

DFGQEMTEEPSYTTKNLMANLGGALGLWSGISVLTVCELIELLVYAILTARQRIAAKKRISQEAAHRAEVAAGNPEPSRS
PVDGEEFEISQLKNENESG

>Deu_Ambulacrar_hemi_Ptyfla_40v0_9_20150316_1g22493_t1_scaffold26776_cov117

MKYVDELTFPAIAICNFNLHRRSAIAKVNLTGLYDLLYTTGNVDFKQYNVSTTASEQFYRESAHEVYSMLYYRSATIHNGI
SDLLEEMYTTYNPDFDKYDIDVTASEQFYRDTAHLFEYTFYAPTWQGNFLSKDDFTPILTDFGVCYTFNAGDGNDTHKV
KNTGRSFGFSVYLNAVEFDYNIGPRQGLGFQVMLYQQGDIPLVADLGFAVSVGEEVLVGVDVTSRYPCLNLTNGFAISQ
ENCEILIWSFTSYTSNSAICNGKTVEVMYENTKRSYTAVTFKIGSNYQRVIYKPNLSHAKYTYQAFANFQAEYFGVPPEYF
EIVWILVLLAGLGLLSYQMWDRVSYFAIYPLNVNVRLNYVEEVTFPAIAICNFNMFSYWNYNIGPRVGAGFQLMLYEQ
GDVPMVSDLGFSVAPGEDVRVGIEVTKVTNLPPPHGVCAEKSLQYYDRYSVNACQLECKTDYVVERCGCRMYYMPGT
VTVLKKQDDRSVGRECNSMRLAAIFLLCTDNIVIVWLLVLLAGLGLLSYQIWDRVSYFVDYPLNVNVKLNYVEEVTFPAIA
ICNFNMFSYVIPHDYGDDGNDIHHFLICHFWLGTSHGLSVNLDVNQWDYNVGPRVGAGFQLMLYEQGDVPLISDLGF
AVAPGEDVRVGIEVTKVVNLPPPHGICGEKSLKYYNRYSVNACQMECKTDHVVEHCGCKAFFMPGNANVCDVKQYY
NCLQYAEFSFTNKTCDCPVPCTQVIYKPSLSHAKYPSDAYATYLESAYGYPREVFTIVWLLVLLAGLGLLSYQIWDRVSYF
VDYPLNVNVKLNYVEELTFPVVAICNFNMFSATWQWRPMELDAVQSILTDFGACYAFNTGEDKYGIRNVTSKGTSHGL
SFYLDVNQWDYNIGPQVGAGFQLMLYEQGDVPIVSDLGFSVAPGEDVRVGIEVTKVINLPPPHGICAEKSLKYYSRYSV
TACQLECLTDYMVEKCGCRAFYMSGNATVCDFEQYFTCGQDAEFAFTNLTCDCPVPCTQVIYRPSLSHAKFPSDAYATY
LGNVTGYPTEVFTINGDGRNLSRIVPR

>Protostome_Lophotroco_annelid_CAC9565522_1__Ofus_G051249_Owenia__fusiformis_

MTFEGRNLKVSIITPKKRWYASFKLSNKVWTHVVVTWEDADGLTLYVNGSARDPVRRWKHEAASPNNQFHFGPPVG
NGSEKDIIQVDEMLMWNVRKTDAEITALFETYTVHNNINIPSCSLDDSSTQTTPRLPPTTTTTTQATTPAETTRPAMSGT
THTEPSTDERITGGSTVATINVTYPALVEDTTGMNESHHIQRRASGEDDVIATDYVTDELDNEVGDELHENDEQKEELSL
NDFLTTTAHNKGDLIYSCRWKGTECSASDFDVTLTDFGVCYTFNLEARSEFAVNESANTSLLADETATKVYQPGTSNGLS
LIINVEQYEYMVGPRTDAGIKVLLHNQSEVPMVTDIGFAVAPGMHSLVAVKRTEFLSLSKPWGTCSEKELDYFDDYTVS
KCMIDCQTRYVVAICHCRAAYMPQKHHNEPRICSLEEYFDCVLPSLQNSTKFMEVNCSCPQPCKKVVYEPRLSYGAISSL
SVNKLLDERNIKTTELRNKYHEVLDIVQHVDEEIWETDQHMFENIINDYNQLVDYTIHFYDKDEDNNQPVNEIVYNVQS
VIKTYVRDLNLTFARHEEFLSIYSRSYKTTNDTLYEVAHDFLELTRLIFESDGPKLVESLPGLTPTDLSGTIRKNVQQAARKG
KDVVDKLLAFREALESGRPLDEQLPDYELSMLPGTIYTSYSSVGGTVDDFLADMWQLLDFYFPNIDEKLNRITSTDSVIPV
ELTEIFNSFNADVLDKSISKLLQSLNDFYNKIHAMKMWYENSLEEIKHENQTFYERFVLNRNTLYDLAIYTKNITHRKETLG
KLYGSYTKSEITKLKLAGGYFSESSINDDILYIENFVSSFKTRVLEELLREITNINEHLTLKYQAILTKLEELQVYYDKKFDYSPKI
LNIWKTVNALGLIDEKNMNDSVLSDKMAEHFRTSVSADIARFDTGLQENERNLIESLNFAKDRMKLYVAASKMDENFV
TTNFLYMDVYFAELSYQKVEQKEAFTWQMLISEIGGFMGLLLGASVLTVFEILDFLLYNGTRKYLNRRGNVNLENGAAR
GNNNDGANRV

>Protostome_Lophotroco_annelid_Pdum_comp411397_c0_seq1_

MTEDKQEAPPPWRRLIDDFTLDTSFHGWRYLHCGTQYLFRRFLWLLVVGFCTVTFGWVVYGRIDKYRKFEKNVNVEV
ESSDSVSFPTLTLCNQNMFRLTKAHEAGVLDLLEFHFKKDLQSRRVTSSKGFANISVTHGNQLSHPSASGYSFKDVSLAV
CSSTCCSPYFASMCRAFQFDFTTKTCDILMYNSSTNGVNITKNENVTYYEVTNCGKIPYESEMRPLSRSEYDISLLNFSIAA
SHSKEDLIQSCTFEGSPCSSDNFTMTLTDFGVCYTFNGLAEDVINTKKTGSSSGLRIVFSIEKDEYLSGPNHDVGVKLFLHE
PNELAMVRDLGVAVDVGKHSYISVKATSLTNQKPPVGKCGSQKLRFSNEYSWFRCMRECETIKTIELSGCRNIYMAPEA
TGVPDYCNIDNYTKNILSDEVKSVAASCNCTEPCQRILYESSITSIGFKVLLLENLLSSDVHSFKAKTRKVKNLKFRVDGEYF
QDIAESVMSVINGYIRFLNETKVLTANFLQQIGPTRALAQKLYTDIYDKDLNTVLFGGITEGQFVVNTLFSPMKTSLFSLLD
QIRVQTNILHMLVWQTMMFRSSDVDEKVMAKASEVIQLLKDVTFGMMVFPSTAYNVAMVANYFRWNVTAYSFFPK

VIIKGDACALTFGSYANLSEKMRSDVQGMADLVNGSSGSAINETEAQVAFGNILRSISSFQEFETAFKSNCLDSFDNYIA
NYVTRRENIKQEVNNKINAISVAYFYDMEIAPVLQDKNNLYQNALRFFNEGNITAYELASSLNQEITTGMSSKIQDFVTK
TALQSSNTMRSTVFDYRNSVVNAYIEMLQELCIYRDYMSPAFVESRISQLAILKGPLVDSDDLSITFTQNTTNSTLLTYMQ
QICTNISVIPLIQAEMSTMVTNSFSKVGRIRMAFDLTLKKEEGVLTDAVKEMSLKMDSITNNYFAVGDTFIKENFGSFVV
FFSELRMEFINQSPAYTASDALSDIGGFMGLLLGASVITVCELVDLFVYNCIRKLLYQKNSNVSSTPVTSLVGMSDQKNAK
KNIYDDILFAQRK

>Deu_Ambulacrar_hemi_Ptyfla_40v0_9_20150316_1g17757_t1_scaffold14980_cov102

DELEQFKTRVEEVIEENSELHMKVEQTGAQEFITPTEWHQLKEQAKLVLEENQLLMEQLDIQQNKHKEALKHQLYEVSK
ISKKLILCESEKSSVEADLSELKHKHSVLKQKYESSLMEGDHKISLQDHYSTVADLKKAVEDLKEKNAQDLENVMMKLQAI
QKEKKSLGLQLTDSKAENKKLEGEIKAMQRAIKKSQQQIVQMQKELEFSENKGAAANHHLNSVLKVAEKTVQERDHYY
KAIEVITAKRLVFPAVSLCNTNKLRKSAIENSEYSDLVAIDTKSFRPYYGWFSCNDSIEMTPMLNKCDGYSDCSDGSDEDE
STCEELFSCDNGEEMINISYKCDGYIDCSDGSDEDLIICEAMKLVIHNYRVWYNLPHHRSNMSELQPRLSIANPSRQMT
MIEPFSLDDDVGKKDVMTHKMLVKDFAASTTAHGLPKIFESRHVLLTLFWTLIFMTAMGVSVWQSAKLIKQYAARDV
NTRIEVVTAKSLPFPAVSICNTNKLRKSAVENSKYSDLLAVDTEIVRPYYASECLDGDFRCNNSGCIKSYLKCDGYDNCRD
KSDEVGCNYGDCGVNQFKCNTGGDDGVCIDRRQMCDRVLDCYGGEDEDRCVCKSKTEFSCSTSGRCISNDKLCNGVS
DCPDGSDESSSKVCTNREYQCDARTCIRSYFRCDGHEDCKTGSDETLCYESFFHMFSCAVGEFDCGISGLDCIKNAHICD
GTPNCNDGKDEHPSICSDNSTAAVNNTDLFVCGNGNAIPISYKCDGTTDCSDGSDEDESTCEGAYFRCTKSGVWIHWS
QYCNKVVDCEDASDETNCQKGRNASHTPTFEEILKFYPSLKEEYSLFEDFMEEYYDDHMFGRVSREDPPDWYGFIAYSK
TADYSDLENVLKLTKDEVKTLGHQKEDFILDCTFDEASCSPSDFLTYQDDEYGNCFGFNHLQGNDTVIYTTKSGSEYDDD
QTVSSYFQNGRHSASSVQTRSATSSAAAASLSPETLKSLIKECLQDLELKVNGISNTTDTILAEQKQLFERVSVVESRCDEF
EKTIQYSSDEVDALRGKCRNLEVAMLN

>Deu_Ambulacrar_hemi_Ptyfla_40v0_9_20150316_1g16000_t1_scaffold12160_cov117

MSTDARLSWNXPPDIPSDEPYPILSIDSGHVDTVKIKEYNDNPKKAWFEEDNKNKAFGLGKEKCVEHDDSIKGMLVDFS
KETTAHGIVQITSAKSNVTRSTWLAIVLAAACLMLLQMTLLLFQFFEYNVSVTVSLVSGKALDFPSVTVCNTNKLRRSAV
RDSNYSEMLMLETTFVQPYYAECGDDQFRCVSGSELGICISNDLLCDKISHCYGGEDETDCGARLAGAEIRIGSHWWSY
NNPICATGIEDHQTQTESFSIRCSPSAVAGRYLSIQVKDNSEDMQLCEVEVMAAACRFRDDFMILTLTSECTDAIVNRY
MYFTDSRYMNLALKQKVSQSSTLAYGSAVRAVDGDKSNNYNERSCTHTQKEYQPWWKVDLGGEYEIHEVIITNRADC
CDIADELPNVALGKPAEQSSTYYTHVAGRAVDGNENFALETGHSCIHSAKEAGAWWRVDLQSIHAVYKIVIINRYDCCS
GRINGAVVKVGYYRDDIEKNVQCGSQIGDDDLQDKTIVKDCLVVILGKYVVLRNHPSREEYLHVCEVKVHAKEFLRGEFV
DELHRRMFLMFDAQEERALTVNSTDVFTDIPLYQCVSKCLYWKRYDCRSLDYNHGSNTCKLHRDTAGSNGMELVKST
GNVYYQKLRESVLIPPDNETACPAEYIRCTSGECVHPYKICDTLVDCADGFDELECLANDPKASNKSTVFYPGWHEPYHA
ITDDLDVYKYFEENIYVHFEYDRVKGEDPPDWMRFRTFSRSPDYTELNKILKLSADEISELGHQAEDFILQCSYAGEVCDA
SADFTVLQDESYGNCFQFNANRNNMRLARGTGPEQGLTLTLFIEQHEYFSIYGQDSATVVSIVPPGDKPFPLDQGFFA
MPGTATSVSLTKSKVERQAQPYGNCSNQAGFDVDQRVMEKYLSDSYSLTACEKACLQTALIRYCGCTDTLNIEMPPCR
ALNKTQVACKQLIYYFFQSDLLPCKCHPSCRANLVRLKVYFETLNYESTSEIPAYTWEDLLSDVGGTLGLYVGLSIITVCEIM
RLLLKVITKFCRLNRKRNSCFPEVLKDSQVRRLE

>Protostome_Lophotroco_annelid_Pdum_comp400938_c0_seq1_

DSVHLRRFLWLILVLSVWGILIFYATDRFIYYFQFNKNVNVEVNYLEKVQFPAVTICNQNTFKITSLADNGWYHIVQTLFT
NTAPKSSSRRCVEGERRCKDASACIPVSYVCDLFPDCGDGSDEEYCDQCPLVRMSGPSDLEKLHPTFVNKNFAMQPNT
LISDRPLYVHESNRNMTFLYYTNNEWRIGEMFGDSQPLAYAINSVGSPERVTNAWYIREKSGYVFRPDFRLTCQQENM
ELLAATDFSSPPGLNLSYLYQATGHTQLDFIQGCKWKGEKCRSNAIQEKLTDMGICYQFNDPEDMKIEQAGPGKGLSLT

LNIEFYEYTKDTSDGVGVKILMHEKSDFPMVAEQGISVSPGLYSFISVETTEVGNLPPPWGLCGETDLTYFDSYSIAKCKV
DCETNLAYDDCGCRDFYQPNDKKGNPPVCQVEKYITCTAVKIEEYRKSGNFCKCPSPCSQVAYVPTTSFAIVLSSNNADE
SEKDALQHKLVRARETSYRIIKSNYLETMRQFHIMFRQMTFLRRFCQISSAALEVEFQDVFDISSRLTSDCAWYRSVVTEV
SKNVTRFLHEPFATYRMAWRQQLINRFISLVNTVEETLYEYQSYVSTPSMSSEEIAEKHYQRLALINRQNKTFTHMYTEV
QDAITLLEDELLQNLEVSVATMPAEMSLRLQESRYKPMQTYVSPNCPGCFDMENTPMTEILRASAEVTMDQVKSVLT
WTRGDLTRLYNTYLRDLTLMTNMGKKEKITFRAMTTEMLKRLQGISDLFQDNIDALQGARNLLHDLLEYTWRGHCQF
ELMIHLDEYQSVHQEMVNLFKEVSDYMEERLTQWGEYRNAYDTRTKKLQNIAQDFPFTNVQTLIDRKETRLETIINKRK
QVLLKIQNHVQNTVYLLMMVLDDDAMATNSALWEKFQAAIPTLIIPESRDDEEITSQPGASVQLSSEEQKYWDDIQEEL
DALANNLDTQTAGLRTITVEALAFFKAYVDGNKISAEFYEKNFLQVELFYKELTERKVSQQAAYSYTNLLSEIGGLMGLLL
GASVLSFMEILDLFFYNSIRKCVDFHRARVHPSPMN

>Deu_Ambulacrar_hemi_Ptyfla_40v0_9_20150316_1g16149_t1_scaffold12353_cov92

MTDQAELEEARLPLNQQPRRVERKIRKPFKETKRRSIGGITRAFALSTSCHGIPRLAASKTVHRRSLWTLVIIIVLSGYVTH
MSELLRTYFEYPVRTRIFKETNGSLQFPSVTICNANKVRLSELTKSSPYGAVVSELMLTMKRGKGIRHSGTAPSKKPKPED
LAPGCLIEQNIAISGTNGTVTTVSANTSAECCAGCSATLGCLAWTFEKSDGICLLMDSLGHRKIANCCDAGELYVNSTQG
IRHSGTAPSKKPKPEDLAPGCLIEQNIAISGTNGTVATFSANTSAECCAGCSATLGCLAWTFEKSDGICRLMDSLGHRKIA
NCCDAGELYVNSSQETVPSYSTSSPPKNVGICAVERNVDRVGFDLLVNATDNKRESAEDCCDACTRTPGCVSWVFYKQ
LEYFGDCNLKFSVSDAKPSPCCDSGIVERGLSTSETVSEKTPGVSTTDEVVTTTTPGDSTTVLNVDTTTPAEVRSHTTEERT
SPTREGVLPTRRTDAIPDYPNPYTYGPFEGYHVDTDYDYLGVDVELEEIEYLRDLFGGVVYDENVFSDFNETINLIKARAEF
EETSAATELRDLLKGLSQDEIFEYGHQGGDMVLQCEYNGMPCNVSTDFMSFQDQQYGNCYTFTIDATAEQFPKAGIN
SGLKLTVFAEQDEYVGPVSPSAGFRVVVHSPQSMASPENEGVSVHPGTSTSLGVNLVSFARMNEPYNYCPDDENLQLF
TEYIYTLKPTNESGDSSCAYRGKPGKSNLSWADVTPIIVTMESMGAHGGADVDDPADSVDDLADVVASVLVSDDDEH
ATGDVSESTSIQRTECPGYKMGVCMRRRTSCPYQHPHVPFPSASKWETHHMNRFRFEAYQFETTNFFSEEEVICYWIV
HNATIVGDIDCAQLPSCPATPAMKTAIARSAIGKPRYCQVLQNHVTKLLQEDVFVKTIKRFAQRYDMITDPIETLLMASA
TDNEVHSRPTDITNILPRLACEALAVGKDSPFGNQVYKTVYQLYVHSVYCSESGCVQLHAFLVKCHLSVDTLENMSHCPI
ESCRSPSFIMHKTYSCGDFRSCKFLSFVYKALKAAVLTYQGIEVDDKVGK

>Deu_Ambulacrar_hemi_Ptyfla_40v0_9_20150316_1g15400_t1_scaffold11115_cov135

MEFSDFCAPKEIKLVTDEQLPFPTVTVCNTNMIRRSAVKDTVHRELLVFDSDLVRPYYVPCYDGDFQCADGKHCIKPFN
QCDGTIHCPDNSDEQNCTYGFCGEEQFRCRSGSESGVCIPEDAECDKKFDCYEGEDEDDCECEGEEDFRCVVSNECIGQ
LSVCDGSFDCRDGSDESPSLCPPPLEILSRYRTTTGSHPVSLTSLAVDGDNDTCYRSDVSWGKHWQVKLDRRYTVYEVRI
THHFVLSSTIPLEGAVIYVGPFAEYTQDTRCTQELSRALVATLTFTVRCDDPINGQFLTIAVDSGDDPTALTLCEVEVLGLD
SLYANVALRKSASQSSQYRDGSAVKAVDGDHDPDYHHRSCMHTDFDYEPWWQVDLGESYVISDVIIANRNDCCSERIL
NAEVQVATRESFNNYTVCGDRISRKTAENTVIMVNCAPHVTGRYIRLQIKERSDYLHVCEVQVYVDVDNLPNVAKGKS
AEQSSTVLSFTADRAVDGDIPEGNNQLTCAVTVVEKNPWWSVDLEKIYRVYRVALSFMDCCRSLNGKQFIVTYNDLLLE
EDLTSLVTRVGYWEINKGEVNALCGTIGDYNADKTQLAVNCNVPMLGKYVSVQHEGAAVSLTLCEVQVHAEEFKRDFS
VQIWRERLYANFDKQNERVLVPSDAQQTETVTQVTIDACIEMCSTAKRYICKSFSYNSDSNTCLLSRENAADEGATLRH
DKHFVYYQNLVLNKYIPPANASSCPDDMWRCGSGECLHKYSVCDVIKDCDDSSDEVNCTNHVDGNRNSEVSDIISRLK
ALFEDLTKNASVFEHFRDNVYVANGFDRVMGEDPPDWTRFKAFSRYADYSDVKPALKLSNAEIAALGHTKEGFIVQCA
FAGHECDMRDYGNCFSFNPGRSDNNDFLFATGPGPSLGLEMILFTEQSEYIALYGQDSGVRVTITPAGALVIPFIDGFTIK
PGTITSAGLTFNVRTRTPKPYGDCLESFPVQGFNTDLKYSTRRKYLLKSLLRKNEKLKDINDMMSARSNLVKLEVYITDM
NYESVNERPVYTWEDLLSDIGGTLGLYVGISVISVCEFIHFCLELCRSSFGQMKSIDPT

>Deutero_Ambulac_Sakowv30039460m

MESSNSNSRNATRLNRPAAYRSRWLQGYDHGNENFYTTVGEGNEMYNTSYSGYTNSQYIRDNGATVTNQPFSISKTG
RVEQRPLPRPDEDTSEDGGGEKKESYSSILKDCGLNTSSHGIPRIFMSDNLLIRVFWILITLAAFCAFFWHSTTLIQDFLER
NVVTKIEVVTEKRLVFPTVTVCNVNKLRKSAIQDSPYRRLLSIDNDNVLPYYVPCIDNDFACGNGELCVKQYLRCDGINH
CRDFSDESGCVYGDCGSTKYKCQSGSDRGFCIAKTKECDRTRNCYDGEDEDDCECKSREFKCVEKGGCIEKARVCDDVV
DCTDGSDELGCGTIANCSAGFLCDNGRLCIPQAFKCDGFADCTDSADEQNCEEDTRVPVPLVGVCPDNQFACDATQC
VPIYWVCDGYPDCSNGYDEQQDCQNMGSDAGSDMGSDMEVDNCATGSDEYFSCNDGTCFQSVYRCDNYADCMD
GSDETNCVCTDFSCDNSTKCISPYWVCDNYNDCQDGTDEPENCTGSSGSGSGSGSGSGPNGSTVGPNDVECSMFAC
DLGVVCLPLMHVCDGTPDCIDGTDENNCTDNCATGSDEYFYCNDGACFQSMYRCDNYADCMDGSDEMNCGTGSSE
LVTGSEMYSNVSTVGPANANCTDFSCDGGTACIEQSWVCDGLSDCEDGTDEHENCSNLSRKRRNPADDPPRQLRKER
SNRNGTEDICNGFQCSNGECIDSYSECDTVAECTDGSDEHEHCIGAVVENKHVFEQNWHVPYEELTTKMSHYNDFRK
RYYRYNGFYRVKGENPPDWNGFRTFSSTPDYSDLEDVLKLEDDTISVLGHQAEDFILQCTFDQRECSASDFSVFQDNRY
GNCFQFPGKNDTFRISRTGSKYGLKLTLFLEQDEYISLYGRETGARVAISPHKVHAYPEDEGVTIRPGAVTSFGLREHVLTR
MPDPYGNCTHGNSELDESRMLRGKSYTVLGCQKACVLRQMLKYCYCVDTFEIDAPRCQLLNKTQDMCRQLIYYFHQE
DLLPCVCPQKCSENLASLEIYFEELNYEQVSEYPAYGVFKLFSDIGGTLGLYIGVIMVTVCEAIDLVYQLIRHKLRR

>Protos_Platyhelmin_Macrostomum_lignano_A0A267DD42_A0A267DD42_9PLAT_Uncharacterized_protein__Fragment__OS_Macrostomum_lignano_OX_282301_GN_BOX15_Mlig026586g1_PE_3_SV_1

HRNFWQSGSPAAIAGMADTEKQSAADASGTAEAEAGGFGIANSFREFAQYTTAHGLGRFAAQSTAFGRTIWIAFLLGT
YGGFAYHLFTLIDRFTSTPLLTSLATSLPYEFPDVYICPKNPIAMSKAYEPFKLKDDMPKLRGLKKDCNTTCQARLSYFKES
DVYLSTPMYAFARMVYSVEAALMPKLNNKGIAGNTSECRFTLDGTEYTGDKSSVPVFGNSLRLTLPPSNLTQPNYVTEL
PCMPWNEAARLLAEAGLKNMSDKILSYNFSNNDARCRTPPGLPLEEFDNFTTYLINLDKDPICYQDSASDRLFVSCFGG
QIPTMNVEKCTECCFKNNFKYYGLQYYDCFCSNRIVPGNTADSSNCKYSCPGQAADADETCGGDGLYMSVKQIRPTAD
LPVFASYGGFLADYLIGPYCFYNDTHNGLALSYCNIPFCSASGTYDCVVRSSGDVAEKQSAKYTGNLNYTNSGRECMVW
DDVKTAIAKRTLSLANSDLISWNFPFRYYSIDQMIILTTDYAISNRTNYCRAAHVLYDATYYDETGTNRVGSSRSTLISSLVP
GCFYVNEEDSEIHFETCAIESCVETSPSVPGIVRNNIGTAELARQFGPSLDVSLIDCDFYGWQCTANHFQEVLHPESGVCY
KFMKEPFITGLSLPELKEASLAVQLFTDATDSDGIGPEQAQDASKAILNELGDSSRAGNRAFDVVIVPNGTFPWISQAFG
VVKSQKSEFKVRLDRTTRLSKPSLECSKAAPPVTYVLDYWKPAGLPSNVTALMETTFWAVENAENWTAKIDLMKQLVIS
YNGMKSSTKQKVTFQTSTQEDCLNEYVLRRFAENCRCLPTFLPIPEQLFAKYPICFNSTDIENLYKTDQCFIQQKKRVKTY
RVEARDHCMPPCETRKYTVSQNSFPWPNVRRQVSEGFLRNLAKTYGYPNRVKSLRGPGVIYSTYLRYRSLSENITELMN
FDYQLLTLQDQLAMVSVSSAQMFGEDILEEEAYPIKNLLADMGGILGLWCGVSILTVCELIELSVYVILTLKQFCAGDKQR
RGSSDRGASGHDGPGAAYAVPELPTAPPSMKDDANNGGEVADEDGGVESSRF

>Deutero_Ambulac_Sakowv30043178m

MFCGSQAELEVGAPDRQSISGETEVVTHGSKPSVRHSHTDQSNHEEISLETEIPGEARKNNKKNKIMELLEQFASSTTAH
GINRVSSSEKPFRKFMWVMVVLVAFIFFTSHASALLIKYFEFEVTVKLKVTKNHTIDFPSVTICNMNKLRKSAIRNTAYSDI
AKFDDYFPIIYKGDCSESEFECHDGTCIPLDQKCDRVFDCFDDSDEADCPIVTCGTDEWTCNSLHCIPMAAFCNGVRDC
YDDSDEPNNVDCRYPLQQCSDEEFRCYDGQCIKRSKRCYKLSSQTACSKGEHLTNCDDYSCSYFKCNDGLCISQYHICNG
IPNCDDEEDEDEDMCAAYTCDAFKCNNNICKPENVVCNGNNDCPDGEDENIDMCRNRTCNGVKCSNFNCVPWYW
VCSGTNECPDGEDEGIDICGNRTCSEFECDDGKCVDEKEVCNGLMYCADGKDENFDICLNRTCHGFRCNTGLCIPEDW
ICDEDDDCLNGDDEDIDICRNQTCQGFQCNNGPCLDEDVNWDCDGDIDCPDGDDEDADMCIHQTCLTGYLPCESDG
ECIEVRKWCNGRTDCLNKQDENFSICQIPWHPFGSAQVKFSAYPWSDFNDIDNTSGEPESFSQKAKPLELHVLQKHTG
TIHCVSDNCITKVSGSSKNAICQHINCFIENIPDGISQCRSTQFIHSDLWCDGYGDCNDGSDERYCNHTCPDDKFQCKGG
WCVDIDNLCNGWNDCGDKSDEENCTELCPAQYQCASGVFSGGNTPAKCIPLYEKCDKFVDCSGGDDEENCDYGSDV
FAESEWFTQFLDITHDLELFANFKEKYYTNISINMIQNGTQLSMKHIILASAYNPAVDIRDVLSIPVEDIAKYGHQQEDMI

LQCQYLTHECQSSDFIEFQNDEYGNCFTFNPSKENRELYTSSKSGKRYGLTVILFVEQNEYLTLFSEGVGFVLSVHNPKNQ
PFPEDESISVIPGYETSIGISVTETNKLQYPWDNCTEEETTPYGHYPYSILACEKECLNMVLLSKCGCVDEITLSAFNKCNIH
NQTQGNRYSYRLTASQSIWPSENYTVIGLFADIGGALGLWIGISVVTLFEIAELTSQIFTVLCTCHRNVGSQ

>Deutero_Cephalo_237150R_t1__name__chr_scaffold6_start_401717_end_415020_strand_pro_len_1
123_

MADDDGNTSSLGGLWKNFRENSSAHGIPRTSTATSMVRRVMWSLLFLGACAYFIFQFHGLMVKYFSYPVVTNVEILFE
PEVTFPAVAVCNLNAVRRSKLNLSRSESEDVVAPQEKRRKRREGDNNPSTSPEPFGTTTDYEHDTTTSPTGSETTLTTTLE
EHSVSRMEEMTTTTTISVVTSAPTSNAPVVTTGTPDPGTTATDQDTTGTATETRGSRSTAPEQTTLPSNTMMGTAPTT
HSSTADSITEMPTDALGRSSAETTVSNAVSGTLTTSSYPTVTTIPSVSSVRWDASSQDTVPPNTSQGTTATSDDLAGTTS
QLYNTTMPQLTTNVSRQTGDHSNLGNFINKPRIVRYSTRDNRSRRHFNKRHSSIQLFIRDQCSMDHYSNHNGIRHATR
DHRTNINKKQFTIKYFRGECNLGRFFITKHDQCTMEHFNHNGVKSVKGGKRAMGHIGTTHLIVSHVIKNYLDVIHVITNY
PDVRHVITNHHDVRHVITNYLNVSDPKPPAGEILAVDEESLPLSPSYEKAQNFSALLASLNTTVRHQMGHDKNDFIQDC
TFNGRVCTPEEDFATILDETYGNCFIFNSGRTRGGAFWTAKQPGPLHGLQLTLFLDPDEYLADLTPAAGARLLIHQYTQY
PFAGEEGISLPPGYTVSIGLSQIQLYLVLCLLGGFGVGSGTVLDMGARLLNHQYTQYPFAGEEGISLPPGYTVSIGLSQVLI
HQYTQYPFAGEEGISLPPGYTVSIGLSQYTQYPFAGEEGISLPPGYTVSIGLSQMLIHQYTQYPFAGEEGISLPPGYTVSIGL
SQYTQYPFAGEEGISLPPGYTVSIGLSQTLLERKGGKYGPCTTGEDKGFYRLKNGSHIRLFPDIYTYSVKLCEKTCLQQQIV
ERCGCADPYIKIPDNERACHPRDPCRTDFKKLYLNGSLQCDCPEACSETVFTSTVGFGEWPAPLYAETFIDDMKDKKGT
GFGNLGTAAEIREGFIQVTVYFENMNIEHIYETPSYEVENLLGDIGGQLGLWMGISIMTVIELAEFLTDVVLFLLYGRRRR
AADSPGDGSELGAERGAAAPGHANGPPGASPAAMPPRREVILLPREFPGTEIVPNRSRVVPKYPIMKGSWRSTANTAF
TEV

>Deu_Ambulacrar_hemi_Ptyfla_40v0_9_20150316_1g26019_t1_scaffold49916_cov105

MNSLSVIEVVTAKSLPFPAVSICNTNKLRKSAVENSKYTDLVAVDTEIVRPYYASECLDGDFRCNNSGCIKSYLKCDGYDN
CRDKSDEVGCNYGDCGVNPFKCNTGGDDGVCIDRRQMCDRVLDCYGGEDEDRCVCKSKTEFSCSTSGRCISNDKLCN
GVSDCPDGSDEISCVGFGNYCPEDYEKCDNDRKCIDPKWKCDGDNDCTDRTDEENCGDEAPSCAVGEFDCGISGLDCI
KNAHICDGTPNCNDGKDEHPSICSDNSTAAVNNTSKSSTFSRLISVPLFRFRITFIFIRFCPDLFVCGNGNAIPLSYKCDGYS
DCSDGSDEDESTCEGLIFTCADHSSKINIAKKCDKFKDCTDGSDEAECGDDTMFNLTTGTYFRCTKSGVWINWSQYCNK
VVDCEDASDETNCQKGRNASHTPTFEEILKFYPSLKEEYSLFEDFMEEYYDDHMFGRVSREDPPDWYGFIAYSKTADYS
DLENVLKLTKDEVKTLGHQKEDFILDCTFDEASCSARDFITYQDDEYGNCFGFNHLQGNDTVIYTTKSGSEYDVCKQFM
YYLLSRDKLDCSCNQACAKNLINLEIYFDELNYENIEEIPAVTIEVVTAKSLPFPAVSICNTNKLRKSAVENSKYTDLVAVDT
EIVRPYYASECLDGDFRCNNSGCIKSYLKCDGYDNCRDKSDEVGCNYGDCGVNQFKCNTGGDDGVCIDRRHMCDRVL
DCYGGEDEDRCVCKSKTEFSCSTSGRCISNDKLCDGVSDCPDGSDEISCVGFGNYCPEDYEKCDNDRKCIDPKWKCDGD
NDCTDRTDEENCGDEAPSCAVGEFDCGISGLDCIKNAHICDGTPNCNDGKDEHPSICSDNSTAAVNNTNLFVCGNGNT
IPLSYKCDGYSDCSDGSDEDESTCEGTYFRCTKSGVWIHWSQYCNKVVDCEDASDETNCQKGLKLILYTEQDEYISLFGR
YSGARVVIKPPGFPAIPASEGFTVSPGTISSVGIREKRIKRKPDPYGNCRTEEKFQSIYGDKYSQMICEETCVQAKMFGNC
GCVDTMLVNATRCMLLNREQDVCKQFMYYLLSRDKLDCGCNQACAKNLINLEVYFDELNYENIEEIPAVTQIQLNVFRT
SDKRLVFSTPHQNI

>Protostome_Lophotroco_annelid_Pdum_Contig10257_

MADANHYDATKVAEKESSTEPWKTLLFDFTQNTTLHGIRYIFEPTKWLLRRIVWFLSVSAALTGFLLINYFNVKQYRAHP
KKVNVEVSVKDNLTFPSVTICNYNFHRFSKVVEMGLYDTFYDVYSPDAKQSIGDDTYSYSFCLPTEIEVGLSICPSQNNSF
ECLVQAPVYHDISSVACERLCSLVHSKTCNGFWYQRSKRNCYIIKQIITADDSSESESCNYEYHKRNRKIDQLHFTCKFDD
NSCKWLSLEIPTESDLEFQEPFIISEAKNPPTDFSTQSERGSYLYLFETNKRPGFHATASTEVFRGNDVCLEFFYTFIGKGVT

ELEISFIGLDNQPISSTNLTGNFDSPIWQRYFAEVPAENVRLMLTAKRPNGKLSSGLAIDDFSLDMCSNKPECLRDRDGR
SYSGSVAVTQGGAPCYNSTCRNLNYEKGPWCYTSSDLEEFDYCNITHCECAFDEVQCSDSKICLPAKFKCDGRRDCDDS
SDEYQICDAERSVSCNFHSKFACGYLVGPLYTSMDVKYVSGINNTDELGPKTDAEGSRLGTFVIMDTQGSNEGDRVSY
WSPVFEHNNTHHCLKFSMRAVGDEMSVFNSLRMYLVDADFIFGSRVHLKTFDLHGTNNPDWQEVTVFVPVIDGHPY
YRFIWEFTMGFPYRSSLALDTISISRCDKPVPGYVNTNAFDTHTSFMCNKFGPNCSTNQKSSSAHANWKWVPKQLLPH
ITDGSPRNGDCGPVRVTPNGYFETPLYPDPVLEDGTFCLVHIVMLNQSSISLNFPGVEGSGKIKITTRQLNSSQEVLLSNS
SKLVELNGASEVYVYLEYDGIGLAASFRGYYNVTHQANEDYVLAAEYTGPLSNVKHQSNIYSDVMLFREPHCLSLVSYVK
TIDIQVHLIGRQSIEIKGSRNKLGKPWFPSLFNINLPPGKYNILISLVLQRALPGIHAVVEDMKVVKGRCQDQDTTGSATC
NFSAPDQCGFLDVSSSATRWMRDVVDWPTVPPRSTDQAPAMVLHHNRRSSSTLVANLLTPKFDITVAGCLAFTVSAT
DKTVIYVSLLESSQHLSDLSSGYIGDVHVAGSIVQRATVVLNSRNHYALFSVRKGNTNSIMIDDVQVTDAPCDSLARKKR
SITNTAIGNSSKLTCNYTKQNHTL

>Deu_Ambulacrar_hemi_Ptyfla_40v0_9_20150316_1g20484_t1_scaffold20820_cov96

MVRRYRSLTTFEFPKKRITAKREYVAPRTNVGFRLMVHGQDEIPNVASYGLSISPGTASTIGLTSKLVSNLGSPYGNCVEN
VENNLKYFDGAYSLSKCWMECETDYAVEQCGCRYFYMPDDFALKVDPCNHCEEPCVQTKYETKLSYDSYPSSTYSYLLT
ATGRWPCNEALAFYGIIHSSNDLMEITANMKNFVLEHYYTVEANGQNISAFISSPEFHAVIVKALTEGLFIDILNDLTFEVL
KYTLTLIYKNGTDDVGSDFYAETKETLQQKYMNLSVNPWTGHNTTMWNMFASANHTTDSIQYLIDSLFQDYYLPLFDE
TFLYLYSELKSFRDNKTDEFVNEGPGDICKAYMQENFSKVTVYFEDLNVNSIIQQPGYHSFNLICDIGGSLGLFFGASIVTF
VEIIDFFIVQFWLMKISGNKVFWLLIVLCALGGLLFQVSTLIMSFADRPINVNIKFEEPNERPDFPAITICNNNMFRFNNV
KYLLGGMSLFYVYFPYLYAIFSSNGEAISEFGDVRVPEDFNYGEGHTQLTFMEESAHDVEDIVRRCSFGGLPCGTENFTK
VITNYGICYKFNSKEQGYVNKVRNAGDIGGSLGLFFGASIVTFVEIIDFFVVQFWLKKVSGDKNSILPLDTVADETEFSDD
RKKQENPIPAGQDDDNSEKDGKTWIGIFKTFTQEAGVSGVKFVGSANVSFGRRLNNIKNLLGGSSIFKFSFPFLYALFSSN
GEEISEYNDFRYTERENYGEGHTQLTFMEESAHDVEDIVRRCSFGGLPCGTENFTKVITNYGICYTFNSNEQGYVNKVRN
AGQQYGLQLLLFVNQREYVAPRTNVGFRLMVHGQDEIPNIASYGLSISPGTASTIGLKSELVLVRMSRYRYRLIICFVNIA
GNSTYCSPYALQNCYFIAMDDFALKVDPCKHCEEPCVQTKYEIKLSYDSYPSSIYSRLLSDTGRLPCNEALAYHAITHSSDD
LMGIATNMENFVLEHYYSLEANGQNISAFIPSPEFHAVILKALTEGPFTDVLNDLTFEVIKYTLTLIYKNGTDAMDSDFYVE
MKVMLQQKYMNLSTYPWLFYIEIMGNMFASANHTSDTIQYLIDTLFQDYYLPLFDETFLYLYSELKSWRDNKIDEFVNE
LLGEICKAYMQ

>Deutero_Cephalo_155310R_t1__name__chr_scaffold30_start_1521843_end_1527439_strand_pro_le
n_1150_

MIDSRLLEVIEETAEERARKMVEKYLKERDEKIKEQRKKDIKTMTKDFCATTTAHGVSRVAEADTTRSRIIWTVILITCITLF
VYQSAILIKSYFEYPVNVDIKVVDNKKLTFPSVTVCNNNRLRKSKLSGSRHGSLLELDERTERDVRDYLGQVPVDMEIQG
VVIGTEPEDGTFDGYNEDFEVAYLSDENSDASSVCRVRSKQLCTLAQILRHSKDDNIMSGWSYYNRDGWAAMLSSQC
TAADQQDTCDVCYNDRILLRRIEDIPDKINSTFCCHPDSYVKLTNQTFETRQEALDSGACANLVGYVEGEACTASQLAAA
YKNGEHEYGEEVLFCGHVDGNGYNFQHLPRPVITRHVAILPRVISDGRNKTLRVEFLGCEESDCILSATHANSSRRITIDPS
SIICSSNDCVSNVTTYRGTVRSAGNAVCQDWTCPNRHFDFCLEENFCRSAPGGASTDLVCPSDTRQANLTSDPCQSIAR
CPISENEDEPVFPTRHKFYQQTYAAQYCEERGKQLCSLEQLVLLNTFKDVKATGWGWIRNHGMEAMFEEDCTTSPSEF
CFNVYRYRVSVAVSNVAYCCDDSLFELTEQKHRNHSSAEKGCRDRGMQLCLPSQLMAVHEASIRRDSYPEDIAWINM
ADKVTLLNDTCGCLVGERCFDNKIPLFEIDRSKSYKALCCEESFFQSEDKFSQQSEAFTVCAKGAKQPCSRLQLNDLYND
GVRNDGFSWFVDPNAVASLQDQCNTSDHNCKDGVTYKDSISSNDTAFHVYCCNPESGTSPQNQSIPRKADETNHFN
GCNNPLGMESGVIVRDQITASSFLHQRTSNHARLNSRSSWKMEENDTRPWIQVDLLDSYILAGVITQGGGKAGGFVK
TFSLSFSIDGDQWLTYREDETIPTVFVGNKDGDTPNHVYFSRAVGTRYVRLYPLLWGRKGGLRLGIVGCRSSSCTMSTN
TVGQTVQFYHHHQAMCAEPECIVGRGLNYRGKQNVTRSGRPCQKWNSHDPQRHCTTPSLYPDADLSENYCRNIPGH

PSKAPWWFTEERDWEYCHVERCEVGCVDLNLNAVGGDNDWAGFLLSSNTDDYTDITDFSMATREEIAEMGHQKED
LIVQCTFDKERCSLDDFKVTQNAKYGNCFTFNHGEDGVVRNTTKVGAEYG

>Cnidar_HydraVulSc4wPfr_1269_g30727_t2

MKRLSDKVVSIIRETNEAAATKSQKEDNQRSTFLDIPLLKTSSSTSLLENNPRLNVYTPDVPHQSSLYLNPLFSYTQDNNSI
CIRENKRNLLCESNRQLNNPEQLSDTDTDSEVEVLKSFKFSSRRNTYSKLINEKIEYFQRKQSLSLIKNAAKKKLKEIAEERL
NAKEIFSRYIEASTLHGYAYVCSDTFYIRRVLWAVLMLLGGIYFVFKLKVGIIEYFQYPFSTLSTEDFPEVLTFPAISICPINNF
NISRVQNSILKQLYDADRLPLDKNWIDPGFDIPGNELADILKRSSLQIDKMHFYCDWISRNTTHPYIGPNPCGIENFTEYF
NFKDQLCFTLNSGKKGFNLLSVDHEGLTYGYELGLDVHTNEIISSYPYNGIKLIIHNQQESPVMSDGFVISPGDRTFIKMA
TVEKKSLQPPYSTECGSVELKYHSVYNRRACLLELLTEYLGLKCGCRSLYMPDKFLNLYDKHFPNKIIRIKTKGLLNPWFTKS
LLKSSKRKQKLYIRFINSKKERDKHNYLEHKRLNERTKKIARTTYFKNKLLELKGNCKKTWDAIKEILGKQKIHNKALPNVIT
VDGTQISDVKLVATKFNEFFVNIGPNLASKIKHTPAHFLSYLSKSDKSLAFKKLTSKELDDSIKKLNKNKATGYDAISESIIKD
CYELIKDVLLDIFNKSLKTGVFPDKMKVAIITPIFKSGDESLVTNYRPISVLPVFSKILERIMYNRIFAFLYDNNLLYWNQYGF
RSNYLTEHALMQLVEVISEASSHNKFTLGIFLDLSKAFDTVNHQILLEKLKYYGISHEYHKWFTSYLTDRKQFISLNDGFQT
NKLKISCGVLQGSILGPLLFLIYVNDLYKASNTMSTIMFADDTTLFYSHSNIISLLQVANNELEQISNWLFANSLFLNTDKTK
FSLFHSKRKFQSIPKSLPMLYINNINIERSFTNKLLGILIDKNLSWRAHINHISNKISRSIGILYKTRRILDKSALTQLYFAFINSY
LSYGNIVWGSTNKSKLDCLLKQQNHAARIIHFQNRFTNAKPLLTSMRALDIYELNIFKITQFMFQHQSKKTPKVFENIFHK
TLNRYNTRSTGRFKKPFRSSKLTSFAISYRGPMLWNHFTSLSGKFKEANTSQKFKKNIIELIANHDSLHDFF

>Proto_ecdy_PANarthopod_Tcas_tr_A0A139WB42_A0A139WB42_TRICA_RNA_helicase_OS_Tribolium_
castaneum_OX_7070_GN_TcasGA2_TC031326_PE_3_SV_1

MATVPGFANPDQTVEINDSTEEKVKKKGSKKGGGFQCMGLSYNVLKGITKRGYQQPTPIQRKTIPLVLEGRDVVAMAR
TGSGKTACFLIPMFEKLKIRSAKAGARALILSPTRELALQTLKFIKELGRFTGLKSAVILGGDSMDDQFSAIHGNPDIIVATP
GRFLHVCIEMELKLNSVEYVVFDEADRLFEMGLGEQLTEIVNRLPDARQTVLFSATLPKVLVEFAKAGLNDPVLLRLDVE
SKIPEELQLWYIIVRPEERLAALLVLLKTHIDNKQQTVVFAATRHHVEYIHMILDRVGVSNTYIYSNLDPSARKINAAKFST
GKVKVLVVTDVAARGIDIPQLDNVINYNFPAKSKLFVHRVGRCARAGRSGTAYSLVTPDEYPYLLDLHLFLGRPLSIKTRT
ENEGSVGKIPQALLEEQHSALLTLHENNIDLVSMKKVTENAYLQYVRSRPAASSDSNRRVKELAFASCGIHPIFGDYKEYV
EEERENLLEKMKSYRPQGTIFEICGKNKSQEYQIMKQKRSFHKEKITSFHQNQEQKENEKQLLNHKDETTLAKSSSEDID
DAFDRVITPKKRKLENLYKRNKKQKTKDENFIPYLPADRHTEEGLAINNFHNEASKVALDLTGDSAETMRKSVQAKKWD
RKKKKMVAVPDKRSGKIKTESGVWIPATYKSNRYAQWKEKSKIGQSNSDDENDETGPSGPKLHTEAKTHWARHNKKL
QQKQRKSELKTTDQILKARKIAEKKKNRQKKKGKKFDKANKMVVLNIPEVPKSEHTENYFSEFSNNTTIHGFKYFVDRGI
SVFEKIWWVLAFCLCAYVCTSMVMSNWVKWDENPVFLSVSTKPMKHWEVPFPAITICPENKIWKTKYNYTYYKEKFK
NRTLTLADYTFNGHFSHKNVLNWDYDNNYAKVNKIGYPFRTTKITDSLTVQLNLDLNNIDKNCDSSQGYKVYAHHPAE
MPSKYVKINNEKKYRFEIKPEAIISDSSLIDYNPQRRKCFYSYERPLKFYKIYTRRNCETECLANFTLKSCGCIPYYMPFKEKS
DLSECNCLPSCTSIKYDMDVSTEESGTEIRQMSNVMTDLTFVFNDVEFIPMTRNELFGNADFLANLGGLLGLFCGFSALS
FFEIVYYLTLRWICRICGLWTRILASLGNISY

>Protostome_Lophotroco_annelid_Pdum_comp419518_c0_seq7_

DVHTSIIRCRSNLTDGRCSGRQTVIRGVSSMVCEYLCHGPVYAAHCNGFWYNRTSLLCIISSVKLKNKITDFVGPCKKNSI
EYYEKQLILETPSFICSFDANECEEHLEYTSRYRVYELFRVIKASEEINDFSNLTTNRGRYAGLFETARRPGFLSEVGTKMLN
FTNSCLEFYYRFLGTGNSTLRIQVFGENKIDEDIEEIRGIGHQHWKRFFTRLPDGPGKVSLTFRRAVEPVTSGLIIDNFGINS
CDFYKSNCLRDRKGESFNGTIAVTKSGYNCISQDKIPLLVNTPYCRNPDNDSQPWCFTDNSTVFWEYCDIPRCSCHEGE
FLCKSTGTCVPESFLCDGLADCEGREDEGKVCEKKKQFNCHFDDDFLCGYVTPFRESISWQRKNAREIYVENKPKTDPSR
QEDGYFMFVDTQADEEGKSTSFWSPPFSLYNDSHACISFHMRSHSEYSMETKLKIYLFSENFIATHTLVKEYDLTVGNSA

KWKKQRLKLPLGDPNIKYRLAWWFTMGLPGHSNIAIDETFFYRPKYCYWPDQEPVKGHAAKESFTCDFSDPDNCELTF
FNNHAKMKWQWAPGDVADYAFDNKSTDCTLSHARKMSFRTLGFPAKRSFLMQCIVGIQGPPGSKIKLRFFNGNFTS
QNCSHSRDFMYMIDGINDTWSQFCEFEPQDDDAVFELDSELVEIRAASLTSGLSFDGEYEILDYGNRSALIFFDLLGDTR
KSVGYIFTPALHLKEVHCLTLDILSEGIVRIVGNGYDYTFTLFTKTPSNLLRFSIPVGFYTIGLANEVTGKAGTITVIRDMKIEK
GWCPSLYYEGDVNCLFESHDICGYQDMSPEMTKWHKSEGGGMVIRSTSSDKKSEGRERANLVSPVFSADKASCIRIRV
TSERDTRLQVSVFTSTDVFDSTDAEYFEIAVHGKKDQKFNILITSDAKRVMFSYEPKEKDVLHLQSVTLLSDYQACSDRSY
VMRMQGPLDNAVASKFQSYAEKLGDMKASSDNAPTPASETTTEKPLVSEEKKIGTHSRRNFTEYFDVGDLNEKTAHQL
EDLIFSCKFDNEECDYENDFVEEKTKYGVCYTFNSARNGSLKSTSIPGSKHGLRLVLNIESYENMKGLKSDDGVKIHIHDP
NDIPYIQYVGKAAAVSHHTYFEISKQDVR

>Deutero_Ambulac_Apla_gbr29_1_t1

MLLKVLTLWLFSSYCVAQEEPKGFSTCCGGCFQGPAGTPGIPGIPGNPGGQGPPGPKGNSGMGLSGPKGDTGVQGQ
KGDQGEPGIQGLPGKLGPAGRKGDIGEGQKGVQGVHGPPGEQGQKGEPGDLTVVTPTLPSAIAFSAYRSSSVTGTNG
DVIIFNNILTNLGNDYSLQSGVFTCSRPGAYFFTVTIQRMASSSDPFVKLKKNGQLVFSIYNTHDNYYHMSSNSAVLVLA
AGDRVWLEFGFGGIYSDSDRNNSQVTYHGAWGAPPIPKKPPVLPSWEEACQPAAKRKRTLGDILQDFAENTTAHGVP
TIARSKRTVTKVFWLAVFLAMLTVFCWQGITVLVDYLGFPSTTKVKLVTNPELTFPSVTICNMNRLRRSQLVGSRFEGLIR
LDGGLTDDSGQTDYSWFFDWDSAWYEHYMDTYGNASGAGQASTGSNLGTGGPTLQSDWSVMYGDGGVDPAGIF
TQDGTDTFTTAASDNMMQVETTMGAMTDFPEAPSTVGIAYTGLAVSTKSSSSAQETPTADHGTLDEGLFEAGTGTGI
DVTALDMTTAHSPPQTNAAVSDSNVSPGSSSEEYPVTDQSATSMGMQATTEEEGMITTEIPESEPFESLASSSTAFEHD
ASPSTLTIESGETTTSKESDSSIVSVTTPLSQATTGEDSVSPPDVVSLASSASSEGVWSPSTVASSGVGKGTTAASVDFSPS
NFPVTSAPTSSTGTESEAPPARTTGTITGSAATQQLEEKTTTAGGDSRRKKRSLRIIRDTSMDDGIRGIIGEFGSWVDQE
WNNEEGGAQFDFQYDQYFNWTGVNSDDDWQQFYDRSTSPDYSDFLDIINPTREELKTLGHQAKDTILQCTFDKKLCS
FKDFHQFQNKEYGNCFTFNHEEELQATKTTKNSGAQYGLHLTLFLEMPEYVGILSPEAGIRVSIHSQTTPPFPEDDGITAS
PGMATNIGLRQVYLERLGKPYNTEDCQPYTATTYSRKSCLKECVQEEMFKRCRCVSDILLGNKTKCSFLNAWQQKCKQL
VEYLGEEKKLNCDCPIACNETTYTKAVSSSYWPSERYEEHLQRRLNENPNSAFAARMATSEERTRVYFEELNYQSILESPL
YTFPNLLGNVGGLLGLYIGVSFLTLAEVIALIFHIIKHVCRKTICFKKVSPTPAVA

>Deutero_Ambulac_Spurpu_018814

MSVPIPNFDVPTLVLGHQPRDIHPPDNHPRQLPFPCEFNYDGLGNNSWKYPESTGNAVFGNTHYPFPASITYNELKEP
HTFTPRRRSKPRSHPLSACLTADKIEKMASGFCRDDTCEEDGCNRRSKLVYSVSERRRLCQGCASEKATGPTQQSDAA
WTCRKHCEPIKFFCVQHNEVICQSCATVAHQKPCDLKDVDDVVTENREKLDDILPSIKSRVPVLQKHMKTNLVTSNDIH
DHFDTVETQIRAHVDVEKEAEKNKAKEKNLAITEAAERKIREIHKERDEQLKHSMEELEFREQEVDKKFNILLGELNEIRG
AVAKEMGGIIEKSKYVLEKLNESLEGIEHVLMSDQELCVRTKGLLRAANDVLQTDTETTSLQRVATTALEVKFFKRPRAKV
GAKNRIDGFRGGWTLVDTIDVPQSVELPGLVGCVGEGAVSIRDKQTGDLYLTNLKTKTTDKIAKGTVDTIVSCVSIDDKT
TLCGKVRKGAFSDCIGLYDADWQPINAPDGSQTQELVLNTKRVFVDVDSTGMILASDFMASDIHIISPSDLTRGFQESIV
HTVPVKDKSTEGFRCLSSGDIVVKSGYGGLAVIDRSGEVKNFIQDYEWGCAVFSVDSLTDAIYVVHKDSEKMMYAVDQ
VCATSGGVFRKILEYPMSDSKRNTSTQRCAVTPAGILITFNGDRNLDELETENASRGSVRRIAVKSADEADEVPRKTAVT
TVSETVTNSAVTTAFQYPTYRDRVSWLAVIYKSVPDAVGLAGLKYAFNSAEVRWRRLLWTLLVLGGMSTCCYQVVDRSI
HFASHPKSLDVSINEVSTIAFPTIAICSYNRFRASAVNGTDFGAFLKDGYLNPTTLNYSQYFDTLSSSIFDDVLFNYTHLLED
SILKLYWNRLPIYPATKYFSLKMTDWGPCFIFNDADNGLDGLVLESSGRATAGLVISVNVHQDEYFYDPWTYLAAGLHV
AIYPRGTQPFIDDIGFSVAPGTSTLVGLDLEENVFLPPPYGDCENKTLNYFDYYGTLECKLECAGSFVASQCGCKVMNLP
GEFANNVTCDCTPPCATRNYVTTLSQAAYPATFYAAPLASIYNETAEYFRENICRLNFYFKEMNVVTKTQQRDYTFFSLL
CDVGGSLGLWLGGSILTLFEIIDLFGHATYVYSRTSPPK

>Protostome_Lophotroco_annelid_CAC9565515_1__Ofus_G051248_partial_Owenia__fusiformis_

MMDKVEKHPKESEESWGDLLHTFTQNSTFHGIRYITETSPYILRKLVWVLIVLAAICGFSYVVVVRCTVYYSWEKTVNVE
VTYTENLTFPSVTICNENAFRVSVAAKINKVELVRKHFGNEKQKEKDSTNDNTTTLVNVTTSAPVTTTPVGEKEHHPTSK
NITSNATVYLNMDSLKSKTEIVGKGIIGFALRDPEIVDGIVNKALYLNGLDQHVEFNNQAVDSVCLKELDDCKLGLTLSLW
IKLGFKEASGSQILHYISTGGHKIGNTGIAMYQRYDRLVVYIRTHKKEWHVTTSVEKNVWNMVAFTWSKERDLTMYIN
GSKADHGQYYPLKAVEIDEDAMEDGLTLGGPNDLLLKNEMFSTDLGEAYMDDLYIWDTYMNGSDIVKVFDGYKGFLK
EDDVKPENPTEPEESQPEPTVFYHMSMDYVWGGSVLGRGIEASMDFDPWIPPPIVETVIENKTDIEIFVEQCFNGTELN
VTIQVLGHDENQTKYVDNVISCIDGSIYLVRHRNKSRIECILINQTVTTTSHIPVTPTEGNDTTMTTSTPLNTTSEYIANTTD
LNDMSNFTVSTNDTRNLTRFMEDIENTPGFMNNMTNTSTAIVSNTTDNYTDYLMNTTSFINSSIPNGNISSINSTHNES
DVYVPLTEEVEVCNNVTFEEEYHVEVECGGNSSELLCPRPVVIEIVDPCEDFDIESSYDPGYDGVLGKSFYMSNFSQRLDF
GFKKDRCFGNIQMCNNGFSYRFWLRDDTHPCNTPRILYTSHYSLNDDSNVPPQGVFVFLDLNEVSVHVIQDWRYWNT
SIGLNPDSWNHLFITWSSELGLNVSVNITGKPGNQHVSNICPGVLESFPVNITNITEGVRFYGPTAIDEFMFWDTIVNKK
RYAGTATVQYFLPMDDQVEDQIYGIGLNATVYKGAQTAPGKLQNSLYLNGINQWASLGNTSGECLGSLDLCEDGFTLA
FWIKKGYKHPIEKYSMFYLSSGAQSPRAQGITVFQQSDKLNVHLKTKSRLYEASIDALKDDEWSHITITWILSHGLRLYHN
GDLKHEDRTWKLLRPSNYTKDVDFVIGRPNDEHKHFGDAFIDDLNFWNIQQDETFIKDLTKSKDIKHIYVSMDRVEDG
MLVGVGMVGKIVEKKKNTMATATVKGILGMAIEFKGGRGYVDFGSMQGECI

>Deutero_Cephalo_155300R_t1__name__chr_scaffold30_start_1500686_end_1518328_strand_pro_le
n_1170_

MRKVLAVVTEIDNLKMADIETKGQEAKEFNKTEDLDVRILKLIDIRAKQHAAKEVAVCLAAHEKMMKERAKTDIKAMTK
QFCSTTTAHGVSKIVDADSKTARVLWVTILLACLGVFLYQAGILIKTYFDYPVVVEIKVVENNKLTFPSVTVCNNNRLRRSK
VPGTRHGALLSLNTQTSRGLKEFVPPPIPLPAPTRPGGGPHLVAIEGVLHTISEILIHHDHAIALRDHDHASSICGKNPMQ
LCTTSQLRKESKSSTYGINAWGYFSDKNFAIHPGQCGKDGDTCTDCIAGRFPLKPADQVHVTTVFCCPPTIAISKPFTDHS
GDNVTANDMPSSPTPKQAIDQCKTVRQVRRGRPCSASQLLQLYKEGVRFDEWGWFVSPLQPHTQIRLNGSCRGEPST
SCYENTLPVTTTVQNAGVFCCSPEVCIQSDCTTKVPSCSNPLGMESGKIWDRQIRSSSAFTDKLAHHGRLNAPSFWQP
DDPNPRFEVDFETFIVLTGVIIQLDSQSSGLEGYMLEYAWDGLEWTRAEPGKSTYIGLKRRVVNRQPHPYGNCSLSTKTS
PLYGGLYTYETCKRSCLQEALIEECGCSDELLAINSTICSTLNESQECCRQRPGQCIEVERNQTGRASTLPTMTGQNEEKN
LETRSKERDKKEELDVRLLKMIDIRVKQHVAKEAAVCLRAQEKMAKERAKTDIKEMTKQFCSKTTAHGVSRVVNADSKT
GKFLWVMILLMCLTVFLYQATIMVKTYFDYPVVVEIKVVENNKLTFPSVTVCNNNRLRVSKVPGTRHQDLVEINEKISA
WIPDFLATSSPEPDGGPPVHIKPGIVNLRSNIIILQSQSDYAQAKIVCKKESKQLCTTSQLKEEAKKQHFPLSAGWGFFSH
ENLAVRIGSCEKDGDTCTDCIAGSFPLRPANQVNQTHAFCCPVTVTMSSPVGSNSMVPRRVIIQLDSHGSGMEGYVLK
YSQNGTSWTRYVDEFGGDVILCGTVDDDYYNIQHLPKAIAAMRISIQPSEDMNMFRGCNNPLGMEAGTIAADQITAK
PEMGSTENAPAHARLNGMSSWKMQSTADGWIQVDLLQQKDPGVKVTVHIPGSTPLPEVNALSAEPGKSTYIGLKRR
VVNRQPHPYGNCSLSTKTSSLYGGLYTYETYGGGKVTGRLKYLIAPVASKPPRIDPFHAH

>Protostome_Lophotroco_annelid_Pdum_comp413503_c0_seq1_

QVQRANCTDLKELHLSLMSSNNTEVDLEPKWYEVMSSEKEENVEKSWKELLLDYSQATTLHGVPYTTGSPEKIRRAVW
VILVLAAFSTFCYLFVIRLQYYFSYPTSVSVEVTYEDELQFPAVTICNQNEFRLTKLQEKGLFDLFQEKYGNQERQASPLKAT
CSEGFSKVREGCLSRNHSVAYMKTYREVTLSGCKNLCQTLDSKTCGAILFIPLERSCIFQPVVPIEENGEGCRPIEVHRRHR
CPANIKSTCDYDGSECHVIQEKEEDSSDWLEVEAHVIEPKQMDHTTATVRGRLLYLPESLSITDVNAMLYTQYSNTNGL
CLELFYSTRGQNLASLTVGYRGEDYVSRILHSSSWMSSSWIRIFAHLPTGLNQILIEGKKPANKYAGLLVDSLTIASCTDFD
KTCMRYPGGSGYTGNTIKTPKGSCIPWVNVPEAWHFRFPEHILASGNKCRSPTSTLSKYKTMDRPWCFTDNVTYSVCN
VSYCECQANFFACEDGRCLPKKMLCDGIVDCPLGKDENQYCVNKRRVTCDFSDNYLCGYNGSKRVEQGAFWSYKTGA
SSIWKSSPERYMKIEIGEIGDPTGFLITPKTTLTSPSCLTFLYHVGNSNKLQIKMADFQEKLSGKVLWWSQNEAQATASV
TIPAHPNGAYLIFQGTVYNTSEMVATADIKLTEGSCPQVTQNGRCSSNEFQCDGKCLATSKLCNGQADCFGNIDEEESQ

CDAKKKAACDFKDDFLCGYLHSSTLPNFQWSQGLLVDFPSPNQTIENQTFALSASISPIDSKTVNPYSMAFLKSPKLELTT
PSCLEFGFSVVSDLTIFLQPIPGTPGQQKRVWHSSMFSGNTWSKGWANLPAGKYFLLFYHQFGFDVKLRASNNRSVTF
TTAISNLNIKNGKCDITASAAKLECSFDSEDQCGYQDMSADVPYWTRVKRMGYYFMAAKLTHESSTNPAVLLSKHVTIT
NKTCVGLDFFCYRCTNNDTIIKLTFITSSGQSSVVSFSPAKNNSFVVRATEMNPGSVKLNISVTGTGVIFLNKVFLKDADC
NSLNKDSSFKLAKLFEKWSYLSFTAVPSTEEYINSEFAKDFNGYDSNFAVESQTVGALFATLGSVVDALHLADIGKVLDST
LGNLTGGLMSTGGVLPPSVTNVSLTNTIGGLTNTVGGLTNNSLTNSLGGLT

>Protostome_Lophotroco_annelid_Pdum_comp414816_c1_seq13_

IGRQITPFSTVTVSFNFFFSDGRKMSFFIDLVYTLGIPTKSKFSLFLIMLFPLISICEPEAVSHSVLEKVEAKSNLACVDFSNSS
SCPLTVTTTSANVKCSLAMPNFYGSGSFPIEGTFQSPKYPADFQVHTEEEYVVANLKSIENSTIFLLFDGFKLLPYSIDVCSG
HIVVYTSNDNVQFDQIQYCIEPLLIDEESSSLLINTHFVKIAASSVAGSSYPVFRGSYKLKMDDRGDHSHQDFSLLCKAQL
GFSFLDSPHSITIATDVLAFETHQCLLYNVVLHEKADIKIIMAGKQQVEIYHGNDTQRQIGIPRSRYSIKFLVGFPNTIRTGN
VIEMKYITVRPGHCFDQENPQYNKYLEDLAKKVSSVSKITTLEEMESTKTVNISLYGINPQEESAKQKTHRRRDYSMYFG
VDEVFEKTAHQLDDLIFTCSFNNLPCDLEKDFKKQKTENGICYTFNPGGDTPLKNITTAGSTHGLRLVLNSEAYEVKEGFT
KVDSGVKIHVHDQNEIPVVHMVGLAAPLSHQTYFAINLKEFISLGHPHGNCGIMDLDLFPNVSYSRFRCNQQCFTKHW
EAECGCRDHYMLGDSDYCTFQKYLKCLRNIAEDTALECDCPTACHIVSFASQSSLASLTKVGVAEMLLEKSKIEKIASKYEK
LLNFESRMIPKYFLKMDDKLTDMHTAIISLRNFMTDNEKLVSQVQAALTKIYNVLRDDLLRVNVILEKVLESEECHNSTAK
SRLKVLQRIVDELDLVTFLLEMKTYDRARMDTLIAGIAENVDVLANVFERHQCLDHHTTDYVEMRSNFPGYEDRTDELL
NISRELRQILEGGKDVSYIRHEVKEMSSYFQGLLLKFKDDFQLDSHLKSALELPKNIEGFVPNMKRTIKVRMKVSYALEFLR
FIFRRQHHFYKKANFSKRLYKIVRSEFPRIKKTSLSFLHWLENEVMPVLMNEATDRGPKYLQHMYMEALQMYANVSM
HLGANIQEKIISAVASMKIWQEPRIDRTNKELIIYAREKEETEEAMKLPLKYLLENQAERQLHATMDGFFSSITDEIDGLAH
EFVDNFQRFEEAFSALNHSLQDIVKKSIIDEEFLLKNFLRLDVFYDELTHEIVEQTPAYELSQFFSELGGYLGLLVGASVITIC
EFLDLIFHYVYVRFIQRKETNTKTQKLDGDPV

>Proto_ecdy_PANarthopod__XP_023231888_1_phospholipase_D1_like__partial__Centruroides_sculpt
uratus_

MGFTYHATTYFSMFFSYPTTVEVKVENTGILEFPAVTICNNNKFLKTLYCQNYPCETETINVEEIMWHLNKSIKMKLGHK
KDDLFLKCKLKGRNELESDDCLNSDEELESSTLPYFSYTRIYKQPFNLSGTHLDLLLPNYPLKIKFLTVEREIETHLLHRDIFSIR
VRHGNFQWIVKRSFNEIEKLHNSLQKYSLGNCATSMPSLQARKSSIIRYENPVTRREKLQQYMRAVLIDPNLRNHRNTM
EFLQITPLSFIHILGHQGKECLSRVNCIGYSENTFLQLGSYITGHCTRWRTFWIIARENFISLINPFNGKIEYVMLIDTKFKVH
YGFLRTGFHSHIVIKNLSMRLSIDLENRRTCKEWIGFFQEIKDGSAKDFIKRNKFKSYAPIRYCTQSCWFVDGASYFDCVA
NALESATEDIFIAGWWVSPEFHLKRPSDESEWWRLDNVLKRKAEQGIKIFVLLYKEVEGTLSIGSAYSETVLMQLHSNIK
VLRHPTLSDLNIWLWAHHEKIIIIDQLYAFVGGLDICYGRWDNREHKLTDFGTDTASHKKDTSSKYLDSDSHKTLTKAVA
ESFLPEEISNSEISEILDYRKKYQITSKESPKWEIGETEIKSQRVKRFELPFHKKPINNKCPLQDLKSKKLWLGKQYTNFVYQ
DLHRVHKPFEDQIDRNIIPRMPWHDVSCLVTGSAARDVARHFIQRWNNIKYYKLRNNDAYPWLLPKAYVDLTTIKPFP
QCPKLYSVNSQVCRSASLWSAGINTCEESIQNAYIYCITNAKYYIYIENQFFITHPPGDNYIYNGIWEAIYNRILQALSENQT
FRIYVMIPLMPAFEGDIISEKSIILRSVTDNLYSSICKGPTSLLQRLCELVPDPTEYINFFSLRNYGFMNGKYVSEMVYVHSK
LLIADDNTAIIGSANINDRSLLGRRDSEIAIVFNDYETVSSIMNGAEYKAGRFNINLRKRIFSEHLGLPENDPLLNDPISDFFY
KQVWLKTAVTNTALYEKVFQCVPTNETTTYKELMEYTNRPRLAVTDIKLAEMLLQKIKGYLILFPLYFLSKESLKPSGRWH
LYYIPDHGNCFTFNAQWNSMEKEIIYAYDSDMWNPPTELILDINVQYDEYLDEERSAAVLVTFHTPDSYPDPTSETVTAR
AGHQYNFGMRQTTTILLPLPYQTNCTDYSNLPWA

>Protostome_Lophotroco_annelid_Pdum_comp419641_c0_seq3_

EVYSVTLIRTSKPLRCPVGNFSCNSTGKCVVQSDVCSGKENCPELYSVMHCEYKNITCDFNSWGPSKCGFTLSTTNYNFK
WAPEVVEAAGGIFKTKTSQGGSCGRLKYKELSGSLWSPEFPGIYRPNLTCNYSIVNQYSQPKKIVLTLLYYEISVSRNCLGD
YLQIYDGDTLLAKKCGMNNSVETYEAIDNMTLVFKTDGMYGDKGFYANYHMTDVPNGAVSSTQNIERSLVARFTSFE
DSFVATYMTSPLLVLPSPACLKFRYVYDCDCDFFVTFHYGRIFKYNIIEKPQRNWTTEYVELPNGTYYVTFGAKRRNLASN
AIETQIENIVVTSDYDACNVVTSSGLSEGYTFSSLAAVRNSTENTVFQDTRGQSLKPKEIHMESFFDTLENITYLDFPQSTE
NYFTIAERMAPEIDNFILYCTFSNGTCKESGVKIGTRFSYTEICVEIKVQEYNTTSSGPYSGLTVVIDMHNYDNMKTVAD
MGSRINLFDTRYSNVPSDAGFSITSGLKINAELSLGSFTYLASSKQGYSSCRSKNKGKMYLQDDCLHKEWVSDVYKQCN
CNMDDKKTDGMKNCTTFEYFNCSKTVWPDMYSNCPIECNFLSYKQSLSYSALSPALLENEQMSDEKRKKILDRLKTALD
IEGRVDPRIFADGMAHISVLSDTLVMFIEVAENQLSNVRIVIHDALASFLKYVKQDVQKVKELTDQLRSSYETEYHVKTDV
FDNQITFISGLCMMYMHVSKNRPANDTRKLLLGDKMTTENIDNFKGIMKGMKESIIGAPMQTDPYSPGAFVKNAEHL
ANCTTTFDVVLAGKNASDICESISPIQLRRCAISYEMALEEAEEAVQNTVDAVEMVLAEDIEIILAREQASYRDSLIQLKDI
QKGLAALIVDLQTHNQTVQDVAGKIGKEYLLEIETKLQEIVALADDEHAKTVRNTIQNFVSRTLNEYHRIIDSCVDLAAFV
DTNKCTVEGVLRSLEIFSRPQLTLQSAILVKNAFLNSEREWPKSLSLRAFTIDTNPMYQTKIKSIMDDILKDSMAFFNSMY
SELHTCKIKAASYLKAVEKVMQPYQDNELDEEFIRKNIAQVNIFFGSLELEYVEQLRSYTLTNFFSDVGGYLGLLVGASVLS
VCEIMDVLLHLCIQHHNKRKKIRDDEKLKDTENQMDSYEAYDAAVRKTYESES

>Deutero_Cephalo_163300R_t1__name__chr_scaffold32_start_2777739_end_2781445_strand_pro_le
n_1184_

MWLALVLTAAGMTIWQGSRLVKKYMSYPVQVKVGIVVNNKLIFPSVTVCNQNRLRKSRLNGSRFQPLIEIDDETIGSVD
FLELQGLDEPLDGEDVAPTNAPTTSAPDTANRSRRSAHWLEMESMNRLHEKRRHMVKRKNRATSTCPTAPVSGTDIV
CLDEKFLMENASEGCAQIGMQLCSYHQLRTAHARGIRDPLPLDRSDSWRFFSRPGWDVTLQNECGPRYENVVDECYD
GMFPHTSRAGTEGFLYCCTKEYIVTDETFTDHQGAKQACDDQGFQLCTPQQLKKIFTDGFRGETWGFVSDPTKKAHLS
PCGCRPKKMCYQLTYGYVPRTNRNMPAYCCQQTYALLEVDSPQQNRAEETCSSKNFHLCPIQRLVDVHKAGIQSDRW
GWVKHPGRIALLGKPCDDLQSNDTECYQQRVTHLDDQTSSNLSHCCSPMIAISNDRFQDEEDASNSCDRQGLQLCTLP
QLVTGYRRGTPSDDYGWARDGTYQGIVTKCSSDANCSNARTVRPGDSFPAYCCQYVEVKITIPPRTQCTNMIRGCSEPL
GMEDGRISNHQITASSTMGRNGTLPYQGRLNGWGAWWADVISDSNLGSNEWFQVDLLKPTYVTGLMTQGVRGGLI
ITYKLFYSNDSHGPWMVYRDKSSNREELFKGVTRVGETSHQHFAYPILTRYVRLKPHFLSNKYATLRMELVGCDPETCA
DPRSTDARGVVDVSLDCQGRDCYVGNGENYRGTKRTTESGRGCQQWTRHDPHPHCNITSDTYPGACLVLDYCRNPD
GRPGGPWCFTKDPDVPWEPCGVGKCTLHENEVLDKAAWRGYIEKSNTSDNSDLSEILPPATRQELRQLGHQRNDFIL
QCSFDKQPCSPEDFITSQNAKYGNCFTFNSGHNGVVRTTTRSGTSYGLKLTLNAESAEYLGLFGQRVGARVTLHPVNST
AFPDTEGIDAPPGTASSVSVRQVVITRTPWPYGQTDCTSNDDDYGSLYSGKIYTLRMCENTCLQCAIINRCRCATDMIDL
TRKKAISECSPLFHNSTVPVCNIAKSRVQACVRGLWKRFQRGELDCKCNQACRDETFDLSLSSSKWPSNQFVPYILRDLH
MIQDVRRNNLPRNMKEVKDNLVRVEISYKELNYESISEEWNYKEENLLGDMGGLLGFYIGVSIITICEFVDYVLDLFGLLC
RKIKRSLKRKK

>Cnidar_HydraVulSc4wPfr_1269_g30727_t3

MKRLSDKVVSIIRETNEAAATKSQKEDNQRSTFLDIPLLKTSSSTSLLENNPRLNVYTPDVPHQSSLYLNPLFSYTQDNNSI
CIRENKRNLLCESNRQLNNPEQLSDTDTDSEVEVLKSFKFSSRRNTYSKLINEKIEYFQRKQSLSLIKNAAKKKLKEIAEERL
NAKEIFSRYIEASTLHGYAYVCSDTFYIRRVLWAVLMLLGGIYFVFKLKVGIIEYFQYPFSTLSTEDFPEVLTFPAISICPINNF
NISRVQNSILKQLYDADRLPLDKNWIDPGFDIPGNELADILKRSSLQIDKMHFYCDWISRNTTHPYIGPNPCGIENFTEYF
NFKDQLCFTLNSGKKGFNLLSVDHEGLTYGYELGLDVHTNEIISSYPYNGIKLIIHNQQESPVMSDGFVISPGDRTFIKMA
TVEYKYQIYSVELGSSSIFRPLSKRYSSNFTSLTNTECWFLKKSLQPPYSTECGSVELKYHSVYNRRACLLELLTEYLGLKCGC
RSLYMPDKFLNLYDKHFPNKIIRIKTKGLLNPWFTKSLLKSSKRKQKLYIRFINSKKERDKHNYLEHKRLNERTKKIARTTYF
KNKLLELKGNCKKTWDAIKEILGKQKIHNKALPNVITVDGTQISDVKLVATKFNEFFVNIGPNLASKIKHTPAHFLSYLSKS

DKSLAFKKLTSKELDDSIKKLNKNKATGYDAISESIIKDCYELIKDVLLDIFNKSLKTGVFPDKMKVAIITPIFKSGDESLVTNY
RPISVLPVFSKILERIMYNRIFAFLYDNNLLYWNQYGFRSNYLTEHALMQLVEVISEASSHNKFTLGIFLDLSKAFDTVNHQ
ILLEKLKYYGISHEYHKWFTSYLTDRKQFISLNDGFQTNKLKISCGVLQGSILGPLLFLIYVNDLYKASNTMSTIMFADDTTL
FYSHSNIISLLQVANNELEQISNWLFANSLFLNTDKTKFSLFHSKRKFQSIPKSLPMLYINNINIERSFTNKLLGILIDKNLSW
RAHINHISNKISRSIGILYKTRRILDKSALTQLYFAFINSYLSYGNIVWGSTNKSKLDCLLKQQNHAARIIHFQNRFTNAKPLL
TSMRALDIYELNIFKITQFMFQHQSKKTPKVFENIFHKTLNRYNTRSTGRFKKPFRSSKLTSFAISYRGPMLWNHFTSLSG
KFKEANTSQKFKKNIIELIANHDSLHDFF

>Deutero_Ambulac_Sakowv30034138m

MDNWYLKKKNPHYTNEEVDLSGHSITTNNLEERNQQLVSGGTHDIQSEKSMKEVVEEFGLSTTTHGVSHIVSATNRCR
RLTWMIILFVAFSVLLLQLALLVKQYFEYNVNVQIAVVSSTELMFPSVTVCNTNKLRRSAVRDTNYSDLLAVDTDLVLPYY
VPCMDGDFNCTNGIHCIKQYLVCDGIIHCEDKSDENDCFYGECGKDEFHCDNGSEQGACIPLNLVCDKILDCYDGQDE
DKCECDVMTDYRCILSLECISKVKTCDGEYDCRDGSDESNTVCETPLTNIAPMAPSSLQHATYADPELAYDGNTSTCSM
TYTIYEPFWSIDLGSNALVYDVTITHNGNYEYASLTNAEIRIGGDNHFLENELCTARLLPTQTSSLTFTVECDDPLYGRYLSI
QVMESTQSLILCEVEIMAVVNSLSNVAYGKPVEQSSQYRDGSAIKAVDGDTSSDYSHHSCMHTQFEYEPWWRVDVGY
QCEIQDIVITNRLDCCGNRLIGAEVRVGISKSLNESNTLCGIVTRDNIDEIDIAINCGDNVVGRYVSIQILNRTDYLHVCEV
QVFGKVAVDTLSNVAFNKSAYMSSVWFGNTADKAVDGNTYPVLDSGTCIHSAEESEPWWRVDLGGIYFVYKVTLYNR
LDCCEKRLAGAAVRVGYYEDAQQNLRCGQQIIPQITSWREIHRECSTPILGRFVSVQIEGVDSGFLHICEVEVFADELLID
PIEKIKQRALLGLFDKQQFRRLISDTGKVYNDMSLEKCLSKCYFQKAFVCKSFDYIQDMYVCTIYSVDATYENSSLMIDGS
ANYYQRLVHGENSSHCPPDMWQCGSGECLHTYRVCDVIIDCLDSSDEEGCTFKENRTRGPEHKHLSLWMDVFKNITE
DVLVYQRFFKSVYRDNLFDRVKGEDPPDWTRFKTFSSTPDHSDLRHVLKLTEAEILAYGHQMDDFIIQCNYNSEKCEPS
DFVAFQNDFYGNCFQFNAGSNQTAKRAMGTGSSYGLQLTLFTEQEDYLSIYGQDSGARVAILPRGILSFPVDEGITVKP
GTITSIALKEKVNVRMPWPYGNCTNGQTEQTVYDQQNNTLYSILSCEKACSENALIHYCGCSEYLFTDAPRCQLLNETQ
EQTSKDVEMATFQVKERERYAVSQWEDVLSDIGGTFGLYVGISIISFCEFVSFVVVLCRQYRKKNKDTVGQWNQKQSR
VREHAVIT

>Deu_Ambulacrar_hemi_Ptyfla_40v0_9_20150316_1g31524_t1_scaffold335965_cov147

MLVNLDDEPAAVRHVSDQFWATKSSASDRHHHDEKKSDANKALKTHRKVIKDFIGSTTAHGLPRIFEDRHFILTLFWGII
FFTAMSVSMWQGYRLIKQYLLRDVTVRIEVVTAPSLMFPSVSICNTNKLRKSAVADSPHRELLAVDTEIVRPYFGNCSNG
KMKCMSGSDMGVCIDLEYRCDRVVDCYDGEDENQCVCKDKKEFQCTTSGRCVDISDLCDGNYDCPDSSDEMNCNDS
GMYCAEDYFKCDNDRNCIPPSWKCDGANDCFDQTDEHDCGTVVPGVCATGEFECDFLDCYSNSIICDGEVQCDDGT
DEDSAICGKYDDTGTSTGGTDHFTCEDDPWSMFPAVKCDGMADCLDGSDEMGCDDTTDILLSYPEITTEAPTFLCSAD
KVELEWSQRCNKVVDCVDASDEIDCPRGRNASYTQDPQKLLEKFSHLSGDISLFEDFVENYYDDHLFGRVLSEDPPDWY
GFIAYSKTADYSDLEGVLKLKQEEVETLGHQIEDFILECQFDAYLISQNRDFRVYQDEKYGNCFQFNYADTEKLVTTKAGV
NYGLKMTLYTEQNEYISTYGRDSGARVVIKTPGVPSIPSSEGFTILPGMISSVGIRENRIKRQPKPYGDCREESSYESLFGN
KYSETACEETCIQMKMLEYCGCVNTMMMEETRCMLLNRTQGVENLIRLEVFFDELNYESITEIPAYTALLTLIMVQLNDI
ATLVNYFNFVTFMVELTGVLGLLWWRYRKPDLPRPLKKSEMGHKVTRLLVLPFVLMLYLTTVSTAEAARHRKCKRDPIQ
LTEDQIYSLQCNQTNIRKLENLPLSYLSRMPPIVTLTRQQERRIDAQYGHMFTTIDEMPSLPSRYKRAGQESQEDYARAC
EVEQSYNPIHLAFNMAGKMVQNIQKSEMGHTTRLLVLPFVLMLYLTTVSNAEAARHRKCKRDPIQLTEDQIYSLQCNQ
TNTRKLENLPLSDLSRMPPIVTLTRQQERRIDAQYGHMFTTIDEMPSLPSRYKREGQESQEDYVRACEVERSYNPIHLAV
NMAGKMVQNIQKSEMGHTTRLLVLPFVLALYLTMISTAQAARHRKCKRGPIKLTNDQIYSIQCNQTFISQLKNRPPSELS
RMSPIVTLTRDQERRIDAQHGDMFTTIDEMPSAPTRYKREGQGSKDGFTRVCEAERSYDPIHLAINMDEETVQMIQVI
HQIPEFMSVYIVRRRYCPSNTEVDDVRFI

>Protostome_Lophotroco_annelid_Pdum_Contig7281_

MDELQVYYLKEPRKADKMPDETKVNKVKPLQQYNSFFRTRTPVVPAFEEKKVSPAWGDLIADFFDVTSLHGIRYCTKV
NGVVLIRRVLWAILVIGLMVISIYYLADRVEKYLKFEQSISVEENYVESMPFPAITICNQNLFSLTEAAKGNWYNVAKDLST
ASSTSTSGGSSSNCANGVQCTSGSRACISQSQQCDGFPDCSLYEDERNCSRCVRLVISGTATSPPSGYNDIIRTTFVLQSN
THFSTKPAYMSEKPDSPLAIYYDGDTSKWVISRAFGKNIIATASQATASPQEVTTAWKFVAGNVDVNLNFTCAEHPLFY
NVSEKKQNDDSITLENLFKDAGSKLQKMLVDCKWKGQVCKTTDFVETVTDMGVCYTFGKNNRSVQGPGPGSGLAVT
MNIAHYDYTKSGASGVQILMHEPSDTPVMSGQSLAVSPGLISLVALERAQIKNLAPPWGNCTKDKLKYYDTFTDNTCKL
ECENIYVRDACDCIDYYMPKKDSKPVCTAAQYLNCVRPKLTEKKKDTKFCVCGLPCENTVFVPSLSFATYNADPEALDSK
AEDVVQTVTAANEEARKVDEDKYRETARIINTGYLGLTNFQNFLLALEGQLNDSKTTLQDVSVLAGADCHTLSGGHKNII
AEIGASLERAINTFSVYYETSTYDSIIKFIEIVKNLTDPYDSVELDYLHSRKKELHFQQENSNISVDYVSMSLDNITVTINMDY
NTMNLSELIDMNITLPNTTTFATNFQVRVNSSVLYAQNSSVTFPDEQFWNHTTDIWIRFATLRSATQRAMASVQLTVN
KLVKQLEDRINAVAFADFVLKVDLVRFIQPEMTSSCVWCYDSNGNDFVQILKTALLNVVGNIKPKLRVMESSLYLIYNDT
LKSMTTGSPSMSSITTRLVTLATESESLITKLESDMMKKLKEGMNFIQNLLQLFDLKCLTVSAMTRALDDNHIKTYTILEEII
TSVDEERKHFYQESNLFTIGRMNLSTLSSSLDRTGKTDLVAARVSSLEDVLLDRKILMEGFVRKVNNVYFWSMTMWKY
SYLANVSQFWKRFSITNPDSSTTSSAAIPQIGDASASDLLIMKGTQSSQTFWDDLLQKVITMVENVKLNYDLFTTTRNKV
QAFFVAFLESNKIGASFYSSNYLKLDVYYKALTDKKVLQKKAYELPALLSEVGGLMGLLVGASLITVFEYIDLFLYNGIVKYR
EKRDDDKKEKKSPPPN

>Deu_Ambulacrar_hemi_Ptyfla_40v0_9_20150316_1g16647_t1_scaffold13124_cov152

MPSINNWTPSNLSTTEDQQHRKKTCYSIFALFTQKAGLTGISHICEPSASIHRRLFWCVIVLLALAGLLYQVSLMIHSFINR
PINVDIEYVTPKEKPDFPAVTICNNNMISCDFGGIACGSENLTKVVTNYGVCYTFNAGDDGRIHKIKNLGLPYVDCVEDV
KNTLKYFDSAYSLSKCWMECETDYAVEQCGCRYFYMPGNETFCSPQVLDRCYFDKMDEFALLNDPCNCKEPCTQIQYA
TKLSYSRLESDDAFHEAMSVVNSFIKERYYNITFNGETVNEFYPSTSFHDTMADKLVSKVFTDTSEAIFYVLNMNILKRRQ
GEDDGINSPLYFGTKTLLASFFMDYKIAYWQPMVTIMEQEFLINDLSQSTAHNITTTLLTQYYIPLYEEFFDVTYVEVRELR
DARIFTADDSDLRDVCQDFTLLFWSIIVLLALSGLLHQVITLIMSFMERPINANIKFDEPDTRPEFPAVTICNNNMFRLVNL
TKFLGGASVFNFLCPYLYTLFSSDGERIPEFTEFRIPDNESANYGEGQTQLTFMEQSAHSVEDIVRRCSFGGVACGADNF
TKIITNYGICYTFNSHKQGYVNQVRNAGQKYGLQLLLYANEREYVAPHTNVGFRLMVHGQDEIPDIANHALAISPGLSTT
IGLKTIQVSNLGFPYGHCVEDVKNNLKYFDGAYSLSKCWMECETDYAVEKCGCRYYYMPDEFALKADICNHCEEPCMN
TKYDTKMSYDSYPSSVYATLLAFRTKWPCDESLEFYINSRSLQERRGIEADFRTYIDEAYYRTKPNGQNISDFLPDAEFHVL
MVTKLVERVFTDIATDLRKELFLHTLAFIIMNETDGLNSEFYNKIKDSLIEKRNLAKVTIYFEDLKINSIIQQPEYHSSNLIFGS
IKNIYVTLAQPIRYILEYAGIEYEEKLYEQGPVGTNDDERIRADLANEIAKDIRTMIGYNAYNPKWLTYADFVMFELFDQFL
KVDPGLLDKYKNLQSIVARNCNRKDHSKAVLAMPVLLGYWKVRGVAQPIRYILEYAGIEYEEKLYEQGPAPDYSRDDWL
KEKYNLGLTFPNVPYLFDGDIKLTETNVIIRYLSSKANLLGTSEDERRRADLANEVARDIRTRMGVAAYNPNWEEFKPDLL
KYADEMFQSFSDFLDENEWFAGQNLTYADFVMYELFYQFLKVDTSLLDKYRNLQEFKDRFEAIPSIAAYMQTDRCKSFP
LYNKQATFGGVLLD

>Deu_Ambulacrar_hemi_Ptyfla_40v0_9_20150316_1g25772_t1_scaffold47285_cov141

MDTISRQCILKSSVCNGQPDCFDGSDEIIGTVCLYPKSVCPRDHFRCRNGECVHYTRRCNGQQDCSDRSDEDGCGNNE
CPENFYRCRYHGDCVLAAAVCDGKHHCADRTDELGCSLIPLPTFCPPGFFTCDGGRCIQMIDVCNGLDDCASVANPLD
RSDERDCENFTCHDDYFQCADGVCVHWLERCDYSQDCHDNSDEEGCDANVCGDAYFRCDDSRCIEMVSRCDGRPD
CGSDEDGCENNVCPAAFFTCSNGQCIPVTDQCNFGQDCNDGSDEVNCTKHVEDVFSDPKWKTSYHDITTDDKIFDEFI
HEVYVNRNFDFVRSENPPSWNQFLLASSTPDFTDLQDVLKLTPEELYRLGHQAEDLVLQCTYDQNKCNFSEFYMIQHD
KYGNCFTFNHGMDNAPIRKVTKAGADYGLKLTLFTEQNEYIGLFGQQSGVRVVIHHPAAKPFPEDEGVYVEPGAVTAIA
IKMRKRTRMGPPHGRCMSDQQNIDSAYEYTALECQKRCLQNYLYEKCGCVDVLTKDKPRCSVLDKTQDACKQLMYYF
YQASLLTCDCQQPCRQLVSLIMHYFEYPVNINMGVATDTSLQFPAVTICNNNKVRKSAILESIYKELLFIDDAGSRPGFLT

YIGECFEGEYSCPNGRCIPEKLVCDFYNDCGDNADEENCSFRVCPENFFRCTYWGECIPKAAFCDGKVHCGDRTDEFPG
SCPEGTPFPNGCYSERQFACDDGQCIDVLRRCNGIPECIDHSDEVGCNSSTYCPPGLFTCSGGRCIDMVDVCNTYDNFT
CFDGYFKCADGKCVPWDKRCDHIEECQDGSDEIGCQNNVCGDGYFRCNDGRCIDMLYRCDEQAHCGSDEDDCENSD
CPNEYFRCRSGQCVVLQMRCNFQVDCVDGSDEYNCTGLEPPPANCDSDEFKCVSLEKCVNASWVCNGVDDCGDNS
DESECTSIANCKIDEFLCQGGDETGDFCIPRLWLCDRINQCGDGSDELNCTYGVENVFSDPKWKLFYHDITVDDNMYK
DFISNVYINRNFDFVRSENPPSWNQFLLASSTPDFTDLQDVLKLTPEELSRLGHQAEDLVLQCTYDQKNCNYSEFYAFQH
DKYGNCFTFNHGMNNDTVRKASKAGADYGLKLTLFTEQNEYIGLFGQQSGVRVVIHHQMTKPFPEDDGVGVKPGAV
TSIAIKMITQLLSDTGGSLGLYVGLSVITLIEFIEYLVLMFGAMRRRRKVTPKPNTTQVKVIRVKSNE

>Protostome_Lophotroco_annelid_Pdum_comp422880_c0_seq1_

MGSPVTMDNFVLASRRDAVDWALARENYQKRRINTIANGHVKVSQEPPPEETAEESDPWPTLVKTFTQETSLHGIRYL
TVPSRFVVRKILWALLLLGLFAVFLYYLVERVKQFLKYEINVSVAENYVESTPFPAITICNQNPFSLTTSATQGWYLVAKRLS
NPIAAAIAKRNCTSNEFRCDTGTCIAKAKECDSFPDCVDGSDEKKCNPCPRLTAIPGVIGIAILAGLENTTFAHQSETYYNK
KPVYISEIPSRPYFIYFDEETELWTLAVGYGKDVIGTSSQSSADSPQDITEMWSLSTNQSVRIKFQCANAIITPDTNKGKFD
GSLKLNYLYNEGSQNYSDLIMSCKWKGQPCSKGAFKTAVTDMGICYTFKIPDLSVERPGPGFALTVDVNIAYYDYTLSGS
SGIQILMHDPNDVPIMGGQSLAISPGLFSMVALERTVVENLEPPWGNCAKKKLKYYSDYTQNTCKLDCENEVAREKCG
CRHNYMPTATKNDVDYCTVDQHAKCLRPELEKAAKDPKWCPCGTPCINTVYVPSLSFATLATDAGANNDQNDQTAIL
SRRLANAREKAHRVDSDKYRKTTRIFNVAYVALMKLENFVNAVEGQFSHGTETVNDVYILAGQDCNKLSKRHSQVITDI
GSTLMDALTSFSNFYQASTFANLVQFRDIMRKLNFEYSPEERSYLQARKEELMFRQEQASVNINQASMRIKNVPIYVRM
DYSRMNLSDLVAMNVTLPNGTFFRTDFHIGVSSGQLNAAKASVVLPDDNFWNHTTEIWIRYTILRTALELSTSSLRLVSS
DLLRRLQEDVKNITTRGFLVKPNLVKLIQPLLRAECSWCSNDKDETIMTILKQDVTDFVKKTIQPKLDKIQIDLVKVYNDFL
KAMASNDKKLQDDIFNTRLPAVIDTAKSLSHDMRQGMIPKVTKAMEFTAELLRQLEPRCDDRIVTKAKEIQDNHAKGV
AILKEVNKYVREVKDEFYNNTVKYTLVELTLSDMARSTDFAGLSNYLQPRMASLQDVLRDRKIILSSFESILDNIYFWFMA
FWDNNLLEDKSQYWRKFNISNPGYGTPSKTPAPGDNAAYDKFVMKPGSASESFWKDLGNRLTSMVNNVQSHINGLS
RTTAEAQNFFNAYLEANKISPSFYSKNFLKLDVYYKAITNKTIVQQKAYEYPALLSELGGLLGLLLGASCITLIEFLDLLWYNC
CVKCCEKNDSALERHRDPENPNSSPGRWSSKPTTNGASR

>Protostome_Lophotroco_annelid_CAC9667542_1__Ofus_G151389_partial_Owenia__fusiformis_

LSSGMASFCRKSISLFFEYDTPKIVHIRSKKVGVINRLIQLAIIGYIIGWAIVWKKGYQETDSVESGVTTKLKGVAYTNSSLY
NITNNSSGIFDRIWDVADYVNPPQENNAFFVMTNMVITPSQAQGECPEDPSMGVLCDGPQNCSAGDPVINGNGVM
TGACSDPYFSPVTNSTVKSCIIKAWCPVEIDLLPVPEKAILAESKDFTVLIKNNIQFPLFKVKRRNIPEAYNNDTYLKRCRW
NAEKDPYCPIFTLESMTDAAGENYEEMAVKGGVMNIIIRWDCNLDYDPSECVPKYEFRRLDDKDAKISKGWNFRYAYY
YEDTKQKTQRTLVKAYGIRWIISVHGTAGKFNFVPLLLNFGSGIALLSLATIVCDIITLYVLKMGTFYKEKKYFNVTGDDAF
EYEALPDHDDDESSSGTKAVQRKPVNSSASNYGTTGEGAQ

>Protostome_Lophotroco_annelid_Pdum_comp414453_c0_seq1_

DHVTVKMADSAKVVVVQSGFKKAVSAFFEYDTPKIVHIKSKKVGVINRILQLGILAYIIGYAIVYQKGYQDFDTVESVALTKL
KGVTYTNFNKYIPGISNRTWDVADYTIPPEEVDAYFVMTNVVVTPNQTQGVCPEDPDMPDVKCENSTESECPEGKPV
VNGNGVRTGNCVPSREEGIKTCEIYAWCDVEVDELPDPNNPVLAASKDFTVLIKNWIEFPKFHKKRRNIIDTVNNKTYL
QNCRYDANDELNRFCPIFTLGTIVKEAGVQNYSDIANNGGVIAINIKWDCNLDYDISNCRPRYSFSRLDDPEAQIAKGW
NFRYANYYYNDGVYTRDLIKAVGIRFVLKVTGQAGKFSVVPLLLNLGSGVGLLAIATVLCDIIVLYLLKGGEFYLEKKYLFVT
AKDAFKLGENNEENEDGVENPRRTAECPYDESESSEPKLLLNMLDDNPVRPSTQNNDPSAESGQEEPNTVVHRGRPG
QYERFE

>Proto_ecdy_PANarthopod__XP_023231978_1_P2X_purinoceptor_4_like__Centruroides_sculpturatus_

MAYELLKSIIFEYSTVKFVNIKSKTVGILNRFIQLVILGYVIGYAIVYNKGYQSFGGLESSVTTKLKGVAFVNNSNDDFIPHV
DPSLYKRIWDIPDIIVPPSDNGAFFVTTNIEITRQNLSTCPEDPSIKDAICNIHNNTCTAGHHYMIGNGVATGECVPIYNST
KANYTCQIEAWCPVEYDEKPLKNGTALFPDTKYFTVLIKNSVDFPTFNIKSNTMLILGQLGAKNEISFLSQQSRVLCIIFVS
HMPCRKGGVVAIIINWDCNFDFNEDECMPTYSFRRLDDPFVKISPGWNFRYAHYYSDTERTLYKAYGIKFEILVGGRGG
KFNIVPLLMNVGAGLGLFAVVSFLSL

>Protostome_Lophotroco_brch_Lingula_anatina_comp134000_c0_seq1_p1_comp134000_c0__comp13
4000_c0_seq1_p1__ORF_type_complete_len_450___score_54_63_comp134000_c0_seq1_139_1488_

MTEKARHVIPGCRILISIIVVIDINKQGRMSCATCCGYLKGALVSFFEYDTPKIVHIKSKKVGLLNRTVQLAVILYIVIWVIVIK
KGYQDTDIPISAATTKLKGVDFTNITKVPSIGARIWDVADYVIPPQEKDAFFVMTNMIITPNQTQGVCGEDPELAPCTD
DSECTKDMPVPVGSGVMTGVCNKATNSCEIHAWCPVEIDEPPLKDSPVLTDSKDFTVLIKNYVYFPKFNVRRRNIPDWI
KKDKYLSKCRWSRDKDPYCPIFRLEDLISAAGEDYLVMAYKGGVIDININWDCNLDYAETNCRPTYTISRVDKNETVSPG
WNFRYADFYVVDGVNTRTLYKAYGIRFIITITGKAGKFNIVPMLLNIGAGLGLLTLSTVLSDIVVLYFLKYRSFYREKKYQNV
DEMDVSYQVLSPELEGKDFGEPDPTNSVRHRPLGSNANNDSET

>Protostome_Lophotroco_brch_Lingula_anatina_comp134000_c0_seq10_p1_comp134000_c0__comp1
34000_c0_seq10_p1__ORF_type_complete_len_429___score_46_60_comp134000_c0_seq10_139_142
5_

MTEKARHVIPGCRILISIIVVIDINKQGRMSCATCCGYLKGALVSFFEYDTPKIVHIKSKKVGLLNRTVQLAVILYIVIWVIVIK
KGYQDTDIPISAATTKLKGVDFTNITKVPSIGARIWDVADYVIPPQEKDAFFVMTNMIITPNQTQGVCGEDPELAPCTD
DSECTKDMPVPVGSGVMTGVCNKATNSCEIHAWCPVEIDEPPLKDSPVLTDSKDFTVLIKNYVYFPKFNVRRRNIPDWI
KKDKYLSKCRWSRDKDPYCPIFRLEDLISAAGEDYLVMAYKGGVIDININWDCNLDYAETNCRPTYTISRVDKNETVSPG
WNFRYADFYVVDGVNTRTLYKAYGIRFIITITGKAGKFNIVPMLLNIGAGLGLLTLSTVLSDIVVLYFLKYRSFYREKKYQNV
DEMDVSYQVCMENISYRQIISN

>Protostome_Lophotroco_brch_Lingula_anatina_comp134000_c0_seq16_p1_comp134000_c0__comp1
34000_c0_seq16_p1__ORF_type_complete_len_254___score_36_11_comp134000_c0_seq16_946_170
7_

MKHLFYSASYLCFSRKDSPVLTDSKDFTVLIKNYVYFPKFNVRRRNIPDWIKKDKYLSKCRWSRDKDPYCPIFRLEDLISAA
GEDYLVMAYKGGVIDININWDCNLDYAETNCRPTYTISRVDKNETVSPGWNFRYADFYVVDGVNTRTLYKAYGIRFIITI
TGKAGKFNIVPMLLNIGAGLGLLTLSTVLSDIVVLYFLKYRSFYREKKYQNVDEMDVSYQELEGKDFGEPDPTNSVRHRP
LGSNANNDSET

>Protostome_Lophotroco_brch_Lingula_anatina_comp151342_c0_seq2_p1_comp151342_c0__comp15
1342_c0_seq2_p1__ORF_type_complete_len_388___score_19_28_comp151342_c0_seq2_930_2093_

MCLTLDDVKIAARGLFEYETPKYVVIKNNVVGTINFTIQLTILLYTAFYALWWERGYQETEVPVSAVTTKLKGIEYTNISSV
GPRIWDVAEYVVPPLEKDAFFLMTNMVVTPNQKLGKCDEDPEVAPCKRDSDCSAGQTTFGSNGRAVSGVMTGRCIP
SSRNSSMTSCEVFAWCPVENDVPLLKGRPVLPGVENCTVLIKNYIVFPKFGVRRRTIPEDYAEKHLKTCRWNKDEEKYCP
TFRIGDIINSIGENFTELAYKGGVININIQWDCNLDYDPKYCLPLFTMRRVDEKTDIAPGLNFRHAYHYVEHGQPARTLIK
AWGIRFVITVSGKGGKFSIAPTLRNIGAGLALMAMASVVCDIVLRIWYRSSFEKEKYNEAKIMRLECC

>Deutero_Cephalo_000580R_t1__name__chr_scaffold1_start_696012_end_705258_strand_pro_len_4
54_

MCPRGGRFAAEGVFATERRTPEERECQHGGRPSICQQIVQVVFEYDTPRIVHIKNKKVGLINRVIQLGIIGYIIGWVFVYK
KGYQDFGDIVSSVSTKLKGVAYPNRSDIGDLNRVWDIAGLVVPPQQRDGFFVMTNMIITPDQHTTTCAEAPWIGHNV
SCTDTDTRNCPAGQVELTGNGMFTGNCANFSSTVQTCEIQGWCPVEQQDNPPSPAVLEDAKEFTVLLKNTISFPLFNF
TKRNILDDYDPDYLKNCRFNNNDPVDKYCPIIKLDTMVKEANQNFTALAEQGGIVGISITWDCNLDYSGSHCLPKYSFSR
LDKADSKVSPGYNFRYARYYRENGTDYRTLIKAYGIRFDVVVSGRAGKFNIIPTTMNIGSGLALLGVATILCDIVVLYLLKK
RYFYQEKKYMMVDDDDDLRLFAKPQSPPGKLSGKYEQLEEDKTVNLHRNEQNYGS

>Deuterostome_chordata_Ggallus_tr_R4GJZ2_R4GJZ2_CHICK_ATP_receptor_OS_Gallus_gallus_OX_9031_GN_P2RX6_PE_3_SV_3

MEDVDCSVGNPVVHGNGIKTGRCVMFNSTHSTCEIFGWCPVENDTLPRKPLLAEAENFTLFIKNTVNFTKFNFSKGNT
LQTNDPTYFKTCMYDPFFSPSCPVFRIRDMVEAAGETFGALALMGGSISVRIEWDCDLDWPAAHCQPRYSFILLDRRY
NFRTASYYWDSKRQHYRSLLKLYGIRFDISVHGQAGKFGIIPAAVSFGTGIAFLGTATIICDMVLLYLDAKADLYWKEKFE
EAKPPKGKKAAAAEAQRAGQDLGDAQQCALE

>Deuterostome_chordata_Ggallus_tr_F1NC81_F1NC81_CHICK_P2X_purinoceptor_OS_Gallus_gallus_OX_9031_GN_P2RX4_PE_3_SV_1

MAACCGAVRGFLFEYDTPRIVLIRSRKVGLINRAVQLAILAYVIGWVFLWEKGYQETDSVVSSVTTKVKGVTMTNTSAL
GSRIWDVADYVIPPQEKNAVFVMTNMIFTLNQSQSHCPELPDDNTECNNNSSCVPGYVSTHSNGIQTGACIPYNSSIKT
CEVFAWCPVEDDYHIPNPAFLQGAENFTILVKNNIWYPKFNFSKRNILPTFSSSYLKNCIYDAQTDPFCPIFRLGKIVEAAG
QNFQEMAVEGGVMGLQINWDCNLDRAASHCVPKYSFRRLDNKDSANTISPGYNFRFAKYYKDSSGIETRTLIKAYGIRL
DIIVFGKAGKFDVIPTMINIGSGLALFGVATVLCDIVVLYCMKKRYFYREKKYKYVEDYELGTSET

>Deuterostome_chordata_Ggallus_tr_Q8AWC0_Q8AWC0_CHICK_P2X_purinoceptor_OS_Gallus_gallus_OX_9031_GN_P2X1_PE_2_SV_1

MAVGQRCMDKLSTFLFEYDTPRMVLVRNKKVGLTFRLIQLVVLAYIIGWVFLYEKGYQTQDSIVSSVSVKLKGLTMTNE
SSLGPHIWDVVDYVFPPQGDSSFVVMTNFIITPGQKQGTCPELPDAGLCRQDSDCSEGTYNRQGQGLMTGKCVNYNS
SMKTCEIFGWCPAEVDYHVPNPALLPEVEKFTIFIKNSITFPKFKVSRRNIVESVTAQYLKECTYHKVTDSLCPMFELGYVV
KESGQNFSMLAVKGGVVGITIDWNCDLDWPIRYCKPIYQFHGLYNDNSNVSPGFNFRYAKYYREDGMEKRTLYKVFGI
RFDILVNGKAGKFDIIPTMTTIGSGIGIFGVASVLCDLLLLHFLQGRDYYKQKKFKYAEQEPSKSNKKEKELDNAP

>Deuterostome_chordata_Ggallus_tr_E1C6P3_E1C6P3_CHICK_P2X_purinoceptor_OS_Gallus_gallus_OX_9031_GN_P2RX7_PE_1_SV_3

LNFQSLKNSQETLTMVAWGWMKDVFNYESPKLIRFPSVGLVCVKWFIYGVIAVYICYTLIVHKRYQEKEELTSSVRVTLK
GVAHVDRIWDAAEYTIPTQTRDSFFVMTNIIRTENQIQKTCPEYPTAKAICSSDKSCAKGIVDVHSNGVQTGKCVHYNIT
HKTCEIKAWCPVQGEERPPVPAVLRSSEDFTVFIKNNIHFPTFNYTVQNISPKLNTSCKFNKVTAPLCPIFRLGDILQEAKE
NFSEMAVKGGIIAIEIKWDCDLDSWSYYCSPEYSFRRLDDKTRTQYPGFSIRFARHYKLPDGTEQRTLFKAYGIRFDVLVF
GMGGQFKLIELFTFIGSTIAYFGLAVTIIEMCFHLYNCSSCCKIQVCENVIRKKYETVLMPEQVILVSYVDKPHITLIKMPLR
TSLQNAEGSIFEDHPVKSYDPRTCCSHKSNEKHGAAQSELRPLTQSSSSTNCPKWCCCGRCQVAQKHHEQLCCRKKEG
QCITTTYWFAQLVLSRDTLNKALLYEDPFLDLTGHSSNSQLRRIAYKQYIHWRFGSFELEDRAIIPSCCRRLIRSTYPKENG
NYTGFNLE

>Deuterostome_chordata_Ggallus_tr_A0A1L1RYK7_A0A1L1RYK7_CHICK_P2X_purinoceptor_OS_Gallus_gallus_OX_9031_GN_P2RX2_PE_3_SV_2

MPICCSPFGLRETQHRCPEQIPSSCWERTNAALCPHRSGKRGAHQETPPGAAGLLNAGRVASDCCTTIAGTRRSYTDA
GLGAHSGHRESLTRGNNHGLQRALLPLPARLRAGGSGPCAAEPPPAATPQPAGSAAGPGRAGRSAAAMGPGCRRAV
SSYSSPQVVVVRDGRLGTAYRALQLLVLLYFIGYVFIVQKGYQERETGPESSVITKVKGVTQSLSKVWDVGEYVAPPEGG
SSFSILTRVEVSAAQAMGTCHEGPSAACHSEQDCVSGAMDASHHGVRTGRCVPGGSGRSCEVLAWCPLHGGSSSESL
AEMAAQFTILIKNHVRFPRFGFSKANIQAAESHYLKSCTFNATSALYCPIFRLGFLAEQAGEDFAVLAEKGGVIGVIISWD
CNLDLPDTECNPRYSFRRLDPKGALASPGYNYRFAKYYSWNGTCTRVLTKAYGIRVDVIVQGQAGKFSLIPTVITLATALT
SVGLGSFLCDWVLLCCMDKERRYSSRKFEQVPLG

>Deuterostome_chordata_Ggallus_tr_Q9IAD2_Q9IAD2_CHICK_P2X_purinoceptor_OS_Gallus_gallus_OX
_9031_GN_P2RX5_PE_2_SV_1

MGQVSWKGLFLSLFDYKTEKYVIAKNKKVGILYRVVQLSILAYLVGWVFVVKKGYQDTDTSLQSSVITKLKGVAFTNTSE
LGERLWDVADYVIPPQGENVFFVMTNLIVTPNQRQTTCPESVNIPDALCHQDEDCPEGQAVVAGNGVKTGRCLKDRD
SIRGSCEVLAWCPVEKRSKPKKPLLASAENFTVFIKNSIRFPKFKFSKMNVLATNNESYLKTCHYSMEHPYCPIFLLGNIVR
WTGNNFQEMALEGGVIGIQIEWNCDLDQAPSECNPHYSFSRLDNKFAEKSVSSGYNFRFAKYYRDAKGIEYRTLFKAYG
IRFDVMVNGKAGKFNIIPTIINIGSGLALMGAGAFFCDLVLLYLIKKSNFYRGKKYEEVKSSSRKSLTSPTLNGSQSPDQLG
GL

>Deutero_Chord_Hsap_sp_O15547_P2RX6_HUMAN_P2X_purinoceptor_6_OS_Homo_sapiens__Human
__OX_9606_GN_P2RX6_PE_1_SV_2

MCPQLAGAGSMGSPGATTGWGLLDYKTEKYVMTRNWRVGALQRLLQFGIVVYVVGWALLAKKGYQERDLEPQFSII
TKLKGVSVTQIKELGNRLWDVADFVKPPQGENVFFLVTNFLVTPAQVQGRCPEHPSVPLANCWVDEDCPEGEGGTHS
HGVKTGQCVVFNGTHRTCEIWSWCPVESGVVPSRPLLAQAQNFTLFIKNTVTFSKFNFSKSNALETWDPTYFKHCRYE
PQFSPYCPVFRIGDLVAKAGGTFEDLALLGGSVGIRVHWDCDLDTGDSGCWPHYSFQLQEKSYNFRTATHWWEQPG
VEARTLLKLYGIRFDILVTGQAGKFGLIPTAVTLGTGAAWLGVVTFFCDLLLLYVDREAHFYWRTKYEEAKAPKATANSV
WRELALASQARLAECLRRSSAPAPTATAAGSQTQTPGWPCPSSDTHLPTHSGSL

>Deutero_Chord_Hsap_sp_P51575_P2RX1_HUMAN_P2X_purinoceptor_1_OS_Homo_sapiens__Human
__OX_9606_GN_P2RX1_PE_1_SV_1

MARRFQEELAAFLFEYDTPRMVLVRNKKVGVIFRLIQLVVLVYVIGWVFLYEKGYQTSSGLISSVSVKLKGLAVTQLPGLG
PQVWDVADYVFPAQGDNSFVVMTNFIVTPKQTQGYCAEHPEGGICKEDSGCTPGKAKRKAQGIRTGKCVAFNDTVK
TCEIFGWCPVEVDDDIPRPALLREAENFTLFIKNSISFPRFKVNRRNLVEEVNAAHMKTCLFHKTLHPLCPVFQLGYVVQE
SGQNFSTLAEKGGVVGITIDWHCDLDWHVRHCRPIYEFHGLYEEKNLSPGFNFRFARHFVENGTNYRHLFKVFGIRFDIL
VDGKAGKFDIIPTMTTIGSGIGIFGVATVLCDLLLLHILPKRHYYKQKKFKYAEDMGPGAAERDLAATSSTLGLQENMRTS

>Deutero_Chord_Hsap_sp_P56373_P2RX3_HUMAN_P2X_purinoceptor_3_OS_Homo_sapiens__Human
__OX_9606_GN_P2RX3_PE_1_SV_2

MNCISDFFTYETTKSVVVKSWTIGIINRVVQLLIISYFVGWVFLHEKAYQVRDTAIESSVVTKVKGSGLYANRVMDVSDY
VTPPQGTSVFVIITKMIVTENQMQGFCPESEEKYRCVSDSQCGPERLPGGGILTGRCVNYSSVLRTCEIQGWCPTEVDT
VETPIMMEAENFTIFIKNSIRFPLFNFEKGNLLPNLTARDMKTCRFHPDKDPFCPILRVGDVVKFAGQDFAKLARTGGVL
GIKIGWVCDLDKAWDQCIPKYSFTRLDSVSEKSSVSPGYNFRFAKYYKMENGSEYRTLLKAFGIRFDVLVYGNAGKFNIIP
TIISSVAAFTSVGVGTVLCDIILLNFLKGADQYKAKKFEEVNETTLKIAALTNPVYPSDQTTAEKQSTDSGAFSIGH

>Deutero_Chord_Hsap_sp_Q93086_P2RX5_HUMAN_P2X_purinoceptor_5_OS_Homo_sapiens__Human
__OX_9606_GN_P2RX5_PE_2_SV_4

MGQAGCKGLCLSLFDYKTEKYVIAKNKKVGLLYRLLQASILAYLVVWVFLIKKGYQDVDTSLQSAVITKVKGVAFTNTSDL
GQRIWDVADYVIPAQGENVFFVVTNLIVTPNQRQNVCAENEGIPDGACSKDSDCHAGEAVTAGNGVKTGRCLRRENL
ARGTCEIFAWCPLETSSRPEEPFLKEAEDFTIFIKNHIRFPKFNFSKSNVMDVKDRSFLKSCHFGPKNHYCPIFRLGSVIRW
AGSDFQDIALEGGVIGINIEWNCDLDKAASECHPHYSFSRLDNKLSKSVSSGYNFRFARYYRDAAGVEFRTLMKAYGIRF
DVMVNGKGAFFCDLVLIYLIKKREFYRDKKYEEVRGLEDSSQEAEDEASGLGLSEQLTSGPGLLGMPEQQELQEPPEAKR
GSSSQKGNGSVCPQLLEPHRST

>Deutero_Chord_Hsap_sp_Q99571_P2RX4_HUMAN_P2X_purinoceptor_4_OS_Homo_sapiens__Human
__OX_9606_GN_P2RX4_PE_1_SV_2

MAGCCAALAAFLFEYDTPRIVLIRSRKVGLMNRAVQLLILAYVIGWVFVWEKGYQETDSVVSSVTTKVKGVAVTNTSKL
GFRIWDVADYVIPAQEENSLFVMTNVILTMNQTQGLCPEIPDATTVCKSDASCTAGSAGTHSNGVSTGRCVAFNGSVK
TCEVAAWCPVEDDTHVPQPAFLKAAENFTLLVKNNIWYPKFNFSKRNILPNITTTYLKSCIYDAKTDPFCPIFRLGKIVEN
AGHSFQDMAVEGGIMGIQVNWDCNLDRAASLCLPRYSFRRLDTRDVEHNVSPGYNFRFAKYYRDLAGNEQRTLIKAY
GIRFDIIVFGKAGKFDIIPTMINIGSGLALLGMATVLCDIIVLYCMKKRLYYREKKYKYVEDYEQGLASELDQ

>Deutero_Chord_Hsap_sp_Q99572_P2RX7_HUMAN_P2X_purinoceptor_7_OS_Homo_sapiens__Human
__OX_9606_GN_P2RX7_PE_1_SV_4

MPACCSCSDVFQYETNKVTRIQSMNYGTIKWFFHVIIFSYVCFALVSDKLYQRKEPVISSVHTKVKGIAEVKEEIVENGVK
KLVHSVFDTADYTFPLQGNSFFVMTNFLKTEGQEQRLCPEYPTRRTLCSSDRGCKKGWMDPQSKGIQTGRCVVYEGN
QKTCEVSAWCPIEAVEEAPRPALLNSAENFTVLIKNNIDFPGHNYTTRNILPGLNITCTFHKTQNPQCPIFRLGDIFRETG
DNFSDVAIQGGIMGIEIYWDCNLDRWFHHCRPKYSFRRLDDKTTNVSLYPGYNFRYAKYYKENNVEKRTLIKVFGIRFDI
LVFGTGGKFDIIQLVVYIGSTLSYFGLAAVFIDFLIDTYSSNCCRSHIYPWCKCCQPCVVNEYYYRKKCESIVEPKPTLKYVSF
VDESHIRMVNQQLLGRSLQDVKGQEVPRPAMDFTDLSRLPLALHDTPPIPGQPEEIQLLRKEATPRSRDSPVWCQCGS
CLPSQLPESHRCLEELCCRKKPGACITTSELFRKLVLSRHVLQFLLLYQEPLLALDVDSTNSRLRHCAYRCYATWRFGSQD
MADFAILPSCCRWRIRKEFPKSEGQYSGFKSPY

>Deutero_Chord_Hsap_sp_Q9UBL9_P2RX2_HUMAN_P2X_purinoceptor_2_OS_Homo_sapiens__Human
__OX_9606_GN_P2RX2_PE_1_SV_1

MAAAQPKYPAGATARRLARGCWSALWDYETPKVIVVRNRRLGVLYRAVQLLILLYFVWYVFIVQKSYQESETGPESSIIT
KVKGITTSEHKVWDVEEYVKPPEGGSVFSIITRVEATHSQTQGTCPESIRVHNATCLSDADCVAGELDMLGNGLRTGRC
VPYYQGPSKTCEVFGWCPVEDGASVSQFLGTMAPNFTILIKNSIHYPKFHFSKGNIADRTDGYLKRCTFHEASDLYCPIFK
LGFIVEKAGESFTELAHKGGVIGVIINWDCDLDLPASECNPKYSFRRLDPKHVPASSGYNFRFAKYYKINGTTTRTLIKAYG
IRIDVIVHGQAGKFSLIPTIINLATALTSVGVGSFLCDWILLTFMNKNKVYSHKKFDKVCTPSHPSGSWPVTLARVLGQAP
PEPGHRSEDQHPSPPSGQEGQQGAECGPAFPPLRPCPISAPSEQMVDTPASEPAQASTPTDPKGLAQL

>Deuterostome_chordata_PetMar_tr_S4R4R7_S4R4R7_PETMA_Uncharacterized_protein_OS_Petromyz
on_marinus_OX_7757_PE_3_SV_1

GEKTGRCVPFDKNVSTCEVNAWCPVEVEGKNTGKRLLRSAENFTILIKNNIRFPKFNVSKGNLPENMDKNFLKSCHHHP
KHNNLCPIFSLRDILRSAGQDFKSIGINGGGRQRRGSCSVEV

>Deuterostome_chordata_PetMar_tr_S4RS70_S4RS70_PETMA_P2X_purinoceptor_OS_Petromyzon_ma
rinus_OX_7757_PE_3_SV_1

LAMQGCGALVCSAFEYKTEKLMIARDRSVGVVYRLMQLVILAYIIGWVFVVKKGYQETDPDPETSVITKVKGVALTNST
EGPGQRLWDTADYVIPPQGESRFFVVTNLVSTPRQRQGLCAENPNVPGARCQEDGDCPAGRTFNTGNGICTGRCVPT

NGTARVCEVHAWCPVENSPTPREAVLREAENFTVFIKVSVNFTRFNVHRRNVLEEGDSSYLKSCEFHPERSPLCPVFRLG
ELAVRAGASFSDMATRGGVIAVHLAWDCDLDRDLSACAAQYRFSRLDSERSRVAAGHNFRFARYSWQDGAETRSLLK
VYGVRFDVMVSGKAGKFSIVPTLLNLGAGLALLGAGAFLCDLFLLYASKESPQYRQRKFERSEEDGPVPPQQDAVRARS
SSTPGTANSS

>Deuterostome_chordata_PetMar_tr_S4RU33_S4RU33_PETMA_ATP_receptor_OS_Petromyzon_marinu
s_OX_7757_PE_3_SV_1

MCQTKRRGLPCNIFFYPRELVIGIENKYCGPIHMVVRIVILGYGARYVVGFEHAYREYEDVTSAVRVKVKGTAKVDIPNE
GVRTFDSADYVVPDQEPNTFFVMTNFIVTKNQTQVTTCCNPEAMKFSVSPCQNDGDCTRDASKHLGSGIQTGRCIRY
DATCKSCEVMGWCPVERRTSPPQPPLLPATENATVYIQNHVYFIKNNKERENFPLNLEGKDLSSCRYHPRDDPSCPIFRL
GDITAMAGHSFSSMSVQGGIIAIVISWDCKLSEVECFPAYSFHRLDRNYTTKRGYNFRSLVYKSMETDVQTRTLVKAFGI
RFNILVTGQ

>Deuterostome_chordata_PetMar_tr_S4RR49_S4RR49_PETMA_ATP_receptor_OS_Petromyzon_marinu
s_OX_7757_PE_3_SV_1

QVRSYQDVLSKFYLNYTITKKARIKSRKAAILTFLVFVMLVISYTGWVIYTHGYQEFDGVVSSSTAKVKGATEVTGMVMD
AADYVISPEEPNSFFVMTNFIETPNQKQGACAEDPDVAPCSSDTDCRAGVTDPEGEGVFTGLCVPINNTTKSCEVNAW
CPLPKDSNPPKHNILGSNASYLKSCRYHRQSDPECPIFILGDIVSEAGEDFKELAIEGGVFSIMIEWNCNLDLNEQACKPN
YKFRRLDKENRNKTISPGYNFRFPKYYKDAKGIETRTLVKAYGIRFMVEVIGQAGKASLLRSLISLGSGVGLLGMVWVICD
LIVLFCCRKSRIYREQLYE

>sp_Q64663_P2RX7_RAT_P2X_purinoceptor_7_OS_Rattus_norvegicus_OX_10116_GN_P2rx7_PE_1_SV
_1

MPACCSWNDVFQYETNKVTRIQSVNYGTIKWILHMTVFSYVSFALMSDKLYQRKEPLISSVHTKVKGVAEVTENVTEG
GVTKLVHGIFDTADYTLPLQGNSFFVMTNYLKSEGQEQKLCPEYPSRGKQCHSDQGCIKGWMDPQSKGIQTGRCIPYD
QKRKTCEIFAWCPAEEGKEAPRPALLRSAENFTVLIKNNIDFPGHNYTTRNILPGMNISCTFHKTWNPQCPIFRLGDIFQ
EIGENFTEVAVQGGIMGIEIYWDCNLDSWSHRCQPKYSFRRLDDKYTNESLFPGYNFRYAKYYKENGMEKRTLIKAFGV
RFDILVFGTGGKFDIIQLVVYIGSTLSYFGLATVCIDLIINTYASTCCRSRVYPSCKCCEPCAVNEYYYRKKCEPIVEPKPTLKY
VSFVDEPHIWMVDQQLLGKSLQDVKGQEVPRPQTDFLELSRLSLSLHHSPPIPGQPEEMQLLQIEAVPRSRDSPDWCQ
CGNCLPSQLPENRRALEELCCRRKPGQCITTSELFSKIVLSREALQLLLLYQEPLLALEGEAINSKLRHCAYRSYATWRFVSQ
DMADFAILPSCCRWKIRKEFPKTQGQYSGFKYPY

>sp_P51578_P2RX5_RAT_P2X_purinoceptor_5_OS_Rattus_norvegicus_OX_10116_GN_P2rx5_PE_2_SV_
1

MGQAAWKGFVLSLFDYKTAKFVVAKSKKVGLLYRVLQLIILLYLLIWVFLIKKSYQDIDTSLQSAVVTKVKGVAYTNTTML
GERLWDVADFVIPSQGENVFFVVTNLIVTPNQRQGICAEREGIPDGECSEDDDCHAGESVVAGHGLKTGRCLRVGNST
RGTCEIFAWCPVETKSMPTDPLLKDAESFTISIKNFIRFPKFNFSKANVLETDNKHFLKTCHFSSTNLYCPIFRLGSIVRWA
GADFQDIALKGGVIGIYIEWDCDLDKAASKCNPHYYFNRLDNKHTHSISSGYNFRFARYYRDPNGVEFRDLMKAYGIRF
DVIVNGKAGKFSIIPTVINIGSGLALMGAGAFFCDLVLIYLIRKSEFYRDKKFEKVRGQKEDANVEVEANEMEQERPEDEP
LERVRQDEQSQELAQSGRKQNSNCQVLLEPARFGLRENAIVNVKQSQILHPVKT

>sp_P49653_P2RX2_RAT_P2X_purinoceptor_2_OS_Rattus_norvegicus_OX_10116_GN_P2rx2_PE_1_SV_
1

MVRRLARGCWSAFWDYETPKVIVVRNRRLGFVHRMVQLLILLYFVWYVFIVQKSYQDSETGPESSIITKVKGITMSEDK
VWDVEEYVKPPEGGSVVSIITRIEVTPSQTLGTCPESMRVHSSTCHSDDDCIAGQLDMQGNGIRTGHCVPYYHGDSKT
CEVSAWCPVEDGTSDNHFLGKMAPNFTILIKNSIHYPKFKFSKGNIASQKSDYLKHCTFDQDSDPYCPIFRLGFIVEKAGE
NFTELAHKGGVIGVIINWNCDLDLSESECNPKYSFRRLDPKYDPASSGYNFRFAKYYKINGTTTTRTLIKAYGIRIDVIVHG
QAGKFSLIPTIINLATALTSIGVGSFLCDWILLTFMNKNKLYSHKKFDKVRTPKHPSSRWPVTLALVLGQIPPPPSHYSQD
QPPSPPSGEGPTLGEGAELPLAVQSPRPCSISALTEQVVDTLGQHMGQRPPVPEPSQQDSTSTDPKGLAQL

>sp_P49654_P2RX3_RAT_P2X_purinoceptor_3_OS_Rattus_norvegicus_OX_10116_GN_P2rx3_PE_1_SV_1

MNCISDFFTYETTKSVVVKSWTIGIINRAVQLLIISYFVGWVFLHEKAYQVRDTAIESSVVTKVKGFGRYANRVMDVSDY
VTPPQGTSVFVIITKMIVTENQMQGFCPENEEKYRCVSDSQCGPERFPGGGILTGRCVNYSSVLRTCEIQGWCPTEVDT
VEMPIMMEAENFTIFIKNSIRFPLFNFEKGNLLPNLTDKDIKRCRFHPEKAPFCPILRVGDVVKFAGQDFAKLARTGGVLG
IKIGWVCDLDKAWDQCIPKYSFTRLDGVSEKSSVSPGYNFRFAKYYKMENGSEYRTLLKAFGIRFDVLVYGNAGKFNIIPT
IISSVAAFTSVGVGTVLCDIILLNFLKGADHYKARKFEEVTETTLKGTASTNPVFASDQATVEKQSTDSGAYSIGH

>sp_P51577_P2RX4_RAT_P2X_purinoceptor_4_OS_Rattus_norvegicus_OX_10116_GN_P2rx4_PE_1_SV_1

MAGCCSVLGSFLFEYDTPRIVLIRSRKVGLMNRAVQLLILAYVIGWVFVWEKGYQETDSVVSSVTTKAKGVAVTNTSQL
GFRIWDVADYVIPAQEENSLFIMTNMIVTVNQTQSTCPEIPDKTSICNSDADCTPGSVDTHSSGVATGRCVPFNESVKT
CEVAAWCPVENDVGVPTPAFLKAAENFTLLVKNNIWYPKFNFSKRNILPNITTSYLKSCIYNAQTDPFCPIFRLGTIVEDA
GHSFQEMAVEGGIMGIQIKWDCNLDRAASLCLPRYSFRRLDTRDLEHNVSPGYNFRFAKYYRDLAGKEQRTLTKAYGIR
FDIIVFGKAGKFDIIPTMINVGSGLALLGVATVLCDVIVLYCMKKKYYYRDKKYKYVEDYEQGLSGEMNQ

>sp_P51579_P2RX6_RAT_P2X_purinoceptor_6_OS_Rattus_norvegicus_OX_10116_GN_P2rx6_PE_1_SV_2

MQLQPAGTGSMASAVAAALVSWGFLDYKTEKYVMTRNCWVGISQRLLQLGVVVYVIGWALLAKKGYQEWDMDPQ
ISVITKLKGVSVTQVKELEKRLWDVADFVRPSQGENVFFLVTNFLVTPAQVQGRCPEHPSVPLANCWADEDCPEGEMG
TYSHGIKTGQCVAFNGTHRTCEIWSWCPVESSAVPRKPLLAQAKNFTLFIKNTVTFNKFNFSRTNALDTWDNTYFKYCL
YDSLSSPYCPVFRIGDLVAMTGGDFEDLALLGGAVGINIHWDCNLDTKGSDCSPQYSFQLQERGYNFRTANYWWAAS
GVESRSLLKLYGIRFDILVTGQAGKFALIPTAITVGTGAAWLGMVTFLCDLLLLYVDREAGFYWRTKYEEARAPKATTNSA

>sp_P47824_P2RX1_RAT_P2X_purinoceptor_1_OS_Rattus_norvegicus_OX_10116_GN_P2rx1_PE_2_SV_1

MARRLQDELSAFFFEYDTPRMVLVRNKKVGVIFRLIQLVVLVYVIGWVFVYEKGYQTSSDLISSVSVKLKGLAVTQLQGL
GPQVWDVADYVFPAHGDSSFVVMTNFIVTPQQTQGHCAENPEGGICQDDSGCTPGKAERKAQGIRTGNCVPFNGT
VKTCEIFGWCPVEVDDKIPSPALLREAENFTLFIKNSISFPRFKVNRRNLVEEVNGTYMKKCLYHKIQHPLCPVFNLGYVV
RESGQDFRSLAEKGGVVGITIDWKCDLDWHVRHCKPIYQFHGLYGEKNLSPGFNFRFARHFVQNGTNRRHLFKVFGIH
FDILVDGKAGKFDIIPTMTTIGSGIGIFGVATVLCDLLLLHILPKRHYYKQKKFKYAEDMGPGEGEHDPVATSSTLGLQEN
MRTS

>Cnidar_AlaAla_c43961_g1_i1_p1__GENE_c43961_g1_i1__c43961_g1_i1_p1__ORF__type_5prime_partial__len_433__score_67_69__c43961_g1_i1_182_1480_

KNPCLKVCSKAGNLMFSYDTLKIVKINNKKVGLLNRTIQLAIIGYVIGYAIVWKKGYQETDNAPVTSVVTKLKGTSLVNFT
SFSSTMLRGMHIFDPADYVVPPQENNAFFVMTNMILTPNQTRGECPEDPKFKDNHCKSDSDCPAMTAVVNGNGIRT

GRCVPSDRKEGLKVCEIYAWCPVEKDILPMPGHGFKDGDPLLDEAENFTVLIKNSISFPKFSFTKRNILNENNSRYLKSCN
YDKTTDNLCPIFKVGDITKEAGTDFKSLAYLGGNIGIIISWNCNLDLSESLCTPEYSFRRIDDPDAPIAKGYNFRFPQYYVED
GAQYRLLYKAYGIKFDVLVKGKAGKFSIVPLLLNLGSGLALLGIATVWCDLVVLYCLNKRNVYKSHKYHQVEDADIQGNK
SVDSDTEETELHDDDIDENRFKYKRLSDEEG

>Cnidar_HydraVulSc4wPfr_308_g5501_t1

MSTILFGAIFGVFVEWDCDLDEVPSKCIPAYKIKRLDNPNDRLASGFNYRVANYSFVNNKQYRTLTKTYGLKFHIIVTGKG
RKKDFYTTIVFLGATLALLSKVAVFFEWIVLFVCSCKKNKEKFDKCKYKTFNEENVCLKITNNKSLSVCSFNCRSFKNSVLQ
RNMFNEYDFNLLQEHWLLPYELGIFNNVHNEFYGTGLSAVNILFDIMIGRPYGGTTILYRKDFAKSIHPNKL

>Cnidar_HydraVulSc4wPfr_1483_g15774_t1

MKSVRTGRCVQSDRPNISYRVCEIYGWCPTETDILPMPGYNFSASVPLLDDAKEFTVLLKNEVIFPKFKVQRRNIVKNNN
SNYLKTCFHNHDTDPLCPIFKIKDVIQECNDDFEQVAYLVIYGILVNWDCNLDLSLDKCTPTYSTRRLDDPDAPISPGFNF
SLIKSIKAYISIQNRYYSDITNPDIYET

>Cnidar_NemVecNVEC200_004811_1_1_protein_AED_0_22_eAED_0_22_QI_0_1_0_91_1_1_1_12_585
_431

IRCRSKMAPSCGNCCACVGSVLFEYDTEKIVHIKNKKIGIINRVIQLVIIAYIIGFVIVYKKGYQEFESPYSSVTTKLKGVALTN
LTGANLSYYGREFLWDSADLVIPPEEEGAVFMMTNMIITPEQSQGKCPEDPSVPGARCTSDADCPPLQSTSTSHGVNT
GNCVPSTHNMTVKTCELYAWCPIENDVTPMPGFNLSYGMPLLDAAKNFTLLIKNSVQFPKFSESRRNIIQKNDSHLKSC
NYHPTSDPLCPVFRLGTMAELAGVNFEKLAYKGGVMAIVITWNCNFDPLAYHCEPKYSFRRLDDDESPIAPGYNFRFPR
YYIKNETLYRTIVKAYGIRFVVTVYGKGGKFSVIPLFLNIGSGLALLGIATVLCDVFVLYVLRKKYFYRDKKYLNVDDSDNEM
SLFQKADKVHSKRGYSQLSDNEGEMMK

>Cnidar_NemVecNVEC200_005314_1_1_protein_AED_0_28_eAED_0_28_QI_121_1_1_1_0_91_0_84_1
3_1162_417

MAACCGKVIDSLLEYNTSKIVHIKSKKVGLINRLIQLAIIGYIIGYVIIYKKGYQEVEKPYSSVTTKLKGTSFTNLTGNESFPLY
GGPFVWDQADYVIPPEENDAFFVATNMVITPDQRQGTCPESPDIANVKCTRDENCTRMEETSYGHGFQTGKCVPSD
QEKGIKVCEVFAWCPVELDITPMPQFPSLKDNKPLLDARNFTVLIKNTVQFPSFGVSKRNIAPDADTSTLKSCQYDKDTS
PLCPIFRLETITKEAGQDFNKLAYKGGVIGIIITWNCDFDSLLYSCKPVYSFRRLDNANAPIAKGYNFRHANYYVKNDTLHR
TLFKAYGIRFVVTVTGQGGKFNVVPLFLNLGSGLALLSIATVLCDVVVLYVLRRKEFYREKKYQYVVENSSDEALVNRKPS
YVKTETGYQNLIS

>Cnidar_Polpod_Hydrif_GBGH01011573_1__p1__GENE_GBGH01011573_1_GBGH01011573_1__p1__O
RF__type_internal__len_104__score_26_67__GBGH01011573_1__1_309_

DSSKTRTHPTIKVDIHNYAMNFTVLLKNSVVFKALGRQERNIAPSMNASYLKKCRYRPGDINDSHCPIFELKTIVQEANAS
FQDLTEKGGVIAIYVDWDCNYD

>Cnidar_Polpod_Hydrif_GBGH01023749_1__p2__GENE_GBGH01023749_1_GBGH01023749_1__p2__O
RF__type_5prime_partial__len_103__score_27_61__GBGH01023749_1__1_309_

PGFNFRFSKYYSCGQEHCRALYKAIGVRFDVITRARGRKFSIIPTLLNVGSGIGLLGVAAVLCDLVVVYLSKHKMKYRRIKF
DQVAESEDELLRDGHPNNID

>Cnidar_Rhopilema_esculentum_TR57293_c0_g1_i1_p1__Rhopilema_esculentum_TR57293_c0_g1__R hopilema_esculentum_TR57293_c0_g1_i1_p1__ORF__type_complete__len_441__score_66_70__Rhopi lema_esculentum_TR57293_c0_g1_i1_588_1871_

MVAKEGIGTKILDTLLSYETIKIVTIKNKKVGFLHRLVQLAILGYVIGYAIIWKKGYQEFDQSLSAVTTKLKGVALANFSTFES PLLNGMHIFDSADYVIPPQENNEFFVMTNMIITPNQTRGRCPEDPRFEDVYCQRDSDCREGEAVANGNGVKTGKCVQ SDQNITAYVCQIYGWCPVELDVLPMPGHGFKKGDPLLQSAVNFTVLIKNQVTFPKFGFARRNIISQNNTTYLKSCNYHH ESDPFCPIFKLGTIVSDAGYDFNDIAYKGGIVAIQIKWECNLDVSEDKCRPQYSFKRIDDPHAPIASGFNFRFAQYYVENN KQHRILYKAYGIKFEVLVNGRAGKFSPVPLFLNLGSGLALLSIATIWCDIIVLYCLKDRQVYRSHKYHKVVDGDEPTDTDD DGDLNKGANVSEFAYQKVEDENDPD

>Cnidar_Rhopilema_esculentum_TR95815_c0_g1_i1_p1__Rhopilema_esculentum_TR95815_c0_g1__R hopilema_esculentum_TR95815_c0_g1_i1_p1__ORF__type_5prime_partial__len_410__score_37_93__ Rhopilema_esculentum_TR95815_c0_g1_i1_850_2079_

SLVPPRIIGIPTRNPQTKKRPSGLSPCDFSFSATYLLKMSMFHYVTYKYVVIKDKRLGFAYYAVAALILIYTFVQIVLNKAYL QFDHSPSGNFRIITNSPPLHGHEGDLKLENEYCCTNNNIAESKDNRCKPCRTYDGLALNWPVESKAVTIATFIKERWEET GHKQEDMEDYIVLREDSYFTRNPEHVLFKIEHSISALESSEDPLALSHRKMNGWLMSASGEERKLNGGDVHKADKIRLIE FLRAAGIQSLDEKTDALNSRKRSFRENGVVLRVLIHYANSANVLFGTTVPEYRYIVRRIPYSDYKVKQEIPVIEENRFESVH NMSMVQRRVVRKRFCVRVEFIQAGKIGQFSMLNLAIHLVSGFGLLSVTGLIIDTLALYVLTSRVVYSKYIYEYSPDLDKNIK QN

>Cnidaria_Myxozoa_T_bryosalmonae_contig_2149_32_1636

MDSKIFNETTRNKESDSISNLNSQLIHQSEKKDNNEIITSEISIINPPLLSSQSSILNNKIKLSEKSDILYEEFHSLNDFSNSLFN GNQLNERINEKSIDNKRKISKNKLAFQKKIIRIFKKFNLAAFISFFIALIQLASACFIIWFLLIKNNTFQLVDLGISTFSIKLINSN TKGRQDSLENLINQFDTENQPNYLFDWNGGGKFPILINSIITVNQTYGTCPEDPSIQNAICYSDSDCSIINLYGNGEFNGK CVKCMEKVTDLKSDYGCCEIIGWCPVQTASTSFSYINIAPLSDPIHSLRKNSVLIKNLVDFRVFNRIKNNIPDYLLGNKAYV CRYNKSSEMYANCPNFLLEDIVDWAGMTYESLQNSGGSINILITWKCNFDLNETCFPTYSFDNYMETNNESSSDINSKF SRYYMINGTFYRDLLTTRLLNIIITTKAEANGFTVISLMLAMGQCFGLILMIVTFFDYLNKNINKSIKIIKKRQQKKKKQKKL NKKDKSTDMEKDLKFNNTIKVVNPEFLLKKNGYYDF

>Cnidaria_Myxozoa_T_bryosalmonae_contig_3145_1_1110

MGNTLSYETNQVISIGDKYVGFLYRLVQLTVIAYVIGYVMVYNGGYQEVDNGVCHTFGKIKGISRLFPNKKLKNMDDKY NILDSVDIVMPSYGADKYFIMTMFGLTTDQTISSCSSNLSCNSSDSCPFNVSNKELNGIYNGKCKENLCQVDGWCPIENE NITKFYSIDVKNYSLLVKNSVKFNKFNRNEVNIFDSLGKDFLESCRYNNKNADMKHCPIFVFSDILDILQIKNNELSMGVNI EISINWNCNYDSNSPCKPEYKISRVDISSKVAPGYNFRCAEKYHGANNQHFRDLYKCYGIGLQLTCISKAVKFSIIPLLLNIG SGLGLLGLATIMCDIVVFYISPNRQRYRKLIVKSLEDSLSDGSDSI

>Deutero_Ambulac_Apla_gbr53_10_t1

MIKNSISFPAFNFSKRNIRETAEKAYLRGCMYDPVNDPLCPIFKLGSIVEMANDTFENMAYKGGVVTIDIQWNCNLDRH RDECLPKYRFFRSDDQHAQIAAGFNFRYSKYYKSNGSEHRTLTKAYGILFNIKITGVAGQFHFIPLMLNVASGFALLSLAT VMCDIVVLYLLKKRNIYKECKYQNVNPVTDDGYEVPALRTFSKADGNSAIFPLRRISYDGNSMDVGKRYATAYLDSSAD NDRNS

>Deutero_Ambulac_Apla_gbr53_13_t1

MTSPEENVPGVGSLLISWALEYDTQKIVHIKNKTIGFLNRLIQLAIVAYVIGYAIIYKKAYQSTDSVISSVTTKLKGVSYTNIS
GLPDLNIKDTTPYDRIWDVSDYVVPPQQSNAFFVMTNMIITPNQQRGQCPEDPRVPNVGCTPAKESQDCVKGEPVIN
GNGVRTGKCVPYYYPNGTETYTCEIEAWCPVEYDVNPTQSEPVLEDAKRFTVLIKNTATYPKFHFSKRNILESSNKSYLSG
CMYNNETDPFCPIFKLGDIVAATGEDFNKLAYSGGVIGISINWDCNLDYSWKDCTPLYTFSRIDNSEAKLAKGYNFRFSN
YYQNETGIDTRTLFKAYGILFEVIVTGTAGKFDFVPLMLNIGTGLALLTMSTVACDIIVCYIMKGHEYYKKKKYLTVKPDAE
VNVDYEPLDNASVCSKETNEE

>Deutero_Ambulac_Spurpu_004826

MSMENKSFAMTEVIDIQHESSSGTMSGVGTIISSAVASFFEYDTPKIVHIKSKKVGLINRLLQLVIIGYIIGWVLVYKKGYQ
STDEVMSAVTTKMKGVAYTNITGEPNLNVKDTTPYNRIWDVSDYVIPPQQTNAFFVMTNMIITPGQSKSECPEDPKIP
NAECSGPEDCQADMPIINGNGIMTGRCVNYTYPDGNVTLTCEIRAWCPVEFDEIPV

>Deutero_Ambulac_Spurpu_013578

MSLSGKITTISCQIDLGAKEPGAVVGIVIHWDCNLDKSLDNCRPSYTFRRLDNRNALIAPGYNFRYSINYKENDTDYRTQT
KAYGILFQVIVSGEAGKFDVVPLILNIASGLALLSLASIMADICVLYLLKKRFLYRQKKFQLVDDIPYEDADDGDRSE

>Deutero_Ambulac_Spurpu_019489

MHMKSTPRQIYPDLRLVSNSIGQKWTSAILEEAENFTVLIKNSISFPKFGFVKRNILDTTDPMYLRSCRYSRLKDPLCPIFR
LGTIVAETGQSFRQMAIKGGVITIDIQWNCNLDLSYKLCLPKYRFIRADEQDAKIAGGFNFRFGKFYRINDTEYRDLTKAY
GILFQVKITGVAGKFDIVPLMLNFASGVALLSLATVMCDVVVLYLLKKRKYYKEAKFQNVASNENGPQKTYEEVAALCEQ
ALVTSLPITQLNNNHQTPPGMGGSNHSLRDEITLPTTNETKKPTSL

>Deutero_Ambulac_Spurpu_021054

MQNLKGAVVGIIIHWDCNLDYSLDNCRPSYTFRRLDNRNALIAPGYNFRYSINYKENDTDYRTQTKAYGILFQVIVSGEA
GKFDVVPLILNIASGLALLSLVSSRFSNQK

>Deutero_Ambulac_Sakowv30020762m

MAPQWCRFVIDAALEYDTPKIVHIKSKKVGIINRLIQLVIISYIVGYVIVYQKGYQEFDDVVSTATTKVKGVSYTNLSNVDP
ANLTVDDTTPYDRLWDVADYVIPPQMHSSFFVMTNMIITVDQTQGKCPEDPGMFGANCTVDADCHEGEAIVAGNG
VATGKCIDNPGYNNTKTCEILAWCPVERDIKPGPRKDKPVLYEAANFTVLIKNSVSFPKFDFQKRNILDSSNASFLRDCRY
DSKDADLQFCPIFRLGDIVKEAKVSAGEEVDFGTMGFEGGVMGIFINWDCNLDHAAEECKPHYSFRRLDNPEAKIAQG
FNFRFAYKYKINATDEYRTLTKAYGILYEVIVTGQAGKFSVIPLFLNIGSGLALLGLATVLCDIVVLYVLKKREYYKENKYLYV
DDLEDVPSEKYRNLDKE

>Deutero_Ambulac_Sakowv30015735m

MIVHKDYQVTDSVESITFIKVKGASYTNLSTIDPSYIAVKDTTPYDRVWDEADYVIPPQMKNSFFVMTNMLITEGQTQG
KFAEPPLSMGHGECIDHPDYPGKKTCEIRTWCPVENDQLP

>Deutero_Ambulac_Sakowv30015736m

MQSSFFVMTNMIITEDQTQGECAEVGHGEFFKHLVFHLKCIEHPDYPGTNTCEIRTWCPVEKDQLPRKDRAVLYEASN
FTVLIKNAIAFPKFGFKKHNILESADYTFLEDCLYTRKMNEIHCVLYSDLVILLGRLEGGSIGIIIDWRCNLDIPHAYDKCKPE
YRFRRIDREDAKIAPGYHFKFSYQYRNNNIQYRTLTKAYGILFEIVVEGEAGQFSWINLVLAVGALFGLLGL

>Deutero_Ambulac_Sakowv30015730m

GGSIIDWICNLDESYDKCKPTYRFRRIDRKDVKVSSGYNFRFSHQYRNNGITYRTLTKAYGILFEIVVEGEVGKFSVTKCVL
AIGASWGALGI

>Deutero_Ambulac_Sakowv30028261m

MVITEDQTQGECAEDPSISSAICVNDSDCSIPSPVRGHGPRTGRCIDHPGTTLEYPGNKTCEVRTWCPVEKDQLPRTDR
AVLYEAANFTVLIKNAITFPKFDFRKRNILQTDNSTFLEKCLYNEEDESYSMCPIFRLGDIVEYAGHDFEKIAYTGGSIGIVID
WTCNLDISHDKCKPKYKFLRIYREDAKIAPGFNFRFSYKYRNNGITYRTLTKAYGILFEIVVEGEAGKCSATKILLLIGATIGAI
SIVSGIFTCIMRASNKQFDNASNEIFEDKKWSCRNRFDEIRQSFKEGFVKLKSCKCFSGEARGDDSNSGEASGDDNKRK
GNN

>Deu_Ambulacrar_hemi_Ptyfla_40v0_9_20150316_1g10203_t1_scaffold5807_cov86

MVQYAKSFNFHFLTPMQTXIVYMTVTVYSVRITHLKLGDGLEIRRNGLKSSIFQLSLSHANANVVGVNRYVIILSKGYQEF
DDVVSTVTTKVKGVSYINLSALDPAIVTVDDTAPYDKLWDVADYIIPPQMQNSFFIMTNMIITVDQTQATCAEDPAIFG
ANCTVNDTCKKGEPIFNGNGVMTGKCIDNPEYEGTRSCEILAWCPVEKNIKPGPRLDKAVLHEAANFTVLLKNSISFPKF
GFSKRNILESSNEAFLKECIYNSRDPDLQFCPKFRLGDIVKETGEDFEKMALEGGVMGIVIDWDCDLDKSGDKCKPHYRF
RRLDNPDAKTAKGFNFRFAYKYKINETGEYRKLTKAYGILYEVVVTGKAGKFNIVPLFLNIGSGVGLLTVLKSRYFYELGEV
NFPAFADCFPKTNRDPDIFGANCTVDDTCKKGEPTINGNGVMTGNCIDNPAYEGTKSCEILAWCPVEKDIKPGPRLDKP
VLYEAANFTVLLKNSISFPKFGITKRNILEGSNETFLKECHYNSRDPDLQLCPKFRLGDISKEAGEDFEKLALEGGVMGIIIE
WNCNLDKSVDKCKPHYRFRRLKDSSAKSAKGFNFRFAYKYKINETGEYRTLTKAYGILYEVIVTGKAGKFSIIPLLVKIGSGL
SLLTVANIICDFVVLKILKKRKYYRKHKYLYVKEEAGEYEELSENDMPLKPINQNGNKIIQTDEILDLSKPMSDEKP

>Deu_Ambulacrar_hemi_Ptyfla_40v0_9_20150316_1g11467_t1_scaffold6869_cov168

MVEYIDSVDDIAEFTTFQKFQVFLLFMSMLPYSSVTFVNTFYFASTNHYCKTNENETYQPDSDWKHCVIPLREMGSTFT
SDSCYRNEISHTGKIPVNKSALCSFNKTAVIPCDKGWIHDRSNFANTQVMEFNLVCDDEWKKQLPKSLTGLALMIGSFL
YGQLADMLMEESPRWLFQKGDEQQAKKILQRMNKMNCNSEDDVRKTMTNHVDEKVKRHNNKKEEKTKYPLYDLM
RTPGLREWLLIVCCNGFSTTLMYYGLAYNADYFSSKTHLLFMMINFVDIPAIFLISLTLHYFPRRWQFIFTMLISGVTWLF
CAFVAATSVNMAPKWCEFIVAAALEYDTPKIVHIKSKKVGFINRLIQLAIIAYIVGYVIVLSKGYQEFDNVISTVTTKVKGVS
YTNLSALDPAVVTVDDTTPYDRLWDVADYIIPPQMQNSFFIMTNMILTVDQTQGTCAEGGVMGIFIEWNCNLDKSGD
ECKPHYKFRRLDNPTAKIAKGFNFRFAYKYKINETDEYRTLTKAYGILYEVVVTGEAGKFSVIPLFLNIGSGLALLSLATIMC
DIVVLYILKKREYYREHKYLYVDDLDDAAGEYRVTHTASALVIIFRNYSYTTENTTRSNIRNKNLRASLALWREYQTGKFKI
GNIPVNVTSTLMFTFRNLREVYRVHHS

>Protostome_Lophotroco_molsk_AplCal_NP_001191558_1_purinergic_ATP_P2X4_receptor_subunit_1_
_Aplysia_californica_

MAPPQVMKSAARSALDVFFEYDTPRIVHIRSKKVGLFNRFLQLSILAYIIGYAIVYKKGYQEFENVQSAVTTKVKGIAFSN
GSIPGIGTRTWDVADYVIPPQENDAFFVMTNVVVTPGQTQTGCAEDPGVTGAICQTNADCEQVKGTILPSGSGPVTG
ECVPSDVPGSKEKVCQIFGWCPLENDKMQGDSPVLENAKNFTVFIKNNIEFPRFGVRQTKPSKQRKRHVLKKCQYDGA
SSRDKFCPIFRLETITEGAGVAFENISKEGGVMQIVIDWTCNLDYSVEDCVPEYTFRRLDKGDYSVSRGFNFRFADRYSVF
NESTGLQLYRNLYKAYGVRFLVTVQGKAGKFSIVPLLLNIGSGMALLGVATIICDIMVLYVLKAKNFYRDKKYLDVKGQD
AFEVLEEEAGPGEQMSEGSTNSVSRRNLTASDGGAAP

>Protostome_Lophotroco_molsk_AplCal_XP_012942306_1_purinergic_ATP_P2X4_receptor_subunit_1_
isoform_X1__Aplysia_californica_

MAPPQVMKSAARSALGVFFEYDTPRIVHIRSKKVGLFNRFLQLSILAYIIGYAIVYKKGYQEFENVQSAVTTKVKGIAFSN
GSIPGIGTRTWDVADYVIPPQENDAFFVMTNVVVTPGQTQTGCAEDPGVTGAICQTNADCEQVKGTILPSGSGPVTG
ECVPSDVPGSKEKVCQIFGWCPLENDKMQGDSPVLENAKNFTVFIKNNIEFPRFGVSRRNILGFYNDTALKTCRWKRG
DPKLKFCPIFVLDDIARDAGVTFENMMMEGGVMQIVIDWTCNLDYSVEDCVPEYTFRRLDKGDYSVSRGFNFRFADR
YSVFNESTGLQLYRNLYKAYGVRFLVTVQGKAGKFSIVPLLLNIGSGMALLGVATIICDIMVLYVLKAKNFYRDKKYLDVK
GQDAFEVLEEEAGPGEQMSEGSTNSVSRRNLTASDGGAAP

>Protostome_Lophotroco_molsk_OctBim_tr_A0A0L8GLW5_A0A0L8GLW5_OCTBM_ATP_receptor_OS_
Octopus_bimaculoides_OX_37653_GN_OCBIM_22031653mg_PE_3_SV_1

MAYCDFILSLIFEYDTPKIVHIHSKKVGIINRLLQLIIIGYIVGYVIVYKKGYQEFDFVVGSISTKVKGIALTNRTILSDTPGVW
DVADYVVPPQENGAVFVMTNAIITYNQTQRCPQDVASGDFCTTDADCPAETMIPSGDGFRTGRCVNSSQNGTKVCEI
YAWCPFENDTLGEPLLKSAANFTIFVKNNVHFPKFKVSRRNIIGFENNRQLSKCRYDPRNATNRFCPIFTIQDILDLVKGD
DSTESILKMGASIQVNIVWNCNLDFSLDECLPEYRFTRLDRSNVVSEGFNFRFAKYFQNENQSMRTLIKAYGIQFLITIQG
RAGKFSIVPLLLNLGSGLALLSLASVVADIIMLYVLKSKNFYREKKYQNVDDNVYSAMKEDTPITTYGSNKDQRSASYH

>Placozoa_Hoilungia_hongkongensis__1_g03619_t2

MATLEESQGYFGVLEYNTAKIVHIKSRKIGILYRLVQLIIIGYVIGYAIIYKKGYQTFDQVQSSVTSKVKGVAFTNLSHSDPAI
GTRIWDTADYVVPAEENDATFVITNMIITPRQKNGICAETPSNKDAHCTKDSQCPQGVPVKEGHGIRTGKCDLSTKTCE
IYAWCPIENDHLKYPYPVLEDAKDFTLLIKNSIQFPKFKFRKRNIANVSSDFLKSCRYNPNDPMSKLCPIFTIGSIVNATGES
FDSFAYLGGVIAILIQWNCNLDFSDRVCLPEYSFQRLDNPNVTLSPGYNFRFPTYYQVDDLQYRTLTKAYGIRFIFIVFGQA
GKFNIVPLFLNLGSGLALLSIATVLCDIIVLYFIKGRMYYYGHKYEVIEEADDESEPFIQPVEYKGL

>Placozoa_Hoilungia_hongkongensis__1_g05782_t1

MENGCNKFANLVFDYNTPKIIHIKSKFVGFINRLIQLAIVGYIIGYVLVYKKGYQEFDTAQSSATSKVKGVAFVNYTKDPNI
GTRVWDPADYVVPPEENGAFFVMTNMIITKNQTNSVCPEDIYIKGANCTSSADCIPGEHLYLGHGVNTGECVPASPDS
DEMTCEIYSWCPLEYDHLRTKYPFLAEAVNFTVLIKNSIAFPKFKVRRSNIDAVKNSEDLKHCMFDPVHDKLCPIFKLGTII
KSAGEDFNKIAYEGGVVAIILDWDCNLDFDIKNCQPKLTFRRLDDPHAPTAVGYNFRYPIYYEVNYKLHRDLIKAYGIRFV
FIIIGRAGKFSVVPLLLNIGSGLALLAIASVISDVVILYILKKRQYYRSKKYQTVSPFDSDDPLLEDA

>Placozoa_TriAdh_evg1623395__type_protein_aalen_411_18%_partial5_utrbad_clen_6520_strand___
offs_6518_5283___evgclass_maina2_okay_match_evg35589_pct_99_100_

DLSMSGRADASRHNPSCMMEDESSGCFGVLEYNTAKIVHIKSKKIGILYRLVQLIIVGYVIGYAIVYKKGYQIFDQVQSSV
TSKVKGVAFTNFSQSDPSIGTRIWDTADYVVPAEENDATFVITNMIITSNQTNGICSETHKNHDAHCTKDMDCPKGIPI
KDGHGVRTGKCDLSTKTCKIYAWCPIETDQLKYPFPILEDAKDFTLLIKNSIQFPKFQYRKRNIANVSSDFLKSCRYSPNNA
MSKLCPIFTIDSIVNATGESFNPIAYLGGVIAILIRWDCDLDFGDSKCLPEYTFQRLDNPNVTLSPGYNFRFPIYYQIDNTQ
HRTLTKAYGIRFIFIVFGQAGKFNIVPLLLNLGSGLALLSIATVLCDIIVLYFIRGRMYYYGHKYEVIEEADEESESFIQPVENK
RL

>Placozoa_TriAdh_evg374767__type_protein_aalen_393_54%_complete_utrpoor_clen_2177_strand__
_offs_2057_876___evgclass_maina2_okay_match_evg591226_pct_99_100_

MENCCNRFADLALDYNTPKIIHIKSKVVGFINRFIQLAIVGYIIGYVLVYKKGYQEFDTAQNSVTSKVKGVAFVNYTKDPNI
GTRVWDPADYIIPPEENGAFFVMTNMIITKNQTNTVCPEDMSIRGANCTDSTDCIPGKHLYLGHGVNTGECVPVSPDS
DEMTCEIYSWCPLEYDHLRTNYPFLGEAANFTVLIKNSIAFPKFNVRRSNIDAVKNSEDLKHCMYDPVHDALCPILKLGTI

VNSAEQDFNKIAYKGGVIAIILNWDCNLDFDIRNCQPKLTFRRLDDPHAPTAVGYNFRYPIYYEVNNKLHRDLIKAYGIRF
VFIIVGRAGRFSVVPLLLNIGSGLALLAIASVISDVVILYVLKRRQYYRSKKYQTVSPFDDDSPLLQEDVE

>Protos_Platyhelmin_Macrostomum_lignano_A0A267G6J9_A0A267G6J9_9PLAT_ATP_receptor__Fragment__OS_Macrostomum_lignano_OX_282301_GN_BOX15_Mlig018619g3_PE_3_SV_1

NVLDYYNSSQLQNCVYNPDTDMYCPVFRIGDILKLAGIDNFTKIATVGGVVSITVNWDCNLDWDASYCNPTYRFRRLD
DENTKIAKGWNFRYANYYRINDTDHRTLIKAYGLRFVVYVTGRAGRFNVIPLTMNLGSGLALLVLRRSCAT

>Protos_Platyhelmin_Macrostomum_lignano_A0A267EI98_A0A267EI98_9PLAT_ATP_receptor_OS_Macrostomum_lignano_OX_282301_GN_BOX15_Mlig018619g1_PE_3_SV_1

MTSNARSNIAVKTVDRGFGALFRYETSKVVQISSIKVSVIYRLIQLLIIAYVIGWVVVYKKGYQDFDDVVSGVTTKVKGVQ
YTNLTGTSIGARIWDVADYVVPPQENNAFFITTNAVVTRDQTQGACDEDPSVAPCFSDADCPAGKPQMLGNGVNTG
RCVPSSQAPNKTSCEILAWCPVEVDQLPAGGTMFPEAVNFTVLIKDSIEFPLFGVKRRNVLDYYNSSQLQNCVYNPDTD
MYCPVFRIGDILKLAGIDNFTKIATVGGVVSITVNWDCNLDWDASYCNPTYRFRRLDDENTKIAKGWNFRYANYYRIND
TDHRTLIKAYGLRFVVYVTGRAGRFNVIPLTMNLGSGLALLGIATVLCDMIVLYLHHKKVIYRGAKYETLEDDDAYRLMN
EEDDGLLSETSQDARQRVRASAQALDPPGGNYGAIE

>Protos_Platyhelmin_Macrostomum_lignano_A0A267GC16_A0A267GC16_9PLAT_ATP_receptor_OS_Macrostomum_lignano_OX_282301_GN_BOX15_Mlig019017g2_PE_3_SV_1

MCSLRSFFSRMLAGFFEYRLQKVVVIRSVKFGLFYRSVQLSVIVYFLFYIIWHKKGYQVMETPEGGVVFKVKGLDFVNRS
GDPDYGARVWDSADYIIPPIMNGAIFITTNARVTDKQAMSECPESVGHCHQDGDCPAGSISSQGRRTGRCSPDTRTCI
SYGWCPAPSQDAFNLPVGSVLTRARNFTVQLKNSVDFRAFDKKRRNILDWMTSEYLNTCLYDKAHPNLTYCPVFRLGD
ILAYSGTTSEEVYKIGGAFGIDISWQCNFDLPEEHCLPKYHFRELDIGKGTDIEGFSVQYAYNYRSEGQDRRMLVHAYGIK
IFVTVSGEGGRFDFYRLSMNLGSGLALFGLSSMLCDVILLRIASQKQRASLQRARVDYCDVGRRRRRSAGALRRKTRETG
TSRKKKTKMSSRRTTRQRRRSWRWSTSRAGPARMSTRRAPGRRRAAPAADGATGSRGGVDIVSDANVTVS

>Protos_Platyhelmin_Macrostomum_lignano_A0A267EZC4_A0A267EZC4_9PLAT_ATP_receptor_OS_Macrostomum_lignano_OX_282301_GN_BOX15_Mlig014818g1_PE_3_SV_1

MASSVKKFLSATLFEYETQKVIAISSYKIGLIYRGIQMLIVTYVIGWVMLREKGYQSFETVVSGVTTKLKGLEYTNFTEIPSIG
DRIWDVADWAVPPQMNSAFFVTTNVVITKNQSQTTCEEDKDNKWHAKCSSDSDCIKGHVHGLGWGVRTGRCLNST
REEGLRICEIYGWCPTEQDVLPLGEGKILMTRVQNFTVLIKNSVEFPKFGVKRRNILSNMGKNYLRGCNYRPDHEMDKY
CPIFRLGDIIKYGQAISPETAKQVWITGGMVAIDIHWECNFDYDEEECKPDYKFRNLEDGHVGRGWNFRHAYHYVEDG
VHRRTLVKAYGIQFFISVTGRGGKFDVLNFTMNLGSGLALLTLATVLCDIIVLNIMRKRKKYRQAKIDKVKRKQIERARSA
RGGGAEDENQSAAAAAAERPGGADADADDADLESAPSFRLHVGARIPGNGAANFGSLVGGVSSFTDEPPRAFVPRD
GQSPIPYMQPSGSEAASESGRERGWRAGSGELAGELSD

>Protos_Platyhelmin_Macrostomum_lignano_A0A267G670_A0A267G670_9PLAT_ATP_receptor__Fragment__OS_Macrostomum_lignano_OX_282301_GN_BOX15_Mlig019017g3_PE_3_SV_1

RLAGSQSPARMCSLRSFFSRMLAGFFEYRLQKVVVIRSVKFGLFYRSVQLSVIVYFLFYIIWHKKGYQVIETPEGGVVFKV
KGLDFVNRSGDPDYGARVWDSADYIIPPIMNGAIFITTNARVTDKQAMSECPESVGHCHQDGDCLAGSISSQGRRTGR
CSPDTRTCISYGWCPAPSQDAFNLPVGSVLTRARNFTVQLKNSVDFRAFDKKRRNILDWMTSEYLNTCLYDKAHPNLTY
CPVFRLGDILAYSGTTSEEVYKIGGAFGIDISWQCNFDLPEEHCLPKYHFRELDIGKGTDIEGFSVQYAYNYRSEGQDRRM
LVHAYGIKIFVTVSGEGGRFDFYRLSMNLGSGLALFGLSSMLCDVILLRIASQKQRASLQRARVDYCDVGRRRRRSAGGT
KKEDERDGNESKEEDQDEFEEDDEAAPEELEMEYQSGRSSEDVDEESAGEEARSASRRRCHREQRRRRHRQRCQRHR

QLNQGSSPVLRQWQQQQQPERSRTPPSPLLPQFAAAGVASKALPYPPPPPPLCLQEPLLARQPPRPHLLADQRQPPPP
YSTGPARLVVGCNGCERCSNSGRRARRLGPRYAESSFCGDDFNDGATTGSGDAGVADFGAYRPPRIVRRYYGDSSQSP
IARETLI

>Protos_Platyhelmin_Macrostomum_lignano_A0A267DPT3_A0A267DPT3_9PLAT_ATP_receptor__Fragment__OS_Macrostomum_lignano_OX_282301_GN_BOX15_Mlig027460g2_PE_3_SV_1

VYKKGYQEFDNAVSGVTSKVKGVQYTNFTGSSIGARIWDVADYVIPPQENNAFFIMTNAVVTADQRQGRCDEDPLVA
PCSTDSDCPAGLAQMLGSGVNTGRCVNSTQRPGVRACEVLAWCPVEVDWLPAASLLPNVANFTVLIKNSIEFPKFKVK
RRNIASGMPEEDYLRFCRWRPDHPLLKFCPIFTVRDMLRHAKVSDNIFIKGGVMAMNIHWDCNLDWAESHCTPEYSF
TRLDDPAAGISKGWNFRYANYYRQNGSDYRVLIKAYGLRFIVYVTGRAGRFNIIPLTMNLGSGLALLGLATILCDVIVLHL
LRKRSAYREAKFQQMEDEDAYKAMNDEEAEESELAAGSAGAAAAAVGSSAKPDYGPLGVTGHGLSVCLSVCLSVCLSV
CLSVCLSVCLSVCLSVCLSVCLSVCLSVCLCLSVCLSVCLSVCLSVCLSVGLSFCLSVCLSVCLSVFLCAPGHSSSQYILTGWR
YLLKILAT

>Protos_Platyhelmin_schman_Q65A65_Q65A65_SCHMA_ATP_receptor_OS_Schistosoma_mansoni_OX_6183_GN_schP2X_PE_2_SV_1

MVKGIAVLFEYETPKLVQISNIKIGVTQRLLQLVILIYVVCWVMIYEKGYQENDIAKSAVTTKVKGVGFTNFSHIPGIGMR
SWDVADYIVPPLENNALFVITNLVKTERQSLSKCQESSWVPEAACYKDSDCKPYFISHLGNGAHTGKCIIKPGNDIGSCEI
YSWCPLENDTLPLGRKSFLFPMVYNYTLLIKNDINFEKFGIHRRNIQNWASKKFLRTCLYNKTDPENRFCPIFQFGTIFEEA
NVDQSIFISGGVIGIDIDWKCDLDWDVQYCNPTYSFRRLDDAHAKIASGFNFRYAHFYSENGTNYRDLIKAYGIRFVIHV
SGEAGKFHLLPLTMNIGSGLALLGLAPTVCDIIALNLLRSRDIYQRAKFETIAEEQAHLSSRRADKAQRKKYGLSKRNTETI
NENSDENGKIIQWKDDQEFTVSDTQTIHSST

>Protos_Platyhelmin_schman_A0A5K4F386_A0A5K4F386_SCHMA_ATP_receptor_OS_Schistosoma_mansoni_OX_6183_PE_3_SV_1

MGFLRSLAEGLLEYETQKVVQIHSKKIGVTFRLIQLGIILYVVLWVMIYEKGYQSFDQAVSGVTAKLKGIAFANVTNNPSL
GATVWDAADYVIPPQQNSAFFVMTNLIYTPGQTLGNCEESHDVFGNSCHNDSQCHSGQLVQRGSGIQTGRCVNSTR
QSNLRVCEIYGWCPTEHDVIPRPHLLASAENFTVILKNSIEFPRYRIKRRNILKWMSNLFLKTCLYDPNDDELKYCPIFRLG
DILKYSDPVNKDIWETGGMVSIHIEWNCNLDFDEEKCLPKYIFRRLDDFQSNVAKGWNFRFSQHYIDGGVRKRNLIKAY
GIQFFITVYGTGGKFNILTFSMNLGSGLALLGIATVLCDMIILNTTRKRGVYRKAKIDYIAEKRRKQQEEKLKSKNKENDFD
TMQIGDNKKFEWKLHPTNLHPDESNTDKEQIYGQNSLLICGPVSHFMWMDNNKCQKRVIDKSHLSTSKNEQLCLNQP
SFLGFHYTPSTSSSSSQVVGIKKQSKYENHSESMLPNHLNGCFTSIPYRNKNNISLYEKNTIPMNDTTYSINKHSKYVQCN
LKSTNMLTFTKESNLNTTYNEMELFKTISTLPKCCTSFIKPTIHS

>Protos_Platyhelmin_schman_A0A5K4F3P5_A0A5K4F3P5_SCHMA_ATP_receptor_OS_Schistosoma_mansoni_OX_6183_PE_3_SV_1

MSQTNGLFDILFKGLVDYETQKVVQIHSKKIGVTFRLIQLGIILYVVLWVMIYEKGYQSFDQAVSGVTAKLKGVAFANVT
NNPSLGATVWDAADYVIPPQQNSAFFVMTNLIYTPGQTLGNCEESHDVFGNSCHNDSQCHSGQLVQRGSGIQTGRC
VNSTRQSNLRVCEIYGWCPTEHDVIPRPHLLASAENFTVMIKNNIEFPHFRVKRRNILNWMSRLFLKTCLYNPNDDKFK
YCPVFRLGDILKYSDPVNKDIWETGGMVSIHIEWNCNLDFDEEKCLPKYIFRRLDDFQSNVAKGWNFRFSQHYIDGGVR
KRNLIKAYGIQFFITVYGTGGKFNILTFSMNLGSGLALLGIATMLCDLIVLNITEKRDIYRKAKIDLIAEKINIHQSSEKFNIHD
NTYPVVKVTKSDKKRHLIIRRHHEFVRTPDSIRRSSFQSIKQFINERSPLSFRKLNVTPINFIPKIKSKQNHISLIQSNLLSKAQ
IPGENLIPINHHAQSRQNVTSINNKTDNNISNASISSNNPHSLISDLSDRTTFQKSSFSLNPPSTTFNINTTNKQQYWRQQ
RLLYE

>Protos_Platyhelmin_schman_A0A5K4FD36_A0A5K4FD36_SCHMA_Uncharacterized_protein_OS_Schistosoma_mansoni_OX_6183_PE_4_SV_1

MPRVVHIRKPGLAILHRLIQFGILTYFIIYVMWLNKGYQTFDHPISGVAVKVNGIAWKQIFKYQNHDTNGIIVLDNGDYL
LQPTQNSGFYLITTIKHFVIQRMTTCSESDKLPDALCNHDIDCPIGYQAGYKPYGPDGIVPNENNIIIDESGHGIFTGKCLI
HSKKCQIYGWCPTTDDYEYELYKQRHYKLNLIKPPNILEGYYNYFKDLNFLDNKRQLNERTQHLIDPFFEIINFTIFIKNSIEF
PLFNIKLSNILKWMTNNYLKNCLYNSKHSIDKYCPRFRLYDIFQLIGIHSNRLLKYGGVITITIQWECNLDYNIKYCLPNYKFL
ELESVHKVRKIKTNNITYTKISNNNNYTYKVYESYNNDDDDIIDDNEEVITIHEGSSLRSTAYYGYDTEFIKQRRRILIHVNGI
QFIIRVNGIAGKFNLLSFTMKFGSGLALLSISTIITDLILFYFTYNRKKYKKVICDDKTINYLSN

>Porifera_AmpQue_tr_A0A1X7VID9_A0A1X7VID9_AMPQE_ATP_receptor_OS_Amphimedon_queenslandica__Sponge__OX_400682_GN_100636400_PE_3_SV_1

MSAASACRWLGSTTVSVLFEYDTPKFVHIRNKKVGLLNRLVQLAIVGYIIGYGIVWENGAQDKEPVQSVVTTKVKGVSV
LNATRQFMTDCNEWPILDHVDLIVPAQEPNSFFVTTNLWATCNQKYGVCPELSTIPDAVCLTNDSCPPGIIYRTGNGVT
TGECTGFNETCQIRGWCPVEDENDTLTDEGPIIDTKDFTVLVKNSISFPKLDSSYRVRNIPERANNKSYLSGCQFDPDSE
MGLFCPIFSLKQIVDMINSSDGYYESLATKGAVVGLEISWDCNLDRSPTHCVPKYSARRLDNPNAQISNGFNFRYPRYYR
DQNGDDVRDVWKVYGIKFEIVVTGEGRKFSFTTLVLALGSTIALLAIATTVTDVIALYLLKKGTYYREKKYQKVAKEDEESV
TLKALNHSQEKSPLLREKRKE

>Porifera_EpMuel_Em0004g75a

MNLCRSAWGTLASFAFDYDTPKIVRIQNKKVGALYRLFQLGIMGFMIGYSIIYSKGYQDKDNVFGGVTMKLKGVEFYNS
SQEPFLQNCQGSTGAQVYDPADYVVPPQEPNSFFVMTNMWVTCDQSYGFCGEDPSVQGSNCTGPSDCVKQNTIKG
NGPRTGVCNNETHTCEVYAWCPTEKENVSYTDMGPKINAEGFTVLIKNSIQFPNLAPNTLRKNYISNYDLSKCGIWSHT
TITSSDLPAGDIVAMIDPPQNFTELALREQIGVVLDDCFGTCDLDSVWL

>Porifera_EpMuel_Em0004g76a

MRPKEKCEWAWSVFIAATFEYDTPKIVHIRNKKVGVLYRLFQLCIMGFMIGYSIIYSKGYQDTQTVIGAVTTKLKGVQYY
NVTGGVPFLQNCVATNEPLVLDPADYVIPPEQPSSFFVMTNMWVTCNQTYGTCVEQDPSLASKCSGPADCTPQSTIK
GDGPRTGTCNNQTHTCDVYAWCPTEYENTNFTNMGPKINAESFTVLIKNTIQFPNLDPNFFKKNYISNGDLANCTNSLI
GNNYCPIFKLADIVNMISEEPKNFSNLATKGAVVVLTITWNCDLDMSLDHCNPRYSARRLDTTPLDTASNSGGFNFRFA
KYWYDSNGVQYRTLIKAYGISFVVELVGQGGKFSFTALIFKLGSTIALLGIAAVLSDMVVLYVVQKRHLYHQAKYQVVND
EDDSVAGDDSHYHSVHPLLNK

>Porifera_EpMuel_Em0004g80a

MTNMWVTCNQTYGTCVEQDPSLASKCSGPADCTPQSTIKGDGVLCLGTGMLDVWSLRCNMGPKINAENFTVLIKNT
IQFPNLDSNFFKKNYISNGDLANCTNSLIGNNYCPIFKLADIVNMISQEPKNFSNLATKGAVVVLMITWNCDLDMSLDH
CNPRYSARSHMGRINHYISNLFK

>Porifera_EpMuel_m_22014_g_22014__ORF_g_22014_m_22014_type_5prime_partial_len_370___comp51346_c1_seq1_3_1112_

NSSRRVIYSRSKRGIISAMLNYVTYKYVVLKDFRLGILYYVLVASIVLYTLTEIFFNKGYLHFDTRPQGTIKIVISDNLSNETFR
PASLPYCQGNAACKFIDPMDLSWPVESRAITIATYIKDRDQLLEEDTNSPEDEYVLKREQDYFTQGAEYIVMKVDHAVV
AHDSSGRERLAASQRKLKGYLVDLNNRPVVYLSPGEGKPDKMSLQQLLQAAGVHSLDEQSDAVSARGRSFRERGCVLR

VTIFYENWYSTWFGTSDIHYTYHVRRVPFMDYSVREVQSPGHRDSTRREEGQERLFRKRYAVRVEFIQAGSIGMFSLTT
LLFQLVSGVGLLTLAVTIIDTLTLYILPARHMYRQYVYDESPNIKKLE

>Porifera_EpMuel_m_190834_g_190834__ORF_g_190834_m_190834_type_complete_len_609___com
p67432_c1_seq1_118_1944_

MMAQKDTSTLLELEMEAEATAGALAGSNETKKTDAGAEQTWKAFVNNSVDSFNAVSVAPHSMTVKNYGIGILSSGM
KLIVLGISLYALFGFFEYQEQYPVQGIAVSSLKGVSVYVLNGSEPFLQNCETNDMNRLVFDTKEFGIIDDYDSFFVTTNM
WLTCNQSVGICGEDERSVPNGQAYCLNNTNCTDGKLLRNGNGPQTGTCNLQTKTCNVLAWCPVEAERKESNLKPKID
AANFTVLVKNTVSFPNSVSNYARSNIYGGDEHWLKNCTYDKDLSPGCPIFTLGQIVSMISPTENFTELSITGAAIAFNIEW
NCDLDETWLQPTCTPVYSVAREDRQNGVSAGYHFRSTRNRYETNGTQYRTLVKAYGIRVIVRVNGLGGKASLIAVARW
LGTSLAYFTSATGFLTLLIVYGYLNYYIQVQELIKSSKIYNCCGCKGKIGNSILYGLISGFCCCCILLPIICDCGKDKSESNNPED
EKKDENKDENKDYPTERILKLLAGAINVIWILVLVIFMVIPVLTMFMIVVLASILSLVVLCCLRYYKVNTTKRLTKVLNTVLS
YEIHDTHAAAILKRKEGIQKYRDAKEKEKGCAPNHVVITFDDAPDEN

>Porifera_EpMuel_m_233174_g_233174__ORF_g_233174_m_233174_type_complete_len_411___com
p68168_c0_seq2_35_1267_

MGPKEKCEWAWSVFVAATFEYDTPKIVHIQNKKVGVLYRLFQLCIMGFMIGYSIIYSKGYQDKDNVFGEVTMKLKGVE
FYNSSQEPFLQNCQGSTGAQVYDPADYVVPPQEPNSFFVMTNMWVTCDQSYGFCGEDPSVQGSNCTGPSDCVKQN
TIKGNGPRTGVCNNETHTCEVYAWCPTEKENVSYTDMGPKINAEGFTVLIKNSIQFPNLAPNTLRKNYISNYDLSKCGI
WSHDNYYCPIFQLEDIVAMIDPPQNFTELATQGAVVVLTIAWNCDLDRSGCDPVYDARRLDTSAGDSTSGGFNFRFAK
YWYEQGIQYRTLIKAYGILFVVELVGQGGKFSITTLVLKLGSTVALLGIAAVLSDVVVLYVVRKHDMYYGRKYLMVKEYD
QEQGEGGEEEKVPFLQHEVQ

>Porifera_OscCar_comp8618_c0_seq2_p1__comp8618_c0__comp8618_c0_seq2_p1__ORF__type_5pri
me_partial__len_413__score_64_89__comp8618_c0_seq2_3_1241_

GRSAVFSMGSACRRCFSATSGALLEYDTSKLVHIRNQKIGLIYRLVQLVVIVYIIGYVIVIQKGYQETDTVVSSVTTKLKGIA
FVEDEPGAGEPAVWDVADYVVPPEQTDAVFVTTNAIITTNQSYGVCNEDENVVDGSCESDSDCPAGVPVRNGNGVR
NGTCDLDVKTCFISAWCPTENDTNRLSPLLGDYPLNFTLLIKNNIQFQKFNFTKSNILASSDRAFLQNCTYDSEDPERRFC
PIFRLGHMFEEAAEDYSLMALKGGVMGVFIKWDCNLDNSATDCKPEYKFKRLDNAAAPIAPGYSFRYPIYYMVDNVM
HRQLIKAYGIRFQILVTGTGGKFSVVPLFLNIGSGIALLGISTLICDLLVLYVVKRRKYYKDNKFQMIEDEEALFSEKSKMSS
DPENLQYSKISD

>Porifera_Syc_Cil_scpid72722_scgid16433_P2X__purinoceptor__4_ATP__receptor_Purinergic__recepto
r

MSGVCGKIGSTLFEYETLKVVHIRSKKVGFINRLIQLVVICYVIGYAIVYEKGYQDTDLVTSGVTAKLKGVAYTDLNSDPAI
GERIWDMADYHIPPQQSSAFYVMTNMHVTPFQKNGTCAEKPSIKSCTNGNDCAAGELVPNGDGRMTGNCVQYNN
TEPVGRMSCQIMGWCPVEDEDPTKSRGAVLEQDARNFTVFIKNSIEFKKFTYKARNINDDVNNRTYLEKCNYHPDTDA
FCPIFRLEDIFRYAGILEQWPQATKDGAVMAIDILWNCNLDSLQFKCAPEYKFRRVDNPAVKLSPGFNFRFGKYYKLQED
GQVVQYRDLTKSYGVLFLINVEGEAGKFSPVPLFINLGTGLALLSVATVLCDFITLNLLKKRTFYREKKYEDVDVDDGFTILK
RSHDDSTSIG

>Porifera_Syc_Cil_scpid13851_scgid33070_P2X__purinoceptor__4_ATP__receptor_Purinergic__recepto
r

MGFAATISSVLLEYDTLKVVHIKSKKVGIINRLIQVVIIAYIIGYVIVYKKGYQDTEEVEGAVTAKLKGAAYPLNAGLASRVW
DAADYHIPPQQTGAFFVMTNMIITPDQRNMTCPESLKNAKCYNASDCPSGQMVPAGNGLMTGRCIPYDSYDSDTRH
DVYEEDFTNGSAVNKTCEIIGWCPLEKDVTPSPARLGNESMEFTVLIKNSIIFRKFDVRRRNIPSTSTEKWLKKCRYSPDH
PQDKYCPIIKLYNIFKYAGVADQWDDAAKNGAVISIDINWSCNLDHDEDECNPKYTFRRLDNPDVALSPGFNFRYSQYH
SDVNGEKTRTLIKAYGILFDIKISGEAGKFSPVPLLVNLGSGLALLSVATIMCDFVVLHFLKKRQFYHEKKFEDVEVQDDLL
PEGLLSPSERDSLIQKH

>Protost_ecdy_cyclo_PriCa_ecdy_XP_014678562_1__PREDICTED_P2X__purinoceptor__4_like_Priapulu
s__caudatus_

MGVLSYARRGVAVFFEYDTPRIVHIRSKKVGVINRIIQFLIVAYIVGYVIIYSKGYQKFGGIDSAVTTKVKGTIYTNVSGLDP
LYNNRIWDVGDYIQPAQENGAFFVMTNVIITPNQTQGECAEDPAIMPCDTADNCTAGEQTPNGNGVMTGECVSYPP
EPTLRSCEIHAWCPVELDELPLEEALLKNSSVFTVLLKNSVEFPQFGIKRRNILNTMDSEFLHKCRYDPNNATAKLCPVFEL
GVLASSAGQDYAKMAIKGGVMEIVITWDCNFDRDVHECISAIQLLPSRRPNDELAKGYNFRYAHYYYTGDVQTRVLYK
AYGIRFMISVTGQAGKFNIIPLATNIGAGLALLGIATICCDVVVLYLLNKRDFYYEKKYLTVKDQDAFLLGEEDNGPPDAGS
TR

>Protostome_Lophotroco_RotSoc__A0A817XFJ5_A0A817XFJ5_9BILA_Hypothetical_protein_OS_Rotaria
_socialis_OX_392032_GN_FME351_LOCUS6022_PE_4_SV_1

MTFQIKDIFHSLLFDVFLTYSTVKLVKVPHTYIAVLNRALQAVIFAYIIGYIVLLKQGYQQFQEPRGTSVIKVKGIAKVTGNN
ASFYTNDQTKYIWDTPEYENPAIENNAFFIATHQTVTPGQTQQTCPTALADKLFCNDTATDKCKTDQPTPNTFGYFTGK
CVVSRENSTLKVCEMNGWCPEELSASMDYIMDGDDLENFTIFLKTMVTFTLFKKNLRNIQDNTDFSCRFDGTPRTADC
PIIRVGYILDQLNTNKTALLLAGGLIEIRQDWTCNFDRHPKRCIPKYEFSLLQSGDDKQSPGINYRTAQKYRVNDTDYRTL
SKVYGLRFVVAITGKGGQFNIVNLFIAIGSGIGFMVIAGIVCDAILMYIHKSREKYKQGKFDVCEVDDEDPTATQNVTEA

>Protostome_Lophotroco_RotSoc__A0A821JA93_A0A821JA93_9BILA_Hypothetical_protein_OS_Rotaria
_socialis_OX_392032_GN_FME351_LOCUS14379_PE_4_SV_1

MQNRRDGVLMVKPNLPLKPQENFTFFKFMLSNVTCYETEKVVRFEYKWIGLLSLIVKLTIAISCLYLMFQRDSYQIFDRSP
ISAVTIKVKQSNICSNNDSIIRYALQYNCSNSLYDVNDFVKPATENGAVSITTRMIEIKHILQPCNTKNVKKNQTLYNASIH
QCDHQARCILPPFYRSQNGTMLNKQTLCWFEALPRFERLNYQALDYVIFIKHFVEFAQLGVVRHNLLPNLLTSDYLRTCE
YDPYEHPFCPKFRILKILQIIEKNSSEYKSIFFHGSLIEIRIAWKCNLDRNLKYCEPTYEFQRLDIVPYSKNPYDSGSNFLTSHY
FFQPNSREVYRMHTHIYNLHIIISVSGEAGRFDLFQTTTSIGSFLGIFGTGSIVCDFIAAFVINFKKVKYDD

>Protostome_Lophotroco_RotSoc__A0A818KSR3_A0A818KSR3_9BILA_Hypothetical_protein_OS_Rotari
a_socialis_OX_392032_GN_FME351_LOCUS19776_PE_4_SV_1

MRLTSFVKDRNSIGKSVTNVLTGFFSEYETQKLVLIHSVKYVGLRRIIQIIVLIYFVIYFLIYEKGYQKQSTAIVSSVALKVKGIG
YVRTPENKTIIMDVADYIIPASENNAIFVMTTFIETDQRRSTCAESSQLKQAKCRSNSDCVTKSFIAGMNGLWTGRCLFT
SKPTVIIGAENTTNSRTGLCEYEGWCPPEDDAVSPMYIRGPLGFRIFIKNFIEFPAFGVKHKNMVADLKPCVFHPTDDKD
CPIFPLDYIVNQAENDSNERDLMLQYGGVIGVKLNWICNLDRNIKMCKPQYSFARLDVPYREKPFSVGFNFRYASTWK
HERGQFRTLTKVYGLRFIIATSGKAGKFDFITLSLNIGSLVGIFGLATFVCDIIILNLSKRAKLYRKHICETVHFRTRFSSAIPLM
KMDSKGSSSNRLNPRRKDETTLLNELITDSTKIREKTVGMRSLIVNDMNHLHCSPLV

>Protostome_Lophotroco_RotSoc__A0A818UWD6_A0A818UWD6_9BILA_Hypothetical_protein_OS_Rot
aria_socialis_OX_392032_GN_FME351_LOCUS27929_PE_4_SV_1

MATRDSSAKLHCEPLREYCSFSNVKSVVRGFVYEYETPKIVTIHSISIALMCRLIQLLILIYGVIYLMIHKKGYQETDTSIISSIT
LKVKGIGYNQTGQNKTLVIDGADYIVPPQENNALFLMTNFIRTDQEHGKCPESPDEKMSSCTDNDYCLANQNFQKAN
GKWTGICLIKNNSFVNASETRQGRCELEGWCPVENDRNIPFPIRDALNFTIFVKNFIEFPRFKVIRKNIQEDAAYLKRCHY
DQGEHKGCPIFRVGTLLDIVESDPFEQLKMLQLGGVIRLKIHWNCNLDKSLNLCKPEYSFGRLDGPYKEQSFSHGFNFRF
ASHWKHHNRSYRTLTKAYGLRFIISVSGLAGRFDVITLTLNIGSLIGILGLATFICDIVALYVHRQSDVYRQQKFQDVDLHF
QQIESLKTLPTDSSTEQTRRRYLNHIDDTVLNNHTADNGDATQNVETASKTEFQLEEVNQRSLYDNLQPTPKQISQQST
PQGAFINIQRPKSGSTSSNGSSLLSSIKQKYATRRSQLPHLSSEENQLYDKSNIFFVDEDTPELNERNELRGRSDENKQLSN
SSSTNHGLSVRPTSTVETLL

>Protostome_Lophotroco_RotSoc__A0A821EE34_A0A821EE34_9BILA_Hypothetical_protein_OS_Rotari
a_socialis_OX_392032_GN_FME351_LOCUS29067_PE_4_SV_1

MDAVPSSDRRCKVSVFFKELFYGFVTEYETPKIVAIHSYTMTILLRILQIVLLLYSIFYLFLNEKGYQKQDSSIISSVTLKVKGI
GHSRTANNETFTFDGSDYIVPPSENNAIFIMTNFIQTDQVRTKCRESPSVSEAICTKHSDCQNKPYNPNINGRWTGRCI
MRPKVNQFNAENTTKNQTTGLCELQGWCPVQDDRQVATFIEEVGNFTIFIKNFIEFPAFHVKHKNMVEHLKPCIFHPV
EEKDCPIFRLDYILNEAEKDDHERGLMLRLGGVIRVKIDWDCNLDRSIKLCKPVYSFARLDTKFQEEKFSLGFNFRFSSNW
KYNRRYLRALTKAYGLRLIICVSGKAGKFDFITLTLNTGSLIGIFGMATFFCDIILLNLTKKASLYRDFVYQRVNIDASRSNSV
SQRIQNRNENVGAIFENPLTNEEKFSLPIWTSETFVSFADKRCTTSTYDFVNSTDSSRRQPSLGLTNILK

>Protostome_Lophotroco_RotSoc__A0A820SYJ6_A0A820SYJ6_9BILA_Hypothetical_protein_OS_Rotaria
_socialis_OX_392032_GN_FME351_LOCUS13480_PE_4_SV_1

MSETTCAKIKRHCQPFTHAFGLTHMKSALISMVTEYETPKIITIHSYSIALTCRFVQLVIVFYAFAYLMWYKRGYQDRDPS
LISSVTLKVTGIGRFDKNNTITLDNADFIIPPQENNALFIMTNFIRTDQQRKRCIEGFDIRDAACANDTHCQALGPYPPKS
NGRWTGKCRQPEGRCEIEGWCPVENDNDKPEPMMGSLNFTIFVKNFVEFTRFGIARTNIFHDSVAITQCLYHPVHDQL
CPVFRVGDLLNIVEPDTDERHKMLISGGVVRIKIDWMCNLDLGNEKCKPIYSFGRLDSRSRDEQFSFGFNFRYASHWKV
KNRSHRTLTKAFGLRLIVTVNGEAGRFDFLVLTLNIGSMIGVLGLATFICDIVALYFCKQGSIYRKQKFQTVKLASTASLSTI
QKHLTLNDELDKNNQPRNKRMNSHLPAGDEC

>Protostome_Lophotroco_RotSoc__A0A818ESC7_A0A818ESC7_9BILA_Hypothetical_protein_OS_Rotari
a_socialis_OX_392032_GN_FME351_LOCUS14282_PE_4_SV_1

MAIKMNSHPVSFSSIITKYTPTTRHRVSSPLKHIWTSVTHYDTPKVVRFESKWIASLSLFVKLSLAISCLYLMCQRSSYQIFD
RSPISAVTIKIKPSPYCLRNASILRYFPKYNCSQSSYDVNDFITPATENSAISITTRMLETEHVLRYCTDKTSRRRQNRSSSFR
LVSLHTCNERARCILPPLYRQEYENETIANKSKLCWFKLSTITVKRNYQALDYILFIKHFVEFTQLGLVRANLIPNKITKEYID
TCEYDPIHHPLCPKFRLSKILEMIETDPHEYEAMFYYGSLIEIKISWKCNLDKKLEYCQPFYKFQRMDIKPYKSNPYDPGSN
FLTSRYFFRPNDGELHRLHTNIYNLHILVSVTGEVGKFDLFQTTTSIGSFLGIFGTGTIVCDFIAAFFTNFKTVKYDP

>Protostome_Lophotroco_RotSoc__A0A820CYY0_A0A820CYY0_9BILA_Hypothetical_protein_OS_Rotari
a_socialis_OX_392032_GN_FME351_LOCUS24632_PE_4_SV_1

MSGEAPVITVQCLSLREYFSASNMKSIFRSFVYEYETPKIVTIHSISIALMCRLIQVLILIYGIAYLMIHKKGYQETNTSIISSITL
KVKGIGYNQTGPNETLVIDGADYIVPPQENNALFLMTNFIRTDQEHRRCAESPDEKESACVNHSQCQLNKEFRKANGK
WTGRCLINPKTVQNKSLSRIGRCELEGWCPVENDRDIPNPIRDALNFTIFVKNFIEFPRFKVIRKNIQQNASYLKKCNYDQ
ANHKICPIFRVSTILDIVEPDPVEQMNMLKYGGVIRVKIDWKCNLDKLLEHCLPEYTFGRLDGPYKEETFSQGFNFRFASH
WKHQGQSFRTLTKAFGLRFIISVSGHAGRFDVITLSLNIGSLIGILGLATFICDIVALYVHRQSDVYRQQKFQDVDLNLLRL
ETLNSLTEGMINEVAKRKQQALLVGHVVENNDAISITDSPHKEKFRRNKEQRPHDLHDQLTTSSPSVVHRTPAVSSELLC

NDYHQRPMAFANSLTNSPSPSSVKHRRSITTRSPQLTSLPKGNHEQIDMIDSINTDFTDEDSPIVSGHKQLNEFRSNLNIKSSSLTTTGQENLKRPPPKLETFL

>Protostome_Lophotroco_RotSoc__A0A818Y2B5_A0A818Y2B5_9BILA_Hypothetical_protein_OS_Rotaria_socialis_OX_392032_GN_FME351_LOCUS6006_PE_4_SV_1

MNSIPFSKNNARIRRNAINGLTGFFTEYETSKAVLIHSVKYAGLLRIIQIIILIYSVAYLLIYQKGYQKQSTAIISSVTLKVKGIGYVRTPENKTIIMDVADYIIPASENNAIFVMTTFIETDQRRSTCAESSQLKQAKCRSNSDCVTELFIADMNGLWTGRCLFTSKPTVIIGAENTTNSRTGLCEYEGWCPPEDDAVSPMFIRGALDFRIFIKNFIEFPVFGVKHKNMVADLKPCVFHPTDDKDCPIFPLDYIVNQAENDSNERDLMLQYGGVIGVKLNWICNLDRNIKMCKPQYSFARLDVPYREKPFSVGFNFRYASTWKHERGQFRTLTKVYGLRFIIATSGKAGKFDFITLSLNIGSLVGIFGLATFVCDIIILNLSKKADMYRMHMIEKIHVRNPYGNQQQASDNSSEQNDEIHSNTNSNIKTELEYGTPGNSIASLEKVVTPRSNSLTISPLS

>Protostome_Lophotroco_RotSoc__A0A820S223_A0A820S223_9BILA_Hypothetical_protein_OS_Rotaria_socialis_OX_392032_GN_FME351_LOCUS1308_PE_4_SV_1

MGSSMKRFFSSALTDTFYTYSTVKIVQVPHTIIASLYRLLQFAIVVYIACYIVWYKKGYQEFQEPRATSIIKVKGLTKVFGNATSGYGNSSSQHLWDTAEYQVPPMENNAFFIATRQIITFDQEEGVCPSALDDKLFCNSTQLDACPKDKQTPNTFGYFTGKCVASLEDYNITVCEIHAWCPEELSNSTEYIVDSNDLANFTVYIKTAVTFTAFNINLRNVRDSTNFSCRYDKTNDRRCPIFRIGDILDSLKTNKTALLHEGGLIEIRQDWTCNFDFDKNSCFPTVAFSVLQSGDDKQSPGINYRAAQKYRIDGVDRRTLSKIYGLRFVVSITGTGGQFNIVNLFVAIGSGIGFMIIADIVCDAIFMYIHKSRERYREGKFSVCEVSATQAYQDSDSSVRLRTI

>Proto_ecdy_PANarthopod_tard_tardig_Hduj_A0A1W0WA52_A0A1W0WA52_HYPDU_ATP_receptor_OS_Hypsibius_dujardini_OX_232323_GN_BV898_13640_PE_3_SV_1

MQLQSVGKNILGRLTDSAESLKAFLFNYETPRVITVHNYRVGLIYRCVQLCIVTYLLGYVFMWEKSYQEFASVESSVTTKVKGILQGTVIGQKNRKDVWDAADFVFPAQETNAFFIATTVIHTVNQTQSECAEDPNVSRNGIICNGRRDCARGDVAVNGVKTGRCVPWNSTVKTCEISAWCPLEVDEPPSAPLLSSYESLTVLVKNHVAFPKFGVKRRNILDSINQTYLETCRFHPSNHPFCPVFSIKDIVKLAGEDFMETATRGAIIGFGIQWNCDLDFNIDYCRPVYAFRRLDDKDDQIARGWNFRYSRRSVDHNGKDRRDLIKLWGIRFVFNVSGRAGRFSAVNFLLTIGSGIGLLGLASLVCEFLLLTCVRPPHRNDLIQQKFCKV

>Proto_ecdy_PANarthopod_tard_tardig_Hduj_A0A1W0WGF5_A0A1W0WGF5_HYPDU_ATP_receptor_OS_Hypsibius_dujardini_OX_232323_GN_BV898_11521_PE_3_SV_1

MCSMTLSSFHVESSSSDGKPSASQFRALGAAARNAFFKYDTPRLVSVHSVSVGLLNRVVQISIIAYVVGFALIYEKGYQQTDTLESTVTTKVKGLVVASGSGHEWISRVWDTADYVVPPLENGAFFVTTNAVVTPNQQHGTCPESPIIANGKCLNHTECPVDVPVISGNGIRTGICNNLTSTCDIFGWCPTEVDILPEEPVLKGTENLTVLVKNYIVFPRFHTKRQSISLTANSSRYFATCLYHPETDPLCPVFRLGDIVSFSGERYEELVHLGAIIGIIIEWNCNLDYDIALCLPKYTFRRIDDRLDETSKGWNFRYADYWTDAAGEERRTLMKVYGIRFVFYVSGQAGKFSIVPFLTNIGAGLALLHVVTFFCDFFLLHVVKFKQQYSDHKFETVSGYPTVKTTDSVTIAADLAMK

>Proto_ecdy_PANarthopod_tard_tardig_Hduj_A0A1W0WB09_A0A1W0WB09_HYPDU_ATP_receptor_OS_Hypsibius_dujardini_OX_232323_GN_BV898_13334_PE_3_SV_1

MAPWESAHAVSDKCLQRTGAFLFEYETPRVVTLRSYRVGSIYRIVQLCIIAYLIGYVFEVWDAVDFVVPPQESGAFFVATSILYTGNQSQSTCPEDANVGKKGIHCLVDSECKESEPTPYGNGIKTGRCVDWDANTRTCEIAGWCPAETDETPNFLSGFESLTVLVKHHVAFAKYGVKRRNILETVNQTYLHDCRFHPTTDPLCPIFSISDIVEFAQEDITQIAVRGGIIGFNIDWDCDLDFDVKYCVPQYSFRRLDNKNDKIAKGWNFRHAYYHFDPSGKSRRDLVKLWGIRFVFNVNGRAGKFSAVNFAMNLGSGL

GLLGIASLICEFVLFNCMSNKSELLAQKFTKLSDPRSHRPLTRIPDSPSVARVTAVNHPNQMMFNNGNCYDSQSLYGNY
DVRRPPLVMWDEV

>Proto_ecdy_PANarthopod_tard_tardig_Hduj_A0A1W0WRS8_A0A1W0WRS8_HYPDU_ATP_receptor_O
S_Hypsibius_dujardini_OX_232323_GN_BV898_08022_PE_3_SV_1

MAWGTKVRGGLGRFFEYDTPKIVHIRSLRVGILYRLVQLTIIAYIVGYVIVWKKGYQLTENVESTVTSKVKGVVLTNVSLF
GEDNPTERVWDTADYVVPPQENGAFFVTTNVIVTPNQQQGICPEDVSSHQLLPSCSKDADCPRLSPSANGVRSGTCN
TTTGSCNVYAWCPVERDQEPALPVLNGTEAFTVLIKNQIIFPKFGIRRRNILDSFQTNSTYLASCHYDPVNDPYCPVFVLA
DIIRLAGENYSAIAVKGAVIGFVIEWDCNLDLPVNWCLPKYSFRRIDNANDVSAKGWNFRYANYWTDTESQNHTVQR
RTLVKAYGIRFIFYVYGVAGKFNMVPFFTNLGAGLALLGLAVIFCDIAILYCSKKRNFYKENKYYEVKDDDAFLIEESQQSS
TNRSPSYSDFQ

>Proto_ecdy_PANarthopod_tard_tardig_Hduj_A0A1W0XB51_A0A1W0XB51_HYPDU_ATP_receptor_OS_
Hypsibius_dujardini_OX_232323_GN_BV898_01713_PE_3_SV_1

MTNFTNTNEQEKLSSYAKRKALDFFFEYETPKVVKVYSTRLGAINRTVQFLVIGYVIGYIFIYAKGYQAFEKAESTTIAKVK
GMIKTNFSASTGANNVWDATDLENDAVFITTNFIQTPSQQSGKCPEDTNNKFARCRTDSGCTAGEQVLLGNGFKTGQ
CVQATGTCEIDGWCPAEVDDKPEPPIFMEAENFTIMVKNYVTFPEFDVKRRNIPDDMKGAYINTCRYHPVSDPYCPIFT
VQRLVELAHENFSVVARTGAIFGFIIEWDCNLDFSVEYCLPKYQIRRIDSVEEKIAKGWNFRHADYWNAPDGTRRRTLYK
FYGIKFVFMVYGKAGRFSPEELAMNLGSGIGLLGIATILCDFLTDFLLRTDAFGKSKFERIRDTKKKGPPTLKTMLENAKPP
GAVMVIENEPDADPDASTLDDDSSSSDEDGSTRRGLPEQTPSGDSGFGVARPEKHFLLDDPVKPVHFQRTRVTSL

>Proto_ecdy_PANarthopod_tard_tardig_Hduj_A0A1W0WAZ9_A0A1W0WAZ9_HYPDU_ATP_receptor__
Fragment__OS_Hypsibius_dujardini_OX_232323_GN_BV898_13326_PE_3_SV_1

MVSFGGDMIDTEGVPTGGPITFSDMMCSRGANFFFAYQTPKVVNIRSKRIGAMYRLVQAAIIAYVIGYVFVWEKGYQE
TDRVESSVVTKVKGIIRTKPPYPLGLQIWDSVDFITPAQENNAFFIATNVLATPNQQQATCPEDKNVGIGGVRCASDED
CPPDTQIVNGNGFRTGTCSNVTGTCEILAWCPTEIDVRPVPPVLAGSENFTILVKNHIMFPKFGVKRRNILEDMNATYLA
NCRYHPVQHPYCPIFVLGDVLKLAGESYSSIAERGGVMGFVIEWNCNLDLSVNSCLPVYHFRRMDNKDDKLAKGWNF
RYANYWADENHVERRTVVKAYGIRFVFNVYGKAGRFSPVVFAMNLGSGLGLLGIATILCDFVLLNCLKNRQTFQEFKFL
RMESKYLRRDSGKRGSTSGAGESISDSESPRQHNPNNRRPSASPNQRYRPYSNNQVQTHIDDVTHSSPSLLLHPRSIVV
MNGGPENGNGHIAYSESSSVQLTPHLWNIWKMSEASAEEPRNFAATVFTLLFSYPTPKVVNIRIKRIGLMYRLVQVAIIS
YVIGYVFIWKKGYQETDRVMSSVVTKVKGVVRADLSHPSLGGHIWDSVDLLVPSQENNAFFITTNLLVTPDQTQGNCP
EDPNIDVGRVSCVTDSECVAGGAVTNGNGFRTGICRTEVGTCEVSAWCPTERDVRPKPPVLTGSENFTVLVKNHIMFP
KFGVKRRNIPDDMNATYLASCRYHPTRDPFCPIFIIGDILRLAGETYDAISDRGGIVAFIIEWYCNLDVSESQCKPVYSFRR
MDTTDEKIAKGWNFRFGHYWTDSTALIGGLFNWHT

>Proto_ecdy_PANarthopod_tard_tardig_Hduj_A0A1W0WB33_A0A1W0WB33_HYPDU_ATP_receptor_O
S_Hypsibius_dujardini_OX_232323_GN_BV898_13333_PE_3_SV_1

MVAASQALSQAVSRFFQYNTPRMVIVRSRFIGIIYRLVQLPILLYLIGYIFIYKKAYQDFETAESSVLSKLKGVLQARDFTHTI
WDSTDISILPQETTSFFIPTNYLFTPNQTRRTCPEDPHLGTPGVFCMTDRDCPAGEPTTYGNGEKTGKCVKLGETISTCEI
QGWCPTEVDIPPREILLTGFESITVLIKNHIAFAKFGVKRRNILFHDEQRMMDCHILQLAGENITQIAARGGIIGFNINWN
CDLDYDVERCIPQYSFKRLDNHNDVLAKGWNFRQANYWHDVRGEKKRDLIKLYGIRVVFSVKGRAGKFSAVMFAMNL
GASVGLLGIASMICDFALIRIFGSDESLQAQKYLRLDNLRRPRLASRANVCENYYYYPANGCGNPPTILANEV

>Proto_ecdy_PANarthopod_tardig_Rvar__A0A1D1VFE8_A0A1D1VFE8_RAMVA_ATP_receptor_OS_Ramazzottius_varieornatus_OX_947166_GN_RvY_11229_PE_3_SV_1

MVRPARLMRSAVGRFFEYDTPKIVHIQSLHIGITYRFVQLVIISYVIGWVLVYKKGYQETDNVESSVTSKVKGVVLTNFTLDPNATADEEKLRVWDTADYVIPPQENGAFFVTTNIIMSPNQEQGICPELGASCSTGFDCPSGSKSANGVRTGTCNNATSTCDIRAWCPVEKDVEPQSPVLQGTEDFTVLVKNQIFFPKFRVRRRNILDGFAKNASYLDSCLYHPSADPFCPIFRLGDIIALAQENYSTIAVKGAVIGFVIEWECNLDLSVESCLPKYSFRRIDNADDVIAKGWNFRYANYWTDTVGRQSVQRRTLIKAYGIRFVFYVYGVAGKFNIVPLATNLGAGLALLGLAVIFCDIVILYCSKRRQFYSDNKYLRVQDEDAVLFEEDDRSGHSGENFFS

>Proto_ecdy_PANarthopod_tardig_Rvar__A0A1D1VAA6_A0A1D1VAA6_RAMVA_Uncharacterized_protein_OS_Ramazzottius_varieornatus_OX_947166_GN_RvY_08057_PE_4_SV_1

MGILSYLSSFLFQYQTPRVVSIRSKRIGATYRIIQLLLIIYLIGYVFLYEKNYQSREAVTSSVVTKVKGTLVRIDNGSTEIWDSSDFVVPAQVSPTSRPLGACGPSCLYMIASPGEQCFLHCHKFDIYRKPDPMSLSRGS

>Proto_ecdy_PANarthopod_tardig_Rvar__A0A1D1V786_A0A1D1V786_RAMVA_Uncharacterized_protein_OS_Ramazzottius_varieornatus_OX_947166_GN_RvY_08057_PE_4_SV_1

MGILSYLSSFLFQYQTPRVVSIRSKRIGATYRIIQLLLIIYLIGYVFLYEKNYQSREAVTSSVVTKVKGTLVRIDNGSTEIWDSSDFVVPAQESNAFFIATNLIYTGNQTQCLCPEDRNVGVSGII

>Proto_ecdy_PANarthopod_tardig_Rvar__A0A1D1VC07_A0A1D1VC07_RAMVA_ATP_receptor_OS_Ramazzottius_varieornatus_OX_947166_GN_RvY_07865_1_PE_3_SV_1

MFPKFNVKRRNILDHMNGSYLANCRYHPVSDPLCPIFVLKDLLTLAGEDYKAISKRGGVMGFVIEWNCNLDMSVENCLPVYSFRRMDNKDDKLAKGWNFRYAHYWTDQNGIQRRTMLKAHGIRFVFNVYGRAGRFSAVVFAMNLGSGIGLLGIATILCDFVLLHCLKNRDTFSEFKFVRLESKALRRGSRAPPTLRDEQSDMDGDTISALSPGRKLLLAQNQVYRPYNNDQIQTAIDDVDCKDAVFSPRTFKTPE

>Proto_ecdy_PANarthopod_tardig_Rvar__A0A1D1V3R0_A0A1D1V3R0_RAMVA_ATP_receptor_OS_Ramazzottius_varieornatus_OX_947166_GN_RvY_07861_1_PE_3_SV_1

MDSTTPNPGAKDGKLGVFCRRVMNTLKHLLSEYETPRLVILRSYPVGIIYRTVQLLIVAYLIGYVFIWTKSYQSFDTAESSVVTKIKGILYRNIAEGSAVIRKEVWDGADFVVPVQETNAFFITTSLLYTANQSQTICPEDPNVGIEGIHCSQDQDCVPGESTPYGNGLRTGRCVPSTSNAELSVCEINGWCPTEFDQTPKSPVLHGFDSLTVLVKNHIAFPKYGLKRRNILDSFDDKYLKSCRFHHVSDPLCPVFKIVDIVSFANESFHQLAVRGGVIAFNIHWDCDLDLSVELCLPNYHFRRLDNEKDKIAKGWNFRHAYYYQDMHGNMRRDLIKSWGIRFVFNVQGVAGKFSAVNFWMNLGSGIGLLGIASLVCEFILMKCIQPTDSLMRQKFLQLKDIKEPASQSSGLFYRNNRNDASFLDGAAVCDNCTLQEGAPLAYIDEFGSSHS

>Proto_ecdy_PANarthopod_tardig_Rvar__A0A1D1V0X0_A0A1D1V0X0_RAMVA_ATP_receptor_OS_Ramazzottius_varieornatus_OX_947166_GN_RvY_06095_1_PE_3_SV_1

MWYEKMSELRTAPGLKKFVKKHTLNFLFEYETPKVVKVYSTRLGLINRTVQLLVLSYVIGYVFIHQKGYQQFEQAESNTVAKVKGMAITNFSQESGTNNIWDQTDLVIPAETDAFFITTNFIQTPSQKTSKCPEDRNVGFFAACKSDSDCPAGISINLGNGFMTGKCVNSTGTCEIEGWCPTENDYRPEPPLFLSSENFTVLIKNYISFPEFNVKRRNIPDWMTTKHIAECRYHPEKDPYCPIFVVGDMVKAAKENFSSVARTGAVFAFIIHWDCNLDYSEEHCLPKYSFRRMDSPNEKIAKGWNFRHADYWNDPNGVRRRTLYKYYGIRFVFMVYGKGGKFDPETFAMNLGSGIGLLGIATVLCEFLMDFLLKTDAFRQTKVSRIRDDGHKQPTLE

RMLGKHAAALGGQPINPEMEKLDNGPTRFSSKDHSRNHLTPEDSDDEIAGLSPHEIEALLNRRQSKGAEELRNSQYSLPLMPAKERTKESHL

>Proto_ecdy_PANarthopod_tardig_Rvar__A0A1D1V8N9_A0A1D1V8N9_RAMVA_Uncharacterized_protein_OS_Ramazzottius_varieornatus_OX_947166_GN_RvY_07864_1_PE_4_SV_1

MALTGSTATFKKIWDTGTSYLFAYETPKVVNIQSKRIGAMHRLVQIGIIAYVIGYVFIWNKGYQDVDHVESSVVTKVKGVLRSRTPYPMDVEIWDSVDFIVPSQENNAFFIATNVIATRNQRQGRCPEVHTDASELTARNACAWSICVLLCQSIN

>Proto_ecdy_PANarthopod_tardig_Rvar__A0A1D1V4M3_A0A1D1V4M3_RAMVA_Uncharacterized_protein_OS_Ramazzottius_varieornatus_OX_947166_GN_RvY_06346_PE_4_SV_1

MIPYKYSVREIVATILNSAGVRYQKAMQSLARKLSKGIGRPLARFTNSFFFQYETPRLLTMRTRHIGIMYRFIQLLLLVYVIAYVFIVMKSYQTFDYVNSSVVTKVKGVLSYNNSGVRYIWDSTDFVVPAQESGAFFIATNVLMTRNQSHGICPEDINVGSAGIECTNDGQCPANQPTLYGNGIRTGRCVKQSGTCEIAGWCPTEIDSTPRTATLTGHESITILVKNHIVFPNFGVKRRNILKHFSQEYLQSCRFHPEKDPICPIFNLLDIVRFANESMETLAIRVSFRR

>Proto_ecdy_PANarthopod_tardig_Rvar__A0A1D1UZ05_A0A1D1UZ05_RAMVA_ATP_receptor_OS_Ramazzottius_varieornatus_OX_947166_GN_RvY_06544_1_PE_3_SV_1

MLRSHYNYTGQRYSVAITRELYWRNILDRFSQEYLQNCRYHPENEPLCPIFRLEDIVHYTNDSFERMAVKGGIIGFHIDWNCDLDYGVEACLPRYFFRRLDSEDDKIAKGWNFRTAYYYYDQNGTFHRDLAKYWGIRFVFLVAGKAGKFSTVHFILHVGSSLGLLGLASAMCEIFLSYHRSRRVNDLKFEYLAKKKPMKAQRRYSNMSAPQLQWDVINQNGHGTYAPDHRLQVRSASASPNRIRHTSVTPVLDRRIVRRSYGSKKSMFSLGNRNGGNPVWI

>Proto_ecdy_PANarthopod_tardig_Rvar__A0A1D1UY75_A0A1D1UY75_RAMVA_ATP_receptor_OS_Ramazzottius_varieornatus_OX_947166_GN_RvY_06346_PE_3_SV_1

METLAIRGGIIGFHIDWHCNLDHGLEACVPSYSFRRLDNSNDKIAKGWNFRHAYYFNDGSGVNHRDVIKYWGIRFVFLVGGQAGKFSLVDFALHLGSSLGLLGLASAMCEIVLSYYASKEVQRLKMERVDEIKQGKDCRPSKLSSPSYSADLLSGHLVSLQNGTGYRYRKDSISTDSVHNTVVTPISQRHLLTAHASSKSLHSNSLHAVRNETIILPVWL

>Proto_ecdy_PANarthopod_tardig_Rvar__A0A1D1V4E8_A0A1D1V4E8_RAMVA_Uncharacterized_protein_OS_Ramazzottius_varieornatus_OX_947166_GN_RvY_08057_PE_4_SV_1

MGILSYLSSFLFQYQTPRVVSIRSKRIGATYRIIQLLLIIYLIGYVFLYEKNYQSREAVTSSVVTKVKGTLVRIDNGSTEIWDSSDFVVPAQDRNVGVSGII

>outgro_CafRoe_KAA0149765_1_hypothetical_protein_FNF29_05775__Cafeteria_roenbergensis_

MDLDNVFAYSTPRFVKIRDRRLGMLNTLFKFFIFVYVVVYNVILLMGYKVQGDVVGTARLQPRAPDLKYLRNSSRHRYCMIDGKNRTYTSSDGLIPDLPAFECRYLDTFDTVYPPVEAQALFISTRVNDTLQQLPPNCTDLPRPSCRYETTSVETYFVPGTELVTLLIDHTMSVPSLDFSKSSLDMTGRIEYTNGTEVNPCDAYTRRGYLCPTFANGAGFEITFGTQDAADIIALDTLFQAADVRSLDERAGNIGKFTSETFRYAGIVLIVTVNYDNYFSYDENKVQYILTSKLVPDADFKAEEINPGPGIDETTRRIIDRHGVRIVITQGGKIGRFAVSQLLIVLVTSMGLLAVSSSIVDGVAMNVLKMRHIYAQYRTVTSVDFSDIAAIKQSDLNRFVAEDLVNPRPAIFEQVGQGAGGVTSGEGQPRQMTSAPAPAKQAGYYGGMGPASGLVDEVDDVDSDEGSGTTGTSVRSPEIVAWGENPAAASSAPSVRMKGWKRGFPSSAAV

>outgro_choano_Mbre_tr_A9URI5_A9URI5_MONBE_ATP_receptor_OS_Monosiga_brevicollis__Choanoflagellate__OX_81824_GN_MONBRDRAFT_31366_PE_3_SV_1

MAASGFWGSIQQGIYSMLEYDTLKTVHIRSKKVGLIFRLLQITILAYVVGYGIIYQKGYQEVDSAVSTVLTKVKGIAITCDN
TDISSMNDCVPGDLRVWDTPDYIKPAQENDAFFVVSNSIQTSKQTQRAEGWDEDPAAPVTGSASAFNCTSDADCPRF
ATSRNGALTGECNTTTERCRIYGWGPVESKDEDDRATTDGLFYARHMPAVKNFTVYIKNTVFFQRFGAKFGSTDESDK
VDVYTCTWSPTGLERHCPIFKIDTILNEAGITDFENQAMRNGALITIQVNYDCNLDSSAHTCSPTYKFTRLDTKSDLSAGY
NFRFANYQIDPPARDLYKVYGLRFVFVVSGTAGRFSMVPLLVALGSGLGLLGLATVIADLLVTKCIRNANVYYGLKYQVV
DEEDIDRAGIDHIPIGDQKLERQPLLP

>outgro_choano_Sros_tr_F2U6L3_F2U6L3_SALR5_ATP_receptor_OS_Salpingoeca_rosetta__strain_ATC
C_50818_BSB_021__OX_946362_GN_PTSG_04102_PE_3_SV_1

MAGGEGKFSCTQAALSVLEYDTLKTVHIKSKKVGLIFRILQLVILGYIIGYAIIWQKGYQAADQAVSTVYSKVKGVAVTCN
DLNVTSIRNCSDSDIRVWDTADYVIPPQESNAAFIVTNSVQTSNQTQQPGGWPEDPDASTKGSQRAYTCEKNADCEA
VRFSPSRNGAVSGKCDLATKRCMVYGWGPVELSKDDDASQVLGYPRQMPQVKNFTMFIKNTIFFPHFQRKFGNTGL
GNSSASYLKSCLWNESTDRYCPVFRIEDILSAAGVKSFEQDVMMAGAVITVQIRYDCNLDKNVETCAPEYRFTRIDEPN
NTLSNGFNFRFARYAIGEVPSRDLYKVYGLRFVFIVSGTAGKFDFVPLLVTFGSGLGLLSLATIVADLLVTKCLRNAEFYYD
RKYEIVNEDEVEGEAINVTIRQKPEEDTPLLH

>outgro_Filast_tr_A0A0D2VGJ3_A0A0D2VGJ3_CAPO3_Uncharacterized_protein_OS_Capsaspora_owcza
rzaki__strain_ATCC_30864__OX_595528_GN_CAOG_000560_PE_4_SV_1

MASVVAAFKRFIATDPGEAHMGSRLDTLPSSSFPLPSSLSLSPPSLGQVAHACFCGCLVLGGGPTPPCLLDQVFIVFAEFP
AFFFFFFYLSLSLCSFPHMQISLIYNLSHCFCFLFFSRSSLLHSQPDLLFNYKTQRIVRINDRRLGLIYYAFMLAILGYVIGYTIIY
NKRYLLLDDSPTGSVRMSLAKPTKVNSNQPYCLQNQPTVNDYTNYMCNYYDDLLVVYPRTEQTALLAASRITIQNNTL
NCDFAADPNCDFTPETPTIVYVGDIEHFTLMIDHTTYSTNLDIQANAQSLEGKLLDLHGNTMKLTGPNVVGQIGQLDIF
ELQTLLLAAGIATLDQDSGALNTKTNNSSLRYTGLVLLVLIEYDNVHSFDTSDIRYTISVRSVKNTQFKSVETRTSTTDQSL
QTWDRHGIRILIQQKGQMGKFDFQTLLINVVSGFGLLAVATLVVDVLAVSVMPEREIYSKYKYDDTLDFSDIRDGITKAE
DDQLISIKTRKSAAQLVGVLDSASQPLLDESSTV

>outgro_Filast_tr_A0A0D2WGT6_A0A0D2WGT6_CAPO3_ATP_receptor_OS_Capsaspora_owczarzaki__st
rain_ATCC_30864__OX_595528_GN_CAOG_000277_PE_3_SV_1

MSESKPTLRSRVATTLGEGFRAVVAYETAKVVEVRSLKLGFFNRFVQLAILGYVIGYTIVYQKGYQTQAPLQSTLTTKVKN
VGFACQSTDSAGGRYGGRGGAAIGVTTCPIPGVDPPGNTELSLWDSTDYVIPANEHDALFILTNSIVTPEQKQTTCAED
PQYAQACTDDSECTKYSSTRNGKRIMTGRCVNSTCEIYAWAPLELDSLSPQSGSVIEQAGNFTVFIKNNVRFAKFGISRT
NVDIGNTSAAYLANCRWNKETDYYCPIFKLGDLVEASGWKIEDMLVLGGVLGVNLDWTCNFDNDESECRPVFSISRID
DPTSTSPGYNFRYADYYTLPSPTDPSISVPHRDLIKAYGIRLIFVVSGEGGKFDVAALTMSLGSGLALLGLATLLCDFFMTH
FFAKKDYYREKKFELVDDLPPLDQTFSRLVGGPRRSEVAVPVPSTSTSVGSVQGGNTASEKSPLLQEDQRHTWS

>Filast_tunicaraptor_GIQG01028472_1__p1__GENE_GIQG01028472_1_GIQG01028472_1__p1__ORF__
type_complete__len_419__score_88_71__GIQG01028472_1__70_1326_

MGNCCAALKAMDLDKVFAYSTVKIVKIQDRRLGITHKLFLLGIFIYVFGWTLIHNKRYLLRETPVGTVRMSLQKPAHPSN
PEDLPYCLTADHGSTNGFENWPCRYWDETLVEYPTTEEYAMHVASRITAYNYTLPADCYDWSRPSCAFESNGKNDTFF
IADIENYTFMMDHTMYAVNLGIQRNAQNLSGELVDTDGHVMTGSNIQVGVRDKLDIMSIDALLKAAGIDSLDSASGA
SNTKHDTDRSSKRFAGIILLVIIEYSNVNTFHTSHISYTVKVRHIKNTEFKGEQLQYAKYPGDFLKKDRHGIRIIIQQSGTIGE
FDFQTMLLSLVSGLGLMAVATVVVDFAALSIMPQREAYKDYKYLVTADFSDVRDGTVTIDSYDTDYKPKSSYAGKGKDE
ESTPILENRGGRAGSVNRSV

>ClCha_Protostome_Lophotroco_annelid_CAC9470383_1__Ofus_G012392_Owenia__fusiformis_

MEDSQANGENCAINCTRDSIGRHDNPEPAGHNSNGTDDTSLSQRHDNNSDGIDLDRIDEIRETHLLNDSDMNNSCQE
RIMERDTTESNLCCFTCQRNRKSKRRRRLKNNCCGLRLCSTRRQNRDDESLDTSDGDRGDADEVDIYAVDREPVRKQL
CRACSTFTYILLAIIAIVVTYSMIANLISSLNNPVRSIQFQQVKHYEAPGIAIYPGGAKFLACGHYYNNVVTSKVRLGQLDLL
NRNCSYVNMTYSDPHNTNISRDVLVFRGPTNFVDREAVFFHFMLEDETRDFFTVSYFVFQQWDRFNVSSDKVERLLNY
HRYVPTYALSAELRMWIKLSMKETHVSDGDMHTEFKVEYSVVRFIDRRPEDDKKRHLFFIFFEWKNDLIEESYMMYTQT
PWSVGAGLCAVLLTLHRGIQIAQLSFKRVQKDKRRRTIRRRTIQEVKEAKRSGRRVPPQLLRGQSAPAFNH

>ClCha_Protostome_Lophotroco_annelid_Pdum_Contig13931_

MPGNGLSDANGANDENLNCSNNVTERTPIKPSLKSSNSNSVDGFRKKRNALNRQQSQVSFDPEDNVTDSSTCLSHNV
EDTKDPAKYIVAPDSSSIGSRSNRSWSEVCCRLPKCFSSRFCDRCCCKKEKPDDPLELQFQLHDDPFRKKCSRACLTFVYV
LLAIIAVVVTYSMIADLVKSMNNPVRSIHYHKVKEYDPPGIAVYPGGAKFLACGHYYNNVVTSKVRTSDLDLVNLNCSYT
RLRYDDPHYIGVERDVIIVRGPTSFEKREAVFIHFELEDITRDFFTVSYFVFPEFDRFNASEISAQITELSDLHRIVPTFALSSG
LRMWIKLIMSESYLQDGSKHTEYRVEYSVVKFWDRRPEDMRGRHLFFIFFEWKNNIIEQSYLIHTTTPWSIGAGLCGVFL
TLHRFFQLAQTSYQRVQKDKQRRSIRKRTLRELRQKSIEHNSKAIQKHL

>ClCha_Protostome_Lophotroco_annelid_Pdum_comp388816_c0_seq1_

FGLIISISCVFGGFVVIYKSVDDVVDAYGHNVRSVSFVKVNRYKPPGIIIYPTNNTFVACGHYFNNIVGSKVRLSELDLQRK
QCTYEHVTYTDPVDHVTLREAIVFRGPRKVKNVEALFVHFQGTNEGRDFVTVSYFVISFWSHFKKSQDHSQYLHDLHGT
VPTFAISAGLRTWVKLSLREVYFIGIDQPARNFLIEASIVKYVGTNSTGPVRNFFIFFEWKNNLIERSRELRGMTPWSVAG
SMCGLILAMYKASKSAHKMLRARLKDVKTDINCIQEAKRLRETNGKNCRVFCVVGSNTL

>ClCha_Protostome_Lophotroco_annelid_Pdum_comp388816_c0_seq3_

FGLIISISCVFGGFVVIYKSVDDVVDAYGHNVRSVSFVKVNRYKPPVRLSELDLQRKQCTYEHVTYTDPVDHVTLREAIVF
RGPRKVKNVEALFVHFQGTNEGRDFVTVSYFVISFWSHFKKSQDHSQYLHDLHGTVPTFAISAGLRTWVKLSLREVYFI
GIDQPARNFLIEASIVKYVGTNSTGPVRNFFIFFEWKNNLIERSRELRGMTPWSVAGSMCGLILAMYKASKSAHKMLRA
RLKDVKTDINCIQEAKRLRETNGKNCRVFCVVGSNTL

>ClCha_Deuterostome_chordata_Ggallus_tr_E1C5B3_E1C5B3_CHICK_Proton_activated_chloride_chann
el_OS_Gallus_gallus_OX_9031_GN_TMEM206_PE_3_SV_3

MLRPELSASYTELSEDVEQVSENPTDEREGEMGVDEDTSSAILPDSELGTSAPSLRFSKTCLKNVFSVILLFIYLLLMAVAV
FLVYQTISDFREKLKHPVMSVSYKEVSLYDAPGIAFYPGKARLLSCKHHYYDHILPLINPGQPGDIECTTQRINYTDPFTNQ
TMKYALVVQGPRDVKKKELVFLQFHLNGTDQDFSAIDYLLFSSFQEFINSPEKAEFMKDCESSYSSWKFSGGFRTWVK
MSLVKTKEEDGRETVEFKQETSVVNYIDRRTKLRSDQLFFVVFEWKDPFIQTVQDIITANPWNTIALLCVEGKTPTTKQS
GLPSPKECLNRQSTLEFYYIIFHLKCIDLNIKKK

>ClCha_Deutero_Chord_Hsap_sp_Q9H813_PACC1_HUMAN_Proton_activated_chloride_channel_OS_H
omo_sapiens__Human__OX_9606_GN_PACC1_PE_1_SV_1

MIRQERSTSYQELSEELVQVVENSELADEQDKETVRVQGPGILPGLDSESASSSIRFSKACLKNVFSVLLIFIYLLLMAVAVF
LVYRTITDFREKLKHPVMSVSYKEVDRYDAPGIALYPGQAQLLSCKHHYEVIPPLTSPGQPGDMNCTTQRINYTDPFSN
QTVKSALIVQGPREVKKRELVFLQFRLNKSSEDFSAIDYLLFSSFQEFLQSPNRVGFMQACESAYSSWKFSGGFRTWVK
MSLVKTKEEDGREAVEFRQETSVVNYIDQRPAAKKSAQLFFVVFEWKDPFIQKVQDIVTANPWNTIALLCGAFLALFKA
AEFAKLSIKWMIKIRKRYLKRRGQATSHIS

>ClCha_Deuterostome_chordata_PetMar_tr_S4RG85_S4RG85_PETMA_Proton_activated_chloride_channel_OS_Petromyzon_marinus_OX_7757_PE_3_SV_1

MRVSKLCLKNLVSVLLVVFYLVLSAVASFVAYQSISDFQHQAQHPAMSVSYKAVERYDAPGIALYLGSAELLSCRHYFHSGVSVMERPGQPGDVRCNIFTVAYPDPYHNTSQKSALVVRGPERVQEKEQIFLHFRLNGSTQQFSAIDYLLFSSYFQVVWESTNRPSFMMQMERTQSMWTFSGGFRTWVKMSLVRTKTDRPSEEFTLESSVVKYNDERPESERSDELFFIVFEWKDALIQEVQGIVTANPWSVAAVLCGVLLLIFKVADFATSSVRWMIRVRKRNRRHRVQKMNNMA

>ClCha_sp_Q66H28_PACC1_RAT_Proton_activated_chloride_channel_OS_Rattus_norvegicus_OX_10116_GN_Pacc1_PE_1_SV_1

MIRQDLSTSYQELSEELEQVVENSEQADERDKELVQVQGPGVVPGVDNESASSSIRFSKACLKNVFSVLLILIYLLLMAVAVFLVYQTITDFREKLKHPVMSVSYKEVDRYDAPGIALYPGQAQLLSCKHHYEVIPPLASPGQPGDRNCTTQRINYTHPFSNHTMQSALIVQGPQEVKKRELVFLQFRLNQSDEDFSAIDYLLFSSFREFMQSPDKAGFMQACESAYSSWKFSGGFRTWVKMSLVKTKEEDGREAVEFRQETSVVNYIDQRPAAEKSTQLFFVVFEWKDPFIQKVQDIITANPWNTIALLCGAFLALFKAAEFAKLSVKWMIKIRKRYLKRRGQATNHIS

>ClCha_Cteno_Ple_Bac_comp47550_c0_seq1_p2__comp47550_c0__comp47550_c0_seq1_p2__ORF__type_complete__len_294__score_22_16__comp47550_c0_seq1_1621_2502_

MVMAGDSIQYDYDYVSESDCIDPSRQCYQVISKGKVLQTDSACVFCCWKCGEIFKAFILPTFYSLVVAGCIVYCYQIVSNYINSVRHPIETLSYKHADKYEAPGIAVFLYDNASFVSCEKSYAIPEHKVERGKRRKAGGVGSLLRGSPDTSPCMFSAISNYESVMSRTNTSMIVFQGPSNVARREYMKIHFKVDQEKADPDIGFLEFFPFANFSEWSVEYNSSQQYLQTREKIQDIWPLAVGFITYSRLAIRKFVPHRINGKFNPLVGSTVAAPPLPHAAALLFQYLQNLSCQ

>ClCha_Cteno_Ple_Bac_comp47550_c0_seq10_p1__comp47550_c0__comp47550_c0_seq10_p1__ORF__type_complete__len_312__score_28_09__comp47550_c0_seq10_1427_2362_

MVMAGDSIQYDYDYVSESDCIDPSRQCYQVISKGKVLQTDSACVFCCWKCGEIFKAFILPTFYSLVVAGCIVYCYQIVSNYINSVRHPIETLSYKHADKYEAPGIAVFLYDNASFVSCEKSYAIPEHKVERGKRRKAGGVGSLLRGSPDTSPCMFSAISNYESVMSRTNTSMIVFQGPSNVARREYMKIHFKVDQEKADPDIGFLEFFPFANFSEWSVEYNSSQQYLQTREKIQDIWPLAVGFITYSRLAIRKFVPHRINDTGWPDSGLEKTLFNPVNDIVKYLPADAPSNYTPRYIDGGGSQFCIALTEN

>ClCha_Cteno_Ple_Bac_comp47550_c0_seq26_p2__comp47550_c0__comp47550_c0_seq26_p2__ORF__type_complete__len_376__score_33_48__comp47550_c0_seq26_196_1323_

MVMAGDSIQYDYDYVSESDCIDPSRQCYQVISKGKVLQTDSACVFCCWKCGEIFKAFILPTFYSLVVAGCIVYCYQIVSNYINSVRHPIETLSYKHADKYEAPGIAVFLYDNASFVSCEKSYAIPEHKVERGKRRKAGGVGSLLRGSPDTSPCMFSAISNYESVMSRTNTSMIVFQGPSNVARREYMKIHFKVDQEKADPDIGFLEFFPFANFSEWSVEYNSSQQYLQTREKIQDIWPLAVGFITYSRLAIRKFVPHRINDTGWPDSGLEKTLFNPVNDIVKYLPADAPSNYTPRYENMYAIFEWRDTEYEVIEHVETQNIWTTVGLMVTAILTILRMFDLTKVIVRRQKVRYGQSRLSFLSLCSCCDPQVPENE

>ClCha_Cteno_Puk_Fal_comp26698_c0_seq2__p1__GENE_comp26698_c0_seq2_comp26698_c0_seq2__p1__ORF__type_5prime_partial__len_386__score_29_53__comp26698_c0_seq2__2_1159_

FKRINLSGMPENGIEYDYDYVSESDCIDPRRKCYQVISKGRVLQTDSSCVFCCWKCGEVFKAFVLPTFYSLVVAGCIVYCYQIVSNYINSVRHPIETLSYKHTDEYEAPGIAVFMYDNATFVSCEKSYAIPEKWGRGQRRRGGGASSLLRPSPKFESTPCIFSAISEYESVMSRTNTSMIVFQGPSNVARREFMTIHFKVDQDKADPDIGFLEFFPFANFSEWSLVYNSSQEYLRTREKVQDI

WPLAVGFITYSRLAIRKFVPHRINDTGWPDGSVEKTLFNPVNDIVKFLPPDAPSNYTPRYENMFAIFEWRDTEYEVIEHV
ETQNIWTTVGLMVTAVLTILRMFDLTKVIVRRQRVRYGQSRLNFLSLCSCCEPNEDQTGSVQ

>ClCha_Ctenoph_Hormiphora_californiensis_evg142601__p1_GENE_evg142601_evg142601__p1__ORF_type_complete_len_420___score_24_84_evg142601__75_1334_

MPLNDIDYDYVSEADCRDPRAKSYQVISKGKVLQTDSGCVFCCWKCGEVVKAFVLPTFYSLVIAGHSHTNIPSTNYSLPC
IYHNVDHRFRLQSYPYSFSVFVSPNSVRAIAGCIVYCYQIVSNYINSVRHPIETLSYKHADEYEAPGIAVFMYDNASFVSCE
KSYAIPEHEVVRGRQRRAGGVSGLLRARPSSDPCRFAAISDYESVMSRTNTSIIVFQGPSNVARRENMKIHFKVDQEKA
DPDIGFLEFFPFANFSEWSTVYNSSMTYLQTREKIQDIWPLAVGFITYSRLAIRKFVPHRVNDTEWPNSGSEKTLFNPVN
DIVKYLPPDAPSNYTPRYENMYAIFEWRDTEYEVIEHVETQNIWTTVGLMVTAVLTILRMFDLTKVIVRRQRVRYGQSR
LSFFTICTCCEPQDQETIN

>ClCha_Deutero_Ambulac_Sakowv30028634m

MDTTGKCWRCPCCINNGNSDEDNPVDYHGDTVNLIQASNESVTPQCKSGKSVLGDIMKAALTLFYVTLFSTAGIVAYQ
TISDLMESRKHPVASIQYNETDMYLPPGIVIYSTDVNAKFLSCGKYFYDLVDPPDINPNHKNCTNEEVVYPDPYFPERAEL
QHKTLQAAKTQTTVPFNSISSARAPVERRSKNRSNKEGFIRQVDIDNPTWTFSGGMLTWIRMSLMETYYLNGSFNSVF
DVSESMVKYNDIRPEEERVNELFFVLYEWRDKYIQQNHIVNTMSVWNTVGVLCGVFLTLFKAGEFVTKAIQRMLKARR
NKKLQEQQRNNSRRQMIT

>ClCha_Deu_Ambulacrar_hemi_Ptyfla_40v0_9_20150316_1g8542_t1_scaffold4379_cov149

MEMKKIADKERENDTGSFYDSIQTKDEENLLNASIDSVTPQCKDPKHLFSHFVKGFLAVFYLSLVAVAGIVAYNTINVYVE
SQKRPVTSITYKEMQVYPAPGIVIYSSDINAVFLSCGHYYNDIVEPPTLDPNHKNCTFVDISYPNPYFPERFRKAVVFRGPT
KLQKKEILFINFSLNLTARNFSGIEYLLFPNWYGFWNR

>ClCha_Porifera_AmpQue_tr_A0A1X7VNE5_A0A1X7VNE5_AMPQE_Proton_activated_chloride_channel_OS_Amphimedon_queenslandica__Sponge__OX_400682_GN_105316455_PE_3_SV_1

MADRSSIHFVPASLPPQLERGRVREDSWKNEMNSFARGRPVSEADAEWLEEDHLQLLHSEEEREGGYGSITEDFPKEE
PAPLRERLSQNRSRRKIKKHHRRLDANPFTSLKRLVIGEDEDEDSSDEEPPTMNPVYRRSGKDLAKLILLPTLYFALMIVAC
VFAFYTIEGVVLSYNNKVQSVQLVEVKDRYPPIGIVILPQFSNYSFCQYRYYDDVSPDPEKPPERCNKYKLPETCTYFNVSF
NSTALVNVTRFAMVFKGPTLVDCKESLLLHFSINTTSRQFSAVEYILFENWNYFESLKENERKGFLASLEQYRDIYTFPAGF
RTWVKMSYTIRTDYRATTNKTDFNIIDNYASFNDPGLDGMYPMEVVFEWKDRYYDYVRDIISTTAWSAFGSICGVFITL
VKAGEFCRMWMRRIRRDKEKKMLHLKKLEEEHEKELEAYKERQKEKRENKLRKSMETTKSYHIQKN

>Protostome_Lophotroco_brch_Lingula_XP_023930484_1_ASIC_channel_Lingula_anatina

MSEKMVRRRSSAFEVQMETLEENKTKESTESKASKESGSVSEGLEQFGEDTDFHGLKRVFRRDYTLLRRTFWLFFFLAGL
TGFIANTVDRLSYYLQYPHSSELDVMYDMELEFPAVTICNMNSYRLSALTDVDILHFGEKLHILDENRRLIHPEYYNQSW
VDWVHGINWTELAANDDEENFDVLEFIRRTGHQLEDMILLCSWKGQHCGPENFTSVFTHFGICYTFNADHSYKYVSRK
AGAGNGLKLYINIEEEEYLTSDVLAGQDAGLKMVIHAQEEPPFVKELGFGVLPGDHHFIAIQKKYVHNLVPPWGNCYEG
KLKYYSHYSVPACRIECETDTIVKECGCKLAEMPGNTSVCLGHMYMGCAYPALEEVEHSDLCSCQNPCDMTTYRRDISS
VRLRDTTLDIIAENNPQVKRETLSRDNLLVLNIYYEELCYETIRQIKAYSIPALLSDIGGQMGLFIGASVLTLLHVIEAVGAVV
GGKFLKTTKQGRVSTTRVQSLK

>Lopho_Phor_aus_TRINITY_DN315771_c0_g2_i2_p1_T

MNSEVNKRNLKGRAKEFCETTSAHALGQTVRSGKVKAIFWSLVFLGSLAGCVWNIIHVVESYAGFGFSVKSKLEMEPSS
LKFPSVTICNLNPISFSRQARFVGDFERSTGLDEKDCDNEFYRNETNRPLVLDLYRWSELPQFWFEYNENVTDMYGHSK
KDFIVDCVFSKKLCEDNFYVTTDPNLYNCYTLEPNKYNSDLHAGVGIEAGLSLTLFVENTRVQNAYTGSFSIDSYQTGNV
GVKVAMHVSGSHPNPNARGVIAEVGKSTDFILRTVNRTALGPPYPSPCNPKKTIESSHRQELQYEENLCFASCLQNAIAK
KCGCVSINPVAVPIADKFGIDLPFCGSYPCNATKVTENYECVHDVTRRFLLKNDPNCRDCTKPCNEVSYEATKSQSKWPS
ELHQSEFIGWLASKDNTALYNQLVRNTSDDKISRFIENNFLRINIYFGDFYVRKDTETLNMDWFDLLSSVGGAFGFWVGI
SVVTGVEVLELLLDCIVIFLNKPSRKRNQKDLRNTTGNNDVEVYNN

>Lopho_Phor_aus_TRINITY_DN291806_c2_g12_i3_p1_T

MAEVELSCVNDVHFGLEMSNGVNPKTHLENMAKEEEEKQSLKEFLKEWAENAACDGVPQIVLRKSWVKRILWLLIVL
GLTAYTIEEVYGVFDEFFTYPVNTNINYTYSNQLTFPAITLCNMNPIRKSMLSEAGTTFDDIFGSTNGTFPGRKKREIGPQ
EETAPSGSQQRGKEKMTFINQFKRQWATLSYPKRIQLGHQLPNFVVTCSLGGQSCNKFFTYSTGTFGNCFVFNSGLRN
TTVQSISKTGPLFGLTMEIFLEESEYLPEVTEQTGARLHINHQKLMPFPENAGYNLAPGYLTTIGLKRVEISRLLGYESNCTK
VTTAEAEGYSIYSKRFGVSYSREACHNTCYQRKVIEICRCGDADYLFDYSFFKSLLPDTVKSIKPCLSATEEACVSRVLKLYA
SNNITCDCPIPCSESDYQATVTISPWPSKLYEPTLRSSYSSLNRNDILNTTLSRNLLKVEIYYDTLNYQYIYESPKYTNLSLISD
LGGQIGLWLGASLFLLLQVLELFIDVIIFGCGRLFKSGSRVSSR

>Lopho_Phor_aus_TRINITY_DN297387_c12_g1_i2_p1_T

MSGRRSAWNDYHSESVWDEQRAGTQNGDKHGEDDCKDEDDQPRSSWQIIKDCGNDYLDVSSMHGLAFFVGSRYH
PLRRLLWFALFTASFTWLMIVVTTAFDNLQRKPVLTKVKVKYEDNARFPTVTICNLNKYRKSYVEKHHPHSEKLLRYIYPV
LREDEHIHLDWSDPKYAWSLNRSSFKEFALKAAPQMMDNIKIVDFGAHWLTISNEFSTIETDFGMCFQFNPNGTWYS
QLSGEMHGLAIVLDVQQDDYYFGDSFSTGFKIALHRHDEVPFPNQFSFGVSPGQELLVGMTMTEVKSLPAPYEDGTCV
DTEAGDFINPLKYFKTYKYNSCRKECQTVFSVGQCGCKPAFDPGPTRTCIGEEYRCYRLATEEYAKNHSIERECNCRLPCH
ERGYVYQLSSLKFPAIQYEAYFKNTHNISLQYARENLMLVRIYFEKMNYELVEQVLGYDGMNFFADFGGNSGLCIGASLL
TVAELLELIAVIVFNLIFRRRKKTSPEKQ

>Lopho_Phor_aus_TRINITY_DN307417_c2_g3_i8_p1_T

MASEPETIPLNGLNHNLRKRRVQNYIVIDLVNNGEQDEQENGVYKSRFECFGEDTDFHGLKYISRGENSGVKRLLWLLLI
IASLIYLVIEIYGRVAYYLTFPHVTKVDVEFKSDLDFPAFTVCNMNKYRAYSMTDMDLFHMGEILDIVDENMNLHHHSH
YKESFVKRVQNINKTEIRLKNESFDLEEFVRRTGHQARDLILYCKWKDELCTGEANFTHVFTHLGNCYTFNSGKKKVAAR
KAGAGNGLTLYLNIEEHTYLENKDAADEGLKVMVHSQMEPPFIRELGFGIMPEQHHYIAIRKTEITNLPHPYGKCVDTN
DSRRHPDFEHYTIPGCRIQCETKNVLETCGCRLPEMPKMPDHEAKICTPPHQYKCAVKELDRISESDVCICQTPCHIEDYR
FTHSSAKLRPETIEKLRELQGDKLVPANLTSNNVAVINVFFEALSMEKIEQRVAYPWPSLLGDIGGQMGLFIGASVLTIVH
AIEFCMNEFARGVKDRARKKNEKSASLRSNNNANRDFQQDPTLPTVLKEESL

>Lopho_Phor_aus_TRINITY_DN307786_c0_g2_i2_p1_T

MSKGVKTSGGVWATNVFNSGQHDNTLYRQHVEAKQDITADIDQPNTGLSIVKDGIQKYAEDSSLHGVQYFGGAKYNP
VRRLIWFLMFVGSFTFLVINVSDAYQTLMRKPVQTKLSVEYTKTARFPTVTLCNFNKFRGIYVWTQPALMEILRYLYPLSE
EDRKLQLNWEDPKYAPFINGSTYLDFAKIAGHQLRDSIYFARFKGMNLDIQKEFKQVVTDFGLCFQFNSNGTWNTSRS
GTNNGLWIQMNAEHYMYFYGDSNSAGFKVALHNFDEEPLVNELGFAISPGQESFVSAQIKEIKSLPPPYEGGKCKNTTK
ENFVNHLAYYETYSMTGCRRECRTNYTVKRCGCKYIYDPGPARVCDAPEMGCYRQADGEYTQDESIEDDCDCFVPCHE
IQYEHRLSTSMYPGNHIIYGLQAQYNVTKEIARENFLELRIFFEKLNYDLVEQVPSYDAMNFFTDLGGNLGLCIGASLLTA
AELLEHIGITFFQVLQKFIGRKQTHPKKIQREGVF

>Lopho_Phor_aus_TRINITY_DN307786_c0_g5_i2_p1_T

MPKGFKSSNDIWATKVTPIAPYGQKESDNKDEGAPESGTVLSIVKEGVTKYAEETSCHGVQYFGGAKHNPGRRIVWFL
MFVGSFTWLVINVNNAYQTLMRNPVQTKLHVEYTQTAKFPTVTFCNFNKFRSPYVLAQPGLIDILRYVYPVTEEDREQ
QFDWTDPLYAPLVNGSQYLEFAKIAGHQIRDSIYVARFKGVNLDIEKEFKRVVTDFGLCFQFNSNGTWNTSRSGTNNGL
WLEMNVEHNLYIYGDSNSAGYKVALHNHDEEPLVNELGFAISPGQETFVSVQIKEIKSLPLPYEGGKCKNTTKEDFVNHL
ENYKPYSLTGCRRECRANYTKQKCGCKYIYDPGPARTCNAPELGCYRNADEEYTRNASIENDCNCFVPCHEMQYEHRLS
TSMYPGNHVIQRLMDDFNITKELARENFLELRIFFEKLNYD

>Lopho_Phor_aus_TRINITY_DN312240_c0_g5_i3_p1_T

MDNFEMAWDNQQTRTQQGVKSGREDCKDEELSRSSWQIFKDCVTDYLDVCSMHGLAFFLGSMYHPLRRILWFLLFT
ASFVWLMIVVATAYDNLKRKPVQTKMKVKYADNASFPTVTICNLNKYRKKYVEENYPDSEEVVRFMHPVVPEDSTANI
DWSEPRHAWSLKESNILHLARDGAHEIHKSFISAQFGGHELPVAKPNFVTVETDSGVCFQFNPKGTWYSELSGKTFGL
MVMLNADQENYYFGDSFSVGFKIALHRHDEEPFVDEFGFVVSPGQEVLVPMVLNEVISLPEPYEGGTCVDTKVKDFTN
PLEYYSPYNYNSCRKECQTRYSVEQCGCKLAFDPGSSRICVGDEYACYRLSTQNYAKNASIERSCNCRMPCHQVGYDYR
LSSAMFPGLQYRSYLANAHNLSLDYARQNLVVVRLYFEKLNYQLVEQVLSYDGMNFFADLGGNSGLCIGASLLTLAELIE
LIGVIIFNMIFRRREKTTPEKNNAK

>Lopho_Phor_aus_TRINITY_DN316400_c0_g1_i8_p1_T

MADTGQKHYSYLTGNDYENVPIHALTYKALWRGFQENTSGHGFSRIYLTPGTVKKALWTLVCIGGFTAWGVHTTTLLV
NFFYYHVDVTVQILTAKNLSYPAVTICNLNQIKSSKISDVLKLDDLYALAAQTVDDGDGNLGQRDQYSEYVDILRGAVAD
MDEAAKSDAGHQRGDFILGCTYKGYECYGSDVMEFFDPIYGNCFTFNGADGSPQKDYMRAHSTGPESGLTLMLNIEQ
SEYLPFTQSAGVRLVVHNDTVMPFPADEGINLEPGKLTMVGMRQIQIIRKGNPHGVCIDPTSYNGDRNLYEDKFRVRY
TYQACKKTCYQKAITERCKCFDPHYPTTVNVSDTLKACRYGTIEQDECSENITQDFADGNLRCDCPLSCSESAYALTISSS
QWPSDNYKTALFTKLTSLNNSNIDAILTNAADKDIYRRNFLKVAVYFEDLNYEEIEQKAAYTGSSLISDLGGSIGLWIGWS
VITIVELFEFLSDVLILGCSRKSSKRAVGHSSDTPVKSIK

>CAE7949834_1_ASIC1_Symbiodinium_KB8

MAPQTPPDVAGSAWWRPAASLAELPSQAAGFANRRDMAVPLRQRRLERQADGAAVALVAGQPEGFQPLQEFSDN
VTVRELRFVTASTTTGALLSAVSVCKRWLWLVLLLGALVALVVIVTDRVQYFLQNPTATVVKWESRAPVPFPAVTVCNE
NGFQQSRVAQYGLSTLNNSVLELMTQPVNSFILRCEFAGLECFADNFSVLLDAQFGACYTFHGLRALTIDGEEFNASGP
NPLTSARTGPDYGLRLVLDIEQAEYVGYSPRTSVRQASGADQGPGTEPDSSGSAQTSPQTSPQTSPQTSPQTSPQMSA
QASAQASPQTSSSSATASSSPPPAAASSSPQASSPAGASPTRTPSPSQPARRLLAPRRVQEDPGPGSARDARAPPDNRN
STVEALFKGPESAGIRVIIHDPSVLANPTSGFGVGPGSASDIAFERVDRSRLFPPWGTCNEAWHSESGATDPAGYAKTA
CERDCYYASVADVCECLLLPWPTTMEERPSLYVCSSEEKACIATQLSRFQAGELGCSASCPDRCLETLYTATAGRQLWPS
VASASTILDVVRSAKQDTSITPEYLAQNMMSVRVFPPTTEYQVVTTVRAYSFESMLGEVGGNTGLWAGMSALTIVEILE
FVMLALSAWLCSRGGCCRRGGKRAPRTNVEHGTELHEGEAIELVAPQADEESSAPAVAPQAAAAASGRAREAWRE
