## Supplementary material for "Function and phylogeny support the independent evolution of acid-sensing ion channels in the Placozoa": File S3

>NeNaC1_tr_A7RI82_A7RI82_NEMVEPredictedproteinOS_NematostellavectensisOX_45351GN_v1g1974
95PE_3SV_1

MKENDKDEDGGDVKIIIKGSDDKAPQGAGSIYAFCNHCKKDISVVYIKSENNGKELSVAESLTNKWPRSEFPENDYDKCL
KDNDYGGCKKLLDGRKKRAKVKELWENFLGGCTLHGFHYCFAGNPPLRRLIWSLLLLGAFAMFFEKCTESFINFFDYPFT
TTTLLVYDKRLPFPAISMCNYNDARMSKMNGTLMNEIFVASKLEGRNTSHLQSQLTGELMQRTLKEAAHRLPDMIKEC
SWQKHGKCSWKNFTSFKSADGDTCYTFNSGRKDPILSMSNVGEENGLRLVIDTQHSEYYYDVKNAGFKVILHDQGETP
VKMQGLSVSPGFTSYMELKRTKVTNLPFPYKTMCGMPELKYFNSYSKSKCFLDKLTQYVVTLCGCRDWFMPVNGITGI
DIGNPVRNLSKLGKARYQCVTTKQRHPACGKLGHTSSSCDYFDGKRGSRVSELQAFFVPTYSVLLISAYFEENKLDQCPV
ACNSVEYSAQLSYARFPANNYAKMLAKEYGLKGSDEENRQYLRDNLIEIKIYYEDLTYFDVQQVPSYDLYSLLGDVGGQI
GLFLGASLLTVVEYLDLLGMVAYTSFKYRN

>NeNaC2_tr_A7RGQ6_A7RGQ6_NEMVEPredictedproteinOS_NematostellavectensisOX_45351GN_v1g80
340PE_3SV_1

MSLDICDAYIQQETDIGKLYFVNTHILDAAITGCLSLYDLKLIAAVMSCPVAQRHFETEEDKKDEDEDDRAEDPVDENPD
DTITVSQMWQDFLHTLTLHGFRFVFERGPTIRKVLWLAILLFAVGMLMMQSKKSIQKYFDHPITTSVQVEFLEEIQFPAV
TICNFNLFPYYLINGTIGEKVMSILAPQKYIDNKEEVLFARSPIPNFLNYARKRRRSTGGTIVTDDMLQSEKDFGELDEKFD
FAEFVRTHGHRIDHMIKKCRWKSQPCGPENFTAVITEFGLCYTFNSGMKGHPLLKVQRAGVDYALRLQLSVQQDQYY
GSLRDSSGFKVMVHDQEEPPLINELGIAIQPGTHTFCGLRKEEMHNLPAPFKTACRDMQLEGFKKYTKSACLLKCRADY
VMKMCKCRSYDLKGPAPPCQPREVKNCVWPAMEIFRNESINCECPVPCEITKYQTQLSYAQTPAKHFSEVLARRKHIN
KDVMRHYLRDNFLELDVYFEEMQVTLIQQRQAYDQESLFGDIGGQVGLFLGASILTVLEFLDLLWRILIHKFKKRKNRKV
RNV

>NeNaC3_tr_A7S2F2_A7S2F2_NEMVEPredictedproteinOS_NematostellavectensisOX_45351GN_v1g205
746PE_3SV_1

MLKLPCSEEMRRNTDPKETALEQINQFLQETTAHGFGRLGATAGSKWRLYWVMFCLAAYCVFVWQLVGLVNQYNSK
PIKTRTQLKHAQKLDFPVVTICNMNVLRASRLPPKLRTKFDQIINNTNKTSPRNSSNSRNAFVDPQDLSFEETKKFEILHA
VTAHDDYRELVSAAHQLEDILLSCNFNGVNCRNSKDPTIPTYWTQTWNDNFGNCYMFNSVKTHNGEKVDLYSSSVPG
ESNGLTLQLNLEQNEYLEGITEVAGMKVTISDQGVLPFPGQQGIRIMPGQSTGIQMTKLQTRRIDPFKNRSCESSNEMS
DKNLFFGYNMTYSIMACKYSCLNARKIERCGCTNYNTPEMQKRNIPLCNRLNSTIIDCLNKAYDTFEDGSCDRECPPSCS
EVSFDLTVSSARWPARSFEKTKLKTLQTEYGINMTKEEMFENIAQVHVYYGELDYLLVQETLAYTFMSLLSDIGGQMGM
WIGISALTCAELIELVCVILANMSNRSKKIVHINSFRPEKLGGVETIEKGGVKRIEQEVQGGVGTIEQ

>NeNaC4_tr_A7SA77_A7SA77_NEMVEPredictedproteinOS_NematostellavectensisOX_45351GN_v1g209
148PE_3SV_1

MTRLESITKKDAWWPDPKEAVRNKDNEEHATEPEQHLSKQADSAKNTLIGNFASYTTLHGLHFLFQPIPRARKIIWGIM
LVIATVGLSIQIGNGLVKVYSYNIFTAKTFVRPESLPFPAITICNQNMLRKTAILGSAGQRYLDQLDELRARFIGEDMNVTS
ERINMDAIVQKSGHQLEKMVYECRFVRDKCDNKSIKVFTSEKRGLCYTFNSGVNGTPLLNVSHSGSEYSLSLRLDAEPTE
YYGPYSYDGTGFKIAVHDQKVLPDIEEDAYDISPGFYTKITMKRAQTLFLPSPYSSGCGSRKLQFSQEYSYKNCIRECHTQL
MIGRCRCRAIGMTLQGRVLMSRTRNKCSKGPLTFFLYPENDTTPYCTTREVQDCIMRVHERSSRQRCDCPVPCEMTTF
TSRISLAYFPSQHVWETFLPFLTTAYDLNLTGLSPDELDMKAEHFVRKRFAMVKIFYETLRTEMYHQRPAYEMSDLFADV
GGNMGLFLGCSMLTIAEFIDLAIMLLITKFWRRRTVDHAQARDQT

>NeNaC5_tr_A7SA83_A7SA83_NEMVEPredictedproteinOS_NematostellavectensisOX_45351GN_v1g209
152PE_3SV_1

MRVHSQRQSEMTTVKVANHENPSESVEVPRESLLEAFSSYTTLHGFHFALSSTNRVRQIIWILLLILTSVVLIYQLAYSTQR
VLEYASMVQVETRNEDSITFPAISICSNNMMQKSKILGKDAQRYLDLLDQKKEDQWDAISQSFSPDFDIEKAVHQYGLN
LSLAMKSCHYGRYLKCNPSHFTTFKEFRYGLCYTFNSGKRESAFISHDTGPTSGLSITLDAQPEEYYSLYSSTGTGFRVIVH
DQSEFPWVEKHGWEIPPGFSTNVRLARKEISSLESPYNSNCSRDTNYASQSYCLVQCYSDMVVKRCGCHMLGMTEET
GHTPWCSPQQIKACVYTTSRRFQPNMCSCPVRCSRVEFDVQLSSLYYPPDNFWETISNERSLTIYANDTKKFQEWYRRR
VIQLNVFYKELTTEVRKEKEAYTISDLAGDFGGNMGLFLGCSILTIAEFIDLLVVYLVHRHKKRTAKIQ

>NeNaC6_tr_A7RL38_A7RL38_NEMVEPredictedproteinOS_NematostellavectensisOX_45351GN_v1g239
048PE_3SV_1

MESFNQNTKTDIKDCKEDEKREPQPTMLQDFAGYTTLHGFHFLIHPGSPFRRFIWLLMLLGCWVALFYQLYNSVDRLLA
HRIVMSRGVEQPDEIDFPAITICNQNIMRMSKINGTEAQKYLDELDDVFRKDIKKENVSYDAETFVNKYGHDWENMFE
NIPYSCMFQRFFICSAKNFTSFLSFTRGLCYTYNSGVNRSYVQRVSEAGRNNRLEFHLEAHPEEYYGPYSYEGIGFKIAVH
DQSYVPNMDQEGYDITAGFYTNVRVKRYKEKSLPHPYKTNCGERKLEYYERYSRSACILECQAREFVRKTKCRIIGFPPIK
VIRDVPFCSVLKIETGLTYMYYNWNTNQCDCPKPCVEISYSAQMSLLQYPTPSLVREIRKSFNDTEDYINNMRANSVIVSI
FYETLLTDVFEEKNDYDISRFGSDLGGNLGLFLGCSLLTLVEFFDLGIRWCLGRKDKVQRS

>NeNaC7_tr_A7SH56_A7SH56_NEMVEPredictedproteinOS_NematostellavectensisOX_45351GN_v1g212
215PE_3SV_1

MTSANDDLTNDSRGETNRVGQVRPLWRDFLSRTTLHGTQYACVTKPLIRRVTWLLLLLGMVGYFGYLFYGNLKRYHSH
PVEVTVEIETPNDGIGFPAVSVCTNNKYMKSKINMLRNHSYFHKLGLDIPECAALQNVSGNMTCGQALMCAIVGKYGY
IINERCKWALEKIRKIINDSDYAFDIEKFTLKYGHDIKALLTPRFCTFRGKPCNEEDFVPVITSTSLCWTFNSGFRGSHGNPA
PRKQVTFSGVDFGLTVLLNTRVDENTIGTSSEGVRAVVHEPGEYFSVDHGVNVMPGAHAAILVHAQKTTTLPLPYKSNC
TESKPGLRLYSMEGCVALCASQELTRRCGCRPVGLPYVDAASVCSFKHETCAMDTFGSFDAQRCMCNNACHRTMYN
AKVSYARFPDQYIIRFIQETTSYNYSAEYFRRNLVLVQVGMESLSYEHHRQVPAFPVESLLGAVGGHLGLLLGCSVLTVFE
FIDFFIVALASMLRTNVTPNMDRETKNL

>NeNaC8_same_NeNaC13_tr_A7SJB5_A7SJB5_NEMVEPredictedproteinOS_NematostellavectensisOX_4
5351GN_v1g213150PE_3SV_1

METLKPHPSVRAIFKDFSDRTSCHGIGQIGGSQSVMWRVSWLMIFLAGLGMVLYQGLTLLDTYLNKPTATAVDITYSEV
TNFPSVTICNMNMIKKSQLQHFPQVKRLVDTFNNMTSSNSSLNSSAFMDTNREKAEKDRLSLNTKNSKNKVILNNVNI
DMQRYIEDKIVQYLSMSDTTKLMKAGHVFRELVFRCVWNGFVCNKGDFLKYWRPFWHWRYGNCYTFNQGVDVNG
TELPSLASSKPGPMYGLTLDLFIDQEQYIIPLSQEAGVKVLLSDQRNIPFPFTHGFTVQPGVSASAGIRQLVVKRIDPFSNG
SCYSGNGLEANSIYHKYKGMRYSVQGCMSSCLANNQFKVCNCTEGKFRAKGRPCMTEPEVKCLNNISKKYENGSLGCS
KSCPQPCTHYSFRRTISQTQWSDSYEKTFQRLVRKSDRGFANKMNDASILRKNFLRVKLYYEELNRETITYSLSYPVENLL
GDVGGQLGLWIGVSVITCAEFLKLLVDLAWYLASKMSGKTKTQVQDLNMQ

>NeNaC9_tr_A7SST7_A7SST7_NEMVEPredictedproteinOS_NematostellavectensisOX_45351GN_v1g216
915PE_3SV_1

MDPPEERNKEDAREKEHKEENETWNMIREFAGYTTLHGFHFLVDSSSRWRRRIWFLLLLFCCAMFIYQFVISVNRLRAY
KVVFSRGAEQPGEVDFPAVTICNKNMLRKSLILNTSAQIYLDEQDADFKRSLKQSNISFDAEEFVKKYGHNITNMLNRAK
GCTFKMLYPCSGENFTSFFSFTRGWCYTFNAGANYIQRVSVAGRETRLKLYLDAKSHEYYGPFSYDGVGFKIAVHDQND
VPDMDNGGYDISPGYLTTISVKRFKEVSLPPPFPTKCGSRTLEHYERYSTKGCEYECFAKDFVRKFKCKTLGMAPIKEIRN
ASFCPVSVVSAALMDFHKNWHLELCDCPKPCETVNYNAQLSTAHYPAPSLLDELSKFPLEGFVNKSKEEYVKFVRDNIV
VVEVFYETLLTDVLKEERDYDFNMFASDLGGILGLYLGTSLLTIAEFLDLGIRWCLRRRSDRSQVRSQAWVK

>NeNaC10_tr_A7S8S3_A7S8S3_NEMVEPredictedproteinOS_NematostellavectensisOX_45351GN_v1g20
8535PE_3SV_1

MNLFQTKVKPLSRDKAKSDSEKEIEEQRKTLIKNFSSYTTLHGFHFLLDSSPMPRRVLWTALVGFGLVVFFIQLVMSYGKL
RARESILAKGVERPMNVLYPAVTICNQNMMRKSRITGTAAQRYLDQLDHIKASLSRVNRTNERFETEEMVRLYGHNIT
DMLWECNFMNKPCSHKDFAMRYTSYSRGLCYTFNAGANGSPIGQATTSGTRTSLSLRLNAESDEYYGPFSYDATGFKL
AVHDQNEIPNMDEDAFDISPGFLTNIRIRREKEINLPSPYRSECGSRDLSNAPKYSMSGCIYECYSKIIADKCKCRVLGMAL
NGLNVTQFCTSDEIINCVSTTYLELNPAMCDCPKPCRALHYKIQLSLAYFPSDHLWDSIFPVLLNFTDTTGKSQDEVLLQI
QEALRKQIAQVQIYYETLLTDVLEEKPAYGISEFGSDVGGNMGLFLGCSLLTFCEFIDLVFMLCLHRHRLRKEAKERQARA
AIQDD

>NeNaC11_tr_A7SJJ5_A7SJJ5_NEMVEPredictedproteinOS_NematostellavectensisOX_45351GN_v1g213
238PE_3SV_1

MKKDAFETSNQTNFSVFDENEDIRQKRKKLISEFSGYTTLHGLHFLIDSGSLFRKVFWMILLLVMFTCFFIQLVESYKRLKE
YGSNLSKGVESPEEVTFPAITICNQNMMRKSLVMGTDAQKYLDGQDIMKIKLGAAQVSNESFEVDKMVREKGHLLES
MLFECSFAGTTCTPENFTTSLSFTRGLCYTFNSGTNNTPVFTARAADIRMAFSAMLFSQPEEYYGPFSHRATGFKIAIHD
QSETPDIDLESYDLSPGFATNIRLIREKAKYLPAPYSSNCSSKRGIDGGTYSETGCLTRCYNNLMTSQCQCKILGHESDYKN
ITGFCSTYQLKACVYEAWMVLRPQNCDCPKPCTSLKYKAQISTSYFPSESLWESLIPFLGQSSLPFVNLTGKTLEQATAEA
QVNVRKSVCMVNVFFETLVTDILEEKPSYDLTMFGADLGGTMGLFLGCSILTICEFIDLVIILVANGWRKGKARVIDVKEK
PEARP

>Nenac12_XP_032238530_1acid_sensingionchannel2isoformX1_Nematostellavectensis_

MSYNCKVEDSSDINSNNEYRGGSKTRQLIKEFSGYTTLHGFHFLVDSYSVTRRVVWTCFIVISLGFLLYQLVNGIKNYNDR
GIIMSRSVEEPNEVDFPAVTICNQNIMKKSLIIGTDAQRYLDEMNYIKADLGLVNSTNERLDAEDFVRKYGHTLGEMMY
GCEFKDRRCTAQDFIVSTSFMRGLCYTFNSGRDNSSVRRIATPGRLESLILRLNAQPEEYYGAYSYENVGFVLAVHDQAE
PPDMELNAYDIPPGFTTNLRIRRFKENSLPEPYPTKCGSRNLSLYKRYSRKACMQECYARLIITHCGCRLLGMPPMKDVIE
APFCTSKEYLDCQLMMPVLLKPSKCDCPKRCQHIHYSVQPSLAHYPSKSVIKELLPSLNMTEVNSTTERINEVNRIIREGH
AIIRVFYETLRTEIIKEKPQYTLATLTSDMGGSMGLFLGCSVLTICEFIDLFIQICLERRKRNEVINQK

>Nenac14_XP_032226792_1degenerinunc_8_Nematostellavectensis_

MSRSGPSVRALLRDFSDRTSCHGIGQINGSHSPTWRIFWLLTFLAGLGMVLFQCITLLGIYLDKPTATSVDVTYDEVTNF
PAVTICNLNMIKKKNLANFTQSKKIFDDFEAFVSSNSSMDSSAFLGSKMETVLKDRLSMDSNDGSNSVSLDDTSMDTEL
YVEDMLIRHMAMVDDKDLIEAGHEFDELVFRCVWNGFTCNKGGFMKFWRRFWHWRYGNCYIFNQGVDENGTLLA
HLTSSKPGPMYGLTLDLFIDQEQYLIPLSQEAGVKVLLSDQRNVPFPFTDGFSVQPGVSASVGIRKLVINRIDPFNNGSCY
SGDGLEKENIYSKYKSLKYSVQGCMSSCLANSEFSICNCTEGKFRVKGRPCMSESEVKCLNTVNKMYEKGTLGCTRKCP
QPCSHFSFRRTISQSQWSESYEQTFQKMVMKSDKGFDKKMRDASVLRQNFLRVKLFYEELNTEAITYSRSYTTESFLGD
VGGQLGLWIGVSVITCAEFFKLLIDVVWYLARKVHGGPKKTVRDLNMN

>Nenac15_XP_032229849_1degenerindel_1_Nematostellavectensis_

MADQPRPSVRALVRDFVDRTTCHGIGQINGSQSPLWRVFWVVTFVAGLGMVVHQGVTLFGTFLDRPTSTTIDMTYA
PAMDFPAVTICNLNAIRKDHLSQFPDADVLLKGFSEASTPSVTVFLGKDVESFIKQHSEAGPNVTLDPELAFKDAMVEIF
AQSELKKLQMAGHGFEELVLGCTWNNIKCNKGDFLKYWRPFWNFRYGNCYTFNQGMSEKGVAIKPLTLSNTGPNYG
LTLDLFINQEQYIAPYTQEAGVRILLSDQNQIPFPDSDGFTVSPSSSSAVGIKKIFITRIDPFNNGSCYKVTKGLEEGSIYKSV

FSHDMGYSVQGCMNSCLANKQREMCNCTEGRFDMMTGLICQQIEAWRCLNRVNMMYQEGGLKCLEKCPQPCTQ
NVFSRSMSHAHWAEEYKKALSKFFPNNGNGSSVDPNELISHNILRVKIYFEELNMETITYKRNYPVESFLGDVGGQLGL
WIGVSVITCAEFAKLLIDLVLCVAKKMNRSDKVQSVCIGRGN

>Nenac16_XP_032229236_1degenerinmec_10isoformX1_Nematostellavectensis_

MKAMAKSGNYTINRNVRLNEEQEAKKKNVLKESKEAIKEFSFETTAHGFARAASSYSRLARSLWILLLLVAFSFCFYHNSV
SIGKYFNFPAKTDVRFSDAWFSIKYPAITLCNINNLKKSSFFVKTVVKELNKTRDAFGDTSNKRRENLERVLYFMSGYKEE
QVKDFGHQAKDMIIACNAPGRWAQYPCDYRNFTWFYDQIHGNCYTFNAARNQSEPVYIEDAGHRHGLHLTLFVEAE
EYWSYLSDSLGFVISIHDREQLPFPAETGYFISPGMATSVALQSKVLKRLPWPYDHKECETRRGIPGVTPSNTIYSEEGCR
KSCVARMMLDKCKCVSLLYRLKPSCNPFNKTQGSCIEDVLRSRSKESMSSCNCPPACREDVFTTLMSSSVWPTRSYMS
NVLNTLPQGSQAQKVVTNLSSARENMAHVMVYFPTLMQESIVQKPAYQIESLLGDIGGQMGLFIGISVLTLVEFIALLW
DLIAIVILKNKKNTSAPDGEKP

>Nenac17_XP_032233690_1acid_sensingionchannel3_likeisoformX1_Nematostellavectensis_

MPRSPKEVLKDFGETTTAHGIGKLVGSKHPIRKTVWLVLLCVCFGYVIYQIKELCGQYKEHQITVMSHAERKKEIPFPGVT
VCNLNPVRKSKIIEHNLTELLKPSGPNAIGQVEMRNKLMSMVMGMNATTRTELGHQEKDFIFGCYWLGQPCNTSFEQ
IFTPEYGNCFTFNSGKLHDLAVTEAGDTEGLSLLLNTEQWDYMGAVSPSIGAIVDVHHPDDIEVPAVRGITVSPGQATSI
GLVLGSMYRLDSPYSKTGCKSIDTPTIYKNKIYSAEACYRGCLIAKSLEKCGCVESRMQVGLTAQTCDPFNKTLRKCIEEVV
DFNSCSCPPACREMKFEFSASTSVWPSEPYWTFLKEAYDSRNTSRSFASFPDSRGNLLAVNIFFKDMSITIKTETFSYEFS
NLVADVGGQLGLFVGMSVLSIMEYWELLIDLGTALLRRDKTMRVAVASKPEPSTALHRKDKVCCRGLKDRT

>Nenac18_XP_032233718_1degenerindeg_1_Nematostellavectensis_

MSQNEEKCRQTVSGLIWDFFGQTTAHGFGKIANAKSRLRRMFWVCLVMAAFGMFSLQIYDLLMIYLSRPTGTQIWM
KHAPSIAFPAVTLCNFNIVRRSQITSELAGSFENFITPNISDVDDARSLRKRRQSVDGCSCSCYLKCRNFHDPHRNTPEPA
TTCAPVPTTIPTTATTTEPTTVPTTIPTTIPTTLPTTEPTTAPTTPEPTTLEPTDPPTTEAPTTLPPGVTPAPTEKPLTIEEKKE
QAFTGEVSQPDPATLDKQTLKTEELVLSLATKPEDMLMEVGHQFNDMVLSCTYRGIPCTNYTENLWSRFWHYRYGNC
YVFNSGKDWQGEPKPILRSNKPGPASGLTLELNAEQHEYVGQLSHEAGMRVLISTQGEMPSPLEKGISVSPGFSTATGI
RLTNILRADPFNNKSCLSSDDIDPENLYRKRYNVSYSRTTCMDSCLAHKQVDMCGCLEYRFPNIDNRTVCDILDIDVIRCL
NKVQSLYQDNKLGCSEKCRPPCNEEVFKLSISSARWPTDDYEAHFLDELKEEKGFVLPANADARAYFVKLQIFYEELNHE
VIEEYRSYELVNFVSDIGGQLGLWIGISALTASEFIELLIVIALHLLRKVTRIGLIDNSAEQPADTNPCYLDDITLTGDLKERRG
ALTESNTSLTVRYLAEVARKRNKKGHKEKREASWHIDEEAFPCYRDQATIHPNVESNTSLTVRYLDDIVQKKSKHRKRSK
GDTSMVQKGDGHRGDLSGRRGELTESSGSLTVRYLAEVASKRKR

>Nenac19_XP_001623796_2amiloride_sensitivesodiumchannelsubunitalpha_Nematostellavectensis_

MDSHSNETKKGEKDQTRCQSFTALLREFCGYTSFGGLGRVTDSPYLLFRITWFFLFMGAVAYASTQLYSLYEKYQDRPV
NTVISLKHSEKLSFPSVAVCNFNILRQSVMQDTLSNVLNMVKEQTPRCVRLRNCSTTSHNVSTTSSPRPEDSDPEVTKEL
EEMGFENPEQLSDEYRLEELVSITIAKLSPSDLKPMGHQFDDMVYKCTWRGVDCKANYTQGKYWTSFWHYRFGNCFV
FNPSIGWNRNTRSRLHIYRTGPLNGLQLQLNIQQEEYIDQITKRAGAMIQVFPFGQMPFPEERGTLVAPGFSTDIGLEK
MDIKRVDPHNNGSCVTDEASINTENIFKQKFNTSYSKTACVLSCLAYHQMHECKCMEYRFPTDSVCDTLNSTIVECLAK
VKFKLSRNELGCSTACGPACREEAYRLTSSFAVWPSQRYQNSIIDNLFDGIFNYSRSWFRDNYLEVNIFFKELNYFIEESVA
YELVNFLSDIGGQLGLWVGVSVMTGAEVVELALLIFFKVFCRKPRKVTVNSEGDPEPNVLSNSQLQLESIDNYKYAP

>Nenac20_XP_032233720_1amiloride_sensitivesodiumchannelsubunitbeta_Nematostellavectensis_

MEEKKEQAFTGEVSQPDPATLDKQTLKTEELVLSLATKPEDMLMEAGHQFYDMVLSCTYRGIPCTNYTENLWSRFWH
YRYGNCYVFNSGKDWQGEPKPILRSNKPGPASGLTLELNAEQHEYVGQLSHEAGMRVLISTQGEMPSPLEKGISVSPGF
STTTGIRMTNILRADPFNNKSCLSSDDIDPENLYRKRYNVSYSRTTCMDSCLAHKQVDMCGCLEYRFPNIDNRTVCDILD
IDVIRCLNKVQSLYQDNKLGCSEKCRPPCNEEVFKLSISSARWPTDDYEAHFLDELKEDKGFVLPANADARAYFVKLQIFY
EELNHEVIEEYRSYELVNFVSDIGGQLGLWIGISALTASELVELLIVIALHLLRKVTRTGLIDNSAEQPADTNPRYLDDITLTG
DLGERRGALTDSNTSLTVRYLGEIARKRKMKGHKEKRETSWHIDNSADQPADTNPHYLDNITFTGDLGERRGALTDSNT
SLTVRYLAEVARKRNKKGHKEKREASWHIDEEAFPCYRDQATIHPNVESNTSLTVRYLDDIVQKKSKHRKRSKGDTSMV
QKGDGHRGDLSGRRGELTESSGSLTVRYLAEVASKRKR

>Nenac21_XP_032233957_1degenerindeg_1isoformX1_Nematostellavectensis_

MGKEKKKIVTKIKDFLGYTTAHGFARLVESKNPVQRAFWVLACLGAFVAFGLQVHKLFSKFNRKPVDTYITVKHEMNLE
FPAVTICNLNPIRKKHLPESLREYFKYSLGDSSTKEPTSKPTTSSLLGKAFSSVDSLLTKPALLSNKTLPSLVNSSLDTLGGAL
NDSLSSVNKLKDSGESLVNSTLGLVRGLLSRRRRGLLGDVLDVDGVLPEKEKDGGKPGLLGNLLDGVLPEKEKDGGKRK
DGERPAPEELCESARRRKYLYWRIAETPTETLIQSGHQFLDMVQECSFAGVSCLGYNWTQFWNHLYGNCFVFNTAMI
GSNRERNNISYTSSIPGPSDGLLLRLRIEGEEYLGNFSTVSGVRVHISDRHEMPFPGDKGFSVSPGFETSVGMRKVFIHRE
DPFGNNSCISDVGKPVVRPDYYMENFGVNYSTIACRSSCVANLQKTMYGCVESAYAITDELRKIVCNALNTTVFRCIVKA
NETAEDSDGKSCLTKCAIRCDENIYKLTISSAVFPSPSYQNISSQDLKSDSGNEEQIRANLLSLRIFFEEHNYETIEQRKSYELI
DFLSDIGGQMGMFVGLSVLTCAELIELLALIIMHLFNRDHKRGAVGADTVTPVIELKKKPLVRLPRVE

>Nenac22_XP_001636193_2acid_sensingionchannel4_B_Nematostellavectensis_

MKQSANIEQDELATVVLEDFKSNTTLHGLPHALNSTRNWRKALWVCLTLGSAAALVTQLTENWETLFSYEIVKFSTPKL
HENLTFPAVTICNENSLRKSKVINTSLQEVYEYLRNKTIGNVTDEVLYYPLGMTKMFGEDGHDMRDMLLTASWNGHLV
TSQDFQPYFYSKYGKCFVFNGSQNKTTPARTTTLKGMGSGFEALLDIQPEEYMDLSPNEASSAIGLVVNIHDQHQDPAII
DTGVSVAPGSRVDFAIKLTQQKRLPFPYPSKCEKVEKARLPLTDRVYSVSRCEEECRAKEMEDKCRCRFLDLANDKDVPY
CTPRNTSCIMAVSNTHKKCSCPTPCEENIYEVRTSAAYHPTDYFLKQAARLYYNVTNLTTEASIRDFEQFYRKRAVYIRVYF
ESLTSDLIEEKATYELPSFLSDIGGSVGLFLGCSFLTACEFLELFAVTVASLFMKFFRRNNNNVRDFRA

>Nenac23_XP_001625049_2acid_sensingionchannel3_Nematostellavectensis_

MSNAPDYRNFIEALHQTLNNPNNRVAVDNRGYALESPLDFYKNNWVGDGESVDQKKAQEVDPWNKDEKLTISQHFA
SLCSSTTMHGISNVFDPSSATLRKGIWSVVFLCSFIFCAYEIGNNVRYYLTKPVNTVFKIDYVDEIKFPAVTICNNNPIRKS
WAATTPYLPVIMAYNANPGEEAIPINWDAYNWTGFGFDKLMSSAAHLASEMIHTCKWKGIYCSAANFTLDATFLGGC
YTFNIDQRLMVTGTGMANALHLVLNIQQNEYIGNVRSGAGFRLLFREKHEPPSTDRFVIALQPGTQTLIPLTMKKLISLP
EPFGVCQEKNNLKMFDKYSVTACEFECRARLGGKLCGCREMYPSSIKTEIPVCLPKAYRDCLNPLLVEISVNNLCRGCKN
PCNKVTFVPRMSYSQYPANHIADSMAMSMNTTRDFVRDNFLEVEIYFEDIMVEIIEQQEAFSLTSLVGIIGGTLGVFIGA
SIITVSEFMEFLILIPFRR

>Nenac24_XP_032222096_1acid_sensingionchannel1B_Nematostellavectensis_

MALCGCCASKPSPTNSASEDESTGPESKEEILLANQNGNASTYENIRKRSAAKRDKKHAEYGKRKMREVFAYYINHCTL
HGFHYIFETKSLFRKIAWFSVLAIAGGFFFKEIKTSTTQYFKYPFSIMSTVEYPRTLVFPAVTICDFHDIRQTLVSNAKGEQAI
NISSSEKLETLARKTYKSFNETLISCSLRRGVRGARPFNIHDFKVFFTAKGQTCYTFNAAMDGKKLLEVDNVGPKFGLEIYL
NAQHWWFKDDVRESGFRFILHQQDDLPLTREGFRVSSGYVTYVDMRLEKVKNLPYPFSSDCDGRGLDLYPKYSRNNC
YMESLTKYILQQCKCRAWFMADVINTSTCSIKEALDCMWPAWEDFDNAYNYTCPVDCEERVYKTRLSSALFLPQKLLPL
TKKYKFLMRPKGIPNDTEGAVDFILENYSVINLFFDELRLDTIQQTPAYGFFRLVGDVGGQLGLVLGASVITIVEIIDLVIMY
SIYWIKSKTVPRAATS

>Nenac25_XP_032221585_1acid_sensingionchannel2_Nematostellavectensis_

MISAMKETWREFASNTTLHGLRYAVGVCDIRRRSLWILCLLACAASYLYMVIISFGTYIDRPIRTEVSHEFPKDGLDFPVV
TICNLNYFVKSKIDTGYEDEAFYTRNLNMSVCDIIRGVSGNLTCGQALLCAYETFGSSVVDGCDDVIRSRVIAAVNSSNRP
FDTEEFMTRYGHDFAPMLMGYCLFSVNEKCDINDWSPHITPGGMCYTFSKANQSKVYFLGGEGGLSIILDAQISEHTFG
DFSTGFRVILSARGTYINRARGFNVFPGSHALVAVTPKKFERLPAPYRTNCSDKFLPGYGKYTKDACYTQCINNATMTDC
GCRLPSQHAKFNNLPKCSIQDQKCRAAASARVRSTLCDCTVPCKEQIYEPRISYSKFPDITITKILTNHFKLNKSASYLRDSL
VFLQIGFEELAYLVDRQAPSYGPGNLFGDLGGNMGLMLGCSILTLIEFIDFLWIALKSSLNKSKRTAINDCGFSQGTGTSN
QDANNVGLDMHVKT

>NeNaC26_XP_032233942_1amiloride_sensitivesodiumchannelsubunitgammaisoformX1_Nematostella
vectensis_

MRVLISTQGEMPSPLEKGISVSPGFSTTTGIRMTNILRADPFNNKSCLSSDDIDPENLYRKRYNVSYSRTTCMDSCLAHK
QVDMCGCLEYRFPNIDNRTVCDILDIDVIRCLNKVQSLYQDNKLGCSEKCRPPCNEEVFKLSISSARWPTDDYEAHFLDE
LKEDKGFVLPANADARAYFVKLQIFYEELNHEVIEEYRSYELVNFVSDIGGQLGLWIGISALTASELVELLIVIALHLLRKVTR
TGLIDNSAEQPADTNPRYLDDITLTGDLGERRGALTDSNTSLTVRYLGEIARKRKMKGHKEKREASWHIDNSADQPADT
NPHYLHDITFTGDLGERRAALTDSNTSLTVRYLGEIARKRNKKGHKEKRKASWHIDEEAFPCYRDQATFHPNVESNTSLT
VRYLDDIVQKKSKHRKRSKGDTSMVQKGDGHRRDLSVRRGRLTESSGSLTVRYLAEVASKRKR

>NeNaC27_XP_032225907_1acid_sensingionchannel3isoformX1_Nematostellavectensis_

MSDTSWKDFLGNTSLHGVRYLTENSFIRRVIWLLVVLAAFCGFGYQLYKSITFYYEWPTVTAMTTEWDEELDFPAVTIC
NLNTLSKENFVKARKRYKSDATTEQLEKEVETFTFFMSAPLNENAKSKLLNDFFENSTFYRERGNFVNKSLNDFSHSIDGI
LSIKWLRPCQFSGEDCDRKNFSSFVDLNFGLCHTFNSGKKEENALKVESLGPNNGLRLRLNVQQEDHVSNFYWLPAGF
KVLVHDRKEYPIMEQTGFALQPGTHTFCSTKLKKYKNLPTPYNTKCGENITDFNRYFSVNYSMSTCGKQCLHDYGIKKC
GCQPIYPITKGVPLCALNDTFECIYPTYYGKFPREAAECFKKYCPVPCDYTKYETKLSYASAISNALARSLPTHLTESKDVAD
AFSDIQNMTEQERKRFFRENVATLDVYFEELSYDKIEQKPSFDSWSLIGNIGGYLGLFLGMSVLTVMEFLDLLVIKFLNRIS
GTKTRVTTIQVGS

>NenaC28_XP_032224488_1acid_sensingionchannel3_Nematostellavectensis_

MADTSWKGFLCNTSLHGARYLTENNFLRRAFWLFVVLAGFAGFIYQSFLSITSFYSWPVSTVMMTEWHKEMNFPAVT
ICNFNPISKSRYAQNYGNLFNLSTQESNNIINGIIFLMSAPRDSSQKSKFSQFSLENYTQRGPMLGKSLRSYAHSIDGMLS
LKWLEPCKYSGKECGKNSFSTFEHLRYGSCHTFNSDGQFGLKVDSIGPISGLQLRLNVEEKDHVSNLYGLQAGFKVLVHD
PREHPMIEETGFALQPGTHTFCSVRMKKVRKLMITKDSKYVNLRAPYRTECGENITDFNRYFNVNYTMAICSKQCLHDY
GIKKCGCQPIFEITKDVPLCSLNDTFECIYPTYNGPYPREAAECFKKSCPVPCYYTKYETKLSYASAISNAIARDIPTYLLPNG
SEFDDIEKMTEGERKTFFRENVASLDVYFEELSYDLIEQKPSFDRWSLIAMIGGYLGLFLGMSLLTVLEFFDLLVMKFLNRI
SSNKKGVNTVTVEPQESQNAFAFKQGPTV

>Proto_ecdy_PANarthopod__XP_023244170_1_amiloride_sensitive_sodium_channel_subunit_gamma_
like__Centruroides_sculpturatus__759

MRKLNLEAQSQNSSLCGLCRSFAYRSSAHGVQRIASSQDNARRLMWSVVFLFAIAGCGFHSVYLILTYLSYPRMTITEEI
HADHIDFPALTVCNLNPLKKSSIREHLLETHQSVSEFYNPDKYDDDDEIEERGICFRSENEFLKKSKTMDLSDVWMSVIAT
KDSLMKCGHQAQDLITQCTYNARNCFNDSSTIVLLDQYPSPRYGLCHTIVVDKEPLRKVKKTGLSLGLRLTLNIEREDYLD
LVSPEFGARLLVHPKGTYPTLQGGGVVLQPGTKTYVGVRMRKIERLPAPYRGCYDNFQSSQLIHFLRKQDQSQAPFAVI

HNVYTYEYCQTLCRDIHLLKNCGCVEEVTPVNNKFCDPCNTTQARCRTRFYRSFTRANVDHDCQKFCLPACTDVRYDLT
VSRSEWPNVRHQNYVLKRWPNLRQSLTVFLTNKTLKNGSDELDVDNLRKNFLRVHVYIQEMNYLSVRDIPAYTLPQLF
ADLGGCLGLYIGVSAITIVEVLEHMASVMAFLYKKRKEYFPKSNSVKKSEIKKRRTNFRLSERSHVQKYSKSKDCKRGNH

>Proto_ecdy_PANarthopod__XP_023215424_1_amiloride_sensitive_sodium_channel_subunit_gamma_like__Centruroides_sculpturatus__870

MRPYIKDPDIHSINTLCGLCQAFAYRSSAHGIPRIASSQNRFRRFMWIIVFLVAVAGFAYHSIYLILTYLSYPRMTTTEEIHA
DQIDFPYVTVCNLNPLKKSYFADYLAEISTSTPRSTEDTQSDDLLEFLTSPSSDDHGDFGGGGICFRSVEEFLQRSRTMDLS
DMWMSVIATKENLSKFGHQSKDLVVQCTYNARNCFNTSYSIIDVDSYPSPRYGLCHTIIVRQESLRKVIKTGSALGLRLTL
NIEREEYLDLISPEYGARLLVHPQGTFPTLQRGGVILQPGTKTYVSVRMRKIERLPEPYRGCSDDFQKTLLAKYLKMTGQE
FLLHHQVYTYEYCQTLCREAHLLNQCGCLEELALEGNKSCDPCNGTQARCRSGFYRSFTRASSTHECQKLCKPACTDIRY
DLTVSQSEWPNVQQQQYALKRWPNLNKRLGQSLSSSNSNNTTDTKEEIQINRKYFRKNFLRVHVYIQEMNYLSVKDIP
GYTLPQLFADLGGCLGLYIGVSAITVVEIIEHVFSVFAFLYVKRKRNSQTRDPIHRMAISSRKSAVVTRHVPTQFPEPYSIQ
KRDLAKNIKHLRAYFPRDTHRRTGGIAETPGNCKTLYPINEQLFRRYPHLTDDSLDSKW

>Proto_ecdy_PANarthopod__XP_023225242_1_amiloride_sensitive_sodium_channel_subunit_gamma_like_isoform_X1__Centruroides_sculpturatus__519

MGFCLFHSIAYLREYFQYPTVLNVQVNTESKLDFPAVTVCNYNRIRSNALDSFCCTSGLLLKASSLCHVLNVTCSQTNNEE
ASSKRLKTLIGSPPLSASTAFATCPRNRTNNTVSYEGSIMKSFSNAYVSLPLSKKIQMGHQVQKFVHQCQFMGQPCDHL
NFTTFHTFLYGNCFTFNSGQNGKDILRATTDGSMSGLHLELNLETDEYIFDLTNNIGARLIIHPSNVKPLPENEGIAITSELE
TSISIEQVNIHRSKPPYPDNCIDYPNDGEGFLYTRNTCLRDCFQQLSQNKCSCADPTWPLPRNSTSCDLRDVTQVCCLDD
VRQLLRNDENICICPLPCCEIQYKLSVSSTKWNPLKSKINKEMYSSILPSDIAKVRIYFQTLDHVVLKSHPKYQVQDIFSNLG
GQMGMWLGISLMTLLHYFETILGSVFCKQKQTIHQSPIGINP

>Proto_ecdy_PANarthopod__XP_023225243_1_amiloride_sensitive_sodium_channel_subunit_gamma_like_isoform_X2__Centruroides_sculpturatus__470

MGFCLFHSIAYLREYFQYPTVLNVQVNTESKLDFPAVTVCNYNRIRSNALDSFCCTSGLLLKASSLCHVLNVTCSQTNNEE
ASSKRLKTLIGSPPLSASTAFATCPRNRTNNTVSYEGSIMKSFSNAYVSLPLSKKIQMGHQVQKFVHQCQFMGQPCDHL
NFTTFHTFLYGNCFTFNSGQNGKDILRATTDGSMSGLHLELNLETDEYIFDLTNNIGARLIIHPSNVKPLPENEGIAITSELE
TSISIEQVNIHRSKPPYPDNCIDYPNDGEGFLYTRNTCLRDCFQQLSQNKCSCADPTWPLPRNSTSCDLRDVTQVCCLDD
VRQLLRNDENICICPLPCCEIQYKLSVSSTKWNPLKSKVQDIFSNLGGQMGMWLGISLMTLLHYFETILGSVFCKQKQTI
HQSPIGINP

>Proto_ecdy_PANarthopod__XP_023240338_1_acid_sensing_ion_channel_4_like__Centruroides_sculpturatus__568

MKSYEKRSNTKPPYIIAWENSSKNKKKLSRKKCSLSGFTYSSYKFLTNFFEYPVVVNLEVENEGELFFPAVTICNSNRMRV
SELHKDVCKSKRKSCEYLRIIHESSSDIENKPEFVFKFATCSNSSKREMTLYHQKLLAFTNTFSRFEDWKIKKMSHQIEDMI
EECTFDGEKCDASSFTPLESFTYGSCYTFNGRWMNSRGQEDMGRRKRKKFIDDGNEKLISRSTNPFSGLSLTLNVQIEEY
LSITSKMGVHVIIHSPDERPYPEETGIDISTGFDTAFAIQENLYRRMSKPFKDKCVLYGKDDFKKIKSQKECLSLCTRDLIKK
ECGCLDPTLKFYRNVTFCDLTKEEDACCMMKMRKVNISYCQCPLPCHEITYDVSLYSTIWPSESEFKLRSLKNLSKQITYE
EYRKGHLQMNVFFDRMERFIYNQHPVYEKDEFFSHLGGQLSLWLGLSLLTLFEYLEKFSLFCIQVYKNFSRK

>Proto_ecdy_PANarthopod__XP_023226921_1_amiloride_sensitive_sodium_channel_subunit_alpha_like__Centruroides_sculpturatus__577

MQEPSEKDKNTSIFEKFLGSSSVIGLSQITKSRSIVRKLLWLAVLVTGLTFCAIESHKFMREFYKYPVVVNLEIENKGALEFP
AVTVCNLNSVRKSEYQKRFVKTITIESAFDSPFLPSAARGFFIPFRKFTSQCDAVEQKSRNENNQNDFYNTYFSLNKTVRG
SLGHQKENFIMSSSFNVKQISLFDNFQMVVKPRHGPCYTFNGRNTTSQNMNVSSVRVSRNIGTLSGLTLVLNLELEEYF
DKTKSVGARIVIHSPDEFPDTEGEGINILPGVETSIAINKQTTKRLPFPYKDRCKDYKKQTIRNVSILNEKDCIIHCLQDLNIQ
RCNCTDPTLIFKKKRQKCDLENNEEVCCLDKVFEEMSSLNFETCDCRVSCSITNYEMTTSMSLWPSPEEFQYNYQYYGF
NYSSFRNTYAKLNVYYATLEETIYAQRPVYQNNEWYSHLGGQLGLWLGISLPAIFDCIETIVLLIHHAIIKYITKN

>Proto_ecdy_PANarthopod__XP_023226919_1_acid_sensing_ion_channel_4_like__Centruroides_sculp
turatus__804

MRELNNKSKEDGGENEEEFSIQTFFGSSAVIGLPQIASSQHILRKLLWCAVLITGITFCALESYKFMTEFYKYPVVINLEIEN
KGILEFPAVTICNINRVRKSEYHVINPENVNNRTTFINSTPRDSYCSRERLVDERNQAEFYSKYFFLNKTVRNKIGHQYHKF
IIMKSLRNMNFASHFFTSITKPNYGACYTFNSKINTSYKDVPMSTHIGPKSGLAMTLNLELEEYLDDVKTVGARVVIHSRE
EFPDTEGEGINIPPGMETSLAITKKSIRRLKSPYKDKCRDYKGEYYANKGPAILNQKDCRIYCLQQKNLKTCKCTDPLQVFT
ANKKCNLKNLNKMCCLDNVFDKMIHSQYAECDCPLECLTTQYEPIISSTAWPAPENFKRLELQNYSEFRKTYSKVNIFYS
TLEETIYVQRPAYENSEWYSHLGGQLGLWLGISLPAIFDCIESIILLLCYICLPKYRNKKKFKNDFETIVKPVNGPCYTFNGR
NTTSENMNVSSVRLSRNIGTLSGLTLVLNLELEEYIYKTQSVGARIVIHSSDEXVTNFYICSSVTSKWLSNQDMGRVIHLM
VETRLVKI

>PanArt_PPK14_NP_609017_2_pickpocket_14__Drosophila_melanogaster__67_OUT_153

MFVRSTEKETRVVADRIRRQDQNPLAPVNTKSEIQRAWTLLIDSYISRSHIHGLYLLFLPSMRRRMRVLWALALICACTV
LFHVSYLLGDRYHNKQFQTIVAHAHASIHHIAFPVVIICNKNRLNWSRLPEIKSLYNITPSQDELFDRILTAYDGFSFHKFN
AFDSLLGESLDELNHLNFTEIVIQMSWRCDEILRDCHWQTASRDCCKLFRPRRLPLGYCLAFNELEKRRGTETGINTGLLL
RLLLREGQHAPGNSGLKGFWLTVVESSVWFGFPIEVVPHSRTNVAVTAVYHYFDESTLSLPSSWRHCVMDYEEESEHF
RTLEGQKYMLENCQAECQQRYLLRYCNCTVDLFYPPSNYPACRLKDLPCLAAHNHLLQNFEQPGEHPYVHREESGLVC
ECLHNCKSLTLLTDMRKSVQQPWLQPNSSAIESMWLNVYFKKPSMLVYKTNLIYTWVDLIVSFGGICQLCLGCSIISLIEF
VFFALYKVPQLYWERFSNEHRSNK

>PanArt_PPK19_NP_651708_2_pickpocket_19__Drosophila_melanogaster__72_PUT_175

MLLYTKELVIPRPRPGLLRFRNNPRGIKFREKLCNSFAHSNIHGMQHVFGEQHLWQRCLWLAIVLGAVITGFSLYTVLM
HRHSEQLLVSLIETTQLPVYHIDFPAVAVCPWNHFNWQRAPSAFIRFLPRHPNAELRETFRQLLASMDIMNFSNFNRIRI
LTKRNLTGISYLKMTDLMNFMTYRCDELFVADSCVFDETPYDCCKLFVREQTVKGQCLVFNSMISENSRKKHLINQFYP
HKLSTAGEDSGLKFTINASYSFMNNIDALTPFGMNLMIKEPRQWSNEMMYHLYPDTENFVAVHPLVTETSPNTYEMS
PKKRRCYFDDEKNPTFQNTSLTYNRENCLVVCLHLVVWKTCQCSLPAFLPPIDGVPECGINDAQCLGNNSDIFTYVKMG
DQEKYINDSRQGHFCDCPDNCNSRLYEMSLNVRKLDYPKNSTDQLIKAQVYYGQRVMTKIITKLKYTNIDLLANFGGIISL
YIGASVMSFIELLFVLGKLMWGFIRDARIKLKEYTK

>PanArt_PPK21_NP_651704_2_pickpocket_21__isoform_A__Drosophila_melanogaster__74_OUT_236

MLYPLELPRARRPLYRDYGGKSGLIQTQRDARKNSRLGKLWHFMLPYLKDYAAESSVHGIRYLADPKMRNYLNIRVIWL
LILLTTSIGAIVVYVDLNELYQTVRIQTTIKNTMLPIFRIPFPSIGLCPRNRLNWKILETEAVDHFLGANVSAAQKDLFVKFFT
AAGDPHLSRLNEMSNFFGNKTLTDELHMLDHLDLREVYKFIQFRCQDLFHTCRWRGNPVNCCEVIEYQFTEAGLCFVF
NTEISPASRQKAREDKYYPLRTPHYGEGSGLDLFLRLNRSFIRPGKRGINVMIKQPQQWSDVVRHVPHEAHTRISITPRF
TVTDERTRTVTPEIRRCIFGDEVDNPHYKNFPDFEYWVGNCRSRCHQEHVLNLCKCSPSIFFPISDKDNFTACKASDFKCL

YDNRFTFSIERHPEEDDFVKNPFKESMICDCFTSCSQLVFDRVFTTTTLDNNETDTEAGTMRLDIFYQSGWFIQYQTNM
RFTFVELLASFGGIIGLFLGASLLSAFELAYYFSIGLYLYIHGKRKLRKPEPGVLTIQFGQRKITPIKF

>PanArt_PPK7_NP_609016_2_pickpocket_7__Drosophila_melanogaster__60_OUT_279

MTLVYFPPSKLQQQQQPSRSSRLAQQLAQSSWQLALRFGKRTTIHGLDRLLSAKASRWERFVWLCTFVSAFLGAVYVC
LILSARYNAAHFQTVVDSTRFPVYRIPFPVITICNRNRLNWQRLAEAKSRFLANGSNSAQQELFELIVGTYDDAYFGHFQS
FERLRNQPTELLNYVNFSQVVDFMTWRCNELLAECLWRHHAYDCCEIFSKRRSKNGLCWAFNSLETEEGRRMQLLDP
MWPWRTGSAGPMSALSVRVLIQPAKHWPGHRETNAMKGIDVMVTEPFVWHNNPFFVAANTETTMEIEPVIYFYDN
DTRGVRSDQRQCVFDDEHNSKDFKSLQGYVYMIENCQSECHQEYLVRYCNCTMDLLFPPGQYRSCRAQDLLCLAEHN
DLLIYSHNPGEKEFVRNQFQGMSCKCFRNCYSLNYISDVRPAFLPPDVYANNSYVDLDVHFRFETIMVYRTSLVFGWVD
LMVSFGGIAGLFLGCSLISGMELAYFLCIEVPAFGLDGLRRRWKARRQMDLGVTVPTPTLNFQQTTPSQLMENYIMQL
KAEKAQQQKANFQNWHRITFAQKHVIGK

>PanArt_PPK20_NP_651705_2_pickpocket_20__isoform_A__Drosophila_melanogaster__73_OUT_291

MAKGDNSVKPTEAAGFGDNERCLADLLKVHFRSYCEKSTIHCVRYLYDSHLHNLERFVVYPLFISTFNFKLYIIIVFRIIWS
MLLIISIGLSFFFYLLLSERFVSQKLQTVVHDPQFPVFLVPFPAVGICTDNRINWNKLEAAKEQFLPTNASVELVESFTVLVS
RMETLRFGSYLSSLAELEDDNLEAVGFVNLTELAIFMTLQCSDIMVPKSCLWRSSSFNCCEYFVLEKTEFGFCLVFNSEVS
PRSKAIKQKEGNNFYPRHNAKAGQSTGLNFDLILNESFRRPDSQANNNVYVSICQAPDQLNNVVYSITQNTETYVTVRP
GLTWTDNTTRSIPPERRNCLFADEQGELDANDSAKNFGKPFQLSNCLNRCHESYLIQLCNCSLPIFFLYNHRVPDCNAVS
LRCLARHNDIFSYDKRRDEDALFSATKLGMTCSCLVDCYLLDYYTSTTTLPLSAHKLPKDPHQKLFRVDVHYQVETTPLYR
TSLEFTIIDLIANLGGIFGLCLGASMVSAFELIYYLTVGLAMHLYDHQYYGVLFKHLKAKWVNLKGYLRNEVGHLAENPAA
HKDTNDRKLRHPFNYRKNVW

>Protost_ecdy_cyclo_nema_Cele_tr_P91103_P91103_CAEEL_DEgenerin_Linked_to_Mechanosensation
_OS_Caenorhabditis_elegans_OX_6239_GN_delm_2_PE_3_SV_1_819_308

MIPTISKPKTNQSTKSARKSSNESPYPSVTFRPNSSLSHLIIEVPVAQLKKFKHAGETVSVERETQHFCETTTMHGPKRIFQ
GKRWATLFWLIMVCTSIGLLITQVCILASNYLSKPVVSDVSFLINEEGIQFPQITICNFTPIRKTFVNEMNKTGQISPNMIN
YIMHWFTEVPILIGSSNWQLLHEGNKDLQEYQKNNPNFTVQGFFIDAGFSCSDIFKLCSFQGETFDCCSISTPVLTPLGKC
YTLDLLSSTKPSMHKQTEPGIQAGLAITLDAHLEEQFDGSNGMDALFTNSFVNGFQYFVHPPNTIPHLSSDEFTVTPNSV
AYTAISSERFELLPTNKWGNCTEHYPSGIKSDLPYLTGNCVSLCKAKFFMENCGCTPAVYNNERNLKECTPFETLACVNN
HNGSNNATGKLEFKLPRCAQCAQQCNNLIYRAFNSQGNQFSARAFEWFRTNNSNWTMAHIKTNFQIIHIFYQDMSN
TEYNQVQDASISDLLSNIGGNMGMFLGMSLITITEICLYFSKIIWLGISKKRREYMYSKRVHEKTHKKEIVETVEKMKVIES
KRKLSSPIGFENQISWNSNDNVEFRINLNNLEREFEAVGSRSVLI

>Cyclo_ACD4_NP_505230_1_Uncharacterized_protein_CELE_F28A12_1__Caenorhabditis_elegans__44_
313

MNRKRKLSCFVSVKFPVDSVKKLRKTEGVGSVYQETQHFSTITTVNGPRRIFYGKRSAQIFWILVVISILAFLVYQIVILIQY
FYSKPTLSQINFITNEGAVYFPSVTVCNLNPVKTSFIKKLNSSGDLSEELLNYLLATKTNSMYMFNNANIFELKRAHLNALV
YLANHPDFEIVNFLNSAQFDCDELFETCFYGGKQFNCCKYMTQSITSLGRCWELNLRNETDAWLTKKGRSGTSPKTGLQ
IIANARQSEQFINFHYSSFQENGFRYFIHPPHVSPDLAAEGITVSPSRVVNSAIKTVLHDLLNHENWGNCTSSWPEHYNT
NLPYSSSACQALCVSNYFKKLCGCSPYSYNIDNNTQVCLPYEEVICMMEKMTKSDSNGTVSLDFPFCAECHLACQKTSYS
SYTSYGDGFNYNSMKWLTRETNRSASYIRQNIAIINIHFMELFYTSYSQVKATTILNTFNKIFGLNGLWFGMSVVSLTELIL
YFTKISWIAVSSKRRQYLFEKKMSEKRKERNIEEAVQEAEISRSRRSSAANLKFLDIESLDDEDYWYRSSSQLSDSHLDNVI
QLAIDFEKPLQRPSAISLPRISECCEDFEEEDEDENNDLGIIIKL

>Protost_ecdy_cyclo_nema_Cele_tr_O45402_O45402_CAEEL_DEgenerin_Linked_to_Mechanosensation_OS_Caenorhabditis_elegans_OX_6239_GN_delm_1_PE_3_SV_2_846_322

MNSPPISPYHVDFRAGSSKESLSPSTPSPMYPKCNFRYNDSRTQMIIEVPMAQLKKFQKAEGTISVKRETQHFCETTTM
HGPKRIFKGKRIFTKLFWLIMVLGSLGLLIYQCWILSASYLSKPIVSQVSFLIPEDGMEFPSVTVCNFTPIRKSYIEAMNRTG
DVSPDMINYLMNWFTEIPILLGNTDQQSLNKGNEELREYQSIHPNFTVDQFFMDASFDCSDLMKLCSFQGESFDCCSL
ASPTLTPVGKCFTLDLSKSPMNKQTEPGIRAGLSLTLDANLDEQFDSSTGIDALLPNSFVNGFRYFVHPKDTIPNLASDEY
TVSPNSVAYSAISKFRYVLLPPNDWGNCTEDYPYGIQSNLSYTSGNCLSLCKAKYFMNQCGCTPALYNIGNNFQECTPFE
TYNCVNNSLSLLNEETGKMEFQPPSCTRCGQQCDSQVYRADNSNGNQFSAGAFNYFNSKNSSWSVDYMKSNFQMIR
IFYRDMSYTEYNQVQDASITDLLSAIGGNMGMFLGGSVITIVELFLFFSKVFWIGFSKTRRNYLYSKRANEKAHKKEVVET
VEKMKVIASQGNLTSLSDATCATPPLKLSSPNKKVEFRINFEDLASQLDFDSSECKGKSEKSSNLQKY

>Cyclo_ACD1_NP_491295_2_Uncharacterized_protein_CELE_C24G7_2__Caenorhabditis_elegans__41_331

MEPTLSPNYRNEAFEHDDSYLVNFVAGSSSGESSTPPPSFVPKCNFRYNQSRSQMIIEVPVAQLKKLRKLEGTVSIKRET
QHFCETTTMHGPKRIFQGKRWATLFWLIMVSCSLGLLITQVFILASEYLSKPTVSDVSFLINEDGMDFPLITICNLNPIRKT
YVNEINKTGEVSPPMINYMMKWFTEIPTLIGGADRPTLHEGNEELKLYMKNHLNFTVDSFFMNSGFSCPDIFKLCSFQG
EIFDCCTLSTEVLTPLGKCFTLDLSSSTKASMHKQTEPGIQAGLAITLDAHLEEQFDGSNGMDALFTNSFVNGFRYFVHP
PNTIPHLSSDEFTVSPNTVAFSAISSDRYVLLPTHQWGNCTENFPDGIQSNLSYSSGNCLSLCKAKFYMENCGCTPALYNI
ENNLKECTPYETTTCLDNILAKPNKETGKIEFQTPNCKACAQQCNSLVYRAYNSYGSQFSAGAFHYLKSINPEWTDGHM
RANFQMINIFYRDMSYTEYNQVQDASVTQLLSDIGGNMGMFLGMSVITITEICLFFSKMFWLGFSKKRRDYMYSKRV
NEKTHEREVCETVEKMKAIASQGNLSSIAGTTSAKNSIPNDNVEFRINLKDLADQLDSDSGYSQNQPDSRNASFQKY

>Cyclo_FLR1_NP_510243_1_Uncharacterized_protein_CELE_F02D10_5__Caenorhabditis_elegans__53_334

METETESERIYLQLYDYETKEFSGLTTYHGLVRIYNSNTWPSRIFWVVVVLSCLSLFMIHSGYLLLGYHAKPTLFQTNTIVP
MNGLLFPEVTICNLNPLNTTKLEELNISKSTWTYIFGYFDEITTSEHKSTKLGEEQFLEIMNNYQELTKQEFNVKNFLKSVS
KSCEETFISCSFGREKLHNCCEHVTTEMTEVGVCFRLSNVNKKYRQWYSGNGFGWEFVLNGNNEIDDHADSLDFEPDR
GFLIMVHESEKYPKINSYGVAVSPDSQLHAAISMKNISLLDKANWGSCSKGWNRNDTDVPYTATHCEIDCKLRKVRNLC
GCSPLAYSARESGSNDTICTPYQIQQCFRKVRGLDNRWEDECDCPSECNMLEFDVTNSYSDLDGRSRGLSSSKVESDIS
HVSLYFSHVAYERIEQQKQLQTADLLSNIAGSMGLFLGMSTVTLLEIFIYLFKSVWGTVNSTRQQQFVDAVAEEEKERSE
SIVIIQNGRNDDMDDQKPSRFPGADRKLSGNSIHIHLDRRNSRMIRGGDLSAASRGSVSIPSQLLSPLSRHNRQSISYGQ
LGRKVSAIGIPLQPNHDTVESGTSLMPPKSPIRRCSTSTTPSMLTRKLSFASQQSDPAQPAHQSRKVSTSSIFKSQLI

>Cyclo_ACD2_NP_001309477_1_Uncharacterized_protein_CELE_C24G7_4__Caenorhabditis_elegans__42_364

MHLEDGPSTKPPDFENEKTQETSLSGEEFENNSTLGTMRDAKSWAAANKQFANQLVIQVPVNSFKNGKKIKGVGSAF
RETKHFSSTTTMHGPKRIFYGKGVARAFWMLIVGLALAMLCFQIFILLQMYFSKPTLSQVSFIVNEGGMDFPAVTVCNF
NPIKKSYVRELNVSGDLTGETLEYLLQTNMDAMFLFSNLDRHNLKETHDEAETYFQNHTDFQIIKFLRTAGYDCGEMFM
TCYFGGRRFDCCKYMKQKVTSLGKCWELDLRNLAPEWMRKQISPGSEAGLQIVVDAQLEEELKGENDDAKAIFSDIYE
NGFRYFIHPPGTNAQLTSEGISVSPSRTVYSAIKTVTHNLLNRGNWGNCSENWPEGYNTFLSYSASACRALCIAQFFNDT
CGCAPFTYNVDGRKKICAPYESITCMDNHMLKKVNGTDYLELPDCEECHMECQSTSYTSYNSYGDGFNRGSLEWLKKIS
NKSETHIKNNVAVINIFFLEMFYTSYSQVQATSLTEILSDIGGNMGMFLGMSVITITELSLFFSKIFWIMVSKRRRQYMYS

KKTHEKEKEHQLDEAVKEFQERRSRRNSRENISALGHYSNRITPVDDFQTKFGYKNAFSEGNMSNSSLDSVMELKFDIN
ELRRQLNQPSTDGIARIRLPHQTSRQNSTENYYSSSPPIFTIEPMSRKQSKTSLPSSLSPR

>Cyclo_ACD5_NP_491196_3_Uncharacterized_protein_CELE_T28F2_7__Caenorhabditis_elegans__45_3
70

MRRVRNLSLLYNDGPMGRFADQENPVENRNKKETVHFQSGSYDDDMSNSPSSSCSTVGDMPNIKPSASKGSFLSELK
PFSKRASQLIVDVPVAHLRKIKNTEGVSSITRESEHFSNTTTLHGPKRIYNGKGWSCVFWVFIWISSMIMLLTQVTSLISM
YISKPTVSQVSFLLSEGGMQFPRVTVCSFNPIKRTTVEALNSTKDLSDDLLDYLMMFNSDAMTLYGRADAASLHSGDNV
FKHYVSSHPNFTADNFFMDAGFSCGDMFKMCSFGGRRFDCCKYATPIFSDLGKCFTLNLQGSDKSWMKMQTEPGIA
AGLQIILDSHLEEQFDSETDGVTPVFSSAFENGFRFYIHSSEEIPFLASEGIAVSPDSVVYSALSSSKYILLSSNAWGNCSDS
WPRGYDYSFPYTSAMCSTMCKAQYFQNLCGCSPSIYNHLNRFNDCTPYETFICMDTKMKKVVNQSFNIEMPTCEECK
VECKSQVYHSFNSYGKGLSRGALMWLTKQGKQETWTIPHMKLNFQVVNVFFRDMSYTEYIQKRGMSLTELLSDIGGN
MGMFMGMSVFTIIELFLFLSKIGWIGFSRKRRDYMYSKKKNEEMHEKELEDVVTGFKLFRHRKSGKDMSHLREKIKGLS
MHRVTSEQLNVCKLAWENEPDIERRLASVTRQNSALKEHKDYKQPTILPFDLKDIKDQITRGRAASMFRSRRSRSETAP
AVIHEA

>Porifera_Syc_Cil_scpid47944_scgid21366_Acid_sensing__ion__channel__3_Amiloride_sensitive__catio
n__channel__3_Dorsal__root__ASIC_800

MDTSATDQANDSSSSNNSAELNADSIFVDVSAHTDKVTPKKRKPIWPLSRFLHGKPDSSTAYEYHVDDDSDFPLKEYTR
NGGGHAGNGGTSNGHIGQQKQTFEQTDEADTPAHHSGIDWRERKAMALKFTYTYVYLAACHGIARAMQNKASVAR
RLFWLLAFSGAVSMFFYQISLLVKQFTDYEFQVDVEFSFNRQLAFPAVTFCNLNPMNVKLLNCSRFHDECLQLAGANQS
ADGQLIKTDDAIGYAHEYNFSGEPITRAEYLQYGFLPEDMIITCTWNGEPCSARNFTISQNPFYGNCFTLNGDPKNPLQT
NYPGREAGLALELYMNPDEGIGGETSTSSGVRMSLHQPGVKPFPDEDGFSLATGEATFIGVRQLEVSRLGSPYSNCTQE
NDNWKSLYPGYSYSRTACVQSCLAQEMFDRCGCVEIFLVDKPICNNANVNKEQADCAAGVRRDYRLDRLNCDLCYNP
CVDRIYLHTLSKATWPAKAYQYSFRETLKELNFSDGNSEDSNAEEIYREKFLRAVVYFEELNYQRIVQKPAYPAENLFADF
GGQMGLWAGLSFLAFVELFDYVGQVLLIALGLKKVVVLP

>Porifera_Syc_Cil_scpid49206_scgid6888_Amiloride_sensitive__sodium__channel__subunit__alpha_Al
pha_NaCH_Epithelial__Na_channel__subunit__alpha_Nonvoltage_gated__sodium__channel__1__subu
nit__alpha_SCNEA_882

MDVDQDALIRAEADNLFEELASTPEHRRPLPHMPSSSQFKRQTSLSFSTSQSSSLQRWYGEKNADEVASEAKEELDDDL
VGDMPPNWESKKTKRRTSAQSAWSTSSSSSKGSRESNVWSRDSLFSFGPKETPQITDRDVADIADEGAVKGLQRKRV
GFIDFTMRFVLDTAGHGMPRIVEKDIWLIRRIIWGCIVLGAAVGFIIQTHALVSKYLDKDFKVDVKYKFNKEIGFPAVTLC
NLNPLNRSKLECSRFDTSVHKGALVCQQTASGNHTTVQVTLAQILQARINALLKSDLQQAKERAFQYNFTHQEAMTAA
EIAKYGYDLSKLMVSCTFRGQPCGIENFTRFESARYGNCYTFNNNISTQIQISQPGPAFGLAIDLAYDPLEEIGLLSHGAGF
QVAVHEPRSKVFPEEDGMSISPATATSIGIRQAVVERLGEPYGNCTNITGYDWGNLYSGYKYTRTICLKECFAQAMLDA
CGCVSENILVGKRLCLPATLNKTEADCRAEIEELYRIYKLPCLQCFNPCKERIYRTTVSTASWPSFEYQREFKQLLKQKGFQ
VDHFDVVRFRDQYARVEVFYEELNFENIVQKPAYEPENLLADMGGQLGLWLGFSVLAILEIIEYLIYMLLVALGKVKLPDE
EDYLKARDD

>Porifera_Syc_Cil_scpid40922_scgid7672_Amiloride_sensitive__sodium__channel__subunit__alpha_Al
pha_NaCH_Epithelial__Na_channel__subunit__alpha_Nonvoltage_gated__sodium__channel__1__subu
nit__alpha_SCNEA_933

MSAELRRDSARLWVEAYLEDLTSHGLSRITATASYLPGIVRLLWGVVFAAAFAFALQQCSILVAQYRKNDVDVSVDFSF
NKEVDFPAVTICNLNALRRQEVLKTRLHEHVNVTASPTTVSAAPVTVTTAAPTTTPTTPYVSTVNPRDFEANVTLPSTAT
RTVHTTVFVEPGTEAPLITTATLGETEPTLATQTTGMPNIAPTTAGDGSGGSRTTPRPHATTASGGGTHAPATGPPVPG
GGTTESSQGVCIIAGVPVTLTLGLPCSVMKLLLGEGAAAGGPANQDGSAITRAPTVAGFIPDDFFTSSTTPDTKTASDAV
LEIADSFEGGNTSLFAHVETERNRYNLQKVDLTSHQVNYQYDEAPVVREEISVLGHQIEDMLFRCRYDSEGCSPSNFTQF
QNGRYGNCYTFNGDEARVLRTERVGPTYGLLMELDIEQSLNYIGQLTQSAGIRMVIHRQGEQAFPEYDGFDLAPGFKT
YVGIRQVVIERQGRPYGNCQEGNPQEELASSNATTYTKWGCLRECLAEHMLRECKCVDATYIDKDARLCHAYNHPEQA
NCLAEVDRQFKTDLICQDCEDPCRSVIFRKSISSERWPSKQYEPALFARLAEENRTIDDDVVTAKIRDNYIQAVVYFEEFN
FQTVTQRPALEIVDLLSYIGGTIGLFVGVSALSILEFLDIIVNLVYFFAKWLYKRVRTRLLGDDVFDGDERELQRRTSDVKA
GLYY

>Cyclo_tr_K7H9J0_K7H9J0_CAEJA_ASIC1_OS_Caenorhabditis_japonica_tested

MLCQSNVNVIANIIMKEEVEVEVEAKKSCCNTLGDCVYTEKPKKKTISCLCATPIKMCVR

IDPPQTNATTLDDRLVKFWHIQPSTTVSPTIKRKEERDKAYGYTGVKDRIALRAKAMENI

IFAVDALTEEQKWKISYNKSDFIVKCSFNGRECNVKHDFVEYLDPTYGACFTYGQKLGNI

TNERSGPAYGLRLEVFVNVTEYLPTTEAAGVRLTVHATDEQPFPDTLGFSAPTGFVSSFG

IKLKSMVRLPAPYGDCVKEGKTEDFIYTKKAYNTEGCQRSCIQKHLSATCGCGDPRFPPY

RESKNCPVDDPYKRECIKSEMHVATRDSKKLGCSCKQPCNQDVYSVSYSASRWPAIAGDL

SGCPLGMSAHHCLGYKREQGSMIEVYFEQLNYESLLESEAYGWSNLLSDFGGQLGLWMGV

SVITIGEVACFIFEMFISIFSAKRVKRRPARKSFSSSLRCSTDYNLNKDGFNLDN

>Lopho_Phor_aus_Phoronis_g6004_t1_TESTED

MKRQGEMEGPDGLLTNFARSTSAHGLARIPGTSQPMQRAVWALLVTALAAALISALAIVVSVYLQYQYTEFAKKVARP
NIKFPVITACNKVPYGLLQTEQTVNEFFRSIGLDEPTCQSEISPGLNDPIIYRRISDAVRPRVLYEYNSKVMQVLGQNLDDF
MVTCAYQGPPCFIYSNFSTSLFKDPYHYNCVTLKVPDEVQVKVNGRGAMELVFFVGDNKTSLQGKHVDKLVTDGTVGI
KLIIHAEGMHPKWAKTLDVAPGHLANIAVRPREIYRLKAPYASDCIENPPDILAASTNTSYTYSKELCTLKCFNKLVADAC
DCIPEPVVVDKFGLDYPYCGESPCNYSYILDKIKCHTNLTERLHEGMLSNQCPECKLPCEELTYDTDMHLSKWPSKFTSKT
ITKYILKNHVIPRNIENDSSALDSYVQQNFLKVVIYLEDLLTVEQTEQESMNLAQLTSSVGGAMGFFLGISIVTVFEFIDLLV
KSVQSLFKKKSVDKVLPFHSKGS

>Lopho_Phor_aus_Phoronis_g5063_t1_TESTED

MNSEVKNWSVKDRMKEFCESTSAHALGQTVSSGEVKAIFWSLVFLSALAGCVWNIVHVVESYTSFGFSVKSKLEMEPS
SLKFPSVTICNLNPISFSRQARFAGDFERSTGLDDGDCDNEFHRDETYRPVVLDLYRWSELPQFWFEYNENVTEMFGHS
KKDFIVDCVFQREGGCENNFSVTKDPNLYNCYTLEPNKNEDLLEIGFAAGLSLTLFVENRRIRNAFTGNYALDSYHTGIVG
VKVAIHVSGSHPNPNTRGVIAEVGKSTDFILRTVNRTALGPPYPSPCNPKKTIESSHRKELQYEENLCFASCLQNAIAKKC
GCVSINPMAVPIADKFGIDLPFCGSYPCNATKVTENYECVHDVIGRFLLKNDPNCRNCTKPCNEVSFEVTKFQSKWPSEL

HQSEFIRWLANKDNTALYNQLVRNISDDNISRFIENNFLRINIYFGDFYVRKDIETLDMDWFDLLSSVGGAFGFWVGISV
VTGVEVLELLLDCIVLFLNRRIKKRNQIDLRNTTGDDDVVTGS

>Macrostomum_Mlig049925_g2_Protos_Platyhelmin_Macrostomum__TESTED_A0A267GWB9_A0A267
GWB9_9PLAT_Uncharacterized_protein_OS_Macrostomum_lignano_OX_282301_GN_BOX15_Mlig0037
20g1_PE_3_SV_1

MQDRLRTVTGAAQEVAGITRTHGLLHIFLSRGSARRLFWLLLFVAALIGCSVHLAKLVQKFTERSVESQLKLRSERAQFP
DVTICNFKPASASLLSIINFLEFHSKNEFKFEVFDYFEIFWGNFDNWFKEDTQKRPQESDESYDNRLEKVCKKVLDVMWV
SGDTRDISQLDETMLLSCRYNSEPCSHKNFSLVQTSRFWNCYTFHPDPRQSSSGPGGRGAELDMVLFTDSNEKYPDLYE
IYPNGALPDGCITRKFLRSKAVNRLLQNNEPQSAGLRIFIHEEKSFPMSETEFVDVASATSTSIKLRPVHNRLMSKPSRRCS
EPPIETINYVRHFSNASLNVVTKAYSKSVSDFVVEAQQSILHSNCGCYSHLLPFSVNTSDLCYFAPPQEWITPSDAVLKRID
CHDHWFEHAQRQSEELAKEFQHRMWCQFTQQRPWQQSRRWPPFSAIQEIWENLMVPQVRHGVEFLPEDGKEHHI
FRNSTKTRQEIQLYLLARGSDYCRFNFLRKFNFIDRLLGKDYKKKIIESATSCVSRSELAKELAAVSISLQSPAADAYEEKYKY
HWTEALSEIGGTLGLWLGISVVSTFELLEFIYILVQKCRRNDDDSDTEE

>Macrostomum_Mlig051885_g1_Protos_Platyhelmin_Macrostomum_lignano_TESTED

MPQYLLNHDSPARRQLESNMLDLMVALRRVYAKGGDIFGTQKYFHWYNLLQHYWASVDTAHLGHNLSLSMVACFY
KKAPCRPADFKLVQNSHYWNCWSFRPKDRSVQGTGPNDGLNIILYTSTMPLDDSSQPDHVVEYPPPKHRSTLRQLDT
VFGKAGMQVSSGVRLLVHEPGTYPHVYWEGADVGNGWSADLRFKMKRNVYVNRTGHNCVENYGHTDYWASEAE
GIQRFRKRGQDCVVRKWQEHLMRKCHCQSTFLPTQDRSALCHYLRAAGVSAQNISAPFGAYKFQQLKSFNDVENQVR
TELYDQFASECGTVQACEQSSYTYSIASVPWPAHTDLEAFIHTFVSPKFHRARYQGQQLLDQIVRFHRHPDGSRDFEKG
WSVSHRFVRQNFVKLQVFAEESKSTLIHESPSYSFVEVLSELGGIGGLWVGMSLVTFVELFEFLAILGIKCTQVASSYLHSR
WTRRAGQMANGACPGSANHLRTPACTRSAQSPDEITSGPCVVLEEASSVVDEDTELVGIFLCPEASRNEHLHDGNSSR
LSRGQAMQHVSVVGS

>Macrostomum_Mlig041003_g1_Protos_Platyhelmin_Macrostomum_lignano_TESTED

MKNTLHMMRCGNLMAAAMAPGLGPCATNIGLNNLQQQQQQQGAQEEQQGNPNAPGNGGGSRIPRRREVMLR
GYINHFTSGTTAHGLNRVNAEQSWVRMVWVGIVLAASFGAIVHTTKQVKTYLQYPVSSTNRQEPNSFEFPDLTFCDPL
NKRFFHKNVISYAREVDVYDEAHMIYGLVYSHFRTDSDWRKERRAIELIYNALEHLLNITDVNIRPMDVVLYCTFDGEPC
SHTDFHVFYHRLYTNCFTFKPKQRSLRSSGMDRGLFLLLYVPSADVNLKLDSLARLASGLENNGIRFQIHQRDTIPHPLEY
GIMAPTGTLTAVGLEQVRTSLADTPTNPCVRDTKLRLFDNQFNFMRQYDRQLFDCIKHQYMREVRKQCGCTLESHVIA
EEGFDPREVVFCHDLISAKNATRVGALTKLYNELKNVTNTAGLHGHIKLGLLMRQNDAELNASLDRMICSDSVSESAAR
PERCYDSCNYNKYEYSLSQCPWPEDSFEVRNSQIQMVKIAQMLEDYKLRYGSANPETALAGLMRKSSLNISSCLTHDTQ
KMQVPDKVDCHRVQLFVRRSVVQLRVYPETLTVRHTIEERSYELVNLCSELGGILGLWIGFSIVTLFEFAELFIICASYYWY
LLISKLPSIRKSRHPLRIPKPHPVGLRRQIRRDIELISNSGQGNSSSQQASPSLLATGRPACTNNRAPLSSSSFANHHHHHQ
LRLDGNSQETRPLRDGMGGS

>Deutero_Cephalo_088140F_t1_XP_019621273_1_PREDICTED__acid_sensing_ion_channel_2_like_isoform_X1__Branchiostoma_belcheri_

MHVKLTCSVELDCCPCCDCGQCCCGSSTAGSEPRDAGSELGDGTDAGEQTYYRKQAQTISDFAAGSTLHGLPHIFPDA
PLSIRQVAWALAFIGSLSVLLYQCSDRVKFYFQYPHITKLDMVLAERLDFPAITICNMNMFRWKQFTQNDLWHMGKGL
NILDENNNLRCSEYATPGDMEALKNKANFDSFKPSPFSIMEFANRTGHQIESFLLDCKWKNFTCGPEYFTPYRGLLWSIT
TVGSQNGCLSGYLCVYLSVYLSGYLSGYQSGYLCVYLSVYLSVYLSGTRHSGSIPASSAFPGASLALGKKRSAADVTSSRQI
PDEHVDTVDVFTRYGKCYTFNSGSPNQPVLKTLKGGIGNGVEFFLDVQQEDYMPAWGESDEVTFEVGFKIQLHTQEEP
PFIHELGFGVGPGMQYYVSTQEQRLNSDLLLSAEPTLLLPPPMCKAGRNQVELLFGWTKYHETITYLPAPWGQCKAEN
DLTFDFAKYTTSACRIDCETKFVVSQCGCKMVHMPGNFPICTPDIYVECADQALDFLVKSDNKKCVCDTPCNTTRYNLF
MSHVKFPSEQAVKYLARKYGKPEDYFRGVGLKYGKPEDYFRKNTLVMNIFFEALNYETIEQQKAYEVASLLGDIGGQMG
LFIGASILTILELFDYLYEVLKDKCTERRHQPRSESNAVSVNLEDCKRDNSRAPLS

>Deutero_Cephalo_XP_019621275_1_PREDICTED__acid_sensing_ion_channel_1_like_isoform_X3__Branchiostoma_belcheri_

MEGGSQNHDVPRQSDINVFAASASMHGLAHIFTEGKFTVRRILWAGVFCGCVAVLLVQSVDRVQYYLSNPHATKLDEI
TAVNGLNFPAVTICNMNSFRFSQVNQQDLFYAGPDILDLIDHTLPRPKLKISEEFLNHMNENPSDHQEMRQKMERIRE
LTNF

DGFVPGKFSMSDFYDRTGHQIEDMLLDCKYKGEPCSAKNFTTVFTRYGKCYTFNSGSPNQPVLKTLKGGIGNGVEFFLD
VQQEDYMPAWGESDEVTFEVGFKIQLHTQEEPPFIHELGFGVGPGMQYYVSTQEQRITYLPAPWGQCKAENDLTFDF
AKYT

TSACRIDCETKFVVSQCGCKMVHMPGNFPICTPDIYVECADQALDFLVKSDNKKCVCDTPCNTTRYNLFMSHVKFPSE
QAVKYLARKYGKPEDYFRKNTLVLNVFFEALNYETIEQQKAYEVASLLGDIGGQMGLFIGASILTILELFDYLYEVLKDKCT

ERRHQPRSESNAVSVNLEDCKRDNSRAPLS

>Deutero_Cephalo_XP_019621276_1_PREDICTED__acid_sensing_ion_channel_1_like_isoform_X4__Branchiostoma_belcheri_

MSSYEENEPKPSDINVFAGNATMHGISHIFTEGSFSFRRFLWAAAFLICFGMVSYNGYDRVIYYLSYPRVTKLDELEAQM
MNFPAITMCNLNPFRFSKLTKDDLWQVGHALEILDENKKLAHSEFVDPEHLSTLRRLAEFGPGYVPKNKDSFNMKEFY
NR

TAHQIQDMVLECSYRGTKCSWANFTVVFTRYGKCYTFNSGSPNQPVLKTLKGGIGNGVEFFLDVQQEDYMPAWGES
DEVTFEVGFKIQLHTQEEPPFIHELGFGVGPGMQYYVSTQEQRITYLPAPWGQCKAENDLTFDFAKYTTSACRIDCETKF
VVSQ

CGCKMVHMPGNFPICTPDIYVECADQALDFLVKSDNKKCVCDTPCNTTRYNLFMSHVKFPSEQAVKYLARKYGKPEDY
FRKNTLVLNVFFEALNYETIEQQKAYEVASLLGDIGGQMGLFIGASILTILELFDYLYEVLKDKCTERRHQPRSESNAVSVN

LEDCKRDNSRAPLS

>XP_019623542_1_PREDICTED__acid_sensing_ion_channel_2_like__Branchiostoma_belcheri_

MAAWDPFSDELEDIIETVELRVRTDVTISFPALTRARSTQGKSCCRASPPPSSRSQPIPTRSRVEVSTVAASIAMELANRK
DMGKEDSFTPLALAATPDVREVKPVQCVPTSTVGTGYTSYRKPSRFRRFALDTDIHGMKHLIKRGPIRRTAWVLMLLLSL
GLMSFECMESLIYYFEFHHVTKVELQYTNNMTFPAVTICNMNKYRRSALGMRDLATVGPYIGIVDRQQFTLQQEHLYP
KAWVDKVKNENLKALSKMTTGQQNFAVFVERVGHQKEDMIVNCEWRGSPCSAKNFTHTFTHLGNCYTFNGMTLSR
KVLSSSKPGEGNGLKVTINIENDEYMATNDVAGDAMDAGIKVMVHPQHEPPFVKELGFAVSPGFHTFVAIRKEVITTLP
APYGNCQQRGGQKYYPDYSLSACRIECETEHVVHECGCRLVEMPGEHVPVCTPERYQCAYEKLGIVDRQQFTLQQEHL
YPKAWVDKVKNENLKALSKMTTGHQNFAVFVERVGHQKEDMIVNCEWRGSPCSAKNFTHTFTHLGNCYTFNGMTL
SRKVLSSSKPGEGNGLKVTINIENDEYMATNDVAGDAMDAGIKVMVHPQHEPPFVKELGFAVSPGFHTFVAIRKEVITT
LPAPYGNCQERGGQKYYPDYSLSACRIECETEHVVHECGCRLVEMPVEYVREDFSLLIHYTKTLNLDYLFISLFYSDNIAVL
NVYYEALNYESIVQIPAIETEGLLGDLGGQMGLFLGASFLTLVEVFEYLMDELYGRVLKKCCRCKATASAHTTSPSADVLS
VNINGSATQLPQTSRA

>Porifera_AmpQue_tr_A0A1X7TFM5_A0A1X7TFM5_AMPQE_Uncharacterized_protein_OS_Amphimedo
n_queenslandica__Sponge__OX_400682_GN_Aqu2_1_13197_PE_3_SV_1_370_0

MCFSQKKKEGCLVLIGLSIKRFGDKPTASTITVVSSHESGLPFPAVTICNLNLKKNDSDFLLNTTYQAMNFLYNPDKSYQF
NSTKKKNHLLNTCTAPHSNSIQNATIWDIVNPDRAVNELIQYCGFLHGADSDVVLCKDLFEPVLTSAGICFTFNGTNKLA
NSTGIRYGLKLVLDVQKKERPSFNGKLGVKLVVHDGRDIARPNLYGLTVPPGIAVDVGVRKVITRDETNEAKCIDGMNL
PFFPSDKFEYSQFACRANAVAENIAKRSKCNCVVQPDRPPGFYTSTPNCTFSKACCLLQEQYTFHPLKKSTVLCHATFNI
MSILLVFPVFLMDNTCNY

>Porifera_AqNaC9_Aqu2_1_25935_001_PorifAmpQue_89_89

MTKKEFKWSDQYFNDFVETTTINGVFHIFRGRSKTRRLLWGLLFLISFVSCTIVLGFSFKRYSEKPTVSAINVISEENGMPF
PSVTVCNRNFYKYPNISNETDTLIHHLFHSSGFLQNINRTQECKIDEDTPDFKLQDLLIPKENNFIYYCALSYQAESEVMLC
KDMFFPTLTPAGICYTFNGVRSKTRALNMTTTGKRYGLRLILNIEQETHLAFDGVAGVQVIVHESNDIPRPNLHGIGVPP
GQNVDIGVKRAISSDETDQDACIENNEKDLPFLPGVVYSQYACRQNELYESLADKNLCNCIIDPFELDNATCFLHDLCCL
QQQFTEHDRNSSCRPPCKYTYFDIINSYSSFPEGHALTEIKNSINTSDDYVKKNFLSVSVFLQALETRETTTRYSYGVVELL
GELGGNLGLFLGISIISVMELLVLVVDELKKCVCPKKVKQKFKKFDEKLKCIPDCVEQEEKNEIQALTNF

>Porifera_AqNaC6_Aqu2_1_26214_001_PorifAmpQue_86_93

MKNAGRFSWSDKYFSDFVETTTINGVIHVFRGRSKIRQIMWGLLLISSFIAVCVVIGFNIKEYVNKPTASTILVSPSVKNGL
PFPAVTICNLNVYTPTGEDKEPDFSSVIHSLFNSQDLVNQTSLFDECSEVINSSDTSDAFDCEVWDLLLPDDKRFIYQCSFS
HDADSEIVSCRDMFYPVLTPAGICYTFNGIRSKMPVPMAKDIGVRHGLNLILNIEQDSHPTFHGLTGVKVIVHDRNDISR
PNLYGISVAPGQNVAISVERKVYIDKTKERDCTNDERELGFFPSTVYSQFACKENALYQHLADESVCGCVPNPYRPSTGP
YINTPNCTLHSLCCLLQEYFDYRVSSATCPLPCQFSMYDYKSSYSSFPNGHALKSIAKSLNLSQDAVKNNFLSVQVYLESLE
THEYVTKYSKTLTGLFGDIGGLIGLFLGMSIISIIEVLVLILDELKKLLCIKKFRKKVKKVDDMLTHFLPDVK

>Porifera_AqNaC11_Aqu2_1_26219_001_PorifAmpQue_91_101

MTKKGFKWSDQYFNDFVETTTINGVFHIFRGRTKTRRLLWGLLFFISFVSCTIVLGFSFKRYSEKPTVSAINVILGENGIPFP
SVTVCNQNFYKDLNVSNETNALIHHLFHSGGFLHDINRTQQCKIDEDRHDFELQDLLIPKENNFIHYCAFSHQAESEIML
CKDKFFPTLTPAGICYTFNGVRSKTRAPKMTTTGKRYGLRLILNIEQETHPVFDGVAGVQVIVHESNDIPRPNLHGVGVS
PGQNVDIGVKRAIMFDETDQDECIKDNGKDLPFLPGVVYSQYACRQNELYESLADKNLCNCIIDPFELGNATCSSHDLC

CLQQQFTEHDRNSSCRPPCKYTYYDIMNSYSSFPEGHALTEIKNSINASEDYIKKNFLSASVFLQALETRETTTRYSYGVVE
LLGELGGNLGLFLGISIISVMELLVLVIDEVKKCVCPKKVKQKFEKFDDKLKCIPDCVEQEEKDENTSGPSAELNKIA

>Porifera_AqNaC2_Aqu2_1_09805_001_PorifAmpQue_82_141

MAEDSHSKSSEEKTVKCHLHDRYLKEFLDDNTIGGINHIFRGRSKVRRLLWALIFIGSIVACITLISISFQTFLEKPTASTITVI
TQDDEGVSFPSVTICNLNLERNESDMVADTGYLLMNHIFNPDENFHLTGLNSSFLLNSCNAISDSFPASFRNTTLWNSQ
HPQETLDKLIHFCGFVSGINSAVIPCKDAFKPVLTSAGICYTFNGSNNRIHSTGVRYGLKLILNIQQEERPSFNGKSGVKLIV
HDGRDIARPNLYGIDVAPAHAVDVGVRRKASKDETNEADCIDSKELPFFSEYRYSQFACRQNAIVENLATSCDCSIHPDR
PSSGPYSSTPHCTFDKGCCVLEQYQTFNPELACPLPCYFPYYEHTASYSSFPNGRYLNYLVEETNMSVDYIKDNFLSINVF
VDDLQLTTTITKYTFGVAELLGEIGGQMGLFLGISIISIVEVVVLFLDELKRLFCTKKMREKMQDIENAIELPEIEGTDVEEDI
DNKV

>Porifera_AqNaC7_Aqu2_1_26220_001_PorifAmpQue_87_142

MQPAKGYTLTDQYFHDFVETTRISGIKHIFRGRSKIRRIMWALFFISSFVGCTLVVGKNIARYIEKPTASSIKVIPHDNGMR
FPSVTICNINIYKDPNVSAVSKETYSLIYYLFNSDTKEYNSTQECIDNATEYYKKHDIWSSLLPKENNFIHDCSFSYETGVISC
KDMFYPVLTPAGICYTFNDFKMNEMLIPNITSGMKYGLKLVLNIEEERYPAFEGKTGAQIIVHERNDIPRPNLAGISVPPG
QNIDIGFIKAIKNDKTDSRDCINDGEKKLDFLPDVVYSKFACQENELYERLASNCNCTINPYGLDVTNTSNCSLSNLCCMR
QQYSKYNRNSSCKSPCKYTFYNLKNSYSSFPRGRTLTEISKTVKMNKSAIRDNFLSVHVFLLDLETKETYSHNSFDISELLGE
LGGTMGLFLGINILAIVEVIILILDEIKKYLCPKKCKQKLNKIENCIPECALSRGRNDTLLTDPVSTPEKENHETPYTTAVDIDN

>Porifera_AqNaC1_Aqu2_1_34376_001_PorifAmpQue_81_145

MPSKDKGENDPQENTGVRNRKEKSCQNPQFTKWAESSTIHGVDHIFLGKSKVRRVVWAVILLLAIGGCLYGIIDRSIYFA
SKPTATTVTADINEDGIPFFPAVTICNLSPISRQYADQHNLTSLLSYILFTDSNTHSKGFISSNCQANLEQITDTTITLKDVFRD
GARNSSFILACHYGSARNKMDCANMTLRTTLTPRGLCYTFNGDPASPPLLVRSVGERFGLRMIFNISQSDYTHSINGDA
GIRVSVHTRDEKPDPLLKGISVPPRSHASIALYPIRSISKPEITRCAPTDTQLSYFPGLSYTTSGCQANEHFERSQAQCGCVD
VAESATNDCTVEDICCLYDEGTSTDTLNSTCLPSCNNMIFSSSVSYSQYPSDVTVSLLTSFQSQSAESIDDNILALNIYFGSL
HTIVTSTYYTYLWSGLLADIGGQLALFVGASVISFMELVLLCFDETKCSGVFIRKKIKKKHEEHELEERDDKESLNGGKIEDK
QANTNV

>Porifera_AqNaC10_Aqu2_1_36244_001_PorifAmpQue_90_167

MDEGNDPVNGEKSKCKNCTCTCTDRYLREFLEDNTIAGISHVFKGQSKVRRSFWALIFIAAIISCIALISLSIQIFLNKPTAST
INVITLNGVPFPSVTICNINFEKNEALPLLSRTYNLMNQLFNADESFHSNSMNASSVISNCRNTVPRNNSDATFEATIWNI
QRPARSKRRLIHYCGFVTGVNSNVSNCKSLFQPVLTPAGICFNYNGSSNTIHSTGTRYGMKLVLNIEQDLRPSYNGKAG
VILSVHDGKDVARPNVNGINVAPGQAIDVGVNLKEYIDETKEANCTAGQDLVFFEDYDYSQYACAQDALIKQIAKPNVC
NCTLLPRRPSNGEYDHTPNCTFPTSCCLLNEYKTINTEEQCPLPCRYRYYDYTTSYASYPNSFILEDLMRDENVTEDYLRK
NYLSINIYINDLRYSVVTTSYTFGVAGLLGDIGGQLGLFIGVSIITFFEVLILCLDELKRICCPDFVINKCKNMKKKGESSDIVA
GERETRRQIQRKAWSNDSL

>Porifera_AqNaC13_Aqu2_1_13200_001_PorifAmpQue_93_173

MCSDGAGHAYDKVSKYFKKKITIAGLSHVFPERTKKDDRTRSGLTNPEKIIMVIWALFFTGCVLGCLVLIGLSTKRFVDKP
TASTITVVSHDKTGLPFFPAVTICNLNLKKNDSDFLLNTTYQAIISLYNADKSSQFTSNKKIDHLLNTCTASLSNSIQNATIWD
IVSPEMAVNEFIHYCGFLHGADSDVVMCKDLFEPVLTSAGICFTFNGTNKLANSTGRRYGLKLVLNIQQEERPSFSGKLG
VKLVIHDGKDIARPSLYGISVPPGFAVDVGVRKMATRDETSEAKCIDDMNLPFFPSDKFQYSQFACRENAVAENIARGS
KCNCVIQPDRSPGLYASTPNCTFSKACCLLQEHYRFHPEEIDCPLPCHFEYYEHTASYSSFPNGQYLQLLMEELNMSAEY

VKDNFLSIDVFFDDFQVTTTTTKYTYGIEALLGEIGGLLGLFIGVNIINFFELLVLSGDGLGMLCRRAGRSCRRALEKIKKRK
NERKEEPMKMDTQPGSSRSGNGHA

>Porifera_AqNaC8_Aqu2_1_34361_001_PorifAmpQue_88_177

MAAEKKDVEMASEKEDMQMMETKKKKGRGCHLNDPYLKEFLDDNTIAGMNHIFKGQSKIKRLVWALIFIGSMIACITL
ISISFRRFINKPTSSTITVVTENTKSGVEFPAVTICNHNLEYNLSDYIIRNTYLLMNYLFNADENFHLTGSNSSSVIKQCQALV
GDAPSEILNATIWNIQNPSKAIDELIHYCGFIEGVNGEVEPCKDAFKPVLTSAGICFTLNGSDHGIHSTGIRYGLKLVLNVQ
QEKRPSFNGKSGIKLVIHNGGDIARPNLYGISVPPGRAIDVGVRRKATKDETSEAGCIDDMDLPFFPKDKFDYSQFACRE
NAIMERIAKRSSCNCVIHPDRPSTGAYSSTPNCTMSNACCLLHEYNTFHPESSSEECPLPCYFSYYEWTASYSSFPNGRYL
DRLVNTLNKSADYIKNNFLSVNVFLDDIQLTTTITQYTYGPEALLGEIGGQLGLFIGVSIITFFEVLVLCVDELKRLCFKLQIIP
EEKINNLESRITLPEVEEDQTEN

>Porifera_AqNaC4_Aqu2_1_26218_001_PorifAmpQue_84_192

MPTLVRNAHVWTDQYFSDFIDTTTINGVFQIFRGRSKIRQIFWGLLFIGSFIGCIVTFGYSFRNFAQKPTASTIKVITQAQT
GLAFPAVTICNLNIYQNPNYSNVLSSEMYALIQYLFETDDIFNEFNITSECKDLIDNASEDYGKESLYDMLLPKDYSSNLIYD
CTFRDDALGDAMSCKDQFYPVLTPGGICYTFNGVRSEMIAPVIKSIGIKYGLKLVLNIEQETHPTFDGRTGVKVIIHERNDI
PRPNLYGINVSPGQNIDIGVSRSFFIDETDQDKCNNDIEGNFPFLPNIVYSQFACRINQLYERLSQQRNCGCLPIPYRPES
GPYTNTPNCTLGNLCCLLREFPIADASSTCQLPCNYSVYEYRDSYSSFPNGRALTEIARKVNMSKTDVKENFLSVNVFFEA
LHTTESITQYTYGAVDLLGELGGNMGLFLGISIISIMEVIMLILDEIKHLCPKKVKKKFDNIDDKLRNYIPDIAPSQTDTPNAL
EAGTEEVHVEDPSPSEADIKSNAEAIEES

>Porifera_AqNaC12_Aqu2_1_34365_001_PorifAmpQue_92_215

MNPADETDAEKVSGSRCKSCQLKDSYFNDLDKTTIAGLNHVFMKDSSKIRRLIWALFFIGCILGCLVLIGLSINRFVDKPT
ASTITVVSNDETGIFFPAVTICNLNLKKNDSDVLLNTTYQVMNFLYNSDESFQFTGSNTMSLGNTCTAPLSNSIQNATIW
DIVNPDRAVDELIHYCGFLHGADSDVVMCEDLFEPVLTSAGICYTFNGTNKLANSTGIRYGLKLILNIQQKERPSFNGKSG
VKLVIHDGRDISRPNLYGISVPPGHAIDVGVHKMATQDETSQANCIKSMNLPFFPSDKFDYSQFACRANAVAENIARRS
KCNCVIQPDRPPGLYASTPNCTFGKACCLLQEHYKFNPEEINCPLPCHFQYYEHTASYSSFPNGQYLQRLMEASNMSAE
DIKDNFLSINVFIDDFQVTTTTTQYTYGIEALLGEIGGLLGLFIGVSIITFFELLVLCVDELKRLCCSQAIIKRMKRIEETALVPV
VESAEGSLSNEEDAPEEVTSQPEAESNKTSPSSRDANSDSNEGKCIKIEEIKL

>Porifera_AqNaC3_Aqu2_1_34363_001_PorifAmpQue_83_251

MNGKDKAKEPLKPADSERDEKGKGTCHLSDPYLSDFIDDNTIAGINKIFRGKSNLRRLIWAIIFIGSLIVCTVMLSFSIKRFI
DKPTASTITIVSNTEQGIAFPAVTFCNLNLERNSSNFLLRSTYQLMNYLYNADENFHLSGLNDSYVLQLCDNVVRSSSEDI
LNATIWNIQNPSKTIDELIHYCGFIEGANSEVKPCKDAFKPILTSAGICFTFNGSDNRIHSTGIRYGLKLILNIQQKERPSFN
GKSGVKLVIHNGRDIARPNLYGISVPPGHAIDVGVRKKAVEDDTNEAQCIHGMNLPFYPSDKFDYSQLACRENALAENI
AHSSKCSCAIDRPSTGPYASTPNCTFSKACCLLKEHYEFNAEEADCPSPCHFEYYEYTSSYSTFPNGLYLDTLVNKTNMSV
NEIRENFLSVNVFVDDLHTTTTITQYTYGVEALLGEVGGQLGLFIGVSIISFFEVLILCIDELKRLCCRGSVKRTMKKLEKMIR
LPEIDSGETDKSNEIELNSVEIISCDEERSIKDLSPCDEDNKAVLLPIEDKSNEVGLQSVKTTEV

>Porifera_AqNaC5_Aqu2_1_09804_001_PorifAmpQue_85_255

MNKGKAGEPLNSLKAADSENDEKGKGRCHLNDPYLSDFIDDNTIAGINKIFRGKSNLRRLTWAIIFIGSLIVCTVMLSFSIK
RFIDKPTASTITIVSNTERGIPFPAVTFCNLNLERNSSNFLLRSTYQLMNYLYNADENFHLSGLNDSYVLQLCDNVVRSSSE
DILNATIWNIQNPSKTIDELIHYCGFIEGPNSEVKPCKDAFKPILTSAGICFTFNGSDNRIHSTGIRYGLKLILNIQQEKRPSF
NGKSGVKLVIHNGRDIARPNLYGISVPPGHAIDVGVRKMAVEDDTNEAQCIHEMNLPFFPSDKYDYSQLACRENAIAE

NIAQSSKCNCVIGRPSTGPYASTPNCTFSKACCLLKEHYEFNPEEADCPSPCHFEYYEQTSSYSSFPNGFYLDTLVNKTNM
SVNEIRENFLSVNVFVGDLHTTTTITQYTYGIEALLGEIGGQLGLFIGVSIITFFEVLILCIDELKRLCCRGSVKRRMKKLEKM
VRLPEIDSGETDKSNEIELNSVKIIQCDKERSIKDLSPCDEDNKAVLLPIEDKSNEVGLQSIKTTEV

>Porifera_AqNaC14_Aqu2_1_20433_001_PorifAmpQue_94_260

MAGNDPQNPKAPVADKTSRSPTVTSGPPSPETNRPPALEPPRSGVDREDTSTVDITPVYDKVSEYVREKITIAGLGHVFP
NPEKKKGVHSLSTCEKTMMVVWAVFFTGCVLGCLVLIGLSFKRFVDKPTASTITVVSHDKAGLPFPAVTICNLNLKKNDS
DFILNTTYQAIISLYNADKSSQFTSNKKIDHLLNTCTASLSNSIQNATIWDIVSPEMAVNEFIHYCGFLHGADSDVVMCKD
LFEPVLTSAGICFTFNGTNKLANSTGRRYGLKLVLNIQQEERPSFSGKLGVKLVIHDGKDIARPSLYGISVPPGFAVDVGVR
KMATRDETSEAKCIDNMNLPFFPSDKFQYSQFACRENAVAENIARGSKCNCVIQPDRPPGLYASTPNCTFSKACCLLQE
HYTFHPEEIDCPLPCHFEYYEHTASYSSFPHGQYLQVLMEELNMSAEYVKNNFLSIDVFFDDFQVTTTTTKYTYGIETLLG
EIGGLLGLFIGVNIINFFELLVLCMDATKLCNKCFKKEKPNKRTNPVELDNVLVTKGSRNSTKPMNTVVGDTVTVN

>Deutero_Ambulac_Sakowv30000480m_382_2

MLQHYGITLELLQRDLERKFPDGLDVENFTRAIGWQLNDFTLPLCQWRGHKCFPQNFTHTFTRFGNCYTFNGGDEYID
QRIPGAGHGLSVVLNIQQSEYTESLYGNIEAGIKFLVHPKEEPPQIGSQGYALQPGTRAFADIRQVRYISLEPPWGQCDN
NAALDYFTSYSLSGCSLEYLKNLIYKDCGCRPLFIPGNEPVCSLNQSSTCVQQISAEWASNAPELPCTVPCNYTLYPTSLSY
TTFPSTNVAEQFEILLDITLEDMRENYVYLDVYYSQIQYEQYKQTKAMTSSALLSDVGGQFGLFIGISVITFVEIIEYIAKKIT
GKCCRAEKTMKVNSYNGEPHSTRN

>Cnidar_HydraVulSc4wPfr_248_2_g2925_t1_384_4

MGRYFPSQPTPKVEDKIKSPQEQKRRTIKQYLAYAGENISIHGLSHVFDKRENFICRTVWLLITIAAFGYAVQKVYESTLNY
FSYPFSTARMKIHVNQMDFPAVSFCNLNDFRYSATKGTKLDKAILSNDKKISGEEYLNITLKARHKIEEMLVDCEFDGNK
CSVHNFTEFNWNQGELCFTFNSGKFPHSLLKVNGVGMKRSLVLTINVQHYEYYGDELDAGIHLILHDQEETPIKKRGPVI
PPGFTTYIQVEKKTILNLESPYKTKCGSIKLKYFDSYSMHTCWLEQLTDHVYKICKCKDFFMPGDVGGQIGLFVGAGVMS
YFEIIDCLVLIIYTRFFQKFKTINSIRNHSL

>Protostome_Lophotroco_brch_Lingula_anatina_comp131826_c1_seq2_p1_comp131826_c1__comp131826_c1_seq2_p1__ORF_type_5prime_partial_len_373___score_4_70_comp131826_c1_seq2_1_1119__401_8

IDYYWSNPVSVNIELHYVDFIPFPAVTICNNNKLRVLNLYHTGEIDVGYLFTSPTATTTKNLSSYNWTEFITEYGHDAPTM
LNAPGSCEWNAQPCSVDNFTRILTEMGVCYTFNHGDTNLKTELLGSTFGLKLILNAEQYNYLATTHSAGFTVLLHHPDD
VPYIENLGFQVAPGESISVGISQTKVTNLPSPYGNCAEKNLAYYDSYSTLNCIAECTLNFTMDNCGCRAYYMINVPGVPV
CSPTQWWECLDELRANTSAALQACACPVPCEEVHYGYTISHSKMSNAITAALMAGYPGTTEDYFRDNYLILQIFYKELN
FQGIEQQPAYSGLALFSDIGGALGLVLGSTLMTAAEIIDFAVSLCLWVAFRRK

>Deutero_Ambulac_Sakowv30023603m_404_9

MTKDNHPSTENGDTTVRRITRRFALETSAHALPKSAKSSHTYTQCMWVVIFCAACSVYLWQIGDLLSLYLSVPKRTENIL
VSANTLPFPAVTLCNTNKVRRSMIQQSGYSSLLDSLEINTLIYYLKSEDPPDWRAFMSFSRDPSWTPLRKVLKFSKDELKV
LGHQLEDLILECSFDGNECDMEKDFHTFQHDEYGNCFTFNHGMHGVPLRNSTKAGGDYGLRLTMNVEQPEYITFYGK
DAGVLMTIHSPEDTPFPEDHGMVLRTGSKTSIGLYKEDDILIPEPNRECENVDTFYGIYGDYKYTTLACDKSCLNRYIQYY
CGCVDTIMIEGVPRCMILNDTQEVCQQLMNYLYYRGLLVCRCPRRCRQVHKNTMR

>Protostome_Lophotroco_annelid_Pdum_comp418513_c0_seq7__408_10

MDGEIGSKKPDNWTSALYRFSQTTTVDGVKQITEPTPFCARRLVWIALLLGGIALVARQIADSVLYYHSRPVTVNLNIKY
NKTLKFPAVTICNQNNFRATVASTRGQFDLISDIYVRNSSLEDKTYEDDFYSESVAGSMFALKHDKHDLIHSCKWDNDY
NCSADDFITSPTDHGICYSFNSGRDDKPIRYISQTGADSGLKMKLNIEQYEYMQGPNHGAGLKILVHDPSDVPMVKDLG
LAVKPGTHTYMDINIIEVHNLPPPHGHCEEKNLTLFSKYAMSTCLLECKTLHITSLCGCIDYYMPWKTSGGPPFCTIRQFL
ECVKPSLEAFNVADECDCPVPCNMTILNPSITLLANSDFDVEKMLHQNHSELQEKYTYAQEV

>Cnidar_HydraVulSc4wPfr_1152_g30239_t1_417_13

MSDTNIVNKHFITNDSKGWQSPCPKIKLNNQFRKESNEFIHRNGKPYFPVLKSEKEARREKLDETIQKFVDASTCHGFKH
CFNSGSKVRQLIWMMILAASVALLVQKLYESGVKFLERPFSTKTTLTYVDQMEFPAISICNMNDMRNSKYDYYEDIKEA
GIKLIIHDQHETPVRMAGVKLSPGFSATVQIKKKKTLNLKAPYATNCGSKPLKYFDHYSTNTCWLERLTDHVVTSCNCKD
SFMPGHARVCSIPELMNCTFLKWEEFNKLKDIQCPIPCESQEFESSISFARYPSNILADKIAKDMQLPGSVQENREFIRDN
YLRVEIFYEEMSYIQVEQTPSYDLMILLGDIGGQFGLFLGSSIITYVEFFDFFAALIYTKYFRIFKPPKI

>Protostome_Lophotroco_molsk_AplCal_XP_035825598_1_acid_sensing_ion_channel_2__partial__Apl
ysia_californica__419_14

MNNTAHHNSFEPRPFNTQYDNNIPESISMKPYAKDGVTSSSLDPDVISPQKKKGSLTWKEALFDFTQNTTLHGIRFIFM
NDVFILRRLLWLALFLTCSVLMSVQIVERIVFFYSYPVTVNVHVNFNKTLAFPAFTVCNQNAFRASAATDRHLYRLIERLH
SGDTSALLSQSPDPLPSNLSLDELYLTTAHRKEDLIVRCEWQNKPCGPENFTLVLTDHGVCYNFNDNPSEPLWVTSTGA
EYGLKLTLNVEQYEYMPGPHDAAGIKILLHDGKEFPKVAELGLSIPTGTHTYVGIQLLKIQNLPAPHGTCRSERSPYYERYS
PDACQLACLTKYVSEQCHCRHFYMPHIDGSPPVCTLGEYLSCYERIIDQVKDRVRAECDCPVPCDFLIYD

>Protostome_Lophotroco_annelid_CAC9660887_1__Ofus_G111053_partial_Owenia__fusiformis__422_
15

IGCTFKGEQCNRSMFVLQKDSHYHNCYTFNGIGSNNFNKNVSKTDPGSGLSLVVFTDTYTDTLTSDPGVIYNPSDPTSG
NNGIRVIVHSPKTRPAPVDKGFDVPNGFSTTVALRATRRTLLTEPHGDCTTDASNKGTEYRYTTDICYEQCHQDIITQKC
GCKSSHFIPPRNETGFQYCGKLNITNLLSGDDTSKESILEDFTHLECEKSVIDTFWSNVTMVSKCNCRPPCNSTYYIKTTSQ
AVWPNEFTQQSFYKTYVNITDSTARPNILFQGLTPKEIKDRGLIQKNFLRLNIYFEDLQVEETSQEADYPITQVISDIGGN
MGFYVGVSVITLVEFLSLIGALLLIFFKSCIKAGGKLREAKVSDVNIQPAHSLENKLGAQYTYEYKSYNDMISR

>Protostome_Lophotroco_brch_Lingula_anatina_comp132718_c0_seq1_p1_comp132718_c0__comp13
2718_c0_seq1_p1__ORF_type_complete_len_399___score_16_97_comp132718_c0_seq1_93_1289__4
23_16

MDRLYLNGATLYLAALSSKNSRAYAHINLDNIKNLNMTAWYGSGYTFANFTQDKGWLLNERNARCTFRGEDCNITAD
FKHVFTEFGNCYTFNSGDDPSRNLFQDQAGIGNGLRIEINIQQEQYTNLILRGDAPDAGILFHVHNQSEPASVETDGRA
VGPGLHAYAGLTRSDFATLHPPYGQCNETASLEFYPVYTMSGCVVECKRKHILRHCNCRLMEHPGYEPVCGLIRILTCVK

PLLVNLTQDFTGMCSCSVPCSSVVYQTALSYSLVPSESTKVEASNTLGVAVDDARKNHIILDVYFQSLNYQQSEQLPAVE
WTALISDIGGQFGLFMGFSLLTVVEFIEFAIMSVVTLTAAKRSQNKVKGHDTIHGKGDKVTKSSKTGRISMQGDFVQVH
EIF

>Cnidar_AlaAla_c59883_g1_i1_p1__GENE_c59883_g1_i1__c59883_g1_i1_p1__ORF__type_5prime_partial__len_400__score_24_40__c59883_g1_i1_3994_5193__424_17

IWSLLILTALGLLVEKLYESTMHYFSYPFSTTTTIKYAGKMKFPAVSFCNLNDLRMSKLNGTKLHTAILQSKNLLTTLTGEEY
KNTTKNANHRIHDMLDTCQFVHDNCSTFNFTQFFHNQGDRCYTFNSGKNNHPVLEVDNTGVGHSLQLVINIQHYDYY
MDTEKSGLHLILHGQNETPVKMQGVIVSPGFITYVDIRRRKLKNLPYPYKTNCGSKPLHYFSGYSMHLCWLESLTDHVV
EKCNCRDWFMPGSHKVCSLNESMNCMWPEWAAFDKFKMYNCPLPCEIETYSSRLSFAQFPSNSHADVMAKRLNLH
GSPIENRLFIRDNYLKIVIYYSELSYDHMEQVPSYDMMVLLGDIGGQLGLFLGSSILTYLEFFDCLVMLIYSRFFEKIKPTSVP
V

>Protostome_Lophotroco_annelid_CAC9477573_1__Ofus_G013340_Owenia__fusiformis__427_19

MIMPQMRMKLYLLGMTWKTFCFHVQLLAMTNMSQIESFINPYFFKCYTMNPKDFNDSSIEPLVHSTGAINGLSALFFL
DIGTAKRIYNPTSPLGGSSGLLVVIHPPGTQPDPLQNGFNIAPGFSTSVELQNKNRELLGHPWGDCENREKLSDFPYKYD
KSSCERQCQQKFIMDQCGCVTSIQPIPDNLKNMSMCGKMDLKNLNSDSPNISQIAEEIDKLECEHAKKRVLKKDYPTEN
KCDCPRACHNTIYDYTISQSEWPSEGVMLDFYKHAITLRDMQGSGYVHSTMYNLLDSKVNSNNTKENRANQEMIRKN
FLRLNVFFSSLNTEITKQVVEYTFTDLISGVGGGFGVYVGFSIVTICEFCVLFAHLVKAMMNNRKNAVNDEPWAFKRKS
MEIPSSK

>Deu_Ambulacrar_hemi_Ptyfla_40v0_9_20150316_1g5442_t1_scaffold2520_cov137_429_20

MPSEDSLTYKALIQNVLRNSSAHGLPNVQRATSLPMRLFWLLAFFTAVGIFVWMSSGLIAEYLRYDVDVNLQIQFSRDL
TFPAVTICNLNPLRKSKLSEFGHDILQRRPLGRMEGRRRQERQTTERSETSNDTLNDDQERTFEEYNYWDQIPTNYHAS
PSSSWSIIERIEHYISDIPAVQRAEIGHQLKDLVVDCQWNNIKCSPRNFTTFTNVMYGNCFTFNGEHNNVMPLSTHYSG
SVFGLTLILFVEQAEYMDDVIDSPGVRVTIHSQDDTPFPEDSGFDIQPGRATSVGILMGRTQRLPKPYTNCISDNIPTYDT
AFGDIYSVKDYIREFLKSRSSKLRTLMEQEERTGLDLLSNANRDTSKCYSEHHRKRSVATFDPLRFICAALLDSNDRYNLRT
SPDDC

>Deutero_Ambulac_Apla_gbr414_1_t1_430_21

MSDAVGIPGIRYVVSSNFHILRRLLWTVALLGGMAAASYQVIDRATFFFSNPKSVNVVIENAEPMLFPAVTFCNTNRFR
DVNGQLSTHPFGQFLAATYFHGGNGTGFTSTLGGANTTALYLEFAHQMEYGQMFIRGALGGKNITSADFRRVLTDAG
VCFTFNSGLNTELRGQSFPGRAHGLYLFLNVEQYYHYYAYEFSGAGILVAVHDQTEIPDMDSMGFGIPPGRHAIVSVGK
KLTSRKDGGCVEKQLKYFSKYTKSMCLRECVIDLLLQNCGCVEPYMTDKRRYVDMCDCPLPCSTVKFTTSVSYTLFPGNF
YSWLLAEQFSTDEFTLPHDFYQENFVIIQMFFEELTVEHREQHSDYTFFALLCDIGGALGLWLGGSILTIVEILDHFTSTTVI
PGSTSHQRG

>Deutero_Ambulac_Apla_XP_022104029_1_ASIClike2

MGCYKRCYLRQWALTDTDLHGVKHIAGEGGILRRLIWAACFLAALVVFLHQATLTLIHFVEKHHVTKVDI

SYRKKLDFPAVTVCNFNKYRESALTDRDIRNVGYHLGIVDEDHNLINPYLYTEEFRRKMAAVDWSVHDID

DEYNMTEFTNRTGHQLDEMIVECSWRDEPCSPDDFHHIFSHLGNCYTFNHVALATERHSSISAGAANGLK

LTLNIQEEEYTPSNDLHGAAEDAGIKWMLHHPSEPPYVKELGFGAGPGHHTFVAVRHEEVESLPSPYTPC

METSAGFLDHFDHYSLQACRIECETEVVVQRCGCRLVEQPGNAPVCNPAETHECAQAALVHAVAGHGDQA

CDCSSPCSVESYPFTTTNVRLRAKYIERIYSNTTHNFSADYIQNNLVLLSIYYEALNAEVIEQLPEMTVP

SLLAALGGNFGLFLGASVLTIVELLEYVFDELTASCTRRKTSGRISNAREILTVRAAPISVIDTEKELEH

GRQWNR

>Deutero_Ambulac_Sakowv30036990m_432_23

MKSPSNADIIRNVNRRRPTPTSTSFDMSNTEKFTSTSSLSLDGGNVRRRGAVYIFSRNFRQFLSETTLHGARYTANNEYH
VVRRLDVKYNINFDWSLLENTTAILNRTQFELEAAHQKESFIVACNYQGDGVERLCGPNNFTTTFTEFGVCYTFNNDPE
NQLFVTKFGSNSGLRVRLFTDENEYTFGKQTGSGFKVLLYSPGDVPLVQQLGFAVPTGVDALVAIRLEKSINLPPPFPTKC
SNAPLKYYDGNYTYVWCALEKITDFVVSACGCREPYMPGSKRVCNLKESIVCVIPVMDDSTERSVTYCGVGCDVTRFDS
RVTYADFPAKPILEHMSSTTNESSEYFRRNYADISFFAEDMTFKLTKTVPVMTGATLVSNFGGLMGLCLGASLLTAVEFI
DFILFCFCKS

>Deu_Ambulacrar_hemi_Ptyfla_40v0_9_20150316_1g16608_t1_scaffold13089_cov115_434_24

MMADAKPVTFSELLKKQLQTTTAHGLPRIEAADKLVRKLFWAFIVVAGLGMFIWQSSVIITKYYARDTTISIEMKFDTSL
DFPAITICNMNPVKLQSLERDPYLMSLVGYDNDEDDNLTGRRRRRKRNVENIIVIPTDSDAGTANYSTPIRGLSLELYIEEE
EYIPELQQSSGVRVIVHSQGLMPFPEDDGFLAAPGFKTSVGLRQLRLDRQPHPYSECVDTIYGPDNIFKDFYETYYSRKV
AAFFLTFCHNCEAFMYACNGYANAVIFFREFKNGEITYEATVSNAVWPNSAYRNVLLRDLMQTSSEIRYKVEADDTFIS
DNMVKIDIYYNDLNYEAIGENVAYTGGDVISNLGGQVGLWIGVSVMTCFEFFEFLYDVMALFLIKLTRAPQRKRVSTPII
PLRHNNRIDFS

>Deu_Ambulacrar_hemi_Ptyfla_40v0_9_20150316_1g4330_t1_scaffold1875_cov91_435_25

MDPLIDSKHKDSFSEVLTHLLANSSSHGLPNIQRSKHPVGKLFWAVLFLAGVGVLCWQVSVLVATYRKRDVDVTLKMH
SNTSIDFPVVTICNTNALKMSALVDDVYMLADLYNANSAFATIMQAYRNIEHLDSEPEKSTGGTEVNVYREEIVLSNSTS
AFVTGKYSNITTSTYPSGEQKSSTGPEPSSQPMINNITYPTFANKTNINSSSLLMERELEDYIDDEDYYNKMDNRHELQYL
LRQIMAAYGNCYMFHVANSNEHTKPLMINKPGRDNGLTLELFINQDNYIPDITETSGIRVAIHPHRVVPFPDDNGISLSP
GYSTEIGLRMVNIERQPRPYGNCIDANNIPDKYYDNDIYIRRYRAAYSVAACEKSCYQNTLIDRCGCYDADYPPTMDGID
LKPCSILGNSTGK

>Deu_Ambulacrar_hemi_Ptyfla_40v0_9_20150316_1g1885_t1_scaffold848_cov143_437_26

MASTTRKPERTSLGDVRQRFVQRSREYLSRTSLHGASYIIDNNIHPFRRCLWFLLVTGLSLSLLVTLHTEVRRYLEYPVSTIV
RMNYLPRLLFPAVTVCNYNRYRKSVIGGTPEDALMRHLYFTGSIYPTNPDGFDWTTLENSTSTVNRSLFELEAAHPKESLI
ESCRFKGDGFDWPCGPENFSTTFTEFGVCYSFNDDLDNAITTNLFGSHTGLRIRLFTNEDEYTHGPQTGSGFKWEHLPH
PYVTNCSHGQLRYSTLTYSYSACCLEKMTDFVSEKCGCKEIHMPAFGHQNTEFCNVACEEVNYIPHISYANIPSKPLVDD
LWDKKNMSRDFLRDNYADVSVFLEDLNCKQITFVPAITLNSIWTMPTAIDDSRLCSVPLARPNLICTDGVTVVRSSSSTG
LSSLSWLLFTY

>Deutero_Ambulac_Sakowv30036989m_439_27

MSSSELNELTYRRDSIRTDIVVTEGNDGNKSTAGLNTTTAMNTQTAFSAKLSQFMGRTTLHGVAFTVGPNNSWKRRS
VWIFLVIVSVVFLCLCLFSLVGLYLTYPVDTMTSLQFTQSLTFPTVTICNKNRYRKSVINGTAFEEFLWSIYPLAGFGPQLHV
NYNWSLLENDPALQNRTEFELTAAHQLEDTILKCQFVNSDDKHECGRENFTTVFTEHGVCYSFNNGLDNILKATSTGSS

TGLHMLINVQRIEYTIGPRTGIGITVILSEPGADITGENTALSIAPGVEALVAMPMDKYLFLKAPYQTNCCTEGTSYYPVYS
YHGCMNEKMSNEMAEKCNCKEIELPGPVRTCTLQESVECALPLKVSNALTNIDCNVPCNFTLYNPRVTYGHFPSPSIIQR
LSDMYGISEESVK

>Protostome_Lophotroco_brch_Lingula_anatina_comp147337_c2_seq1_p1_comp147337_c2__comp147337_c2_seq1_p1__ORF_type_complete_len_421___score__4_96_comp147337_c2_seq1_410_1672__443_28

MFYKFFEYDVGVKLEITSNSTLKFPAVTVCNENAFRKSALLSSPSKLPTLDAFVTNGTVPTTGFNSSLIDSELGGRSNRATLI
DKTLEDISELTTGEKQALGHQRSDFILDCEYGGYSCNGKTFESFYSTYNGNCFIFNSGWNNSSILLATESGPNFGLSLTFNI
EQSEYVGDLTQTAGVRVLVHDQDVMPFPEDQGFLAAPGEMTFVGIKMVEIDRYGGRYTTCKKTDTFNITENMYQALY
PSVGYSQKACKKTCYQRTVIETCKCALSIYPRVETLFDSTTSVRTCDSLNATDVACTVKVQTQFINGELNCSCIQPCSATSF
SIDTSSAYWPSDEYATEFLSNYNSKSSIVRKIYNTSGSAGVMKNLAKVHIFYKDLNYEYISEYVYYDVSIIGIASCILVVFLLN
YEKRTLCTQLTA

>Deutero_Ambulac_Apla_gbr467_14_t1_444_29

MGIRVKPIEDDISIISFNTDEVKIDKRERDKETTGDSLKTGRAGPQKGSKYHSSRCTKFLATWSSTLPDAVGITGIQYIVSN
QDHILRRFVWLLALFGGFAAASYQTIDRTIYFLSNPKSVDYEIKYISPLEFPAVTVCNYNSYRAYWDGWGSVTSANFTRVI
TDFGVCYTFNAGQAGQELLKQKVAGKGHGLSLMLDAQQYFYFYSSKALQVSAGFVVAIHNQSEVPQVDSLGVGVAPG
TEVRIGLKRKEAINLEPPHGECGSKELKYFSSYSINSCRQECLTDFVLEGCGCVEPYMAERFNSEGVCECPIPCQQTTYTTS
LSFATFPSDFYIQTLVNLYSDFNVTPEYFSSNTLKIEIYYEELSVESMEQQEAYTFFALLCDLGGALGLWLGGSILTFVEILDH
FGHTAFLRGTAFQHS

>Cnidar_NemVecNVEC200_012401_1_1_protein_AED_0_06_eAED_0_06_QI_181_1_1_1_0_72_0_58_12_321_422_446_30

MSRSGPSVRALLRDFSDRTSCHGIGQINGSHSPTWRIFWLLTFLAGLGMVLFQCITLLGIYLDKPTATSVDVTYDEVTNF
PAVTICNLNMIKKKNLANFTQSKKIFDDFEAFVSSNSSMDSSAFLGSKMETVLKDRLSMDSNDGSNSVSLDDTSMDTEL
YVEDMLIRHMAMVDDKDLIKAGHEFDELVFRCVWNGFTCNKGGFMKFWRRFWHWRYGNCYIFNQGVDENGTLLA
HLTSSKPGPMYGLTLDLFIDQEQYLIPLSQEAGVKVLLSDQRNVPFPFTDGFSVQPGVSASVGIRKLVINRIDPFNNGSCY
SGDGLEKDNIYSKHKSLKYSVQGCMSSCLANSEFSICNCTEGKFRVKGRPCMSESEVKCLNTVNKMYEKGTLGCTRKCP
QPCSHFSFRRTISQSQWSESYEQTFQKWS

>Deutero_Ambulac_Spurpu_014120_448_31

MKAPKVDGQEGSWGTAPEREEKSLRTILNSRMENSSAHGVPNIQRSSGPVTKLAWSLVFLTGISVMTWQAVILFQTYF
EWNYSVNLEVRFNRTQSFPAITICNANPIKRSELETRDALFQALFDVHYVPSMPDLPDQQPLPDEQPLPDVPGLSLELFV
QQDEYVEGMTEVAGFRVSVHHPSIMPFPEYNGLLVSPGFATDIGLRVLEVDRLPKPYGDCKADLTQGIEDDIFHQHYNI
TYNRKTCEVSCFQNEVISRCDCFDATYPNSLKVNRTVYPCEYINDVETQCMADIEMEHARDELECNCPLACRETTYLTSV
SSSIWPSDAYESTLFQKMVKYNAEIRRNVVGENASDWTRRNMAKVEIFYDEFNYEHIRQDAAYTISTIMETRYLHPFFIH
TNSIAYISETVHNLKQTFNVLSKSI

>Deutero_Ambulac_Apla_gbr500_4_t1_459_35

MRASEKPADTPRFVCSVLERIGAHGIPNIGRAGTQQRRLAWTGLVLAGLGFLVWQGTLLVTRYLQYDVKVSVGLGYEA
LTTFPAVTVCNLNRIPRTAMHTNSKLAELEGYIEYILETQVYQAWSDEMRYTNQVADFGITTESFDIMADIPYANRSSAG

HQLSRMLLYCTFNGYPCSPLNFTHFYNYFYGNCYTFNFDSRLARRVAQAGPFYGLTMELFVEQTEYVRGLQNSAGLKVF
ISGQNQVPFPEDRGIIVSPGRETSIALHKACEKSCVAQEILEMCHCADPREVEFEETVSAASWPNENYKFALENIITRLSA
DFSSAIEEDDRFIEKNVVKLNVFLSSLQYQYIYQTPDYSQENLISDLGGQVGLWMGFSILTFFEIAELVLDLFVYTRGKSAN
RVSREAKVGEGLVGMSLSPSVNTLTPIG

>Deutero_Ambulac_Apla_gbr243_2_t1_462_36

MEGNRSKLVADFKKFGDQTTFHGLRYVTNNTYHSVRRLIWLAVVLGMTTWLVINIIRAVVYMFQYPETSSISVNYVPNI
TFPAVTLCNFNQYRKDALDARAIKILAKVFGNPALRESIDLDVSDVKLFPTTNQANVTATTVQATHQIEDMLVDCHWRT
EPCSHLNFTQRLTDYGVCYTFNDDLAGLGDVLTIQNPGASNGLHLRLNIQQDLYTFGESTAAGMKVLLHPQGEFPIMKE
FAFSLSPGFETSVAVRKETVTSLKAPYKSDCIDGSLKEFPYKYSVAACQLECRALYVIDKCGCRDIRWPEEFLSRGERCHCP
MACETTTYQSKLSLAYWPAGYLTAELQAKRNLTEDFIRKNYIDVYIYFEEIMYMAIEQKMAYTIDNVQSDIGGYLGLLCG
MSLITLVEWADFILITLYKRCRRLASRRIST

>Deu_Ambulacrar_hemi_Ptyfla_40v0_9_20150316_1g8126_t1_scaffold4191_cov132_466_38

MSSSEFAWDGKKICEHETAKNTLSVIFDELLKNSSSHGLPNIQRSVSAVGKLSWALLFFVAVIMLLIQTVGLLQTYFSYPV
AVSLTMEFDKKLFFPAVTVCNANPVRMSELRAASDYRLRSQFDPSFSPSVKVEVEEVTDSGKPLITEGGSNTTPVGKLKF
SRTTDVETTLAQPTTTENSSEPIMSWDDRVGGDFYRKPSNGFSKANSLSVLMANQSTQTKMTMGHQLSTMLLDCFY
NYKYGNCFTFNSGRNGEALTINKPGPMYGLSLELYIEQIEYMIGTTEAAGVRVLLHHQDEMPFPEDEGFSVAPGQAASV
GVRQMELLRKSAPYGDCKDMENLSQEEIDNNIFMSRFDVGYSVQVCGRNCYQYEVIEECGCFDSNYPIPVSAEGRYDA
CDIVSDEKAITVMNLLPQTLIETYKLLPNMEWNMPLLC

>Deutero_Ambulac_Apla_gbr40_206_t1_468_39

MRVDKLVKILGWEVHGDPNSAAVSIPGEVLDQTLCAEAPREVFHKPTSGEVPKTRQPAWHVCSRMDRVWTKEEKRR
TSHGASLRLKTALKDDTSVVGIKNVVDEKVQTGRKKCAVEKYGPEFVDYIDLAYPVGMMNNKQTPPDFMVFKDLDLK
QFYVDTGHLKDDMILECSWQGQPCGHDDFISTLTDFGMCHTFNSHTKGMAVRRIKHRGSRFGLRLRLNVETDQYMP
GPRDSVGIKVALHHQNDIPKGMSKDLGFALSPGTYNLAEVTMSQITNQPMPFGKCATKQLKHVEMYSVMACILDCQT
DYVVEKCHCKDAFMPESFISSFETNCSCPVPCERTEYSTMLSSGIFPARHVAMALDKELNMSMDYSRNNLLELIVYFEEL
KYTRITQQPAYSLESLLSDIGGSMGLFMGVSILTLFELMDVLLRTVY

>Deu_Ambulacrar_hemi_Ptyfla_40v0_9_20150316_1g28005_t1_scaffold79692_cov99_470_40

MFSESPDLSQLTGVLQLSVDEISELGHQAEDFILQCTYAGEKCDANETTYDMTISQSQWPSHAYLNNLLKLVQVSNKKT
QNMHNLESARANLVRLKIYFETLNHDLTSELPAYTWEDLLSDVGGTLGLYVGFSVITVCEFVTLLFNLCGKSPHLDHTKKE
EQPLHERVSDTNEGTNEFSDKTNVQHDFFKHSDDTGKQHDESVKHVLEDFGRETTAHGIVHITTAKSNVTRSTWIMIV
LVAACVMFVQMTLLLVQYFEYNVNVKVTLVSEKALHFPSVTICNTNKLRHSAIRSSKYSEMLMLERNFVSPYYAPCIEGD
FQCKNGIHCIKPYLICDGVNHCGDMSDEHNCAYPECGPDQFRCDSGSELGVCIPNDKFCDRISHCYGAEDETSCGECSE
DEFQCITSRECVASLKTCDQLFDCRDGSDESASICGKQ

>Deu_Ambulacrar_hemi_Ptyfla_40v0_9_20150316_1g5424_t1_scaffold2509_cov138_473_42

MACDDVDKEFFTGTTIHGVGRIVERSRLVFKILWFFIVLVAFGFCAWQITDRFVRFFEFNVNTEITLKFESSIEFPAITICNF
NRYYSSAITASDQAIVNQLLYAMDYDYDYSDSEYSSFDWDNWYSNLSLAADFDYANFTRRVGMQLDDTTLLSCEWRG
RSSSCSAREEYTESASGNVQAGVKFLVHDKNTPPLMDSQGSAVAPGTYAFVAVQQVRTVSLSKPYGSCVSSSTLQYYDE
YSLSACLIEWRLELIVQECGCKPFRYPGPARDCTVTESATCAKTTLGRIKNGDFGQSSCYVPCNMTSYPTSVSYAQYPSQS
VAPEMASFYNVSESYGRNNLIYLDIYYDEINYEELKQTEAMTPSALVSDIGGQLGFFLGASIITLCEFIEYLIMKCHGACRTK
SKKVWDSNDEKEKKVDVAANGDTELNFGPRFAFS

>Deutero_Ambulac_Spurpu_024200_477_43

MAQQAKEAWYDAPSKPVTAFKSLDKESATPIEQPSFNMSKEEENSVLFLVGNRMASTGAHGIPNIQRASNIWRRLIWL
LLFTGGVTVVVWQLTTIINNFFAYEVDVNIDFVRQRAIKFPAITVCNLNPVPLSKADVLWNIASNEEDATSLRDFLPAQK
PSGVMNATQNQNDAGASMRGYQDWVNVDFNQLNTRVSNLEVAQSIIGSTAYENRSETGHDLDDMLLDCNFENYPC
APDNFTHFYNYMYGNCYTFNSGQTGKPLEVSTVGPLYGLSLELYIQQSEYISDLQPSAGLRLLVHDQNEMPFPEDRGINL
APGAHTSVGLSLVYLERKESPYTNCSLEFKPGNVFKEQFDHLEYSRSIYSLISDLGGQIGLWIGVSILTIFEFIELIYDLIKLICLK
LMDPRMKKKRSSQSQTKQSNVMALDGQQGRTNSLFADR

>Deutero_Ambulac_Sakowv30044075m_476_44

MPTTLRASPIKDIVNIHSLQERYQELTQDTRLFDEFRVNYYQGNHFYQSDLANNDTPDWKRYLTYSSSTDYSDLRDVLTL
NMSDVFKYGHQFEDFVLQCAFNENKCDQSDFSTFQNDRYGNCFTFNNVRNDALTRRADKMGSQYGLKLTLYIEQDEY
IPLYGQEAGVRILVHPSDITPFPEDDGITVAPGRKASIALREELHSSVAHPYSNCENNPQIDSIFGSQYNYSVQACRKTYLH
QKIQQKCNCRDTLWLSKDRPACKITNTAQEVCRRLINFLDRRGVFENNCKEPCRKTTFKKTVSQSMWPSNKHLGGLLK
VLRPINTKIRDKVVDEKSARDNLVLIEVFFEVPNYRSILNTPAYTVDQLFADIGGLMGLYVGISLITVVEFIELVYNLYNYFSL
NREEKLAKRAKKMESKNMNHKYDPIPCNLPQSMIGTSV

>Placozoa_HhoNaC67_TR9228_c0_g1_i2_m_20240__Hoilungia_hongkongensis__30_45

MSQEKFNHEERFATSTSYHGMAHIYDGQTSKQTRSFWLVLTLIATGACISQCVIIILNASQLPTRIVTKTVLQNSSIFPSVTI
CNTNDYDRNSLSSSDIQHLSTIVDFVYGYDIEPHRFEAAISNLSIKFGKGFNYELFLRRAGHKLENMLLSCTWMGRPCSIE
DFVNTTTAAGSCFTFNPGSTNTSIKNQTASGNANGLRLILNIEQYKYYPALYTPGQPDAGLRYSLHYYKDPPNLIAESYYA
APGFHTYVPMTLKREKRLKKPWGDCEELSLKRYQFYSRYACLNELSAKYASESCNCSYQDNEAPNPCDGVQFLSCIIPTA
GYYRTQLSQNLSVCPIACETFSFATEISQSSIASQAFNSAIDSFINVSGLISDAQSKYWLPQSYTNVDFIRDNIVYLDIYYRKL
DMTQILQLEDTGFSKVLSTVCIYLQITFTQ

>Deutero_Urochord_tr_F6ULW5_F6ULW5_CIOIN_Uncharacterized_protein_OS_CioInt_481_47

VMSSLSDLIYEFAQRTTCHGLSQILQTKRKTIYRCLWFMMVSGLCGGATWQLTQAVQEYLSYKTSTDIQEMYTSELPFP
AVTICNINSFSHLVKKFLNVLSDKLFSVWPSPEFEMIAFNNFDNWWRREVELYYEKLNERGIDKFYLSDEILLKGWTLNET
TLSHCTFRQKPCSIQNFTQVVTPMGLCYTFTSNESMYQNVPGSGHGLVVHLNIDQKRYSEHPKYGNPNAGVKVHVHN
HNDPSEVQNYGIGVPSGFGAHIVIRKTERILMYKPWGVCSEKTKTLLHHDHYSTAGCLRECRAEYVYSKCGCKPYWALN
MANVPECDALNILNCSGSADAVFSTKFSLRTCECVEPCNFVVYDSSTSYSTFPSIEVGEFFALHAGKNLRKYRENHLMLDI
FYDSLSYVRMKQSKAVTEISLMSDMGGLLGLWLGISLVTVSEFVQLFF

>Deutero_Ambulac_Sakowv30035231m_482_48

MDASNSSRNIMETVFQTTNDVHPSSDEDMSHQRNCAPRCRDRLREFTQETTLHGIRFTTDSKSNIYRRLMWVLFVSLG
TAGVVITLKDSAIRFIAQPVTTVVEIHHPNAVPFPAVTVCNYNKYRQSVLAGTWAEDLLYKMYGSPVQGEIGEVDWSNY
TDVITNLNRTAFEMEAAHQIEDMLIACTWSSETACGPENFTNVMTNYGVCYTFNGDVKNGLTVKNRGSMFGLYMVL
DIEQHEYMPGVSKGAGVKIMTHTQTDVPLVNEVSMSFGPGMDSTIALRLHQTKLLRCTNQPLQYFDVYSRANCEMES
FTLAATKSCTCREPYMPEESELSCESACQEVIYDSTISYGSYPSLQILKKIQLMRNITEAEIRENYLAISVFIEDLTVLSTEEEYS
YTYDAFLGDIGGQFGLCLGASLLTLLEFFDFLMMSLCGACRRWYNRSRKQ

>Cnidar_NemVecNVEC200_006432_1_1_protein_AED_0_27_eAED_0_27_QI_414_1_1_1_1_1_6_1486_
449_486_51

MIMAAAKKLLDGRKKRAKVKELWENFLGGCTLHGFHYCFAGNPPLRRLIWSLLLLGAFAMFFEKCTESFINFFDYPFTTT
TLLVYDKRLPFPAISMCNYNDARMSKMNGTLMNEIFVASKLEGRNTSHLQSQLTGELMQRTLKEAAHRLPDMIKECS
WQKHGKCSWKNFTSFKSADGDTCYTFNSGRKDPILSMSNVGEENGLRLVIDTQHSEYYYDVKNAGFKVILHDQGETPV
KMQGLSVSPGFTSYMELKRTKVTNLPFPYKTMCGMPELKYFNSYSKSKCFLDKLTQYVVTLCGCRDWFMPGQGKIPVC
DYETAASCMWKAWAYFEENKLDQCPVACNSVEYSAQLSYARFPANNYAKMLAKEYGLKGSDEENRQYLRDNLIEIKIY
YEDLTYFDVQQVPSYDLYSLLGDVGGQIGLFLGASLLTVVEYLDLLGMVAYTSFKYRN

>Cnidaria_HyNaC9_NP_001296667_1_acidsensing_ion_channel_1like__Hydra_vulgaris__31_52

MSEEDEKKIKSPHEQRNDRVKEHVAHLIKNVSFHGLSYVADKRNNYFRRAIWFLITVGAFIYAVEKVYESTVNYFSYPFKT
ARMKIYVNELNFPAVSFCNLNDFLFSKLNGTKLDESILYPDDPEKNNVSEIEISNITSDATIRLDQMLVDCEFEGKKCTHEN
FTDFWTMQGELCFTFNSGKNSHSLLKVSGVGVLRSLKLTINVQHYEYYRDEMAGGIHLMIHGQDEEPVKMQGQIVSP
GYSFYVKVEKKTIMNLEKPYKTECGTVKLKYFDRYSMHTCWLEQLTDYVNKMCHCKDFFMPGNIPYCSLPELQNCTWI
EWAKFNKDKMYKCPLPCKIDLYGVSLSRALFPTTQYSSILAEQFRKQPHVLSIVHNITDELLFMRDNLLRFIIYYDDLSYEVL
EQKPSYETLVWLGDIGGQIGLFIGAGVMSYFEFLDCLAIVIYTRFFQKFTSS

>Deutero_Ambulac_Spurpu_015893_493_53

MGSDDLQNETVIGSVNTFGNETTVHGVRYMIARGHFLFRLCWLAIIVLAFVLFVIQAFNVYGDYASSPYSTKIDIVQQTF
KAFPAVTVCNANRIRRSMLYDTNYEGLIDIDDNNRSESEVEDGEAERSEEIWRKVNGLYDWWGFYSASVADDFSDLIN
VVNPSQEQLMGYGHQLEKFLIQCTYDQQPCNMSSDFRVWQSRYYGNCFTFNFGMMETAEAITTSKTGALNGLHLTL
WVEEDEYMGLLSPSRGAKVTIHPQNTLPFPEDEGINVGTGMATSIGIREEVIERISGMTNCVEEDGAGTNFTVTRTNTIY
TVAYCLKLCFQQSLINICGCVDGILLDYTHCDVLNSTQRACVSMVHQLYINNGLYCNCPLPCERNLIRLQIYYETLNNEVV
EQVPRYTIPNILGNWGGLMGLFTGMSFISVFEVIFLLLRVSKISCTKLFFPNRVQPVKT

>Protostome_Lophotroco_brch_Lingula_anatina_comp141843_c0_seq1_p1_comp141843_c0__comp14
1843_c0_seq1_p1__ORF_type_complete_len_456___score_27_90_comp141843_c0_seq1_2150_3517_
_494_54

MNRNRVTPTDIPKGKIGDEATKDSEIDQTSSTQEKLATFAGEVSILGLKQSTNPKYSPIRRLIWLCFVLIGFGFLIFHLKNRF
DYYLSNPATLNMEVVPNDTLVFPAITICNSSPLKASNLAANNESELGNIFSPFVNVDWSQYNLSHINWTDFHMRNGFQ
LEELLFWCTWKKTNCTKHDFTPIITEIGLCYQFNAGGEEQLHLETQGSGYGFGMTLLAKQRDYVSYAYPAGFLIIVHSPG
DHPDINLGTLISSGGFTGVSLKLKRLINLPPPHGKCGEKPLVYYNKYSKANCHAECFHNFTLTKCDCRLTHFPRVPGYREC
SPEDSRVCVDNLKLIITESVYLRECGCKTPCDSFKYDVSLSSARLSETAMRTTAAILRTTQEEISQDFLSVFIYYSEMSYEEVV
QQKAYSALALLADIGGALGLVLGSTLMTLAEVVDFFFGLCFWKLFGVEM

>Cnidaria_HyNaC10_NP_001296656_1_acidsensing_ion_channel_1like__Hydra_vulgaris__39_56

MAKVEDKIKSPQEQKRRTIKQYLAYAGENISIHGLSHVFDKRENFICRTVWLLITIAAFGYAVQKVYESTLNYFSYPFSTAR
MKIHVNQMDFPAVSFCNLNDFRYSATKGTKLDKAILSNDKKSISGEEYLNITLKARHKIEEMLVDCEFDGNKCSVHNFTE
FNWNQGELCFTFNSGKFPHSLLKVNGVGMKRSLVLTINVQHYEYYGDELDAGIHLILHDQEETPIKKRGPVIPPGFTTYI
QVEKKTILNLESPYKTKCGSIKLKYFDSYSMHTCWLEQLTDHVYKICKCKDFFMPGEIKVCSFEVLENCTWKEWAKFDNY
KLHTCPLPCKIDSYEMSLSRALFPTARYASSLANRFRKLPHVLNTVKSISNDLLFMRENLLRVVIYYNDLSYELLEQKPNYD
TLVWLGDVGGQIGLFVGAGVMSYFEIIDCLVLIIYTRFFQKFKTINSIRNHSL

>Cnidar_Polpod_Hydrif_GBGH01017297_1__p1__GENE_GBGH01017297_1_GBGH01017297_1__p1__O
RF__type_complete__len_459__score_52_80__GBGH01017297_1__223_1599__498_58

MQVSRGAKPNTLVEVSSKKQTAPTTLVRKHLSNFFGQSTIHGLHYCFDGNLPMRRVVWCCLVLGALALALQKISESATE
YLSYPISTVNTIRYEETMVFPAVTLCNMNDWRLSVTVNTSLFYTVTSFYNTGEKKEVSAEELLATQRRANHRLENMLLSC
KWKGQSCGLQNFSEYFTVQGERCYTFNSGRDNHSTLYAETASIIDSLELILNVEYYDYFIDNFVSGIRIILHSQEELPVTSQ
GYVLTAGDKTYIKVKKTKARNLPYPYKTNCSSIPLKYFNKYTADMCWLEQLTDHVTSKCGCRDVCMPGNASVCPLSVM
DACGWAEWDMFNIGKLAKCPLPCEVDKYSALMSHAHMAPSIVSTTLTESLRIERPTMAHETFVGENVIRAVIYYEDLSY
VLIEQEPTYTLMKLLGDIGGNLGLLVGASVLSYMEIIDCIILIVITLIKNSRSSSKKQKS

>Protostome_Lophotroco_brch_Lingula_anatina_comp138624_c0_seq6_p1_comp138624_c0__comp13
8624_c0_seq6_p1__ORF_type_complete_len_459___score_26_12_comp138624_c0_seq6_318_1694__
502_62

MISPSRLPRKTDNSSDQETPVIEKFAERVSCQALQEIVSRESGYIRRLIWLVCFLGGLILMAFQVYERVTFYAGHPVRFNT
YTVTPLSVMFPAVTICNYNAISKMRARELDLLEEFERYFPLKVGSENFTGLGDDWFDWDSVLNISKPFVKECTFRGISCS
MKNFTPVYTDMGICYTFNSGYSGHRLTSDSSGRLFGLQLTLDVDQRNYFRGFGGYGAGFKVRVHARDPPMMSNMGF
SVGAGMHTLVGLKMINISNLDTSQWGKCNRDIKLKHHVVYSQSACRLECLYEYMEKKCQCVRFLKFKHHQGPRARLC
DPEQLFNCYTPNYNLFIQEKGQCSHTCPVACESIKYEVSLSQAPVADYHAYNSATTTNETKVNKYRKNFVQLDIFFQEISY
ERREQYQAYNAFTLGCDIGGALGLILGASVLTLFEVGDFIILMILKGKWRRQTQVTSVSVHKK

>Deutero_Ambulac_Spurpu_009765_505_63

MGDLPRDTKTSPEKMADYSEKAYGRPDYDESPKEGTRKIVNEFVDNATTHGIPRVLNASRPWQSRLFWCVVTLIFAGV
FLFQGSKLVQSYIARPTTTKISLITKSRLEFPAVTICNLNMLRRSMLKGDEYDYSGIEDIEDPNDWEALYNLSKGSDYNNFK
NFVKPTKEELRTLGHKAKDFILQCSFDTEACSYENFTVIQNAEYGNCFVFNNAHKLKRGKRTTTSRTGSQYGLQLTLMVE
QPEYIGILSPNSGVKVAIKDPRIYAFPEDDGIEAAPGFATSIGLTKTSISRLPEPYGNCIRKHDSFYAPEKYDFSQRSCLKLCL
QETLNETCSCITDVLIDGTMCEVTNREQGNCRNSVFKDFLKNKLNCDCSNACRSRSSSNSSSSSSSSSSSSSRSSSSSSSSSIS
SIFSSISSGSRSSRSSSIGSSSRGSSSSSSSSSSSSNIRSSIIVVAVVEVVQ

>Deutero_Ambulac_Apla_gbr40_182_t1_508_68

MILDSEARPGSDRGAGNYRHGISLTMMTEATAGGTSPSGTASTTEQPSEECENSSLKERIATFGDETTFHGVRFVTNHQ
LHFLRRLLWFCIVGGMTGYLIYGVSVSFITFYQRPVTSVVKINYVRQIPFPTVTFCNYNQWKKSRVPPEFYDAVRSLNPA
NKKPVDMTYLKAHDVTNLTDKLINDTHQIEDMLLECTFKEANCSAENFTQIITDFGVCYSFNDDPSNVHYVHQSGSQN
ALFMKITVEQEEYIFGENNAAGIKVVNLKYPYESNCTDRGNVKYSAEYTVPLCQYEHKIETVKAKCGCKDMRYPANFTSE
DKSFNCPVPCDLTIYDTRMSFAYYPGQHHLEEIMQQNNLSENSIRKNVLDLRIFFEELSFQKIEEIPSYTFYSLQSSIGGYM
GLLIGASLITLFEFLDFILVTTTSCLLKHSVLARPRGNRVRGAARQRDRSSHGEAKPCFIKK

>Deutero_Ambulac_Sakowv30004505m_514_69

MDVVVPLSQMDNTDHQTKKAEDDTASQSQEPEKPSRLTLAVIYAQNTTAHGLANTVNAGTLHKRLFWILTFIICMVTL
VVQAQSVIRLYLSYPKAVSVEIKSRSKLDFPAVTVCNLNMARVSRWNGTRFQELLDVDRMQFDIDEWFSENLGSSDAP
EPDTSTQADVPINATSSGDAPVNTTSSGDAPVNTTSSGDAPVNTTSSGARKRRAVVAKHPVTDLLGQGTLSEFLKTLGP
GLQSSWIRQYAKENVITRPKRNSGNDSETSETDGSDDTDNGNSEGSEMGSERGSMGFYASDDYDDFSSNAFMDYGL

GGTNPNDWSTFYTKSSTDDFSDLRAALRMTQAEIEEYGERAEDFIVQCSFDRSSCSSKDFTVSQNDVYGNCFTFNGITG
NKTRRTTKAGEQYGLKLTLYVDADEYVGLFTQQEGARVVVHPGSLYPVPESVGLTAGVGQSLSIGLKKVKPKY

>Protostome_Lophotroco_annelid_CAC9474682_1__Ofus_G012955_partial_Owenia__fusiformis__516_70

MACGCCCTKSQRKLDYEFATSTSCHGLGRIADSSRKPQRIFWILVTLFCFGYMVSQIVDRFDHFMQMKQTTELEVMM
DQTSLEFPAVTICNYNKYRLSAMTESDAIILSNSALLMSYLQWGVPLDEEHVKFAQELEYNITQLYPDGFDMGNLTRRV
GWQMDDTTLLWCRIAGKQCTFEDFSLVQTSYGNCYTFNRYPREPNVYKQTVHGTGGGLRIVLDINQSDYTEVFAFGG
NSEAGIKFLVHPQYEPPSVESEGYAVSPGFRAYAGIRYDELQNAVSPWGKCDPEAKLKFYEKYSFEGCLSECREGKLLERC
GCKPYSIPGNIPNCLPTKAVTCANEILGAIEENPKEECPQCTVACEEVEYPVSLSYATFPNHNIAKELFEIFDVTEDYIRDNI
VELDIYFRDLYRHLTKMSPAVTPSGLLSDLGGQMGLFVGMSILTLFEFLEFTLRRLYLGCFRRNAED

>Cnidar_Rhopilema_esculentum_TR103737_c1_g2_i1_p1__Rhopilema_esculentum_TR103737_c1_g2__Rhopilema_esculentum_TR103737_c1_g2_i1_p1__ORF__type_complete__len_466__score_59_67__Rhopilema_esculentum_TR103737_c1_g2_i1_277_1674__517_71

MWRIINWRKSTAGEYAEARRETEVRYPGIPEKAFHMEQRRKRKRNAEHFNTFISDSTLHGLHFCFDKKHLARRIIWTLILI
SAFGYVGEKIYASLLQFFQYPMSTTTTLVYDEPPIKLPAISICNTNDFKRSIVVTTKLGDILRNKTDNITGSEYSSLIKLGNHR
LVDMLVTCRQSHKTCNVKDFVEYWDNSGTVRCHTFNSGKTGPIMTINRTGTTEAIEMVINIQQYDYNDFDTAGIKLIIH
GQDQTPIKTAGLLLAPGTVTNIEISKRKRVNLQHPYKTNCGSLKLKAFDSYSEHLCWLDQLTSYVVDKCGCVDAFMPGK
HKVCDFKTVGCVWNNWERFDIEQKYQCPIPCVVEEFIAKQSSAMFPSNAAADNLAIDLNLSGTAMENRLFIRENFLKFK
VFFGEYSYELVEQVPSYDLKVLLGDIGGQLGLFLGSSLLTYVEFFDCLLMIIYTKFFEVLLPR

>Cnidaria_HyNaC4_CAL36113_1_hydra_Na_channel_4__Hydra_vulgaris__37_72

MNFEELAKIVVEAVQENNNQPINTNKIKTPRELRNNKTKEYVSEMIDNSSFHGISYIAGKENHFIRRTIWLLITMTAFGYA
AQKVYESTVNYFSFPISTTQMRIYVNEIDFPAVSFCNFNEFRLSKMDGTKVDQAILNPKLQGLVTAEEYRNVTFGAMFDL
KEMLVDCEFNGIPCSDENFTMFSWMQGERCFTFNSGKFPHKLLKIGGAGMKRSLKITINIIHYEYYKDEMDAAIHMIVH
GQQDTPLKMRGPTLSPGFTTYIQLEKKMIINLEAPYKTKCGSLKLKYFDSYSLDTCWLEQLTDHVHSVCKCKDFFMPGDI
PICSLNDAMSCMWPEWARFDQLKMQNCPLPCVIDSYKLSLSRALFPSPQYADSLTNSLQQPPHIINSIKNISDKVQFAR
DNLLRFVIYYDDLSYELIEQKPSYDTLVWLGDIGGQIGLLIGAGAMSYFEFIDCLFLVIYNQFFKTYR

>Cnidaria_HyNaC8_NP_001296668_1_acidsensing_ion_channel_5like__Hydra_vulgaris__36_73

MLELKCEISKRRRKRYQNAISYLPGANHDTVDELRNQSINDYLSDMAVNSSFHGINYICDSTYKVRRIIWIVVTLTAMLYA
MREVYESTRKYLNYPVSTVRMKIYVDDLEFPAVSLCNLNDVRMSIVNGTSFDNALIDQNAQNISADDALMIAREARHN
LEDMLLECKFNGRSCSAKNFSEFNWMQGDRCFTINSGKPGHSRLSVKGTGIKRNLELILNLQHYEYYRDEMESGIHFILH
SQEETPVRMRGPIVSPGFTTYFRINKIKTKNLKYPYKTRCGSLNLKFFKGYSKQLCWLDQLTDYVNSKCGCKDFFMPGNI
SICTFSTAFGCMWPAWEEFEKKKISNCPLPCDADTYFGQTVSRALFPSNEYKKSFIKNLMHIQQFRDVYGNKKKELQFM
RDNLLRIVLYYDDLSYELLEQKPSYDLLSWLGDIGGQIGLFVGSSAMSYFEFLDCLIMIIYAKYFKQYK

>Cnidaria_HyNaC2_CAL36110_1_hydra_Na_channel_2__Hydra_vulgaris__32_75

MKARFARRMEKIVEERMTVKQIFNRYVQTSTLHGFRFIFMDTFIVRRVLWTILTLTMATIFFKELRNSINLFYEYPFTTTST
IQYEPSLTFPAISVCNLNHFLLSKIKKSKLKPLYDQGRLPFDNNLENPGFDIQGEELYSILKTSSQSIDEIFLSCEWKSRDTAK
NGVPNPCKPNNFTVYSGLYGQSCYTFNPGVSGYPLLSLSETGVNMGFKLELDLKSKDSLQGIQEIGAIVIVHHQQETPVL

QAGFVVSPGFQTFVEIKVRQTENLPPPYATKCGSKPLKNYQIYRQSSCFLEQLGDAIETKCKCKSSFMAGRNIPYCSLRQT
VTCLMPTIYDFDRKTNNNCPVDCETIQYLSSLSYARFISNVTYLSKNAEQSSYIRKLKNSMSPKKLQKYIEENIVAVQFFYQ
EMKKEKVKQEPSYDFYKLIGDVGGQLGLLLGASVLTLVEFVDLFIFTLYHQLLRLSKKKS

>Cnidaria_HyNaC3_CAL36112_1_hydra_Na_channel_3__Hydra_vulgaris__40_76

MLNFQDIAQITVEAIQEKNELPNEEEKNTQHPRDLRNKKIKEHISYMIDNSSFHGLSYIFDKRHSIRRTIWFFITIAAFVYA
MQKVYESTMNYFSYPFYTARMRMYVNQINFPAVSFCNLNDIRFSVMNGTIVDDAIINNNQEANITGEEMRTFIQAAR
HTLKEMLVDCDFEGKKCSYENFTEFSWMQGESCFTFNSGKPPHTLLKVNGAGINRSLKLTINVQHYEYYRDKMDAGIH
LILHGQDETPVKMRGLHLPPGFTSYIQIEKKTIINLEAPYKTKCGSVELKYFDSYSMHTCWLEQLTDHVYKVCKCKDVFM
PGDIPICSFDEAFNCMWPQWETFDKLKLYHCPLPCKIDSYAVSLSRALFPTARYASSLANELRKHQHVAMNLKSNTDEL
AFMRENLLRLVIYYDDLSYELLEQKPNYDTLVWLGDVGGQIGLFIGAGVMSYFEFIDCLAMILYTRFFEKFTTK

>Deu_Ambulacrar_hemi_Ptyfla_40v0_9_20150316_1g18733_t1_scaffold17031_cov102_519_77

MDPLIDSKHKDSFSEVLTHLLANSSSHGLPNIQRSKHPVGKLFWALLFLAGVGVLCWQVSVLVATYRKRDVDVTLKMH
SNTSIEFPVVTICNTNALKLSALVDDVYMLADLYNANSAFATIMQAYRNIEHLDSEPEKSTGGAEVNDYREEIVLSNSTSA
FEAGGYSNITTSTYPSGEQKSSTGPEPSSQPMINNITYPTFANKTNINSSSLSMKTELKDYMDDEDYYNKMDNRHELQY
LLRQIMAGLTLELFVNQDNYIPDITETSGIRVAIHPHRVVPFPEDNGISLSPGYSTEIGLRMVNIERQPRPYGNCIDANNIP
DKYYDNDIYIRRYRAAYSVAEFDLCSDIGGQLGLWIGVSVITLCEMFEFFASISRLFYRRLSVCVFNFGLFDSWINNNSAIY
HHPARNNCNGGQHPNNHTTTNNPDTNDCFDDHYPCNNPTICYNCSLNIQGTFDNPTFQRYDSFNLVSP

>Protost_ecdy_cyclo_PriCa_ecdy_XP_014670949_1__PREDICTED_acid_sensing__ion__channel__1_like
_Priapulus__caudatus__521_79

MADAKIDSQERDTCPADRKPPESFKQQLQQFSHGVSATGLRYVFDDDVSIYRRIIWGVLVACGLLYSFYQIGKSIHVYAS
RPVAVSIDLSYRDELEFPVVTFCNENVIKKSAMPQQGFEDYVRVIRMLVGGVSSPLDRNMTEIDALLDGLDVQLLQEVL
ETGMYHQDEIMRQCWWRGQPCEQKYVTSVYTDMGLCFSFNANGSSPLIARDTSGHQSGLRLLLDAQQHEYVASREG
TVGFRVMLHSPYETPVLQARGFKVGPGMHILTSVSKSVTHRLGSPYSQCRQVAGDASEHYSYYACLQECENAFVEGVC
GCHNLQGRNVARVCTEKTELQCLMRLMQVSSAFDRRHCKCPTQCRDIDYTPSTSFSPFANYLSAMMINATADYVTNN
LLLLDIYFTELEENVITSVPAYTFVALMADIGGSLSLFLGSGLFTIFEVVDFAIVAANKKRVARNGECTDPEQTAMKSTKR

>Deutero_Ambulac_Sakowv30036986m_524_81

MNTLGATVSPEEASSTAKIYAMTGIKEGKDDIGQGDRSRKHGCTCEYTKERIRKFTQETSMHGVRYTGMTDIHSVRRII
WTVLVLWSLGWLTYFLYTSVVKYRAFNVNTLLSMKYTDVLLFPAVTVCNYNRYRKSYIQGTTFAEWQQTLYHPINLIHG
STTINWTTDAAIEELAKNRTYLELSAAHQKESMILECKLGRASSGCNAVDNFTTTLTDYGVCYTYNNALSNHMLVTESGS
RHGLSLTLNLEQEEYTYGQNYGAGFKILVHFPGDVPLVTDFGTAVSPGTEVFMPVRLVVEKNKESPYRTNCTNVSLEYYA
HYTKSNCIMEKVTQMVVSKCGCREPYMPGNVRVCNYQEKEECVQIAVNAANCPTACDIETFQTSSSFATYPSYHMND
YLEQTYNMTIANIRENVAVLHVFYEELSFMSIEQSAAYNGMDFLADFGGQMGLCVGASLLTVCEAFEFVFIGIWKRLCR

>Deutero_Urochord_tr_F6VJJ2_F6VJJ2_CIOIN_Uncharacterized_protein_OS_CioInt_525_82

NNHTEQNGLSVLKRLNNRIRNNLRMKPKLGLRTKQKKLAVTFSEETSCHGILQIKNNAKIWLKVFWALLVLAAFSVLIW
QIIDRINAYNKHETNTQISEEYVQSVHFPAVTICNSNRILYEAIQRIKKYFYFSSDEERLAMLTLSKAFDTGVPQNASVVQN
LGLASNFSLREFMMAKGWRLDVGGALAFCGVSKTEVCSHVSNSLGLCWTLNAQRRQSIPGYPHGIHLIFITHQSDSTED
PISGNSETGIKFQVHSQDEPPQVESLGHSVGVGLWASVQMHKQEKSNMYSPWGACRPSNENLRFYSKYTLNACLKQC

YDTLIQQTCNCRGPLMEYNYEDVVECTPIQVLNCTAPVRYAIESSYTPQKCGCYQPCNEVKYQTTLSYSSFNPKHSEFFEK
GMNLSSGYLSENSVGISVYFDALSVTKHVESKAITESALLSDIGGQLGLWVGVSVVTLFELLQFLTSNIYCILIYL

>Cnidaria_HyNaC11_NP_001296594_1_acidsensing_ion_channel_1like__Hydra_vulgaris__34_85

MLNFKDIAQITVEAIQETNEVTNKDEKNIQTIQDLRNKKIREHISYMIDNSSFHGLSYIFDKRHSVRRTIWFFITIAAFAYA
MQKVYESTMNYFSYPFYTVRMRMYVNQIDFPAISFCNLNDIKFSAMNGTIVDDAVVTQNHEANITGEEYRSYNQAAR
HTLNEMLVDCDFEGKKCSHKNFTEFSWMQGESCFTFNSGKPPHTLLKVKGAGINRSLKLTINVQHYDYYRDKMDSGIR
LILHGQDETPVKMSGLTVPPGFTTYIQIEKKTIINLEAPYKTKCGSVKLKYFDSYSMHTCWLEQLTDYVYKTCNCKDYFMP
GDIPICSFDLLYNCAWPEWETFDKQKLYQCPLPCKIDSYEVSLSRALFPTGLYASSLANDLRKYQQVPIALKSKTDELIFMR
ENLLRLVIYYDDLAYELVEQKPSYNTLLWLGDVGGQIGLFIGAGVMSYFEFIDCLAMVIYTRFFEKISLKNPTTV

>Deutero_Ambulac_Apla_gbr173_7_t1_528_86

MWPGAAGVVRRLIWAVVTLAALGFFLYQTALSFISFFEWNHVTKVDINYADELEFPAVTLCNFNKYRESAWTERDLKN
VGTHLGIIDEDHNLINPHLYTDEFKQNLAAVNWTALEVEEDYDMTEFTNRTGHQLDEMIVECTWKGSPCNSQDFHHV
FSHLGNCYTFNHVELDQDRHSTISAGAANGLKLTLNIQEEEYTPSNDLDGAAEDAGIKWMLHHPSEPPYVKELGFGAG
PGHHTFVATRHEQISSLPFPYTMCEEDGSLKYFDHYSIQACRIECETDLVVDQCGCRLVEQPGTAPVCNPAESHECAHV
VLVQAVTGETESSCDCKSPCELEVYPFSTTNVKLRANFIETIYSNSTTHNFTAEYIENNLVLLSIYFEALNHEVIEQLPEMTV
FTILATLGGNFGLFLGGSVLTVMQLLEYLFDEVTASCASKGSSNRVGDSRANLTGKEKDVENAEDGISIIDMGKDKSKDR

>Deutero_Cephalo_194160R_t1__name__chr_scaffold41_start_1865858_end_1873272_strand_pro_le
n_477__530_88

MSPGEEKESPEKEFAGNTTLHGLNRIVIAPSGYFRVLWVVAILGSYSGFAYMFSSMIIDYFSYRTITDTKLEFAEDLTFPAV
TICNMNKFDESKLKIIDFSYLSLFLYGVQLDVDAILASPGAYPDETVNSTLEGVTVGDLLLEKGFNLQWPRMPLCKWKGN
PCTQQDFNHSFGYYGNCYTFNADANSPLKQTIQGSGNGFVAFVDLKVNDYTESYYASGNSEVGLKLLLHDQNEPPMM
DTQGIALAPGSHAFIAAKRTLYENHVPPWGVCQDLQLEYYDTYTLNGCYLECRSKHVVRNCSCRPYDLPGTAPSCDPNT
MFSCVRAVLDQVTRGELECNCPVPCSMTAYSMSVSYAGWPNKLTGEYISPLWGSTGEYLELNGVMFSVYYEKLNYQKI
TELKAMDGGQLASNIGGMMGLFIGASLLTVLEVWEYLWQRVLGLLGKQRRPRVIEVKSPPDDMERGIRNVAFPMENT
KIN

>Protostome_Lophotroco_brch_Lingula_anatina_comp134404_c2_seq5_p1_comp134404_c2__comp13
4404_c2_seq5_p1__ORF_type_complete_len_479___score_28_84_comp134404_c2_seq5_272_1708__
536_92

MSSVHPFSFEEKGTEFIKQYNNMHGDFSNKSPHRGTNITKRKPDQPKDETIPSLKDEISDFANNVSILGMKQVANPKFS
PVRRLLWLAFVLCGSGFLIFHLWNRISYFLENPASVDIKFHHNDSIHFPTVTICNNNRFKLPYLRANGHLEQANYFDMFN
FNLTNIEQYNWTDFHTRNQYKMADLMPSFCLFNQVICTDADFSMIETEMGSCIQFNAEGELKSVSRAGSLSGLQMWL
FSHQDDYLNFTTKAGYTVLMHERGEFPHMIGLGLKVSPGESVDIAMKQRRRTNLPPPHGQCKERTLKYFSKYTKLNCD
AECSFIAMQACGCRLFHQPKTGNQTIDGQRTCNLKDTMFGCFNISKEPMAGYSSCACMEACQSTLYTHSISHTRMSEV

FIKRLIAMYNNTDTSFFRENIIMLNIFYSDISVEEVVQQEAYSALTLFADIGGALGLVLGSTLMTAAEIVDFFLGVSLWKLFRIRV

>Ctenoph_Hormiphora_californiensis_evg1166627__p1_GENE_evg1166627

MQPNGRPKDYVFTDIDQPLSANGFYKRIPAPPRPYRPTPAFPIYDNLLDREDVMSSNCPSTIPGVYPGVEAFTPYSKYKQDESAFDMTGRGERTSYYPYLEHYQPSYPRPARPKTTANPTTTTANHRAEPNGKIKYPPNHHNRDTDSSGSPTEMDDRDGEECEEFMPPPKRKKKKKRRANRSLENRKNLRLRRRKKAALEQRREDMEPDEMESFKEFASTVTADYLAEMFAKASNGRRAIWVTLWLASFVYAWYNIALSIGVYLSKPTATKLNFYEAPPNGVAFPTVTICNFNKYNKSFFEREGDSVNAQLRDFLVETTPIWDHKERLAKDQWLSKYPSIMNLELKQVLTAATHRLEDTVEYCRFGGKPCLDSQYRETSIPSEAFTKIYTEHGLCYSFNQDGKLNMTRTGAQYGLSLRLHVAQDDYFKTQDVAGFKVLLHNWYEPPMIEEYGFALRPGSESYVRIRLQKFRDTERPLGYCDPLLEQDYGPGSYTIAKCVMANRASLTIKKCGCIPFYMDVARSAFAVPPKYPSSTPVCNLYQEWDCVEPLLEDIALGTLVIDGEPIGDPDCPSPCNYLTYSYTISQSEYPSQSVEQEILDNVLGARGKSWNIEKIRDNYLKVHIYFEELSTLEMERHPSYLISNLIGDVGGQLGLLLGMNICSLVQFTDYIIRFSFFNGIMKLVRISQRNSKRRAKL

>Cteno_MneLey_comp19805_c0_seq2__p1__GENE_comp19805_c0_seq2__comp19805_c0_seq2__p1___ORF__type_complete__len_802___score_191_14__comp19805_c0_seq2__85_2490_

MSNRVITTDLRLACQTALTPNTMPETKIKIGLTRGLVDGGEIFAQFSLSSTAHYQHLLTYISAICSREKVLAIYWFNRHDREFVLIHDNASMDYCLVQSGLHEILYIITETGGKPAGYNSKHKPEPIQLKTSSPSSGPSPDTANLERSSDEDRPGGTVQREDHIPLRKMKKSASLEWSYDNEQPPSVQEISSRDESAFDMTGRGERPTYYPYLEHYQPSYPARPKPHRTPTRNKRPEPKVNNKYHTHSDQSGSLSDSDDDRNAKEVDQFMEPEPPKRKKKAKRRANRAALEQKRQEIEMETFKEFASTVTADYLAEMCSKASNGRRLIWVSLWLASFVYAWYNIALSIGVYMSKPTATKLNFYEASSDGVTFPTVTICNFNKFNKSYFDTDDITTKKGQLKDFLKITTPIWGHKQELDSASWDTKYDKIVQQSMKDALQTASHSLEGTVEFCKFSGKPCLPADASPGDVADVEDADPTAAFNPVFTEHGLCYSFNQNGNLTMSRTGALYGLSLRLHVAQDDYFETQDVAGFKVLLHNSYEPPMIEEYGFALRPGSETYVRIRLQKDVYPQYKDTPRPLGYCDPYLEDQFSTNYTISVCVMQYRAINMLKVCGCITFYMDRPDIVLSEGKPYMKPKICNLKEERTCADNVLEGLALGTLKIDGEKVPEPDCPSPCVYTTYTYTVSQSEYPSSAVKDKVVEEVNRQKGTSPGEDDASSWTIERIRDNYLKVHIYFEELSTLEMERHPSYLISNLIGDVGGQLGLLLGMNICSLVQFTDYIVRFSFFKGFMSLVRLYKARRRNK

>Deutero_Ambulac_Apla_gbr256_12_t1_540_96

MDPRVIIETTPRNGFNERRVMGKDNLDELNIIKNDRNRHTQDDDANRTFAERFQDFGNSTTLHGISYVINLNYKKQRRLLWLLLVLVMSIWLVYSIVEAFITYFNYPMTSAISIHYMDTLTFPAVTLCNFNQFRRSALTPGQVMVMNEVYGPNPPESIDFGPFNEITNYDFSEDHLANISHHLDRMLVKCKWRGTQPCTGQNFTRRFTDHGVCFTFNDPKDASHRLEVSNAGRLDGLFLRLFAETDEYTFGENSAAGFRILLHPQGLMPLVKELGISVSPGFESSISIRQNVFKSLPWPFESNCTSSKSKNSTTYSVPTCKYECKVQYVVERCGCRDYRWTGSAPLCPPEQHFNCIDKYKAEYQKQEKYRDCHCPVSCELKTYDSRVSFGYWPSSYVSENLPLNSTSEEYIRKNYLDVYIFFEELVYLQIEQKSAYDFAALQSGVGGNMGLLCGMSLITVVEWADFIVISLLRKFKARRI

>Protostome_Lophotroco_annelid_Pdum_Contig8016__542_97

MSKVTHYPMDDLGKSPRPLVNDMGQGHRVRKSSSMQRVLSDYSEHSSMHGVWIVANPDYPKLRRLIWMLALITTTTVFVYAFIQQMTMFFQYKVTTVLSVEEQATLNFPAVTLCNYNQFRKSKAEQKPGVMGVLRDLFSVDAIIERQKSKLRMSAQSLVSQDINIINELVALAHTLKDMVRHCQWCGKTIPCSDYFSQTLTEVGVCYTFNSHDVRKKIPPLEVHDAGTDCGLKLRLNIEEYEYYYGEGSSAGIQMLIHDPEDVPLVRQHGFSAAPGTSVMAAIRLTKETYAQSPYGDCVDAKHPSFENPLRYHEIYRYSACKLEHRVDYIYNACGCVMYYNPVGSRSVCKTVEEVLCFKDAKENHTRYEMQAMRHACPVTCQSDNYEARLS

WAAFPAPQILYDLKKNFNMSGDQVRKNLLQVTVYFEDLGIHKKEQQPAYTGNSLFADIGGMMGLCIGASLLTIVELLEL
GAQLMLSAGKR

>Protostome_Lophotroco_brch_Lingula_anatina_comp149479_c1_seq4_p1_comp149479_c1__comp14
9479_c1_seq4_p1__ORF_type_complete_len_482___score_29_72_comp149479_c1_seq4_182_1627__
543_98

MGKGEGESLETEFASGTTCHGFNRLAVAKKWPHRAFWTVAILWGFGMTTWMISSRIEAYFKYETSTEIRVEEKYALEFP
AVTFCNFNKYRRSAMTALDWYIYANSKLAEKYAFEADYDPEATAALRAAFEGFEANLTATYPGGFDFGNFTRRIGWEL
NNETMPLCLFRMKQCFPENFTHAFQSEYGNCYTFNSHHDEDSLSLQQTRPGAQQGLYILFDIQQAEYTELLNLNGALG
VGIAFQIHSWYERPNVAQHGLGAGVGTAALVSMHQNELQNLAHPWGNCSNDRTLKHFPRYSRSGCEVECYLEKVLEL
CGCKPFSYPGNAVNCNPVDQVLCADPTIFKVASNFQKECGARCPEACQIIEYPPTVSYAPFPNYYMAQELATMYGTNA
NYASENLVGLSIYYKDLNNRLTKKNPAMTPSGLLSDIGGQLGLFIGMSTLTLMEFGEYIIFKIIQCCRKRKPKTSQSKGVAN
NVKVNESGM

>Deutero_Ambulac_Apla_gbr54_23_t1_545_100

MVREDADQSPRPSLEREFAESTSLHGIAKVFHASRLLVRVFWVIIMLTCLCVCVWQISDRFHRFLQYKANTEISVEYVGH
LPFPAVTICNFNRYRSGALAPGDEDALAYIWELSNYQNDYDYEGESNSTDSSYLDSLVDFNYTEFTLRAGFRMDNVTLLA
CSWRGKRNSCSAEDFSHVFTSYGNCWTFNSGKTGPVLTESYSGSGNGVSLLVDIQQSEYTETETIEAGLKIQVHDQVTP
PTVESSGLALSPGVHAFMAVRRQQYLNLEHPYGKCDASRTLKRYDNYTLEGCNIECRANAVWEKCRCRLIQHPGNEVE
CTPEEVRFCAREAIDGLIGGDTESCDCPVPCNFTAFTTSLSTAFLPNEKILPILLLLLNNISTSDERNVSRSISEQQEYIRKNW
VAIDTFYEDINYQKFEQTEAITPSALISDIGGQLGLFLGASFITLAEIFSYLSRKIQRLIHGPTRRPEKQRQMSHSVETLA

>Deutero_Ambulac_Apla_gbr658_1_t1_550_104

MNRPCDGTDHNFDMETDDLRDGRERRRPSVTRRKRGFKRKKSKDTCKNILAARIQQFGDDTTFHGLRYVTNSTYHTFR
RFVWLLVVTGMTALLIFDIIRAIMLMFEYPEMSSISVQYPPNITFPALTLCNINMLRKDALDASEVQLLSDIFANPATPPD
TNALVDSKLFNSSSNEELMGRMIDASYRIEDMLVKCRWVSETCSYRNFTRSITDHGVCYTFNDPLSDHEALIVRHPGSR
HGLFMRLNVQQYLYTYGESTAAGLKVLLHAQGEYPMIKDFAFSLSPGFETSIVVRQKVMINQKAPYKSNCTDGNWGR
HRFKYSAASCKYACKILYVTRKCGCRDYRWSIRDIPICSLEKQVHCIYRYEDEFVLSQENQCHCPIACKAITYGRETSQAYW
PASFLSKELQAEMNVTEDFIRENYLDVYIYYDEIMYTRIEQIGAYTVDNLQSDIGGYLGLLCGMSLMTVVELLDFILVALCS
RLKR

>Deutero_Cephalo_298690R_t1__name__chr_scaffold94_start_933624_end_940952_strand_pro_len_
484__551_105

MITSDEKTPEREFMDNTTLHGFNKIHHPNRAIRTVWIVAMLAAYAGFIYMAVNMAQTYFSYDTVCDVKLSFESKMTFP
AVTICNYNRIDSHKLTWEEWSILSTLLYGRAMDVPTLQAFLGPTFTDRNRTTNGTVTNATTMDMAAAVRMKGFDISP
ARMYLCSFAGDSCTELNYTHSFSNYGNCYTFNADAANKLNQTIPGSTMGFQIVVNVMADSYTETMAVGGHEEVGLKF
LIHDQNEPPMVETHGLAIKPGIHSFVSVKRTEYYNHVPPWGDCHDKALTYFDAYTLPGCNAECRGKHVQDRCGCKAYY
LPGSAPPCAADVLFTCVQEVLAEVNEGKLTCDCPLPCTMVKYSPTLSFSAWPNQLTEDYYSRTYNLSRGYLTLFMLFPRD
NVVALTVYYEDFSYMKISQLKAMDSGQLGCNMGGMMGLFIGASVLTLVELTEYLVARLSISWGKKKRPLHVQPAPDA
EMSDKKQPPDSLVFHK

>Deuterostome_chordata_PetMar_tr_S4RS34_S4RS34_PETMA_Acid_sensing_ion_channel_subunit_fam
ily_member_5_OS_Petromyzon_marinus_OX_7757_PE_3_SV_1_552_106

STGSPAGPTDSAKETIAFSSSTSLHGVARATSAQSRAQRAFWMMVVVAGVAMAGWQVALRLAAYYAWPTSTSLTLH
YVRDLSFPAVTFCNYNRYRRSSLHGTTDLLAALSILSAGIKTPHPEYLSGFLWRITLRGYSSPPITQTYSARNQDLSTTILEID
DTYCENSCTPQDFQHVFTEYGNCYTFNGGFSEEQRSLKQKTAGAGKGLQVLFDVQQKSHLWDYWENPELGFTGVGIR
LVVHDPDEPPQVEALGLSIGVGTQAYVSVQHTKSINQQYPWGDCDPSKKLQSPAPYSMARCLQECEADLVERICGCKP
FNFPGDIKECSLYQHYHCARDALVIFLAQGMCTGGSHLSQCPVPCLHKNFPSSVSYATFPSLQAAQRYERSLNYSLVYM
RENLLSIDVAYTDLNYALIQQQKTLPESVLLSDIGGQLGLFVGASVITIIEVLEFVARSAARALNAAASFLSNRAATCTKAAL
SPPVQG

>Protostome_Lophotroco_brch_Lingula_anatina_comp131676_c0_seq1_p1_comp131676_c0__comp131676_c0_seq1_p1__ORF_type_complete_len_484___score_23_39_comp131676_c0_seq1_234_1685__554_107

MSQVDPFSVDYRTSERKQAFTDDGLISFNVYNRGKQKQNSRDIVKQFKAEKDGNDEFSLSSELSSFANDVSILGMKQIV
NPKFSLFRRATWLCFVLAGTGFLIYHLWNRLDYYFSNPASVNIRVNYDGSLLFPTVTICNNNVFLKSKLAANNETYIGDLF
YPSFDSYPNWSAYNFTGFNWTDFYYRNGYSVMDVFGGFCFWKSSPCQANVLKPVITEHGVCYQFNSGDEGKRSDVE
GSLYGLMISFMTKQEEYLVGSYIAGYTILLHNSKDHPNVVTQGFQVAPGGSANVAIKQKYVKNLPQPHGQCADKQLAY
FPHYSKINCQAECLHNKTIEKCGCRLVYYPEVPGLRECSPIDVNTCWGPLLTNVSQIMKNCACTEACETFYFEYSVTTGLV
SDVSAQAAAVFLNTTQTYVEKNYLAVYLYYKEMSYEEVIQQEAYSTLTLLADIGGALGLILGSTLMTVAEILDFFLGLSIWK
FFGVKL

>Placozoa_TadNaC4_MK547545_14_111

MAKEDLSMEEKFASTTSCHGISHVYQNDSGRLPRMIWLILTIAATAVCISQCIIIIYDASQLPTRITFSDVSANSTVFPSITIC
NTDNSPRDALPESDLKYLSILLHTQSTENAYNETQVQEAARYFIKKYGDKFDYENYIRTSSAKLENMLLRCQWMNRPCSL
SDFTSIVTDYGNCFTFNPGTKDLPLKNQTVPGEVYGLRLAFNIGQYKYYPDLLDRNRPDAGIRFTIHYHKEPPNLLAKSIIA
PVGSHTYVSFTRTYHKKLEKPWGECGSRKLLFHEFYSHVACIDEVSAIYASTLCNCSIIGAFGPYGVCDSVKFITCIVPILAK
ARVQANENIAACPVACETYTYPTVISYGNLALIPISSLITNYLNVSGIIRHAQSLDWIKDPNYSTANFVRDNILYLDIYYSDL
HTNAIEQKRATGFAEVLSNIGGQMGLFIGASIITIAEILQYLIRIFYNKSFNKPADKSKEDLNLEQNRKSSSV

>Cnidar_HydraVulSc4wPfr_824_g11302_t1_557_112

MLNFKDIAQITVEAIQETNEVTNKDEKNIQTIQDLRNKKIREHISYMIDNSSFHGLSYIFDKRHSVRRTIWFFITIAAFAYA
MQKVYESTMNYFSYPFYTVRMRMYVNQIDFPAISFCNLNDIKFSAMNGTIVDDAVVTQNHEANITGEEYRSYNQAAR
HTLNEMLVDCDFEGKKCSHKNFTEFSWMQGESCFTFNSGKPPHTLLKVKGAGINRSLKLTINVQHYDYYRDKMDSGIR
LILHGQDETPVKMSGLTVPPGFTTYIQIEKKTIINLEAPYKTKCGSVKLKYFDSYSMHTCWLEQLTDYVYKTCNCKDYFMP
GDIPICSFDLLYNCAWPEWETFDKQKLYQCPLPCKIDSYEVSLSRALFPTGLYASSLANDLRKYQQVPIALKSKTDELIFMR
ENLLRLVIYYDDLAYELVEQKPSYNTLLWLGDVGGQIGLFIGAGVMSYFEFIDCLAMVNYKVLHITQVANIIIFFTDVVASF
VTMV

>Cnidar_Rhopilema_esculentum_TR103127_c0_g2_i1_p1__Rhopilema_esculentum_TR103127_c0_g2__Rhopilema_esculentum_TR103127_c0_g2_i1_p1__ORF__type_complete__len_487__score_56_61__Rhopilema_esculentum_TR103127_c0_g2_i1_686_2146__558_113

MYSHSESVREKEVVKSVDWIFDFGDEFSNRDIYRKVMTHMADAELLDLRKEKEVRRQKVKQHFDDLVSNSTLHGLHFC
FDKQHFFRRIFWTLLILVALGLLVQKLYESTTYFFSYPFSTTTTVKYVNQMVFPAVSICNLNDFRLSVMNGTKLHKAIQER
GNILALDGKEYTDTIRKANHRLEDMLQGCTMNNKICTHKNFTQFFHNQGDRCFTFNSGMNGQPLITVNNTGLSQALT
FLINIEHHDYYVDSQHSGIHLILHGQDETPVKMQGVILSPGFVSYIEIKKRKVKNLPHPYETNCGKLKLKYFRSYSKHLCWL
EKLTDHVVNLCGCKDWFMPGDHRVCTLNESMNCMWIHWSQFDLTQNYSCPLPCIIDSYTPTLSFARFPANNLADKM

AKRLNISGTKAEQRGWIRDNFLKVVIYYGDLNYEYMEQIPSYDLMVLLGDIGGQLGLFLGSSVLTYIEFFDFFAMVIYTRF
FEVFKRSPSRV

>Cnidar_Rhopilema_esculentum_TR52437_c0_g1_i1_p1__Rhopilema_esculentum_TR52437_c0_g1__R
hopilema_esculentum_TR52437_c0_g1_i1_p1__ORF__type_complete__len_487__score_65_71__Rhopi
lema_esculentum_TR52437_c0_g1_i1_68_1528__559_114

MEKGEESKDEKPPENAVRNMLFDFASVTTAHGISRFVTSGPIYARISWFFIWLAVMIGFIYMIIQLVLLYTSRPVSTSISISF
EESLTFPGVTICNLNVFPASSMKNVRVKFPNMTIINNMFKKQFASQSSLDVGDLEMDDVSSIQENTLTAINNLPIADRLK
NGQEFSSLITYCRWAGFVCNKGVFLRDYWHKTWNWKYGNCFTFNGKMKNINGSSIAPLESTSPGTSGGLTLNIDVNR
DEYLKGIAMETGVRVIIQDQDVFAEPTEHGFSAASGYSVSVGLRKEMIKREDVSGTGSCYDTSKANTKYFQQTCIAICKA
GTQKRRCNCTSLQYRTYFGSKQCESLAESSCLLEVLSNFTNKIFDCSSRCPPSCSDIAFSKTIATSKLYPEGYLEIHKTKNLTA
DYVEKNIINLQIHYADQTVTTISTGQYYTFDNLVSDVGGQLGLWIGVSAVTCVEFFSLIWNLLLYCICRKKEKRVQDENAP
C

>Deutero_Cephalo_293250R_t1__name__chr_scaffold9_start_4819346_end_4829052_strand_pro_len
_487__560_115

MAACCSCQDGALDHEYASNTSLHGPGNIINAKRPAHRAVWVVLFLAAFGVAVWQISERFVAYFSYNTVTSVKVEFKDE
LDFPAVTICNFNKFQLSKVTPSELNYITEVLDLSTGFEGDDTGELGFDYDDYEEEGFGPEEDDYSLNVSAIPSNFDLADLTL
NAGFVLDETTLQDCRWRGKRCYADNFTHAFTSYGNCWTFNSNDDKILRQTIPGSGNGLYLVIDVQQEQYTEKPPSGNS
DAGLKFLVHPLAEPPKIDSQGTAVQPGTHAYASIQNILYKNEIPPWGTCDPTWRLDNYDTYTKTGCLLECRAEWVKKDC
GCRTVSMPGNATYCSPTQLTQCVKTVVGKLADGRYPCECPTPCVANTFPTTVSYAAWPSISAQDYYTNLFNHTAEYLQ
RNFVVMDLYYAQLNYQEVIQTRQYTVGSFLGDFGGQLGLFLGASVITIAEFIEYIVMKVTQPCSSRGKRRTDTEMTSLGI
DTKSNQSSPLS

>Deutero_Cephalo_091790R_t1__name__chr_scaffold19_start_3687569_end_3692166_strand_pro_le
n_488__562_116

MGDEQNSLDHEFANYTTCHGAARIANAKNKPHMALWTLIFLAAFGLCTWQIVDRFNNYFAYRTGTEINVQLESELTFP
AVTICDFNRIRGSGVTENDLAYLGALYQIAALVSDPVALRNLSESIDWDAFNASAGLVEDFPGFVRRNGFVLDNVTLAEC
SWRGSRCYAENFTHVFTEYGNCWTFNSEKNSPLRQTSPGAGNGLKLILDIQTHEYTEDPFIGNLEYGLVFQVHDQYEPP
RPELVGTAVAPGSHTYASTFQTHVKNQQAPWGQCDPRKDQNFLKYFDKYTLAGCLLECRAEIVVRACNCRPLYYPATF
LSETNEFDFCGCTVPCEYTSYSSQVSYAGYPSLKAAEFYETTFNYSRGYMSTNVVLLDVYYHELRSTSWEQFKAVEESGLI
SDVGGQLGFFIGCSVITLWEFLEYLALKFSAFFLFSSKPSSKTATKDGVKVDNAMVFENTAAGDAEKTEITELSTIRTTSNG
EKSAVLEMT

>Deutero_Ambulac_Apla_gbr81_93_t1_564_118

MENTAQKELNTAKNKQKPNLEYEFATTTTLHGISRVAESNGARAKTLWVLVLLIAGCFYVVVTVDRVRAYLKFDANTNV
IVEFNDTLEFPAVTICNYNRFMNNKISEDDTKYVRHLLELNEDLVDYEEDFNYTHIDDLFGEGFNYTQFALTTGFTLNESVI
RCDWKGVAYSCNALNFSSYYSPSYGQCYTFNTRGDQHSHEQSQQGIGNGLQLIVDIGQSEYTETFKSGHLEAGLKFAIH
PRDELPLIGTMGLSAAPGFHTYASMRRVRHINLPEPWGVCGTGETIAVTGDSRKYSRNTCLRICRRDAIIEVCGCQPFGY
ERAVEPDSSELVPLCDIDRFYCVAHALERYRASFSALDCTCPVACEYITYETTLSMAKYPSQSVINDILSGFPDDEASKAGA
VDAEYLDENLVYLDVYFEELSTITYKQVEALPFTALVGDMGGQLGLFLGASIITGAELLDYAIRRLKLLILKVKSAKKIRSGL
MRS

>Protostome_Lophotroco_annelid_Pdum_comp404514_c0_seq1__567_119

MDTKKEEASAFALIAAMANNTGMHGLPNVGRSQHIVRKVFWMLIFLAGAGMLIWQVYEALAQYYSYPTKTSIIPKVY
NKLNLPAVTICNVNPVRKSLLYQLDALTDPSDRNYENFQTAMNALNVTEKQKIGHQLKDMMLYCNYNGYTCEVGNFT
TFYNSYLGNCFTFNGGNRPTVDGLGKTGPYYGLDIEFYIEQDEYEPEYSEYAGMKVVIHNPKEMSFPEDQGLFVSPGAK
SQVSLKKIENTRADPPHGECRTFNEEETKARNAYKDSYDYLEYTSKACGKTCYQQYVMSKCGCYDTSYPSTGSAFENITS
GKTFSSCKSLNSTYTDCNTAVYYNYTDNIISCANMCPPACTEVLYDMTTSNLAWPSVNALSEIVKKLEDKSTALNTTIQN
MTEIEKESFVRKNVVSISVFYGSLEVTKIKTEPSFGLVDLLFAIGGSVGLWLGLSVISVIEIVELLYDISSILVCCCFKTKPNGN
NPKTIQVDPQS

>Deutero_Ambulac_Apla_gbr54_19_t1_572_121

MKDSSSQKQSDLDQEFASTTTLHGIARIFDSTRLPVRLLWLSILLGCLGVCVWQITDRFQRYLLHEATTAVSVEYVGDLD
FPAVTICNFNRYRSSALTEDDKAHLEDLIEYADYDYDTYDDEEPDARRTAQSQGETTNFSFSEVTLRTGFQMDEETLLDC
KWRGKRNSCNGQNFTHVFTSFGNCWTFNSGETQDEKEISVLNQIQPGSGNGLTMVINIQQAEYTEPVQNGNLEAGL
KVLVHDQETPPSVDSEGFAIAPGVHAFVGLRKIEYENLEYPWGECDKSRRLLHYDKYTLPGCDIECRAERIYERCQCKLVR
HPGDETECSPLQVKKCATPVLAKLKTGEVEGCGCPVPCNYSEFRTSLSMATLPSNNLLVDLWKLYGGDDSVNYTFDETL
RYIRDNWIFLDVYYESLNFEKYVQSEAITLSALISDIGGQMGLFLGASFITITEILHYLGRKAGLWVTSRSRPHANQSKVRP
GNETVAISDLSSPH

>Deutero_Ambulac_Sakowv30024946m_573_122

MDSMNDVKDADRKSQSSIVHNFMSSTTAHGLPRAFEGRSLWLSLFWVLLFCSAIAVSIWQIGLLVKHFIARDVSVKTEI
VTAPQMPFPSVSICNTNKLRKSAVSESAYSELLTVDTGIAEPYFGDCLEGDFQCTNGGCVKTYLKCDGIDNCGDKSDESG
CVYGDCDAGKIKCVSGSPTGACIDEKFKCDREFNCYEGEDENDCVCKRKEFKCGDTGRCIDGTLQCDESIDCNDGSDEE
ECGAECPSDHLECNGLCIPPEWLCDNLQDCPDGTDEEDCGDPEIQICSPFEFACDLYQCIPIYWECDGVPDCINGKDED
NCPLQAAPQSFGFVTCSEDEYQCDLFTCIPLNQKCNGIVDCALEGDEDDCPEMTGWMTTQGSNKTATYQPVAKDLLKI
YTALSYDITLYEDFVDNHYHDHQFGRVKSEDPPDWPGFITYSSTPDYSDLENVLKLRADEIAELGHQLEDFVLECSYDERI
CDLNMLSFTSLYFNFS

>Placozoa_Hoilungia_hongkongensis__1_g02560_t1_574_123

MSQEKFNHEERFATSTSYHGMAHIYDGQTSKQTRSLWLVLTLIATGACISQCVIIILNASQLPTRIVTKTVLQNSSIFPSVTI
CNTNDYDRNSLSSSDIQHLSTIVDFVYGYDIEPHRFEAAVNSLSIKFGKGFNYELFLRRAGHKLENMLLSCTWMGRPCSIE
DFVNTTTAAGSCFTFNPGSTNTSIKNQTASGNANGLRLILNIEQYKYYPALYTPGQPDAGLRYSLHYYKDPPNLIAESYYA
APGFHTYVPMTLKREKRLKKPWGDCEELSLKRYQFYSRYACLNELSAKHASESCNCSYQDNEALNPCDGVQFLSCIIPTA
GYYRTQLSQNLSVCPIACETFSFATEISQSSIASQAFNSAIDSFINVSGLISDAQSKYWLPQSYTNIDFIRDNIVYLDIYYRKL
DMTQILQLEDTGFSKVLSEIGGQMGFFIGASVLTVCEIVQYLFEKCFTSNLKSGTKKNTTNVVASDNHIFALERTTGFTPR
DD

>Placozoa_TadNaC6_MK547547_16_124

MMAKGKIDHDERFATTTSYHGVAHIYDSNNSKKTKSIWIILVILATAICISQCVIIIYNASLLPTRMVIRKRLMNSSVFPSVT
ICNTNDFDYTGLPANDLNHLSSIVNAIYGFTPASAADDAIQYFVRKHGDDFQIDNYTRMAGHKLKNMLLSCTWMGEP
CTVNDFTNIISNGGSCFTFNPGTNAIPLKNQTVSGNINGLRLILNVEQYKYYSPLFSPQAPDAGIRFTINYYKQPPNFISKP
YYAPTGFHTYVPITLHRDKRLTKPWGECGELLLKDHSYYSRDACLTEYASELAALTCNCSSSGQTATNSCNGAKFLTCIIPS

SFVIRMSLSQNLSVCPIACETYSYPTEISQSSLGTFAFSRVLDSVINISTILGKAKDEHWIPPSLPYSVSEFIRDNIVYLDIYYSD
LRVTETEQQEDTGFSKVLSEIGGQLGLCIGASVITLCEIIQYLIGKFFTSEKKTNLNKRQNTTSPLFYNRNSPDADATNPDM

>Placozoa_TadNaC7_MK547548_17_125

MDEEKISEEERFATTTSYHGWAHIYDGNNGRMTKIIWMILVILATAACISQCIIIIYNASKLPTRMVIRKQLMNTSIFPSVT
LCNTNDYDRSALPAADLYHQSALVKAIYGSGDPNKTNAAINYFDKKFGGNFQFENLTRVAGHKLSNMLLSCTWMEQP
CYATDFVNIITDGGSCFTFNPGTGNLSLKNETVSGNSNGLRLILNVEQYKYYSGIFALEQPDAGIRFTTHYYKQTPNFISKS
YYAPTGFHTYVPITLQHDKRLKKPWGQCGEEILVYNSFYSRDACLSEYAGRLASYICNCTFEPQIASNPCNGTQFLTCITPL
ASKFRQDLSQNLSICPIACETYSYPTEISQSALAALAFASSLDPLINVSGIIANAKANNWIAPNQSYFAADFIRDNIVYLDIYY
SDLHLTQTLQEEDTGFSKIISELGGQLGICIGASALTICEIVQYLIKKYFASNKKSKSDKSSTNTISLGHDKKPFETTTVTNIDL

>Deutero_Ambulac_Apla_gbr256_11_t1_576_126

MDVWTVEKSLNRIDESTSADSIKMKGWRKDEGGVNGDGRKRSVHTQTESDRRNADNCRQIFNRRKWSEFGTVTTLH
GMRYITQPNHSFTRKVIWAIILLSFMGVLIFVVIDNTVRYFNYNVTSIVTVKYVNDLTFPAVTICNYNLLRKSYSEQAFDED
LLEAFKIISPGGGQNSVNFTALKDWNITEDLIKGAHQKEDMIIECTWRSTRNCTADNFTQVLTDFGVCYTFNNPKDPND
ALTVQQTGYDSGLYLRLNLEQYDYFYGSSKGAGFKIMLHKQGEFPFVKQLGFAVSPGFETLVTLKFTQLVNLETPYVSEC
VVAEDIQGFKYTVETCQALCKTKYILEGCGCRTTDLPAGPGAFMADLDARCNCSVPCFREVFTPRISEVFWPAEHITKAL
QDAFNSSESFLRKNIIDMVIFFEELNYEEIRQIAAYTFTDLLSDIGGYMGLMVGASLITFLEFIDFLVLGAIEYCQKHNMFKR
RGKINISQSHLHR

>Deutero_Ambulac_Apla_gbr54_18_t1_578_128

MTGSASGKTTDLDREFAANTSLHGVARFFDSTRLPVRLLWLSILLGCLGVCVWQITDRFQRYLLHEATTAVSVEYVGDL
DFPAVTICNFNRYRKSALTEADVAKLRAYQRYIDYDYDFYDYDSSVGGTPPPDDISDFRFTNFTLRTGFQMDEDTLLGCL
WQSNRNSCTAKNFTHVFTPFGNCWTFNSGEDGEKDVPILKQTQPGSDNGLVMTIDILQREYTQHLQSGYVEAGLKILV
HDQKTPPPIDSEGSAIAPGVHAFVGVRKIEYSNLEPPWGKCDKSRRLTYYDKYTLSGCVIECRARKIDEECKCRLFSHPGN
AVECTPTQVKDCAIPVIVKLRSGERAGCGCPLPCNYSEFRTSLSMATLPSNNLRDEIWEELRQEGINDTLPETYIRDNWIL
LDVYYETLNYEKYVQSEAITPSALISDIGGQLGLFLGASFITVTEVLHYLGCKAGRWMTTTCIKVDGKPTHESAKEVVMEE
GQQTSGSKDSTLT

>Deutero_Ambulac_Spurpu_015892_583_129

MGNIEKEKLNELDDDTVVGAVRVFSNETTLHGIRYLIGRGHLLIRLCWLAIVVLAFALLVMQAQIVYQDFRSSPYSTKIDI
VSEVVKTFPAVTVCNDNKIRRSQLYNTKYEGLIDIDDGGRSTEGRSSTRDKSSDNVWASVKDDYDWQGFYSASSADDF
SDFINVVNPTKAELNDYGHTLEDFVIQCTYNQKSCNLSADFTVSQNRHYGNCFTFNFGDGDDEPPRVTSKTGALSGLHL
SLMVDEEEYMGLLAPNRGAMVVIHAPNTVPFPEDDGINLETGSAISIGIREEVIQRLPKFSQCVVDRKIGTNTSISGTNYT
VRCCLKICYQQSLLERCSCVDDILLDYRQCSVTNKSEGDHGGHGSPVYHLSIDLLYLECNVPPAYRGLKFKKGKRDVRVG
KRVLNHSMVYFRGKTYEFGANSLVLHEGGVGRTNPTSCPTVRKTTAGRSLCTEYQATELAVRYHQDHNFNLFSKNCH
MFAEWLINRLLNNKCTI

>Protostome_Lophotroco_annelid_Pdum_Contig6895__580_131

MTRVGEPSRDPPVVIASQGGANKPTATKNPEEETLRKILDDYVEHSSMHGIWKAAKTSPFSRAVWTVVLLCMICLCLFT
FVSQLLFFLQFKTNTLVTVIEQPFLPFPAVTVCNYNSFRKSKALEKPSITKVLEQLFSLDRIFHINVYTTSTAEQSDDVDILKE

LIDLAHTLEGMVVYCFWCGMRVDCKQHFVQTMTDVGVCYTFNSKNTTGDTPMLVRDAGSDCGLKLILNIQRHEYFYG
QGDSTGVKVLVHDPKEVPLVGQQGFSAAPGTTVIASIKLQQELYASSPYGDCVDVSKRDFKNPLQYHDEYTYTGCKME
HRVNYIYSSCHCVQYFHPVGGREICTTFDQVECSRKAQENHTMYQLEQLSQDCPVTCEYNTFDEKLSWASFPAQHILD
DLQTTFNMSAQAISDNLVQLHIYYENLAVHRQEQTPAYTVGALFADTGGEMGLCIGASLLTVVELFELCGHLAAFSWKK
ITRHMFYKKPRSSLSKPI

>Deutero_Ambulac_Apla_gbr256_13_t1_582_133

MYAWTEGTKTPNLLGDQESIFSIGSPDFTVRSTIINMEDKETKGNNCRSQTCSADVDRRSFADRFREFANSTTFHGISNV
TNSKHNGIRRLFWSLIVIGVSSWLVTGITQTVIEFFKYPVTSAISINYVDSMTFPAVTICNFNQFRRSVIPDDQVDFINKLY
GLDPEAPTDLNISDFEITGLTRDQYEDLFAAMSHQLETMLSECVWRSSETCTVQNFTRRFTDHGICFTFNDPKNGSEILQ
VNSAGTRNGLFLRLYADTEEYMFSENTAAGFRMLLHPQGVAPLVKELGFSVSPGFESSISVRQTLVESLPQPYESNCTNS
TLRYSDTYTVPSCRFECKVEYVVDKCGCRDYRWPGPAPICNPQEQFKCVYPQEAEFLRQNVECSCPVSCETKTYESRISL
GYWPAGHVTEFVEAHMNLTEEYVRKNYLDVYIFFEELMYLKIEQKEAYSTSSLQSNIGGYMGLLCGMSLITVLEWADFT
LVSIIRRFRSRKE

>Deutero_Ambulac_Spurpu_006217_591_135

MAYDERMNATSVRHTTTNSTMDGRPPKSEVYAMVKPMLENTAAHGIPNIVRAESTPRRVAWSFLFVVALCCFIGLSG
NLIRKYYSFDFNVNVEVLFEPSINFPAVTLCNMNPFRQSSVVNASLELTEILGLENDPMSKWNSSTNALYGDWENLEFSS
LDAQTEILISAIQVVGNMSYEQRFDIGHDLGDMLLSCSFHGLPCAPAMETGRTSNFHRSMLKLRSSYRQSRSLATCLTN
NGLTSGMISVICYSVVRSMVYRVLQRLAIELFIDQEEYISTLQSSAGIRIIIHDPNEMPFPEDSGSTLAPGRQTSIGLTKVAV
HQLDYPYTTCTNDFPNDNIFQERFPHTIYSVLACEKNCLFKYIKQECNCADARYRYNETIETCDFDLNEAKIRTKNQFLKQ
FGQTMLIGTNVVKVIVYFSSLEYENFYQTADYTVYDLISALGGQVGLWIGVSVLTVFEFVELLYDILKFGCSKLTKLGKRTIF
AKDSLEMGPHRT

>sp_Q9R0W5_ASIC5_RAT_Acid_sensing_ion_channel_5_OS_Rattus_norvegicus_OX_10116_GN_Asic5_P
E_1_SV_1_586_137

MEHTEKSKGPAEKGLLGKIRRYLSKRPLPSPTDRKKFDHDFAISTSFHGIHNIAQNQNKVRKVIWLSVVLGSVSLLVWQIY
SRLVNYFMWPTTTSIEVQYVEKIEFPAVTFCNLNRFQTEAVSRFGIIFFLWDIVSKVLRLQEISGNNTGSPEALDFVASHR
NFSITEFVKNNGFYLNHDTLVHCEFFGKTCDPKDFKHVFTEYGNCFTFNYGENVQSKNKVSVSGRGLKLLLDVHQEEFT
DNPVPGFADAGVIFVIHSPKKEPQFDGLGLSSPVGMHARVTIRQLKTIHQEYPWGECNPDIKLRNFTTYSTYGCLKECKA
KHIQRLCGCLPFLLPGNGVECDLLKYYNCVSPILDHIERKGLCTMGTHNSSCPVPCEETEYPATIAYSTFPSQRATKFLAKK
LNQSQEYIRENLVNIEINYSDLNYKITQQQKAVSVPELLADVGGQLGLFCGASLITIIEIIEYLFTSFYWVFIFFLLKILEMIQR
TSPPQTV

>Cnidar_Rhopilema_esculentum_TR90022_c1_g5_i1_p1__Rhopilema_esculentum_TR90022_c1_g5__R
hopilema_esculentum_TR90022_c1_g5_i1_p1__ORF__type_complete__len_497__score_69_36__Rhopi
lema_esculentum_TR90022_c1_g5_i1_126_1616__588_139

MDNEGDEHLSSSTDESAFDEEYCDFQQEEVFKGIEWILDCDDLTESEYLRYITRFLAEMEDHARRLDKQTRQRKIHIYKR
SLISKSTVHGLRNCFDRKSRVRRFIWTILLLTAVGLLMQKLYECTMHFISHPFSTTTTVRYEDSMRFPAVSICNLNDMRTS

VMKGTKLDMILKGDRNVSLSGDEYRNTIRKANHELPDMLYKCKILDKECSVQDFIQFNKDQGDRCFTFNHGHEGQQIL
FFNKTGPTHALELILNIEEYEYYPGTDYSGIQLILHGQDETPVKMLGVMLSPGFITYVQVKRRKIKNLPPPYKSNCGSKKLK
YFSERYSKHLCWLETLTDHVIEECGCKEWFMPGGYRVCSLNESKECMWPRWAEFDKYMKYDCPLPCVMDTYTPSLSL
SQFLLDRDVDEIAQDPRFRGAKEMNRKTIRGTFVKAVIFYGELSYEYSEQVPSYDKLDFLADIGGQLGLFLGSSALTYMEF
IDCLVMIIYTKYISVKP

>Protostome_Lophotroco_annelid_CAC9633462_1__Ofus_G091394_Owenia__fusiformis__592_143

MEVMTETKEKQTDPVIEHSFSTRLKEFGEDVEFHGVRHLFRDNSIVTKVTWILLFLCGCSWLTYQIHDRIIYYYQFPHITKI
DKLYVPSLDFPTITICNINTFRRHKLNEYDLLHYGTSLNILDENWTLLHPEHYDKAFDDWVYSINWTDVEIHDRSINHSME
EMYERAGHQIEDMLIYCKWKQQECSVANFTLINTHYGRCYQFNSGKDGVKHQSFKGGKANGLKLYLNVEELEYLNYM
EASDLGFKILAHDQDEPPLIQELGFGVTTGNHYFIALETERVTSLPDPYGNCEEDHKLDHYDHYSIPACRIECETLIVEEKCS
CRLVEMHGDHGIRVCTAEEYHDCALPTLESITESDTCVCQNPCELTQFQHSISSVKLRESAVEMIHGHTSSNINLTEFKSV
EFIEKNLLVVNLFFDSLNYQYIEQTVAYPGVSLLSDVGGQMGLCIGASILTVLHLVQFAIGEIIKKFQKRENKEVNTNVIHV
ASANPVDDKL

>Deutero_Ambulac_Spurpu_015310_593_144

MGCRSGRCHLRKWAADTSDLHGLKHIAGSGNLFRRLVWLCLFLTALGFCIYECYLVFVGFLSFNHVTQVDVLYSTEVEFP
AVTVCNINKYRESAFTEDDIKNVGVHLGIIDEDHNLLLPELYTDEFRDFIDSVNWTVVEEDPDYNMTDFTIRTGHQKED
MILSCLWKEEPCEESDFQHKLTHLGNCYSFNLQGANSDEADWKHSYAAGAANGLQLILNMETPEYTPTNDLDGGAM
DAGLRIMFHYPTEPPYPKELGFAVAAGDHSFISMRHEKITSLPDPYSECESDGADIITHFDHYSLQACRIECETLVVVAEC
GCRLPEMPGDDDVCGPADLHECAHPTLVEFITGNLDSAESCECFSPCEVENYPFTLSTSRLRLTYLEALFANTSTNFTAAY
IGENIAVVSMYYEALNFETIDMLPEYTVATLLAVLGGNLGLFLGASFLTLAQLGEYCFDEVIGWCICTPKKEDDDKNDGG
NKVEPVSTVTAMDAWGAQKL

>Cnidar_Rhopilema_esculentum_TR40535_c0_g1_i1_p1__Rhopilema_esculentum_TR40535_c0_g1__R
hopilema_esculentum_TR40535_c0_g1_i1_p1__ORF__type_complete__len_501__score_55_33__Rhopi
lema_esculentum_TR40535_c0_g1_i1_597_2099__597_147

MDSKTEMATLEREWTEQYLEYYVKKYLEEYEESTSDTSEGSDYSMMDWDFQKKGLLERERLWLKAKRRKKITKHFNT
MVKYSTFHGLQFCFKKESPLRRTVWCLLLLICNGLLVQKMFESTQHYLEHPFSTTKSVKYKESLTFPAVSLCNLNDMRHS
RMVGTKLHKLIIERQQNISKQLSGNEYMNTIRQANHRLENMLYSCTMDGVKCTSEDFSLFYHKQGDKCFTFNSGRPRY
KLVKTNRIGPQHALELTINIEFWDYYDDAVQSGIHLILHGQEETPVTMEGFRISPGFITYAEVKKTQRRNLPSPYKTQCGS
LKLKYFKGYSRNLCWLEQLTDDVVGHCGCKDWFMPGDYKVCSLTELEMCLWSRWVDFDKYKKYSCPLPCVIDSFTTKT
SFSRFPTHDGSESMAKELNLNGTSRDNHKFLSDNFLKVVIYYGDLSYEYLEQKPSYDLLVFLGDIGGQIGLFVGASVLTFFE
FIDCFLMCIHARFFEIFRPRETQL

>Deu_Ambulacrar_hemi_Ptyfla_40v0_9_20150316_1g17738_t1_scaffold14937_cov156_599_148

MPSAWCYTTNRDVGYELCDLKIDRKECVTDHVLNDFNVSFAPLEDLEYDQQYEQMTEAQCAAMCLEYSAFVCMFFDY
SVQNHTCYLSSNSSDQSQPSTVQQDIVRYIRILDDTEVEILHLKFSRYPNKKMIEGGLQRKVIAGSISECITACIEMDIFKCK
SLDFSPSQSTCNLYEKPSANVGGLMKEDISYIHLTRQGEYYDHRTVTEQFLDISGITPEILNDFENGYYRDNGFSGVVKGE
VPPDWYGFKTFSSTPDYSDLRSVLKLTGEEIAKYGHQAEDFILQCTYDEKTCSHSDFSTFQDDHYGNCFKFNHGLNGTV
VRDANRQGALSGLRLTLFLEQNEYISIYGRNAGARVNINPPNDTVLPQDEGITIMPGTVTSIGIREKHTFRKTKPYGKCKD

GTNEDNLYGPGNTYKVLIEDFLSDIGGTLGLYIGLSVITVAEFIELVASVIRYMYCSQKDSNPRTAESENKDRERRFSGAYAYPSFGRVDINDTKVAKL

>Deuterostome_chordata_Ggallus_tr_F1P0L7_F1P0L7_CHICK_Uncharacterized_protein_OS_Gallus_gallus_OX_9031_GN_ASIC5_PE_3_SV_3_600_149

KTNKSHLISEFSIIFHCLSLDKMLPRLEDRRKYRQEFASSTSFHGVYNIVQTQTRTRRVLWLLVVTGCLGIVIWQICSRFNYYFSWPTTTTVVVQHVENVKFPAVTFCNLNRFQAHAVSNLQIIFFLWNIVSGIVQKFAMEDKYFHELNGFLLGNQNFSIKEFTRENGFYLNGSTLLECEFFGKTCHPEDFEHIFTEYGNCFTFNYNDLPARRVSLSGRGLHLLFDVQQEQFTDDPALGYTDAGITFVIHSPKEIPRFDGLGLLTPVGMHAQVSIQQLKSIIQEYPWGECKPDIKLQYQDTYSTNGCLKECKAWYIQDWCGCLPFILPGNGIECDLMKYYNCVYPAIHDIEVKGLCTVGTHNSTCPAPCEETHYPTTVTYSSFGGENAIKYISAKLKKSPEYIRQNLVIIDIKYDDLNYRITQQQKALTISELLADVGGQLGLFCGASMITIIEVLEYIFTNFFWMCLFLLLKAPEIPRWNNPSHDQPTHVEKNKGIQEC

>Cnidar_Rhopilema_esculentum_TR106708_c0_g1_i1_p1__Rhopilema_esculentum_TR106708_c0_g1__Rhopilema_esculentum_TR106708_c0_g1_i1_p1__ORF__type_complete__len_504__score_95_58__Rhopilema_esculentum_TR106708_c0_g1_i1_88_1599__601_150

MMKSSEKKENNRNEQGSVKKIFDDFANSTTAHGIQHLVSSKKTLGKAVWLIVCLAALTTNLFLCAQLVKKYLSKPVSTTVGVRHEQTMIFPSVMVCNHNMLRKSEFQKAKRRKLINFKQDASMMSLYKEMMLKAIGMANQASNTSSVNISVNTDIIERRCLLEYLSNIEKPQLKGLGHQFHNFVVRCDWKGIDCKSGMLENRWYKVWNHIYGNCFIFNYGYNMERKKQEPLRLEDVGMQNGLTLDLNIELYEYDEELTKETGVRVLLADQGVFPLPSIQGFSVPPGVHASIGIKKVEAVRVDPFNNGSCRRSDSLDEDGDKYTMQRCTQRCHWQRQLKKCNCTGLRLSSPNRTCLTVEEHMCKAEVSTEMDALGINKICSTKCPPPCFEAKFSHSVTYSTLYTAALLKLKTSIFRNDDDYRKNFVRLQFFFESDIVEFIETYTSYEMEQLLSDIGGQLGLWVGVSFVTCVEIFLFLYSFSYHVLKKVWCTSFKVHNNKQTQEN

>Deutero_Cephalo_292990F_t1__name__chr_scaffold9_start_4371418_end_4385499_strand_pro_len_504__602_151

MRNVVGHKIGGTVACQDSNSGPLGSESTTLPLRHTTPQNLFLNPSFMATCCGDDSVEREYAAETSLHGVGKIAGARHRGVRLLWAVLFLGMFGVATWQITERFVAYFQFDTVTNMKVEFRDVLDFPTVTICNFNKYRDSQITEEEEQYVSTLLEGSTGFSDYDYDADYYDDDGDGEYQYSYDWNNTGIPDDFNLAEFTLRAGFDIHTSLKHCLWRGQTCNSDNFTHIFTSFGNCWMFNAEGRMNQTISGVGNGLQVAIDIQQDEYTENAPTGNLDAGIRFIVHSPSEPPKVDTQGISVGPGIHAYASISKIEFRNEIPPWGQCDPGRALQYYSGYTKTGCLLECRADHVADECGCRTVSMPGTLDYCQPAVVTGCVKTTIAELKTGKRSCNCPTPCSATAYPATLSYGGWPSSSTMDYFTDLLNKTEQEIKDNIVLLDVYYQQLNLQTVQQRRAISTNALLGDLGGQLGLFLGASVITIIEFLEFLVKKGTSCCFRHNNRVKT

>Cnidaria_HyNaC7_CDG50528_1_Hydra_sodium_channel_7__Hydra_vulgaris__33_152

MRIRLKERMRKIVQERLTVKQIFKRYIESSTLHGFCYVCMDTFLGRRLIWAVLMILGAIYFIFKLRYGIKEYFDYPFSTLSTVEYVDDLLFPAVSVCATNSYIASQVYTNQLNTMYKEGRLPLDNNQSIPEYNIPGDELVKTLKNSSLTIESLLKYCDWIMQDTSHPLVTPNNCGALNFTSYFNYKGEQCHTLNSGAKGHELLKVSDVGISHGYELVFDLQTNEVIKNYQLSGMRIVIHDQVFPPQLVDGFFISPGFKTYIKLGITQSQSLPPPYSTECGQKKLKYYAIYSQRLCLLETLTDFTGDLCGCRDVFMPENGLPFCSLQELYSCMYPAKESFSEFTMRKECPSDCEERTFSYELSEARYLHNPPIGLSLSRLDKLQLDKSLPSEAHLSKLAKSLTPKELDAYIESNIISVILFFGDTRIDYHEQEATNDFFQFLGNMGGEFGLMLGASLLTFVEFIDLFIVLLYHQMLRLYNLRKIPDIFASKRNRNKEKTKKCENV

>Chordat_hASIC5_NP_059115_1_acidsensing_ion_channel_5__Homo_sapiens__7_154

MEQTEKSKVYAENGLLEKIKLCLSKKPLPSPTERKKFDHDFAISTSFHGIHNIVQNRSKIRRVLWLVVVLGSVSLVTWQIYI
RLLNYFTWPTTTSIEVQYVEKMEFPAVTFCNLNRFQTDAVAKFGVIFFLWHIVSKVLHLQEITANSTGSREATDFAASHQ
NFSIVEFIRNKGFYLNNSTLLDCEFFGKPCSPKDFAHVFTEYGNCFTFNHGETLQAKRKVSVSGRGLSLLFNVNQEAFTD
NPALGFVDAGIIFVIHSPKKVPQFDGLGLLSPVGMHARVTIRQVKTVHQEYPWGECNPNIKLQNFSSYSTSGCLKECKA
QHIKKQCGCVPFLLPGYGIECDLQKYFSCVSPVLDHIEFKDLCTVGTHNSSCPVSCEEIEYPATISYSSFPSQKALKYLSKKL
NQSRKYIRENLVKIEINYSDLNYKITQQQKAVSVSELLADLGGQLGLFCGASLITIIEIIEYLFTNFYWICIFFLLKISEMTQWT
PPPQNHLGNKNRIEEC

>Deutero_Ambulac_Apla_gbr54_16_t1_606_155

MTASAPDTETNLEEEFVSNTTLHGIARVAHNSRTIFRLAWLAIMLVFLGVCIAQITDRLQRFFMYKANTAISVEYVGDLD
FPAVTICNFNRYRWSELTSSDLRSLNYILEATDYDYDSYGEASSSSLSDTYPLDDDNASSAFNYTEFTLRAGFLMDDLTLLS
CEWRGKTNSCSAANFSHVFTSFGNCWTFNSGSDGGPLLKEIQAGAGNGLRVQIDIQQNEYTEPLSGNREAGLKVLVH
DQQTPPMMDSQGFAIQPGVHAFVATRRQQFLNLEPPYGKCDASRTLNRFPNYTLEGCNIECRTEKIIERCNCRLVRHP
GTEVECTPMQTSECARDVLSKLRSGEIPACDCPVPCNYTTFSTSLSTATLPNQRATQLLVDQYDDLAQYSSNISDTTNTG
SASSAGQSNPILLDSTYMQNNWILLDVFYENINFQRYEQSEAITPSALISDIGGQLGLFLGASFITLTEILSYLGRKLGSILAS
RRPRRAATSPDTSPGWDMNGVKIT

>Protostome_Lophotroco_annelid_Pdum_Contig1978__608_157

FHSINRMHMMAEIGGDGNPAMSFLTRLKTLTENTGMDVLPHVGRARSKVKRLFWTLLLFVGTTMMISGVAKSFQRF
YSRPVETHLIQRVSESLPFPAITICNENPVRFSAVERAYPDFLKKLQNSSDKTFWVREELTSFLNSLPLEERINLGHNLTTML
LDCQQKGITCNESNFESMNSLYHGNCYTYKGMGDNLASNIGPHTGIGITLYIEESEYIAGVADGTGFKVVIHNGSDLPFP
EEEGFSVTPGFITFVALTKREVERAEHPYGVCHQEEDVPIDDIQYRTIGCRRRCLLRRQIAKCGCGLATFPYTEQTSNITDD
FNPPQCNTAEEKECSENVQQKYMDGKLNCNVGCPPRCKEVYHTYELSYTGWPPVCSRQLIMKIIERNTHTPLQLANKC
GKKYENIRYQTLKLVFFFNTMETTITKTASAYTCQDVLSNIGGQMGLWLGLSLCTLCEFLEFFFECFHILLRYLCKKNITTV
QSPQFGMRDHSHQNANNSSPKDKQ

>Deu_Ambulacrar_hemi_Ptyfla_40v0_9_20150316_1g15975_t1_scaffold12140_cov134_611_159

MKPRENSEDIHQEKSPSFSVQFKNTTGSSVIGLFFGDADSKKGTVQNLDNKYENLDENAAIQDFTQKATISGMKYVGD
KKSRIPRRCFWLLVVLTGAGFLTYQIIRSVNLYRSFPVNVDVKIKYAQDLPFPAITICNMNLLRKKEMEKHNEKNVAGVLK
LLYPLVTEIIEDSDALHNFTLSSVNNLTELLLRYGHRKEDMIIWCEWKGRRCSYENFTSTLTDYGLCHTFNSGSNGLPSLRV
DNTGARYALTLILNAEQWEYGRGPSQSAGFKIAIHSQYDIPMVSDLGFAVAPGTSAFIGLKMTKVSSFPEPYGTCKSRKL
KYHGNYTMSKCQLECLIDFIVEKCGCKRPYMPVAVDAPYCSPEEEYLCCLPKLDEYASHRQMCDCPVPCERTLYSASLSY
AQFPSDFMAVEYSKLLNKSDPQYPRRNGAKLNIFFEELSYEEISHRAAYELFTLFSDIGGSMGLLLGASFVTVVEIVDFACV
NIYRRWRRKTSSKIYPKTSSNDR

>Deu_Ambulacrar_hemi_Ptyfla_40v0_9_20150316_1g14166_t1_scaffold9674_cov125_616_162

MNKARARKIKMTGKKKRGLRSIISKFAKNTTAHGVANISNSTSTFGRTVWSVVCVTAFSLFLWQSSELLKDYLEYGIKIKF
DVVSVPKLSFPTVTVCNTNKIRKSEILKSEHRRILVVDEDVVHPYYALQAFKSTYNKRLSPKDAVLKTNTSLGLCALSCMHF
SEFTCLSFDYDSQRRECYINPGSKDTLVRFTLTNDDDYDYYERIYRGMECYTLDDGTDYRGHVSLSSWGERCMNWSSL
DPSKFKYTPSTAPGKGLGNHNYCRNPNDEPGTWCYVYRFNSNVPVAARCTVNPPKYGCLSQKQWEKEIVESDTFRDS
KPIPAIAVNCSADPSGQGYRGNKSTTVSGKQCQNCDFYMFQDDQYGNCFKFNDGEKEPVIRQADRQGALSGLTLTLFL

EQNEYISIYGRDAGARVNINPLNETVFPQDEGITIMPGTVTSIGIRELDDILSDIGGTLGLYIGLSIITVAEFIEFVAAILRYLY
QSRRIKKLRTVDSDIKEKKRRRKCKF

>Deutero_Ambulac_Apla_gbr129_26_t1_617_163

MADEKSGENDARSLRALFKNLMGNTSAHGLPNIDRAGNPFRRCFWSVLFVWALGMFLWQCYELVYAYFQWEVDVN
INIQYKTQIDFPAVTICNLNPVKASSLSNSRELEILLGSLDSETSGSTDGSWTAVTAVTPKVTSDQLSVNSTDGELFGGTAS
SVTKLLPATEAGLTATGDTHTLKASSERDQLTPTSTDASSPSSGVSSASPAGSASIHVVGGTQMATNAVTAGVGIPTKEV
ERRKRDTSTNSERQYQNWDNIDYTAKESDVDLFDQVADAIAKTEYGKRLNFGHSIDDMLLACSYKGYPCGPTNFSYFH
NYLYGNCYTFNAGMNSDIQTSSKPGPLYGLILELNLEETEYIPSIQQASGARVVIDQQYRMPFPEDAGINVAPGVLTSIGI
RKVEIDRKSDPYTNCTERDSDDSQTVFSDFFGANYSLEGFGLISNLGGQVGLWIGISMCTVFEFVELLYDVCKILLLKVIFC
SHAKRATAPSASDLKLDEKPSKPRLALNN

>Deutero_Ambulac_Sakowv30038064m_622_164

MDNTEKRECDKANNGFPKFKKSTSLLALFFRDLSEQRSQKDRDSQDSSDEEENGDEKKVEDESVIGDFAQKSTISGMKY
VSDKKSRIPRRCFWLLVVLTGAGFLVYQIVRSVNLYLDYPVNVDVKITYANDLPFPAITICNMNLLRRLELEHYEGHDAIEF
LELLYPLVTEKTSPVPENIIQNFTLTTVANLTDLLMRYGHQKENMVILCHWKSSACSSVNFTTTITDYGLCHTFNSGLEGT
APLRVDNTGARYGLTLYVDIEQGEYGRGPRESAGLKIAIHNQRDIPLVSDLGFAVSPGTHTLIGLRMTKISSLPNPYGLCK
SKKLQFYGDYTMSKCQLECLTHFIVGRCGCKQAYMPGQVRHCSPQDVYNCYLPNLGEYAGHKDMCECPVPCERTIYSS
RLSYAQFPSDFAANEYSKLLNNSDSDYIRRNGLKLEIFYEELSFKEVTHQVEYELFKLFSDIGGSMGLLLGASFVTIVEIFDFA
CLNLYRRVRSKWRPRITPIETKS

>Deutero_Cephalo_032830F_t2__name__chr_scaffold117_start_653459_end_666145_strand_pro_len
_509__623_166

MPKIKTPMSSDEGHDWPESPEYVFATTTSFHGIGRIATAPNIPLRVLWTLATLASYIGFFILMVEMLQSYFTHGTVTDVT
LEFVPEVRFPAVTICNLNKFEDKALTVEEKHYLSYILYGVQEDIDVIAGAPVNNKYTPNSTDANVTTETPRTEAPHTHGSF
SVDSITREKGFRLGNDMHVCTWMGQTCTELDFTHSFSTYGNCYTFNASPENPINQTIPGSGYGLSVMIDIKSHLYTENP
LLPGGTADVGLKLLVHDQNEPPKMDTQGIAVAPGSHAYIAIRQIQYQNHIPPWGVCQDLQLEYYDTYTLTGCQLECRS
KYVVQNCTCRPIYLPGDAPYCEPADVAHCVRPVEAAVTSGALPCDCPVPCSLMTYRTSSSYAKWPNNKADVVYTTAFG
LSPGYMADNSVVFDVYYEELNYEIISQLKAMDSGQLASDLGGQMGLFIGASILTLLEIFEYLGLRLIAFIQRRKVRTRRVHT
SAYPMPDTLSVNQAAKNKESKEKSLMNLKE

>Protostome_Lophotroco_annelid_Pdum_comp414620_c0_seq3__625_168

MGCKEWFTMGKSNPPLTYTEVTNQFADNTNCNGISSIKRSNHVVRKAFWTLLVLAGAGLTIKLLVDVFTKFFAYPVSIS
VNVTYESTVQFPAVTLCNVNMIRKSQIGLMTGSSALDKLTFTVGGNPDISAWATTEGRSILRHSFVDGWSQLNDTVKK
QQGHQLEEMLLDCSYNGDVCGTNNFTHYINPNFGNCYTFNSNWENQTIQYSHRAGPFYGLMLELFIEQDEYIPELTEV
AGVIVSIHHQDVMPFPEDDGVLIPPDMSSNIAITAETITRKAKPYPSDCLNATFKEAAHYSAYSELHPVRYTHTACYRTCY
QAHLFSECGCGDPVMPLEGQALYNLVTDKQNKRTVCTSQDEITCELKVYKQYSNNELECNCSVACEDHIYQTAVTSLR
WPSANYKNLYYGHLATRNPHLATIMQSYVTNGTNPGEHLVKLLIYFNEFNFNYIEEYASYPSSSLLSDLGGAIGFYLGISVI
AAFEFIEYIFDLIFIGCKKMKNRKANEVKSID

>Protostome_Lophotroco_annelid_Pdum_comp405972_c0_seq2__629_171

MMSGESSIKPILGSMAENTAMHGLPNVKRSISTARKIFWSIIVLAGVAMILYEVIIATIKYYQYPVQTTISINSAIVHQFPV
VTICNQNSIRRSALEDAGSDIYMLLNSTGSRQTGSSIVSLRGNILKYYSQLTPQVRMDIGHQMETMMLDCAYGAMECN
TTYFEQFSNYYYGNCYSFNNGLNYGIKTASTVGPDHAFQIVLFIQQDEYISQLSEGAGVRVIVHNRTHMPFPENEGFSAP
PGQITYVGLSRREIKRASPPYGDCSENHLTMNSYTDFLTGVDYSKMACKKSCLQYTIIETCSCADTSLVTNSTALNTFFKNI
EPCRNLTQDACISTVTQKYYNNELNCSTGCSPECFEVQHERTLSYSLWPGDNALNDTLTLLSENTNAAQLMKNMNDDE
KESFLRKNVVKLAVYFESLDYDYIATSSSYSSSDLAASYGGQMGLWLGVSVCTLFEVVEMLWEMGLVISQKMNSRKNK
TGNMETTNSDGKKMPSDKAVIVKKDSWTASY

>Placozoa_TadNaC11_XP_002114391_1_hypothetical_protein_TRIADDRAFT_58144__Trichoplax_adhaerens__21_176

MGSKSENPRKVSNGQTDIVSFDLEENFATSTTCHGVAHIFETKGSRRSMWFGITLVSTIACLVQCCIIVYDASLYPTRVNI
KVKYNTQSLFPSVTLCNTNLIASIDNTREELKYAMLLYRHYYAKNQITHQELQSAENYFNRVYGKNFSLEEYVARYRFKAE
DMIVSCQWGEEPCGSANFSQVITDYGTCYTFNSGQKGYPLLYQTISGSSHGLRLAINVQQYVYPTLPLSFLTPDAGIRLS
VHHYNEIANMGSRGVFVPPGMHGYIAITDTHVLSNLGPPWGECGQKNLQYFRYYSRSACRREFEANLAAQQCNCSYQ
QATVNTENKKTLNSFCPYPCQQTEYPIILSYAGIATNAIIDSSSHNGKLANLLDSVKNDPSKYPNYTSEAFIRENLIYLDVYF
RELITSTTTESKATGYAQVLSDVGGQLGLFIGASIITLCEIITYLCDRCERKKREKREMRIHRQSQMIRDIMQVGVEKKDLQ
TETKPNKNNDFVFNTSEDGQETSSTNPNV

>Cnidar_AlaAla_c46701_g1_i1_p1__GENE_c46701_g1_i1__c46701_g1_i1_p1__ORF__type_complete__len_515__score_83_16__c46701_g1_i1_223_1767__638_178

MDIKEKDLPENVKAVLKQFASSTSAHGLSHIASAKYSYQKLLWAFVLLLNLFMFFFMITRLITLYTSKPVSTKVEFTFEKEL
AFPAVTICNQNMFRRSKLTPSDHARLRPSETQITSTMVEKMQKKFQDELKRQGHEGTHGGSGGTTGATSGGSNSTST
DNMVTVDSSHLMRMSILESVNLMEEQEKMGKGHSFEDLIKFCRWDGVACHKGFMKQYWYRTWDWYYGNCYTFNY
GKTGNGTSLKRLKATGSGSENGLFLEIDIQNKEYFPELTLESGVRLLLTEQEVNLFPLQEGFSIPPAVSAMVGVKMSTAIR
ADPFNNGSCRANEYADGGFRYTVQNCVQNCTALAQESKCNCTLLKYAAVNSTRVTCHTEAEKYCADEVVKSLKNGTN
DCKANCPQPCSESYYSKSMSYSRLYGEGYQKDIAIRDKSNPKEKDIMENYIRLLVFFEDQNVHKIRTAVYYEVENLLSDIG
GQLGLWIGVSMVTMVELILFIYKFVEYLMEKKKAGKVIHVAH

>Deu_Ambulacrar_hemi_Ptyfla_40v0_9_20150316_1g17281_t1_scaffold14250_cov87_639_179

MLIYSYFSWQDDRPGDPYPLVSLNPTTLNGNGEGQDQNRNLRKRQGNNTDERTDEISSLDKSAGETNGEYEDSVKHM
LGDFGRETTAHGIVKITTAKSNVTRSVWSAIVLLAASVMLVQMTLLLVQYFEYNVNVKVTQVSEKALEFPPVTICNTNKL
RKSAVRASKYSEILLLEQSFVPPYYTPCIEGDFLCQDGIHCIKPYLVCDGVNNCGDMSDEDNCTYRECGPNQFRCDSGSN
LGICLTKDRVCDRIKDCYEGEDETDCGSERIYGAVVRIGNSDVIVNNTQCGEIIISDNIDSNAAVEVGYALRGRFVSIQLEN
RTDYLSLCEVEVIGEASRILGAVIRIGYYKEEITQNVPCGGQITETDISNKTIIRDCIIPILGRYVIVQNDPYREDYLHFCEVKV
SAKEFVPSEFTDPLKTRKFVMFEPQEEKTLPVNSTDVFSNLLLNKCLSECLNWKRYECHSFDFHFDSKTCRIHRQKAGMD
GLHVVHSPGTVYYQKLRMSGMKNRYSVLIVT

>Placozoa_TadNaC9_MK547550_19_180

METANNDIQSTNLEKDFANATSYHGFGQFYKPQTTHYRRYFWMLLTVVATSACILQSSRIVIAAFQYPTKITSKVRYSNS
SVFPAVTICNSNYLDKTKMDPDILAVVNSVFSNYNRSALDLVNLGLTPEIIAARFGQHADYGFFARAFGFKLKDIILECTYH
NFNCLNSFTEVVEPNFGLCYQFNAGILQRGGNGNGGNNSSFHRQVGRGPSFGLQLVLDAEQYKYSTLDFFYRYQAGFII
AIHDQFETSSMTINEISVGVGRYTSISLHQTLIEKYLKPRWGECDDQELAFYRKYTREACKQEKESLYVMNLCKCRFKEVS
ESTVNYTICTTFDTITCVIPQLDLASRNYRISNSCKIACARKLFPKTISSSSIGTLAYGNILNKRLDLSTKLADMQGLGLLPSSY

TVQDYIRDNIVQLDVFFSDLAHTTVQQEQDITPEEVLSNIGGQLGLFIGVSMLTVCEIVFYIFDKMQYWPQRRNRDNRQ
TTPVDFEMKKSKVNNNANTIGVMGYGAET

>Protostome_Lophotroco_brch_Lingula_anatina_comp136986_c0_seq2_p1_comp136986_c0__comp136986_c0_seq2_p1__ORF_type_complete_len_518___score_46_16_comp136986_c0_seq2_175_1728__644_183

MTEACEKMVLQTIIVEPMDKCKGLEKDKVSQDITMEEEESLEREFATGTTAHGFSRAAETKSIFRRSAWILLILGGLGMA
TWMISLRVINYFQYDTSTEITLTFEPRMAFPAVTICNFNRYMRRNLNAEDYAIISLTEDVYSGYDSGYSYGQYYYNYYYDY
SGDGESSPSTTTQPTNSNFNESWANISASYPNGFDFGNFTLEKGWKLDNISVLECTFKGIKCNLKKDFIHTFSTYGNCYTY
NPSLNSNRSRYQDQPGLGNGLHLTIDVEQSEYTESLPDGSAQTGVIFQVHPQWEPPLVESKGLGASPGFHAFGALKRQ
ETVNIEPPWGKCNKSLTLLHFDQYTFSGCIIECKLQKIETECGCKPVQYPGTYRICDPAEMMYCVKPLLKTINQAFSDHCH
MCTVPCNSTQYNVQLSYTTIPNNNIEAAFATKYNLPTGSNYIKDNIVALDIYYEELNLETMEQKKAVEESGLLSDIGGQLG
LFMGFSALTFLEFFEYIILKFRRITQRKKRIKPLA

>Protostome_Lophotroco_brch_Lingula_XP_023930484_1_ASIC_channel_Lingula_anatina

MSEKMVRRRSSAFEVQMETLEENKTKESTESKASKESGSVSEGLEQFGEDTDFHGLKRVFRRDYTLLRRT

FWLFFFLAGLTGFIANTVDRLSYYLQYPHSSELDVMYDMELEFPAVTICNMNSYRLSALTDVDILHFGEK

LHILDENRRLIHPEYYNQSWVDWVHGINWTELAANDDEENFDVLEFIRRTGHQLEDMILLCSWKGQHCGP

ENFTSVFTHFGICYTFNADHSYKYVSRKAGAGNGLKLYINIEEEEYLTSDVLAGQDAGLKMVIHAQEEPP

FVKELGFGVLPGDHHFIAIQKKYVHNLVPPWGNCYEGKLKYYSHYSVPACRIECETDTIVKECGCKLAEM

PGNTSVCLGHMYMGCAYPALEEVEHSDLCSCQNPCDMTTYRRDISSVRLRDTTLDIIAENNPQVKRETLS

RDNLLVLNIYYEELCYETIRQIKAYSIPALLSDIGGQMGLFIGASVLTLLHVIEAVGAVVGGKFLKTTKQ

GRVSTTRVQSLK

>Deutero_Ambulac_Apla_gbr54_20_t1_645_184

MTHEASRQRHGKSSLEREFAESTSLHGIAKVFHASRLLIRFVWVAIMLTCLCVCVWQISDRFHRFLQYKANTEISVEYTR
DLDFPAVTICNFNRYRSSALTESDKTYLPYIIWANDYDYDSFGDSQGGVEYWDMVADDGRDNDTHFNYTDFTLRAGFL
MDDITLRQCEWRRKTKSCSPANFSHVFTSFGNCWTFNSGKTGPILKETQAGSGNGVRMLIDIQQSEYTETVDGNIPAG
LKVLIHDQATPPFVESSGLAISPGVYAFIGVRKQQYLNLEHPYGKCNASKTLTRFSHYTREGCKIECRARAILEKCNCRLVR
HPGNETECSPQQTSGCAMSTLGALVGGEDNPCDCPVPCNYTTFTTSLSTASIPSNSVAAHLQGLTETSYSSDYSYSSFSV
PYPAQLDLFEEGYIPKNVILLDVFYEDINFERYEQSEAITPSALISDIGGQLGLFLGASFITLAEILSYFGKKIDFWIQRVLGKH
RRLLERQQDNRHGRDVPLKSVRIDNGNHRTLWADV

>Deuterostome_chordata_PetMar_tr_S4RU13_S4RU13_PETMA_Uncharacterized_protein_OS_Petromyzon_marinus_OX_7757_PE_4_SV_1_653_189

KAAHAAFVGRAKIHGVRHFLWPRHAVTRRILWLLAFMAALGLLLAWSADRIQHLLSRPAHTRVSTSWVRGLPFPAVTF
CNNNPVRFPRLTKSDLYVAGMWLGLLTHAESQQPQATFAATAANLTANPHMMAGLSEPRRQWFQRMSDFSRFLPP

REAPGISQGLLDRLGHPLDEMVLSCRFCGQDCKPQDFITVYTRYGKCYTFNSGKGGRPLLTTVKGGMGNGLEIMLDIQ
QEEYLPVWGENDETSFEAGIKVQIHSQDEPPLIDQLGFGVSPGFQTFVSCQEQRLIYLPPPWGDCKSTPISSDFFETYSIT
ACRIDCETRYLVENCNCRMVHMPGDAPYCTPEQYTECARPVLDVLVKRDNDFCLCETPCNMTRYGKELSMVKIPSKAS
AKYLAKKYNKTEAYIAENVLVLDIFFEALNYETIEQKKAYEVAGLLETGDIGGQMGLFIGASILTILEIFDYVYELIRDKILFYF
RAKKKQKTESKDTGPCEALRGHSESAGYAASMLPPPHHHHQHHAL

>Placozoa_HhoNaC9_TR3133_c0_g1_i7_m_7432__Hoilungia_hongkongensis__26_190

MNVKVDDSQNIEKPNLEKEFAEITSFHGFNQFYRPKTSTYRRYFWILLTLAATAVCIVQSSRILATAYTFPTKISSKIRFKDK
AIFPAVTICNVNLFQRTKVDPNSKSVMNVIYGSTTKVNLTQLGISTAQLDAAYGAHADLADFAEFYGYELKNAMIQCKF
HNQDCSENFTEVFDPNFGRCFQFNAGQIARADNNTANATASQSTKTNDTLNGPSFYEQEGRGSRFGLQLLLNAEQYK
YTRLDFTNRFEAGFLIAVHDQYEPHGMASNGISVGVGAYTGIVLQENLNMKYLKPRWGECDHQVLKLFTHYTREACKY
EKEAFYVMELCKCRLKSVPESAINLTLCTTTDSLTCTIPQLERASRNYSLSRSCRIECYRKLYPKTISTSTLASLAYANYLDSTF
QISSQISLLQSYGLVPTPYTPIQFVSDNIVEVDVYFADLIDRIVEQEPDASVEVVLSNIGGQLGLLVGVSLLTVCELIFYVFDK
IKYWLNGRSPRTSMTAQHDSDEKNTAELTTVNGW

>Placozoa_HhoNaC3_TR31058_c1_g1_i2_m_67045__Hoilungia_hongkongensis__25_191

MESSPSRPRKPSIDFSTLNTTISTTGHGSQEDDDSQQHRSSLNISYQHEEEFAKSTSCHGISHVFDDSSSNMRRLLWMIL
TVSMTVICLIQCIDRVLYLASNPTRIVLEYNMPHQIRFPAVTICNYNRFRNSTINQSNRHRLPHLLRLMSPWHTQSTARQ
LNSRVQAINTTHDRQDDLKLLAQVGEQADIFIKYCIWNNQPDSCSAENFTLAYTEYGNCFTFNGDLVGNATLYQKHAG
RSHGLRLVMNIEADEYTTMNPEPDIGLKFRIHEPDEPPDIEARGVAVPAGYHAFARLKYNEAIFLKEPFGKCNSRPLQYY
HHYTRTGCKLECKTNRSIEICGCRTLYMPGSAPICTDKQMNQCFIHNLDKIENGECYCPTHCIWSSYDPKMTFAIMPTET
ITREEVIALGISADELQANTVINESLTNHLRKNFIFLDIYMEKLKYRTAKQDLSYTFNALLSDIGGQLGLFIGASILTMLEIGE
YLMTKILSLCKTLHRPTKQSRNETADFIQTSNLKNNPVPV

>Placozoa_TadNaC3_MK547544_13_193

MDRNFQQRPRKPSVDFSTITTITTSTKDQSQDSDFFSQRQSPLNISYQHEEDFAKRTSCHGISHVFDDETPKLRRLSWVF
LTLAMATICIIQCADRMIYLISNPTKIVIKNEIPPRIRFPTVTICNFNRFRNSTINESNHYRLQYLLRLMNAPQAPSKRSEFSF
RNDLVNYSRNTNDDLDLLSKVGIQRQILIRYCNWDNQPHSCSAENFTLTYTKYGNCFLFNGQSADNEALYQKRIGRSHG
LRLAFNVEADQYTTMNPEPDVGIKFRIHEPDEPADIEAQGVAIPPGYHAYVRLKYNEAKFLRKPIGKCDSRPLQYYDRYT
RSGCKLECKTNKSIEVCGCRALYMPGSAPFCTAHQINQCMTNNIEEINNSSCYCPTACLWTSYDPAVSYSMTPTETITSE
EAPLFGLNLDEVDANRVLNSSISRYMRKNYIFLDIFFERLRYNTAIQELGYTFNAFLSDIGGQLGLFIGASVLTMLEIGEYFL
TKLKNLCKIIKRPTSQNEYEIAPASPNTTFEENPIPV

>Deu_Ambulacrar_hemi_Ptyfla_40v0_9_20150316_1g15265_t1_scaffold10904_cov157_656_194

MSSTVYCDRYSDGALQFDLQPVATATLAKPKTRKSFAVTSFRDRGYHSKVEVIKEEEKKGDRESDSRSVGQKDTCDDCC
CCRTFYSERFRQFSEDTTLHGLRYAVGPHVGTSRRIFWTLLMIGMIVSSCIYLYLCWVKFLAFPVKTVVSMNYKTEISFPAI
TICNYNHYRKSYVNGTRFEDWVRQRYPTFNFSDDEPPVELDEDFLRINRTWFEISAAHDKWNMIYQCTFGRSKIPCDAS
NFTLTLTDFGVCYTFNGGKESEDNVLHISNGGSEHGLRLRLFVEQFEYTYSENSAAGFKILAHRQNDVPMIRNFGFAIPP
GTESLIGLKMIQEHNLPAPYVTRCSNEKLEYFEVYTHSNCVRELNTNAVVENCGCREPYMPGSARVCNYNESVKCAVPV
IETLHTDKDCPVACSNIGYETRVSYASFPGRHIVKDLEDRLNLTQYDIRENIADLKIYFEEISYEEIRQEKGYAFPDLIGDSGG
ELGLCLGASLLTMCEIVEFFLMWIWRKMTRFQRGVGPNLKR

>Protostome_Lophotroco_annelid_Pdum_comp415688_c0_seq3__659_196

MKYTKCLKTGKSNEPLTYTEVTNNFADNTNCHAIGSIKRSESALRKSFWIILFLTGAALTIHQICEIFETYFSYPVSINVNVTY
EKSLVFPTIAICNYNMVRKSQLGLMKGSEALDLLTYTVGGDPDMSQWENSEGRAELRHKFLNSWSQLNESSKAAQGH
QIEEMLLDCSYNEKRCYPDDFTHFFNPKFGNCYMFNSNWKNQNVQVSHQPGPFYGLKLELFIDQDEYIPELTESAGVVI
AIHDQDVMPFPEEDGVLITPDAQSNIAIVVEKIQRMQSPYPSDCLNVSRKEAYDYSAYSELYPVRYTHHACYKTCLQAHL
FKECNCGDPEIPMEGVALYNLVDDKENIRTVCTKDQQACMEKVHKKYLNGELKCNCSVACIDTVFEKTVSTLQWPNAN
YEDLYYSRLATRNPQMKSVLKSHSQSGTNPGTNLVKLLIYYSEFNFKAIEEYASYPVSSLLSDLGGSIGFYLGISIIAAFELIEY
LFDLFLVTCQKSRKKDDNRNAKSSPESRILNDLENEREERQ

>Protostome_Lophotroco_annelid_Pdum_comp408485_c0_seq2__664_198

MSTSKEQQDGVAVCMDSCAEDSSTQCIPQIIRAKNRLRQLTWVLITVTLVTAWIWHSCFTIGKYLKFDVNVGTKIIPSRQ
LANPAVTVCNKNAYHGVKADEFLTEMGQRTKSALKNNLENASLYGPSWFLAWVEALKSNKNVYKIPSSAQFLQMFSE
FTDIYGPTYGVDYSNFTMKYEQFVQRCLYAGAPCALKGVHYQVVFNWLYGNCFTFTVDKNATGTGPLYGLQLSLFIDQ
EGYSELTTGAGVQLVVHDESQMPFPDEDGISVPPGQSAFVGIRLHEVQRLGGAFGDCFDEAKQRNSLDYFRKYSYSTSY
SLRTCERSCFQRNMMSQCQCLDNRYPPYTVKHIESSKYNNCITQYLSNFNFSNGLPEDLGCKDRLEDMFNNVELGCDI
DCNIPCNTTKYEVMVSYAKWPSEKAMNETKAFMSLKEAPEMSDEEFRRNFLEVTVFFKTLDYVITSESKAVTILQTISDL
GGNMGLYIGASVFTFFELTQLVLDFMNWLCGKKEKKKNATGEAQEGHERYEVKL

>Protostome_Lophotroco_brch_Lingula_anatina_comp147337_c2_seq2_p1_comp147337_c2__comp14
7337_c2_seq2_p1__ORF_type_complete_len_527___score_4_70_comp147337_c2_seq2_294_1874__6
65_199

MTTKTKFFRYDVPSDFDEENDTLTYYVLWKGLKERTTSHIIPQVTNAKGIFKHFFWILITLATFVGLGYHLYTMFYKFFEYD
VGVKLEITSNSTLKFPAVTVCNENAFRKSALLSSPSKLPTLDAFVTNGTVPTTGFNSSLIDSELGGRSNRATLIDKTLEDISEL
TTGEKQALGHQRSDFILDCEYGGYSCNGKTFESFYSTYNGNCFIFNSGWNNSSILRATESGPNFGLSLTFNIEQSEYVGDL
TQTAGVRVLVHDQDVMPFPEDQGFLAAPGEMTFVGIKMVEIDRYGGRYTTCKKTDTFNITENMYQALYPSVGYSQKA
CKKTCYQRTVIETCKCALSIYPRVETLFDSTTSVRTCDSLNATDVACTVKVQTQFINGELNCSCIQPCSATSFSIDTSSAYW
PSDEYATEFLSNYNSKSSIVRKIYNTSGSAGVMKNLAKVHIFYKDLNYEYISEYVYYDSIQFLSDFGGALGLWLGWSVVTF
FEFFELFMDVLAFSCRCGRKRKVSRSSVNDITLKEKASF

>Cyclo_DEL4_NP_492230_2_DEgenerin_Like__Caenorhabditis_elegans__47_200

MGVFWTGLKYVFTDFSCWTSTHGVPHIGMANARWLRAFWILVVVVSIALFIWQFITLLTNYLSFSVNTETTLQFAERTF
PTVTICHLNPWKLSETKSVDPDMSALIDAYNSDSSSAQFGLPASLTADRQQQASKWTLMYSERLNEKQYNDAADIAYS
YDDMVVSCTYNAKTCNITDFNDFYNPSYGNCLQFNTDGMYSSSRAGPLYGLRMVMRTDQDTYLPWTEASGVIIDIHM
QDEIPYPDVFGYFAPPGTASSLGVSYVQTTRLSKPYGSCTTKTKLKTTHYTGTYTVEACFRSCMQEKIIASCGCYYPAYSH
ASNTTQYVSCDNGVQTLSNLNCVDLINSADSTEFDVLTDCDCPQPCEIDSYGVTVSTAQWPSDSYVPTECNPGGPSGP
WDASGESCLDWYKANTILIEIYYERMNFQVLTESPAYTFVNFISDVGGQVGLFLGMSIISAIEYLVLIFLVFFYCCTHKSRR
AEIEQLEMDIKKAKDDVDQVAEKRKKHQKANAELYEMDTAHDIVPPKPHSND

>Deuterostome_chordata_Ggallus_sp_Q1XA76_ASIC1_CHICK_Acid_sensing_ion_channel_1_OS_Gallus_
gallus_OX_9031_GN_ASIC1_PE_1_SV_1_667_201

MMDLKVDEEEVDSGQPVSIQAFASSSTLHGISHIFSYERLSLKRVVWALCFMGSLALLALVCTNRIQYYFLYPHVTKLDEV
AATRLTFPAVTFCNLNEFRFSRVTKNDLYHAGELLALLNNRYEIPDTQTADEKQLEILQDKANFRNFKPKPFNMLEFYDR
AGHDIREMLLSCFFRGEQCSPEDFKVVFTRYGKCYTFNAGQDGKPRLITMKGGTGNGLEIMLDIQQDEYLPVWGETDE
TSFEAGIKVQIHSQDEPPLIDQLGFGVAPGFQTFVSCQEQRLIYLPPPWGDCKATTGDSEFYDTYSITACRIDCETRYLVE
NCNCRMVHMPGDAPYCTPEQYKECADPALDFLVEKDNEYCVCEMPCNVTRYGKELSMVKIPSKASAKYLAKKYNKSE

QYIGENILVLDIFFEALNYETIEQKKAYEVAGLLGDIGGQMGLFIGASILTVLELFDYAYEVIKHRLCRRGKCRKNHKRNNT
DKGVALSMDDVKRHNPCESLRGHPAGMTYAANILPHHPARGTFEDFTC

>Deu_Ambulacrar_hemi_Ptyfla_40v0_9_20150316_1g2678_t1_scaffold1096_cov89_668_202

MSCTVLDENNNCPDEYRECGSGECVYQYAICDTIHNCADGSDEVNCTTTTKGENISLRTVTEQFLDISGVTPEILSEFENG
YYKDNGFSGVVKGEVPPDWYGFKTFSSTPDYSDLRRVLKLTREEIAKYGHQAEDFILQCTYDEKSCSPSEFLTFQDDHYG
NCFKFNHGRGGTVLRDAKRQGALYGLRLTLFLEQNEYISIYGRNAGTRVNISPLDASVLPQDEGITIMPGTVTSIGIREIPV
DNSLDICFKHRNRTVIAGFRARLYIGLSVITVAEFIELVASIIQYMYCGQRDMKSRVKSANEETKEHFSGAVAFPGFRSFD
VDVAHSTKKFRIVSGKIFHRSVKYGTIVNLTLYNHMAEDYTTMHWHGIYQIRTPWSDGVPYITECPIPPGGVYVYAFAAI
PAENPVDQPTFDSGCGIPHDFYGESGTITSMGYDGTTLYENNAKCEFRITASEGKKIVLNFVAFDLEEGSDFLSVNGIGG
ESIAEYTGSNIPGPLATSSSQVIVHFNTNESGQAKGFHADWEAV

>Deutero_Chord_Hsap_sp_P78348_ASIC1_HUMAN_Acid_sensing_ion_channel_1_OS_Homo_sapiens__
Human__OX_9606_GN_ASIC1_PE_1_SV_3_669_203

MELKAEEEEVGGVQPVSIQAFASSSTLHGLAHIFSYERLSLKRALWALCFLGSLAVLLCVCTERVQYYFHYHHVTKLDEVA
ASQLTFPAVTLCNLNEFRFSQVSKNDLYHAGELLALLNNRYEIPDTQMADEKQLEILQDKANFRSFKPKPFNMREFYDR
AGHDIRDMLLSCHFRGEVCSAEDFKVVFTRYGKCYTFNSGRDGRPRLKTMKGGTGNGLEIMLDIQQDEYLPVWGETD
ETSFEAGIKVQIHSQDEPPFIDQLGFGVAPGFQTFVACQEQRLIYLPPPWGTCKAVTMDSDLDFFDSYSITACRIDCETRY
LVENCNCRMVHMPGDAPYCTPEQYKECADPALDFLVEKDQEYCVCEMPCNLTRYGKELSMVKIPSKASAKYLAKKFNK
SEQYIGENILVLDIFFEVLNYETIEQKKAYEIAGLLGDIGGQMGLFIGASILTVLELFDYAYEVIKHKLCRRGKCQKEAKRSSA
DKGVALSLDDVKRHNPCESLRGHPAGMTYAANILPHHPARGTFEDFTC

>Chordat_hASIC3_NP_004760_1_acidsensing_ion_channel_3_isoform_a__Homo_sapiens__5_204

MKPTSGPEEARRPASDIRVFASNCSMHGLGHVFGPGSLSLRRGMWAAAVVLSVATFLYQVAERVRYYREFHHQTALD
ERESHRLIFPAVTLCNINPLRRSRLTPNDLHWAGSALLGLDPAEHAAFLRALGRPPAPPGFMPSPTFDMAQLYARAGHS
LDDMLLDCRFRGQPCGPENFTTIFTRMGKCYTFNSGADGAELLTTTRGGMGNGLDIMLDVQQEEYLPVWRDNEETPF
EVGIRVQIHSQEEPPIIDQLGLGVSPGYQTFVSCQQQQLSFLPPPWGDCSSASLNPNYEPEPSDPLGSPSPSPSPPYTLM
GCRLACETRYVARKCGCRMVYMPGDVPVCSPQQYKNCAHPAIDAMLRKDSCACPNPCASTRYAKELSMVRIPSRAAA
RFLARKLNRSEAYIAENVLALDIFFEALNYETVEQKKAYEMSELLGDIGGQMGLFIGASLLTILEILDYLCEVFRDKVLGYF
WNRQHSQRHSSTNLLQEGLGSHRTQVPHLSLGPRPPTPPCAVTKTLSASHRTCYLVTQL

>Deu_Ambulacrar_hemi_Ptyfla_40v0_9_20150316_1g19294_t1_scaffold17970_cov134_672_205

MFHTEASYAFQRKKRIIPIIVEKDYRPDGWLGLLTLPELYFDFTSDEQLEKNFDELVRQIGDHGNLNDDGKDELHISLQRS
KEGLKHHQLPLKSGAYNVERGKAYQKCQTNDNSEEDESDVDDKQQRVDSRTCGDGQERSQATLKHVLADFGKGTTA
HGIVHITEAETSVSRTVWLAIVVAAAGIMVIQMTLLLVQYFEFGVTVKVSLVSEKMLEFPSVTVCNTNKLRKSAVRNSIY
NQMLILESNFVPAYYECEEPNFRCLASSECINNLKVCDGTYDCRDGSDESGSICEPKLKNVARGKATSLTGPILENRDSNL
AVDGYTYTCVQTDWIWEPLWSVDLGKNYLVYKLKITHMATYDPKLVNAEIRVGLAPKAGPDNPICVKSIREHETQTETF
TVRCHPVPVAGRHVFIQVKGKTEDMRLCEVEVLATDIAEELENVAYGKPAEQSSTYYEHVASRAVDGNENFALETGHSC
IHSAKEAGAWWRVDLLNVYSVYKVVMINRYDCCRERASFVVGYSVLQVLSSLSYP

>Placozoa_HhoNaC11_TR28363_c2_g1_i3_m_60033__Hoilungia_hongkongensis__23_208

MDTKQQHSTNEDSSDSASVDLEEDFATTTSCHGVGHIFTNAGSRRSMWFTITLICSVACICQCFIIVYDASFFPTRVNIKL
EYNIESLFPSVTICNTNLIASLSIGDVEYKNALTLYRYYYINDNRTFGELDEAEKFFTRIYGKNFTIHDYIVKYRFRIEDMIISCQ
WDSNPCGPQNFSQVVTDYGICYTFNSGQGQHPLLYQRASGSFHGLKLILDVQQYLYPTYPISIQAPDAGIRLSIHSFNEIP
NMDSQGVFVPPGAHSYVSITSTHLFTSLKPPWGQCGDKKLKYFSHYSRSACRREFEVDIAADMCNCSYHHYHHYNHQ
SKKPICSTRQLFECVLPKLTNGEGKVNFNRYCPHPCQQSDYPMVLSYAGIANYATLQMAHNNSVLANLMRKQKLANA
TAQSFIRENLVYLDVYFRELITSLTIESKATGYAQVLSDIGGQLGLFVGASIITLCEIITYFCDRCKRDKREKQTKKNNALNM
TNKPKNKESAIKRGNSSSIIRNDSYENSENKKPPASINGTRNNPGVPV

>sp_O35240_ASIC3_RAT_Acid_sensing_ion_channel_3_OS_Rattus_norvegicus_OX_10116_GN_Asic3_PE_1_SV_1_675_209

MKPRSGLEEAQRRQASDIRVFASSCTMHGLGHIFGPGGLTLRRGLWATAVLLSLAAFLYQVAERVRYYGEFHHKTTLDE
RESHQLTFPAVTLCNINPLRRSRLTPNDLHWAGTALLGLDPAEHAAYLRALGQPPAPPGFMPSPTFDMAQLYARAGHS
LEDMLLDCRYRGQPCGPENFTVIFTRMGQCYTFNSGAHGAELLTTPKGGAGNGLEIMLDVQQEEYLPIWKDMEETPF
EVGIRVQIHSQDEPPAIDQLGFGAAPGHQTFVSCQQQQLSFLPPPWGDCNTASLDPDDFDPEPSDPLGSPRPRPSPPY
SLIGCRLACESRYVARKCGCRMMHMPGNSPVCSPQQYKDCASPALDAMLRKDTCVCPNPCATTRYAKELSMVRIPSR
ASARYLARKYNRSESYITENVLVLDIFFEALNYEAVEQKAAYEVSELLGDIGGQMGLFIGASLLTILEILDYLCEVFQDRVLG
YFWNRRSAQKRSGNTLLQEELNGHRTHVPHLSLGPRPPTTPCAVTKTLSASHRTCYLVTRL

>Deutero_Ambulac_Apla_gbr160_4_t1_678_211

MMYRNRSADEAHLNNSTLSGWPTIGSVYSLPTTINEGKVTATKTPAGGKNGAVGERLGGFGHASTFHGLNYVMNNSL
SRNRRLVWFIVVLGMTTLLVVNTIQAILVLLRHPETSAISLNYVPRITFPAVTVCNINLLRNSALDESVIPTLAAIYLEDRSDS
TGFDFGPVDALYHYPNGSDSLETIFNASHQIEDMLFRCRWRHEKCSAANFTRRLTDHGVCYTFNDPADERDALQVQN
PGSSNGLFMRLNIEQDLYTYGESTSAGIKVLLHPQGEFPIVKEFAFSLTPGFETSIAVRKHMVRSLRAPYKSNCTESTLREF
PFKYSVAACHFECLVHFVVEHCGCRDYRWPAVKAPICSVEKQVTCVYHYEDEFVWDSGNCSCPMSCETTTYESKISQA
QWPAKYFDRDLQEKYNLSHQYLRDNYVDVYIFMEEIMFMEIEQQEAYTLEHFEGDIGGYMGLLCGMSLITVVEWVDFI
IITIYKRFIGSKWMCSRKTRVMSSDDAGELETFGEEPSYAWRARRGASLLGLAQLSLAGT

>Protostome_Lophotroco_annelid_Pdum_comp409581_c1_seq8__679_212

MFVIRQKPTDHQNETSEKGIIAEFCGSTTGHGYYKIYSARHQPLARFFWIFLTFFAMVCFIVHATSLGLRYVSYPLRSDQIT
QSEEVHFPDITICNIYPYDVATWVLNDLEPIKKYRSELDCLRANHSSIGNEKQWNTLKSITGIYANMKPEEFSQLGHKKED
LIKDCVYGGKPCNMDHWKLMPHPKLVNCYTFHPKHEGLVYRQKSLSATVYIQSDDISQQCLHSSNYQIGSAHGLRLSIH
PPGTAPRKKHAMNLPPGVLVLLPLTNTKSTRPGHPYSDCVTDRNLVALKEDLNYSLTACSSLCQQQLVQRRCNCSSTGE
LIPDRNDPDLCKKVIPSDLDLTFKKMTCEDEADEEAAEMDTVTECGCTWPCELWSYKVVPSTAVWPTESATLDILKTYV
SATNLEKCLEKEAGPMPPELKTIDHSSPIPAEYHSYLYKANRWIKGHFLRINIYMRDPRVVEYRYQAAYGLQEFLSDLGGS
LGLWMGFSILTIVELFELIPRFLIRFSNLTTTTQPTGGAVHNYSKENASISKISH

>Deutero_Ambulac_Sakowv30041176m_681_213

MTDNKLETPIRGTFCGLVQNQMSNTTAHGVPRIESSTNLLRRLYWVVVFLGGVSLFLWQCSTIISKFYDRETTVNIDMK
FKPMLSFPAVTICNLNPVKSPDATPPTLQDNSTANPIEAITNKEFTMPYKATTDTGTTADYSRNWESKHVAFNEKREGF
KLREKVQNYMASKDYEDRSAYGHSLNDLLFDCTWKGYSCSPSNFTSFFNHMYGNCYTFNSGSLEETLMTSKAGPLYGL
NLELYIEEKEYISDLQESSGVRIVIHDQNEMPFPEDDGFMAAPGFLTSVGLRMIKVTRQPDPYSTCKTDADADATTNIFS
ELFSTGYTRKACEKSCFSWAVTEHCMCTDIIYKYHSYETVCNSTDPDIVECQEMVSLLNDNGELSCNVTCAQPCEQRLYE
ATVSNVVWPNEQYKSALTEELFKFSDDVKQKLNDNNNFISENMVKIDIYYNELNYEYIQEQIAYDNGDVVSDLGGQVG
LWLGVSVITCCEFIEFMIDVCILLWKKLTKASNLKVTPSKLKETKGFSEEKSGMPKEFQMVNT

>Protostome_Lophotroco_annelid_CAC9667732_1__Ofus_G151678_Owenia__fusiformis__684_214

MKKLLEDFAGNTTAHGWGNVIKSTTKIGKAVWIVITSGCCIVALYLIINVIIRYGQFDTKDRVRVTKDDAVFPSVTICPLSP
VSRKGVVNYGKDVKDNFELSAFNNLMNTLEQYDLNISNLLDASQEVQMLYNQSFKRLNTNSGQYENFQLSQLLRYTM
THEDFIPSCKYQGNKCKEEPYRSGHHAHFNCFTYNGPSSNIPGLTSEIAGPTAGLSLLLFLDTDEDFSTSFDPDYPTSGSQ
GVRVVIHDRGTLPDPENDGFDIKAGNSVNVALSAKRRELMIPPWGECVDHNDIRYDVWKYSTKACTSICLQRMIYKTC
GCILGSLPVDDQTKNAEFCANIQAVEWLKEDPNVTILRQDLEQLNCSMQEYTWTDVLAQEGCECKENCSNNKYDITISE
SQWPMKGIEFNFYCKLIVKLDNYENSSIYDVFHQRISQQCSNNSMENIRVVNKQVGDDLREQFIRLNVYFKDLDTKITT
QTEDFTLGSMVSEIGGSLGLLVGMSVISMTEAFVLWYGIIVGLLTKFKRKSKVETLKDKMKN

>Protostome_Lophotroco_annelid_CAC9485189_1__Ofus_G014735_Owenia__fusiformis__686_216

MKRIVNKFASNTSGHGWGMVGCTNSKLGKLLWVAITLGSTIAAIIWVLTIVQTYAKFETKDRVELRADVDIVFPSVTVCL
LYGFSPKKIRATLTDPNSGYVLHNSFLKTIYTNHVVLTQMTNESILENIVGQARTNRGYYENLGPSEISSLGHNIGSLIPYC
MYQESKCTEADFRKLVHHEYKNCYTFNGEDVNLTKPIITEVGAQSGLSFTLYLDHKEGARVTYEKSRAMTVASGARVIIH
EKGTLPDPDSKGFDVEPGHLINVALSVDKRELMKPPWGECADHTTLHSTDFKYTRNTCKRKCMEKMIEKQCGCWKDL
LPVDLNDTRDDDLQACGKMNREEWLKTTRANVTVLKEDIARIKCQNEKYSWSEVYEAVVGCECKLNCTYLSYNIETSQS
VWPMKGSELDFYCIGFSYDTGYNGSSIYDEFHDKIGVHCSDHFEFENVSKNWGDELRANFLRINVYFKDTETKYIIEEEN
FPLVSMISEIGGVLGLFIGVSLITILEVFVLCSSIIYTLLGRNLEETRYMKKKICQSLKPI

>sp_Q9JHS6_ASIC4_RAT_Acid_sensing_ion_channel_4_OS_Rattus_norvegicus_OX_10116_GN_Asic4_PE
_2_SV_1_687_217

MPIEIVCKIKFAEEDAKPKEKEAGDEQSLLGAAQGPAAPRDLATFASTSTLHGLGRACGPGPHGLRRTLWVLALLTSLAA
FLYQAASLARGYLTRPHLVAMDPAAPAPVAGFPAVTLCNINRFRHSALSDADIFHLANLTGLPPKDRDGHRAAGLRYPE
PDMVDILNRTGHQLADMLKSCNFSGHHCSASNFSVVYTRYGKCYTFNADPQSSLPSRAGGMGSGLEIMLDIQQEEYLP
IWRETNETSFEAGIRVQIHSQEEPPYIHQLGFGVSPGFQTFVSCQKQRLTYLPQPWGNCRAESKLREPELQGYSAYSVSA
CRLRCEKEAVLQRCHCRMVHMPGNETICPPNIYIECADHTLDSLGGGSEGPCFCPTPCNLTRYGKEISMVKIPNRGSAR
YLARKYNRNETYIRENFLVLDVFFEALTSEAMEQRAAYGLSALLGDLGGQMGLFIGASILTLLEILDYIYEVSWDRLKRVW
RRPKTPLRTSTGGISTLGLQELKEQSPCPNRGRAEGGGASNLLPNHHHPHGPPGSLFEDFAC

>Deutero_Ambulac_Spurpu_015894_690_219

MRGKEKDTLGKSLTDFGNETTIHGVQYIVNKDNIIYRLCWVAICIGFLVVFIIQGSDIITDFLQWPYSTKIDIVGRPSLTFPA
VTVCNANMMRRSQLVDTRFEGLIALDGGVSGADYDYSWWFSSDFVNGFEQSSSSVQRKKRSPPASIDYDMFSWYDP
SDVSDFQYYADNWDTVTSGDDWTGFYMSSRADDYSDFVDVVNPTRAELTDYGHQLEDFVLQCTFDRRQCNISEDFY
VWQNRYYGNCFTFNSALSPANQTRTTGKTGGLSGLQLTLFVEQPEYLGILSPHTGAKVSIHHPDVFPFPEDDALELSTG
QETSIGIRQEYIKRLGGFYTNCTIDGDDTNFTSSEYIYSKAACKKICYQLHLSQFCGCVDNQFFDDYPTKCDVLNATQQW
CRQYVEDRYLDDELACVCPSPCAETKYVKTASSLLWPSERYEEHLLRRLNASSHSNAVRILASSELSSKNLIRVKIFYEDLNY
EFVEMVPVYTIPSVLGSVGGLMGLYIGMSFISVFEVLSLLLRICRIGFAKLFFNGERVLPIKT

>Protostome_Lophotroco_annelid_CAC9669404_1__Ofus_G17801_Owenia__fusiformis__689_220

MTDIVETGAETNKQRYSEEHLYPSSVVETGHQDVNDFDKNRQEENKSSIEIDQQEEHTNEFSCDVQKQESTTRNDMHL
REGVLIDTLLGQESHSNKDHVENKLELSVENKIPDDPNISASTSGAESSHNKPAIMATQGEQMKSFRRKISSSLAESAEM
TSIHGSKRILKAKGKLSTIKWTIIFLGTVSMAVSLIASVVVKFYSYPVQPVEDGISVTTQPLAFPAVTLCPFIPYYMRDINPK
VRGKLKNFTEKLWARVHMLDCSYKVIFQPDENYDANTIRKIDKYERGCLPLQDIFQLRHKLISSGWLVENLPIDYETFATP
IKVEDFILECTYNNKPCESQINITRTNTVYGQCFTLTVNEPNDHVSEIGPDKGIALILYTGRANPLSSVNSNKEVRDGFTTIL

STPSDGVQVVLHKPGTMPRPYNEGFYICPGRFSSVEITQETRTRLLPPYGECTDAGYITKTTYGYSYDVCVDQCIQERIAK
SCGCISPMFITPTKHNFMKTPFCGNIQGRIWERTRGGGPNFRSEKKTSAKFWAP

>Placozoa_HhoNaC5_TR3219_c3_g1_i1_m_8097__Hoilungia_hongkongensis__27_221

MNNSPDDSDDISLASFDLEEHFANVTSCHGMIHIFDHKTSFLRRYVWGIATFVALTSCIIGCIQLLQNLLSHPTNISVKIHH
TNKMLFPAVTLCNFNQISRTLISAKDIRHLNTVLSAYNHDHTIPKSVLKETEEYFKSKIHGHFDFSQYNKEIGLQKEDLILSC
TWNGQKCGPKNFTRIFTSYGNCYTFNGDSSKQALLQQHGRGAAHGLSLILNIEQYKYTPELMLGAPNVGIRCSIHYYKSL
PRMESQGIAIPPGAHAYAAIPGTEITKYQKKPWGQCGYKKLKYYKYYSMSACLHEDETLYAESICHCRDPRLPGKEKVCS
PTQMLECLIPAMARYREQGKNLSSCPEVCERVEYVPQISYAKIPSKIMAEEIALKYGLHNVRQQAISGGGLIDNSTSVKQF
IRDNLVFLDVFFKNLYNTTTTQGRGSSFLQFLSNVGGQIGLFIGGSFLSVLEIVEYVFDKLAQQRSRYRRKTGQNEKAEWK
KMYASSMIVHKDSDDCRETNQPPPYEGTLIQSDDPDSTEGFLNGPKYVHVHKVGKE

>Protostome_Lophotroco_annelid_CAC9674716_1__Ofus_G26148_Owenia__fusiformis__693_222

MAKTEKQFNPKEILREFSENATPHGPGLIANSKTTVGKLISVCILLTCTGAAVVNIGFQLNSFFKFQYKDTIQVDNVVIEM
PSVTICPTLGFSRMKYISYNFSGETSVNIWSAIPINYQLLKDHSHENFSQFEDLFITGSIALDNMLTEDRLDVGHDLDGLIIK
CRFAGRDCNTSHFGQFLHPELNHCYTFNGKDVNMDIDTRIKASTTDAGLHLKIFLDAFIPEILPYGSYFNEDVITSGNIGL
RVTVHPKDTITFPKATGFDIPPGYASTITLKQHKIERLGLPYSECTNRKLLEGTKYAYTQAGCQAQCFQKNLMKTCGCVSS
WHPIPPNNTLKICTKLDSENIYKEMVYFMNLSTSRGNMDILSKDFTHLGCLTKTVNEKMETPLNCEECLPACKEISYSKSI
SQAYWPDEVIQGFILKMILQSPERNTRGQQMIQKYINFSEIERNHYQITNRTHADTIRKNMVSLSISFGDLSTEVTKQKV
EYEGAQLLCDIGGALGLYIGVSFISLCEVGKLLMRLLRYAFCRTSTSRIEATSAEN

>Deuterostome_chordata_PetMar_tr_S4RG90_S4RG90_PETMA_Uncharacterized_protein_OS_Petromy
zon_marinus_OX_7757_PE_4_SV_1_695_223

MPINIICTIPLPLEEDSKLGIKERGKSGQDLKDMHSIEEDGQPQASAITEFASESTLHGINHIFAPGEFSLRRAIWACAFLAS
LGMFLYQSADRIVFYLEHHHVTQLDEQESHLMLFPAITLCNFNSFRRSKVTPVDKHYAGPILGIPDNDPDLATVDVRLLR
AWSRYSTYEFYDRAGHTMDEMLLRCRYRDRECGPTNFTTVFTRYGKCYMFNSGKGGRPLLTTLKGGTGNGLEMMLDI
QQDEYLPVWGETDEISLEAGVKVQIHSQDEPPFVHQLGFGVAPGFQTFVSCQEQRLIYLPSPWGDCKADPINSDFFDT
YSITACRIDCETRYVVENCNCRMVHMPGDAPYCTPEQYKECADPALDFLVEKDNNYCVCDTPCNTTRYGKELSMVKIPS
KASAKYLAKKYNKTEDYIRENVLVLDIFFEAALNYETIEQKKAYEVAGLLGDIGGQMGLFIGASILTILEIFDYLYEVLKDKVC
KCRKKHNKKRSENMVSLSCTINKIHLYDTPCETLRSHHSQDVHFNSSILPHHTVRGTFEDFAC

>Protostome_Lophotroco_annelid_CAC9640639_1__Ofus_G092493_partial_Owenia__fusiformis__697_
224

VSYGYLKKNISLVRLFFPEFSMTYNMKKVVEEFADHTTAHGWGNISQRSSTLGKTVWIIFTLGCTCLAIASIIGTISKYFEYE
TKDIVELRDEPLEFPSVTVCPLNPIAHTHQLMYRQNIYYEIVDYHNLTNNILPTERKTPQHNTLYKDYINQLVSYQGYYEN
AMQILVYSHNKDDFITSCTYMENMCNRSDITTSLHHEHYNCFTYSTSRSLPDFAKTTVGPNSGLSMILYLDVSKLSMTLY
DPNYPTSGSNGARVVIHERGSLPDPEKDGSDIEPGHSVNAALSVNRRELMKKPWGNCVDYSALDAGGFVHTMNSCIK
RCQQKRVYEECGCVKASLPLEPDLENAQFCGKLDVKEWLKNEPNITILNEDLNRLECQKSAWRPCKTCAKNCTYHTYDV
SLSQSEWPTDGAVKSFYERWIMQQPNYENSTLYHRFHREMKDLKDIYSVTETTKSAIKNNFVRLNVYFKESETKVTTKA
QAFDLANLIAETGGFLGFYVGISVISIAEVMILLYNLLNFLRNKLIQQKYNNTNNAGVNIKQNGK

>Placozoa_TadNaC5_MK547546_15_225

MTTNPSDDSDDSNSLASFDLEEHFANVTSCHGMIHIFDHKTSFLRRYVWGIATFAAFTACIIGCINLLHNLLSHPTNISVKI
HHTNKMLFPTVTLCNFNQFSRTLISHKDIRHLDTVLSAYNHDHTIPKEDLKEAEDYFKSKIHGHFDFSQYNQELGLQKED

LILSCTWNGEKCGSKNFSRVFTSYGNCYTFNGGSANHPLLNQHGRGAAHGLSLILNIEQYKYTPELMVGAPNVGIRCSIH
YYKSLPRMESQGIAIPPGAHAYAAIPGTEVTNYQKKPWGQCGEKKLKYYKYYSMSACLHEDETLFAETTCQCRDPRLPG
NEKACTPIQMLECLIPAMSRYREQDRNLSSCPEVCERVEYVPQVSYAKIPAKIMAEEIASKYNLHQVHKKAIIEGGIVDNS
TSVKQFIRDNLVFLDIFFKNLYNTTTTQARDASFSQFLSNVGGQIGLFIGGSFLTLLEIVEYIFDKLAQQRSRYRHKTSQIEK
AEWKKIYASSMIVPKTSYHNGNANHPPPYERTLVQMDEYDSTEGFLNGPKYVHVHKQCKE

>Protost_ecdy_cyclo_nema_Cele_sp_Q22851_ASIC2_CAEEL_Degenerin_like_protein_asic_2_OS_Caenorhabditis_elegans_OX_6239_GN_asic_2_PE_3_SV_1_699_226

MRGGGFVQIFKDFSNWSTVAVVPHVANANNKISRIFWIAIFLFVLGMFAYELYILIAKFFSYPATVNTEILFEKQIFPVVTV
CNMNPYKYSVVKSNSAFSSVNTLMTTYSDATVGTFSTDKWGLYADNDETYDLDSRAADALVLEANLISDTAKVPALYTY
ADLIQDCSFAGIPCSESDFTKFIDPVYGACYSFNEDASLNYSVSREGIQFGLKLMLTVTQTKTNGNTDSLPTTKLAGARIG
VNSRGSSPGLDSNGIDAGVGYESAVSVSLTQNVRAKKPYGTCVDREPDSSDYYKDFIYTLETCFNGCKQRDTIAKCQCA
NPRLALGSTDTACQPIKADLDCLQTLKGNQTSSTPNIDLLVECNCNPPCDESTYTPTVSLAQFPSTSYYVATSSTAGVGSC
SSTNSKFSSKSDCQKWYNNNGMIIQVFLETLSYELYTETAGYTVSNVINDLGGQAGLWLGLSVISVVEMTGLMLVMGA
FCVTGGAIKMAPDDDEIENDHRIKDVEDVKKEIDHLEKKHGEMESGSDGEVDDIENKGDEEKKKK

>Protostome_Lophotroco_annelid_CAC9618795_1__Ofus_G081188_Owenia__fusiformis__700_227

MRTLIGEFAGNTSAHGWGKASQTTSKLAKAAWIFTSSACSIVALFLIIKVVIKYCEFNTKDSIRVSKDEAVFPSVTVCPLTPI
SRKGMATFRKELAENNSAIDSMLYYNNFLYVLGQYNLNISSLQNASEELQLLYNQSYKRLHTNAGYYENYQQSERLRYTI
KQKDFIPSCKYQSEVCSEELYHSGHHEHFNCFTFNGPLSNISSLDVDSGPSAGLSLILYLDIDLEHSSFYDPDYPTSGSQGV
RVVIHEQGTWPDPENDGFDIKAGNSVNVALSSERRELMGQPWGDCADHNDIKYDRWKYSTKACLLTCIQRLGFKRCG
CIRGSLPVDDKTKDVEFCANIQSREWLKSDPNVTLLRQDLNQLNCSLQEYSLSDILAQEGCECKENCKIDKYDIMLSDSE
WPMKGIEFHFYCDMIMKLGNYNENSSIYQAFHDRVASNCSSLAYDVERKKIINKDIGKDLREQFLRLNVYFKDSDTKVT
TQTEDFSIGSMVSEIGGSLGLLIGMSVISISEVFILCLGILAILIRKSKAQTKVETFKDKMEYP

>Protostome_Lophotroco_annelid_CAC9667655_1__Ofus_G151630_Owenia__fusiformis__703_229

MKRILKDFSSNTTAHGWGNIHQRTTTLSKVIWIIICLGCTAVAIWQVITIVMRYGLFETKDRIKVEVGEIVFPSVTVCPLIP
VPHDGNLKFERDMKKGNIDVEPFLNFRLVFHSETFYINSNVSHNLHSEKLEFYNVFAEQIYSNQGFFENFKFMSDYTHQ
QGNFIPVCSYQGKPCLRSDFHTLDHPEYNSCFTFNGANKTINNPITKITGPKGGLSLVLYLDVGQLSMIYTPSYPTSGSQG
VRVVIHEKGTLPDPENDGFDIEPGHSINAALSVNERRLMKPPWGECNDQNPELKGGFTYTKKHCQLSCLQKFFYRTCG
CIKSSLPIDEETVYKEYCLKLHAKEWAKLTTDQYNKTTIVSDMENFRCQNREFLSSDVEVQDGCQCLEKCSYQTYDMTLS
QSEWPMKGVEIEFYCRTIMSMDSYINSSIYDRFHDIFPEYCDHSKDEDLLTKIQKETLRENFIRLNVYFKDLETKITNQVED
FTISSMISEIGGSLGFFVGMSIITIAEIILLCWNITCLLGKKFVISPFEQKIGSDMKHASTNTS

>Deutero_Ambulac_Apla_gbr361_5_t1_704_230

MYHQESGSQEGLHLAKRKVSPMTVEDISAEDGAAPAAPPDPAASAKQTSLLYVVSHRLAESGAHGIPNIQRANSNFRR
VAWTLLFLAGFGMFIYQGCDLIIKFYKWPYNVNIEVRTPKRVEFPAVTWCNLNPIRKDALADFTDLRSLLEGGPSSCGTA
VNATFGGYPDDWESADIDYKLEDCTSQTVDAAIDIVGAMPYGERAVTGHRLNDMLLMCTFQQKPCSPRNFTSFYNSR
YGNCFTFNSASRGVAPLKVTRTGPSYGLSMELYIQQDRYITGVETGAGIRIMVHNQTEMAFPEDIGANIAPGTESFLGLY
RAMVNRLSDPYGDCATDFTQDNIFKEQFKRLEYTKEACEKDCFFRAVLLDCNCAYINYRYNDSQAPCNSSVEAVKTCIEK
VENNFTSGELRCGHCHKSCNETRYQISFSNSKWPNREYMETVRKNIQESNKETISLIKNNSYFIEDNVLRVNVYFDSLNY
DDIFQTAAYTLGSLVSDLGGQVGLWIGVSVLTLFEFLELMIDIFVLISTRCRAGGENRRNGKTGQENSRPQA

>Placozoa_TadNaC2_MK547543_12_231

MDTHPWQRPRSKSINLEKKFAERTTCHGLGHIVDQDVPKVRRYLWSIVTLAASVGCMIGCIQLLQYVLSFPTNIDIEIIHQ
DSLIFPAVTICNFNQLTKTALSEEDNRHLQTFLRIYHRKGAINKKDLELATSYFRNKTGVDFHLQNLTQDLGHRKDDIIVSC
LWDDQVCGPENFTTIFTIYGNCYTFNSGAKKEGMLSQRGKGSAHGLRLVLNIEQYKYSGKLSYGSPDAGIRFAVHSTAD
LPEMDAEGMSIPPGMHAYASIPGADVIEGLPKPWGQCGSQKLKYFDHYSVSSCRREKEIDFILQRCGCVEPHHARNLTP
CSPEIMLECVLPLMSSSNPVSVGINASVCPVACVQTEYNVEVSYALIPSQVVVNDISDQYNVTKIIENARNQRLNLTMSK
LEFIRENFAFLDVYYKDLYLFKTIQKQASGFVAFLSDIGGQLGLFVGGSFLTMFEFFEYIYDKCFQQTKRSKREISRRMQSIR
EKRRPESMASSTSVRGNLSTNVHNFKVNESISSHNVRSNGIKSKFRRSNSEHLANIRISPSSID

>Protostome_Lophotroco_annelid_CAC9628722_1__Ofus_G09545_Owenia__fusiformis__706_232

MSLTLLLDHPVDDRGTISHYMSLITTHGCQQIYQTKSKLIKICWTIFLLASVCVATYFVSVAISKYYKYRTHEETSLKDTYVD
FPAVTVCRLDDDGVSRLFSEHYTPDKGDEEIIKQEMNILMKIIDIDLDFENEDVSYLHLLLLYAEYAPRLLDRQSIPQAFPAI
AEYLASDYDRFIVSCQFKEHDCNKTHFVEKKNGQYFNCFTFNHTDTIKKEGPTSGLSLVFFLDTLKEIELQEISLGDSSLRFT
DSGIKVAIHEPNTIPDIINDGFIVAPGYSTNIALKQSVFEKMKSPWGECDKSRIKINDITNENTDILYNRETCMKICQQRYI
QGKCGCLNVMLPVPNDLNKEKYCFYIDKDNMLQNPNWTQVFHEELKHVDCESKHLLDLPDSYVGQYKCDCPLSCHYK
TYNSLISQANWPPQNGVHRFIKKYRNEQPTLLNGTDIYKAYEENNATGDLYDMVQNNFLRLNVYHRSLTVQTTTQVAA
YTVQDLVSEVGGIIGICVGMSVLSILELFQLVAIVIRSSGKKADRRGMSLKDGDMELNEEGV

>Deutero_Ambulac_Spurpu_015895_708_233

MDTEEKDTVKKSVTDFGNETTIHGLQFVVNKKNIIYRLCWLGICTTFLVVFLIQGNVILKDFLRWPYSTKIDIVGRPNLAFP
AVTVCNANMMRRSQIEGSRFEDLVNLDGGVEGADYDYSWWFSSAYRNWYASSSASSYGQSSDQNSNGRSSSSSQSD
SASSSDGQSSSSNYDPSSSEESSSASSGESSGVTSDGQSSSSSYSPSSSEESSSASSGESSGGTSDGQSSSSNDGPSSSEEP
ASSSPGGSADGQSSSTNYGPPPSGSVSSFGLPNSEFPAPNWVNEWSWYDPSFFADFEFSENGWDGVSGGEDWQGF
YEASKADDYSDFLNVINPTKEELEQYGHQLEDFIVQCTFDRRPCNISRDFHVWQNRHYGNCFTFNTELSPDNKNILTGK
TGGLNGLHLTLFVEQPEYLGVLSHQTGAKVTIHHPNEYPFPEDNALSLGTGQETSIGIRQEYIKRLGGYYTNCTSDGKDT
NFTSTTELSYSSVACKKICYQLHLSQLCKCVDDQFYDGFPTKCDVLNMTHQICRKFVEDLFLDDKLPCSCPPPCT

>Deuterostome_chordata_PetMar_tr_S4RTA3_S4RTA3_PETMA_Sodium_channel_epithelial_1_alpha_subunit_OS_Petromyzon_marinus_OX_7757_PE_4_SV_1_707_234

PPTVLEFWRGSVTDFYDSYDEMFEFFCDNTTIHGTIRLVCSKRNKLKTAFWSLLFTVTVILFYYTSALVFLQYYSYTVAVT
MGLMFQQSTFPAITVCSLNPYRYEVVQSSLSQLDSMTGQALQQLYGYQPPATKAAGTAAAPGIRLDTGVVLERTGPDT
VGFKLCNATGGDCFYQSYGSGVQAVTEWYTFQYVNIMSQVPSYIKQSDDANIEDFIFSCMFSGMPCSDSEYSRFHHPT
YGNCYTFNSANSSKLWQASKPGRDYGLSLILRTEQNDYIPFLSTVAGARIMVHDQESPPFMEEGGFDMRPGFETSLGIR
MLEATRMPDPYGNCTEDGSNVPVLNLYSSAYTVQVPECSCFQLALVEACGCGYYFYPLPPNASYCSYNNTAWAGHCY
YKLYRQFISDELGCVDKCAQPCTTKRFAVTPGYAAWPDSSSEKWIFNLLSLQNNYSVTTVRNDVAKLNVYFRELNMKTI
SESAATNVIWLLSNIGSQWSLWFGSSVLSWLEVGELGIDCCIMVFVLAYRRRRSRAERRRARGTGDSEAAPPPPPSF

>Placozoa_HhoNaC1_TR8933_c0_g1_i1_m_19478__Hoilungia_hongkongensis__29_235

MSKAFDDVRLTSYEIDEVSSISYEDQIDSDDVFESAESAMPTSHHTEHLNISTKDKFEFSDDSNYDDRFALQTSCNGIIHIF
GRGGRIRHGIWFILTFTMIIFCISTCIQRYDYLISRPTSTVINYTVSDRLKFPAVSVCNFNRFRFSSLEYDDWHRIGYLINLFT
STDSNDIFSGLNGKTGEEWNDYLKNISIELYDNITFDITQFLNVKSNQAKIFIKHCTWNNGRQPCSINNFTRIYTDYGSCF
TFNAGVKAPILYQDRAGSRHGLKLILNIEEEEYTHLNPDPDIGIKFRVHDQDEPADINAEGIAVPPGYHAYTKLEYTKSDFL
KPPWGNCGEVKLKYFKSYNRASCHLECLADSYKGQCSCRTPYMPGPFPICSPEHIKKCISKYTGKNRSRNVTCHCPNDC

QIKSFHPQVTYAEIPIQHITAATAHRYGINELELEFLEYNNISINDYLRDNYVFLDLFYDDLSYTTFKENKAYDENQFISDIGG
QLGLFVGGSFLTFWEIFEWSQIKAFLVIRKIIHEYKKGRRRTRKRFNSTPEDTERLL

>Placozoa_HhoNaC2_TR15906_c0_g1_i7_m_30554__Hoilungia_hongkongensis__22_237

MDTWAKPRKESINLEKKFAERTTCHGLGHIVDQDVPKPRRYWWSVVTLAASIGCLIGCIQLLQYVLSYPTNIDIEIEHQD
SLVFPAVTICNFNQLTKTSLTEEDNRHLETLLRMYHHKGSVDKKDLEIASSYFRNKTGTDFHLQSLTDDLGHRKNDIIISCL
WDGQVCGPENFTTTFTIYGNCYTFNSGKPNHSLVSQQSKGSAFGLRLVLNIEQYKYSGKLSYGSPDAGLRFAVHSVDDL
PEMDAEGMSIPPGMHAYASIPGVDIIQGLTKPWGKCGQRKLKYHDHYSVPSCKRELEIDYILNECSCQQPHHPGNSTH
CPPFLMFDCVLPALGNFRASNMESIASKCPVACIEMEYNADVSYSLIPSQVAADEISQKYNVSTIIENAQKQGINITSTNIE
FIRQNLAFLDVYYKDLYTSKTIQKEVGGFVSFLSDIGGQLGLFVGGSFLTIFEIGEYIYDKCFQQTKRKQRAISRRVQTFREK
RRTASMRQSPIHANLSGNLSNLKPNDDIKSSRNVRANGIKSKFTRSPHITSPTNPISPIRTDKNGIR

>Placozoa_HhoNaC8_TR7882_c1_g1_i1_m_16465__Hoilungia_hongkongensis__28_238

MSQSSDPDSGKVTSEDSSGEHPLQPRKVSLLMSSEKSHEDDSRNLLTLDFRRRPSIEHDQNFPFSSTVHGIPHIFDEKYG
TRTILWILLVATAVVGCFAFIIIQIVRYCQFHSTTKTSLIYAKQLEFPAITICNYNSFRRSALTANDLVHMAYLVKAYGLDDG
MLRKLLTKKEKEKLIEYWKKYDQTHTKKFNYQLFVERVGYHASDMIKSCHFGVEECGPKNFSNVLTTYGNCITFNVGGV
GKPLHQKFPGSNHGLKLLVNIQEYEYTGTYRTDRLDIGIKFVVHEKNYPPDVLQQGKAVGPGSHAYASVKYRTISNLPAP
YGKCSSRVLPFYAQYTFAGCHICCKTEYINDKCGCRAPDMPGENTIPVCSPQVMLQCVRPELVHFQSEEDKVCDCPIPC
FTGHYDTTVSYAKIPNPQMAKSLAVTMNKTEFIHETHGLVDPNISPTLYISQNYLLLNIFFDDLYYEETTSLPVTTFSSLLGN
IGGQLGLFVGASLLTIAELVEYGFYHSRGAIRRYQQKRNQRMSDIAKQTSCAEKEPLVENGKASLPS

>Deu_Ambulacrar_hemi_Ptyfla_40v0_9_20150316_1g20853_t1_scaffold21886_cov93_711_240

MSKEGLKHRQLPLKSGAYNVERGKANQQCQTDDNSEEDDSDVDDKQQRVDSRTCGDGQERSQATFKNVLADFGKG
TTAHGIVHITEAETSVSRTVWLAMVVAAAGIMVIQMTLLLVQYFEYGVTVKTFFRGINICTTIFVIPIEFLRDDFKDIMKRR
MFSMFDRQVDRSLSVNSTDILPSIPLYKCFESCLRWKRYSCRSFDYNRTTLTCRLFKESAGNSGGEVVKAIGTDFYQNLRT
RIPDTVFKPGWQMPYQSIIDDPDIYKYFKENVFINPGYHRVKGEDPPDWLRFKSFSKSPDYTELADVLKVNTNEIAAMG
HQAEDFILQCTYGGEVCHPRSDFVITQDEVYGNCFHFNADPNKDTRFTRGTGTAHGLSLVLFTEQYEYLGIYGQDSATV
VSIVPRNLRPFPVDHGFFAMPGTATSVSLTESKIHRQNQPYGNCTDENSNLHFTDNMKKDSSDKMYSIVDNLLKLVKV
KNQKTRNLNDLESARSNLVRLKVYFESLNYESTSEHPAYTVRILVCFDDVTVSSLGEFRSVCLRLKVNFDFVSLLLTF

>Deutero_Ambulac_Apla_gbr412_20_t1_713_242

MGSGDQKETFLGALNTLLETSSAHGLPNIHRSKSFLSKAAWTLLFFAGVTALTIQVISLVTTYLKFEYTVSLDVRFDRSLNF
PAVTVCNINPVRKSKLERASDGFRELFDVNFAPPRPEDQTGAVNPGDNTDMSPGSTAGGTMKQAGQPLTTQGLAGA
GMGQGKGQGGSGMPTAGATMPIAIPTGITTEGPETTAGIEYSNSTSSVNNGTTEATILAWEERQTGDFFKKRDDDYLK
EQRLISHLANLTETQRMDMGHLLEDMLLDCQWQGYPCSPANFTSFYHYKFGNCYTFNSFRFGLTLELFLEQEEFMPEIT
EAAGFRLVVHDRDTMPFPEDDGISISPGSKTAITARVVTIERLGNPYGNCTKKHDEPDDGNIFRDRYGVSYSLKACERSC
YQKEVISQCGCFDPHYPNTLNDTVYPCDIDNDEAQSCMSDLEEKYKQGTLTCYCFQQCNKNVAKIEVYFQEFNFEYIKQ
SPAHTIPSLMSDIGGQLGLWLGLSILTVFELFEHCGTFLAVIVSKLCREGGNLKSSSSKVQKIKIFPTEDPGITLQNY

>Protostome_Lophotroco_annelid_CAC9661516_1__Ofus_G111721_Owenia__fusiformis__714_243

MKRTISDFASNTTAHGWGNVTQRTTKLGRTIWIGLCLGCTGVAIWQVITIILRYGTFDTRDRIVVKNGDIVFPSVTVCPL
VGMPASGYDQLFQDMREGKQETIPLMRLTELLQNLTAYVDNNISQSAQSEKLEFYEEFHTQLYSFQGFFDNSKKSIQDY
SHKQDNFIPVCVYQGKSCLKSDFTTLDHHEHYTCFTFNGGNTTIAKPSTKISGPKGGLSLTLYLDAVEGSLIYNPNYATSGS
QGVRVVIHEKGTLPDPENDGFDIEPGHSVNAALSVNERQLMKAPWGECADHDSEAYGSFKYTKKHCLLKCLQKTIYET
CGCVQSSLPVDDDMAHTRYCLKLHLNEWVKLQSNEYNLTIIQNDLEQLRCQNNFSLSDFVGQDGCQCREKCLYQSYE
MTLSQSVWPLEGVEIDLYCRSLMNMDNYINSSIYNSFHDVFPEYCGHAKSEEENKILRDTYGIKLRKNVVRLNVYFKDLD
TKITQQAEDFTINSMISEIGGSFGFFIGMSVITIAEVIILCLNLAQVIRKQMVISPTHQEKEMARKNAWVETWPKQ

>Deutero_Ambulac_Sakowv30044073m_715_244

MNSTRELSHGTLTVESMDASPKFKKRMKRFTRRDIVNEFSRNTRCHGLPRIISAKTLPSRFVWSLVFFAALGAFVFQATK
LIKLYLNYDVTVTIEDETIASLMFPAITICNTNKLRFSEIEKSEHAVLLRTDPNHPNSLHRSLSYQGPCLKGDFECSDGIHCIK
PHLHCDGYIHCWDGLSDEVNCTYPPCGRDQFKCNGPGYYGICIDSDKRCDGLPHCLSGEDEAYCNECKSGFKCDDNDD
GQGGKCVQEEQRCDRFEDCKDGQDESKCDSVYADPCDKNLSATSMQENLYSPLYPKAYPVEKICTVTIIAPPNKSILITFL
VFDIEHNSFCDYDYLKIQDNMNSDLNVKLCGGLRPPPWISSSNVVTVTFRSDTEYTARGFHLVYEAVNSTQYTMRWEV
GPWSNCSKRCGGGVQTRAVMCNGASQTGEVTNLDECKAIGSRPVSTKVCKQEVCESTCQSRLTHCCQVIQSKGYPDQ
YTNGQNCYTNIINEGGCINITFSDFPLEQGGSCGDFVELSDHNKPTLYRRICDMETGDSNPTWGSFSGNVSVTYR

>Protostome_Lophotroco_annelid_CAC9674854_1__Ofus_G26524_partial_Owenia__fusiformis__718_2
45

IQEMTVKSLLSDFSKTTTGHGWGNIHERKSMCTKVAWLIFCIASTLASVGHVTSIIYRYSLFETKDRVLVQRQDLLFPSVTI
CPLTQMSKQGILELTQDSINGEDTYLSIFALQTLLADMVKIEQNSTLLMEASSERQQLYEKFFNRLYTYNGFYENVPTVTK
YGHKKHDFISSCMYQGKPCSDEDVHSVIHSHYYKCFTFNDINTSLSNPYSNTTGPTGGLSLQLFLDTDENLYAIYNPEYPT
SGSQGVRVVIHEKGTLPDPENEGFDIEQGHSANVALSVTRRELMKPPWGDCTEHPQLDEDGFTFNTRVCQMRCLQK
MVYKSCGCKRGSLPMYPDVENVQFCNKIQYNEWIKKNPNITALLYDLEKLQCGMTEYNWDVVLDQDDCQCQDNCSF
HTYDMTLSQSQWPKKGVELDFYCRKIIHLPNYSNSSVYKAFHDQIEQLCTDIYSNASTTKWNEVKQEVGDRLRENFIRL
NVYFKDLETKIIKQEPDFALGSMISEIGGSLGIFIGVSIITIVEVFILCLSIIRVLSQKSKVMSIQDPNGVNHEKQWP

>Protostome_Lophotroco_annelid_CAC9539641_1__Ofus_G033690_Owenia__fusiformis__722_247

MEGNKIRETFVEFTENTSAHGCGQIPTAKFKLTKIFWGLTFLGCFTWATYSTGNLIARYLSYPTKDVVSIDFSEIQFPAVTL
CSLQPLTLSGMLNLQENPLPPDHPTSILLYVMLNFKIIDKNIGGLGKRYQRQLSRIVSNQFIYENSDEKNTEGFIHDLRSFL
LGCKYNSKTCDLEKLFITYQDGQFFNCFTFNSELLVPKAFIVEKTDPSSGLSLVIFVDALSSAPAHYTIYNPDDPTSGKMGV
RAIIHSPGTRPMPFEKGFDIPTGFSTSVALQGSIRNLMSEPHGNCTTETLIPGTNYTYSSDTCVQECKQEKLIKECGCKSSL
LTASADKPHVPYCGMFNMTNILRQMHYPTDDEDITMAVRDLERIECEGNILRLFAYPAEINKCACKEACSSMKYTKTIS
QSVWPNDNNQNIFYETYVNVTDHTLRPNILFKQQNITEIINNDLIHKNFLRLNIYFESLQVETTSEVEDYPLSQLISDIGGN
MGFYVGISVITLLEFLSLTGTVLLYFFKDRLASCGQSKTTKVSQINIQHGDSKLDHYTDGFKDKY

>Protostome_Lophotroco_annelid_CAC9657917_1__Ofus_G102514_Owenia__fusiformis__723_248

MGISKMDGKETKKKAKDLVTEFADDATAHGFGLIKRSGNKYSKILAIAVVLTCATFAVKNIWEQIKRYYDYHYIDTISIEER
AIEMPSVSICTALPYPMTIVFQEDIILRAPSLYRYASLTSYGYLLKNLLETSVSDFKLQDGYDYIFENVLFSQQMGNAHVPD
QEMAYLFHNINNLVIQCRFAGKPCNSSFFTEFRHPSFGRCFTFNGAGLKMDQDMTIRSTDPKSGLHFKVFLDGYKPIPF
PNGMFYDNKDPLGGIVGLHVTIHQPGSVPFPIEDGFEIPPGYATYVTLKRKKRTRLGKPWNSCEDQLFLNGTDFAYTKA
GCVSQCYQEKIMHMCGCVSYDFIIPKNNTMVFCEHLDLDNMFKLIQRQYGSRNLGNISILLQDLDKLKCLVNIKNMRLP

CTECLPPCVETDYEHVQSQAYWPGDMTLGYFARSIYERSDIDLNENNAWKFIETHIINYTTLAAKEFQISNITQSAMMH
KNFASLIVYFKEMTTEVTKQEADYNVSQLICDIGGALGIYIGASLISLYELGKLLVDLCHIVLHRNKNQVEEIKT

>Protostome_Lophotroco_annelid_CAC9662966_1__Ofus_G12887_Owenia__fusiformis__724_249

MNKLITDFASSTSAHGWGTIGQRGSKCGKLLWVIFTIACQIAAVVWVAIILTRYLQYKTMDRIKMTNDYQVKFPSVTVC
PLNPTPISMLGQWLLDYKKPDSELLKLNDYYALLFKLSGYLVEGIEIVDEEMQSLASKYNQLDSHRGYLENIPYLWKYGHE
QNNFIPGCMYKKKTCVNESFERIHHTAFQNCYTYNGATLNDMYVTSQGMEGGLSLIIFLDMGSEGLSLYNPYHSDSGS
SGARVIIHEKGTMPDPDNDGFNVEPGLSINVALSVNRRELMKQPWGQCKDEFSLGLGKFKYSKNACRQKCRKDLICEA
CGCVYETLPMLNVTLARDEVLCGKINTDEWLKNNPNITLLRNDLDRLECLEKYLNYTWREISEEQGCECRENCTTHSYNI
ETSQALWPIEGSESGFYCRQIMMLTPNYINTTIYERFHERVSDCCSNKTRGKDIGDELRKNFIRLNVYFKDLDTRVIIQDQ
DFSFWSMLCEVGGILGFFIGVSIISIVEFTLLFLNIFNYVTFKRSKINNHVCSGNDDIDHKRKKTEHSTGLTFYQEHK

>Protostome_Lophotroco_annelid_CAC9672326_1__Ofus_G21693_Owenia__fusiformis__725_250

MPADDENEEKPKENNSGNARSILSNFASSTTAHGCSQINQSQSSVAKCVWLLIFIGCTIGAIICITALLQKYFAFPTKDVIV
LKTGAIAAPSISVCSLNPIPSSFRTRLSGIAPDNKVNTVLRALDLASERVANTSFGLHEKLSESYLRLNSYIWIYENAPGENE
TLQFAHELDDLVVSCMYQGIPCDKSTVKIFTNPYFYKCYTFNAMDYNDTYNGPFIHSTGTKSGISMLFFLDTHDSGETLY
NPYCPLGGNTGVRVVIHQPGTIPDPFNDGFDVAPGYSTNVGITKRKRELLGRPWGDCERREKLDDGLPFLYDRRSCKLL
CLQAFVAERCGCVTSYLPISENMKNITRCGQMDLENFNSASPNLTTLEHELDRLSCELKSVYVILDDQDDVDCDCPQSCE
RDIYDYTLSQSEWPSGGVQLHFYRHIAHLKGEAYFNSSMYHLLDDKVNTNDSMVNMKNQRMIQQNFLRVNTFFTSM
DTEIVKQVEEYPFTDMISGVGGGFGVYVGFSIVTMCEFCVLFVHLLRAMIMDREKRKVKSAELHSSKVIPTKTAW

>Protostome_Lophotroco_annelid_Pdum_comp402494_c0_seq1__729_252

MDNNADFNKNRNCRNVRPQEKLMYLSHRLTEETTAHGIPHIARANGFWRSLIWIIISLVMLVTWIYHSQYTISQYLKME
VNVKLEFKTAQNLPFPAMTVCNKNAFKNTQGMWLMERMIMRFNTPEGKNASYPPWLISYLKYLATFNGTDEPEAGT
VQTSNPTQLGEEMMQLLYDSIIKYGPEYNLTVDDLTYSFKDFVLTCDYERTPCEKRGVKINRIFNWLHGHCYVLNAEKAT
LLSTQTGPRYGLKVALNIDQDEYWEVAEQAGVRVLIHDPDQMPFPEDDGYNIPPGLAASLGVRLVKQSRLGGSYTPCF
NEKEQNEKHDIFRAHYGWTKYSLKTCMISCYQREVEARCQCSDIRYPAYKKGGGKCVSCTENMTMSFGAEMKTHNPN
ECLRFVQEMYRNQSLNCDKECLVPCSQNSYVSSVSYSGWPSTKSLETVTQNLKKIPSAAKLLDESIANIYKNFVEVTLYYE
DFSYNAVSERPAIRLTECLSDLGGNMGMYVGASLFTFFEFFQYLIDVFLWSCVWCCKSLTRNGDGHLKELFAATKVRKI
QVKPEH

>Deutero_Ambulac_Spurpu_018446_731_253

MDAPKVNCEESSWGTAPEREEKSLRTLLNSRMENSSAHGIPNIQRSSGLVTKLAWSLIFLAGIGVMTWQAVILFQTYFE
WKYSVDIEMRFNRTQSFPAITICNTNPVKRSELETRDASFRRAFDVHYVSSTPDQQPLPDVPDGVTPSMLNNSDSGNE
TQGGSTNEQVDMSEAVNDWRSRIAIPKFYRMKSEDYGKKRIRVVTLANETLEERVSLGHKLDDMLLDCSWKGIPCSPE
NFTKFYDSILGNCYTFNSGKNGEQLTTNRPGSTHGLTLELFVQQDEYVEGMTEEAGFRVSIHHPSKMPFPQFNGLLVSP
GFATNIGLRKLEVDRLPKPYGDCEADLSKNIEDDIYHQHYSITYNRKTCEVSCFQNEVISRCDCFDATYPNSLKVNHTVYP
CEYINDVETQCIADIEMEHARDELECNCPLACRETTYLTGASSSIWPSDAYESTLIEKMLKYNAEIRGHVVGENASDWTR
RNMAKVEIFYDEFNYEYIRQDPAYTIPDLLSDIGGQLGLWLGLSIITIFEFFEGAWLVLAFFCSRSGNKTNRSEIDPGTKTT

>Placozoa_TadNaC8_MK547549_18_259

MSQSSDHDSNKTASDESSTDAHPNSQKVPILSHQDEVDQDNPEPRNRFDLDFRWRPSIEHDENFPFSASFHGIEHIYEG
RYGTRKILWILLVAATMIACFVFIFIQIAHYSAFHTTTKSTLVYEKQLAFPAVTICNYNSFRRSAVTANDLIHMAYLVKAYRL
NQGVVSDFIGEKERQKLINYWKKYDATHAKKFNYQLFVERVGYHASQMIKSCHFRGLKCGPKNFSNVLTSYGNCITFN
GPKLESNPKLYQKNPGAHQGLELLINIQEYEYTGSWHSDRPDIGIKFVIHERHYPPDVTSQGKAVGPGSHAYASVKYKTI
SNLPSPYGHCGSKKLAFYKKYTYAGCQISCKTEYVQKKCGCRAPDMPGQNIIPVCSPQKMIECVSPNLEKLITINDKVCIC
PIPCHIVHFDTTISYAKIPNPQMAKDLTEKINKTAFQIETHGLVDPDIDPTLYISQNYILLNVFFDDLYYEKTVSTPVYTFTSL
LGNIGGQLGLFVGASVLTLVEIIEFGFYRSRGVIRRSDWKQNLRKSISRSREVTATEEKEPLCSVENGDTQLSK

>Protostome_Lophotroco_annelid_CAC9522438_1__Ofus_G031129_Owenia__fusiformis__736_261

MKRIINEFTSNTSGHGWGMINQTSSKFGKALWVAITLLSTIAAIIWVSTIIVRYVKYETIDRVEAKVDEDIIFPSVTVCPLYG
ISSMKIAEVYSDPNSDYISLNNFLSSSYALVDELAQITNETYLSIMTNTILVARSNRGYYANLGVSGVSTIGHEFGDFIPYCL
YQEKECSAADFQKLVHHEYKNCYTFNGGDINISKPIISSTGAQKGLSLTLYLENSENAYVAYNVNRVMTAASGARVIIHEK
GTLPDPDNEGFDVEPGHLISVALSANKRQLLKQPWGECAEHNTLHSTDYKYTRNTCRIKCIEKVIQRQCGCRYDLLPVDI
NATTELDIQPCGKFNIKEWLKATEANTTVLKEDLAKVICSKEEHQWTALAKDIPSCECEYNCTYVDYEIETSQSVWPMRG
SELDFFCQLSYLNISGSFFERFQNSIGIHCNSFMEYINVAQNHGDELRANVLRVNVYFKTLEVKYTIQDEGFSLVSMISEIG
GVLGIFVGVSIITILEIFVLCSGIVHVIINSKKVSSEINVKEIKSTSKSKLEESSNHKPDDSYYDQFKKY

>Protostome_Lophotroco_annelid_CAC9477543_1__Ofus_G013338_Owenia__fusiformis__739_262

MDTKKENMSPNESVSEKKKDSSIKSILSTFASSTTAHGCSRINESQSARGKLIWMAIFVACFIGATVCITSLLQKYFKFPTK
DVIEVSVDPISFPSVSFCSLNPVPSSYQRKKSRMKPGNRFLDVAATYNNLEAHLFNESFPLYDRMSPNRLRIKSYKWMYD
NAPNEDETLAIGHDLEDILLSCSIAGYDCNISQTQSFTSPFYFKCYTLNAKDFNDSSIEPLVHSTGARSGLSAIFFLDTHDKA
KGIYNPSCPIGGSSGLHVVIHPPGTQPDPINEGFDIVPGFATNIGLQKKSRELLGYPWGDCENRESLSELPYKYDKLSCER
QCQQKFITDQCGCVTSFQPIPDDLKNMSMCGKMDLKNWNSASPNMSKIAEEMDTIECEFGKFWVLKKDYPEENKCE
CPSACHNDIYDYTISQSEWPSGGIKLDFYKHVMHMRNMRGSGYINSTMYDLLDSKLNTNDTDENHKNQGMVRQNFL
RLNIFFTSLNTEIVKQVEEYPFTDMISGVGGGFGVYVGFSFVTICEFCVLFAHLLKAIMNHRNKAVEDIPSGNIITVKEYKM

>Protostome_Lophotroco_annelid_CAC9510306_1__Ofus_G024263_Owenia__fusiformis__740_263

MKQILNEFASNTSGHGWGMVGRTTNAFAKTLWIVITVLSTIAAIIWVATIVGRYIKYETKDKVELKADVDIVFPSVTVCPL
HGYSNDKMNDPKLVSSFLLQSRLIMAVYPRIKNITQMTNETGDVILELVHRIRSHRGYYENMGKRETSALSHEIINLVPYC
LYQESRCTPEHFQKLVHHEYMNCYTFNGKDINLTKPVISSSGAQNGLSLTLFLEQKDDTYAPYDRSRAITATAGARVIIHE
KGTLPDPDNDGFNVEPGHLVNVALSVNKRELMKPPWGECADHNTLQSTDFKYSRNACKSKCLEKMIQQECGCKKDLL
PVELETRDDDIQACGKFNREEWLKVPNITEANITILKEDVTRFKCGEKEYAWSKISEDIPGCKCEYNCSYLTYDIETSQSLW
PMKGSEMDFFCGGLAYEDTYVGSTIYDEFHDKIGMYCLNYFGQYANISKLWGDKLRDNFLRVNVYLKEPVTKYTVEEV
DFTVITMVSEIGGVLGIFIGVSIITILEVFVLCNGIVYTLYRAKKTSSNIDVTEVIVTQSKNSENLSDPKTKYYQSKLSK

>Protostome_Lophotroco_annelid_CAC9574840_1__Ofus_G053378_Owenia__fusiformis__743_264

MMKEQNKMDGKSIQERVKEFIETTSAHGCGQIPTAKFKLTKFLWGLTFLSFFTFATVSSGILIAKYFTYPTKDVVSIEFGNI
EFPAVTFCSLQPITITSLIDFRQNPPPPGHSVSNMIYVMQNIRNIIIDRDIGGLGQWYLKHFDRFLSNQFIFENGEENDME
QFIHDLRSFKLGCKFQGQVCNGTSFVLHKDGGFHNCFTFNGAGSDLSNVNIVRTDPASGLSLVLFMDAYAKGKSVSAYT
IYNPDDPTSGQTGVRAVIHSPRTRPMPFDKGFDIPTGLSTSVALKETKRELMTQPHNNCTNDEFNQGTNYSYSVDTCYE
QCQQDLLIEECGCKSSLLIPPSFDKSDLEYCGKVNITNMMLMMNQNDDITQSIAEKELENLDCEEGVMYSFGNLAILEQ

CACKDHCQTINYVKTISQAVWPNDNAQEAFYDTYVNRTDQTLRPNKLFKGYNTSEIINNDMIHKNFLRLNIYFESLYVET
TSEVEDYPAGTLISEIGGNMGFYVGISIITLMEIFCITGAIILYCFKDWLIKCGHMNKTKVSHVNVKPVELDGDSTEKIKDKY

>Protostome_Lophotroco_annelid_CAC9674171_1__Ofus_G24722_Owenia__fusiformis__745_266

MDGKNMIATVKEFTENTSAHGCGQIPTSKFKLTKFLWGLIFVGFFMFATISSVNLITIYFTFPTKDVVSVNFEDIEFPAVTF
CTQQPVTATALIDFIDNQSHRDDAIGKLVLTMGSFPGMLVKPNIGGLGDWYFKYSYRFLSNQFVFENLDKKDLKTIQHD
LKSLKISCKFKGHPCKDSAFVVHQDGGFQHCFTFNGAGAQLNDTKVIKADPTSGLSLILFVDAYTTKRDLSELTLYNPFDP
TSGQSGVRVIIHSPKTRPMPFEKGFDIPTGFSTSVALKETKRELMTEPHGNCTMAKFNGGTNYAYSEDTCLEQCKQKILI
ENCRCMSSLLPTPSGELKPQYCGKVNVTNMFTMMYGKLDATMRGIAVKELEMLDCETKLMESLGNSEVLNKCECKKP
CVHTNYVKMLSQAVWPSDYNQKNFLAESINMSDPTQRATILFEGLDIGQITKNESEMIQKNFLRFNVYFESLQVKTTSQ
VEDFPASTLISEIGGNMGFYVGISIITLMEMLTLTGAIILYCFKDLIIKCGQSNTTKVPDVNIKQIKSYDNHMDEDKENINAE
IHG

>Protostome_Lophotroco_annelid_CAC9602553_1__Ofus_G07376_Owenia__fusiformis__749_267

MEKLKEQLKHFAESTTAHGLARIPTSRTKIVKFIWIIIILGCGITSFVFLYQSFAKYLAFKPKDVVSISREITVKFPSVTVCPLYP
IATGKNFNIYDYLDNNSTYVYKLNQIISGFYEKFDEPEDPLYAKYIRHKNRVVAWQWIYENNPNITNASHSWPDLVPICS
FKGKPCKAEHIEKFVDPNYYNCYTFNGMNSSTLNGENHTSTEDEMIVESIGPTSGLSLIMFLDLHEEQRSMFNPMIPTS
GSAGIRVVIHETGTVPDPTANGFDIPPGFSTNVPLRLIQRNHMEEPWGTCDKKADPYLKGSRYTYSRSSCKRQCVQGLL
KAKCGCISSLWPINNDADDSKYCGTFNIDEWIKQESNRSILEEDLLRLDCEEDLLSHLTEISEFIQSPDCQSNPDCHDCDCR
PACEHYKYYQDISMSYWPMESAQMTFYESLMTMPGYNDSTIYKLLHKPHDDLGTADNSSQVNKDLIRKNFLRLNIYYK
TIETEVISMYAEFTSGELVAEVGGTLGIFLGVSFVTLCEVVGLLSNLISSLLCNKNRKIKDTHDLSEKISKYDANDVKIVNQ

>Protostome_Lophotroco_annelid_CAC9607166_1__Ofus_G071156_Owenia__fusiformis__751_268

MGFKKDNTFKDIVSEFANDTATNGVPNIARAGSIPRRIIWTIIVLVAAGWMFTQLAQSFITYYKRPHSTLLTETFSSKIYFP
AVTICNINPVRESQIYLSNSTEIEALLRPTSQKEGSERFSQLTELQRSMQTLPTSSLQAMGHLPETMIMSCTYAGEDCSYS
DFTMFTNYKYGNCITYNAGPDIVTSSQAGAQHGLSLELFVEENEYLTLTDTVGFKVSIERQNKALFPEETGIFVPVGARTA
LSLKRQEIIRLPDPYSSDCWNLTANQSIYDNAYSDETEPSPKNYTRLACLKTCYQLNLITQCACLSHKIKTRGTAYDKANV
NLEELDFCNLTDSCYLKVKDDYEDGSLDCNCNSECWELAYSATISQTSWPAPKYLDDLLRTYAAKNDVIRGLLESPSSRKK
RDVGNGTETQNGNGTVNGNVTNNGNETVNENIGVEEAPIEIEPGTDRYFAKLDVYFDELNFIRIEETIAYTEWNLFSDL
GGQFGFWLGFSIVSIFEIIEFLIDVSIFLVYKSLNREKLKAKVGDSATELEGRPVSSGPQVFVTHGGETKFGLDENVKAGVI

>Protostome_Lophotroco_annelid_CAC9672069_1__Ofus_G21330_Owenia__fusiformis__752_269

MKQTLNDFASNTTAHGWGNIHQRTTTLSKAVWIILCLGCTGVAIWQVITIVIRYGLFETKDRIQVEEGDITFPSVTVCPL
VGIPKSENSKLLKDLYNGKIDADTYAEMLAFDTQIVYSSIYLDSNISQNAYREKLEFHNVFSKQLFSFQGYIENFNIFRDYSH
KQEEFIPICRYQGKPCLKSDFQSLDHHEHNKCYTFNGANNTISNPSTNITGPKGGLSLVLFVDASEASQVYNPDYPTSGS
QGVRVVIHEKGTFPDPENDGFDIEPGHSVNAALSVNQRQLLKPPWGKCANYDAEVYGGFTYRKKLCQLSCTQRFIYKT
CGCIKSSLPVDEVTAHKEYCLKIHPQEWLKLNSSQYNLTIIQNDLEKLRCQNREYPSSDILAQDGCQCREKCMYHTYDMT
LSQAEWPMRGVEITFYCRLIMTADNYINSSIYDTFHNISEYCDFAKPENESKILRERYGDTLRQNFIRLNVYFKDLEMKVIK
QAEDFTINSMISEIGGSLGFFVGMSIITIAEIFLLCWNITGVLKNRVENRVVSTPHQENKTDQKSALERANRSNAWEDEK
SIKW

>Protostome_Lophotroco_annelid_Pdum_comp402143_c0_seq3__753_270

MYENSYIIFPLKMSPIKPKTTIAFTEKDVIKKPTEKEILVKEFLDETTLHGVTHIAKAKGPATTFLWAFITLIMIIACILHSKESV
AKFLEYKANTEIHILSKRQLAFPAVTLCNKNPFKKEQFYKFFNAVLSGMKNARANHYHNASLYGPEWLLRAERMYPGHE
PFPEILLRNPEYGDLILQSMSEMVEEYGHLYGYTIDNITYSQGDLLLTCLYMQKPCNTSNVEVREFISWLYGKCFKIVPKK
DCTRIGPRFGLQLTLNIDQDSYLVTSTYAGVQVEIHSPDTMPFPEDKGIAISPGQAGFIAVKVSKHRKLNGKFGDCFDEE
QQEKAQDVFRETHDWVEYGQKNCERTCFQREMIQKCKCADIRFPRQGYSVNFPWRRAPACLLDKLYSFGLVYKSLEV
PKNKSSDCYEKVINMFHANNLSCHGEDNCHQPCKETYFDSSISLSDWPSEASTEYVVDGLRNKSKSADKVIKAGIANISK
NFLELNVFYTVLSLERITQTISVEVTEMLSDLGSNIGMYLGASLFAIFELFKIFFDTTLLSCKRPNVKKKKTKADKEQIANDA
A

>Protostome_Lophotroco_annelid_Pdum_comp413248_c0_seq11__754_271

MESTEDSGKSAGDTVGEKTDSCNSEIPFYLSDTPPPVYEMPEKDKKRTYGSVTYNYLDTTSAHALPHLISQKTKLQKFIW
FLIFWAAMGYSFYQLRGIVVEFRKYQVTVKTVVRHRTVADFPAVTICNENKLKKNKISGTPFQSILKLENKIMKKNDDAY
ASDDGDYSDDDSGYDSQSYDSDVQEFQQQEDEDGDYDTLGSDDLFVGLRGEHDYEKLFNLSRTTDYSDVINMLRPNT
TELDLYGHQEKDFILQCTFDGENCKDSHFTKFYNEAFGNCYTFNGMNQNNTVVRTSKFGSKFGLKLSLNVEADEYLGLF
THQIGAKISIHPSNMTPFPEDSAVSAPVGKLTDIALSSNLLEGVQSPYETNCSTGEGLYLFYPGDYTAQNCMHSCLRKALL
EDCHCVETIDVGPRETKCKISNVTQVICRLNVYKKFKNNKYPCQQGCNEPCHQVTYNILASYSEWPTEVYRALLKDQLD
KRNISADGYFLEGNILRANIYFKTLSYSYQGAFPTYTWETLLANIGGTWGLFIGFSVCTFLEAAEYFLELISLVFRSCCCKSKE
KAISPKK

>Protostome_Lophotroco_annelid_Pdum_comp419533_c0_seq2__757_272

MVSTKEMVLAGKFGREDAALENMGIATLIGSMAENTGMHGLPNISRSKSLFRKLFWLIIFLAGVAMMTWQVYEAFKK
FQSHPTETSIEPKSNTTLPFPAVTFCNLNPIKKSKLSDLGDIESLLNSNSNSSSEWTPSGTAQPSSFFDWDAQGDAAYANE
DAEFTLREKILEGLYSLTVAEKQAAGHQLNDILLSCTYQGYACGPKNFSTFYNSLFGNCYTFNGGEMNTAAKAGKVGPF
YSLSLELYIEQTEYIPGLADAAGMRVVVHNPNAMPFPEDEGFSVAPGELSYVGLHRVEFTRSKPPHGECQKFSENETLAR
NAWKQKYNFLEYTRRSCAKTCYQQYVMSICGCSDRDFPNEGTPFENITKGQNFSACNSEDEAMEECQQQVYTNFTEN
ILNCSKTCPPPCSEATHEITQSHARWPTEAKANDLVNQVLQKSSELFDALINLDAEERTETVRENVLKILVYFKSLEYTTITT
KPSFGIVDLLASIGGQVGLWLGLSVITLFEIVELFFDAWAFAFCKCLETVPKKKSKNLIQVTPRSEENEIGTSPSEIHFNHVN
EPNQKIVW

>Deu_Ambulacrar_hemi_Ptyfla_40v0_9_20150316_1g16271_t1_scaffold12531_cov113_758_273

MCDAWKGLKTLSGLGKAKDKIGGSTLAEQLSMANELNKFYCRFDKFDFSNVIGDIRSDLEQRVGEELFEITDEQITIVTKT
SLTFPTVTICNTNKVRRSAIADSSHGDVLVIDDAITLPYYGPCMEGDFMCDDGLLCIKPFLKCDGVRNCIWDGSDELGCE
YGPCMEGDFMCDDGLLCIKPFLKCDGVRNCIWDGSDELGCEYGTCGHNHFRCANGSQYGFCIEKHLYCDKKKDCYDG
EDEVGCECQRDEFQCQDKRGCISKKQRCNVHFDCDDKSDEKDCPAKDCGDALWCAADDYCILNEFICDGYKDCSDGL
DERNCTAEIPTCYGYSCDDGLKCISFTKRCDGQQDCADNLDEKDCPVAPVYTDMYMTERMGVLLSPDYPMNYPNNIF
HRYSVMMLPTERTESTIRVKFTFEDFNIEYSENCTNDWLQSCKNLCLHSTIVKCGCSQTMNIQSAPPCSILNKTQGNEQ
GFIILLKVSVSPHPADWLYECSYWQSLNLSNVDIHYDELTYQEIREEPAYPIESLFSDIGGSLGLYIGLSVITVFEFFEFVVEAL
RVCLRRENGGRHS

>Protostome_Lophotroco_annelid_CAC9650466_1__Ofus_G101010_Owenia__fusiformis__759_274

MDGVGKDEKNSDEDKDSSLRDILGNFASSTTAHGCERIDKSTSTIARVIWATIFIGCTIAAIVCIAALLEKYFQYPSKDVYE
MNSDKIEFPSISFCSLKPIVSSYKKRVAKMEPGNPYMEIDKAIKNIDGVLDYLSQENNTQNNLYQRLKEMYYLFASYTSM
YIHATDETRSLGHQLEDLLLSCKVNMMSCVNMPDVKIKQFDNALYWNCYTFNWKDYNTTGESEPYVEMVGGLSGFQ
ATFFLDNIDTLESYYNPMCPIDGSVGAKLIIHRPGSRPDPLREGLEVPVGFSTNVAVSKKKRELIGQPWGDCENREKLDG

APYRYDQANCERGCQQKEIVKKCGCVSPWLPVSDEQKNISWCGKFNLDNFNSASPNLTILDKDLIPFECHMMMTKMT
PNVTECDCPPACENHMYEYKTSQSQWLLGGNRLNFYQSVVNRMGDSYINSSMYHLLDAALNSNDSAENFKNRNLVS
QNFLRVNIFFDSMHTDLIRRVEEYTFTDMISGVGGGFGVYVGFSIVTMCEFVVLFASLVTALIQRYGLRQNKVAMESAP
THVHIIHVKHANNHLADVD

>Cnidar_Polpod_Hydrif_GBGH01000761_1__p1__GENE_GBGH01000761_1_GBGH01000761_1__p1__ORF__type_complete__len_571__score_27_43__GBGH01000761_1__228_1940__760_275

MSTNNGPVMRPESHQDKLAVPSSAKRSALMDKALGSNARNSGCEKGVNLSALKDGPIPLRVAQKDPTQKFRKIGLQA
MIAKRIERYAEERLSPKQIFKRFTESSTLHGFRYIFTAGTIVRRFSWFVLCVTMTSLFLSELQKLIMLYSEYPFTTTTTLESVQ
FHEFPSISICNTNSFRKSISLSSGIDHLVFDGSSGSSTKPTSNTTSDVIDGNVLFEKLNASSHQIEDMLIYCSFIDAFEQGKHV
HSDMECGPHNYTAYTNLAGALCYRFNPGPTFGVPLLSVTNTGILHGLKVYVKLQTDDYSPQVQEAGLQLVLHDHDENP
LSFSPFVVPPGFQTYVEMKKQQVLNLPPPYKTHCGSRTLNISTTYRRSMCYIQKLSEFVLERCGCKGPFMEKSVPYQVPL
CNSSQYKDCYRPTINSFDPKTAGDQCPVDCIRSNYMYTLSYGRFIADPKIGTYPSMTAKRNINLARLNMSYQDQRQYIR
DNFVAFVIYFSDMTVENISQERSYDIYKLMGDIGGQLGLMLGASVLTVIEFIDLIFFIVYTKISRIIKRRQVNRRFPTPAESKT
TTGV

>Deutero_Ambulac_Spurpu_014519_764_277

MTTGYTFEWYDCLLTQATLLLGRVKRDSLFLMRTDKDLPHLNSVNAVLIDYSGSTSAHGIPRIITSRSVKSRLFWSFVTLV
CLGAFLWQGSLLLLDFSKYPYTTQIDVVARTELQFPAVTVCNMNKMKRSAMVHTRFQSLIQADLGVNGGDADYSWW
FDWSSQWSQEEEEQEEQEEQELASASAPDTGSSSSDWSSDAREHYSEQAPALVTGSSTSSSDSSSDAQEHYSEQYEW
WDPGWDTDGFDFHEYDWTNVSDDDDWEGFYKQSTSDDFSDLLEVVNPTREELKVMGHQAEDFILQCTFDRHQCNY
TDFHQFQNKYYGNCFTFNREVGNSTRARSTGKTGAQYGLHLTLFTEQPEYVGLFAQEAGVRVAIHPPNVFPFPEDDGV
VASTGQATDIGMRQSYFNRLPHPHGNCTEGTRTIFMSEEYAYTTRACVKSCVQQQLFDNCGCVTDIIMNETMCGARN
KSQQVCRQAIEHFHEDQSSCNCPIACDRNLARVRIYFEELNFEQMIQKPKYTIESLLGGIGGLLGLYIGFSVITICEVGVLVV
DLVKFLLRKAYNRREKIVPIELKC

>Protostome_Lophotroco_annelid_CAC9661806_1__Ofus_G112085_Owenia__fusiformis__763_278

MHRLVCSQKIGSFTFDDANGYAITVNIKLYFVIPIEFLGTHARADTEEDIPKGKKTPMSYYYLLGNDYQNCSDHEVRPMG
LMRGLAENTSAHGITEIYYSKGFFKITCWVLLTLAAIAVMILHMTTLFIDFYSYETTMSISVQNKRSLPFPAVTICNVNPIRA
SRLSQSTILQSTIDGRDNITGATRDDKQTGLGAMSNEFLIEELIEETIADIDTDTKIAMGHQYSDFILDCQYDGYYCDEGEF
LAFYNYKYGNCFTFNSGSNSDVFLSGRAGPLHGLKLVLDIEAAEYIGDMSPAYGVRVLVHSQGDMPFPEDQGITIGPGQ
ATVIGTNMLNIQLAGDKYSDCTNSSFSNVNINVYEEHYPGTSYTQTACMKTCYQSHLLSGCQCGDPSVPLDGKAFPSAF
HTEPATSCNSDNQAQVTCEENTYASYVNGSLSCTCYPECNQTTFEAHVSSTLWPTDQYISRLIQDLSPKLNQYTNDSAG
LRKNVAHLQVYYEELNLQMIQEQPSYSTVQFASDVGGTVGLYVGASLLTAFEFGEFFLDLLVYFIRKPFHKNKVSEVKME
QTKSNVQFS

>Protostome_Lophotroco_annelid_Pdum_Contig13621__765_280

MAGKRVLKSKSNIAHLFWLLVCLAAIAMFCMQMAEVLQRYFSYPKKVTVEVISAQVPFPAVSLCNMRNMDVHVLNTL
NRKFIEDHSPINHINNSDNDFVREYMKLSAKYGPLWYKYQGKYPLAFQEVFSRTTYSANIPQHIVSSAAVQLDEFVVSCY
FGEYHCNTTNDFTTFFDPYYFNCFTYNMADPDQASSSEGIENGWSSVLYSGSGILDRNKDIRVLPGLHESRSAVSANEG
VRVVIHPPGTIPFPLTEGFDVPPGFSASFGIRPRQIKRIGPPHGNCITSNPFGDSSSQYRVLSCQKMCLQRYIINSCGCQDK
TLPDVPDLKATPCRNATDFPDSCMTDASDECLKQFFHLYDRIQCVRSTKDRFMKNSSLLTDCGCFPQCDEISYDVSYSLS

KWPASGYEGDAAFFDIFYIEGFRERFLNTPKYNMVMSYFHEDTREKTMQDFARLNVYIADSNVIITQEMEDYTTTQLVS
DIGGQLGLWIGISIITLAEVLELFIDTCRMLFSKRRIMVRNARRKETDRYRSHSGIAGLVPIKKTNCNNIFMSPLDRSETHA
MILRRQNGDSSSIR

>Chordat_hASIC1_NP_064423_2_acidsensing_ion_channel_1_isoform_a__Homo_sapiens__3_281

MELKAEEEEVGGVQPVSIQAFASSSTLHGLAHIFSYERLSLKRALWALCFLGSLAVLLCVCTERVQYYFHYHHVTKLDEVA
ASQLTFPAVTLCNLNEFRFSQVSKNDLYHAGELLALLNNRYEIPDTQMADEKQLEILQDKANFRSFKPKPFNMREFYDR
AGHDIRDMLLSCHFRGEVCSAEDFKVVFTRYGKCYTFNSGRDGRPRLKTMKGGTGNGLEIMLDIQQDEYLPVWGETD
ETSFEAGIKVQIHSQDEPPFIDQLGFGVAPGFQTFVACQEQRLIYLPPPWGTCKAVTMDSDLDFFDSYSITACRIDCETRY
LVENCNCRMVHMPGDAPYCTPEQYKECADPALDFLVEKDQEYCVCEMPCNLTRYGKELSMVKIPSKASAKYLAKKFNK
SEQYIGENILVLDIFFEVLNYETIEQKKAYEIAGLLGELLMTPVPFSCHGHGVAPYHPKAGCSLLSHEGPPPQRPFPKPCCL
GDIGGQMGLFIGASILTVLELFDYAYEVIKHKLCRRGKCQKEAKRSSADKGVALSLDDVKRHNPCESLRGHPAGMTYAA
NILPHHPARGTFEDFTC

>Deu_Ambulacrar_hemi_Ptyfla_40v0_9_20150316_1g8763_t1_scaffold4594_cov140_769_283

YGVMAGKSLTEFGQETSIGGVKYVTDVSSRNLRRFFWFLVVCTALGALTFQIVNICTAYIARPVSVNREYIPMGQMKFP
AVTICNYNQIRKTGLLEAFGYSHGDGVIKTAHAIMSPSPETMDQDVNLVGLNANYDFEYLMTTVGHKKDEMIMRCKW
PGRQCSADNFTTTYARERYGLTVYLDVESQEYVDNWQNFVGFRVIVHDQDDKPNMKDKGFNVAPGTYTAAALKKLS
VMHLFAMSYSFTIAMCQRWYRFLPPTAKFTDVQGLCSHCKTPCYEESYGEHLSYALFPSGPSVEELAEKAEKPCPKAMA
LYYKEKITAYVFASLSMANLSKILITTKDKLLDAIFLGETHISIDRIINDITASAPEGGLLPSVDVTDSPIENDQLTVGKLYEILT
EALQSYYDQIYSEISIPLSSNLYRYIHSERQSSVEVGGDIYRKVYETMINSSASARRLKKIAPDLAVQGTFRAALNVTDSYGA
LVNDTKYALSDAFTDIFDLIYTTVLDTFEENRITALENHTMYQICMEYMSMCPLDRLPGPSHVFQAFGGGNGNDTERSA
IAPKKLILEVGEPMMLL

>Deutero_Urochord_tr_F6SR69_F6SR69_CIOIN_Uncharacterized_protein_OS_CioInt_771_285

MHSIHNRDVRVSSLSAHFEPVENCVVHSDDVTVADDVSYTSSSMSSDLESMKPSNIHVFSSSSTLHGFNHIFALHHSSLR
RFAWTVAFLTSLSILLYQSSNRAIYYLSYPHVTRLDEVAAGNLTFPAVTICNLNEFQFSKITKNDMYHVGELLGFLDKNYAL
CCNTIVPERQTLNFLNIQRSSFRHFNPRGTFSMKEFYERVGHNLSDFLLACNYRGRFCNADNFTTVYTRYGKCYTFNSGV
KQPPLKTLKGGVDNGLELLLDTQQNEYMPVWKETDEITIEAGFKVQIHVQSEPPFIHELGFGISPGFQTLVSTQEQRITFL
PNPWGKCRNGFDKSEHYHFPNYSISACRISCETLYVYLHCNCRMIHMPGKERYCTPEEYKTCADKALDFLVMTDDQLC
VCETPCEVIRYNLEMSTLQLPSTQASSYLAYKYQVSEDYVRKNFAKLNIFFEALNYETIEQKVAYEIPGLFGDIGGQMGLFI
GASILTILELVDYFYEVVKDRTWGRKLRKNTDRSPTREEANNKTVSVMVREQAIFPVITNLFLGVVCCLYIYIILVVYKCCN
VPHILTTESGS

>Cnidar_NemVecNVEC200_011692_1_1_protein_AED_0_06_eAED_0_06_QI_233_1_1_1_0_91_0_84_1
3_1484_578_773_286

MLKLPCNKEMLRNTDPKETALEQINQFLQETTAHGFGRLGATAGSKWRIYWVMFCLAAYCVFAWQLVGLVNQYNSK
PIKTRTQLKHAQKLDFPVVTICNMNVLRASRLPPKLRTKFDEIINNTKKTSSRKSSNSRNAFVDPQDLSFEETKKIEILHAVT
THDNYRELVSAAHQLEDILLSCNFNGVNCRNSNDPTIPTSWTQTWNDNFGNCYMFNPAQTHNGEKVDPYSSSIPGES
NGLTLQLNIEQNEYLEGITEVAGIKVSISDQGVLPFPGQQGIRIMPGQSTGIQMTKLQTRRIDPFKNRSCENSNEMSDK
NLFFGYNMRYSKMACKYSCLNAKTIERCGCTNYNTPELQKRNISLCNRLNNAIIDCLNKAYDTFEDGSCDRECPPSCSEV
SFDLTISSAKWPAKSYEKTVLKTLQTEYGINMTKEEMFENIAQVHVYYGELDYLLVQETLAYTFMSLLSDIGGQMGMWI

GISALTCAELVELVCVILANMSNRSKKIVHISSNIGCFRPAKLGGVEAIEKVGGVKTIEQEVQGGVGIIEQEVQGAVGTIEQ
EVQGGVGTIEQEMQGRVGTIEQ

>Deutero_Ambulac_Sakowv30035415m_780_288

MIADYDNMVEDYENMIADCDSIVADYDNMVADYDSMVADYDNMIADYNNNLVVFKELGECGSKANRYRIMASSYS
FYGDRYSDGALQFSQPIPHPCKTKTRNTIALSSFSDGRFRSKVNSIKEDEEKKVQLQKDPPTATSKGDVNACGDCCCCRV
FYSDRFREFSSDTTLHGLRYAVAEGVEPWRRFLWTVLLVIAVSATAVYMYLCWYKFFSFPVNTVVTLTYKSKLRFPAITIC
NYNQYRKSVVKGTRFEEWVRQQYPLFNFSDDDVQVELDESILEANRTWFDQMAAHSKWSMIYQCTFGSVKIPCSASN
FTETFTDFGVCYTFNGADREGGALLVGHSGSEHGLRMRLFVNQSEYTFGEHCGAGFKILPHPQDEVPLVRNFGFAVSP
GTEALVGLKMIQEHNLPAPYASQCSNETLQYFAKYARSNCVRERQTDAVVAKCGCREPYMPSDARVCNYNETRVCVR
PELERIQVQCPMACESSTYAPRVSYAMFPGRHIVEDLQMRFNLSYVDIRENVVDVKIYFEEISLEEIYQEDGYPISELIGELF
YNIDDARAQDNDVINITVRIIRYDDCSIPH

>Placozoa_HhoNaC10_TR2949_c0_g1_i6_m_5640__Hoilungia_hongkongensis__24_290

MPTKHLKVVSQHNGGISARKKRLLAITITKGFEDCGAQGISNMARAQTTQARIIWAILTIVAVAFCIVMAVDLIDKYYRFE
YDVQLEISFKPQLDFPVVTICNLNPIKKSEMLTRPAFKPLFAKDFPDDPIAPLPTTAGSTMSGSGTGGTGTATKEKTGTGT
ATRSGTSTPTGTPKPRVRRGIAGVNDDIERPTPPANLTFDDLDASSENYNLFSEINAELFNKLSNQKMYDIGTKASEFIAK
CTFKQKPCAASNFTFSFNYLYGNCFSFNTGTLRRQKIESVNYPGPLYGLQLYLDINKGEYISREVPAAGVRVAITPQGIRP
NPADEGFDVAPGSLTSIGLKMRNITRLSKPYNPEGCLNNPPRNSSLTLYENGTAKYSYKGCIKSCIASLQYNTCGCIAPRYS
FTASTKGFTICKLSNKTETNCQEKLEREFINGKIKCGCVRACNEKSFEATISQAQMPTEVNLDSNYELLEYFKRSVTHLHLT
RKNVTTFLRENLVSLNIYYDELNFETIKQTPRYSSVDLASDIGGVLGLWIGVSVLTVFEFAEILMDTLLIIFGKEKIASRRNTIT
EFKAQLEANGFA

>Protostome_Lophotroco_annelid_Pdum_comp411104_c0_seq3__791_292

MEVIKEFGDTTTMHGVQKIAKAQYKRLRVFWIVAVICAIGMFIFQLYLLISQYLKWPTRTTMELSRDSIRFPDVSVCNMR
NIDVDVLYSLVKQFSLNKHPFELLTWTNITRASPEFEVKFLQLTGEYYLFYVYHYFAYPEVFSTLFSRSNLVGNFGKDTMIK
GAVPPWELTLRCSWQGAPCYSSNFTTFYDTYYSNCVTFKAPVEKIGSEGAESGLSLVALVGSGMIDWANISAKKVLIPGL
QENNYPLAGDGGLRVVIHPPGSHPLPSIEGYDVPPGFSGSFALKVTNNSLLGKPYGNCSTDYVEQDGSSYRVLNCLRKC
MQKQIIEKCKCVDARLPFSDDMIVNTDFCGKVAPLTAACKMSSSVSCMGPLFEAANNITCAKQVAGEMISKDTLVSDC
NCFPPCKETLYDLTYGLAKWPADHETDFVYQELLLQDRFFEKLNASNTSALKLDLFHQYFQFQNREKNLKEFAKINVYFA
DLTITKTKQVPDYNLIDLMSDIGGNMGLWLGMSLLTWTEFLQLALDLCIHAKSANSNKTKVQKMKNVTKVSVLPADD
AKYDFDYKKRTTSRENLIKQAMETDGLYYINR

>Lopho_annelid_Pdum_MGIC_AWC68057_1_MIPgated_ion_channel__Platynereis_dumerilii__8_293

MALRALMQEFAGGTTMHGIPKAIRSRSISARIFWSIVCICAATMFCVQFAQLISKFYAFPKKVTIEIVPAMVPFPAISLCN
MRNLDIMVLNTLNSIFKNATDPLTWTNITEDPFINAYMMTVAKYHPMFVRNDTDMKIFQTILTRTLIATNVDRHLVQK
AGVPFKEFIVTCRYGGLACNRSEEFTQFFDPYYYNCFTYTAPELMYADTTLAEGLENGWSTVVLTGSGMLDQNDELRIIP
GTHEKFSPMASNEGVRLVIHPPHTEPFPHTEGFDVPPGFSVSLGVKARLNLRIGPPHGNCSHIDPFGQGKSREYRLISCQ
KKCLQREIVKECGCKEISLPNHEKYDNLKYCTQDDDLPDSCSVGATPECFERLYQVYDRFLCVQNTTARLTRNMTFAGQ
CKCFPPCREVSYDVTYSLSKWPAESFDGEEAYVDIFETEAYPVRFMGPDDYKKFELYANYFDMSNRKRAMKDFARLNV
YIADSNVLKTEESQDYTQSQLLSDIGGQLGLWVGISVITLAEVLELIIDLCKFIASNHGPYSKGRTFNKRNNKYSAPNDEPV
PNCRSCRLYGQMNGTIPLTAVPEPMDPSHMV

>Deutero_Cephalo_107760F_t1__name__chr_scaffold21_start_1172790_end_1185891_strand_pro_len_591__793_295

MSPEDDKESPEKAFAGNTSLHGFNRIASSPGGSPRVLWVIAILGSYTGFGYMFGSMMASYFKYDTITDTTLEFADELLFPAVTICNMNKFDSDKLQVADWSYLSLFLYGIQADIPTILALGVPLDETVNSTMTGVKIEDLVLAKGFNVSPNRMSLCWWRGTNCSERNFTHSFGHYGNCYTFNADADNPLKQTMPGAVNGFMAFVDIKEDKYTESFLVSGNAEVGLKLLVHDPREPPMMDTQGIALAPGNHAFIAIKQILYENHVPPWGVCKDLQLEYYDTYTLNGCYLECRSKHLVRNCSCRPYDLPGTAPSCDPRTMFTCVRAVLAQVITGDLKCDCPVPCRMTSYSTSLSFAGFPNKHTREYLSPLLGMEPSYMGDNGVVFSVFYEKLNYQKIRQLKAMEEGQLASNIGGMMGLFLGASVLSLLEVCEYLLKRPLGFLGRTRHAKVVHVQPQETKNDVIQHVPVLRGISQQTHKGIPLPAAGDGAKLHGMFSFVTYSIRFRDNGVVFSVFYEKLNYQKIRQLKAMEEGQLASNIGGMMGLFLGASVLSLLEVCEYLLKRPLGFLGRTRHAKVVHVQPQEPKSATALHK

>Protostome_Lophotroco_annelid_Pdum_comp414793_c0_seq1__795_296

MGKLQGLMDDYAQSMTAHGVGRISRASSFKAKIFWAVIWTGMIAMFFLQATTLMRRFFAYPKSVRLEMANQPVPFPAVSLCNLRPLDVFMFKDAIFKDNVTMRWIPGNYSYTGRSGASDFERFLIKYGNVYMSYRSYIEVLLEKEQVDQDWFWEARRALFSRLTLAANFNEKAAQQGGIQADQFIAQCEYAGKKCSFENFTYFLDPAYFNCYTFDMNQMGWESQTIYEGPDHGLSVVLFIPSLPSTDLSPEAYRLIAFTQELSFGGEGIRLVIHEQNTVPYPLTDGLDIPRGVSASIGVQLNHNMRLSPPYGNCTDKNTLEGTLVNYTYTMASCKKTCLQGLIADTCGCTDIGLPLSENFEEGSCSRFDALPTECQQKGDIRMNLDKCKTFFEPWFERNLCKRTVRTNISKNLSAWNLCKCFPRCYDVAYVTSTSQSDWPTIESSVYLLQDILTAGGFASKFPEEKAKQYFGPLLGAETPTFQEARDHILAKNWLRINLYISDTSVVKIEETEDYGISQLISDIGGQLGLWIGVSIISIIEIFDLIYQICKYLLDPERQQKRHRGSTIRETKQNGSNDLSYAQSNGNPLRF

>Protostome_Lophotroco_annelid_Pdum_comp418040_c0_seq1__796_297

MMVLTLLLNLRFKSKMALRDVLGNFAVSTTMHGVPKVINAKSAMARLFWSIVCLAAGSMFCLQMSEVLTRYYSYPKKVTVEVVPTPVPFPSISICNMRNLDVHILNTLNAAFLRDDNPINHINKSDNAFINSYMYNVGRLAHLFWKYQDAHPEVFQEVFSRTTFSANIPEDIIALAAVQLDGFVVNCHYAGHRCNKSRDFLRFFDPYYFSCFTYKAPEPSELDDSLSEGIENGWSSILLSGSGMLDKNDEIRMLPGLHEWRSAVSASEGVRVVIHPPNTEPYPFTEGYDVPPGFSASFGIKPRRNIRIGPPHGNCTNKNPFDQATERYRLISCQKMCLQSAIIKSCNCSDVGLPRLNHMDVPLCRSSESFPESCFDDATDECLEALMNMSSLIKCVRVTKAKLTKNTTQMEECQCFPPCDEVAYDVSYSLSKWPASGYEGDAAYFDVFGIEGFNERFNNSETKGKYELFAKYFNVSNREETMKNFARLNVYIADSNVVKTQESADYTRNQLVSDIGGQLGLWVGISVITLTEVLELLSDLVRFFTSSYSRISRHDKTYRNNNMNHHHNIIPLKRPMLHDHDLQFRMENTTL

>Protostome_Lophotroco_annelid_CAC9486619_1__Ofus_G015124_Owenia__fusiformis__802_301

MHSESNNKLEFPPISGYDFEVTTTPKEKRYSEFDFTATTPRSKGLDGIDEKNESPKRKEITPEITPLFREFCDNASVHGVNHLRTERVKVKRWVWTIFVILALGFNLFHCSLLIDKYLGFPSEETQFVDQSWIEFPSVTICNINAMSRITRKKMLEDNSTLLYKWHDYVNNRFEALVSDLADGYSLEGSNLLDTLELLYNRVHLPTGYLENIGEEALEVGHKLNDLVVDCTFGITKCHAVNFTSFFESTYYNCYTFNGGNVSTSHNNLITRTTGPQEGLSIIMYLESDNGNINENGSYLTMSKLNNAAGARVMIHAPNTMPSPTDQGFDIPPGFSSSVGVSVSTRTRLGEPYAKCSKREINIGTDYLYSDNICLKLCQQRYVMGNCNCISSLLPFDRSTDLYFCGHMDYNNNASFFENMACESRVLEEFVHNEDVRRECGCHPPCHNYIYNYQTSQSYWPLEYYQADFYDLYILSDPNKENLKAYQNLKHHNVPQLIEKGLIRKNFLRLNVYLKDLTIVQHIEKRSYAIENLFSDVGGTFGLWAGMSILTICEFSELFFRVMAILVSKFLSIFGPMSTRAVVEDSSSNSHTPVRRT

>Cnidar_Rhopilema_esculentum_TR55888_c0_g2_i1_p1__Rhopilema_esculentum_TR55888_c0_g2__Rhopilema_esculentum_TR55888_c0_g2_i1_p1__ORF__type_complete__len_604__score_83_96__Rhopilema_esculentum_TR55888_c0_g2_i1_896_2707__809_303

MAATNTRGLFLRAQRHIMDRELSKDENEEEPSQPYSSADESKESLLLQNFGMKYGHAFKVKPKKTEPEQKNGNTELPQ
RKVSPEIETNKPAQVTVNPAFKNVALDRFRRAGMKARLVRRVNKIVEDRLTAKQIFARFTKESTLHGFRFIFTKTFYIRRFI
WFVITITMAAMFLKELTDSINLYFQHPFSTTSTIEYVNRLTFPAISFCNLNDFRFSKINGSDLQDVFMHEKGKFYLHRNSSF
DLDGKKLGERLEDASHRITDMFMKCVWLFSQTAAGQPVPCNYTNITTYYGLNGQTCYTFNPGDRGHRLLTLNETGLFH
AFELQLDLETHEYLKDIQEGGVRVHIHDQNETPFSSAGFAVPPGFKTFVSLNVQKIENLPPPYTTQCGSKKLKYYSSYKKS
KCYLEELSKFVTKKCKCRGRFMPDASYPLCSLKQTVECLRPTIRTFDHRAGGACPEDCLSTQYNAQLSYARYVSRPPIGTS
GITIPERFQKANLTNSQLEAVIRDNIVYLIFFFPELRVENVKQTVTYDFYKLVGDVGGQLGLMLGASVLTLVEFIDLIAFIIYH
QILRLTRLKKKKKEETEQLKLREMNGNGAREGRFINF

>Protostome_Lophotroco_annelid_CAC9670605_1__Ofus_G181373_Owenia__fusiformis__810_304

MVYCSSSIIKLNMAKRIHDVVQRFASGTTIHGVPKLIRAQTIPGKIFWSLICLAALGVFIFELALLLQKFFGYPKSVDVKVVQ
EPVAFPSVTVCSVHAMDPFVIHKIYKLRQEQMEHRMRRKPWVISKNVSDFEKWYFDTFLKKSMSYVLEPSYLEGLIDRH
DLKTEFFPSLSSRLTLAANLNKSITFPGVEPAELIAFCRYARQECSFEHFKTVFDPYYFNCFTFNASVIAGSLKTLAEGIENAL
SIVMYIPKLHNEIKIGTKVKLPGIVEHDIRDPLAGSGGVRVVIHPPDTQPHPATEGFDIMPGYSVSIGVKTTENTRLGRPY
GNCTETTKGVFSDKASYRYTMTSCRKKCLQNLLSREVKDGCGCLDVVLPTFPEVNGVDFCAKFDDIPRKCIFGPFARRNE
SCKELKARWLKRMICMRDIETSGSHEHIAITENCNCHPPCKDLSYEFFYSNSLWPGQQHKQDVYRDLFISRRFERRFQE
GQRTYYFGAENAQALNVTEQEKTVDRFARLNVYLYDTNVVKITETKDYTGIQLISDVGGQLGLWLGISIITLTEVFELIADI
VHLCLKRKRKQSIREGNCTPAEITALNRRNGSTTV

>Deu_Ambulacrar_hemi_Ptyfla_40v0_9_20150316_1g24013_t1_scaffold33951_cov84_813_305

MARKDTDERSFRQLFVELVQNSSAHGIPNAGRAKTKFRTVVWSIIFVIGVAGFLFQFSELFIRFIDFPVTTNVDVTSNRSL
VFPAVTICNQNPVRVSALEYASDPKLRENFDDTYEPSTSSQAPQDVTGNTAFTSPDIASAPPIGNPPPPQGRKRRATDYL
ERQGNKQPPSPDSYNGPSAGLAQVSLLTELMSKVSIEERIELGHQAQDFIVNCTWQGERCSSDNFTVFNNPVYGNCFT
FNGKDVADEDVLLTNFPGPLFGLSLQLFIEQGEYLVDYTEIAGVRIVIHDRDKMPFPEDTGYTVPPGDSTSLGVEITRKPY
PYSNCVKKYDSAYSNGYSELYGVGYSVQRHDLRERSISGSVAGKDFNGLTLQLNIEQDDYLEEYSETAGVRVLIHAQDVA
LMPEDDGFDVNPGVLTSIGIRRTEITRLGDPYSNCVSKTGDIGQYSNVFTDMFNANYSQETCMKACYLTRVAERCDCY
DGRYPKPNSTEDLDFCNTHNNANYRCVKNIEEEVRTNSTSFRCDSPQPCEQVTYTTEFSTAYWPARISQRYVNDWLTQ
RNRELHRMVMKQEDLNEDLVRLNILKLRVYYRDLNYETITQSPSYTDAM

>Deutero_Ambulac_Spurpu_025881_821_307

MDTEEKDTVKKSVTDFGNETTIHGLQFVVNKKNIIYRLCWLGICTTFLVVFLIQGNVILKDFLRWPYSTKIDIVGRPNLAFP
AVTVCNANMMRRSQIEGSRFEDLVNLDGGVEGADYDYSWWFSSAYRNWYASSSASSYGQSSDQNSNGRSSSSSQSD
SATSSDGQSSSSNYDPSSSEESSSASSVESSGGTSDGQSSSSNDGPSSSEEPASSSPGRSADGHSSSTNYGPATVRVSIVF
WVYNSEFPAPNSXVNEWSWYDPSFFADFEFSENGWDGVSGEKTGRASMRPPKPMTTATFLTSSTRRRKSWKSTVTS
WKISLCSVPSIVDHVISGLHLTLFVEQPEYLGVLSHQTGAKVTIHHPNEYPFPEDNALSLGTGQETSIGIRQEYIKRLGGYY
TNCTSDGKDTNFSSTTELSYSSEACKKICYQLHLSQLCKCVDDQFFDGFPTKCDVLNMTHQICRKFVEDLFLDDKLPCSCP

PPCTEYKYVRTPSSSLWPSERYEEHLLRRLNGSANENLIRVLQSNELSRKNLIRLKIFYEDLNYEVVEMVPVYTIPSVLGSIG
GLMGLYIGMSFISVFEVLFLVLRLIKIALIRIYSRINRVQPYPVKNV

>Placozoa_TriAdh_evg1307556__type_protein_aalen_609_92__partial3_clen_1981_strand___offs_155_1981___evgclass_main_okay_match_evg1307557_pct_99_100__822_309

MSQSSYHDSNKTASDESSTDAHPNSQKVPILSHQDEVNQDNPEPRNRFDLDFRWRPSIEHDENFPFSASFHGIEHIYEG
RYGTRKILWILLVAATMIACFVFIFIQIAHYSAFHTTTKSTLVYEKQLAFPAVTICNYNSFRRSAVTANDLIHMAYLVKAYRL
NQGVVSDFIGEKERQKLINYWKKYDATHAKKFNYQLFVERVGYHASQMIKSCHFRGLKCGPKNFSNVLTSYGNCITFN
GPKLESNPKLYQKNPGAHQGLELLINIQEYEYTGSWHSDRPDIGIKFVIHERHYPPDVTSQGKAVGPGSHAYASVKYKTI
SNLPSPYGHCGSKKLAFYKKYTYAGCQISCKTEYVQKKCGCRAPDMPGQNVIPVCSPQKMIECVSPNLEKLRTINDKVCI
YTYAGCQISCKTEYVQKKCGCRAPDMPGQNIIPVCSPQKMIECVSPNLEKLITINDKVCICPIPCHIVHFDTTISYAKIPNP
QMAKDLTEKINKTAFQIETHGLVDPDIDPTLYISQNYILLNVFFDDLYYEKTVSTPVYTFTSLLGNIGGQLGLFVGASVLTLV
EIIEFGFYRSRGVIRRSDWKQNLRKSISRSREVTATEEKEPLCS

>Protostome_Lophotroco_annelid_CAC9670604_1__Ofus_G181372_Owenia__fusiformis__823_310

MVRMFDEVVQHFASGTTIHGVPKLLKAQTIQGKIFWSVICLSALGVFIFELVLLLSKYFEHPKSVDIKIVQEPVAFPSVSVC
SVYAIDPFVIHEIYKLKTGTQKDSAMATDDVGYRFTRDMSGELDDTQFGVENAFIKWYLDDFIEKSSDFMGKQENRLAI
KRMLGSEAEEKYRTKFLPSLASRLTLTANMLHEHIAFPGIKPKEFIALCRYARKECSYKHFTTVFDPYYYDCFTFNASIIAGA
RKTLAEGIENALSLVMYLPELHRDLKFNKMKTHLPGILGHDLRDPLSGSGGVRVVIHAPGTHPYPATDGFDVMPGSSVS
IGVKTTENIRLGWPYGNCQKEEDDTIQIQQKTNTYKYTMTSCREKCIQSNLLMNKPKGCECLDVNLPTFTEVEHGEYCA
RFDDVPKHCQNKTECAELKARWLKRMNCMHDKEANNNSLVGECNCHPPCEDVDYDFFYSSTPWPDANYVNGVYRD
LFISHKFQDNFDNAQKRTAYFGDKIESDLDNEGRIKKTVGRLARLNVYLYDTNVVKITENEDYTGIQLISDVGGQLGLWL
GVSIITLTEVFELIAHIIHIVCKRDDKRDVDVQHEELDYQRLPIPEIPHNNE

>Deutero_Ambulac_Apla_gbr361_4_t1_824_311

MYRSEAASPERWQAAASDQHKYRPELNNHNSDGWTSAENGQQAATDSNAVNTPFQPAAMSRRTSRGFKRGESLIG
DDRRRDWDRFPQQEPRHLQQQKPGMTDEERLAKTSVAQSLVLSLLENTGAHGIPNIKRAPNNVRKVFWTILFFFGLG
MFIWQTSWLLERYYAFGVDVKMEISSDRMVEFPAVTICNLNPIKSDFWEELDERGIPIPPSLTRPNTGYGPDDSEEEFD
DAFEMLGDAIELVTAFKYKDRRTAGHLLRDMLVRCVFQGRPCHARDFTYFFHYLYGNCYIFNAGPEPNATTPRLFVTKA
GPLYGLTLELYIEQDQYIEDIQPAAGARVVVHARHNMPFPDDEGVSVSPGQETFIGFKRTNFTRLPHPYSNCTVVENAN
ATIFNLPLPHVRYSQRACENNCYYQNLTRVCGCADVRFRYDDDVPLCLNSTQEQCLEGVERMYLAGMLDCMCSPPCR
ELQYDIVTGQARWPNEKFKLFLDDNIGNFSDDLYQDIDSGGDPDFLEKNLLKVNVYYDKLEYTTIYQDVAYDHIALLSDM
GGNVGLWIGVSVLTVFEFIELLYDLGKLIFYRVINKNATRKTGKVTSSDVDLQHVSNLKIASEA

>Protostome_Lophotroco_annelid_CAC9484202_1__Ofus_G014375_Owenia__fusiformis__827_312

MPQKYYDKNMKLPGVLQVQPTAEGYEEYRDVGKDNTKDKTPQNYYNYLLQEKDKNLLNDEVKLTHLYRGLVEHTSSL
GITQISYSKGPVKIGFWVALTLAMSVLTIWNVSTVFMEYLKFDVDMTIVVEQRATLDFPAITVCNMNSIRLSKYLSNPML
NATLNGDPGSDGANATATSTTTTTTTTTTTVAPAITTAASKGNGKNSKTKGGTNTDKGNGKTDKKPINGNGNKQTND
NLGEVDPMFAFQERVSEMLANMGESEKVPMGHQREDFIIDCKYNGYECFIDQFATFYNPKHGNCYIFNAGWNNSQS
RFTSSRPGPFYGLQLTFNIEQSEYIGKLTSTAGVRVQVHHQNVMPFPEDEGINIVPGQSTSVGIQMLNLQKQGGKYSDC
FDDDVRNERMNVYEEYYNVKYSNPACMKTCFQRHLLESCGCADKQYPMTGVAFNNANATVETCNSNDISQSRCIQNI
TDQHIGGKLRCSCYSACNETLYKTEVSSVLWPADAYKDDLLANLALRSPSASAVVSDADTDAYKRNFAHLQIYYSEFNFQ
SIVENIAYDEGALASDLGGAFGLWLGASILTICEYIDFIMDVCVWALRKLKKKMTVNKVDKHPS

>Protostome_Lophotroco_annelid_Pdum_comp408841_c0_seq5__833_314

MAHPLHRTMYLGASHTYGADMRNHYHNHSPRPLKPVRPNRKTMDVHRVFREFADSTSMHGVPRIINARSLAARVF
WSITCICAFGIFLWQCIILLQRFYSYPKKVNVEVVQRPVRFPSVSFCNTDHLDLVVVKRLEEMLLESDNITYGNDTHFMKF
KEAYQAFWDSSSFFFQPYLQHVKPYEKAMTHMLAAYSRLGLIANLGVDLASQGGIEMRDFIVNCRFMGEPCDIEKSFV
KFFDPYFFNCFTFDPSTILSSKTTRLQGAEYGLTIILFTGSAGQLTKKSEFEYVIPGMEEADGVLASGRGARVLVHSPGTAP
RPASSGFDAPPEFSVSLGVRARENVRIDKPWGNCSFGNMDSKFKYTLEDCQNTCLQRNIMQKCGCIDNKISIPSYTKGL
PFCLTLPTIPYTCYDMPLPEKCAKIMDEWTFRMDCRKEVYENLTMKDPDAMDNCGCFPPCNDIIYEASYSLSTIPEQTEE
NSAFYSIISNFLSSLAEPKKKLLNDLYKLNKKDGRIRGYIGRINVFIADSNVVKTTEAPDYEAIRLISDIGGQLGLWIGISVMT
LFEVMQLTCDICRFLSASGRQKSRDRRRARPTRVDRVADRDVEIHFEVGDKLTAV

>Placozoa_TadNaC10_MK547551_20_315

MPTKQLKVMSQHNGGISVRKKRLLAITVSRGFEDCGAQGISNMARAQTTQARILWAILTIAAISLCTVMAVDLIQKYYR
FEYDVTLKISFKPQLDFPVITICNLNPMLRSEMIKRPIFKPLYGRDFESVNTSQTSPLVNNSSLSNNTVGQSDSNSSTPLPT
NEVPQDATAAAAATTVAANSQGGTAAGSTKESGKTATPTGTKTSLPRVKKSLSNDVNENIDRPTPPSGVQFENLDKSH
ENYHLFSEISAELFNRLTDQRMYELGTQASNFIAKCTFKQKPCAANNFSISFNYLYGNCFSFNTGNSRGKRIESVNYPGPL
FGLQLYLDINKNEYISREVPTAGVRIAVTPQGVRPNPEDEGFDVPPGALTSIGLKMRNISRLSRPYNKEGCLKHPPNNKTL
NIYENGSDRYSYKGCIKSCIASLQNKTCGCIAARYSFTSSTKGLKVCSLKNETEINCQSRLQNRFIAGKINCGCVRACNEQS
FEATTSQAQFPSEVNLDNNNDLLEYFKRDLTKFQLTRKTVTPFLRENLVAVNIYYEELNFETIEQTPRYSEIDLASDIGGVL
GLWIGISVLTVFEFMEILVDSLLIIFGKEKIASRRNTITEFKAQLQASGLA

>Deu_Ambulacrar_hemi_Ptyfla_40v0_9_20150316_1g34414_t1_scaffold1171760_cov98_837_316

MTLYPPVSEVPKIHNFPQFTWENTLKHLNVNQNVRNNVNQAEIPQRLDKHGNCGHAEPMERSQTVNDEYHVGNTR
SVAAQADEDISRSPTRQKIREKSNTTNAARLPNWTIDKQGHELKYSEELWDRASLRTITTSRARTLCDSGRLRGHISLLNR
VDNSNRKLMWLAVFSIAVAVFAFQAYELVALFLNYDVSVNIEIGTATSLAFPAITICNTNKLRLSEIEKSEQHQDLAKTDPE
HPDSVHRLLSYEATCRDGVTCDFENDGRTFTCVPRKNQCDRFPDCEDGTDEKNCDLNDQCPTHTVYIWNSPEVLFSVN
YPSNYDDDYRCSWILSAYSSYCILLSFLDLDIEPSEDCSKDYLEISDVGNLSIKMRYCGSRIPPSWRSSSNTVKIEFVSNDKIT
GKGFRMEYFSVVCSSSQISDVTSWSECSVSCGWGVRKRNIKCSVVHEESLSNSDTGSGYQPHYGSGESPDSNDHDDED
YLSDAESMPSSDMVNHYNRTNVTYLNPNDDMLXLKLTLFIEQNEYIPLYGQEAGVRVLINPQDITPFPEDEAITVAPGLK
TSIGIRKDCHKSMLQQYIRHYCDCVDTLHLKGRYCNIANQEEGKRMNMFFVVINHLLFDNTCSN

>Protostome_Lophotroco_annelid_Pdum_comp417306_c0_seq28__839_317

MMNPEKVPPPMTPVSAFSDPPQYNDSFKSVTLEYLDTTTAHGLPSVITKRRKIQKVLWFLIFWALMGYAVYQLFGIVNE
FNEYQVTVKTLKKHRNLAEFPAVTICNENKLKKSKLGGTNYASIIELQQKYSEEVFGPSAPTTSNTAGSSLPTNVANSTAL
QNVGTASLLNTTTNGTSSTNLTNTISNAVSNLTNNGTNVPNLSNTISNALTNLTTNSTSLTNLTNTISNAMSNLTTNGTS
LTNLTNTISNALSNLTTNATGLTNLTNTISNALSNLTANGTSLTNLTNTISNAVSNLTSPLTNSTGNTNQTGSTGLNLPSIP
LGRKKRQILSGNTLSPRVPPGESGTNNLLSMELKGDHDYETLFASSKTSDFSDVVSMLRPSMTDLDKYGHQFDDFVLM
CTFDGENCTSDDFVRVYNEVYGNCFTFNRQTNGSTVRTTTKFGSQYGLKLSLNIEKDEYLGLFSHQYGARVTVHSNNIT
AFPQDNSVSIPVGYYASVAVDTEVMDSKERPYETNCTHGKSADLFYEGLYSTDNCLNSCLRKMVKEVCSCVETVLVTGN
ESRCSISNTTEVACRLEVYANFRDNAYSCQHECNEPCLEVTYRTSVSYANWPTDGFNALVGRQL

>Deu_Ambulacrar_hemi_Ptyfla_40v0_9_20150316_1g14838_t1_scaffold10372_cov150_840_318

MEMKDNNYDDANSRKNNNVSPDSISAAIARQSYIDSPETPGVDIDSTPSGRIRIWAAQISDIHGVKHIVGERSRLRKLL
WALVVLASFGILLQQCIKAAINYGEFHHVTKVDVEYVQHMPFPAITICNFNKYRKSAITPADMVHVGKPLGLMDEDGN

LNSDLFSEEFVRQMKDLDWDSEEKKRFNYTEFTYRVGSQVKETIVECTWNGHKCTEHDFVKVFTHYGICFAFNKYHRD
TEARHAGKPGADNGLRVVLNAQTSEHLPTADLEDSFINVGFKLMIHPPTEPPYPKELGFAVGPGSHIFLAITRQEIKRLSK
PYGECDMKSVGSKYFDHYSMSACRIECETALLLEMCGCRLVEQPGNGPVCTPKIVKECAHVKLLEYIEGHIEFDCPCHIP
CDSEVYSVTPSSSRLKPDRSGKSPAMSNYTQEYIDSNVLVLTIFYEELNFETITQLPETSLVGLLGQLGGNMGLFLGASILTL
IQIIEYFVDECIHCFRPMAPKKPKRTYRGEDKDVNTPLSVQHWQGSHPVRNTTLMIVQHDCEIVSVEHFIPLYEILTGSAN
TGLFVFPRCLRTRAANMLRFTGPVWRQRLSFELLRNAENQFITFLRHDAWYSCLTYKMSSKLVR

>Deuterostome_chordata_Ggallus_tr_A0A3Q2UMV1_A0A3Q2UMV1_CHICK_Uncharacterized_protein_
OS_Gallus_gallus_OX_9031_GN_ASIC4_PE_3_SV_1_844_320

MPGQPAWGSVTLPQPPAGLGEEFSIPKSRLGKACVGTSPLLSLPSPVTAQEGACHIRLWEGAEQKAPGRETASGARSE
AGHAGSVLTTAMPLPLSCCPDGEALAPAQGGFLRATRIPGLHYMGTRPQSCLRRLLWGLAFLASAGLLATGATDRLHH
LLSRPVLTRARLTRVPQLRFPAVTLCNPNRARFLQLTKPDLYSVGQWLGLSREDRSLVPELLAMLGDEQRRWLTRLANY
SRFLPPRRSERTMQSFFHRLSHQIEDMLVECRFQGKRCGPQHFTPVYTRYGKCYTFNGDRRNPRVTRQGGMGNGLEI
MLDIQQEEYLPIWRETNETSFEAGIRVQIHSQDEPPYIHQLGFGVSPGFQTFVSCQEQRLTYLPQPWGNCRASVQGEQ
MLPGYDTYSIAACRLQCEKEAVVRSCHCRMVHMPGNESICSPNVYIECADHTLDAAVEDSQERCSCPTPCNLTRYGKEI
SMVRIPNKGSARYLARKYNKNETYIRENFLVLDIFFEALNYEAIEQKKAYDLAGLLGDIGGQMGLFIGASILTILEILDYIYEV
IRDRVSRVLRHSKPPLKKPSGSIATLGLEELKDQSPCETLGRHVEGTYNAGILPNHHHRHHYPHQGVFEDFAC

>Protostome_Lophotroco_molsk_OctBim_tr_A0A0L8FW45_A0A0L8FW45_OCTBM_Uncharacterized_pr
otein_OS_Octopus_bimaculoides_OX_37653_GN_OCBIM_22006431mg_PE_3_SV_1_847_323

MKVTGFDKLGFNSNQLRTMRQQLRRRNYNALSIITELAAESNAHGLAKIATSTQTPRKVLWALMVIVGFTAATLQLSLL
VRKYLEFQVVEVSKMKDGMDVEFPAITVCNIAPISLTKTKELLSSDTSELTQWLRFIDKYNFGAQTDRMFTVQSLYENLA
DEAQLLGRDLNDFLIHCQYNQEICNIQNFTRYFDGHYFNCYTFNSGHQTGTSLLTHATGPQSGLSLILSLDNDDPPVGGY
GSYNIKSNIEHSAGVRVVVHPPSTMPSPVDHGFDVPPGYSSSVGLKTIMHARLSNPYGNCQNSRLQNSSKYIHTVFSCL
ELCKQRIVMSTCGCRSSNLPEMTSANFTFCGLVENKWNWKDMMKNPENYNITKINLTELACEERVLKEFSINRAYESSC
NCFQPCQETTYQKSISLSYWPLEFNQLSALETFYGDKINETFLHDAYKMLKYLSNLGSVPTPESPPPEEINITVDDNNNST
NSRMHSPALLSQLLANSTEAPKPAATRDDFIEHVSAKLNSSDKEKQIRASNMIRQNLLRLNVYLEDLSIIEFRQMPAYELA
DLFADIGGTLGLWMGISVLTIMELIELFTRLLMLIFSSEKKIPNADPVTNGMLDHDCDCQKSSMESPF

>Cnidar_HydraVulSc4wPfr_147_g8550_t1_848_324

MLKDQASYYQIINEEVDSNEASFPSISNNNKTFDRRFSSENCKIEKSKHSSNLIGSAKWAALTKKNAEQLLLKSENTKPVID
SAYDLNLTELDTITDISAYKKDNSPVKGNIEPPAVKPGVLMKRTNTIPKQADKWGTIKERTSLIVKPTDEFTKKVMPSNDK
PLTTTPKLLKVVTAVTAAKTATTKFREASKMRIRLKERMRKIVQERLTVKQIFKRYIESSTLHGFCYVCMDTFLGRRLIWAV
LMILGAIYFIFKLRYGIKEYFDYPFSTLSTVEYVDDLLFPAVSVCATNSYIASQVYTNQLNTMYKEGRLPLDNNQSIPEYNIP
GDELVKTLKNSSLTIESLLKYCDWIMQDTSHPLVTPNNCGALNFTSYFNYKGEQCHTLNSGAKGHELLKVSDVGISHGYE
LVFDLQTNEVIKNYQLSGMRIVIHDQVFPPQLVDGFFISPGFKTYIKLGITQSQSLPPPYSTECGQKKLKYYAIYSQRLCLLE
TLTDFTGDLCGCRDVFMPENGLPFCSLQELYSXSFSEFTMRKECPSDCEERTFSYELSEARYLHNPPIGLSLSRLDKSNIISV
ILFFGDTRIDYHEQEATNDFFQFLGNMGALMAGVGLKLSFYTQDLSSYSAFYKS

>Protostome_Lophotroco_annelid_Pdum_comp417306_c0_seq24__852_325

MMNPEKVPPPMTPVSAFSDPPQYNDSFKSVTLEYLDTTTAHGLPSVITKRRKIQKVLWFLIFWALMGYAVYQLFGIVNE
FNEYQVTVKTLKKHRNLAEFPAVTICNENKLKKSKLGGTNYASIIELQQKYSEEVFGPSAPTTSNTAGSSLPTNVFNSTAL

QNASPPVVGTGSLLNTTTNGTSSTNLTNTISNAVSNLTNNVSNLTSPLTNSTGNTNQTGSTGLNLPSIPLGRKKRQILSG
NTLSPRVPPGESGTNNLLSMELKGDHDYETLFASSKTSDFSDVVSMLRPSMTDLDKYGHQFDDFVLMCTFDGENCTSD
DFVRVYNEVYGNCFTFNRQTNGSTVRTTTKFGSQYGLKLSLNIEKDEYLGLFSHQYGARVTVHSNNITAFPQDNSVSIPV
GYYASVAVDTEVMDSKERPYETNCTHGKSADLFYEGLYSTDNCLNSCLRKMVKEVCSCVETVLVTGNESRCSISNTTEV
ACRLEVYANFRDNAYSCQHECNEPCLEVTYRTSVSYANWPTDGFNALVGRQLVGKMARNEIKEYIENNILRVNIFFRTL
TYSHQQVFPTYTWETLLSNIGGTWGLFVGFSVCTFLEAAEYFLELICVCCGLSSKKKKEKVAPEGFNYAKGLS

>Deutero_Ambulac_Spurpu_025728_853_326

MEVPLDEEEKPLSPEKPSSPEKPKAVLEKPKALVREMTNDILVKSWRTVVQSFCKSTTMHGMSRVIDSTRAIFRLAWLLT
VLTFLCVLMWQAYRLVDEYRGNPTTTTIQMVTNNKLSFPAVTVCNMNRLRRSKLVGTRFEPLLDIDRHVTILNLGLDDG
EIEVDDHAEPSPSLTPATTTSTITTTTASQVTTKSTSVPPNDEDLVSSTTVEGLEMSSTKDTPLVVDVDNPDARRRKRSVN
WQTTNKRLSSLRQMNNVPAPAIPLIGAPVENDDSNWGYLLQLSESDDFQDFIGSVNPSKGELRKLGHQAEEFILQCSF
NQHYCDYRNFTTTHNSQYGNCFTFNRPAENNSVLQTGKIGSRFGLHLTLFTDQSEYIGLLSHQSGVRVAIHEPNARPFP
EDEGITASTGTLTSIGLRLRNITRLSGRYSDCRKKNRGDPSEFTHDFDYSLRACLNACYQGKLRYECGCVNDVMGNTTICS
TLDRRQEACQRRVDQMAIDDKLGCPCVSPCRKNLVRVEIFYEKLNYEAYTQMPKYTFGSLLGGIGGIMGFFAGMSLITV
FELCGFIIQLLGLLCGRLTTPMPEEAATEGGEPAPPHPESRIGRVGRRFTKVFSHHPHAHGHGGSARARQTDL

>Deutero_Ambulac_Spurpu_020954_856_327

MKAPKVDGEKSSRGTTPEREEKSLRTILNSRMENSSAHGVPNIQRSSGLVTKLAWSLIFLAGIGVMIWQVVTIFQAYYE
WNYSVIIEVKFNRTQSFPAITLCNANPMRKSKLKTKNASFQETFDVNYVPSMPDLPEQQPQPDVPALAPTCPVGLGHY
RPADQQPLPDMPDGVTPSMLNNSDSGNETQGGSTNEQVDMSEAVKDWKSRIASPSFYVMESVDYEKRRIRVDALA
NETLEERVSLGHKLDDMLLDCSWKGKPCSPENFTKFYDSQLGNCYTFNSGQNGEQLTTTRPGSKYVNGPIDCLVLKPGL
SLELFVQQDEYVEGMTEEASFRVSVHHPSIMPFPADDGVLVSPGFATAIAFIKLELDRLPKPYGDCKADLSTDIEDDIYHQ
HYNITYNMKTCEVSCFQNEVISRCDCFDATYPNSLKVNRTVYPCEYNNRVACPYCQQVLHLIIIKKLKSWESFFSHFCLRF
EITDQFFAHSTLFEKMVEYNAEIRRYVNRENTAEWTRRNMAKVEIFYDEFNYEHIRQEPAYMVCKKTRQIASYDVHKPP
PPPPHSENGRTPCVLPKCPFRNKRKRSTRIDFTHRTFSEKGGAEKGSYTKEKGRAVLTLHTGLLLKKGGGRLEHAYAYPK

>Deutero_Ambulac_Sakowv30039242m_855_330

MSHIGVDNLVSPGSSNCKEKQQTFSSILGSCFVQSSAHGLPNISRAGNAPRRSLWIISLGGTVAIFCLISATLIVRYFDYDV
NVSVAMQFSRELTFPAVTICNLNPLRKSKTQMDGPSDPGQQRAEEAERNSQTSQGTNIKTETTQHPNHIAVSEDIQRN
PFTQVPTTTKIYENRQYSRFTHQTYHVEQDQQSIYTHENNTDGQSLDVTNSDWNGRTLDDTVTNSEENGRPPPETSTG
GAPELVNVLLNETNDTEAEVYDDATGDTEAEVDDDATGDTEAEVDDDETSDTEAKVDDDETGDTEAEVDDDSEASE
WDYFDWDDLDTKTEFYNEPTQYWNKSASLNQWLASMPTQQKEEWGHQLKDFLLDCQWNDVMCSPKNFTKFVSSR
YGNCYTFNSGANNSTVVKTNYAGPYYGLTLELFIEQNEYLDDYPDFAGVRVAIHSQKTMPFPEDDGFNVEPGRVTSVGI
RRTCMKSCYQKRVAERCRCLDGRYPPPIVVIHRTLNACFIGANMAYQYDVMSYLFQDYVNVQLKKKSQQLKLMIEEEE
KGGNEFVRENIVKLQVFYQDLNYEGITQSIAYTEESLASDLGGQVGLWIGISFLTLIEFIELVYDLFKLWIRRLCCMATKNT
GVTNVI

>Protostome_Lophotroco_annelid_Pdum_Contig8214__858_332

GTALQICQHTKMAKKDKGLWKEFSQNTTIHGLNKVAEENGLTFRRLFWITLILGGTALFAYQVVERSLYYIKWPVSVNY
KFVYQSEMPFPTVTICNQNAFKLSTSVQSGLHDVITSIFSKRNSIPKYSRNNISSELEDIFGNSSSLKTDDVWQLLAHAKED
MIYSCSWQEDPCSPDNFTTLLTDHGVCFQFNSASNSHPPLKVTQSGSQQGLSLVMNVEQYEYMTGPHDSVGLKVLLH

DSSHVAMVRDLGEALPTGSHAFVAVTLSKIQNLKHPYGECVNNVSLDYYDTFSLSTCRRDCETRKTVKDCGCRDAYMP
SYTDGTPQVCTLQEYSQCIQANKDQQSGIAGDVCSSNCPIPCLTTSYKSSTTFSTAFENYADDIDKESYWNRTRQHYILA
REISQKVNQGLRRKELEILSRLETEWQVITQIINEVKETVKHSKRFKADILDEMEPRTQFHTFWGLEKIKFVVDYDYVRIW
DVMEERTMAYATIGFYQTAHLYRNMVDKFLLEPELTVEKVAIFTSTMRDLDVRLQLTMSTQINLTQVCDAFANGTLRW
KYLTSPKRDNFVKVNIFMKDLYLEDMESNEAYSLIALLSDIGGSMGLFIGGSVMTILEVLDILVIACMRKSKKKNETNET

>Deuterostome_chordata_Ggallus_sp_Q92075_SCNNA_CHICK_Amiloride_sensitive_sodium_channel_subunit_alpha_OS_Gallus_gallus_OX_9031_GN_SCNN1A_PE_2_SV_1_860_333

MGTASRGGSVKAEKMPEGEKTRQCKQETEQQQKEDEREGLIEFYGSYQDVFQFFCSNTTIHGAIRLVCSKKNKMKTAF
WSVLFILTFGLMYWQFGILYREYFSYPVNLNLNLNSDRLTFPAVTLCTLNPYRYSAIRKKLDELDQITHQTLLDLYDYNMS
LARSDGSAQFSHRRTSRSLLHHVQRHPLRRQKRDNLVSLPENSPSVDKNDWKIGFVLCSENNEDCFHQTYSSGVDAVR
EWYSFHYINILAQMPDAKDLDESDFENFIYACRFNEATCDKANYTHFHHPLYGNCYTFNDNSSSLWTSSLPGINNGLSL
VVRTEQNDFIPLLSTVTGARVMVHDQNEPAFMDDGGFNVRPGIETSISMRKEMTERLGGSYSDCTEDGSDVPVQNLY
SSRYTEQVCIRSCFQLNMVKRCSCAYYFYPLPDGAEYCDYTKHVAWGYCYYKLLAEFKADVLGCFHKCRKPCKMTEYQL
SAGYSRWPSAVSEDWVFYMLSQQNKYNITSKRNGVAKVNIFFEEWNYKTNGESPAFTVVTLLSQLGNQWSLWFGSS
VLSVMELAELILDFTVITFILAFRWFRSKQWHSSPAPPNSHDNTAFQDEASGLDAPHRFTVEAVVTTLPSYNSLEPCGP
SKDGETGLE

>sp_P37090_SCNNB_RAT_Amiloride_sensitive_sodium_channel_subunit_beta_OS_Rattus_norvegicus_OX_10116_GN_Scnn1b_PE_1_SV_2_861_335

MPVKKYLLKCLHRLQKGPGYTYKELLVWYCNNTNTHGPKRIICEGPKKKAMWFLLTLLFACLVCWQWGVFIQTYLSWE
VSVSLSMGFKTMNFPAVTVCNSSPFQYSKVKHLLKDLYKLMEAVLDKILAPKSSHTNTTSTLNFTIWNHTPLVLIDERNP
DHPVVLNLFGDSHNSSNPAPGSTCNAQGCKVAMRLCSANGTVCTFRNFTSATQAVTEWYILQATNIFSQVLPQDLVG
MGYAPDRIILACLFGTEPCSHRNFTPIFYPDYGNCYIFNWGMTEKALPSANPGTEFGLKLILDIGQEDYVPFLASTAGARL
MLHEQRTYPFIREEGIYAMAGTETSIGVLLDKLQGKGEPYSPCTMNGSDVAIQNLYSDYNTTYSIQACLHSCFQDHMIH
NCSCGHYLYPLPAGEKYCNNRDFPDWAYCYLSLQMSVVQRETCLSMCKESCNDTQYKMTISMADWPSEASEDWILH
VLSQERDQSSNITLSRKGIVKLNIYFQEFNYRTIEESPANNIVWLLSNLGGQFGFWMGGSVLCLIEFGEIIIDFIWITVIKLV
ASCKGLRRRRPQAPYTGPPPTVAELVEAHTNFGFQPDTTSCRPNAEVYPDQQTLPIPGTPPPNYDSLRLQPLDTMESDS
EVEAI

>Chordat_hENaCbeta_NP_000327_2_amiloridesensitive_sodium_channel_subunit_beta__Homo_sapiens__1_336

MHVKKYLLKGLHRLQKGPGYTYKELLVWYCDNTNTHGPKRIICEGPKKKAMWFLLTLLFAALVCWQWGIFIRTYLSWE
VSVSLSVGFKTMDFPAVTICNASPFKYSKIKHLLKDLDELMEAVLERILAPELSHANATRNLNFSIWNHTPLVLIDERNPH
HPMVLDLFGDNHNGLTSSSASEKICNAHGCKMAMRLCSLNRTQCTFRNFTSATQALTEWYILQATNIFAQVPQQELVE
MSYPGEQMILACLFGAEPCNYRNFTSIFYPHYGNCYIFNWGMTEKALPSANPGTEFGLKLILDIGQEDYVPFLASTAGVR
LMLHEQRSYPFIRDEGIYAMSGTETSIGVLVDKLQRMGEPYSPCTVNGSEVPVQNFYSDYNTTYSIQACLRSCFQDHMI
RNCNCGHYLYPLPRGEKYCNNRDFPDWAHCYSDLQMSVAQRETCIGMCKESCNDTQYKMTISMADWPSEASEDWI
FHVLSQERDQSTNITLSRKGIVKLNIYFQEFNYRTIEESAANNIVWLLSNLGGQFGFWMGGSVLCLIEFGEIIIDFVWITIIK
LVALAKSLRQRRAQASYAGPPPTVAELVEAHTNFGFQPDTAPRSPNTGPYPSEQALPIPGTPPPNYDSLRLQPLDVIESD
SEGDAI

>Deuterostome_chordata_Ggallus_tr_A0A3Q2U1X6_A0A3Q2U1X6_CHICK_Uncharacterized_protein_OS_Gallus_gallus_OX_9031_GN_SCNN1D_PE_4_SV_1_862_337

MEQEAAREEEERKEGLIEFYDSFKDMFEFFCKNTTIHGTIRLVCSSSNKMKTAFWTLLLLASFGMLYWQFALMFSQYW
DYPVVLTMSMHSEPKMFPAITICNLDPYRFDLVSEHLAQLDRMAEKSVTVLYGINTSASLFHVNEKSIHVRDLPSTGNH
NGSSFKLSQKFSLLRTTEFNNRTGKRQSLVGFRQCNATGGNCFYKTYSSGMDAILEWYRFHYMNIMSQQPVIINISDHE
EKIEDMVYSCQYDGEPCRPSDYVHFHHPVFGSCYTFNSKGTDPFWTATKPGIPYGLSLILRAEQKDHIPLLSTVAGVKVM
IHNHNQTPFLEHEGFDIRPGIATTIGIQQDKVNRLGGNYGKCTTDGSDVKVKLLYNSYTLQACLHSCFQHIMVQKCGCG
YYYYPLPPGAEYCNYNKQPAWGHCFYQLYSRLRNHHLNCFDQCPKPCRESLYKVSAGTAKWPSRKSQDWIRQALRHQ
NGYNSTSNRKDIAKVTIYYKQLNYQSVNESPLLSDNLLLSSMGSQWSLWFGSSVLSVVEMLELLIDTLVLSLLFCYQRFRS
KTLNVARTPSIPSVSLTLESYRVVQEAGNGTAPAHGHTSGVPMAVANSSDPHPAQLSSKAIPEHCPDVVLNGFRYMKD
SSLGGEINH

>Deuterostome_chordata_PetMar_tr_S4RK61_S4RK61_PETMA_Sodium_channel_epithelial_1_gamma_
subunit_OS_Petromyzon_marinus_OX_7757_PE_4_SV_1_863_338

MASEGDSKRVLHRVKDTLKIDGPDPSITDLLDFYLNNTNMHGMRRIAVSKGPIKKTIWIVFSLIAVAMVFWQGIQLIQSF
YSIAVSVTINYQKLPFPAITVCSLNPYKYNQSQALLEKLDRNTAVALHNIGIAVTNLSAAKRDDEPLPIPLVWLDTTVTNQT
VVTDVISGKFHVVPGKVEMRSYFSQNYQSSEPLIAIEVCGEERKCIYNAFTSAIDAVIQWYRLHFINIMAIVPEKDKDKLG
YSADEFIIDCLFSGTVCDPSTSFKKLQHPILGNCFTFNDGSDGKSLDIASAGIDYGLHMVLNTRQDNSLPYLAMGAGAKI
GIHLQNTTPFIEAVGIDIPPAMESSLGLRVNDVQKLGDPYSDCTMDGSDIDVKSLYDSPYSVQTCQNSCFQWEMIKSCG
CANYEQPLPEGSRFCNYDNNPGWEYCYYRLYDMYIKEELKCIQVCRQICSETEHEVTLSLADWPSKASKGWLLRALSKE
QGLPANDTLKPSDIAIVNIYFKDLTQKTISESPASSIVTLLSNLGGLLGLWLSCSMLCVVEVLEIFCVDFPWILLKKLLTTCSS
AFASLIRGPDPAPSVHFPVQLPVGGSPAEDPPTFHTAMQCPREPIPMPNTPPPQYNTLRLRQIAGYVPDGGSDGE

>Protos_Platyhelmin_Macrostomum_lignano_A0A267GUI2_A0A267GUI2_9PLAT_Uncharacterized_prot
ein__Fragment__OS_Macrostomum_lignano_OX_282301_GN_BOX15_Mlig004814g1_PE_3_SV_1_866_
340

CSNFLPNTMTTEKKTPTDEEHKEPDAKTGNSFLELASSWGSNIGMHGIPNITRSSSVAKKFLWTLLLLAGLCLTVVQIKSI
VDKYYSYPIAVARGAEINLPREFPAVTVCNQSPARKSKVQSATSTSTSTTETSNATANSTSTTTMSPAGYSNVTSSTPTST
MNSSTIIAANSTPAPSKRKKRSAGSGYKDEEDSYSGLTSSMYSSGTTRNLSTMDSKTVPDSVRMTYMFLDFYNTLDDDV
KTEIGYQISDMLVDCSMGTSYCSVSNFTRFLHPMYGNCYTFNAAVTNSSRVSVKQQGPLFGLTLTLYIDQSDYVSTVAQ
SAGAVVVLHEPTTQPFPEESGIRVSPGRETYIGMKQTYKKLLGSPYSDNCAIDYKTANKKYNMYRSLDEWALPKVNYTK
MVCLKTCIQRKTESRCNCSSPKLPPSNITRTPALPICKYSYNSTSKSVSQEAECLHAVQESEFETCRSGCQSQCEEMQYDA
SISMAAWPSMGYESDAFHQVMMYNPIVRTQMDGHKTDDSKADFISQNMVKLIVFFADPETRTEISTKGYEITDLLSD
MGGQVGLWLGLSVLTLFELIEMLLDFVVLAASKAALRQPKLPRDSKQPPTENLDKKLSMEMKNSIRFNKQQNPAMWV
DEKHRQP

>Protostome_Lophotroco_annelid_Pdum_Contig18637__869_342

MAGQYIKDPIKRDINKGHTLIICHEFATETSAHGMSHIIRARGTCRKLFWLVVTLGLIAIWLLQSQDTIRKYLKNEVNVQV
EIKASRELPFPAVTICNKNPYKGDDTIRFIELMNHTASLAISRNQSLEDIAPSWFVPWVRSLTGSDDGTFETGISSAKFGE
QMLQLLAELVEVYGEYHNFSLAELSVDPDNFILNCMYERVHCDMRNDIEMTSIFNWLYGGCIVYIIKQNTTRTGPRFGL
ELTLNIEQSDYMDLTTQAGALVLVHPPEEMPFPEDDGISIPPGQAALIGVKVKRTERLGGKFGDCAHANKLPNFTNYFQ
AENPWTNYSLKACQRSCFQWNVRQMCQCLDVRFPPFDEYKNFTSCVNKYFSTFNFSQGFPESVGCLDEVTKKFNAHQ
LGCDKLCHVPCKQTYYEPFVSYAAWPSDAALNVTLENLKGSSASADEILEVGHDNFRKNFLQLTVYFQDLNFEFITEKEA
VELTQMLSDLGGNTGMYVGASLFTMIEFVELFGDLWLWVCCHRMRRKRPKYAPPTEEDLDLYFITPNPSKQNNNHTN

NYQNRQGHAPNQQNRHGHANNKQNRQPHPNKHDRRHSKNNGRAPGRGQDNNGYHFNYNGHLERPRKSLPRTPL
PMDMAEVQDTMVGLY

>Deutero_Ambulac_Apla_gbr2_220_t1_870_343

MMSKNTMTTGIPAGYDDAFVVMPSESIPLPSYHNKAPTSVEAKAAEPEKENHPVTCGSIVTNFADSTTAHGVARIANA
SSCFASFLWLVILCVAFGGFFFQGTNLVLDFFSWPYGTTIDIITNTSVDFPAVTVCNMNRLRRSKLPGTRFEGVIAIDGGIS
GGDNDYSWFFEWSSAGDFYNQFVSASAGGGSSGGGGGSSAGGGNSSSVGGGSSAGGGNSSSVGGGSSAGGGGSS
AGGGGSSAGGGGGSSAAGSSYSYPPASSNYYYWETEFGDDFNFEYDNYYDFGTVTGESDWDGFLANSKSEDFSDIINVA
NPTQDEMDELGHQAEDFILQCTFDRRKCNYTDFYKFQNSHYGNCFTFNHGRNETVRTTSKSGFQYGLHLTLFIEQPEYV
GLFSPESGVRVSINHWQTTPHPEDSGITATTGQATSIALRKNFIKRLGGWYSNCTYGTETNFTSDFFTYSPLTCKKQCMQ
RNLKDRCDCVTDLLLDGEKCSYLNSTQQRCRQLVEALYEEDKLDCLCPVACEENVFKTSASVSIWPSERYEEHLYSRLSSK
NAVAARMLQDVETTRKNLARVRIYFEELNYQKVEQIPQWTIESILGAVGGLMGLYVGISSITLMEIIVFVFSLLKQCCKSVI
CLNKVQPSN

>Chordat_hASIC4_NP_878267_2_acidsensing_ion_channel_4_isoform_2__Homo_sapiens__6_344

MLSGAAGAARRGGAALAPSLTRSLAGTHAGADSCAGADKGSHKETIEERDKRQQRQQRQRQHQGCGAAGSGSDSP
TSGPHPVPVLFPLALSLEEQPLPPLPLGRAPGLLAREGQGREALASPSSRGQMPIEIVCKIKFAEEDAKPKEKEAGDEQSLL
GAVAPGAAPRDLATFASTSTLHGLGRACGPGPHGLRRTLWALALLTSLAAFLYQAAGLARGYLTRPHLVAMDPAAPAP
VAGFPAVTLCNINRFRHSALSDADIFHLANLTGLPPKDRDGHRAAGLRYPEPDMVDILNRTGHQLADMLKSCNFSGHH
CSASNFSVVYTRYGKCYTFNADPRSSLPSRAGGMGSGLEIMLDIQQEEYLPIWRETNETSFEAGIRVQIHSQEEPPYIHQL
GFGVSPGFQTFVSCQEQRLTYLPQPWGNCRAESELREPELQGYSAYSVSACRLRCEKEAVLQRCHCRMVHMPDSLGG
GPEGPCFCPTPCNLTRYGKEISMVRIPNRGSARYLARKYNRNETYIRENFLVLDVFFEALTSEAMEQRAAYGLSALLGDL
GGQMGLFIGASILTLLEILDYIYEVSWDRLKRVWRRPKTPLRTSTGGISTLGLQELKEQSPCPSRGRVEGGGVSSLLPNHH
HPHGPPGGLFEDFAC

>Chordat_hENaCgamma_NP_001030_2_amiloridesensitive_sodium_channel_subunit_gamma__Homo_
sapiens__2_346

MAPGEKIKAKIKKNLPVTGPQAPTIKELMRWYCLNTNTHGCRRIVVSRGRLRRLLWIGFTLTAVALILWQCALLVFSFYT
VSVSIKVHFRKLDFPAVTICNINPYKYSTVRHLLADLEQETREALKSLYGFPESRKRREAESWNSVSEGKQPRFSHRIPLLIF
DQDEKGKARDFFTGRKRKVGGSIIHKASNVMHIESKQVVGFQLCSNDTSDCATYTFSSGINAIQEWYKLHYMNIMAQV
PLEKKINMSYSAEELLVTCFFDGVSCDARNFTLFHHPMHGNCYTFNNRENETILSTSMGGSEYGLQVILYINEEEYNPFLV
SSTGAKVIIHRQDEYPFVEDVGTEIETAMVTSIGMHLTESFKLSEPYSQCTEDGSDVPIRNIYNAAYSLQICLHSCFQTKM
VEKCGCAQYSQPLPPAANYCNYQQHPNWMYCYYQLHRAFVQEELGCQSVCKEACSFKEWTLTTSLAQWPSVVSEK
WLLPVLTWDQGRQVNKKLNKTDLAKLLIFYKDLNQRSIMESPANSIEMLLSNFGGQLGLWMSCSVVCVIEIIEVFFIDFF
SIIARRQWQKAKEWWAWKQAPPCPEAPRSPQGQDNPALDIDDDLPTFNSALHLPPALGTQVPGTPPPKYNTLRLERA
FSNQLTDTQMLDEL

>Deuterostome_chordata_Ggallus_tr_F1NW62_F1NW62_CHICK_Uncharacterized_protein_OS_Gallus_g
allus_OX_9031_GN_SCNN1G_PE_4_SV_2_875_347

MAPGKITARIKKTLPVRGPQAPTLRELMRWYCLNTNTHGCRRIVVSRGRLRRFIWILLTLSAVGLILWQCAELLLNYYSAS
VSVTVQFQKLPFPAVTICNINPYKYSSMKDYLSELDKETKKALETFYGFSEGKTKVRRAAGDWNGTESLFFRHVPLLRFEN
SFRAATDLRSGRKRKVEGSVFHKDSSIVNSGDSNDIIGFQLCDANNSSECALYTFSSGVNAIQEWYKLHYMNIMAQIPLE

TKEELSYSADDLLLTCFFDGLSCDKRHFTRFHHPLHGNCYTFNSGENGTVLSTSTGGSEYGLQVVLYIDEADYNPFLVTST
GAKIIVHDQDEYPFIEDIGTEIETAAATSIGMHFTRSRKLSKPYSDCTETGADIPVENLYNKSYSLQICLHSCFQKAMVESC
GCAQYAQPLPNGAEYCNYKKNPNWMYCYYRLHEKFVKEQLGCQQICKDACSFKEWALTTSIAQWPSTVSEDWMLRV
LSWDKGQKINKKLNKTDLANLMVFYKDLNERFISENPANTLVILLSNFGGQLGLWMSCSVVCVIEIIEVFFIDSFSIVMRR
QWQKAKKWWNHRKRDETGKPPEVGDAEQQGHDNPACSDEDLPTFNTALRLPLPQEGHPPRTPPPNYSTLRLETAFT
EQLPDTLEAGQH

>sp_P37091_SCNNG_RAT_Amiloride_sensitive_sodium_channel_subunit_gamma_OS_Rattus_norvegicus_OX_10116_GN_Scnn1g_PE_1_SV_2_877_348

MAPGEKIKAKIKKNLPVRGPQAPTIKDLMHWYCMNTNTHGCRRIVVSRGRLRRLLWIAFTLTAVALIIWQCALLVFSFY
TVSVSIKVHFQKLDFPAVTICNINPYKYSAVSDLLTDLDSETKQALLSLYGVKESRKRREAGSMPSTLEGTPPRFFKLIPLLV
FNENEKGKARDFFTGRKRKISGKIIHKASNVMHVHESKKLVGFQLCSNDTSDCATYTFSSGINAIQEWYKLHYMNIMAQ
VPLEKKINMSYSAEELLVTCFFDGMSCDARNFTLFHHPMYGNCYTFNNKENATILSTSMGGSEYGLQVILYINEDEYNPF
LVSSTGAKVLIHQQNEYPFIEDVGMEIETAMSTSIGMHLTESFKLSEPYSQCTEDGSDVPVTNIYNAAYSLQICLYSCFQT
KMVEKCGCAQYSQPLPPAANYCNYQQHPNWMYCYYQLYQAFVREELGCQSVCKQSCSFKEWTLTTSLAQWPSEASE
KWLLNVLTWDQSQQINKKLNKTDLAKLLIFYKDLNQRSIMESPANSIEMLLSNFGGQLGLWMSCSVVCVIEIIEVFFIDFF
SIIARRQWHKAKDWWARRQTPPSTETPSSRQGQDNPALDTDDDLPTFTSAMRLPPAPGSTVPGTPPPRYNTLRLDRA
FSSQLTDTQLTNEL

>Deuterostome_chordata_PetMar_tr_S4RY81_S4RY81_PETMA_Sodium_channel_epithelial_1_beta_subunit_OS_Petromyzon_marinus_OX_7757_GN_SCNN1B_PE_4_SV_1_879_349

MKIRKYLTRSLHRLQKGPVASVSELLYWYCMNTNTHGCKRIVVYGKKKRVLWFLITIIMLGVVLWQWVLLFQAYLSYGV
SVSVNMGFQRMNFPAVTVCNLNAYRYSSMKDKIKDLEAYTRVALQTLYNYTDSSTPSAYDTSYAVGPWQEIPLVLIDRR
DPNRTVVTEVMCSRAAVGIETHTVDNRVFHIGFCKSALGVCCDSAGDKCFYSEYLSGMTAVKQWFHFNLLSLLGNLST
EEKSNLSSSGDELIRSCLFSSDTCSATNFTTFFHPMYGNCFIFNWGENETVMQVSNPGVEYGLKLVLSIDQDEYIPFLTTIA
GAVIMVHDQNTYPFLSDLGVFVKTGVETSVGIEVGQLQRQGAPYSDCTMDGTDLPITNLYNGTAYSVQACLRSCFQTK
MIEMCGCGYYLYPLPPGEKYCQNQNFTGWRYCYYKLYEQFVEEDMDCYTICKQPCIESEYKMSISMSDWPSQSSEDWI
FHILSKERKHNVSRIFNRKQDIIKLNLFFQEFNSMTISESPAQTIVTLLSNLGGQFGFWMGGSVLCIIEFIEIIIDCVWIGMIK
ASNDVRERRKTSRKPRYSDEPPTLSSIVQGQGNSGFEMEERGPPGEAPEANGSAAQPPAAAAEQQPDVPGTPPPHYD
TLRISKTELHDEINSDDDGEFV

>Cyclo_DEL1_sp_Q19038_1_DEL1_CAEEL_RecName__Full_Degenerin_del1_46_351

MARKYIDILKKSKMMLFQDVGKSFEDDSPCKEEAPKTQIQHSVRDFCEQTTFHGVNMIFTTSLYWVRFLWVVVSLVCIC
LCMYSFSHVKDKYDRKEKIVNVELVFESAPFPAITVCNLNPFKNHLARSVPEISETLDAFHQAVVYSNDATMDELSGRGR
RSLNDGPSFKYLQYEPVYSDCSCVPGRQECIAQTSAPRTLENACICNYDRHDGSAWPCYSAQTWEKSICPECNDIGFCN
VPNTTGSGNIPCYCQLEMGYCVFQPESRVRRIWEFQGNKIPEKGSPLRKEYMEQLTQLGYGNMTDQVAITTQAKEKMI
LKMSGLHPQRRAALGYGKSELIKMCSFNGQQCNIDTEFKLHIDPSFGNCYTFNANPEKKLASSRAGPSYGLRLMMFVN
SSDYLPTTEATGVRIAIHGKEECPFPDTFGYSAPTGVISSFGISLRNINRLPQPYGNCLQKDNPQSRSIYKGYKYEPEGCFRS
CYQYRIIAKCGCADPRYPKPWKRSAWCDSTNTTTLNCLTTEGAKLSTKENQKHCKCIQPCQQDQYTTTYSAAKWPSGSI
QTSCDNHSKDCNSYLREHAAMIEIYYEQMSYEILRESESYSWFNLMADMGGQAGLFLGASIMSVIEFLFFAVRTLGIACK
PRRWRQKTELLRAEELNDAEKGVSTNNN

>Protostome_Lophotroco_brch_Lingula_anatina_comp148540_c0_seq7_p1_comp148540_c0__comp14
8540_c0_seq7_p1__ORF_type_complete_len_666___score_77_65_comp148540_c0_seq7_79_2076__8
87_353

MSTGAQTMYPSEKNQFDFFKNSLVVDHNSQDPEKSAREESLIALAYDWASNSPVNGIPNVVRSQNLIKRFFWLVALLG
CFGVMGYQTWELVAKFYRFPVDTTLTYTHSKEVYFPAVTVCNVNPLRRSMLSSAGDDVYALLGPPKSQMGSSPGGAQ
APAGPAGSFASQSQTTQAAAGPPGPASSSITTQAPAAPGPKGSGSGGTTQAPSASQTTAGAPAGGGGGTTQAPAGP
PAPPSGRKRRSTGTNSTSTSRRKPGQMNERFEMRKRFGNIWAKLNYTTRETVGHELDTMIISCTFNGVTCSSANFTRF
NNHQFGNCYTFNSGWDSSIPVETSSNAGPLYGLSLEMYIEQSEYIGDLSESAGVRVQIHSQRSMAFPEDEGFNIAPGYL
TSMAMTRVEITRRPHPYPSKCRNFTTEESKQKSVFTSVHNVDYSVSGCMKTCYQRYVISECACGDPAYPFSLDLEAFSDL
KNNINGTTAESVDPCVSDADDECVAGIKRRFADDDLSCDCPLTCVDVEYSGVPSLAKWPSKQYMSTLHATLASAGAHL
DNIINGKNGMAAEPGDNLLKLEVYFQELNFQKISENVAYDVFALLADIGGQIGFWVGLSIMALFEVVELILDVFRLLLFRV
GSFCFKKQQKRPPSRQVKVQPASEENVLTPRLEKVRFS

>Deuterostome_chordata_Ggallus_tr_F1NE95_F1NE95_CHICK_Uncharacterized_protein_OS_Gallus_gall
us_OX_9031_GN_SCNN1B_PE_4_SV_4_889_354

MNLKRYFVRALHRLQKGPGYTYKELLVWYCDNTNTHGPKRIIKEGPKKKVMWFFLTLLFASLVFWQWGILINTYLSYNV
TSSLSIGFKTMKFPAVTVCNANPFKYSEVRPLLKELDKLIEAALERILQPTHGDPISPLLLNNSNATEGLDLDLWNQIPLVLI
DEQDKDNPVIVEIFETNQSAAGNQTAAPPAPANVTSEEKKYKLAVKLCSHQGSNNCTYRNFTSAAQAVTEWYILQSTSI
LSKVPLQERIRMGYQAEDMILACLYGAEPCNYKNFTQIYHPDHGNCYIFNWGMDKEALNSSNPGAEFGLKLILDISQQ
DYIPYLSSAAGARLMLHQQKSFPFLKDQGIYAMAGTETSIGVLVDELERMGYPYSDCTANGSDVPVKNLYSEYNTSYSIQ
ACLRSCFQNHMTEICGCGHYMFPLPEGVTYCNNEDNPGWAYCYSSLRSSIRHRQICIDSCKETCNDTQYKMTISMAD
WPSEASEDWIFHILSYERDMSTNVTLDRNGIIKLNIYFQEYNYRTISESAATTIVWLLSSLGGQFGFWMGGSVLCLIEFGE
IIIDSLWITVINIISWCKGLKQKRVRARYPDTPPTVSELVEAHTNLGFQHEEAGTETQGEALPPEPGTPPPNYDSLRVQPS
HNPGTDSDIECEEQRPAANHHGDASVWAE

>Protostome_Lophotroco_annelid_CAC9584386_1__Ofus_G061163_Owenia__fusiformis__891_355

MNEKDDLDNMTVIDTKEEEIKGASDAKTVTDEFMGSFGAHGLPRVWSSASPVKKIVWLVLFLAASGYCLNIIIRVGIKFG
TFPSGINSKVYPRSIVDFPAVTICNLNMLKATSISNNNVEKYINLLSVQSEAEASMNSWITWALNYTDSVTTSFGPTTDST
SRTNSSNAGTTDHTTVADRRKRAVEEFPFNEDLLEDELLLNLRADHVNDLTRLIHMPEVVKMATLRLLKKRHGFDMRE
QRSAPPIPHRQKRDNQAAETNSSDYYDDNFYYYDNYDYSYYDDYGFGDVSENDFRTLLTKSKTDDYQDLMGVLKPTKT
ELELYGHSAQDMIVSCTFDSKKCNYTLFKTFQNSYYGNCFTFNYDDGNSTTQEVVFNTTKKGSRFGLKLTLNIERQEYIGL
FAHGSGVRVAVHPRNATPFPEDYGISAPTGWETAIGVRENRVTRLTKPFESNCSHGNHDPSETYTGLYTLLECNQRCLQ
KQIRSKCKCLDEIMDDQTWESVYASDSATSMLNPCSILNSTEEKCRQRMYYLSQTNRLGCDCPLPCLELGFEESLHSSE
WPSAQYYPLMMQKLTDSNIISEGQLKIMNDLKTTRENFVRLHVYYKDMILESLEEFEAYTVDSLIGDIGGILGLFIGMSVL
TIAEAVEYLIDLCLLRCKTNNKRNSVVALK

>Chordat_hENaCalpha_NP_001029_1_amiloridesensitive_sodium_channel_subunit_alpha_isoform_1__
Homo_sapiens__0_356

MEGNKLEEQDSSPPQSTPGLMKGNKREEQGLGPEPAAPQQPTAEEEALIEFHRSYRELFEFFCNNTTIHGAIRLVCSQH
NRMKTAFWAVLWLCTFGMMYWQFGLLFGEYFSYPVSLNINLNSDKLVFPAVTICTLNPYRYPEIKEELEELDRITEQTLF
DLYKYSSFTTLVAGSRSRRDLRGTLPHPLQRLRVPPPPHGARRARSVASSLRDNNPQVDWKDWKIGFQLCNQNKSDCF
YQTYSSGVDAVREWYRFHYINILSRLPETLPSLEEDTLGNFIFACRFNQVSCNQANYSHFHHPMYGNCYTFNDKNNSNL
WMSSMPGINNGLSLMLRAEQNDFIPLLSTVTGARVMVHGQDEPAFMDDGGFNLRPGVETSISMRKETLDRLGGDYG

DCTKNGSDVPVENLYPSKYTQQVCIHSCFQESMIKECGCAYIFYPRPQNVEYCDYRKHSSWGYCYYKLQVDFSSDHLGC
FTKCRKPCSVTSYQLSAGYSRWPSVTSQEWVFQMLSRQNNYTVNNKRNGVAKVNIFFKELNYKTNSESPSVTMVTLLS
NLGSQWSLWFGSSVLSVVEMAELVFDLLVIMFLMLLRRFRSRYWSPGRGGRGAQEVASTLASSPPSHFCPHPMSLSLS
QPGPAPSPALTAPPPAYATLGPRPSPGGSAGASSSTCPLGGP

>Deutero_Ambulac_Sakowv30002567m_893_357

MADISVSPEVNKQALRGWSAQRKKDIRNDGYKTWKKFSQETSLAGVKFVGDDSTIYSRRIIWLLIVLAGAAGFVYQVYR
MVDTFAEMPVTVNYRYTYPEYQRSFPAITLCNNNLFSYVSMEKYYGILGITEYLFLLFSFIPSKGLEEMAGFIPLEEVPENF
GDNLTAFEYMDMHNHDIKDMLISCDFGGIYCDAHNFTSTVTNGGICYTFNGGQGNEDLLKLTGTGLTHGLNIVLDANK
NDYLAPSDTVGFQFAVHDQKDIPNIRDKGIGIPTGMHSRIALSTTAISNLGSPHSDCVTDHTRTLNYFPGEYTESKCLME
CEEDFAVRQCECRYYYMPAYAFRTNANQNFILENFTVHTSTICDCPERCNTISYNWKLSYNTYPTNVISIFKADMGYPPC
DRAVGNYVFFATTANESYTSGALPLIDAIINNTISAQDLTRLSPSRGIPDSVFRDVTYWAALEWMIDLNYEICFNSLFYILY
VQKGNYSFYDGKETSYHNQTIALINEKYSSQEKLDQIFTLVLKNNNISLLKEEFVNVSRVFYENYYLQILRQALEVTYGYFW
VERNIVIDYLFRYRRHSDICEKFMRENFAKVSIFFEDLKFENITQKADYEPFQLVCDFGGSLGLFFGASLISFLEIFDFLLSWT
FARFKKKNQMNPDEELSETPGTIVK

>Deu_Ambulacrar_hemi_Ptyfla_40v0_9_20150316_1g10161_t1_scaffold5763_cov99_895_358

MWSSNNTVQPNGLFKDTTGEKNGTELTDFGSNLRLTREPGRMDHGAAGKRPGCMGILLSGFADDCGISGMKFLAGK
SVVPLRRIVWLLVLLTGIGLLSYQILDRVSYFAIYPLNVNVQLNYVEEVTFPAIAICNFNMYSLQYTNELNLTGLFYEMYSTY
NPNFTNYDIELTASEQFYRDAAQQPSAMFYRATWQWRPMEIDAVQSILTDFGACYAFNTGNDKYGIRKVRNKGTSHG
LSVYLDVNQWDYNIGPRVGAGFQLMLYEQGDVPMVSDLGFSVAPGEDVRVGIEVTKVTNLPPPHGVCAEKSLQYYDR
YSVNACQLECKTDYVVERCGCRMYYMPGNATVCDLQQFSFCVQTTLWYMNQTCDCPVPCTQVMYKPSLSHAKYPS
DAYATYLENTYGFPKDVFSLTYTNEVNLTGILYEMYGTFNPNFDNYDIEITTSEQFYRDAAQQPSSMFYTATWQWRPM
ELDAVQSILTDFGACYAFNTGNDKYGIRKVRSKGTSHGLSVNLDVNQWDYNVGPRVGAGFQLMLYEQGDVPMISDL
GFAVAPGEDVRVGIEVIKVVNLPPPHGICGEKSLKYYNRYSVNACQMECKTDHVVEHCGCKMFFMPGNANICDVKHY
YNCLQYAEFSFTNVTCDCPVPCTQVIYKPSLSHAKYPSDAYAMHLENAYGFPKEVFS

>Protostome_Lophotroco_annelid_CAC9662587_1__Ofus_G12132_Owenia__fusiformis__896_359

MSEKNVEPHDASVKDLTQGFADATSLHGLPRVYSSKSLTRKIIWGLVFFGCFVGFCWQVTALTLNYFEYPISVTTEVKTR
LKVDFPAVTVCNMNMLKKSLLMGTRFENLTKVDKRLSDIYGTSATSNSQTTPPAEMPQDPENEDPLSGADFNEGSTTE
MDTEDDGTSTIEAEDTTDGITYVTTEDPYDETIDTTQEDIATEFITTEASAGDATTDYDYLTTETSRKKRSVGKRRPKENE
HKLPEQLHKQMKPLDGSAHKHAHPYSNNAKGESRRRKRQADSYYDDNYEDDGDEYDSDPYDDYYDNYYDNGDDYYP
EERFEFEDTGFSEIEPNDYFGFLQNSQTQDLSDLVGIVVPTTDEMDSYGHTFEDFVLQCSFDKKNCSNSDWKKLYNSQY
GNCYTWNFGYNNTIKSTSRFGSRYGLRLTLNAQADEYIGLLSHTVGARVTVHSHNVMPFPEDQGVSASVGRKTGIGVQ
MQNIKRKPHPFPTNCTYGTHLQSNYEGDYSVLSCMMSCLQNKIKTNCKCVDKIVHNQTACDITNTTQEKCRQRMYHL
FDEAKLGCDCPQACEELVYGTTISGSEWPSNQYSPYLLQKLEGGSPVSDIKQNLVRVHVYFETLAVHTIEEVPSYTWDNL
LADIGGTMGMFIGISICTAFEIVELLMEIGKLVVGKMTNKNKVTSLK

>Placozoa_TadNaC1_XP_002114386_1_hypothetical_protein_TRIADDRAFT_58138__Trichoplax_adhaer
ens__11_360

MNEELNEGSEKPITSSLYTKNFKVKTGKEFKDDLLTPKDQEDYSVFEKIELTTPSSPNAKDSKFEIEKEYQYHQEDRSVFKA
VKQPIPSLSCIESSKVKRDKVFENVLLTPGDDQEDYPAVEKVKHPMLSLACAESSKVKADKEFEDIILTPYDDNDDSSVFEE
ANQTTVSASYAENLHISTKDKFEFSDDSNYDDRFALQTSCNGIIRIFGRGGRVRHAVWFLLTLTMTILCIITCVQRYDYLFT
YPTNTAINYTVSKKLKFPAVSVCNFNRFRFSSLEYGDWHRIGYLVNLFTTTDNNQIFTGLNGKTGKEWNDYLKNISFELY

DNITFDITQFLNVKSNQAEVFIKHCTWNDGRQPCSIENFTRIYTDYGSCFTFNAGVDAPILYQKRPGSRYGLKLILNIEEEE
YTHLNPDPDIGIKFRVHNQFEPPDINAEGIAVPPGYHAYTKLLYTESDFLKPPWGNCGQKKLKYFKSYSRASCQLECLADS
YRRRCDCRTPYMPGSSPICSPEHIKKCVTKYLGLSTPENFTCNCPNDCRIKSFNPHVTYAEIPLRQTSRFAAHRYGLNELEI
DFLKYNNISMGEYIRDNYVFLDLFYDDLSYTTFKEKKAYDVNQFISDIGGQLGLFLGGSFLTFWEIFEWSQIKSFLVIRKIIH
EYKKGRRRTRRRFNSTPEDTERLL

>AcFaNaC_XP_012938733_1_PREDICTED__FMRFamideactivated_amiloridesensitive_sodium_channel_isoform_X1__Aplysia_californica__10_362

MWGRGKRQRNKNYPSGSGGGGGAFRSPAMRNDNELEGFVSILHTSGDNYVPIRDSSADHMKYTSVSAKSGMVPEH
RYTMVRSRHHGRHHHHHSYQEYNTQRSAISLIAELGSESNAHGLAKIVTSRDTKRKVIWALMVIIGFTAATLQLSLLVRK
YLQFQVVELSEIKDSMPVEYPSVTICNIEPISLRKIRKAYNKNESQNLKDWLNFTQTFHFKDMSFMNSIRAFYENLGSDAK
KISHDLRDLLIHCRFNREECTTENFTSSFDGNYFNCFTFNGGQLRDQLQMHATGPENGLSLIISIEKDEPLPGTYGVYNFE
NNILHSAGVRVVVHAPGSMPSPVDHGFDIPPGYSSSVGLKALLHTRLSEPYGNCTEDSLEGIQTYRNTFFACLQLCKQRR
LIRECKCKSSALPDLSVENITFCGVIPDWKDIRRNVTGEYKMNQTIPTISLACEARVQKQLNNDRSYETECGCYQPCSETS
YLKSVSLSYWPLEFYQLSALERFFSQKNPTDQQHFMKIAQDFLSRLAHPQQQALARNNSHDKDILTTSYSLSEKEMAKE
ASDLIRQNLLRLNIYLEDLSVVEYRQLPAYGLADLFADIGGTLGLWMGISVLTIMELMELIIRLFALIFNAEREVPKAPVHSS
NNGGGGGGDGQHNFANGDVEHERDTHFPDLGSSDFDFRRGGGIGAESPV

>Deu_Ambulacrar_hemi_Ptyfla_40v0_9_20150316_1g2128_t1_scaffold965_cov108_905_363

MTNSFEEIDKFSHISPDIVPNQRLDVIVTHVSPSIIMVTLDIHSDYDHGHTALPSMYTLDIEPDIGDAFPINITVDKESETEY
TFSENELGCFIGRSCQDLLNPCDGNPCGEHGLCFRDSQSCFQYACVCNECHAGKNCQIHVNDPCQFFSPCRNGGLCIPS
PDSCMEYTCECSQCFTGKFCEQVPCEPNDFQCNSGIHCVKKYLVCDGIYHCPDTSDEFGCDYHSYECSRNQIKCETGGP
SGICIDDSKQCDGVEDCYQGGDEANCDTYGEKNTHICTGFQCDDGLCHPLEDRCNYFTDCNDATDEEDCKFRDCQPW
EFSCDNHRCIDADSACDWKDDCGDNSDEVACDYRECTESEFKCVSNSQCIAGWKRCNSYSECEDQSDELECTTPGNS
HPGREMLFRGNWTNIYANVTTNQTYFDDFQYHYFTDPGFTRVKREKPPDLNGFVTFSSTPDFSDVRRVLKLTADEIEEY
GHQAKDFILQXLKLTLFVEQDEYLGVFGQSAGVRVTVHPNNQLPWPEDVGMTAKTGAATSFAIKQTCKKRCVQDYMI
RYCRCTDTFDSDHAQCPILNILQEACRQIIHYFYQKALLACDCPPLCKKNLALVSVYYETLSSDLVKESPGYGGEDLTSDLG
GLLGLYIGVSVITTIECVIFVKGVIVIAFRSMRDNNKDSRDITKDEIDESASDDSR

>Deutero_Ambulac_Spurpu_018442_909_366

MDKSRRPGASYQSLCYRASPIIAKENSFFPPEYDDIHPMKTDSVKAVLNEYSGVTTAHGVPRIITSKSILSKLFWACVTLAA
LGAFLWQGSLLLFDYKGHPYTTQIDVVTQTEVQFPAVTVCNMNKMRRSAMVGTRFESLIEADGGAMGGDVDYSWW
FDWSSEWWLKYEDSSSSREESSKEDIGPSAEGSVISGTSQSVVDMSGDSATDGSSFNTMANFTDIPDGNSSTYPSSTPQ
NQEYSTQSDVDTASATSDGTESFVDEEWSSTDEVPPLRKRRSGLRLIPDPTTGKLDQRRFVRRKRQTNTGPSSSDPSPS
DSSEDPERFDWWEPEWETNMFDYQAYDWGDVTSDNDWEGFYRQSTADDFSDLLDVINPTREELEIMGHQAEDFIL
QCTFDRRPCNYTNFRQFQNKYYGNCFTFNQEVGNSSNARRTGQTGAQYGLHLTLFTEQPEYVGLFAQEAGVRVAIHP
PNVFPFPEDDGVVASTGQATNIGIRQSYFERLTKPHGNCSDGTQTNFTSEEYTYTTRACVKSCVQQHIFNKCGCVTDIM
MNDTICSPRNRTEQMCRQAIEQFFHEGQLECFCPIACNDLNSLDHVITFQTHLKSRLTNEKATRILQNIEQTRKNLARVR
IYFEELNYEQMIQKPQYTFESLLGGIGGLLGLYIGFSVITICEVGVLVLDIVKYLFRKAYSHDRVVPVDFKT

>Protostome_Lophotroco_brch_Lingula_anatina_comp148540_c0_seq4_p1_comp148540_c0__comp148540_c0_seq4_p1__ORF_type_complete_len_699___score_91_82_comp148540_c0_seq4_170_2266__907_367

MQQDKYEQEQRTTSQRSLLNSQKNGGTGFDSEQSENGKYSSCSKTASFEGGGLFVRQAQDQLPSAAEQQDETSMLLLCKNWGDNSPINGVPNIARSTSGVGKVTWTLLLLMCFGVMGYQTWELVAKFYRFPVDTTLTYTHSKEVYFPAVTVCNVNPLRRSMLSSAGDDVYALLGPPKSQMGSSPGGAQAPAGPAGSFASQSQTTQAAAGPPGPASSSITTQAPAAPGPKGSGSGGTTQAPSASQTTAGAPAGGGGGGTTQAPAGPPAPPSGRKRRSTGTNSTSTSRRKPGQMNERFEMRKRFGNIWAKLNYTTRETVGHELDTMIISCTFNGVTCSSANFTRFNNHQFGNCYTFNSGWDSSIPVETSSNAGPLYGLSLEMYIEQSEYIGDLSESAGVRVQIHSQRSMAFPEDEGFNIAPGYLTSMAMTRVEITRRPHPYPSKCRNFTTEESKQKSVFTSVHNVDYSVSGCMKTCYQRYVISECACGDPAYPFSLDLEAFSDLKNNINGTTAESVDPCVSDADDECVAGIKRRFADDDLSCDCPLTCVDVEYSGVPSLAKWPSKQYMSTLHATLASAGAHLDNIINGKNGMAAEPGDNLLKLEVYFQELNFQKISENVAYDVFALLADIGGQIGFWVGLSIMALFEVVELILDVFRLLLFRVGSFCFKKQQKRPPSRQVKVQPASEENVLTPRLEKVRFS

>sp_P37089_SCNNA_RAT_Amiloride_sensitive_sodium_channel_subunit_alpha_OS_Rattus_norvegicus_OX_10116_GN_Scnn1a_PE_1_SV_2_908_368

MLDHTRAPELNIDLDLHASNSPKGSMKGNQFKEQDPCPPQPMQGLGKGDKREEQGLGPEPSAPRQPTEEEEALIEFHRSYRELFQFFCNNTTIHGAIRLVCSKHNRMKTAFWAVLWLCTFGMMYWQFALLFEEYLSYPVSLNINLNSDKLVFPAVTVCTLNPYRYTEIKEELEELDRITEQTLFDLYKYNSSYTRQAGARRRSSRDLLGAFPHPLQRLRTPPPPYSGRTARSGSSSVRDNNPQVDRKDWKIGFQLCNQNKSDCFYQTYSSGVDAVREWYRFHYINILSRLSDTSPALEEEALGNFIFTCRFNQAPCNQANYSKFHHPMYGNCYTFNDKNNSNLWMSSMPGVNNGLSLTLRTEQNDFIPLLSTVTGARVMVHGQDEPAFMDDGGFNLRPGVETSISMRKEALDSLGGNYGDCTENGSDVPVKNLYPSKYTQQVCIHSCFQENMIKKCGCAYIFYPKPKGVEFCDYRKQSSWGYCYYKLQGAFSLDSLGCFSKCRKPCSVINYKLSAGYSRWPSVKSQDWIFEMLSLQNNYTINNKRNGVAKLNIFFKELNYKTNSESPSVTMVSLLSNLGSQWSLWFGSSVLSVVEMAELIFDLLVITLLMLLRRFRSRYWSPGRGARGAREVASTPASSFPSRFCPHPTSPPPSLPQQGMTPPLALTAPPPAYATLGPSAPPLDSAAPDCSACALAAL

>Protostome_Lophotroco_annelid_CAC9509085_1__Ofus_G024052_Owenia__fusiformis__920_371

MAKQVECKKNVIALFKATLHTSPKITQQSVLTDGYVIIVQNKDDENQIAIEMKEMIKCDKDTFAPSVKQDTKCIGNGYVKADSSIDSYTLYDLENVVSVITDVQGDKEGSINPKVLNEQTDDSSQNGTQTVKKSLKVQIRSALEEFSNTTSIHGPKRILKANGKYSTVGWTFVFIGVIILTVTLVTGVVVKYYSYPVEDGISITTPPLGFPAVTLCPLIPFDIDQILVDLENEAIVTPDHVYDLIRYFFLRSEHFTPDIECISDVNCEVEADDNILQSWLELSSWLHRKLNPTYRWQVENIPLNETIVSSMLPTKTEEFILECTYNERQCDEQIEVTRILTAYGKCFTLNIKNQAEHINEIGPNKGITLILYTGKHNPIQGIIGAEFSTMGFTSTYSQPDDGVQVVIHNPGTMPQPFREGFHVTPGRFSSIKLTKTERTRLLPPYGECTDENYLFNSEFKYSYEMCVEQCLQERIMQECGCVSPMYIIPRNHNFNQTQYCGNISSVLSDKMLINAMFFRNSNNKLIKALKGVIDNLKCEKKLSKIGKELCQCKRPCKENTNIHLINTIPWPEKSVVNTGGQRGFVKFKKKHKNVSKLIKQLIKYTGNLHSLNDSIDDMIRDFLTHIDHVNMSKPEYNRDSLAIDVVETNFMKLNIHFESLDVRTVYEERTYTYADVMKDIGNVFGFYLGMSAVSVAEGVYIIGVMIKLIISRILC

>Protostome_Lophotroco_annelid_CAC9657286_1__Ofus_G102358_Owenia__fusiformis__922_373

MVDKKTFSSRCDRTSPQQMITILTYDGDQGKCAKAYVMTSDDEDFIALRPLAYQRGAKPKFVVKNAEDIDIDKLSDDSDFDDKSLKDIAVEFANETTLHGVPKIIKDRSCGTRLFWSCVCLSALCMFLIQASTLMGKYLDYKKKVDIEVLDDPAPFPSISLCNIRHIDAILFEEIQGYSQQHYMKVLEVMEASKNPNYVRNKTVEEDKPFYWTSLETSINVQGYLSTKPELVKELGFDNIN

DLTEIMFTRETPAANIDKDKVTQFGIEAIEFIITCVYSGDRCNYSDFKRFFHPFFYNCFTFNTSSFLTDNSTTGRSFSLIAFLG
KMFSKESTKLDRGDTFGDPVYNSDGLRIVIHSSNSEPDPIQDGFNIPAGFSASIGVKATQYERIDYPYGNCSEQGSLSMD
NGIYDYTLISCQNLCLQNEIIEHCQCIDIALPIPENVNVSFCQDIENPPIFDCLEDKANEWKCNDAIRRSWNKFKCMRSVK
GRVSITQLAKQKCRCYPPCHEITYGVFHSLTSWPSMDQTIPTMEKVITGDFMQRFTTDHERNLVWENYFPNYSNDTDL
VKLYLETDNLTVQHDLTKVCARFLKDFSHVYVYIADDNVVKITESEFYSGVQLVSDIGGQLGLWVGISVVTLAELLQLCAT
LCGFLCKNKHRRKLKKEFERRSMRKKQRGASVRTKRYNHHHHRPRSRSLSNTRQNGTPTRNHLLKNQNHHVNV

>Cyclo_MEC10_NP_509438_1_Degenerin_mec10__Caenorhabditis_elegans__51_375

MNRNPRMSKFQPNPRSRSRFQDETDLRSLRSFKTDFSNYLASDTNFLNVAEIMTSYAYGESNNAHEKEIQCDLLTENG
GIEIDPTRLSYRERIRWHLQQFCYKTSSHGIPMLGQAPNSLYRAAWVFLLLICAIQFINQAVAVIQKYQKMDKITDIQLKF
DTAPFPAITLCNLNPYKDSVIRSHDSISKILGVFKSVMKKAGDSSSEALEEEEETEYDMNGITIQAKRKKRGAGEKGTFEPA
NSACECDEEDGSNECEERSTEKPSGDNDMCICAFDRQTNDAWPCHRKEQWTNTTCQTCDEHYLCSKKAKKGTKRSEL
KKEPCICESKGLFCIKHEHAAMVLNLWEYFGDSEDFSEISTEEREALGFGNMTDEVAIVTKAKENIIFAMSALSEEQRILM
SQAKHNLIHKCSFNGKPCDIDQDFELVADPTFGNCFVFNHDREIFKSSVRAGPQYGLRVMLFVNASDYLPTSEAVGIRLT
IHDKDDFPFPDTFGYSAPTGYISSFGMRMKKMSRLPAPYGDCVEDGATSNYIYKGYAYSTEGCYRTCFQELIIDRCGCSD
PRFPSIGGVQPCQVFNKNHRECLEKHTHQIGEIHGSFKCRCQQPCNQTIYTTSYSEAIWPSQALNISLGQCEKEAEECNE
EYKENAAMLEVFYEALNFEVLSESEAYGIVKMMADFGGHLGLWSGVSVMTCCEFVCLAFELIYMAIAHHINQQRIRRR
ENAANEY

>Deu_Ambulacrar_hemi_Ptyfla_40v0_9_20150316_1g9562_t1_scaffold5200_cov88_926_376

MMLYSRSNLNFSKDYLLAELRLSSTLYVDNDDSKRFAVFESRQDDRSDNPFPLLSVNSPYVDDTEKAEQPTDRRVSDTN
EGTKELSDKSNVQHEFFKLADKTTEEHDESVKHVLEDFGRETTAHGIVHITNATSSVTRSTWIIIVLVAACAMLVQMTLLL
IQYFEYNVHVKVTLVSEKALGFPSVTICNTNKLRHSAIRSSKYSEMLMLERNFVPPYYTPCIEGDFTCRNGIHCIRPFLVCD
GVNQCGDMSDEYDCIYHSKYWSLAFKQKASQSSTFAYGSAMKAVDGDKSNNYNERTCTHTQKEYQPWWKVDLGGE
YEIDKVIITNRADCCDDRLSGAVVRVGNDDVIENNERCGQTVTKGDINQDGEVTVECVLQGRFVSVQLENKTDYLRLCE
VQVFGEAFISSDNQTECPDDYIRCTSGECVHPYKICDVIVDCADGFDEMECVNSESNANGAQIDHFQNCSEEERQPHK
PKLCPTDITSDDKGQAKKVAEQLANITSEADDFEENDVTLSVDVIDNILESGVNGTEIEVAEDVLMSVDNLLHVDHEVLV
ASQQNENTASRLIESVEKLSLIVQFINDSMDSGNQSVTIETENIVLILAEINFESFAGLNFWFSSGKAVDINGTNAGFSPVI
AASIRDLRITNLRDPVRIQIPRNNQNIYSNDIDETHQKSLSIISYIGCGISLIASAITLVSVIVYRRYKSVPRVGLCRQKGKDFL
TDC

>Protostome_Lophotroco_annelid_CAC9663096_1__Ofus_G121090_Owenia__fusiformis__927_377

MAPPCELSDDEEIVFKSGVHQQQSNHEYQIEIEMKEKNELKNDSSVPSIEQEIACQNDECIEPELIMNRCIRDSKEIEDIVP
QKLVIQGDTCESTNVNLLAEQREDCYQYTQSFKAQISSTLAGFSNTTSIHGPKRILKARGRTSRAGWTFVFIGVVSLAVYL
IAGVVVKYYSYPIEDGISIKTQSLAFPSVTLCPLIPNDDFLVMSDLEAYWQSNQGDKYLLYTLKAYKYLIGKDRLVSDIRTDF
PKNESSFSNALIQSWLDLSSWLSEKLSPVSRWQVENFPLNDININVSEIGPTKGLSSDDINVAGMDPVKEQDFILECVYN
DKPCDEQFNVTKVTTAYGQCFTLNISESAEDVNEIGPNKGLTLILYTGRYNPIPIRPNFYMPTMGFSSTYAQPSDGVQIV
VHNPGTMPQPYREGFHVTPGRLTSVKITKTERTRLVPPYGDCTDKDYLVNSEFRYSYEMCVEQCLQERIIQKCGCVSPM
YIIPADHNFSLIQYCGNISKIFGFSENKNCIHSNSWMTRGSMCLYGTPQLRLRMINLANDIIDRLKCEQYWSKSSIQSCKC
RRPCKENTYIHLINTIPWPEKSVVSNDGQRGFLKFRQKHKNIDKTFKLFERELRLYTYLWDHQFEVFEDCFDHNITDGHK
QYNINSLIENIVDTNFLKLNIHIESLDVRTVYEERTYTYDNVMKDIGNVFGFYLGMSAVSVVEGVYIIGVLIKLIIYRMMCR
RN

>Protost_ecdy_cyclo_nema__A0A0V1BKZ2_A0A0V1BKZ2_TRISP_Degenerin_mec_10_OS_Trichinella_spiralis_OX_6334_GN_mec_10_PE_3_SV_1_932_380

MSNRVTFQRDTTTKRRFKNLLLNFSDKTTAHGVPRIGMATTKREGAFWTFVCVTSLTVFLFQLSLLIARFLRFETTTQVEL
KFETSLFPVVTLCNLNPYRNSLLKQVDQMKLMFDAFDNKPLSTPKTSTLLARKRRASSADEKQYRYTPLYARCSCPNEES
DCVPFTSAPLEDGEEICICYFDHVTKSPWPCYSEKQWRSEWCSRCSPNGNCLQILKKRQLSLYEDVKQCVCHSNGKTCL
VADKNKPERIWDPAYRDVRLDRYTTSTTTTTTTTTTTTTTTTTTTTSTATTTTTPIPTTPSEVQEYVDALGFEGITDETAK
VSRAQENTLFIMAELPPETKKKVSHQKKDFLLQCSFNGRQCDIENDFELLMDPVYGNCYMFNSNSTINRTTSRAGPIYG
LSLVIYVDADDYLPTTKASGVRLLVHSQEEYPFPDTNGYNAPTGLLSSFGIRMKKIERLPQPYGDCIKEGKTKDYIYKDQIY
SLEGCYRSCFQMEMIKSCGCGDPRFPVPEGRRHCRVKERKARECLENVIQKSGGLHGSFSGKCDCRQPCVQYVYDMSF
SAAKWPTPSIELDDCNDTAEECIRKLKKNALSLEIYFEQLNYEVLKELQAYQWVNLMADFGGQLGLWMGVSVITIIEVLV
LIYEVIKICCTRRKKPTMNKHHRRSIYDDSTVEMTIKIHPPNQNNPKDSIATAHSENDDECSQNSYAMENHLHRHDAEIP
NSVITFCAHCSLNIKQ

>Deutero_Ambulac_Sakowv30041168m_935_382

MSVSRRKILLTSDVEEESESECHGRSFEDGEIRLVDMSLLNGALEKEICQFCKVGNISIVERRRHGLGSCIRSKLSQPSFPSS
RKKGAVFLINKKAAIGMRAIGKGTVRSGETVKLLGVTSTSNGSALSPEYPTNDFYLAGMENVSLRRESKQVALRKLFGR
MVNNSTAHGIPNAARAESLPRRLFWSVLFMVAVGMFLWQFSDQLVRFIDRPVNVKLNISFSRELTFPAVTVCNQNPIR
QSAIDFNQEPQSSPPTQNLNQTLNDTALDEDLGNHGGTNNNTNSPPYNTRRRRQTAQTSGDQSGQESKNDPPHPDG
DNGPSAGNRDMIRITEILTALPTSERITLGHQADMFIVNCTWQGEQCYASNFTTFSNSMYGNCFTINGPSGYIDPYWTT
SFSGPIYGLSLQLFIEQDEYIKGYTAIAGARIVIHDQDKMPFPEDNGFTIAPGAATSIGIRKVLISREPDPYSDCVEEGDSDY
TNIYMDSYDVGYSVQACMKACYQSEVINNCGCADPLYPFPYGVYEACKTSNSTQMYCKALVEYDYSTDGLECDCPQAC
SDVGFTTETSSAMWPANSYKETLLNYLKGSNKKLESLVGDDLKNGTNSVSENLLRVDIYYQELNYEVIEQEPAYLFSDLGS
DFGGLIGLWIGVSILTCFEFLELIFDFCHVFCSKGALAVNRTTDISAVSGGDIFFRDKYGNDPALTLNDRSSNKPRMNINN
RTTYVPETNAQSLYIPQYSMKIYEPGLLNTS

>Deu_Ambulacrar_hemi_Ptyfla_40v0_9_20150316_1g7150_t1_scaffold3457_cov142_936_383

MELDELRYDPPKRPSSPPAGAVDRDVGDTGFLALLYQFGDTVSAAGLPRVFTERSSLWSRLIYAIVFVASFGTGVYFSVKI
VQEFLEYPVIVTTAFTAESRMDFPSVTICNTNRVRYSAIIESKHSDLEYLLTDRSLGGLYYPPCLEDDMLCDTNECLRRFKV
CDGIRDCDGGEDEVNCYYGDCGYRQHRCETGGDHGYCINHNDYVCNGEQDCYDNSDESDCDNVFFGLNSYVDDGD
CLGFRCTNSSDCISQTEVCDGTYDCSDGSDELNCTSAFTCMTGMVKCATIDYCIYPSWVCDNYADCSDRSDEAPLATCP
SSSAARKRRSSSSGVSPTSSSGCYWCSAKYGSDNTTLQCFPLWYECDGYEDCVGGTDESNCDNINYHCPDNFFACTDG
ECIHEQFECDGWDDCASGSDEDNCEEPTRSSNSSTISDLFQDGWYSKYLHITDEEWFYEDFEDNYLQEKYNVGFERVSS
EDPPDWHGFATFSATGDYTDLEDVVKLTKAEMADLGHQNEDFILQCTFDQTKCHYEDFEVIQDDKYGNCFTWNHHS
RNMSLRSTGPGSTHGLKLTLFTEQSEYISIYGQDSGVRVSIHEMGTTPFPEDDGFTVAPGKATGVGLKQLIINREDPPHG
ECSNSTSFESMYGDGYSVAYHLLKTIHSINDKTKDINDFEQAQSNLVRLEVYFEEMTYETMSETPKYEGVYNLLAEIGGVI
GIYVGLSFITVVEFVEFFVAVVRYLFRRIKRSMKVEALK

>Deu_Ambulacrar_hemi_Ptyfla_40v0_9_20150316_1g8260_t1_scaffold4268_cov134_944_386

MEMQAQNGTTERNLRMASMEASEDPDQKNCSILVTNFGQSTTCHGLQQILIGDSPFRKLIWLAVFLTAMGVFTFQAY
ELVSLFLTYDVRINMEVGTASSLPFPAVTICNTNKLRLSEIERSVHHELTKTDPEHPDSIHRQLSYETPCGSEEFNCGQAGF
HGICISNEKRCDGVRDCFDGRDEMNCTVCYGGRQCDVGLDSHASKCIPEDNICDRFPDCIDGADEESCVLTDQCEMG
VLFARSDPSVLYSRNYPKKYDTDYTCELRLTAETGFDGLQSSDCISLAFLELDIERGKNCQRDYIECSKTCGGGVRKRNVTC
SVIESDDPSMDFADMSLDSGENFDNPLVEGSNGKMVVARGTRGHYDSPDTGDYFADEGDVSRRCTSAGSSPERVESC
NNTPCSGGPGGIQILDKLFEKYGHISDDKKIYENFELNYYKSNNFDRVQTFPDPNWKGEFSFVKTTSPDYSDLQKVLHLD
RDEVARFGHQAEDFILQCSFDEVYCDHSNFYRFENDVYGNCFTFNSLQQNNQTTVTSSRPGSRYGLKLTLFIEQDEYIPL
YGQEAGVRVLIHPQDITPFPEDEAITVAPGLKTSIGIRMDTIKSLSEPYTNCSNDEDFESVYGEGYKYSDMGGVIKVLRPIN
EKVRNIVVDEQSARIYILQGGVIKVLRPINEKVRNIVVDEQSARENLVRLAVYYEELNYQKITKLPAYTIEELLADLGGILGLY
IGMSLITAFEVLELFLNALRHLAKKFCKPKPEDDGPTYL

>Cyclo_MEC4_NP_510712_2_Degenerin_mec4__Caenorhabditis_elegans__52_388

MSWMQNLKNYQHLRDPSEYMSQVYGDPLAYLQETTKFVTEREYYEDFGYGECFNSTESEVQCELITGEFDPKLLPYDK
RLAWHFKEFCYKTSAHGIPMIGEAPNVYYRAVWVVLFLGCMIMLYLNAQSVLDKYNRNEKIVDIQLKFDTAPFPAITLC
NLNPYKASLATSVDLVKRTLSAFDGAMGKAGGNKDHEEEREVVTEPPTTPAPTTKPARRRGKRDLSGAFFEPGFARCLC
GSQGSSEQEDKDEEKEEELLETTTKKVFNINDADEEWDGMEEYDNEHYENYDVEATTGMNMMEECQSERTKFDEPT
GFDDRCICAFDRSTHDAWPCFLNGTWETTECDTCNEHAFCTKDNKTAKGHRSPCICAPSRFCVAYNGKTPPIEIWTYLQ
GGTPTEDPNFLEAMGFQGMTDEVAIVTKAKENIMFAMATLSMQDRERLSTTKRELVHKCSFNGKACDIEADFLTHIDP
AFGSCFTFNHNRTVNLTSIRAGPMYGLRMLVYVNASDYMPTTEATGVRLTIHDKEDFPFPDTFGYSAPTGYVSSFGLRL
RKMSRLPAPYGDCVPDGKTSDYIYSNYEYSVEGCYRSCFQQLVLKECRCGDPRFPVPENARHCDAADPIARKCLDARM
NDLGGLHGSFRCRCQQPCRQSIYSVTYSPAKWPSLSLQIQLGSCNGTAVECNKHYKENGAMVEVFYEQLNFEMLTESE
AYGFVNLLADFGGQLGLWCGISFLTCCEFVFLFLETAYMSAEHNYSLYKKKKAEKAKKIASGSF

>Protostome_Lophotroco_annelid_CAC9521433_1__Ofus_G03942_Owenia__fusiformis__951_390

MIMDHPMINTGTMADGENVTKTVKADVEINADNTNIEGTENVAQTGKVDDEKNTDDVNRADIDRVTQAVNVDESN
DIEMGTFKRIGNTGVVIELNQNVGIVPVDEPNIERATEDMITEHTQNNEHIVNERMAIDELISDTPIVTSQTQHHDIIAD
ALEFKYNSLKNQISKFALITSIQSLLQVRNAKSVLGRLIWFAITLFVCSLAIIGVHEITRKYLDHPSEDILYSREELVTFPSVTIC
GVKPVVFTDEHLKKKLSSRANTGDTINTWIYHDYRLKMLESRAFSDEREKDHERINAVMRRIVSEERYIENLNEWRDYRL
RMYDDLILECRFKGKPCGKTDFKQVMLGRYGNCFTFQPNSKDDTNILSEPGPDFGLNLMLFTNSFSPLDEFTDSDTFNYI
KLIGSSDSAILNEFNSDFSTASDGIRLSIHAPGTMPDPEKDGIDLSTGTFSSIGLSQDVRTLLGPPHGNCTEERLNTHTTYTY
TQNMCLRQCQQIETIKRCKCMNPKLPLPPLTEKPHKIHYCGYEMRQEYLKEKEREMTYGNVFPLENVQFDWKIFIENFK
LYECEKEVTELFSREGEKCGCKRECKSTAYYYLKHSIPWPNERYYKGLKHRNFISFLRNHPKADQFLNKTYWSGNRLNRL
GICMSGDNIFSSYSSAVVKGAYPYLTCPCNIANSTDQPDRCAPIINWYRFINKFVETNFLKVNVFYSTLNVNRIAAEPSYP
FNKWLSDFGGIIGMNMGMSVVSLVELGYLMVSLAFILYCKHKRDN

>Protostome_Lophotroco_annelid_CAC9668492_1__Ofus_G161027_Owenia__fusiformis__952_391

MNTCAKIEEQIKVSVIGYAEEHVLNESHEAETSGDTKRDTNLEVDCVSHDELELSIEKTNDGDDNYGHINKETSEQCLNI
DCDARDEPQVRQENKSCNISKRKAMCSIEDDHDDPIVDEAQTAKENENYDVCKGTTSYKTDTEDDNEDAIINEASSDK
NNDHNTNDDIEDPFDSWIWKYNQLKERLSYMASITSIQGCINIRNSQTMFGRIIWFSIALALGFTTVIKVSNIVVQYLSYP
SADALYSREEDVTFPSITICGARPIVFSENFHRSCMNSNCMPKTMDVQDGRLNQGILLLMNESGYESQTETLDFLRKRLL
SPERYVENVDDFPYSRLRYLKKNDLILACTFQGKECPDALKFKEVKFGRYGNCVTIDSNNTMKESGPDSGLSLILFSNSYS

PLDEFQKAMNLFIYLVSGVSEFDSTHATSSDGIRLAIHAPGTIAEPEHDGIDLSPGTFNTIGLTQDVRTLLEPPYGDCTKKIF
NEKTTYKYTKDLCLDKCMQEEMVAKCGCMTPKAQASLGKNGYDVSYCGRYYNTIKPKGYPQYLKKPNWRRRLRSQLL
QFLRRLKCESDVLRLYSRDSSNMCTSKCKTECTKIVYDPLTHATPWPNEKYYKQLNQRPFITFMKSKDNVAKLLEYWFD
VRRSVLDSYCELLHGDLRKNLLKRRVTGHVFKPLFADVNFPFVTCSENITAHINRYNIINKVIEKNFLKVNIFFRSLNVNRIA
AEPNYPLNKFLSEFGGIIGMYLGMSAVSLVELGYIMVAMVFILLFKYKR

>Cyclo_UNC8_NP_501138_1_Degenerin_unc8__Caenorhabditis_elegans__56_392

MSPLLTWNLICVSSRWYTILCLKNKVKFWLGTRLVHEPESMESRSSPYIRPSPYAGGVHPHFEEEDDRSKLHASALYSER
RTSSRKSLRSQKIDYHTTTIKSLWFDFCARTSSHGIPYVATSSFFGRYVWAALFMCMLMAFLLQTYWTMSEYLQYRTIIE
MQLQFEAAAFPAATVCNLNAFKYSELTQYEEIKEGFDYWERVINARMMSDSMKPGGDILEAISVRKKRSKSRDQLLFPI
DDEDLEGAVYQPVFVRCTCMNMEQCVPNRNPLEVNASICMCFEDVTRGLIWPCYPTSVWTVKKCSGCSISNTCPDPD
GPNASKQIAKHNSPLPCLCQSISHHCMVHPKDEIRWWNPNNYTVYSVTEPPTTEITETEEAFGLSDLKDAGAITTQTKE
NLIFLVAALPRETRRNLSYTLNEFVLRCSFNSKDCSMERDFKLHVDPEYGNCYTFNFNDSVELKNSRAGPMYGLRLLLNV
HQSDYMPTTEAAGVRLVVHEQDQEPFPDTFGYSAPTGFISSFGLKTKELHRLSAPWGNCSDTFRPVPYIYNEHYSPEGC
HRNCFQLKVLEICGCGDPRFPLPSEEHRHCNAKSKIDRQCLSNLTSDSGGYHHLHEQCECRQPCHEKVFETAYSASAWP
SQNFKIGTDCPAVSDIFNDTEACTEYYRQNTAYIEIYYEQLNFESLKETAGYTLVNLFSDFGGNIGLWIGFSVITFAEFAELF
CEICKLMYFKGIVYVQKKMQGKEYTSSSLMHIDFLQRSPKKSQPGEDEVSTNESTKELMSK

>Protost_ecdy_cyclo_nema_Cele_sp_P24585_DEG1_CAEEL_Degenerin_deg_1_OS_Caenorhabditis_elegans_OX_6239_GN_deg_1_PE_1_SV_2_955_393

MSNHHSKTKKTSMLGREDYIYSHDITNKNKKEKLNGASKNNDYNQDDDDETMKSKMMDFCDKTTAHGAKRVLIARN
SFSKLMWGLIIFSFLLMFAYQASKLIFKFSAHEKITDISLKFDDVEFPAITFCNLNPYKKSLVMMVPSIRDTMDVYDNAKT
HSKSEGEKKKPKVSRKQHSDASQQMVRELFAKEIEEGMVELKKSNKTLQSQNKSGRRRSQRSIENRRYEAIEAHCKCVG
NIGMECIRFESPPRDPSSKCICTYDRDMEVAWPCFNISVWYDHECPLCHDDGYCESTLPSGTTSSDKWPCMCRNRGDT
SERDDTPYCIGKAGVGKIEIRKLWLENNMTTTSTTTTTTTTPPPTTTSTTTTTTTTPPPTTTARPNQRAIVSNPETIKAMGF
QGMTDGVAMLTRAKENLMFTMAALSDKQRIALSQSKHEFIEMCSFNGKECDIDEDFRLHVDPEFGNCFTFNYDVNN
NYTSSRAGPMYGIRVLLFVNTSDYMSTSESSGVRLAIHPPTEYPFPDTFGYSAPVGFASSFGIKKKVMQRLPAPYGECVE
TKKVVDRNYIYAGYDYHPEGCHRSCFQNGLIDDCSCGDPRFPVPEGYRHCSAFNATARTCLEKNIGSVGDFHHITQKM
DKCVCKQSCEEIIHEVTFSCSKWPSGATDLGDCDGMTESECEQYYRLNAAMIEVFYEQLNYELLQESEAYGLVNLIADFG
GHLGLWLGFSVITVMEVCVLLVDMISLFFKSRHEEKLLRQSTKRKDVPEDKRQITVGSGRKSDAFVSI

>Deutero_Cephalo_154560F_t1__name__chr_scaffold30_start_582601_end_590481_strand_pro_len_784__956_394

MRLFLELLRTLYKEAPPIDRGQENLIYPAKMTATSDIDLRVLERIDSRLLRVIEETAEDRAQKIVEKCLKERDEKMKEQRKK
DIKTMTKEFCASTTAHGVNRVAEADTTRSRIIWTVILITCVALFVYQSAILIKSYFEYPVNVDIKVVDNKKLTFPSVTVCNN
NRLRKSQLPGSRHGSLLELDEQTARDVSDYLWEKQIGEIDIDKDMVIDLESGYPGLGSGFESELGLTNTFLSDESIDASTE
CRLQSKQLCTLAQLLREPRNDGIHTEWSYFSREGWAVMISHPCTGTEQDTCDQCYNDRILLKRIEDIPYKINATFCCPPD
SYIALTDHTPNHARLNSRSYWKMDEYDATPWIQVDLRDSHVLAGVITQGGGDAGGYVTSFSLSFSVDGNQWQTDNE
TGERDWEYCHVERCEVGCVDLDLNAVAGENDWGEFLLSSNTDDYTDITDFSMATREEIAEMGHQKEDLILQCTFDKE
RCSLDDFKVTQNAKYGNCFTFNHGEDDIVRNTTKVGAEYGLKLTLSIESNEYVGLFGQDIGAKVTIHSVGSTPFPEGNAI
NTKPGESTFVSLKRSSIKRRPYPYGDCTSHQERDALYGGRYTYETCQHSCLQAALLHECGCSDELIAINSTLCSVLNKTQEC
CRQDVRSRHEDGNLSCDCRQSCNEDSYALWLSSSLWPSDSYAWYVLENIHTRSQAQNLPLNPDELRQNLARIHVYFRD
LNYELITENPTYTEESLLSGLGGLLGLYVGLSVITVFEFINLLVDLFKAAFNKEVKDTANPGHNI

>Deutero_Ambulac_Apla_gbr29_2_t1_957_395

MEYGTTVVEGQRLTRRVLVTGSLDYLQRTGEEDKLSQEKKRGALRQAVTTFADTTTAHGLPLIINGTYTVSRIAWSVIWL
IAFGLAILQSTLLLVEYFQVRPLTTTIELITKTNLDFPAVTVCNMNRMRRSKLVGTRFEGLIAIDGGVQGEDYDYSWFFDW
SSLEWFDRWDSRGEPSSSPDGNSLQDSSPVNSEPDDSTAGGIGSSESIPGVGGESDVTPQGSGGATVGGSSVDGTDGS
DSVPGAGDDDDGMSSDSSTSGPSHVASNGEGDNPSTAGDSIDEMSSIDNESSLPGGSSNTPSHDVEKEEASSTAPPHS
LNTHASTSSGIFPVTQVNSPDSSSSQTTTDPPGSETVAPGQERDSDGASSSFNPGEESDTSILDSERRRRSVSSRQERQR
RTLSSSTEDFWWESDDFEYDSYYNWDGVTDENDWRGFYDRSTADDFSDLIDIANPTREELEVLGHQAQDFILQCTFDK
RNCSYRDFRVFQNTDYGNCFTFNHGTGNSKVRTTSRFGAKYGLHVTLFIEQPEYLGIFSPESGVRLTVHGQRTMPLPED
NGITAAAGLATSVGMRQDFISRQPFPYGNCSDSFDTRFNVTAGDYDYSLSACTKFCLQSEMLSRCGCVTDILIYSALKCS
YLNTTQQTCRRAVENLYEDNKLQSCKCFSACRETTFKLGVSSSRWPSERYEEHLYSRISATNEKAARLMQEVEQTRKNLV
RLRIYFEELNYQSIIQTPKYSVETLLGSLGGLFGLFIGFSVITIFEVVLFLLKLLKIFFFWGQPCHRREIKPVIS

>Deu_Ambulacrar_hemi_Ptyfla_40v0_9_20150316_1g15264_t1_scaffold10904_cov157_958_396

MSLMIGTRFQNLHENNCDTRTIVKIESCIQYSCGVGIYHISMHRMVAKLDKINQRKTVNLSNSRVKPKYIYDRRRVAERR
NMIESSLGNNSRVAPETPEISKKTHAGSTEGVSADRWNMIESGLSNNSQVAPETSEMSEKAHAGSSSMVSMNDMRE
NKPVGCCRTCCCGRSEKFREFTQETTLHGIKYTTDTNISSYRRLLWFILLLAMVASFTTTLRESAMRFWSNPVNTVVSYRE
MESMPFPAVTICNYNVYRKSVVKESVLGQYLDAMYGNPFSPETDENTTLPEPPANYFANKTQIMYQSAHRIEDMLKEC
KLSSDVYCSAENFTKVITNFGVCYTFNGEVGNALKVRARGQQNGFFVSIDIQQNEYYYGPNVGAGLKNKLLKCDEEQLD
YFDEYSTANCEMEKRTKAVVKDCQCKQPHMPNKNEYGCSVPCDDVIYTGTLSFAAFPGPHVVKRLEAKYNQTEVQMR
LLWIALMLFTIILLIFAVKDSAIRYSTDPVTTVTTFQHHTEVTFPAITLCNYNQYRKSVIGGTWFEDFLKELFTAPAPQRRQI
NGLNDYEDRLRNMNRTEFEIKAAHRIEDMLLECVWSSTSNCDENNFTLVLTDFGVCYTFNGDPNNILLVRNRGSLYGL
QVLLNIQQIEYTVGRKQEEFSCKMPCETVSHASTLSFGSFPGVQVSEQLKIRFNMTEKSIEQNFVSLNIYVEDLMVLVTSE
VMAYTYDDFVGDLGGQIGLFLGASLLTVFEFAEFFVTCVVRRCRRLRAKFRAINKKKNKALESASSQKNKPECAGYKS

>Protostome_Lophotroco_annelid_CAC9599174_1__Ofus_G063499_Owenia__fusiformis__959_397

MQRVADDPRHNMIKGPGNNVADINAASCAAGINEGLGPNEANEHIYADVVVEIKINTLEEPSKKEPLHAMATASSNP
ASTEDRANGTIDSNEISNAESKGRNVAFNLPLTAGIEETRNVSAKDQPNDSMRKKVSSTLSEYAEMASVNGPKRIVRAK
SRPSMVAWSVIFLGVVCMAVYLIVGVVVKYYTYPIQDGITIKTQPLAFPAVTLCPFIPNSMSDLKKALRSKEAKLPFANLL
GDKITETMIRYHISAVHTRNMYEFSINPETFGLLKFNGTDINMKKIFQLREALRSSGWLVENVPINDFSRNLLKQIKSKDFI
MECTYNDKACDDLIDISISNTVYGRCFTLNVNESIGYVQEVGPDKGLVLLLYTGRHNPLQRIGMGNGGYEFESKYAEPSD
GVKVVVHKPGTMPRPYSDGFFVTPGRLAAVGITQSERTRLLPPHGECTDAVFNTNTTYKYTYEICVDQCIQERIKENCGC
VSPMFPVPKDHDFKRIQYCGNISDIFKSYNDNETSAELSMFSVLLEYTERLHASEKGDFEHYIDTILSTPELRKQLANTVDE
IIARLQCENHWMISDKDSCKCKRQCKENEYTHLINTIPWPEKSVVDTGGHRGFVKFFQKHKHFETLMHNLVGLTREIGD
LHFYRQSQIENMLRGEKSEFRNTAEALVALILTTRFDFMNELFLYTENFQNVYGISSFTIATYLDKFIDTNFLKLNIHFESLD
VRVVYEERTYTYASVIKDIGNVFGFYLGMSVFSVMEAVFMICLLIKLVICQMVRSEENDDDNIGEQHIKGTDDID

>Deutero_Chord_Hsap_sp_P51172_SCNND_HUMAN_Amiloride_sensitive_sodium_channel_subunit_delta_OS_Homo_sapiens__Human__OX_9606_GN_SCNN1D_PE_1_SV_3_962_399

MRAVLSQKTTPLPRYLWPGHLSGPRRLTWSWCSDHRTPTCRELGSPHPTPCTGPARGWPRRGGGPCGFTSAGHVLC
GYPLCLLSGPIQGCGTGLGDSSMAFLSRTSPVAAASFQSRQEARGSILLQSCQLPPQWLSTEAWTGEWKQPHGGALTS
RSPGPVAPQRPCHLKGWQHRPTQHNAACKQGQAAAQTPPRPGPPSAPPPPPKEGHQEGLVELPASFRELLTFFCTNA
TIHGAIRLVCSRGNRLKTTSWGLLSLGALVALCWQLGLLFERHWHRPVLMAVSVHSERKLLPLVTLCDGNPRRPSPVLR
HLELLDEFARENIDSLYNVNLSKGRAALSATVPRHEPPFHLDREIRLQRLSHSGSRVRVGFRLCNSTGGDCFYRGYTSGVA

AVQDWYHFHYVDILALLPAAWEDSHGSQDGHFVLSCSYDGLDCQARQFRTFHHPTYGSCYTVDGVWTAQRPGITHG
VGLVLRVEQQPHLPLLSTLAGIRVMVHGRNHTPFLGHHSFSVRPGTEATISIREDEVHRLGSPYGHCTAGGEGVEVELLH
NTSYTRQACLVSCFQQLMVETCSCGYYLHPLPAGAEYCSSARHPAWGHCFYRLYQDLETHRLPCTSRCPRPCRESAFKL
STGTSRWPSAKSAGWTLATLGEQGLPHQSHRQRSSLAKINIVYQELNYRSVEEAPVYSVPQLLSAMGSLCSLWFGASVL
SLLELLELLLDASALTLVLGGRRLRRAWFSWPRASPASGASSIKPEASQMPPPAGGTSDDPEPSGPHLPRVMLPGVLAG
VSAEESWAGPQPLETLDT

>Deu_Ambulacrar_hemi_Ptyfla_40v0_9_20150316_1g21852_t1_scaffold24782_cov84_964_400

MRFDDVDRPTWRRSASMHVSFVKEASFSTQHIVISVRQSVPFARVVHTGTRCTVAICGAISHFSKIPNTNMQAYDADV
IENVTLEVCASVCFRYEPWECMSFDYHETFRVCRINPGKADVFVRLKNGKSSYDHFAKTTYGIECYAELDSSDYRGHVSV
TRNGNICQPWNVQVPHVHYLYGRXXXXXXXXXXXXXXXXXXXXXXXXXXDPNIRFELCDVGQPRVGCLSKVEPSARECF
TSANGDDYRGTVSTTESGYICQPWTSQLPYSHRYTPEIYPNAGLGDHNYCRNPDGWSATWCYTLDPLVRWESCDVGS
PQTTCQNTDMPIHEFFRTTNQTQRGFDSMTGNVTVEECASSCLNSRKFTCLSFDFSNEDGICYHSDANESVSKPSTVSG
ILVDHYDRRFEDEERLLLDFWKFPDYRIPSDHVALLTNVTAHGCAAKCLDEITYKCQSFDYKPSLAKCYLYDKPFLSVIADT
PEKDISYIHYELKVASSNGCPSDYRQCGSGECIHQYSFCDTIINCADESDEDNCYGNHRAIPDMDDIPGVDWHDHPDKA
TLMADFLSDYYHDNGFASVKAEVPPDWYGFKTFSSTPDYSDLQSVLKLSRDELRRYGHQREDFVLQCTYNEEDCVQNS
DFYTFQDEKYGNCFKFNHGRGNETVKMATKQGATFGLKLTLCLEQDEYIGIYGHDAGVRVAIEPSKYVAFPLDAGITVQ
PGTVTSIGLREIRVSKKPHPFGNCTLDDTQDSIFGSHYQYSVLREDLLSDIGGTLGLYIGLSVITVAEFIELLIVVIKFVGRRRG
ALRTAKLSPSS

>Cnidar_HydraVulSc4wPfr_2074_g13182_t1_968_403

MYLQFVNHFTTSSGKVWYIKTSISCNSKNIIYFLSCNRCDGKTTYIGKTNSLSFLKKMSSIARFANIAKVSSRHVDDVSKKE
SEPLIEKKEEEIDTLLKNKDAIFNNETKHENKKSFKRWSLLKEHPNYSQIINEVVEVPDFQETDTNKELISQGTDTKEVPNF
QETSNKEITNFQGTESKNDEASDKIVLPADNQSVDSEVTLNLSGNTKWAALTKKNAKKIILESDKLKQPVLSLSSDSTSQT
DNKILNNKNSLGKSKWNAVRERTNSIKNTKDDVNSNEKPSLPKLLQVVNTATATKTAASKFLEASQMRVNLKARMRKI
VQERLTVKQIFKRYVESSTLHGFCYVCGDTFLVRRVLWALLMILGAIYFIIKLRYGIEEYLNYPFSTLSTVDYVHELFFPAMS
LCATNSYLASMVSKNQLKLLYDEGRLPLDNNQTNPSYNMSGNELVKAIQESSLSIESMLAFCDWIQQDTNNPDIPPNPC
GPQNFTTYLNYKGEQCYTLNSGLEGHKLLKVDTVGLTYGYELIFDLQTNKVIKNNQFSGMRVVIHNQDVPPQLADGFIIP
PGFKTFVKMGTVQSKSLPPPYSTECGTKKLKYYNTYSQRFCLLETLTDFTGKLCGCRDVFMPENGLPFCSLKELYSCMYP
AKESFSEFTMRKECPADCEERTYPYELSEARFIHNPPIGLSIQGLANLQHKKHLEEELYLSKLAQSMTSEELDAYVEDNIVS
VIFFFGDTRIDYNEQEATNDFFQFLGNMGGEFGLMLGASLLTFVEFVDLFIFLIYHQMLRLHTLKKVPDIFGRKRSRINEYI
KKREKTRV

>Protost_ecdy_cyclo_nema_Cele_sp_O01635_ASIC1_CAEEL_Degenerin_like_protein_asic_1_OS_Caenorhabditis_elegans_OX_6239_GN_asic_1_PE_3_SV_2_970_405

MGKNSLKRALELDVVDFAEHTSAHGIPRAYVSTGWRRYMWLLCFLFCLSCFGHQAYLIVERFNRNDIIVGVEIKFEEIKFPAVTICNMNPYKNSAARELGAIRNAIEAFELAIDKSDGNAHSKRKKRSANSKMVPIDLLCKEEHGMFTAHDYGHVECTCVTFEDMSKVGDTDDDEIFWNCHQRKDWTHKICHLAEGSNQLKTCKCFEDTCVSDEVTKQLVWPLQLSKNGTKLCISPESSGPRYCASAQKFQVSTCSNCDWLGKCEESDDMDLEEEIDSKTCICHHGNCFQIKGNVKKRKRRTPERKVHERLLSRYEGLLAVYSHCNCTKQHGCVSTSVPDMDLENSNKTCLCFYNKKNEQIWPCYKEPEWEERKCSRCNTMGDCVYSDKPKKQTISCLCATPIKMCVRIDPPQTNDTSLDDRVVKFWDIQPSTTMSPIVKKKEERDKAYGYTGVKDRIALRAKAMENMIFAVDALTEEEKWKISYNKSDFIMKCSFNGRECNVKHDFVEYLDPTYGACFTYGQKLGNNTNERSGPAYGLRLEVFVNVTEYLPTTEAAGVRLTVHATDEQPFPDTLGFSAPTGFVSSFGIKLKSMVRLPAPYGDCVREGKTEDFIYTQKAYNTEGCQRSCIQKHLSKTCGCGDPRFPPYRESKNCPVDDPYKRECIKNEMHVATRDSKKLGCSCKQPCNQDVYSVSYSASRWPAIAGDLSGCPLGMAAHHCLNYKREQGSMIEVYFEQLNYESLLESEAYGWSNLLSDFGGQLGLWMGVSVITIGEVACFFFEVFISLISSNRTKRRPARKSFSSSLRCSTDYNLNKDGFNLDN

>Protostome_Lophotroco_annelid_CAC9662918_1__Ofus_G12800_Owenia__fusiformis__973_408

METKISTNDIKIQQENEPTTFKGLLTDFANNTTLHGLPRVITSAQLIRKFVWFCIFFGSFGYFSFQVAELINDYYKWPVLIRSEMKLRTFMEVPAVTICNMNMLRRSKIEGTMFNSLVELDNDTANEIIEKQDTTENEINNQNQIQNSDKTSRKKRNINMIQKSIQRPMSKENNALKGHENKFEISRVKTKSMESVNNILAFEHEGSGSTKIEHTFFDGSLPENEADDPHSLNDEYIALEDGHSVEPIDDTNDNGRRVRREVAGGSGGEATDSYVSDPTDTTTDSYDTYTDPTDTTTDSYDTNTDPTDTTTDSYDTNTDPTDTTTDSFDTNTNPTDTTIDSYDTNTDPTDTTTDSYDTNTDPTDTTTDSYDTNTDPTDTIEDSFDTNTDPVSTNTYPIYTDSYPTDPNADPMDFDTYPTESYTDPNFQSSDYYESYEYSEYESSEQEMDQESGEDPGYDVPFQFEQSKLMGIKDPNNYAEILRLAEAEDLSDIFGLASPTVEQLDEYGHNLSDLVALCTFDGENCNRSSDHWLETYNKYYGKCYTWNSVINKPRGERPLVTTNFGSRYGLRLTLNVERDEYTGLFSPTYGTRVAIHDSSIISYPENDGITASVGNEMVIRLRLKKYERETEPYPSDCVKDDDLVKQFGGYTVIGCMRECLEERLVKRCKCLDRIVEKDEHRCSYFNRTEELCRQRVYYAYNSGKLGCKCRARCRENKYDISTTVSTWPSKHHDYYLLNRLKERGIQTLEEIEDNILRVHIYFEDFNVETIKEVPAYSVISLLSNIGGTMGMFVGLSICTLGEFLELIWEMIMLLRRKISRRHNMVNASSA

>Deu_Ambulacrar_hemi_Ptyfla_40v0_9_20150316_1g11917_t1_scaffold7296_cov167_974_409

MYGRHARYFERGERTKYGGYAEGLAKPTDTDIDRQIRNQTFSVSRSVNMYSEPTLDSPQRTDEKASKRSIIAWKVVKEYGLNSTSHGIPRIFGAKSRVLGFFWTVLTLTAFGAFLWQGSELIMQFKRFDVITKVEVVTEKRLVFPSVTVCNVNKLRKSAIASSSYKKMLIVDEQIVLPYYAPCIPGDFACANRIYCVKEYLRCDGINHCKDFSDEDGCVYELHKSYVMKTGGHKLKTCQRLATKSTYFNDNKLRNQRCSLRMPFMTVCEEQAVMRSKSPVTDNRQIPVKPGNCGENQFKCINGSDFGFCISKKYECDRKSHCYDGADEDNCRACGSYDQFTCDDGVCFPAWYQCDNYEDCADGSDERDCDCTFSCDNGTLCIPDFWICDGVQDCMDKSDEPDYCNYDAIPDSGDETSQMLFPSDDSDVLTMPSDEEGDLFSFSLLHDDAMSVASPSNEDLTTEAENQTLKYNCEGFRCFDGQCVELYSDCDTIVHCEDSSDEDDCIPSDDRYGNCYTFPGSRNNEKPLYVSRSGQSHGLKLTLFTEQDEYISIYGRQAGIRVTITAPENSESIHTQDGITIQPGTETHIGLREKNLLRTIHSLNNKTLNINDKYSTRGDRTQYGGFTEGLAKPTDTDIELPIRDHTLSVSCSTNMYSEPTLDSPQRTGEKVSKRSTIAWKVVKEYGLNTTSHGIPKIVGAKNKVLGFFWTVLTLTAFGAFLWHGSELIREFQRFDVITKIEVVTEKRLVFPSVTICNVNKLRKSAIASSPYKEMLIIDEQIALPYPAPCIPGDFVCANGIYCVKQYLRCDGIDHCKDFSDEDGCVYGELKE

>Deutero_Ambulac_Sakowv30020102m_976_411

MEMQNSQVTEKDDTQRKKDDKESYKALVNHFISTTNAHGLPKAFEGRGIFVSSFWAIVFLAALTGAALQISELLVQYLE
WEVKIKMKVVSESYLQFPGISICNTNKLRKSAIEKSEHKGLNGVDDDIVFPYYDEGGCMANDFACNNTGCIKNYLHCDG
YDNCRDRSDEMGCKYGRCGNTEFRCPSGSDMGVCIKLKFKCDRHLDCYDGEDENECVCKEKKEFSCLDNGRCVPVNK
LCDGNNDCADGSDEKECSEDGNYCPSDYFKCDNGRTCIPAIFKCDGGADCSDETDEMSCPTQNPGCADDQFHCDESY
CIDNSWLCDGEYDCMDGTDELEMNCITSSGTNSTNTNPCQDWEFYCGDDKCINGDWECDTESDCPGGQDEVNCLT
TRAPTSTEVGSFMCELSGKLVERQYVCNNVVDCDDASDEYNCHNDASPTYIPDAEELLRNYSTLSTDLSLFEDFVSRYYID
NMFGRVVRENPPDWPGFIAYSSSPDYSDLAYVLKLRTDEMHKLGHQLNDFVLHCTFDSVKCDMEKDFLTFYDDRYGN
CFQYNYKHDHDSEQLLSTKTGPRYGLKLTLFTEQDEYISVYGHDSGARVVIHPSYMRAMPWSEGFTIAPGKIAFIGIKETR
VDRQPAPYGECATNIYQETVYGKYYKESTCEESCIQDRMMEYCGCVDTMLRNATRCMLLNRTQDTCRQLIYYSIQQKL
LGCDCPQSCKERYYETSLSQSLWPSNTYLKHLLKQIHAENPKTLNINNLETSRQNLIRLELYYDDLNFQQITEKPEVSEEAL
LSSIGGSLGLYCGFSFLTIVEFIQFAFDLVKLTYVRVFLRRLKPVAVSA

>Protostome_Lophotroco_annelid_CAC9671100_1__Ofus_G19963_Owenia__fusiformis__983_412

MANKTSPEEVDTQHDKDHPTFKQLIIDFADNSTLHGLPRSLRNSVRVFRRLIWLGIFAGCFGYFIFQVIELIKDYNKWPILI
KSEIRLRTFMDVPAVTICNVNMLRKSKIEGTTFDSLVDLDRKNFANGNTRSTTVHKKATPKPRNKRDGKVEETWERSRP
PLNSNTDVKAGRVNIRQILRGYRPQVGIGSKENGVSHSVNTLRDVKPDDPGPYNENGLQGEGIGENNYETFDNRPIEAY
NAVNDESPSRDDGLLRQHDTADDVANDKIRIKRTLDTKPSAPGATATDPTTMFDQTTTTISGQIPTKSDPVTTISDQTTT
ISGQIPTKSDPVTTISDQTTTISGQTPTKSDPVTTISDQTTTISGQTPTKSDPVTTISDQTTTISGQIPTKLDQTTTIFDQTAT
PGQIATNSDQTTTVTSVPICTCTIPVAATSAHTQTSPPGSTTVSNDTFVESDTTSSYTGNNGWSNNSSLGQGHDPNTYD
RYGPRMDTGYTAPFTFKESPFLKIKNRNNFGQILNSSTAEDLSDIFGLATPTVQQLDMYGHRISDLAALCTFDMENCKRT
SGHWLETYNKYYGKCYTWNSAINKPEGERPLVTTNFGSRYGLRLTLNVEPDEYTGLLSPTYGVRVSVHDNSVVSYPESD
GLTVSVGNEMVFRLRKKKYERQPNPYPSNCMDGWMRPSKNRNRGEYTVLGCMMECLQQRLFEICGCFDRIVDEGEH
RCNYFNRTEEFCRQKVYHAYNTGRLGCRCPAKCRESTYEVSTSVSAWPSKHHDHFIRERLNKINNISLHDIKNTTVRVHV
YFEDFNEETVIEVPVYTVSGILSNIGGTMGMFIGLSLCTIGELLELMLDVAILLYKKLSTRFNIRRVGTAVT

>Deu_Ambulacrar_hemi_Ptyfla_40v0_9_20150316_1g26495_t1_scaffold55703_cov149_984_413

MVSVVMVVMRQPLSRNLSQKITPYVLESLSVVYLGEIDITTRQRLLNVLFPGETGISMDRKIIDNILVSTPGGGLLSPNLQ
VIDLPVENNQLTVGQLYEILHNALQSYYNLIYAEISIPLSSNLYRYIHSETQSSVDVGSDIYRKVYEATMNRNTSAWHLRTT
ASHYAVEIAFRQALNGDHPYGALMNATMHAVSDAFPDIFDLIYTTVLDTFEENRLTALENITAYQICMEYMRGAGYRG
WSRNRYNIFFAISNVEEAMYVHTCKTTVLAITCRKTGLLEAFGYSHGDDVIKTAHAILSPSTETMDQDVNLVGVNANYD
FEYLMTTVGHKKDEMIMRCKWPGHQCSVDNFTTTYASDSICVTYNLGERGDIVEQVNGGERYGLTVYLDVESHEYVN
SRQHYVGFRVIVHDQDDIPNTKDRGFNVAPGAYTAAALKKSSSSYLKEPVGHTDCIDSPRGTLHYFHGNYSLAKCKVEC
ETDYAIKQCQCRYYTMPGDAPICNVMQLHNCYVPKMAKFTDVQGVCSYCKQPCYEESFTEHLSYALFPSGPLVEELAKK
AEKPCPKAMALHYKKKITPYVFASLSIVNLDEIHITTQDKVLDIIFPGETNISVHRIVENITASMPEAGLLPLIDVHDSPVKIH
QLTVGQLSEILRKALQSYYEQIYAEITIPLSLNLYRYIHSEGQSSVDIGSDRYRKVFEATMNSSASARLLRTIAPDLVVNGEF
RAALNMSYSYGALVNTTKHAVSNEFMDIFDMIYTTILDTFEENRITALENRTMYQICMEYMRKNTLELRVYFNEMREEK
MTQQQHYEPFNLVCDIGGTLGLFFGASLLSVIEIIDFLLFRRCNKVESQKNESSIALENRGSEKSATSPNEEHKT

>Cyclo_UNC105_NP_001122595_1_Degenerinlike_protein_unc105__Caenorhabditis_elegans__55_414

MAEDRIKSKLRRPASIESTMSSRTKPRHKPSPMSILMPHLMVGESFRKYRPHGLRNIRMNGHLDWNQLRKSFEKQSTF
HGISHAATADGKWRWFWYTAFTICLLALLIQIFFLISKYRQYGKTVDLDLKFENAPFPSITICNLNPYKKSAIQSNPNTKA
MMEAYSRRIGSGDKTEGIAAALSATGGLHAKVRRAKRKAKGKPRLRDRRYHQAFAQCLCDIEQLTGDRKGSCFAAFKG
KIEIDTNNTAGFMNLHTSRCLCQLDTVSKALWPCFPYSSWKEKLCSECVDNTGHCPMRFYKGNELYENIKEQVDLCLCH
KEYNHCVSTRDDGIILEISPNDELNDLDIGKKIASQLSAQQEKQAEVTTTEAPTVTQALGFEELTDDIAITSQAQENLMFA

VGEMSEKAKESMSYELDELVLKCSFNQKDCQMDRDFTLHYDNTFGNCYTFNYNRTAEVASHRAGANYGLRVLLYANV
SEYLPTTEAVGFRITVHDKHIVPFPDAFGYSAPTGFMSSFGVRMKQFIRLEPPYGHCRHGGEDAATFVYTGFQYSVEAC
HRSCAQKVIVEACGCADPMYPVAEMFGNNTKPCQAVNMDQRECLRNTTLWLGELYSKGKEAIIPDCYCHQPCQETN
YEVTYSSARWPSGSAKVMECLPGDFLCLEKYRKNAAMVQIFYEELNYETMQESPAYTLTSVLADLGGLTGLWIGASVVS
LLEIVTLIVFATQAYVRKRKGSISAQSHHSVPVHRASRVSLNTLHKSSTTQSVKLSVMDIRSIKSIHSNHSSKSKQSILIEDLP
PAIQEQSDDEEETTESSRTNGSCRYLAPGEDLPCLCKYHPDGSIRIMKALCPVHGYMVRRNYDYSVSNSEEEDAEDEVH
REPEPFYSAPYEHRKK

>Lopho_Phor_aus_TRINITY_DN315771_c0_g2_i2_p1_T

MNSEVNKRNLKGRAKEFCETTSAHALGQTVRSGKVKAIFWSLVFLGSLAGCVWNIIHVVE

SYAGFGFSVKSKLEMEPSSLKFPSVTICNLNPISFSRQARFVGDFERSTGLDEKDCDNEF

YRNETNRPLVLDLYRWSELPQFWFEYNENVTDMYGHSKKDFIVDCVFSKKLCEDNFYVTT

DPNLYNCYTLEPNKYNSDLHAGVGIEAGLSLTLFVENTRVQNAYTGSFSIDSYQTGNVGV

KVAMHVSGSHPNPNARGVIAEVGKSTDFILRTVNRTALGPPYPSPCNPKKTIESSHRQEL

QYEENLCFASCLQNAIAKKCGCVSINPVAVPIADKFGIDLPFCGSYPCNATKVTENYECV

HDVTRRFLLKNDPNCRDCTKPCNEVSYEATKSQSKWPSELHQSEFIGWLASKDNTALYNQ

LVRNTSDDKISRFIENNFLRINIYFGDFYVRKDTETLNMDWFDLLSSVGGAFGFWVGISV

VTGVEVLELLLDCIVIFLNKPSRKRNQKDLRNTTGNNDVEVYNN

>Lopho_Phor_aus_TRINITY_DN291806_c2_g12_i3_p1_T

MAEVELSCVNDVHFGLEMSNGVNPKTHLENMAKEEEEKQSLKEFLKEWAENAACDGVPQI

VLRKSWVKRILWLLIVLGLTAYTIEEVYGVFDEFFTYPVNTNINYTYSNQLTFPAITLCN

MNPIRKSMLSEAGTTFDDIFGSTNGTFPGRKKREIGPQEETAPSGSQQRGKEKMTFINQF

KRQWATLSYPKRIQLGHQLPNFVVTCSLGGQSCNKFFTYSTGTFGNCFVFNSGLRNTTVQ

SISKTGPLFGLTMEIFLEESEYLPEVTEQTGARLHINHQKLMPFPENAGYNLAPGYLTTI

GLKRVEISRLLGYESNCTKVTTAEAEGYSIYSKRFGVSYSREACHNTCYQRKVIEICRCG

DADYLFDYSFFKSLLPDTVKSIKPCLSATEEACVSRVLKLYASNNITCDCPIPCSESDYQ

ATVTISPWPSKLYEPTLRSSYSSLNRNDILNTTLSRNLLKVEIYYDTLNYQYIYESPKYT

NLSLISDLGGQIGLWLGASLFLLLQVLELFIDVIIFGCGRLFKSGSRVSSR

>Lopho_Phor_aus_TRINITY_DN297387_c12_g1_i2_p1_T

MSGRRSAWNDYHSESVWDEQRAGTQNGDKHGEDDCKDEDDQPRSSWQIIKDCGNDYLDVS

SMHGLAFFVGSRYHPLRRLLWFALFTASFTWLMIVVTTAFDNLQRKPVLTKVKVKYEDNA

RFPTVTICNLNKYRKSYVEKHHPHSEKLLRYIYPVLREDEHIHLDWSDPKYAWSLNRSSF

KEFALKAAPQMMDNIKIVDFGAHWLTISNEFSTIETDFGMCFQFNPNGTWYSQLSGEMHG

LAIVLDVQQDDYYFGDSFSTGFKIALHRHDEVPFPNQFSFGVSPGQELLVGMTMTEVKSL

PAPYEDGTCVDTEAGDFINPLKYFKTYKYNSCRKECQTVFSVGQCGCKPAFDPGPTRTCI

GEEYRCYRLATEEYAKNHSIERECNCRLPCHERGYVYQLSSLKFPAIQYEAYFKNTHNIS

LQYARENLMLVRIYFEKMNYELVEQVLGYDGMNFFADFGGNSGLCIGASLLTVAELLELI

AVIVFNLIFRRRKKTSPEKQ

>Lopho_Phor_aus_TRINITY_DN307417_c2_g3_i8_p1_T

MASEPETIPLNGLNHNLRKRRVQNYIVIDLVNNGEQDEQENGVYKSRFECFGEDTDFHGL

KYISRGENSGVKRLLWLLLIIASLIYLVIEIYGRVAYYLTFPHVTKVDVEFKSDLDFPAF

TVCNMNKYRAYSMTDMDLFHMGEILDIVDENMNLHHHSHYKESFVKRVQNINKTEIRLKN

ESFDLEEFVRRTGHQARDLILYCKWKDELCTGEANFTHVFTHLGNCYTFNSGKKKVAARK

AGAGNGLTLYLNIEEHTYLENKDAADEGLKVMVHSQMEPPFIRELGFGIMPEQHHYIAIR

KTEITNLPHPYGKCVDTNDSRRHPDFEHYTIPGCRIQCETKNVLETCGCRLPEMPKMPDH

EAKICTPPHQYKCAVKELDRISESDVCICQTPCHIEDYRFTHSSAKLRPETIEKLRELQG

DKLVPANLTSNNVAVINVFFEALSMEKIEQRVAYPWPSLLGDIGGQMGLFIGASVLTIVH

AIEFCMNEFARGVKDRARKKNEKSASLRSNNNANRDFQQDPTLPTVLKEESL

>Lopho_Phor_aus_TRINITY_DN307786_c0_g2_i2_p1_T

MSKGVKTSGGVWATNVFNSGQHDNTLYRQHVEAKQDITADIDQPNTGLSIVKDGIQKYAE

DSSLHGVQYFGGAKYNPVRRLIWFLMFVGSFTFLVINVSDAYQTLMRKPVQTKLSVEYTK

TARFPTVTLCNFNKFRGIYVWTQPALMEILRYLYPLSEEDRKLQLNWEDPKYAPFINGST

YLDFAKIAGHQLRDSIYFARFKGMNLDIQKEFKQVVTDFGLCFQFNSNGTWNTSRSGTNN

GLWIQMNAEHYMYFYGDSNSAGFKVALHNFDEEPLVNELGFAISPGQESFVSAQIKEIKS
LPPPYEGGKCKNTTKENFVNHLAYYETYSMTGCRRECRTNYTVKRCGCKYIYDPGPARVC
DAPEMGCYRQADGEYTQDESIEDDCDCFVPCHEIQYEHRLSTSMYPGNHIIYGLQAQYNV
TKEIARENFLELRIFFEKLNYDLVEQVPSYDAMNFFTDLGGNLGLCIGASLLTAAELLEH
IGITFFQVLQKFIGRKQTHPKKIQREGVF
>Lopho_Phor_aus_TRINITY_DN307786_c0_g5_i2_p1_T
MPKGFKSSNDIWATKVTPIAPYGQKESDNKDEGAPESGTVLSIVKEGVTKYAEETSCHGV
QYFGGAKHNPGRRIVWFLMFVGSFTWLVINVNNAYQTLMRNPVQTKLHVEYTQTAKFPTV
TFCNFNKFRSPYVLAQPGLIDILRYVYPVTEEDREQQFDWTDPLYAPLVNGSQYLEFAKI
AGHQIRDSIYVARFKGVNLDIEKEFKRVVTDFGLCFQFNSNGTWNTSRSGTNNGLWLEMN
VEHNLYIYGDSNSAGYKVALHNHDEEPLVNELGFAISPGQETFVSVQIKEIKSLPLPYEG
GKCKNTTKEDFVNHLENYKPYSLTGCRRECRANYTKQKCGCKYIYDPGPARTCNAPELGC
YRNADEEYTRNASIENDCNCFVPCHEMQYEHRLSTSMYPGNHVIQRLMDDFNITKELARE
NFLELRIFFEKLNYD
>Lopho_Phor_aus_TRINITY_DN312240_c0_g5_i3_p1_T
MDNFEMAWDNQQTRTQQGVKSGREDCKDEELSRSSWQIFKDCVTDYLDVCSMHGLAFFLG
SMYHPLRRILWFLLFTASFVWLMIVVATAYDNLKRKPVQTKMKVKYADNASFPTVTICNL
NKYRKKYVEENYPDSEEVVRFMHPVVPEDSTANIDWSEPRHAWSLKESNILHLARDGAHE
IHKSFISAQFGGHELPVAKPNFVTVETDSGVCFQFNPKGTWYSELSGKTFGLMVMLNADQ
ENYYFGDSFSVGFKIALHRHDEEPFVDEFGFVVSPGQEVLVPMVLNEVISLPEPYEGGTC
VDTKVKDFTNPLEYYSPYNYNSCRKECQTRYSVEQCGCKLAFDPGSSRICVGDEYACYRL
STQNYAKNASIERSCNCRMPCHQVGYDYRLSSAMFPGLQYRSYLANAHNLSLDYARQNLV
VVRLYFEKLNYQLVEQVLSYDGMNFFADLGGNSGLCIGASLLTLAELIELIGVIIFNMIF
RRREKTTPEKNNAK
>Lopho_Phor_aus_TRINITY_DN316400_c0_g1_i8_p1_T
MADTGQKHYSYLTGNDYENVPIHALTYKALWRGFQENTSGHGFSRIYLTPGTVKKALWTL
VCIGGFTAWGVHTTTLLVNFFYYHVDVTVQILTAKNLSYPAVTICNLNQIKSSKISDVLK
LDDLYALAAQTVDDGDGNLGQRDQYSEYVDILRGAVADMDEAAKSDAGHQRGDFILGCTY
KGYECYGSDVMEFFDPIYGNCFTFNGADGSPQKDYMRAHSTGPESGLTLMLNIEQSEYLP

FTQSAGVRLVVHNDTVMPFPADEGINLEPGKLTMVGMRQIQIIRKGNPHGVCIDPTSYNG

DRNLYEDKFRVRYTYQACKKTCYQKAITERCKCFDPHYPTTVNVSDTLKACRYGTIEQDE

CSENITQDFADGNLRCDCPLSCSESAYALTISSSQWPSDNYKTALFTKLTSLNNSNIDAI

LTNAADKDIYRRNFLKVAVYFEDLNYEEIEQKAAYTGSSLISDLGGSIGLWIGWSVITIV

ELFEFLSDVLILGCSRKSSKRAVGHSSDTPVKSIK

>CAE7949834_1_ASIC1_Symbiodinium_KB8

MAPQTPPDVAGSAWWRPAASLAELPSQAAGFANRRDMAVPLRQRRLERQADGAAVALVAGQPEGFQPLQE

FSDNVTVRELRFVTASTTTGALLSAVSVCKRWLWLVLLLGALVALVVIVTDRVQYFLQNPTATVVKWESR

APVPFPAVTVCNENGFQQSRVAQYGLSTLNNSVLELMTQPVNSFILRCEFAGLECFADNFSVLLDAQFGA

CYTFHGLRALTIDGEEFNASGPNPLTSARTGPDYGLRLVLDIEQAEYVGYSPRTSVRQASGADQGPGTEP

DSSGSAQTSPQTSPQTSPQTSPQTSPQMSAQASAQASPQTSSSSATASSSPPPAAASSSPQASSPAGASP

TRTPSPSQPARRLLAPRRVQEDPGPGSARDARAPPDNRNSTVEALFKGPESAGIRVIIHDPSVLANPTSG

FGVGPGSASDIAFERVDRSRLFPPWGTCNEAWHSESGATDPAGYAKTACERDCYYASVADVCECLLLPWP

TTMEERPSLYVCSSEEKACIATQLSRFQAGELGCSASCPDRCLETLYTATAGRQLWPSVASASTILDVVR

SAKQDTSITPEYLAQNMMSVRVFPPTTEYQVVTTVRAYSFESMLGEVGGNTGLWAGMSALTIVEILEFVM

LALSAWLCSRGGCCRRGGKRAPRTNVEHGTELHEGEAIELVAPQADEESSAPAVAPQAAAAASGRAREAW

RE

>Filast_tunicaraptor_GIQG01049703_1__p1__GENE_GIQG01049703_1__GIQG01049703_1__p1___ORF__type_complete__len_874____score_117_73__GIQG01049703_1__2484_5063___979

MTGAASDERLEKLQAQLDVLAAQLRAARKEADNGGEYDDAALGSPAEDPYGAGSGESCLSMLTGCMKESTVHGIPHA
FAFNRTIWRRAMWVTYFTAAMCGLLAFCVGRVRYYYERPFSSSVSLEYAPNLAFPAVSVCLTNGYKNSGYVAWFRHA
AAEAVREHVPGAAAFDNAEVQQHLESLLFYGDTLASPNGASGGTSVARRVRQSPAMRKVRAEKVAAAAAVAAAAAR
VAREAEDGGACPRPLSDTPGDVAFAELIALLHDSGLASSALSFNVDDTVETACAAWADAYYEGEETPTVVSYGESAVNF

EPMCAKYVLLGDDPVYHEFKTALTAAGAPAHNSTAAWSDALAAEGVASSQNALVLPLRCEVIASLSELGVAPPTPVPSS
TGAAPTPTARAAAPSATPSAGSAHISTLIHPAVLSAKESALIAFISDELDEEELEKMAYTAEEVILGCLWKGGALPCSHRNF
TTFFANEWGNCFTFNNPQTLRNDVLHSAAPGPKGGLVVTLFIHQDEYEALPAGSITEDNFAALAIEGASVKLLMHDQR
DHPEMDSALDVAPGRYHLAAMRKLVVNNLGQPYSDCQEDVAAALTSTDGVVLSTIPGRTRYNPKTCFKECTVREIFKE
CACLPPTSRVARGKRVDRCLSDEQVACQDAVEGRLSNDPNLQTTCACVKPCESVTFPVSLSAAQWPSEVNRDVFLADI
EGVLGMQVPPTFFDVNIVNLNVFYSDLNFERIEQQGVMTVGSLLAEIGGMLGLFVGMSVLTLAEFIEFLFFGSRRQVRK
CKQGGDPIGPKNGVPSDACGGADRVDVVLEEPNAMNMQGLPLSKEATAAGAGRGADPVRILKLQRADSVNSVHSAV

>Filast_tunicaraptor_GIQG01084220_1__p1__GENE_GIQG01084220_1__GIQG01084220_1__p1___ORF
__type_5prime_partial__len_641___score_99_41__GIQG01084220_1__3_1925__864

DGDGDGDNDGDGDGDGDGDGAGDGEGEGGGNGDGDDDAVALYRRFRSSLGAAGYDVTYVARDVGFQVAGLCAS
TAAALDAGQPTPAYVMWNNATVEGGLADVNWAPLCDKYVPLYNDPLYLHVMDLATKVTETTDSADLANLMDAVNS
ADVQESSNVVLGALVADINAFLTDDPFLHPRSDGLAPIGSLDWASTQTEEQVARAERLSTEAVSLDAAKLWTFEASEFII
NCRWKNGALPCSAANWTTFYVEEYGSCFTFNGPKEDAAVLASGSPGPNGGLTLTLYIDQPEHFTVEAGVLDDDNLDIL
LTPGAGVKILMHDQKDSPEMDGATDLAPGRYYSLSMQKIIIDNLGPPYGDCREELGVGTRYSIRSCFRECFYRRVADEC
GCLPIDSPSTRASTSLRCTTVALYDCQQSVESSLNRQNVPDLCGCPKPCKAVLFPPSMSFQQWPTKKDKAIFLADVRRIV
GGDVDETLFEENFLTANIFFQDLNFQQIEQSPAYSVGNLMAEIGGMCGLFAGISILTVMEFVEMLFLGGWTLRKLRRNA
AASDANAHATNGHHQQPQQLPQTGRTDTIEVELAERQPTKMAEMPAVPVVGKLKLTGGSDSDCDDDDDCRPAALR
LRSSLSRGGSLASVASMQSEV

>Filast_tunicaraptor_GIQG01095491_1__p1__GENE_GIQG01095491_1__GIQG01095491_1__p1___ORF
__type_complete__len_581___score_92_90__GIQG01095491_1__198_1940__777

MTARGNNWSLELDEANPGNAPEAVGAHREGMGRARNHDKPFDEDEETKMFENAYGCMPLTRFFMHTTTIHGLPR
VFEKGHHWFRTALWAALFCASFGMFVYVAQFRFDEFYAYNTSTSVDISFQPRVQFPTVTICNSNMYRRSALSYDVLEAL
WDDEGLTNQEIVEFVAVTINELNSDSSLPTLTREQLFAIAVDELGKEGEDDQELITDDLIELLADRYHFDDDDFYDLLLEKY
LRSNRNGGTDAQVLASLFSDEQIGEIGHQLDDLVAFGYAKFKGESIDIATEFERTISHPYGNCYSWNSNGNYVTTRPGPD
LAFDLIINLEQQEYLSSSLSGTGKGIDDVSIIPAGLRVHIHDKSEPPFPDAGLTVAPGTQAFISLQQQTITKMTPPHGECVE
RTDTAAYSVESCFKGCFEKAVFTQCGCRYPAAHAVSRTVGTCSTSDFTDCVDDIEDAFSTDVASFSCGCDDPCNRITYRK
NLSHGRWPSVAVSGIISDAYNISTREVGDSFLRVTIYYEELNVIAMTQEEKYSMSALGAELGGLLGLFLGVSVMTVFEFCE
YMTFFFRPRMYRKKSVKVTPQS

>outgro_CafRoe_KAA0150573_1_hypothetical_protein_FNF29_05148__Cafeteria_roenbergensis__747

MAEGTVKGQRRWWGPRLAARWASSTSAHGFGRIAHYCRQPGSCLRLAGCVWFSSVVAFLVLLVVFILDRLAFYRTWP
VTTSFALKQERPIQFPAVTVCSNVFAFDSRIALVNFTDVIEHPTDAALSRLGAQPDELVIMCVFGGRKCSGLHAEEWIHP
AFGLCLTLNGGQRPFRVGNTIYDAASLPSSFSTVGEYLSAREERVDYPPLEASEIAGLSNSPKQLSVKAPRADDGLTLYLG
VWQNEYTAVASSADEIGIGWSNEPVTGVFNAESAGVRIAVHDRLKTPDMDTATMLHPGKETHVTYRKRVVTSQPPS
WGDCSPRFWNETAGQPLPSGNYSVAGCELVCAAKLVEAECGCLTIPTPRGFVPANPSMRACPLTMECPVVQQEKVLL
GQAECPDCVKDCATTEYPLTTSTLSWPAAPSQQVFLDLFDQFSQIGIGASALENISTAEDGLQRLQSNVLLARIYPSSATY
EEVTSNRAYSITSLLSDLGGQAGLWGGISILTAIEFVEFIGLGFMACFGVACARNAFTEPGDEDELTPDDQATTHTSPIGS
AHALQSKAATDAA

>outgro_CafRoe_KAA0157466_1_hypothetical_protein_FNF29_00042__Cafeteria_roenbergensis__674

MASSKNAPGSMAITGADEGFQPLQEFSDNVTAHGVRKVMYLSKERSCRKWVWVALLVGALIALVVIVSERIGYFMEN
PTSTVVVWENIEPVPFPAVTVCNENGFQQSRVAEYGLDTLNNSALLMMVQPVGNLIVRCEFAGQECFADNFSVLLDA
QFGACYTFHGLQPVTVGSEVYDANNGNRLTSARTGPDYGLRLMLDIEQSEYVGYTPKTSVQPPQQDVKAAPDNRNST
VEALFEGPESAGIRVIIHDSSVLANPTSGIGVAPGTASDLAFVRADRSRLFAPWGTCNQAWHSESQGTDPTGYAKSACE
RDCYYASVADVCECLLLPWPESLSDRPSLFVCSVEEKTCIAEQLSKFQSGKLGCSENCPDRCLETLYTATVGSQLWPSSSS
ADTILSLVRDTRQDNTISAEYLTENMMSVRVFAPSSEFQVITTVRAYTFESMLGEVGGNTGLWAGMSALTLVEILEFVFL
ALSAWLCAGCRCCSRDRAVRTVGGSELHAEMSPGSARPGPDAAASRAAFDDRELAAKASP

>outgro_CafRoe_KAA0153446_1_hypothetical_protein_FNF29_03263__Cafeteria_roenbergensis__865

MSSESRQPEGFQPLQEFSDNVTAHGVGKVFYLSKERSCRRWLWLVLLLGALVALVVIVTDRVQYFLQNPTATVVKWES
RAPVPFPAVTVCNENGFQQSRVAQYGLSTLNNSVLELMTQPVNNFILRCEFAGLECFADNFSVLLDAQFGACYTFHGL
RALTIDGEEFNASGPNPLTSARTGPDYGLRLVLDIEQAEYVGYSPRTSVRQASGADQGPGTEPDSSGSAQTSPQTSPQT
SPQTSPQTSPQTSAQASAQASPQTSSSSATASSSPPPAAASSSPQASSPAGASPTRTPSPSQPARRLLAPRRVQEDPGP
GSASDARAPPDNRNSTVEALFKGPESAGIRVIIHDPSVLANPTSGFGVGPGSASDIAFERVDRSRLFPPWGTCNEAWHS
ESGATDPAGYAKTACERDCYYASVADVCECLLLPWPTTMEERPSLYVCSSEEKACIATQLSRFQAGELGCSASCPDRCLE
TLYTATAGRQLWPSVASASTILDVVRSAKQDTSITPEYLAQNMMSVRVFPPTTEYQVVTTVRAYSFESMLGEVGGNTG
LWAGMSALTIVEILEFVMLALSAWLCSRGGCCRRGGKRAPRTNVEHGTELHEGEAIELVAPQADEESSAPAVAPQAAA
AASGRAREAWRE

>Cteno_MneLey_comp601992_c0_seq1__p1__GENE_comp601992_c0_seq1__comp601992_c0_seq1__p1___ORF__type_complete__len_212____score_42_61__comp601992_c0_seq1__142_726___205

MLNKTCGCLNSTPPLNDVFNYKECTIKQVATCSLKAYYQYVLDFADVKSDYEECGCDVVACEYTLFRTTVTATPLPDTEIE
HKLKAWNSPRHFDRFQRFSNYTAEDLAKNMARIEFFFGSSSITKVEEIISYDFDNLLGDIGGVMGLFLGASIFTTMEFCTL
VVNLLSRLWQWKIQPWLRSETRDRKIISETNL

>Cteno_Ple_Bac_comp46220_c0_seq11_p1__comp46220_c0__comp46220_c0_seq11_p1___ORF__type_complete__len_202___score_27_04__comp46220_c0_seq11_1766_2371__211

MLCSDVLKDNSFTREASVNGNCFRVNPEGKLKGKGGEYGRLRLMFFADLNDYTAPAKNQAQYGYTVAFHDHETYSSTI
PSGFFMSPGSIYKVDLGLVREFREPPPAGLCDSTLTTNTYGRYEENSCIAECRDNAIIRACGCVHVVPPHPDNNPLVEYLP
CTLEQWAKCGMREYKKWGESNSNRHFKARKILSKLLETQKT

>Cteno_Ple_Bac_comp46220_c0_seq48_p1__comp46220_c0__comp46220_c0_seq48_p1___ORF__type_3prime_partial__len_242___score_38_17__comp46220_c0_seq48_1236_1958__259

MIQTLFLFMWLASIGYAVYVIASSTMSFIGKPTGTKFQVIASQLDHQENVNLQSAVQFPTISVCSHNQVSLKWSSERPG
LEKLLEKFDKWNPVLAANIDWETPQFNKWRNATYEEVIKKGGPDNRTFIQCEHGIMLCSDVLKDNSFTREASVNGNCF
RVNPEGKLKGKGGEYGRLRLMFFADLNDYTAPAKNQAQYGYTVAFHDHETYSSTIPSGFFMSPGSIYKVDLGLVREFRE
PPPAG

>Ctenoph_Hormiphora_californiensis_evg190294__p1_GENE_evg190294__evg190294__p1__ORF_type_internal_len_253____score_30_09_evg190294__2_757___271

LMAEYSPANRRVSDGGRSSATSRSDTPSSWVKGFQLFAETTTAHGVNQIFTGSNRAVRLFYLLAWIACVIYTMYMIITSI
VTFINKPTGTKYEVIVNDLGGLKPQSIKFPTVSICSHNKVKKSYLRKEGNEPIRDLWSKLDMYDQAEADKINWTNIGTASI
QHMTLEDIISEGGPDDDRLLSCTQRNKNCQDVLGKNESYSVRENTPSGKCWRINPEGKIFGKSGDWGKMQLIVFADL
QDYSKRSADQPQYG

>Ctenoph_Hormiphora_californiensis_evg1639__p1_GENE_evg1639__evg1639__p1__ORF_type_internal_len_264____score_48_35_evg1639__2_790___291

HGVKYVFHSNHRVIQALFLIMWFVSIGYAVYVIASSTISFIGKPTGTKFQVIASQLDRNNVNLQSAVQFPTISVCSHNQVS
AKWSRQRPGLEKILERFDTWSPGITEAINWDRPEFNEWRNSTFETIIKEGGPQDSTFIQCEQGIQLCEEVLGAKYVTREP
SVNGNCFRVNPEGQLRGKGGDYGRLRLMFFADLNDYTVPAKNQAQYGYTVAFHDHETYSSTIPSGFYMSPGSIYKVDL
GLVREFREPPPAGLCDSTLTYNTY

>Ctenoph_Hormiphora_californiensis_evg16615__p1_GENE_evg16615__evg16615__p1__ORF_type_5prime_partial_len_265____score_62_43_evg16615__102_896___294

AFHDSVYHGVTLNSGYYMSPGSIYKVDLSLKKETREPPPGGNCNASLTMTTYGAYSEGSCIQECKDKDLMKRCGCINM
VPPQNYLNYSSCTVRQWAECGLSAYLNFSERYRDPNLTDDICPCNSPCSETHYEAQLSSSSLSPAYTENLFASKYIKGYLSK
VSQPQYGIIYNSSDDILKNVMVLEILFSSMVTSEIQQVVTYTSANLLGDIGGVLGLFCGASVFTIVEFLQLFLSSIQKHCLTD
LCSSDEDTVEAADDKRGDKEL

>Cteno_Puk_Fal_comp26850_c0_seq10__p1__GENE_comp26850_c0_seq10__comp26850_c0_seq10__p1___ORF__type_5prime_partial__len_320____score_71_31__comp26850_c0_seq10__394_1353___349

RRVRGHEVIPEPEVNFFTREVSLTGNCFRFNPNGVLRGKGGDYGKMQLMFFTDLNEYSDLTRREPQYGYTVVFHDHES
YSSTIPSGFWLSPGSIYKADLSLSQEFREGPPAGTCDPTRVNNTYGRYEENSCIAQCRDDALMAECGCVHVAPPLPENEP
GKDYRGCTLQEWATCGLRAYKEWVKKYSDVNRVSMGCNCPPTCFETEYKAQISSSSLSKFYAESKVGRLPPGYNTTQD
VMENLLIIDILFTSMQVSEIREIVTYGWANFLGDVGGVLGLFLGASIFTIMEFLQFFVLAIWDALCGLPGRKKAPNHDTVS
LT

>Cteno_Ple_Bac_comp48643_c0_seq3_p1__comp48643_c0__comp48643_c0_seq3_p1___ORF__type_5prime_partial__len_344___score_41_94__comp48643_c0_seq3_3_1034__377

LSSLYFRVPKFTLFQEVSLSGSCYRINPKGSLLTQVGDYGVLKLTFFTDLNDYSKFSKKDPMYGYTVVFHDNGTYSSTMTS
GFYMSPGNVYKADMRKYKEFNIGPPYGRCNPHLKNNVYGTYSSSSCIARCRDQYVKDMCGCIQILPPHPEYEPELDFRG
CTLKEWVECGARATLNFTDRFVDLKEEAMCECTPACMETWYDATISSSRLSKFYANDKAISIDGQPAQLARIVDGYGLN
HGIEETYNISKETVYENLMVLQVLFTSIQESNYKEYVKYDYTQLIGDVGGQMGMLLGASLFTIIEFVHFFIKVAWRYCVKK
ATRRAGEAAERPAYEMHRDQVEE

>Cteno_MneLey_comp13853_c0_seq1__p1__GENE_comp13853_c0_seq1__comp13853_c0_seq1__p1___ORF__type_3prime_partial__len_479____score_87_11__comp13853_c0_seq1__3_1436___533

MLLNNDTKPPDVPKKQSFFEDFKEFTTETTSHGIKYVFHAAYKWVRVIYLILWLAAVGYTVSIVFISCKKYFSMETGTKIEE
KHTPKHSSNEVAITHPTITVCPNNLISKSYLAKHPNLDELMTKLVTFNENETPSLFDDPKYAGYGDMTYHDLMWQGRP
RVGLLQCTYFAHKCGAVDPFNSEDYETKTYHWDKTYNGSNHKFYEWEVSLSGSCYRINPRGNLWAKVGDYGEIKLTFF

ADLNDYSSASKEEPEYGYLVTFTDSGTYSSTATSGFFMSPGNVYKADLRKYKEENLGPPAGKCDIESKTNVYGAYSTNSC
NKLCRDQHVMEKCGCVHILPPHPENDPDTKFVGCTLKQWAECGLRESTKFTEEFVNMNDDKVHCEHCNPACMEIWY
EATISSSRLSTFYAEAKVSELDKLITGYQLNEGNKTTNYTIDKDTVYENLMVLKVMFTSIQESTIKEYYKYDYTQLIGDVGG
QM

>Ctenoph_Hormiphora_californiensis_evg120240__p1_GENE_evg120240__evg120240__p1__ORF_type
_complete_len_484____score_100_73_evg120240__313_1764___548

MGWLADFKEFSDGTTAHGVKWIFHPSVHGNFKIVKLIFFIVWLAVIGYSLFTIYNAVHNYVNKPTGTKFQVIAEGPEQE
KPASIQFPTISVCSHNLIKQSYFDSNTGLEELWSDLDKWDPDVTEKLNFTDSKFARYRNMTYQEIIRAGGPDNTTFVHCR
HFINKCEDAIPNPEENFYVREASLSGNCFRINPNGVLHGKGGDYGKMQLLFFADLKEYSTMTRSEPQYGYTVVFHDHES
YSSTMSSGFWLSPGSIYKVDLSLSQEFREGPPAGTCDPTRVNNTYGRYEENSCIAQCRDDALMEKCGCVHISPPLPHNE
PGKDYRGCTLEEWATCGLRAYKEWVVKYSDVNRISMGCDCPPTCFETEYKAQVSSSKLNRFYAEEKANMGLLPPGYN
GTQDVMDNLLIIDILFTSMQVSEIREIVTYGWGNFLGDVGGVLGLFLGASIFTIMEFIQFFILAICNACCPGLTSRSKANSP
VDTYALS

>Ctenoph_Hormiphora_californiensis_evg197715__p1_GENE_evg197715__evg197715__p1__ORF_type
_complete_len_492____score_113_33_evg197715__97_1572___570

MSGSIQGELRNFADNTSAHGVRLIFHGHHVAVKVAFLIFWVSAVTFCVFRASLSIRKYCQQETTSRYEMFPASPTAVMD
FPTITACNLNMIRKSYLKANPGLMHFWKLASIHNFQALAELDWESEELSPYQNLTYYELLKEAQMYNDTFISCTQGRMR
YCVDTVMGADYYQRDIEFVSGSCFRINPYGTLKGKSGDYGVLTMMLLADRDEYIYNSSNVGWVMAIHESERYGASIDN
GIKISPGYTYFVNLDMMNIKNHRKHCEKSVNMISGYGRYDQTTCLLDCRDRQLYEKCGCLMTVPPNAKQAGLNYLPCT
LKQVSQCALRTYVKYVKNYADVRAKSTTADCDCFVTACSYTLYTTSVTATPIPDLELQFKVDSWKNQARFSRFRNYTYED
LGKNMAVIEVYFGTSSITSIEEVVAYDFDNLLGDIGGVMGLFLGASMFTAMEFVTLIISLSLRIWRTHFPTKDSSQDHMIP
HHSLKLPRASVSEVNL

>Cteno_Ple_Bac_comp47842_c0_seq1_p1__comp47842_c0__comp47842_c0_seq1_p1___ORF__type_
complete__len_492___score_104_93__comp47842_c0_seq1_138_1613__571

MTWIRDFRDFAGDTSAHGVKNIFEGPTKLLKLIFLICWIICSVYATYVIMNSIITFINRPTGTKFTFLVENEQRKLGKPAVVP
MPTISVCSHNKVKKSFLEKKGNEELKKIYTILDKYDLGEAAELAENITGEVADMKYEDLLEEGSISNDRILMCQQRSTNCF
QLPAFQNSEWVTRENSITGTCFRINPKGTLNALMGDYGALQLRFWADIQDYSESSKNNPYHGFTVAFHDKDSYGSSLS
SGYLMSPGSYYKVDLMMRKESREPPPSGHCDDSQPFTTYGNYSEGACVMECKDRVLNDTCGCVNVVPPINSLNYTSC
TVKQWASCGLKTYLDWYKKYTNPDTESELPSETMCRCHLPCTETNYEAKVSSSDIPMSYAEKMFSNPAIQGWMTGISD
PELDINYNTAKDIKDNMMVLEVMFSSSQISQVKEFVKYTTNNLLGDIGGVLGLFLGASLFTILEFFQFIIMSIAKYCCMWT
PGGRKKLEASEMS

>Cteno_MneLey_comp15203_c0_seq1__p1__GENE_comp15203_c0_seq1__comp15203_c0_seq1__p1_
__ORF__type_complete__len_500___score_87_69__comp15203_c0_seq1__52_1551__595

MKKVTEMSWVKDFREFANDTSAHGVKYIFEGRYKLVKLLFLVTWLGFSIYACHVIITSIVRFVEKPTSTKYEVIQDDSEGR
PERIEFPTISVCSMNKVRKSYLEAPENEAIREYYEVVDKYNVSLVKDLAKRFKNSDDPLHSIKDMTYEELIRNGGPNPDRLL
KCTQRAKYCHELPAFNGRDVSVMENSMTGNCWRVNPEGRLMGKMGDYGAMKLMFWADVQDYSARTADVENQG
FVVAFHDNSTYGSTMTAGFLMSPGTYYKADLRRKQEIRNRDKIESCNASLTENTYGAYNEGSCALECKDEALHKACGCT
NVVPPLNNGKYKSCTLEQWVDCGLAVYKEWFHNFTDTDRADQLCPCQIQCEEVRYEAQISSSSISPAFAEKLFPSVQPII
SQPQYGYSNPDFNILYNTTQDILDNVMVLEVLFTSMRTNEIKEIISYGLSNLLGDIGGVLGLFLGASLFTILEVFQFVFFSISK
YCFDKGQPKHDSLTNGDKLSI

>Cteno_Puk_Fal_comp20088_c0_seq2__p1__GENE_comp20088_c0_seq2__comp20088_c0_seq2__p1___ORF__type_internal__len_505___score_74_57__comp20088_c0_seq2__2_1513__604

AEFTTAHGVNLIFVGGHKALRFLYFVIWLLCVLYAIYIILQAAVAFISRPTGTTEQLVVETDHGQDIQFVKFPTITVCSTNIM
KRSFTETDKSISALWYLISRLKVMDPQYVSHVLEHPKRFFQSYCDYHGVTFEHLKNYTVKEMFERGGPDDNRLVYCKQR
NIDCQDLTESYFVRETSSDGMCWRVNPNGSMYGRSGEWGRMELFFFADIQDYTSTSPVYGYKVKFHDSEYCGEGITG
AHFMSPGFVYKVDLGLTKRSWIPKPIGNCDANAKKSTYGKNSEESCLKECEDRYMNKVCGCVDLLPPHNHHNYTACTI
GQYIGCGFREMVHFRMGLRDASRRHGTGEDGCSCSHSCDETHYDVKLLSTSALSPVWADDLSRDKWIAEAVANFSQP
SFNISYNVSSDLLKNIVILQVEFSTVFTKEFKQIVTYSYSNLLGDIGGVLGLFLGASLFSVLEFFQFTLTSLHKHFCGREDKPK
TLPETEKLSSNLAFSPAIRSTPVLCV

>Cteno_MneLey_comp16014_c0_seq2__p1__GENE_comp16014_c0_seq2__comp16014_c0_seq2__p1___ORF__type_5prime_partial__len_507____score_115_90__comp16014_c0_seq2__352_1872___610

PTRKNRRIKCPRCRQHFVFNKVRKDGIMGWLQDFKEFSDGTTAHGVKYIFHPSVHGNFKIVRILFLVVWLVAIAYSLFVI
YNAVGNYVGKPTGTKFQVIASNPYKEKPASIQFPTISVCSHNLVTKSYFKSNDGLEELWSELDQWNPDTAENIDFNEGS
PAAKYKDWTYEKIIAEGGPANWTFLQCEHFINLCQEVIPQPDFFERETTLTGNCFRINPNGKLRGKGGDYGKMQLMFF
ADLNEYSDLTRREPQYGYTVVFHDHESYSSTIPSGFWLSPGSIYKVDLSLGKEFREPPPAGSCDPTRVNNTYGRYEENSCI
AQCRDDVLMEKCGCVHVSPPHPRNVPDKYYRGCTLEEWATCGLRAYKEWVVEYSNVNKVKTECNCPATCTEIAYKAQ
LSSSQLSKFYAEEKAAKLPPGYENAQDVLENLLIIDILFTSMQISEIREIVTYGWGNFLGDVGGVLGLFLGASMFTIIEFIQFI
VCAVLKGCCGLNDKESRSHSPLYNENL

>Ctenoph_Hormiphora_californiensis_evg1138204__p1_GENE_evg1138204__evg1138204__p1__ORF_type_complete_len_508____score_90_51_evg1138204__158_1681___615

MGITQMTWIKDFREFAGDTSAHGVKNIFEGPTKILKLLFLITWIVCSVYATYVIFNAIVTFINKPTGTKFQVTVESTEREEG
KPAVIKFPTISVCSYNKVKKSFLRAPGNELLRRVYLTLNKYDLDEAAELAAEFEDPDSEVAQASVGDMKYEDLLADGSPDS
NRFLMCSQRAKNCDKLAVFKGNASEGIPPGHFVRENSISGMCWRVNPNGKLKGLMGDYGALELRFWADIQDYSEST
KNMATHGFIVAFHDNGTFGSTLSSGYLMSPGTYYKVDLRLRKEIRKPPPGGNCNASLTDTSYGVYTEGSCIMECKDRWL
YERCGCVNVVPPINKDNYESCNLRQWASCGLKAYLEWYDRYTDTDTANPICTCSMPCEETNYEAKVSSSDISQAYADH
MFTTKGVAGFLNSISDPSLNISYDSANDIRENMMVVEVLFSSAQVSEIREVVTYTANNLLGDIGGVLGLFLGASLFTILEFF
QFIIISVAKHCCNWIPGGGKKKYESAEMS

>Cteno_Puk_Fal_comp27454_c0_seq1__p1__GENE_comp27454_c0_seq1__comp27454_c0_seq1__p1___ORF__type_complete__len_520____score_91_49__comp27454_c0_seq1__400_1959___649

MSRYGERYPPSSGLQLFPDPDKEEEGISPVQQPCRCTCPCNTQNTRDTGSQMDVKTISTSSGKRATFYDRWFDFSDGT
SLHGFKFIFSGGRVTRAAFAVFMVLFVFKLGWQLRESVISYMEHPTSSKISVSNVEGPALVTFPTISICSHNLVSQSYLDD
NPGLDFLFEMIDTWQPEVVSQIDFTDPQFAPYADFKYSDILRDGGPPSINVLQCSHFTNDCTPDLQSEKSMERELSMRG
NCYRINPKGQLMGKGGDYGKLSIKLFSDLNEYSATGKKLAQVGYSVQFHDHEQYSGAIPNSYWMSPGLIYKADLSLTRE
TRLTPPAGRCNDTVLYNSYGRNEENSCLAQCRDDAMMRACGCVLVPPPYDPTSYTPCTLQQWSECGLTAYNGWYQD
YVDVSRTEEMCYCYPSCREVFYTVSMSSSLLSRFYARDHVQFLPKGYNGTNDVLNNLVVLDILFVNTQELEIEQVVAYG
WTNFLGDVGGVLGLFLGASAFSLIEFFFFVGHLFWTYVLCIWREEHQ

>Cteno_Puk_Fal_comp26884_c0_seq1__p1__GENE_comp26884_c0_seq1__comp26884_c0_seq1__p1___ORF__type_5prime_partial__len_531___score_77_18__comp26884_c0_seq1__1_1593__671

QRRVRGFFICIYPSCVCFEARNVKRTAHSTGTSTIGRRMTWIKDFRDFAGDTSAHGVKNIFEGPTKLLKLIFLICWIVCSVY
ATYVIMVSIITFMQRPTGTKYTFLVENENRKLGKSAIVTMPTISVCSHNKVSKSFLKTEGNEILSEVYTTLDKYDKEDAEEL
AKRFDNSSDPLSEVADYLYEDLLADGSIPDDRLLLCQQRAKQCFKLPAFAKTPFFTRENSITGKCFRINPMGTLSALMGD
YGALKLYFWADVQDYSEASSNSATHGFTVAFHDSDTYGSTLSSGYLMSPGSYYKVDLRLRKEVREPPPSGHCDKTREFT
SYGNYSEGACVMECKDKTLNETCGCVNVVPPINDLGYRSCTLKEWATCGLKTYLDWYDAYTDTDTAQTTCKCHLPCQE
TNYEAKVSSSSISTAYSEKMYSNPDVVNWMKKISKPELGINYSTAEDIRENIMALEVMFSSSQVSQVEEFVKYTSTNLLG
DIGGVLGLFLGASLFTILEFFQFIIMSIAKYCCQWTPDGRKKLDGHEMS

>Ctenoph_Hormiphora_californiensis_evg1163616__p1_GENE_evg1163616__evg1163616__p1__ORF_type_5prime_partial_len_545____score_81_53_evg1163616__2_1636___694

FKRINLSSILLSPSHILPNDQYAMLLSNGAPQGETKRKSFYEDFKEFTTETTSHGIQYVFNAAYKWVRLIYLVLWLSAVLYT
MYIVCISVGKYISMETGTKIEEMVTPRPANLNQPAITHPTVTICPHNIVSKAFLNKNEGLEELFAELHKWEDDSAWNTSA
SYHKYANWTYHDVMWKGRPRVGLLQCEHYRHECKDLDFTNRSVPEYHWDREYDASNHLHYDWEVSLSGSCYRVNP
QGSLFTQVGDYGELKLFFFTDLNDYSQFAKDEAQYGYTVTFSDNGTYSSTMTSGFYMSPGNVYKADMRKYREENIGPP
YGRCNPHHKTNVYGTYSLNSCVAQCRDSYVMERCGCVQILPPHPEWNQDGVEMVGCTLRQWMDCGAKAQKEFTD
RFVNLYAEEKLCHCSPACMEIWYEATISSSRLSKFYAKKKATDRLPGIISYYNLSAHIDEPYTIDTETVYENLMVLQVIFTSIQ
ESNYKEYIKYDYTQLIGDVGGQMGMLLGASLFTIIEFVHFFVRVAWRYCLRKRGARAAEEQPAYEMHRDQE

>Ctenoph_Hormiphora_californiensis_evg156929__p1_GENE_evg156929__evg156929__p1__ORF_type_complete_len_561____score_102_52_evg156929__369_2051___738

MPYSEIPGPAEPDGAADQGYAGTSRTPPQAHSSGLQLFPDPDRDVEVPVQVCQCTCSCNRATPGGEGPGELASAAGD
GRMEVRTVSQTSSRGEKGSSKFYRRWDEFSDGTTMHGIKFIFSGSRLTRLAFFLFMLLFVFKLAWQLRESLANYMEYPT
SSKITVSNVEGPALVTFPTVSICSHNLVGQSYLDSKPGLDDLFERVERWDAEVASSINFSDPTFAPYADLPYETVLREGGP
PDSNVLQCSHFTNDCLGNSSDKYMERELSLRGLCYRINPAGKLMGKGGDYGKLSIKLFSDLDEYSATGKKLAQVGYSVQ
FHDHEQYSGAIPNSYWLSPGLIYKADLSLTRETRLPPPAGRCNDTLLYNSYGRHEENSCLSQCRDDAIMAKCGCVLVSPP
NPHYAPSTNYTPCTLKQWTECGLAAYSNWYQQYVNVSRTEELCYCYPSCREVLYSVSLSSSHLSRFYARDHVQYLPPGY
NGTRDVLNNLVVLDILFVNTQELEIEQVLSYGWTNFLGDVGGVLGLFLGASAFTLIEFVLFLGQLFWHYVLCAWRDDHE
LQIRDL

>Cteno_MneLey_comp614729_c0_seq1__p1__GENE_comp614729_c0_seq1__comp614729_c0_seq1__p1___ORF__type_internal__len_184___score_30_14__comp614729_c0_seq1__1_549__193

NFISLSYERVRNLKQPYGDCRDDAMPLYRNYSISGCERQCFTEIMIKECNCSSYYLKAMPGGKDVECNLYQEKNCVKELI
RRYHAGEFINSSECRCPDPCVSESYLATLSTAEYPAKNHGEALQRYNEANRGPNDVVQDVTFYRENLMKVHIYFEILSIKE
VEKVGSYKWSNLMGDVGGQLGL

>Cteno_MneLey_comp1086195_c0_seq1__p1__GENE_comp1086195_c0_seq1__comp1086195_c0_seq1__p1___ORF__type_internal__len_190____score_29_76__comp1086195_c0_seq1__2_568___202

KLLWAVLFLGFFVYSFQCCLKSFNTYFSMPTSTKYNMYFEPNKLLRFPSITICNMNPHNTAYLDKEENNPMKAYLLSESR
VPWDTSNLTPDVSETDYRNFNVKKYYRQAGQKLNDFISEVVFFEEEIETSDAFREQFTTHGMCWTFNADGSRQVNRT
GMDFGLSFKLNINQSSYPNFVTTAGVKVMFHN

>Cteno_Puk_Fal_comp26472_c0_seq1__p1__GENE_comp26472_c0_seq1__comp26472_c0_seq1__p1___ORF__type_5prime_partial__len_221____score_43_26__comp26472_c0_seq1__674_1336___234

NAEYGDPVLEQDYDPDGYTIAKCVMNNRARYTIQKCNCIPFYMEVPRDLSSTSPAVCNLYQEWDCVEPLLEDLALGTLT
MGGQLVQDPDCPSPCKYLTYSYTISQSEYPSQSVEDQVLSRVGASRGAGWNIERIRDNYLKVHIYFEELSTLEMERHPSY
MIANLIGDVGGQLGLLLGMNICSLVQFTDYIVRFSFFKGIMKLVRMSKRNNKRSVGGRSRL

>Cteno_MneLey_comp860380_c0_seq1__p1__GENE_comp860380_c0_seq1__comp860380_c0_seq1__
p1___ORF__type_internal__len_249___score_49_23__comp860380_c0_seq1__1_744__269

FPEINTELGICYTFNADGKKRVKRTGNDFGATFTININQSEYAETQESAGIKVMLHHFKEPPLIKEYGLAIPPGSEAYISTRT
LQLNNLREPYFNCTDRKLRTTDYYSVVACTLECHAKILDEECQCKQSHMPLENITICSLYTHRMCTENAIKKIHNGSYADR
ISDCWCIDACKDTYYEYSLSRAQYASINAAKKILEVNRNVSEYSTISDLRENFIRLHVYFETLAIENVEKVPAYTTMNLLGD
VG

>Cteno_Ple_Bac_comp46643_c0_seq4_p1__comp46643_c0__comp46643_c0_seq4_p1___ORF__type_
5prime_partial__len_272___score_55_68__comp46643_c0_seq4_3_818__300

DDYFRTQDVAGFKVLLHNSYEPPMIEEYGFALRPGSESYVRIRLQKFRDTERPLGYCDPVLEQDYQPAGYTIAKCVMNN
RARYTIQKCNCIPFYMEVPRDLSSISPKVCNLYQEWDCVEPLLEEIALGTLTMGGELIQDPDCPSPCNYVTYSYTISQSEYP
SQSVEAEVKSKVVGARGAGWNIDRIRDNYLKVHIYFEELSTLEMERHPSYMISNLIGDVGGQLGLLLGMNICSLVQFTD
YIIRFSFFNGIMKLVRMSHRNNKRAAGRSRL

>Ctenoph_Hormiphora_californiensis_evg16414__p1_GENE_evg16414__evg16414__p1__ORF_type_int
ernal_len_297____score_76_10_evg16414__3_890___327

APVKKKNEFRPYPAPYHVPPQQESASAYQSVNEYQAPSERERNPVKRSQSDSSTAGSFRAAPWLNGQTKILAPKYPVM
SSVAATPTNSLKMRSMGGGGGRTVLPPPPTWHDSLLGEDRPDQVIQDREEETTVKSSGDKEPTKSESFADFTGDMTL
NYLRYVWNSPEYIRKLIWAVLFLGFFIYSFSCCYRSIAHYLARPTSTKYNMFYEPNKQLRFPAITICNMNPHNKTYLDQDE
QIAMRSYLRKTWKTPWAKDLPEAKEELMQNFNVKGYYRNSGQRVNDFIRKAVFIEEEQVV

>Ctenoph_Hormiphora_californiensis_evg15900__p1_GENE_evg15900__evg15900__p1__ORF_type_5p
rime_partial_len_299____score_42_58_evg15900__2_898___328

EEQVVEENFVERFTTHGLCWTFNAKGDKFVNRTGMDFGLSFSMDINQGGYPDFVRTAGVKVMFHNYYEPALIDEYG
MALSPGTENFISLSYERIRNLDSPYGNCTKDPSPPFANYSVSGCERICFTRALEGDCNCTSYYLPVGEKKECTLFDEDVCVE
KVIKRYHAGLYSDRPYCQCQDACVSENYIASLSTAEYPSKNYGEALEAHNKETLNDTVRDVTFYRENYIKVHIYFEILSIKEV
EKVASYVWSNLMGDVGGQLGLFLGANIVTIFEFVDFFLRVFWYRLLTPYIFKRFRGN

>Cteno_Puk_Fal_comp27722_c0_seq1__p1__GENE_comp27722_c0_seq1__comp27722_c0_seq1__p1_
__ORF__type_5prime_partial__len_329____score_62_50__comp27722_c0_seq1__161_1147___358

FQYFNVKNYYRDSGQRLENFIERPVFLEEPIDVSEAFVEKFTTHGLCWTFNDKGDKFVNRTGMDFGLSFNMNINQDGY
PDFITTAGVKVMFHNFYEPALIDEYGMALAPGTENYISLSYERIRNLQSPYGNCTKDSSNLPFANYSVSGCERMCFTLALK
EECECTSYYLPPSNVKECTLFDEQECVEKVIKKYHAGEYADKPFCKCQDACVSENYIATLSTAEYPSKNYGEAIERENKKN
ANGSDVRDILFYRENMLKVHIYFEILSIKEVEKVKSYIWSNLMGDVGGQLGLFLGANIVTIFEFVDFFLRVIWYRFLTPYVF
KRFQRH

>Cteno_Ple_Bac_comp47578_c1_seq3_p1__comp47578_c1__comp47578_c1_seq3_p1___ORF__type_
complete__len_384____score_68_54__comp47578_c1_seq3_141_1292___410

MELNTSPGGMKKLVTDVKDLAILMNYRNFKEGVYPGVEAFTPYSKYKQDESAFDMTGRGERTSYYPYLEHYQPSYPRP
PRGGKPPPPTGAPVPNNHRTSDQNGKIKYPPNHRDTGSPTELEDRDGEDCEEFMPPPKRKKKKKRRANRAALEQRRE

EMEPDEMESFKEFASTVTADYLAEMFAKASNGRRAIWVSLWLASFVYAWYNIALSVGVYLSKPTATKLNFYEAPPGGV
AFPTVTICNFNKYNKSFFEREGDSVNSQLRDFLVATTPIWDHKERLDQDQWTSKYPNIMSLNLKELYANGSHLITNTVEF
CRFGGKPCKDTANRETSIPSEAFTKVYTEHGLCYAFNLDGKLNMTRTGAQYGLSLRLHVAQDDYFRTQVT

>Cteno_Ple_Bac_comp47578_c1_seq5_p1__comp47578_c1__comp47578_c1_seq5_p1___ORF__type_
5prime_partial__len_459____score_71_82__comp47578_c1_seq5_141_1517___500

TDVKDLAILMNYRNFKEVQLVVPATLSVPAPCMHQNGRPKDYIFTEIEQPLSANGFFKRVPVPNRTTGPGFHGRGGPP
PREPAFPIYDNLLDRDDVLSSNCPSTVPGVYPGVEAFTPYSKYKQDESAFDMTGRGERTSYYPYLEHYQPSYPRPPRGGK
PPPPTGAPVPNNHRTSDQNGKIKYPPNHRDTGSPTELEDRDGEDCEEFMPPPKRKKKKKRRANRAALEQRREEMEPD
EMESFKEFASTVTADYLAEMFAKASNGRRAIWVSLWLASFVYAWYNIALSVGVYLSKPTATKLNFYEAPPGGVAFPTVT
ICNFNKYNKSFFEREGDSVNSQLRDFLVATTPIWDHKERLDQDQWTSKYPNIMSLNLKELYANGSHLITNTVEFCRFGG
KPCKDTANRETSIPSEAFTKVYTEHGLCYAFNLDGKLNMTRTGAQYGLSLRLHVAQDDYFRTQVT
