## Supplementary material for "Function and phylogeny support the independent evolution of acid-sensing ion channels in the Placozoa": FIle S4

>NeNaC1_tr_A7RI82_A7RI82_NEMVEPredictedproteinOS_NematostellavectensisOX_4535
1GN_v1g197495PE_3SV_1
------------------------------------------------------------
----------M-------------------------------------------------
------------------------------------------------------------
------------------------------------------------------------
-----------------------------------------------------------K
ENDKDEDGGDVKIIIK--------------------------------------------
-----------------------------------GSDDKAPQGAG-S----------IY
AFCNHCKKDISVVYIKSENN-------------GKELSVAESLTNK---------WPR--
------------------------------------------------------------
-----------SEFPENDYDKCLKDNDYGGC--------------------------KKL
L----------------------DGRKK--------------------------------
--------R----------------------A---------KV-----K-----------
-EL--WENFL-GGCTLHGFHYCF---------A-GN--------------PP---LR---
--RLIWS----LLLLGAFAMFFEKCTESFI-NFFD----------Y---PFTTTT----L
-L-VY-D----------K-RLPF-----PAI-----------------------SMCNYN
DA------RMS-----KMNGTL------------------M-----------------N-
---------EI----------------------------------FV-------------
------------------------------------------------------------
------------------------------------------------------------
------------------------------------------------------------
------------------------------------------------------------
------------------------------------------------------------
------------------------------------------------------------
------------------------------------------------------------
------------------------------------------------------------
------------------------------------------------------------
------------------------------------------------------------
------------------------------------------------------------
------------------------------------------ASKL--------------
------------------------E-GRNTSHLQSQ-------------------LTGE-
---------------------------------LMQRTL---------------------
------------------------------------------------KEA----AHRLP
--DMI------KECS----WQ-K----H---------GK---C-----------S-W---
----------------------------------------------K-------------
------------------------------------------------------------
------------------------------------------------------------
-----------N----------F-------T-----------SFK-S-ADGD-------T
CYTFN-------SG-------------RK-D-------------------------PI--
-------LSMSNV------GE--------EN-------------GLRLVIDTQHS-----
----------E-----------------Y------------Y-Y-D--------------
------------------------------------------------------------
------------------------------------------------------------
---------VKNAGFKVILH-DQGETP---VKMQGLSVSPGFTSYMELKRT------K--
-----VTNL-----------PFP-YK-TM-C--GMP--------E-------LK----YF
------------------N-SYSKSKCFLDKLTQYVV----TL--CGCRDWF----M---
------------P-V--NGITGIDI-GNPVRNLSKLGKARYQ------------------
C----VTTKQRH-----P---ACGKLG---------------------------------
------------------------------------HTS---SSCDYFDGKRGSELQ---
A-FFV-PTYSVLLIS-------AYFE-E-------NK---------------------LD
------------QCP-VACNSVE---YSAQ-LSY------------------ARF-PA--
-------NNY------AKM-L-----------AK--------------------------
------------------------------------------------EY----------
------------------------------------------------------------
-G---------------L------------------KGS---------------------
------------------------DE-ENR------------------------------

-----Q-------YL-----------R------DNLIEIKIYYEDLTYFDVQ-QVPSYDL
YSL-----------------LGDVGGQIGLFLGASLLTVVEYLDLLGMV-----------
--------AYTS--------F----------K----------------------------
------------------------------------------------------------
------------------------------------------------------------
------------------------------------------------------------
------------------------------------------------------------
------------------------------------------------------------
------------------------------------------------------------
------------------------------------------------------------
-----YRN
>NeNaC2_tr_A7RGQ6_A7RGQ6_NEMVEPredictedproteinOS_NematostellavectensisOX_4535
1GN_v1g80340PE_3SV_1
------------------------------------------------------------
----------MSLDI-------------------------------C-------------
------------------------------------------------------------
------------------------------------------------------------
----------------------------------------------------------DA
YIQQETDIGKLYFV-----NTHILDAAITGC------------YDLKLIA----------
------------------------------------AVMSCPV-----------------
-------AQRHFETEEDKKD-------------EDEDDRAE-------------------
------------------------------------------------------------
---------------------------------------------------------DPV
D----------------------ENPDDT-------------------------------
-------------------------------I---------TV-----S-----------
-QM--WQDFL-HTLTLHGFRFVF---------E-RG--------------PT---IR---
--KVLWL----AILLFAVGMLMMQSKKSIQ-KYFD----------H---PITTSV----Q
-V-EF-L----------E-EIQF-----PAV-----------------------TICNFN
LF------PYY-----LINGTI-----------------------------------GE-
---------KV------------M-SILA------------PQKYID----NK-------
---------------------------------EE-------VLFAR-------------
------------------------------------------------------------
------------------------------------------------------------
------------------------------------------------------------
------------------------------------------------------------
------------------------------------------------------------
------------------------------------------------------------
-------------------------SPIPN------------------------------
------------------------------------------------------------
------------------------------------------------------------
------------------------------------------------------------
----FLNYARKR------------------------------------------------
------------------------RRSTGGTIVTDDMLQ---SEKDFGEL-----DEKF-
---------------------------------DFAEFV---------------------
------------------------------------------------RTH----GHRID
--HMI------KKCR----WK-S----Q----------P---C-----------G-P---
----------------------------------------------E-------------
------------------------------------------------------------
------------------------------------------------------------
-----------N----------F-------T-----------AVI-T-EFG--------L
CYTFN-------SG-------------MK-G--------------------H----PL--
-------LKVQRA------GV--------DY-------------ALRLQLSVQQD-----
----------Q-----------------Y------------Y-G-S---LR---------
------------------------------------------------------------
------------------------------------------------------------
----------DSSGFKVMVH-DQEEPP--LINELGIAIQPGTHTFCGLRKE------E--
-----MHNL-----------PAP-FK-TA-C--RDM--------Q-------LE----GF

------------------K-KYTKSACLLKCRADYVM----KM--CKCRSYD----L---
------------K-G---------------------------------------------
----------PA-----P---PCQP-----------------------------------
------------------------------------REV---KNCVW---------P---
A-M-------------------EIFR-N-------ES--------------------INC
------------ECP-VPCEITK---YQTQ-LSY------------------AQT-PA--
-------KHF------SEV-L-----------AR--------------------------
------------------------------------------------RK------HI--
------------------------------------------------------------
-N----------------------------------------------------------
------------------------KD-VMR------------------------------
-----H-------YL-----------R------DNFLELDVYFEEMQVTLIQ-QRQAYDQ
ESL-----------------FGDIGGQVGLFLGASILTVLEFLDLLWRI-----------
--------LIHK--------F----------K-------KR-------------------
KN------------------------------RK------------------VR------
------------------------------------------------------------
------------------------------------------------------------
------------------------------------------------------------
------------------------------------------------------------
------------------------------------------------------------
------------------------------------------------------------
------NV
>NeNaC3_tr_A7S2F2_A7S2F2_NEMVEPredictedproteinOS_NematostellavectensisOX_4535
1GN_v1g205746PE_3SV_1
------------------------------------------------------------
----------M-------------------------------------------------
------------------------------------------------------------
------------------------------------------------------------
------------------------------------------------------------
------------------------------------------------------------
------------------------------------------------------------
------------------------------------------------------------
------------------------------------------------------------
------------------------------------------------------------
--------------------LKLPCSEEM-------------------------------
--RRNTDPK----------------------E---------TA-----L-----------
-EQ--INQFL-QETTAHGFGRLG---------A-TA--------------GS---KW---
--RLYWV----MFCLAAYCVFVWQLVGLVN-QYNS----------K---PIKTRT----Q
-L-KH-A----------Q-KLDF-----PVV-----------------------TICNMN
VL------RAS-----RLPPKLR-----------------------------------T-
---------KF-------------------------------DQIIN-------------
---------------------------------NT-------------------------
------------------------------------------------------------
------------------------------------------------------------
-----------------NKTSPRN-------------------SSNSRNAFV--------
------------------------------------------------------------
------------------------------------------------------------
------------------------------------------------------------
-------------------------DP---------------------------------
------------------------------------------------------------
------------------------------------------------------------
------------------------------------------------------------
------------------------------------------QDLSFEE-----------
-------------------------------------------------------TKKFE
---------------------------------ILHAVT---------------------
------------------------AHD----------------DYRELVSA----AHQLE
--DIL------LSCN----FN-G----V----------N---C-----------R-N---

----------------------------------------------S-------------
------------------------------------------------------------
------------------------------------------------------------
-----------KDPTIP---TYW-------T-----------QTW-NDNFG--------N
CYMFN-------SV-------------KT-H-NGEK---------------V----DL--
-------YSSSVP------GE--------SN-------------GLTLQLNLEQN-----
----------E-----------------Y------------L-E-G---IT---------
------------------------------------------------------------
------------------------------------------------------------
----------EVAGMKVTIS-DQGVLP--FPGQQGIRIMPGQSTGIQMTKL------Q--
-----TRRI------------DP-FKNRS-C--ESS--------NE------MSD-KNLF
----FG---YN-------M-TYSIMACKYSCLNARKI----ER--CGCTNYN----T---
------------P-E---------------------------------------------
------MQKRNI-----P---LCNRL----------N-----------------------
------------------------------------STI---IDCLN---------K---
A-Y-------------------DTFE-D------------------------------GSC
----------DRECP-PSCSEVS---FDLT-VSS------------------ARW-PA--
-------RSF------EKT-K-----------LK--------------------------
------------------------------------------------TL-Q--------
----------------------------------TEYG--------I-------------
-N----------------------------------------------------------
--------------------------MTK-------------------------------
-----E-------EM-----------F------ENIAQVHVYYGELDYLLVQ-ETLAYTF
MSL-----------------LSDIGGQMGMWIGISALTCAELIELVCVI-----------
--------LANM--------S----------N-------RS-------------------
KKIVHINS-----------F------------RPEKLGG---------VETIEK------
---------------------------------------GGVKRIEQEVQGGV-------
------------------------------------------------------------
------------------------------------------------------------
------------------------------------------------------------
------------------------------------------------------------
------------------------------------------------------------
---GTIEQ
>NeNaC4_tr_A7SA77_A7SA77_NEMVEPredictedproteinOS_NematostellavectensisOX_4535
1GN_v1g209148PE_3SV_1
------------------------------------------------------------
----------M-------------------------------------------------
------------------------------------------------------------
------------------------------------------------------------
------------------------------------------------------------
------------------------------------------------------------
------------------------------------------------------------
--------------TRLESI-------------TKKDA----------------WWP---
------------------------------------------------------------
---------------------------------------------------------DPK
EAV----------------RNKDNEEHATEPE----------------------------
-QHLSKQAD----------------------S---------AK-----N-----------
-TL--IGNFA-SYTTLHGLHFLF---------Q-PI--------------PR---AR---
--KIIWG----IMLVIATVGLSIQIGNGLV-KVYS----------Y---NIFTAK----T
-F-VR-P----------E-SLPF-----PAI-----------------------TICNQN
ML------RKT-----AILGSA------------------G-----------------Q-
-------------------------------------------RYLD----QL-------
---------------------------------DE-------LRARFIG-----------
------------------------------------------------------------
------------------------------------------------------------
------------------------------------------------------------
------------------------------------------------------------

------------------------------------------------------------
------------------------------------------------------------
------------------------------------------------------------
------------------------------------------------------------
------------------------------------------------------------
------------------------------------------------------------
------------------------------------------------------------
--------------------------------------------EDMNVT-----SERI-
---------------------------------NMDAIV---------------------
------------------------------------------------QKS----GHQLE
--KMV------YECR----FV-R----D----------K---C-----------D-N---
----------------------------------------------K-------------
------------------------------------------------------------
------------------------------------------------------------
-----------S----------I-------K-----------VFT-SEKRG--------L
CYTFN-------SG-------------VN-G--------------------T----PL--
-------LNVSHS------GS--------EY-------------SLSLRLDAEPT-----
----------E-----------------Y------------Y-G-P---YS---------
------------------------------------------------------------
------------------------------------------------------------
------Y---DGTGFKIAVH-DQKVLP--DIEEDAYDISPGFYTKITMKRA------Q--
-----TLFL-----------PSP-YS-SG-C--GSR--------K-------LQ----FS
------------------Q-EYSYKNCIRECHTQLMI----GR--CRCRAIG----MTLQ
GRVLMRNKCSKGP----LT-----------------------------------------
--FFLYPENDTT-----P---YCTT-----------------------------------
------------------------------------REV---QDCIM---------R---
V-H-------------------ERSS-R-----------------------------QRC
------------DCP-VPCEMTT---FTSR-ISL------------------AYF-PS--
-------QHV------WET-F-----------LP--------------------------
------------------------------------------------FLTT--------
---------------------------------------------------------AYD
LN---------------L------------------TGL---------------------
-------------------SPDELDM-KAE------------------------------
-----H-------FV-----------R------KRFAMVKIFYETLRTEMYH-QRPAYEM
SDL-----------------FADVGGNMGLFLGCSMLTIAEFIDLAIML-----------
--------LITK--------F----------W-------RR-------------------
RTVDHA--------------------------QA-------------------R------
------------------------------------------------------------
------------------------------------------------------------
------------------------------------------------------------
------------------------------------------------------------
------------------------------------------------------------
------------------------------------------------------------
-----DQT
>NeNaC5_tr_A7SA83_A7SA83_NEMVEPredictedproteinOS_NematostellavectensisOX_4535
1GN_v1g209152PE_3SV_1
------------------------------------------------------------
----------M-------------------------------------------------
------------------------------------------------------------
------------------------------------------------------------
------------------------------------------------------------
------------------------------------------------------------
------------------------------------------------------------
------------RVHSQRQS-------------EMTTVKVANHE----------------
------------------------------------------------------------
------------------------------------------------------------
-----------------------NPSESV-------------------------------

-------------------------------EV--------PR-----E-----------
-SL--LEAFS-SYTTLHGFHFAL---------S-ST--------------NR---VR---
--QIIWI----LLLILTSVVLIYQLAYSTQ-RVLE----------Y---ASMVQV----E
-T-RN-E----------D-SITF-----PAI-----------------------SICSNN
MM------QKS-----KILGKD------------------A-----------------Q-
-------------------------------------------RYLD----LL-------
---------------------------------DQ-------KKEDQW------------
------------------------------------------------------------
------------------------------------------------------------
-----------------D------------------------------------------
------------------------------------------------------------
------------------------------------------------------------
------------------------------------------------------------
------------------------------------------------------------
------------------------------------------------------------
------------------------------------------------------------
------------------------------------------------------------
------------------------------------------------------------
--------------------------------------------AISQSF-----SPDF-
---------------------------------DIEKAV---------------------
------------------------------------------------HQY----GLNLS
--LAM------KSCH----YG-R----Y---------LK---C-----------N-P---
----------------------------------------------S-------------
------------------------------------------------------------
------------------------------------------------------------
-----------H----------F-------T-----------TFK-EFRYG--------L
CYTFN-------SG-------------KR-E-------------------------SA--
-------FISHDT------GP--------TS-------------GLSITLDAQPE-----
----------E-----------------Y------------Y-S-L---YS---------
------------------------------------------------------------
------------------------------------------------------------
---------STGTGFRVIVH-DQSEFP--WVEKHGWEIPPGFSTNVRLARK------E--
-----ISSL-----------ESP-YN-SN-C--SRD--------T---------------
------------------N-YASQSYCLVQCYSDMVV----KR--CGCHMLG----M---
------------T-E---------------------------------------------
-------ETGHT-----P---WCSP-----------------------------------
------------------------------------QQI---KACVY---------T---
T-S-------------------RRFQ-P-----------------------------NMC
------------SCP-VRCSRVE---FDVQ-LSS------------------LYY-PP--
-------DNF------WET-I-----------SNER------------------------
------------------------------------------------SL----------
---------------------------------------------------------TIY
AN----------------------------------------------------------
-----------------------DTK-KFQ------------------------------
-----E-------WY-----------R------RRVIQLNVFYKELTTEVRK-EKEAYTI
SDL-----------------AGDFGGNMGLFLGCSILTIAEFIDLLVVY-----------
--------LVHR--------H----------K-------KR-------------------
TA------------------------------KI-------------------Q------
------------------------------------------------------------
------------------------------------------------------------
------------------------------------------------------------
------------------------------------------------------------
------------------------------------------------------------
------------------------------------------------------------
--------
>NeNaC6_tr_A7RL38_A7RL38_NEMVEPredictedproteinOS_NematostellavectensisOX_4535
1GN_v1g239048PE_3SV_1

```
------------------------------------------------------------
----------M-------------------------------------------------
------------------------------------------------------------
------------------------------------------------------------
------------------------------------------------------------
------------------------------------------------------------
------------------------------------------------------------
--------------ESFNQN-------------TKTDIKDC-------------------
------------------------------------------------------------
------------------------------------------------------------
------------------------------------------------------------
---KEDEKR----------------------E---------PQ-----P-----------
-TM--LQDFA-GYTTLHGFHFLI---------H-PG--------------SP---FR---
--RFIWL----LMLLGCWVALFYQLYNSVD-RLLA----------H---RIVMSR----G
-V-EQ-P----------D-EIDF-----PAI-----------------------TICNQN
IM------RMS-----KINGTE------------------A-----------------Q-
-------------------------------------------KYLD----EL-------
---------------------------------DD-------VFR---------------
------------------------------------------------------------
------------------------------------------------------------
------------------------------------------------------------
------------------------------------------------------------
------------------------------------------------------------
------------------------------------------------------------
------------------------------------------------------------
------------------------------------------------------------
------------------------------------------------------------
------------------------------------------------------------
------------------------------------------------------------
--------------------------------------------KDIKKE-----NVSY-
---------------------------------DAETFV---------------------
------------------------------------------------NKY----GHDWE
--NMF--ENIPYSCM----FQ-RF---F----------I---C-----------S-A---
----------------------------------------------K-------------
------------------------------------------------------------
------------------------------------------------------------
-----------N----------F-------T-----------SFL-SFTRG--------L
CYTYN-------SG-------------VN-R--------------------S----YV--
-------QRVSEA------GR--------NN-------------RLEFHLEAHPE-----
----------E-----------------Y------------Y-G-P---YS---------
------------------------------------------------------------
------------------------------------------------------------
---------YEGIGFKIAVH-DQSYVP--NMDQEGYDITAGFYTNVRVKRY------K--
-----EKSL-----------PHP-YK-TN-C--GER--------K-------LE----YY
------------------E-RYSRSACILECQAREFV----RK--TKCRIIG----F---
------------P-P--IK-----------------------------------------
-------VIRDV-----P---FCSV-----------------------------------
------------------------------------LKI---ETGLT---------Y---
M-Y-------------------YNWN-T-----------------------------NQC
------------DCP-KPCVEIS---YSAQ-MSL------------------LQY-PT--
-------PSL------VRE-I-----------RK--------------------------
------------------------------------------------SF----------
------------------------------------------------------------
-N----------------------------------------------------------
---------------------------DTE------------------------------
-----D-------YI---NNM-----R------ANSVIVSIFYETLLTDVFE-EKNDYDI
SRF-----------------GSDLGGNLGLFLGCSLLTLVEFFDLGIRW-----------
```

```
--------CLGR-------------------KD------KV-------------------
QR----------------------------------------------------------
------------------------------------------------------------
------------------------------------------------------------
------------------------------------------------------------
------------------------------------------------------------
------------------------------------------------------------
------------------------------------------------------------
-------S
>NeNaC7_tr_A7SH56_A7SH56_NEMVEPredictedproteinOS_NematostellavectensisOX_4535
1GN_v1g212215PE_3SV_1
------------------------------------------------------------
----------M-------------------------------------------------
------------------------------------------------------------
------------------------------------------------------------
------------------------------------------------------------
------------------------------------------------------------
------------------------------------------------------------
------------------------------------------------------------
------------------------------------------------------------
------------------------------------------------------------
-----------------------TSANDDLTND---------------------------
-SRGETNRV----------------------G---------QV-----R-----------
-PL--WRDFL-SRTTLHGTQYAC---------V-TK--------------PL---IR---
--RVTWL----LLLLGMVGYFGYLFYGNLK-RYHS----------H---PVEVTV----E
-I-ETPN----------D-GIGF-----PAV-----------------------SVCTNN
KY------MKS-----KINMLR------------------------------------N-
------------------------------------------HSYFH----KLG------
----L-------------------------------------------------------
------------------------------------------------------------
------------------------------------------------------------
-----------------D------------------------------------------
------------------------------------------------------------
------------------------------------------------------------
------------------------------------------------------------
-------------------------IPECAALQNV-------------------------
------------------------------------------------------------
------------------------------------------------------------
------------------------------------------------------------
------------------------------------------------------------
-------------------------------------------------------SGNMT
CGQALMCAIVGKY-GYII-----NE-RCKWALEKIRKII---------------------
----------------------NDSDY-------------AFDIEKFTLKY----GHDIK
--ALL----TPRFCT----FR-G----K----------P---C-----------N-E---
----------------------------------------------E-------------
------------------------------------------------------------
------------------------------------------------------------
-----------D----------F-------V-----------PVI-T-STS--------L
CWTFN-------SG-------------FR-G-SHG--------------NPA----PR--
-------KQVTFS------GV--------DF-------------GLTVLLNTRVD-----
----------E-----------------N------------T---I---GT---------
------------------------------------------------------------
------------------------------------------------------------
----------SSEGVRAVVH-EPGEY---FSVDHGVNVMPGAHAAILVHAQ------K--
-----TTTL-----------PLP-YK-SN-C--TES--------KPG-----LR------
--------------------LYSMEGCVALCASQELT----RR--CGCRPVG----L---
------------P-Y--VD-----------------------------------------
```

----------AA-----S---VCSF-----------------------------------
------------------------------------KH----ETCAM---------D---
T-F-------------------GSFD-A-----------------------------QRC
------------MCN-NACHRTM---YNAK-VSY------------------ARF-PD--
-------QYI------IRF-I-----------QETT------------------------
------------------------------------------------SY----------
------------------------------------------------------------
-N----------------------------------------------------------
---------------------------YSA------------------------------
-----E-------YF-----------R------RNLVLVQVGMESLSYEHHR-QVPAFPV
ESL-----------------LGAVGGHLGLLLGCSVLTVFEFIDFFIVA-----------
--------LASM--------L----------R-------TN-------------------
VTPNMD--------------------------RE--------------------------
------------------------------------------------------------
------------------------------------------------------------
------------------------------------------------------------
------------------------------------------------------------
------------------------------------------------------------
------------------------------------------------------------
----TKNL
>NeNaC8_same_NeNaC13_tr_A7SJB5_A7SJB5_NEMVEPredictedproteinOS_Nematostellavec
tensisOX_45351GN_v1g213150PE_3SV_1
------------------------------------------------------------
----------M-------------------------------------------------
------------------------------------------------------------
------------------------------------------------------------
------------------------------------------------------------
------------------------------------------------------------
------------------------------------------------------------
------------------------------------------------------------
------------------------------------------------------------
------------------------------------------------------------
------------------------------------------------------------
---ETLKPH----------------------P---------SV-----R-----------
-AI--FKDFS-DRTSCHGIGQIG---------G-SQ--------------SV---MW---
--RVSWL----MIFLAGLGMVLYQGLTLLD-TYLN----------K---PTATAV----D
-I-TY-S----------E-VTNF-----PSV-----------------------TICNMN
MI------KKS-----QLQHFP--------------------------------------
---------QV-------------------------------KRLVD-------------
------------------------------TF-NN-------------------------
------------------------------------------------------------
------------------------------------------------------------
---------MTSSNSSLNSS--A------------------FMDTNREKAEK--------
------------------------------------------------------------
------------------------------------------------------------
------------------------------------------------------------
------------------------------------------------------------
------------------------------------------------------------
------------------------------------------------------------
------------------------------------------------------------
--DRLSLNTKNS------------------------------KNKV--------------
------------------------------------------ILNNVNI------DMQRY
------------------------------IEDKIVQYL---------------------
------------------------SMS----------------DTTKLMKA----GHVFR
--ELV------FRCV----WN-G----F----------V---C-----------N-K---
----------------------------------------------G-------------
------------------------------------------------------------

```
------------------------------------------------------------
-----------D----------F---LKYWR-----------PFW-HWRYG--------N
CYTFN-------QG-------------VD-V-NGTE---------------L----PS--
-------LASSKP------GP--------MY-------------GLTLDLFIDQE-----
----------Q-----------------Y------------I-I-P---LS---------
------------------------------------------------------------
------------------------------------------------------------
----------QEAGVKVLLS-DQRNIP--FPFTHGFTVQPGVSASAGIRQL------V--
-----VKRI------------DP-FSNGS-C--YSG--------NG------LEA-NSIY
----HKY--KG-------M-RYSVQGCMSSCLANNQF----KV--CNCTEGK----F---
------------R-A---------------------------------------------
----------KG-----R---PCMT-----------------------------------
------------------------------------EPE---VKCLN---------N---
I-S-------------------KKYE-N-------GS--------L------------GC
----------SKSCP-QPCTHYS---FRRT-ISQ------------------TQW-S---
-------DSY------EKT-F-----------QR--------------------------
------------------------------------------------LVR---------
----------------------------------KS------------------------
-D-------------RGF------------------ANKM--------------------
---------------------------NDA------------------------------
-----S-------IL-----------R------KNFLRVKLYYEELNRETIT-YSLSYPV
ENL-----------------LGDVGGQLGLWIGVSVITCAEFLKLLVDL-----------
--------AWYL--------A----------S-------KM-------------------
SG------------------------------KT-------------------K------
-----------------------------------------TQVQD--------------
------------------------------------------------------------
------------------------------------------------------------
------------------------------------------------------------
------------------------------------------------------------
------------------------------------------------------------
----LNMQ
>NeNaC9_tr_A7SST7_A7SST7_NEMVEPredictedproteinOS_NematostellavectensisOX_4535
1GN_v1g216915PE_3SV_1
------------------------------------------------------------
----------M-------------------------------------------------
------------------------------------------------------------
------------------------------------------------------------
------------------------------------------------------------
------------------------------------------------------------
------------------------------------------------------------
--------------DPPEER-------------NKEDAR---------------------
------------------------------------------------------------
------------------------------------------------------------
------------------------------------------------------------
--EKEHKEE----------------------N---------ET-----------------
WNM--IREFA-GYTTLHGFHFLV---------D-SS--------------SR---WR---
--RRIWF----LLLLFCCAMFIYQFVISVN-RLRA----------Y---KVVFSR----G
-A-EQ-P----------G-EVDF-----PAV-----------------------TICNKN
ML------RKS-----LIILNTS-----------------A-----------------Q-
-------------------------------------------IYLD----EQ-------
---------------------------------DA-------------------------
------------------------------------------------------------
------------------------------------------------------------
-----------------DFK----------------------------------------
------------------------------------------------------------
------------------------------------------------------------
------------------------------------------------------------
```

------------------------------------------------------------
------------------------------------------------------------
------------------------------------------------------------
------------------------------------------------------------
------------------------------------------------------------
--------------------------------------------RSLKQS-----NISF-
---------------------------------DAEEFV---------------------
------------------------------------------------KKY----GHNIT
--NML---NRAKGCT----FK-ML---Y----------P---C-----------S-G---
----------------------------------------------E-------------
------------------------------------------------------------
------------------------------------------------------------
-----------N----------F-------T-----------SFF-SFTRG--------W
CYTFN-------AG-------------AN---------------------------YI--
-------QRVSVA------GR--------ET-------------RLKLYLDAKSH-----
----------E-----------------Y------------Y-G-P---FS---------
------------------------------------------------------------
------------------------------------------------------------
---------YDGVGFKIAVH-DQNDVP--DMDNGGYDISPGYLTTISVKRF------K--
-----EVSL-----------PPP-FP-TK-C--GSR--------T-------LE----HY
------------------E-RYSTKGCEYECFAKDFV----RK--FKCKTLG----M---
-----------AP----IK-----------------------------------------
-------EIRNA-----S---FCPV-----------------------------------
------------------------------------SVV---SAALM---------D---
F-H-------------------KNWH-------------------L-----------ELC
------------DCP-KPCETVN---YNAQ-LST------------------AHY-PA--
-------PSL------LDE-L-----------SK-------------F------------
------------------------------------------------PL----------
---------------------------------------------------------EGF
VN----------------------------------------------------------
----------------------KSKE-EYV------------------------------
-----K-------FV-----------R------DNIVVVEVFYETLLTDVLK-EERDYDF
NMF-----------------ASDLGGILGLYLGTSLLTIAEFLDLGIRW-----------
--------CLRR-------------------RS------DR-------------------
SQV-----------------------------RS--------------------------
------------------------------------------------------------
------------------------------------------------------------
------------------------------------------------------------
------------------------------------------------------------
------------------------------------------------------------
------------------------------------------------------------
---QAWVK
>NeNaC10_tr_A7S8S3_A7S8S3_NEMVEPredictedproteinOS_NematostellavectensisOX_453
51GN_v1g208535PE_3SV_1
------------------------------------------------------------
----------M-------------------------------------------------
------------------------------------------------------------
------------------------------------------------------------
------------------------------------------------------------
------------------------------------------------------------
------------------------------------------------------------
--------------NLFQTK-------------VKPLSR---------------------
------------------------------------------------------------
---------------------------------------------------------DKA
K-----------------------------------------------------------
-SDSEKEIE----------------------E---------QR-----K-----------
-TL--IKNFS-SYTTLHGFHFLL---------D-SS--------------PM---PR---

--RVLWT----ALVGFGLVVFFIQLVMSYG-KLRA----------R---ESILAK----G
-V-ER-P----------M-NVLY-----PAV-----------------------TICNQN
MM------RKS-----RITGTA------------------A-----------------Q-
-------------------------------------------RYLD----QL-------
---------------------------------DH-------IKASL-------------
------------------------------------------------------------
------------------------------------------------------------
------------------------------------------------------------
------------------------------------------------------------
------------------------------------------------------------
------------------------------------------------------------
------------------------------------------------------------
------------------------------------------------------------
------------------------------------------------------------
------------------------------------------------------------
------------------------------------------------------------
--------------------------------------------SRVNRT-----NERF-
---------------------------------ETEEMV---------------------
------------------------------------------------RLY----GHNIT
--DML------WECN----FM-N----K----------P---C-----------S-H---
----------------------------------------------K-------------
------------------------------------------------------------
------------------------------------------------------------
-----------D----------F-------A-----------MRYTSYSRG--------L
CYTFN-------AG-------------AN-G--------------------S----PI--
-------GQATTS------GT--------RT-------------SLSLRLNAESD-----
----------E-----------------Y------------Y-G-P---FS---------
------------------------------------------------------------
------------------------------------------------------------
---------YDATGFKLAVH-DQNEIP--NMDEDAFDISPGFLTNIRIRRE------K--
-----EINL-----------PSP-YR-SE-C--GSR--------D-------LS----NA
------------------P-KYSMSGCIYECYSKIIA----DK--CKCRVLG----M---
------------A----LN-----------------------------------------
-------GLNVT-----Q---FCTS-----------------------------------
------------------------------------DEI---INCVS---------T---
T-Y-------------------LELN-P-----------------------------AMC
------------DCP-KPCRALH---YKIQ-LSL------------------AYF-PS--
-------DHL------WDS-I-----------FP--------------------------
------------------------------------------------VL-L--------
------------------------------------------------------------
-N------------FTDT------------------TGK---------------------
-------------------SQDEVLL-QIQ------------------------------
-----E-------AL-----------R------KQIAQVQIYYETLLTDVLE-EKPAYGI
SEF-----------------GSDVGGNMGLFLGCSLLTFCEFIDLVFML-----------
--------CLHR--------H----------R-------LR-------------------
KEAKE---------------------------RQ------------------AR------
------------------------------------------------------------
------------------------------------------------------------
------------------------------------------------------------
------------------------------------------------------------
------------------------------------------------------------
------------------------------------------------------------
--AAIQDD
>NeNaC11_tr_A7SJJ5_A7SJJ5_NEMVEPredictedproteinOS_NematostellavectensisOX_453
51GN_v1g213238PE_3SV_1
------------------------------------------------------------
----------M-------------------------------------------------

```
------------------------------------------------------------
------------------------------------------------------------
------------------------------------------------------------
------------------------------------------------------------
------------------------------------------------------------
------------KKDAFETS-------------NQTNFSV--------------------
------------------------------------------------------------
------------------------------------------------------------
------------------------------------------------------------
-FDENEDIR----------------------Q---------KR-----K-----------
-KL--ISEFS-GYTTLHGLHFLI----------D-SG--------------SL---FR---
--KVFWM----ILLLVMFTCFFIQLVESYK-RLKE----------Y---GSNLSK----G
-V-ES-P----------E-EVTF-----PAI-----------------------TICNQN
MM------RKS-----LVMGTD------------------A-----------------Q-
-------------------------------------------KYLD----GQ-------
---------------------------------DI-------MKIKL-------------
------------------------------------------------------------
------------------------------------------------------------
------------------------------------------------------------
------------------------------------------------------------
------------------------------------------------------------
------------------------------------------------------------
------------------------------------------------------------
------------------------------------------------------------
------------------------------------------------------------
------------------------------------------------------------
------------------------------------------------------------
--------------------------------------------GAAQVS-----NESF-
---------------------------------EVDKMV---------------------
------------------------------------------------REK----GHLLE
--SML------FECS----FA-G----T----------T---C-----------T-P---
----------------------------------------------E-------------
------------------------------------------------------------
------------------------------------------------------------
-----------N----------F-------T-----------TSL-SFTRG--------L
CYTFN-------SG-------------TN-N--------------------T----PV--
-------FTARAA------DI--------RM-------------AFSAMLFSQPE-----
----------E-----------------Y------------Y-G-P---FS---------
------------------------------------------------------------
------------------------------------------------------------
---------HRATGFKIAIH-DQSETP--DIDLESYDLSPGFATNIRLIRE------K--
-----AKYL-----------PAP-YS-SN-C--SSK--------RG------IDG-----
------------------G-TYSETGCLTRCYNNLMT----SQ--CQCKILG----H---
------------E-S---------------------------------------------
-------DYKNI-----TG--FCST-----------------------------------
------------------------------------YQL---KACVY---------E---
A-W-------------------MVLR-P-----------------------------QNC
------------DCP-KPCTSLK---YKAQ-IST------------------SYF-PS--
-------ESL------WES-L-----------IP--------------------------
------------------------------------------------FL----------
--G-------------------------------QSS--------------------LPF
VN---------------L------------------TGK---------------------
-------------------TLEQATA-EAQ------------------------------
-----V-------NV-----------R------KSVCMVNVFFETLVTDILE-EKPSYDL
TMF-----------------GADLGGTMGLFLGCSILTICEFIDLVIIL-----------
--------VANG--------W----------R-------KG-------------------
KA------------------------------RV--------------------------
```

------------------------------------------------------------
------------------------------------------------------------
------------------------------------------------------------
------------------------------------------------------------
------------------------------------------------------------
------------------------------------------I---------------DV
KEKPEARP
>Nenac12_XP_032238530_1acid_sensingionchannel2isoformX1_Nematostellavectensis
_
------------------------------------------------------------
----------M-------------------------------------------------
------------------------------------------------------------
------------------------------------------------------------
------------------------------------------------------------
------------------------------------------------------------
------------------------------------------------------------
------------------------------------------------------------
------------------------------------------------------------
------------------------------------------------------------
-------------------SYNCKVEDSSDIN----------------------------
-SNNEYRGG----------------------S---------KT-----R-----------
-QL--IKEFS-GYTTLHGFHFLV---------D-SY--------------SV---TR---
--RVVWT----CFIVISLGFLLYQLVNGIK-NYND----------R---GIIMSR----S
-V-EE-P----------N-EVDF-----PAV-----------------------TICNQN
IM------KKS-----LIIGTD------------------A-----------------Q-
-------------------------------------------RYLD----EM-------
---------------------------------NY-------IKADL-------------
------------------------------------------------------------
------------------------------------------------------------
------------------------------------------------------------
------------------------------------------------------------
------------------------------------------------------------
------------------------------------------------------------
------------------------------------------------------------
------------------------------------------------------------
------------------------------------------------------------
------------------------------------------------------------
------------------------------------------------------------
--------------------------------------------GLVNST-----NERL-
---------------------------------DAEDFV---------------------
------------------------------------------------RKY----GHTLG
--EMM------YGCE----FK-D----R----------R---C-----------T-A---
----------------------------------------------Q-------------
------------------------------------------------------------
------------------------------------------------------------
-----------D----------F-------I-----------VST-SFMRG--------L
CYTFN-------SG-------------RD-N--------------------S----SV--
-------RRIATP------GR--------LE-------------SLILRLNAQPE-----
----------E-----------------Y------------Y-G-A---YS---------
------------------------------------------------------------
------------------------------------------------------------
---------YENVGFVLAVH-DQAEPP--DMELNAYDIPPGFTTNLRIRRF------K--
-----ENSL-----------PEP-YP-TK-C--GSR--------N-------LS----LY
------------------K-RYSRKACMQECYARLII----TH--CGCRLLG----M---
------------P-P--MK-----------------------------------------
-------DVIEA-----P---FCTS-----------------------------------
------------------------------------KEY---LDCQL---------M---

```
M-P-------------------VLLK-P-----------------------------SKC
------------DCP-KRCQHIH---YSVQ-PSL------------------AHY-PS--
-------KSV------IKE-L-----------LP--------------------------
------------------------------------------------SL----------
------------------------------------------------------------
-N---------------M------------------TEV---------------------
------------------NSTTERIN-EVN------------------------------
-----R-------II-----------R------EGHAIIRVFYETLRTEIIK-EKPQYTL
ATL-----------------TSDMGGSMGLFLGCSVLTICEFIDLFIQI-----------
--------CLER---------------------------RK-------------------
--------------------------------RN--------------------------
------------------------------------------------------------
------------------------------------------------------------
------------------------------------------------------------
------------------------------------------------------------
------------------------------------------------------------
------------------------------------------------------------
--EVINQK
>Nenac14_XP_032226792_1degenerinunc_8_Nematostellavectensis_
------------------------------------------------------------
----------M-------------------------------------------------
------------------------------------------------------------
------------------------------------------------------------
------------------------------------------------------------
------------------------------------------------------------
------------------------------------------------------------
------------------------------------------------------------
------------------------------------------------------------
------------------------------------------------------------
------------------------------------------------------------
-----SRSG----------------------P---------SV-----R-----------
-AL--LRDFS-DRTSCHGIGQIN---------G-SH--------------SP---TW---
--RIFWL----LTFLAGLGMVLFQCITLLG-IYLD----------K---PTATSV----D
-V-TY-D----------E-VTNF-----PAV-----------------------TICNLN
MI------KKK-----NLANFT--------------------------------------
---------QS-------------------------------KKIFD-------------
------------------------------DF-EA-------------------------
------------------------------------------------------------
------------------------------------------------------------
---------FVSSNSSMDSS--A------------------FLGSKMETVLK--------
------------------------------------------------------------
------------------------------------------------------------
------------------------------------------------------------
------------------------------------------------------------
------------------------------------------------------------
------------------------------------------------------------
------------------------------------------------------------
--DRLSMDSNDG------------------------------SNSV--------------
------------------------------------------SLDDTSM------DTELY
------------------------------VEDMLIRHM---------------------
------------------------AMV----------------DDKDLIEA----GHEFD
--ELV------FRCV----WN-G----F----------T---C-----------N-K---
----------------------------------------------G-------------
------------------------------------------------------------
------------------------------------------------------------
-----------G----------F---MKFWR-----------RFW-HWRYG--------N
CYIFN-------QG-------------VD-E-NGTL---------------L----AH--
```

-------LTSSKP------GP--------MY-------------GLTLDLFIDQE-----
----------Q-----------------Y------------L-I-P---LS---------
------------------------------------------------------------
------------------------------------------------------------
----------QEAGVKVLLS-DQRNVP--FPFTDGFSVQPGVSASVGIRKL------V--
-----INRI------------DP-FNNGS-C--YSG--------DG------LEK-ENIY
----SKY--KS-------L-KYSVQGCMSSCLANSEF----SI--CNCTEGK----F---
------------R-V---------------------------------------------
----------KG-----R---PCMS-----------------------------------
------------------------------------ESE---VKCLN---------T---
V-N-------------------KMYE-K-------GT--------L------------GC
----------TRKCP-QPCSHFS---FRRT-ISQ------------------SQW-S---
-------ESY------EQT-F-----------QK--------------------------
------------------------------------------------MVM---------
----------------------------------KS------------------------
-D-------------KGF------------------DKKM--------------------
---------------------------RDA------------------------------
-----S-------VL-----------R------QNFLRVKLFYEELNTEAIT-YSRSYTT
ESF-----------------LGDVGGQLGLWIGVSVITCAEFFKLLIDV-----------
--------VWYL--------A----------R-------KV-------------------
HG------------------------------GP-------------------K------
-----------------------------------------KTVRD--------------
------------------------------------------------------------
------------------------------------------------------------
------------------------------------------------------------
------------------------------------------------------------
------------------------------------------------------------
----LNMN
>Nenac15_XP_032229849_1degenerindel_1_Nematostellavectensis_
------------------------------------------------------------
----------M-------------------------------------------------
------------------------------------------------------------
------------------------------------------------------------
------------------------------------------------------------
------------------------------------------------------------
------------------------------------------------------------
------------------------------------------------------------
------------------------------------------------------------
------------------------------------------------------------
------------------------------------------------------------
----ADQPR----------------------P---------SV-----R-----------
-AL--VRDFV-DRTTCHGIGQIN---------G-SQ--------------SP---LW---
--RVFWV----VTFVAGLGMVVHQGVTLFG-TFLD----------R---PTSTTI----D
-M-TY-A----------P-AMDF-----PAV-----------------------TICNLN
AI------RKD-----HLSQFP--------------------------------------
---------DA-------------------------------DVLLK-------------
------------------------------GF-SE-------------------------
------------------------------------------------------------
------------------------------------------------------------
--------A-----STPSVT--V------------------FLGKDVESFIK--------
------------------------------------------------------------
------------------------------------------------------------
------------------------------------------------------------
------------------------------------------------------------
------------------------------------------------------------
------------------------------------------------------------
------------------------------------------------------------

QH------SEAG------------------------------------------------
--------------------------------------------PNVTL------DPELA
------------------------------FKDAMVEIF---------------------
------------------------AQS----------------ELKKLQMA----GHGFE
--ELV------LGCT----WN-N----I----------K---C-----------N-K---
----------------------------------------------G-------------
------------------------------------------------------------
------------------------------------------------------------
-----------D----------F---LKYWR-----------PFW-NFRYG--------N
CYTFN-------QG-------------MS-E-KGVA---------------I----KP--
-------LTLSNT------GP--------NY-------------GLTLDLFINQE-----
----------Q-----------------Y------------I-A-P---YT---------
------------------------------------------------------------
------------------------------------------------------------
----------QEAGVRILLS-DQNQIP--FPDSDGFTVSPSSSSAVGIKKI------F--
-----ITRI------------DP-FNNGS-C--YKV-T------KG------LEE-GSIY
----KSVFSHD-------M-GYSVQGCMNSCLANKQR----EM--CNCTEGR----F---
------------D-M---------------------------------------------
---------MTG-----L---ICQQ-----------------------------------
------------------------------------IEA---WRCLN---------R---
V-N-------------------MMYQ-E-------GG--------L------------KC
----------LEKCP-QPCTQNV---FSRS-MSH------------------AHW--A--
-------EEY------KKA-L-----------SK--------------------------
------------------------------------------------FF-P--------
----------------------------------NNG-----------------------
-N----------------------------------------------------------
----------------------GSSV-DPN------------------------------
-----E-------LI-----------S------HNILRVKIYFEELNMETIT-YKRNYPV
ESF-----------------LGDVGGQLGLWIGVSVITCAEFAKLLIDL-----------
--------VLCV--------A----------K-------KM-------------------
NR------------------------------SD-------------------K------
-----------------------------------------VQSVC--------------
------------------------------------------------------------
------------------------------------------------------------
------------------------------------------------------------
------------------------------------------------------------
------------------------------------------------------------
---IGRGN
>Nenac16_XP_032229236_1degenerinmec_10isoformX1_Nematostellavectensis_
------------------------------------------------------------
----------M-------------------------------------------------
------------------------------------------------------------
------------------------------------------------------------
------------------------------------------------------------
------------------------------------------------------------
------------------------------------------------------------
------------------------------------------------------------
------------------------------------------------------------
------------------------------------------------------------
-----------------------KAMAKSGNYTINRNVR--------------------L
NEEQEAKKKNVL-------------------K---------ES-----K-----------
-EA--IKEFS-FETTAHGFARAA---------S-SY--------------SR---LA---
--RSLWI----LLLLVAFSFCFYHNSVSIG-KYFN----------F---PAKTDV----R
-F-SD-AW--------F--SIKY-----PAI-----------------------TLCNIN
NL------KKS-----SFFVKT------------------VV----------------K-
---------EL----------------------------NKTRDAFG-------------
---------------------------------DT-------------------------

------------------------------------------------------------
------------------------------------------------------------
------------------------------------------------------------
------------------------------------------------------------
------------------------------------------------------------
------------------------------------------------------------
------------------------------------------------------------
------------------------------------------------------------
------------------------------------------------------------
------------------------------------------------------------
------------------------------------------SNKR--------------
-------------------------------------------------------RENLE
---------------------------------RVLYFM---------------------
------------------------SGY----------------KEEQVKDF----GHQAK
--DMI------IACNAPGRWA-Q----Y----------P---C-----------D-Y---
----------------------------------------------R-------------
------------------------------------------------------------
------------------------------------------------------------
-----------N----------F-------T-----------WFY-DQIHG--------N
CYTFN-------AA-------------RN-Q--------------------S----EP--
-------VYIEDA------GH--------RH-------------GLHLTLFVEAE-----
----------E-----------------Y------------W-S-Y---LS---------
------------------------------------------------------------
------------------------------------------------------------
----------DSLGFVISIH-DREQLP--FPAETGYFISPGMATSVALQSK------V--
-----LKRL-----------PWP-YDHKE-C--ETR--------RG------IPG-----
-------------VTPSNT-IYSEEGCRKSCVARMML----DK--CKCVSLL----YR--
----------LKP-----------------------------------------------
---------------------SCNPF----------N-----------------------
------------------------------------KTQ---GSCIE---------D---
V-L-------------------RSRS-K-------ES--------M-----------SSC
------------NCP-PACREDV---FTTL-MSS------------------SVW-PT--
-------RSY------MSN-V-----------LN--------------------------
------------------------------------------------TL--P-------
------------------------------------------------------------
QG-------------SQA------------------QKV------------V--------
---------------------------TNL------------------------------
-----S-------SA-----------R------ENMAHVMVYFPTLMQESIV-QKPAYQI
ESL-----------------LGDIGGQMGLFIGISVLTLVEFIALLWDL-----------
--------IAIV--------I----------L-------KN-------------------
KK------------------------------NT--------------------------
------------------------------------------------------------
------------------------------------------------------------
------------------------------------------------------------
------------------------------------------------------------
-------------------------------SAP--------------------------
------------------------------------------------------------
---DGEKP
>Nenac17_XP_032233690_1acid_sensingionchannel3_likeisoformX1_Nematostellavect
ensis_
------------------------------------------------------------
----------M-------------------------------------------------
------------------------------------------------------------
------------------------------------------------------------
------------------------------------------------------------
------------------------------------------------------------
-----------------------------------------P------------------

------------------------------------------------------------
------------------------------------------------------------
------------------------------------------------------------
------------------------------------------------------------
-------------------------------R---------SP-----K-----------
-EV--LKDFG-ETTTAHGIGKLV---------G-SK--------------HP---IR---
--KTVWL----VLLCVCFGYVIYQIKELCG-QYKE----------H---QITVMS----H
-A-ER-K----------K-EIPF-----PGV-----------------------TVCNLN
PV------RKS-----KIIEHN------------------L-------------------
------------------------------------------TELLK-------------
------------------------------------------------------------
------------------------------------------------------------
----------------------P-------------------------------------
-----------------S------------------------------------------
------------------------------------------------------------
------------------------------------------------------------
------------------------------------------------------------
-------------------------GPNA-------------------------------
------------------------------------------------------------
------------------------------------------------------------
------------------------------------------------------------
------------------------------------------------------------
-------------------------------------------------------IGQVE
------------------------------MRNKLMSMV---------------------
------------------------MGM----------------NATTRTEL----GHQEK
--DFI------FGCY----WL-G----Q----------P---C-----------N-----
----------------------------------------------T-------------
------------------------------------------------------------
------------------------------------------------------------
-----------S----------F-------E-----------QIF-TPEYG--------N
CFTFN-------SG-------------KL-H-D---------------------------
-------LAVTEA------GD--------TE-------------GLSLLLNTEQW-----
----------D-----------------Y------------M-G-A---VS---------
------------------------------------------------------------
------------------------------------------------------------
----------PSIGAIVDVH-HPDDIE--VPAVRGITVSPGQATSIGLVLG------S--
-----MYRL-----------DSP-YSKTG-C-------------KS------IDT-PTIY
----KNK-------------IYSAEACYRGCLIAKSL----EK--CGCVESR----M---
------------Q-V---------------------------------------------
--------GLTA-----Q---TCDPF----------N-----------------------
------------------------------------KTL---RKCIE---------E---
V-V------------------------D-----------------F-----------NSC
------------SCP-PACREMK---FEFS-AST------------------SVW-PS--
-------EPY------WTF-L-----------KE--------------------------
------------------------------------------------AY----------
---------------------------------DSR------------------------
-N----------------------------------------------------------
---------------------------TSR------------------------------
-----S-------FASFPD------SR------GNLLAVNIFFKDMSITIKT-ETFSYEF
SNL-----------------VADVGGQLGLFVGMSVLSIMEYWELLIDL-----------
--------GTAL--------L----------R-------RD-------------------
KTM-----------------------------RV--------------------------
-----------------------------------------AVASKPE------------
------------------------------------------------PSTA--------
------------------------------------------------------------
------------------------------------------------------------
------------------------------------------------------------

----------------------------------------------------LHRKDKVC
CRGLKDRT
>Nenac18_XP_032233718_1degenerindeg_1_Nematostellavectensis_
------------------------------------------------------------
----------M-------------------------------------------------
------------------------------------------------------------
------------------------------------------------------------
------------------------------------------------------------
------------------------------------------------------------
------------------------------------------------------------
------------------------------------------------------------
------------------------------------------------------------
------------------------------------------------------------
------------------------------------------------------------
-SQNEEKCR----------------------Q---------TV-----S-----------
-GL--IWDFF-GQTTAHGFGKIA---------N-AK--------------SR---LR---
--RMFWV----CLVMAAFGMFSLQIYDLLM-IYLS----------R---PTGTQI----W
-M-KH-A----------P-SIAF-----PAV-----------------------TLCNFN
IV------RRS-----QITSEL--------A-------GSF-----------------EN
FI-----TPNI-------------------------SDVDDARSLRK----RRQSVD---
----------GCSCSCY---------LKCRNFHDPHR-----------------------
------------------------------------------------------------
------------------------------------------------------------
-----------------NTPEPATTCAP-------VPTTIPTTATTTEPTTVPT------
------------------------------------------------------------
------------------------------------------------------------
-------------------------------------------------------TIPT-
----------------TIPTTLPTTEPTTAPTTPEPTT----------------------
------------------------------------------------------------
--------LEPTDPPTTEAPTTLP------------------------------------
-------------------------------------------------------PGVTP
APTEKPLTIEEK------------------K-----------EQAFTGE-----------
----------------------------------------VSQPDPATL------DKQTL
------------------------------KTEELVLSL---------------------
------------------------ATK----------------PEDMLMEV----GHQFN
--DMV------LSCT----YR-G----I----------P---C-----------T-NYT-
----------------------------------------------E-------------
------------------------------------------------------------
------------------------------------------------------------
-----------N---------LW-------S-----------RFW-HYRYG--------N
CYVFN-------SG-------------KD-W-QGEP---------------K----PI--
-------LRSNKP------GP--------AS-------------GLTLELNAEQH-----
----------E-----------------Y------------V-G-Q---LS---------
------------------------------------------------------------
------------------------------------------------------------
----------HEAGMRVLIS-TQGEMP--SPLEKGISVSPGFSTATGIRLT------N--
-----ILRA------------DP-FNNKS-C--LSS--------DD------IDP-ENLY
----RKR--YN-------V-SYSRTTCMDSCLAHKQV----DM--CGCLEYR----F---
------------P-N---------------------------------------------
--------IDNR-----T---VCDIL----------D-----------------------
------------------------------------IDV---IRCLN---------K---
V-Q-------------------SLYQ-D-------NK--------L------------GC
----------SEKCR-PPCNEEV---FKLS-ISS------------------ARW-PT--
-------DDY------EAH-F-----------LD--------------------------
------------------------------------------------EL-K--------
----------------------------------EE------------------------
-K-------------GFVL-----------------------------------------

---------------------------PAN------------------------------
-----A-------DA-----------R------AYFVKLQIFYEELNHEVIE-EYRSYEL
VNF-----------------VSDIGGQLGLWIGISALTASEFIELLIVI-----------
--------ALHL--------L----------R-------KV-------------------
TRIGLIDNSAEQPADTNPCYLDDITLTGDLKERR--------------------------
---------------------------------------GALTESNTSLTV---------
-----------------RYLAEVARKRNKKGH----------------KEKREAS-----
------------------------------------------------------------
------------------------------------------WHIDEEAFPCYR------
---------------------------DQATIHPNVESNTSLTVRYLDDIVQKKSKHRKR
SKGDTSMVQKGDGHRGDLSGRRG-----------ELTE-----------SSGSLTVRYLA
EVASKRKR
>Nenac19_XP_001623796_2amiloride_sensitivesodiumchannelsubunitalpha_Nematostellavectensis_
------------------------------------------------------------
----------M-------------------------------------------------
------------------------------------------------------------
------------------------------------------------------------
------------------------------------------------------------
------------------------------------------------------------
------------------------------------------------------------
------------------------------------------------------------
------------------------------------------------------------
------------------------------------------------------------
-----------------------DSHSNE-----------------------------TK
KGEKDQTRC----------------------Q---------SF-----T-----------
-AL--LREFC-GYTSFGGLGRVT---------D-SP--------------YL---LF---
--RITWF----FLFMGAVAYASTQLYSLYE-KYQD----------R---PVNTVI----S
-L-KH-S----------E-KLSF-----PSV-----------------------AVCNFN
IL------RQS-----VMQDTL------------------------------------S-
---------NV-------------------------------LNMVK-------------
---------------------------------EQ-------------------------
------------------------------------------------------------
------------------------------------------------------------
------------------------------------------------------------
------------------------------------------------------------
------------------------------------------------------------
------------------------------------------------------------
-------------------------TPRCVRLR---------------------------
------------------------------------------------------------
------------------------------------------------------------
--------------------------------------------------------NCST
TSHNVSTTSSPRP-----------------------------EDSDPEV-----------
---------------------TKELEEMGFENPEQL-------------------SDEYR
------------------------------LEELVSITI---------------------
------------------------AKL----------------SPSDLKPM----GHQFD
--DMV------YKCT----WR-G----V----------D---C-----------K-A---
------------------------------------------------------------
------------------------------------------------------------
------------------------------------------------------------
-----------N----------YT-QGKYWT-----------SFW-HYRFG--------N
CFVFN-------PS-------------IG-W-NRNT---------------R----SR--
-------LHIYRT------GP--------LN-------------GLQLQLNIQQE-----
----------E-----------------Y------------I-D-Q---IT---------
------------------------------------------------------------
------------------------------------------------------------
----------KRAGAMIQVF-PFGQMP--FPEERGTLVAPGFSTDIGLEKM------D--

-----IKRV------------DP-HNNGS-C--VTD--------EAS-----INT-ENIF
----KQK--FN-------T-SYSKTACVLSCLAYHQM----HE--CKCMEYR----F---
------------P-T--DS-----------------------------------------
---------------------VCDTL----------N-----------------------
------------------------------------STI---VECLA---------K---
V-K-------------------FKLS-R-------NE--------L------------GC
----------STACG-PACREEA---YRLT-SSF------------------AVW-PS--
-------QRY------QNS-I-----------ID--------------------------
------------------------------------------------NLFD--------
------------------------------------G----------------------I
FN----------------------------------------------------------
---------------------------YSR------------------------------
-----S-------WF-----------R------DNYLEVNIFFKELNY-FIE-ESVAYEL
VNF-----------------LSDIGGQLGLWVGVSVMTGAEVVELALLI-----------
--------FFKV--------F----------C-------RK-------------------
PR------------------------------KV--------------------------
----------------------------------------TVNSEGDPEP----------
---------------------------------------------NVLSNSQ--------
------------------------------------------------------------
----------------L-------------------------------------------
------------------------------------------------------------
--------------------------------------------------------QLES
IDNYKYAP
>Nenac20_XP_032233720_1amiloride_sensitivesodiumchannelsubunitbeta_Nematostel
lavectensis_
------------------------------------------------------------
------------------------------------------------------------
------------------------------------------------------------
------------------------------------------------------------
------------------------------------------------------------
------------------------------------------------------------
------------------------------------------------------------
------------------------------------------------------------
------------------------------------------------------------
------------------------------------------------------------
------------------------------------------------------------
------------------------------------------------------------
------------------------------------------------------------
------------------------------------------------------------
------------------------------------------------------------
------------------------------------------------------------
------------------------------------------------------------
------------------------------------------------------------
------------------------------------------------------------
------------------------------------------------------------
------------------------------------------------------------
------------------------------------------------------------
------------------------------------------------------------
------------------------------------------------------------
------------------------------------------------------------
------------------------------------------------------------
------------------------------------------------------------
------------------------------------------------------------
--------MEEK------------------K-----------EQAFTGE-----------
----------------------------------------VSQPDPATL------DKQTL
------------------------------KTEELVLSL---------------------
------------------------ATK----------------PEDMLMEA----GHQFY

--DMV------LSCT----YR-G----I----------P---C-----------T-NYT-
----------------------------------------------E-------------
------------------------------------------------------------
------------------------------------------------------------
-----------N---------LW-------S-----------RFW-HYRYG--------N
CYVFN-------SG-------------KD-W-QGEP---------------K----PI--
-------LRSNKP------GP--------AS-------------GLTLELNAEQH-----
----------E-----------------Y------------V-G-Q---LS---------
------------------------------------------------------------
------------------------------------------------------------
----------HEAGMRVLIS-TQGEMP--SPLEKGISVSPGFSTTTGIRMT------N--
-----ILRA------------DP-FNNKS-C--LSS--------DD------IDP-ENLY
----RKR--YN-------V-SYSRTTCMDSCLAHKQV----DM--CGCLEYR----F---
------------P-N---------------------------------------------
--------IDNR-----T---VCDIL----------D-----------------------
------------------------------------IDV---IRCLN---------K---
V-Q-------------------SLYQ-D-------NK--------L------------GC
----------SEKCR-PPCNEEV---FKLS-ISS------------------ARW-PT--
-------DDY------EAH-F-----------LD--------------------------
------------------------------------------------EL-K--------
----------------------------------ED------------------------
-K-------------GFVL-----------------------------------------
---------------------------PAN------------------------------
-----A-------DA-----------R------AYFVKLQIFYEELNHEVIE-EYRSYEL
VNF-----------------VSDIGGQLGLWIGISALTASELVELLIVI-----------
--------ALHL--------L----------R-------KV-------------------
TRTGLIDNSAEQPADTNPRYLDDITLTGDLGERRGALTDSNTSLTVRYLGEIARKRKMKG
HKEKRETSWHIDNSADQPADTNPHYLDNITFTGDLGERRGALTDSNTSLTV---------
-----------------RYLAEVARKRNKKGH----------------KEKREAS-----
------------------------------------------------------------
------------------------------------------WHIDEEAFPCYR------
---------------------------DQATIHPNVESNTSLTVRYLDDIVQKKSKHRKR
SKGDTSMVQKGDGHRGDLSGRRG-----------ELTE-----------SSGSLTVRYLA
EVASKRKR
>Nenac21_XP_032233957_1degenerindeg_1isoformX1_Nematostellavectensis_
------------------------------------------------------------
----------M-------------------------------------------------
------------------------------------------------------------
------------------------------------------------------------
------------------------------------------------------------
------------------------------------------------------------
------------------------------------------------------------
------------------------------------------------------------
------------------------------------------------------------
------------------------------------------------------------
------------------------------------------------------------
-----GKEK----------------------K---------KI-----V-----------
-TK--IKDFL-GYTTAHGFARLV---------E-SK--------------NP---VQ---
--RAFWV----LACLGAFVAFGLQVHKLFS-KFNR----------K---PVDTYI----T
-V-KH-E----------M-NLEF-----PAV-----------------------TICNLN
PI------RKK-----HLPESLR-----------------------------------EY
FKY------SL--------GDSSTKEPTS---------KPTTSSLLG----K-AFSSV--
---------------------------------DS-------------------------
------------------------------------------------------------
------------------------------------------------------------
------------------------------------------------------------
------------------------------------------------------------

```
------------------------------------------------------------
------------------------------------------------------------
----------------------LLTKPALLSN----------------------------
------------------------------------------------------------
------------------------------------------------------------
------------------------------------------------------------
--KTLPSLVNSSLD----------------------TLGGALNDSLSSVNK---------
-----------------------------------------LKDSGESL-----VNSTLG
LVRGLLSRRRRGLLGDVLDLPKEKDGGKPGLLGNLLDGVLPEKEKDGGKRKDGERPAPEE
LC---------ESARRRKYLYWRIAET----------------PTETLIQS----GHQFL
--DMV------QECS----FA-G----V----------S---C-----------L-G---
----------------------------------------------Y-------------
------------------------------------------------------------
------------------------------------------------------------
-----------N----------W-------T-----------QFW-NHLYG--------N
CFVFN-------TA-------------MI-G-SNRE-------------RNN----IS--
-------YTSSIP------GP--------SD-------------GLLLRLRIEGE-----
----------E-----------------Y------------L-G-N---FS---------
------------------------------------------------------------
------------------------------------------------------------
----------TVSGVRVHIS-DRHEMP--FPGDKGFSVSPGFETSVGMRKV------F--
-----IHRE------------DP-FGNNS-C--ISDVG------KP------VVR-PDYY
----MEN--FG-------V-NYSTIACRSSCVANLQK----TM--YGCVESA----Y---
------------A-I--TD-----------------------------------------
----------EL-----RK-IVCNAL----------N-----------------------
------------------------------------TTV---FRCIV---------K---
A-N-------------------ETAE-D-------SD-------------------GKSC
----------LTKCA-IRCDENI---YKLT-ISS------------------AVF-PS--
-------PSY------QNI-S-----------SQ--------------------------
------------------------------------------------DL-K--------
----------------------------------SD------------------------
-S----------------------------------------------------------
---------------------------GNE------------------------------
-----E-------QI-----------R------ANLLSLRIFFEEHNYETIE-QRKSYEL
IDF-----------------LSDIGGQMGMFVGLSVLTCAELIELLALI-----------
--------IMHL--------F----------N-------RD-------------------
HK------------------------------RG-------------------A------
-----------------------------------------VGADTVT------------
------------------PVIELKKKPLVR------------------------------
------------------------------------------------------------
------------------------------------------------------------
------------------------------------------------------------
------------------------------------------------------------
---LPRVE
>Nenac22_XP_001636193_2acid_sensingionchannel4_B_Nematostellavectensis_
------------------------------------------------------------
----------M-------------------------------------------------
------------------------------------------------------------
------------------------------------------------------------
------------------------------------------------------------
------------------------------------------------------------
------------------------------------------------------------
------------------------------------------------------------
------------------------------------------------------------
------------------------------------------------------------
-----------------------KQSANI-------------------------------
------EQD----------------------E---------LA-----T-----------
```

```
-VV--LEDFK-SNTTLHGLPHAL---------N-ST--------------RN---WR---
--KALWV----CLTLGSAAALVTQLTENWE-TLFS----------Y---EIVKFS----T
-P-KL-H----------E-NLTF-----PAV-----------------------TICNEN
SL------RKS-----KVINTS------------------L-----------------Q-
---------EV------------Y-EYL-------------RNKTIG----VT-------
---------------------------------DE-------VLYY--------------
------------------------------------------------------------
------------------------------------------------------------
------------------------------------------------------------
------------------------------------------------------------
------------------------------------------------------------
------------------------------------------------------------
--------------------------PL--------------------------------
------------------------------------------------------------
------------------------------------------------------------
------------------------------------------------------------
------------------------------------------------------------
------------------------------------------------------------
---------------------------------GMTKMF---------------------
------------------------------------------------GED----GHDMR
--DML------LTAS----WN-G----H----------L---V-----------T-S---
----------------------------------------------Q-------------
------------------------------------------------------------
------------------------------------------------------------
-----------D----------F-------Q-----------PYF-YSKYG--------K
CFVFN-------GS-------------QN-K--------------------T---TPA--
-------RTTTLK------GM--------GS-------------GFEALLDIQPE-----
----------E-----------------Y------------M-DLS---PN---------
------------------------------------------------------------
------------------------------------------------------------
-------EASSAIGLVVNIH-DQHQDP--AIIDTGVSVAPGSRVDFAIKLT------Q--
-----QKRL-----------PFP-YP-SK-C--EKV--------EKA----RLPL-----
----TD------------R-VYSVSRCEEECRAKEME----DK--CRCRFLD----L---
------------A-N--DK-----------------------------------------
----------DV-----P---YCTP-----------------------------------
------------------------------------RNT----SCIM---------A---
V-S-------------------NTHK--------------------------------KC
------------SCP-TPCEENI---YEVR-TSA------------------AYH-PT--
-------DYF------LKQ-A-----------AR--------------------------
------------------------------------------------LY----------
------------------------------------------------------------
YN---------------V------------------TNL---------------------
---------------------------TTEASIR--------------------------
--DFEQ-------FY-----------R------KRAVYIRVYFESLTSDLIE-EKATYEL
PSF-----------------LSDIGGSVGLFLGCSFLTACEFLELFAVT-----------
--------VASL--------F----------M-------KF-------------------
FR------------------------------RN-------------------N------
-----------------------------------------NNVRD--------------
------------------------------------------------------------
------------------------------------------------------------
------------------------------------------------------------
------------------------------------------------------------
------------------------------------------------------------
-----FRA
>Nenac23_XP_001625049_2acid_sensingionchannel3_Nematostellavectensis_
------------------------------------------------------------
----------M-------------------------------------------------
```

```
------------------------------------------------------------
------------------------------------------------------------
------------------------------------------------------------
------------------------------------------------------------
--------------------------------------SNAPDYRNF-------------
-------------IEALHQTL------------NNPNNRVAVDNRGYAL-----ESPL--
--DFYKNNW------------------------------------------VGD------
------------------------------------------G--------------ESV
DQ---------------------KKAQEV-------------------------------
-DPWNKDEK----------------------L---------TI-----S-----------
-QH--FASLC-SSTTMHGISNVF---------D-PS-------------SAT---LR---
--KGIWS----VVFLCSFIFCAYEIGNNVR-YYLT----------K---PVNTVF----K
-I-DY-V----------D-EIKF-----PAV-----------------------TICNNN
PI------RKS-----WAATTP------------------YL------------------
---------PV-------------------------------IMAYN-------------
------------------------------------------------------------
------------------------------------------------------------
------------------------------------------------------------
------------------------------------------------------------
------------------------------------------------------------
------------------------------------------------------------
------------------------------------------------------------
------------------------ANPGEE------------------------------
------------------------------------------------------------
------------------------------------------------------------
------------------------------------------------------------
------------------------------------------------------------
------------------------AIPINWDAYN---------------------WTGF-
---------------------------------GFDKLM---------------------
------------------------------------------------SSA----AHLAS
--EMI------HTCK----WK-G----I----------Y---C-----------S-A---
----------------------------------------------A-------------
------------------------------------------------------------
------------------------------------------------------------
-----------N----------F-------T-----------LDA-T-FLG--------G
CYTFN-------ID-------------QR-------------------------------
-------LMVTGT------GM--------AN-------------ALHLVLNIQQN-----
----------E-----------------Y------------I-G-N---VR---------
------------------------------------------------------------
------------------------------------------------------------
----------SGAGFRLLFR-EKHEPP--STDRFVIALQPGTQTLIPLTMK------K--
-----LISL-----------PEP-F--GV-C--QEK-N------N-------LK----MF
------------------D-KYSVTACEFECRARLGG----KL--CGCREMY--------
------------PSS---------------------------------------------
-------IKTEI-----P---VCLP-----------------------------------
------------------------------------KAY---RDCLN---------P---
L-L-------------------VEIS-V-------NN---------------------LC
R-----------GCK-NPCNKVT---FVPR-MSY------------------SQY-PA--
-------NHI------ADS-M-----------AM--------------------------
------------------------------------------------SM----------
------------------------------------------------------------
-N----------------------------------------------------------
---------------------------TTR------------------------------
-----D-------FV-----------R------DNFLEVEIYFEDIMVEIIE-QQEAFSL
TSL-----------------VGIIGGTLGVFIGASIITVSEFMEFLILI-----------
--------PFRR------------------------------------------------
------------------------------------------------------------
```

------------------------------------------------------------
------------------------------------------------------------
------------------------------------------------------------
------------------------------------------------------------
------------------------------------------------------------
------------------------------------------------------------
--------
>Nenac24_XP_032222096_1acid_sensingionchannel1B_Nematostellavectensis_
------------------------------------------------------------
----------M---------------------------------ALC-----GCC-----
------------------------------------------------------------
------------------------------------------------------------
------------------------------------------------------ASKPSP
TNSASEDESTG-------------------------------------------------
-----------------------------------------PE-----------------
----SKEEILLANQNGNAST-------------YENIRKRSA------------------
------------------------------------------------------------
---------------------------------------------------------AKR
D----------------------KKHAEY-------------------------------
-------GK----------------------R---------KM-----R-----------
-EV--FAYYI-NHCTLHGFHYIF---------E-TK--------------SL---FR---
--KIAWF----SVLAIAGGFFFKEIKTSTT-QYFK----------Y---PFSIMS----T
-V-EY-P----------R-TLVF-----PAV-----------------------TICDFH
DI------RQT-----LVSNAK-----------------------------------GE-
-------------------------------------------QAIN-------------
------------------------------------------------------------
------------------------------------------------------------
------------------------------------------------------------
------------------------------------------------------------
------------------------------------------------------------
------------------------------------------------------------
------------------------------------------------------------
------------------------------------------------------------
------------------------------------------------------------
------------------------------------------------------------
------------------------------------------------------------
------------------------------------------------------------
-------------------------------------------------------ISSSE
---------------------------------KLETLA---------------------
------------------------------------------------RKT----YKSFN
--ETL------ISCS----LRRGVRGAR----------P---F-----------N-I---
----------------------------------------------H-------------
------------------------------------------------------------
------------------------------------------------------------
-----------D----------F-------K-----------VFF-T-AKGQ-------T
CYTFN-------AA-------------MD-G--------------------K----KL--
-------LEVDNV------GP--------KF-------------GLEIYLNAQHW-----
----------W-----------------F------------K-D-D--------------
------------------------------------------------------------
------------------------------------------------------------
---------VRESGFRFILH-QQDDLP---LTREGFRVSSGYVTYVDMRLE------K--
-----VKNL-----------PYP-FS-SD-C--DGR--------G-------LD----LY
------------------P-KYSRNNCYMESLTKYIL----QQ--CKCRAWF----M---
------------A-DVI-------------------------------------------
----------NT-----S---TCSI-----------------------------------
------------------------------------KEA---LDCMW---------P---
A-W-------------------EDFD-N-------AY---------------------NY

```
------------TCP-VDCEERV---YKTR-LSS------------------ALFLPQ--
-------KLL--P-------L------T----KK-------------Y------------
------------------------------------------------KFLMRP------
---------------------------------------------------------KGI
PN----------------------------------------------------------
-----------------------DTE-GAV------------------------------
-----D-------FI-----------L------ENYSVINLFFDELRLDTIQ-QTPAYGF
FRL-----------------VGDVGGQLGLVLGASVITIVEIIDLVIMY-----------
--------SIYW--------I----------K-------SK-------------------
TV-----------------------------PRA--------------------------
------------------------------------------------------------
------------------------------------------------------------
------------------------------------------------------------
------------------------------------------------------------
------------------------------------------------------------
------------------------------------------------------------
-----ATS
>Nenac25_XP_032221585_1acid_sensingionchannel2_Nematostellavectensis_
------------------------------------------------------------
----------M-------------------------------------------------
------------------------------------------------------------
------------------------------------------------------------
------------------------------------------------------------
------------------------------------------------------------
------------------------------------------------------------
------------------------------------------------------------
------------------------------------------------------------
------------------------------------------------------------
------------------------------------------------------------
-------------------I-----------S---------AM-----K-----------
-ET--WREFA-SNTTLHGLRYAV---------G-VC--------------DI---RR---
--RSLWI----LCLLACAASYLYMVIISFG-TYID----------R---PIRTEV----S
-H-EFPK----------D-GLDF-----PVV-----------------------TICNLN
YF------VKS-----KIDTGY------------------------------------ED
------------------------------------------EAFYT----R--------
---NL----------------------------NMSV-----------------------
------------------------------------------------------------
------------------------------------------------------------
-C---------------DIIRGV-------------------------------------
------------------------------------------------------------
------------------------------------------------------------
------------------------------------------------------------
------------------------------------------------------------
------------------------------------------------------------
------------------------------------------------------------
------------------------------------------------------------
------------------------------------------------------------
-------------------------------------------------------SGNLT
CGQALLCAY-ETF-GSSVV--D---GCDDVIRSRVIAAV---------------------
----------------------NSSNRP-------------FDTEEFMTRY----GHDFA
--PML--M---GYCL----FSVN----E----------K---C-----------D-I---
----------------------------------------------N-------------
------------------------------------------------------------
------------------------------------------------------------
-----------D----------W-------S-----------PHI-T-PGG--------M
CYTFS-------KA-------------NQ-------------------------------
-------SKVYFL------GG--------EG-------------GLSIILDAQIS-----
```

----------E-----------------H------------T-------FG----D----
------------------------------------------------------------
------------------------------------------------------------
----------FSTGFRVILS-ARGTY---INRARGFNVFPGSHALVAVTPK------K--
-----FERL-----------PAP-YR-TN-C--SDK-F--------------LPG-----
-----------------YG-KYTKDACYTQCINNATM----TD--CGCRLPS----Q---
------------------------------------------------------------
-----HAKFNNL-----P---KCSIQ----------D-----------------------
------------------------------------------QKCRA---------A---
A-S-------------------ARVR-S-----------------------------TLC
------------DCT-VPCKEQI---YEPR-ISY------------------SKF-PD--
-------ITI------TKI-L-----------TN--------------------------
------------------------------------------------HF----------
-----------------------------------------------------------K
LN----------------------------------------------------------
---------------------------KSA------------------------------
-----S-------YL-----------R------DSLVFLQIGFEELAYLVDR-QAPSYGP
GNL-----------------FGDLGGNMGLMLGCSILTLIEFIDFLWIA-----------
--------LKSS--------L----------N-------KS-------------------
KRTAI---------------------------ND---------------CGFSQGTG---
------------------------------------------------------------
------------------------------------------------TSNQDA------
------------------------------------------------------------
------------------------------------------------------------
------------------------------------------------------------
---------------------------------------------------------NNV
GLDMHVKT
>NeNaC26_XP_032233942_1amiloride_sensitivesodiumchannelsubunitgammaisoformX1_
Nematostellavectensis_
------------------------------------------------------------
------------------------------------------------------------
------------------------------------------------------------
------------------------------------------------------------
------------------------------------------------------------
------------------------------------------------------------
------------------------------------------------------------
------------------------------------------------------------
------------------------------------------------------------
------------------------------------------------------------
------------------------------------------------------------
------------------------------------------------------------
------------------------------------------------------------
------------------------------------------------------------
------------------------------------------------------------
------------------------------------------------------------
------------------------------------------------------------
------------------------------------------------------------
------------------------------------------------------------
------------------------------------------------------------
------------------------------------------------------------
------------------------------------------------------------
------------------------------------------------------------
------------------------------------------------------------
------------------------------------------------------------
------------------------------------------------------------
------------------------------------------------------------
------------------------------------------------------------

------------------------------------------------------------
------------------------------------------------------------
------------------------------------------------------------
------------------------------------------------------------
------------------------------------------------------------
------------------------------------------------------------
------------------------------------------------------------
------------------------------------------------------------
------------------------------------------------------------
------------------------------------------------------------
------------------------------------------------------------
------------------------------------------------------------
------------------------------------------------------------
------------------------------------------------------------
--------------MRVLIS-TQGEMP--SPLEKGISVSPGFSTTTGIRMT------N--
-----ILRA------------DP-FNNKS-C--LSS--------DD------IDP-ENLY
----RKR--YN-------V-SYSRTTCMDSCLAHKQV----DM--CGCLEYR----F---
------------P-N---------------------------------------------
--------IDNR-----T---VCDIL----------D-----------------------
------------------------------------IDV---IRCLN---------K---
V-Q-------------------SLYQ-D-------NK--------L------------GC
----------SEKCR-PPCNEEV---FKLS-ISS------------------ARW-PT--
-------DDY------EAH-F-----------LD--------------------------
------------------------------------------------EL-K--------
----------------------------------ED------------------------
-K-------------GFVL-----------------------------------------
---------------------------PAN------------------------------
-----A-------DA-----------R------AYFVKLQIFYEELNHEVIE-EYRSYEL
VNF-----------------VSDIGGQLGLWIGISALTASELVELLIVI-----------
--------ALHL--------L----------R-------KV-------------------
TRTGLIDNSAEQPADTNPRYLDDITLTGDLGERRGALTDSNTSLTVRYLGEIARKRKMKG
HKEKREASWHIDNSADQPADTNPHYLHDITFTGDLGERRAALTDSNTSLTV---------
-----------------RYLGEIARKRNKKGH----------------KEKRKAS-----
------------------------------------------------------------
------------------------------------------WHIDEEAFPCYR------
---------------------------DQATFHPNVESNTSLTVRYLDDIVQKKSKHRKR
SKGDTSMVQKGDGHRRDLSVRRG-----------RLTE-----------SSGSLTVRYLA
EVASKRKR
>NeNaC27_XP_032225907_1acid_sensingionchannel3isoformX1_Nematostellavectensis
_
------------------------------------------------------------
----------M-------------------------------------------------
------------------------------------------------------------
------------------------------------------------------------
------------------------------------------------------------
------------------------------------------------------------
------------------------------------------------------------
------------------------------------------------------------
------------------------------------------------------------
------------------------------------------------------------
------------------------------------------------------------
------------------------------------------S-----D-----------
-TS--WKDFL-GNTSLHGVRYLT---------E-NS-------------F-----IR---
--RVIWL----LVVLAAFCGFGYQLYKSIT-FYYE----------W---PTVTAM----T
-T-EW-D----------E-ELDF-----PAV-----------------------TICNLN
TL------SKE-----NFVKARKRYK----------SDATT-----------------E-
---------QL------------E-KEV-------------ETFTFFM------------

---------------------------------SA-------------------------
------------------------------------------------------------
------------------------------------------------------------
------------------------------------------------------------
------------------------------------------------------------
------------------------------------------------------------
------------------------------------------------------------
--------------------------P---------------------------------
------------------------------------------------------------
------------------------------------------------------------
------------------------------------------------------------
--------LNE-------------------------------------------------
---------------------------NAKSKLLNDFFE---NS--TFYR-----ERGN-
---------------------------------FVNKSL---------------------
------------------------------------------------NDF----SHSID
--GILS-IKWLRPCQ----FS-G----E----------D---C-----------D-R---
----------------------------------------------K-------------
------------------------------------------------------------
------------------------------------------------------------
-----------N----------F-------S-----------SFV-DLNFG--------L
CHTFN-------SG-------------KK-E--------------------E----NA--
-------LKVESL------GP--------NN-------------GLRLRLNVQQE-----
----------D-----------------H------------V-S-N---FY---------
------------------------------------------------------------
------------------------------------------------------------
-------W--LPAGFKVLVH-DRKEYP--IMEQTGFALQPGTHTFCSTKLK------K--
-----YKNL-----------PTP-YN-TK-C--GEN--------I-------TDF-NRYF
------------------SVNYSMSTCGKQCLHDYGI----KK--CGCQPIY----P---
------------I-T--K------------------------------------------
----------GV-----P---LCAL-----------------------------------
------------------------------------NDT---FECIY---------P---
T-YY------------------GKFP-R-------EA--------A------------EC
F---------KKYCP-VPCDYTK---YETK-LSY------------------ASA-IS--
-------NAL------ARS-L-----------PT--------------------------
------------------------------------------------HL-T--------
----------------------------------ESKD-------V---ADAFSDIQ---
-N----------------------------------------------------------
---------------------------MTEQERK--------------------------
-----R-------FF-----------R------ENVATLDVYFEELSYDKIE-QKPSFDS
WSL-----------------IGNIGGYLGLFLGMSVLTVMEFLDLLVIK-----------
--------FLNR--------I----------S--------GT------------------
KT------------------------------RV-------------------TTIQ---
-----------------------------------------VGS----------------
------------------------------------------------------------
------------------------------------------------------------
------------------------------------------------------------
------------------------------------------------------------
------------------------------------------------------------
--------
>NenaC28_XP_032224488_1acid_sensingionchannel3_Nematostellavectensis_
------------------------------------------------------------
----------M-------------------------------------------------
------------------------------------------------------------
------------------------------------------------------------
------------------------------------------------------------
------------------------------------------------------------
------------------------------------------------------------

```
------------------------------------------------------------
------------------------------------------------------------
------------------------------------------------------------
------------------------------------------------------------
------------------------------------------A-----D-----------
-TS--WKGFL-CNTSLHGARYLT---------E-NN-------------F-----LR---
--RAFWL----FVVLAGFAGFIYQSFLSIT-SFYS----------W---PVSTVM----M
-T-EW-H----------K-EMNF-----PAV-----------------------TICNFN
PI------SKS-----RYAQNYGNL-----------FNLST-----------------Q-
---------ES------------N-NII-------------NGIIFLM------------
---------------------------------SA-------------------------
------------------------------------------------------------
------------------------------------------------------------
------------------------------------------------------------
------------------------------------------------------------
------------------------------------------------------------
------------------------------------------------------------
--------------------------P---------------------------------
------------------------------------------------------------
------------------------------------------------------------
------------------------------------------------------------
--------RDS-------------------------------------------------
---------------------------SQKSKFSQFSLE---N-----YT-----QRGP-
---------------------------------MLGKSL---------------------
------------------------------------------------RSY----AHSID
--GMLS-LKWLEPCK----YS-G----K----------E---C-----------G-K---
----------------------------------------------N-------------
------------------------------------------------------------
------------------------------------------------------------
-----------S----------F-------S-----------TFE-HLRYG--------S
CHTFN-------S---------------D-G--------------------Q----FG--
-------LKVDSI------GP--------IS-------------GLQLRLNVEEK-----
----------D-----------------H------------V-S-N---LY---------
------------------------------------------------------------
------------------------------------------------------------
-------G--LQAGFKVLVH-DPREHP--MIEETGFALQPGTHTFCSVRMKMITKDSK--
-----YVNL-----------RAP-YR-TE-C--GEN--------I-------TDF-NRYF
------------------NVNYTMAICSKQCLHDYGI----KK--CGCQPIF----E---
------------I-T--K------------------------------------------
----------DV-----P---LCSL-----------------------------------
------------------------------------NDT---FECIY---------P---
T-YN------------------GPYP-R-------EA--------A------------EC
F---------KKSCP-VPCYYTK---YETK-LSY------------------ASA-IS--
-------NAI------ARD-I-----------PT--------------------------
------------------------------------------------YL-L--------
----------------------------------PN-------------GSEFDDIE---
-K----------------------------------------------------------
---------------------------MTEGERK--------------------------
-----T-------FF-----------R------ENVASLDVYFEELSYDLIE-QKPSFDR
WSL-----------------IAMIGGYLGLFLGMSLLTVLEFFDLLVMK-----------
--------FLNR--------I----------S-------SN-------------------
KK------------------------------GV-------------------NTVT---
-----------------------------------------VEPQE--------------
-----------------------SQNA---------------------FAFK--------
---------QG-------------------------------------------------
------------------------------------------------------------
------------------------------------------------------------
```

------------------------------------------------------------
-----PTV
>Proto_ecdy_PANarthopod__XP_023244170_1_amiloride_sensitive_sodium_channel_subunit_gamma_like__Centruroides_sculpturatus__759
------------------------------------------------------------
----------M-------------------------------------------------
------------------------------------------------------------
------------------------------------------------------------
------------------------------------------------------------
------------------------------------------------------------
------------------------------------------------------------
------------------------------------------------------------
------------------------------------------------------------
---------------------------------------------------------RKL
N-----------------------------------------------------------
--LEAQSQN----------------------S---------SL-----C-----------
-GL--CRSFA-YRSSAHGVQRIA---------S-SQ--------------DN---AR---
--RLMWS----VVFLFAIAGCGFHSVYLIL-TYLS----------Y---PRMTIT----E
-E-IH-A----------D-HIDF-----PAL-----------------------TVCNLN
PL------KKS-----SIREHL------------------L-----------------ET
-------HQSV-------------------------------SEFYN-------------
------------------------------------------------------------
------------------------------------------------------------
------------------------------------------------------------
------------------------------------------------------------
------------------------------------------------------------
------------------------------------------------------------
------------------------------------------------------------
--------------------------P---------------------------------
------------------------------------------------------------
------------------------------------------------------------
------------------------------------------------------------
------DKYDDD------------------------------DEIEE-------------
----------------------RGICFRSENEFLK--------------------KSKTM
---------------------------------DLSDVW---------------------
----------------------MSVIA----------------TKDSLMKC----GHQAQ
--DLI------TQCT----YN-A----R----------N---C-----------F-N---
----------------------------------------------D-------------
------------------------------------------------------------
------------------------------------------------------------
-----------S----------S-------TI------VLLDQYP-SPRYG--------L
CHTIV-------VD-------------KE---------------------------PL--
-------RKVKKT------GL--------SL-------------GLRLTLNIERE-----
----------D-----------------Y------------L-D-L---VS---------
------------------------------------------------------------
------------------------------------------------------------
----------PEFGARLLVH-PKGTYP--TLQGGGVVLQPGTKTYVGVRMR------K--
-----IERL-----------PAP-Y--RG-C--YDN----FQS-SQ----------LIHF
----LRKQDQSQPFAVIHN-VYTYEYCQTLCRDIHLL----KN--CGCVEEV----T---
------------P-V---------------------------------------------
----------NN-----K---FCDPC----------N-----------------------
------------------------------------TTQ---ARCRT---------R---
F-Y-------------------RSFT-R-------AN--------VD----------HDC
----------QKFCL-PACTDVR---YDLT-VSR------------------SEW-PN--
-------VRH------QNY-V-----------LK--------------------------
------------------------------------------------RW----------
-----P------------------------------------------------------

```
-N-----------LRQSL------------------TVF-LTNKTLKNG-----------
-----------------------SDE-LDV------------------------------
-----D-------NL-----------R------KNFLRVHVYIQEMNYLSVR-DIPAYTL
PQL-----------------FADLGGCLGLYIGVSAITIVEVLEHMASV-----------
--------MAFL--------Y----------K-------KR-------------------
KEYFP---------------------------KS-------------------N------
------------------------------------------------------------
-----------------------SVKK---------------------SEIK--------
--------KRRTNFRL-------------------SERSHVQKYSKSK------------
------------------------------------------------------------
------------------------------------------------------------
------------------------------------------------------------
-DCKRGNH
>Proto_ecdy_PANarthopod__XP_023215424_1_amiloride_sensitive_sodium_channel_su
bunit_gamma_like__Centruroides_sculpturatus__870
------------------------------------------------------------
----------M-------------------------------------------------
------------------------------------------------------------
------------------------------------------------------------
------------------------------------------------------------
------------------------------------------------------------
------------------------------------------------------------
------------------------------------------------------------
------------------------------------------------------------
------------------------------------------------------------
----------------------------------------------------------RP
YIKDPDIHS-------------I--------N---------TL-----C-----------
-GL--CQAFA-YRSSAHGIPRIA---------S-SQ--------------NR---FR---
--RFMWI----IVFLVAVAGFAYHSIYLIL-TYLS----------Y---PRMTTT----E
-E-IH-A----------D-QIDF-----PYV-----------------------TVCNLN
PL------KKS-----YFADYL----------------AEIST---------STPRSTED
TQ-----SDDL-------------------------------LEFLT-------------
------------------------------------------------------------
------------------------------------------------------------
------------------------------------------------------------
------------------------------------------------------------
------------------------------------------------------------
------------------------------------------------------------
------------------------------------------------------------
-------------------------SPS--------------------------------
------------------------------------------------------------
------------------------------------------------------------
------------------------------------------------------------
--------SDDH------------------------------GDFGG-------------
----------------------GGICFRSVEEFLQ--------------------RSRTM
---------------------------------DLSDMW---------------------
----------------------MSVIA----------------TKENLSKF----GHQSK
--DLV------VQCT----YN-A----R----------N---C-----------F-N---
----------------------------------------------T-------------
------------------------------------------------------------
------------------------------------------------------------
-----------S----------Y-------SI------IDVDSYP-SPRYG--------L
CHTII-------VR-------------QE---------------------------SL--
-------RKVIKT------GS--------AL-------------GLRLTLNIERE-----
----------E-----------------Y------------L-D-L---IS---------
------------------------------------------------------------
------------------------------------------------------------
```

```
----------PEYGARLLVH-PQGTFP--TLQRGGVILQPGTKTYVSVRMR------K--
-----IERL-----------PEP-Y--RG-C--SDD-F------QKTL--------LAKY
----LKMTGQE--FLLHHQ-VYTYEYCQTLCREAHLL----NQ--CGCLEELA---L---
------------------------------------------------------------
---------EGN-----K---SCDPC----------N-----------------------
------------------------------------GTQ---ARCRS---------G---
F-Y-------------------RSFT-R-------AS--------ST----------HEC
----------QKLCK-PACTDIR---YDLT-VSQ------------------SEW-PN--
-------VQQ------QQY-A-----------LK--------------------------
------------------------------------------------RW----------
-----P------------------------------------------------------
-N-----------LNKRL------------------GQS-LSSSNSNNT-----------
------------------TDTKEEIQ-INR------------------------------
-----K-------YF-----------R------KNFLRVHVYIQEMNYLSVK-DIPGYTL
PQL-----------------FADLGGCLGLYIGVSAITVVEIIEHVFSV-----------
--------FAFL--------Y----------V-------KR-------------------
KRNSQT--------------------------RD----------------PIHR------
------------------------------------------------------------
--------------------MAISSRK---------------------SAVV--------
--------TRHVPTQF-------------------PEPYSIQKRDLAKNIKHLRAY----
-------------------------------------------------FPRDTHRR---
--------------------------TGGIAETPGNCKTLYPIN----------------
-------------------------------------------------EQLFRRYPHLT
DDSLDSKW
>Proto_ecdy_PANarthopod__XP_023225242_1_amiloride_sensitive_sodium_channel_su
bunit_gamma_like_isoform_X1__Centruroides_sculpturatus__519
------------------------------------------------------------
----------M-------------------------------------------------
------------------------------------------------------------
------------------------------------------------------------
------------------------------------------------------------
------------------------------------------------------------
------------------------------------------------------------
------------------------------------------------------------
------------------------------------------------------------
------------------------------------------G-----------------
------------------------------------------------------------
------------------------------------------------------------
------------------------------------------------------------
-------------------FCLFHSIAYLR-EYFQ----------Y---PTVLNV----Q
-V-NT-E----------S-KLDF-----PAV-----------------------TVCNYN
RI------RSN-----ALDS--------------------FC--------------CTSG
LLLK---ASSL-------------------------------CHVLN-------------
--------------------------VTCSQT-NN-------------------------
------------------------------------------------------------
------------------------------------------------------------
-----------------EEA------------------------SSKRLKTL--------
------------------------------------------------------------
------------------------------------------------------------
------------------------------------------------------------
-----------------------IGSPPLSA-----------------------------
------------------------------------------------------------
------------------------------------------------------------
------------------------------------------------------------
-STAFATCPRNRT-----------------------------------------------
-------------------------------------------------------NNTVS
--------------------------YEGSIMKSFSNAY---------------------
```

```
------------------------VSL----------------PLSKKIQM----GHQVQ
--KFV------HQCQ----FM-G----Q----------P---C-----------D-H---
----------------------------------------------L-------------
------------------------------------------------------------
------------------------------------------------------------
-----------N----------F-------T-----------TFH-TFLYG--------N
CFTFN-------SG-------------QN-G--------------------K----DI--
-------LRATTD------GS--------MS-------------GLHLELNLETD-----
----------E-----------------Y------------I-F-D---LT---------
------------------------------------------------------------
------------------------------------------------------------
----------NNIGARLIIH-PSNVKP--LPENEGIAITSELETSISIEQV------N--
-----IHRS-----------KPP-YP-DN-C--IDY-P------ND----------GEGF
--------------------LYTRNTCLRDCFQQLSQ----NK--CSCADPT----W---
------------P-L--PR-----------------------------------------
----------NS-----T---SCDLR----------D-----------------------
------------------------------------VTQ---VCCLD---------D---
V-R-------------------QLLR-N-------DE--------------------NIC
------------ICP-LPCCEIQ---YKLS-VSS------------------TKWNPL--
-------KSK----------I---------------------------------------
------------------------------------------------------------
------------------------------------------------------------
-N----------------------------------------------------------
------------------------KE-MYS------------------------------
-----S-------IL-----------P------SDIAKVRIYFQTLDHVVLK-SHPKYQV
QDI-----------------FSNLGGQMGMWLGISLMTLLHYFETILGS-----------
--------VFCK---------------------------QK-------------------
QTI-------H---------------------QS--------------------------
------------------------------------------------------------
------------------------------------------------------------
------------------------------------------------------------
------------------------------------------------------------
---------------------------------P--------------------------
------------------------------------------------------------
---IGINP
>Proto_ecdy_PANarthopod__XP_023225243_1_amiloride_sensitive_sodium_channel_su
bunit_gamma_like_isoform_X2__Centruroides_sculpturatus__470
------------------------------------------------------------
----------M-------------------------------------------------
------------------------------------------------------------
------------------------------------------------------------
------------------------------------------------------------
------------------------------------------------------------
------------------------------------------------------------
------------------------------------------------------------
------------------------------------------------------------
------------------------------------------G-----------------
------------------------------------------------------------
------------------------------------------------------------
------------------------------------------------------------
-------------------FCLFHSIAYLR-EYFQ----------Y---PTVLNV----Q
-V-NT-E----------S-KLDF-----PAV-----------------------TVCNYN
RI------RSN-----ALDS--------------------FC--------------CTSG
LLLK---ASSL-------------------------------CHVLN-------------
--------------------------VTCSQT-NN-------------------------
------------------------------------------------------------
------------------------------------------------------------
```

-----------------EEA------------------------SSKRLKTL--------
------------------------------------------------------------
------------------------------------------------------------
------------------------------------------------------------
-----------------------IGSPPLSA-----------------------------
------------------------------------------------------------
------------------------------------------------------------
------------------------------------------------------------
-STAFATCPRNRT-----------------------------------------------
-------------------------------------------------------NNTVS
--------------------------YEGSIMKSFSNAY---------------------
------------------------VSL----------------PLSKKIQM----GHQVQ
--KFV------HQCQ----FM-G----Q----------P---C-----------D-H---
----------------------------------------------L-------------
------------------------------------------------------------
------------------------------------------------------------
-----------N----------F-------T-----------TFH-TFLYG--------N
CFTFN-------SG-------------QN-G--------------------K----DI--
-------LRATTD------GS--------MS-------------GLHLELNLETD-----
----------E-----------------Y------------I-F-D---LT---------
------------------------------------------------------------
------------------------------------------------------------
----------NNIGARLIIH-PSNVKP--LPENEGIAITSELETSISIEQV------N--
-----IHRS-----------KPP-YP-DN-C--IDY-P------ND----------GEGF
--------------------LYTRNTCLRDCFQQLSQ----NK--CSCADPT----W---
------------P-L--PR-----------------------------------------
----------NS-----T---SCDLR----------D-----------------------
------------------------------------VTQ---VCCLD---------D---
V-R-------------------QLLR-N-------DE--------------------NIC
------------ICP-LPCCEIQ---YKLS-VSS------------------TKWNPL--
-------KSK--------------------------------------------------
------------------------------------------------------------
------------------------------------------------------------
------------------------------------------------------------
------------------------------------------------------------
-----------------------------------------------------------V
QDI-----------------FSNLGGQMGMWLGISLMTLLHYFETILGS-----------
--------VFCK---------------------------QK-------------------
QTI-------H---------------------QS--------------------------
------------------------------------------------------------
------------------------------------------------------------
------------------------------------------------------------
------------------------------------------------------------
---------------------------------P--------------------------
------------------------------------------------------------
---IGINP
>Proto_ecdy_PANarthopod__XP_023240338_1_acid_sensing_ion_channel_4_like__Cent
ruroides_sculpturatus__568
------------------------------------------------------------
----------M-------------------------------------------------
------------------------------------------------------------
------------------------------------------------------------
------------------------------------------------------------
------------------------------------------------------------
------------------------------------------------------------
------------------------------------------------------------
------------------------------------------------------------

------------------------------------------------------------
------------------------------------------------------------
-------------------------------K---------SY-----------------
-----------EKRSNTKPPYII------AWEN-SS--------------KN---KK---
--KLSRK-------KCSLSGFTYSSYKFLT-NFFE----------Y---PVVVNL----E
-V-EN-E----------G-ELFF-----PAV-----------------------TICNSN
RM------RVS-----ELHKDV----------C-------------------------KS
KRK----SCEY-------------------------------LRIIH-------------
---------------------------------ESSS-----------------------
------------------------------------------------------------
------------------------------------------------------------
-----------------DIEN---------------------------------------
------------------------------------------------------------
------------------------------------------------------------
------------------------------------------------------------
-------------------------KPEFVFK----------------------------
------------------------------------------------------------
------------------------------------------------------------
------------------------------------------------------------
----FATCSNSS------------------------------------------------
-------------------------------------------------------KREMT
--------------------------LYHQKLLAFTNTF---------------------
------------------------SRF----------------EDWKIKKM----SHQIE
--DMI------EECT----FD-G----E----------K---C-----------D-A---
----------------------------------------------S-------------
------------------------------------------------------------
------------------------------------------------------------
-----------S----------F-------T-----------PLE-SFTYG--------S
CYTFN-------GR-------------WM-N-SRGQEDMGRRKRKKFIDDGN----EK--
-------LISRST------NP--------FS-------------GLSLTLNVQIE-----
----------E-----------------Y------------L-S-----IT---------
------------------------------------------------------------
------------------------------------------------------------
----------SKMGVHVIIH-SPDERP--YPEETGIDISTGFDTAFAIQEN------L--
-----YRRM-----------SKP-F--KDKC--VLY-G------------------KDDF
----K-------------K-IKSQKECLSLCTRDLIK----KE--CGCLDPT----L---
------------K-F---------------------------------------------
--------YRNV-----T---FCDLT----------K-----------------------
------------------------------------EED---ACCMM---------K---
M---------------------RKVN-------------------I-----------SYC
------------QCP-LPCHEIT---YDVS-LYS------------------TIW-PS--
-------ESE----------F----------KLR--------------------------
------------------------------------------------SL----------
------------------------------------------------------------
KN----------------------------------------------------------
----------------------LSKQ-ITY------------------------------
-----E-------EY-----------R------KGHLQMNVFFDRMERFIYN-QHPVYEK
DEF-----------------FSHLGGQLSLWLGLSLLTLFEYLEKFSLF-----------
--------CIQV--------Y------------------KNFS-----------------
--------------------------------RK--------------------------
------------------------------------------------------------
------------------------------------------------------------
------------------------------------------------------------
------------------------------------------------------------
------------------------------------------------------------
------------------------------------------------------------
--------

>Proto_ecdy_PANarthopod__XP_023226921_1_amiloride_sensitive_sodium_channel_subunit_alpha_like__Centruroides_sculpturatus__577
------------------------------------------------------------
----------M-------------------------------------------------
------------------------------------------------------------
------------------------------------------------------------
------------------------------------------------------------
------------------------------------------------------------
------------------------------------------------------------
------------------------------------------------------------
------------------------------------------------------------
------------------------------------------------------------
------------------------------------------------------------
---QEPSEK----------------------D---------KN-----T-----------
-SI--FEKFL-GSSSVIGLSQIT---------K-SR--------------SI---VR---
--KLLWL----AVLVTGLTFCAIESHKFMR-EFYK----------Y---PVVVNL----E
-I-EN-K----------G-ALEF-----PAV-----------------------TVCNLN
SV------RKS-----EYQKRF------------------V-----------------K-
---------TI-----------------------------TIESAFD-------------
------------------------------------------------------------
------------------------------------------------------------
------------------------------------------------------------
------------------------------------------------------------
------------------------------------------------------------
------------------------------------------------------------
------------------------------------------------------------
-------------------------SPFL-------------------------------
------------------------------------------------------------
------------------------------------------------------------
------------------------------------------------------------
------------------------------------------PSAARGF----F------
----------------------------------------IPFRKFTSQCD--AVEQKSR
---------------------------NENNQNDFYNTY---------------------
------------------------FSL----------------NKTVRGSL----GHQKE
--NFI------MSSS----FN-V----K----------Q---I-----------S-LF--
----------------------------------------------D-------------
------------------------------------------------------------
------------------------------------------------------------
-----------N----------F-------Q-----------MVV-KPRHG--------P
CYTFN-------GR-------------NT-T-SQNM-------------NVS----SV--
-------RVSRNI------GT--------LS-------------GLTLVLNLELE-----
----------E-----------------Y------------F-D-----KT---------
------------------------------------------------------------
------------------------------------------------------------
----------KSVGARIVIH-SPDEFP--DTEGEGINILPGVETSIAINKQ------T--
-----TKRL-----------PFP-Y--KDRC--KDY--------K-----------KQTI
----RNV-----------S-ILNEKDCIIHCLQDLNI----QR--CNCTDPT----L---
------------I-F--KK-----------------------------------------
----------KR-----Q---KCDLE----------N-----------------------
------------------------------------NEE---VCCLD---------K---
V-F-------------------EEMS-S-------LN--------F-----------ETC
------------DCR-VSCSITN---YEMT-TSM------------------SLW-PS--
-------PEE------FQY------------NYQ--------------------------
------------------------------------------------YY----------
------------------------------------------------------------
-G----------------------------------------------------------
---------------------------FNY------------------------------

-----S-------SF-----------R------NTYAKLNVYYATLEETIYA-QRPVYQN
NEW-----------------YSHLGGQLGLWLGISLPAIFDCIETIVLL-----------
--------IHHA--------I----------I-------KY-------------------
IT------------------------------KN--------------------------
------------------------------------------------------------
------------------------------------------------------------
------------------------------------------------------------
------------------------------------------------------------
------------------------------------------------------------
------------------------------------------------------------
--------
>Proto_ecdy_PANarthopod__XP_023226919_1_acid_sensing_ion_channel_4_like__Cent
ruroides_sculpturatus__804
------------------------------------------------------------
----------M-------------------------------------------------
------------------------------------------------------------
------------------------------------------------------------
------------------------------------------------------------
------------------------------------------------------------
------------------------------------------------------------
------------------------------------------------------------
------------------------------------------------------------
------------------------------------------------------------
-----------------------------------------------------------R
ELNNKSKED-----------------GGENEE---------EF-----S-----------
-----IQTFF-GSSAVIGLPQIA---------S-SQ--------------HI---LR---
--KLLWC----AVLITGITFCALESYKFMT-EFYK----------Y---PVVINL----E
-I-EN-K----------G-ILEF-----PAV-----------------------TICNIN
RV------RKS-----EYHVINP-----------------------------------E-
---------NV----------------------------NNRTTFIN-------------
------------------------------------------------------------
------------------------------------------------------------
------------------------------------------------------------
------------------------------------------------------------
------------------------------------------------------------
------------------------------------------------------------
------------------------------------------------------------
------------------------STP---------------------------------
------------------------------------------------------------
------------------------------------------------------------
------------------------------------------------------------
------------------------------------------RDSYCS------------
-------------------------------------------------------RERLV
---------------------------DERNQAEFYSKY---------------------
------------------------FFL----------------NKTVRNKI----GHQYH
--KFI------IMKS----LR-N----M----------N---F-----------A-S---
----------------------------------------------H-------------
------------------------------------------------------------
------------------------------------------------------------
-----------F----------F-------T-----------SIT-KPNYG--------A
CYTFN-------SK-------------IN-T-SY-----------------K----DV--
-------PMSTHI------GP--------KS-------------GLAMTLNLELE-----
----------E-----------------Y------------L-D-D--------------
------------------------------------------------------------
------------------------------------------------------------
---------VKTVGARVVIH-SREEFP--DTEGEGINIPPGMETSLAITKK------S--
-----IRRL-----------KSP-Y--KDKC--RDY--------K-----------GEYY

----ANK---------GPA-ILNQKDCRIYCLQQKNL----KT--CKCTDPL----Q---
------------V-F--TA-----------------------------------------
---------N-------K---KCNLK----------N-----------------------
------------------------------------LNK---MCCLD---------N---
V-F-------------------DKMI-H-------SQ--------Y-----------AEC
------------DCP-LECLTTQ---YEPI-ISS------------------TAW-PA--
-------PEN------FKR-L---------------------------------------
------------------------------------------------EL----------
------------------------------------------------------------
------------------------------------------------------------
---------------------------QNY------------------------------
-----S-------EF-----------R------KTYSKVNIFYSTLEETIYV-QRPAYEN
SEW-----------------YSHLGGQLGLWLGISLPAIFDCIESIILL-----------
--------LCYI--------C----------LP------KYRNK----------------
KKF-----------------------------KN-----D--------FETIVKPVNGPC
Y----------------------------TFNGNTTSENMNVSSVRLSRNIGTLSGLTL-
------------VLNLELEEYIYKTQSVGARI-------------VIHSSDEXVTNFY--
-------ICSSVTSKWLS------------------------------------------
------------------------------------------------------------
------------------------------------------------------------
-------------------------------------------------NQDMGRVIHLM
VETRLVKI
>PanArt_PPK14_NP_609017_2_pickpocket_14__Drosophila_melanogaster__67_OUT_153
------------------------------------------------------------
----------MFVRSTE-------------------------------------------
------------------------------------------------------------
------------------------------------------------------------
------------------------------------------------------------
------------------------------------------------------------
------------------------------------------------------------
------------------------------------------------------------
------------------------------------------------------------
--------------------------------------------------KETRVVADRI
R--------------------RQDQNPLAPVN----------------------------
---TKSEIQ----------------------R---------AW-----T-----------
-LL--IDSYI-SRSHIHGLYLLF---------L-PS-------------MRR---RM---
--RVLWA----LALICACTVLFHVSYLLGD-RYHN----------K---QFQTIVAH--A
HASIH--------------HIAF-----PVV-----------------------IICNKN
RL------NWS-----RLPEIKSLY----------NITPSQ-----------------D-
---------ELF------------------------------DRILT----AY-------
------------------------------------------------------------
------------------------------------------------------------
------------------------------------------------------------
-----------------D------------------------------------------
------------------------------------------------------------
------------------------------------------------------------
------------------------------------------------------------
------------------------------------------------------------
------------------------------------------------------------
------------------------------------------------------------
------------------------------------------------------------
--GFSFHKFNAFDS----------------------------------------------
-----------------------------------------------------LLGESLD
ELNHL----------------------------NFTEIV---------------------
------------------------------------------------IQM----SWRCD
--EIL------RDCH----WQ-T----A----------SR-DCC----------------
------------------------------------------------------------

------------------------------------------------------------
------------------------------------------------------------
-----------K---------LF-------------------RPR-RLPLG--------Y
CLAFN-------EL-------------EK-------------------------------
-------RRGTET------GI--------NT-------------GLLLRLLLREG-----
----------Q-----------------H------------A-P-G---NS---------
------------------------------------------------------------
------------------------------------------------------------
----------GLKGFWLTVV-ESSVWF-GFP----IEVVPHSRTNVAVTAVYHYFDES--
-----TLSL-----------PSS-W--RH-C--VMD-Y------EEE---------SEHF
----RTL---------EGQ-KYMLENCQAECQQRYLL----RY--CNCTVDL----F---
---------Y--P----PS-----------------------------------------
----------NY-----P---ACRL-----------------------------------
------------------------------------KDL----PCLA---------AHNH
L-LQN-----------FEQPGEHPYV-H------REE--------SG----------LVC
------------ECL-HNCKSLT---LLTD-MRK------------------SVQQPW--
--------------------LQ--------------------------------------
------------------------------------------------------------
------------------------------------------------------------
PN----------------------------------------------------------
------------------------------------------------------------
-----SS------AI-------------------ESMWLNVYFKKPSMLVYK-TNLIYTW
VDL-----------------IVSFGGICQLCLGCSIISLIEFVFFALYK-----------
--------VPQL--------Y------W---E-------RF-------------------
SNE-------H---------------------RS--------------------------
------------------------------------------------------------
------------------------------------------------------------
------------------------------------------------------------
------------------------------------------------------------
------------------------------------------------------------
------------------------------------------------------------
------NK
>PanArt_PPK19_NP_651708_2_pickpocket_19__Drosophila_melanogaster__72_PUT_175
------------------------------------------------------------
----------MLLYTKEL------------------------------------------
------------------------------------------------------------
------------------------------------------------------------
------------------------------------------------------------
------------------------------------------------------------
---------------------------------------VIPRPR---------------
--------------------------------------------------------P---
G-------------------------------------------------LL--------
------------------------------------------------------------
---------------------RFRNNPRG-------------------------------
--------------------------------I--------KF-----R-----------
-EK--LCNSF-AHSNIHGMQHVF---------G-EQ--------------HL---WQ---
--RCLWL----AIVLGAVITGFSLYTVLMH-RHSE----------Q---LLVSLIET--T
QLPVY--------------HIDF-----PAV-----------------------AVCPWN
HF------NWQ-----RAPSAFIRFL-------PRHPNAEL-----------------R-
---------ETFR------------------------------QLLA----SM-------
------------------------------------------------------------
------------------------------------------------------------
------------------------------------------------------------
-----------------D------------------------------------------
------------------------------------------------------------
------------------------------------------------------------
------------------------------------------------------------

------------------------------------------------------------
------------------------------------------------------------
------------------------------------------------------------
------------------------------------------------------------
--IMNFSNFNRIRI----------------------------------------------
-----------------------------------------------------LTKRNLT
GISYL----------------------------KMTDLM---------------------
------------------------------------------------NFM----TYRCD
--ELF----VADSCV----FD-E----T----------PY-DCC----------------
------------------------------------------------------------
------------------------------------------------------------
------------------------------------------------------------
-----------K---------LF-------------------VRE-QTVKG--------Q
CLVFN-------SM-------------IS-E-NSR---------------KK----HL--
-INQFYPHKLSTA------GE--------DS-------------GLKFTINASYS-----
----------------------------F------------M-N-N---IDAL-------
------------------------------------------------------------
------------------------------------------------------------
----------TPFGMNLMIK-EPRQWS----NEMMYHLYPDTENFVAVHPLVTETSPN--
-----TYEM-----------SPK-K--RR-CYFDDE--------K-----------NPTF
----QNT----------SL-TYNRENCLVVCLHLVVW----KT--CQCSLPA----F---
---------L--PPI---------------------------------------------
---------DGV-----P---ECGI-----------------------------------
------------------------------------NDA----QCLG---------NNSD
I-FTY-----------VKMGDQEKYI-N------DSR--------QG----------HFC
------------DCP-DNCNSRL---YEMS-LNV----------------RKLDY-PK--
------------------------------------------------------------
------------------------------------------------------------
------------------------------------------------------------
-N----------------------------------------------------------
------------------------------------------------------------
-----S--------------------T------DQLIKAQVYYGQRVMTKII-TKLKYTN
IDL-----------------LANFGGIISLYIGASVMSFIELLFVLGKL-----------
--------MWGF--------I------------------RD-------------------
ARIKL---------------------------KE-----------------YTK------
------------------------------------------------------------
------------------------------------------------------------
------------------------------------------------------------
------------------------------------------------------------
------------------------------------------------------------
------------------------------------------------------------
--------
>PanArt_PPK21_NP_651704_2_pickpocket_21__isoform_A__Drosophila_melanogaster__
74_OUT_236
------------------------------------------------------------
----------M-------------------------------------------------
------------------------------------------------------------
------------------------------------------------------------
------------------------------------------------------------
------------------------------------------------------------
-----------------------------------LYPLELPRARR--------------
--------------------------------------------------------PL--
------------------------------------------------------------
------------------------------------------------------------
-------------------YRDYGGKSGL------------------------------I
QTQRDARKN----------------------S---------RL----GK-----LWHFML
-PY--LKDYA-AESSVHGIRYLA---------D-PK-------------------MRNYL

```
NIRVIWL----LILLTTSIGAIVVYVDLNE-LYQT----------V---RIQTTIKN--T
MLPIF--------------RIPF-----PSI-----------------------GLCPRN
RL------NWK-----ILETEAVDHFL------GANVSAAQ-----------------K-
---------DLF------------------------------VKFFT----AAG------
------------------------------------------------------------
------------------------------------------------------------
------------------------------------------------------------
-----------------D------------------------------------------
------------------------------------------------------------
------------------------------------------------------------
------------------------------------------------------------
--------------------------P---------------------------------
------------------------------------------------------------
------------------------------------------------------------
------------------------------------------------------------
----HLSRLNEMSN----------------FFG---------NKTL--------------
-------------------------------------------------------TDELH
MLDHL----------------------------DLREVY---------------------
------------------------------------------------KFI----QFRCQ
--DLF------HTCR----WR-G----N----------PV-NCC----------------
------------------------------------------------------------
------------------------------------------------------------
------------------------------------------------------------
-----------E---------VI-------------------EYQ-FTEAG--------L
CFVFN-------TE-------------IS-P-ASR---------------QK----AR--
-EDKYYPLRTPHY------GE--------GS-------------GLDLFLRLNRS-----
----------------------------F------------IRP----------------
------------------------------------------------------------
------------------------------------------------------------
----------GKRGINVMIK-QPQQWS-----DVVRHVPHEAHTRISITPRFTVTDER--
-----TRTV-----------TPE-I--RR-CIFGDE----------------VD--NPHY
----KNF---------PDF-EYWVGNCRSRCHQEHVL----NL--CKCSPSI----F---
---------F--P-I--SD-----------------------------------------
--------KDNF-----T---ACKAS----------D-----------------------
------------------------------------------FKCLY---------DNRF
T-FSI-----------ERHPEEDDFVKNPF----KES--------------------MIC
------------DCF-TSCSQLV---FDRV-FTT------------------TTL-DN--
------------------------------------------------------------
------------------------------------------------------------
------------------------------------------------------------
-N----------------------------------------------------------
------------------------------------------------------------
-----ET------DT-----------E------AGTMRLDIFYQSGWFIQYQ-TNMRFTF
VEL-----------------LASFGGIIGLFLGASLLSAFELAYYFSIG-----------
--------LYLY--------IHG--------K-------RK-------------------
--------------------L-----------RK--------------------------
------------------------------------------------------------
------------------------------------------------------------
------------------------------------------------------------
------------------------------------------------------------
-------------------------------PEPG-------------------------
----------------------------------------------------VLTIQFGQ
RKITPIKF
>PanArt_PPK7_NP_609016_2_pickpocket_7__Drosophila_melanogaster__60_OUT_279
------------------------------------------------------------
----------M-------------------------------------------------
------------------------------------------------------------
```

```
------------------------------------------------------------
------------------------------------------------------------
------------------------------------------------------------
------------------------------------------------------------
------------------------------------------------------------
------------------------------------------------------------
--------------------------------------------------TLVYFPPSKL
Q--------------------QQQQPSRSSRL----------------------------
---AQQLAQ----------------------S---------SW-----------------
-QL--ALRFG-KRTTIHGLDRLL---------S-AK-------------ASR---WE---
--RFVWL----CTFVSAFLGAVYVCLILSA-RYNA----------A---HFQTVVDS--T
RFPVY--------------RIPF-----PVI-----------------------TICNRN
RL------NWQ-----RLAEAKSRFL-------ANGSNSAQ-----------------Q-
---------ELFE------------------------------LIVG----TY-------
------------------------------------------------------------
------------------------------------------------------------
------------------------------------------------------------
-----------------D------------------------------------------
------------------------------------------------------------
------------------------------------------------------------
------------------------------------------------------------
------------------------------------------------------------
------------------------------------------------------------
------------------------------------------------------------
------------------------------------------------------------
--DAYFGHFQSFER----------------------------------------------
-----------------------------------------------------LRNQPTE
LLNYV----------------------------NFSQVV---------------------
------------------------------------------------DFM----TWRCN
--ELL------AECL----WR-H----H----------AY-DCC----------------
------------------------------------------------------------
------------------------------------------------------------
------------------------------------------------------------
-----------E---------IF-------------------SKR-RSKNG--------L
CWAFN-------SL-------------ET-E-EGR---------------RM----QL--
-LDPMWPWRTGSA------GP--------MS-------------ALSVRVLIQPA-----
----------K-----------------H------------W-P-GHRETN---------
------------------------------------------------------------
------------------------------------------------------------
----------AMKGIDVMVT-EPFVWH-----NNPFFVAANTETTMEIEPVIYFYDND--
-----TRGV-----------RSD-Q--RQ-CVFDDE--------HN----------SKDF
----KSL---------QGY-VYMIENCQSECHQEYLV----RY--CNCTMDL----L---
---------F--P----PG-----------------------------------------
----------QY-----R---SCRA-----------------------------------
------------------------------------QDL----LCLA---------EHND
L-LIY-----------SHNPGEKEFV-R-------NQ-------FQG----------MSC
------------KCF-RNCYSLN---YISD-VRP------------------AFL-PP--
-------DVY--------------------------------------------------
------------------------------------------------------------
------------------------------------------------------------
AN----------------------------------------------------------
------------------------------------------------------------
---------------------------------NSYVDLDVHFRFETIMVYR-TSLVFGW
VDL-----------------MVSFGGIAGLFLGCSLISGMELAYFLCIE-----------
--------VPAFG-------L-------DGLR-------RRWK-----------------
AR------------------------------RQMDLG-----------VTVPTPT----
------------------------------------LNFQQTTPSQLMEN----------
```

```
------------------YIMQLKAEK---------------------AQQQK-------
------------------------------------------------------------
------------------------------------------------------------
------------------------------------------------------------
-------------------------------------------------ANFQNWHRITF
AQKHVIGK
>PanArt_PPK20_NP_651705_2_pickpocket_20__isoform_A__Drosophila_melanogaster__
73_OUT_291
------------------------------------------------------------
----------M-------------------------------------------------
------------------------------------------------------------
------------------------------------------------------------
------------------------------------------------------------
------------------------------------------------------------
---------------------------------AKGDNSVKPTEAA--------------
------------------------------------------------------------
------------------------------------------------------------
------------------------------------------------------------
---------------------GFGDNERC-------------------------------
---------L--------------------AD---------LL-----K-----------
-VH--FRSYC-EKSTIHCVRYLY---------D-SHLHNLERFVVYPLFISTFN-FKLYI
VFRIIWS----MLLIISIGLSFFFYLLLSE-RFVS----------Q---KLQTVVHDP-Q
-FPVF--------------LVPF-----PAV-----------------------GICTDN
RI------NWN-----KLEAAKEQFL-------PTNASVELV----------------E-
---------SF-------------------------------TVLVS----RM-------
------------------------------------------------------------
------------------------------------------------------------
------------------------------------------------------------
-----------------E------------------------------------------
------------------------------------------------------------
------------------------------------------------------------
------------------------------------------------------------
------------------------------TLR---------------------------
------------------------------------------------------------
------------------------------------------------------------
------------------------------------------------------------
-FGSYLSSLAELE-----------------------------------------------
-------------------------------------------------------DDNLE
AVGFV----------------------------NLTELA---------------------
------------------------------------------------IFM----TLQCS
--DIM----VPKSCL----WR-S----S----------SF-NCC----------------
------------------------------------------------------------
------------------------------------------------------------
------------------------------------------------------------
-----------E---------YF-------------------VLE-KTEFG--------F
CLVFN-------SE-------------VS-P-RSKA-------------IKQ----KE--
-GNNFYPRHNAKA------GQ--------ST-------------GLNFDLILNES-----
----------------------------F--------RR----P-D---SQ---------
------------------------------------------------------------
------------------------------------------------------------
----------ANNNVYVSIC----QAPD-QLNNVVYSITQNTETYVTVRPGLTWTDNT--
-----TRSI-----------PPE-R--RN-CLFADE--------QGE-----LDA-NDSA
----KNF----------GK-PFQLSNCLNRCHESYLI----QL--CNCSLPI----F---
------------------------------------------------------------
-----FLYNHRV-----P---DCNA-----------------------------------
------------------------------------VSL----RCLA---------RHND
I-FSY-----------DKRRDEDALF-S------ATK--------LG----------MTC
```

```
------------SCL-VDCYLLD---YYTS-TTTL---------------------PL--
-------SAHK---------L-----------PK--------------------------
------------------------------------------------------------
------------------------------------------------------------
------------------------------------------------------------
------------------------------------------------------------
-----D-------------------PH------QKLFRVDVHYQVETTPLYR-TSLEFTI
IDL-----------------IANLGGIFGLCLGASMVSAFELIYYLTVG-----------
--------LAMH--------L------YDH-QYYG-VLFKHLK-----------------
AKWVNLKGYL----------------------RN--------------------------
------------------------------------------------------------
------------------EVGHLAENP---------------------AAHK--------
------------------------------------------------------------
------------------------------------------------------------
------------------------------------------------------------
--------------------------------------------------DTNDRKLRHP
FNYRKNVW
>Protost_ecdy_cyclo_nema_Cele_tr_P91103_P91103_CAEEL_DEgenerin_Linked_to_Mech
anosensation_OS_Caenorhabditis_elegans_OX_6239_GN_delm_2_PE_3_SV_1_819_308
------------------------------------------------------------
----------M-------------------------------------------------
------------------------------------------------------------
------------------------------------------------------------
------------------------------------------------------------
------------------------------------------------------------
----------------------------------------IPTISK--------------
---------PKTNQSTKSAR------------KSSNESPYPS------------------
------------------------------------------------------------
--------------------VTFRPNSSLSH-------------------LIIEVPVAQL
K--------------------KFKHAGET-------------------------------
-------------------------------V---------SV-----E-----------
-RE--TQHFC-ETTTMHGPKRIF---------Q-GK--------------R----WA---
--TLFWL----IMVCTSIGLLITQVCILAS-NYLS----------K---PVVSDV----S
FL-IN-E----------E-GIQF-----PQI-----------------------TICNFT
PI------RKT-----FVNEMN-----------------------------------KTG
---------QI------------SPNMINYIMHW----FTEVPILIG-------------
---------------------------------SS-------------------------
------------------------------------------------------------
------------------------------------------------------------
-----------------N------------------------------------------
------------------------------------------------------------
------------------------------------------------------------
------------------------------------------------------------
------------------------------------------------------------
------------------------------------------------------------
------------------------------------------------------------
------------------------------------------------------------
------------------------------------------------------------
-------------------------------------------------------WQLLH
EGNK-----------------------------DLQEYQ---------------------
---------------------KNNPNF---------------TVQGFFIDA----GFSCS
--DIF------KLCS----FQ-G----E----------TF-DCC----------------
------------------------------------------------------------
------------------------------------------------------------
------------------------------------------------------------
-----------S----------I-------S-----------TPV-LTPLG--------K
CYTLD-------LL-------------SS-T-------------------KP----SM--
```

-------HKQTEP------GI--------QA-------------GLAITLDAHLE-----
----------E-----------------Q------------F-D-G---SN---------
------------------------------------------------------------
------------------------------------------------------------
GMDALF-TNSFVNGFQYFVH-PPNTIP--HLSSDEFTVTPNSVAYTAISSE------R--
-----FELL-----------PTNKW--GN-C--TEH-Y------------------PSGI
----KSD-----------L-PYLTGNCVSLCKAKFFM----EN--CGCTPAV----Y---
------------N-N--ER-----------------------------------------
----------NL-----K---ECTP-----------------------------------
------------------------------------FET---LACVN---------N---
H---------------------NGSN-NA-----TGK--------LEF-------KLPRC
A-----------QCA-QQCNNLI---YRAF-NSQ------------------GNQ-FS--
-------ARA------FEW-F-----------RT--------------------------
------------------------------------------------------------
------------------------------------------------------------
NN----------------------------------------------------------
------------------------SN-WTM------------------------------
-----A-------HI-----------K------TNFQIIHIFYQDMSNTEYN-QVQDASI
SDL-----------------LSNIGGNMGMFLGMSLITITEICLYFSKI-----------
--------IWLG--------I----------S-------KK-------------------
RREYMYS-------------------------KRVHEK------------THKKEIV---
---------------------------------------ETVEKMKVIESKRKL------
SSPIG-------------------------------------------FENQ--------
--------ISWNSNDNV-------------------------------------------
----EFRINLNNLEREFEAVGSRSVL----------------------------------
------------------------------------------------------------
------------------------------------------------------------
-------I
>Cyclo_ACD4_NP_505230_1_Uncharacterized_protein_CELE_F28A12_1__Caenorhabditis
_elegans__44_313
------------------------------------------------------------
------------------------------------------------------------
------------------------------------------------------------
------------------------------------------------------------
------------------------------------------------------------
------------------------------------------------------------
------------------------------------------------------------
------------------------------------------------------------
------------------------------------------------------------
---------------------MNRKRKLSCF-------------------VSVKFPVDSV
K--------------------KLRKTEGV-------------------------------
-------------------------------G---------SV-----Y-----------
-QE--TQHFS-TITTVNGPRRIF---------Y-GK--------------R----SA---
--QIFWI----LVVISILAFLVYQIVILIQ-YFYS----------K---PTLSQI----N
FI-TN-E----------G-AVYF-----PSV-----------------------TVCNLN
PV------KTS-----FIKKLN-----------------------------------SSG
---------DL------------SEELLNYLLAT----KTNSMYMFN-------------
---------------------------------NA-------------------------
------------------------------------------------------------
------------------------------------------------------------
-----------------N------------------------------------------
------------------------------------------------------------
------------------------------------------------------------
------------------------------------------------------------
------------------------------------------------------------
------------------------------------------------------------
------------------------------------------------------------

------------------------------------------------------------
------------------------------------------------------------
-------------------------------------------------------IFELK
RAHL-----------------------------NALVYL---------------------
---------------------ANHPDF---------------EIVNFLNSA----QFDCD
--ELF------ETCF----YG-G----K----------QF-NCC----------------
------------------------------------------------------------
------------------------------------------------------------
------------------------------------------------------------
-----------K----------Y-------M-----------TQS-ITSLG--------R
CWELN-------LR-------------NE-T-------------------DA----WL--
-------TKKGRS------GT------SPKT-------------GLQIIANARQS-----
----------E-----------------Q------------F-I-N---FH---------
------------------------------------------------------------
------------------------------------------------------------
-----Y-SSFQENGFRYFIH-PPHVSP--DLAAEGITVSPSRVVNSAIKTV------L--
-----HDLL-----------NHENW--GN-C--TSS-W------------------PEHY
----NTN-----------L-PYSSSACQALCVSNYFK----KL--CGCSPYS----Y---
------------N-I--DN-----------------------------------------
----------NT-----Q---VCLP-----------------------------------
------------------------------------YEE---VICMM---------E---
K-M-------------------TKSDSN-------GT--------VSL-------DFPFC
A-----------ECH-LACQKTS---YSSY-TSY------------------GDG-FN--
-------YNS------MKW-L-----------TR--------------------------
------------------------------------------------------------
-----------------------------------------------------------E
TN----------------------------------------------------------
---------------------------RSA------------------------------
-----S-------YI-----------R------QNIAIINIHFMELFYTSYS-QVKATTI
LNT-----------------FNKIFGLNGLWFGMSVVSLTELILYFTKI-----------
--------SWIA--------V----------S-------SK-------------------
RRQYLFE-------------------------KKMSEK------------RKERNIE---
---------------------------------------EAVQEAEISRSRRSS-----A
ANLKFLD---------------IESLDD---EDYW----YR-------SSSQ--------
-------LSDSHLDNVI-------------------------------------------
----QLAID---FEKPLQRPSAISLPRI--------------------------------
-------------------------------SECCEDFEEEDEDENNDLGII--------
---------------------------------------------IKL------------
--------
>Protost_ecdy_cyclo_nema_Cele_tr_O45402_O45402_CAEEL_DEgenerin_Linked_to_Mechanosensation_OS_Caenorhabditis_elegans_OX_6239_GN_delm_1_PE_3_SV_2_846_322
------------------------------------------------------------
----------M-------------------------------------------------
------------------------------------------------------------
------------------------------------------------------------
------------------------------------------------------------
------------------------------------------------------------
---------------------------------------NSPPISP--------------
---------YHVDFRAGSSK-------------ESLSPSTPSPMY-----------P---
------------------------------------------------------------
-------------------KCNFRYNDSRTQ-------------------MIIEVPMAQL
K--------------------KFQKAEGT-------------------------------
-------------------------------I---------SV-----K-----------
-RE--TQHFC-ETTTMHGPKRIF---------K-GK--------------RI---FT---
--KLFWL----IMVLGSLGLLIYQCWILSA-SYLS----------K---PIVSQV----S
FL-IP-E----------D-GMEF-----PSV-----------------------TVCNFT
PI------RKS-----YIEAMN-----------------------------------RTG

```
---------DV------------SPDMINYLMNW----FTEIPILLG-------------
---------------------------------NT-------------------------
------------------------------------------------------------
------------------------------------------------------------
-----------------D------------------------------------------
------------------------------------------------------------
------------------------------------------------------------
------------------------------------------------------------
------------------------------------------------------------
------------------------------------------------------------
------------------------------------------------------------
------------------------------------------------------------
------------------------------------------------------------
-------------------------------------------------------QQSLN
KGNE-----------------------------ELREYQ---------------------
---------------------SIHPNF---------------TVDQFFMDA----SFDCS
--DLM------KLCS----FQ-G----E----------SF-DCC----------------
------------------------------------------------------------
------------------------------------------------------------
------------------------------------------------------------
-----------S----------L-------A-----------SPT-LTPVG--------K
CFTLD-------LS------------------------------------KS----PM--
-------NKQTEP------GI--------RA-------------GLSLTLDANLD-----
----------E-----------------Q------------F-D-S---ST---------
------------------------------------------------------------
------------------------------------------------------------
GIDALL-PNSFVNGFRYFVH-PKDTIP--NLASDEYTVSPNSVAYSAISKF------R--
-----YVLL-----------PPNDW--GN-C--TED-Y------------------PYGI
----QSN-----------L-SYTSGNCLSLCKAKYFM----NQ--CGCTPAL----Y---
------------N-I--GN-----------------------------------------
----------NF-----Q---ECTP-----------------------------------
------------------------------------FET---YNCVN---------N---
S-L-------------------SLLN-EE-----TGK--------MEF-------QPPSC
T-----------RCG-QQCDSQV---YRAD-NSN------------------GNQ-FS--
-------AGA------FNY-F-----------NS--------------------------
------------------------------------------------------------
------------------------------------------------------------
KN----------------------------------------------------------
------------------------SS-WSV------------------------------
-----D-------YM-----------K------SNFQMIRIFYRDMSYTEYN-QVQDASI
TDL-----------------LSAIGGNMGMFLGGSVITIVELFLFFSKV-----------
--------FWIG--------F----------S-------KT-------------------
RRNYLYS-------------------------KRANEK------------AHKKEVV---
---------------------------------------ETVEKMKVIASQGNL------
TSLSD-------------------------------------------ATCATPP-----
-------LKLSSPNKKV-------------------------------------------
----EFRINFEDLASQLDFDSSECKGK---------------------------------
------------------------------------------------------------
----------------------------------------------------------SE
KSSNLQKY
>Cyclo_ACD1_NP_491295_2_Uncharacterized_protein_CELE_C24G7_2__Caenorhabditis_
elegans__41_331
------------------------------------------------------------
----------M-------------------------------------------------
------------------------------------------------------------
------------------------------------------------------------
------------------------------------------------------------
```

```
------------------------------------------------------------
----------------------------------------EPTLSPN-----------YR
NEAFEHDDSYLVNFVAGS---------------SSGESSTPPPSFV----------P---
------------------------------------------------------------
-------------------KCNFRYNQSRSQ-------------------MIIEVPVAQL
K--------------------KLRKLEGT-------------------------------
-------------------------------V---------SI-----K-----------
-RE--TQHFC-ETTTMHGPKRIF---------Q-GK--------------R----WA---
--TLFWL----IMVSCSLGLLITQVFILAS-EYLS----------K---PTVSDV----S
FL-IN-E----------D-GMDF-----PLI-----------------------TICNLN
PI------RKT-----YVNEIN-----------------------------------KTG
---------EV------------SPPMINYMMKW----FTEIPTLIG-------------
---------------------------------GA-------------------------
------------------------------------------------------------
------------------------------------------------------------
-----------------D------------------------------------------
------------------------------------------------------------
------------------------------------------------------------
------------------------------------------------------------
------------------------------------------------------------
------------------------------------------------------------
------------------------------------------------------------
------------------------------------------------------------
------------------------------------------------------------
-------------------------------------------------------RPTLH
EGNE-----------------------------ELKLYM---------------------
---------------------KNHLNF---------------TVDSFFMNS----GFSCP
--DIF------KLCS----FQ-G----E----------IF-DCC----------------
------------------------------------------------------------
------------------------------------------------------------
------------------------------------------------------------
-----------T----------L-------S-----------TEV-LTPLG--------K
CFTLD-------LS-------------SS-T-------------------KA----SM--
-------HKQTEP------GI--------QA-------------GLAITLDAHLE-----
----------E-----------------Q------------F-D-G---SN---------
------------------------------------------------------------
------------------------------------------------------------
GMDALF-TNSFVNGFRYFVH-PPNTIP--HLSSDEFTVSPNTVAFSAISSD------R--
-----YVLL-----------PTHQW--GN-C--TEN-F------------------PDGI
----QSN-----------L-SYSSGNCLSLCKAKFYM----EN--CGCTPAL----Y---
------------N-I--EN-----------------------------------------
----------NL-----K---ECTP-----------------------------------
------------------------------------YET---TTCLD---------N---
I-L-------------------AKPN-KE-----TGK--------IEF-------QTPNC
K-----------ACA-QQCNSLV---YRAY-NSY------------------GSQ-FS--
-------AGA------FHY-L-----------KS--------------------------
------------------------------------------------------------
------------------------------------------------------------
IN----------------------------------------------------------
------------------------PE-WTD------------------------------
-----G-------HM-----------R------ANFQMINIFYRDMSYTEYN-QVQDASV
TQL-----------------LSDIGGNMGMFLGMSVITITEICLFFSKM-----------
--------FWLG--------F----------S-------KK-------------------
RRDYMYS-------------------------KRVNEK------------THEREVC---
---------------------------------------ETVEKMKAIASQGNL------
SSIAG-------------------------------------------TTSA--------
--------KNSIPNDNV-------------------------------------------
```

----EFRINLKDLADQLDSDSGYSQNQP--------------------------------
------------------------------------------------------------
----------------------------------------------------------DS
RNASFQKY
>Cyclo_FLR1_NP_510243_1_Uncharacterized_protein_CELE_F02D10_5__Caenorhabditis_elegans__53_334
------------------------------------------------------------
----------M-------------------------------------------------
------------------------------------------------------------
------------------------------------------------------------
------------------------------------------------------------
------------------------------------------------------------
------------------------------------------------------------
------------------------------------------------------------
------------------------------------------------------------
------------------------------------------------------------
--------------------ETETESERIYL-----------------------------
-------------------------------Q---------LY-----D-----------
-YE--TKEFS-GLTTYHGLVRIY---------N-SN--------------TW---PS---
--RIFWV----VVVLSCLSLFMIHSGYLLL-GYHA----------K---PTLFQT----N
TI-VP-M----------N-GLLF-----PEV-----------------------TICNLN
PL------NTT-----KLEELN------------------I-----------------SK
---------STW------------------------------TYIFG-------------
------------------------------------------------------------
--------------------------------------------YF--------------
------------------------------------------------------------
-----------------DEITT--------------------------------------
------------------------------------------------------------
------------------------------------------------------------
------------------------------------------------------------
------------------------------------------------------------
------------------------------------------------------------
------------------------------------------------------------
------------------------------------------------------------
------------------------------------------SEHKSTKL----------
------------------------------------------------------GEEQFL
EIMN-----------------------------NYQELT---------------------
-----------------------KQEF---------------NVKNFLKSV----SKSCE
--ETF------ISCS----FG-R----E----------KLHNCC----------------
----------------------------------------------E-------------
------------------------------------------------------------
------------------------------------------------------------
-----------H------------------V-----------TTE-MTEVG--------V
CFRLS-------NV-------------NK-K-----------------------------
-------YRQWYS------GN--------GF-------------GWEFVLNGNNE----I
D---------D-----------------HA---------DSL-D-----FE---------
------------------------------------------------------------
------------------------------------------------------------
----------PDRGFLIMVH-ESEKYP--KINSYGVAVSPDSQLHAAISMK------N--
-----ISLL-----------DKANW--GS-C--SKG-W------N-------RNDT----
-----------------DV-PYTATHCEIDCKLRKVR----NL--CGCSPLA----Y---
------------S-A--RE-----------------------------------------
-------SGSND-----T---ICTP-----------------------------------
------------------------------------YQI---QQCFR---------K---
V-RGL----------------DNRWE-------------------------------DEC
------------DCP-SECNMLE---FDVT-NSY------------------SDL-DG--
-------RSR--------------------------------------------------

```
------------------------------------------------------------
------------------------------------------------------------
-G----------------------------------------------------------
---------------------------LSS------------------------------
-----S-------KV-----------E------SDISHVSLYFSHVAYERIE-QQKQLQT
ADL-----------------LSNIAGSMGLFLGMSTVTLLEIFIYLFKS-----------
--------VWGT--------V----------NST-----RQ-------------------
QQFVDAVAEE-------------------EKERS--ES-----------IVIIQN--G--
-------------------------------------RNDDMDDQKPSRFPGAD------
RKLSG-------------NSIHIHLDRRNS-----------IRGGDLSAASRG-------
--------SVSIPSQLLSPLSRHNRQSISYGQ--------GRKVSAIGIPLQPNHDTVES
GTSPIRRCSTSTTPSMLTRKLSFASQQS--------------------------------
--------------------------------DPAQ------------------------
-------------------------------------------------PAHQSRKVSTS
SIFKSQLI
>Cyclo_ACD2_NP_001309477_1_Uncharacterized_protein_CELE_C24G7_4__Caenorhabdit
is_elegans__42_364
------------------------------------------------------------
----------M-------------------------------------------------
------------------------------------------------------------
------------------------------------------------------------
------------------------------------------------------------
------------------------------------------------------------
------------------------------------HLEDGPSTKPPD------------
---FENEKTQETSLSGEE---------------FENNSTLGTMRDAK-------------
------------------------------------------------------------
-------------------SWAAANKQFANQ-------------------LVIQVPVNSF
K--------------------NGKKIKGV-------------------------------
-------------------------------G---------SA-----F-----------
-RE--TKHFS-STTTMHGPKRIF---------Y-GK---------------G---VA---
--RAFWM----LIVGLALAMLCFQIFILLQ-MYFS----------K---PTLSQV----S
FI-VN-E----------G-GMDF-----PAV-----------------------TVCNFN
PI------KKS-----YVRELN-----------------------------------VSG
---------DL------------TGETLEYLLQT----NMDAMFLFS-------------
---------------------------------NL-------------------------
------------------------------------------------------------
------------------------------------------------------------
-----------------D------------------------------------------
------------------------------------------------------------
------------------------------------------------------------
------------------------------------------------------------
------------------------------------------------------------
------------------------------------------------------------
------------------------------------------------------------
------------------------------------------------------------
------------------------------------------------------------
-------------------------------------------------------RHNLK
ETHD-----------------------------EAETYF---------------------
---------------------QNHTDF---------------QIIKFLRTA----GYDCG
--EMF------MTCY----FG-G----R----------RF-DCC----------------
------------------------------------------------------------
------------------------------------------------------------
------------------------------------------------------------
-----------K----------Y-------M-----------KQK-VTSLG--------K
CWELD-------LR-------------NL-A-------------------PE----WM--
-------RKQISP------GS--------EA-------------GLQIVVDAQLE-----
----------E-----------------E------------L-K-G---EN---------
```

------------------------------------------------------------
-----------------------------------------------------------D
DAKAIF-SDIYENGFRYFIH-PPGTNA--QLTSEGISVSPSRTVYSAIKTV------T--
-----HNLL-----------NRGNW--GN-C--SEN-W------------------PEGY
----NTF-----------L-SYSASACRALCIAQFFN----DT--CGCAPFT----Y---
------------N-V--DG-----------------------------------------
----------RK-----K---ICAP-----------------------------------
------------------------------------YES---ITCMD---------N---
H-M-------------------LKKV-N-------GT--------DYL-------ELPDC
E-----------ECH-MECQSTS---YTSY-NSY------------------GDG-FN--
-------RGS------LEW-L-----------KK--------------------------
------------------------------------------------------------
-----------------------------------------------------------I
SN----------------------------------------------------------
---------------------------KSE------------------------------
-----T-------HI-----------K------NNVAVINIFFLEMFYTSYS-QVQATSL
TEI-----------------LSDIGGNMGMFLGMSVITITELSLFFSKI-----------
--------FWIM--------V----------S-------KR-------------------
RRQYMYS-------------------------KKTHEK------------EKEHQLD---
---------------------------------------EAVKEFQERRSRRNSR-----
ENISALGH----------YSNRITPVDDFQ-TKFG----YK----NAFSEGN--------
-------MSNSSLDSVM-------------------------------------------
----ELKFDINELRRQLNQPSTDGIARI--------------------------------
-------------------------------RLPHQTSRQNSTENYYSSSPP--------
---------------------------------------------IFTIEPMSRKQSKTS
LPSSLSPR
>Cyclo_ACD5_NP_491196_3_Uncharacterized_protein_CELE_T28F2_7__Caenorhabditis_
elegans__45_370
------------------------------------------------------------
----------M-------------------------------------------------
------------------------------------------------------------
------------------------------------------------------------
------------------------------------------------------------
------------------------------------------------------------
----------------------------RRVRNLSLLYNDGPMGRFAD----------QE
NPVENRNKKETVHFQSGSYDD------DMSNSPSSSCSTVGDMPNIK---------PSA-
--------------------------------------------------SKGSF-----
------------------LSELKPFSKRASQ-------------------LIVDVPVAHL
R--------------------KIKNTEGV-------------------------------
-------------------------------S---------SI-----T-----------
-RE--SEHFS-NTTTLHGPKRIY---------N-GK-------------G-----WS---
--CVFWV----FIWISSMIMLLTQVTSLIS-MYIS----------K---PTVSQV----S
FL-LS-E----------G-GMQF-----PRV-----------------------TVCSFN
PI------KRT-----TVEALN-----------------------------------STK
---------DL------------SDDLLDYLMMF----NSDAMTLYG-------------
---------------------------------RA-------------------------
------------------------------------------------------------
------------------------------------------------------------
-----------------D------------------------------------------
------------------------------------------------------------
------------------------------------------------------------
------------------------------------------------------------
------------------------------------------------------------
------------------------------------------------------------
------------------------------------------------------------
------------------------------------------------------------
------------------------------------------------------------

```
-------------------------------------------------------AASLH
SGDN-----------------------------VFKHYV---------------------
---------------------SSHPNF---------------TADNFFMDA----GFSCG
--DMF------KMCS----FG-G----R----------RF-DCC----------------
------------------------------------------------------------
------------------------------------------------------------
------------------------------------------------------------
-----------K----------Y-------A-----------TPI-FSDLG--------K
CFTLN-------LQ-------------GS-D-------------------KS----WM--
-------KMQTEP------GI--------AA-------------GLQIILDSHLE-----
----------E-----------------Q------------F-D-S---ET---------
------------------------------------------------------------
-----------------------------------------------------------D
GVTPVF-SSAFENGFRFYIH-SSEEIP--FLASEGIAVSPDSVVYSALSSS------K--
-----YILL-----------SSNAW--GN-C--SDS-W------------------PRGY
----DYS-----------F-PYTSAMCSTMCKAQYFQ----NL--CGCSPSI----Y---
------------N-H--LN-----------------------------------------
----------RF-----N---DCTP-----------------------------------
------------------------------------YET---FICMD---------T---
K-M-------------------KKVV-N-------QS--------FNI-------EMPTC
E-----------ECK-VECKSQV---YHSF-NSY------------------GKG-LS--
-------RGA------LMW-L-----------TK--------------------------
------------------------------------------------------------
------------------------------------------------------------
QG----------------------------------------------------------
----------------------KQET-WTI------------------------------
-----P-------HM-----------K------LNFQVVNVFFRDMSYTEYI-QKRGMSL
TEL-----------------LSDIGGNMGMFMGMSVFTIIELFLFLSKI-----------
--------GWIG--------F----------S-------RK-------------------
RRDYMYS-------------------------KKKNEE------------MHEKELE---
---------------------------------------DVVTGFKLFRHRKSGKDMSLR
EKIKGL------------SMHRVTSEQLNVCKLAWEPDIER----RLASVTR--------
--------QNSALKEHKDYKQP--------------------------------------
---TILPFDLKDIKDQITRGRAASMFR---------------------------------
------------------------------------------------------------
----------------------------------------------------SRRSRSET
APAVIHEA
>Porifera_Syc_Cil_scpid47944_scgid21366_Acid_sensing__ion__channel__3_Amilori
de_sensitive__cation__channel__3_Dorsal__root__ASIC_800
------------------------------------------------------------
----------M-------------------------------------------------
------------------------------------------------------------
------------------------------------------------------------
---------------------------------------------------------DTS
ATDQANDSSSSNN---S--AELNADSIFV-------------------------------
DV--SAH-----------------------------TDKVTPKKRK------------PI
WPLSRFLHGKPDSSTAYEYH-------------VDDDSDFPLKEYTR-N-----G-----
------------------------------------GGH------------AGN------
----------------------------------------G-G-------TSNGHIGQQK
Q--------------------TFEQTDEADTPAHHS------------------------
-GIDWRERK----------------------AM--------AL-----------------
-KF--TYTYV-YLAACHGIARAM---------Q-NK-------------ASV---AR---
--RLFWL----LAFSGAVSMFFYQISLLVK-QFTD----------Y---EFQVDV----E
-F-SF-N----------R-QLAF-----PAV-----------------------TFCNLN
PM------NVK-----LLNCSR------------------FH----------------DE
CL-------QLAG----------ANQSAD-------------GQLIK-------------
------------------------------------------------------------
```

------------------------------------------------------------
----------------T-------------------------------------------
------------------------------------------------------------
------------------------------------------------------------
------------------------------------------------------------
------------------------------------------------------------
------------------------------------------------------------
------------------------------------------------------------
------------------------------------------------------------
------------------------------------------------------------
------------------------------------------DDAIGYAH----------
-------------------------------------------------------EYNFS
------------------------------------------------------------
-----------------------GEPI----------------TRAEYLQY----GFLPE
--DMI------ITCT----WN-G----E----------P---C-----------S-A---
----------------------------------------------R-------------
------------------------------------------------------------
------------------------------------------------------------
-----------N----------F-------T-----------ISQ-NPFYG--------N
CFTLN-------GD-------------PK-N--------------------------P--
-------LQTNYP------GR--------EA-------------GLALELYMNPD-----
----------E-----------------G------------IGG-E---TS---------
------------------------------------------------------------
------------------------------------------------------------
----------TSSGVRMSLH-QPGVKP--FPDEDGFSLATGEATFIGVRQL------E--
-----VSRL-----------GSP-Y--SN-C--TQE-N------DNW---------KSLY
----PGY-------------SYSRTACVQSCLAQEMF----DR--CGCVEIF----L---
------------V-----------------------------------------------
----------DK-----P---ICNNAN--------VN-----------------------
------------------------------------KEQ---ADCAA---------G---
V-R-------------------RDYR-L-------DR--------L------------NC
D-----------LCY-NPCVDRI---YLHT-LSK------------------ATW-PA--
-------KAY------QYS-F-----------RE--------------------------
------------------------------------------------TL-K--------
----------------------------------EL------------------------
-N-------------F--------------------------------------------
------------------SDGNSEDS-NAE------------------------------
-----E-------IY-----------R------EKFLRAVVYFEELNYQRIV-QKPAYPA
ENL-----------------FADFGGQMGLWAGLSFLAFVELFDYVGQV-----------
--------LLIA--------L----------G---L---KK-------------------
------------------------------------------------------------
------------------------------------------------------------
------------------------------------------------------------
------------------------------------------------------------
------------------------------------------------------------
------------------------------------------------------------
------------------------------------------------------------
---VVVLP
>Porifera_Syc_Cil_scpid49206_scgid6888_Amiloride_sensitive__sodium__channel__subunit__alpha_Alpha_NaCH_Epithelial__Na_channel__subunit__alpha_Nonvoltage_gated__sodium__channel__1__subunit__alpha_SCNEA_882
------------------------------------------------------------
----------MDVDQDALIR----------------------------------------
------------------------------------------------------------
-------------------------AEADNLFEELA------------------------
--------------STPEHR----------RPLP-HM---PSS--------SQFKRQTSL
SFSTSQSSSLQ------RYGEKNADEV---------------------------------

-------ASEAKEELDDDL------------------VGDMPP------------------
------------NWESKKTK-------------RRTSAQSAWSTSSSS------------
------------------------------------------------------------
-------------------SKGSRESNVWSRDSLF---SFGP---------------KET
P--------------------QITDRDVADIADEGA------------------------
-VKGLQRKR----------------------V---------GF-----I-----------
-DF--TMRFV-LDTAGHGMPRIV---------E-KD-------------IWL---IR---
--RIIWG----CIVLGAAVGFIIQTHALVS-KYLD----------K---DFKVDV----K
-Y-KF-N----------K-EIGF-----PAV-----------------------TLCNLN
PL------NRS-----KLECSR------------------F-----------------DT
SVH----KGALVCQQT----ASGNHTTVQ----------VTLAQILQ-------------
------------------------------------------------------------
------------------------------------------------------------
----------------A-------------------------------------------
------------------------------------------------------------
------------------------------------------------------------
------------------------------------------------------------
------------------------------------------------------------
------------------------------------------------------------
------------------------------------------------------------
------------------------------------------------------------
------------------------------------------------------------
--RINALLKSDL------------------------------QQAK--------------
-------------------------------------------------------ERAFQ
YN-----F-------------------------THQEAM---------------------
-------------------------------------------TAAEIAKY----GYDLS
--KLM------VSCT----FR-G----Q----------P---C-----------G-I---
----------------------------------------------E-------------
------------------------------------------------------------
------------------------------------------------------------
-----------N----------F-------T-----------RFE-SARYG--------N
CYTFN-------NN-------------IS-T-------------------------Q---
-------IQISQP------GP--------AF-------------GLAIDLAYDPL-----
----------E-----------------E------------I-G-L---LS---------
------------------------------------------------------------
------------------------------------------------------------
----------HGAGFQVAVH-EPRSKV--FPEEDGMSISPATATSIGIRQA------V--
-----VERL-----------GEP-Y--GN-C--TNI--------TG------YDW-GNLY
----SGY-------------KYTRTICLKECFAQAML----DA--CGCVSEN----I---
------------L-V---------------------------------------------
----------GK-----R---LCLPAT--------LN-----------------------
------------------------------------KTE---ADCRA---------E---
I-E-------------------ELYR-I-------YK--------L------------PC
L-----------QCF-NPCKERI---YRTT-VST------------------ASW-PS--
-------FEY------QRE-F-----------KQ--------------------------
------------------------------------------------LL-K--------
----------------------------------QK------------------------
-G-------------FQV------------------------------------------
------------------------DH-FDV------------------------------
-----V-------RF-----------R------DQYARVEVFYEELNFENIV-QKPAYEP
ENL-----------------LADMGGQLGLWLGFSVLAILEIIEYLIYM-----------
--------LLVA--------L----------G-------KV-------------------
KLPDEEDYL-----------------------KA-------------------R------
------------------------------------------------------------
------------------------------------------------------------
------------------------------------------------------------
------------------------------------------------------------

------------------------------------------------------------
------------------------------------------------------------
------DD
>Porifera_Syc_Cil_scpid40922_scgid7672_Amiloride_sensitive__sodium__channel__
subunit__alpha_Alpha_NaCH_Epithelial__Na_channel__subunit__alpha_Nonvoltage_g
ated__sodium__channel__1__subunit__alpha_SCNEA_933
------------------------------------------------------------
----------M-------------------------------------------------
------------------------------------------------------------
------------------------------------------------------------
------------------------------------------------------------
------------------------------------------------------------
------------------------------------------------------------
------------------------------------------------------------
------------------------------------------------------------
------------------------------------------------------------
------------------------------------------------------------
---SAELRR----------------------D---------SA-----------------
-RLW-VEAYL-EDLTSHGLSRIT---------A-TA--------------SYLPGIV---
--RLLWG----VVFAAAFAFALQQCSILVA-QYRK----------N---DVDVSV----D
-F-SF-N----------K-EVDF-----PAV-----------------------TICNLN
AL------RRQ-----EVLKTR------------------LH----------------E-
-------HVNVTASPT---TVSAAPVTVTTAAPTT-TPTTPYVSTVN-------------
------PR-------------------------DFEA-----------------------
------------------------------------------------------------
------------------------------------------------------------
-----------------NVT-------------------LPSTATRTVHTTV----FVEP
G-----------------------------------------------------------
------------------------------------------------------------
-----------------------------------------------------TE-APLI
-----------------TTATLGETEPTLATQTTGM-----------------PNIAPTT
AGD-----------------------------------------------GSG-------
------------------------------------------------------------
--------------------------------------------------------GSRT
TPRPHATTASGGGTHAPATG--------------PPVPGGGTTESSQGVCIIALTLGL--
-------PCSVMKLLLGEGAAAGGPANQDGSAITRAPTVFIPDDFFTSSTTPLEIADSFE
GGNTSLFAHVE-----------TERNRYNLQKVDLTSHQVN-------------------
-------------------YQYDEAPV----------------VREEISVL----GHQIE
--DML------FRCR----YD-S----E----------G---C-----------S-P---
----------------------------------------------S-------------
------------------------------------------------------------
------------------------------------------------------------
-----------N----------F-------T-----------QFQ-NGRYG--------N
CYTFN-------GD-------------EA---------------------------RV--
-------LRTERV------GP--------TY-------------GLLMELDIEQSL----
----------N-----------------Y------------I-G-Q---LT---------
------------------------------------------------------------
------------------------------------------------------------
----------QSAGIRMVIH-RQGEQA--FPEYDGFDLAPGFKTYVGIRQV------V--
-----IERQ-----------GRP-Y--GN-C--QEG--------NPQE---ELAS-----
---------SN------AT-TYTKWGCLRECLAEHML----RE--CKCVDAT----Y---
------------------------------------------------------------
-------IDKDA-----R---LCHAY----------N-----------------------
---------------------------H--------PEQ---ANCLA---------E---
V-D-------------------RQFK-T-------DL---------------------IC
Q-----------DCE-DPCRSVI---FRKS-ISS------------------ERW-PS--
-------KQY------EPA-L-----------FA--------------------------

```
------------------------------------------------RL-----A----
----------------------------------EE------------------------
-N-------------RTI------------------------------------------
-----------------------DDD-VVT------------------------------
-----A-------KI-----------R------DNYIQAVVYFEEFNFQTVT-QRPALEI
VDL-----------------LSYIGGTIGLFVGVSALSILEFLDIIVNL-----------
--------VYFF--------A----------KWLY----KRVR-----------------
TRLLGDDVFDGDE-------------------RE----------------LQRRT-----
------------------------------------------------------------
------------------------------------------------------------
------------------------------------------------------------
------------------------------------------------------------
------------------------------------------------------------
-----------------------------------------------------------S
DVKAGLYY
>Cyclo_tr_K7H9J0_K7H9J0_CAEJA_ASIC1_OS_Caenorhabditis_japonica_tested
------------------------------------------------------------
------------------------------------------------------------
------------------------------------------------------------
------------------------------------------------------------
------------------------------------------------------------
------------------------------------------------------------
------------------------------------------------------------
------------------------------------------------------------
------------------------------------------------------------
------------------------------------------------------------
------------------------------------------------------------
------------------------------------------------------------
------------------------------------------------------------
------------------------------------------------------------
------------------------------------------------------------
------------------------------------------------------------
------------------------------------------------------------
------------------------------------------------------------
------------------------------------------------------------
------------------------------------------------------------
------------------------------------------------------------
------------------------------------------------------------
------------------------------------------------------------
------------------------------------------------------------
-------------MLCQSNVNVIANIIMKEEVEVE-----------------VEAKKSCC
NTL-GDCVYTEK-------P-KKKTISCLCATPIKM-----------CVR----------
----IDP-PQTNATTLDDRLVK--------------------------------------
-----------------------------------------FWHIQPSTT----------
----VSPTIKRK------------------------------EERD--------------
-----------------------------------------KAYGYTGV------KDRIA
--------------------------LRAKAMENIIFAV---------------------
------------------------DAL----------------TEEQKWKI----SYNKS
--DFI------VKCS----FN-G----R----------E---C-----------N-VK--
----------------------------------------------H-------------
------------------------------------------------------------
------------------------------------------------------------
-----------D----------F-------V-----------EYL-DPTYG--------A
CFTYG-------QK-------------LG-N-----------------------------
-------ITNERS------GP--------AY-------------GLRLEVFVNVT-----
----------E-----------------Y------------L-P-----TT---------
------------------------------------------------------------
```

------------------------------------------------------------
----------EAAGVRLTVH-ATDEQP--FPDTLGFSAPTGFVSSFGIKLK------S--
-----MVRL-----------PAP-Y--GD-C--VKE-G------K-------TED--FIY
----TKK-------------AYNTEGCQRSCIQKHLS----AT--CGCGDPR----F---
------------P-P--YR-----------------------------------------
----------ES-----K---NCPVD----------D-----------------------
------------------------------------PYK---RECIK---------S---
E-M-------------------HVAT-R-------DS--------KK----------LGC
------------SCK-QPCNQDV---YSVS-YSA------------------SRW-PA--
-------IAG------DLS-----------------------------------------
------------------------------------------------------------
------------------------------------------------------------
-G----------------------------------------------------------
------------------CPLGMSAH-HCL------------------------------
-----G-------YK-----------R------EQGSMIEVYFEQLNYESLL-ESEAYGW
SNL-----------------LSDFGGQLGLWMGVSVITIGEVACFIFEM-----------
--------FISI--------F----------S-------AK-------------------
RV------------------------------KR-------------------R------
-----------------------------------------PARKSFS------------
------------------------------------------------SSLR--------
------------------------------------------------------------
------------------------------------------------------------
------------------------------------------------------------
----------------------------------------------------CSTDYNLN
KDGFNLDN
>Lopho_Phor_aus_Phoronis_g6004_t1_TESTED
------------------------------------------------------------
----------M-------------------------------------------------
------------------------------------------------------------
------------------------------------------------------------
------------------------------------------------------------
------------------------------------------------------------
------------------------------------------------------------
------------------------------------------------------------
------------------------------------------------------------
------------------------------------------------------------
-----------------------KRQGEM-------------------------------
-------------------------------E---------GP-----D-----------
-GL--LTNFA-RSTSAHGLARIP---------G-TS--------------QP---MQ---
--RAVWA----LLVTALAAALISALAIVVS-VYLQ----------Y---QYTEFA----K
-K-VA-R----------P-NIKF-----PVI-----------------------TACNKV
PY------GLL-----QTEQTV--------------------------------------
------------------------------------------NEFFR----SIG------
----L----------------------------DE-------------------------
------------------------------------------------------------
------------------------------------------------------------
------------------------------------------------------------
------------------------------------------------------------
------------------------------------------------------------
------------------------------------------------------------
--------------------------PTC-------------------------------
------------------------------------------------------------
------------------------------------------------------------
------------------------------------------------------------
------------------------------------------------------------
-------------------------------------------------------QSEIS
PGLND----------PIIY---R----------RISDAV---------------RP----

```
---------------------RVLYEY----------------NSKVMQVL----GQNLD
--DFM------VTCA----YQ-G----P----------P---C-----------F-IY--
----------------------------------------------S-------------
------------------------------------------------------------
------------------------------------------------------------
-----------N----------FS------T-S---------LFK-DPYHY--------N
CVTLK-------VP-------------DE-------------------------------
-------VQVKVN------GR---------------G-------AMELVFFVGDN-----
----------K-------TSLQGK----H------------V-D-K-LVTD---------
------------------------------------------------------------
------------------------------------------------------------
----------GTVGIKLIIH-AEGMHP---KWAKTLDVAPGHLANIAVRPR------E--
-----IYRL-----------KAP-YA-SD-C--IEN--------PPDI----LAASNTSY
--------------------TYSKELCTLKCFNKLVA----DA--CDCIPEP----VV--
--------------V--DK-----------------------------------------
-------FGLDY-----P---YCGESP--------CNYS---------------------
----------------------------------YILDK---IKCHT---------N---
L-T-------------------ERLH-E-------GM--------LS----------NQC
P-----------ECK-LPCEELT---YDTD-MHL------------------SKW-PS--
-------KFT------SKT-I-----------TK--------------------------
------------------------------------------------YILK--------
------------------------------------------------------------
-N-------------HVIP-----------------------------------------
------------------RNIENDSS-ALD------------------------------
-----S-------YV-----------Q------QNFLKVVIYLEDLLTVEQT-EQESMNL
AQL-----------------TSSVGGAMGFFLGISIVTVFEFIDLLVKS-----------
--------VQSL--------F----------K-------KK-------------------
SVD-----------------------------KV--------------------------
------------------------------------------------------------
------------------------------------------------------------
------------------------------------------------------------
------------------------------------------------------------
--------------------------------LP--------------------------
-----------------------------------------F------------------
---HSKGS
>Lopho_Phor_aus_Phoronis_g5063_t1_TESTED
------------------------------------------------------------
----------M-------------------------------------------------
------------------------------------------------------------
------------------------------------------------------------
------------------------------------------------------------
------------------------------------------------------------
------------------------------------------------------------
------------------------------------------------------------
------------------------------------------------------------
------------------------------------------------------------
------------------------------------------------------------
---NSEVKN----------------------W---------SV-----K-----------
-DR--MKEFC-ESTSAHALGQTV---------S-SGE------------------VK---
--AIFWS----LVFLSALAGCVWNIVHVVE-SYTS----------F---GFSVKS----K
-L-EM-E---------PS-SLKF-----PSV-----------------------TICNLN
PI------SFS-----RQARFA--------GD--------F-----------------ER
ST-------GL-------------------------------------------------
---------------------------------DD-------------------------
----------------------------------------G-------------------
------------------------------------------------------------
-----------------D------------------------------------------
```

------------------------------------------------------------
------------------------------------------------------------
------------------------------------------------------------
------------------------------------------------------------
-----------------------------------------------C------------
------------------------------------------------------------
------------------------------------------------------------
------------------------------------------------------------
-------------------------------------------------------DNEFH
RDETYR---------PVV--------LDLYRWSELPQFW---------------------
------------------------FEY----------------NENVTEMF----GHSKK
--DFI------VDCV----FQ-R----E----------G--GC-----------E-----
----------------------------------------------N-------------
------------------------------------------------------------
------------------------------------------------------------
-----------N----------F-------S-----------VTK-DPNLY--------N
CYTLE-------PN-------------KN-------------------------------
-------EDLLEI------GF--------AA-------------GLSLTLFVENR-----
----------RIRNA-------------F------------T-G-N---YA---------
------------------------------------------------------------
------------------------------------------------------------
-LDS---YHTGIVGVKVAIH-VSGSHP--NPNTRGVIAEVGKSTDFILRTV------N--
-----RTAL-----------GPP-YP-SP-C--NPK--------KTI---------ESSH
----RKE-----------L-QYEENLCFASCLQNAIA----KK--CGCVSINP---M---
----------AVP-I--AD-----------------------------------------
------KFGIDL-----P---FCGSYP--------CNAT---------------------
----------------------------------KVTEN---YECVH---------D---
V-I-------------------GRFLLK-------ND--------------------PNC
R-----------NCT-KPCNEVS---FEVT-KFQ------------------SKW-PS--
-------ELH------QSE-F-----------IR--------------------------
------------------------------------------------WLANKD------
------------------------------------------------------------
-N-------------TAL------------------YNQ---------------------
------------------LVRNISDD-NIS------------------------------
-----R-------FI-----------E------NNFLRINIYFGDFYVRKDI-ETLDMDW
FDL-----------------LSSVGGAFGFWVGISVVTGVEVLELLLDC-----------
--------IVLF--------L----------N-------RR-------------------
IKK-----------------------------RN-------------------Q------
---------------------------------------IDLRNTT--------------
------------------------------------------------GDDD--------
------------------------------------------------------------
------------------------------------------------------------
------------------------------------------------------------
------------------------------------------------------------
---VVTGS
>Macrostomum_Mlig049925_g2_Protos_Platyhelmin_Macrostomum__TESTED_A0A267GWB9_
A0A267GWB9_9PLAT_Uncharacterized_protein_OS_Macrostomum_lignano_OX_282301_GN_
BOX15_Mlig003720g1_PE_3_SV_1
------------------------------------------------------------
----------M-------------------------------------------------
------------------------------------------------------------
------------------------------------------------------------
------------------------------------------------------------
------------------------------------------------------------
------------------------------------------------------------
------------------------------------------------------------
------------------------------------------------------------

------------------------------------------------------------
------------------------------------------------------------
-----QDRL----------------------R---------TV-----T-----------
-GA--AQEVA-GITRTHGLLHIF---------L-SR--------------GS---AR---
--RLFWL----LLFVAALIGCSVHLAKLVQ-KFTE----------R---SVESQL----K
-L-RS-E------------RAQF-----PDV-----------------------TICNFK
PA------SAS-----LLSIIN------------------FL----------------EF
HS-----KNEF------------------------------KFEVFD----YFEIFW---
--------------------GNF----------DN-------------------------
--------------------------------------------WFKE------------
------------------------------------------------------------
-----------------DTQK---------------------------------------
------------------------------------------------------------
------------------------------------------------------------
------------------------------------------------------------
-------------------------RPQES------------------------------
------------------------------------------------------------
------------------------------------------------------------
------------------------------------------------------------
------------------------------------------DESY--------------
-------------------------------------------------------DNRLE
KVCK-----------------------------KVLDVM---------------------
------------------------------------------WVSGDTRDI----SQLDE
--TML------LSCR----YN-S----E----------P---C-----------S-H---
----------------------------------------------K-------------
------------------------------------------------------------
------------------------------------------------------------
-----------N----------F-------S-----------LVQ-TSRFW--------N
CYTFH-------PD-------------PR-------------------------------
-------QSSSGPG-----GR--------GA-------------ELDMVLFTDSN-----
----------EKYPDL------------YEIYP-----NGAL-P-D-GCIT--RKF----
------------------------------------------------------------
--------------------------------------------------------LRSK
AVNRLLNNEPQSAGLRIFIH-EEKSFP--MSETEFVDVASATSTSIKLRPV------H--
-----NRLM-----------SKP-S--RR-C--SEP-----------P----IET-INYV
----RHFSNASLNVVTKAY-SKSVSDFVVEAQQSILH----SN--CGCYSHL----L---
------------P-F--SV-----------------------------------------
----------NT-----S--DLCYFAPPQEW-ITPSD-----------------------
----------------------------------AVLKR---IDCHDHWFEHA-QRQSEE
L----A----------------KEFQ---------HR-----------------------
----------------MWCQFTQQRPWQQS----------------------RRW-PP--
-------FSA------IQE-I-----------WE--------------------------
------------------------------------------------NLMVPQVRH---
----------------------------GVEFLPEDGKEH-----------------HIF
RNSTK--------TRQEIQLYLDCRFNFLRKLGKD-YKK---------------------
------------------KIIESATSCVSR------------------------------
-----S-------EL-----------A------KELAAVSISLQSPAADAYE-EKYKYHW
TEA-----------------LSEIGGTLGLWLGISVVSTFELLEFIYIL-----------
--------VQKC-------------------R----------------------------
--------------------------------RN--------------------------
------------------------------------------------------------
------------------------------------------------------------
------------------------------------------------------------
------------------------------------------------------------
------------------------------------------------------------
------------------------------------------------------------
DDDSDTEE

```
>Macrostomum_Mlig051885_g1_Protos_Platyhelmin_Macrostomum_lignano_TESTED
------------------------------------------------------------
----------M-------------------------------------------------
------------------------------------------------------------
------------------------------------------------------------
------------------------------------------------------------
------------------------------------------------------------
------------------------------------------------------------
--------------------------------------------------------P---
------------------------------------------------------------
------------------------------------------------------------
------------------------------------------------------------
------------------------------------------------------------
------------------------------------------------------------
-------------------------------QYLLN---------H-------------D
-----------------S----------PARR----QLE-------------------SN
ML----------------------------------------------------------
---------DL---------------------------MVALRRVYA----K--------
------------------------------------------------------------
------------------------------------------------------------
-------------------------------------------------GG---------
-----------------D-------------------------------------IFG--
------------------------------------------------------------
------------------------------------------------------------
------------------------------------------------------------
------------------------------------------------------------
------------------------------------------------------------
------------------------------------------------------------
------------------------------------------------------------
------------------------------------------------------------
-------------------------------------------------------TQKYF
--------H----Y-------------------NLLQHY---------------------
------------------------------------------WASVDTAHL----GHNLS
--LSM------VACF----YK-K----A----------P---C-----------R-P---
----------------------------------------------A-------------
------------------------------------------------------------
------------------------------------------------------------
-----------D----------F-------K-----------LVQ-NSHYW--------N
CWSFR-------PK----------------D-----------------------------
-------RSVQGT------GP--------ND-------------GLNIILYTSTM---PL
DDSSQPDHVVE-----------------Y---PPSTLQLDTVFG-K-AGMQ---------
------------------------------------------------------------
------------------------------------------------------------
----------VSSGVRLLVH-EPGTYP--HVYWEGADVGNGWSADLRFKMK------R--
-----NVYV-----------NRT-G--HN-C--VEN-YGH----------------TDYW
----ASEA-----EGIQRF-RKRGQDCVVRKWQEHLM----RK--CHCQSTF----L---
------------P-T--QD-----------------------------------------
----------RS-----A---LCHYLRAAG--VSAQNIS---------------------
--------APFGA---------------------YKFQQ---LKSFN---------D---
V-E-------------------NQVRTELY-----DQ--------FA----------SEC
------------GTV-QACEQSS---YTYS-IAS------------------VPW-PA--
-------HTD------LEA-F-----------IH--------------------------
------------------------------------------------TFVSPKFHR---
---------------------------------------------------------ARY
QG-------------QQLL-----------------DQI---------------------
--------VRFHRHPDGSRDFEKGWS-VSH------------------------------
-----R-------FV-----------R------QNFVKLQVFAEESKSTLIH-ESPSYSF
```

VEV-----------------LSELGGIGGLWVGMSLVTFVELFEFLAILG----------
--------IKCTQV-ASS-YLHS--------RWT-----RR-----A-------------
GQMANGACPGSAN-----------------HLRT---------------PACTR------
---------------------------------------SAQSPDEITSGPCVVL-----
----------------EEASSVVDEDTELVG----------IFLCPEASRNEHLHDG---
------------------------------------------------------------
------------------------------------------------------------
------------------------------------------------------------
-------------------------------------------------NSSRLSRGQAM
QHVSVVGS
>Macrostomum_Mlig041003_g1_Protos_Platyhelmin_Macrostomum_lignano_TESTED
MKNTLH------------------------------------------------------
----------M---------------------------------MRC-------------
------------------------------------------------------------
--------------------------GN--------------------------------
------------------------------------------------------------
------------------------------------------------------------
----------------------------------LMAAAMAPG-----------------
------------------------------------------------------------
---------------------------------------------------LGP------
--------------------------------------------------CATNIGLNNL
QQQQQ--------------QQGAQEEQQGNPNAPGNG----------------------G
GSRIPRRRE----------------------V---------ML-----R-----------
-GY--INHFT-SGTTAHGLNRVN-----------AE--------------QS---WV---
--RMVWV----GIVLAASFGAIVHTTKQVK-TYLQ----------Y---PVSSTN----R
---QE-P----------N-SFEF-----PDL-----------------------TFCDLN
KR----FFHKN-----VISYAR------------------------------------E-
--V------DV---------------------------YDEAHMIYG----LVYSHFRT-
---------------------------------DS-------------------------
------------------------------------------------------------
------------------------------------------------------------
-----------------D--------------------------------------WRKE
------------------------------------------------------------
------------------------------------------------------------
------------------------------------------------------------
------------------------------------------------------------
------------------------------------------------------------
------------------------------------------------------------
------------------------------------------------------------
------------------------------------------------------------
-------------------------------------------------------RRAIE
LIYN-----------------------------ALEHLL---------------------
----------------------------------------------NITDV----NIRPM
--DVV------LYCT----FD-G----E----------P---C-----------S-H---
----------------------------------------------T-------------
------------------------------------------------------------
------------------------------------------------------------
-----------D----------F-------H-----------VFY-HRLYT--------N
CFTFK-------PK----------------Q-----------------------------
-------RSLRSS------GM--------DR-------------GLFLLLYVPSA-----
----------D-----------------VNL--K-------L-D-SLARLA---------
------------------------------------------------------------
------------------------------------------------------------
-------SGLENNGIRFQIH-QRDTIP--HPLEYGIMAPTGTLTAVGLEQV------R--
-----TSLA-----------DTP-T--NP-C--VRD--------TK------LRLFDNQF
----NFM---------RQY-DRQLFDCIKHQYMREVR----KQ--CGCTLESHVIGF---
--------D---P-----------------------------------------------

---------REV--------VFCHDLISAK------N-----------------------
------------------------------------ATR---VGALT---------K---
L-Y-------------------NELK-N----V-TNTAMRQNDAELNAS-----LDRMIC
SDSVSESAARPERCY-DSCNYNK---YEYS-LSQ------------------CPW-PE--
-------DSFEV----RNSQIQM---------VKIA------------------------
------------------------------------------------QMLEDYKLR---
---------------------------------------------------------YGS
AN-----------PETALAGLM--------------RKSSLN------------------
---------------------------ISS-------CLTHDTQKMQVPDKV--------
-----DCH-RVQLFV-----------R------RSVVQLRVYPETLTVRHTI-EERSYEL
VNL-----------------CSELGGILGLWIGFSIVTLFEFAELFIIC-----------
--------ASYYWY-L----L----I-----S-------KL-------------PSI---
RKS-----------------------------RH--------------PLRIPKPHPVG-
------------------------------------LRRQIRRDIELISNSGQGN-----
------------------------------------------------SSSQQASP----
------------------------------------------------------------
-------------------------------------------SLLATGRPACTNN----
-------------------------------RAPLSSSSFANHHHHHQL-----------
-------------------------------------------------RLDGNSQETRP
LRDGMGGS
>Deutero_Cephalo_088140F_t1_XP_019621273_1_PREDICTED__acid_sensing_ion_channe
l_2_like_isoform_X1__Branchiostoma_belcheri_
------------------------------------------------------------
----------M-------------------------------------------------
------------------------------------------------------------
------------------------------------------------------------
------------------------------------------------------------
HVKLTCSVELDCC------------------------------------PCGQCCCGSST
AGSEPRDAGSEL------------------------------------------------
--------------GDGTDA-------------GEQTYY---------------------
------------------------------------------------------------
------------------------------------------------------------
------------------------------------------------------------
-------------------------------R---------KQ-----A-----------
-QT--ISDFA-AGSTLHGLPHIF---------P-DA-------------PLS---IR---
--QVAWA----LAFIGSLSVLLYQCSDRVK-FYFQ----------Y---PHITKL----D
-M-VL-A----------E-RLDF-----PAI-----------------------TICNMN
MF------RWK-----QFTQND------------------L-----------------W-
---------HM------------G-KGL---------------NILD----E--------
---------------------------------NN-------------------------
------------------------------------------------------------
------------------------------------------------------------
-------------NLRCS-E-YA-------------------------------------
------------------------------------------------------------
------------------------------------------------------------
------------------------------------------------------------
-------------------------TPGD-------------------------------
------------------------------------------------------------
------------------------------------------------------------
------------------------------------------------------------
------------------------------------------MEA---------------
------------------------LK--NKANFD----------SFKP--------SPF-
---------------------------------SIMEFA---------------------
------------------------------------------------NRT----GHQIE
--SFL------LDCK----WK-N----F----------T---C-----------G-P---
----------------------------------------------E-------------
------------------------------------------------------------

------------------------------------------------------------
-----------Y----------F-------TPYRAADVVDTVDVF-T-RYG--------K
CYTFN-------SG-------------SP-N--------------------Q----PV--
-------LKTLKG------GI--------GN-------------GVEFFLDVQQE-----
----------D-----------------Y------------M-P-AWGESD---------
------------------------------------------------------------
------------------------------------------------------------
-------EVTFEVGFKIQLH-TQEEPP--FIHELGFGVGPGMQYYVSTQEQ------RLD
LWTKTITYL-----------PAP-W--GQ-C--KAE-N------D-------LTF---DF
------------------A-KYTTSACRIDCETKFVV----SQ--CGCKMVH----M---
------------P-G---------------------------------------------
----------NF-----P---ICTP-----------------------------------
------------------------------------DIY---VECAD---------Q---
A-L-------------------DFLV-K-------SD--------N-----------KKC
------------VCD-TPCNTTR---YNLF-MSH------------------VKF-PS--
-------EQA------VKY-L-----------AR--------------------------
----------------------------------KYGKPEDYFRGVGLKY----------
------------------------------------------------------------
-G----------------------------------------------------------
---------------------------KPE------------------------------
-----D-------YF-----------R------KNTLVMNIFFEALNYETIE-QQKAYEV
ASL-----------------LGDIGGQMGLFIGASILTILELFDYLYEV-----------
--------LKDK--------C----------T-------ER-------------------
RH------------------------------QP--------------------------
------------------------------------------------------------
-------------------------RS---------------------ESNA--------
-----------VS-----------------------------------------------
----------------VNLEDCKRD------------------------NSRAPLS----
------------------------------------------------------------
------------------------------------------------------------
--------
>Deutero_Cephalo_XP_019621275_1_PREDICTED__acid_sensing_ion_channel_1_like_is
oform_X3__Branchiostoma_belcheri_
------------------------------------------------------------
----------M-------------------------------------------------
------------------------------------------------------------
------------------------------------------------------------
------------------------------------------------------------
------------------------------------------------------------
------------------------------------------------------------
------------------EG-------------GSQNHD---------------------
------------------------------------------------------------
------------------------------------------------------------
------------------------------------------------------------
-------------------------------V---------PR-----Q-----------
-SD--INVFA-ASASMHGLAHIF---------T-EG-------------KFT---VR---
--RILWA----GVFCGCVAVLLVQSVDRVQ-YYLS----------N---PHATKL----D
-E-IT-A---------VN-GLNF-----PAV-----------------------TICNMN
SF------RFS-----QVNQQD------------------L-----------------F-
---------YA------------GPDIL---------------DLID----H--------
---------------------------------TL-------PRPK--------------
------------------------------------------------------------
------------------------------------------------------------
-------------LKISE-E-FLN------------------------------------
------------------------------------------------------------
------------------------------------------------------------
------------------------------------------------------------

----------------HMNE-----NPSDH------------------------------
------------------------------------------------------------
------------------------------------------------------------
------------------------------------------------------------
------QEMRQK------------------------------MER---------------
------------------------IR--ELTNFD----------GFVP--------GKF-
---------------------------------SMSDFY---------------------
------------------------------------------------DRT----GHQIE
--DML------LDCK----YK-G----E----------P---C-----------S-A---
----------------------------------------------K-------------
------------------------------------------------------------
------------------------------------------------------------
-----------N----------F-------T-----------TVF-T-RYG--------K
CYTFN-------SG-------------SP-N--------------------Q----PV--
-------LKTLKG------GI--------GN-------------GVEFFLDVQQE-----
----------D-----------------Y------------M-P-AWGESD---------
------------------------------------------------------------
------------------------------------------------------------
-------EVTFEVGFKIQLH-TQEEPP--FIHELGFGVGPGMQYYVSTQEQ------R--
-----ITYL-----------PAP-W--GQ-C--KAE-N------D-------LTF---DF
------------------A-KYTTSACRIDCETKFVV----SQ--CGCKMVH----M---
------------P-G---------------------------------------------
----------NF-----P---ICTP-----------------------------------
------------------------------------DIY---VECAD---------Q---
A-L-------------------DFLV-K-------SD--------N-----------KKC
------------VCD-TPCNTTR---YNLF-MSH------------------VKF-PS--
-------EQA------VKY-L-----------AR--------------------------
------------------------------------------------KY----------
------------------------------------------------------------
-G----------------------------------------------------------
---------------------------KPE------------------------------
-----D-------YF-----------R------KNTLVLNVFFEALNYETIE-QQKAYEV
ASL-----------------LGDIGGQMGLFIGASILTILELFDYLYEV-----------
--------LKDK--------C----------T-------ER-------------------
RH------------------------------QP--------------------------
------------------------------------------------------------
-------------------------RS---------------------ESNA--------
-----------VS-----------------------------------------------
----------------VNLEDCKRD------------------------NSRAPLS----
------------------------------------------------------------
------------------------------------------------------------
--------
>Deutero_Cephalo_XP_019621276_1_PREDICTED__acid_sensing_ion_channel_1_like_isoform_X4__Branchiostoma_belcheri_
------------------------------------------------------------
----------M-------------------------------------------------
------------------------------------------------------------
------------------------------------------------------------
------------------------------------------------------------
------------------------------------------------------------
------------------------------------------------------------
------------------SS-------------YEEN-----------------------
------------------------------------------------------------
------------------------------------------------------------
------------------------------------------------------------
-------------------------------E---------PK-----P-----------
-SD--INVFA-GNATMHGISHIF---------T-EG-------------SFS---FR---

--RFLWA----AAFLICFGMVSYNGYDRVI-YYLS----------Y---PRVTKL----D
-E-LE-A----------Q-MMNF-----PAI-----------------------TMCNLN
PF------RFS-----KLTKDD------------------L-----------------W-
---------QV------------G-HAL---------------EILD----E--------
---------------------------------NK-------------------------
------------------------------------------------------------
------------------------------------------------------------
-------------KLAHS-E-FV-------------------------------------
------------------------------------------------------------
------------------------------------------------------------
------------------------------------------------------------
------------------------------------------------------------
------------------------------------------------------------
------------------------------------------------------------
------------------------------------------------------------
--------DPEH------------------------------LST---------------
------------------------LR--RLAEFGP---------GYVPKN-----KDSF-
---------------------------------NMKEFY---------------------
------------------------------------------------NRT----AHQIQ
--DMV------LECS----YR-G----T----------K---C-----------S-W---
----------------------------------------------A-------------
------------------------------------------------------------
------------------------------------------------------------
-----------N----------F-------T-----------VVF-T-RYG--------K
CYTFN-------SG-------------SP-N--------------------Q----PV--
-------LKTLKG------GI--------GN-------------GVEFFLDVQQE-----
----------D-----------------Y------------M-P-AWGESD---------
------------------------------------------------------------
------------------------------------------------------------
-------EVTFEVGFKIQLH-TQEEPP--FIHELGFGVGPGMQYYVSTQEQ------R--
-----ITYL-----------PAP-W--GQ-C--KAE-N------D-------LTF---DF
------------------A-KYTTSACRIDCETKFVV----SQ--CGCKMVH----M---
------------P-G---------------------------------------------
----------NF-----P---ICTP-----------------------------------
------------------------------------DIY---VECAD---------Q---
A-L-------------------DFLV-K-------SD--------N-----------KKC
------------VCD-TPCNTTR---YNLF-MSH------------------VKF-PS--
-------EQA------VKY-L-----------AR--------------------------
------------------------------------------------KY----------
------------------------------------------------------------
-G----------------------------------------------------------
---------------------------KPE------------------------------
-----D-------YF-----------R------KNTLVLNVFFEALNYETIE-QQKAYEV
ASL-----------------LGDIGGQMGLFIGASILTILELFDYLYEV-----------
--------LKDK--------C----------T-------ER-------------------
RH------------------------------QP--------------------------
------------------------------------------------------------
-------------------------RS---------------------ESNA--------
-----------VS-----------------------------------------------
----------------VNLEDCKRD------------------------NSRAPLS----
------------------------------------------------------------
------------------------------------------------------------
--------
>XP_019623542_1_PREDICTED__acid_sensing_ion_channel_2_like__Branchiostoma_bel
cheri_
------------------------------------------------------------
----------M-------------------------------------------------

```
--------------------------------AAWDPFSD--------------ELED-I
-IETVELRVR----------------------TDVTISFP-------ALTRARSTQGKS-
-----------CCRASPPP-----SSRSQPIPTRS---------------RVEVS-----
--TVAASIAMELA---NR-KDMGKED--------------------------------SF
TPLALAATPDVREVKPV-----------------Q----CVP------------------
----------TSTV-GTGYT-------------SYRK-----------------------
------------------------------------------------------------
------------------------------------------------------------
------------------------------------------------------------
------------------------------------------------P-----------
-SR--FRRFA-LDTDIHGMKHLI------------K-------------RGP---IR---
--RTAWV----LMLLLSLGLMSFECMESLI-YYFE----------F---HHVTKV----E
-L-QY-T----------N-NMTF-----PAV-----------------------TICNMN
KY------RRS-----ALGMRD------------------L-----------------A-
---------TV------------G-PYI---------------GIVD----R--------
---------------------------------QQ-------------------------
------------------------------------------------------------
------------------------------------------------------------
------------FTLQQE-H-LY-------------------------------------
------------------------------------------------------------
------------------------------------------------------------
------------------------------------------------------------
------------------------------------------------------------
------------------------------------------------------------
------------------------------------------------------------
------------------------------------------------------------
--------PKAW------------------------------VDK---------------
------------------------VKNENLKALS------------K-MT-----TGQQ-
---------------------------------NFAVFV---------------------
------------------------------------------------ERV----GHQKE
--DMI------VNCE----WR-G----S----------P---C-----------S-A---
----------------------------------------------K-------------
------------------------------------------------------------
------------------------------------------------------------
-----------N----------F-------T-----------HTF-T-HLG--------N
CYTFN-------GM-------------TL-S--------------------R----KV--
-------LSSSKP------GE--------GN-------------GLKVTINIEND-----
----------E-----------------Y------------M-A-T-NDVA---------
------------------------------------------------------------
------------------------------------------------------------
-------GDAMDAGIKVMVH-PQHEPP--FVKELGFAVSPGFHTFVAIRKE------V--
-----ITTL-----------PAP-Y--GN-C--QQR----G---G-------QK----YY
------------------P-DYSLSACRIECETEHVV----HE--CGCRLVE----M---
------------P-G--EH-----------------------------------------
-----------V-----P---VCTP-----------------------------------
-------------------------------------ER---YQCAY---------E---
K-LGIVDRQQFTLQQVENLKALSKMT-T-------GH--------Q--NFVERVGEDMIV
------------NCEGSPCSAKN---FTHT-FTHLGTINIENAGDAMDKETFVAI-RK--
-------EVI------TTL-P-----------APYGN--CQERGGQKY------YPDYSL
SACRIE----ETEHVVHCRLVEMPVEYVRE----DFSLLIHYTKTLNLDY----------
------------------------------------------------------------
-L----------------------------------------------------------
---------------------------FIS------------------------------
-----L-------FY-----------S------DNIAVLNVYYEALNYESIV-QIPAIET
EGL-----------------LGDLGGQMGLFLGASFLTLVEVFEYLMDE-----------
--------LYGR--------V----------L-------KK----------------CC-
RC------------------------------KA-------------------T------
```

-----------------------------------------ASAHT--------------
-----------------------T-SP---------------------SADV--------
-----------LS-----------------------------------------------
----------------VNINGSATQ------------------------LPQTSR-----
------------------------------------------------------------
------------------------------------------------------------
-------A
>Porifera_AmpQue_tr_A0A1X7TFM5_A0A1X7TFM5_AMPQE_Uncharacterized_protein_OS_Am
phimedon_queenslandica__Sponge__OX_400682_GN_Aqu2_1_13197_PE_3_SV_1_370_0
------------------------------------------------------------
------------------------------------------------------------
------------------------------------------------------------
------------------------------------------------------------
------------------------------------------------------------
------------------------------------------------------------
------------------------------------------------------------
------------------------------------------------------------
------------------------------------------------------------
------------------------------------------------------------
------------------------------------------------------------
------------------------------------------------------------
------------------------------------------------------------
------M----CFSQKKKEGCLVLIGLSIK-RFGD----------K---PTASTI----T
-V-VS-SH--------ES-GLPF-----PAV-----------------------TICNLN
LK------KNDS--D-FLLNTT------------------Y-------------------
---------QA------------M------------------NFLYN-------------
------------------------------P--DK-------S-----------------
------------------------------------------------------------
------------------------------------------------------------
-----------YQFNS-TKKK---------------------------------------
------------------------------------------------------------
------------------------------------------------------------
------------------------------------------------------------
------------------------------------------------------------
------------------------------------------------------------
------------------------------------------------------------
------------------------------------------------------------
-----NHLLNTC------------------------------TAPH--------------
-------------------------------------------------------SNSIQ
NA-------------------------------TIWDIV---------------------
-----------------------------------------------------NPDRAVN
--ELI------QYCG----FLHGA-DSD--------VVL---C-----------------
----------------------------------------------K-------------
------------------------------------------------------------
------------------------------------------------------------
-----------DL---------F-------E-----------PVL-T-SAG--------I
CFTFN-------GT-------------NK-------------------------------
-------L-ANST------GI--------RY-------------GLKLVLDVQKK-----
----------E-----------------R--------------P-S---FN---------
------------------------------------------------------------
------------------------------------------------------------
----------GKLGVKLVVH-DGRDIA--RPNLYGLTVPPGIAVDVGVRKV------I--
-----TRDE-----------TNE----AK-C--IDG---M----N-------LP----FF
----PS---DK-------F-EYSQFACRANAVAENIA----KRSKCNCVVQP----D---
-----------RP-P---G-----------------------------------------
-------FYTST-----P---NCTF-----------------------------------
------------------------------------SKA----CCLL---------Q---

E-Q-------------------YTFH-P-----------------------------LKK
------------STV-L-CHATFNFLMDNT-CNY--------------------------
------------------------------------------------------------
------------------------------------------------------------
------------------------------------------------------------
------------------------------------------------------------
------------------------------------------------------------
------------------------------------------------------------
------------------------------------------------------------
------------------------------------------------------------
------------------------------------------------------------
------------------------------------------------------------
------------------------------------------------------------
------------------------------------------------------------
------------------------------------------------------------
------------------------------------------------------------
------------------------------------------------------------
--------
>Porifera_AqNaC9_Aqu2_1_25935_001_PorifAmpQue_89_89
------------------------------------------------------------
----------M-------------------------------------------------
------------------------------------------------------------
------------------------------------------------------------
------------------------------------------------------------
------------------------------------------------------------
------------------------------------------------------------
------------------------------------------------------------
------------------------------------------------------------
------------------------------------------------------------
------------------------TKKEF-------------------------------
-------------------------------K---------WS-----D-----------
-QY--FNDFV-ETTTINGVFHIF---------R-GR--------------SK---TR---
--RLLWG----LLFLISFVSCTIVLGFSFK-RYSE----------K---PTVSAI----N
-V-IS-E---------EN-GMPF-----PSV-----------------------TVCNRN
FY------KYP-----NISNET------------------------------------D-
---------TL------------I------------------HHLFH-------------
------------------------------S--SG-------F-----------------
------------------------------------------------------------
------------------------------------------------------------
------------------------------------------------------------
------------------------------------------------------------
------------------------------------------------------------
------------------------------------------------------------
------------------------------------------------------------
------------------------------------------------------------
------------------------------------------------------------
------------------------------------------------------------
--LQNINRTQEC------------------------------------------------
--------------------------------------------KIDED------TPDF-
---------------------------------KLQDLL---------------------
-------------------------------------------------------IPKEN
--NFI------YYCA----LSYQA-ESE--------VML---C-----------------
----------------------------------------------K-------------
------------------------------------------------------------
------------------------------------------------------------
-----------DM---------F-------F-----------PTL-T-PAG--------I
CYTFN-------GV-------------RS-K--------------------T----RA--

-------LNMTTT------GK--------RY-------------GLRLILNIEQE-----
----------T-----------------H------------L---A---FD---------
------------------------------------------------------------
------------------------------------------------------------
----------GVAGVQVIVH-ESNDIP--RPNLHGIGVPPGQNVDIGVKRA------I--
-----SSDE-----------TDQ----DA-C--IEN-NEK----D-------LP----FL
----PG------------V-VYSQYACRQNELYESLA----DKNLCNCIIDP----F---
-----------EL-----------------------------------------------
----------DN-----A---TCFL-----------------------------------
------------------------------------HDL----CCLQ---------Q---
Q-F-------------------TEHD-------------------------------RNS
------------SCR-PPCKYTY---FDII-NSY------------------SSF-PE--
-------GHA------LTE-I-----------KN--------------------------
------------------------------------------------SI----------
------------------------------------------------------------
-N----------------------------------------------------------
---------------------------TSD------------------------------
-----D-------YV-----------K------KNFLSVSVFLQALETRETT-TRYSYGV
VEL-----------------LGELGGNLGLFLGISIISVMELLVLVVDE-----------
--------LKKC--------V----------------CPKKV----------------K-
QKF-----------------------------KK--------------------------
----------------------------------------FDEKLK--------------
--------------------CIPDCVE---------------------QEEK--------
------------------------------------------------------------
------------------------------------------------------------
------------------------------------------------------------
-----------------------------------------------------NEIQALT
NF------
>Porifera_AqNaC6_Aqu2_1_26214_001_PorifAmpQue_86_93
------------------------------------------------------------
----------M-------------------------------------------------
------------------------------------------------------------
------------------------------------------------------------
------------------------------------------------------------
------------------------------------------------------------
------------------------------------------------------------
------------------------------------------------------------
------------------------------------------------------------
------------------------------------------------------------
-----------------------KNAGRF-------------------------------
-------------------------------S---------WS-----D-----------
-KY--FSDFV-ETTTINGVIHVF---------R-GR--------------SK---IR---
--QIMWG----LLLISSFIAVCVVIGFNIK-EYVN----------K---PTASTI----L
-V-SP-SV--------KN-GLPF-----PAV-----------------------TICNLN
VY------TPT-----GEDKEP-----------------DF-----------------S-
---------SV------------I------------------HSLFN-------------
------------------------------S--QD-------L-----------------
------------------------------------------------------------
------------------------------------------------------------
------------------------------------------------------------
------------------------------------------------------------
------------------------------------------------------------
------------------------------------------------------------
------------------------------------------------------------
------------------------------------------------------------
------------------------------------------------------------
------------------------------------------------------------

--VNQTSLFDEC------------------------------SE----------------
--------------------------------------------VINSSD-----TSDAF
DC-------------------------------EVWDLL---------------------
-------------------------------------------------------LPDDK
--RFI------YQCS----FSHDA-DSE--------IVS---C-----------------
----------------------------------------------R-------------
------------------------------------------------------------
------------------------------------------------------------
-----------DM---------F-------Y-----------PVL-T-PAG--------I
CYTFN-------GI-------------RS-K--------------------M----PV--
-------PMAKDI------GV--------RH-------------GLNLILNIEQD-----
----------S-----------------H--------------P-T---FH---------
------------------------------------------------------------
------------------------------------------------------------
----------GLTGVKVIVH-DRNDIS--RPNLYGISVAPGQNVAISVERK------V--
-----YIDK-----------TKE----RD-C--TND--ER----E-------LG----FF
----PS------------T-VYSQFACKENALYQHLA----DESVCGCVPNP----Y---
-----------RP-S--TG-----------------------------------------
-------PYINT-----P---NCTL-----------------------------------
------------------------------------HSL----CCLL---------Q---
E-Y-------------------FDYR-------------------V-----------SSA
------------TCP-LPCQFSM---YDYK-SSY------------------SSF-PN--
-------GHA------LKS-I-----------AK--------------------------
------------------------------------------------SL----------
------------------------------------------------------------
-N----------------------------------------------------------
---------------------------LSQ------------------------------
-----D-------AV-----------K------NNFLSVQVYLESLETHEYV-TKYSKTL
TGL-----------------FGDIGGLIGLFLGMSIISIIEVLVLILDE-----------
--------LKKL--------L----------------CIKKF----------------R-
KKV-----------------------------KK--------------------------
----------------------------------------VDDMLT--------------
------------H-------FLPDVK----------------------------------
------------------------------------------------------------
------------------------------------------------------------
------------------------------------------------------------
------------------------------------------------------------
--------
>Porifera_AqNaC11_Aqu2_1_26219_001_PorifAmpQue_91_101
------------------------------------------------------------
----------M-------------------------------------------------
------------------------------------------------------------
------------------------------------------------------------
------------------------------------------------------------
------------------------------------------------------------
------------------------------------------------------------
------------------------------------------------------------
------------------------------------------------------------
------------------------------------------------------------
------------------------TKKGF-------------------------------
-------------------------------K---------WS-----D-----------
-QY--FNDFV-ETTTINGVFHIF---------R-GR--------------TK---TR---
--RLLWG----LLFFISFVSCTIVLGFSFK-RYSE----------K---PTVSAI----N
-V-IL-G---------EN-GIPF-----PSV-----------------------TVCNQN
FY------KDL-----NVSNET------------------------------------N-
---------AL------------I------------------HHLFH-------------
------------------------------S--GG-------F-----------------

------------------------------------------------------------
------------------------------------------------------------
------------------------------------------------------------
------------------------------------------------------------
------------------------------------------------------------
------------------------------------------------------------
------------------------------------------------------------
------------------------------------------------------------
------------------------------------------------------------
------------------------------------------------------------
--LHDINRTQQC------------------------------------------------
--------------------------------------------KIDED------RHDF-
---------------------------------ELQDLL---------------------
-------------------------------------------------------IPKEN
--NFI------HYCA----FSHQA-ESE--------IML---C-----------------
----------------------------------------------K-------------
------------------------------------------------------------
------------------------------------------------------------
-----------DK---------F-------F-----------PTL-T-PAG--------I
CYTFN-------GV-------------RS-K--------------------T----RA--
-------PKMTTT------GK--------RY-------------GLRLILNIEQE-----
----------T-----------------H--------------P-V---FD---------
------------------------------------------------------------
------------------------------------------------------------
----------GVAGVQVIVH-ESNDIP--RPNLHGVGVSPGQNVDIGVKRA------I--
-----MFDE-----------TDQ----DE-C--IKD-NGK----D-------LP----FL
----PG------------V-VYSQYACRQNELYESLA----DKNLCNCIIDP----F---
-----------EL-----------------------------------------------
----------GN-----A---TCSS-----------------------------------
------------------------------------HDL----CCLQ---------Q---
Q-F-------------------TEHD-------------------------------RNS
------------SCR-PPCKYTY---YDIM-NSY------------------SSF-PE--
-------GHA------LTE-I-----------KN--------------------------
------------------------------------------------SI----------
------------------------------------------------------------
-N----------------------------------------------------------
---------------------------ASE------------------------------
-----D-------YI-----------K------KNFLSASVFLQALETRETT-TRYSYGV
VEL-----------------LGELGGNLGLFLGISIISVMELLVLVIDE-----------
--------VKKC--------V----------------CPKKV----------------K-
QKF-----------------------------EK--------------------------
----------------------------------------FDDKLK--------------
--------------------CIPDCVE---------------------QEEK--------
------------------------------------------------------------
------------------------------------------------------------
------------------------------------------------------------
-----------------------------------------------------DENTSGP
SAELNKIA
>Porifera_AqNaC2_Aqu2_1_09805_001_PorifAmpQue_82_141
------------------------------------------------------------
----------M-------------------------------------------------
------------------------------------------------------------
------------------------------------------------------------
------------------------------------------------------------
--AEDS------------------------------------------------------
------------------------------------------------------------
------------------------------------------------------------

```
------------------------------------------------------------
-------------------------------------------------------HSKSS
E----------------------EKTVKC-------------------------------
-------------------------------H---------LH-----D-----------
-RY--LKEFL-DDNTIGGINHIF---------R-GR--------------SK---VR---
--RLLWA----LIFIGSIVACITLISISFQ-TFLE----------K---PTASTI----T
-V-IT-QD--------DE-GVSF-----PSV-----------------------TICNLN
LE------RNES--D-MVADTG------------------Y-------------------
---------LL------------M------------------NHIFN-------------
------------------------------P--DE-------N-----------------
------------------------------------------------------------
------------------------------------------------------------
-----------FHLTG--------------------------------------------
------------------------------------------------------------
------------------------------------------------------------
------------------------------------------------------------
------------------------------------------------------------
------------------------------------------------------------
------------------------------------------------------------
------------------------------------------------------------
--LNSSFLLNSC------------------------------NAIS--------------
----------------------------------------------DSF------PASFR
NT-------------------------------TLWNSQ---------------------
-----------------------------------------------------HPQETLD
--KLI------HFCG----FVSGI-NSA--------VIP---C-----------------
----------------------------------------------K-------------
------------------------------------------------------------
------------------------------------------------------------
-----------DA---------F-------K-----------PVL-T-SAG--------I
CYTFN-------GS-------------NN-------------------------------
--------RIHST------GV--------RY-------------GLKLILNIQQE-----
----------E-----------------R--------------P-S---FN---------
------------------------------------------------------------
------------------------------------------------------------
----------GKSGVKLIVH-DGRDIA--RPNLYGIDVAPAHAVDVGVRRK------A--
-----SKDE-----------TNE----AD-C--IDS---K----E-------LP----FF
---------SE-------Y-RYSQFACRQNAIVENLA----TS--CDCSIHP----D---
-----------RP-S--SG-----------------------------------------
-------PYSST-----P---HCTF-----------------------------------
------------------------------------DKG----CCVL---------E---
Q-Y-------------------QTFN-P------------------------------EL
------------ACP-LPCYFPY---YEHT-ASY------------------SSF-PN--
-------GRY------LNY-L-----------VE--------------------------
------------------------------------------------ET----------
------------------------------------------------------------
-N----------------------------------------------------------
---------------------------MSV------------------------------
-----D-------YI-----------K------DNFLSINVFVDDLQLTTTI-TKYTFGV
AEL-----------------LGEIGGQMGLFLGISIISIVEVVVLFLDE-----------
--------LKRL--------F----------------CTKKM----------------R-
EKM-----------------------------QD--------------------------
----------------------------------------IENAIE--------------
---------------------LPEIEG---------------------TDVE--------
------------------------------------------------------------
------------------------------------------------------------
------------------------------------------------------------
------------------------------------------------------------
```

-EDIDNKV
>Porifera_AqNaC7_Aqu2_1_26220_001_PorifAmpQue_87_142
------------------------------------------------------------
----------M-------------------------------------------------
------------------------------------------------------------
------------------------------------------------------------
------------------------------------------------------------
------------------------------------------------------------
------------------------------------------------------------
------------------------------------------------------------
------------------------------------------------------------
------------------------------------------------------------
-----------------------QPAKGY-------------------------------
-------------------------------T---------LT-----D-----------
-QY--FHDFV-ETTRISGIKHIF---------R-GR--------------SK---IR---
--RIMWA----LFFISSFVGCTLVVGKNIA-RYIE----------K---PTASSI----K
-V-IP-H---------DN-GMRF-----PSV-----------------------TICNIN
IY------KDPNV-S-AVSKET------------------Y-------------------
---------SL------------I------------------YYLFN-------------
------------------------------S--DT-------------------------
------------------------------------------------------------
------------------------------------------------------------
------------------------------------------------------------
------------------------------------------------------------
------------------------------------------------------------
------------------------------------------------------------
------------------------------------------------------------
------------------------------------------------------------
------------------------------------------------------------
------------------------------------------------------------
---KEYNSTQEC------------------------------------------------
---------------------------------------------IDNA------TEYYK
KH-------------------------------DIWSSL---------------------
-------------------------------------------------------LPKEN
--NFI------HDCS----FSYET---G--------VIS---C-----------------
----------------------------------------------K-------------
------------------------------------------------------------
------------------------------------------------------------
-----------DM---------F-------Y-----------PVL-T-PAG--------I
CYTFN-------DF-------------KM-N--------------------E----ML--
-------IPNITS------GM--------KY-------------GLKLVLNIEEE-----
----------R-----------------Y--------------P-A---FE---------
------------------------------------------------------------
------------------------------------------------------------
----------GKTGAQIIVH-ERNDIP--RPNLAGISVPPGQNIDIGFIKA------I--
-----KNDK-----------TDS----RD-C--IND-GEK----K-------LD----FL
----PD------------V-VYSKFACQENELYERLA----SN--CNCTINP----Y---
-----------GL-----------------------------------------------
-------DVTNT-----S---NCSL-----------------------------------
------------------------------------SNL----CCMR---------Q---
Q-Y-------------------SKYN-------------------------------RNS
------------SCK-SPCKYTF---YNLK-NSY------------------SSF-PR--
-------GRT------LTE-I-----------SK--------------------------
------------------------------------------------TV----------
------------------------------------------------------------
-K----------------------------------------------------------
---------------------------MNK------------------------------

-----S-------AI-----------R------DNFLSVHVFLLDLETKETY-SHNSFDI
SEL-----------------LGELGGTMGLFLGINILAIVEVIILILDE-----------
--------IKKY--------L----------------CPKKC----------------K-
QKL-----------------------------NK--------------------------
----------------------------------------IEN-----------------
--------------------CIPECAL---------------------SRGR--------
--------NDTLL-----------------------------------------------
------------------------------------------------------------
-------------------------------TDPVS------------------------
-------------------------------------------------TPEKENHETPY
TTAVDIDN
>Porifera_AqNaC1_Aqu2_1_34376_001_PorifAmpQue_81_145
------------------------------------------------------------
----------M-------------------------------------------------
------------------------------------------------------------
------------------------------------------------------------
------------------------------------------------------------
------------------------------------------------------------
-----------------------------------------PSK----------------
------------------------------------------------------------
------------------------------------------------------------
---------------------------------------------------------DKG
E----------------------NDPQENTGV----------------------------
---RNRKEK----------------------S---------CQ-----N-----------
-PQ--FTKWA-ESSTIHGVDHIF---------L-GK--------------SK---VR---
--RVVWA----VILLLAIGGCLYGIIDRSI-YFAS----------K---PTATTV----T
-A-DINE----------D-GIPF-----PAV-----------------------TICNLS
PI------SRQ-----YADQHN------------------L-----------------T-
---------SLLS-----------------------------YILFT-------------
------------------------------------------------------------
------------------------------------------------------------
------------------------------------------------------------
-----------------D------------------------------------------
------------------------------------------------------------
------------------------------------------------------------
------------------------------------------------------------
------------------------------------------------------------
------------------------------------------------------------
------------------------------------------------------------
------------------------------------------------------------
------------------------------------------SNTH--------------
-------------------------------------------------------SKGFI
SSNCQA---------------------------NLEQIT---------------------
-----------------------DTTI----------------TLKDVFRD----GARNS
--SFI------LACH----YG-SA-RNK---------MD---C-----------A-N---
----------------------------------------------M-------------
------------------------------------------------------------
------------------------------------------------------------
-----------T----------L-------R-----------TTL-T-PRG--------L
CYTFN-------GD-------------PA-S-------------------------PP--
-------LLVRSV------GE--------RF-------------GLRMIFNISQS-----
----------D-----------------Y------------T-H-S---IN---------
------------------------------------------------------------
------------------------------------------------------------
----------GDAGIRVSVH-TRDEKP--DPLLKGISVPPRSHASIALYPI------RS-
----I---------------SKPEI--TR-CAPTDT--------Q-----------LSYF
----PGL-------------SYTTSGCQANEHFERSQ----AQ--CGCVDVA----E---

------------S-----------------------------------------------
----------AT-----N---DCTV-----------------------------------
------------------------------------EDI----CCLY---------D---
----------------------EGTS-T-------DT--------L------------NS
------------TCL-PSCNNMI---FSSS-VSY------------------SQY-PS--
-------DVT------VSL-L-----------TS--------------------------
------------------------------------------------------------
-----------------------------------------------------------F
QS----------------------------------------------------------
---------------------------QSA------------------------------
-----E-------SI-----------D------DNILALNIYFGSLHTIVTS-TYYTYLW
SGL-----------------LADIGGQLALFVGASVISFMELVLLCFDE-----------
--------TKCS------GVFI---------R-------KKI------------------
KK------------------------------KH-------------------EEHEL--
---------------------------------------EERDDKESL------------
------------------NGGKIEDKQ---------------------------------
------------------------------------------------------------
------------------------------------------------------------
------------------------------------------------------------
------------------------------------------------------------
---ANTNV
>Porifera_AqNaC10_Aqu2_1_36244_001_PorifAmpQue_90_167
------------------------------------------------------------
----------M-------------------------------------------------
------------------------------------------------------------
------------------------------------------------------------
------------------------------------------------------------
------------------------------------------------------------
------------------------------------DEGNDPV-----------------
------------------------------------------------------------
------------------------------------------------------------
-------------------------------------------------------NGEKS
K----------------------CKNCTC-------------------------------
-------------------------------T---------CT-----D-----------
-RY--LREFL-EDNTIAGISHVF---------K-GQ--------------SK---VR---
--RSFWA----LIFIAAIISCIALISLSIQ-IFLN----------K---PTASTI----N
-V-IT-L----------N-GVPF-----PSV-----------------------TICNIN
FE------KNEA--L-PLLSRT------------------Y-------------------
---------NL------------M------------------NQLFN-------------
------------------------------A--DE-------S-----------------
------------------------------------------------------------
------------------------------------------------------------
-----------FHSNS--------------------------------------------
------------------------------------------------------------
------------------------------------------------------------
------------------------------------------------------------
------------------------------------------------------------
------------------------------------------------------------
------------------------------------------------------------
------------------------------------------------------------
--MNASSVISNC------------------------------RNTVP-------------
----------------------------------------------RNN------SDATF
EA-------------------------------TIWNIQ---------------------
-----------------------------------------------------RPARSKR
--RLI------HYCG----FVTGV-NSN--------VSN---C-----------------
----------------------------------------------K-------------
------------------------------------------------------------

------------------------------------------------------------
-----------SL---------F-------Q-----------PVL-T-PAG--------I
CFNYN-------GS-------------SN-------------------------------
--------TIHST------GT--------RY-------------GMKLVLNIEQD-----
----------L-----------------R--------------P-S---YN---------
------------------------------------------------------------
------------------------------------------------------------
----------GKAGVILSVH-DGKDVA--RPNVNGINVAPGQAIDVGVNLK------E--
-----YIDE-----------TKE----AN-C--TAG---Q----D-------LV----FF
----ED------------Y-DYSQYACAQDALIKQIA----KPNVCNCTLLP----R---
-----------RP-S--NG-----------------------------------------
-------EYDHT-----P---NCTF-----------------------------------
------------------------------------PTS----CCLL---------N---
E-Y-------------------KTIN-------------------------------TEE
------------QCP-LPCRYRY---YDYT-TSY------------------ASY-PN--
-------SFI------LED-L-----------MR--------------------------
------------------------------------------------DE----------
------------------------------------------------------------
-N----------------------------------------------------------
---------------------------VTE------------------------------
-----D-------YL-----------R------KNYLSINIYINDLRYSVVT-TSYTFGV
AGL-----------------LGDIGGQLGLFIGVSIITFFEVLILCLDE-----------
--------LKRI--------C----------------CPDFV----------------I-
NKC-----------------------------KN--------------------------
----------------------------------------MKKKGE--------------
---------------------SSDIVA---------------------GERE--------
------------------------------------------------------------
------------------------------------------------------------
------------------------------------------------------------
-----------------------------------------------------TRRQIQR
KAWSNDSL
>Porifera_AqNaC13_Aqu2_1_13200_001_PorifAmpQue_93_173
------------------------------------------------------------
----------M-------------------------------------------------
------------------------------------------------------------
------------------------------------------------------------
------------------------------------------------------------
------------------------------------------------------------
------------------------------------------------------------
------------------------------------------------------------
------------------------------------------------------------
------------------------------------------------------------
-----------------------CSDGAG-------------------------------
-------------------------------H---------AY-----D-----------
-KV--SKYFK-KKITIAGLSHVF---------P-ERTKKDDRTRSGLTNPEK---II---
--MVIWA----LFFTGCVLGCLVLIGLSTK-RFVD----------K---PTASTI----T
-V-VS-HD--------KT-GLPF-----PAV-----------------------TICNLN
LK------KNDS--D-FLLNTT------------------Y-------------------
---------QA------------I------------------ISLYN-------------
------------------------------A--DK-------S-----------------
------------------------------------------------------------
------------------------------------------------------------
-----------SQFTS-NKKI---------------------------------------
------------------------------------------------------------
------------------------------------------------------------
------------------------------------------------------------
------------------------------------------------------------

------------------------------------------------------------
------------------------------------------------------------
------------------------------------------------------------
-----DHLLNTC------------------------------TASL--------------
-------------------------------------------------------SNSIQ
NA-------------------------------TIWDIV---------------------
-----------------------------------------------------SPEMAVN
--EFI------HYCG----FLHGA-DSD--------VVM---C-----------------
----------------------------------------------K-------------
------------------------------------------------------------
------------------------------------------------------------
-----------DL---------F-------E-----------PVL-T-SAG--------I
CFTFN-------GT-------------NK-------------------------------
--------LANST------GR--------RY-------------GLKLVLNIQQE-----
----------E-----------------R--------------P-S---FS---------
------------------------------------------------------------
------------------------------------------------------------
----------GKLGVKLVIH-DGKDIA--RPSLYGISVPPGFAVDVGVRKM------A--
-----TRDE-----------TSE----AK-C--IDD---M----N-------LP----FF
----PS---DK-------F-QYSQFACRENAVAENIA----RGSKCNCVIQP----D---
-----------RS----PG-----------------------------------------
-------LYAST-----P---NCTF-----------------------------------
------------------------------------SKA----CCLL---------Q---
E-H-------------------YRFH-P-----------------------------EEI
------------DCP-LPCHFEY---YEHT-ASY------------------SSF-PN--
-------GQY------LQL-L-----------ME--------------------------
------------------------------------------------EL----------
------------------------------------------------------------
-N----------------------------------------------------------
---------------------------MSA------------------------------
-----E-------YV-----------K------DNFLSIDVFFDDFQVTTTT-TKYTYGI
EAL-----------------LGEIGGLLGLFIGVNIINFFELLVLSGDG-----------
--------LGML--------C----------RRAGRSCRRALEKI-------------K-
KRK-----------------------------NE--------------------------
---------------------------------------RKEEPMK--------------
---------------------MDTQPG---------------------SSRS--------
--------GNGH------------------------------------------------
------------------------------------------------------------
------------------------------------------------------------
------------------------------------------------------------
-------A
>Porifera_AqNaC8_Aqu2_1_34361_001_PorifAmpQue_88_177
------------------------------------------------------------
----------M-------------------------------------------------
------------------------------------------------------------
------------------------------------------------------------
------------------------------------------------------------
--AAEKKDVEM-------------------------------------------------
-----------------------------------ASEKEDMQ-----------------
------------------------------------------------------------
------------------------------------------------------------
-------------------------------------------------------MMETK
K----------------------KKGRGC-------------------------------
-------------------------------H---------LN-----D-----------
-PY--LKEFL-DDNTIAGMNHIF---------K-GQ--------------SK---IK---
--RLVWA----LIFIGSMIACITLISISFR-RFIN----------K---PTSSTI----T
-V-VT-EN-------TKS-GVEF-----PAV-----------------------TICNHN

```
LE------YNLS--D-YIIRNT------------------Y-------------------
---------LL------------M------------------NYLFN-------------
------------------------------A--DE-------N-----------------
------------------------------------------------------------
------------------------------------------------------------
-----------FHLTG--------------------------------------------
------------------------------------------------------------
------------------------------------------------------------
------------------------------------------------------------
------------------------------------------------------------
------------------------------------------------------------
------------------------------------------------------------
------------------------------------------------------------
--SNSSSVIKQC------------------------------QALV--------------
----------------------------------------------GDA------PSEIL
NA-------------------------------TIWNIQ---------------------
-----------------------------------------------------NPSKAID
--ELI------HYCG----FIEGV-NGE--------VEP---C-----------------
----------------------------------------------K-------------
------------------------------------------------------------
------------------------------------------------------------
-----------DA---------F-------K-----------PVL-T-SAG--------I
CFTLN-------GS-------------DH-------------------------------
--------GIHST------GI--------RY-------------GLKLVLNVQQE-----
----------K-----------------R--------------P-S---FN---------
------------------------------------------------------------
------------------------------------------------------------
----------GKSGIKLVIH-NGGDIA--RPNLYGISVPPGRAIDVGVRRK------A--
-----TKDE-----------TSE----AG-C--IDD---M----D-------LP----FF
----PK---DK-------F-DYSQFACRENAIMERIA----KRSSCNCVIHP----D---
-----------RP-S--TG-----------------------------------------
-------AYSST-----P---NCTM-----------------------------------
------------------------------------SNA----CCLL---------H---
E-Y-------------------NTFH-P-----------------------------ESS
----------SEECP-LPCYFSY---YEWT-ASY------------------SSF-PN--
-------GRY------LDR-L-----------VN--------------------------
------------------------------------------------TL----------
------------------------------------------------------------
-N----------------------------------------------------------
---------------------------KSA------------------------------
-----D-------YI-----------K------NNFLSVNVFLDDIQLTTTI-TQYTYGP
EAL-----------------LGEIGGQLGLFIGVSIITFFEVLVLCVDE-----------
--------LKRL--------C----------------FKLQII--------------PE-
EKI-----------------------------NN--------------------------
----------------------------------------LESRIT--------------
---------------------LPEVEE---------------------DQTE--------
------------------------------------------------------------
------------------------------------------------------------
------------------------------------------------------------
------------------------------------------------------------
-------N
>Porifera_AqNaC4_Aqu2_1_26218_001_PorifAmpQue_84_192
------------------------------------------------------------
----------M-------------------------------------------------
------------------------------------------------------------
------------------------------------------------------------
------------------------------------------------------------
```

```
------------------------------------------------------------
-----------------------------------------PT-----------------
------------------------------------------------------------
------------------------------------------------------------
------------------------------------------------------------
-----------------------LVRNAH-------------------------------
-------------------------------V---------WT-----D-----------
-QY--FSDFI-DTTTINGVFQIF---------R-GR--------------SK---IR---
--QIFWG----LLFIGSFIGCIVTFGYSFR-NFAQ----------K---PTASTI----K
-V-IT-QA--------QT-GLAF-----PAV-----------------------TICNLN
IY------QNPNYSN-VLSSEM------------------Y-------------------
---------AL------------I------------------QYLFE-------------
------------------------------T--DD-------I-----------------
------------------------------------------------------------
------------------------------------------------------------
------------------------------------------------------------
------------------------------------------------------------
------------------------------------------------------------
------------------------------------------------------------
------------------------------------------------------------
------------------------------------------------------------
------------------------------------------------------------
------------------------------------------------------------
--FNEFNITSEC------------------------------KD----------------
--------------------------------------------LIDNA------SEDYG
KE-------------------------------SLYDML---------------------
-----------------------------------------------------LPKDYSS
--NLI------YDCT----FRDDA-LGD--------AMS---C-----------------
----------------------------------------------K-------------
------------------------------------------------------------
------------------------------------------------------------
-----------DQ---------F-------Y-----------PVL-T-PGG--------I
CYTFN-------GV-------------RS-E--------------------M----IA--
-------PVIKSI------GI--------KY-------------GLKLVLNIEQE-----
----------T-----------------H--------------P-T---FD---------
------------------------------------------------------------
------------------------------------------------------------
----------GRTGVKVIIH-ERNDIP--RPNLYGINVSPGQNIDIGVSRS------F--
-----FIDE-----------TDQ----DK-C--NND-IEG----N-------FP----FL
----PN------------I-VYSQFACRINQLYERLS----QQRNCGCLPIP----Y---
-----------RP-E--SG-----------------------------------------
-------PYTNT-----P---NCTL-----------------------------------
------------------------------------GNL----CCLL---------R---
E-F-------------------PIAD-------------------------------ASS
------------TCQ-LPCNYSV---YEYR-DSY------------------SSF-PN--
-------GRA------LTE-I-----------AR--------------------------
------------------------------------------------KV----------
------------------------------------------------------------
-N----------------------------------------------------------
---------------------------MSK------------------------------
-----T-------DV-----------K------ENFLSVNVFFEALHTTESI-TQYTYGA
VDL-----------------LGELGGNMGLFLGISIISIMEVIMLILDE-----------
--------IKHL-------------------------CPKKV----------------K-
KKF-----------------------------DN--------------------------
----------------------------------------IDDKLR--------------
------------N-------YIPDIAP---------------------SQTD--------
--------TPNALEAGTEEVHV--------------------------------------
```

------------------------------------------------------------
-------------------------------EDPS-------------------------
----------------------------------------------------PSEADIKS
NAEAIEES
>Porifera_AqNaC12_Aqu2_1_34365_001_PorifAmpQue_92_215
------------------------------------------------------------
----------M-------------------------------------------------
------------------------------------------------------------
------------------------------------------------------------
------------------------------------------------------------
------------------------------------------------------------
----------------------------------------NPA-----------------
------------------------------------------------------------
------------------------------------------------------------
----------------------------------------------------DETDAEKV
S---------------------GSRCKSC-------------------------------
-------------------------------Q---------LK-----D-----------
-SY--FNDL--DKTTIAGLNHVF--------MK-DS--------------SK---IR---
--RLIWA----LFFIGCILGCLVLIGLSIN-RFVD----------K---PTASTI----T
-V-VS-ND--------ET-GIFF-----PAV-----------------------TICNLN
LK------KNDS--D-VLLNTT------------------Y-------------------
---------QV------------M------------------NFLYN-------------
------------------------------S--DE-------S-----------------
------------------------------------------------------------
------------------------------------------------------------
-----------FQFTG--------------------------------------------
------------------------------------------------------------
------------------------------------------------------------
------------------------------------------------------------
------------------------------------------------------------
------------------------------------------------------------
------------------------------------------------------------
------------------------------------------------------------
--SNTMSLGNTC------------------------------TAPL--------------
-------------------------------------------------------SNSIQ
NA-------------------------------TIWDIV---------------------
-----------------------------------------------------NPDRAVD
--ELI------HYCG----FLHGA-DSD--------VVM---C-----------------
----------------------------------------------E-------------
------------------------------------------------------------
------------------------------------------------------------
-----------DL---------F-------E-----------PVL-T-SAG--------I
CYTFN-------GT-------------NK-------------------------------
--------LANST------GI--------RY-------------GLKLILNIQQK-----
----------E-----------------R--------------P-S---FN---------
------------------------------------------------------------
------------------------------------------------------------
----------GKSGVKLVIH-DGRDIS--RPNLYGISVPPGHAIDVGVHKM------A--
-----TQDE-----------TSQ----AN-C--IKS---M----N-------LP----FF
----PS---DK-------F-DYSQFACRANAVAENIA----RRSKCNCVIQP----D---
-----------RP----PG-----------------------------------------
-------LYAST-----P---NCTF-----------------------------------
------------------------------------GKA----CCLL---------Q---
E-H-------------------YKFN-P-----------------------------EEI
------------NCP-LPCHFQY---YEHT-ASY------------------SSF-PN--
-------GQY------LQR-L-----------ME--------------------------
------------------------------------------------AS----------

------------------------------------------------------------
-N----------------------------------------------------------
---------------------------MSA------------------------------
-----E-------DI-----------K------DNFLSINVFIDDFQVTTTT-TQYTYGI
EAL-----------------LGEIGGLLGLFIGVSIITFFELLVLCVDE-----------
--------LKRL--------C----------------CSQAI----------------I-
KRM-----------------------------KR--------------------------
----------------------------------------IEETAL--------------
---------------------VPVVES---------------------AEGS--------
--------LSNEEDAPEE-------------------------VTSQPEAES--------
-----------NKTSPSSRDA---------------------------------------
------------------------------------------------------------
---------------------------------------------------NSDSNEGKC
IKIEEIKL
>Porifera_AqNaC3_Aqu2_1_34363_001_PorifAmpQue_83_251
------------------------------------------------------------
----------M-------------------------------------------------
------------------------------------------------------------
------------------------------------------------------------
------------------------------------------------------------
--NGKDKAKEP-------------------------------------------------
---------------------------------------LKPA-----------------
------------------------------------------------------------
------------------------------------------------------------
-------------------------------------------------------DSERD
E----------------------KGKGTC-------------------------------
-------------------------------H---------LS-----D-----------
-PY--LSDFI-DDNTIAGINKIF---------R-GK--------------SN---LR---
--RLIWA----IIFIGSLIVCTVMLSFSIK-RFID----------K---PTASTI----T
-I-VS-NT--------EQ-GIAF-----PAV-----------------------TFCNLN
LE------RNSS--N-FLLRST------------------Y-------------------
---------QL------------M------------------NYLYN-------------
------------------------------A--DE-------N-----------------
------------------------------------------------------------
------------------------------------------------------------
-----------FHLSG--------------------------------------------
------------------------------------------------------------
------------------------------------------------------------
------------------------------------------------------------
------------------------------------------------------------
------------------------------------------------------------
------------------------------------------------------------
------------------------------------------------------------
--LNDSYVLQLC------------------------------DNVV--------------
----------------------------------------------RSS------SEDIL
NA-------------------------------TIWNIQ---------------------
-----------------------------------------------------NPSKTID
--ELI------HYCG----FIEGA-NSE--------VKP---C-----------------
----------------------------------------------K-------------
------------------------------------------------------------
------------------------------------------------------------
-----------DA---------F-------K-----------PIL-T-SAG--------I
CFTFN-------GS-------------DN-------------------------------
--------RIHST------GI--------RY-------------GLKLILNIQQK-----
----------E-----------------R--------------P-S---FN---------
------------------------------------------------------------
------------------------------------------------------------

```
----------GKSGVKLVIH-NGRDIA--RPNLYGISVPPGHAIDVGVRKK------A--
-----VEDD-----------TNE----AQ-C--IHG---M----N-------LP----FY
----PS---DK-------F-DYSQLACRENALAENIA----HSSKCSCAI------D---
-----------RP-S--TG-----------------------------------------
-------PYAST-----P---NCTF-----------------------------------
------------------------------------SKA----CCLL---------K---
E-H-------------------YEFN-A-----------------------------EEA
------------DCP-SPCHFEY---YEYT-SSY------------------STF-PN--
-------GLY------LDT-L-----------VN--------------------------
------------------------------------------------KT----------
------------------------------------------------------------
-N----------------------------------------------------------
---------------------------MSV------------------------------
-----N-------EI-----------R------ENFLSVNVFVDDLHTTTTI-TQYTYGV
EAL-----------------LGEVGGQLGLFIGVSIISFFEVLILCIDE-----------
--------LKRL--------C----------------CRGSV----------------K-
RTM-----------------------------KK--------------------------
----------------------------------------LEKMIR--------------
---------------------LPEIDS---------------------GETD--------
--------KSNEIELNSVEI-----------------------ISCDEERSI--------
-----------KDLSPCDEDNKAV------------------------------------
-------------------------------LLPI-------------------------
---------------------------------------------------EDKSNEVGL
QSVKTTEV
>Porifera_AqNaC5_Aqu2_1_09804_001_PorifAmpQue_85_255
------------------------------------------------------------
----------M-------------------------------------------------
------------------------------------------------------------
------------------------------------------------------------
------------------------------------------------------------
--N-KGKAGEP-------------------------------------------------
------------------------------------LNSLKAA-----------------
------------------------------------------------------------
------------------------------------------------------------
-------------------------------------------------------DSEND
E----------------------KGKGRC-------------------------------
-------------------------------H---------LN-----D-----------
-PY--LSDFI-DDNTIAGINKIF---------R-GK--------------SN---LR---
--RLTWA----IIFIGSLIVCTVMLSFSIK-RFID----------K---PTASTI----T
-I-VS-NT--------ER-GIPF-----PAV-----------------------TFCNLN
LE------RNSS--N-FLLRST------------------Y-------------------
---------QL------------M------------------NYLYN-------------
------------------------------A--DE-------N-----------------
------------------------------------------------------------
------------------------------------------------------------
-----------FHLSG--------------------------------------------
------------------------------------------------------------
------------------------------------------------------------
------------------------------------------------------------
------------------------------------------------------------
------------------------------------------------------------
------------------------------------------------------------
------------------------------------------------------------
--LNDSYVLQLC------------------------------DNVV--------------
----------------------------------------------RSS------SEDIL
NA-------------------------------TIWNIQ---------------------
-----------------------------------------------------NPSKTID
```

--ELI------HYCG----FIEGP-NSE--------VKP---C-----------------
----------------------------------------------K-------------
------------------------------------------------------------
------------------------------------------------------------
-----------DA---------F-------K-----------PIL-T-SAG--------I
CFTFN-------GS-------------DN-------------------------------
--------RIHST------GI--------RY-------------GLKLILNIQQE-----
----------K-----------------R--------------P-S---FN---------
------------------------------------------------------------
------------------------------------------------------------
----------GKSGVKLVIH-NGRDIA--RPNLYGISVPPGHAIDVGVRKM------A--
-----VEDD-----------TNE----AQ-C--IHE---M----N-------LP----FF
----PS---DK-------Y-DYSQLACRENAIAENIA----QSSKCNCVI------G---
-----------RP-S--TG-----------------------------------------
-------PYAST-----P---NCTF-----------------------------------
------------------------------------SKA----CCLL---------K---
E-H-------------------YEFN-P-----------------------------EEA
------------DCP-SPCHFEY---YEQT-SSY------------------SSF-PN--
-------GFY------LDT-L-----------VN--------------------------
------------------------------------------------KT----------
------------------------------------------------------------
-N----------------------------------------------------------
---------------------------MSV------------------------------
-----N-------EI-----------R------ENFLSVNVFVGDLHTTTTI-TQYTYGI
EAL-----------------LGEIGGQLGLFIGVSIITFFEVLILCIDE-----------
--------LKRL--------C----------------CRGSV----------------K-
RRM-----------------------------KK--------------------------
----------------------------------------LEKMVR--------------
---------------------LPEIDS---------------------GETD--------
--------KSNEIELNSVKI-----------------------IQCDKERSI--------
-----------KDLSPCDEDNKAV------------------------------------
-------------------------------LLPI-------------------------
---------------------------------------------------EDKSNEVGL
QSIKTTEV
>Porifera_AqNaC14_Aqu2_1_20433_001_PorifAmpQue_94_260
------------------------------------------------------------
----------M-------------------------------------------------
------------------------------------------------------------
------------------------------------------------------------
------------------------------------------------------------
--AGNDPQNPKAP-----------------------------------------------
---------------------------------VADKTSRSPTVTS--------------
------------------------------GPPSPETNRPPALE------------P---
------------------------------------------------------------
--------------------------------------------------PRSGVDREDT
-----------------------STVDIT-------------------------------
-------------------------------P---------VY-----D-----------
-KV--SEYVR-EKITIAGLGHVF---------P-NPEKK--KGVHSLSTCEK---TM---
--MVVWA----VFFTGCVLGCLVLIGLSFK-RFVD----------K---PTASTI----T
-V-VS-HD--------KA-GLPF-----PAV-----------------------TICNLN
LK------KNDS--D-FILNTT------------------Y-------------------
---------QA------------I------------------ISLYN-------------
------------------------------A--DK-------S-----------------
------------------------------------------------------------
------------------------------------------------------------
-----------SQFTS-NKKI---------------------------------------
------------------------------------------------------------

------------------------------------------------------------
------------------------------------------------------------
------------------------------------------------------------
------------------------------------------------------------
------------------------------------------------------------
------------------------------------------------------------
-----DHLLNTC------------------------------TASL--------------
-------------------------------------------------------SNSIQ
NA-------------------------------TIWDIV---------------------
-----------------------------------------------------SPEMAVN
--EFI------HYCG----FLHGA-DSD--------VVM---C-----------------
----------------------------------------------K-------------
------------------------------------------------------------
------------------------------------------------------------
-----------DL---------F-------E-----------PVL-T-SAG--------I
CFTFN-------GT-------------NK-------------------------------
--------LANST------GR--------RY-------------GLKLVLNIQQE-----
----------E-----------------R--------------P-S---FS---------
------------------------------------------------------------
------------------------------------------------------------
----------GKLGVKLVIH-DGKDIA--RPSLYGISVPPGFAVDVGVRKM------A--
-----TRDE-----------TSE----AK-C--IDN---M----N-------LP----FF
----PS---DK-------F-QYSQFACRENAVAENIA----RGSKCNCVIQP----D---
-----------RP----PG-----------------------------------------
-------LYAST-----P---NCTF-----------------------------------
------------------------------------SKA----CCLL---------Q---
E-H-------------------YTFH-P-----------------------------EEI
------------DCP-LPCHFEY---YEHT-ASY------------------SSF-PH--
-------GQY------LQV-L-----------ME--------------------------
------------------------------------------------EL----------
------------------------------------------------------------
-N----------------------------------------------------------
---------------------------MSA------------------------------
-----E-------YV-----------K------NNFLSIDVFFDDFQVTTTT-TKYTYGI
ETL-----------------LGEIGGLLGLFIGVNIINFFELLVLCMDA-----------
--------TK----------L----------------CNKCF----------------K-
KEK-----------------------------PN--------------------------
---------------------------------------KRTNPVE--------------
---------------------LDNVLV---------------------TKGS--------
--------RNST------------------------------------------------
------------------------------------------------------------
--------------------------------KPMN------------------------
----------------------------------------------------------TV
VGDTVTVN
>Deutero_Ambulac_Sakowv30000480m_382_2
------------------------------------------------------------
----------M-------------------------------------------------
------------------------------------------------------------
------------------------------------------------------------
------------------------------------------------------------
------------------------------------------------------------
------------------------------------------------------------
------------------------------------------------------------
------------------------------------------------------------
------------------------------------------------------------
------------------------------------------------------------
------------------------------------------------------------

-----LQHYG--------------------------------------------------
-----------ITL-----------------ELLQ----------R-------------D
-L-----------------ERKF-------------------------------------
------------------------------------------------------------
------------------------------------------------------------
------------------------------------------------------------
------------------------------------------------------------
------------------------------------------------------------
------------------------------------------------------------
------------------------------------------------------------
------------------------------------------------------------
------------------------------------------------------------
------------------------------------------------------------
------------------------------------------------------------
------------------------------------------------------------
------------------------------------------------------------
------------------------------------------------------------
-------------------------------------------------------PDGL-
---------------------------------DVENFT---------------------
------------------------------------------------RAI----GWQLN
-DFTL------PLCQ----WR-G----H----------K---C-----------F-P---
----------------------------------------------Q-------------
------------------------------------------------------------
------------------------------------------------------------
-----------N----------F-------T-----------HTF-T-RFG--------N
CYTFN-------GG-------------DE-----------------------------Y-
-------IDQRIP------GA--------GH-------------GLSVVLNIQQS-----
----------E-----------------Y------------T-E-S--------------
------------------------------------------------------------
------------------------------------------------------------
------LYGNIEAGIKFLVH-PKEEPP--QIGSQGYALQPGTRAFADIRQV------R--
-----YISL-----------EPP-W--GQ-C--DNN-A------A-------LD----YF
------------------T-SYSLSGCSLEYLKNLIY----KD--CGCRPLF----I---
------------P-G---------------------------------------------
----------NE-----P---VCSL-----------------------------------
------------------------------------NQS---STCVQ---------Q---
I-S-------------------AEWA-S-------NA--------------------PEL
------------PCT-VPCNYTL---YPTS-LSY------------------TTF-PS--
-------TNV------AEQ-F-----------EI--------------------------
------------------------------------------------LL----------
------------------------------------------------------------
-D----------------------------------------------------------
---------------------------ITL------------------------------
-----E-------DM-----------R------ENYVYLDVYYSQIQYEQYK-QTKAMTS
SAL-----------------LSDVGGQFGLFIGISVITFVEIIEYIAKK-----------
--------ITGK--------C----------C-------RA-------------------
EK------------------------------TM-------------------K------
-----------------------------------------VNSYN--------------
------------------------------------------------------------
------------------------------------------------------------
------------------------------------------------------------
------------------------------------------------------------
--------G---------------------------------------------------
-EPHSTRN
>Cnidar_HydraVulSc4wPfr_248_2_g2925_t1_384_4
------------------------------------------------------------
----------M-------------------------------------------------

```
----------------------------------GRYF----------------------
------------------------------------------------------------
------------------------------------------------------------
------------------------------------------------------------
------------------------------------------------------------
------------------------------------------------------------
------------------------------------------------------------
---------------------PSQPTPKVED--------------------------KIK
S----------------------PQEQK--------------------------------
--------R----------------------R---------TI-----K-----------
-QY--LAYAG-ENISIHGLSHVF---------D-KR-------------ENF---IC---
--RTVWL----LITIAAFGYAVQKVYESTL-NYFS----------Y---PFSTAR----M
-K-IH-V----------N-QMDF-----PAV-----------------------SFCNLN
DF------RYS-----ATKGTK------------------L-----------------D-
---------KA----------------------------------IL-------------
------------------------------------------------------------
------------------------------------------------------------
------------------------------------------------------------
------------------------------------------------------------
------------------------------------------------------------
------------------------------------------------------------
------------------------------------------------------------
------------------------------------------------------------
------------------------------------------------------------
------------------------------------------------------------
------------------------------------------------------------
------------------------------------------SNDKK-------------
-------------------------------------------------------ISGE-
---------------------------------EYLNIT---------------------
------------------------------------------------LKA----RHKIE
--EML------VDCE----FD-G----N----------K---C-----------S-V---
----------------------------------------------H-------------
------------------------------------------------------------
------------------------------------------------------------
-----------N----------F-------T-----------EFN-W-NQGE-------L
CFTFN-------SG-------------KF-P--------------------H----SL--
-------LKVNGV------GM--------KR-------------SLVLTINVQHY-----
----------E-----------------Y------------Y-G-D--------------
------------------------------------------------------------
------------------------------------------------------------
---------ELDAGIHLILH-DQEETP---IKKRGPVIPPGFTTYIQVEKK------T--
-----ILNL-----------ESP-YK-TK-C--GSI--------K-------LK----YF
------------------D-SYSMHTCWLEQLTDHVY----KI--CKCKDFF----M---
------------P-----------------------------------------------
------------------------------------------------------------
------------------------------------------------------------
------------------------------------------------------------
------------------------------------------------------------
------------------------------------------------------------
------------------------------------------------------------
------------------------------------------------------------
------------------------------------------------------------
------------------------------------------------------------
------------------------------------------------------------
---------------------GDVGGQIGLFVGAGVMSYFEIIDCLVLI-----------
--------IYTR--------F----------F-------QK-------------------
FK------------------------------TI-------------------N------
```

------------------------------------------------------------
------------------------------------------------------------
------------------------------------------------------------
------------------------------------------------------------
------------------------------------------------------------
------------------------------------------------------------
-SIRNHSL
>Protostome_Lophotroco_brch_Lingula_anatina_comp131826_c1_seq2_p1_comp131826_c1__comp131826_c1_seq2_p1__ORF_type_5prime_partial_len_373___score_4_70_comp131826_c1_seq2_1_1119__401_8
------------------------------------------------------------
------------------------------------------------------------
------------------------------------------------------------
------------------------------------------------------------
------------------------------------------------------------
------------------------------------------------------------
------------------------------------------------------------
------------------------------------------------------------
------------------------------------------------------------
------------------------------------------------------------
------------------------------------------------------------
------------------------------------------------------------
------------------------------------------------------------
----------------------------ID-YYWS----------N---PVSVNI----E
-L-HY-V----------D-FIPF-----PAV-----------------------TICNNN
KL------RVL-----NLYHTG------------------------------------E-
-----I---DV-------------------------------GYLFT-------------
------------------------------------------------------------
------------------------------------------------------------
------------------------------------------------------------
------------------------------------------------------------
------------------------------------------------------------
------------------------------------------------------------
------------------------------------------------------------
-------------------------SPTATTTKN--------------------------
------------------------------------------------------------
------------------------------------------------------------
------------------------------------------------------------
------------------------------------------------------------
-------------------------------------------------------LSSY-
---------------------------------NWTEFI---------------------
------------------------------------------------TEY----GHDAP
--TMLN---APGSCE----WN-A----Q----------P---C-----------S-V---
----------------------------------------------D-------------
------------------------------------------------------------
------------------------------------------------------------
-----------N----------F-------T-----------RIL-T-EMG--------V
CYTFN-------HG-------------DT-N-----------------------------
-------LKTELL------GS--------TF-------------GLKLILNAEQY-----
----------N-----------------Y------------L-A-----TT---------
------------------------------------------------------------
------------------------------------------------------------
----------HSAGFTVLLH-HPDDVP--YIENLGFQVAPGESISVGISQT------K--
-----VTNL-----------PSP-Y--GN-C--AEK--------N-------LA----YY
------------------D-SYSTLNCIAECTLNFTM----DN--CGCRAYY----M---
---------INVP-----------------------------------------------
----------GV-----P---VCSP-----------------------------------

------------------------------------TQW---WECLD---------E---
L-R-------------------ANTS-A-------AL--------------------QAC
------------ACP-VPCEEVH---YGYT-ISH------------------SKM-SN--
-------AIT------AAL-M-----------AG--------------------------
------------------------------------------------YP----------
------------------------------------------------------------
-G----------------------------------------------------------
---------------------------TTE------------------------------
-----D-------YF-----------R------DNYLILQIFYKELNFQGIE-QQPAYSG
LAL-----------------FSDIGGALGLVLGSTLMTAAEIIDFAVSL-----------
--------CLWV--------A----------F-------RR-------------------
-----------------------------------------------------K------
------------------------------------------------------------
------------------------------------------------------------
------------------------------------------------------------
------------------------------------------------------------
------------------------------------------------------------
------------------------------------------------------------
--------
>Deutero_Ambulac_Sakowv30023603m_404_9
------------------------------------------------------------
----------M-------------------------------------------------
------------------------------------------------------------
------------------------------------------------------------
------------------------------------------------------------
------------------------------------------------------------
------------------------------------------------------------
------------------------------------------------------------
------------------------------------------------------------
------------------------------------------------------------
-----------------------TKDNHP-------------------------------
---STENGD----------------------T---------TV-----R-----------
-RI--TRRFA-LETSAHALPKSA---------K-SS--------------HT---YT---
--QCMWV----VIFCAACSVYLWQIGDLLS-LYLS----------V---PKRTEN----I
-L-VS-A----------N-TLPF-----PAV-----------------------TLCNTN
KV------RRS-----MIQQSG------------------Y-----------------S-
---------SLLDS-------------------------LEINTLIY-------------
------------------------------------------------------------
------------------------------------------------------------
------------------------------------------------------------
------------------------------------------------------------
------------------------------------------------------------
------------------------------------------------------------
------------------------------------------------------------
------------------------------------------------------------
------------------------------------------------------------
------------------------------------------------------------
------------------------------------------------------------
-----------------------------------------------------Y------
---------------------LKSEDPPDWRAFMSF-----------SR------DPSWT
---------------------------------PLRKVL---------------------
-------------------------KF----------------SKDELKVL----GHQLE
--DLI------LECS----FD-G----N----------E---C-----------D-ME--
----------------------------------------------K-------------
------------------------------------------------------------
------------------------------------------------------------
-----------D----------F-------H-----------TFQ-HDEYG--------N

CFTFN-------HG-------------MH-G--------------------V----PL--
-------RNSTKA------GG--------DY-------------GLRLTMNVEQP-----
----------E-----------------Y------------I-T-F---YG---------
------------------------------------------------------------
------------------------------------------------------------
----------KDAGVLMTIH-SPEDTP--FPEDHGMVLRTGSKTSIGLYKE------D--
-----DILI-----------PEP-N--RE-C--ENV--------DTF---------YGIY
----GDY-------------KYTTLACDKSCLNRYIQ----YY--CGCVDTI----M---
------------I-----------------------------------------------
---------EGV-----P---RCMIL----------N-----------------------
------------------------------------DTQ---EVCQQ---------L---
M-N-------------------YLYY-R-------GL--------L------------VC
------------RCP-RRCRQVH---KNTM------------------------------
------------------------------------------------------------
------------------------------------------------------------
------------------------------------------------------------
------------------------------------------------------------
------------------------------------------------------------
--------------------------R---------------------------------
------------------------------------------------------------
------------------------------------------------------------
------------------------------------------------------------
------------------------------------------------------------
------------------------------------------------------------
------------------------------------------------------------
------------------------------------------------------------
------------------------------------------------------------
------------------------------------------------------------
--------
>Protostome_Lophotroco_annelid_Pdum_comp418513_c0_seq7__408_10
------------------------------------------------------------
----------M-------------------------------------------------
------------------------------------------------------------
------------------------------------------------------------
------------------------------------------------------------
------------------------------------------------------------
------------------------------------------------------------
------------------------------------------------------------
------------------------------------------------------------
------------------DGE---------------------------------------
-----------------------IGSKKP-------------------------------
-------------------------------D---------NW-----T-----------
-SA--LYRFS-QTTTVDGVKQIT---------E-PT-------------PFC---AR---
--RLVWI----ALLLGGIALVARQIADSVL-YYHS----------R---PVTVNL----N
-I-KY-N----------K-TLKF-----PAV-----------------------TICNQN
NF------RAT-----VASTRGQ-----------------F-------------------
---------DL------------I------------------SDIYV-----R-------
---------------------------------NS-------SL----------------
------------------------------------------------------------
------------------------------------------------------------
------------------------------------------------------------
------------------------------------------------------------
------------------------------------------------------------
------------------------------------------------------------
------------------------------------------------------------
------------------------------------------------------------
------------------------------------------------------------

------------------------------------------------------------
------------------------------------------EDKTY-------------
-------------------------------------------------------EDDFY
S-------------------------------ESVAGSM---------------------
------------------------------------------------FAL----KHDKH
--DLI------HSCK----WD-N----D---------YN---C-----------S-A---
----------------------------------------------D-------------
------------------------------------------------------------
------------------------------------------------------------
-----------D----------F-------I-----------TSP-T-DHG--------I
CYSFN-------SG-------------RD-D--------------------K----PI--
-------RYISQT------GA--------DS-------------GLKMKLNIEQY-----
----------E-----------------Y------------M-Q-G---PN---------
------------------------------------------------------------
------------------------------------------------------------
----------HGAGLKILVH-DPSDVP--MVKDLGLAVKPGTHTYMDINII------E--
-----VHNL-----------PPP-H--GH-C--EEK--------N-------LT----LF
------------------S-KYAMSTCLLECKTLHIT----SL--CGCIDYY----M---
------------PWK---------------------------------------------
-------TSGGP-----P---FCTI-----------------------------------
------------------------------------RQF---LECVK---------P---
S-L-------------------EAFN-V-------AD---------------------EC
------------DCP-VPCNMTI---LNPS-ITL------------------LAN-SD--
-------FDV------EKM-L-----------HQ--------------------------
------------------------------------------------------------
------------------------------------------------------------
-NH---------------------------------------------------------
------------------------------------------------------------
-----S-------EL-----------Q------E----------KYTYA-----------
QEV---------------------------------------------------------
------------------------------------------------------------
------------------------------------------------------------
------------------------------------------------------------
------------------------------------------------------------
------------------------------------------------------------
------------------------------------------------------------
------------------------------------------------------------
------------------------------------------------------------
--------
>Cnidar_HydraVulSc4wPfr_1152_g30239_t1_417_13
------------------------------------------------------------
----------M-------------------------------------------------
------------------------------------------------------------
------------------------------------------------------------
------------------------------------------------------------
-SDTN-IVN-------------------------------------------------KH
FITNDSKGWQSPC-----------------------------------------------
-------PKIKLN-NQFRKE-------------SNEFIH---------------------
------------------------------------------------------------
-----------------RNGKPYFP--------------------------------VLK
S----------------------EKEAR--------------------------------
--------R----------------------E---------KL-----D-----------
-ET--IQKFV-DASTCHGFKHCF---------N-SG--------------SK---VR---
--QLIWM----MILAASVALLVQKLYESGV-KFLE----------R---PFSTKT----T
-L-TY-V----------D-QMEF-----PAI-----------------------SICNMN
DM------RNS-----K-------------------------------------------
------------------------------------------------------------

------------------------------------------------------------
------------------------------------------------------------
------------------------------------------------------------
------------------------------------------------------------
------------------------------------------------------------
------------------------------------------------------------
------------------------------------------------------------
------------------------------------------------------------
------------------------------------------------------------
------------------------------------------------------------
------------------------------------------------------------
------------------------------------------------------------
------------------------------------------------------------
------------------------------------------------------------
------------------------------------------------------------
------------------------------------------------------------
------------------------------------------------------------
------------------------------------------------------------
------------------------------------------------------------
------------------------------------------------------------
------------------------------------------------------------
------------------------------------------------------Y-----
----------D-----------------Y------------Y-E-D--------------
------------------------------------------------------------
------------------------------------------------------------
---------IKEAGIKLIIH-DQHETP---VRMAGVKLSPGFSATVQIKKK------K--
-----TLNL-----------KAP-YA-TN-C--GSK--------P-------LK----YF
------------------D-HYSTNTCWLERLTDHVV----TS--CNCKDSF----M---
------------P-G--H------------------------------------------
-----------A-----R---VCSI-----------------------------------
------------------------------------PEL---MNCTF---------L---
K-W-------------------EEFN-K-------LK---------------------DI
------------QCP-IPCESQE---FESS-ISF------------------ARY-PS--
-------NIL------ADK-I-----------AK--------------------------
------------------------------------------------DM----------
------------------------------------------------------------
-Q---------------L------------------PGS---------------------
------------------------VQ-ENR------------------------------
-----E-------FI-----------R------DNYLRVEIFYEEMSYIQVE-QTPSYDL
MIL-----------------LGDIGGQFGLFLGSSIITYVEFFDFFAAL-----------
--------IYTK--------Y----------F-------RI-------------------
FK------------------------------PP--------------------------
------------------------------------------------------------
------------------------------------------------------------
------------------------------------------------------------
------------------------------------------------------------
------------------------------------------------------------
------------------------------------------------------------
------KI
>Protostome_Lophotroco_molsk_AplCal_XP_035825598_1_acid_sensing_ion_channel_2
__partial__Aplysia_californica__419_14
------------------------------------------------------------
----------M-------------------------------------------------
------------------------------------------------------------
------------------------------------------------------------
------------------------------------------------------------
-NNTAHHNSFE-------------------------------------------------

```
-----------------------------------------P-----------------R
PFNTQYDNNIPESISMKPYA-------------KDGVTSSSLD-------------P---
------------------------------------------------------------
---------------------------------------------------------DVI
SP---------------------QKKKGS-------------------------------
-------------------------------L---------TW-----K-----------
-EA--LFDFT-QNTTLHGIRFIF---------M-ND-------------VFI---LR---
--RLLWL----ALFLTCSVLMSVQIVERIV-FFYS----------Y---PVTVNV----H
-V-NF-N----------K-TLAF-----PAF-----------------------TVCNQN
AF------RAS-----AATDRH------------------L-----------------Y-
---------RL------------I------------------ERLHS-----G-------
---------------------------------DT-------SALLSQS-----------
------------------------------------------------------------
----------------------P-------------------------------------
-----------------D------------------------------------------
------------------------------------------------------------
------------------------------------------------------------
------------------------------------------------------------
--------------------------PL--------------------------------
------------------------------------------------------------
------------------------------------------------------------
------------------------------------------------------------
------------------------------------------------------------
-------------------------------------------------------PSNL-
---------------------------------SLDELY---------------------
------------------------------------------------LTT----AHRKE
--DLI------VRCE----WQ-N----K----------P---C-----------G-P---
----------------------------------------------E-------------
------------------------------------------------------------
------------------------------------------------------------
-----------N----------F-------T-----------LVL-T-DHG--------V
CYNFN-------DN-------------PS---------------------------EP--
-------LWVTST------GA--------EY-------------GLKLTLNVEQY-----
----------E-----------------Y------------M-P-G---PH---------
------------------------------------------------------------
------------------------------------------------------------
----------DAAGIKILLH-DGKEFP--KVAELGLSIPTGTHTYVGIQLL------K--
-----IQNL-----------PAP-H--GT-C--RSE--------R-------SP----YY
------------------E-RYSPDACQLACLTKYVS----EQ--CHCRHFY----M---
------------P-H---------------------------------------------
-------IDGSP-----P---VCTL-----------------------------------
------------------------------------GEY---LSCYE---------R---
I-I-------------------DQVK-D-------RV--------R-----------AEC
------------DCP-VPCDFLI---YD--------------------------------
------------------------------------------------------------
------------------------------------------------------------
------------------------------------------------------------
------------------------------------------------------------
------------------------------------------------------------
------------------------------------------------------------
------------------------------------------------------------
------------------------------------------------------------
------------------------------------------------------------
------------------------------------------------------------
------------------------------------------------------------
------------------------------------------------------------
------------------------------------------------------------
```

------------------------------------------------------------
------------------------------------------------------------
--------
>Protostome_Lophotroco_annelid_CAC9660887_1__Ofus_G111053_partial_Owenia__fus
iformis__422_15
------------------------------------------------------------
------------------------------------------------------------
------------------------------------------------------------
------------------------------------------------------------
------------------------------------------------------------
------------------------------------------------------------
------------------------------------------------------------
------------------------------------------------------------
------------------------------------------------------------
------------------------------------------------------------
------------------------------------------------------------
------------------------------------------------------------
------------------------------------------------------------
------------------------------------------------------------
------------------------------------------------------------
------------------------------------------------------------
------------------------------------------------------------
------------------------------------------------------------
------------------------------------------------------------
------------------------------------------------------------
------------------------------------------------------------
------------------------------------------------------------
------------------------------------------------------------
------------------------------------------------------------
------------------------------------------------------------
------------------------------------------------------------
------------------------------------------------------------
------------------------------------------------------------
------------------------------------------------------------
------------------------------------------------------------
------------------------------------------------------------
------------------------------------------------------------
-----------IGCT----FK-G----E----------Q---C-----------N-R---
----------------------------------------------S-------------
------------------------------------------------------------
------------------------------------------------------------
-----------M----------F-------V-----------LQK-DSHYH--------N
CYTFN-------GI-------------GS-N-NF-----------------------N--
-------KNVSKT------DP--------GS-------------GLSLVVFTDTY-----
----------TDTL---TSDPGVI----YN-------------P-S-DPTS---------
------------------------------------------------------------
------------------------------------------------------------
----------GNNGIRVIVH-SPKTRP--APVDKGFDVPNGFSTTVALRAT------R--
-----RTLL-----------TEP-H--GD-C--TTD--------AS------NKG--TEY
--------------------RYTTDICYEQCHQDIIT----QK--CGCKSSH----F---
------------I-P--PR-----------------------------------------
-----NE--TGF-----Q---YCGKL----------NIT-NLLSG--------DDT----
---------------------------SKESIL-EDFTH---LECEK---------S---
V-I-------------------DTFW-S-------NV-------TMV----------SKC
------------NCR-PPCNSTY---YIKT-TSQ------------------AVW-PN--
-------EFT------QQS-F-----------YK--------------------------
------------------------------------------------TYVN--------

-------------------------------ITDSTA-----------------------
RP-------------NIL------------------F-----------------------
------------------QGLTPKEI-KDR------------------------------
-----G-------LI-----------Q------KNFLRLNIYFEDLQVEETS-QEADYPI
TQV-----------------ISDIGGNMGFYVGVSVITLVEFLSLIGAL-----------
--------LLIF--------F----------K-------SC-------------------
---------------------------------------------------IKA------
-----------------------------------------GGKLREA------------
------------------KVSDVNIQP---------------------AHSL--------
--------ENKLGAQYTYEYK---------------------------------------
------------------------------------------------------------
------------------------------------------------------------
------------------------------------------------------------
SYNDMISR
>Protostome_Lophotroco_brch_Lingula_anatina_comp132718_c0_seq1_p1_comp132718_c0__comp132718_c0_seq1_p1__ORF_type_complete_len_399___score_16_97_comp132718_c0_seq1_93_1289__423_16
------------------------------------------------------------
----------M-------------------------------------------------
------------------------------------------------------------
------------------------------------------------------------
------------------------------------------------------------
------------------------------------------------------------
------------------------------------------------------------
------------------------------------------------------------
------------------------------------------------------------
------------------------------------------------------------
------------------------------------------------------------
------------------------------------------------------------
-----DRLYL----------------------N-GA------------------------
-----------TLYLAALS----------------------------------------S
------K----------N-SRAY-----AHI-----------------------NLDN--
------------------------------------------------------------
------------------------------------------------------------
------------------------------------------------------------
------------------------------------------------------------
------------------------------------------------------------
------------------------------------------------------------
------------------------------------------------------------
------------------------------------------------------------
------------------------------------------------------------
------------------------------------------------------------
------------------------------------------------------------
------------------------------------------------------------
------------------------------------------------------------
------------------------------------------------------------
------------------------IKNLNMTAWY---------------------GSGY-
---------------------------------TFANFT---------------------
------------------------------------------------QDK----GWLLN
--ERN------ARCT----FR-G----E----------D---C-----------NIT---
----------------------------------------------A-------------
------------------------------------------------------------
------------------------------------------------------------
-----------D----------F-------K-----------HVF-T-EFG--------N
CYTFN-------SG-------------DD-P--------------------S----RN--
-------LFQDQA------GI--------GN-------------GLRIEINIQQE-----
----------Q-----------------Y------------T-N-L---IL---------

------------------------------------------------------------
------------------------------------------------------------
------RGDAPDAGILFHVH-NQSEPA--SVETDGRAVGPGLHAYAGLTRS------D--
-----FATL-----------HPP-Y--GQ-C--NET-A------S-------LE----FY
------------------P-VYTMSGCVVECKRKHIL----RH--CNCRLME----H---
------------P-G---------------------------------------------
----------YE-----P---VCGL-----------------------------------
------------------------------------IRI---LTCVK---------P---
L-L-------------------VNLT-Q-------DF--------T-----------GMC
------------SCS-VPCSSVV---YQTA-LSY------------------SLV-PS--
-------EST------KVE-A-----------SN--------------------------
------------------------------------------------TL----------
------------------------------------------------------------
-G----------------------------------------------------------
---------------------------VAV------------------------------
-----D-------DA-----------R------KNHIILDVYFQSLNYQQSE-QLPAVEW
TAL-----------------ISDIGGQFGLFMGFSLLTVVEFIEFAIMS-----------
--------VVTL--------T----------A-------AK-------------------
RS------------------------------QN-------------------K------
-----------------------------------------VKGHDTIHG----------
--------------------KGDKVTK---------------------SSKTGR------
------------------------------------------------------------
------------------------------------------------------------
------------------------------------------------------------
------------------------------------------------------ISMQGD
FVQVHEIF
>Cnidar_AlaAla_c59883_g1_i1_p1__GENE_c59883_g1_i1__c59883_g1_i1_p1__ORF__type
_5prime_partial__len_400__score_24_40__c59883_g1_i1_3994_5193__424_17
------------------------------------------------------------
------------------------------------------------------------
------------------------------------------------------------
------------------------------------------------------------
------------------------------------------------------------
------------------------------------------------------------
------------------------------------------------------------
------------------------------------------------------------
------------------------------------------------------------
------------------------------------------------------------
------------------------------------------------------------
------------------------------------------------------------
------------------------------------------------------------
----IWS----LLILTALGLLVEKLYESTM-HYFS----------Y---PFSTTT----T
-I-KY-A----------G-KMKF-----PAV-----------------------SFCNLN
DL------RMS-----KLNGTK------------------L-----------------H-
---------TA----------------------------------IL-------------
------------------------------------------------------------
------------------------------------------------------------
------------------------------------------------------------
------------------------------------------------------------
------------------------------------------------------------
------------------------------------------------------------
------------------------------------------------------------
------------------------------------------------------------
------------------------------------------------------------
------------------------------------------------------------
------------------------------------------------------------
------------------------------------------QSKN--------------

--------------------------------LLTT-------------------LTGE-
---------------------------------EYKNTT---------------------
------------------------------------------------KNA----NHRIH
--DML------DTCQ----FV-H----D----------N---C-----------S-T---
----------------------------------------------F-------------
------------------------------------------------------------
------------------------------------------------------------
-----------N----------F-------T-----------QFF-H-NQGD-------R
CYTFN-------SG-------------KN-N--------------------H----PV--
-------LEVDNT------GV--------GH-------------SLQLVINIQHY-----
----------D-----------------Y------------Y-M-D--------------
------------------------------------------------------------
------------------------------------------------------------
---------TEKSGLHLILH-GQNETP---VKMQGVIVSPGFITYVDIRRR------K--
-----LKNL-----------PYP-YK-TN-C--GSK--------P-------LH----YF
------------------S-GYSMHLCWLESLTDHVV----EK--CNCRDWF----M---
------------P-G---------------------------------------------
----------SH-----K---VCSL-----------------------------------
------------------------------------NES---MNCMW---------P---
E-W-------------------AAFD-K-------FK---------------------MY
------------NCP-LPCEIET---YSSR-LSF------------------AQF-PS--
-------NSH------ADV-M----------AK--------------------------
------------------------------------------------RL----------
------------------------------------------------------------
-N---------------L------------------HGS---------------------
------------------------PI-ENR------------------------------
-----L-------FI-----------R------DNYLKIVIYYSELSYDHME-QVPSYDM
MVL-----------------LGDIGGQLGLFLGSSILTYLEFFDCLVML-----------
--------IYSR--------F----------F-------EK-------------------
IK------------------------------PT-------------------S------
------------------------------------------------------------
------------------------------------------------------------
------------------------------------------------------------
------------------------------------------------------------
------------------------------------------------------------
------------------------------------------------------------
-----VPV
>Protostome_Lophotroco_annelid_CAC9477573_1__Ofus_G013340_Owenia__fusiformis_
_427_19
------------------------------------------------------------
------------------------------------------------------------
------------------------------------------------------------
------------------------------------------------------------
------------------------------------------------------------
------------------------------------------------------------
------------------------------------------------------------
------------------------------------------------------------
------------------------------------------------------------
------------------------------------------------------------
------------------------------------------------------------
------------------------------------------------------------
------------------------------------------------------------
------------------------------------------------------------
------------------------------------------------------------
------------------------------------------------------------
------------------------------------------------------------
------------------------------------------------------------

------------------------------------------------------------
------------------------------------------------------------
------------------------------------------------------------
------------------------------------------------------------
------------------------------------------------------------
------------------------------------------------------------
------------------------------------------------------------
------------------------------------------------------------
------------------------------------------------------------
------------------------------------------------------------
------------------------------------------------------------
---------------------------------------------MIMP------QMRMK
---------------------------------LY-------------------------
-------------------------------------------------LL----GMTWK
--TFC------FHVQ----LL-A----M--------------T-----------N-M---
----------------------------------------------S-------------
------------------------------------------------------------
------------------------------------------------------------
-----------Q----------I-------E-----------SFI-NPYFF--------K
CYTMN-------PK-------------DF-N-DS-----------------S----IE--
-------PLVHST------GA--------IN-------------GLSALFFLDI------
----------G--------TAKRI----YN-------------P-T-SPLG---------
------------------------------------------------------------
------------------------------------------------------------
----------GSSGLLVVIH-PPGTQP--DPLQNGFNIAPGFSTSVELQNK------N--
-----RELL-----------GHP-W--GD-C--ENR--------EK------LSD--FPY
--------------------KYDKSSCERQCQQKFIM----DQ--CGCVTSI----Q---
------------P-I--PD-----------------------------------------
-----NL--KNM-----S---MCGKM----------DLK-NLNSD------SP-------
---------------------------NISQIA-EEIDK---LECEH---------A---
K-K-------------------RVLK-K-------DY-------PTE----------NKC
------------DCP-RACHNTI---YDYT-ISQ------------------SEW-PS--
-------EGV------MLD-F-----------YK--------------------------
------------------------------------------------HAITLRD-----
-------------------------------MQGSGY-----------------------
VH-------------STM------------------YNL-LDSKVN--------------
------------------SNNTKENR-ANQ------------------------------
-----E-------MI-----------R------KNFLRLNVFFSSLNTEITK-QVVEYTF
TDL-----------------ISGVGGGFGVYVGFSIVTICEFCVLFAHL-----------
--------VKAM--------M----------N-------NR-------------------
KNAVNDEPWAFK--------------------RK--------------------------
------------------------------------------------------------
------------------SMEIPSSK----------------------------------
------------------------------------------------------------
------------------------------------------------------------
------------------------------------------------------------
------------------------------------------------------------
--------
>Deu_Ambulacrar_hemi_Ptyfla_40v0_9_20150316_1g5442_t1_scaffold2520_cov137_429
_20
------------------------------------------------------------
----------M-------------------------------------------------
------------------------------------------------------------
------------------------------------------------------------
------------------------------------------------------------
------------------------------------------------------------
------------------------------------------------------------

------------------------------------------------------------
------------------------------------------------------------
------------------------------------------------------------
------------------------------------------------------------
----PSEDS----------------------L---------TY-----K-----------
-AL--IQNVL-RNSSAHGLPNVQ---------R-AT--------------SL---PM---
--RLFWL----LAFFTAVGIFVWMSSGLIA-EYLR----------Y---DVDVNL----Q
-I-QF-S----------R-DLTF-----PAV-----------------------TICNLN
PL------RKS-----KLSEFG--------------------------------------
------------------------------------------HDILQ----R--------
------------------------------------------------------------
------------------------------------------------------------
------------------------------------------------------------
------------------------------------------------------------
------------------------------------------------------------
------------------------------------------------------------
------------------------------------------------------------
-------------------------RPLGRMEGRRRQ-----------------------
------------------------------------------------------------
------------------------------------------------------------
----------------------------------------------------------ER
QTTERSETSNDTLN----------------------------DDQERTF-----------
------------------------EEYNYWDQI--------PTNYHASP------SSSWS
------------------------------IIERIEHYI---------------------
------------------------SDI----------------PAVQRAEI----GHQLK
--DLV------VDCQ----WN-N----I----------K---C-----------S-P---
----------------------------------------------R-------------
------------------------------------------------------------
------------------------------------------------------------
-----------N----------F-------T-----------TFT-NVMYG--------N
CFTFN-------GE-------------HN-N--------------------V----MP--
-------LSTHYS------GS--------VF-------------GLTLILFVEQA-----
----------E-----------------Y------------M-D-D---VI---------
------------------------------------------------------------
------------------------------------------------------------
----------DSPGVRVTIH-SQDDTP--FPEDSGFDIQPGRATSVGILMG------R--
-----TQRL-----------PKP-Y--TN-C--ISD-N--------------IPTYDTAF
----GD--------------IYSVKDYIREFLKSRSS----KLR--TLMEQEERTGL---
------------------------------------------------------------
-----------------D---LLSNA----------N-----------------------
------------------------------------RDT---SKCYS---------E---
------------------------------------------------------------
--------------------------HHRK-RSV------------------ATFDPLRF
--------------------I-------------C-------------------------
-----------------------------------------------AAL----------
---------------------------------LDS------------------------
-N----------------------------------------------------------
------------------------------------------------------------
-----D--------------------R------------------------------YN-
------------------------------------------------------------
--------------------L---------------------------------------
--------------------------------RT--------------------------
------------------------------------------------------------
------------------------------------------------------------
------------------------------------------------------------
------------------------------------------------------------
--------------------------------SPD-------------------------

-----------------------------------------------------------D
C-------
>Deutero_Ambulac_Apla_gbr414_1_t1_430_21
------------------------------------------------------------
------------------------------------------------------------
------------------------------------------------------------
------------------------------------------------------------
------------------------------------------------------------
------------------------------------------------------------
------------------------------------------------------------
------------------------------------------------------------
------------------------------------------------------------
------------------------------------------------------------
------------------------------------------------------------
------------------------------------------------------------
--------MS-DAVGIPGIRYVV---------S-SN-------------FHI---LR---
--RLLWT----VALLGGMAAASYQVIDRAT-FFFS----------N---PKSVNV----V
-I-EN-A----------E-PMLF-----PAV-----------------------TFCNTN
RF------RDVN---GQLSTHP------------------F----------------G--
---------QF------------L-----------------AATYFH-------------
------------------------------------------------------------
------------------------------------------------------------
------------------------------------------------------------
------------------------------------------------------------
------------------------------------------------------------
------------------------------------------------------------
------------------------------------------------------------
------------------------------------------------------------
------------------------------------------------------------
------------------------------------------------------------
------------------------------------------------------------
------------------------------------------------------------
--------------------------GGNGTGFTST-------------------LGGA-
---------------------------------NTTALY---------------------
------------------------------------------------LEF----AHQME
YGQMF------IRGA----LG-G----K----------N---I-----------T-S---
----------------------------------------------A-------------
------------------------------------------------------------
------------------------------------------------------------
-----------D----------F-------R-----------RVL-T-DAG--------V
CFTFN-------SG--------------L-N--------------------T----EL--
-------RGQSFP------GR--------AH-------------GLYLFLNVEQY-----
----------Y-----------------H------------Y-Y-A---YE---------
------------------------------------------------------------
------------------------------------------------------------
-------F--SGAGILVAVH-DQTEIP--DMDSMGFGIPPGRHAIVSVGKK------L--
-----TSRK-----------D------GG-C--VEK--------Q-------LK----YF
------------------S-KYTKSMCLRECVIDLLL----QN--CGCVEPY----M---
------------T-D---------------------------------------------
----------KR-----R------------------------------------------
------------------------------------------------------------
--------------------------------------------YV-----------DMC
------------DCP-LPCSTVK---FTTS-VSY------------------TLF-PG--
-------NFY------SWL-L-----------AE--------------------------
------------------------------------------------QF----------
------------------------------------------------------------
-S----------------------------------------------------------

---------------------------TDE------------------------------
FTLPHD-------FY-----------Q------ENFVIIQMFFEELTVEHRE-QHSDYTF
FAL-----------------LCDIGGALGLWLGGSILTIVEILDHFTST-----------
--------TVIPG------------------S-------TS-------------------
HQ------------------------------RG--------------------------
------------------------------------------------------------
------------------------------------------------------------
------------------------------------------------------------
------------------------------------------------------------
------------------------------------------------------------
------------------------------------------------------------
--------
>Deutero_Ambulac_Apla_XP_022104029_1_ASIClike2
------------------------------------------------------------
----------M-------------------------------------------------
------------------------------------------------------------
------------------------------------------------------------
------------------------------------------------------------
------------------------------------------------------------
------------------------------------------------------------
------------------------------------------------------------
------------------------------------------------------------
------------------------------------------------------------
------------------------------------------------------------
-----------------------------GCY---------------KR-----------
-CY--LRQWALTDTDLHGVKHIA---------G-EG--------------GI---LR---
--RLIWA----ACFLAALVVFLHQATLTLI-HFVE----------K---HHVTKV----D
-I-SY-R----------K-KLDF-----PAV-----------------------TVCNFN
KY------RES-----ALTDRD------------------I-----------------R-
---------NV------------G-YHL---------------GIVD----E--------
---------------------------------DH-------NLIN--------------
------------------------------------------------------------
------------------------------------------------------------
------------------------------------------------------------
------------------------------------------------------------
------------------------------------------------------------
------------------------------------------------------------
--------------------------PYLY------------------------------
------------------------------------------------------------
------------------------------------------------------------
------------------------------------------------------------
--------TEEF------------------------------RRK---------------
------------------------MAAVDWSVHDI--------------------DDEY-
---------------------------------NMTEFT---------------------
------------------------------------------------NRT----GHQLD
--EMI------VECS----WR-D----E----------P---C-----------S-P---
----------------------------------------------D-------------
------------------------------------------------------------
------------------------------------------------------------
-----------D----------F-------H-----------HIF-S-HLG--------N
CYTFN-------HV-------------AL-A--------------------T----ER--
-------HSSISA------GA--------AN-------------GLKLTLNIQEE-----
----------E-----------------Y------------T-P-S-NDLH---------
------------------------------------------------------------
------------------------------------------------------------
-------GAAEDAGIKWMLH-HPSEPP--YVKELGFGAGPGHHTFVAVRHE------E--
-----VESL-----------PSP-Y--TP-C--MET-S-AG---F-------LD----HF

```
------------------D-HYSLQACRIECETEVVV----QR--CGCRLVE----Q---
------------P-G---------------------------------------------
----------NA-----P---VCNP-----------------------------------
------------------------------------AET---HECAQ---------A---
A-L-------------------VHAV-A-------GH--------GD----------QAC
------------DCS-SPCSVES---YPFT-TTN------------------VRL-RA--
-------KYI------ERI-Y-----------SN--------------------------
-----------------------------------------------TTH----------
------------------------------------------------------------
-N----------------------------------------------------------
---------------------------FSA------------------------------
-----D-------YI-----------Q------NNLVLLSIYYEALNAEVIE-QLPEMTV
PSL-----------------LAALGGNFGLFLGASVLTIVELLEYVFDE-----------
--------LTAS--------C----------T-------RR-------------------
KT------------------------------SG-------------------R------
-----------------------------------------ISNAREI------------
--------------------LTVRAAP---------------------------------
------------------------------------------------------------
----------------ISVIDTEKEL----------------------------------
------------------------------------------------------------
------------------------------------------------------------
EHGRQWNR
>Deutero_Ambulac_Sakowv30036990m_432_23
------------------------------------------------------------
----------M-------------------------------------------------
------------------------------------------------------------
------------------------------------------------------------
------------------------------------------------------------
-KSPSNADIIR-------------------------------------------------
------------------------------------------------------------
------------NVNRRRPT-------------PTSTSFDM-------------------
------------------------------------------------------------
---------------------------------------------------------SNT
EKF--------------------TSTSSLSLD----------------------------
---GGNVRR----------------------R--------GAVYIF--S-----------
-RN--FRQFL-SETTLHGARYTA---------N-NE-------------YHV---VR---
--RL-------------------------------------------------------D
-V-KY--------------NINF--------D--------------------WSLLENTT
AI--------------------------------------L-------------------
------------------------------------------------------------
------------------------------------------------------------
------------------------------------------------------------
------------------------------------------------------------
------------------------------------------------------------
------------------------------------------------------------
------------------------------------------------------------
------------------------------------------------------------
------------------------------------------------------------
------------------------------------------------------------
------------------------------------------------------------
------------------------------------------------------------
------------------------------------------------------------
------------------------------------------------------------
---------------------------------NRTQFE---------------------
------------------------------------------------LEA----AHQKE
--SFI------VACN----YQ-G----D---G---VERL---C-----------G-P---
----------------------------------------------N-------------
```

------------------------------------------------------------
------------------------------------------------------------
-----------N----------F-------T-----------TTF-T-EFG--------V
CYTFN-------ND-------------PE-N--------------------Q--------
-------LFVTKF------GS--------NS-------------GLRVRLFTDEN-----
----------E-----------------Y------------T-F-G---KQ---------
------------------------------------------------------------
------------------------------------------------------------
----------TGSGFKVLLY-SPGDVP--LVQQLGFAVPTGVDALVAIRLE------K--
-----SINL-----------PPP-FP-TK-C--SNA--------P-------LK----YY
------------------DGNYTYVWCALEKITDFVV----SA--CGCREPY----M---
------------P-G---------------------------------------------
----------SK-----R---VCNL-----------------------------------
------------------------------------KES---IVCVI---------P---
V-M-------------------DDST-E-------RS--------V-------------T
------------YCG-VGCDVTR---FDSR-VTY------------------ADF-PA--
-------KPI------LEH-M-----------SS--------------------------
------------------------------------------------TT----------
------------------------------------------------------------
-N----------------------------------------------------------
---------------------------ESS------------------------------
-----E-------YF-----------R------RNYADISFFAEDMTFKLTK-TVPVMTG
ATL-----------------VSNFGGLMGLCLGASLLTAVEFIDFILFC-----------
--------FCKS------------------------------------------------
------------------------------------------------------------
------------------------------------------------------------
------------------------------------------------------------
------------------------------------------------------------
------------------------------------------------------------
------------------------------------------------------------
------------------------------------------------------------
--------
>Deu_Ambulacrar_hemi_Ptyfla_40v0_9_20150316_1g16608_t1_scaffold13089_cov115_4
34_24
------------------------------------------------------------
----------M-------------------------------------------------
------------------------------------------------------------
------------------------------------------------------------
------------------------------------------------------------
------------------------------------------------------------
------------------------------------------------------------
------------------------------------------------------------
------------------------------------------------------------
------------------------------------------------------------
------------------------------------------------------------
---MADAKP----------------------V---------TF-----S-----------
-EL--LKKQL-QTTTAHGLPRIE---------A-AD--------------KL---VR---
--KLFWA----FIVVAGLGMFIWQSSVIIT-KYYA----------R---DTTISI----E
-M-KF-D----------T-SLDF-----PAI-----------------------TICNMN
PV------KLQ-----SLERDP--------------------------------------
---------YL-------------------------------MSLVG----Y--------
---------------------------------DN-------------------------
------------------------------------------------------------
------------------------------------------------------------
-----------------D--E---------------------------------------
------------------------------------------------------------
------------------------------------------------------------

------------------------------------------------------------
------------------------------------------------------------
------------------------------------------------------------
------------------------------------------------------------
------------------------------------------------------------
------------------------------------------------------------
-------------------------------------------------------DDNLT
------------------------------------------------------------
-------------------------------------------------------GRRRR
---------------------------R----------K---------------------
----------------------------------------------R-------------
------------------------------------------------------------
------------------------------------------------------------
-----------N----------V-------E----------------------------N
IIVIP----------------------TD----------------------S----DA--
-------GTANYS------TP--------IR-------------GLSLELYIEEE-----
----------E-----------------Y------------I-P-E---LQ---------
------------------------------------------------------------
------------------------------------------------------------
----------QSSGVRVIVH-SQGLMP--FPEDDGFLAAPGFKTSVGLRQL------R--
-----LDRQ-----------PHP-Y--SE-C--VDT-I--------------YGP-DNIF
----KDF--YE-------T-YYSRK------VAAFFL----TF--CNCEAFM----Y---
------------------------------------------------------------
---------------------ACNGYA---------------------------------
------------------------------------NAV---------------------
IFF-------------------REFK-N--------------------------------
-------------------GEIT---YEAT-VSN------------------AVW-PN--
-------SAY------RNV-L-----------LR--------------------------
------------------------------------------------DL-M--------
----------------------------------QT------------------------
-S-------------SEI------------------RYKVE-------------------
---------------------------ADD------------------------------
-----T-------FI-----------S------DNMVKIDIYYNDLNYEAIG-ENVAYTG
GDV-----------------ISNLGGQVGLWIGVSVMTCFEFFEFLYDV-----------
--------MALF--------L----------I-------KL-------------------
TRAPQ---------------------------RK-------------------R------
----------------------------------------------VSTP----------
------------------------------------------------------------
------------------------------------------------------------
------------------------------------------------------------
-------------------------------IIP--------------------------
----------------------------------------------------------LR
HNNRIDFS
>Deu_Ambulacrar_hemi_Ptyfla_40v0_9_20150316_1g4330_t1_scaffold1875_cov91_435_
25
------------------------------------------------------------
----------M-------------------------------------------------
------------------------------------------------------------
------------------------------------------------------------
------------------------------------------------------------
--DP--------------------------------------------------------
------------------------------------------------------------
------------------------------------------------------------
------------------------------------------------------------
------------------------------------------------------------
------------------------------------------------------------
--LIDSKHK----------------------D---------SF-----S-----------

-EV--LTHLL-ANSSSHGLPNIQ---------R-SK--------------HP---VG---
--KLFWA----VLFLAGVGVLCWQVSVLVA-TYRK----------R---DVDVTL----K
-M-HS-N----------T-SIDF-----PVV-----------------------TICNTN
AL------KMS-----ALVDDV------------------Y-------------------
---------ML-------------------------------ADLYN----A--------
---------------------------------NS-------A-----------------
------------------------------------------------------------
---------------------------------FAT-----------IMQA---------
-------YRNIEHLDS-E-------------------------PEKSTGGT-EVNVYREE
------------------------------------------------------------
------------------------------------------------------------
------------------------------------------------------------
--------------------------------------------------IVL----SNS
TSAFVTGKY---------------------------------------------------
-----------SNIT----TST--------------------------------------
------------------------------------------------------------
-----YPSGEQK------------------------------SSTGPE------------
---------------------PSSQPMINNITYPTFAN------KTNIN------SSSLL
------------------------------MERELEDYI---------------------
------------------------DDE----------------DYYNKMDN----RHELQ
--YLL-------------------------------------------------------
----------------------------------------------R-------------
------------------------------------------------------------
------------------------------------------------------------
-----------Q----------I-----------------------MAAYG--------N
CYMFH-------VA-------------NS-N------------------EHT----KP--
-------LMINKP------GR--------DN-------------GLTLELFINQD-----
----------N-----------------Y------------I-P-D---IT---------
------------------------------------------------------------
------------------------------------------------------------
----------ETSGIRVAIH-PHRVVP--FPDDNGISLSPGYSTEIGLRMV------N--
-----IERQ-----------PRP-Y--GN-C--IDA--NNI-PDK-------YYD-NDIY
----IRR--YR-------A-AYSVAACEKSCYQNTLI----DR--CGCYDAD----Y---
------------P-P--TM-----------------------------------------
-------DGIDL-----K---PCSI-----------------------------------
------------------------------------------------------------
------------------------------------------------------------
------------------------------------------------------------
------------------------------------------------------------
------------------------------------------------------------
------------------------------------------------------------
------------------------------------------------------------
------------------------------------------------------------
------------------------------------------------------------
-----------------------LGNSTG-------------------------------
-------------------------------K----------------------------
------------------------------------------------------------
------------------------------------------------------------
------------------------------------------------------------
------------------------------------------------------------
------------------------------------------------------------
------------------------------------------------------------
------------------------------------------------------------
--------
>Deu_Ambulacrar_hemi_Ptyfla_40v0_9_20150316_1g1885_t1_scaffold848_cov143_437_
26
------------------------------------------------------------

----------M-------------------------------------------------
------------------------------------------------------------
------------------------------------------------------------
------------------------------------------------------------
------------------------------------------------------------
------------------------------------------------------------
------------------------------------------------------------
------------------------------------------------------------
---------------------------------------------------------AST
T----------------------RKPERT-------------------------------
---SLGDVR----------------------Q---------RF-----V-----------
-QR--SREYL-SRTSLHGASYII---------D-NN-------------IHP---FR---
--RCLWF----LLVTGLSLSLLVTLHTEVR-RYLE----------Y---PVSTIV----R
-M-NY-L----------P-RLLF-----PAV-----------------------TVCNYN
RY------RKS-----VIGGTP-----------------------------------ED-
---------AL------------------------------MRHLYF-------------
------------------------------------------------------------
------------------------------------------------------------
------------------------------------------------------------
------------------------------------------------------------
------------------------------------------------------------
------------------------------------------------------------
------------------------------------------------------------
---------------------TGSIYPTN-------------------------------
------------------------------------------------------------
------------------------------------------------------------
------------------------------------------------------------
------------------------------------------------------------
------------------------PDGFDWTTLENS-------------------TSTV-
---------------------------------NRSLFE---------------------
------------------------------------------------LEA----AHPKE
--SLI------ESCR----FK-G----D------GFDWP---C-----------G-P---
----------------------------------------------E-------------
------------------------------------------------------------
------------------------------------------------------------
-----------N----------F-------S-----------TTF-T-EFG--------V
CYSFN-------DD-------------LD-N--------------------A--------
-------ITTNLF------GS--------HT-------------GLRIRLFTNED-----
----------E-----------------Y------------T-H-G---PQ---------
------------------------------------------------------------
------------------------------------------------------------
----------TGSGFK--------------------------------------------
-----WEHL-----------PHP-YV-TN-C--SHG--------Q-------LR----YS
------------------TLTYSYSACCLEKMTDFVS----EK--CGCKEIH----M---
------------P-AF--------------------------------------------
-------GHQNT-----E------------------------------------------
------------------------------------------------------------
------------------------------------------------------------
------------FCN-VACEEVN---YIPH-ISY------------------ANI-PS--
-------KPL------VDD-L-----------WD--------------------------
------------------------------------------------KK----------
------------------------------------------------------------
-N----------------------------------------------------------
---------------------------MSR------------------------------
-----D-------FL-----------R------DNYADVSVFLEDLNCKQIT-FVPAITL
NSI---W-------------------TMPTAIDDSRLCSVPLARPNLIC-----------
-----TDGVTVV------------------------------------------------

--------------------------------RS--------------------------
------------------------------------------------------------
------------------------------------------------SSSTGL------
------------------------------------------------------------
------------------------------------------------------------
------------------------------------------------------------
----------------------------------------------------------SS
LSWLLFTY
>Deutero_Ambulac_Sakowv30036989m_439_27
------------------------------------------------------------
----------M-------------------------------------------------
---S--------------------------------------------------------
------------------------------------------------------------
------------------------------------------------------------
------------------------------------------------------------
------------------------------------------------------------
---------------------------------SSELNELTYRR----------------
------------------------------------------------------------
---------------------------------------------------------DSI
RTDIV--------------VTEGNDGNKSTAGLN--------------------------
-TTTAMNTQ----------------------T---------AF-----S-----------
-AK--LSQFM-GRTTLHGVAFTV---------G-PN-------------NSW---KR---
--RSVWI----FLVIVSVVFLCLCLFSLVG-LYLT----------Y---PVDTMT----S
-L-QF-T----------Q-SLTF-----PTV-----------------------TICNKN
RY------RKS-----VINGTA------------------F-----------------E-
---------EF------------L------------------WSIYP-------------
------------------------------------------------------------
------------------------------------------------------------
------------------------------------------------------------
------------------------------------------------------------
------------------------------------------------------------
------------------------------------------------------------
------------------------------------------------------------
---------------------LAGFGPQL-------------------------------
------------------------------------------------------------
------------------------------------------------------------
------------------------------------------------------------
------------------------------------------------------------
------------------------HVNYNWSLLENDPAL--Q------------------
---------------------------------NRTEFE---------------------
------------------------------------------------LTA----AHQLE
--DTI------LKCQ----FV-NSDDKH----------E---C-----------G-R---
----------------------------------------------E-------------
------------------------------------------------------------
------------------------------------------------------------
-----------N----------F-------T-----------TVF-T-EHG--------V
CYSFN-------NG-------------LD---------------------------NI--
-------LKATST------GS--------ST-------------GLHMLINVQRI-----
----------E-----------------Y------------T-I-G---PR---------
------------------------------------------------------------
------------------------------------------------------------
----------TGIGITVILS-EPGADI--TGENTALSIAPGVEALVAMPMD------K--
-----YLFL-----------KAP-YQ-TN-C--CTE--------G-------TS----YY
------------------P-VYSYHGCMNEKMSNEMA----EK--CNCKEIE----L---
------------P-G---------------------------------------------
----------PV-----R---TCTL-----------------------------------
------------------------------------QES---VECAL---------P---

L-K------------V-----------S-------NA--------L-----------TNI
------------DCN-VPCNFTL---YNPR-VTY------------------GHF-PS--
-------PSI------IQR-L-----------SD--------------------------
------------------------------------------------MY----------
------------------------------------------------------------
-G----------------------------------------------------------
---------------------------ISE------------------------------
-----E-------SV-----------K---------------------------------
------------------------------------------------------------
------------------------------------------------------------
------------------------------------------------------------
------------------------------------------------------------
------------------------------------------------------------
------------------------------------------------------------
------------------------------------------------------------
------------------------------------------------------------
------------------------------------------------------------
--------
>Protostome_Lophotroco_brch_Lingula_anatina_comp147337_c2_seq1_p1_comp147337_
c2__comp147337_c2_seq1_p1__ORF_type_complete_len_421___score__4_96_comp147337
_c2_seq1_410_1672__443_28
------------------------------------------------------------
------------------------------------------------------------
------------------------------------------------------------
------------------------------------------------------------
------------------------------------------------------------
------------------------------------------------------------
------------------------------------------------------------
------------------------------------------------------------
------------------------------------------------------------
------------------------------------------------------------
------------------------------------------------------------
------------------------------------------------------------
------------------------------------------------------------
---------------------------MFY-KFFE----------Y---DVGVKL----E
-I-TS-N----------S-TLKF-----PAV-----------------------TVCNEN
AF------RKS-----ALLSSP------------------------------------S-
---------KL----------------------------PTLDAFVT-------------
------------------------------------------------------------
------------------------------------------------------------
------------------------------------------------------------
-----------------NGT----------------------------------------
------------------------------------------------------------
------------------------------------------------------------
------------------------------------------------------------
-------------------------VPTTGF-----------------------------
------------------------------------------------------------
------------------------------------------------------------
------------------------------------------------------------
------------------------------------------NSSL--------------
-----------------------------------------IDSELGGR------SNRAT
------------------------------LIDKTLEDI---------------------
------------------------SEL----------------TTGEKQAL----GHQRS
--DFI------LDCE----YG-G----Y----------S---C-----------N-G---
----------------------------------------------K-------------
------------------------------------------------------------
------------------------------------------------------------

-----------T----------F-------E-----------SFY-STYNG--------N
CFIFN-------SG-------------WN-N--------------------S----SI--
-------LLATES------GP--------NF-------------GLSLTFNIEQS-----
----------E-----------------Y------------V-G-D---LT---------
------------------------------------------------------------
------------------------------------------------------------
----------QTAGVRVLVH-DQDVMP--FPEDQGFLAAPGEMTFVGIKMV------E--
-----IDRY-----------GGR-Y--TT-C--KKT--------DTF-----NIT-ENMY
----QAL--YP------SV-GYSQKACKKTCYQRTVI----ET--CKCALSI----Y---
------------P-R--VE-------------------TLFD------------------
-------STTSV-----R---TCDSL----------N-----------------------
------------------------------------ATD---VACTV---------K---
V-Q-------------------TQFI-N-------GE--------L------------NC
------------SCI-QPCSATS---FSID-TSS------------------AYW-PS--
-------DEY------ATE-F-----------LS--------------------------
------------------------------------------------NY-N--------
----------------------------------SK------------------------
-S-------------SIV------------------RKI---------------------
-----------------------YNT-SGS------------------------------
-----A-------GV-----------M------KNLAKVHIFYKDLNYEYIS-EYVYYD-
--------------------VSIIGIASCI-----LVVFLLNYEKRTLC-----------
--------TQLT--------A---------------------------------------
------------------------------------------------------------
------------------------------------------------------------
------------------------------------------------------------
------------------------------------------------------------
------------------------------------------------------------
------------------------------------------------------------
------------------------------------------------------------
--------
>Deutero_Ambulac_Apla_gbr467_14_t1_444_29
------------------------------------------------------------
----------M--------------------------------G----------------
------------------------------------------------------------
------------------------------------------------------------
----------------------------------------------------------IR
VKPIEDDISII-------------------------------------------------
------------------------------------------------------------
--SFNTDEVKIDKRERDKET-------------TGDSLKTGRAG------------P---
------------------------------------------------------------
------------------------------------------------------------
-----------------------QKGSKY-------------------------------
-----HSSR----------CT----------K---------FL-----A-----------
-TW--SSTLP-DAVGITGIQYIV---------S-NQ-------------DHI---LR---
--RFVWL----LALFGGFAAASYQTIDRTI-YFLS----------N---PKSVDY----E
-I-KY-I----------S-PLEF-----PAV-----------------------TVCNYN
SY------RAY-------------------------------------------------
------------------------------------------------------------
------------------------------------------------------------
------------------------------------------------------------
------------------------------------------------------------
------------------------------------------------------------
------------------------------------------------------------
------------------------------------------------------------
------------------------------------------------------------
------------------------------------------------------------
------------------------------------------------------------

```
------------------------------------------------------------
------------------------------------------------------------
------------------------------------------------------------
------------------------------------------------------------
------------------------------------------------------------
------------------------------------------------------------
-------------------WD-G---WG----------S---V-----------T-S---
----------------------------------------------A-------------
------------------------------------------------------------
------------------------------------------------------------
-----------N----------F-------T-----------RVI-T-DFG--------V
CYTFN-------AG-------------QA-G--------------------Q----EL--
-------LKQKVA------GK--------GH-------------GLSLMLDAQQY-----
----------F-----------------Y------------F-Y-S---SK---------
------------------------------------------------------------
------------------------------------------------------------
-------ALQVSAGFVVAIH-NQSEVP--QVDSLGVGVAPGTEVRIGLKRK------E--
-----AINL-----------EPP-H--GE-C--GSK--------E-------LK----YF
------------------S-SYSINSCRQECLTDFVL----EG--CGCVEPY----M---
------------A-----------------------------------------------
------------------------------------------------------------
------------------------------------------------------------
----------------------ERFN-S----------------------------EGVC
------------ECP-IPCQQTT---YTTS-LSF------------------ATF-PS--
-------DFY------IQT-L-----------VN--------------------------
---------------------------------------------LYSDF----------
------------------------------------------------------------
-N----------------------------------------------------------
---------------------------VTP------------------------------
-----E-------YF-----------S------SNTLKIEIYYEELSVESME-QQEAYTF
FAL-----------------LCDLGGALGLWLGGSILTFVEILDHFGHT-----------
--------AFLRG------------------T-------AF-------------------
QH----------------------------------------------------------
------------------------------------------------------------
------------------------------------------------------------
------------------------------------------------------------
------------------------------------------------------------
------------------------------------------------------------
------------------------------------------------------------
-------S
>Cnidar_NemVecNVEC200_012401_1_1_protein_AED_0_06_eAED_0_06_QI_181_1_1_1_0_72
_0_58_12_321_422_446_30
------------------------------------------------------------
----------M-------------------------------------------------
------------------------------------------------------------
------------------------------------------------------------
------------------------------------------------------------
------------------------------------------------------------
------------------------------------------------------------
------------------------------------------------------------
------------------------------------------------------------
------------------------------------------------------------
------------------------------------------------------------
-----SRSG----------------------P---------SV-----R-----------
-AL--LRDFS-DRTSCHGIGQIN---------G-SH--------------SP---TW---
--RIFWL----LTFLAGLGMVLFQCITLLG-IYLD----------K---PTATSV----D
-V-TY-D----------E-VTNF-----PAV-----------------------TICNLN
```

MI------KKK-----NLANFT--------------------------------------
---------QS-------------------------------KKIFD-------------
------------------------------DF-EA-------------------------
------------------------------------------------------------
------------------------------------------------------------
---------FVSSNSSMDSS--A------------------FLGSKMETVLK--------
------------------------------------------------------------
------------------------------------------------------------
------------------------------------------------------------
------------------------------------------------------------
------------------------------------------------------------
------------------------------------------------------------
------------------------------------------------------------
--DRLSMDSNDG------------------------------SNSV--------------
------------------------------------------SLDDTSM------DTELY
------------------------------VEDMLIRHM---------------------
------------------------AMV----------------DDKDLIKA----GHEFD
--ELV------FRCV----WN-G----F----------T---C-----------N-K---
----------------------------------------------G-------------
------------------------------------------------------------
------------------------------------------------------------
-----------G----------F---MKFWR-----------RFW-HWRYG--------N
CYIFN-------QG-------------VD-E-NGTL---------------L----AH--
-------LTSSKP------GP--------MY-------------GLTLDLFIDQE-----
----------Q-----------------Y------------L-I-P---LS---------
------------------------------------------------------------
------------------------------------------------------------
----------QEAGVKVLLS-DQRNVP--FPFTDGFSVQPGVSASVGIRKL------V--
-----INRI------------DP-FNNGS-C--YSG--------DG------LEK-DNIY
----SKH--KS-------L-KYSVQGCMSSCLANSEF----SI--CNCTEGK----F---
------------R-V---------------------------------------------
----------KG-----R---PCMS-----------------------------------
------------------------------------ESE---VKCLN---------T---
V-N-------------------KMYE-K-------GT--------L------------GC
----------TRKCP-QPCSHFS---FRRT-ISQ------------------SQW-S---
-------ESY------EQT-F-----------QK--------------------------
------------------------------------------------WS----------
------------------------------------------------------------
------------------------------------------------------------
------------------------------------------------------------
------------------------------------------------------------
------------------------------------------------------------
------------------------------------------------------------
------------------------------------------------------------
------------------------------------------------------------
------------------------------------------------------------
------------------------------------------------------------
------------------------------------------------------------
------------------------------------------------------------
------------------------------------------------------------
--------
>Deutero_Ambulac_Spurpu_014120_448_31
------------------------------------------------------------
----------M-------------------------------------------------
------------------------------------------------------------
------------------------------------------------------------
------------------------------------------------------------

```
------------------------------------------------------------
------------------------------------------------------------
------------------------------------------------------------
------------------------------------------------------------
------------------------------------------------------------
-----------------------KAPKVDGQEG--------------------------S
WGTAPEREE----------------------K---------SL-----R-----------
-TI--LNSRM-ENSSAHGVPNIQ---------R-SS--------------GP---VT---
--KLAWS----LVFLTGISVMTWQAVILFQ-TYFE----------W---NYSVNL----E
-V-RF-N----------R-TQSF-----PAI-----------------------TICNAN
PI------KRS-----ELETRD--------A-----------------------------
---------LF-------------------------------QALFD----V-HYVPSMP
DLP------------------------------DQ-------------------------
------------------------------------------------------------
---------------------QPLP-----------------------------------
-----------------D------------------------------------------
------------------------------------------------------------
------------------------------------------------------------
------------------------------------------------------------
------------------------------------------------------------
------------------------------------------------------------
------------------------------------------------------------
------------------------------------------------------------
------------------------------------------------------------
------------------------------------------------------------
------------------------------------------------------------
------------------------------------------------------------
------------------------------------------------------------
------------------------------------------------------------
------------------------------------------------------------
------------------------------------------------------------
------------------------------------------------------------
--------------------------------------------------------EQ--
-------PLPDVP-------------------------------GLSLELFVQQD-----
----------E-----------------Y------------V-E-G---MT---------
------------------------------------------------------------
------------------------------------------------------------
----------EVAGFRVSVH-HPSIMP--FPEYNGLLVSPGFATDIGLRVL------E--
-----VDRL-----------PKP-Y--GD-C--KAD--------LTQ-----GIE-DDIF
----HQH--YN-------I-TYNRKTCEVSCFQNEVI----SR--CDCFDAT----Y---
------------P-N--SL-----------------------------------------
------KVNRTV-----Y---PCEYI----------N-----------------------
------------------------------------DVE---TQCMA---------D---
I-E-------------------MEHA-R-------DE--------L------------EC
------------NCP-LACRETT---YLTS-VSS------------------SIW-PS--
-------DAY------EST-L-----------FQ--------------------------
------------------------------------------------KM-V--------
----------------------------------KY------------------------
-N-------------AEI------------------RRN---------------------
----------------------VVGE-NAS------------------------------
-----D-------WT-----------R------RNMAKVEIFYDEFNYEHIR-QDAAYTI
STI-----------------METRYLHPFFIHTNSIAYISETVHNLKQT-----------
--------FNVL--------S----------K-------SI-------------------
------------------------------------------------------------
------------------------------------------------------------
------------------------------------------------------------
------------------------------------------------------------
```

```
------------------------------------------------------------
------------------------------------------------------------
------------------------------------------------------------
--------
>Deutero_Ambulac_Apla_gbr500_4_t1_459_35
------------------------------------------------------------
----------M-------------------------------------------------
------------------------------------------------------------
------------------------------------------------------------
------------------------------------------------------------
------------------------------------------------------------
------------------------------------------------------------
------------------------------------------------------------
------------------------------------------------------------
------------------------------------------------------------
------------------------------------------------------------
---RASEKP----------------------A---------DT-----P-----------
-RF--VCSVL-ERIGAHGIPNIG---------R-AG--------------TQ---QR---
--RLAWT----GLVLAGLGFLVWQGTLLVT-RYLQ----------Y---DVKVSV----G
-L-GY-E----------A-LTTF-----PAV-----------------------TVCNLN
RI------PRT-----AMHTNS--------------------------------------
---------KL-------------------------------AELEG----YIEYIL---
---------------------------------ET-------------------------
------------------------------------------------------------
------------------------------------------------------------
------------------------------------------------------------
------------------------------------------------------------
------------------------------------------------------------
------------------------------------------------------------
------------------------------------------------------------
------------------------------------------------------------
------------------------------------------------------------
------------------------------------------------------------
------------------------------------------------------------
------------------------QVYQAWS----------DEMRYTNQ------VADFG
------------------------------ITTESFDIM---------------------
------------------------ADI----------------PYANRSSA----GHQLS
--RML------LYCT----FN-G----Y----------P---C-----------S-P---
----------------------------------------------L-------------
------------------------------------------------------------
------------------------------------------------------------
-----------N----------F-------T-----------HFY-NYFYG--------N
CYTFN-------FD-------------SR---------------------------LA--
-------RRVAQA------GP--------FY-------------GLTMELFVEQT-----
----------E-----------------Y------------V-R-G---LQ---------
------------------------------------------------------------
------------------------------------------------------------
----------NSAGLKVFIS-GQNQVP--FPEDRGIIVSPGRETSIALHKA---------
------------------------------------------------------------
--------------------------CEKSCVAQEIL----EM--CHCADPR--------
------------------------------------------------------------
------------------------------------------------------------
------------------------------------------------------------
------------------------------------------------------------
--------------------EVE---FEET-VSA------------------ASW-PN--
-------ENY------KFA-L-----------EN--------------------------
------------------------------------------------II-T--------
```

----------------------------------RL------------------------
-S-------------ADF------------------SSAIE-------------------
---------------------------EDD------------------------------
-----R-------FI-----------E------KNVVKLNVFLSSLQYQYIY-QTPDYSQ
ENL-----------------ISDLGGQVGLWMGFSILTFFEIAELVLDL-----------
--------FVYT-------------------RG------KS-------------------
AN------------------------------RV--------------------------
-----------------------------------------SREAKVGEGLVGMS-----
------------------------------------------------------------
------------------------------------------------------------
------------------------------------------------------------
-------------------------------LSP--------------------------
-----------------------------------------------------------S
VNTLTPIG
>Deutero_Ambulac_Apla_gbr243_2_t1_462_36
------------------------------------------------------------
----------M-------------------------------------------------
------------------------------------------------------------
------------------------------------------------------------
------------------------------------------------------------
------------------------------------------------------------
------------------------------------------------------------
------------------------------------------------------------
------------------------------------------------------------
------------------------------------------------------------
------------------------------------------------------------
-----EGNR----------------------S---------KL-----V-----------
-AD--FKKFG-DQTTFHGLRYVT---------N-NT-------------YHS---VR---
--RLIWL----AVVLGMTTWLVINIIRAVV-YMFQ----------Y---PETSSI----S
-V-NY-V----------P-NITF-----PAV-----------------------TLCNFN
QY------RKD-----ALDARA------------------I-----------------K-
---------IL-------------------------------AKVFG-------------
------------------------------------------------------------
------------------------------------------------------------
------------------------------------------------------------
------------------------------------------------------------
------------------------------------------------------------
------------------------------------------------------------
------------------------------------------------------------
-------------------------NPALR------------------------------
------------------------------------------------------------
------------------------------------------------------------
------------------------------------------------------------
------------------------------------------------------------
-----------------------ESIDLDVSDVKLFPT-----------------TNQA-
---------------------------------NVTATT---------------------
------------------------------------------------VQA----THQIE
--DML------VDCH----WR-T----E----------P---C-----------S-H---
----------------------------------------------L-------------
------------------------------------------------------------
------------------------------------------------------------
-----------N----------F-------T-----------QRL-T-DYG--------V
CYTFN-------DD-------------LA-G-------------------LG----DV--
-------LTIQNP------GA--------SN-------------GLHLRLNIQQD-----
----------L-----------------Y------------T-F-G---ES---------
------------------------------------------------------------
------------------------------------------------------------

----------TAAGMKVLLH-PQGEFP--IMKEFAFSLSPGFETSVAVRKE------T--
-----VTSL-----------KAP-YK-SD-C--IDG--------S-------LK----EF
------------------PYKYSVAACQLECRALYVI----DK--CGCRDIR----W---
------------P-----------------------------------------------
------------------------------------------------------------
------------------------------------------------------------
----------------------EEFL-S-------RG--------------------ERC
------------HCP-MACETTT---YQSK-LSL------------------AYW-PA--
-------GYL------TAE-L-----------QA--------------------------
------------------------------------------------KR----------
------------------------------------------------------------
-N----------------------------------------------------------
---------------------------LTE------------------------------
-----D-------FI-----------R------KNYIDVYIYFEEIMYMAIE-QKMAYTI
DNV-----------------QSDIGGYLGLLCGMSLITLVEWADFILIT-----------
--------LYKR--------C----------R-------RL-------------------
AS------------------------------RR--------------------------
------------------------------------------------------------
------------------------------------------------------------
------------------------------------------------------------
------------------------------------------------------------
------------------------------------------------------------
------------------------------------------------------------
-----IST
>Deu_Ambulacrar_hemi_Ptyfla_40v0_9_20150316_1g8126_t1_scaffold4191_cov132_466
_38
------------------------------------------------------------
----------M-------------------------------------------------
------------------------------------------------------------
------------------------------------------------------------
------------------------------------------------------------
------------------------------------------------------------
------------------------------------------------------------
------------------------------------------------------------
------------------------------------------------------------
------------------------------------------------------------
-----------------------SSSEFA---------------------------WDGK
KICEHETAK----------------------N---------TL-----S-----------
-VI--FDELL-KNSSSHGLPNIQ--------R-SV---------------SA---VG---
--KLSWA----LLFFVAVIMLLIQTVGLLQ-TYFS----------Y---PVAVSL----T
-M-EF-D----------K-KLFF-----PAV-----------------------TVCNAN
PV------RMS-----ELRAAS------------------------------------DY
---------RL------------------------------RSQFD-----P-SFSPSVK
--------------------------VEVEEVTDS-------------------------
------------------------------------------------------------
--------------------GKPLIT-----------------------EGGS-------
-----------------NTT---------------------PVGKLKFSRTTDV------
------------------------------------------------------------
------------------------------------------------------------
------------------------------------------------------------
--------------------ETTLAQPTTTENSS--------------------------
------------------------------------------------------------
------------------------------------------------------------
------------------------------------------------------------
------------------------------------------------------------
------------------------EPIMSWDDR-------VGGDFYRKP------SNGFS
------------------------------KANSLSVLM---------------------

------------------------ANQ----------------STQTKMTM----GHQLS
--TML------LDCF---------------------------------------------
------------------------------------------------------------
------------------------------------------------------------
------------------------------------------------------------
----------------------Y-----------------------NYKYG--------N
CFTFN-------SG-------------RN----------------------G----EA--
-------LTINKP------GP--------MY-------------GLSLELYIEQI-----
----------E-----------------Y------------M-I-G---TT---------
------------------------------------------------------------
------------------------------------------------------------
----------EAAGVRVLLH-HQDEMP--FPEDEGFSVAPGQAASVGVRQM------E--
-----LLRK-----------SAP-Y--GD-C--KDM--------ENLSQ--EEID-NNIF
----MSR--FD-------V-GYSVQVCGRNCYQYEVI----EE--CGCFDSN----Y---
------------P-I--PV-----------------------------------------
------SAEGRY-----D---ACDIV----------S-----------------------
------------------------------------DEK---AITVM---------N---
L-LP------------------QTLI---------ET-----------------------
--------------------------YKLL--------------------------PN--
--------------------M---------------------------------------
------------------------------------------------EW----------
------------------------------------------------------------
-N----------------------------------------------------------
------------------------------------------------------------
------------------------------------------------------------
------------------------------------------------------------
--------MPLL--------C---------------------------------------
------------------------------------------------------------
------------------------------------------------------------
------------------------------------------------------------
------------------------------------------------------------
------------------------------------------------------------
------------------------------------------------------------
------------------------------------------------------------
--------
>Deutero_Ambulac_Apla_gbr40_206_t1_468_39
------------------------------------------------------------
----------M---------------------RVDKLVKI----L---------------
---------------------------------GWE------------------------
------------------------------------------------------------
----------------------------------------------------------VH
GDPNSAAVSI------------------------------------------------PG
EVLDQTLCAEAPREVF--------------------------------------------
----------------HKPT-------------SGEVPKTR-------------------
-----QPAWHVCSRM---------------------------------------------
---------------------------------------------------------DRV
W----------------------TKEEK--------------------------------
--------R----------------------R---------TS---HGA-----------
-SLRLKTALK-DDTSVVGIKNVV---------D-EK-------------VQT---GR---
--K---------------------------------------------------------
-----------------K--------------------------------------CAVE
KY------GPE-----FVDYID------------------L-----------------A-
-------------------------YPV---------------GMMN-------------
---------------------------------NK-------QT----------------
------------------------------------------------------------
------------------------------------------------------------
------------------------------------------------------------

------------------------------------------------------------
------------------------------------------------------------
------------------------------------------------------------
--------------------------P---------------------------------
------------------------------------------------------------
------------------------------------------------------------
------------------------------------------------------------
------------------------------------------------------------
---------------------------------------------PDFMV-----FKDL-
---------------------------------DLKQFY---------------------
------------------------------------------------VDT----GHLKD
--DMI------LECS----WQ-G----Q----------P---C-----------G-H---
----------------------------------------------D-------------
------------------------------------------------------------
------------------------------------------------------------
-----------D----------F-------I-----------STL-T-DFG--------M
CHTFN-------SH-------------TK-G--------------------M----AV--
-------RRIKHR------GS--------RF-------------GLRLRLNVETD-----
----------Q-----------------Y------------M-P-G---PR---------
------------------------------------------------------------
------------------------------------------------------------
----------DSVGIKVALH-HQNDIPKGMSKDLGFALSPGTYNLAEVTMS------Q--
-----ITNQ-----------PMP-F--GK-C--ATK--------Q-------LK----HV
------------------E-MYSVMACILDCQTDYVV----EK--CHCKDAF----M---
------------P-----------------------------------------------
------------------------------------------------------------
------------------------------------------------------------
----------------------ESFI-S-------SF--------ET-----------NC
------------SCP-VPCERTE---YSTM-LSS------------------GIF-PA--
-------RHV------AMA-L-----------DK--------------------------
------------------------------------------------EL----------
------------------------------------------------------------
-N----------------------------------------------------------
---------------------------MSM------------------------------
-----D-------YS-----------R------NNLLELIVYFEELKYTRIT-QQPAYSL
ESL-----------------LSDIGGSMGLFMGVSILTLFELMDVLLRT-----------
--------VY--------------------------------------------------
------------------------------------------------------------
------------------------------------------------------------
------------------------------------------------------------
------------------------------------------------------------
------------------------------------------------------------
------------------------------------------------------------
------------------------------------------------------------
--------
>Deu_Ambulacrar_hemi_Ptyfla_40v0_9_20150316_1g28005_t1_scaffold79692_cov99_47
0_40
------------------------------------------------------------
----------MFSESPDLSQLT-----GVLQLSVDEISELGHFILQCTYAGEKCDANETN
LESARANLVRLKI-YFETLNHDL-TSE--LPAYTWEDLLS--------------------
---------------------------DVGGTLGLYVG--------------------FS
VITVCE--------------------------FVTLLFNLCGKSPHLDHTKKEEQPLHER
VSDTNEG-----------------------------------------------------
------------------------------------------------------------
---------------------------------TNEFS----------------------
------------------------------------------------------------
-----------------------DKTNV-------------------------QHDFFKH

S----------------------DDTGK--------------------------------
------QHD----------------------E---------SV-----K-----------
-HV--LEDFG-RETTAHGIVHIT---------T-AK--------------SN---VT---
--RSTWI----MIVLVAACVMFVQMTLLLV-QYFE----------Y---NVNVKV----T
-L-VS-E----------K-ALHF-----PSV-----------------------TICNTN
KL------RHS-----AIRSSK------------------Y-----------------S-
---------EML---------MLERNFVS--------------PYYA-------------
---------------------------PCIEG-DF-------------------------
------------------------------------------------------------
-----------QCK-----------------------------------NGIH-------
-CIKPYLICDGVNHCG-D-M------------------------SDEHNCA-----YP-E
------------------------------------------------------------
------------------------------------------------------------
-----------------CGPD------------------------Q--------------
------------------------------------------------------------
------------------------------------------------------------
------------------------------------------------------------
------------------------------------------F-----------------
----RCDSGSEL---GVC---IPNDKFC---------------DRISHC----Y------
--------GA--------------EDETSCG-----------------------------
------------------------------------------------------------
------------------------------------------------------------
------------------------------------------------------------
-------------ECSEDEFQCITSRECVASLKTCDQLFDCRDGSDESASICGKQ-----
------------------------------------------------------------
------------------------------------------------------------
------------------------------------------------------------
------------------------------------------------------------
------------------------------------------------------------
------------------------------------------------------------
------------------------------------------------------------
------------------------------------------------------------
------------------------------------------------------------
------------------------------------------------------------
------------------------------------------------------------
------------------------------------------------------------
------------------------------------------------------------
------------------------------------------------------------
------------------------------------------------------------
------------------------------------------------------------
------------------------------------------------------------
------------------------------------------------------------
------------------------------------------------------------
------------------------------------------------------------
------------------------------------------------------------
------------------------------------------------------------
------------------------------------------------------------
------------------------------------------------------------
------------------------------------------------------------
------------------------------------------------------------
------------------------------------------------------------
------------------------------------------------------------
------------------------------------------------------------
------------------------------------------------------------
------------------------------------------------------------
--------

>Deu_Ambulacrar_hemi_Ptyfla_40v0_9_20150316_1g5424_t1_scaffold2509_cov138_473
_42
------------------------------------------------------------
----------M-------------------------------------------------
------------------------------------------------------------
------------------------------------------------------------
------------------------------------------------------------
------------------------------------------------------------
------------------------------------------------------------
------------------------------------------------------------
------------------------------------------------------------
------------------------------------------------------------
------------------------------------------------------------
-----------------------------------------AC-----D-----------
-DV--DKEFF-TGTTIHGVGRIV---------E-RS--------------RL---VF---
--KILWF----FIVLVAFGFCAWQITDRFV-RFFE----------F---NVNTEI----T
-L-KF-E----------S-SIEF-----PAI-----------------------TICNFN
RY------YSS-----AITASD------------------Q-----------------A-
---------IV------------N-QLL---------------YAMD----Y--------
---------------------------------DY-------DYSDS-------------
------------------------------------------------------------
------------------------------------------------------------
-----------------E------------------------------------------
------------------------------------------------------------
------------------------------------------------------------
------------------------------------------------------------
------------------------------------------------------------
------------------------------------------------------------
------------------------------------------------------------
------------------------------------------------------------
------------------------------------------------------------
------------------------YSSFDWDNWYSNLSL----------------AADF-
---------------------------------DYANFT---------------------
------------------------------------------------RRV----GMQLD
-DTTL------LSCE----WR-G----R--------SSS---C-----------S-A---
----------------------------------------------R-------------
------------------------------------------------------------
------------------------------------------------------------
-----------E------------------------------------------------
------------------------------------------------------------
------------------------------------------------------------
----------E-----------------Y------------T-E-S--------------
------------------------------------------------------------
------------------------------------------------------------
------ASGNVQAGVKFLVH-DKNTPP--LMDSQGSAVAPGTYAFVAVQQV------R--
-----TVSL-----------SKP-Y--GS-C--VSS-S------T-------LQ----YY
------------------D-EYSLSACLIEWRLELIV----QE--CGCKPFR----Y---
------------P-G---------------------------------------------
----------PA-----R---DCTV-----------------------------------
------------------------------------TES---ATCAK---------T---
T-L-------------------GRIK-N-------GD--------F-----------GQS
------------SCY-VPCNMTS---YPTS-VSY------------------AQY-PS--
-------QSV------APE-M-----------AS--------------------------
------------------------------------------------FY----------
------------------------------------------------------------
-N----------------------------------------------------------
---------------------------VSE------------------------------

```
-----S-------YG-----------R------NNLIYLDIYYDEINYEELK-QTEAMTP
SAL-----------------VSDIGGQLGFFLGASIITLCEFIEYLIMK-----------
--------CHGA--------C----------R-------TK-------------------
SK------------------------------KV--------------------------
--------W--------------------------------DSNDE--------------
--------------------KEKKVDVA-------------------ANGDTEL------
------------------------------------------------------------
------------------------------------------------------------
------------------------------------------------------------
-----------------------------------------------------------N
FGPRFAFS
>Deutero_Ambulac_Spurpu_024200_477_43
------------------------------------------------------------
----------M-AQ----------------------------------------------
------------------------------------------------------------
------------------------------------------------------------
------------------------------------------------------------
------------------------------------------------------------
-----------------------------QAKEAWYDAPSKPVTA---------------
------------------------------------------------------------
------------------------------------------------------------
------------------------------------------------------------
-------------------FKSLDKESATPIEQ--------------------------P
SFNMSKEEE----------------------N---------SV-----L-----------
-FL--VGNRM-ASTGAHGIPNIQ---------R-AS--------------NI---WR---
--RLIWL----LLFTGGVTVVVWQLTTIIN-NFFA----------Y---EVDVNI----D
-F-VR-Q----------R-AIKF-----PAI-----------------------TVCNLN
PV------PLS-----KADVLW-----------------NIAS-------------NEED
AT-------SL-------------------------------RDFLP----A--------
------------------------------------------------------------
------------------------------------------------------------
------------------------------------------------------------
------------------------------------------------------------
------------------------------------------------------------
------------------------------------------------------------
------------------------------------------------------------
------------------------QKPSGVMNATQNQ-----------------------
------------------------------------------------------------
------------------------------------------------------------
------------------------------------------------------------
------------------------------------------NDAGASM-----------
------------------------RGYQDW-----------VNVDFNQL------NTRVS
------------------------------NLEVAQSII---------------------
------------------------GST----------------AYENRSET----GHDLD
--DML------LDCN----FE-N----Y----------P---C-----------A-P---
----------------------------------------------D-------------
------------------------------------------------------------
------------------------------------------------------------
-----------N----------F-------T-----------HFY-NYMYG--------N
CYTFN-------SG-------------QT----------------------G----KP--
-------LEVSTV------GP--------LY-------------GLSLELYIQQS-----
----------E-----------------Y------------I-S-D---LQ---------
------------------------------------------------------------
------------------------------------------------------------
----------PSAGLRLLVH-DQNEMP--FPEDRGINLAPGAHTSVGLSLV------Y--
-----LERK-----------ESP-Y--TN-C--SLE----------------FKP-GNVF
----KEQ--FD------HL-EYS-------------------------------------
```

------------------------------------------------------------
------------------------------------------------------------
------------------------------------------------------------
------------------------------------------------------------
------------------------------------------------------------
------------------------------------------------------------
------------------------------------------------------------
------------------------------------------------------------
------------------------------------------------------------
------------------------------------------------------------
--------------------------R------S-------------------------I
YSL-----------------ISDLGGQIGLWIGVSILTIFEFIELIYDL-----------
--------IKLI--------C----------L-------KL-------------------
MDPRM---------------------------KK-------------------KR-----
---------------------------------------SSQSQTKQSNVM---------
------------------------------------------------------------
------------------------------------------------------------
------------------------------------------------------------
------------------------------------------------------------
----------------------------------------------------ALDGQQGR
TNSLFADR
>Deutero_Ambulac_Sakowv30044075m_476_44
------------------------------------------------------------
----------M-------------------------------------------------
------------------------------------------------------------
------------------------------------------------------------
------------------------------------------------------------
------------------------------------------------------------
------------------------------------------------------------
------------------------------------------------------------
------------------------------------------------------------
------------------------------------------------------------
------------------------------------------------------------
------------------------------------------------------------
------------------------------------------------------------
------------------------------------------------------------
------------------------------------------------------------
------------------------------------------------------------
------------------------------------------------------------
------------------------------------------------------------
------------------------------------------------------------
------------------------------------------------------------
------------------------------------------------------------
------------------------------------------------------------
------------------------------------------------------------
------------------------------------------------------------
------------------------------------------------------------
------------------------------------------------------------
------------------------------------------------------------
-------------------------------------------PTTLRASPIKDIVNIHS
LQERYQELTQDTRLFDEF----RVNY--------------Y-QGNH-------FY-----
-------QSDL-----------ANNDTPDWKRYLTY-----------SS------STDYS
---------------------------------DLRDVL---------------------
-------------------------TL----------------NMSDVFKY----GHQFE
--DFV------LQCA----FN-E----N----------K---C-----------D-Q---
----------------------------------------------S-------------
------------------------------------------------------------

------------------------------------------------------------
-----------D----------F-------S-----------TFQ-NDRYG--------N
CFTFN-------NV-------------RN-D--------------------A----LT--
-------RRADKM------GS--------QY-------------GLKLTLYIEQD-----
----------E-----------------Y------------I-P-L---YG---------
------------------------------------------------------------
------------------------------------------------------------
----------QEAGVRILVH-PSDITP--FPEDDGITVAPGRKASIALREE------L--
-----HSSV-----------AHP-Y--SN-C--ENN--------PQ------I---DSIF
----GSQ-----------Y-NYSVQACRKTYLHQKIQ----QK--CNCRDTL----W---
------------L----SK-----------------------------------------
----------DR-----P---ACKIT----------N-----------------------
------------------------------------TAQ---EVCRR---------L---
I-N-------------------FLDR-R-------GV--------F------------EN
------------NCK-EPCRKTT---FKKT-VSQ------------------SMW-PS--
-------NKH------LGG-L-----------LK--------------------------
------------------------------------------------VL-R--------
----------------------------------PI------------------------
-N-------------TKI------------------RDKV--------------------
---------------------------VDE------------------------------
-----K-------SA-----------R------DNLVLIEVFFEVPNYRSIL-NTPAYTV
DQL-----------------FADIGGLMGLYVGISLITVVEFIELVYNLYNYFSL-----
------NREEKL--------A----------K-------RA-------------------
KKM-----------------------------ES-------------------K------
----------------------------------------NMNHKYDP------------
------------------------------------------------------------
------------------------------------------------------------
------------------------------------------------------------
------------------------------------------------------------
------------------------------------------------------IPCNLP
QSMIGTSV
>Placozoa_HhoNaC67_TR9228_c0_g1_i2_m_20240__Hoilungia_hongkongensis__30_45
------------------------------------------------------------
----------M-------------------------------------------------
------------------------------------------------------------
------------------------------------------------------------
------------------------------------------------------------
------------------------------------------------------------
------------------------------------------------------------
------------------------------------------------------------
------------------------------------------------------------
------------------------------------------------------------
------------------------------------------------------------
----------------------------S--Q---------EK-----F-----------
-NH--EERFA-TSTSYHGMAHIY---------D-GQ-------------TSK---QT---
--RSFWL----VLTLIATGACISQCVIIIL-NASQ----------L---PTRIVT----K
-T-VL-Q----------N-SSIF-----PSV-----------------------TICNTN
DY------DRN-----SLSSSD------------------I-----------------Q-
---------HL------------S-TIV--------------DFVYG-------------
------------------------------------------------------------
------------------------------------------------------------
-----------------Y-DI---------------------------------------
-----------------E------------------------------------------
------------------------------------------------------------
------------------------------------------------------------
------------------------------------------------------------
------------------------------------------------------------

------------------------------------------------------------
------------------------------------------------------------
------------------------------------------------------------
--------PHRF------------------------------EAA---------------
------------------------ISNLSI------------------KF-----GKGF-
---------------------------------NYELFL---------------------
------------------------------------------------RRA----GHKLE
--NML------LSCT----WM-G----R----------P---C-----------S-I---
----------------------------------------------E-------------
------------------------------------------------------------
------------------------------------------------------------
-----------D----------F-------V-----------NTT-T-AAG--------S
CFTFN-------PG-------------ST-N--------------------T----SI--
-------KNQTAS------GN--------AN-------------GLRLILNIEQY-----
----------K-----------------Y------------Y-P-A--LYT---------
------------------------------------------------------------
------------------------------------------------------------
-------PGQPDAGLRYSLH-YYKDPP--NLIAESYYAAPGFHTYVPMTLK------R--
-----EKRL-----------KKP-W--GD-C--EEL--------S-------LK----RY
------------------Q-FYSRYACLNELSAKYAS----ES--CNCSYQD----N---
------------E-A---------------------------------------------
----------------PN---PCDG-----------------------------------
------------------------------------VQF---LSCII---------P---
T-A-------------------GYYR-T-------QL--------SQ----------NLS
------------VCP-IACETFS---FATE-ISQ------------------SSI-AS--
-------QAF------NSA-I-----------DS--------------------------
------------------------------------------------FI----------
------------------------------------------------------------
-N---------------V------------------SGLISDAQSKYW------------
-------------------LP--QSY-TNV------------------------------
-----D-------FI-----------R------DNIVYLDIYYRKLDMTQIL-QLEDTGF
SKV-----------------LS---------------TVCIYLQITFTQ-----------
------------------------------------------------------------
------------------------------------------------------------
------------------------------------------------------------
------------------------------------------------------------
------------------------------------------------------------
------------------------------------------------------------
------------------------------------------------------------
------------------------------------------------------------
--------
>Deutero_Urochord_tr_F6ULW5_F6ULW5_CIOIN_Uncharacterized_protein_OS_CioInt_48
1_47
---------------------------V--------------------------------
----------M-------------------------------------------------
------------------------------------------------------------
------------------------------------------------------------
------------------------------------------------------------
------------------------------------------------------------
------------------------------------------------------------
------------------------------------------------------------
------------------------------------------------------------
------------------------------------------------------------
------------------------------------------------------------
-------------------------------S---------SL-----S-----------
-DL--IYEFA-QRTTCHGLSQIL---------Q-TK-------------RKT---IY---
--RCLWF----MMVSGLCGGATWQLTQAVQ-EYLS----------Y---KTSTDI----Q

-E-MY-T----------S-ELPF-----PAV-----------------------TICNIN
SF-------------SHLVKKF------------------L-----------------N-
---------VL------------------------------SDKLFS--V----------
------------------------------------------------------------
------------------------------------------------------------
---------------------WP-------------------------------------
------------------------------------------------------------
------------------------------------------------------------
------------------------------------------------------------
------------------------------------------------------------
-------------------------SPEFEMIA---------------------------
------------------------------------------------------------
------------------------------------------------------------
------------------------------------------------------------
--------FNNF------------------------------DNWWRR------------
------------------------EVELYYEKLNERG------------------IDKF-
---------------------------------YLSDEI---------------------
------------------------------------------------LLK----GWTLN
-ETTL------SHCT----FR-Q----K----------P---C-----------S-I---
----------------------------------------------Q-------------
------------------------------------------------------------
------------------------------------------------------------
-----------N----------F-------T-----------QVV-T-PMG--------L
CYTFT-------SN-------------ES-------------------------------
-------MYQNVP------GS--------GH-------------GLVVHLNIDQK-----
----------R-----------------Y------------S-E-H---PK---------
------------------------------------------------------------
------------------------------------------------------------
-------YGNPNAGVKVHVH-NHNDPS--EVQNYGIGVPSGFGAHIVIRKT------E--
-----RILM-----------YKP-W--GV-C--SEK-T------KT------LL----HH
------------------D-HYSTAGCLRECRAEYVY----SK--CGCKPYWAL-NM---
--------------A---------------------------------------------
----------NV-----P---ECDA-----------------------------------
------------------------------------LNI---LNCSG---------S---
A-D-------------------AVFS-T-------KF--------SL----------RTC
------------ECV-EPCNFVV---YDSS-TSY------------------STF-PS--
-------IEV------GEF-F-----------AL--------------------------
------------------------------------------------HA----------
------------------------------------------------------------
-G----------------------------------------------------------
---------------------------KNL------------------------------
-----R-------KY-----------R------ENHLMLDIFYDSLSYVRMK-QSKAVTE
ISL-----------------MSDMGGLLGLWLGISLVTVSEFVQLFF-------------
------------------------------------------------------------
------------------------------------------------------------
------------------------------------------------------------
------------------------------------------------------------
------------------------------------------------------------
------------------------------------------------------------
------------------------------------------------------------
------------------------------------------------------------
--------
>Deutero_Ambulac_Sakowv30035231m_482_48
------------------------------------------------------------
----------M-------------------------------------------------
------------------------------------------------------------
------------------------------------------------------------

```
-----------------------------------------------------------D
ASNSSRNIM---------------------------------------------------
------------------------------------------------------------
--------------ETVFQT-------------TNDVHPSSD------------------
------------------------------------------------------------
---------------------------------------------------------EDM
S----------------------HQRNCA-------------------------------
-------------------------------P---------RC-----R-----------
-DR--LREFT-QETTLHGIRFTT---------D-SK-------------SNI---YR---
--RLMWV----LFVSLGTAGVVITLKDSAI-RFIA----------Q---PVTTVV----E
-I-HH-P----------N-AVPF-----PAV-----------------------TVCNYN
KY------RQS-----VLAGTW------------------A-----------------E-
---------DL------------L------------------YKMYG-------------
------------------------------------------------------------
------------------------------------------------------------
------------------------------------------------------------
------------------------------------------------------------
------------------------------------------------------------
------------------------------------------------------------
------------------------------------------------------------
-------------------------SP---------------------------------
------------------------------------------------------------
------------------------------------------------------------
------------------------------------------------------------
--------VQGE------------------------------------------------
------------------------IGEVDWSNYTDV-------------------ITNL-
---------------------------------NRTAFE---------------------
------------------------------------------------MEA----AHQIE
--DML------IACT----WS-S----E---------TA---C-----------G-P---
----------------------------------------------E-------------
------------------------------------------------------------
------------------------------------------------------------
-----------N----------F-------T-----------NVM-T-NYG--------V
CYTFN-------GD-------------VK-N--------------------G--------
-------LTVKNR------GS--------MF-------------GLYMVLDIEQH-----
----------E-----------------Y------------M-P-G---VS---------
------------------------------------------------------------
------------------------------------------------------------
----------KGAGVKIMTH-TQTDVP--LVNEVSMSFGPGMDSTIALRLH------Q--
-----TKLLR--------------------C--TNQ--------P-------LQ----YF
------------------D-VYSRANCEMESFTLAAT----KS--CTCREPY----M---
------------P-----------------------------------------------
------------------------------------------------------------
------------------------------------EES---------------------
----------------------------------------------------------EL
------------SCE-SACQEVI---YDST-ISY------------------GSY-PS--
-------LQI------LKK-I-----------QL--------------------------
------------------------------------------------MR----------
------------------------------------------------------------
-N----------------------------------------------------------
---------------------------ITE------------------------------
-----A-------EI-----------R------ENYLAISVFIEDLTVLSTE-EEYSYTY
DAF-----------------LGDIGGQFGLCLGASLLTLLEFFDFLMMS-----------
--------LCGA--------C----------R-------RW-------------------
YN------------------------------RS-------------------R------
------------------------------------------------------------
------------------------------------------------------------
```

------------------------------------------------------------
------------------------------------------------------------
------------------------------------------------------------
------------------------------------------------------------
------KQ
>Cnidar_NemVecNVEC200_006432_1_1_protein_AED_0_27_eAED_0_27_QI_414_1_1_1_1_1_
6_1486_449_486_51
------------------------------------------------------------
----------M-------------------------------------------------
------------------------------------------------------------
------------------------------------------------------------
------------------------------------------------------------
------------------------------------------------------------
------------------------------------------------------------
------------------------------------------------------------
------------------------------------------------------------
--------------------------IMAAA--------------------------KKL
L----------------------DGRKK--------------------------------
--------R----------------------A---------KV-----K-----------
-EL--WENFL-GGCTLHGFHYCF---------A-GN--------------PP---LR---
--RLIWS----LLLLGAFAMFFEKCTESFI-NFFD----------Y---PFTTTT----L
-L-VY-D----------K-RLPF-----PAI-----------------------SMCNYN
DA------RMS-----KMNGTL------------------M-----------------N-
---------EI----------------------------------FV-------------
------------------------------------------------------------
------------------------------------------------------------
------------------------------------------------------------
------------------------------------------------------------
------------------------------------------------------------
------------------------------------------------------------
------------------------------------------------------------
------------------------------------------------------------
------------------------------------------------------------
------------------------------------------------------------
------------------------------------------------------------
------------------------------------------ASKL--------------
------------------------E-GRNTSHLQSQ-------------------LTGE-
---------------------------------LMQRTL---------------------
------------------------------------------------KEA----AHRLP
--DMI------KECS----WQ-K----H---------GK---C-----------S-W---
----------------------------------------------K-------------
------------------------------------------------------------
------------------------------------------------------------
-----------N----------F-------T-----------SFK-S-ADGD-------T
CYTFN-------SG-------------RK-D-------------------------PI--
-------LSMSNV------GE--------EN-------------GLRLVIDTQHS-----
----------E-----------------Y------------Y-Y-D--------------
------------------------------------------------------------
------------------------------------------------------------
---------VKNAGFKVILH-DQGETP---VKMQGLSVSPGFTSYMELKRT------K--
-----VTNL-----------PFP-YK-TM-C--GMP--------E-------LK----YF
------------------N-SYSKSKCFLDKLTQYVV----TL--CGCRDWF----M---
------------P-G--QG-----------------------------------------
----------KI-----P---VCDY-----------------------------------
------------------------------------ETA---ASCMW---------K---
A-W-------------------AYFE-E-------NK---------------------LD
------------QCP-VACNSVE---YSAQ-LSY------------------ARF-PA--

-------NNY------AKM-L-----------AK--------------------------
------------------------------------------------EY----------
------------------------------------------------------------
-G---------------L------------------KGS---------------------
------------------------DE-ENR------------------------------
-----Q-------YL-----------R------DNLIEIKIYYEDLTYFDVQ-QVPSYDL
YSL-----------------LGDVGGQIGLFLGASLLTVVEYLDLLGMV-----------
--------AYTS--------F----------K----------------------------
------------------------------------------------------------
------------------------------------------------------------
------------------------------------------------------------
------------------------------------------------------------
------------------------------------------------------------
------------------------------------------------------------
------------------------------------------------------------
-----YRN
>Cnidaria_HyNaC9_NP_001296667_1_acidsensing_ion_channel_1like__Hydra_vulgaris
__31_52
------------------------------------------------------------
----------M-------------------------------------------------
------------------------------------------------------------
------------------------------------------------------------
------------------------------------------------------------
------------------------------------------------------------
------------------------------------------------------------
------------------------------------------------------------
------------------------------------------------------------
-------------------------SEEDEK--------------------------KIK
S----------------------PHEQR--------------------------------
--------N----------------------D---------RV-----K-----------
-EH--VAHLI-KNVSFHGLSYVA---------D-KR-------------NNY---FR---
--RAIWF----LITVGAFIYAVEKVYESTV-NYFS----------Y---PFKTAR----M
-K-IY-V----------N-ELNF-----PAV-----------------------SFCNLN
DF------LFS-----KLNGTK------------------L-----------------D-
---------ES----------------------------------IL-------------
------------------------------------------------------------
------------------------------------------------------------
------------------------------------------------------------
------------------------------------------------------------
------------------------------------------------------------
------------------------------------------------------------
------------------------------------------------------------
------------------------------------------------------------
------------------------------------------------------------
------------------------------------------------------------
------------------------------------------------------------
------------------------------------------YPDDPE------------
---------------------------------KNN-------------------VSEI-
---------------------------------EISNIT---------------------
------------------------------------------------SDA----TIRLD
--QML------VDCE----FE-G----K----------K---C-----------T-H---
----------------------------------------------E-------------
------------------------------------------------------------
------------------------------------------------------------
-----------N----------F-------T-----------DFW-T-MQGE-------L
CFTFN-------SG-------------KN-S--------------------H----SL--
-------LKVSGV------GV--------LR-------------SLKLTINVQHY-----

```
----------E-----------------Y------------Y-R-D--------------
------------------------------------------------------------
------------------------------------------------------------
---------EMAGGIHLMIH-GQDEEP---VKMQGQIVSPGYSFYVKVEKK------T--
-----IMNL-----------EKP-YK-TE-C--GTV--------K-------LK----YF
------------------D-RYSMHTCWLEQLTDYVN----KM--CHCKDFF----M---
------------P-G---------------------------------------------
----------NI-----P---YCSL-----------------------------------
------------------------------------PEL---QNCTW---------I---
E-W-------------------AKFN-K-------DK---------------------MY
------------KCP-LPCKIDL---YGVS-LSR------------------ALF-PT--
-------TQY------SSI-L-----------AE--------------------------
------------------------------------------------QFRKQP--H---
-----------------------------------------------------------V
LS---------------I------------------VHN---------------------
------------------------IT-DEL------------------------------
-----L-------FM-----------R------DNLLRFIIYYDDLSYEVLE-QKPSYET
LVW-----------------LGDIGGQIGLFIGAGVMSYFEFLDCLAIV-----------
--------IYTR--------F----------F-------QK-------------------
FT------------------------------SS--------------------------
------------------------------------------------------------
------------------------------------------------------------
------------------------------------------------------------
------------------------------------------------------------
------------------------------------------------------------
------------------------------------------------------------
--------
&gt;Deutero_Ambulac_Spurpu_015893_493_53
------------------------------------------------------------
----------M-------------------------------------------------
------------------------------------------------------------
------------------------------------------------------------
------------------------------------------------------------
------------------------------------------------------------
------------------------------------------------------------
------------------------------------------------------------
------------------------------------------------------------
------------------------------------------------------------
--------------------G---------------------------------------
---SDDLQN----------------------E---------TV-----I-----------
-GS--VNTFG-NETTVHGVRYMI---------A-RG--------------HF---LF---
--RLCWL----AIIVLAFVLFVIQAFNVYG-DYAS----------S---PYSTKI----D
-I-VQ-Q----------T-FKAF-----PAV-----------------------TVCNAN
RI------RRS-----MLYDTN------------------Y-----------------E-
---------GLIDI----------------------------------------------
---------------------------------DD-------------------------
------------------------------------------------------------
------------------------------------------------------------
--------NNRSESEVEDGEAER-------------------------------------
------------------------------------------------------------
------------------------------------------------------------
------------------------------------------------------------
------------------------------------------------------------
------------------------------------------------------------
------------------------------------------------------------
------------------------------------------------------------
------------------------------------------SEEI-------W------
```

--------RKV-------------NGLYDWWGFYSA-----------SV------ADDFS
---------------------------------DLINVV---------------------
-------------------------NP----------------SQEQLMGY----GHQLE
--KFL------IQCT----YD-Q----Q----------P---C-----------N-MS--
----------------------------------------------S-------------
------------------------------------------------------------
------------------------------------------------------------
-----------D----------F-------R-----------VWQ-SRYYG--------N
CFTFN-------FG-------------MM-E-------------------TA----EA--
-------ITTSKT------GA--------LN-------------GLHLTLWVEED-----
----------E-----------------Y------------M-G-L---LS---------
------------------------------------------------------------
------------------------------------------------------------
----------PSRGAKVTIH-PQNTLP--FPEDEGINVGTGMATSIGIREE------V--
-----IERI-----------SGM----TN-C--VEE--------DGAG--------TNFT
----VTR---------TNT-IYTVAYCLKLCFQQSLI----NI--CGCVDGI----L---
------------L-----------------------------------------------
----------DY-----T---HCDVL----------N-----------------------
------------------------------------STQ---RACVS---------M---
V-H-------------------QLYI-N-------NG--------L------------YC
------------NCP-LPCE----------------------------------------
------------------------------------------------------------
------------------------------------------------------------
------------------------------------------------------------
------------------------------------------------------------
------------------------------------------------------------
---------------------------------RNLIRLQIYYETLNNEVVE-QVPRYTI
PNI-----------------LGNWGGLMGLFTGMSFISVFEVIFLLLRV-----------
--------SKIS--------C----------T-------KLFF-----------------
--------------------------------PN-------------------R------
-----------------------------------------VQPVK--------------
------------------------------------------------------------
------------------------------------------------------------
------------------------------------------------------------
------------------------------------------------------------
------------------------------------------------------------
-------T
>Protostome_Lophotroco_brch_Lingula_anatina_comp141843_c0_seq1_p1_comp141843_
c0__comp141843_c0_seq1_p1__ORF_type_complete_len_456___score_27_90_comp141843
_c0_seq1_2150_3517__494_54
------------------------------------------------------------
----------M-------------------------------------------------
------------------------------------------------------------
------------------------------------------------------------
------------------------------------------------------------
------------------------------------------------------------
------------------------------------------------------------
--------------NRNRVT-------------PTDIPKGKIGDE---------------
------------------------------------------------------------
---------------------------------------------------------ATK
D----------------------SEIDQT-------------------------------
-------------------------------S---------ST-----Q-----------
-EK--LATFA-GEVSILGLKQST---------N-PK-------------YSP---IR---
--RLIWL----CFVLIGFGFLIFHLKNRFD-YYLS----------N---PATLNM----E
-V-VP-N----------D-TLVF-----PAI-----------------------TICNSS
PL------KAS-----NLAANN------------------E-----------------S-
---------EL-------------------------------GNIFS-------------

```
------------------------------------------------------------
------------------------------------------------------------
------------------------------------------------------------
------------------------------------------------------------
------------------------------------------------------------
------------------------------------------------------------
------------------------------------------------------------
--------------------------P---------------------------------
------------------------------------------------------------
------------------------------------------------------------
------------------------------------------------------------
------------------------------------------------------------
------------------------FVNVDWSQYN---------------------LSHI-
---------------------------------NWTDFH---------------------
------------------------------------------------MRN----GFQLE
--ELL------FWCT----WK-K----T----------N---C-----------T-K---
----------------------------------------------H-------------
------------------------------------------------------------
------------------------------------------------------------
-----------D----------F-------T-----------PII-T-EIG--------L
CYQFN-------AG-------------GE-E--------------------Q--------
-------LHLETQ------GS--------GY-------------GFGMTLLAKQR-----
----------D-----------------Y------------V-S-----YA---------
------------------------------------------------------------
------------------------------------------------------------
----------YPAGFLIIVH-SPGDHP--DI-NLGTLISSGGFTGVSLKLK------R--
-----LINL-----------PPP-H--GK-C--GEK--------P-------LV----YY
------------------N-KYSKANCHAECFHNFTL----TK--CDCRLTH----F---
------------P-R---------------------------------------------
--------VPGY-----R---ECSP-----------------------------------
------------------------------------EDS---RVCVD---------N---
L-K-------------------LIIT-E-------SV--------YL----------REC
------------GCK-TPCDSFK---YDVS-LSS------------------ARL-SE--
-------TAM------RTT-------------AA--------------------------
------------------------------------------------IL----------
------------------------------------------------------------
-R----------------------------------------------------------
---------------------------TTQ------------------------------
-----E-------EI-----------S------QDFLSVFIYYSEMSYEEVV-QQKAYSA
LAL-----------------LADIGGALGLVLGSTLMTLAEVVDFFFGL-----------
--------CFWK--------L----------F-------GV-------------------
EM----------------------------------------------------------
------------------------------------------------------------
------------------------------------------------------------
------------------------------------------------------------
------------------------------------------------------------
------------------------------------------------------------
------------------------------------------------------------
--------
>Cnidaria_HyNaC10_NP_001296656_1_acidsensing_ion_channel_1like__Hydra_vulgari
s__39_56
------------------------------------------------------------
----------M-------------------------------------------------
----------------------------------A-------------------------
------------------------------------------------------------
------------------------------------------------------------
------------------------------------------------------------
```

```
------------------------------------------------------------
------------------------------------------------------------
------------------------------------------------------------
---------------------------KVED--------------------------KIK
S----------------------PQEQK--------------------------------
--------R----------------------R---------TI-----K-----------
-QY--LAYAG-ENISIHGLSHVF---------D-KR-------------ENF---IC---
--RTVWL----LITIAAFGYAVQKVYESTL-NYFS----------Y---PFSTAR----M
-K-IH-V----------N-QMDF-----PAV-----------------------SFCNLN
DF------RYS-----ATKGTK------------------L-----------------D-
---------KA----------------------------------IL-------------
------------------------------------------------------------
------------------------------------------------------------
------------------------------------------------------------
------------------------------------------------------------
------------------------------------------------------------
------------------------------------------------------------
------------------------------------------------------------
------------------------------------------------------------
------------------------------------------------------------
------------------------------------------------------------
------------------------------------------------------------
------------------------------------------SNDKKS------------
-------------------------------------------------------ISGE-
---------------------------------EYLNIT---------------------
------------------------------------------------LKA----RHKIE
--EML------VDCE----FD-G----N----------K---C-----------S-V---
----------------------------------------------H-------------
------------------------------------------------------------
------------------------------------------------------------
-----------N----------F-------T-----------EFN-W-NQGE-------L
CFTFN-------SG-------------KF-P--------------------H----SL--
-------LKVNGV------GM--------KR-------------SLVLTINVQHY-----
----------E-----------------Y------------Y-G-D--------------
------------------------------------------------------------
------------------------------------------------------------
---------ELDAGIHLILH-DQEETP---IKKRGPVIPPGFTTYIQVEKK------T--
-----ILNL-----------ESP-YK-TK-C--GSI--------K-------LK----YF
------------------D-SYSMHTCWLEQLTDHVY----KI--CKCKDFF----M---
------------P-G---------------------------------------------
----------EI-----K---VCSF-----------------------------------
------------------------------------EVL---ENCTW---------K---
E-W-------------------AKFD-N-------YK---------------------LH
------------TCP-LPCKIDS---YEMS-LSR------------------ALF-PT--
-------ARY------ASS-L-----------AN--------------------------
------------------------------------------------RFRKLP--H---
-----------------------------------------------------------V
LN---------------T------------------VKS---------------------
------------------------IS-NDL------------------------------
-----L-------FM-----------R------ENLLRVVIYYNDLSYELLE-QKPNYDT
LVW-----------------LGDVGGQIGLFVGAGVMSYFEIIDCLVLI-----------
--------IYTR--------F----------F-------QK-------------------
FK------------------------------TI-------------------N------
------------------------------------------------------------
------------------------------------------------------------
------------------------------------------------------------
------------------------------------------------------------
```

```
------------------------------------------------------------
------------------------------------------------------------
-SIRNHSL
>Cnidar_Polpod_Hydrif_GBGH01017297_1__p1__GENE_GBGH01017297_1_GBGH01017297_1_
_p1__ORF__type_complete__len_459__score_52_80__GBGH01017297_1__223_1599__498_
58
------------------------------------------------------------
----------M-------------------------------------------------
------------------------------------------------------------
------------------------------------------------------------
------------------------------------------------------------
------------------------------------------------------------
------------------------------------------------------------
------------------------------------------------------------
------------------------------------------------------------
--------------QVSRGAKPNTLVEVSSK-----------------------------
-----------------------KQTAPT-------------------------------
-------------------------------T---------LV-----R-----------
-KH--LSNFF-GQSTIHGLHYCF---------D-GN--------------LP---MR---
--RVVWC----CLVLGALALALQKISESAT-EYLS----------Y---PISTVN----T
-I-RY-E----------E-TMVF-----PAV-----------------------TLCNMN
DW------RLS-----VTVNTS------------------L-----------------F-
---------YT------------------------------VTSFYN-------------
------------------------------------------------------------
------------------------------------------------------------
------------------------------------------------------------
------------------------------------------------------------
------------------------------------------------------------
------------------------------------------------------------
------------------------------------------------------------
------------------------------------------------------------
------------------------------------------------------------
------------------------------------------------------------
------------------------------------------------------------
------------------------------------------------------------
------------------------------TGEKKE-------------------VSAE-
---------------------------------ELLATQ---------------------
------------------------------------------------RRA----NHRLE
--NML------LSCK----WK-G----Q----------S---C-----------G-L---
----------------------------------------------Q-------------
------------------------------------------------------------
------------------------------------------------------------
-----------N----------F-------S-----------EYF-T-VQGE-------R
CYTFN-------SG-------------RD-N--------------------H----ST--
-------LYAETA------SI--------ID-------------SLELILNVEYY-----
----------D-----------------Y------------F-I-D--------------
------------------------------------------------------------
------------------------------------------------------------
---------NFVSGIRIILH-SQEELP---VTSQGYVLTAGDKTYIKVKKT------K--
-----ARNL-----------PYP-YK-TN-C--SSI--------P-------LK----YF
------------------N-KYTADMCWLEQLTDHVT----SK--CGCRDVC----M---
------------P-G---------------------------------------------
----------NA-----S---VCPL-----------------------------------
------------------------------------SVM---DACGW---------A---
E-W-------------------DMFN-I-------GK---------------------LA
------------KCP-LPCEVDK---YSAL-MSH------------------AHM-AP--
-------SIV------STT-L-----------TE--------------------------
```

------------------------------------------------SL----------
------------------------------------------------------------
-R---------------I------------------ERP---------------------
------------------------TM-AHE------------------------------
-----T-------FV-----------G------ENVIRAVIYYEDLSYVLIE-QEPTYTL
MKL-----------------LGDIGGNLGLLVGASVLSYMEIIDCIILI-----------
--------VITL--------I----------K-------NS-------------------
RS------------------------------SS-------------------K------
------------------------------------------------------------
------------------------------------------------------------
------------------------------------------------------------
------------------------------------------------------------
------------------------------------------------------------
------------------------------------------------------------
----KQKS
>Protostome_Lophotroco_brch_Lingula_anatina_comp138624_c0_seq6_p1_comp138624_
c0__comp138624_c0_seq6_p1__ORF_type_complete_len_459___score_26_12_comp138624
_c0_seq6_318_1694__502_62
------------------------------------------------------------
----------M-------------------------------------------------
------------------------------------------------------------
------------------------------------------------------------
------------------------------------------------------------
------------------------------------------------------------
------------------------------------------------------------
------------------------------------------------------------
------------------------------------------------------------
------------------------------------------------------------
-----------------------ISPSRL-------------------------------
-PRKTDNSS----------------------D---------QE-----T-----------
-PV--IEKFA-ERVSCQALQEIV---------S-RE-------------SGY---IR---
--RLIWL----VCFLGGLILMAFQVYERVT-FYAG----------H---PVRFNT----Y
-T-VT-P----------L-SVMF-----PAV-----------------------TICNYN
AI------SKM-----RARELD------------------LL----------------E-
---------EF-------------------------------ERYFPL---KVG------
------------------------------------------------------------
------------------------------------------------------------
------------------------------------------------------------
------------------------------------------------------------
------------------------------------------------------------
------------------------------------------------------------
------------------------------------------------------------
------------------------------------------------------------
------------------------------------------------------------
------------------------------------------------------------
------------------------------------------------------------
------------------------------------------------------------
-------------------------------------------------------SENFT
GLGDDWF--------------------------DWDSVL---------------------
-------------------------NIS--------------------------------
-KPFV------KECT----FR-G----I----------S---C-----------S-M---
----------------------------------------------K-------------
------------------------------------------------------------
------------------------------------------------------------
-----------N----------F-------T-----------PVY-T-DMG--------I
CYTFN-------SG-------------YS-G-------------------------HR--
-------LTSDSS------GR--------LF-------------GLQLTLDVDQR-----

----------N-----------------Y------------F-R-G---FG---------
------------------------------------------------------------
------------------------------------------------------------
---------GYGAGFKVRVH-ARD-PP--MMSNMGFSVGAGMHTLVGLKMI------N--
-----ISNL-----------DTSQW--GK-C--NRD-I------K-------LK----HH
------------------V-VYSQSACRLECLYEYME----KK--CQCVRFLK---F---
-------KHHQGP-----------------------------------------------
----------RA-----R---LCDP-----------------------------------
------------------------------------EQL---FNCYT---------P---
N-Y-------------------NLFI-Q-------EK--------------------GQC
----------SHTCP-VACESIK---YEVS-LSQ------------------APV-AD--
-------YHA------YNS-A-----------TT--------------------------
------------------------------------------------------------
------------------------------------------------------------
TN--------------------E-------------------------------------
---------------------------TKV------------------------------
-----N-------KY-----------R------KNFVQLDIFFQEISYERRE-QYQAYNA
FTL-----------------GCDIGGALGLILGASVLTLFEVGDFIILM-----------
--------ILKG-------------------KW------RR-------------------
QT------------------------------QV--------------------------
------------------------------------------------------------
------------------------------------------------------------
------------------------------------------------------------
------------------------------------------------------------
------------------------------------------------------------
------------------------------------------------------------
TSVSVHKK
>Deutero_Ambulac_Spurpu_009765_505_63
------------------------------------------------------------
----------MG------------------------------------------------
------------------------------------------------------------
------------------------------------------------------------
------------------------------------------------------------
--DLPRDTKTS-------------------------------------------------
-----------------------------------------P------------------
------------------------------------------------------------
------------------------------------------------------------
---------------------------------------------------------EKM
A--------------------DYSEKAYGRPD----------------------------
---YDESPK----------------------E---------GT-----R-----------
-KI--VNEFV-DNATTHGIPRVL---------N-AS--------------RPW--QS---
--RLFWC----VVTLIFAGVFLFQGSKLVQ-SYIA----------R---PTTTKI----S
-L-IT-K----------S-RLEF-----PAV-----------------------TICNLN
ML------RRS-----MLKGDE------------------Y-----------------DY
--------SGI-------------------------------------------------
--E------------------------------DI-------------------------
------------------------------------------------------------
------------------------------------------------------------
------------------------------------------------------------
------------------------------------------------------------
------------------------------------------------------------
------------------------------------------------------------
------------------------------------------------------------
------------------------------------------------------------
------------------------------------------------------------
------------------------------------------------------------
------------------------------------------------------------

```
------------------------EDPNDWEALYNL-----------SK------GSDYN
---------------------------------NFKNFV---------------------
-------------------------KP----------------TKEELRTL----GHKAK
--DFI------LQCS----FD-T----E----------A---C-----------S-Y---
----------------------------------------------E-------------
------------------------------------------------------------
------------------------------------------------------------
-----------N----------F-------T-----------VIQ-NAEYG--------N
CFVFN-------NA-------------HKLK-------------------RG----KR--
-------TTTSRT------GS--------QY-------------GLQLTLMVEQP-----
----------E-----------------Y------------I-G-I---LS---------
------------------------------------------------------------
------------------------------------------------------------
----------PNSGVKVAIK-DPRIYA--FPEDDGIEAAPGFATSIGLTKT------S--
-----ISRL-----------PEP-Y--GN-C--IRK-H------------------DSFY
----APE----------KY-DFSQRSCLKLCLQETLN----ET--CSCITDV----L---
------------I-----------------------------------------------
----------DG-----T---MCEVT----------N-----------------------
------------------------------------REQ---GNCRN---------S---
V-F-------------------KDFL-K-------NK--------L------------NC
------------DCS-NACRSRS---SSNS-SSS------------------SSS-SS--
-------SSS------SRS-----------------------------------------
------------------------------------------------------------
------------------------------------------------------------
------------------------------------------------------------
------------------------------------------------------------
-----S--------------------S------SSSSSISSIFSSISSGSRSSGSSSRGS
SSS-----------------SSSSSSSSNIRSSIIVVAVVEVVQ----------------
------------------------------------------------------------
------------------------------------------------------------
------------------------------------------------------------
------------------------------------------------------------
------------------------------------------------------------
------------------------------------------------------------
------------------------------------------------------------
------------------------------------------------------------
--------
>Deutero_Ambulac_Apla_gbr40_182_t1_508_68
------------------------------------------------------------
----------M-------------------------------------------------
------------------------------------------------------------
------------------------------------------------------------
-----------------------------------------------------------I
LDSEARPGSDR-------------------------------------------------
------------------------------------------------------------
--GAGNYRHGISLTMMTEAT-------------AGGTSPSGTA-----------------
------------------------------------------------------------
---------------------------------------------------------STT
E----------------------QPSEEC-------------------------------
-------EN----------------------S---------SL-----K-----------
-ER--IATFG-DETTFHGVRFVT---------N-HQ-------------LHF---LR---
--RLLWF----CIVGGMTGYLIYGVSVSFI-TFYQ----------R---PVTSVV----K
-I-NY-V----------R-QIPF-----PTV-----------------------TFCNYN
QW------KKS-----RVPPEF------------------Y-----------------D-
---------AV--------------------------------RSLN-------------
------------------------------------------------------------
------------------------------------------------------------
```

------------------------------------------------------------
------------------------------------------------------------
------------------------------------------------------------
------------------------------------------------------------
------------------------------------------------------------
--------------------------PAN-------------------------------
------------------------------------------------------------
------------------------------------------------------------
------------------------------------------------------------
------------------------------------------------------------
------------------------KKPVDMTYLK---------------------AHDVT
---------------------------------NLTDKL---------------------
------------------------------------------------IND----THQIE
--DML------LECT----FK-E----A----------N---C-----------S-A---
----------------------------------------------E-------------
------------------------------------------------------------
------------------------------------------------------------
-----------N----------F-------T-----------QII-T-DFG--------V
CYSFN-------DD-------------PS---------------------------NV--
-------HYVHQS------GS--------QN-------------ALFMKITVEQE-----
----------E-----------------Y------------I-F-G---EN---------
------------------------------------------------------------
------------------------------------------------------------
----------NAAGIKVV------------------------------------------
-------NL-----------KYP-YE-SN-C--TDR-G------N-------VK----YS
------------------A-EYTVPLCQYEHKIETVK----AK--CGCKDMR----Y---
------------P-----------------------------------------------
------------------------------------------------------------
------------------------------------------------------------
----------------------ANFT-S-------ED--------------------KSF
------------NCP-VPCDLTI---YDTR-MSF------------------AYY-PG--
-------QHH------LEE-I-----------MQ--------------------------
------------------------------------------------QN----------
------------------------------------------------------------
-N----------------------------------------------------------
---------------------------LSE------------------------------
-----N-------SI-----------R------KNVLDLRIFFEELSFQKIE-EIPSYTF
YSL-----------------QSSIGGYMGLLIGASLITLFEFLDFILVT-----------
--------TTSC--------L----------L-------KH-------------------
SVLAR--------------------------PRG------------------NR------
-----------------------------------------VRGAARQRD----------
------------------------------------------------RSSH--------
------------------------------------------------------------
------------------------------------------------------------
------------------------------------------------------------
----------------------------------------------------------GE
AKPCFIKK
>Deutero_Ambulac_Sakowv30004505m_514_69
------------------------------------------------------------
----------M-------------------------------------------------
------------------------------------------------------------
------------------------------------------------------------
------------------------------------------------------------
------------------------------------------------------------
-------------------------------------DVVVPLSQM--------------
------------------------------------------------------------
------------------------------------------------------------

```
---------------------------------------------------------DNT
DHQTK------------------KAEDDT-------------------------------
ASQSQEPEK----------------------P---------SR-----L-----------
-TL--AVIYA-QNTTAHGLANTV---------N-AG--------------TL---HK---
--RLFWI----LTFIICMVTLVVQAQSVIR-LYLS----------Y---PKAVSV----E
-I-KS-R----------S-KLDF-----PAV-----------------------TVCNLN
MA------RVS-----RWNGTR------------------F-----------------Q-
---------ELLDVDR---M------------------QFDIDEWFS----E--------
-----------------NLGSS-----------DA-------------------------
------------------------------------------------------------
----------------------PEP-----------------------------------
-----------------DTSTQAD---------------VPINATSSGDAPV--------
------------------------------------------------------------
-------------NTTSSGDAPVNT---------TSSGDAPVNTTSSGARKRRAVV----
-----------------------------------------------------AK-HPVT
DG--------------TLSEFLKTLGPGLQSS----------------------------
---------------------------------------WI-------------------
------RQYAKENVITRPKRNSGNDSETSETDGSD-------------------------
--------------------------------------------------------DTDN
GNSEGSEMGSERGSMGFY---ASDDY----------------DDFSSNA----FM-----
------------------DYGLGGTNPNDWSTFYTK-----------SS------TDDFS
---------------------------------DLRAAL---------------------
-------------------------RM----------------TQAEIEEY----GERAE
--DFI------VQCS----FD-R----S----------S---C-----------S-S---
----------------------------------------------K-------------
------------------------------------------------------------
------------------------------------------------------------
-----------D----------F-------T-----------VSQ-NDVYG--------N
CFTFN-------GI-------------TG-N-------------------------KT--
-------RRTTKA------GE--------QY-------------GLKLTLYVDAD-----
----------E-----------------Y------------V-G-L---FT---------
------------------------------------------------------------
------------------------------------------------------------
----------QQEGARVVVH-PGSLYP--VPESVGLTAGVGQSLSIGLKKV------K--
----------------------PKY-----------------------------------
------------------------------------------------------------
------------------------------------------------------------
------------------------------------------------------------
------------------------------------------------------------
------------------------------------------------------------
------------------------------------------------------------
------------------------------------------------------------
------------------------------------------------------------
------------------------------------------------------------
------------------------------------------------------------
------------------------------------------------------------
------------------------------------------------------------
------------------------------------------------------------
------------------------------------------------------------
------------------------------------------------------------
------------------------------------------------------------
------------------------------------------------------------
------------------------------------------------------------
------------------------------------------------------------
------------------------------------------------------------
------------------------------------------------------------
--------
```

>Protostome_Lophotroco_annelid_CAC9474682_1__Ofus_G012955_partial_Owenia__fus
iformis__516_70
------------------------------------------------------------
----------M-------------------------------------------------
------------------------------------------------------------
------------------------------------------------------------
------------------------------------------------------------
------------------------------------------------------------
------------------------------------------------------------
----------------ACGC-------------CCT------------------------
------------------------------------------------------------
------------------------------------------------------------
------------------------------------------------------------
-------------------------------K---------SQ-----R-----------
-KL--DYEFA-TSTSCHGLGRIA---------D-SS--------------RK---PQ---
--RIFWI----LVTLFCFGYMVSQIVDRFD-HFMQ----------M---KQTTEL----E
-V-MMDQ----------T-SLEF-----PAV-----------------------TICNYN
KY------RLS-----AMTESD------------------A-----------------I-
---------IL------------S-NSA---------------LLMS----YL-------
---------------------------------QWG-----V------------------
------------------------------------------------------------
------------------------------------------------------------
------------------------------------------------------------
------------------------------------------------------------
------------------------------------------------------------
------------------------------------------------------------
--------------------------PL--------------------------------
------------------------------------------------------------
------------------------------------------------------------
------------------------------------------------------------
--------DEEHVK---------------------------FAQE---------------
-------------------------LEYNITQLY---------------------PDGF-
---------------------------------DMGNLT---------------------
------------------------------------------------RRV----GWQMD
-DTTL------LWCR----IA-G----K----------Q---C-----------T-F---
----------------------------------------------E-------------
------------------------------------------------------------
------------------------------------------------------------
-----------D----------F-------S-----------LVQ-T-SYG--------N
CYTFN-------RY-------------PR-E--------------------P----NV--
-------YKQTVH------GT--------GG-------------GLRIVLDINQS-----
----------D-----------------Y------------T-E-V---FA---------
------------------------------------------------------------
------------------------------------------------------------
------FGGNSEAGIKFLVH-PQYEPP--SVESEGYAVSPGFRAYAGIRYD------E--
-----LQNA-----------VSP-W--GK-C--DPE-A------K-------LK----FY
------------------E-KYSFEGCLSECREGKLL----ER--CGCKPYS----I---
------------P-G---------------------------------------------
----------NI-----P---NCLP-----------------------------------
------------------------------------TKA---VTCAN---------E---
I-L-------------------GAIE-E-------NP--------K-----------EEC
-----------PQCT-VACEEVE---YPVS-LSY------------------ATF-PN--
-------HNI------AKE-L-----------FE--------------------------
------------------------------------------------IF----------
------------------------------------------------------------
-D----------------------------------------------------------
---------------------------VTE------------------------------

-----D-------YI-----------R------DNIVELDIYFRDLYRHLTK-MSPAVTP
SGL-----------------LSDLGGQMGLFVGMSILTLFEFLEFTLRR-----------
--------LYLG--------C----------F-------RR-------------------
NA------------------------------ED--------------------------
------------------------------------------------------------
------------------------------------------------------------
------------------------------------------------------------
------------------------------------------------------------
------------------------------------------------------------
------------------------------------------------------------
--------
>Cnidar_Rhopilema_esculentum_TR103737_c1_g2_i1_p1__Rhopilema_esculentum_TR103737_c1_g2__Rhopilema_esculentum_TR103737_c1_g2_i1_p1__ORF__type_complete__len_466__score_59_67__Rhopilema_esculentum_TR103737_c1_g2_i1_277_1674__517_71
------------------------------------------------------------
----------M-------------------------------------------------
----------------------------------WRII----------------------
------------------------------------------------------------
------------------------------------------------------------
------------------------------------------------------------
------------------------------------------------------------
--------------NWRKST-------------AGEYAEAR-------------------
------------------------------------------------------------
-----------------RETEVRYPG-IPEK--------------------------AFH
M----------------------EQRRK--------------------------------
--------R----------------------K---------RN-----A-----------
-EH--FNTFI-SDSTLHGLHFCF---------D-KK--------------HL---AR---
--RIIWT----LILISAFGYVGEKIYASLL-QFFQ----------Y---PMSTTT----T
-L-VY-D----------EPPIKL-----PAI-----------------------SICNTN
DF------KRS-----IVVTTK------------------L-----------------G-
---------DI----------------------------------LR-------------
------------------------------------------------------------
------------------------------------------------------------
------------------------------------------------------------
------------------------------------------------------------
------------------------------------------------------------
------------------------------------------------------------
------------------------------------------------------------
------------------------------------------------------------
------------------------------------------------------------
------------------------------------------------------------
------------------------------------------------------------
------------------------------------------NKTDN-------------
-------------------------------------------------------ITGS-
---------------------------------EYSSLI---------------------
------------------------------------------------KLG----NHRLV
--DML------VTCR----QS-H----K----------T---C-----------N-V---
----------------------------------------------K-------------
------------------------------------------------------------
------------------------------------------------------------
-----------D----------F-------V-----------EYW-D-NSGT------VR
CHTFN-------SG-------------KT-G-------------------------PI--
-------MTINRT------GT--------TE-------------AIEMVINIQQY-----
----------D-----------------Y--------------N-D--------------
------------------------------------------------------------
------------------------------------------------------------
---------FDTAGIKLIIH-GQDQTP---IKTAGLLLAPGTVTNIEISKR------K--

```
-----RVNL-----------QHP-YK-TN-C--GSL--------K-------LK----AF
------------------D-SYSEHLCWLDQLTSYVV----DK--CGCVDAF----M---
------------P-G---------------------------------------------
----------KH-----K---VCDF-----------------------------------
------------------------------------KTV----GCVW---------N---
N-W-------------------ERFD-I-------EQ---------------------KY
------------QCP-IPCVVEE---FIAK-QSS------------------AMF-PS--
-------NAA------ADN-L-----------AI--------------------------
------------------------------------------------DL----------
------------------------------------------------------------
-N---------------L------------------SGT---------------------
------------------------AM-ENR------------------------------
-----L-------FI-----------R------ENFLKFKVFFGEYSYELVE-QVPSYDL
KVL-----------------LGDIGGQLGLFLGSSLLTYVEFFDCLLMI-----------
--------IYTK--------F----------F-------EV-------------------
LL------------------------------PR--------------------------
------------------------------------------------------------
------------------------------------------------------------
------------------------------------------------------------
------------------------------------------------------------
------------------------------------------------------------
------------------------------------------------------------
--------
>Cnidaria_HyNaC4_CAL36113_1_hydra_Na_channel_4__Hydra_vulgaris__37_72
------------------------------------------------------------
----------M-------------------------------------------------
------------------------------------------------------------
------------------------------------------------------------
------------------------------------------------------------
------------------------------------------------------------
------------------------------------------------------------
-------------------N-------------FEELAKIV-------------------
------------------------------------------------------------
---------------VEAVQENNNQP-INTN--------------------------KIK
T----------------------PRELR--------------------------------
--------N----------------------N---------KT-----K-----------
-EY--VSEMI-DNSSFHGISYIA---------G-KE-------------NHF---IR---
--RTIWL----LITMTAFGYAAQKVYESTV-NYFS----------F---PISTTQ----M
-R-IY-V----------N-EIDF-----PAV-----------------------SFCNFN
EF------RLS-----KMDGTK------------------V-----------------D-
---------QA----------------------------------IL-------------
------------------------------------------------------------
------------------------------------------------------------
------------------------------------------------------------
------------------------------------------------------------
------------------------------------------------------------
------------------------------------------------------------
------------------------------------------------------------
------------------------------------------------------------
------------------------------------------------------------
------------------------------------------------------------
------------------------------------------------------------
------------------------------------------NPKL--------------
---------------------------------QGL-------------------VTAE-
---------------------------------EYRNVT---------------------
------------------------------------------------FGA----MFDLK
--EML------VDCE----FN-G----I----------P---C-----------S-D---
```

```
----------------------------------------------E-------------
------------------------------------------------------------
------------------------------------------------------------
-----------N----------F-------T-----------MFS-W-MQGE-------R
CFTFN-------SG-------------KF-P--------------------H----KL--
-------LKIGGA------GM--------KR-------------SLKITINIIHY-----
----------E-----------------Y------------Y-K-D--------------
------------------------------------------------------------
------------------------------------------------------------
---------EMDAAIHMIVH-GQQDTP---LKMRGPTLSPGFTTYIQLEKK------M--
-----IINL-----------EAP-YK-TK-C--GSL--------K-------LK----YF
------------------D-SYSLDTCWLEQLTDHVH----SV--CKCKDFF----M---
------------P-G---------------------------------------------
----------DI-----P---ICSL-----------------------------------
------------------------------------NDA---MSCMW---------P---
E-W-------------------ARFD-Q-------LK---------------------MQ
------------NCP-LPCVIDS---YKLS-LSR------------------ALF-PS--
-------PQY------ADS-L-----------TN--------------------------
------------------------------------------------SLQQPP--H---
-----------------------------------------------------------I
IN---------------S------------------IKN---------------------
------------------------IS-DKV------------------------------
-----Q-------FA-----------R------DNLLRFVIYYDDLSYELIE-QKPSYDT
LVW-----------------LGDIGGQIGLLIGAGAMSYFEFIDCLFLV-----------
--------IYNQ--------F----------F-------KT-------------------
YR----------------------------------------------------------
------------------------------------------------------------
------------------------------------------------------------
------------------------------------------------------------
------------------------------------------------------------
------------------------------------------------------------
------------------------------------------------------------
--------
>Cnidaria_HyNaC8_NP_001296668_1_acidsensing_ion_channel_5like__Hydra_vulgaris
__36_73
------------------------------------------------------------
----------M-------------------------------LELKC-------------
------------------------------------------------------------
------------------------------------------------------------
------------------------------------------------------------
------------------------------------------------------------
------------------------------------------------------------
-----------------------------------EISKRR-------------------
------------------------------------------------------------
-----------------RKRYQNAISYLPGA--------------------------NHD
T----------------------VDELR--------------------------------
--------N----------------------Q---------SI-----N-----------
-DY--LSDMA-VNSSFHGINYIC---------D-ST--------------YK---VR---
--RIIWI----VVTLTAMLYAMREVYESTR-KYLN----------Y---PVSTVR----M
-K-IY-V----------D-DLEF-----PAV-----------------------SLCNLN
DV------RMS-----IVNGTS------------------F-----------------D-
---------NA----------------------------------LI-------------
------------------------------------------------------------
------------------------------------------------------------
------------------------------------------------------------
------------------------------------------------------------
------------------------------------------------------------
```

------------------------------------------------------------
------------------------------------------------------------
------------------------------------------------------------
------------------------------------------------------------
------------------------------------------------------------
------------------------------------------------------------
------------------------------------------DQNAQN------------
-------------------------------------------------------ISAD-
---------------------------------DALMIA---------------------
------------------------------------------------REA----RHNLE
--DML------LECK----FN-G----R----------S---C-----------S-A---
----------------------------------------------K-------------
------------------------------------------------------------
------------------------------------------------------------
-----------N----------F-------S-----------EFN-W-MQGD-------R
CFTIN-------SG-------------KP-G--------------------H----SR--
-------LSVKGT------GI--------KR-------------NLELILNLQHY-----
----------E-----------------Y------------Y-R-D--------------
------------------------------------------------------------
------------------------------------------------------------
---------EMESGIHFILH-SQEETP---VRMRGPIVSPGFTTYFRINKI------K--
-----TKNL-----------KYP-YK-TR-C--GSL--------N-------LK----FF
------------------K-GYSKQLCWLDQLTDYVN----SK--CGCKDFF----M---
------------P-G---------------------------------------------
----------NI-----S---ICTF-----------------------------------
------------------------------------STA---FGCMW---------P---
A-W-------------------EEFE-K-------KK---------------------IS
------------NCP-LPCDADTY--FGQT-VSR------------------ALF-PS--
-------NEY------KKS-F-----------IK--------------------------
------------------------------------------------NLMHIQ--Q---
-----------------------------------------------------------F
RD---------------V------------------YGN---------------------
------------------------KK-KEL------------------------------
-----Q-------FM-----------R------DNLLRIVLYYDDLSYELLE-QKPSYDL
LSW-----------------LGDIGGQIGLFVGSSAMSYFEFLDCLIMI-----------
--------IYAK--------Y----------F-------KQ-------------------
YK----------------------------------------------------------
------------------------------------------------------------
------------------------------------------------------------
------------------------------------------------------------
------------------------------------------------------------
------------------------------------------------------------
------------------------------------------------------------
--------
>Cnidaria_HyNaC2_CAL36110_1_hydra_Na_channel_2__Hydra_vulgaris__32_75
------------------------------------------------------------
----------M-------------------------------------------------
------------------------------------------------------------
------------------------------------------------------------
------------------------------------------------------------
------------------------------------------------------------
------------------------------------------------------------
------------------------------------------------------------
------------------------------------------------------------
--------------------------KARFA--------------------------RRM
E----------------------KIVEE--------------------------------
--------R----------------------M---------TV-----K-----------

-QI--FNRYV-QTSTLHGFRFIF---------M-DT--------------FI---VR---
--RVLWT----ILTLTMATIFFKELRNSIN-LFYE----------Y---PFTTTS----T
-I-QY-E----------P-SLTF-----PAI-----------------------SVCNLN
HF------LLS-----KIKKSK------------------L-----------------K-
---------PL----------------------------------YD-------------
------------------------------------------------------------
------------------------------------------------------------
------------------------------------------------------------
------------------------------------------------------------
------------------------------------------------------------
------------------------------------------------------------
------------------------------------------------------------
------------------------------------------------------------
------------------------------------------------------------
------------------------------------------------------------
------------------------------------------------------------
------------------------------------------QGRL--------------
------------------------PFDNNLENPGFD-------------------IQGE-
---------------------------------ELYSIL---------------------
------------------------------------------------KTS----SQSID
--EIF------LSCE----WK-S----RDTAKNGV-PNP---C-----------K-P---
----------------------------------------------N-------------
------------------------------------------------------------
------------------------------------------------------------
-----------N----------F-------T-----------VYS-G-LYGQ-------S
CYTFN-------PG-------------VS-G--------------------Y----PL--
-------LSLSET------GV--------NM-------------GFKLELDLKSK-----
----------D-----------------S------------L-Q-G--------------
------------------------------------------------------------
------------------------------------------------------------
---------IQEIGAIVIVH-HQQETP---VLQAGFVVSPGFQTFVEIKVR------Q--
-----TENL-----------PPP-YA-TK-C--GSK--------P-------LK----NY
------------------Q-IYRQSSCFLEQLGDAIE----TK--CKCKSSF----M---
------------A-G---------------------------------------------
---------RNI-----P---YCSL-----------------------------------
------------------------------------RQT---VTCLM---------P---
T-I-------------------YDFD-R-------KT---------------------NN
------------NCP-VDCETIQ---YLSS-LSY------------------ARF-IS--
-------NVT---Y------L-----------SK--------------------------
------------------------------------------------NAEQSS--YI--
---------------------------------------------------------RKL
KN----------------------------------SMS---------------------
------------------------PK-KLQ------------------------------
-----K-------YI-----------E------ENIVAVQFFYQEMKKEKVK-QEPSYDF
YKL-----------------IGDVGGQLGLLLGASVLTLVEFVDLFIFT-----------
--------LYHQ--------L----------L-------RL-------------------
SK------------------------------KK-------------------S------
------------------------------------------------------------
------------------------------------------------------------
------------------------------------------------------------
------------------------------------------------------------
------------------------------------------------------------
------------------------------------------------------------
--------
>Cnidaria_HyNaC3_CAL36112_1_hydra_Na_channel_3__Hydra_vulgaris__40_76
------------------------------------------------------------
----------M-------------------------------------------------

```
------------------------------------------------------------
------------------------------------------------------------
------------------------------------------------------------
------------------------------------------------------------
------------------------------------------------------------
------------------LN-------------FQDIAQIT-------------------
------------------------------------------------------------
---------------VEAIQEKNELPNEEEK--------------------------NTQ
H----------------------PRDLR--------------------------------
--------N----------------------K---------KI-----K-----------
-EH--ISYMI-DNSSFHGLSYIF---------D-KR--------------HS---IR---
--RTIWF----FITIAAFVYAMQKVYESTM-NYFS----------Y---PFYTAR----M
-R-MY-V----------N-QINF-----PAV-----------------------SFCNLN
DI------RFS-----VMNGTI------------------V-----------------D-
---------DA----------------------------------II-------------
------------------------------------------------------------
------------------------------------------------------------
------------------------------------------------------------
------------------------------------------------------------
------------------------------------------------------------
------------------------------------------------------------
------------------------------------------------------------
------------------------------------------------------------
------------------------------------------------------------
------------------------------------------------------------
------------------------------------------------------------
------------------------------------------NNNQ--------------
---------------------------------EAN-------------------ITGE-
---------------------------------EMRTFI---------------------
------------------------------------------------QAA----RHTLK
--EML------VDCD----FE-G----K----------K---C-----------S-Y---
----------------------------------------------E-------------
------------------------------------------------------------
------------------------------------------------------------
-----------N----------F-------T-----------EFS-W-MQGE-------S
CFTFN-------SG-------------KP-P--------------------H----TL--
-------LKVNGA------GI--------NR-------------SLKLTINVQHY-----
----------E-----------------Y------------Y-R-D--------------
------------------------------------------------------------
------------------------------------------------------------
---------KMDAGIHLILH-GQDETP---VKMRGLHLPPGFTSYIQIEKK------T--
-----IINL-----------EAP-YK-TK-C--GSV--------E-------LK----YF
------------------D-SYSMHTCWLEQLTDHVY----KV--CKCKDVF----M---
------------P-G---------------------------------------------
----------DI-----P---ICSF-----------------------------------
------------------------------------DEA---FNCMW---------P---
Q-W-------------------ETFD-K-------LK---------------------LY
------------HCP-LPCKIDS---YAVS-LSR------------------ALF-PT--
-------ARY------ASS-L-----------AN--------------------------
------------------------------------------------ELRKHQ--H---
-----------------------------------------------------------V
AM---------------N------------------LKS---------------------
------------------------NT-DEL------------------------------
-----A-------FM-----------R------ENLLRLVIYYDDLSYELLE-QKPNYDT
LVW-----------------LGDVGGQIGLFIGAGVMSYFEFIDCLAMI-----------
--------LYTR--------F----------F-------EK-------------------
FT------------------------------TK--------------------------
```

------------------------------------------------------------
------------------------------------------------------------
------------------------------------------------------------
------------------------------------------------------------
------------------------------------------------------------
------------------------------------------------------------
--------
>Deu_Ambulacrar_hemi_Ptyfla_40v0_9_20150316_1g18733_t1_scaffold17031_cov102_5
19_77
------------------------------------------------------------
----------M-------------------------------------------------
------------------------------------------------------------
------------------------------------------------------------
------------------------------------------------------------
------------------------------------------------------------
------------------------------------------------------------
------------------------------------------------------------
------------------------------------------------------------
------------------------------------------------------------
------------------------------------------------------------
DPLIDSKHK----------------------D---------SF-----S-----------
-EV--LTHLL-ANSSSHGLPNIQ---------R-SK--------------HP---VG---
--KLFWA----LLFLAGVGVLCWQVSVLVA-TYRK----------R---DVDVTL----K
-M-HS-N----------T-SIEF-----PVV-----------------------TICNTN
AL------KLS-----ALVDDV------------------Y-------------------
---------ML-------------------------------ADLYN----A--------
---------------------------------NS-------A-----------------
------------------------------------------------------------
---------------------------------FAT-----------IMQA---------
-------YRNIEHLDS-E-------------------------PEKSTGGA-EVNDYREE
------------------------------------------------------------
------------------------------------------------------------
------------------------------------------------------------
--------------------------------------------------IVL----SNS
TSAFEAGGY---------------------------------------------------
-----------SNIT----TST--------------------------------------
------------------------------------------------------------
-----YPSGEQK------------------------------SSTGPE------------
---------------------PSSQPMINNITYPTFAN------KTNIN------SSSLS
------------------------------MKTELKDYM---------------------
------------------------DDE----------------DYYNKMDN----RHELQ
--YLL-------------------------------------------------------
----------------------------------------------R-------------
------------------------------------------------------------
------------------------------------------------------------
-----------Q----------I-------------------------------------
------------------------------------------------------------
-----------------------------MA-------------GLTLELFVNQD-----
----------N-----------------Y------------I-P-D---IT---------
------------------------------------------------------------
------------------------------------------------------------
----------ETSGIRVAIH-PHRVVP--FPEDNGISLSPGYSTEIGLRMV------N--
-----IERQ-----------PRP-Y--GN-C--IDA-N-NI-PDK-------YYD-NDIY
----IRR--YR-------A-AYS-------------------------------------
------------------------------------------------------------
------------------------------------------------------------
------------------------------------------------------------

```
V-----------------------------------------------------------
----------------------------------------------------AEF-----
------------------------------------------------------------
------------------------------------------------------------
------------------------------------------------------------
------------------------------------------------------------
------------------------------------------------------------
------------------------------------------------------------
-DL-----------------CSDIGGQLGLWIGVSVITLCEMFEFFASI-----------
--------SRLF--------Y----------R-------RLSVCV--FNFGLFDSWINNN
SAIYHHPA------------------------RN--------------NCNGGQH-----
-------------------------------------PNNHTTTNNPDTNDCFDD-----
------------------------------------------------------------
------------------------------------------------------------
------------------------------------------------HYPC--------
-------------------------------NNPTICYNCS--LNIQGT-----------
--------------------------------------------F-------DNPTFQRY
DSFNLVSP
>Protost_ecdy_cyclo_PriCa_ecdy_XP_014670949_1__PREDICTED_acid_sensing__ion__c
hannel__1_like_Priapulus__caudatus__521_79
------------------------------------------------------------
----------M-------------------------------------------------
------------------------------------------------------------
------------------------------------------------------------
------------------------------------------------------------
------------------------------------------------------------
------------------------------------------------------------
------------------------------------------------------------
------------------------------------------------------------
-------------------------------------------------------ADAKI
D----------------------SQERDT-------------------------------
-CPADRKPP----------------------E---------SF-----K-----------
-QQ--LQQFS-HGVSATGLRYVF---------D-DD-------------VSI---YR---
--RIIWG----VLVACGLLYSFYQIGKSIH-VYAS----------R---PVAVSI----D
-L-SY-R----------D-ELEF-----PVV-----------------------TFCNEN
VI------KKS-----AMPQQG------------------F-----------------E-
---------DYV-------------RVI--------------RMLVG-------------
------------------------------------------------------------
------------------------------------------------------------
------------------------------------------------------------
------------------------------------------------------------
------------------------------------------------------------
------------------------------------------------------------
------------------------------------------------------------
----------------------GVSSPL--------------------------------
------------------------------------------------------------
------------------------------------------------------------
------------------------------------------------------------
------------------------------------------------------------
-------------------------------------------------------DRNMT
---------------------------------EIDALL---------------------
--------------------DGLDVQL----------------LQEVLETG----MYHQD
--EIM------RQCW----WR-G----Q----------P---C-----------E-Q---
----------------------------------------------K-------------
------------------------------------------------------------
------------------------------------------------------------
----------------------YV------T-----------SVY-T-DMG--------L
```

```
CFSFN-------AN-------------GS-S-------------------------PL--
-------IARDTS------GH--------QS-------------GLRLLLDAQQH-----
----------E-----------------Y------------V-A-S---RE---------
------------------------------------------------------------
------------------------------------------------------------
----------GTVGFRVMLH-SPYETP--VLQARGFKVGPGMHILTSVSKS------V--
-----THRL-----------GSP-Y--SQ-C--RQV--------A-----------GDAS
------------------E-HYSYYACLQECENAFVE----GV--CGCHNLQ--------
--------------G---------------------------------------------
---------RNVA----R---VCTE-----------------------------------
------------------------------------KTE---LQCLM---------R---
L-M-------------------QVSS-A------FDR--------------------RHC
------------KCP-TQCRDID---YTPS-TSF------------------SPF-----
-------ANY------LSA-M---------------------------------------
------------------------------------------------MI----------
------------------------------------------------------------
-N----------------------------------------------------------
---------------------------ATA------------------------------
-----D-------YV-----------T------NNLLLLDIYFTELEENVIT-SVPAYTF
VAL-----------------MADIGGSLSLFLGSGLFTIFEVVDFAIVA-----------
--------ANKK---------------------------RV-------------------
-----------------A--------------RN--------------------------
------------------------------------------------------------
------------------------------------------------------------
------------------------------------------------------------
------------------------------------------------------------
----------------------------GECTDP--------------------------
----------------------------------------------------------EQ
TAMKSTKR
>Deutero_Ambulac_Sakowv30036986m_524_81
------------------------------------------------------------
----------M-------------------------------------------------
------------------------------------------------------------
------------------------------------------------------------
------------------------------------------------------------
-NTLGATVSPE-------------------------------------------------
------------------------------------------------------------
---EASSTAKIYAMTGIKEG-------------KDDIGQG--------------------
------------------------------------------------------------
---------------------------------------------------------DRS
R----------------------KHGCTC-------------------------------
-------------------------------E---------YT-----K-----------
-ER--IRKFT-QETSMHGVRYTG---------M-TD-------------IHS---VR---
--RIIWT----VLVLWSLGWLTYFLYTSVV-KYRA----------F---NVNTLL----S
-M-KY-T----------D-VLLF-----PAV-----------------------TVCNYN
RY------RKS-----YIQGTT------------------F-----------------A-
---------EW------------Q------------------QTLYH-------------
------------------------------------------------------------
------------------------------------------------------------
------------------------------------------------------------
------------------------------------------------------------
------------------------------------------------------------
------------------------------------------------------------
------------------------------------------------------------
--------------------------PINLIHG---------------------------
------------------------------------------------------------
------------------------------------------------------------
```

------------------------------------------------------------
------------------------------------------------------------
------------------------STTINWTTDAA--------------------IEELA
K--------------------------------NRTYLE---------------------
------------------------------------------------LSA----AHQKE
--SMI------LECK----LG-R----A--------SSG---C-----------N-AV--
----------------------------------------------D-------------
------------------------------------------------------------
------------------------------------------------------------
-----------N----------F-------T-----------TTL-T-DYG--------V
CYTYN-------NA-------------LS-N--------------------H--------
-------MLVTES------GS--------RH-------------GLSLTLNLEQE-----
----------E-----------------Y------------T-Y-G---QN---------
------------------------------------------------------------
------------------------------------------------------------
----------YGAGFKILVH-FPGDVP--LVTDFGTAVSPGTEVFMPVRLV------V--
-----EKNK-----------ESP-YR-TN-C--TNV--------S-------LE----YY
------------------A-HYTKSNCIMEKVTQMVV----SK--CGCREPY----M---
------------P-G---------------------------------------------
----------NV-----R---VCNY-----------------------------------
------------------------------------QEK---EECVQ---------I---
A-V------------------------------------------------------NAA
------------NCP-TACDIET---FQTS-SSF------------------ATY-PS--
-------YHM------NDY-L-----------EQ--------------------------
------------------------------------------------TY----------
------------------------------------------------------------
-N----------------------------------------------------------
---------------------------MTI------------------------------
-----A-------NI-----------R------ENVAVLHVFYEELSFMSIE-QSAAYNG
MDF-----------------LADFGGQMGLCVGASLLTVCEAFEFVFIG-----------
--------IWKR--------L----------C----------------------------
-----------------------------------------------------R------
------------------------------------------------------------
------------------------------------------------------------
------------------------------------------------------------
------------------------------------------------------------
------------------------------------------------------------
------------------------------------------------------------
--------
>Deutero_Urochord_tr_F6VJJ2_F6VJJ2_CIOIN_Uncharacterized_protein_OS_CioInt_52
5_82
NNHTEQNGLS--VLK--RLNNRIRNN-LR-------------------------------
----------M-------------------------------------------------
------------------------------------------------------------
------------------------------------------------------------
------------------------------------------------------------
------------------------------------------------------------
------------------------------------------------------------
------------------------------------------------------------
------------------------------------------------------------
------------------------------------------------------------
------------------------------------------------------------
--KPKLGLR----------------------T---------KQ-----K-----------
-KL--AVTFS-EETSCHGILQIK---------N-NA--------------KI---WL---
--KVFWA----LLVLAAFSVLIWQIIDRIN-AYNK----------H---ETNTQI----S
-E-EY-V----------Q-SVHF-----PAV-----------------------TICNSN
RI------LYE-----AIQRIKKYFY--------------FSS-------------DEE-

---------RL--------------AML------------TLSKAFD-------------
------------------------------------------------------------
------------------------------------------------------------
------------------------------------------------------------
------------------------------------------------------------
------------------------------------------------------------
------------------------------------------------------------
------------------------------------------------------------
-----------------------TGVPQ--------------------------------
------------------------------------------------------------
------------------------------------------------------------
------------------------------------------------------------
------------------------------------------------------------
----------------------------NASVVQNLGL-----------------ASNF-
---------------------------------SLREFM---------------------
------------------------------------------------MAK----GWRLD
VGGAL------AFCG----VS-KT---E-----------V--C-----------------
------------------------------------------------------------
------------------------------------------------------------
------------------------------------------------------------
------------------------------S-----------HVS-N-SLG--------L
CWTLN-------AQ----------------------------------------------
-------RRQSIP------GY--------PH-------------GIHLIFITHQS-----
----------D-----------------S------------T-E-D---PI---------
------------------------------------------------------------
------------------------------------------------------------
-------SGNSETGIKFQVH-SQDEPP--QVESLGHSVGVGLWASVQMHKQ------E--
-----KSNM-----------YSP-W--GA-C--RPS-N-----EN-------LR----FY
------------------S-KYTLNACLKQCYDTLIQ----QT--CNCRGPL----M---
------------E-Y---------------------------------------------
-------NYEDV-----V---ECTP-----------------------------------
------------------------------------IQV---LNCTA---------P---
V-R-------------------YAIE-S-------SY--------TP----------QKC
------------GCY-QPCNEVK---YQTT-LSY------------------SSFNPK--
---------------HSEF-F-----------EK--------------------------
------------------------------------------------GM----------
------------------------------------------------------------
-N----------------------------------------------------------
---------------------------LSS------------------------------
-----G-------YL-----------S------ENSVGISVYFDALSVTKHV-ESKAITE
SAL-----------------LSDIGGQLGLWVGVSVVTLFELLQFLTSN-----------
--------IYCI--------L----------I-------YL-------------------
------------------------------------------------------------
------------------------------------------------------------
------------------------------------------------------------
------------------------------------------------------------
------------------------------------------------------------
------------------------------------------------------------
------------------------------------------------------------
--------
>Cnidaria_HyNaC11_NP_001296594_1_acidsensing_ion_channel_1like__Hydra_vulgari
s__34_85
------------------------------------------------------------
----------M-------------------------------------------------
------------------------------------------------------------
------------------------------------------------------------
------------------------------------------------------------

```
------------------------------------------------------------
------------------------------------------------------------
------------------LN-------------FKDIAQIT-------------------
------------------------------------------------------------
---------------VEAIQETNEVTNKDEK--------------------------NIQ
T----------------------IQDLR--------------------------------
--------N----------------------K---------KI-----R-----------
-EH--ISYMI-DNSSFHGLSYIF---------D-KR--------------HS---VR---
--RTIWF----FITIAAFAYAMQKVYESTM-NYFS----------Y---PFYTVR----M
-R-MY-V----------N-QIDF-----PAI-----------------------SFCNLN
DI------KFS-----AMNGTI------------------V-----------------D-
---------DA----------------------------------VV-------------
------------------------------------------------------------
------------------------------------------------------------
------------------------------------------------------------
------------------------------------------------------------
------------------------------------------------------------
------------------------------------------------------------
------------------------------------------------------------
------------------------------------------------------------
------------------------------------------------------------
------------------------------------------------------------
------------------------------------------------------------
------------------------------------------TQNH--------------
---------------------------------EAN-------------------ITGE-
---------------------------------EYRSYN---------------------
------------------------------------------------QAA----RHTLN
--EML------VDCD----FE-G----K----------K---C-----------S-H---
----------------------------------------------K-------------
------------------------------------------------------------
------------------------------------------------------------
-----------N----------F-------T-----------EFS-W-MQGE-------S
CFTFN-------SG-------------KP-P--------------------H----TL--
-------LKVKGA------GI--------NR-------------SLKLTINVQHY-----
----------D-----------------Y------------Y-R-D--------------
------------------------------------------------------------
------------------------------------------------------------
---------KMDSGIRLILH-GQDETP---VKMSGLTVPPGFTTYIQIEKK------T--
-----IINL-----------EAP-YK-TK-C--GSV--------K-------LK----YF
------------------D-SYSMHTCWLEQLTDYVY----KT--CNCKDYF----M---
------------P-G---------------------------------------------
----------DI-----P---ICSF-----------------------------------
------------------------------------DLL---YNCAW---------P---
E-W-------------------ETFD-K-------QK---------------------LY
------------QCP-LPCKIDS---YEVS-LSR------------------ALF-PT--
-------GLY------ASS-L-----------AN--------------------------
------------------------------------------------DLRKYQ--Q---
-----------------------------------------------------------V
PI---------------A------------------LKS---------------------
------------------------KT-DEL------------------------------
-----I-------FM-----------R------ENLLRLVIYYDDLAYELVE-QKPSYNT
LLW-----------------LGDVGGQIGLFIGAGVMSYFEFIDCLAMV-----------
--------IYTR--------F----------F-------EK-------------------
IS------------------------------LK--------------------------
------------------------------------------------------------
------------------------------------------------------------
------------------------------------------------------------
```

------------------------------------------------------------
------------------------------------------------------------
------------------------------------------------------------
---NPTTV
>Deutero_Ambulac_Apla_gbr173_7_t1_528_86
------------------------------------------------------------
------------------------------------------------------------
------------------------------------------------------------
------------------------------------------------------------
------------------------------------------------------------
------------------------------------------------------------
------------------------------------------------------------
------------------------------------------------------------
------------------------------------------------------------
------------------------------------------------------------
------------------------------------------------------------
------------------------------------------------------------
--------------------MWP---------G-AA--------------GV---VR---
--RLIWA----VVTLAALGFFLYQTALSFI-SFFE----------W---NHVTKV----D
-I-NY-A----------D-ELEF-----PAV-----------------------TLCNFN
KY------RES-----AWTERD------------------L-----------------K-
---------NV------------G-THL---------------GIID----E--------
---------------------------------DH-------NLIN--------------
------------------------------------------------------------
------------------------------------------------------------
------------------------------------------------------------
------------------------------------------------------------
------------------------------------------------------------
------------------------------------------------------------
--------------------------PHLY------------------------------
------------------------------------------------------------
------------------------------------------------------------
------------------------------------------------------------
--------TDEF------------------------------KQN---------------
------------------------LAAVNWTALEV--------------------EEDY-
---------------------------------DMTEFT---------------------
------------------------------------------------NRT----GHQLD
--EMI------VECT----WK-G----S----------P---C-----------N-S---
----------------------------------------------Q-------------
------------------------------------------------------------
------------------------------------------------------------
-----------D----------F-------H-----------HVF-S-HLG--------N
CYTFN-------HV-------------EL-D--------------------Q----DR--
-------HSTISA------GA--------AN-------------GLKLTLNIQEE-----
----------E-----------------Y------------T-P-S-NDLD---------
------------------------------------------------------------
------------------------------------------------------------
-------GAAEDAGIKWMLH-HPSEPP--YVKELGFGAGPGHHTFVATRHE------Q--
-----ISSL-----------PFP-Y--TM-C--EED----G---S-------LK----YF
------------------D-HYSIQACRIECETDLVV----DQ--CGCRLVE----Q---
------------P-G---------------------------------------------
----------TA-----P---VCNP-----------------------------------
------------------------------------AES---HECAH---------V---
V-L-------------------VQAV-T-------GE--------TE----------SSC
------------DCK-SPCELEV---YPFS-TTN------------------VKL-RA--
-------NFI------ETI-Y-----------SN--------------------------
----------------------------------------------STTH----------

------------------------------------------------------------
-N----------------------------------------------------------
---------------------------FTA------------------------------
-----E-------YI-----------E------NNLVLLSIYFEALNHEVIE-QLPEMTV
FTI-----------------LATLGGNFGLFLGGSVLTVMQLLEYLFDE-----------
--------VTAS--------C----------A-------SK-------------------
GS------------------------------SN-------------------R------
-----------------------------------------VGDSRAN------------
--------------------LTGKEKDVE------------------NAEDG--------
------------------------------------------------------------
----------------ISIIDMGKD-----------------------------------
------------------------------------------------------------
------------------------------------------------------------
---KSKDR
>Deutero_Cephalo_194160R_t1__name__chr_scaffold41_start_1865858_end_1873272_s
trand_pro_len_477__530_88
------------------------------------------------------------
----------M-------------------------------------------------
------------------------------------------------------------
------------------------------------------------------------
------------------------------------------------------------
------------------------------------------------------------
------------------------------------------------------------
------------------------------------------------------------
------------------------------------------------------------
------------------------------------------------------------
------------------------------------------------------------
------SPG----------------------E---------EK-----E-----------
-SP--EKEFA-GNTTLHGLNRIV---------I-AP--------------SG---YF---
--RVLWV----VAILGSYSGFAYMFSSMII-DYFS----------Y---RTITDT----K
-L-EF-A----------E-DLTF-----PAV-----------------------TICNMN
KF------DES-----KLKIID------------------F-----------------S-
---------YL------------S-LFL---------------YGVQ----L--------
---------------------------------DV-------DAIL--------------
------------------------------------------------------------
------------------------------------------------------------
-----------------ASPGA--------------------------------------
------------------------------------------------------------
------------------------------------------------------------
------------------------------------------------------------
-------------------------YP---------------------------------
------------------------------------------------------------
------------------------------------------------------------
------------------------------------------------------------
------------------------------------------------------------
------------------------DETVNST------------------------LEGV-
---------------------------------TVGDLL---------------------
------------------------------------------------LEK----GFNLQ
-WPRM------PLCK----WK-G----N----------P---C-----------T-Q---
----------------------------------------------Q-------------
------------------------------------------------------------
------------------------------------------------------------
-----------D----------F-------N----------HSF-G-YYG---------N
CYTFN-------AD-------------AN-S--------------------P--------
-------LKQTIQ------GS--------GN-------------GFVAFVDLKVN-----
----------D-----------------Y------------T-E-S--YYA---------
------------------------------------------------------------

------------------------------------------------------------
-------SGNSEVGLKLLLH-DQNEPP--MMDTQGIALAPGSHAFIAAKRT------L--
-----YENH-----------VPP-W--GV-C--QDL--------Q-------LE----YY
------------------D-TYTLNGCYLECRSKHVV----RN--CSCRPYD----L---
------------P-G---------------------------------------------
----------TA-----P---SCDP-----------------------------------
------------------------------------NTM---FSCVR---------A---
V-L-------------------DQVT-R-------GE--------L------------EC
------------NCP-VPCSMTA---YSMS-VSY------------------AGW-PN--
-------KLT------GEY-I-----------SP--------------------------
------------------------------------------------LW----------
------------------------------------------------------------
-G----------------------------------------------------------
---------------------------STG------------------------------
-----E-------YL-----------E------LNGVMFSVYYEKLNYQKIT-ELKAMDG
GQL-----------------ASNIGGMMGLFIGASLLTVLEVWEYLWQR-----------
--------VLGL--------L----------G-------KQ-------------------
RR------------------------------PR--------------------------
-----------------------------------------VIEVK--------------
-------------------------SP---------------------PDDMER-GIRNV
AFP---MENTKIN-----------------------------------------------
------------------------------------------------------------
------------------------------------------------------------
------------------------------------------------------------
--------
>Protostome_Lophotroco_brch_Lingula_anatina_comp134404_c2_seq5_p1_comp134404_
c2__comp134404_c2_seq5_p1__ORF_type_complete_len_479___score_28_84_comp134404
_c2_seq5_272_1708__536_92
------------------------------------------------------------
----------M-------------------------------------------------
------------------------------------------------------------
------------------------------------------------------------
------------------------------------------------------------
--SSVHPFSFEEKGTE--------------------------------------------
------------------------------------------------------------
-FIKQYNNMHGDFSNKSPHR-------------GTNITKR--------------------
------------------------------------------------------------
---------------------------------------------------------KPD
Q----------------------PKDETI-------------------------------
-------------------------------P---------SL-----K-----------
-DE--ISDFA-NNVSILGMKQVA---------N-PK-------------FSP---VR---
--RLLWL----AFVLCGSGFLIFHLWNRIS-YFLE----------N---PASVDI----K
-F-HH-N----------D-SIHF-----PTV-----------------------TICNNN
RF------KLP-----YLRANG--------H---------L-------------------
---------EQ-------------------------------ANYFD-------------
------------------------------------------------------------
------------------------------------------------------------
------------------------------------------------------------
------------------------------------------------------------
------------------------------------------------------------
------------------------------------------------------------
------------------------------------------------------------
------------------------------------------------------------
------------------------------------------------------------
------------------------------------------------------------
------------------------------------------------------------
------------------------------------------------------------

------------------------MFNFNLTN-----------------------IEQY-
---------------------------------NWTDFH---------------------
------------------------------------------------TRN----QYKMA
--DLM-----PSFCL----FN-Q----V----------I---C-----------T-D---
----------------------------------------------A-------------
------------------------------------------------------------
------------------------------------------------------------
-----------D----------F-------S-----------MIE-T-EMG--------S
CIQFN-------AE----------------G-------------------------EL--
-------KSVSRA------GS--------LS-------------GLQMWLFSHQD-----
----------D-----------------Y------------L-N-----FT---------
------------------------------------------------------------
------------------------------------------------------------
----------TKAGYTVLMH-ERGEFP--HMIGLGLKVSPGESVDIAMKQR------R--
-----RTNL-----------PPP-H--GQ-C--KER--------T-------LK----YF
------------------S-KYTKLNCDAEC-SFIAM----QA--CGCRLFH----Q---
------------P-K--TGNQ---------------------------------------
-------TIDGQ-----R---TCNL-----------------------------------
------------------------------------KDT--MFGCFN-------------
ISK-------------------EPMA---------GY--------------------SSC
------------ACM-EACQSTL---YTHS-ISH------------------TRM-SE--
-------VFI------KRL-I-----------AM--------------------------
------------------------------------------------YN----------
------------------------------------------------------------
-N----------------------------------------------------------
---------------------------TDT------------------------------
-----S-------FF-----------R------ENIIMLNIFYSDISVEEVV-QQEAYSA
LTL-----------------FADIGGALGLVLGSTLMTAAEIVDFFLGV-----------
--------SLWK--------L----------F-------RI-------------------
RV----------------------------------------------------------
------------------------------------------------------------
------------------------------------------------------------
------------------------------------------------------------
------------------------------------------------------------
------------------------------------------------------------
------------------------------------------------------------
--------
>Ctenoph_Hormiphora_californiensis_evg1166627__p1_GENE_evg1166627
------------------------------------------------------------
----------M------------Q------------------------------------
------------------------------------------------------------
----------------------------------------------------PNGRPKD-
-YVFTDI-------DQPLS----ANGFYKRIPAPP---RPYRP-----------------
--TPAFPIYDN--------L-----------LDRED---VMSS----------NCPSTIP
GVY-----PGVEAFTP--------------------------------------------
---------------YSKYK-------------QDESAFDMTGRGERTS-----YYPYL-
--EHYQPSYPRPARPKTTANPTTTT--AN-HR-AEPNGKIKYPPNHRDTDSSGSPTEMDD
RDGEECEEFM--PPPKRKKKKKRRANRSLENRKNL---------------RLRRRKKAAL
E----------------------QRREDM-------------------------------
--------E----------------------P---------DE-----M-----------
-ES--FKEFA-STVTADYLAEMF-----------AK-------------ASN---GR---
--RAIWV----TLWLASFVYAWYNIALSIG-VYLS----------K---PTATKL----N
-F-YE-AP--------PN-GVAF-----PTV-----------------------TICNFN
KY------NKS-----FFEREG------------------D-----------------SV
N------A-QL------------R-DFL---------------VETT-------------
------------------------------------------------------------
------------------------------------------------------------

------------------------------------------------------------
------------------------------------------------------------
------------------------------------------------------------
------------------------------------------------------------
------------------------------------------------------------
--------------------------PIWDH-----------------------------
------------------------------------------------------------
------------------------------------------------------------
------------------------------------------------------------
--------KERL------------------------------AKD---------------
----------------------QWLSKYPS-------------------------IMNL-
---------------------------------ELKQVL---------------------
------------------------------------------------TAA----THRLE
--DTV------EYCR----FG-G----K----------P---CL-DSQYRETSIP-S---
----------------------------------------------E-------------
------------------------------------------------------------
------------------------------------------------------------
-----------A----------F-------T-----------KIY-T-EHG--------L
CYSFN-------QD-------------GK-------------------------------
-------LNMTRT------GA--------QY-------------GLSLRLHVAQD-----
----------D-----------------Y------------F-K-T--------------
------------------------------------------------------------
------------------------------------------------------------
---------QDVAGFKVLLH-NWYEPP--MIEEYGFALRPGSESYVRIRLQ------K--
-----FRDT-----------ERP-L--GY-C--DPL----------------LEQ---DY
----------------GPG-SYTIAKCVMANRASLTI----KK--CGCIPFY----MDVA
RSAFA-----VPP-K---------------------------------------------
-------YPSST-----P---VCNL-----------------------------------
------------------------------------YQE---WDCVE---------P---
L-L-------------------EDIA-L-------GT--------LVID-GEPI---GDP
------------DCP-SPCNYLT---YSYT-ISQ------------------SEY-PS--
-------QSV------EQE-I-----------LD--------------------------
------------------------------------------------NV----------
------------------------------------------------------------
-LGARG------------------------------------------------------
------------------------KS-WNI------------------------------
-----E-------KI-----------R------DNYLKVHIYFEELSTLEME-RHPSYLI
SNL-----------------IGDVGGQLGLLLGMNICSLVQFTDYIIRFSFFN-------
-------GIMKL--------V------------------RI-------------------
SQ------------------------------RN-------------------S------
-----------------------------------------KR-----------------
------------------------------------------------------------
------------------------------------------------------------
------------------------------------------------------------
------------------------------------------------------------
------------------------------------------------------------
----RAKL
>Cteno_MneLey_comp19805_c0_seq2__p1__GENE_comp19805_c0_seq2__comp19805_c0_seq
2__p1___ORF__type_complete__len_802___score_191_14__comp19805_c0_seq2__85_249
0_
------------------------------------------------------------
----------MSNRVITTDLRLCQTAL---TPNTMPETKI--------------------
-----------KIGLTRGLV------DGGEIFAQFSLSST-AHYQHLLTYISAI------
CSREKVLAIY-WFN-RHDREFVLI-------HDNASMDY----------CLVQSGLHEI-
LYIITET------GGKPAG----YNSKHKPEPIQL---KTSSP------------SSGPS
PDTANLERSSDEDRPGGT-------------VQREDHI----------------------
-----------------------------------------PLRKM--------------

```
------KKSASLEWSYDNEQ----PPSVQEISSRDESAFDMTGRGERPT-----YYPYL-
--EHYQPSYPARPKPHRTPTRNKRPEPNNYHTHSDQSGSL------SDSD--------DD
RNAKEVDQFMEPEPPKRKKKAKRRANR------------------------------AAL
E----------------------QKRQEI-------------------------------
-------------------------------E----------------M-----------
-ET--FKEFA-STVTADYLAEMC-----------SK-------------ASN---GR---
--RLIWV----SLWLASFVYAWYNIALSIG-VYMS----------K---PTATKL----N
-F-YE-AS--------SD-GVTF-----PTV-----------------------TICNFN
KF------NKS-----YFDTDD------------------I-----------------TT
K------KGQL------------K-DFL---------------KITT-------------
------------------------------------------------------------
------------------------------------------------------------
------------------------------------------------------------
------------------------------------------------------------
------------------------------------------------------------
------------------------------------------------------------
------------------------------------------------------------
--------------------------PIWGH-----------------------------
------------------------------------------------------------
------------------------------------------------------------
------------------------------------------------------------
--------KQEL------------------------------DSA---------------
----------------------SWDTKYDK-------------------------IVQQ-
---------------------------------SMKDAL---------------------
------------------------------------------------QTA----SHSLE
--GTV------EFCK----FS-G----K----------P---CLPDVADVEDADP-T---
----------------------------------------------A-------------
------------------------------------------------------------
------------------------------------------------------------
-----------A----------F-------N-----------PVF-T-EHG--------L
CYSFN-------QN-------------GN-------------------------------
-------LTMSRT------GA--------LY-------------GLSLRLHVAQD-----
----------D-----------------Y------------F-E-T--------------
------------------------------------------------------------
------------------------------------------------------------
---------QDVAGFKVLLH-NSYEPP--MIEEYGFALRPGSETYVRIRLQ------K-D
VYPQ-YKDT-----------PRP-L--GY-C--DPY----------------LED---QF
-----------------ST-NYTISVCVMQYRAINML----KV--CGCITFY----MDRP
DIVLSEGKPYMKP-----------------------------------------------
-----------------K---ICNL-----------------------------------
------------------------------------KEE---RTCAD---------N---
V-L-------------------EGLA-L-------GT--------LKID-GEKV---PEP
------------DCP-SPCVYTT---YTYT-VSQ------------------SEY-PS--
-------SAV------KDK-V-----------VE--------------------------
------------------------------------------------EV----------
------------------------------------------------------------
-NRQKG------------------------------------------------------
----------------TSPGEDDASS-WTI------------------------------
-----E-------RI-----------R------DNYLKVHIYFEELSTLEME-RHPSYLI
SNL-----------------IGDVGGQLGLLLGMNICSLVQFTDYIVRFSFFK-------
-------GFMSL--------V------------------RLYK-----------------
AR------------------------------RR--------------------------
------------------------------------------------------------
------------------------------------------------------------
------------------------------------------------------------
------------------------------------------------------------
------------------------------------------------------------
```

```
------------------------------------------------------------
------NK
>Deutero_Ambulac_Apla_gbr256_12_t1_540_96
------------------------------------------------------------
----------M----------------------------------------------D--
------------------------------------------------------------
------------------------------------------------------------
--------------------------------------------------------PRVI
IETTPRNGFNERRV----------------------------------------------
-----------------------------------M------------------------
----------------GKDN-------------LDELNIIKNDR----------------
------------------------------------------------------------
---------------------------------------------------------NRH
T----------------------QDDDA--------------------------------
--------N----------------------R---------TF-----A-----------
-ER--FQDFG-NSTTLHGISYVI---------N-LN-------------YKK---QR---
--RLLWL----LLVLVMSIWLVYSIVEAFI-TYFN----------Y---PMTSAI----S
-I-HY-M----------D-TLTF-----PAV-----------------------TLCNFN
QF------RRS-----ALTPGQ------------------V-----------------M-
---------VM-------------------------------NEVYG-------------
------------------------------------------------------------
------------------------------------------------------------
------------------------------------------------------------
------------------------------------------------------------
------------------------------------------------------------
------------------------------------------------------------
------------------------------------------------------------
--------------------------PNP-------------------------------
------------------------------------------------------------
------------------------------------------------------------
------------------------------------------------------------
------------------------------------------------------------
------------------------PESIDFGPFNE--------------------ITNYD
---------------------------------FSEDHL---------------------
------------------------------------------------ANI----SHHLD
--RML------VKCK----WR-G---TQ----------P---C-----------T-G---
----------------------------------------------Q-------------
------------------------------------------------------------
------------------------------------------------------------
-----------N----------F-------T-----------RRF-T-DHG--------V
CFTFN-------DP-------------KD-A--------------------S----HR--
-------LEVSNA------GR--------LD-------------GLFLRLFAETD-----
----------E-----------------Y------------T-F-G---EN---------
------------------------------------------------------------
------------------------------------------------------------
----------SAAGFRILLH-PQGLMP--LVKELGISVSPGFESSISIRQN------V--
-----FKSL-----------PWP-FE-SN-C--TSS--------K-------SK----NS
------------------T-TYSVPTCKYECKVQYVV----ER--CGCRDYR----W---
------------T-G---------------------------------------------
----------SA-----P---LCPP-----------------------------------
------------------------------------EQH---FNCID---------K---
Y-K-------------------AEYQ-K-------QE--------KY----------RDC
------------HCP-VSCELKT---YDSR-VSF------------------GYW-PS--
-------SYV------SEN-L-----------PL--------------------------
------------------------------------------------------------
------------------------------------------------------------
-NS---------------------------------------------------------
```

---------------------------TSE------------------------------
-----E-------YI-----------R------KNYLDVYIFFEELVYLQIE-QKSAYDF
AAL-----------------QSGVGGNMGLLCGMSLITVVEWADFIVIS-----------
--------LLRK--------F----------K-------AR-------------------
RI----------------------------------------------------------
------------------------------------------------------------
------------------------------------------------------------
------------------------------------------------------------
------------------------------------------------------------
------------------------------------------------------------
------------------------------------------------------------
--------
>Protostome_Lophotroco_annelid_Pdum_Contig8016__542_97
------------------------------------------------------------
----------M-------------------------------------------------
------------------------------------------------------------
------------------------------------------------------------
------------------------------------------------------------
------------------------------------------------------------
------------------------------------------------------------
-------------------------------------SKVT-------------HYPM--
------------------------------------------------------------
---------------------------------------------------------DDL
G----------------------KSPRPLVNDM---------------------------
GQGHRVRKS----------------------S---------SM-----Q-----------
-RV--LSDYS-EHSSMHGVWIVA---------N-PD-------------YPK---LR---
--RLIWM----LALITTTTVFVYAFIQQMT-MFFQ----------Y---KVTTVL----S
-V-EE-Q----------A-TLNF-----PAV-----------------------TLCNYN
QF------RKS-----KAEQKP--------G---------V-----------------M-
---------GV------------------------------LRDLFS--V----------
---------------------------------DA-------------------------
------------------------------------------------------------
------------------------------------------------------------
------------------------------------------------------------
------------------------------------------------------------
------------------------------------------------------------
------------------------------------------------------------
------------------------------------------------------------
------------------------------------------------------------
------------------------------------------------------------
------------------------------------------------------------
------------------------------------------------------------
------------------------IIERQKSKLRMSAQ--------------SLVSQDI-
---------------------------------NIINEL---------------------
------------------------------------------------VAL----AHTLK
--DMV------RHCQ----WC-G----K--------TIP---C-----------S-----
------------------------------------------------------------
------------------------------------------------------------
------------------------------------------------------------
-----------D---------YF-------S-----------QTL-T-EVG--------V
CYTFN-------SH-------------DV-R--------------------K--KIPP--
-------LEVHDA------GT--------DC-------------GLKLRLNIEEY-----
----------E-----------------Y------------Y-Y-G---EG---------
------------------------------------------------------------
------------------------------------------------------------
----------SSAGIQMLIH-DPEDVP--LVRQHGFSAAPGTSVMAAIRLT------K--
-----ETYA-----------QSP-Y--GD-C--VDA--------KHPSFENPLR----YH

------------------E-IYRYSACKLEHRVDYIY----NA--CGCVMYY----N---
------------PVG---------------------------------------------
----------SR-----S---VCKT-----------------------------------
------------------------------------VEE---VLCFK---------D---
A-K-------------------ENHT-R-------YE--------MQ---------AMRH
------------ACP-VTCQSDN---YEAR-LSW------------------AAF-PA--
-------PQI------LYD-L-----------KK--------------------------
------------------------------------------------NF----------
------------------------------------------------------------
-N----------------------------------------------------------
---------------------------MSG------------------------------
-----D-------QV-----------R------KNLLQVTVYFEDLGIHKKE-QQPAYTG
NSL-----------------FADIGGMMGLCIGASLLTIVELLELGAQL-----------
--------MLSA--------G----------K----------------------------
--------------------------------R---------------------------
------------------------------------------------------------
------------------------------------------------------------
------------------------------------------------------------
------------------------------------------------------------
------------------------------------------------------------
------------------------------------------------------------
--------
>Protostome_Lophotroco_brch_Lingula_anatina_comp149479_c1_seq4_p1_comp149479_
c1__comp149479_c1_seq4_p1__ORF_type_complete_len_482___score_29_72_comp149479
_c1_seq4_182_1627__543_98
------------------------------------------------------------
----------M-------------------------------------------------
------------------------------------------------------------
------------------------------------------------------------
------------------------------------------------------------
------------------------------------------------------------
------------------------------------------------------------
------------------------------------------------------------
------------------------------------------------------------
------------------------------------------------------------
------------------------------------------------------------
-------GK----------------------G---------EG-----E-----------
-SL--ETEFA-SGTTCHGFNRLA---------V-AK---------------KW---PH---
--RAFWT----VAILWGFGMTTWMISSRIE-AYFK----------Y---ETSTEI----R
-V-EE-K----------Y-ALEF-----PAV-----------------------TFCNFN
KY------RRS-----AMTALD------------------W-------------------
---------YI-----------YANSKLA------------EKYAFE----A--------
---------------------------------DY-------------------------
------------------------------------------------------------
------------------------------------------------------------
-----------------D------------------------------------------
------------------------------------------------------------
------------------------------------------------------------
------------------------------------------------------------
--------------------------PEA-------------------------------
------------------------------------------------------------
------------------------------------------------------------
------------------------------------------------------------
----TAALRAAF------------------------------------------------
-----------------------EGFEANLTATY---------------------PGGF-
---------------------------------DFGNFT---------------------
------------------------------------------------RRI----GWELN

-NETM------PLCL----FR-M----K----------Q---C-----------F-P---
----------------------------------------------E-------------
------------------------------------------------------------
------------------------------------------------------------
-----------N----------F-------T-----------HAFQS-EYG--------N
CYTFN-------SH-------------HD-E--------------------D----SLS-
-------LQQTRP------GA--------QQ-------------GLYILFDIQQA-----
----------E-----------------Y------------T-E-L---LN---------
------------------------------------------------------------
------------------------------------------------------------
------LNGALGVGIAFQIH-SWYERP--NVAQHGLGAGVGTAALVSMHQN------E--
-----LQNL-----------AHP-W--GN-C--SND-R------T-------LK----HF
------------------P-RYSRSGCEVECYLEKVL----EL--CGCKPFS----Y---
------------P-G---------------------------------------------
----------NA-----V---NCNP-----------------------------------
------------------------------------VDQ---VLCAD---------P---
T-I-------------------FKVA-S-------NF--------Q-----------KEC
----------GARCP-EACQIIE---YPPT-VSY------------------APF-PN--
-------YYM------AQE-L-----------AT--------------------------
------------------------------------------------MY----------
------------------------------------------------------------
-G----------------------------------------------------------
---------------------------TNA------------------------------
-----N-------YA-----------S------ENLVGLSIYYKDLNNRLTK-KNPAMTP
SGL-----------------LSDIGGQLGLFIGMSTLTLMEFGEYIIFK-----------
--------IIQC--------C----------R-------KR-------------------
KP------------------------------KT-------------------S------
-----------------------------------------QSKGV--------------
------------------------------------------------ANNV--------
------------------------------------------------------------
------------------------------------------------------------
------------------------------------------------------------
------------------------------------------------------------
-KVNESGM
>Deutero_Ambulac_Apla_gbr54_23_t1_545_100
------------------------------------------------------------
----------M-------------------------------------------------
------------------------------------------------------------
------------------------------------------------------------
------------------------------------------------------------
------------------------------------------------------------
------------------------------------------------------------
----------------VRED-------------ADQ------------------------
------------------------------------------------------------
------------------------------------------------------------
------------------------------------------------------------
-------------------------------S---------PR-----P-----------
-SL--EREFA-ESTSLHGIAKVF---------H-AS--------------RL---LV---
--RVFWV----IIMLTCLCVCWQISDRFH-RFLQ-----------Y---KANTEI----S
-V-EY-V----------G-HLPF-----PAV-----------------------TICNFN
RY------RSG-----ALAPGD------------------E-----------------D-
---------AL------------A-YIW---------------ELSN----YQN------
---------------------------------DY-------DYEG--------------
------------------------------------------------------------
------------------------------------------------------------
-----------------E------------------------------------------
------------------------------------------------------------

```
------------------------------------------------------------
------------------------------------------------------------
------------------------------------------------------------
------------------------------------------------------------
------------------------------------------------------------
------------------------------------------------------------
----SNSTDSSY------------------------------LDS---------------
-------------------------------------------------------LVDF-
---------------------------------NYTEFT---------------------
------------------------------------------------LRA----GFRMD
-NVTL------LACS----WR-G----K--------RNS---C-----------S-A---
----------------------------------------------E-------------
------------------------------------------------------------
------------------------------------------------------------
-----------D----------F-------S-----------HVF-T-SYG--------N
CWTFN-------SG-------------KT-G-------------------------PV--
-------LTESYS------GS--------GN-------------GVSLLVDIQQS-----
----------E-----------------Y------------T-E----------------
------------------------------------------------------------
------------------------------------------------------------
-------TETIEAGLKIQVH-DQVTPP--TVESSGLALSPGVHAFMAVRRQ------Q--
-----YLNL-----------EHP-Y--GK-C--DAS-R------T-------LK----RY
------------------D-NYTLEGCNIECRANAVW----EK--CRCRLIQ----H---
------------P-G---------------------------------------------
----------NE-----V---ECTP-----------------------------------
------------------------------------EEV---RFCAR---------E---
A-I-------------------DGLI-G-------GD--------T-----------ESC
------------DCP-VPCNFTA---FTTS-LST------------------AFL-PN--
-------EKI------LPI-L-----------LL--------------------------
------------------------------------------------LL----------
------------------------------------------------------------
-NNIST------------------------------SDERNVSRSIS-------------
---------------------------EQQ------------------------------
-----E-------YI-----------R------KNWVAIDTFYEDINYQKFE-QTEAITP
SAL-----------------ISDIGGQLGLFLGASFITLAEIFSYLSRK-----------
--------IQRL--------I----------H-------GP-------------------
TR------------------------------RP--------------------------
-----------------------------------------EKQRQ--------------
------------------------MSH---------------------SVET--------
------------------------------------------------------------
------------------------------------------------------------
------------------------------------------------------------
------------------------------------------------------------
------LA
>Deutero_Ambulac_Apla_gbr658_1_t1_550_104
------------------------------------------------------------
----------M-------------------------------------------------
------------------------------------------------------------
------------------------------------------------------------
------------------------------------------------------------
--NRPCDGTDH-------------------------------------------------
------------------------------------------------------------
----NFDMETDDLRDGRERR-------------RPSVTRRKRGF----------------
------------------------------------------------------------
---------------------------------------------------------KRK
K----------------------SKDTC--------------------------------
--------K----------------------N---------IL-----A-----------
```

-AR--IQQFG-DDTTFHGLRYVT---------N-ST-------------YHT---FR---
--RFVWL----LVVTGMTALLIFDIIRAIM-LMFE----------Y---PEMSSI----S
-V-QY-P----------P-NITF-----PAL-----------------------TLCNIN
ML------RKD-----ALDASE------------------V-----------------Q-
---------LL-------------------------------SDIFA-------------
------------------------------------------------------------
------------------------------------------------------------
------------------------------------------------------------
------------------------------------------------------------
------------------------------------------------------------
------------------------------------------------------------
------------------------------------------------------------
-------------------------NPATP------------------------------
------------------------------------------------------------
------------------------------------------------------------
------------------------------------------------------------
------------------------------------------PDT---------------
------------------------NALVDSKLFNS--------------------SSNE-
---------------------------------ELMGRM---------------------
------------------------------------------------IDA----SYRIE
--DML------VKCR----WV-S----E----------T---C-----------S-Y---
----------------------------------------------R-------------
------------------------------------------------------------
------------------------------------------------------------
-----------N----------F-------T-----------RSI-T-DHG--------V
CYTFN-------DP--------------LS-D-------------------H----EA--
-------LIVRHP------GS--------RH-------------GLFMRLNVQQY-----
----------L-----------------Y------------T-Y-G---ES---------
------------------------------------------------------------
------------------------------------------------------------
----------TAAGLKVLLH-AQGEYP--MIKDFAFSLSPGFETSIVVRQK------V--
-----MINQ-----------KAP-YK-SN-C--TDG--------N-------WG----RH
------------------RFKYSAASCKYACKILYVT----RK--CGCRDYR----W---
------------S-I---------------------------------------------
---------RDI-----P---ICSL-----------------------------------
------------------------------------EKQ---VHCIY---------R---
Y-E-------------------DEFVLS-------QE--------------------NQC
------------HCP-IACKAIT---YGRE-TSQ------------------AYW-PA--
-------SFL------SKE-L-----------QA--------------------------
------------------------------------------------EM----------
------------------------------------------------------------
-N----------------------------------------------------------
---------------------------VTE------------------------------
-----D-------FI-----------R------ENYLDVYIYYDEIMYTRIE-QIGAYTV
DNL-----------------QSDIGGYLGLLCGMSLMTVVELLDFILVA-----------
--------LCSR--------L----------K----------------------------
-----------------------------------------------------R------
------------------------------------------------------------
------------------------------------------------------------
------------------------------------------------------------
------------------------------------------------------------
------------------------------------------------------------
------------------------------------------------------------
--------
>Deutero_Cephalo_298690R_t1__name__chr_scaffold94_start_933624_end_940952_strand_pro_len_484__551_105
------------------------------------------------------------

```
----------M-------------------------------------------------
------------------------------------------------------------
------------------------------------------------------------
------------------------------------------------------------
------------------------------------------------------------
------------------------------------------------------------
------------------------------------------------------------
------------------------------------------------------------
------------------------------------------------------------
------------------------------------------------------------
-----------IT------------------S---------DE-----K-----------
-TP--EREFM-DNTTLHGFNKIH---------H-PN-------------R-----AI---
--RTVWI----VAMLAAYAGFIYMAVNMAQ-TYFS----------Y---DTVCDV----K
-L-SF-E----------S-KMTF-----PAV-----------------------TICNYN
RI------DSH-----KLTWEE------------------W-----------------S-
---------IL------------S-TLL---------------YGRA----M--------
---------------------------------DV-------PTLQ--------------
------------------------------------------------------------
------------------------------------------------------------
-----------------AFLG---------------------------------------
------------------------------------------------------------
------------------------------------------------------------
------------------------------------------------------------
--------------------------PTF-------------------------------
------------------------------------------------------------
------------------------------------------------------------
------------------------------------------------------------
------------------------------------------TDR---------------
------------------------NRTTNGTVTN---------------------ATTM-
---------------------------------DMAAAV---------------------
------------------------------------------------RMK----GFDIS
-PARM------YLCS----FA-G----D----------S---C-----------T-E---
----------------------------------------------L-------------
------------------------------------------------------------
------------------------------------------------------------
-----------N----------Y-------T-----------HSF-S-NYG--------N
CYTFN-------AD-------------AA-N--------------------K--------
-------LNQTIP------GS--------TM-------------GFQIVVNVMAD-----
----------S-----------------Y------------T-E-T---MA---------
------------------------------------------------------------
------------------------------------------------------------
------VGGHEEVGLKFLIH-DQNEPP--MVETHGLAIKPGIHSFVSVKRT------E--
-----YYNH-----------VPP-W--GD-C--HDK--------A-------LT----YF
------------------D-AYTLPGCNAECRGKHVQ----DR--CGCKAYY----L---
------------P-G---------------------------------------------
----------SA-----P---PCAA-----------------------------------
------------------------------------DVL---FTCVQ---------E---
V-L-------------------AEVN-E-------GK--------L------------TC
------------DCP-LPCTMVK---YSPT-LSF------------------SAW-PN--
-------QLT------EDY-Y-----------SR--------------------------
------------------------------------------------TY----------
------------------------------------------------------------
-N----------------------------------------------------------
---------------------------LSR------------------------------
-----G-------YLTL----FMLFPR------DNVVALTVYYEDFSYMKIS-QLKAMDS
GQL-----------------GCNMGGMMGLFIGASVLTLVELTEYLVAR-----------
--------LSIS--------W----------G-------KK-------------------
```

```
KR------------------------------PL-------------------H------
-----------------------------------------VQPAPDAEM----------
---------------------SDKKQP---------------------------------
------------------------------------------------------------
------------------------------------------------------------
---------------------------------P--------------------------
------------------------------------------------------------
-DSLVFHK
>Deuterostome_chordata_PetMar_tr_S4RS34_S4RS34_PETMA_Acid_sensing_ion_channel
_subunit_family_member_5_OS_Petromyzon_marinus_OX_7757_PE_3_SV_1_552_106
------------------------------------------------------------
------------------------------------------------------------
------------------------------------------------------------
------------------------------------------------------------
------------------------------------------------------------
------------------------------------------------------------
------------------------------------------------------------
------------------------------------------------------------
------------------------------------------------------------
------------------------------------------------------------
-----------------------STGSPA-------------------------------
-----------------------------GPT---------DS-----A-----------
-KE--TIAFS-SSTSLHGVARAT---------S-AQ--------------SR---AQ---
--RAFWM----MVVVAGVAMAGWQVALRLA-AYYA----------W---PTSTSL----T
-L-HY-V----------R-DLSF-----PAV-----------------------TFCNYN
RY------RRS-----SLHGTT-----------------DLL----------------A-
---------AL--------------------------------SILS-------------
------------------------------------------------------------
------------------------------------------------------------
-----------------------A------------------------------------
-----------GIKTPH-------------------------------------------
------------------------------------------------------------
------------------------------------------------------------
------------------------------------------------------------
--------------------------PEY-------------------------------
------------------------------------------------------------
------------------------------------------------------------
------------------------------------------------------------
--------LSGFL-----------------------------------------------
--------------------------------------------------W-RITLRGYS
S-------------------------------PPITQTY---------------------
------------------------------------------------SAR----NQDLS
--TTI------LEIDD--TYC-E----N----------S---C-----------T-P---
----------------------------------------------Q-------------
------------------------------------------------------------
------------------------------------------------------------
-----------D----------F-------Q-----------HVF-T-EYG--------N
CYTFN-------GGF------------SE-E--------------------Q----RS--
-------LKQKTA------GA--------GK-------------GLQVLFDVQQKS----
-------HLWD-----------------Y------------W-E-N---PE---------
------------------------------------------------------------
------------------------------------------------------------
-------LGFTGVGIRLVVH-DPDEPP--QVEALGLSIGVGTQAYVSVQHT------K--
-----SINQ-----------QYP-W--GD-C--DPS-K------K-------LQ----SP
------------------A-PYSMARCLQECEADLVE----RI--CGCKPFN----F---
------------P-G---------------------------------------------
----------DI-----K---ECSL-----------------------------------
```

```
------------------------------------YQH---YHCAR---------D---
A-L-------------------VIFL-A-------QG---------------------MC
TGGS-----HLSQCP-VPCLHKN---FPSS-VSY------------------ATF-PS--
-------LQA------AQR-Y-----------ER--------------------------
------------------------------------------------SL----------
------------------------------------------------------------
-N----------------------------------------------------------
---------------------------YSL------------------------------
-----V-------YM-----------R------ENLLSIDVAYTDLNYALIQ-QQKTLPE
SVL-----------------LSDIGGQLGLFVGASVITIIEVLEFVARS-----------
--------AARA--------L----------N-------AA-------------------
ASFLS--------------------------NRA---------------ATCTKA-----
------------------------------------------------------------
-----------------------ALSP---------------------------------
------------------------------------------------------------
------------------------------------------------------------
---------------------------------P--------------------------
------------------------------------------------------------
-----VQG
>Protostome_Lophotroco_brch_Lingula_anatina_comp131676_c0_seq1_p1_comp131676_
c0__comp131676_c0_seq1_p1__ORF_type_complete_len_484___score_23_39_comp131676
_c0_seq1_234_1685__554_107
------------------------------------------------------------
----------M-------------------------------------------------
------------------------------------------------------------
------------------------------------------------------------
------------------------------------------------------------
--SQVDPFSVDYRTSE--RKQAFTDDGLI-------------------------------
------------------------------------------------------------
-------SFNVYNRGKQKQN-------------SRDIVKQ--------------------
------------------------------------------------------------
---------------------------------------------------------FKA
E----------------------KDGNDE-------------------------------
-------------------------------F---------SL-----S-----------
-SE--LSSFA-NDVSILGMKQIV---------N-PK-------------FSL---FR---
--RATWL----CFVLAGTGFLIYHLWNRLD-YYFS----------N---PASVNI----R
-V-NY-D----------G-SLLF-----PTV-----------------------TICNNN
VF------LKS-----KLAANN------------------E-----------------T-
---------YI-------------------------------GDLFY-------------
------------------------------------------------------------
------------------------------------------------------------
------------------------------------------------------------
------------------------------------------------------------
------------------------------------------------------------
------------------------------------------------------------
------------------------------------------------------------
--------------------------PSF-------------------------------
------------------------------------------------------------
------------------------------------------------------------
------------------------------------------------------------
------------------------------------------------------------
------------------------DSYPNWSAYN---------------------FTGF-
---------------------------------NWTDFY---------------------
------------------------------------------------YRN----GYSVM
--DVFG-----GFCF----WK-S----S----------P---C-----------Q-A---
----------------------------------------------N-------------
------------------------------------------------------------
```

------------------------------------------------------------
-----------V----------L-------K-----------PVI-T-EHG--------V
CYQFN-------SG-------------DE-G-----------------------------
-------KRSDVE------GS--------LY-------------GLMISFMTKQE-----
----------E-----------------Y------------L-V-G--------------
------------------------------------------------------------
------------------------------------------------------------
---------SYIAGYTILLH-NSKDHP--NVVTQGFQVAPGGSANVAIKQK------Y--
-----VKNL-----------PQP-H--GQ-C--ADK--------Q-------LA----YF
------------------P-HYSKINCQAECLHNKTI----EK--CGCRLVY----Y---
------------P-E---------------------------------------------
--------VPGL-----R---ECSP-----------------------------------
------------------------------------IDV---NTCWG---------P---
L-L-------------------TNVS-Q-----------------IM----------KNC
------------ACT-EACETFY---FEYS-VTT------------------GLV-SD--
--------------VSAQA-A-----------AV--------------------------
------------------------------------------------FL----------
------------------------------------------------------------
-N----------------------------------------------------------
---------------------------TTQ------------------------------
-----T-------YV-----------E------KNYLAVYLYYKEMSYEEVI-QQEAYST
LTL-----------------LADIGGALGLILGSTLMTVAEILDFFLGL-----------
--------SIWK--------F----------F-------GV-------------------
KL----------------------------------------------------------
------------------------------------------------------------
------------------------------------------------------------
------------------------------------------------------------
------------------------------------------------------------
------------------------------------------------------------
------------------------------------------------------------
--------
>Placozoa_TadNaC4_MK547545_14_111
------------------------------------------------------------
----------M-------------------------------------------------
------------------------------------------------------------
------------------------------------------------------------
------------------------------------------------------------
------------------------------------------------------------
------------------------------------------------------------
------------------------------------------------------------
------------------------------------------------------------
------------------------------------------------------------
------------------------------------------------------------
----------------------------A--K---------ED-----L-----------
-SM--EEKFA-STTSCHGISHVY---------Q-ND-------------SGR---LP---
--RMIWL----ILTIAATAVCISQCIIIIY-DASQ----------L---PTRITF----S
-D-VS-A----------N-STVF-----PSI-----------------------TICNTD
NS------PRD-----ALPESD------------------L-----------------K-
---------YL------------S-ILLH---------TQSTENAYN-------------
------------------------------------------------------------
------------------------------------------------------------
------------------------------------------------------------
------------------------------------------------------------
------------------------------------------------------------
------------------------------------------------------------
------------------------------------------------------------
------------------------------------------------------------

------------------------------------------------------------
------------------------------------------------------------
------------------------------------------------------------
--------ETQV------------------------------QEA---------------
------------------------ARYFIK------------------KY-----GDKF-
---------------------------------DYENYI---------------------
------------------------------------------------RTS----SAKLE
--NML------LRCQ----WM-N----R----------P---C-----------S-L---
----------------------------------------------S-------------
------------------------------------------------------------
------------------------------------------------------------
-----------D----------F-------T-----------SIV-T-DYG--------N
CFTFN-------PG-------------TK-D--------------------L----PL--
-------KNQTVP------GE--------VY-------------GLRLAFNIGQY-----
----------K-----------------Y------------Y-P-D--LLD---------
------------------------------------------------------------
------------------------------------------------------------
-------RNRPDAGIRFTIH-YHKEPP--NLLAKSIIAPVGSHTYVSFTRT------Y--
-----HKKL-----------EKP-W--GE-C--GSR--------K-------LL----FH
------------------E-FYSHVACIDEVSAIYAS----TL--CNCSIIG----A---
------------F-G---------------------------------------------
----------PY-----G---VCDS-----------------------------------
------------------------------------VKF---ITCIV---------P---
I-L-------------------AKAR-V-------QA--------NE----------NIA
------------ACP-VACETYT---YPTV-ISY------------------GNL-AL--
-------IPI------SSL-I-----------TN--------------------------
------------------------------------------------YL----------
------------------------------------------------------------
-N---------------V------------------SGIIRHAQSLDW------------
-------------------IK-DPNY-STA------------------------------
-----N-------FV-----------R------DNILYLDIYYSDLHTNAIE-QKRATGF
AEV-----------------LSNIGGQMGLFIGASIITIAEILQYLIRI-----------
--------FYNK--------S----------F-------NK-------------------
PA------------------------------DK-------------------S------
-----------------------------------------KEDLNLEQN----------
------------------------------------------------------------
------------------------------------------------------------
------------------------------------------------------------
------------------------------------------------------------
------------------------------------------------------------
--RKSSSV
>Cnidar_HydraVulSc4wPfr_824_g11302_t1_557_112
------------------------------------------------------------
----------M-------------------------------------------------
------------------------------------------------------------
------------------------------------------------------------
------------------------------------------------------------
------------------------------------------------------------
------------------------------------------------------------
------------------LN-------------FKDIAQIT-------------------
------------------------------------------------------------
---------------VEAIQETNEVTNKDEK--------------------------NIQ
T----------------------IQDLR--------------------------------
--------N----------------------K---------KI-----R-----------
-EH--ISYMI-DNSSFHGLSYIF---------D-KR--------------HS---VR---
--RTIWF----FITIAAFAYAMQKVYESTM-NYFS----------Y---PFYTVR----M
-R-MY-V----------N-QIDF-----PAI-----------------------SFCNLN

DI------KFS-----AMNGTI------------------V-----------------D-
---------DA----------------------------------VV-------------
------------------------------------------------------------
------------------------------------------------------------
------------------------------------------------------------
------------------------------------------------------------
------------------------------------------------------------
------------------------------------------------------------
------------------------------------------------------------
------------------------------------------------------------
------------------------------------------------------------
------------------------------------------------------------
------------------------------------------------------------
------------------------------------------TQNH--------------
---------------------------------EAN-------------------ITGE-
---------------------------------EYRSYN---------------------
------------------------------------------------QAA----RHTLN
--EML------VDCD----FE-G----K----------K---C-----------S-H---
----------------------------------------------K-------------
------------------------------------------------------------
------------------------------------------------------------
-----------N----------F-------T-----------EFS-W-MQGE-------S
CFTFN-------SG-------------KP-P--------------------H----TL--
-------LKVKGA------GI--------NR-------------SLKLTINVQHY-----
----------D-----------------Y------------Y-R-D--------------
------------------------------------------------------------
------------------------------------------------------------
---------KMDSGIRLILH-GQDETP---VKMSGLTVPPGFTTYIQIEKK------T--
-----IINL-----------EAP-YK-TK-C--GSV--------K-------LK----YF
------------------D-SYSMHTCWLEQLTDYVY----KT--CNCKDYF----M---
------------P-G---------------------------------------------
----------DI-----P---ICSF-----------------------------------
------------------------------------DLL---YNCAW---------P---
E-W-------------------ETFD-K-------QK---------------------LY
------------QCP-LPCKIDS---YEVS-LSR------------------ALF-PT--
-------GLY------ASS-L-----------AN--------------------------
------------------------------------------------DLRKYQ--Q---
-----------------------------------------------------------V
PI---------------A------------------LKS---------------------
------------------------KT-DEL------------------------------
-----I-------FM-----------R------ENLLRLVIYYDDLAYELVE-QKPSYNT
LLW-----------------LGDVGGQIGLFIGAGVMSYFEFIDCLAMV-----------
--------NYKVLHTANIIIF----------F-------TD-------------------
VV------------------------------AS--------------------------
------------------------------------------------------------
------------------------------------------------------------
------------------------------------------------------------
------------------------------------------------------------
------------------------------------------------------------
------------------------------------------------------------
---FVTMV
>Cnidar_Rhopilema_esculentum_TR103127_c0_g2_i1_p1__Rhopilema_esculentum_TR103
127_c0_g2__Rhopilema_esculentum_TR103127_c0_g2_i1_p1__ORF__type_complete__len
_487__score_56_61__Rhopilema_esculentum_TR103127_c0_g2_i1_686_2146__558_113
------------------------------------------------------------
----------M-------------------------------------------------
------------------------------------------------------------

```
------------------------------------------------------------
------------------------------------------------------------
-YSHSESVR---------------------------------------------------
------------------------------------------------------------
------EKEVVKSVDWIFDF-------------GDEFSNRD-------------------
------------------------------------------------------------
---------------IYRKVMTHMAD-AELL--------------------------DLR
K----------------------EKEVR--------------------------------
--------R----------------------Q---------KV-----K-----------
-QH--FDDLV-SNSTLHGLHFCF---------D-KQ--------------HF---FR---
--RIFWT----LLILVALGLLVQKLYESTT-YFFS----------Y---PFSTTT----T
-V-KY-V----------N-QMVF-----PAV-----------------------SICNLN
DF------RLS-----VMNGTK------------------L-----------------H-
---------KA----------------------------------IQ-------------
------------------------------------------------------------
------------------------------------------------------------
------------------------------------------------------------
------------------------------------------------------------
------------------------------------------------------------
------------------------------------------------------------
------------------------------------------------------------
------------------------------------------------------------
------------------------------------------------------------
------------------------------------------------------------
------------------------------------------------------------
------------------------------------------ERGNILA-----------
-------------------------------------------------------LDGK-
---------------------------------EYTDTI---------------------
------------------------------------------------RKA----NHRLE
--DML------QGCT----MN-N----K----------I---C-----------T-H---
----------------------------------------------K-------------
------------------------------------------------------------
------------------------------------------------------------
-----------N----------F-------T-----------QFF-H-NQGD-------R
CFTFN-------SG-------------MN-G--------------------Q----PL--
-------ITVNNT------GL--------SQ-------------ALTFLINIEHH-----
----------D-----------------Y------------Y-V-D--------------
------------------------------------------------------------
------------------------------------------------------------
---------SQHSGIHLILH-GQDETP---VKMQGVILSPGFVSYIEIKKR------K--
-----VKNL-----------PHP-YE-TN-C--GKL--------K-------LK----YF
------------------R-SYSKHLCWLEKLTDHVV----NL--CGCKDWF----M---
------------P-G---------------------------------------------
----------DH-----R---VCTL-----------------------------------
------------------------------------NES---MNCMW---------I---
H-W-------------------SQFD-L-------TQ---------------------NY
------------SCP-LPCIIDS---YTPT-LSF------------------ARF-PA--
-------NNL------ADK-M-----------AK--------------------------
------------------------------------------------RL----------
------------------------------------------------------------
-N---------------I------------------SGT---------------------
------------------------KA-EQR------------------------------
-----G-------WI-----------R------DNFLKVVIYYGDLNYEYME-QIPSYDL
MVL-----------------LGDIGGQLGLFLGSSVLTYIEFFDFFAMV-----------
--------IYTR--------F----------F-------EV-------------------
FK------------------------------RS-------------------P------
------------------------------------------------------------
```

------------------------------------------------------------
------------------------------------------------------------
------------------------------------------------------------
------------------------------------------------------------
------------------------------------------------------------
-----SRV
>Cnidar_Rhopilema_esculentum_TR52437_c0_g1_i1_p1__Rhopilema_esculentum_TR5243
7_c0_g1__Rhopilema_esculentum_TR52437_c0_g1_i1_p1__ORF__type_complete__len_48
7__score_65_71__Rhopilema_esculentum_TR52437_c0_g1_i1_68_1528__559_114
------------------------------------------------------------
----------M-------------------------------------------------
------------------------------------------------------------
------------------------------------------------------------
------------------------------------------------------------
------------------------------------------------------------
------------------------------------------------------------
------------------------------------------------------------
------------------------------------------------------------
------------------------------------------------------------
-----------------------EKGEES-------------------------------
--KDEKPPE----------------------N---------AV-----R-----------
-NM--LFDFA-SVTTAHGISRFV---------T-SG--------------PI---YA---
--RISWF----FIWLAVMIGFIYMIIQLVL-LYTS----------R---PVSTSI----S
-I-SF-E----------E-SLTF-----PGV-----------------------TICNLN
VF------PAS-----SMKNVR------------------V-----------------K-
-------FPNM----------------------------TIINNMFK-------------
---------------------------------KQFA-----------------------
------------------------------------------------------------
------------------------------------------------------------
------------------------------------------------------------
------------------------------------------------------------
------------------------------------------------------------
------------------------------------------------------------
------------------------------------------------------------
------------------------------------------------------------
------------------------------------------------------------
------------------------------------------------------------
------------------------------------------SQSS--------------
-----------------------------------------LDVGDLEM------DDVSS
------------------------------IQENTLTAI---------------------
------------------------NNL----------------PIADRLKN----GQEFS
--SLI------TYCR----WA-G----F----------V---C-----------N-K---
----------------------------------------------G-------------
------------------------------------------------------------
------------------------------------------------------------
-----------V----------F--LRDYWH-----------KTW-NWKYG--------N
CFTFN-------GK-------------MK-NINGSS---------------I----AP--
-------LESTSP------GT--------SG-------------GLTLNIDVNRD-----
----------E-----------------Y------------L-K-G---IA---------
------------------------------------------------------------
------------------------------------------------------------
----------METGVRVIIQ-DQDVFA--EPTEHGFSAASGYSVSVGLRKE------M--
-----IKRE-----------DVSGT--GS-C--YDT--------SK--------------
---------AN-------T-KYFQQTCIAICKAGTQK----RR--CNCTSLQ----Y---
------------R-T---------------------------------------------
--------YFGS-----K---QCES-----------------------------------
------------------------------------LAE---SSCLL---------E---

V-L-------------------SNFT-N-------KI--------F------------DC
----------SSRCP-PSCSDIA---FSKT-IAT------------------SKLYP---
-------EGY----------L---------------------------------------
------------------------------------------------EIHK--------
----------------------------------TK------------------------
-N----------------------------------------------------------
---------------------------LTA------------------------------
-----D-------YV-----------E------KNIINLQIHYADQTVTTIS-TGQYYTF
DNL-----------------VSDVGGQLGLWIGVSAVTCVEFFSLIWNL-----------
--------LLYC--------I----------C-------RK-------------------
KE------------------------------KR--------------------------
------------------------------------------------------------
------------------------------------------------------------
------------------------------------------------------------
------------------------------------------------------------
------------------------------------------------------------
------------------------------------------------------------
VQDENAPC
>Deutero_Cephalo_293250R_t1__name__chr_scaffold9_start_4819346_end_4829052_st
rand_pro_len_487__560_115
------------------------------------------------------------
----------M-------------------------------------------------
------------------------------------------------------------
------------------------------------------------------------
------------------------------------------------------------
------------------------------------------------------------
------------------------------------------------------------
----------------AACC-------------SC-------------------------
------------------------------------------------------------
------------------------------------------------------------
------------------------------------------------------------
-----------------------------------------QD-----G-----------
-AL--DHEYA-SNTSLHGPGNII---------N-AK--------------RP---AH---
--RAVWV----VLFLAAFGVAVWQISERFV-AYFS----------Y---NTVTSV----K
-V-EF-K----------D-ELDF-----PAV-----------------------TICNFN
KF------QLS-----KVTPSE------------------L-----------------N-
---------YI------------T-EVLDLSTGFE-G---DDTGELG----F--------
---------------------------------DY-------DDYEE-------------
------------------------------------------------------------
------------------------------------------------------------
-----------------EGF----------------------------------------
------------------------------------------------------------
------------------------------------------------------------
------------------------------------------------------------
-------------------------GPE--------------------------------
------------------------------------------------------------
------------------------------------------------------------
------------------------------------------------------------
------------------------------------------------------------
------------------------EDDYSLNVSAI--------------------PSNF-
---------------------------------DLADLT---------------------
------------------------------------------------LNA----GFVLD
-ETTL------QDCR----WR-G----K----------R---C-----------Y-A---
----------------------------------------------D-------------
------------------------------------------------------------
------------------------------------------------------------
-----------N----------F-------T-----------HAF-T-SYG--------N

```
CWTFN-------SN-------------DD---------------------------KI--
-------LRQTIP------GS--------GN-------------GLYLVIDVQQE-----
----------Q-----------------Y------------T-E-K---PP---------
------------------------------------------------------------
------------------------------------------------------------
-------SGNSDAGLKFLVH-PLAEPP--KIDSQGTAVQPGTHAYASIQNI------L--
-----YKNE-----------IPP-W--GT-C--DPT-W------R-------LD----NY
------------------D-TYTKTGCLLECRAEWVK----KD--CGCRTVS----M---
------------P-G---------------------------------------------
----------NA-----T---YCSP-----------------------------------
------------------------------------TQL---TQCVK---------T---
V-V-------------------GKLA-D-------GR--------------------YPC
------------ECP-TPCVANT---FPTT-VSY------------------AAW-PS--
-------ISA------QDY-Y-----------TN--------------------------
------------------------------------------------LF----------
------------------------------------------------------------
-N----------------------------------------------------------
---------------------------HTA------------------------------
-----E-------YL-----------Q------RNFVVMDLYYAQLNYQEVI-QTRQYTV
GSF-----------------LGDFGGQLGLFLGASVITIAEFIEYIVMK-----------
--------VTQP--------C----------S-------SR-------------------
GK------------------------------RR-------------------T------
-----------------------------------------DTEMTSL------------
------------------------------------------------GIDT--------
------------------------------------------------------------
------------------------------------------------------------
------------------------------------------------------------
-----------------------------------------------------------K
SNQSSPLS
>Deutero_Cephalo_091790R_t1__name__chr_scaffold19_start_3687569_end_3692166_s
trand_pro_len_488__562_116
------------------------------------------------------------
----------M-------------------------------------------------
------------------------------------------------------------
------------------------------------------------------------
------------------------------------------------------------
------------------------------------------------------------
------------------------------------------------------------
------------------------------------------------------------
------------------------------------------------------------
------------------------------------------G-----------------
------------------------------------------------------------
-------------------------------D---------EQ-----N-----------
-SL--DHEFA-NYTTCHGAARIA---------N-AK--------------NK---PH---
--MALWT----LIFLAAFGLCTWQIVDRFN-NYFA----------Y---RTGTEI----N
-V-QL-E----------S-ELTF-----PAV-----------------------TICDFN
RI------RGS-----GVTEND------------------L-----------------A-
---------YL------------G-ALY---------------QIAA----LVS------
------------------------------------------------------------
------------------------------------------------------------
------------------------------------------------------------
-----------------D------------------------------------------
------------------------------------------------------------
------------------------------------------------------------
------------------------------------------------------------
--------------------------PVA-------------------------------
------------------------------------------------------------
```

------------------------------------------------------------
------------------------------------------------------------
--------LRNL------------------------------------------------
------------------------SESIDWDAFNA--------------------SAGLV
E--------------------------------DFPGFV---------------------
------------------------------------------------RRN----GFVLD
-NVTL------AECS----WR-G----S----------R---C-----------Y-A---
----------------------------------------------E-------------
------------------------------------------------------------
------------------------------------------------------------
-----------N----------F-------T-----------HVF-T-EYG--------N
CWTFN-------SE--------------KN-S-------------------P--------
-------LRQTSP------GA--------GN-------------GLKLILDIQTH-----
----------E-----------------Y------------T-E-D---PF---------
------------------------------------------------------------
------------------------------------------------------------
-------IGNLEYGLVFQVH-DQYEPP--RPELVGTAVAPGSHTYASTFQT------H--
-----VKNQ-----------QAP-W--GQ-C--DPR--------KDQNF---LK----YF
------------------D-KYTLAGCLLECRAEIVV----RA--CNCRPLY----Y---
------------P-A---------------------------------------------
----------TF------------------------------------------------
------------------------------------------------------------
--L-------------------SETN-E-----------------F-----------DFC
------------GCT-VPCEYTS---YSSQ-VSY------------------AGY-PS--
-------LKA------AEF-Y-----------ET--------------------------
------------------------------------------------TF----------
------------------------------------------------------------
-N----------------------------------------------------------
---------------------------YSR------------------------------
-----G-------YM-----------S------TNVVLLDVYYHELRSTSWE-QFKAVEE
SGL-----------------ISDVGGQLGFFIGCSVITLWEFLEYLALK-----------
--------FSAF--------F----------L-------FS-------------------
SK------------------------------PS-------------------S------
-----------------------------------------KTATK--------------
--------------------DGVKVDNAMVFE-------------NTAAGDAEK------
------TEITELS-----------------------------------------------
------------------------------------------------------------
------------------------------------------------------------
---------------------------------------------------TIRTTSNGE
KSAVLEMT
>Deutero_Ambulac_Apla_gbr81_93_t1_564_118
------------------------------------------------------------
----------M-------------------------------------------------
------------------------------------------------------------
------------------------------------------------------------
------------------------------------------------------------
------------------------------------------------------------
------------------------------------------------------------
----------------ENTA-------------QKELNT---------------------
------------------------------------------------------------
------------------------------------------------------------
------------------------------------------------------------
------AKN----------------------K---------QK-----P-----------
-NL--EYEFA-TTTTLHGISRVA---------E-SN--------------GA---RA---
--KTLWV----LVLLIAGCFYVVVTVDRVR-AYLK----------F---DANTNV----I
-V-EF-N----------D-TLEF-----PAV-----------------------TICNYN
RF------MNN-----KISEDD------------------T-----------------K-

```
---------YV------------R-HLL---------------ELNE-------------
------------------------------------------------------------
------------------------------------------------------------
------------------------------------------------------------
-----------------D------------------------------------------
------------------------------------------------------------
------------------------------------------------------------
------------------------------------------------------------
---------------------L--------------------------------------
------------------------------------------------------------
------------------------------------------------------------
------------------------------------------------------------
------------------------------------------VDY---------------
------------------------EEDFNYTHIDDL------------------FGEGF-
---------------------------------NYTQFA---------------------
------------------------------------------------LTT----GFTLN
--ESV------IRCD----WK-GV--AY----------S---C-----------N-A---
----------------------------------------------L-------------
------------------------------------------------------------
------------------------------------------------------------
-----------N----------F-------S-----------SYY-SPSYG--------Q
CYTFN-------TR-------------GD-Q-------------------------HS--
-------HEQSQQ------GI--------GN-------------GLQLIVDIGQS-----
----------E-----------------Y------------T-E-T---FK---------
------------------------------------------------------------
------------------------------------------------------------
-------SGHLEAGLKFAIH-PRDELP--LIGTMGLSAAPGFHTYASMRRV------R--
-----HINL-----------PEP-W--GV-C--GTG-------ET-------IAV-TGDS
------------------R-KYSRNTCLRICRRDAII----EV--CGCQPFG----YERA
----------VEPDS--SE-----------------------------------------
----------LV-----P---LCDI-----------------------------------
------------------------------------DRF----YCVA---------H---
A-L-------------------ERYR-A-------SF--------SA----------LDC
------------TCP-VACEYIT---YETT-LSM------------------AKY-PS--
-------QSV------IND-I-----------LS--------------------------
------------------------------------------------GF----------
-----P------------------------------------------------------
-D-------------DEA------------------SKA---------------------
------------------------GA-VDA------------------------------
-----E-------YL-----------D------ENLVYLDVYFEELSTITYK-QVEALPF
TAL-----------------VGDMGGQLGLFLGASIITGAELLDYAIRR-----------
--------LKLL--------I----------L-------KV-------------------
KS------------------------------AK-------------------K------
-----------------------------------------IRSGL--------------
------------------------------------------------------------
------------------------------------------------------------
------------------------------------------------------------
------------------------------------------------------------
------------------------------------------------------------
-----MRS
>Protostome_Lophotroco_annelid_Pdum_comp404514_c0_seq1__567_119
------------------------------------------------------------
----------M-------------------------------------------------
------------------------------------------------------------
------------------------------------------------------------
------------------------------------------------------------
------------------------------------------------------------
```

------------------------------------------------------------
------------------------------------------------------------
------------------------------------------------------------
------------------------------------------------------------
------------------------------------------------------------
---DTKKEE----------------------A---------SA-----F-----------
-AL--IAAMA-NNTGMHGLPNVG---------R-SQ--------------HI---VR---
--KVFWM----LIFLAGAGMLIWQVYEALA-QYYS----------Y---PTKTSI----I
-P-KV-Y----------N-KLNL-----PAV-----------------------TICNVN
PV------RKS-----LLYQLD--------------------------------------
---------AL-------------------------------------------------
------------------------------------------------------------
------------------------------------------------------------
------------------------------------------------------------
------------------------------------------------------------
------------------------------------------------------------
------------------------------------------------------------
------------------------------------------------------------
------------------------TDPS--------------------------------
------------------------------------------------------------
------------------------------------------------------------
------------------------------------------------------------
------------------------------------------------------------
-------------------------------------------------------DRNYE
---------------------------------NFQTAM---------------------
------------------------NAL----------------NVTEKQKI----GHQLK
--DMM------LYCN----YN-G----Y----------T---C-----------E-V---
----------------------------------------------G-------------
------------------------------------------------------------
------------------------------------------------------------
-----------N----------F-------T-----------TFY-NSYLG--------N
CFTFN-------GG-------------NR----------------------P----TV--
-------DGLGKT------GP--------YY-------------GLDIEFYIEQD-----
----------E-----------------Y------------E-P-E---YS---------
------------------------------------------------------------
------------------------------------------------------------
----------EYAGMKVVIH-NPKEMS--FPEDQGLFVSPGAKSQVSLKKI------E--
-----NTRA-----------DPP-H--GE-C--RTF--------NEEE----TKA-RNAY
----KDS--YD------YL-EYTSKACGKTCYQQYVM----SK--CGCYDTS----Y---
------------P-S--TG-------------------SAFEN-----------------
-----ITSGKTF-----S---SCKSL----------N-----------------------
------------------------------------STY---TDCNT---------A---
V-Y-------------------YNYT-D-------NI--------I------------SC
----------ANMCP-PACTEVL---YDMT-TSN------------------LAW-PS--
-------VNA------LSE-I-----------VK--------------------------
------------------------------------------------KL-E--------
----------------------------------DK------------------------
-S-------------TAL------------------NTTIQN------------------
----------------------MTEI-EKE------------------------------
-----S-------FV-----------R------KNVVSISVFYGSLEVTKIK-TEPSFGL
VDL-----------------LFAIGGSVGLWLGLSVISVIEIVELLYDI-----------
--------SSIL--------V----------C----CCFKT-------------------
--------------------------------KP--------------------------
-------------------------------------NGNNPKTIQVD------------
------------------------------------------------------------
------------------------------------------------------------
------------------------------------------------------------

```
---------------------------------P--------------------------
------------------------------------------------------------
------QS
>Deutero_Ambulac_Apla_gbr54_19_t1_572_121
------------------------------------------------------------
----------M-------------------------------------------------
------------------------------------------------------------
------------------------------------------------------------
------------------------------------------------------------
------------------------------------------------------------
------------------------------------------------------------
----------------KDSS-------------SQ-------------------------
------------------------------------------------------------
------------------------------------------------------------
------------------------------------------------------------
-----------------------------------------KQ-----S-----------
-DL--DQEFA-STTTLHGIARIF---------D-ST--------------RL---PV---
--RLLWL----SILLGCLGVCVWQITDRFQ-RYLL----------H---EATTAV----S
-V-EY-V----------G-DLDF-----PAV-----------------------TICNFN
RY------RSS-----ALTEDD------------------K-----------------A-
---------HL------------E-DLI---------------EYAD----Y--------
---------------------------------DY-------DTYD--------------
------------------------------------------------------------
------------------------------------------------------------
-----------------DEE----------------------------------------
------------------------------------------------------------
------------------------------------------------------------
------------------------------------------------------------
--------------------------P---------------------------------
------------------------------------------------------------
------------------------------------------------------------
------------------------------------------------------------
----DARRTAQS------------------------------QGE---------------
-------------------------------------------------------TTNF-
---------------------------------SFSEVT---------------------
------------------------------------------------LRT----GFQMD
-EETL------LDCK----WR-G----K--------RNS---C-----------N-G---
----------------------------------------------Q-------------
------------------------------------------------------------
------------------------------------------------------------
-----------N----------F-------T-----------HVF-T-SFG--------N
CWTFN-------SG-------------ET-Q--------------------DEKEISV--
-------LNQIQP------GS--------GN-------------GLTMVINIQQA-----
----------E-----------------Y------------T-E-P---VQ---------
------------------------------------------------------------
------------------------------------------------------------
-------NGNLEAGLKVLVH-DQETPP--SVDSEGFAIAPGVHAFVGLRKI------E--
-----YENL-----------EYP-W--GE-C--DKS-R------R-------LL----HY
------------------D-KYTLPGCDIECRAERIY----ER--CQCKLVR----H---
------------P-G---------------------------------------------
----------DE-----T---ECSP-----------------------------------
------------------------------------LQV---KKCAT---------P---
V-L-------------------AKLK-T-------GE--------V-----------EGC
------------GCP-VPCNYSE---FRTS-LSM------------------ATL-PS--
-------NNL------LVD-L-----------WK--------------------------
------------------------------------------------LY----------
------------------------------------------------------------
```

```
-GG-----------DDSV------------------NYT-----------F---------
--------------D------------ETL------------------------------
-----R-------YI-----------R------DNWIFLDVYYESLNFEKYV-QSEAITL
SAL-----------------ISDIGGQMGLFLGASFITITEILHYLGRK-----------
--------AGLW--------V----------T-------SR-------------------
SR------------------------------PH--------------------------
-----------------------------------------ANQSK--------------
------------------------VRP---------------------GNET--------
------------------------------------------------------------
--------------VAISD-----------------------------------------
------------------------------------------------------------
------------------------------------------------------------
---LSSPH
>Deutero_Ambulac_Sakowv30024946m_573_122
------------------------------------------------------------
----------M-------------------------------------------------
------------------------------------------------------------
------------------------------------------------------------
------------------------------------------------------------
--DSMNDV----------------------------------------------------
------------------------------------------------------------
------------------------------------------------------------
------------------------------------------------------------
------------------------------------------------------------
-----------------------KDAD---------------------------------
-------RK--------------------------------SQ-----S-----------
-SI--VHNFM-SSTTAHGLPRAF---------E-GR--------------SL---WL---
--SLFWV----LLFCSAIAVSIWQIGLLVK-HFIA----------R---DVSVKT----E
-I-VT-A----------P-QMPF-----PSV-----------------------SICNTN
KL------RKS-----AVSESA------------------Y-----------------S-
---------EL----------------------------------LT-------------
------------------------------V--DT-------GIAEPYF-----------
------------------------------------------------------------
----------GDCL----------------EGDFQCTN-----------GG---------
-CVKTYLKCDGIDNCG-D-K------------------------SDESGCV-----YGDC
D--------AGKIKCVSGSPTGACIDEK-FKCDREFNCYEGEDENDCVCK----------
---RKEFKCG------DTGRCIDGT----------LQCDESIDCNDGSD-------EE--
--EC---GAECPSDHLECNG---------------------------LCIPP--------
----EWLCDNLQDC------------P--------DGTDE------C-----GDPEIQIC
SPFEFAC----------------DLYQCI-------PIYWECDGVPDCINGKD-------
----------EDNCPLQAAPQSFGFVTCSEDEYQCDLFTCIPLNQKCNGIVDCALEGDED
DC---------------PEMTGWMTTQG------------------SNKTA-TYQPVAKD
LLKIYTALSYDITLYEDF----VDNH--------------Y-HDHQ-------F------
--------GRV-----------KSEDPPDWPGFITY-----------SS------TPDYS
---------------------------------DLEN-V---------------------
------------------------LKL----------------RADEIAEL----GHQLE
--DFV------LECS----YD-E----R----------I---C-----------D-L---
------------------------------------------------------------
------------------------------------------------------------
------------------------------------------------------------
-----------N----------M-------L-----------SF---------------T
SLYFN-------FS----------------------------------------------
------------------------------------------------------------
------------------------------------------------------------
------------------------------------------------------------
------------------------------------------------------------
------------------------------------------------------------
```

------------------------------------------------------------
------------------------------------------------------------
------------------------------------------------------------
------------------------------------------------------------
------------------------------------------------------------
------------------------------------------------------------
------------------------------------------------------------
------------------------------------------------------------
------------------------------------------------------------
------------------------------------------------------------
------------------------------------------------------------
------------------------------------------------------------
------------------------------------------------------------
------------------------------------------------------------
------------------------------------------------------------
------------------------------------------------------------
------------------------------------------------------------
------------------------------------------------------------
------------------------------------------------------------
------------------------------------------------------------
------------------------------------------------------------
------------------------------------------------------------
--------
>Placozoa_Hoilungia_hongkongensis__1_g02560_t1_574_123
------------------------------------------------------------
----------M-------------------------------------------------
------------------------------------------------------------
------------------------------------------------------------
------------------------------------------------------------
------------------------------------------------------------
------------------------------------------------------------
------------------------------------------------------------
------------------------------------------------------------
------------------------------------------------------------
------------------------------------------------------------
----------------------------S--Q---------EK-----F-----------
-NH--EERFA-TSTSYHGMAHIY---------D-GQ-------------TSK---QT---
--RSLWL----VLTLIATGACISQCVIIIL-NASQ----------L---PTRIVT----K
-T-VL-Q----------N-SSIF-----PSV-----------------------TICNTN
DY------DRN-----SLSSSD------------------I-----------------Q-
---------HL------------S-TIV--------------DFVYG-------------
------------------------------------------------------------
------------------------------------------------------------
-----------------Y-DI---------------------------------------
-----------------E------------------------------------------
------------------------------------------------------------
------------------------------------------------------------
------------------------------------------------------------
------------------------------------------------------------
------------------------------------------------------------
------------------------------------------------------------
------------------------------------------------------------
--------PHRF------------------------------EAA---------------
------------------------VNSLSI------------------KF-----GKGF-
---------------------------------NYELFL---------------------
------------------------------------------------RRA----GHKLE
--NML------LSCT----WM-G----R----------P---C-----------S-I---

----------------------------------------------E-------------
------------------------------------------------------------
------------------------------------------------------------
-----------D----------F-------V-----------NTT-T-AAG--------S
CFTFN-------PG-------------ST-N--------------------T----SI--
-------KNQTAS------GN--------AN-------------GLRLILNIEQY-----
----------K-----------------Y------------Y-P-A--LYT---------
------------------------------------------------------------
------------------------------------------------------------
-------PGQPDAGLRYSLH-YYKDPP--NLIAESYYAAPGFHTYVPMTLK------R--
-----EKRL-----------KKP-W--GD-C--EEL--------S-------LK----RY
------------------Q-FYSRYACLNELSAKHAS----ES--CNCSYQD----N---
------------E-A---------------------------------------------
----------------LN---PCDG-----------------------------------
------------------------------------VQF---LSCII---------P---
T-A-------------------GYYR-T-------QL--------SQ----------NLS
------------VCP-IACETFS---FATE-ISQ------------------SSI-AS--
-------QAF------NSA-I-----------DS--------------------------
------------------------------------------------FI----------
------------------------------------------------------------
-N---------------V------------------SGLISDAQSKYW------------
-------------------LP--QSY-TNI------------------------------
-----D-------FI-----------R------DNIVYLDIYYRKLDMTQIL-QLEDTGF
SKV-----------------LSEIGGQMGFFIGASVLTVCEIVQYLFEK-----------
--------CFTS--------N----------L-------KS-------------------
GT------------------------------KK-------------------N------
-----------------------------------------TTNVVASDN----------
-----------------------HIFA--------------------LERTT--------
------------------------------------------------------------
------------------------------------------------------------
------------------------------------------------------------
------------------------------------------------------------
-GFTPRDD
>Placozoa_TadNaC6_MK547547_16_124
---------------------------M--------------------------------
----------M-------------------------------------------------
------------------------------------------------------------
------------------------------------------------------------
------------------------------------------------------------
------------------------------------------------------------
------------------------------------------------------------
------------------------------------------------------------
------------------------------------------------------------
------------------------------------------------------------
------------------------------------------------------------
----------------------------A--K---------GK-----I-----------
-DH--DERFA-TTTSYHGVAHIY---------D-SN-------------NSK---KT---
--KSIWI----ILVILATAICISQCVIIIY-NASL----------L---PTRMVI----R
-K-RL-M----------N-SSVF-----PSV-----------------------TICNTN
DF------DYT-----GLPAND------------------L-----------------N-
---------HL------------S-SIV--------------NAIYG-------------
------------------------------------------------------------
------------------------------------------------------------
-----------------F-T----------------------------------------
------------------------------------------------------------
------------------------------------------------------------
------------------------------------------------------------

------------------------------------------------------------
--------------------------P---------------------------------
------------------------------------------------------------
------------------------------------------------------------
------------------------------------------------------------
--------ASAA------------------------------DDA---------------
------------------------IQYFVR------------------KH-----GDDF-
---------------------------------QIDNYT---------------------
------------------------------------------------RMA----GHKLK
--NML------LSCT----WM-G----E----------P---C-----------T-V---
----------------------------------------------N-------------
------------------------------------------------------------
------------------------------------------------------------
-----------D----------F-------T-----------NII-S-NGG--------S
CFTFN-------PG-------------TN-A--------------------I----PL--
-------KNQTVS------GN--------IN-------------GLRLILNVEQY-----
----------K-----------------Y------------Y-S-P--LFS---------
------------------------------------------------------------
------------------------------------------------------------
-------PQAPDAGIRFTIN-YYKQPP--NFISKPYYAPTGFHTYVPITLH------R--
-----DKRL-----------TKP-W--GE-C--GEL--------L-------LK----DH
------------------S-YYSRDACLTEYASELAA----LT--CNCSSSG----Q---
------------T-A---------------------------------------------
----------------TN---SCNG-----------------------------------
------------------------------------AKF---LTCII---------P---
S-S-------------------FVIR-M-------SL--------SQ----------NLS
------------VCP-IACETYS---YPTE-ISQ------------------SSL-GT--
-------FAF------SRV-L-----------DS--------------------------
------------------------------------------------VI----------
------------------------------------------------------------
-N---------------I------------------STILGKAKDEHW------------
-------------------IPPSLPY-SVS------------------------------
-----E-------FI-----------R------DNIVYLDIYYSDLRVTETE-QQEDTGF
SKV-----------------LSEIGGQLGLCIGASVITLCEIIQYLIGK-----------
--------FFTS--------E----------K-------KT-------------------
NL------------------------------NK-------------------R------
-----------------------------------------QNTTSPLFY----------
-----------------------NRNS--------------------PDADA--------
------------------------------------------------------------
------------------------------------------------------------
------------------------------------------------------------
------------------------------------------------------------
---TNPDM
>Placozoa_TadNaC7_MK547548_17_125
------------------------------------------------------------
----------M-------------------------------------------------
------------------------------------------------------------
------------------------------------------------------------
------------------------------------------------------------
------------------------------------------------------------
------------------------------------------------------------
------------------------------------------------------------
------------------------------------------------------------
------------------------------------------------------------
------------------------------------------------------------
----------------------------D--E---------EK-----I-----------
-SE--EERFA-TTTSYHGWAHIY---------D-GN-------------NGR---MT---

--KIIWM----ILVILATAACISQCIIIIY-NASK----------L---PTRMVI----R
-K-QL-M----------N-TSIF-----PSV-----------------------TLCNTN
DY------DRS-----ALPAAD------------------L-----------------Y-
---------HQ------------S-ALV--------------KAIYG-------------
------------------------------------------------------------
------------------------------------------------------------
-----------------S-G----------------------------------------
-----------------D------------------------------------------
------------------------------------------------------------
------------------------------------------------------------
------------------------------------------------------------
------------------------------------------------------------
------------------------------------------------------------
------------------------------------------------------------
------------------------------------------------------------
--------PNKT------------------------------NAA---------------
------------------------INYFDK------------------KF-----GGNF-
---------------------------------QFENLT---------------------
------------------------------------------------RVA----GHKLS
--NML------LSCT----WM-E----Q----------P---C-----------Y-A---
----------------------------------------------T-------------
------------------------------------------------------------
------------------------------------------------------------
-----------D----------F-------V-----------NII-T-DGG--------S
CFTFN-------PG-------------TG-N--------------------L----SL--
-------KNETVS------GN--------SN-------------GLRLILNVEQY-----
----------K-----------------Y------------Y-S-G--IFA---------
------------------------------------------------------------
------------------------------------------------------------
-------LEQPDAGIRFTTH-YYKQTP--NFISKSYYAPTGFHTYVPITLQ------H--
-----DKRL-----------KKP-W--GQ-C--GEE--------I-------LV----YN
------------------S-FYSRDACLSEYAGRLAS----YI--CNCTFEP----Q---
------------I-A---------------------------------------------
----------------SN---PCNG-----------------------------------
------------------------------------TQF---LTCIT---------P---
L-A-------------------SKFR-Q-------DL--------SQ----------NLS
------------ICP-IACETYS---YPTE-ISQ------------------SAL-AA--
-------LAF------ASS-L-----------DP--------------------------
------------------------------------------------LI----------
------------------------------------------------------------
-N---------------V------------------SGIIANAKANNW------------
-------------------IAPNQSY-FAA------------------------------
-----D-------FI-----------R------DNIVYLDIYYSDLHLTQTL-QEEDTGF
SKI-----------------ISELGGQLGICIGASALTICEIVQYLIKK-----------
--------YFAS--------N----------K-------KS-------------------
KS------------------------------DK-------------------S------
-----------------------------------------STNTISLGH----------
-----------------------DKKP--------------------FETTT--------
------------------------------------------------------------
------------------------------------------------------------
------------------------------------------------------------
------------------------------------------------------------
--VTNIDL
>Deutero_Ambulac_Apla_gbr256_11_t1_576_126
------------------------------------------------------------
----------M-------------------------------------------------
------------------------------------------------------------

```
------------------------------------------------------------
-----------------------------------------------------------D
VWTVEKSLNRI-------------------------------------------------
------------------------------------------------------------
----------------DEST-------------SADSIKMKGWRKDEGGV----------
---------------------------------------------------NGD------
------------------------------------------G--------------RKR
SVH--------------------TQTESD-------------------------------
-RRNADNCR----------------------Q---------IF----NR-----------
-RK--WSEFG-TVTTLHGMRYIT---------Q-PN-------------HSF---TR---
--KVIWA----IILLSFMGVLIFVVIDNTV-RYFN----------Y---NVTSIV----T
-V-KY-V----------N-DLTF-----PAV-----------------------TICNYN
LL------RKS-----YSEQAF-----------------------------------DE-
---------DL-------------------------------LEAFK-------------
------------------------------------------------------------
------------------------------------------------------------
------------------------------------------------------------
------------------------------------------------------------
------------------------------------------------------------
------------------------------------------------------------
------------------------------------------------------------
-----------------------IISPGGG------------------------------
------------------------------------------------------------
------------------------------------------------------------
------------------------------------------------------------
------------------------------------------------------------
------------------------QNSVNFTA-----------------------LKDW-
---------------------------------NITEDL---------------------
------------------------------------------------IKG----AHQKE
--DMI------IECT----WR-S---TR----------N---C-----------T-A---
----------------------------------------------D-------------
------------------------------------------------------------
------------------------------------------------------------
-----------N----------F-------T-----------QVL-T-DFG--------V
CYTFN-------NP-------------KD-P--------------------N----DA--
-------LTVQQT------GY--------DS-------------GLYLRLNLEQY-----
----------D-----------------Y------------F-Y-G---SS---------
------------------------------------------------------------
------------------------------------------------------------
----------KGAGFKIMLH-KQGEFP--FVKQLGFAVSPGFETLVTLKFT------Q--
-----LVNL-----------ETP-YV-SE-C--VVA--------ED------IQ----GF
--------------------KYTVETCQALCKTKYIL----EG--CGCRTTD----L---
------------P-AG--------------------------------------------
-----------------P------------------------------------------
------------------------------------------------------------
----------------------GAFM-A-------DL-------------------DARC
------------NCS-VPCFREV---FTPR-ISE------------------VFW-PA--
-------EHI------TKA-L-----------QD--------------------------
------------------------------------------------AF----------
------------------------------------------------------------
-N----------------------------------------------------------
---------------------------SSE------------------------------
-----S-------FL-----------R------KNIIDMVIFFEELNYEEIR-QIAAYTF
TDL-----------------LSDIGGYMGLMVGASLITFLEFIDFLVLG-----------
--------AIEY--------C----------Q-------KH-------------------
NMFK----------------------------RR------------------GK------
------------------------------------------------------------
```

------------------------------------------------------------
------------------------------------------------------------
------------------------------------------------------------
------------------------------------------------------------
----------------------------------------------------------IN
ISQSHLHR
>Deutero_Ambulac_Apla_gbr54_18_t1_578_128
------------------------------------------------------------
----------M-------------------------------------------------
------------------------------------------------------------
------------------------------------------------------------
------------------------------------------------------------
------------------------------------------------------------
------------------------------------------------------------
----------------TGSA-------------SG-------------------------
------------------------------------------------------------
------------------------------------------------------------
------------------------------------------------------------
-----------------------------------------KT-----T-----------
-DL--DREFA-ANTSLHGVARFF---------D-ST--------------RL---PV---
--RLLWL----SILLGCLGVCVWQITDRFQ-RYLL----------H---EATTAV----S
-V-EY-V----------G-DLDF-----PAV-----------------------TICNFN
RY------RKS-----ALTEAD------------------V-----------------A-
---------KL------------R-AYQ---------------RYID----Y--------
---------------------------------DY-------DFYDY-------------
------------------------------------------------------------
------------------------------------------------------------
-----------------D------------------------------------------
------------------------------------------------------------
------------------------------------------------------------
------------------------------------------------------------
------------------------------------------------------------
------------------------------------------------------------
------------------------------------------------------------
------------------------------------------------------------
----SSVGGTPP------------------------------PDD---------------
-------------------------------------------------------ISDF-
---------------------------------RFTNFT---------------------
------------------------------------------------LRT----GFQMD
-EDTL------LGCL----WQ-S----N--------RNS---C-----------T-A---
----------------------------------------------K-------------
------------------------------------------------------------
------------------------------------------------------------
-----------N----------F-------T-----------HVF-T-PFG--------N
CWTFN-------SG-------------ED-G--------------------E-KDVPI--
-------LKQTQP------GS--------DN-------------GLVMTIDILQR-----
----------E-----------------Y------------T-Q-H---LQ---------
------------------------------------------------------------
------------------------------------------------------------
-------SGYVEAGLKILVH-DQKTPP--PIDSEGSAIAPGVHAFVGVRKI------E--
-----YSNL-----------EPP-W--GK-C--DKS-R------R-------LT----YY
------------------D-KYTLSGCVIECRARKID----EE--CKCRLFS----H---
------------P-G---------------------------------------------
----------NA-----V---ECTP-----------------------------------
------------------------------------TQV---KDCAI---------P---
V-I-------------------VKLR-S-------GE--------R-----------AGC
------------GCP-LPCNYSE---FRTS-LSM------------------ATL-PS--

```
-------NNL------RDE-I-----------WE--------------------------
------------------------------------------------EL----------
------------------------------------------------------------
-R------------QEGI------------------NDT---------------------
---------------------------LPE------------------------------
-----T-------YI-----------R------DNWILLDVYYETLNYEKYV-QSEAITP
SAL-----------------ISDIGGQLGLFLGASFITVTEVLHYLGCK-----------
--------AGRW--------M----------T-------TTCIKV-----------DGKP
TH------------------------------ES--------------------------
-----------------------------------------AKEVV--------------
------------------------MEE---------------------GQQT--------
------------------------------------------------------------
--------------SGSKD-----------------------------------------
------------------------------------------------------------
------------------------------------------------------------
----STLT
>Deutero_Ambulac_Spurpu_015892_583_129
------------------------------------------------------------
----------M-------------------------------------------------
------------------------------------------------------------
------------------------------------------------------------
------------------------------------------------------------
------------------------------------------------------------
------------------------------------------------------------
------------------------------------------------------------
------------------------------------------------------------
------------------------------------------------------------
---------------------GNIEKEKL-------------------------------
----NELDD----------------------D---------TV-----V-----------
-GA--VRVFS-NETTLHGIRYLI---------G-RG--------------HL---LI---
--RLCWL----AIVVLAFALLVMQAQIVYQ-DFRS----------S---PYSTKI----D
-I-VS-E----------V-VKTF-----PAV-----------------------TVCNDN
KI------RRS-----QLYNTK------------------Y-----------------E-
---------GLIDI----------------------------------------------
---------------------------------DD-------------------------
------------------------------------------------------GG----
------------------------------------------------------------
-------RSTEGRSSTRDKS----------------------------------------
------------------------------------------------------------
------------------------------------------------------------
------------------------------------------------------------
------------------------------------------------------------
------------------------------------------------------------
------------------------------------------------------------
------------------------------------------------------------
------------------------------------------SDNV-------W------
--------ASV-------------KDDYDWQGFYSA-----------SS------ADDFS
---------------------------------DFINVV---------------------
-------------------------NP----------------TKAELNDY----GHTLE
--DFV------IQCT----YN-Q----K----------S---C-----------N-LS--
----------------------------------------------A-------------
------------------------------------------------------------
------------------------------------------------------------
-----------D----------F-------T-----------VSQ-NRHYG--------N
CFTFN-------FG-------------DG-D-------------------DE----PP--
-------RVTSKT------GA--------LS-------------GLHLSLMVDEE-----
----------E-----------------Y------------M-G-L---LA---------
```

------------------------------------------------------------
------------------------------------------------------------
----------PNRGAMVVIH-APNTVP--FPEDDGINLETGSAISIGIREE------V--
-----IQRL-----------PK--F--SQ-C--VVD--------RKIG--------TNTS
----ISG-----------T-NYTVRCCLKICYQQSLL----ER--CSCVDDI----L---
------------L-----------------------------------------------
----------DY-----R---QCSVT----------N-----------------------
------------------------------------KSE----------GDHGG--HGSP
V-Y-------------------HLSI-D----------------LLY----------LEC
------------NVP-PAYRGLK---FKKG------------------------------
-------KRD----------V---------------------------------------
------------------------------------------------RV----------
------------------------------------------------------------
-G----------------------------------------------------------
------------------------------------------------------------
-----K--------------------R-------VLNHSMVYFRGKTYE--------FGA
NSL-----------------VLHEGGVRKTTAGRSLCTEYQATELAVRYHQDHN------
--------------------F------------------NLFS-----------------
--------------------------------KN---------CHMFAEWLINRLLN---
------------------------------------------------------------
------------------------------------------------------------
------------------------------------------------------------
------------------------------------------------------------
------------------------------------------------------------
------------------------------------------------------------
---NKCTI
>Protostome_Lophotroco_annelid_Pdum_Contig6895__580_131
------------------------------------------------------------
----------M-------------------------------------------------
------------------------------------------------------------
------------------------------------------------------------
------------------------------------------------------------
------------------------------------------------------------
------------------------------------TRVGEPS-----------------
------------------------------------------------------------
------------------------------------------------------------
------------------------------------------------------------
-----------------------RDPPVVIASQGGAN----------------------K
PTATKNPEE----------------------E---------TL-----R-----------
-KI--LDDYV-EHSSMHGIWKAA---------K----------------TSP---FS---
--RAVWT----VVLLCMICLCLFTFVSQLL-FFLQ----------F---KTNTLV----T
-V-IE-Q----------P-FLPF-----PAV-----------------------TVCNYN
SF------RKS-----KALEKP--------S---------I-----------------T-
---------KV-----------------------------LEQLFS--L----------
---------------------------------DR-------------------------
------------------------------------------------------------
------------------------------------------------------------
------------------------------------------------------------
------------------------------------------------------------
------------------------------------------------------------
------------------------------------------------------------
------------------------------------------------------------
------------------------------------------------------------
------------------------------------------------------------
------------------------------------------------------------
------------------------------------------------------------
------------------------IFHINVYTTSTAEQ-----------------SDDV-

---------------------------------DILKEL---------------------
------------------------------------------------IDL----AHTLE
--GMV------VYCF----WC-G----M--------RVD---C-----------K-----
----------------------------------------------Q-------------
------------------------------------------------------------
------------------------------------------------------------
-----------H----------F-------V-----------QTM-T-DVG--------V
CYTFN-------SK-------------NTTG--------------------D----TP--
-------MLVRDA------GS--------DC-------------GLKLILNIQRH-----
----------E-----------------Y------------F-Y-G---QG---------
------------------------------------------------------------
------------------------------------------------------------
----------DSTGVKVLVH-DPKEVP--LVGQQGFSAAPGTTVIASIKLQ------Q--
-----ELYA-----------SSP-Y--GD-C--VDV--------SKRDFKNPLQ----YH
------------------D-EYTYTGCKMEHRVNYIY----SS--CHCVQYF----H---
------------PVG---------------------------------------------
----------GR-----E---ICTT-----------------------------------
------------------------------------FDQ---VECSR---------K---
A-Q-------------------ENHT-M-------YQ--------LE---------QLSQ
------------DCP-VTCEYNT---FDEK-LSW------------------ASF-PA--
-------QHI------LDD-L-----------QT--------------------------
------------------------------------------------TF----------
------------------------------------------------------------
-N----------------------------------------------------------
---------------------------MSA------------------------------
-----Q-------AI-----------S------DNLVQLHIYYENLAVHRQE-QTPAYTV
GAL-----------------FADTGGEMGLCIGASLLTVVELFELCGHL-----------
--------AAFS--------W----------K-------KI-------------------
TRHMFY--------------------------KK--------------------------
------------------------------------------------------------
------------------------------------------------------------
------------------------------------------------------------
------------------------------------------------------------
---------------------------------P--------------------------
------------------------------------------------------------
RSSLSKPI
>Deutero_Ambulac_Apla_gbr256_13_t1_582_133
------------------------------------------------------------
----------M----------------------------------------------Y--
------------------------------------------------------------
------------------------------------------------------------
--------------------------------------------------------AWTE
GTKTPNLLGDQESI----------------------------------------------
------------------------------------FSIGSP------------------
---DFTVRSTIINMEDKETK-------------GNNCR----------------------
------------------------------------------------------------
---------------------------------------------------------SQT
C----------------------SADVD--------------------------------
--------R----------------------R---------SF-----A-----------
-DR--FREFA-NSTTFHGISNVT---------N-SK-------------HNG---IR---
--RLFWS----LIVIGVSSWLVTGITQTVI-EFFK----------Y---PVTSAI----S
-I-NY-V----------D-SMTF-----PAV-----------------------TICNFN
QF------RRS-----VIPDDQ------------------V-----------------D-
---------FI-------------------------------NKLYG-------------
------------------------------------------------------------
------------------------------------------------------------
------------------------------------------------------------

------------------------------------------------------------
------------------------------------------------------------
------------------------------------------------------------
------------------------------------------------------------
------------------------LDPEA-------------------------------
------------------------------------------------------------
------------------------------------------------------------
------------------------------------------------------------
------------------------------------------------------------
------------------------PTDLNISDFEITG------------------LTRD-
---------------------------------QYEDLF---------------------
------------------------------------------------AAM----SHQLE
--TML------SECV----WR-S---SE----------T---C-----------T-V---
----------------------------------------------Q-------------
------------------------------------------------------------
------------------------------------------------------------
-----------N----------F-------T-----------RRF-T-DHG--------I
CFTFN-------DP-------------KN-G--------------------S----EI--
-------LQVNSA------GT--------RN-------------GLFLRLYADTE-----
----------E-----------------Y------------M-F-S---EN---------
------------------------------------------------------------
------------------------------------------------------------
----------TAAGFRMLLH-PQGVAP--LVKELGFSVSPGFESSISVRQT------L--
-----VESL-----------PQP-YE-SN-C--TNS--------T-------LR----YS
------------------D-TYTVPSCRFECKVEYVV----DK--CGCRDYR----W---
------------P-G---------------------------------------------
----------PA-----P---ICNP-----------------------------------
------------------------------------QEQ---FKCVY---------P---
Q-E-------------------AEFL-R-------QN--------V------------EC
------------SCP-VSCETKT---YESR-ISL------------------GYW-PA--
-------GHV------TEF-V-----------EA--------------------------
------------------------------------------------HM----------
------------------------------------------------------------
-N----------------------------------------------------------
---------------------------LTE------------------------------
-----E-------YV-----------R------KNYLDVYIFFEELMYLKIE-QKEAYST
SSL-----------------QSNIGGYMGLLCGMSLITVLEWADFTLVS-----------
--------IIRR--------F----------R-------SR-------------------
KE----------------------------------------------------------
------------------------------------------------------------
------------------------------------------------------------
------------------------------------------------------------
------------------------------------------------------------
------------------------------------------------------------
------------------------------------------------------------
--------
>Deutero_Ambulac_Spurpu_006217_591_135
------------------------------------------------------------
----------M-------------------------------------------------
------------------------------------------------------------
------------------------------------------------------------
------------------------------------------------------------
------------------------------------------------------------
------------------------------------------------------------
------------------------------------------------------------
------------------------------------------------------------
------------------------------------------------------------

-----------------------AYDERMNATSVRHTTT--------------------N
STMDGRPPK----------------------S---------EV-----Y-----------
-AM--VKPML-ENTAAHGIPNIV---------R-AE--------------ST---PR---
--RVAWS----FLFVVALCCFIGLSGNLIR-KYYS----------F---DFNVNV----E
-V-LF-E----------P-SINF-----PAV-----------------------TLCNMN
PF------RQS-----SVVNAS------------------L-------------------
---------EL-------------------------------TEILG----L--------
---------------------------------EN-------------------------
------------------------------------------------------------
------------------------------------------------------------
-----------------D------------------------------------------
------------------------------------------------------------
------------------------------------------------------------
------------------------------------------------------------
--------------------------PMSKW-----------------------------
------------------------------------------------------------
------------------------------------------------------------
------------------------------------------------------------
------------------------------------------NSSTN-------------
------------------------ALYGDWENL-----------EFSSL------DAQTE
------------------------------ILISAIQVV---------------------
------------------------GNM----------------SYEQRFDI----GHDLG
--DML------LSCS----FH-G----L----------P---C-----------A-PAME
TG------------------------------------------RTS-------------
------------------------------------------------------------
------------------------------------------------------------
-----------N----------FHRMLKLRS-----------SYRQSRSLA--------T
CLTNN-------GL-------------TS-G-MISV-------------ICY----SV--
-------VRSMVY------RV--------LQ-------------RLAIELFIDQE-----
----------E-----------------Y------------I-S-T---LQ---------
------------------------------------------------------------
------------------------------------------------------------
----------SSAGIRIIIH-DPNEMP--FPEDSGSTLAPGRQTSIGLTKV------A--
-----VHQL-----------DYP-Y--TT-C--TND----------------FPN-DNIF
----QER--FP------HT-IYSVLACEKNCLFKYIK----QE--CNCADAR----Y---
------------R-Y---------------------------------------------
--------NETI-----E---TCDFDL---------N-----------------------
------------------------------------EAK---------------------
IRT------------------KNQFL---------KQ-----------------------
--------------------------FGQT------------------------------
--------------------M---------------------------------------
------------------------------------------------------------
------------------------------------------------------------
------------------------------------------------------------
------------------------------------------------------------
-------------LI-----------G------TNVVKVIVYFSSLEYENFY-QTADYTV
YDL-----------------ISALGGQVGLWIGVSVLTVFEFVELLYDI-----------
--------LKFG--------C----------S-------KL-------------------
TKLGK---------------------------RT----------------IFAKD-----
------------------------------------------------------------
------------------------------------------------------------
------------------------------------------------------------
------------------------------------------------------------
------------------------------------------------------------
-----------------------------------------------------------S
LEMGPHRT

>sp_Q9R0W5_ASIC5_RAT_Acid_sensing_ion_channel_5_OS_Rattus_norvegicus_OX_10116_GN_Asic5_PE_1_SV_1_586_137
MEHTEKSKGP--AEK--GLLGKIRRY----------------------------------
----------L-------------------------------------------------
------------------------------------------------------------
------------------------------------------------------------
------------------------------------------------------------
------------------------------------------------------------
------------------------------------------------------------
---------------SKRPL-------------PSPT-----------------------
------------------------------------------------------------
------------------------------------------------------------
------------------------------------------------------------
-----------------------------------------DR-----K-----------
-KF--DHDFA-ISTSFHGIHNIA---------Q-NQ--------------NK---VR---
--KVIWL----SVVLGSVSLLVWQIYSRLV-NYFM----------W---PTTTSI----E
-V-QY-V----------E-KIEF-----PAV-----------------------TFCNLN
RF------QTE-----AVSRFG------------------I-----------------I-
---------FF------------L-WDI--VSKV--------LRLQE----I--------
---------------------------------SG-------------------------
------------------------------------------------------------
------------------------------------------------------------
-------------NNTGS------------------------------------------
------------------------------------------------------------
------------------------------------------------------------
------------------------------------------------------------
--------------------------PE--------------------------------
------------------------------------------------------------
------------------------------------------------------------
------------------------------------------------------------
--------ALDF------------------------------VAS---------------
-------------------------------------------------------HRNF-
---------------------------------SITEFV---------------------
------------------------------------------------KNN----GFYLN
-HDTL------VHCE----FF-G----K----------T---C-----------D-P---
----------------------------------------------K-------------
------------------------------------------------------------
------------------------------------------------------------
-----------D----------F-------K-----------HVF-T-EYG--------N
CFTFN-------YG-------------EN-V--------------------Q--------
-------SKNKVS------VS--------GR-------------GLKLLLDVHQE-----
----------E-----------------F------------T-D-N---PV---------
------------------------------------------------------------
------------------------------------------------------------
-------PGFADAGVIFVIH-SPKKEP--QFDGLGLSSPVGMHARVTIRQL------K--
-----TIHQ-----------EYP-W--GE-C--NPD-I------K-------LR----NF
------------------T-TYSTYGCLKECKAKHIQ----RL--CGCLPFL----L---
------------P-G---------------------------------------------
----------NG-----V---ECDL-----------------------------------
------------------------------------LKY---YNCVS---------P---
I-L-------------------DHIE-R-------KG---------------------LC
TMGT-----HNSSCP-VPCEETE---YPAT-IAY------------------STF-PS--
-------QRA------TKF-L-----------AK--------------------------
------------------------------------------------KL----------
------------------------------------------------------------
-N----------------------------------------------------------
---------------------------QSQ------------------------------

-----E-------YI-----------R------ENLVNIEINYSDLNYKITQ-QQKAVSV
PEL-----------------LADVGGQLGLFCGASLITIIEIIEYLFTS-----------
--------FYWV--------F----------I-------FF-------------------
LL------------------------------KI-------------------L------
-----------------------------------------EMIQR--------------
------------------------TSP---------------------PQTV--------
------------------------------------------------------------
------------------------------------------------------------
------------------------------------------------------------
------------------------------------------------------------
--------
>Cnidar_Rhopilema_esculentum_TR90022_c1_g5_i1_p1__Rhopilema_esculentum_TR9002
2_c1_g5__Rhopilema_esculentum_TR90022_c1_g5_i1_p1__ORF__type_complete__len_49
7__score_69_36__Rhopilema_esculentum_TR90022_c1_g5_i1_126_1616__588_139
------------------------------------------------------------
----------M-------------------------------------------------
------------------------------------------------------------
------------------------------------------------------------
------------------------------------------------------------
-DNEGDEHLSS-------------------------------------------------
-STDESAFDEEYCD----------------------------------------------
----FQQEEVFKGIEWI-LD-------------CDDLTESE-------------------
------------------------------------------------------------
---------------YLRYITRFLAE-MEDH--------------------------ARR
L----------------------DKQTR--------------------------------
--------Q----------------------R---------KI-----H-----------
-IY--KRSLI-SKSTVHGLRNCF---------D-RK--------------SR---VR---
--RFIWT----ILLLTAVGLLMQKLYECTM-HFIS----------H---PFSTTT----T
-V-RY-E----------D-SMRF-----PAV-----------------------SICNLN
DM------RTS-----VMKGTK------------------L-----------------D-
---------MI----------------------------------LK-------------
------------------------------------------------------------
------------------------------------------------------------
------------------------------------------------------------
------------------------------------------------------------
------------------------------------------------------------
------------------------------------------------------------
------------------------------------------------------------
------------------------------------------------------------
------------------------------------------------------------
------------------------------------------------------------
------------------------------------------------------------
------------------------------------------GDRNVS------------
-------------------------------------------------------LSGD-
---------------------------------EYRNTI---------------------
------------------------------------------------RKA----NHELP
--DML------YKCK----IL-D----K----------E---C-----------S-V---
----------------------------------------------Q-------------
------------------------------------------------------------
------------------------------------------------------------
-----------D----------F-------I-----------QFN-K-DQGD-------R
CFTFN-------HG-------------HE-G--------------------Q----QI--
-------LFFNKT------GP--------TH-------------ALELILNIEEY-----
----------E-----------------Y------------Y-P-G--------------
------------------------------------------------------------
------------------------------------------------------------
---------TDYSGIQLILH-GQDETP---VKMLGVMLSPGFITYVQVKRR------K--

-----IKNL-----------PPP-YK-SN-C--GSK--------K-------LK----YF
-----------------SE-RYSKHLCWLETLTDHVI----EE--CGCKEWF----M---
------------P-G---------------------------------------------
----------GY-----R---VCSL-----------------------------------
------------------------------------NES---KECMW---------P---
R-W-------------------AEFD-K-------YM---------------------KY
------------DCP-LPCVMDT---YTPS-LSL------------------SQF-LL--
-------DRD------VDE-I-----------AQ--------------------------
------------------------------------------------DP----------
------------------------------------------------------------
-R---------------F------------------RGA---------------------
------------------------KE-MNR------------------------------
-----K-------TI-----------R------GTFVKAVIFYGELSYEYSE-QVPSYDK
LDF-----------------LADIGGQLGLFLGSSALTYMEFIDCLVMI-----------
--------IYTK--------Y----------I-------SV-------------------
KP----------------------------------------------------------
------------------------------------------------------------
------------------------------------------------------------
------------------------------------------------------------
------------------------------------------------------------
------------------------------------------------------------
------------------------------------------------------------
--------
>Protostome_Lophotroco_annelid_CAC9633462_1__Ofus_G091394_Owenia__fusiformis_
_592_143
------------------------------------------------------------
----------M-------------------------------------------------
------------------------------------------------------------
------------------------------------------------------------
------------------------------------------------------------
------------------------------------------------------------
------------------------------------------------------------
------------------------------------------------------------
------------------------------------------------------------
---------------------------------------------------------EVM
TE---------------------TKEKQTDPVIE--------------------------
-------------------------------H---------SF-----S-----------
-TR--LKEFG-EDVEFHGVRHLF---------R-DN--------------SI---VT---
--KVTWI----LLFLCGCSWLTYQIHDRII-YYYQ----------F---PHITKI----D
-K-LY-V----------P-SLDF-----PTI-----------------------TICNIN
TF------RRH-----KLNEYD------------------L-----------------L-
---------HY------------G-TSL---------------NILD----E--------
---------------------------------NW-------TLLH--------------
------------------------------------------------------------
------------------------------------------------------------
------------------------------------------------------------
------------------------------------------------------------
------------------------------------------------------------
------------------------------------------------------------
--------------------------PEHY------------------------------
------------------------------------------------------------
------------------------------------------------------------
------------------------------------------------------------
--------DKAF------------------------------DDW---------------
------------------------VYSINWTDVEIHDR-----------------SINH-
---------------------------------SMEEMY---------------------
------------------------------------------------ERA----GHQIE

--DML------IYCK----WK-Q----Q----------E---C-----------S-V---
----------------------------------------------A-------------
------------------------------------------------------------
------------------------------------------------------------
-----------N----------F-------T-----------LIN-T-HYG--------R
CYQFN-------SG-------------KD-G--------------------VK-------
-------HQSFKG------GK--------AN-------------GLKLYLNVEEL-----
----------E-----------------Y------------L-N-Y--------------
------------------------------------------------------------
------------------------------------------------------------
-------MEASDLGFKILAH-DQDEPP--LIQELGFGVTTGNHYFIALETE------R--
-----VTSL-----------PDP-Y--GN-C--EED-H------K-------LD----HY
------------------D-HYSIPACRIECETLIVE----EK--CSCRLVE----M---
------------H-G--DH-----------------------------------------
----------GI-----R---VCTA-----------------------------------
------------------------------------EEY---HDCAL---------P---
T-L-------------------ESIT-E-------SD---------------------TC
------------VCQ-NPCELTQ---FQHS-ISS------------------VKL-RE--
-------SAV------EMI--HGHT-------SS--------------------------
------------------------------------------------NI----------
------------------------------------------------------------
-N----------------------------------------------------------
----------------------LTEF-KSV------------------------------
-----E-------FI-----------E------KNLLVVNLFFDSLNYQYIE-QTVAYPG
VSL-----------------LSDVGGQMGLCIGASILTVLHLVQFAIGE-----------
--------IIKK--------F----------Q-------KR-------------------
EN------------------------------KE------------------VN------
-----------------------------------------TNVIHVA------------
------------------------------------------------------------
------------------------------------------------------------
------------------------------------------------------------
------------------------------------------------------------
-----------------------------------------------------------S
ANPVDDKL
>Deutero_Ambulac_Spurpu_015310_593_144
------------------------------------------------------------
----------M-------------------------------------------------
------------------------------------------------------------
------------------------------------------------------------
------------------------------------------------------------
------------------------------------------------------------
------------------------------------------------------------
------------------------------------------------------------
------------------------------------------------------------
------------------------------------------------------------
------------------------------------------------------------
-----------------------------GCR---------SG-----R-----------
-CH--LRKWAADTSDLHGLKHIA---------G-SG--------------NL---FR---
--RLVWL----CLFLTALGFCIYECYLVFV-GFLS----------F---NHVTQV----D
-V-LY-S----------T-EVEF-----PAV-----------------------TVCNIN
KY------RES-----AFTEDD------------------I-----------------K-
---------NV------------G-VHL---------------GIID----E--------
---------------------------------DH-------NLLL--------------
------------------------------------------------------------
------------------------------------------------------------
------------------------------------------------------------
------------------------------------------------------------

------------------------------------------------------------
------------------------------------------------------------
--------------------------PELY------------------------------
------------------------------------------------------------
------------------------------------------------------------
------------------------------------------------------------
--------TDEF------------------------------RDF---------------
------------------------IDSVNWTVVEE--------------------DPDY-
---------------------------------NMTDFT---------------------
------------------------------------------------IRT----GHQKE
--DMI------LSCL----WK-E----E----------P---C-----------E-E---
----------------------------------------------S-------------
------------------------------------------------------------
------------------------------------------------------------
-----------D----------F-------Q-----------HKL-T-HLG--------N
CYSFN-------LQ-------------GA-N--------------------SD-EADW--
-------KHSYAA------GA--------AN-------------GLQLILNMETP-----
----------E-----------------Y------------T-P-T-NDLD---------
------------------------------------------------------------
------------------------------------------------------------
-------GGAMDAGLRIMFH-YPTEPP--YPKELGFAVAAGDHSFISMRHE------K--
-----ITSL-----------PDP-Y--SE-C--ESD-G------ADI-----IT----HF
------------------D-HYSLQACRIECETLVVV----AE--CGCRLPE----M---
------------P-G---------------------------------------------
----------DD-----D---VCGP-----------------------------------
------------------------------------ADL---HECAH---------P---
T-L-------------------VEFI-T-------GN--------LDS--------AESC
------------ECF-SPCEVEN---YPFT-LST------------------SRL-RL--
-------TYL------EAL-F-----------AN--------------------------
-----------------------------------------------TST----------
------------------------------------------------------------
-N----------------------------------------------------------
---------------------------FTA------------------------------
-----A-------YI-----------G------ENIAVVSMYYEALNFETID-MLPEYTV
ATL-----------------LAVLGGNLGLFLGASFLTLAQLGEYCFDE-----------
--------VIGW--------C----------I-------CT-------------------
PK------------------------------KE-------------------D------
-----------------------------------------DDKND--------------
--------------------GGNKVEPV--------------------STVT--------
------------------------------------------------------------
------------------------------------------------------------
------------------------------------------------------------
----------------------------------------------------------AM
DAWGAQKL
>Cnidar_Rhopilema_esculentum_TR40535_c0_g1_i1_p1__Rhopilema_esculentum_TR40535_c0_g1__Rhopilema_esculentum_TR40535_c0_g1_i1_p1__ORF__type_complete__len_501__score_55_33__Rhopilema_esculentum_TR40535_c0_g1_i1_597_2099__597_147
------------------------------------------------------------
----------M-------------------------------------------------
------------------------------------------------------------
------------------------------------------------------------
------------------------------------------------------------
-DSKTEMATLE-------------------------------------------------
-----REWTEQYL-----------------------------------------------
------EYYVKKYLEEYEES-------------TSDTSEGS-------------------
------------------------------------------------------------
--------------DYSMMDWDFQKKGLLER--------------------------ERL

W----------------------LKAKR--------------------------------
--------R----------------------K---------KI-----T-----------
-KH--FNTMV-KYSTFHGLQFCF---------K-KE--------------SP---LR---
--RTVWC----LLLLICNGLLVQKMFESTQ-HYLE----------H---PFSTTK----S
-V-KY-K----------E-SLTF-----PAV-----------------------SLCNLN
DM------RHS-----RMVGTK------------------L-----------------H-
---------KL----------------------------------II-------------
------------------------------------------------------------
------------------------------------------------------------
------------------------------------------------------------
------------------------------------------------------------
------------------------------------------------------------
------------------------------------------------------------
------------------------------------------------------------
------------------------------------------------------------
------------------------------------------------------------
------------------------------------------------------------
------------------------------------------------------------
------------------------------------------ERQQ--------------
-------------------------------NISKQ-------------------LSGN-
---------------------------------EYMNTI---------------------
------------------------------------------------RQA----NHRLE
--NML------YSCT----MD-G----V----------K---C-----------T-S---
----------------------------------------------E-------------
------------------------------------------------------------
------------------------------------------------------------
-----------D----------F-------S-----------LFY-H-KQGD-------K
CFTFN-------SG-------------RP-R--------------------Y----KL--
-------VKTNRI------GP--------QH-------------ALELTINIEFW-----
----------D-----------------Y------------Y-D-D--------------
------------------------------------------------------------
------------------------------------------------------------
---------AVQSGIHLILH-GQEETP---VTMEGFRISPGFITYAEVKKT------Q--
-----RRNL-----------PSP-YK-TQ-C--GSL--------K-------LK----YF
------------------K-GYSRNLCWLEQLTDDVV----GH--CGCKDWF----M---
------------P-G---------------------------------------------
----------DY-----K---VCSL-----------------------------------
------------------------------------TEL---EMCLW---------S---
R-W-------------------VDFD-K-------YK---------------------KY
------------SCP-LPCVIDS---FTTK-TSF------------------SRF-PT--
-------HDG------SES-M-----------AK--------------------------
------------------------------------------------EL----------
------------------------------------------------------------
-N---------------L------------------NGT---------------------
------------------------SR-DNH------------------------------
-----K-------FL-----------S------DNFLKVVIYYGDLSYEYLE-QKPSYDL
LVF-----------------LGDIGGQIGLFVGASVLTFFEFIDCFLMC-----------
--------IHAR--------F----------F-------EI-------------------
FR------------------------------PR-------------------E------
------------------------------------------------------------
------------------------------------------------------------
------------------------------------------------------------
------------------------------------------------------------
------------------------------------------------------------
------------------------------------------------------------
-----TQL

>Deu_Ambulacrar_hemi_Ptyfla_40v0_9_20150316_1g17738_t1_scaffold14937_cov156_5
99_148
------------------------------------------------------------
----------M-------------------------------------------------
------------------------------------------------------------
------------------------------------------------------------
------------------------------------------------------------
------------------------------------------------------------
------------------------------------------------------------
------------------------------------------------------------
------------------------------------------------------------
------------------------------------------------------------
------------------------------------------------------------
------------------------------------------------------------
------------------------------------------------------------
------------------------------------------------------------
----------------------------PSA-----------------------WCYTTN
RD------VGY-----ELCD----------------------------------------
-----------L---------KIDRKECV-----TDHVLNDFNVSFA----PLEDLEYDQ
QYEQMTEA--QCAAMCLE----Y-SAFVCMFF-DY-------SVQNHTCYLSSNSSDQSQ
PSTVQQDIV--RYIRILDDTEVE-------------ILHLKF-SRYPNKKMIEG-GLQRK
VIAGSISECITAC--------------------IEMD--------IFKCKSLD-------
-FSPSQSTCNLYEKPS-ANVGG-------------------LMKEDISYIHL--------
------------------------------------------------------------
---------------TRQGEYYDHRT----------------------------------
------------------------------------------------------------
------------------VTEQFLDISGI-------------------------------
------------------------------------------------------------
------------------------------------------------------------
------------------------------------------------------------
----TPEILNDF----------ENGYYR--------------DNGFSGV-----------
---------------------VKGEVPPDWYGF-----------------------KTFS
STP------------------------------DYSDLR---------------------
----------------------SVLKL----------------TGEEIAKY----GHQAE
--DFI------LQCT----YD-E----K----------T---C-----------S-H---
----------------------------------------------S-------------
------------------------------------------------------------
------------------------------------------------------------
-----------D----------F-------S-----------TFQ-DDHYG--------N
CFKFN-------HG-------------LN-G--------------------T----VV--
-------RDANRQ------GA--------LS-------------GLRLTLFLEQN-----
----------E-----------------Y------------I-S-I---YG---------
------------------------------------------------------------
------------------------------------------------------------
----------RNAGARVNIN-PPNDTV--LPQDEGITIMPGTVTSIGIREK------H--
-----TFRK-----------TKP-Y--GK-C--KDG--------TNE---------DNLY
----GPGN------------TYK-------------------------------------
------------------------------------------------------------
------------------------------------------------------------
------------------------------------------------------------
V-----------------------------------------------------------
------------------------------------------------------------
--------------------L---------------------------------------
------------------------------------------------------------
------------------------------------------------------------
------------------------------------------------------------
------------------------------------------------------------

-----------------------------------------------------------I
EDF-----------------LSDIGGTLGLYIGLSVITVAEFIELVASV-----------
--------IRYM--------YCS--------Q-------KD-------------------
SNP-----------------------------RT------------------AE------
---------------------------------------SENKDRERRFSGAYA------
----------------YPSFGRVDIND---------------------TKVAKL------
------------------------------------------------------------
------------------------------------------------------------
------------------------------------------------------------
------------------------------------------------------------
--------
>Deuterostome_chordata_Ggallus_tr_F1P0L7_F1P0L7_CHICK_Uncharacterized_protein
_OS_Gallus_gallus_OX_9031_GN_ASIC5_PE_3_SV_3_600_149
--KTNKSHLI--SEF--SIIFHCLSL---------------------DK-----------
----------M-------------------------------------------------
------------------------------------------------------------
------------------------------------------------------------
------------------------------------------------------------
------------------------------------------------------------
------------------------------------------------------------
-------------------------------------------------------LPRL-
------------------------------------------------------------
------------------------------------------------------------
------------------------------------------------------------
-------------------------------E---------DR-----R-----------
-KY--RQEFA-SSTSFHGVYNIV---------Q-TQ--------------TR---TR---
--RVLWL----LVVTGCLGIVIWQICSRFN-YYFS----------W---PTTTTV----V
-V-QH-V----------E-NVKF-----PAV-----------------------TFCNLN
RF------QAH-----AVSNLQ------------------I-----------------I-
---------FF------------L-WNI--VSGIV------QKFAME-------------
---------------------------------DK-------------------------
------------------------------------------------------------
------------------------------------------------------------
------------------------------------------------------------
------------------------------------------------------------
------------------------------------------------------------
------------------------------------------------------------
------------------------------------------------------------
------------------------------------------------------------
------------------------------------------------------------
------------------------------------------------------------
----YFHELNGF------------------------------------------------
----------------------------------------------------LLGNQNF-
---------------------------------SIKEFT---------------------
------------------------------------------------REN----GFYLN
-GSTL------LECE----FF-G----K----------T---C-----------H-P---
----------------------------------------------E-------------
------------------------------------------------------------
------------------------------------------------------------
-----------D----------F-------E-----------HIF-T-EYG--------N
CFTFN-------YN-------------DL-------------------------------
-------PARRVS------LS--------GR-------------GLHLLFDVQQE-----
----------Q-----------------F------------T-D-D---PA---------
------------------------------------------------------------
------------------------------------------------------------
-------LGYTDAGITFVIH-SPKEIP--RFDGLGLLTPVGMHAQVSIQQL------K--
-----SIIQ-----------EYP-W--GE-C--KPD-I------K-------LQ----YQ

------------------D-TYSTNGCLKECKAWYIQ----DW--CGCLPFI----L---
------------P-G---------------------------------------------
----------NG-----I---ECDL-----------------------------------
------------------------------------MKY---YNCVY---------P---
A-I-------------------HDIE-V-------KG---------------------LC
TVGT-----HNSTCP-APCEETH---YPTT-VTY------------------SSF-GG--
-------ENA------IKY-I-----------SA--------------------------
------------------------------------------------KL----------
------------------------------------------------------------
-K----------------------------------------------------------
---------------------------KSP------------------------------
-----E-------YI-----------R------QNLVIIDIKYDDLNYRITQ-QQKALTI
SEL-----------------LADVGGQLGLFCGASMITIIEVLEYIFTN-----------
--------FFWM--------C----------L-------FL-------------------
LL------------------------------KA-------------------P------
-----------------------------------------EIPRW--------------
------------------------NNP---------------------SHDQ--------
------------------------------------------------------------
------------------------------------------------------------
---------------------------------P--------------------------
--------------------------------------------------------THVE
KNKGIQEC
>Cnidar_Rhopilema_esculentum_TR106708_c0_g1_i1_p1__Rhopilema_esculentum_TR106
708_c0_g1__Rhopilema_esculentum_TR106708_c0_g1_i1_p1__ORF__type_complete__len
_504__score_95_58__Rhopilema_esculentum_TR106708_c0_g1_i1_88_1599__601_150
------------------------------------------------------------
----------M-------------------------------------------------
------------------------------------------------------------
------------------------------------------------------------
------------------------------------------------------------
------------------------------------------------------------
------------------------------------------------------------
------------------------------------------------------------
------------------------------------------------------------
------------------------------------------------------------
-----------------------MKSSEK-------------------------------
-KENNRNEQ----------------------G---------SV-----K-----------
-KI--FDDFA-NSTTAHGIQHLV---------S-SK--------------KT---LG---
--KAVWL----IVCLAALTTNLFLCAQLVK-KYLS----------K---PVSTTV----G
-V-RH-E----------Q-TMIF-----PSV-----------------------MVCNHN
ML------RKS-----EFQKAKR-----------------------------------R-
---------KLINFK--------------Q--------DASMMSLYK-------------
---------------------------------EM-------------------------
------------------------------------------------------------
------------------------------------------------------------
------------------------------------------------------------
------------------------------------------------------------
------------------------------------------------------------
------------------------------------------------------------
------------------------------------------------------------
------------------------------------------------------------
------------------------------------------------------------
------------------------------------------------------------
MLKAIGMANQAS------------------------------NTSSVNISV---------
-------------------------------------------------------NTDII
------------------------------ERRCLLEYL---------------------
------------------------SNI----------------EKPQLKGL----GHQFH

```
--NFV------VRCD----WK-G----I----------D---C-----------K-S---
-GML------------------------------------------E-------------
------------------------------------------------------------
------------------------------------------------------------
-----------N---------RW-------Y-----------KVW-NHIYG--------N
CFIFN-------YG-------------YN-M-ERKK---------------Q----EP--
-------LRLEDV------GM--------QN-------------GLTLDLNIELY-----
----------E-----------------Y------------D-E-E---LT---------
------------------------------------------------------------
------------------------------------------------------------
----------KETGVRVLLA-DQGVFP--LPSIQGFSVPPGVHASIGIKKV------E--
-----AVRV------------DP-FNNGS-C--RRS--------DS------LDE-----
----------------DGD-KYTMQRCTQRCHWQRQL----KK--CNCTGLR----L---
------------S-S--PN-----------------------------------------
-----------------R---TCLT-----------------------------------
------------------------------------VEE---HMCKA---------E---
V-S-------------------TEMD-A-----------------LGI--------NKIC
----------STKCP-PPCFEAK---FSHS-VTY------------------STL-YT--
-------AAL----------L---------------------------------------
------------------------------------------------KL----------
------------------------------------------------------------
-K-------------TSIF-----------------------------------------
---------------------------RND------------------------------
-----D-------DY-----------R------KNFVRLQFFFESDIVEFIE-TYTSYEM
EQL-----------------LSDIGGQLGLWVGVSFVTCVEIFLFLYSF-----------
--------SYHV--------L----------K-------KVWC-----------------
TSF-----------------------------KV--------------------------
------------------------------------------------------------
------------------------------------------------------------
------------------------------------------------------------
------------------------------------------------------------
------------------------------------------------------------
-----------------------------------------------------------H
NNKQTQEN
>Deutero_Cephalo_292990F_t1__name__chr_scaffold9_start_4371418_end_4385499_st
rand_pro_len_504__602_151
------------------------------------------------------------
----------M-------------------------------------------------
------------------------------------------------------------
------------------------------------------------------------
--------------------------------------------------------RNVV
GHKIGGTVACQ-------------------------------------------------
------------------------------------DSNSGPL-----------------
---------------GSEST----------------------------------TLPLR-
---------------------------------------------H--------------
------------------------------------------------------------
-----------------------TTPQNLFLNP---------------------------
-------------------------------SFMATCC---GD-----D-----------
-SV--EREYA-AETSLHGVGKIA---------G-AR--------------HR---GV---
--RLLWA----VLFLGMFGVATWQITERFV-AYFQ----------F---DTVTNM----K
-V-EF-R----------D-VLDF-----PTV-----------------------TICNFN
KY------RDS-----QITEEE------------------E-----------------Q-
---------YV------------S-TLLEGSTGF---------SDYD----Y--------
---------------------------------DA-------DYYDD-------------
------------------------------------------------------------
------------------------------------------------------------
-----------------DG-----------------------------------------
```

```
------------------------------------------------------------
------------------------------------------------------------
------------------------------------------------------------
------------------------------------------------------------
------------------------------------------------------------
------------------------------------------------------------
------------------------------------------------------------
--------DGEY------------------------------------------------
------------------------QYSYDWNNTGI--------------------PDDF-
---------------------------------NLAEFT---------------------
------------------------------------------------LRA----GFDIH
--TSL------KHCL----WR-G----Q----------T---C-----------N-S---
----------------------------------------------D-------------
------------------------------------------------------------
------------------------------------------------------------
-----------N----------F-------T-----------HIF-T-SFG--------N
CWMFN-------AE-------------GR-------------------------------
-------MNQTIS------GV--------GN-------------GLQVAIDIQQD-----
----------E-----------------Y------------T-E-N---AP---------
------------------------------------------------------------
------------------------------------------------------------
-------TGNLDAGIRFIVH-SPSEPP--KVDTQGISVGPGIHAYASISKI------E--
-----FRNE-----------IPP-W--GQ-C--DPG-R------A-------LQ----YY
------------------S-GYTKTGCLLECRADHVA----DE--CGCRTVS----M---
------------P-G---------------------------------------------
----------TL-----D---YCQP-----------------------------------
------------------------------------AVV---TGCVK---------T---
T-I-------------------AELK-T-------GK--------------------RSC
------------NCP-TPCSATA---YPAT-LSY------------------GGW-PS--
-------SST------MDY-F-----------TD--------------------------
------------------------------------------------LL----------
------------------------------------------------------------
-N----------------------------------------------------------
---------------------------KTE------------------------------
-----Q-------EI-----------K------DNIVLLDVYYQQLNLQTVQ-QRRAIST
NAL-----------------LGDLGGQLGLFLGASVITIIEFLEFLVKK-----------
--------GTSC--------C----------F-------RH-------------------
NN------------------------------RV-------------------K------
------------------------------------------------------------
------------------------------------------------------------
------------------------------------------------------------
------------------------------------------------------------
------------------------------------------------------------
------------------------------------------------------------
-------T
>Cnidaria_HyNaC7_CDG50528_1_Hydra_sodium_channel_7__Hydra_vulgaris__33_152
------------------------------------------------------------
----------M-------------------------------------------------
------------------------------------------------------------
------------------------------------------------------------
------------------------------------------------------------
------------------------------------------------------------
------------------------------------------------------------
------------------------------------------------------------
------------------------------------------------------------
--------------------------RIRLK--------------------------ERM
R----------------------KIVQE--------------------------------
```

```
--------R----------------------L---------TV-----K-----------
-QI--FKRYI-ESSTLHGFCYVC---------M-DT--------------FL---GR---
--RLIWA----VLMILGAIYFIFKLRYGIK-EYFD----------Y---PFSTLS----T
-V-EY-V----------D-DLLF-----PAV-----------------------SVCATN
SY------IAS-----QVYTNQ------------------L-----------------N-
---------TM----------------------------------YK-------------
------------------------------------------------------------
------------------------------------------------------------
------------------------------------------------------------
------------------------------------------------------------
------------------------------------------------------------
------------------------------------------------------------
------------------------------------------------------------
------------------------------------------------------------
------------------------------------------------------------
------------------------------------------------------------
------------------------------------------------------------
------------------------------------------EGRL--------------
------------------------PLDNNQSIPEYN-------------------IPGD-
---------------------------------ELVKTL---------------------
------------------------------------------------KNS----SLTIE
--SLL------KYCD----WI-M----QDTSHPLVTPNN---C-----------G-A---
----------------------------------------------L-------------
------------------------------------------------------------
------------------------------------------------------------
-----------N----------F-------T-----------SYF-N-YKGE-------Q
CHTLN-------SG-------------AK-G--------------------H----EL--
-------LKVSDV------GI--------SH-------------GYELVFDLQTN-----
----------E-----------------V------------I-K-N--------------
------------------------------------------------------------
------------------------------------------------------------
---------YQLSGMRIVIH-DQVFPP---QLVDGFFISPGFKTYIKLGIT------Q--
-----SQSL-----------PPP-YS-TE-C--GQK--------K-------LK----YY
------------------A-IYSQRLCLLETLTDFTG----DL--CGCRDVF----M---
------------P-E--N------------------------------------------
----------GL-----P---FCSL-----------------------------------
------------------------------------QEL---YSCMY---------P---
A-K-------------------ESFS-E-------FT--------------------MRK
------------ECP-SDCEERT---FSYE-LSE------------------ARY-LH--
-------NPP------IGL-SLSRLDK--LQLDK--------------------------
------------------------------------------------SLPSEA--H---
-----------------------------------------------------------L
SK---------------LA----------------KSLT---------------------
------------------------PK-ELD------------------------------
-----A-------YI-----------E------SNIISVILFFGDTRIDYHE-QEATNDF
FQF-----------------LGNMGGEFGLMLGASLLTFVEFIDLFIVL-----------
--------LYHQ--------M----------L-------RL-------------------
YN------------------------------LR-------------------KIPDI--
--------------------------------------FASKRNRNKEKT----------
-----------------------KKCE---------------------------------
------------------------------------------------------------
------------------------------------------------------------
------------------------------------------------------------
------------------------------------------------------------
------NV
>Chordat_hASIC5_NP_059115_1_acidsensing_ion_channel_5__Homo_sapiens__7_154
MEQTEKSKVY--AEN--GLLEKIKLC----------------------------------
```

```
----------L-------------------------------------------------
------------------------------------------------------------
------------------------------------------------------------
------------------------------------------------------------
------------------------------------------------------------
------------------------------------------------------------
---------------SKKPL-------------PSPT-----------------------
------------------------------------------------------------
------------------------------------------------------------
------------------------------------------------------------
-----------------------------------------ER-----K-----------
-KF--DHDFA-ISTSFHGIHNIV---------Q-NR--------------SK---IR---
--RVLWL----VVVLGSVSLVTWQIYIRLL-NYFT----------W---PTTTSI----E
-V-QY-V----------E-KMEF-----PAV-----------------------TFCNLN
RF------QTD-----AVAKFG------------------V-----------------I-
---------FF------------L-WHI--VSKV--------LHLQE----I--------
---------------------------------TA-------------------------
------------------------------------------------------------
------------------------------------------------------------
-------------NSTGS------------------------------------------
------------------------------------------------------------
------------------------------------------------------------
------------------------------------------------------------
--------------------------RE--------------------------------
------------------------------------------------------------
------------------------------------------------------------
------------------------------------------------------------
--------ATDF------------------------------AAS---------------
-------------------------------------------------------HQNF-
---------------------------------SIVEFI---------------------
------------------------------------------------RNK----GFYLN
-NSTL------LDCE----FF-G----K----------P---C-----------S-P---
----------------------------------------------K-------------
------------------------------------------------------------
------------------------------------------------------------
-----------D----------F-------A-----------HVF-T-EYG--------N
CFTFN-------HG-------------ET-L--------------------Q--------
-------AKRKVS------VS--------GR-------------GLSLLFNVNQE-----
----------A-----------------F------------T-D-N---PA---------
------------------------------------------------------------
------------------------------------------------------------
-------LGFVDAGIIFVIH-SPKKVP--QFDGLGLLSPVGMHARVTIRQV------K--
-----TVHQ-----------EYP-W--GE-C--NPN-I------K-------LQ----NF
------------------S-SYSTSGCLKECKAQHIK----KQ--CGCVPFL----L---
------------P-G---------------------------------------------
----------YG-----I---ECDL-----------------------------------
------------------------------------QKY---FSCVS---------P---
V-L-------------------DHIE-F-------KD---------------------LC
TVGT-----HNSSCP-VSCEEIE---YPAT-ISY------------------SSF-PS--
-------QKA------LKY-L-----------SK--------------------------
------------------------------------------------KL----------
------------------------------------------------------------
-N----------------------------------------------------------
---------------------------QSR------------------------------
-----K-------YI-----------R------ENLVKIEINYSDLNYKITQ-QQKAVSV
SEL-----------------LADLGGQLGLFCGASLITIIEIIEYLFTN-----------
--------FYWI--------C----------I-------FF-------------------
```

```
LL------------------------------KI-------------------S------
-----------------------------------------EMTQW--------------
------------------------TPP---------------------PQNH--------
------------------------------------------------------------
------------------------------------------------------------
------------------------------------------------------------
----------------------------------------------------------LG
NKNRIEEC
>Deutero_Ambulac_Apla_gbr54_16_t1_606_155
------------------------------------------------------------
----------M-------------------------------------------------
------------------------------------------------------------
------------------------------------------------------------
------------------------------------------------------------
------------------------------------------------------------
------------------------------------------------------------
----------------TASA-------------PD-------------------------
------------------------------------------------------------
------------------------------------------------------------
------------------------------------------------------------
-----------------------------------------TE-----T-----------
-NL--EEEFV-SNTTLHGIARVA---------H-NS--------------RT---IF---
--RLAWL----AIMLVFLGVCIAQITDRLQ-RFFM----------Y---KANTAI----S
-V-EY-V----------G-DLDF-----PAV-----------------------TICNFN
RY------RWS-----ELTSSD------------------L-----------------R-
---------SL------------N-YIL---------------EATD----Y--------
---------------------------------DY-------DSYG--------------
------------------------------------------------------------
------------------------------------------------------------
-----------------EAS----------------------------------------
------------------------------------------------------------
------------------------------------------------------------
------------------------------------------------------------
------------------------------------------------------------
------------------------------------------------------------
------------------------------------------------------------
------------------------------------------------------------
--------SSSL------------------------------SDT---------------
-------------------------YPLDDDNA----------------------SSAF-
---------------------------------NYTEFT---------------------
------------------------------------------------LRA----GFLMD
-DLTL------LSCE----WR-G----K--------TNS---C-----------S-A---
----------------------------------------------A-------------
------------------------------------------------------------
------------------------------------------------------------
-----------N----------F-------S-----------HVF-T-SFG--------N
CWTFN-------SG-------------SD-G--------------------G----PL--
-------LKEIQA------GA--------GN-------------GLRVQIDIQQN-----
----------E-----------------Y------------T-E-P--------------
------------------------------------------------------------
------------------------------------------------------------
------LSGNREAGLKVLVH-DQQTPP--MMDSQGFAIQPGVHAFVATRRQ------Q--
-----FLNL-----------EPP-Y--GK-C--DAS-R------T-------LN----RF
------------------P-NYTLEGCNIECRTEKII----ER--CNCRLVR----H---
------------P-G---------------------------------------------
----------TE-----V---ECTP-----------------------------------
------------------------------------MQT---SECAR---------D---
```

```
V-L-------------------SKLR-S-------GE--------I-----------PAC
------------DCP-VPCNYTT---FSTS-LST------------------ATL-PN--
-------QRA------TQL-L-----------VD--------------------------
------------------------------------------------QY----------
------------------------------------------------------------
-DDLAQ-------YSSNI------------------SDTTNTGSASSA------------
-------------------GQSNPIL-LDS------------------------------
-----T-------YM-----------Q------NNWILLDVFYENINFQRYE-QSEAITP
SAL-----------------ISDIGGQLGLFLGASFITLTEILSYLGRK-----------
--------LGSI--------L----------A-------SR-------------------
RP------------------------------RR-------------------A------
-----------------------------------------ATSPD--------------
------------------------TSP---------------------GWDM--------
------------------------------------------------------------
--------------NGVKI-----------------------------------------
------------------------------------------------------------
------------------------------------------------------------
-------T
>Protostome_Lophotroco_annelid_Pdum_Contig1978__608_157
FHSINRMHM---------------------------------------------------
----------M-------------------------------------------------
------------------------------------------------------------
------------------------------------------------------------
------------------------------------------------------------
------------------------------------------------------------
------------------------------------------------------------
------------------------------------------------------------
------------------------------------------------------------
------------------------------------------------------------
-----------------------------------------------------------A
EIGGDGNPA----------------------M---------SF-----L-----------
-TR--LKTLT-ENTGMDVLPHVG---------R-AR--------------SK---VK---
--RLFWT----LLLFVGTTMMISGVAKSFQ-RFYS----------R---PVETHL----I
-Q-RV-S----------E-SLPF-----PAI-----------------------TICNEN
PV------RFS-----AVERAYP-----------------------------------D-
---------FL-------------------------------KKLQN-------------
---------------------------------SS-------------------------
------------------------------------------------------------
------------------------------------------------------------
------------------------------------------------------------
------------------------------------------------------------
------------------------------------------------------------
------------------------------------------------------------
------------------------------------------------------------
------------------------------------------------------------
------------------------------------------------------------
------------------------------------------------------------
------------------------------------------------------------
-------------------------------------------------------DKTFW
------------------------------VREELTSFL---------------------
------------------------NSL----------------PLEERINL----GHNLT
--TML------LDCQ----QK-G----I----------T---C-----------N-E---
----------------------------------------------S-------------
------------------------------------------------------------
------------------------------------------------------------
-----------N----------F-------E-----------SMN-SLYHG--------N
CYTYK-------GM-------------GD-------------------------------
```

-------NLASNI------GP--------HT-------------GIGITLYIEES-----
----------E-----------------Y------------I-A-G---VA---------
------------------------------------------------------------
------------------------------------------------------------
----------DGTGFKVVIH-NGSDLP--FPEEEGFSVTPGFITFVALTKR------E--
-----VERA-----------EHP-Y--GV-C--HQE--------EDVP----IDD-----
------------------I-QYRTIGCRRRCLLRRQI----AK--CGCGLAT----F---
------------P-Y--TEQT---------------------------------------
-----SNITDDFNP---P---QCNT-----------------------------------
------------------------------------AEE---KECSE---------N---
V-Q-------------------QKYM-D-------GK--------L------------NC
----------NVGCP-PRCKEVY---HTYE-LSY------------------TGW-PP--
-------VCS------RQL-I-----------MK--------------------------
------------------------------------------------II----------
----------------------------------ER------------------------
-N-------------THT------------------PLQLANKCG---------------
---------------------------KKY------------------------------
-----E-------NI-----------R------YQTLKLVFFFNTMETTITK-TASAYTC
QDV-----------------LSNIGGQMGLWLGLSLCTLCEFLEFFFEC-----------
--------FHIL--------L----------R-------YL-------------------
CK------------------------------KN--------------------------
-----------------------------------------ITTVQ--------------
------------------------------------------------------------
------------------------------------------------------------
------------------------------------------------------------
--------------------------------SPQ-------------------------
--------------------------------------------F----GMRDHSHQNAN
NSSPKDKQ
>Deu_Ambulacrar_hemi_Ptyfla_40v0_9_20150316_1g15975_t1_scaffold12140_cov134_6
11_159
------------------------------------------------------------
----------M-------------------------------------------------
------------------------------------------------------------
------------------------------------------------------------
-----------------------------------------------------------K
PRENSEDIHQE-------------------------------------------------
------------------------------------------------------------
-------KSPSFSVQFKNTT--------------GSSVIGLFFG----------------
------------------------------------------------------------
---------------------------------------------------------DAD
S----------------------KKGTVQNLD----------------------------
----NKYEN----------------------L---------DE-----N-----------
-AA--IQDFT-QKATISGMKYVG---------D-KK-------------SRI---PR---
--RCFWL----LVVLTGAGFLTYQIIRSVN-LYRS----------F---PVNVDV----K
-I-KY-A----------Q-DLPF-----PAI-----------------------TICNMN
LL------RKK-----EMEKHN------------------E-----------------K-
---------NV------------A-GVL---------------KLLY-------------
------------------------------------------------------------
------------------------------------------------------------
------------------------------------------------------------
------------------------------------------------------------
------------------------------------------------------------
------------------------------------------------------------
------------------------------------------------------------
--------------------------PLVT------------------------------
------------------------------------------------------------
------------------------------------------------------------

```
------------------------------------------------------------
------------------------------------------------------------
------------------------EIIEDSDALHNFTL-----------------SSVN-
---------------------------------NLTELL---------------------
------------------------------------------------LRY----GHRKE
--DMI------IWCE----WK-G----R----------R---C-----------S-Y---
----------------------------------------------E-------------
------------------------------------------------------------
------------------------------------------------------------
-----------N----------F-------T-----------STL-T-DYG--------L
CHTFN-------SG-------------SN-G--------------------L----PS--
-------LRVDNT------GA--------RY-------------ALTLILNAEQW-----
----------E-----------------Y------------G-R-G---PS---------
------------------------------------------------------------
------------------------------------------------------------
----------QSAGFKIAIH-SQYDIP--MVSDLGFAVAPGTSAFIGLKMT------K--
-----VSSF-----------PEP-Y--GT-C--KSR--------K-------LK----YH
------------------G-NYTMSKCQLECLIDFIV----EK--CGCKRPY----M---
------------P-V--AV-----------------------------------------
----------DA-----P---YCSP-----------------------------------
------------------------------------EEE---YLCCL---------P---
K-L-------------------DEYA-S-------HR--------------------QMC
------------DCP-VPCERTL---YSAS-LSY------------------AQF-PS--
-------DFM------AVE-Y-----------SK--------------------------
------------------------------------------------LL----------
------------------------------------------------------------
-N----------------------------------------------------------
---------------------------KSD-----------------------------P
-----Q-------YP-----------R------RNGAKLNIFFEELSYEEIS-HRAAYEL
FTL-----------------FSDIGGSMGLLLGASFVTVVEIVDFACVN-----------
--------IYRR--------W----------R-------RK-------------------
TS------------------------------SK--------------------------
-----------------------------------------IYPKT--------------
------------------------------------------------------------
------------------------------------------------------------
------------------------------------------------------------
------------------------------------------------------------
------------------------------------------------------------
---SSNDR
>Deu_Ambulacrar_hemi_Ptyfla_40v0_9_20150316_1g14166_t1_scaffold9674_cov125_61
6_162
------------------------------------------------------------
----------M-------------------------------------------------
------------------------------------------------------------
------------------------------------------------------------
------------------------------------------------------------
------------------------------------------------------------
------------------------------------------------------------
------------------------------------------------------------
------------------------------------------------------------
------------------------------------------------------------
------------------------NKARARK-----------------------------
-IKMTGKKK----------------------R---------GL-----R-----------
-SI--ISKFA-KNTTAHGVANIS---------N-ST--------------ST---FG---
--RTVWS----VVCVTAFSLFLWQSSELLK-DYLE----------Y---GIKIKF----D
-V-VS-V----------P-KLSF-----PTV-----------------------TVCNTN
KI------RKS-----EILKSE------------------H-----------------R-
```

---------RIL---------VVDEDVVH-----PYYALQAFKSTYN----KRLSPKDAV
LKTNTSLG--LCALSCMH----F-SEFTCLSF-DY-------DSQRRECYINPGSKDT--
-------LV--RFTLTNDDD-------------------------YDYYERIYR-G----
----------MEC--------------------YTLD------------DGTD-------
-YRGHVSLSSWGERCM-NW-SS-------------------LDPSKFKYTPS--------
------------------------------------------------------------
---------------TAPGKGLGNHN----------------------------------
------------------------------------------------------------
------------------YCRNPNDEPGT-------------------------------
------------------------------------------------------------
------------------------------------------------------------
------------------------------------------------------------
----WCYV-----------------YRF--------------NSNVPVA-----------
---------------------ARCTVNPPKYGCLSQKQ-W---------------EKEIV
ESD------------------------------TFRDSK---------------------
----------------------PIPAI----------------AVNCSADP----SGQGY
--RGN------KSTT----VS-G----K----------Q---C-----------Q-N---
----------------------------------------------C-------------
------------------------------------------------------------
------------------------------------------------------------
-----------D----------F-------Y-----------MFQ-DDQYG--------N
CFKFN-------DG-------------EK-E--------------------P----VI--
-------RQADRQ------GA--------LS-------------GLTLTLFLEQN-----
----------E-----------------Y------------I-S-I---YG---------
------------------------------------------------------------
------------------------------------------------------------
----------RDAGARVNIN-PLNETV--FPQDEGITIMPGTVTSIGIREL---------
------------------------------------------------------------
------------------------------------------------------------
------------------------------------------------------------
------------------------------------------------------------
------------------------------------------------------------
------------------------------------------------------------
------------------------------------------------------------
------------------------------------------------------------
------------------------------------------------------------
------------------------------------------------------------
------------------------------------------------------------
------------------------------------------------------------
------------------------------------------------------------
DDI-----------------LSDIGGTLGLYIGLSIITVAEFIEFVAAI-----------
--------LRYL--------YQS--------R-------RI-------------------
KKL-----------------------------RT------------------VD------
---------------------------------------SDIKEKKRRRKCKF-------
------------------------------------------------------------
------------------------------------------------------------
------------------------------------------------------------
------------------------------------------------------------
------------------------------------------------------------
--------
>Deutero_Ambulac_Apla_gbr129_26_t1_617_163
------------------------------------------------------------
----------M-------------------------------------------------
------------------------------------------------------------
------------------------------------------------------------
------------------------------------------------------------
------------------------------------------------------------

```
------------------------------------------------------------
------------------------------------------------------------
------------------------------------------------------------
------------------------------------------------------------
-----------------------------------------------------------A
DEKSGENDA----------------------R---------SL-----R-----------
-AL--FKNLM-GNTSAHGLPNID---------R-AG--------------NP---FR---
--RCFWS----VLFVWALGMFLWQCYELVY-AYFQ----------W---EVDVNI----N
-I-QY-K----------T-QIDF-----PAV-----------------------TICNLN
PV------KAS-----SLSNSR--------------------------------------
---------EL-------------------------------EILLG----S--------
-------L-------------------------DSETS---GSTDGSWTAVT--------
------------------------------------------------------------
-------------------AVTPKVT----------------------------------
-----------SDQLSVNSTDGELFGGT-------ASSVTKLLPATEAGLTAT-------
G-----------------------------------------------------------
------------------------------------------------------------
------------------------------------------------------------
-----------DTHTLKASSERDQLTPTSTDASSP-------------------------
------------------------------------------------------------
------------------------------------------------------------
------------------------------------------------------------
SSGVSSASPAGSASIHVVGGQMATNAVTAGGIPTKEVERRKRDTSTNSE-----------
------------------------RQYQNWDNI-----------DYTAK------ESDVD
------------------------------LFDQVADAI---------------------
------------------------AKT----------------EYGKRLNF----GHSID
--DML------LACS----YK-G----Y----------P---C-----------G-P---
----------------------------------------------T-------------
------------------------------------------------------------
------------------------------------------------------------
-----------N----------F-------S-----------YFH-NYLYG--------N
CYTFN-------AG-------------MN----------------------S----DI--
-------QTSSKP------GP--------LY-------------GLILELNLEET-----
----------E-----------------Y------------I-P-S---IQ---------
------------------------------------------------------------
------------------------------------------------------------
----------QASGARVVID-QQYRMP--FPEDAGINVAPGVLTSIGIRKV------E--
-----IDRK-----------SDP-Y--TN-C--TER--------DS------DDS-QTVF
----SDF--FG-------A-NYSL------------------------------------
------------------------------------------------------------
------------------------------------------------------------
------------------------------------------------------------
------------------------------------------------------------
------------------------------------------------------------
------------------------------------------------------------
------------------------------------------------------------
------------------------------------------------------------
------------------------------------------------------------
------------------------------------------------------------
---------------------------------E-------------------------G
FGL-----------------ISNLGGQVGLWIGISMCTVFEFVELLYDV-----------
--------CKIL--------L----------L-------KVIFC--------------SH
AK------------------------------RA--------------------T-----
--------------------------------------APSASDLKLD------------
------------------------------------------------------------
------------------------------------------------------------
------------------------------------------------------------
```

-------------------------------EKP--------------------------
-----------------------------------------------------------S
KPRLALNN
>Deutero_Ambulac_Sakowv30038064m_622_164
------------------------------------------------------------
----------M-------------------------------------------------
------------------------------------------------------------
------------------------------------------------------------
------------------------------------------------------------
--DNTEKRECD-------------------------------------------------
------------------------------------------------------------
-------KANNGFPKFKKST-------SLLALFFRDLSEQRSQK----------------
------------------------------------------------------------
---------------------------------------------------------DRD
S----------------------QDSSDEEEN----------------------------
----GDEKK----------------------V---------ED-----E-----------
-SV--IGDFA-QKSTISGMKYVS---------D-KK-------------SRI---PR---
--RCFWL----LVVLTGAGFLVYQIVRSVN-LYLD----------Y---PVNVDV----K
-I-TY-A----------N-DLPF-----PAI-----------------------TICNMN
LL------RRL-----ELEHYE-----------------------------------GH-
---------DA------------I-EFL---------------ELLY-------------
------------------------------------------------------------
------------------------------------------------------------
------------------------------------------------------------
------------------------------------------------------------
------------------------------------------------------------
------------------------------------------------------------
------------------------------------------------------------
--------------------------PLVTEK----------------------------
------------------------------------------------------------
------------------------------------------------------------
------------------------------------------------------------
------------------------------------------------------------
------------------------TSPVPENIIQNFTL-----------------TTVA-
---------------------------------NLTDLL---------------------
------------------------------------------------MRY----GHQKE
--NMV------ILCH----WK-S----S----------A---C-----------S-S---
----------------------------------------------V-------------
------------------------------------------------------------
------------------------------------------------------------
-----------N----------F-------T-----------TTI-T-DYG--------L
CHTFN-------SG-------------LE-G--------------------T----AP--
-------LRVDNT------GA--------RY-------------GLTLYVDIEQG-----
----------E-----------------Y------------G-R-G---PR---------
------------------------------------------------------------
------------------------------------------------------------
----------ESAGLKIAIH-NQRDIP--LVSDLGFAVSPGTHTLIGLRMT------K--
-----ISSL-----------PNP-Y--GL-C--KSK--------K-------LQ----FY
------------------G-DYTMSKCQLECLTHFIV----GR--CGCKQAY----M---
------------P-G---------------------------------------------
----------QV-----R---HCSP-----------------------------------
------------------------------------QDV---YNCYL---------P---
N-L-------------------GEYA-G-------HK--------------------DMC
------------ECP-VPCERTI---YSSR-LSY------------------AQF-PS--
-------DFA------ANE-Y-----------SK--------------------------
------------------------------------------------LL----------
------------------------------------------------------------

-N----------------------------------------------------------
---------------------------NSD-----------------------------S
-----D-------YI-----------R------RNGLKLEIFYEELSFKEVT-HQVEYEL
FKL-----------------FSDIGGSMGLLLGASFVTIVEIFDFACLN-----------
--------LYRR--------V----------R-------SK-------------------
WR------------------------------PR--------------------------
-----------------------------------------ITPIE--------------
------------------------------------------------------------
------------------------------------------------------------
------------------------------------------------------------
------------------------------------------------------------
------------------------------------------------------------
-----TKS
>Deutero_Cephalo_032830F_t2__name__chr_scaffold117_start_653459_end_666145_st
rand_pro_len_509__623_166
------------------------------------------------------------
----------M-------------------------------------------------
------------------------------------------------------------
------------------------------------------------------------
------------------------------------------------------------
------------------------------------------------------------
------------------------------------------------------------
--------------------------------------------------------P---
------------------------------------------------------------
------------------------------------------------------------
-----------------------KIKTPMS------------------------------
-------------------------------S---------DEGHDWPE-----------
-SP--EYVFA-TTTSFHGIGRIA---------T-AP--------------NI---PL---
--RVLWT----LATLASYIGFFILMVEMLQ-SYFT----------H---GTVTDV----T
-L-EF-V----------P-EVRF-----PAV-----------------------TICNLN
KF------EDK-----ALTVEE------------------K-----------------H-
---------YL------------S-YIL---------------YGVQ----E--------
---------------------------------DI-------DVIA--------------
------------------------------------------------------------
------------------------------------------------------------
-----------------G------------------------------------------
------------------------------------------------------------
------------------------------------------------------------
------------------------------------------------------------
-------------------------APV--------------------------------
------------------------------------------------------------
------------------------------------------------------------
------------------------------------------------------------
--------NNKY------------------------------TPN---------------
------------------------STDANVTTETPR--------TEAPHT-----HGSF-
---------------------------------SVDSIT---------------------
------------------------------------------------REK----GFRLG
--NDM------HVCT----WM-G----Q----------T---C-----------T-E---
----------------------------------------------L-------------
------------------------------------------------------------
------------------------------------------------------------
-----------D----------F-------T-----------HSF-S-TYG--------N
CYTFN-------AS-------------PE-N--------------------P--------
-------INQTIP------GS--------GY-------------GLSVMIDIKSH-----
----------L-----------------Y------------T-E-N-PLLP---------
------------------------------------------------------------
------------------------------------------------------------

-------GGTADVGLKLLVH-DQNEPP--KMDTQGIAVAPGSHAYIAIRQI------Q--
-----YQNH-----------IPP-W--GV-C--QDL--------Q-------LE----YY
------------------D-TYTLTGCQLECRSKYVV----QN--CTCRPIY----L---
------------P-G---------------------------------------------
----------DA-----P---YCEP-----------------------------------
------------------------------------ADV---AHCVR---------P---
V-E-------------------AAVT-S-------GA--------L------------PC
------------DCP-VPCSLMT---YRTS-SSY------------------AKW-PN--
-------NKA------DVV-Y-----------TT--------------------------
------------------------------------------------AF----------
------------------------------------------------------------
-G----------------------------------------------------------
---------------------------LSP------------------------------
-----G-------YM-----------A------DNSVVFDVYYEELNYEIIS-QLKAMDS
GQL-----------------ASDLGGQMGLFIGASILTLLEIFEYLGLR-----------
--------LIAF--------I----------Q-------RR-------------------
KV------------------------------RT-------------------R------
-----------------------------------------RVHTS--------------
------------------------AYP---------------------MPDT--------
------------------------------------------------------------
----------------LSVNQAAKNKE---------------------------------
------------------------------------------------------------
---------------------------------------------------------SKE
KSLMNLKE
>Protostome_Lophotroco_annelid_Pdum_comp414620_c0_seq3__625_168
------------------------------------------------------------
----------M-------------------------------------------------
------------------------------------------------------------
------------------------------------------------------------
------------------------------------------------------------
------------------------------------------------------------
------------------------------------------------------------
------------------------------------------------------------
------------------------------------------------------------
------------------------------------------------------------
-----------------------GCKEWF-------------------------------
-TMGKSNPP----------------------L---------TY-----T-----------
-EV--TNQFA-DNTNCNGISSIK---------R-SN--------------HV---VR---
--KAFWT----LLVLAGAGLTIKLLVDVFT-KFFA----------Y---PVSISV----N
-V-TY-E----------S-TVQF-----PAV-----------------------TLCNVN
MI------RKS-----QIGLMT--------G------SSAL-----------------D-
---------KL-------------------------------TFTVG----G--------
---------------------------------NP-------------------------
------------------------------------------------------------
------------------------------------------------------------
-----------------DIS----------------------------------------
------------------------------------------------------------
------------------------------------------------------------
------------------------------------------------------------
------------------------------------------------------------
------------------------------------------------------------
------------------------------------------------------------
------------------------------------------------------------
------------------------------------------------------------
----------------------------AWATT----------------------EGRSI
------------------------------LRHSFVDGW---------------------
------------------------SQL----------------NDTVKKQQ----GHQLE

--EML------LDCS----YN-G----D----------V---C-----------G-T---
----------------------------------------------N-------------
------------------------------------------------------------
------------------------------------------------------------
-----------N----------F-------T-----------HYI-NPNFG--------N
CYTFN-------SN-------------WE-N--------------------Q----TI--
-------QYSHRA------GP--------FY-------------GLMLELFIEQD-----
----------E-----------------Y------------I-P-E---LT---------
------------------------------------------------------------
------------------------------------------------------------
----------EVAGVIVSIH-HQDVMP--FPEDDGVLIPPDMSSNIAITAE------T--
-----ITRK-----------AKP-YP-SD-C--LNA--------TFKE----AAH-YSAY
----SEL--HP-------V-RYTHTACYRTCYQAHLF----SE--CGCGDPV----M---
------------P-L--EG-------------------QALYNL----------------
-----VTDKQNKR----T---VCTS-----------------------------------
------------------------------------QDE---ITCEL---------K---
V-Y-------------------KQYS-N-------NE--------L------------EC
------------NCS-VACEDHI---YQTA-VTS------------------LRW-PS--
-------ANY------KNL-Y-----------YG--------------------------
------------------------------------------------HL-A--------
----------------------------------TR------------------------
-N-------------PHL------------------ATIMQSYV----------------
---------------------------TNG------------------------------
-----T-------NP-----------G------EHLVKLLIYFNEFNFNYIE-EYASYPS
SSL-----------------LSDLGGAIGFYLGISVIAAFEFIEYIFDL-----------
--------IFIG--------C----------K-------KM-------------------
KN------------------------------RK--------------------------
----------------------------------------ANEVKSID------------
------------------------------------------------------------
------------------------------------------------------------
------------------------------------------------------------
------------------------------------------------------------
------------------------------------------------------------
--------
>Protostome_Lophotroco_annelid_Pdum_comp405972_c0_seq2__629_171
------------------------------------------------------------
----------M-------------------------------------------------
------------------------------------------------------------
------------------------------------------------------------
------------------------------------------------------------
------------------------------------------------------------
------------------------------------------------------------
------------------------------------------------------------
------------------------------------------------------------
------------------------------------------------------------
------------------------------------------------------------
-----MSGE----------------------S---------SI-----K-----------
-PI--LGSMA-ENTAMHGLPNVK---------R-SI--------------ST---AR---
--KIFWS----IIVLAGVAMILYEVIIATI-KYYQ----------Y---PVQTTI----S
-I-NS-A----------I-VHQF-----PVV-----------------------TICNQN
SI------RRS-----ALEDAG------------------------------------S-
---------DI-------------------------------YMLLN----ST-------
---------------------------------GSRQTG---------------------
------------------------------------------------------------
------------------------------------------------------------
------------------------------------------------------------
------------------------------------------------------------

------------------------------------------------------------
------------------------------------------------------------
------------------------------------------------------------
------------------------------------------------------------
------------------------------------------------------------
------------------------------------------------------------
------------------------------------------------------------
-------------------------------------------------------SSIVS
------------------------------LRGNILKYY---------------------
------------------------SQL----------------TPQVRMDI----GHQME
--TMM------LDCA----YG-A----M----------E---C-----------N-T---
----------------------------------------------T-------------
------------------------------------------------------------
------------------------------------------------------------
-----------Y----------F-------E-----------QFS-NYYYG--------N
CYSFN-------NG-------------LN----------------------Y----GI--
-------KTASTV------GP--------DH-------------AFQIVLFIQQD-----
----------E-----------------Y------------I-S-Q---LS---------
------------------------------------------------------------
------------------------------------------------------------
----------EGAGVRVIVH-NRTHMP--FPENEGFSAPPGQITYVGLSRR------E--
-----IKRA-----------SPP-Y--GD-C--SEN----------------HLT-MNSY
----TDF--LT------GV-DYSKMACKKSCLQYTII----ET--CSCADTS----L---
------------V-T--NS-------------------TALN------------------
------TFFKNI-----E---PCRN-----------------------------------
------------------------------------LTQ---DACIS---------T---
V-T-------------------QKYY-N-------NE--------L------------NC
----------STGCS-PECFEVQ---HERT-LSY------------------SLW-PG--
-------DNA------LND-T-----------LT--------------------------
------------------------------------------------LL----------
----------------------------------SE------------------------
-N-------------TNA------------------AQLMKN------------------
----------------------MNDD-EKE------------------------------
-----S-------FL-----------R------KNVVKLAVYFESLDYDYIA-TSSSYSS
SDL-----------------AASYGGQMGLWLGVSVCTLFEVVEMLWEM-----------
--------GLVI--------S----------Q-------KM-------------------
NS------------------------------RK------------------NKT-----
---------------------------------------GNMETTNSDGK----------
------------------------------------------------------------
------------------------------------------------------------
------------------------------------------------------------
-------------------------------KMP--------------------------
----------------------------------------------------SDKAVIVK
KDSWTASY
>Placozoa_TadNaC11_XP_002114391_1_hypothetical_protein_TRIADDRAFT_58144__Tric
hoplax_adhaerens__21_176
------------------------------------------------------------
----------M-------------------------------------------------
------------------------------------------------------------
------------------------------------------------------------
------------------------------------------------------------
------------------------------------------------------------
------------------------------------------------------------
----------------GSKS-------------ENP------------------------
------------------------------------------------------------
---------------------------------------------------------RKV
S----------------------NGQTDI-------------------------------

-----------------------------------------VS-----F-----------
-DL--EENFA-TSTTCHGVAHIF---------E-TK------------G------SR---
--RSMWF----GITLVSTIACLVQCCIIVY-DASL----------Y---PTRVNI----K
-V-KY-N----------T-QSLF-----PSV-----------------------TLCNTN
LI------ASI-----DNTREE------------------L-----------------K-
---------YA------------M-LLY---------------RHYY----A--------
---------------------------------KN-------QI----------------
------------------------------------------------------------
------------------------------------------------------------
-----------------T------------------------------------------
------------------------------------------------------------
------------------------------------------------------------
------------------------------------------------------------
------------------------------------------------------------
------------------------------------------------------------
------------------------------------------------------------
------------------------------------------------------------
--------HQEL------------------------------QSA---------------
------------------------ENYFNR------------------VY-----GKNF-
---------------------------------SLEEYV---------------------
------------------------------------------------ARY----RFKAE
--DMI------VSCQ----WG-E----E----------P---C-----------G-S---
----------------------------------------------A-------------
------------------------------------------------------------
------------------------------------------------------------
-----------N----------F-------S-----------QVI-T-DYG--------T
CYTFN-------SG-------------QK-G--------------------Y----PL--
-------LYQTIS------GS--------SH-------------GLRLAINVQQY-----
----------V-----------------Y------------P-T-L--PLS---------
------------------------------------------------------------
------------------------------------------------------------
-------FLTPDAGIRLSVH-HYNEIA--NMGSRGVFVPPGMHGYIAITDT------HV-
-----LSNL-----------GPP-W--GE-C--GQK--------N-------LQ----YF
------------------R-YYSRSACRREFEANLAA----QQ--CNCSYQQ--------
------------------------------------------------------------
------------------------------------------------------------
------------------------------------------------------------
A-T-------------------VNTE-N-------KK--------T-----------LNS
------------FCP-YPCQQTE---YPII-LSY------------------AGI-AT--
-------NAI------IDS-S-----------SH--------------------------
------------------------------------------------------------
------------------------------------------------------------
-N-------------GKL------------------ANLLDSVKNDPSK-----------
----------------------YPNY-TSE------------------------------
-----A-------FI-----------R------ENLIYLDVYFRELITSTTT-ESKATGY
AQV-----------------LSDVGGQLGLFIGASIITLCEIITYLCDR-----------
--------CERK--------K----------R-------EK-------------------
REMRI--------------------------HRQ---------------SQMIRD-----
------------------------------------------------------------
-----------------------IMQV---------------------GVEK--------
---------KDLQ-----------------------------------------------
----------------TETK-------------------------------PNKNNDFVF
-----------------NTSEDG-------------QET---------------------
------------------------------------------------------------
-SSTNPNV
>Cnidar_AlaAla_c46701_g1_i1_p1__GENE_c46701_g1_i1__c46701_g1_i1_p1__ORF__type
_complete__len_515__score_83_16__c46701_g1_i1_223_1767__638_178

------------------------------------------------------------
----------M-------------------------------------------------
------------------------------------------------------------
------------------------------------------------------------
------------------------------------------------------------
------------------------------------------------------------
------------------------------------------------------------
------------------------------------------------------------
------------------------------------------------------------
------------------------------------------------------------
------------------------------------------------------------
-DIKEKDLP----------------------E---------NV-----K-----------
-AV--LKQFA-SSTSAHGLSHIA---------S-AK--------------YS---YQ---
--KLLWA----FVLLLNLFMFFFMITRLIT-LYTS----------K---PVSTKV----E
-F-TF-E----------K-ELAF-----PAV-----------------------TICNQN
MF------RRS-----KLTPSD--------------------------------------
-------HARLRPSE-----------------------TQITSTMVE----KMQ------
-----------------------------KKFQDELKRQGH-------------------
------------------------------------------------------------
------------------------------------------------------------
-----------------EGTHGGSG-----------------------------------
------------------------------------------------------------
------------------------------------------------------------
------------------------------------------------------------
------------------------------------------------------------
------------------------------------------------------------
------------------------------------------------------------
------------------------------------------------------------
--GTTGATSGGS------------------------------NSTS--------------
------------------------------------------TDNMVTV------DSSHL
------------------------------MRMSILESV---------------------
------------------------NLM----------------EEQEKMGK----GHSFE
--DLI------KFCR----WD-G----V----------A---C-----------H-K---
----------------------------------------------G-------------
------------------------------------------------------------
------------------------------------------------------------
----------------------F--MKQYWY-----------RTW-DWYYG--------N
CYTFN-------YG-------------KT-G-NGTS---------------L----KR--
-------LKATGS------GS--------EN-------------GLFLEIDIQNK-----
----------E-----------------Y------------F-P-E---LT---------
------------------------------------------------------------
------------------------------------------------------------
----------LESGVRLLLT-EQEVNL--FPLQEGFSIPPAVSAMVGVKMS------T--
-----AIRA------------DP-FNNGS-C--RAN--------EYA---------DGGF
--------------------RYTVQNCVQNCTALAQE----SK--CNCTLLK----Y---
------------A-A--VN-----------------------------------------
----------ST-----R--VTCHT-----------------------------------
------------------------------------EAE---KYCAD---------E---
V-V-------------------KSLK-N-------GT--------------------NDC
----------KANCP-QPCSESY---YSKS-MSY------------------SRL-YG--
-------EGY------QKD-I---------------------------------------
------------------------------------------------AIRD--------
----------------------------------KS------------------------
-N----------------------------------------------------------
---------------------------PKE------------------------------
-----K-------DI-----------M------ENYIRLLVFFEDQNVHKIR-TAVYYEV
ENL-----------------LSDIGGQLGLWIGVSMVTMVELILFIYKF-----------

--------VEYL--------M----------E-------KK-------------------
KAG-----------------------------KV--------------------------
------------------------------------------------------------
------------------------------------------------------------
------------------------------------------------------------
------------------------------------------------------------
------------------------------------------------------------
------------------------------------------------------------
---IHVAH
>Deu_Ambulacrar_hemi_Ptyfla_40v0_9_20150316_1g17281_t1_scaffold14250_cov87_63
9_179
------------------------------------------------------------
----------M-------------------------------------------------
------------------------------LIYSYFSWQD--------------------
---------------------------DRPGDPYPLVS--------------------LN
PTTL----------------------------------NGNGEGQDQNRNLRKRQG----
-NNTDER-----------------------------------------------------
------------------------------------------------------------
---------------------------------TDEIS----------------------
------------------------------------------------------------
-------------------------------------------------------SLDKS
A----------------------GETNG--------------------------------
------EYE----------------------D---------SV-----K-----------
-HM--LGDFG-RETTAHGIVKIT---------T-AK--------------SN---VT---
--RSVWS----AIVLLAASVMLVQMTLLLV-QYFE----------Y---NVNVKV----T
-Q-VS-E----------K-ALEF-----PPV-----------------------TICNTN
KL------RKS-----AVRASK------------------Y-----------------S-
---------EIL---------LLEQSFVP--------------PYYT-------------
---------------------------PCIEG-DF-------------------------
------------------------------------------------------------
-----------LCQ-----------------------------------DGIH-------
-CIKPYLVCDGVNNCG-D-M------------------------SDEDNCT-----YR-E
------------------------------------------------------------
------------------------------------------------------------
-----------------CGPN------------------------Q--------------
------------------------------------------------------------
------------------------------------------------------------
------------------------------------------------------------
------------------------------------------F-----------------
----RCDSGSNL---GIC---LTKDRVC---------------DRIKDC----YEGEDET
DCGSERIYGAVVR---IG-NSDVIVNNTQCGE-------------------------III
SDNI-----------------------------DSNA-----------------------
-----------------------AVEV----------------GY---ALR----GRFVS
--IQL------ENRT----DY-L----S----------L---CE----------V-EVIG
EASRIRVPCGGQITETDISNKT-IIRDCIIPILGRYVIVQ-NDPYRE-------------
----------DYLHFC---E--VKVS--------------AKEFVPSE------FTDPLK
T----------RKFVMF-EPQEEKTLPVN---------------------STDVFSNLLL
NKCLSECLNWKR----------Y-------E-----------CHS-FD------------
-------------F-------------HF-D--------------------S----KT--
-------CRIHRQ------KA--------GM-------------DGLHVVHSPG------
----------T-----------------V------------Y-Y-Q---K----------
------------------------------------------------------------
------------------------------------------------------------
--------------LRMSGMKNRYSVL--IVT----------------------------
------------------------------------------------------------
------------------------------------------------------------
------------------------------------------------------------

------------------------------------------------------------
------------------------------------------------------------
------------------------------------------------------------
------------------------------------------------------------
------------------------------------------------------------
------------------------------------------------------------
------------------------------------------------------------
------------------------------------------------------------
------------------------------------------------------------
------------------------------------------------------------
------------------------------------------------------------
------------------------------------------------------------
------------------------------------------------------------
------------------------------------------------------------
------------------------------------------------------------
------------------------------------------------------------
------------------------------------------------------------
------------------------------------------------------------
------------------------------------------------------------
--------
>Placozoa_TadNaC9_MK547550_19_180
------------------------------------------------------------
----------M-------------------------------------------------
------------------------------------------------------------
------------------------------------------------------------
------------------------------------------------------------
------------------------------------------------------------
------------------------------------------------------------
------------------------------------------------------------
------------------------------------------------------------
------------------------------------------------------------
-----------------------ETANND--------------I----------------
-----------------------------------------QS-----T-----------
-NL--EKDFA-NATSYHGFGQFY---------K-PQ-------------TTH---YR---
--RYFWM----LLTVVATSACILQSSRIVI-AAFQ----------Y---PTKITS----K
-V-RY-S----------N-SSVF-----PAV-----------------------TICNSN
YL------DKT-----KMDPDI------------------L-----------------A-
---------VV-------------------------------NSVFS-------------
---------------------------------NY-------------------------
------------------------------------------------------------
------------------------------------------------------------
-----------------N------------------------------------------
------------------------------------------------------------
------------------------------------------------------------
------------------------------------------------------------
------------------------------------------------------------
------------------------------------------------------------
------------------------------------------------------------
------------------------------------------------------------
------------------------------------------------------------
------------------------RSALDLVNLGLTPE------IIAARF-----GQHA-
---------------------------------DYGFFA---------------------
------------------------------------------------RAF----GFKLK
--DII------LECT----YH-N----F----------N---C-----------L-----
----------------------------------------------N-------------
------------------------------------------------------------
------------------------------------------------------------

-----------S----------F-------T-----------EVV-EPNFG--------L
CYQFN-------AGILQRGGNGN---GGN-N--------------------S----SF--
-------HRQVGR------GP--------SF-------------GLQLVLDAEQY-----
----------K-----------------Y------------S-T-----LD---------
------------------------------------------------------------
------------------------------------------------------------
------FFYRYQAGFIIAIH-DQFETS--SMTINEISVGVGRYTSISLHQT------L--
-----IEKY-----------LKPRW--GE-C--DDQ--------E-------LA----FY
------------------R-KYTREACKQEKESLYVM----NL--CKCRFKE----V---
------------S-E--ST-----------------------------------------
---------VNY-----T---ICTT-----------------------------------
------------------------------------FDT---ITCVI---------P---
Q-L-------------------DLAS-R-------NY-------------------RISN
------------SCK-IACARKL---FPKT-ISS------------------SSI-GT--
-------LAY------GNI-L-----------NK--------------------------
------------------------------------------------RL----------
------------------------------------------------------------
-D---------------L------------------STKLADMQGLGLL-----------
----------------------PSSY-TVQ------------------------------
-----D-------YI-----------R------DNIVQLDVFFSDLAHTTVQ-QEQDITP
EEV-----------------LSNIGGQLGLFIGVSMLTVCEIVFYIFDK-----------
--------MQYW--P----------------Q-------RR-------------------
NR------------------------------DN-------------------R------
-----------------------------------------QTTPVDFEM----------
--------------------KKSKVNN---------------------NANT--------
------------------------------------------------------------
------------------------------------------------------------
------------------------------------------------------------
----------------------------------------------------------IG
VMGYGAET
>Protostome_Lophotroco_brch_Lingula_anatina_comp136986_c0_seq2_p1_comp136986_
c0__comp136986_c0_seq2_p1__ORF_type_complete_len_518___score_46_16_comp136986
_c0_seq2_175_1728__644_183
------------------------------------------------------------
----------M-------------------------------------------------
------------------------------------------------------------
------------------------------------------------------------
------------------------------------------------------------
------------------------------------------------------------
------------------------------------------------------------
----------------TEAC-------------EKMVLQTIIVE------------PM--
------------------------------------------------------------
---------------------------------------------------------DKC
KG---------------------LEKDKV-------------------------------
---SQDITM----------------------E---------EE-----E-----------
-SL--EREFA-TGTTAHGFSRAA---------E-TK--------------SI---FR---
--RSAWI----LLILGGLGMATWMISLRVI-NYFQ----------Y---DTSTEI----T
-L-TF-E----------P-RMAF-----PAV-----------------------TICNFN
RY------MRR-----NLNAED------------------Y-----------------A-
---------II------------S-LTEDVYSGYD-SGYSYGQYYYN----YYY------
---------------------------------DYSG-----------------------
------------------------------------------------------------
------------------------------------------------------------
-----------------DGES---------------------------------------
------------------------------------------------------------
------------------------------------------------------------
------------------------------------------------------------

-------------------------SPSTT------------------------------
------------------------------------------------------------
------------------------------------------------------------
------------------------------------------------------------
----TQPTNSNF------------------------------------------------
-----------------------NESWANISASY---------------------PNGF-
---------------------------------DFGNFT---------------------
------------------------------------------------LEK----GWKLD
-NISV------LECT----FK-G----I----------K---C-----------NLK---
----------------------------------------------K-------------
------------------------------------------------------------
------------------------------------------------------------
-----------D----------F-------I-----------HTF-S-TYG--------N
CYTYN-------PS-------------LN-S--------------------N----RS--
-------RYQDQP------GL--------GN-------------GLHLTIDVEQS-----
----------E-----------------Y------------T-E-S---LP---------
------------------------------------------------------------
------------------------------------------------------------
-------DGSAQTGVIFQVH-PQWEPP--LVESKGLGASPGFHAFGALKRQ------E--
-----TVNI-----------EPP-W--GK-C--NKS-L------T-------LL----HF
------------------D-QYTFSGCIIECKLQKIE----TE--CGCKPVQ----Y---
------------P-G---------------------------------------------
----------TY-----R---ICDP-----------------------------------
------------------------------------AEM---MYCVK---------P---
L-L-------------------KTIN-Q-------AF--------S-----------DHC
H-----------MCT-VPCNSTQ---YNVQ-LSY------------------TTI-PN--
-------NNI------EAA-F-----------AT--------------------------
------------------------------------------------KY----------
------------------------------------------------------------
-N---------------LP-----------------------------------------
---------------------------TGS------------------------------
-----N-------YI-----------K------DNIVALDIYYEELNLETME-QKKAVEE
SGL-----------------LSDIGGQLGLFMGFSALTFLEFFEYIILK-----------
--------FRRI--------T----------Q-------RK-------------------
KRI-----------------------------KP--------------------------
------------------------------------------------------------
------------------------------------------------------------
------------------------------------------------------------
------------------------------------------------------------
------------------------------------------------------------
------------------------------------------------------------
------LA
>Protostome_Lophotroco_brch_Lingula_XP_023930484_1_ASIC_channel_Lingula_anati
na
------------------------------------------------------------
----------M-------------------------------------------------
------------------------------------------------------------
------------------------------------------------------------
------------------------------------------------------SEKMVR
RRSSAFEVQME-------------------------------------------------
-TLEENKTKESTES----------------------------------------------
------------------------------------------------------------
------------------------------------------------------------
------------------------------------------------------------
-----------------------KASKES-------------------------------
-------------------------------G---------SV-----S-----------
-EG--LEQFG-EDTDFHGLKRVF---------R-RD-------------YTL---LR---

--RTFWL----FFFLAGLTGFIANTVDRLS-YYLQ----------Y---PHSSEL----D
-V-MY-D----------M-ELEF-----PAV-----------------------TICNMN
SY------RLS-----ALTDVD------------------I-----------------L-
---------HF------------G-EKL---------------HILD----E--------
---------------------------------NR-------RLIH--------------
------------------------------------------------------------
------------------------------------------------------------
------------------------------------------------------------
------------------------------------------------------------
------------------------------------------------------------
------------------------------------------------------------
--------------------------PEYY------------------------------
------------------------------------------------------------
------------------------------------------------------------
------------------------------------------------------------
--------NQSW------------------------------VDW---------------
------------------------VHGINWTELAANDD-----------------EENF-
---------------------------------DVLEFI---------------------
------------------------------------------------RRT----GHQLE
--DMI------LLCS----WK-G----Q----------H---C-----------G-P---
----------------------------------------------E-------------
------------------------------------------------------------
------------------------------------------------------------
-----------N----------F-------T-----------SVF-T-HFG--------I
CYTFN-------AD-------------HS-Y--------------------K--------
-------YVSRKA------GA--------GN-------------GLKLYINIEEE-----
----------E-----------------Y------------L-T-S---DV---------
------------------------------------------------------------
------------------------------------------------------------
-------LAGQDAGLKMVIH-AQEEPP--FVKELGFGVLPGDHHFIAIQKK------Y--
-----VHNL-----------VPP-W--GN-C--YEG--------K-------LK----YY
------------------S-HYSVPACRIECETDTIV----KE--CGCKLAE----M---
------------P-G---------------------------------------------
----------NT-----S---VCLG-----------------------------------
------------------------------------HMY---MGCAY---------P---
A-L-------------------EEVE-H-------SD---------------------LC
------------SCQ-NPCDMTT---YRRD-ISS------------------VRL-RD--
-------TTL------DII-A-----------EN--------------------------
------------------------------------------------------------
------------------------------------------------------------
-N----------------------------------------------------------
-----------------------PQV-KRE------------------------------
-----T-------LS-----------R------DNLLVLNIYYEELCYETIR-QIKAYSI
PAL-----------------LSDIGGQMGLFIGASVLTLLHVIEAVGAV-----------
--------VGGK--------F----------L-------KT-------------------
TK------------------------------QG-------------------R------
-----------------------------------------VSTTR--------------
------------------------------------------------------------
------------------------------------------------------------
------------------------------------------------------------
------------------------------------------------------------
------------------------------------------------------------
---VQSLK
>Deutero_Ambulac_Apla_gbr54_20_t1_645_184
------------------------------------------------------------
----------M-------------------------------------------------
------------------------------------------------------------

```
------------------------------------------------------------
------------------------------------------------------------
------------------------------------------------------------
------------------------------------------------------------
----------------THEA-------------SRQ------------------------
------------------------------------------------------------
------------------------------------------------------------
------------------------------------------------------------
-------------------------------R------H--GK-----S-----------
-SL--EREFA-ESTSLHGIAKVF---------H-AS--------------RL---LI---
--RFVWV----AIMLTCLCVCWQISDRFH-RFLQ-----------Y---KANTEI----S
-V-EY-T----------R-DLDF-----PAV-----------------------TICNFN
RY------RSS-----ALTESD------------------K-----------------T-
---------YL------------P-YII---------------WAND----Y--------
---------------------------------DY-------DSFG--------------
------------------------------------------------------------
------------------------------------------------------------
-----------------DSQG---------------------------------------
------------------------------------------------------------
------------------------------------------------------------
------------------------------------------------------------
------------------------------------------------------------
------------------------------------------------------------
------------------------------------------------------------
------------------------------------------------------------
--------GVEY------------------------------WDMVADDGRDN-------
-------------------------------------------------------DTHF-
---------------------------------NYTDFT---------------------
------------------------------------------------LRA----GFLMD
-DITL------RQCE----WR-R----K--------TKS---C-----------S-P---
----------------------------------------------A-------------
------------------------------------------------------------
------------------------------------------------------------
-----------N----------F-------S-----------HVF-T-SFG--------N
CWTFN-------SG-------------KT-G-------------------------PI--
-------LKETQA------GS--------GN-------------GVRMLIDIQQS-----
----------E-----------------Y------------T-E-T--------------
------------------------------------------------------------
------------------------------------------------------------
------VDGNIPAGLKVLIH-DQATPP--FVESSGLAISPGVYAFIGVRKQ------Q--
-----YLNL-----------EHP-Y--GK-C--NAS-K------T-------LT----RF
------------------S-HYTREGCKIECRARAIL----EK--CNCRLVR----H---
------------P-G---------------------------------------------
----------NE-----T---ECSP-----------------------------------
------------------------------------QQT---SGCAM---------S---
T-L-------------------GALV-G-------GE--------D-----------NPC
------------DCP-VPCNYTT---FTTS-LST------------------ASI-PS--
-------NSV------AAH-L-----------QG--------------------------
------------------------------------------------LT----------
------------------------------------------------------------
-ETSYS------------------------------SDYSYSSFSVP-------------
-------------------YPAQLDL-FEE------------------------------
-----G-------YI-----------P------KNVILLDVFYEDINFERYE-QSEAITP
SAL-----------------ISDIGGQLGLFLGASFITLAEILSYFGKK-----------
--------IDFW--------I----------Q-------RV---------------LGK-
HR------------------------------RL--------------------------
--------L--------------------------------ERQQD--------------
```

------------------------NRH---------------------GRDVPL------
------------------------------------------------------------
--------------KSVRID----------------------------------------
------------------------------------------------------------
---------------------------------------------------------NGN
HRTLWADV
>Deuterostome_chordata_PetMar_tr_S4RU13_S4RU13_PETMA_Uncharacterized_protein_
OS_Petromyzon_marinus_OX_7757_PE_4_SV_1_653_189
------------------------------------------------------------
------------------------------------------------------------
------------------------------------------------------------
------------------------------------------------------------
------------------------------------------------------------
------------------------------------------------------------
------------------------------------------------------------
------------------------------------------------------------
------------------------------------------------------------
------------------------------------------------------------
------------------------------------------------------------
------------------------------------------------K-----------
-AA--HAAFV-GRAKIHGVRHFL---------W-PR-------------HAV---TR---
--RILWL----LAFMAALGLLLAWSADRIQ-HLLS----------R---PAHTRV----S
-T-SW-V----------R-GLPF-----PAV-----------------------TFCNNN
PV------RFP-----RLTKSD------------------L-----------------Y-
---------VA------------G-MWL---------------GLLT----H--------
---------------------------------AE-------S-----------------
------------------------------------------------------------
------------------------------------------------------------
-------------QQPQA-T-FAATA----------------------------------
------------------------------------------------------------
------------------------------------------------------------
------------------------------------------------------------
---------------ANLTA-----NPHMM------------------------------
------------------------------------------------------------
------------------------------------------------------------
------------------------------------------------------------
-----AGLSEPR------------------------------RQW---------------
------------------------FQ--RMSDFS----------RFLPPR-----EAPG-
---------------------------------ISQGLL---------------------
------------------------------------------------DRL----GHPLD
--EMV------LSCR----FC-G----Q----------D---C-----------K-P---
---------------------------------------------Q--------------
------------------------------------------------------------
------------------------------------------------------------
-----------D----------F-------I-----------TVY-T-RYG--------K
CYTFN-------SG-------------KG-G--------------------R----PL--
-------LTTVKG------GM--------GN-------------GLEIMLDIQQE-----
----------E-----------------Y------------L-P-VWGEND---------
------------------------------------------------------------
------------------------------------------------------------
-------ETSFEAGIKVQIH-SQDEPP--LIDQLGFGVSPGFQTFVSCQEQ------R--
-----LIYL-----------PPP-W--GD-C--KST-PISS---D-------------FF
------------------E-TYSITACRIDCETRYLV----EN--CNCRMVH----M---
------------P-G---------------------------------------------
----------DA-----P---YCTP-----------------------------------
------------------------------------EQY---TECAR---------P---
V-L-------------------DVLV-K-------RD--------N-----------DFC

```
------------LCE-TPCNMTR---YGKE-LSM------------------VKI-PS--
-------KAS------AKY-L-----------AK--------------------------
------------------------------------------------KY----------
------------------------------------------------------------
-N----------------------------------------------------------
---------------------------KTE------------------------------
-----A-------YI-----------A------ENVLVLDIFFEALNYETIE-QKKAYEV
AGLLE---------------TGDIGGQMGLFIGASILTILEIFDYVYEL-----------
--------IRDK--------I----------L-------FY----------------FR-
AK------------------------------KK-------------------Q------
------------------------------------------------------------
-----------------------KTES---------------------KDTG--------
------------------------------------------------------------
--------------------------------------------------PCEALRGHSE
------------------------SAGYAASMLPPPHHHHQHH-----------------
---------------------AL-------------------------------------
--------
>Placozoa_HhoNaC9_TR3133_c0_g1_i7_m_7432__Hoilungia_hongkongensis__26_190
------------------------------------------------------------
----------M-------------------------------------------------
------------------------------------------------------------
------------------------------------------------------------
------------------------------------------------------------
------------------------------------------------------------
------------------------------------------------------------
------------------------------------------------------------
------------------------------------------------------------
---------------------------------------------------------NVK
V----------------------DDSQNI-------------------------------
-----------------------------------------EK-----P-----------
-NL--EKEFA-EITSFHGFNQFY---------R-PK-------------TST---YR---
--RYFWI----LLTLAATAVCIVQSSRILA-TAYT----------F---PTKISS----K
-I-RF-K----------D-KAIF-----PAV-----------------------TICNVN
LF------QRT-----KVDPNS------------------K-----------------S-
---------VM-------------------------------NVIYG-------------
------------------------------------------------------------
------------------------------------------------------------
------------------------------------------------------------
-----------------S------------------------------------------
------------------------------------------------------------
------------------------------------------------------------
------------------------------------------------------------
------------------------------------------------------------
------------------------------------------------------------
------------------------------------------------------------
------------------------------------------------------------
------------------------------------------------------------
------------------------TTKVNLTQLGISTA------QLDAAY-----GAHA-
---------------------------------DLADFA---------------------
------------------------------------------------EFY----GYELK
--NAM------IQCK----FH-N----Q----------D---C-----------S-----
----------------------------------------------E-------------
------------------------------------------------------------
------------------------------------------------------------
-----------N----------F-------T-----------EVF-DPNFG--------R
CFQFN-------AGQIAR--------ADN-N--------------------TANGPSF--
-------YEQEGR------GS--------RF-------------GLQLLLNAEQY-----
```

----------K-----------------Y------------T-R-----LD---------
------------------------------------------------------------
------------------------------------------------------------
------FTNRFEAGFLIAVH-DQYEPH--GMASNGISVGVGAYTGIVLQEN------L--
-----NMKY-----------LKPRW--GE-C--DHQ--------V-------LK----LF
------------------T-HYTREACKYEKEAFYVM----EL--CKCRLKS----V---
------------P-E--SA-----------------------------------------
---------INL-----T---LCTT-----------------------------------
------------------------------------TDS---LTCTI---------P---
Q-L-------------------ERAS-R-------NY--------S-----------LSR
------------SCR-IECYRKL---YPKT-IST------------------STL-AS--
-------LAY------ANY-L-----------DS--------------------------
------------------------------------------------TF----------
------------------------------------------------------------
-Q---------------I------------------SSQISLLQSYGLV-----------
----------------------PTPY-TPI------------------------------
-----Q-------FV-----------S------DNIVEVDVYFADLIDRIVE-QEPDASV
EVV-----------------LSNIGGQLGLLVGVSLLTVCELIFYVFDK-----------
--------IKYW--------L----------N-------GR-------------------
SP------------------------------RT-------------------S------
-----------------------------------------MTAQHDSD-----------
------------------------------------------------EKNT--------
------------------------------------------------------------
------------------------------------------------------------
------------------------------------------------------------
------------------------------------------------------------
-----------------------------------------------------------A
ELTTVNGW
>Placozoa_HhoNaC3_TR31058_c1_g1_i2_m_67045__Hoilungia_hongkongensis__25_191
------------------------------------------------------------
----------M-------------------------------------------------
------------------------------------------------------------
------------------------------------------------------------
------------------------------------------------------------
------------------------------------------------------------
------------------------------------------------------------
-----------------ESS-------------PSRPRK---------------------
----------------------------------------------------------PS
IDFSTLNTTI-----------STTGHGSQED--------------------------DDS
Q----------------------QHRSSL-------------------------------
-------------------------------N---------IS-----Y-----------
-QH--EEEFA-KSTSCHGISHVF---------D-DS-------------SSN---MR---
--RLLWM----ILTVSMTVICLIQCIDRVL-YLAS----------N---PTRIVL----E
-Y-NM-P----------H-QIRF-----PAV-----------------------TICNYN
RF------RNS-----TINQSN------------------R-----------------H-
---------RL------------P-HLL---------------RLMS-------------
------------------------------------------------------------
------------------------------------------------------------
------------------------------------------------------------
------------------------------------------------------------
------------------------------------------------------------
------------------------------------------------------------
------------------------------------------------------------
--------------------------PWH-------------------------------
------------------------------------------------------------
------------------------------------------------------------
------------------------------------------------------------
----TQSTARQL------------------------------NSR---------------

------------------------VQAINTT------------------------HDRQ-
---------------------------------DDLKLL---------------------
------------------------------------------------AQV----GEQAD
--IFI------KYCI----WN-N----Q--------PDS---C-----------S-A---
----------------------------------------------E-------------
------------------------------------------------------------
------------------------------------------------------------
-----------N----------F-------T-----------LAY-T-EYG--------N
CFTFN-------GD-------------LV-G--------------------N----AT--
-------LYQKHA------GR--------SH-------------GLRLVMNIEAD-----
----------E-----------------Y------------T-T-M--------------
------------------------------------------------------------
------------------------------------------------------------
-------NPEPDIGLKFRIH-EPDEPP--DIEARGVAVPAGYHAFARLKYN------E--
-----AIFL-----------KEP-F--GK-C--NSR--------P-------LQ----YY
------------------H-HYTRTGCKLECKTNRSI----EI--CGCRTLY----M---
------------P-G---------------------------------------------
----------SA-----P---ICTD-----------------------------------
------------------------------------KQM---NQCFI---------H---
N-L-------------------DKIE-N-----------------------------GEC
------------YCP-THCIWSS---YDPK-MTF------------------AIM-PT--
-------ETI------TRE-E-----------VI--------------------------
------------------------------------------------AL----------
------------------------------------------------------------
-G---------------I------------------SADELQANT---------------
----------------------VINE-SLT------------------------------
-----N-------HL-----------R------KNFIFLDIYMEKLKYRTAK-QDLSYTF
NAL-----------------LSDIGGQLGLFIGASILTMLEIGEYLMTK-----------
--------ILSL--------C----------K-------TL-------------------
HR------------------------------PT-------------------K------
-----------------------------------------QSRNE--------------
-----------------------TADF---------------------IQTS--------
---------NLKN-----------------------------------------------
----------------NPVP----------------------------------------
------------------------------------------------------------
------------------------------------------------------------
-------V
>Placozoa_TadNaC3_MK547544_13_193
------------------------------------------------------------
----------M-------------------------------------------------
------------------------------------------------------------
------------------------------------------------------------
------------------------------------------------------------
------------------------------------------------------------
------------------------------------------------------------
----------------DRNF-------------QQRPRK---------------------
----------------------------------------------------------PS
VDFSTITTIT-----------TSTKDQSQDS--------------------------DFF
S----------------------QRQSPL-------------------------------
-------------------------------N---------IS-----Y-----------
-QH--EEDFA-KRTSCHGISHVF---------D-DE-------------TPK---LR---
--RLSWV----FLTLAMATICIIQCADRMI-YLIS----------N---PTKIVI----K
-N-EI-P----------P-RIRF-----PTV-----------------------TICNFN
RF------RNS-----TINESN------------------H-----------------Y-
---------RL------------Q-YLL---------------RLMN-------------
------------------------------------------------------------
------------------------------------------------------------

------------------------------------------------------------
------------------------------------------------------------
------------------------------------------------------------
------------------------------------------------------------
------------------------------------------------------------
-------------------------APQ--------------------------------
------------------------------------------------------------
------------------------------------------------------------
------------------------------------------------------------
----APSKRSEF------------------------------SFR---------------
------------------------NDLVNYS------------------------RNTN-
---------------------------------DDLDLL---------------------
------------------------------------------------SKV----GIQRQ
--ILI------RYCN----WD-N----Q--------PHS---C-----------S-A---
----------------------------------------------E-------------
------------------------------------------------------------
------------------------------------------------------------
------------------------------------------------------------
-----------N----------F-------T-----------LTY-T-KYG--------N
CFLFN-------GQ-------------SA-D--------------------N----EA--
-------LYQKRI------GR--------SH-------------GLRLAFNVEAD-----
----------Q-----------------Y------------T-T-M--------------
------------------------------------------------------------
------------------------------------------------------------
-------NPEPDVGIKFRIH-EPDEPA--DIEAQGVAIPPGYHAYVRLKYN------E--
-----AKFL-----------RKP-I--GK-C--DSR--------P-------LQ----YY
------------------D-RYTRSGCKLECKTNKSI----EV--CGCRALY----M---
------------P-G---------------------------------------------
----------SA-----P---FCTA-----------------------------------
------------------------------------HQI---NQCMT---------N---
N-I-------------------EEIN-N-----------------------------SSC
------------YCP-TACLWTS---YDPA-VSY------------------SMT-PT--
-------ETI------TSE-E-----------AP--------------------------
------------------------------------------------LF----------
------------------------------------------------------------
-G---------------L------------------NLDEVDANR---------------
----------------------VLNS-SIS------------------------------
-----R-------YM-----------R------KNYIFLDIFFERLRYNTAI-QELGYTF
NAF-----------------LSDIGGQLGLFIGASVLTMLEIGEYFLTK-----------
--------LKNL--------C----------K-------II-------------------
KR------------------------------PT-------------------S------
-----------------------------------------QNEYE--------------
-----------------------IAPA---------------------SPNT--------
---------TFEE-----------------------------------------------
----------------NPIP----------------------------------------
------------------------------------------------------------
------------------------------------------------------------
-------V
>Deu_Ambulacrar_hemi_Ptyfla_40v0_9_20150316_1g15265_t1_scaffold10904_cov157_6
56_194
------------------------------------------------------------
----------MSSTV-------Y-------------------------------------
------------------------------------------------------------
------------------------------------------------------------
------------------------------------------------------------
-------------------CDRYSDGALQ--------------FDLQPVATATLAKPKTR
KSF-AVTSFRDRGY----------------------------------------------
-------HSKV-EVIKEEEK-------------KGDRESDS-------------------

------------------------------------------------------------
-------------------------RSVGQK--------------------------DTC
D----------------------DCCCC--------------------------------
--------R----------------------T---------FY-----S-----------
-ER--FRQFS-EDTTLHGLRYAV---------G-PH-------------VGT---SR---
--RIFWT----LLMIGMIVSSCIYLYLCWV-KFLA----------F---PVKTVV----S
-M-NY-K----------T-EISF-----PAI-----------------------TICNYN
HY------RKS-----YVNGTR------------------F-----------------E-
---------DW------------V------------------RQRY--------------
------------------------------------------------------------
------------------------------------------------------------
------------------------------------------------------------
------------------------------------------------------------
------------------------------------------------------------
------------------------------------------------------------
------------------------------------------------------------
--------------------------P---------------------------------
------------------------------------------------------------
------------------------------------------------------------
------------------------------------------------------------
--------TFNF------------------------------SDD---------------
---------------------------EPPVELDED-------------------FLRI-
---------------------------------NRTWFE---------------------
------------------------------------------------ISA----AHDKW
--NMI------YQCT----FG-R----S--------KIP---C-----------D-A---
----------------------------------------------S-------------
------------------------------------------------------------
------------------------------------------------------------
-----------N----------F-------T-----------LTL-T-DFG--------V
CYTFN-------GG-------------KE-S--------------------ED---NV--
-------LHISNG------GS--------EH-------------GLRLRLFVEQF-----
----------E-----------------Y------------T-Y-S---EN---------
------------------------------------------------------------
------------------------------------------------------------
----------SAAGFKILAH-RQNDVP--MIRNFGFAIPPGTESLIGLKMI------Q--
-----EHNL-----------PAP-YV-TR-C--SNE--------K-------LE----YF
------------------E-VYTHSNCVRELNTNAVV----EN--CGCREPY----M---
------------P-G---------------------------------------------
----------SA-----R---VCNY-----------------------------------
------------------------------------NES---VKCAV---------P---
V-I-------------------ETLH-T-------DK-----------------------
------------DCP-VACSNIG---YETR-VSY------------------ASF-PG--
-------RHI------VKD-L-----------ED--------------------------
------------------------------------------------RL----------
------------------------------------------------------------
-N----------------------------------------------------------
---------------------------LTQ------------------------------
-----Y-------DI-----------R------ENIADLKIYFEEISYEEIR-QEKGYAF
PDL-----------------IGDSGGELGLCLGASLLTMCEIVEFFLMW-----------
--------IWRK--------M----------T-------RF-------------------
QRGVG---------------------------PN--------------------------
------------------------------------------------------------
------------------------------------------------------------
------------------------------------------------------------
------------------------------------------------------------
------------------------------------------------------------
------------------------------------------------------------

-----LKR
>Protostome_Lophotroco_annelid_Pdum_comp415688_c0_seq3__659_196
------------------------------------------------------------
----------M-------------------------------------------------
------------------------------------------------------------
------------------------------------------------------------
------------------------------------------------------------
------------------------------------------------------------
------------------------------------------------------------
------------------------------------------------------------
------------------------------------------------------------
------------------------------------------------------------
-----------------------KYTKCL-------------------------------
-KTGKSNEP----------------------L---------TY-----T-----------
-EV--TNNFA-DNTNCHAIGSIK---------R-SE--------------SA---LR---
--KSFWI----ILFLTGAALTIHQICEIFE-TYFS----------Y---PVSINV----N
-V-TY-E----------K-SLVF-----PTI-----------------------AICNYN
MV------RKS-----QLGLMK--------G------SEAL-----------------D-
---------LL-------------------------------TYTVG----G--------
---------------------------------DP-------------------------
------------------------------------------------------------
------------------------------------------------------------
-----------------DMS----------------------------------------
------------------------------------------------------------
------------------------------------------------------------
------------------------------------------------------------
------------------------------------------------------------
------------------------------------------------------------
------------------------------------------------------------
------------------------------------------------------------
------------------------------------------------------------
----------------------------QWENS----------------------EGRAE
------------------------------LRHKFLNSW---------------------
------------------------SQL----------------NESSKAAQ----GHQIE
--EML------LDCS----YN-E----K----------R---C-----------Y-P---
----------------------------------------------D-------------
------------------------------------------------------------
------------------------------------------------------------
-----------D----------F-------T-----------HFF-NPKFG--------N
CYMFN-------SN-------------WK-N--------------------Q----NV--
-------QVSHQP------GP--------FY-------------GLKLELFIDQD-----
----------E-----------------Y------------I-P-E---LT---------
------------------------------------------------------------
------------------------------------------------------------
----------ESAGVVIAIH-DQDVMP--FPEEDGVLITPDAQSNIAIVVE------K--
-----IQRM-----------QSP-YP-SD-C--LNV--------SRKE----AYD-YSAY
----SEL--YP-------V-RYTHHACYKTCLQAHLF----KE--CNCGDPE----I---
------------P-M--EG-------------------VALYNL----------------
-----VDDKENIR----T---VCT------------------------------------
------------------------------------KDQ---QACME---------K---
V-H-------------------KKYL-N-------GE--------L------------KC
------------NCS-VACIDTV---FEKT-VST------------------LQW-PN--
-------ANY------EDL-Y-----------YS--------------------------
------------------------------------------------RL-A--------
----------------------------------TR------------------------
-N-------------PQM------------------KSVLKSHS----------------
---------------------------QSG------------------------------

-----T-------NP-----------G------TNLVKLLIYYSEFNFKAIE-EYASYPV
SSL-----------------LSDLGGSIGFYLGISIIAAFELIEYLFDL-----------
--------FLVT--------C----------Q-------KS-------------------
RK------------------------------KD--------------------------
----------------------------------------DNRNAKSS------------
------------------------------------------------------------
------------------------------------------------------------
------------------------------------------------------------
---------------------------------P--------------------------
----------------------------------------------------ESRILNDL
ENEREERQ
>Protostome_Lophotroco_annelid_Pdum_comp408485_c0_seq2__664_198
------------------------------------------------------------
----------M-------------------------------------------------
------------------------------------------------------------
------------------------------------------------------------
------------------------------------------------------------
------------------------------------------------------------
------------------------------------------------------------
------------------------------------------------------------
------------------------------------------------------------
------------------------------------------------------------
------------------------------------------------------------
--STSKEQQ----------------------D---------GV-----A-----------
-VC--MDSCA-EDSSTQCIPQII---------R-AK--------------NR---LR---
--QLTWV----LITVTLVTAWIWHSCFTIG-KYLK----------F---DVNVGT----K
-I-IP-S----------R-QLAN-----PAV-----------------------TVCNKN
AY------HGV-----KADEFL------------------------------------T-
---------EM---------GQRTKSALKN--------NLENASLYG-------------
------------------------------------------------------------
------------------------------------------------------------
------------------------------------------------------------
------------------------------------------------------------
------------------------------------------------------------
------------------------------------------------------------
------------------------------------------------------------
--------------------------PSWFLAWVEALKS---------------------
------------------------------------------------------------
------------------------------------------------------------
------------------------------------------------------------
------------------------------------------NKNVYKIPS---------
-------------------------------------------------------SAQFL
Q-----------------------------MFSEFTDIY---------------------
------------------------GPT----------------YGVDYSNF----TMKYE
--QFV------QRCL----YA-G----A----------P---C-----------A-L---
----------------------------------------------KG------------
------------------------------------------------------------
------------------------------------------------------------
----------VH----------Y-------Q-----------VVF-NWLYG--------N
CFTFT-------VD----------------------------------------------
-------KNATGT------GP--------LY-------------GLQLSLFIDQE-----
----------G-----------------Y------------S-E-----LT---------
------------------------------------------------------------
------------------------------------------------------------
----------TGAGVQLVVH-DESQMP--FPDEDGISVPPGQSAFVGIRLH------E--
-----VQRL-----------GGA-F--GD-C--FDE-A------KQ------RNS-LDYF
----RKYS-YS-------T-SYSLRTCERSCFQRNMM----SQ--CQCLDNR----Y---

------------P-P--YTVK---------------------------------------
-----HIESSKY-----N---NCITQY-----LSNFNFS---------------------
---------------------------------NGLPED---LGCKD---------R---
L-E-------------------DMFN-N-------VE--------L------------GC
----------DIDCN-IPCNTTK---YEVM-VSY------------------AKW-PS--
-------EKA------MNE-T-----------KA--------------------------
------------------------------------------------FM----------
------------------------------------------------------------
-S----------------------------------------------------------
--------------------LKEAPE-MSD------------------------------
-----E-------EF-----------R------RNFLEVTVFFKTLDYVITS-ESKAVTI
LQT-----------------ISDLGGNMGLYIGASVFTFFELTQLVLDF-----------
--------MNWL--------C----------G-------KK-------------------
EK------------------------------KK--------------------------
---------------------------------------NATGEAQ--------------
------------------------------------------------------------
------------------------------------------------------------
------------------------------------------------------------
------------------------------------------------------------
----------------------------------------------------------EG
HERYEVKL
>Protostome_Lophotroco_brch_Lingula_anatina_comp147337_c2_seq2_p1_comp147337_
c2__comp147337_c2_seq2_p1__ORF_type_complete_len_527___score_4_70_comp147337_
c2_seq2_294_1874__665_199
------------------------------------------------------------
----------M-------------------------------------------------
------------------------------------------------------------
------------------------------------------------------------
------------------------------------------------------------
------------------------------------------------------------
------------------------------------------------------------
------------------------------------------------------------
------------------------------------------------------------
------------------------------------------------------------
-----------------------TTKTKFFRYDV-------------------------P
SDFDEENDT----------------------L---------TY-----Y-----------
-VL--WKGLK-ERTTSHIIPQVT---------N-AK--------------GI---FK---
--HFFWI----LITLATFVGLGYHLYTMFY-KFFE----------Y---DVGVKL----E
-I-TS-N----------S-TLKF-----PAV-----------------------TVCNEN
AF------RKS-----ALLSSP------------------------------------S-
---------KL----------------------------PTLDAFVT-------------
------------------------------------------------------------
------------------------------------------------------------
------------------------------------------------------------
-----------------NGT----------------------------------------
------------------------------------------------------------
------------------------------------------------------------
------------------------------------------------------------
-------------------------VPTTGF-----------------------------
------------------------------------------------------------
------------------------------------------------------------
------------------------------------------------------------
------------------------------------------NSSL--------------
-----------------------------------------IDSELGGR------SNRAT
------------------------------LIDKTLEDI---------------------
------------------------SEL----------------TTGEKQAL----GHQRS
--DFI------LDCE----YG-G----Y----------S---C-----------N-G---

----------------------------------------------K-------------
------------------------------------------------------------
------------------------------------------------------------
-----------T----------F-------E-----------SFY-STYNG--------N
CFIFN-------SG-------------WN-N--------------------S----SI--
-------LRATES------GP--------NF-------------GLSLTFNIEQS-----
----------E-----------------Y------------V-G-D---LT---------
------------------------------------------------------------
------------------------------------------------------------
----------QTAGVRVLVH-DQDVMP--FPEDQGFLAAPGEMTFVGIKMV------E--
-----IDRY-----------GGR-Y--TT-C--KKT--------DTF-----NIT-ENMY
----QAL--YP------SV-GYSQKACKKTCYQRTVI----ET--CKCALSI----Y---
------------P-R--VE--------------------TLFD------------------
-------STTSV-----R---TCDSL----------N-----------------------
------------------------------------ATD---VACTV---------K---
V-Q-------------------TQFI-N-------GE--------L------------NC
------------SCI-QPCSATS---FSID-TSS------------------AYW-PS--
-------DEY------ATE-F-----------LS--------------------------
------------------------------------------------NY-N--------
----------------------------------SK------------------------
-S-------------SIV------------------RKI---------------------
-----------------------YNT-SGS------------------------------
-----A-------GV-----------M------KNLAKVHIFYKDLNYEYIS-EYVYYDS
IQF-----------------LSDFGGALGLWLGWSVVTFFEFFELFMDV-----------
--------LAFS--------C----------R-----CGRK-------------------
--------------------------------RK-----------------VSR------
---------------------------------------SSVNDITLK------------
------------------------------------------------------------
------------------------------------------------------------
------------------------------------------------------------
------------------------------------------------------------
------------------------------------------------------------
---EKASF
>Cyclo_DEL4_NP_492230_2_DEgenerin_Like__Caenorhabditis_elegans__47_200
------------------------------------------------------------
----------M-------------------------------------------------
-------------GV------------------FW-------------------------
------------------------------------------------------------
------------------------------------------------------------
------------------------------------------------------------
------------------------------------------------------------
------------------------------------------------------------
------------------------------------------------------------
------------------------------------------------------------
------------------------------------------------------------
-------------------------------T---------GL-----K-----------
-YV--FTDFS-CWTSTHGVPHIG---------M-AN--------------AR---WL---
--RAFWI----LVVVVSIALFIWQFITLLT-NYLS----------F---SVNTET----T
-L-QF-A------------ERTF-----PTV-----------------------TICHLN
PW------KLS-----ETKSVD--------------------------------------
--------PDM-------------------------------SALID-------------
------------------------------AY-NS-------------------------
------------------------------------------------------------
------------------------------------------------------------
-----------------D--------------------------SSSAQFGL--------
------------------------------------------------------------
------------------------------------------------------------

------------------------------------------------------------
--------------------------P---------------------------------
------------------------------------------------------------
------------------------------------------------------------
------------------------------------------------------------
-----ASLTADR------------------------------------------------
-------------------------------------------------------QQQAS
KWTL-----------------------------MYSERL---------------------
------------------------NEK----------------QYNDAADI----AYSYD
--DMV------VSCT----YN-A----K----------T---C-----------N-I---
----------------------------------------------T-------------
------------------------------------------------------------
------------------------------------------------------------
-----------D----------F-------N-----------DFY-NPSYG--------N
CLQFN-------TD-------------GM-------------------------------
-------YSSSRA------GP--------LY-------------GLRMVMRTDQD-----
----------T-----------------Y------------L-P-----WT---------
------------------------------------------------------------
------------------------------------------------------------
----------EASGVIIDIH-MQDEIP--YPDVFGYFAPPGTASSLGVSYV------Q--
-----TTRL-----------SKP-Y--GS-C--TTK-T------K-------LKT--THY
----T-------------G-TYTVEACFRSCMQEKII----AS--CGCYYPA----Y---
------------S-H--AS-----------------------------------------
----------NT-----TQYVSCDNGV---------------------------------
----------------------------------QTLSN---LNCVDL------------
I-N-------------------SADS-T-----EFDV--------L-----------TDC
------------DCP-QPCEIDS---YGVT-VST------------------AQW-PS--
-------DSY------VPT-----------------------------------------
------------------------------------------------EC----------
------------------------------------------------------------
-N------------------------------------------------------PGG-
----------------PSGPWDASGE-SCL------------------------------
-----D-------WY-----------K------ANTILIEIYYERMNFQVLT-ESPAYTF
VNF-----------------ISDVGGQVGLFLGMSIISAIEYLVLIFLV-----------
--------FFYC--------C----------T-------HK-------------------
SR------------------------------RA-------------------E------
-----------------------------------------IEQLE--------------
-----M-----------------DIKK---------------------AKDD--------
--------VDQVAEKRK------------------------KHQKANAELYEMD------
------------------------------------------------------------
------------------------------------------------------------
------------------------------------------------------TAHDIV
PPKPHSND
>Deuterostome_chordata_Ggallus_sp_Q1XA76_ASIC1_CHICK_Acid_sensing_ion_channel
_1_OS_Gallus_gallus_OX_9031_GN_ASIC1_PE_1_SV_1_667_201
------------------------------------------------------------
----------M-------------------------------------------------
------------------------------------------------------------
------------------------------------------------------------
------------------------------------------------------------
------------------------------------------------------------
------------------------------------------------------------
---------------MDLKV-------------DEEEVD---------------------
------------------------------------------------------------
------------------------------------------------------------
------------------------------------------------------------
-------------------------------S---------GQ-----P-----------

-VS--IQAFA-SSSTLHGISHIF---------S-YE-------------RLS---LK---
--RVVWA----LCFMGSLALLALVCTNRIQ-YYFL----------Y---PHVTKL----D
-E-VA-A----------T-RLTF-----PAV-----------------------TFCNLN
EF------RFS-----RVTKND------------------L-----------------Y-
---------HA------------G-ELL---------------ALLN----N--------
---------------------------------RY-------------------------
------------------------------------------------------------
------------------------------------------------------------
-------------EIPDT-Q-TA--------------------------------------
------------------------------------------------------------
------------------------------------------------------------
------------------------------------------------------------
------------------------------------------------------------
------------------------------------------------------------
------------------------------------------------------------
------------------------------------------------------------
--------DEKQ------------------------------LEI---------------
------------------------LQ--DKANFR----------NFK--------PKPF-
---------------------------------NMLEFY---------------------
------------------------------------------------DRA----GHDIR
--EML------LSCF----FR-G----E----------Q---C-----------S-P---
----------------------------------------------E-------------
------------------------------------------------------------
------------------------------------------------------------
-----------D----------F-------K-----------VVF-T-RYG--------K
CYTFN-------AG-------------QD-G--------------------K----PR--
-------LITMKG------GT--------GN-------------GLEIMLDIQQD-----
----------E-----------------Y------------L-P-VWGETD---------
------------------------------------------------------------
------------------------------------------------------------
-------ETSFEAGIKVQIH-SQDEPP--LIDQLGFGVAPGFQTFVSCQEQ------R--
-----LIYL-----------PPP-W--GD-C--KAT-TGDS------------E----FY
------------------D-TYSITACRIDCETRYLV----EN--CNCRMVH----M---
------------P-G---------------------------------------------
----------DA-----P---YCTP-----------------------------------
------------------------------------EQY---KECAD---------P---
A-L-------------------DFLV-E-------KD--------N-----------EYC
------------VCE-MPCNVTR---YGKE-LSM------------------VKI-PS--
-------KAS------AKY-L-----------AK--------------------------
------------------------------------------------KY----------
------------------------------------------------------------
-N----------------------------------------------------------
---------------------------KSE------------------------------
-----Q-------YI-----------G------ENILVLDIFFEALNYETIE-QKKAYEV
AGL-----------------LGDIGGQMGLFIGASILTVLELFDYAYEV-----------
--------IKHR--------L----------C-------RR----------------GK-
CR------------------------------KN-------------------H------
------------------------------------------------------------
-----------------------KRNN---------------------TDKG--------
-----------VA-----------------------------------------------
----------------LSMDDVKRH------------------------NPCESLRGHPA
------------------------GMTYAANILP-------HH---------PA------
---------RGT-----FEDFTC-------------------------------------
--------
>Deu_Ambulacrar_hemi_Ptyfla_40v0_9_20150316_1g2678_t1_scaffold1096_cov89_668_
202
------------------------------------------------------------

```
------------------------------------------------------------
------------------------------------------------------------
------------------------------------------------------------
------------------------------------------------------------
------------------------------------------------------------
------------------------------------------------------------
------------------------------------------------------------
------------------------------------------------------------
------------------------------------------------------------
------------------------------------------------------------
------------------------------------------------------------
------------------------------------------------------------
------------------------------------------------------------
------------------------------------------------------------
------------------------------------------------------------
------------------------------------------------------------
------------------------------------------------------------
------------------------------------------------------------
----------MSC--------------------TVLDENNNCPDEYRECGSGE-------
-CVYQYAICDTIHNCA-DG-------------------------SDEVNCTT--------
------------------------------------------------------------
---------------TTKGENISLRT----------------------------------
------------------------------------------------------------
------------------VTEQFLDISGV-------------------------------
------------------------------------------------------------
------------------------------------------------------------
------------------------------------------------------------
----TPEILSEF----------ENGYYK--------------DNGFSGV-----------
---------------------VKGEVPPDWYGF-----------------------KTFS
STP------------------------------DYSDLR---------------------
----------------------RVLKL----------------TREEIAKY----GHQAE
--DFI------LQCT----YD-E----K----------S---C-----------S-P---
----------------------------------------------S-------------
------------------------------------------------------------
------------------------------------------------------------
-----------E----------F-------L-----------TFQ-DDHYG--------N
CFKFN-------HG-------------RG-G--------------------T----VL--
-------RDAKRQ------GA--------LY-------------GLRLTLFLEQN-----
----------E-----------------Y------------I-S-I---YG---------
------------------------------------------------------------
------------------------------------------------------------
----------RNAGTRVNIS-PLDASV--LPQDEGITIMPGTVTSIGIREI------P--
-----VDNS-----------LDI-C--FK-H--RNR--------T---------------
------------------------------------------------------------
------------------------------------------------------------
------------------------------------------------------------
------------------------------------------------------------
------------------------------------------------------------
------------------------------------------------------------
--------------------V---------------------------------------
------------------------------------------------------------
------------------------------------------------------------
------------------------------------------------------------
------------------------------------------------------------
-----------------------------------------------------------I
AGF-----------------RAR------LYIGLSVITVAEFIELVASI-----------
--------IQYM--------YCG--------Q-------RD-------------------
```

```
MKS-----------------------------RV-------------------K------
---------------------------------------SANEETKEHFSGAVA------
----------------FPGFRSFDVDV---------------------AHSTKKF-----
------------------------RIVSGKIF----------------------------
------RSVKYGTIVNLTLYNHMAEDYTT-MHWHG--IYQ-IRTPWSDGVPYITECPIPP
GGVYVYAFAAIPAENPVDQPT-----FDSGCGIPHDFYGESGTITS--------------
------MGYDGTTL---YENNAKCEFRVLNFVAFDLEEGSDFIQVIVH-FNTNESGQAKG
FHADWEAV
>Deutero_Chord_Hsap_sp_P78348_ASIC1_HUMAN_Acid_sensing_ion_channel_1_OS_Homo_
sapiens__Human__OX_9606_GN_ASIC1_PE_1_SV_3_669_203
------------------------------------------------------------
----------M-------------------------------------------------
------------------------------------------------------------
------------------------------------------------------------
------------------------------------------------------------
------------------------------------------------------------
------------------------------------------------------------
----------------ELKA-------------EEEEVG---------------------
------------------------------------------------------------
------------------------------------------------------------
------------------------------------------------------------
-------------------------------G---------VQ-----P-----------
-VS--IQAFA-SSSTLHGLAHIF---------S-YE-------------RLS---LK---
--RALWA----LCFLGSLAVLLCVCTERVQ-YYFH----------Y---HHVTKL----D
-E-VA-A----------S-QLTF-----PAV-----------------------TLCNLN
EF------RFS-----QVSKND------------------L-----------------Y-
---------HA------------G-ELL---------------ALLN----N--------
---------------------------------RY-------------------------
------------------------------------------------------------
------------------------------------------------------------
-------------EIPDT-Q-MA-------------------------------------
------------------------------------------------------------
------------------------------------------------------------
------------------------------------------------------------
------------------------------------------------------------
------------------------------------------------------------
------------------------------------------------------------
------------------------------------------------------------
--------DEKQ------------------------------LEI---------------
------------------------LQ--DKANFR----------SFK--------PKPF-
---------------------------------NMREFY---------------------
------------------------------------------------DRA----GHDIR
--DML------LSCH----FR-G----E----------V---C-----------S-A---
----------------------------------------------E-------------
------------------------------------------------------------
------------------------------------------------------------
-----------D----------F-------K-----------VVF-T-RYG--------K
CYTFN-------SG-------------RD-G--------------------R----PR--
-------LKTMKG------GT--------GN-------------GLEIMLDIQQD-----
----------E-----------------Y------------L-P-VWGETD---------
------------------------------------------------------------
------------------------------------------------------------
-------ETSFEAGIKVQIH-SQDEPP--FIDQLGFGVAPGFQTFVACQEQ------R--
-----LIYL-----------PPP-W--GT-C--KAV-TMDS---D-------LD----FF
------------------D-SYSITACRIDCETRYLV----EN--CNCRMVH----M---
------------P-G---------------------------------------------
----------DA-----P---YCTP-----------------------------------
```

```
------------------------------------EQY---KECAD---------P---
A-L-------------------DFLV-E-------KD--------Q-----------EYC
------------VCE-MPCNLTR---YGKE-LSM------------------VKI-PS--
-------KAS------AKY-L-----------AK--------------------------
------------------------------------------------KF----------
------------------------------------------------------------
-N----------------------------------------------------------
---------------------------KSE------------------------------
-----Q-------YI-----------G------ENILVLDIFFEVLNYETIE-QKKAYEI
AGL-----------------LGDIGGQMGLFIGASILTVLELFDYAYEV-----------
--------IKHK--------L----------C-------RR----------------GK-
CQ------------------------------KE-------------------A------
------------------------------------------------------------
-----------------------KRSS---------------------ADKG--------
-----------VA-----------------------------------------------
----------------LSLDDVKRH------------------------NPCESLRGHPA
------------------------GMTYAANILP-------HH---------PA------
---------RGT-----FEDFTC-------------------------------------
--------
>Chordat_hASIC3_NP_004760_1_acidsensing_ion_channel_3_isoform_a__Homo_sapiens
__5_204
------------------------------------------------------------
----------M-------------------------------------------------
------------------------------------------------------------
------------------------------------------------------------
------------------------------------------------------------
------------------------------------------------------------
------------------------------------------------------------
----------------KPTS-------------GPEEA----------------------
------------------------------------------------------------
------------------------------------------------------------
------------------------------------------------------------
-------------------------------R---------RP-----A-----------
-SD--IRVFA-SNCSMHGLGHVF---------G-PG-------------SLS---LR---
--RGMWA----AAVVLSVATFLYQVAERVR-YYRE----------F---HHQTAL----D
-E-RE-S----------H-RLIF-----PAV-----------------------TLCNIN
PL------RRS-----RLTPND------------------L-----------------H-
---------WA------------G-SAL---------------LGLD-------------
------------------------------------------------------------
------------------------------------------------------------
------------------------------------------------------------
------------------------------------------------------------
------------------------------------------------------------
------------------------------------------------------------
------------------------------------------------------------
------------------------------------------------------------
------------------------------------------------------------
------------------------------------------------------------
------------------------------------------------------------
--------PAEH------------------------------AAF---------------
------------------------LRALGRPPAPPG---------FMP-------SPTF-
---------------------------------DMAQLY---------------------
------------------------------------------------ARA----GHSLD
--DML------LDCR----FR-G----Q----------P---C-----------G-P---
----------------------------------------------E-------------
------------------------------------------------------------
------------------------------------------------------------
```

-----------N----------F-------T-----------TIF-T-RMG--------K
CYTFN-------SG-------------AD-G--------------------A----EL--
-------LTTTRG------GM--------GN-------------GLDIMLDVQQE-----
----------E-----------------Y------------L-P-VWRDNE---------
------------------------------------------------------------
------------------------------------------------------------
-------ETPFEVGIRVQIH-SQEEPP--IIDQLGLGVSPGYQTFVSCQQQ------Q--
-----LSFL-----------PPP-W--GD-C--SSA-SLNP-----------------NY
EPEPSDPLGSP--SPSPSP-PYTLMGCRLACETRYVA----RK--CGCRMVY----M---
------------P-G---------------------------------------------
----------DV-----P---VCSP-----------------------------------
------------------------------------QQY---KNCAH---------P---
A-I-------------------DAML-R-------KD---------------------SC
------------ACP-NPCASTR---YAKE-LSM------------------VRI-PS--
-------RAA------ARF-L-----------AR--------------------------
------------------------------------------------KL----------
------------------------------------------------------------
-N----------------------------------------------------------
---------------------------RSE------------------------------
-----A-------YI-----------A------ENVLALDIFFEALNYETVE-QKKAYEM
SEL-----------------LGDIGGQMGLFIGASLLTILEILDYLCEV-----------
--------FRDK--------V----------L-------GY----------------FW-
NR------------------------------QH-------------------S------
------------------------------------------------------------
-----------------------QRHS---------------------STNL--------
-----------LQ-----------------------------------------------
----------------EGLGSHRTQ------------------------VPHLSLGPRPP
------------------TP-------------P---------CAVTKTLSASH------
---------RTC-----YLVTQL-------------------------------------
--------
>Deu_Ambulacrar_hemi_Ptyfla_40v0_9_20150316_1g19294_t1_scaffold17970_cov134_6
72_205
------------------------------------------------------------
----------MF-----------------------------------------------H
TEASYAFQRKKRI-IPIIVEKDY-RPDGWLGLLTLPELYF--------------------
---------------------------DFTSDEQLEKN--------------------FD
E-----------------------------------LVRQIGDHGNLNDDGKDE--LHIS
LQRSKEGLKHHQ------------------------------------------------
----------------------------------------LP------------------
--LKSGAYNVERGKAYQKCQ-------------TNDNSE---------------------
------------------------------------------------------------
----------------------EDESDVDDK---------------------QQRVDSRT
C----------------------GDGQE--------------------------------
------RSQ----------------------A---------TL-----K-----------
-HV--LADFG-KGTTAHGIVHIT---------E-AE--------------TS---VS---
--RTVWL----AIVVAAAGIMVIQMTLLLV-QYFE----------F---GVTVKV----S
-L-VS-E----------K-MLEF-----PSV-----------------------TVCNTN
KL------RKS-----AVRNSI------------------Y-----------------N-
---------QML---------ILESNFVP--------------AYYE-------------
----------------------------CEEP-NF-------------------------
------------------------------------------------------------
-----------RCL-----------------------------------ASSE-------
-CINNLKVCDGTYDCR-D-G------------------------SDESGSI-----CE-P
K-----------------------------------------------------------
--------LKNVAR--GKATSLT---------------GPILENRDSNLAVDGYTY----
----------------TCVQT------------------------DW-------------
-----------------------IWEP---------------------------------

------------------------------------------------------------
------------------------------------------------------------
-----------------------------------------LW-----------------
----SVDLGKNYLVYKLK---ITHMATY--------------DPKLV-------------
--------NAEIR---VGLAPKAGPDNPICVK-------------------------SIR
EHE------------------------------TQTE-----------------------
-----------------------TFTV----------------RCHPVPVA----GRHVF
--IQV------KGKT----ED-M----R----------L---CE----------V-EVLA
------------------------------------------------------------
-------------------------------------------------------TDIAE
E----------LENVAYGKPAEQSS-----------------------------------
--------------------------------------------T-YY------------
-------------E-------------HV-A--------------------S----RA--
-------VDGNEN------FA--------LE-------------TGHSCIHSAKE-----
----------A-----------------G------------A-W-W---R----------
------------------------------------------------------------
------------------------------------------------------------
--------------VDLLNVYSVYKVV--MIN----------------------------
------------------------------------------------------------
------------------------------------------------------------
------------------------------------------------------------
------------------------------------------------------------
------------------------------------------------------------
------------------------------------------------------------
------------------------------------------------------------
------------------------------------------------------------
------------------------------------------------------------
------------------------------------------------------------
------------------------------------------------------------
------------------------------------------------------------
------------------------------------------------------------
------------------------------------------------------------
------------------------------------------------------------
------------------------------------------------------------
------------------------------------------------------------
------------------------------------------------------------
----------RYDCCR-ERASF----VVGY------------------------------
--------------SVLQVLSSLSY-------------------------P---------
------------------------------------------------------------
------------------------------------------------------------
--------
>Placozoa_HhoNaC11_TR28363_c2_g1_i3_m_60033__Hoilungia_hongkongensis__23_208
------------------------------------------------------------
----------M-------------------------------------------------
------------------------------------------------------------
------------------------------------------------------------
------------------------------------------------------------
------------------------------------------------------------
------------------------------------------------------------
----------------DTKQ-------------QHSTN----------------------
------------------------------------------------------------
------------------------------------------------------------
-----------------------EDSSD--------------------------------
-------------------------------S---------AS-----V-----------
-DL--EEDFA-TTTSCHGVGHIF---------T-NA------------G------SR---
--RSMWF----TITLICSVACICQCFIIVY-DASF----------F---PTRVNI----K
-L-EY-N----------I-ESLF-----PSV-----------------------TICNTN

LI------ASL-----SIGDVE------------------Y-----------------K-
---------NA------------L-TLY---------------RYYY----I--------
---------------------------------ND-------NR----------------
------------------------------------------------------------
------------------------------------------------------------
-----------------T------------------------------------------
------------------------------------------------------------
------------------------------------------------------------
------------------------------------------------------------
------------------------------------------------------------
------------------------------------------------------------
------------------------------------------------------------
------------------------------------------------------------
--------FGEL------------------------------DEA---------------
------------------------EKFFTR------------------IY-----GKNF-
---------------------------------TIHDYI---------------------
------------------------------------------------VKY----RFRIE
--DMI------ISCQ----WD-S----N----------P---C-----------G-P---
----------------------------------------------Q-------------
------------------------------------------------------------
------------------------------------------------------------
-----------N----------F-------S-----------QVV-T-DYG--------I
CYTFN-------SG-------------QG-Q--------------------H----PL--
-------LYQRAS------GS--------FH-------------GLKLILDVQQY-----
----------L-----------------Y------------P-T-Y--PIS---------
------------------------------------------------------------
------------------------------------------------------------
-------IQAPDAGIRLSIH-SFNEIP--NMDSQGVFVPPGAHSYVSITST------HL-
-----FTSL-----------KPP-W--GQ-C--GDK--------K-------LK----YF
------------------S-HYSRSACRREFEVDIAA----DM--CNCSYHH----YHHY
----------NHQ-S---------------------------------------------
----------KK-----P---ICST-----------------------------------
------------------------------------RQL---FECVL---------P---
K-L-------------------TNGE-G-------KV--------N-----------FNR
------------YCP-HPCQQSD---YPMV-LSY------------------AGI-AN--
-------YAT------LQM-A-----------HN--------------------------
------------------------------------------------------------
------------------------------------------------------------
-N-------------SVL------------------ANLMRKQK----------------
----------------------LANA-TAQ------------------------------
-----S-------FI-----------R------ENLVYLDVYFRELITSLTI-ESKATGY
AQV-----------------LSDIGGQLGLFVGASIITLCEIITYFCDR-----------
--------CKRD--------K----------R-------EK-------------------
QT------------------------------KK-------------------NN-----
------------------------------------------------------------
-----------------------ALNM---------------------TNKP--------
---------KNKE-----------------------------------------------
----------------SAIK-------------------------------RGNSSSIIR
-----------------NDSYEN-------------SENKKPP-----------------
------------------------------------------------------ASINGT
RNNPGVPV
>sp_O35240_ASIC3_RAT_Acid_sensing_ion_channel_3_OS_Rattus_norvegicus_OX_10116
_GN_Asic3_PE_1_SV_1_675_209
------------------------------------------------------------
----------M-------------------------------------------------
------------------------------------------------------------
------------------------------------------------------------

```
------------------------------------------------------------
------------------------------------------------------------
------------------------------------------------------------
----------------KPRS-------------GLEEAQ---------------------
------------------------------------------------------------
------------------------------------------------------------
------------------------------------------------------------
-------------------------------R---------RQ-----A-----------
-SD--IRVFA-SSCTMHGLGHIF---------G-PG-------------GLT---LR---
--RGLWA----TAVLLSLAAFLYQVAERVR-YYGE----------F---HHKTTL----D
-E-RE-S----------H-QLTF-----PAV-----------------------TLCNIN
PL------RRS-----RLTPND------------------L-----------------H-
---------WA------------G-TAL---------------LGLD-------------
------------------------------------------------------------
------------------------------------------------------------
------------------------------------------------------------
------------------------------------------------------------
------------------------------------------------------------
------------------------------------------------------------
------------------------------------------------------------
------------------------------------------------------------
------------------------------------------------------------
------------------------------------------------------------
------------------------------------------------------------
--------PAEH------------------------------AAY---------------
------------------------LRALGQPPAPPG---------FMP-------SPTF-
---------------------------------DMAQLY---------------------
------------------------------------------------ARA----GHSLE
--DML------LDCR----YR-G----Q----------P---C-----------G-P---
----------------------------------------------E-------------
------------------------------------------------------------
------------------------------------------------------------
-----------N----------F-------T-----------VIF-T-RMG--------Q
CYTFN-------SG-------------AH-G--------------------A----EL--
-------LTTPKG------GA--------GN-------------GLEIMLDVQQE-----
----------E-----------------Y------------L-P-IWKDME---------
------------------------------------------------------------
------------------------------------------------------------
-------ETPFEVGIRVQIH-SQDEPP--AIDQLGFGAAPGHQTFVSCQQQ------Q--
-----LSFL-----------PPP-W--GD-C--NTA-SLDP---D-------------DF
DPEPSDPLGSP--RPRPSP-PYSLIGCRLACESRYVA----RK--CGCRMMH----M---
------------P-G---------------------------------------------
----------NS-----P---VCSP-----------------------------------
------------------------------------QQY---KDCAS---------P---
A-L-------------------DAML-R-------KD---------------------TC
------------VCP-NPCATTR---YAKE-LSM------------------VRI-PS--
-------RAS------ARY-L-----------AR--------------------------
------------------------------------------------KY----------
------------------------------------------------------------
-N----------------------------------------------------------
---------------------------RSE------------------------------
-----S-------YI-----------T------ENVLVLDIFFEALNYEAVE-QKAAYEV
SEL-----------------LGDIGGQMGLFIGASLLTILEILDYLCEV-----------
--------FQDR--------V----------L-------GY----------------FW-
NR------------------------------RS-------------------A------
------------------------------------------------------------
-----------------------QKRS---------------------GNTL--------
```

-----------LQ-----------------------------------------------
----------------EELNGHRTH------------------------VPHLSLGPRPP
------------------TT-------------P---------CAVTKTLSASH------
---------RTC-----YLVTRL-------------------------------------
--------
>Deutero_Ambulac_Apla_gbr160_4_t1_678_211
------------------------------------------------------------
----------M-------------------------------------------------
------------------------------------------------------------
------------------------------------------------------------
-----------------------------------------------------------M
YRNRSADEAHL-------------------------------------------------
------------------------------------------------------------
----NNSTLSGWPTIGSVYS-------------LPTTINEGKVT----------------
------------------------------------------------------------
---------------------------------------------------------ATK
T-----------------------------------------------------------
---PAGGKN----------------------G---------AV-----G-----------
-ER--LGGFG-HASTFHGLNYVM---------N-NS-------------LSR---NR---
--RLVWF----IVVLGMTTLLVVNTIQAIL-VLLR----------H---PETSAI----S
-L-NY-V----------P-RITF-----PAV-----------------------TVCNIN
LL------RNS-----ALDESV------------------I-----------------P-
---------TL-------------------------------AAIYL-------------
------------------------------------------------------------
------------------------------------------------------------
------------------------------------------------------------
------------------------------------------------------------
------------------------------------------------------------
------------------------------------------------------------
------------------------------------------------------------
------------------------------------------------------------
------------------------------------------------------------
------------------------------------------------------------
------------------------------------------------------------
------------------------------------------EDRS--------------
-----------------------DSTGFDFGPVDALYHY----------------PNGS-
---------------------------------DSLETI---------------------
------------------------------------------------FNA----SHQIE
--DML------FRCR----WR-H----E----------K---C-----------S-A---
----------------------------------------------A-------------
------------------------------------------------------------
------------------------------------------------------------
-----------N----------F-------T----------RRL-T-DHG---------V
CYTFN-------DP-------------AD-E--------------------R----DA--
-------LQVQNP------GS--------SN-------------GLFMRLNIEQD-----
----------L-----------------Y------------T-Y-G---ES---------
------------------------------------------------------------
------------------------------------------------------------
----------TSAGIKVLLH-PQGEFP--IVKEFAFSLTPGFETSIAVRKH------M--
-----VRSL-----------RAP-YK-SN-C--TES--------T-------LR----EF
------------------PFKYSVAACHFECLVHFVV----EH--CGCRDYR----W---
------------P-A---------------------------------------------
---------VKA-----P---ICSV-----------------------------------
------------------------------------EKQ---VTCVY---------H---
Y-E-------------------DEFV-W-------DS--------------------GNC
------------SCP-MSCETTT---YESK-ISQ------------------AQW-PA--
-------KYF------DRD-L-----------QE--------------------------

------------------------------------------------KY----------
------------------------------------------------------------
-N----------------------------------------------------------
---------------------------LSH------------------------------
-----Q-------YL-----------R------DNYVDVYIFMEEIMFMEIE-QQEAYTL
EHF-----------------EGDIGGYMGLLCGMSLITVVEWVDFIIIT-----------
--------IYKR--------F----------I-------GS-------------------
KWMCS---------------------------RK------------------TR------
-----------------------------------------VMSSDDAGEL---------
--------------------ETFGEEPSYAW-----------------RARRGA------
------------------------------------------------------------
------------------------------------------------------------
------------------------------------------------------------
-------------------------------------------------------SLLGL
AQLSLAGT
>Protostome_Lophotroco_annelid_Pdum_comp409581_c1_seq8__679_212
------------------------------------------------------------
----------MFV-----------------------------------------------
------------------------------------------------------------
------------------------------------------------------------
------------------------------------------------------------
------------------------------------------------------------
------------------------------------------------------------
------------------------------------------------------------
------------------------------------------------------------
------------------------------------------------------------
----------------------------------------------------------IR
QKPTDHQNE----------------------T---------SE-----K-----------
-GI--IAEFC-GSTTGHGYYKIY---------S-AR-------------HQP---LA---
--RFFWI----FLTFFAMVCFIVHATSLGL-RYVS----------Y---PLRSDQ-----
-I-TQ-S----------E-EVHF-----PDI-----------------------TICNIY
PY------DVA-----TWVLND------------------L-----------------EP
IKK---YRSEL-------------------------DCLRANHSSIG-------------
------------------------------------------------------------
------------------------------------------------------------
------------------------------------------------------------
------------------------------------------------------------
------------------------------------------------------------
------------------------------------------------------------
------------------------------------------------------------
------------------------------------------------------------
------------------------------------------------------------
------------------------------------------------------------
------------------------------------------------------------
------------------------------------------NEKQ--------------
-------------------------------------------------------WNTLK
---------------------------------SITGIY---------------------
------------------------ANM----------------KPEEFSQL----GHKKE
--DLI------KDCV----YG-G----K----------P---C-----------N-M---
----------------------------------------------D-------------
------------------------------------------------------------
------------------------------------------------------------
-----------H----------W-------K-----------LMP-HPKLV--------N
CYTFH-------PK-------------HE-------------------------------
-------------------GL-VYR----QK-------------SLSATVYIQSD-----
----------D-------ISQQCLHSSNY----------------Q---IG---------
------------------------------------------------------------

```
------------------------------------------------------------
----------SAHGLRLSIH-PPGTAP---RKKHAMNLPPGVLVLLPLTNT------K--
-----STRP-----------GHP-Y--SD-C--VTD--------RNLVA---LKE-----
-----------------DL-NYSLTACSSLCQQQLVQ----RR--CNCSSTGE---L---
---------I--P-----------------------------------------------
--------DRND-----P--DLCKKVI-------PSDLD---------------------
----------------------------------LTFKK---MTCED---------E---
A-D-------------------EEAA-E------MDT-------------------VTEC
------------GCT-WPCELWS---YKVV-PST------------------AVW-PT--
-------ESA------TLD-I-----------LK--------------------------
------------------------------------------------TYVSAT------
------------------------------------------------------------
-NLEKCLEKEAGPMPPEL------------------KTI---------------------
-----------DHSSPIPAEYHSYLY-KAN------------------------------
-----R-------WI-----------K------GHFLRINIYMRDPRVVEYR-YQAAYGL
QEF-----------------LSDLGGSLGLWMGFSILTIVELFELIPRF-----------
--------LIRF----SN--L----------T-------TT-------------------
TQPTGGAVHNYS--------------------KE--------------------------
------------------------------------------------------------
------------------------------------------------------------
------------------------------------------------------------
------------------------------------------------------------
------------------------------------------------------------
-----------------------------------------------------------N
ASISKISH
>Deutero_Ambulac_Sakowv30041176m_681_213
------------------------------------------------------------
----------M-------------------------------------------------
------------------------------------------------------------
------------------------------------------------------------
------------------------------------------------------------
------------------------------------------------------------
------------------------------------------------------------
------------------------------------------------------------
------------------------------------------------------------
------------------------------------------------------------
-----------------------------------------------------------T
DNKLETPIR----------------------G---------TF-----C-----------
-GL--VQNQM-SNTTAHGVPRIE---------S-ST--------------NL---LR---
--RLYWV----VVFLGGVSLFLWQCSTIIS-KFYD----------R---ETTVNI----D
-M-KF-K----------P-MLSF-----PAV-----------------------TICNLN
PV------KSP-----DATPPT------------------LQ----------------DN
ST-----ANPI-------------------------------EAITN----K--------
------------------------------------------------------------
------------------------------------------------------------
------------------------------------------------------------
-----------------EFT----------------------------------------
------------------------------------------------------------
------------------------------------------------------------
------------------------------------------------------------
-------------------------MPYKAT-----------------------------
------------------------------------------------------------
------------------------------------------------------------
------------------------------------------------------------
------------------------------------------TDTGTTA-----------
------------------------DYSRNWES---------KHVAFNEK------REGFK
------------------------------LREKVQNYM---------------------
```

------------------------ASK----------------DYEDRSAY----GHSLN
--DLL------FDCT----WK-G----Y----------S---C-----------S-P---
----------------------------------------------S-------------
------------------------------------------------------------
------------------------------------------------------------
-----------N----------F-------T-----------SFF-NHMYG--------N
CYTFN-------SG-------------SL----------------------E----ET--
-------LMTSKA------GP--------LY-------------GLNLELYIEEK-----
----------E-----------------Y------------I-S-D---LQ---------
------------------------------------------------------------
------------------------------------------------------------
----------ESSGVRIVIH-DQNEMP--FPEDDGFMAAPGFLTSVGLRMI------K--
-----VTRQ-----------PDP-Y--ST-C--KTD--------ADA-----DAT-TNIF
----SEL--FS-------T-GYTRKACEKSCFSWAVT----EH--CMCTDII----Y---
------------K-Y--HSYE---------------------------------------
-----------------T---VCNST----------D-----------------------
------------------------------------PDI---VECQE---------M---
V-S-------------------LLND-N-------GE--------L------------SC
----------NVTCA-QPCEQRL---YEAT-VSN------------------VVW-PN--
-------EQY------KSA-L-----------TE--------------------------
------------------------------------------------EL-F--------
----------------------------------KF------------------------
-S-------------DDV------------------KQKLN-------------------
---------------------------DNN------------------------------
-----N-------FI-----------S------ENMVKIDIYYNELNYEYIQ-EQIAYDN
GDV-----------------VSDLGGQVGLWLGVSVITCCEFIEFMIDV-----------
--------CILL--------W----------K-------KL-------------------
TKASNL--------------------------KV--------------------------
----------------------------------------TPSKLKETKGF---------
------------------------------------------------SEEK--------
------------------------------------------------------------
------------------------------------------------------------
------------------------------SGMP--------------------------
------------------------------------------------------------
KEFQMVNT
>Protostome_Lophotroco_annelid_CAC9667732_1__Ofus_G151678_Owenia__fusiformis_
_684_214
------------------------------------------------------------
------------------------------------------------------------
------------------------------------------------------------
------------------------------------------------------------
------------------------------------------------------------
------------------------------------------------------------
------------------------------------------------------------
------------------------------------------------------------
------------------------------------------------------------
------------------------------------------------------------
------------------------------------------------------------
------------------------------------------M-----K-----------
-KL--LEDFA-GNTTAHGWGNVI---------K-ST--------------TK---IG---
--KAVWI----VITSGCCIVALYLIINVII-RYGQ----------F---DTKDRV----R
-V-TK-D------------DAVF-----PSV-----------------------TICPLS
PV------SRK-----GVVNYGKDVK---------DNF----------------------
---------EL----------------------------SAFNNLMN----TLEQYD---
------------------------------------------------------------
------------------------------------------------------------
------------------------------------------------------------

------------------------------------------------------------
------------------------------------------------------------
------------------------------------------------------------
------------------------------------------------------------
------------------------------------------------------------
------------------------------------------------------------
------------------------------------------------------------
------------------------------------------------------------
--LNISNLLDAS------------------------------QEVQ--------------
-------------------------------------------MLYNQS------FKRLN
---------------------------------TNSGQY---------------------
------------------------ENF----------------QLSQLLRY----TMTHE
--DFI------PSCK----YQ-G----N----------K---C-----------K-E---
----------------------------------------------E-------------
------------------------------------------------------------
------------------------------------------------------------
-----------P----------Y-------R-----------SGH-HAH-F--------N
CFTYN-------GP-------------SS-N--------------------I----PG--
-------LTSEIA------GP--------TA-------------GLSLLLFLDTD-----
----------E--------DFSTS----FD-------------P-D-YPTS---------
------------------------------------------------------------
------------------------------------------------------------
----------GSQGVRVVIH-DRGTLP--DPENDGFDIKAGNSVNVALSAK------R--
-----RELM-----------IPP-W--GE-C--VDH--------ND------IRY--DVW
--------------------KYSTKACTSICLQRMIY----KT--CGCILGS----L---
------------P-V--DD-----------------------------------------
-----QT--KNA-----E---FCANI----------QAV-EWLKE------DP-------
---------------------------NVTILR-QDLEQ---LNCSM---------Q---
-----------------------EYTWT-------DV-------LAQ----------EGC
------------ECK-ENCSNNK---YDIT-ISE------------------SQW-PM--
-------KGI------EFN-F-----------YC--------------------------
------------------------------------------------KLIV--------
-------------------------------KL-DNY-----------------------
EN-------------SSI------------------YDV-FHQRISQQC-----------
-----------SNNS--MENIRVVNK-QVG------------------------------
-----D-------DL-----------R------EQFIRLNVYFKDLDTKITT-QTEDFTL
GSM-----------------VSEIGGSLGLLVGMSVISMTEAFVLWYGI-----------
--------IVGL--------L----------T-------KF-------------------
KR------------------------------KS----------------KVET------
------------------------------------------------------------
----------------------LKDKM---------------------KN----------
------------------------------------------------------------
------------------------------------------------------------
------------------------------------------------------------
------------------------------------------------------------
--------
>Protostome_Lophotroco_annelid_CAC9485189_1__Ofus_G014735_Owenia__fusiformis_
_686_216
------------------------------------------------------------
------------------------------------------------------------
------------------------------------------------------------
------------------------------------------------------------
------------------------------------------------------------
------------------------------------------------------------
------------------------------------------------------------
------------------------------------------------------------
------------------------------------------------------------

------------------------------------------------------------
------------------------------------------------------------
------------------------------------------M-----K-----------
-RI--VNKFA-SNTSGHGWGMVG---------C-TN--------------SK---LG---
--KLLWV----AITLGSTIAAIIWVLTIVQ-TYAK----------F---ETKDRV----E
-L-RA-D----------V-DIVF-----PSV-----------------------TVCLLY
GF------SPK-----KIRATLTDPN---------SGY----------------------
---------------------------------------VLHNSFLK----TI--Y----
------------------------------------------------------------
------------------------------------------------------------
------------------------------------------------------------
------------------------------------------------------------
------------------------------------------------------------
------------------------------------------------------------
------------------------------------------------------------
------------------------------------------------------------
------------------------------------------------------------
------------------------------------------------------------
------------------------------------------------------------
--TNHVVLTQMT------------------------------NES---------------
--------------------------------------------ILENI------VGQAR
---------------------------------TNRGYY---------------------
------------------------ENL----------------GPSEISSL----GHNIG
--SLI------PYCM----YQ-E----S----------K---C-----------T-E---
----------------------------------------------A-------------
------------------------------------------------------------
------------------------------------------------------------
-----------D----------F-------R-----------KLV-HHEYK--------N
CYTFN-------GE-------------DV-N--------------------L----TK--
-------PIITEV------GA--------QS-------------GLSFTLYLDHK-----
----------E--------GARVT----YE-------------K-S-RAMT---------
------------------------------------------------------------
------------------------------------------------------------
----------VASGARVIIH-EKGTLP--DPDSKGFDVEPGHLINVALSVD------K--
-----RELM-----------KPP-W--GE-C--ADH--------TT------LHS--TDF
--------------------KYTRNTCKRKCMEKMIE----KQ--CGCWKDL----L---
------------P-V--DL-----------------------------------------
----NDTRDDDL-----Q---ACGKM----------NRE-EWLKT--------TRA----
---------------------------NVTVLK-EDIAR---IKCQN---------E---
-----------------------KYSWS-------EV-------YEA---------VVGC
------------ECK-LNCTYLS---YNIE-TSQ------------------SVW-PM--
-------KGS------ELD-F-----------YC--------------------------
------------------------------------------------IGFS--------
-------------------------------YD-TGY-----------------------
NG-------------SSI------------------YDE-FHDKIGVHC-----------
-----------SDHF----EFENVSK-NWG------------------------------
-----D-------EL-----------R------ANFLRINVYFKDTETKYII-EEENFPL
VSM-----------------ISEIGGVLGLFIGVSLITILEVFVLCSSI-----------
--------IYTL--------L---------------------------------------
------------------------------------------------------------
------------------------------------------------------------
-------------------------GR---------------------NLEE--------
--------TRY-------------------------------------------------
------------------------------------------------------------
---------------------------------MKKKI----------------------
------------------------------------------------------------
-CQSLKPI

>sp_Q9JHS6_ASIC4_RAT_Acid_sensing_ion_channel_4_OS_Rattus_norvegicus_OX_10116
_GN_Asic4_PE_2_SV_1_687_217
------------------------------------------------------------
----------M-------------------------------------------------
------------------------------------------------------------
------------------------------------------------------------
------------------------------------------------------------
PIEIVCKIKFA-------------------------------------------------
---EEDAK----------------------------------------------------
--------------PKEKEA-------------GDEQSLLGAAQ------------G---
------------------------------------------------------------
------------------------------------------------------------
------------------------------------------------------------
-------------------------------P---------AA-----P-----------
-RD--LATFA-STSTLHGLGRAC---------G-PG-------------PHG---LR---
--RTLWV----LALLTSLAAFLYQAASLAR-GYLT----------R---PHLVAM----D
-P-AA-P----------APVAGF-----PAV-----------------------TLCNIN
RF------RHS-----ALSDAD------------------I-----------------F-
---------HL------------A-NLT---------------GLPP----K--------
---------------------------------DR-------------------------
------------------------------------------------------------
------------------------------------------------------------
-----------------DGH----------------------------------------
------------------------------------------------------------
------------------------------------------------------------
------------------------------------------------------------
------------------------------------------------------------
------------------------------------------------------------
------------------------------------------------------------
------------------------------------------------------------
------------------------------------------------------------
--------------------------------------------RAAGLR-----YPEP-
---------------------------------DMVDIL---------------------
------------------------------------------------NRT----GHQLA
--DML------KSCN----FS-G----H----------H---C-----------S-A---
----------------------------------------------S-------------
------------------------------------------------------------
------------------------------------------------------------
-----------N----------F-------S-----------VVY-T-RYG--------K
CYTFN-------AD-------------PQ---------------------------SS--
-------LPSRAG------GM--------GS-------------GLEIMLDIQQE-----
----------E-----------------Y------------L-P-IWRETN---------
------------------------------------------------------------
------------------------------------------------------------
-------ETSFEAGIRVQIH-SQEEPP--YIHQLGFGVSPGFQTFVSCQKQ------R--
-----LTYL-----------PQP-W--GN-C--RAE-SKLR-EPE-------LQ----GY
------------------S-AYSVSACRLRCEKEAVL----QR--CHCRMVH----M---
------------P-G---------------------------------------------
----------NE-----T---ICPP-----------------------------------
------------------------------------NIY---IECAD---------H---
T-L-------------------DSLG-G-------GS--------E-----------GPC
------------FCP-TPCNLTR---YGKE-ISM------------------VKI-PN--
-------RGS------ARY-L-----------AR--------------------------
------------------------------------------------KY----------
------------------------------------------------------------
-N----------------------------------------------------------
---------------------------RNE------------------------------

```
-----T-------YI-----------R------ENFLVLDVFFEALTSEAME-QRAAYGL
SAL-----------------LGDLGGQMGLFIGASILTLLEILDYIYEV-----------
--------SWDR--------L----------K-------RV----------------WR-
RP------------------------------KT-------------------P------
------------------------------------------------------------
-----------------------LRTS---------------------TGGI--------
-----------ST-----------------------------------------------
----------------LGLQELKEQ------------------------SPCPNRGR---
-----------------------AEGGGASNLLP--NHHHPHG---------PP------
---------GSL-----FEDFAC-------------------------------------
--------
>Deutero_Ambulac_Spurpu_015894_690_219
------------------------------------------------------------
----------M-------------------------------------------------
------------------------------------------------------------
------------------------------------------------------------
------------------------------------------------------------
------------------------------------------------------------
------------------------------------------------------------
------------------------------------------------------------
------------------------------------------------------------
------------------------------------------------------------
------------------------------------------------------------
----RGKEK----------------------D---------TL-----G-----------
-KS--LTDFG-NETTIHGVQYIV---------N-KD--------------NI---IY---
--RLCWV----AICIGFLVVFIIQGSDIIT-DFLQ----------W---PYSTKI----D
-I-VG-R----------P-SLTF-----PAV-----------------------TVCNAN
MM------RRS-----QLVDTR------------------F-----------------E-
---------GLIAL----------------------------------------------
---------------------------------DG-------GVSGADYDYSWWFSSDFV
-------------------------------------------NGFEQ------------
---------SSSSV----------------------------------------------
------------------------------------------------------------
------------------------------------------------------------
------------------------------------------------------------
------------------------------------------------------------
--------------------QRKKRSPPASI-----------------------------
------------------------------------------------------------
------------------------------------------------------------
------------------------------------------------------------
----------DYDMFSWY----DPSDVS---------DFQY-YADN-------W------
--------DTV-------------TSGDDWTGFYMS-----------SR------ADDYS
---------------------------------DFVDVV---------------------
-------------------------NP----------------TRAELTDY----GHQLE
--DFV------LQCT----FD-R----R----------Q---C-----------N-IS--
----------------------------------------------E-------------
------------------------------------------------------------
------------------------------------------------------------
-----------D----------F-------Y-----------VWQ-NRYYG--------N
CFTFN-------SA-------------LS-P-------------------AN----QT--
-------RTTGKT------GG--------LS-------------GLQLTLFVEQP-----
----------E-----------------Y------------L-G-I---LS---------
------------------------------------------------------------
------------------------------------------------------------
----------PHTGAKVSIH-HPDVFP--FPEDDALELSTGQETSIGIRQE------Y--
-----IKRL-----------GGF-Y--TN-C--TID-G------DD----------TNFT
----SSE-----------Y-IYSKAACKKICYQLHLS----QF--CGCVDNQ----F---
```

------------F-----------------------------------------------
----------DDYP---T---KCDVL----------N-----------------------
------------------------------------ATQ---QWCRQ---------Y---
V-E-------------------DRYL-D-------DE--------L------------AC
------------VCP-SPCAETK---YVKT-ASS------------------LLW-PS--
-------ERY------EEH-L-----------LR--------------------------
------------------------------------------------RL-N--------
----------------------------------ASS-----------------------
-H-------------SNA------------------VRIL--------------------
---------------------------ASS------------------------------
-----E-------LS-----------S------KNLIRVKIFYEDLNYEFVE-MVPVYTI
PSV-----------------LGSVGGLMGLYIGMSFISVFEVLSLLLRI-----------
--------CRIG--------F----------A-------KLFFN----------------
--------------------------------GE-------------------R------
-----------------------------------------VLPIK--------------
------------------------------------------------------------
------------------------------------------------------------
------------------------------------------------------------
------------------------------------------------------------
------------------------------------------------------------
-------T
>Protostome_Lophotroco_annelid_CAC9669404_1__Ofus_G17801_Owenia__fusiformis__
689_220
MTDIVETG-------AETNKQRYSEEHLYPSSVVETGHQDVNDF---D------KNRQE-
-----ENKSSI-------------------------------------------------
------------------------------------------------------------
------------------------------------------------------------
------------------------------------------------------------
--EIDQQEEHT--------NEF--------------------------------------
------SCDVQKQESTTRN-----------------DMHLREGVLIDTLL----------
-G------------QESHSN-------------KDHVENKLELS----------------
----------------------------------VENK----------------------
------------IPDDPNISASTSGAESSHNKPA--------------------IMATQG
E----------------------QMK----------------------------------
------SFR----------------------R---------KI-----S-----------
-SS--LAESA-EMTSIHGSKRIL---------K-AK--------------GK---LS---
--TIKWT----IIFLGTVSMAVSLIASVVV-KFYS----------YPVQPVEDGI----S
-V-TT------------Q-PLAF-----PAV-----------------------TLCPFI
PY------YMR-----DINPKVRGK-----------------------------------
----------------------LKNFTE---------------KLWA----RVH------
------------------------------ML-DC-------SYKVIF------------
------------------------------------------------------------
------------------------------------------------------------
-----------------QPD------------------------ENYDAN----------
------------------------------------------------------------
------------------------------------------------------------
------------------------------------------------------------
------------------------------------------------------------
------------------------------------------------------------
------------------------------------------------------------
------------------------------------------------------------
----TIRKIDKY------------------------------ERGC--------------
---------------------------------------L-PLQDIFQL------RHKLI
---------------------------------SSGWLV---------------------
------------------------ENLPID---------------YETFAT----PIKVE
--DFI------LECT----YN-N----K----------P---C-----------E-S---
----------------------------------------------Q-------------

------------------------------------------------------------
------------------------------------------------------------
-----------I----------N-------I-----------TRT-NTVYG--------Q
CFTLT-------VN-------------EP----------------------------N--
-------DHVSEI------GP--------DK-------------GIALILYTGRA-----
----------NPLSSVNSNKEVRD----GF-------------T-T-ILST---------
------------------------------------------------------------
------------------------------------------------------------
----------PSDGVQVVLH-KPGTMP--RPYNEGFYICPGRFSSVEITQE------T--
-----RTRL-----------LPP-Y--GE-C--TDA--------GY------ITK--TTY
--------------------GYSYDVCVDQCIQERIA----KS--CGCISPM----F---
------------I-T--PT-----------------------------------------
-----KHNFMKT-----P---FCGNIQ---------GRI---------------------
---WERT---RG-------GGPNFRS-----------EK---KTSAK---------F---
W------------------------------------------------AP---------
------------------------------------------------------------
------------------------------------------------------------
------------------------------------------------------------
------------------------------------------------------------
------------------------------------------------------------
------------------------------------------------------------
------------------------------------------------------------
------------------------------------------------------------
------------------------------------------------------------
------------------------------------------------------------
------------------------------------------------------------
------------------------------------------------------------
------------------------------------------------------------
------------------------------------------------------------
------------------------------------------------------------
------------------------------------------------------------
--------
>Placozoa_HhoNaC5_TR3219_c3_g1_i1_m_8097__Hoilungia_hongkongensis__27_221
------------------------------------------------------------
----------M-------------------------------------------------
------------------------------------------------------------
------------------------------------------------------------
------------------------------------------------------------
------------------------------------------------------------
------------------------------------------------------------
----------------NNSP--------------DDSDDI--------------------
------------------------------------------------------------
------------------------------------------------------------
------------------------------------------------------------
-------------------------------SL--------AS-----F-----------
-DL--EEHFA-NVTSCHGMIHIF---------D-HK-------------TSF---LR---
--RYVWG----IATFVALTSCIIGCIQLLQ-NLLS----------H---PTNISV----K
-I-HH-T----------N-KMLF-----PAV-----------------------TLCNFN
QI------SRT-----LISAKD------------------I-----------------R-
---------HL------------N-TVL---------------SAYN----H--------
---------------------------------DH-------TI----------------
------------------------------------------------------------
------------------------------------------------------------
------------------------------------------------------------
------------------------------------------------------------
------------------------------------------------------------
------------------------------------------------------------

--------------------------P---------------------------------
------------------------------------------------------------
------------------------------------------------------------
------------------------------------------------------------
--------KSVL------------------------------KET---------------
------------------------EEYFKS------------------KI-----HGHF-
---------------------------------DFSQYN---------------------
------------------------------------------------KEI----GLQKE
--DLI------LSCT----WN-G----Q----------K---C-----------G-P---
----------------------------------------------K-------------
------------------------------------------------------------
------------------------------------------------------------
-----------N----------F-------T-----------RIF-T-SYG--------N
CYTFN-------GD-------------SS-K--------------------Q----AL--
-------LQQHGR------GA--------AH-------------GLSLILNIEQY-----
----------K-----------------Y------------T-P-E---LM---------
------------------------------------------------------------
------------------------------------------------------------
-------LGAPNVGIRCSIH-YYKSLP--RMESQGIAIPPGAHAYAAIPGT------E--
-----ITKYQ----------KKP-W--GQ-C--GYK--------K-------LK----YY
------------------K-YYSMSACLHEDETLYAE----SI--CHCRDPR----L---
------------P-G---------------------------------------------
----------KE-----K---VCSP-----------------------------------
------------------------------------TQM---LECLI---------P---
A-M-------------------ARYR-E-------QG--------K-----------NLS
------------SCP-EVCERVE---YVPQ-ISY------------------AKI-PS--
-------KIM------AEE-I-----------AL--------------------------
------------------------------------------------KY----------
------------------------------------------------------------
-G---------------L------------------HNVRQQAISGGG------------
-------------------LI-DNST-SVK------------------------------
-----Q-------FI-----------R------DNLVFLDVFFKNLYNTTTT-QGRGSSF
LQF-----------------LSNVGGQIGLFIGGSFLSVLEIVEYVFDK-----------
--------LAQQ--------R----------S-------RY-------------------
RR------------------------------KT-------------------G------
-----------------------------------------QNEKAEWK-----------
--------------------KMYASSM---------------------IVHK----DSDD
C--------RETNQPPPYEG----------------------------------------
--------------TLIQSDDPDSTE----------------------------------
------------------------------------------------------------
----------GF-----LNGPKY------------------------------------V
HVHKVGKE
>Protostome_Lophotroco_annelid_CAC9674716_1__Ofus_G26148_Owenia__fusiformis__
693_222
------------------------------------------------------------
----------M-------------------------------------------------
------------------------------------------------------------
------------------------------------------------------------
------------------------------------------------------------
------------------------------------------------------------
------------------------------------------------------------
------------------------------------------------------------
------------------------------------------------------------
------------------------------------------------------------
------------------------------------------------------------
---AKTEKQ-------------F------------------NP-----K-----------
-EI--LREFS-ENATPHGPGLIA---------N-SK--------------TT---VG---

--KLISV----CILLTCTGAAVVNIGFQLN-SFFK----------F---QYKDTI----Q
-V-DN------------V-VIEM-----PSV-----------------------TICPTL
GF------SRM-----KYISYN------------FSGETSV-------------------
---------NIWSA------------------------IPINYQLLK-------------
---------------------------------DH-------------------------
------------------------------------------------------------
------------------------------------------------------------
------------------------------------------------------------
------------------------------------------------------------
------------------------------------------------------------
------------------------------------------------------------
------------------------------------------------------------
------------------------------------------------------------
------------------------------------------------------------
------------------------------------------------------------
------------------------------------------------------------
-----------------------------------------SHENFSQF------EDLFI
---------------------------------TGSIAL---------------------
------------------------DNM----------------LTEDRLDV----GHDLD
--GLI------IKCR----FA-G----R----------D---C-----------N-T---
----------------------------------------------S-------------
------------------------------------------------------------
------------------------------------------------------------
-----------H----------F-------G-----------QFL-HPELN--------H
CYTFN-------GK-------------DV-N--------------------M----DID-
-------TRIKAS------TT--------DA-------------GLHLKIFLDAF-----
--------IPE------ILPYGSY----FN-------------E-D-VITS---------
------------------------------------------------------------
------------------------------------------------------------
----------GNIGLRVTVH-PKDTIT--FPKATGFDIPPGYASTITLKQH------K--
-----IERL-----------GLP-Y--SE-C--TNR--------KL------LEG--TKY
--------------------AYTQAGCQAQCFQKNLM----KT--CGCVSSW----H---
------------P-I--PP-----------------------------------------
--------NNTL-----K---ICTKL----------DSE-NIYKEMVYMNLSTSRG----
---------------------------NMDILS-KDFTH---LGCLT---------K---
T-V-------------------NEKM-E-------TP--------------------LNC
E-----------ECL-PACKEIS---YSKS-ISQ------------------AYW-PD--
-------EVI------QGF-I-----------LK--------------------------
------------------------------------------------MILQSPERNT--
----------------------------RGQQMIQKY-----------------------
IN------------FSEI------------------------------------------
----------------ERNHYQITNR-THA------------------------------
-----D-------TI-----------R------KNMVSLSISFGDLSTEVTK-QKVEYEG
AQL-----------------LCDIGGALGLYIGVSFISLCEVGKLLMRL-----------
--------LRYA--------F----------C-------RT-------------------
STS-----------------------------RI--------------------------
------------------------------------------------------------
------------------------------------------------------------
------------------------------------------------------------
------------------------------------------------------------
------------------------------------------------------------
------------------------------------------------------------
-EATSAEN
>Deuterostome_chordata_PetMar_tr_S4RG90_S4RG90_PETMA_Uncharacterized_protein_
OS_Petromyzon_marinus_OX_7757_PE_4_SV_1_695_223
------------------------------------------------------------
----------M-------------------------------------------------

```
------------------------------------------------------------
------------------------------------------------------------
------------------------------------------------------------
PINIICTIPLP-------------------------------------------------
--LEEDSK----------------------------------------------------
--------------LGIKER-------------GKSGQDLKDMH----------------
------------------------------------------------------------
------------------------------------------------------------
------------------------------------------------------------
---SIEEDG----------------------Q---------PQ-----A-----------
-SA--ITEFA-SESTLHGINHIF---------A-PG-------------EFS---LR---
--RAIWA----CAFLASLGMFLYQSADRIV-FYLE----------H---HHVTQL----D
-E-QE-S----------H-LMLF-----PAI-----------------------TLCNFN
SF------RRS-----KVTPVD------------------K-----------------H-
---------YA------------G-PIL---------------GIPD----N--------
---------------------------------DP-------------------------
------------------------------------------------------------
------------------------------------------------------------
-----------------DLA-TV-------------------------------------
------------------------------------------------------------
------------------------------------------------------------
------------------------------------------------------------
------------------------------------------------------------
------------------------------------------------------------
------------------------------------------------------------
------------------------------------------------------------
--------DVRL------------------------------LRA---------------
-------------------------------------------------------WSRY-
---------------------------------STYEFY---------------------
------------------------------------------------DRA----GHTMD
--EML------LRCR----YR-D----R----------E---C-----------G-P---
----------------------------------------------T-------------
------------------------------------------------------------
------------------------------------------------------------
-----------N----------F-------T-----------TVF-T-RYG--------K
CYMFN-------SG-------------KG-G--------------------R----PL--
-------LTTLKG------GT--------GN-------------GLEMMLDIQQD-----
----------E-----------------Y------------L-P-VWGETD---------
------------------------------------------------------------
------------------------------------------------------------
-------EISLEAGVKVQIH-SQDEPP--FVHQLGFGVAPGFQTFVSCQEQ------R--
-----LIYL-----------PSP-W--GD-C--KAD-PINS---D-------------FF
------------------D-TYSITACRIDCETRYVV----EN--CNCRMVH----M---
------------P-G---------------------------------------------
----------DA-----P---YCTP-----------------------------------
------------------------------------EQY---KECAD---------P---
A-L-------------------DFLV-E-------KD--------N-----------NYC
------------VCD-TPCNTTR---YGKE-LSM------------------VKI-PS--
-------KAS------AKY-L-----------AK--------------------------
------------------------------------------------KY----------
------------------------------------------------------------
-N----------------------------------------------------------
---------------------------KTE------------------------------
-----D-------YI-----------R------ENVLVLDIFFEALNYETIE-QKKAYEV
AGL-----------------LGDIGGQMGLFIGASILTILEIFDYLYEV-----------
--------LKDK--------V----------C-------KC-------------------
RK------------------------------KH--------------------------
```

------------------------------------------------------------
-----------------------NKKR---------------------SENM--------
-----------VS-----------------------------------------------
----------------LSCTINKIHLYD---------------------TPCETLRSHHS
-----------------------QDVHFNSSILP-------HH---------TV------
---------RGT-----FEDFAC-------------------------------------
--------
>Protostome_Lophotroco_annelid_CAC9640639_1__Ofus_G092493_partial_Owenia__fus
iformis__697_224
------------------------------------------------------------
----------V-------------------------------------------------
------------------------------------------------------------
------------------------------------------------------------
------------------------------------------------------------
------------------------------------------------------------
------------------------------------------------------------
------------------------------------------------------------
------------------------------------------------------------
------------------------------------------------------------
------------------SYGYLKKNISL-----------------------------VR
LFFPEFSMT----------------------Y---------NM-----K-----------
-KV--VEEFA-DHTTAHGWGNIS---------Q-RS--------------ST---LG---
--KTVWI----IFTLGCTCLAIASIIGTIS-KYFE----------Y---ETKDIV----E
-L-RD-E------------PLEF-----PSV-----------------------TVCPLN
PI------AHT-----HQLMYRQNIYY---------------------------------
---------EI----------------------------VDYHNLTN-------------
------------------------------------------------------------
------------------------------------------------------------
------------------------------------------------------------
------------------------------------------------------------
------------------------------------------------------------
------------------------------------------------------------
------------------------------------------------------------
------------------------------------------------------------
------------------------------------------------------------
------------------------------------------------------------
------------------------------------------------------------
---NILPTERKT------------------------------PQHN--------------
-------------------------------------------TLYKDY------INQLV
---------------------------------SYQGYY---------------------
------------------------ENA------------------MQILVY----SHNKD
--DFI------TSCT----YM-E----N----------M---C-----------N-R---
----------------------------------------------S-------------
------------------------------------------------------------
------------------------------------------------------------
-----------D----------I-------T-----------TSL-HHEHY--------N
CFTYS-------TS-------------RS----------------------L----PD--
-------FAKTTV------GP--------NS-------------GLSMILYLDVS-----
----------K--------LSMTL----YD-------------P-N-YPTS---------
------------------------------------------------------------
------------------------------------------------------------
----------GSNGARVVIH-ERGSLP--DPEKDGSDIEPGHSVNAALSVN------R--
-----RELM-----------KKP-W--GN-C--VDY--------SA------LDA--GGF
--------------------VHTMNSCIKRCQQKRVY----EE--CGCVKAS----L---
------------P-L--EP-----------------------------------------
-----DL--ENA-----Q---FCGKL----------DVK-EWLKN------EP-------
---------------------------NITILN-EDLNR---LECQK---------S---

```
A-W------------------------------------------------------RPC
K-----------TCA-KNCTYHT---YDVS-LSQ------------------SEW-PT--
-------DGA------VKS-F-----------YE--------------------------
------------------------------------------------RWIMQQ------
----------------------------------PNY-----------------------
EN-------------STL------------------YHR-FHREMK--------------
----------------DLKDIYSVTE-TTK------------------------------
-----S-------AI-----------K------NNFVRLNVYFKESETKVTT-KAQAFDL
ANL-----------------IAETGGFLGFYVGISVISIAEVMILLYNL-----------
--------LNFL--------R----------N-------KL-------------------
IQ------------------------------QK----------------YNNT------
------------------------------------------------------------
------------------NNAGVNIKQ---------------------------------
------------------------------------------------------------
------------------------------------------------------------
------------------------------------------------------------
------------------------------------------------------------
-----NGK
>Placozoa_TadNaC5_MK547546_15_225
------------------------------------------------------------
----------M-------------------------------------------------
------------------------------------------------------------
------------------------------------------------------------
------------------------------------------------------------
------------------------------------------------------------
------------------------------------------------------------
----------------TTNP-------------SDDSDDSN-------------------
------------------------------------------------------------
------------------------------------------------------------
------------------------------------------------------------
-------------------------------SL--------AS-----F-----------
-DL--EEHFA-NVTSCHGMIHIF---------D-HK-------------TSF---LR---
--RYVWG----IATFAAFTACIIGCINLLH-NLLS----------H---PTNISV----K
-I-HH-T----------N-KMLF-----PTV-----------------------TLCNFN
QF------SRT-----LISHKD------------------I-----------------R-
---------HL------------D-TVL---------------SAYN----H--------
---------------------------------DH-------TI----------------
------------------------------------------------------------
------------------------------------------------------------
------------------------------------------------------------
------------------------------------------------------------
------------------------------------------------------------
------------------------------------------------------------
--------------------------P---------------------------------
------------------------------------------------------------
------------------------------------------------------------
------------------------------------------------------------
--------KEDL------------------------------KEA---------------
------------------------EDYFKS------------------KI-----HGHF-
---------------------------------DFSQYN---------------------
------------------------------------------------QEL----GLQKE
--DLI------LSCT----WN-G----E----------K---C-----------G-S---
----------------------------------------------K-------------
------------------------------------------------------------
------------------------------------------------------------
-----------N----------F-------S-----------RVF-T-SYG--------N
CYTFN-------GG-------------SA-N--------------------H----PL--
```

-------LNQHGR------GA--------AH-------------GLSLILNIEQY-----
----------K-----------------Y------------T-P-E---LM---------
------------------------------------------------------------
------------------------------------------------------------
-------VGAPNVGIRCSIH-YYKSLP--RMESQGIAIPPGAHAYAAIPGT------E--
-----VTNYQ----------KKP-W--GQ-C--GEK--------K-------LK----YY
------------------K-YYSMSACLHEDETLFAE----TT--CQCRDPR----L---
------------P-G---------------------------------------------
----------NE-----K---ACTP-----------------------------------
------------------------------------IQM---LECLI---------P---
A-M-------------------SRYR-E-------QD--------R-----------NLS
------------SCP-EVCERVE---YVPQ-VSY------------------AKI-PA--
-------KIM------AEE-I-----------AS--------------------------
------------------------------------------------KY----------
------------------------------------------------------------
-N---------------L------------------HQVHKKAIIEGG------------
-------------------IV-DNST-SVK------------------------------
-----Q-------FI-----------R------DNLVFLDIFFKNLYNTTTT-QARDASF
SQF-----------------LSNVGGQIGLFIGGSFLTLLEIVEYIFDK-----------
--------LAQQ--------R----------S-------RY-------------------
RH------------------------------KT-------------------S------
-----------------------------------------QIEKAEWK-----------
--------------------KIYASSM---------------------IVPK----TSYH
N--------GNANHPPPYER----------------------------------------
--------------TLVQMDEYDSTE----------------------------------
------------------------------------------------------------
----------GF-----LNGPKY------------------------------------V
HVHKQCKE
>Protost_ecdy_cyclo_nema_Cele_sp_Q22851_ASIC2_CAEEL_Degenerin_like_protein_asic_2_OS_Caenorhabditis_elegans_OX_6239_GN_asic_2_PE_3_SV_1_699_226
------------------------------------------------------------
----------M-------------------------------------------------
------------------------------------------------------------
------------------------------------------------------------
------------------------------------------------------------
------------------------------------------------------------
------------------------------------------------------------
------------------------------------------------------------
------------------------------------------------------------
------------------------------------------------------------
------------------------------------------------------------
-------RG----------------------G---------GF-----V-----------
-QI--FKDFS-NWSTVAVVPHVA---------N-AN--------------NK---IS---
--RIFWI----AIFLFVLGMFAYELYILIA-KFFS----------Y---PATVNT----E
-I-LF-E------------KQIF-----PVV-----------------------TVCNMN
PY------KYS-----VVKSNS--------A---------F-----------------S-
---------SV--------------NTL--------------MTTYS-------------
------------------------------------------------------------
------------------------------------------------------------
------------------------------------------------------------
-----------------D-A----------------------------------------
------------------------------------------------------------
------------------------------------------------------------
------------------------------------------------------------
------------------------------------------------------------
------------------------------------------------------------
------------------------------------------------------------

------------------------------------------------------------
---TVGTFSTDKW--GLY---AD------------------NDETYD------------
------------------------------------------------------LDSRAA
---------------------------------DALVLE--------------------
------------------------ANL--------------ISDTAKVPA----LYTYA
--DLI------QDCS----FA-G----I----------P---C-----------S-E---
-----------------------------------------------S------------
------------------------------------------------------------
------------------------------------------------------------
-----------D----------F-------T-----------KFI-DPVYG--------A
CYSFN-------ED-------------ASLN-----------------------------
-------YSVSRE------GI--------QF-------------GLKLMLTVTQT-----
----------K------------------TNGNT------DSL-P-----TT--------
------------------------------------------------------------
------------------------------------------------------------
----------KLAGARIGVN-SRGSSP--GLDSNGIDAGVGYESAVSVSLT------Q--
-----NVRA-----------KKP-Y--GT-C--VDR--------E-------PDS-SDYY
----KDF-------------IYTLETCFNGCKQRDTI----AK--CQCANPR----L---
------------A-L--GS-----------------------------------------
----------TD-----T---ACQPI----------------------------------
------------------------------------KAD---LDCLQ---------T---
L-K-------------------GNQT-S-------ST--------PNI---DLL---VEC
------------NCN-PPCDEST---YTPT-VSL------------------AQF-PS--
-------TSY------YVA-T-----------SS--------------------------
------------------------------------------------TA----------
----------------------------GVGSCSST------------------------
-N----------------------------------------------------------
-------------------SKFSSKS-DCQ------------------------------
-----K-------WY-----------N------NNGMIIQVFLETLSYELYT-ETAGYTV
SNV-----------------INDLGGQAGLWLGLSVISVVEMTGLMLVM-----------
--------GAFC--------V----------TG---GAIKMAP--------------DD-
DEI-----------------------------EN-----------------DHR------
-----------------------------------------IKDVEDVKK----------
------------------EIDHLEKKH---------------------GEME--------
--------SGSDGEVD--------------------------------------------
------------------------------------------------------------
------------------------------------------------------------
-------------------------------------------------------DIENK
GDEEKKKK
>Protostome_Lophotroco_annelid_CAC9618795_1__Ofus_G081188_Owenia__fusiformis_
_700_227
------------------------------------------------------------
------------------------------------------------------------
------------------------------------------------------------
------------------------------------------------------------
------------------------------------------------------------
------------------------------------------------------------
------------------------------------------------------------
------------------------------------------------------------
------------------------------------------------------------
------------------------------------------------------------
------------------------------------------------------------
-----------------------------------------M-----R------------
-TL--IGEFA-GNTSAHGWGKAS---------Q-TT--------------SK---LA---
--KAAWI----FTSSACSIVALFLIIKVVI-KYCE----------F---NTKDSI----R
-V-SK-D------------EAVF-----PSV-----------------------TVCPLT
PI------SRK-----GMATFRKELA---------ENNSAI-------------------

--------DSM----------------------------LYYNNFLY----VLGQYN---
------------------------------------------------------------
------------------------------------------------------------
------------------------------------------------------------
------------------------------------------------------------
------------------------------------------------------------
------------------------------------------------------------
------------------------------------------------------------
------------------------------------------------------------
------------------------------------------------------------
------------------------------------------------------------
------------------------------------------------------------
--LNISSLQNAS------------------------------EELQ--------------
-------------------------------------------LLYNQS------YKRLH
---------------------------------TNAGYY---------------------
------------------------ENY----------------QQSERLRY----TIKQK
--DFI------PSCK----YQ-S----E----------V---C-----------S-E---
----------------------------------------------E-------------
------------------------------------------------------------
------------------------------------------------------------
-----------L----------Y-------H-----------SGH-HEH-F--------N
CFTFN-------GP-------------LS-N--------------------I----SS--
-------LDVD-S------GP--------SA-------------GLSLILYLDID-----
----------L--------EHSSF----YD-------------P-D-YPTS---------
------------------------------------------------------------
------------------------------------------------------------
----------GSQGVRVVIH-EQGTWP--DPENDGFDIKAGNSVNVALSSE------R--
-----RELM-----------GQP-W--GD-C--ADH--------ND------IKY--DRW
--------------------KYSTKACLLTCIQRLGF----KR--CGCIRGS----L---
------------P-V--DD-----------------------------------------
-----KT--KDV-----E---FCANI----------QSR-EWLKS------DP-------
---------------------------NVTLLR-QDLNQ---LNCSL---------Q---
-----------------------EYSLS-------DI-------LAQ----------EGC
------------ECK-ENCKIDK---YDIM-LSD------------------SEW-PM--
-------KGI------EFH-F-----------YC--------------------------
------------------------------------------------DMIM--------
-------------------------------KL-GNY----------------------N
EN-------------SSI------------------YQA-FHDRVASNC-----------
-----------SSLAYDVERKKIINK-DIG------------------------------
-----K-------DL-----------R------EQFLRLNVYFKDSDTKVTT-QTEDFSI
GSM-----------------VSEIGGSLGLLIGMSVISISEVFILCLGI-----------
--------LAIL--------I----------R-------KS-------------------
KA------------------------------QT----------------KVET------
------------------------------------------------------------
----------------------FKDKM---------------------EYP---------
------------------------------------------------------------
------------------------------------------------------------
------------------------------------------------------------
------------------------------------------------------------
--------
>Protostome_Lophotroco_annelid_CAC9667655_1__Ofus_G151630_Owenia__fusiformis_
_703_229
------------------------------------------------------------
------------------------------------------------------------
------------------------------------------------------------
------------------------------------------------------------
------------------------------------------------------------

------------------------------------------------------------
------------------------------------------------------------
------------------------------------------------------------
------------------------------------------------------------
------------------------------------------------------------
------------------------------------------------------------
------------------------------------------M-----K-----------
-RI--LKDFS-SNTTAHGWGNIH---------Q-RT--------------TT---LS---
--KVIWI----IICLGCTAVAIWQVITIVM-RYGL----------F---ETKDRI----K
-V-EV-G------------EIVF-----PSV-----------------------TVCPLI
PV------PHD-----GNLKFERDMK---------KGNIDV-------------------
--------EPF----------------------------LNFRLVFH----SETFY----
------------------------------------------------------------
------------------------------------------------------------
------------------------------------------------------------
------------------------------------------------------------
------------------------------------------------------------
------------------------------------------------------------
------------------------------------------------------------
------------------------------------------------------------
------------------------------------------------------------
------------------------------------------------------------
------------------------------------------------------------
--INSNVSHNLH------------------------------SEKL--------------
-------------------------------------------EFYNVF------AEQIY
---------------------------------SNQGFF---------------------
------------------------ENF------------------KFMSDY----THQQG
--NFI------PVCS----YQ-G----K----------P---C-----------L-R---
----------------------------------------------S-------------
------------------------------------------------------------
------------------------------------------------------------
-----------D----------F-------H-----------TLD-HPEYN--------S
CFTFN-------GA-------------NK-T--------------------I----NN--
-------PITKIT------GP--------KG-------------GLSLVLYLDVG-----
----------Q---------LSMI----YT-------------P-S-YPTS---------
------------------------------------------------------------
------------------------------------------------------------
----------GSQGVRVVIH-EKGTLP--DPENDGFDIEPGHSINAALSVN------E--
-----RRLM-----------KPP-W--GE-C--NDQ--------NP------ELK--GGF
--------------------TYTKKHCQLSCLQKFFY----RT--CGCIKSS----L---
------------P-I--DE-----------------------------------------
-----ET--VYK-----E---YCLKL----------HAK-EWAKL------TTDQY----
---------------------------NKTTIV-SDMEN---FRCQN---------R---
-----------------------EFLSS-------DV-------EVQ----------DGC
------------QCL-EKCSYQT---YDMT-LSQ------------------SEW-PM--
-------KGV------EIE-F-----------YC--------------------------
------------------------------------------------RTIM--------
-------------------------------SM-DSY-----------------------
IN-------------SSI------------------YDR-FHDIFPEYC-----------
-----------DHSK----DEDLLTK-IQK------------------------------
-----E-------TL-----------R------ENFIRLNVYFKDLETKITN-QVEDFTI
SSM-----------------ISEIGGSLGFFVGMSIITIAEIILLCWNI-----------
--------TCLL--------G----------K-------KF-------------------
VI------------------------------SP----------------FEQK------
------------------------------------------------------------
------------------I--GSDMKH---------------------ASTN--------
--------TS--------------------------------------------------

------------------------------------------------------------
------------------------------------------------------------
------------------------------------------------------------
--------
>Deutero_Ambulac_Apla_gbr361_5_t1_704_230
------------------------------------------------------------
----------M-------------------------------------------------
------------------------------------------------------------
------------------------------------------------------------
------------------------------------------------------------
--YHQESGSQE-------------------------------------------------
------------------------------------------------------------
------------------------------------------------------------
------------------------------------------------------------
------------------------------------------GL---------HLAKRKV
SPMTV---------------EDISAEDGAAPAA--------------------------P
PDPAASAKQ----------------------T---------SL-----L-----------
-YV--VSHRL-AESGAHGIPNIQ---------R-AN--------------SN---FR---
--RVAWT----LLFLAGFGMFIYQGCDLII-KFYK----------W---PYNVNI----E
-V-RT-P----------K-RVEF-----PAV-----------------------TWCNLN
PI------RKD-----ALADFT--------------------------------------
---------DL-------------------------------RSLLE----G--------
------------------------------------------------------------
------------------------------------------------------------
------------------------------------------------------------
------------------------------------------------------------
------------------------------------------------------------
------------------------------------------------------------
------------------------------------------------------------
-------------------------GP---------------------------------
------------------------------------------------------------
------------------------------------------------------------
------------------------------------------------------------
--SSCGTAVNATFG----------------------------------------------
------------------------GYPDDWES---------ADIDYKLE------DCTSQ
------------------------------TVDAAIDIV---------------------
------------------------GAM----------------PYGERAVT----GHRLN
--DML------LMCT----FQ-Q----K----------P---C-----------S-P---
----------------------------------------------R-------------
------------------------------------------------------------
------------------------------------------------------------
-----------N----------F-------T-----------SFY-NSRYG--------N
CFTFN-------SA-------------SR-G--------------------V----AP--
-------LKVTRT------GP--------SY-------------GLSMELYIQQD-----
----------R-----------------Y------------I-T-G---VE---------
------------------------------------------------------------
------------------------------------------------------------
----------TGAGIRIMVH-NQTEMA--FPEDIGANIAPGTESFLGLYRA------M--
-----VNRL-----------SDP-Y--GD-C--ATD----------------FTQ-DNIF
----KEQ--FK------RL-EYTKEACEKDCFFRAVL----LD--CNCAYIN----Y---
------------R-Y---------------------------------------------
---------NDSQ-----A---PCNSSV--------------------------------
------------------------------------EAV---KTCIE---------K---
V-E-------------------NNFT-S-------GE--------L------------RC
G-----------HCH-KSCNETR---YQIS-FSN------------------SKW-PN--
-------REY------MET-V-----------RK--------------------------
------------------------------------------------NI-Q--------

----------------------------------ES------------------------
-N-------------KET------------------ISLIK-------------------
---------------------------NNS------------------------------
-----Y-------FI-----------E------DNVLRVNVYFDSLNYDDIF-QTAAYTL
GSL-----------------VSDLGGQVGLWIGVSVLTLFEFLELMIDI-----------
--------FVLI--------S----------T-------RC-------------------
RAGGENR-------------------------RN------------------GK------
------------------------------------------------------------
------------------------------------------------------------
------------------------------------------------------------
------------------------------------------------------------
------------------------------------------------------------
----------------------------------------------------------TG
QENSRPQA
>Placozoa_TadNaC2_MK547543_12_231
------------------------------------------------------------
----------M-------------------------------------------------
------------------------------------------------------------
------------------------------------------------------------
------------------------------------------------------------
------------------------------------------------------------
------------------------------------------------------------
------------------------------------------------------------
------------------------------------------------------------
---------------------------------------------------------DTH
P-----------------------------------------------------------
----WQRPR----------------------S---------KS-----I-----------
-NL--EKKFA-ERTTCHGLGHIV---------D-QD-------------VPK---VR---
--RYLWS----IVTLAASVGCMIGCIQLLQ-YVLS----------F---PTNIDI----E
-I-IH-Q----------D-SLIF-----PAV-----------------------TICNFN
QL------TKT-----ALSEED------------------N-----------------R-
---------HL------------Q-TFL---------------RIYH----R--------
---------------------------------KG-------AI----------------
------------------------------------------------------------
------------------------------------------------------------
-----------------N------------------------------------------
------------------------------------------------------------
------------------------------------------------------------
------------------------------------------------------------
------------------------------------------------------------
------------------------------------------------------------
------------------------------------------------------------
------------------------------------------------------------
--------KKDL------------------------------ELA---------------
------------------------TSYFRN------------------KT-----GVDF-
---------------------------------HLQNLT---------------------
------------------------------------------------QDL----GHRKD
--DII------VSCL----WD-D----Q----------V---C-----------G-P---
----------------------------------------------E-------------
------------------------------------------------------------
------------------------------------------------------------
-----------N----------F-------T-----------TIF-T-IYG--------N
CYTFN-------SG-------------AK-K--------------------E----GM--
-------LSQRGK------GS--------AH-------------GLRLVLNIEQY-----
----------K-----------------Y------------S-G-K---LS---------
------------------------------------------------------------
------------------------------------------------------------

```
-------YGSPDAGIRFAVH-STADLP--EMDAEGMSIPPGMHAYASIPGA------DV-
-----IEGL-----------PKP-W--GQ-C--GSQ--------K-------LK----YF
------------------D-HYSVSSCRREKEIDFIL----QR--CGCVEPH----H---
------------A-R---------------------------------------------
----------NL-----T---PCSP-----------------------------------
------------------------------------EIM---LECVL---------P---
L-M-------------------SSSN-P-------VS--------VGI---------NAS
------------VCP-VACVQTE---YNVE-VSY------------------ALI-PS--
-------QVV------VND-I-----------SD--------------------------
------------------------------------------------QY----------
------------------------------------------------------------
-N---------------V------------------TKIIENARNQRL------------
----------------------NLTM-SKL------------------------------
-----E-------FI-----------R------ENFAFLDVYYKDLYLFKTI-QKQASGF
VAF-----------------LSDIGGQLGLFVGGSFLTMFEFFEYIYDK-----------
--------CFQQ--------T-----------K-------RS------------------
KR------------------------------EI------------------S------
-----------------------------------------RRMQSIRE-----------
--------------------KRRPESM---------------------ASST--------
---VRGNLSTNVHNFKVNESI---------------------------------------
------------SSHNVRSNGIKSKFRR--------------------------------
------------------------------------------------------------
---------------------------------------------------SNSEHLANI
RISPSSID
>Protostome_Lophotroco_annelid_CAC9628722_1__Ofus_G09545_Owenia__fusiformis__
706_232
------------------------------------------------------------
----------M-------------------------------------------------
------------------------------------------------------------
------------------------------------------------------------
------------------------------------------------------------
--SLTLLL----------------------------------------------------
------------------------------------------------------------
------------------------------------------------------------
------------------------------------------------------------
------------------------------------------------------------
------------------------------------------------------------
------DHP-----------------------V---------DD-----R----------
-GT--ISHYM-SLITTHGCQQIY---------Q-TK--------------SK---LI---
--KICWT----IFLLASVCVATYFVSVAIS-KYYK----------Y---RTHEET----S
-L-KD------------T-YVDF-----PAV-----------------------TVCRLD
DD------GVS-----RLFSEHYTPD---------KGDEEI-------------------
-----------------------IKQEM---------------NILM----KIIDIDLD-
------------------------------------------------------------
------------------------------------------------------------
------------------------------------------------------------
------------------------------------------------------------
------------------------------------------------------------
------------------------------------------------------------
------------------------------------------------------------
------------------------------------------------------------
------------------------------------------------------------
------------------------------------------------------------
------------------------------------------------------------
--------FENE--------------------------------DVSYL-----------
----------------------------------------H-LLLLYAEY------APRL-
--------------------------------LDRQSIP---------------------
```

```
------------------------QAF-----------------PAIAEYL----ASDYD
--RFI------VSCQ----FK-E----H----------D---C-----------N-K---
----------------------------------------------T-------------
------------------------------------------------------------
------------------------------------------------------------
-----------H----------F-------V-----------EKK-NGQYF--------N
CFTFN-------HT-------------D--------------------------------
--------TIKKE------GP--------TS-------------GLSLVFFLDTL-----
----------KEIE-----LQEIS----LG-------------D-S-SLRF---------
------------------------------------------------------------
------------------------------------------------------------
----------TDSGIKVAIH-EPNTIP--DIINDGFIVAPGYSTNIALKQS------V--
-----FEKM-----------KSP-W--GE-C--DKS--------RIKINDITNEN--TDI
--------------------LYNRETCMKICQQRYIQ----GK--CGCLNVM----L---
------------P-V--PN-----------------------------------------
-----DL--NKE-----K---YCFYI----------DKD-NMLQN------PN-------
---------------------------WTQVFH-EELKH---VDCES---------K---
H-L-------------------LDLP-D-------SY--------------VGQ---YKC
------------DCP-LSCHYKT---YNSL-ISQ------------------ANW-PP--
-------QNG------VHR-F-----------IK--------------------------
------------------------------------------------KYRN--------
-------------------------------EQPTLL-----------------------
NG-------------TDI------------------YKA---------------------
------------------YEENNATG-DLY------------------------------
-----D-------MV-----------Q------NNFLRLNVYHRSLTVQTTT-QVAAYTV
QDL-----------------VSEVGGIIGICVGMSVLSILELFQLVAIV-----------
--------IRSS--------G----------K-------KA-------------------
DR------------------------------R---------------------------
------------------------------------------------------------
---------------------GMSLKD---------------------GDME--------
--------LNEEG-----------------------------------------------
------------------------------------------------------------
------------------------------------------------------------
------------------------------------------------------------
-------V
>Deutero_Ambulac_Spurpu_015895_708_233
------------------------------------------------------------
----------M-------------------------------------------------
------------------------------------------------------------
------------------------------------------------------------
------------------------------------------------------------
------------------------------------------------------------
------------------------------------------------------------
------------------------------------------------------------
------------------------------------------------------------
------------------------------------------------------------
------------------------------------------------------------
----DTEEK----------------------D---------TV-----K-----------
-KS--VTDFG-NETTIHGLQFVV---------N-KK--------------NI---IY---
--RLCWL----GICTTFLVVFLIQGNVILK-DFLR----------W---PYSTKI----D
-I-VG-R----------P-NLAF-----PAV-----------------------TVCNAN
MM------RRS-----QIEGSR------------------F-----------------E-
---------DLVNL----------------------------------------------
---------------------------------DG-------GVEGADYDYSWWFSSAYR
-------------------------------------------NWYAS------------
---------SSASS----------------------------------------------
-----YGQSSDQNSNGRSSSSSQSDS---------------ASSSDGQSSSSN---YD--
```

---------PSSSEESSSASSGESSGVTS----------DGQSSS---------------
-----------SSYSPSSSEESSSA----------SSGESSGGTSDGQSSSSND------
--------------------------------------------------------GP--
----------------SSSEEPASSSPGGSADGQSSSTN---------------------
------------------------------------------------------------
------------------------------------------------------YGPPPS
GSVSSFGLPNSEFPAP--------------------------------------------
---------NWVNEWSWY----DPSFFA---------DFEF-SENG-------W------
--------DGV-------------SGGEDWQGFYEA-----------SK------ADDYS
---------------------------------DFLNVI---------------------
-------------------------NP----------------TKEELEQY----GHQLE
--DFI------VQCT----FD-R----R----------P---C-----------N-IS--
----------------------------------------------R-------------
------------------------------------------------------------
------------------------------------------------------------
-----------D----------F-------H-----------VWQ-NRHYG--------N
CFTFN-------TE-------------LS-P-------------------DN----KN--
-------ILTGKT------GG--------LN-------------GLHLTLFVEQP-----
----------E-----------------Y------------L-G-V---LS---------
------------------------------------------------------------
------------------------------------------------------------
----------HQTGAKVTIH-HPNEYP--FPEDNALSLGTGQETSIGIRQE------Y--
-----IKRL-----------GGY-Y--TN-C--TSD-G------KD----------TNFT
---STTE-----------L-SYSSVACKKICYQLHLS----QL--CKCVDDQ----F---
------------Y-----------------------------------------------
----------DGFP---T---KCDVL----------N-----------------------
------------------------------------MTH---QICRK---------F---
V-E-------------------DLFL-D-------DK--------L------------PC
------------SCP-PPCT----------------------------------------
------------------------------------------------------------
------------------------------------------------------------
------------------------------------------------------------
------------------------------------------------------------
------------------------------------------------------------
------------------------------------------------------------
------------------------------------------------------------
------------------------------------------------------------
------------------------------------------------------------
------------------------------------------------------------
------------------------------------------------------------
------------------------------------------------------------
------------------------------------------------------------
------------------------------------------------------------
------------------------------------------------------------
--------
>Deuterostome_chordata_PetMar_tr_S4RTA3_S4RTA3_PETMA_Sodium_channel_epithelia
l_1_alpha_subunit_OS_Petromyzon_marinus_OX_7757_PE_4_SV_1_707_234
------------------------------------------------------------
------------------------------------------------------------
------------------------------------------------------------
------------------------------------------------------------
------------------------------------------------------------
------------------------------------------------------------
------------------------------------------------------------
------------------------------------------------------------
------------------------------------------------------------
------------------------------------------------------------

-----------------------------------------------------------P
PTVLEFWRGSVT----DFY------------D---------SY-----D-----------
-EM--FEFFC-DNTTIHGTIRLV---------C-SK-------------RNK---LK---
--TAFWS----LLFTVTVILFYYTSALVFL-QYYS----------Y---TVAVTM----G
-L-MF------------Q-QSTF-----PAI-----------------------TVCSLN
PY------RYE-----VVQSSL----------------SQL-----------------D-
---------SM------------TGQAL--------------QQLYG----Y--------
---------------------------------QP-------------------------
------------------------------------------------------------
------------------------------------------------------------
----------PATKAAGTAA----------------------------------------
------------------------------------------------------------
------------------------------------------------------------
------------------------------------------------------------
-------------------------APGIRLDT-------------------GVVLERTG
PD----------------------------------------------------------
-------------------TVG--------------------------------------
------------------------------------------FKLC--------------
--------NATG------------------------------GDC---------------
--------------------------------------------FYQSY------GSGVQ
AV-------------------------TEWYTFQYVNIM---------------------
------------------------SQV----------------PSYIKQSD----DANIE
--DFI------FSCM----FS-G----M----------P---C-----------S-D---
----------------------------------------------S-------------
------------------------------------------------------------
------------------------------------------------------------
-----------E----------Y-------S-----------RFH-HPTYG--------N
CYTFN-------SA-------------NS-S-------------------------KL--
-------WQASKP------GR--------DY-------------GLSLILRTEQN-----
----------D-----------------Y------------I-P-F---LS---------
------------------------------------------------------------
------------------------------------------------------------
----------TVAGARIMVH-DQESPP--FMEEGGFDMRPGFETSLGIRML------E--
-----ATRM-----------PDP-Y--GN-C--TED-G------SN------VPV-LNLY
----S-------------S-AYTVQVPECSCFQLALV----EA--CGCGYYF----Y---
------------P-----------------------------------------------
-------LPPNA-----S---YCSY-----------------------------------
------------------------------------NNTAWAGHCYY---------K---
L-Y-------------------RQFI-S-------DE--------L------------GC
----------VDKCA-QPCTTKR---FAVT-PGY------------------AAW-PD--
-------SSS------EKW-I-----------FN--------------------------
------------------------------------------------LL-S--------
----------------------------------LQN-----------------------
-N----------------------------------------------------------
---------------------------YSV------------------------------
-----T-------TV-----------R------NDVAKLNVYFRELNMKTIS-ESAATNV
IWL-----------------LSNIGSQWSLWFGSSVLSWLEVGELGIDC-----------
--------CIMV--------F----------V-------LAY------------------
RRR-----------------------------RS------------RAERRRARGT----
------------------------------------------------------------
------------------------------------------------GDSEAAPP----
------------------------------------------------------------
------------------------------------------------------------
---------------------------------PPPSF----------------------
------------------------------------------------------------
--------
>Placozoa_HhoNaC1_TR8933_c0_g1_i1_m_19478__Hoilungia_hongkongensis__29_235

```
------------------------------------------------------------
----------M-------------------------------------------------
------------------------------------------------------------
------------------------------------------------------------
------------------------------------------------------------
------------------------------------------------------------
------------------------------------------------------------
------------------------------------------------------------
------------------------------------------------------------
-------------------------SKAFDDVRLTSYE---------IDEVSSISYEDQI
DSDD-----------------VFESAESAMPTSHHTEH----------------------
-LNISTKDK------------F-------EFS---------DD-----S-----------
-NY--DDRFA-LQTSCNGIIHIF---------G-RG--------------GR---IR---
--HGIWF----ILTFTMIIFCISTCIQRYD-YLIS----------R---PTSTVI----N
-Y-TV-S----------D-RLKF-----PAV-----------------------SVCNFN
RF------RFS-----SLEYDD------------------W-----------------H-
---------RI------------G-YLI---------------NLFT----ST-------
---------------------------------DS-------------------------
------------------------------------------------------------
------------------------------------------------------------
-----------------N------------------------------------------
------------------------------------------------------------
------------------------------------------------------------
------------------------------------------------------------
--------------------DI-----------FS-------------------------
------------------------------------------------------------
------------------------------------------------------------
------------------------------------------------------------
--GLNGKTGEEW------------------------------NDY---------------
------------------------LKNISIELYD---------------------NITF-
---------------------------------DITQFL---------------------
------------------------------------------------NVK----SNQAK
--IFI------KHCT----WN-N--GRQ----------P---C-----------S-I---
----------------------------------------------N-------------
------------------------------------------------------------
------------------------------------------------------------
-----------N----------F-------T-----------RIY-T-DYG--------S
CFTFN-------AG-------------VK-A-------------------------PI--
-------LYQDRA------GS--------RH-------------GLKLILNIEEE-----
----------E-----------------Y------------T-H-L--------------
------------------------------------------------------------
------------------------------------------------------------
-------NPDPDIGIKFRVH-DQDEPA--DINAEGIAVPPGYHAYTKLEYT------K--
-----SDFL-----------KPP-W--GN-C--GEV--------K-------LK----YF
------------------K-SYNRASCHLECLADSYK----GQ--CSCRTPY----M---
------------P-G---------------------------------------------
----------PF-----P---ICSP-----------------------------------
------------------------------------EHI---KKCIS---------K---
Y-T-------------------GKNR-S-------RN--------------------VTC
------------HCP-NDCQIKS---FHPQ-VTY------------------AEI-PI--
-------QHI------TAA-T-----------AH--------------------------
------------------------------------------------RY----------
------------------------------------------------------------
-G---------------I------------------NELELEFLE---------------
----------------------YNNI-SIN------------------------------
-----D-------YL-----------R------DNYVFLDLFYDDLSYTTFK-ENKAYDE
NQF-----------------ISDIGGQLGLFVGGSFLTFWEIFEWSQIK-----------
```

--------AFLV--------I----------R-------KIIHEY------------KKG
RR------------------------------RT-------------------R------
------------------------------------------------------------
--------------------KRFNSTP---------------------------------
------------------------------------------------------------
------------------------------------------------------------
------------------------------------------------------------
------------------------------------------------------------
-EDTERLL
>Placozoa_HhoNaC2_TR15906_c0_g1_i7_m_30554__Hoilungia_hongkongensis__22_237
------------------------------------------------------------
----------M-------------------------------------------------
------------------------------------------------------------
------------------------------------------------------------
------------------------------------------------------------
------------------------------------------------------------
------------------------------------------------------------
------------------------------------------------------------
------------------------------------------------------------
---------------------------------------------------------DT-
------------------------------------------------------------
----WAKPR----------------------K---------ES-----I-----------
-NL--EKKFA-ERTTCHGLGHIV---------D-QD-------------VPK---PR---
--RYWWS----VVTLAASIGCLIGCIQLLQ-YVLS----------Y---PTNIDI----E
-I-EH-Q----------D-SLVF-----PAV-----------------------TICNFN
QL------TKT-----SLTEED------------------N-----------------R-
---------HL------------E-TLL---------------RMYH----H--------
---------------------------------KG-------SV----------------
------------------------------------------------------------
------------------------------------------------------------
-----------------D------------------------------------------
------------------------------------------------------------
------------------------------------------------------------
------------------------------------------------------------
------------------------------------------------------------
------------------------------------------------------------
------------------------------------------------------------
------------------------------------------------------------
--------KKDL------------------------------EIA---------------
------------------------SSYFRN------------------KT-----GTDF-
---------------------------------HLQSLT---------------------
------------------------------------------------DDL----GHRKN
--DII------ISCL----WD-G----Q----------V---C-----------G-P---
----------------------------------------------E-------------
------------------------------------------------------------
------------------------------------------------------------
-----------N----------F-------T-----------TTF-T-IYG--------N
CYTFN-------SG-------------KP-N--------------------H----SL--
-------VSQQSK------GS--------AF-------------GLRLVLNIEQY-----
----------K-----------------Y------------S-G-K---LS---------
------------------------------------------------------------
------------------------------------------------------------
-------YGSPDAGLRFAVH-SVDDLP--EMDAEGMSIPPGMHAYASIPGV------DI-
-----IQGL-----------TKP-W--GK-C--GQR--------K-------LK----YH
------------------D-HYSVPSCKRELEIDYIL----NE--CSCQQPH----H---
------------P-G---------------------------------------------
----------NS-----T---HCPP-----------------------------------

------------------------------------FLM---FDCVL---------P---
A-L-------------------GNFR-A-------SN--------MES---------IAS
------------KCP-VACIEME---YNAD-VSY------------------SLI-PS--
-------QVA------ADE-I-----------SQ--------------------------
------------------------------------------------KY----------
------------------------------------------------------------
-N---------------V------------------STIIENAQKQGI------------
----------------------NITS-TNI------------------------------
-----E-------FI-----------R------QNLAFLDVYYKDLYTSKTI-QKEVGGF
VSF-----------------LSDIGGQLGLFVGGSFLTIFEIGEYIYDK-----------
--------CFQQ--------T----------K-------RK-------------------
QR------------------------------AI-------------------S------
-----------------------------------------RRVQTFRE-----------
--------------------KRRTASM---------------------RQSP--------
---IHANLSGNLSNLKPNDDI---------------------------------------
-----------KSSRNVRANGIKSKFTR--------------------------------
--------------------------------SPH-------------------------
-------------------------------------------------ITSPTNPISPI
RTDKNGIR
>Placozoa_HhoNaC8_TR7882_c1_g1_i1_m_16465__Hoilungia_hongkongensis__28_238
------------------------------------------------------------
----------M-------------------------------------------------
------------------------------------------------------------
------------------------------------------------------------
------------------------------------------------------------
------------------------------------------------------------
------------------------------------------------------------
----------------SQSS-------------DPDSGK---------------------
----------------------------------------------------------VT
SEDSSGEHPLQ----PRKVSLLMSSEKSHED-----------------------------
-----------------------DSRNLLTLD-FRR------------------------
-------------------------------R---------PS-----I-----------
-EH--DQNFP-FSSTVHGIPHIF---------D-EK--------------YG---TR---
--TILWI----LLVATAVVGCFAFIIIQIV-RYCQ----------F---HSTTKT----S
-L-IY-A----------K-QLEF-----PAI-----------------------TICNYN
SF------RRS-----ALTAND------------------L-----------------V-
---------HM------------A-YLV---------------KAYG----L--------
---------------------------------DD-------------------------
------------------------------------------------------------
------------------------------------------------------------
-------------GMLRK---LLT------------------------------------
------------------------------------------------------------
------------------------------------------------------------
------------------------------------------------------------
------------------------------------------------------------
------------------------------------------------------------
------------------------------------------------------------
------------------------------------------------------------
--------KKEK------------------------------EKL---------------
------------------------IEYWKK---------------YDQTH-----TKKF-
---------------------------------NYQLFV---------------------
------------------------------------------------ERV----GYHAS
--DMI------KSCH----FG-V----E----------E---C-----------G-P---
----------------------------------------------K-------------
------------------------------------------------------------
------------------------------------------------------------
-----------N----------F-------S-----------NVL-T-TYG--------N

```
CITFN-------VG-------------GV-G-------------------------KP--
-------LHQKFP------GS--------NH-------------GLKLLVNIQEY-----
----------E-----------------Y------------T-G-T---YR---------
------------------------------------------------------------
------------------------------------------------------------
-------TDRLDIGIKFVVH-EKNYPP--DVLQQGKAVGPGSHAYASVKYR------T--
-----ISNL-----------PAP-Y--GK-C--SSR--------V-------LP----FY
------------------A-QYTFAGCHICCKTEYIN----DK--CGCRAPD----M---
------------P-G--EN-----------------------------------------
----------TI-----P---VCSP-----------------------------------
------------------------------------QVM---LQCVR---------P---
E-L-------------------VHFQ-S-------EE--------D-----------KVC
------------DCP-IPCFTGH---YDTT-VSY------------------AKI-PN--
-------PQM------AKS-L-----------AV--------------------------
------------------------------------------------TM----------
------------------------------------------------------------
-N---------------K------------------TEFIHETHG---------------
-------------------LV-DPNI-SPT------------------------------
-----L-------YI-----------S------QNYLLLNIFFDDLYYEETT-SLPVTTF
SSL-----------------LGNIGGQLGLFVGASLLTIAELVEYGFYH-----------
--------SRGA--------I----------R--------RY------------------
QQ------------------------------KR-------------------N------
-----------------------------------------QRMSD--------------
------------------------------------------------IAKQ--------
---------TSCA-----------------------------------------------
----------------EKEPLVE----------NG-------------------------
------------------------------------------------------------
------------------------------------------------------------
--KASLPS
>Deu_Ambulacrar_hemi_Ptyfla_40v0_9_20150316_1g20853_t1_scaffold21886_cov93_71
1_240
------------------------------------------------------------
----------M-------------------------------------------------
------------------------------------------------------------
------------------------------------------------------------
------------------------------------------------------------
---SKEGLKHRQ------------------------------------------------
----------------------------------------LP------------------
--LKSGAYNVERGKANQQCQ-------------TDDNSE---------------------
------------------------------------------------------------
----------------------EDDSDVDDK---------------------QQRVDSRT
C----------------------GDGQE--------------------------------
------RSQ----------------------A---------TF-----K-----------
-NV--LADFG-KGTTAHGIVHIT---------E-AE--------------TS---VS---
--RTVWL----AMVVAAAGIMVIQMTLLLV-QYFE----------Y---GVT--------
-V-----------------KTFF-----RGI-----------------------NICTTI
FVIPIEFLRDD--F-KDIMK--------------RRMFSMF-----------------DR
QV-----DRSL---------SVNSTDILP------------SIPLYK-------------
-----------CFESCLR----W-KRYSCRSF-DY-------NRTT--------------
------------------------------------------------------------
----------LTCR-L-------------------FKESAGN-------SGGEV------
-----------VKAIGTD-----------------------F--------------YQN-
------------------------------------------------------------
------------------------------------LR----------------------
-------------------------------------------------TR---------
--------------IP---DT--VFKP---------------------------------
------------------------------------------------------------
```

------------------------------------------------------------
-----------------------------------------GWQMPYQSI----------
--IDDPDIYKYFK-----------------------------ENVFINP-----------
-----------------GYHRVKGEDPPDWLR-----------------------FKSFS
KSP------------------------------DYTELA---------------------
----------------------DVLKV----------------NTNEIAAM----GHQAE
--DFI------LQCT----YG-G----E----------V---C-----------H-P---
-----R----------------------------------------S-------------
------------------------------------------------------------
------------------------------------------------------------
-----------D----------F-------V-----------ITQ-DEVYG--------N
CFHFN-------AD-------------PN----------------------K----DT--
-------RFTRGT------GT--------AH-------------GLSLVLFTEQY-----
----------E-----------------Y------------L-G-I---YG---------
------------------------------------------------------------
------------------------------------------------------------
----------QDSATVVSIV-PRNLRP--FPVDHGFFAMPGTATSVSLTES------K--
-----IHRQ-----------NQP-Y--GN-C--TDE--------NS----------NLHF
----TDNM---------------------------------KK-----------------
------------------------------------------------------------
----------DS-----S------------------------------------------
----------------------------------------------D---------K---
M-----------------------------------------------------------
--------------------------YSIV--------------------------DN--
--------------------L-----------LKLV------------------------
------------------------------------------------KV----------
------------------------------------------------------------
KN-------------QKT------------------RNL---------------------
---------------------------NDL------------------------------
-----E-------SA-----------R------SNLVRLKVYFESLNYESTS-EHPAYTV
RILVCF----------DDVTVSSLGEFRSVCLRLKV--NFDFVSLLLTF-----------
------------------------------------------------------------
------------------------------------------------------------
------------------------------------------------------------
------------------------------------------------------------
------------------------------------------------------------
------------------------------------------------------------
------------------------------------------------------------
------------------------------------------------------------
--------
>Deutero_Ambulac_Apla_gbr412_20_t1_713_242
------------------------------------------------------------
----------M-------------------------------------------------
------------------------------------------------------------
------------------------------------------------------------
------------------------------------------------------------
------------------------------------------------------------
------------------------------------------------------------
------------------------------------------------------------
------------------------------------------------------------
------------------------------------------------------------
------------------------------------------------------------
---GSGDQK----------------------E---------TF-----L-----------
-GA--LNTLL-ETSSAHGLPNIH---------R-SK--------------SF---LS---
--KAAWT----LLFFAGVTALTIQVISLVT-TYLK----------F---EYTVSL----D
-V-RF-D----------R-SLNF-----PAV-----------------------TVCNIN
PV------RKS-----KLERAS------------------------------------D-

---------GF-------------------------------RELFD----V-NFAPPRP
E--------------------------------DQ--T---GA-----------------
------------------------------------------------------------
--------------------VNPGDN----------T-----------------------
-----------------DMSPGSTAGG--------------TMKQAGQPLTTQ-------
GLA---------------------------------------------------------
------------------------------------------------------------
--------------------------------------------------------GAGM
G---------------QGKGQGGSGMPTAGATM------------------------PIA
IP----------------------------------------------------------
------------------------------------------------------------
------------------------------------------------------------
-TGITTEGPETTAGIEY-------------------------SNSTSSVNNG--------
---------------------TTEATILAWEERQ-------TGDFFKKR------DDDYL
------------------------------KEQRLISHL---------------------
------------------------ANL----------------TETQRMDM----GHLLE
--DML------LDCQ----WQ-G----Y----------P---C-----------S-P---
----------------------------------------------A-------------
------------------------------------------------------------
------------------------------------------------------------
-----------N----------F-------T-----------SFY-HYKFG--------N
CYTFN-------SF----------------------------------------------
-----------------------------RF-------------GLTLELFLEQE-----
----------E-----------------F------------M-P-E---IT---------
------------------------------------------------------------
------------------------------------------------------------
----------EAAGFRLVVH-DRDTMP--FPEDDGISISPGSKTAITARVV------T--
-----IERL-----------GNP-Y--GN-C--TKK--------HDE-----PDD-GNIF
----RDR--YG-------V-SYSLKACERSCYQKEVI----SQ--CGCFDPH----Y---
------------P-N--TL-----------------------------------------
--------NDTV-----Y---PCDID----------N-----------------------
------------------------------------DEA---QSCMS---------D---
L-E-------------------EKYK-Q-------GT--------L------------TC
------------YCF-QQC-----------------------------------------
------------------------------------------------------------
------------------------------------------------------------
------------------------------------------------------------
-N----------------------------------------------------------
------------------------------------------------------------
---------------------------------KNVAKIEVYFQEFNFEYIK-QSPAHTI
PSL-----------------MSDIGGQLGLWLGLSILTVFELFEHCGTF-----------
--------LAVI--------V----------S-------KL-------------------
CREGGNL-------------------------KS----------------SSSK------
-----------------------------------------VQKIKIF------------
------------------------------------------------PTED--------
------------------------------------------------------------
------------------------------------------------------------
---------------------------------P--------------------------
------------------------------------------------------------
-GITLQNY
>Protostome_Lophotroco_annelid_CAC9661516_1__Ofus_G111721_Owenia__fusiformis_
_714_243
------------------------------------------------------------
------------------------------------------------------------
------------------------------------------------------------
------------------------------------------------------------
------------------------------------------------------------

```
------------------------------------------------------------
------------------------------------------------------------
------------------------------------------------------------
------------------------------------------------------------
------------------------------------------------------------
------------------------------------------------------------
------------------------------------------M-----K-----------
-RT--ISDFA-SNTTAHGWGNVT---------Q-RT--------------TK---LG---
--RTIWI----GLCLGCTGVAIWQVITIIL-RYGT----------F---DTRDRI----V
-V-KN-G------------DIVF-----PSV-----------------------TVCPLV
GM------PAS-----GYDQLFQDMR---------EGKQET-------------------
--------IPL----------------------------MRLTELLQ----NLTAY----
------------------------------------------------------------
------------------------------------------------------------
------------------------------------------------------------
------------------------------------------------------------
------------------------------------------------------------
------------------------------------------------------------
------------------------------------------------------------
------------------------------------------------------------
------------------------------------------------------------
------------------------------------------------------------
------------------------------------------------------------
--VDNNISQSAQ------------------------------SEKL--------------
-------------------------------------------EFYEEF------HTQLY
---------------------------------SFQGFF---------------------
------------------------DNS-----------------KKSIQDY----SHKQD
--NFI------PVCV----YQ-G----K----------S---C-----------L-K---
----------------------------------------------S-------------
------------------------------------------------------------
------------------------------------------------------------
-----------D----------F-------T-----------TLD-HHEHY--------T
CFTFN-------GG-------------NT-T--------------------I----AK--
-------PSTKIS------GP--------KG-------------GLSLTLYLDAV-----
----------E---------GSLI----YN-------------P-N-YATS---------
------------------------------------------------------------
------------------------------------------------------------
----------GSQGVRVVIH-EKGTLP--DPENDGFDIEPGHSVNAALSVN------E--
-----RQLM-----------KAP-W--GE-C--ADH--------DS------EAY--GSF
--------------------KYTKKHCLLKCLQKTIY----ET--CGCVQSS----L---
------------P-V--DD-----------------------------------------
-----DM--AHT-----R---YCLKL----------HLN-EWVKL------QSNEY----
---------------------------NLTIIQ-NDLEQ---LRCQN-------------
-----------------------NFSLS-------DF-------VGQ----------DGC
------------QCR-EKCLYQS---YEMT-LSQ------------------SVW-PL--
-------EGV------EID-L-----------YC--------------------------
------------------------------------------------RSLM--------
-------------------------------NM-DNY-----------------------
IN-------------SSI------------------YNS-FHDVFPEYC-----------
-----------GHAK-SEEENKILRD-TYG------------------------------
-----I-------KL-----------R------KNVVRLNVYFKDLDTKITQ-QAEDFTI
NSM-----------------ISEIGGSFGFFIGMSVITIAEVIILCLNL-----------
--------AQVI--------R----------K-------QM-------------------
VI------------------------------SP----------------THQE------
------------------------------------------------------------
------------------K--EMARKN---------------------AWVE--------
------------------------------------------------------------
```

```
------------------------------------------------------------
------------------------------------------------------------
------------------------------------------------------------
---TWPKQ
>Deutero_Ambulac_Sakowv30044073m_715_244
------------------------------------------------------------
----------MNS-----------------------------------------------
------------------------------------------------------------
------------------------------------------------------------
------------------------------------------------------------
------------------------------------------------------------
------------------------------------------------------------
---------------------------------TRELS----------------------
------------------------------------------------------------
-----------------------HGTLT-------------------------VESMDAS
P----------------------KFKKR--------------------------------
------MKR----------------------F---------TR-----R-----------
-DI--VNEFS-RNTRCHGLPRII---------S-AK--------------TL---PS---
--RFVWS----LVFFAALGAFVFQATKLIK-LYLN----------Y---DVTVTI----E
-D-ET-I----------A-SLMF-----PAI-----------------------TICNTN
KL------RFS-----EIEKSE------------------H-----------------A-
---------VLL---------RTDPNHPNSLHR--------SLSYQG-------------
---------------------------PCLKG-DF-------------------------
------------------------------------------------------------
-----------ECS-----------------------------------DGIH-------
-CIKPHLHCDGYIHCW-DGL------------------------SDEVNCT-----YP-P
------------------------------------------------------------
------------------------------------------------------------
-----------------CGRD------------------------Q--------------
------------------------------------------------------------
------------------------------------------------------------
------------------------------------------------------------
------------------------------------------F-----------------
----KCNGPGYY---GIC---IDSDKRC---------------DGLPHC----LS-----
--------GE---------------DEAYCNE----------------------------
------------------------------------------------------------
------------------------------------------------------------
------------------------------------------CK----------------
-----------------SGFKCDDNGKCVQEEQRCDRFEDCKDGQDESKCDSVYADCDKN
NLYSPLYPKAYPVEKICTVTPNEHNSFCDYDYLKIQDNMNSDLWISSSNVVTVTFRSDTE
YTARGFHLVYTMRWEVGPWSNCSKRCGGGVQTRAVMCNGASQTGDECKAIGSRP-VSTKV
CKQEVCE-----------------------------------------------------
-------------S-------------TC-Q--------------------S----RL--
-------THCCQV------IQ--------SK-------------GYPDQYTNGQN-----
----------C-----------------Y------------T-N-I---IN---------
------------------------------------------------------------
------------------------------------------------------------
----------EGGCINITFSDFPLEQG--GSCGDFVELSDHNKPTLYRRIC------D--
-----METGDSNPTWGSFSGNV--------------------------------------
------------------------------------------------------------
------------------------------------------------------------
------------------------------------------------------------
------------------------------------------------------------
------------------------------------------------------------
------------------------------------------------------------
------------------------------------------------------------
------------------------------------------------------------
```

------------------------------------------------------------
------------------------------------------------------------
------------------------------------------------------------
------------------------------------------------------------
------------------------------------------------------------
------------------------------------------------------------
------------------------------------------------------------
------------------------------------------------------------
------------------------------------------------------------
------------------------------------------------------------
---------------------SVTYR----------------------------------
------------------------------------------------------------
------------------------------------------------------------
--------
>Protostome_Lophotroco_annelid_CAC9674854_1__Ofus_G26524_partial_Owenia__fusi
formis__718_245
------------------------------------------------------------
----------I-------------------------------------------------
------------------------------------------------------------
------------------------------------------------------------
------------------------------------------------------------
------------------------------------------------------------
------------------------------------------------------------
------------------------------------------------------------
------------------------------------------------------------
------------------------------------------------------------
------------------------------------------------------------
-------QE----------------------M---------TV-----K-----------
-SL--LSDFS-KTTTGHGWGNIH---------E-RK--------------SM---CT---
--KVAWL----IFCIASTLASVGHVTSIIY-RYSL----------F---ETKDRV----L
-V-QR-Q------------DLLF-----PSV-----------------------TICPLT
QM------SKQ-----GILELTQDSI---------NGEDTY-------------------
--------LSI----------------------------FALQTLLA----DMVKIE---
------------------------------------------------------------
------------------------------------------------------------
------------------------------------------------------------
------------------------------------------------------------
------------------------------------------------------------
------------------------------------------------------------
------------------------------------------------------------
------------------------------------------------------------
------------------------------------------------------------
------------------------------------------------------------
------------------------------------------------------------
--QNSTLLMEAS------------------------------SERQ--------------
-------------------------------------------QLYEKF------FNRLY
---------------------------------TYNGFY---------------------
------------------------EN------------------VPTVTKY----GHKKH
--DFI------SSCM----YQ-G----K----------P---C-----------S-D---
----------------------------------------------E-------------
------------------------------------------------------------
------------------------------------------------------------
-----------D----------V-------H-----------SVI-HSHYY--------K
CFTFN-------DI-------------NT-S--------------------L----SN--
-------PYSNTT------GP--------TG-------------GLSLQLFLDTD-----
----------E--------NLYAI----YN-------------P-E-YPTS---------
------------------------------------------------------------

------------------------------------------------------------
----------GSQGVRVVIH-EKGTLP--DPENEGFDIEQGHSANVALSVT------R--
-----RELM-----------KPP-W--GD-C--TEH--------PQ------LDE--DGF
--------------------TFNTRVCQMRCLQKMVY----KS--CGCKRGS----L---
------------P-M--YP-----------------------------------------
-----DV--ENV-----Q---FCNKI----------QYN-EWIKK------NP-------
---------------------------NITALL-YDLEK---LQCGM---------T---
-----------------------EYNWD-------VV-------LDQ----------DDC
------------QCQ-DNCSFHT---YDMT-LSQ------------------SQW-PK--
-------KGV------ELD-F-----------YC--------------------------
------------------------------------------------RKII--------
-------------------------------HL-PNY-----------------------
SN-------------SSV------------------YKA-FHDQIEQLCTD---------
---------IYSNAS--TTKWNEVKQ-EVG------------------------------
-----D-------RL-----------R------ENFIRLNVYFKDLETKIIK-QEPDFAL
GSM-----------------ISEIGGSLGIFIGVSIITIVEVFILCLSI-----------
--------IRVL--------S----------Q-------KS-------------------
KV------------------------------MS----------------IQDP------
------------------------------------------------------------
------------------N--GVNHEK---------------------QWP---------
------------------------------------------------------------
------------------------------------------------------------
------------------------------------------------------------
------------------------------------------------------------
--------
>Protostome_Lophotroco_annelid_CAC9539641_1__Ofus_G033690_Owenia__fusiformis_
_722_247
------------------------------------------------------------
----------M-------------------------------------------------
------------------------------------------------------------
------------------------------------------------------------
------------------------------------------------------------
------------------------------------------------------------
------------------------------------------------------------
------------------------------------------------------------
------------------------------------------------------------
------------------------------------------------------------
------------------------------------------------------------
-----------------------------EGN---------KI-----R-----------
-ET--FVEFT-ENTSAHGCGQIP---------T-AK--------------FK---LT---
--KIFWG----LTFLGCFTWATYSTGNLIA-RYLS----------Y---PTKDVV----S
-I-DF-S------------EIQF-----PAV-----------------------TLCSLQ
PL------TLS-----GMLNLQENPL---------PPDHPTS------------------
---------IL-------------------------------LYVML----NFK-III--
------------------------------------------------------------
------------------------------------------------------------
------------------------------------------------------------
------------------------------------------------------------
------------------------------------------------------------
------------------------------------------------------------
------------------------------------------------------------
------------------------------------------------------------
------------------------------------------------------------
------------------------------------------------------------
------------------------------------------------------------
------------------------------------------DKNIGG------------
-----------------------------------------LGKRYQRQ------LSRIV

---------------------------------SNQFIY---------------------
------------------------ENS----------------DEKNTEGF----IHDLR
--SFL------LGCK----YN-S----K----------T---C-----------D-L---
----------------------------------------------E-------------
------------------------------------------------------------
------------------------------------------------------------
------------------------------------------------------------
-----------K---------LF-------I-----------TYQ-DGQFF--------N
CFTFN-------SE-------------LL-V--------------------P----KA--
-------FIVEKT------DP--------SS-------------GLSLVIFVDAL-----
----------SSAP-----AHYTI----YN-------------P-D-DPTS---------
------------------------------------------------------------
------------------------------------------------------------
----------GKMGVRAIIH-SPGTRP--MPFEKGFDIPTGFSTSVALQGS------I--
-----RNLM-----------SEP-H--GN-C--TTE--------TL------IPG--TNY
--------------------TYSSDTCVQECKQEKLI----KE--CGCKSSL----L---
------------T-A--SA-----------------------------------------
-----DK--PHV-----P---YCGMF----------NMT-NILRQMH---YPTDDE----
---------------------------DITMAV-RDLER---IECEG---------N---
I-L-------------------RLFA---------YP-------AEI----------NKC
------------ACK-EACSSMK---YTKT-ISQ------------------SVW-PN--
-------DNN------QNI-F-----------YE--------------------------
------------------------------------------------TYVN--------
-------------------------------VTDHTL-----------------------
RP-------------NIL------------------F-----------------------
------------------KQQNITEI-INN------------------------------
-----D-------LI-----------H------KNFLRLNIYFESLQVETTS-EVEDYPL
SQL-----------------ISDIGGNMGFYVGISVITLLEFLSLTGTV-----------
--------LLYF--------F----------K-------DR-------------------
---------------------------------------------------LAS------
-----------------------------------------CGQSKTT------------
------------------KVSQINIQH---------------------GDSK--------
--------LDHYTDGFKDKY----------------------------------------
------------------------------------------------------------
------------------------------------------------------------
------------------------------------------------------------
--------
>Protostome_Lophotroco_annelid_CAC9657917_1__Ofus_G102514_Owenia__fusiformis_
_723_248
------------------------------------------------------------
----------M-------------------------------------------------
------------------------------------------------------------
------------------------------------------------------------
------------------------------------------------------------
------------------------------------------------------------
------------------------------------------------------------
------------------------------------------------------------
------------------------------------------------------------
------------------------------------------G-----------------
-----------------------------------------------------------I
SKMDGKETK----------------------K---------KA-----K-----------
-DL--VTEFA-DDATAHGFGLIK---------R-SG--------------NK---YS---
--KILAI----AVVLTCATFAVKNIWEQIK-RYYD----------Y---HYIDTI----S
-I-EE------------R-AIEM-----PSV-----------------------SICTAL
PY------PMT-----IVFQ--------------------------------------E-
---------DIILR---------APSLYRY------ASLTSYGYLLK----NLLETSVS-
---------------------------------DF-------------------------
------------------------------------------------------------

------------------------------------------------------------
------------------------------------------------------------
------------------------------------------------------------
------------------------------------------------------------
------------------------------------------------------------
------------------------------------------------------------
------------------------------------------------------------
------------------------------------------------------------
------------------------------------------------------------
------------------------------------------------------------
---------------------------------------K-LQDGYDYIF-----ENVLF
---------------------------------SQQMGN---------------------
------------------------AHV----------------PDQEMAYL----FHNIN
--NLV------IQCR----FA-G----K----------P---C-----------N-S---
----------------------------------------------S-------------
------------------------------------------------------------
------------------------------------------------------------
-----------F----------F-------T-----------EFR-HPSFG--------R
CFTFN-------GA-------------GL-K--------------------M----DQD-
-------MTIRST------DP--------KS-------------GLHFKVFLDGY-----
----------KPIP----FPNGMF----YD-------------N-K-DPLG---------
------------------------------------------------------------
------------------------------------------------------------
----------GIVGLHVTIH-QPGSVP--FPIEDGFEIPPGYATYVTLKRK------K--
-----RTRL-----------GKP-W--NS-C--EDQ--------LF------LNG--TDF
--------------------AYTKAGCVSQCYQEKIM----HM--CGCVSYD----F---
------------I-I--PK-----------------------------------------
--------NNTM-----V---FCEHL----------DLD-NMFKLIQRQYGSRNLG----
---------------------------NISILL-QDLDK---LKCLV---------N---
I---------------------KNMR-------------------------------LPC
T-----------ECL-PPCVETD---YEHV-QSQ------------------AYW-PG--
-------DMT------LGY-F-----------AR--------------------------
------------------------------------------------SIYERSDIDL--
-N------------------------ENNAWKFIETH----------------------I
IN------------YTTL------------------------------------------
----------------AAKEFQISNI-TQS------------------------------
-----A-------MM-----------H------KNFASLIVYFKEMTTEVTK-QEADYNV
SQL-----------------ICDIGGALGIYIGASLISLYELGKLLVDL-----------
--------CHIV--------L----------H-------RN-------------------
KN------------------------------QV--------------------------
------------------------------------------------------------
------------------------------------------------------------
------------------------------------------------------------
------------------------------------------------------------
------------------------------------------------------------
------------------------------------------------------------
---EEIKT
>Protostome_Lophotroco_annelid_CAC9662966_1__Ofus_G12887_Owenia__fusiformis__
724_249
------------------------------------------------------------
------------------------------------------------------------
------------------------------------------------------------
------------------------------------------------------------
------------------------------------------------------------
------------------------------------------------------------
------------------------------------------------------------
------------------------------------------------------------

------------------------------------------------------------
------------------------------------------------------------
------------------------------------------------------------
------------------------------------------M-----N-----------
-KL--ITDFA-SSTSAHGWGTIG---------Q-RG--------------SK---CG---
--KLLWV----IFTIACQIAAVVWVAIILT-RYLQ----------Y---KTMDRI----K
-M-TN-D----------Y-QVKF-----PSV-----------------------TVCPLN
PT------PIS-----MLGQWLLDYK---------KPDSEL-------------------
--------LKL----------------------------NDYYALLF----KLSGY----
------------------------------------------------------------
------------------------------------------------------------
------------------------------------------------------------
------------------------------------------------------------
------------------------------------------------------------
------------------------------------------------------------
------------------------------------------------------------
------------------------------------------------------------
------------------------------------------------------------
------------------------------------------------------------
------------------------------------------------------------
----LVEGIEIV------------------------------DEEM--------------
-------------------------------------------QSLASK------YNQLD
---------------------------------SHRGYL---------------------
------------------------ENI------------------PYLWKY----GHEQN
--NFI------PGCM----YK-K----K----------T---C-----------V-N---
----------------------------------------------E-------------
------------------------------------------------------------
------------------------------------------------------------
-----------S----------F-------E-----------RIH-HTAFQ--------N
CYTYN-------GA----------------T--------------------L----ND--
-------MYVTSQ------GM--------EG-------------GLSLIIFLDMG-----
----------S--------EGLSL----YN-------------P-Y-HSDS---------
------------------------------------------------------------
------------------------------------------------------------
----------GSSGARVIIH-EKGTMP--DPDNDGFNVEPGLSINVALSVN------R--
-----RELM-----------KQP-W--GQ-C--KDE--------FS------LGL--GKF
--------------------KYSKNACRQKCRKDLIC----EA--CGCVYET----L---
------------P-M--LN-----------------------------------------
-----VTLARDE-----V---LCGKI----------NTD-EWLKN------NP-------
---------------------------NITLLR-NDLDR---LECLE---------K---
Y---------------------LNYTWR-------EI-------SEE----------QGC
------------ECR-ENCTTHS---YNIE-TSQ------------------ALW-PI--
-------EGS------ESG-F-----------YC--------------------------
------------------------------------------------RQIM--------
-------------------------------MLTPNY-----------------------
IN-------------TTI------------------YER-FHERVSDCC-----------
-------------------SNKTRGK-DIG------------------------------
-----D-------EL-----------R------KNFIRLNVYFKDLDTRVII-QDQDFSF
WSM-----------------LCEVGGILGFFIGVSIISIVEFTLLFLNI-----------
--------FNYV--------T----------F-------KRS------------------
KI------------------------------NN----------------HVCS------
------------------------------------------------------------
------------------GNDDIDHKR---------------------KKTE--------
--------HST-------------------------------------------------
------------------------------------------------------------
---------------------------------GLTFY----------------------
------------------------------------------------------------

----QEHK
>Protostome_Lophotroco_annelid_CAC9672326_1__Ofus_G21693_Owenia__fusiformis__725_250
------------------------------------------------------------
----------M-------------------------------------------------
------------------------------------------------------------
------------------------------------------------------------
------------------------------------------------------------
------------------------------------------------------------
------------------------------------------------------------
--------------------------------------------------------PAD-
------------------------------------------------------------
------------------------------------------------------------
-----------------------DENEEK-------------------------------
---PKENNS----------------------G---------NA-----R-----------
-SI--LSNFA-SSTTAHGCSQIN---------Q-SQ--------------SS---VA---
--KCVWL----LIFIGCTIGAIICITALLQ-KYFA----------F---PTKDVI----V
-L-KT-G------------AIAA-----PSI-----------------------SVCSLN
PI------PSS-----FRTRLSGIAP--------DNKVNTV-------------------
------------------------------------------LRALD----LASERVA--
------------------------------------------------------------
------------------------------------------------------------
------------------------------------------------------------
------------------------------------------------------------
------------------------------------------------------------
------------------------------------------------------------
------------------------------------------------------------
------------------------------------------------------------
------------------------------------------------------------
------------------------------------------------------------
------------------------------------------------------------
------------------------------------------NTSFG-------------
-----------------------------------------LHEKLSES------YLRLN
---------------------------------SYIWIY---------------------
------------------------ENA---------------PGENETLQF----AHELD
--DLV------VSCM----YQ-G----I----------P---C-----------D-K---
----------------------------------------------S-------------
------------------------------------------------------------
------------------------------------------------------------
-----------T----------V-------K-----------IFT-NPYFY--------K
CYTFN-------AM-------------DY-N-DT-----------------Y----NG--
-------PFIHST------GT--------KS-------------GISMLFFLDTH-----
----------D--------SGETL----YN-------------P-Y-CPLG---------
------------------------------------------------------------
------------------------------------------------------------
----------GNTGVRVVIH-QPGTIP--DPFNDGFDVAPGYSTNVGITKR------K--
-----RELL-----------GRP-W--GD-C--ERR--------EK------LDD-GLPF
--------------------LYDRRSCKLLCLQAFVA----ER--CGCVTSY----L---
------------P-I--SE-----------------------------------------
-----NM--KNI-----T---RCGQM----------DLE-NFNSA------SP-------
---------------------------NLTTLE-HELDR---LSCEL---------K---
S-V-------------------YVIL-D-------DQ--------DD----------VDC
------------DCP-QSCERDI---YDYT-LSQ------------------SEW-PS--
-------GGV------QLH-F-----------YR--------------------------
------------------------------------------------HIAH--------
-------------------------------LKGEAY-----------------------
FN-------------SSM------------------YHL-LDDKVN--------------

```
------------------TNDSMVNM-KNQ------------------------------
-----R-------MI-----------Q------QNFLRVNTFFTSMDTEIVK-QVEEYPF
TDM-----------------ISGVGGGFGVYVGFSIVTMCEFCVLFVHL-----------
--------LRAM--------I----------M-------DR-------------------
EK------------------------------RK--------------------------
-----------------------------------------VKSAELH------------
------------------SSKVIPTKT---------------------------------
------------------------------------------------------------
------------------------------------------------------------
------------------------------------------------------------
------------------------------------------------------------
------AW
>Protostome_Lophotroco_annelid_Pdum_comp402494_c0_seq1__729_252
------------------------------------------------------------
----------M-------------------------------------------------
------------------------------------------------------------
------------------------------------------------------------
------------------------------------------------------------
------------------------------------------------------------
------------------------------------------------------------
------------------------------------------------------------
------------------------------------------------------------
---------------------------------------------------------DNN
A--------------------DFNKNRNC-------------------------------
---RNVRPQ----------------------E---------KL-----M-----------
-YL--SHRLT-EETTAHGIPHIA---------R-AN--------------GF---WR---
--SLIWI----IISLVMLVTWIYHSQYTIS-QYLK----------M---EVNVKL----E
-F-KT-A----------Q-NLPF-----PAM-----------------------TVCNKN
AF------KNT-----QGMW--------------------LM----------------E-
---------RM-------------------------------IMRFN-------------
------------------------------------------------------------
------------------------------------------------------------
------------------------------------TP-----------EGK--------
-----------------NAS----------------------------------------
------------------------------------------------------------
------------------------------------------------------------
------------------------------------------------------------
-------------------------YPPWLI-----------------------------
------------------------------------------------------------
------------------------------------------------------------
------------------------------------------------------------
SYLKYLATFNGT------------------------------DEPEAGTVQ---------
-------------------------------------------------------TSNPT
Q-----------------------------LGEEMMQLL---------------------
-----------------------YDSII--------KYGP--EYNLTVDDL----TYSFK
--DFV------LTCD----YE-R----T----------P---C-----------E-K---
----------------------------------------------RG------------
------------------------------------------------------------
------------------------------------------------------------
----------VK----------I-------N-----------RIF-NWLHG--------H
CYVLN-------AE-------------KA-T-----------------------------
-------LLSTQT------GP--------RY-------------GLKVALNIDQD-----
----------E-----------------Y------------W-E-----VA---------
------------------------------------------------------------
------------------------------------------------------------
----------EQAGVRVLIH-DPDQMP--FPEDDGYNIPPGLAASLGVRLV------K--
-----QSRL-----------GGS-Y--TP-C--FNE-K------EQ------NEK-HDIF
```

----RAH--YG------WT-KYSLKTCMISCYQREVE----AR--CQCSDIR----Y---
------------P-A--YKKGGG-------------------------------------
-----------------KC-VSCTENM-----TMSFGAE---------------------
---------------------------------MKTHNP---NECLR---------F---
V-Q-------------------EMYR-N-------QS--------L------------NC
----------DKECL-VPCSQNS---YVSS-VSY------------------SGW-PS--
-------TKS------LET-V-----------TQ--------------------------
------------------------------------------------NL----------
------------------------------------------------------------
-KK-----------IPSA------------------AKL---------------------
------------------------LD-ESI------------------------------
-----A-------NI-----------Y------KNFVEVTLYYEDFSYNAVS-ERPAIRL
TEC-----------------LSDLGGNMGMYVGASLFTFFEFFQYLIDV-----------
--------FLWS--------C----------V-W---CCKSL------------------
TRNGDGH------------------------LKELF--------------AATK------
-----------------------------------------VRKIQ--------------
------------------------------------------------------------
------------------------------------------------------------
------------------------------------------------------------
------------------------------------------------------------
------------------------------------------------------------
---VKPEH
>Deutero_Ambulac_Spurpu_018446_731_253
------------------------------------------------------------
----------M-------------------------------------------------
------------------------------------------------------------
------------------------------------------------------------
------------------------------------------------------------
------------------------------------------------------------
------------------------------------------------------------
------------------------------------------------------------
------------------------------------------------------------
------------------------------------------------------------
-----------------------DAPKVNCEES--------------------------S
WGTAPEREE----------------------K---------SL-----R-----------
-TL--LNSRM-ENSSAHGIPNIQ---------R-SS--------------GL---VT---
--KLAWS----LIFLAGIGVMTWQAVILFQ-TYFE----------W---KYSVDI----E
-M-RF-N----------R-TQSF-----PAI-----------------------TICNTN
PV------KRS-----ELETRD--------A-----------------------------
---------SF-------------------------------RRAFD----V-HYVSSTP
---------------------------------DQ-------------------------
------------------------------------------------------------
---------------------QPLP-----------------------------------
-----------------DVPDGV-------------------------------------
------------------------------------------------------------
------------------------------------------------------------
------------------------------------------------------------
-------------------------TPSM-------------------------------
------------------------------------------------------------
------------------------------------------------------------
------------------------------------------------------------
--LNNSDSGNETQG-------------------------GSTNEQVDMS-----------
------------------------EAVNDWRSRIA------IPKFYRMK------SEDYG
------------------------------KKRIRVVTL---------------------
------------------------ANE----------------TLEERVSL----GHKLD
--DML------LDCS----WK-G----I----------P---C-----------S-P---
----------------------------------------------E-------------

```
------------------------------------------------------------
------------------------------------------------------------
-----------N----------F-------T-----------KFY-DSILG--------N
CYTFN-------SG-------------KN----------------------G----EQ--
-------LTTNRP------GS--------TH-------------GLTLELFVQQD-----
----------E-----------------Y------------V-E-G---MT---------
------------------------------------------------------------
------------------------------------------------------------
----------EEAGFRVSIH-HPSKMP--FPQFNGLLVSPGFATNIGLRKL------E--
-----VDRL-----------PKP-Y--GD-C--EAD--------LSK-----NIE-DDIY
----HQH--YS-------I-TYNRKTCEVSCFQNEVI----SR--CDCFDAT----Y---
------------P-N--SL-----------------------------------------
------KVNHTV-----Y---PCEYI----------N-----------------------
------------------------------------DVE---TQCIA---------D---
I-E-------------------MEHA-R-------DE--------L------------EC
------------NCP-LACRETT---YLTG-ASS------------------SIW-PS--
-------DAY------EST-L-----------IE--------------------------
------------------------------------------------KM-L--------
----------------------------------KY------------------------
-N-------------AEI------------------RGH---------------------
----------------------VVGE-NAS------------------------------
-----D-------WT-----------R------RNMAKVEIFYDEFNYEYIR-QDPAYTI
PDL-----------------LSDIGGQLGLWLGLSIITIFEFFEGAWLV-----------
--------LAFF--------C----------S-------RS-------------------
GN------------------------------KT-------------------N------
-----------------------------------------RSEIDPG------------
------------------------------------------------------------
------------------------------------------------------------
------------------------------------------------------------
------------------------------------------------------------
------------------------------------------------------------
----TKTT
>Placozoa_TadNaC8_MK547549_18_259
------------------------------------------------------------
----------M-------------------------------------------------
------------------------------------------------------------
------------------------------------------------------------
------------------------------------------------------------
------------------------------------------------------------
------------------------------------------------------------
----------------SQSS-------------DHDSNK---------------------
----------------------------------------------------------TA
SDESSTDAHPN----SQKVPILSHQDEVDQD-----------------------------
---------------------NPEPRNRFDLD-FRW------------------------
-------------------------------R---------PS-----I-----------
-EH--DENFP-FSASFHGIEHIY---------E-GR--------------YG---TR---
--KILWI----LLVAATMIACFVFIFIQIA-HYSA----------F---HTTTKS----T
-L-VY-E----------K-QLAF-----PAV-----------------------TICNYN
SF------RRS-----AVTAND------------------L-----------------I-
---------HM------------A-YLV---------------KAYR----L--------
---------------------------------NQ-------------------------
------------------------------------------------------------
------------------------------------------------------------
-------------GVVSD---FIG------------------------------------
------------------------------------------------------------
------------------------------------------------------------
------------------------------------------------------------
```

------------------------------------------------------------
------------------------------------------------------------
------------------------------------------------------------
------------------------------------------------------------
--------EKER------------------------------QKL---------------
------------------------INYWKK---------------YDATH-----AKKF-
---------------------------------NYQLFV---------------------
------------------------------------------------ERV----GYHAS
--QMI------KSCH----FR-G----L----------K---C-----------G-P---
----------------------------------------------K-------------
------------------------------------------------------------
------------------------------------------------------------
-----------N----------F-------S-----------NVL-T-SYG--------N
CITFN-------GP-------------KLES--------------------N----PK--
-------LYQKNP------GA--------HQ-------------GLELLINIQEY-----
----------E-----------------Y------------T-G-S---WH---------
------------------------------------------------------------
------------------------------------------------------------
-------SDRPDIGIKFVIH-ERHYPP--DVTSQGKAVGPGSHAYASVKYK------T--
-----ISNL-----------PSP-Y--GH-C--GSK--------K-------LA----FY
------------------K-KYTYAGCQISCKTEYVQ----KK--CGCRAPD----M---
------------P-G---------------------------------------------
---------QNII----P---VCSP-----------------------------------
------------------------------------QKM---IECVS---------P---
N-L-------------------EKLI-TI------ND--------------------KVC
------------ICP-IPCHIVH---FDTT-ISY------------------AKI-PN--
-------PQM------AKD-L-----------TE--------------------------
------------------------------------------------KI----------
------------------------------------------------------------
-N---------------K------------------TAFQIETHG---------------
-------------------LV-DPDI-DPT------------------------------
-----L-------YI-----------S------QNYILLNVFFDDLYYEKTV-STPVYTF
TSL-----------------LGNIGGQLGLFVGASVLTLVEIIEFGFYR-----------
--------SRGV--------I----------R-------RS-------------------
DW------------------------------KQ-------------------N------
-----------------------------------------LRKSISR------------
------------------------------------------------SREV--------
---------TATE-----------------------------------------------
----------------EKEPLCS-------------------------------------
------------------------------------------------------------
----------------------------------------------------------VE
NGDTQLSK
>Protostome_Lophotroco_annelid_CAC9522438_1__Ofus_G031129_Owenia__fusiformis_
_736_261
------------------------------------------------------------
----------M-------------------------------------------------
------------------------------------------------------------
------------------------------------------------------------
------------------------------------------------------------
------------------------------------------------------------
------------------------------------------------------------
------------------------------------------------------------
------------------------------------------------------------
------------------------------------------------------------
------------------------------------------------------------
------------------------------------------------K-----------
-RI--INEFT-SNTSGHGWGMIN---------Q-TS--------------SK---FG---

--KALWV----AITLLSTIAAIIWVSTIIV-RYVK----------Y---ETIDRV----E
-A-KV-D----------E-DIIF-----PSV-----------------------TVCPLY
GI------SSM-----KIAEVYSDPN---------SDY----------------------
---------------------------------------ISLNNFLS----SS--Y----
------------------------------------------------------------
------------------------------------------------------------
------------------------------------------------------------
------------------------------------------------------------
------------------------------------------------------------
------------------------------------------------------------
------------------------------------------------------------
------------------------------------------------------------
------------------------------------------------------------
------------------------------------------------------------
------------------------------------------------------------
--ALVDELAQIT------------------------------NETY--------------
------L------------------------------------SIMTNT------ILVAR
---------------------------------SNRGYY---------------------
------------------------ANL----------------GVSGVSTI----GHEFG
--DFI------PYCL----YQ-E----K----------E---C-----------S-A---
----------------------------------------------A-------------
------------------------------------------------------------
------------------------------------------------------------
-----------D----------F-------Q-----------KLV-HHEYK--------N
CYTFN-------GG-------------DI-N--------------------I----SK--
-------PIISST------GA--------QK-------------GLSLTLYLENS-----
----------E--------NAYVA----YN-------------V-N-RVMT---------
------------------------------------------------------------
------------------------------------------------------------
----------AASGARVIIH-EKGTLP--DPDNEGFDVEPGHLISVALSAN------K--
-----RQLL-----------KQP-W--GE-C--AEH--------NT------LHS--TDY
--------------------KYTRNTCRIKCIEKVIQ----RQ--CGCRYDL----L---
------------P-V--DI-----------------------------------------
----NATTELDI-----Q---PCGKF----------NIK-EWLKA--------TEA----
---------------------------NTTVLK-EDLAK---VICSK---------E---
-----------------------EHQWT-------AL-------AKD---------IPSC
------------ECE-YNCTYVD---YEIE-TSQ------------------SVW-PM--
-------RGS------ELD-F-----------FC--------------------------
------------------------------------------------QL----------
-----------------------------------SY-----------------------
LN-----------ISGSF------------------FER-FQNSIGIHC-----------
------------NSF---MEYINVAQ-NHG------------------------------
-----D-------EL-----------R------ANVLRVNVYFKTLEVKYTI-QDEGFSL
VSM-----------------ISEIGGVLGIFVGVSIITILEIFVLCSGI-----------
--------VHVI--------I----------N-------SK-------------------
KV------------------------------SS----------------EINV------
------------------------------------------------------------
------------------KEIKSTSKS---------------------KLEE--------
--------SSN-------------------------------------------------
------------------------------------------------------------
-------------------------------HKPDDSY----------------------
------------------------------------------------------------
-YDQFKKY
>Protostome_Lophotroco_annelid_CAC9477543_1__Ofus_G013338_Owenia__fusiformis_
_739_262
------------------------------------------------------------
----------M-------------------------------------------------

```
------------------------------------------------------------
------------------------------------------------------------
------------------------------------------------------------
------------------------------------------------------------
------------------------------------------------------------
--------------------------------------------------------DTK-
------------------------------------------------------------
------------------------------------------------------------
-------------------KENMSPNESV-------------------------------
---SEKKKD----------------------S---------SI-----K-----------
-SI--LSTFA-SSTTAHGCSRIN---------E-SQ--------------SA---RG---
--KLIWM----AIFVACFIGATVCITSLLQ-KYFK----------F---PTKDVI----E
-V-SV-D------------PISF-----PSV-----------------------SFCSLN
PV------PSS-----YQRKKSRMKP--------GNRFLDV-------------------
------------------------------------------AATYN----NLEAHLF--
------------------------------------------------------------
------------------------------------------------------------
------------------------------------------------------------
------------------------------------------------------------
------------------------------------------------------------
------------------------------------------------------------
------------------------------------------------------------
------------------------------------------------------------
------------------------------------------------------------
------------------------------------------------------------
------------------------------------------------------------
------------------------------------------NESFP-------------
-----------------------------------------LYDRMSPN------RLRIK
---------------------------------SYKWMY---------------------
------------------------DNA---------------PNEDETLAI----GHDLE
--DIL------LSCS----IA-G----Y----------D---C-----------N-I---
----------------------------------------------S-------------
------------------------------------------------------------
------------------------------------------------------------
-----------Q----------T-------Q-----------SFT-SPFYF--------K
CYTLN-------AK-------------DF-N-DS-----------------S----IE--
-------PLVHST------GA--------RS-------------GLSAIFFLDTH-----
----------D--------KAKGI----YN-------------P-S-CPIG---------
------------------------------------------------------------
------------------------------------------------------------
----------GSSGLHVVIH-PPGTQP--DPINEGFDIVPGFATNIGLQKK------S--
-----RELL-----------GYP-W--GD-C--ENR--------ES------LSE--LPY
--------------------KYDKLSCERQCQQKFIT----DQ--CGCVTSF----Q---
------------P-I--PD-----------------------------------------
-----DL--KNM-----S---MCGKM----------DLK-NWNSA------SP-------
---------------------------NMSKIA-EEMDT---IECEF---------G---
K-F-------------------WVLK-K-------DY-------PEE----------NKC
------------ECP-SACHNDI---YDYT-ISQ------------------SEW-PS--
-------GGI------KLD-F-----------YK--------------------------
------------------------------------------------HVMHMRN-----
-------------------------------MRGSGY-----------------------
IN-------------STM------------------YDL-LDSKLN--------------
------------------TNDTDENH-KNQ------------------------------
-----G-------MV-----------R------QNFLRLNIFFTSLNTEIVK-QVEEYPF
TDM-----------------ISGVGGGFGVYVGFSFVTICEFCVLFAHL-----------
--------LKAI--------M----------N-------HR-------------------
NKAVEDIP----------------------------------------------------
```

------------------------------------------------------------
------------------SGNIITVKE---------------------------------
------------------------------------------------------------
------------------------------------------------------------
------------------------------------------------------------
------------------------------------------------------------
-----YKM
>Protostome_Lophotroco_annelid_CAC9510306_1__Ofus_G024263_Owenia__fusiformis_
_740_263
------------------------------------------------------------
------------------------------------------------------------
------------------------------------------------------------
------------------------------------------------------------
------------------------------------------------------------
------------------------------------------------------------
------------------------------------------------------------
------------------------------------------------------------
------------------------------------------------------------
------------------------------------------------------------
------------------------------------------------------------
------------------------------------------M-----K-----------
-QI--LNEFA-SNTSGHGWGMVG---------R-TT--------------NA---FA---
--KTLWI----VITVLSTIAAIIWVATIVG-RYIK----------Y---ETKDKV----E
-L-KA-D----------V-DIVF-----PSV-----------------------TVCPLH
GY------SND-----KMN----DPKL-------VSSF----------------------
---------------------------------------LLQSRLIM----AV--Y----
------------------------------------------------------------
------------------------------------------------------------
------------------------------------------------------------
------------------------------------------------------------
------------------------------------------------------------
------------------------------------------------------------
------------------------------------------------------------
------------------------------------------------------------
------------------------------------------------------------
------------------------------------------------------------
------------------------------------------------------------
--PRIKNITQMT------------------------------NETG--------------
-------------------------------------------DVILEL------VHRIR
---------------------------------SHRGYY---------------------
------------------------ENM----------------GKRETSAL----SHEII
--NLV------PYCL----YQ-E----S----------R---C-----------T-P---
----------------------------------------------E-------------
------------------------------------------------------------
------------------------------------------------------------
-----------H----------F-------Q-----------KLV-HHEYM--------N
CYTFN-------GK-------------DI-N--------------------L----TK--
-------PVISSS------GA--------QN-------------GLSLTLFLEQK-----
----------D--------DTYAP----YD-------------R-S-RAIT---------
------------------------------------------------------------
------------------------------------------------------------
----------ATAGARVIIH-EKGTLP--DPDNDGFNVEPGHLVNVALSVN------K--
-----RELM-----------KPP-W--GE-C--ADH--------NT------LQS--TDF
--------------------KYSRNACKSKCLEKMIQ----QE--CGCKKDL----L---
------------P-V--EL-----------------------------------------
-----ETRDDDI-----Q---ACGKF----------NRE-EWLKVP-----NITEA----
---------------------------NITILK-EDVTR---FKCGE---------K---

```
-----------------------EYAWS-------KI-------SED---------IPGC
------------KCE-YNCSYLT---YDIE-TSQ------------------SLW-PM--
-------KGS------EMD-F-----------FC--------------------------
------------------------------------------------GGLA--------
-------------------------------YE-DTY-----------------------
VG-------------STI------------------YDE-FHDKIGMYC-----------
-----------LNYF---GQYANISK-LWG------------------------------
-----D-------KL-----------R------DNFLRVNVYLKEPVTKYTV-EEVDFTV
ITM-----------------VSEIGGVLGIFIGVSIITILEVFVLCNGI-----------
--------VYTL--------Y----------R-------AK-------------------
KT------------------------------SS----------------NIDV------
------------------------------------------------------------
------------------TEVIVTQSK---------------------NSEN--------
--------LSD-------------------------------------------------
------------------------------------------------------------
---------------------------------PKTKY----------------------
------------------------------------------------------------
-YQSKLSK
>Protostome_Lophotroco_annelid_CAC9574840_1__Ofus_G053378_Owenia__fusiformis_
_743_264
------------------------------------------------------------
----------M-------------------------------------------------
------------------------------------------------------------
------------------------------------------------------------
------------------------------------------------------------
------------------------------------------------------------
------------------------------------------------------------
------------------------------------------------------------
------------------------------------------------------------
------------------------------------------------------------
------------------------------------------------------------
---MKEQNKM-------------------DGK---------SI-----Q-----------
-ER--VKEFI-ETTSAHGCGQIP---------T-AK--------------FK---LT---
--KFLWG----LTFLSFFTFATVSSGILIA-KYFT----------Y---PTKDVV----S
-I-EF-G------------NIEF-----PAV-----------------------TFCSLQ
PI------TIT-----SLIDFRQNPP---------PPGHSVS------------------
---------NM-------------------------------IYVMQ----NIRNIII--
------------------------------------------------------------
------------------------------------------------------------
------------------------------------------------------------
------------------------------------------------------------
------------------------------------------------------------
------------------------------------------------------------
------------------------------------------------------------
------------------------------------------------------------
------------------------------------------------------------
------------------------------------------------------------
------------------------------------------------------------
------------------------------------------DRDIGG------------
-----------------------------------------LGQWYLKH------FDRFL
---------------------------------SNQFIF---------------------
------------------------ENG----------------EENDMEQF----IHDLR
--SFK------LGCK----FQ-G----Q----------V---C-----------N-G---
----------------------------------------------T-------------
------------------------------------------------------------
------------------------------------------------------------
-----------S----------F-------V-----------LHK-DGGFH--------N
```

CFTFN-------GA-------------GS-D--------------------L----SN--
-------VNIVRT------DP--------AS-------------GLSLVLFMDAY-----
----------AKGK---SVSAYTI----YN-------------P-D-DPTS---------
------------------------------------------------------------
------------------------------------------------------------
----------GQTGVRAVIH-SPRTRP--MPFDKGFDIPTGLSTSVALKET------K--
-----RELM-----------TQP-H--NN-C--TND--------EF------NQG--TNY
--------------------SYSVDTCYEQCQQDLLI----EE--CGCKSSL----L---
-----------IP----PS-----------------------------------------
----FDK--SDL-----E---YCGKV----------NIT-NMMLMMN----QNDDI----
---------------------------TQSIAE-KELEN---LDCEE---------G---
V-M-------------------YSFG----------NL------AIL----------EQC
------------ACK-DHCQTIN---YVKT-ISQ------------------AVW-PN--
-------DNA------QEA-F-----------YD--------------------------
------------------------------------------------TYVN--------
-------------------------------RTDQTL-----------------------
RP-------------NKL------------------F-----------------------
------------------KGYNTSEI-INN------------------------------
-----D-------MI-----------H------KNFLRLNIYFESLYVETTS-EVEDYPA
GTL-----------------ISEIGGNMGFYVGISIITLMEIFCITGAI-----------
--------ILYC--------F----------K-------DW-------------------
---------------------------------------------------LIK------
-----------------------------------------CGHMNKT------------
------------------KVSHVNVKP---------------------VELD--------
--------GDSTEKI-KDKY----------------------------------------
------------------------------------------------------------
------------------------------------------------------------
------------------------------------------------------------
--------
>Protostome_Lophotroco_annelid_CAC9674171_1__Ofus_G24722_Owenia__fusiformis__
745_266
------------------------------------------------------------
----------M-------------------------------------------------
------------------------------------------------------------
------------------------------------------------------------
------------------------------------------------------------
------------------------------------------------------------
------------------------------------------------------------
------------------------------------------------------------
------------------------------------------------------------
------------------------------------------------------------
------------------------------------------------------------
-----------------------------DGK---------NM-----I-----------
-AT--VKEFT-ENTSAHGCGQIP---------T-SK--------------FK---LT---
--KFLWG----LIFVGFFMFATISSVNLIT-IYFT----------F---PTKDVV----S
-V-NF-E------------DIEF-----PAV-----------------------TFCTQQ
PV------TAT-----ALIDFIDNQS---------HRDDAIG------------------
---------KL-------------------------------VLTMG----SFPGMLV--
------------------------------------------------------------
------------------------------------------------------------
------------------------------------------------------------
------------------------------------------------------------
------------------------------------------------------------
------------------------------------------------------------
------------------------------------------------------------
------------------------------------------------------------
------------------------------------------------------------

------------------------------------------------------------
------------------------------------------------------------
------------------------------------------KPNIGG------------
-----------------------------------------LGDWYFKY------SYRFL
---------------------------------SNQFVF---------------------
------------------------ENL----------------DKKDLKTI----QHDLK
--SLK------ISCK----FK-G----H----------P---C-----------K-D---
----------------------------------------------S-------------
------------------------------------------------------------
------------------------------------------------------------
-----------A----------F-------V-----------VHQ-DGGFQ--------H
CFTFN-------GA-------------GA-Q--------------------L----ND--
-------TKVIKA------DP--------TS-------------GLSLILFVDAY-----
----------TTKR---DLSELTL----YN-------------P-F-DPTS---------
------------------------------------------------------------
------------------------------------------------------------
----------GQSGVRVIIH-SPKTRP--MPFEKGFDIPTGFSTSVALKET------K--
-----RELM-----------TEP-H--GN-C--TMA--------KF------NGG--TNY
--------------------AYSEDTCLEQCKQKILI----EN--CRCMSSL----L---
------------P-T--PS-----------------------------------------
-----GE--LKP-----Q---YCGKV----------NVT-NMFTMMY---GKLDAT----
---------------------------MRGIAV-KELEM---LDCET---------K---
L-M-------------------ESLG---------NS-------EVL----------NKC
------------ECK-KPCVHTN---YVKM-LSQ------------------AVW-PS--
-------DYN------QKN-F-----------LA--------------------------
------------------------------------------------ESIN--------
-------------------------------MSDPTQ-----------------------
RA-------------TIL------------------F-----------------------
------------------EGLDIGQI-TKNES----------------------------
-----E-------MI-----------Q------KNFLRFNVYFESLQVKTTS-QVEDFPA
STL-----------------ISEIGGNMGFYVGISIITLMEMLTLTGAI-----------
--------ILYC--------F----------K-------DL-------------------
---------------------------------------------------IIK------
-----------------------------------------CGQSNTT------------
------------------KVPDVNIKQ---------------------IKSY--------
--------DNHMDED-KENI----------------------------------------
------------------------------------------------------------
------------------------------------------------------------
------------------------------------------------------------
--NAEIHG
>Protostome_Lophotroco_annelid_CAC9602553_1__Ofus_G07376_Owenia__fusiformis__
749_267
------------------------------------------------------------
----------M-------------------------------------------------
------------------------------------------------------------
------------------------------------------------------------
------------------------------------------------------------
------------------------------------------------------------
------------------------------------------------------------
------------------------------------------------------------
------------------------------------------------------------
------------------------------------------------------------
------------------------------------------------------------
-------------------------------E---------KL-----K-----------
-EQ--LKHFA-ESTTAHGLARIP---------T-SR--------------TK---IV---
--KFIWI----IIILGCGITSFVFLYQSFA-KYLA----------F---KPKDVV----S
-I-SR-E----------I-TVKF-----PSV-----------------------TVCPLY

PI------ATG-----KNFNIYDYLD---------NNSTYVY------------------
---------KL-------------------------------NQIIS----GFYEKF---
---------------------------------DE-------------------------
------------------------------------------------------------
----------------------PE------------------------------------
-----------------D------------------------------------------
------------------------------------------------------------
------------------------------------------------------------
------------------------------------------------------------
--------------------------P---------------------------------
------------------------------------------------------------
------------------------------------------------------------
------------------------------------------------------------
------------------------------------------------------------
-----------------------------------------LYAKYIRH------KNRVV
---------------------------------AWQWIY---------------------
------------------------ENN------------------PNITNA----SHSWP
--DLV------PICS----FK-G----K----------P---C-----------K-A---
----------------------------------------------E-------------
------------------------------------------------------------
------------------------------------------------------------
-----------H----------I-------E-----------KFV-DPNYY--------N
CYTFN-------GM-------------NS-S-TLNGENHT---------STE----DE--
-------MIVESI------GP--------TS-------------GLSLIMFLDLH-----
----------E--------EQRSM----FN-------------P-M-IPTS---------
------------------------------------------------------------
------------------------------------------------------------
----------GSAGIRVVIH-ETGTVP--DPTANGFDIPPGFSTNVPLRLI------Q--
-----RNHM-----------EEP-W--GT-C--DKK--------ADPY----LKG--SRY
--------------------TYSRSSCKRQCVQGLLK----AK--CGCISSL----W---
------------P-I--NN-----------------------------------------
-----DA--DDS-----K---YCGTF----------NID-EWIKQ------ES-------
---------------------------NRSILE-EDLLR---LDCEE---------D---
L-L-------------------SHLT-E------ISEFIQSPDNPDC----------HDC
------------DCR-PACEHYK---YYQD-ISM------------------SYW-PM--
-------ESA------QMT-F-----------YE--------------------------
------------------------------------------------SLMTM-------
----------------------------------PGY-----------------------
-N------------DSTI------------------YKL-LHKPHD--------------
----------------DLGTADNSSQ-VNK------------------------------
-----D-------LI-----------R------KNFLRLNIYYKTIETEVIS-MYAEFTS
GEL-----------------VAEVGGTLGIFLGVSFVTLCEVVGLLSNL-----------
--------ISSL--------L----------C-------NK-------------------
NR------------------------------KI-------------------K------
------------------------------------------------------------
------------------DTHDLSEKI---------------------SKYD--------
------------------------------------------------------------
------------------------------------------------------------
------------------------------------------------------------
-----------------------------------------------------------A
NDVKIVNQ
>Protostome_Lophotroco_annelid_CAC9607166_1__Ofus_G071156_Owenia__fusiformis_
_751_268
------------------------------------------------------------
----------M-------------------------------------------------
------------------------------------------------------------
------------------------------------------------------------

```
------------------------------------------------------------
------------------------------------------------------------
------------------------------------------------------------
------------------------------------------------------------
------------------------------------------------------------
------------------------------------------------------------
------------------------------------------------------------
----GFKKD----------------------N---------TF-----K-----------
-DI--VSEFA-NDTATNGVPNIA---------R-AG--------------SI---PR---
--RIIWT----IIVLVAAGWMFTQLAQSFI-TYYK----------R---PHSTLL----T
-E-TF-S----------S-KIYF-----PAV-----------------------TICNIN
PV------RES-----QIYLSN------------------------------------ST
---------EI-------------------------------EALLR-------------
------------------------------------------------------------
------------------------------------------------------------
------------------------------------------------------------
------------------------------------------------------------
------------------------------------------------------------
------------------------------------------------------------
------------------------------------------------------------
--------------------------PTSQKEG---------------------------
------------------------------------------------------------
------------------------------------------------------------
------------------------------------------------------------
------------------------------------------------------------
-------------------------------------------------------SERFS
------------------------------QLTELQRSM---------------------
------------------------QTL----------------PTSSLQAM----GHLPE
--TMI------MSCT----YA-G----E----------D---C-----------S-Y---
----------------------------------------------S-------------
------------------------------------------------------------
------------------------------------------------------------
-----------D----------F-------T-----------MFT-NYKYG--------N
CITYN-------AG-------------------------------------P----DI--
-------VTSSQA------GA--------QH-------------GLSLELFVEEN-----
----------E-----------------Y------------L-T-----LT---------
------------------------------------------------------------
------------------------------------------------------------
----------DTVGFKVSIE-RQNKAL--FPEETGIFVPVGARTALSLKRQ------E--
-----IIRL-----------PDP-YS-SD-C--WNL--------TANQ----SIY-DNAY
----SDE--TEP----SPK-NYTRLACLKTCYQLNLI----TQ--CACLSHK----I---
------------K-T--RG-------------------TAYDKAN---------------
------VNLEEL-----D---FCNLT----------------------------------
------------------------------------------DSCYL---------K---
V-K-------------------DDYE-D-------GS--------L------------DC
------------NCN-SECWELA---YSAT-ISQ------------------TSW-PA--
-------PKY------LDD-L-----------LR--------------------------
------------------------------------------------TY-A--------
----------------------------------AK------------------------
-N-------------DVI-----------------RGLLESPSSRKKRDVGN-------
---------GTETQNGNGTVNGNVTN-NGN------------------------------
-----E-------TV-----------NENGT-DRYFAKLDVYFDELNFIRIE-ETIAYTE
WNL-----------------FSDLGGQFGFWLGFSIVSIFEIIEFLIDV-----------
--------SIFL--------V----------Y-------KS-------------------
LN------------------------------RE-------------------K------
-----------------------------------------LKAKVGD------------
------------------------------------------------SATE----LE--
```

-----GRPVSS-------------------------------------------------
------------------------------------------------------------
--------------------------------GPQ-------------------------
----------------------------------------------VFVTHGGETKFGLD
ENVKAGVI
>Protostome_Lophotroco_annelid_CAC9672069_1__Ofus_G21330_Owenia__fusiformis__752_269
------------------------------------------------------------
------------------------------------------------------------
------------------------------------------------------------
------------------------------------------------------------
------------------------------------------------------------
------------------------------------------------------------
------------------------------------------------------------
------------------------------------------------------------
------------------------------------------------------------
------------------------------------------------------------
------------------------------------------------------------
------------------------------------------M-----K-----------
-QT--LNDFA-SNTTAHGWGNIH---------Q-RT--------------TT---LS---
--KAVWI----ILCLGCTGVAIWQVITIVI-RYGL----------F---ETKDRI----Q
-V-EE-G------------DITF-----PSV-----------------------TVCPLV
GI------PKS-----ENSKLLKDLY---------NGKIDA-----------------DT
-------YAEM----------------------------LAFDTQIV----YSSIY----
------------------------------------------------------------
------------------------------------------------------------
------------------------------------------------------------
------------------------------------------------------------
------------------------------------------------------------
------------------------------------------------------------
------------------------------------------------------------
------------------------------------------------------------
------------------------------------------------------------
------------------------------------------------------------
------------------------------------------------------------
--LDSNISQNAY------------------------------REKL--------------
-------------------------------------------EFHNVF------SKQLF
---------------------------------SFQGYI---------------------
------------------------ENF------------------NIFRDY----SHKQE
--EFI------PICR----YQ-G----K----------P---C-----------L-K---
----------------------------------------------S-------------
------------------------------------------------------------
------------------------------------------------------------
-----------D----------F-------Q-----------SLD-HHEHN--------K
CYTFN-------GA-------------NN-T--------------------I----SN--
-------PSTNIT------GP--------KG-------------GLSLVLFVDAS-----
----------E---------ASQV----YN-------------P-D-YPTS---------
------------------------------------------------------------
------------------------------------------------------------
----------GSQGVRVVIH-EKGTFP--DPENDGFDIEPGHSVNAALSVN------Q--
-----RQLL-----------KPP-W--GK-C--ANY--------DA------EVY--GGF
--------------------TYRKKLCQLSCTQRFIY----KT--CGCIKSS----L---
------------P-V--DE-----------------------------------------
-----VT--AHK-----E---YCLKI----------HPQ-EWLKL------NSSQY----
---------------------------NLTIIQ-NDLEK---LRCQN---------R---
-----------------------EYPSS-------DI-------LAQ----------DGC
------------QCR-EKCMYHT---YDMT-LSQ------------------AEW-PM--

```
-------RGV------EIT-F-----------YC--------------------------
------------------------------------------------RLIM--------
-------------------------------TA-DNY-----------------------
IN-------------SSI------------------YDT-FHNI-SEYC-----------
-----------DFAK-PENESKILRE-RYG------------------------------
-----D-------TL-----------R------QNFIRLNVYFKDLEMKVIK-QAEDFTI
NSM-----------------ISEIGGSLGFFVGMSIITIAEIFLLCWNI-----------
--------TGVL--------K----------N-------RVENR----------------
VV------------------------------ST----------------PHQE------
------------------------------------------------------------
------------------N--KTDQKS---------------------ALER--------
--------ANR-------------------------------------------------
------------------------------------------------------------
------------------------------------------------------------
--------------------------------------------------------SNAW
EDEKSIKW
>Protostome_Lophotroco_annelid_Pdum_comp402143_c0_seq3__753_270
------------------------------------------------------------
----------M-------------------------------------------------
------------------------------------------------------------
------------------------------------------------------------
------------------------------------------------------------
------------------------------------------------------------
---------------------------------YENSYIIFPLKM---------------
------------------------------------------------------------
------------------------------------------------------------
---------------------------------------------------------SPI
KPKTT---------------IAFTEKDVI-------------------------------
------KKP----------------------T---------EK-----E-----------
-IL--VKEFL-DETTLHGVTHIA---------K-AK--------------GP---AT---
--TFLWA----FITLIMIIACILHSKESVA-KFLE----------Y---KANTEI----H
-I-LS-K----------R-QLAF-----PAV-----------------------TLCNKN
PF------KKE-----QFYK--------------------FF----------------N-
---------AV---------LSGMKNARAN--------HYHNASLYG-------------
------------------------------------------------------------
------------------------------------------------------------
------------------------------------------------------------
------------------------------------------------------------
------------------------------------------------------------
------------------------------------------------------------
------------------------------------------------------------
--------------------------PEWLLRAERMYPG---------------------
------------------------------------------------------------
------------------------------------------------------------
------------------------------------------------------------
------------------------------------------HEPFPEILLR--------
-------------------------------------------------------NPEYG
DL----------------------------ILQSMSEMV---------------------
---------------------EEYGHL----------------YGYTIDNI----TYSQG
--DLL------LTCL----YM-Q----K----------P---C-----------N-T---
----------------------------------------------S-------------
------------------------------------------------------------
------------------------------------------------------------
-----------N----------V-------EV---------REFI-SWLYG--------K
CFKIV-------PK----------------------------------------------
-------KDCTRI------GP--------RF-------------GLQLTLNIDQD-----
----------S-----------------Y------------L-V-----TS---------
```

```
------------------------------------------------------------
------------------------------------------------------------
----------TYAGVQVEIH-SPDTMP--FPEDKGIAISPGQAGFIAVKVS------K--
-----HRKL-----------NGK-F--GD-C--FDE--------EQQ-----EKA-QDVF
----RET--HD------WV-EYGQKNCERTCFQREMI----QK--CKCADIR----F---
------------P-R--QG-----------------------------------------
--YSVNFPWRRA-----P---ACLLDK-----LYSFGLV---------------------
----------------------------YKSLEVPKNKS---SDCYE---------K---
V-I-------------------NMFH-A-------NN--------L------------SC
H--------GEDNCH-QPCKETY---FDSS-ISL------------------SDW-PS--
-------EAS------TEY-V-----------VD--------------------------
------------------------------------------------GL----------
------------------------------------------------------------
RNK-----------SKSA------------------DKVIK-------------------
---------------------------AGI------------------------------
-----A-------NI-----------S------KNFLELNVFYTVLSLERIT-QTISVEV
TEM-----------------LSDLGSNIGMYLGASLFAIFELFKIFFDT-----------
--------TLLS--------C----------K-------RP-------------------
NV------------------------------KK-------------------K------
-----------------------------------------KTKADKEQI----------
------------------------------------------------------------
------------------------------------------------------------
------------------------------------------------------------
------------------------------------------------------------
------------------------------------------------------------
---ANDAA
>Protostome_Lophotroco_annelid_Pdum_comp413248_c0_seq11__754_271
------------------------------------------------------------
----------M-------------------------------------------------
------------------------------------------------------------
------------------------------------------------------------
------------------------------------------------------------
--ESTEDSGKS-------------------------------------------------
------------------------------------------------------------
------------------------------------------------------------
---------------------------------------------------AG-------
---------------------------------------------------------DTV
G----------------------EKTDSCNSEIPFYLSDTPPP-----------VY----
-EMPEKDKK----------------------R---------TY-----G-----------
-SV--TYNYL-DTTSAHALPHLI---------S-QK--------------TK---LQ---
--KFIWF----LIFWAAMGYSFYQLRGIVV-EFRK----------Y---QVTVKT----V
-V-RH-R----------T-VADF-----PAV-----------------------TICNEN
KL------KKN-----KISGTP------------------F-----------------Q-
---------SILKLE---------------------------NKIMK----K--------
---------------------------------ND-------------------------
------------------------------------------------------------
------------------------------------------------------------
-----------------D-----------------------AYASDDGDYSD--------
---------------DDSGYDSQSYDSD--------------------------------
------------------------------------------------------------
------------------------------------------------------------
------------------------------------------------------------
------------------------------------------------------------
------------------------------------------------------------
------------------------------------------------------------
-VQEFQQQEDEDGDYDTLG-----------------------SDDL-------F------
---------------------VGLRGEHDYEKLFNL-----------SR------TTDYS
```

---------------------------------DVINML---------------------
-------------------------RP----------------NTTELDLY----GHQEK
--DFI------LQCT----FD-G----E----------N---C-----------K-D---
----------------------------------------------S-------------
------------------------------------------------------------
------------------------------------------------------------
-----------H----------F-------T-----------KFY-NEAFG--------N
CYTFN-------GM-------------NQ-N--------------------N----TV--
-------VRTSKF------GS--------KF-------------GLKLSLNVEAD-----
----------E-----------------Y------------L-G-L---FT---------
------------------------------------------------------------
------------------------------------------------------------
----------HQIGAKISIH-PSNMTP--FPEDSAVSAPVGKLTDIALSSN------L--
-----LEGV-----------QSP-YE-TN-C--STG--------EGL---------YLFY
----P-------------G-DYTAQNCMHSCLRKALL----ED--CHCVETID---V--G
------------P-----------------------------------------------
----------RE-----T---KCKIS----------N-----------------------
------------------------------------VTQ---VICRL---------N---
V-Y-------------------KKFK-N-------NK--------Y------------PC
----------QQGCN-EPCHQVT---YNIL-ASY------------------SEW-PT--
-------EVY------RAL-L-----------KD--------------------------
------------------------------------------------QL-D--------
----------------------------------KR------------------------
-N----------------------------------------------------------
---------------------------ISA------------------------------
-----DG-----YFL-----------E------GNILRANIYFKTLSYSYQG-AFPTYTW
ETL-----------------LANIGGTWGLFIGFSVCTFLEAAEYFLEL-----------
--------ISLV--------F----------R-------SCCC-----------------
--------------------------------KS-------------------KE-----
---------------------------------------KAISPKK--------------
------------------------------------------------------------
------------------------------------------------------------
------------------------------------------------------------
------------------------------------------------------------
------------------------------------------------------------
--------
>Protostome_Lophotroco_annelid_Pdum_comp419533_c0_seq2__757_272
------------------------------------------------------------
----------M-------------------------------------------------
------------------------------------------------------------
------------------------------------------------------------
------------------------------------------------------------
------------------------------------------------------------
------------------------------------------------------------
------------------------------------------------------------
------------------------------------------------------------
------------------------------------------------------------
-----------------------VSTKEMVLAGK-------------------------F
GREDAALEN----------------------M---------GI-----A-----------
-TL--IGSMA-ENTGMHGLPNIS---------R-SK--------------SL---FR---
--KLFWL----IIFLAGVAMMTWQVYEAFK-KFQS----------H---PTETSI----E
-P-KS-N----------T-TLPF-----PAV-----------------------TFCNLN
PI------KKS-----KLSDLG--------------------------------------
---------DI-------------------------------ESLLN----S--------
---------------------------------NS-------------------------
------------------------------------------------------------
------------------------------------------------------------

```
-----------------NSS----------------------------------------
------------------------------------------------------------
------------------------------------------------------------
------------------------------------------------------------
----------------------SEWTPSGTAQP---------------------------
------------------------------------------------------------
------------------------------------------------------------
------------------------------------------------------------
------------------------------------------------------------
------------------------SSFFDWDAQ--------GDAAYANE------DAEFT
------------------------------LREKILEGL---------------------
------------------------YSL----------------TVAEKQAA----GHQLN
--DIL------LSCT----YQ-G----Y----------A---C-----------G-P---
----------------------------------------------K-------------
------------------------------------------------------------
------------------------------------------------------------
-----------N----------F-------S-----------TFY-NSLFG--------N
CYTFN-------GG-------------EM----------------------N----TA--
-------AKAGKV------GP--------FY-------------SLSLELYIEQT-----
----------E-----------------Y------------I-P-G---LA---------
------------------------------------------------------------
------------------------------------------------------------
----------DAAGMRVVVH-NPNAMP--FPEDEGFSVAPGELSYVGLHRV------E--
-----FTRS-----------KPP-H--GE-C--QKF--------SENE----TLA-RNAW
----KQK--YN------FL-EYTRRSCAKTCYQQYVM----SI--CGCSDRD----F---
------------P-N--EG-------------------TPFEN-----------------
-----ITKGQNF-----S---ACNSE----------D-----------------------
------------------------------------EAM---EECQQ---------Q---
V-Y-------------------TNFT-E-------NI--------L------------NC
----------SKTCP-PPCSEAT---HEIT-QSH------------------ARW-PT--
-------EAK------AND-L-----------VN--------------------------
------------------------------------------------QV-L--------
----------------------------------QK------------------------
-S-------------SEL------------------FDALIN------------------
----------------------LDAE-ERT------------------------------
-----E-------TV-----------R------ENVLKILVYFKSLEYTTIT-TKPSFGI
VDL-----------------LASIGGQVGLWLGLSVITLFEIVELFFDA-----------
--------WAFA--------F----------C-------KC-------------------
LETVP---------------------------KK-------------------K------
----------------------------------------SKNLIQVT------------
------------------------------------------------PRSEENE-----
------------------------------------------------------------
-----------------------------------------------I------------
------------------------------GTSPS-------------------------
----------------------------------------------------EIHFNHVN
EPNQKIVW
>Deu_Ambulacrar_hemi_Ptyfla_40v0_9_20150316_1g16271_t1_scaffold12531_cov113_7
58_273
------------------------------------------------------------
----------M-----------------------------------C-------------
------------------------------------------------------------
------------------------------------------------------------
------------------------------------------------------------
------------------------------------------------------------
------------------------------------------------------------
------------------------------------------------------------
------------------------------------------------------------
```

```
------------------------------------------------------------
------------------------------------------------------------
-------------------------------D---------AW-----K-----------
-GL----------KTLSGLGKAKDKIGGS---TLAE------------QLSMANELN---
--K-FYCRFDKFDFSNVIGDIRSDLEQRVGEELFE--------------ITDEQI----T
-I-VT-K----------T-SLTF-----PTV-----------------------TICNTN
KV------RRS-----AIADSS------------------------------------HG
---------DVL---------VIDDAIT--------------LPYYG-------------
---------------------------PCMEG-DF-------------------------
------------------------------------------------------------
-----------MCD-----------------------------------DGLL-------
-CIKPFLKCDGVRNCIWD-G------------------------SDELGCE-----YG--
------------------------------------------------------------
------------------------------------------------------------
------------------------------------------------------------
--------------------------PCM-E-----------------------------
------------------------------------------------------------
------------------------------------------------------------
----------------------------------------GDFMC---------------
------------------------------------------DDGLL-------------
------------------------------------------------------------
---------------------------------CIKPFL---------------------
---------------------------K-------------C------------------
--DGV------RNCI----WD-G--SDE-------LGCEYGTC-----------G-H---
------------------------------------------------------------
------------------------------------------------------------
------------------------------------------------------------
-----------N---------HF-------R---CAN---------GSQYGFLYCDKKKD
CYDGE-------VGCEQR---------DEFQCQDKRGCISKK-------QRC----NVHF
DCD----DKSDEKDPAKDCGDALW-CAADDYCILNEGDGLSCDDGLKCISFTKRCD-GQQ
DCADNLD---EKDCPV---------APVY------------T-D---MYMT---------
------------------------------------------------------------
------------------------------------------------------------
----------ERMGVLLSPD-YPMNYPNNIFHRYSVMMLPTERTESTIRVKF-----T--
-----FEDF-----------NIE-YS-EN-C--TND-W----------------------
-----------------------LQSCKNLCLHSTIV---K----CGCSQTMN---I---
--------Q-SAP-----------------------------------------------
---------------------PCSIL----------N-----------------------
------------------------------------KTQ---------G-------N---
---------------------------E--------------------------------
-------------------QGFI---ILLK-VSV------------------SPH-PADW
--------------------L-----------YE--C-----------------------
------------------------------------------------------------
------------------------------------------------------------
------------------------------------------------------------
------------------------------------------------------------
-----S-------YW-----------Q------SNLSNVDIHYDELTYQEIR-EEPAYPI
ESL-----------------FSDIGGSLGLYIGLSVITVFEFFEFVVEA-----------
--------LRVC--------L----------R-------RE-------------------
NGG-----------------------------RH--------------------------
------------------------------------------------------------
------------------------------------------------------------
------------------------------------------------------------
------------------------------------------------------------
------------------------------------------------------------
------------------------------------------------------------
-------S
```

```
>Protostome_Lophotroco_annelid_CAC9650466_1__Ofus_G101010_Owenia__fusiformis_
_759_274
------------------------------------------------------------
----------M-------------------------------------------------
------------------------------------------------------------
------------------------------------------------------------
------------------------------------------------------------
------------------------------------------------------------
------------------------------------------------------------
--------------------------------------------------------DGV-
------------------------------------------------------------
------------------------------------------------------------
-----------------------GKDEKN-------------------------------
---SDEDKD----------------------S---------SL-----R-----------
-DI--LGNFA-SSTTAHGCERID---------K-ST--------------ST---IA---
--RVIWA----TIFIGCTIAAIVCIAALLE-KYFQ----------Y---PSKDVY----E
-M-NS-D------------KIEF-----PSI-----------------------SFCSLK
PI------VSS-----YKKRVAKMEP--------GNPYMEI-------------------
------------------------------------------DKAIK----NIDGVL---
------------------------------------------------------------
------------------------------------------------------------
------------------------------------------------------------
-----------------D------------------------------------------
------------------------------------------------------------
------------------------------------------------------------
------------------------------------------------------------
------------------------------------------------------------
------------------------------------------------------------
------------------------------------------------------------
------------------------------------------------------------
-Y-----LSQEN-----------------------------NTQNN--------------
-----------------------------------------LYQRLKEM------YYLFA
---------------------------------SYTSMY---------------------
---------------------------I--------------HATDETRSL----GHQLE
--DLL------LSCK----VN-M----M----------S---CV----------N-M---
----------------------------------------------P-------------
------------------------------------------------------------
------------------------------------------------------------
-----------DV--------KI-------K-----------QFD-NALYW--------N
CYTFN-------WK-------------DY-N-TT----------------GE----SE--
-------PYVEMV------GG--------LS-------------GFQATFFLDNI-----
----------D--------TLESY----YN-------------P-M-CPID---------
------------------------------------------------------------
------------------------------------------------------------
----------GSVGAKLIIH-RPGSRP--DPLREGLEVPVGFSTNVAVSKK------K--
-----RELI-----------GQP-W--GD-C--ENR--------EK------LDG--APY
--------------------RYDQANCERGCQQKEIV----KK--CGCVSPW----L---
------------P-V--SD-----------------------------------------
-----EQ--KNI-----S---WCGKF----------NLD-NFNSA------SP-------
---------------------------NLTILD-KDLIP---FECHM-------------
M-M-------------------TKMT------------------PNV----------TEC
------------DCP-PACENHM---YEYK-TSQ------------------SQW-LL--
-------GGN------RLN-F-----------YQ--------------------------
------------------------------------------------SVVN--------
-------------------------------RMGDSY-----------------------
IN-------------SSM------------------YHL-LDAALN--------------
------------------SNDSAENF-KNR------------------------------
```

-----N-------LV-----------S------QNFLRVNIFFDSMHTDLIR-RVEEYTF
TDM-----------------ISGVGGGFGVYVGFSIVTMCEFVVLFASL-----------
--------VTAL--------I----------Q-------RY-------------------
------------------------------GLRQ------------------NKV-----
----------------------------------------AMESAPT-------------
------------------HVHIIHVKH---------------------ANNH--------
------------------------------------------------------------
------------------------------------------------------------
------------------------------------------------------------
------------------------------------------------------------
---LADVD
>Cnidar_Polpod_Hydrif_GBGH01000761_1__p1__GENE_GBGH01000761_1_GBGH01000761_1__p1__ORF__type_complete__len_571__score_27_43__GBGH01000761_1__228_1940__760_275
------------------------------------------------------------
----------M-------------------------------------------------
------------------------------------------------------------
------------------------------------------------------------
------------------------------------------------------------
------------------------------------------------------------
------------------------------------STNNGPVMR---------------
--PESHQDKLAVPSSAKRSAL-------MDKALGSNARNSGCEKGVNLSALKDGPIPLRV
------------------------------------------------------------
------------AQKDPTQKFRKIGLQAMIA--------------------------KRI
E----------------------RYAEE--------------------------------
--------R----------------------L---------SP-----K-----------
-QI--FKRFT-ESSTLHGFRYIF---------T-AG--------------TI---VR---
--RFSWF----VLCVTMTSLFLSELQKLIM-LYSE----------Y---PFTTTT----T
-L-ES-V----------Q-FHEF-----PSI-----------------------SICNTN
SF------RKS-----ISLSSG------------------I-----------------D-
---------HL---------------------------------VFD-------------
------------------------------------------------------------
------------------------------------------------------------
------------------------------------------------------------
------------------------------------------------------------
------------------------------------------------------------
------------------------------------------------------------
------------------------------------------------------------
------------------------------------------------------------
------------------------------------------------------------
------------------------------------------------------------
------------------------------------------------------------
------------------------------------------GSSG--------------
------------------------SSTKPTSNTTSDV------------------IDGN-
---------------------------------VLFEKL---------------------
------------------------------------------------NAS----SHQIE
--DML------IYCS----FI-DA-FEQ--GKHVHSDME---C-----------G-P---
----------------------------------------------H-------------
------------------------------------------------------------
------------------------------------------------------------
-----------N----------Y-------T-----------AYT-N-LAGA-------L
CYRFN-------PG-------------PTFG--------------------V----PL--
-------LSVTNT------GI--------LH-------------GLKVYVKLQTD-----
----------D-----------------Y------------S-P-Q--------------
------------------------------------------------------------
------------------------------------------------------------
---------VQEAGLQLVLH-DHDENP---LSFSPFVVPPGFQTYVEMKKQ------Q--

-----VLNL-----------PPP-YK-TH-C--GSR--------T-------LN----IS
------------------T-TYRRSMCYIQKLSEFVL----ER--CGCKGPF----MEK-
----------SVP-Y---------------------------------------------
----------QV-----P---LCNS-----------------------------------
------------------------------------SQY---KDCYR---------P---
T-I-------------------NSFD-P-------KT--------------------AGD
------------QCP-VDCIRSN---YMYT-LSY------------------GRF-IA--
-------DPK------IGT-YPSMT-------AK----------------R---------
------------------------------------------------NI----------
------------------------------------------------------------
-N------------LARL------------------NMS---------------------
------------------------YQ-DQR------------------------------
-----Q-------YI-----------R------DNFVAFVIYFSDMTVENIS-QERSYDI
YKL-----------------MGDIGGQLGLMLGASVLTVIEFIDLIFFI-----------
--------VYTK--------I----------S-------RI-------------------
IK------------------------------RR-------------------Q------
-----------------------------------------VNRRFPT------------
------------------------------------------------------------
------------------------------------------------------------
------------------------------------------------------------
---------------------------------P--------------------------
-----------------------------------------------------------A
ESKTTTGV
>Deutero_Ambulac_Spurpu_014519_764_277
------------------------------------------------------------
----------M-------------------------------------------------
------------------------------------------------------------
------------------------------------------------------------
------------------------------------------------------------
--TTGYTFEWYDCLLT--------QAT---------------------------------
------------------------------------------------------------
------------------------------------------------------------
------------------------------------------------------------
---------------------------------------------------LLLGRVKRD
S-------------------LFLMRTDKDLP-----------------------------
-------HL----------------------N---------SV-----N-----------
-AV--LIDYS-GSTSAHGIPRII---------T-SR--------------SV---KS---
--RLFWS----FVTLVCLGAFLWQGSLLLL-DFSK----------Y---PYTTQI----D
-V-VA-R----------T-ELQF-----PAV-----------------------TVCNMN
KM------KRS-----AMVHTR------------------F-----------------Q-
---------SL----------------------------------IQ-------------
------------------------------A--DL-------GVNGGDADYSWWFDWSSQ
------------W-----------------------------------------------
------------------------------------------------------------
---------SQEEEEQEE------------------------------------------
------------------------------------------------------------
----------------QEEQELASA----------SAPDTGSSSSDWSS---DA------
------------------------------------------------------------
------------------REHYSEQAPAL-------------------------------
------------------------------------------------------------
-------------VTGSSTSSS--------------------------------------
------------------------------------------------------------
-DSSSDAQEHYSEQYEWW----DPGWDTD--------GFDF-HEYD-------W------
--------TNV-------------SDDDDWEGFYKQ-----------ST------SDDFS
---------------------------------DLLEVV---------------------
-------------------------NP----------------TREELKVM----GHQAE
--DFI------LQCT----FD-R----H----------Q---C-----------N-Y---

----------------------------------------------T-------------
------------------------------------------------------------
------------------------------------------------------------
-----------D----------F-------H-----------QFQ-NKYYG--------N
CFTFN-------RE-------------VG-N-S------------------T----RA--
-------RSTGKT------GA--------QY-------------GLHLTLFTEQP-----
----------E-----------------Y------------V-G-L---FA---------
------------------------------------------------------------
------------------------------------------------------------
----------QEAGVRVAIH-PPNVFP--FPEDDGVVASTGQATDIGMRQS------Y--
-----FNRL-----------PHP-H--GN-C--TEG--------TR----------TIFM
----SEE-----------Y-AYTTRACVKSCVQQQLF----DN--CGCVTDI----I---
------------M-----------------------------------------------
----------NE-----T---MCGAR----------N-----------------------
------------------------------------KSQ---QVCRQ---------A---
I---------------------EHFH-E-------DQ--------S------------SC
------------NCP-IACD----------------------------------------
------------------------------------------------------------
------------------------------------------------------------
------------------------------------------------------------
------------------------------------------------------------
------------------------------------------------------------
---------------------------------RNLARVRIYFEELNFEQMI-QKPKYTI
ESL-----------------LGGIGGLLGLYIGFSVITICEVGVLVVDL-----------
--------VKFL--------L----------R-------KAY------------------
NR------------------------------RE-------------------K------
-----------------------------------------IVPIE--------------
------------------------------------------------------------
------------------------------------------------------------
------------------------------------------------------------
------------------------------------------------------------
------------------------------------------------------------
-----LKC
>Protostome_Lophotroco_annelid_CAC9661806_1__Ofus_G112085_Owenia__fusiformis_
_763_278
------------------------------------------------------------
----------M-------------------------------HRLVC-------------
------------------------------------------------------------
------------------------------------------------------------
------------------------------------------------------------
------------------------------------------------------------
------------------------------------------------------------
---------------SQKIG---------SFTFDDANGYAITVNIKLYF-----VIPIE-
--------------------------------------------------FLGTHAR---
------------------------------------------------------------
----------------ADTEEDIPKGKKTPMSYY----------YLLG-----------N
DYQNCSDHE----------------------V---------RP-----M-----------
-GL--MRGLA-ENTSAHGITEIY---------Y-SK--------------GF---FK---
--ITCWV----LLTLAAIAVMILHMTTLFI-DFYS----------Y---ETTMSI----S
-V-QN-K----------R-SLPF-----PAV-----------------------TICNVN
PI------RAS-----RLSQST--------------------------------------
---------IL-------------------------------QSTID----GR-------
---------------------------------DN-------------------------
------------------------------------------------------------
------------------------------------------------------------
------------------------------------------------------------
------------------------------------------------------------

```
------------------------------------------------------------
------------------------------------------------------------
------------------------------------------------------------
------------------------------------------------------------
------------------------------------------------------------
------------------------------------------------------------
---ITGATRDDK------------------------------------------------
------------------------------------------QTGLGAM------SNEFL
------------------------------IEELIEETI---------------------
------------------------ADI----------------DTDTKIAM----GHQYS
--DFI------LDCQ----YD-G----Y----------Y---C-----------D-E---
----------------------------------------------G-------------
------------------------------------------------------------
------------------------------------------------------------
-----------E----------F-------L-----------AFY-NYKYG--------N
CFTFN-------SG-------------SN----------------------S----DV--
-------FLSGRA------GP--------LH-------------GLKLVLDIEAA-----
----------E-----------------Y------------I-G-D---MS---------
------------------------------------------------------------
------------------------------------------------------------
----------PAYGVRVLVH-SQGDMP--FPEDQGITIGPGQATVIGTNML------N--
-----IQLA-----------GDK-Y--SD-C--TNS--------SFS-----NVN-INVY
----EEH--YP------GT-SYTQTACMKTCYQSHLL----SG--CQCGDPS----V---
------------P-L--DG-------------------KAFPS-----------------
-----AFHTEPA-----T---SCNSD----------N-----------------------
------------------------------------QAQ---VTCEE---------N---
T-Y-------------------ASYV-N-------GS--------L------------SC
------------TCY-PECNQTT---FEAH-VSS------------------TLW-PT--
-------DQY------ISR-L-----------IQ--------------------------
------------------------------------------------DL----------
------------------------------------------------------------
-S-------------PKL------------------NQYT--------------------
---------------------------NDS------------------------------
-----A-------GL-----------R------KNVAHLQVYYEELNLQMIQ-EQPSYST
VQF-----------------ASDVGGTVGLYVGASLLTAFEFGEFFLDL-----------
--------LVYF--------I----------R-------KP-------------------
FH------------------------------KN-------------------K------
-----------------------------------------VSEVKME------------
------------------------------------------------------------
------------------------------------------------------------
------------------------------------------------------------
------------------------------------------------------------
-----------------------------------------------------------Q
TKSNVQFS
>Protostome_Lophotroco_annelid_Pdum_Contig13621__765_280
------------------------------------------------------------
----------M-------------------------------------------------
------------------------------------------------------------
------------------------------------------------------------
------------------------------------------------------------
------------------------------------------------------------
------------------------------------------------------------
------------------------------------------------------------
------------------------------------------------------------
------------------------------------------------------------
------------------------------------------------------------
------------------------------------------------------------
```

-----------------AGKRVL---------K-SK--------------SN---IA---
--HLFWL----LVCLAAIAMFCMQMAEVLQ-RYFS----------Y---PKKVTV----E
-V-IS------------A-QVPF-----PAV-----------------------SLCNMR
NM------DVH-----VLNTLNRKFI---------EDHSPI-NH----------INNSDN
---------DF---------------------------VREYMKLSA----KYGPLWYK-
------------------------------------------------------------
------------------------------------------------------------
------------------------------------------------------------
------------------------------------------------------------
------------------------------------------------------------
------------------------------------------------------------
------------------------------------------------------------
------------------------------------------------------------
------------------------------------------------------------
------------------------------------------------------------
------------------------------------------------------------
------------------------------------------------------------
-----------------------------------------YQGKYPLA------FQEVF
---------------------------------SRTTYS---------------------
------------------------ANI----------------PQHIVSSA----AVQLD
--EFV------VSCY----FG-E----Y----------H---C-----------N-TT--
----------------------------------------------N-------------
------------------------------------------------------------
------------------------------------------------------------
-----------D----------F-------T-----------TFF-DPYYF--------N
CFTYN-------MA-------------DP-D-----------------------------
-------QASSSE------GI--------EN-------------GWSSVLYSGSG---IL
DRNK------D------IRVLPGL----HE-------------S-R-SAVS---------
------------------------------------------------------------
------------------------------------------------------------
----------ANEGVRVVIH-PPGTIP--FPLTEGFDVPPGFSASFGIRPR------Q--
-----IKRI-----------GPP-H--GN-C--ITS--------NP------FGD--SSS
--------------------QYRVLSCQKMCLQRYII----NS--CGCQDKT----L---
------------P-D--VP-----------------------------------------
-----DL---KA-----T---PCRNAT---------DFPDSCMT-------DASDE----
---------------------------CLKQFF-HLYDR---IQCVR---------S---
T-K-------------------DRFM-K-------NS-------SLL----------TDC
------------GCF-PQCDEIS---YDVS-YSL------------------SKW-PA--
-------SGY------EGDAA-----------FF--------------------------
------------------------------------------------DIFYIE------
-------------------------------GFRERF-----------------------
LN-------------TPK------------------------------------------
----------------YNMVMSYFHE-DTR------------------------------
-----E-------KT-----------M------QDFARLNVYIADSNVIITQ-EMEDYTT
TQL-----------------VSDIGGQLGLWIGISIITLAEVLELFIDT-----------
--------CRML--------F----------S-------KR-------------------
RIMV----------------------------RN---------------ARRKE------
-----------------------------------------TDRYRSHSGIAGLVPIK-K
TNCNNIFMSPLD--RSETHAMILRRQN---------------------------------
------------------------------------------------------------
------------------------------------------------------------
------------------------------------------------------------
------------------------------------------------------------
-GDSSSIR
>Chordat_hASIC1_NP_064423_2_acidsensing_ion_channel_1_isoform_a__Homo_sapiens
__3_281
------------------------------------------------------------

```
----------M-------------------------------------------------
------------------------------------------------------------
------------------------------------------------------------
------------------------------------------------------------
------------------------------------------------------------
------------------------------------------------------------
----------------ELKA-------------EEEEVG---------------------
------------------------------------------------------------
------------------------------------------------------------
------------------------------------------------------------
-------------------------------G---------VQ-----P-----------
-VS--IQAFA-SSSTLHGLAHIF---------S-YE-------------RLS---LK---
--RALWA----LCFLGSLAVLLCVCTERVQ-YYFH----------Y---HHVTKL----D
-E-VA-A----------S-QLTF-----PAV-----------------------TLCNLN
EF------RFS-----QVSKND------------------L-----------------Y-
---------HA------------G-ELL---------------ALLN----N--------
---------------------------------RY-------------------------
------------------------------------------------------------
------------------------------------------------------------
-------------EIPDT-Q-MA-------------------------------------
------------------------------------------------------------
------------------------------------------------------------
------------------------------------------------------------
------------------------------------------------------------
------------------------------------------------------------
------------------------------------------------------------
------------------------------------------------------------
--------DEKQ------------------------------LEI---------------
------------------------LQ--DKANFR----------SFK--------PKPF-
---------------------------------NMREFY---------------------
------------------------------------------------DRA----GHDIR
--DML------LSCH----FR-G----E----------V---C-----------S-A---
----------------------------------------------E-------------
------------------------------------------------------------
------------------------------------------------------------
-----------D----------F-------K-----------VVF-T-RYG--------K
CYTFN-------SG-------------RD-G--------------------R----PR--
-------LKTMKG------GT--------GN-------------GLEIMLDIQQD-----
----------E-----------------Y------------L-P-VWGETD---------
------------------------------------------------------------
------------------------------------------------------------
-------ETSFEAGIKVQIH-SQDEPP--FIDQLGFGVAPGFQTFVACQEQ------R--
-----LIYL-----------PPP-W--GT-C--KAV-TMDS---D-------LD----FF
------------------D-SYSITACRIDCETRYLV----EN--CNCRMVH----M---
------------P-G---------------------------------------------
----------DA-----P---YCTP-----------------------------------
------------------------------------EQY---KECAD---------P---
A-L-------------------DFLV-E-------KD--------Q-----------EYC
------------VCE-MPCNLTR---YGKE-LSM-----------------VKI-PS--
-------KAS------AKY-L-----------AK--------------------------
------------------------------------------------KF----------
------------------------------------------------------------
-N----------------------------------------------------------
---------------------------KSE------------------------------
-----Q-------YI-----------G------ENILVLDIFFEVLNYETIE-QKKAYEI
AGLLGEMTPVPFSCHGKPCCLGDIGGQMGLFIGASILTVLELFDYAYEV-----------
--------IKHK--------L----------C-------RR----------------GK-
```

```
CQ------------------------------KE-------------------A------
------------------------------------------------------------
-----------------------KRSS---------------------ADKG--------
-----------VA-----------------------------------------------
----------------LSLDDVKRH------------------------NPCESLRGHPA
------------------------GMTYAANILP-------HH---------PA------
---------RGT-----FEDFTC-------------------------------------
--------
>Deu_Ambulacrar_hemi_Ptyfla_40v0_9_20150316_1g8763_t1_scaffold4594_cov140_769
_283
------------------------------------------------------------
------------------------------------------------------------
------------------------------------------------------------
------------------------------------------------------------
------------------------------------------------------------
------------------------------------------------------------
------------------------------------------------------------
------------------------------------------------------------
------------------------------------------------------------
------------------------------------------------------------
------------------------------------------------------------
---------------------------------------YGVM----AG-----------
-KS--LTEFG-QETSIGGVKYVT---------DVSS--------------RN---LR---
--RFFWF----LVVCTALGALTFQIVNICT-AYIA----------R---PVSVNR----E
-Y-IP-M----------G-QMKF-----PAV-----------------------TICNYN
QI------RKT-----GLLEAF--------G---------YSHG--------------D-
---------GV---------------------------IKTAHAIMS-------------
---------------------------PSPETMDQ-------------------------
------------------------------------------------------------
------------------------------------------------------------
-----------------D------------------------------------------
------------------------------------------------------------
---------------------------V--------------------------------
------------------------------------------------------------
------------------------------------------------------------
------------------------------------------------------------
------------------------------------------------------------
------------------------------------------------------------
------------------------------------------------------------
------------------------------------------------------------
---------------------------------NLVGLN---------------------
------------------------ANY---------------DFEYLMTTV----GHKKD
--EMI------MRCK----WP-G----R----------Q---C-----------S-A---
----------------------------------------------D-------------
------------------------------------------------------------
------------------------------------------------------------
-----------N----------F-------T-----------TTY-AR------------
------------------------------------------------------------
--------------------E--------RY-------------GLTVYLDVESQ-----
----------E-----------------Y------------V-D-N---WQ---------
------------------------------------------------------------
------------------------------------------------------------
----------NFVGFRVIVH-DQDDKP--NMKDKGFNVAPGTYTAAALKKL------S--
-----------------------------------------------V--------MHLF
-----------------AM-SYS-----------FTI----AM--CQ-RWYR--------
------------------------------------------------------------
------------------------------------------------------------
```

---------------------------------------------FL--------PP---
T---------------------AKFT-D-----------------VQ----------GLC
S-----------HCK-TPCYEES---YGEH-LSY------------------ALF-PS--
-------GPS------VEE-L-----------AEKAEKPCPKAMALYYKEKITAYVFASL
SMANLSKILITTK-----------DKLLD-------AIFLGETHISIDRIINDITASAPE
GG-LLPSVDVTDSPIENDQLTVGKL-YEILTEALQSYYDQIYS-EI---------SIPLS
SNLYRYIH---SERQSSVE------------VGGDIYRKVYETMINSSASARRLKKIAPD
LAVQGTFRAALNVTDSYGALVNDTKYALSDAFTDIFDLIYTTVLDTFEENRITALENHTM
YQICME-------YM-----------S------MCPLDRLPGPSHVFQANDT-ERSAIAP
KKL-----------------ILEVGEPMMLL-----------------------------
------------------------------------------------------------
------------------------------------------------------------
------------------------------------------------------------
------------------------------------------------------------
------------------------------------------------------------
------------------------------------------------------------
------------------------------------------------------------
------------------------------------------------------------
--------
>Deutero_Urochord_tr_F6SR69_F6SR69_CIOIN_Uncharacterized_protein_OS_CioInt_77
1_285
------------------------------------------------------------
----------M-------------------------------------------------
------------------------------------------------------------
------------------------------------------------------------
------------------------------------------------------------
HSIHNRDVRVS----------------------------------------------SLS
AHFEPVENCVVHSD----------------------------------------------
------DVTVADDVSYTSSS-------------MSSDLE---------------------
------------------------------------------------------------
------------------------------------------------------------
------------------------------------------------------------
-------------------------------S---------MK-----P-----------
-SN--IHVFS-SSSTLHGFNHIF---------A-LH-------------HSS---LR---
--RFAWT----VAFLTSLSILLYQSSNRAI-YYLS----------Y---PHVTRL----D
-E-VA-A----------G-NLTF-----PAV-----------------------TICNLN
EF------QFS-----KITKND------------------M-----------------Y-
---------HV------------G-ELL---------------GFLD----K--------
---------------------------------NY-------------------------
------------------------------------------------------------
------------------------------------------------------------
-------------ALCCN-T-IV-------------------------------------
------------------------------------------------------------
------------------------------------------------------------
------------------------------------------------------------
------------------------------------------------------------
------------------------------------------------------------
------------------------------------------------------------
------------------------------------------------------------
--------PERQ------------------------------TLN---------------
------------------------FLNIQRSSFR----------HFNP-------RGTF-
---------------------------------SMKEFY---------------------
------------------------------------------------ERV----GHNLS
--DFL------LACN----YR-G----R----------F---C-----------N-A---
----------------------------------------------D-------------
------------------------------------------------------------
------------------------------------------------------------

-----------N----------F-------T-----------TVY-T-RYG--------K
CYTFN-------SG-------------VK----------------------Q----PP--
-------LKTLKG------GV--------DN-------------GLELLLDTQQN-----
----------E-----------------Y------------M-P-VWKETD---------
------------------------------------------------------------
------------------------------------------------------------
-------EITIEAGFKVQIH-VQSEPP--FIHELGFGISPGFQTLVSTQEQ------R--
-----ITFL-----------PNP-W--GK-C--RNG-FDKS---E-----------HYHF
------------------P-NYSISACRISCETLYVY----LH--CNCRMIH----M---
------------P-G---------------------------------------------
----------KE-----R---YCTP-----------------------------------
------------------------------------EEY---KTCAD---------K---
A-L-------------------DFLV-M-------TD--------D-----------QLC
------------VCE-TPCEVIR---YNLE-MST------------------LQL-PS--
-------TQA------SSY-L-----------AY--------------------------
------------------------------------------------KY----------
------------------------------------------------------------
-Q----------------------------------------------------------
---------------------------VSE------------------------------
-----D-------YV-----------R------KNFAKLNIFFEALNYETIE-QKVAYEI
PGL-----------------FGDIGGQMGLFIGASILTILELVDYFYEV-----------
--------VKDR--------T----------W-------GR-------------------
KL------------------------------RK-------------------NT-----
------------------------------------------------------------
-----------------------DRSPTRE------------------EANN--------
-----------KT-----------------------------------------------
----------------VSV-----------------------------------------
-------------------------MVREQAIFP--------------------------
-------VITNL-----FLGVVCC-------------------LYIYIILVVYKCCNVPH
ILTTESGS
>Cnidar_NemVecNVEC200_011692_1_1_protein_AED_0_06_eAED_0_06_QI_233_1_1_1_0_91_0_84_13_1484_578_773_286
------------------------------------------------------------
----------M-------------------------------------------------
------------------------------------------------------------
------------------------------------------------------------
------------------------------------------------------------
------------------------------------------------------------
------------------------------------------------------------
------------------------------------------------------------
------------------------------------------------------------
------------------------------------------------------------
--------------------LKLPCNKEM-------------------------------
--LRNTDPK----------------------E---------TA-----L-----------
-EQ--INQFL-QETTAHGFGRLG---------A-TA--------------GS---KW---
--RIYWV----MFCLAAYCVFAWQLVGLVN-QYNS----------K---PIKTRT----Q
-L-KH-A----------Q-KLDF-----PVV-----------------------TICNMN
VL------RAS-----RLPPKLR-----------------------------------T-
---------KF-------------------------------DEIIN-------------
---------------------------------NT-------------------------
------------------------------------------------------------
------------------------------------------------------------
-----------------KKTSSRK-------------------SSNSRNAFV--------
------------------------------------------------------------
------------------------------------------------------------
------------------------------------------------------------
-------------------------DP---------------------------------

------------------------------------------------------------
------------------------------------------------------------
------------------------------------------------------------
------------------------------------------QDLSFEE-----------
-------------------------------------------------------TKKIE
---------------------------------ILHAVT---------------------
------------------------THD----------------NYRELVSA----AHQLE
--DIL------LSCN----FN-G----V----------N---C-----------R-N---
----------------------------------------------S-------------
------------------------------------------------------------
------------------------------------------------------------
-----------NDPTIP---TSW-------T-----------QTW-NDNFG--------N
CYMFN-------PA-------------QT-H-NGEK---------------V----DP--
-------YSSSIP------GE--------SN-------------GLTLQLNIEQN-----
----------E-----------------Y------------L-E-G---IT---------
------------------------------------------------------------
------------------------------------------------------------
----------EVAGIKVSIS-DQGVLP--FPGQQGIRIMPGQSTGIQMTKL------Q--
-----TRRI------------DP-FKNRS-C--ENS--------NE------MSD-KNLF
----FG---YN-------M-RYSKMACKYSCLNAKTI----ER--CGCTNYN----T---
------------P-E---------------------------------------------
------LQKRNI-----S---LCNRL----------N-----------------------
------------------------------------NAI---IDCLN---------K---
A-Y-------------------DTFE-D-----------------------------GSC
----------DRECP-PSCSEVS---FDLT-ISS------------------AKW-PA--
-------KSY------EKT-V-----------LK--------------------------
------------------------------------------------TL-Q--------
----------------------------------TEYG--------I-------------
-N----------------------------------------------------------
---------------------------MTK------------------------------
-----E-------EM-----------F------ENIAQVHVYYGELDYLLVQ-ETLAYTF
MSL-----------------LSDIGGQMGMWIGISALTCAELVELVCVI-----------
--------LANM--------S----------N-------RS-------------------
KKIVHISSNIG-------CF------------RPAKLGG---------VEAIEKV-----
---------------------------------------GGVKTIEQEVQGGV-------
------------------GIIEQEVQGAVG---------------TIEQEVQGGV-----
------------------------------------------------------------
------------------------------------------------------------
------------------------------------------------------------
----------------------------------------------------GTIEQEMQ
GRVGTIEQ
>Deutero_Ambulac_Sakowv30035415m_780_288
------------------------------------------------------------
----------MIADYDNMVED-YENMI-----------------ADCDSI----------
-----------------------V-----------------ADYDNMVAD----------
-------------------------------YDSMVADYD-------N------------
-----------MIADYNNN--LVVFKELGE------------------------GSKANR
YRIMASSYSFY--------GDRYSDGALQ--------------FS-QPIPHPC--KTKTR
NTI-ALSSFSDGRF----------------------------------------------
-------RSKVNSIKEDEEK-------------KVQLQKDP-------------------
------------------------------------------------------------
----------------------PTATSKGDV--------------------------NAC
G----------------------DCCCC--------------------------------
--------R----------------------V---------FY-----S-----------
-DR--FREFS-SDTTLHGLRYAV---------A-EG-------------VEP---WR---
--RFLWT----VLLVIAVSATAVYMYLCWY-KFFS----------F---PVNTVV----T
-L-TY-K----------S-KLRF-----PAI-----------------------TICNYN

```
QY------RKS-----VVKGTR------------------F-----------------E-
---------EW------------V------------------RQQY--------------
------------------------------------------------------------
------------------------------------------------------------
------------------------------------------------------------
------------------------------------------------------------
------------------------------------------------------------
------------------------------------------------------------
------------------------------------------------------------
--------------------------P---------------------------------
------------------------------------------------------------
------------------------------------------------------------
------------------------------------------------------------
--------LFNF------------------------------SDD---------------
---------------------------DVQVELDES-------------------ILEA-
---------------------------------NRTWFD---------------------
------------------------------------------------QMA----AHSKW
--SMI------YQCT----FG-S----V--------KIP---C-----------S-A---
----------------------------------------------S-------------
------------------------------------------------------------
------------------------------------------------------------
-----------N----------F-------T-----------ETF-T-DFG--------V
CYTFN-------GA-------------DR-E--------------------G----GA--
-------LLVGHS------GS--------EH-------------GLRMRLFVNQS-----
----------E-----------------Y------------T-F-G---EH---------
------------------------------------------------------------
------------------------------------------------------------
----------CGAGFKILPH-PQDEVP--LVRNFGFAVSPGTEALVGLKMI------Q--
-----EHNL-----------PAP-YA-SQ-C--SNE--------T-------LQ----YF
------------------A-KYARSNCVRERQTDAVV----AK--CGCREPY----M---
------------P-S---------------------------------------------
----------DA-----R---VCNY-----------------------------------
------------------------------------NET---RVCVR---------P---
E-L-------------------ERIQ-V--------------------------------
------------QCP-MACESST---YAPR-VSY------------------AMF-PG--
-------RHI------VED-L-----------QM--------------------------
------------------------------------------------RF----------
------------------------------------------------------------
-N----------------------------------------------------------
---------------------------LSY------------------------------
-----V-------DI-----------R------ENVVDVKIYFEEISLEEIY-QEDGYPI
SEL-----------------I----GELFYNIDDARAQDNDVINITVRI-----------
--------------------I------------------RY-------------------
DD----------------------------------------------------------
------------------------------------------------------------
------------------------------------------------------------
------------------------------------------------------------
------------------------------------------------------------
------------------------------------------------------------
------------------------------------------------------------
---CSIPH
>Placozoa_HhoNaC10_TR2949_c0_g1_i6_m_5640__Hoilungia_hongkongensis__24_290
------------------------------------------------------------
----------M-------------------------------------------------
------------------------------------------------------------
------------------------------------------------------------
------------------------------------------------------------
```

```
------------------------------------------------------------
------------------------------------------------------------
------------------------------------------------------------
------------------------------------------------------------
------------------------------------------------------------
-----------------------PTKHLKVVSQHN-------------------------
-GGISARKK----------------------R---------LL-----------------
-AIT-ITKGF-EDCGAQGISNMA---------R-AQ--------------TT---QA---
--RIIWA----ILTIVAVAFCIVMAVDLID-KYYR----------F---EYDVQL----E
-I-SF-K----------P-QLDF-----PVV-----------------------TICNLN
PI------KKS-----EMLTRP--------A---------F-----------------K-
---------PLFAKDFP-----------------------------D-------------
---------------------------------DP-------------------------
------------------------------------------------------------
------------IAP---L---PT------------------------------------
---------TAGSTMSGSGTGGT------------------GTATKEKTGTG--------
------------------------------------------------------------
------------------------------------------------------------
------------------------------------------------------------
-----------------TATRSGTSTPTGTPK----------------------------
------------------------------------------------------------
------------------------------------------------------------
------------------------------------------------------------
-PRVRRG-IAGV------------------------------NDDIERP-----------
----------------------TPPANLTFDDLDAS-------------------SENYN
--------L---------------------FSEINAELF---------------------
------------------------NKL----------------SNQKMYDI----GTKAS
--EFI------AKCT----FK-Q----K----------P---C-----------A-A---
----------------------------------------------S-------------
------------------------------------------------------------
------------------------------------------------------------
-----------N----------F-------T-----------FSF-NYLYG--------N
CFSFN-------TG-------------TL-R-------------------RQ----KI--
-------ESVNYP------GP--------LY-------------GLQLYLDINKG-----
----------E-----------------Y------------I-S-R---EV---------
------------------------------------------------------------
------------------------------------------------------------
----------PAAGVRVAIT-PQGIRP--NPADEGFDVAPGSLTSIGLKMR------N--
-----ITRL-----------SKP-YNPEG-C--LNN--------PPR-----NSS-LTLY
----ENG----------TA-KYSYKGCIKSCIASLQY----NT--CGCIAPR----Y---
------------S-F--TA-----------------------------------------
-------STKGF-----T---ICKLS----------N-----------------------
------------------------------------KTE---TNCQE---------K---
L-E-------------------REFI-N-------GK--------I------------KC
------------GCV-RACNEKS---FEAT-ISQ------------------AQM-PT--
-------EVN------LDS------------NYE--------------------------
------------------------------------------------LL----------
----------------------------------EYF-----------------------
-K-------------RSV------------------THL---------------------
---------------------HLTRK-NVT------------------------------
-----T-------FL-----------R------ENLVSLNIYYDELNFETIK-QTPRYSS
VDL-----------------ASDIGGVLGLWIGVSVLTVFEFAEILMDT-----------
--------LLII--------F----------G-------KEKIA----------------
SR------------------------------RN-------------------T------
-----------------------------------------ITEFK--------------
------------------------------------------------AQLE--------
------------------------------------------------------------
```

------------------------------------------------------------
------------------------------------------------------------
------------------------------------------------------------
---ANGFA
>Protostome_Lophotroco_annelid_Pdum_comp411104_c0_seq3__791_292
------------------------------------------------------------
----------M-------------------------------------------------
------------------------------------------------------------
------------------------------------------------------------
------------------------------------------------------------
------------------------------------------------------------
------------------------------------------------------------
------------------------------------------------------------
------------------------------------------------------------
------------------------------------------------------------
------------------------------------------------------------
------------------------------------------------------------
-EV--IKEFG-DTTTMHGVQKIA---------K-AQ--------------YK---RL---
--RVFWI----VAVICAIGMFIFQLYLLIS-QYLK----------W---PTRTTM----E
-L-SR------------D-SIRF-----PDV-----------------------SVCNMR
NI------DVD-----VLYSLVKQFSL-------NKHPFELLTW----------TNITRA
S-------PEF---------------------------EVKFLQLTG----EYYLFYVY-
------------------------------------------------------------
------------------------------------------------------------
------------------------------------------------------------
------------------------------------------------------------
------------------------------------------------------------
------------------------------------------------------------
------------------------------------------------------------
------------------------------------------------------------
------------------------------------------------------------
------------------------------------------------------------
------------------------------------------------------------
------------------------------------------------------------
-----------------------------------------HYFAYPEV------FSTLF
---------------------------------SRSNLV---------------------
------------------------GNF----------------GKDTMIKG----AVPPW
--ELT------LRCS----WQ-G----A----------P---C-----------Y-S---
----------------------------------------------S-------------
------------------------------------------------------------
------------------------------------------------------------
-----------N----------F-------T-----------TFY-DTYYS--------N
CVTFK-------AP------------V---------------------------------
-------EKIGSE------GA--------ES-------------GLSLVALVGSG---MI
DWANISA---K------KVLIPGLQENNY--------------P-----LA---------
------------------------------------------------------------
------------------------------------------------------------
----------GDGGLRVVIH-PPGSHP--LPSIEGYDVPPGFSGSFALKVT------N--
-----NSLL-----------GKP-Y--GN-C--STD-Y--------------VEQ-----
----------------DGS-SYRVLNCLRKCMQKQII----EK--CKCVDAR----L---
------------P-F--SD-----------------------------------------
-----DMI-VNT-----D---FCGKVA---------PLTAACKM-------SSSVS----
---------------------------CMGPLF-EAANN---ITCAK---------Q---
V-A-------------------GEMI-S-------KD-------TLV----------SDC
------------NCF-PPCKETL---YDLT-YGL------------------AKW-PA--
-------DHE------TDF-V-----------YQ--------------------------
------------------------------------------------ELLLQD------

-------------------------------RFFEKLN---------------------A
SN-------------TSA------------------LKL---------------------
-----------------DLFHQYFQF-QNR------------------------------
-----E--------------------K----NLKEFAKINVYFADLTITKTK-QVPDYNL
IDL-----------------MSDIGGNMGLWLGMSLLTWTEFLQLALDL-----------
--------CIHA---------------------------KSAN-----------------
SN------------------------------KT----------------KVQKM-----
---------------------------------------KNVTKVSVLPADD--------
------------------AKYDFDYKKR--------------------TTSREN------
------------------------------------------------------------
------------------------------------------------------------
------------------------------------------------------------
----------------------------------------------------LIKQAMET
DGLYYINR
>Lopho_annelid_Pdum_MGIC_AWC68057_1_MIPgated_ion_channel__Platynereis_dumeril
ii__8_293
------------------------------------------------------------
----------M-------------------------------------------------
------------------------------------------------------------
------------------------------------------------------------
------------------------------------------------------------
------------------------------------------------------------
------------------------------------------------------------
------------------------------------------------------------
------------------------------------------------------------
------------------------------------------------------------
------------------------------------------------------------
-----------------------------------------AL-----R-----------
-AL--MQEFA-GGTTMHGIPKAI---------R-SR--------------SI---SA---
--RIFWS----IVCICAATMFCVQFAQLIS-KFYA----------F---PKKVTI----E
-I-VP------------A-MVPF-----PAI-----------------------SLCNMR
NL------DIM-----VLNTLNSIFK---------NATDPL-TW----------TNITED
---------PF---------------------------INAYMMTVA----KYHPMFVRN
------------------------------------------------------------
------------------------------------------------------------
------------------------------------------------------------
------------------------------------------------------------
------------------------------------------------------------
------------------------------------------------------------
------------------------------------------------------------
------------------------------------------------------------
------------------------------------------------------------
------------------------------------------------------------
------------------------------------------------------------
------------------------------------------------------------
-------------------------------------------DTDMKI------FQTIL
---------------------------------TRTLIA---------------------
------------------------TNV----------------DRHLVQKA----GVPFK
--EFI------VTCR----YG-G----L----------A---C-----------N-RS--
----------------------------------------------E-------------
------------------------------------------------------------
------------------------------------------------------------
-----------E----------F-------T-----------QFF-DPYYY--------N
CFTYT-------AP-------------EL-M-------------------------YA--
-------DTTLAE------GL--------EN-------------GWSTVVLTGSG---ML
DQND------E------LRIIPGT----HE-------------K-F-SPMA---------
------------------------------------------------------------

------------------------------------------------------------
----------SNEGVRLVIH-PPHTEP--FPHTEGFDVPPGFSVSLGVKAR------L--
-----NLRI-----------GPP-H--GN-C--SHI--------DP------FGQ-GKSR
--------------------EYRLISCQKKCLQREIV----KE--CGCKEIS----L---
------------P-N--HE-----------------------------------------
-----KY--DNL-----K---YCTQDD---------DLPDSCSV-------GATPE----
---------------------------CFERLY-QVYDR---FLCVQ---------N---
T-T-------------------ARLT-R-------NM-------TFA----------GQC
------------KCF-PPCREVS---YDVT-YSL------------------SKW-PA--
-------ESF------DGEEA-----------YV--------------------------
------------------------------------------------DIFETE------
-------------------------------AYPVRF-----------------------
MG-------------PDD------------------YKK---------------------
----------------FELYANYFDM-SNR------------------------------
-----K-------RA-----------M------KDFARLNVYIADSNVLKTE-ESQDYTQ
SQL-----------------LSDIGGQLGLWVGISVITLAEVLELIIDL-----------
--------CKFI--------A----------S-------NH-------------------
GP------------------------------YSKGRT-------------FNK------
-----------------------------------------RNNKYSAPND---EPV---
PNCRS--------------CRLYGQMN---------------------GTIPLT------
------------------------------------------------------------
------------------------------------------------------------
------------------------------------------------------------
----------------------------------------------------AVPEPMD-
---PSHMV
>Deutero_Cephalo_107760F_t1__name__chr_scaffold21_start_1172790_end_1185891_s
trand_pro_len_591__793_295
------------------------------------------------------------
----------M-------------------------------------------------
------------------------------------------------------------
------------------------------------------------------------
------------------------------------------------------------
------------------------------------------------------------
------------------------------------------------------------
------------------------------------------------------------
------------------------------------------------------------
------------------------------------------------------------
------------------------------------------------------------
------SPE----------------------D---------DK-----E-----------
-SP--EKAFA-GNTSLHGFNRIA---------S-SP--------------GG---SP---
--RVLWV----IAILGSYTGFGYMFGSMMA-SYFK----------Y---DTITDT----T
-L-EF-A----------D-ELLF-----PAV-----------------------TICNMN
KF------DSD-----KLQVAD------------------W-----------------S-
---------YL------------S-LFL---------------YGIQ----A--------
---------------------------------DI-------PTIL--------------
------------------------------------------------------------
------------------------------------------------------------
-----------------ALG----------------------------------------
------------------------------------------------------------
------------------------------------------------------------
------------------------------------------------------------
-------------------------VPL--------------------------------
------------------------------------------------------------
------------------------------------------------------------
------------------------------------------------------------
------------------------------------------------------------
------------------------DETVNST------------------------MTGV-

---------------------------------KIEDLV---------------------
------------------------------------------------LAK----GFNVS
-PNRM------SLCW----WR-G----T----------N---C-----------S-E---
----------------------------------------------R-------------
------------------------------------------------------------
------------------------------------------------------------
-----------N----------F-------T-----------HSF-G-HYG--------N
CYTFN-------AD-------------AD-N--------------------P--------
-------LKQTMP------GA--------VN-------------GFMAFVDIKED-----
----------K-----------------Y------------T-E-S--FLV---------
------------------------------------------------------------
------------------------------------------------------------
-------SGNAEVGLKLLVH-DPREPP--MMDTQGIALAPGNHAFIAIKQI------L--
-----YENH-----------VPP-W--GV-C--KDL--------Q-------LE----YY
------------------D-TYTLNGCYLECRSKHLV----RN--CSCRPYD----L---
------------P-G---------------------------------------------
----------TA-----P---SCDP-----------------------------------
------------------------------------RTM---FTCVR---------A---
V-L-------------------AQVI-T-------GD--------L------------KC
------------DCP-VPCRMTS---YSTS-LSF------------------AGF-PN--
-------KHT------REY-L-----------SP--------------------------
------------------------------------------------LL----------
------------------------------------------------------------
-G----------------------------------------------------------
---------------------------MEP------------------------------
-----S-------YM-----------G------DNGVVFSVFYEKLNYQKIR-QLKAMEE
GQL-----------------ASNIGGMMGLFLGASVLSLLEVCEYLLKR-----------
--------PLGF--------L----------G-------RT-------------------
RH------------------------------AK--------------------------
-----------------------------------------VVHVQ--------------
--------------------------P---------------------QETKND-VIQHV
PVL---RGISQQTHKGIPLPAAGDGKLHGMFSFVT--------------------YSIRF
RDNGVVFSVFYEKLNYQKIRQLKAMEEGQLASNIGGMLSLLEVCEYLLKRPLGFLGRTRH
--------------------------AKVVHVQP--------------------------
---------------------------------------------------------QEP
KSATALHK
>Protostome_Lophotroco_annelid_Pdum_comp414793_c0_seq1__795_296
------------------------------------------------------------
----------M-------------------------------------------------
------------------------------------------------------------
------------------------------------------------------------
------------------------------------------------------------
------------------------------------------------------------
------------------------------------------------------------
------------------------------------------------------------
------------------------------------------------------------
------------------------------------------------------------
------------------------------------------------------------
-------------------------------G---------KL-----Q-----------
-GL--MDDYA-QSMTAHGVGRIS---------R-AS--------------SF---KA---
--KIFWA----VIWTGMIAMFFLQATTLMR-RFFA----------Y---PKSVRL----E
-M-AN------------Q-PVPF-----PAV-----------------------SLCNLR
PL------DVF-----MFKDAI------------FKDNVTM-RWIPGNYSYTGRSGASD-
-------FERF------------LIKYG---------------NVYM----SYRSY----
------------------------------------------------------------
------------------------------------------------------------
------------------------------------------------------------

------------------------------------------------------------
------------------------------------------------------------
------------------------------------------------------------
------------------------------------------------------------
------------------------------------------------------------
------------------------------------------------------------
------------------------------------------------------------
------------------------------------------------------------
----IEVLLEKE------------------------------------------------
---------------------------------------QVDQDWFWEA------RRALF
---------------------------------SRLTLA---------------------
------------------------ANF----------------NEKAAQQG----GIQAD
--QFI------AQCE----YA-G----K----------K---C-----------S-F---
----------------------------------------------E-------------
------------------------------------------------------------
------------------------------------------------------------
-----------N----------F-------T-----------YFL-DPAYF--------N
CYTFD-------MN-------------QM-G-------------------------WE--
-------SQTIYE------GP--------DH-------------GLSVVLFIPSL--PST
DLSP------E----------------AYRL----------I-AFT-QELS---------
------------------------------------------------------------
------------------------------------------------------------
---------FGGEGIRLVIH-EQNTVP--YPLTDGLDIPRGVSASIGVQLN------H--
-----NMRL-----------SPP-Y--GN-C--TDK--------NT------LEGTLVNY
--------------------TYTMASCKKTCLQGLIA----DT--CGCTDIG----L---
------------P-L--SE-----------------------------------------
-----NF--EEG---------SCSRFD---------ALPTECQQ-------KGDIRMN--
------------------------LDKCKTFFE-PWFER---NLCKR---------T---
V-R-------------------TNIS-K-------NL-------SAW----------NLC
------------KCF-PRCYDVA---YVTS-TSQ------------------SDW-PT--
-------IES------SVY-L-----------LQ--------------------------
------------------------------------------------DILTAG------
-------------------------------GFASKFPE---------------------
EK----------------------------------AKQYFGPLLG--------------
------------------AETPTFQE-ARD------------------------------
-----H-------IL-----------A------KNWLRINLYISDTSVVKIE-ETEDYGI
SQL-----------------ISDIGGQLGLWIGVSIISIIEIFDLIYQI-----------
--------CKYL--------L----------DPE-----RQ-------------------
QK------------------------------RH-------------------RG-----
---------------------------------------STIRETKQN------------
------------------------------------------------GSND--------
------------------------------------------------------------
------------------------------------------------------------
------------------------------------------------------------
-----------------------------------------------------L--SYAQ
SNGNPLRF
>Protostome_Lophotroco_annelid_Pdum_comp418040_c0_seq1__796_297
------------------------------------------------------------
----------MMVLTLL-------------------------------------------
------------------------------------------------------------
------------------------------------------------------------
------------------------------------------------------------
------------------------------------------------------------
------------------------------------------------------------
------------------------------------------------------------
------------------------------------------------------------
------------------------------------------------------------

------------------------------------------------------------
-LNLRFKSK----------------------M---------AL-----R-----------
-DV--LGNFA-VSTTMHGVPKVI---------N-AK--------------SA---MA---
--RLFWS----IVCLAAGSMFCLQMSEVLT-RYYS----------Y---PKKVTV----E
-V-VP------------T-PVPF-----PSI-----------------------SICNMR
NL------DVH-----ILNTLNAAFL---------RDDNPI-NH----------INKSDN
---------AF---------------------------INSYMYNVG----RLAHLFWK-
------------------------------------------------------------
------------------------------------------------------------
------------------------------------------------------------
------------------------------------------------------------
------------------------------------------------------------
------------------------------------------------------------
------------------------------------------------------------
------------------------------------------------------------
------------------------------------------------------------
------------------------------------------------------------
------------------------------------------------------------
------------------------------------------------------------
-----------------------------------------YQDAHPEV------FQEVF
---------------------------------SRTTFS---------------------
------------------------ANI----------------PEDIIALA----AVQLD
--GFV------VNCH----YA-G----H----------R---C-----------N-KS--
----------------------------------------------R-------------
------------------------------------------------------------
------------------------------------------------------------
-----------D----------F-------L----------RFF-DPYYF---------S
CFTYK-------AP-------------EP-S-------------------------EL--
-------DDSLSE------GI--------EN-------------GWSSILLSGSG---ML
DKND------E------IRMLPGL----HEW--------------R-SAVS---------
------------------------------------------------------------
------------------------------------------------------------
----------ASEGVRVVIH-PPNTEP--YPFTEGYDVPPGFSASFGIKPR------R--
-----NIRI-----------GPP-H--GN-C--TNK--------NP------FDQ--ATE
--------------------RYRLISCQKMCLQSAII----KS--CNCSDVG----L---
------------P-R--LN-----------------------------------------
-----HM---DV-----P---LCRSSE---------SFPESCFD-------DATDE----
---------------------------CLEALM-NMSSL---IKCVR---------V---
T-K-------------------AKLT-K-------NT-------TQM----------EEC
------------QCF-PPCDEVA---YDVS-YSL------------------SKW-PA--
-------SGY------EGDAA-----------YF--------------------------
------------------------------------------------DVFGIE------
-------------------------------GFNERF-----------------------
NN-------------SET------------------KGK---------------------
----------------YELFAKYFNV-SNR------------------------------
-----E-------ET-----------M------KNFARLNVYIADSNVVKTQ-ESADYTR
NQL-----------------VSDIGGQLGLWVGISVITLTEVLELLSDL-----------
--------VRFF--------T----------S-------SY-------------------
SRIS----------------------------RH---------------DKTYR------
-----------------------------------------NNNMNHHHNI---IPLK-R
PMLHD-------------HDLQFRMEN---------------------------------
------------------------------------------------------------
------------------------------------------------------------
------------------------------------------------------------
------------------------------------------------------------
-----TTL

>Protostome_Lophotroco_annelid_CAC9486619_1__Ofus_G015124_Owenia__fusiformis_
_802_301
------------------------------------------------------------
----------M-------------------------------------------------
------------------------------------------------------------
------------------------------------------------------------
------------------------------------------------------------
HSESNNKLEFPPISG---Y-----------------------------------------
----------------------------------DFEVTTTPKEKR--------------
------------------------------------------------------------
------------------------------------------------------------
-----------------------------------------------YSEFDFTATTPRS
K---GL-------------DGIDEKNESP-------------------------------
---KRKEIT----------------------P---------EI-----T-----------
-PL--FREFC-DNASVHGVNHLR---------T-ER--------------VK---VK---
--RWVWT----IFVILALGFNLFHCSLLID-KYLG----------F---PSEETQ----F
-V-DQ------------S-WIEF-----PSV-----------------------TICNIN
AM------SRI-----TRKKMLEDNS----------------------------------
---------TLLYK---------------------------WHDYVN----NRFEALVS-
---------------------------------DLA------------------------
------------------------------------------------------------
------------------------------------------------------------
-----------------DGYS---------------------------------------
------------------------------------------------------------
---------------------------L--------------------------------
------------------------------------------------------------
------------------------------------------------------------
------------------------------------------------------------
------------------------------------------------------------
------------------------------------------------------------
------------------------------------------EGSN--------------
-----------------------------------------LLDTLELL------YNRVH
---------------------------------LPTGYL---------------------
------------------------ENI-----------------GEEALEV----GHKLN
--DLV------VDCT----FG-I----T----------K---C-----------H-A---
----------------------------------------------V-------------
------------------------------------------------------------
------------------------------------------------------------
-----------N----------F-------T-----------SFF-ESTYY--------N
CYTFN--------GG------------NV-S-TS-----------------H----NN--
-------LITRTT------GP--------QE-------------GLSIIMYLESD-----
----------N-----GNINENGS----Y------------L-T-M-SKLN---------
------------------------------------------------------------
------------------------------------------------------------
----------NAAGARVMIH-APNTMP--SPTDQGFDIPPGFSSSVGVSVS------T--
-----RTRL-----------GEP-Y--AK-C--SKR--------EINI--------GTDY
--------------------LYSDNICLKLCQQRYVM----GN--CNCISSL----L---
------------P-F--DR-----------------------------------------
--------STDL-----Y---FCGHM----------DYN-NNASF---------------
------------------------------------FEN---MACES---------R---
V-L-------------------EEFV-H-------NE-------DVR----------REC
------------GCH-PPCHNYI---YNYQ-TSQ------------------SYW-PL--
-------EYY------QAD-F-----------YD--------------------------
------------------------------------------------LYILSDPNK---
------------------------------------------------------------
EN-------------LKA------------------YQN---------------------
-----------------LKHHNVPQL-IEK------------------------------

-----G-------LI-----------R------KNFLRLNVYLKDLTIVQHI-EKRSYAI
ENL-----------------FSDVGGTFGLWAGMSILTICEFSELFFRV-----------
--------MAIL--------V----------S-------KF-------------------
LSIFGPMST-----------------------RA----------------VVED------
------------------------------------------------------------
------------------------------------------------SSSN--------
------------------------------------------------------------
------------------------------------------------------------
------------------------------------------------------------
------------------------------------------------------------
SHTPVRRT
>Cnidar_Rhopilema_esculentum_TR55888_c0_g2_i1_p1__Rhopilema_esculentum_TR5588
8_c0_g2__Rhopilema_esculentum_TR55888_c0_g2_i1_p1__ORF__type_complete__len_60
4__score_83_96__Rhopilema_esculentum_TR55888_c0_g2_i1_896_2707__809_303
------------------------------------------------------------
----------M-------------------------------------------------
------------------------------------------------------------
------------------------------------------------------------
-------------AATNTR-GLFLRAQ---RHIMDRE-----------------LSKDEN
EEEPSQPYSSA--------------------DESKESLLLQNF---------GMKYGHAF
KVKPKKTEPEQKNGNTELP--------QRKVS---------PE-----------------
-IETNKPAQVTVNPAFKNVA-------------LDRFRRAG-------------------
------------------------------------------------------------
-------------------------MKARLV--------------------------RRV
N----------------------KIVED--------------------------------
--------R----------------------L---------TA-----K-----------
-QI--FARFT-KESTLHGFRFIF---------T-KT--------------FY---IR---
--RFIWF----VITITMAAMFLKELTDSIN-LYFQ----------H---PFSTTS----T
-I-EY-V----------N-RLTF-----PAI-----------------------SFCNLN
DF------RFS-----KINGSD------------------L-----------------Q-
---------DV----------------------------------FM-------------
------------------------------------------------------------
------------------------------------------------------------
------------------------------------------------------------
------------------------------------------------------------
------------------------------------------------------------
------------------------------------------------------------
------------------------------------------------------------
------------------------------------------------------------
------------------------------------------------------------
------------------------------------------------------------
------------------------------------------------------------
------------------------------------------HEKG--------------
------------------------KFYLH-RNSSFD-------------------LDGK-
---------------------------------KLGERL---------------------
------------------------------------------------EDA----SHRIT
--DMF------MKCV----WL-F----SQTAA-GQ-PVP---C-----------N-Y---
----------------------------------------------T-------------
------------------------------------------------------------
------------------------------------------------------------
-----------N----------I-------T-----------TYY-G-LNGQ-------T
CYTFN-------PG-------------DR-G--------------------H----RL--
-------LTLNET------GL--------FH-------------AFELQLDLETH-----
----------E-----------------Y------------L-K-D--------------
------------------------------------------------------------
------------------------------------------------------------
---------IQEGGVRVHIH-DQNETP---FSSAGFAVPPGFKTFVSLNVQ------K--

-----IENL-----------PPP-YT-TQ-C--GSK--------K-------LK----YY
------------------S-SYKKSKCYLEELSKFVT----KK--CKCRGRF----M---
------------P-D---------------------------------------------
---------ASY-----P---LCSL-----------------------------------
------------------------------------KQT---VECLR---------P---
T-I-------------------RTFD-H-------RA---------------------GG
------------ACP-EDCLSTQ---YNAQ-LSY------------------ARY-VS--
-------RPP------IGT-S---------------------------------------
------------------------------------------------GITIPE------
---------------------------------------------------------RFQ
KA----------------------------------NLT---------------------
------------------------NS-QLE------------------------------
-----A-------VI-----------R------DNIVYLIFFFPELRVENVK-QTVTYDF
YKL-----------------VGDVGGQLGLMLGASVLTLVEFIDLIAFI-----------
--------IYHQ--------I----------L-------RL-------------------
TR------------------------------LK-------------------K------
-----------------------------------------KKKEETEQL----------
-----------------------KLRE---------------------MNGN--------
------------------------------------------------------------
----------------------------------G-------------------------
------------------------------------------------------------
-----------------------------------------------------------A
REGRFINF
>Protostome_Lophotroco_annelid_CAC9670605_1__Ofus_G181373_Owenia__fusiformis_
_810_304
------------------------------------------------------------
----------MVYCSS--------------------------------------------
------------------------------------------------------------
------------------------------------------------------------
------------------------------------------------------------
------------------------------------------------------------
------------------------------------------------------------
------------------------------------------------------------
------------------------------------------------------------
------------------------------------------------------------
------------------------------------------------------------
-SIIKLNMA----------------------K---------RI-----H-----------
-DV--VQRFA-SGTTIHGVPKLI---------R-AQ--------------TI---PG---
--KIFWS----LICLAALGVFIFELALLLQ-KFFG----------Y---PKSVDV----K
-V-VQ------------E-PVAF-----PSV-----------------------TVCSVH
AM------DPF-----VIHKIYKLRQ------------EQM-EHRMRRKPWVISKNVSD-
----------------------FEKWYF--------------DTFLK----KSMSY----
------------------------------------------------------------
------------------------------------------------------------
------------------------------------------------------------
------------------------------------------------------------
------------------------------------------------------------
------------------------------------------------------------
------------------------------------------------------------
------------------------------------------------------------
------------------------------------------------------------
------------------------------------------------------------
------------------------------------------------------------
--VLEPSYLEGL------------------------------------------------
---------------------------------------IDRHDLKTEF------FPSLS
---------------------------------SRLTLA---------------------
------------------------ANL-----------------NKSITFP----GVEPA

--ELI------AFCR----YA-R----Q----------E---C-----------S-F---
----------------------------------------------E-------------
------------------------------------------------------------
------------------------------------------------------------
-----------H----------F-------K-----------TVF-DPYYF--------N
CFTFN-------AS-------------VI-A-------------------------GS--
-------LKTLAE------GI--------EN-------------ALSIVMYIPKLHN---
----------EIKIG-TKVKLPGIVE--HD-----------I---R-DPLA---------
------------------------------------------------------------
------------------------------------------------------------
----------GSGGVRVVIH-PPDTQP--HPATEGFDIMPGYSVSIGVKTT------E--
-----NTRL-----------GRP-Y--GN-C--TET--------TKGV----FSD-KASY
--------------------RYTMTSCRKKCLQNLLSREVKDG--CGCLDVV----L---
------------P-T--FP-----------------------------------------
-----EV--NGV-----D---FCAKFD---------DIPRKCIFGPF---ARRNES----
---------------------------CKELKA-RWLKR---MICMR---------D---
I-E-------------------TSGS-H-------EH------IAIT----------ENC
------------NCH-PPCKDLS---YEFF-YSN------------------SLW-PG--
-------QQH------KQD-V-----------YR--------------------------
------------------------------------------------DLFISR------
-------------------------------RFERRF-----------------------
QE----------------G----------------QRTYYFGA-----------------
------------------ENAQALNV-TEQ------------------------------
-----E-------KT-----------V------DRFARLNVYLYDTNVVKIT-ETKDYTG
IQL-----------------ISDVGGQLGLWLGISIITLTEVFELIADI-----------
--------VHLC--------L----------K-------RK-------------------
RK------------------------------QS--------------------------
-----------------------------------------IREGNCTPA----------
------------------EITALNRRN---------------------------------
------------------------------------------------------------
------------------------------------------------------------
------------------------------------------------------------
------------------------------------------------------------
---GSTTV
>Deu_Ambulacrar_hemi_Ptyfla_40v0_9_20150316_1g24013_t1_scaffold33951_cov84_81
3_305
------------------------------------------------------------
----------M-------------------------------------------------
------------------------------------------------------------
------------------------------------------------------------
------------------------------------------------------------
------------------------------------------------------------
------------------------------------------------------------
------------------------------------------------------------
------------------------------------------------------------
------------------------------------------------------------
------------------------------------------------------------
--ARKDTDE----------------------R---------SF-----R-----------
-QL--FVELV-QNSSAHGIPNAG---------R-AK--------------TK---FR---
--TVVWS----IIFVIGVAGFLFQFSELFI-RFID----------F---PVTTNV----D
-V-TS-N----------R-SLVF-----PAV-----------------------TICNQN
PV------RVS-----ALEYAS------------------------------------DP
---------KL-------------------------------RENFD-------------
---------------------------------DT-------------------------
------------------------------------------------------------
--------------------YEP-------------------------------------
---------STSSQAPQDVTGNTAF-----------------------------------

------------------------------------------------------------
------------------------------------------------------------
------------------------------------------------------------
------------------------TSPDIASAPPI-------------------------
------------------------------------------------------------
------------------------------------------------------------
--------------------------------------------------------GNPP
PPQGRKRRATDYLE----------------------------RQGNKQP-----------
----------------------------------------PSPDSYNGP------SAGLA
------------------------------QVSLLTELM---------------------
------------------------SKV----------------SIEERIEL----GHQAQ
--DFI------VNCT----WQ-G----E----------R---C-----------S-S---
----------------------------------------------D-------------
------------------------------------------------------------
------------------------------------------------------------
-----------N----------F-------T-----------VFN-NPVYG--------N
CFTFN-------GK-------------DV-A-------------------DE----DV--
-------LLTNFP------GP--------LF-------------GLSLQLFIEQG-----
----------E-----------------Y------------L-V-D---YT---------
------------------------------------------------------------
------------------------------------------------------------
----------EIAGVRIVIH-DRDKMPALMPEDDGFDVNPGVLTSIGIRRT------E--
-----ITRL-----------GDP-Y--SN-C--VSK--------TGDI----GQY-SNVF
----TDM--FN-------A-NYSQETCMKACYLTRVA----ER--CDCYDGR----Y---
------------P-K--PN-----------------------------------------
-------STEDL-----D---FCNTH----------N-----------------------
------------------------------------NAN---YRCVK---------N---
I-E-------------------EEVR-T-------NS--------TS----------FRC
------------DSP-QPCEQVT---YTTE-FST------------------AYW-PA--
-------RIS------QRY-V-----------ND--------------------------
------------------------------------------------WL-T--------
----------------------------------QR------------------------
-N-------------REL------------------HRMVM-------------------
----------------------KQED-LNE------------------------------
-----D-------LV-----------R------LNILKLRVYYRDLNYETIT-QSPSYTD
AM----------------------------------------------------------
------------------------------------------------------------
------------------------------------------------------------
------------------------------------------------------------
------------------------------------------------------------
------------------------------------------------------------
------------------------------------------------------------
------------------------------------------------------------
------------------------------------------------------------
--------
>Deutero_Ambulac_Spurpu_025881_821_307
------------------------------------------------------------
----------M-------------------------------------------------
------------------------------------------------------------
------------------------------------------------------------
------------------------------------------------------------
------------------------------------------------------------
------------------------------------------------------------
------------------------------------------------------------
------------------------------------------------------------
------------------------------------------------------------
------------------------------------------------------------

```
----DTEEK----------------------D---------TV-----K-----------
-KS--VTDFG-NETTIHGLQFVV---------N-KK--------------NI---IY---
--RLCWL----GICTTFLVVFLIQGNVILK-DFLR----------W---PYSTKI----D
-I-VG-R----------P-NLAF-----PAV-----------------------TVCNAN
MM------RRS-----QIEGSR------------------F-----------------E-
---------DLVNL----------------------------------------------
---------------------------------DG-------GVEGADYDYSWWFSSAYR
-------------------------------------------NWYAS------------
---------SSASS----------------------------------------------
-----YGQSSDQNSNGRSSSSSQSDS---------------ATSSDGQSSSSN---YD--
------------------------------------------------------------
---------------PSSSEESSSA----------SSVESSGGTSDGQSSSSND------
--------------------------------------------------------GP--
----------------SSSEEPASSSPGRSADGHSSSTN---------------------
------------------------------------------------------------
------------------------------------------------------YGPATV
RVSIVFWVYNSEFPAP--------------------------------------------
--------NSXVNEWSWY----DPSFFA---------DFEF-SENG-------W------
--------DGV-------------SGEKTGRASMRP-----------PK------PMTTA
---------------------------------TFLT-----------------------
------------------------------------------------------------
------------------------------------------------------S-ST--
----------------------------------------------R-------------
------------------------------------------------------------
------------------------------------------------------------
-----------R----------R-------K-----------SWK-STVTS--------W
KISLC-------SV-------------PS-I-------------------VD----HV--
-------I----------------------S-------------GLHLTLFVEQP-----
----------E-----------------Y------------L-G-V---LS---------
------------------------------------------------------------
------------------------------------------------------------
----------HQTGAKVTIH-HPNEYP--FPEDNALSLGTGQETSIGIRQE------Y--
-----IKRL-----------GGY-Y--TN-C--TSD-G------KD----------TNFS
---STTE-----------L-SYSSEACKKICYQLHLS----QL--CKCVDDQ----F---
------------F-----------------------------------------------
----------DGFP---T---KCDVL----------N-----------------------
------------------------------------MTH---QICRK---------F---
V-E-------------------DLFL-D-------DK--------L------------PC
------------SCP-PPCTEYK---YVRT-PSS------------------SLW-PS--
-------ERY------EEH-L-----------LR--------------------------
------------------------------------------------RL-N--------
----------------------------------GSA-----------------------
-N-------------ENL------------------IRVL--------------------
---------------------------QSN------------------------------
-----E-------LS-----------R------KNLIRLKIFYEDLNYEVVE-MVPVYTI
PSV-----------------LGSIGGLMGLYIGMSFISVFEVLFLVLRL-----------
--------IKIA--------L----------I-------RIYS-----------------
--------------------------------RI------------------NR------
-----------------------------------------VQPYP--------------
------------------------------------------------------------
------------------------------------------------------------
------------------------------------------------------------
------------------------------------------------------------
------------------------------------------------------------
----VKNV
```

>Placozoa_TriAdh_evg1307556__type_protein_aalen_609_92__partial3_clen_1981_st
rand___offs_155_1981___evgclass_main_okay_match_evg1307557_pct_99_100__822_30
9
------------------------------------------------------------
----------M-------------------------------------------------
------------------------------------------------------------
------------------------------------------------------------
------------------------------------------------------------
------------------------------------------------------------
------------------------------------------------------------
----------------SQSS-------------YHDSNK---------------------
----------------------------------------------------------TA
SDESSTDAHPN----SQKVPILSHQDEVNQD-----------------------------
---------------------NPEPRNRFDLD-FRW------------------------
-------------------------------R---------PS-----I-----------
-EH--DENFP-FSASFHGIEHIY---------E-GR--------------YG---TR---
--KILWI----LLVAATMIACFVFIFIQIA-HYSA----------F---HTTTKS----T
-L-VY-E----------K-QLAF-----PAV-----------------------TICNYN
SF------RRS-----AVTAND------------------L-----------------I-
---------HM------------A-YLV---------------KAYR----L--------
---------------------------------NQ-------------------------
------------------------------------------------------------
------------------------------------------------------------
-------------GVVSD---FIG------------------------------------
------------------------------------------------------------
------------------------------------------------------------
------------------------------------------------------------
------------------------------------------------------------
------------------------------------------------------------
------------------------------------------------------------
------------------------------------------------------------
--------EKER------------------------------QKL---------------
------------------------INYWKK---------------YDATH-----AKKF-
---------------------------------NYQLFV---------------------
------------------------------------------------ERV----GYHAS
--QMI------KSCH----FR-G----L----------K---C-----------G-P---
----------------------------------------------K-------------
------------------------------------------------------------
------------------------------------------------------------
-----------N----------F-------S-----------NVL-T-SYG--------N
CITFN-------GP-------------KLES--------------------N----PK--
-------LYQKNP------GA--------HQ-------------GLELLINIQEY-----
----------E-----------------Y------------T-G-S---WH---------
------------------------------------------------------------
------------------------------------------------------------
-------SDRPDIGIKFVIH-ERHYPP--DVTSQGKAVGPGSHAYASVKYK------T--
-----ISNL-----------PSP-Y--GH-C--GSK--------K-------LA----FY
------------------K-KYTYAGCQISCKTEYVQ----KK--CGCRAPD----M---
------------P-G--QNVIPVCSP-------QKMIECVSPNLE-KLRTINDKVCIYTY
CCRAPDMPGQNII----P---VCSP-----------------------------------
------------------------------------QKM---IECVS---------P---
N-L-------------------EKLI-TI------ND--------------------KVC
------------ICP-IPCHIVH---FDDT-ISY------------------AKI-PN--
-------PQM------AKD-L-----------TE--------------------------
------------------------------------------------KI----------
------------------------------------------------------------
-N---------------K------------------TAFQIETHG---------------

-------------------LV-DPDI-DPT------------------------------
-----L-------YI-----------S------QNYILLNVFFDDLYYEKTV-STPVYTF
TSL-----------------LGNIGGQLGLFVGASVLTLVEIIEFGFYR-----------
--------SRGV--------I----------R-------RS-------------------
DW------------------------------KQ-------------------N------
-----------------------------------------LRKSISR------------
------------------------------------------------SREV--------
---------TATE-----------------------------------------------
----------------EKEPLCS-------------------------------------
------------------------------------------------------------
------------------------------------------------------------
--------
>Protostome_Lophotroco_annelid_CAC9670604_1__Ofus_G181372_Owenia__fusiformis_
_823_310
------------------------------------------------------------
----------M-------------------------------------------------
-----------------------V------------------------------------
------------------------------------------------------------
------------------------------------------------------------
------------------------------------------------------------
------------------------------------------------------------
------------------------------------------------------------
------------------------------------------------------------
------------------------------------------------------------
------------------------------------------------------------
-------------------------------R---------MF-----D-----------
-EV--VQHFA-SGTTIHGVPKLL---------K-AQ--------------TI---QG---
--KIFWS----VICLSALGVFIFELVLLLS-KYFE----------H---PKSVDI----K
-I-VQ------------E-PVAF-----PSV-----------------------SVCSVY
AI------DPF-----VIHEIYKLKTG-------TQKDSAMATYRFTRD---MSGELDDT
---------QF------GVENAFIKWYL--------------DDFIE----KSSDFMGKQ
---------------------------------EN-------------------------
------------------------------------------------------------
------------------------------------------------------------
------------------------------------------------------------
------------------------------------------------------------
------------------------------------------------------------
------------------------------------------------------------
------------------------------------------------------------
------------------------------------------------------------
------------------------------------------------------------
------------------------------------------------------------
--RLAIKRMLGS------------------------------------------------
---------------------------------------EAEEKYRTKF------LPSLA
---------------------------------SRLTLT---------------------
------------------------ANM----------------LHEHIAFP----GIKPK
--EFI------ALCR----YA-R----K----------E---C-----------S-Y---
----------------------------------------------K-------------
------------------------------------------------------------
------------------------------------------------------------
-----------H----------F-------T-----------TVF-DPYYY--------D
CFTFN-------AS-------------II-A-------------------------GA--
-------RKTLAE------GI--------EN-------------ALSLVMYLPELHR---
----------DLKFNKMKTHLPGILG--HD-----------L---R-DPLS---------
------------------------------------------------------------
------------------------------------------------------------
----------GSGGVRVVIH-APGTHP--YPATDGFDVMPGSSVSIGVKTT------E--

-----NIRL-----------GWP-Y--GN-C--QKE--------EDDTIQ--IQQKTNTY
--------------------KYTMTSCREKCIQSNLLMNKPKG--CECLDVN----L---
------------P-T--FT-----------------------------------------
-----EV--EHG-----E---YCARFD---------DVPKHC---------QNKTE----
---------------------------CAELKA-RWLKR---MNCMH---------D---
----------------------KEAN---------NN-------SLV----------GEC
------------NCH-PPCEDVD---YDFF-YSS------------------TPW-PD--
-------ANY------VNG-V-----------YR--------------------------
------------------------------------------------DLFISH------
-------------------------------KFQDNF-----------------------
DN-------------AQK------------------RTAYFGD-----------------
-----------------KIESDLDNE-GRI------------------------------
-----K-------KT-----------V------GRLARLNVYLYDTNVVKIT-ENEDYTG
IQL-----------------ISDVGGQLGLWLGVSIITLTEVFELIAHI-----------
--------IHIV--------C----------K-------RD-------------------
DK------------------------------RD--------------------------
-------------------------------------VDVQHEELDYQRL----------
------------------PIPEIPHNNE--------------------------------
------------------------------------------------------------
------------------------------------------------------------
------------------------------------------------------------
------------------------------------------------------------
--------
>Deutero_Ambulac_Apla_gbr361_4_t1_824_311
------------------------------------------------------------
----------M-------------------------------------------------
------------------------------------------------------------
------------------------------------------------------------
---------------------------------------------------YRSEAASPE
RWQAAASDQHK-------------------------------------------------
----------------------------------Y-----RPE--------L--------
------NNHNSDGWTSAENG-------------QQAATDSNAVNTPFQ--------P---
------------------------------------------------------AA----
------------MSRRTSRGFKRGESLIGDDRR-----RDWD-----RFPQQEPRHLQQQ
K-------------------PGMTDEERL-------------------------------
------AKT----------------------S---------VA-----Q-----------
-SL--VLSLL-ENTGAHGIPNIK---------R-AP--------------NN---VR---
--KVFWT----ILFFFGLGMFIWQTSWLLE-RYYA----------F---GVDVKM----E
-I-SS-D----------R-MVEF-----PAV-----------------------TICNLN
PI------KSD-----FWEELD------------------------------------ER
---------GI----------------------------PIPPSLTR-------------
------------------------------------------------------------
------------------------------------------------------------
------------------------------------------------------------
------------------------------------------------------------
------------------------------------------------------------
------------------------------------------------------------
------------------------------------------------------------
--------------------------P---------------------------------
------------------------------------------------------------
------------------------------------------------------------
------------------------------------------------------------
---NTGYGPDDS------------------------------EEEF--------------
-------------------------------------------------------DDAFE
------------------------------MLGDAIELV---------------------
------------------------TAF----------------KYKDRRTA----GHLLR
--DML------VRCV----FQ-G----R----------P---C-----------H-A---

----------------------------------------------R-------------
------------------------------------------------------------
------------------------------------------------------------
-----------D----------F-------T-----------YFF-HYLYG--------N
CYIFN-------AG-------------PE-P-NAT----------------T----PR--
-------LFVTKA------GP--------LY-------------GLTLELYIEQD-----
----------Q-----------------Y------------I-E-D---IQ---------
------------------------------------------------------------
------------------------------------------------------------
----------PAAGARVVVH-ARHNMP--FPDDEGVSVSPGQETFIGFKRT------N--
-----FTRL-----------PHP-Y--SN-C--TVV--------E-------NAN-ATIF
----NLP--LP------HV-RYSQRACENNCYYQNLT----RV--CGCADVR----F---
------------R-Y---------------------------------------------
--------DDDV-----P---LCLN-----------------------------------
------------------------------------STQ---EQCLE---------G---
V-E-------------------RMYL-A-------GM--------L------------DC
------------MCS-PPCRELQ---YDIV-TGQ------------------ARW-PN--
-------EKF------KLF-L-----------DD--------------------------
------------------------------------------------NI-G--------
----------------------------------NF------------------------
-S-------------DDL------------------YQDIDSG-----------------
---------------------------GDP------------------------------
-----D-------FL-----------E------KNLLKVNVYYDKLEYTTIY-QDVAYDH
IAL-----------------LSDMGGNVGLWIGVSVLTVFEFIELLYDL-----------
--------GKLI--------F----------Y-------RV-------------------
IN------------------------------KN----------------ATRK------
------------------------------------------------------------
-----------------------TGKV---------------------TSSD--------
------------------------------------------------------------
------------------------------------------------------------
------------------------------------------------------------
-----------------------------------------------------VDLQHVS
NLKIASEA
>Protostome_Lophotroco_annelid_CAC9484202_1__Ofus_G014375_Owenia__fusiformis_
_827_312
------------------------------------------------------------
----------M-------------------------------------------------
------------------------------------------------------------
------------------------------------------------------------
------------------------------------------------------------
------------------------------------------------------------
-----------------------------------------PQKYYDKNMKL--------
--------------------------------------------------------P---
------------------------------------------------------------
------------------------------------------GVLQV---QPTAEGYEEY
RDVGKD-----------------NTKDKTPQNYYN---------YLLQ-----------E
KDKNLLNDE----------------------V---------KL-----T-----------
-HL--YRGLV-EHTSSLGITQIS---------Y-SK--------------GP---VK---
--IGFWV----ALTLAMSVLTIWNVSTVFM-EYLK----------F---DVDMTI----V
-V-EQ-R----------A-TLDF-----PAI-----------------------TVCNMN
SI------RLS-----KYLSNP--------------------------------------
---------ML-------------------------------NATLN----G--------
---------------------------------DP------GS-----------------
------------------------------------------------------------
------------------------------------------------------------
-----------------DGANATA-----------------TSTTTTTTTTT--------
------------------------------------------------------------

```
------------------------------------------------------------
------------------------------------------------------------
----------------------TTVAPAITTAASK-------------------------
------------------------------------------------------------
------------------------------------------------------------
--------------------------------------------------------GNGK
NSKTKGGTNTDKGN--------------------GKTDKKPINGNGNKQ-----------
-----------------------------------------TNDNLGEV------DPMFA
------------------------------FQERVSEML---------------------
------------------------ANM----------------GESEKVPM----GHQRE
--DFI------IDCK----YN-G----Y----------E---C-----------F-I---
----------------------------------------------D-------------
------------------------------------------------------------
------------------------------------------------------------
-----------Q----------F-------A-----------TFY-NPKHG--------N
CYIFN-------AG-------------WN-N-------------------SQ----SR--
-------FTSSRP------GP--------FY-------------GLQLTFNIEQS-----
----------E-----------------Y------------I-G-K---LT---------
------------------------------------------------------------
------------------------------------------------------------
----------STAGVRVQVH-HQNVMP--FPEDEGINIVPGQSTSVGIQML------N--
-----LQKQ-----------GGK-Y--SD-C--FDD--------DVR-----NER-MNVY
----EEY--YN-------V-KYSNPACMKTCFQRHLL----ES--CGCADKQ----Y---
------------P-M--TG-------------------VAFN------------------
------NANATV-----E---TCNSN----------D-----------------------
------------------------------------ISQ---SRCIQ---------N---
I-T-------------------DQHI-G-------GK--------L------------RC
------------SCY-SACNETL---YKTE-VSS------------------VLW-PA--
-------DAY------KDD-L-----------LA--------------------------
------------------------------------------------NL-A--------
----------------------------------LR------------------------
-S-------------PSA------------------SAVVSD------------------
---------------------------ADT------------------------------
-----D-------AY-----------K------RNFAHLQIYYSEFNFQSIV-ENIAYDE
GAL-----------------ASDLGGAFGLWLGASILTICEYIDFIMDV-----------
--------CVWA--------L----------R-------KL-------------------
KK------------------------------KM--------------------------
----------------------------------------TVNKVDKH------------
------------------------------------------------------------
------------------------------------------------------------
------------------------------------------------------------
---------------------------------P--------------------------
------------------------------------------------------------
-------S
>Protostome_Lophotroco_annelid_Pdum_comp408841_c0_seq5__833_314
------------------------------------------------------------
----------M-------------------------------------------------
------------------------------------------------------------
------------------------------------------------------------
------------------------------------------------------------
------------------------------------------------------------
---------------------------------------AHPLHRTMYLGA---------
------------------------------------------------------------
------------------------------------------------------------
------------------------------------------------------------
-----------------------SHTYGADMRNHYHNHSP----------------RPLK
P-VRPNRKT----------------------M---------DV-----H-----------
```

```
-RV--FREFA-DSTSMHGVPRII---------N-AR--------------SL---AA---
--RVFWS----ITCICAFGIFLWQCIILLQ-RFYS----------Y---PKKVNV----E
-V-VQ------------R-PVRF-----PSV-----------------------SFCNTD
HL------DLV-----VVKRLEEMLL--------ESDNITYGN---------------DT
H------FMKF---------------------------KEAYQAFWD----SSSFFF--Q
------------------------------------------------------------
------------------------------------------------------------
------------------------------------------------------------
------------------------------------------------------------
------------------------------------------------------------
------------------------------------------------------------
------------------------------------------------------------
--------------------------PYL-------------------------------
------------------------------------------------------------
------------------------------------------------------------
------------------------------------------------------------
------------------------------------------------------------
----------------------------------------QHVKPYEKAM-----THMLA
A-------------------------------YSRLGLI---------------------
------------------------ANL----------------GVDLASQG----GIEMR
--DFI------VNCR----FM-G----E----------P---C-----------DIE---
----------------------------------------------K-------------
------------------------------------------------------------
------------------------------------------------------------
-----------S----------F-------V-----------KFF-DPYFF--------N
CFTFD-------PS-------------TI-L-------------------------SS--
-------KTTRLQ------GA--------EY-------------GLTIILFTGSA--GQL
TKKSEF----E-----------------YVIP--------GMEEAD-GVLA---------
------------------------------------------------------------
------------------------------------------------------------
----------SGRGARVLVH-SPGTAP--RPASSGFDAPPEFSVSLGVRAR------E--
-----NVRI-----------DKP-W--GN-C--SFG--------NM----------DSKF
--------------------KYTLEDCQNTCLQRNIM----QK--CGCIDNK----I---
------------S-I--PS-----------------------------------------
-----YT--KGL-----P---FCLTLP---------TIPYTCYD-------MPLPEK---
---------------------------CAKIMD-EWTFR---MDCRK---------E---
V-Y-------------------ENLTMK-------DP-------DAM----------DNC
------------GCF-PPCNDII---YEAS-YSL------------------STI-PE--
-------QTE------ENS-A-----------FY--------------------------
------------------------------------------------SIIS--------
-------------------------------NFLSSLAEPKKK----------------L
LN--------------DL------------------YKL---------------------
---------------------------NKK------------------------------
-----DG------RI-----------R------GYIGRINVFIADSNVVKTT-EAPDYEA
IRL-----------------ISDIGGQLGLWIGISVMTLFEVMQLTCDI-----------
--------CRFL--------S----------A-------SG-------------------
RQ------------------------------KSRDR-------------RRAR------
-----------------------------------------PTRVDRV------------
------------------------------------------------ADRD--------
------------------------------------------------------------
------------------------------------------------------------
------------------------------------------------------------
------------------------------------------------------VEIHFE
VGDKLTAV
>Placozoa_TadNaC10_MK547551_20_315
------------------------------------------------------------
----------M-------------------------------------------------
```

```
------------------------------------------------------------
------------------------------------------------------------
------------------------------------------------------------
------------------------------------------------------------
------------------------------------------------------------
------------------------------------------------------------
------------------------------------------------------------
------------------------------------------------------------
-----------------------PTKQLKVMSQHN-------------------------
-GGISVRKK----------------------R---------LL-----------------
-AIT-VSRGF-EDCGAQGISNMA---------R-AQ--------------TT---QA---
--RILWA----ILTIAAISLCTVMAVDLIQ-KYYR----------F---EYDVTL----K
-I-SF-K----------P-QLDF-----PVI-----------------------TICNLN
PM------LRS-----EMIKRP--------I---------F-----------------K-
---------PLYGRDFESVNTSQTSPLVN-------------NSSLS-------------
---------------------------------NN-------TVGQSDS-----------
------------------------------------------------------------
---------NSS-TP---L---PTN-----------------------------------
EVPQDATAAAAATTVAANSQGGT------------------AAGSTKESGK---------
------------------------------------------------------------
------------------------------------------------------------
------------------------------------------------------------
-----------------------TATPTGTKTS---------------------------
------------------------------------------------------------
------------------------------------------------------------
------------------------------------------------------------
LPRVKKSLSNDV------------------------------NENIDRP-----------
----------------------TPPSGVQFENLDKS-------------------HENYH
--------L---------------------FSEISAELF---------------------
------------------------NRL----------------TDQRMYEL----GTQAS
--NFI------AKCT----FK-Q----K----------P---C-----------A-A---
----------------------------------------------N-------------
------------------------------------------------------------
------------------------------------------------------------
-----------N----------F-------S-----------ISF-NYLYG--------N
CFSFN-------TG-------------NS-R-------------------GK----RI--
-------ESVNYP------GP--------LF-------------GLQLYLDINKN-----
----------E-----------------Y------------I-S-R---EV---------
------------------------------------------------------------
------------------------------------------------------------
----------PTAGVRIAVT-PQGVRP--NPEDEGFDVPPGALTSIGLKMR------N--
-----ISRL-----------SRP-YNKEG-C--LKH--------PPN-----NKT-LNIY
----ENG----------SD-RYSYKGCIKSCIASLQN----KT--CGCIAAR----Y---
------------S-F--TS-----------------------------------------
-------STKGL-----K---VCSLK----------N-----------------------
------------------------------------ETE---INCQS---------R---
L-Q-------------------NRFI-A-------GK--------I------------NC
------------GCV-RACNEQS---FEAT-TSQ------------------AQF-PS--
-------EVN------LDN------------NND--------------------------
------------------------------------------------LL----------
----------------------------------EYF-----------------------
-K------------RDL-------------------TKF---------------------
---------------------QLTRK-TVT------------------------------
-----P-------FL-----------R------ENLVAVNIYYEELNFETIE-QTPRYSE
IDL-----------------ASDIGGVLGLWIGISVLTVFEFMEILVDS-----------
--------LLII--------F----------G-------KEKIA----------------
SR------------------------------RN-------------------T------
```

-----------------------------------------ITEFK--------------
------------------------------------------------AQLQ--------
------------------------------------------------------------
------------------------------------------------------------
------------------------------------------------------------
------------------------------------------------------------
---ASGLA
>Deu_Ambulacrar_hemi_Ptyfla_40v0_9_20150316_1g34414_t1_scaffold1171760_cov98_
837_316
------------------------------------------------------------
----------M-------------------------------------------------
------------------------------------------------------------
-----------------------------TLYPPVS------------------------
--------------EVP----------------------------KIHNFPQFTWENTLK
HLNVNQNVRNN-------------------------------------------------
-----------------------------------VNQAEIPQRLDKHGNC---------
-------------------------------GHAEPMERSQTVNDEYH------------
---------------------------------------------------VGN------
------------------------------------------------TRSVAAQADEDI
SRSPT--------------RQKIREKSNTTNAARLPNW----------------------
-TIDKQGHE--------LKYSE---------ELWDR----ASL-----R-----------
-TITRARTLC-DSGRLRGHISLLN--------R-VD--------------NS---NR---
--KLMWL----AVFSIAVAVFAFQAYELVA-LFLN----------Y---DVSVNI----E
-I-GT-A----------T-SLAF-----PAI-----------------------TICNTN
KL------RLS-----EIEKSE------------------------------------Q-
-------HQDLAKTDP---E-----------------HPDSVHRLLS----Y--------
------------EATCR----------------DG-------------------------
------------------------------------------------------------
----------VTCD--------------------------------FENDGRTF------
TCVPRKNQCDRFPDCE-D-------------------------GTDEKNCDL--------
------------------------------------------------------------
------------------------------------------------------------
------------------------------------------------------------
----------------------NDQCPTHTVYI--------------WN-----------
------------------------------------------------------------
------------------------------------------------------------
----------------------------------------------------------SP
EVLFSVNYPSNY------------------------------DDDYRCS-----------
----------------------------------------WILSAYSSYCIP---SEDCS
K--------------------------------DYLEIS---------------------
----------------------DVGNLSIKMRYCGSRIPPSWRSSSNTVKIETGKGFRME
ISDVTSWSECSVSCG----WG--V--RK-------RNIK---C-----------S-VV--
--------------------------------------------HEE-SL----------
------------------------------------------------------------
------------------------------------------------------------
----------SNSD----------------T-G---------SGY-QPHYG---------
------------SG-------------ESPD-SNDH-------------DDE----DY--
-------LSDAESM----PSSDMYNRTNVTYLNPND-DML----XLKLTLFIEQN-----
----------E-----------------Y------------I-P-L---YG---------
------------------------------------------------------------
------------------------------------------------------------
----------QEAGVRVLIN-PQDITP--FPEDEAITVAPGLKTSIGIRKD---------
------------------------------------------------------------
--------------------------CHKSMLQQYIR----HY--CDCVDTLH---L---
-----------------KG-----------------------------------------
-----------------R---YCNIA----------N-----------------------
------------------------------------QEE---------------------

```
-----------------------------------GK-----------------------
------------------------------------------------------------
------------------------------------------------------------
------------------------------------------------------------
------------------------------------------------------------
------------------------------------------------------------
------------------------------------------------------------
--------------------------R-----------MNMFFV--------V------I
NHL-----------------LFD-------------------------------------
------------NT------C---------------------------------------
--------------------------------SN--------------------------
------------------------------------------------------------
------------------------------------------------------------
------------------------------------------------------------
------------------------------------------------------------
------------------------------------------------------------
------------------------------------------------------------
--------
>Protostome_Lophotroco_annelid_Pdum_comp417306_c0_seq28__839_317
------------------------------------------------------------
----------M-------------------------------------------------
------------------------------------------------------------
------------------------------------------------------------
------------------------------------------------------------
------------------------------------------------------------
------------------------------------------------------------
------------------------------------------------------------
------------------------------------------------------------
------------------------------------------------------------
-----------------------MNPEKVPPPMTPVSA----------------------
-FSDPPQYN----------------------D---------SF-----K-----------
-SV--TLEYL-DTTTAHGLPSVI---------T-KR--------------RK---IQ---
--KVLWF----LIFWALMGYAVYQLFGIVN-EFNE----------Y---QVTVKT-----
-LKKH-R----------N-LAEF-----PAV-----------------------TICNEN
KL------KKS-----KLGGTN------------------Y-----------------A-
---------SIIEL---------QQKYS--------------EEVFG----P-SAPTTSN
TAG---------------------SSLPTNVA-NS-------TALQN-------------
------------------------------------------------------------
-------------------------------------------------VGTA-------
---------SLLNTTT-N-------------------------GTSSTNLTN--------
---------------TISNAVSNLTNNG-TNVPNLSNTISNALTN---------------
-------LTTNSTSLTNLTNTISNAM--------SNLTTNGTSLTNLTNTISNA------
-LSNLTTNATGLTNLTNTISNALSNLT--------------------------AN-GTSL
TNLTNTISNAVSNLT----------SPLT--------------------NSTGNTNQTGS
TGL---------------------NLPSI-------PL----------------------
---------GRKKRQILSGNTLSPRVPPGE------------------------------
------------------------------------------------------------
------------------------------------------SGTNN-------------
------------------LLSMELKGDHDYETLFAS-----------SK------TSDFS
---------------------------------DVVSML---------------------
-------------------------RP----------------SMTDLDKY----GHQFD
--DFV------LMCT----FD-G----E----------N---C-----------T-S---
----------------------------------------------D-------------
------------------------------------------------------------
------------------------------------------------------------
-----------D----------F-------V-----------RVY-NEVYG--------N
CFTFN-------RQ-------------TN-G--------------------S----TV--
```

-------RTTTKF------GS--------QY-------------GLKLSLNIEKD-----
----------E-----------------Y------------L-G-L---FS---------
------------------------------------------------------------
------------------------------------------------------------
----------HQYGARVTVH-SNNITA--FPQDNSVSIPVGYYASVAVDTE------V--
-----MDSK-----------ERP-YE-TN-C--THG--------KSA---------DLFY
----E-------------G-LYSTDNCLNSCLRKMVK----EV--CSCVETV----L---
------------V-T---------------------------------------------
---------GNE-----S---RCSIS----------N-----------------------
------------------------------------TTE---VACRL---------E---
V-Y-------------------ANFR-D-------NA--------Y------------SC
----------QHECN-EPCLEVT---YRTS-VSY------------------ANW-PT--
-------DGF------NAL-V-----------GR--------------------------
------------------------------------------------QL----------
------------------------------------------------------------
------------------------------------------------------------
------------------------------------------------------------
------------------------------------------------------------
------------------------------------------------------------
------------------------------------------------------------
------------------------------------------------------------
------------------------------------------------------------
------------------------------------------------------------
------------------------------------------------------------
------------------------------------------------------------
------------------------------------------------------------
------------------------------------------------------------
--------
>Deu_Ambulacrar_hemi_Ptyfla_40v0_9_20150316_1g14838_t1_scaffold10372_cov150_840_318
------------------------------------------------------------
----------M-------------------------------------------------
------------------------------------------------------------
------------------------------------------------------------
------------------------------------------------------------
------------------------------------------------------------
------------------------------------------------------------
--------------EMKDNN-------------YDDANSRKNNNVS----------P---
------------------------------------------------------------
---------------------------------------------------------DSI
SAAIA-------------RQSYIDSPETPGVDI---------------------------
-------------------------------D---------ST----PS-----------
-GR--IRIWAAQISDIHGVKHIV---------G-ER--------------SR---LR---
--KLLWA----LVVLASFGILLQQCIKAAI-NYGE----------F---HHVTKV----D
-V-EY-V----------Q-HMPF-----PAI-----------------------TICNFN
KY------RKS-----AITPAD------------------M-----------------V-
---------HV------------G-KPL---------------GLMD----E--------
---------------------------------DG-------NL----------------
------------------------------------------------------------
------------------------------------------------------------
------------------------------------------------------------
------------------------------------------------------------
------------------------------------------------------------
------------------------------------------------------------
-------------------------NSDLF------------------------------
------------------------------------------------------------
------------------------------------------------------------

------------------------------------------------------------
--------SEEF------------------------------VRQ---------------
------------------------MKDLDWDSEE---------------------KKRF-
---------------------------------NYTEFT---------------------
------------------------------------------------YRV----GSQVK
--ETI------VECT----WN-G----H----------K---C-----------T-E---
----------------------------------------------H-------------
------------------------------------------------------------
------------------------------------------------------------
-----------D----------F-------V-----------KVF-T-HYG--------I
CFAFN-------KY-------------HR-D--------------------T----EA--
-------RHAGKP------GA--------DN-------------GLRVVLNAQTS-----
----------E-----------------H------------L-P-T-ADLE---------
------------------------------------------------------------
------------------------------------------------------------
-------DSFINVGFKLMIH-PPTEPP--YPKELGFAVGPGSHIFLAITRQ------E--
-----IKRL-----------SKP-Y--GE-C--DMK-SV-----G-------SK----YF
------------------D-HYSMSACRIECETALLL----EM--CGCRLVE----Q---
------------P-G---------------------------------------------
----------NG-----P---VCTP-----------------------------------
------------------------------------KIV---KECAH---------V---
K-L-------------------LEYI-E-------GH--------IE----------FDC
------------PCH-IPCDSEV---YSVT-PSS------------------SRL-KP--
-------DRS------GKS---------P---AM--------------------------
------------------------------------------------------------
------------------------------------------------------------
SN----------------------------------------------------------
---------------------------YTQ------------------------------
-----E-------YI-----------D------SNVLVLTIFYEELNFETIT-QLPETSL
VGL-----------------LGQLGGNMGLFLGASILTLIQIIEYFVDE-----------
--------CIHC--------F----------R-------PM-------------------
AP------------------------------KK-------------------P------
-----------------------------------------KRTYR--------------
------------------------------------------------GEDKDVN-----
------------------------------------------------------------
--------------TPLSVQHWQGSHP----------------------VRNTTLMIVQH
D----------------------CEIVSVEHFIP-----------LYEILTGSAN-----
---------TGL-----FVFPRCLRTRAANMLSFELLRNAEN-QFITFLRHDAWYSCLTY
KMSSKLVR
>Deuterostome_chordata_Ggallus_tr_A0A3Q2UMV1_A0A3Q2UMV1_CHICK_Uncharacterized
_protein_OS_Gallus_gallus_OX_9031_GN_ASIC4_PE_3_SV_1_844_320
------------------------------------------------------------
----------M-------------------------------------------------
------------------------------------------------------------
--------------------------PGQPAWGSVTLPQP-PAGLGEEFSIPKSRLGKA-
-----------CVGTSPLL--------SLPSPVTA-------------------------
-QEGACHIRLW-------------------------------------------------
EGAEQKAPGRETAS----------------------------------------------
--------------GARSEA-------------GHAGSVLTTAM------------PL--
------------------------------------------------------------
------------------------------------------------------------
------------------------------------------------------------
-PLSCCPDG----------------------E---------AL-----A-----------
-PA--QGGFL-RATRIPGLHYMG---------T-RP-------------QSC---LR---
--RLLWG----LAFLASAGLLATGATDRLH-HLLS----------R---PVLTRA----R
-L-TR-V----------P-QLRF-----PAV-----------------------TLCNPN
RA------RFL-----QLTKPD------------------L-----------------Y-

---------SV------------G-QWL---------------GLSR----E--------
---------------------------------DR-------SLV---------------
------------------------------------------------------------
------------------------------------------------------------
------------------------------------------------------------
------------------------------------------------------------
------------------------------------------------------------
------------------------------------------------------------
--------------------------PELL------------------------------
------------------------------------------------------------
------------------------------------------------------------
------------------------------------------------------------
-----AMLGDEQ-----------------------------RRW----------------
------------------------LT--RLANYS----------RFLPPR-----RSER-
---------------------------------TMQSFF---------------------
------------------------------------------------HRL----SHQIE
--DML------VECR----FQ-G----K----------R---C-----------G-P---
----------------------------------------------Q-------------
------------------------------------------------------------
------------------------------------------------------------
-----------H----------F-------T-----------PVY-T-RYG--------K
CYTFN-------GD-------------RR-N--------------------P--------
-------RVTRQG------GM--------GN-------------GLEIMLDIQQE-----
----------E-----------------Y------------L-P-IWRETN---------
------------------------------------------------------------
------------------------------------------------------------
-------ETSFEAGIRVQIH-SQDEPP--YIHQLGFGVSPGFQTFVSCQEQ------R--
-----LTYL-----------PQP-W--GN-C--RAS-VQGE-Q-M-------LP----GY
------------------D-TYSIAACRLQCEKEAVV----RS--CHCRMVH----M---
------------P-G---------------------------------------------
----------NE-----S---ICSP-----------------------------------
------------------------------------NVY---IECAD---------H---
T-L-------------------DAAV-E-------DS--------Q-----------ERC
------------SCP-TPCNLTR---YGKE-ISM------------------VRI-PN--
-------KGS------ARY-L-----------AR--------------------------
------------------------------------------------KY----------
------------------------------------------------------------
-N----------------------------------------------------------
---------------------------KNE------------------------------
-----T-------YI-----------R------ENFLVLDIFFEALNYEAIE-QKKAYDL
AGL-----------------LGDIGGQMGLFIGASILTILEILDYIYEV-----------
--------IRDR--------V----------S-------RV----------------LR-
HS------------------------------KP-------------------P------
------------------------------------------------------------
-----------------------LKKP---------------------SGSI--------
-----------AT-----------------------------------------------
----------------LGLEELKDQ------------------------SPCETLGRHVE
-------------------------GTYNAGILP--NHHHRHH--------YPH------
---------QGV-----FEDFAC-------------------------------------
--------
>Protostome_Lophotroco_molsk_OctBim_tr_A0A0L8FW45_A0A0L8FW45_OCTBM_Uncharacte
rized_protein_OS_Octopus_bimaculoides_OX_37653_GN_OCBIM_22006431mg_PE_3_SV_1_
847_323
-----------------------------------------------------------M
KVTGFD-KLGF-------------------------------------------------
------------------------------------------------------------
------------------------------------------------------------

------------------------------------------------------------
--NSNQLRTM--------------------------------------------------
------------------------------------------------------------
------------------------------------------------------------
------------------------------------------------------------
----------------------------------------------------------RQ
Q-------------------LRRRNY----------------------------------
-----------------------------------------NA-----L-----------
-SI--ITELA-AESNAHGLAKIA---------T-ST--------------QT---PR---
--KVLWA----LMVIVGFTAATLQLSLLVR-KYLE----------F---QVVEVS----K
-M-KD-G----------M-DVEF-----PAI-----------------------TVCNIA
PI------SLT-----KTKELLSSDT---------SE-----------------------
-----------------------LTQWL---------------RFID----KYN------
------------------------------------------------------------
------------------------------------------------------------
------------------------------------------------------------
------------------------------------------------------------
------------------------------------------------------------
------------------------------------------------------------
------------------------------------------------------------
------------------------------------------------------------
------------------------------------------------------------
------------------------------------------------------------
------------------------------------------------------------
------------------------------------------------------------
---------------------------------------------FGAQ------TDRMF
---------------------------------TVQSLY---------------------
------------------------ENL-----------------ADEAQLL----GRDLN
--DFL------IHCQ----YN-Q----E----------I---C-----------N-I---
----------------------------------------------Q-------------
------------------------------------------------------------
------------------------------------------------------------
-----------N----------F-------T-----------RYF-DGHYF--------N
CYTFN-------SG-------------HQ-T-GT-----------------------S--
-------LLTHAT------GP--------QS-------------GLSLILSLDND-----
----------DPPV-----GGYGS----YN-------------I-K-SNIE---------
------------------------------------------------------------
------------------------------------------------------------
----------HSAGVRVVVH-PPSTMP--SPVDHGFDVPPGYSSSVGLKTI------M--
-----HARL-----------SNP-Y--GN-C--QNS--------RL------QNS--SKY
--------------------IHTVFSCLELCKQRIVM----ST--CGCRSSN----L---
------------P-E--MT-----------------------------------------
--------SANF-----T---FCGLVENK------WNWK-DMMKN------PE-------
---------------------------NYNITK-INLTE---LACEE---------R---
V-L-------------------KEFS-I-------NR-------AYE----------SSC
------------NCF-QPCQETT---YQKS-ISL------------------SYW-PL--
-------EFN------QLS-A-----------LE--------------------------
------------------------------------------------TFYGDKINETFL
HDYMLKYLSNLGSVPTPESPPPEEI---NITVDDNNN----------------------S
TN-------------SRM------------------HSPALLSQLLANSTEAPKPAATRD
DFI------EHVSAK-LNSSDKEKQI-RAS------------------------------
-----N-------MI-----------R------QNLLRLNVYLEDLSIIEFR-QMPAYEL
ADL-----------------FADIGGTLGLWMGISVLTIMELIELFTRL-----------
--------LMLI--------F----------S-------SE-------------------
KK-------------------------------------------------IPN------
-----------------------------------------ADPV---------------
------------------TNGMLDHDC---------------------DCQK--------

--------SSM-------------------------------------------------
------------------------------------------------------------
------------------------------------------------------------
------------------------------------------------------------
----ESPF
>Cnidar_HydraVulSc4wPfr_147_g8550_t1_848_324
------------------------------------------------------------
----------M-------------------------------------------------
-------------------------------------LKDQASYYQIINE----EVDSN-
-----------EASFPSISN-------NNKTFDRRFSSENC-----------KIEKSKHS
SNLIGSAKWAA-LTKKNAE-QLLLKSENTK-PVIDSAY----------------DLNLTE
LDTITDISAYK-------KDNSPVKGNIEP-PAVKPGVLMKRT---------NTIPKQA-
---DKWGTIKERTSLIVKP--------TDEFTKKVMPSNDKPL-----------------
-TTTPKLLKVVTAVTAAKTA-------------TTKFREAS-------------------
------------------------------------------------------------
------------------------KMRIRLK--------------------------ERM
R----------------------KIVQE--------------------------------
--------R----------------------L---------TV-----K-----------
-QI--FKRYI-ESSTLHGFCYVC---------M-DT--------------FL---GR---
--RLIWA----VLMILGAIYFIFKLRYGIK-EYFD----------Y---PFSTLS----T
-V-EY-V----------D-DLLF-----PAV-----------------------SVCATN
SY------IAS-----QVYTNQ------------------L-----------------N-
---------TM----------------------------------YK-------------
------------------------------------------------------------
------------------------------------------------------------
------------------------------------------------------------
------------------------------------------------------------
------------------------------------------------------------
------------------------------------------------------------
------------------------------------------------------------
------------------------------------------------------------
------------------------------------------------------------
------------------------------------------------------------
------------------------------------------------------------
------------------------------------------EGRL--------------
------------------------PLDNNQSIPEYN-------------------IPGD-
---------------------------------ELVKTL---------------------
------------------------------------------------KNS----SLTIE
--SLL------KYCD----WI-M----QDTSHPLVTPNN---C-----------G-A---
----------------------------------------------L-------------
------------------------------------------------------------
------------------------------------------------------------
-----------N----------F-------T-----------SYF-N-YKGE-------Q
CHTLN-------SG-------------AK-G--------------------H----EL--
-------LKVSDV------GI--------SH-------------GYELVFDLQTN-----
----------E-----------------V------------I-K-N--------------
------------------------------------------------------------
------------------------------------------------------------
---------YQLSGMRIVIH-DQVFPP---QLVDGFFISPGFKTYIKLGIT------Q--
-----SQSL-----------PPP-YS-TE-C--GQK--------K-------LK----YY
------------------A-IYSQRLCLLETLTDFTG----DL--CGCRDVF----M---
------------P-E--N------------------------------------------
----------GL-----P---FCSL-----------------------------------
------------------------------------QEL---YS----------------
----------------------XSFS-E-------FT--------------------MRK
------------ECP-SDCEERT---FSYE-LSE------------------ARY-LH--
-------NPP------IGL-SLSRLD----------------------------------

```
------------------------------------------------------------
------------------------------------------------------------
------------------------------------------------------------
------------------------------------------------------------
--------------------------K------SNIISVILFFGDTRIDYHE-QEATNDF
FQF-----------------LGNMGA---LMAGVGLKLSFYTQDLSSYS-----------
--------AFYK--------S---------------------------------------
------------------------------------------------------------
------------------------------------------------------------
------------------------------------------------------------
------------------------------------------------------------
------------------------------------------------------------
------------------------------------------------------------
------------------------------------------------------------
--------
>Protostome_Lophotroco_annelid_Pdum_comp417306_c0_seq24__852_325
------------------------------------------------------------
----------M-------------------------------------------------
------------------------------------------------------------
------------------------------------------------------------
------------------------------------------------------------
------------------------------------------------------------
------------------------------------------------------------
------------------------------------------------------------
------------------------------------------------------------
------------------------------------------------------------
-----------------------MNPEKVPPPMTPVSA----------------------
-FSDPPQYN----------------------D---------SF-----K-----------
-SV--TLEYL-DTTTAHGLPSVI---------T-KR--------------RK---IQ---
--KVLWF----LIFWALMGYAVYQLFGIVN-EFNE----------Y---QVTVKT-----
-LKKH-R----------N-LAEF-----PAV-----------------------TICNEN
KL------KKS-----KLGGTN------------------Y-----------------A-
---------SIIEL---------QQKYS--------------EEVFG----P-SAPTTSN
TAG---------------------SSLPTNVF-NS-------TALQNAS-----------
------------------------------------------------------------
--------------------------------------P--------PVVGTG-------
---------SLLNTTT-N-------------------------GTSSTNLTN--------
---------------TISNAVSNLTNN---------------------------------
------------------------------------------------------------
------------------------------------------------------------
----------VSNLT----------SPLT--------------------NSTGNTNQTGS
TGL---------------------NLPSI-------PL----------------------
---------GRKKRQILSGNTLSPRVPPGE------------------------------
------------------------------------------------------------
------------------------------------------SGTNN-------------
------------------LLSMELKGDHDYETLFAS-----------SK------TSDFS
---------------------------------DVVSML---------------------
-------------------------RP----------------SMTDLDKY----GHQFD
--DFV------LMCT----FD-G----E----------N---C-----------T-S---
----------------------------------------------D-------------
------------------------------------------------------------
------------------------------------------------------------
-----------D----------F-------V-----------RVY-NEVYG--------N
CFTFN-------RQ-------------TN-G--------------------S----TV--
-------RTTTKF------GS--------QY-------------GLKLSLNIEKD-----
----------E-----------------Y------------L-G-L---FS---------
------------------------------------------------------------
```

```
------------------------------------------------------------
----------HQYGARVTVH-SNNITA--FPQDNSVSIPVGYYASVAVDTE------V--
-----MDSK-----------ERP-YE-TN-C--THG--------KSA---------DLFY
----E-------------G-LYSTDNCLNSCLRKMVK----EV--CSCVETV----L---
------------V-T---------------------------------------------
---------GNE-----S---RCSIS----------N-----------------------
------------------------------------TTE---VACRL---------E---
V-Y-------------------ANFR-D-------NA--------Y------------SC
----------QHECN-EPCLEVT---YRTS-VSY------------------ANW-PT--
-------DGF------NAL-V-----------GR--------------------------
------------------------------------------------QLVG--------
----------------------------------KMA-----------------------
RN----------------------------------------------------------
---------------------------EIK------------------------------
-----E-------YI-----------E------NNILRVNIFFRTLTYSHQQ-VFPTYTW
ETL-----------------LSNIGGTWGLFVGFSVCTFLEAAEYFLEL-----------
--------ICVC--------C----------G-------LS-------------------
SK------------------------------KK-------------------K------
---------------------------------------EKVAPEGFNY-----------
------------------------------------------------------------
------------------------------------------------------------
------------------------------------------------------------
------------------------------------------------------------
------------------------------------------------------------
---AKGLS
>Deutero_Ambulac_Spurpu_025728_853_326
------------------------------------------------------------
----------M-------------------------------------------------
------------------------------------------------------------
------------------------------------------------------------
------------------------------------------------------------
--EVPLDEEEK-------------------------------------------------
--------------------------------------PLSPEK----------------
--------------------------------------------------------P---
------------------------------------------------------------
------------------------------------------------------------
-----------------------SSPEKPKAVLEKP-----------------------K
ALVREMTNDILV-------------------K---------SW-----R-----------
-TV--VQSFC-KSTTMHGMSRVI---------D-ST--------------RA---IF---
--RLAWL----LTVLTFLCVLMWQAYRLVD-EYRG----------N---PTTTTI----Q
-M-VT-N----------N-KLSF-----PAV-----------------------TVCNMN
RL------RRS-----KLVGTR------------------F-----------------E-
---------PLLDIDRH-------------------------VTILNLGL----------
---------------------------------DD-------------------------
-------------------------------------------------------G----
---------EIEVD----------------------------------------------
-----------------DHAEPSPSL---------------TPATTTSTITT--------
------------------------------------------------------------
---------------TTASQVTTKS----------TSVPPNDEDLVSSTTVEGL------
------------------------------------------------------------
------------------EMSSTKDTPLV-------------------------------
------------------------------------------------------------
--------VDVDNPDARRRKRS--------------------------------------
------------------------------------------------------------
------------------------------------VNWQTTNKRLSSLRQ---------
-------MNNVPAPAIPLIGAPVENDDSNWGYLLQL-----------SE------SDDFQ
---------------------------------DFIGSV---------------------
```

```
-------------------------NP----------------SKGELRKL----GHQAE
--EFI------LQCS----FN-Q----H----------Y---C-----------D-Y---
----------------------------------------------R-------------
------------------------------------------------------------
------------------------------------------------------------
-----------N----------F-------T-----------TTH-NSQYG--------N
CFTFN-------RP-------------AE-N--------------------N----SV--
-------LQTGKI------GS--------RF-------------GLHLTLFTDQS-----
----------E-----------------Y------------I-G-L---LS---------
------------------------------------------------------------
------------------------------------------------------------
----------HQSGVRVAIH-EPNARP--FPEDEGITASTGTLTSIGLRLR------N--
-----ITRL-----------SGR-Y--SD-C--RKK--------NRGD--------PSEF
----THD-----------F-DYSLRACLNACYQGKLR----YE--CGCVNDV----M---
------------------------------------------------------------
---------GNT-----T---ICSTL----------D-----------------------
------------------------------------RRQ---EACQR---------R---
V-D-------------------QMAI-D-------DK--------L------------GC
------------PCV-SPC-----------------------------------------
------------------------------------------------------------
------------------------------------------------------------
------------------------------------------------------------
------------------------------------------------------------
------------------------------------------------------------
--------------------------R------KNLVRVEIFYEKLNYEAYT-QMPKYTF
GSL-----------------LGGIGGIMGFFAGMSLITVFELCGFIIQL-----------
--------LGLL--------C----------G-------RL-------------------
TTPMPEEAATEGG-EPAPPHPESRIGRVG---RR-----------------FTKVFS---
------------------------------------------------------------
------------------------------------------------------------
------------------------------------------------------------
------------------------------------------------------------
-------------------------------HHP--------------------------
----------------------------------------------------HAHGHGGS
ARARQTDL
>Deutero_Ambulac_Spurpu_020954_856_327
------------------------------------------------------------
----------M-------------------------------------------------
------------------------------------------------------------
------------------------------------------------------------
------------------------------------------------------------
------------------------------------------------------------
------------------------------------------------------------
------------------------------------------------------------
------------------------------------------------------------
------------------------------------------------------------
-----------------------KAPKVDGEKS--------------------------S
RGTTPEREE----------------------K---------SL-----R-----------
-TI--LNSRM-ENSSAHGVPNIQ---------R-SS--------------GL---VT---
--KLAWS----LIFLAGIGVMIWQVVTIFQ-AYYE----------W---NYSVII----E
-V-KF-N----------R-TQSF-----PAI-----------------------TLCNAN
PM------RKS-----KLKTKN--------A-----------------------------
---------SF-------------------------------QETFD----VN-YVPSMP
DLPEQQPQDVALAPTCPVGLGHYRPA-------DQ-------------------------
------------------------------------------------------------
---------------------QPLP-----------------------------------
-----------------DMPDGV-------------------------------------
```

------------------------------------------------------------
------------------------------------------------------------
------------------------------------------------------------
-------------------------TPSM-------------------------------
------------------------------------------------------------
------------------------------------------------------------
------------------------------------------------------------
--LNNSDSGNETQG-------------------------GSTNEQVDMS-----------
------------------------EAVKDWKSRIA------SPSFYVME------SVDYE
------------------------------KRRIRVDAL---------------------
------------------------ANE----------------TLEERVSL----GHKLD
--DML------LDCS----WK-G----K----------P---C-----------S-P---
----------------------------------------------E-------------
------------------------------------------------------------
------------------------------------------------------------
-----------N----------F-------T-----------KFY-DSQLG--------N
CYTFN-------SG-------------QN----------------------G----EQ--
-------LTTTRP------GS--------KYVNGPI-DCLVLKPGLSLELFVQQD-----
----------E-----------------Y------------V-E-G---MT---------
------------------------------------------------------------
------------------------------------------------------------
----------EEASFRVSVH-HPSIMP--FPADDGVLVSPGFATAIAFIKL------E--
-----LDRL-----------PKP-Y--GD-C--KAD--------LST-----DIE-DDIY
----HQH--YN-------I-TYNMKTCEVSCFQNEVI----SR--CDCFDAT----Y---
------------P-N--SL-----------------------------------------
------KVNRTV-----Y---PCEYN----------N-----------------------
------------------------------------RVA---------------------
-----------------------------------------------------------C
----------------PYCQQVL---HLII-IKK-----------------LKSW-ESFF
CLRFEITDQFF----AHST-L-----------FE--------------------------
------------------------------------------------KM-V--------
----------------------------------EY------------------------
-N-------------AEI------------------RRY---------------------
----------------------VNRE-NTA------------------------------
-----E-------WT-----------R------RNMAKVEIFYDEFNYEHIR-QEPAYMV
CKK-----------------TRQIAS--------------------YDVHK----PPPPP
PHSENGRTPCVLP------KCP--------FR----------------------------
NK------------------------------RK-------------------R------
-----------------------------------------STRIDFTHR----------
---------------------TFSEKGGAEK--------------GSYTKEKGRAV----
-----------LT-----------------------------------------------
------------------------------------------------------------
------------------------------------------------------------
----------------------------------------------LHTGLLLKKGGGRL
EHAYAYPK
>Deutero_Ambulac_Sakowv30039242m_855_330
------------------------------------------------------------
----------M-------------------------------------------------
------------------------------------------------------------
------------------------------------------------------------
------------------------------------------------------------
------------------------------------------------------------
------------------------------------------------------------
------------------------------------------------------------
------------------------------------------------------------
------------------------------------------------------------
-----------------------SHIGVDNLVS--------------------------P

```
GSSNCKEKQ----------------------Q---------TF-----S-----------
-SI--LGSCF-VQSSAHGLPNIS---------R-AG--------------NA---PR---
--RSLWI----ISLGGTVAIFCLISATLIV-RYFD----------Y---DVNVSV----A
-M-QF-S----------R-ELTF-----PAV-----------------------TICNLN
PL------RKS-----KTQMDGPSDPGQQRAEEQTSQGTNIKT---------------ET
TQHPNIVSEDI------------QRNPFTQ--------VPTTTKIYE----NRQYSRFTH
QTYHV---EQ-----------------------DQQSIYTH-------------------
------------------------------------------------------------
---------------------------------------------ENNTDGQSL------
-----------------DVTNSDWNGR----------TLDDTVTNSEENGRPP-------
------------------------------------------------------------
------------------------------------------------------------
------------------------------------------------------------
------------------PETSTGGAPELVNVLLNETNDTEAEVYD---DATGDTEAEVD
DDA---------------------------------------------------------
------------------------------------------------------------
--------------------------------------------------------TGDT
EAEVDDDETSDTEA------KVDDDE-------TGDTEAEVDDDSEASE-----------
------------------------WDYFDWDDLDT------KTEFYNEP------TQYWN
------------------------------KSASLNQWL---------------------
------------------------ASM----------------PTQQKEEW----GHQLK
--DFL------LDCQ----WN-D----V----------M---C-----------S-P---
----------------------------------------------K-------------
------------------------------------------------------------
------------------------------------------------------------
-----------N----------F-------T----------KFV-SSRYG---------N
CYTFN-------SG-------------AN-N--------------------S----TV--
-------VKTNYA------GP--------YY-------------GLTLELFIEQN-----
----------E-----------------Y------------L-D-D---YP---------
------------------------------------------------------------
------------------------------------------------------------
----------DFAGVRVAIH-SQKTMP--FPEDDGFNVEPGRVTSVGIRRT---------
------------------------------------------------------------
--------------------------CMKSCYQKRVA----ER--CRCLDGR----Y---
------------P-P--PI-------------------VVIH------------------
------------------------------------------------------------
------------------------------------RTL---NACFI---------G---
A--------------------------N-----------------MA-------------
--------------------------YQYDVMSYLF------------------------
-------QDY----------V---------------------------------------
--------------------------------NV--------------QL-K--------
----------------------------------KK------------------------
-S-------------QQL------------------KLMIE-------------------
----------------------EEEK-GGN------------------------------
-----E-------FV-----------R------ENIVKLQVFYQDLNYEGIT-QSIAYTE
ESL-----------------ASDLGGQVGLWIGISFLTLIEFIELVYDL-----------
--------FKLW--------I----------R-------RL-------------------
CCMAT---------------------------KN--------------------------
------------------------------------------------------------
------------------------------------------------------------
------------------------------------------------------------
------------------------------------------------------------
------------------------------------------------------------
------------------------------------------------------------
-TGVTNVI
>Protostome_Lophotroco_annelid_Pdum_Contig8214__858_332
--------------------------G--TA-----------------------------
```

----------L---------------------------------QIC-------------
------------------------------------------------------------
------------------------------------------------------------
------------------------------------------------------------
------------------------------------------------------------
------------------------------------------------------------
------------------------------------------------------------
------------------------------------------------------------
------------------------------------------------------------
-----------------------QHTKMA-------------------------------
-------------------------------K---------KD-----K-----------
-GL--WKEFS-QNTTIHGLNKVA---------E-EN-------------GLT---FR---
--RLFWI----TLILGGTALFAYQVVERSL-YYIK----------W---PVSVNY----K
-F-VY-Q----------S-EMPF-----PTV-----------------------TICNQN
AF------KLS-----TSVQSG------------------L-----------------H-
---------DV------------I------------------TSIFS----KR-------
---------------------------------NS-------------------------
------------------------------------------------------------
------------------------------------------------------------
------------------------------------------------------------
------------------------------------------------------------
------------------------------------------------------------
------------------------------------------------------------
-------------------------IPKYS------------------------------
------------------------------------------------------------
------------------------------------------------------------
------------------------------------------------------------
----RNNISSEL------------------------------EDIFGN------------
-------------------------------------------------------SSSL-
---------------------------------KTDDVW---------------------
------------------------------------------------QLL----AHAKE
--DMI------YSCS----WQ-E----D----------P---C-----------S-P---
----------------------------------------------D-------------
------------------------------------------------------------
------------------------------------------------------------
-----------N----------F-------T-----------TLL-T-DHG--------V
CFQFN-------SA-------------SN-S--------------------H----PP--
-------LKVTQS------GS--------QQ-------------GLSLVMNVEQY-----
----------E-----------------Y------------M-T-G---PH---------
------------------------------------------------------------
------------------------------------------------------------
----------DSVGLKVLLH-DSSHVA--MVRDLGEALPTGSHAFVAVTLS------K--
-----IQNL-----------KHP-Y--GE-C--VNN-V------S-------LD----YY
------------------D-TFSLSTCRRDCETRKTV----KD--CGCRDAY----M---
------------P-SY--------------------------------------------
-------TDGTP-----Q---VCTL-----------------------------------
------------------------------------QEY---SQCIQ---------A---
N-K-------------------DQQS-GI------AG--------------------DVC
----------SSNCP-IPCLTTS---YKSS-TTF--STAFENYADDIDKE--SYWNRTR-
-------QHYIL----ARE-I-----------SQKVNQ----------------------
------EILLETEWQVITQIINEVKETVKHSKDI--------------KFVVDYD-YV--
------R----------------------IWDVMEERTMAYATIG-------FYQTAHLY
RNMVDKFLLEPELTVEKVA-I--------------FTSTMRDLDVR---L----------
----------------------QLTM-STQINLTQV----------------------C-
-----DAFNLRWKYLTSP-------KR------DNFVKVNIFMKDLYLEDME-SNEAYSL
IAL-----------------LSDIGGSMGLFIGGSVMTILEVLDILVIA-----------
--------CMRK--------S----------K-------KK-------------------

NE------------------------------TN--------------------------
------------------------------------------------------------
------------------------------------------------------------
------------------------------------------------------------
------------------------------------------------------------
------------------------------------------------------------
------------------------------------------------------------
------ET
>Deuterostome_chordata_Ggallus_sp_Q92075_SCNNA_CHICK_Amiloride_sensitive_sodium_channel_subunit_alpha_OS_Gallus_gallus_OX_9031_GN_SCNN1A_PE_2_SV_1_860_333
------------------------------------------------------------
----------M-------------------------------------------------
------------------------------------------------------------
------------------------------------------------------------
------------------------------------------------------------
------------------------------------------------------------
------------------------------------------------------------
---------------GTASR----------------------------------------
------------------------------------------------------------
-----------------------------------------GG----------SVKAEKM
-----------------------PEGEKTRQCK-----------------------QETE
QQQKEDEREGLI----EFY------------G---------SY-----Q-----------
-DV--FQFFC-SNTTIHGAIRLV---------C-SK-------------KNK---MK---
--TAFWS----VLFILTFGLMYWQFGILYR-EYFS----------Y---PVNLNL----N
-L-NS------------D-RLTF-----PAV-----------------------TLCTLN
PY------RYS-----AIRKKL----------------DEL-----------------D-
---------QI------------THQTL--------------LDLYD----Y--------
---------------------------------NM-------------------------
------------------------------------------------------------
------------------------------------------------------------
----------SLARSDGSAQFSHRR------------------TSRSLLHHVQ-------
------------------------------------------------------------
------------------------------------------------------------
------------------------------------------------------------
------------------------RHPLRRQK--------------------RDNLVSLP
ENSP--------------------------------------------------------
------------SVDKNDWKIG--------------------------------------
------------------------------------------FVLC--------------
--------SENN------------------------------EDC---------------
--------------------------------------------FHQTY------SSGVD
AV-------------------------REWYSFHYINIL---------------------
------------------------AQMP-----------------DAKDLD----ESDFE
--NFI------YACR----FN-E----A----------T---C-----------D-K---
----------------------------------------------A-------------
------------------------------------------------------------
------------------------------------------------------------
-----------N----------Y-------T-----------HFH-HPLYG--------N
CYTFN-------DN-------------SS---------------------------SL--
-------WTSSLP------GI--------NN-------------GLSLVVRTEQN-----
----------D-----------------F------------I-P-L---LS---------
------------------------------------------------------------
------------------------------------------------------------
----------TVTGARVMVH-DQNEPA--FMDDGGFNVRPGIETSISMRKE------M--
-----TERL-----------GGS-Y--SD-C--TED-G------SD------VPV-QNLY
----S-------------S-RYTEQVCIRSCFQLNMV----KR--CSCAYYF----Y---
------------P-----------------------------------------------
-------LPDGA-----E---YCDY-----------------------------------

------------------------------------TKHVAWGYCYY---------K---
L-L-------------------AEFK-A-------DV--------L------------GC
----------FHKCR-KPCKMTE---YQLS-AGY------------------SRW-PS--
-------AVS------EDW-V-----------FY--------------------------
------------------------------------------------ML-S--------
----------------------------------QQ------------------------
-NK---------------------------------------------------------
---------------------------YNI------------------------------
-----T-------SK-----------R------NGVAKVNIFFEEWNYKTNG-ESPAFTV
VTL-----------------LSQLGNQWSLWFGSSVLSVMELAELILDF-----------
--------TVIT--------F----------I-------LAF------------------
RWF-----------------------------RS------------K-QWHSSPAPP---
------------------------------------------------------------
----------------------PNSHDNTAF-----------------QDEA--------
------------------------------------------------------------
--------------SGLDAP----------------------------------------
-----------------------HRFTVEAVVTTLPSYNSLEPCGP--------------
-----------------------------------------------------------S
KDGETGLE
>sp_P37090_SCNNB_RAT_Amiloride_sensitive_sodium_channel_subunit_beta_OS_Rattu
s_norvegicus_OX_10116_GN_Scnn1b_PE_1_SV_2_861_335
------------------------------------------------------------
----------M-------------------------------------------------
------------------------------------------------------------
------------------------------------------------------------
------------------------------------------------------------
------------------------------------------------------------
------------------------------------------------------------
------------------------------------------------------------
------------------------------------------------------------
------------------------------------------------------------
-----------------------PVKKYLLKCL---------------------------
-HRLQKGPG----------------------Y---------TY-----K-----------
-EL--LVWYC-NNTNTHGPKRII-----------CE--------------GP---KK---
--KAMWF----LLTLLFACLVCWQWGVFIQ-TYLS----------W---EVSVSL----S
-M-GF------------K-TMNF-----PAV-----------------------TVCNSS
PF------QYS-----KVKHLL----------------KDL-----------------Y-
---------KL------------MEAVL--------------DKILA----PKS------
------------------------------------------------------------
------------------------------------------------------------
------------------------------------------------------------
-----------------SHTNTTS------------------------------------
------------------------------------------------------------
------------------------------------------------------------
------------------------------------------------------------
----------------TLNFTIWNHTPLVLIDERNPDHPVVLNL---F-GDSHN--SSNP
APGS--------------------------------------------------------
------------TCNAQGCKVA--------------------------------------
------------------------------------------MRLC--------------
--------SANG------------------------------TVC---------------
--------------------------------------------TFRNF------TSATQ
AV-------------------------TEWYILQATNIF---------------------
------------------------SQV----------------LPQDLVGM----GYAPD
--RII------LACL----FG-T----E----------P---C-----------S-H---
----------------------------------------------R-------------
------------------------------------------------------------
------------------------------------------------------------

-----------N----------F-------T-----------PIF-YPDYG--------N
CYIFN-------WG-------------MT-E-------------------------KA--
-------LPSANP------GT--------EF-------------GLKLILDIGQE-----
----------D-----------------Y------------V-P-F---LA---------
------------------------------------------------------------
------------------------------------------------------------
----------STAGARLMLH-EQRTYP--FIREEGIYAMAGTETSIGVLLD------K--
-----LQGK-----------GEP-Y--SP-C--TMN-G------SD------VAI-QNLY
----SDY----------NT-TYSIQACLHSCFQDHMI----HN--CSCGHYL----Y---
------------P-----------------------------------------------
-------LPAGE-----K---YCNN-----------------------------------
------------------------------------RDFPDWAYCYL---------S---
L-Q-------------------MSVV-Q-------RE---------------------TC
----------LSMCK-ESCNDTQ---YKMT-ISM------------------ADW-PS--
-------EAS------EDW-I-----------LH--------------------------
------------------------------------------------VL-S--------
----------------------------------QERD----------------------
QS----------------------------------------------------------
---------------------------SNI------------------------------
-----T-------LS-----------R------KGIVKLNIYFQEFNYRTIE-ESPANNI
VWL-----------------LSNLGGQFGFWMGGSVLCLIEFGEIIIDF-----------
--------IWIT--------V----------I-------KLVASC---------------
KGL-----------------------------RR----------------RRPQAPY---
--------------------------------------TGPPPTVAEL------------
----------------------VEAHTNFGF-----------------QPDTTS------
------------------------------------------------------------
-----------CRPNAEVYP---DQQTL--------------------------------
----------------------------PIPGTPPPNYDSLRLQPLDT------------
----------------------------------------------------------ME
SDSEVEAI
>Chordat_hENaCbeta_NP_000327_2_amiloridesensitive_sodium_channel_subunit_beta
__Homo_sapiens__1_336
------------------------------------------------------------
----------M-------------------------------------------------
------------------------------------------------------------
------------------------------------------------------------
------------------------------------------------------------
------------------------------------------------------------
------------------------------------------------------------
------------------------------------------------------------
------------------------------------------------------------
------------------------------------------------------------
-----------------------HVKKYLLKGL---------------------------
-HRLQKGPG----------------------Y---------TY-----K-----------
-EL--LVWYC-DNTNTHGPKRII-----------CE--------------GP---KK---
--KAMWF----LLTLLFAALVCWQWGIFIR-TYLS----------W---EVSVSL----S
-V-GF------------K-TMDF-----PAV-----------------------TICNAS
PF------KYS-----KIKHLL----------------KDL-----------------D-
---------EL------------MEAVL--------------ERILA----PEL------
------------------------------------------------------------
------------------------------------------------------------
------------------------------------------------------------
-----------------SHANATR------------------------------------
------------------------------------------------------------
------------------------------------------------------------
------------------------------------------------------------
----------------NLNFSIWNHTPLVLIDERNPHHPMVLDL---F-GDNHNGLTSSS

ASEK--------------------------------------------------------
------------ICNAHGCKMA--------------------------------------
------------------------------------------MRLC--------------
--------SLNR------------------------------TQC---------------
--------------------------------------------TFRNF------TSATQ
AL-------------------------TEWYILQATNIF---------------------
------------------------AQV----------------PQQELVEM----SYPGE
--QMI------LACL----FG-A----E----------P---C-----------N-Y---
----------------------------------------------R-------------
------------------------------------------------------------
------------------------------------------------------------
-----------N----------F-------T-----------SIF-YPHYG--------N
CYIFN-------WG-------------MT-E-------------------------KA--
-------LPSANP------GT--------EF-------------GLKLILDIGQE-----
----------D-----------------Y------------V-P-F---LA---------
------------------------------------------------------------
------------------------------------------------------------
----------STAGVRLMLH-EQRSYP--FIRDEGIYAMSGTETSIGVLVD------K--
-----LQRM-----------GEP-Y--SP-C--TVN-G------SE------VPV-QNFY
----SDY----------NT-TYSIQACLRSCFQDHMI----RN--CNCGHYL----Y---
------------P-----------------------------------------------
-------LPRGE-----K---YCNN-----------------------------------
------------------------------------RDFPDWAHCYS---------D---
L-Q-------------------MSVA-Q-------RE---------------------TC
----------IGMCK-ESCNDTQ---YKMT-ISM------------------ADW-PS--
-------EAS------EDW-I-----------FH--------------------------
------------------------------------------------VL-S--------
----------------------------------QERD----------------------
QS----------------------------------------------------------
---------------------------TNI------------------------------
-----T-------LS-----------R------KGIVKLNIYFQEFNYRTIE-ESAANNI
VWL-----------------LSNLGGQFGFWMGGSVLCLIEFGEIIIDF-----------
--------VWIT--------I----------I-------KLVALA---------------
KSL-----------------------------RQ----------------RRAQASY---
--------------------------------------AGPPPTVAEL------------
----------------------VEAHTNFGF-----------------QPDTAP------
------------------------------------------------------------
-----------RSPNTGPYP---SEQAL--------------------------------
----------------------------PIPGTPPPNYDSLRLQPLDV------------
----------------------------------------------------------IE
SDSEGDAI
>Deuterostome_chordata_Ggallus_tr_A0A3Q2U1X6_A0A3Q2U1X6_CHICK_Uncharacterized
_protein_OS_Gallus_gallus_OX_9031_GN_SCNN1D_PE_4_SV_1_862_337
------------------------------------------------------------
----------M-------------------------------------------------
------------------------------------------------------------
------------------------------------------------------------
------------------------------------------------------------
------------------------------------------------------------
------------------------------------------------------------
------------------------------------------------------------
------------------------------------------------------------
------------------------------------------------------------
--------------------------------------------------------EQEA
AREEEERKEGLI----EFY------------D---------SF-----K-----------
-DM--FEFFC-KNTTIHGTIRLV---------C-SS-------------SNK---MK---
--TAFWT----LLLLASFGMLYWQFALMFS-QYWD----------Y---PVVLTM----S

-M-HS------------E-PKMF-----PAI-----------------------TICNLD
PY------RFD-----LVSEHL----------------AQL-----------------D-
---------RM------------AEKSV--------------TVLYG----I--------
---------------------------------NT-------------------------
------------------------------------------------------------
------------------------------------------------------------
----------SASLFHVNEKSIHVR-----------------------------------
------------------------------------------------------------
------------------------------------------------------------
------------------------------------------------------------
------------------------DLPSTGNHNGSS-----------FKLSQKFSLLRTT
E-----------------------------------------------------------
---------FNNRTGKRQSLVG--------------------------------------
------------------------------------------FRQC--------------
--------NATG------------------------------GNC---------------
--------------------------------------------FYKTY------SSGMD
AI-------------------------LEWYRFHYMNIM---------------------
------------------------SQQP---------------VIINISDH----EEKIE
--DMV------YSCQ----YD-G----E----------P---C-----------R-P---
----------------------------------------------S-------------
------------------------------------------------------------
------------------------------------------------------------
-----------D----------Y-------V-----------HFH-HPVFG--------S
CYTFN-------SK-------------GT-D-------------------------PF--
-------WTATKP------GI--------PY-------------GLSLILRAEQK-----
----------D-----------------H------------I-P-L---LS---------
------------------------------------------------------------
------------------------------------------------------------
----------TVAGVKVMIH-NHNQTP--FLEHEGFDIRPGIATTIGIQQD------K--
-----VNRL-----------GGN-Y--GK-C--TTD-G------SD------VKV-KLLY
----N---------------SYTLQACLHSCFQHIMV----QK--CGCGYYY----Y---
------------P-----------------------------------------------
-------LPPGA-----E---YCNY-----------------------------------
------------------------------------NKQPAWGHCFY---------Q---
L-Y-------------------SRLR-N-------HH--------L------------NC
----------FDQCP-KPCRESL---YKVS-AGT------------------AKW-PS--
-------RKS------QDW-I-----------RQ--------------------------
------------------------------------------------AL-R--------
----------------------------------HQ------------------------
-NG---------------------------------------------------------
---------------------------YNS------------------------------
-----T-------SN-----------R------KDIAKVTIYYKQLNYQSVN-ESPLLSD
NLL-----------------LSSMGSQWSLWFGSSVLSVVEMLELLIDT-----------
--------LVLS--------L----------L-------FCY------------------
QRF-----------------------------RS------------K-TLNVART-----
---------------------------------------PSIPSVSLTLES---------
------------Y-------RVVQEAGNGTA--------------PAHGHTSGVPMAV--
------------------------------------------------------------
------------------------------------------------------------
----------------------------ANSSDPHPAQLSSKAIPEH-----CPD-----
------VVL------------------------------------------NGFRYMKDS
SLGGEINH
>Deuterostome_chordata_PetMar_tr_S4RK61_S4RK61_PETMA_Sodium_channel_epithelia
l_1_gamma_subunit_OS_Petromyzon_marinus_OX_7757_PE_4_SV_1_863_338
------------------------------------------------------------
----------M-------------------------------------------------
------------------------------------------------------------

```
------------------------------------------------------------
------------------------------------------------------------
------------------------------------------------------------
------------------------------------------------------------
------------------------------------------------------------
------------------------------------------------------------
------------------------------------------------------------
-----------------------ASEGDSKRVLHRV----------------------K-
DTLKIDGPD----------------------P---------SI-----T-----------
-DL--LDFYL-NNTNMHGMRRIA---------V-SK--------------GP---IK---
--KTIWI----VFSLIAVAMVFWQGIQLIQ-SFYS----------I---AVSVTI----N
-Y---------------Q-KLPF-----PAI-----------------------TVCSLN
PY------KYN-----QSQALL----------------EKL-----------------D-
---------RN------------TAVALH----------NIGIAVTN-------------
------------------------------------------------------------
------------------------------------------------------------
------------------------------------------------------------
-----------LSAAKRD------------------------------------------
------------------------------------------------------------
------------------------------------------------------------
------------------------------------------------------------
--------------------DEPLPIPLVWLDTTVTNQTVVTDV---ISGKFHVVPGKVE
MRSY--------------------------------------------------------
---------FSQNYQSSEPLIA--------------------------------------
------------------------------------------IEVCGEE-----------
------------------------------------------RKC---------------
--------------------------------------------IYNAF------TSAID
AV-------------------------IQWYRLHFINIM---------------------
------------------------AIV----------------PEKDKDKL----GYSAD
--EFI------IDCL----FS-G----T----------V---C-----------DPS---
----------------------------------------------T-------------
------------------------------------------------------------
------------------------------------------------------------
-----------S----------F-------K-----------KLQ-HPILG--------N
CFTFN-------DG-------------SD-G-------------------------KS--
-------LDIASA------GI--------DY-------------GLHMVLNTRQD-----
----------N-----------------S------------L-P-Y---LA---------
------------------------------------------------------------
------------------------------------------------------------
----------MGAGAKIGIH-LQNTTP--FIEAVGIDIPPAMESSLGLRVN------D--
-----VQKL-----------GDP-Y--SD-C--TMD-G------SD------IDV-KSLY
----D-------------S-PYSVQTCQNSCFQWEMI----KS--CGCANYE----Q---
------------P-----------------------------------------------
-------LPEGS-----R---FCNY-----------------------------------
------------------------------------DNNPGWEYCYY---------R---
L-Y-------------------DMYI-K-------EE--------L------------KC
----------IQVCR-QICSETE---HEVT-LSL------------------ADW-PS--
-------KAS------KGW-L-----------LR--------------------------
------------------------------------------------AL-S--------
----------------------------------KEQ-----------------------
-G-------------------------------LP-------------------------
---------------------------AND------------------------------
-----T-------LK-----------P------SDIAIVNIYFKDLTQKTIS-ESPASSI
VTL-----------------LSNLGGLLGLWLSCSMLCVVEVLEIFCVDF----------
--------PWIL--------L----------K-------KLL------------------
TTC-----------------------------SS-------------AFASLIRGPD---
--------------------------------------PAPSVHFPVQLPV---------
```

--------------------GG--------------------------SPAEDPP-----
------------------------------------------------------------
-----------TFHTAMQCP----REPI--------------------------------
----------------------------PMPNTPPPQYNTLRLRQIAGY-----------
------------------------------------------------------VP----
-DGGSDGE
>Protos_Platyhelmin_Macrostomum_lignano_A0A267GUI2_A0A267GUI2_9PLAT_Uncharact
erized_protein__Fragment__OS_Macrostomum_lignano_OX_282301_GN_BOX15_Mlig00481
4g1_PE_3_SV_1_866_340
CSNFLPNT----------------------------------------------------
----------M-------------------------------------------------
------------------------------------------------------------
------------------------------------------------------------
------------------------------------------------------------
------------------------------------------------------------
------------------------------------------------------------
------------------------------------------------------------
------------------------------------------------------------
------------------------------------------------------------
-----------------------TTEKKTPTDE--------------------------E
HKEPDAKTG----------------------N---------SF-----L-----------
-EL--ASSWG-SNIGMHGIPNIT---------R-SS--------------SV---AK---
--KFLWT----LLLLAGLCLTVVQIKSIVD-KYYS----------Y---PIAVAR----G
-A-EI-N----------L-PREF-----PAV-----------------------TVCNQS
PA------RKS-----KVQSAT------------------------------------ST
---------STSTTE--------TSNATA---------NSTSTTTMS-------------
------------------------------------------------------------
------------------------------------------------------------
----------------------P--------------------------AGYS-------
-----------------NVT----------------------------------------
------------------------------------------------------------
------------------------------------------------------------
------------------------------------------------------------
-----------------------SSTPTSTMNSSTIIAA---------------------
------------------------------------------------------------
------------------------------------------------------------
-----------------------------------------------NSTP----APSKR
KKRSAGSGYKDE------------------------------EDSYSGL-----------
-----------------------------------------TSSMYSSGT-----TRNLS
TMDSKT-------------VP-----DSVRMTYMFLDFY---------------------
------------------------NTL----------------DDDVKTEI----GYQIS
--DML------VDCS----MG-T----S----------Y---C-----------S-V---
----------------------------------------------S-------------
------------------------------------------------------------
------------------------------------------------------------
-----------N----------F-------T-----------RFL-HPMYG--------N
CYTFN-------AA-------------VT-N--------------------S----SR--
-------VSVKQQ------GP--------LF-------------GLTLTLYIDQS-----
----------D-----------------Y------------V-S-T---VA---------
------------------------------------------------------------
------------------------------------------------------------
----------QSAGAVVVLH-EPTTQP--FPEESGIRVSPGRETYIGMKQT------Y--
-----KKLL-----------GSP-Y--SDNC--AID-Y------KTA-----NKK-YNMY
----RSLDEWA----LPKV-NYTKMVCLKTCIQRKTE----SR--CNCSSPK----L---
------------P-P--SN-----------------------------------------
-----ITRTPAL-----P---ICKYSY---------NST---------------------
-----------------SK---------------SVSQE---AECLH---------A---

V-Q------------------ESEFE--------------------------------TC
----------RSGCQ-SQCEEMQ---YDAS-ISM------------------AAW-PS--
-------MGY------ESD-A-----------FH--------------------------
------------------------------------------------QVMM--------
-------------------------Y----------------------------------
-N-------------PIV------------------RTQMDG------------------
---------------------HKTDD-SKA------------------------------
-----D-------FI-----------S------QNMVKLIVFFADPETRTEI-STKGYEI
TDL-----------------LSDMGGQVGLWLGLSVLTLFELIEMLLDF-----------
--------VVLA--------A----------S-------KAAL-----------------
--------------------------------RQ---------------PKLPRDSK---
--------------------------------------QPPTENLDKKLS----------
----------------------MEMKNSIRF-----------------NKQQ--------
------------------------------------------------------------
------------------------------------------------------------
--------------------------------NPA-------------------------
----------------------------------------------------------MW
VDEKHRQP
>Protostome_Lophotroco_annelid_Pdum_Contig18637__869_342
------------------------------------------------------------
----------M-------------------------------------------------
------------------------------------------------------------
------------------------------------------------------------
------------------------------------------------------------
------------------------------------------------------------
------------------------------------------------------------
------------------------------------------------------------
------------------------------------------------------------
------------------------------------------------------------
-----------------------AGQYIKDP-----------------------------
-IKRDINKG----------------------H---------TL-----------------
-II--CHEFA-TETSAHGMSHII---------R-AR--------------GT---CR---
--KLFWL----VVTLGLIAIWLLQSQDTIR-KYLK----------N---EVNVQV----E
-I-KA-S----------R-ELPF-----PAV-----------------------TICNKN
PY------KGD-----DTIR--------------------FI----------------E-
---------LM---------NHTASLAISR--------N-QSLEDIA-------------
------------------------------------------------------------
------------------------------------------------------------
------------------------------------------------------------
------------------------------------------------------------
------------------------------------------------------------
------------------------------------------------------------
------------------------------------------------------------
--------------------------PSW-------------------------------
------------------------------------------------------------
------------------------------------------------------------
------------------------------------------------------------
-FVPWVRSLTGS------------------------------DDGTFET-----------
------------------------------------------GISSAKF------GEQML
Q-----------------------------LLAELVEVY---------------------
------------------------GEY----------------HNFSLAEL----SVDPD
--NFI------LNCM----YE-R----V----------H---C-----------D-M---
----------------------------------------------R-------------
------------------------------------------------------------
------------------------------------------------------------
-----------ND-------IEM-------T-----------SIF-NWLYG--------G
CIVYI-------IK----------------------------------------------

-------QNTTRT------GP--------RF-------------GLELTLNIEQS-----
----------D-----------------Y------------M-D-----LT---------
------------------------------------------------------------
------------------------------------------------------------
----------TQAGALVLVH-PPEEMP--FPEDDGISIPPGQAALIGVKVK------R--
-----TERL-----------GGK-F--GD-C--AHA-N------K-------LPNFTNYF
----QAE--NP------WT-NYSLKACQRSCFQWNVR----QM--CQCLDVR----F---
------------P-P--FD-----------------------------------------
-------EYKNF-----T---SCVNKY-----FSTFNFS---------------------
---------------------------------QGFPES---VGCLD---------E---
V-T-------------------KKFN-A-------HQ--------L------------GC
----------DKLCH-VPCKQTY---YEPF-VSY------------------AAW-PS--
-------DAA------LNV-T-----------LE--------------------------
------------------------------------------------NL----------
------------------------------------------------------------
KGS-----------SASA------------------DEI---------------------
------------------------LE-VGH------------------------------
-----D-------NF-----------R------KNFLQLTVYFQDLNFEFIT-EKEAVEL
TQM-----------------LSDLGGNTGMYVGASLFTMIEFVELFGDL-----------
--------WLWV--------C----------C-------HR-------------------
MR------------------------------RK-------------------RP-----
---------------------------------------KYAPPTEEDL-----------
------------------DLYFITPNP---------------------SKQNNNH-----
--------TNNYQNRQGHAPNQQNR--------------HGHANNKQNRQPHPNKH----
--------DRRHSKNNGRAPGRGQDNNGYHFNYNGHL------ERPRKSLPRT-------
-------------------------------PLP--------------------------
------------------------------------------------------MDMAEV
QDTMVGLY
>Deutero_Ambulac_Apla_gbr2_220_t1_870_343
------------------------------------------------------------
----------M-------------------------------------------------
------------------------------------------------------------
------------------------------------------------------------
------------------------------------------------------------
--MSKNTMTTGIPAGY--------DDAF--------------------------------
--------------------------------------VVMPSESI--------------
--------------------------------------------------------PLP-
------------------------------------------------------------
------------------------------------------------------------
-------------------SYHNKAPTSVEAK---------------------------A
AEPEKENHP----------------------V---------TC-----G-----------
-SI--VTNFA-DSTTAHGVARIA---------N-AS--------------SC---FA---
--SFLWL----VILCVAFGGFFFQGTNLVL-DFFS----------W---PYGTTI----D
-I-IT-N----------T-SVDF-----PAV-----------------------TVCNMN
RL------RRS-----KLPGTR------------------F-----------------E-
---------GVIAI----------------------------------------------
---------------------------------DG-------GISGGDNDYSWFFEWSSA
------------------------------------------GDFYNQF-----------
------------------------------------------------------------
------VSASAGGGSSGGGGGSS------------------AGGGNSSSVGGG-------
--------------SSAGGGNSSSVGGGS-------------------------------
-----------------------------------SAGGGGSSAGGGGSSAGGG------
-------------------------------------------G----------------
----------------SSAAGSSYSYPPASSN----------------------------
------------------------------------------------------------
------------------------------------------------------------
------------------------------------------------------------

--------------YYYW----ETEFGDD-------FNFEYDNYYD-------F------
--------GTV-------------TGESDWDGFLAN-----------SK------SEDFS
---------------------------------DIINVA---------------------
-------------------------NP----------------TQDEMDEL----GHQAE
--DFI------LQCT----FD-R----R----------K---C-----------N-Y---
----------------------------------------------T-------------
------------------------------------------------------------
------------------------------------------------------------
-----------D----------F-------Y-----------KFQ-NSHYG--------N
CFTFN-------HG-------------RN----------------------E----TV--
-------RTTSKS------GF--------QY-------------GLHLTLFIEQP-----
----------E-----------------Y------------V-G-L---FS---------
------------------------------------------------------------
------------------------------------------------------------
----------PESGVRVSIN-HWQTTP--HPEDSGITATTGQATSIALRKN------F--
-----IKRL-----------GGW-Y--SN-C--TYG--------TE----------TNFT
----SDF-----------F-TYSPLTCKKQCMQRNLK----DR--CDCVTDL----L---
------------L-----------------------------------------------
----------DG-----E---KCSYL----------N-----------------------
------------------------------------STQ---QRCRQ---------L---
V-E-------------------ALYE-E-------DK--------L------------DC
------------LCP-VACEENV---FKTS-ASV------------------SIW-PS--
-------ERY------EEH-L-----------YS--------------------------
------------------------------------------------RL-S--------
----------------------------------SK------------------------
-N-------------AVA------------------ARML--------------------
---------------------------QDV------------------------------
-----E-------TT-----------R------KNLARVRIYFEELNYQKVE-QIPQWTI
ESI-----------------LGAVGGLMGLYVGISSITLMEIIVFVFSL-----------
--------LKQC--------C----------K-------SVIC-----------------
--------------------------------LN-------------------K------
-----------------------------------------VQPSN--------------
------------------------------------------------------------
------------------------------------------------------------
------------------------------------------------------------
------------------------------------------------------------
------------------------------------------------------------
--------
>Chordat_hASIC4_NP_878267_2_acidsensing_ion_channel_4_isoform_2__Homo_sapiens
__6_344
------------------------------------------------------------
----------M-----------LSGAAGLAPSLTRSLAGTHAGADSC-------------
--------------------------------AGADKGSH--------------------
-KETIEERDKRQQRQQRQRQHQGCGAAGSGSDSPTSGPHPVPVLFPLALSLEEQPLPPL-
-----------PLGRAPGLLREGQGREALASPSSR----------------------GQM
PIEIVCKIKFA-------------------------------------------------
---EEDAK----------------------------------------------------
--------------PKEKEA-------------GDEQSLLGAVA------------P---
------------------------------------------------------------
------------------------------------------------------------
------------------------------------------------------------
-------------------------------G---------AA-----P-----------
-RD--LATFA-STSTLHGLGRAC--------G-PG--------------PHG---LR---
--RTLWA----LALLTSLAAFLYQAAGLAR-GYLT----------R---PHLVAM----D
-P-AA-P----------APVAGF-----PAV-----------------------TLCNIN
RF------RHS-----ALSDAD------------------I-----------------F-
---------HL------------A-NLT---------------GLPP----K--------

---------------------------------DR-------------------------
------------------------------------------------------------
------------------------------------------------------------
-----------------DGH----------------------------------------
------------------------------------------------------------
------------------------------------------------------------
------------------------------------------------------------
------------------------------------------------------------
------------------------------------------------------------
------------------------------------------------------------
------------------------------------------------------------
------------------------------------------------------------
--------------------------------------------RAAGLR-----YPEP-
---------------------------------DMVDIL---------------------
------------------------------------------------NRT----GHQLA
--DML------KSCN----FS-G----H----------H---C-----------S-A---
----------------------------------------------S-------------
------------------------------------------------------------
------------------------------------------------------------
-----------N----------F-------S-----------VVY-T-RYG--------K
CYTFN-------AD-------------PR---------------------------SS--
-------LPSRAG------GM--------GS-------------GLEIMLDIQQE-----
----------E-----------------Y------------L-P-IWRETN---------
------------------------------------------------------------
------------------------------------------------------------
-------ETSFEAGIRVQIH-SQEEPP--YIHQLGFGVSPGFQTFVSCQEQ------R--
-----LTYL-----------PQP-W--GN-C--RAE-SELR-EPE-------LQ----GY
------------------S-AYSVSACRLRCEKEAVL----QR--CHCRMVH----M---
------------P-----------------------------------------------
------------------------------------------------------------
------------------------------------------------------------
----------------------DSLG-G-------GP--------E-----------GPC
------------FCP-TPCNLTR---YGKE-ISM------------------VRI-PN--
-------RGS------ARY-L-----------AR--------------------------
------------------------------------------------KY----------
------------------------------------------------------------
-N----------------------------------------------------------
---------------------------RNE------------------------------
-----T-------YI-----------R------ENFLVLDVFFEALTSEAME-QRAAYGL
SAL-----------------LGDLGGQMGLFIGASILTLLEILDYIYEV-----------
--------SWDR--------L----------K-------RV----------------WR-
RP------------------------------KT-------------------P------
------------------------------------------------------------
-----------------------LRTS---------------------TGGI--------
-----------ST-----------------------------------------------
----------------LGLQELKEQ------------------------SPCPSRGR---
-----------------------VEGGGVSSLLP--NHHHPHG---------PP------
---------GGL-----FEDFAC-------------------------------------
--------
>Chordat_hENaCgamma_NP_001030_2_amiloridesensitive_sodium_channel_subunit_gamma__Homo_sapiens__2_346
------------------------------------------------------------
----------M-------------------------------------------------
------------------------------------------------------------
------------------------------------------------------------
------------------------------------------------------------
------------------------------------------------------------

```
------------------------------------------------------------
------------------------------------------------------------
------------------------------------------------------------
------------------------------------------------------------
-----------------------APGEKIKAKIK-------------------------K
NLPVTGPQA----------------------P---------TI-----K-----------
-EL--MRWYC-LNTNTHGCRRIV---------V-SR--------------GR---LR---
--RLLWI----GFTLTAVALILWQCALLVF-SFYT----------V---SVSIKV----H
-F---------------R-KLDF-----PAV-----------------------TICNIN
PY------KYS-----TVRHLL----------------ADL-----------------E-
---------QE------------TREAL--------------KSLYG----FP-------
---------------------------------ES-------------------------
------------------------------------------------------------
------------------------------------------------------------
----------RKRREAESWNSVSE------------------------------------
------------------------------------------------------------
------------------------------------------------------------
------------------------------------------------------------
----------------GKQPRFSHRIPLLIFDQDEKG--KARDF---FTGRKRKVGGSII
HKAS--------------------------------------------------------
---------NVMHI-ESKQVVG--------------------------------------
------------------------------------------FQLC--------------
--------SNDT------------------------------SDC---------------
--------------------------------------------ATYTF------SSGIN
AI-------------------------QEWYKLHYMNIM---------------------
------------------------AQV----------------PLEKKINM----SYSAE
--ELL------VTCF----FD-G----V----------S---C-----------D-A---
----------------------------------------------R-------------
------------------------------------------------------------
------------------------------------------------------------
-----------N----------F-------T-----------LFH-HPMHG--------N
CYTFN-------NR-------------EN-E-------------------------TI--
-------LSTSMG------GS--------EY-------------GLQVILYINEE-----
----------E-----------------Y------------N-P-F---LV---------
------------------------------------------------------------
------------------------------------------------------------
----------SSTGAKVIIH-RQDEYP--FVEDVGTEIETAMVTSIGMHLT------E--
-----SFKL-----------SEP-Y--SQ-C--TED-G------SD------VPI-RNIY
----N-------------A-AYSLQICLHSCFQTKMV----EK--CGCAQYS----Q---
------------P-----------------------------------------------
-------LPPAA-----N---YCNY-----------------------------------
------------------------------------QQHPNWMYCYY---------Q---
L-H-------------------RAFV-Q-------EE--------L------------GC
----------QSVCK-EACSFKE---WTLT-TSL------------------AQW-PS--
-------VVS------EKW-L-----------LP--------------------------
------------------------------------------------VL-T--------
----------------------------------WDQG----------------------
RQ----------------------------------------------------------
---------------------------VNK------------------------------
-----K-------LN-----------K------TDLAKLLIFYKDLNQRSIM-ESPANSI
EML-----------------LSNFGGQLGLWMSCSVVCVIEIIEVFFIDF----------
--------FSII--------A----------R-------RQW------------------
QKA-----------------------------KE------------W-WAWKQAPPC---
--------------------------------------PEAPRS----------------
----------------------PQGQDNPAL-----------------DIDDDLP-----
------------------------------------------------------------
-----------TFNSALHLP---PALGT--------------------------------
```

----------------------------QVPGTPPPKYNTLRL-----------------
---------------------------------------------------ERAFSNQLT
DTQMLDEL
>Deuterostome_chordata_Ggallus_tr_F1NW62_F1NW62_CHICK_Uncharacterized_protein
_OS_Gallus_gallus_OX_9031_GN_SCNN1G_PE_4_SV_2_875_347
------------------------------------------------------------
----------M-------------------------------------------------
------------------------------------------------------------
------------------------------------------------------------
------------------------------------------------------------
------------------------------------------------------------
------------------------------------------------------------
------------------------------------------------------------
------------------------------------------------------------
------------------------------------------------------------
-----------------------APGKIT-ARIK-------------------------K
TLPVRGPQA----------------------P---------TL-----R-----------
-EL--MRWYC-LNTNTHGCRRIV---------V-SR--------------GR---LR---
--RFIWI----LLTLSAVGLILWQCAELLL-NYYS----------A---SVSVTV----Q
-F---------------Q-KLPF-----PAV-----------------------TICNIN
PY------KYS-----SMKDYL----------------SEL-----------------D-
---------KE------------TKKAL--------------ETFYG----FS-------
---------------------------------EGK------------------------
------------------------------------------------------------
------------------------------------------------------------
---------TKVRRAAGDWN----------------------------------------
------------------------------------------------------------
------------------------------------------------------------
------------------------------------------------------------
----------------GTESLFFRHVPLLRFENSFR---AATDL---RSGRKRKVEGSVF
HKDS--------------------------------------------------------
---------SIVNSGDSNDIIG--------------------------------------
------------------------------------------FQLCD-------------
--------ANNS------------------------------SEC---------------
--------------------------------------------ALYTF------SSGVN
AI-------------------------QEWYKLHYMNIM---------------------
------------------------AQI----------------PLETKEEL----SYSAD
--DLL------LTCF----FD-G----L----------S---C-----------D-K---
----------------------------------------------R-------------
------------------------------------------------------------
------------------------------------------------------------
-----------H----------F-------T-----------RFH-HPLHG--------N
CYTFN-------SG-------------EN-G-------------------------TV--
-------LSTSTG------GS--------EY-------------GLQVVLYIDEA-----
----------D-----------------Y------------N-P-F---LV---------
------------------------------------------------------------
------------------------------------------------------------
----------TSTGAKIIVH-DQDEYP--FIEDIGTEIETAAATSIGMHFT------R--
-----SRKL-----------SKP-Y--SD-C--TET-G------AD------IPV-ENLY
----N-------------K-SYSLQICLHSCFQKAMV----ES--CGCAQYA----Q---
------------P-----------------------------------------------
-------LPNGA-----E---YCNY-----------------------------------
------------------------------------KKNPNWMYCYY---------R---
L-H-------------------EKFV-K-------EQ--------L------------GC
----------QQICK-DACSFKE---WALT-TSI------------------AQW-PS--
-------TVS------EDW-M-----------LR--------------------------
------------------------------------------------VL-S--------

----------------------------------WDKG----------------------
QK----------------------------------------------------------
---------------------------INK------------------------------
-----K-------LN-----------K------TDLANLMVFYKDLNERFIS-ENPANTL
VIL-----------------LSNFGGQLGLWMSCSVVCVIEIIEVFFIDS----------
--------FSIV--------M----------R-------RQW------------------
QKA-----------------------------KK------------W-WNHRKRDET---
--------------------------------------GKPPEVGDAE------------
----------------------QQGHDNPA------------------CSDEDLP-----
------------------------------------------------------------
-----------TFNTALRLP---LPQEG--------------------------------
----------------------------HPPRTPPPNYSTLRL-----------------
---------------------------------------------------ETAFTEQLP
DTLEAGQH
>sp_P37091_SCNNG_RAT_Amiloride_sensitive_sodium_channel_subunit_gamma_OS_Ratt
us_norvegicus_OX_10116_GN_Scnn1g_PE_1_SV_2_877_348
------------------------------------------------------------
----------M-------------------------------------------------
------------------------------------------------------------
------------------------------------------------------------
------------------------------------------------------------
------------------------------------------------------------
------------------------------------------------------------
------------------------------------------------------------
------------------------------------------------------------
------------------------------------------------------------
-----------------------APGEKIKAKIK-------------------------K
NLPVRGPQA----------------------P---------TI-----K-----------
-DL--MHWYC-MNTNTHGCRRIV---------V-SR--------------GR---LR---
--RLLWI----AFTLTAVALIIWQCALLVF-SFYT----------V---SVSIKV----H
-F---------------Q-KLDF-----PAV-----------------------TICNIN
PY------KYS-----AVSDLL----------------TDL-----------------D-
---------SE------------TKQAL--------------LSLYG----VK-------
---------------------------------ES-------------------------
------------------------------------------------------------
------------------------------------------------------------
----------RKRREAGSMPSTLE------------------------------------
------------------------------------------------------------
------------------------------------------------------------
------------------------------------------------------------
----------------GTPPRFFKLIPLLVFNENEKG--KARDF---FTGRKRKISGKII
HKAS--------------------------------------------------------
---------NVMHVHESKKLVG--------------------------------------
------------------------------------------FQLC--------------
--------SNDT------------------------------SDC---------------
--------------------------------------------ATYTF------SSGIN
AI-------------------------QEWYKLHYMNIM---------------------
------------------------AQV----------------PLEKKINM----SYSAE
--ELL------VTCF----FD-G----M----------S---C-----------D-A---
----------------------------------------------R-------------
------------------------------------------------------------
------------------------------------------------------------
-----------N----------F-------T-----------LFH-HPMYG--------N
CYTFN-------NK-------------EN-A-------------------------TI--
-------LSTSMG------GS--------EY-------------GLQVILYINED-----
----------E-----------------Y------------N-P-F---LV---------
------------------------------------------------------------

------------------------------------------------------------
----------SSTGAKVLIH-QQNEYP--FIEDVGMEIETAMSTSIGMHLT------E--
-----SFKL-----------SEP-Y--SQ-C--TED-G------SD------VPV-TNIY
----N-------------A-AYSLQICLYSCFQTKMV----EK--CGCAQYS----Q---
------------P-----------------------------------------------
-------LPPAA-----N---YCNY-----------------------------------
------------------------------------QQHPNWMYCYY---------Q---
L-Y-------------------QAFV-R-------EE--------L------------GC
----------QSVCK-QSCSFKE---WTLT-TSL------------------AQW-PS--
-------EAS------EKW-L-----------LN--------------------------
------------------------------------------------VL-T--------
----------------------------------WDQS----------------------
QQ----------------------------------------------------------
---------------------------INK------------------------------
-----K-------LN-----------K------TDLAKLLIFYKDLNQRSIM-ESPANSI
EML-----------------LSNFGGQLGLWMSCSVVCVIEIIEVFFIDF----------
--------FSII--------A----------R-------RQW------------------
HKA-----------------------------KD------------W-WARRQTPPS---
--------------------------------------TETPSS----------------
----------------------RQGQDNPAL-----------------DTDDDLP-----
------------------------------------------------------------
-----------TFTSAMRLP---PAPGS--------------------------------
----------------------------TVPGTPPPRYNTLRL-----------------
---------------------------------------------------DRAFSSQLT
DTQLTNEL
>Deuterostome_chordata_PetMar_tr_S4RY81_S4RY81_PETMA_Sodium_channel_epithelia
l_1_beta_subunit_OS_Petromyzon_marinus_OX_7757_GN_SCNN1B_PE_4_SV_1_879_349
------------------------------------------------------------
----------M-------------------------------------------------
------------------------------------------------------------
------------------------------------------------------------
------------------------------------------------------------
------------------------------------------------------------
------------------------------------------------------------
------------------------------------------------------------
------------------------------------------------------------
------------------------------------------------------------
-----------------------KIRKYLTRSL---------------------------
-HRLQKGPV----------------------A---------SV-----S-----------
-EL--LYWYC-MNTNTHGCKRIV---------V----------------YGK---KK---
--RVLWF----LITIIMLGVVLWQWVLLFQ-AYLS----------Y---GVSVSV----N
-M-GF------------Q-RMNF-----PAV-----------------------TVCNLN
AY------RYS-----SMKDKI----------------KDL-----------------E-
---------AY------------TRVAL--------------QTLYN----YT-------
---------------------------------DS-------------------------
------------------------------------------------------------
------------------------------------------------------------
-----------STPSAYDTSYAVGP-----------------------------------
------------------------------------------------------------
------------------------------------------------------------
------------------------------------------------------------
----------------------WQEIPLVLIDRRDPNRTVVTEV-----MCSRAAVGIET
HT----------------------------------------------------------
--------VDNRVFHIGFCKSA--------------------------------------
------------------------------------------LGVCC-------------
--------DSAG------------------------------DKC---------------
--------------------------------------------FYSEY------LSGMT

AV-------------------------KQWFHFNLLSLL---------------------
------------------------GNL----------------STEEKSNL----SSSGD
--ELI------RSCL----FS-S----D----------T---C-----------S-A---
----------------------------------------------T-------------
------------------------------------------------------------
------------------------------------------------------------
-----------N----------F-------T-----------TFF-HPMYG--------N
CFIFN-------WG-------------EN-E-------------------------TV--
-------MQVSNP------GV--------EY-------------GLKLVLSIDQD-----
----------E-----------------Y------------I-P-F---LT---------
------------------------------------------------------------
------------------------------------------------------------
----------TIAGAVIMVH-DQNTYP--FLSDLGVFVKTGVETSVGIEVG------Q--
-----LQRQ-----------GAP-Y--SD-C--TMD-G------TD------LPI-TNLY
----N------------GT-AYSVQACLRSCFQTKMI----EM--CGCGYYL----Y---
------------P-----------------------------------------------
-------LPPGE-----K---YCQN-----------------------------------
------------------------------------QNFTGWRYCYY---------K---
L-Y-------------------EQFV-E-------ED--------M------------DC
----------YTICK-QPCIESE---YKMS-ISM------------------SDW-PS--
-------QSS------EDW-I-----------FH--------------------------
------------------------------------------------IL-S--------
----------------------------------KERK----------------------
HN----------------------------------------------------------
---------------------------VSR------------------------------
--------I----FN----------RK------QDIIKLNLFFQEFNSMTIS-ESPAQTI
VTL-----------------LSNLGGQFGFWMGGSVLCIIEFIEIIIDC-----------
--------VWIG--------M----------I-------KASNDV---------------
RER-----------------------------RK----------------TSRKPRY---
--------------------------------------SDEPPTLSSI------------
----------------------VQGQGNSGF-----------------EMEERGPPGE--
------------------------------------------------------------
--------APEANGSAAQPPAAAAEQQP--------------------------------
----------------------------DVPGTPPPHYDTLRIS----------------
---------------------------------------------------KTELHDEIN
SDDDGEFV
>Cyclo_DEL1_sp_Q19038_1_DEL1_CAEEL_RecName__Full_Degenerin_del1_46_351
------------------------------------------------------------
----------M-------------------------------------------------
-------------------------------ARKY-------------------------
------------------------------------------------------------
------------------------------------------------------------
------------------------------------------------------------
------------------------------------------------------------
------------------------------------------------------------
------------------------------------------------------------
------------------------------------------------------------
-------------------IDILKKSKMMLFQDVGKSFE-------------------DD
SPCKEEAPK----------------------T---------QI-----Q-----------
-HS--VRDFC-EQTTFHGVNMIF---------T-TS--------------LY---WV---
--RFLWV----VVSLVCICLCMYSFSHVKD-KYDR----------K---EKIVNV----E
-L-VF-E------------SAPF-----PAI-----------------------TVCNLN
PF------KNH-----LARSVP--------------------------------------
---------EI-------------------------------SETLD-------------
------------------------------AF-HQAV-----VYSN--------------
------------------------------------------------------------
------------------------------------------------------------

-----------------DATMDEL------------------------------------
------------------------------------------------------------
---------SGRGRRSLNDG--------------PSFKYLQYEP----------------
-----------VYSDCSCVPG------------------------RQECIAQTSA--P--
-------RTLENACICNYDRHDGSAWPCYSAQTWE------------------KSICPEC
NDI-GFCNVPNTTGSG--------NIPCYCQLEMGY-----------CVFQP--------
----------------ESRVRR--------------------------------------
-----------------------------------------IWEFQGNKIP---------
--EKGSPLRKEY------------------------------MEQL--------------
-----------------------------------------TQLGYGNM------TDQVA
--------------------------ITTQAKEKMILKM---------------------
------------------------SGL----------------HPQRRAAL----GYGKS
--ELI------KMCS----FN-G----Q----------Q---C-----------N-ID--
----------------------------------------------T-------------
------------------------------------------------------------
------------------------------------------------------------
-----------E----------F-------K-----------LHI-DPSFG--------N
CYTFN-------AN-------------PE-K-K---------------------------
-------LASSRA------GP--------SY-------------GLRLMMFVNSS-----
----------D-----------------Y------------L-P-----TT---------
------------------------------------------------------------
------------------------------------------------------------
----------EATGVRIAIH-GKEECP--FPDTFGYSAPTGVISSFGISLR------N--
-----INRL-----------PQP-Y--GN-C--LQK--------DN------PQS-RSIY
----KGY-------------KYEPEGCFRSCYQYRII----AK--CGCADPR----Y---
------------P-K--PW-----------------------------------------
---------KRS-----A---WCDST----------N-----------------------
------------------------------------TTT---LNCLT---------T---
E-G-------------------AKLS-TKE-----NQ--------------------KHC
------------KCI-QPCQQDQ---YTTT-YSA------------------AKW-PS--
-------GSI------QTS-----------------------------------------
------------------------------------------------------------
------------------------------------------------------------
------------------------------------------------------------
--------------------CDNHSK-DCN------------------------------
-----S-------YL-----------R------EHAAMIEIYYEQMSYEILR-ESESYSW
FNL-----------------MADMGGQAGLFLGASIMSVIEFLFFAVRT-----------
--------LGIA--------C----------KP------RR--------------W----
--------------------------------RQ-------------------KT-----
---------------------------------------ELLRAEELNDA----------
------------------------------------------------------------
------------------------------------------------------------
------------------------------------------------------------
------------------------------------------------------------
-----------------------------------------------------------E
KGVSTNNN
>Protostome_Lophotroco_brch_Lingula_anatina_comp148540_c0_seq7_p1_comp148540_c0__comp148540_c0_seq7_p1__ORF_type_complete_len_666___score_77_65_comp148540_c0_seq7_79_2076__887_353
------------------------------------------------------------
----------M-------------------------------------------------
------------------------------------------------------------
------------------------------------------------------------
------------------------------------------------------------
------------------------------------------------------------
------------------------------------------------------------
------------------------------------------------------------

------------------------------------------------------------
------------------------------------------------------------
-----------------------STGAQTMYPSEKN---QFD--FFKNSLVVDHNSQD--
--PEKSARE----------------------E---------SL-----I-----------
-AL--AYDWA-SNSPVNGIPNVV---------R-SQ--------------NL---IK---
--RFFWL----VALLGCFGVMGYQTWELVA-KFYR----------F---PVDTTL----T
-Y-TH-S----------K-EVYF-----PAV-----------------------TVCNVN
PL------RRS-----MLSSAG------------------------------------D-
---------DV-------------------------------YALLG----P--------
------------------------P--------KSQM----GSSPGG-------------
------------------------------------------------------------
-------------------AQAPAGP----------------------------------
---------AGSFASQSQTTQAAAG----------------PPGPASSSITTQ-------
APA---------------------------------------------------------
------------------------------------------------------------
--------------------------------------------------------APGP
K---------------GSGSGGTTQAPSASQTTAGAPAG-------GGGG---TTQAPAG
PPAP--------------------------------------------------------
------------------------------------------------------------
-------------------------------------------------------PSGRK
R-RSTGTNSTST------------------------------------------------
-----------------------------------------SRRKPGQM------NERFE
------------------------------MRKRFGNIW---------------------
------------------------AKL----------------NYTTRETV----GHELD
--TMI------ISCT----FN-G----V----------T---C-----------S-S---
----------------------------------------------A-------------
------------------------------------------------------------
------------------------------------------------------------
-----------N----------F-------T-----------RFN-NHQFG--------N
CYTFN-------SG-------------WD-S-------------------SI----PV--
-------ETSSNA------GP--------LY-------------GLSLEMYIEQS-----
----------E-----------------Y------------I-G-D---LS---------
------------------------------------------------------------
------------------------------------------------------------
----------ESAGVRVQIH-SQRSMA--FPEDEGFNIAPGYLTSMAMTRV------E--
-----ITRR-----------PHP-YP-SK-C--RNF--------TTEE----SKQ-KSVF
----TSV--HN-------V-DYSVSGCMKTCYQRYVI----SE--CACGDPA----Y---
------------P-F--SLD-----------------LEAFSDLKN--------------
--NINGTTAESV-----D---PCVS-----------------------------------
------------------------------------DAD---DECVA---------G---
I-K-------------------RRFA-D-------DD--------L------------SC
------------DCP-LTCVDVE---YSGV-PSL------------------AKW-PS--
-------KQY------MST-L-----------HA--------------------------
------------------------------------------------TL-A--------
----------------------------------SA------------------------
-G-------------AHL------------------DNIINGK-----------------
----------------------N----GMA------------------------------
-----A-------EP-----------G------DNLLKLEVYFQELNFQKIS-ENVAYDV
FAL-----------------LADIGGQIGFWVGLSIMALFEVVELILDV-----------
--------FRLL--------L----------F-------RV-------------------
GSFCF---------------------------KK----------------QQKR------
---------------------------------------PPSRQVKVQ------------
------------------------------------------------PASEENV-----
------------------------------------------------------------
------------------------------------------------------------
-------------------------------LTP--------------------------
------------------------------------------------------------

```
RLEKVRFS
>Deuterostome_chordata_Ggallus_tr_F1NE95_F1NE95_CHICK_Uncharacterized_protein
_OS_Gallus_gallus_OX_9031_GN_SCNN1B_PE_4_SV_4_889_354
------------------------------------------------------------
----------M-------------------------------------------------
------------------------------------------------------------
------------------------------------------------------------
------------------------------------------------------------
------------------------------------------------------------
------------------------------------------------------------
------------------------------------------------------------
------------------------------------------------------------
------------------------------------------------------------
-----------------------NLKRYFVRAL---------------------------
-HRLQKGPG----------------------Y---------TY-----K-----------
-EL--LVWYC-DNTNTHGPKRII-----------KE--------------GP---KK---
--KVMWF----FLTLLFASLVFWQWGILIN-TYLS----------Y---NVTSSL----S
-I-GF------------K-TMKF-----PAV-----------------------TVCNAN
PF------KYS-----EVRPLL----------------KEL-----------------D-
---------KL------------IEAAL--------------ERILQ----PTHG-----
---------------------------------DP-------------------------
------------------------------------------------------------
------------------------------------------------------------
-----------ISPLLLNNSNATE------------------------------------
------------------------------------------------------------
------------------------------------------------------------
------------------------------------------------------------
----------------GLDLDLWNQIPLVLIDEQDKDNPVIVEI---F-ETNQSAAGNQT
AAPPA-------------------------------------------------------
--------PANVTSEEKKYKLA--------------------------------------
------------------------------------------VKLC--------------
-------SHQGS------------------------------NNC---------------
--------------------------------------------TYRNF------TSAAQ
AV-------------------------TEWYILQSTSIL---------------------
------------------------SKV----------------PLQERIRM----GYQAE
--DMI------LACL----YG-A----E----------P---C-----------N-Y---
----------------------------------------------K-------------
------------------------------------------------------------
------------------------------------------------------------
-----------N----------F-------T-----------QIY-HPDHG--------N
CYIFN-------WG-------------MD-K-------------------------EA--
-------LNSSNP------GA--------EF-------------GLKLILDISQQ-----
----------D-----------------Y------------I-P-Y---LS---------
------------------------------------------------------------
------------------------------------------------------------
----------SAAGARLMLH-QQKSFP--FLKDQGIYAMAGTETSIGVLVD------E--
-----LERM-----------GYP-Y--SD-C--TAN-G------SD------VPV-KNLY
----SEY----------NT-SYSIQACLRSCFQNHMT----EI--CGCGHYM----F---
------------P-----------------------------------------------
-------LPEGV-----T---YCNN-----------------------------------
------------------------------------EDNPGWAYCYS---------S---
L-R-------------------SSIR-H-------RQ---------------------IC
----------IDSCK-ETCNDTQ---YKMT-ISM------------------ADW-PS--
-------EAS------EDW-I-----------FH--------------------------
------------------------------------------------IL-S--------
----------------------------------YERD----------------------
MS----------------------------------------------------------
```

---------------------------TNV------------------------------
-----T-------LD-----------R------NGIIKLNIYFQEYNYRTIS-ESAATTI
VWL-----------------LSSLGGQFGFWMGGSVLCLIEFGEIIIDS-----------
--------LWIT--------V----------I-------NIISWC---------------
KGL-----------------------------KQ----------------KRVRARY---
--------------------------------------PDTPPTVSEL------------
----------------------VEAHTNLGF-----------------QHEEAG------
------------------------------------------------------------
-----------TETQGEALP----------------------------------------
----------------------------PEPGTPPPNYDSLRVQPSHNPGTDSDIE----
-------------------------------------------------CEEQRPAANHH
GDASVWAE
>Protostome_Lophotroco_annelid_CAC9584386_1__Ofus_G061163_Owenia__fusiformis_
_891_355
------------------------------------------------------------
----------M-------------------------------------------------
------------------------------------------------------------
------------------------------------------------------------
------------------------------------------------------------
------------------------------------------------------------
------------------------------------------------------------
------------------------------------------------------------
------------------------------------------------------------
------------------------------------------------------------
-----------------------NEKDDLDNMT---------------------------
-VIDTKEEEI------------------KGAS---------DA-----K-----------
-TV--TDEFM-GSFGAHGLPRVW---------S-SA--------------SP---VK---
--KIVWL----VLFLAASGYCLNIIIRVGI-KFGT----------F---PSGINS----K
-V-YP-R----------S-IVDF-----PAV-----------------------TICNLN
ML------KAT-----SISNNN------------------V-----------------E-
---------KY-------------------------------INLLS-------------
------------------------------------------VQSEAEASMNSWITWALN
------------------------------------------------------------
---------YT-------------------------------------------------
-----------------DSVTTSF-----------------GPTTDSTSRTN--------
------------------------------------------------------------
-----------------------------------SSNAGTTDHTTVADRRKRA------
-VEEFPFNEDLLEDEL--------------------------------------------
--LLNLRADHVNDLT------RLIHMPEVVK-----------------------------
------------------------------------------------------------
--MATLRLLKKRHGFDMREQRSAPPIPHRQKRD---------------------------
------------------------------------------------------------
-NQAAETNSSDYYDDNFYY---YDNY-----------DYSYYDDYG-------F------
----------------------GDVSENDFRTLLTK-----------SK------TDDYQ
---------------------------------DLMGVL---------------------
-------------------------KP----------------TKTELELY----GHSAQ
--DMI------VSCT----FD-S----K----------K---C-----------N-Y---
----------------------------------------------T-------------
------------------------------------------------------------
------------------------------------------------------------
-----------L----------F-------K-----------TFQ-NSYYG--------N
CFTFN-------YD-------------DG-N-STT---------------QE----VV--
-------FNTTKK------GS--------RF-------------GLKLTLNIERQ-----
----------E-----------------Y------------I-G-L---FA---------
------------------------------------------------------------
------------------------------------------------------------
----------HGSGVRVAVH-PRNATP--FPEDYGISAPTGWETAIGVREN------R--

-----VTRL-----------TKP-FE-SN-C--SHG--------NH------DP--SETY
----T-------------G-LYTLLECNQRCLQKQIR----SK--CKCLDEI----M---
------------D-D--QT-----------------------------------------
---WESVYASDSATSMLN---PCSIL----------N-----------------------
------------------------------------STE---EKCRQ---------R---
M-Y-------------------YLSQ-T-------NR--------L------------GC
------------DCP-LPCLELG---FEES-LHS------------------SEW-PS--
-------AQY------YPL-M-----------MQ--------------------------
------------------------------------------------KL-T--------
----------------------------------DS------------------------
-N-I---------ISEGQ------------------LKI------------M--------
---------------------------NDL------------------------------
-----K-------TT-----------R------ENFVRLHVYYKDMILESLE-EFEAYTV
DSL-----------------IGDIGGILGLFIGMSVLTIAEAVEYLIDL-----------
--------CLLR--------C----------K-------TN-------------------
NK------------------------------RN-------------------S------
-----------------------------------------VVALK--------------
------------------------------------------------------------
------------------------------------------------------------
------------------------------------------------------------
------------------------------------------------------------
------------------------------------------------------------
--------
>Chordat_hENaCalpha_NP_001029_1_amiloridesensitive_sodium_channel_subunit_alp
ha_isoform_1__Homo_sapiens__0_356
------------------------------------------------------------
----------M-------------------------------------------------
------------------------------------------------------------
------------------------------------------------------------
------------------------------------------------------------
------------------------------------------------------------
------------------------------------------------------------
------------------------------------------------------------
------------------------------------------------------------
-------------------------------------------------------EGNKL
EEQDSS-------PPQ--STPGLMKGNKREEQGLGP--------------------EPAA
PQQPTAEEEALI----EFH------------R---------SY-----R-----------
-EL--FEFFC-NNTTIHGAIRLV---------C-SQ-------------HNR---MK---
--TAFWA----VLWLCTFGMMYWQFGLLFG-EYFS----------Y---PVSLNI----N
-L-NS------------D-KLVF-----PAV-----------------------TICTLN
PY------RYP-----EIKEEL----------------EEL-----------------D-
---------RI------------TEQTL--------------FDLYK----Y--------
---------------------------------SS-------------------------
------------------------------------------------------------
------------------------------------------------------------
----------FTTLVAGSRSR------------------------RDLRGTLP-------
------------------------------------------------------------
------------------------------------------------------------
------------------------------------------------------------
-------------------------HPLQRLRVPPPPH---------GARRARSVASSLR
DNNP--------------------------------------------------------
------------QVDWKDWKIG--------------------------------------
------------------------------------------FQLC--------------
--------NQNK------------------------------SDC---------------
--------------------------------------------FYQTY------SSGVD
AV-------------------------REWYRFHYINIL---------------------
------------------------SRL----------------PETLPSLE----EDTLG

--NFI------FACR----FN-Q----V----------S---C-----------N-Q---
----------------------------------------------A-------------
------------------------------------------------------------
------------------------------------------------------------
-----------N----------Y-------S-----------HFH-HPMYG--------N
CYTFN-------DK-------------NN-S-------------------------NL--
-------WMSSMP------GI--------NN-------------GLSLMLRAEQN-----
----------D-----------------F------------I-P-L---LS---------
------------------------------------------------------------
------------------------------------------------------------
----------TVTGARVMVH-GQDEPA--FMDDGGFNLRPGVETSISMRKE------T--
-----LDRL-----------GGD-Y--GD-C--TKN-G------SD------VPV-ENLY
----P-------------S-KYTQQVCIHSCFQESMI----KE--CGCAYIF----Y---
------------P-----------------------------------------------
-------RPQNV-----E---YCDY-----------------------------------
------------------------------------RKHSSWGYCYY---------K---
L-Q-------------------VDFS-S-------DH--------L------------GC
----------FTKCR-KPCSVTS---YQLS-AGY------------------SRW-PS--
-------VTS------QEW-V-----------FQ--------------------------
------------------------------------------------ML-S--------
----------------------------------RQN-----------------------
-N----------------------------------------------------------
---------------------------YTV------------------------------
-----N-------NK-----------R------NGVAKVNIFFKELNYKTNS-ESPSVTM
VTL-----------------LSNLGSQWSLWFGSSVLSVVEMAELVFDL-----------
--------LVIM--------F----------L-------MLL------------------
RRF-----------------------------RS------------R-YWSPGRG-----
--------------------------------------GRGAQEVASTLASS--------
------------------------------------------------PPSHFCPHPM--
------------------------------------------------------------
-----------S--LSLSQP----------------------------------------
-----------------------GPAPSPALTAPPPAYATLGPRPS--------------
---------------------------------------------------PGGSAGASS
STCPLGGP
>Deutero_Ambulac_Sakowv30002567m_893_357
------------------------------------------------------------
----------M-------------------------------------------------
------------------------------------------------------------
------------------------------------------------------------
------------------------------------------------------------
------------------------------------------------------------
-----------------------------------ADISVSPE-----------------
------------------------------------------------------------
------------------------------------------------------------
------------------------------------------------------------
-----------------------VNKQAL-------------------------------
-RGWSAQRK----------------------K---------DI-----RND-----GY--
-KT--WKKFS-QETSLAGVKFVG---------D-DS--------------TIY--SR---
--RIIWL----LIVLAGAAGFVYQVYRMVD-TFAE----------M---PVTVNY----R
-Y-TYPE----------Y-QRSF-----PAI-----------------------TLCNNN
LF------SYV-----SMEKYY--------GI--------L--------------GITE-
---------YL-------------------------------FLLFS-------------
------------------------------------------------------------
------------------------------------------------------------
------------------------------------------------------------
------------------------------------------------------------
------------------------------------------------------------

```
---------------------------F--------------------------------
------------------------------------------------------------
-------------------------IP---------------------------------
------------------------------------------------------------
------------------------------------------------------------
------------------------------------------------------------
------------------------------------------------------------
-------------------------------------------------------SKGLE
---------------------------------EMAGFI----------------P----
-------------LEEV--PENFGDNL---------------TAFEYMDMH----NHDIK
--DML------ISCD----FG-G----I----------Y---C-----------D-A---
----------------------------------------------H-------------
------------------------------------------------------------
------------------------------------------------------------
-----------N----------F-------T-----------STV-T-NGG--------I
CYTFN-------GG-------------QG-N--------------------E----DL--
-------LKLTGT------GL--------TH-------------GLNIVLDANKN-----
----------D-----------------Y------------L-A-----PS---------
------------------------------------------------------------
------------------------------------------------------------
----------DTVGFQFAVH-DQKDIP--NIRDKGIGIPTGMHSRIALSTT------A--
-----ISNL-----------GSP-H--SD-C--VTD-H--------------TRT-LNYF
-----------------PG-EYTESKCLMECEEDFAV----RQ--CECRYYY----M---
------------P-A--YA-----------------------------------------
---F-------------R-----------------TN-----------------------
------------------------------------ANQ-----------------N--F
I-L-------------------ENFTVH-------TS--------------------TIC
------------DCP-ERCNTIS---YNWK-LSY------------------NTY-PTNV
ISIFKANESYTSPLISAQD-LTRLSPSTYWAALE-------WMIDLNYE-----ICFNSL
-------------F---------YILYVQKGN---Y------------SFYDGKETSYHN
Q---------------------------TIALINEKYSSQEKLDQI---------FTLVL
KN-------------NNISL---LKEEFVNV-----SRVFYENYYLQILRQALEVTYGY-
------FWVERNIVIDYLFRYRRHSD-ICE------------------------------
-----K-------FM-----------R------ENFAKVSIFFEDLKFENIT-QKADYEP
FQL-----------------VCDFGGSLGLFFGASLISFLEIFDFLLSW-----------
--------TFAR--------F----------K-------KK-------------------
NQMNPD--------------------------EE--------------------------
------------------------------------------------------------
------------------------------------------------------------
------------------------------------------------------------
------------------------------------------------------------
------------------------------------------------------------
----------------------------------------------------------LS
ETPGTIVK
>Deu_Ambulacrar_hemi_Ptyfla_40v0_9_20150316_1g10161_t1_scaffold5763_cov99_895
_358
------------------------------------------------------------
----------M-------------------------------------------------
------------------------------------------------------------
------------------------------------------------------------
------------------------------------------------------------
-WSSNNTVQ---------------------------------------------------
-----------------------------------------P------------------
-------NGLFKDTTGEKNG-------------TELTDFGSN------------------
------------------------------------------------------------
----------------------LRLTREPGR--------------------------MDH
G----------------------AAGKR--------------------------------
```

--------P----------------------G---------CM-----G-----------
-IL--LSGFA-DDCGISGMKFLA---------G-KS-------------VVP---LR---
--RIVWL----LVLLTGIGLLSYQILDRVS-YFAI----------Y---PLNVNV----Q
-L-NY-V----------E-EVTF-----PAI-----------------------AICNFN
MY------SLQ-----YTNELN------------------L-----------------T-
---------GL------------F------------------YEMYS-------------
------------------------------------------------------------
------------------------------------------------------------
------------------------------------------------------------
------------------------------------------------------------
------------------------------------------------------------
------------------------------------------------------------
------------------------------------------------------------
------------------------------------------------------------
------------------------------------------------------------
------------------------------------------------------------
------------------------------------------------------------
------------------------------------------------------------
------------------------TYNPNFTNYDIELT----------------------
---------------------------------ASEQFY---------------------
------------------------------------------------RDA----AQQPS
--AMF------YRAT----WQ-W----R----------P---M-----------E-I---
----------------------------------------------D-------------
------------------------------------------------------------
------------------------------------------------------------
-----------A----------V-------Q-----------SIL-T-DFG--------A
CYAFN-------TG-------------ND-K--------------------Y----GI--
-------RKVRNK------GT--------SH-------------GLSVYLDVNQW-----
----------D-----------------Y------------N-I-G---PR---------
------------------------------------------------------------
------------------------------------------------------------
----------VGAGFQLMLY-EQGDVP--MVSDLGFSVAPGEDVRVGIEVT------K--
-----VTNL-----------PPP-H--GV-C--AEK--------S-------LQ----YY
------------------D-RYSVNACQLECKTDYVV----ER--CGCRMYY----M---
------------P-G--N------------------------------------------
-----------A-----T---VCDL-----------------------------------
------------------------------------QQF---SFCVQ---------T---
T-L-------------------WYMN-Q------------------------------TC
------------DCP-VPCTQVM---YKPS-LSH------------------AKY-PS--
-------DAY------ATY-L-----------EN--------------------------
------------------------------------------------TY----------
------------------------------------------------------------
-G----------------------------------------------------------
---------------------------FPK------------------------------
-----DV--FSLTYT---NEVGILY-EMYGTNFDNY-DIEITTSEQFYRDAA-QQPSSMF
YTAT--WQWRPMELDAVQSILTDFGACYAFNTGNDKYGIRKVRSKGTSHGLSVNLNVGPR
VGAGFQLMLYEQ---GDVPMISDLGFAPGEDV-------RVGIEVIKVNLPPPHGICGE-
KSL-----------------------------KY--------------------------
-----------------------------------------YNRYS------VNA-----
-------------C---------QMEC---------------------KTDH----VVEH
CG-----MPGNANICDVKHYY---------------------------------------
--------------NCLQYAEFSFTNV-----------------TCDCPVPCT-------
---QVIYKPSLSHAKYPSDAYAMLE---NAYGFP--------------------------
------------------------------------------------------------
---KEVFS
>Protostome_Lophotroco_annelid_CAC9662587_1__Ofus_G12132_Owenia__fusiformis__
896_359

```
------------------------------------------------------------
----------M-------------------------------------------------
------------------------------------------------------------
------------------------------------------------------------
------------------------------------------------------------
------------------------------------------------------------
------------------------------------------------------------
------------------------------------------------------------
------------------------------------------------------------
------------------------------------------------------------
-----------------------SEKNVE-------------------------------
------PHD----------------------A---------SV-----K-----------
-DL--TQGFA-DATSLHGLPRVY---------S-SK--------------SL---TR---
--KIIWG----LVFFGCFVGFCWQVTALTL-NYFE----------Y---PISVTT----E
-V-KT-R----------L-KVDF-----PAV-----------------------TVCNMN
ML------KKS-----LLMGTR------------------F-----------------E-
---------NLTKV------------------------DKRLSDIYG----T-SATS---
---------------------------------NS-------------------------
------------------------------------------------------------
-----------QTT--------P-------PAEMPQDPENED-----PLSGADF------
---------NEGSTTEMD-------------------------TEDDGTSTI--------
------------------------------------------------------------
-------------EAEDTTDGITYV----------TTEDPYDETIDTTQEDIATEF---I
TTEASAGDATTDYDYLTTETSRKKRSVGK---------------RR--------------
---------------PK---ENEHKLPEQLHK----------------------------
------------------------------------------------------------
---------QMKPLDGSAHKHAHPYSNNAKGESR--------------------------
-------------------RRKRQADSYYDDNYEDDGD-------EYDSDP------YDD
YYDNYYDNGDDYYPEERF---------------------EF-EDTG-------F------
----------------------SEIEPNDYFGFLQN-----------SQ------TQDLS
---------------------------------DLVGIV---------------------
-------------------------VP----------------TTDEMDSY----GHTFE
--DFV------LQCS----FD-K----K----------N---C-----------S-N---
----------------------------------------------S-------------
------------------------------------------------------------
------------------------------------------------------------
-----------D----------W-------K-----------KLY-NSQYG--------N
CYTWN-------FG-------------YN-N-------------------------TI--
-------KSTSRF------GS--------RY-------------GLRLTLNAQAD-----
----------E-----------------Y------------I-G-L---LS---------
------------------------------------------------------------
------------------------------------------------------------
----------HTVGARVTVH-SHNVMP--FPEDQGVSASVGRKTGIGVQMQ------N--
-----IKRK-----------PHP-FP-TN-C--TYG--------TH------L---QSNY
----E-------------G-DYSVLSCMMSCLQNKIK----TN--CKCVDKI----V---
------------------------------------------------------------
---------HNQ-----T---ACDIT----------N-----------------------
------------------------------------TTQ---EKCRQ---------R---
M-Y-------------------HLFD-E-------AK--------L------------GC
------------DCP-QACEELV---YGTT-ISG------------------SEW-PS--
-------NQY------SPY-L-----------LQ--------------------------
------------------------------------------------KL-E--------
------------------------------------------------------------
GG----------------------------------------------------------
---------------------------SPV------------------------------
-----S-------DI-----------K------QNLVRVHVYFETLAVHTIE-EVPSYTW
DNL-----------------LADIGGTMGMFIGISICTAFEIVELLMEI-----------
```

--------GKLV--------V----------G-------KM-------------------
TN------------------------------KN-------------------K------
-----------------------------------------VTSLK--------------
------------------------------------------------------------
------------------------------------------------------------
------------------------------------------------------------
------------------------------------------------------------
------------------------------------------------------------
--------
>Placozoa_TadNaC1_XP_002114386_1_hypothetical_protein_TRIADDRAFT_58138__Trichoplax_adhaerens__11_360
------------------------------------------------------------
----------M-------------------------------------------------
------------------------------------------------------------
------------------------------------------------------------
--------------------------------------------------------NEEL
NEGSEKPITSSLYTKNFK-------------------------------------VKTGK
EFKDDLLTPKDQEDYSVFEK-------------IELTTPSSPNAKD--------------
---SKFEIEKEYQYHQEDRSVFKQPIPSLSCIESSKVKRDKVFENVLLT-------P---
----------------------------------------------------GD----DQ
EDYPAVEKVKHPLSLACAESSKVKADKEFEDIILTPY--------------------DDN
DDSS-----------------VFEEANQTTVSASYAEN----------------------
-LHISTKDK------------F-------EFS---------DD-----S-----------
-NY--DDRFA-LQTSCNGIIRIF---------G-RG--------------GR---VR---
--HAVWF----LLTLTMTILCIITCVQRYD-YLFT----------Y---PTNTAI----N
-Y-TV-S----------K-KLKF-----PAV-----------------------SVCNFN
RF------RFS-----SLEYGD------------------W-----------------H-
---------RI------------G-YLV---------------NLFT----TT-------
---------------------------------DN-------------------------
------------------------------------------------------------
------------------------------------------------------------
-----------------N------------------------------------------
------------------------------------------------------------
------------------------------------------------------------
------------------------------------------------------------
--------------------QI-----------FT-------------------------
------------------------------------------------------------
------------------------------------------------------------
------------------------------------------------------------
--GLNGKTGKEW------------------------------NDY---------------
------------------------LKNISFELYD---------------------NITF-
---------------------------------DITQFL---------------------
------------------------------------------------NVK----SNQAE
--VFI------KHCT----WN-D--GRQ----------P---C-----------S-I---
----------------------------------------------E-------------
------------------------------------------------------------
------------------------------------------------------------
-----------N----------F-------T-----------RIY-T-DYG--------S
CFTFN-------AG-------------VD-A-------------------------PI--
-------LYQKRP------GS--------RY-------------GLKLILNIEEE-----
----------E-----------------Y------------T-H-L--------------
------------------------------------------------------------
------------------------------------------------------------
-------NPDPDIGIKFRVH-NQFEPP--DINAEGIAVPPGYHAYTKLLYT------E--
-----SDFL-----------KPP-W--GN-C--GQK--------K-------LK----YF
------------------K-SYSRASCQLECLADSYR----RR--CDCRTPY----M---
------------P-G---------------------------------------------

```
----------SS-----P---ICSP-----------------------------------
------------------------------------EHI---KKCVT---------K---
Y-L-------------------GLST-P-------EN--------------------FTC
------------NCP-NDCRIKS---FNPH-VTY------------------AEI-PL--
-------RQT------SRF-A-----------AH--------------------------
------------------------------------------------RY----------
------------------------------------------------------------
-G---------------L------------------NELEIDFLK---------------
----------------------YNNI-SMG------------------------------
-----E-------YI-----------R------DNYVFLDLFYDDLSYTTFK-EKKAYDV
NQF-----------------ISDIGGQLGLFLGGSFLTFWEIFEWSQIK-----------
--------SFLV--------I----------R-------KIIHEY------------KKG
RR------------------------------RT-------------------R------
------------------------------------------------------------
--------------------RRFNSTP---------------------------------
------------------------------------------------------------
------------------------------------------------------------
------------------------------------------------------------
------------------------------------------------------------
-EDTERLL
>AcFaNaC_XP_012938733_1_PREDICTED__FMRFamideactivated_amiloridesensitive_sodi
um_channel_isoform_X1__Aplysia_californica__10_362
MWGRGKRQRNKYAFRSPAMR-NDNELEGFVSILHTS----------GDNYVPIRDSSADM
KYTSVSAKSGM-------------------------------------------------
------------------------------------------------------------
------------------------------------------------------------
------------------------------------------------------------
--VPEHRYTMV-------------------------------------------------
------------------------------------------------------------
------------------------------------------------------------
------------------------------------------------------------
-----------------------------------------------------R---SRH
H-------------------GRHHHHHSY-------------------------------
---QEYNTQ----------------------R---------SA-----I-----------
-SL--IAELG-SESNAHGLAKIV---------T-SR--------------DT---KR---
--KVIWA----LMVIIGFTAATLQLSLLVR-KYLQ----------F---QVVELS----E
-I-KD-S----------M-PVEY-----PSV-----------------------TICNIE
PI------SLR-----KIRKAYNKNE---------SQN----------------------
-----------------------LKDWL---------------NFTQ----TFH------
------------------------------------------------------------
------------------------------------------------------------
------------------------------------------------------------
------------------------------------------------------------
------------------------------------------------------------
------------------------------------------------------------
------------------------------------------------------------
------------------------------------------------------------
------------------------------------------------------------
------------------------------------------------------------
------------------------------------------------------------
------------------------------------------------------------
---------------------------------------------F-KD------MSFMN
---------------------------------SIRAFY---------------------
------------------------ENL-----------------GSDAKKI----SHDLR
--DLL------IHCR----FN-R----E----------E---C-----------T-T---
----------------------------------------------E-------------
------------------------------------------------------------
```

------------------------------------------------------------
-----------N----------F-------T-----------SSF-DGNYF--------N
CFTFN-------GG--------------Q-L-RD-----------------------Q--
-------LQMHAT------GP--------EN-------------GLSLIISIEKD-----
----------EPLP-----GTYGV----YN-------------F-E-NNIL---------
------------------------------------------------------------
------------------------------------------------------------
----------HSAGVRVVVH-APGSMP--SPVDHGFDIPPGYSSSVGLKAL------L--
-----HTRL-----------SEP-Y--GN-C--TED--------SL------EGI--QTY
--------------------RNTFFACLQLCKQRRLI----RE--CKCKSSA----L---
------------P-D--LS-----------------------------------------
--------VENI-----T---FCGVIP---------DWK-DIRRNV-----TG-------
---------------------------EYKMNQTIPTIS---LACEA---------R---
V-Q-------------------KQLN-N-------DR-------SYE----------TEC
------------GCY-QPCSETS---YLKS-VSL------------------SYW-PL--
-------EFY------QLS-A-----------LE--------------------------
------------------------------------------------RFFSQK------
------------------NPTDQQH---FMKIAQD-------------------------
----------------------------------------FLSRLAHPQQQALARNNSHD
KDI------LTTSYS-L--SEKEMAK-EAS------------------------------
-----D-------LI-----------R------QNLLRLNIYLEDLSVVEYR-QLPAYGL
ADL-----------------FADIGGTLGLWMGISVLTIMELMELIIRL-----------
--------FALI--------F----------N-------AE-------------------
RE-------------------------------------------------VPK------
-----------------------------------------A-PVHSSNNGGG-------
------GGGDG---QHNFANGDVEHER---------------------DTHF--------
--------PDLGSSDF-DFRRG--------------------------------------
------------------------------------------------------------
------------------------------------------------------------
-----------------------------------------------------------G
GIGAESPV
>Deu_Ambulacrar_hemi_Ptyfla_40v0_9_20150316_1g2128_t1_scaffold965_cov108_905_
363
------------------------------------------------------------
----------M-------------------------------------------------
------------------------------------------------------------
-------------------------T----------------------------------
------------------------------------------------------------
------------------------------------------------------------
------------------------------------------------------------
------------------------------------------------------------
------------------------------------------------------------
------------------------------------------------------------
------------------------------------------------------------
-------------------------------N---------SF-----------------
-EE--IDKFS------HISPDIVP----------NQ------------------------
--RLDVI-------VTHVSPSIIMVTLDIHSDYDH-GHTALPSM-YTLFPINITV---DK
---ES-E----------T-EYTFSENLNPCD-GNPC---GEHGLCRDSQSFQYACVCN--
----ECHAGKN----CQIHVND-----PCQ----------FF----------------SP
CR-----NGGL--------CIPSPDSCMEY--------TCECSQCFT-------------
--------------------GKFCEQVPCEPN-DF-------------------------
------------------------------------------------------------
-----------QCN-----------------------------------SGIH-------
-CVKKYLVCDGIYHCP-D-------------------------TSDEFGCDYH--SYE--
------------------------------------------------------------
------------------------------------------------------------
-----------------CSRN------------------------QIKC-----------

```
----------------------ETGGPSGI------------------------------
-----------------------------------------------CI-----------
------------------------------------------------------------
------------------------------------------------------------
------------------------------------------DDSKQCD-----------
------------------------------------------------------------
---------------------------------GVEDCY---------------------
-------------------QGGDEANC---------------DTYGEKNTH-ICTGFQCD
--DGLCHYF--TDCNDATKFR-DC---Q----------PWEDC-GD-------NSDEVAC
D---YR-------ECTESEFKCVSNSQCIAGWKRCNSYSECEDQSDELECTT--------
--------------------PG--------------------------------------
------------------------------------------------------------
-----------NSH--PGREMLFRN----WT-NIYANVTTNQTYFDDFQYH-YFTDDLNG
FVTFS-------ST-------------PDFS------------------DVR----RV--
-------LKLTAD-EIEEYGH-----QAKDF-ILQ---------XLKLTLFVEQD-----
----------E-----------------Y------------L-G-V---FG---------
------------------------------------------------------------
------------------------------------------------------------
----------QSAGVRVTVH-PNNQLP--WPEDVGMTAKTGAATSFAIKQT---------
------------------------------------------------------------
--------------------------CKKRCVQDYMI----RY--CRCTDTF--------
------------------------------------------------------------
--------DSDH-----A---QCPIL----------N-----------------------
------------------------------------ILQ---EACRQI------------
I-H-------------------YFYQ-K-------AL--------------------LAC
------------DCP-PLC-----------------------------------------
------------------------------------------------------------
------------------------------------------------------------
------------------------------------------------------------
------------------------------------------------------------
------------------------------------------------------------
--------------------------K------KNLALVSVYYETLSSDLVK-ESPGYGG
EDL-----------------TSDLGGLLGLYIGVSVITTIECVIFVKGV-----------
--------IVIA--------F----------R-------SM-------------------
RDNNKDS-------------------------RD-------------------------I
------------------------------------------------------------
------------------------------------------------TKDE--------
------------------------------------------------------------
------------------------------------------------------------
------------------------------------------------------------
----------------------------------------------------------ID
ESASDDSR
>Deutero_Ambulac_Spurpu_018442_909_366
------------------------------------------------------------
----------M-------------------------------------------------
------------------------------------------------------------
------------------------------------------------------------
------------------------------------------------------------
-DKSRRPGASYQSLCY--------RAS---------------------------------
------------------------------------------------------------
--------------------------------------------------------PII-
------------------------------------------------------------
--------------------------------------------------------AKEN
S-------------------FFPPEYDDI-------------------------------
----HPMKT----------------------D---------SV-----K-----------
-AV--LNEYS-GVTTAHGVPRII---------T-SK--------------SI---LS---
--KLFWA----CVTLAALGAFLWQGSLLLF-DYKG----------H---PYTTQI----D
```

-V-VT-Q----------T-EVQF-----PAV-----------------------TVCNMN
KM------RRS-----AMVGTR------------------F-----------------E-
---------SLIEA----------------------------------------------
---------------------------------DG-------GAMGGDVDYSWWFDWSSE
------------W-------------------------------WLKY------------
---------EDSSS----------------------------------------------
---------SREESSKEDIGPSAEGSV--------------ISGTSQSVVDMS-------
--------G----DSATDGSSFNTMANFT-------DIPDGNSST---------------
-----------YPSSTPQNQEYSTQ----------SDVDTASATSDGTESFVDE------
------------------------------------------------------------
------------------EWSSTDEVPPLRKRR---------------------------
-----------------------SGLRLI-------P-----DPTTGKLDQRR------F
--------VRRKRQTNTGPSSS--------------------------------------
------------------------------------------------------------
-DPSPSDSSEDPERFDWW----EPEWETN--------MFDY-QAYD-------W------
--------GDV-------------TSDNDWEGFYRQ-----------ST------ADDFS
---------------------------------DLLDVI---------------------
-------------------------NP----------------TREELEIM----GHQAE
--DFI------LQCT----FD-R----R----------P---C-----------N-Y---
----------------------------------------------T-------------
------------------------------------------------------------
------------------------------------------------------------
-----------N----------F-------R-----------QFQ-NKYYG--------N
CFTFN-------QE-------------VG-N-------------------SS----NA--
-------RRTGQT------GA--------QY-------------GLHLTLFTEQP-----
----------E-----------------Y------------V-G-L---FA---------
------------------------------------------------------------
------------------------------------------------------------
----------QEAGVRVAIH-PPNVFP--FPEDDGVVASTGQATNIGIRQS------Y--
-----FERL-----------TKP-H--GN-C--SDG--------TQ----------TNFT
----SEE-----------Y-TYTTRACVKSCVQQHIF----NK--CGCVTDI----M---
------------M-----------------------------------------------
----------ND-----T---ICSPR----------N-----------------------
------------------------------------RTE---QMCRQ---------A---
I-E-------------------QFFH-E-------GQ--------L------------EC
------------FCP-IACNDLNS--LDHV-ITF--------------------------
----------------QTH-L-----------KS--------------------------
------------------------------------------------RL----------
------------------------------------------------------------
TN-------------EKA------------------TRIL--------------------
---------------------------QNI------------------------------
-----E-------QT-----------R------KNLARVRIYFEELNYEQMI-QKPQYTF
ESL-----------------LGGIGGLLGLYIGFSVITICEVGVLVLDI-----------
--------VKYL--------F----------R-------KAYS-----------------
--------------------------------HD-------------------R------
-----------------------------------------VVPVD--------------
------------------------------------------------------------
------------------------------------------------------------
------------------------------------------------------------
------------------------------------------------------------
------------------------------------------------------------
-----FKT
>Protostome_Lophotroco_brch_Lingula_anatina_comp148540_c0_seq4_p1_comp148540_
c0__comp148540_c0_seq4_p1__ORF_type_complete_len_699___score_91_82_comp148540
_c0_seq4_170_2266__907_367
------------------------------------------------------------
----------M-------------------------------------------------

------------------------------------------------------------
------------------------------------------------------------
------------------------------------------------------------
------------------------------------------------------------
------------------------------------------------------------
------------------------------------------------------------
------------------------------------------------------------
-------------------------------------------------------QQDKY
E------------------QEQRTTSQRSLLNSQKNGGTGFDSTSFEGGGLFVRQAQDQL
PSAAEQQDE----------------------T---------SM-----L-----------
-LL--CKNWG-DNSPINGVPNIA---------R-ST--------------SG---VG---
--KVTWT----LLLLMCFGVMGYQTWELVA-KFYR----------F---PVDTTL----T
-Y-TH-S----------K-EVYF-----PAV-----------------------TVCNVN
PL------RRS-----MLSSAG------------------------------------D-
---------DV-------------------------------YALLG----P--------
------------------------P--------KSQM----GSSPGG-------------
------------------------------------------------------------
-------------------AQAPAGP----------------------------------
---------AGSFASQSQTTQAAAG----------------PPGPASSSITTQ-------
APA---------------------------------------------------------
------------------------------------------------------------
--------------------------------------------------------APGP
K---------------GSGSGGTTQAPSASQTTAGAPAG-------GGGG---TTQAPAG
PPAP--------------------------------------------------------
------------------------------------------------------------
-------------------------------------------------------PSGRK
R-RSTGTNSTST------------------------------------------------
-----------------------------------------SRRKPGQM------NERFE
------------------------------MRKRFGNIW---------------------
------------------------AKL----------------NYTTRETV----GHELD
--TMI------ISCT----FN-G----V----------T---C-----------S-S---
----------------------------------------------A-------------
------------------------------------------------------------
------------------------------------------------------------
-----------N----------F-------T-----------RFN-NHQFG--------N
CYTFN-------SG-------------WD-S-------------------SI----PV--
-------ETSSNA------GP--------LY-------------GLSLEMYIEQS-----
----------E-----------------Y------------I-G-D---LS---------
------------------------------------------------------------
------------------------------------------------------------
----------ESAGVRVQIH-SQRSMA--FPEDEGFNIAPGYLTSMAMTRV------E--
-----ITRR-----------PHP-YP-SK-C--RNF--------TTEE----SKQ-KSVF
----TSV--HN-------V-DYSVSGCMKTCYQRYVI----SE--CACGDPA----Y---
------------P-F--SLD-----------------LEAFSDLKN--------------
--NINGTTAESV-----D---PCVS-----------------------------------
------------------------------------DAD---DECVA---------G---
I-K-------------------RRFA-D-------DD--------L------------SC
------------DCP-LTCVDVE---YSGV-PSL------------------AKW-PS--
-------KQY------MST-L-----------HA--------------------------
------------------------------------------------TL-A--------
----------------------------------SA------------------------
-G-------------AHL------------------DNIINGK-----------------
----------------------N----GMA------------------------------
-----A-------EP-----------G------DNLLKLEVYFQELNFQKIS-ENVAYDV
FAL-----------------LADIGGQIGFWVGLSIMALFEVVELILDV-----------
--------FRLL--------L----------F-------RV-------------------
GSFCF---------------------------KK----------------QQKR------

```
---------------------------------------PPSRQVKVQ------------
------------------------------------------------PASEENV-----
------------------------------------------------------------
------------------------------------------------------------
-------------------------------LTP--------------------------
------------------------------------------------------------
RLEKVRFS
>sp_P37089_SCNNA_RAT_Amiloride_sensitive_sodium_channel_subunit_alpha_OS_Ratt
us_norvegicus_OX_10116_GN_Scnn1a_PE_1_SV_2_908_368
------------------------------------------------------------
----------M-------------------------------------------------
------------------------------------------------------------
------------------------------------------------------------
------------------------------------------------------------
------------------------------------------------------------
-----------------------------------LDHTRAPELNIDL------------
------------DLHASNSP----------------------------------------
------------------------------------------------------------
-----------------------------------------KG----------SMKGNQF
KEQDPC-------PPQ--PMQGLGKGDKREEQGLGP--------------------EPSA
PRQPTEEEEALI----EFH------------R---------SY-----R-----------
-EL--FQFFC-NNTTIHGAIRLV---------C-SK-------------HNR---MK---
--TAFWA----VLWLCTFGMMYWQFALLFE-EYLS----------Y---PVSLNI----N
-L-NS------------D-KLVF-----PAV-----------------------TVCTLN
PY------RYT-----EIKEEL----------------EEL-----------------D-
---------RI------------TEQTL--------------FDLYK----Y--------
---------------------------------NS-------------------------
------------------------------------------------------------
------------------------------------------------------------
----------SYTRQAGARRR----------------------SSRDLLGAFP-------
------------------------------------------------------------
------------------------------------------------------------
------------------------------------------------------------
-------------------------HPLQRLRTPPPPY---------SGRTARSGSSSVR
DNNP--------------------------------------------------------
------------QVDRKDWKIG--------------------------------------
------------------------------------------FQLC--------------
--------NQNK------------------------------SDC---------------
--------------------------------------------FYQTY------SSGVD
AV-------------------------REWYRFHYINIL---------------------
------------------------SRL----------------SDTSPALE----EEALG
--NFI------FTCR----FN-Q----A----------P---C-----------N-Q---
----------------------------------------------A-------------
------------------------------------------------------------
------------------------------------------------------------
-----------N----------Y-------S-----------KFH-HPMYG--------N
CYTFN-------DK-------------NN-S-------------------------NL--
-------WMSSMP------GV--------NN-------------GLSLTLRTEQN-----
----------D-----------------F------------I-P-L---LS---------
------------------------------------------------------------
------------------------------------------------------------
----------TVTGARVMVH-GQDEPA--FMDDGGFNLRPGVETSISMRKE------A--
-----LDSL-----------GGN-Y--GD-C--TEN-G------SD------VPV-KNLY
----P-------------S-KYTQQVCIHSCFQENMI----KK--CGCAYIF----Y---
------------P-----------------------------------------------
-------KPKGV-----E---FCDY-----------------------------------
------------------------------------RKQSSWGYCYY---------K---
```

L-Q-------------------GAFS-L-------DS--------L------------GC
----------FSKCR-KPCSVIN---YKLS-AGY------------------SRW-PS--
-------VKS------QDW-I-----------FE--------------------------
------------------------------------------------ML-S--------
----------------------------------LQN-----------------------
-N----------------------------------------------------------
-------------------------YTI--------------------------------
-----N-------NK-----------R------NGVAKLNIFFKELNYKTNS-ESPSVTM
VSL-----------------LSNLGSQWSLWFGSSVLSVVEMAELIFDL-----------
--------LVIT--------L----------L-------MLL------------------
RRF-----------------------------RS------------R-YWSPGRG-----
--------------------------------------ARGAREVASTPASS--------
------------------------------------------------FPSRFCPHPT--
------------------------------------------------------------
----------SPPPSLPQQ-----------------------------------------
----------------------GMTPPLALTAPPPAYATLGPSAP---------------
---------------------------------------------------PLDSAAPDC
SACALAAL
>Protostome_Lophotroco_annelid_CAC9509085_1__Ofus_G024052_Owenia__fusiformis_
_920_371
M--------------AKQVECKKNVIALFKATLHTS-PKITQQSVLTDGYVIIVQNKDD-
-----ENQIAI-------------------------------------------------
------------------------------------------------------------
------------------------------------------------------------
------------------------------------------------------------
--EMKEMIKCD--------KDT--------------------------------------
------FAPSVKQDTKCIG-----------------NGYVKADSSIDS------------
--------------YTLYDL-------------ENVVSVI--TD----------------
----------------------------------VQGD----------------------
------------KEGSINPKVLNEQTDDSS------------------------QNGTQ
T----------------------VKK----------------------------------
------SLK----------------------V---------QI-----R-----------
-SA--LEEFS-NTTSIHGPKRIL---------K-AN--------------GK---YS---
--TVGWT----FVFIGVIILTVTLVTGVVV-KYYS----------Y---PVEDGI----S
-I-TT------------P-PLGF-----PAV-----------------------TLCPLI
PF------DID-----QILVDLENEAIV-------TPDH---------------------
-----------------------VYDLI----------------RYFF----LRSEHFTPD
I----------------------------ECIS-DV-------NC---------------
------------------------------------------------------------
------------------------------------------------------------
-----------------EVE------------------------ADDN------------
------------------------------------------------------------
------------------------------------------------------------
------------------------------------------------------------
------------------------------------------------------------
------------------------------------------------------------
------------------------------------------------------------
------------------------------------------------------------
------------------------------------------------------------
-----------------------------------------ILQSWLEL------SSWLH
RK----------------------------LNPTYRWQV---------------------
------------------------ENIPLN-------------ETIVSSML----PTKTE
--EFI------LECT----YN-E----R----------Q---C-----------D-E---
------------------------------------------------Q-----------
------------------------------------------------------------
------------------------------------------------------------
-----------I----------E-------V-----------TRI-LTAYG--------K

CFTLN-------IK-------------NQ----------------------------A--
-------EHINEI------GP--------NK-------------GITLILYTGKH-----
----------NPIQGIIGAEFSTM----GF-------------T-S-TYSQ---------
------------------------------------------------------------
------------------------------------------------------------
----------PDDGVQVVIH-NPGTMP--QPFREGFHVTPGRFSSIKLTKT------E--
-----RTRL-----------LPP-Y--GE-C--TDE--------NY------LFN--SEF
--------------------KYSYEMCVEQCLQERIM----QE--CGCVSPM----Y---
------------I-I--PR-----------------------------------------
-----NHNFNQT-----Q---YCGNIS---------SVL-------------SDKMLINA
MFF-------RNS-----------NNKLIKALK-GVIDN---LKCEK---------K---
L------------------------------------------------SKIGK---ELC
------------QCK-RPCKENT---NIHL-INT------------------IPW-PE--
-------KSV------VNT-G-----------GQ--------------------------
------------------------------------------------RGFVKFKKK---
------------------HKNVS--------KLIKQLIKYTG------------------
NL-------------HSL------------------NDS-IDDMIRDFLTHI--------
-----------DHVN-MSKPEYNRDS-LAI------------------------------
-----D-------VV-----------E------TNFMKLNIHFESLDVRTVY-EERTYTY
ADV-----------------MKDIGNVFGFYLGMSAVSVAEGVYIIGVM-----------
--------IKLI--------I----------S-------RI-------------------
LC----------------------------------------------------------
------------------------------------------------------------
------------------------------------------------------------
------------------------------------------------------------
------------------------------------------------------------
------------------------------------------------------------
------------------------------------------------------------
--------
>Protostome_Lophotroco_annelid_CAC9657286_1__Ofus_G102358_Owenia__fusiformis_
_922_373
------------------------------------------------------------
----------M------VDKKT--------------------FSSRC-------------
------------------------------------------------------------
------------------------------------------------------------
------------------------------------------------------------
------------------------------------------------------------
-------------------------------------DRTSPQQMITILTYDGDQGKC--
---------------------------AKAYVMTSDDEDFIALR------------PLAY
Q--------------------------------------------------RGAKP----
------------------------------------------------------------
-------------------KFVVKNAEDIDIDK---------------------------
LSDDSDFDD----------------------K---------SL-----K-----------
-DI--AVEFA-NETTLHGVPKII---------K-DR--------------SC---GT---
--RLFWS----CVCLSALCMFLIQASTLMG-KYLD----------Y---KKKVDI----E
-V-LD------------D-PAPF-----PSI-----------------------SLCNIR
HI------DAI-----LFEEIQGYSQ---------QHYMKVL------------------
---------EVMEA-------SKNPNYVR-------------NKTVE----EDKPFYWTS
L--------------------------------ETSI-----------------------
------------------------------------------------------------
------------------------------------------------------------
-----------------NVQG-----------------------------------YL--
------------------------------------------------------------
------------------------------------------------------------
------------------------------------------------------------
-----------------------STKPELVKE----------------------------
------------------------------------------------------------

------------------------------------------------------------
------------------------------------------------------------
-L----------------------------------------------------------
-----------------------------------------GFDNINDL------TEIMF
---------------------------------TRETPA---------------------
------------------------ANI----------------DKDKVTQF----GIEAI
--EFI------ITCV----YS-G----D----------R---C-----------N-Y---
----------------------------------------------S-------------
------------------------------------------------------------
------------------------------------------------------------
-----------D----------F-------K-----------RFF-HPFFY--------N
CFTFN-------TS-------------SF-L-----------------------------
-------TDNSTT------GR-----------------------SFSLIAFLGKM-----
----FSK---E------STKLDRGDT--F----------------G-DPVY---------
------------------------------------------------------------
------------------------------------------------------------
----------NSDGLRIVIH-SSNSEP--DPIQDGFNIPAGFSASIGVKAT------Q--
-----YERI-----------DYP-Y--GN-C--SEQ--------GS------LSMDNGIY
--------------------DYTLISCQNLCLQNEII----EH--CQCIDIA----L---
------------P-I--PE-----------------------------------------
-----NV---NV-----S---FCQDIE---------NPPFDCLE------DKANEWK---
---------------------------CNDAIR-RSWNK---FKCMR---------S---
V-K-------------------GRVS--------ITQ-------LAK----------QKC
------------RCY-PPCHEIT---YGVF-HSL------------------TSW-PS--
-------MDQ------TIP-T-----------ME--------------------------
------------------------------------------------KVIT--G-----
-------------------------------DFMQRFTTDHERNLVWEN------YFPNY
SN------------DTDL------------------------------------------
----------------VKLYLETDNL-TVQ------------------------------
----HD-L----TKVC---------AR----FLKDFSHVYVYIADDNVVKIT-ESEFYSG
VQL-----------------VSDIGGQLGLWVGISVVTLAELLQLCATL-----------
--------CGFL--------C----------K-------NK-------------------
HR------------------------------RKLKK-------------EFER------
---------------------------------------RSMRKKQRGASV---------
------------------RTKRYNHHHHRPRS-------------RSLSNTRQNG-----
------------------------------------------------------------
------------------------------------------------------------
--------------------------------TP--------------------------
-----------------------------------------------------TRNHLLK
NQNHHVNV
>Cyclo_MEC10_NP_509438_1_Degenerin_mec10__Caenorhabditis_elegans__51_375
------------------------------------------------------------
----------M-------------------------------------------------
------------------------------------------------------------
------------------------------------------------------------
-------------NRNP-------------------------------------------
------------------------------------------------------------
-----------------------------------RMSKFQPNPRSRS------------
---RFQDETDLRSLRSFKTDF--------SNYLASDTNFLNVAEI---------------
------------------------------------------------------------
--------------------------------------------------MTSYAYGESN
N----------------------AHEKEIQCDLLTENGG-------------------IE
IDPTRLSYR----------------------E---------RI-----R-----------
-WH--LQQFC-YKTSSHGIPMLG---------Q-AP--------------NS---LY---
--RAAWV----FLLLICAIQFINQAVAVIQ-KYQK----------M---DKITDI----Q
-L-KF-D------------TAPF-----PAI-----------------------TLCNLN
PY------KDS-----VIRSHD--------------------------------------

---------SI-------------------------------SKILG-------------
------------------------------VF-KSVM-----KKAG--------------
------------------------------------------------------------
------------------------------------------------------------
-----------------DSSSEALEE----------------------------------
---------EEETEYDMNGI----------------------------------------
-------TIQAKRKKRGAGE------------------KGTFEP----------------
-----------ANSACECDEE---------------------DG-SNECEERSTE-KP--
-------SGDNDMCICAFDRQTNDAWPCHRKEQWT------------------NTTCQTC
DEH-YLCSKKAKKGTK---RSELKKEPCICESKGLF-----------CIKHEH-------
----------------AAMVLN--------------------------------------
-----------------------------------------LWEYFGDS-----------
--EDFSEISTEE------------------------------------------------
----------------------------------------REALGFGNM------TDEVA
--------------------------IVTKAKENIIFAM---------------------
------------------------SAL----------------SEEQRILM----SQAKH
--NLI------HKCS----FN-G----K----------P---C-----------D-ID--
----------------------------------------------Q-------------
------------------------------------------------------------
------------------------------------------------------------
-----------D----------F-------E-----------LVA-DPTFG--------N
CFVFN-------HD-------------RE---------------------------IF--
-------KSSVRA------GP--------QY-------------GLRVMLFVNAS-----
----------D-----------------Y------------L-P-----TS---------
------------------------------------------------------------
------------------------------------------------------------
----------EAVGIRLTIH-DKDDFP--FPDTFGYSAPTGYISSFGMRMK------K--
-----MSRL-----------PAP-Y--GD-C--VED-G------A-------TS--NYIY
----KGY-------------AYSTEGCYRTCFQELII----DR--CGCSDPR----F---
------------P-S--IG-----------------------------------------
----------GV-----Q---PCQVF----------N-----------------------
------------------------------------KNH---RECLE---------K---
H-T-------------------HQIG-EIH-----GS--------------------FKC
------------RCQ-QPCNQTI---YTTS-YSE------------------AIW-PS--
-------QAL------NIS-L---------------------------------------
------------------------------------------------------------
------------------------------------------------------------
-G----------------------------------------------------------
-------------------QCEKEAE-ECN------------------------------
-----E-------EY-----------K------ENAAMLEVFYEALNFEVLS-ESEAYGI
VKM-----------------MADFGGHLGLWSGVSVMTCCEFVCLAFEL-----------
--------IYMA--------I----------A-------HHINQ----------------
QRI-----------------------------RR--------------------------
------------------------------------------------------------
------------------------------------------------------------
------------------------------------------------------------
------------------------------------------------------------
------------------------------------------------------------
------------------------------------------------------------
RENAANEY
>Deu_Ambulacrar_hemi_Ptyfla_40v0_9_20150316_1g9562_t1_scaffold5200_cov88_926_
376
------------------------------------------------------------
----------M-------------------------------------------------
MLYSRSNLNFSKD-Y----------------------LLA--------------------
---------------------------ELRLSSTLYVDNDD--------------SKRFA
VFESRQDD-RSD------------------NPFPLLSVN----SPYVDDTEKAEQPTDRR

VSDTNEG-----------------------------------------------------
------------------------------------------------------------
---------------------------------TKELS----------------------
------------------------------------------------------------
-----------------------DKSNV-------------------------QHEFFKL
A----------------------DKTTE--------------------------------
------EHD----------------------E---------SV-----K-----------
-HV--LEDFG-RETTAHGIVHIT---------N-AT--------------SS---VT---
--RSTWI----IIVLVAACAMLVQMTLLLI-QYFE----------Y---NVHVKV----T
-L-VS-E----------K-ALGF-----PSV-----------------------TICNTN
KL------RHS-----AIRSSK------------------Y-----------------S-
---------EML---------MLERNFVP--------------PYYT-------------
---------------------------PCIEG-DF-------------------------
------------------------------------------------------------
-----------TCR-----------------------------------NGIH-------
-CIRPFLVCDGVNQCG-D-M------------------------SDEYDCI-----YH-S
K--YWSL-----------------------------------------------------
-------AFKQKAS--QSSTFAY---------------GSAMKAVDGDKSNNYNER----
----------------TCTHT------------------------QK-------------
-----------------------EYQP---------------------------------
------------------------------------------------------------
------------------------------------------------------------
-----------------------------------------WW-----------------
----KVDLGGEYEIDKVI---ITNRADCC-------------DDRLS-------------
--------GAVVR---VG-NDDVIENNERCGQ-------------------------TVT
KGDI-----------------------------NQDG-----------------------
-----------------------EVTV----------------EC---VLQ----GRFVS
--VQL------ENKT----DY-L----R----------L---CE----------V-QVFG
EAF---ISSDNQTECPDDYIRC-TSGECVHPYKICDVIVDCADGFDEMECVNSES-NANG
AQ-------IDHFQNCSEEE--RQPH--------------KPKLCPTD------ITSDDK
G----------QAKKVA-EQLANITSEADD------------FEENDVTLSVDVIDNILE
SGVNGTEIEVAE----------D-------V-----------LMS-VDN-----------
------------LL-------------HV-D--------------------H----EV--
-------LVASQQ------NE--------NT-------------ASRLIESVEKL-----
----------S-----------------L------------I-V-Q---F----------
------------------------------------------------------------
------------------------------------------------------------
--------------INDSMDSGNQSVT--IETE------------------------N--
-----IVLILAEINFESFAGLNF-W--FS-S--GKA----------------V-------
------------------------------------------------------------
------------------------------------------------------------
------------------------------------------------------------
------------------------------------------------------------
------------------------------------------------------------
------------------------------------------------------------
------------------------------------------------------------
------------------------------------------------------------
------------------------------------------------------------
------------------------------------------------------------
------------------------------------------------------------
-----------------------------------------DINGTN-------------
AGF-----------------SPV----IAASIRDLRI-----------------------
-------------------------------TNL-----RDPVRIQ------------IP
RNN-----------------------------Q---------------------------
------------------------------------------------------------
-------------------------N------------------------IY--------
--------SNDIDETH-QKSLS----IISYI-GCG---------------ISLIA-----

```
--------------SAITLVSVIVYRRY-------------------KSVPRVGLC----
------------------------------------------------------------
-------RQKG----KDF-----------------LTD----------------------
-------C
>Protostome_Lophotroco_annelid_CAC9663096_1__Ofus_G121090_Owenia__fusiformis_
_927_377
M--------------APPCELSDDEEIVFKSGVHQ-------------------QQSNH-
-----EYQIEI-------------------------------------------------
------------------------------------------------------------
------------------------------------------------------------
------------------------------------------------------------
--EMKEKNELK--------NDS--------------------------------------
------SVPSIEQEIACQN-----------------DECIEPELIMNRC-----------
-------------IRDSKEI-------------EDIVPQK--LV----------------
----------------------------------IQGD----------------------
------------TCESTNVNLLAEQREDCY-------------------------Q----
-----------------------YTQ----------------------------------
------SFK----------------------A---------QI-----S-----------
-ST--LAGFS-NTTSIHGPKRIL---------K-AR--------------GR---TS---
--RAGWT----FVFIGVVSLAVYLIAGVVV-KYYS----------Y---PIEDGI----S
-I-KT------------Q-SLAF-----PSV-----------------------TLCPLI
PN------DDF-----LVMSDL--EAYW-------QSNQ---------------------
-----------------------GDKYL---------------LYTL----KAYKYLIGK
D---------------------------RLVS-DI-------RT----------------
------------------------------------------------------------
------------------------------------------------------------
-----------------DFP------------------------KNESSF----------
------------------------------------------------------------
------------------------------------------------------------
------------------------------------------------------------
------------------------------------------------------------
------------------------------------------------------------
------------------------------------------------------------
------------------------------------------------------------
-----------S------------------------------N-----------------
---------------------------------------A-LIQSWLDL------SSWLS
EK----------------------------LSPVSRWQV---------------------
------------------------ENFPLNININVSEIGPSSDDINVAGMD----PVKEQ
--DFI------LECV----YN-D----K----------P---C-----------D-E---
----------------------------------------------Q-------------
------------------------------------------------------------
------------------------------------------------------------
-----------F----------N-------V-----------TKV-TTAYG--------Q
CFTLN-------IS-------------ES----------------------------A--
-------EDVNEI------GP--------NK-------------GLTLILYTGRY-----
----------NPIPIRPNFYMPTM----GF-------------S-S-TYAQ---------
------------------------------------------------------------
------------------------------------------------------------
----------PSDGVQIVVH-NPGTMP--QPYREGFHVTPGRLTSVKITKT------E--
-----RTRL-----------VPP-Y--GD-C--TDK--------DY------LVN--SEF
--------------------RYSYEMCVEQCLQERII----QK--CGCVSPM----Y---
------------I-I--PA-----------------------------------------
-----DHNFSLI-----Q---YCGNIS---------KIF--GFSE------NKNCIHSNS
WMT-------RGSMCLY--GTPQLRLRMINLAN-DIIDR---LKCEQ---------Y---
W------------------------------------------------SKSSI---QSC
------------KCR-RPCKENT---YIHL-INT------------------IPW-PE--
-------KSV------VSN-D-----------GQ--------------------------
```

------------------------------------------------RGFLKFRQK---
------------------HKNID--------KTFKLFERELR------------------
LY-------------TYL------------------WDH-QFEVFEDCFDH---------
------------NIT-DGHKQYNINS-LIE------------------------------
-----N-------IV-----------D------TNFLKLNIHIESLDVRTVY-EERTYTY
DNV-----------------MKDIGNVFGFYLGMSAVSVVEGVYIIGVL-----------
--------IKLI--------I----------Y-------RM-------------------
MC------------------------------RR--------------------------
------------------------------------------------------------
------------------------------------------------------------
------------------------------------------------------------
------------------------------------------------------------
------------------------------------------------------------
------------------------------------------------------------
-------N
>Protost_ecdy_cyclo_nema__A0A0V1BKZ2_A0A0V1BKZ2_TRISP_Degenerin_mec_10_OS_Trichinella_spiralis_OX_6334_GN_mec_10_PE_3_SV_1_932_380
------------------------------------------------------------
----------M-------------------------------------------------
------------------------------------------------------------
------------------------------------------------------------
------------------------------------------------------------
------------------------------------------------------------
------------------------------------------------------------
------------------------------------------------------------
------------------------------------------------------------
--------------------------------------------------------SNRV
T-----------------------------------------------------------
-FQRDTTTK----------------------R---------RF-----K-----------
-NL--LLNFS-DKTTAHGVPRIG---------M-AT--------------TK---RE---
--GAFWT----FVCVTSLTVFLFQLSLLIA-RFLR----------F---ETTTQV----E
-L-KF-E------------TSLF-----PVV-----------------------TLCNLN
PY------RNS-----LLKQVD--------------------------------------
---------QM-------------------------------KLMFD-------------
------------------------------AF-DN-------------------------
------------------------------------------------------------
------------------------------------------------------------
-----------------------------------------KPLSTPKTSTL--------
------------------------------------------------------------
--------LARKRRASSADE-----------------KQYRYTP----------------
-----------LYARCSCPNE------------------------ESDCVPFTSA--P--
------LEDGEEICICYFDHVTKSPWPCYSEKQWR------------------SEWCSRC
SPN-GNCLQIILKKRQLSLY---EDVKQCVCHSNGKT----------CLVADKNKPERIY
RDVRLDR-YTTSTTTTTTTTTT--------------------------------------
-----------------------------------------TTTTTTTTTTTSTAT---T
TTTPIPTTPSEV------------------------------QEYV--------------
-----------------------------------------DALGFEGI------TDETA
--------------------------KVSRAQENTLFIM---------------------
------------------------AEL----------------PPETKKKV----SHQKK
--DFL------LQCS----FN-G----R----------Q---C-----------D-IE--
----------------------------------------------N-------------
------------------------------------------------------------
------------------------------------------------------------
-----------D----------F-------E-----------LLM-DPVYG--------N
CYMFN-------SN-------------STIN-----------------------------
-------RTTSRA------GP--------IY-------------GLSLVIYVDAD-----
----------D-----------------Y------------L-P-----TT---------

------------------------------------------------------------
------------------------------------------------------------
----------KASGVRLLVH-SQEEYP--FPDTNGYNAPTGLLSSFGIRMK------K--
-----IERL-----------PQP-Y--GD-C--IKE-G------K-------TK--DYIY
----KDQ-------------IYSLEGCYRSCFQMEMI----KS--CGCGDPR----F---
------------P-V--PE-----------------------------------------
----------GR-----R---HCRVK----------E-----------------------
------------------------------------RKA---RECLE---------N---
V-I-------------------QKSG-GLH-----GS--------FS----------GKC
------------DCR-QPCVQYV---YDMS-FSA------------------AKW-PT--
-------PSIE---------L---------------------------------------
------------------------------------------------------------
------------------------------------------------------------
-D----------------------------------------------------------
-------------------DCNDTAE-ECI------------------------------
-----R-------KL-----------K------KNALSLEIYFEQLNYEVLK-ELQAYQW
VNL-----------------MADFGGQLGLWMGVSVITIIEVLVLIYEV-----------
--------IKIC--------C----------T-------RR-------------------
KKPTMN--------------------------KH-----------------HRRSI----
---------------------------------------YDDSTVEMTIK----------
----------------IHPPNQNNPKDSIA---------------TAHSENDDE------
--------CSQNSYAMENHLHR--------------------------------------
------------------------------------------------------------
----------------------------HDAEIP--------------------------
----------------------------------------------------NSVITFCA
HCSLNIKQ
>Deutero_Ambulac_Sakowv30041168m_935_382
------------------------------------------------------------
----------MSVS----------------------------------------------
------------------------------------------------------------
------------------------------------------------------------
---------------------------------------------------RRKILLTSD
VEEESESECHGRSFED---GEIRLVDMSL-------------------------------
--LNGALEKEICQFCKVGNISIVERHGLGSCIRSKLSQPSFPSSRKKGAVFLI-------
---------------NKKAA----------------------------------------
---------------------------------------------------IGMRAI---
----------------------------------------GKGTV---------RSGETV
KLLGV---------------TSTSNGSALSPEYPTNDF------YLAGME---------N
VSLRRESKQ----------------------V---------AL-----R-----------
-KL--FGRMV-NNSTAHGIPNAA---------R-AE--------------SL---PR---
--RLFWS----VLFMVAVGMFLWQFSDQLV-RFID----------R---PVNVKL----N
-I-SF-S----------R-ELTF-----PAV-----------------------TVCNQN
PI------RQS-----AIDFNQ------------------------------------EP
QSSPP--TQNL-------------------------------NQTLN-------------
---------------------------------DTALDE---------------------
------------------------------------------------------------
------------------------------------------------------------
-----------------DLGN---------------------------------------
------------------------------------------------------------
------------------------------------------------------------
------------------------------------------------------------
----------------HGGTNNNTNSPPYNTRRR--------------------------
------------------------------------------------------------
------------------------------------------------------------
------------------------------------------------------------
--RQTAQTSGDQSG----------------------------QESKNDP-----------
----------------------------------------PHPDGDNGP------SAGNR

```
------------------------------DMIRITEIL---------------------
------------------------TAL----------------PTSERITL----GHQAD
--MFI------VNCT----WQ-G----E----------Q---C-----------Y-A---
----------------------------------------------S-------------
------------------------------------------------------------
------------------------------------------------------------
-----------N----------F-------T-----------TFS-NSMYG--------N
CFTIN-------GP-------------SG-Y-------------------ID----PY--
-------WTTSFS------GP--------IY-------------GLSLQLFIEQD-----
----------E-----------------Y------------I-K-G---YT---------
------------------------------------------------------------
------------------------------------------------------------
----------AIAGARIVIH-DQDKMP--FPEDNGFTIAPGAATSIGIRKV------L--
-----ISRE-----------PDP-Y--SD-C--VEE--------GD------SDY-TNIY
----MDS--YD-------V-GYSVQACMKACYQSEVI----NN--CGCADPL----Y---
------------P-F--PY-------------------GVY-------------------
-----------------E---ACKTS----------N-----------------------
------------------------------------STQ---MYCKA---------L---
V-E-------------------YDYS-T-------DG--------L------------EC
------------DCP-QACSDVG---FTTE-TSS------------------AMW-PA--
-------NSY------KET-L-----------LN--------------------------
------------------------------------------------YL-K--------
----------------------------------GS------------------------
-N-------------KKL------------------ESLVG-------------------
----------------------DDLK-NGT------------------------------
-----N-------SV-----------S------ENLLRVDIYYQELNYEVIE-QEPAYLF
SDL-----------------GSDFGGLIGLWIGVSILTCFEFLELIFDF-----------
--------CHVF--------C----------S-------KG-------------------
ALAVN---------------------------RT--------------------------
---------------------------------------TDISAVSGGDIF---------
------------------FRDKYGNDPALTL--------------NDRSSNKPRMNI---
------------------------------------------------------------
----------------------------------------------------NN------
----------------------------RTTYVPETNAQSLYI-----------------
----------------------------------------------------PQYSMKIY
EPGLLNTS
>Deu_Ambulacrar_hemi_Ptyfla_40v0_9_20150316_1g7150_t1_scaffold3457_cov142_936
_383
------------------------------------------------------------
----------M-------------------------------------------------
------------------------------------------------------------
---------------------------EL-------------------------------
------------------------------------------------------------
------------------------------------------------------------
------------------------------------------------------------
------------------------------------------------------------
------------------------------------------------------------
---------------------------------------------------------DEL
RYDPP------------------KRPSSPPAG----------------------------
-AVDRDVGD----------------------T---------GF----L------------
-AL--LYQFG-DTVSAAGLPRVF---------T-ER-------------SSL---WS---
--RLIYA----IVFVASFGTGVYFSVKIVQ-EFLE----------Y---PVIVTT----A
-F-TA-E----------S-RMDF-----PSV-----------------------TICNTN
RV------RYS-----AIIESK------------------H-----------------S-
---------DL-------------------------------EYLLT-------------
---------------------------------DR-------SLGGLYY-----------
------------------------------------------------------------
```

----------------------PPC----LEDDMLCDTN---------------------
ECLRRFKVCDGIRDC--D------G------------------GEDEVNCY-----YGDC
G--------YRQHRCETGGDHGYCINHNDYVCNGEQDCYDNSDESDCDNV-------FFG
LNSYVDDGDCLGFRCTNSSDCISQT----------EVCDGTYDCSDGSDELNCTSA--F-
---------TCMTGMVKCATID-------------------------YCI----------
--YPSWVCDNYADCSDRSDEAPLATCPSSSAARKR--------------RSSSSGVSPTS
SSGCYWCSAK--------YGSDNTTLQCF-------PLWYECDGYEDCVGGT--------
---------DESNCDNINYHCPDNFFACTDGE-------CIHEQFECDGWDDCASGSDED
NC--------------------------------------EEPTRSSNSSTISDL-FQDG
WYSKYLHITDEEWFYEDF----EDNYL---------------QEKYNVG----F------
--------ERV-----------SSEDPPDWHGFATF-----------SA------TGDYT
---------------------------------DLEDVV---------------------
-------------------------KL----------------TKAEMADL----GHQNE
--DFI------LQCT----FD-Q----T----------K---C-----------H-Y---
----------------------------------------------E-------------
------------------------------------------------------------
------------------------------------------------------------
-----------D----------F-------E-----------VIQ-DDKYG--------N
CFTWN-------HH-------------SR-N-------------------------MS--
-------LRSTGP------GS--------TH-------------GLKLTLFTEQS-----
----------E-----------------Y------------I-S-I---YG---------
------------------------------------------------------------
------------------------------------------------------------
----------QDSGVRVSIH-EMGTTP--FPEDDGFTVAPGKATGVGLKQL------I--
-----INRE-----------DPP-H--GE-C--SNS--------TSF---------ESMY
----GD--------------GYS-------------------------------------
------------------------------------------------------------
------------------------------------------------------------
------------------------------------------------------------
------------------------------------------------------------
-------------------------------VAY-----------------H--------
--------------------L-----------LK--------------------------
------------------------------------------------TI----------
--------------------------------H-SI------------------------
-N-------------DKT------------------KDI---------------------
---------------------------NDF------------------------------
-----E-------QA-----------Q------SNLVRLEVYFEEMTYETMS-ETPKYEV
YNL-----------------LAEIGGVIGIYVGLSFITVVEFVEFFVAV-----------
--------VRYL--------F----------R-------RI-------------------
KR------------------------------SM-------------------K------
-----------------------------------------VEALK--------------
------------------------------------------------------------
------------------------------------------------------------
------------------------------------------------------------
------------------------------------------------------------
------------------------------------------------------------
--------
>Deu_Ambulacrar_hemi_Ptyfla_40v0_9_20150316_1g8260_t1_scaffold4268_cov134_944
_386
------------------------------------------------------------
----------M-------------------------------------------------
------------------------------------------------------------
------------------------------------------------------------
------------------------------------------------------------
--EMQAQNGTT-------------------------------------------------
------------------------------------------------------------
------------------------------------------------------------

```
------------------------------------------------------------
---------------------------------------------------------ERN
LRMA-----------------SMEASEDP-------------------------------
-------DQ----------------------K---------NC-----S-----------
-IL--VTNFG-QSTTCHGLQQIL---------I-GD--------------SP---FR---
--KLIWL----AVFLTAMGVFTFQAYELVS-LFLT----------Y---DVRINM----E
-V-GT-A----------S-SLPF-----PAV-----------------------TICNTN
KL------RLS-----EIERSV------------------H-----------------H-
---------ELTKT---------DPEHPD-----------SIHRQLS-------------
------------------------------------------------------------
------------Y-----------------------------------------------
---------ETPCG----------------SEEFNCGQ-----------AGFHG------
ICISNEKRCDGVRDCF-D-------------------------GRDEMNCTVC---YG--
---------GRQCDVGLDSHASKCIPED-NICDRFPDCIDGADEESCVLTDQCEMGVLFA
RSDPSVLYSRNYPKKYDTDYTCELR----------LTAETGFDGLQSSDCISLAFL----
-ELDIERGKNCQRDYIECSKTCGGGVR-K---------------RNVTCSVIESD-DPSM
D-FADMSLDSGENFD----------NPLV------EGSNGKMVVAR---GTRGHYDSPDT
--------------------------------------------------GDY-------
---------FADEGDVSRRCTSAGSSPERV------------------------------
---------------------------------------ESCNNTPCSGGP-GGIQILDK
LFEKYGHISDDKKIYENF----ELNY--------------Y-KSNN-------F------
--------DRV-----------QTFPDPNWKGEFSFV---------KTT------SPDYS
---------------------------------DLQKVL---------------------
-------------------------HL----------------DRDEVARF----GHQAE
--DFI------LQCS----FD-E----V----------Y---C-----------D-H---
----------------------------------------------S-------------
------------------------------------------------------------
------------------------------------------------------------
-----------N----------F-------Y-----------RFE-NDVYG--------N
CFTFN-------SL------------QQN-N--------------------Q----TT--
-------VTSSRP------GS--------RY-------------GLKLTLFIEQD-----
----------E-----------------Y------------I-P-L---YG---------
------------------------------------------------------------
------------------------------------------------------------
----------QEAGVRVLIH-PQDITP--FPEDEAITVAPGLKTSIGIRMD------T--
-----IKSL-----------SEP-Y--TN-C--SND--------ED------F---ESVY
----GEG-----------Y-KYS-------------------------------------
------------------------------------------------------------
------------------------------------------------------------
------------------------------------------------------------
------------------------DM-G-------GV--------I------------K-
-------------VL-RPINEKV---RNIV-VDE------------------QSA-RI--
-------YIL------QGG-V-----------IK--------------------------
------------------------------------------------VL-R--------
----------------------------------PI------------------------
-N-------------EKV------------------RNIV--------------------
---------------------------VDE------------------------------
-----Q-------SA-----------R------ENLVRLAVYYEELNYQKIT-KLPAYTI
EEL-----------------LADLGGILGLYIGMSLITAFEVLELF--------L-----
------NALRHL--------A----------K-------KF-------------------
CKP-----------------------------KP-------------------E------
----------------------------------------DDGPTYL-------------
------------------------------------------------------------
------------------------------------------------------------
------------------------------------------------------------
------------------------------------------------------------
------------------------------------------------------------
```

```
--------
>Cyclo_MEC4_NP_510712_2_Degenerin_mec4__Caenorhabditis_elegans__52_388
------------------------------------------------------------
----------M-------------------------------------------------
---------------------------------SW-------------------------
------------------------------------------------------------
---------------------------------------------------MQNLKNYQH
LRDPSEYMSQVYG-----------------------------------------------
----------------------------------------DP------------------
-----------------------------LAYLQETTKFVTEREY---------------
------------------------------------------------------------
--------------------------------------------------YEDFGYGECF
N----------------------STESEVQCELITG-----------------------E
FDPKLLPYD----------------------K---------RL-----A-----------
-WH--FKEFC-YKTSAHGIPMIG---------E-AP--------------NV---YY---
--RAVWV----VLFLGCMIMLYLNAQSVLD-KYNR----------N---EKIVDI----Q
-L-KF-D------------TAPF-----PAI-----------------------TLCNLN
PY------KAS-----LATSVD--------------------------------------
---------LV-------------------------------KRTLS-------------
------------------------------AF-DGAM-----GKAGGNK-----------
------------------------------------------------------------
------------------------------------------------------------
-----------------DHEEER-------------------------------------
---------EVVTEPPTTPA----------------------------------------
-------PTTKPARRRGKRD----------------LSGAFFEP----------------
-----------GFARCLCGSQGSSEQEDELLETTTKKVFEEWDGMMEECQSERTKDEP--
-------TGFDDRCICAFDRSTHDAWPCFLNGTWE------------------TTECDTC
NEH-AFCTKDNK-------TAKGHRSPCICA-PSRF-----------CVAYNG-------
----------------KTPPIE--------------------------------------
-----------------------------------------IWTYLQGGTPT--------
-----------------------------------------EDPN--------F------
----------------------------------------LEAMGFQGM------TDEVA
--------------------------IVTKAKENIMFAM---------------------
------------------------ATL----------------SMQDRERL----STTKR
--ELV------HKCS----FN-G----K----------A---C-----------D-IE--
----------------------------------------------A-------------
------------------------------------------------------------
------------------------------------------------------------
-----------D----------F-------L-----------THI-DPAFG--------S
CFTFN--------HN------------RTVN-----------------------------
-------LTSIRA------GP--------MY-------------GLRMLVYVNAS-----
----------D-----------------Y------------M-P-----TT---------
------------------------------------------------------------
------------------------------------------------------------
----------EATGVRLTIH-DKEDFP--FPDTFGYSAPTGYVSSFGLRLR------K--
-----MSRL-----------PAP-Y--GD-C--VPD-G------K-------TS--DYIY
----SNY-------------EYSVEGCYRSCFQQLVL----KE--CRCGDPR----F---
------------P-V--PE-----------------------------------------
----------NA-----R---HCDAA----------D-----------------------
------------------------------------PIA---RKCLD---------A---
R-M-------------------NDLG-GLH-----GS--------------------FRC
------------RCQ-QPCRQSI---YSVT-YSP------------------AKW-PS--
-------LSL------QIQ-L---------------------------------------
------------------------------------------------------------
------------------------------------------------------------
-G----------------------------------------------------------
-------------------SCNGTAV-ECN------------------------------
```

-----K-------HY-----------K------ENGAMVEVFYEQLNFEMLT-ESEAYGF
VNL-----------------LADFGGQLGLWCGISFLTCCEFVFLFLET-----------
--------AYMS--------A----------E-------HNYSLY---------------
KK------------------------------KK-----------------AEK------
------------------------------------------------------------
------------------------------------------------------------
------------------------------------------------------------
------------------------------------------------------------
------------------------------------------------------------
-----------------------------------------------------------A
KKIASGSF
>Protostome_Lophotroco_annelid_CAC9521433_1__Ofus_G03942_Owenia__fusiformis__
951_390
M----------I------------------------------------------------
----------M-------------------------------------------------
------------------------------------------------------------
------------------------------------------------------------
------------------------------------------------DHPMINTGTMAD
GENVTKTVKAD--------VEINADNT---------------------------------
------------------------------------------------------------
-NIEGTENVAQTGKVDDEKN-------------TDDVNRADIDRVTQAVNV---------
----------------------------------DESNDIE----------MGTFK----
-----------------RIGNTGVVIELNQNV----------GIV-----PVDEPNIERA
TEDMI--------------TEHTQNNEHIVNERMAIDELI----------------SDTP
IVTSQTQHHDIIADALEFKY-----------N---------SL-----K-----------
-NQ--ISKFA-LITSIQSLLQVR---------N-AK--------------SV---LG---
--RLIWF----AITLFVCSLAIIGVHEITR-KYLD----------H---PSEDIL-----
-Y-SR-E----------E-LVTF-----PSV-----------------------TICGVK
PVVFTDEHLKK-----KLSSRA-------------NTGDTINTWIYH-----------D-
------YRLKMLE-----------------------------SRAFS-------------
---------------------------------DEREK----------------------
------------------------------------------------------------
------------------------------------------------------------
-----------------D------------------------------------------
------------------------------------------------------------
------------------------------------------------------------
------------------------------------------------------------
------------------------------------------------------------
------------------------------------------------------------
------------------------------------------------------------
------------------------------------------------------------
------------------------------------------------------------
------------------------------------------HERINAV------MRRIV
---------------------------------SEERYI---------------------
------------------------ENL------------------NEWRDYR---LRMYD
--DLI------LECR----FK-G----K----------P---C-----------G-K---
----------------------------------------------T-------------
------------------------------------------------------------
------------------------------------------------------------
-----------D----------F-------K-----------QVM-LGRYG--------N
CFTFQ-------PN-------------SK-D-------------------------DT--
-------NILSEP------GP--------DF-------------GLNLMLFTNSF--SPL
D---------E------FTDSDTF---NY-----IKL----IGSSDSAILNEFNS-----
------------------------------------------------------------
------------------------------------------------------------
------DFSTASDGIRLSIH-APGTMP--DPEKDGIDLSTGTFSSIGLSQD------V--
-----RTLL-----------GPP-H--GN-C--TEE--------R-------LNT-HTTY

```
--------------------TYTQNMCLRQCQQIETI----KR--CKCMNPK----L---
------------P-L--PP-----------------------------------------
----LTEKPHKI-----H---YCGYEM----R-------QEYLKE------KEREMTYGN
-------VFPLENVQF-----------DWKIFI-ENFKL---YECEK---------E---
V-T-------------------ELFS-R-------EG--------------------EKC
------------GCK-RECKSTA---YYYL-KHS------------------IPW-PN--
-------ERY------YKG-L-----------KH--------------------------
------------------------RNFI--------------------SFLRNH------
-------------------PKADQ-------FLNKTY------------------WSGNR
LNRLGI-----CMSGDNI------------------FSSYSSAVVKGAYPY---------
----------------LTCPCNIANS-TDQ------------------PDRCAPIINW--
YRFINK-------FV-----------E------TNFLKVNVFYSTLNVNRIA-AEPSYPF
NKW-----------------LSDFGGIIGMNMGMSVVSLVELGYLMVSL-----------
--------AFIL--------Y----------C-------KH-------------------
KR------------------------------DN--------------------------
------------------------------------------------------------
------------------------------------------------------------
------------------------------------------------------------
------------------------------------------------------------
------------------------------------------------------------
------------------------------------------------------------
--------
>Protostome_Lophotroco_annelid_CAC9668492_1__Ofus_G161027_Owenia__fusiformis_
_952_391
------------------------------------------------------------
----------M---------------------------------NTC-------------
------------------------------------------------------------
----------------------------AKIEEQIKVSVIG-------------------
--------------------------------------------YA-----EEHVLNESH
EAETSGDTKRD--------TNLEVDCVS-H------------------------------
---DELELSIEKTNDGDDNYGHINKETSEQCLNIDCDARDEPQVRQENKSCNI-------
-----SKRKAMCSIEDDHDDPIVD-----EAQTAKENENYDVCK----------------
------------------------------------------------------------
-------------------GTTSYKTDTEDD--------------------NEDAIINEA
S----------------------SDKNNDHNT---NDDIE----------------DPFD
---SWIWKY----------------------N---------QL-----K-----------
-ER--LSYMA-SITSIQGCINIR---------N-SQ--------------TM---FG---
--RIIWF----SIALALGFTTVIKVSNIVV-QYLS----------Y---PSADAL-----
-Y-SR-E----------E-DVTF-----PSI-----------------------TICGAR
PIVFSENFHRS-----CMNSNC------------MPKTMDVQ----------------DG
---------RL------------NQGIL---------------LLMN-------------
------------------------------------------------------------
------------------------------------------------------------
------------------------------------------------------------
------------------------------------------------------------
------------------------------------------------------------
------------------------------------------------------------
------------------------------------------------------------
------------------------------------------------------------
------------------------------------------------------------
------------------------------------------------------------
------------------------------------------------------------
-------------------------------------------ESGYES-----------
----------------------------------------QT-ETLDFL------RKRLL
---------------------------------SPERYV---------------------
------------------------ENV----------------DDFPYSRLR---YLKKN
--DLI------LACT----FQ-G----K----------E---CP----------D-A---
```

```
----------------------------------------------L-------------
------------------------------------------------------------
------------------------------------------------------------
-----------K----------F-------K-----------EVK-FGRYG--------N
CVTID-------SN----------------------------------------------
-------NTMKES------GP--------DS-------------GLSLILFSNSY--SPL
D---------E------FQKAMNL-FI-Y------------LVS-G---VSEFDS-----
------------------------------------------------------------
------------------------------------------------------------
------THATSSDGIRLAIH-APGTIA--EPEHDGIDLSPGTFNTIGLTQD------V--
-----RTLL-----------EPP-Y--GD-C--TKK-I--------------FNE-KTTY
--------------------KYTKDLCLDKCMQEEMV----AK--CGCMTPK----A---
------------Q-A--SL-----------------------------------------
-----GKNGYDV-----S---YCGRYY---------NTI---------------------
--------KPKGYPQYL--KKPNWRRRLRSQLL-QFLRR---LKCES---------D---
V-L-------------------RLYS-R-------DS-------------------SNMC
----------TSKCK-TECTKIV---YDPL-THA------------------TPW-PN--
-------EKY------YKQ-L-----------NQ--------------------------
------------------------RPFI--------------------TFMKSK------
-------------------DNVAKLYFDVRRSVLDSYCELLH--------------GDLR
KNLLK------RRVTGHV------------------FKPLFADVNFPFVTC---------
------------------------SENITA-----------------H-------INR--
YNIINK-------VI-----------E------KNFLKVNIFFRSLNVNRIA-AEPNYPL
NKF-----------------LSEFGGIIGMYLGMSAVSLVELGYIMVAM-----------
--------VFIL--------L----------F-------KY-------------------
KR----------------------------------------------------------
------------------------------------------------------------
------------------------------------------------------------
------------------------------------------------------------
------------------------------------------------------------
------------------------------------------------------------
------------------------------------------------------------
--------
>Cyclo_UNC8_NP_501138_1_Degenerin_unc8__Caenorhabditis_elegans__56_392
------------------------------------------------------------
----------MSPLLTWNLI--------------------------CVSS----------
---------------------------------RWYT-----------------------
-----------------------------------------------ILCL----KNKVK
FWL---------------------------------------------------GTRLVH
EPESMESRSSPY------------------------------------------------
---------------------------------------IRPS-----------------
-------------------------------------PYAGGVH------------PHF-
------------------------------------------------------------
-------------------------EEEDDR----------------SKLHASALYSERR
T----------------------SSRKSLRSQ----------------------------
---KIDYHT----------------------T---------TI-----K-----------
-SL--WFDFC-ARTSSHGIPYVA---------T-SS-------------F-----FG---
--RYVWA----ALFMCMLMAFLLQTYWTMS-EYLQ----------Y---RTIIEM----Q
-L-QF-E------------AAAF-----PAA-----------------------TVCNLN
AF------KYS-----ELTQYE--------------------------------------
---------EI-------------------------------KEGFD-------------
------------------------------YW-ERVI-----NARM--------------
------------------------------------------------------------
------------------------------------------------------------
-------MSDSMKPGG-D------------------------------------------
------------------------------------------------------------
---ILEAISVRKKRSKSRDQ-------LLFPIDDEDLEGAVYQP----------------
```

-----------VFVRCTCMN-------------------------MEQCVPNRN---P--
------LEVNASICMCFEDVTRGLIWPCYPTSVWT------------------VKKCSGC
SIS-NTCPDPDGPNASKQIAKHNSPLPCLCQSISHH-----------CM-----------
----VHPK--------------E-------------------------------------
---------------------------------------IRWWNPNNYTVY---------
--SVTEPPTTEI------------------------------TETE--------------
-----------------------------------------EAFGLSDL------KDAGA
--------------------------ITTQTKENLIFLV---------------------
------------------------AAL----------------PRETRRNL----SYTLN
--EFV------LRCS----FN-S----K----------D---C-----------S-ME--
----------------------------------------------R-------------
------------------------------------------------------------
------------------------------------------------------------
-----------D----------F-------K-----------LHV-DPEYG--------N
CYTFN-------FN-------------DS-V-E---------------------------
-------LKNSRA------GP--------MY-------------GLRLLLNVHQS-----
----------D-----------------Y------------M-P-----TT---------
------------------------------------------------------------
------------------------------------------------------------
----------EAAGVRLVVH-EQDQEP--FPDTFGYSAPTGFISSFGLKTK------E--
-----LHRL-----------SAP-W--GN-C--SDT-F------RPV---------PYIY
----NE--------------HYSPEGCHRNCFQLKVL----EI--CGCGDPR----F---
------------P-L--PS-----------------------------------------
---------EEH-----R---HCNAK----------S-----------------------
------------------------------------KID---RQCLS---------N---
L-T-------------------SDSG-GYHH--------------LH----------EQC
------------ECR-QPCHEKV---FETA-YSA------------------SAW-PS--
-------QNF------KIG-------------TD-----CP-------------------
------------------------------------------------AV----------
---------------------------------------------------------SDI
FN----------------------------------------------------------
-----------------------DTE-ACT------------------------------
-----E-------YY-----------R------QNTAYIEIYYEQLNFESLK-ETAGYTL
VNL-----------------FSDFGGNIGLWIGFSVITFAEFAELFCEI-----------
--------CKLM--------Y----------F-------KGIVYV---------------
--------------------------------QK-------------------K------
-----------------------------------------MQGKEYTSSS---------
--------------LMHIDFLQRSPKKSQP------------------GEDE--------
--------VSTN------------------------------------------------
------------------------------------------------------------
------------------------------------------------------------
------------------------------------------------------------
-----------------------------------------------------------E
STKELMSK
>Protost_ecdy_cyclo_nema_Cele_sp_P24585_DEG1_CAEEL_Degenerin_deg_1_OS_Caenorhabditis_elegans_OX_6239_GN_deg_1_PE_1_SV_2_955_393
------------------------------------------------------------
----------M-------------------------------------------------
------------------------------------------------------------
------------------------------------------------------------
------------------------------------------------------------
------------------------------------------------------------
------------------------------------------------------------
---------------------------------SNHHSKTKKTSM---------------
---------------------------------------------------LG-------
--------------------------------------------------REDYIYSHDI
T----------------------NKNKKEKLNGASK-----------------------N
NDYNQDDDD----------------------E---------TM-----K-----------

-SK--MMDFC-DKTTAHGAKRVL---------I-AR--------------NS---FS---
--KLMWG----LIIFSFLLMFAYQASKLIF-KFSA----------H---EKITDI----S
-L-KF-D------------DVEF-----PAI-----------------------TFCNLN
PY------KKS-----LVMMVP--------------------------------------
---------SI-------------------------------RDTMD-------------
------------------------------VY-DNAKTH---SKSEGEK-----------
------------------------------------------------------------
--------------------KKPKVS-----------------------------RKQHS
-----------------DASQQMVRELFAKEIEEG------MVELKKSNKTLQ-------
------------------------------------------------------------
-------SQNKSGRRRSQRS----------------IENRRYEA----------------
-----------IEAHCKCVGN-----------------------IGMECIRFESP--P--
-------RDPSSKCICTYDRDMEVAWPCFNISVWY------------------DHECPLC
HDD-GYCESTLPSGTT-----SSDKWPCMCRNRGDTS---ERDDTPYCIGKAG------V
GKIEIRKLWLENNMTTTSTTTT--------------------------------------
-----------------------------------------TTTTPPPTTTSTTTT---T
TTTPPPTTTARPNQRAIVSN----------------------PETI--------------
-----------------------------------------KAMGFQGM------TDGVA
--------------------------MLTRAKENLMFTM---------------------
------------------------AAL----------------SDKQRIAL----SQSKH
--EFI------EMCS----FN-G----K----------E---C-----------D-ID--
----------------------------------------------E-------------
------------------------------------------------------------
------------------------------------------------------------
-----------D----------F-------R-----------LHV-DPEFG--------N
CFTFN-------YD-------------VN-N--------------------N--------
-------YTSSRA------GP--------MY-------------GIRVLLFVNTS-----
----------D-----------------Y------------M-S-----TS---------
------------------------------------------------------------
------------------------------------------------------------
----------ESSGVRLAIH-PPTEYP--FPDTFGYSAPVGFASSFGIKKK------V--
-----MQRL-----------PAP-Y--GE-C--VET--------KK------VVDRNYIY
----AGY-------------DYHPEGCHRSCFQNGLI----DD--CSCGDPR----F---
------------P-V--PE-----------------------------------------
----------GY-----R---HCSAF----------N-----------------------
------------------------------------ATA---RTCLE---------K---
N-I-------------------GSVG-DFHHIT--QK--------M-----------DKC
------------VCK-QSCEEII---HEVT-FSC------------------SKW-PS--
-------GATD---------L---------------------------------------
------------------------------------------------------------
------------------------------------------------------------
-G----------------------------------------------------------
-------------------DCDGMTESECE------------------------------
-----Q-------YY-----------R------LNAAMIEVFYEQLNYELLQ-ESEAYGL
VNL-----------------IADFGGHLGLWLGFSVITVMEVCVLLVDM-----------
--------ISLF--------F----------K-------SR-------------------
HEEKLL--------------------------RQ---------------STKRKDV----
------------------------------------------------------------
------------------------------------------------PEDK--------
------------------------------------------------------------
------------------------------------------------------------
------------------------------------------------------------
---------------------------------------------------RQITVGSGR
KSDAFVSI
>Deutero_Cephalo_154560F_t1__name__chr_scaffold30_start_582601_end_590481_str
and_pro_len_784__956_394
------------------------------------------------------------

```
----------M-RLFLELL-----------------------------------------
------------------------------------------------------------
------------------------------------------------------------
--------------------------------------------------------RTLY
KEAPPIDRGQE--------------NL---------------------------------
---------------------------------------IYPAKMT--------------
--------------------------------ATSDIDLRVLERIDSR------------
------------------------------------------------------------
-----------------------------------------------LLRVIEETAEDRA
QKIVE---------------KCLKERDEK-------------------------------
---MKEQRK----------------------K---------DI-----K-----------
-TM--TKEFC-ASTTAHGVNRVA---------E-AD--------------TT---RS---
--RIIWT----VILITCVALFVYQSAILIK-SYFE----------Y---PVNVDI----K
-V-VD-N----------K-KLTF-----PSV-----------------------TVCNNN
RL------RKS-----QLPGSR------------------H-----------------G-
---------SLLEL----------------------------------------------
---------------------------------DE-------QTARDVSDYLW-------
------------------------------------------------------------
---------EKQIG------------EIDIDKDMVIDLESG-Y--PGLGSGFES------
---ELGLTNTFLSDESIDASTE-------------------CRLQSKQLCTL--------
---------------------AQL------------------------------------
-------------LREPRNDGIHTEWSYFSRWAVMISHPCTGTEQDTCDQCYND------
------------------------------------------------------------
----RILLKRIEDIPYKIN--ATFCCPPDSY-----------------------------
------------------------------------------------------------
----------------IALTDHTPNHARLNSRSYWKMD----------------------
---------------------------------------------EYDATPWIQV-DLRD
SHVLAGVITQGGGDAGGYVTSFSLSFSVDG------NQWQTDNETGERDWE--YCHVERC
EVGCVDLLNAV-------------AGENDWGEFLLS-----------SN------TDDYT
---------------------------------DITDFS---------------------
-------------------------MA----------------TREEIAEM----GHQKE
--DLI------LQCT----FD-K----E----------R---C-----------S-L---
----------------------------------------------D-------------
------------------------------------------------------------
------------------------------------------------------------
-----------D----------F-------K-----------VTQ-NAKYG--------N
CFTFN-------HG-------------ED---------------------D----IV--
-------RNTTKV------GA--------EY-------------GLKLTLSIESN-----
----------E-----------------Y------------V-G-L---FG---------
------------------------------------------------------------
------------------------------------------------------------
----------QDIGAKVTIH-SVGSTP--FPEGNAINTKPGESTFVSLKRS------S--
-----IKRR-----------PYP-Y--GD-C--TSH--------QER---------DALY
----G-------------G-RYTYETCQHSCLQAALL----HE--CGCSDEL----I---
------------A-I---------------------------------------------
----------NS-----T---LCSVL----------N-----------------------
------------------------------------KTQ---ECCRQ---------D---
V-R-------------------SRHE-D-------GN--------L------------SC
------------DCR-QSCNEDS---YALW-LSS------------------SLW-PS--
-------DSY------AWY-V-----------LE--------------------------
------------------------------------------------NI-H--------
----------------------------------TR---------------------SQA
QN----------------------------------------------------------
------------------------LP-LNP------------------------------
-----D-------EL-----------R------QNLARIHVYFRDLNYELIT-ENPTYTE
ESL-----------------LSGLGGLLGLYVGLSVITVFEFINLLVDL-----------
--------FKAA--------F----------N-------KE-------------------
```

```
VK------------------------------DT--------------------------
-----------------------------------------ANPGH--------------
------------------------------------------------------------
------------------------------------------------------------
------------------------------------------------------------
------------------------------------------------------------
------------------------------------------------------------
------NI
>Deutero_Ambulac_Apla_gbr29_2_t1_957_395
------------------------------------------------------------
----------M-------------------------------------------------
------------------------------------------------------------
------------------------------------------------------------
------------------------------------------------------------
--EYGTTVVEGQRLTR--------------------------------------------
------------------------------------------------------------
------------------------------------------------------------
------------------------------------------------------------
-------------------------------------------------RVLVTGSLDYL
Q--------------------RTGEEDKL-------------------------------
---SQEKKR----------------------G---------AL-----R-----------
-QA--VTTFA-DTTTAHGLPLII---------N-GT--------------YT---VS---
--RIAWS----VIWLIAFGLAILQSTLLLV-EYFQ---------VR---PLTTTI----E
-L-IT-K----------T-NLDF-----PAV-----------------------TVCNMN
RM------RRS-----KLVGTR------------------F-----------------E-
---------GLIAI----------------------------------------------
---------------------------------DG-------GVQGEDYDYSWFFDWSSL
-------------------------------------------EWFDRWDS---RG----
---------EPSSS--------P--------------------------DGNSLQ-----
-------DSSPVNSEPDDSTAG-------------------GIGSSESIPGVG-------
----------GESDVTPQGSGGATVGGSS---------VDGTDGS---------------
-----------------------------------DSVPGAGDDDDGMSS----------
------------------------------------------------------------
--------------------DSSTSGPSHVASN-------------------GEGDNPST
AGDSI------------------DEMSSI-------------DNESSLPGGSSNTPSHD-
--------VEKEEASSTAPPHSLNTHASTSSGIFPVTQ---------------VNSPDSS
SSQTTTDPPGSETVAPGQERDSDGASSSFNPGEESDTSILDSERRRRSVSS-----RQER
QRRTLSSSTEDF----WW----ESD------------DFEYDSYYN-------W------
--------DGV-------------TDENDWRGFYDR-----------ST------ADDFS
---------------------------------DLIDIA---------------------
-------------------------NP----------------TREELEVL----GHQAQ
--DFI------LQCT----FD-K----R----------N---C-----------S-Y---
----------------------------------------------R-------------
------------------------------------------------------------
------------------------------------------------------------
-----------D----------F-------R-----------VFQ-NTDYG--------N
CFTFN-------HG-------------TG-N--------------------S----KV--
-------RTTSRF------GA--------KY-------------GLHVTLFIEQP-----
----------E-----------------Y------------L-G-I---FS---------
------------------------------------------------------------
------------------------------------------------------------
----------PESGVRLTVH-GQRTMP--LPEDNGITAAAGLATSVGMRQD------F--
-----ISRQ-----------PFP-Y--GN-C--SDS-F------DTR---------FNVT
----AGD-----------Y-DYSLSACTKFCLQSEML----SR--CGCVTDI----L---
------------I-Y---------------------------------------------
----------SA-----L---KCSYL----------N-----------------------
------------------------------------TTQ---QTCRR---------A---
```

```
V-E-------------------NLYE-D-------NK--------L-----------QSC
------------KCF-SACRETT---FKLG-VSS------------------SRW-PS--
-------ERY------EEH-L-----------YS--------------------------
------------------------------------------------RI-S--------
----------------------------------AT------------------------
-N-------------EKA------------------ARLM--------------------
---------------------------QEV------------------------------
-----E-------QT-----------R------KNLVRLRIYFEELNYQSII-QTPKYSV
ETL-----------------LGSLGGLFGLFIGFSVITIFEVVLFLLKL-----------
--------LKIF--------F-----FWG--QPC--------------------------
HR------------------------------RE--------------------------
-----------------------------------------IKPVI--------------
------------------------------------------------------------
------------------------------------------------------------
------------------------------------------------------------
------------------------------------------------------------
------------------------------------------------------------
-------S
>Deu_Ambulacrar_hemi_Ptyfla_40v0_9_20150316_1g15264_t1_scaffold10904_cov157_9
58_396
------------------------------------------------------------
----------MSLMIGTRFQNLHEN-----NCDTRTIVKI----ESCIQY--SCG-----
------------VG----------------------------------------------
------------------------------------------------------------
----------------------------------------------IYHISMHRMVAKLD
KINQRKTVNLS-------------------------------------------------
------------------------------------NSRVKPK-----------------
--YIYDRRRVAERRNMIESS-------------LGNNSRVAPET------------P---
------------------------------------------------------------
--------------EISKKTHAGSTEGVSADR----WN------------MIESGLSNNS
QVAP----------------ETSEMSEKAHAGSSSMVS----------------------
-MNDMRENK--------VGCC----------R---------TCCGR--S-----------
-EK--FREFT-QETTLHGIKYTT---------D-TN-------------ISS---YR---
--RLLWF----ILLLAMVASFTTTLRESAM-RFWS----------N---PVNTVV----S
-Y-RE-M----------E-SMPF-----PAV-----------------------TICNYN
VY------RKS-----VVKESV------------------L-----------------G-
---------QY------------L------------------DAMYG-------------
------------------------------------------------------------
------------------------------------------------------------
------------------------------------------------------------
-----------------N------------------------------------------
------------------------------------------------------------
------------------------------------------------------------
------------------------------------------------------------
--------------------------PFSP------------------------------
------------------------------------------------------------
------------------------------------------------------------
------------------------------------------------------------
------------------------------------------------------------
------------------------ETDENTTLPEP--------------------PANYF
A--------------------------------NKTQIM---------------------
------------------------------------------------YQS----AHRIE
--DML------KECK----LS-S----D---------VY---C-----------S-A---
----------------------------------------------E-------------
------------------------------------------------------------
------------------------------------------------------------
-----------N----------F-------T-----------KVI-T-NFG--------V
```

CYTFN-------GE-------------VG-N--------------------A--------
-------LKVRAR------GQ--------QN-------------GFFVSIDIQQN-----
----------E-----------------Y------------Y-Y-G---PN---------
------------------------------------------------------------
------------------------------------------------------------
----------VGAGLKNKLL-------------------------------------K--
------------------------------C--DEE--------Q-------LD----YF
------------------D-EYSTANCEMEKRTKAVV----KD--CQCKQPH----M---
------------P-N--KNEYGCSVPAFPGPHVVKRLEAKYNQTEVQMRLLWIALMLFTI
-ILLIFAVKDSAIRVTFPAITLCNYNQYRKSRINGLNDYEDRLRNM-----NRTEFEI--
------------------------KAAH------RIEDM--LLECVW---------S---
S-T-------------SNCD-ENNFT-----------L------VLT--------DFGVC
YTFNGDRKQEEFSCK-MPCETVS---HAST-LSF------------------GSF-PG--
-------VQV------SEQ-L-----------KI--------------------------
------------------------------------------------RF----------
------------------------------------------------------------
-N----------------------------------------------------------
---------------------------MTE------------------------------
-----K-------SI-----------E------QNFVSLNIYVEDLMVLVTS-EVMAYTY
DDF-----------------VGDLGGQIGLFLGASLLTVFEFAEFFVTC-----------
--------VVRR--------C----------R-------RL-------------------
RA------------------------------KF-------------------RA-----
-----------------------------------------INKKNKALE----------
-----------------------SASS---------------------QKNK--------
------------------------------------------------------------
------------------------------------------------------------
---------------------------------P--------------------------
------------------------------------------------------------
-ECAGYKS
>Protostome_Lophotroco_annelid_CAC9599174_1__Ofus_G063499_Owenia__fusiformis_
_959_397
MQRVADDPRHN-MIKGPGNNVADINAASCAAGINEGGPNEANEHIYAD------------
--------VVV-------------------------------------------------
------------------------------------------------------------
------------------------------------------------------------
------------------------------------------------------------
--EIK-----I--------NTL--------------------------------------
------EEPSKKEPLHAMA-----------------TASSNPASTEDRAN----------
-G-----------TIDSNEI-------------SNAES----------------------
------------------------------------KG----------------------
------------RNVAFNLPLTAGIEETRN-------------------------VSAKD
Q----------------------PND----------------------------------
------SMR----------------------K---------KV-----S-----------
-ST--LSEYA-EMASVNGPKRIV---------R-AK--------------SR---PS---
--MVAWS----VIFLGVVCMAVYLIVGVVV-KYYT----------Y---PIQDGI----T
-I-KT------------Q-PLAF-----PAV-----------------------TLCPFI
PN------SMS-----DLKKALRSKEAK-------LPFANL-------------------
----------------------LGDKIT---------------ETMI----RYH------
------------------------------IS-AV-------HTRNMY------------
------------------------------------------------------------
------------------------------------------------------------
-----------------EFS------------------------INPETF----------
------------------------------------------------------------
------------------------------------------------------------
------------------------------------------------------------
------------------------------------------------------------
------------------------------------------------------------

------------------------------------------------------------
------------------------------------------------------------
----GLLKFNGT------------------------------D-----------------
---------------------------------------I-NMKKIFQL------REALR
---------------------------------SSGWLV---------------------
------------------------ENVPIN-------------DFSRNLLK----QIKSK
--DFI------MECT----YN-D----K----------A---C-----------D-D---
----------------------------------------------L-------------
------------------------------------------------------------
------------------------------------------------------------
-----------I----------D-------I-----------SIS-NTVYG--------R
CFTLN-------VN-------------ES----------------------------I--
-------GYVQEV------GP--------DK-------------GLVLLLYTGRH-----
----------NPLQ-RIGMGNGGY----EF-------------E-S-KYAE---------
------------------------------------------------------------
------------------------------------------------------------
----------PSDGVKVVVH-KPGTMP--RPYSDGFFVTPGRLAAVGITQS------E--
-----RTRL-----------LPP-H--GE-C--TDA--------VF------NTN--TTY
--------------------KYTYEICVDQCIQERIK----EN--CGCVSPM----F---
------------P-V--PK-----------------------------------------
-----DHDFKRI-----Q---YCGNIS---------DIF-KSYND------NETSAELSM
FSVTERLASEKGDFEHYDTSTPELRKQLANTVD-EIIAR---LQCEN---------H---
W------------------------------------------------MISDK---DSC
------------KCK-RQCKENE---YTHL-INT------------------IPW-PE--
-------KSV------VDT-G-----------GH--------------------------
------------------------------------------------RGFVKFFQK---
------------------HKHFE--------TLMHNLVGLTREIGDIENFRNTAEALVAL
IL-------------TTR------------------FDF-MNELFLYTENF---------
------------QNV-YGISSFTIAT-YLD------------------------------
-----K-------FI-----------D------TNFLKLNIHFESLDVRVVY-EERTYTY
ASV-----------------IKDIGNVFGFYLGMSVFSVMEAVFMICLL-----------
--------IKLV--------I----------C-------QM-------------------
VR------------------------------SEENDD----------------------
------------------------------------------------------------
--------DNI---------GEQHIKG---------------------TDD---------
------------------------------------------------------------
------------------------------------------------------------
------------------------------------------------------------
------------------------------------------------------------
------ID
>Deutero_Chord_Hsap_sp_P51172_SCNND_HUMAN_Amiloride_sensitive_sodium_channel_
subunit_delta_OS_Homo_sapiens__Human__OX_9606_GN_SCNN1D_PE_1_SV_3_962_399
------------------------------------------------------------
----------M-------------------------------------------------
----------------------------------------RA------------------
-------------------------VLS--------------------------------
------------QKTTPLPR----------YLWPGHL---SGP-----------------
------------------------------------------------------------
----------------------------RRLTWSWCSDHRTPTCRELGSPHPTPCTPCPL
CLLSGPIQGCGTGLGDSSMAFL---------SRTSPVAAASFQSRQEARGSQSCQLPPQ-
WL---------------------------STEAW-----------------TGEWK----
--------------------------------------QPHGG-------ALTSRSPGPV
APQRPCHLKWQHRPTQ--HNAACKQGQAAAQTPPRPGPP-----------------SAPP
PPPKEGHQEGLV----ELP------------A---------SF-----R-----------
-EL--LTFFC-TNATIHGAIRLV---------C-SR-------------GNR---LK---
--TTSWG----LLSLGALVALCWQLGLLFE-RHWH----------R---PVLMAV----S
-V-HS------------E-RKLL-----PLV-----------------------TLCDGN

```
PR------RPS-----PVLRHL----------------ELL-----------------D-
---------EF------------ARENI--------------DSLYN----V--------
---------------------------------NL-------------------------
------------------------------------------------------------
------------------------------------------------------------
----------SKGRAALSAT----------------------------------------
------------------------------------------------------------
------------------------------------------------------------
------------------------------------------------------------
---------------------VPRHEPPFHLDR-------------------EIRLQRLS
HSG---------------------------------------------------------
----------------SRVRVG--------------------------------------
------------------------------------------FRLC--------------
--------NSTG------------------------------GDC---------------
--------------------------------------------FYRGY------TSGVA
AV-------------------------QDWYHFHYVDIL---------------------
------------------------ALL----------------PAAWEDSH----GSQDG
--HFV------LSCS----YD-G----L----------D---C-----------Q-A---
----------------------------------------------R-------------
------------------------------------------------------------
------------------------------------------------------------
-----------Q----------F-------R-----------TFH-HPTYG--------S
CYTVD-------GV----------------------------------------------
-------WTAQRP------GI--------TH-------------GVGLVLRVEQQ-----
----------P-----------------H------------L-P-L---LS---------
------------------------------------------------------------
------------------------------------------------------------
----------TLAGIRVMVH-GRNHTP--FLGHHSFSVRPGTEATISIRED------E--
-----VHRL-----------GSP-Y--GH-C--TAG-G------EG------VEV-ELLH
----N-------------T-SYTRQACLVSCFQQLMV----ET--CSCGYYL----H---
------------P-----------------------------------------------
-------LPAGA-----E---YCSS-----------------------------------
------------------------------------ARHPAWGHCFY---------R---
L-Y-------------------QDLE-T-------HR--------L------------PC
----------TSRCP-RPCRESA---FKLS-TGT------------------SRW-PS--
-------AKS------AGW-T-----------LA--------------------------
------------------------------------------------TL-G--------
----------------------------------EQ------------------------
-G-------------------------------LP-------------------------
---------------------------HQS------------------------------
-----H-------RQ-----------R------SSLAKINIVYQELNYRSVE-EAPVYSV
PQL-----------------LSAMGSLCSLWFGASVLSLLELLELLLDA-----------
--------SALT--------L----------V------------------------LGG-
RRL-----------------------------RR------------A-WFSWPRASP---
--------------------------------------ASGASSIK--------------
-----------P------------------------------------EASQMPPPAG--
------------------------------------------------------------
------------------------------------------------------------
----------------------------GTSDDPEPSGPHLPRVMLPGVLA---------
-------------------------------------------------GVSAEESWAGP
QPLETLDT
>Deu_Ambulacrar_hemi_Ptyfla_40v0_9_20150316_1g21852_t1_scaffold24782_cov84_96
4_400
------------------------------------------------------------
----------M-------------------------------------------------
------------------------------------------------------------
------------------------------------------------------------
```

```
------------------------------------------------------------
------------------------------------------------------------
------------------------------------------------------------
------------------------------------------------------------
------------------------------------------------------------
------------------------------------------------------------
------------------------------------------------------------
---------------------RFDDVDRP-------------------------------
----TWRRS----------------------A---------SM-----H-----------
-VS--FVKEA-SFSTQHIVISVR---------Q-SV--------------P----FA---
--RVVHTRCTVAICISHFSCFRYEPWECMSFDYHETGKSSYDHFTY---GIEIRF----E
-L-CD-V--------GQP-RVGCLSKVEPSARGYICQLPGDHNYCRNPDGWSATWCYTLD
PL------VRW-----ESCD----------------------------------------
-----------V---------GSPQTTCQN----TDMPIHEFFRTTN----QTQR-GFDS
MTGNVTVE--ECASSCLN----S-RKFTCLSF-DF-------SNEDGICYHSDANESVSK
PSTVSGILV-DHYDRRFEDEER---------------LLLDF-WKFPDYRIPSDVALLTN
VTAHG---CAAKCLDLAKIADTPEKDISYIHYELKVASSNGCPSDYRQCGSGE-------
-CIHQYSFCDTIINCA-DE-------------------------SDEDNCYG--------
------------------------------------------------------------
---------------NHRA-----------------------------------------
------------------------------------------------------------
-------------------IPDMDDIPGVDWH----------------------------
------------------------------------------------------------
------------------------------------------------------------
------------------------------------------------------------
DHPDKATLMADF----------LSDYYH--------------DNGFAS------------
---------------------VKAEVPPDWYGF-----------------------KTFS
STP------------------------------DYSDLQ---------------------
----------------------SVLKL----------------SRDELRRY----GHQRE
--DFV------LQCT----YN-E----E----------D---CV----------Q-N---
----------------------------------------------S-------------
------------------------------------------------------------
------------------------------------------------------------
-----------D----------F-------Y-----------TFQ-DEKYG--------N
CFKFN-------HG-------------RG-N--------------------E----TV--
-------KMATKQ------GA--------TF-------------GLKLTLCLEQD-----
----------E-----------------Y------------I-G-I---YG---------
------------------------------------------------------------
------------------------------------------------------------
----------HDAGVRVAIE-PSKYVA--FPLDAGITVQPGTVTSIGLREI------R--
-----VSKK-----------PHP-F--GN-C--TLD--------DTQ---------DSIF
----GSHY------------QYS-------------------------------------
------------------------------------------------------------
------------------------------------------------------------
------------------------------------------------------------
V-----------------------------------------------------------
------------------------------------------------------------
--------------------L---------------------------------------
------------------------------------------------------------
------------------------------------------------------------
------------------------------------------------------------
------------------------------------------------------------
-----------------------------------------------------------R
EDL-----------------LSDIGGTLGLYIGLSVITVAEFIELLIVV-----------
--------IKFV----------G--------R-------RR-------------------
GAL-----------------------------RT------------------AK------
---------------------------------------LSPSS----------------
------------------------------------------------------------
```

------------------------------------------------------------
------------------------------------------------------------
------------------------------------------------------------
------------------------------------------------------------
--------
>Cnidar_HydraVulSc4wPfr_2074_g13182_t1_968_403
------------------------------------------------------------
----------MYLQFVNHFTT-----------SSGKVWYIK-TSISCNSKNLSCNRCDG-
--KTT------YIGKTNSLSKDAFKHENKKSFKRWSLLKEHPNYSQIINEVVEVETDTNK
GTDTKEVPNFQETSNKEITNFQGTESKNDEASDKIVLPADN-----------QSVDSEVT
LNLSGNTKWAA-LTKKNAK-KIILESDKLKQPVL--------------------------
--SLSSDSTSQ-------TDNK----------------ILNNK---------NSLGK---
---SKWNAVRERTNSIKNT--------KDDVN-----SNEKP------------------
--SLPKLLQVVNTATATKTA-------------ASKFLEAS-------------------
------------------------------------------------------------
------------------------QMRVNLK--------------------------ARM
R----------------------KIVQE--------------------------------
--------R----------------------L---------TV-----K-----------
-QI--FKRYV-ESSTLHGFCYVC---------G-DT--------------FL---VR---
--RVLWA----LLMILGAIYFIIKLRYGIE-EYLN----------Y---PFSTLS----T
-V-DY-V----------H-ELFF-----PAM-----------------------SLCATN
SY------LAS-----MVSKNQ------------------L-----------------K-
---------LL----------------------------------YD-------------
------------------------------------------------------------
------------------------------------------------------------
------------------------------------------------------------
------------------------------------------------------------
------------------------------------------------------------
------------------------------------------------------------
------------------------------------------------------------
------------------------------------------------------------
------------------------------------------------------------
------------------------------------------------------------
------------------------------------------------------------
------------------------------------------EGRL--------------
------------------------PLDNNQTNPSYN-------------------MSGN-
---------------------------------ELVKAI---------------------
------------------------------------------------QES----SLSIE
--SML------AFCD----WI-Q----QDTNNPDIPPNP---C-----------G-P---
----------------------------------------------Q-------------
------------------------------------------------------------
------------------------------------------------------------
-----------N----------F-------T-----------TYL-N-YKGE-------Q
CYTLN-------SG-------------LE-G--------------------H----KL--
-------LKVDTV------GL--------TY-------------GYELIFDLQTN-----
----------K-----------------V------------I-K-N--------------
------------------------------------------------------------
------------------------------------------------------------
---------NQFSGMRVVIH-NQDVPP---QLADGFIIPPGFKTFVKMGTV------Q--
-----SKSL-----------PPP-YS-TE-C--GTK--------K-------LK----YY
------------------N-TYSQRFCLLETLTDFTG----KL--CGCRDVF----M---
------------P-E--N------------------------------------------
----------GL-----P---FCSL-----------------------------------
------------------------------------KEL---YSCMY---------P---
A-K-------------------ESFS-E-------FT--------------------MRK
------------ECP-ADCEERT---YPYE-LSE------------------ARF-IH--
-------NPP------IGL-SIQGLAN--LQHKK--------------------------

------------------------------------------------HLEEEL--Y---
-----------------------------------------------------------L
SK---------------LA----------------QSMT---------------------
------------------------SE-ELD------------------------------
-----A-------YV-----------E------DNIVSVIFFFGDTRIDYNE-QEATNDF
FQF-----------------LGNMGGEFGLMLGASLLTFVEFVDLFIFL-----------
--------IYHQ--------M----------L-------RL-------------------
HT------------------------------LK-------------------KVPDI--
--------------------------------------FGRKRSRINEYI----------
-----------------------KKRE---------------------------------
------------------------------------------------------------
------------------------------------------------------------
------------------------------------------------------------
------------------------------------------------------------
----KTRV
>Protost_ecdy_cyclo_nema_Cele_sp_O01635_ASIC1_CAEEL_Degenerin_like_protein_as
ic_1_OS_Caenorhabditis_elegans_OX_6239_GN_asic_1_PE_3_SV_2_970_405
------------------------------------------------------------
----------M-------------------------------------------------
------------------------------------------------------------
------------------------------------------------------------
------------------------------------------------------------
------------------------------------------------------------
------------------------------------------------------------
------------------------------------------------------------
------------------------------------------------------------
------------------------------------------------------------
------------------------------------------------------------
---GKNSLK----------------------R---------AL-----------------
-ELD-VVDFA-EHTSAHGIPRAYV----------ST------------------GWR---
--RYMWL----LCFLFCLSCFGHQAYLIVE-RFNR----------N---DIIVGV----E
-I-KF-E------------EIKF-----PAV-----------------------TICNMN
PY------KNS-----AARE--------------------LG------------------
---------AI-------------------------------RNAIE-------------
------------------------------AF-ELAI-----DKSDGNAH----------
------------------------------------------------------------
-------------------SKRKKRSSKMVPIDLLCKEEHGMF--TAHDYGHVEC-----
TCV----TFEDMSKVG-DTDDDEIFWNCHQRLAEGCKCFEDTCVSDEVTKQLV---WPLQ
LSK----NGKLCISPESSGPRYCASAQKFSNCDWLGKCEESDDMDLEEEIDSKTCICHHG
NCFQIKGNVKKRKRRTPERKVHERL--L-----------SRYEG----------------
--------LLAVYSHCNCTK-------------------------QHGCVSTSV---PDM
D-----LENSNKTCLCFYNKKNEQIWPCYKEPEWE------------------ERKCSRC
NTM-GDCVYSDK-------P-KKQTISCLCATPIKM-----------CVR----------
----IDP-PQTNDTSLDDRVVK--------------------------------------
-----------------------------------------FWDIQPSTT----------
----MSPIVKKK------------------------------EERD--------------
-----------------------------------------KAYGYTGV------KDRIA
--------------------------LRAKAMENMIFAV---------------------
------------------------DAL----------------TEEEKWKI----SYNKS
--DFI------MKCS----FN-G----R----------E---C-----------N-VK--
----------------------------------------------H-------------
------------------------------------------------------------
------------------------------------------------------------
-----------D----------F-------V-----------EYL-DPTYG--------A
CFTYG-------QK-------------LG-N-----------------------------
-------NTNERS------GP--------AY-------------GLRLEVFVNVT-----
----------E-----------------Y------------L-P-----TT---------

------------------------------------------------------------
------------------------------------------------------------
----------EAAGVRLTVH-ATDEQP--FPDTLGFSAPTGFVSSFGIKLK------S--
-----MVRL-----------PAP-Y--GD-C--VRE-G------K-------TED--FIY
----TQK-------------AYNTEGCQRSCIQKHLS----KT--CGCGDPR----F---
------------P-P--YR-----------------------------------------
----------ES-----K---NCPVD----------D-----------------------
------------------------------------PYK---RECIK---------N---
E-M-------------------HVAT-R-------DS--------KK----------LGC
------------SCK-QPCNQDV---YSVS-YSA------------------SRW-PA--
-------IAG------DLS-----------------------------------------
------------------------------------------------------------
------------------------------------------------------------
-G----------------------------------------------------------
------------------CPLGMAAH-HCL------------------------------
-----N-------YK-----------R------EQGSMIEVYFEQLNYESLL-ESEAYGW
SNL-----------------LSDFGGQLGLWMGVSVITIGEVACFFFEV-----------
--------FISL--------I----------S-------SN-------------------
RT------------------------------KR-------------------R------
-----------------------------------------PARKSFS------------
------------------------------------------------SSLR--------
------------------------------------------------------------
------------------------------------------------------------
------------------------------------------------------------
----------------------------------------------------CSTDYNLN
KDGFNLDN
>Protostome_Lophotroco_annelid_CAC9662918_1__Ofus_G12800_Owenia__fusiformis__
973_408
------------------------------------------------------------
----------M-------------------------------------------------
------------------------------------------------------------
------------------------------------------------------------
------------------------------------------------------------
------------------------------------------------------------
------------------------------------------------------------
------------------------------------------------------------
------------------------------------------------------------
------------------------------------------------------------
-----------------------ETKISTNDI----------------------------
-KIQQENEP----------------------T---------TF-----K-----------
-GL--LTDFA-NNTTLHGLPRVI---------T-SA--------------QL---IR---
--KFVWF----CIFFGSFGYFSFQVAELIN-DYYK----------W---PVLIRS----E
-M-KL-R----------T-FMEV-----PAV-----------------------TICNMN
ML------RRS-----KIEGTM------------------F-----------------N-
---------SLVELD-----------------------NDTANEIIE----K-Q------
---------------------------------DTTENEINNQIQNSDKTSRKKRNINMI
QKSIQRPMSKNNALKGHENKFEISRVKTKSMESVNNIL------AFEH------EGSGST
KIEHTFSLPENEADD-------P----HSLNDEYIALE-----------DGHSV------
---------EPIDDTN-DNGRRVR------------------------------------
-------------REVAGGSGGEATD---SYVSDPTDTTTDSYDTYTDPT----------
-------DTTTDSYDTNTDPTDTTT----------DSYDTNTDPTDTTTDSFDT------
----NTNPTDTTIDSYDTNT----------------------------------------
-------------------------DPTDTTTD----------------SYDTNTDPTDT
TTDSYDT---------------------------------NTDPTDTIEDS---------
---------FDTNTDPVSTNT-YPIYTDS-------------------------YPTDPN
ADPMDFDTYPTESYTDPNFQSSDYYESYEYSEYESS------------------------
EQEMDQESGEDP------------GYDV---------PFQF-EQSK-------L------

---------------------MGIKDPNNYAEILRL-----------AE------AEDLS
---------------------------------DIFGLA---------------------
-------------------------SP----------------TVEQLDEY----GHNLS
--DLV------ALCT----FD-G----E----------N---C-----------N-R---
----------------------------------------------SS------------
------------------------------------------------------------
------------------------------------------------------------
-----------D---------HW-------L-----------ETY-NKYYG--------K
CYTWN-------SV-------------IN-K-PR----------------GE----RP--
-------LVTTNF------GS--------RY-------------GLRLTLNVERD-----
----------E-----------------Y------------T-G-L---FS---------
------------------------------------------------------------
------------------------------------------------------------
----------PTYGTRVAIH-DSSIIS--YPENDGITASVGNEMVIRLRLK------K--
-----YERE-----------TEP-YP-SD-C--VKD--------DD------L---VKQF
------------------G-GYTVIGCMRECLEERLV----KR--CKCLDRI----V---
------------E-----------------------------------------------
---------KDE-----H---RCSYF----------N-----------------------
------------------------------------RTE---ELCRQ---------R---
V-Y-------------------YAYN-S-------GK--------L------------GC
------------KCR-ARCRENK---YDIS-TTV------------------STW-PS--
-------KHH------DYY-L-----------LN--------------------------
------------------------------------------------RL-K--------
----------------------------------ER------------------------
-GI---------------------------------------------------------
---------------------------QTL------------------------------
-----E-------EI-----------E------DNILRVHIYFEDFNVETIK-EVPAYSV
ISL-----------------LSNIGGTMGMFVGLSICTLGEFLELIWEM-----------
--------IMLL--------R----------R-------KI-------------------
SR------------------------------RH-------------------N------
-----------------------------------------MVNAS--------------
------------------------------------------------------------
------------------------------------------------------------
------------------------------------------------------------
------------------------------------------------------------
------------------------------------------------------------
------SA
>Deu_Ambulacrar_hemi_Ptyfla_40v0_9_20150316_1g11917_t1_scaffold7296_cov167_97
4_409
------------------------------------------------------------
----------MYG-----------------------------------------------
------------------------------------------------------------
------------------------------------------------------------
-----------------------------------------------RHARYFERGERTK
YGGYAEGLA---------------------------------------------------
------------------------------------------------------------
---KPTDTDIDRQIRNQ----------------TFSVS----------------------
------------------------------------------------------------
-----------------------RSVNM----------------------------YSEP
T----------------------LDSPQ--------------------------------
------RTD----------------------E---------KA-----SKR--SIIAW--
-KV--VKEYG-LNSTSHGIPRIF---------G-AK--------------SR---VL---
--GFFWT----VLTLTAFGAFLWQGSELIM-QFKR----------F---DVITKV----E
-V-VT-E----------K-RLVF-----PSV-----------------------TVCNVN
KL------RKS-----AIASSS------------------Y-----------------K-
---------KML---------IVDEQIVL--------------PYYA-------------
---------------------------PCIPG-DF-------------------------

------------------------------------------------------------
-----------ACA-----------------------------------NRIY-------
-CVKEYLRCDGINHCK-D-F------------------------SDEDGCV-----YE-L
HKSYVMKGG---------------------------------------------------
------HKLKTCQRLATKSTYFN---------------DNKLRNQRCSLRMPFMTVCEMR
SKSPVTDNRQIPVKPGNCGEN------------------------Q--------------
------------------------------------------------------------
------------------------------------------------------------
------------------------------------------------------------
------------------------------------------F-----------------
----KCINGSDF---GFC---ISKKYEC---------------DRKSHC----YD-----
--------GA---------------DEDNC------------------------------
------------------------------------------------------------
------------------------------------------------------------
---------------------------R----------A---CG----------------
---------------SYDQFTC-DDGVCFPAWYQCDNYEDCADGSDERDCDCTFS-CDNG
TLCIPDFWICDGVQDCMDKS--DEPDYCNYDAIPDSGDETSQMLFPSDDSDVLTMPSDEE
GDLFSFSLLHDDAMSVASPSNEDLTTEAENQTLKYNCEGFRCFDGQCVELYSDC-DTIVH
CEDSSDE----D----------D-------C-----------IPS-DDRYG--------N
CYTFP-------GS-------------RN-N--------------------E----KP--
-------LYVSRS------GQ--------SH-------------GLKLTLFTEQD-----
----------E-----------------Y------------I-S-I---YG---------
------------------------------------------------------------
------------------------------------------------------------
----------RQAGIRVTITAPENSES--IHTQDGITIQPGTETHIGLREK------N--
-----LLRTIHSLNNKTLNINDK-Y--ST-R--GDR----------------T-------
------------------------------------------------------------
------------------------------------------------------------
------------------------------------------------------------
------------------------------------------------------------
------------------------------------------------------------
------------------------------------------------------------
------------------------------------------------------------
------------------------------------------------------------
------------------------------------------------------------
------------------------------------------------------------
------------------------------------------------------------
-----------------------------------------QYGGFT-------------
EGL-----------------AKPTDTDIELPIRDHTLSVSC-------------------
--------------------S----------TNM-----YSEPTLD------------SP
QRT-----------------------------GE-------------------K------
----------------------------------------VSKRST--------------
-----------IWKVV----KEYGLN------------------------TT--------
--------SHGIPKIV-GAKNK----VLGFF-WTV---------------LTLTAFGAFL
W-HGSELIREFQRFDVITKIEVVTEKRL--------------------VFPSVTICNVNK
LRKSAIASSPYKEMLIIDEQIAL--PYPAPC-IPGDFVCANGIYC------VKQY-----
------LRCDGIDHCKDFSDEDGCVYG-------ELKE----------------------
--------
>Deutero_Ambulac_Sakowv30020102m_976_411
------------------------------------------------------------
----------M-------------------------------------------------
------------------------------------------------------------
------------------------------------------------------------
------------------------------------------------------------
--EMQNS-----------------------------------------------------
------------------------------------------------------------
------------------------------------------------------------

```
------------------------------------------------------------
------------------------------------------------------------
---------------------QVTEKDDT-------------------------------
--QRKKDDK----------------------E---------SY-----K-----------
-AL--VNHFI-STTNAHGLPKAF---------E-GR--------------GI---FV---
--SSFWA----IVFLAALTGAALQISELLV-QYLE----------W---EVKIKM----K
-V-VS-E----------S-YLQF-----PGI-----------------------SICNTN
KL------RKS-----AIEKSE------------------H-----------------K-
---------GLNGV----------------------------------------------
---------------------------------DD-------DIVFPYYD----------
------------------------------------------------------------
---------EGGCM----------------ANDFACNN-----------TG---------
-CIKNYLHCDGYDNCR-D-------------------------RSDEMGCK-----YGRC
G--------NTEFRCPSGSDMGVCIKLK-FKCDRHLDCYDGEDENECVCKE---------
---KKEFSCL------DNGRCVPVN----------KLCDGNNDCADGSDE----------
-KECSEDGNYCPSDYFKCDNGR-------------------------TCIPA--------
----IFKCDGGADCS---DETDEMSCPTQN---------------------------PGC
ADDQFHC----------------DESYCI-------DNSWLCDGEYDCMDGTD-------
--------ELEMNCITSSGTNSTNTNPCQDWEFYCGDDKCINGDWECDTESDCPGGQDEV
NCLTTRAPTSTEVGSFMCELSGKLVEQYVCNNVVDCDDASDEYNCHNDASP-TYIPDAEE
LLRNYSTLSTDLSLFEDF----VSRY--------------Y-IDNM-------F------
--------GRV-----------VRENPPDWPGFIAY-----------SS------SPDYS
---------------------------------DLAYVL---------------------
-------------------------KL----------------RTDEMHKL----GHQLN
--DFV------LHCT----FD-S----V----------K---C-----------D-ME--
----------------------------------------------K-------------
------------------------------------------------------------
------------------------------------------------------------
-----------D----------F-------L-----------TFY-DDRYG--------N
CFQYN-------YK-------------HD-H-------------------DS----EQ--
-------LLSTKT------GP--------RY-------------GLKLTLFTEQD-----
----------E-----------------Y------------I-S-V---YG---------
------------------------------------------------------------
------------------------------------------------------------
----------HDSGARVVIH-PSYMRA--MPWSEGFTIAPGKIAFIGIKET------R--
-----VDRQ-----------PAP-Y--GE-C--ATN----------------IYQ-ETVY
----G-------------K-YYKESTCEESCIQDRMM----EY--CGCVDTM----L---
------------------------------------------------------------
---------RNA-----T---RCMLL----------N-----------------------
------------------------------------RTQ---DTCRQ---------L---
I-Y-------------------YSIQ-Q-------KL--------L------------GC
------------DCP-QSCKERY---YETS-LSQ------------------SLW-PS--
-------NTY------LKH-L-----------LK--------------------------
------------------------------------------------QI-H--------
----------------------------------AE------------------------
-N-------------PKT------------------LNI---------------------
---------------------------NNL------------------------------
-----E-------TS-----------R------QNLIRLELYYDDLNFQQIT-EKPEVSE
EAL-----------------LSSIGGSLGLYCGFSFLTIVEFIQFAFDL-----------
--------VKLT--------Y----------V-------RVFL-----------------
--------------------------------RR--------------------------
------------------------------------------------------------
------------------------------------------------------------
---------------------------------------------------------L--
------------------------------------------------------------
------------------------------------------------------------
------------------------------------------------------------
```

-KPVAVSA
>Protostome_Lophotroco_annelid_CAC9671100_1__Ofus_G19963_Owenia__fusiformis__
983_412
------------------------------------------------------------
----------M-------------------------------------------------
------------------------------------------------------------
------------------------------------------------------------
------------------------------------------------------------
------------------------------------------------------------
------------------------------------------------------------
------------------------------------------------------------
------------------------------------------------------------
------------------------------------------------------------
-----------------------ANKTSPEEV----------------------------
-DTQHDKDH----------------------P---------TF-----K-----------
-QL--IIDFA-DNSTLHGLPRSLR--------N-SV--------------RV---FR---
--RLIWL----GIFAGCFGYFIFQVIELIK-DYNK----------W---PILIKS----E
-I-RL-R----------T-FMDV-----PAV-----------------------TICNVN
ML------RKS-----KIEGTT------------------F-----------------D-
---------SLVDLD--------RKNFAN---------GNTRSTTVH----KKATPKPRN
K--------------------R-----------DG-------KVEETWERSRPPLNSNTD
VKAGRVNIRQ--ILRGYRPQVGIGSKENGVSHSVNTLRDVKPENGLQG------EGIG--
---------ENNYETF---DNRPIEAYNAVNDESPSRD-----------DGLLR------
-------QHDTADDVANDKIRIKRTLD----------TKPSAPGATATDPTT--------
------------------------------MFDQTTTTISGQIPTKSDPV----------
-------TTISDQTTTISGQIPTKS----------DPVTTISDQTTTISGQTPT------
----KSDPVTTISDQTTTIS----------------------------------------
------------------GQTPTKSDPVTTISD-----------------------QTTT
ISGQIPT---------------------------------KLDQTTTIFDQTA-------
-----TPGQIATNSDQTTTVTSVPICTCT-------------------------IPVAAT
SAHTQTSPPGSTTVSNDTFVESDTTSSYTGNNGWSN------------------------
--NSSLGQGHDPNTYDRYGPRMDTGYTA---------PFTF-KESP-------F------
---------------------LKIKNRNNFGQILNS-----------ST------AEDLS
---------------------------------DIFGLA---------------------
-------------------------TP----------------TVQQLDMY----GHRIS
--DLA------ALCT----FD-M----E----------N---C-----------K-R---
----------------------------------------------T-------------
------------------------------------------------------------
------------------------------------------------------------
-----------S--------GHW-------L-----------ETY-NKYYG--------K
CYTWN-------SA-------------IN-K-PE----------------GE----RP--
-------LVTTNF------GS--------RY-------------GLRLTLNVEPD-----
----------E-----------------Y------------T-G-L---LS---------
------------------------------------------------------------
------------------------------------------------------------
----------PTYGVRVSVH-DNSVVS--YPESDGLTVSVGNEMVFRLRKK------K--
-----YERQ-----------PNP-YP-SN-C--MDG-W--------------MR--PSKN
----RNR-----------G-EYTVLGCMMECLQQRLF----EI--CGCFDRI----V---
------------D-----------------------------------------------
---------EGE-----H---RCNYF----------N-----------------------
------------------------------------RTE---EFCRQ---------K---
V-Y-------------------HAYN-T-------GR--------L------------GC
------------RCP-AKCREST---YEVS-TSV------------------SAW-PS--
-------KHH------DHF-I-----------RE--------------------------
------------------------------------------------RL-N--------
----------------------------------KIN-----------------------
-N----------------------------------------------------------

```
---------------------------ISL------------------------------
-----H-------DI-----------K------NTTVRVHVYFEDFNEETVI-EVPVYTV
SGI-----------------LSNIGGTMGMFIGLSLCTIGELLELMLDV-----------
--------AILL--------Y----------K-------KL-------------------
ST------------------------------RF-------------------N------
-----------------------------------------IRRVG--------------
------------------------------------------------------------
------------------------------------------------------------
------------------------------------------------------------
------------------------------------------------------------
------------------------------------------------------------
----TAVT
>Deu_Ambulacrar_hemi_Ptyfla_40v0_9_20150316_1g26495_t1_scaffold55703_cov149_9
84_413
------------------------------------------------------------
----------M-------------------------------------------------
------------------------------------------------------------
------------------------------------------------------------
--------------------------------------------------------VSVV
MVVMRQPLSRN-------------------------------------------------
-----------------------------------LSQKITPYVLESLSVVYL-------
-------------------------------GEIDITTRQRLLNVLF---------PGE-
---------------------------------------------------TGI------
----------------------SMDRKIIDNILVS---TPGGGLL-SPNLQVIDLPVENN
QLTVGQLYEILHNALQAERYIHSETQSSVDVGSDIYRKV-------------------YE
ATMNRNTSAWHLRTTASHYA---V-------EIAFRQAHYGAL----MN-----------
-AT--MHAVS-DAPDIFDLIYTT---------VLDT--------------FE---EN---
--RL-----------TALENIT--AYQICM-EYMRG--AGY----R---GWSRNRYNIAM
-Y-VH-T----------C-KTTV-----LAI-----------------------T-----
-C------RKT-----GLLEAF--------G---------YSHG--------------D-
---------DV---------------------------IKTAHAILS-------------
---------------------------PSTETMDQ-------------------------
------------------------------------------------------------
------------------------------------------------------------
-----------------D------------------------------------------
------------------------------------------------------------
---------------------------V--------------------------------
------------------------------------------------------------
------------------------------------------------------------
------------------------------------------------------------
------------------------------------------------------------
------------------------------------------------------------
------------------------------------------------------------
------------------------------------------------------------
---------------------------------NLVGVN---------------------
------------------------ANY---------------DFEYLMTTV----GHKKD
--EMI------MRCK----WP-G----H----------Q---C-----------S-V---
----------------------------------------------D-------------
------------------------------------------------------------
------------------------------------------------------------
-----------N----------F-------T-----------TTY-ASD---------SI
CVTYN-------LG-------------ER-G-------------------------DI--
-------VEQVNG------GE--------RY-------------GLTVYLDVESH-----
----------E-----------------Y------------V-N-S---RQ---------
------------------------------------------------------------
------------------------------------------------------------
----------HYVGFRVIVH-DQDDIP--NTKDRGFNVAPGAYTAAALKKS------S--
```

-----SSYL-----------KEP-VGHTD-C--IDS--------PRGT--------LHYF
-----------------HG-NYSLAKCKVECETDYAI----KQ--CQCRYYT----M---
------------P-G---------------------------------------------
----------DA-----P---ICNV-----------------------------------
------------------------------------MQL---HNCYV--------PK---
M---------------------AKFT-D-----------------VQ----------GVC
S-----------YCK-QPCYEES---FTEH-LSY------------------ALF-PS--
-------GPL------VEE-L-----------AKKAEKPCPKAMALHYKKKITPYVFASL
SIVNLDEIHITTQ-----------DKVLD-------IIFPGETNISVHRIVENITASMPE
AG-LLPLIDVHDSPVKIHQLTVGQL-SEILRKALQSYYEQIYA-EI---------TIPLS
LNLYRYIH---SEGQSSVD------------IGSDRYRKVFEATMNSSASARLLRTIAPD
LVVNGEFRAALNMSYSYGALVNTTKHAVSNEFMDIFDMIYTTILDTFEENRITALENRTM
YQICME-------YM-----------R------KNTLELRVYFNEMREEKMT-QQQHYEP
FNL-----------------VCDIGGTLGLFFGASLLSVIEIIDFLLFR-----------
-------------------------------R-----CNKV-------------------
ESQ-----------------------------KN------------ESSIALENR-----
------------------------------------------------------------
------------------------------------------------GSEK--------
------------------------------------------------------------
------------------------------------------------------------
------------------------------------------------------------
---------------------------------------------------------SAT
SPNEEHKT
>Cyclo_UNC105_NP_001122595_1_Degenerinlike_protein_unc105__Caenorhabditis_ele
gans__55_414
------------------------------------------------------------
----------M-------------------------------------------------
------------------------------------------------------------
------------------------------------------------------------
------------------------------------------------------------
------------------------------------------------------------
------------------------------------------------------------
------------------------------------------------------------
------------------------------------------------------------
-------------------------------------------------------AEDRI
K-------------------SKLRRPASIESTMS--------------------------
-SRTKPRHKP----------SPSLMPHLMVGE---------SF----RKYRPIRMNGHLD
WQL--RKSFE-KQSTFHGISHAA---------T-AD--------------GKW-------
--RWFWY----TAFTICLLALLIQIFFLIS-KYRQ----------Y---GKTVDL----D
-L-KF-E------------NAPF-----PSI-----------------------TICNLN
PY------KKS-----AIQSNP--------------------------------------
---------NT-------------------------------KAMME-------------
------------------------------AY-SR-------------------------
------------------------------------------------------------
----------------------------------------------RIGSG---------
-----------------DKTEGIA---------------AALSATGGLHAKV--------
------------------------------------------------------------
----------RRAKRKAKGK--------------PRLRDRRYHQ----------------
-----------AFAQCLCDIEQ-------LTGDRKGSCFAAFKG-KIEIDTNNTA-----
----GFMNLHTSRCLCQLDTVSKALWPCFPYSSWK------------------EKLCSEC
VDNTGHCPMRFYKGNELYENIKEQVDLCLCHKEYNH-----------CVSTRDDGII---
--LEISPNDELNDLDIGKKIAS--------------------------------------
-----------------------------------------QLSAQ--------------
QEKQAEVTTTEA----------------------------------PTV-----------
----------------------------------------TQALGFEEL------TDDIA
--------------------------ITSQAQENLMFAV---------------------
------------------------GEM----------------SEKAKESM----SYELD

--ELV------LKCS----FN-Q----K----------D---C-----------Q-MD--
----------------------------------------------R-------------
------------------------------------------------------------
------------------------------------------------------------
-----------D----------F-------T-----------LHY-DNTFG--------N
CYTFN-------YN-------------RT-A-E---------------------------
-------VASHRA------GA--------NY-------------GLRVLLYANVS-----
----------E-----------------Y------------L-P-----TT---------
------------------------------------------------------------
------------------------------------------------------------
----------EAVGFRITVH-DKHIVP--FPDAFGYSAPTGFMSSFGVRMK------Q--
-----FIRL-----------EPP-Y--GH-C--RHG-G------EDAA--------TFVY
----TGF-------------QYSVEACHRSCAQKVIV----EA--CGCADPM----Y---
------------P-V--AE-----------------------------------------
------MFGNNT-----K---PCQAV----------N-----------------------
------------------------------------MDQ---RECLR---------N---
T-TLWL---------------GELYS-K-------GKE------AII----------PDC
------------YCH-QPCQETN---YEVT-YSS------------------ARW-PS--
-------GSAK---------V-----------ME--------------------------
------------------------------------------------CL----------
------------------------------------------------------------
------------------------------------------------------------
----------------------PGDF-LCL------------------------------
-----E-------KY-----------R------KNAAMVQIFYEELNYETMQ-ESPAYTL
TSV-----------------LADLGGLTGLWIGASVVSLLEIVTLIVFA-----------
--------TQAY--------V----------R-------KR-------------------
KGSISAQSHHSVP----------------VH-RASRVS----------LNTLHKS-----
---------------------------------------STTQSVKLSVMDIRSI-----
--------------KSIHSNHSSKSKQSILIE---DLPPAI----QEQSDDEEE------
--------TTESSRTNGSCRYL--------------------------------------
--------------------------------------------APGEDLPCLCKY----
---------------HPDGSIRI-----MKALCPVHGYMVRRNY----------------
----------------DYS--------VSNS---EEEDAED--------EVHREPEPFYS
APYEHRKK
>Lopho_Phor_aus_TRINITY_DN315771_c0_g2_i2_p1_T
------------------------------------------------------------
----------M-------------------------------------------------
------------------------------------------------------------
------------------------------------------------------------
------------------------------------------------------------
------------------------------------------------------------
------------------------------------------------------------
------------------------------------------------------------
------------------------------------------------------------
------------------------------------------------------------
------------------------------------------------------------
---NSEVNK----------------------R---------NL-----K-----------
-GR--AKEFC-ETTSAHALGQTV--------R-SGK-------------------VK---
--AIFWS----LVFLGSLAGCVWNIIHVVE-SYAG----------F---GFSVKS----K
-L-EM-E---------PS-SLKF-----PSV-----------------------TICNLN
PI------SFS-----RQARFV--------GD--------F-----------------ER
ST-------GL-------------------------------------------------
---------------------------------DE-------------------------
----------------------------------------K-------------------
------------------------------------------------------------
-----------------D------------------------------------------
------------------------------------------------------------

------------------------------------------------------------
------------------------------------------------------------
------------------------------------------------------------
-----------------------------------------------C------------
------------------------------------------------------------
------------------------------------------------------------
------------------------------------------------------------
-------------------------------------------------------DNEFY
RNETNR---------PLV--------LDLYRWSELPQFW---------------------
------------------------FEY----------------NENVTDMY----GHSKK
--DFI------VDCV----FS-K----K----------L---C-----------E-----
----------------------------------------------D-------------
------------------------------------------------------------
------------------------------------------------------------
-----------N----------F-------Y-----------VTT-DPNLY--------N
CYTLE-------PN-------------KY-N-S---------------------------
-------DLHAGV------GI--------EA-------------GLSLTLFVENT-----
----------RVQNA-------------Y------------T-G-S---FS---------
------------------------------------------------------------
------------------------------------------------------------
-IDS---YQTGNVGVKVAMH-VSGSHP--NPNARGVIAEVGKSTDFILRTV------N--
-----RTAL-----------GPP-YP-SP-C--NPK--------KTI---------ESSH
----RQE-----------L-QYEENLCFASCLQNAIA----KK--CGCVSINP---V---
----------AVP-I--AD-----------------------------------------
------KFGIDL-----P---FCGSYP--------CNAT---------------------
----------------------------------KVTEN---YECVH---------D---
V-T-------------------RRFLLK-------ND--------------------PNC
R-----------DCT-KPCNEVS---YEAT-KSQ------------------SKW-PS--
-------ELH------QSE-F-----------IG--------------------------
------------------------------------------------WLASKD------
------------------------------------------------------------
-N-------------TAL------------------YNQ---------------------
------------------LVRNTSDD-KIS------------------------------
-----R-------FI-----------E------NNFLRINIYFGDFYVRKDT-ETLNMDW
FDL-----------------LSSVGGAFGFWVGISVVTGVEVLELLLDC-----------
--------IVIF--------L----------N-------KP-------------------
SRK-----------------------------RN-------------------Q------
---------------------------------------KDLRNTT--------------
------------------------------------------------GNND--------
------------------------------------------------------------
------------------------------------------------------------
------------------------------------------------------------
------------------------------------------------------------
--VEVYNN
>Lopho_Phor_aus_TRINITY_DN291806_c2_g12_i3_p1_T
------------------------------------------------------------
----------M-------------------------------------------------
------------------------------------------------------------
------------------------------------------------------------
------------------------------------------------------------
------------------------------------------------------------
------------------------------------------------------------
------------------------------------------------------------
------------------------------------------------------------
------------------------------------------------------------
-----------------------AEVELSCVNDVHFGLE-----MSNGVN----PKTHLE
NMAKEEEEK----------------------Q----------SL-----K----------

-EF--LKEWA-ENAACDGVPQIV---------L-RK--------------SW---VK---
--RILWL----LIVLGLTAYTIEEVYGVFD-EFFT----------Y---PVNTNI----N
-Y-TY-S----------N-QLTF-----PAI-----------------------TLCNMN
PI------RKS-----MLSEAG------------------------------------T-
---------TF-------------------------------DDIFG-------------
---------------------------------ST-------------------------
------------------------------------------------------------
------------------------------------------------------------
-----------------NGT---------------------FPGRKKREIGPQ-------
------------------------------------------------------------
------------------------------------------------------------
------------------------------------------------------------
----------------------EETAPSGSQQRG--------------------------
------------------------------------------------------------
------------------------------------------------------------
------------------------------------------------------------
------------------------------------------------------------
-------------------------------------------------------KEKMT
------------------------------FINQFKRQW---------------------
------------------------ATL----------------SYPKRIQL----GHQLP
--NFV------VTCS----LG-G----Q----------S---C-----------N-----
------------------------------------------------------------
------------------------------------------------------------
------------------------------------------------------------
-----------K----------F-------F-----------TYS-TGTFG--------N
CFVFN-------SG-------------LR-N--------------------T----TV--
-------QSISKT------GP--------LF-------------GLTMEIFLEES-----
----------E-----------------Y------------L-P-E---VT---------
------------------------------------------------------------
------------------------------------------------------------
----------EQTGARLHIN-HQKLMP--FPENAGYNLAPGYLTTIGLKRV------E--
-----ISRL-----------LGY-E--SN-C--TKV--------TTAE----AEG-YSIY
----SKR--FG-------V-SYSREACHNTCYQRKVI----EI--CRCGDAD----Y---
------------L-F--DY-------------------SFFKSLL---------------
-----PDTVKSI-----K---PCLS-----------------------------------
------------------------------------ATE---EACVS---------R---
V-L-------------------KLYA-S-------NN--------I------------TC
------------DCP-IPCSESD---YQAT-VTI------------------SPW-PS--
-------KLY------EPT-L-----------RS--------------------------
------------------------------------------------SY-S--------
----------------------------------SL------------------------
-N----------------------------------------------------------
-----------------------RND-ILN------------------------------
-----T-------TL-----------S------RNLLKVEIYYDTLNYQYIY-ESPKYTN
LSL-----------------ISDLGGQIGLWLGASLFLLLQVLELFIDV-----------
--------IIFG--------C----------G-------RL-------------------
FKSGS---------------------------RV--------------------------
------------------------------------------------------------
------------------------------------------------------------
------------------------------------------------------------
------------------------------------------------------------
------------------------------------------------------------
------------------------------------------------------------
-----SSR
>Lopho_Phor_aus_TRINITY_DN297387_c12_g1_i2_p1_T
------------------------------------------------------------
----------M-------------------------------------------------

```
------------------------------------------------------------
------------------------------------------------------------
----------------------------------------------------------SG
RRSAWNDYHSE-------------------------------------------------
------------------------------------------------------------
-----------SVWDEQRAG-------------TQNGDKHGEDDCK--------------
------------------------------------------------------------
---------------------------------------------------------DED
D----------------------QPRSSW-------------------------------
-------------------------------Q---------II-----K-----------
-DC--GNDYL-DVSSMHGLAFFV---------G-SR-------------YHP---LR---
--RLLWF----ALFTASFTWLMIVVTTAFD-NLQR----------K---PVLTKV----K
-V-KY-E----------D-NARF-----PTV-----------------------TICNLN
KY------RKS-----YVEKHH----------------PHS-----------------E-
---------KL-------------------------------LRYIY-------------
------------------------------------------------------------
------------------------------------------------------------
------------------------------------------------------------
------------------------------------------------------------
------------------------------------------------------------
------------------------------------------------------------
------------------------------------------------------------
--------------------------PVLR------------------------------
------------------------------------------------------------
------------------------------------------------------------
------------------------------------------------------------
------------------------------------------EDE---------------
------------------------HIHLDWSDPKYAWS-----------------LNRS-
---------------------------------SFKEFA---------------------
------------------------------------------------LKA----APQMM
--DNI------KIVD----FG-A----H--------WLT---I-----------S-----
----------------------------------------------N-------------
------------------------------------------------------------
------------------------------------------------------------
-----------E----------F-------S-----------TIE-T-DFG--------M
CFQFN-------PN-------------GT-------------------------------
-------WYSQLS------GE--------MH-------------GLAIVLDVQQD-----
----------D-----------------Y------------Y-F-G---DS---------
------------------------------------------------------------
------------------------------------------------------------
----------FSTGFKIALH-RHDEVP--FPNQFSFGVSPGQELLVGMTMT------E--
-----VKSL-----------PAP-YEDGT-C--VDT-EAGDFINP-------LK----YF
------------------K-TYKYNSCRKECQTVFSV----GQ--CGCKPAF----D---
------------P-G---------------------------------------------
----------PT-----R---TCIG-----------------------------------
------------------------------------EEY----RCYR---------L---
A-T-------------------EEYA-K-------NH--------SI---------EREC
------------NCR-LPCHERG---YVYQ-LSS------------------LKF-PA--
-------IQY------EAY-F-----------KN--------------------------
------------------------------------------------TH----------
------------------------------------------------------------
-N----------------------------------------------------------
---------------------------ISL------------------------------
-----Q-------YA-----------R------ENLMLVRIYFEKMNYELVE-QVLGYDG
MNF-----------------FADFGGNSGLCIGASLLTVAELLELIAVI-----------
--------VFNL--------I----------F-------RR-------------------
RK------------------------------KT-------------------S------
```

-----------------------------------------PEKQ---------------
------------------------------------------------------------
------------------------------------------------------------
------------------------------------------------------------
------------------------------------------------------------
------------------------------------------------------------
--------
>Lopho_Phor_aus_TRINITY_DN307417_c2_g3_i8_p1_T
------------------------------------------------------------
----------M-------------------------------------------------
------------------------------------------------------------
------------------------------------------------------------
------------------------------------------------------------
-ASEPETIPLN-------------------------------------------------
-GLNHNLRKRRVQNYIVI------------------------------------------
------------------------------------------------------------
------------------------------------------------------------
---------------------------------------------------------DLV
NNGE-------------------QDEQEN-------------------------------
-------------------------------G---------VY-----K-----------
-SR--FECFG-EDTDFHGLKYIS---------R-GE-------------NSG---VK---
--RLLWL----LLIIASLIYLVIEIYGRVA-YYLT----------F---PHVTKV----D
-V-EF-K----------S-DLDF-----PAF-----------------------TVCNMN
KY------RAY-----SMTDMD------------------L-----------------F-
---------HM------------G-EIL---------------DIVD----E--------
---------------------------------NM-------NLHH--------------
------------------------------------------------------------
------------------------------------------------------------
------------------------------------------------------------
------------------------------------------------------------
------------------------------------------------------------
------------------------------------------------------------
--------------------------HSHY------------------------------
------------------------------------------------------------
------------------------------------------------------------
------------------------------------------------------------
--------KESF------------------------------VKR---------------
------------------------VQNINKTEIRLK-------------------NESF-
---------------------------------DLEEFV---------------------
------------------------------------------------RRT----GHQAR
--DLI------LYCK----WK-D----E----------L---C-----------T-G---
----------------------------------------------EA------------
------------------------------------------------------------
------------------------------------------------------------
-----------N----------F-------T-----------HVF-T-HLG--------N
CYTFN-------SG-------------KK-K-----------------------------
-------VAARKA------GA--------GN-------------GLTLYLNIEEH-----
----------T-----------------Y------------L-E-N--------------
------------------------------------------------------------
------------------------------------------------------------
-------KDAADEGLKVMVH-SQMEPP--FIRELGFGIMPEQHHYIAIRKT------E--
-----ITNL-----------PHP-Y--GK-C--VDT-NDSR---R-------HP----DF
------------------E-HYTIPGCRIQCETKNVL----ET--CGCRLPE----M---
------------P-KMPDH-----------------------------------------
----------EA-----K---ICTP-----------------------------------
------------------------------------PHQ---YKCAV---------K---
E-L-------------------DRIS-E-------SD---------------------VC

------------ICQ-TPCHIED---YRFT-HSS------------------AKL-RP--
-------ETI------EKL---------R-------------------------------
------------------------------------------------EL----------
------------------------------------------------------------
QG-------------DKL------------------------------------------
---------------------------VPA------------------------------
-----N-------LT-----------S------NNVAVINVFFEALSMEKIE-QRVAYPW
PSL-----------------LGDIGGQMGLFIGASVLTIVHAIEFCMNE-----------
--------FARG--------V----------K-------DR-------------------
AR------------------------------KK-------------------N------
-----------------------------------------EKSASLRSNNN--------
------------------ANRDFQQDPT--------------------------------
------------------------------------------------------------
------------------------------------------------------------
--------------------------------LP--------------------------
------------------------------------------------------------
TVLKEESL
>Lopho_Phor_aus_TRINITY_DN307786_c0_g2_i2_p1_T
------------------------------------------------------------
----------M-------------------------------------------------
------------------------------------------------------------
------------------------------------------------------------
----------------------------------------------------------SK
GVKTSGGVWAT-------------------------------------------------
------------------------------------------------------------
--------NVFNSGQHDNTL-------------YRQHVEAKQDIT---------------
------------------------------------------------------------
---------------------------------------------------------ADI
D----------------------QPNTGL-------------------------------
-------------------------------S---------IV-----K-----------
-DG--IQKYA-EDSSLHGVQYFG----------G-AK------------YNP---VR---
--RLIWF----LMFVGSFTFLVINVSDAYQ-TLMR----------K---PVQTKL----S
-V-EY-T----------K-TARF-----PTV-----------------------TLCNFN
KF------RGI-----YVWTQP-------------A----L-----------------M-
---------EI-------------------------------LRYLY-------------
------------------------------------------------------------
------------------------------------------------------------
------------------------------------------------------------
------------------------------------------------------------
------------------------------------------------------------
------------------------------------------------------------
------------------------------------------------------------
--------------------------PLSE------------------------------
------------------------------------------------------------
------------------------------------------------------------
------------------------------------------------------------
------------------------------------------EDR---------------
------------------------KLQLNWEDPKYAPF-----------------INGS-
---------------------------------TYLDFA---------------------
------------------------------------------------KIA----GHQLR
--DSI------YFAR----FK-G----M----------N---L-----------DIQ---
----------------------------------------------K-------------
------------------------------------------------------------
------------------------------------------------------------
-----------E----------F-------K-----------QVV-T-DFG--------L
CFQFN-------SN-------------GT-------------------------------
-------WNTSRS------GT--------NN-------------GLWIQMNAEHY-----

```
----------M-----------------Y------------F-Y-G---DS---------
------------------------------------------------------------
------------------------------------------------------------
----------NSAGFKVALH-NFDEEP--LVNELGFAISPGQESFVSAQIK------E--
-----IKSL-----------PPP-YEGGK-C--KNT-TKENFVNH-------LA----YY
------------------E-TYSMTGCRRECRTNYTV----KR--CGCKYIY----D---
------------P-G---------------------------------------------
----------PA-----R---VCDA-----------------------------------
------------------------------------PEM----GCYR---------Q---
A-D-------------------GEYT-Q-------DE--------SI---------EDDC
------------DCF-VPCHEIQ---YEHR-LST------------------SMY-PG--
-------NHI------IYG-L-----------QA--------------------------
------------------------------------------------QY----------
------------------------------------------------------------
-N----------------------------------------------------------
---------------------------VTK------------------------------
-----E-------IA-----------R------ENFLELRIFFEKLNYDLVE-QVPSYDA
MNF-----------------FTDLGGNLGLCIGASLLTAAELLEHIGIT-----------
--------FFQV--------L----------Q-------KFIG-----------------
RK------------------------------QT-------------------H------
-----------------------------------------PKKIQ--------------
------------------------------------------------------------
------------------------------------------------------------
------------------------------------------------------------
------------------------------------------------------------
------------------------------------------------------------
---REGVF
>Lopho_Phor_aus_TRINITY_DN307786_c0_g5_i2_p1_T
------------------------------------------------------------
----------M-------------------------------------------------
------------------------------------------------------------
------------------------------------------------------------
----------------------------------------------------------PK
GFKSSNDIWAT-------------------------------------------------
------------------------------------------------------------
--------KVTPIAPYGQ----------------------KESDN---------------
------------------------------------------------------------
---------------------------------------------------------KDE
G----------------------APESGTVL-----------------------------
-------------------------------S---------IV-----K-----------
-EG--VTKYA-EETSCHGVQYFG---------G-AK-------------HNP---GR---
--RIVWF----LMFVGSFTWLVINVNNAYQ-TLMR----------N---PVQTKL----H
-V-EY-T----------Q-TAKF-----PTV-----------------------TFCNFN
KF------RSP-----YVLAQP-------------G----L-----------------I-
---------DI-------------------------------LRYVY-------------
------------------------------------------------------------
------------------------------------------------------------
------------------------------------------------------------
------------------------------------------------------------
------------------------------------------------------------
------------------------------------------------------------
------------------------------------------------------------
--------------------------PVTE------------------------------
------------------------------------------------------------
------------------------------------------------------------
------------------------------------------------------------
------------------------------------------EDR---------------
```

------------------------EQQFDWTDPLYAPL-----------------VNGS-
---------------------------------QYLEFA---------------------
------------------------------------------------KIA----GHQIR
--DSI------YVAR----FK-G----V----------N---L-----------DIE---
----------------------------------------------K-------------
------------------------------------------------------------
------------------------------------------------------------
-----------E----------F-------K-----------RVV-T-DFG--------L
CFQFN-------SN-------------GT-------------------------------
-------WNTSRS------GT--------NN-------------GLWLEMNVEHN-----
----------L-----------------Y------------I-Y-G---DS---------
------------------------------------------------------------
------------------------------------------------------------
----------NSAGYKVALH-NHDEEP--LVNELGFAISPGQETFVSVQIK------E--
-----IKSL-----------PLP-YEGGK-C--KNT-TKEDFVNH-------LE----NY
------------------K-PYSLTGCRRECRANYTK----QK--CGCKYIY----D---
------------P-G---------------------------------------------
----------PA-----R---TCNA-----------------------------------
------------------------------------PEL----GCYR---------N---
A-D-------------------EEYT-R-------NA--------SI---------ENDC
------------NCF-VPCHEMQ---YEHR-LST------------------SMY-PG--
-------NHV------IQR-L-----------MD--------------------------
------------------------------------------------DF----------
------------------------------------------------------------
-N----------------------------------------------------------
---------------------------ITK------------------------------
-----E-------LA-----------R------ENFLELRIFFEKLNYD-----------
------------------------------------------------------------
------------------------------------------------------------
------------------------------------------------------------
------------------------------------------------------------
------------------------------------------------------------
------------------------------------------------------------
------------------------------------------------------------
------------------------------------------------------------
------------------------------------------------------------
--------
>Lopho_Phor_aus_TRINITY_DN312240_c0_g5_i3_p1_T
------------------------------------------------------------
----------M-------------------------------------------------
------------------------------------------------------------
------------------------------------------------------------
------------------------------------------------------------
------------------------------------------------------------
------------------------------------------------------------
-------DNFEMAWDNQQTR-------------TQQGVKSGREDC---------------
------------------------------------------------------------
---------------------------------------------------------KDE
E----------------------LSRSSW-------------------------------
-------------------------------Q---------IF-----K-----------
-DC--VTDYL-DVCSMHGLAFFL---------G-SM-------------YHP---LR---
--RILWF----LLFTASFVWLMIVVATAYD-NLKR----------K---PVQTKM----K
-V-KY-A----------D-NASF-----PTV-----------------------TICNLN
KY------RKK-----YVEENY----------------PDS-----------------E-
---------EV-------------------------------VRFMH-------------
------------------------------------------------------------
------------------------------------------------------------

------------------------------------------------------------
------------------------------------------------------------
------------------------------------------------------------
------------------------------------------------------------
------------------------------------------------------------
--------------------------PVVP------------------------------
------------------------------------------------------------
------------------------------------------------------------
------------------------------------------------------------
------------------------------------------EDS---------------
------------------------TANIDWSEPRHAWS-----------------LKES-
---------------------------------NILHLA---------------------
------------------------------------------------RDG----AHEIH
--KSF------ISAQ----FG-G----H--------ELP---V-----------A-K---
----------------------------------------------P-------------
------------------------------------------------------------
------------------------------------------------------------
-----------N----------F-------V-----------TVE-T-DSG--------V
CFQFN-------PK-------------GT-------------------------------
-------WYSELS------GK--------TF-------------GLMVMLNADQE-----
----------N-----------------Y------------Y-F-G---DS---------
------------------------------------------------------------
------------------------------------------------------------
----------FSVGFKIALH-RHDEEP--FVDEFGFVVSPGQEVLVPMVLN------E--
-----VISL-----------PEP-YEGGT-C--VDT-KVKDFTNP-------LE----YY
------------------S-PYNYNSCRKECQTRYSV----EQ--CGCKLAF----D---
------------P-G---------------------------------------------
----------SS-----R---ICVG-----------------------------------
------------------------------------DEY----ACYR---------L---
S-T-------------------QNYA-K-------NA--------SI---------ERSC
------------NCR-MPCHQVG---YDYR-LSS------------------AMF-PG--
-------LQY------RSY-L-----------AN--------------------------
------------------------------------------------AH----------
------------------------------------------------------------
-N----------------------------------------------------------
---------------------------LSL------------------------------
-----D-------YA-----------R------QNLVVVRLYFEKLNYQLVE-QVLSYDG
MNF-----------------FADLGGNSGLCIGASLLTLAELIELIGVI-----------
--------IFNM--------I----------F-------RR-------------------
RE------------------------------KT-------------------T------
-----------------------------------------PEKNN--------------
------------------------------------------------------------
------------------------------------------------------------
------------------------------------------------------------
------------------------------------------------------------
------------------------------------------------------------
------AK
>Lopho_Phor_aus_TRINITY_DN316400_c0_g1_i8_p1_T
------------------------------------------------------------
----------M-------------------------------------------------
------------------------------------------------------------
------------------------------------------------------------
------------------------------------------------------------
------------------------------------------------------------
------------------------------------------------------------
------------------------------------------------------------
------------------------------------------------------------

------------------------------------------------------------
-----------------------ADTGQKHYSYLTG-----------------------N
DYENVPIHA----------------------L---------TY-----K-----------
-AL--WRGFQ-ENTSGHGFSRIY---------L-TP--------------GT---VK---
--KALWT----LVCIGGFTAWGVHTTTLLV-NFFY----------Y---HVDVTV----Q
-I-LT-A----------K-NLSY-----PAV-----------------------TICNLN
QI------KSS-----KISDVL--------------------------------------
---------KL-------------------------------DDLYA-------------
------------------------------------------------------------
------------------------------------------------------------
------------------------------------------------------------
------------------------------------------------------------
------------------------------------------------------------
------------------------------------------------------------
------------------------------------------------------------
------------------------------------------------------------
------------------------------------------------------------
------------------------------------------------------------
------------------------------------------------------------
---LAAQTVDDG------------------------------------------------
------------------------------------------DGNLGQR------DQYSE
------------------------------YVDILRGAV---------------------
------------------------ADM----------------DEAAKSDA----GHQRG
--DFI------LGCT----YK-G----Y----------E---C-----------Y-G---
----------------------------------------------S-------------
------------------------------------------------------------
------------------------------------------------------------
-----------D----------V-------M-----------EFF-DPIYG--------N
CFTFN-------GA-------------DG-S------------------PQK----DY--
-------MRAHST------GP--------ES-------------GLTLMLNIEQS-----
----------E-----------------Y------------L-P-----FT---------
------------------------------------------------------------
------------------------------------------------------------
----------QSAGVRLVVH-NDTVMP--FPADEGINLEPGKLTMVGMRQI------Q--
-----IIRK-----------GNP-H--GV-C--IDP--------TSY-----NGD-RNLY
----EDK--FR-------V-RYTYQACKKTCYQKAIT----ER--CKCFDPH----Y---
------------P-T--TV-----------------------------------------
------NVSDTL-----K---ACRYG----------T-----------------------
------------------------------------IEQ---DECSE---------N---
I-T-------------------QDFA-D-------GN--------L------------RC
------------DCP-LSCSESA---YALT-ISS------------------SQW-PS--
-------DNY------KTA-L-----------FT--------------------------
------------------------------------------------KL-T--------
----------------------------------SL------------------------
-N------------NSNI------------------DAI---------------------
----------------------LTNA-ADK------------------------------
-----D-------IY-----------R------RNFLKVAVYFEDLNYEEIE-QKAAYTG
SSL-----------------ISDLGGSIGLWIGWSVITIVELFEFLSDV-----------
--------LILG--------C----------S-------RK-------------------
SS------------------------------KR-------------------A------
-----------------------------------------VGHSSDT------------
------------------------------------------------------------
------------------------------------------------------------
------------------------------------------------------------
---------------------------------P--------------------------
------------------------------------------------------------
---VKSIK

>CAE7949834_1_ASIC1_Symbiodinium_KB8
------------------------------------------------------------
----------M-------------------------------------------------
------------------------------------------------------------
------------------------------------------------------------
------------------------------------------------------------
------------------------------------------------------------
----------------------------------------APQTPPDV------------
------------------------------------------------------------
---------------------------------------------------AGSAWW---
-------------------RPAASLAELPSQ---------------------AAGFANRR
DMAVPL------------RQRRLERQADGAAV----------------------------
---ALVAGQ---------------------PE---------GF-----------------
-QP--LQEFS-DNVTVRELRFVTASTTTGALLS-AV--------------SV---CK---
--RWLWL----VLLLGALVALVVIVTDRVQ-YFLQ----------N---PTATVV----K
-W-ES-R----------A-PVPF-----PAV-----------------------TVCNEN
GF------QQS-----RVAQYG------------------L------------------S-
---------TL-------------------------------------------------
------------------------------------------------------------
------------------------------------------------------------
------------------------------------------------------------
------------------------------------------------------------
------------------------------------------------------------
------------------------------------------------------------
------------------------------------------------------------
------------------------------------------------------------
------------------------------------------------------------
------------------------------------------------------------
------------------------------------------------------------
------------------------------------------------------------
------------------------------------------------------------
------------------------------------------------------------
-------------------------------------------NNSVLELM----TQPVN
--SFI------LRCE----FA-G----L----------E---C-----------F-A---
----------------------------------------------D-------------
------------------------------------------------------------
------------------------------------------------------------
-----------N----------F-------S-----------VLL-DAQFG--------A
CYTFHGLRALTIDG----E----EFN-AS-G--------------------P----NP--
-------LTSART------GP--------DY-------------GLRLVLDIEQA-----
----------E-----------------Y------------V-G-----YSPRTSVRQAS
GADQGPGTEPDSSGSAQTSPQTSPQTSPQTSPQTSPQMSAQASAQASPQTSSSSATASSS
PPPAAASSSPQASSPAGASPTRTPSPSQPARRLLAPRRVQEDPGPGSARDARAPPDNRNS
TVEALF-KGPESAGIRVIIH-DPSVLA--NP-TSGFGVGPGSASDIAFERV------D--
-----RSRL-----------FPP-W--GT-C--NEA-----------W---HSES-----
-------------GATDPA-GYAKTACERDCYYASVA----DV--CECLLLP----W---
------------P-T--TMEERP-------------------------------------
----------SL--------YVCSS-----------------------------------
------------------------------------EEK----ACIA---------T---
Q-L-------------------SRFQ-A-------GE--------L------------GC
----------SASCP-DRCLETL---YTAT-AGR------------------QLW-PS--
-------VAS------AST-I-----------LD--------------------------
------------------------------------------------VVRSAK------
------------------------------------------------------------
QD----------------------------------------------------------
------------------------TS-ITP------------------------------
-----E-------YL-----------A------QNMMSVRVFPPTTEYQVVT-TVRAYSF

ESM-----------------LGEVGGNTGLWAGMSALTIVEILEFVMLA-----------
--------LSAW--------LCSRG---GCCR-------RG-----------------GK
RAP-----------------------------RT--------------------------
---------------------------------------NVEHGTELHEGEAI-------
------------------ELVAPQADE---------------------ESSA--------
--------PAVAPQAAA-------------------------------------------
------------------------------------------------------------
------------------------------------------------------------
--------------------------------------------------------AASG
RAREAWRE
>Filast_tunicaraptor_GIQG01049703_1__p1__GENE_GIQG01049703_1__GIQG01049703_1_
_p1___ORF__type_complete__len_874____score_117_73__GIQG01049703_1__2484_5063_
__979
------------------------------------------------------------
----------M-------------------------------------------------
------------------------------------------------------------
------------------------------------------------------------
------------------------------------------------------------
------------------------------------------------------------
------------------------------------------------------------
------------------------------------------------------------
---------------------------------------------------TG-------
--------------------------------------------------AASDERLEKL
QAQLDVL-----------AAQLRAARKEADNGGEYD-----------------------D
AALGSPAED---------------PYGAGSGE---------SC--------L--------
-SM--LTGCM-KESTVHGIPHAF---------AFNR--------------TI---WR---
--RAMWV----TYFTAAMCGLLAFCVGRVR-YYYE----------R---PFSSSV----S
-L-EY-A----------P-NLAF-----PAV-----------------------SVCLTN
GY------KNSGYAAEAVREHV-------------PGAAAF-----------------D-
-------NAEV-------------------------QQHLESLLFYG-------------
---------------------------------DTLA----S------------------
------------------------------------------------------------
----------------------P----------------------NGASGGTSVARKVRA
AAAAVAAAAARVAREAEDGGA-C-----------------PRPLSDTPGDVA----FA--
------------------------------------------------------------
--------------------------ELI-----ALLHDSGLASSALSFNVDDTVE----
---------------TACAAW---------------------------------------
----------------ADAYYEGEETPTVVSY----------------------------
------------------------------------------------------------
------------------------------------------------------------
----------------------------------------------------------GE
SAVNFEPMCAKY------------------------VLLG--DDPVY-HE---FKTA---
------------------LTAAGAPAHNSTAAWSDALA-----------------AEGVA
SSQNA---------------------LVLPLRCEVIASLSEVAPPTPVPSSTGAAPTPTA
RAISTLIHPAVLSAKESALIAFISDEL----------------DEEELEKM----AYTAE
--EVI------LGCL----WK-G----G--------ALP---C------------S-H---
----------------------------------------------R-------------
------------------------------------------------------------
------------------------------------------------------------
-----------N----------F-------T-----------TFF-ANEWG--------N
CFTFN-------NP-------------QTLR--------------------N----DV--
-------LHSAAP------GP--------KG-------------GLVVTLFIHQD-----
----------E-----------------Y----------EAL-P-A-GSIT---------
------------------------------------------------------------
----------------------------------------------------------ED
NFAA---LAIEGASVKLLMH-DQRDHP---EMDSALDVAPGRYHLAAMRKL------V--
-----VNNL-----------GQP-Y--SD-C--QED----------------VAAALTST

```
----DGVVLSTIP---GRT-RYNPKTCFKECTVREIF----KE--CACLPPT----S---
------------R-V--AR-----------------------------------------
--------GKRV-----D---RCLS-----------------------------------
------------------------------------DEQ---VACQD---------A---
V-E-------------------GRLS-N-------DP-------NLQ----------TTC
------------ACV-KPCESVT---FPVS-LSA------------------AQW-PS--
-------EVN------RDV-F-----------LA--------------------------
------------------------------------------------DI----------
------------------------------------------------------------
---------------EGV-------------LGMQ-------------------------
---------------------------VPP------------------------------
-----T-------FF-----------D------VNIVNLNVFYSDLNFERIE-QQGVMTV
GSL-----------------LAEIGGMLGLFVGMSVLTLAEFIEFLFFG-----------
--------SRRQ--------V----------R-------KC-------------------
KQGGDPIG-P----------------------KN----------------GVPSD-----
-------------------------------------ACGGADRVDVVLEEPNA------
MNMQGLP---------------LSKEATAA------------------GAGRGADP----
------------------------------------------------------------
-------------VRILKLQRA--------------------------------------
------------------------------------------------------------
----------------------------------------------------------DS
VNSVHSAV
>Filast_tunicaraptor_GIQG01084220_1__p1__GENE_GIQG01084220_1__GIQG01084220_1_
_p1___ORF__type_5prime_partial__len_641___score_99_41__GIQG01084220_1__3_1925
__864
------------------------------------------------------------
------------------------------------------------------------
------------------------------------------------------------
------------------------------------------------------------
------------------------------------------------------------
------------------------------------------------------------
------------------------------------------------------------
------------------------------------------------------------
------------------------------------------------------------
------------------------------------------------------------
-----------------------------DGDGDGD-----------------------N
DGDGDGDGD---------------GDGAGDGE----------------------------
------------------------------------------------------------
------------------------------------------------------------
------------------------------------------------------------
------------------------------------------------------------
------------------------------------------------------------
------------------------------------------------------------
------------------------------------------------------------
----------------------------------------------GEGGGNG-------
-----------------DG--------------------------DDDAVAL----YR--
------------------------------------------------------------
--------------------------RFR-----SSLGAAGYDVTYVARDVGFQVA----
---------------GLCAST---------------------------------------
----------------AAALDAGQPTPAYVMWNNATV-----------------------
------------------------------------------------------------
------------------------------------------------------------
--------------------------------------------------------EGGL
ADVNWAPLCDKY------------------------VPLY--NDPLYLHV---MDLA---
------------------TKVTETTDSADLANLMDAVN-----------------SADVQ
ESSNV---------------------VLGALVADINAFLTD--DPFLHPRSDGLAP----
--IGSLDWASTQTEEQVARAERLSTEA----------------VSLDAAKLW---TFEAS
```

--EFI------INCR----WK-N----G--------ALP---C-----------S-A---
----------------------------------------------A-------------
------------------------------------------------------------
------------------------------------------------------------
-----------N----------W-------T-----------TFY-VEEYG--------S
CFTFN-------GP-------------KE-D--------------------A----AV--
-------LASGSP------GP--------NG-------------GLTLTLYIDQP-----
----------E-----------------H----------FTV-E-A-GVLD---------
------------------------------------------------------------
----------------------------------------------------------DD
NLDI---LLTPGAGVKILMH-DQKDSP---EMDGATDLAPGRYYSLSMQKI------I--
-----IDNL-----------GPP-Y--GD-C--REE----------------LGVG----
------------------T-RYSIRSCFRECFYRRVA----DE--CGCLPID----S---
------------P-S--TR-----------------------------------------
--------ASTS-----L---RCTT-----------------------------------
------------------------------------VAL---YDCQQ---------S---
V-E-------------------SSL--N-------RQ-------NVP----------DLC
------------GCP-KPCKAVL---FPPS-MSF------------------QQW-PT--
-------KKD------KAI-F-----------LA--------------------------
------------------------------------------------DV----------
------------------------------------------------------------
---------------RRI-------------VGGD-------------------------
---------------------------VDE------------------------------
-----T-------LF-----------E------ENFLTANIFFQDLNFQQIE-QSPAYSV
GNL-----------------MAEIGGMCGLFAGISILTVMEFVEMLFLGG----------
-------WTLRK--------L----------R-------RN-------------------
AAASDANAHA----------------------TN----------------GHHQQPQQL-
------------------------------------PQTGRTDTIEVELAERQP------
TKMAEMP-----------AVPVVGKLKLTG------------------GSDSDCDD----
------------------------------------------------------------
--------DDDCRPAALRLRSSL-------------------------------------
------------------------------------------------------------
----------------------------------------------------SRGGSLAS
VASMQSEV
>Filast_tunicaraptor_GIQG01095491_1__p1__GENE_GIQG01095491_1__GIQG01095491_1_
_p1___ORF__type_complete__len_581___score_92_90__GIQG01095491_1__198_1940__77
7
------------------------------------------------------------
----------M-------------------------------------------------
------------------------------------------------------------
------------------------------------------------------------
------------------------------------------------------------
------------------------------------------------------------
------------------------------------------------------------
-----TARGNNWSLELDEAN-------------PGNAPEAVGAH----------------
------------------------------------------------------------
-------------------REGMGRARNHDK-----------------------------
---------------------PFDEDEET-------------------------------
---KMFENA----------------------Y---------GC-----M-----------
-PL--TRFFM-HTTTIHGLPRVF---------E-KG-------------HHW---FR---
--TALWA----ALFCASFGMFVYVAQFRFD-EFYA----------Y---NTSTSV----D
-I-SF-Q----------P-RVQF-----PTV-----------------------TICNSN
MY------RRS-----ALSYDV------------------L-----------------E-
---------ALWDDE-----GLTNQEIV---------------EFVA-------------
--------------------------VTINEL-NS-------------------------
------------------------------------------------------------
------------------------------------------------------------

-----------------DSS----------------------------------------
------------------------------------------------------------
---------------------------L--------------------------------
------------------------------------------------------------
--------------------------P----------------------TLTREQL----
------------------------------------------------------------
------------------------------------------------------------
------------------------------------------FA----------------
--IAVDELGKEG------------------------------EDDQELI-----------
-------------------------------------------------------TDDLI
ELLA-------------------DR--YHFDDDDFYDLLLE-----KYLRSN--------
--------------RNGGTDAQVLASL---------------FSDEQIGEI----GHQLD
--DLVA-F---GYAK----FK-G----E----------S---I-----------DIA---
----------------------------------------------T-------------
------------------------------------------------------------
------------------------------------------------------------
-----------E----------F-------E-----------RTI-SHPYG--------N
CYSWN-------SN-------------GN-------------------------------
-------YVTTRP------GP--------DL-------------AFDLIINLEQQ-----
----------E-----------------Y------------L-S-S--SLS---------
------------------------------------------------------------
--------------------------------------------------------GTGK
GID---DVSIIPAGLRVHIH-DKSEPP--FP-DAGLTVAPGTQAFISLQQQ------T--
-----ITKM-----------TPP-H--GE-C--VER----------------TDT-----
------------------A-AYSVESCFKGCFEKAVF----TQ--CGCRYPA----A---
-----------HA-V--SR-----------------------------------------
----------TV-----G---TCST-----------------------------------
------------------------------------SDF---TDCVD---------D---
I-E-------------------DAFS-T-------DV--------AS----------FSC
------------GCD-DPCNRIT---YRKN-LSH------------------GRW-PS--
-------VAV------SGI-I-----------SD--------------------------
------------------------------------------------AY----------
------------------------------------------------------------
-N----------------------------------------------------------
---------------------------IST------------------------------
-----R-------EV-----------G------DSFLRVTIYYEELNVIAMT-QEEKYSM
SAL-----------------GAELGGLLGLFLGVSVMTVFEFCEYMTFF-----------
--------FRPR--------M----------Y-------RK-------------------
KSV-----------------------------KV-----------------TPQ------
------------------------------------------------------------
------------------------------------------------------------
------------------------------------------------------------
------------------------------------------------------------
------------------------------------------------------------
------------------------------------------------------------
-------S
>outgro_CafRoe_KAA0150573_1_hypothetical_protein_FNF29_05148__Cafeteria_roenb
ergensis__747
------------------------------------------------------------
----------M-------------------------------------------------
-------------A----------------------------------------------
------------------------------------------------------------
------------------------------------------------------------
------------------------------------------------------------
------------------------------------------------------------
------------------------------------------------------------
------------------------------------------------------------

------------------------------------------------------------
------------------------------------------------------------
-------------------------------E--------GTV-----KGQ-RRWWGP--
-RL--AARWA-SSTSAHGFGRIA------HYCR-QP--------------GSC--LRL--
-AGCVWF----SSVVAFLVLLVVFILDRLA-FYRT----------W---PVTTSF----A
-L-KQ-E----------R-PIQF-----PAV-----------------------TVCS-N
VFA-----FDS-----RIAL----------------------------------------
------------------------------------------------------------
------------------------------------------------------------
------------------------------------------------------------
------------------------------------------------------------
------------------------------------------------------------
------------------------------------------------------------
------------------------------------------------------------
------------------------------------------------------------
------------------------------------------------------------
------------------------------------------------------------
------------------------------------------------------------
------------------------------------------------------------
------------------------------------------------------------
------------------------------------------------------------
-------------------------V-------NFTDVI---------------------
------------------------EHP----------------TDAALSRL----GAQPD
--ELV------IMCV----FG-G----R----------K---C-----------S-GL--
------------------------------------------------------------
------------------------------------------------------------
------------------------------------------------------------
-----------H----------A-------E-----------EWI-HPAFG--------L
CLTLN-------GG--QRPFRGNIYDPSSFS-TVGEYLSAREERVDYPLEAS----EIAG
LSNSPKQLSVKAP------RA--------DD-------------GLTLYLGVWQN-----
----------E-----------------Y------------T-A-V---ASSADEI----
----GIG-----------------------------------------------------
--------------------------------------------------------WSNE
PVTGV--FNAESAGVRIAVH-DRLKTP---DMDTATMLHPGKETHVTYRKR------V--
-----VTSQ-----------PPS-W--GD-C--SPRFW-----NETAGQ--PLPS-----
------------------G-NYSVAGCELVCAAKLVE----AE--CGCLTIP----T---
------------P-RGVPA-----------------------------------------
--------NPSM-----R---ACPLT----------------------------------
------------------------------------------MECPV-------------
VQQ-------------------EKVL--------LGQ--------------------AEC
P-----------DCV-KDCATTE---YPLT-TST------------------LSW-PA--
-------APS------QQV-F-----------LD--------------------------
------------------------------------------------LF----------
----------------------------------DQFSQ----IGI---------GASAL
EN----------------------------------------------------------
---------------------------IST-----------------AEDGL--------
-----Q-------RL-----------Q------SNVLLARIYPSSATYEEVT-SNRAYSI
TSL-----------------LSDLGGQAGLWGGISILTAIEFVEFIGLG-----------
--------FMAC--------F---GV---ACA----------------------------
--------------------------------RN----------------AFTEP-----
---------------------------------------GDEDELTPDDQ----------
------------------------------------------------ATTH--------
------------------------------------------------------------
----------------------------------------T-------------------
--------------------------------SP--------------------------
----------------------------------------------------IGSAHALQ
SKAATDAA

>outgro_CafRoe_KAA0157466_1_hypothetical_protein_FNF29_00042__Cafeteria_roenb
ergensis__674
------------------------------------------------------------
----------M-------------------------------------------------
------------------------------------------------------------
------------------------------------------------------------
------------------------------------------------------------
------------------------------------------------------------
------------------------------------------------------------
------------------------------------------------------------
------------------------------------------------------------
------------------------------------------------------------
-----------------------ASSKNAPGS----------------------------
---MAITGA---------------------DE---------GF-----------------
-QP--LQEFS-DNVTAHGVRKVM-------YLS-KE--------------RS---CR---
--KWVWV----ALLVGALIALVVIVSERIG-YFME----------N---PTSTVV----V
-W-EN-I----------E-PVPF-----PAV-----------------------TVCNEN
GF------QQS-----RVAEYG------------------L-----------------D-
---------TL-------------------------------------------------
------------------------------------------------------------
------------------------------------------------------------
------------------------------------------------------------
------------------------------------------------------------
------------------------------------------------------------
------------------------------------------------------------
------------------------------------------------------------
------------------------------------------------------------
------------------------------------------------------------
------------------------------------------------------------
------------------------------------------------------------
------------------------------------------------------------
------------------------------------------------------------
------------------------------------------------------------
-------------------------------------------NNSALLMM----VQPVG
--NLI------VRCE----FA-G----Q----------E---C-----------F-A---
----------------------------------------------D-------------
------------------------------------------------------------
------------------------------------------------------------
-----------N----------F-------S-----------VLL-DAQFG--------A
CYTFHGLQPVTVGS----E----VYD-AN-N--------------------G----NR--
-------LTSART------GP--------DY-------------GLRLMLDIEQS-----
----------E-----------------Y------------V-G-----YTPKTSVQ---
------------------------------------------------------------
-PP--------------------------------------------QQDVKAAPDNRNS
TVEALF-EGPESAGIRVIIH-DSSVLA--NP-TSGIGVAPGTASDLAFVRA------D--
-----RSRL-----------FAP-W--GT-C--NQA-----------W---HSES-----
-------------QGTDPT-GYAKSACERDCYYASVA----DV--CECLLLP----W---
------------P-E--SLSDRP-------------------------------------
----------SL--------FVCSV-----------------------------------
------------------------------------EEK----TCIA---------E---
Q-L-------------------SKFQ-S-------GK--------L------------GC
----------SENCP-DRCLETL---YTAT-VGS------------------QLW-PS--
-------SSS------ADT-I-----------LS--------------------------
------------------------------------------------LVRDTR------
------------------------------------------------------------
QD----------------------------------------------------------
------------------------NT-ISA------------------------------

-----E-------YL-----------T------ENMMSVRVFAPSSEFQVIT-TVRAYTF
ESM-----------------LGEVGGNTGLWAGMSALTLVEILEFVFLA-----------
--------LSAW--------LCAGC---RCCS-------RD-------------------
RAV-----------------------------RT--------------------------
-----------------------------------------VGGSELHAEMSP-------
------------------GSARPGPDA---------------------AASR--------
--------AAFDDRELA-------------------------------------------
------------------------------------------------------------
------------------------------------------------------------
------------------------------------------------------------
---AKASP
>outgro_CafRoe_KAA0153446_1_hypothetical_protein_FNF29_03263__Cafeteria_roenb
ergensis__865
------------------------------------------------------------
----------M-------------------------------------------------
------------------------------------------------------------
------------------------------------------------------------
------------------------------------------------------------
------------------------------------------------------------
------------------------------------------------------------
------------------------------------------------------------
------------------------------------------------------------
------------------------------------------------------------
------------------------------------------------------------
---SSESRQ---------------------PE---------GF-----------------
-QP--LQEFS-DNVTAHGVGKVF-------YLS-KE--------------RS---CR---
--RWLWL----VLLLGALVALVVIVTDRVQ-YFLQ----------N---PTATVV----K
-W-ES-R----------A-PVPF-----PAV-----------------------TVCNEN
GF------QQS-----RVAQYG------------------L-----------------S-
---------TL-------------------------------------------------
------------------------------------------------------------
------------------------------------------------------------
------------------------------------------------------------
------------------------------------------------------------
------------------------------------------------------------
------------------------------------------------------------
------------------------------------------------------------
------------------------------------------------------------
------------------------------------------------------------
------------------------------------------------------------
------------------------------------------------------------
------------------------------------------------------------
------------------------------------------------------------
------------------------------------------------------------
-------------------------------------------NNSVLELM----TQPVN
--NFI------LRCE----FA-G----L----------E---C-----------F-A---
----------------------------------------------D-------------
------------------------------------------------------------
------------------------------------------------------------
-----------N----------F-------S-----------VLL-DAQFG--------A
CYTFHGLRALTIDG----E----EFN-AS-G--------------------P----NP--
-------LTSART------GP--------DY-------------GLRLVLDIEQA-----
----------E-----------------Y------------V-G-----YSPRTSVRQAS
GADQGPGTEPDSSGSAQTSPQTSPQTSPQTSPQTSPQTSAQASAQASPQTSSSSATASSS
PPPAAASSSPQASSPAGASPTRTPSPSQPARRLLAPRRVQEDPGPGSASDARAPPDNRNS
TVEALF-KGPESAGIRVIIH-DPSVLA--NP-TSGFGVGPGSASDIAFERV------D--
-----RSRL-----------FPP-W--GT-C--NEA-----------W---HSES-----

```
-------------GATDPA-GYAKTACERDCYYASVA----DV--CECLLLP----W---
------------P-T--TMEERP-------------------------------------
----------SL--------YVCSS-----------------------------------
------------------------------------EEK----ACIA---------T---
Q-L-------------------SRFQ-A-------GE--------L------------GC
----------SASCP-DRCLETL---YTAT-AGR------------------QLW-PS--
-------VAS------AST-I-----------LD--------------------------
------------------------------------------------VVRSAK------
------------------------------------------------------------
QD----------------------------------------------------------
------------------------TS-ITP------------------------------
-----E-------YL-----------A------QNMMSVRVFPPTTEYQVVT-TVRAYSF
ESM-----------------LGEVGGNTGLWAGMSALTIVEILEFVMLA-----------
--------LSAW--------LCSRG---GCCR-------RG-----------------GK
RAP-----------------------------RT--------------------------
---------------------------------------NVEHGTELHEGEAI-------
------------------ELVAPQADE---------------------ESSA--------
--------PAVAPQAAA-------------------------------------------
------------------------------------------------------------
------------------------------------------------------------
--------------------------------------------------------AASG
RAREAWRE
>Cteno_MneLey_comp601992_c0_seq1__p1__GENE_comp601992_c0_seq1__comp601992_c0_
seq1__p1___ORF__type_complete__len_212____score_42_61__comp601992_c0_seq1__14
2_726___205
------------------------------------------------------------
------------------------------------------------------------
------------------------------------------------------------
------------------------------------------------------------
------------------------------------------------------------
------------------------------------------------------------
------------------------------------------------------------
------------------------------------------------------------
------------------------------------------------------------
------------------------------------------------------------
------------------------------------------------------------
------------------------------------------------------------
------------------------------------------------------------
------------------------------------------------------------
------------------------------------------------------------
------------------------------------------------------------
------------------------------------------------------------
------------------------------------------------------------
------------------------------------------------------------
------------------------------------------------------------
------------------------------------------------------------
------------------------------------------------------------
------------------------------------------------------------
------------------------------------------------------------
------------------------------------------------------------
------------------------------------------------------------
------------------------------------------------------------
------------------------------------------------------------
------------------------------------------------------------
------------------------------------------------------------
------------------------------------------------------------
------------------------------------------------------------
```

------------------------------------------------------------
------------------------------------------------------------
------------------------------------------------------------
------------------------------------------------------------
------------------------------------------------------------
------------------------------------------------------------
------------------------------------------------------------
------------------------------------------------------------
------------------------------------------------------------
------------------------------------------------------------
------------------------------------------------------------
------------------------------------------------------------
----------------------------------MLN----KT--CGCLNST----P---
------------P-----L-----------------------------------------
------NDVFNY-----K---ECTI-----------------------------------
------------------------------------KQV---ATCSL---------K---
A-Y-------------------YQYV-LDFADV--KS--------DY----------EEC
------------GCDVVACEYTL---FRTT-VTA------------------TPL-PD--
-------TEI------EHK-L-----------KA--------------------------
----------------------------------------WNSPRHFDRF----------
---------------------------------------------------------QRF
SN----------------------------------------------------------
---------------------------YTA------------------------------
-----E-------DL-----------A------KNMARIEFFFGSSSITKVE-EIISYDF
DNL-----------------LGDIGGVMGLFLGASIFTTMEFCTLVVNL-----------
--------LSRL--------W----------QW------KI-------------------
QPW----------L------------------RS--------------------------
------------------------------------------------------------
-------------------------------------------------ETR--------
------------------------------------------------------------
------------------------------------------------------------
------------------------------------------------------------
----------------------------------------------------------DR
KIISETNL
>Cteno_Ple_Bac_comp46220_c0_seq11_p1__comp46220_c0__comp46220_c0_seq11_p1___O
RF__type_complete__len_202___score_27_04__comp46220_c0_seq11_1766_2371__211
------------------------------------------------------------
------------------------------------------------------------
------------------------------------------------------------
------------------------------------------------------------
------------------------------------------------------------
------------------------------------------------------------
------------------------------------------------------------
------------------------------------------------------------
------------------------------------------------------------
------------------------------------------------------------
------------------------------------------------------------
------------------------------------------------------------
------------------------------------------------------------
------------------------------------------------------------
------------------------------------------------------------
------------------------------------------------------------
------------------------------------------------------------
------------------------------------------------------------
------------------------------------------------------------
------------------------------------------------------------
------------------------------------------------------------

------------------------------------------------------------
------------------------------------------------------------
------------------------------------------------------------
------------------------------------------------------------
------------------------------------------------------------
------------------------------------------------------------
------------------------------------------------------------
------------------------------------------------------------
------------------------------------------------------------
------------------------------------------------------------
------------------------------------------------------------
---------------------------M----------L---C-----------S-DVLK
D-----------------------------------------------------------
------------------------------------------------------------
------------------------------------------------------------
-----------N---------SF-------T-----------REA-S-VNG--------N
CFRVN-------PE-------------GK-------------------------------
--------LKGKG------GE--------YG-------------RLRLMFFADLN-----
----------D-----------------Y------------T-A-----PA---------
------------------------------------------------------------
------------------------------------------------------------
-------KNQAQYGYTVAFH-DHETYS--STIPSGFFMSPGSIYKVDLGLV------R--
-----EFRE-----------PPP-A--GL-C--DST----------------LTT--NTY
------------------G-RYEENSCIAECRDNAII----RA--CGCVHVV----P---
------------P-H--PD-----------------------------------------
-----NNPLVEY-----L---PCTL-----------------------------------
------------------------------------EQW---AKCGM---------R---
E-Y-------------------KKWG-E--SNS--NR--------HF----------KAR
------------KIL-SKLLETQ---KT--------------------------------
------------------------------------------------------------
------------------------------------------------------------
------------------------------------------------------------
------------------------------------------------------------
------------------------------------------------------------
------------------------------------------------------------
------------------------------------------------------------
------------------------------------------------------------
------------------------------------------------------------
------------------------------------------------------------
------------------------------------------------------------
------------------------------------------------------------
------------------------------------------------------------
------------------------------------------------------------
------------------------------------------------------------
--------
>Cteno_Ple_Bac_comp46220_c0_seq48_p1__comp46220_c0__comp46220_c0_seq48_p1___O
RF__type_3prime_partial__len_242___score_38_17__comp46220_c0_seq48_1236_1958_
_259
------------------------------------------------------------
------------------------------------------------------------
------------------------------------------------------------
------------------------------------------------------------
------------------------------------------------------------
------------------------------------------------------------
------------------------------------------------------------
------------------------------------------------------------
------------------------------------------------------------

```
------------------------------------------------------------
------------------------------------------------------------
------------------------------------------------------------
-------------------------------------------------------MI---
--QTLFL----FMWLASIGYAVYVIASSTM-SFIG----------K---PTGTKF----Q
-V-IA-SQLDQENVNLQS-AVQF-----PTI-----------------------SVCSHN
QV------SLK-----WSSE-----------------RPGL-----------------E-
---------KL-------------------------------LEKFD-------------
------------------------------KW-NP-------VLAANI------------
------------------------------------------------------------
------------------------------------------------------------
-----------------D-------------------------WET--------------
------------------------------------------------------------
------------------------------------------------------------
------------------------------------------------------------
--------------------------P---------------------------------
------------------------------------------------------------
------------------------------------------------------------
------------------------------------------------------------
------------------------------------------------------------
------------------------------------------------------------
---------------------------------QFNKWR---------------------
-------------------------NA----------------TYEEVIKK----GGPDN
--RTF------IQCE----HG-I----M----------L---C-----------S-DVLK
D-----------------------------------------------------------
------------------------------------------------------------
------------------------------------------------------------
-----------N---------SF-------T-----------REA-S-VNG--------N
CFRVN-------PE-------------GK-------------------------------
--------LKGKG------GE--------YG-------------RLRLMFFADLN-----
----------D-----------------Y------------T-A-----PA---------
------------------------------------------------------------
------------------------------------------------------------
-------KNQAQYGYTVAFH-DHETYS--STIPSGFFMSPGSIYKVDLGLV------R--
-----EFRE-----------PPP-A--G--------------------------------
------------------------------------------------------------
------------------------------------------------------------
------------------------------------------------------------
------------------------------------------------------------
------------------------------------------------------------
------------------------------------------------------------
------------------------------------------------------------
------------------------------------------------------------
------------------------------------------------------------
------------------------------------------------------------
------------------------------------------------------------
------------------------------------------------------------
------------------------------------------------------------
------------------------------------------------------------
------------------------------------------------------------
------------------------------------------------------------
------------------------------------------------------------
------------------------------------------------------------
------------------------------------------------------------
------------------------------------------------------------
------------------------------------------------------------
--------
```

>Ctenoph_Hormiphora_californiensis_evg190294__p1_GENE_evg190294__evg190294__p
1__ORF_type_internal_len_253____score_30_09_evg190294__2_757___271
-------L----------------------------------------------------
----------M-------------------------------------------------
------------------------------------------------------------
------------------------------------------------------------
------------------------------------------------------------
------------------------------------------------------------
------------------------------------------------------------
------------------------------------------------------------
------------------------------------------------------------
------------------------------------------------------AEYSPA
NR-------------------RVSDGGRSSAT----------------------------
---SRSDTP----------------------S---------SW-----V-----------
-KG--FQLFA-ETTTAHGVNQIF---------T-GS--------------NR---AV---
--RLFYL----LAWIACVIYTMYMIITSIV-TFIN----------K---PTGTKY----E
-V-IV-NDLGGL---KPQ-SIKF-----PTV-----------------------SICSHN
KV------KKS-----YLRKEG---------------NEPI-----------------R-
---------DL-------------------------------WSKLD-------------
------------------------------MY-DQ-------AEADKI------------
------------------------------------------------------------
------------------------------------------------------------
-----------------N-------------------------WTN--------------
------------------------------------------------------------
---------------------------I--------------------------------
------------------------------------------------------------
------------------------------------------------------------
------------------------------------------------------------
------------------------------------------------------------
------------------------------------------------------------
------------------------------------------------------------
------------------------------------------------------------
---------------------------------GTASIQ---------------------
-------------------------HM----------------TLEDIISE----GGPDD
--DRL------LSCT----QR-N----K----------N---C-----------Q-DVLG
--------------------------------------------KNE-------------
------------------------------------------------------------
------------------------------------------------------------
-----------S----------Y-------SV----------REN-T-PSG--------K
CWRIN-------PE-------------GK-------------------------------
--------IFGKS------GD--------WG-------------KMQLIVFADLQ-----
----------D-----------------Y------------S-K-----RS---------
------------------------------------------------------------
------------------------------------------------------------
-------ADQPQYG----------------------------------------------
------------------------------------------------------------
------------------------------------------------------------
------------------------------------------------------------
------------------------------------------------------------
------------------------------------------------------------
------------------------------------------------------------
------------------------------------------------------------
------------------------------------------------------------
------------------------------------------------------------
------------------------------------------------------------
------------------------------------------------------------
------------------------------------------------------------

------------------------------------------------------------
------------------------------------------------------------
------------------------------------------------------------
------------------------------------------------------------
------------------------------------------------------------
------------------------------------------------------------
------------------------------------------------------------
------------------------------------------------------------
------------------------------------------------------------
------------------------------------------------------------
--------
>Ctenoph_Hormiphora_californiensis_evg1639__p1_GENE_evg1639__evg1639__p1__ORF
_type_internal_len_264____score_48_35_evg1639__2_790___291
------------------------------------------------------------
------------------------------------------------------------
------------------------------------------------------------
------------------------------------------------------------
------------------------------------------------------------
------------------------------------------------------------
------------------------------------------------------------
------------------------------------------------------------
------------------------------------------------------------
------------------------------------------------------------
------------------------------------------------------------
------------------------------------------------------------
----------------HGVKYVF---------H-SN--------------HR---VI---
--QALFL----IMWFVSIGYAVYVIASSTI-SFIG----------K---PTGTKF----Q
-V-IA-SQLDRNNVNLQS-AVQF-----PTI-----------------------SVCSHN
QV------SAK-----WSRQ-----------------RPGL-----------------E-
---------KI-------------------------------LERFD-------------
------------------------------TW-SP-------GITEAI------------
------------------------------------------------------------
------------------------------------------------------------
-----------------N-------------------------WDR--------------
------------------------------------------------------------
------------------------------------------------------------
------------------------------------------------------------
--------------------------P---------------------------------
------------------------------------------------------------
------------------------------------------------------------
------------------------------------------------------------
------------------------------------------------------------
------------------------------------------------------------
---------------------------------EFNEWR---------------------
-------------------------NS----------------TFETIIKE----GGPQD
--STF------IQCE----QG-I----Q----------L---C-----------E-EVLG
A-----------------------------------------------------------
------------------------------------------------------------
------------------------------------------------------------
-----------K---------YV-------T-----------REP-S-VNG--------N
CFRVN-------PE-------------GQ-------------------------------
--------LRGKG------GD--------YG-------------RLRLMFFADLN-----
----------D-----------------Y------------T-V-----PA---------
------------------------------------------------------------
------------------------------------------------------------
-------KNQAQYGYTVAFH-DHETYS--STIPSGFYMSPGSIYKVDLGLV------R--
-----EFRE-----------PPP-A--GL-C--DST----------------LTY--NTY

------------------------------------------------------------
------------------------------------------------------------
------------------------------------------------------------
------------------------------------------------------------
------------------------------------------------------------
------------------------------------------------------------
------------------------------------------------------------
------------------------------------------------------------
------------------------------------------------------------
------------------------------------------------------------
------------------------------------------------------------
------------------------------------------------------------
------------------------------------------------------------
------------------------------------------------------------
------------------------------------------------------------
------------------------------------------------------------
------------------------------------------------------------
------------------------------------------------------------
------------------------------------------------------------
------------------------------------------------------------
------------------------------------------------------------
--------
>Ctenoph_Hormiphora_californiensis_evg16615__p1_GENE_evg16615__evg16615__p1__
ORF_type_5prime_partial_len_265____score_62_43_evg16615__102_896___294
------------------------------------------------------------
------------------------------------------------------------
------------------------------------------------------------
------------------------------------------------------------
------------------------------------------------------------
------------------------------------------------------------
------------------------------------------------------------
------------------------------------------------------------
------------------------------------------------------------
------------------------------------------------------------
------------------------------------------------------------
------------------------------------------------------------
------------------------------------------------------------
------------------------------------------------------------
------------------------------------------------------------
------------------------------------------------------------
------------------------------------------------------------
------------------------------------------------------------
------------------------------------------------------------
------------------------------------------------------------
------------------------------------------------------------
------------------------------------------------------------
------------------------------------------------------------
------------------------------------------------------------
------------------------------------------------------------
------------------------------------------------------------
------------------------------------------------------------
------------------------------------------------------------
------------------------------------------------------------
------------------------------------------------------------
------------------------------------------------------------
------------------------------------------------------------
------------------------------------------------------------

------------------------------------------------------------
------------------------------------------------------------
------------------------------------------------------------
------------------------------------------------------------
------------------------------------------------------------
------------------------------------------------------------
------------------------------------------------------------
------------------------------------------------------------
------------------------------------------------------------
-----------------AFH-DSVYHG--VTLNSGYYMSPGSIYKVDLSLK------K--
-----ETRE-----------PPP-G--GN-C--NAS----------------LTM--TTY
------------------G-AYSEGSCIQECKDKDLM----KR--CGCINMV----P---
------------P-Q--NY-----------------------------------------
---------LNY-----S---SCTV-----------------------------------
------------------------------------RQW---AECGL---------S---
A-Y-------------------LNFS-ERYRDP--NL--------TD----------DIC
------------PCN-SPCSETH---YEAQ-LSS------------------SSL-SP--
-------AYT------ENL-F-----------AS--------------------------
-------------------------KYIK-G---------YLSKVSQPQY----------
---------------------------------------------------------GII
YN----------------------------------------------------------
----------------------------SS------------------------------
-----D-------DI-----------L------KNVMVLEILFSSMVTSEIQ-QVVTYTS
ANL-----------------LGDIGGVLGLFCGASVFTIVEFLQLFLSS-----------
--------IQKH--------C----------L-------TD-------------------
LCSSD---------------------------ED--------------------------
----------------------------------------TVEAADDKRG----------
------------------------------------------------------------
------------------------------------------------------------
------------------------------------------------------------
------------------------------------------------------------
------------------------------------------------------------
----DKEL
>Cteno_Puk_Fal_comp26850_c0_seq10__p1__GENE_comp26850_c0_seq10__comp26850_c0_
seq10__p1___ORF__type_5prime_partial__len_320____score_71_31__comp26850_c0_se
q10__394_1353___349
------------------------------------------------------------
------------------------------------------------------------
------------------------------------------------------------
------------------------------------------------------------
------------------------------------------------------------
------------------------------------------------------------
------------------------------------------------------------
------------------------------------------------------------
------------------------------------------------------------
------------------------------------------------------------
------------------------------------------------------------
------------------------------------------------------------
------------------------------------------------------------
------------------------------------------------------------
------------------------------------------------------------
------------------------------------------------------------
------------------------------------------------------------
------------------------------------------------------------
------------------------------------------------------------
------------------------------------------------------------
------------------------------------------------------------

```
------------------------------------------------------------
------------------------------------------------------------
------------------------------------------------------------
------------------------------------------------------------
------------------------------------------------------------
------------------------------------------------------------
------------------------------------------------------------
------------------------------------------------------------
------------------------------------------------------------
------------------------------------------------------------
------------------------------------------------------------
--------------------R-R----V----------R---G-----------H-EVIP
E-------------------------------------------PEV-------------
------------------------------------------------------------
------------------------------------------------------------
-----------N---------FF-------T-----------REV-S-LTG--------N
CFRFN-------PN-------------GV-------------------------------
--------LRGKG------GD--------YG-------------KMQLMFFTDLN-----
----------E-----------------Y------------S-D-----LT---------
------------------------------------------------------------
------------------------------------------------------------
-------RREPQYGYTVVFH-DHESYS--STIPSGFWLSPGSIYKADLSLS------Q--
-----EFRE-----------GPP-A--GT-C--DPT----------------RVN--NTY
------------------G-RYEENSCIAQCRDDALM----AE--CGCVHVA----P---
------------P-L--PE-----------------------------------------
-----NEPGKDY-----R---GCTL-----------------------------------
------------------------------------QEW---ATCGL---------R---
A-Y-------------------KEWV-KKYSDV--NR--------VS----------MGC
------------NCP-PTCFETE---YKAQ-ISS------------------SSL-SK--
-------FYA------ESK-----------------------------------------
----------------------------------------------VGRL----------
---------------------------------------------------------PPG
YN----------------------------------------------------------
----------------------------TT------------------------------
-----Q-------DV-----------M------ENLLIIDILFTSMQVSEIR-EIVTYGW
ANF-----------------LGDVGGVLGLFLGASIFTIMEFLQFFVLA-----------
--------IWDA--------L----------C-------GL-------------------
PG------------------------------RK-------------------K------
------------------------------------------------------------
------------------------------------------------APNH--------
------------------------------------------------------------
------------------------------------------------------------
------------------------------------------------------------
------------------------------------------------------------
--DTVSLT
>Cteno_Ple_Bac_comp48643_c0_seq3_p1__comp48643_c0__comp48643_c0_seq3_p1___ORF
__type_5prime_partial__len_344___score_41_94__comp48643_c0_seq3_3_1034__377
------------------------------------------------------------
------------------------------------------------------------
------------------------------------------------------------
------------------------------------------------------------
------------------------------------------------------------
------------------------------------------------------------
------------------------------------------------------------
------------------------------------------------------------
------------------------------------------------------------
------------------------------------------------------------
```

------------------------------------------------------------
------------------------------------------------------------
------------------------------------------------------------
------------------------------------------------------------
------------------------------------------------------------
------------------------------------------------------------
------------------------------------------------------------
------------------------------------------------------------
------------------------------------------------------------
------------------------------------------------------------
------------------------------------------------------------
------------------------------------------------------------
------------------------------------------------------------
------------------------------------------------------------
------------------------------------------------------------
------------------------------------------------------------
------------------------------------------------------------
------------------------------------------------------------
------------------------------------------------------------
------------------------------------------------------------
------------------------------------------------------------
------------------------------------------------------------
------------------------------------------------------------
-------------------------------------------------------L----
------------------------------------------------------------
------------------------------------------------------------
-----------SSLYFRVPKFTL-------F-----------QEV-S-LSG--------S
CYRIN-------PK-------------GS-------------------------------
--------LLTQV------GD--------YG-------------VLKLTFFTDLN-----
----------D-----------------Y------------S-K-----FS---------
------------------------------------------------------------
------------------------------------------------------------
-------KKDPMYGYTVVFH-DNGTYS--STMTSGFYMSPGNVYKADMRKY------K--
-----EFNI-----------GPP-Y--GR-C--NPH----------------LKN--NVY
------------------G-TYSSSSCIARCRDQYVK----DM--CGCIQIL----P---
------------P-H--PE-----------------------------------------
-----YEPELDF-----R---GCTL-----------------------------------
------------------------------------KEW---VECGA---------R---
A-T-------------------LNFT-DRFVDL---K--------EE----------AMC
------------ECT-PACMETW---YDAT-ISS------------------SRL-SK--
-------FYA------NDK-A-----------ISIDGQP------------------AQL
------------------------ARIVD-G-----------YGLNH-GI----------
---------------------------------------------------------EET
YN----------------------------------------------------------
---------------------------ISK------------------------------
-----E-------TV-----------Y------ENLMVLQVLFTSIQESNYK-EYVKYDY
TQL-----------------IGDVGGQMGMLLGASLFTIIEFVHFFIKV-----------
--------AWRY--------C----------V-------KK-------------------
ATR-----------------------------RA---------------GEAAE------
-----------------------------------------RPAYEMHRD----------
------------------------------------------------------------
------------------------------------------------------------
------------------------------------------------------------
------------------------------------------------------------
------------------------------------------------------------
----QVEE

>Cteno_MneLey_comp13853_c0_seq1__p1__GENE_comp13853_c0_seq1__comp13853_c0_seq
1__p1___ORF__type_3prime_partial__len_479____score_87_11__comp13853_c0_seq1__
3_1436___533
------------------------------------------------------------
----------M-------------------------------------------------
------------------------------------------------------------
------------------------------------------------------------
------------------------------------------------------------
------------------------------------------------------------
------------------------------------------------------------
------------------------------------------------------------
------------------------------------------------------------
------------------------------------------------------------
---------------------LLNNDTKP-------------------------------
---PDVPKK----------------------Q---------SF-----F-----------
-ED--FKEFT-TETTSHGIKYVF---------H-AA--------------YK---WV---
--RVIYL----ILWLAAVGYTVSIVFISCK-KYFS----------M---ETGTKI----E
-E-KH-TPKHSS---NEV-AITH-----PTI-----------------------TVCPNN
LI------SKS-----YLAK-----------------HPNL-----------------D-
---------EL-------------------------------MTKLV-------------
------------------------------TF-NE-------NETPSLF-----------
------------------------------------------------------------
------------------------------------------------------------
-----------------D------------------------------------------
------------------------------------------------------------
------------------------------------------------------------
------------------------------------------------------------
-------------------------DP---------------------------------
------------------------------------------------------------
------------------------------------------------------------
------------------------------------------------------------
------------------------------------------------------------
------------------------------------------------------------
---------------------------------KYAGYG---------------------
-------------------------DM----------------TYHDLMWQ----GRPR-
--VGL------LQCT----YF-A----H----------K---C-----------G-AVDP
F-------------------------------------------NSEDYETKTYHW----
------------------------------------------------------------
------------------------------------------------------------
-----------DKTYNGSNHKFY-------E-----------WEV-S-LSG--------S
CYRIN-------PR-------------GN-------------------------------
--------LWAKV------GD--------YG-------------EIKLTFFADLN-----
----------D-----------------Y------------S-S-----AS---------
------------------------------------------------------------
------------------------------------------------------------
-------KEEPEYGYLVTFT-DSGTYS--STATSGFFMSPGNVYKADLRKY------K--
-----EENL-----------GPP-A--GK-C--DIE----------------SKT--NVY
------------------G-AYSTNSCNKLCRDQHVM----EK--CGCVHIL----P---
------------P-H--PE-----------------------------------------
-----NDPDTKF-----V---GCTL-----------------------------------
------------------------------------KQW---AECGL---------R---
E-S-------------------TKFT-EEFVNM--ND--------DK----------VHC
E-----------HCN-PACMEIW---YEAT-ISS------------------SRL-ST--
-------FYA------EAK-V-----------SE-------------------------L
------------------------DKLIT-G-----------YQLNEGNK----------
---------------------------------------------------------TTN
YT----------------------------------------------------------

---------------------------IDK------------------------------
-----D-------TV-----------Y------ENLMVLKVMFTSIQESTIK-EYYKYDY
TQL-----------------IGDVGGQM--------------------------------
------------------------------------------------------------
------------------------------------------------------------
------------------------------------------------------------
------------------------------------------------------------
------------------------------------------------------------
------------------------------------------------------------
------------------------------------------------------------
------------------------------------------------------------
--------
>Ctenoph_Hormiphora_californiensis_evg120240__p1_GENE_evg120240__evg120240__p
1__ORF_type_complete_len_484____score_100_73_evg120240__313_1764___548
------------------------------------------------------------
----------M-------------------------------------------------
------------------------------------------------------------
------------------------------------------------------------
------------------------------------------------------------
------------------------------------------------------------
------------------------------------------------------------
------------------------------------------------------------
------------------------------------------------------------
------------------------------------------------------------
------------------------------------------------------------
-----------------------------------------GW-----L-----------
-AD--FKEFS-DGTTAHGVKWIF---------H-PSVH----------GNFK---IV---
--KLIFF----IVWLAVIGYSLFTIYNAVH-NYVN----------K---PTGTKF----Q
-V-IA-EGPEQE---KPA-SIQF-----PTI-----------------------SVCSHN
LI------KQS-----YFDS-----------------NTGL-----------------E-
---------EL-------------------------------WSDLD-------------
------------------------------KW-DP-------DVTEKL------------
------------------------------------------------------------
------------------------------------------------------------
-----------------N-------------------------FTD--------------
------------------------------------------------------------
------------------------------------------------------------
------------------------------------------------------------
--------------------------S---------------------------------
------------------------------------------------------------
------------------------------------------------------------
------------------------------------------------------------
------------------------------------------------------------
------------------------------------------------------------
---------------------------------KFARYR---------------------
-------------------------NM----------------TYQEIIRA----GGPDN
--TTF------VHCR----HF-I----N----------K---C-----------E-DAIP
N-------------------------------------------PEE-------------
------------------------------------------------------------
------------------------------------------------------------
-----------N---------FY-------V-----------REA-S-LSG--------N
CFRIN-------PN-------------GV-------------------------------
--------LHGKG------GD--------YG-------------KMQLLFFADLK-----
----------E-----------------Y------------S-T-----MT---------
------------------------------------------------------------
------------------------------------------------------------
-------RSEPQYGYTVVFH-DHESYS--STMSSGFWLSPGSIYKVDLSLS------Q--

-----EFRE-----------GPP-A--GT-C--DPT----------------RVN--NTY
------------------G-RYEENSCIAQCRDDALM----EK--CGCVHIS----P---
------------P-L--PH-----------------------------------------
-----NEPGKDY-----R---GCTL-----------------------------------
------------------------------------EEW---ATCGL---------R---
A-Y-------------------KEWV-VKYSDV--NR--------IS----------MGC
------------DCP-PTCFETE---YKAQ-VSS------------------SKL-NR--
-------FYA------EEK-A---------------------------------------
---------------------------------------------NMGLL----------
---------------------------------------------------------PPG
YN----------------------------------------------------------
----------------------------GT------------------------------
-----Q-------DV-----------M------DNLLIIDILFTSMQVSEIR-EIVTYGW
GNF-----------------LGDVGGVLGLFLGASIFTIMEFIQFFILA-----------
--------ICNA--------C----------CP------GL-------------------
TS------------------------------RS-------------------K------
------------------------------------------------------------
------------------------------------------------ANSP--------
------------------------------------------------------------
------------------------------------------------------------
------------------------------------------------------------
------------------------------------------------------------
-VDTYALS
>Ctenoph_Hormiphora_californiensis_evg197715__p1_GENE_evg197715__evg197715__p
1__ORF_type_complete_len_492____score_113_33_evg197715__97_1572___570
------------------------------------------------------------
----------M-------------------------------------------------
------------------------------------------------------------
------------------------------------------------------------
------------------------------------------------------------
------------------------------------------------------------
------------------------------------------------------------
------------------------------------------------------------
------------------------------------------------------------
------------------------------------------------------------
------------------------------------------------------------
-------------------------------S--------GSI-----Q-----------
-GE--LRNFA-DNTSAHGVRLIF---------H-GH--------------HV---AV---
--KVAFL----IFWVSAVTFCVFRASLSIR-KYCQ----------Q---ETTSRY----E
-M-FP-ASPT-------A-VMDF-----PTI-----------------------TACNLN
MI------RKS-----YLKA-----------------NPGLM------------------
---------HF-------------------------------WKLAS-------------
------------------------------IH-NF-------QALAEL------------
------------------------------------------------------------
------------------------------------------------------------
-----------------D-------------------------WESE-------------
------------------------------------------------------------
------------------------------------------------------------
------------------------------------------------------------
------------------------------------------------------------
------------------------------------------------------------
------------------------------------------------------------
------------------------------------------------------------
------------------------------------------------------------
------------------------------------------------------------
---------------------------------ELSPYQ---------------------
-------------------------NL----------------TYYELLKE----AQMYN

--DTF------ISCTQ------G----R-------M--RY--CV----------D-TVMG
----------------------------------------------A-------------
------------------------------------------------------------
------------------------------------------------------------
-----------D---------YY-------Q-----------RDI-EFVSG--------S
CFRIN-------PY-------------GT-------------------------------
--------LKGKS------GD--------YG-------------VLTMMLLADRD-----
----------E-----------------Y------------I-Y-----NS---------
------------------------------------------------------------
------------------------------------------------------------
----------SNVGWVMAIH-ESERYG--ASIDNGIKISPGYTYFVNLDMM------N--
-----IKNH---------------R--KH-C--EKS----------------VNM-ISGY
------------------G-RYDQTTCLLDCRDRQLY----EK--CGCLMTV----P---
------------P-N--AK-----------------------------------------
------QAGLNY-----L---PCTL-----------------------------------
------------------------------------KQV---SQCAL---------R---
T-Y-------------------VKYV-KNYADVRAKS--------TT----------ADC
------------DCFVTACSYTL---YTTS-VTA------------------TPI-PD--
-------LEL------QFK-V-----------DS--------------------------
----------------------------------------W---KNQARF----------
---------------------------------------------------------SRF
RN----------------------------------------------------------
---------------------------YTY------------------------------
-----E-------DL-----------G------KNMAVIEVYFGTSSITSIE-EVVAYDF
DNL-----------------LGDIGGVMGLFLGASMFTAMEFVTLIISL-----------
--------SLRI--------W----------RT------HF-------------------
PT------------------------------KD----S---------------------
------------------------------------------------------------
------------------------------------------------SQDHM-------
------------------------------------------------------------
------------------------------------------------------------
--------------------------------IP--------------------------
----------------------------------------------------HHSLKLPR
ASVSEVNL
>Cteno_Ple_Bac_comp47842_c0_seq1_p1__comp47842_c0__comp47842_c0_seq1_p1___ORF
__type_complete__len_492___score_104_93__comp47842_c0_seq1_138_1613__571
------------------------------------------------------------
------------------------------------------------------------
------------------------------------------------------------
------------------------------------------------------------
------------------------------------------------------------
------------------------------------------------------------
------------------------------------------------------------
------------------------------------------------------------
------------------------------------------------------------
------------------------------------------------------------
------------------------------------------------------------
-------------------------------M---------TW-----I-----------
-RD--FRDFA-GDTSAHGVKNIF---------E-GP--------------TK---LL---
--KLIFL----ICWIICSVYATYVIMNSII-TFIN----------R---PTGTKF----T
-F-LV-ENEQRK-LGKPA-VVPM-----PTI-----------------------SVCSHN
KV------KKS-----FLEKKG---------------NEEL-----------------K-
---------KI-------------------------------YTILD-------------
------------------------------KY-DL-------GEAAEL------------
------------------------------------------------------------
------------------------------------------------------------
------------------------------------------------------------

```
------------------------------------------------------------
------------------------------------------------------------
------------------------------------------------------------
------------------------------------------------------------
------------------------------------------------------------
------------------------------------------------------------
------------------------------------------------------------
------------------------------------------------------------
-------------------------------------------------------AENIT
-----------------------------------GEVA---------------------
-------------------------DM----------------KYEDLLEE----GSISN
--DRI------LMCQ----QR-S----T----------N---C-----------F-QLPA
F-------------------------------------------QNS-------------
------------------------------------------------------------
------------------------------------------------------------
-----------E---------WV-------T-----------REN-S-ITG--------T
CFRIN-------PK-------------GT-------------------------------
--------LNALM------GD--------YG-------------ALQLRFWADIQ-----
----------D-----------------Y------------S-E-----SS---------
------------------------------------------------------------
------------------------------------------------------------
-------KNNPYHGFTVAFH-DKDSYG--SSLSSGYLMSPGSYYKVDLMMR------K--
-----ESRE-----------PPP-S--GH-C--DDS----------------QPF--TTY
------------------G-NYSEGACVMECKDRVLN----DT--CGCVNVV----P---
------------P-I--NS-----------------------------------------
---------LNY-----T---SCTV-----------------------------------
------------------------------------KQW---ASCGL---------K---
T-Y-------------------LDWY-KKYTNP--DTE---SELPSE----------TMC
------------RCH-LPCTETN---YEAK-VSS------------------SDI-PM--
-------SYA------EKM-F-----------SN--------------------------
-------------------------PAIQ-G---------WMTGISDPEL----------
---------------------------------------------------------DIN
YN----------------------------------------------------------
----------------------------TA------------------------------
-----K-------DI-----------K------DNMMVLEVMFSSSQISQVK-EFVKYTT
NNL-----------------LGDIGGVLGLFLGASLFTILEFFQFIIMS-----------
--------IAKY--------C----------C-------MW-------------------
TP-GG---------------------------RK--------------------------
----------------------------------------KLEASEMS------------
------------------------------------------------------------
------------------------------------------------------------
------------------------------------------------------------
------------------------------------------------------------
------------------------------------------------------------
--------
>Cteno_MneLey_comp15203_c0_seq1__p1__GENE_comp15203_c0_seq1__comp15203_c0_seq
1__p1___ORF__type_complete__len_500___score_87_69__comp15203_c0_seq1__52_1551
__595
------------------------------------------------------------
----------M-------------------------------------------------
------------------------------------------------------------
------------------------------------------------------------
------------------------------------------------------------
------------------------------------------------------------
------------------------------------------------------------
------------------------------------------------------------
------------------------------------------------------------
```

------------------------------------------------------------
------------------------------------------------------------
----KKVTE----------------------M---------SW-----V-----------
-KD--FREFA-NDTSAHGVKYIF---------E-GR--------------YK---LV---
--KLLFL----VTWLGFSIYACHVIITSIV-RFVE----------K---PTSTKY----E
-V-IQ-DDSE----GRPE-RIEF-----PTI-----------------------SVCSMN
KV------RKS-----YLEAPE---------------NEAI-----------------R-
---------EY-------------------------------YEVVD-------------
------------------------------KY-NV-------SLVKDL------------
------------------------------------------------------------
------------------------------------------------------------
------------------------------------------------------------
------------------------------------------------------------
------------------------------------------------------------
------------------------------------------------------------
------------------------------------------------------------
------------------------------------------------------------
------------------------------------------------------------
------------------------------------------------------------
------------------------------------------------------------
-------------------------------------------------------AKRFK
NSDD-----------------------------PLHSIK---------------------
-------------------------DM----------------TYEELIRN----GGPNP
--DRL------LKCT----QR-A----K----------Y---C-----------H-ELPA
F-------------------------------------------NGR-------------
------------------------------------------------------------
------------------------------------------------------------
-----------D---------VS-------V-----------MEN-S-MTG--------N
CWRVN-------PE-------------GR-------------------------------
--------LMGKM------GD--------YG-------------AMKLMFWADVQ-----
----------D-----------------Y------------S-A-----RT---------
------------------------------------------------------------
------------------------------------------------------------
-------ADVENQGFVVAFH-DNSTYG--STMTAGFLMSPGTYYKADLRRK------Q--
-----EIRN-----------RDK-I--ES-C--NAS----------------LTE--NTY
------------------G-AYNEGSCALECKDEALH----KA--CGCTNVV----P---
------------P-L--NN-----------------------------------------
---------GKY-----K---SCTL-----------------------------------
------------------------------------EQW---VDCGL---------A---
V-Y-------------------KEWF-HNFTDT--DR--------AD----------QLC
------------PCQ-IQCEEVR---YEAQ-ISS------------------SSI-SP--
-------AFA------EKL-F-----------PSV-------------------------
------------------------QPIIS-Q---------PQYGYSNPDF----------
---------------------------------------------------------NIL
YN----------------------------------------------------------
----------------------------TT------------------------------
-----Q-------DI-----------L------DNVMVLEVLFTSMRTNEIK-EIISYGL
SNL-----------------LGDIGGVLGLFLGASLFTILEVFQFVFFS-----------
--------ISKY--------C----------FD------KG-------------------
QP------------------------------KH--------------------D-----
---------------------------------------SLTNGDKLS------------
------------------------------------------------------------
------------------------------------------------------------
------------------------------------------------------------
------------------------------------------------------------
------------------------------------------------------------
-------I

>Cteno_Puk_Fal_comp20088_c0_seq2__p1__GENE_comp20088_c0_seq2__comp20088_c0_se
q2__p1___ORF__type_internal__len_505___score_74_57__comp20088_c0_seq2__2_1513
__604
------------------------------------------------------------
------------------------------------------------------------
------------------------------------------------------------
------------------------------------------------------------
------------------------------------------------------------
------------------------------------------------------------
------------------------------------------------------------
------------------------------------------------------------
------------------------------------------------------------
------------------------------------------------------------
------------------------------------------------------------
------------------------------------------------------------
---------A-EFTTAHGVNLIF---------V-GG--------------HK---AL---
--RFLYF----VIWLLCVLYAIYIILQAAV-AFIS----------R---PTGTTE----Q
-L-VV-ETDHGQ---DIQ-FVKF-----PTI-----------------------TVCSTN
IM------KRS-----FTET-----------------DKSI-----------------S-
---------AL-------------------------------WYLIS----RL-------
-----------------------------KVM-DP-------QYVSHVL-----------
------------------------------------------------------------
------------------------------------------------------------
-----------------E------------------------------------------
------------------------------------------------------------
------------------------------------------------------------
------------------------------------------------------------
-------------------------HP---------------------------------
------------------------------------------------------------
------------------------------------------------------------
------------------------------------------------------------
------------------------------------------------------------
-------------------------------------------------------KRFFQ
SYCDYHGV-------------------------TFEHLK---------------------
-------------------------NY----------------TVKEMFER----GGPDD
--NRL------VYCK----QR-N----I----------D---C-----------Q-DLT-
----------------------------------------------E-------------
------------------------------------------------------------
------------------------------------------------------------
-----------S---------YF-------V-----------RET-S-SDG--------M
CWRVN-------PN-------------GS-------------------------------
--------MYGRS------GE--------WG-------------RMELFFFADIQ-----
----------D-----------------Y------------T-S-----TS---------
------------------------------------------------------------
------------------------------------------------------------
----------PVYGYKVKFH-DSEYCG--EGITGAHFMSPGFVYKVDLGLT------K--
-----RSWI-----------PKP-I--GN-C--DAN----------------AKK--STY
------------------G-KNSEESCLKECEDRYMN----KV--CGCVDLL----P---
------------P-H--NH-----------------------------------------
---------HNY-----T---ACTI-----------------------------------
------------------------------------GQY---IGCGF---------R---
E-M-------------------VHFR-MGLRDA--SR----RHGTGE----------DGC
------------SCS-HSCDETH---YDVKLLST------------------SAL-SP--
-------VWA------DDL-------------SR--------------------------
------------------------DKWIA-E---------AVANFSQPSF----------
---------------------------------------------------------NIS
YN----------------------------------------------------------

```
---------------------------VSS------------------------------
-------------DL-----------L------KNIVILQVEFSTVFTKEFK-QIVTYSY
SNL-----------------LGDIGGVLGLFLGASLFSVLEFFQFTLTS-----------
--------LHKH--------F----------CG------RE-------------------
DKP-----------------------------KT--------------------------
----------------------------------------LPETEKLSSNLAF-------
------------------------------------------------------------
------------------------------------------------------------
------------------------------------------------------------
--------------------------------SP--------------------------
----------------------------------------------------------AI
RSTPVLCV
>Cteno_MneLey_comp16014_c0_seq2__p1__GENE_comp16014_c0_seq2__comp16014_c0_seq
2__p1___ORF__type_5prime_partial__len_507____score_115_90__comp16014_c0_seq2_
_352_1872___610
PTRKNRRIKC------PRCRQHFVFNKVRKDGI---------------------------
----------M-------------------------------------------------
------------------------------------------------------------
------------------------------------------------------------
------------------------------------------------------------
------------------------------------------------------------
------------------------------------------------------------
------------------------------------------------------------
------------------------------------------------------------
------------------------------------------------------------
------------------------------------------------------------
-----------------------------------------GW-----L-----------
-QD--FKEFS-DGTTAHGVKYIF---------H-PSVH----------GNFK---IV---
--RILFL----VVWLVAIAYSLFVIYNAVG-NYVG----------K---PTGTKF----Q
-V-IA-SNPYKE---KPA-SIQF-----PTI-----------------------SVCSHN
LV------TKS-----YFKS-----------------NDGL-----------------E-
---------EL-------------------------------WSELD-------------
------------------------------QW-NP-------DTAENI------------
------------------------------------------------------------
------------------------------------------------------------
-----------------D-------------------------FNE--------------
------------------------------------------------------------
------------------------------------------------------------
------------------------------------------------------------
-------------------------GS---------------------------------
------------------------------------------------------------
------------------------------------------------------------
------------------------------------------------------------
------------------------------------------------------------
------------------------------------------------------------
---------------------------------PAAKYK---------------------
-------------------------DW----------------TYEKIIAE----GGPAN
--WTF------LQCE----HF-I----N----------L---C-----------Q-EVIP
Q-------------------------------------------P---------------
------------------------------------------------------------
------------------------------------------------------------
-----------D---------FF-------E-----------RET-T-LTG--------N
CFRIN-------PN-------------GK-------------------------------
--------LRGKG------GD--------YG-------------KMQLMFFADLN-----
----------E-----------------Y------------S-D-----LT---------
------------------------------------------------------------
------------------------------------------------------------
```

-------RREPQYGYTVVFH-DHESYS--STIPSGFWLSPGSIYKVDLSLG------K--
-----EFRE-----------PPP-A--GS-C--DPT----------------RVN--NTY
------------------G-RYEENSCIAQCRDDVLM----EK--CGCVHVS----P---
------------P-H--PR-----------------------------------------
-----NVPDKYY-----R---GCTL-----------------------------------
------------------------------------EEW---ATCGL---------R---
A-Y-------------------KEWV-VEYSNV--NK--------VK----------TEC
------------NCP-ATCTEIA---YKAQ-LSS------------------SQL-SK--
-------FYA------EEK-----------------------------------------
----------------------------------------------AAKL----------
---------------------------------------------------------PPG
YE----------------------------------------------------------
----------------------------NA------------------------------
-----Q-------DV-----------L------ENLLIIDILFTSMQISEIR-EIVTYGW
GNF-----------------LGDVGGVLGLFLGASMFTIIEFIQFIVCA-----------
--------VLKG--------C----------C-------GL----------------N--
DK------------------------------ES-------------------R------
------------------------------------------------------------
------------------------------------------------SHSP--------
------------------------------------------------------------
------------------------------------------------------------
------------------------------------------------------------
------------------------------------------------------------
--LYNENL
>Ctenoph_Hormiphora_californiensis_evg1138204__p1_GENE_evg1138204__evg1138204
__p1__ORF_type_complete_len_508____score_90_51_evg1138204__158_1681___615
------------------------------------------------------------
----------M-------------------------------------------------
------------------------------------------------------------
------------------------------------------------------------
------------------------------------------------------------
------------------------------------------------------------
------------------------------------------------------------
------------------------------------------------------------
------------------------------------------------------------
------------------------------------------------------------
------------------------------------------------------------
-----GITQ----------------------M---------TW-----I-----------
-KD--FREFA-GDTSAHGVKNIF---------E-GP--------------TK---IL---
--KLLFL----ITWIVCSVYATYVIFNAIV-TFIN----------K---PTGTKF----Q
-V-TV-ESTERE-EGKPA-VIKF-----PTI-----------------------SVCSYN
KV------KKS-----FLRAPG---------------NELL-----------------R-
---------RV------------------------------YLTLN-------------
------------------------------KY-DL-------DEAAEL------------
------------------------------------------------------------
------------------------------------------------------------
------------------------------------------------------------
------------------------------------------------------------
------------------------------------------------------------
------------------------------------------------------------
------------------------------------------------------------
------------------------------------------------------------
------------------------------------------------------------
------------------------------------------------------------
------------------------------------------------------------
-------------------------------------------------------AAEFE
DPDSEV---------------------------AQASVG---------------------

```
-------------------------DM----------------KYEDLLAD----GSPDS
--NRF------LMCS----QR-A----K----------N---C-----------D-KLAV
F-------------------------------------------KGNAS-----------
------------------------------------------------------------
------------------------------------------------------------
-----------E----GIPPGHF-------V-----------REN-S-ISG--------M
CWRVN-------PN-------------GK-------------------------------
--------LKGLM------GD--------YG-------------ALELRFWADIQ-----
----------D-----------------Y------------S-E-----ST---------
------------------------------------------------------------
------------------------------------------------------------
-------KNMATHGFIVAFH-DNGTFG--STLSSGYLMSPGTYYKVDLRLR------K--
-----EIRK-----------PPP-G--GN-C--NAS----------------LTD--TSY
------------------G-VYTEGSCIMECKDRWLY----ER--CGCVNVV----P---
------------P-I--NK-----------------------------------------
---------DNY-----E---SCNL-----------------------------------
------------------------------------RQW---ASCGL---------K---
A-Y-------------------LEWY-DRYTDT--DT--------AN----------PIC
------------TCS-MPCEETN---YEAK-VSS------------------SDI-SQ--
-------AYA------DHM-F-----------TT--------------------------
-------------------------KGVA-G---------FLNSISDPSL----------
---------------------------------------------------------NIS
YD----------------------------------------------------------
----------------------------SA------------------------------
-----N-------DI-----------R------ENMMVVEVLFSSAQVSEIR-EVVTYTA
NNL-----------------LGDIGGVLGLFLGASLFTILEFFQFIIIS-----------
--------VAKH--------C----------C-------NW-------------------
IPGGG---------------------------KK--------------------------
----------------------------------------KYESAEMS------------
------------------------------------------------------------
------------------------------------------------------------
------------------------------------------------------------
------------------------------------------------------------
------------------------------------------------------------
--------
>Cteno_Puk_Fal_comp27454_c0_seq1__p1__GENE_comp27454_c0_seq1__comp27454_c0_se
q1__p1___ORF__type_complete__len_520____score_91_49__comp27454_c0_seq1__400_1
959___649
------------------------------------------------------------
----------MSR-YGE-------------------------------------------
------------------------------------------------------------
------------------------------------------------------------
------------------------------------------------------------
------------------------------------------------------------
--------------------------------------RYPP---SSGLQLF--------
-------------------------------PDPDKEEEG-------I-------SPVQQ
------------------------------------------------------------
---------------PCRCTCPCNTQNTRDT---------G----------------SQM
DV-----------------KTISTSSGK--------------------------------
--------R----------------------A---------TF-----Y-----------
-DR--WFDFS-DGTSLHGFKFIF---------S-GG---------------R---VT---
--RAAFA----VFMVLFVFKLGWQLRESVI-SYME----------H---PTSSKI----S
-V---------SNVEGPA-LVTF-----PTI-----------------------SICSHN
LV------SQS-----YLDD-----------------NPGL-----------------D-
---------FL-------------------------------FEMID-------------
------------------------------TW-QP-------EVVSQI------------
------------------------------------------------------------
```

------------------------------------------------------------
-----------------D-------------------------FTD--------------
------------------------------------------------------------
------------------------------------------------------------
------------------------------------------------------------
--------------------------P---------------------------------
------------------------------------------------------------
------------------------------------------------------------
------------------------------------------------------------
------------------------------------------------------------
------------------------------------------------------------
---------------------------------QFAPYA---------------------
-------------------------DF----------------KYSDILRD----GGPPS
--INV------LQCS----HF-T----N----------D---C-----------T-PDL-
--------------------------------------------QSE-------------
------------------------------------------------------------
------------------------------------------------------------
-----------K---------SM-------E-----------REL-S-MRG--------N
CYRIN-------PK-------------GQ-------------------------------
--------LMGKG------GD--------YG-------------KLSIKLFSDLN-----
----------E-----------------Y------------S-A-----TG---------
------------------------------------------------------------
------------------------------------------------------------
-------KKLAQVGYSVQFH-DHEQYS--GAIPNSYWMSPGLIYKADLSLT------R--
-----ETRL-----------TPP-A--GR-C--NDT----------------VLY--NSY
------------------G-RNEENSCLAQCRDDAMM----RA--CGCVLVP----P---
------------P-Y--DP-----------------------------------------
---------TSY-----T---PCTL-----------------------------------
------------------------------------QQW---SECGL---------T---
A-Y-------------------NGWY-QDYVDV--SR--------TE----------EMC
------------YCY-PSCREVF---YTVS-MSS------------------SLL-SR--
-------FYA------RDH-----------------------------------------
----------------------------------------------VQFL----------
---------------------------------------------------------PKG
YN----------------------------------------------------------
----------------------------GT------------------------------
-----N-------DV-----------L------NNLVVLDILFVNTQELEIE-QVVAYGW
TNF-----------------LGDVGGVLGLFLGASAFSLIEFFFFVGHL-----------
--------FWTY--------V----------L-------CI----------------W--
RE------------------------------EH-------------------Q------
------------------------------------------------------------
------------------------------------------------------------
------------------------------------------------------------
------------------------------------------------------------
------------------------------------------------------------
------------------------------------------------------------
--------
>Cteno_Puk_Fal_comp26884_c0_seq1__p1__GENE_comp26884_c0_seq1__comp26884_c0_se
q1__p1___ORF__type_5prime_partial__len_531___score_77_18__comp26884_c0_seq1__
1_1593__671
QRRVRGFFICI----YPSCVCFEARNVKRTAHSTGT------------------------
----------S-------------------------------------------------
------------------------------------------------------------
------------------------------------------------------------
------------------------------------------------------------
------------------------------------------------------------
------------------------------------------------------------

```
------------------------------------------------------------
------------------------------------------------------------
------------------------------------------------------------
------------------------------------------------------------
----TIGRR----------------------M---------TW-----I-----------
-KD--FRDFA-GDTSAHGVKNIF---------E-GP--------------TK---LL---
--KLIFL----ICWIVCSVYATYVIMVSII-TFMQ----------R---PTGTKY----T
-F-LV-ENENRK-LGKSA-IVTM-----PTI-----------------------SVCSHN
KV------SKS-----FLKTEG---------------NEIL-----------------S-
---------EV-------------------------------YTTLD-------------
------------------------------KY-DK-------EDAEEL------------
------------------------------------------------------------
------------------------------------------------------------
------------------------------------------------------------
------------------------------------------------------------
------------------------------------------------------------
------------------------------------------------------------
------------------------------------------------------------
------------------------------------------------------------
------------------------------------------------------------
------------------------------------------------------------
------------------------------------------------------------
-------------------------------------------------------AKRFD
NSSD-----------------------------PLSEVA---------------------
-------------------------DY----------------LYEDLLAD----GSIPD
--DRL------LLCQ----QR-A----K----------Q---C-----------F-KLPA
F-------------------------------------------AKT-------------
------------------------------------------------------------
------------------------------------------------------------
-----------P---------FF-------T-----------REN-S-ITG--------K
CFRIN-------PM-------------GT-------------------------------
--------LSALM------GD--------YG-------------ALKLYFWADVQ-----
----------D-----------------Y------------S-E-----AS---------
------------------------------------------------------------
------------------------------------------------------------
-------SNSATHGFTVAFH-DSDTYG--STLSSGYLMSPGSYYKVDLRLR------K--
-----EVRE-----------PPP-S--GH-C--DKT----------------REF--TSY
------------------G-NYSEGACVMECKDKTLN----ET--CGCVNVV----P---
------------P-I--ND-----------------------------------------
---------LGY-----R---SCTL-----------------------------------
------------------------------------KEW---ATCGL---------K---
T-Y-------------------LDWY-DAYTDT--DT--------AQ----------TTC
------------KCH-LPCQETN---YEAK-VSS------------------SSI-ST--
-------AYS------EKM-Y-----------SN--------------------------
-------------------------PDVV-N---------WMKKISKPEL----------
---------------------------------------------------------GIN
YS----------------------------------------------------------
----------------------------TA------------------------------
-----E-------DI-----------R------ENIMALEVMFSSSQVSQVE-EFVKYTS
TNL-----------------LGDIGGVLGLFLGASLFTILEFFQFIIMS-----------
--------IAKY--------C----------C-------QW-------------------
TP-DG---------------------------RK--------------------------
----------------------------------------KLDGHEMS------------
------------------------------------------------------------
------------------------------------------------------------
------------------------------------------------------------
------------------------------------------------------------
```

------------------------------------------------------------
--------
>Ctenoph_Hormiphora_californiensis_evg1163616__p1_GENE_evg1163616__evg1163616
__p1__ORF_type_5prime_partial_len_545____score_81_53_evg1163616__2_1636___694
FKRINLSSILL----SPSHILPNDQYA---------------------------------
----------M-------------------------------------------------
------------------------------------------------------------
------------------------------------------------------------
------------------------------------------------------------
------------------------------------------------------------
------------------------------------------------------------
------------------------------------------------------------
------------------------------------------------------------
------------------------------------------------------------
---------------------LLSNGAP--------------------------------
---QGETKR----------------------K---------SF-----Y-----------
-ED--FKEFT-TETTSHGIQYVF---------N-AA--------------YK---WV---
--RLIYL----VLWLSAVLYTMYIVCISVG-KYIS----------M---ETGTKI----E
-E-MV-TPRPAN--LNQP-AITH-----PTV-----------------------TICPHN
IV------SKA-----FLNK-----------------NEGL-----------------E-
---------EL-------------------------------FAELH-------------
------------------------------KW-ED-------DSAWNT------------
------------------------------------------------------------
------------------------------------------------------------
------------------------------------------------------------
------------------------------------------------------------
------------------------------------------------------------
------------------------------------------------------------
-------------------------SA---------------------------------
------------------------------------------------------------
------------------------------------------------------------
------------------------------------------------------------
------------------------------------------------------------
------------------------------------------------------------
---------------------------------SYHKYA---------------------
-------------------------NW----------------TYHDVMWK----GRPR-
--VGL------LQCE----HY-R----H----------E---C-----------K-DLDF
T-------------------------------------------NRSVPE---YHW----
------------------------------------------------------------
------------------------------------------------------------
-----------DREYDASNHLHY-------D-----------WEV-S-LSG--------S
CYRVN-------PQ-------------GS-------------------------------
--------LFTQV------GD--------YG-------------ELKLFFFTDLN-----
----------D-----------------Y------------S-Q-----FA---------
------------------------------------------------------------
------------------------------------------------------------
-------KDEAQYGYTVTFS-DNGTYS--STMTSGFYMSPGNVYKADMRKY------R--
-----EENI-----------GPP-Y--GR-C--NPH----------------HKT--NVY
------------------G-TYSLNSCVAQCRDSYVM----ER--CGCVQIL----P---
------------P-H--PE-----------------------------------------
----WNQDGVEM-----V---GCTL-----------------------------------
------------------------------------RQW---MDCGA---------K---
A-Q-------------------KEFT-DRFVNL--YA--------EE----------KLC
------------HCS-PACMEIW---YEAT-ISS------------------SRL-SK--
-------FYA------KKK-A-----------TD------------------------RL
------------------------PGIIS-Y-----------YNLSA-HI----------
---------------------------------------------------------DEP

YT----------------------------------------------------------
---------------------------IDT------------------------------
-----E-------TV-----------Y------ENLMVLQVIFTSIQESNYK-EYIKYDY
TQL-----------------IGDVGGQMGMLLGASLFTIIEFVHFFVRV-----------
--------AWRY--------C----------L-------RK-------------------
RGA-----------------------------RA---------------AE--E------
-----------------------------------------QPAYEMHRD----------
------------------------------------------------------------
------------------------------------------------------------
------------------------------------------------------------
------------------------------------------------------------
------------------------------------------------------------
------QE
>Ctenoph_Hormiphora_californiensis_evg156929__p1_GENE_evg156929__evg156929__p
1__ORF_type_complete_len_561____score_102_52_evg156929__369_2051___738
------------------------------------------------------------
----------MP--YSEI------------------------------------------
------------------------------------------------------------
------------------------------------------------------------
------------------------------------------------------------
------------------PGPAEPDGAADQG-----------------------------
---------------------------------YAGTSRTPPQAHSSGLQLF--------
-------------------------------PDPDRDVE----------------VPVQV
------------------------------------------------------------
----------------CQCTCSCNRATPGGE---------GPG------ELASAAGDGRM
EV-----------------RTVSQTSSR--------------------------------
----GEKGS----------------------S---------KF-----Y-----------
-RR--WDEFS-DGTTMHGIKFIF---------S-GS---------------R---LT---
--RLAFF----LFMLLFVFKLAWQLRESLA-NYME----------Y---PTSSKI----T
-V---------SNVEGPA-LVTF-----PTV-----------------------SICSHN
LV------GQS-----YLDS-----------------KPGL-----------------D-
---------DL-------------------------------FERVE-------------
------------------------------RW-DA-------EVASSI------------
------------------------------------------------------------
------------------------------------------------------------
-----------------N-------------------------FSD--------------
------------------------------------------------------------
------------------------------------------------------------
------------------------------------------------------------
--------------------------P---------------------------------
------------------------------------------------------------
------------------------------------------------------------
------------------------------------------------------------
------------------------------------------------------------
------------------------------------------------------------
---------------------------------TFAPYA---------------------
-------------------------DL----------------PYETVLRE----GGPPD
--SNV------LQCS----HF-T----N----------D---C-----------L-GN--
--------------------------------------------SSD-------------
------------------------------------------------------------
------------------------------------------------------------
-----------K---------YM-------E-----------REL-S-LRG--------L
CYRIN-------PA-------------GK-------------------------------
--------LMGKG------GD--------YG-------------KLSIKLFSDLD-----
----------E-----------------Y------------S-A-----TG---------
------------------------------------------------------------
------------------------------------------------------------

-------KKLAQVGYSVQFH-DHEQYS--GAIPNSYWLSPGLIYKADLSLT------R--
-----ETRL-----------PPP-A--GR-C--NDT----------------LLY--NSY
------------------G-RHEENSCLSQCRDDAIM----AK--CGCVLVS----P---
------------P-N--PH-----------------------------------------
-----YAPSTNY-----T---PCTL-----------------------------------
------------------------------------KQW---TECGL---------A---
A-Y-------------------SNWY-QQYVNV--SR--------TE----------ELC
------------YCY-PSCREVL---YSVS-LSS------------------SHL-SR--
-------FYA------RDH-----------------------------------------
----------------------------------------------VQYL----------
---------------------------------------------------------PPG
YN----------------------------------------------------------
----------------------------GT------------------------------
-----R-------DV-----------L------NNLVVLDILFVNTQELEIE-QVLSYGW
TNF-----------------LGDVGGVLGLFLGASAFTLIEFVLFLGQL-----------
--------FWHY--------V----------L-------CA----------------W--
RD------------------------------DH-------------------E------
------------------------------------------------------------
------------------------------------------------------------
------------------------------------------------------------
------------------------------------------------------------
------------------------------------------------------------
------------------------------------------------------------
--LQIRDL
>Cteno_MneLey_comp614729_c0_seq1__p1__GENE_comp614729_c0_seq1__comp614729_c0_
seq1__p1___ORF__type_internal__len_184___score_30_14__comp614729_c0_seq1__1_5
49__193
------------------------------------------------------------
------------------------------------------------------------
------------------------------------------------------------
------------------------------------------------------------
------------------------------------------------------------
------------------------------------------------------------
------------------------------------------------------------
------------------------------------------------------------
------------------------------------------------------------
------------------------------------------------------------
------------------------------------------------------------
------------------------------------------------------------
------------------------------------------------------------
------------------------------------------------------------
------------------------------------------------------------
------------------------------------------------------------
------------------------------------------------------------
------------------------------------------------------------
------------------------------------------------------------
------------------------------------------------------------
------------------------------------------------------------
------------------------------------------------------------
------------------------------------------------------------
------------------------------------------------------------
------------------------------------------------------------
------------------------------------------------------------
------------------------------------------------------------
------------------------------------------------------------
------------------------------------------------------------
------------------------------------------------------------

------------------------------------------------------------
------------------------------------------------------------
------------------------------------------------------------
------------------------------------------------------------
------------------------------------------------------------
------------------------------------------------------------
------------------------------------------------------------
------------------------------------------------------------
------------------------------------------------------------
------------------------------------------------------------
------------------------------------------------------------
------------------------------------------------------------
-------------------------------------------NFISLSYE------R--
-----VRNL-----------KQP-Y--GD-C--RDD--------A-------MP----LY
------------------R-NYSISGCERQCFTEIMI----KE--CNCSSYY----LK--
----------AMP-G---------------------------------------------
---------GKD-----V---ECNL-----------------------------------
------------------------------------YQE---KNCVK---------E---
L-I-------------------RRYH-A-------GE--------FI--------NSSEC
------------RCP-DPCVSES---YLAT-LST------------------AEY-PA--
-------KNH------GEA-L-----------QR--------------------------
------------------------------------------------YN----------
----------------------------------EANRG---------------------
PN----------------------------------------------------------
-----------------------DVV-QDV------------------------------
-----T-------FY-----------R------ENLMKVHIYFEILSIKEVE-KVGSYKW
SNL-----------------MGDVGGQLGL------------------------------
------------------------------------------------------------
------------------------------------------------------------
------------------------------------------------------------
------------------------------------------------------------
------------------------------------------------------------
------------------------------------------------------------
------------------------------------------------------------
------------------------------------------------------------
--------
>Cteno_MneLey_comp1086195_c0_seq1__p1__GENE_comp1086195_c0_seq1__comp1086195_
c0_seq1__p1___ORF__type_internal__len_190____score_29_76__comp1086195_c0_seq1
__2_568___202
------------------------------------------------------------
------------------------------------------------------------
------------------------------------------------------------
------------------------------------------------------------
------------------------------------------------------------
------------------------------------------------------------
------------------------------------------------------------
------------------------------------------------------------
------------------------------------------------------------
------------------------------------------------------------
------------------------------------------------------------
------------------------------------------------------------
------------------------------------------------------------
--KLLWA----VLFLGFFVYSFQCCLKSFN-TYFS----------M---PTSTKY----N
-M-YF-EP--------NK-LLRF-----PSI-----------------------TICNMN
PH------NTA-----YLDKEE-----------------------------------NN-
---------PM------------------------------KAYLLS-------------
------------------------------------------------------------

------------------------------------------------------------
------------------------------------------------------------
-----------------ESR----------------------------------------
------------------------------------------------------------
------------------------------------------------------------
------------------------------------------------------------
-------------------------VPWD-------------------------------
------------------------------------------------------------
------------------------------------------------------------
------------------------------------------------------------
----TSNLTPDV------------------------------SETD--------------
-------------------------------------------------------YRNF-
---------------------------------NVKKYY---------------------
------------------------------------------------RQA----GQKLN
--DFI------SEVV----FF-E----E----------E---I-----------E-T---
----------------------------------------------S-------------
------------------------------------------------------------
------------------------------------------------------------
-----------DA---------F-------R-----------EQF-T-THG--------M
CWTFN-------AD-------------GS-------------------------------
-------RQVNRT------GM--------DF-------------GLSFKLNINQS-----
----------S-----------------Y--------------P-N---FV---------
------------------------------------------------------------
------------------------------------------------------------
----------TTAGVKVMFH----------------------------------------
------------------------------------------------------------
------------------------------------------------------------
------------------------------------------------------------
------------------------------------------------------------
------------------------------------------------------------
------------------------------------------------------------
------------------------------------------------------------
------------------------------------------------------------
------------------------------------------------------------
------------------------------------------------------------
------------------------------------------------------------
------------------------------------------------------------
----------------------------------N-------------------------
------------------------------------------------------------
------------------------------------------------------------
------------------------------------------------------------
------------------------------------------------------------
------------------------------------------------------------
------------------------------------------------------------
------------------------------------------------------------
------------------------------------------------------------
------------------------------------------------------------
--------
>Cteno_Puk_Fal_comp26472_c0_seq1__p1__GENE_comp26472_c0_seq1__comp26472_c0_se
q1__p1___ORF__type_5prime_partial__len_221____score_43_26__comp26472_c0_seq1_
_674_1336___234
------------------------------------------------------------
------------------------------------------------------------
------------------------------------------------------------
------------------------------------------------------------
------------------------------------------------------------
------------------------------------------------------------

```
------------------------------------------------------------
------------------------------------------------------------
------------------------------------------------------------
------------------------------------------------------------
------------------------------------------------------------
------------------------------------------------------------
------------------------------------------------------------
------------------------------------------------------------
------------------------------------------------------------
------------------------------------------------------------
------------------------------------------------------------
------------------------------------------------------------
------------------------------------------------------------
------------------------------------------------------------
------------------------------------------------------------
------------------------------------------------------------
------------------------------------------------------------
------------------------------------------------------------
------------------------------------------------------------
------------------------------------------------------------
------------------------------------------------------------
------------------------------------------------------------
------------------------------------------------------------
------------------------------------------------------------
------------------------------------------------------------
------------------------------------------------------------
------------------------------------------------------------
------------------------------------------------------------
------------------------------------------------------------
------------------------------------------------------------
------------------------------------------------------------
------------------------------------------------------------
------------------------------------------------------------
------------------------------------------------------------
------------------------------------------------------------
------------------------------------------------------------
------------------------------------------------------------
----------------------N-A--EY-G--DPV----------------LEQ---DY
----------------DPD-GYTIAKCVMNNRARYTI----QK--CNCIPFY----MEVP
RDLSS-----TSP-A---------------------------------------------
---------------------VCNL-----------------------------------
------------------------------------YQE---WDCVE---------P---
L-L-------------------EDLA-L-------GT--------LTMG-GQLV---QDP
------------DCP-SPCKYLT---YSYT-ISQ------------------SEY-PS--
-------QSV------EDQ-V-----------LS--------------------------
------------------------------------------------RV----------
------------------------------------------------------------
-GASRG------------------------------------------------------
------------------------AG-WNI------------------------------
-----E-------RI-----------R------DNYLKVHIYFEELSTLEME-RHPSYMI
ANL-----------------IGDVGGQLGLLLGMNICSLVQFTDYIVRFSFFK-------
-------GIMKL--------V------------------RM-------------------
SK------------------------------RN-------------------N------
-----------------------------------------KRSVG--------------
------------------------------------------------------------
------------------------------------------------------------
------------------------------------------------------------
```

------------------------------------------------------------
------------------------------------------------------------
---GRSRL
>Cteno_MneLey_comp860380_c0_seq1__p1__GENE_comp860380_c0_seq1__comp860380_c0_
seq1__p1___ORF__type_internal__len_249___score_49_23__comp860380_c0_seq1__1_7
44__269
------------------------------------------------------------
------------------------------------------------------------
------------------------------------------------------------
------------------------------------------------------------
------------------------------------------------------------
------------------------------------------------------------
------------------------------------------------------------
------------------------------------------------------------
------------------------------------------------------------
------------------------------------------------------------
------------------------------------------------------------
------------------------------------------------------------
------------------------------------------------------------
------------------------------------------------------------
------------------------------------------------------------
------------------------------------------------------------
------------------------------------------------------------
------------------------------------------------------------
------------------------------------------------------------
------------------------------------------------------------
------------------------------------------------------------
------------------------------------------------------------
------------------------------------------------------------
------------------------------------------------------------
------------------------------------------------------------
------------------------------------------------------------
------------------------------------------------------------
------------------------------------------------------------
------------------------------------------------------------
------------------------------------------------------------
------------------------------------------------------------
------------------------------------------------------------
------------------------------------------------------------
------------------------------------------------------------
------------------------------------------------------------
------------------------------------------------------------
----------------------F-------P-----------EIN-T-ELG--------I
CYTFN-------AD-------------GK-------------------------------
-------KRVKRT------GN--------DF-------------GATFTININQS-----
----------E-----------------Y------------A-E-----TQ---------
------------------------------------------------------------
------------------------------------------------------------
----------ESAGIKVMLH-HFKEPP--LIKEYGLAIPPGSEAYISTRTL------Q--
-----LNNL-----------REP-Y--FN-C--TDR--------K-------LR----TT
------------------D-YYSVVACTLECHAKILD----EE--CQCKQSH----M---
------------P-L---------------------------------------------
---------ENI-----T---ICSL-----------------------------------
------------------------------------YTH---RMCTE---------N---
A-I-------------------KKIH-N-------GS--------YA----DRI---SDC
------------WCI-DACKDTY---YEYS-LSR------------------AQY-AS--
-------INA------AKK-I------------LE--------------------------

```
------------------------------------------------VN----------
------------------------------------------------------------
RN----------------------------------------------------------
----------------------VSEY-STI------------------------------
-----S-------DL-----------R------ENFIRLHVYFETLAIENVE-KVPAYTT
MNL-----------------LGDVG-----------------------------------
------------------------------------------------------------
------------------------------------------------------------
------------------------------------------------------------
------------------------------------------------------------
------------------------------------------------------------
------------------------------------------------------------
------------------------------------------------------------
------------------------------------------------------------
--------
>Cteno_Ple_Bac_comp46643_c0_seq4_p1__comp46643_c0__comp46643_c0_seq4_p1___ORF
__type_5prime_partial__len_272___score_55_68__comp46643_c0_seq4_3_818__300
------------------------------------------------------------
------------------------------------------------------------
------------------------------------------------------------
------------------------------------------------------------
------------------------------------------------------------
------------------------------------------------------------
------------------------------------------------------------
------------------------------------------------------------
------------------------------------------------------------
------------------------------------------------------------
------------------------------------------------------------
------------------------------------------------------------
------------------------------------------------------------
------------------------------------------------------------
------------------------------------------------------------
------------------------------------------------------------
------------------------------------------------------------
------------------------------------------------------------
------------------------------------------------------------
------------------------------------------------------------
------------------------------------------------------------
------------------------------------------------------------
------------------------------------------------------------
------------------------------------------------------------
------------------------------------------------------------
------------------------------------------------------------
------------------------------------------------------------
------------------------------------------------------------
------------------------------------------------------------
------------------------------------------------------------
------------------------------------------------------------
------------------------------------------------------------
------------------------------------------------------------
------------------------------------------------------------
------------------------------------------------------------
------------------------------------------------------------
------------------------------------------------------------
------------------------------------------------------------
------------------------------------------------------D-----
----------D-----------------Y------------F-R-T--------------
```

------------------------------------------------------------
------------------------------------------------------------
---------QDVAGFKVLLH-NSYEPP--MIEEYGFALRPGSESYVRIRLQ------K--
-----FRDT-----------ERP-L--GY-C--DPV----------------LEQ---DY
----------------QPA-GYTIAKCVMNNRARYTI----QK--CNCIPFY----MEVP
RDLSS-----ISP-K---------------------------------------------
---------------------VCNL-----------------------------------
------------------------------------YQE---WDCVE---------P---
L-L-------------------EEIA-L-------GT--------LTMG-GELI---QDP
------------DCP-SPCNYVT---YSYT-ISQ------------------SEY-PS--
-------QSV------EAE-V-----------KS--------------------------
------------------------------------------------KV----------
------------------------------------------------------------
-VGARG------------------------------------------------------
------------------------AG-WNI------------------------------
-----D-------RI-----------R------DNYLKVHIYFEELSTLEME-RHPSYMI
SNL-----------------IGDVGGQLGLLLGMNICSLVQFTDYIIRFSFFN-------
-------GIMKL--------V------------------RM-------------------
SH------------------------------RN-------------------N------
-----------------------------------------KRA-A--------------
------------------------------------------------------------
------------------------------------------------------------
------------------------------------------------------------
------------------------------------------------------------
------------------------------------------------------------
---GRSRL
>Ctenoph_Hormiphora_californiensis_evg16414__p1_GENE_evg16414__evg16414__p1__ORF_type_internal_len_297____score_76_10_evg16414__3_890___327
---------------AP-------------------------------------------
----------V-------------------------------------------------
--------------------K---------------------------------------
------------------------------------------------------------
-----------------------KKNEFRPYPAPYHV--------------------PPQ
QESASAYQSVN-------------------------------------------------
-------------------------------------EYQAPSER---------------
------------ERNPVKRS-------------QSDSSTAGSFRAA----------PWLN
GQKILAPKYPVMSSV---------------------------------------------
------------------AATPTNSLKMRSM-------GGGGGRTVPPTWHDSLLGEDRP
DQVI-----------------QDREEETT-------------------------------
-VKSSGDKE----------------------P---------TK-----S-----------
-ES--FADFT-GDMTLNYLRYVW---------N-SP--------------EY---IR---
--KLIWA----VLFLGFFIYSFSCCYRSIA-HYLA----------R---PTSTKY----N
-M-FY-EP--------NK-QLRF-----PAI-----------------------TICNMN
PH------NKT-----YLDQDE------------------------------------Q-
-I-------AM-------------------------------RSYLR-------------
------------------------------------------------------------
------------------------------------------------------------
----------------------K-------------------------------------
-----------------TWK----------------------------------------
------------------------------------------------------------
------------------------------------------------------------
------------------------------------------------------------
-------------------------TPW--------------------------------
------------------------------------------------------------
------------------------------------------------------------
------------------------------------------------------------
-----AKDLPEA------------------------------KEEL--------------

-------------------------------------------------------MQNF-
---------------------------------NVKGYY---------------------
------------------------------------------------RNS----GQRVN
--DFI------RKAV----FI-E----E----------E---Q-----------V-V---
------------------------------------------------------------
------------------------------------------------------------
------------------------------------------------------------
------------------------------------------------------------
------------------------------------------------------------
------------------------------------------------------------
------------------------------------------------------------
------------------------------------------------------------
------------------------------------------------------------
------------------------------------------------------------
------------------------------------------------------------
------------------------------------------------------------
------------------------------------------------------------
------------------------------------------------------------
------------------------------------------------------------
------------------------------------------------------------
------------------------------------------------------------
------------------------------------------------------------
------------------------------------------------------------
------------------------------------------------------------
------------------------------------------------------------
------------------------------------------------------------
------------------------------------------------------------
------------------------------------------------------------
------------------------------------------------------------
------------------------------------------------------------
------------------------------------------------------------
------------------------------------------------------------
------------------------------------------------------------
------------------------------------------------------------
------------------------------------------------------------
------------------------------------------------------------
--------
>Ctenoph_Hormiphora_californiensis_evg15900__p1_GENE_evg15900__evg15900__p1__
ORF_type_5prime_partial_len_299____score_42_58_evg15900__2_898___328
------------------------------------------------------------
------------------------------------------------------------
------------------------------------------------------------
------------------------------------------------------------
------------------------------------------------------------
------------------------------------------------------------
------------------------------------------------------------
------------------------------------------------------------
------------------------------------------------------------
------------------------------------------------------------
------------------------------------------------------------
------------------------------------------------------------
------------------------------------------------------------
------------------------------------------------------------
------------------------------------------------------------
------------------------------------------------------------
------------------------------------------------------------
------------------------------------------------------------

------------------------------------------------------------
------------------------------------------------------------
------------------------------------------------------------
------------------------------------------------------------
------------------------------------------------------------
------------------------------------------------------------
------------------------------------------------------------
------------------------------------------------------------
------------------------------------------------------------
------------------------------------------------------------
------------------------------------------------------------
------------------------------------------------------------
------------------------------------------------------------
------------------------------------------------------------
----------------------E----E----------Q---V-----------V-E---
----------------------------------------------E-------------
------------------------------------------------------------
------------------------------------------------------------
-----------N----------F-------V-----------ERF-T-THG--------L
CWTFN-------AK-------------GD-------------------------------
-------KFVNRT------GM--------DF-------------GLSFSMDINQG-----
----------G-----------------Y------------P-D-----FV---------
------------------------------------------------------------
------------------------------------------------------------
----------RTAGVKVMFH-NYYEPA--LIDEYGMALSPGTENFISLSYE------R--
-----IRNL-----------DSP-Y--GN-C--TKD--------P-------SP----PF
------------------A-NYSVSGCERICFTRALE----GD--CNCTSYY----L---
------------P-V---------------------------------------------
---------GEK-----K---ECTL-----------------------------------
------------------------------------FDE---DVCVE---------K---
V-I-------------------KRYH-A-------GL--------YS--------DRPYC
------------QCQ-DACVSEN---YIAS-LST------------------AEY-PS--
-------KNY------GEA-L-----------EA--------------------------
------------------------------------------------HN----------
----------------------------------KETL----------------------
-N----------------------------------------------------------
-----------------------DTV-RDV------------------------------
-----T-------FY-----------R------ENYIKVHIYFEILSIKEVE-KVASYVW
SNL-----------------MGDVGGQLGLFLGANIVTIFEFVDFFLRV-----------
--------FWYR--------L----------LTPY--IFKRF------------------
--------------------------------RG--------------------------
------------------------------------------------------------
------------------------------------------------------------
------------------------------------------------------------
------------------------------------------------------------
------------------------------------------------------------
------------------------------------------------------------
-------N
>Cteno_Puk_Fal_comp27722_c0_seq1__p1__GENE_comp27722_c0_seq1__comp27722_c0_se
q1__p1___ORF__type_5prime_partial__len_329____score_62_50__comp27722_c0_seq1_
_161_1147___358
------------------------------------------------------------
------------------------------------------------------------
------------------------------------------------------------
------------------------------------------------------------
------------------------------------------------------------
------------------------------------------------------------

------------------------------------------------------------
------------------------------------------------------------
------------------------------------------------------------
------------------------------------------------------------
------------------------------------------------------------
------------------------------------------------------------
------------------------------------------------------------
---------------------------------------------F--------------
---QY-----------------F-------------------------------------
------------------------------------------------------------
------------------------------------------------------------
------------------------------------------------------------
------------------------------------------------------------
------------------------------------------------------------
------------------------------------------------------------
------------------------------------------------------------
------------------------------------------------------------
------------------------------------------------------------
------------------------------------------------------------
------------------------------------------------------------
------------------------------------------------------------
------------------------------------------------------------
------------------------------------------------------------
------------------------------------------------------------
---------------------------------NVKNYY---------------------
------------------------------------------------RDS----GQRLE
--NFI------ERPV----FL-E----E----------P---I-----------DVS---
----------------------------------------------E-------------
------------------------------------------------------------
------------------------------------------------------------
-----------A----------F-------V-----------EKF-T-THG--------L
CWTFN-------DK-------------GD-------------------------------
-------KFVNRT------GM--------DF-------------GLSFNMNINQD-----
----------G-----------------Y------------P-D-----FI---------
------------------------------------------------------------
------------------------------------------------------------
----------TTAGVKVMFH-NFYEPA--LIDEYGMALAPGTENYISLSYE------R--
-----IRNL-----------QSP-Y--GN-C--TKD--------S-------SNL---PF
------------------A-NYSVSGCERMCFTLALK----EE--CECTSYY----L---
------------P-P---------------------------------------------
---------SNV-----K---ECTL-----------------------------------
------------------------------------FDE---QECVE---------K---
V-I-------------------KKYH-A-------GE--------YA--------DKPFC
------------KCQ-DACVSEN---YIAT-LST------------------AEY-PS--
-------KNY------GEA-I-----------ER--------------------------
------------------------------------------------EN----------
----------------------------------KKNA----------------------
-N----------------------------------------------------------
----------------------GSDV-RDI------------------------------
-----L-------FY-----------R------ENMLKVHIYFEILSIKEVE-KVKSYIW
SNL-----------------MGDVGGQLGLFLGANIVTIFEFVDFFLRV-----------
--------IWYR--------F----------LTPY--VFKRF------------------
--------------------------------QR--------------------------
------------------------------------------------------------
------------------------------------------------------------
------------------------------------------------------------
------------------------------------------------------------

------------------------------------------------------------
------------------------------------------------------------
-------H
>Cteno_Ple_Bac_comp47578_c1_seq3_p1__comp47578_c1__comp47578_c1_seq3_p1___ORF
__type_complete__len_384____score_68_54__comp47578_c1_seq3_141_1292___410
------------------------------------------------------------
----------M------------EL-------NTSP------------------------
-------------GGMKKLV----------------------------------------
-TDVKDLAIL--MNYRNFKE----------------------------------------
------------------------------------------------------------
------------------------------------------------------------
GVY-----PGVEAFTP--------------------------------------------
---------------YSKYK-------------QDESAFDMTGRGERTS-----YYPYL-
--EHYQPSYPRPPRGGKPPPPTGAPVPNN-HRTSDQNGKIKYPPNHRDT---GSPTELED
RDGEDCEEFM--PPPKRKKKKKRRANR------------------------------AAL
E----------------------QRREEM-------------------------------
--------E----------------------P---------DE-----M-----------
-ES--FKEFA-STVTADYLAEMF-----------AK-------------ASN---GR---
--RAIWV----SLWLASFVYAWYNIALSVG-VYLS----------K---PTATKL----N
-F-YE-AP--------PG-GVAF-----PTV-----------------------TICNFN
KY------NKS-----FFEREG------------------D-----------------SV
N------S-QL------------R-DFL---------------VATTPI-W---------
---------------------------------DH-------------------------
------------------------------------------------------------
------------------------------------------------------------
------------------------------------------------------------
------------------------------------------------------------
------------------------------------------------------------
------------------------------------------------------------
------------------------------------------------------------
------------------------------------------------------------
------------------------------------------------------------
------------------------------------------------------------
--------KERL------------------------------DQD---------------
----------------------QWTSKYPN-------------------------IMSL-
---------------------------------NLKELY---------------------
------------------------------------------------ANG----SHLIT
--NTV------EFCR----FG-G----K----------P---CK-DTANRETSIP-S---
----------------------------------------------E-------------
------------------------------------------------------------
------------------------------------------------------------
-----------A----------F-------T-----------KVY-T-EHG--------L
CYAFN-------LD-------------GK-------------------------------
-------LNMTRT------GA--------QY-------------GLSLRLHVAQD-----
----------D-----------------Y------------F-R-T--------------
------------------------------------------------------------
------------------------------------------------------------
---------QVT------------------------------------------------
------------------------------------------------------------
------------------------------------------------------------
------------------------------------------------------------
------------------------------------------------------------
------------------------------------------------------------
------------------------------------------------------------
------------------------------------------------------------
------------------------------------------------------------
------------------------------------------------------------

```
------------------------------------------------------------
------------------------------------------------------------
------------------------------------------------------------
------------------------------------------------------------
------------------------------------------------------------
------------------------------------------------------------
------------------------------------------------------------
------------------------------------------------------------
------------------------------------------------------------
------------------------------------------------------------
------------------------------------------------------------
------------------------------------------------------------
------------------------------------------------------------
--------
>Cteno_Ple_Bac_comp47578_c1_seq5_p1__comp47578_c1__comp47578_c1_seq5_p1___ORF
__type_5prime_partial__len_459____score_71_82__comp47578_c1_seq5_141_1517___5
00
------------------------------------------------------------
------------------------------------------------------------
------------------------------------------------------------
-TDVKDLAIL--MNYRNFKEVQLVVP--------ATLSVP-------APCMHQNGRPKD-
-YIFTEI-------EQPLS----ANGFFKRVPVPN---RTTGPGFH--------GRGGPP
PREPAFPIYDN--------L-----------LDRDD---VLSS----------NCPSTVP
GVY-----PGVEAFTP--------------------------------------------
---------------YSKYK-------------QDESAFDMTGRGERTS-----YYPYL-
--EHYQPSYPRPPRGGKPPPPTGAPVPNN-HRTSDQNGKIKYPPNHRDT---GSPTELED
RDGEDCEEFM--PPPKRKKKKKRRANR------------------------------AAL
E----------------------QRREEM-------------------------------
--------E----------------------P---------DE-----M-----------
-ES--FKEFA-STVTADYLAEMF-----------AK-------------ASN---GR---
--RAIWV----SLWLASFVYAWYNIALSVG-VYLS----------K---PTATKL----N
-F-YE-AP--------PG-GVAF-----PTV-----------------------TICNFN
KY------NKS-----FFEREG------------------D-----------------SV
N------S-QL------------R-DFL---------------VATTPI-W---------
---------------------------------DH-------------------------
------------------------------------------------------------
------------------------------------------------------------
------------------------------------------------------------
------------------------------------------------------------
------------------------------------------------------------
------------------------------------------------------------
------------------------------------------------------------
------------------------------------------------------------
------------------------------------------------------------
------------------------------------------------------------
--------KERL------------------------------DQD---------------
----------------------QWTSKYPN-------------------------IMSL-
---------------------------------NLKELY---------------------
------------------------------------------------ANG----SHLIT
--NTV------EFCR----FG-G----K----------P---CK-DTANRETSIP-S---
----------------------------------------------E-------------
------------------------------------------------------------
------------------------------------------------------------
-----------A----------F-------T-----------KVY-T-EHG--------L
CYAFN-------LD-------------GK-------------------------------
-------LNMTRT------GA--------QY-------------GLSLRLHVAQD-----
----------D-----------------Y------------F-R-T--------------
```

```
------------------------------------------------------------
------------------------------------------------------------
---------QVT------------------------------------------------
------------------------------------------------------------
------------------------------------------------------------
------------------------------------------------------------
------------------------------------------------------------
------------------------------------------------------------
------------------------------------------------------------
------------------------------------------------------------
------------------------------------------------------------
------------------------------------------------------------
------------------------------------------------------------
------------------------------------------------------------
------------------------------------------------------------
------------------------------------------------------------
------------------------------------------------------------
------------------------------------------------------------
------------------------------------------------------------
------------------------------------------------------------
------------------------------------------------------------
------------------------------------------------------------
------------------------------------------------------------
------------------------------------------------------------
------------------------------------------------------------
--------
```
