## Supplementary material for "Function and phylogeny support the independent evolution of acid-sensing ion channels in the Placozoa": Table S1

**Table S1. Y2H hits for TadNaC2 and TadNaC10**

TadNaC2

| Sequence ID | Number of | SwissProt | Accession | E-value | Percent identity (%) |
|---|---|---|---|---|---|
| Tricho_evg1230856 | 7 | Filamin-C (Homo sapiens) | Q14315 | 6.30E-101 | 27.1 |
| Tricho_evg958972 | 5 | 40S ribosomal protein S20 (Rattus norvegicus) | P60868 | 7.30E-50 | 76.7 |
| Tricho_evg108179 | 3 | Carboxypeptidase D (Homo sapiens) | O75976 | 3.70E-134 | 52.6 |
| Tricho_evg1238756 | 2 | U5 small nuclear ribonucleoprotein 200 kDa helicase (Homo sapiens) | O75643 | 0 | 67.9 |
| Tricho_evg1767418 | 1 | Elongin-B (Rattus norvegicus) | P62870 | 1.70E-39 | 54.3 |
| Tricho_evg1030571 | 1 | Serine/threonine-protein kinase PRP4 homolog (Mus musculus) | Q61136 | 0 | 46.7 |
| Tricho_evg800323 | 1 | Y-box-binding protein 3 (Homo sapiens) | P16989 | 5.70E-52 | 44 |
| Tricho_evg1089899 | 1 | Lysine--tRNA ligase (Mus musculus) | Q99MN1 | 0 | 66.5 |
| Tricho_evg1144705 | 1 | Lysosomal acid glucosylceramidase (Sus scrofa) | Q70KH2 | 4.80E-160 | 47.6 |
| Tricho_evg1806985 | 1 | Contactin-6 (Rattus norvegicus) | P97528 | 3.00E-58 | 27.2 |
| Tricho_evg1037342 | 1 | Protein VAC14 homolog (Homo sapiens) | Q08AM6 | 0 | 43.3 |

TadNaC10

| Sequence ID | Number of | SwissProt | Accession | E-value | Percent Identity (%) |
|---|---|---|---|---|---|
| Tricho_evg1230856 | 1 | Filamin-C (Homo sapiens) | Q14315 | 6.30E-101 | 27.1 |
| Tricho_evg1238756 | 1 | U5 small nuclear ribonucleoprotein 200 kDa helicase (Homo sapiens) | O75643 | 0 | 67.9 |
