## Supplementary material for "Function and phylogeny support the independent evolution of acid-sensing ion channels in the Placozoa": Table S2

| Clade | Phylum | Species | Source | Accession/link | | Completeness | Singles | Doubles | Fragmented | Missing | BUSCOS |
|---|---|---|---|---|---|---|---|---|---|---|---|
| Ambulacraria | Echinodermata | *Acanthaster planci* | | https://marinegenomics.oist.jp/cots/viewer/download?project_id=46 | | 91.4 | 91.4 | 0 | 5.9 | 2.7 | n:255 |
| Ambulacraria | Echinodermata | *Strongylocentrotus purpuratus* | UniProt | UP000007110 | | 86.3 | 80 | 6.3 | 11 | 2.7 | n:255 |
| Ambulacraria | Hemichordata | *Saccoglossus kowalevskii* | | https://groups.oist.jp/molgenu/hemichordate-genomes | | 94.1 | 91.4 | 2.7 | 3.5 | 2.4 | n:255 |
| Ambulacraria | Hemichordata | *Ptychodera flava* | | https://groups.oist.jp/molgenu/hemichordate-genomes | | 53 | 51.4 | 1.6 | 25.1 | 21.9 | n:255 |
| Chordata | Cephalochordata | *Branchiostoma belcheri* | | http://genome.bucm.edu.cn/lancelet/download_data.php | | 96.9 | 85.1 | 11.8 | 2.4 | 0.7 | n:255 |
| Chordata | Urochordata | *Ciona intestinalis* | UniProt | UP000008144 | | 85.5 | 85.5 | 0 | 7.1 | 7.4 | n:255 |
| Chordata | Craniata | *Gallus gallus* | UniProt | UP000000539 | | 96.1 | 86.3 | 9.8 | 2 | 1.9 | n:255 |
| Chordata | Craniata | *Homo sapiens* | UniProt | UP000005640 | | 100 | 97.6 | 2.4 | 0 | 0 | n:255 |
| Chordata | Craniata | *Rattus norvegicus* | UniProt | UP000002494 | | 100 | 98 | 2 | 0 | 0 | n:255 |
| Chordata | Craniata | *Petromyzon marinus* | UniProt | UP000245300 | | 54.5 | 52.9 | 1.6 | 9.8 | 35.7 | n:255 |
| Cnidaria | Cubozoa | *Alatina Alata* | https://bitbucket.org/caseywdunn/cnidaria2014-assemblies/src/master/Cnidaria_Transcriptomes/ | | | 96.5 | 76.5 | 20 | 2.4 | 1.1 | n:255 |
| Cnidaria | Hydrozoa | *Hydra vulgaris* | https://research.nhgri.nih.gov/hydra/ | | | 86.6 | 82.7 | 3.9 | 10.2 | 3.2 | n:255 |
| Cnidaria | Hexacoralia | *Nematostella vectensis* | http://metazoa.ensembl.org/species.html | | | 99.2 | 92.5 | 6.7 | 0.8 | 0 | n:255 |
| Cnidaria | Polypodium | *Polypodium hydriforme* | https://www.ncbi.nlm.nih.gov/nuccore/671211599 | | | 78.4 | 13.7 | 64.7 | 14.5 | 7.1 | n:255 |
| Cnidaria | Scyphozoa | *Rhopilema_esculentum* | https://academic.oup.com/gigascience/article/9/4/giaa036/5823175 | | | 98.1 | 62.4 | 35.7 | 1.2 | 0.7 | n:255 |
| Ctenophora | Lobata | *Mnemiopsis leidyi* | https://research.nhgri.nih.gov/mnemiopsis/download/download.cgi?dl=transcript | | Trinity assembly | 89.8 | 83.5 | 6.3 | 5.1 | 5.1 | n:255 |
| Ctenophora | Cydippida | *Pleurobrachia_bachei* | https://neurobase.rc.ufl.edu/pleurobrachia/download | | | 79.6 | 30.2 | 49.4 | 13.7 | 6.7 | n:255 |
| Ctenophora | Cydippida | *Pukia_falcata* | https://neurobase.rc.ufl.edu/ | | | 71 | 35.3 | 35.7 | 20.8 | 8.2 | n:255 |
| Ctenophora | Cydippida | *Hormiphora californiensis* | JGI portal (Project accession: PRJNA281977, runs accession: SRR1992642) | | Trinity assembly | 94.9 | 39.6 | 55.3 | 1.6 | 3.5 | 255 |
| Protostomia | Ecdysozoa | *Centruroides sculpturatus* | PRJNA281977, runs accession: SRR1992642 | https://www.ncbi.nlm.nih.gov/genome/?term=centruroides+sculpturatus | | 89.1 | 67.5 | 21.6 | 9 | 1.9 | n:255 |
| Protostomia | Ecdysozoa | *Drosophila melanogaster* | UniProt | UP000000803 | | 100 | 99.6 | 0.4 | 0 | 0 | n:255 |
| Protostomia | Ecdysozoa | *Priapulus caudatus* | NCBI | PRJNA20497 | | 89 | 73.3 | 15.7 | 7.8 | 3.2 | n:255 |
| Protostomia | Ecdysozoa | *Ramazzottius variernatus* | UniProt | UP000186922 | | 92.5 | 88.2 | 4.3 | 3.1 | 4.4 | 225 |
| Protostomia | Ecdysozoa | *Hypsibius dujardini* | UniProt | UP000192578 | | 76.9 | 74.9 | 2 | 4.7 | 18.4 | 225 |
| Protostomia | Ecdysozoa | *Tribolium castaneum* | UniProt | UP000007266 | | 97.3 | 96.9 | 0.4 | 2 | 0.7 | n:255 |
| Protostomia | Ecdysozoa | *Trichinella spiralis* | UniProt | UP000054776□ | | 87.1 | 85.5 | 1.6 | 2 | 10.9 | 255 |
| Protostomia | Ecdysozoa | *Caenorhabditis elegans* | UniProt | UP000001940 | | 97.7 | 96.9 | 0.8 | 0.8 | 1.5 | 255 |
| Protostomia | Spiralia | *Phoronis australis* | NCBI | https://www.ncbi.nlm.nih.gov/assembly/GCA_002633005.1/ | | 96 | 73.3 | 22.7 | 2.4 | 1.6 | 255 |
| Protostomia | Spiralia | *Aplysia californica* | UniProt | UP000694888 | | 98.4 | 75.3 | 23.1 | 0.8 | 0.8 | 255 |
| Protostomia | Spiralia | *Lingula_anatina* | https://marinegenomics.oist.jp/lan_v2/viewer/info?project_id=50 | | | 94.9 | 72.2 | 22.7 | 3.9 | 1.2 | n:255 |
| Protostomia | Spiralia | *Octopus bimaculoides* | UniProt | UP000053454 | | 87.9 | 87.5 | 0.4 | 9.4 | 2.7 | n:255 |
| Protostomia | Spiralia | *Owenia fusiformis* | NCBI | https://www.ncbi.nlm.nih.gov/assembly/GCA_903813345.2 | | 93.7 | 84.3 | 9.4 | 2 | 4.3 | n:255 |
| Protostomia | Spiralia | *Platynereis dumerilii* | PdumBase | http://pdumbase.gdcb.iastate.edu/ | | 90.6 | 87.5 | 3.1 | 5.5 | 3.9 | n:255 |
| Protostomia | platyhelminthes | *Schistosoma mansoni* | NCBI | UP000008854 | | 95.3 | 78.4 | 16.9 | 2 | 2.7 | 255 |
| Protostomia | platyhelminthes | *Macrostomum lignano* | NCBI | UP000215902 | | 91.8 | 12.2 | 79.6 | 3.1 | 5.1 | 255 |
| Protostomia | Spiralia | *Rotaria socialis* | UniProt | | | 95.7 | 79.2 | 16.5 | 2 | 2.3 | 255 |
| Placozoa | Trichoplacidae | *Hoilungia hongkongensis* | | https://bitbucket.org/molpalmuc/hoilungia-genome/src/master/sequences/Hhon_TRINITY_transdecoder_proteins.fasta.gz | | 92.9 | 89 | 3.9 | 2.4 | 4.7 | n:255 |
| Placozoa | Trichoplacidae | *Trichoplax adhaerens* | https://www.researchsquare.com/article/rs-5181/v1 | | | 92.9 | 92.5 | 0.4 | 4.7 | 2.4 | n:255 |
| Porifera | Demospongiae | *Amphimedon queenslandica* | UniProt | UP000007879 | | 92.5 | 88.2 | 4.3 | 7.1 | 0.4 | n:255 |
| Porifera | Demospongiae | *Ephydatia muelleri* | | https://spaces.facsci.ualberta.ca/ephybase/ | | 97.3 | 81.2 | 16.1 | 1.6 | 1.1 | n:255 |
| Porifera | Homoscleromorpł | *Oscarella carmela* | COMPAGEN | | | 95.3 | 87.1 | 8.2 | 2.7 | 2 | n:255 |
| Porifera | Calcarea | *Sycon ciliatum* | COMPAGEN | | | 97.3 | 91 | 6.3 | 2 | 0.7 | n:255 |
| Choanoflagellatea | Salpingoecidae | *Monosiga_brevicolis* | UniProt | UP000001357 | | 78.8 | 78.8 | 0 | 7.5 | 13.7 | n:255 |
| Choanoflagellatea | Salpingoecidae | *Salpingoeca rosetta* | UniProt | UP000007799 | | 83.1 | 83.1 | 0 | 6.7 | 10.2 | n:255 |
| Filasterea | | *Capsaspora owczarzaki* | UniProt | UP000008743 | | 93.7 | 92.9 | 0.8 | 2.4 | 3.9 | n:255 |
| Filasterea-related "pl | position debated | *Tunicaraptor unikontum* | NCBI | PRJNA638967 | | 74.1 | 64.7 | 9.4 | 16.5 | 9.4 | n:255 |
| Alveolata | Symbiodinium | Symbyodinium sp_KB8 | NCBI | https://www.ncbi.nlm.nih.gov/assembly/GCA_905221625.1/#/def | | 63.4 | 44.3 | 20 | 12.2 | 23.5 | n:255 |
| Stramenopiles | Cafeteriaceae | Cafeteria roenbergensis | NCBI | https://www.ncbi.nlm.nih.gov/bioproject/PRJNA552725/ | | 67.1 | 65.9 | 1.2 | 9 | 23.9 | 255 |
