## Supplementary material for "Function and phylogeny support the independent evolution of acid-sensing ion channels in the Placozoa": Table S3

**Table S3: Primers used for cloning TadNaC2 constructs**

| Primer Name | DNA Sequence (5' to 3') |
|---|---|
| TadNaC2_F1 | CTTTCGACACTGCTGTACAGCTC |
| TadNaC2_F2 | ATTATACTCGAGGCCGCCACCATGGATACTCATCCTTGGCAAAGGCC |
| TadNaC2_R1 | GATGATAATCTAAACATCGAACTACGTAGC |
| TadNaC2_R2 | ATTATAGGATCCAACCTAGTCAATACTGGATGGACTAATTCG |
| TadNaC10_F1 | ACGACACTGAAACAGCTTTCGCG |
| TadNaC10_F2 | ATTATACTCGAGGCCGCCACCATGCCAACCAAGCAGTTAAAGGTTATGAG |
| TadNaC10_R1 | GTTGATAATGAAAGTATCGGTCTTGCGACG |
| TadNaC10_R2 | ATTATAGGATCCACTTTATGCAAGTCCAGAAGCTTGCAACTG |
| TadNaC2_F70A_F | TTGCAGTATGTTTTATCCGCCCCAACGAATATTGATATC |
| TadNaC2_F70A_R | GATATCAATATTCGTTGGGGCGGATAAAACATACTGCAA |
| TadNaC2_F70H_F | TTGCAGTATGTTTTATCCCACCCAACGAATATTGATATC |
| TadNaC2_F70H_R | GATATCAATATTCGTTGGGTGGGATAAAACATACTGCAA |
| TadNaC2_D75A_F | CCTTTCCAACGAATATTGCCATCGAAATTATACACCAAG |
| TadNaC2_D75A_R | CTTGGTGTATAATTTCGATGGCAATATTCGTTGGAAAGG |
| TadNaC2_E77A_F | CCAACGAATATTGATATCGCCATTATACACCAAGATAGC |
| TadNaC2_E77A_R | GCTATCTTGGTGTATAATGGCGATATCAATATTCGTTGG |
| TadNaC2_H80A_F | ATTGATATCGAAATTATAGCCCAAGATAGCTTAATATTC |
| TadNaC2_H80A_R | GAATATTAAGCTATCTTGGGCTATAATTTCGATATCAAT |
| TadNaC2_E104A_F | CCAAAACAGCATTATCGGCCGAAGATAATCGACACTTGC |
| TadNaC2_E104A_R | GCAAGTGTCGATTATCTTCGGCCGATAATGCTGTTTTGG |
| TadNaC2_E105A_F | CCAAAACAGCATTATCGGAAGCCGATAATCGACACTTGC |
| TadNaC2_E105A_R | GCAAGTGTCGATTATCGGCTTCCGATAATGCTGTTTTGG |
| TadNaC2_E104-5A_F | CCAAAACAGCATTATCGGCCGCCGATAATCGACACTTGC |
| TadNaC2_E104-5A_R | GCAAGTGTCGATTATCGGCGGCCGATAATGCTGTTTTGG |
| TadNaC2_H109A_F | TCGGAAGAAGATAATCGAGCCTTGCAGACATTTCTTAGA |
| TadNaC2_H109A_R | TCTAAGAAATGTCTGCAAGGCTCGATTATCTTCTTCCGA |
| TadNaC2E104-5-9AF | CCAAAACAGCATTATCGGCCGCCGATAATCGAGCCTTGCAGACATTTCTTAGA |
| TadNaC2E104-5-9AR | TCTAAGAAATGTCTGCAAGGCTCGATTATCGGCGGCCGATAATGCTGTTTTGG |
| TadNaC2_R201A_F | GAAGGTATGTTATCTCAGGCCGGGAAAGGCTCTGCACAC |
| TadNaC2_R201A_R | GTGTGCAGAGCCTTTCCCGGCCTGAGATAACATACCTTC |
| TadNaC2_K203E_F | ATGTTATCTCAGCGTGGGGAGGGCTCTGCACACGGACTACG |
| TadNaC2_K203E_R | CGTAGTCCGTGTGCAGAGCCCTCCCCACGCTGAGATAACAT |
| TadNaC2_K203Del_F | ATGTTATCTCAGCGTGGGGGCTCTGCACACGGACTACG |
| TadNaC2_K203Del_R | CGTAGTCCGTGTGCAGAGCCCCCACGCTGAGATAACAT |
| TadNaC2_pGBK_F | AATTTAGAATTCCGAATGCAAAGTATACGAG |
| TadNaC2_pGBK_R | AATTTAGTCGACCTAGTCAATACTGGATG |
| TadNaC10_pGBK_F | AATTTAGAATTCGAGCAAACACCACGCTACTC |
| TadNaC10_pGBK_R | AATTTAGTCGACTTATGCAAGTCCAGAAGC |
